## Additional file 2 for "Discovery of 20 novel ribosomal leaders in bacteria and archaea"

### Supplementary File 1: alignments and information on downstream genes and taxonomy
### Discovery of 20 novel ribosomal leaders in bacteria and archaea

Iris Eckert & Zasha Weinberg

Note: the presentation and explanatory text of this supplementary data on novel RNA motifs follows the pattern of a presentation of previously found conserved RNA motifs (Weinberg, *et al.*, 2010).

12.2 Gene contexts . . . 1024
12.3 Conserved domains . . . 1214
12.4 Multiple-sequence alignment . . . 1216

**13 L31-Corynebacteriaceae** **1252**
13.1 Taxa . . . 1252
13.2 Gene contexts . . . 1255
13.3 Conserved domains . . . 1262
13.4 Multiple-sequence alignment . . . 1263

**14 S4-Bacteroidia** **1267**
14.1 Taxa . . . 1267
14.2 Gene contexts . . . 1268
14.3 Conserved domains . . . 1283
14.4 Multiple-sequence alignment . . . 1283

**15 S4-Flavobacteria** **1297**
15.1 Taxa . . . 1297
15.2 Gene contexts . . . 1299
15.3 Conserved domains . . . 1324
15.4 Multiple-sequence alignment . . . 1325

**16 S4-Clostridia** **1337**
16.1 Taxa . . . 1337
16.2 Gene contexts . . . 1338
16.3 Conserved domains . . . 1351
16.4 Multiple-sequence alignment . . . 1352

**17 S4-Fusobacteriales** **1383**
17.1 Taxa . . . 1383
17.2 Gene contexts . . . 1385
17.3 Conserved domains . . . 1402
17.4 Multiple-sequence alignment . . . 1403

**18 S6-S18-Chlorobi** **1408**
18.1 Taxa . . . 1408
18.2 Gene contexts . . . 1408
18.3 Conserved domains . . . 1410
18.4 Multiple-sequence alignment . . . 1411

**19 S10-Clostridia 1412**
19.1 Taxa . . . 1412
19.2 Gene contexts . . . 1414
19.3 Conserved domains . . . 1461
19.4 Multiple-sequence alignment . . . 1462

**20 S15-Halobacteria 1499**
20.1 Taxa . . . 1499
20.2 Gene contexts . . . 1501
20.3 Conserved domains . . . 1508
20.4 Multiple-sequence alignment . . . 1509

**21 S15-Methanobacteria 1516**
21.1 Taxa . . . 1516
21.2 Gene contexts . . . 1517
21.3 Conserved domains . . . 1522
21.4 Multiple-sequence alignment . . . 1523

**22 S15-Flavobacteria 1531**
22.1 Taxa . . . 1531
22.2 Notes . . . 1535
22.3 Gene contexts . . . 1536
22.4 Conserved domains . . . 1586
22.5 Multiple-sequence alignment . . . 1586

**23 S16-Flavobacteria 1681**
23.1 Taxa . . . 1681
23.2 Gene contexts . . . 1682
23.3 Conserved domains . . . 1686
23.4 Multiple-sequence alignment . . . 1687

### 1 L2-Alphaproteobacteria

#### 1.1 Taxa

The taxonomy of each organism containing a putative L2-Alphaproteobacteria RNA is listed, with abbreviations identifying each hit (e.g., “Eco-1-1” and “Eco-1-2” might hypothetically represent two distinct RNAs in *E. coli*). The abbreviations will be used to identify each individual L2-Alphaproteobacteria RNA in Sections 1.2 and 1.4:.

| abbrev. of hits | taxonomy of species |
|---|---|
| Aat-1-1 | Bacteria Proteobacteria Alphaproteobacteria Rhodobacterales Rhodobacteraceae *Actibacterium atlanticum* |
| Amu-1-1 | Bacteria Proteobacteria Alphaproteobacteria Rhodobacterales Rhodobacteraceae *Actibacterium mucosum* KCTC 23349 |
| Aat-2-1 | Bacteria Proteobacteria Alphaproteobacteria Rhodobacterales Rhodobacteraceae *Aestuariivita atlantica* |
| Abo-1-1 | Bacteria Proteobacteria Alphaproteobacteria Rhodobacterales Rhodobacteraceae *Aestuariivita boseongensis* |

| | |
|---|---|
| Cba-1-1 | Bacteria Proteobacteria Alphaproteobacteria Rhodobacterales Rhodobacteraceae *Celeribacter baekdonensis* B30 |
| Csp-1-1 | Bacteria Proteobacteria Alphaproteobacteria Rhodobacterales Rhodobacteraceae *Citreicella sp.* 357 |
| Csp-2-1 | Bacteria Proteobacteria Alphaproteobacteria Rhodobacterales Rhodobacteraceae *Citreicella sp.* SE45 |
| Dsp-1-1 | Bacteria Proteobacteria Alphaproteobacteria Rhodobacterales Rhodobacteraceae *Defluviimonas sp.* 20V17 |
| Dsh-1-1 | Bacteria Proteobacteria Alphaproteobacteria Rhodobacterales Rhodobacteraceae *Dinoroseobacter shibae* DFL 12 |
| Gne-1-1 | Bacteria Proteobacteria Alphaproteobacteria Rhodobacterales Rhodobacteraceae *Gemmobacter nectariphilus* DSM 15620 |
| Hma-1-1 | Bacteria Proteobacteria Alphaproteobacteria Rhodobacterales Rhodobacteraceae *Haematobacter massiliensis* |
| Hmi-1-1 | Bacteria Proteobacteria Alphaproteobacteria Rhodobacterales Rhodobacteraceae *Haematobacter missouriensis* |
| Hna-1-1 | Bacteria Proteobacteria Alphaproteobacteria Rhodobacterales Rhodobacteraceae *Halocynthiibacter namhaensis* |
| Hsp-1-1 | Bacteria Proteobacteria Alphaproteobacteria Rhodobacterales Rhodobacteraceae *Halocynthiibacter sp.* PAMC 20958 |
| Jaq-1-1 | Bacteria Proteobacteria Alphaproteobacteria Rhodobacterales Rhodobacteraceae *Jannaschia aquimarina* |
| Jsp-1-1 | Bacteria Proteobacteria Alphaproteobacteria Rhodobacterales Rhodobacteraceae *Jannaschia sp.* CCS1 |
| Kvu-1-1 | Bacteria Proteobacteria Alphaproteobacteria Rhodobacterales Rhodobacteraceae *Ketogulonicigenium vulgare* WSH-001 |
| Kvu-2-1 | Bacteria Proteobacteria Alphaproteobacteria Rhodobacterales Rhodobacteraceae *Ketogulonicigenium vulgare* Y25 |
| Laq-1-1 | Bacteria Proteobacteria Alphaproteobacteria Rhodobacterales Rhodobacteraceae *Leisingera aquimarina* DSM 24565 |
| Lca-1-1 | Bacteria Proteobacteria Alphaproteobacteria Rhodobacterales Rhodobacteraceae *Leisingera caerulea* DSM 24564 |
| Lda-1-1 | Bacteria Proteobacteria Alphaproteobacteria Rhodobacterales Rhodobacteraceae *Leisingera daeponensis* DSM 23529 |
| Lme-1-1 | Bacteria Proteobacteria Alphaproteobacteria Rhodobacterales Rhodobacteraceae *Leisingera methylohalidivorans* DSM 14336 |
| Lsp-1-1 | Bacteria Proteobacteria Alphaproteobacteria Rhodobacterales Rhodobacteraceae *Leisingera sp.* ANG-DT |
| Lsp-2-1 | Bacteria Proteobacteria Alphaproteobacteria Rhodobacterales Rhodobacteraceae *Leisingera sp.* ANG-M1 |
| Lsp-3-1 | Bacteria Proteobacteria Alphaproteobacteria Rhodobacterales Rhodobacteraceae *Leisingera sp.* ANG-M6 |
| Lsp-4-1 | Bacteria Proteobacteria Alphaproteobacteria Rhodobacterales Rhodobacteraceae *Leisingera sp.* ANG-M7 |
| Lsp-5-1 | Bacteria Proteobacteria Alphaproteobacteria Rhodobacterales Rhodobacteraceae *Leisingera sp.* ANG-S |
| Lsp-6-1 | Bacteria Proteobacteria Alphaproteobacteria Rhodobacterales Rhodobacteraceae *Leisingera sp.* ANG-S3 |
| Lsp-7-1 | Bacteria Proteobacteria Alphaproteobacteria Rhodobacterales Rhodobacteraceae *Leisingera sp.* ANG-S5 |
| Lsp-8-1 | Bacteria Proteobacteria Alphaproteobacteria Rhodobacterales Rhodobacteraceae *Leisingera sp.* ANG-Vp |
| Lsp-9-1 | Bacteria Proteobacteria Alphaproteobacteria Rhodobacterales Rhodobacteraceae *Leisingera sp.* ANG1 |
| Lci-1-1 | Bacteria Proteobacteria Alphaproteobacteria Rhodobacterales Rhodobacteraceae *Loktanella cinnabarina* LL-001 |
| Lho-1-1 | Bacteria Proteobacteria Alphaproteobacteria Rhodobacterales Rhodobacteraceae *Loktanella hongkongensis* DSM 17492 |
| Lsp-10-1 | Bacteria Proteobacteria Alphaproteobacteria Rhodobacterales Rhodobacteraceae *Loktanella sp.* S4079 |
| Lve-1-1 | Bacteria Proteobacteria Alphaproteobacteria Rhodobacterales Rhodobacteraceae *Loktanella vestfoldensis* DSM 16212 |
| Lve-2-1 | Bacteria Proteobacteria Alphaproteobacteria Rhodobacterales Rhodobacteraceae *Loktanella vestfoldensis* SKA53 |
| Mal-1-1 | Bacteria Proteobacteria Alphaproteobacteria Rhodobacterales Rhodobacteraceae *Mameliella alba* |
| Msp-1-1 | Bacteria Proteobacteria Alphaproteobacteria Rhodobacterales Rhodobacteraceae *Maribius sp.* MOLA 401 |
| Mal-2-1 | Bacteria Proteobacteria Alphaproteobacteria Rhodobacterales Rhodobacteraceae *Marinovum algicola* DG 898 |
| Nit-1-1 to Nit-1-2 | Bacteria Proteobacteria Alphaproteobacteria Rhodobacterales Rhodobacteraceae *Nautella italica* |
| Nsp-1-1 | Bacteria Proteobacteria Alphaproteobacteria Rhodobacterales Rhodobacteraceae *Nautella sp.* ECSMB14104 |
| Nig-1-1 | Bacteria Proteobacteria Alphaproteobacteria Rhodobacterales Rhodobacteraceae *Nereida ignava* |
| Oin-1-1 | Bacteria Proteobacteria Alphaproteobacteria Rhodobacterales Rhodobacteraceae *Oceanibulbus indolifex* HEL-45 |
| Oba-1-1 | Bacteria Proteobacteria Alphaproteobacteria Rhodobacterales Rhodobacteraceae *Oceanicola batsensis* HTCC2597 |
| Ogr-1-1 | Bacteria Proteobacteria Alphaproteobacteria Rhodobacterales Rhodobacteraceae *Oceanicola granulosus* HTCC2516 |
| Osp-1-1 | Bacteria Proteobacteria Alphaproteobacteria Rhodobacterales Rhodobacteraceae *Oceanicola sp.* HL-35 |
| Osp-2-1 | Bacteria Proteobacteria Alphaproteobacteria Rhodobacterales Rhodobacteraceae *Oceanicola sp.* MCTG156(1a) |
| Osp-3-1 | Bacteria Proteobacteria Alphaproteobacteria Rhodobacterales Rhodobacteraceae *Oceanicola sp.* S124 |
| Ogu-1-1 | Bacteria Proteobacteria Alphaproteobacteria Rhodobacterales Rhodobacteraceae *Oceaniovalibus guishaninsula* JLT2003 |
| Oan-1-1 | Bacteria Proteobacteria Alphaproteobacteria Rhodobacterales Rhodobacteraceae *Octadecabacter antarcticus* 307 |
| Oar-1-1 | Bacteria Proteobacteria Alphaproteobacteria Rhodobacterales Rhodobacteraceae *Octadecabacter arcticus* 238 |
| Ote-1-1 | Bacteria Proteobacteria Alphaproteobacteria Rhodobacterales Rhodobacteraceae *Octadecabacter temperatus* |
| Pde-1-1 | Bacteria Proteobacteria Alphaproteobacteria Rhodobacterales Rhodobacteraceae *Paracoccus denitrificans* PD1222 |
| Pga-1-1 | Bacteria Proteobacteria Alphaproteobacteria Rhodobacterales Rhodobacteraceae *Phaeobacter gallaeciensis* 2.10 |
| Pga-2-1 | Bacteria Proteobacteria Alphaproteobacteria Rhodobacterales Rhodobacteraceae *Phaeobacter gallaeciensis* DSM 17395 |
| Pga-3-1 | Bacteria Proteobacteria Alphaproteobacteria Rhodobacterales Rhodobacteraceae *Phaeobacter gallaeciensis* DSM 26640 |
| Pin-1-1 | Bacteria Proteobacteria Alphaproteobacteria Rhodobacterales Rhodobacteraceae *Phaeobacter inhibens* DSM 16374 |
| Psp-1-1 | Bacteria Proteobacteria Alphaproteobacteria Rhodobacterales Rhodobacteraceae *Phaeobacter sp.* S26 |
| Psp-2-1 | Bacteria Proteobacteria Alphaproteobacteria Rhodobacterales Rhodobacteraceae *Phaeobacter sp.* S60 |
| Pte-1-1 | Bacteria Proteobacteria Alphaproteobacteria Rhodobacterales Rhodobacteraceae *Planktomarina temperata* RCA23 |
| Pxi-1-1 | Bacteria Proteobacteria Alphaproteobacteria Rhodobacterales Rhodobacteraceae *Pseudodonghicola xiamenensis* DSM 18339 |
| Pat-1-1 | Bacteria Proteobacteria Alphaproteobacteria Rhodobacterales Rhodobacteraceae *Pseudooceanicola atlanticus* |
| Pna-1-1 | Bacteria Proteobacteria Alphaproteobacteria Rhodobacterales Rhodobacteraceae *Pseudooceanicola nanhaiensis* DSM 18065 |
| Par-1-1 | Bacteria Proteobacteria Alphaproteobacteria Rhodobacterales Rhodobacteraceae *Pseudophaeobacter arcticus* DSM 23566 |
| Pan-1-1 | Bacteria Proteobacteria Alphaproteobacteria Rhodobacterales Rhodobacteraceae *Pseudorhodobacter antarcticus* |
| Paq-1-1 | Bacteria Proteobacteria Alphaproteobacteria Rhodobacterales Rhodobacteraceae *Pseudorhodobacter aquimaris* |
| Pfe-1-1 | Bacteria Proteobacteria Alphaproteobacteria Rhodobacterales Rhodobacteraceae *Pseudorhodobacter ferrugineus* |
| Pfe-2-1 | Bacteria Proteobacteria Alphaproteobacteria Rhodobacterales Rhodobacteraceae *Pseudorhodobacter ferrugineus* DSM 5888 |
| Pwa-1-1 | Bacteria Proteobacteria Alphaproteobacteria Rhodobacterales Rhodobacteraceae *Pseudorhodobacter wandonensis* |

| | |
|---|---|
| Psp-3-1 | Bacteria Proteobacteria Alphaproteobacteria Rhodobacterales Rhodobacteraceae *Puniceibacterium sp.* IMCC21224 |
| CRh-1-1 | Bacteria Proteobacteria Alphaproteobacteria Rhodobacterales Rhodobacteraceae Rhodobacter *Candidatus* Rhodobacter lobularis |
| Rca-1-1 | Bacteria Proteobacteria Alphaproteobacteria Rhodobacterales Rhodobacteraceae *Rhodobacter capsulatus* B6 |
| Rca-2-1 | Bacteria Proteobacteria Alphaproteobacteria Rhodobacterales Rhodobacteraceae *Rhodobacter capsulatus* DE442 |
| Rca-3-1 | Bacteria Proteobacteria Alphaproteobacteria Rhodobacterales Rhodobacteraceae *Rhodobacter capsulatus* R121 |
| Rca-4-1 | Bacteria Proteobacteria Alphaproteobacteria Rhodobacterales Rhodobacteraceae *Rhodobacter capsulatus* SB 1003 |
| Rca-5-1 | Bacteria Proteobacteria Alphaproteobacteria Rhodobacterales Rhodobacteraceae *Rhodobacter capsulatus* Y262 |
| Rca-6-1 | Bacteria Proteobacteria Alphaproteobacteria Rhodobacterales Rhodobacteraceae *Rhodobacter capsulatus* YW1 |
| Rca-7-1 | Bacteria Proteobacteria Alphaproteobacteria Rhodobacterales Rhodobacteraceae *Rhodobacter capsulatus* YW2 |
| Rsp-1-1 | Bacteria Proteobacteria Alphaproteobacteria Rhodobacterales Rhodobacteraceae *Rhodobacter sp.* AKP1 |
| Rsp-2-1 | Bacteria Proteobacteria Alphaproteobacteria Rhodobacterales Rhodobacteraceae *Rhodobacter sp.* SW2 |
| Rsp-3-1 | Bacteria Proteobacteria Alphaproteobacteria Rhodobacterales Rhodobacteraceae *Rhodobacter sphaeroides* 2.4.1 |
| Rsp-4-1 | Bacteria Proteobacteria Alphaproteobacteria Rhodobacterales Rhodobacteraceae *Rhodobacter sphaeroides* ATCC 17025 |
| Rsp-5-1 | Bacteria Proteobacteria Alphaproteobacteria Rhodobacterales Rhodobacteraceae *Rhodobacter sphaeroides* ATCC 17029 |
| Rsp-6-1 | Bacteria Proteobacteria Alphaproteobacteria Rhodobacterales Rhodobacteraceae *Rhodobacter sphaeroides* KD131 |
| Rsp-7-1 | Bacteria Proteobacteria Alphaproteobacteria Rhodobacterales Rhodobacteraceae *Rhodobacter sphaeroides* WS8N |
| Rba-1-1 | Bacteria Proteobacteria Alphaproteobacteria Rhodobacterales *Rhodobacteraceae bacterium* HIMB11 |
| Rba-2-1 | Bacteria Proteobacteria Alphaproteobacteria Rhodobacterales *Rhodobacteraceae bacterium* KLH11 |
| Rba-3-1 | Bacteria Proteobacteria Alphaproteobacteria Rhodobacterales *Rhodobacteraceae bacterium* PD-2 |
| Rel-1-1 | Bacteria Proteobacteria Alphaproteobacteria Rhodobacterales Rhodobacteraceae *Roseibacterium elongatum* DSM 19469 |
| Rat-1-1 | Bacteria Proteobacteria Alphaproteobacteria Rhodobacterales Rhodobacteraceae *Roseivivax atlanticus* |
| Rha-1-1 | Bacteria Proteobacteria Alphaproteobacteria Rhodobacterales Rhodobacteraceae *Roseivivax halodurans* JCM 10272 |
| Ris-1-1 | Bacteria Proteobacteria Alphaproteobacteria Rhodobacterales Rhodobacteraceae *Roseivivax isoporae* LMG 25204 |
| Rde-1-1 | Bacteria Proteobacteria Alphaproteobacteria Rhodobacterales Rhodobacteraceae *Roseobacter denitrificans* OCh 114 |
| Rli-1-1 | Bacteria Proteobacteria Alphaproteobacteria Rhodobacterales Rhodobacteraceae *Roseobacter litoralis* Och 149 |
| Rsp-8-1 | Bacteria Proteobacteria Alphaproteobacteria Rhodobacterales Rhodobacteraceae *Roseobacter sp.* AzwK-3b |
| Rsp-9-1 | Bacteria Proteobacteria Alphaproteobacteria Rhodobacterales Rhodobacteraceae *Roseobacter sp.* CCS2 |
| Rsp-10-1 | Bacteria Proteobacteria Alphaproteobacteria Rhodobacterales Rhodobacteraceae *Roseobacter sp.* GAI101 |
| Rsp-11-1 | Bacteria Proteobacteria Alphaproteobacteria Rhodobacterales Rhodobacteraceae *Roseobacter sp.* MED193 |
| Rsp-12-1 | Bacteria Proteobacteria Alphaproteobacteria Rhodobacterales Rhodobacteraceae *Roseobacter sp.* SK209-2-6 |
| Rmu-1-1 | Bacteria Proteobacteria Alphaproteobacteria Rhodobacterales Rhodobacteraceae *Roseovarius mucosus* DSM 17069 |
| Rnu-1-1 | Bacteria Proteobacteria Alphaproteobacteria Rhodobacterales Rhodobacteraceae *Roseovarius nubinhibens* ISM |
| Rsp-13-1 | Bacteria Proteobacteria Alphaproteobacteria Rhodobacterales Rhodobacteraceae *Roseovarius sp.* 217 |
| Rsp-14-1 | Bacteria Proteobacteria Alphaproteobacteria Rhodobacterales Rhodobacteraceae *Roseovarius sp.* HTCC2601 |
| Rsp-15-1 | Bacteria Proteobacteria Alphaproteobacteria Rhodobacterales Rhodobacteraceae *Roseovarius sp.* MCTG156(2b) |
| Rsp-16-1 | Bacteria Proteobacteria Alphaproteobacteria Rhodobacterales Rhodobacteraceae *Roseovarius sp.* TM1035 |
| Rto-1-1 | Bacteria Proteobacteria Alphaproteobacteria Rhodobacterales Rhodobacteraceae *Roseovarius tolerans* |
| Rme-1-1 | Bacteria Proteobacteria Alphaproteobacteria Rhodobacterales Rhodobacteraceae *Rubellimicrobium mesophilum* DSM 19309 |
| Rha-2-1 | Bacteria Proteobacteria Alphaproteobacteria Rhodobacterales Rhodobacteraceae *Ruegeria halocynthiae* |
| Rmo-1-1 | Bacteria Proteobacteria Alphaproteobacteria Rhodobacterales Rhodobacteraceae *Ruegeria mobilis* |
| Rmo-2-1 | Bacteria Proteobacteria Alphaproteobacteria Rhodobacterales Rhodobacteraceae *Ruegeria mobilis* F1926 |
| Rpo-1-1 | Bacteria Proteobacteria Alphaproteobacteria Rhodobacterales Rhodobacteraceae *Ruegeria pomeroyi* DSS-3 |
| Rsp-17-1 | Bacteria Proteobacteria Alphaproteobacteria Rhodobacterales Rhodobacteraceae *Ruegeria sp.* 6PALISEP08 |
| Rsp-18-1 | Bacteria Proteobacteria Alphaproteobacteria Rhodobacterales Rhodobacteraceae *Ruegeria sp.* ANG-R |
| Rsp-19-1 | Bacteria Proteobacteria Alphaproteobacteria Rhodobacterales Rhodobacteraceae *Ruegeria sp.* ANG-S4 |
| Rsp-20-1 | Bacteria Proteobacteria Alphaproteobacteria Rhodobacterales Rhodobacteraceae *Ruegeria sp.* R11 |
| Rsp-21-1 | Bacteria Proteobacteria Alphaproteobacteria Rhodobacterales Rhodobacteraceae *Ruegeria sp.* TM1040 |
| Rsp-22-1 | Bacteria Proteobacteria Alphaproteobacteria Rhodobacterales Rhodobacteraceae *Ruegeria sp.* TW15 |
| Sla-1-1 | Bacteria Proteobacteria Alphaproteobacteria Rhodobacterales Rhodobacteraceae Ruegeria *Silicibacter lacuscaerulensis* ITI-1157 |
| Ssp-1-1 | Bacteria Proteobacteria Alphaproteobacteria Rhodobacterales Rhodobacteraceae Ruegeria *Silicibacter sp.* TrichCH4B |
| Sst-1-1 | Bacteria Proteobacteria Alphaproteobacteria Rhodobacterales Rhodobacteraceae *Sagittula stellata* E-37 |
| Smu-1-1 | Bacteria Proteobacteria Alphaproteobacteria Rhodobacterales Rhodobacteraceae *Salipiger mucosus* DSM 16094 |
| Sna-1-1 | Bacteria Proteobacteria Alphaproteobacteria Rhodobacterales Rhodobacteraceae *Sedimentitalea nanhaiensis* DSM 24252 |
| Sqi-1-1 | Bacteria Proteobacteria Alphaproteobacteria Rhodobacterales Rhodobacteraceae *Sediminimonas qiaohouensis* DSM 21189 |
| Sdo-1-1 to Sdo-1-2 | Bacteria Proteobacteria Alphaproteobacteria Rhodobacterales Rhodobacteraceae *Sulfitobacter donghicola* DSW-25 = KCTC 12864 = JCM 14565 |
| Sge-1-1 | Bacteria Proteobacteria Alphaproteobacteria Rhodobacterales Rhodobacteraceae *Sulfitobacter geojensis* |
| Sgu-1-1 | Bacteria Proteobacteria Alphaproteobacteria Rhodobacterales Rhodobacteraceae *Sulfitobacter guttiformis* KCTC 32187 |
| Sme-1-1 | Bacteria Proteobacteria Alphaproteobacteria Rhodobacterales Rhodobacteraceae *Sulfitobacter mediterraneus* |
| Sme-2-1 | Bacteria Proteobacteria Alphaproteobacteria Rhodobacterales Rhodobacteraceae *Sulfitobacter mediterraneus* KCTC 32188 |
| Sno-1-1 | Bacteria Proteobacteria Alphaproteobacteria Rhodobacterales Rhodobacteraceae *Sulfitobacter noctilucae* |
| Sno-2-1 | Bacteria Proteobacteria Alphaproteobacteria Rhodobacterales Rhodobacteraceae *Sulfitobacter noctilucicola* |
| Spo-1-1 | Bacteria Proteobacteria Alphaproteobacteria Rhodobacterales Rhodobacteraceae *Sulfitobacter pontiacus* 3SOLIMAR09 |
| Sps-1-1 | Bacteria Proteobacteria Alphaproteobacteria Rhodobacterales Rhodobacteraceae *Sulfitobacter pseudonitzschiae* |
| Ssp-2-1 | Bacteria Proteobacteria Alphaproteobacteria Rhodobacterales Rhodobacteraceae *Sulfitobacter sp.* 20_GPM-1509m |
| Ssp-3-1 | Bacteria Proteobacteria Alphaproteobacteria Rhodobacterales Rhodobacteraceae *Sulfitobacter sp.* CB2047 |

| | |
|---|---|
| Ssp-4-1 | Bacteria Proteobacteria Alphaproteobacteria Rhodobacterales Rhodobacteraceae *Sulfitobacter sp.* EE-36 |
| Ssp-5-1 | Bacteria Proteobacteria Alphaproteobacteria Rhodobacterales Rhodobacteraceae *Sulfitobacter sp.* NAS-14.1 |
| Tsp-1-1 | Bacteria Proteobacteria Alphaproteobacteria Rhodobacterales Rhodobacteraceae *Tateyamaria sp.* ANG-S1 |
| Tar-1-1 | Bacteria Proteobacteria Alphaproteobacteria Rhodobacterales Rhodobacteraceae *Thalassobacter arenae* DSM 19593 |
| Tsp-2-1 | Bacteria Proteobacteria Alphaproteobacteria Rhodobacterales Rhodobacteraceae *Thalassobacter sp.* 16PALIMAR09 |
| Tst-1-1 | Bacteria Proteobacteria Alphaproteobacteria Rhodobacterales Rhodobacteraceae *Thalassobacter stenotrophicus* |
| Tsp-3-1 | Bacteria Proteobacteria Alphaproteobacteria Rhodobacterales Rhodobacteraceae Thalassobium *Thalassiobium sp.* R2A62 |
| Tat-1-1 | Bacteria Proteobacteria Alphaproteobacteria Rhodobacterales Rhodobacteraceae *Thioclava atlantica* |
| Tda-1-1 | Bacteria Proteobacteria Alphaproteobacteria Rhodobacterales Rhodobacteraceae *Thioclava dalianensis* |
| Tin-1-1 | Bacteria Proteobacteria Alphaproteobacteria Rhodobacterales Rhodobacteraceae *Thioclava indica* |
| Tpa-1-1 | Bacteria Proteobacteria Alphaproteobacteria Rhodobacterales Rhodobacteraceae *Thioclava pacifica* DSM 10166 |
| Wma-1-1 | Bacteria Proteobacteria Alphaproteobacteria Rhodobacterales Rhodobacteraceae *Wenxinia marina* DSM 24838 |
| Rba-4-1 | Bacteria Proteobacteria Alphaproteobacteria *Rhodobacterales bacterium* HTCC2150 |
| Rba-5-1 | Bacteria Proteobacteria Alphaproteobacteria *Rhodobacterales bacterium* HTCC2255 |
| Rba-6-1 | Bacteria Proteobacteria Alphaproteobacteria *Rhodobacterales bacterium* HTCC2654 |
| Rba-7-1 | Bacteria Proteobacteria Alphaproteobacteria *Rhodobacterales bacterium* Y4I |
| env-1 to env-656 | environmental samples |

#### 1.2 Gene contexts

Each L2-Alphaproteobacteria RNA (indicated by "RNA→") is listed. For each hit, the downstream genes predicted to reside in a regulated operon are listed. If the nearest downstream gene is encoding in the opposite strand (and therefore presumed to not be a part of a regulated operon), then that gene is still depicted. Some environmental sequences and some RefSeq entries lack gene annotations, and so no genes are available for such sequences. The direction of each gene is indicated with an arrow (→), and each predicted conserved domain in the gene is named. Conserved domains associated with more than one L2-Alphaproteobacteria RNA are assigned a color; other domains are gray. Information about these conserved domains is given in Section 1.3. The accession of the sequence containing each L2-Alphaproteobacteria RNA is given in the column named "Seq. accession". Accessions beginning with "NC_", "NS_", "NW_" or "NZ_" are contained in RefSeq. Other accession refer to environmental samples. Nucleotide coordinates are given for the 5′ and 3′ boundaries of each L2-Alphaproteobacteria RNA. If the 5′ coordinate is greater than the 3′ coordinate, the RNA is present on the reverse-complement strand of the containing genomic DNA sequence. Each hit is denoted by an abbreviation (like "Eco-1-1") that refers to a taxonomy given in Section 1.1.

| **abbrev.** | **Seq. accession** | | **5′ at** | **3′ at** | **genes** |
|---|---|---|---|---|---|
| env-1 | SL_3KL_010_SED_10464608 | + | 263 | 326 | **RNA**→ RplB (COG0090)rplB (PRK09374)→ |
| Dsh-1-1 | NC_009952.1 | + | 285364 | 285406 | **RNA**→ RplB (COG0090)rplB (PRK09374)→ rpsS (PRK00357)RpsS (COG0185)→ rplV (PRK00565)Ribosomal_L22 (pfam00237)→ RpsC (COG0092)rpsC (PRK00310)→ RplP (COG0197)rplP (PRK09203)→ |
| Pde-1-1 | NC_008686.1 | + | 746785 | 746857 | **RNA**→ RplB (COG0090)rplB (PRK09374)→ rpsS (PRK00357)RpsS (COG0185)→ rplV (PRK00565)Ribosomal_L22 (pfam00237)→ RpsC (COG0092)rpsC (PRK00310)→ RplP (COG0197)rplP (PRK09203)→ |
| env-2 | CEVH01071484.1 | - | 2499 | 2437 | **RNA**→ RplB (COG0090)rplB (PRK09374)→ rpsS (PRK00357)RpsS (COG0185)→ rplV (PRK00565)Ribosomal_L22 (pfam00237)→ RpsC (COG0092)rpsC (PRK00310)→ rplP (PRK09203)Ribosomal_L16 (pfam00252)→ |
| Aat-2-1 | NZ_AQQZ01000010.1 | + | 3052 | 3115 | **RNA**→ RplB (COG0090)rplB (PRK09374)→ rpsS (PRK00357)RpsS (COG0185)→ rplV (PRK00565)Ribosomal_L22 (pfam00237)→ RpsC (COG0092)rpsC (PRK00310)→ RplP (COG0197)rplP (PRK09203)→ |
| Rsp-2-1 | NZ_ACYY01000029.1 | - | 40278 | 40238 | **RNA**→ RplB (COG0090)rplB (PRK09374)→ RplB (COG0090)rplB (PRK09374)→ rpsS (PRK00357)RpsS (COG0185)→ rpsS (PRK00357)RpsS (COG0185)→ rplV (PRK00565)Ribosomal_L22 (pfam00237)→ rplV (PRK00565)Ribosomal_L22 (pfam00237)→ RpsC (COG0092)rpsC (PRK00310)→ RpsC (COG0092)rpsC (PRK00310)→ RplP (COG0197)rplP (PRK09203)→ RplP (COG0197)rplP (PRK09203)→ |
| env-3 | JGI20154J14316_10014726 | + | 1330 | 1394 | **RNA**→ RplB (COG0090)rplB (PRK09374)→ rpsS (PRK00357)RpsS (COG0185)→ rplV (PRK00565)Ribosomal_L22 (pfam00237)→ RpsC (COG0092)rpsC (PRK00310)→ RplP (COG0197)rplP (PRK09203)→ GIY-YIG_SF (cd00719)GIYc (smart00465)→ |

| | | | | | |
|---|---|---|---|---|---|
| Amu-1-1 | NZ_JFKE01000006.1 | - | 218957 | 218896 | **RNA**→ RplB (COG0090)rplB (PRK09374)→ rpsS (PRK00357)RpsS (COG0185)→ rplV (PRK00565)Ribosomal_L22 (pfam00237)→ RpsC (COG0092)rpsC (PRK00310)→ RplP (COG0197)rplP (PRK09203)→ |
| env-4 | AglaG_F499EG101CWTC2 | - | 345 | 276 | **RNA**→ rplB (PRK09374)Ribosomal_L2 (pfam00181)→ |
| env-5 | AglaG_contig17173 | - | 368 | 299 | **RNA**→ rplB (PRK09374)Ribosomal_L2 (pfam00181)→ |
| Rat-1-1 | NZ_AQQW01000010.1 | + | 2615 | 2692 | **RNA**→ RplB (COG0090)rplB (PRK09374)→ rpsS (PRK00357)RpsS (COG0185)→ rplV (PRK00565)Ribosomal_L22 (pfam00237)→ RpsC (COG0092)rpsC (PRK00310)→ RplP (COG0197)rplP (PRK09203)→ hypo→ 2OG-FeII_Oxy_5 (pfam13759)→ |
| env-6 | JGI24724J26744_10003266 | - | 8748 | 8659 | **RNA**→ RplB (COG0090)rplB (PRK09374)→ rpsS (PRK00357)RpsS (COG0185)→ rplV (PRK00565)Ribosomal_L22 (pfam00237)→ RpsC (COG0092)rpsC (PRK00310)→ RplP (COG0197)rplP (PRK09203)→ |
| env-7 | JGI24025J20009_10035938 | - | 1770 | 1681 | **RNA**→ RplB (COG0090)rplB (PRK09374)→ rpsS (PRK00357)RpsS (COG0185)→ rplV (PRK00565)Ribosomal_L22 (pfam00237)→ rpsC (PRK00310)30S_S3_KH (cd02412)→ |
| env-8 | GX8B9KJ01CH3VN | - | 334 | 271 | **RNA**→ rplB (PRK09374)Ribosomal_L2 (pfam00181)→ |
| env-9 | JGI12109J20283_12362431 | + | 37 | 102 | **RNA**→ |
| Tst-1-1 | NZ_JGVS01000007.1 | + | 4252 | 4317 | **RNA**→ RplB (COG0090)rplB (PRK09374)→ rpsS (PRK00357)RpsS (COG0185)→ rplV (PRK00565)Ribosomal_L22 (pfam00237)→ RpsC (COG0092)rpsC (PRK00310)→ RplP (COG0197)rplP (PRK09203)→ |
| Tsp-2-1 | NZ_JHAK01000011.1 | - | 52173 | 52108 | **RNA**→ RplB (COG0090)rplB (PRK09374)→ rpsS (PRK00357)RpsS (COG0185)→ rplV (PRK00565)Ribosomal_L22 (pfam00237)→ RpsC (COG0092)rpsC (PRK00310)→ RplP (COG0197)rplP (PRK09203)→ |
| env-10 | P_2C_Liq_1_UnCty_1004676 | - | 3400 | 3335 | **RNA**→ RplB (COG0090)rplB (PRK09374)→ rpsS (PRK00357)RpsS (COG0185)→ rplV (PRK00565)Ribosomal_L22 (pfam00237)→ RpsC (COG0092)rpsC (PRK00310)→ RplP (COG0197)rplP (PRK09203)→ |
| env-11 | P_2C_Liq_2_UnCty_1000207 | - | 15707 | 15642 | **RNA**→ RplB (COG0090)rplB (PRK09374)→ rpsS (PRK00357)RpsS (COG0185)→ rplV (PRK00565)Ribosomal_L22 (pfam00237)→ RpsC (COG0092)rpsC (PRK00310)→ RplP (COG0197)rplP (PRK09203)→ |
| env-12 | P_2C_Liq_2_UnCty_1009999 | + | 1288 | 1353 | **RNA**→ RplB (COG0090)rplB (PRK09374)→ |
| env-13 | ATLU01000110.1 | + | 1285 | 1375 | **RNA**→ RplB (COG0090)rplB (PRK09374)→ rpsS (PRK00357)RpsS (COG0185)→ rplV (PRK00565)Ribosomal_L22 (pfam00237)→ RpsC (COG0092)rpsC (PRK00310)→ RplP (COG0197)rplP (PRK09203)→ |
| Rca-7-1 | NZ_AYPZ01000027.1 | + | 120952 | 121013 | **RNA**→ RplB (COG0090)rplB (PRK09374)→ rpsS (PRK00357)RpsS (COG0185)→ rplV (PRK00565)Ribosomal_L22 (pfam00237)→ RpsC (COG0092)rpsC (PRK00310)→ RplP (COG0197)rplP (PRK09203)→ |
| Rca-4-1 | NC_014034.1 | + | 342742 | 342803 | **RNA**→ RplB (COG0090)rplB (PRK09374)→ rpsS (PRK00357)RpsS (COG0185)→ rplV (PRK00565)Ribosomal_L22 (pfam00237)→ RpsC (COG0092)rpsC (PRK00310)→ RplP (COG0197)rplP (PRK09203)→ |
| Rca-3-1 | NZ_AYQC01000005.1 | + | 120809 | 120870 | **RNA**→ RplB (COG0090)rplB (PRK09374)→ rpsS (PRK00357)RpsS (COG0185)→ rplV (PRK00565)Ribosomal_L22 (pfam00237)→ RpsC (COG0092)rpsC (PRK00310)→ RplP (COG0197)rplP (PRK09203)→ |
| Rca-5-1 | NZ_AYQB01000006.1 | + | 120842 | 120903 | **RNA**→ RplB (COG0090)rplB (PRK09374)→ rpsS (PRK00357)RpsS (COG0185)→ rplV (PRK00565)Ribosomal_L22 (pfam00237)→ RpsC (COG0092)rpsC (PRK00310)→ RplP (COG0197)rplP (PRK09203)→ |
| Rca-2-1 | NZ_AYPR01000005.1 | + | 120776 | 120837 | **RNA**→ RplB (COG0090)rplB (PRK09374)→ rpsS (PRK00357)RpsS (COG0185)→ rplV (PRK00565)Ribosomal_L22 (pfam00237)→ RpsC (COG0092)rpsC (PRK00310)→ RplP (COG0197)rplP (PRK09203)→ |
| Rca-1-1 | NZ_AYQA01000038.1 | + | 120869 | 120930 | **RNA**→ RplB (COG0090)rplB (PRK09374)→ rpsS (PRK00357)RpsS (COG0185)→ rplV (PRK00565)Ribosomal_L22 (pfam00237)→ RpsC (COG0092)rpsC (PRK00310)→ RplP (COG0197)rplP (PRK09203)→ |
| Rsp-4-1 | NC_009428.1 | - | 2571487 | 2571448 | **RNA**→ RplB (COG0090)rplB (PRK09374)→ rpsS (PRK00357)RpsS (COG0185)→ rplV (PRK00565)Ribosomal_L22 (pfam00237)→ RpsC (COG0092)rpsC (PRK00310)→ RplP (COG0197)rplP (PRK09203)→ |
| Rsp-5-1 | NC_009049.1 | + | 387361 | 387400 | **RNA**→ RplB (COG0090)rplB (PRK09374)→ rpsS (PRK00357)RpsS (COG0185)→ rplV (PRK00565)Ribosomal_L22 (pfam00237)→ RpsC (COG0092)rpsC (PRK00310)→ RplP (COG0197)rplP (PRK09203)→ |

| | | | | | |
|---|---|---|---|---|---|
| Rsp-6-1 | NC_011963.1 | + | 19443 | 19482 | **RNA**→ RplB (COG0090)rplB (PRK09374)→ rpsS (PRK00357)RpsS (COG0185)→ rplV (PRK00565)Ribosomal_L22 (pfam00237)→ RpsC (COG0092)rpsC (PRK00310)→ RplP (COG0197)rplP (PRK09203)→ |
| Rsp-7-1 | NZ_CM001161.1 | + | 2879562 | 2879601 | **RNA**→ RplB (COG0090)rplB (PRK09374)→ rpsS (PRK00357)RpsS (COG0185)→ rplV (PRK00565)Ribosomal_L22 (pfam00237)→ RpsC (COG0092)rpsC (PRK00310)→ RplP (COG0197)rplP (PRK09203)→ |
| Rsp-3-1 | NC_007493.2 | + | 309766 | 309805 | **RNA**→ RplB (COG0090)rplB (PRK09374)→ rpsS (PRK00357)RpsS (COG0185)→ rplV (PRK00565)Ribosomal_L22 (pfam00237)→ RpsC (COG0092)rpsC (PRK00310)→ RplP (COG0197)rplP (PRK09203)→ |
| env-14 | JQGR01000094.1 | + | 4628 | 4725 | **RNA**→ RplB (COG0090)rplB (PRK09374)→ rpsS (PRK00357)RpsS (COG0185)→ rplV (PRK00565)Ribosomal_L22 (pfam00237)→ RpsC (COG0092)rpsC (PRK00310)→ RplP (COG0197)rplP (PRK09203)→ |
| env-15 | 4455295.3.316264 | + | 69 | 130 | **RNA**→ RplB (COG0090)rplB (PRK09374)→ |
| env-16 | AYRF01032015.1 | + | 99 | 189 | **RNA**→ ←hypo |
| env-17 | 3300000052_005679 | + | 305 | 368 | **RNA**→ RplB (COG0090)rplB (PRK09374)→ rpsS (PRK00357)RpsS (COG0185)→ rplV (PRK00565)Ribosomal_L22 (pfam00237)→ RpsC (COG0092)rpsC (PRK00310)→ rplP (PRK09203)Ribosomal_L16 (pfam00252)→ |
| Sqi-1-1 | NZ_AUIJ01000004.1 | - | 73400 | 73337 | **RNA**→ RplB (COG0090)rplB (PRK09374)→ rpsS (PRK00357)RpsS (COG0185)→ rplV (PRK00565)Ribosomal_L22 (pfam00237)→ RpsC (COG0092)rpsC (PRK00310)→ RplP (COG0197)rplP (PRK09203)→ PRK00306 (PRK00306)Ribosomal_L29 (pfam00831)→ RpsQ (COG0186)rpsQ (PRK05610)→ rplN (PRK05483)Ribosomal_L14 (pfam00238)→ RplX (COG0198)rplX (PRK00004)→ rplE (PRK00010)RplE (COG0094)→ rpsN (PRK08881)Ribosomal_S14 (pfam00253)→ rpsH (PRK00136)Ribosomal_S8 (pfam00410)→ RplF (COG0097)rplF (PRK05498)→ RplR (COG0256)rplR (PRK05593)→ RpsE (COG0098)rpsE (PRK00550)→ Ribosomal_L30 (cd01658)rpmD (PRK05611)→ |
| env-18 | AYRH01014365.1 | - | 242 | 152 | **RNA**→ RplB (COG0090)rplB (PRK09374)→ |
| env-19 | JCVI_READ_1105499881630 | - | 824 | 763 | **RNA**→ RplB (COG0090)rplB (PRK09374)→ |
| env-20 | JCVI_SCAF_1101668060777 | + | 269 | 332 | **RNA**→ RplB (COG0090)rplB (PRK09374)→ rpsS (PRK00357)RpsS (COG0185)→ |
| Hsp-1-1 | NZ_JWIE01000061.1 | + | 80663 | 80737 | **RNA**→ RplB (COG0090)rplB (PRK09374)→ rpsS (PRK00357)RpsS (COG0185)→ rplV (PRK00565)Ribosomal_L22 (pfam00237)→ RpsC (COG0092)rpsC (PRK00310)→ RplP (COG0197)rplP (PRK09203)→ |
| Hna-1-1 | NZ_JWIF01000023.1 | - | 41827 | 41752 | **RNA**→ RplB (COG0090)rplB (PRK09374)→ rpsS (PRK00357)RpsS (COG0185)→ rplV (PRK00565)Ribosomal_L22 (pfam00237)→ RpsC (COG0092)rpsC (PRK00310)→ RplP (COG0197)rplP (PRK09203)→ |
| Rba-6-1 | NZ_CH902578.1 | - | 2097076 | 2096975 | **RNA**→ RplB (COG0090)rplB (PRK09374)→ rpsS (PRK00357)RpsS (COG0185)→ rplV (PRK00565)Ribosomal_L22 (pfam00237)→ RpsC (COG0092)rpsC (PRK00310)→ RplP (COG0197)rplP (PRK09203)→ |
| env-21 | CENL01121776.1 | + | 1754 | 1825 | **RNA**→ RplB (COG0090)rplB (PRK09374)→ rpsS (PRK00357)RpsS (COG0185)→ rplV (PRK00565)Ribosomal_L22 (pfam00237)→ RpsC (COG0092)rpsC (PRK00310)→ RplP (COG0197)rplP (PRK09203)→ |
| Cba-1-1 | NZ_AMRK01000018.1 | + | 2718 | 2789 | **RNA**→ RplB (COG0090)rplB (PRK09374)→ RplB (COG0090)rplB (PRK09374)→ rpsS (PRK00357)RpsS (COG0185)→ rpsS (PRK00357)RpsS (COG0185)→ rplV (PRK00565)Ribosomal_L22 (pfam00237)→ rplV (PRK00565)Ribosomal_L22 (pfam00237)→ RpsC (COG0092)rpsC (PRK00310)→ RpsC (COG0092)rpsC (PRK00310)→ RplP (COG0197)rplP (PRK09203)→ RplP (COG0197)rplP (PRK09203)→ |
| Rsp-1-1 | NZ_ANFS01000015.1 | + | 2126 | 2188 | **RNA**→ RplB (COG0090)rplB (PRK09374)→ rpsS (PRK00357)RpsS (COG0185)→ rplV (PRK00565)Ribosomal_L22 (pfam00237)→ RpsC (COG0092)rpsC (PRK00310)→ RplP (COG0197)rplP (PRK09203)→ |
| env-22 | JGI24724J26744_10214823 | + | 124 | 189 | **RNA**→ rplB (PRK09374)Ribosomal_L2 (pfam00181)→ |
| env-23 | NCBI_READ_1112637934486 | + | 240 | 306 | **RNA**→ RplB (COG0090)rplB (PRK09374)→ |
| env-24 | CESE01117110.1 | - | 1184 | 1118 | **RNA**→ RplB (COG0090)rplB (PRK09374)→ rpsS (PRK00357)RpsS (COG0185)→ |
| env-25 | CEST01016851.1 | + | 2161 | 2227 | **RNA**→ RplB (COG0090)rplB (PRK09374)→ rpsS (PRK00357)RpsS (COG0185)→ |
| env-26 | 2236876009_187088 | + | 204 | 270 | **RNA**→ RplB (COG0090)rplB (PRK09374)→ |
| env-27 | DelMOSum2010_c10002940 | - | 12329 | 12263 | **RNA**→ RplB (COG0090)rplB (PRK09374)→ rpsS (PRK00357)RpsS (COG0185)→ rplV (PRK00565)Ribosomal_L22 (pfam00237)→ RpsC (COG0092)rpsC (PRK00310)→ RplP (COG0197)rplP (PRK09203)→ |

| | | | | | |
|---|---|---|---|---|---|
| env-28 | JGI20152J14361_10100075 | - | 149 | 83 | **RNA**→ RplB (COG0090)rplB (PRK09374)→ |
| env-29 | JGI20151J14362_10028856 | + | 2610 | 2676 | **RNA**→ RplB (COG0090)rplB (PRK09374)→ |
| env-30 | JGI20154J14316_10002806 | + | 4976 | 5042 | **RNA**→ RplB (COG0090)rplB (PRK09374)→ rpsS (PRK00357)RpsS (COG0185)→ rplV (PRK00565)Ribosomal_L22 (pfam00237)→ RpsC (COG0092)rpsC (PRK00310)→ RplP (COG0197)rplP (PRK09203)→ |
| env-31 | JGI20160J14292_10200265 | + | 405 | 471 | **RNA**→ RplB (COG0090)rplB (PRK09374)→ |
| env-32 | JGI20153J14295_10377309 | - | 117 | 51 | **RNA**→ hypo→ |
| env-33 | JGI20153J14318_10377309 | - | 117 | 51 | **RNA**→ hypo→ |
| env-34 | JGI20157J14317_10000107 | - | 35008 | 34942 | **RNA**→ RplB (COG0090)rplB (PRK09374)→ rpsS (PRK00357)RpsS (COG0185)→ rplV (PRK00565)Ribosomal_L22 (pfam00237)→ RpsC (COG0092)rpsC (PRK00310)→ RplP (COG0197)rplP (PRK09203)→ |
| env-35 | CEUP01061385.1 | - | 473 | 377 | **RNA**→ rplB (PRK09374)Ribosomal_L2 (pfam00181)→ |
| Csp-1-1 | NZ_AJKJ01000048.1 | + | 3267 | 3349 | **RNA**→ RplB (COG0090)rplB (PRK09374)→ RplB (COG0090)rplB (PRK09374)→ rpsS (PRK00357)RpsS (COG0185)→ rpsS (PRK00357)RpsS (COG0185)→ rplV (PRK00565)Ribosomal_L22 (pfam00237)→ rplV (PRK00565)Ribosomal_L22 (pfam00237)→ RpsC (COG0092)rpsC (PRK00310)→ RpsC (COG0092)rpsC (PRK00310)→ RplP (COG0197)rplP (PRK09203)→ RplP (COG0197)rplP (PRK09203)→ hypo→ hypo→ |
| Pat-1-1 | NZ_AQQX01000010.1 | - | 63842 | 63774 | **RNA**→ RplB (COG0090)rplB (PRK09374)→ rpsS (PRK00357)RpsS (COG0185)→ rplV (PRK00565)Ribosomal_L22 (pfam00237)→ RpsC (COG0092)rpsC (PRK00310)→ RplP (COG0197)rplP (PRK09203)→ |
| Rba-3-1 | NZ_KN714899.1 | - | 63316 | 63257 | **RNA**→ RplB (COG0090)rplB (PRK09374)→ rpsS (PRK00357)RpsS (COG0185)→ rplV (PRK00565)Ribosomal_L22 (pfam00237)→ RpsC (COG0092)rpsC (PRK00310)→ RplP (COG0197)rplP (PRK09203)→ |
| env-36 | JGI24025J20009_10174089 | + | 339 | 403 | **RNA**→ RplB (COG0090)rplB (PRK09374)→ |
| env-37 | CEQG01114380.1 | - | 89 | 29 | **RNA**→ |
| Rca-6-1 | NZ_AYPY01000003.1 | - | 128998 | 128937 | **RNA**→ RplB (COG0090)rplB (PRK09374)→ rpsS (PRK00357)RpsS (COG0185)→ rplV (PRK00565)Ribosomal_L22 (pfam00237)→ RpsC (COG0092)rpsC (PRK00310)→ RplP (COG0197)rplP (PRK09203)→ |
| env-38 | JGI24723J26617_10335773 | - | 212 | 147 | **RNA**→ RplB (COG0090)rplB (PRK09374)→ |
| env-39 | JGI24724J26744_10397626 | + | 94 | 159 | **RNA**→ rplB (PRK09374)Ribosomal_L2 (pfam00181)→ |
| env-40 | JGI24723J26617_10265371 | + | 184 | 249 | **RNA**→ RplB (COG0090)rplB (PRK09374)→ |
| Oba-1-1 | NZ_CH724131.1 | - | 1639797 | 1639719 | **RNA**→ RplB (COG0090)rplB (PRK09374)→ rpsS (PRK00357)RpsS (COG0185)→ rplV (PRK00565)Ribosomal_L22 (pfam00237)→ RpsC (COG0092)rpsC (PRK00310)→ RplP (COG0197)rplP (PRK09203)→ |
| env-41 | JGI12210J13797_10911808 | - | 154 | 79 | **RNA**→ RplB (COG0090)rplB (PRK09374)→ |
| env-42 | SA_S1_NOR08_45m_c10088539 | - | 163 | 103 | **RNA**→ RplB (COG0090)rplB (PRK09374)→ |
| env-43 | SA_S1_NOR05_45m_c10012273 | + | 805 | 865 | **RNA**→ RplB (COG0090)rplB (PRK09374)→ rpsS (PRK00357)RpsS (COG0185)→ rplV (PRK00565)Ribosomal_L22 (pfam00237)→ KH (smart00322)30S_S3_KH (cd02412)→ |
| env-44 | SA_S2_NOR15_50m_c10022560 | - | 1787 | 1727 | **RNA**→ RplB (COG0090)rplB (PRK09374)→ rpsS (PRK00357)Ribosomal_S19 (pfam00203)→ rplV (PRK00565)Ribosomal_L22 (pfam00237)→ rpsC (PRK00310)30S_S3_KH (cd02412)→ |
| env-45 | KGI_S2_ANT05_2345m_c1008115 | + | 360 | 420 | **RNA**→ RplB (COG0090)rplB (PRK09374)→ rpsS (PRK00357)RpsS (COG0185)→ rplV (PRK00565)Ribosomal_L22 (pfam00237)→ RpsC (COG0092)rpsC (PRK00310)→ |
| env-46 | KGI_S1_ANT02_95m_c10175114 | - | 173 | 113 | **RNA**→ RplB (COG0090)rplB (PRK09374)→ |
| env-47 | SA_S2_NOR13_50m_c1107468 | - | 300 | 240 | **RNA**→ RplB (COG0090)rplB (PRK09374)→ |
| env-48 | KGI_S1_ANT03_95m_c1050218 | + | 27 | 87 | **RNA**→ rplB (PRK09374)Ribosomal_L2 (pfam00181)→ |
| env-49 | JGI24723J26617_10562187 | - | 219 | 156 | **RNA**→ RplB (COG0090)rplB (PRK09374)→ |
| env-50 | JGI24724J26744_10017014 | - | 875 | 812 | **RNA**→ RplB (COG0090)rplB (PRK09374)→ |
| env-51 | JGI24025J20009_10010867 | - | 4900 | 4837 | **RNA**→ RplB (COG0090)rplB (PRK09374)→ rpsS (PRK00357)RpsS (COG0185)→ rplV (PRK00565)Ribosomal_L22 (pfam00237)→ RpsC (COG0092)rpsC (PRK00310)→ RplP (COG0197)rplP (PRK09203)→ |
| env-52 | KGI_S1_ANT01_95m_c10003918 | + | 2539 | 2601 | **RNA**→ RplB (COG0090)rplB (PRK09374)→ rpsS (PRK00357)RpsS (COG0185)→ rplV (PRK00565)Ribosomal_L22 (pfam00237)→ RpsC (COG0092)rpsC (PRK00310)→ RplP (COG0197)rplP (PRK09203)→ |
| env-53 | KGI_S2_ANT05_2345m_c1031605 | - | 159 | 97 | **RNA**→ RplB (COG0090)rplB (PRK09374)→ |
| env-54 | KGI_S1_ANT02_95m_c10012913 | + | 2664 | 2726 | **RNA**→ rplB (PRK09374)Ribosomal_L2 (pfam00181)→ |

| | | | | | |
|---|---|---|---|---|---|
| env-55 | AGBJ01004298.1 | - | 284 | 223 | **RNA**→ ←hypo |
| env-56 | JGI24025J20009_10035212 | - | 340 | 279 | **RNA**→ rplB (PRK09374)Ribosomal_L2 (pfam00181)→ |
| env-57 | LGVE01157751.1 | + | 1200 | 1261 | **RNA**→ rplB (PRK09374)Ribosomal_L2 (pfam00181)→ |
| env-58 | LGVF01211244.1 | - | 1626 | 1565 | **RNA**→ RplB (COG0090)rplB (PRK09374)→ rpsS (PRK00357)Ribosomal_S19 (pfam00203)→ rplV (PRK00565)→ |
| | | | | | rplV (PRK00565)Ribosomal_L22 (pfam00237)→ rpsC (PRK00310)30S_S3_KH (cd02412)→ |
| env-59 | LGVD01030433.1 | + | 391 | 452 | **RNA**→ rplB (PRK09374)Ribosomal_L2 (pfam00181)→ |
| env-60 | SA_S1_NOR08_45m_c10143553 | + | 557 | 618 | **RNA**→ RplB (COG0090)rplB (PRK09374)→ |
| env-61 | SA_S2_NOR15_50m_c10181039 | - | 134 | 73 | **RNA**→ RplB (COG0090)rplB (PRK09374)→ |
| env-62 | SA_S1_NOR02_45m_c1048433 | - | 379 | 318 | **RNA**→ rplB (PRK09374)Ribosomal_L2 (pfam00181)→ |
| env-63 | SA_S2_NOR18_50m_1043735 | + | 332 | 393 | **RNA**→ rplB (PRK09374)Ribosomal_L2 (pfam00181)→ |
| env-64 | LGOV01019582.1 | + | 657 | 718 | **RNA**→ RplB (COG0090)rplB (PRK09374)→ |
| Rli-1-1 | NC_015730.1 | - | 3449426 | 3449359 | **RNA**→ RplB (COG0090)rplB (PRK09374)→ rpsS (PRK00357)RpsS (COG0185)→ |
| | | | | | rplV (PRK00565)Ribosomal_L22 (pfam00237)→ RpsC (COG0092)rpsC (PRK00310)→ |
| | | | | | RplP (COG0197)rplP (PRK09203)→ |
| env-65 | JGI12109J20283_10879218 | - | 201 | 135 | **RNA**→ RplB (COG0090)rplB (PRK09374)→ |
| Rde-1-1 | NC_008209.1 | + | 1346820 | 1346887 | **RNA**→ RplB (COG0090)rplB (PRK09374)→ rpsS (PRK00357)RpsS (COG0185)→ |
| | | | | | rplV (PRK00565)Ribosomal_L22 (pfam00237)→ RpsC (COG0092)rpsC (PRK00310)→ |
| | | | | | RplP (COG0197)rplP (PRK09203)→ |
| env-66 | JGI11958J13698_1070723 | - | 120 | 52 | **RNA**→ ←hypo |
| env-67 | CAM_READ_0234679589 | - | 364 | 296 | **RNA**→ rplB (PRK09374)Ribosomal_L2 (pfam00181)→ |
| env-68 | CAM_READ_0234796905 | - | 364 | 296 | **RNA**→ rplB (PRK09374)Ribosomal_L2 (pfam00181)→ |
| Rsp-15-1 | NZ_JQLS01000008.1 | + | 336402 | 336479 | **RNA**→ RplB (COG0090)rplB (PRK09374)→ rpsS (PRK00357)RpsS (COG0185)→ |
| | | | | | rplV (PRK00565)Ribosomal_L22 (pfam00237)→ RpsC (COG0092)rpsC (PRK00310)→ |
| | | | | | RplP (COG0197)rplP (PRK09203)→ |
| env-69 | CESQ01192690.1 | - | 546 | 470 | **RNA**→ rplB (PRK09374)Ribosomal_L2 (pfam00181)→ |
| Rsp-14-1 | NZ_DS022276.1 | - | 423367 | 423284 | **RNA**→ RplB (COG0090)rplB (PRK09374)→ rpsS (PRK00357)RpsS (COG0185)→ |
| | | | | | rplV (PRK00565)Ribosomal_L22 (pfam00237)→ RpsC (COG0092)rpsC (PRK00310)→ |
| | | | | | RplP (COG0197)rplP (PRK09203)→ |
| env-70 | CETR01181359.1 | - | 335 | 252 | **RNA**→ RplB (COG0090)rplB (PRK09374)→ |
| env-71 | JGI11800J13271_1007151 | + | 1158 | 1250 | **RNA**→ |
| Csp-2-1 | NZ_GG704599.1 | + | 399690 | 399782 | **RNA**→ RplB (COG0090)rplB (PRK09374)→ RplB (COG0090)rplB (PRK09374)→ |
| | | | | | rpsS (PRK00357)RpsS (COG0185)→ rpsS (PRK00357)RpsS (COG0185)→ |
| | | | | | rplV (PRK00565)Ribosomal_L22 (pfam00237)→ rplV (PRK00565)Ribosomal_L22 (pfam00237)→ |
| | | | | | RpsC (COG0092)rpsC (PRK00310)→ RpsC (COG0092)rpsC (PRK00310)→ RplP (COG0197)rplP (PRK09203)→ |
| | | | | | RplP (COG0197)rplP (PRK09203)→ |
| env-72 | P_2C_Liq_1_UnCty_1008935 | - | 2463 | 2386 | **RNA**→ RplB (COG0090)rplB (PRK09374)→ rpsS (PRK00357)RpsS (COG0185)→ |
| | | | | | rplV (PRK00565)Ribosomal_L22 (pfam00237)→ RpsC (COG0092)rpsC (PRK00310)→ |
| | | | | | rplP (PRK09203)Ribosomal_L16 (pfam00252)→ |
| env-73 | P_2C_Liq_2_UnCty_1004597 | + | 433 | 510 | **RNA**→ RplB (COG0090)rplB (PRK09374)→ rpsS (PRK00357)RpsS (COG0185)→ |
| | | | | | rplV (PRK00565)Ribosomal_L22 (pfam00237)→ RpsC (COG0092)rpsC (PRK00310)→ |
| | | | | | rplP (PRK09203)Ribosomal_L16 (pfam00252)→ |
| Nig-1-1 | NZ_CVPC01000014.1 | - | 4708 | 4631 | **RNA**→ RplB (COG0090)rplB (PRK09374)→ rpsS (PRK00357)RpsS (COG0185)→ |
| | | | | | rplV (PRK00565)Ribosomal_L22 (pfam00237)→ RpsC (COG0092)rpsC (PRK00310)→ |
| | | | | | RplP (COG0197)rplP (PRK09203)→ |
| env-74 | JGI12273J12029_10524331 | + | 216 | 286 | **RNA**→ |
| env-75 | JGI24025J20009_10000374 | - | 31294 | 31182 | **RNA**→ RplB (COG0090)rplB (PRK09374)→ rpsS (PRK00357)RpsS (COG0185)→ |
| | | | | | rplV (PRK00565)Ribosomal_L22 (pfam00237)→ RpsC (COG0092)rpsC (PRK00310)→ |
| | | | | | RplP (COG0197)rplP (PRK09203)→ |
| env-76 | JGI24025J20009_10010033 | + | 1689 | 1756 | **RNA**→ RplB (COG0090)rplB (PRK09374)→ rpsS (PRK00357)RpsS (COG0185)→ |
| | | | | | rplV (PRK00565)Ribosomal_L22 (pfam00237)→ RpsC (COG0092)rpsC (PRK00310)→ |
| | | | | | RplP (COG0197)rplP (PRK09203)→ |
| Tsp-1-1 | NZ_JWLL01000009.1 | - | 233099 | 233024 | **RNA**→ RplB (COG0090)rplB (PRK09374)→ rpsS (PRK00357)RpsS (COG0185)→ |
| | | | | | rplV (PRK00565)Ribosomal_L22 (pfam00237)→ RpsC (COG0092)rpsC (PRK00310)→ |
| | | | | | RplP (COG0197)rplP (PRK09203)→ |
| env-77 | 2236876003_101132 | + | 388 | 448 | **RNA**→ RplB (COG0090)rplB (PRK09374)→ |

| | | | | | |
|---|---|---|---|---|---|
| env-78 | PML_READ_00355505 | + | 143 | 210 | **RNA**→ |
| env-79 | JGI20152J14361_10000074 | + | 12924 | 12991 | **RNA**→ RplB (COG0090)rplB (PRK09374)→ rpsS (PRK00357)RpsS (COG0185)→ rplV (PRK00565)Ribosomal_L22 (pfam00237)→ RpsC (COG0092)rpsC (PRK00310)→ RplP (COG0197)rplP (PRK09203)→ hypo→ |
| env-80 | JGI20154J14316_10160884 | + | 228 | 295 | **RNA**→ rplB (PRK09374)Ribosomal_L2 (pfam00181)→ |
| env-81 | JGI20153J14295_10001123 | + | 7148 | 7215 | **RNA**→ RplB (COG0090)rplB (PRK09374)→ rpsS (PRK00357)RpsS (COG0185)→ rplV (PRK00565)Ribosomal_L22 (pfam00237)→ RpsC (COG0092)rpsC (PRK00310)→ RplP (COG0197)rplP (PRK09203)→ hypo→ |
| env-82 | JGI20153J14318_10001123 | + | 7148 | 7215 | **RNA**→ RplB (COG0090)rplB (PRK09374)→ rpsS (PRK00357)RpsS (COG0185)→ rplV (PRK00565)Ribosomal_L22 (pfam00237)→ RpsC (COG0092)rpsC (PRK00310)→ RplP (COG0197)rplP (PRK09203)→ hypo→ |
| Lve-2-1 | NZ_CH672414.1 | + | 2669496 | 2669565 | **RNA**→ RplB (COG0090)rplB (PRK09374)→ rpsS (PRK00357)RpsS (COG0185)→ rplV (PRK00565)Ribosomal_L22 (pfam00237)→ RpsC (COG0092)rpsC (PRK00310)→ RplP (COG0197)rplP (PRK09203)→ |
| Lve-1-1 | NZ_KB907973.1 | + | 38196 | 38265 | **RNA**→ RplB (COG0090)rplB (PRK09374)→ rpsS (PRK00357)RpsS (COG0185)→ rplV (PRK00565)Ribosomal_L22 (pfam00237)→ RpsC (COG0092)rpsC (PRK00310)→ RplP (COG0197)rplP (PRK09203)→ |
| env-83 | JGI12210J13797_10455319 | + | 174 | 245 | **RNA**→ |
| env-84 | JGI12210J13797_10023293 | - | 122 | 51 | **RNA**→ ←hypo |
| Ogr-1-1 | NZ_CH724107.1 | - | 1761097 | 1761017 | **RNA**→ RplB (COG0090)rplB (PRK09374)→ rpsS (PRK00357)RpsS (COG0185)→ RplV (COG0091)rplV (PRK00565)→ RpsC (COG0092)rpsC (PRK00310)→ RplP (COG0197)rplP (PRK09203)→ |
| env-85 | JCVI_SCAF_1096627353865 | + | 440 | 504 | **RNA**→ RplB (COG0090)rplB (PRK09374)→ Zw10 (pfam06248)→ rpsS (PRK00357)RpsS (COG0185)→ hypo→ rplV (PRK00565)Ribosomal_L22 (pfam00237)→ RpsC (COG0092)rpsC (PRK00310)→ |
| env-86 | PR_CR_10_Liq_1_inCR_1002101 | + | 720 | 784 | **RNA**→ RplB (COG0090)rplB (PRK09374)→ rpsS (PRK00357)RpsS (COG0185)→ rplV (PRK00565)Ribosomal_L22 (pfam00237)→ RpsC (COG0092)rpsC (PRK00310)→ RplP (COG0197)rplP (PRK09203)→ |
| env-87 | PR_CR_10_Liq_3_inCR_1034197 | - | 612 | 548 | **RNA**→ RplB (COG0090)rplB (PRK09374)→ |
| Lsp-10-1 | NZ_JXYE01000005.1 | - | 18000 | 17935 | **RNA**→ RplB (COG0090)rplB (PRK09374)→ rpsS (PRK00357)RpsS (COG0185)→ rplV (PRK00565)Ribosomal_L22 (pfam00237)→ RpsC (COG0092)rpsC (PRK00310)→ RplP (COG0197)rplP (PRK09203)→ |
| env-88 | Antartic2_1216799 | + | 177 | 241 | **RNA**→ RplB (COG0090)rplB (PRK09374)→ |
| env-89 | JGI25920J50251_10282450 | - | 274 | 210 | **RNA**→ ←hypo |
| Aat-1-1 | NZ_AQQY01000002.1 | + | 4134 | 4211 | **RNA**→ RplB (COG0090)rplB (PRK09374)→ rpsS (PRK00357)RpsS (COG0185)→ rplV (PRK00565)Ribosomal_L22 (pfam00237)→ RpsC (COG0092)rpsC (PRK00310)→ RplP (COG0197)rplP (PRK09203)→ |
| env-90 | HLSNC_1002228 | - | 3054 | 2987 | **RNA**→ RplB (COG0090)rplB (PRK09374)→ rpsS (PRK00357)RpsS (COG0185)→ rplV (PRK00565)Ribosomal_L22 (pfam00237)→ RpsC (COG0092)rpsC (PRK00310)→ RplP (COG0197)rplP (PRK09203)→ hypo→ |
| env-91 | KVRMV2_106490902 | + | 59 | 134 | **RNA**→ RplB (COG0090)rplB (PRK09374)→ |
| env-92 | GYQO9XB02HJOI1 | - | 85 | 21 | **RNA**→ ←hypo |
| env-93 | 4455295.3.416243 | + | 77 | 139 | **RNA**→ rplB (PRK09374)Ribosomal_L2 (pfam00181)→ |
| env-94 | JGI20152J14361_10000792 | - | 16131 | 16066 | **RNA**→ RplB (COG0090)rplB (PRK09374)→ rpsS (PRK00357)RpsS (COG0185)→ rplV (PRK00565)Ribosomal_L22 (pfam00237)→ RpsC (COG0092)rpsC (PRK00310)→ RplP (COG0197)rplP (PRK09203)→ |
| env-95 | JGI20152J14361_10021758 | + | 662 | 727 | **RNA**→ RplB (COG0090)rplB (PRK09374)→ rpsS (PRK00357)RpsS (COG0185)→ rplV (PRK00565)Ribosomal_L22 (pfam00237)→ |
| Pna-1-1 | NZ_JHZF01000011.1 | - | 1853666 | 1853592 | **RNA**→ RplB (COG0090)rplB (PRK09374)→ rpsS (PRK00357)RpsS (COG0185)→ rplV (PRK00565)Ribosomal_L22 (pfam00237)→ RpsC (COG0092)rpsC (PRK00310)→ RplP (COG0197)rplP (PRK09203)→ |
| env-96 | C687J26610_1065482 | - | 330 | 262 | **RNA**→ rplB (PRK09374)Ribosomal_L2 (pfam00181)→ |
| Sna-1-1 | NZ_KI421509.1 | - | 2029881 | 2029817 | **RNA**→ RplB (COG0090)rplB (PRK09374)→ rpsS (PRK00357)RpsS (COG0185)→ rplV (PRK00565)Ribosomal_L22 (pfam00237)→ RpsC (COG0092)rpsC (PRK00310)→ RplP (COG0197)rplP (PRK09203)→ TIGR02466 (TIGR02466)→ |
| env-97 | GYQO9XB02GIUBI | + | 54 | 123 | **RNA**→ |
| env-98 | DCrCPGB_c116271 | + | 75 | 144 | **RNA**→ rplB (PRK09374)Ribosomal_L2 (pfam00181)→ |

| | | | | | |
|---|---|---|---|---|---|
| env-99 | KGI_S2_ANT04_2345m_c1008211 | - | 1538 | 1465 | **RNA**→ RplB (COG0090)rplB (PRK09374)→ rpsS (PRK00357)RpsS (COG0185)→ rplV (PRK00565)Ribosomal_L22 (pfam00237)→ |
| env-100 | KGI_S1_ANT02_95m_c10000174 | + | 24541 | 24614 | **RNA**→ RplB (COG0090)rplB (PRK09374)→ rpsS (PRK00357)RpsS (COG0185)→ rplV (PRK00565)Ribosomal_L22 (pfam00237)→ RpsC (COG0092)rpsC (PRK00310)→ RplP (COG0197)rplP (PRK09203)→ |
| env-101 | KGI_S1_ANT01_95m_c10002670 | + | 4835 | 4908 | **RNA**→ RplB (COG0090)rplB (PRK09374)→ rpsS (PRK00357)RpsS (COG0185)→ rplV (PRK00565)Ribosomal_L22 (pfam00237)→ RpsC (COG0092)rpsC (PRK00310)→ RplP (COG0197)rplP (PRK09203)→ |
| env-102 | KGI_S2_ANT05_2345m_c1002025 | - | 4049 | 3976 | **RNA**→ RplB (COG0090)rplB (PRK09374)→ rpsS (PRK00357)RpsS (COG0185)→ rplV (PRK00565)Ribosomal_L22 (pfam00237)→ RpsC (COG0092)rpsC (PRK00310)→ RplP (COG0197)rplP (PRK09203)→ |
| env-103 | SR_TP_S13sol_1015872 | - | 193 | 120 | **RNA**→ RplB (COG0090)rplB (PRK09374)→ |
| env-104 | HLSNC_1000102 | + | 4292 | 4365 | **RNA**→ RplB (COG0090)rplB (PRK09374)→ rpsS (PRK00357)RpsS (COG0185)→ rplV (PRK00565)Ribosomal_L22 (pfam00237)→ RpsC (COG0092)rpsC (PRK00310)→ RplP (COG0197)rplP (PRK09203)→ |
| Jsp-1-1 | NC_007802.1 | + | 574447 | 574519 | **RNA**→ RplB (COG0090)rplB (PRK09374)→ rpsS (PRK00357)RpsS (COG0185)→ rplV (PRK00565)Ribosomal_L22 (pfam00237)→ RpsC (COG0092)rpsC (PRK00310)→ RplP (COG0197)rplP (PRK09203)→ |
| env-105 | KVRMV2_101198589 | + | 2281 | 2349 | **RNA**→ rplB (PRK09374)Ribosomal_L2 (pfam00181)→ |
| env-106 | PR_CR_10_Liq_1_inCR_1005216 | - | 2908 | 2842 | **RNA**→ RplB (COG0090)rplB (PRK09374)→ rpsS (PRK00357)RpsS (COG0185)→ rplV (PRK00565)Ribosomal_L22 (pfam00237)→ RpsC (COG0092)rpsC (PRK00310)→ RplP (COG0197)rplP (PRK09203)→ |
| env-107 | PR_CR_10_Liq_3_inCR_1001525 | - | 3425 | 3359 | **RNA**→ RplB (COG0090)rplB (PRK09374)→ rpsS (PRK00357)RpsS (COG0185)→ rplV (PRK00565)Ribosomal_L22 (pfam00237)→ RpsC (COG0092)rpsC (PRK00310)→ RplP (COG0197)rplP (PRK09203)→ |
| env-108 | GSLSAAI_contig03181 | + | 626 | 692 | **RNA**→ |
| env-109 | ElkS_mat_CD2A_1005801 | - | 1817 | 1752 | **RNA**→ RplB (COG0090)rplB (PRK09374)→ rpsS (PRK00357)RpsS (COG0185)→ rplV (PRK00565)Ribosomal_L22 (pfam00237)→ rpsC (PRK00310)30S_S3_KH (cd02412)→ |
| env-110 | ElkS_mat_MD6A_1013394 | + | 164 | 229 | **RNA**→ RplB (COG0090)rplB (PRK09374)→ rpsS (PRK00357)RpsS (COG0185)→ rplV (PRK00565)Ribosomal_L22 (pfam00237)→ rpsC (PRK00310)30S_S3_KH (cd02412)→ |
| env-111 | JGI12210J13797_11132003 | + | 2467 | 2532 | **RNA**→ RplB (COG0090)rplB (PRK09374)→ rpsS (PRK00357)RpsS (COG0185)→ rplV (PRK00565)Ribosomal_L22 (pfam00237)→ RpsC (COG0092)rpsC (PRK00310)→ RplP (COG0197)rplP (PRK09203)→ |
| env-112 | JGI12210J13797_11132005 | - | 3162 | 3097 | **RNA**→ RplB (COG0090)rplB (PRK09374)→ rpsS (PRK00357)RpsS (COG0185)→ rplV (PRK00565)Ribosomal_L22 (pfam00237)→ RpsC (COG0092)rpsC (PRK00310)→ RplP (COG0197)rplP (PRK09203)→ |
| env-113 | JGI11876J14442_10003279 | - | 3251 | 3186 | **RNA**→ RplB (COG0090)rplB (PRK09374)→ rpsS (PRK00357)RpsS (COG0185)→ rplV (PRK00565)Ribosomal_L22 (pfam00237)→ RpsC (COG0092)rpsC (PRK00310)→ RplP (COG0197)rplP (PRK09203)→ |
| env-114 | JGI12109J20283_10578297 | + | 2553 | 2618 | **RNA**→ rplB (PRK09374)Ribosomal_L2 (pfam00181)→ |
| env-115 | JGI12109J20283_10578298 | - | 338 | 273 | **RNA**→ rplB (PRK09374)Ribosomal_L2 (pfam00181)→ |
| env-116 | JGI12210J13797_11558909 | + | 43 | 103 | **RNA**→ ←hypo |
| env-117 | JGI12210J13797_11938347 | - | 70 | 10 | **RNA**→ ←hypo |
| env-118 | CENR01018324.1 | + | 1102 | 1170 | **RNA**→ RplB (COG0090)rplB (PRK09374)→ |
| env-119 | CENV01010391.1 | - | 2234 | 2166 | **RNA**→ RplB (COG0090)rplB (PRK09374)→ rpsS (PRK00357)RpsS (COG0185)→ RplV (COG0091)rplV (PRK00565)→ RpsC (COG0092)rpsC (PRK00310)→ |
| env-120 | CERO01013343.1 | + | 295 | 363 | **RNA**→ RplB (COG0090)rplB (PRK09374)→ |
| env-121 | JGI24724J26744_10870797 | + | 32 | 99 | **RNA**→ RplB (COG0090)rplB (PRK09374)→ |
| env-122 | JGI12210J13797_10838428 | - | 914 | 850 | **RNA**→ RplB (COG0090)rplB (PRK09374)→ |
| env-123 | JGI12210J13797_10838429 | + | 87 | 151 | **RNA**→ RplB (COG0090)rplB (PRK09374)→ |
| env-124 | JGI20152J14361_10179686 | + | 269 | 339 | **RNA**→ ←hypo |
| env-125 | SA_S1_NOR08_45m_c10004046 | + | 4200 | 4267 | **RNA**→ RplB (COG0090)rplB (PRK09374)→ rpsS (PRK00357)RpsS (COG0185)→ rplV (PRK00565)Ribosomal_L22 (pfam00237)→ RpsC (COG0092)rpsC (PRK00310)→ RplP (COG0197)rplP (PRK09203)→ |

| | | | | | |
|---|---|---|---|---|---|
| env-126 | SA_S2_NOR15_50m_c10018770 | + | 306 | 373 | **RNA**→ RplB (COG0090)rplB (PRK09374)→ rpsS (PRK00357)RpsS (COG0185)→ rplV (PRK00565)Ribosomal_L22 (pfam00237)→ RpsC (COG0092)rpsC (PRK00310)→ RplP (COG0197)rplP (PRK09203)→ |
| env-127 | SA_S1_NOR05_45m_c10315404 | + | 255 | 322 | **RNA**→ |
| Ote-1-1 | NZ_CP012160.1 | + | 252086 | 252153 | **RNA**→ RplB (COG0090)rplB (PRK09374)→ rpsS (PRK00357)RpsS (COG0185)→ rplV (PRK00565)Ribosomal_L22 (pfam00237)→ RpsC (COG0092)rpsC (PRK00310)→ RplP (COG0197)rplP (PRK09203)→ |
| Oan-1-1 | NC_020911.1 | - | 4346201 | 4346135 | **RNA**→ RplB (COG0090)rplB (PRK09374)→ rpsS (PRK00357)RpsS (COG0185)→ rplV (PRK00565)Ribosomal_L22 (pfam00237)→ RpsC (COG0092)rpsC (PRK00310)→ RplP (COG0197)rplP (PRK09203)→ |
| Oar-1-1 | NC_020908.1 | + | 3931342 | 3931406 | **RNA**→ RplB (COG0090)rplB (PRK09374)→ rpsS (PRK00357)RpsS (COG0185)→ rplV (PRK00565)Ribosomal_L22 (pfam00237)→ RpsC (COG0092)rpsC (PRK00310)→ RplP (COG0197)rplP (PRK09203)→ |
| env-128 | KGI_S1_ANT01_95m_c10350259 | - | 325 | 258 | **RNA**→ rplB (PRK09374)Ribosomal_L2 (pfam00181)→ |
| env-129 | SA_S1_NOR08_45m_c10019190 | + | 380 | 447 | **RNA**→ RplB (COG0090)rplB (PRK09374)→ rpsS (PRK00357)RpsS (COG0185)→ rplV (PRK00565)Ribosomal_L22 (pfam00237)→ RpsC (COG0092)rpsC (PRK00310)→ rplP (PRK09203)Ribosomal_L16 (pfam00252)→ |
| env-130 | KGI_S2_ANT04_2345m_c1106538 | - | 368 | 301 | **RNA**→ rplB (PRK09374)Ribosomal_L2 (pfam00181)→ |
| env-131 | KGI_S2_ANT05_2345m_c1147958 | + | 34 | 101 | **RNA**→ rplB (PRK09374)Ribosomal_L2 (pfam00181)→ |
| env-132 | KGI_S1_ANT03_95m_c1027476 | + | 53 | 120 | **RNA**→ RplB (COG0090)rplB (PRK09374)→ |
| env-133 | KGI_S1_ANT02_95m_c10267034 | - | 323 | 256 | **RNA**→ RplB (COG0090)rplB (PRK09374)→ |
| env-134 | SA_S2_NOR18_50m_1258724 | + | 183 | 250 | **RNA**→ |
| env-135 | SA_S1_NOR05_45m_c10100002 | + | 326 | 393 | **RNA**→ rplB (PRK09374)Ribosomal_L2 (pfam00181)→ |
| Rsp-9-1 | NZ_AAYB01000008.1 | + | 62353 | 62420 | **RNA**→ RplB (COG0090)rplB (PRK09374)→ RplB (COG0090)rplB (PRK09374)→ RplB (COG0090)rplB (PRK09374)→ RplB (COG0090)rplB (PRK09374)→ RplB (COG0090)rplB (PRK09374)→ rpsS (PRK00357)RpsS (COG0185)→ rpsS (PRK00357)RpsS (COG0185)→ rplV (PRK00565)Ribosomal_L22 (pfam00237)→ rplV (PRK00565)Ribosomal_L22 (pfam00237)→ RpsC (COG0092)rpsC (PRK00310)→ RpsC (COG0092)rpsC (PRK00310)→ RplP (COG0197)rplP (PRK09203)→ RplP (COG0197)rplP (PRK09203)→ |
| env-136 | JGI24724J26744_10208358 | + | 330 | 396 | **RNA**→ RplB (COG0090)rplB (PRK09374)→ |
| env-137 | JGI24723J26617_10144364 | - | 304 | 238 | **RNA**→ RplB (COG0090)rplB (PRK09374)→ |
| Pan-1-1 | NZ_LGHU01000123.1 | + | 98116 | 98182 | **RNA**→ RplB (COG0090)rplB (PRK09374)→ rpsS (PRK00357)RpsS (COG0185)→ rplV (PRK00565)Ribosomal_L22 (pfam00237)→ RpsC (COG0092)rpsC (PRK00310)→ RplP (COG0197)rplP (PRK09203)→ |
| Pfe-1-1 | NZ_LGHV01000017.1 | - | 8115 | 8049 | **RNA**→ RplB (COG0090)rplB (PRK09374)→ rpsS (PRK00357)RpsS (COG0185)→ rplV (PRK00565)Ribosomal_L22 (pfam00237)→ RpsC (COG0092)rpsC (PRK00310)→ RplP (COG0197)rplP (PRK09203)→ |
| Pfe-2-1 | NZ_ATVN01000032.1 | - | 7903 | 7837 | **RNA**→ RplB (COG0090)rplB (PRK09374)→ rpsS (PRK00357)RpsS (COG0185)→ rplV (PRK00565)Ribosomal_L22 (pfam00237)→ RpsC (COG0092)rpsC (PRK00310)→ RplP (COG0197)rplP (PRK09203)→ |
| Lci-1-1 | NZ_BATB01000009.1 | - | 87402 | 87331 | **RNA**→ RplB (COG0090)rplB (PRK09374)→ rpsS (PRK00357)RpsS (COG0185)→ rplV (PRK00565)Ribosomal_L22 (pfam00237)→ RpsC (COG0092)rpsC (PRK00310)→ RpsC (COG0092)rpsC (PRK00310)→ hypo→ RplP (COG0197)rplP (PRK09203)→ |
| env-138 | JGI24724J26744_10487725 | + | 233 | 347 | **RNA**→ ←hypo |
| Abo-1-1 | NZ_JXYH01000044.1 | - | 31116 | 31026 | **RNA**→ RplB (COG0090)rplB (PRK09374)→ rpsS (PRK00357)RpsS (COG0185)→ rplV (PRK00565)Ribosomal_L22 (pfam00237)→ RpsC (COG0092)rpsC (PRK00310)→ RplP (COG0197)rplP (PRK09203)→ |
| Rha-1-1 | NZ_JALZ01000013.1 | + | 2420 | 2528 | **RNA**→ RplB (COG0090)rplB (PRK09374)→ rpsS (PRK00357)RpsS (COG0185)→ rplV (PRK00565)Ribosomal_L22 (pfam00237)→ RpsC (COG0092)rpsC (PRK00310)→ RplP (COG0197)rplP (PRK09203)→ hypo→ 2OG-FeII_Oxy_5 (pfam13759)→ |
| Mal-1-1 | NZ_JSUQ01000002.1 | - | 319288 | 319208 | **RNA**→ RplB (COG0090)rplB (PRK09374)→ rpsS (PRK00357)RpsS (COG0185)→ rplV (PRK00565)Ribosomal_L22 (pfam00237)→ RpsC (COG0092)rpsC (PRK00310)→ RplP (COG0197)rplP (PRK09203)→ |
| env-139 | JGI12075J13256_1001549 | + | 1682 | 1762 | **RNA**→ RplB (COG0090)rplB (PRK09374)→ rpsS (PRK00357)RpsS (COG0185)→ rplV (PRK00565)Ribosomal_L22 (pfam00237)→ |
| env-140 | JGI12112J13074_1001921 | - | 771 | 691 | **RNA**→ RplB (COG0090)rplB (PRK09374)→ |

| | | | | | |
|---|---|---|---|---|---|
| env-141 | JGI24724J26744_10066135 | - | 1234 | 1166 | **RNA**→ RplB (COG0090)rplB (PRK09374)→ rpsS (PRK00357)RpsS (COG0185)→ |
| env-142 | JGI24723J26617_10483647 | + | 129 | 197 | **RNA**→ RplB (COG0090)rplB (PRK09374)→ |
| env-143 | JCVI_SCAF_1101667320527 | - | 473 | 406 | **RNA**→ rplB (PRK09374)Ribosomal_L2 (pfam00181)→ |
| env-144 | APMI01028553.1 | - | 2484 | 2421 | **RNA**→ RplB (COG0090)rplB (PRK09374)→ rpsS (PRK00357)RpsS (COG0185)→<br>rplV (PRK00565)Ribosomal_L22 (pfam00237)→ RpsC (COG0092)rpsC (PRK00310)→<br>rplP (PRK09203)Ribosomal_L16 (pfam00252)→ |
| env-145 | JGI24724J26744_10205793 | - | 600 | 536 | **RNA**→ RplB (COG0090)rplB (PRK09374)→ |
| env-146 | JGI25321J39296_1078939 | - | 191 | 127 | **RNA**→ RplB (COG0090)rplB (PRK09374)→ |
| env-147 | JGI25321J50212_10013221 | - | 2584 | 2520 | **RNA**→ RplB (COG0090)rplB (PRK09374)→ rpsS (PRK00357)RpsS (COG0185)→<br>rplV (PRK00565)Ribosomal_L22 (pfam00237)→ RpsC (COG0092)rpsC (PRK00310)→<br>RplP (COG0197)rplP (PRK09203)→ |
| env-148 | JGI24025J20009_10017180 | + | 442 | 544 | **RNA**→ RplB (COG0090)rplB (PRK09374)→ rpsS (PRK00357)RpsS (COG0185)→<br>rplV (PRK00565)Ribosomal_L22 (pfam00237)→ RpsC (COG0092)rpsC (PRK00310)→<br>RplP (COG0197)rplP (PRK09203)→ |
| env-149 | ATLU01001954.1 | + | 1098 | 1219 | **RNA**→ rplB (PRK09374)Ribosomal_L2 (pfam00181)→ |
| env-150 | JGI12210J13797_10050283 | + | 46 | 137 | **RNA**→ RplB (COG0090)rplB (PRK09374)→ |
| env-151 | JGI12210J13797_10199298 | - | 104 | 13 | **RNA**→ |
| env-152 | JGI11876J14442_10101235 | - | 251 | 160 | **RNA**→ RplB (COG0090)rplB (PRK09374)→ |
| env-153 | JGI12109J20283_10360159 | - | 6078 | 5987 | **RNA**→ RplB (COG0090)rplB (PRK09374)→ rpsS (PRK00357)RpsS (COG0185)→<br>rplV (PRK00565)Ribosomal_L22 (pfam00237)→ RpsC (COG0092)rpsC (PRK00310)→<br>RplP (COG0197)rplP (PRK09203)→ |
| env-154 | JGI12109J20283_10360162 | + | 86 | 177 | **RNA**→ RplB (COG0090)rplB (PRK09374)→ rpsS (PRK00357)RpsS (COG0185)→<br>rplV (PRK00565)Ribosomal_L22 (pfam00237)→ rpsC (PRK00310)30S_S3_KH (cd02412)→ |
| env-155 | ElkS_mat_MD6A_1245876 | + | 17 | 108 | **RNA**→ RplB (COG0090)rplB (PRK09374)→ |
| env-156 | 3300001605_10196433 | + | 760 | 822 | **RNA**→ rplB (PRK09374)Ribosomal_L2 (pfam00181)→ |
| env-157 | JGI24723J26617_10002649 | - | 2300 | 2237 | **RNA**→ RplB (COG0090)rplB (PRK09374)→ rpsS (PRK00357)RpsS (COG0185)→<br>rplV (PRK00565)Ribosomal_L22 (pfam00237)→ RpsC (COG0092)rpsC (PRK00310)→ |
| env-158 | SA_S1_NOR05_45m_c10038069 | - | 1091 | 1029 | **RNA**→ RplB (COG0090)rplB (PRK09374)→ rpsS (PRK00357)Ribosomal_S19 (pfam00203)→ |
| env-159 | SA_S1_NOR08_45m_c10005006 | - | 2877 | 2815 | **RNA**→ RplB (COG0090)rplB (PRK09374)→ rpsS (PRK00357)RpsS (COG0185)→<br>rplV (PRK00565)Ribosomal_L22 (pfam00237)→ RpsC (COG0092)rpsC (PRK00310)→<br>RplP (COG0197)rplP (PRK09203)→ |
| env-160 | SA_S2_NOR15_50m_c10038301 | + | 1693 | 1755 | **RNA**→ rplB (PRK09374)Ribosomal_L2 (pfam00181)→ |
| env-161 | SA_S1_NOR02_45m_c1051392 | - | 97 | 35 | **RNA**→ |
| env-162 | KGI_S1_ANT01_95m_c10509536 | - | 192 | 130 | **RNA**→ RplB (COG0090)rplB (PRK09374)→ |
| Kvu-1-1 | NC_017384.1 | - | 2210425 | 2210363 | **RNA**→ RplB (COG0090)rplB (PRK09374)→ rpsS (PRK00357)RpsS (COG0185)→<br>rplV (PRK00565)Ribosomal_L22 (pfam00237)→ RpsC (COG0092)rpsC (PRK00310)→<br>RplP (COG0197)rplP (PRK09203)→ |
| Kvu-2-1 | NC_014625.1 | - | 2496662 | 2496600 | **RNA**→ RplB (COG0090)rplB (PRK09374)→ rpsS (PRK00357)RpsS (COG0185)→ |
| env-163 | JGI24723J26617_10010575 | - | 238 | 176 | **RNA**→ RplB (COG0090)rplB (PRK09374)→ |
| Lho-1-1 | NZ_KB822996.1 | + | 454151 | 454214 | **RNA**→ RplB (COG0090)rplB (PRK09374)→ rpsS (PRK00357)RpsS (COG0185)→<br>rplV (PRK00565)Ribosomal_L22 (pfam00237)→ RpsC (COG0092)rpsC (PRK00310)→<br>RplP (COG0197)rplP (PRK09203)→ |
| CRh-1-1 | NZ_LFTY01000002.1 | + | 138295 | 138372 | **RNA**→ RplB (COG0090)rplB (PRK09374)→ rpsS (PRK00357)RpsS (COG0185)→<br>rplV (PRK00565)Ribosomal_L22 (pfam00237)→ RpsC (COG0092)rpsC (PRK00310)→<br>RplP (COG0197)rplP (PRK09203)→ |
| env-164 | CETY01316910.1 | + | 295 | 359 | **RNA**→ rplB (PRK09374)Ribosomal_L2 (pfam00181)→ |
| env-165 | SI34jun09_120m_1114364 | + | 319 | 383 | **RNA**→ |
| Ogu-1-1 | NZ_AMGO01000054.1 | + | 2304 | 2368 | **RNA**→ RplB (COG0090)rplB (PRK09374)→ RplB (COG0090)rplB (PRK09374)→<br>rpsS (PRK00357)RpsS (COG0185)→ rpsS (PRK00357)RpsS (COG0185)→<br>rplV (PRK00565)Ribosomal_L22 (pfam00237)→ rplV (PRK00565)Ribosomal_L22 (pfam00237)→<br>RpsC (COG0092)rpsC (PRK00310)→ RpsC (COG0092)rpsC (PRK00310)→ RplP (COG0197)rplP (PRK09203)→<br>RplP (COG0197)rplP (PRK09203)→ |
| env-166 | _GX4IIKE02G7TTM | + | 176 | 245 | **RNA**→ RplB (COG0090)rplB (PRK09374)→ |
| Paq-1-1 | NZ_LGHS01000007.1 | - | 43447 | 43382 | **RNA**→ RplB (COG0090)rplB (PRK09374)→ rpsS (PRK00357)RpsS (COG0185)→<br>rplV (PRK00565)Ribosomal_L22 (pfam00237)→ RpsC (COG0092)rpsC (PRK00310)→<br>RplP (COG0197)rplP (PRK09203)→ |

| | | | | | |
|---|---|---|---|---|---|
| env-167 | CERE01101430.1 | + | 2524 | 2585 | RNA→ RplB (COG0090)rplB (PRK09374)→ rpsS (PRK00357)RpsS (COG0185)→ rplV (PRK00565)Ribosomal_L22 (pfam00237)→ RpsC (COG0092)rpsC (PRK00310)→ |
| env-168 | ATLU01000020.1 | + | 4112 | 4189 | RNA→ RplB (COG0090)rplB (PRK09374)→ rpsS (PRK00357)RpsS (COG0185)→ rplV (PRK00565)Ribosomal_L22 (pfam00237)→ RpsC (COG0092)rpsC (PRK00310)→ RplP (COG0197)rplP (PRK09203)→ |
| env-169 | CERH01017500.1 | - | 305 | 240 | RNA→ RplB (COG0090)rplB (PRK09374)→ |
| Rnu-1-1 | NZ_CH724156.1 | - | 347220 | 347155 | RNA→ RplB (COG0090)rplB (PRK09374)→ rpsS (PRK00357)RpsS (COG0185)→ RplV (COG0091)rplV (PRK00565)→ RpsC (COG0092)rpsC (PRK00310)→ |
| env-170 | CEQZ01016289.1 | - | 1777 | 1712 | RNA→ RplB (COG0090)rplB (PRK09374)→ rpsS (PRK00357)RpsS (COG0185)→ rplV (PRK00565)Ribosomal_L22 (pfam00237)→ rpsC (PRK00310)30S_S3_KH (cd02412)→ |
| env-171 | KVRMV2_100592709 | + | 54 | 115 | RNA→ RplB (COG0090)rplB (PRK09374)→ |
| env-172 | 2236876011_238237 | + | 256 | 328 | RNA→ ←hypo |
| Rba-4-1 | NZ_AAXZ01000001.1 | + | 629695 | 629767 | RNA→ RplB (COG0090)rplB (PRK09374)→ RplB (COG0090)rplB (PRK09374)→ rpsS (PRK00357)RpsS (COG0185)→ rpsS (PRK00357)RpsS (COG0185)→ rplV (PRK00565)Ribosomal_L22 (pfam00237)→ rplV (PRK00565)Ribosomal_L22 (pfam00237)→ RpsC (COG0092)rpsC (PRK00310)→ RpsC (COG0092)rpsC (PRK00310)→ RplP (COG0197)rplP (PRK09203)→ RplP (COG0197)rplP (PRK09203)→ |
| env-173 | JCVI_SCAF_1096628174387 | + | 532 | 596 | RNA→ RplB (COG0090)rplB (PRK09374)→ rpsC (PRK00310)Ribosomal_S3_C (pfam00189)→ RplP (COG0197)rplP (PRK09203)→ |
| Sps-1-1 | NZ_JAMD01000003.1 | - | 509244 | 509179 | RNA→ RplB (COG0090)rplB (PRK09374)→ rpsS (PRK00357)RpsS (COG0185)→ rplV (PRK00565)Ribosomal_L22 (pfam00237)→ RpsC (COG0092)rpsC (PRK00310)→ RplP (COG0197)rplP (PRK09203)→ |
| Ssp-2-1 | NZ_JIBC01000009.1 | + | 270672 | 270737 | RNA→ RplB (COG0090)rplB (PRK09374)→ rpsS (PRK00357)RpsS (COG0185)→ rplV (PRK00565)Ribosomal_L22 (pfam00237)→ RpsC (COG0092)rpsC (PRK00310)→ RplP (COG0197)rplP (PRK09203)→ |
| env-174 | JGI24723J26617_10018613 | + | 746 | 824 | RNA→ RplB (COG0090)rplB (PRK09374)→ rpsS (PRK00357)RpsS (COG0185)→ rplV (PRK00565)Ribosomal_L22 (pfam00237)→ RpsC (COG0092)rpsC (PRK00310)→ rplP (PRK09203)Ribosomal_L16 (pfam00252)→ |
| Osp-1-1 | NZ_JAFT01000005.1 | - | 653626 | 653563 | RNA→ RplB (COG0090)rplB (PRK09374)→ rpsS (PRK00357)RpsS (COG0185)→ rplV (PRK00565)Ribosomal_L22 (pfam00237)→ RpsC (COG0092)rpsC (PRK00310)→ RplP (COG0197)rplP (PRK09203)→ |
| Sst-1-1 | NZ_AAYA01000001.1 | + | 200930 | 200993 | RNA→ RplB (COG0090)rplB (PRK09374)→ RplB (COG0090)rplB (PRK09374)→ rpsS (PRK00357)RpsS (COG0185)→ rpsS (PRK00357)RpsS (COG0185)→ rplV (PRK00565)Ribosomal_L22 (pfam00237)→ rplV (PRK00565)Ribosomal_L22 (pfam00237)→ RpsC (COG0092)rpsC (PRK00310)→ RpsC (COG0092)rpsC (PRK00310)→ RplP (COG0197)rplP (PRK09203)→ RplP (COG0197)rplP (PRK09203)→ |
| env-175 | JCVI_READ_1103242074397 | + | 704 | 765 | RNA→ |
| env-176 | JGI24724J26744_10049941 | + | 488 | 552 | RNA→ RplB (COG0090)rplB (PRK09374)→ rpsS (PRK00357)RpsS (COG0185)→ hypo→ |
| Rsp-16-1 | NZ_ABCL01000006.1 | + | 350229 | 350292 | RNA→ RplB (COG0090)rplB (PRK09374)→ RplB (COG0090)rplB (PRK09374)→ rpsS (PRK00357)RpsS (COG0185)→ rpsS (PRK00357)RpsS (COG0185)→ rplV (PRK00565)Ribosomal_L22 (pfam00237)→ rplV (PRK00565)Ribosomal_L22 (pfam00237)→ RpsC (COG0092)rpsC (PRK00310)→ RpsC (COG0092)rpsC (PRK00310)→ RplP (COG0197)rplP (PRK09203)→ RplP (COG0197)rplP (PRK09203)→ |
| env-177 | JGI24724J26744_10021376 | + | 90 | 155 | RNA→ RplB (COG0090)rplB (PRK09374)→ rpsS (PRK00357)RpsS (COG0185)→ rplV (PRK00565)Ribosomal_L22 (pfam00237)→ RpsC (COG0092)rpsC (PRK00310)→ RplP (COG0197)rplP (PRK09203)→ |
| env-178 | 3300001592_10000630 | + | 2376 | 2441 | RNA→ RplB (COG0090)rplB (PRK09374)→ rpsS (PRK00357)RpsS (COG0185)→ rplV (PRK00565)Ribosomal_L22 (pfam00237)→ KH (smart00322)30S_S3_KH (cd02412)→ rpsC (PRK00310)Ribosomal_S3_C (pfam00189)→ RplP (COG0197)rplP (PRK09203)→ |
| env-179 | JGI25317J35169_1006515 | + | 816 | 879 | RNA→ RplB (COG0090)rplB (PRK09374)→ rpsS (PRK00357)RpsS (COG0185)→ rplV (PRK00565)Ribosomal_L22 (pfam00237)→ RpsC (COG0092)rpsC (PRK00310)→ RplP (COG0197)rplP (PRK09203)→ |
| env-180 | JGI25319J35699_1002819 | + | 816 | 879 | RNA→ RplB (COG0090)rplB (PRK09374)→ rpsS (PRK00357)RpsS (COG0185)→ rplV (PRK00565)Ribosomal_L22 (pfam00237)→ RpsC (COG0092)rpsC (PRK00310)→ RplP (COG0197)rplP (PRK09203)→ |

| | | | | | |
|---|---|---|---|---|---|
| env-181 | JGI25322J35698_1003836 | + | 3368 | 3431 | **RNA**→ RplB (COG0090)rplB (PRK09374)→ rpsS (PRK00357)RpsS (COG0185)→ rplV (PRK00565)Ribosomal_L22 (pfam00237)→ RpsC (COG0092)rpsC (PRK00310)→ RplP (COG0197)rplP (PRK09203)→ |
| env-182 | JGI25318J35512_1004712 | + | 816 | 879 | **RNA**→ RplB (COG0090)rplB (PRK09374)→ rpsS (PRK00357)RpsS (COG0185)→ rplV (PRK00565)Ribosomal_L22 (pfam00237)→ RpsC (COG0092)rpsC (PRK00310)→ RplP (COG0197)rplP (PRK09203)→ |
| env-183 | JGI25321J39296_1001477 | - | 10112 | 10049 | **RNA**→ RplB (COG0090)rplB (PRK09374)→ rpsS (PRK00357)RpsS (COG0185)→ rplV (PRK00565)Ribosomal_L22 (pfam00237)→ RpsC (COG0092)rpsC (PRK00310)→ RplP (COG0197)rplP (PRK09203)→ |
| env-184 | JGI25320J50211_1001516 | - | 2747 | 2684 | **RNA**→ RplB (COG0090)rplB (PRK09374)→ rpsS (PRK00357)RpsS (COG0185)→ rplV (PRK00565)Ribosomal_L22 (pfam00237)→ RpsC (COG0092)rpsC (PRK00310)→ RplP (COG0197)rplP (PRK09203)→ |
| env-185 | JGI25321J50212_10005715 | - | 4273 | 4210 | **RNA**→ RplB (COG0090)rplB (PRK09374)→ rpsS (PRK00357)RpsS (COG0185)→ rplV (PRK00565)Ribosomal_L22 (pfam00237)→ RpsC (COG0092)rpsC (PRK00310)→ RplP (COG0197)rplP (PRK09203)→ |
| env-186 | JGI24724J26744_10053916 | + | 237 | 300 | **RNA**→ RplB (COG0090)rplB (PRK09374)→ rpsS (PRK00357)RpsS (COG0185)→ rplV (PRK00565)Ribosomal_L22 (pfam00237)→ |
| Rsp-13-1 | NZ_CH902584.1 | + | 1991396 | 1991459 | **RNA**→ RplB (COG0090)rplB (PRK09374)→ rpsS (PRK00357)RpsS (COG0185)→ rplV (PRK00565)Ribosomal_L22 (pfam00237)→ RpsC (COG0092)rpsC (PRK00310)→ RplP (COG0197)rplP (PRK09203)→ |
| env-187 | CERE01077594.1 | + | 40 | 103 | **RNA**→ rplB (PRK09374)Ribosomal_L2 (pfam00181)→ |
| Rmu-1-1 | NZ_KN293980.1 | + | 258018 | 258081 | **RNA**→ RplB (COG0090)rplB (PRK09374)→ rpsS (PRK00357)RpsS (COG0185)→ rplV (PRK00565)Ribosomal_L22 (pfam00237)→ RpsC (COG0092)rpsC (PRK00310)→ RplP (COG0197)rplP (PRK09203)→ |
| env-188 | GSLSAS_GLMUJHB01ANH3P | - | 111 | 36 | **RNA**→ ←hypo |
| env-189 | PR_CR_10_Liq_1_inCR_1002975 | - | 4609 | 4543 | **RNA**→ RplB (COG0090)rplB (PRK09374)→ rpsS (PRK00357)RpsS (COG0185)→ rplV (PRK00565)Ribosomal_L22 (pfam00237)→ RpsC (COG0092)rpsC (PRK00310)→ RplP (COG0197)rplP (PRK09203)→ |
| env-190 | GSLSAS_contig20752 | + | 91 | 157 | **RNA**→ rplB (PRK09374)Ribosomal_L2 (pfam00181)→ |
| env-191 | GSLSAAI_GFP0ZZ301AT1AJ | + | 118 | 184 | **RNA**→ RplB (COG0090)rplB (PRK09374)→ |
| env-192 | P_2C_Liq_2_UnCty_1112442 | + | 202 | 268 | **RNA**→ RplB (COG0090)rplB (PRK09374)→ |
| env-193 | JCVI_SCAF_1101668363300 | - | 793 | 725 | **RNA**→ RplB (COG0090)rplB (PRK09374)→ |
| env-194 | CESM01031007.1 | + | 3318 | 3386 | **RNA**→ RplB (COG0090)rplB (PRK09374)→ rpsS (PRK00357)RpsS (COG0185)→ rplV (PRK00565)Ribosomal_L22 (pfam00237)→ rpsC (PRK00310)30S_S3_KH (cd02412)→ |
| env-195 | LPaug09P1610m_c1075736 | - | 346 | 278 | **RNA**→ rplB (PRK09374)Ribosomal_L2 (pfam00181)→ |
| env-196 | CENZ01075739.1 | - | 473 | 405 | **RNA**→ rplB (PRK09374)Ribosomal_L2 (pfam00181)→ |
| env-197 | JCVI_SCAF_1096627291439 | + | 336 | 404 | **RNA**→ RplB (COG0090)rplB (PRK09374)→ rpsS (PRK00357)RpsS (COG0185)→ |
| env-198 | CERM01028656.1 | + | 736 | 804 | **RNA**→ RplB (COG0090)rplB (PRK09374)→ |
| env-199 | CEWO01045537.1 | + | 245 | 313 | **RNA**→ rplB (PRK09374)Ribosomal_L2 (pfam00181)→ |
| env-200 | CEVX01073106.1 | + | 997 | 1065 | **RNA**→ RplB (COG0090)rplB (PRK09374)→ rpsS (PRK00357)RpsS (COG0185)→ rplV (PRK00565)Ribosomal_L22 (pfam00237)→ RpsC (COG0092)rpsC (PRK00310)→ RplP (COG0197)rplP (PRK09203)→ |
| env-201 | CEUB01117696.1 | + | 96 | 164 | **RNA**→ rplB (PRK09374)Ribosomal_L2 (pfam00181)→ |
| env-202 | CEUG01069584.1 | - | 338 | 270 | **RNA**→ rplB (PRK09374)Ribosomal_L2 (pfam00181)→ |
| env-203 | CEVF01073172.1 | - | 937 | 869 | **RNA**→ RplB (COG0090)rplB (PRK09374)→ |
| env-204 | CEQW01075709.1 | + | 117 | 185 | **RNA**→ rplB (PRK09374)Ribosomal_L2 (pfam00181)→ |
| env-205 | CESW01144144.1 | - | 3772 | 3704 | **RNA**→ RplB (COG0090)rplB (PRK09374)→ rpsS (PRK00357)RpsS (COG0185)→ rplV (PRK00565)Ribosomal_L22 (pfam00237)→ RpsC (COG0092)rpsC (PRK00310)→ RplP (COG0197)rplP (PRK09203)→ |
| env-206 | CERG01046692.1 | - | 1707 | 1639 | **RNA**→ RplB (COG0090)rplB (PRK09374)→ rpsS (PRK00357)RpsS (COG0185)→ rplV (PRK00565)Ribosomal_L22 (pfam00237)→ rpsC (PRK00310)30S_S3_KH (cd02412)→ |
| env-207 | CENJ01096940.1 | + | 1807 | 1875 | **RNA**→ RplB (COG0090)rplB (PRK09374)→ rpsS (PRK00357)RpsS (COG0185)→ rplV (PRK00565)Ribosomal_L22 (pfam00237)→ RpsC (COG0092)rpsC (PRK00310)→ RplP (COG0197)rplP (PRK09203)→ |
| env-208 | CESN01179177.1 | - | 635 | 567 | **RNA**→ RplB (COG0090)rplB (PRK09374)→ |

| | | | | | |
|---|---|---|---|---|---|
| env-209 | CESP01157305.1 | - | 3522 | 3454 | **RNA**→ RplB (COG0090)rplB (PRK09374)→ rpsS (PRK00357)RpsS (COG0185)→ rplV (PRK00565)Ribosomal_L22 (pfam00237)→ RpsC (COG0092)rpsC (PRK00310)→ RplP (COG0197)rplP (PRK09203)→ |
| env-210 | CEVU01049360.1 | + | 90 | 158 | **RNA**→ RplB (COG0090)rplB (PRK09374)→ rpsS (PRK00357)RpsS (COG0185)→ rplV (PRK00565)Ribosomal_L22 (pfam00237)→ RpsC (COG0092)rpsC (PRK00310)→ RplP (COG0197)rplP (PRK09203)→ |
| env-211 | CEVX01093179.1 | + | 444 | 512 | **RNA**→ |
| env-212 | CEWK01264123.1 | + | 1805 | 1873 | **RNA**→ ←hypo |
| env-213 | CENP01218092.1 | - | 190 | 122 | **RNA**→ ←hypo |
| env-214 | CEVP01021979.1 | + | 1007 | 1075 | **RNA**→ ←hypo |
| env-215 | JCVI_READ_1103242821391 | + | 471 | 539 | **RNA**→ RplB (COG0090)rplB (PRK09374)→ |
| env-216 | JCVI_SCAF_1101668667046 | + | 950 | 1018 | **RNA**→ rplB (PRK09374)Ribosomal_L2 (pfam00181)→ |
| env-217 | CETT01105554.1 | + | 279 | 347 | **RNA**→ rplB (PRK09374)Ribosomal_L2 (pfam00181)→ |
| env-218 | CEUU01047001.1 | + | 1170 | 1238 | **RNA**→ ←hypo |
| env-219 | CEPA01032629.1 | - | 488 | 420 | **RNA**→ rplB (PRK09374)Ribosomal_L2 (pfam00181)→ |
| env-220 | JGI24820J26691_1001055 | - | 5955 | 5887 | **RNA**→ RplB (COG0090)rplB (PRK09374)→ rpsS (PRK00357)RpsS (COG0185)→ rplV (PRK00565)Ribosomal_L22 (pfam00237)→ RpsC (COG0092)rpsC (PRK00310)→ RplP (COG0197)rplP (PRK09203)→ |
| env-221 | CEVE01064005.1 | + | 149 | 217 | **RNA**→ RplB (COG0090)rplB (PRK09374)→ rpsS (PRK00357)RpsS (COG0185)→ rplV (PRK00565)Ribosomal_L22 (pfam00237)→ rpsC (PRK00310)30S_S3_KH (cd02412)→ |
| env-222 | JCVI_SCAF_1096627794593 | - | 1348 | 1280 | **RNA**→ rplB (PRK09374)Ribosomal_L2 (pfam00181)→ |
| env-223 | JCVI_SCAF_1096627334841 | + | 695 | 763 | **RNA**→ RplB (COG0090)rplB (PRK09374)→ rpsS (PRK00357)RpsS (COG0185)→ rplV (PRK00565)Ribosomal_L22 (pfam00237)→ |
| env-224 | JCVI_SCAF_1101668673697 | + | 999 | 1067 | **RNA**→ rplB (PRK09374)Ribosomal_L2 (pfam00181)→ |
| env-225 | CEUQ01027845.1 | - | 417 | 349 | **RNA**→ rplB (PRK09374)Ribosomal_L2 (pfam00181)→ |
| env-226 | CEUK01022496.1 | + | 194 | 262 | **RNA**→ rplB (PRK09374)Ribosomal_L2 (pfam00181)→ |
| env-227 | JCVI_SCAF_1101668339934 | - | 213 | 145 | **RNA**→ |
| env-228 | 2236876001_088379 | + | 363 | 434 | **RNA**→ RplB (COG0090)rplB (PRK09374)→ |
| env-229 | JCVI_READ_1103769037343 | + | 211 | 279 | **RNA**→ RplB (COG0090)rplB (PRK09374)→ |
| env-230 | JCVI_READ_1103769614257 | + | 956 | 1021 | **RNA**→ ←hypo |
| env-231 | CEWK01092565.1 | + | 421 | 489 | **RNA**→ |
| env-232 | CENQ01040580.1 | + | 79 | 147 | **RNA**→ rplB (PRK09374)Ribosomal_L2 (pfam00181)→ |
| env-233 | CEVN01064966.1 | - | 373 | 305 | **RNA**→ rplB (PRK09374)Ribosomal_L2 (pfam00181)→ |
| env-234 | CEPL01019978.1 | + | 61 | 129 | **RNA**→ hypo→ rplB (PRK09374)Ribosomal_L2 (pfam00181)→ |
| env-235 | CEPU01001285.1 | + | 124 | 192 | **RNA**→ RplB (COG0090)rplB (PRK09374)→ |
| env-236 | JCVI_READ_1103769096640 | - | 624 | 556 | **RNA**→ RplB (COG0090)rplB (PRK09374)→ |
| env-237 | CENO01039587.1 | - | 4337 | 4269 | **RNA**→ RplB (COG0090)rplB (PRK09374)→ rpsS (PRK00357)RpsS (COG0185)→ rplV (PRK00565)Ribosomal_L22 (pfam00237)→ RpsC (COG0092)rpsC (PRK00310)→ RplP (COG0197)rplP (PRK09203)→ |
| env-238 | CENW01212150.1 | - | 201 | 133 | **RNA**→ |
| env-239 | CEOJ01093429.1 | - | 167 | 99 | **RNA**→ |
| env-240 | CEOS01129456.1 | + | 419 | 487 | **RNA**→ RplB (COG0090)rplB (PRK09374)→ |
| env-241 | CEOV01114312.1 | - | 112 | 44 | **RNA**→ |
| env-242 | CEOW01046858.1 | + | 2221 | 2289 | **RNA**→ RplB (COG0090)rplB (PRK09374)→ rpsS (PRK00357)RpsS (COG0185)→ rplV (PRK00565)Ribosomal_L22 (pfam00237)→ |
| env-243 | CEPB01039107.1 | + | 1192 | 1260 | **RNA**→ RplB (COG0090)rplB (PRK09374)→ rpsS (PRK00357)RpsS (COG0185)→ rplV (PRK00565)Ribosomal_L22 (pfam00237)→ RpsC (COG0092)rpsC (PRK00310)→ RplP (COG0197)rplP (PRK09203)→ |
| env-244 | CEPC01014332.1 | + | 2318 | 2386 | **RNA**→ RplB (COG0090)rplB (PRK09374)→ rpsS (PRK00357)RpsS (COG0185)→ |
| env-245 | CEVG01269443.1 | + | 380 | 448 | **RNA**→ rplB (PRK09374)Ribosomal_L2 (pfam00181)→ |
| env-246 | CEVY01099477.1 | + | 2626 | 2694 | **RNA**→ RplB (COG0090)rplB (PRK09374)→ rpsS (PRK00357)RpsS (COG0185)→ rplV (PRK00565)Ribosomal_L22 (pfam00237)→ RpsC (COG0092)rpsC (PRK00310)→ RplP (COG0197)rplP (PRK09203)→ |
| env-247 | CEQD01048550.1 | - | 1102 | 1034 | **RNA**→ RplB (COG0090)rplB (PRK09374)→ rpsS (PRK00357)RpsS (COG0185)→ |

| | | | | | |
|---|---|---|---|---|---|
| env-248 | CEOC01035558.1 | + | 378 | 446 | **RNA**→ rplB (PRK09374)Ribosomal_L2 (pfam00181)→ |
| env-249 | CEQM01116065.1 | + | 207 | 275 | **RNA**→ rplB (PRK09374)Ribosomal_L2 (pfam00181)→ |
| env-250 | CEWA01125852.1 | - | 329 | 261 | **RNA**→ rplB (PRK09374)Ribosomal_L2 (pfam00181)→ |
| env-251 | CEWO01164779.1 | - | 401 | 333 | **RNA**→ rplB (PRK09374)Ribosomal_L2 (pfam00181)→ |
| env-252 | CENG01071547.1 | - | 1142 | 1074 | **RNA**→ RplB (COG0090)rplB (PRK09374)→ rpsS (PRK00357)Ribosomal_S19 (pfam00203)→ |
| env-253 | CENJ01189867.1 | - | 96 | 28 | **RNA**→ |
| env-254 | CENU01109952.1 | + | 428 | 496 | **RNA**→ rplB (PRK09374)Ribosomal_L2 (pfam00181)→ |
| env-255 | CEVW01291667.1 | - | 97 | 29 | **RNA**→ |
| env-256 | CAM_READ_0234711973 | + | 389 | 459 | **RNA**→ ←hypo |
| env-257 | JCVI_SCAF_1096626302357 | - | 446 | 378 | **RNA**→ rplB (PRK09374)Ribosomal_L2 (pfam00181)→ |
| env-258 | CESA01208526.1 | + | 115 | 183 | **RNA**→ RplB (COG0090)rplB (PRK09374)→ |
| env-259 | JGI24723J26617_10000490 | - | 34468 | 34404 | **RNA**→ RplB (COG0090)rplB (PRK09374)→ rpsS (PRK00357)RpsS (COG0185)→ |
| | | | | | rplV (PRK00565)Ribosomal_L22 (pfam00237)→ RpsC (COG0092)rpsC (PRK00310)→ |
| | | | | | RplP (COG0197)rplP (PRK09203)→ |
| Rsp-8-1 | NZ_ABCR01000011.1 | + | 18945 | 19019 | **RNA**→ RplB (COG0090)rplB (PRK09374)→ RplB (COG0090)rplB (PRK09374)→ |
| | | | | | rpsS (PRK00357)RpsS (COG0185)→ rpsS (PRK00357)RpsS (COG0185)→ |
| | | | | | rplV (PRK00565)Ribosomal_L22 (pfam00237)→ rplV (PRK00565)Ribosomal_L22 (pfam00237)→ |
| | | | | | RpsC (COG0092)rpsC (PRK00310)→ RpsC (COG0092)rpsC (PRK00310)→ RplP (COG0197)rplP (PRK09203)→ |
| | | | | | RplP (COG0197)rplP (PRK09203)→ |
| env-260 | CENP01298067.1 | - | 497 | 434 | **RNA**→ rplB (PRK09374)Ribosomal_L2 (pfam00181)→ |
| env-261 | SI34jun09_10m_1176238 | - | 203 | 140 | **RNA**→ RplB (COG0090)rplB (PRK09374)→ |
| env-262 | CESO01033516.1 | - | 891 | 828 | **RNA**→ RplB (COG0090)rplB (PRK09374)→ |
| env-263 | 2236876002_126045 | - | 189 | 126 | **RNA**→ RplB (COG0090)rplB (PRK09374)→ |
| env-264 | JGI20151J14362_10250213 | - | 362 | 299 | **RNA**→ rplB (PRK09374)Ribosomal_L2 (pfam00181)→ |
| env-265 | CENZ01260479.1 | + | 344 | 407 | **RNA**→ RplB (COG0090)rplB (PRK09374)→ |
| env-266 | 2236876002_106467 | - | 167 | 104 | **RNA**→ RplB (COG0090)rplB (PRK09374)→ |
| env-267 | CENT01014152.1 | + | 496 | 559 | **RNA**→ RplB (COG0090)rplB (PRK09374)→ rpsS (PRK00357)RpsS (COG0185)→ |
| | | | | | rplV (PRK00565)Ribosomal_L22 (pfam00237)→ RpsC (COG0092)rpsC (PRK00310)→ |
| | | | | | RplP (COG0197)rplP (PRK09203)→ |
| env-268 | CERM01014689.1 | - | 2838 | 2775 | **RNA**→ RplB (COG0090)rplB (PRK09374)→ rpsS (PRK00357)RpsS (COG0185)→ |
| | | | | | rplV (PRK00565)Ribosomal_L22 (pfam00237)→ RpsC (COG0092)rpsC (PRK00310)→ |
| | | | | | RplP (COG0197)rplP (PRK09203)→ |
| env-269 | CENN01049021.1 | + | 1200 | 1263 | **RNA**→ rplB (PRK09374)Ribosomal_L2 (pfam00181)→ |
| env-270 | CESS01003967.1 | - | 3048 | 2985 | **RNA**→ RplB (COG0090)rplB (PRK09374)→ rpsS (PRK00357)RpsS (COG0185)→ |
| | | | | | rplV (PRK00565)Ribosomal_L22 (pfam00237)→ RpsC (COG0092)rpsC (PRK00310)→ |
| | | | | | RplP (COG0197)rplP (PRK09203)→ |
| env-271 | JGI20158J14315_10236147 | - | 342 | 279 | **RNA**→ rplB (PRK09374)Ribosomal_L2 (pfam00181)→ |
| env-272 | DelMOWin2010_c10431137 | + | 54 | 117 | **RNA**→ RplB (COG0090)rplB (PRK09374)→ |
| env-273 | JCVI_SCAF_1096628356459 | - | 1073 | 1010 | **RNA**→ rpsS (PRK00357)RpsS (COG0185)→ rplV (PRK00565)Ribosomal_L22 (pfam00237)→ |
| env-274 | JCVI_READ_1103769101289 | - | 925 | 866 | **RNA**→ RplB (COG0090)rplB (PRK09374)→ |
| env-275 | JCVI_SCAF_1096627271074 | + | 1225 | 1289 | **RNA**→ RplB (COG0090)rplB (PRK09374)→ rpsS (PRK00357)RpsS (COG0185)→ |
| env-276 | JCVI_READ_1108799738541 | - | 650 | 586 | **RNA**→ RplB (COG0090)rplB (PRK09374)→ |
| env-277 | JCVI_SCAF_1096627976978 | + | 67 | 130 | **RNA**→ RplB (COG0090)rplB (PRK09374)→ |
| env-278 | CEWK01197169.1 | - | 120 | 57 | **RNA**→ |
| env-279 | CESM01101386.1 | - | 277 | 213 | **RNA**→ ←hypo |
| env-280 | CEUH01138205.1 | + | 837 | 901 | **RNA**→ |
| env-281 | CEWG01043024.1 | + | 1807 | 1871 | **RNA**→ RplB (COG0090)rplB (PRK09374)→ |
| env-282 | CENF01097607.1 | + | 586 | 650 | **RNA**→ |
| env-283 | CEUG01060018.1 | - | 123 | 59 | **RNA**→ |
| env-284 | CEQW01039727.1 | - | 82 | 18 | **RNA**→ |
| env-285 | JGI24820J26691_1324391 | + | 37 | 101 | **RNA**→ ←hypo |
| env-286 | CESM01340684.1 | - | 97 | 33 | **RNA**→ |
| env-287 | JCVI_SCAF_1096627811062 | - | 972 | 908 | **RNA**→ RplB (COG0090)rplB (PRK09374)→ |
| env-288 | JGI24820J26691_1087074 | + | 510 | 574 | **RNA**→ hypo→ |

| | | | | | |
|---|---|---|---|---|---|
| env-289 | JCVI_SCAF_1096627081604 | - | 1775 | 1711 | **RNA**→ RplB (COG0090)rplB (PRK09374)→ rpsS (PRK00357)RpsS (COG0185)→ rplV (PRK00565)Ribosomal_L22 (pfam00237)→ rpsC (PRK00310)30S_S3_KH (cd02412)→ |
| env-290 | JCVI_SCAF_1096627977260 | + | 343 | 407 | **RNA**→ rplB (PRK09374)Ribosomal_L2 (pfam00181)→ |
| env-291 | CEPJ01112005.1 | + | 720 | 784 | **RNA**→ rplB (PRK09374)Ribosomal_L2 (pfam00181)→ |
| env-292 | CAM_READ_0232770725 | - | 286 | 222 | **RNA**→ RplB (COG0090)rplB (PRK09374)→ |
| env-293 | CAM_READ_0232288387 | - | 248 | 184 | **RNA**→ hypo→ |
| env-294 | JCVI_READ_1108800192449 | + | 288 | 352 | **RNA**→ rplB (PRK09374)Ribosomal_L2 (pfam00181)→ |
| env-295 | JCVI_SCAF_1096627092934 | + | 576 | 640 | **RNA**→ RplB (COG0090)rplB (PRK09374)→ |
| env-296 | JCVI_SCAF_1096627099260 | + | 352 | 416 | **RNA**→ RplB (COG0090)rplB (PRK09374)→ rpsS (PRK00357)RpsS (COG0185)→ rplV (PRK00565)Ribosomal_L22 (pfam00237)→ rpsC (PRK00310)30S_S3_KH (cd02412)→ |
| env-297 | JGI24820J26691_1028973 | + | 1055 | 1119 | **RNA**→ |
| env-298 | CEUV01003215.1 | + | 768 | 832 | **RNA**→ |
| env-299 | CEUQ01156669.1 | - | 85 | 21 | **RNA**→ |
| env-300 | JCVI_SCAF_1101667251435 | - | 166 | 102 | **RNA**→ hypo→ |
| env-301 | CENF01068770.1 | - | 88 | 24 | **RNA**→ |
| env-302 | CEOM01237289.1 | + | 465 | 529 | **RNA**→ |
| env-303 | CERW01142606.1 | + | 975 | 1039 | **RNA**→ |
| env-304 | CEVS01036929.1 | - | 88 | 24 | **RNA**→ |
| env-305 | CEVU01131676.1 | + | 1938 | 2002 | **RNA**→ |
| env-306 | CEVX01017747.1 | - | 2836 | 2772 | **RNA**→ RplB (COG0090)rplB (PRK09374)→ rpsS (PRK00357)RpsS (COG0185)→ rplV (PRK00565)Ribosomal_L22 (pfam00237)→ RpsC (COG0092)rpsC (PRK00310)→ RplP (COG0197)rplP (PRK09203)→ |
| env-307 | CEWO01083412.1 | + | 3433 | 3497 | **RNA**→ RplB (COG0090)rplB (PRK09374)→ |
| env-308 | JCVI_SCAF_1096627863534 | - | 717 | 653 | **RNA**→ RplB (COG0090)rplB (PRK09374)→ |
| env-309 | JCVI_SCAF_1101668429250 | + | 407 | 471 | **RNA**→ RplB (COG0090)rplB (PRK09374)→ |
| env-310 | JCVI_SCAF_1101668671537 | - | 1477 | 1413 | **RNA**→ rplB (PRK09374)Ribosomal_L2 (pfam00181)→ rplB (PRK09374)Ribosomal_L2_C (pfam03947)→ rpsS (PRK00357)RpsS (COG0185)→ rplV (PRK00565)Ribosomal_L22 (pfam00237)→ |
| env-311 | CETT01040907.1 | - | 3530 | 3466 | **RNA**→ RplB (COG0090)rplB (PRK09374)→ rpsS (PRK00357)RpsS (COG0185)→ rplV (PRK00565)Ribosomal_L22 (pfam00237)→ RpsC (COG0092)rpsC (PRK00310)→ RplP (COG0197)rplP (PRK09203)→ |
| env-312 | JCVI_READ_1103242680220 | - | 329 | 265 | **RNA**→ RplB (COG0090)rplB (PRK09374)→ |
| env-313 | JCVI_SCAF_1101668723498 | + | 490 | 554 | **RNA**→ RplB (COG0090)rplB (PRK09374)→ |
| env-314 | JCVI_READ_1108800250675 | - | 813 | 749 | **RNA**→ RplB (COG0090)rplB (PRK09374)→ |
| env-315 | JCVI_SCAF_1101667072466 | - | 232 | 168 | **RNA**→ |
| env-316 | JCVI_READ_1104230004824 | + | 69 | 133 | **RNA**→ RplB (COG0090)rplB (PRK09374)→ |
| env-317 | CEPU01102359.1 | - | 315 | 251 | **RNA**→ RplB (COG0090)rplB (PRK09374)→ |
| env-318 | JCVI_READ_1104230151507 | - | 139 | 75 | **RNA**→ |
| env-319 | JCVI_SCAF_1096626924486 | - | 857 | 793 | **RNA**→ RplB (COG0090)rplB (PRK09374)→ |
| env-320 | CEQD01002240.1 | + | 207 | 271 | **RNA**→ RplB (COG0090)rplB (PRK09374)→ |
| env-321 | JCVI_SCAF_1101668260309 | - | 1155 | 1091 | **RNA**→ RplB (COG0090)rplB (PRK09374)→ rpsS (PRK00357)RpsS (COG0185)→ |
| env-322 | CEWK01132454.1 | + | 1102 | 1166 | **RNA**→ RplB (COG0090)rplB (PRK09374)→ rpsS (PRK00357)RpsS (COG0185)→ rplV (PRK00565)Ribosomal_L22 (pfam00237)→ RpsC (COG0092)rpsC (PRK00310)→ RplP (COG0197)rplP (PRK09203)→ |
| env-323 | CETB01173647.1 | + | 105 | 169 | **RNA**→ rplB (PRK09374)Ribosomal_L2 (pfam00181)→ |
| env-324 | CENY01053602.1 | + | 393 | 457 | **RNA**→ rplB (PRK09374)Ribosomal_L2 (pfam00181)→ |
| env-325 | JCVI_READ_1108799747928 | + | 140 | 204 | **RNA**→ RplB (COG0090)rplB (PRK09374)→ |
| env-326 | JCVI_SCAF_1096627704005 | - | 1325 | 1261 | **RNA**→ RplB (COG0090)rplB (PRK09374)→ RplB (COG0090)rplB (PRK09374)→ rpsS (PRK00357)RpsS (COG0185)→ |
| env-327 | JCVI_SCAF_1101668170846 | - | 710 | 646 | **RNA**→ RplB (COG0090)rplB (PRK09374)→ |
| env-328 | CEVY01136175.1 | + | 444 | 508 | **RNA**→ |
| env-329 | CAM_READ_0072915149 | + | 365 | 429 | **RNA**→ |
| env-330 | JCVI_READ_1103180852614 | - | 134 | 70 | **RNA**→ |
| env-331 | CETS01117025.1 | + | 341 | 405 | **RNA**→ |
| env-332 | CEUJ01061937.1 | - | 283 | 219 | **RNA**→ RplB (COG0090)rplB (PRK09374)→ |

| | | | | | |
|---|---|---|---|---|---|
| env-333 | CEVE01027800.1 | - | 84 | 20 | **RNA**→ |
| env-334 | CEWJ01004911.1 | - | 89 | 25 | **RNA**→ |
| env-335 | CEWH01205319.1 | + | 544 | 608 | **RNA**→ |
| env-336 | JCVI_SCAF_1096627835679 | - | 1455 | 1391 | **RNA**→ RplB (COG0090)rplB (PRK09374)→ RplB (COG0090)rplB (PRK09374)→ rpsS (PRK00357)RpsS (COG0185)→ rplV (PRK00565)Ribosomal_L22 (pfam00237)→ |
| env-337 | JCVI_SCAF_1096627978139 | - | 713 | 649 | **RNA**→ RplB (COG0090)rplB (PRK09374)→ rpsS (PRK00357)RpsS (COG0185)→ |
| env-338 | JCVI_SCAF_1096628267383 | + | 720 | 784 | **RNA**→ rplV (PRK00565)Ribosomal_L22 (pfam00237)→ RpsC (COG0092)rpsC (PRK00310)→ |
| env-339 | JCVI_SCAF_1096627226171 | - | 356 | 292 | **RNA**→ rplB (PRK09374)Ribosomal_L2 (pfam00181)→ |
| env-340 | CESB01055415.1 | + | 1248 | 1312 | **RNA**→ RplB (COG0090)rplB (PRK09374)→ |
| env-341 | JCVI_SCAF_1101668333168 | + | 132 | 196 | **RNA**→ RplB (COG0090)rplB (PRK09374)→ rpsS (PRK00357)RpsS (COG0185)→ rplV (PRK00565)Ribosomal_L22 (pfam00237)→ |
| env-342 | CEVZ01074731.1 | + | 606 | 670 | **RNA**→ |
| env-343 | JCVI_READ_1108829670529 | - | 825 | 762 | **RNA**→ RplB (COG0090)rplB (PRK09374)→ |
| env-344 | JCVI_SCAF_1101668663947 | - | 95 | 31 | **RNA**→ |
| env-345 | JCVI_READ_1103242633429 | + | 739 | 803 | **RNA**→ |
| env-346 | JCVI_SCAF_1101668342543 | - | 593 | 529 | **RNA**→ RplB (COG0090)rplB (PRK09374)→ |
| env-347 | CENF01127799.1 | + | 676 | 740 | **RNA**→ |
| env-348 | CEQO01040664.1 | - | 501 | 437 | **RNA**→ rplB (PRK09374)Ribosomal_L2 (pfam00181)→ |
| env-349 | CEQW01046424.1 | + | 1679 | 1743 | **RNA**→ ←hypo |
| env-350 | CEQY01100769.1 | - | 387 | 323 | **RNA**→ rplB (PRK09374)Ribosomal_L2 (pfam00181)→ |
| env-351 | CERR01085567.1 | + | 474 | 538 | **RNA**→ |
| env-352 | CETI01093028.1 | - | 91 | 27 | **RNA**→ |
| env-353 | CEVU01103598.1 | - | 85 | 21 | **RNA**→ |
| env-354 | CEVX01107346.1 | + | 461 | 525 | **RNA**→ |
| env-355 | CEWB01023124.1 | + | 120 | 184 | **RNA**→ rplB (PRK09374)Ribosomal_L2 (pfam00181)→ |
| env-356 | JCVI_SCAF_1096627339121 | - | 1737 | 1673 | **RNA**→ RplB (COG0090)rplB (PRK09374)→ rpsS (PRK00357)RpsS (COG0185)→ rplV (PRK00565)Ribosomal_L22 (pfam00237)→ rpsC (PRK00310)30S_S3_KH (cd02412)→ |
| env-357 | JCVI_SCAF_1101667362029 | + | 16 | 80 | **RNA**→ rplB (PRK09374)Ribosomal_L2 (pfam00181)→ |
| env-358 | CETP01093824.1 | + | 77 | 141 | **RNA**→ RplB (COG0090)rplB (PRK09374)→ |
| env-359 | CEUU01096110.1 | - | 95 | 31 | **RNA**→ |
| env-360 | JCVI_SCAF_1101668661533 | - | 618 | 555 | **RNA**→ RplB (COG0090)rplB (PRK09374)→ |
| env-361 | JCVI_SCAF_1101668691223 | - | 229 | 166 | **RNA**→ RplB (COG0090)rplB (PRK09374)→ |
| Psp-3-1 | NZ_LDPY01000001.1 | + | 1628103 | 1628171 | **RNA**→ RplB (COG0090)rplB (PRK09374)→ rpsS (PRK00357)RpsS (COG0185)→ rplV (PRK00565)Ribosomal_L22 (pfam00237)→ RpsC (COG0092)rpsC (PRK00310)→ RplP (COG0197)rplP (PRK09203)→ |
| Hmi-1-1 | NZ_JFGS01000025.1 | - | 34255 | 34184 | **RNA**→ RplB (COG0090)rplB (PRK09374)→ rpsS (PRK00357)RpsS (COG0185)→ rplV (PRK00565)Ribosomal_L22 (pfam00237)→ RpsC (COG0092)rpsC (PRK00310)→ RplP (COG0197)rplP (PRK09203)→ MhpC (COG0596)Abhydrolase_6 (pfam12697)→ MhpC (COG0596)Abhydrolase_6 (pfam12697)→ |
| Hma-1-1 | NZ_JGYG01000018.1 | - | 57450 | 57378 | **RNA**→ RplB (COG0090)rplB (PRK09374)→ rpsS (PRK00357)RpsS (COG0185)→ rplV (PRK00565)Ribosomal_L22 (pfam00237)→ RpsC (COG0092)rpsC (PRK00310)→ RplP (COG0197)rplP (PRK09203)→ MhpC (COG0596)Abhydrolase_6 (pfam12697)→ |
| Laq-1-1 | NZ_KI421504.1 | + | 1151633 | 1151741 | **RNA**→ RplB (COG0090)rplB (PRK09374)→ rpsS (PRK00357)RpsS (COG0185)→ rplV (PRK00565)Ribosomal_L22 (pfam00237)→ RpsC (COG0092)rpsC (PRK00310)→ RplP (COG0197)rplP (PRK09203)→ |
| env-362 | JGI24723J26617_10003411 | - | 7982 | 7888 | **RNA**→ RplB (COG0090)rplB (PRK09374)→ rpsS (PRK00357)RpsS (COG0185)→ rplV (PRK00565)Ribosomal_L22 (pfam00237)→ RpsC (COG0092)rpsC (PRK00310)→ RplP (COG0197)rplP (PRK09203)→ |
| Rpo-1-1 | NC_003911.12 | + | 487501 | 487565 | **RNA**→ RplB (COG0090)rplB (PRK09374)→ rpsS (PRK00357)RpsS (COG0185)→ rplV (PRK00565)Ribosomal_L22 (pfam00237)→ RpsC (COG0092)rpsC (PRK00310)→ RplP (COG0197)rplP (PRK09203)→ |
| env-363 | CETY01001211.1 | - | 2221 | 2155 | **RNA**→ RplB (COG0090)rplB (PRK09374)→ rpsS (PRK00357)RpsS (COG0185)→ rplV (PRK00565)Ribosomal_L22 (pfam00237)→ RpsC (COG0092)rpsC (PRK00310)→ |
| env-364 | JGI24023J19991_10525132 | - | 195 | 105 | **RNA**→ RplB (COG0090)rplB (PRK09374)→ |

| | | | | | |
|---|---|---|---|---|---|
| Sla-1-1 | NZ_GG704596.1 | + | 1508433 | 1508499 | **RNA**→ RplB (COG0090)rplB (PRK09374)→ RplB (COG0090)rplB (PRK09374)→ rpsS (PRK00357)RpsS (COG0185)→ rpsS (PRK00357)RpsS (COG0185)→ rplV (PRK00565)Ribosomal_L22 (pfam00237)→ rplV (PRK00565)Ribosomal_L22 (pfam00237)→ RpsC (COG0092)rpsC (PRK00310)→ RpsC (COG0092)rpsC (PRK00310)→ RplP (COG0197)rplP (PRK09203)→ RplP (COG0197)rplP (PRK09203)→ |
| Rha-2-1 | NZ_JQEZ01000002.1 | - | 890631 | 890567 | **RNA**→ RplB (COG0090)rplB (PRK09374)→ rpsS (PRK00357)RpsS (COG0185)→ rplV (PRK00565)Ribosomal_L22 (pfam00237)→ RpsC (COG0092)rpsC (PRK00310)→ RplP (COG0197)rplP (PRK09203)→ |
| Rsp-17-1 | NZ_LGXZ01000018.1 | - | 15524 | 15460 | **RNA**→ RplB (COG0090)rplB (PRK09374)→ rpsS (PRK00357)RpsS (COG0185)→ rplV (PRK00565)Ribosomal_L22 (pfam00237)→ RpsC (COG0092)rpsC (PRK00310)→ RplP (COG0197)rplP (PRK09203)→ |
| Rsp-18-1 | NZ_JWLJ01000003.1 | + | 9098 | 9163 | **RNA**→ RplB (COG0090)rplB (PRK09374)→ rpsS (PRK00357)RpsS (COG0185)→ rplV (PRK00565)Ribosomal_L22 (pfam00237)→ RpsC (COG0092)rpsC (PRK00310)→ RplP (COG0197)rplP (PRK09203)→ |
| Rsp-19-1 | NZ_JWLK01000001.1 | + | 450619 | 450684 | **RNA**→ RplB (COG0090)rplB (PRK09374)→ rpsS (PRK00357)RpsS (COG0185)→ rplV (PRK00565)Ribosomal_L22 (pfam00237)→ RpsC (COG0092)rpsC (PRK00310)→ RplP (COG0197)rplP (PRK09203)→ |
| Rsp-22-1 | NZ_AEYW01000001.1 | + | 6172 | 6237 | **RNA**→ RplB (COG0090)rplB (PRK09374)→ RplB (COG0090)rplB (PRK09374)→ rpsS (PRK00357)RpsS (COG0185)→ rpsS (PRK00357)RpsS (COG0185)→ rplV (PRK00565)Ribosomal_L22 (pfam00237)→ rplV (PRK00565)Ribosomal_L22 (pfam00237)→ RpsC (COG0092)rpsC (PRK00310)→ RpsC (COG0092)rpsC (PRK00310)→ RplP (COG0197)rplP (PRK09203)→ RplP (COG0197)rplP (PRK09203)→ |
| Rba-2-1 | NZ_DS999531.1 | + | 702885 | 702950 | **RNA**→ RplB (COG0090)rplB (PRK09374)→ RplB (COG0090)rplB (PRK09374)→ rpsS (PRK00357)RpsS (COG0185)→ rpsS (PRK00357)RpsS (COG0185)→ rplV (PRK00565)Ribosomal_L22 (pfam00237)→ rplV (PRK00565)Ribosomal_L22 (pfam00237)→ RpsC (COG0092)rpsC (PRK00310)→ RpsC (COG0092)rpsC (PRK00310)→ RplP (COG0197)rplP (PRK09203)→ RplP (COG0197)rplP (PRK09203)→ |
| env-365 | KGI_S1_ANT01_95m_c10129328 | + | 275 | 337 | **RNA**→ rplB (PRK09374)Ribosomal_L2 (pfam00181)→ |
| env-366 | KGI_S1_ANT02_95m_c10007513 | + | 664 | 726 | **RNA**→ RplB (COG0090)rplB (PRK09374)→ rpsS (PRK00357)RpsS (COG0185)→ rplV (PRK00565)Ribosomal_L22 (pfam00237)→ RpsC (COG0092)rpsC (PRK00310)→ RplP (COG0197)rplP (PRK09203)→ hypo→ M20_Acy1 (cd03886)amidohydrolases (TIGR01891)→ |
| env-367 | KGI_S2_ANT04_2345m_c1167548 | + | 316 | 378 | **RNA**→ ←hypo |
| env-368 | JGI24723J26617_10802083 | + | 118 | 180 | **RNA**→ rplW (PRK05738)→ |
| env-369 | JGI12109J20283_10872926 | - | 175 | 77 | **RNA**→ hypo→ |
| env-370 | JGI12109J20283_10006735 | - | 753 | 682 | **RNA**→ RplB (COG0090)rplB (PRK09374)→ |
| env-371 | JGI12109J20283_10006736 | + | 78 | 149 | **RNA**→ rplB (PRK09374)Ribosomal_L2 (pfam00181)→ |
| env-372 | HLSNC_1001521 | - | 9879 | 9807 | **RNA**→ RplB (COG0090)rplB (PRK09374)→ rpsS (PRK00357)RpsS (COG0185)→ rplV (PRK00565)Ribosomal_L22 (pfam00237)→ RpsC (COG0092)rpsC (PRK00310)→ RplP (COG0197)rplP (PRK09203)→ QdoI (COG1917)DMSP_lyase (pfam16867)→ 2OG-FeII_Oxy_5 (pfam13759)→ |
| env-373 | KGI_S2_ANT06_2345m_c1009878 | - | 2821 | 2753 | **RNA**→ RplB (COG0090)rplB (PRK09374)→ rpsS (PRK00357)RpsS (COG0185)→ rplV (PRK00565)Ribosomal_L22 (pfam00237)→ RpsC (COG0092)rpsC (PRK00310)→ RplP (COG0197)rplP (PRK09203)→ |
| env-374 | KGI_S2_ANT04_2345m_c1010869 | - | 2778 | 2711 | **RNA**→ RplB (COG0090)rplB (PRK09374)→ rpsS (PRK00357)RpsS (COG0185)→ rplV (PRK00565)Ribosomal_L22 (pfam00237)→ RpsC (COG0092)rpsC (PRK00310)→ RplP (COG0197)rplP (PRK09203)→ |
| env-375 | JGI20153J14295_10082023 | - | 314 | 252 | **RNA**→ RplB (COG0090)rplB (PRK09374)→ |
| env-376 | JGI20153J14318_10082023 | - | 314 | 252 | **RNA**→ RplB (COG0090)rplB (PRK09374)→ |
| env-377 | JGI20159J14440_10153261 | + | 97 | 159 | **RNA**→ RplB (COG0090)rplB (PRK09374)→ |
| env-378 | AMFJ01022780.1 | - | 134 | 67 | **RNA**→ RplB (COG0090)rplB (PRK09374)→ |
| env-379 | ACD54_3 | - | 1770 | 1703 | **RNA**→ RplB (COG0090)rplB (PRK09374)→ rpsS (PRK00357)RpsS (COG0185)→ rplV (PRK00565)Ribosomal_L22 (pfam00237)→ rpsC (PRK00310)30S_S3_KH (cd02412)→ |
| env-380 | CESJ01069642.1 | - | 121 | 57 | **RNA**→ |
| env-381 | JGI12109J20283_10580390 | - | 247 | 150 | **RNA**→ RplB (COG0090)rplB (PRK09374)→ |
| Osp-2-1 | NZ_JQMY01000001.1 | + | 973809 | 973875 | **RNA**→ RplB (COG0090)rplB (PRK09374)→ rpsS (PRK00357)RpsS (COG0185)→ rplV (PRK00565)Ribosomal_L22 (pfam00237)→ RpsC (COG0092)rpsC (PRK00310)→ RplP (COG0197)rplP (PRK09203)→ |

| | | | | | |
|---|---|---|---|---|---|
| env-382 | JGI12396J12026_1059309 | + | 985 | 1055 | **RNA**→ RplB (COG0090)rplB (PRK09374)→ |
| env-383 | JGI12396J12026_1059311 | - | 316 | 246 | **RNA**→ RplB (COG0090)rplB (PRK09374)→ |
| env-384 | JGI12467J12023_1025468 | + | 360 | 430 | **RNA**→ RplB (COG0090)rplB (PRK09374)→ |
| env-385 | JGI12273J12029_10120939 | + | 462 | 532 | **RNA**→ RplB (COG0090)rplB (PRK09374)→ |
| env-386 | JGI12467J12023_1167671 | + | 157 | 226 | **RNA**→ RplB (COG0090)rplB (PRK09374)→ |
| env-387 | JGI12273J12029_10075303 | + | 308 | 377 | **RNA**→ RplB (COG0090)rplB (PRK09374)→ |
| env-388 | JGI12273J12029_10163269 | + | 383 | 452 | **RNA**→ |
| Rme-1-1 | NZ_KK088521.1 | - | 243248 | 243181 | **RNA**→ hypo→ hypo→ rpsS (PRK00357)RpsS (COG0185)→ rplV (PRK00565)Ribosomal_L22 (pfam00237)→ RpsC (COG0092)rpsC (PRK00310)→ RplP (COG0197)rplP (PRK09203)→ PRK00306 (PRK00306)Ribosomal_L29 (pfam00831)→ rpsQ (PRK05610)Ribosomal_S17 (pfam00366)→ rplN (PRK05483)Ribosomal_L14 (pfam00238)→ RplX (COG0198)rplX (PRK00004)→ rplE (PRK00010)RplE (COG0094)→ rpsN (PRK08881)Ribosomal_S14 (pfam00253)→ rpsH (PRK00136)Ribosomal_S8 (pfam00410)→ RplF (COG0097)rplF (PRK05498)→ hypo→ rplR (PRK05593)Ribosomal_L18p (pfam00861)→ |
| env-389 | JGI12467J12023_1134564 | + | 240 | 309 | **RNA**→ |
| env-390 | JGI12273J12029_10074910 | + | 717 | 786 | **RNA**→ RplB (COG0090)rplB (PRK09374)→ |
| env-391 | JGI12273J12029_10159171 | + | 74 | 143 | **RNA**→ RplB (COG0090)rplB (PRK09374)→ |
| env-392 | CESO01257586.1 | + | 654 | 720 | **RNA**→ |
| env-393 | CEVX01283897.1 | - | 753 | 687 | **RNA**→ RplB (COG0090)rplB (PRK09374)→ |
| env-394 | CESM01223547.1 | + | 62 | 128 | **RNA**→ RplB (COG0090)rplB (PRK09374)→ rpsS (PRK00357)Ribosomal_S19 (pfam00203)→ |
| env-395 | JGI26524J50256_1001720 | + | 3021 | 3111 | **RNA**→ RplB (COG0090)rplB (PRK09374)→ rpsS (PRK00357)RpsS (COG0185)→ rplV (PRK00565)Ribosomal_L22 (pfam00237)→ RpsC (COG0092)rpsC (PRK00310)→ RplP (COG0197)rplP (PRK09203)→ |
| env-396 | 3300001592_10003260 | + | 2614 | 2692 | **RNA**→ RplB (COG0090)rplB (PRK09374)→ rpsS (PRK00357)RpsS (COG0185)→ rplV (PRK00565)Ribosomal_L22 (pfam00237)→ RpsC (COG0092)rpsC (PRK00310)→ RplP (COG0197)rplP (PRK09203)→ |
| Pxi-1-1 | NZ_AUBS01000014.1 | + | 30482 | 30560 | **RNA**→ RplB (COG0090)rplB (PRK09374)→ rpsS (PRK00357)RpsS (COG0185)→ rplV (PRK00565)Ribosomal_L22 (pfam00237)→ RpsC (COG0092)rpsC (PRK00310)→ RplP (COG0197)rplP (PRK09203)→ |
| Rmo-1-1 | NZ_JXYG01000033.1 | - | 76643 | 76542 | **RNA**→ RplB (COG0090)rplB (PRK09374)→ rpsS (PRK00357)RpsS (COG0185)→ rplV (PRK00565)Ribosomal_L22 (pfam00237)→ RpsC (COG0092)rpsC (PRK00310)→ RplP (COG0197)rplP (PRK09203)→ |
| Rmo-2-1 | NZ_AQCH01000135.1 | - | 22727 | 22626 | **RNA**→ RplB (COG0090)rplB (PRK09374)→ rpsS (PRK00357)RpsS (COG0185)→ rplV (PRK00565)Ribosomal_L22 (pfam00237)→ RpsC (COG0092)rpsC (PRK00310)→ RplP (COG0197)rplP (PRK09203)→ |
| Rsp-21-1 | NC_008044.1 | + | 268831 | 268932 | **RNA**→ RplB (COG0090)rplB (PRK09374)→ rpsS (PRK00357)RpsS (COG0185)→ rplV (PRK00565)Ribosomal_L22 (pfam00237)→ RpsC (COG0092)rpsC (PRK00310)→ RplP (COG0197)rplP (PRK09203)→ |
| Ssp-1-1 | NZ_GG703517.1 | + | 54267 | 54368 | **RNA**→ RplB (COG0090)rplB (PRK09374)→ RplB (COG0090)rplB (PRK09374)→ rpsS (PRK00357)RpsS (COG0185)→ rpsS (PRK00357)RpsS (COG0185)→ rplV (PRK00565)Ribosomal_L22 (pfam00237)→ rplV (PRK00565)Ribosomal_L22 (pfam00237)→ RpsC (COG0092)rpsC (PRK00310)→ RpsC (COG0092)rpsC (PRK00310)→ RplP (COG0197)rplP (PRK09203)→ RplP (COG0197)rplP (PRK09203)→ |
| Psp-1-1 | NZ_JSWK01000016.1 | - | 260767 | 260659 | **RNA**→ RplB (COG0090)rplB (PRK09374)→ rpsS (PRK00357)RpsS (COG0185)→ rplV (PRK00565)Ribosomal_L22 (pfam00237)→ RpsC (COG0092)rpsC (PRK00310)→ RplP (COG0197)rplP (PRK09203)→ |
| Pga-1-1 | NC_018286.1 | - | 3196082 | 3195974 | **RNA**→ RplB (COG0090)rplB (PRK09374)→ rpsS (PRK00357)RpsS (COG0185)→ rplV (PRK00565)Ribosomal_L22 (pfam00237)→ RpsC (COG0092)rpsC (PRK00310)→ RplP (COG0197)rplP (PRK09203)→ |
| Pin-1-1 | NZ_KI421498.1 | + | 3089651 | 3089759 | **RNA**→ RplB (COG0090)rplB (PRK09374)→ rpsS (PRK00357)RpsS (COG0185)→ rplV (PRK00565)Ribosomal_L22 (pfam00237)→ RpsC (COG0092)rpsC (PRK00310)→ RplP (COG0197)rplP (PRK09203)→ |
| Psp-2-1 | NZ_JSWJ01000014.1 | - | 97388 | 97280 | **RNA**→ RplB (COG0090)rplB (PRK09374)→ rpsS (PRK00357)RpsS (COG0185)→ rplV (PRK00565)Ribosomal_L22 (pfam00237)→ RpsC (COG0092)rpsC (PRK00310)→ RplP (COG0197)rplP (PRK09203)→ |

| | | | | | |
|---|---|---|---|---|---|
| Rsp-11-1 | NZ_CH902583.1 | + | 2863415 | 2863523 | **RNA**→ RplB (COG0090)rplB (PRK09374)→ rpsS (PRK00357)RpsS (COG0185)→ rplV (PRK00565)Ribosomal_L22 (pfam00237)→ RpsC (COG0092)rpsC (PRK00310)→ RplP (COG0197)rplP (PRK09203)→ |
| Par-1-1 | NZ_KI421507.1 | + | 92373 | 92482 | **RNA**→ RplB (COG0090)rplB (PRK09374)→ rpsS (PRK00357)RpsS (COG0185)→ rplV (PRK00565)Ribosomal_L22 (pfam00237)→ RpsC (COG0092)rpsC (PRK00310)→ RplP (COG0197)rplP (PRK09203)→ |
| env-397 | CAM_READ_0074161329 | - | 292 | 213 | **RNA**→ RplB (COG0090)rplB (PRK09374)→ |
| env-398 | JGI24025J20009_10459630 | + | 162 | 240 | **RNA**→ |
| Rsp-12-1 | NZ_AAYC01000008.1 | - | 172899 | 172821 | **RNA**→ RplB (COG0090)rplB (PRK09374)→ RplB (COG0090)rplB (PRK09374)→ rpsS (PRK00357)RpsS (COG0185)→ rpsS (PRK00357)RpsS (COG0185)→ rplV (PRK00565)Ribosomal_L22 (pfam00237)→ rplV (PRK00565)Ribosomal_L22 (pfam00237)→ RpsC (COG0092)rpsC (PRK00310)→ RpsC (COG0092)rpsC (PRK00310)→ RplP (COG0197)rplP (PRK09203)→ RplP (COG0197)rplP (PRK09203)→ |
| Lsp-7-1 | NZ_JWLH01000015.1 | - | 13431 | 13353 | **RNA**→ RplB (COG0090)rplB (PRK09374)→ rpsS (PRK00357)RpsS (COG0185)→ rplV (PRK00565)Ribosomal_L22 (pfam00237)→ RpsC (COG0092)rpsC (PRK00310)→ RplP (COG0197)rplP (PRK09203)→ |
| Lsp-1-1 | NZ_JWLE01000039.1 | - | 14129 | 14051 | **RNA**→ RplB (COG0090)rplB (PRK09374)→ rpsS (PRK00357)RpsS (COG0185)→ rplV (PRK00565)Ribosomal_L22 (pfam00237)→ RpsC (COG0092)rpsC (PRK00310)→ RplP (COG0197)rplP (PRK09203)→ |
| Lsp-3-1 | NZ_JWLG01000013.1 | + | 28657 | 28735 | **RNA**→ RplB (COG0090)rplB (PRK09374)→ rpsS (PRK00357)RpsS (COG0185)→ rplV (PRK00565)Ribosomal_L22 (pfam00237)→ RpsC (COG0092)rpsC (PRK00310)→ RplP (COG0197)rplP (PRK09203)→ |
| Lsp-4-1 | NZ_JWLI01000007.1 | - | 341160 | 341082 | **RNA**→ RplB (COG0090)rplB (PRK09374)→ rpsS (PRK00357)RpsS (COG0185)→ rplV (PRK00565)Ribosomal_L22 (pfam00237)→ RpsC (COG0092)rpsC (PRK00310)→ RplP (COG0197)rplP (PRK09203)→ |
| Lsp-6-1 | NZ_JWLF01000005.1 | - | 489868 | 489790 | **RNA**→ RplB (COG0090)rplB (PRK09374)→ rpsS (PRK00357)RpsS (COG0185)→ rplV (PRK00565)Ribosomal_L22 (pfam00237)→ RpsC (COG0092)rpsC (PRK00310)→ RplP (COG0197)rplP (PRK09203)→ |
| Lsp-9-1 | NZ_AFCF02000006.1 | + | 28882 | 28960 | **RNA**→ RplB (COG0090)rplB (PRK09374)→ rpsS (PRK00357)RpsS (COG0185)→ rplV (PRK00565)Ribosomal_L22 (pfam00237)→ RpsC (COG0092)rpsC (PRK00310)→ RplP (COG0197)rplP (PRK09203)→ |
| Lsp-5-1 | NZ_JWLM01000001.1 | + | 28726 | 28804 | **RNA**→ RplB (COG0090)rplB (PRK09374)→ rpsS (PRK00357)RpsS (COG0185)→ rplV (PRK00565)Ribosomal_L22 (pfam00237)→ RpsC (COG0092)rpsC (PRK00310)→ RplP (COG0197)rplP (PRK09203)→ |
| Pga-2-1 | NC_018290.1 | + | 154778 | 154856 | **RNA**→ RplB (COG0090)rplB (PRK09374)→ rpsS (PRK00357)RpsS (COG0185)→ rplV (PRK00565)Ribosomal_L22 (pfam00237)→ RpsC (COG0092)rpsC (PRK00310)→ RplP (COG0197)rplP (PRK09203)→ |
| Nsp-1-1 | NZ_LAUK01000016.1 | - | 42619 | 42541 | **RNA**→ RplB (COG0090)rplB (PRK09374)→ rpsS (PRK00357)RpsS (COG0185)→ rplV (PRK00565)Ribosomal_L22 (pfam00237)→ RpsC (COG0092)rpsC (PRK00310)→ RplP (COG0197)rplP (PRK09203)→ |
| Nit-1-1 | NZ_CVRL01000015.1 | - | 39753 | 39675 | **RNA**→ RplB (COG0090)rplB (PRK09374)→ rpsS (PRK00357)RpsS (COG0185)→ rplV (PRK00565)Ribosomal_L22 (pfam00237)→ RpsC (COG0092)rpsC (PRK00310)→ RplP (COG0197)rplP (PRK09203)→ |
| Nit-1-2 | NZ_CVRM01000022.1 | + | 261598 | 261676 | **RNA**→ RplB (COG0090)rplB (PRK09374)→ rpsS (PRK00357)RpsS (COG0185)→ rplV (PRK00565)Ribosomal_L22 (pfam00237)→ RpsC (COG0092)rpsC (PRK00310)→ RplP (COG0197)rplP (PRK09203)→ |
| Rsp-20-1 | NZ_DS999054.1 | + | 3164594 | 3164672 | **RNA**→ RplB (COG0090)rplB (PRK09374)→ RplB (COG0090)rplB (PRK09374)→ rpsS (PRK00357)RpsS (COG0185)→ rpsS (PRK00357)RpsS (COG0185)→ rplV (PRK00565)Ribosomal_L22 (pfam00237)→ rplV (PRK00565)Ribosomal_L22 (pfam00237)→ RpsC (COG0092)rpsC (PRK00310)→ RpsC (COG0092)rpsC (PRK00310)→ RplP (COG0197)rplP (PRK09203)→ RplP (COG0197)rplP (PRK09203)→ |
| Lsp-2-1 | NZ_JWLC01000007.1 | - | 265099 | 265021 | **RNA**→ RplB (COG0090)rplB (PRK09374)→ rpsS (PRK00357)RpsS (COG0185)→ rplV (PRK00565)Ribosomal_L22 (pfam00237)→ RpsC (COG0092)rpsC (PRK00310)→ RplP (COG0197)rplP (PRK09203)→ |

| | | | | | |
|---|---|---|---|---|---|
| Rba-7-1 | NZ_DS995281.1 | - | 3127551 | 3127473 | **RNA**→ RplB (COG0090)rplB (PRK09374)→ RplB (COG0090)rplB (PRK09374)→ rpsS (PRK00357)RpsS (COG0185)→ rpsS (PRK00357)RpsS (COG0185)→ rplV (PRK00565)Ribosomal_L22 (pfam00237)→ rplV (PRK00565)Ribosomal_L22 (pfam00237)→ RpsC (COG0092)rpsC (PRK00310)→ RpsC (COG0092)rpsC (PRK00310)→ RplP (COG0197)rplP (PRK09203)→ RplP (COG0197)rplP (PRK09203)→ |
| Lca-1-1 | NZ_KI421513.1 | - | 919197 | 919119 | **RNA**→ RplB (COG0090)rplB (PRK09374)→ rpsS (PRK00357)RpsS (COG0185)→ rplV (PRK00565)Ribosomal_L22 (pfam00237)→ RpsC (COG0092)rpsC (PRK00310)→ RplP (COG0197)rplP (PRK09203)→ |
| Lda-1-1 | NZ_KI421500.1 | - | 2486523 | 2486445 | **RNA**→ RplB (COG0090)rplB (PRK09374)→ rpsS (PRK00357)RpsS (COG0185)→ rplV (PRK00565)Ribosomal_L22 (pfam00237)→ RpsC (COG0092)rpsC (PRK00310)→ RplP (COG0197)rplP (PRK09203)→ |
| Lsp-8-1 | NZ_JWLD01000017.1 | - | 20237 | 20159 | **RNA**→ RplB (COG0090)rplB (PRK09374)→ rpsS (PRK00357)RpsS (COG0185)→ rplV (PRK00565)Ribosomal_L22 (pfam00237)→ RpsC (COG0092)rpsC (PRK00310)→ RplP (COG0197)rplP (PRK09203)→ |
| env-399 | 3300001605_10201626 | + | 440 | 500 | **RNA**→ RplB (COG0090)rplB (PRK09374)→ |
| env-400 | 3300001239_10025476 | + | 68 | 128 | **RNA**→ rplB (PRK09374)Ribosomal_L2 (pfam00181)→ |
| env-401 | 3300001605_10465888 | + | 68 | 128 | **RNA**→ RplB (COG0090)rplB (PRK09374)→ |
| env-402 | 3300001605_10229328 | - | 253 | 194 | **RNA**→ RplB (COG0090)rplB (PRK09374)→ |
| env-403 | Antartic2_1036878 | - | 110 | 44 | **RNA**→ ←hypo |
| env-404 | CESI01066207.1 | + | 305 | 371 | **RNA**→ rplB (PRK09374)Ribosomal_L2 (pfam00181)→ |
| env-405 | JCVI_SCAF_1096627291198 | + | 567 | 632 | **RNA**→ RplB (COG0090)rplB (PRK09374)→ |
| env-406 | DelMOSum2010_c10000597 | + | 3357 | 3422 | **RNA**→ RplB (COG0090)rplB (PRK09374)→ rpsS (PRK00357)RpsS (COG0185)→ rplV (PRK00565)Ribosomal_L22 (pfam00237)→ RpsC (COG0092)rpsC (PRK00310)→ RplP (COG0197)rplP (PRK09203)→ |
| env-407 | DelMOSum2011_c10059128 | + | 607 | 672 | **RNA**→ RplB (COG0090)rplB (PRK09374)→ |
| env-408 | JGI20156J14371_10007799 | - | 4716 | 4651 | **RNA**→ RplB (COG0090)rplB (PRK09374)→ rpsS (PRK00357)RpsS (COG0185)→ rplV (PRK00565)Ribosomal_L22 (pfam00237)→ RpsC (COG0092)rpsC (PRK00310)→ RplP (COG0197)rplP (PRK09203)→ |
| env-409 | JGI20154J14316_10019027 | + | 1070 | 1135 | **RNA**→ RplB (COG0090)rplB (PRK09374)→ rpsS (PRK00357)RpsS (COG0185)→ rplV (PRK00565)Ribosomal_L22 (pfam00237)→ RpsC (COG0092)rpsC (PRK00310)→ RplP (COG0197)rplP (PRK09203)→ |
| env-410 | JGI20160J14292_10005213 | + | 2291 | 2356 | **RNA**→ RplB (COG0090)rplB (PRK09374)→ rpsS (PRK00357)RpsS (COG0185)→ rplV (PRK00565)Ribosomal_L22 (pfam00237)→ RpsC (COG0092)rpsC (PRK00310)→ RplP (COG0197)rplP (PRK09203)→ |
| env-411 | JGI20157J14317_10008243 | - | 4807 | 4742 | **RNA**→ RplB (COG0090)rplB (PRK09374)→ rpsS (PRK00357)RpsS (COG0185)→ rplV (PRK00565)Ribosomal_L22 (pfam00237)→ RpsC (COG0092)rpsC (PRK00310)→ RplP (COG0197)rplP (PRK09203)→ |
| env-412 | JGI20159J14440_10014743 | - | 561 | 496 | **RNA**→ RplB (COG0090)rplB (PRK09374)→ |
| env-413 | JGI20155J14468_10115856 | + | 624 | 689 | **RNA**→ RplB (COG0090)rplB (PRK09374)→ |
| env-414 | JGI20153J14295_10306322 | - | 286 | 219 | **RNA**→ RplB (COG0090)rplB (PRK09374)→ |
| env-415 | JGI20153J14318_10306322 | - | 286 | 219 | **RNA**→ RplB (COG0090)rplB (PRK09374)→ |
| Rel-1-1 | NZ_CP004372.1 | + | 1532613 | 1532674 | **RNA**→ RplB (COG0090)rplB (PRK09374)→ rpsS (PRK00357)RpsS (COG0185)→ rplV (PRK00565)Ribosomal_L22 (pfam00237)→ RpsC (COG0092)rpsC (PRK00310)→ RplP (COG0197)rplP (PRK09203)→ |
| env-416 | JGI12210J13797_11047960 | - | 109 | 48 | **RNA**→ |
| env-417 | SA_S1_NOR08_45m_c10022254 | + | 16 | 83 | **RNA**→ RplB (COG0090)rplB (PRK09374)→ rpsS (PRK00357)RpsS (COG0185)→ rplV (PRK00565)Ribosomal_L22 (pfam00237)→ RpsC (COG0092)rpsC (PRK00310)→ RplP (COG0197)rplP (PRK09203)→ |
| env-418 | 3300001605_10068995 | - | 547 | 488 | **RNA**→ rplB (PRK09374)Ribosomal_L2 (pfam00181)→ |
| env-419 | JGI24768J34885_10007107 | + | 613 | 672 | **RNA**→ RplB (COG0090)rplB (PRK09374)→ rpsS (PRK00357)RpsS (COG0185)→ rplV (PRK00565)Ribosomal_L22 (pfam00237)→ RpsC (COG0092)rpsC (PRK00310)→ RplP (COG0197)rplP (PRK09203)→ |
| env-420 | JGI25925J51416_10384036 | + | 89 | 148 | **RNA**→ RplB (COG0090)rplB (PRK09374)→ |
| env-421 | 3300001592_10004376 | - | 13045 | 12987 | **RNA**→ RplB (COG0090)rplB (PRK09374)→ rpsS (PRK00357)Ribosomal_S19 (pfam00203)→ rplV (PRK00565)Ribosomal_L22 (pfam00237)→ RpsC (COG0092)rpsC (PRK00310)→ RplP (COG0197)rplP (PRK09203)→ |

| | | | | | |
|---|---|---|---|---|---|
| Gne-1-1 | NZ_KE386949.1 | - | 572600 | 572542 | **RNA**→ RplB (COG0090)rplB (PRK09374)→ rpsS (PRK00357)RpsS (COG0185)→ rplV (PRK00565)Ribosomal_L22 (pfam00237)→ RpsC (COG0092)rpsC (PRK00310)→ RplP (COG0197)rplP (PRK09203)→ |
| env-422 | KVRMV2_101642803 | + | 262 | 356 | **RNA**→ rplB (PRK09374)Ribosomal_L2 (pfam00181)→ |
| env-423 | JGI24724J26744_10668225 | - | 103 | 15 | **RNA**→ |
| env-424 | JGI24723J26617_10025028 | + | 221 | 309 | **RNA**→ RplB (COG0090)rplB (PRK09374)→ rpsS (PRK00357)RpsS (COG0185)→ rplV (PRK00565)Ribosomal_L22 (pfam00237)→ RpsC (COG0092)rpsC (PRK00310)→ |
| env-425 | JGI24723J26617_10031254 | + | 1462 | 1550 | **RNA**→ RplB (COG0090)rplB (PRK09374)→ |
| env-426 | JGI24025J20009_10003199 | + | 1288 | 1376 | **RNA**→ RplB (COG0090)rplB (PRK09374)→ rpsS (PRK00357)RpsS (COG0185)→ rplV (PRK00565)Ribosomal_L22 (pfam00237)→ RpsC (COG0092)rpsC (PRK00310)→ RplP (COG0197)rplP (PRK09203)→ |
| Msp-1-1 | NZ_JQEY01000002.1 | + | 278853 | 278919 | **RNA**→ RplB (COG0090)rplB (PRK09374)→ rpsS (PRK00357)RpsS (COG0185)→ rplV (PRK00565)Ribosomal_L22 (pfam00237)→ RpsC (COG0092)rpsC (PRK00310)→ RplP (COG0197)rplP (PRK09203)→ |
| Sno-1-1 | NZ_JASC01000003.1 | + | 4659 | 4742 | **RNA**→ RplB (COG0090)rplB (PRK09374)→ rpsS (PRK00357)RpsS (COG0185)→ rplV (PRK00565)Ribosomal_L22 (pfam00237)→ RpsC (COG0092)rpsC (PRK00310)→ RplP (COG0197)rplP (PRK09203)→ |
| Sno-2-1 | NZ_JASD01000008.1 | - | 392212 | 392129 | **RNA**→ RplB (COG0090)rplB (PRK09374)→ rpsS (PRK00357)RpsS (COG0185)→ rplV (PRK00565)Ribosomal_L22 (pfam00237)→ RpsC (COG0092)rpsC (PRK00310)→ RplP (COG0197)rplP (PRK09203)→ |
| env-427 | CEWO01366965.1 | + | 1207 | 1273 | **RNA**→ rplB (PRK09374)Ribosomal_L2 (pfam00181)→ |
| env-428 | wf2_AHAA5765_x1 | - | 329 | 263 | **RNA**→ |
| Tin-1-1 | NZ_AUNB01000021.1 | + | 2941 | 3007 | **RNA**→ RplB (COG0090)rplB (PRK09374)→ rpsS (PRK00357)RpsS (COG0185)→ rplV (PRK00565)Ribosomal_L22 (pfam00237)→ RpsC (COG0092)rpsC (PRK00310)→ RplP (COG0197)rplP (PRK09203)→ |
| Pwa-1-1 | NZ_LGHT01000042.1 | - | 148727 | 148661 | **RNA**→ RplB (COG0090)rplB (PRK09374)→ rpsS (PRK00357)RpsS (COG0185)→ rplV (PRK00565)Ribosomal_L22 (pfam00237)→ RpsC (COG0092)rpsC (PRK00310)→ RplP (COG0197)rplP (PRK09203)→ |
| env-429 | 3300001605_10795549 | + | 347 | 413 | **RNA**→ ←hypo |
| env-430 | F3OLC3J02I91T9 | - | 208 | 142 | **RNA**→ hypo→ |
| env-431 | 3300001605_10634355 | - | 145 | 79 | **RNA**→ |
| env-432 | TB_FS06_10DRAFT_1000581 | + | 16462 | 16528 | **RNA**→ RplB (COG0090)rplB (PRK09374)→ rpsS (PRK00357)RpsS (COG0185)→ rplV (PRK00565)Ribosomal_L22 (pfam00237)→ RpsC (COG0092)rpsC (PRK00310)→ |
| env-433 | 3300000052_245369 | + | 133 | 199 | **RNA**→ |
| Osp-3-1 | NZ_AFPM01000017.1 | + | 4012 | 4166 | **RNA**→ ←hypo |
| env-434 | JGI20160J14292_10238378 | + | 75 | 140 | **RNA**→ hypo→ rplB (PRK09374)Ribosomal_L2 (pfam00181)→ |
| env-435 | GYQO9XB02IIOI9 | - | 83 | 18 | **RNA**→ |
| env-436 | JGI24724J26744_10484319 | - | 357 | 266 | **RNA**→ rplB (PRK09374)Ribosomal_L2 (pfam00181)→ |
| env-437 | 3300001592_10003340 | - | 10494 | 10403 | **RNA**→ RplB (COG0090)rplB (PRK09374)→ rpsS (PRK00357)RpsS (COG0185)→ rplV (PRK00565)Ribosomal_L22 (pfam00237)→ RpsC (COG0092)rpsC (PRK00310)→ RplP (COG0197)rplP (PRK09203)→ |
| env-438 | GSLSAS_GLMUJHB01D1RM5 | - | 167 | 75 | **RNA**→ RplB (COG0090)rplB (PRK09374)→ |
| env-439 | JGI24025J20009_10404831 | + | 123 | 211 | **RNA**→ ←hypo |
| Rsp-10-1 | NZ_DS999213.1 | + | 3307378 | 3307453 | **RNA**→ RplB (COG0090)rplB (PRK09374)→ RplB (COG0090)rplB (PRK09374)→ RplB (COG0090)rplB (PRK09374)→ RplB (COG0090)rplB (PRK09374)→ RplB (COG0090)rplB (PRK09374)→ rpsS (PRK00357)RpsS (COG0185)→ rpsS (PRK00357)RpsS (COG0185)→ rpsS (PRK00357)RpsS (COG0185)→ rplV (PRK00565)Ribosomal_L22 (pfam00237)→ rplV (PRK00565)Ribosomal_L22 (pfam00237)→ RpsC (COG0092)rpsC (PRK00310)→ RpsC (COG0092)rpsC (PRK00310)→ RplP (COG0197)rplP (PRK09203)→ RplP (COG0197)rplP (PRK09203)→ |
| Tar-1-1 | NZ_KE557314.1 | - | 63321 | 63252 | **RNA**→ RplB (COG0090)rplB (PRK09374)→ RplB (COG0090)rplB (PRK09374)→ RplB (COG0090)rplB (PRK09374)→ rpsS (PRK00357)RpsS (COG0185)→ rplV (PRK00565)Ribosomal_L22 (pfam00237)→ RpsC (COG0092)rpsC (PRK00310)→ RplP (COG0197)rplP (PRK09203)→ |
| env-440 | JGI12109J20283_11967054 | + | 55 | 116 | **RNA**→ hypo→ |

| | | | | | |
|---|---|---|---|---|---|
| Sme-2-1 | NZ_JASH01000010.1 | - | 261103 | 261034 | **RNA**→ RplB (COG0090)rplB (PRK09374)→ rpsS (PRK00357)RpsS (COG0185)→ rplV (PRK00565)Ribosomal_L22 (pfam00237)→ RpsC (COG0092)rpsC (PRK00310)→ RplP (COG0197)rplP (PRK09203)→ |
| Sme-1-1 | NZ_JEMU01000021.1 | + | 4343 | 4412 | **RNA**→ RplB (COG0090)rplB (PRK09374)→ rpsS (PRK00357)RpsS (COG0185)→ rplV (PRK00565)Ribosomal_L22 (pfam00237)→ RpsC (COG0092)rpsC (PRK00310)→ RplP (COG0197)rplP (PRK09203)→ |
| Tsp-3-1 | NZ_GG697169.2 | + | 2896550 | 2896617 | **RNA**→ RplB (COG0090)rplB (PRK09374)→ RplB (COG0090)rplB (PRK09374)→ RplB (COG0090)rplB (PRK09374)→ rpsS (PRK00357)RpsS (COG0185)→ rpsS (PRK00357)RpsS (COG0185)→ rplV (PRK00565)Ribosomal_L22 (pfam00237)→ rplV (PRK00565)Ribosomal_L22 (pfam00237)→ RpsC (COG0092)rpsC (PRK00310)→ RpsC (COG0092)rpsC (PRK00310)→ RplP (COG0197)rplP (PRK09203)→ RplP (COG0197)rplP (PRK09203)→ |
| env-441 | P_1C_Liq_3_UnCty_1043905 | - | 143 | 47 | **RNA**→ |
| env-442 | P_2C_Liq_1_UnCty_1002030 | + | 5317 | 5413 | **RNA**→ RplB (COG0090)rplB (PRK09374)→ rpsS (PRK00357)RpsS (COG0185)→ rplV (PRK00565)Ribosomal_L22 (pfam00237)→ rpsC (PRK00310)30S_S3_KH (cd02412)→ |
| env-443 | P_2C_Liq_2_UnCty_1008969 | + | 248 | 344 | **RNA**→ RplB (COG0090)rplB (PRK09374)→ rpsS (PRK00357)RpsS (COG0185)→ rplV (PRK00565)Ribosomal_L22 (pfam00237)→ rpsC (PRK00310)30S_S3_KH (cd02412)→ |
| env-444 | P_2C_Liq_2_UnCty_1056800 | + | 368 | 464 | **RNA**→ RplB (COG0090)rplB (PRK09374)→ |
| env-445 | JGI12210J13797_10174589 | - | 302 | 232 | **RNA**→ RplB (COG0090)rplB (PRK09374)→ |
| env-446 | JGI12210J13797_10174590 | + | 188 | 258 | **RNA**→ RplB (COG0090)rplB (PRK09374)→ |
| Rto-1-1 | NZ_LGVV01000034.1 | - | 33402 | 33310 | **RNA**→ RplB (COG0090)rplB (PRK09374)→ rpsS (PRK00357)RpsS (COG0185)→ RplV (COG0091)rplV (PRK00565)→ RpsC (COG0092)rpsC (PRK00310)→ RplP (COG0197)rplP (PRK09203)→ |
| Ris-1-1 | NZ_JAME01000033.1 | - | 54293 | 54185 | **RNA**→ RplB (COG0090)rplB (PRK09374)→ rpsS (PRK00357)RpsS (COG0185)→ rplV (PRK00565)Ribosomal_L22 (pfam00237)→ RpsC (COG0092)rpsC (PRK00310)→ RplP (COG0197)rplP (PRK09203)→ |
| env-447 | KGI_S1_ANT02_95m_c10357205 | + | 47 | 140 | **RNA**→ RplB (COG0090)rplB (PRK09374)→ |
| env-448 | LADL01003569.1 | + | 6160 | 6249 | **RNA**→ RplB (COG0090)rplB (PRK09374)→ rpsS (PRK00357)RpsS (COG0185)→ RplV (COG0091)rplV (PRK00565)→ RpsC (COG0092)rpsC (PRK00310)→ RplP (COG0197)rplP (PRK09203)→ |
| env-449 | 3300001239_10287636 | - | 266 | 177 | **RNA**→ RplB (COG0090)rplB (PRK09374)→ |
| env-450 | 3300001605_10004081 | - | 3779 | 3690 | **RNA**→ RplB (COG0090)rplB (PRK09374)→ rpsS (PRK00357)RpsS (COG0185)→ RplV (COG0091)rplV (PRK00565)→ RpsC (COG0092)rpsC (PRK00310)→ RplP (COG0197)rplP (PRK09203)→ |
| env-451 | JGI25317J35169_1001357 | - | 3662 | 3573 | **RNA**→ RplB (COG0090)rplB (PRK09374)→ rpsS (PRK00357)RpsS (COG0185)→ RplV (COG0091)rplV (PRK00565)→ RpsC (COG0092)rpsC (PRK00310)→ RplP (COG0197)rplP (PRK09203)→ |
| env-452 | JGI25319J35699_1000784 | - | 3894 | 3805 | **RNA**→ RplB (COG0090)rplB (PRK09374)→ rpsS (PRK00357)RpsS (COG0185)→ RplV (COG0091)rplV (PRK00565)→ RpsC (COG0092)rpsC (PRK00310)→ RplP (COG0197)rplP (PRK09203)→ |
| env-453 | JGI25322J35698_1004507 | - | 4384 | 4295 | **RNA**→ RplB (COG0090)rplB (PRK09374)→ rpsS (PRK00357)RpsS (COG0185)→ RplV (COG0091)rplV (PRK00565)→ RpsC (COG0092)rpsC (PRK00310)→ RplP (COG0197)rplP (PRK09203)→ |
| env-454 | JGI25318J35512_1000721 | - | 3894 | 3805 | **RNA**→ RplB (COG0090)rplB (PRK09374)→ rpsS (PRK00357)RpsS (COG0185)→ RplV (COG0091)rplV (PRK00565)→ RpsC (COG0092)rpsC (PRK00310)→ RplP (COG0197)rplP (PRK09203)→ |
| env-455 | JGI25321J39296_1001405 | + | 10390 | 10479 | **RNA**→ RplB (COG0090)rplB (PRK09374)→ rpsS (PRK00357)RpsS (COG0185)→ RplV (COG0091)rplV (PRK00565)→ RpsC (COG0092)rpsC (PRK00310)→ RplP (COG0197)rplP (PRK09203)→ |
| env-456 | JGI25320J50211_1000655 | + | 15085 | 15174 | **RNA**→ RplB (COG0090)rplB (PRK09374)→ rpsS (PRK00357)RpsS (COG0185)→ RplV (COG0091)rplV (PRK00565)→ RpsC (COG0092)rpsC (PRK00310)→ RplP (COG0197)rplP (PRK09203)→ |
| env-457 | JGI25321J50212_10002409 | + | 10390 | 10479 | **RNA**→ RplB (COG0090)rplB (PRK09374)→ rpsS (PRK00357)RpsS (COG0185)→ RplV (COG0091)rplV (PRK00565)→ RpsC (COG0092)rpsC (PRK00310)→ RplP (COG0197)rplP (PRK09203)→ |
| Tpa-1-1 | NZ_AUND01000002.1 | + | 2906 | 2974 | **RNA**→ RplB (COG0090)rplB (PRK09374)→ rpsS (PRK00357)RpsS (COG0185)→ rplV (PRK00565)Ribosomal_L22 (pfam00237)→ RpsC (COG0092)rpsC (PRK00310)→ RplP (COG0197)rplP (PRK09203)→ hypo→ |
| env-458 | CEVN01086405.1 | - | 332 | 264 | **RNA**→ rplB (PRK09374)Ribosomal_L2 (pfam00181)→ |
| Tat-1-1 | NZ_AQRC01000003.1 | - | 337439 | 337371 | **RNA**→ RplB (COG0090)rplB (PRK09374)→ rpsS (PRK00357)RpsS (COG0185)→ rplV (PRK00565)Ribosomal_L22 (pfam00237)→ RpsC (COG0092)rpsC (PRK00310)→ RplP (COG0197)rplP (PRK09203)→ |
| env-459 | TFO_HY__Contig_1825 | + | 253 | 321 | **RNA**→ RplB (COG0090)rplB (PRK09374)→ rpsS (PRK00357)RpsS (COG0185)→ rplV (PRK00565)Ribosomal_L22 (pfam00237)→ KH (smart00322)30S_S3_KH (cd02412)→ |
| env-460 | CESE01105678.1 | + | 551 | 617 | **RNA**→ |
| env-461 | CESI01080217.1 | - | 93 | 27 | **RNA**→ |
| env-462 | CESU01020394.1 | - | 93 | 27 | **RNA**→ |
| env-463 | CESJ01170158.1 | - | 78 | 12 | **RNA**→ |

| | | | | | |
|---|---|---|---|---|---|
| env-464 | CEST01002662.1 | - | 308 | 242 | **RNA**→ RplB (COG0090)rplB (PRK09374)→ |
| env-465 | CESI01225209.1 | - | 91 | 25 | **RNA**→ |
| env-466 | CESU01161643.1 | - | 91 | 25 | **RNA**→ |
| env-467 | DelMOSum2010_c10465078 | - | 126 | 60 | **RNA**→ RplB (COG0090)rplB (PRK09374)→ |
| env-468 | JGI20157J14317_10424860 | + | 203 | 269 | **RNA**→ RplB (COG0090)rplB (PRK09374)→ |
| env-469 | JGI20152J14361_10060366 | + | 794 | 860 | **RNA**→ RplB (COG0090)rplB (PRK09374)→ |
| env-470 | JGI20151J14362_10458567 | + | 170 | 236 | **RNA**→ RplB (COG0090)rplB (PRK09374)→ |
| env-471 | JGI20157J14317_10486189 | + | 185 | 251 | **RNA**→ |
| env-472 | CERM01031828.1 | - | 1160 | 1094 | **RNA**→ RplB (COG0090)rplB (PRK09374)→ rpsS (PRK00357)Ribosomal_S19 (pfam00203)→ |
| env-473 | JGI20152J14361_10019476 | - | 938 | 873 | **RNA**→ rplB (PRK09374)Ribosomal_L2 (pfam00181)→ rplB (PRK09374)Ribosomal_L2_C (pfam03947)→ |
| env-474 | JGI20157J14317_10015572 | - | 3473 | 3408 | **RNA**→ RplB (COG0090)rplB (PRK09374)→ rpsS (PRK00357)RpsS (COG0185)→ rplV (PRK00565)Ribosomal_L22 (pfam00237)→ RpsC (COG0092)rpsC (PRK00310)→ RplP (COG0197)rplP (PRK09203)→ |
| env-475 | 2236876002_042693 | - | 416 | 351 | **RNA**→ rplB (PRK09374)Ribosomal_L2 (pfam00181)→ |
| env-476 | JGI20158J14315_10019288 | + | 625 | 691 | **RNA**→ rplB (PRK09374)Ribosomal_L2 (pfam00181)→ rplB (PRK09374)Ribosomal_L2_C (pfam03947)→ rpsS (PRK00357)RpsS (COG0185)→ rplV (PRK00565)Ribosomal_L22 (pfam00237)→ RpsC (COG0092)rpsC (PRK00310)→ RplP (COG0197)rplP (PRK09203)→ |
| env-477 | P_2C_Liq_1_UnCty_1012554 | + | 141 | 207 | **RNA**→ RplB (COG0090)rplB (PRK09374)→ rpsS (PRK00357)RpsS (COG0185)→ rplV (PRK00565)Ribosomal_L22 (pfam00237)→ rpsC (PRK00310)30S_S3_KH (cd02412)→ |
| env-478 | P_2C_Liq_2_UnCty_1004631 | + | 159 | 225 | **RNA**→ RplB (COG0090)rplB (PRK09374)→ rpsS (PRK00357)RpsS (COG0185)→ rplV (PRK00565)Ribosomal_L22 (pfam00237)→ RpsC (COG0092)rpsC (PRK00310)→ RplP (COG0197)rplP (PRK09203)→ |
| env-479 | JGI20151J14362_10307003 | + | 248 | 314 | **RNA**→ hypo→ |
| env-480 | JGI20154J14316_10012324 | + | 1879 | 1945 | **RNA**→ RplB (COG0090)rplB (PRK09374)→ rpsS (PRK00357)RpsS (COG0185)→ rplV (PRK00565)Ribosomal_L22 (pfam00237)→ KH (smart00322)30S_S3_KH (cd02412)→ rpsC (PRK00310)Ribosomal_S3_C (pfam00189)→ RplP (COG0197)rplP (PRK09203)→ |
| env-481 | JGI20160J14292_10120340 | - | 142 | 76 | **RNA**→ |
| env-482 | JCVI_SCAF_1101668062396 | - | 504 | 438 | **RNA**→ rplB (PRK09374)Ribosomal_L2 (pfam00181)→ |
| env-483 | JGI20156J14371_10029263 | - | 2057 | 1991 | **RNA**→ RplB (COG0090)rplB (PRK09374)→ rpsS (PRK00357)RpsS (COG0185)→ rplV (PRK00565)Ribosomal_L22 (pfam00237)→ RpsC (COG0092)rpsC (PRK00310)→ |
| env-484 | JCVI_SCAF_1101667021862 | + | 537 | 603 | **RNA**→ rplB (PRK09374)Ribosomal_L2 (pfam00181)→ |
| env-485 | JCVI_SCAF_1096626884361 | - | 4210 | 4144 | **RNA**→ RplB (COG0090)rplB (PRK09374)→ rpsS (PRK00357)RpsS (COG0185)→ rplV (PRK00565)Ribosomal_L22 (pfam00237)→ RpsC (COG0092)rpsC (PRK00310)→ RplP (COG0197)rplP (PRK09203)→ |
| env-486 | JCVI_SCAF_1096627292964 | - | 129 | 63 | **RNA**→ RplB (COG0090)rplB (PRK09374)→ |
| env-487 | DelMOSum2010_c10287456 | - | 395 | 329 | **RNA**→ rplB (PRK09374)Ribosomal_L2 (pfam00181)→ |
| env-488 | DelMOSum2011_c10012372 | - | 3519 | 3453 | **RNA**→ RplB (COG0090)rplB (PRK09374)→ rpsS (PRK00357)RpsS (COG0185)→ rplV (PRK00565)Ribosomal_L22 (pfam00237)→ RpsC (COG0092)rpsC (PRK00310)→ RplP (COG0197)rplP (PRK09203)→ |
| env-489 | JGI20153J14295_10009487 | - | 3477 | 3411 | **RNA**→ RplB (COG0090)rplB (PRK09374)→ rpsS (PRK00357)RpsS (COG0185)→ rplV (PRK00565)Ribosomal_L22 (pfam00237)→ RpsC (COG0092)rpsC (PRK00310)→ RplP (COG0197)rplP (PRK09203)→ |
| env-490 | JGI20153J14318_10009487 | - | 3477 | 3411 | **RNA**→ RplB (COG0090)rplB (PRK09374)→ rpsS (PRK00357)RpsS (COG0185)→ rplV (PRK00565)Ribosomal_L22 (pfam00237)→ RpsC (COG0092)rpsC (PRK00310)→ RplP (COG0197)rplP (PRK09203)→ |
| env-491 | JGI20159J14440_10009403 | + | 2075 | 2141 | **RNA**→ RplB (COG0090)rplB (PRK09374)→ rpsS (PRK00357)RpsS (COG0185)→ rplV (PRK00565)Ribosomal_L22 (pfam00237)→ RpsC (COG0092)rpsC (PRK00310)→ RplP (COG0197)rplP (PRK09203)→ |
| env-492 | JGI20155J14468_10011527 | + | 2004 | 2070 | **RNA**→ RplB (COG0090)rplB (PRK09374)→ rpsS (PRK00357)RpsS (COG0185)→ rplV (PRK00565)Ribosomal_L22 (pfam00237)→ RpsC (COG0092)rpsC (PRK00310)→ RplP (COG0197)rplP (PRK09203)→ |
| Pte-1-1 | NZ_CP003984.1 | + | 2755361 | 2755427 | **RNA**→ RplB (COG0090)rplB (PRK09374)→ rpsS (PRK00357)RpsS (COG0185)→ rplV (PRK00565)Ribosomal_L22 (pfam00237)→ RpsC (COG0092)rpsC (PRK00310)→ RplP (COG0197)rplP (PRK09203)→ |
| env-493 | JCVI_SCAF_1101668090558 | + | 847 | 913 | **RNA**→ RplB (COG0090)rplB (PRK09374)→ |
| env-494 | DelMOSum2010_c10054733 | - | 257 | 191 | **RNA**→ RplB (COG0090)rplB (PRK09374)→ |

| | | | | | |
|---|---|---|---|---|---|
| env-495 | JGI20151J14362_10072490 | - | 659 | 593 | RNA→ RplB (COG0090)rplB (PRK09374)→ |
| env-496 | JGI20160J14292_10100804 | - | 278 | 212 | RNA→ ←hypo |
| env-497 | JGI24724J26744_10002738 | - | 11069 | 10947 | RNA→ RplB (COG0090)rplB (PRK09374)→ rpsS (PRK00357)RpsS (COG0185)→ |
| | | | | | rplV (PRK00565)Ribosomal_L22 (pfam00237)→ RpsC (COG0092)rpsC (PRK00310)→ |
| | | | | | RplP (COG0197)rplP (PRK09203)→ |
| env-498 | JGI24723J26617_10030085 | - | 1778 | 1656 | RNA→ RplB (COG0090)rplB (PRK09374)→ rpsS (PRK00357)RpsS (COG0185)→ |
| | | | | | rplV (PRK00565)Ribosomal_L22 (pfam00237)→ rpsC (PRK00310)30S_S3_KH (cd02412)→ |
| env-499 | JGI24723J26617_10138499 | + | 494 | 616 | RNA→ |
| env-500 | P_2C_Liq_1_UnCty_1025942 | + | 228 | 318 | RNA→ RplB (COG0090)rplB (PRK09374)→ |
| env-501 | P_2C_Liq_2_UnCty_1020991 | + | 163 | 253 | RNA→ RplB (COG0090)rplB (PRK09374)→ |
| env-502 | JCVI_SCAF_1096627363432 | - | 1000 | 917 | RNA→ RplB (COG0090)rplB (PRK09374)→ |
| env-503 | P_1C_Liq_1_UnCty_1060565 | + | 89 | 180 | RNA→ RplB (COG0090)rplB (PRK09374)→ |
| env-504 | P_2C_Liq_1_UnCty_1025617 | + | 238 | 329 | RNA→ RplB (COG0090)rplB (PRK09374)→ |
| env-505 | P_2C_Liq_2_UnCty_1030521 | + | 237 | 327 | RNA→ RplB (COG0090)rplB (PRK09374)→ |
| Smu-1-1 | NZ_KE557273.1 | + | 282336 | 282429 | RNA→ RplB (COG0090)rplB (PRK09374)→ rpsS (PRK00357)RpsS (COG0185)→ |
| | | | | | rplV (PRK00565)Ribosomal_L22 (pfam00237)→ RpsC (COG0092)rpsC (PRK00310)→ |
| | | | | | RplP (COG0197)rplP (PRK09203)→ |
| env-506 | JGI20154J14316_10000278 | - | 10091 | 10028 | RNA→ RplB (COG0090)rplB (PRK09374)→ rpsS (PRK00357)RpsS (COG0185)→ |
| | | | | | rplV (PRK00565)Ribosomal_L22 (pfam00237)→ RpsC (COG0092)rpsC (PRK00310)→ |
| | | | | | RplP (COG0197)rplP (PRK09203)→ PRK00306 (PRK00306)Ribosomal_L29 (pfam00831)→ |
| | | | | | rpsQ (PRK05610)Ribosomal_S17 (pfam00366)→ rplN (PRK05483)Ribosomal_L14 (pfam00238)→ |
| | | | | | RplX (COG0198)rplX (PRK00004)→ rplE (PRK00010)RplE (COG0094)→ |
| | | | | | rpsN (PRK08881)Ribosomal_S14 (pfam00253)→ rpsH (PRK00136)Ribosomal_S8 (pfam00410)→ |
| | | | | | RplF (COG0097)rplF (PRK05498)→ rplR (PRK05593)Ribosomal_L18p (pfam00861)→ |
| | | | | | RpsE (COG0098)rpsE (PRK00550)→ Ribosomal_L30 (cd01658)rpmD (PRK05611)→ |
| env-507 | CESJ01079969.1 | + | 363 | 426 | RNA→ RplB (COG0090)rplB (PRK09374)→ |
| env-508 | DelMOSum2011_c10084907 | - | 321 | 258 | RNA→ rplB (PRK09374)Ribosomal_L2 (pfam00181)→ |
| env-509 | JGI20160J14292_10062109 | + | 335 | 398 | RNA→ RplB (COG0090)rplB (PRK09374)→ rpsS (PRK00357)RpsS (COG0185)→ |
| env-510 | CESI01081701.1 | - | 1672 | 1609 | RNA→ RplB (COG0090)rplB (PRK09374)→ rpsS (PRK00357)RpsS (COG0185)→ |
| | | | | | rplV (PRK00565)Ribosomal_L22 (pfam00237)→ rpsC (PRK00310)30S_S3_KH (cd02412)→ |
| env-511 | CESU01035091.1 | - | 8481 | 8418 | RNA→ RplB (COG0090)rplB (PRK09374)→ rpsS (PRK00357)RpsS (COG0185)→ |
| | | | | | rplV (PRK00565)Ribosomal_L22 (pfam00237)→ RpsC (COG0092)rpsC (PRK00310)→ |
| | | | | | RplP (COG0197)rplP (PRK09203)→ PRK00306 (PRK00306)Ribosomal_L29 (pfam00831)→ |
| | | | | | rpsQ (PRK05610)Ribosomal_S17 (pfam00366)→ rplN (PRK05483)Ribosomal_L14 (pfam00238)→ |
| | | | | | RplX (COG0198)rplX (PRK00004)→ rplE (PRK00010)RplE (COG0094)→ |
| | | | | | rpsN (PRK08881)Ribosomal_S14 (pfam00253)→ rpsH (PRK00136)Ribosomal_S8 (pfam00410)→ |
| | | | | | RplF (COG0097)rplF (PRK05498)→ rplR (PRK05593)Ribosomal_L18p (pfam00861)→ |
| | | | | | RpsE (COG0098)rpsE (PRK00550)→ Ribosomal_L30 (cd01658)rpmD (PRK05611)→ |
| env-512 | JGI20151J14362_10229901 | + | 58 | 122 | RNA→ rplB (PRK09374)Ribosomal_L2 (pfam00181)→ |
| env-513 | 2236876002_225186 | + | 183 | 247 | RNA→ RplB (COG0090)rplB (PRK09374)→ |
| env-514 | 2236876010_0411961 | + | 127 | 191 | RNA→ |
| env-515 | CERM01014667.1 | + | 479 | 543 | RNA→ RplB (COG0090)rplB (PRK09374)→ rpsS (PRK00357)RpsS (COG0185)→ |
| | | | | | rplV (PRK00565)Ribosomal_L22 (pfam00237)→ RpsC (COG0092)rpsC (PRK00310)→ |
| | | | | | RplP (COG0197)rplP (PRK09203)→ PRK00306 (PRK00306)Ribosomal_L29 (pfam00831)→ |
| | | | | | rpsQ (PRK05610)Ribosomal_S17 (pfam00366)→ rplN (PRK05483)Ribosomal_L14 (pfam00238)→ |
| | | | | | RplX (COG0198)rplX (PRK00004)→ rplE (PRK00010)RplE (COG0094)→ |
| | | | | | rpsN (PRK08881)Ribosomal_S14 (pfam00253)→ rpsH (PRK00136)Ribosomal_S8 (pfam00410)→ |
| | | | | | RplF (COG0097)rplF (PRK05498)→ rplR (PRK05593)Ribosomal_L18p (pfam00861)→ |
| | | | | | RpsE (COG0098)rpsE (PRK00550)→ Ribosomal_L30 (cd01658)rpmD (PRK05611)→ |
| env-516 | 2236876009_129660 | + | 384 | 447 | RNA→ |
| env-517 | JGI20160J14292_10014797 | + | 4611 | 4675 | RNA→ |
| env-518 | JGI20157J14317_10244667 | + | 41 | 105 | RNA→ rplB (PRK09374)Ribosomal_L2 (pfam00181)→ |
| env-519 | JCVI_READ_1105499001019 | - | 595 | 532 | RNA→ rplB (PRK09374)Ribosomal_L2 (pfam00181)→ |
| env-520 | 2236876010_0225648 | + | 55 | 119 | RNA→ RplB (COG0090)rplB (PRK09374)→ |
| env-521 | 2236876011_173305 | + | 341 | 406 | RNA→ |

| | | | | | |
|---|---|---|---|---|---|
| env-522 | P_2C_Liq_2_UnCty_1069081 | + | 299 | 389 | RNA→ RplB (COG0090)rplB (PRK09374)→ |
| env-523 | P_2C_Liq_1_UnCty_1043093 | - | 165 | 75 | RNA→ RplB (COG0090)rplB (PRK09374)→ |
| env-524 | KGI_S2_ANT04_2345m_c1261754 | + | 93 | 130 | RNA→ RplB (COG0090)rplB (PRK09374)→ |
| env-525 | JGI24723J26617_10273187 | - | 116 | 79 | RNA→ |
| env-526 | CESU01002408.1 | + | 477 | 543 | RNA→ |
| Tda-1-1 | NZ_JHEH01000004.1 | + | 3299 | 3366 | RNA→ RplB (COG0090)rplB (PRK09374)→ rpsS (PRK00357)RpsS (COG0185)→ rplV (PRK00565)Ribosomal_L22 (pfam00237)→ RpsC (COG0092)rpsC (PRK00310)→ RplP (COG0197)rplP (PRK09203)→ |
| Dsp-1-1 | NZ_AYXI01000047.1 | + | 2274 | 2353 | RNA→ RplB (COG0090)rplB (PRK09374)→ rpsS (PRK00357)RpsS (COG0185)→ rplV (PRK00565)Ribosomal_L22 (pfam00237)→ RpsC (COG0092)rpsC (PRK00310)→ RplP (COG0197)rplP (PRK09203)→ |
| env-527 | JGI24770J26754_10000998 | - | 16793 | 16732 | RNA→ RplB (COG0090)rplB (PRK09374)→ rpsS (PRK00357)RpsS (COG0185)→ rplV (PRK00565)Ribosomal_L22 (pfam00237)→ RpsC (COG0092)rpsC (PRK00310)→ RplP (COG0197)rplP (PRK09203)→ 2OG-FeII_Oxy_5 (pfam13759)→ |
| Jaq-1-1 | NZ_JYFE01000080.1 | - | 61937 | 61870 | RNA→ RplB (COG0090)rplB (PRK09374)→ rpsS (PRK00357)RpsS (COG0185)→ rplV (PRK00565)Ribosomal_L22 (pfam00237)→ RpsC (COG0092)rpsC (PRK00310)→ RplP (COG0197)rplP (PRK09203)→ |
| Sge-1-1 | NZ_JASE01000005.1 | - | 3545549 | 3545479 | RNA→ RplB (COG0090)rplB (PRK09374)→ rpsS (PRK00357)RpsS (COG0185)→ rplV (PRK00565)Ribosomal_L22 (pfam00237)→ RpsC (COG0092)rpsC (PRK00310)→ RplP (COG0197)rplP (PRK09203)→ |
| env-528 | CESE01069575.1 | - | 1653 | 1587 | RNA→ RplB (COG0090)rplB (PRK09374)→ rpsS (PRK00357)RpsS (COG0185)→ rplV (PRK00565)Ribosomal_L22 (pfam00237)→ rpsC (PRK00310)30S_S3_KH (cd02412)→ |
| env-529 | CEST01031231.1 | - | 1873 | 1807 | RNA→ RplB (COG0090)rplB (PRK09374)→ rpsS (PRK00357)RpsS (COG0185)→ rplV (PRK00565)Ribosomal_L22 (pfam00237)→ KH (smart00322)30S_S3_KH (cd02412)→ |
| env-530 | 2236876001_338482 | - | 82 | 18 | RNA→ hypo→ |
| env-531 | 2236876009_222696 | - | 140 | 75 | RNA→ RplB (COG0090)rplB (PRK09374)→ |
| env-532 | JGI20152J14361_10053033 | + | 949 | 1014 | RNA→ |
| env-533 | 2236876001_438614 | - | 371 | 305 | RNA→ rplB (PRK09374)Ribosomal_L2 (pfam00181)→ |
| env-534 | SI34jun09_10m_1003973 | - | 1624 | 1559 | RNA→ RplB (COG0090)rplB (PRK09374)→ rpsS (PRK00357)RpsS (COG0185)→ rplV (PRK00565)Ribosomal_L22 (pfam00237)→ |
| env-535 | CESU01069477.1 | - | 82 | 17 | RNA→ |
| env-536 | DelMOSum2010_c10041356 | - | 259 | 194 | RNA→ RplB (COG0090)rplB (PRK09374)→ |
| env-537 | DelMOSum2011_c10032717 | - | 140 | 75 | RNA→ |
| env-538 | SI60aug11_100m_c1018162 | - | 144 | 79 | RNA→ |
| env-539 | SI47jul10_120m_c1099469 | + | 199 | 264 | RNA→ RplB (COG0090)rplB (PRK09374)→ |
| env-540 | SI34jun09_135m_1040685 | - | 788 | 723 | RNA→ RplB (COG0090)rplB (PRK09374)→ |
| env-541 | SI34jun09_100m_1002546 | - | 626 | 561 | RNA→ RplB (COG0090)rplB (PRK09374)→ |
| env-542 | JGI20151J14362_10030512 | + | 2508 | 2573 | RNA→ |
| env-543 | CESI01072935.1 | + | 1105 | 1170 | RNA→ |
| env-544 | PML_READ_01030272 | - | 197 | 132 | RNA→ RplB (COG0090)rplB (PRK09374)→ |
| env-545 | F10SI0310a_Sequence0000021673 | - | 388 | 323 | RNA→ rplB (PRK09374)Ribosomal_L2 (pfam00181)→ |
| env-546 | 2236876003_146267 | + | 345 | 409 | RNA→ RplB (COG0090)rplB (PRK09374)→ |
| env-547 | 2236876010_0560637 | + | 290 | 354 | RNA→ ←hypo |
| env-548 | 2236876010_0169121 | - | 311 | 246 | RNA→ RplB (COG0090)rplB (PRK09374)→ |
| env-549 | 2236876009_267955 | - | 333 | 268 | RNA→ rplB (PRK09374)Ribosomal_L2 (pfam00181)→ |
| env-550 | 2236876009_052156 | - | 273 | 208 | RNA→ RplB (COG0090)rplB (PRK09374)→ |
| env-551 | 2236876010_0456997 | - | 181 | 116 | RNA→ RplB (COG0090)rplB (PRK09374)→ |
| env-552 | DelMOSum2011_c10406745 | - | 105 | 40 | RNA→ ←hypo |
| env-553 | SI54feb11_100m_c1113376 | - | 189 | 124 | RNA→ RplB (COG0090)rplB (PRK09374)→ |
| env-554 | SI48aug10_10m_c1000222 | + | 2332 | 2397 | RNA→ RplB (COG0090)rplB (PRK09374)→ rpsS (PRK00357)RpsS (COG0185)→ rplV (PRK00565)Ribosomal_L22 (pfam00237)→ RpsC (COG0092)rpsC (PRK00310)→ RplP (COG0197)rplP (PRK09203)→ |
| env-555 | SI34jun09_120m_1016761 | - | 1571 | 1506 | RNA→ RplB (COG0090)rplB (PRK09374)→ rpsS (PRK00357)RpsS (COG0185)→ rplV (PRK00565)Ribosomal_L22 (pfam00237)→ |
| env-556 | SI48aug10_100m_1019662 | - | 473 | 408 | RNA→ rplB (PRK09374)Ribosomal_L2 (pfam00181)→ |

| | | | | | |
|---|---|---|---|---|---|
| env-557 | JGI20156J14371_10036454 | - | 128 | 63 | **RNA**→ |
| env-558 | JGI20154J14316_10048488 | + | 1717 | 1782 | **RNA**→ |
| env-559 | JGI20160J14292_10049921 | - | 944 | 879 | **RNA**→ RplB (COG0090)rplB (PRK09374)→ |
| env-560 | JGI20153J14295_10106585 | + | 637 | 702 | **RNA**→ |
| env-561 | JGI20153J14318_10106585 | + | 637 | 702 | **RNA**→ |
| env-562 | JGI20157J14317_10081233 | + | 1167 | 1232 | **RNA**→ |
| env-563 | JGI20158J14315_10394374 | + | 254 | 319 | **RNA**→ rplW (PRK05738)Ribosomal_L23 (pfam00276)→ |
| env-564 | CERI01006995.1 | - | 80 | 15 | **RNA**→ |
| env-565 | CESO01025210.1 | - | 5544 | 5479 | **RNA**→ RplB (COG0090)rplB (PRK09374)→ rpsS (PRK00357)RpsS (COG0185)→ rplV (PRK00565)Ribosomal_L22 (pfam00237)→ RpsC (COG0092)rpsC (PRK00310)→ RplP (COG0197)rplP (PRK09203)→ |
| env-566 | F10SI03100a_Sequence0000087588 | - | 597 | 532 | **RNA**→ RplB (COG0090)rplB (PRK09374)→ |
| Rba-5-1 | NZ_DS022282.1 | - | 1056719 | 1056654 | **RNA**→ RplB (COG0090)rplB (PRK09374)→ rpsS (PRK00357)RpsS (COG0185)→ rplV (PRK00565)Ribosomal_L22 (pfam00237)→ RpsC (COG0092)rpsC (PRK00310)→ RplP (COG0197)rplP (PRK09203)→ |
| env-567 | JCVI_SCAF_1101667005364 | + | 568 | 633 | **RNA**→ rplB (PRK09374)Ribosomal_L2 (pfam00181)→ |
| env-568 | DelMOSum2010_c10035518 | + | 2424 | 2489 | **RNA**→ RplB (COG0090)rplB (PRK09374)→ |
| env-569 | DelMOSum2011_c10035518 | + | 2007 | 2072 | **RNA**→ |
| env-570 | JGI20152J14361_10025542 | + | 1917 | 1982 | **RNA**→ |
| env-571 | JGI20153J14295_10035283 | - | 131 | 66 | **RNA**→ |
| env-572 | JGI20153J14318_10035283 | - | 131 | 66 | **RNA**→ |
| env-573 | JGI20157J14317_10045573 | + | 1946 | 2011 | **RNA**→ |
| env-574 | CESI01143925.1 | + | 1396 | 1461 | **RNA**→ |
| env-575 | CESU01042376.1 | - | 78 | 13 | **RNA**→ |
| env-576 | DelMOWin2010_c10019689 | - | 113 | 48 | **RNA**→ ←hypo |
| env-577 | JGI20152J14361_10025431 | - | 148 | 83 | **RNA**→ |
| env-578 | JGI20160J14292_10037631 | - | 132 | 67 | **RNA**→ |
| env-579 | JGI20151J14362_10009967 | - | 5544 | 5479 | **RNA**→ RplB (COG0090)rplB (PRK09374)→ rpsS (PRK00357)RpsS (COG0185)→ rplV (PRK00565)Ribosomal_L22 (pfam00237)→ RpsC (COG0092)rpsC (PRK00310)→ |
| env-580 | JGI20157J14317_10012557 | + | 1945 | 2010 | **RNA**→ RplB (COG0090)rplB (PRK09374)→ rpsS (PRK00357)RpsS (COG0185)→ rplV (PRK00565)Ribosomal_L22 (pfam00237)→ RpsC (COG0092)rpsC (PRK00310)→ RplP (COG0197)rplP (PRK09203)→ |
| env-581 | CERM01019450.1 | + | 628 | 693 | **RNA**→ |
| env-582 | 2236876001_009312 | - | 254 | 189 | **RNA**→ RplB (COG0090)rplB (PRK09374)→ |
| env-583 | JGI20156J14371_10029870 | + | 2238 | 2303 | **RNA**→ rplB (PRK09374)Ribosomal_L2 (pfam00181)→ |
| env-584 | JGI20154J14316_10019470 | - | 1818 | 1753 | **RNA**→ RplB (COG0090)rplB (PRK09374)→ rpsS (PRK00357)RpsS (COG0185)→ rplV (PRK00565)Ribosomal_L22 (pfam00237)→ rpsC (PRK00310)30S_S3_KH (cd02412)→ |
| env-585 | JGI20153J14295_10035952 | + | 1907 | 1972 | **RNA**→ |
| env-586 | JGI20153J14318_10035952 | + | 1907 | 1972 | **RNA**→ |
| env-587 | JGI20159J14440_10007957 | + | 3434 | 3499 | **RNA**→ RplB (COG0090)rplB (PRK09374)→ rpsS (PRK00357)RpsS (COG0185)→ rplV (PRK00565)Ribosomal_L22 (pfam00237)→ RpsC (COG0092)rpsC (PRK00310)→ rplP (PRK09203)Ribosomal_L16 (pfam00252)→ |
| env-588 | JGI20155J14468_10004488 | + | 2462 | 2527 | **RNA**→ RplB (COG0090)rplB (PRK09374)→ rpsS (PRK00357)RpsS (COG0185)→ rplV (PRK00565)Ribosomal_L22 (pfam00237)→ RpsC (COG0092)rpsC (PRK00310)→ RplP (COG0197)rplP (PRK09203)→ |
| env-589 | JGI20158J14315_10007229 | + | 2960 | 3025 | **RNA**→ RplB (COG0090)rplB (PRK09374)→ rpsS (PRK00357)RpsS (COG0185)→ rplV (PRK00565)Ribosomal_L22 (pfam00237)→ RpsC (COG0092)rpsC (PRK00310)→ RplP (COG0197)rplP (PRK09203)→ |
| Sgu-1-1 | NZ_JASG01000004.1 | - | 2771313 | 2771244 | **RNA**→ RplB (COG0090)rplB (PRK09374)→ rpsS (PRK00357)RpsS (COG0185)→ rplV (PRK00565)Ribosomal_L22 (pfam00237)→ RpsC (COG0092)rpsC (PRK00310)→ RplP (COG0197)rplP (PRK09203)→ |
| env-590 | CENJ01071080.1 | + | 446 | 521 | **RNA**→ RplB (COG0090)rplB (PRK09374)→ rpsS (PRK00357)RpsS (COG0185)→ |
| env-591 | CENR01032483.1 | + | 411 | 486 | **RNA**→ |
| env-592 | JGI12120J13076_100262 | + | 774 | 849 | **RNA**→ RplB (COG0090)rplB (PRK09374)→ rpsS (PRK00357)RpsS (COG0185)→ rplV (PRK00565)Ribosomal_L22 (pfam00237)→ rpsC (PRK00310)30S_S3_KH (cd02412)→ |

| | | | | | |
|---|---|---|---|---|---|
| env-593 | JGI11929J13085_109680 | + | 171 | 246 | **RNA**→ RplB (COG0090)rplB (PRK09374)→ |
| env-594 | JGI12207J13218_1020216 | + | 89 | 164 | **RNA**→ rplB (PRK09374)Ribosomal_L2 (pfam00181)→ |
| env-595 | CEOE01076289.1 | - | 558 | 483 | **RNA**→ rplB (PRK09374)Ribosomal_L2 (pfam00181)→ |
| env-596 | LAZR01000132.1 | - | 92425 | 92350 | **RNA**→ RplB (COG0090)rplB (PRK09374)→ rpsS (PRK00357)RpsS (COG0185)→ rplV (PRK00565)Ribosomal_L22 (pfam00237)→ RpsC (COG0092)rpsC (PRK00310)→ RplP (COG0197)rplP (PRK09203)→ |
| env-597 | CEWQ01009550.1 | + | 346 | 421 | **RNA**→ RplB (COG0090)rplB (PRK09374)→ |
| Ssp-3-1 | NZ_JPOY01000011.1 | - | 2334383 | 2334308 | **RNA**→ RplB (COG0090)rplB (PRK09374)→ rpsS (PRK00357)RpsS (COG0185)→ rplV (PRK00565)Ribosomal_L22 (pfam00237)→ RpsC (COG0092)rpsC (PRK00310)→ RplP (COG0197)rplP (PRK09203)→ |
| Ssp-4-1 | NZ_CH959310.1 | - | 345269 | 345194 | **RNA**→ RplB (COG0090)rplB (PRK09374)→ rpsS (PRK00357)RpsS (COG0185)→ rplV (PRK00565)Ribosomal_L22 (pfam00237)→ RpsC (COG0092)rpsC (PRK00310)→ RplP (COG0197)rplP (PRK09203)→ |
| env-598 | CEQX01294317.1 | - | 107 | 32 | **RNA**→ |
| env-599 | CESO01088717.1 | + | 87 | 162 | **RNA**→ rplB (PRK09374)Ribosomal_L2 (pfam00181)→ |
| Ssp-5-1 | NZ_CH959312.1 | - | 929126 | 929051 | **RNA**→ RplB (COG0090)rplB (PRK09374)→ rpsS (PRK00357)RpsS (COG0185)→ rplV (PRK00565)Ribosomal_L22 (pfam00237)→ RpsC (COG0092)rpsC (PRK00310)→ RplP (COG0197)rplP (PRK09203)→ |
| Spo-1-1 | NZ_AXZR01000005.1 | + | 3608 | 3683 | **RNA**→ RplB (COG0090)rplB (PRK09374)→ rpsS (PRK00357)RpsS (COG0185)→ rplV (PRK00565)Ribosomal_L22 (pfam00237)→ RpsC (COG0092)rpsC (PRK00310)→ RplP (COG0197)rplP (PRK09203)→ |
| env-600 | CEQZ01007430.1 | - | 103 | 28 | **RNA**→ |
| env-601 | CERD01103759.1 | + | 510 | 585 | **RNA**→ RplB (COG0090)rplB (PRK09374)→ rpsS (PRK00357)RpsS (COG0185)→ rplV (PRK00565)Ribosomal_L22 (pfam00237)→ RpsC (COG0092)rpsC (PRK00310)→ |
| env-602 | CERE01033137.1 | + | 1304 | 1379 | **RNA**→ RplB (COG0090)rplB (PRK09374)→ rpsS (PRK00357)RpsS (COG0185)→ rplV (PRK00565)Ribosomal_L22 (pfam00237)→ RpsC (COG0092)rpsC (PRK00310)→ RplP (COG0197)rplP (PRK09203)→ |
| env-603 | CERJ01019869.1 | - | 717 | 642 | **RNA**→ RplB (COG0090)rplB (PRK09374)→ |
| env-604 | CESK01030715.1 | - | 2698 | 2623 | **RNA**→ RplB (COG0090)rplB (PRK09374)→ rpsS (PRK00357)RpsS (COG0185)→ rplV (PRK00565)Ribosomal_L22 (pfam00237)→ RpsC (COG0092)rpsC (PRK00310)→ RplP (COG0197)rplP (PRK09203)→ |
| env-605 | CEOG01072571.1 | - | 582 | 507 | **RNA**→ RplB (COG0090)rplB (PRK09374)→ |
| env-606 | JGI11755J13075_100039 | + | 3752 | 3827 | **RNA**→ RplB (COG0090)rplB (PRK09374)→ rpsS (PRK00357)RpsS (COG0185)→ rplV (PRK00565)Ribosomal_L22 (pfam00237)→ RpsC (COG0092)rpsC (PRK00310)→ RplP (COG0197)rplP (PRK09203)→ |
| env-607 | JGI11949J13268_1000009 | + | 3765 | 3840 | **RNA**→ RplB (COG0090)rplB (PRK09374)→ rpsS (PRK00357)RpsS (COG0185)→ rplV (PRK00565)Ribosomal_L22 (pfam00237)→ RpsC (COG0092)rpsC (PRK00310)→ RplP (COG0197)rplP (PRK09203)→ |
| env-608 | CEQG01048819.1 | - | 2905 | 2830 | **RNA**→ RplB (COG0090)rplB (PRK09374)→ rpsS (PRK00357)RpsS (COG0185)→ rplV (PRK00565)Ribosomal_L22 (pfam00237)→ RpsC (COG0092)rpsC (PRK00310)→ RplP (COG0197)rplP (PRK09203)→ |
| env-609 | CEVJ01067098.1 | + | 321 | 396 | **RNA**→ RplB (COG0090)rplB (PRK09374)→ |
| env-610 | CAM_READ_0079652197 | + | 1 | 76 | **RNA**→ |
| env-611 | SL_1KL_011_SED_10748645 | + | 106 | 170 | **RNA**→ RplB (COG0090)rplB (PRK09374)→ |
| env-612 | SL_9KL_010_SED_10060232 | - | 1582 | 1518 | **RNA**→ RplB (COG0090)rplB (PRK09374)→ rpsS (PRK00357)RpsS (COG0185)→ rplV (PRK00565)Ribosomal_L22 (pfam00237)→ |
| env-613 | JGI24025J20009_10228543 | - | 185 | 112 | **RNA**→ RplB (COG0090)rplB (PRK09374)→ |
| env-614 | JGI24724J26744_10018762 | + | 3389 | 3462 | **RNA**→ |
| Oin-1-1 | NZ_ABID01000001.1 | + | 1417271 | 1417344 | **RNA**→ RplB (COG0090)rplB (PRK09374)→ RplB (COG0090)rplB (PRK09374)→ rpsS (PRK00357)RpsS (COG0185)→ rpsS (PRK00357)RpsS (COG0185)→ rplV (PRK00565)Ribosomal_L22 (pfam00237)→ rplV (PRK00565)Ribosomal_L22 (pfam00237)→ RpsC (COG0092)rpsC (PRK00310)→ RpsC (COG0092)rpsC (PRK00310)→ RplP (COG0197)rplP (PRK09203)→ RplP (COG0197)rplP (PRK09203)→ |
| env-615 | LAZR01017474.1 | + | 457 | 530 | **RNA**→ RplB (COG0090)rplB (PRK09374)→ rpsS (PRK00357)RpsS (COG0185)→ rplV (PRK00565)Ribosomal_L22 (pfam00237)→ RpsC (COG0092)rpsC (PRK00310)→ |
| env-616 | JGI24724J26744_10081951 | + | 450 | 522 | **RNA**→ RplB (COG0090)rplB (PRK09374)→ |

| | | | | | |
|---|---|---|---|---|---|
| env-617 | ABEF01021384.1 | - | 622 | 549 | **RNA**→ RplB (COG0090)rplB (PRK09374)→ |
| env-618 | CEQG01019720.1 | + | 5395 | 5468 | **RNA**→ RplB (COG0090)rplB (PRK09374)→ rpsS (PRK00357)RpsS (COG0185)→ rplV (PRK00565)Ribosomal_L22 (pfam00237)→ RpsC (COG0092)rpsC (PRK00310)→ RplP (COG0197)rplP (PRK09203)→ |
| env-619 | CEQZ01013875.1 | - | 101 | 28 | **RNA**→ |
| env-620 | CESC01001606.1 | - | 1753 | 1680 | **RNA**→ RplB (COG0090)rplB (PRK09374)→ rpsS (PRK00357)RpsS (COG0185)→ rplV (PRK00565)Ribosomal_L22 (pfam00237)→ rpsC (PRK00310)30S_S3_KH (cd02412)→ |
| env-621 | CEVI01215114.1 | - | 178 | 105 | **RNA**→ RplB (COG0090)rplB (PRK09374)→ |
| env-622 | CEWF01005413.1 | + | 2525 | 2598 | **RNA**→ RplB (COG0090)rplB (PRK09374)→ rpsS (PRK00357)RpsS (COG0185)→ rplV (PRK00565)Ribosomal_L22 (pfam00237)→ rpsC (PRK00310)30S_S3_KH (cd02412)→ |
| env-623 | CERO01018958.1 | - | 2363 | 2290 | **RNA**→ RplB (COG0090)rplB (PRK09374)→ rpsS (PRK00357)RpsS (COG0185)→ rplV (PRK00565)Ribosomal_L22 (pfam00237)→ RpsC (COG0092)rpsC (PRK00310)→ |
| env-624 | CEVA01074518.1 | + | 66 | 139 | **RNA**→ rplB (PRK09374)Ribosomal_L2 (pfam00181)→ |
| env-625 | JGI24724J26744_10034861 | + | 2147 | 2220 | **RNA**→ |
| env-626 | JGI24723J26617_10013494 | + | 1334 | 1407 | **RNA**→ RplB (COG0090)rplB (PRK09374)→ rpsS (PRK00357)RpsS (COG0185)→ rplV (PRK00565)Ribosomal_L22 (pfam00237)→ RpsC (COG0092)rpsC (PRK00310)→ RplP (COG0197)rplP (PRK09203)→ |
| env-627 | LAZR01008751.1 | + | 1162 | 1237 | **RNA**→ RplB (COG0090)rplB (PRK09374)→ rpsS (PRK00357)RpsS (COG0185)→ rplV (PRK00565)Ribosomal_L22 (pfam00237)→ RpsC (COG0092)rpsC (PRK00310)→ RplP (COG0197)rplP (PRK09203)→ |
| env-628 | JGI24724J26744_10022772 | + | 2527 | 2568 | **RNA**→ RplB (COG0090)rplB (PRK09374)→ |
| env-629 | JGI24724J26744_10281221 | + | 457 | 498 | **RNA**→ ←hypo |
| Sdo-1-1 | NZ_JASF01000005.1 | + | 1880338 | 1880379 | **RNA**→ RplB (COG0090)rplB (PRK09374)→ rpsS (PRK00357)RpsS (COG0185)→ rplV (PRK00565)Ribosomal_L22 (pfam00237)→ RpsC (COG0092)rpsC (PRK00310)→ RplP (COG0197)rplP (PRK09203)→ |
| Sdo-1-2 | NZ_JAMC01000005.1 | + | 6216 | 6257 | **RNA**→ RplB (COG0090)rplB (PRK09374)→ rpsS (PRK00357)RpsS (COG0185)→ rplV (PRK00565)Ribosomal_L22 (pfam00237)→ RpsC (COG0092)rpsC (PRK00310)→ RplP (COG0197)rplP (PRK09203)→ |
| env-630 | KVRMV2_105402183 | - | 158 | 94 | **RNA**→ RplB (COG0090)rplB (PRK09374)→ |
| env-631 | KVRMV2_100053128 | + | 1810 | 1873 | **RNA**→ RplB (COG0090)rplB (PRK09374)→ rpsS (PRK00357)RpsS (COG0185)→ rplV (PRK00565)Ribosomal_L22 (pfam00237)→ |
| env-632 | KVRMV2_100511851 | + | 614 | 679 | **RNA**→ RplB (COG0090)rplB (PRK09374)→ rpsS (PRK00357)RpsS (COG0185)→ |
| env-633 | 3300001592_10006696 | - | 11253 | 11192 | **RNA**→ RplB (COG0090)rplB (PRK09374)→ rpsS (PRK00357)RpsS (COG0185)→ rplV (PRK00565)Ribosomal_L22 (pfam00237)→ RpsC (COG0092)rpsC (PRK00310)→ RplP (COG0197)rplP (PRK09203)→ |
| env-634 | JGI24770J26754_10633113 | - | 235 | 176 | **RNA**→ RplB (COG0090)rplB (PRK09374)→ |
| env-635 | JGI24770J26754_10084963 | - | 885 | 824 | **RNA**→ RplB (COG0090)rplB (PRK09374)→ |
| env-636 | APMI01044789.1 | + | 283 | 344 | **RNA**→ RplB (COG0090)rplB (PRK09374)→ |
| env-637 | JGI24770J26754_10316505 | + | 277 | 338 | **RNA**→ RplB (COG0090)rplB (PRK09374)→ |
| Lme-1-1 | NC_023135.1 | + | 198601 | 198709 | **RNA**→ RplB (COG0090)rplB (PRK09374)→ rpsS (PRK00357)RpsS (COG0185)→ rplV (PRK00565)Ribosomal_L22 (pfam00237)→ hypo→ RpsC (COG0092)rpsC (PRK00310)→ RplP (COG0197)rplP (PRK09203)→ |
| env-638 | CAM_READ_0234721741 | - | 220 | 152 | **RNA**→ RplB (COG0090)rplB (PRK09374)→ |
| env-639 | 84546017 | - | 387 | 322 | **RNA**→ rplB (PRK09374)Ribosomal_L2 (pfam00181)→ |
| Rba-1-1 | NZ_AVDB01000002.1 | + | 2929 | 2994 | **RNA**→ RplB (COG0090)rplB (PRK09374)→ rpsS (PRK00357)RpsS (COG0185)→ rplV (PRK00565)Ribosomal_L22 (pfam00237)→ RpsC (COG0092)rpsC (PRK00310)→ RplP (COG0197)rplP (PRK09203)→ |
| env-640 | 84518656 | - | 774 | 709 | **RNA**→ RplB (COG0090)rplB (PRK09374)→ |
| env-641 | CAM_READ_0234724723 | + | 190 | 256 | **RNA**→ rplB (PRK09374)Ribosomal_L2 (pfam00181)→ |
| env-642 | CAM_READ_0235767883 | + | 190 | 256 | **RNA**→ RplB (COG0090)rplB (PRK09374)→ |
| env-643 | JCVI_SCAF_1101667302592 | - | 664 | 599 | **RNA**→ RplB (COG0090)rplB (PRK09374)→ |
| env-644 | CAM_READ_0236589595 | - | 346 | 280 | **RNA**→ rplB (PRK09374)Ribosomal_L2 (pfam00181)→ |
| env-645 | 84547844 | + | 884 | 948 | **RNA**→ |
| env-646 | CAM_READ_0236055591 | - | 90 | 24 | **RNA**→ hypo→ |
| env-647 | CAM_READ_0236222577 | - | 217 | 152 | **RNA**→ RplB (COG0090)rplB (PRK09374)→ |

| | | | | | |
|---|---|---|---|---|---|
| env-648 | CAM_READ_0235239301 | + | 303 | 368 | **RNA**→ |
| env-649 | JCVI_READ_1101854610991 | + | 341 | 406 | **RNA**→ rplB (PRK09374)Ribosomal_L2 (pfam00181)→ |
| env-650 | CAM_READ_0235244455 | + | 234 | 302 | **RNA**→ |
| Wma-1-1 | NZ_KB902287.1 | - | 74480 | 74410 | **RNA**→ RplB (COG0090)rplB (PRK09374)→ rpsS (PRK00357)RpsS (COG0185)→ rplV (PRK00565)Ribosomal_L22 (pfam00237)→ RpsC (COG0092)rpsC (PRK00310)→ RplP (COG0197)rplP (PRK09203)→ |
| env-651 | JCVI_READ_1108839274073 | + | 792 | 857 | **RNA**→ |
| env-652 | JGI11829J13087_100089 | + | 1021 | 1083 | **RNA**→ RplB (COG0090)rplB (PRK09374)→ rpsS (PRK00357)RpsS (COG0185)→ rplV (PRK00565)Ribosomal_L22 (pfam00237)→ RpsC (COG0092)rpsC (PRK00310)→ RplP (COG0197)rplP (PRK09203)→ hypo→ |
| Mal-2-1 | NZ_CP010855.1 | - | 3005866 | 3005804 | **RNA**→ RplB (COG0090)rplB (PRK09374)→ rpsS (PRK00357)RpsS (COG0185)→ rplV (PRK00565)Ribosomal_L22 (pfam00237)→ RpsC (COG0092)rpsC (PRK00310)→ RplP (COG0197)rplP (PRK09203)→ |
| env-653 | JGI12120J13076_116087 | + | 62 | 124 | **RNA**→ RplB (COG0090)rplB (PRK09374)→ |
| env-654 | JGI11929J13085_103365 | - | 387 | 325 | **RNA**→ rplB (PRK09374)Ribosomal_L2 (pfam00181)→ |
| env-655 | JGI11819J13099_125412 | + | 134 | 196 | **RNA**→ |
| env-656 | JGI12210J13797_10226873 | - | 121 | 59 | **RNA**→ hypo→ |
| Pga-3-1 | NC_023137.1 | + | 301203 | 301281 | **RNA**→ RplB (COG0090)rplB (PRK09374)→ rpsS (PRK00357)RpsS (COG0185)→ rplV (PRK00565)Ribosomal_L22 (pfam00237)→ RpsC (COG0092)rpsC (PRK00310)→ RplP (COG0197)rplP (PRK09203)→ |

#### 1.3 Conserved domains

Conserved domains found in protein-coding genes listed in Section 1.2 are shown below, with the first sentence in their description from the Conserved Domain Database (if any). Conserved domains associated with more than one L2-Alphaproteobacteria RNA are assigned a color, while others are shown in gray. The number in parentheses after the colored domain name is the number of occurrences in Section 1.2.

cd00719 (1) GIY-YIG nuclease domain superfamily.
cd01658 (4) Ribosomal protein L30, which is found in eukaryotes and prokaryotes but not in archaea, is one of the smallest ribosomal proteins with a molecular mass of about 7kDa.
cd02412 (29) K homology RNA-binding (KH) domain of the prokaryotic 30S small ribosomal subunit protein S3.
cd03886 (1) M20 Peptidase Aminoacylase 1 family.
COG0090 (567) Ribosomal protein L2 [Translation, ribosomal structure and biogenesis]
COG0091 (13) Ribosomal protein L22 [Translation, ribosomal structure and biogenesis]
COG0092 (310) Ribosomal protein S3 [Translation, ribosomal structure and biogenesis]
COG0094 (5) Ribosomal protein L5 [Translation, ribosomal structure and biogenesis]
COG0097 (5) Ribosomal protein L6P/L9E [Translation, ribosomal structure and biogenesis]
COG0098 (4) Ribosomal protein S5 [Translation, ribosomal structure and biogenesis]
COG0185 (365) Ribosomal protein S19 [Translation, ribosomal structure and biogenesis]
COG0186 (1) Ribosomal protein S17 [Translation, ribosomal structure and biogenesis]
COG0197 (289) Ribosomal protein L16/L10AE [Translation, ribosomal structure and biogenesis]
COG0198 (5) Ribosomal protein L24 [Translation, ribosomal structure and biogenesis]
COG0256 (1) Ribosomal protein L18 [Translation, ribosomal structure and biogenesis]
COG0596 (3) Pimeloyl-ACP methyl ester carboxylesterase [Coenzyme transport and metabolism, General function prediction only]
COG1917 (1) Cupin domain protein related to quercetin dioxygenase [General function prediction only]
pfam00181 (107) Ribosomal Proteins L2, RNA binding domain.
pfam00189 (3) Ribosomal protein S3, C-terminal domain.
pfam00203 (7) Ribosomal protein S19.
pfam00237 (339) Ribosomal protein L22p/L17e.
pfam00238 (5) Ribosomal protein L14p/L23e.
pfam00252 (8) Ribosomal protein L16p/L10e.
pfam00253 (5) Ribosomal protein S14p/S29e.
pfam00276 (1) Ribosomal protein L23.
pfam00366 (4) Ribosomal protein S17.
pfam00410 (5) Ribosomal protein S8.
pfam00831 (5) Ribosomal L29 protein.
pfam00861 (4) Ribosomal L18 of archaea, bacteria, mitoch.
pfam03947 (3) Ribosomal Proteins L2, C-terminal domain.
pfam06248 (1) Centromere/kinetochore Zw10.
pfam12697 (3) Alpha/beta hydrolase family.
pfam13759 (4) Putative 2OG-Fe(II) oxygenase.
pfam16867 (1) Dimethlysulfonioproprionate lyase.
PRK00004 (5) 50S ribosomal protein L24; Reviewed
PRK00010 (5) 50S ribosomal protein L5; Validated
PRK00136 (5) 30S ribosomal protein S8; Validated
PRK00306 (5) 50S ribosomal protein L29; Reviewed
PRK00310 (337) 30S ribosomal protein S3; Reviewed
PRK00357 (372) 30S ribosomal protein S19; Reviewed

PRK00550 (4) 30S ribosomal protein S5; Validated
PRK00565 (353) 50S ribosomal protein L22; Reviewed
PRK05483 (5) 50S ribosomal protein L14; Validated
PRK05498 (5) 50S ribosomal protein L6; Validated
PRK05593 (5) 50S ribosomal protein L18; Reviewed
PRK05610 (5) 30S ribosomal protein S17; Reviewed
PRK05611 (4) 50S ribosomal protein L30; Reviewed
PRK05738 (2) 50S ribosomal protein L23; Reviewed
PRK08881 (5) 30S ribosomal protein S14; Reviewed
PRK09203 (297) 50S ribosomal protein L16; Reviewed
PRK09374 (677) 50S ribosomal protein L2; Validated
smart00322 (5) K homology RNA-binding domain.
smart00465 (1) GIY-YIG type nucleases (URI domain).
TIGR01891 (1) amidohydrolase. [Protein fate, Degradation of proteins, peptides, and glycopeptides]
TIGR02466 (1) conserved hypothetical protein. [Hypothetical proteins, Conserved]

#### 1.4 Multiple-sequence alignment

Each L2-Alphaproteobacteria RNA is denoted by an abbreviation (like "Eco-1-1") that refers to a taxonomy given in Section 1.1. The alignment may include sequences containing the RNA motif, as well as flanking sequence. The L2-Alphaproteobacteria RNA itself is denoted by the line underneath marked 5′ and 3′ on either end. Nucleotides in flanking sequences (i.e., not part of the motif) are written in gray letters. Stems of predicted rho-independent transcription terminators, if any, are shaded yellow. (Note: terminator predictions have not been analyzed manually, and many are likely to be false positives. Terminator predictions are those of the RNIE software.) Nucleotides predicted to function (as DNA) as transcription-factor binding sites, if any, are shaded in green. (But note: these predictions are manually annotated, so they might be under-predicted.) Annotated start codons, if any, are shaded green. (Note: start codons are frequently misannotated, especially in environmental samples.) Nucleotides proposed to basepair as part of the consensus structure are shaded in color when they comprise Watson-Crick or G-U pairs. Otherwise they are shaded gray. Conserved stems are also indicated at the bottom of the alignment by angle brackets, where matching $<$ and $>$ denote base-paired columns. Below these angle brackets, the symbol "2" denotes base pairs exhibiting covariation according to the statistically well-founded R-scape method. "1" denotes base pairs exhibiting covariation according to R2R's simplistic method. "0" denotes base pairs that are not observed to mutate and "?" denotes base pairs that have a significant frequency of non-canonical nucleotides for Watson-Crick or G-U pairs ($>5\%$). Below these base pair annotation is the consensus sequence: "R" = "A" or "G", "Y" = "C" or "U", red nucleotides: nucleotide identity conserved more than 97% of the time, black nucleotides: 90%, gray nucleotides: 75%, red circle (•): nucleotide is present 97% of the time, black circle (•): 90%, gray circle (•): 75%, white circle (◦): 50%. All percentages of sequences just described (e.g. 97% conserved) assume that sequences have been weighted by the GSC algorithm implemented by the Infernal software package.

Duplicate sequences: the following putative homologs are not shown in the alignment because their sequences are identical to a homolog already shown: Cba-1-1, Gne-1-1, Kvu-2-1, Lca-1-1, Lsp-5-1, Lsp-9-1, Nig-1-1, Nit-1-1, Nit-1-2, Oin-1-1, Pfe-2-1, Rba-5-1, Rca-1-1, Rca-2-1, Rca-3-1, Rca-5-1, Rmo-2-1, Rnu-1-1, Rsp-20-1, Rsp-3-1, Rsp-6-1, Sdo-1-2, Spo-1-1, Ssp-1-1, Ssp-3-1, Tsp-2-1, env-100, env-102, env-107, env-11, env-110, env-111, env-112, env-113, env-114, env-115, env-12, env-123, env-126, env-129, env-130, env-131, env-132, env-133, env-134, env-137, env-140, env-142, env-147, env-152, env-153, env-154, env-159, env-160, env-161, env-180, env-181, env-182, env-183, env-184, env-185, env-198, env-199, env-208, env-209, env-210, env-211, env-212, env-238, env-239, env-240, env-241, env-242, env-243, env-244, env-245, env-246, env-249, env-250, env-251, env-253, env-267, env-268, env-270, env-28, env-281, env-283, env-29, env-292, env-294, env-30, env-302, env-303, env-304, env-305, env-306, env-307, env-31, env-318, env-32, env-328, env-33, env-333, env-334, env-34, env-348, env-349, env-350, env-351, env-352, env-353, env-354, env-355, env-366, env-368, env-376, env-377, env-383, env-384, env-385, env-387, env-390, env-393, env-401, env-406, env-408, env-409, env-410, env-411, env-412, env-415, env-420, env-424, env-425, env-44, env-442, env-443, env-444, env-446, env-449, env-45, env-450, env-451, env-452, env-453, env-454, env-455, env-456, env-457, env-46, env-461, env-462, env-466, env-468, env-470, env-471, env-474, env-478, env-479, env-48, env-480, env-481, env-485, env-486, env-487, env-488, env-489, env-490, env-491, env-492, env-496, env-498, env-499, env-501, env-504, env-509, env-510, env-511, env-529, env-53, env-532, env-535, env-537, env-538, env-539, env-54, env-540, env-541, env-542, env-543, env-544, env-545, env-547, env-550, env-551, env-552, env-553, env-554, env-555, env-556, env-557, env-558, env-559, env-560, env-561, env-562, env-563, env-564, env-565, env-566, env-568, env-569, env-570, env-571, env-572, env-573, env-574, env-575, env-577, env-578, env-58, env-580, env-581, env-584, env-585, env-586, env-587, env-588, env-589, env-593, env-594, env-599, env-601, env-602, env-603, env-604, env-607, env-608, env-609, env-61, env-610, env-619, env-62, env-620, env-621, env-622, env-63, env-642, env-654, env-655, env-73, env-80, env-81, env-82, env-87, env-95

alignment positions 1 · · · 180

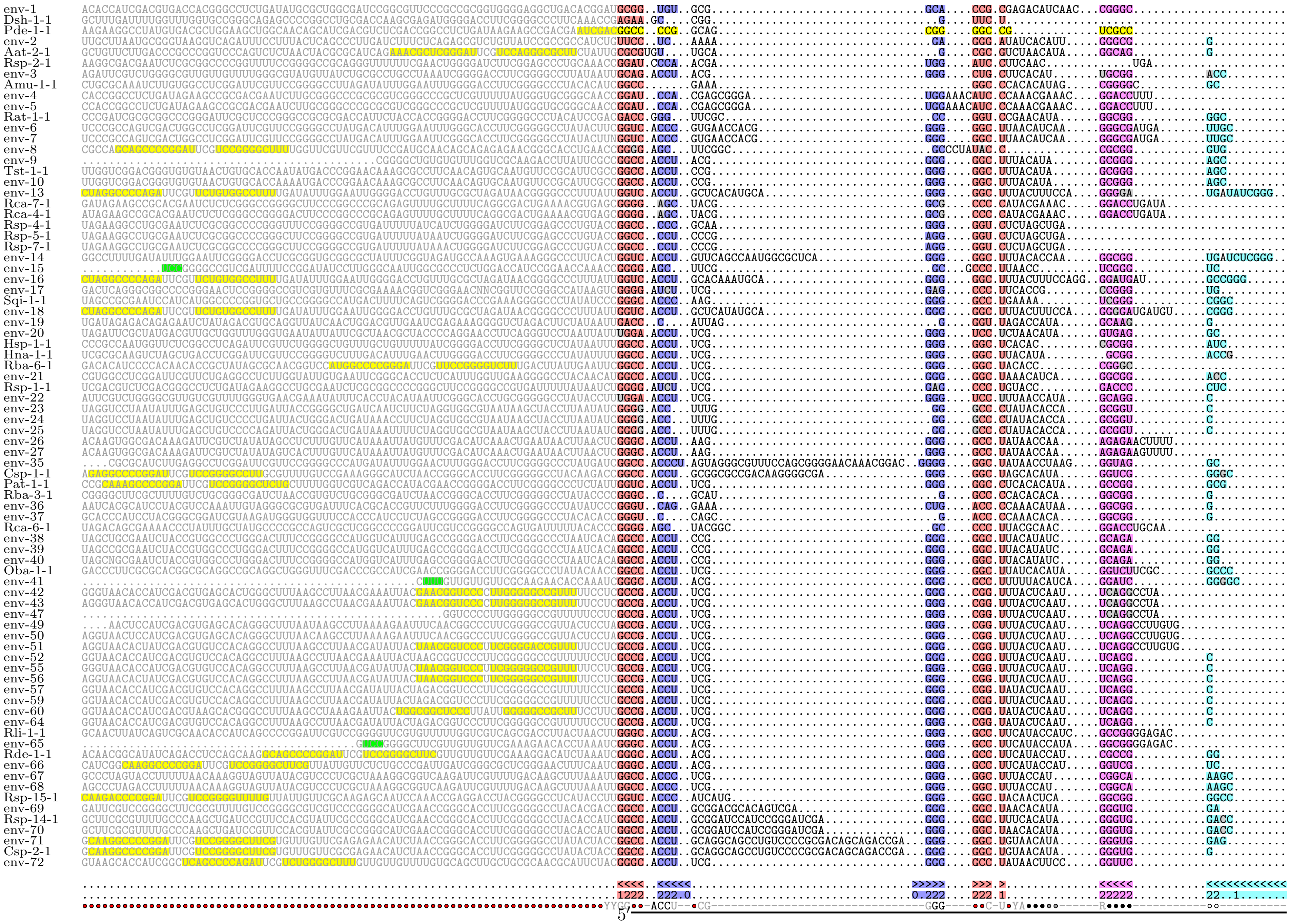

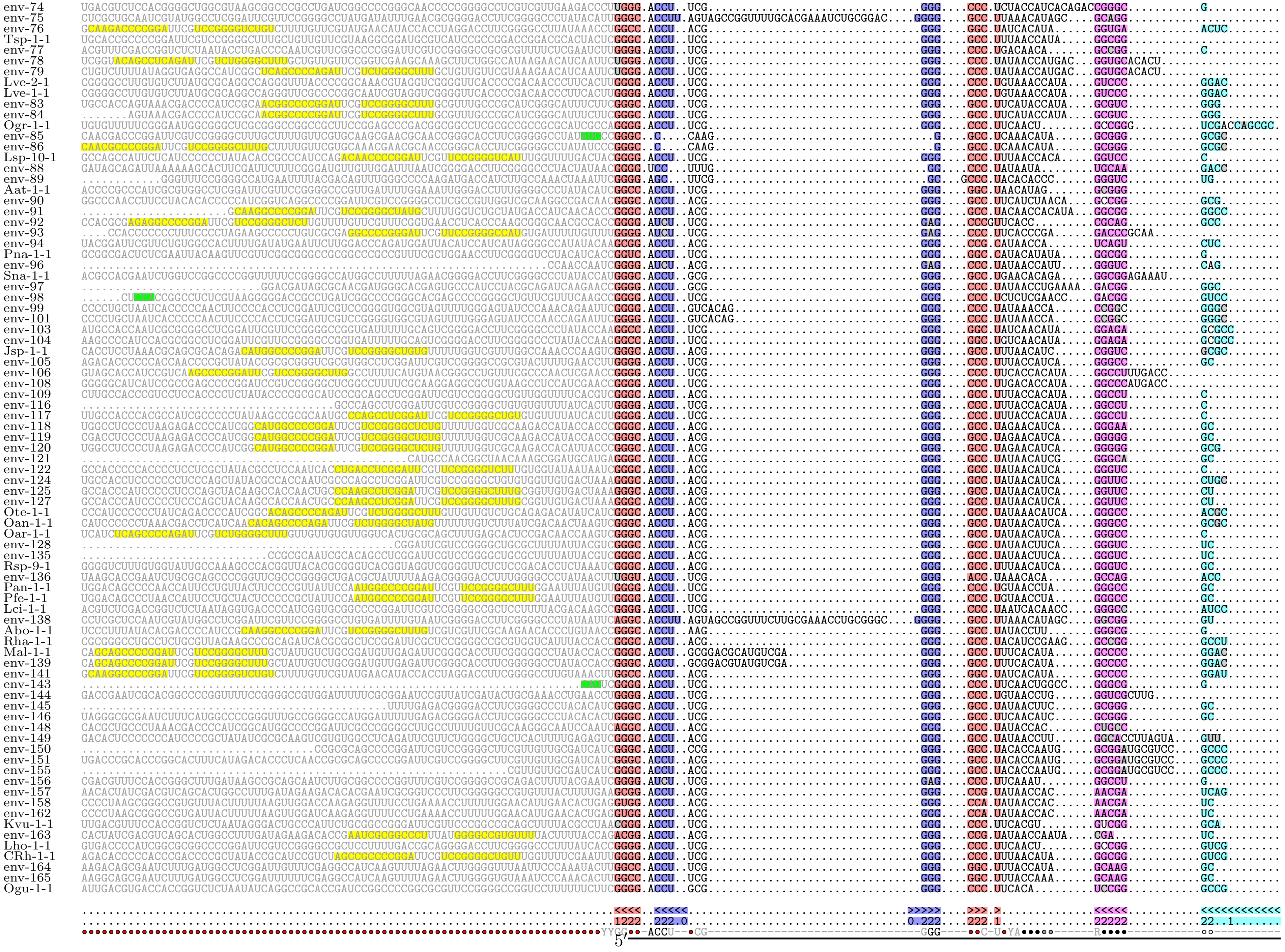
env-74
env-75
env-76
Tsp-1-1
env-77
env-78
env-79
Lve-2-1
Lve-1-1
env-83
env-84
Ogr-1-1
env-85
env-86
Lsp-10-1
env-88
env-89
Aat-1-1
env-90
env-91
env-92
env-93
env-94
Pna-1-1
env-96
Sna-1-1
env-97
env-98
env-99
env-101
env-103
env-104
Jsp-1-1
env-105
env-106
env-108
env-109
env-116
env-117
env-118
env-119
env-120
env-121
env-122
env-124
env-125
env-127
Ote-1-1
Oan-1-1
Oar-1-1
env-128
env-135
Rsp-9-1
env-136
Pan-1-1
Pfe-1-1
Lci-1-1
env-138
Abo-1-1
Rha-1-1
Mal-1-1
env-139
env-141
env-143
env-144
env-145
env-146
env-148
env-149
env-150
env-151
env-155
env-156
env-157
env-158
env-162
Kvu-1-1
env-163
Lho-1-1
CRh-1-1
env-164
env-165
Ogu-1-1
5′

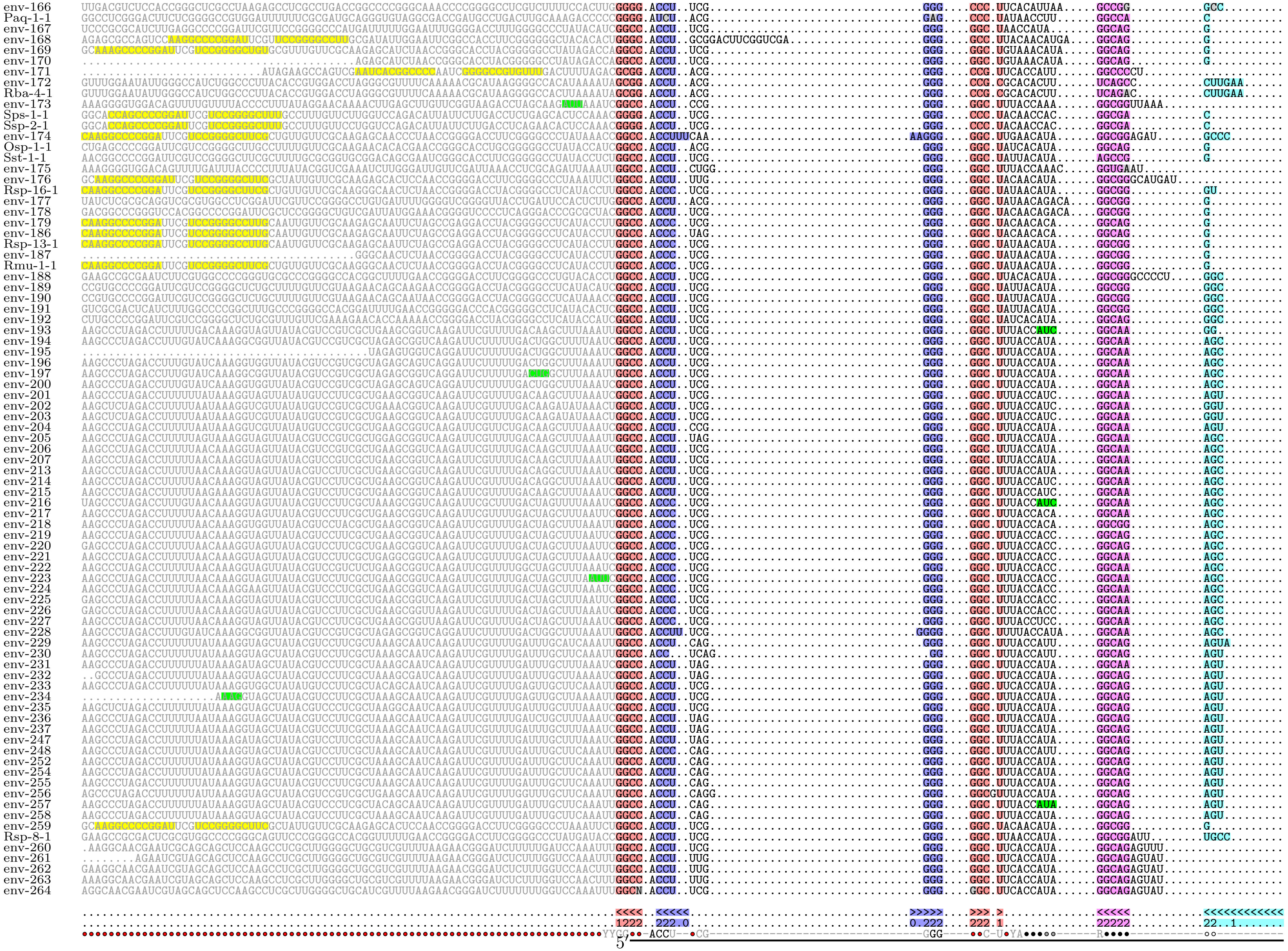
env-166
Paq-1-1
env-167
env-168
env-169
env-170
env-171
env-172
Rba-4-1
env-173
Sps-1-1
Ssp-2-1
env-174
Osp-1-1
Sst-1-1
env-175
env-176
Rsp-16-1
env-177
env-178
env-179
env-186
Rsp-13-1
env-187
Rmu-1-1
env-188
env-189
env-190
env-191
env-192
env-193
env-194
env-195
env-196
env-197
env-200
env-201
env-202
env-203
env-204
env-205
env-206
env-207
env-213
env-214
env-215
env-216
env-217
env-218
env-219
env-220
env-221
env-222
env-223
env-224
env-225
env-226
env-227
env-228
env-229
env-230
env-231
env-232
env-233
env-234
env-235
env-236
env-237
env-247
env-248
env-252
env-254
env-255
env-256
env-257
env-258
env-259
Rsp-8-1
env-260
env-261
env-262
env-263
env-264
5′

```
env-265   GAAGGCAAUGAAUCGUAGCAGCUCCAAGCCUCGCUUGGGGCUGCGUCGUUUUAAGAACGGGAUCUUUUUGGUCCAAAUUUGGCC.ACCU..UCG...............................GGG....GGC.UUCACCAUA.....GGCAGAGUAU.................
env-266   AGGCAACGAAUCGUAGCAGCUCCAAGCCUCGCUUGGGGCUGCGUCGUUUUAAGAACGGGAUCUUUUUUGGUCCAAAUUUGGCC.ACCU..UCG...............................GGG....GGC.UUCACCAUA.....GGCAGAGUAU.................
env-269   GAAGGCAACGAAUCGCAGCAGCUCCAAGCCUCGCUUGGGGCUGCGUCGUUUUAAGAACGGGAUCUUUUUGAUCCAAAUUUGGCC.ACCU..UCG...............................GGG....GGC.UUCACCAUA.....GGCAGAGUAU.................
env-271   GAAGGCAACGAAUCGUAGCAGCUCCAAGCCUCGCUUGGGGCUGCGUCGUUUUAAGAACGGGAUCUUUUUGGUCCAAAUUUGGCC.ACCU..UCG...............................GGG....GGC.UUCACCAUA.....GGCAGAGUAU.................
env-272   ............................CUCGCUUGGGGCUGCCUCGUUUUAAGAACGGGAUCUUUUUUGGUCCAAAUUUGGCC.ACCU..UCG...............................GGG....GGC.UUCACCAUA.....GGCAGAGUAU.................
env-273   AAAGGCAACGAAUCGCAGCAGCUCCAAGCCUCGCUUGGGGCUGCGUCGUUUUAAGAACGGGAUCUUUUUGAUCCAAAUUUGGCC.ACCU..UCG...............................GGG....GGC.UUCACCAUA.....GGCAGAGUUU.................
env-274   UGCACUACAAAAAUUAGGGGUGGACAGUUUGGUUACCCUUAUACGACGAAUCUGGGUGUAAAGAAUAUCGCCUAACAACCGGCC..CC...UUCG..............................GG....GGC.UUACCAAA.......GGCGGUAGU..................
env-275   AAGGGGUGGACAGUUGUGAUUUACCCUUUAUAGGAACCAAAUCUUGCUGCUUCUAAAAAAGUAUCGCAAGAUUAAAUUGGCC.ACCU..CCG...............................GGG....GGC.UUUACCAAA.....GGUGGUUAAA.................
env-276   UUAAGGGGUGGACACUUUCGCUUUACCCCUUUAUACGACCGAAAUCUUGGGUGGUAAAAGUUUAUCGCCUAAACAACCGGCC.ACCU..UCG...............................GGG....GGC.UUUACCAAA.....GGCGG...........UU.........
env-277   ..............UUCUGAUUUACCCCUUUAUACGAUCGAAAUCUUGAGAUCGUCGAUAAGAUCUCUCAAGAGUAAUUUGGCC.ACCU..UCG...............................GGG....GGC.UUUACCAAA.....GGCAGUAAUA.................
env-278   AAGGGGUGGACAGUUCUGAUUUACCCCUUUAUACGAUCGAAAUCUUGGAUACCUCGAUAAGAGCUUCUAAGGUUAAAAUGGCC.ACCU..UCG...............................GGG....GGC.UUUACCAAA.....GGCGUUAAA..................
env-279   AAGGGGUGGACAGUUCUGAAUUACCCCUUUAUACGAUCGCAAUCUUGGAACUUUGAAUAAAAGAUUGCAAGAUUAAAUUGGCC.ACCU..UCG...............................GGG....GGC.UUUACCGAA.....GGCGAUAAAA.................
env-280   AAGGGGUGGACAGUUCUGAAUUACCCCUUUAUACGAUCGCAAUCUUAGGAACUUUGAAUAAAAGAUUGCAAGAUUAAAUUGGCC.ACCU..UCG...............................GGG....GGC.UUUACCGAA.....GGCGAUAAAU.................
env-282   AAGGGGUGGACAGUUCUGAAUUACCCCUUUAUACGAUCGCAAUCUUAGGAACUUUGGAUAAAAGAUUGCAAGAUUAAAUUGGCC.ACCU..UCG...............................GGG....GGC.UUUACCGAA.....GGCGAUAAAA.................
env-284   AAGGGGUGGACAGUUCUGAAUUACCCCUUUAUACGAUCGCAAUCUUAGGAAUUUGGAUAAAAGAUUGCAAGAUUAAAUUGGCC.ACCU..UCG...............................GGG....GGC.UUUACCGAA.....GGCGAUAAAA.................
env-285   ..............................................UUAGGAACUUUGGAUAAAAGAUUUCAAGAUUAAAUUGGCC.ACCU..UCG...............................GGG....GGC.UUUACCGAA.....GGCGAUAAAA.................
env-286   AAGGGGUGGACAGUUUUAGUUUGCCCCAUUAUAUGAUCGCAAUCUUGGAGUAGCUUUAGUAAUGGCUAACUAGAUUAACUUGGCC.ACCU..CCG...............................GGG....GGC.UUUACCACA.....GGCGCUAACA.................
env-287   AAAGGGGUGGACAGUUCUGAUUUACCCCUUUAUACGAUCGAAAUCUUGAGAACUUCGAUAAAGGUUCGAAAGAAUAAACUGGCC.ACCU..GUG...............................GGG....GGC.UUUACCAUA.....GGCGGUGAAU.................
env-288   AUGGGGUGGACAGUUCUGAUUUACCACAUUAUACGAUCCAAAACCUGCAGAAUCUAGAAAUUUGGCUGCAGUUUAAAAUGGCC.ACCU..UCG...............................GGG....GGC.UUUACCGAA.....GGCGGUAUUA.................
env-289   AAAGGGGUGGACAGUUCUGAUUUACCCCUUUAUAUGAUCGAAAUCUUGAGAACUUAGAUACACGUUCGCAAGAAUAAACUGGCC.ACCU..GCG...............................GGG....GGC.UUUACCAUA.....GGCAUUGAAA.................
env-290   AGGGGCUGGACAGUUUUGGUUUACACCUUUAUAUGAACGCAAUCUUGUGAUUGUUCGAUAGAACCUUGCAAAAUUAAAUUGGCC.ACCU..CUG...............................GGG....GGC.UUUACCAAA.....GGCGGUAAAU.................
env-291   AAGGGGCUGGACAGUUUUGAUUUACCCCUUUAUACGAAUGCAAUCUUGGAUAUGUUCGAUGGAACCUUCAAGAUUAAAUUGGCC.ACCU..CAG...............................GGG....GGC.UUUACCAAA.....GGCGGUAAAU.................
env-293   AAGGGGCUGGACAGUUUUGAUUUACCCCUUUAUACGAAUGCAAUCUUGGAUAUGUUCGAUGGAACCUUCAAGAUUAAAUUGGCC.ACCU..AAG...............................GGG....GGC.UUUACCAAA.....GGCGGUAAAU.................
env-295   AAGGGGUGGACAGUUCUGAUUUACCCCUUUAUACGAUCGAAAUCUUGAGAACGUCGAUAAGAGUUCGCAAGAAUAAACUGGCC.ACCU..GCG...............................GGG....GGC.UUUACCAUA.....GGCGGUAAGA.................
env-296   AAGGGGUGGACAGUUCUGAUUUACCCCUUUAUACGAUCGAAAUCUUGUGAGCUUGGAUAAAAGCGCGCAAGAAUAAACUGGCC.ACCU..GCG...............................GGG....GGC.UUUACCAUA.....GGCGGUAAGA.................
env-297   AUGGGGUGGACAGUUUUUAUUUACCACUUUAUAUGAUCCAAAUCUUGUAGAUUCUAGGUAAGAGUUUGCAAGUGUAAAAUGGCC.ACCU..UCG...............................GGG....GGC.UUUACCAUA.....GGCGGCUAAU.................
env-298   AUGGGGUGGACAGUUUUUAUUUACCACUUUAUAUGAUCCAAAUCUUGUAGAUUCUAGGUAAGAGUUUGCAAGUGUAAAAUGGCC.ACCU..UCG...............................GGG....GGC.UUUACCAAA.....GGCGGCAAAU.................
env-299   AUGGGGUGGACAGUUUUUAUUUACCACUUUAUAUGAUCCAAAUCUUGUAGAUUCUAGGUAAGAGUUCGCAAGUGUAAAAUGGCU.ACCU..UCG...............................GGG....GGC.UUUACCAAA.....GGCGGCAAAU.................
env-300   GGGGUGGACAGUUCUCUGAUUUACCCCUUUAUACGGUCGAAAUCUUGAGAACUUGGAUAAAAGUUCGCAAGAUAAACUGGCC.ACCU..GCG...............................GGG....GGC.UUUACCAUA.....GGCAGUAAAA.................
env-301   AAGGGGUGGACAGUUUUAAUUUACCCCUUUAUACGAUCGCAAUCUUGGGUUCUCUAAAAAAGAGAUCACAAGGUCAACCUGGCC.ACCU..UCG...............................GGG....GGC.UUUACCAAA.....AGCGGUAAAA.................
env-308   AAAGGGGUGGACAGUUCUGAUUUACCCCUUUAUACGAUCGAAAUCUUGAGAUCUUUGAUAAAAGGUCUCAAGAAGAAACCGGCC.ACCU..UCG...............................GGG....GGC.UUUACCAUA.....GGCGGUAAAA.................
env-309   AAAGGGGUGGACAGUUCUGAUUUACCCCUUUAUACGAUCGCAAUCUUAAGAACUUCGAUAAAAGUUCGCAAGAAUAAACUGGCC.ACCU..UCG...............................GGG....GGC.UUUACCAUA.....GGCGGUAAAA.................
env-310   ...............UUUGAUUUACCCCUUUAUACGAUCCAAAUCUCGAGAACUUCGAUAAAAGUUCGAAAGAAUAAACUGGCC.ACCU..UCG...............................GGG....GGC.UUUACCAUA.....GGCGGUAAAA.................
env-311   .....................................................CUUCGAUAAAAGUUCGCAAGAAUAAACCGGCC.ACCU..UCG...............................GGG....GGC.UUUACCAUA.....GGCGGUAAAA.................
env-312   AAAGGGGUGGACAGUUCUGAUUUACCCCUUUAUACGAUCGAAAUCUUGUGAACUCCGUUAAAAGUUCGCAAGAAUAAACCGGCC.ACCU..UCG...............................GGG....GGC.UUUACCAUA.....GGCGGUAAAA.................
env-313   AAAGGGGUGGACAGUUCUGAUUUACCCCUUUAUACGAUCCAAAUCUUGAGAUCUUCGAUAAAAGUUCGCAAGAAUAAACCGGCC.ACCU..UCG...............................GGG....GGC.UUUACCAUA.....GGCGGUAAAA.................
env-314   AAAUUUAAGGGGUGGACAGUUUGCUUUACCCUUUAUACGACGAAAUCUUGGGUGGUAAAUGGUUUAUCGCUAAACGCAUCGGCC.ACUU..UCG...............................GGG....GGC.UUUACCAAA.....GGCGGUUAGU.................
env-315   UUAAGGGGUGGACAGUUUUGCUUUACCCCUUUAUAUGACCGAAAUCUUGGGUGGUAAAAGUUUUAUCAGCCAAACAAACCGGCC.ACCC..UCG...............................GGG....GGC.UUUACCAAA.....GGCGGUUAGU.................
env-316   ............UAUAGGCGGAGAUACCCUUUAUACGACCAAAAUCUUGGGUGGUAAAAGAAUAUCGCCUAAACAACCGGCC.ACCU..UCG...............................GGG....GGC.UUUACCAAA.....GGCGGUUAGU.................
env-317   UUAAGGGGUGGACACUUUAGGUUUACCCCUUUAUACGACCGAAAUCUUGGGUGGUAAAAGUUUUAUCAGCCAAACAAACCGGCC.ACCU..UCG...............................GGG....GGC.UUUACCAAA.....GGCGGUUAGU.................
env-319   UUAAGGGGUGGACAGUUUUGCUUUACCCCUUUAUACGACCGAAAUCUUGGGUGGUAAAAGUUUAUCGCCUAAACACAUCGGCC.ACCU..UCG...............................GGG....GGC.UUUACCAAA.....GGCGGUUAGU.................
env-320   UUAAGGGGUGGACAGUUUUGCUUUACCCCUUUAUACGACCGAAAUCUUGGGUGGUAAAAGAUUUAUCGCCUAAACAAUCUGGCC.ACCU..UCG...............................GGG....GGC.UUUACCAAA.....GGCGGUUAGU.................
env-321   UUAAGGGGUGGACAGUUUCGCUUUACCCCUUUAUACGACCGAAAUCUUGGGCGGUAAAAUAAAAUAUCGCUUAAAUAAACCGGCC.ACCU..UCG...............................GGG....GGC.UUUACCAAA.....GGCGGUUAGU.................
env-322   UUAAGGGGUGGACAGUUUUGCUUUACCCCUUUAUACGACCGAGAUCUUGGGCGGUAAAAGAUUUAUCGCCUAAACAACCGGCC.ACCU..UCG...............................GGG....GGC.UUUACCAAA.....GGCGGUUAGU.................
env-323   AAGGGGUGGACAGUUUUGAUUUACCCCUUUAUACGGUCGAAAUCUUGGGAUUGUUCGAUCAAACCUCGCAAGAUUAAAUUGGCC.ACCU..GCG...............................GGG....GGC.UUUACCAAA.....GGCGGUAAGU.................
env-324   UUAAGGGGUGGACAGUUUUGCUUUACCCCUUUAUACGACCGAAAUCUUGGGUGGUAAAAGAUUUAUCGCCUAAACAAUCUGGCC.ACCU..UCG...............................GGG....GGC.UUUACCAAA.....GGCGGUAAGU.................
env-325   UUAAGGGGUGGACAGUUUUGCUUUACCCCAUUAUACGACCGAAAUCUUGGGUGGUAAAAGAUUUAUCGCCUAAACAAUCUGGCC.ACCU..UCG...............................GGG....GGC.UUUACCAAA.....GGCGGUAAGU.................
env-326   UUAGGGGGUGGACAGUUUUGCUUUACCCCUUUAUAUGACCGAAAUCUUGGGUGGUAAAAGAUUUAUUGCCUAAAAAAUCUGGCC.ACCU..UCG...............................GGG....GGC.UUUACCAAA.....GGCGGUAAGU.................
env-327   UUAAGGGGUGGACAGUUUUGCUUUACCCCUUUAUACGACCGAAAUCUUGGGUGGUAAAAGAUUUAUCGCCUAAACAACCGGCC.ACCU..UCG...............................GGG....GGC.UUUACCAAA.....GGCGGUAAGU.................
env-329   UUAAGGGGUGGACAGUUUUGCUUUACCCCUUUAUACGACCAAAAUCUUGGGUGGUAAAAAAUUUAUCGCCUAAACAACCGGCC.ACCU..UCG...............................GGG....GGC.UUUACCAAA.....GGCGGUAAGU.................
env-330   UUAAGGGGUGGACAGUUUUGCUUUACCCCUUUAUACGACCAAAAUCUUGGGUGGUAAAAAAUUUAUCGCCUAAAUAAUCUGGCC.ACCU..UCG...............................GGG....GGC.UUUACCAAA.....GGCGGUAAGU.................
env-331   AUGGGGUGGACAGUUUUUAUUUACCACUUUAUAUGAUCCAAAACUUGUAGAUUCUAAUUAAGACUUUGCAAGUGUAAAAUGGCC.ACCU..UCG...............................GGG....GGC.UUUACCAAA.....GGCGGUAAAU.................
env-332   AUGGGGUGGACAGUUUUUAUUUACCACUUUAUAUGAUCCAAAUCUUGUAGAUUCUAGGUAAGAGUUUGCAAGUGUAAAAUGGCC.ACCU..UCG...............................GGG....GGC.UUUACCAAA.....GGCGGUAAAU.................
env-335   AUGGGGUGGACAGUUUUUAUUUACCACUUUAUAUGAUCCAAAUCUUGCAGAUUCUAGAUAAGAGUUUGCAGGUGUAAAAUGGCC.ACCU..UCG...............................GGG....GGC.UUUACCAAA.....GGCGGUAAAU.................
env-336   AAAGGGGUGGACAGUUCUGAUUUACCCCUUUAUACGAUCGCAAUCUUAAGAACUUCGAUAAAAGUUCGCAAGAAUAAACUGGCC.ACCU..UCG...............................GGG....GGC.UUUACCAUA.....GGCGGUAAAA.................
env-337   AAAGGGGUGGACAGUUCUGAUUUACCCCUUUAUACGAUCGAGAUCUUGAAGACUUUUAUCAAAGUUCGCAAGAAUAAACCGGCC.ACCU..UCG...............................GGG....GGC.UUUACCAAA.....GGCGGUAAAA.................
env-338   AAGGGGUGGACAGUUCUGGAUUACCCCUUUAUACGAGCGCAAUCUUGGGAAUUUUGGAAAAGAGUUCGAAAGAUAAAAACGGCC.ACCU..UCG...............................GGG....GGC.UUUACCAAA.....GGCGGUAAAA.................
env-339   AAGGGGUGGACAGUUCUGGAUUACCCCUCUAUAUGAGCGAAAUCUUGGGACUUUUGGAAAAGAGUUUAAAAGAUAAAAACGGCC.ACCU..UCG...............................GGG....GGC.UUUACCAAA.....GGCGGUAAAA.................
env-340   AAAGGGGUGGACAGUUCUGAUUUACCCCUUUAUACGAUCGAAAUCUUGAGAACUUCGAUAAAAGUUCGCAAGAAUAAACUGGCC.ACCU..UCG...............................GGG....GGC.UUUACCAAA.....GGCGGUAAAA.................
env-341   AAAGGGGUGGACAGUUCUGAUUUAGCCCUUUAUACGAUCGAAAUCUUGAAAGCUUUAAUCAAAGUUCGCAAGAGUAAACUGGCC.ACCU..UCG...............................GGG....GGC.UUUACCAAA.....GGCGGUAAAA.................
env-342   AGGGGGUGGACAGUUCUGAGUUACCCCUUUAUACGAUCGAAAUCUUGGGGACUCUCUAUAUGGGUUUGUAAGAUUAAAUCGGCC.ACCU..UCG...............................GGG....GGC.UUUACCAAA.....GGCGGUAAAA.................
env-343   UUUAAGGGGUGGACAGUUUUGCUUUACCCCUUUAUACGACCGAAAUCUUGGGUGGUAAAAGAUUAUCGCCUAAACAACCUGGCC.ACU...UCGG..............................GG....GGC.UUUACCAAA.....GGCGGUAAGU.................
env-344   UAAGGGGUGGACAGUCUGCUUUACCCCUUUAUACGACCAAAAAACUUGGGUGGUAAAAGAAUUAUCGCCUAAACAACCGGCC.ACCU..UCG...............................GGG....GGC.UUUACCAAA.....GGCGGCAAGU.................
env-345   AAGGGGUGGACAGUUUUGAUUUACCCCUUUAUAUGAUCAAAAUCUUAGGAUCGUUAGACAAGACAUCCAAAGAUUAAAUCGGCC.ACCU..CCG...............................GGG....GGC.UUUACCAGA.....GGCGGUAAAA.................
env-346   AAAGGGGUGGACAGUUCUGAUUUACCCCUUUAUACGAUCGAAAUCUUGAGAACUUCGAUAAAAGUUCGCAAGAAUAAACUGGCC.ACCU..GCG...............................GGG....GGC.UUUACCAGA.....GGCGGUAAAA.................
env-347   AAAGGGGUGGACAGUUCUGAUUUACCCCUUUAUACGAUCGAAAUCUUGAGAACUUCGAUAAAAGUUCGCAAGAAUAAACUGGCC.ACCU..GCG...............................GGG....GGC.UUUACCAUA.....GGCGGUAAAA.................
env-356   AAAGGGGUGGACAGUUCUGAUUUACCCCUUUAUAUGAUCGAAAUCUUGAGAACUUCGAUAAAAGUUCGCAAGAAUAAACUGGCC.ACCU..GCG...............................GGG....GGC.UUUACCAUA.....GGCGGUAAAA.................
env-357   ...................................................................UUGGCUAAACAACCUGGCC.ACCC..UCG...............................GGG....GGC.UUUACCAAA.....GGCGGAAAGU.................
env-358   ....GGUGGACAGUUCUGAUUUACCACUUUAUAUGAUCCCAACCUUGUAGAAUCUAUAACUUAGGCUGCAAGUUUAAAACGGCC.ACCU..UCG...............................GGG....GGC.UUUACCAUA.....GGCGGUUAUA.................
env-359   AUGGGGUGGACAGUUCUGAUUUACCACUUUAUAUGAUCCCAACCUUGUAGAAUCUAUAACUUAGGCUGCAAGUUUAAAACGGCC.ACCU..UCG...............................GGG....GGC.UUUACCAUA.....GGCGGUUAUA.................
env-360   AAAGGGGUGGACAGUUGUGAAUUACCCCUUUAUACGAGCGAAACCUUGAGAACUUUUAUAAAAUUUCUCAAGAGUAAACUGGCC.ACCU..UCG...............................GGG....GGC.UUUACCAAA.....GGCGGUAAA..................
env-361   AAAGGGGUGGACAGUUCUGAUUUACCCCUUUAUACGAUCGAAAUCUUGAGAGCUUCGAUAGAUUUCUCAAGAUUUAACUGGCC.ACCU..UCG...............................GGG....GGC.UUUACCAGA.....GGCGAUAAA..................
Psp-3-1   GGCUUGCCUUUUUUGUUCGGCGCGGGUUUGCCCGCACUUCAGGACAGCGAAAAUCGGGCACCUUCGGGGGCCUAUACUCAGGCC.ACCU..UCG...............................GGG....GGC.UUUAACAUA.....GGCGG...........GGGC.......
Hmi-1-1   GAAUCUCGUGGCCCCGGACUCUUCCGGGGCCACGCGCUUUCAGCAGAAGCGUAGCUUUCAUUGCCGAAAGUGCACUCAGCGGGG.AUCU..UCG...............................GAG....CCC.UAUAACCCCC.....GGCGGAC.........GC.........
Hma-1-1   GAAUCUCGUGGCCCCGGAUUCUUCCGGGGCCACGCGCUUUCAGCAGAGACGUGGCUUUCUUGCCGGAAGUGCACUCAGCGGGG.AUCU..UCG...............................GAG....CCC.UAUAACCUCC.....GGGCCAC.........GGC........
Laq-1-1   UGCAGACCCCGGAUUCGUUCGGGGUUGUUACGUUUGUCAGGAUGUUUGAACUUUGGGACCUUCGGGGCCCUAUAUCUCAGGCA.ACCCU.AGUAGGGCGUCUCAUCGUGGGACAACCGGGC...GGGG....GGC.UUACACAUA.....GCGGGU..........CCAAAG.....
env-362   CAUCCCCCCUAAACGAGCCUAUCUGUGUGGCCCCGGAUUCGUCCGGGGCUUCGCAUUCGUUCGUAAGAACACCCUAAGCCGGGC.ACCU..ACG...............................GGG....GCC.UAUAAACAU......GGUGG...........G..........
Rpo-1-1   ACGCCACCAAUCUUGCGCCCUGGGAUUCGUCCGGGGCUCGUGAUUUUGCAGCCGGGGACCUUCGGGGCCCUAUACCAUGGGG.ACCU..ACG...............................GGG....GCC.UCAAUACAUA.....GGCGG...........G..........
env-363   ..............UUCUUUAUUUACCCAUAUAUUUUGGGAUAAGUGUUAUUGGUAGCUAUUUAAGCUACUUUAAUUCAGGGG.ACCU..CUG...............................GGG....CCC.UAUAAACCA.......GCGG...........UCU........
env-364   AUGUCUCAACGGGACUCUGAUAGUUAGCCAGGACUCUGGUCUGGCCCUGAUAUCAAUCAGGGCCUAGGCCGUUUAAGCUGGGG.ACCU..UUG...............................GGG....CCC.UAAUUCAGA.....AGCGAGCUAA.......UCGUU.UCGC..
Sla-1-1   ACGCUCCCAAUCUUGCGGCCUCGGAUUCGUUCCGGGGCUUCUGAUAUUUGCAGCCGGGGACCUUCGGGGCCCUAUAACCUGGGG.ACCU..ACG...............................GGG....GCC.UCUCACAUA.....GCGGG...........GAUU.......
Rha-2-1   AUGCACCCAAUCUUGCGGCCUCGGAUUCGUUCCGGGGCCUCUGAUAUUUGAAACCGGGGACCUUCGGGGCCCUAUACCUUGGGG.ACCU..ACG...............................GGG....GCC.UAUCACAUA.....GCGGG...........G..........
Rsp-17-1  AUGCACCCAAUCUUGCGGCCUCGGAUUCGUUCCGGGGCCUAAGAUAUUUGAAACCGGGGACCUUCGGGGCCCUAUACCUUGGCC.ACCU..ACG...............................GGG....GCC.UAUCACAUA.....GCGGG...........G..........

          ..................................................................................<<<<..<<<<<..............................>>>>>....>>>.>..............<<<<<...........<<<<<<<<<<<<
          ..................................................................................1222..222.0..............................0.222....222.1..............22222...........22..1.......
          ..................................................................................YYGG••-ACCU--•CG----------------------------GGG----••C-U•YA••••○○------R••••-----------○○---------
                                                                                            57
```

```
Rsp-18-1   AUGCUCCCAAUCUUGCGGCCUCGGAUUCGUUCCGGGGCCUCUGAUAUUUGUAACCGGGGACCUUCGGGGCCCUAUACCCUUGGC.ACCU..ACG.............................GGG....GCC.UAUCACAUA.....GCGGG...........G..........
Rsp-19-1   ACGCUCCCAAUCUUGCGGCCUCGGAUUCGUUCCGGGGCUUCUGAUAUUUGUAACCGGGGACCUUCGGGGCCCUAUACCCUUGGC.ACCU..ACG.............................GGG....GCC.UAUCACAUA.....GCAGG...........G..........
Rsp-22-1   ACGCUCCCAAUCUUGCGGCCUCGGAUUCGUUCCGGGGCCUCUGAUAUUUGUAACUGGGGACCUUCGGGGCCCUAUACCCUUGGC.ACCU..ACG.............................GGG....GCC.UAUCACAUA.....GCGGG...........G..........
Rba-2-1    AUGCACCCAAUCUUGCGGCCUCGGAUUCGUUCCGGGGCCUCUGAUAUUUGUAACCGGGGACCUUCGGGGCCCUUCACCUCGGGC.ACCU..ACG.............................GGG....GCC.UAUCACAUA.....GCGGG...........GC.........
env-365    CUUUAAGGGGCCGUGUUAUACUUUUACCAGACGGACUUAUAGGUUUUCCUGAAAACCUUUUUGGAUCGUUAAACACAAAAGCGG.ACCU..ACG.............................GGG....CCG.UAUAACCAAUA....CGAUC......................
env-367    ACUAUCGACGUCAGCACUGGCCUUUGAUAGAAGACACCGAAUCACGGCUCCAUCGGGCCGUGUUUACUUUUUAAGCAAGCGG.ACCU..ACG.............................GGG....CCG.UAUAACCAAUA....CGAUC......................
env-369    ..CUUUCCAAUCGGCUGGCCUCGGAUUCGUCCGGGGCCUGGUUUUUGGUUGCUCCCGGGCGGCCGCCUGCCGAGACGAUCGGGG.ACCU..UCG.............................GGG....CCC.UUCACAUCACAGACAGCCGG...........GCCCA..UACCC
env-370    .............................................................................AUCGAUCUGGG.ACCU..UCG.............................GGG....CCC.UUCACACAAGAGAGAGCCGG...........GU.........
env-371    ...GGCCCCGGAUUCGUCCGGGGCUUGGUUUUUGUCGGCUGACGGGCGGGUGGCCGUCCGGCGCGUCGGGCAUAUCGAUCUGGG.ACCU..UCG.............................GGG....CCC.UUCACACAAGAGAGAGCCGG...........GU.........
env-372    GGCUGGCCCCGGAUUCGUCCGGGGCUUCGCUUUUGGUCGACGGAUGGCGGGACAAGUCCCGCCCCUACGGCGCCAUAACCCGGGG.ACCU..CAG.............................GGG....CCC.UACACACCACAGACAGCCGG...........GC.........
env-373    ........................................CCUUUCUCGGAUUCGUUCCGGGGCCGUUUGGUUUAUUCUUUGGG.ACCU..UCG.............................GGG....CCC.UUCACUUCA.....GGCAC...........CCU........
env-374    ..........................................................................UAUUUUUUGGG.ACCU..UCG.............................GGG....CCC.UUCACUUCA.....GGCAC...........CUC........
env-375    UAGACGUUUCUACAGGUCUUUGAUAGAAGCAGCGAAUCAAGCGGCCCCAGCCUGUCUGGGGCCGUGGACUUUUUAAAAUUGGGG.AUCU..UCG.............................GAG....CCC.UUUAACCUC.....GGGUU...........C..........
env-378    ..........CGGUUCUCCGGGGCCGUGGAAUUUUUACGAAUCGGGUUGCUACGGUCCCACCAAGGCAGCCCAUAUCUCGGGG.AUCU..ACG.............................GAG....CCC.UAUAACCUC.....GGGUC...........C..........
env-379    NNNNNNNNNNNCGGUUCUCCGGGGCCGUGGAAUUUUUACGAAUCGGGUUGCUACGGUCCCACCAAGGCAGCCCAUAUCUCGGGG.AUCU..ACG.............................GAG....CCC.UAUAACCUC.....GGGUC...........C..........
env-380    GAUACCUGAUUCUAUCGGCGCUCCGAAUCUUUUUGGGCGUUGUGGGACUUCAACGGGGACCUUCGGGGCCCUAUCAUAACGGGG.ACCU..UUG.............................GGG....CCC.UAUAAUA.......UGCAA...........GAC........
env-381    .......................CGCUUUUGGUCGCUCGGUGGUGGCAGAUUGCCCACCCUACGGCGCCAGAUAACCCGGGG.ACCU..CAG.............................GGG....CCC.UACACACAUAGACA.GCCGG...........GCCC.......
Osp-2-1    AGACACCCACCCCAACCCUGUUACAUCCCCCCAAUCUUGCGGCCUCGGAUUCGUUCCGGGGCCACUUGAUUUUGAGCUCAGGG.ACCU..UCG.............................GGG....CCC.UUCACACAUA.....GGGUC...........G..........
env-382    UGACGUGUCCACGGGCCUCGCGUAAGGGGCUCGCCUGACCGGCCCCGGGGCACCCCCGGGGCUUCGUCGUUCCGAAACCGGGG.ACCU..ACG.............................GGG....CCC.UAUCCACCAAGA...GACGC...........GGCG.......
env-386    GACGUGUCCACGGGCCUCGCGUAAGGGGCCGCCUGACACCGGCCCCGGGGGCACCCCCGGGGCUUCGUCAUUCCAAGCCGGGG.ACCU..ACG.............................GGG....CCC.UAUCGAUCACA....GACGG...........GGCG.......
env-388    CAUUGACGUCACCACCGGGCUCGCCUAAGAGCCCCACUGACCGGCCCCGGGGGAACCCCGGGGCCUCGUCUUUUCUCUCGGGG.ACCU..UCG.............................GGG....CCC.UUCACCAAAC.....CGAGGC..........GCCG.......
Rme-1-1    UGACGUCUCCACGGGUCUCGCGUAAGGGACCGCACCUGACCGGCCCCGGGCGCAACCCCGGGGCCUCGUCAUUCCAAGCCGGGG.ACCU..UCG.............................GGG....CCC.UUCAUCUAACG....GACGC...........ACC........
env-389    UGACGUCUCCACCGGUCUCGCGUAAGGACCGCACCUGACCGGCCCCGGGUAACACCCGGGGCCUCGUCGUUUCACCACCGGGG.ACCU..UCG.............................GGG....CCC.UUCAUAACACC....GACGG...........GCCG.......
env-391    ......CUCUACCGGUCUCGCGUAAGGGACCGCACUUGACUGGCCCCGGGACAACCCCGGGGCCUCGUCUUUCUCACCGGGG.ACCU..GCG.............................GGG....CCC.UUCAGAUAACG....GACGG...........ACCG.......
env-392    ....................AUUUCCCCAUGUAUUUUGGGAUAAAUGUUAUGGGUAGCUAUUUAAGCUACUUUAAUUCAGGGG.ACCU..CUG.............................GGG....CCC.UAUAAACCA......GCGAU...........CU.........
env-394    ...................UUAUUUGCCCAUAUAUUUUGGGAUAAAUGUUAUGGGUAGCUAUUUAAGCUACUUUAAUUCAGGGG.ACCU..CUG.............................GGG....CCC.UAUAAACCA......GCGAU...........CU.........
env-395    AUAGAAGCCGCUACGAAUCGCGCGGCCCCGGAUUUCCGGGGCCGCCGGUCUUUUGCGGAAUCUGAUUUCGCAUGUCACCGGGG.AUCU..UCG.............................GAG....CCC.UUCAACC.........GCGCGAAA........GCGACCUGAAAG
env-396    ACGCCACCAAUCCUGCAGGCCCCGGAUUCGUUCCGGGGCUGCUUGAUUUUGUAACCGGGGACCUUCGGGGCCCUUCAAUGGGG.ACCU..UCG.............................GGG....GGC.UUUAACAUA......GCGGA...........CCC........
Pxi-1-1    CACGCCACCAAUCCUGCGGCCUCGGAUUCGUUCCGGGGCCCAAGAUUUUUGUAACCGGGGACCUUCGGGGCCCUUCAACUGGGC.ACCU..UCG.............................GGG....GGC.UUUAACAUA......GCGGG...........UCC........
Rmo-1-1    UCCUGACAGGCCUCGGAUUCGUUCCGGGGCCGUUAUCUUUGUCGGGACAUUCGACCCAGGGACCUUCGGGGCCCUUCAUUGGUC.ACCU..GAACCCGGCGUGGAUGCGCCAAGGCA........GGG....GGC.UUUAACAUA......GCGGU...........GCC........
Rsp-21-1   UCCUGACAGGCCUCGGAUUCGUUCCGGGGCCGUUAUCUUUGUCGGGACAUUCGACCCAGGGACCUUCGGGGCCCUUCAUCGGUC.ACCU..GAACCCGGCGCGGAUGCGCCAAGGCA........GGG....GGC.UUUAACAUA......GCUGG...........GCC........
Psp-1-1    CUGACGACCCCGGAUUCGCUCCGGGGUUAUUACGUUUGUCGGGAUGUUCGAACCUGGGGACCUUCGGGGCCCUAUAAUCUGGCC.ACCCU.AGUAGGGCGUCUCAUCUUGGGACAACCGGGC...GGGG....GGC.UUACACAUA......GCUGG...........ACC........
Pga-1-1    CUGACGACCCCGGAUUCGCUCCGGGGUUAUUACGUUUGUCAGGAUGUUCGAACCUGGGGACCUUCGGGGCCCUAUAAUCUGGCC.ACCCU.AGUAGGGCGUCUCAUCGUGGGACAACCGGGC...GGGG....GGC.UUACACAUA......GCGGG...........CCC........
Pin-1-1    CUGACGACCCCGGAUUCGCUCCGGGGUUAUUACGUUUGUCGGGAUGUUCGAACCUGGGGACCUUCGGGGCCCUAUAAUCUGGCC.ACCCU.AGUAGGGCGUCUCAUUGUGGGACAACCGGGC...GGGG....GGC.UUACACAUA......GCGGG...........CCC........
Psp-2-1    CUGACGACCCCGGAUUCGCUCCGGGGUUAUUACGUUUGUCAGGAUGUUCGAACCUGGGGACCUUCGGGGCCCUAUAAUCUGGCC.ACCCU.AGUAGGGCGUCUCAUCUUGGGACAACCGGGC...GGGG....GGC.UUACACAUA......GCGGA...........CCC........
Rsp-11-1   CUACUGGCCUCAGAUUCGUUCUGGGGCCGUUAUCUUUGUGGGAAACACAUGAGCUGGGGACCUUCGGGGCCCUAAAGACUGGCC.ACCU..UAGUAGUCAUUGUUCCAUAGGGGCAGGCGGAC...GGG...GGGG.UUACACAUA......GCGGG...........UCC........
Par-1-1    CCUACUGGCCUCAGAUUCGUUCUGGGGCCGUAUUCUUUGUGGGAUGAUUUGAGCUGGGGACCUUCGGGGCCCUAAAGACUGGCC.ACCU..UAGUAGUCAUUGUCCCAUGGGGGCUCGGCGGAC..GGG...GGGG.UUACACAUA......GCGGG...........UCC........
env-397    CGCACGCAUCGCAUUCGGCCUCAGAUUCGUUCUGGGGCCGGUUGCUGUUUACCGGGGACCUGAAAAGGGCCCUAUACAUUUGGC.ACCU..ACG.............................GGG....GCC.UUUAACAUA......GCUGG...........UCC........
env-398    CGUGCACACCCCGGAUUCGUUCCGGGGUUGUUACGUUUGUCAGGAUGCAUGAACUCGGGGACCUUAGGGGCCCUAUAACUCAGGC.ACCU..ACG.............................GGG....GCC.UUAAACAUA......GCUGG...........GCC........
Rsp-12-1   CCUGUAGACCCCGGAUUCGCUCCGGGGUCGUUACGUUUGUCAGGAUGUAUCACUUGGGGACCUUCGGGGCCCUAUAUCCCAGGC.ACCU..ACG.............................GGG....GCC.UUAAACAUA......GCUGG...........GCC........
Lsp-7-1    UGAAGACCCCGGAUUCGUUUCCGGGGUCGUUAUGUUUGUCAGGACAUACGAACUCGGGGACCUUCGGGGCCCUAUAACACAGGC.ACCU..ACG.............................GGG....GCC.UUCAACAUA......GCCGG...........GCC........
Lsp-1-1    UGAAGACCCCGGAUUCGUUUCCGGGGUCGUUAUGUUUGUCAGGACACACGAACUCGGGGACCUUCGGGGCCCUAUAACACAGGC.ACCU..ACG.............................GGG....GCC.UUCAACAUA......GCCGG...........GCC........
Lsp-3-1    UGAAGACCCCGGAUUCGUUUCCGGGGUCGUUAUGUUUGUCAGGACAUACGAACUCGGGGACCUUCGGGGCCCUAUAACACAGGC.ACCU..ACG.............................GGG....GCC.UUCAACAUA......GCUGG...........GCC........
Lsp-4-1    UGCAGACCCCGGAUUCGUUUCCGGGGUCGUUAUGUUUGUCAGGACAUACGAACUCGGGGACCUUCGGGGCCCUAUAUCUCAGGC.ACCU..ACG.............................GGG....GCC.UUCAACAUA......GCUGG...........GCC........
Lsp-6-1    UGAAGACCCCGGAUUCGUUUCCGGGGUCGUUAUGUUUGUCAGGACACACGAACUCGGGGACCUUCGGGGCCCUAUAACACAGGC.ACCU..ACG.............................GGG....GCC.UUCAACAUA......GCUGG...........GCC........
Pga-2-1    CUGAAGACCCCGGAUUCGUUCCGGGGUCGUUACGUUUGUCGGGAUGUUCGAACUCGGGGACCUUUGGGGCCCUAUAUCACUGGC.ACCU..ACG.............................GGG....GCC.UUAAACAUA......GCUGG...........ACU........
Nsp-1-1    CUGAAGACCCCGGAUUCGUUCCGGGGUCGUUACGUUUGUCGGGAUGUUCGAACUCGGGGACCUUUGGGGCCCUAUAUCACUGGC.ACCU..ACG.............................GGG....GCC.UUAAACAUA......GCUGG...........ACC........
Lsp-2-1    CUGAAGGCCCCGGAUUCGCUCCGGGGUCGUUACGUUUGUCAGGAUGUGUGAACUCGGGGACCUUCGGGGCCCUAAAACUUUGGC.ACCU..ACG.............................GGG....GCC.UUAAACAUA......GCUGG...........GCC........
Rba-7-1    CUGCAGACCCCGGAUUCGUUCCGGGGUCGUUACGUUUGUCAGGAUGUUCGAACCUGGGGACCUUCGGGGCCCUAUAACUCAGGC.ACCU..ACG.............................GGG....GCC.UUAAACAUA......GCGGG...........UCC........
Lda-1-1    CUGCAGACCCCGGAUUCGUUCCGGGGUCGUUACGUUUGUCAGGAUGUUCGAACUCGGGGACCUUCGGGGCCCUAUAAUCUCAGGC.ACCU..CCG.............................GGG....GCC.UUAAACAUA......GCGGG...........UCC........
Lsp-8-1    UCCUGAAGGCCCCGGAUUCGCUCCGGGGUCGUUACGUUUGUCAGGAUGUUGAACUCGGGGACCUUCGGGGCCCUACAUCAGGC.ACCU..ACG.............................GGG....GCC.UUAAACAUA......GCGGG...........CCC........
env-399    UUUGAUAGAAGGCACCGAAUCUCGCGGCCCCGGGUUUCCGGGGCCGUGUGNUUUGUNCGGCAAGNGCCGGAUUUAACCUUGGGG.ACCU..UCG.............................GGG....CCC.UGCACCUC.......GGGCC......................
env-400    ............ACCGAAUCUCGCGGCCCCGGGUUUCCGGGGCCGUGUGUUUUGUCCGGCAAGUGCCGGAUUUAACCUUGGGG.ACCU..UCG.............................GGG....CCC.UGCACCUC.......GGGCC......................
env-402    UUGAUAGAAGGCCAGCGAAUCUGCGGCCCCGGGUUUCCGGGGCCGUGUGUUUUGUCCGACAUGCGUCGGAUACCAGUCGGGG.ACCU..UCG.............................GGG....CCC.UACACCC........GGGCC......................
env-403    ...............................UUGGGGCUAAUUAAUCAUAUAGGUGGCGUUAUGAGCCACCUUUAAACUGGGG.ACCU..UUG.............................GGG....CCC.UACACACC.......GGCG............GUCU.......
env-404    CUAGGGCCUAAUCUUUGAUUUGUCUCAAAUUUUUGGGGCUAAUUAAUCAUAUAGGUGGCUUUAUGAGCCACCUUUAACACUGGGG.ACCU..UUG.............................GGG....CCC.UACACACC.......GGCG............GUCU.......
env-405    CUAGAGCCUAAUCUUUAAUUUAUCUCAAAUUUUUGAGGCUAAUUAAUUAUAUAGGUGGCUUUACGAGCCAUCUUAACACUGGGG.ACCU..UUG.............................GGG....CCC.UAAACAUC.......GGCG............GUCU.......
env-407    CUAGAGCCUAAUCUUUAAUUUAUCUCAAAUUUUUGAGGCUAAUUAAUUAUAUAGGUGGCUUUACGAGCCAUCUUAACACUGGGG.ACCU..UUG.............................GGG....CCC.UAAACACC.......GGCG............GUCU.......
env-413    CUAGAGCCUAAUCUUUAAUUUAUCUCAAAUUUUUGAGGCUAAUUAAUUAUAUAGAUGGCUUUACAAGCCAUCUUAACACUGGGG.ACCU..UUG.............................GGG....CCC.UAAACACC.......GGCG............GUCU.......
env-414    ........................GCUGGCCCCGGAUUCGUCCGGGGCUUUGCCUUUGCUCAUUCGAGCAGACAAUCUCGGGG.ACCU..ACG.............................GGG....GCC.UAAACCCAA......GCCGG...........G..........
Rel-1-1    GACGUGUCGACCGGUCUUUAAUAGACGGCGACAUCGCUCGGCCCCGGACCAUCCGGGGCCGUCGUUUUUAGAAAGAAUCGGGG.ACCU..UCG.............................GGG....CCC.UUCACCACC......GGGGC......................
env-416    ACGUGUCGACCGGUCUUUAAUAGACGGCGGCAUCGCGCCGGCCCCGGAUAUCUCCGGGGCCGUCGUUUUUAGAAAGAAUCGGGG.ACCU..UCG.............................GGG....CCC.UUCACAACC......GGGGC......................
env-417    ...........................................................CGGGGCCAUAUAUACGCGG.ACCU..ACG.............................GGG....CCG.CAUAACCU.......UUCAAAC.........CCC........
env-418    GAAUCUCGCGGCCCCGGGUUUUCCGGGGCCGUGGAAUUUUAUCGAAUCGGCCUUUGAUCCUGCAUCAAUGUGCUGAACCGGGG.ACCU..UCG.............................GGG....CCC.UUAACCC........GGGUC......................
env-419    GAAUCUCGCGGCCCCGGUUCAUCCGGGGCCGUGGAAUUUUAUCGAAUCGGCCUUUGAUCUUGCAUCAAAGCGCCAGAACCGGGG.ACCU..UCG.............................GGG....CCC.UUAACCC........GGGUC......................
env-421    CAGCGAAUCUCGCGGCCCCGGGUUUCCGGGGCCGUGGAUUUUUCCGGUUUCCGCCUUGCGCGGGGGCGGAACCUGAACCGGGG.ACCU..UCG.............................GGG....CCC.UAUAACCC.......GGGUC......................
env-422    CGGGCCUGACCCGGGACCUUGCCGCCAGAGGCCCCGGAUCAAGUCCGGGGCGUGAUGGGUGGGGAGGCGAGCGAAGAGCCGGGG.ACCU..AUG.............................GGG....CCC.UAUAACAUGA.....GGUGG...........GUU........
env-423    AUAGACACCACCCCCAUCCCUGUUAGCUCGCACCAGUCGCGGCCUCAGAUUCGUUUUGGGGCCGGUACUUGUUUAACCGGGG.ACCU..UCG.............................GGG....CCC.UAUACCAAC......GGUGG...........GCU........
env-426    GAACCCCCUAAACGAACCCAUCCGCAAGGCCCCGGAUUCGUCCGGGGCUUCGCUGUUGUUCGCAAGAACAGACCAAAACCGGGG.ACCU..UCG.............................GGG....CCC.UAUACCAAC......GGUGG...........GUU........
Msp-1-1    GAUUGACGUCACGACGGGUCUGUAAUAGCUGACCCGUUCUGCGGCCCCGGACCACUUUGGGGCCGCGUAUCUCACUGAUCCGGGG.ACCU..ACG.............................GGG....CCC.UUAAAUUGC......GGCGG...........G..........
Sno-1-1    AUCCCCCCUAAACGACCCUAUCGGCAUGGCCCCAGAUUCGUCUGGGGUUUUGUUAUUGCUGCGCAUGCAGCAGAUAAAUCGGGG.ACCU..UCG.............................GGG....CCC.UAUAACCAUC.....GGCAG......................
Sno-2-1    AUCCCCCCUAAACGACCCCAUCGGCAUGGCCCCAGAUUCGUCUGGGGUUUUGUUAUUGCUGCGCAUGCAGCAGAUAAAUCGGGG.ACCU..UCG.............................GGG....CCC.UAUAACCAUC.....GGCAG......................
env-427    UAAUAAUAGGACUGUUCUUUAAUUUUCCCAUGUAUUUUGGGAUAAAUGUUAUGGGUAGCUAUUUAAGCUACUUUAAUUCAGGGG.ACCU..CUG.............................GGG....CCC.UAUAAACCAG.....CGAUC......................
env-428    GACAGCCCCGCGCAACCCUGCUAGCUACGCGCCAGUCUCGAGGCCUCGGAUUCGUUCCGGGGCCUGUGACUUUUGCAUUCGGGG.ACCU..UCG.............................GGG....CCC.UACACAUAA......GUAGG...........CGGG.......
Tin-1-1    GACACCUCCCACAUCCCUGCUAGCUACGCGCCAGUCUCGUGGCCUCGGAUUCGUUCCGGGGCCUGUGACUUUUGCAUUCGGGG.ACCU..UCG.............................GGG....CCC.UACACAUAA......GCAGG...........CGGG.......
Pwa-1-1    GGACGCGGCGGGGGUUUUGACCUAAGACGCCCCAUGUCAAACGGCCUCGGAUUCGCUCCGGGGCCUUGAUAUUGUUUUCGGGG.ACCU..UCG.............................GGG....CCC.UAUAAUCC.......GGGUC...........UGU........
env-429    GGGAUUCGUUCCGGGGCUUUGCCUUUGUUCUGCCUGCAAAGGUGGAUGAACCACCGGGGACCUUCGGGGCCCUAUAAGCCGGGG.ACCU..UCG.............................GGG....CCC.UAUAAUUC.......GGGUC...........UAGC.......
env-430    CCCGGGUUUCCGGGGUCCGGUGGAUUUUUUGUAAGAAUCGGUCUGAUGUGCAGGUUCCUCUGCCGAACAGACAAACCACCGGGG.ACCU..UCG.............................GGG....CCC.UAUAACUC.......GGGUC...........UGG........
env-431    GAUUUUUUGCAGAAUCAGUCUGUUUUGUGAGGUUUCACAGACAGAUUAUACCACCGGGGACCUUCGGGGCCCUAUAAACCGGGG.ACCU..UCG.............................GGG....CCC.UAUAAUUC.......GGGUU...........UGU........
env-432    CGGCCCCGGUUUUCCGGGGCCGCGGGUUUUUUCUGAAAUGGUCUGCCAUGCUGGGGGUUCCAGCCGACAGAUCGAACACCGGGG.ACCU..UCG.............................GGG....CCC.UAUAACUC.......GGGUC...........UGU........
env-433    GAUUUUUUGCAGAAUCGGUCUGUUUUGUGGGCUUCCACGCGCAGAUCAAACCACCGGGGACCUUCGGGGCCCUAUAAGCCGGGG.ACCU..UCG.............................GGG....CCC.UAUAAUUC.......GGGUC......................
Osp-3-1    CACCCCUCCACCCCCCUGCUAUACGCGCCCAAUCCUGACGGCCCCGGAUUCGCUCCGGGGCCUUCGGGUGUUUGACGCCUGGGG.ACCU..UCG.............................GGG....CCC.UGUAACAUC......CGGGUAGUCUGUCAGCGGCGG......
env-434    ......UUUCUAUCAGCCGACAACGCUAUCCAGCCAGAUUCGUCUGGGAUGGAGGGGUAUGGGGAAGCCCAAUCUAAACGGGG.ACCU..UCG.............................GGG....CCC.UAUAACACC......AGUGG...........CCC........

           .........................................................................<<<<..<<<<<..............................>>>>>....>>>.>..............<<<<<...........<<<<<<<<<<<
           .........................................................................1222..222.0..............................0.222....222.1..............22222...........22..1.......
           ●●●●●●●●●●●●●●●●●●●●●●●●●●●●●●●●●●●●●●●●●●●●●●●●●●●●●●●●●●●●●●●●●●●●●●YYGG●●−ACCU−−●CG−−−−−−−−−−−−−−−−−−−−−−−−−−−−−−GGG−−−−●●C−U●YA●●●●○○−−−−−−R●●●●−−−−−−−−−−−○○−−−−−−−−−
                                                                                57
```

```
env-435   AUAGAGCGCGACAUCCUUCCGGCCCUGGGUUCGCCCCGGGGCCGCGAUCUUUCCCGGGCCUUGUGUCCGGGCACUGAGCCGGGG.ACCU..GCG..............................GGG....CCC.UACACCACGACA...GGGGC......................
env-436   ..........................................................GGGGCCAUGGCCGUUUAAGCCGGGG.ACCU..UCG..............................GGG....CCC.UAUACCAAGG....GGUGG...........GUU........
env-437   UCAUCUGCGCCGCCCCGGUCAUUGCCGGGGCUGCGUGUUUUUCGGGGAAUGGCGGGCAUAUGCGCGACACCCCGCGAGCCGGGG.ACCU..UCG..............................GGG....CCC.UAUAACAGC.....GGUGG...........GCC........
env-438   UGCUCUCGCCUAACAGGACCUGUCCGCACGGUCCCGGAUUCGUCCGGGGCCGUGCUGUUGUUCGCAAGAACAUCCAAACCCGGG.ACCUU.GCG.............................AGGG....CCC.UUUACCCGA.....ACCGGGCUCUAA....CCCGGUC.....
env-439   AUAGACACCCACCCCCAUCCCUGUUAGCUCGCACCAGUCGCGGCCUCAGAUUCGUUUUUGGGGCCGGUACUGUUUAACCGGGG.ACCU..UCG..............................GGG....CCC.UAUACCAAC.....GGUGG...........GUUU.......
Rsp-10-1  UGUGUUGUUUUGCGCAUCGGUGGGUUUCACUCGCCUUGCGCUGGUCCGGCAAAGGGAUUCUUUGCCAGACGUCUUACAUUGGGG.ACCU..UCG..............................GGG....CCC.UAUAACCAU.....CUCGG...........CCUC.......
Tar-1-1   GCAUAGGCCCCGGAUUCGUCCGGGGCUUUGCUGUUGUUCCCGGUUCUGCCCAGCAGAUGCCGAAUGGGACACUUCAAGUCGGGG.ACCU..UCG..............................GGG....CCC.UAUAAACACA....AGCGG...........CGG........
env-440   ...........................UUUCCGGAUUCGUCCGGGGCCUUGCAUUUGUUCGCACAGCCGGACAGUUCAGCCGGGG.ACCU..UCG..............................GGG....CCC.UUUACCAC......CGGGGUUGACC.................
Sme-2-1   AUCGGCACAGCCCCAGAUUCGUCUGGGGCUUUGUUGUUGUUUGCAUUGCGGGGCGAUAGCCCAUCAAGCAGACAUCCACCGGGG.ACCU..UCG..............................GGG....CCC.UAUAACCAU.....UGUCA...........CCU........
Sme-1-1   AUCGGCACAGCCCCAGAUUCGUCUGGGGCUUUGUUGUUGUUUGUUCUGUGGGGCCAUAGCCCAACAAGCAGACAUCCACCGGGG.ACCU..UCG..............................GGG....CCC.UAUAACCAU.....UGCCA...........CCU........
Tsp-3-1   UGACCCACGCCCCCCAUACACCUACACACCCCCAUCUACACCGCCCCAGAUUCGUCUGGGGUGUUUGUAGUUUUGAAAUACGGG.ACCU..UCG..............................GGG....CCC.UACAACCAU.....GGCGC...........CU.........
env-441   GCGCUUUAUUACCAGACCCUAUCGGAAAGGCCCCGGAUUCGUCCGGGGCUGCGUGUUUUUGCCCUUUGGCCAUUUCAGCCGGGC.ACCU..UCG..............................GGG....GCC.UAUACCC.......GGUGGGCAGAUU....GCCC.......
env-445   UCCAAACCCCCUAAACGACCCCAUCGGCUGGCCCCGGAUUCGUCCGGGGCUUCGCUUUUGCUCGCUAGAGCAUCGAAUUCGGGG.ACCU..UCG..............................GGG....CCC.UAUACACACAGACA.GCCGG...........GC.........
Rto-1-1   CACUCCCCUAUCAGAGCCCAUCCGCAAGGCCCCGGAUUCGUCCGGGGCUUCGCUGUUGUUCGCAAGGGCAACGAAGAGCCGGGG.ACCU..UCG..............................GGG....CCC.UAUAACCACG.....GUGGGAUUGAAA....CCC........
Ris-1-1   CUGCCUGCCGCUAUCGGCGGAGUGGGGGCCCGGCGAUGACGCGGAACGCGCGCUCGGCGCGGGUCGCACCCCACCACCGGGG.ACCU..UCG..............................GGG....CCC.UACAUCC.......GAUGG...........CCG........
env-447   ...................................GAUUCGUCCGGGGCUUUGCUAUUGUUCGCAAGAACACCAAUAAGCCGGGG.ACCU..AAG..............................GGG....CCC.UAUAACACG.....GUGGGUUUU.......ACCC.......
env-448   GCGCUCCCCUAACAGACCCCAUCCGCAAGGCCCCGGAUUCGUCCGGGGCUUCGCUGUUGUUCGCAAGAGCAAACCGAGCCGGGG.ACCU..UCG..............................GGG....CCC.UUUAACCAAA.....GGUGGGUCC.......GC.........
Tpa-1-1   AGACUCGCCCACCCAUCCCUGCUACAUCGCGCCAGUCUCGAGGCCUCGGAUUCGUUCCGGGGCCCGUGAUUUUGAAUUCGGGG.ACCU..UCG..............................GGG....CCC.UAUACACACAGACA.GGCGG...........G..........
env-458   AGACUCACCCCCACAUCCCUGCUACAUCGCGCCAGUCUCGCGGCCUCGGAUUCGUUCCGGGGCCCGUGAUUUUGAAUUCGGGG.ACCU..UCG..............................GGG....CCC.UAUACACACAGACA.GGCGG...........G..........
Tat-1-1   AGACUCGUCCCCGCAACCCUGCUACAUCGCGCCAAUCUCGAGGCCUCGGAUUCGUUCCGGGGCCCGUGAUUUUGAAUUCGGGG.ACCU..UCG..............................GGG....CCC.UAUACACACAGACA.GGCGG...........G..........
env-459   GACAGGCCCCGGCAACCCUGCUACACGCCCCCCAUCGCGAGGGCCCGGGAUUCGUUCCGGGCCCCGUGAUGUUUGAACGUGGGG.ACCU..UCG..............................GGG....CCC.UAAACACACAGACA.GGCGG...........G..........
env-460   CACCCCCUGUUAUCAGCCAACAACGUCACACCCCAGAUUCGUCUGGGGUGGUGACGUUCGGGGAAACCCAAUUAGAACGGGG.ACCU..UCG..............................GGG....CCC.UAAACACCA.....GCGGU...........CU.........
env-463   GCCCCCCUGUUAUCAGCCAACAACGCUGCACACCCCAGAUUCGUCUGGGGUGGUGUCGUUUGGGGAAACCCAAUUUAACGGGG.ACCU..UCG..............................GGG....CCC.UAAACACCA.....GCAGC...........CC.........
env-464   CACCCCCUGUUAUCAGCCAACAACGUCACACCCCAGAUUCGUCUGGGGUGGUGACGUUCGGGGAAACCCAAUUAGAACGGGG.ACCU..UCG..............................GGG....CCC.UAAACACCA.....GCAGU...........CU.........
env-465   UCCCCACUGCUAUCAGCAGAAACGCUAUACACCCCAGAUUCGUCUGGGGUGGUAUCGUUUGGGGAAACUCAAUUAAAGCGGGG.ACCU..UCG..............................GGG....CCC.UAAACACCA.....GCGGC...........CC.........
env-467   CGCCCCCUGCUAUCAGCCUUCAUCGCCACUCACCCCAGAUUCGUCUGGGGUGGUGUCGUUUGGGGAAACCCAAAUUCAUCGGGG.ACCU..ACG..............................GGG....GCC.UAAACACCA.....GCGGC...........CC.........
env-469   CGCCCCCUGCUAUCAGCCCUCAUCGUCACUCACCCCAGAUUCGUCUGGGGUGGUGUCGUUUGGGGAAACCCAAAUUCAUCGGGG.ACCU..ACG..............................GGG....GCC.UAAACACCA.....GCGGC...........CC.........
env-472   GCCCCCUGCUAUCAGCGCCACAUCGUCAGUCACCCCAGAUUCGUCUGGGGUGGUGUCGUUUGGGGAUACCCAAAUUCAUCGGGG.ACCU..ACG..............................GGG....GCC.UAAACACCA.....GCGGC...........CC.........
env-473   CGCCCCCUGCUAUCAGCCCUCAUCGUCAUUCACCCCAGAUUCGUCUGGGGUGGUGUCGUUUGGGGGAACCCAAAUUCAUCGGGG.ACCU..ACG..............................GGG....GCC.UAAACACCA.....GCGGC...........CC.........
env-475   ..................UUUAUCGUCAUUCACCCCAGAUUCGUCUGGGGUGGUGUCGUUUGGGGGAACCCAAAUUCAUCGGGG.ACCU..ACG..............................GGG....GCC.UAAACACCA.....GCGGC...........CU.........
env-476   CACCCCCUGCUAUUAGCCCUCAUCGUCAUUCACCCCAGAUUCGUCUGGGGUGGUGUCGUUUGGGGGAACCCAAAUUCAUCGGGG.ACCU..ACG..............................GGG....GCC.UAAACACCA.....GCGGC...........CC.........
env-477   CGCCCCCUGCUAUCAGCCCUCAUCGUCAUUCACCCCAGAUUCGUCUGGGGUGGUGUCGUUUGGGGGAACCCAAAUUCAUCGGGG.ACCU..ACG..............................GGG....GCC.UAAACACCA.....GCGGC...........CC.........
env-482   CGCCCCCUGCUAUCAGCCCUCAUCGUCAUUCACCCCAGAUUCGUCUGGGGUGGUGUCGUUUGGGGGAACCCAAAUUCAUCGGGG.ACCU..ACG..............................GGG....GCC.UAAACACCA.....GCGGC...........CC.........
env-483   CACCCCCUGCUAUCAGCCCUCAUCGUCAUUCACCCCAGAUUCGUCUGGGGUGGUGUCGUUUGGGGGAACCCAAAUUCAUCGGGG.ACCU..ACG..............................GGG....GCC.UAAACACCA.....GCGGC...........CC.........
env-484   CGCCCCCUGCUAUCAGCCCUCAUCGUCAUUCACCCCAGAUUCGUCUGGGGUGGUGUCGUUUGGGGGAACCCAAAUUCAUCGGGG.ACCU..ACG..............................GGG....GCC.UAAACACCA.....GCGGC...........CC.........
Pte-1-1   CGCCCCCUGCUAUCAGCGCUCAUCGUCAUUCACCCCAGAUUCGUCUGGGGUGGUGUCGUUUGGGGGAACCCAAAUUCAUCGGGG.ACCU..ACG..............................GGG....GCC.UAAACACCA.....GCGGC...........CC.........
env-493   CGCCCCCUGCUAUCAGCCCUCAUCGUUAUUCACCCCAGAUUCGUCUGGGGUAUUGUCUUUUGGGGGAACCCAAAUUCAUCGGGG.ACCU..ACG..............................GGG....GCC.UAAACACCA.....GCGGC...........CC.........
env-494   CGCCCCCUGCUAUCAGCCCUCAUCGUCAUUCACCCCAGAUUCGUCUGGGGUGGUGUCGUUUGGGGAAACCCAAAUUAAUAGGGG.ACCU..ACG..............................GGG....GCC.UAAACACCA.....GCGGC...........CC.........
env-495   GCCCCCUGCUAUCAGCGCCACAUCGUAAGUCACCCCAGAUUCGUCUGGGGUGGUGUCGUUUGGGGAUACCCAAAUUCAUCGGGG.ACCU..ACG..............................GGG....GCC.UAAACACCA.....GCGGC...........CC.........
env-497   GUGUUCCCCUAAACGACCCCAUCGGCAAAGCCCCGGAUUCGUCCGGGGCUGUGCUUUUCGUUCGCAGGAACGCACGAACCGGGG.ACCU..UCG..............................GGG....CCC.UAUACCAC.......UGGUCACCCGAUCCU.CUGCCUUUGUGU
env-500   UAGACAGCCCCAAGCUUCCCUACUAUACGAGCCAGAUCGCGUGGCCUCAGAUUCGUCUGGGGCCCGUUAUAUUUUUCGCUGGGG.ACCU..UCG..............................GGG....CCC.UAAAAGAC.......CGAGC......................
env-502   GUGGCGCCUUUUCCUACACGACCCCAUCCGCGAGAGGCCCCGGAUUCGUCCGGGGCUAUGUCUUUGCGGGCACGAACACCGGGG.ACCU..ACG..............................GGG....GCC.UAUACCCG........GCGCAUC.........GC.........
env-503   UAGACAGCCCCGAGCUUCCCCUCUAUACGAGCCAAGUCGCGUGGCCUCAGAUUCGUCUGGGGCCCGUUAUAUUUUUCGCUGGGG.ACCU..UCG..............................GGG....CCC.UAAAAGAC.......CGAGCAAAACAAA...UGCUCGAAACA.
env-505   UAGACAGCCCCGAGCUUCCCCUCUAUACGAGCCAAGUCGCGUGGCCUCAGAUUCGUCUGGGGCCCGUUAUAUUUUUCGCUGGGG.ACCU..UCG..............................GGG....CCC.UAAAAGAC.......CGAGCAAAACAAA...GCUCGAAACA..
Smu-1-1   CGGCGCCGGAUUCGUUCGGCGCCGCGUGAUCUUUGGCGGAAACGCCGUCCGGGGUGCCCGUUUCGGGCACGCCGUUCCACCGGGG.ACCU..UCG..............................GGG....CCC.UACGACCGAG.....GCGGG...........AUC........
env-506   AUAGCACGCCUCAUCAACGCUCCGAAUCUGUUCGGGGCGUUGUGUUUUAAGUCUGGGGCCUUCGGGCCCUAUUACAACGGGG.ACCU..UCG..............................GGG....CCC.UAUCAUC........GCGGC...........GC.........
env-507   AUAGCACGCUUCAUCAACGCUCCGAAUCUGUUCGGGGCGUUGUGUUUUAAGUCUGGGACCUUCGGGCCCUAUUACAACGGGG.ACCU..UCG..............................GGG....CCC.UAUCAUC........GCGGC...........GC.........
env-508   AUAGCACGCCUCAUCAACGCUCCGAAUCUGUUCGGGGCGUUGUGUUUUAAGUCUGGGACCUUCGGGCCCUAUUACAACGGGG.ACCU..UCG..............................GGG....CCC.UAUCAUC........GCGGC...........GC.........
env-512   ........................CCGAAUCUGUUCGGAGCGUUGUUGUUUUAAGUCGGGGACCUUCGGGCCCUAUUAUCUCGGGG.ACCU..UCG..............................GGG....CCC.UAUCAAUC.......GCGGC...........GC.........
env-513   GAUAACACGCCUCAUCAACGCUCCGAAUCUGUUCGGAGCGUUGUUGUUUUAAGUCGGGGACCUUCGGGCCCUAUUAUCUCGGGG.ACCU..UCG..............................GGG....CCC.UAUCAAUC.......GCGGC...........GC.........
env-514   GAUAGCACGCCUCGUCAACGCUCCGAAUCUGUUCGGAGCGUUGUUGUUUUAAGUCGGGGACCUUCGGGCCCUAUUAUCUCGGGG.ACCU..UCG..............................GGG....CCC.UAUCAAUC.......GCGGC...........GC.........
env-515   GAUAGCACGCCUCAUCAACGCUCCGAAUCUGUUCGGAGCGUUGUUGUUUUAAGUCGGGGACCUUCGGGCCCUAUUAUCUCGGGG.ACCU..UCG..............................GGG....CCC.UAUCAAUC.......GCGGC...........GC.........
env-516   AUAGCACGCCUCAUCAACGCUCCGAAUCUGUUCGGAGCGUUUGUGUUUUAAGUCGGGGACCUUCGGGCCCUAUUAUCUCGGGG.ACCU..UCG..............................GGG....CCC.UAUCAAUC.......GCGGC...........GC.........
env-517   GAUAGCACGCCUCAUCAACGCUCCGAAUCUGUUCGGAGCGUUGUUGUUUUAAGUCGGGGACCUUCGGGCCCUAUUAUCUCGGGG.ACCU..UCG..............................GGG....CCC.UAUCAAUC.......GCGGC...........GC.........
env-518   .........................................CAACGCUCCGAAUCUGUUCGGAGCGUUGUUGUUUUAAGUCGGGG.ACCU..UCG..............................GGG....CCC.UAUCAAUC.......GCGGC...........GC.........
env-519   ACCAUCGACGUGAACACCGGACUGUGAUAGCACGCCUCAUCAACGCUCUGAAUCUGUUCGGAGCGUUGUGUUUUAAGUCUUGGGG.ACC...UUCG.............................GG....CCC.UAUCAAUC.......GCGGC...........GC.........
env-520   ..........................AUCUGUUCGGAGCGUUGUUGUUUUAAGUCGGGGACCUUCGGGCCCUAUUAUCUCGGGG.ACCU..UCG..............................GGG....CCC.UAUCAAUC.......GCGGC...........GC.........
env-521   UAGCACGCCUCAUCAACGCUCCGAAUCUGUUCGGAGCGUUGUUGUUUUAAGUCGGGGACCUUCGGGCCCUAUUAUCUCGGGG.ACCU..UCGG.............................GGG....CCC.UAUCAAUC.......GCGGC...........GC.........
env-522   UGACGCCCCUAGCGGUAUCCCUUAAACGGCACCAACCGCGCAGCCCCGGAUUCGUCCGGGGCUUCGUUGUUGUUAAACCGGGG.ACCU..UCG..............................GGG....CCC.UAUACCCAU......GGCGGAUAC.......GUC........
env-523   UGACGCCCCUAGCGGUAUCCCUUAAACGGCACCAACCGCGCAGCCCCGGAUUCGUCCGGGGCUUCGUUGUUGAGUUAACCGGGG.ACCU..UCG..............................GGG....CCC.UAUACCCAU......GGCGGAUAC.......GUC........
env-524   AUAGACGUCUCUACAGGCCUCUGAUAGAAGCCAACGAAUCUACGGCCCCGGCAGAUUCGGGGCCGUACACUUUUCAACCAGGG.ACCU..ACG..............................GGG....CCC.UA..................AU.......................
env-525   CUAUCGACGUAUCUACGGGCCUCUGAUAAAGGGGUCCGAAUCUGCGGCCCUGGUAUCCGGGGCCGUCUACUUUUCAAACAGGG.ACCU..UCG..............................GGG....CCC.UA...................A.......................
env-526   CCCCCACUGCUAUCAGCCCACAACGCCAUGUACCCCAGAUUCGUCUGGGGUGGUGUCGUUCUUGGGGAAACCCAAUUAACGGGG.ACCU..CCG..............................GGG....CCC.UAAACACCG......GCAGC...........CC.........
Tda-1-1   GACAGCCCCCGCCAACCCUGCUAGCUACGCGCCAGUCUCGCGGCCUCGGAUUCGUUCCGGGGCCGGUGAUUUUUGCAUUCGGGG.ACCU..UAG..............................GGG....CCC.UAUACCAAUCAUA..GGCGG...........G..........
Dsp-1-1   ACGUGACCACCGGCCUCUGAUAGAAGGCCGCGAAUCUUUCGGCCCCGGGUUCGCCUCGGGGCCGGUGAUUUUCAAGCUUCGGGG.ACCU..UCG..............................GGG....CCC.UAUACCAACCAUA..GGCGG...........G..........
env-527   GGCGGCGAAUCUCGCGGCCCCGGGUUCCCUGGGGCCGUCUGAUUUUUGAAGGCGAGGACCUCCGGGGCCUCAUGCAACCGGGG.ACCU..ACG..............................GGG....CCC.CUUAACCAAC.....GGACCCUCU.......................
Jaq-1-1   GCCCCCAUCCGCAAGGCCCCGGAUUCGUCCGGGGCUUUGUCGUCGCGCCGGCCCGGCAGGUGGGGGCGGCAUCCAAGCCGGGG.ACCU..ACG..............................GGG....CCC.UAUACCCAC......GGCGG...........G..........
Sge-1-1   UGUCUUUUGUCGCAUGUCGGGAUCAAUCCCGCGCUACGGCAAACCGGAAAGCGGGAUUUCCCGCCGCACCAAAUCCAUCGGGG.ACCU..UCG..............................GGG....CCC.UAUAACCACA.....GGCGG...........GGC........
env-528   UGUUUAUCUGAUACUUGGCACAGACAAUAAGUAGCUGGAUUCGUCCCAAAGCUGUUGAGAUUAAAAAUUAUUAAGAAUCGCGG.ACCU..ACG..............................GGG....CCG.CAUAACCUU......UCGGG...........CCCC.......
env-530   CUCGUUAUUUAUCUGAUACUUGGCACGGACUAUGCUUCUCAAGAUUCGUUCAGAGAAGCAUUUUUAUUGAUUAAUUUAUUGGCGG.ACCU..ACG..............................GGG....CCG.CAUAACCUU......UCAGG...........CCC........
env-531   UCGUUAUUUAUCUGAUACUUGGCACAGACUAUGCUUCUCAAGAUUCGUUCAGAGAAGCAUUUUUAUUGAUUAAUUUAUUCGCGG.ACCU..ACG..............................GGG....CCG.CAUAACCUU......UCAGG...........CCC........
env-533   .................................UUCUCAAGAUUCGUUCAGAGAAGCUAUUUUUAUUGAUUAAUUUAUUGGCGG.ACCU..ACG..............................GGG....CCG.CAUAACCUU......UCAGG...........CCC........
env-534   UCGUUAUUUAUCUGAUACUUGGCACGGACUAUGCUUCUCAAGAUUCGUUCAGAGAAGCAUUUUUAUUGAAUAAUUUAUUCGCGG.ACCU..ACG..............................GGG....CCG.CAUAACCUU......UCAGG...........CCC........
env-536   UCGUUAUUUAUCUGAUACUUGGCACAGACUAUGCUUCUCAAGAUUCGUUCAGAGAAGCAUUUUUAUUGAAUAAUUUAUUCGCGG.ACCU..ACG..............................GGG....CCG.CAUAACCUU......UCAGG...........CCC........
env-546   UCGUUAUUUAUCUGAUACUUGGCACGGACUAUGCUUCUCAAGAUUCGUUCAGAGAAGCAUUUUUAUUGAUUAAUUUAUUGGCGG.ACCU..ACG..............................GGG....CCG.CAUAACCUU......UCAGG...........CCC........
env-548   ..............................AUGCUUCUCAAGAUUCGUUCAGAGAAGCAUUUUUAUUGAAUAAUUUAUUCGCGG.ACCU..ACG..............................GGG....CCG.CAUAACCUU......UCAGG...........CCC........
env-549   UCGUUAUUUAUCUGAUACUUGGCACGGACUAUGCUUCUCAAGAUUCGUUCAGAGAAGCAUUUUUAUUGAUUAAUUUAUUGGCGG.ACCU..ACG..............................GGG....CCG.CAUAACCUU......UCAGG...........CCC........
env-567   UAGUUAUUUAUCUGAUACUUGGCACAGAUAGUGCUUCUCAAGAUUCGAUCAGAGAAGCAUUUUUAUUGAAUAAUAUUUUCGCGG.ACCU..ACG..............................GGG....CCG.CAUAACCUU......UCAGA...........CCC........
env-576   CCAGUAAUUUAUCUGUUACUUGGCACAGACUUUGCUUCUCAAGAUUCGUAAAGAGGAGCAUUUUAUUGAGUAACUCAUUCGCGG.ACCU..ACG..............................GGG....CCG.CAUAACCUU......UCAGG...........CCU........
env-579   CCAGUAAUUUAUCUGUUACUUGGCACAGACUUUGCUUCUCAAGAUUCGUAAAGAGGAGCAUUUUAUUGAGUAACUCAUUCGCGG.ACCU..ACG..............................GGG....CCG.CAUAACCUU......UCAGA...........CCU........
env-582   CCAGUAAUUUAUCUGUUACUUGGCACAGACUUUGCUUCUCCAGAUUCGUAAAGAGGAGCAUUUUAUUGAGUAACUUAUUCGCGG.ACCU..ACG..............................GGG....CCG.CAUAACCUU......UCAGA...........CCU........
env-583   CCAGUAAUUUAUCUGUUACUUGGCACAGACUUUGCUUCUCAAGAUUCGUAAAGAGGGGCAUUUUAUUGAGUAACUCAUUCGCGG.ACCU..ACG..............................GGG....CCG.CAUAACCUU......UCAGA...........CCU........
Sgu-1-1   CCUAAAGAACCCUAUCGGCACAGCCCCAGAUUCGUCUGGGGCUUCGUCUUUGUGCAAGGCUCAAAUCCUGCACACUAUCUUGGG.ACCU..UCG..............................GGG....CCC.UCUAACCAC......AGCGG...........ACC........

          .....................................................................................&lt;&lt;&lt;&lt;..&lt;&lt;&lt;&lt;&lt;.............................&gt;&gt;&gt;&gt;&gt;....&gt;&gt;&gt;.&gt;.............&lt;&lt;&lt;&lt;&lt;...........&lt;&lt;&lt;&lt;&lt;&lt;&lt;&lt;&lt;&lt;&lt;&lt;
          .....................................................................................1222..222.0.............................0.222....222.1.............22222...........22..1........
          ●●●●●●●●●●●●●●●●●●●●●●●●●●●●●●●●●●●●●●●●●●●●●●●●●●●●●●●●●●●●●●●●●●●●●●●●●●●●●●●●●●●●●YYGG●●-ACCU--●CG-------------------------------GGG----●●C-U●YA●●●●○------R●●●●-----------○○----------
                                                                                     57
```

```
env-590   GCUGUGUCGUUUUGCGCAUUGGUGAGUUCUAGCCACGCGCAACGUCGGGCAAAGGGGUUCUUUGUCUGGCAUUUCAAAUUGGGG.ACCU..UCG..............................GGG....CCC.UACAACCAU.....CUCGG...........CGCC........
env-591   GCUGUGUCGUUUUGCGCAUUGGUGAGUUCUAGCCACGCGCGAGGUUGGGCAAAGGGGUUCUUUGUCUGGCAUUUCAAAUUGGGG.ACCU..UCG..............................GGG....CCC.UACAACCAU.....CUCGG...........CACC........
env-592   GCUGUGUCGUUUUGCGCAUUGGUGAGUUCUAGCCACGCGCGAGGUUGGGCAAAGGGGUUCUUUGUCUGGCAUUUCAAAUUGGGG.ACCU..UCG..............................GGG....CCC.UACAACCAU.....CUCGG...........CACC........
env-595   ..................UUGGUGAGUUCUAGCCAAGCGCGAGGUUGGGCAAAGGGGUUCUUUGUCCGGCAUUUCAAAUUGGGG.ACCU..UCG..............................GGG....CCC.UACAACCAU.....CUCGG...........CACC........
env-596   GCUGUGUCGUUUUGCGCGUUGGUAGGUUCUAGCCGCACGCAACGUCCGGCAAAGGGGUUCUUUGUCUGGCAUUUCAAAUUGGGG.ACCU..UCG..............................GGG....CCC.UACAACCAU.....CUCGG...........CACC........
env-597   GCUGUGUCGUUUUGCGCAUAGGUGAGUUCUAGCCACGCGCGAGGUUGGGCAAAGGGGUUCUUUGUCUGGCAUUUCAAAUUGGGG.ACCU..UCG..............................GGG....CCC.UACAACCAU.....CUCGG...........CACC........
Ssp-4-1   GCUGUGUCGUUUUGCGCAUGGGUGAGUUCUAGCCAAGCGCGAGGUUGGGCAAAGGGGUUCUUUGUCUGGCAUUUCAAAUUGGGG.ACCU..UCG..............................GGG....CCC.UACAACCAU.....CUCGG...........CACC........
env-598   GCUGUGUCGUUCUGCGCAUUGGUGAGUUCUAGCCACGCGCGAGGUUGGGCAAAGGGGUUCUUUGUCUGGCAUUUCAAAUUGGGG.ACCU..UCG..............................GGG....CCC.UACAACCAU.....CUCGG...........CACC........
Ssp-5-1   GCUGUGUCGUUUUGCGCAUUGGUGAGUUCUAGCCACGCGCGAGGUUGGGCAAAGGGGUUCUUUGUCUGGCAUUUCAAAUUGGGG.ACCU..UCG..............................GGG....CCC.UACAACCAU.....CUCGG...........CACC........
env-600   GCUGUGUCGUUUUGCGCAUUGGUGAGUUCUAGCCAAGCGCGAGGUUGGGCAAAGGGGUUCUUUGUCUGGCAUUUCAAAUUGGGG.ACCU..UCG..............................GGG....CCC.UACAACCAU.....CUCGG...........CACC........
env-605   GCUGUGUCGUUCUGCGCAUUGGUGAGUUCUAGCCACGCGCGAGGUUGGGCAAAGGGGUUCUUUGUCUGGCAUUUCAAAUUGGGG.ACCU..UCG..............................GGG....CCC.UACAACCAU.....CUCGG...........CACC........
env-606   ................................................................................GGGG.ACCU..UCG..............................GGG....CCC.UACAACCAU.....CUCGG...........CACC........
env-611   UAUCGACGUGACCACGGGCCUUUGAUAUAGGCAUCACGACUCAGCGGCCCCGACCCUCUCGGGGCCGCUGGUUAUUUGGCGCGG.ACCC..ACG..............................GGG....CCG.CAUCACGACA....ACCGGACCUU.....................
env-612   ...................................................CCCGACGCUCUCGGGGCCGCUGGUUAUUUGGCGCGG.ACCU..ACG..............................GGG....CCG.CAUCACGACA....ACCGGACCUU.....................
env-613   GGGGUUUGGCUAUUUGAACGAUACCGGUGGAUCGGAUGCUUAGCUCGCAAGAGAUAAUCGCCCCAUCUGCCACUCCUACAGGGG.ACCU..UCG..............................GGG....CCC.UAUAACCAU.....CGCGCCUACAUU....UU..........
env-614   GGGGUUUGGCUAUUUGAACGAUCCCGGUGGAUCGGAUGCUUACCUCGAGAGAGAUAAUCGCCCCAUUCGCCACUCCUACAGGGG.ACCU..UCG..............................GGG....CCC.UACAACCAA.....CGCGCUUUCAU.....CUC.........
env-615   GGGGUUUGGCUAUUUGAACGAUCCCGGUGGAUCGGAUGCUUACCUCGAGAGAGAUAAUCGCCCCAUUCGCCACUCCUACAGGGG.ACCU..UCG..............................GGG....CCC.UACAACCAA.....CGCGCUUUCAU.....CUC.........
env-616   CGGGGUUUGGCUAUUUGAACGAUCCCGGUGGUUGGGAUGCCUGCCUCGCAAGAGAUUUAAGCCCACCCACCACUCCUACAGGGG.ACCU..UCG..............................GGG....CCC.UACAACCAA.....CGCGCUUUC.......AUCU........
env-617   CGGGGUUUGGCUAUUUGAACGAUCCCGGUGGUUGGGAUGCCUGCCUCGCAAGAGAUUUAAGCCCACCCACCACUCCUACAGGGG.ACCU..UCG..............................GGG....CCC.UACAACCAA.....CGCGCUUUC.......AUCU........
env-618   CGGGGUUUGGCUAUUUGAACGAUCCCGGUGGUUGGGAUGCCUGCCUCGCAAGAGAUUUAAGCCCAUCCACCACUCCUACAGGGG.ACCU..UCG..............................GGG....CCC.UACAACCAA.....CGCGCUUUC.......AUCA........
env-623   .................................................................ACCCACCACUCCUACAGGGG.ACCU..UCG..............................GGG....CCC.UACAACCAA.....CGCGCUUUC.......AUCA........
env-624   ...............UGAACGAUCCCGGUGGUUGGGAUGCCUGCCUCGCAAGAGAUUUAAACCCAUCCACCACUCCCACAGGGG.ACCU..UCG..............................GGG....CCC.UAUAACCAA.....CGCGCUUUCACC....UUG.........
env-625   GGGGUUUGGCUAUUUGAACGAUCCCGGUGGAUCGGAUGCUUAACUCGCAAGAGAUAAUCGCCCCAUCUGCCACUCCUACAGGGG.ACCU..UCG..............................GGG....CCC.UAUAACCAA.....CGCGCCUUCGUC....UCG.........
env-626   GGGGUUUGGCUAUUUGAACGAUCCCGGUGGAUCGGAUGCUUAGCUCGCAAGAGAUAAUCGCCCCAUCUGCCACUCCUACAGGGG.ACCU..UCG..............................GGG....CCC.UAUAACCAA.....CGCGCCUUCGUC....UCG.........
env-627   UUGUUUUGCGAUACGGUCAAUCACACUGACCCGACGCGCACUGAUGGGGCUGACGAUAUUCCGCCCUUAUCAUUAUAUUUUGGG.ACCU..UCG..............................GGG....CCC.UAUAACCAU.....CUCGG...........CGUC........
env-628   AGAUUCGUCUGGGGCUUCGUCUUUGUGCACGGCUCAAACCUUGCACGCUACGUUGGGACCUUCGGGGCCCUCUAACCACAGCGG.ACCU..ACG..............................GGG....CCG.CAUA..........................................
env-629   AUUCGUCUGGGGCUUCGUCUUUGUGCAUGGGUCAAACCCUGUGCGCGCUACUUUGGGACCUUCGGGGCCCUCUAACCACAGCGG.ACCU..ACG..............................GGG....CCG.CAUA..........................................
Sdo-1-1   CACAGCCCCAGAUUCGUCUGGGGCUUUGUUGUUGUUCGCAAGAACAUAUCUACUGGGACCUUCGGGGCCCUCUAACCACAGCGG.ACCU..ACG..............................GGG....CCG.CAUA..........................................
env-630   ....................GCUAUACCGCGCGGGUGUUUGGCCUCAGAUUCGUUCUGGGGCUGAAUUCCUUUUACCAAAUGCGG.ACCU..ACG..............................GGG....CCG.UUCACCAUA.....GGGGC...........C...........
env-631   GCCACCCCCCNAAACCCCCGCUAUACCGCGCGGGUGUUUGGCCUCAGANUCGUUCUGGGGCCGAAUUCCUUUUACCAAAUGCGG.ACCU..ACG..............................GGG....ACG.UUCACCAUA.....GGGGC...........C...........
env-632   UUCGUUCUGGGGCCUGAUACGUUUUACCAUUAGNGGACCCAAGNGGUUUGAACAAACCUCCGCCGGAACCUAAACACCAUGCGG.ACCU..ACG..............................GGG....CCG.UUCACAAUA.....GGGGC...........C...........
env-633   CACGAGUCUCGCGGCCCCGGGUAAUCGGGGCCGCGUGCCUUUUCGGGAAUCGUGCUUCGGCAUUUUCCCGAAUUCACGACGGGG.ACCU..GCG..............................GGG....CCC.UAUAUCGAAAC....GGACCGAAA...................
env-634   CACCAGCAGCCCCGGAUUCGUCCGGGGCUCUUGGUUUGUUUGUCCCUGUGCAUUCAAGUAUCGGGCUGGACAGUCAAGCCGGGG.ACCU..UCG..............................GGG....CCC.UUCAUCUAAC.....GGACCUAU....................
env-635   AACACGAUCGACGUUCAGUCGGGCCUGUGACGAUCUGGUCCCGCGGGCCGUGAAGGCACGCGGGAUCAGCUUUAGAAGACGGGG.ACCU..UCG..............................GGG....CCC.UUCAUCCACCAC...GGACCAUG....................
env-636   AACACGAUCGACGUGCAGUCGGGCCUGUGACGAUCUGAUCCCGCGCGCCGUGAAGGCACGCGGGAUCAGCUUUAGAAGACGGGG.ACCU..UCG..............................GGG....CCC.UUCACCAACCAC...GGACCAUG....................
env-637   GCGAAUCUCGCGGCCUCGGGAAUACCGGGGCCGUGAAUUUUUGGGUGGGGCGCAUCAGCAAGUCACUCAAACGACACGGCGGGG.ACCU..UCG..............................GGG....CCC.UAUAAAUCCAUC...GGACCUAU....................
Lme-1-1   CUGAAGACCCCGGAUUCGUUCCGGGGUCGUUACGUUUGUCAGGAUGUUCGAACUUUGGGACCUUCGGGGCCCUAUAAAUCGGCC.ACCUU.AGUAGCUUGUUCCAAACGGAACCUGCGGGC....GGGG....GGC.UUUCAACAUA.....GCGGG...........UCC.........
env-638   ..................................UUUCCGUUCGGUUUCGGGCCCUUCGUUGAAUUAUUUUAAUAAACGUCGGCCUACCG..UUCG.............................GGG....GGC.UUUACCAUG.....GGCGG...........G...........
env-639   CACCCACGCCAUUCACCACUAUACCGCAGCGAUCUUGUAUGGCCUUAGAUUCGUCUGGGGCCUCUUGAUGUUUAUAAACUGGCC.ACCU..UCG..............................GGG....GGC.UUUACCAUG.....GGCGGGGAGAC.................
Rba-1-1   CACCCACGUCAUUCACCACUAUACCGCAGCGAUCUUGUAUGGCCUUAGAUUCGUCUGGGGCCUCUCGAUGUUUAUAAACUGGCC.ACCU..UCG..............................GGG....GGC.UUUACCAUG.....GGCGGGGAGAC.................
env-640   CACCCAAGCCAUUCACCACUAUACCGCAGCAAUCUUGCAUGGCCCUAGAUUCGUCUUGGGCCUCUUGAUAUUUAUAAACUGGCC.ACCU..UCG..............................GGG....GGC.UUUACCAUG.....GGCGG...........G...........
env-641   CACCCACGCCAUUCACCACUAUACCGCAACGAUCUUGUAUGGCCCUGGAUUCGUCUAGGGCCACUUGAUGUUUAUAAACUGGCC.ACCU..UCG..............................GGG...GGGC.UUUACCAUG.....GGCGG...........G...........
env-643   CACCCACGCCAUUCACCACUAUACGACAGCAAUCUUGUAUGGCCCUAGAUUCGUCUAGGGCCUACUGAUAUUUAUGAACUGGCC.ACCU..UCG..............................GGG....GGC.UUUACCAUG.....GGCGG...........G...........
env-644   CACCCAAGCCAUUCACCACUAUACCGCAGCAAUCUUGCAUGGCCCUAGAUUCGUCUUGGGCCUCUUGAUAUUUAUAAACUGGCC.ACCU..UCGG.............................GGG....GGC.UUUACCAUG.....GGCGG...........G...........
env-645   CACCCAAGCCAUUCACCACUAUACCGCAGCAAUCUUGCAUGGCCCUAGAUUCGUCUUGGGCCUCUUGAUAUUUAUAAACUGGCC.ACCU..UCG..............................GGG....GGC.UUUACCAUG.....GGCGG...........G...........
env-646   CAACCAAGCCAUUCACCACUAUACCGCAGCAAUCUUGCAUGGCCCUAGAUUCGUCUAGGGCCUCUUGAUGUUUAUAAACUGGCC.ACCU..UCGG.............................GGG....GGC.UUUACCAUG.....GGCGG...........G...........
env-647   CACCCACGCCAUUCACCACUAUACCGCAACGAUCUUGUAUGGCCCUGGAUUCGUCUAGGGCCACUUGAUGUUUAUAAACUGGCC.ACCU..UCG..............................GGG....GGC.UUUACCAUG.....GGCGG...........G...........
env-648   CACCCAAGCCAUUCACCAUUAUACCGCAGCAAUCUUGUACGGCCCUAGAUUCGUCUAGGGCCUCUUGAUGUUUAUAAACUGGCC.ACCU..UCG..............................GGG....GGC.UUUACCAUG.....GGCGG...........G...........
env-649   CAGCCAAGCCAUUCACCACUAUACCGCAGCAAUCUUGUAUGGCCCUAGAUUCGUCUAGGGCUUCUUGAUGUUUAUAAACUGGCC.ACCU..UCG..............................GGG....GGC.UUUACCAUG.....GGCGG...........G...........
env-650   CACCCAAGCCAUUCACCACUAUACCGCAGCAAUCUUGCAUGGCCCUAGAUUCGUCUUGGGCCUCUUGAUAUUUAUAAACUGGCCAACCU..UCGG.............................GGG....GGC.UUUUACCAUG....GGCGG...........G...........
Wma-1-1   CCUCUAGUAGGGGGCUCCAUCCGAGCGGCCCCGGAGCGAUCCGGGGCUUCUCUUUACCCGGCGGGCCGAUUGCCCGUCACUGGG.ACCU..UCG..............................GGG....CCC.UUCACUGACA....GCCGG...........GCGC........
env-651   AAAGCCCUAGACCUUUUUAAUAAAGAUAGCUAUACGUCCUUCGCUAAAGCAAUCAAGAUUCGUUUGAUUUGCUUUAAAUCGGCC.ACCU..UAU..............................GGG....GGC.UUUACCAUA.....GCAGA...........GUA.........
env-652   GGGGGCCCCGGGGCAAGCCCGGGGCGCGAUGGGCGGGGCAGGUGGCGCACGAACCGGGGACCUUCGGGGCCCUAUACUCUGGCU..C....GCAA..............................G....AGC.CCAAACAUA.....GCGGG...........GC..........
Mal-2-1   AGGGGCCCCGGGGCAAGCCCGGGGCGCGAUGGGCGGGGCAGGUGGCGCACGAACCGGGGACCUUCGGGGCCCUAUACUCUGGCU..C....GCAA..............................G....AGC.CCAAACAUA.....GCGGG...........GC..........
env-653   ...................CGGGGCGCGAUGGGCGGGGCAGGUGGCGCACGAACCGGGGACCUUCGGGGCCCUAUACUCUGGCU..C....GCAA..............................G....AGC.CCAAACAUA.....GCGGG...........GC..........
env-656   UAACCCCCAACCGCGCAGCCCCGGAUUCGUCCGGGGCUUCGUUGUUGUUCUAACCGGGGACCUUCGGGGCCCUAUACCAAGGCU..C....CCAG..............................G....AGC.CCAAACAUA.....GCGGG...........UCG.........
Pga-3-1   CUGACGACCCCGGAUUCGCUCCGGGGUUAUUACGUUUGUCAGGAUGCACGAACUCGGGGACCUUUGGGGCCCUAUAUCACUGGC.ACCU..ACG..............................GGG....GCC.UUAAACAUA.....GCUGG...........ACC.........
          ................................................................................<<<<..<<<<<...........................>>>>>....>>>.>...............<<<<<...........<<<<<<<<<<<<
          ................................................................................1222..222.0...........................0.222....222.1...............22222...........22..1.......
          ●●●●●●●●●●●●●●●●●●●●●●●●●●●●●●●●●●●●●●●●●●●●●●●●●●●●●●●●●●●●●●●●●●●●●●●●●●●●●●●YYGG●●-ACCU--●CG-----------------------------GGG----●●C-U●YA●●●○○------R●●●●------------○○---------
                                                                                5′
```

alignment positions 181 · · · 343

```
env-1     ................CUGAG...........................................................................................G.CCCG......AAAGC.AGACGGA.AGACAGA.AAG.CAUG
Dsh-1-1   ................ACCCGGCACGCAAC.......................................................................................................AACGGA.AGACAGA.AAG.CAUG
Pde-1-1   ................UUUUGACUGAAACGGACCACCUCA..........................................................................G.GUCC......AAAGC..AACGGA.AGACAGA.AAG.CAUG
env-2     ................GGGA.............................................................C...............................C.GUCCG..U..U.UGC.UAACGGA.AGACAGA.AAG.CAUG
Aat-2-1   ................GCAA.............................................................C...............................C.UGCCG..U..U.UGCUAAACGGA.AGACAGA.AAG.CAUG
Rsp-2-1   ...............................................................................................................................CGGA.AGACAGA.AAG.CAUG
env-3     ................UACG............................................................GGG.............................C.CGCCA..CA.U.....ACACGGA.AGACAGU.AAG.AAUG
Amu-1-1   ................AAA..............................................................GC.............................C.CUCG...CA.C.AGCUAAACGGA.AGACAGA.AAG.CAUG
env-4     ..................................................................................................................G.GUCC.....AAAGC..AACGGA.AGACAGA.GAG.CAUG
env-5     ..................................................................................................................G.GUCC.....AAAGC..AACGGA.AGACAGA.GAG.CAUG
Rat-1-1   ................GCCCAUCCGGGGCA..................................................GCU.............................U.CGCCG.UC..U..GC.AAACGGA.AGACAGA.AAG.CAUG
env-6     ................GGGUAAACUCGCA..................................................GCAA.............................C.GCUC.....A.AGCUAAACGGA.AGACAGA.AAG.CAUG
env-7     ................GGGUAAACUCGCA..................................................GCAA.............................C.GCUC.....A.AGCUAAACGGA.AGACAGA.AAG.CAUG
env-8     ................GCAA............................................................CAC.............................C.CGUG...C..A.AGC.AAACGGA.AGACAGA.GAA.CAUG
env-9     ................AACU............................................................GCU.............................C.UCGC...C..C.AGA.CAACGGA.AGACAGU.AAA.CAUG
Tst-1-1   ................GCAG............................................................GCU.............................C.UCGC...C..C.AGA.CAACGGA.AGACAGU.AAA.CAUG

          <<<<<<<<<<<<<<<<.....................................................>>>>>>>>>>>>>>>>>>>>>>>>>>>>.............................>.>>>>...........................................
          ....1.000.00.000.....................................................000.00.000.1..........1..22.............................2.2222...........................................
          ----------------○○○○---------------------------------------------------------------○○-----------------------------●-●●●Y○--○--○-○GC-○AACGGA-AGACAGA-AAG-CAUG
                                                                                                                                                                      3′
```

```
env-10     ................GCAG...........................................................GCU.....................C.UCGC...C..C.AGA.CAACGGA.AGACAGU.AAA.CAUG
env-13     ................UAAA....................................................CUCGGUG.CAU.....................C.ACCU.....G.AGCUAAACGGA.AGACAGA.AAG.CAUG
Rca-7-1    .........................................................................................................G.GUCCA....A.AGC..AACGGA.AGACAGA.GAG.CAUG
Rca-4-1    .........................................................................................................G.GUCCA....A.AGC..AACGGA.AGACAGA.GAG.CAUG
Rsp-4-1    .....................................................................................................................................CGGA.AGACAGG.AAG.CAUG
Rsp-5-1    .....................................................................................................................................CGGA.AGACAGG.AAG.CAUG
Rsp-7-1    .....................................................................................................................................CGGA.AGACAGG.AAG.CAUG
env-14     ................CGAA....................................................CUCGGGG.CA......................U.CGCU.....UAUGCUAAACGGA.AGACAGA.AAG.CAUG
env-15     ................GCAA...........................................................GA......................C.CUGGA....U.AGC.AAGCGGA.AGACAGU.AAU.CAUG
env-16     ................UAAA........................................................CUCGGUGCAU...................C.ACCUG......AGCUAAACGGA.AGACAGA.AAG.CAUG
env-17     ................GCGA...........................................................CA......................C.CCGUG..C...AAGC.AAACGGA.AGACAGG.AAA.CAUG
Sqi-1-1    ................ACG...........................................................GCCG......................C.CUCA.....U.AGC.AAACGGA.AGACAGA.AAG.CAUG
env-18     ................AAAA.........................................................CUCGG.....................UGCAUCG..CCUG.AGCUAAACGGA.AGACAGA.AAG.CAUG
env-19     ................AUCAG...........................................................C......................G.UUGCA..U..U.AGA.AAACGGA.AGACAGA.AAG.CAUG
env-20     ................GAU............................................................GC......................C.UCGCA..CU....GA.AAACGAA.AGACAGU.AAG.AAUG
Hsp-1-1    ................ACGCCAGUAGGGCAG...............................................GGU......................U.CGCU.....U.AGCUAAACGGA.AGACAGA.AAG.CAUG
Hna-1-1    ................UGACCAUUCGGGUU................................................GGGU......................U.CGC.A..C....AGCUAAACGGA.AGACAGA.AAG.CAUG
Rba-6-1    ................ACCGCUCAGCGAGCAAAGCCCACACACAGACAGGCGGGGCACGCACCC.....................................C.GCUG...C..AAAGC..AACGGA.AGACAGA.AAG.CAUG
env-21     ................AGUUACAG......................................................GAU......................C.CGCC...U..C.AGCUAGACGGA.AGACAGA.AAG.AAUG
Rsp-1-1    ................CC............................................................GAG......................G.GGUCU..C..U.AGC.UGACGGA.AGACAGG.AAG.CAUG
env-22     ................CAAUC...........................................................G......................C.CUGCG..UUUAA....AAGCGGA.AGACAGU.AAG.AAUG
env-23     ................UUAAUC..........................................................G......................G.CCGCG..CU.A.AGC.AAACGGA.AGACAGA.AAG.CAUG
env-24     ................UUAAUC..........................................................G......................G.CCGCG..CU.A.AGC.AAACGGA.AGACAGA.AAG.CAUG
env-25     ................UUAAUC..........................................................G......................G.CCGCG..CU.A.AGC.AAACGGA.AGACAGA.AAG.CAUG
env-26     .........................................................................................................U.CUCUU.AAUAU.AGC.AAACGGA.AGACAGC.AAU.CAUG
env-27     .........................................................................................................U.CUCUU.AAUAU.AGC.AAACGGA.AGACAGC.AAU.CAUG
env-35     ................AAUC...........................................................GC......................C.UGCCU..CACA.....AAACGGA.AGACAGA..AG.CAUG
Csp-1-1    .............................................................................GACC......................C.GACCG.UCU....GU.AAACGGA.AGACAGA.AAG.CAUG
Pat-1-1    ................CUCA...........................................................CGC......................C.UGGCA..C..U..GCUAAACGGA.AGACAGA.AAG.CAUG
Rba-3-1    ................GCAA.............................................................C......................C.CGCCG....GAAGC.UUACGGA.AGACAGA.AAG.CAUG
env-36     ................GUAA.............................................................C......................C.CGUC.....UAAGCUAAACGGA.AGACAGA.AAG.CAUG
env-37     ................ACCA.............................................................C......................C.CGCC...UUUU.AGC.UAACGGA.AGACAGA.AAG.CAUG
Rca-6-1    .........................................................................................................G.GUCCA......AAGC..AACGGA.AGACAGA.GAG.CAUG
env-38     ................CACA............................................................CC......................U.CUGCA....U.UGCUAAACGGA.AGACAGA.AAG.CAUG
env-39     ................CACA............................................................CC......................U.CUGCA....U.UGCUAAACGGA.AGACAGA.AAG.CAUG
env-40     ................CACA............................................................CC......................U.CUGCA....U.UGCUAAACGGA.AGACAGA.AAG.CAUG
Oba-1-1    ................GGAG..........................................................GGGCAGG...................G.GAUCG..C..U.AGCUAAACGGA.AGACAGA.AAG.CAUG
env-41     ................CGC..........................................................GCGCCUUC...................G.GUCCG..C..CAAGC.AAACGGA.AGA.AGC.UAA.CAUG
env-42     .........................................................................................................G.CCUGA.ACA.C......AACGGA.AGACAGA.UAA.AAUG
env-43     .........................................................................................................G.CCUGA.ACA.C......AACGGA.AGACAGA.UAA.AAUG
env-47     .........................................................................................................G.CCUGA.ACA.C......AACGGA.AGACAGA.UAA.AAUG
env-49     .........................................................................................................C.CUGA...CA...UAC..AACGGA.AGACAGU.UAA.CAUG
env-50     .........................................................................................................C.CUGA...CA...UAC..AACGGA.AGACAGU.UAA.CAUG
env-51     .........................................................................................................C.CUGA...CA...UAC..AACGGA.AGACAGU.AAA.CAUG
env-52     ................GCAA.............................................................G......................U.CUGAA......AUAC..AACGGA.AGACAGA.UAA.AAUG
env-55     ................CUA..............................................................G......................C.CUGAA......AUAC..AACGGA.AGACAGA.GAA.CAUG
env-56     ................CUA..............................................................G......................C.CUGAA......AUAC..AACGGA.AGACAGA.GAA.CAUG
env-57     ................CUA..............................................................G......................C.CUGAA......AUAC..AACGGA.AGACAGA.GAA.CAUG
env-59     ................CUA..............................................................G......................C.CUGAA......AUAC..AACGGA.AGACAGA.UAA.AAUG
env-60     ................CUA..............................................................G......................C.CUGAA......AUAC..AACGGA.AGACAGA.UAA.AAUG
env-64     ................CUA..............................................................G......................C.CUGAA......AUAC..AACGGA.AGACAGA.UAA.AAUG
Rli-1-1    .........................................................................................................C.CGGCG..CA.U.AGCAAAACGGA.AGACAGA.CAA.CAUG
env-65     .........................................................................................................C.CGCUG..U..UAAGC.AAACGGA.AGACAGA.CAA.CAUG
Rde-1-1    ................GCAA............................................................CC......................C.GGCG...CA.U.AGCAAAACGGA.AGACAGA.CAA.CAUG
env-66     ................AUCGC...........................................................GA......................C.GACCG.AC....AGCAAAACGGA.AGACAGA.CAA.CAUG
env-67     ................AUU...........................................................GCUU......................U.GUCG...CU.UAAGC.AAACGGA.AGACAGA.AAG.CAUG
env-68     ................AUU...........................................................GCUU......................U.GUCG...CU.UAAGC.AAACGGA.AGACAGA.AAG.CAUG
Rsp-15-1   ................AUUUCGGU......................................................GGUU......................CACGUCA.A...U.AGC.AAACGGA.AGACAGA.AAG.CAUG
env-69     ................UCC.............................................................UC......................C.ACCCG..U..C.UGC.AAACGGA.AGACAGA..AG.CAUG
Rsp-14-1   ................C.............................................................GUUU......................C.ACCCG..UUC..UGC.AAACGGA.AGACAGA..AG.CAUG
env-70     ................C.............................................................GUUU......................C.ACCCG..UUC..UGC.AAACGGA.AGACAGA..AG.CAUG
env-71     ................U..............................................................CUC......................C.ACCCG..UC...UGC.AAACGGA.AGACAGA..AG.CAUG
Csp-2-1    ................AUUCU............................................................C......................C.ACCCG..U..C.UGC.AAACGGA.AGACAGA..AG.CAUG
env-72     ................UGAUGGCGAAAGCCGAAU.......................................................................G.AGCCG..C..AAAGC.AAAUGGA.AGACAGA.AAG.CAUG
env-74     ................GCAA.............................................................C......................G.CCUGG..C..GAAGC.AGACGGA.AGACAGG.AAG.CAUG
env-75     ................AAAACCUGCGCGCGGCGGGGAGA..................................................................C.CCGC...CG.G.AGCUAAACGGA.AGACAGA.AAG.CAUG
env-76     ................AA............................................................GGGU......................U.CACCA..C..U..GCUAAACGGA.AGACAGA.AAG.CAUG
Tsp-1-1    ................GGGUGAUCGCACCCA..........................................................................U.CGUCG..UU.U.AGCAAAACGGA.AGACAGU.AA..CAUG
env-77     ................UAACC............................................................G......................C.CCGC......AAAGC..AACGGA.AGACAGA..AA.CAUG
env-78     .........................................................................................................G.CACC...CGAUAAGCAAAACGGA.AGACAUC..AA.CAUG
env-79     .........................................................................................................G.CACC...CGAUAAGCAAAACGGA.AGACAUC..AA.CAUG
Lve-2-1    ................CU............................................................GUCC......................G.GGACG..C..CAAGC.AAACGGA.AGA.AGC.UAA.CAUG
Lve-1-1    ................CU............................................................GUCC......................G.GGACG..C..CAAGC.AAACGGA.AGA.AGC.AAA.CAUG
env-83     ................AGGA...........................................................CCC......................G.ACGUG..UU.U.AGCUAAACGGA.AGACAGC.AA..CAUG
env-84     ................AGGA...........................................................CCC......................G.ACGUG..UU.U.AGCUAAACGGA.AGACAGC.AA..CAUG
Ogr-1-1    ................CAC.....................................................GCGCCG..CGAGAU...................C.CG.C......AAAGC..AACGGAAAGACAGU.AAA.CAUG
env-85     ................CCA...........................................................ACGC......................C.UCGCA..C..U..GCUAAACGGA.AGACAGA.AAG.CAUG
env-86     ................CCA...........................................................ACGC......................C.UCGCA..C..U..GCUAAACGGA.AGACAGA.AAG.CAUG
Lsp-10-1   ................AUCU.............................................................G......................G.GACCG.ACA.G.AGC.AAACGGA.AGA.AGC.UAA.CAUG
env-88     ................CAAC..........................................................UGUU......................U.UGCA...CAUA.....AAACGGA.AGACAGA.AAG.CAAG
env-89     ................AAACCU..........................................................CA......................G.ACCC...C.....CC.UAACGGA.AGACAGA.GAG.CAUG
Aat-1-1    ................GAACCGCCCUGCGGGUCA.......................................................................C.CCCCG..CAGU.UGCUAAACGGA.AGACAGA.AAG.CAUG
env-90     ................CGAAC..........................................................CGC......................C.CG.C......AAAGC..AACGGA.AGACAGA.AAG.CAUG

           <<<<<<<<<<<<<<<<.......................................................>>>>>>>>>>>>>>>>>>>>>>>>>>>>.......................>.>>>>....................................
           ....1.000.00.000.......................................................000.00.000.1..........1..22.......................2.2222....................................
           ----------------oooo---------------------------------------------------------------------------oo-----------------------●-●●●Yo--o--●-●GC-●AACGGA-AGACAGA-AAG-CAUG
                                                                                                                                                                       3′
```

```
env-91    ................AAAGA..........................GGCC..........UGCGUCG..UU...UGCUAAACGGA.AGACAGA.CAA.CAUG
env-92    ................GCAA............................GGU..........C.UGUG...C..A.AGC.AAACGGA.AGACAGA.GAA.CAUG
env-93    .................................................................G.GGUCG..C..GAAGC.AAACGGA.AGACAGA.GAA.CAUG
env-94    ................CACAA...........................GAG..........G.CUGAG..CU.A.....AAACGGA.AGACAGA.AAG.AAAA
Pna-1-1   ................CCGGCCUGGCCGCA....................C..........C.CGCCG..C..U.AGC.AAACGGA.AGACAGA.AAG.CAUG
env-96    ................ACCAU...........................C.G..........G.ACCCA..U..C.AGC.UGACGGA.AGACAGA.AAG.CAUG
Sna-1-1   .................................................................C.CGCCG....U.AGCUUAACGGA.AGACAGA.AAG.CAUG
env-97    ................GAAU............................GCC..........C.CG.C......AAAGC.AAACGGA.AGACAGU.AAG.CAUG
env-98    ................CAA............................GGAC..........C.CG.CG.....A.AGCACAACGGA.AGACAGG.AAG.CAUG
env-99    ................AAAU...........................UCCC............GUCGG..C..CAAGC.AAACGGA.AGACAGA.AAG.CAUG
env-101   ................AAAU...........................UCCC............GUCGG..C..CAAGC.AAACGGA.AGACAGA.AAG.CAUG
env-103   ................GCAAG.........................GGC.C..........U.CUUCG..UU...UGCUAAACGGA.AGACAGA.AAG.CAUG
env-104   ................GCAAG.........................GGC.C..........U.CUUCG..UU...UGCUAAACGGA.AGACAGA.AAG.CAUG
Jsp-1-1   ................GAAACAU........................GCGG..........G.GCCG...UC.A.AGC.AAACGGA.AGACAGU.GAA.CAUG
env-105   ................GCAG.............................GC..........G.GUCCG..CA.CAAGC.AAACGGA.AGACAGU.AAA.CAUG
env-106   .................................................................A.GGCCG..C..AAAGC..AACGGAAAGACAGU.AAA.CAUG
env-108   .................................................................G.GGCCG..C..AAAGC..AACGGAAAGUCAGU.AAA.CAUG
env-109   ................GCAA..............................G..........A.GGCCG..C..CAAGC.AAACGGA.AGA.AGC.UAA.CAUG
env-116   ................GCAA..............................G..........A.GGCCG..C..CAAGC.AAACGGA.AGA.............
env-117   ................GCAA..............................G..........A.GGCCG..C..CAAGC.AAACGGA.AGA.............
env-118   ................GAAA.............................GC..........U.UCCCG..CA.CAAGC.AAACGGA.AGACAGU.UAA.CAUG
env-119   ................GCAG.............................GC..........C.UCCCG..CA.CAAGC.AAACGGA.AGACAGU.UAA.CAUG
env-120   ................CA..............................CGC..........C.UCCCG..CA.CAAGC.AAACGGA.AGACAGU.UAA.CAUG
env-121   ................GCAA.............................GC..........C.GCCCA..CA.C.AGA.AAACGGA.AGACAGU.UAA.CAUG
env-122   ................GCAA..............................G..........G.GCCCG..C..CAAGC.AAACGGA.AGA.AGC.UAA.CAUG
env-124   ................ACG............................UCAG..........G.GACCG..C..CAAGC.AAACGGA.AGA.AGCUAAU.CAUG
env-125   ................AUCU.............................AG..........G.GACCG..C..CAAGC.AAACGGA.AGAAGCU.AAU.CAUG
env-127   ................AAAU.............................AG..........G.GACCG..C..CAAGC.AAACGGA.AGAAGCU.AAU.CAUG
Ote-1-1   ...............................................GAGU..........G.GUCCG..C..CAAGC.AAACGGA.AGU.AGC.AAA.CAUG
Oan-1-1   ...............................................GAGC..........G.GUCCG..C..CAAGC.AAACGGA.AGU.AGC.AAA.CAUG
Oar-1-1   ................AUAU..............................G..........G.GUCCG..C..CAAGC.AAACGGA.AGU.AGC.AAA.CAUG
env-128   ................CUCGC............................GA..........G.GCCCG..C..CAAGC.AAACGGA.AGA.AGC.AAA.CAUG
env-135   ................CUCGC............................GA..........G.GCCCG..C..CAAGC.AAACGGA.AGA.AGC.AAA.CAUG
Rsp-9-1   ................GCAA.............................GU..........G.GCCCG..CC.A.AGC.AAACGGA.AGAAGCU.AA..CAUG
env-136   ................AAAU............................GGU..........U.UGGCA....U.AGCUAAACGGA.AGACAGA.AAG.CAUG
Pan-1-1   ................AAACU............................GC..........G.GUCCU..C..A.AGC.UAGCGGA.AGACAGA.GAG.CAUG
Pfe-1-1   ................AAACU............................GC..........G.GUCCU..C..A.AGC.UAGCGGA.AGACAGA.GAG.CAUG
Lci-1-1   ................AUUCG..........................GGAU..........C.GCCCG..C..AAAGC..AACGGA.AGACACA..AG.CAUG
env-138   ................AAACCUGCCGCCAGUCGGGGAA...........AC..........C.CGAC...U..U.AGCUAAACGGA.AGACAGA.AAG.CAUG
Abo-1-1   ................CAUUGCCGUCCUCACAUAUUGGUUGAACU......C..........U.GUUCG.ACC.G.AGCUAAACGGA.AGACAGA.AAG.CAUG
Rha-1-1   ................CGCGCCGCGGCCGACCACAUAGGCGGACGCCCCGACCGGGGGC....GGGU..........CGCUGC........UGA.CAACGGA.AGACAGA.AAG.CAUG
Mal-1-1   ................ACU............................UUCC..........G.GGGCG..C..U.AGC.UAACGGA.AGACAGA.AAG.CAUG
env-139   ................CCU............................UUCC..........G.GGGCG..C..U.AGC.UAACGGA.AGACAGA.AAG.CAUG
env-141   ................CAU............................GUCC..........G.GGGCA..C..U..GCUAAACGGA.AGACAGA.AAG.CAUG
env-143   ................GACA..............................C..........C.GUCCG..C..AAAGC..AACGGAAAGACAGAGAAA.CAUG
env-144   .................................................................C.GACCC.ACC.U.AGC.UGACGGA.AGACAGA.GAA.CAUG
env-145   ................UUU..............................GC..........C.CUGCA.AC.....GCACAACGGA.AGACAGA.AAG.CAUG
env-146   ................UCU..............................GC..........C.CUGC......UACGCACAACGGA.AGACAGA.AAG.CAUG
env-148   ..................ACCUUCGGGGGGCUUACAUCGCGGGGGAUGGCCAUCGGGCCGAACUC..........GCACUG...C..U.AG..AAACGGA.AGACAGU.AAU.CAUG
env-149   ................GGCUUUGAGAUCAAAACCAGCGAGCAGGGGCCUUCAUCUACGCAGAGAGCACGC....UCU............UUGC.......AUGC.ACACGGA.AGACAGA.AAG.CAUG
env-150   ................AAACAUAGCCCCGGGCAAAAUCCG.......GGGC..........A.CUGC......U......AACGGA.AGACAGA.AAG.CAUG
env-151   ................AAACAUAGCCCCGGGCAAAAUCCG.......GGGC..........A.CUGC......U......AACGGA.AGACAGA.AAG.CAUG
env-155   ................AAACAUAGCCCCGGGCAAAAUCCG.......GGGC..........A.CUGC......U......AACGGA.AGACAGA.AAG.CAUG
env-156   ................CAACGU............................C..........A.GGCC...CC.C.UAC..AACGGA.AGACAGA.AAG.CAUG
env-157   ...............................................CUGA..........U.CGUU...UUUA.....AAACGGA.AGACAGA.AAG.AAUG
env-158   ................AAUU.............................GA..........U.CGUU...UA.A.....AAACGGA.AGACAGA.AAG.AAUG
env-162   ................AAUU.............................GA..........U.CGUU...UA.A.....AAACGGA.AGACAGA.AAG.AAUG
Kvu-1-1   ................UGAC............................UGC..........C.CG.C......AAAGC.AAACGGA.AGACAGA..AG.CAUG
env-163   ................AACU.............................GAU.........C.GUUUA.....A.....AAACGGA.AGACAGA.AAG.AAUG
Lho-1-1   ................ACA............................CGAC..........C.CG.C......AAAGC..AACGGA.AGACACA.AAG.CAUG
CRh-1-1   ................GCUUUCUAACA....................UGGC..........C.CGCCG.UC..U.AGC.UAACGGA.AGAGAGC.AAG.CAUG
env-164   ................AUU..............................GC..........C.UUGCG.AU....AGC.AAACGGA.AGACAGA.AAG.CAUG
env-165   ................AUU..............................GC..........C.UUGCG.AU....AGC.AAACGGA.AGACAGA.AAG.CAUG
Ogu-1-1   ................ACC............................UGGC..........C.CGCA...C..A.AGC.AGACGGA.AGACAGA.AAG.CAUG
env-166   ................UCAGG...........................GUC..........G.CGGCA..C..A.AGC.AGACGGA.AGACAGA.AAG.CAUG
Paq-1-1   ................AUUUU.............................G..........U.GGCC...CU.UAAGC.UGACGGA.AGACAGA.GAG.CAUG
env-167   ................GUAA..............................C..........C.UGCCG..C....AGC.UAACGGA.AGACAGA.AAG.CAUG
env-168   ................GCAA..............................C..........C.UGCCG..C..AAAGC..AACGGA.AGACAGA.AAG.CAUG
env-169   ................CAAGA.............................C..........C.UGCCG..U..U.AGC.UAACGGA.AGACAGA.AAG.CAUG
env-170   ................CAAGA.............................C..........C.UGCCG..U..U.AGC.UAACGGA.AGACAGA.AAG.CAUG
env-171   .................................................................G.GGUCAAAC..A.AGC.AAACGGA.AGACAGC.AAG..AUG
env-172   ................UUA..........................UUCAAG..........G.CUGAG..CA...UG..AAACGGA.AGACAGA.AAG.CAUG
Rba-4-1   ................AUA..........................UUCAAG..........G.CUGAG..CA...UG..AAACGGA.AGACAGA.AAG.CAUG
env-173   .................................................................C.UGCCG....ACAUAGCAAACGGA.AGACAGA.AAG.CAUG
Sps-1-1   ................UUCG..............................G..........U.CGCUG.AC..A.AGCAAAACGGA.AGACAGC.AAA.CAUG
Ssp-2-1   ................UUCG..............................G..........U.CGCUG.AC..A.AGCAAAACGGA.AGACAGC.AAA.CAUG
env-174   ................GAA............................GGGCCU........U.CGCCG..U..U.AGC.AAACGGA.AGACAGA.AAG.CAUG
Osp-1-1   ................AAUA..............................C..........C.UGCCG..UU...UGC.UAACGGA.AGACAGA.AAG.CAUG
Sst-1-1   ................GCAA..............................C..........C.GGCUG..C..U.AGC.UAACGGA.AGACAGA.AAG.CAUG
env-175   .................................................................C.CGCCC.AUA.U.AGC.AAACGG..AGACAGA..AG.CAUG
env-176   .................................................................C.CGCUG..C..A.AGC.AAACGGA.AGACAGA.AAG.CAUG
Rsp-16-1  ................UU...............................AC..........C.UGCCG..U..A.AGC.AAACGGA.AGACAGA.AAG.CAUG
env-177   ................GUAA..............................C..........C.CGCCG..C..AAAGC..AACGGA.AGACAGA.AAG.CAUG

          <<<<<<<<<<<<<<<<..................>>>>>>>>>>>>>>>>>>>>>>>>>>>>>..........>.>>>>...................................
          ....1.000.00.000..................000.00.000.1..........1..22..........2.2222...................................
          ----------------oooo------------------------------oo----------●-●●●Yo--o--o-oGC-oAACGGA-AGACAGA-AAG-CAUG
                                                                                                            3′
```

```
env-178    ................GCAA.............................................................................C.......................C.CGCCG..C..AAAGC..AACGGA.AGACAGA.AAG.CAUG
env-179    ................UUAU.............................................................................C.......................C.UGCCG.....UAAGC.AAACGGA.AGACAGA.AAG.CAUG
env-186    ................UUAU.............................................................................C.......................C.UGCCG.....UAAGC.AAACGGA.AGACAGA.AAG.CAUG
Rsp-13-1   ................GUAU.............................................................................C.......................C.CGCCG.....UAAGC.AAACGGA.AGACAGA.AAG.CAUG
env-187    ................GCAU.............................................................................C.......................C.CGCCG.....UAAGC.AAACGGA.AGACAGA.AAG.CAUG
Rmu-1-1    ................GCAU.............................................................................C.......................C.CGCCG.....UAAGC.AAACGGA.AGACAGA.AAG.CAUG
env-188    ................AACCAG.........................................................................GCC.......................C.CGCCG..C..U.AGC.AAACGGA.AGACAGA.AAG.CAUG
env-189    ................AGG............................................................................GCC.......................C.UGCCG..U..U.AGC.AAACGGA.AGACAGA.AAG.CAUG
env-190    ................AGG............................................................................GCC.......................C.UGCCG..U..U.AGC.AAACGGA.AGACAGA.AAG.CAUG
env-191    ................AGG............................................................................GCC.......................C.UGCCG..C..U.AGC.AGACGGA.AGACAGA.AAG.CAUG
env-192    ................AGG............................................................................GCC.......................U.UGCCG.....UAAGC.AAACGGA.AGACAGA.AAG.CAUG
env-193    ................GAUCG...........................................................................CU.......................U.UGCCG..CA.UAAGC.AAACGGA.AGACAGA.AAG.CAUG
env-194    ................UGU............................................................................GCU.......................U.UGCCG..CA.UAAGC.AAACGGA.AGACACA.AAG.CAUG
env-195    ................GAU............................................................................GCU.......................U.UGCCG..CA.UAAGC.AAACGGA.AGACACA.AAG.CAUG
env-196    ................GAU............................................................................GCU.......................U.UGCCG..CA.CAAGC.AAACGGA.AGACACA.AAG.CAUG
env-197    ................GAU............................................................................GCU.......................U.UGCCG..CA.CAAGC.AAACGGA.AGACACA.AAG.CAUG
env-200    ................GAU............................................................................GCU.......................U.UGCCG..CA.CAAGC.AAACGGA.AGACACA.AAG.CAUG
env-201    ................AUC............................................................................GCU.......................U.UGCCA..CA.UAAGC.AAACGGA.AGACAGA.AAG.CAUG
env-202    ................AUC............................................................................GCU.......................U.UGCCG..CA.UAAGC.AAACGGA.AGACAGA.AAG.CAUG
env-203    ................AUC............................................................................GCU.......................U.UGCCG..CA.UAAGC.AAACGGA.AGACAGA.AAG.CAUG
env-204    ................AUC............................................................................GCU.......................U.UGCCG..CA.UAAGC.AAACGGA.AGACAGA.AAG.CAUG
env-205    ................AUC............................................................................GCU.......................U.UGCCG..CU.UAAGC.AAACGGA.AGACAGA.AAG.CAUG
env-206    ................AUC............................................................................GCU.......................U.UGCCG..CU.UAAGC.AAACGGA.AGACAGA.AAG.CAUG
env-207    ................AUC............................................................................GCU.......................U.UGCCG..CU.UAAGC.AAACGGA.AGACAGA.AAG.CAUG
env-213    ................AUC............................................................................GCU.......................U.UGCCG..CU.UAAGC.AAACGGA.AGACAGA.AAG.CAUG
env-214    ................AUC............................................................................GCU.......................U.UGCCG..CU.UAAGC.AAACGGA.AGACAGA.AAG.CAUG
env-215    ................AUC............................................................................GCU.......................U.UGCCG..CU.UAAGC.AAACGGA.AGACAGA.AAG.CAUG
env-216    ................UUU............................................................................GCU.......................U.UGUCG..CU.UAAGC.AAACGGA.AGACAGA.AAG.CAUG
env-217    ................AUU............................................................................GCU.......................U.UGUCG..UU.UAAGC.AAACGGA.AGACAGA.AAG.CAUG
env-218    ................AUU............................................................................GCU.......................U.UGUCG..CU.UAAGC.AAACGGA.AGACAGA.AAG.CAUG
env-219    ................AUU............................................................................GCU.......................U.UGUCG..CU.UAAGC.AAACGGA.AGACAGA.AAG.CAUG
env-220    ................AUU............................................................................GCU.......................U.UGUCG..CU.UAAGC.AAACGGA.AGACAGA.AAG.CAUG
env-221    ................AUU............................................................................GCU.......................U.UGUCG..CU.UAAGC.AAACGGA.AGACAGA.AAG.CAUG
env-222    ................AUU............................................................................GCU.......................U.UGUCG..CU.UAAGC.AAACGGA.AGACAGA.AAG.CAUG
env-223    ................AUU............................................................................GCU.......................U.UGUCG..CU.UAAGC.AAACGGA.AGACAGA.AAG.CAUG
env-224    ................AUU............................................................................GCU.......................U.UGUCG..CU.UAAGC.AAACGGA.AGACAGA.AAG.CAUG
env-225    ................AUU............................................................................GCU.......................U.UGUCG..CU.UAAGC.AAACGGA.AGACAGA.AAG.CAUG
env-226    ................AUU............................................................................GCU.......................U.UGUCG..CU.UAAGC.AAACGGA.AGACAGA.AAG.CAUG
env-227    ................AUU............................................................................GCU.......................U.UGUCG..CU.UAAGC.AAACGGA.AGACAGA.AAG.CAUG
env-228    ................GUU............................................................................GCU.......................U.UGCCG..CA.UAAGC.AAACGGA.AGACACA.AAG.CAUG
env-229    ................U.............................................................................UACU.......................U.UGCCG..UACA.AGC.AAACGGA.AGACAGA.AAG.CAUG
env-230    ................AUU............................................................................ACU.......................U.UGCCG..UA.CAAGC..AACGG..AGACAGA.AAG.CAUG
env-231    ................AAU............................................................................ACU.......................U.UGCCA..UA.CAAGC.AAACGGA.AGACAGA.AAG.CAUG
env-232    ................AAA............................................................................ACU.......................U.UGCCG..UA.CAAGC.AAACGGA.AGACAGA.AAG.CAUG
env-233    ................AAU............................................................................ACU.......................U.UGUCG..UA.CAAGC.AAACGGA.AGACAGA.AAG.CAUG
env-234    ................AAU............................................................................ACU.......................U.UGUCG..UA.CAAGC.AAACGGA.AGACAGA.AAG.CAUG
env-235    ................AAU............................................................................ACU.......................U.UGCCG..UA.CAAGC.AAACGGA.AGACAGA.AAG.CAUG
env-236    ................AAU............................................................................ACU.......................U.UGCCG..UA.CAAGC.AAACGGA.AGACAGA.AAG.CAUG
env-237    ................AAU............................................................................ACU.......................U.UGCCG..UA.CAAGC.AAACGGA.AGACAGA.AAG.CAUG
env-247    ................AAU............................................................................ACU.......................U.UGCCG..UA.CAAGC.AAACGGA.AGACAGA.AAG.CAUG
env-248    ................AUU............................................................................ACU.......................U.UGCCG..UA.CAAGC.AAACGGA.AGACAGA.AAG.CAUG
env-252    ................AUU............................................................................ACU.......................U.UGUC...UUACAAGC.AAACGGA.AGACAGA.AAG.CAUG
env-254    ................AUU............................................................................ACU.......................U.UGUCG..UA.AAAGC.AAACGGA.AGACAGA.AAG.CAUG
env-255    ................ACU............................................................................ACU.......................U.UGUCG..UA.CAAGC.AAACGGA.AGACAGA.AAG.CAUG
env-256    ................AUU............................................................................ACU.......................U.UGCCG..UA.CAAGC.AAACGGA.AGACAGA.AAG.CAUG
env-257    ................AUU............................................................................ACU.......................U.UGUCG..UA.CAAGC.AAACGGA.AGACAGA.AAG.CAUG
env-258    ................AUU............................................................................ACU.......................U.UGUCG..UA.CAAGC.AAACGGA.AGACAGA.AAG.CAUG
env-259    ................CAUGA............................................................................C.......................C.CGCUG..C..A.AGC.AAACGGA.AGACAGA.AAG.CAUG
Rsp-8-1    ................UCAACG........................................................................GGCG.......................C.CGCCG..U..U.AGC.AAACGGA.AGACAGA.AAG.CAUG
env-260    ...........................................................................................................................C.UGCCG..C..UAAGC.AAACGGA.AGACAGA.AAG.CAUG
env-261    ...........................................................................................................................C.UGCCG..C..UAAGC.AAACGGA.AGACAGA.AAG.CAUG
env-262    ...........................................................................................................................C.UGCCG..C..UAAGC.AAACGGA.AGACAGA.AAG.CAUG
env-263    ...........................................................................................................................C.UGCCG..C..UAAGC.AAACGGA.AGACAGA.AAG.CAUG
env-264    ...........................................................................................................................C.UGCCG..C..UAAGC.AAACGGA.AGACAGA.AAG.CAUG
env-265    ...........................................................................................................................C.UGCCG..C..UAAGC.AAACGGA.AGACAGA.AAG.CAUG
env-266    ...........................................................................................................................C.UGCCG..C..UAAGC.AAACGGA.AGACAGA.AAG.CAUG
env-269    ...........................................................................................................................C.UGCCG..C..UAAGC.AAACGGA.AGACAGA.AAG.CAUG
env-271    ...........................................................................................................................C.UGCCG..C..UAAGC.AAACGGA.AGACAGA.AAG.CAUG
env-272    ...........................................................................................................................C.UGCCG..C..UAAGC.AAACGGA.AGACAGA.AAG.CAUG
env-273    ...........................................................................................................................C.UGCCG..CU.C.AGC.AAACGGA.AGACAGA.AAG.CAUG
env-274    ...........................................................................................................................C.CGCCG.ACA.U.AGC.AAACGG..AGACAGA.AAG.CAUG
env-275    ...........................................................................................................................C.CACCG.GCA.U.AGC.AAACGGA.AGACAGA.AAG.CAUG
env-276    ................A..............................................................................GA.......................C.CGCCG.ACA.U.AGC.AAACGGA.AGACAGA.AAG.CAUG
env-277    .............................................................................................................................CGCCA.ACA.U.AGC.AGACGGA.AGACAGA.AAG.CAUG
env-278    ...........................................................................................................................A.UGCCG.ACA.U.AGC.AAACGGA.AGACAGA.AAG.CAUG
env-279    ...........................................................................................................................C.CGCCG.ACA.C.AGC.AAACGGA.AGACAGA.AAG.CAUG
env-280    ...........................................................................................................................U.CGCCG.ACA.U.AGC.AAACGGA.AGACAGA.AAG.CAUG
env-282    ...........................................................................................................................U.CGCCG.ACA.U.AGC.AAACGGA.AGACAGA.AAG.CAUG
env-284    ...........................................................................................................................U.CGCCG.ACA.U.AGC.AAACGGA.AGACAGA.AAG.CAUG
env-285    ...........................................................................................................................U.CGCCG.ACA.U.AGC.AAACGGA.AGACAGA.AAG.CAUG
env-286    ...........................................................................................................................C.CGCCG.ACA.U.AGC.AAACGGA.AGACAGA.AAG.CAUG
env-287    ...........................................................................................................................C.CGCCG.ACA.C.AGC.AAACGGA.AGACAGA.AAG.CAUG
env-288    ...........................................................................................................................C.CGCCG.ACA.U.AGC.AAACGGA.AGACAGA.AAG.CAUG

           <<<<<<<<<<<<<<<<......................................................>>>>>>>>>>>>>>>>>>>>>>>>>>>>.......................>.>>>>.....................................
           ....1.000.00.000......................................................000.00.000.1...........1..22.......................2.2222.....................................
           ----------------○○○○----------------------------------------------------------------------------○○-----------------------●-●●●Y○--○--○-○GC-○AACGGA-AGACAGA-AAG-CAUG
                                                                                                                                                                          3'
```

```
env-289   .........................................................................................................................C.UGCCG.AUA.U.AGC.AAACGGA.AGACAGA.AAG.CAUG
env-290   .........................................................................................................................C.CGCCG.ACA.U.AGC.AAACGGA.AGACAGA.AAG.CAUG
env-291   .........................................................................................................................C.CGCCG.ACA.U.AGC.AAACGGA.AGACAGA.AAG.CAUG
env-293   .........................................................................................................................C.CGCCG.ACA.U.AGC.AAACGGA.AGACAGA.AAG.CAUG
env-295   .........................................................................................................................C.CGCCG.ACA.U.AGC.AAACGGA.AGACAGA.AAG.CAUG
env-296   .........................................................................................................................C.CGCCG.ACA.U.AGC.AAACGGA.AGACAGA.AAG.CAUG
env-297   .........................................................................................................................C.CGUCG.ACA.U.AGC.AAACGGA.AGACAGA.AAG.CAUG
env-298   .........................................................................................................................C.CGUCG.ACA.U.AGC.AAACGGA.AGACAGA.AAG.CAUG
env-299   .........................................................................................................................C.CGCCG.ACA.U.AGC.AAACGGA.AGACAGA.AAG.CAUG
env-300   .........................................................................................................................C.UGCCG.ACA.U.AGC.AAACGGA.AGACAGA.AAG.CAUG
env-301   .........................................................................................................................C.CGCUG.ACA.U.AGC.AAACGGA.AGACAGA.AAG.CAUG
env-308   .........................................................................................................................C.UGCCG.ACG.U.AGC.AAACGGA.AGACAGA.AAG.CAUG
env-309   .........................................................................................................................C.UGCCG.ACA.U.AGC.AAACGGA.AGACAGA.AAG.CAUG
env-310   .........................................................................................................................C.UGCCG.ACA.U.AGC.AAACGGA.AGACAGA.AAG.CAUG
env-311   .........................................................................................................................C.UGCCG.ACA.U.AGC.AAACGGA.AGACAGA.AAG.CAUG
env-312   .........................................................................................................................C.UGCCG.ACA.U.AGC.AAACGGA.AGACAGA.AAG.CAUG
env-313   .........................................................................................................................C.UGCCG.ACA.U.AGC.AAACGGA.AGACAGA.AAG.CAUG
env-314   .........................................................................................................................C.CGCCG.ACA.U.AGC.AAACGGA.AGACAGA.AAG.CAUG
env-315   .........................................................................................................................C.CGCCG.ACA.U.AGC.AAACGGA.AGACAGA.AAG.CAUG
env-316   .........................................................................................................................C.CGCCG.ACA.U.AGC.AAACGGA.AGACAGA.AAG.CAUG
env-317   .........................................................................................................................C.CGCCG.ACA.U.AGC.AAACGGA.AGACAGA.AAG.CAUG
env-319   .........................................................................................................................C.CGCCG.ACA.U.AGC.AAACGGA.AGACAGA.AAG.CAUG
env-320   .........................................................................................................................C.CGCCG.ACA.U.AGC.AAACGGA.AGACAGA.AAG.CAUG
env-321   .........................................................................................................................C.CGCCG.ACA.U.AGC.AAACGGA.AGACAGA.AAG.CAUG
env-322   .........................................................................................................................C.CGCCG.ACA.U.AGC.AAACGGA.AGACAGA.AAG.CAUG
env-323   .........................................................................................................................C.CGCCG.ACA.U.AGC.AAACGGA.AGACAGA.AAG.CAUG
env-324   .........................................................................................................................C.CGCCG.ACA.U.AGC.AAACGGA.AGACAGA.AAG.CAUG
env-325   .........................................................................................................................C.CGCCG.ACA.U.AGC.AAACGGA.AGACAGA.AAG.CAUG
env-326   .........................................................................................................................C.CGCCG.ACA.U.AGC.AAACGGA.AGACAGA.AAG.CAUG
env-327   .........................................................................................................................C.CGCCG.ACA.U.AGC.AAACGGA.AGACAGA.AAG.CAUG
env-329   .........................................................................................................................C.CGCCG.ACA.U.AGC.AAACGGA.AGACAGA.AAG.CAUG
env-330   .........................................................................................................................C.CGCCG.ACA.U.AGC.AAACGGA.AGACAGA.AAG.CAUG
env-331   .........................................................................................................................C.CGCCG.ACA.U.AGC.AAACGGA.AGACAGA.AAG.CAUG
env-332   .........................................................................................................................C.CGCCG.ACA.U.AGC.AAACGGA.AGACAGA.AAG.CAUG
env-335   .........................................................................................................................C.CGCCG.ACA.U.AGC.AAACGGA.AGACAGA.AAG.CAUG
env-336   .........................................................................................................................C.CGCCG.ACA.U.AGC.AAACGGA.AGACAGA.AAG.CAUG
env-337   .........................................................................................................................C.CGCCG.ACA.U.AGC.AAACGGA.AGACAGA.AAG.CAUG
env-338   .........................................................................................................................C.CGCCG.ACA.U.AGC.AAACGGA.AGACAGA.AAG.CAUG
env-339   .........................................................................................................................C.CGCCG.ACA.U.AGC.AAACGGA.AGACAGA.AAG.CAUG
env-340   .........................................................................................................................C.CGCCG.ACA.U.AGC.AAACGGA.AGACAGA.AAG.CAUG
env-341   .........................................................................................................................C.CGCCG.ACA.U.AGC.AAACGGA.AGACAGA.AAG.CAUG
env-342   .........................................................................................................................C.CGCCG.ACA.U.AGC.AAACGGA.AGACAGA.AAG.CAUG
env-343   .........................................................................................................................C.CGCCG.ACA.U.AGC.AAACGGA.AGACAGA.AAG.CAUG
env-344   .........................................................................................................................C.CGCCG.ACA.C.AGC.AAACGGA.AGACAGA.AAG.CAUG
env-345   .........................................................................................................................C.CGCCG.ACA.U.AGC.AAACGGA.AGACAGA.AAG.CAUG
env-346   .........................................................................................................................C.CGCCG.ACA.C.AGC.AAACGGA.AGACAGA.AAG.CAUG
env-347   .........................................................................................................................C.CGCCG.ACA.C.AGC.AAACGGA.AGACAGA.AAG.CAUG
env-356   .........................................................................................................................C.CGCCG.ACA.C.AGC.AAACGGA.AGACAGA.AAG.CAUG
env-357   .........................................................................................................................C.CGCCG.ACA.U.AGC.AAACGGA.AGACAGA.AAG.CAUG
env-358   .........................................................................................................................C.CGCCA.ACA.U.AGC.AAACGGA.AGACAGA.AAG.CAUG
env-359   .........................................................................................................................C.CGCCA.ACA.U.AGC.AAACGGA.AGACAGA.AAG.CAUG
env-360   .........................................................................................................................A.CGCCA.AAA.U.AGC.AGACGGA.AGACAGA.AAG.CAUG
env-361   .........................................................................................................................A.CGCCA.ACA.U.AGC.AGACGGA.AGACAGA.AAG.CAUG
Psp-3-1   ................CUG...........................................................................UUCC.......................C.UGCCG.UC..U..GC.UAACGGA.AGACAGA.AAG.CAUG
Hmi-1-1   ................AUUCG...........................................................................GCCG.....................C.GCCCG..C..A.AGC.AGACGGA.AGACAGA.AAG.CAUG
Hma-1-1   ................AUUCG..........................................................................GCC.......................GCGCCCG..C...AAGC.AGACGGA.AGACAGA.AAG.CAUG
Laq-1-1   ................ACAAAAAC....................................................................CUAAGGG......................C.CUGCA.......CGCUAAACGGA.AGACAGA.AAG.CAUG
env-362   ................AUCAAAUCCCACCCUACAUAGGUGGGGCGAAAUC...............................................C.......................C.UGCCG..UU...UGCUAAACGGA.AGACAGA.AAG.CAUG
Rpo-1-1   ................GGAA.............................................................................C.......................C.CGCCG..U....UGCUAAACGGA.AGACAGA.AAG.CAUG
env-363   ................UAAU...........................................................................GGG.......................U.CGC.G..C..UAAGC.AAACGGA.AGACAGA.AAG.CAUG
env-364   ................AACAUA..................................................................GCGAUGGGGAAAC....................U.CAUC...C..U.UUCUAAACGGA.AGACAGA.AAG.CAUG
Sla-1-1   ..............................................................................................GAUC.......................C.UCGCG..CU...UGCUAAACGGA.AGACAGA.AAG.CAUG
Rha-2-1   ................AUGA.............................................................................C.......................C.CCGCA..CU...UGCUAAACGGA.AGACAGA.AAG.CAUG
Rsp-17-1  ................AAAU.............................................................................C.......................C.CCGCA..CU...UGCUAAACGGA.AGACAGA.AAG.CAUG
Rsp-18-1  ................AAAUC............................................................................C.......................C.CUGCA..CU...UGCUAAACGGA.AGACAGA.AAG.CAUG
Rsp-19-1  ................AAACC............................................................................C.......................C.CUGCA..CU...UGCUAAACGGA.AGACAGA.AAG.CAUG
Rsp-22-1  ................AAACC............................................................................C.......................C.CUGCA..CU...UGCUAAACGGA.AGACAGA.AAG.CAUG
Rba-2-1   ................AGC.............................................................................GC.......................C.UCGCA..CU...UGCUAAACGGA.AGACAGA.AAG.CAUG
env-365   ................AUUU.....................................................................................................G.AUCGU..UUAA.....AAACGGA.AGACAGA.AAG.AAUG
env-367   ................AAUUGAU..................................................................................................C.GUUU....A.A.....AAACGGA.AGACAGA.AAG.AAUG
env-369   GGCGC...........AAACCAGA..........................................................GAG.CGGGCCUUGGGU.......................C.CGACG.........C.AAACGGA.AGACAGA.AAG.CAUG
env-370   ................CCAU............................................................................AC.......................C.CGGCG..C..AAACC..AACGGA.AGACAGA.AAG.CAUG
env-371   ................CCAU............................................................................AC.......................C.CGGCG..C..AAACC..AACGGA.AGACAGA.AAG.CAUG
env-372   ................CCAU............................................................................AC.......................C.CGGCG..CG.A.AGC.AAACGGA.AGACAGA.AAG.CAUG
env-373   ................GCAA...........................................................................GGG.......................G.CGCCG..C..CAAGC.AAACGGA.AGACAGA.AA..CAUG
env-374   ................GCAA...........................................................................GGG.......................G.CGCCG..C..CAAGC.AAACGGA.AGACAGA..AA.CAUG
env-375   ................UACG.............................................................................G.......................A.ACCC...C..U.AGA.UGACGGA.AGACAGA.AAG.CAUG
env-378   ................AAGUUAUC.........................................................................G.......................G.GCCCG..C..U.AGC.UGACGGA.AGACAGA.AAG.CAUG
env-379   ................AAGUUAUC.........................................................................G.......................G.GCCCG..C..U.AGC.UGACGGA.AGACAGA.AAG.CAUG
env-380   ................UUUUU..........................................................................GUC.......................U.UGCA...CG.U..AC..AACGGA.AGACAGA.AAG.CAUG
env-381   ................AUACCCGGCGCGACCCAAAGAGCGGACCUC................................................UGGU.......................C.CGACG..C..A.....AAACGGA.AGACAGA.AAG.CAUG
Osp-2-1   ................AAAG.............................................................................C.......................G.ACCCG..UA.U.AGCUAAACGGA.AGAGAGC.AAG.CAUG
env-382   ................GCAA..........................................................................CGCU.......................G.CG.CG.....A.AGC.AGACGGA.AGACAGA.AAG.CAUG

          <<<<<<<<<<<<<<<<......................................................>>>>>>>>>>>>>>>>>>>>>>>>>>>>.......................>.>>>>....................................
          ....1.000.00.000......................................................000.00.000.1...........1..22.......................2.2222....................................
          ----------------●●●○----------------------------------------------------------------------------○○-----------------------●-●●●Y○--○--●-●GC-●AACGGA-AGACAGA-AAG-CAUG
                                                                                                                                                                     3′
```

```
env-386   ................GCAA...........................................................................CGUC.......................C.CG.CG.....A.AGC.AGACGGA.AGACAGA.AAG.CAUG
env-388   ................GCAA...........................................................................CGGU.......................G.CGCG.....A.AGC.AGACGGA.AGACAGA.AAG.CAUG
Rme-1-1   ................GUAC............................................................................GGU.......................G.CG.CG.....A.AGC.AGACGGA.AGACAGA.AAG.CAUG
env-389   ................GCAA...........................................................................CGGU.......................C.CG.CG.....A.AGC.AGACGGA.AGACAGA.AAG.CAUG
env-391   ................GCAA...........................................................................CGGU.......................C.CG.CG.....A.AGC.AGACGGA.AGACAGA.AAG.CAUG
env-392   ................UAAU.............................................................................GG.......................G.UCGCG..C..UAAGC.AAACGGA.AGACAGA.AAG.CAUG
env-394   ................UAAU.............................................................................GG.......................G.UCGCG..C..UAAGC.AAACGGA.AGACAGA.AAG.CAUG
env-395   CC..............GCAA.................................................................GGACCGAGGGGCGU.......................U.AUGCG..C..A.AGC.UAACGGA.AGACAGA.AAG.CAUG
env-396   ................GCAGGCAAGCGCCGAG................................................................GGG.......................U.CCGUG.....U..GCUAGACGGA.AGACAGA.AAG.CAUG
Pxi-1-1   ................AAAGACAAGAACCUAA................................................................GGA.......................C.CCGUG.....U..GCUAGACGGA.AGACAGA.AAG.CAUG
Rmo-1-1   ................AAAGACAAGAACCUAA................................................................GGC.......................A.CUGCA.......CGCUAAACGGA.AGACAGA.AAG.CAUG
Rsp-21-1  ................AAAGAUAAUAACCUAA................................................................GGC.......................U.CGGCA..C.....GCUAAACGGA.AGACAGA.AAG.CAUG
Psp-1-1   ................GAAGACAAAAACCUAA................................................................GGU.......................C.UGGCA.......CGCUAAACGGA.AGACAGA.AAG.CAUG
Pga-1-1   ................AAAGACAAAAACCUAA................................................................GGG.......................U.CUGCA.......CGCUAAACGGA.AGACAGA.AAG.CAUG
Pin-1-1   ................AAAGACAAAAACCUAA................................................................GGG.......................U.CUGCA.......CGCUAAACGGA.AGACAGA.AAG.CAUG
Psp-2-1   ................AAAGACAAAAACCUAA................................................................GGG.......................U.CUGCA.......CGCUAAACGGA.AGACAGA.AAG.CAUG
Rsp-11-1  ................AAAGACAAAAACCUAG................................................................GGA.......................U.CCGCA.......CGCUAAACGGA.AGACAGA.AAG.CAUG
Par-1-1   ................AAAGACAAUAACCUAA................................................................GGG.......................U.CCGCA.......CGCUAAACGGA.AGACAGA.AAG.CAUG
env-397   ................AAAGACAAGAACCUAA................................................................GGA.......................C.CA.CG..C..A.CGCUAAACGGA.AGACAGA.AAG.CAUG
env-398   ................AAAGAAAAUAACCUAA................................................................GGC.......................U.CGGC......A.AGCUAAACGGA.AGACAGA.AAG.CAUG
Rsp-12-1  ................AAAGACAACAACCUAA................................................................GGC.......................U.CGGCA..C.....GCUAAACGGA.AGACAGA.AAG.CAUG
Lsp-7-1   ................AAAGACAAGAACCUAA................................................................GGC.......................C.UGGCA.......CGCUAAACGGA.AGACAGA.AAG.CAUG
Lsp-1-1   ................AAAGACAAGAACCUAA................................................................GGC.......................C.UGGCA.......CGCUAAACGGA.AGACAGA.AAG.CAUG
Lsp-3-1   ................AAAGACAAGAACCUAA................................................................GGC.......................C.UGGCA.......CGCUAAACGGA.AGACAGA.AAG.CAUG
Lsp-4-1   ................AAAGACAAGAACCUAA................................................................GGC.......................C.UGGCA.......CGCUAAACGGA.AGACAGA.AAG.CAUG
Lsp-6-1   ................AAAGACAAGAACCUAA................................................................GGC.......................C.UGGCA.......CGCUAAACGGA.AGACAGA.AAG.CAUG
Pga-2-1   ................AAAGACAAAAACCUAA................................................................GGU.......................C.UGGCA.......CGCUAAACGGA.AGACAGA.AAG.CAUG
Nsp-1-1   ................AAAGACAAAAACCUAA................................................................GGU.......................C.UGGCA.......CGCUAAACGGA.AGACAGA.AAG.CAUG
Lsp-2-1   ................AAAGACAAUAACCUAA................................................................GGC.......................C.UGGCA.......CGCUAAACGGA.AGACAGA.AAG.CAUG
Rba-7-1   ................AAAGACAAGAACCUAA................................................................GGG.......................C.CUGCA.......CGCUAAACGGA.AGACAGA.AAG.CAUG
Lda-1-1   ................AAAGACAAGAACCUAA................................................................GGG.......................C.CUGCA.......CGCUAAACGGA.AGACAGA.AAG.CAUG
Lsp-8-1   ................AAAGACAAUAACCUAA................................................................GGG.......................U.CUGCA.......CGCUAAACGGA.AGACAGA.AAG.CAUG
env-399   ................CAAG......................................................................................................G.GCCC...C..NAAAC.AGACGGA.AGACAGA.AAC.CAUG
env-400   ................CAAG......................................................................................................G.GCCC...C..CAAAC.AGACGGA.AGACAGA.AAC.CAUG
env-402   ................UUUG......................................................................................................G.GCCC...CU.G.AAC.AGACGGA.AGACAGA.AAC.CAUG
env-403   ................UAUUG..........................................................................AGAC.......................A.CGCU......A.AGC.AAACGGA.AGACAGA.AAG.CAUG
env-404   ................UAUUG..........................................................................AGAC.......................A.CGCU......A.AGC.AAACGGA.AGACAGA.AAG.CAUG
env-405   ................AUAA...........................................................................AGAU.......................A.CGCU......A.AGC.AAACGGA.AGACAGA.AAG.CAUG
env-407   ................AUAA...........................................................................AGAU.......................A.CGCU......A.AGC.AAACGGA.AGACAGA.AAG.CAUG
env-413   ................AUAA...........................................................................AGAC.......................A.CGCU......A.AGC.AAACGGA.AGACAGA.AAG.CAUG
env-414   ................GCAA..............................................................................C.......................C.CGGCAGACA.U.AGCAAAACGGA.AGACAGC.AAU.CAUG
Rel-1-1   ................UGAC......................................................................................................C.CCCCG..C..AAAGC..AACGGA.AGACAGA.AAG.CAUG
env-416   ................UGAC......................................................................................................C.CCCCG..C..AAAGC..UACGGA.AGACAGA.AAG.CAUG
env-417   ................ACAU............................................................................GGGU......................U.UGAAG..CU.A.....AAACGGA.AGACAGA.AAGUAAUG
env-418   ................GCAA......................................................................................................G.GCCCG.AU....AGC.UAACGGA.AGACAGA.GAA.CAUG
env-419   ................GUAA......................................................................................................G.GCCCG.AU....AGC.UAACGGA.AGACAGA.GAA.CAUG
env-421   ................GCAA......................................................................................................G.ACCCG..C....AGC.AGACGGA.AGACAGA..AA.CAUG
env-422   ................UCACCCACCCUACACAUAGGCGAAUUCAAG..................................................GAU.......................U.CGCCA..C..U..GCUAAACGGA.AGACAGA.AAG.CAUG
env-423   ................UCACCCACCCUACAUAAGACGGGGA.......................................................GAC.......................C.CGUC......UAAGCUAAACGGA.AGACAGA.AAG.CAUG
env-426   ................UCACCCACCCUACAUAAGACGGGGA.......................................................GAC.......................C.CGUC......UAAGCUAAACGGA.AGACAGA.AAG.CAUG
Msp-1-1   ................ACCGA.............................................................................C.......................C.CGUCG..CCACAAUC..AACGGA.AGACAGA.AAG.CAUG
Sno-1-1   ................ACACCGGCCUUCAAAAGCCAUCU...................................................................................C.UGCCG..CA.C.AGCUAAACGGA.AGACAGA.CAA.CAUG
Sno-2-1   ................ACACCGGCCUUCAAAAGCCAUCU...................................................................................C.UGCCG..CA.C.AGCUAAACGGA.AGACAGA.CAA.CAUG
env-427   ................UUAAUG....................................................................................................G.GUCG...CACUAAGC.AAACGGA.AGACAGA.AAG.CAUG
env-428   ................GCAA...........................................................................CCCG.......................U.CU.C......A.AGC.UAACGGA.AGACAGA.AAG.CAUG
Tin-1-1   ................GCAA...........................................................................CCCG.......................U.CU.C......A.AGC.UAACGGA.AGACAGA.AAG.CAUG
Pwa-1-1   ................AUCA............................................................................ACA.......................G.ACCC...C..CAAGC.UAACGGA.AGACAGA.GAG.CAUG
env-429   ................AC.............................................................................GCCA.......................G.ACCCU..C..U.AGC.UGACGGA.AGACAGA.AAG.CAUG
env-430   ................AACU............................................................................CCA.......................G.ACCCU..C..U.AGC.UGACGGA.AGACAGA.AAG.CAUG
env-431   ................AACA............................................................................ACA.......................G.ACCCU..C..U.AGC.UGACGGA.AGACAGA.AAG.CAUG
env-432   ................AACA............................................................................GCA.......................G.ACCCU..C..U.AGC.UGACGGA.AGACAGA.AAG.CAUG
env-433   ................UGGCACAUCA................................................................................................G.ACCCU..C..U.AGC.UGACGGA.AGACAGA.AAG.CAUG
Osp-3-1   ................UACUUCCAACCGCCCGGCAAAUACCCAAUACAUAGCCGCCUCACGUGGACCC..........................CGGCCCGACUUUUCCGGACCAAACGAGGGCGGCG..C..U.UGCUAAACGGA.AGACAGA.AAG.CAUG
env-434   ................UUCG............................................................................GGG.......................U.UGCCG.....A.AGC.AAACGGA.AGACAGA.AAG.CAUG
env-435   ................CCUCGC....................................................................................................G.CC.CA..C..AAAGC..AACGGA.AGACAGA.AAG.CAUG
env-436   ................CGACCCGCCCUACAUAACCGGCAGGGA.....................................................AAC.......................C.UGUCG.....A.AGCUAAACGGA.AGACAGA.AAG.CAUG
env-437   ................UUACACCCACCCGAUAUAGCGGGCCCAAG...................................................GGC.......................G.CGCA...UU...UGC.AAACGGA.AGACAGA.AAG.CAUG
env-438   ................ACAACACA....................................................................GGCGGGGCGAC...................C.CGUCG.....CAAGC.UAACGGA.AGACAGA.AAG.CAUG
env-439   ................CACCCACCCUACAUAAGACGGGG........................................................AAAC.......................C.CGUC......UAAGCUAAACGGA.AGACAGA.AAG.CAUG
Rsp-10-1  ................AACAUGUU.......................................................................GAAG.......................U.CGGG...CA.CAAGCAAAACGGA.AGACAGA.AAA.CAUG
Tar-1-1   ................GGCGAC..........................................................................CCGU......................U.UGCUG..C....AAC..AACGGA.AGACAGA.AAG.CAUG
env-440   ..........................................................................................................................C.CCCG...C..AAAGC..AACGGA.AGACAGA.AAG.CAUG
Sme-2-1   ................UUCG............................................................................GGG.......................U.GGCA...CUAC.AGCUAAACGGA.AGACAGA.CAA.CAUG
Sme-1-1   ................UUCG............................................................................GGG.......................U.GGCA...CUAC.AGCUAAACGGA.AGACAGA.CAA.CAUG
Tsp-3-1   ................UACG.............................................................................GG.......................G.CGUCG..C..CAAGC.AAACGGA.AGACAGAUAAU.CAUG
env-441   ................ACCCUACACAUAGCGGACCCCACAAA.....................................................GGGU.......................A.CG.CG..UU...UGCUAAACGGA.AGACAGA.AAG.CAUG
env-445   ................CCAU.............................................................................AC.......................C.CGGCG..CA.G.ACC..AACGGA.AGACAGA.AAG.CAUG
Rto-1-1   ................AUCUUACAUAGCGGGCCUUC............................................................GGG.......................C.ACGCG..UU...UGCUAAACGGA.AGACAGA.AAG.CAUG
Ris-1-1   ................GAUCGCUCCGGCCGACACCAGGCGGACAGGCGGGCCCGAGGGCACGC.................................AGG.......................C.CGCCA..U..C.UGA.AAACGGA.AGACAGG.AAG.CAUG
env-447   ................CACCGAUACACAUAGCGGAACCUU.......................................................UGGU.......................U.CUGCA..C..U..GCUAAACGGA.AGACAGA.AAG.CAUG
env-448   ................CCAUCCGACAUACAGGCGGGGCA..........................................................AC.......................C.CGCCG..C..U.AGC.UAACGGA.AGACAGA.AAG.CAUG
Tpa-1-1   ................GUAA..............................................................................C.......................U.CGCCG..C..AAACC..AACGGA.AGACAGA.AAG.CAUG
env-458   ................GCAA..............................................................................C.......................U.CGCCG..C..AAACC..AACGGA.AGACAGA.AAG.CAUG

          <<<<<<<<<<<<<<<<.......................................................>>>>>>>>>>>>>>>>>>>>>>>>>>>>.......................>.>>>>.....................................
          ....1.000.00.000.......................................................000.00.000.1...........1..22.......................2.2222.....................................
          --------------oooo----------------------------------------------------------------------------oo-----------------------●-●●●Yo--o--o-o-oGC-oAACGGA-AGACAGA-AAG-CAUG
                                                                                                                                                                            3′
```

```
Tat-1-1  ................GCAA.............................................................................C........................U.CGCCG..C..A.AGC.UAACGGA.AGACAGA.AAG.CAUG
env-459  ................GCAA.............................................................................C........................C.CGCCG..C..AAAGC..AACGGA.AGACAGA.AAG.CAUG
env-460  ................UUUG............................................................................GG........................A.CUGCA..U..GAAGC.AAACGGA.AGACAGA.AAG.CAUG
env-463  ................UUUG............................................................................GG........................A.CUGCA..U..GAAGC.AAACGGA.AGACAGA.AAG.CAUG
env-464  ................UUUG............................................................................GG........................A.CUGCA..CG.A.AGC.AAACGGA.AGACAGA.AAG.CAUG
env-465  ................UUCG............................................................................GG........................G.UUGCA..CG.A.AGC.AAACGGA.AGACAGA.AAG.CAUG
env-467  ................AUGC............................................................................GG........................G.CUGCA..CG.A.AGC.AAACGGA.AGACAGA.AAG.CAUG
env-469  ................AUGC............................................................................GG........................G.CUGCA..CG.A.AGC.AAACGGA.AGACAGA.AAG.CAUG
env-472  ................UCUU............................................................................GG........................G.CUGCA..CG.A.AGC.AAACGGA.AGACAGA.AAG.CAUG
env-473  ................AAA.............................................................................GG........................G.CUGCA..CG.A.AGC.AAACGGA.AGACAGA.AAG.CAUG
env-475  ................AUAA............................................................................GG........................G.CUGCA..CG.A.AGC..AACGGA.AGACAGA.AAG.CAUG
env-476  ................ACAA............................................................................GG........................G.CUGCA..CG.A.AGC.AAACGGA.AGACAGA.AAG.CAUG
env-477  ................ACAA............................................................................GG........................G.CUGCA..CG.A.AGC.AAACGGA.AGACAGA.AAG.CAUG
env-482  ................AUAA............................................................................GG........................G.CUGCG..CG.A.AGC.AAACGGA.AGACAGA.AAG.CAUG
env-483  ................AUAA............................................................................GG........................G.CUGCA..CG.A.AGC.AAACGGA.AGACAGA.AAG.CAUG
env-484  ................AUAA............................................................................GG........................G.CUGCA..CG.A.AGC.AAACGGA.AGACAGA.AAG.CAUG
Pte-1-1  ................AUAA............................................................................GG........................G.CUGCA..CG.A.AGC.AAACGGA.AGACAGA.AAG.CAUG
env-493  ................AUAA............................................................................GG........................G.CUGCA..CG.A.AGC.AAACGGA.AGACAGA.AAG.CAUG
env-494  ................UUUA............................................................................GG........................G.CUGCA..CG.A.AGC.AAACGGA.AGACAGA.AAG.CAUG
env-495  ................UUUC............................................................................GG........................G.CUGCA..CG.A.AGC.AAACGGA.AGACAGA.AAG.CAUG
env-497  GAUGGACGGGGGCUGAACACA.................................................UAGGCCGCCG.CCG....AC.AGGGCAGG.......................G.GCCG...C....AGCUAAACGGA.AGACAGA.AAG.CAUG
env-500  ................AAAACAAAUGCUCGAAACAUACGGUCGGGGCA..........................................................................A.CCC.G.ACC.U.UGCUUAACGGA.AGACAGA.AAG.CAUG
env-502  ................GCCCAAACAUA.....................................................................GCCCCUCCAAC...............C.UGGGG..CA.G.AGC.UAACGGA.AGACAGA.AAG.CAUG
env-503  ................UAC....................................................................UGGUCGGGGCA........................A.CCCGA..UC.U.UGCUUAACGGA.AGACAGA.AAG.CAUG
env-505  ................UAC.....................................................................UGGUCGGGGCA.......................A.CCCGA..UC.U.UGCUUAACGGA.AGACAGA.AAG.CAUG
Smu-1-1  ................UCCCGCCGACCACAUAGCGGCCCCACGG...................................................GGU........................C.CCGUG..U..C.UGA.AAACGGA.AGACAGA.AAG.CAUG
env-506  ................GCAAA...........................................................................GU........................G.CUGCA..UUUA.....UAACGGA.AGACAGA.AAG.CAUG
env-507  ................GCAAA...........................................................................GC........................G.CUGCA..UUUA.....UAACGGA.AGACAGA.AAG.CAUG
env-508  ................GCAAA...........................................................................GC........................G.CUGCA..UUUA.....UAACGGA.AGACAGA.AAG.CAUG
env-512  ................GUAAA...........................................................................GU........................G.CUGCA.UCU.A.....UAACGGA.AGACAGA.AAG.CAUG
env-513  ................GUAAA...........................................................................GU........................G.CUGCA.UCU.A.....UAACGGA.AGACAGA.AAG.CAUG
env-514  ................GUAAA...........................................................................GU........................G.CUGCA.UCU.A.....UAACGGA.AGACAGA.AAG.CAUG
env-515  ................GUAAA...........................................................................GU........................G.CUGCA.UCU.A.....UAACGGA.AGACAGA.AAG.CAUG
env-516  ................GUAAA...........................................................................GU........................G.CUGCA.UCU.A.....UAACGG..AGACAGA.AAG.CAUG
env-517  ................GUAAA...........................................................................GU........................G.CUGCA..UNUA.....UAACGGA.AGACAGA.AAG.CAUG
env-518  ................GUAAA...........................................................................GU........................G.CUGCA..UUUA.....UAACGGA.AGACAGA.AAG.CAUG
env-519  ................GUAAA...........................................................................GU........................G.CUGCA..UUUA.....UAACGGA.AGACAGA.AAG.CAUG
env-520  ................GUAAA...........................................................................GU........................G.CUGCA..UUUA.....UAACGGA.AGACAGA.AAG.CAUG
env-521  ................GUAAA...........................................................................GU........................G.CUGCA..UUUA.....UAACGGA.AGACAGA.AAG.CAUG
env-522  ................CGCCCAACAUAGCGGGUCGUCAAA.......................................................GAC........................C.UG.CA..CU...UGC.UAACGGA.AGACAGA.AAG.CAUG
env-523  ................CGCCCAACAUAGCGGGUCAUCUAA.......................................................GAC........................C.UGCA...CU...UGC.UAACGGA.AGACAGA.AAG.CAUG
env-524  ..........................................................................................................................C........UAACGGA.AGACAGA.AAG.CAUG
env-525  ..........................................................................................................................UU.C......AACGGA.AGACAGA.AAG.CAUG
env-526  ................UCCG............................................................................GG........................G.CCGCA..C..GAAGC.AAACGGA.AGACAGA.AAG.CAUG
Tda-1-1  ................GCAA.............................................................................C........................U.CGCCG..C..AAAGC..AACGGA.AGACAGA.AAG.CAUG
Dsp-1-1  ................CCGUUCCGAAACGGAC.................................................................C........................C.CGCCG..C..A.AGC.AGACGGA.AGACAGG.AAG.CAUG
env-527  ..........................................................................................................................G.GUCC......AAAGC.AAACGGA.AGACAGA.AAG.CAUG
Jaq-1-1  ................GCGA.............................................................................C........................C.CGUCUGACA.U.AGC.AAACGGA.AGACAGCUGA..CAUG
Sge-1-1  ................AACA...........................................................................UCC........................C.UGCCG..CA.C.AGCUAAACGGA.AGACAGA.CAA.CAUG
env-528  ................ACA...........................................................................UGGG........................U.CUGAG..CU.A.....AAACGGA.AGACAGA.AAG.AAUG
env-530  ................AAAU...........................................................................UGG........................U.CUGA.........GCUAAACGGA.AGACAGA.AAG.AAUG
env-531  ................AAAU...........................................................................UGG........................U.CUGAG..CU.A.....AAACGGA.AGACAGA.AAG.AAUG
env-533  ................AAAU...........................................................................UGG........................U.CUGAG.ACU.A.....AAACGGA.AGACAGA.AAG.AAUG
env-534  ................AAAU...........................................................................UGG........................U.CUGAG..CU.A.....AAACGGA.AGACAGA.AAG.AAUG
env-536  ................AAAU...........................................................................UGG........................U.CUGAG..CU.A.....AAACGGA.AGACAGA.AAG.AAUG
env-546  ................AAAU...........................................................................UGG........................U.CUGAG..CU.A.....AAACGG..AGACAGA.AAG.AAUG
env-548  ................AAAU...........................................................................UGG........................U.CUGAG..CU.A.....AAACGGA.AGACAGA.AAG.AAUG
env-549  ................AAAU...........................................................................UGG........................U.CUGAG..CU.A.....AAACGGA.AGACAGA.AAG.AAUG
env-567  ................UAAU...........................................................................UGG........................U.CUGAG..CU.A.....AAACGGA.AGACAGA.AAG.AAUG
env-576  ................UAAU...........................................................................UGG........................U.CUGAG..CU.A.....AAACGGA.AGACAGA.AAG.AAUG
env-579  ................UAAU...........................................................................UGG........................U.CUGAG..CU.A.....AAACGGA.AGACAGA.AAG.AAUG
env-582  ................UAAU...........................................................................UGG........................U.CUGAG..CU.A.....AAACGGA.AGACAGA.AAG.AAUG
env-583  ................UAAU...........................................................................UGG........................U.CUGAG..CU.A.....AAACGGA.AGACAGA.AAG.AAUG
Sgu-1-1  ................UACG...........................................................................GGG........................C.CGCAG.AC..A.AGCAAAACGGA.AGACAGA.GAA.CAUG
env-590  ................AUCACCAC......................................................................GGGG........................C.CGGGG.AC..A.AGCAAAACGGA.AGACAGA.GAA.CAUG
env-591  ................AUCACGAC......................................................................GGCG........................U.CGGGG.AC..A.AGCAAAACGGA.AGACAGA.GAA.CAUG
env-592  ................AGCACUAC......................................................................GGGG........................U.CGGGG.AC..A.AGCAAAACGGA.AGACAGA.GAA.CAUG
env-595  ................AUCACGAC......................................................................GGGG........................C.CGGGG.AC..A.AGCAAAACGGA.AGACAGA.GAA.CAUG
env-596  ................AUCACGAC......................................................................GGGG........................C.CGGGG.AC..A.AGCAAAACGGA.AGACAGA.GAA.CAUG
env-597  ................AUCACGAC......................................................................GGGG........................C.CGGGG.AC..A.AGCAAAACGGA.AGACAGA.GAA.CAUG
Ssp-4-1  ................AUCACGAC......................................................................GGGG........................C.CGGGG.AC..A.AGCAAAACGGA.AGACAGA.GAA.CAUG
env-598  ................AUCACGAC......................................................................GGGG........................C.CGGGG.AC..A.AGCAAAACGGA.AGACAGA.GAA.CAUG
Ssp-5-1  ................AUCACGAC......................................................................GGGG........................C.CGGGG.AC..A.AGCAAAACGGA.AGACAGA.GAA.CAUG
env-600  ................AUCACGAC......................................................................GGGG........................C.CGGGG.AC..A.AGCAAAACGGA.AGACAGA.GAA.CAUG
env-605  ................AGCACGAC......................................................................GGGG........................C.CGGGG.AC..A.AGCAAAACGGA.AGACAGA.GAA.CAUG
env-606  ................AGCACGAC......................................................................GGGG........................C.CGGGG.AC..A.AGCAAAACGGA.AGACAGA.GAA.CAUG
env-611  ..........................................................................................................................U.UGGUC..CA.G.AGC.ACACGGA.AGACAGA.AAG.CAUG
env-612  ..........................................................................................................................U.UGGUC..CA.G.AGC.ACACGGA.AGACAGA.AAG.CAUG
env-613  ................GAUGU...........................................................................GA........................G.CGUG...CC.U.AGC.AAACGGA.AGACAGA.GAA.CAUG
env-614  ................GAUG...........................................................................GAG........................G.CGUGC..C..A.AGC.AAACGGA.AGACAGA.GAA.CAUG
env-615  ................GGUG...........................................................................GAG........................G.CGUG...C..CAAGC.AAACGGA.AGACAGA.GAA.CAUG
env-616  ................GAU...........................................................................GGAU........................G.CGUG...C..CAAGC.AAACGGA.AGACAGA.GAA.CAUG
env-617  ................CGAU..........................................................................GGAU........................G.CGUG...CC.A.AGC.AAACGGA.AGACAGA.GAA.CAUG

         <<<<<<<<<<<<<<<<......................................................>>>>>>>>>>>>>>>>>>>>>>>>>>>>........................>.>>>>....................................
         ....1.000.00.000......................................................000.00.000.1...........1..22........................2.2222....................................
         ----------------○○○○----------------------------------------------------------------------------○○------------------------●-●●●Y○--○--●-●GC-●AACGGA-AGACAGA-AAG-CAUG
                                                                                                                                                              3′
```

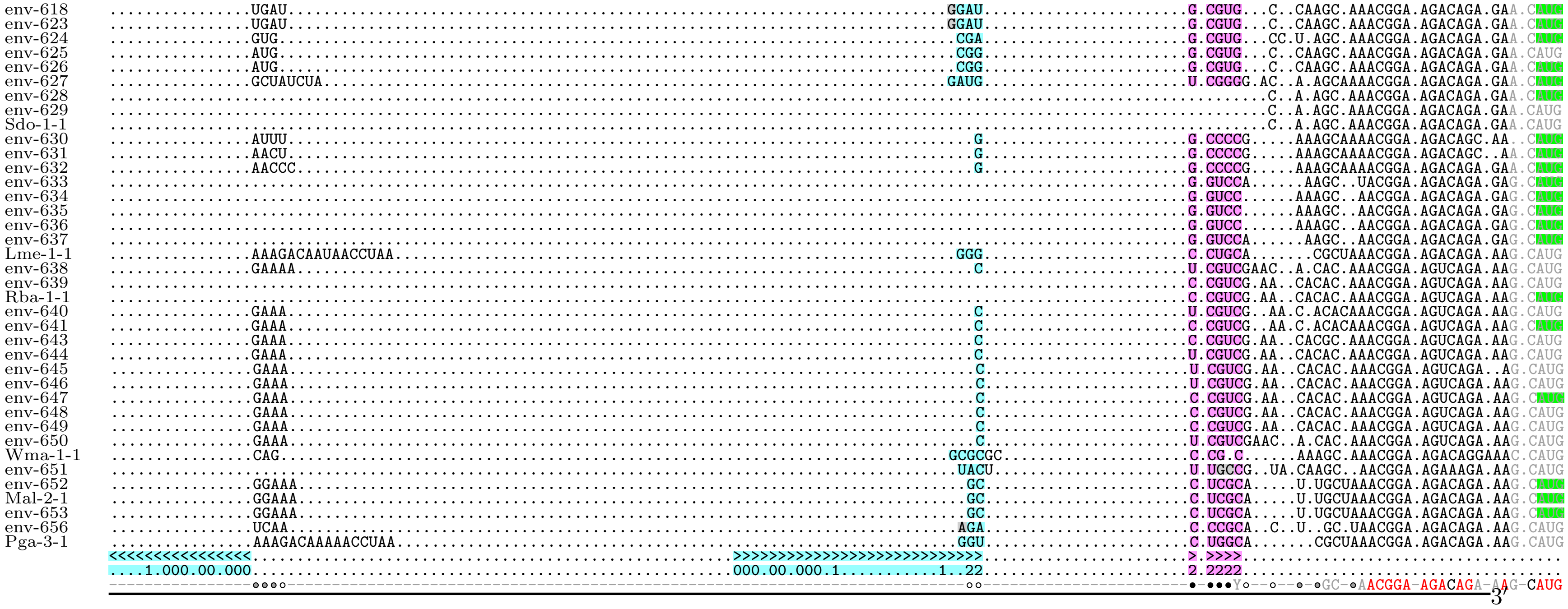

### 2 L4-Archaeoglobi

#### 2.1 Taxa

The taxonomy of each organism containing a putative L4-Archaeoglobi RNA is listed, with abbreviations identifying each hit (e.g., “Eco-1-1” and “Eco-1-2” might hypothetically represent two distinct RNAs in *E. coli*). The abbreviations will be used to identify each individual L4-Archaeoglobi RNA in Sections 2.2 and 2.4:.

| **abbrev. of hits** | **taxonomy of species** |
|---|---|
| har-1-1 | halophilic archaeon DL31 Archaea |
| h3A-1-1 | Archaea Euryarchaeota Halobacteria Halobacteriales haloarchaeon 3A1_DGR |
| Hsa-1-1 | Archaea Euryarchaeota Halobacteria Halobacteriales Haloarculaceae *Halapricum salinum* |
| Hsp-1-1 | Archaea Euryarchaeota Halobacteria Halobacteriales Haloarculaceae *Haloarcula sp.* CBA1115 |
| CHa-1-1 | Archaea Euryarchaeota Halobacteria Halobacteriales Halobacteriaceae *Candidatus* Halobonum tyrrellensis G22 |
| Hci-1-1 | Archaea Euryarchaeota Halobacteria Halobacteriales Halobacteriaceae *Haladaptatus cibarius* D43 |
| Hpa-1-1 to Hpa-1-2 | Archaea Euryarchaeota Halobacteria Halobacteriales Halobacteriaceae *Haladaptatus paucihalophilus* DX253 |
| Hje-1-1 | Archaea Euryarchaeota Halobacteria Halobacteriales Halobacteriaceae *Halalkalicoccus jeotgali* B3 |
| Hsu-1-1 | Archaea Euryarchaeota Halobacteria Halobacteriales Halobacteriaceae *Halanaeroarchaeum sulfurireducens* |
| Hac-1-1 to Hac-1-2 | Archaea Euryarchaeota Halobacteria Halobacteriales Halobacteriaceae *Halarchaeum acidiphilum* MH1-52-1 |
| Ham-1-1 | Archaea Euryarchaeota Halobacteria Halobacteriales Halobacteriaceae *Haloarcula amylolytica* JCM 13557 |
| Har-1-1 | Archaea Euryarchaeota Halobacteria Halobacteriales Halobacteriaceae *Haloarcula argentinensis* DSM 12282 |
| Hca-1-1 | Archaea Euryarchaeota Halobacteria Halobacteriales Halobacteriaceae *Haloarcula californiae* ATCC 33799 |
| Hhi-1-1 | Archaea Euryarchaeota Halobacteria Halobacteriales Halobacteriaceae *Haloarcula hispanica* ATCC 33960 |
| Hhi-2-1 | Archaea Euryarchaeota Halobacteria Halobacteriales Halobacteriaceae *Haloarcula hispanica* N601 |
| Hja-1-1 | Archaea Euryarchaeota Halobacteria Halobacteriales Halobacteriaceae *Haloarcula japonica* DSM 6131 |

| | |
|---|---|
| Hma-1-1 | Archaea Euryarchaeota Halobacteria Halobacteriales Halobacteriaceae *Haloarcula marismortui* ATCC 43049 |
| Hsa-2-1 | Archaea Euryarchaeota Halobacteria Halobacteriales Halobacteriaceae *Haloarcula salaria* |
| Hsi-1-1 | Archaea Euryarchaeota Halobacteria Halobacteriales Halobacteriaceae *Haloarcula sinaiiensis* ATCC 33800 |
| Hva-1-1 | Archaea Euryarchaeota Halobacteria Halobacteriales Halobacteriaceae *Haloarcula vallismortis* ATCC 29715 |
| Hsa-3-1 | Archaea Euryarchaeota Halobacteria Halobacteriales Halobacteriaceae *Halobacterium salinarum* R1 |
| Hsp-2-1 | Archaea Euryarchaeota Halobacteria Halobacteriales Halobacteriaceae *Halobacterium sp.* DL1 |
| Hsp-3-1 | Archaea Euryarchaeota Halobacteria Halobacteriales Halobacteriaceae *Halobacterium sp.* NRC-1 |
| Hla-1-1 to Hla-1-2 | Archaea Euryarchaeota Halobacteria Halobacteriales Halobacteriaceae *Halobiforma lacisalsi* AJ5 |
| Hni-1-1 | Archaea Euryarchaeota Halobacteria Halobacteriales Halobacteriaceae *Halobiforma nitratireducens* JCM 10879 |
| Hha-1-1 to Hha-1-2 | Archaea Euryarchaeota Halobacteria Halobacteriales Halobacteriaceae *Halococcus hamelinensis* 100A6 |
| Hmo-1-1 | Archaea Euryarchaeota Halobacteria Halobacteriales Halobacteriaceae *Halococcus morrhuae* DSM 1307 |
| Hsa-4-1 | Archaea Euryarchaeota Halobacteria Halobacteriales Halobacteriaceae *Halococcus saccharolyticus* DSM 5350 |
| Hsa-5-1 | Archaea Euryarchaeota Halobacteria Halobacteriales Halobacteriaceae *Halococcus salifodinae* DSM 8989 |
| Hth-1-1 | Archaea Euryarchaeota Halobacteria Halobacteriales Halobacteriaceae *Halococcus thailandensis* JCM 13552 |
| Hal-1-1 | Archaea Euryarchaeota Halobacteria Halobacteriales Halobacteriaceae *Haloferax alexandrinus* JCM 10717 |
| Hde-1-1 | Archaea Euryarchaeota Halobacteria Halobacteriales Halobacteriaceae *Haloferax denitrificans* ATCC 35960 |
| Hel-1-1 | Archaea Euryarchaeota Halobacteria Halobacteriales Halobacteriaceae *Haloferax elongans* ATCC BAA-1513 |
| Hgi-1-1 | Archaea Euryarchaeota Halobacteria Halobacteriales Halobacteriaceae *Haloferax gibbonsii* ATCC 33959 |
| Hla-2-1 | Archaea Euryarchaeota Halobacteria Halobacteriales Halobacteriaceae *Haloferax larsenii* JCM 13917 |
| Hlu-1-1 | Archaea Euryarchaeota Halobacteria Halobacteriales Halobacteriaceae *Haloferax lucentense* DSM 14919 |
| Hme-1-1 to Hme-1-2 | Archaea Euryarchaeota Halobacteria Halobacteriales Halobacteriaceae *Haloferax mediterranei* ATCC 33500 |
| Hmu-1-1 | Archaea Euryarchaeota Halobacteria Halobacteriales Halobacteriaceae *Haloferax mucosum* ATCC BAA-1512 |
| Hpr-1-1 | Archaea Euryarchaeota Halobacteria Halobacteriales Halobacteriaceae *Haloferax prahovense* DSM 18310 |
| Hsp-4-1 | Archaea Euryarchaeota Halobacteria Halobacteriales Halobacteriaceae *Haloferax sp.* ATCC BAA-644 |
| Hsp-5-1 | Archaea Euryarchaeota Halobacteria Halobacteriales Halobacteriaceae *Haloferax sp.* ATCC BAA-645 |
| Hsp-6-1 | Archaea Euryarchaeota Halobacteria Halobacteriales Halobacteriaceae *Haloferax sp.* ATCC BAA-646 |
| Hsu-2-1 | Archaea Euryarchaeota Halobacteria Halobacteriales Halobacteriaceae *Haloferax sulfurifontis* ATCC BAA-897 |
| Hvo-1-1 | Archaea Euryarchaeota Halobacteria Halobacteriales Halobacteriaceae *Haloferax volcanii* DS2 |
| Hbo-1-1 | Archaea Euryarchaeota Halobacteria Halobacteriales Halobacteriaceae *Halogeometricum borinquense* DSM 11551 |
| Hsa-6-1 | Archaea Euryarchaeota Halobacteria Halobacteriales Halobacteriaceae *Halogranum salarium* B-1 |
| Hka-1-1 | Archaea Euryarchaeota Halobacteria Halobacteriales Halobacteriaceae *Halomicrobium katesii* DSM 19301 |
| Hmu-2-1 | Archaea Euryarchaeota Halobacteria Halobacteriales Halobacteriaceae *Halomicrobium mukohataei* DSM 12286 |
| Hxa-1-1 | Archaea Euryarchaeota Halobacteria Halobacteriales Halobacteriaceae *Halopiger xanaduensis* SH-6 |
| Hwa-1-1 | Archaea Euryarchaeota Halobacteria Halobacteriales Halobacteriaceae *Haloquadratum walsbyi* C23 |
| Hwa-2-1 | Archaea Euryarchaeota Halobacteria Halobacteriales Halobacteriaceae *Haloquadratum walsbyi* DSM 16790 |
| Hti-1-1 | Archaea Euryarchaeota Halobacteria Halobacteriales Halobacteriaceae *Halorhabdus tiamatea* SARL4B |
| Hut-1-1 | Archaea Euryarchaeota Halobacteria Halobacteriales Halobacteriaceae *Halorhabdus utahensis* DSM 12940 |
| Hai-1-1 | Archaea Euryarchaeota Halobacteria Halobacteriales Halobacteriaceae *Halorubrum aidingense* JCM 13560 |
| Har-2-1 | Archaea Euryarchaeota Halobacteria Halobacteriales Halobacteriaceae *Halorubrum arcis* JCM 13916 |
| Hca-2-1 | Archaea Euryarchaeota Halobacteria Halobacteriales Halobacteriaceae *Halorubrum californiensis* DSM 19288 |
| Hco-1-1 | Archaea Euryarchaeota Halobacteria Halobacteriales Halobacteriaceae *Halorubrum coriense* DSM 10284 |
| Hdi-1-1 | Archaea Euryarchaeota Halobacteria Halobacteriales Halobacteriaceae *Halorubrum distributum* JCM 10118 |
| Hdi-2-1 | Archaea Euryarchaeota Halobacteria Halobacteriales Halobacteriaceae *Halorubrum distributum* JCM 9100 |
| Hho-1-1 | Archaea Euryarchaeota Halobacteria Halobacteriales Halobacteriaceae *Halorubrum hochstenium* ATCC 700873 |
| Hko-1-1 | Archaea Euryarchaeota Halobacteria Halobacteriales Halobacteriaceae *Halorubrum kocurii* JCM 14978 |
| Hla-3-1 | Archaea Euryarchaeota Halobacteria Halobacteriales Halobacteriaceae *Halorubrum lacusprofundi* ATCC 49239 |
| Hli-1-1 | Archaea Euryarchaeota Halobacteria Halobacteriales Halobacteriaceae *Halorubrum lipolyticum* DSM 21995 |
| Hli-2-1 | Archaea Euryarchaeota Halobacteria Halobacteriales Halobacteriaceae *Halorubrum litoreum* JCM 13561 |
| Hsa-7-1 | Archaea Euryarchaeota Halobacteria Halobacteriales Halobacteriaceae *Halorubrum saccharovorum* DSM 1137 |
| Hsp-7-1 | Archaea Euryarchaeota Halobacteria Halobacteriales Halobacteriaceae *Halorubrum sp.* T3 |
| Hte-1-1 | Archaea Euryarchaeota Halobacteria Halobacteriales Halobacteriaceae *Halorubrum tebenquichense* DSM 14210 |
| Hte-2-1 | Archaea Euryarchaeota Halobacteria Halobacteriales Halobacteriaceae *Halorubrum terrestre* JCM 10247 |
| Hpa-2-1 | Archaea Euryarchaeota Halobacteria Halobacteriales Halobacteriaceae *Halosarcina pallida* JCM 14848 |
| Hca-3-1 | Archaea Euryarchaeota Halobacteria Halobacteriales Halobacteriaceae *Halosimplex carlsbadense* 2-9-1 |
| Hli-3-1 | Archaea Euryarchaeota Halobacteria Halobacteriales Halobacteriaceae *Haloterrigena limicola* JCM 13563 |
| Hsa-8-1 | Archaea Euryarchaeota Halobacteria Halobacteriales Halobacteriaceae *Haloterrigena salina* JCM 13891 |
| Hth-2-1 | Archaea Euryarchaeota Halobacteria Halobacteriales Halobacteriaceae *Haloterrigena thermotolerans* DSM 11522 |
| Htu-1-1 | Archaea Euryarchaeota Halobacteria Halobacteriales Halobacteriaceae *Haloterrigena turkmenica* DSM 5511 |
| Has-1-1 | Archaea Euryarchaeota Halobacteria Halobacteriales Halobacteriaceae *Halovivax asiaticus* JCM 14624 |
| Nae-1-1 | Archaea Euryarchaeota Halobacteria Halobacteriales Halobacteriaceae *Natrialba aegyptia* DSM 13077 |
| Nas-1-1 | Archaea Euryarchaeota Halobacteria Halobacteriales Halobacteriaceae *Natrialba asiatica* DSM 12278 |
| Nch-1-1 | Archaea Euryarchaeota Halobacteria Halobacteriales Halobacteriaceae *Natrialba chahannaoensis* JCM 10990 |
| Nhu-1-1 | Archaea Euryarchaeota Halobacteria Halobacteriales Halobacteriaceae *Natrialba hulunbeirensis* JCM 10989 |
| Nma-1-1 | Archaea Euryarchaeota Halobacteria Halobacteriales Halobacteriaceae *Natrialba magadii* ATCC 43099 |
| Nta-1-1 | Archaea Euryarchaeota Halobacteria Halobacteriales Halobacteriaceae *Natrialba taiwanensis* DSM 12281 |

| | |
|---|---|
| Nal-1-1 | Archaea Euryarchaeota Halobacteria Halobacteriales Halobacteriaceae *Natrinema altunense* JCM 12890 |
| Nga-1-1 | Archaea Euryarchaeota Halobacteria Halobacteriales Halobacteriaceae *Natrinema gari* JCM 14663 |
| Npa-1-1 | Archaea Euryarchaeota Halobacteria Halobacteriales Halobacteriaceae *Natrinema pallidum* DSM 3751 |
| Npe-1-1 | Archaea Euryarchaeota Halobacteria Halobacteriales Halobacteriaceae *Natrinema pellirubrum* DSM 15624 |
| Nsp-1-1 | Archaea Euryarchaeota Halobacteria Halobacteriales Halobacteriaceae *Natrinema sp.* J7-2 |
| Nve-1-1 | Archaea Euryarchaeota Halobacteria Halobacteriales Halobacteriaceae *Natrinema versiforme* JCM 10478 |
| Ngr-1-1 | Archaea Euryarchaeota Halobacteria Halobacteriales Halobacteriaceae *Natronobacterium gregoryi* SP2 |
| Nam-1-1 | Archaea Euryarchaeota Halobacteria Halobacteriales Halobacteriaceae *Natronococcus amylolyticus* DSM 10524 |
| Nje-1-1 | Archaea Euryarchaeota Halobacteria Halobacteriales Halobacteriaceae *Natronococcus jeotgali* DSM 18795 |
| Noc-1-1 | Archaea Euryarchaeota Halobacteria Halobacteriales Halobacteriaceae *Natronococcus occultus* SP4 |
| Nin-1-1 | Archaea Euryarchaeota Halobacteria Halobacteriales Halobacteriaceae *Natronolimnobius innermongolicus* JCM 12255 |
| Nmo-1-1 | Archaea Euryarchaeota Halobacteria Halobacteriales Halobacteriaceae *Natronomonas moolapensis* 8.8.11 |
| Nph-1-1 | Archaea Euryarchaeota Halobacteria Halobacteriales Halobacteriaceae *Natronomonas pharaonis* DSM 2160 |
| Nba-1-1 | Archaea Euryarchaeota Halobacteria Halobacteriales Halobacteriaceae *Natronorubrum bangense* JCM 10635 |
| Nsu-1-1 | Archaea Euryarchaeota Halobacteria Halobacteriales Halobacteriaceae *Natronorubrum sulfidifaciens* JCM 14089 |
| Nti-1-1 | Archaea Euryarchaeota Halobacteria Halobacteriales Halobacteriaceae *Natronorubrum tibetense* GA33 |
| Ssp-1-1 | Archaea Euryarchaeota Halobacteria Halobacteriales Halobacteriaceae *Salinarchaeum sp.* Harcht-Bsk1 |
| Hag-1-1 | Archaea Euryarchaeota Halobacteria Halobacteriales Halococcaceae *Halococcus agarilyticus* |
| Hse-1-1 | Archaea Euryarchaeota Halobacteria Halobacteriales Halococcaceae *Halococcus sediminicola* |
| Hru-1-1 | Archaea Euryarchaeota Halobacteria Haloferacales Haloferacaceae *Halobellus rufus* |
| Hal-2-1 | Archaea Euryarchaeota Halobacteria Haloferacales Haloferacaceae *Haloferax alexandrinus* |
| Hgi-2-1 | Archaea Euryarchaeota Halobacteria Haloferacales Haloferacaceae *Haloferax gibbonsii* |
| Hme-1-1 to Hme-1-2 | Archaea Euryarchaeota Halobacteria Haloferacales Haloferacaceae *Haloferax mediterranei* ATCC 33500 |
| Hsp-8-1 | Archaea Euryarchaeota Halobacteria Haloferacales Haloferacaceae *Haloferax sp.* ATB1 |
| Hna-1-1 | Archaea Euryarchaeota Halobacteria Haloferacales Haloferacaceae *Haloplanus natans* DSM 17983 |
| Hru-2-1 | Archaea Euryarchaeota Halobacteria Haloferacales Halorubraceae *Halolamina rubra* |
| Hez-1-1 | Archaea Euryarchaeota Halobacteria Haloferacales Halorubraceae *Halorubrum ezzemoulense* DSM 17463 |
| Hha-2-1 | Archaea Euryarchaeota Halobacteria Haloferacales Halorubraceae *Halorubrum halophilum* |
| Hsa-9-1 | Archaea Euryarchaeota Halobacteria Haloferacales Halorubraceae *Halorubrum saccharovorum* |
| Hsp-9-1 | Archaea Euryarchaeota Halobacteria Haloferacales Halorubraceae *Halorubrum sp.* AJ67 |
| Hsp-10-1 | Archaea Euryarchaeota Halobacteria Haloferacales Halorubraceae *Halorubrum sp.* BV1 |
| Hdj-1-1 | Archaea Euryarchaeota Halobacteria Natrialbales Natrialbaceae *Halopiger djelfimassiliensis* |
| Hgo-1-1 | Archaea Euryarchaeota Halobacteria Natrialbales Natrialbaceae *Halopiger goleimassiliensis* |
| Hsa-10-1 | Archaea Euryarchaeota Halobacteria Natrialbales Natrialbaceae *Halopiger salifodinae* |
| Hla-4-1 | Archaea Euryarchaeota Halobacteria Natrialbales Natrialbaceae *Halostagnicola larsenii* XH-48 |
| Hsp-11-1 | Archaea Euryarchaeota Halobacteria Natrialbales Natrialbaceae *Halostagnicola sp.* A56 |
| Hje-2-1 | Archaea Euryarchaeota Halobacteria Natrialbales Natrialbaceae *Haloterrigena jeotgali* A29 |
| Nal-2-1 | Archaea Euryarchaeota Halobacteria Natrialbales Natrialbaceae *Natrinema altunense* |
| Nsp-2-1 | Archaea Euryarchaeota Halobacteria Natrialbales Natrialbaceae *Natrinema sp.* J7-1 |
| Mco-1-1 | Archaea Euryarchaeota Methanomicrobia Methanocellales Methanocellaceae *Methanocella conradii* HZ254 |
| CMe-1-1 | Archaea Euryarchaeota Methanomicrobia Methanomicrobiales Generaincertae sedis *Candidatus* Methanoregula boonei 6A8 |
| Mpa-1-1 | Archaea Euryarchaeota Methanomicrobia Methanomicrobiales Generaincertae sedis *Methanosphaerula palustris* E1-9c |
| Mbo-1-1 | Archaea Euryarchaeota Methanomicrobia Methanomicrobiales Methanomicrobiaceae *Methanoculleus bourgensis* MS2 |
| Mma-1-1 | Archaea Euryarchaeota Methanomicrobia Methanomicrobiales Methanomicrobiaceae *Methanoculleus marisnigri* JR1 |
| Mse-1-1 | Archaea Euryarchaeota Methanomicrobia Methanomicrobiales Methanomicrobiaceae *Methanoculleus sediminis* |
| Msp-1-1 | Archaea Euryarchaeota Methanomicrobia Methanomicrobiales Methanomicrobiaceae *Methanoculleus sp.* MH98A |
| Mli-1-1 | Archaea Euryarchaeota Methanomicrobia Methanomicrobiales Methanomicrobiaceae *Methanofollis liminatans* DSM 4140 |
| Mpa-2-1 | Archaea Euryarchaeota Methanomicrobia Methanomicrobiales Methanomicrobiaceae *Methanolacinia paynteri* |
| Mmo-1-1 | Archaea Euryarchaeota Methanomicrobia Methanomicrobiales Methanomicrobiaceae *Methanomicrobium mobile* BP |
| Mli-2-1 | Archaea Euryarchaeota Methanomicrobia Methanomicrobiales Methanomicrobiaceae *Methanoplanus limicola* DSM 2279 |
| Mpe-1-1 | Archaea Euryarchaeota Methanomicrobia Methanomicrobiales Methanomicrobiaceae *Methanoplanus petrolearius* DSM 11571 |
| Mta-1-1 | Archaea Euryarchaeota Methanomicrobia Methanomicrobiales Methanoregulaceae *Methanolinea tarda* NOBI-1 |
| Mfo-1-1 | Archaea Euryarchaeota Methanomicrobia Methanomicrobiales Methanoregulaceae *Methanoregula formicica* SMSP |
| Mhu-1-1 | Archaea Euryarchaeota Methanomicrobia Methanomicrobiales Methanospirillaceae *Methanospirillum hungatei* JF-1 |
| Mco-2-1 | Archaea Euryarchaeota Methanomicrobia Methanosarcinales Methanosaetaceae *Methanosaeta concilii* GP6 |
| Mha-1-1 | Archaea Euryarchaeota Methanomicrobia Methanosarcinales Methanosaetaceae *Methanosaeta harundinacea* 6Ac |
| Msh-1-1 | Archaea Euryarchaeota Methanomicrobia Methanosarcinales Methermicoccaceae *Methermicoccus shengliensis* DSM 18856 |
| env-1 to env-513 | environmental samples |

#### 2.2 Gene contexts

Each L4-Archaeoglobi RNA (indicated by "RNA→") is listed. For each hit, the downstream genes predicted to reside in a regulated operon are listed. If the nearest downstream gene is encoding in the opposite strand (and therefore presumed to not be a part of a regulated operon), then that gene is still depicted. Some environmental sequences and some RefSeq entries lack gene annotations, and so no genes are available for such sequences. The direction of each gene is indicated with an arrow (→), and each predicted conserved domain in the gene is named. Conserved domains associated with more than one L4-Archaeoglobi RNA are assigned a color; other domains are gray. Information about these conserved domains is given in Section 2.3. The accession of the sequence containing each L4-Archaeoglobi RNA is given in the column named "Seq. accession". Accessions beginning with "NC_", "NS_", "NW_" or "NZ_" are contained in RefSeq. Other accession refer to environmental samples. Nucleotide coordinates are given for the 5′ and 3′ boundaries of each L4-Archaeoglobi RNA. If the 5′ coordinate is greater than the 3′ coordinate, the RNA is present on the reverse-complement strand of the containing genomic DNA sequence. Each hit is denoted by an abbreviation (like "Eco-1-1") that refers to a taxonomy given in Section 2.1.

| abbrev. | Seq. accession | | 5′ at | 3′ at | genes |
|---|---|---|---|---|---|
| Mli-1-1 | NZ_CM001555.1 | + | 465849 | 465921 | **RNA**→ RplC (COG0087)rpl3p (PRK04231)→ RplC (COG0087)rpl3p (PRK04231)→ RplD (COG0088)rpl4lp (PRK04042)→ RplD (COG0088)rpl4lp (PRK04042)→ RplW (COG0089)PRK14548 (PRK14548)→ RplW (COG0089)PRK14548 (PRK14548)→ RplB (COG0090)rpl2p (PRK09612)→ RplB (COG0090)rpl2p (PRK09612)→ RpsS (COG0185)rps19p (PRK04038)→ RpsS (COG0185)rps19p (PRK04038)→ rpl22p (PRK04223)RplV (COG0091)→ rpl22p (PRK04223)RplV (COG0091)→ RpsC (COG0092)rps3p (PRK04191)→ RpsC (COG0092)rps3p (PRK04191)→ PRK14549 (PRK14549)Ribosomal_L29 (pfam00831)→ PRK14549 (PRK14549)Ribosomal_L29 (pfam00831)→ PRK03879 (PRK03879)UPF0086 (pfam01868)→ PRK03879 (PRK03879)UPF0086 (pfam01868)→ RpsQ (COG0186)rps17p (PRK08572)→ RpsQ (COG0186)rps17p (PRK08572)→ rpl14p (PRK08571)Ribosomal_L14 (pfam00238)→ rpl14p (PRK08571)Ribosomal_L14 (pfam00238)→ rpl24p (PRK01191)KOW_RPL26 (cd06089)→ rpl24p (PRK01191)KOW_RPL26 (cd06089)→ PRK04313 (PRK04313)S4 (cd00165)KOW_RPS4 (cd06087)Ribosomal_S4e (pfam00900)→ PRK04313 (PRK04313)S4 (cd00165)KOW_RPS4 (cd06087)Ribosomal_S4e (pfam00900)→ RplE (COG0094)rpl5p (PRK04219)→ RplE (COG0094)rpl5p (PRK04219)→ RpsN (COG0199)rps14P (PRK05766)→ RpsN (COG0199)rps14P (PRK05766)→ rps8p (PRK04034)RpsH (COG0096)→ rps8p (PRK04034)RpsH (COG0096)→ RplF (COG0097)rpl6p (PRK05518)→ RplF (COG0097)rpl6p (PRK05518)→ rpl32e (PRK08562)Ribosomal_L32e (pfam01655)→ rpl32e (PRK08562)Ribosomal_L32e (pfam01655)→ rpl19e (PRK08570)Ribosomal_L19e (pfam01280)→ rpl19e (PRK08570)Ribosomal_L19e (pfam01280)→ RplR (COG0256)rpl18p (PRK08569)→ RplR (COG0256)rpl18p (PRK08569)→ RpsE (COG0098)rps5p (PRK04044)→ RpsE (COG0098)rps5p (PRK04044)→ |
| env-1 | SakLake120mDRAFT_c0000096 | + | 22741 | 22827 | **RNA**→ RplC (COG0087)rpl3p (PRK04231)→ RplD (COG0088)rpl4lp (PRK04042)→ uL23_arch (TIGR03636)Ribosomal_L23 (pfam00276)→ RplB (COG0090)rpl2p (PRK09612)→ RpsS (COG0185)rps19p (PRK04038)→ rpl22p (PRK04223)RplV (COG0091)→ RpsC (COG0092)rps3p (PRK04191)Spore_IV_A (pfam09547)→ PRK14549 (PRK14549)Ribosomal_L29 (pfam00831)→ PRK03879 (PRK03879)UPF0086 (pfam01868)→ RpsQ (COG0186)rps17p (PRK08572)→ rpl14p (PRK08571)Ribosomal_L14 (pfam00238)→ rpl24p (PRK01191)KOW_RPL26 (cd06089)→ PRK04313 (PRK04313)KOW_RPS4 (cd06087)Ribosomal_S4e (pfam00900)→ RplE (COG0094)rpl5p (PRK04219)→ RpsN (COG0199)rps14P (PRK05766)→ rps8p (PRK04034)RpsH (COG0096)→ RplF (COG0097)rpl6p (PRK05518)→ rpl32e (PRK08562)Ribosomal_L32e (pfam01655)→ rpl19e (PRK08570)Ribosomal_L19e (pfam01280)→ RplR (COG0256)rpl18p (PRK08569)→ RpsE (COG0098)rps5p (PRK04044)→ |
| env-2 | SL_8KL_010_SED_10004849 | - | 1682 | 1589 | **RNA**→ Methyltrn_RNA_3 (pfam02598)→ RplC (COG0087)rpl3p (PRK04231)→ RplD (COG0088)rpl4lp (PRK04042)→ |
| env-3 | SL_8KL_010_SED_10000147 | + | 55887 | 55980 | **RNA**→ MTH1 (COG2106)→ RplC (COG0087)rpl3p (PRK04231)→ |
| env-4 | JGI24732J26686_1113329 | + | 312 | 377 | **RNA**→ |
| env-5 | JGI24731J21663_1001690 | + | 821 | 886 | **RNA**→ hypo→ |
| env-6 | JGI24729J20445_1012742 | - | 868 | 807 | **RNA**→ hypo→ rpl3p (PRK04231)Ribosomal_L3 (pfam00297)→ |
| env-7 | JGI24730J26740_1315709 | + | 120 | 181 | **RNA**→ ←hypo |

| | | | | | |
|---|---|---|---|---|---|
| env-8 | 2014733311 | - | 1671 | 1608 | **RNA**→ RplC (COG0087)rpl3p (PRK04231)→ RplD (COG0088)rpl4lp (PRK04042)→ |
| env-9 | ANASMECb_4085297_C1539 | - | 1671 | 1608 | **RNA**→ RplC (COG0087)rpl3p (PRK04231)→ RplD (COG0088)rpl4lp (PRK04042)→ |
| env-10 | PCEOT_contig04337 | + | 574 | 637 | **RNA**→ rpl3p (PRK04231)Ribosomal_L3 (pfam00297)→ RplC (COG0087)rpl3p (PRK04231)→<br>RplD (COG0088)rpl4lp (PRK04042)→ hypo→ |
| env-11 | JGI12104J13512_1001493 | + | 1210 | 1273 | **RNA**→ RplC (COG0087)rpl3p (PRK04231)→ RplD (COG0088)rpl4lp (PRK04042)→<br>RplW (COG0089)PRK14548 (PRK14548)→ RplB (COG0090)rpl2p (PRK09612)→<br>RpsS (COG0185)rps19p (PRK04038)→ rpl22p (PRK04223)RplV (COG0091)→<br>RpsC (COG0092)rps3p (PRK04191)→ PRK14549 (PRK14549)Ribosomal_L29_HIP (cd00427)→<br>PRK03879 (PRK03879)UPF0086 (pfam01868)→ RpsQ (COG0186)rps17p (PRK08572)→<br>rpl14p (PRK08571)Ribosomal_L14 (pfam00238)→ rpl24p (PRK01191)KOW_RPL26 (cd06089)→<br>PRK04313 (PRK04313)KOW_RPS4 (cd06087)Ribosomal_S4e (pfam00900)→<br>rpl5p (PRK04219)Ribosomal_L5 (pfam00281)Ribosomal_L5_C (pfam00673)→<br>RpsN (COG0199)rps14P (PRK05766)→ rps8p (PRK04034)RpsH (COG0096)→<br>RplF (COG0097)rpl6p (PRK05518)→ rpl32e (PRK08562)Ribosomal_L32_L32e (cd00513)→<br>rpl19e (PRK08570)Ribosomal_L19e (pfam01280)→ RplR (COG0256)rpl18p (PRK08569)→<br>RpsE (COG0098)rps5p (PRK04044)→ Ribosomal_L7_archeal_euk (cd01657)rpl30p (PRK06049)→<br>rpl15p (PRK06419)Ribosomal_L27A (pfam00828)→ SecY (COG0201)PRK08568 (PRK08568)→<br>AdkA (COG2019)PRK04040 (PRK04040)→ COG1422 (COG1422)→ DUF106 (pfam01956)→<br>CmkB (COG1102)PRK04182 (PRK04182)→ |
| env-12 | JGI11944J13513_1000575 | - | 22491 | 22428 | **RNA**→ RplC (COG0087)rpl3p (PRK04231)→ RplD (COG0088)rpl4lp (PRK04042)→<br>RplW (COG0089)PRK14548 (PRK14548)→ RplB (COG0090)rpl2p (PRK09612)→<br>RpsS (COG0185)rps19p (PRK04038)→ rpl22p (PRK04223)RplV (COG0091)→<br>RpsC (COG0092)rps3p (PRK04191)→ PRK14549 (PRK14549)Ribosomal_L29_HIP (cd00427)→<br>PRK03879 (PRK03879)UPF0086 (pfam01868)→ RpsQ (COG0186)rps17p (PRK08572)→<br>rpl14p (PRK08571)Ribosomal_L14 (pfam00238)→ rpl24p (PRK01191)KOW_RPL26 (cd06089)→<br>PRK04313 (PRK04313)KOW_RPS4 (cd06087)Ribosomal_S4e (pfam00900)→<br>rpl5p (PRK04219)Ribosomal_L5 (pfam00281)Ribosomal_L5_C (pfam00673)→<br>RpsN (COG0199)rps14P (PRK05766)→ rps8p (PRK04034)RpsH (COG0096)→<br>uL6_arch (TIGR03653)Ribosomal_L6 (pfam00347)→ rpl6p (PRK05518)Ribosomal_L6 (pfam00347)→<br>rpl32e (PRK08562)Ribosomal_L32_L32e (cd00513)→ rpl19e (PRK08570)Ribosomal_L19e (pfam01280)→<br>RplR (COG0256)rpl18p (PRK08569)→ RpsE (COG0098)rps5p (PRK04044)→ |
| Mhu-1-1 | NC_007796.1 | - | 2515622 | 2515559 | **RNA**→ hypo→ RplC (COG0087)rpl3p (PRK04231)→ RplD (COG0088)rpl4lp (PRK04042)→<br>RplW (COG0089)PRK14548 (PRK14548)→ RplB (COG0090)rpl2p (PRK09612)→<br>RpsS (COG0185)rps19p (PRK04038)→ rpl22p (PRK04223)RplV (COG0091)→<br>RpsC (COG0092)rps3p (PRK04191)→ PRK14549 (PRK14549)Ribosomal_L29_HIP (cd00427)→<br>POP4 (COG1588)PRK03879 (PRK03879)→ RpsQ (COG0186)rps17p (PRK08572)→<br>rpl14p (PRK08571)Ribosomal_L14 (pfam00238)→ rpl24p (PRK01191)KOW_RPL26 (cd06089)→<br>PRK04313 (PRK04313)KOW_RPS4 (cd06087)Ribosomal_S4e (pfam00900)→<br>rpl5p (PRK04219)Ribosomal_L5 (pfam00281)Ribosomal_L5_C (pfam00673)→<br>RpsN (COG0199)rps14P (PRK05766)→ |
| env-13 | 3300001567_10007289 | + | 4411 | 4498 | **RNA**→ hypo→ |
| env-14 | BDMC2_contig11431 | + | 2927 | 3013 | **RNA**→ rpl3p (PRK04231)Ribosomal_L3 (pfam00297)→ RplC (COG0087)rpl3p (PRK04231)→<br>RplD (COG0088)rpl4lp (PRK04042)→ |
| env-15 | APHM01006905.1 | - | 6665 | 6566 | **RNA**→ Methyltrn_RNA_3 (pfam02598)→ RplC (COG0087)rpl3p (PRK04231)→<br>RplD (COG0088)rpl4lp (PRK04042)→ RplW (COG0089)PRK14548 (PRK14548)→<br>RplB (COG0090)rpl2p (PRK09612)→ |
| env-16 | LTJ07As871_J07ABscf098861 | + | 1109424 | 1109523 | **RNA**→ Methyltrn_RNA_3 (pfam02598)→ RplC (COG0087)rpl3p (PRK04231)→<br>RplD (COG0088)rpl4lp (PRK04042)→ RplW (COG0089)PRK14548 (PRK14548)→<br>RplB (COG0090)rpl2p (PRK09612)→ |
| Hwa-1-1 | NC_017459.1 | - | 2240787 | 2240688 | **RNA**→ Methyltrn_RNA_3 (pfam02598)→ RplC (COG0087)rpl3p (PRK04231)→<br>RplD (COG0088)rpl4lp (PRK04042)→ RplW (COG0089)PRK14548 (PRK14548)→<br>RplB (COG0090)rpl2p (PRK09612)→ |
| env-17 | 77514049 | - | 684 | 585 | **RNA**→ Methyltrn_RNA_3 (pfam02598)→ |
| env-18 | FLAS_READ_83861724 | - | 684 | 585 | **RNA**→ ←hypo |

| | | | | | |
|---|---|---|---|---|---|
| env-19 | P_A23_Liq_1_Fmt_1000094 | + | 5234 | 5333 | **RNA**→ Methyltrn_RNA_3 (pfam02598)→ RplC (COG0087)rpl3p (PRK04231)→ RplD (COG0088)rpl4lp (PRK04042)→ RplW (COG0089)PRK14548 (PRK14548)→ RplB (COG0090)rpl2p (PRK09612)→ |
| env-20 | P_A23_Liq_3_Fmt_1000024 | + | 11155 | 11254 | **RNA**→ Methyltrn_RNA_3 (pfam02598)→ RplC (COG0087)rpl3p (PRK04231)→ RplD (COG0088)rpl4lp (PRK04042)→ RplW (COG0089)PRK14548 (PRK14548)→ RplB (COG0090)rpl2p (PRK09612)→ |
| env-21 | P_A23_Liq_2_Fmt_1000003 | - | 57459 | 57360 | **RNA**→ Methyltrn_RNA_3 (pfam02598)→ RplC (COG0087)rpl3p (PRK04231)→ RplD (COG0088)rpl4lp (PRK04042)→ RplW (COG0089)PRK14548 (PRK14548)→ RplB (COG0090)rpl2p (PRK09612)→ |
| env-22 | GSLNARP_contig28763 | + | 955 | 1054 | **RNA**→ Methyltrn_RNA_3 (pfam02598)→ RplC (COG0087)rpl3p (PRK04231)→ rpl3p (PRK04231)Ribosomal_L3 (pfam00297)→ RplD (COG0088)rpl4lp (PRK04042)→ RplW (COG0089)PRK14548 (PRK14548)→ RplB (COG0090)rpl2p (PRK09612)→ |
| Hwa-2-1 | NC_008212.1 | - | 2141989 | 2141890 | **RNA**→ Methyltrn_RNA_3 (pfam02598)→ RplC (COG0087)rpl3p (PRK04231)→ RplD (COG0088)rpl4lp (PRK04042)→ RplW (COG0089)PRK14548 (PRK14548)→ RplB (COG0090)rpl2p (PRK09612)→ |
| env-23 | GSLNARP_contig23680 | + | 166 | 268 | **RNA**→ PRK07233 (PRK07233)Methyltrn_RNA_3 (pfam02598)→ rpl3p (PRK04231)Ribosomal_L3 (pfam00297)→ |
| env-24 | GSLNAS_F7I09SK02F1FOF | + | 46 | 143 | **RNA**→ MTH1 (COG2106)→ |
| env-25 | APHM01021632.1 | - | 3874 | 3775 | **RNA**→ Methyltrn_RNA_3 (pfam02598)→ RplC (COG0087)rpl3p (PRK04231)→ RplD (COG0088)rpl4lp (PRK04042)→ RplW (COG0089)PRK14548 (PRK14548)→ RplB (COG0090)rpl2p (PRK09612)→ RpsS (COG0185)rps19p (PRK04038)→ rpl22p (PRK04223)RplV (COG0091)→ |
| env-26 | LTJ07As871_J07ABscf098869 | - | 558211 | 558112 | **RNA**→ Methyltrn_RNA_3 (pfam02598)→ RplC (COG0087)rpl3p (PRK04231)→ RplD (COG0088)rpl4lp (PRK04042)→ RplW (COG0089)PRK14548 (PRK14548)→ RplB (COG0090)rpl2p (PRK09612)→ RpsS (COG0185)rps19p (PRK04038)→ rpl22p (PRK04223)RplV (COG0091)→ rpl22p (PRK04223)Ribosomal_L22 (pfam00237)→ RpsC (COG0092)rps3p (PRK04191)Astro_capsid_p (pfam12226)→ PRK14549 (PRK14549)Ribosomal_L29_HIP (cd00427)→ PRK03879 (PRK03879)UPF0086 (pfam01868)→ RpsQ (COG0186)rps17p (PRK08572)→ rpl14p (PRK08571)Ribosomal_L14 (pfam00238)→ RplX (COG0198)rpl24p (PRK01191)→ PRK04313 (PRK04313)KOW_RPS4 (cd06087)Ribosomal_S4e (pfam00900)→ |
| env-27 | PR_CR_10_Liq_1_inCR_1027519 | - | 654 | 552 | **RNA**→ Methyltrn_RNA_3 (pfam02598)→ rpl3p (PRK04231)Ribosomal_L3 (pfam00297)→ |
| env-28 | SL_4KL_010_BRINE_10026912 | - | 325 | 226 | **RNA**→ Methyltrn_RNA_3 (pfam02598)→ |
| env-29 | SL_5KL_010_BRINE_10001539 | - | 6337 | 6238 | **RNA**→ Methyltrn_RNA_3 (pfam02598)→ RplC (COG0087)rpl3p (PRK04231)→ RplD (COG0088)rpl4lp (PRK04042)→ RplW (COG0089)PRK14548 (PRK14548)→ RplB (COG0090)rpl2p (PRK09612)→ RpsS (COG0185)rps19p (PRK04038)→ rpl22p (PRK04223)RplV (COG0091)→ RpsC (COG0092)rps3p (PRK04191)→ RpmC (COG0255)PRK14549 (PRK14549)→ PRK03879 (PRK03879)UPF0086 (pfam01868)→ RpsQ (COG0186)rps17p (PRK08572)→ rpl14p (PRK08571)→ |
| env-30 | SL_5KL_010_BRINE_10041088 | - | 903 | 804 | **RNA**→ Methyltrn_RNA_3 (pfam02598)→ rpl3p (PRK04231)Ribosomal_L3 (pfam00297)→ |
| env-31 | P_1C_Sed_2_UnCty_1050957 | - | 389 | 290 | **RNA**→ Methyltrn_RNA_3 (pfam02598)→ Methyltrn_RNA_3 (pfam02598)→ |
| Hut-1-1 | NC_013158.1 | + | 2340327 | 2340426 | **RNA**→ Methyltrn_RNA_3 (pfam02598)→ rpl3p (PRK04231)Ribosomal_L3 (pfam00297)→ RplD (COG0088)rpl4lp (PRK04042)→ RplW (COG0089)PRK14548 (PRK14548)→ RplB (COG0090)rpl2p (PRK09612)→ RpsS (COG0185)rps19p (PRK04038)→ rpl22p (PRK04223)RplV (COG0091)→ RpsC (COG0092)rps3p (PRK04191)→ PRK14549 (PRK14549)Ribosomal_L29 (pfam00831)→ PRK03879 (PRK03879)UPF0086 (pfam01868)→ RpsQ (COG0186)rps17p (PRK08572)→ rpl14p (PRK08571)Ribosomal_L14 (pfam00238)→ RplX (COG0198)rpl24p (PRK01191)→ PRK04313 (PRK04313)KOW_RPS4 (cd06087)Ribosomal_S4e (pfam00900)→ RplE (COG0094)rpl5p (PRK04219)→ RpsN (COG0199)rps14P (PRK05766)→ rps8p (PRK04034)RpsH (COG0096)→ RplF (COG0097)rpl6p (PRK05518)→ NT_POLXc (cd00141)PRK12766 (PRK12766)Rpl32e (COG1717)COG2251 (COG2251)HHH_5 (pfam14520)→ |
| env-32 | GSLSAAI_contig12629 | - | 3841 | 3739 | **RNA**→ Methyltrn_RNA_3 (pfam02598)→ RplC (COG0087)rpl3p (PRK04231)→ rpl4lp (PRK04042)Ribosomal_L4 (pfam00573)→ RplW (COG0089)PRK14548 (PRK14548)→ RplB (COG0090)rpl2p (PRK09612)→ RpsS (COG0185)rps19p (PRK04038)→ rpl22p (PRK04223)RplV (COG0091)→ |
| env-33 | GSLSAAI_contig12562 | - | 1374 | 1272 | **RNA**→ Methyltrn_RNA_3 (pfam02598)→ RplC (COG0087)rpl3p (PRK04231)→ |

| | | | | | |
|---|---|---|---|---|---|
| env-34 | GSLSAAI_contig12554 | - | 3225 | 3123 | **RNA**→ Methyltrn_RNA_3 (pfam02598)→ RplC (COG0087)rpl3p (PRK04231)→<br>RplD (COG0088)rpl4lp (PRK04042)→ RplW (COG0089)PRK14548 (PRK14548)→<br>RplB (COG0090)rpl2p (PRK09612)→ |
| env-35 | GSLSAAI_contig12558 | - | 4430 | 4328 | **RNA**→ Methyltrn_RNA_3 (pfam02598)→ RplC (COG0087)rpl3p (PRK04231)→<br>RplD (COG0088)rpl4lp (PRK04042)→ RplW (COG0089)PRK14548 (PRK14548)→<br>RplB (COG0090)rpl2p (PRK09612)→ RpsS (COG0185)rps19p (PRK04038)→<br>rpl22p (PRK04223)RplV (COG0091)→ KH (smart00322)archeal_30S_S3_KH (cd02411)→ |
| env-36 | SL_6KL_011_BRINE_1000585 | - | 15210 | 15108 | **RNA**→ Methyltrn_RNA_3 (pfam02598)→ RplC (COG0087)rpl3p (PRK04231)→<br>RplD (COG0088)rpl4lp (PRK04042)→ RplW (COG0089)uL23_arch (TIGR03636)→<br>RplB (COG0090)rpl2p (PRK09612)→ RpsS (COG0185)rps19p (PRK04038)→<br>rpl22p (PRK04223)RplV (COG0091)→ RpsC (COG0092)rps3p (PRK04191)→<br>PRK14549 (PRK14549)Ribosomal_L29 (pfam00831)→ POP4 (COG1588)PRK03879 (PRK03879)→<br>RpsQ (COG0186)rps17p (PRK08572)→ rpl14p (PRK08571)Ribosomal_L14 (pfam00238)→<br>rpl24p (PRK01191)KOW_RPL26 (cd06089)→<br>PRK04313 (PRK04313)KOW_RPS4 (cd06087)S4 (pfam01479)Ribosomal_S4e (pfam00900)→<br>RplE (COG0094)rpl5p (PRK04219)→ RpsN (COG0199)rps14P (PRK05766)→<br>rps8p (PRK04034)RpsH (COG0096)→ RplF (COG0097)rpl6p (PRK05518)→<br>Rpl32e (COG1717)rpl32e (PRK08562)HHH_5 (pfam14520)→ |
| env-37 | SL_5KL_010_BRINE_10000390 | + | 6046 | 6145 | **RNA**→ Methyltrn_RNA_3 (pfam02598)→ rpl3p (PRK04231)Ribosomal_L3 (pfam00297)→<br>RplD (COG0088)rpl4lp (PRK04042)→ RplW (COG0089)PRK14548 (PRK14548)→<br>RplB (COG0090)rpl2p (PRK09612)→ RpsS (COG0185)rps19p (PRK04038)→<br>rpl22p (PRK04223)RplV (COG0091)→ RpsC (COG0092)rps3p (PRK04191)→<br>PRK14549 (PRK14549)Ribosomal_L29 (pfam00831)→ PRK03879 (PRK03879)UPF0086 (pfam01868)→<br>RpsQ (COG0186)rps17p (PRK08572)→ rpl14p (PRK08571)Ribosomal_L14 (pfam00238)→<br>RplX (COG0198)rpl24p (PRK01191)→<br>PRK04313 (PRK04313)KOW_RPS4 (cd06087)S4 (pfam01479)Ribosomal_S4e (pfam00900)→<br>RplE (COG0094)rpl5p (PRK04219)→ RpsN (COG0199)rps14P (PRK05766)→<br>rps8p (PRK04034)RpsH (COG0096)→ RplF (COG0097)rpl6p (PRK05518)→<br>ComEA (COG1555)Rpl32e (COG1717)rpl32e (PRK08562)HHH_5 (pfam14520)→<br>rpl19e (PRK08570)Ribosomal_L19e (pfam01280)→ RplR (COG0256)rpl18p (PRK08569)→<br>RpsE (COG0098)rps5p (PRK04044)→ Ribosomal_L7_archeal_euk (cd01657)rpl30p (PRK06049)→<br>rpl15p (PRK06419)Ribosomal_L27A (pfam00828)→ SecY (COG0201)PRK08568 (PRK08568)→ |
| env-38 | SL_4KL_010_BRINE_10108326 | + | 36 | 135 | **RNA**→ Methyltrn_RNA_3 (pfam02598)→ rpl3p (PRK04231)Ribosomal_L3 (pfam00297)→ |
| env-39 | SL_8KL_010_SED_10227597 | - | 345 | 246 | **RNA**→ Methyltrn_RNA_3 (pfam02598)→ |
| env-40 | SS_3KL_010_SOIL_10336605 | - | 246 | 147 | **RNA**→ Methyltrn_RNA_3 (pfam02598)→ |
| env-41 | APHM01002125.1 | + | 421 | 520 | **RNA**→ Methyltrn_RNA_3 (pfam02598)→ RplC (COG0087)rpl3p (PRK04231)→<br>RplD (COG0088)rpl4lp (PRK04042)→ RplW (COG0089)PRK14548 (PRK14548)→<br>rpl2p (PRK09612)Ribosomal_L2 (pfam00181)→ |
| env-42 | LTJ07As871_J07ABscf098859 | - | 2359772 | 2359673 | **RNA**→ Methyltrn_RNA_3 (pfam02598)→ RplC (COG0087)rpl3p (PRK04231)→<br>RplD (COG0088)rpl4lp (PRK04042)→ RplW (COG0089)PRK14548 (PRK14548)→<br>rpl2p (PRK09612)Ribosomal_L2 (pfam00181)→<br>PRK12317 (PRK12317)Translation_factor_III (cd01513)GTPBP_II (cd03694)→ |
| env-43 | JXWU01069486.1 | + | 361 | 460 | **RNA**→ Methyltrn_RNA_3 (pfam02598)→ |
| env-44 | SS_3KL_010_SOIL_10060740 | - | 118 | 19 | **RNA**→ Methyltrn_RNA_3 (pfam02598)→ |
| env-45 | SL_5KL_010_BRINE_10001245 | + | 528 | 630 | **RNA**→ Methyltrn_RNA_3 (pfam02598)→ rpl3p (PRK04231)Ribosomal_L3 (pfam00297)→<br>RplD (COG0088)rpl4lp (PRK04042)→ RplW (COG0089)PRK14548 (PRK14548)→<br>RplB (COG0090)rpl2p (PRK09612)→ RpsS (COG0185)rps19p (PRK04038)→<br>rpl22p (PRK04223)RplV (COG0091)→ RpsC (COG0092)rps3p (PRK04191)→<br>PRK14549 (PRK14549)Ribosomal_L29 (pfam00831)→ PRK03879 (PRK03879)UPF0086 (pfam01868)→<br>RpsQ (COG0186)rps17p (PRK08572)→ rpl14p (PRK08571)Ribosomal_L14 (pfam00238)→<br>rpl24p (PRK01191)KOW_RPL26 (cd06089)→<br>PRK04313 (PRK04313)KOW_RPS4 (cd06087)Ribosomal_S4e (pfam00900)→ RplE (COG0094)rpl5p (PRK04219)→<br>RpsN (COG0199)rps14P (PRK05766)→ rps8p (PRK04034)RpsH (COG0096)→<br>RplF (COG0097)rpl6p (PRK05518)→ PRK12766 (PRK12766)RecA (COG0468)Rpl32e (COG1717)→<br>rpl19e (PRK08570)Ribosomal_L19e (pfam01280)→ RplR (COG0256)rpl18p (PRK08569)→<br>RpsE (COG0098)rps5p (PRK04044)→ Ribosomal_L7_archeal_euk (cd01657)rpl30p (PRK06049)→ |

| | | | | | |
|---|---|---|---|---|---|
| env-46 | SL_6KL_011_BRINE_1001562 | + | 7391 | 7493 | **RNA**→ Methyltrn_RNA_3 (pfam02598)→ rpl3p (PRK04231)Ribosomal_L3 (pfam00297)→<br>RplD (COG0088)rpl4lp (PRK04042)→ RplW (COG0089)PRK14548 (PRK14548)→<br>RplB (COG0090)rpl2p (PRK09612)→ RpsS (COG0185)rps19p (PRK04038)→<br>rpl22p (PRK04223)RplV (COG0091)→ |
| env-47 | SL_4KL_010_BRINE_10002586 | + | 550 | 652 | **RNA**→ Methyltrn_RNA_3 (pfam02598)→ rpl3p (PRK04231)Ribosomal_L3 (pfam00297)→<br>RplD (COG0088)rpl4lp (PRK04042)→ RplW (COG0089)PRK14548 (PRK14548)→<br>RplB (COG0090)rpl2p (PRK09612)→ RpsS (COG0185)rps19p (PRK04038)→<br>rpl22p (PRK04223)RplV (COG0091)→ RpsC (COG0092)rps3p (PRK04191)→<br>PRK14549 (PRK14549)Ribosomal_L29 (pfam00831)→ PRK03879 (PRK03879)UPF0086 (pfam01868)→<br>RpsQ (COG0186)rps17p (PRK08572)→ rpl14p (PRK08571)Ribosomal_L14 (pfam00238)→<br>rpl24p (PRK01191)KOW_RPL26 (cd06089)→<br>PRK04313 (PRK04313)KOW_RPS4 (cd06087)Ribosomal_S4e (pfam00900)→ RplE (COG0094)rpl5p (PRK04219)→<br>RpsN (COG0199)rps14P (PRK05766)→ rps8p (PRK04034)RpsH (COG0096)→<br>RplF (COG0097)rpl6p (PRK05518)→ RecA (COG0468)Rpl32e (COG1717)HHH_5 (pfam14520)→ |
| env-48 | SL_7KL_010_BRINE_10000614 | + | 5277 | 5379 | **RNA**→ Methyltrn_RNA_3 (pfam02598)→ rpl3p (PRK04231)Ribosomal_L3 (pfam00297)→<br>RplD (COG0088)rpl4lp (PRK04042)→ RplW (COG0089)PRK14548 (PRK14548)→<br>RplB (COG0090)rpl2p (PRK09612)→ RpsS (COG0185)rps19p (PRK04038)→<br>rpl22p (PRK04223)RplV (COG0091)→ RpsC (COG0092)rps3p (PRK04191)→<br>PRK14549 (PRK14549)Ribosomal_L29 (pfam00831)→ PRK03879 (PRK03879)UPF0086 (pfam01868)→<br>RpsQ (COG0186)rps17p (PRK08572)→ rpl14p (PRK08571)Ribosomal_L14 (pfam00238)→<br>rpl24p (PRK01191)KOW_RPL26 (cd06089)→<br>PRK04313 (PRK04313)KOW_RPS4 (cd06087)Ribosomal_S4e (pfam00900)→ RplE (COG0094)rpl5p (PRK04219)→<br>RpsN (COG0199)rps14P (PRK05766)→ rps8p (PRK04034)RpsH (COG0096)→<br>RplF (COG0097)rpl6p (PRK05518)→<br>RecA (COG0468)Rpl32e (COG1717)rpl32e (PRK08562)HHH_5 (pfam14520)→<br>rpl19e (PRK08570)Ribosomal_L19e (pfam01280)→ RplR (COG0256)rpl18p (PRK08569)→<br>RpsE (COG0098)rps5p (PRK04044)→ |
| env-49 | GSLNARP_contig28693 | + | 496 | 598 | **RNA**→ Methyltrn_RNA_3 (pfam02598)→ RplC (COG0087)rpl3p (PRK04231)→<br>RplD (COG0088)rpl4lp (PRK04042)→ RplW (COG0089)PRK14548 (PRK14548)→<br>RplB (COG0090)rpl2p (PRK09612)→ RpsS (COG0185)rps19p (PRK04038)→<br>rpl22p (PRK04223)RplV (COG0091)→ RpsC (COG0092)rps3p (PRK04191)→<br>PRK14549 (PRK14549)Ribosomal_L29 (pfam00831)→ PRK03879 (PRK03879)UPF0086 (pfam01868)→<br>RpsQ (COG0186)rps17p (PRK08572)→ rpl14p (PRK08571)Ribosomal_L14 (pfam00238)→<br>rpl24p (PRK01191)KOW_RPL26 (cd06089)→ |
| env-50 | SL_5KL_010_BRINE_10001521 | + | 7795 | 7897 | **RNA**→ Methyltrn_RNA_3 (pfam02598)→ RplC (COG0087)rpl3p (PRK04231)→<br>RplD (COG0088)rpl4lp (PRK04042)→ RplW (COG0089)PRK14548 (PRK14548)→<br>RplB (COG0090)rpl2p (PRK09612)→ rps19p (PRK04038)→ |
| env-51 | SL_6KL_011_BRINE_1000203 | - | 3778 | 3676 | **RNA**→ Methyltrn_RNA_3 (pfam02598)→ RplC (COG0087)rpl3p (PRK04231)→<br>RplD (COG0088)rpl4lp (PRK04042)→ RplW (COG0089)PRK14548 (PRK14548)→<br>RplB (COG0090)rpl2p (PRK09612)→ RpsS (COG0185)rps19p (PRK04038)→ |
| env-52 | SL_5KL_010_SED_10001876 | - | 10224 | 10122 | **RNA**→ Methyltrn_RNA_3 (pfam02598)→ RplC (COG0087)rpl3p (PRK04231)→<br>RplD (COG0088)rpl4lp (PRK04042)→ RplW (COG0089)PRK14548 (PRK14548)→<br>RplB (COG0090)rpl2p (PRK09612)→ RpsS (COG0185)rps19p (PRK04038)→<br>rpl22p (PRK04223)RplV (COG0091)→ RpsC (COG0092)rps3p (PRK04191)→<br>PRK14549 (PRK14549)Ribosomal_L29 (pfam00831)→ PRK03879 (PRK03879)UPF0086 (pfam01868)→<br>RpsQ (COG0186)rps17p (PRK08572)→ rpl14p (PRK08571)Ribosomal_L14 (pfam00238)→<br>rpl24p (PRK01191)KOW_RPL26 (cd06089)→<br>PRK04313 (PRK04313)S4 (cd00165)KOW_RPS4 (cd06087)Ribosomal_S4e (pfam00900)→<br>RplE (COG0094)rpl5p (PRK04219)→ RpsN (COG0199)rps14P (PRK05766)→<br>rps8p (PRK04034)RpsH (COG0096)→ RplF (COG0097)rpl6p (PRK05518)→<br>UvrC (COG0322)rpl32e (PRK08562)Ribosomal_L32e (pfam01655)HHH_5 (pfam14520)→<br>rpl19e (PRK08570)Ribosomal_L19e (pfam01280)→ |
| env-53 | SL_4KL_010_BRINE_10000708 | + | 14067 | 14169 | **RNA**→ Methyltrn_RNA_3 (pfam02598)→ RplC (COG0087)rpl3p (PRK04231)→<br>RplD (COG0088)rpl4lp (PRK04042)→ RplW (COG0089)PRK14548 (PRK14548)→<br>RplB (COG0090)rpl2p (PRK09612)→ rps19p (PRK04038)→ |
| env-54 | P_1C_Liq_1_UnCty_1162688 | - | 122 | 20 | **RNA**→ MTH1 (COG2106)→ |

| | | | | | |
|---|---|---|---|---|---|
| env-55 | SL_6KL_011_BRINE_1072209 | - | 402 | 300 | **RNA**→ Methyltrn_RNA_3 (pfam02598)→ |
| env-56 | SL_5KL_010_BRINE_10188769 | - | 355 | 253 | **RNA**→ Methyltrn_RNA_3 (pfam02598)→ |
| env-57 | SL_4KL_010_BRINE_10162798 | + | 316 | 418 | **RNA**→ Methyltrn_RNA_3 (pfam02598)→ |
| env-58 | SL_6KL_011_BRINE_1009000 | + | 854 | 956 | **RNA**→ Methyltrn_RNA_3 (pfam02598)→ RplC (COG0087)rpl3p (PRK04231)→ RplD (COG0088)rpl4lp (PRK04042)→ RplW (COG0089)PRK14548 (PRK14548)→ RplB (COG0090)rpl2p (PRK09612)→ rps19p (PRK04038)→ |
| env-59 | ADL20m3uS_contig12843 | - | 170 | 68 | **RNA**→ Methyltrn_RNA_3 (pfam02598)→ |
| env-60 | 2140918027-12578 | + | 373 | 475 | **RNA**→ |
| env-61 | 2140918027-03821 | + | 1792 | 1894 | **RNA**→ Methyltrn_RNA_3 (pfam02598)→ |
| env-62 | ADL20m3uS_contig10848 | - | 468 | 366 | **RNA**→ Methyltrn_RNA_3 (pfam02598)→ |
| env-63 | ADL5mRS1u_contig00538 | + | 967 | 1069 | **RNA**→ Methyltrn_RNA_3 (pfam02598)→ rpl3p (PRK04231)Ribosomal_L3 (pfam00297)→ RplC (COG0087)rpl3p (PRK04231)→ RplD (COG0088)rpl4lp (PRK04042)→ RplW (COG0089)PRK14548 (PRK14548)→ RplB (COG0090)rpl2p (PRK09612)→ RpsS (COG0185)rps19p (PRK04038)→ |
| env-64 | GQKUMWE02JS9JS | - | 377 | 275 | **RNA**→ Methyltrn_RNA_3 (pfam02598)→ |
| env-65 | ADL20m3uS_contig10578 | - | 446 | 344 | **RNA**→ Methyltrn_RNA_3 (pfam02598)→ |
| env-66 | 2140918017-36 | - | 14923 | 14821 | **RNA**→ Methyltrn_RNA_3 (pfam02598)→ rpl3p (PRK04231)Ribosomal_L3 (pfam00297)→ RplC (COG0087)rpl3p (PRK04231)→ RplD (COG0088)rpl4lp (PRK04042)→ RplW (COG0089)PRK14548 (PRK14548)→ RplB (COG0090)rpl2p (PRK09612)→ RpsS (COG0185)rps19p (PRK04038)→ rpl22p (PRK04223)RplV (COG0091)→ RpsC (COG0092)rps3p (PRK04191)→ PRK14549 (PRK14549)Ribosomal_L29 (pfam00831)→ PRK03879 (PRK03879)UPF0086 (pfam01868)→ RpsQ (COG0186)rps17p (PRK08572)→ rpl14p (PRK08571)Ribosomal_L14 (pfam00238)→ rpl24p (PRK01191)KOW_RPL26 (cd06089)→ PRK04313 (PRK04313)KOW_RPS4 (cd06087)Ribosomal_S4e (pfam00900)→ RplE (COG0094)rpl5p (PRK04219)→ RpsN (COG0199)rps14P (PRK05766)→ rps8p (PRK04034)RpsH (COG0096)→ RplF (COG0097)rpl6p (PRK05518)→ UvrC (COG0322)Rpl32e (COG1717)HHH_5 (pfam14520)→ rpl19e (PRK08570)Ribosomal_L19e (pfam01280)→ |
| env-67 | GRCP39E01CI3EF | - | 306 | 204 | **RNA**→ Methyltrn_RNA_3 (pfam02598)→ |
| env-68 | GRCP39E01CZCI9 | + | 86 | 188 | **RNA**→ Methyltrn_RNA_3 (pfam02598)→ |
| env-69 | 2140918027-00954 | + | 88 | 190 | **RNA**→ Methyltrn_RNA_3 (pfam02598)→ rpl3p (PRK04231)Ribosomal_L3 (pfam00297)→ RplC (COG0087)rpl3p (PRK04231)→ RplD (COG0088)rpl4lp (PRK04042)→ RplW (COG0089)PRK14548 (PRK14548)→ RplB (COG0090)rpl2p (PRK09612)→ RpsS (COG0185)rps19p (PRK04038)→ rpl22p (PRK04223)RplV (COG0091)→ RpsC (COG0092)rps3p (PRK04191)→ PRK14549 (PRK14549)Ribosomal_L29 (pfam00831)→ PRK03879 (PRK03879)UPF0086 (pfam01868)→ |
| env-70 | GQKUMWE02F1GR6 | + | 61 | 163 | **RNA**→ Methyltrn_RNA_3 (pfam02598)→ |
| env-71 | GQKUMWE02JMQQS | + | 64 | 166 | **RNA**→ Methyltrn_RNA_3 (pfam02598)→ |
| env-72 | SL_8KL_010_SED_10008943 | + | 2731 | 2830 | **RNA**→ Methyltrn_RNA_3 (pfam02598)→ RplC (COG0087)rpl3p (PRK04231)→ RplD (COG0088)rpl4lp (PRK04042)→ RplW (COG0089)PRK14548 (PRK14548)→ RplB (COG0090)rpl2p (PRK09612)→ RpsS (COG0185)rps19p (PRK04038)→ rpl22p (PRK04223)RplV (COG0091)→ KH (smart00322)archeal_30S_S3_KH (cd02411)→ |
| env-73 | SL_4KL_010_BRINE_10005953 | + | 2732 | 2831 | **RNA**→ Methyltrn_RNA_3 (pfam02598)→ RplC (COG0087)rpl3p (PRK04231)→ RplD (COG0088)rpl4lp (PRK04042)→ RplW (COG0089)PRK14548 (PRK14548)→ RplB (COG0090)rpl2p (PRK09612)→ RpsS (COG0185)rps19p (PRK04038)→ rpl22p (PRK04223)RplV (COG0091)→ KH (smart00322)archeal_30S_S3_KH (cd02411)→ |
| env-74 | SL_5KL_010_BRINE_10000601 | + | 12793 | 12892 | **RNA**→ Methyltrn_RNA_3 (pfam02598)→ RplC (COG0087)rpl3p (PRK04231)→ rpl4lp (PRK04042)Ribosomal_L4 (pfam00573)→ rpl4lp (PRK04042)→ RplW (COG0089)PRK14548 (PRK14548)→ RplB (COG0090)rpl2p (PRK09612)→ RpsS (COG0185)rps19p (PRK04038)→ rpl22p (PRK04223)RplV (COG0091)→ KH (smart00322)archeal_30S_S3_KH (cd02411)→ |
| env-75 | SL_5KL_010_SED_10018610 | - | 1058 | 959 | **RNA**→ Methyltrn_RNA_3 (pfam02598)→ rpl3p (PRK04231)Ribosomal_L3 (pfam00297)→ |
| env-76 | JXWU01116399.1 | - | 162 | 60 | **RNA**→ Methyltrn_RNA_3 (pfam02598)→ |
| env-77 | GSLNARP_GFPJP1N02IUPJO | - | 257 | 155 | **RNA**→ Methyltrn_RNA_3 (pfam02598)→ |
| env-78 | GSLNARP_GFPJP1N02J3NU6 | - | 222 | 120 | **RNA**→ Methyltrn_RNA_3 (pfam02598)→ |

| | | | | | |
|---|---|---|---|---|---|
| Hsa-1-1 | NZ_BBMO01000001.1 | - | 854034 | 853935 | **RNA**→ Methyltrn_RNA_3 (pfam02598)→ RplC (COG0087)rpl3p (PRK04231)→ RplD (COG0088)rpl4lp (PRK04042)→ RplW (COG0089)PRK14548 (PRK14548)→ RplB (COG0090)rpl2p (PRK09612)→ RpsS (COG0185)rps19p (PRK04038)→ rpl22p (PRK04223)RplV (COG0091)→ RpsC (COG0092)rps3p (PRK04191)→ PRK14549 (PRK14549)Ribosomal_L29 (pfam00831)→ PRK03879 (PRK03879)UPF0086 (pfam01868)→ RpsQ (COG0186)rps17p (PRK08572)→ rpl14p (PRK08571)Ribosomal_L14 (pfam00238)→ rpl24p (PRK01191)KOW_RPL26 (cd06089)→ PRK04313 (PRK04313)KOW_RPS4 (cd06087)S4 (pfam01479)Ribosomal_S4e (pfam00900)→ RplE (COG0094)rpl5p (PRK04219)→ RpsN (COG0199)rps14P (PRK05766)→ rps8p (PRK04034)RpsH (COG0096)→ RplF (COG0097)rpl6p (PRK05518)→ PRK12766 (PRK12766)RecA (COG0468)Rpl32e (COG1717)→ rpl19e (PRK08570)Ribosomal_L19e (pfam01280)→ |
| Hgi-1-1 | NZ_AOLJ01000014.1 | + | 202046 | 202145 | **RNA**→ Methyltrn_RNA_3 (pfam02598)→ rpl3p (PRK04231)Ribosomal_L3 (pfam00297)→ RplD (COG0088)rpl4lp (PRK04042)→ RplW (COG0089)PRK14548 (PRK14548)→ RplB (COG0090)rpl2p (PRK09612)→ RpsS (COG0185)rps19p (PRK04038)→ rpl22p (PRK04223)RplV (COG0091)→ RpsC (COG0092)rps3p (PRK04191)→ PRK14549 (PRK14549)Ribosomal_L29 (pfam00831)→ PRK03879 (PRK03879)UPF0086 (pfam01868)→ RpsQ (COG0186)rps17p (PRK08572)→ rpl14p (PRK08571)Ribosomal_L14 (pfam00238)→ RplX (COG0198)rpl24p (PRK01191)→ PRK04313 (PRK04313)KOW_RPS4 (cd06087)Ribosomal_S4e (pfam00900)→ RplE (COG0094)rpl5p (PRK04219)→ RpsN (COG0199)rps14P (PRK05766)→ rps8p (PRK04034)RpsH (COG0096)→ RplF (COG0097)rpl6p (PRK05518)→ PRK12766 (PRK12766)RecA (COG0468)Rpl32e (COG1717)→ rpl19e (PRK08570)Ribosomal_L19e (pfam01280)→ |
| Hgi-2-1 | NZ_CP011947.1 | - | 2522923 | 2522824 | **RNA**→ Methyltrn_RNA_3 (pfam02598)→ rpl3p (PRK04231)Ribosomal_L3 (pfam00297)→ RplD (COG0088)rpl4lp (PRK04042)→ RplW (COG0089)PRK14548 (PRK14548)→ RplB (COG0090)rpl2p (PRK09612)→ RpsS (COG0185)rps19p (PRK04038)→ rpl22p (PRK04223)RplV (COG0091)→ RpsC (COG0092)rps3p (PRK04191)→ PRK14549 (PRK14549)Ribosomal_L29 (pfam00831)→ PRK03879 (PRK03879)UPF0086 (pfam01868)→ RpsQ (COG0186)rps17p (PRK08572)→ rpl14p (PRK08571)Ribosomal_L14 (pfam00238)→ RplX (COG0198)rpl24p (PRK01191)→ PRK04313 (PRK04313)KOW_RPS4 (cd06087)Ribosomal_S4e (pfam00900)→ RplE (COG0094)rpl5p (PRK04219)→ RpsN (COG0199)rps14P (PRK05766)→ rps8p (PRK04034)RpsH (COG0096)→ RplF (COG0097)rpl6p (PRK05518)→ PRK12766 (PRK12766)RecA (COG0468)Rpl32e (COG1717)→ rpl19e (PRK08570)Ribosomal_L19e (pfam01280)→ |
| Hsu-2-1 | NZ_AOLM01000020.1 | + | 180613 | 180712 | **RNA**→ Methyltrn_RNA_3 (pfam02598)→ rpl3p (PRK04231)Ribosomal_L3 (pfam00297)→ RplD (COG0088)rpl4lp (PRK04042)→ RplW (COG0089)PRK14548 (PRK14548)→ RplB (COG0090)rpl2p (PRK09612)→ RpsS (COG0185)rps19p (PRK04038)→ rpl22p (PRK04223)RplV (COG0091)→ RpsC (COG0092)rps3p (PRK04191)→ PRK14549 (PRK14549)Ribosomal_L29 (pfam00831)→ PRK03879 (PRK03879)UPF0086 (pfam01868)→ RpsQ (COG0186)rps17p (PRK08572)→ rpl14p (PRK08571)Ribosomal_L14 (pfam00238)→ RplX (COG0198)rpl24p (PRK01191)→ PRK04313 (PRK04313)KOW_RPS4 (cd06087)Ribosomal_S4e (pfam00900)→ RplE (COG0094)rpl5p (PRK04219)→ RpsN (COG0199)rps14P (PRK05766)→ rps8p (PRK04034)RpsH (COG0096)→ RplF (COG0097)rpl6p (PRK05518)→ PRK12766 (PRK12766)RecA (COG0468)Rpl32e (COG1717)→ rpl19e (PRK08570)Ribosomal_L19e (pfam01280)→ |
| Hpr-1-1 | NZ_AOLG01000047.1 | + | 231631 | 231730 | **RNA**→ Methyltrn_RNA_3 (pfam02598)→ rpl3p (PRK04231)Ribosomal_L3 (pfam00297)→ RplD (COG0088)rpl4lp (PRK04042)→ RplW (COG0089)PRK14548 (PRK14548)→ RplB (COG0090)rpl2p (PRK09612)→ RpsS (COG0185)rps19p (PRK04038)→ rpl22p (PRK04223)RplV (COG0091)→ RpsC (COG0092)rps3p (PRK04191)→ PRK14549 (PRK14549)Ribosomal_L29 (pfam00831)→ PRK03879 (PRK03879)UPF0086 (pfam01868)→ RpsQ (COG0186)rps17p (PRK08572)→ rpl14p (PRK08571)Ribosomal_L14 (pfam00238)→ RplX (COG0198)rpl24p (PRK01191)→ PRK04313 (PRK04313)KOW_RPS4 (cd06087)Ribosomal_S4e (pfam00900)→ RplE (COG0094)rpl5p (PRK04219)→ RpsN (COG0199)rps14P (PRK05766)→ rps8p (PRK04034)RpsH (COG0096)→ RplF (COG0097)rpl6p (PRK05518)→ PRK12766 (PRK12766)RecA (COG0468)Rpl32e (COG1717)→ rpl19e (PRK08570)Ribosomal_L19e (pfam01280)→ |

| | | | | | |
|---|---|---|---|---|---|
| Hsp-8-1 | NZ_JPES01000061.1 | + | 4380 | 4479 | **RNA**→ Methyltrn_RNA_3 (pfam02598)→ hypo→ rpl3p (PRK04231)Ribosomal_L3 (pfam00297)→ RplD (COG0088)rpl4lp (PRK04042)→ RplW (COG0089)PRK14548 (PRK14548)→ RplB (COG0090)rpl2p (PRK09612)→ RpsS (COG0185)rps19p (PRK04038)→ rpl22p (PRK04223)RplV (COG0091)→ hypo→ RpsC (COG0092)rps3p (PRK04191)→ PRK14549 (PRK14549)Ribosomal_L29 (pfam00831)→ PRK03879 (PRK03879)UPF0086 (pfam01868)→ RpsQ (COG0186)rps17p (PRK08572)→ hypo→ rpl14p (PRK08571)Ribosomal_L14 (pfam00238)→ RplX (COG0198)rpl24p (PRK01191)→ PRK04313 (PRK04313)KOW_RPS4 (cd06087)Ribosomal_S4e (pfam00900)→ RplE (COG0094)rpl5p (PRK04219)→ RpsN (COG0199)rps14P (PRK05766)→ rps8p (PRK04034)RpsH (COG0096)→ RplF (COG0097)rpl6p (PRK05518)→ PRK12766 (PRK12766)RecA (COG0468)Rpl32e (COG1717)→ rpl19e (PRK08570)Ribosomal_L19e (pfam01280)→ RplR (COG0256)rpl18p (PRK08569)→ hypo→ RpsE (COG0098)rps5p (PRK04044)→ Ribosomal_L7_archeal_euk (cd01657)rpl30p (PRK06049)→ rpl15p (PRK06419)Ribosomal_L27A (pfam00828)→ SecY (COG0201)PRK08568 (PRK08568)→ |
| Hde-1-1 | NZ_AOLP01000011.1 | + | 331865 | 331964 | **RNA**→ Methyltrn_RNA_3 (pfam02598)→ rpl3p (PRK04231)Ribosomal_L3 (pfam00297)→ RplD (COG0088)rpl4lp (PRK04042)→ RplW (COG0089)PRK14548 (PRK14548)→ RplB (COG0090)rpl2p (PRK09612)→ RpsS (COG0185)rps19p (PRK04038)→ rpl22p (PRK04223)RplV (COG0091)→ RpsC (COG0092)rps3p (PRK04191)→ PRK14549 (PRK14549)Ribosomal_L29 (pfam00831)→ PRK03879 (PRK03879)UPF0086 (pfam01868)→ RpsQ (COG0186)rps17p (PRK08572)→ rpl14p (PRK08571)Ribosomal_L14 (pfam00238)→ RplX (COG0198)rpl24p (PRK01191)→ PRK04313 (PRK04313)KOW_RPS4 (cd06087)Ribosomal_S4e (pfam00900)→ RplE (COG0094)rpl5p (PRK04219)→ RpsN (COG0199)rps14P (PRK05766)→ rps8p (PRK04034)RpsH (COG0096)→ RplF (COG0097)rpl6p (PRK05518)→ PRK12766 (PRK12766)RecA (COG0468)Rpl32e (COG1717)→ rpl19e (PRK08570)Ribosomal_L19e (pfam01280)→ |
| Hlu-1-1 | NZ_AOLH01000006.1 | + | 206888 | 206987 | **RNA**→ Methyltrn_RNA_3 (pfam02598)→ rpl3p (PRK04231)Ribosomal_L3 (pfam00297)→ RplD (COG0088)rpl4lp (PRK04042)→ RplW (COG0089)PRK14548 (PRK14548)→ RplB (COG0090)rpl2p (PRK09612)→ RpsS (COG0185)rps19p (PRK04038)→ rpl22p (PRK04223)RplV (COG0091)→ RpsC (COG0092)rps3p (PRK04191)→ PRK14549 (PRK14549)Ribosomal_L29 (pfam00831)→ PRK03879 (PRK03879)UPF0086 (pfam01868)→ RpsQ (COG0186)rps17p (PRK08572)→ rpl14p (PRK08571)Ribosomal_L14 (pfam00238)→ RplX (COG0198)rpl24p (PRK01191)→ PRK04313 (PRK04313)KOW_RPS4 (cd06087)Ribosomal_S4e (pfam00900)→ RplE (COG0094)rpl5p (PRK04219)→ RpsN (COG0199)rps14P (PRK05766)→ rps8p (PRK04034)RpsH (COG0096)→ RplF (COG0097)rpl6p (PRK05518)→ PRK12766 (PRK12766)RecA (COG0468)Rpl32e (COG1717)→ rpl19e (PRK08570)Ribosomal_L19e (pfam01280)→ |
| Hal-2-1 | NZ_LK053000.1 | - | 908912 | 908813 | **RNA**→ Methyltrn_RNA_3 (pfam02598)→ rpl3p (PRK04231)Ribosomal_L3 (pfam00297)→ RplD (COG0088)rpl4lp (PRK04042)→ RplW (COG0089)PRK14548 (PRK14548)→ RplB (COG0090)rpl2p (PRK09612)→ RpsS (COG0185)rps19p (PRK04038)→ rpl22p (PRK04223)RplV (COG0091)→ RpsC (COG0092)rps3p (PRK04191)→ PRK14549 (PRK14549)Ribosomal_L29 (pfam00831)→ PRK03879 (PRK03879)UPF0086 (pfam01868)→ RpsQ (COG0186)rps17p (PRK08572)→ rpl14p (PRK08571)Ribosomal_L14 (pfam00238)→ RplX (COG0198)rpl24p (PRK01191)→ PRK04313 (PRK04313)KOW_RPS4 (cd06087)Ribosomal_S4e (pfam00900)→ RplE (COG0094)rpl5p (PRK04219)→ RpsN (COG0199)rps14P (PRK05766)→ rps8p (PRK04034)RpsH (COG0096)→ RplF (COG0097)rpl6p (PRK05518)→ PRK12766 (PRK12766)RecA (COG0468)Rpl32e (COG1717)→ rpl19e (PRK08570)Ribosomal_L19e (pfam01280)→ |

| | | | | | |
|---|---|---|---|---|---|
| Hsp-6-1 | NZ_AOLD01000045.1 | - | 26639 | 26540 | **RNA**→ Methyltrn_RNA_3 (pfam02598)→ rpl3p (PRK04231)Ribosomal_L3 (pfam00297)→<br>RplD (COG0088)rpl4lp (PRK04042)→ RplW (COG0089)PRK14548 (PRK14548)→<br>RplB (COG0090)rpl2p (PRK09612)→ RpsS (COG0185)rps19p (PRK04038)→<br>rpl22p (PRK04223)RplV (COG0091)→ RpsC (COG0092)rps3p (PRK04191)→<br>PRK14549 (PRK14549)Ribosomal_L29 (pfam00831)→ PRK03879 (PRK03879)UPF0086 (pfam01868)→<br>RpsQ (COG0186)rps17p (PRK08572)→ rpl14p (PRK08571)Ribosomal_L14 (pfam00238)→<br>RplX (COG0198)rpl24p (PRK01191)→<br>PRK04313 (PRK04313)KOW_RPS4 (cd06087)Ribosomal_S4e (pfam00900)→ RplE (COG0094)rpl5p (PRK04219)→<br>RpsN (COG0199)rps14P (PRK05766)→ rps8p (PRK04034)RpsH (COG0096)→<br>RplF (COG0097)rpl6p (PRK05518)→ PRK12766 (PRK12766)RecA (COG0468)Rpl32e (COG1717)→<br>rpl19e (PRK08570)Ribosomal_L19e (pfam01280)→ |
| Hsp-4-1 | NZ_AOLF01000024.1 | - | 14963 | 14864 | **RNA**→ Methyltrn_RNA_3 (pfam02598)→ rpl3p (PRK04231)Ribosomal_L3 (pfam00297)→<br>RplD (COG0088)rpl4lp (PRK04042)→ RplW (COG0089)PRK14548 (PRK14548)→<br>RplB (COG0090)rpl2p (PRK09612)→ RpsS (COG0185)rps19p (PRK04038)→<br>rpl22p (PRK04223)RplV (COG0091)→ RpsC (COG0092)rps3p (PRK04191)→<br>PRK14549 (PRK14549)Ribosomal_L29 (pfam00831)→ PRK03879 (PRK03879)UPF0086 (pfam01868)→<br>RpsQ (COG0186)rps17p (PRK08572)→ rpl14p (PRK08571)Ribosomal_L14 (pfam00238)→<br>RplX (COG0198)rpl24p (PRK01191)→<br>PRK04313 (PRK04313)KOW_RPS4 (cd06087)Ribosomal_S4e (pfam00900)→ RplE (COG0094)rpl5p (PRK04219)→<br>RpsN (COG0199)rps14P (PRK05766)→ rps8p (PRK04034)RpsH (COG0096)→<br>RplF (COG0097)rpl6p (PRK05518)→ PRK12766 (PRK12766)RecA (COG0468)Rpl32e (COG1717)→<br>rpl19e (PRK08570)Ribosomal_L19e (pfam01280)→ |
| Hsp-5-1 | NZ_AOLE01000011.1 | - | 83691 | 83592 | **RNA**→ Methyltrn_RNA_3 (pfam02598)→ rpl3p (PRK04231)Ribosomal_L3 (pfam00297)→<br>RplD (COG0088)rpl4lp (PRK04042)→ RplW (COG0089)PRK14548 (PRK14548)→<br>RplB (COG0090)rpl2p (PRK09612)→ RpsS (COG0185)rps19p (PRK04038)→<br>rpl22p (PRK04223)RplV (COG0091)→ RpsC (COG0092)rps3p (PRK04191)→<br>PRK14549 (PRK14549)Ribosomal_L29 (pfam00831)→ PRK03879 (PRK03879)UPF0086 (pfam01868)→<br>RpsQ (COG0186)rps17p (PRK08572)→ rpl14p (PRK08571)Ribosomal_L14 (pfam00238)→<br>RplX (COG0198)rpl24p (PRK01191)→<br>PRK04313 (PRK04313)KOW_RPS4 (cd06087)Ribosomal_S4e (pfam00900)→ RplE (COG0094)rpl5p (PRK04219)→<br>RpsN (COG0199)rps14P (PRK05766)→ rps8p (PRK04034)RpsH (COG0096)→<br>RplF (COG0097)rpl6p (PRK05518)→ PRK12766 (PRK12766)RecA (COG0468)Rpl32e (COG1717)→<br>rpl19e (PRK08570)Ribosomal_L19e (pfam01280)→ |
| Hme-1-1 | NZ_AOLO01000012.1 | - | 67999 | 67900 | **RNA**→ Methyltrn_RNA_3 (pfam02598)→ rpl3p (PRK04231)Ribosomal_L3 (pfam00297)→<br>RplD (COG0088)rpl4lp (PRK04042)→ RplW (COG0089)PRK14548 (PRK14548)→<br>RplB (COG0090)rpl2p (PRK09612)→ RpsS (COG0185)rps19p (PRK04038)→<br>rpl22p (PRK04223)RplV (COG0091)→ RpsC (COG0092)rps3p (PRK04191)→<br>PRK14549 (PRK14549)Ribosomal_L29 (pfam00831)→ PRK03879 (PRK03879)UPF0086 (pfam01868)→<br>RpsQ (COG0186)rps17p (PRK08572)→ rpl14p (PRK08571)Ribosomal_L14 (pfam00238)→<br>RplX (COG0198)rpl24p (PRK01191)→<br>PRK04313 (PRK04313)KOW_RPS4 (cd06087)Ribosomal_S4e (pfam00900)→ RplE (COG0094)rpl5p (PRK04219)→<br>RpsN (COG0199)rps14P (PRK05766)→ rps8p (PRK04034)RpsH (COG0096)→<br>RplF (COG0097)rpl6p (PRK05518)→ PRK12766 (PRK12766)RecA (COG0468)Rpl32e (COG1717)→<br>rpl19e (PRK08570)Ribosomal_L19e (pfam01280)→ |
| Hme-1-2 | NC_017941.2 | + | 2068879 | 2068978 | **RNA**→ Methyltrn_RNA_3 (pfam02598)→ rpl3p (PRK04231)Ribosomal_L3 (pfam00297)→<br>RplD (COG0088)rpl4lp (PRK04042)→ RplW (COG0089)PRK14548 (PRK14548)→<br>RplB (COG0090)rpl2p (PRK09612)→ RpsS (COG0185)rps19p (PRK04038)→<br>rpl22p (PRK04223)RplV (COG0091)→ RpsC (COG0092)rps3p (PRK04191)→<br>PRK14549 (PRK14549)Ribosomal_L29 (pfam00831)→ PRK03879 (PRK03879)UPF0086 (pfam01868)→<br>RpsQ (COG0186)rps17p (PRK08572)→ rpl14p (PRK08571)Ribosomal_L14 (pfam00238)→<br>RplX (COG0198)rpl24p (PRK01191)→<br>PRK04313 (PRK04313)KOW_RPS4 (cd06087)Ribosomal_S4e (pfam00900)→ RplE (COG0094)rpl5p (PRK04219)→<br>RpsN (COG0199)rps14P (PRK05766)→ rps8p (PRK04034)RpsH (COG0096)→<br>RplF (COG0097)rpl6p (PRK05518)→ PRK12766 (PRK12766)RecA (COG0468)Rpl32e (COG1717)→<br>rpl19e (PRK08570)Ribosomal_L19e (pfam01280)→ |

| | | | | | |
|---|---|---|---|---|---|
| Hla-2-1 | NZ_AOLI01000019.1 | - | 425010 | 424911 | **RNA**→ Methyltrn_RNA_3 (pfam02598)→ rpl3p (PRK04231)Ribosomal_L3 (pfam00297)→<br>RplD (COG0088)rpl4lp (PRK04042)→ RplW (COG0089)PRK14548 (PRK14548)→<br>RplB (COG0090)rpl2p (PRK09612)→ RpsS (COG0185)rps19p (PRK04038)→<br>rpl22p (PRK04223)RplV (COG0091)→ RpsC (COG0092)rps3p (PRK04191)→<br>PRK14549 (PRK14549)Ribosomal_L29 (pfam00831)→ PRK03879 (PRK03879)UPF0086 (pfam01868)→<br>RpsQ (COG0186)rps17p (PRK08572)→ rpl14p (PRK08571)Ribosomal_L14 (pfam00238)→<br>RplX (COG0198)rpl24p (PRK01191)→<br>PRK04313 (PRK04313)KOW_RPS4 (cd06087)Ribosomal_S4e (pfam00900)→ RplE (COG0094)rpl5p (PRK04219)→<br>RpsN (COG0199)rps14P (PRK05766)→ rps8p (PRK04034)RpsH (COG0096)→<br>RplF (COG0097)rpl6p (PRK05518)→ PRK12766 (PRK12766)RecA (COG0468)Rpl32e (COG1717)→<br>rpl19e (PRK08570)Ribosomal_L19e (pfam01280)→ |
| Hel-1-1 | NZ_AOLK01000022.1 | + | 66470 | 66569 | **RNA**→ Methyltrn_RNA_3 (pfam02598)→ RplC (COG0087)rpl3p (PRK04231)→<br>RplD (COG0088)rpl4lp (PRK04042)→ RplW (COG0089)PRK14548 (PRK14548)→<br>RplB (COG0090)rpl2p (PRK09612)→ RpsS (COG0185)rps19p (PRK04038)→<br>rpl22p (PRK04223)RplV (COG0091)→ RpsC (COG0092)rps3p (PRK04191)→<br>PRK14549 (PRK14549)Ribosomal_L29 (pfam00831)→ PRK03879 (PRK03879)UPF0086 (pfam01868)→<br>RpsQ (COG0186)rps17p (PRK08572)→ rpl14p (PRK08571)Ribosomal_L14 (pfam00238)→<br>RplX (COG0198)rpl24p (PRK01191)→<br>PRK04313 (PRK04313)KOW_RPS4 (cd06087)Ribosomal_S4e (pfam00900)→ RplE (COG0094)rpl5p (PRK04219)→<br>RpsN (COG0199)rps14P (PRK05766)→ rps8p (PRK04034)RpsH (COG0096)→<br>RplF (COG0097)rpl6p (PRK05518)→ PRK12766 (PRK12766)RecA (COG0468)Rpl32e (COG1717)→<br>rpl19e (PRK08570)Ribosomal_L19e (pfam01280)→ |
| Hmu-1-1 | NZ_AOLN01000013.1 | - | 37008 | 36909 | **RNA**→ Methyltrn_RNA_3 (pfam02598)→ rpl3p (PRK04231)Ribosomal_L3 (pfam00297)→<br>RplD (COG0088)rpl4lp (PRK04042)→ RplW (COG0089)PRK14548 (PRK14548)→<br>RplB (COG0090)rpl2p (PRK09612)→ RpsS (COG0185)rps19p (PRK04038)→<br>rpl22p (PRK04223)RplV (COG0091)→ RpsC (COG0092)rps3p (PRK04191)→<br>PRK14549 (PRK14549)Ribosomal_L29 (pfam00831)→ PRK03879 (PRK03879)UPF0086 (pfam01868)→<br>RpsQ (COG0186)rps17p (PRK08572)→ rpl14p (PRK08571)Ribosomal_L14 (pfam00238)→<br>RplX (COG0198)rpl24p (PRK01191)→<br>PRK04313 (PRK04313)KOW_RPS4 (cd06087)Ribosomal_S4e (pfam00900)→ RplE (COG0094)rpl5p (PRK04219)→<br>RpsN (COG0199)rps14P (PRK05766)→ rps8p (PRK04034)RpsH (COG0096)→<br>RplF (COG0097)rpl6p (PRK05518)→ PRK12766 (PRK12766)RecA (COG0468)Rpl32e (COG1717)→<br>rpl19e (PRK08570)Ribosomal_L19e (pfam01280)→ |
| env-79 | SL_4KL_010_BRINE_10099275 | - | 413 | 314 | **RNA**→ Methyltrn_RNA_3 (pfam02598)→ |
| env-80 | SL_4KL_010_BRINE_10236927 | - | 339 | 240 | **RNA**→ Methyltrn_RNA_3 (pfam02598)→ |
| env-81 | SL_4KL_010_BRINE_10017268 | + | 119 | 218 | **RNA**→ Methyltrn_RNA_3 (pfam02598)→ RplC (COG0087)rpl3p (PRK04231)→<br>RplD (COG0088)rpl4lp (PRK04042)→ RplW (COG0089)PRK14548 (PRK14548)→<br>RplB (COG0090)rpl2p (PRK09612)→ RpsS (COG0185)rps19p (PRK04038)→<br>rpl22p (PRK04223)RplV (COG0091)→ |
| env-82 | SL_4KL_010_BRINE_10000094 | - | 9413 | 9314 | **RNA**→ Methyltrn_RNA_3 (pfam02598)→ RplC (COG0087)rpl3p (PRK04231)→<br>RplD (COG0088)rpl4lp (PRK04042)→ RplW (COG0089)PRK14548 (PRK14548)→<br>RplB (COG0090)rpl2p (PRK09612)→ RpsS (COG0185)rps19p (PRK04038)→<br>rpl22p (PRK04223)RplV (COG0091)→ RpsC (COG0092)rps3p (PRK04191)→<br>PRK14549 (PRK14549)Ribosomal_L29_HIP (cd00427)→ PRK03879 (PRK03879)UPF0086 (pfam01868)→<br>RpsQ (COG0186)rps17p (PRK08572)→ rpl14p (PRK08571)Ribosomal_L14 (pfam00238)→<br>RplX (COG0198)rpl24p (PRK01191)→<br>PRK04313 (PRK04313)S4 (cd00165)KOW_RPS4 (cd06087)Ribosomal_S4e (pfam00900)→<br>RplE (COG0094)rpl5p (PRK04219)→ RpsN (COG0199)rps14P (PRK05766)→<br>rps8p (PRK04034)RpsH (COG0096)→ RplF (COG0097)rpl6p (PRK05518)→<br>COG2251 (COG2251)HHH_5 (pfam14520)→ |
| env-83 | SL_4KL_010_BRINE_10065590 | + | 860 | 959 | **RNA**→ Methyltrn_RNA_3 (pfam02598)→ rpl3p (PRK04231)Ribosomal_L3 (pfam00297)→ |

| | | | | | |
|---|---|---|---|---|---|
| Hla-4-1 | NZ_CP007055.1 | + | 1979362 | 1979461 | **RNA**→ Methyltrn_RNA_3 (pfam02598)→ rpl3p (PRK04231)Ribosomal_L3 (pfam00297)→<br>RplD (COG0088)rpl4lp (PRK04042)→ RplW (COG0089)PRK14548 (PRK14548)→<br>RplB (COG0090)rpl2p (PRK09612)→ RpsS (COG0185)rps19p (PRK04038)→<br>rpl22p (PRK04223)RplV (COG0091)→ RpsC (COG0092)rps3p (PRK04191)→<br>PRK14549 (PRK14549)Ribosomal_L29 (pfam00831)→ PRK03879 (PRK03879)UPF0086 (pfam01868)→<br>RpsQ (COG0186)rps17p (PRK08572)→ rpl14p (PRK08571)Ribosomal_L14 (pfam00238)→<br>rpl24p (PRK01191)KOW_RPL26 (cd06089)→<br>PRK04313 (PRK04313)KOW_RPS4 (cd06087)Ribosomal_S4e (pfam00900)→ RplE (COG0094)rpl5p (PRK04219)→<br>RpsN (COG0199)rps14P (PRK05766)→ rps8p (PRK04034)RpsH (COG0096)→<br>RplF (COG0097)rpl6p (PRK05518)→<br>RecA (COG0468)Rpl32e (COG1717)rpl32e (PRK08562)HHH_5 (pfam14520)→ |
| Ssp-1-1 | NC_021313.1 | + | 2429676 | 2429775 | **RNA**→ Methyltrn_RNA_3 (pfam02598)→ RplC (COG0087)rpl3p (PRK04231)→<br>RplD (COG0088)rpl4lp (PRK04042)→ RplW (COG0089)PRK14548 (PRK14548)→<br>RplB (COG0090)rpl2p (PRK09612)→ RpsS (COG0185)rps19p (PRK04038)→<br>rpl22p (PRK04223)RplV (COG0091)→ RpsC (COG0092)rps3p (PRK04191)→<br>PRK14549 (PRK14549)Ribosomal_L29 (pfam00831)→ PRK03879 (PRK03879)UPF0086 (pfam01868)→ hypo→<br>RpsQ (COG0186)rps17p (PRK08572)→ rpl14p (PRK08571)Ribosomal_L14 (pfam00238)→<br>RplX (COG0198)rpl24p (PRK01191)→<br>PRK04313 (PRK04313)KOW_RPS4 (cd06087)S4 (pfam01479)Ribosomal_S4e (pfam00900)→<br>RplE (COG0094)rpl5p (PRK04219)→ RpsN (COG0199)rps14P (PRK05766)→<br>rps8p (PRK04034)RpsH (COG0096)→ RplF (COG0097)rpl6p (PRK05518)→<br>UvrC (COG0322)Rpl32e (COG1717)rpl32e (PRK08562)HHH_5 (pfam14520)→<br>Rpl32e (COG1717)rpl32e (PRK08562)HHH_5 (pfam14520)→ |
| env-84 | APHM01017236.1 | - | 204 | 105 | **RNA**→ MTH1 (COG2106)→ |
| env-85 | LTJ07As871_J07ABscf096273 | - | 204 | 105 | **RNA**→ MTH1 (COG2106)→ |
| env-86 | SL_4KL_010_BRINE_10002302 | + | 5048 | 5147 | **RNA**→ Methyltrn_RNA_3 (pfam02598)→ rpl3p (PRK04231)Ribosomal_L3 (pfam00297)→<br>RplD (COG0088)rpl4lp (PRK04042)→ RplW (COG0089)PRK14548 (PRK14548)→<br>RplB (COG0090)rpl2p (PRK09612)→ RpsS (COG0185)rps19p (PRK04038)→<br>rpl22p (PRK04223)RplV (COG0091)→ RpsC (COG0092)rps3p (PRK04191)→<br>RpmC (COG0255)PRK14549 (PRK14549)→ PRK03879 (PRK03879)UPF0086 (pfam01868)→ |
| Hsp-11-1 | NZ_JMIP02000014.1 | - | 158997 | 158898 | **RNA**→ Methyltrn_RNA_3 (pfam02598)→ RplC (COG0087)rpl3p (PRK04231)→ hypo→<br>RplD (COG0088)rpl4lp (PRK04042)→ RplW (COG0089)PRK14548 (PRK14548)→ hypo→ hypo→<br>RpsS (COG0185)rps19p (PRK04038)→ rpl22p (PRK04223)RplV (COG0091)→<br>RpsC (COG0092)rps3p (PRK04191)→ PRK14549 (PRK14549)Ribosomal_L29 (pfam00831)→<br>PRK03879 (PRK03879)UPF0086 (pfam01868)→ RpsQ (COG0186)rps17p (PRK08572)→<br>rpl14p (PRK08571)Ribosomal_L14 (pfam00238)→ hypo→ rpl24p (PRK01191)KOW_RPL26 (cd06089)→<br>PRK04313 (PRK04313)KOW_RPS4 (cd06087)Ribosomal_S4e (pfam00900)→ RplE (COG0094)rpl5p (PRK04219)→<br>RpsN (COG0199)rps14P (PRK05766)→ rps8p (PRK04034)RpsH (COG0096)→<br>RplF (COG0097)rpl6p (PRK05518)→ PRK12766 (PRK12766)UvrC (COG0322)Rpl32e (COG1717)→ |
| env-87 | GSLNARP_GFPJP1N02HFKWJ | - | 439 | 343 | **RNA**→ PRK12442 (PRK12442)Methyltrn_RNA_3 (pfam02598)→ |
| env-88 | SL_4KL_010_BRINE_10091194 | + | 755 | 857 | **RNA**→ Methyltrn_RNA_3 (pfam02598)→ |
| env-89 | SL_4KL_010_BRINE_10095573 | - | 111 | 9 | **RNA**→ MTH1 (COG2106)→ |
| Hka-1-1 | NZ_KB905379.1 | + | 273922 | 274024 | **RNA**→ Methyltrn_RNA_3 (pfam02598)→ rpl3p (PRK04231)Ribosomal_L3 (pfam00297)→<br>RplD (COG0088)rpl4lp (PRK04042)→ RplW (COG0089)PRK14548 (PRK14548)→<br>RplB (COG0090)rpl2p (PRK09612)→ RpsS (COG0185)rps19p (PRK04038)→<br>rpl22p (PRK04223)RplV (COG0091)→ RpsC (COG0092)rps3p (PRK04191)→<br>PRK14549 (PRK14549)Ribosomal_L29 (pfam00831)→ |
| Hmu-2-1 | NC_013202.1 | + | 1747213 | 1747315 | **RNA**→ Methyltrn_RNA_3 (pfam02598)→ rpl3p (PRK04231)Ribosomal_L3 (pfam00297)→<br>RplD (COG0088)rpl4lp (PRK04042)→ RplW (COG0089)PRK14548 (PRK14548)→<br>RplB (COG0090)rpl2p (PRK09612)→ RpsS (COG0185)rps19p (PRK04038)→<br>rpl22p (PRK04223)RplV (COG0091)→ RpsC (COG0092)rps3p (PRK04191)→<br>PRK14549 (PRK14549)Ribosomal_L29 (pfam00831)→ |

| | | | | | |
|---|---|---|---|---|---|
| Hca-3-1 | NZ_AOIU01000034.1 | - | 38153 | 38054 | **RNA**→ Methyltrn_RNA_3 (pfam02598)→ rpl3p (PRK04231)Ribosomal_L3 (pfam00297)→<br>RplD (COG0088)rpl4lp (PRK04042)→ PRK14548 (PRK14548)Ribosomal_L23 (pfam00276)→<br>RplB (COG0090)rpl2p (PRK09612)→ RpsS (COG0185)rps19p (PRK04038)→<br>rpl22p (PRK04223)RplV (COG0091)→ RpsC (COG0092)rps3p (PRK04191)→<br>PRK14549 (PRK14549)Ribosomal_L29 (pfam00831)→ PRK03879 (PRK03879)UPF0086 (pfam01868)→<br>RpsQ (COG0186)rps17p (PRK08572)→ rpl14p (PRK08571)Ribosomal_L14 (pfam00238)→<br>RplX (COG0198)rpl24p (PRK01191)→<br>PRK04313 (PRK04313)KOW_RPS4 (cd06087)S4 (pfam01479)Ribosomal_S4e (pfam00900)→<br>RplE (COG0094)rpl5p (PRK04219)→ RpsN (COG0199)rps14P (PRK05766)→<br>rps8p (PRK04034)RpsH (COG0096)→ RplF (COG0097)rpl6p (PRK05518)→<br>Rpl32e (COG1717)rpl32e (PRK08562)HHH_5 (pfam14520)→ rpl19e (PRK08570)Ribosomal_L19e (pfam01280)→ |
| Hsa-5-1 | NZ_AOME01000079.1 | - | 8539 | 8440 | **RNA**→ Methyltrn_RNA_3 (pfam02598)→ RplC (COG0087)rpl3p (PRK04231)→<br>RplD (COG0088)rpl4lp (PRK04042)→ RplW (COG0089)PRK14548 (PRK14548)→<br>RplB (COG0090)rpl2p (PRK09612)→ RpsS (COG0185)rps19p (PRK04038)→<br>rpl22p (PRK04223)RplV (COG0091)→ RpsC (COG0092)rps3p (PRK04191)HHH_5 (pfam14520)→<br>PRK14549 (PRK14549)Ribosomal_L29 (pfam00831)→ PRK03879 (PRK03879)UPF0086 (pfam01868)→<br>RpsQ (COG0186)rps17p (PRK08572)→ rpl14p (PRK08571)Ribosomal_L14 (pfam00238)→<br>RplX (COG0198)rpl24p (PRK01191)→<br>PRK04313 (PRK04313)S4 (cd00165)KOW_RPS4 (cd06087)Ribosomal_S4e (pfam00900)→<br>RplE (COG0094)rpl5p (PRK04219)→ hypo→ hypo→ RpsN (COG0199)rps14P (PRK05766)→ |
| env-90 | GQKUMWE01CQSAM | + | 397 | 496 | **RNA**→ MTH1 (COG2106)→ |
| Hag-1-1 | NZ_BAFM01000007.1 | + | 1659 | 1758 | **RNA**→ Methyltrn_RNA_3 (pfam02598)→ RplC (COG0087)rpl3p (PRK04231)→<br>RplD (COG0088)rpl4lp (PRK04042)→ RplW (COG0089)PRK14548 (PRK14548)→<br>RplB (COG0090)rpl2p (PRK09612)→ RpsS (COG0185)rps19p (PRK04038)→<br>rpl22p (PRK04223)RplV (COG0091)→<br>RpsC (COG0092)UvrC (COG0322)rps3p (PRK04191)HHH_5 (pfam14520)→<br>PRK14549 (PRK14549)Ribosomal_L29 (pfam00831)→ PRK03879 (PRK03879)UPF0086 (pfam01868)→<br>RpsQ (COG0186)rps17p (PRK08572)→ rpl14p (PRK08571)Ribosomal_L14 (pfam00238)→<br>RplX (COG0198)rpl24p (PRK01191)→<br>PRK04313 (PRK04313)S4 (cd00165)KOW_RPS4 (cd06087)Ribosomal_S4e (pfam00900)→<br>RplE (COG0094)rpl5p (PRK04219)→ RpsN (COG0199)rps14P (PRK05766)→<br>rps8p (PRK04034)RpsH (COG0096)→ RplF (COG0097)rpl6p (PRK05518)→<br>RecA (COG0468)Rpl32e (COG1717)rpl32e (PRK08562)HHH_5 (pfam14520)→<br>rpl19e (PRK08570)Ribosomal_L19e (pfam01280)→ RplR (COG0256)rpl18p (PRK08569)→<br>RpsE (COG0098)rps5p (PRK04044)→ Ribosomal_L7_archeal_euk (cd01657)rpl30p (PRK06049)→<br>rpl15p (PRK06419)Ribosomal_L27A (pfam00828)→ SecY (COG0201)PRK08568 (PRK08568)→ |
| Hsa-4-1 | NZ_AOMD01000025.1 | + | 140368 | 140467 | **RNA**→ Methyltrn_RNA_3 (pfam02598)→ RplC (COG0087)rpl3p (PRK04231)→<br>RplD (COG0088)rpl4lp (PRK04042)→ RplW (COG0089)PRK14548 (PRK14548)→<br>RplB (COG0090)rpl2p (PRK09612)→ RpsS (COG0185)rps19p (PRK04038)→<br>rpl22p (PRK04223)RplV (COG0091)→ RpsC (COG0092)rps3p (PRK04191)HHH_5 (pfam14520)→<br>PRK14549 (PRK14549)Ribosomal_L29 (pfam00831)→ PRK03879 (PRK03879)UPF0086 (pfam01868)→<br>RpsQ (COG0186)rps17p (PRK08572)→ rpl14p (PRK08571)Ribosomal_L14 (pfam00238)→<br>rpl24p (PRK01191)KOW_RPL26 (cd06089)→<br>PRK04313 (PRK04313)S4 (cd00165)KOW_RPS4 (cd06087)Ribosomal_S4e (pfam00900)→<br>RplE (COG0094)rpl5p (PRK04219)→ RpsN (COG0199)rps14P (PRK05766)→<br>rps8p (PRK04034)RpsH (COG0096)→ RplF (COG0097)rpl6p (PRK05518)→<br>RecA (COG0468)Rpl32e (COG1717)COG2251 (COG2251)rpl32e (PRK08562)HHH_5 (pfam14520)→ |
| env-91 | GSLNARP_contig29074 | - | 461 | 359 | **RNA**→ Methyltrn_RNA_3 (pfam02598)→ |
| env-92 | P_1C_Liq_1_UnCty_1017446 | + | 565 | 667 | **RNA**→ MTH1 (COG2106)→ |
| env-93 | P_1C_Liq_3_UnCty_1022117 | - | 224 | 122 | **RNA**→ Methyltrn_RNA_3 (pfam02598)→ |
| env-94 | P_1C_Sed_2_UnCty_1128195 | - | 252 | 150 | **RNA**→ Methyltrn_RNA_3 (pfam02598)→ |
| env-95 | P_1C_Liq_2_UnCty_1026371 | - | 311 | 209 | **RNA**→ Methyltrn_RNA_3 (pfam02598)→ |
| env-96 | 77515515 | + | 477 | 579 | **RNA**→ MTH1 (COG2106)→ |
| env-97 | FLAS_READ_83860922 | + | 477 | 579 | **RNA**→ MTH1 (COG2106)→ |
| env-98 | P_A23_Liq_3_Fmt_1016381 | - | 364 | 262 | **RNA**→ MTH1 (COG2106)→ |

| | | | | | |
|---|---|---|---|---|---|
| env-99 | APHM01021628.1 | + | 4390 | 4492 | **RNA**→ Methyltrn_RNA_3 (pfam02598)→ RplC (COG0087)rpl3p (PRK04231)→ RplD (COG0088)rpl4lp (PRK04042)→ RplW (COG0089)PRK14548 (PRK14548)→ RplB (COG0090)rpl2p (PRK09612)→ RpsS (COG0185)rps19p (PRK04038)→ rpl22p (PRK04223)RplV (COG0091)→ RpsC (COG0092)rps3p (PRK04191)→ |
| env-100 | LTJ07As871_J07ABscf098869 | + | 456526 | 456628 | **RNA**→ Methyltrn_RNA_3 (pfam02598)→ RplC (COG0087)rpl3p (PRK04231)→ RplD (COG0088)rpl4lp (PRK04042)→ RplW (COG0089)PRK14548 (PRK14548)→ RplB (COG0090)rpl2p (PRK09612)→ RpsS (COG0185)rps19p (PRK04038)→ rpl22p (PRK04223)RplV (COG0091)→ RpsC (COG0092)rps3p (PRK04191)→ |
| env-101 | P_1C_Sed_2_UnCty_1030622 | + | 624 | 726 | **RNA**→ MTH1 (COG2106)→ |
| env-102 | SL_5KL_010_BRINE_10180849 | - | 369 | 267 | **RNA**→ PRK10575 (PRK10575)Methyltrn_RNA_3 (pfam02598)→ |
| env-103 | SL_6KL_011_BRINE_1082566 | - | 592 | 490 | **RNA**→ PRK10575 (PRK10575)Methyltrn_RNA_3 (pfam02598)→ |
| env-104 | SL_8KL_010_SED_10233400 | + | 212 | 314 | **RNA**→ Methyltrn_RNA_3 (pfam02598)→ |
| env-105 | SL_4KL_010_BRINE_10134291 | + | 227 | 329 | **RNA**→ Methyltrn_RNA_3 (pfam02598)→ |
| env-106 | SL_5KL_010_BRINE_10114435 | + | 29 | 131 | **RNA**→ Methyltrn_RNA_3 (pfam02598)→ |
| env-107 | P_1C_Sed_2_UnCty_1330058 | + | 56 | 155 | **RNA**→ Methyltrn_RNA_3 (pfam02598)→ |
| env-108 | SL_4KL_010_BRINE_10687755 | + | 77 | 176 | **RNA**→ Methyltrn_RNA_3 (pfam02598)→ |
| env-109 | JXWU01090489.1 | + | 605 | 701 | **RNA**→ Methyltrn_RNA_3 (pfam02598)→ |
| CHa-1-1 | NZ_ASGZ01000021.1 | + | 7594 | 7693 | **RNA**→ Methyltrn_RNA_3 (pfam02598)→ RplC (COG0087)rpl3p (PRK04231)→ RplD (COG0088)rpl4lp (PRK04042)→ RplW (COG0089)PRK14548 (PRK14548)→ RplB (COG0090)rpl2p (PRK09612)→ RpsS (COG0185)rps19p (PRK04038)→ rpl22p (PRK04223)RplV (COG0091)→ RpsC (COG0092)rps3p (PRK04191)→ PRK14549 (PRK14549)Ribosomal_L29 (pfam00831)→ PRK03879 (PRK03879)UPF0086 (pfam01868)→ RpsQ (COG0186)rps17p (PRK08572)→ rpl14p (PRK08571)Ribosomal_L14 (pfam00238)→ RplX (COG0198)rpl24p (PRK01191)→ PRK04313 (PRK04313)Ribosomal_S4e (pfam00900)→ RplE (COG0094)rpl5p (PRK04219)→ RpsN (COG0199)rps14P (PRK05766)→ rps8p (PRK04034)RpsH (COG0096)→ RplF (COG0097)rpl6p (PRK05518)→ PRK12766 (PRK12766)RecA (COG0468)Rpl32e (COG1717)→ rpl19e (PRK08570)Ribosomal_L19e (pfam01280)→ RplR (COG0256)rpl18p (PRK08569)→ RpsE (COG0098)rps5p (PRK04044)→ RpsE (COG0098)rps5p (PRK04044)→ Ribosomal_L7_archeal_euk (cd01657)rpl30p (PRK06049)→ rpl15p (PRK06419)Ribosomal_L27A (pfam00828)→ |
| Hpa-2-1 | NZ_AOIV01000008.1 | - | 240007 | 239908 | **RNA**→ Methyltrn_RNA_3 (pfam02598)→ RplC (COG0087)rpl3p (PRK04231)→ RplD (COG0088)rpl4lp (PRK04042)→ RplW (COG0089)PRK14548 (PRK14548)→ RplB (COG0090)rpl2p (PRK09612)→ RpsS (COG0185)rps19p (PRK04038)→ rpl22p (PRK04223)RplV (COG0091)→ RpsC (COG0092)rps3p (PRK04191)→ PRK14549 (PRK14549)Ribosomal_L29 (pfam00831)→ PRK03879 (PRK03879)UPF0086 (pfam01868)→ RpsQ (COG0186)rps17p (PRK08572)→ rpl14p (PRK08571)Ribosomal_L14 (pfam00238)→ RplX (COG0198)rpl24p (PRK01191)→ PRK04313 (PRK04313)KOW_RPS4 (cd06087)Ribosomal_S4e (pfam00900)→ RplE (COG0094)rpl5p (PRK04219)→ RpsN (COG0199)rps14P (PRK05766)→ rps8p (PRK04034)RpsH (COG0096)→ RplF (COG0097)rpl6p (PRK05518)→ PRK12766 (PRK12766)RecA (COG0468)Rpl32e (COG1717)→ rpl19e (PRK08570)Ribosomal_L19e (pfam01280)→ |
| env-110 | SL_5KL_010_BRINE_10000579 | - | 6143 | 6044 | **RNA**→ Methyltrn_RNA_3 (pfam02598)→ RplC (COG0087)rpl3p (PRK04231)→ RplD (COG0088)rpl4lp (PRK04042)→ RplW (COG0089)PRK14548 (PRK14548)→ RplB (COG0090)rpl2p (PRK09612)→ RpsS (COG0185)rps19p (PRK04038)→ rpl22p (PRK04223)RplV (COG0091)→ RpsC (COG0092)rps3p (PRK04191)→ RpmC (COG0255)PRK14549 (PRK14549)→ hypo→ PRK03879 (PRK03879)UPF0086 (pfam01868)→ RpsQ (COG0186)rps17p (PRK08572)→ |
| env-111 | SL_4KL_010_BRINE_10006116 | - | 6219 | 6120 | **RNA**→ Methyltrn_RNA_3 (pfam02598)→ RplC (COG0087)rpl3p (PRK04231)→ RplD (COG0088)rpl4lp (PRK04042)→ RplW (COG0089)PRK14548 (PRK14548)→ RplB (COG0090)rpl2p (PRK09612)→ RpsS (COG0185)rps19p (PRK04038)→ rpl22p (PRK04223)RplV (COG0091)→ RpsC (COG0092)rps3p (PRK04191)→ RpmC (COG0255)PRK14549 (PRK14549)→ hypo→ PRK03879 (PRK03879)UPF0086 (pfam01868)→ RpsQ (COG0186)rps17p (PRK08572)→ |
| env-112 | SL_4KL_010_BRINE_10037372 | + | 1564 | 1663 | **RNA**→ Methyltrn_RNA_3 (pfam02598)→ rpl3p (PRK04231)Ribosomal_L3 (pfam00297)→ |

| | | | | | |
|---|---|---|---|---|---|
| Nhu-1-1 | NZ_AOIM01000008.1 | + | 9157 | 9256 | **RNA**→ Methyltrn_RNA_3 (pfam02598)→ rpl3p (PRK04231)Ribosomal_L3 (pfam00297)→ RplD (COG0088)rpl4lp (PRK04042)→ RplW (COG0089)PRK14548 (PRK14548)→ RplB (COG0090)rpl2p (PRK09612)→ RpsS (COG0185)rps19p (PRK04038)→ rpl22p (PRK04223)RplV (COG0091)→ RpsC (COG0092)rps3p (PRK04191)→ RpmC (COG0255)PRK14549 (PRK14549)→ PRK03879 (PRK03879)UPF0086 (pfam01868)→ RpsQ (COG0186)rps17p (PRK08572)→ rpl14p (PRK08571)Ribosomal_L14 (pfam00238)→ rpl24p (PRK01191)KOW_RPL26 (cd06089)→ PRK04313 (PRK04313)S4 (pfam01479)→ hypo→ |
| env-113 | P_A23_Liq_1_Fmt_1001862 | - | 2728 | 2626 | **RNA**→ MTH1 (COG2106)→ RplC (COG0087)rpl3p (PRK04231)→ RplD (COG0088)rpl4lp (PRK04042)→ RplW (COG0089)PRK14548 (PRK14548)→ RplB (COG0090)rpl2p (PRK09612)→ |
| env-114 | P_A23_Liq_1_Fmt_1006312 | - | 589 | 487 | **RNA**→ Methyltrn_RNA_3 (pfam02598)→ |
| env-115 | P_A23_Liq_2_Fmt_1001778 | - | 147 | 45 | **RNA**→ MTH1 (COG2106)→ |
| env-116 | SL_5KL_010_BRINE_10182593 | - | 355 | 253 | **RNA**→ PRK10575 (PRK10575)Methyltrn_RNA_3 (pfam02598)→ |
| env-117 | SL_6KL_011_BRINE_1122878 | + | 59 | 161 | **RNA**→ Methyltrn_RNA_3 (pfam02598)→ |
| env-118 | SL_5KL_010_SED_10176812 | - | 343 | 241 | **RNA**→ Methyltrn_RNA_3 (pfam02598)→ |
| env-119 | GSLSAAI_GFP0ZZ301BFHUD | + | 260 | 362 | **RNA**→ Methyltrn_RNA_3 (pfam02598)→ |
| env-120 | APHM01000442.1 | + | 2818 | 2917 | **RNA**→ Methyltrn_RNA_3 (pfam02598)→ RplC (COG0087)rpl3p (PRK04231)→ RplD (COG0088)rpl4lp (PRK04042)→ RplW (COG0089)PRK14548 (PRK14548)→ RplB (COG0090)rpl2p (PRK09612)→ rps19p (PRK04038)→ |
| env-121 | LTJ07As871_J07ABscf098878 | - | 600442 | 600343 | **RNA**→ Methyltrn_RNA_3 (pfam02598)→ RplC (COG0087)rpl3p (PRK04231)→ RplD (COG0088)rpl4lp (PRK04042)→ RplW (COG0089)PRK14548 (PRK14548)→ RplB (COG0090)rpl2p (PRK09612)→ |
| env-122 | JXWS01030537.1 | + | 838 | 937 | **RNA**→ Methyltrn_RNA_3 (pfam02598)→ |
| env-123 | P_A23_Liq_1_Fmt_1000580 | + | 837 | 936 | **RNA**→ Methyltrn_RNA_3 (pfam02598)→ RplC (COG0087)rpl3p (PRK04231)→ RplD (COG0088)rpl4lp (PRK04042)→ RplW (COG0089)PRK14548 (PRK14548)→ RplB (COG0090)rpl2p (PRK09612)→ RpsS (COG0185)rps19p (PRK04038)→ rpl22p (PRK04223)RplV (COG0091)→ RpsC (COG0092)rps3p (PRK04191)→ PRK14549 (PRK14549)Ribosomal_L29 (pfam00831)→ PRK03879 (PRK03879)UPF0086 (pfam01868)→ |
| env-124 | P_A23_Liq_3_Fmt_1000785 | - | 1067 | 968 | **RNA**→ Methyltrn_RNA_3 (pfam02598)→ rpl3p (PRK04231)Ribosomal_L3 (pfam00297)→ |
| env-125 | P_A23_Liq_2_Fmt_1000789 | - | 1096 | 997 | **RNA**→ Methyltrn_RNA_3 (pfam02598)→ rpl3p (PRK04231)Ribosomal_L3 (pfam00297)→ |
| env-126 | GSLNARP_GFPJP1N02GVYAL | + | 71 | 170 | **RNA**→ MTH1 (COG2106)→ |
| env-127 | P_1C_Liq_2_UnCty_1129324 | + | 141 | 240 | **RNA**→ MTH1 (COG2106)→ |
| env-128 | JXWS01088076.1 | + | 222 | 321 | **RNA**→ MTH1 (COG2106)→ |
| env-129 | SS_3KL_010_SOIL_10082725 | - | 438 | 339 | **RNA**→ Methyltrn_RNA_3 (pfam02598)→ |
| Hna-1-1 | NZ_KE386573.1 | - | 1301517 | 1301418 | **RNA**→ Methyltrn_RNA_3 (pfam02598)→ rpl3p (PRK04231)Ribosomal_L3 (pfam00297)→ RplD (COG0088)rpl4lp (PRK04042)→ RplW (COG0089)PRK14548 (PRK14548)→ RplB (COG0090)rpl2p (PRK09612)→ RpsS (COG0185)rps19p (PRK04038)→ rpl22p (PRK04223)RplV (COG0091)→ RpsC (COG0092)rps3p (PRK04191)→ PRK14549 (PRK14549)Ribosomal_L29 (pfam00831)→ PRK03879 (PRK03879)UPF0086 (pfam01868)→ RpsQ (COG0186)rps17p (PRK08572)→ rpl14p (PRK08571)Ribosomal_L14 (pfam00238)→ RplX (COG0198)rpl24p (PRK01191)→ PRK04313 (PRK04313)KOW_RPS4 (cd06087)Ribosomal_S4e (pfam00900)→ RplE (COG0094)rpl5p (PRK04219)→ RpsN (COG0199)rps14P (PRK05766)→ rps8p (PRK04034)RpsH (COG0096)→ RplF (COG0097)rpl6p (PRK05518)→ PRK12766 (PRK12766)RecA (COG0468)Rpl32e (COG1717)→ rpl19e (PRK08570)Ribosomal_L19e (pfam01280)→ |
| env-130 | P_1C_Sed_2_UnCty_1009263 | - | 1341 | 1242 | **RNA**→ Methyltrn_RNA_3 (pfam02598)→ RplC (COG0087)rpl3p (PRK04231)→ |
| env-131 | ADL20m3uS_contig11573 | - | 6008 | 5909 | **RNA**→ Methyltrn_RNA_3 (pfam02598)→ rpl3p (PRK04231)Ribosomal_L3 (pfam00297)→ RplC (COG0087)rpl3p (PRK04231)→ RplD (COG0088)rpl4lp (PRK04042)→ RplW (COG0089)PRK14548 (PRK14548)→ RplB (COG0090)rpl2p (PRK09612)→ RpsS (COG0185)rps19p (PRK04038)→ rpl22p (PRK04223)RplV (COG0091)→ RpsC (COG0092)rps3p (PRK04191)→ PRK14549 (PRK14549)Ribosomal_L29 (pfam00831)→ PRK03879 (PRK03879)UPF0086 (pfam01868)→ |

| | | | | | |
|---|---|---|---|---|---|
| env-132 | 2140918017-125 | - | 28013 | 27914 | **RNA**→ Methyltrn_RNA_3 (pfam02598)→ rpl3p (PRK04231)Ribosomal_L3 (pfam00297)→<br>RplC (COG0087)rpl3p (PRK04231)→ RplD (COG0088)rpl4lp (PRK04042)→<br>RplW (COG0089)PRK14548 (PRK14548)→ RplB (COG0090)rpl2p (PRK09612)→<br>RpsS (COG0185)rps19p (PRK04038)→ rpl22p (PRK04223)RplV (COG0091)→<br>RpsC (COG0092)rps3p (PRK04191)→ PRK14549 (PRK14549)Ribosomal_L29 (pfam00831)→<br>PRK03879 (PRK03879)UPF0086 (pfam01868)→ RpsQ (COG0186)rps17p (PRK08572)→<br>rpl14p (PRK08571)Ribosomal_L14 (pfam00238)→ rpl24p (PRK01191)KOW_RPL26 (cd06089)→<br>PRK04313 (PRK04313)KOW_RPS4 (cd06087)Ribosomal_S4e (pfam00900)→<br>rpl5p (PRK04219)Ribosomal_L5 (pfam00281)→ RpsN (COG0199)rps14P (PRK05766)→<br>rps8p (PRK04034)RpsH (COG0096)→ RplF (COG0097)rpl6p (PRK05518)→<br>PRK12766 (PRK12766)RecA (COG0468)Rpl32e (COG1717)→ rpl19e (PRK08570)Ribosomal_L19e (pfam01280)→ |
| env-133 | 2140918027-00843 | + | 673 | 772 | **RNA**→ Methyltrn_RNA_3 (pfam02598)→ rpl3p (PRK04231)Ribosomal_L3 (pfam00297)→<br>RplC (COG0087)rpl3p (PRK04231)→ RplD (COG0088)rpl4lp (PRK04042)→<br>RplW (COG0089)PRK14548 (PRK14548)→ RplB (COG0090)rpl2p (PRK09612)→<br>RpsS (COG0185)rps19p (PRK04038)→ rpl22p (PRK04223)RplV (COG0091)→<br>RpsC (COG0092)rps3p (PRK04191)→ PRK14549 (PRK14549)Ribosomal_L29 (pfam00831)→<br>PRK03879 (PRK03879)UPF0086 (pfam01868)→ |
| har-1-1 | NC_015954.1 | - | 1701293 | 1701194 | **RNA**→ Methyltrn_RNA_3 (pfam02598)→ RplC (COG0087)rpl3p (PRK04231)→<br>RplD (COG0088)rpl4lp (PRK04042)→ RplW (COG0089)PRK14548 (PRK14548)→<br>RplB (COG0090)rpl2p (PRK09612)→ RpsS (COG0185)rps19p (PRK04038)→<br>rpl22p (PRK04223)RplV (COG0091)→ RpsC (COG0092)rps3p (PRK04191)→<br>PRK14549 (PRK14549)Ribosomal_L29 (pfam00831)→ PRK03879 (PRK03879)UPF0086 (pfam01868)→<br>RpsQ (COG0186)rps17p (PRK08572)→ rpl14p (PRK08571)Ribosomal_L14 (pfam00238)→<br>rpl24p (PRK01191)KOW_RPL26 (cd06089)→<br>PRK04313 (PRK04313)KOW_RPS4 (cd06087)Ribosomal_S4e (pfam00900)→<br>rpl5p (PRK04219)Ribosomal_L5 (pfam00281)→ RpsN (COG0199)rps14P (PRK05766)→<br>rps8p (PRK04034)RpsH (COG0096)→ RplF (COG0097)rpl6p (PRK05518)→<br>PRK12766 (PRK12766)RecA (COG0468)Rpl32e (COG1717)→ rpl19e (PRK08570)Ribosomal_L19e (pfam01280)→ |
| env-134 | SS_3KL_010_SOIL_10099808 | - | 212 | 113 | **RNA**→ Methyltrn_RNA_3 (pfam02598)→ |
| env-135 | 2140918017-82247 | + | 63 | 164 | **RNA**→ |
| Nph-1-1 | NC_007426.1 | + | 2369313 | 2369412 | **RNA**→ Methyltrn_RNA_3 (pfam02598)→ rpl3p (PRK04231)Ribosomal_L3 (pfam00297)→<br>RplD (COG0088)rpl4lp (PRK04042)→ RplW (COG0089)PRK14548 (PRK14548)→<br>RplB (COG0090)rpl2p (PRK09612)→ RpsS (COG0185)rps19p (PRK04038)→<br>rpl22p (PRK04223)RplV (COG0091)→ RpsC (COG0092)rps3p (PRK04191)→<br>RpmC (COG0255)PRK14549 (PRK14549)→ PRK03879 (PRK03879)UPF0086 (pfam01868)→ hypo→<br>RpsQ (COG0186)rps17p (PRK08572)→ rpl14p (PRK08571)Ribosomal_L14 (pfam00238)→<br>RplX (COG0198)rpl24p (PRK01191)→<br>PRK04313 (PRK04313)KOW_RPS4 (cd06087)Ribosomal_S4e (pfam00900)→ RplE (COG0094)rpl5p (PRK04219)→<br>RpsN (COG0199)rps14P (PRK05766)→ rps8p (PRK04034)RpsH (COG0096)→<br>RplF (COG0097)rpl6p (PRK05518)→ NT_POLXc (cd00141)PRK12766 (PRK12766)Rpl32e (COG1717)→<br>rpl19e (PRK08570)Ribosomal_L19e (pfam01280)→ RplR (COG0256)rpl18p (PRK08569)→ |
| env-136 | SL_4KL_010_BRINE_10016891 | - | 2655 | 2556 | **RNA**→ Methyltrn_RNA_3 (pfam02598)→ RplC (COG0087)rpl3p (PRK04231)→<br>RplD (COG0088)rpl4lp (PRK04042)→ RplW (COG0089)PRK14548 (PRK14548)→<br>RplB (COG0090)rpl2p (PRK09612)→ |

| | | | | | |
|---|---|---|---|---|---|
| env-137 | SL_6KL_011_BRINE_1000523 | + | 4060 | 4159 | **RNA**→ Methyltrn_RNA_3 (pfam02598)→ RplC (COG0087)rpl3p (PRK04231)→<br>RplD (COG0088)rpl4lp (PRK04042)→ RplW (COG0089)PRK14548 (PRK14548)→<br>RplB (COG0090)rpl2p (PRK09612)→ RpsS (COG0185)rps19p (PRK04038)→<br>rpl22p (PRK04223)RplV (COG0091)→ RpsC (COG0092)rps3p (PRK04191)→<br>RpmC (COG0255)PRK14549 (PRK14549)→ PRK03879 (PRK03879)UPF0086 (pfam01868)→<br>RpsQ (COG0186)rps17p (PRK08572)→ rpl14p (PRK08571)Ribosomal_L14 (pfam00238)→<br>RplX (COG0198)rpl24p (PRK01191)→<br>PRK04313 (PRK04313)KOW_RPS4 (cd06087)Ribosomal_S4e (pfam00900)→ RplE (COG0094)rpl5p (PRK04219)→<br>RpsN (COG0199)rps14P (PRK05766)→ rps8p (PRK04034)RpsH (COG0096)→<br>RplF (COG0097)rpl6p (PRK05518)→ PRK12766 (PRK12766)RecA (COG0468)Rpl32e (COG1717)→<br>rpl19e (PRK08570)Ribosomal_L19e (pfam01280)→ RplR (COG0256)rpl18p (PRK08569)→<br>RpsE (COG0098)rps5p (PRK04044)→ Ribosomal_L7_archeal_euk (cd01657)rpl30p (PRK06049)→<br>rpl15p (PRK06419)Ribosomal_L27A (pfam00828)→ SecY (COG0201)PRK08568 (PRK08568)→ |
| env-138 | P_1C_Liq_1_UnCty_1007975 | + | 1341 | 1443 | **RNA**→ Methyltrn_RNA_3 (pfam02598)→ |
| env-139 | SL_6KL_011_BRINE_1092333 | + | 64 | 166 | **RNA**→ Methyltrn_RNA_3 (pfam02598)→ |
| env-140 | SL_5KL_010_BRINE_10000145 | - | 14973 | 14871 | **RNA**→ MTH1 (COG2106)→ rpl3p (PRK04231)Ribosomal_L3 (pfam00297)→<br>RplD (COG0088)rpl4lp (PRK04042)→ RplW (COG0089)PRK14548 (PRK14548)→<br>RplB (COG0090)rpl2p (PRK09612)→ RpsS (COG0185)rps19p (PRK04038)→<br>rpl22p (PRK04223)RplV (COG0091)→ RpsC (COG0092)rps3p (PRK04191)→<br>PRK14549 (PRK14549)Ribosomal_L29_HIP (cd00427)→ PRK03879 (PRK03879)UPF0086 (pfam01868)→<br>RpsQ (COG0186)rps17p (PRK08572)→ rpl14p (PRK08571)Ribosomal_L14 (pfam00238)→<br>rpl24p (PRK01191)KOW_RPL26 (cd06089)→<br>PRK04313 (PRK04313)KOW_RPS4 (cd06087)Ribosomal_S4e (pfam00900)→ RplE (COG0094)rpl5p (PRK04219)→<br>RpsN (COG0199)rps14P (PRK05766)→ rps8p (PRK04034)RpsH (COG0096)→<br>RplF (COG0097)rpl6p (PRK05518)→ PRK12766 (PRK12766)RecA (COG0468)Rpl32e (COG1717)→<br>rpl19e (PRK08570)Ribosomal_L19e (pfam01280)→ |
| env-141 | SL_6KL_011_BRINE_1010384 | + | 87 | 189 | **RNA**→ Methyltrn_RNA_3 (pfam02598)→ rpl3p (PRK04231)Ribosomal_L3 (pfam00297)→<br>RplD (COG0088)rpl4lp (PRK04042)→ RplW (COG0089)PRK14548 (PRK14548)→<br>RplB (COG0090)rpl2p (PRK09612)→ RpsS (COG0185)rps19p (PRK04038)→<br>rpl22p (PRK04223)RplV (COG0091)→ |
| env-142 | SL_5KL_010_SED_10000993 | - | 17753 | 17651 | **RNA**→ Methyltrn_RNA_3 (pfam02598)→ rpl3p (PRK04231)Ribosomal_L3 (pfam00297)→<br>RplD (COG0088)rpl4lp (PRK04042)→ RplW (COG0089)PRK14548 (PRK14548)→<br>RplB (COG0090)rpl2p (PRK09612)→ RpsS (COG0185)rps19p (PRK04038)→<br>rpl22p (PRK04223)RplV (COG0091)→ RpsC (COG0092)rps3p (PRK04191)→<br>PRK14549 (PRK14549)Ribosomal_L29_HIP (cd00427)→ PRK03879 (PRK03879)UPF0086 (pfam01868)→<br>RpsQ (COG0186)rps17p (PRK08572)→ rpl14p (PRK08571)Ribosomal_L14 (pfam00238)→<br>rpl24p (PRK01191)KOW_RPL26 (cd06089)→<br>PRK04313 (PRK04313)KOW_RPS4 (cd06087)Ribosomal_S4e (pfam00900)→ RplE (COG0094)rpl5p (PRK04219)→<br>RpsN (COG0199)rps14P (PRK05766)→ rps8p (PRK04034)RpsH (COG0096)→<br>RplF (COG0097)rpl6p (PRK05518)→ PRK12766 (PRK12766)RecA (COG0468)Rpl32e (COG1717)→<br>rpl19e (PRK08570)Ribosomal_L19e (pfam01280)→ |
| env-143 | SL_8KL_010_SED_10049127 | - | 1844 | 1742 | **RNA**→ Methyltrn_RNA_3 (pfam02598)→ rpl3p (PRK04231)Ribosomal_L3 (pfam00297)→<br>RplD (COG0088)rpl4lp (PRK04042)→ |
| env-144 | SL_4KL_010_BRINE_10002885 | - | 1066 | 964 | **RNA**→ MTH1 (COG2106)→ rpl3p (PRK04231)Ribosomal_L3 (pfam00297)→ |
| env-145 | SL_6KL_011_BRINE_1040731 | - | 290 | 188 | **RNA**→ Methyltrn_RNA_3 (pfam02598)→ |
| env-146 | SL_6KL_011_BRINE_1090778 | - | 540 | 438 | **RNA**→ Methyltrn_RNA_3 (pfam02598)→ |
| env-147 | SL_6KL_011_BRINE_1020969 | + | 1942 | 2044 | **RNA**→ MTH1 (COG2106)→ |
| env-148 | SL_6KL_011_BRINE_1001075 | - | 545 | 443 | **RNA**→ Methyltrn_RNA_3 (pfam02598)→ |
| Hsp-9-1 | NZ_CBVY010000043.1 | + | 1088 | 1190 | **RNA**→ MTH1 (COG2106)→ RplC (COG0087)rpl3p (PRK04231)→ |

| | | | | | |
|---|---|---|---|---|---|
| Hez-1-1 | NZ_ATXS01000002.1 | + | 57125 | 57227 | **RNA**→ MTH1 (COG2106)→ RplC (COG0087)rpl3p (PRK04231)→ RplD (COG0088)rpl4lp (PRK04042)→<br>RplW (COG0089)PRK14548 (PRK14548)→ RplB (COG0090)rpl2p (PRK09612)→<br>RpsS (COG0185)rps19p (PRK04038)→ rpl22p (PRK04223)RplV (COG0091)→<br>RpsC (COG0092)rps3p (PRK04191)→ RpmC (COG0255)PRK14549 (PRK14549)→<br>PRK03879 (PRK03879)UPF0086 (pfam01868)→ RpsQ (COG0186)rps17p (PRK08572)→<br>rpl14p (PRK08571)Ribosomal_L14 (pfam00238)→ rpl24p (PRK01191)KOW_RPL26 (cd06089)→<br>PRK04313 (PRK04313)KOW_RPS4 (cd06087)Ribosomal_S4e (pfam00900)→ RplE (COG0094)rpl5p (PRK04219)→<br>RpsN (COG0199)rps14P (PRK05766)→ rps8p (PRK04034)RpsH (COG0096)→<br>RplF (COG0097)rpl6p (PRK05518)→ PRK12766 (PRK12766)RecA (COG0468)Rpl32e (COG1717)→<br>rpl19e (PRK08570)Ribosomal_L19e (pfam01280)→ |
| env-149 | P_1C_Liq_1_UnCty_1110322 | + | 40 | 142 | **RNA**→ Methyltrn_RNA_3 (pfam02598)→ |
| env-150 | P_1C_Liq_2_UnCty_1085242 | - | 257 | 155 | **RNA**→ Methyltrn_RNA_3 (pfam02598)→ |
| env-151 | P_1C_Sed_2_UnCty_1018040 | - | 1354 | 1252 | **RNA**→ Methyltrn_RNA_3 (pfam02598)→ RplC (COG0087)rpl3p (PRK04231)→ |
| Hco-1-1 | NZ_AOJL01000060.1 | - | 167010 | 166908 | **RNA**→ MTH1 (COG2106)→ RplC (COG0087)rpl3p (PRK04231)→ RplD (COG0088)rpl4lp (PRK04042)→<br>RplW (COG0089)PRK14548 (PRK14548)→ RplB (COG0090)rpl2p (PRK09612)→<br>RplB (COG0090)rpl2p (PRK09612)→ RpsS (COG0185)rps19p (PRK04038)→<br>rpl22p (PRK04223)RplV (COG0091)→ RpsC (COG0092)rps3p (PRK04191)→<br>RpmC (COG0255)PRK14549 (PRK14549)→ PRK03879 (PRK03879)UPF0086 (pfam01868)→<br>RpsQ (COG0186)rps17p (PRK08572)→ rpl14p (PRK08571)Ribosomal_L14 (pfam00238)→<br>rpl24p (PRK01191)KOW_RPL26 (cd06089)→<br>PRK04313 (PRK04313)KOW_RPS4 (cd06087)Ribosomal_S4e (pfam00900)→<br>rpl5p (PRK04219)Ribosomal_L5 (pfam00281)Ribosomal_L5_C (pfam00673)→ hypo→<br>RpsN (COG0199)rps14P (PRK05766)→ rps8p (PRK04034)RpsH (COG0096)→<br>RplF (COG0097)rpl6p (PRK05518)→ PRK12766 (PRK12766)RecA (COG0468)Rpl32e (COG1717)→<br>rpl19e (PRK08570)Ribosomal_L19e (pfam01280)→ |
| env-152 | P_1C_Liq_3_UnCty_1007582 | - | 296 | 194 | **RNA**→ MTH1 (COG2106)→ |
| env-153 | P_A23_Liq_1_Fmt_1009395 | - | 369 | 267 | **RNA**→ MTH1 (COG2106)→ |
| env-154 | P_A23_Liq_2_Fmt_1002028 | + | 19 | 121 | **RNA**→ Methyltrn_RNA_3 (pfam02598)→ RplC (COG0087)rpl3p (PRK04231)→<br>RplD (COG0088)rpl4lp (PRK04042)→ RplW (COG0089)PRK14548 (PRK14548)→<br>RplB (COG0090)rpl2p (PRK09612)→ RpsS (COG0185)rps19p (PRK04038)→ |
| env-155 | P_A23_Liq_2_Fmt_1023865 | + | 29 | 131 | **RNA**→ Methyltrn_RNA_3 (pfam02598)→ |
| env-156 | P_A23_Liq_3_Fmt_1023393 | + | 38 | 140 | **RNA**→ Methyltrn_RNA_3 (pfam02598)→ |
| Hsp-10-1 | NZ_JQKV01000002.1 | - | 421137 | 421035 | **RNA**→ Methyltrn_RNA_3 (pfam02598)→ RplC (COG0087)rpl3p (PRK04231)→<br>RplD (COG0088)rpl4lp (PRK04042)→ RplW (COG0089)PRK14548 (PRK14548)→<br>RplB (COG0090)rpl2p (PRK09612)→ RpsS (COG0185)rps19p (PRK04038)→<br>rpl22p (PRK04223)RplV (COG0091)→ RpsC (COG0092)rps3p (PRK04191)→<br>PRK14549 (PRK14549)Ribosomal_L29_HIP (cd00427)→ PRK03879 (PRK03879)UPF0086 (pfam01868)→<br>RpsQ (COG0186)rps17p (PRK08572)→ rpl14p (PRK08571)Ribosomal_L14 (pfam00238)→<br>RplX (COG0198)rpl24p (PRK01191)→<br>PRK04313 (PRK04313)KOW_RPS4 (cd06087)Ribosomal_S4e (pfam00900)→ RplE (COG0094)rpl5p (PRK04219)→<br>RpsN (COG0199)rps14P (PRK05766)→ rps8p (PRK04034)RpsH (COG0096)→<br>RplF (COG0097)rpl6p (PRK05518)→ PRK12766 (PRK12766)RecA (COG0468)Rpl32e (COG1717)→<br>rpl19e (PRK08570)Ribosomal_L19e (pfam01280)→ |
| Hli-1-1 | NZ_AOJG01000040.1 | - | 82466 | 82364 | **RNA**→ MTH1 (COG2106)→ rpl3p (PRK04231)Ribosomal_L3 (pfam00297)→<br>RplD (COG0088)rpl4lp (PRK04042)→ RplW (COG0089)PRK14548 (PRK14548)→<br>RplB (COG0090)rpl2p (PRK09612)→ RpsS (COG0185)rps19p (PRK04038)→<br>rpl22p (PRK04223)RplV (COG0091)→ RpsC (COG0092)rps3p (PRK04191)→<br>PRK14549 (PRK14549)Ribosomal_L29_HIP (cd00427)→ PRK03879 (PRK03879)UPF0086 (pfam01868)→<br>RpsQ (COG0186)rps17p (PRK08572)→ rpl14p (PRK08571)Ribosomal_L14 (pfam00238)→<br>rpl24p (PRK01191)KOW_RPL26 (cd06089)→ PRK04313 (PRK04313)Ribosomal_S4e (pfam00900)→<br>RplE (COG0094)rpl5p (PRK04219)→ RpsN (COG0199)rps14P (PRK05766)→<br>rps8p (PRK04034)RpsH (COG0096)→ RplF (COG0097)rpl6p (PRK05518)→<br>PRK12766 (PRK12766)RecA (COG0468)Rpl32e (COG1717)→ rpl19e (PRK08570)Ribosomal_L19e (pfam01280)→ |

| | | | | | |
|---|---|---|---|---|---|
| Hte-1-1 | NZ_AOJD01000064.1 | + | 2215 | 2317 | **RNA**→ MTH1 (COG2106)→ RplC (COG0087)rpl3p (PRK04231)→ RplD (COG0088)rpl4lp (PRK04042)→<br>RplW (COG0089)PRK14548 (PRK14548)→ RplB (COG0090)rpl2p (PRK09612)→<br>RplB (COG0090)rpl2p (PRK09612)→ RpsS (COG0185)rps19p (PRK04038)→<br>rpl22p (PRK04223)RplV (COG0091)→ RpsC (COG0092)rps3p (PRK04191)→<br>PRK14549 (PRK14549)Ribosomal_L29_HIP (cd00427)→ PRK03879 (PRK03879)UPF0086 (pfam01868)→<br>RpsQ (COG0186)rps17p (PRK08572)→ rpl14p (PRK08571)Ribosomal_L14 (pfam00238)→<br>rpl24p (PRK01191)KOW_RPL26 (cd06089)→<br>PRK04313 (PRK04313)KOW_RPS4 (cd06087)Ribosomal_S4e (pfam00900)→ RplE (COG0094)rpl5p (PRK04219)→<br>RpsN (COG0199)rps14P (PRK05766)→ rps8p (PRK04034)RpsH (COG0096)→<br>RplF (COG0097)rpl6p (PRK05518)→ PRK12766 (PRK12766)RecA (COG0468)Rpl32e (COG1717)→<br>rpl19e (PRK08570)Ribosomal_L19e (pfam01280)→ RplR (COG0256)rpl18p (PRK08569)→<br>RpsE (COG0098)rps5p (PRK04044)→ Ribosomal_L7_archeal_euk (cd01657)rpl30p (PRK06049)→<br>rpl15p (PRK06419)Ribosomal_L27A (pfam00828)→ SecY (COG0201)PRK08568 (PRK08568)→ |
| Hko-1-1 | NZ_AOJH01000027.1 | + | 94307 | 94409 | **RNA**→ Methyltrn_RNA_3 (pfam02598)→ rpl3p (PRK04231)Ribosomal_L3 (pfam00297)→<br>RplD (COG0088)rpl4lp (PRK04042)→ RplW (COG0089)PRK14548 (PRK14548)→<br>RplB (COG0090)rpl2p (PRK09612)→ RpsS (COG0185)rps19p (PRK04038)→<br>rpl22p (PRK04223)RplV (COG0091)→ RpsC (COG0092)rps3p (PRK04191)→<br>PRK14549 (PRK14549)Ribosomal_L29_HIP (cd00427)→ PRK03879 (PRK03879)UPF0086 (pfam01868)→<br>RpsQ (COG0186)rps17p (PRK08572)→ rpl14p (PRK08571)Ribosomal_L14 (pfam00238)→<br>rpl24p (PRK01191)KOW_RPL26 (cd06089)→ PRK04313 (PRK04313)Ribosomal_S4e (pfam00900)→<br>RplE (COG0094)rpl5p (PRK04219)→ RpsN (COG0199)rps14P (PRK05766)→<br>rps8p (PRK04034)RpsH (COG0096)→ RplF (COG0097)rpl6p (PRK05518)→<br>PRK12766 (PRK12766)RecA (COG0468)Rpl32e (COG1717)→ rpl19e (PRK08570)Ribosomal_L19e (pfam01280)→ |
| Hli-2-1 | NZ_AOJF01000042.1 | + | 2248 | 2350 | **RNA**→ MTH1 (COG2106)→ RplC (COG0087)rpl3p (PRK04231)→ RplD (COG0088)rpl4lp (PRK04042)→<br>RplW (COG0089)PRK14548 (PRK14548)→ RplB (COG0090)rpl2p (PRK09612)→<br>RplB (COG0090)rpl2p (PRK09612)→ RpsS (COG0185)rps19p (PRK04038)→<br>rpl22p (PRK04223)RplV (COG0091)→ RpsC (COG0092)rps3p (PRK04191)→<br>RpmC (COG0255)PRK14549 (PRK14549)→ PRK03879 (PRK03879)UPF0086 (pfam01868)→<br>RpsQ (COG0186)rps17p (PRK08572)→ rpl14p (PRK08571)Ribosomal_L14 (pfam00238)→<br>rpl24p (PRK01191)KOW_RPL26 (cd06089)→ hypo→<br>PRK04313 (PRK04313)rpl5p (PRK04219)Ribosomal_L5 (pfam00281)Ribosomal_L5_C (pfam00673)Ribosomal_S4e (pfam0090[illegible]<br>rpl5p (PRK04219)Ribosomal_L5 (pfam00281)Ribosomal_L5_C (pfam00673)→<br>RpsN (COG0199)rps14P (PRK05766)→ rps8p (PRK04034)RpsH (COG0096)→<br>RplF (COG0097)rpl6p (PRK05518)→ PRK12766 (PRK12766)RecA (COG0468)Rpl32e (COG1717)→<br>rpl19e (PRK08570)Ribosomal_L19e (pfam01280)→ RplR (COG0256)rpl18p (PRK08569)→<br>RpsE (COG0098)rps5p (PRK04044)→ Ribosomal_L7_archeal_euk (cd01657)rpl30p (PRK06049)→<br>rpl15p (PRK06419)Ribosomal_L27A (pfam00828)→ SecY (COG0201)PRK08568 (PRK08568)→ |
| Hsa-7-1 | NZ_AOJE01000020.1 | - | 44999 | 44897 | **RNA**→ MTH1 (COG2106)→ rpl3p (PRK04231)Ribosomal_L3 (pfam00297)→<br>RplD (COG0088)rpl4lp (PRK04042)→ RplW (COG0089)PRK14548 (PRK14548)→<br>RplB (COG0090)rpl2p (PRK09612)→ RpsS (COG0185)rps19p (PRK04038)→<br>rpl22p (PRK04223)RplV (COG0091)→ RpsC (COG0092)rps3p (PRK04191)→<br>PRK14549 (PRK14549)Ribosomal_L29_HIP (cd00427)→ PRK03879 (PRK03879)UPF0086 (pfam01868)→<br>RpsQ (COG0186)rps17p (PRK08572)→ rpl14p (PRK08571)Ribosomal_L14 (pfam00238)→<br>rpl24p (PRK01191)KOW_RPL26 (cd06089)→ PRK04313 (PRK04313)Ribosomal_S4e (pfam00900)→<br>RplE (COG0094)rpl5p (PRK04219)→ RpsN (COG0199)rps14P (PRK05766)→<br>rps8p (PRK04034)RpsH (COG0096)→ RplF (COG0097)rpl6p (PRK05518)→<br>PRK12766 (PRK12766)RecA (COG0468)Rpl32e (COG1717)→ rpl19e (PRK08570)Ribosomal_L19e (pfam01280)→ |
| Hca-2-1 | NZ_AOJK01000017.1 | + | 53669 | 53771 | **RNA**→ MTH1 (COG2106)→ RplC (COG0087)rpl3p (PRK04231)→ RplD (COG0088)rpl4lp (PRK04042)→<br>RplW (COG0089)PRK14548 (PRK14548)→ RplB (COG0090)rpl2p (PRK09612)→<br>RplB (COG0090)rpl2p (PRK09612)→ RpsS (COG0185)rps19p (PRK04038)→<br>rpl22p (PRK04223)RplV (COG0091)→ RpsC (COG0092)rps3p (PRK04191)→<br>RpmC (COG0255)PRK14549 (PRK14549)→ PRK03879 (PRK03879)UPF0086 (pfam01868)→<br>RpsQ (COG0186)rps17p (PRK08572)→ rpl14p (PRK08571)Ribosomal_L14 (pfam00238)→<br>rpl24p (PRK01191)KOW_RPL26 (cd06089)→ PRK04313 (PRK04313)Ribosomal_S4e (pfam00900)→ |

| | | | | | |
|---|---|---|---|---|---|
| Hsp-7-1 | NZ_JH815330.1 | + | 360566 | 360668 | **RNA**→ MTH1 (COG2106)→ rpl3p (PRK04231)Ribosomal_L3 (pfam00297)→<br>RplC (COG0087)rpl3p (PRK04231)→ RplC (COG0087)rpl3p (PRK04231)→<br>RplD (COG0088)rpl4lp (PRK04042)→ RplW (COG0089)PRK14548 (PRK14548)→<br>RplB (COG0090)rpl2p (PRK09612)→ RplB (COG0090)rpl2p (PRK09612)→<br>RpsS (COG0185)rps19p (PRK04038)→ rpl22p (PRK04223)RplV (COG0091)→<br>RpsC (COG0092)rps3p (PRK04191)→ RpmC (COG0255)PRK14549 (PRK14549)→<br>PRK03879 (PRK03879)UPF0086 (pfam01868)→ RpsQ (COG0186)rps17p (PRK08572)→<br>rpl14p (PRK08571)Ribosomal_L14 (pfam00238)→ rpl24p (PRK01191)KOW_RPL26 (cd06089)→<br>PRK04313 (PRK04313)KOW_RPS4 (cd06087)Ribosomal_S4e (pfam00900)→ RplE (COG0094)rpl5p (PRK04219)→<br>RpsN (COG0199)rps14P (PRK05766)→ rps8p (PRK04034)RpsH (COG0096)→<br>RplF (COG0097)rpl6p (PRK05518)→ PRK12766 (PRK12766)RecA (COG0468)Rpl32e (COG1717)→<br>rpl19e (PRK08570)Ribosomal_L19e (pfam01280)→ |
| Hha-2-1 | NZ_BBJP01000034.1 | + | 23045 | 23147 | **RNA**→ MTH1 (COG2106)→ rpl3p (PRK04231)Ribosomal_L3 (pfam00297)→<br>RplD (COG0088)rpl4lp (PRK04042)→ RplW (COG0089)PRK14548 (PRK14548)→<br>RplB (COG0090)rpl2p (PRK09612)→ RpsS (COG0185)rps19p (PRK04038)→<br>rpl22p (PRK04223)RplV (COG0091)→ RpsC (COG0092)rps3p (PRK04191)→<br>PRK14549 (PRK14549)Ribosomal_L29_HIP (cd00427)→ PRK03879 (PRK03879)UPF0086 (pfam01868)→<br>RpsQ (COG0186)rps17p (PRK08572)→ rpl14p (PRK08571)Ribosomal_L14 (pfam00238)→<br>rpl24p (PRK01191)KOW_RPL26 (cd06089)→ PRK04313 (PRK04313)Ribosomal_S4e (pfam00900)→<br>RplE (COG0094)rpl5p (PRK04219)→ RpsN (COG0199)rps14P (PRK05766)→<br>rps8p (PRK04034)RpsH (COG0096)→ RplF (COG0097)rpl6p (PRK05518)→<br>PRK12766 (PRK12766)RecA (COG0468)Rpl32e (COG1717)→ rpl19e (PRK08570)Ribosomal_L19e (pfam01280)→ |
| Hdi-1-1 | NZ_AOJN01000013.1 | + | 82475 | 82577 | **RNA**→ MTH1 (COG2106)→ RplC (COG0087)rpl3p (PRK04231)→ RplD (COG0088)rpl4lp (PRK04042)→<br>RplW (COG0089)PRK14548 (PRK14548)→ RplB (COG0090)rpl2p (PRK09612)→<br>RplB (COG0090)rpl2p (PRK09612)→ RpsS (COG0185)rps19p (PRK04038)→<br>rpl22p (PRK04223)RplV (COG0091)→ RpsC (COG0092)rps3p (PRK04191)→<br>RpmC (COG0255)PRK14549 (PRK14549)→ PRK03879 (PRK03879)UPF0086 (pfam01868)→<br>RpsQ (COG0186)rps17p (PRK08572)→ rpl14p (PRK08571)Ribosomal_L14 (pfam00238)→<br>rpl24p (PRK01191)KOW_RPL26 (cd06089)→<br>PRK04313 (PRK04313)KOW_RPS4 (cd06087)Ribosomal_S4e (pfam00900)→ RplE (COG0094)rpl5p (PRK04219)→<br>RpsN (COG0199)rps14P (PRK05766)→ rps8p (PRK04034)RpsH (COG0096)→<br>RplF (COG0097)rpl6p (PRK05518)→ PRK12766 (PRK12766)RecA (COG0468)Rpl32e (COG1717)→<br>rpl19e (PRK08570)Ribosomal_L19e (pfam01280)→ |
| Hdi-2-1 | NZ_AOJM01000049.1 | - | 26024 | 25922 | **RNA**→ MTH1 (COG2106)→ RplC (COG0087)rpl3p (PRK04231)→ RplD (COG0088)rpl4lp (PRK04042)→<br>RplW (COG0089)PRK14548 (PRK14548)→ RplB (COG0090)rpl2p (PRK09612)→<br>RplB (COG0090)rpl2p (PRK09612)→ RpsS (COG0185)rps19p (PRK04038)→<br>rpl22p (PRK04223)RplV (COG0091)→ RpsC (COG0092)rps3p (PRK04191)→<br>RpmC (COG0255)PRK14549 (PRK14549)→ PRK03879 (PRK03879)UPF0086 (pfam01868)→<br>RpsQ (COG0186)rps17p (PRK08572)→ rpl14p (PRK08571)Ribosomal_L14 (pfam00238)→<br>rpl24p (PRK01191)KOW_RPL26 (cd06089)→<br>PRK04313 (PRK04313)KOW_RPS4 (cd06087)Ribosomal_S4e (pfam00900)→<br>rpl5p (PRK04219)Ribosomal_L5 (pfam00281)Ribosomal_L5_C (pfam00673)→<br>RpsN (COG0199)rps14P (PRK05766)→ rps8p (PRK04034)RpsH (COG0096)→<br>RplF (COG0097)rpl6p (PRK05518)→ PRK12766 (PRK12766)RecA (COG0468)Rpl32e (COG1717)→<br>rpl19e (PRK08570)Ribosomal_L19e (pfam01280)→ |

| | | | | | |
|---|---|---|---|---|---|
| Hsa-9-1 | NZ_JNFH02000042.1 | + | 4162 | 4264 | **RNA**→ hypo→ hypo→ rpl3p (PRK04231)Ribosomal_L3 (pfam00297)→ RplD (COG0088)rpl4lp (PRK04042)→<br>RplW (COG0089)PRK14548 (PRK14548)→ RplB (COG0090)rpl2p (PRK09612)→<br>RpsS (COG0185)rps19p (PRK04038)→ rpl22p (PRK04223)RplV (COG0091)→ hypo→<br>RpsC (COG0092)rps3p (PRK04191)→ PRK14549 (PRK14549)Ribosomal_L29_HIP (cd00427)→<br>PRK03879 (PRK03879)UPF0086 (pfam01868)→ RpsQ (COG0186)rps17p (PRK08572)→<br>rpl14p (PRK08571)Ribosomal_L14 (pfam00238)→ rpl24p (PRK01191)KOW_RPL26 (cd06089)→<br>PRK04313 (PRK04313)Ribosomal_S4e (pfam00900)→ RplE (COG0094)rpl5p (PRK04219)→<br>RpsN (COG0199)rps14P (PRK05766)→ rps8p (PRK04034)RpsH (COG0096)→<br>RplF (COG0097)rpl6p (PRK05518)→ PRK12766 (PRK12766)RecA (COG0468)Rpl32e (COG1717)→<br>rpl19e (PRK08570)Ribosomal_L19e (pfam01280)→ RplR (COG0256)rpl18p (PRK08569)→<br>RpsE (COG0098)rps5p (PRK04044)→ Ribosomal_L7_archeal_euk (cd01657)rpl30p (PRK06049)→<br>rpl15p (PRK06419)Ribosomal_L27A (pfam00828)→ SecY (COG0201)PRK08568 (PRK08568)→ |
| Har-2-1 | NZ_AOJJ01000075.1 | + | 39843 | 39945 | **RNA**→ MTH1 (COG2106)→ RplC (COG0087)rpl3p (PRK04231)→ RplD (COG0088)rpl4lp (PRK04042)→<br>RplW (COG0089)PRK14548 (PRK14548)→ RplB (COG0090)rpl2p (PRK09612)→<br>RplB (COG0090)rpl2p (PRK09612)→ RpsS (COG0185)rps19p (PRK04038)→<br>rpl22p (PRK04223)RplV (COG0091)→ RpsC (COG0092)rps3p (PRK04191)→<br>RpmC (COG0255)PRK14549 (PRK14549)→ PRK03879 (PRK03879)UPF0086 (pfam01868)→<br>RpsQ (COG0186)rps17p (PRK08572)→ rpl14p (PRK08571)Ribosomal_L14 (pfam00238)→<br>rpl24p (PRK01191)KOW_RPL26 (cd06089)→<br>PRK04313 (PRK04313)KOW_RPS4 (cd06087)Ribosomal_S4e (pfam00900)→ RplE (COG0094)rpl5p (PRK04219)→<br>RpsN (COG0199)rps14P (PRK05766)→ rps8p (PRK04034)RpsH (COG0096)→<br>RplF (COG0097)rpl6p (PRK05518)→ PRK12766 (PRK12766)RecA (COG0468)Rpl32e (COG1717)→<br>rpl19e (PRK08570)Ribosomal_L19e (pfam01280)→ |
| Hte-2-1 | NZ_AOIW01000054.1 | - | 29402 | 29300 | **RNA**→ MTH1 (COG2106)→ RplC (COG0087)rpl3p (PRK04231)→ RplD (COG0088)rpl4lp (PRK04042)→<br>RplW (COG0089)PRK14548 (PRK14548)→ RplB (COG0090)rpl2p (PRK09612)→<br>RplB (COG0090)rpl2p (PRK09612)→ RpsS (COG0185)rps19p (PRK04038)→<br>rpl22p (PRK04223)RplV (COG0091)→ RpsC (COG0092)rps3p (PRK04191)→<br>RpmC (COG0255)PRK14549 (PRK14549)→ PRK03879 (PRK03879)UPF0086 (pfam01868)→<br>RpsQ (COG0186)rps17p (PRK08572)→ rpl14p (PRK08571)Ribosomal_L14 (pfam00238)→<br>rpl24p (PRK01191)KOW_RPL26 (cd06089)→<br>PRK04313 (PRK04313)KOW_RPS4 (cd06087)Ribosomal_S4e (pfam00900)→ RplE (COG0094)rpl5p (PRK04219)→<br>RpsN (COG0199)rps14P (PRK05766)→ rps8p (PRK04034)RpsH (COG0096)→<br>RplF (COG0097)rpl6p (PRK05518)→ PRK12766 (PRK12766)RecA (COG0468)Rpl32e (COG1717)→<br>rpl19e (PRK08570)Ribosomal_L19e (pfam01280)→ |
| env-157 | P_1C_Liq_2_UnCty_1073497 | - | 265 | 163 | **RNA**→ Methyltrn_RNA_3 (pfam02598)→ |
| env-158 | SL_6KL_011_BRINE_1032700 | - | 1548 | 1446 | **RNA**→ Methyltrn_RNA_3 (pfam02598)→ RplC (COG0087)rpl3p (PRK04231)→ |
| env-159 | SL_4KL_010_BRINE_10001141 | + | 89 | 191 | **RNA**→ MTH1 (COG2106)→ RplC (COG0087)rpl3p (PRK04231)→ RplD (COG0088)rpl4lp (PRK04042)→<br>RplW (COG0089)PRK14548 (PRK14548)→ RplB (COG0090)rpl2p (PRK09612)→<br>RpsS (COG0185)rps19p (PRK04038)→ rpl22p (PRK04223)RplV (COG0091)→<br>RpsC (COG0092)rps3p (PRK04191)→ RpmC (COG0255)PRK14549 (PRK14549)→<br>PRK03879 (PRK03879)UPF0086 (pfam01868)→ RpsQ (COG0186)rps17p (PRK08572)→<br>rpl14p (PRK08571)Ribosomal_L14 (pfam00238)→ rpl24p (PRK01191)KOW_RPL26 (cd06089)→<br>PRK04313 (PRK04313)KOW_RPS4 (cd06087)Ribosomal_S4e (pfam00900)→ RplE (COG0094)rpl5p (PRK04219)→<br>RpsN (COG0199)rps14P (PRK05766)→ rps8p (PRK04034)RpsH (COG0096)→<br>RplF (COG0097)rpl6p (PRK05518)→ PRK12766 (PRK12766)RecA (COG0468)Rpl32e (COG1717)→<br>rpl19e (PRK08570)Ribosomal_L19e (pfam01280)→ RplR (COG0256)rpl18p (PRK08569)→<br>RpsE (COG0098)rps5p (PRK04044)→ Ribosomal_L7_archeal_euk (cd01657)rpl30p (PRK06049)→<br>rpl15p (PRK06419)Ribosomal_L27A (pfam00828)→ SecY (COG0201)PRK08568 (PRK08568)→ hypo→ |
| env-160 | SL_6KL_011_BRINE_1001616 | - | 387 | 285 | **RNA**→ MTH1 (COG2106)→ |

| | | | | | |
|---|---|---|---|---|---|
| h3A-1-1 | NZ_KK033115.1 | + | 666108 | 666210 | **RNA**→ Methyltrn_RNA_3 (pfam02598)→ rpl3p (PRK04231)Ribosomal_L3 (pfam00297)→ RplD (COG0088)rpl4lp (PRK04042)→ RplW (COG0089)PRK14548 (PRK14548)→ RplB (COG0090)rpl2p (PRK09612)→ RpsS (COG0185)rps19p (PRK04038)→ rpl22p (PRK04223)RplV (COG0091)→ RpsC (COG0092)rps3p (PRK04191)→ PRK14549 (PRK14549)Ribosomal_L29_HIP (cd00427)→ PRK03879 (PRK03879)UPF0086 (pfam01868)→ RpsQ (COG0186)rps17p (PRK08572)→ rpl14p (PRK08571)Ribosomal_L14 (pfam00238)→ RplX (COG0198)rpl24p (PRK01191)→ PRK04313 (PRK04313)KOW_RPS4 (cd06087)Ribosomal_S4e (pfam00900)→ RplE (COG0094)rpl5p (PRK04219)→ RpsN (COG0199)rps14P (PRK05766)→ rps8p (PRK04034)RpsH (COG0096)→ RplF (COG0097)rpl6p (PRK05518)→ PRK12766 (PRK12766)RecA (COG0468)Rpl32e (COG1717)→ rpl19e (PRK08570)Ribosomal_L19e (pfam01280)→ |
| Hai-1-1 | NZ_AOJI01000026.1 | - | 211502 | 211400 | **RNA**→ MTH1 (COG2106)→ RplC (COG0087)rpl3p (PRK04231)→ RplD (COG0088)rpl4lp (PRK04042)→ RplW (COG0089)PRK14548 (PRK14548)→ RplB (COG0090)rpl2p (PRK09612)→ RpsS (COG0185)rps19p (PRK04038)→ rpl22p (PRK04223)RplV (COG0091)→ RpsC (COG0092)rps3p (PRK04191)→ PRK14549 (PRK14549)Ribosomal_L29_HIP (cd00427)→ PRK03879 (PRK03879)UPF0086 (pfam01868)→ RpsQ (COG0186)rps17p (PRK08572)→ rpl14p (PRK08571)Ribosomal_L14 (pfam00238)→ RplX (COG0198)rpl24p (PRK01191)→ PRK04313 (PRK04313)KOW_RPS4 (cd06087)Ribosomal_S4e (pfam00900)→ RplE (COG0094)rpl5p (PRK04219)→ RpsN (COG0199)rps14P (PRK05766)→ rps8p (PRK04034)RpsH (COG0096)→ RplF (COG0097)rpl6p (PRK05518)→ PRK12766 (PRK12766)RecA (COG0468)Rpl32e (COG1717)→ rpl19e (PRK08570)Ribosomal_L19e (pfam01280)→ |
| env-161 | ADL20m3uS_contig08402 | - | 1754 | 1652 | **RNA**→ Methyltrn_RNA_3 (pfam02598)→ rpl3p (PRK04231)Ribosomal_L3 (pfam00297)→ RplC (COG0087)rpl3p (PRK04231)→ RplD (COG0088)rpl4lp (PRK04042)→ |
| env-162 | 2140918017-4520 | - | 13975 | 13873 | **RNA**→ Methyltrn_RNA_3 (pfam02598)→ rpl3p (PRK04231)Ribosomal_L3 (pfam00297)→ RplC (COG0087)rpl3p (PRK04231)→ RplD (COG0088)rpl4lp (PRK04042)→ RplW (COG0089)PRK14548 (PRK14548)→ |
| env-163 | 2140918027-00057 | - | 14028 | 13926 | **RNA**→ Methyltrn_RNA_3 (pfam02598)→ rpl3p (PRK04231)Ribosomal_L3 (pfam00297)→ RplC (COG0087)rpl3p (PRK04231)→ RplD (COG0088)rpl4lp (PRK04042)→ RplW (COG0089)PRK14548 (PRK14548)→ RplB (COG0090)rpl2p (PRK09612)→ RpsS (COG0185)rps19p (PRK04038)→ rpl22p (PRK04223)RplV (COG0091)→ RpsC (COG0092)rps3p (PRK04191)→ PRK14549 (PRK14549)Ribosomal_L29_HIP (cd00427)→ PRK03879 (PRK03879)UPF0086 (pfam01868)→ RpsQ (COG0186)rps17p (PRK08572)→ rpl14p (PRK08571)Ribosomal_L14 (pfam00238)→ rpl24p (PRK01191)KOW_RPL26 (cd06089)→ PRK04313 (PRK04313)Ribosomal_S4e (pfam00900)→ RplE (COG0094)rpl5p (PRK04219)→ RpsN (COG0199)rps14P (PRK05766)→ rps8p (PRK04034)RpsH (COG0096)→ RplF (COG0097)rpl6p (PRK05518)→ PRK12766 (PRK12766)RecA (COG0468)Rpl32e (COG1717)→ rpl19e (PRK08570)Ribosomal_L19e (pfam01280)→ |
| Hla-3-1 | NC_012029.1 | - | 2428716 | 2428614 | **RNA**→ Methyltrn_RNA_3 (pfam02598)→ rpl3p (PRK04231)Ribosomal_L3 (pfam00297)→ RplD (COG0088)rpl4lp (PRK04042)→ RplW (COG0089)PRK14548 (PRK14548)→ RplB (COG0090)rpl2p (PRK09612)→ RpsS (COG0185)rps19p (PRK04038)→ rpl22p (PRK04223)RplV (COG0091)→ RpsC (COG0092)rps3p (PRK04191)→ PRK14549 (PRK14549)Ribosomal_L29_HIP (cd00427)→ PRK03879 (PRK03879)UPF0086 (pfam01868)→ RpsQ (COG0186)rps17p (PRK08572)→ rpl14p (PRK08571)Ribosomal_L14 (pfam00238)→ rpl24p (PRK01191)KOW_RPL26 (cd06089)→ PRK04313 (PRK04313)Ribosomal_S4e (pfam00900)→ RplE (COG0094)rpl5p (PRK04219)→ RpsN (COG0199)rps14P (PRK05766)→ rps8p (PRK04034)RpsH (COG0096)→ RplF (COG0097)rpl6p (PRK05518)→ PRK12766 (PRK12766)RecA (COG0468)Rpl32e (COG1717)→ rpl19e (PRK08570)Ribosomal_L19e (pfam01280)→ |

| | | | | | |
|---|---|---|---|---|---|
| Hho-1-1 | NZ_AOJO01000045.1 | - | 129551 | 129449 | **RNA**→ MTH1 (COG2106)→ RplC (COG0087)rpl3p (PRK04231)→ RplD (COG0088)rpl4lp (PRK04042)→ RplW (COG0089)PRK14548 (PRK14548)→ RplB (COG0090)rpl2p (PRK09612)→ RpsS (COG0185)rps19p (PRK04038)→ rpl22p (PRK04223)RplV (COG0091)→ RpsC (COG0092)rps3p (PRK04191)→ PRK14549 (PRK14549)Ribosomal_L29_HIP (cd00427)→ PRK03879 (PRK03879)UPF0086 (pfam01868)→ RpsQ (COG0186)rps17p (PRK08572)→ rpl14p (PRK08571)Ribosomal_L14 (pfam00238)→ rpl24p (PRK01191)KOW_RPL26 (cd06089)→ PRK04313 (PRK04313)KOW_RPS4 (cd06087)Ribosomal_S4e (pfam00900)→ RplE (COG0094)rpl5p (PRK04219)→ RpsN (COG0199)rps14P (PRK05766)→ rps8p (PRK04034)RpsH (COG0096)→ RplF (COG0097)rpl6p (PRK05518)→ PRK12766 (PRK12766)RecA (COG0468)Rpl32e (COG1717)→ rpl19e (PRK08570)Ribosomal_L19e (pfam01280)→ |
| env-164 | SL_6KL_011_BRINE_1124153 | + | 36 | 138 | **RNA**→ PHA03252 (PHA03252)Methyltrn_RNA_3 (pfam02598)→ |
| env-165 | P_A23_Liq_2_Fmt_1045640 | + | 57 | 159 | **RNA**→ Methyltrn_RNA_3 (pfam02598)→ |
| env-166 | P_A23_Liq_1_Fmt_1009104 | + | 715 | 817 | **RNA**→ Methyltrn_RNA_3 (pfam02598)→ |
| env-167 | P_1C_Liq_1_UnCty_1002372 | + | 283 | 385 | **RNA**→ Methyltrn_RNA_3 (pfam02598)→ RplC (COG0087)rpl3p (PRK04231)→ RplD (COG0088)rpl4lp (PRK04042)→ RplW (COG0089)PRK14548 (PRK14548)→ RplB (COG0090)rpl2p (PRK09612)→ rps19p (PRK04038)→ |
| env-168 | P_A23_Liq_3_Fmt_1018522 | - | 551 | 449 | **RNA**→ Methyltrn_RNA_3 (pfam02598)→ |
| env-169 | P_2C_Liq_1_UnCty_1117457 | - | 344 | 242 | **RNA**→ Methyltrn_RNA_3 (pfam02598)→ |
| env-170 | P_A23_Liq_1_Fmt_1046280 | - | 249 | 147 | **RNA**→ Methyltrn_RNA_3 (pfam02598)→ |
| env-171 | P_A23_Liq_2_Fmt_1063179 | - | 245 | 143 | **RNA**→ Methyltrn_RNA_3 (pfam02598)→ |
| env-172 | P_2C_Liq_1_UnCty_1157684 | - | 265 | 163 | **RNA**→ Methyltrn_RNA_3 (pfam02598)→ |
| env-173 | SL_8KL_010_SED_10202168 | - | 335 | 236 | **RNA**→ Methyltrn_RNA_3 (pfam02598)→ |
| Hru-1-1 | NZ_BBJO01000003.1 | - | 101302 | 101203 | **RNA**→ Methyltrn_RNA_3 (pfam02598)→ RplC (COG0087)rpl3p (PRK04231)→ RplD (COG0088)rpl4lp (PRK04042)→ RplW (COG0089)PRK14548 (PRK14548)→ hypo→ hypo→ RpsS (COG0185)rps19p (PRK04038)→ rpl22p (PRK04223)RplV (COG0091)→ RpsC (COG0092)rps3p (PRK04191)→ PRK14549 (PRK14549)Ribosomal_L29 (pfam00831)→ PRK03879 (PRK03879)UPF0086 (pfam01868)→ RpsQ (COG0186)rps17p (PRK08572)→ rpl14p (PRK08571)Ribosomal_L14 (pfam00238)→ RplX (COG0198)rpl24p (PRK01191)→ PRK04313 (PRK04313)S4 (cd00165)KOW_RPS4 (cd06087)Ribosomal_S4e (pfam00900)→ RplE (COG0094)rpl5p (PRK04219)→ RpsN (COG0199)rps14P (PRK05766)→ rps8p (PRK04034)RpsH (COG0096)→ RplF (COG0097)rpl6p (PRK05518)→ PRK12766 (PRK12766)RecA (COG0468)Rpl32e (COG1717)→ rpl19e (PRK08570)Ribosomal_L19e (pfam01280)→ |
| env-174 | P_1C_Sed_2_UnCty_1300983 | + | 148 | 247 | **RNA**→ Methyltrn_RNA_3 (pfam02598)→ |
| Hru-2-1 | NZ_BBJN01000060.1 | + | 9207 | 9306 | **RNA**→ Methyltrn_RNA_3 (pfam02598)→ hypo→ rpl3p (PRK04231)Ribosomal_L3 (pfam00297)→ |
| env-175 | GSLNAS_F7I09SK02H8ZT2 | + | 136 | 236 | **RNA**→ Methyltrn_RNA_3 (pfam02598)→ |
| env-176 | SL_6KL_011_BRINE_1005460 | + | 5299 | 5398 | **RNA**→ Methyltrn_RNA_3 (pfam02598)→ |
| env-177 | SL_6KL_011_BRINE_1051712 | - | 689 | 590 | **RNA**→ Methyltrn_RNA_3 (pfam02598)→ rpl3p (PRK04231)Ribosomal_L3 (pfam00297)→ |
| env-178 | SL_4KL_010_BRINE_10003656 | - | 660 | 561 | **RNA**→ Methyltrn_RNA_3 (pfam02598)→ rpl3p (PRK04231)Ribosomal_L3 (pfam00297)→ |
| env-179 | JXWS01003911.1 | - | 291 | 192 | **RNA**→ Methyltrn_RNA_3 (pfam02598)→ |
| env-180 | SL_6KL_011_BRINE_1016959 | - | 122 | 23 | **RNA**→ Methyltrn_RNA_3 (pfam02598)→ |
| Nmo-1-1 | NC_020388.1 | - | 578139 | 578040 | **RNA**→ Methyltrn_RNA_3 (pfam02598)→ rpl3p (PRK04231)Ribosomal_L3 (pfam00297)→ RplD (COG0088)rpl4lp (PRK04042)→ RplW (COG0089)PRK14548 (PRK14548)→ RplB (COG0090)rpl2p (PRK09612)→ RpsS (COG0185)rps19p (PRK04038)→ rpl22p (PRK04223)RplV (COG0091)→ RpsC (COG0092)rps3p (PRK04191)→ PRK14549 (PRK14549)Ribosomal_L29 (pfam00831)→ PRK03879 (PRK03879)UPF0086 (pfam01868)→ RpsQ (COG0186)rps17p (PRK08572)→ rpl14p (PRK08571)Ribosomal_L14 (pfam00238)→ RplX (COG0198)rpl24p (PRK01191)→ PRK04313 (PRK04313)KOW_RPS4 (cd06087)Ribosomal_S4e (pfam00900)→ RplE (COG0094)rpl5p (PRK04219)→ RpsN (COG0199)rps14P (PRK05766)→ rps8p (PRK04034)RpsH (COG0096)→ RplF (COG0097)rpl6p (PRK05518)→ PolY_Pol_IV_kappa (cd03586)PRK12766 (PRK12766)Rpl32e (COG1717)→ rpl19e (PRK08570)Ribosomal_L19e (pfam01280)→ |

| | | | | | |
|---|---|---|---|---|---|
| Hha-1-1 | NZ_AJRK01000444.1 | - | 10548 | 10449 | **RNA**→ Methyltrn_RNA_3 (pfam02598)→ Methyltrn_RNA_3 (pfam02598)→ RplC (COG0087)rpl3p (PRK04231)→<br>RplC (COG0087)rpl3p (PRK04231)→ RplD (COG0088)rpl4lp (PRK04042)→<br>RplD (COG0088)rpl4lp (PRK04042)→ RplW (COG0089)PRK14548 (PRK14548)→<br>RplW (COG0089)PRK14548 (PRK14548)→ RplB (COG0090)rpl2p (PRK09612)→<br>RplB (COG0090)rpl2p (PRK09612)→ RpsS (COG0185)rps19p (PRK04038)→<br>RpsS (COG0185)rps19p (PRK04038)→ rpl22p (PRK04223)RplV (COG0091)→<br>rpl22p (PRK04223)RplV (COG0091)→<br>NT_POLXc (cd00141)RpsC (COG0092)rps3p (PRK04191)HHH_5 (pfam14520)→<br>NT_POLXc (cd00141)RpsC (COG0092)rps3p (PRK04191)HHH_5 (pfam14520)→<br>PRK14549 (PRK14549)Ribosomal_L29 (pfam00831)→ PRK14549 (PRK14549)Ribosomal_L29 (pfam00831)→<br>PRK03879 (PRK03879)UPF0086 (pfam01868)→ PRK03879 (PRK03879)UPF0086 (pfam01868)→<br>RpsQ (COG0186)rps17p (PRK08572)→ RpsQ (COG0186)rps17p (PRK08572)→<br>rpl14p (PRK08571)Ribosomal_L14 (pfam00238)→ rpl14p (PRK08571)Ribosomal_L14 (pfam00238)→<br>RplX (COG0198)rpl24p (PRK01191)→ RplX (COG0198)rpl24p (PRK01191)→<br>PRK04313 (PRK04313)KOW_RPS4 (cd06087)S4 (pfam01479)Ribosomal_S4e (pfam00900)→<br>PRK04313 (PRK04313)KOW_RPS4 (cd06087)S4 (pfam01479)Ribosomal_S4e (pfam00900)→<br>RplE (COG0094)rpl5p (PRK04219)→ RplE (COG0094)rpl5p (PRK04219)→<br>RpsN (COG0199)rps14P (PRK05766)→ RpsN (COG0199)rps14P (PRK05766)→<br>rps8p (PRK04034)RpsH (COG0096)→ rps8p (PRK04034)RpsH (COG0096)→<br>RplF (COG0097)rpl6p (PRK05518)→ RplF (COG0097)rpl6p (PRK05518)→<br>Lig (COG0272)Rpl32e (COG1717)rpl32e (PRK08562)HHH_5 (pfam14520)→<br>Lig (COG0272)Rpl32e (COG1717)rpl32e (PRK08562)HHH_5 (pfam14520)→ |
| Hha-1-2 | NZ_AOMB01000043.1 | - | 108188 | 108089 | **RNA**→ Methyltrn_RNA_3 (pfam02598)→ RplC (COG0087)rpl3p (PRK04231)→<br>RplD (COG0088)rpl4lp (PRK04042)→ RplW (COG0089)PRK14548 (PRK14548)→<br>RplB (COG0090)rpl2p (PRK09612)→ RpsS (COG0185)rps19p (PRK04038)→<br>rpl22p (PRK04223)RplV (COG0091)→<br>NT_POLXc (cd00141)RpsC (COG0092)rps3p (PRK04191)HHH_5 (pfam14520)→<br>PRK14549 (PRK14549)Ribosomal_L29 (pfam00831)→ PRK03879 (PRK03879)UPF0086 (pfam01868)→<br>RpsQ (COG0186)rps17p (PRK08572)→ rpl14p (PRK08571)Ribosomal_L14 (pfam00238)→<br>RplX (COG0198)rpl24p (PRK01191)→<br>PRK04313 (PRK04313)KOW_RPS4 (cd06087)S4 (pfam01479)Ribosomal_S4e (pfam00900)→<br>RplE (COG0094)rpl5p (PRK04219)→ RpsN (COG0199)rps14P (PRK05766)→<br>rps8p (PRK04034)RpsH (COG0096)→ RplF (COG0097)rpl6p (PRK05518)→<br>Lig (COG0272)Rpl32e (COG1717)rpl32e (PRK08562)HHH_5 (pfam14520)→ |
| env-181 | SL_4KL_010_BRINE_10212872 | - | 267 | 168 | **RNA**→ Methyltrn_RNA_3 (pfam02598)→ |

| | | | | | |
|---|---|---|---|---|---|
| Hpa-1-1 | NZ_AEMG01000030.1 | - | 41019 | 40920 | **RNA**→ Methyltrn_RNA_3 (pfam02598)→ Methyltrn_RNA_3 (pfam02598)→<br>rpl3p (PRK04231)Ribosomal_L3 (pfam00297)→ rpl3p (PRK04231)Ribosomal_L3 (pfam00297)→<br>RplD (COG0088)rpl4lp (PRK04042)→ RplD (COG0088)rpl4lp (PRK04042)→<br>RplW (COG0089)PRK14548 (PRK14548)→ RplW (COG0089)PRK14548 (PRK14548)→<br>RplB (COG0090)rpl2p (PRK09612)→ RplB (COG0090)rpl2p (PRK09612)→<br>RpsS (COG0185)rps19p (PRK04038)→ RpsS (COG0185)rps19p (PRK04038)→<br>rpl22p (PRK04223)RplV (COG0091)→ rpl22p (PRK04223)RplV (COG0091)→<br>RpsC (COG0092)rps3p (PRK04191)→ RpsC (COG0092)rps3p (PRK04191)→<br>PRK14549 (PRK14549)Ribosomal_L29 (pfam00831)→ PRK14549 (PRK14549)Ribosomal_L29 (pfam00831)→<br>POP4 (COG1588)PRK03879 (PRK03879)→ POP4 (COG1588)PRK03879 (PRK03879)→<br>RpsQ (COG0186)rps17p (PRK08572)→ RpsQ (COG0186)rps17p (PRK08572)→<br>rpl14p (PRK08571)Ribosomal_L14 (pfam00238)→ rpl14p (PRK08571)Ribosomal_L14 (pfam00238)→<br>RplX (COG0198)rpl24p (PRK01191)→ RplX (COG0198)rpl24p (PRK01191)→<br>PRK04313 (PRK04313)KOW_RPS4 (cd06087)S4 (pfam01479)Ribosomal_S4e (pfam00900)→<br>PRK04313 (PRK04313)KOW_RPS4 (cd06087)S4 (pfam01479)Ribosomal_S4e (pfam00900)→<br>RplE (COG0094)rpl5p (PRK04219)→ RplE (COG0094)rpl5p (PRK04219)→<br>RpsN (COG0199)rps14P (PRK05766)→ RpsN (COG0199)rps14P (PRK05766)→<br>rps8p (PRK04034)RpsH (COG0096)→ rps8p (PRK04034)RpsH (COG0096)→<br>RplF (COG0097)rpl6p (PRK05518)→ RplF (COG0097)rpl6p (PRK05518)→<br>RecA (COG0468)rpl32e (PRK08562)Ribosomal_L32e (pfam01655)HHH_5 (pfam14520)→<br>RecA (COG0468)rpl32e (PRK08562)Ribosomal_L32e (pfam01655)HHH_5 (pfam14520)→<br>rpl19e (PRK08570)Ribosomal_L19e (pfam01280)→ rpl19e (PRK08570)Ribosomal_L19e (pfam01280)→ |
| Hpa-1-2 | NZ_AQXI01000001.1 | - | 1281236 | 1281137 | **RNA**→ Methyltrn_RNA_3 (pfam02598)→ rpl3p (PRK04231)Ribosomal_L3 (pfam00297)→<br>RplC (COG0087)rpl3p (PRK04231)→ rpl3p (PRK04231)Ribosomal_L3 (pfam00297)→<br>RplD (COG0088)rpl4lp (PRK04042)→ RplW (COG0089)PRK14548 (PRK14548)→<br>RplB (COG0090)rpl2p (PRK09612)→ RpsS (COG0185)rps19p (PRK04038)→<br>rpl22p (PRK04223)RplV (COG0091)→ RpsC (COG0092)rps3p (PRK04191)→<br>PRK14549 (PRK14549)Ribosomal_L29 (pfam00831)→ POP4 (COG1588)PRK03879 (PRK03879)→<br>RpsQ (COG0186)rps17p (PRK08572)→ RpsQ (COG0186)rps17p (PRK08572)→<br>rpl14p (PRK08571)Ribosomal_L14 (pfam00238)→ RplX (COG0198)rpl24p (PRK01191)→<br>PRK04313 (PRK04313)KOW_RPS4 (cd06087)S4 (pfam01479)Ribosomal_S4e (pfam00900)→<br>RplE (COG0094)rpl5p (PRK04219)→ RpsN (COG0199)rps14P (PRK05766)→<br>rps8p (PRK04034)RpsH (COG0096)→ RplF (COG0097)rpl6p (PRK05518)→<br>RecA (COG0468)rpl32e (PRK08562)Ribosomal_L32e (pfam01655)HHH_5 (pfam14520)→<br>rpl19e (PRK08570)Ribosomal_L19e (pfam01280)→ |
| Hci-1-1 | NZ_JDTH01000001.1 | + | 807675 | 807774 | **RNA**→ Methyltrn_RNA_3 (pfam02598)→ rpl3p (PRK04231)Ribosomal_L3 (pfam00297)→<br>RplD (COG0088)rpl4lp (PRK04042)→ RplW (COG0089)PRK14548 (PRK14548)→<br>RplB (COG0090)rpl2p (PRK09612)→ RpsS (COG0185)rps19p (PRK04038)→<br>rpl22p (PRK04223)RplV (COG0091)→ RpsC (COG0092)rps3p (PRK04191)→<br>PRK14549 (PRK14549)Ribosomal_L29 (pfam00831)→ POP4 (COG1588)PRK03879 (PRK03879)→<br>RpsQ (COG0186)rps17p (PRK08572)→ rpl14p (PRK08571)Ribosomal_L14 (pfam00238)→<br>rpl24p (PRK01191)KOW_RPL26 (cd06089)→<br>PRK04313 (PRK04313)KOW_RPS4 (cd06087)Ribosomal_S4e (pfam00900)→ RplE (COG0094)rpl5p (PRK04219)→<br>RpsN (COG0199)rps14P (PRK05766)→ rps8p (PRK04034)RpsH (COG0096)→<br>RplF (COG0097)rpl6p (PRK05518)→ UvrC (COG0322)Ribosomal_L32_L32e (cd00513)HHH_5 (pfam14520)→<br>rpl19e (PRK08570)Ribosomal_L19e (pfam01280)→ |
| env-182 | GRANPEH02G2JJD | + | 188 | 287 | **RNA**→ MTH1 (COG2106)→ |
| env-183 | GQKUMWE02FE1GX | - | 166 | 68 | **RNA**→ hypo→ |
| env-184 | GSLNARP_GFPJP1N02H7ZY7 | - | 182 | 83 | **RNA**→ Methyltrn_RNA_3 (pfam02598)→ |
| env-185 | SL_5KL_010_BRINE_10021216 | + | 2341 | 2440 | **RNA**→ Methyltrn_RNA_3 (pfam02598)→ |

| | | | | | |
|---|---|---|---|---|---|
| Hdj-1-1 | NZ_HG315688.1 | - | 94795 | 94696 | **RNA**→ Methyltrn_RNA_3 (pfam02598)→ rpl3p (PRK04231)Ribosomal_L3 (pfam00297)→ |
| | | | | | RplD (COG0088)rpl4lp (PRK04042)→ RplW (COG0089)PRK14548 (PRK14548)→ |
| | | | | | RplB (COG0090)rpl2p (PRK09612)→ RpsS (COG0185)rps19p (PRK04038)→ |
| | | | | | rpl22p (PRK04223)RplV (COG0091)→ RpsC (COG0092)rps3p (PRK04191)→ |
| | | | | | RpmC (COG0255)PRK14549 (PRK14549)→ PRK03879 (PRK03879)UPF0086 (pfam01868)→ |
| | | | | | RpsQ (COG0186)rps17p (PRK08572)→ rpl14p (PRK08571)Ribosomal_L14 (pfam00238)→ |
| | | | | | rpl24p (PRK01191)KOW_RPL26 (cd06089)→ |
| | | | | | PRK04313 (PRK04313)KOW_RPS4 (cd06087)S4 (pfam01479)Ribosomal_S4e (pfam00900)→ |
| | | | | | RplE (COG0094)rpl5p (PRK04219)SDH (cd12188)→ RpsN (COG0199)rps14P (PRK05766)→ |
| | | | | | rps8p (PRK04034)RpsH (COG0096)→ RplF (COG0097)rpl6p (PRK05518)→ |
| | | | | | PRK12766 (PRK12766)RecA (COG0468)Rpl32e (COG1717)→ rpl19e (PRK08570)Ribosomal_L19e (pfam01280)→ |
| Hsa-10-1 | NZ_JROF01000007.1 | - | 139445 | 139346 | **RNA**→ Methyltrn_RNA_3 (pfam02598)→ rpl3p (PRK04231)Ribosomal_L3 (pfam00297)→ |
| | | | | | RplD (COG0088)rpl4lp (PRK04042)→ RplW (COG0089)PRK14548 (PRK14548)→ |
| | | | | | RplB (COG0090)rpl2p (PRK09612)→ RpsS (COG0185)rps19p (PRK04038)→ |
| | | | | | rpl22p (PRK04223)RplV (COG0091)→ RpsC (COG0092)rps3p (PRK04191)→ |
| | | | | | PRK14549 (PRK14549)Ribosomal_L29 (pfam00831)→ PRK03879 (PRK03879)UPF0086 (pfam01868)→ |
| | | | | | RpsQ (COG0186)rps17p (PRK08572)→ rpl14p (PRK08571)Ribosomal_L14 (pfam00238)→ |
| | | | | | rpl24p (PRK01191)KOW_RPL26 (cd06089)→ |
| | | | | | PRK04313 (PRK04313)KOW_RPS4 (cd06087)Ribosomal_S4e (pfam00900)→ RplE (COG0094)rpl5p (PRK04219)→ |
| | | | | | RpsN (COG0199)rps14P (PRK05766)→ rps8p (PRK04034)RpsH (COG0096)→ |
| | | | | | RplF (COG0097)rpl6p (PRK05518)→ PRK12766 (PRK12766)RecA (COG0468)Rpl32e (COG1717)→ |
| | | | | | rpl19e (PRK08570)Ribosomal_L19e (pfam01280)→ |
| Nsu-1-1 | NZ_AOHX01000025.1 | + | 4846 | 4945 | **RNA**→ Methyltrn_RNA_3 (pfam02598)→ rpl3p (PRK04231)Ribosomal_L3 (pfam00297)→ |
| | | | | | RplD (COG0088)rpl4lp (PRK04042)→ RplW (COG0089)PRK14548 (PRK14548)→ |
| | | | | | RplB (COG0090)rpl2p (PRK09612)→ RpsS (COG0185)rps19p (PRK04038)→ |
| | | | | | rpl22p (PRK04223)RplV (COG0091)→ RpsC (COG0092)rps3p (PRK04191)→ |
| | | | | | RpmC (COG0255)PRK14549 (PRK14549)→ PRK03879 (PRK03879)UPF0086 (pfam01868)→ |
| | | | | | RpsQ (COG0186)rps17p (PRK08572)→ rpl14p (PRK08571)Ribosomal_L14 (pfam00238)→ |
| | | | | | rpl24p (PRK01191)KOW_RPL26 (cd06089)→ |
| | | | | | PRK04313 (PRK04313)KOW_RPS4 (cd06087)Ribosomal_S4e (pfam00900)→ RplE (COG0094)rpl5p (PRK04219)→ |
| | | | | | RpsN (COG0199)rps14P (PRK05766)→ rps8p (PRK04034)RpsH (COG0096)→ |
| | | | | | RplF (COG0097)rpl6p (PRK05518)→ |
| | | | | | UvrC (COG0322)Rpl32e (COG1717)rpl32e (PRK08562)HHH_5 (pfam14520)→ |
| | | | | | rpl19e (PRK08570)Ribosomal_L19e (pfam01280)→ RplR (COG0256)rpl18p (PRK08569)→ |
| | | | | | RpsE (COG0098)rps5p (PRK04044)→ Ribosomal_L7_archeal_euk (cd01657)rpl30p (PRK06049)→ |
| | | | | | rpl15p (PRK06419)Ribosomal_L27A (pfam00828)→ SecY (COG0201)PRK08568 (PRK08568)→ |
| Nba-1-1 | NZ_AOHY01000055.1 | + | 4766 | 4865 | **RNA**→ Methyltrn_RNA_3 (pfam02598)→ rpl3p (PRK04231)Ribosomal_L3 (pfam00297)→ |
| | | | | | RplD (COG0088)rpl4lp (PRK04042)→ RplW (COG0089)PRK14548 (PRK14548)→ |
| | | | | | RplB (COG0090)rpl2p (PRK09612)→ RpsS (COG0185)rps19p (PRK04038)→ |
| | | | | | rpl22p (PRK04223)RplV (COG0091)→ RpsC (COG0092)rps3p (PRK04191)→ |
| | | | | | RpmC (COG0255)PRK14549 (PRK14549)→ PRK03879 (PRK03879)UPF0086 (pfam01868)→ |
| | | | | | RpsQ (COG0186)rps17p (PRK08572)→ rpl14p (PRK08571)Ribosomal_L14 (pfam00238)→ |
| | | | | | rpl24p (PRK01191)KOW_RPL26 (cd06089)→ |
| | | | | | PRK04313 (PRK04313)KOW_RPS4 (cd06087)S4 (pfam01479)Ribosomal_S4e (pfam00900)→ |
| | | | | | RplE (COG0094)rpl5p (PRK04219)→ RpsN (COG0199)rps14P (PRK05766)→ |
| | | | | | rps8p (PRK04034)RpsH (COG0096)→ RplF (COG0097)rpl6p (PRK05518)→ |
| | | | | | RecA (COG0468)Rpl32e (COG1717)rpl32e (PRK08562)HHH_5 (pfam14520)→ |
| | | | | | rpl19e (PRK08570)Ribosomal_L19e (pfam01280)→ RplR (COG0256)rpl18p (PRK08569)→ |
| | | | | | RpsE (COG0098)rps5p (PRK04044)→ Ribosomal_L7_archeal_euk (cd01657)rpl30p (PRK06049)→ |
| | | | | | rpl15p (PRK06419)Ribosomal_L27A (pfam00828)→ SecY (COG0201)PRK08568 (PRK08568)→ |
| | | | | | SecY (COG0201)PRK08568 (PRK08568)→ SecY (COG0201)PRK08568 (PRK08568)→ |

| | | | | | |
|---|---|---|---|---|---|
| Nin-1-1 | NZ_AOHZ01000041.1 | - | 187314 | 187215 | **RNA**→ Methyltrn_RNA_3 (pfam02598)→ rpl3p (PRK04231)Ribosomal_L3 (pfam00297)→ RplD (COG0088)rpl4lp (PRK04042)→ RplW (COG0089)PRK14548 (PRK14548)→ RplB (COG0090)rpl2p (PRK09612)→ RpsS (COG0185)rps19p (PRK04038)→ rpl22p (PRK04223)RplV (COG0091)→ RpsC (COG0092)rps3p (PRK04191)→ RpmC (COG0255)PRK14549 (PRK14549)→ PRK03879 (PRK03879)UPF0086 (pfam01868)→ RpsQ (COG0186)rps17p (PRK08572)→ rpl14p (PRK08571)Ribosomal_L14 (pfam00238)→ rpl24p (PRK01191)KOW_RPL26 (cd06089)→ PRK04313 (PRK04313)S4 (cd00165)KOW_RPS4 (cd06087)Ribosomal_S4e (pfam00900)→ RplE (COG0094)rpl5p (PRK04219)→ RpsN (COG0199)rps14P (PRK05766)→ rps8p (PRK04034)RpsH (COG0096)→ RplF (COG0097)rpl6p (PRK05518)→ RecA (COG0468)Rpl32e (COG1717)rpl32e (PRK08562)HHH_5 (pfam14520)→ rpl19e (PRK08570)Ribosomal_L19e (pfam01280)→ |
| env-186 | SL_4KL_010_BRINE_10024882 | - | 1898 | 1799 | **RNA**→ Methyltrn_RNA_3 (pfam02598)→ rpl3p (PRK04231)Ribosomal_L3 (pfam00297)→ RplD (COG0088)rpl4lp (PRK04042)→ |
| env-187 | ADL20m3uS_GFI381O02JC3VL | + | 316 | 416 | **RNA**→ |
| env-188 | SL_6KL_011_BRINE_1000304 | + | 9714 | 9813 | **RNA**→ Methyltrn_RNA_3 (pfam02598)→ rpl3p (PRK04231)Ribosomal_L3 (pfam00297)→ RplD (COG0088)rpl4lp (PRK04042)→ RplW (COG0089)PRK14548 (PRK14548)→ RplB (COG0090)rpl2p (PRK09612)→ RpsS (COG0185)rps19p (PRK04038)→ rpl22p (PRK04223)RplV (COG0091)→ RpsC (COG0092)rps3p (PRK04191)→ PRK14549 (PRK14549)Ribosomal_L29 (pfam00831)→ PRK03879 (PRK03879)UPF0086 (pfam01868)→ RpsQ (COG0186)rps17p (PRK08572)→ rpl14p (PRK08571)Ribosomal_L14 (pfam00238)→ rpl24p (PRK01191)KOW_RPL26 (cd06089)→ PRK04313 (PRK04313)KOW_RPS4 (cd06087)S4 (pfam01479)Ribosomal_S4e (pfam00900)→ RplE (COG0094)rpl5p (PRK04219)→ RpsN (COG0199)rps14P (PRK05766)→ rps8p (PRK04034)RpsH (COG0096)→ RplF (COG0097)rpl6p (PRK05518)→ UvrC (COG0322)Rpl32e (COG1717)rpl32e (PRK08562)HHH_5 (pfam14520)→ rpl19e (PRK08570)Ribosomal_L19e (pfam01280)→ |
| env-189 | SL_8KL_010_SED_10375651 | - | 274 | 175 | **RNA**→ Methyltrn_RNA_3 (pfam02598)→ |
| env-190 | SL_6KL_011_BRINE_1014318 | - | 155 | 56 | **RNA**→ Methyltrn_RNA_3 (pfam02598)→ |
| env-191 | SL_8KL_010_SED_10000480 | + | 6111 | 6210 | **RNA**→ Methyltrn_RNA_3 (pfam02598)→ rpl3p (PRK04231)Ribosomal_L3 (pfam00297)→ RplD (COG0088)rpl4lp (PRK04042)→ RplW (COG0089)PRK14548 (PRK14548)→ RplB (COG0090)rpl2p (PRK09612)→ RpsS (COG0185)rps19p (PRK04038)→ rpl22p (PRK04223)RplV (COG0091)→ RpsC (COG0092)rps3p (PRK04191)→ RpmC (COG0255)PRK14549 (PRK14549)→ PRK03879 (PRK03879)UPF0086 (pfam01868)→ RpsQ (COG0186)rps17p (PRK08572)→ rpl14p (PRK08571)Ribosomal_L14 (pfam00238)→ rpl24p (PRK01191)KOW_RPL26 (cd06089)→ PRK04313 (PRK04313)KOW_RPS4 (cd06087)S4 (pfam01479)Ribosomal_S4e (pfam00900)→ RplE (COG0094)rpl5p (PRK04219)→ RpsN (COG0199)rps14P (PRK05766)→ rps8p (PRK04034)RpsH (COG0096)→ RplF (COG0097)rpl6p (PRK05518)→ PRK12766 (PRK12766)UvrC (COG0322)Rpl32e (COG1717)→ rpl19e (PRK08570)Ribosomal_L19e (pfam01280)→ RplR (COG0256)rpl18p (PRK08569)→ RpsE (COG0098)rps5p (PRK04044)→ Ribosomal_L7_archeal_euk (cd01657)rpl30p (PRK06049)→ rpl15p (PRK06419)Ribosomal_L27A (pfam00828)→ SecY (COG0201)PRK08568 (PRK08568)→ |
| env-192 | SL_4KL_010_BRINE_10005807 | - | 721 | 622 | **RNA**→ Methyltrn_RNA_3 (pfam02598)→ rpl3p (PRK04231)Ribosomal_L3 (pfam00297)→ |

| | | | | | |
|---|---|---|---|---|---|
| Hla-1-1 | NZ_AGFZ01000103.1 | - | 104403 | 104304 | **RNA**→ Methyltrn_RNA_3 (pfam02598)→ Methyltrn_RNA_3 (pfam02598)→ RplC (COG0087)rpl3p (PRK04231)→<br>RplC (COG0087)rpl3p (PRK04231)→ RplD (COG0088)rpl4lp (PRK04042)→<br>RplD (COG0088)rpl4lp (PRK04042)→ RplW (COG0089)PRK14548 (PRK14548)→<br>RplW (COG0089)PRK14548 (PRK14548)→ RplB (COG0090)rpl2p (PRK09612)→<br>RplB (COG0090)rpl2p (PRK09612)→ RpsS (COG0185)rps19p (PRK04038)→<br>RpsS (COG0185)rps19p (PRK04038)→ rpl22p (PRK04223)RplV (COG0091)→<br>rpl22p (PRK04223)RplV (COG0091)→ RpsC (COG0092)rps3p (PRK04191)→<br>RpsC (COG0092)rps3p (PRK04191)→ PRK14549 (PRK14549)Ribosomal_L29 (pfam00831)→<br>PRK14549 (PRK14549)Ribosomal_L29 (pfam00831)→ PRK03879 (PRK03879)UPF0086 (pfam01868)→<br>PRK03879 (PRK03879)UPF0086 (pfam01868)→ RpsQ (COG0186)rps17p (PRK08572)→<br>RpsQ (COG0186)rps17p (PRK08572)→ rpl14p (PRK08571)Ribosomal_L14 (pfam00238)→<br>rpl14p (PRK08571)Ribosomal_L14 (pfam00238)→ rpl24p (PRK01191)KOW_RPL26 (cd06089)→<br>rpl24p (PRK01191)KOW_RPL26 (cd06089)→<br>PRK04313 (PRK04313)S4 (cd00165)KOW_RPS4 (cd06087)Ribosomal_S4e (pfam00900)→<br>PRK04313 (PRK04313)S4 (cd00165)KOW_RPS4 (cd06087)Ribosomal_S4e (pfam00900)→<br>RplE (COG0094)rpl5p (PRK04219)→ RplE (COG0094)rpl5p (PRK04219)→<br>RpsN (COG0199)rps14P (PRK05766)→ RpsN (COG0199)rps14P (PRK05766)→<br>rps8p (PRK04034)RpsH (COG0096)→ rps8p (PRK04034)RpsH (COG0096)→<br>RplF (COG0097)rpl6p (PRK05518)→ RplF (COG0097)rpl6p (PRK05518)→<br>RecA (COG0468)Rpl32e (COG1717)rpl32e (PRK08562)HHH_5 (pfam14520)→<br>RecA (COG0468)Rpl32e (COG1717)rpl32e (PRK08562)HHH_5 (pfam14520)→ |
| Hla-1-2 | NZ_AOLZ01000040.1 | - | 62859 | 62760 | **RNA**→ Methyltrn_RNA_3 (pfam02598)→ RplC (COG0087)rpl3p (PRK04231)→<br>RplD (COG0088)rpl4lp (PRK04042)→ RplW (COG0089)PRK14548 (PRK14548)→<br>RplB (COG0090)rpl2p (PRK09612)→ RpsS (COG0185)rps19p (PRK04038)→<br>rpl22p (PRK04223)RplV (COG0091)→ RpsC (COG0092)rps3p (PRK04191)→<br>PRK14549 (PRK14549)Ribosomal_L29 (pfam00831)→ PRK03879 (PRK03879)UPF0086 (pfam01868)→<br>RpsQ (COG0186)rps17p (PRK08572)→ rpl14p (PRK08571)Ribosomal_L14 (pfam00238)→<br>rpl24p (PRK01191)KOW_RPL26 (cd06089)→<br>PRK04313 (PRK04313)S4 (cd00165)KOW_RPS4 (cd06087)Ribosomal_S4e (pfam00900)→<br>RplE (COG0094)rpl5p (PRK04219)→ RpsN (COG0199)rps14P (PRK05766)→<br>rps8p (PRK04034)RpsH (COG0096)→ RplF (COG0097)rpl6p (PRK05518)→<br>RecA (COG0468)Rpl32e (COG1717)rpl32e (PRK08562)HHH_5 (pfam14520)→ |
| env-193 | SL_5KL_010_BRINE_10000580 | - | 14326 | 14227 | **RNA**→ Methyltrn_RNA_3 (pfam02598)→ rpl3p (PRK04231)Ribosomal_L3 (pfam00297)→<br>RplD (COG0088)rpl4lp (PRK04042)→ RplW (COG0089)PRK14548 (PRK14548)→<br>RplB (COG0090)rpl2p (PRK09612)→ RpsS (COG0185)rps19p (PRK04038)→<br>rpl22p (PRK04223)RplV (COG0091)→ RpsC (COG0092)rps3p (PRK04191)→<br>RpmC (COG0255)PRK14549 (PRK14549)→ PRK03879 (PRK03879)UPF0086 (pfam01868)→<br>RpsQ (COG0186)rps17p (PRK08572)→ rpl14p (PRK08571)Ribosomal_L14 (pfam00238)→<br>RplX (COG0198)rpl24p (PRK01191)→<br>PRK04313 (PRK04313)S4 (cd00165)KOW_RPS4 (cd06087)Ribosomal_S4e (pfam00900)→<br>RplE (COG0094)rpl5p (PRK04219)→ RpsN (COG0199)rps14P (PRK05766)→<br>rps8p (PRK04034)RpsH (COG0096)→ RplF (COG0097)rpl6p (PRK05518)→<br>PRK12766 (PRK12766)UvrC (COG0322)Rpl32e (COG1717)→ |
| env-194 | SL_5KL_010_SED_10042598 | + | 887 | 986 | **RNA**→ Methyltrn_RNA_3 (pfam02598)→ |
| env-195 | SL_4KL_010_BRINE_10006165 | + | 1728 | 1827 | **RNA**→ Methyltrn_RNA_3 (pfam02598)→ rpl3p (PRK04231)Ribosomal_L3 (pfam00297)→<br>RplD (COG0088)rpl4lp (PRK04042)→ RplW (COG0089)PRK14548 (PRK14548)→<br>RplB (COG0090)rpl2p (PRK09612)→ RpsS (COG0185)rps19p (PRK04038)→<br>rpl22p (PRK04223)RplV (COG0091)→ RpsC (COG0092)rps3p (PRK04191)→<br>PRK14549 (PRK14549)Ribosomal_L29_HIP (cd00427)→ |
| Hac-1-1 | NZ_BATA01000095.1 | + | 7771 | 7870 | **RNA**→ MTH1 (COG2106)→ hypo→ hypo→ |

| | | | | | |
|---|---|---|---|---|---|
| Hac-1-2 | NZ_BANO01000013.1 | - | 14331 | 14232 | **RNA**→ Methyltrn_RNA_3 (pfam02598)→ rpl3p (PRK04231)Ribosomal_L3 (pfam00297)→<br>RplC (COG0087)rpl3p (PRK04231)→ RplC (COG0087)rpl3p (PRK04231)→<br>RplD (COG0088)rpl4lp (PRK04042)→ RplW (COG0089)PRK14548 (PRK14548)→<br>RplB (COG0090)rpl2p (PRK09612)→ RpsS (COG0185)rps19p (PRK04038)→<br>rpl22p (PRK04223)RplV (COG0091)→ RpsC (COG0092)rps3p (PRK04191)→<br>PRK14549 (PRK14549)Ribosomal_L29 (pfam00831)→ POP4 (COG1588)PRK03879 (PRK03879)→<br>RpsQ (COG0186)rps17p (PRK08572)→ rps17p (PRK08572)Ribosomal_S17 (pfam00366)→<br>rpl14p (PRK08571)Ribosomal_L14 (pfam00238)→ RplX (COG0198)rpl24p (PRK01191)→<br>PRK04313 (PRK04313)KOW_RPS4 (cd06087)Ribosomal_S4e (pfam00900)→ RplE (COG0094)rpl5p (PRK04219)→<br>RpsN (COG0199)rps14P (PRK05766)→ rps8p (PRK04034)RpsH (COG0096)→<br>RplF (COG0097)rpl6p (PRK05518)→ |
| env-196 | ADL20m3uS_GFI381O02IYW5A | - | 121 | 22 | **RNA**→ MTH1 (COG2106)→ |
| env-197 | 2140918017-63356 | - | 193 | 94 | **RNA**→ hypo→ |
| env-198 | GQKUMWE02GP8XW | - | 475 | 376 | **RNA**→ Methyltrn_RNA_3 (pfam02598)→ |
| Hsp-2-1 | NZ_CP007060.1 | + | 2200475 | 2200574 | **RNA**→ Methyltrn_RNA_3 (pfam02598)→ rpl3p (PRK04231)Ribosomal_L3 (pfam00297)→<br>RplD (COG0088)rpl4lp (PRK04042)→ RplW (COG0089)PRK14548 (PRK14548)→<br>RplB (COG0090)rpl2p (PRK09612)→ RpsS (COG0185)rps19p (PRK04038)→<br>rpl22p (PRK04223)RplV (COG0091)→ RpsC (COG0092)rps3p (PRK04191)→<br>PRK14549 (PRK14549)Ribosomal_L29 (pfam00831)→ PRK03879 (PRK03879)UPF0086 (pfam01868)→<br>RpsQ (COG0186)rps17p (PRK08572)→ rpl14p (PRK08571)Ribosomal_L14 (pfam00238)→<br>RplX (COG0198)rpl24p (PRK01191)→<br>PRK04313 (PRK04313)KOW_RPS4 (cd06087)Ribosomal_S4e (pfam00900)→ RplE (COG0094)rpl5p (PRK04219)→<br>RpsN (COG0199)rps14P (PRK05766)→ rps8p (PRK04034)RpsH (COG0096)→<br>RplF (COG0097)rpl6p (PRK05518)→ PRK12766 (PRK12766)RecA (COG0468)Rpl32e (COG1717)→<br>rpl19e (PRK08570)Ribosomal_L19e (pfam01280)→ |
| env-199 | APHM01006302.1 | - | 1366 | 1273 | **RNA**→ Methyltrn_RNA_3 (pfam02598)→ RplC (COG0087)rpl3p (PRK04231)→ |
| env-200 | LTJ07As871_J07ABscf092781 | + | 25242 | 25335 | **RNA**→ Methyltrn_RNA_3 (pfam02598)→ rpl3p (PRK04231)Ribosomal_L3 (pfam00297)→ hypo→<br>PRK12766 (PRK12766)Rpl32e (COG1717)→ rpl19e (PRK08570)Ribosomal_L19e (pfam01280)→ |
| Hsu-1-1 | NZ_CP008874.1 | - | 1720180 | 1720081 | **RNA**→ Methyltrn_RNA_3 (pfam02598)→ RplC (COG0087)rpl3p (PRK04231)→<br>RplD (COG0088)rpl4lp (PRK04042)→ RplW (COG0089)PRK14548 (PRK14548)→<br>RplB (COG0090)rpl2p (PRK09612)→ RpsS (COG0185)rps19p (PRK04038)→<br>rpl22p (PRK04223)RplV (COG0091)→ RpsC (COG0092)rps3p (PRK04191)→<br>PRK14549 (PRK14549)Ribosomal_L29 (pfam00831)→ PRK03879 (PRK03879)UPF0086 (pfam01868)→<br>RpsQ (COG0186)rps17p (PRK08572)→ rpl14p (PRK08571)Ribosomal_L14 (pfam00238)→<br>RplX (COG0198)rpl24p (PRK01191)→<br>PRK04313 (PRK04313)KOW_RPS4 (cd06087)Ribosomal_S4e (pfam00900)→ RplE (COG0094)rpl5p (PRK04219)→<br>RpsN (COG0199)rps14P (PRK05766)→ rps8p (PRK04034)RpsH (COG0096)→<br>RplF (COG0097)rpl6p (PRK05518)→<br>RecA (COG0468)Rpl32e (COG1717)rpl32e (PRK08562)HHH_5 (pfam14520)→<br>rpl19e (PRK08570)Ribosomal_L19e (pfam01280)→ |
| env-201 | P_A23_Liq_1_Fmt_1021786 | - | 270 | 171 | **RNA**→ Methyltrn_RNA_3 (pfam02598)→ |
| env-202 | P_A23_Liq_2_Fmt_1024942 | - | 260 | 161 | **RNA**→ Methyltrn_RNA_3 (pfam02598)→ |
| env-203 | P_A23_Liq_3_Fmt_1011772 | + | 655 | 754 | **RNA**→ Methyltrn_RNA_3 (pfam02598)→ |
| Hca-1-1 | NZ_AOLS01000002.1 | - | 28213 | 28114 | **RNA**→ Methyltrn_RNA_3 (pfam02598)→ rpl3p (PRK04231)Ribosomal_L3 (pfam00297)→<br>rpl4lp (PRK04042)Ribosomal_L4 (pfam00573)→ RplW (COG0089)PRK14548 (PRK14548)→<br>RplB (COG0090)rpl2p (PRK09612)→ RpsS (COG0185)rps19p (PRK04038)→<br>rpl22p (PRK04223)RplV (COG0091)→ RpsC (COG0092)rps3p (PRK04191)→<br>PRK14549 (PRK14549)Ribosomal_L29 (pfam00831)→ PRK03879 (PRK03879)UPF0086 (pfam01868)→<br>RpsQ (COG0186)rps17p (PRK08572)→ rpl14p (PRK08571)Ribosomal_L14 (pfam00238)→<br>rpl24p (PRK01191)KOW_RPL26 (cd06089)→<br>PRK04313 (PRK04313)KOW_RPS4 (cd06087)S4 (pfam01479)Ribosomal_S4e (pfam00900)→<br>RplE (COG0094)rpl5p (PRK04219)→ RpsN (COG0199)rps14P (PRK05766)→<br>rps8p (PRK04034)RpsH (COG0096)→ RplF (COG0097)rpl6p (PRK05518)→<br>PRK12766 (PRK12766)RecA (COG0468)Rpl32e (COG1717)→ rpl19e (PRK08570)Ribosomal_L19e (pfam01280)→<br>RplR (COG0256)rpl18p (PRK08569)→ RplR (COG0256)rpl18p (PRK08569)→ |

| | | | | | |
|---|---|---|---|---|---|
| Hsi-1-1 | NZ_AOLR01000002.1 | - | 251120 | 251021 | **RNA**→ Methyltrn_RNA_3 (pfam02598)→ rpl3p (PRK04231)Ribosomal_L3 (pfam00297)→<br>rpl4lp (PRK04042)Ribosomal_L4 (pfam00573)→ RplW (COG0089)PRK14548 (PRK14548)→<br>RplB (COG0090)rpl2p (PRK09612)→ RpsS (COG0185)rps19p (PRK04038)→<br>rpl22p (PRK04223)RplV (COG0091)→ RpsC (COG0092)rps3p (PRK04191)→<br>PRK14549 (PRK14549)Ribosomal_L29 (pfam00831)→ POP4 (COG1588)PRK03879 (PRK03879)→<br>RpsQ (COG0186)rps17p (PRK08572)→ rpl14p (PRK08571)Ribosomal_L14 (pfam00238)→<br>rpl24p (PRK01191)KOW_RPL26 (cd06089)→<br>PRK04313 (PRK04313)KOW_RPS4 (cd06087)S4 (pfam01479)Ribosomal_S4e (pfam00900)→<br>RplE (COG0094)rpl5p (PRK04219)→ RpsN (COG0199)rps14P (PRK05766)→<br>rps8p (PRK04034)RpsH (COG0096)→ RplF (COG0097)rpl6p (PRK05518)→<br>PRK12766 (PRK12766)RecA (COG0468)Rpl32e (COG1717)→ rpl19e (PRK08570)Ribosomal_L19e (pfam01280)→<br>RplR (COG0256)rpl18p (PRK08569)→ RplR (COG0256)rpl18p (PRK08569)→ |
| Hma-1-1 | NC_006396.1 | - | 1443199 | 1443100 | **RNA**→ Methyltrn_RNA_3 (pfam02598)→ rpl3p (PRK04231)Ribosomal_L3 (pfam00297)→<br>rpl4lp (PRK04042)Ribosomal_L4 (pfam00573)→ RplW (COG0089)PRK14548 (PRK14548)→<br>RplB (COG0090)rpl2p (PRK09612)→ RpsS (COG0185)rps19p (PRK04038)→<br>rpl22p (PRK04223)RplV (COG0091)→ RpsC (COG0092)rps3p (PRK04191)→<br>PRK14549 (PRK14549)Ribosomal_L29 (pfam00831)→ POP4 (COG1588)PRK03879 (PRK03879)→<br>POP4 (COG1588)PRK03879 (PRK03879)→ PRK03879 (PRK03879)UPF0086 (pfam01868)→<br>RpsQ (COG0186)rps17p (PRK08572)→ rpl14p (PRK08571)Ribosomal_L14 (pfam00238)→<br>rpl24p (PRK01191)KOW_RPL26 (cd06089)→<br>PRK04313 (PRK04313)KOW_RPS4 (cd06087)S4 (pfam01479)Ribosomal_S4e (pfam00900)→<br>RplE (COG0094)rpl5p (PRK04219)→ RpsN (COG0199)rps14P (PRK05766)→<br>rps8p (PRK04034)RpsH (COG0096)→ RplF (COG0097)rpl6p (PRK05518)→<br>PRK12766 (PRK12766)RecA (COG0468)Rpl32e (COG1717)→ rpl19e (PRK08570)Ribosomal_L19e (pfam01280)→<br>RplR (COG0256)rpl18p (PRK08569)→ |
| Hsp-1-1 | NZ_CP010529.1 | - | 696901 | 696802 | **RNA**→ Methyltrn_RNA_3 (pfam02598)→ rpl3p (PRK04231)Ribosomal_L3 (pfam00297)→<br>rpl4lp (PRK04042)Ribosomal_L4 (pfam00573)→ RplW (COG0089)PRK14548 (PRK14548)→<br>RplB (COG0090)rpl2p (PRK09612)→ RpsS (COG0185)rps19p (PRK04038)→<br>rpl22p (PRK04223)RplV (COG0091)→ RpsC (COG0092)rps3p (PRK04191)→<br>PRK14549 (PRK14549)Ribosomal_L29 (pfam00831)→ PRK03879 (PRK03879)UPF0086 (pfam01868)→<br>RpsQ (COG0186)rps17p (PRK08572)→ rpl14p (PRK08571)Ribosomal_L14 (pfam00238)→<br>rpl24p (PRK01191)KOW_RPL26 (cd06089)→<br>PRK04313 (PRK04313)KOW_RPS4 (cd06087)S4 (pfam01479)Ribosomal_S4e (pfam00900)→<br>RplE (COG0094)rpl5p (PRK04219)→ RpsN (COG0199)rps14P (PRK05766)→<br>rps8p (PRK04034)RpsH (COG0096)→ RplF (COG0097)rpl6p (PRK05518)→<br>PRK12766 (PRK12766)RecA (COG0468)Rpl32e (COG1717)→ rpl19e (PRK08570)Ribosomal_L19e (pfam01280)→ |
| Ham-1-1 | NZ_AOLW01000047.1 | + | 239419 | 239518 | **RNA**→ Methyltrn_RNA_3 (pfam02598)→ rpl3p (PRK04231)Ribosomal_L3 (pfam00297)→<br>rpl4lp (PRK04042)Ribosomal_L4 (pfam00573)→ RplW (COG0089)PRK14548 (PRK14548)→ |
| Hhi-2-1 | NC_023013.1 | - | 2093063 | 2092964 | **RNA**→ Methyltrn_RNA_3 (pfam02598)→ rpl3p (PRK04231)Ribosomal_L3 (pfam00297)→<br>rpl4lp (PRK04042)Ribosomal_L4 (pfam00573)→ RplW (COG0089)PRK14548 (PRK14548)→<br>RplB (COG0090)rpl2p (PRK09612)→ RpsS (COG0185)rps19p (PRK04038)→<br>rpl22p (PRK04223)RplV (COG0091)→ RpsC (COG0092)rps3p (PRK04191)→<br>PRK14549 (PRK14549)Ribosomal_L29 (pfam00831)→ PRK03879 (PRK03879)UPF0086 (pfam01868)→<br>RpsQ (COG0186)rps17p (PRK08572)→ rpl14p (PRK08571)Ribosomal_L14 (pfam00238)→<br>rpl24p (PRK01191)KOW_RPL26 (cd06089)→<br>PRK04313 (PRK04313)KOW_RPS4 (cd06087)S4 (pfam01479)Ribosomal_S4e (pfam00900)→<br>RplE (COG0094)rpl5p (PRK04219)→ RpsN (COG0199)rps14P (PRK05766)→<br>rps8p (PRK04034)RpsH (COG0096)→ RplF (COG0097)rpl6p (PRK05518)→<br>PRK12766 (PRK12766)RecA (COG0468)Rpl32e (COG1717)→ rpl19e (PRK08570)Ribosomal_L19e (pfam01280)→ |

| | | | | | |
|---|---|---|---|---|---|
| Hhi-1-1 | NC_015948.1 | - | 2081630 | 2081531 | **RNA**→ Methyltrn_RNA_3 (pfam02598)→ rpl3p (PRK04231)Ribosomal_L3 (pfam00297)→<br>rpl4lp (PRK04042)Ribosomal_L4 (pfam00573)→ RplW (COG0089)PRK14548 (PRK14548)→<br>RplB (COG0090)rpl2p (PRK09612)→ RpsS (COG0185)rps19p (PRK04038)→<br>rpl22p (PRK04223)RplV (COG0091)→ RpsC (COG0092)rps3p (PRK04191)→<br>PRK14549 (PRK14549)Ribosomal_L29 (pfam00831)→ PRK03879 (PRK03879)UPF0086 (pfam01868)→<br>RpsQ (COG0186)rps17p (PRK08572)→ rpl14p (PRK08571)Ribosomal_L14 (pfam00238)→<br>rpl24p (PRK01191)KOW_RPL26 (cd06089)→<br>PRK04313 (PRK04313)KOW_RPS4 (cd06087)S4 (pfam01479)Ribosomal_S4e (pfam00900)→<br>PRK04313 (PRK04313)KOW_RPS4 (cd06087)S4 (pfam01479)Ribosomal_S4e (pfam00900)→<br>RplE (COG0094)rpl5p (PRK04219)→ RpsN (COG0199)rps14P (PRK05766)→<br>rps8p (PRK04034)RpsH (COG0096)→ RplF (COG0097)rpl6p (PRK05518)→<br>PRK12766 (PRK12766)RecA (COG0468)Rpl32e (COG1717)→ rpl19e (PRK08570)Ribosomal_L19e (pfam01280)→ |
| Hsa-2-1 | NZ_JXAO01000052.1 | + | 4509 | 4608 | **RNA**→ hypo→ rpl3p (PRK04231)Ribosomal_L3 (pfam00297)→ rpl4lp (PRK04042)Ribosomal_L4 (pfam00573)→<br>RplW (COG0089)PRK14548 (PRK14548)→ hypo→ RpsS (COG0185)rps19p (PRK04038)→<br>rpl22p (PRK04223)RplV (COG0091)→ RpsC (COG0092)rps3p (PRK04191)→<br>PRK14549 (PRK14549)Ribosomal_L29 (pfam00831)→ POP4 (COG1588)PRK03879 (PRK03879)→<br>RpsQ (COG0186)rps17p (PRK08572)→ rpl14p (PRK08571)Ribosomal_L14 (pfam00238)→<br>rpl24p (PRK01191)KOW_RPL26 (cd06089)→ hypo→ RplE (COG0094)rpl5p (PRK04219)→<br>RpsN (COG0199)rps14P (PRK05766)→ rps8p (PRK04034)RpsH (COG0096)→<br>RplF (COG0097)rpl6p (PRK05518)→ hypo→ rpl19e (PRK08570)Ribosomal_L19e (pfam01280)→ |
| Har-1-1 | NZ_AOLX01000026.1 | + | 231236 | 231335 | **RNA**→ Methyltrn_RNA_3 (pfam02598)→ rpl3p (PRK04231)Ribosomal_L3 (pfam00297)→<br>rpl4lp (PRK04042)Ribosomal_L4 (pfam00573)→ RplW (COG0089)PRK14548 (PRK14548)→ |
| Hva-1-1 | NZ_AOLQ01000042.1 | - | 26191 | 26092 | **RNA**→ Methyltrn_RNA_3 (pfam02598)→ rpl3p (PRK04231)Ribosomal_L3 (pfam00297)→<br>rpl4lp (PRK04042)Ribosomal_L4 (pfam00573)→ RplW (COG0089)PRK14548 (PRK14548)→<br>RplB (COG0090)rpl2p (PRK09612)→ RpsS (COG0185)rps19p (PRK04038)→<br>rpl22p (PRK04223)RplV (COG0091)→ RpsC (COG0092)rps3p (PRK04191)→<br>PRK14549 (PRK14549)Ribosomal_L29 (pfam00831)→ POP4 (COG1588)PRK03879 (PRK03879)→<br>RpsQ (COG0186)rps17p (PRK08572)→ rpl14p (PRK08571)Ribosomal_L14 (pfam00238)→<br>rpl24p (PRK01191)KOW_RPL26 (cd06089)→<br>PRK04313 (PRK04313)KOW_RPS4 (cd06087)S4 (pfam01479)Ribosomal_S4e (pfam00900)→<br>RplE (COG0094)rpl5p (PRK04219)→ RpsN (COG0199)rps14P (PRK05766)→<br>rps8p (PRK04034)RpsH (COG0096)→ RplF (COG0097)rpl6p (PRK05518)→<br>PRK12766 (PRK12766)RecA (COG0468)Rpl32e (COG1717)→ rpl19e (PRK08570)Ribosomal_L19e (pfam01280)→ |
| Hja-1-1 | NZ_AOLY01000009.1 | + | 92110 | 92209 | **RNA**→ Methyltrn_RNA_3 (pfam02598)→ rpl3p (PRK04231)Ribosomal_L3 (pfam00297)→<br>rpl4lp (PRK04042)Ribosomal_L4 (pfam00573)→ RplW (COG0089)PRK14548 (PRK14548)→<br>RplB (COG0090)rpl2p (PRK09612)→ RpsS (COG0185)rps19p (PRK04038)→<br>rpl22p (PRK04223)RplV (COG0091)→ RpsC (COG0092)rps3p (PRK04191)→<br>PRK14549 (PRK14549)Ribosomal_L29 (pfam00831)→ PRK03879 (PRK03879)UPF0086 (pfam01868)→<br>RpsQ (COG0186)rps17p (PRK08572)→ rpl14p (PRK08571)Ribosomal_L14 (pfam00238)→<br>rpl24p (PRK01191)KOW_RPL26 (cd06089)→<br>PRK04313 (PRK04313)KOW_RPS4 (cd06087)S4 (pfam01479)Ribosomal_S4e (pfam00900)→<br>RplE (COG0094)rpl5p (PRK04219)→ RpsN (COG0199)rps14P (PRK05766)→<br>rps8p (PRK04034)RpsH (COG0096)→ RplF (COG0097)rpl6p (PRK05518)→<br>PRK12766 (PRK12766)RecA (COG0468)Rpl32e (COG1717)→ rpl19e (PRK08570)Ribosomal_L19e (pfam01280)→ |
| env-204 | APHM01021165.1 | + | 2893 | 2995 | **RNA**→ Methyltrn_RNA_3 (pfam02598)→ RplC (COG0087)rpl3p (PRK04231)→<br>RplD (COG0088)rpl4lp (PRK04042)→ rpl4lp (PRK04042)→ RplW (COG0089)PRK14548 (PRK14548)→<br>RplB (COG0090)rpl2p (PRK09612)→ RpsS (COG0185)rps19p (PRK04038)→ |
| env-205 | LTJ07As871_J07ABscf098843 | + | 48637 | 48739 | **RNA**→ Methyltrn_RNA_3 (pfam02598)→ RplC (COG0087)rpl3p (PRK04231)→<br>RplD (COG0088)rpl4lp (PRK04042)→ rpl4lp (PRK04042)→ RplW (COG0089)PRK14548 (PRK14548)→<br>RplB (COG0090)rpl2p (PRK09612)→ RpsS (COG0185)rps19p (PRK04038)→ |

| | | | | | |
|---|---|---|---|---|---|
| Hse-1-1 | NZ_BBMP01000022.1 | - | 550285 | 550186 | **RNA**→ Methyltrn_RNA_3 (pfam02598)→ RplC (COG0087)rpl3p (PRK04231)→ |
| | | | | | RplD (COG0088)rpl4lp (PRK04042)→ RplW (COG0089)PRK14548 (PRK14548)→ |
| | | | | | RplB (COG0090)rpl2p (PRK09612)→ RpsS (COG0185)rps19p (PRK04038)→ |
| | | | | | rpl22p (PRK04223)RplV (COG0091)→ |
| | | | | | RpsC (COG0092)MDN1 (COG5271)rps3p (PRK04191)HHH_5 (pfam14520)→ |
| | | | | | PRK14549 (PRK14549)Ribosomal_L29 (pfam00831)→ PRK03879 (PRK03879)UPF0086 (pfam01868)→ |
| | | | | | RpsQ (COG0186)rps17p (PRK08572)→ rpl14p (PRK08571)Ribosomal_L14 (pfam00238)→ |
| | | | | | RplX (COG0198)rpl24p (PRK01191)→ |
| | | | | | PRK04313 (PRK04313)KOW_RPS4 (cd06087)S4 (pfam01479)Ribosomal_S4e (pfam00900)→ |
| | | | | | RplE (COG0094)rpl5p (PRK04219)→ RpsN (COG0199)rps14P (PRK05766)→ |
| | | | | | rps8p (PRK04034)RpsH (COG0096)→ RplF (COG0097)rpl6p (PRK05518)→ |
| | | | | | RecA (COG0468)Rpl32e (COG1717)COG2251 (COG2251)rpl32e (PRK08562)HHH_5 (pfam14520)→ |
| | | | | | RecA (COG0468)rpl32e (PRK08562)Ribosomal_L32_L32e (cd00513)HHH_5 (pfam14520)→ |
| Hth-1-1 | NZ_AOMF01000153.1 | + | 44875 | 44974 | **RNA**→ Methyltrn_RNA_3 (pfam02598)→ RplC (COG0087)rpl3p (PRK04231)→ |
| | | | | | RplD (COG0088)rpl4lp (PRK04042)→ RplW (COG0089)PRK14548 (PRK14548)→ |
| | | | | | RplB (COG0090)rpl2p (PRK09612)→ RpsS (COG0185)rps19p (PRK04038)→ |
| | | | | | rpl22p (PRK04223)RplV (COG0091)→ |
| | | | | | RpsC (COG0092)Lig (COG0272)rps3p (PRK04191)Nop14 (pfam04147)HHH_5 (pfam14520)→ |
| | | | | | PRK14549 (PRK14549)Ribosomal_L29 (pfam00831)→ PRK03879 (PRK03879)UPF0086 (pfam01868)→ |
| | | | | | RpsQ (COG0186)rps17p (PRK08572)→ rpl14p (PRK08571)Ribosomal_L14 (pfam00238)→ |
| | | | | | RplX (COG0198)rpl24p (PRK01191)→ |
| | | | | | PRK04313 (PRK04313)KOW_RPS4 (cd06087)Ribosomal_S4e (pfam00900)→ RplE (COG0094)rpl5p (PRK04219)→ |
| | | | | | RpsN (COG0199)rps14P (PRK05766)→ rps8p (PRK04034)RpsH (COG0096)→ |
| | | | | | RplF (COG0097)rpl6p (PRK05518)→ |
| | | | | | NT_POLXc (cd00141)Rpl32e (COG1717)rpl32e (PRK08562)HHH_5 (pfam14520)→ |
| | | | | | RecA (COG0468)Rpl32e (COG1717)rpl32e (PRK08562)HHH_5 (pfam14520)→ |
| Hmo-1-1 | NZ_AOMC01000109.1 | - | 103550 | 103451 | **RNA**→ Methyltrn_RNA_3 (pfam02598)→ RplC (COG0087)rpl3p (PRK04231)→ |
| | | | | | RplD (COG0088)rpl4lp (PRK04042)→ RplW (COG0089)PRK14548 (PRK14548)→ |
| | | | | | RplB (COG0090)rpl2p (PRK09612)→ RpsS (COG0185)rps19p (PRK04038)→ |
| | | | | | rpl22p (PRK04223)RplV (COG0091)→ |
| | | | | | RpsC (COG0092)UvrC (COG0322)rps3p (PRK04191)HHH_5 (pfam14520)→ |
| | | | | | PRK14549 (PRK14549)Ribosomal_L29 (pfam00831)→ PRK03879 (PRK03879)UPF0086 (pfam01868)→ |
| | | | | | RpsQ (COG0186)rps17p (PRK08572)→ rpl14p (PRK08571)Ribosomal_L14 (pfam00238)→ |
| | | | | | rpl24p (PRK01191)KOW_RPL26 (cd06089)→ |
| | | | | | PRK04313 (PRK04313)KOW_RPS4 (cd06087)Ribosomal_S4e (pfam00900)→ RplE (COG0094)rpl5p (PRK04219)→ |
| | | | | | RpsN (COG0199)rps14P (PRK05766)→ rps8p (PRK04034)RpsH (COG0096)→ |
| | | | | | rpl6p (PRK05518)Ribosomal_L6 (pfam00347)→ |
| env-206 | P_1C_Sed_2_UnCty_1102543 | + | 116 | 215 | **RNA**→ Methyltrn_RNA_3 (pfam02598)→ |
| env-207 | APHM01019633.1 | + | 691 | 784 | **RNA**→ Methyltrn_RNA_3 (pfam02598)→ rpl3p (PRK04231)Ribosomal_L3 (pfam00297)→ |
| env-208 | LTJ07As871_J07ABscf090118 | - | 8900 | 8807 | **RNA**→ Methyltrn_RNA_3 (pfam02598)→ rpl3p (PRK04231)Ribosomal_L3 (pfam00297)→ |
| env-209 | P_1C_Sed_2_UnCty_1164547 | + | 86 | 185 | **RNA**→ Methyltrn_RNA_3 (pfam02598)→ |
| Nta-1-1 | NZ_AOIL01000042.1 | + | 111800 | 111899 | **RNA**→ Methyltrn_RNA_3 (pfam02598)→ rpl3p (PRK04231)Ribosomal_L3 (pfam00297)→ |
| | | | | | RplD (COG0088)rpl4lp (PRK04042)→ RplW (COG0089)PRK14548 (PRK14548)→ |
| | | | | | RplB (COG0090)rpl2p (PRK09612)→ RpsS (COG0185)rps19p (PRK04038)→ |
| | | | | | rpl22p (PRK04223)RplV (COG0091)→ RpsC (COG0092)rps3p (PRK04191)→ |
| | | | | | RpmC (COG0255)PRK14549 (PRK14549)→ PRK03879 (PRK03879)UPF0086 (pfam01868)→ |
| | | | | | RpsQ (COG0186)rps17p (PRK08572)→ rpl14p (PRK08571)Ribosomal_L14 (pfam00238)→ |
| | | | | | rpl24p (PRK01191)KOW_RPL26 (cd06089)→ |
| | | | | | PRK04313 (PRK04313)KOW_RPS4 (cd06087)Ribosomal_S4e (pfam00900)→ RplE (COG0094)rpl5p (PRK04219)→ |
| | | | | | RpsN (COG0199)rps14P (PRK05766)→ rps8p (PRK04034)RpsH (COG0096)→ |
| | | | | | RplF (COG0097)rpl6p (PRK05518)→ PRK12766 (PRK12766)RecA (COG0468)Rpl32e (COG1717)→ |

| | | | | | |
|---|---|---|---|---|---|
| Nas-1-1 | NZ_AOIO01000023.1 | - | 59520 | 59421 | **RNA**→ Methyltrn_RNA_3 (pfam02598)→ rpl3p (PRK04231)Ribosomal_L3 (pfam00297)→<br>RplD (COG0088)rpl4lp (PRK04042)→ RplW (COG0089)PRK14548 (PRK14548)→<br>RplB (COG0090)rpl2p (PRK09612)→ RpsS (COG0185)rps19p (PRK04038)→<br>rpl22p (PRK04223)RplV (COG0091)→ RpsC (COG0092)rps3p (PRK04191)→<br>RpmC (COG0255)PRK14549 (PRK14549)→ PRK03879 (PRK03879)UPF0086 (pfam01868)→<br>RpsQ (COG0186)rps17p (PRK08572)→ rpl14p (PRK08571)Ribosomal_L14 (pfam00238)→<br>rpl24p (PRK01191)KOW_RPL26 (cd06089)→<br>PRK04313 (PRK04313)KOW_RPS4 (cd06087)Ribosomal_S4e (pfam00900)→ RplE (COG0094)rpl5p (PRK04219)→<br>RpsN (COG0199)rps14P (PRK05766)→ rps8p (PRK04034)RpsH (COG0096)→<br>RplF (COG0097)rpl6p (PRK05518)→ PRK12766 (PRK12766)RecA (COG0468)Rpl32e (COG1717)→ |
| Noc-1-1 | NC_019974.1 | - | 3406135 | 3406036 | **RNA**→ Methyltrn_RNA_3 (pfam02598)→ rpl3p (PRK04231)Ribosomal_L3 (pfam00297)→<br>RplD (COG0088)rpl4lp (PRK04042)→ RplW (COG0089)PRK14548 (PRK14548)→<br>RplB (COG0090)rpl2p (PRK09612)→ RpsS (COG0185)rps19p (PRK04038)→<br>rpl22p (PRK04223)RplV (COG0091)→ RpsC (COG0092)rps3p (PRK04191)→<br>PRK14549 (PRK14549)Ribosomal_L29 (pfam00831)→ PRK03879 (PRK03879)UPF0086 (pfam01868)→<br>RpsQ (COG0186)rps17p (PRK08572)→ rpl14p (PRK08571)Ribosomal_L14 (pfam00238)→<br>rpl24p (PRK01191)KOW_RPL26 (cd06089)→<br>PRK04313 (PRK04313)KOW_RPS4 (cd06087)S4 (pfam01479)Ribosomal_S4e (pfam00900)→<br>RplE (COG0094)rpl5p (PRK04219)→ RpsN (COG0199)rps14P (PRK05766)→<br>rps8p (PRK04034)RpsH (COG0096)→ RplF (COG0097)rpl6p (PRK05518)→<br>UvrC (COG0322)Rpl32e (COG1717)rpl32e (PRK08562)HHH_5 (pfam14520)→ |
| Nje-1-1 | NZ_AOIA01000019.1 | + | 36830 | 36929 | **RNA**→ Methyltrn_RNA_3 (pfam02598)→ rpl3p (PRK04231)Ribosomal_L3 (pfam00297)→<br>RplD (COG0088)rpl4lp (PRK04042)→ RplW (COG0089)PRK14548 (PRK14548)→<br>RplB (COG0090)rpl2p (PRK09612)→ RpsS (COG0185)rps19p (PRK04038)→<br>rpl22p (PRK04223)RplV (COG0091)→ RpsC (COG0092)rps3p (PRK04191)→<br>PRK14549 (PRK14549)Ribosomal_L29 (pfam00831)→ PRK03879 (PRK03879)UPF0086 (pfam01868)→<br>RpsQ (COG0186)rps17p (PRK08572)→ rpl14p (PRK08571)Ribosomal_L14 (pfam00238)→<br>rpl24p (PRK01191)KOW_RPL26 (cd06089)→<br>PRK04313 (PRK04313)KOW_RPS4 (cd06087)S4 (pfam01479)Ribosomal_S4e (pfam00900)→<br>RplE (COG0094)rpl5p (PRK04219)→ RpsN (COG0199)rps14P (PRK05766)→<br>rps8p (PRK04034)RpsH (COG0096)→ RplF (COG0097)rpl6p (PRK05518)→<br>RecA (COG0468)Rpl32e (COG1717)HHH_5 (pfam14520)→ |
| Nam-1-1 | NZ_AOIB01000037.1 | - | 63552 | 63453 | **RNA**→ Methyltrn_RNA_3 (pfam02598)→ rpl3p (PRK04231)Ribosomal_L3 (pfam00297)→<br>RplD (COG0088)rpl4lp (PRK04042)→ RplW (COG0089)PRK14548 (PRK14548)→<br>RplB (COG0090)rpl2p (PRK09612)→ RpsS (COG0185)rps19p (PRK04038)→<br>rpl22p (PRK04223)RplV (COG0091)→ RpsC (COG0092)rps3p (PRK04191)→<br>PRK14549 (PRK14549)Ribosomal_L29 (pfam00831)→ PRK03879 (PRK03879)UPF0086 (pfam01868)→<br>RpsQ (COG0186)rps17p (PRK08572)→ rpl14p (PRK08571)Ribosomal_L14 (pfam00238)→<br>rpl24p (PRK01191)KOW_RPL26 (cd06089)→<br>PRK04313 (PRK04313)KOW_RPS4 (cd06087)S4 (pfam01479)Ribosomal_S4e (pfam00900)→<br>RplE (COG0094)rpl5p (PRK04219)→ RpsN (COG0199)rps14P (PRK05766)→<br>rps8p (PRK04034)RpsH (COG0096)→ RplF (COG0097)rpl6p (PRK05518)→<br>UvrC (COG0322)Rpl32e (COG1717)rpl32e (PRK08562)HHH_5 (pfam14520)→<br>rpl19e (PRK08570)Ribosomal_L19e (pfam01280)→ |
| env-210 | SL_5KL_010_BRINE_10002219 | + | 679 | 778 | **RNA**→ Methyltrn_RNA_3 (pfam02598)→ rpl3p (PRK04231)Ribosomal_L3 (pfam00297)→<br>RplD (COG0088)rpl4lp (PRK04042)→ RplW (COG0089)PRK14548 (PRK14548)→<br>RplB (COG0090)rpl2p (PRK09612)→ RpsS (COG0185)rps19p (PRK04038)→<br>rpl22p (PRK04223)RplV (COG0091)→ RpsC (COG0092)rps3p (PRK04191)→<br>PRK14549 (PRK14549)Ribosomal_L29 (pfam00831)→ PRK03879 (PRK03879)UPF0086 (pfam01868)→<br>RpsQ (COG0186)rps17p (PRK08572)→ rpl14p (PRK08571)Ribosomal_L14 (pfam00238)→<br>rpl24p (PRK01191)KOW_RPL26 (cd06089)→<br>PRK04313 (PRK04313)S4 (cd00165)KOW_RPS4 (cd06087)Ribosomal_S4e (pfam00900)→<br>RplE (COG0094)rpl5p (PRK04219)SDH (cd12188)→ RpsN (COG0199)rps14P (PRK05766)→ rps8p (PRK04034)→ |
| env-211 | SL_5KL_010_BRINE_10004382 | + | 5582 | 5681 | **RNA**→ Methyltrn_RNA_3 (pfam02598)→ rpl3p (PRK04231)Ribosomal_L3 (pfam00297)→ |

| | | | | | |
|---|---|---|---|---|---|
| env-212 | SL_6KL_011_BRINE_1000450 | + | 5568 | 5667 | **RNA**→ Methyltrn_RNA_3 (pfam02598)→ rpl3p (PRK04231)Ribosomal_L3 (pfam00297)→<br>RplD (COG0088)rpl4lp (PRK04042)→ RplW (COG0089)PRK14548 (PRK14548)→<br>RplB (COG0090)rpl2p (PRK09612)→ RpsS (COG0185)rps19p (PRK04038)→<br>rpl22p (PRK04223)RplV (COG0091)→ RpsC (COG0092)rps3p (PRK04191)→<br>PRK14549 (PRK14549)Ribosomal_L29 (pfam00831)→ PRK03879 (PRK03879)UPF0086 (pfam01868)→<br>RpsQ (COG0186)rps17p (PRK08572)→ rpl14p (PRK08571)Ribosomal_L14 (pfam00238)→<br>rpl24p (PRK01191)KOW_RPL26 (cd06089)→<br>PRK04313 (PRK04313)S4 (cd00165)KOW_RPS4 (cd06087)Ribosomal_S4e (pfam00900)→<br>RplE (COG0094)rpl5p (PRK04219)SDH (cd12188)→ RpsN (COG0199)rps14P (PRK05766)→<br>rps8p (PRK04034)RpsH (COG0096)→ RplF (COG0097)rpl6p (PRK05518)→<br>RecA (COG0468)Rpl32e (COG1717)rpl32e (PRK08562)HHH_5 (pfam14520)→<br>rpl19e (PRK08570)Ribosomal_L19e (pfam01280)→ RplR (COG0256)rpl18p (PRK08569)→<br>RpsE (COG0098)rps5p (PRK04044)→ Ribosomal_L7_archeal_euk (cd01657)rpl30p (PRK06049)→<br>rpl15p (PRK06419)Ribosomal_L27A (pfam00828)→ Plug_translocon (pfam10559)→ |
| env-213 | SL_4KL_010_BRINE_10000109 | + | 5553 | 5652 | **RNA**→ Methyltrn_RNA_3 (pfam02598)→ rpl3p (PRK04231)Ribosomal_L3 (pfam00297)→<br>RplD (COG0088)rpl4lp (PRK04042)→ RplW (COG0089)PRK14548 (PRK14548)→<br>RplB (COG0090)rpl2p (PRK09612)→ RpsS (COG0185)rps19p (PRK04038)→<br>rpl22p (PRK04223)RplV (COG0091)→ RpsC (COG0092)rps3p (PRK04191)→<br>PRK14549 (PRK14549)Ribosomal_L29 (pfam00831)→ PRK03879 (PRK03879)UPF0086 (pfam01868)→<br>RpsQ (COG0186)rps17p (PRK08572)→ rpl14p (PRK08571)Ribosomal_L14 (pfam00238)→<br>rpl24p (PRK01191)KOW_RPL26 (cd06089)→<br>PRK04313 (PRK04313)S4 (cd00165)KOW_RPS4 (cd06087)Ribosomal_S4e (pfam00900)→<br>RplE (COG0094)rpl5p (PRK04219)SDH (cd12188)→ RpsN (COG0199)rps14P (PRK05766)→<br>rps8p (PRK04034)RpsH (COG0096)→ RplF (COG0097)rpl6p (PRK05518)→<br>RecA (COG0468)Rpl32e (COG1717)rpl32e (PRK08562)HHH_5 (pfam14520)→<br>rpl19e (PRK08570)Ribosomal_L19e (pfam01280)→ RplR (COG0256)rpl18p (PRK08569)→<br>RpsE (COG0098)rps5p (PRK04044)→ Ribosomal_L7_archeal_euk (cd01657)rpl30p (PRK06049)→<br>rpl15p (PRK06419)Ribosomal_L27A (pfam00828)→ SecY (COG0201)PRK08568 (PRK08568)→ |
| Hsp-3-1 | NC_002607.1 | + | 1258523 | 1258622 | **RNA**→ MTH1 (COG2106)→ rpl3p (PRK04231)Ribosomal_L3 (pfam00297)→<br>RplD (COG0088)rpl4lp (PRK04042)→ RplW (COG0089)PRK14548 (PRK14548)→<br>RplB (COG0090)rpl2p (PRK09612)→ RpsS (COG0185)rps19p (PRK04038)→<br>rpl22p (PRK04223)RplV (COG0091)→ RpsC (COG0092)rps3p (PRK04191)→<br>RpmC (COG0255)PRK14549 (PRK14549)→ POP4 (COG1588)PRK03879 (PRK03879)→<br>RpsQ (COG0186)rps17p (PRK08572)→ rpl14p (PRK08571)Ribosomal_L14 (pfam00238)→<br>RplX (COG0198)rpl24p (PRK01191)→<br>PRK04313 (PRK04313)KOW_RPS4 (cd06087)Ribosomal_S4e (pfam00900)→ RplE (COG0094)rpl5p (PRK04219)→<br>RpsN (COG0199)rps14P (PRK05766)→ rps8p (PRK04034)RpsH (COG0096)→<br>rps8p (PRK04034)RpsH (COG0096)→ RplF (COG0097)rpl6p (PRK05518)→<br>PRK12766 (PRK12766)RecA (COG0468)Rpl32e (COG1717)→ rpl19e (PRK08570)Ribosomal_L19e (pfam01280)→ |
| Hsa-3-1 | NC_010364.1 | + | 1247763 | 1247862 | **RNA**→ MTH1 (COG2106)→ rpl3p (PRK04231)Ribosomal_L3 (pfam00297)→<br>RplD (COG0088)rpl4lp (PRK04042)→ RplW (COG0089)PRK14548 (PRK14548)→<br>RplB (COG0090)rpl2p (PRK09612)→ RpsS (COG0185)rps19p (PRK04038)→<br>rpl22p (PRK04223)RplV (COG0091)→ RpsC (COG0092)rps3p (PRK04191)→<br>RpmC (COG0255)PRK14549 (PRK14549)→ POP4 (COG1588)PRK03879 (PRK03879)→<br>RpsQ (COG0186)rps17p (PRK08572)→ rpl14p (PRK08571)Ribosomal_L14 (pfam00238)→<br>RplX (COG0198)rpl24p (PRK01191)→<br>PRK04313 (PRK04313)KOW_RPS4 (cd06087)Ribosomal_S4e (pfam00900)→ RplE (COG0094)rpl5p (PRK04219)→<br>RpsN (COG0199)rps14P (PRK05766)→ rps8p (PRK04034)RpsH (COG0096)→<br>RplF (COG0097)rpl6p (PRK05518)→ PRK12766 (PRK12766)RecA (COG0468)Rpl32e (COG1717)→<br>rpl19e (PRK08570)Ribosomal_L19e (pfam01280)→ |

| | | | | | |
|---|---|---|---|---|---|
| Nch-1-1 | NZ_AOIN01000067.1 | + | 25511 | 25610 | **RNA**→ Methyltrn_RNA_3 (pfam02598)→ rpl3p (PRK04231)Ribosomal_L3 (pfam00297)→<br>RplD (COG0088)rpl4lp (PRK04042)→ RplW (COG0089)PRK14548 (PRK14548)→<br>RplB (COG0090)rpl2p (PRK09612)→ RpsS (COG0185)rps19p (PRK04038)→<br>rpl22p (PRK04223)RplV (COG0091)→ RpsC (COG0092)rps3p (PRK04191)→<br>PRK14549 (PRK14549)Ribosomal_L29 (pfam00831)→ PRK03879 (PRK03879)UPF0086 (pfam01868)→<br>RpsQ (COG0186)rps17p (PRK08572)→ rpl14p (PRK08571)Ribosomal_L14 (pfam00238)→<br>rpl24p (PRK01191)KOW_RPL26 (cd06089)→<br>PRK04313 (PRK04313)KOW_RPS4 (cd06087)S4 (pfam01479)Ribosomal_S4e (pfam00900)→<br>RplE (COG0094)rpl5p (PRK04219)→ RpsN (COG0199)rps14P (PRK05766)→<br>rps8p (PRK04034)RpsH (COG0096)→ RplF (COG0097)rpl6p (PRK05518)→<br>PRK12766 (PRK12766)RecA (COG0468)Rpl32e (COG1717)→ |
| Nae-1-1 | NZ_AOIP01000033.1 | - | 55488 | 55389 | **RNA**→ PRK13493 (PRK13493)Methyltrn_RNA_3 (pfam02598)→ rpl3p (PRK04231)Ribosomal_L3 (pfam00297)→<br>RplD (COG0088)rpl4lp (PRK04042)→ RplW (COG0089)PRK14548 (PRK14548)→<br>RplB (COG0090)rpl2p (PRK09612)→ RpsS (COG0185)rps19p (PRK04038)→<br>rpl22p (PRK04223)RplV (COG0091)→ RpsC (COG0092)rps3p (PRK04191)→<br>RpmC (COG0255)PRK14549 (PRK14549)→ PRK03879 (PRK03879)UPF0086 (pfam01868)→<br>RpsQ (COG0186)rps17p (PRK08572)→ rpl14p (PRK08571)Ribosomal_L14 (pfam00238)→<br>rpl24p (PRK01191)KOW_RPL26 (cd06089)→<br>PRK04313 (PRK04313)KOW_RPS4 (cd06087)Ribosomal_S4e (pfam00900)→ RplE (COG0094)rpl5p (PRK04219)→<br>RpsN (COG0199)rps14P (PRK05766)→ rps8p (PRK04034)RpsH (COG0096)→<br>RplF (COG0097)rpl6p (PRK05518)→ PRK12766 (PRK12766)RecA (COG0468)Rpl32e (COG1717)→ |
| env-214 | SS_3KL_010_SOIL_10059833 | - | 123 | 24 | **RNA**→ Methyltrn_RNA_3 (pfam02598)→ |
| env-215 | SL_3KL_010_SED_10538221 | + | 53 | 152 | **RNA**→ MTH1 (COG2106)→ |
| env-216 | P_1C_Sed_2_UnCty_1120614 | - | 293 | 194 | **RNA**→ Methyltrn_RNA_3 (pfam02598)→ |
| Hni-1-1 | NZ_AOMA01000004.1 | + | 593 | 692 | **RNA**→ Methyltrn_RNA_3 (pfam02598)→ rpl3p (PRK04231)Ribosomal_L3 (pfam00297)→<br>RplD (COG0088)rpl4lp (PRK04042)→ RplW (COG0089)PRK14548 (PRK14548)→<br>RplB (COG0090)rpl2p (PRK09612)→ RpsS (COG0185)rps19p (PRK04038)→<br>rpl22p (PRK04223)RplV (COG0091)→ RpsC (COG0092)rps3p (PRK04191)→<br>RpmC (COG0255)PRK14549 (PRK14549)→ PRK03879 (PRK03879)UPF0086 (pfam01868)→<br>RpsQ (COG0186)rps17p (PRK08572)→ rpl14p (PRK08571)Ribosomal_L14 (pfam00238)→<br>rpl24p (PRK01191)KOW_RPL26 (cd06089)→<br>PRK04313 (PRK04313)KOW_RPS4 (cd06087)Ribosomal_S4e (pfam00900)→ RplE (COG0094)rpl5p (PRK04219)→<br>RpsN (COG0199)rps14P (PRK05766)→ rps8p (PRK04034)RpsH (COG0096)→<br>RplF (COG0097)rpl6p (PRK05518)→ UvrC (COG0322)HHH_5 (pfam14520)→ hypo→ |
| Has-1-1 | NZ_AOIQ01000021.1 | + | 411355 | 411454 | **RNA**→ Methyltrn_RNA_3 (pfam02598)→ RplC (COG0087)rpl3p (PRK04231)→<br>RplD (COG0088)rpl4lp (PRK04042)→ RplW (COG0089)PRK14548 (PRK14548)→<br>RplB (COG0090)rpl2p (PRK09612)→ RpsS (COG0185)rps19p (PRK04038)→<br>rpl22p (PRK04223)RplV (COG0091)→ RpsC (COG0092)rps3p (PRK04191)→<br>RpmC (COG0255)PRK14549 (PRK14549)→ PRK03879 (PRK03879)UPF0086 (pfam01868)→<br>RpsQ (COG0186)rps17p (PRK08572)→ rpl14p (PRK08571)Ribosomal_L14 (pfam00238)→<br>rpl24p (PRK01191)KOW_RPL26 (cd06089)→<br>PRK04313 (PRK04313)KOW_RPS4 (cd06087)Ribosomal_S4e (pfam00900)→ RplE (COG0094)rpl5p (PRK04219)→<br>RpsN (COG0199)rps14P (PRK05766)→ rps8p (PRK04034)RpsH (COG0096)→<br>RplF (COG0097)rpl6p (PRK05518)→<br>UvrC (COG0322)Rpl32e (COG1717)rpl32e (PRK08562)HHH_5 (pfam14520)→<br>rpl19e (PRK08570)Ribosomal_L19e (pfam01280)→ |
| env-217 | 2140918027-11734 | - | 146 | 47 | **RNA**→ Methyltrn_RNA_3 (pfam02598)→ |

| | | | | | |
|---|---|---|---|---|---|
| Nal-2-1 | NZ_JNCS01000001.1 | - | 130682 | 130583 | **RNA**→ Methyltrn_RNA_3 (pfam02598)→ rpl3p (PRK04231)Ribosomal_L3 (pfam00297)→<br>RplD (COG0088)rpl4lp (PRK04042)→ RplW (COG0089)PRK14548 (PRK14548)→<br>RplB (COG0090)rpl2p (PRK09612)→ RpsS (COG0185)rps19p (PRK04038)→<br>rpl22p (PRK04223)RplV (COG0091)→ RpsC (COG0092)rps3p (PRK04191)→<br>RpmC (COG0255)PRK14549 (PRK14549)→ PRK03879 (PRK03879)UPF0086 (pfam01868)→<br>RpsQ (COG0186)rps17p (PRK08572)→ rpl14p (PRK08571)Ribosomal_L14 (pfam00238)→<br>rpl24p (PRK01191)KOW_RPL26 (cd06089)→<br>PRK04313 (PRK04313)KOW_RPS4 (cd06087)Ribosomal_S4e (pfam00900)→ RplE (COG0094)rpl5p (PRK04219)→<br>RpsN (COG0199)rps14P (PRK05766)→ rps8p (PRK04034)RpsH (COG0096)→<br>RplF (COG0097)rpl6p (PRK05518)→ PRK12766 (PRK12766)UvrC (COG0322)Rpl32e (COG1717)→ |
| Nal-1-1 | NZ_AOIK01000025.1 | + | 71137 | 71236 | **RNA**→ Methyltrn_RNA_3 (pfam02598)→ rpl3p (PRK04231)Ribosomal_L3 (pfam00297)→<br>RplD (COG0088)rpl4lp (PRK04042)→ RplW (COG0089)PRK14548 (PRK14548)→<br>RplB (COG0090)rpl2p (PRK09612)→ RpsS (COG0185)rps19p (PRK04038)→<br>rpl22p (PRK04223)RplV (COG0091)→ RpsC (COG0092)rps3p (PRK04191)→<br>RpmC (COG0255)PRK14549 (PRK14549)→ PRK03879 (PRK03879)UPF0086 (pfam01868)→<br>RpsQ (COG0186)rps17p (PRK08572)→ rpl14p (PRK08571)Ribosomal_L14 (pfam00238)→<br>rpl24p (PRK01191)KOW_RPL26 (cd06089)→<br>PRK04313 (PRK04313)KOW_RPS4 (cd06087)Ribosomal_S4e (pfam00900)→ RplE (COG0094)rpl5p (PRK04219)→<br>RpsN (COG0199)rps14P (PRK05766)→ rps8p (PRK04034)RpsH (COG0096)→<br>RplF (COG0097)rpl6p (PRK05518)→ PRK12766 (PRK12766)UvrC (COG0322)Rpl32e (COG1717)→ |
| Npa-1-1 | NZ_AOII01000067.1 | - | 29396 | 29297 | **RNA**→ Methyltrn_RNA_3 (pfam02598)→ rpl3p (PRK04231)Ribosomal_L3 (pfam00297)→<br>RplD (COG0088)rpl4lp (PRK04042)→ RplW (COG0089)PRK14548 (PRK14548)→<br>RplB (COG0090)rpl2p (PRK09612)→ RpsS (COG0185)rps19p (PRK04038)→<br>rpl22p (PRK04223)RplV (COG0091)→ RpsC (COG0092)rps3p (PRK04191)→<br>RpmC (COG0255)PRK14549 (PRK14549)→ PRK03879 (PRK03879)UPF0086 (pfam01868)→<br>RpsQ (COG0186)rps17p (PRK08572)→ rpl14p (PRK08571)Ribosomal_L14 (pfam00238)→<br>rpl24p (PRK01191)KOW_RPL26 (cd06089)→<br>PRK04313 (PRK04313)KOW_RPS4 (cd06087)Ribosomal_S4e (pfam00900)→ RplE (COG0094)rpl5p (PRK04219)→<br>RpsN (COG0199)rps14P (PRK05766)→ rps8p (PRK04034)RpsH (COG0096)→<br>RplF (COG0097)rpl6p (PRK05518)→<br>UvrC (COG0322)Rpl32e (COG1717)rpl32e (PRK08562)HHH_5 (pfam14520)→ |
| Nga-1-1 | NZ_AOIJ01000042.1 | - | 42266 | 42167 | **RNA**→ Methyltrn_RNA_3 (pfam02598)→ rpl3p (PRK04231)Ribosomal_L3 (pfam00297)→<br>RplD (COG0088)rpl4lp (PRK04042)→ RplW (COG0089)PRK14548 (PRK14548)→<br>RplB (COG0090)rpl2p (PRK09612)→ RpsS (COG0185)rps19p (PRK04038)→<br>rpl22p (PRK04223)RplV (COG0091)→ RpsC (COG0092)rps3p (PRK04191)→<br>RpmC (COG0255)PRK14549 (PRK14549)→ PRK03879 (PRK03879)UPF0086 (pfam01868)→<br>RpsQ (COG0186)rps17p (PRK08572)→ rpl14p (PRK08571)Ribosomal_L14 (pfam00238)→<br>rpl24p (PRK01191)KOW_RPL26 (cd06089)→<br>PRK04313 (PRK04313)KOW_RPS4 (cd06087)Ribosomal_S4e (pfam00900)→ RplE (COG0094)rpl5p (PRK04219)→<br>RpsN (COG0199)rps14P (PRK05766)→ rps8p (PRK04034)RpsH (COG0096)→<br>RplF (COG0097)rpl6p (PRK05518)→<br>RecA (COG0468)Rpl32e (COG1717)rpl32e (PRK08562)HHH_5 (pfam14520)→<br>rpl19e (PRK08570)Ribosomal_L19e (pfam01280)→ |
| Nsp-2-1 | NZ_AJVG01000002.1 | - | 354632 | 354533 | **RNA**→ Methyltrn_RNA_3 (pfam02598)→ rpl3p (PRK04231)Ribosomal_L3 (pfam00297)→<br>RplD (COG0088)rpl4lp (PRK04042)→ RplW (COG0089)PRK14548 (PRK14548)→<br>RplB (COG0090)rpl2p (PRK09612)→ RpsS (COG0185)rps19p (PRK04038)→<br>rpl22p (PRK04223)RplV (COG0091)→ RpsC (COG0092)rps3p (PRK04191)→<br>RpmC (COG0255)PRK14549 (PRK14549)→ PRK03879 (PRK03879)UPF0086 (pfam01868)→<br>RpsQ (COG0186)rps17p (PRK08572)→ rpl14p (PRK08571)Ribosomal_L14 (pfam00238)→<br>rpl24p (PRK01191)KOW_RPL26 (cd06089)→<br>PRK04313 (PRK04313)KOW_RPS4 (cd06087)Ribosomal_S4e (pfam00900)→ RplE (COG0094)rpl5p (PRK04219)→<br>RpsN (COG0199)rps14P (PRK05766)→ rps8p (PRK04034)RpsH (COG0096)→<br>RplF (COG0097)rpl6p (PRK05518)→<br>RecA (COG0468)Rpl32e (COG1717)rpl32e (PRK08562)HHH_5 (pfam14520)→<br>rpl19e (PRK08570)Ribosomal_L19e (pfam01280)→ |

| | | | | | |
|---|---|---|---|---|---|
| Nsp-1-1 | NC_018224.1 | - | 1817666 | 1817567 | **RNA**→ Methyltrn_RNA_3 (pfam02598)→ rpl3p (PRK04231)Ribosomal_L3 (pfam00297)→ RplD (COG0088)rpl4lp (PRK04042)→ RplW (COG0089)PRK14548 (PRK14548)→ RplB (COG0090)rpl2p (PRK09612)→ RpsS (COG0185)rps19p (PRK04038)→ rpl22p (PRK04223)RplV (COG0091)→ RpsC (COG0092)rps3p (PRK04191)→ RpmC (COG0255)PRK14549 (PRK14549)→ PRK03879 (PRK03879)UPF0086 (pfam01868)→ RpsQ (COG0186)rps17p (PRK08572)→ rpl14p (PRK08571)Ribosomal_L14 (pfam00238)→ rpl24p (PRK01191)KOW_RPL26 (cd06089)→ PRK04313 (PRK04313)KOW_RPS4 (cd06087)Ribosomal_S4e (pfam00900)→ RplE (COG0094)rpl5p (PRK04219)→ RpsN (COG0199)rps14P (PRK05766)→ rps8p (PRK04034)RpsH (COG0096)→ RplF (COG0097)rpl6p (PRK05518)→ RecA (COG0468)Rpl32e (COG1717)rpl32e (PRK08562)HHH_5 (pfam14520)→ rpl19e (PRK08570)Ribosomal_L19e (pfam01280)→ |
| Htu-1-1 | NC_013743.1 | + | 2295636 | 2295735 | **RNA**→ Methyltrn_RNA_3 (pfam02598)→ rpl3p (PRK04231)Ribosomal_L3 (pfam00297)→ RplD (COG0088)rpl4lp (PRK04042)→ RplW (COG0089)PRK14548 (PRK14548)→ RplB (COG0090)rpl2p (PRK09612)→ RpsS (COG0185)rps19p (PRK04038)→ rpl22p (PRK04223)RplV (COG0091)→ RpsC (COG0092)rps3p (PRK04191)→ RpmC (COG0255)PRK14549 (PRK14549)→ PRK03879 (PRK03879)UPF0086 (pfam01868)→ RpsQ (COG0186)rps17p (PRK08572)→ rpl14p (PRK08571)Ribosomal_L14 (pfam00238)→ rpl24p (PRK01191)KOW_RPL26 (cd06089)→ PRK04313 (PRK04313)S4 (cd00165)KOW_RPS4 (cd06087)Ribosomal_S4e (pfam00900)→ RplE (COG0094)rpl5p (PRK04219)→ RpsN (COG0199)rps14P (PRK05766)→ rps8p (PRK04034)RpsH (COG0096)→ RplF (COG0097)rpl6p (PRK05518)→ PRK12766 (PRK12766)UvrC (COG0322)Rpl32e (COG1717)→ rpl19e (PRK08570)Ribosomal_L19e (pfam01280)→ |
| Hsa-8-1 | NZ_AOIS01000045.1 | - | 15715 | 15616 | **RNA**→ Methyltrn_RNA_3 (pfam02598)→ rpl3p (PRK04231)Ribosomal_L3 (pfam00297)→ RplD (COG0088)rpl4lp (PRK04042)→ RplW (COG0089)PRK14548 (PRK14548)→ RplB (COG0090)rpl2p (PRK09612)→ RpsS (COG0185)rps19p (PRK04038)→ rpl22p (PRK04223)RplV (COG0091)→ RpsC (COG0092)rps3p (PRK04191)→ PRK14549 (PRK14549)Ribosomal_L29 (pfam00831)→ PRK03879 (PRK03879)UPF0086 (pfam01868)→ RpsQ (COG0186)rps17p (PRK08572)→ rpl14p (PRK08571)Ribosomal_L14 (pfam00238)→ rpl24p (PRK01191)KOW_RPL26 (cd06089)→ PRK04313 (PRK04313)S4 (cd00165)KOW_RPS4 (cd06087)Ribosomal_S4e (pfam00900)→ RplE (COG0094)rpl5p (PRK04219)→ RpsN (COG0199)rps14P (PRK05766)→ rps8p (PRK04034)RpsH (COG0096)→ RplF (COG0097)rpl6p (PRK05518)→ PRK12766 (PRK12766)UvrC (COG0322)Rpl32e (COG1717)→ rpl19e (PRK08570)Ribosomal_L19e (pfam01280)→ |
| Hth-2-1 | NZ_AOIR01000063.1 | + | 173689 | 173788 | **RNA**→ Methyltrn_RNA_3 (pfam02598)→ RplC (COG0087)rpl3p (PRK04231)→ RplD (COG0088)rpl4lp (PRK04042)→ RplW (COG0089)PRK14548 (PRK14548)→ RplB (COG0090)rpl2p (PRK09612)→ RpsS (COG0185)rps19p (PRK04038)→ rpl22p (PRK04223)RplV (COG0091)→ RpsC (COG0092)rps3p (PRK04191)→ RpmC (COG0255)PRK14549 (PRK14549)→ PRK03879 (PRK03879)UPF0086 (pfam01868)→ RpsQ (COG0186)rps17p (PRK08572)→ rpl14p (PRK08571)Ribosomal_L14 (pfam00238)→ rpl24p (PRK01191)KOW_RPL26 (cd06089)→ PRK04313 (PRK04313)KOW_RPS4 (cd06087)Ribosomal_S4e (pfam00900)→ RplE (COG0094)rpl5p (PRK04219)→ RpsN (COG0199)rps14P (PRK05766)→ rps8p (PRK04034)RpsH (COG0096)→ RplF (COG0097)rpl6p (PRK05518)→ UvrC (COG0322)Rpl32e (COG1717)rpl32e (PRK08562)HHH_5 (pfam14520)→ rpl19e (PRK08570)Ribosomal_L19e (pfam01280)→ |

| | | | | | |
|---|---|---|---|---|---|
| Hje-2-1 | NZ_JDTG01000014.1 | + | 597395 | 597494 | **RNA**→ Methyltrn_RNA_3 (pfam02598)→ RplC (COG0087)rpl3p (PRK04231)→ |
| | | | | | RplD (COG0088)rpl4lp (PRK04042)→ RplW (COG0089)PRK14548 (PRK14548)→ |
| | | | | | RplB (COG0090)rpl2p (PRK09612)→ RpsS (COG0185)rps19p (PRK04038)→ |
| | | | | | rpl22p (PRK04223)RplV (COG0091)→ RpsC (COG0092)rps3p (PRK04191)→ |
| | | | | | RpmC (COG0255)PRK14549 (PRK14549)→ PRK03879 (PRK03879)UPF0086 (pfam01868)→ |
| | | | | | RpsQ (COG0186)rps17p (PRK08572)→ rpl14p (PRK08571)Ribosomal_L14 (pfam00238)→ |
| | | | | | rpl24p (PRK01191)KOW_RPL26 (cd06089)→ |
| | | | | | PRK04313 (PRK04313)KOW_RPS4 (cd06087)Ribosomal_S4e (pfam00900)→ RplE (COG0094)rpl5p (PRK04219)→ |
| | | | | | RpsN (COG0199)rps14P (PRK05766)→ rps8p (PRK04034)RpsH (COG0096)→ |
| | | | | | RplF (COG0097)rpl6p (PRK05518)→ |
| | | | | | UvrC (COG0322)Rpl32e (COG1717)rpl32e (PRK08562)HHH_5 (pfam14520)→ |
| | | | | | rpl19e (PRK08570)Ribosomal_L19e (pfam01280)→ |
| Nve-1-1 | NZ_AOID01000027.1 | + | 70079 | 70178 | **RNA**→ Methyltrn_RNA_3 (pfam02598)→ rpl3p (PRK04231)Ribosomal_L3 (pfam00297)→ |
| | | | | | RplD (COG0088)rpl4lp (PRK04042)→ RplW (COG0089)PRK14548 (PRK14548)→ |
| | | | | | RplB (COG0090)rpl2p (PRK09612)→ RpsS (COG0185)rps19p (PRK04038)→ |
| | | | | | rpl22p (PRK04223)RplV (COG0091)→ RpsC (COG0092)rps3p (PRK04191)→ |
| | | | | | PRK14549 (PRK14549)Ribosomal_L29 (pfam00831)→ PRK03879 (PRK03879)UPF0086 (pfam01868)→ |
| | | | | | RpsQ (COG0186)rps17p (PRK08572)→ rpl14p (PRK08571)Ribosomal_L14 (pfam00238)→ |
| | | | | | rpl24p (PRK01191)KOW_RPL26 (cd06089)→ |
| | | | | | PRK04313 (PRK04313)KOW_RPS4 (cd06087)Ribosomal_S4e (pfam00900)→ RplE (COG0094)rpl5p (PRK04219)→ |
| | | | | | RpsN (COG0199)rps14P (PRK05766)→ rps8p (PRK04034)RpsH (COG0096)→ |
| | | | | | RplF (COG0097)rpl6p (PRK05518)→ PRK12766 (PRK12766)UvrC (COG0322)Rpl32e (COG1717)→ |
| Hli-3-1 | NZ_AOIT01000030.1 | + | 41883 | 41982 | **RNA**→ Methyltrn_RNA_3 (pfam02598)→ rpl3p (PRK04231)Ribosomal_L3 (pfam00297)→ |
| | | | | | RplD (COG0088)rpl4lp (PRK04042)→ RplW (COG0089)PRK14548 (PRK14548)→ |
| | | | | | RplB (COG0090)rpl2p (PRK09612)→ RpsS (COG0185)rps19p (PRK04038)→ |
| | | | | | rpl22p (PRK04223)RplV (COG0091)→ RpsC (COG0092)rps3p (PRK04191)→ |
| | | | | | RpmC (COG0255)PRK14549 (PRK14549)→ PRK03879 (PRK03879)UPF0086 (pfam01868)→ |
| | | | | | RpsQ (COG0186)rps17p (PRK08572)→ rpl14p (PRK08571)Ribosomal_L14 (pfam00238)→ |
| | | | | | rpl24p (PRK01191)KOW_RPL26 (cd06089)→ |
| | | | | | PRK04313 (PRK04313)S4 (cd00165)KOW_RPS4 (cd06087)Ribosomal_S4e (pfam00900)→ |
| | | | | | RplE (COG0094)rpl5p (PRK04219)→ RpsN (COG0199)rps14P (PRK05766)→ |
| | | | | | rps8p (PRK04034)RpsH (COG0096)→ RplF (COG0097)rpl6p (PRK05518)→ |
| | | | | | RecA (COG0468)rpl32e (PRK08562)Ribosomal_L32e (pfam01655)HHH_5 (pfam14520)→ |
| | | | | | rpl19e (PRK08570)Ribosomal_L19e (pfam01280)→ |
| Hxa-1-1 | NC_015666.1 | - | 345106 | 345007 | **RNA**→ Methyltrn_RNA_3 (pfam02598)→ rpl3p (PRK04231)Ribosomal_L3 (pfam00297)→ |
| | | | | | RplD (COG0088)rpl4lp (PRK04042)→ RplW (COG0089)PRK14548 (PRK14548)→ |
| | | | | | RplB (COG0090)rpl2p (PRK09612)→ RpsS (COG0185)rps19p (PRK04038)→ |
| | | | | | rpl22p (PRK04223)RplV (COG0091)→ RpsC (COG0092)rps3p (PRK04191)→ |
| | | | | | RpmC (COG0255)PRK14549 (PRK14549)→ PRK03879 (PRK03879)UPF0086 (pfam01868)→ |
| | | | | | RpsQ (COG0186)rps17p (PRK08572)→ rpl14p (PRK08571)Ribosomal_L14 (pfam00238)→ |
| | | | | | rpl24p (PRK01191)KOW_RPL26 (cd06089)→ |
| | | | | | PRK04313 (PRK04313)S4 (cd00165)KOW_RPS4 (cd06087)Ribosomal_S4e (pfam00900)→ |
| | | | | | RplE (COG0094)rpl5p (PRK04219)→ RpsN (COG0199)rps14P (PRK05766)→ |
| | | | | | rps8p (PRK04034)RpsH (COG0096)→ RplF (COG0097)rpl6p (PRK05518)→ |
| | | | | | RecA (COG0468)Rpl32e (COG1717)rpl32e (PRK08562)HHH_5 (pfam14520)→ |
| | | | | | rpl19e (PRK08570)Ribosomal_L19e (pfam01280)→ |

| | | | | | |
|---|---|---|---|---|---|
| Hgo-1-1 | NZ_HG315690.1 | + | 2029000 | 2029099 | **RNA**→ Methyltrn_RNA_3 (pfam02598)→ rpl3p (PRK04231)Ribosomal_L3 (pfam00297)→ |
| | | | | | RplD (COG0088)rpl4lp (PRK04042)→ RplW (COG0089)PRK14548 (PRK14548)→ |
| | | | | | RplB (COG0090)rpl2p (PRK09612)→ RpsS (COG0185)rps19p (PRK04038)→ |
| | | | | | rpl22p (PRK04223)RplV (COG0091)→ RpsC (COG0092)rps3p (PRK04191)→ |
| | | | | | RpmC (COG0255)PRK14549 (PRK14549)→ PRK03879 (PRK03879)UPF0086 (pfam01868)→ |
| | | | | | RpsQ (COG0186)rps17p (PRK08572)→ rpl14p (PRK08571)Ribosomal_L14 (pfam00238)→ |
| | | | | | rpl24p (PRK01191)KOW_RPL26 (cd06089)→ |
| | | | | | PRK04313 (PRK04313)KOW_RPS4 (cd06087)Ribosomal_S4e (pfam00900)→ RplE (COG0094)rpl5p (PRK04219)→ |
| | | | | | RpsN (COG0199)rps14P (PRK05766)→ rps8p (PRK04034)RpsH (COG0096)→ |
| | | | | | RplF (COG0097)rpl6p (PRK05518)→ |
| | | | | | UvrC (COG0322)Rpl32e (COG1717)rpl32e (PRK08562)HHH_5 (pfam14520)→ |
| env-218 | SL_5KL_010_BRINE_10298960 | + | 132 | 231 | **RNA**→ Methyltrn_RNA_3 (pfam02598)→ |
| env-219 | SL_4KL_010_BRINE_10234182 | + | 69 | 168 | **RNA**→ MTH1 (COG2106)→ |
| env-220 | GSLNARP_GFPJP1N02GH0H6 | + | 114 | 213 | **RNA**→ MTH1 (COG2106)→ |
| env-221 | SL_5KL_010_BRINE_10027796 | + | 1534 | 1633 | **RNA**→ Methyltrn_RNA_3 (pfam02598)→ |
| Hsa-6-1 | NZ_ALJD01000008.1 | + | 211183 | 211282 | **RNA**→ Methyltrn_RNA_3 (pfam02598)→ Methyltrn_RNA_3 (pfam02598)→ RplC (COG0087)rpl3p (PRK04231)→ |
| | | | | | RplC (COG0087)rpl3p (PRK04231)→ RplD (COG0088)rpl4lp (PRK04042)→ |
| | | | | | RplD (COG0088)rpl4lp (PRK04042)→ RplW (COG0089)PRK14548 (PRK14548)→ |
| | | | | | RplW (COG0089)PRK14548 (PRK14548)→ RplB (COG0090)rpl2p (PRK09612)→ |
| | | | | | RplB (COG0090)rpl2p (PRK09612)→ RpsS (COG0185)rps19p (PRK04038)→ |
| | | | | | RpsS (COG0185)rps19p (PRK04038)→ rpl22p (PRK04223)RplV (COG0091)→ |
| | | | | | rpl22p (PRK04223)RplV (COG0091)→ RpsC (COG0092)rps3p (PRK04191)→ |
| | | | | | RpsC (COG0092)rps3p (PRK04191)→ PRK14549 (PRK14549)Ribosomal_L29 (pfam00831)→ |
| | | | | | PRK14549 (PRK14549)Ribosomal_L29 (pfam00831)→ PRK03879 (PRK03879)UPF0086 (pfam01868)→ |
| | | | | | PRK03879 (PRK03879)UPF0086 (pfam01868)→ RpsQ (COG0186)rps17p (PRK08572)→ |
| | | | | | rps17p (PRK08572)Ribosomal_S17 (pfam00366)→ rps17p (PRK08572)Ribosomal_S17 (pfam00366)→ |
| | | | | | rpl14p (PRK08571)Ribosomal_L14 (pfam00238)→ rpl14p (PRK08571)Ribosomal_L14 (pfam00238)→ |
| | | | | | rpl24p (PRK01191)KOW_RPL26 (cd06089)→ rpl24p (PRK01191)KOW_RPL26 (cd06089)→ |
| | | | | | PRK04313 (PRK04313)KOW_RPS4 (cd06087)Ribosomal_S4e (pfam00900)→ |
| | | | | | PRK04313 (PRK04313)KOW_RPS4 (cd06087)Ribosomal_S4e (pfam00900)→ RplE (COG0094)rpl5p (PRK04219)→ |
| | | | | | RplE (COG0094)rpl5p (PRK04219)→ RpsN (COG0199)rps14P (PRK05766)→ |
| | | | | | RpsN (COG0199)rps14P (PRK05766)→ rps8p (PRK04034)RpsH (COG0096)→ |
| | | | | | rps8p (PRK04034)RpsH (COG0096)→ RplF (COG0097)rpl6p (PRK05518)→ |
| | | | | | RplF (COG0097)rpl6p (PRK05518)→ PRK12766 (PRK12766)RecA (COG0468)Rpl32e (COG1717)→ |
| | | | | | PRK12766 (PRK12766)RecA (COG0468)Rpl32e (COG1717)→ rpl19e (PRK08570)Ribosomal_L19e (pfam01280)→ |
| | | | | | rpl19e (PRK08570)Ribosomal_L19e (pfam01280)→ |
| env-222 | SL_4KL_010_BRINE_10138442 | + | 547 | 646 | **RNA**→ Methyltrn_RNA_3 (pfam02598)→ |
| env-223 | GSLNARP_GFPJP1N02IH3RV | + | 123 | 222 | **RNA**→ MTH1 (COG2106)→ |
| env-224 | GSLNARP_GFPJP1N02G85LW | - | 148 | 49 | **RNA**→ |
| env-225 | CEQN01112068.1 | - | 3026 | 2927 | **RNA**→ Methyltrn_RNA_3 (pfam02598)→ RplC (COG0087)rpl3p (PRK04231)→ |
| | | | | | RplD (COG0088)rpl4lp (PRK04042)→ RplW (COG0089)PRK14548 (PRK14548)→ |
| | | | | | rpl2p (PRK09612)Ribosomal_L2 (pfam00181)Ribosomal_L2_C (pfam03947)→ |
| env-226 | CEUZ01492716.1 | + | 1180 | 1279 | **RNA**→ MTH1 (COG2106)→ |
| env-227 | CEWR01279639.1 | - | 119 | 20 | **RNA**→ Methyltrn_RNA_3 (pfam02598)→ |
| env-228 | APHM01021436.1 | - | 16571 | 16472 | **RNA**→ Methyltrn_RNA_3 (pfam02598)→ RplC (COG0087)rpl3p (PRK04231)→ |
| | | | | | RplD (COG0088)rpl4lp (PRK04042)→ RplW (COG0089)PRK14548 (PRK14548)→ |
| | | | | | RplB (COG0090)rpl2p (PRK09612)→ |
| env-229 | LTJ07As871_J07ABscf098859 | - | 1081824 | 1081725 | **RNA**→ Methyltrn_RNA_3 (pfam02598)→ RplC (COG0087)rpl3p (PRK04231)→ |
| | | | | | RplD (COG0088)rpl4lp (PRK04042)→ RplW (COG0089)PRK14548 (PRK14548)→ |
| | | | | | RplB (COG0090)rpl2p (PRK09612)→ |
| env-230 | JXWS01149363.1 | - | 957 | 858 | **RNA**→ Methyltrn_RNA_3 (pfam02598)→ rpl3p (PRK04231)Ribosomal_L3 (pfam00297)→ |
| env-231 | 3300000032_0007378 | - | 20967 | 20877 | **RNA**→ RplC (COG0087)rpl3p (PRK04231)→ rpl4lp (PRK04042)Ribosomal_L4 (pfam00573)→ |
| env-232 | 3300000568_10935138 | + | 219 | 307 | **RNA**→ AF6_RA_repeat2 (cd01781)→ |
| env-233 | 2232068851 | - | 371 | 281 | **RNA**→ ←hypo |

| | | | | | |
|---|---|---|---|---|---|
| env-234 | 3300000558_10378615 | + | 2773 | 2863 | **RNA**→ RplC (COG0087)rpl3p (PRK04231)→ RplD (COG0088)rpl4lp (PRK04042)→<br>RplW (COG0089)PRK14548 (PRK14548)→ rpl2p (PRK09612)Ribosomal_L2 (pfam00181)→ |
| env-235 | 3300000568_10054768 | + | 16095 | 16185 | **RNA**→ RplC (COG0087)rpl3p (PRK04231)→ RplD (COG0088)rpl4lp (PRK04042)→<br>RplW (COG0089)PRK14548 (PRK14548)→ RplB (COG0090)rpl2p (PRK09612)→<br>RpsS (COG0185)rps19p (PRK04038)→ rpl22p (PRK04223)RplV (COG0091)→<br>RpsC (COG0092)rps3p (PRK04191)→ PRK14549 (PRK14549)Ribosomal_L29 (pfam00831)→<br>PRK03879 (PRK03879)UPF0086 (pfam01868)→ RpsQ (COG0186)rps17p (PRK08572)→<br>rpl14p (PRK08571)Ribosomal_L14 (pfam00238)→ rpl24p (PRK01191)KOW_RPL26 (cd06089)→<br>PRK04313 (PRK04313)KOW_RPS4 (cd06087)Ribosomal_S4e (pfam00900)→ RplE (COG0094)rpl5p (PRK04219)→<br>RpsN (COG0199)rps14P (PRK05766)→ rps8p (PRK04034)RpsH (COG0096)→<br>RplF (COG0097)rpl6p (PRK05518)→ rpl32e (PRK08562)Ribosomal_L32e (pfam01655)→<br>rpl19e (PRK08570)Ribosomal_L19e (pfam01280)→ RplR (COG0256)rpl18p (PRK08569)→<br>RpsE (COG0098)rps5p (PRK04044)→ |
| env-236 | 3300001580_10000682 | + | 16095 | 16185 | **RNA**→ RplC (COG0087)rpl3p (PRK04231)→ RplD (COG0088)rpl4lp (PRK04042)→<br>RplW (COG0089)PRK14548 (PRK14548)→ RplB (COG0090)rpl2p (PRK09612)→<br>RpsS (COG0185)rps19p (PRK04038)→ rpl22p (PRK04223)RplV (COG0091)→<br>RpsC (COG0092)rps3p (PRK04191)→ PRK14549 (PRK14549)Ribosomal_L29 (pfam00831)→<br>PRK03879 (PRK03879)UPF0086 (pfam01868)→ RpsQ (COG0186)rps17p (PRK08572)→<br>rpl14p (PRK08571)Ribosomal_L14 (pfam00238)→ rpl24p (PRK01191)KOW_RPL26 (cd06089)→<br>PRK04313 (PRK04313)KOW_RPS4 (cd06087)Ribosomal_S4e (pfam00900)→ RplE (COG0094)rpl5p (PRK04219)→<br>RpsN (COG0199)rps14P (PRK05766)→ rps8p (PRK04034)RpsH (COG0096)→<br>RplF (COG0097)rpl6p (PRK05518)→ rpl32e (PRK08562)Ribosomal_L32e (pfam01655)→<br>rpl19e (PRK08570)Ribosomal_L19e (pfam01280)→ RplR (COG0256)rpl18p (PRK08569)→<br>RpsE (COG0098)rps5p (PRK04044)→ |
| env-237 | WSSedB2T_c028865 | - | 329 | 244 | **RNA**→ ←hypo |
| env-238 | JGI24020J35080_1001304 | - | 6969 | 6882 | **RNA**→ PRK05883 (PRK05883)Methyltrn_RNA_3 (pfam02598)→ RplC (COG0087)rpl3p (PRK04231)→<br>RplD (COG0088)rpl4lp (PRK04042)→ RplW (COG0089)PRK14548 (PRK14548)→<br>RplB (COG0090)rpl2p (PRK09612)→ RpsS (COG0185)rps19p (PRK04038)→<br>rpl22p (PRK04223)RplV (COG0091)→ RpsC (COG0092)rps3p (PRK04191)→<br>PRK14549 (PRK14549)Ribosomal_L29 (pfam00831)→ POP4 (COG1588)PRK03879 (PRK03879)→<br>RpsQ (COG0186)rps17p (PRK08572)→ rpl14p (PRK08571)Ribosomal_L14 (pfam00238)→<br>rpl24p (PRK01191)KOW_RPL26 (cd06089)→<br>PRK04313 (PRK04313)S4 (cd00165)KOW_RPS4 (cd06087)Ribosomal_S4e (pfam00900)→ |
| env-239 | JGI24019J35510_1000070 | - | 95685 | 95598 | **RNA**→ PRK05883 (PRK05883)Methyltrn_RNA_3 (pfam02598)→ RplC (COG0087)rpl3p (PRK04231)→<br>RplD (COG0088)rpl4lp (PRK04042)→ RplW (COG0089)PRK14548 (PRK14548)→<br>RplB (COG0090)rpl2p (PRK09612)→ RpsS (COG0185)rps19p (PRK04038)→<br>rpl22p (PRK04223)RplV (COG0091)→ RpsC (COG0092)rps3p (PRK04191)→<br>PRK14549 (PRK14549)Ribosomal_L29 (pfam00831)→ POP4 (COG1588)PRK03879 (PRK03879)→<br>RpsQ (COG0186)rps17p (PRK08572)→ rpl14p (PRK08571)Ribosomal_L14 (pfam00238)→<br>rpl24p (PRK01191)KOW_RPL26 (cd06089)→<br>PRK04313 (PRK04313)S4 (cd00165)KOW_RPS4 (cd06087)Ribosomal_S4e (pfam00900)→<br>RplE (COG0094)rpl5p (PRK04219)→ RpsN (COG0199)rps14P (PRK05766)→<br>rps8p (PRK04034)RpsH (COG0096)→ RplF (COG0097)rpl6p (PRK05518)→<br>Rpl32e (COG1717)rpl32e (PRK08562)HHH (pfam00633)HHH_5 (pfam14520)→<br>rpl19e (PRK08570)Ribosomal_L19e (pfam01280)→ RplR (COG0256)rpl18p (PRK08569)→<br>RpsE (COG0098)rps5p (PRK04044)→ |
| env-240 | PBDCA2_contig02310 | + | 406 | 502 | **RNA**→ Methyltrn_RNA_3 (pfam02598)→ |
| env-241 | LGVC01039323.1 | + | 218 | 314 | **RNA**→ lepA (TIGR01393)Methyltrn_RNA_3 (pfam02598)→ |

| | | | | | |
|---|---|---|---|---|---|
| env-242 | JGI24730J26740_1000024 | - | 31005 | 30928 | **RNA**→ RplC (COG0087)rpl3p (PRK04231)→ RplD (COG0088)rpl4lp (PRK04042)→<br>RplW (COG0089)PRK14548 (PRK14548)→ RplB (COG0090)rpl2p (PRK09612)→<br>rps19p (PRK04038)Ribosomal_S19 (pfam00203)→ rpl22p (PRK04223)RplV (COG0091)→<br>RpsC (COG0092)rps3p (PRK04191)→ PRK14549 (PRK14549)Ribosomal_L29_HIP (cd00427)→<br>POP4 (COG1588)PRK03879 (PRK03879)→ RpsQ (COG0186)rps17p (PRK08572)→<br>rpl14p (PRK08571)Ribosomal_L14 (pfam00238)→ rpl24p (PRK01191)KOW_RPL26 (cd06089)→<br>PRK04313 (PRK04313)S4 (cd00165)KOW_RPS4 (cd06087)Ribosomal_S4e (pfam00900)→<br>rpl5p (PRK04219)Ribosomal_L5 (pfam00281)Ribosomal_L5_C (pfam00673)→<br>RpsN (COG0199)rps14P (PRK05766)→ rps8p (PRK04034)RpsH (COG0096)→<br>RplF (COG0097)rpl6p (PRK05518)→ rpl32e (PRK08562)Ribosomal_L32e (pfam01655)→<br>rpl19e (PRK08570)Ribosomal_L19e (pfam01280)→ RplR (COG0256)rpl18p (PRK08569)→<br>RpsE (COG0098)rps5p (PRK04044)→ |
| env-243 | JGI24729J20445_1009152 | + | 2242 | 2319 | **RNA**→ |
| env-244 | 3300001580_10004676 | + | 12012 | 12089 | **RNA**→ RplC (COG0087)rpl3p (PRK04231)→ |
| env-245 | 3300001592_10154490 | - | 853 | 774 | **RNA**→ ←hypo |
| env-246 | 3300000032_0003629 | - | 1709 | 1630 | **RNA**→ RplC (COG0087)rpl3p (PRK04231)→ RplD (COG0088)rpl4lp (PRK04042)→ |
| env-247 | 3300000568_10010498 | - | 796 | 717 | **RNA**→ hypo→ rpl3p (PRK04231)Ribosomal_L3 (pfam00297)→ |
| env-248 | 3300000558_10003870 | - | 1201 | 1122 | **RNA**→ rpl3p (PRK04231)Ribosomal_L3 (pfam00297)→ rpl3p (PRK04231)Ribosomal_L3 (pfam00297)→ |
| env-249 | 3300001592_10087609 | + | 1266 | 1344 | **RNA**→ |
| env-250 | 3300001567_10002847 | + | 2463 | 2541 | **RNA**→ RplC (COG0087)rpl3p (PRK04231)→ RplD (COG0088)rpl4lp (PRK04042)→<br>RplW (COG0089)PRK14548 (PRK14548)→ RplB (COG0090)rpl2p (PRK09612)→<br>RpsS (COG0185)rps19p (PRK04038)→ rpl22p (PRK04223)RplV (COG0091)→<br>RpsC (COG0092)rps3p (PRK04191)→ PRK14549 (PRK14549)Ribosomal_L29 (pfam00831)→<br>PRK03879 (PRK03879)UPF0086 (pfam01868)→ RpsQ (COG0186)rps17p (PRK08572)→<br>rpl14p (PRK08571)Ribosomal_L14 (pfam00238)→ rpl24p (PRK01191)KOW_RPL26 (cd06089)→<br>PRK04313 (PRK04313)S4 (cd00165)KOW_RPS4 (cd06087)Ribosomal_S4e (pfam00900)→<br>RplE (COG0094)rpl5p (PRK04219)→ |
| env-251 | SL_3KL_010_SED_10010201 | - | 3562 | 3491 | **RNA**→ RplC (COG0087)rpl3p (PRK04231)→ RplD (COG0088)rpl4lp (PRK04042)→<br>RplW (COG0089)PRK14548 (PRK14548)→ RplB (COG0090)rpl2p (PRK09612)→<br>RpsS (COG0185)rps19p (PRK04038)→ hypo→ |
| env-252 | SL_3KL_010_SED_10013818 | - | 1092 | 1021 | **RNA**→ rpl3p (PRK04231)Ribosomal_L3 (pfam00297)→ |
| env-253 | SL_3KL_010_SED_10042920 | + | 1511 | 1582 | **RNA**→ RplC (COG0087)rpl3p (PRK04231)→ RplD (COG0088)rpl4lp (PRK04042)→ |
| env-254 | BDMC2_FXAG6RB01ERNKC | - | 187 | 100 | **RNA**→ hypo→ |
| env-255 | JGI24730J26740_1038966 | - | 597 | 511 | **RNA**→ ←hypo |
| env-256 | 3300000568_10019878 | + | 535 | 613 | **RNA**→ hypo→ |
| env-257 | 3300001592_10003410 | - | 16086 | 16008 | **RNA**→ RplC (COG0087)rpl3p (PRK04231)→ RplD (COG0088)rpl4lp (PRK04042)→<br>RplW (COG0089)PRK14548 (PRK14548)→ rpl2p (PRK09612)Ribosomal_L2_C (pfam03947)→<br>RpsS (COG0185)rps19p (PRK04038)→ rpl22p (PRK04223)RplV (COG0091)→<br>RpsC (COG0092)rps3p (PRK04191)→ PRK14549 (PRK14549)Ribosomal_L29 (pfam00831)→<br>PRK03879 (PRK03879)UPF0086 (pfam01868)→ RpsQ (COG0186)rps17p (PRK08572)→<br>rpl14p (PRK08571)Ribosomal_L14 (pfam00238)→ RplX (COG0198)rpl24p (PRK01191)→<br>PRK04313 (PRK04313)KOW_RPS4 (cd06087)Ribosomal_S4e (pfam00900)→ RplE (COG0094)rpl5p (PRK04219)→<br>RpsN (COG0199)rps14P (PRK05766)→ rps8p (PRK04034)RpsH (COG0096)→<br>RplF (COG0097)rpl6p (PRK05518)→ rpl32e (PRK08562)Ribosomal_L32e (pfam01655)→<br>rpl19e (PRK08570)Ribosomal_L19e (pfam01280)→ RplR (COG0256)rpl18p (PRK08569)→<br>RpsE (COG0098)rps5p (PRK04044)→ |

| | | | | | |
|---|---|---|---|---|---|
| env-258 | 3300001567_10001101 | + | 23848 | 23926 | **RNA**→ RplC (COG0087)rpl3p (PRK04231)→ RplD (COG0088)rpl4lp (PRK04042)→ RplW (COG0089)PRK14548 (PRK14548)→ rpl2p (PRK09612)Ribosomal_L2_C (pfam03947)→ RpsS (COG0185)rps19p (PRK04038)→ rpl22p (PRK04223)RplV (COG0091)→ RpsC (COG0092)rps3p (PRK04191)→ PRK14549 (PRK14549)Ribosomal_L29 (pfam00831)→ PRK03879 (PRK03879)UPF0086 (pfam01868)→ RpsQ (COG0186)rps17p (PRK08572)→ rpl14p (PRK08571)Ribosomal_L14 (pfam00238)→ RplX (COG0198)rpl24p (PRK01191)→ PRK04313 (PRK04313)KOW_RPS4 (cd06087)Ribosomal_S4e (pfam00900)→ RplE (COG0094)rpl5p (PRK04219)→ RpsN (COG0199)rps14P (PRK05766)→ rps8p (PRK04034)RpsH (COG0096)→ RplF (COG0097)rpl6p (PRK05518)→ rpl32e (PRK08562)Ribosomal_L32e (pfam01655)→ rpl19e (PRK08570)Ribosomal_L19e (pfam01280)→ RplR (COG0256)rpl18p (PRK08569)→ RpsE (COG0098)rps5p (PRK04044)→ |
| env-259 | 3300000558_10195618 | + | 1540 | 1618 | **RNA**→ hypo→ |
| env-260 | 3300000032_0041450 | - | 12420 | 12342 | **RNA**→ hypo→ |
| env-261 | JGI24713J26584_10001748 | - | 5737 | 5659 | **RNA**→ RplC (COG0087)rpl3p (PRK04231)→ RplD (COG0088)rpl4lp (PRK04042)→ RplW (COG0089)PRK14548 (PRK14548)→ rpl2p (PRK09612)Ribosomal_L2_C (pfam03947)→ RpsS (COG0185)rps19p (PRK04038)→ rpl22p (PRK04223)RplV (COG0091)→ RpsC (COG0092)rps3p (PRK04191)→ PRK14549 (PRK14549)Ribosomal_L29 (pfam00831)→ PRK03879 (PRK03879)UPF0086 (pfam01868)→ RpsQ (COG0186)rps17p (PRK08572)→ rpl14p (PRK08571)Ribosomal_L14 (pfam00238)→ |
| env-262 | JGI24714J26587_10004747 | + | 2188 | 2266 | **RNA**→ RplC (COG0087)rpl3p (PRK04231)→ RplD (COG0088)rpl4lp (PRK04042)→ RplW (COG0089)PRK14548 (PRK14548)→ rpl2p (PRK09612)Ribosomal_L2_C (pfam03947)→ RpsS (COG0185)rps19p (PRK04038)→ rpl22p (PRK04223)RplV (COG0091)→ RpsC (COG0092)rps3p (PRK04191)→ PRK14549 (PRK14549)Ribosomal_L29 (pfam00831)→ PRK03879 (PRK03879)UPF0086 (pfam01868)→ RpsQ (COG0186)rps17p (PRK08572)→ rpl14p (PRK08571)Ribosomal_L14 (pfam00238)→ |
| env-263 | JGI24504J29685_1004785 | + | 209 | 287 | **RNA**→ RplC (COG0087)rpl3p (PRK04231)→ RplD (COG0088)rpl4lp (PRK04042)→ RplW (COG0089)PRK14548 (PRK14548)→ rpl2p (PRK09612)Ribosomal_L2_C (pfam03947)→ RpsS (COG0185)rps19p (PRK04038)→ rpl22p (PRK04223)RplV (COG0091)→ RpsC (COG0092)rps3p (PRK04191)→ PRK14549 (PRK14549)Ribosomal_L29 (pfam00831)→ PRK03879 (PRK03879)UPF0086 (pfam01868)→ RpsQ (COG0186)rps17p (PRK08572)→ rpl14p (PRK08571)Ribosomal_L14 (pfam00238)→ RplX (COG0198)rpl24p (PRK01191)→ PRK04313 (PRK04313)KOW_RPS4 (cd06087)Ribosomal_S4e (pfam00900)→ rpl5p (PRK04219)Ribosomal_L5 (pfam00281)→ |
| env-264 | JGI24505J29691_1002356 | + | 209 | 287 | **RNA**→ RplC (COG0087)rpl3p (PRK04231)→ RplD (COG0088)rpl4lp (PRK04042)→ RplW (COG0089)PRK14548 (PRK14548)→ rpl2p (PRK09612)Ribosomal_L2_C (pfam03947)→ RpsS (COG0185)rps19p (PRK04038)→ rpl22p (PRK04223)RplV (COG0091)→ RpsC (COG0092)rps3p (PRK04191)→ PRK14549 (PRK14549)Ribosomal_L29 (pfam00831)→ PRK03879 (PRK03879)UPF0086 (pfam01868)→ RpsQ (COG0186)rps17p (PRK08572)→ rpl14p (PRK08571)Ribosomal_L14 (pfam00238)→ RplX (COG0198)rpl24p (PRK01191)→ PRK04313 (PRK04313)KOW_RPS4 (cd06087)Ribosomal_S4e (pfam00900)→ RplE (COG0094)rpl5p (PRK04219)→ RpsN (COG0199)rps14P (PRK05766)→ rps8p (PRK04034)RpsH (COG0096)→ RplF (COG0097)rpl6p (PRK05518)→ rpl32e (PRK08562)Ribosomal_L32e (pfam01655)→ rpl19e (PRK08570)Ribosomal_L19e (pfam01280)→ RplR (COG0256)rpl18p (PRK08569)→ RpsE (COG0098)rps5p (PRK04044)→ Ribosomal_L7_archeal_euk (cd01657)rpl30p (PRK06049)→ rpl15p (PRK06419)Ribosomal_L27A (pfam00828)→ |
| env-265 | 3300000507_100004 | - | 18302 | 18224 | **RNA**→ hypo→ RplC (COG0087)rpl3p (PRK04231)→ RplD (COG0088)rpl4lp (PRK04042)→ RplW (COG0089)PRK14548 (PRK14548)→ RplB (COG0090)rpl2p (PRK09612)→ RpsS (COG0185)rps19p (PRK04038)→ rpl22p (PRK04223)RplV (COG0091)→ RpsC (COG0092)rps3p (PRK04191)→ PRK14549 (PRK14549)Ribosomal_L29 (pfam00831)→ PRK03879 (PRK03879)UPF0086 (pfam01868)→ RpsQ (COG0186)rps17p (PRK08572)→ rpl14p (PRK08571)Ribosomal_L14 (pfam00238)→ RplX (COG0198)rpl24p (PRK01191)→ PRK04313 (PRK04313)KOW_RPS4 (cd06087)Ribosomal_S4e (pfam00900)→ RplE (COG0094)rpl5p (PRK04219)→ RpsN (COG0199)rps14P (PRK05766)→ rps8p (PRK04034)RpsH (COG0096)→ RplF (COG0097)rpl6p (PRK05518)→ rpl32e (PRK08562)Ribosomal_L32e (pfam01655)→ rpl19e (PRK08570)Ribosomal_L19e (pfam01280)→ RplR (COG0256)rpl18p (PRK08569)→ RpsE (COG0098)rps5p (PRK04044)→ |

| | | | | | |
|---|---|---|---|---|---|
| env-266 | 3300000558_10023046 | - | 1376 | 1297 | **RNA**→ RplC (COG0087)rpl3p (PRK04231)→ |
| env-267 | 3300001592_10042930 | - | 999 | 920 | **RNA**→ ←hypo |
| env-268 | 3300000568_10983650 | + | 160 | 239 | **RNA**→ hypo→ |
| env-269 | 3300000568_10052832 | + | 512 | 591 | **RNA**→ hypo→ |
| env-270 | 3300000032_0016008 | + | 347 | 426 | **RNA**→ hypo→ |
| env-271 | JGI24711J26586_10020110 | - | 584 | 506 | **RNA**→ rpl3p (PRK04231)Ribosomal_L3 (pfam00297)→ |
| env-272 | JGI24708J26588_10012893 | + | 2107 | 2185 | **RNA**→ RplC (COG0087)rpl3p (PRK04231)→ RplD (COG0088)rpl4lp (PRK04042)→ RplW (COG0089)PRK14548 (PRK14548)→ |
| env-273 | JGI24710J26742_10008143 | + | 5280 | 5358 | **RNA**→ RplC (COG0087)rpl3p (PRK04231)→ RplD (COG0088)rpl4lp (PRK04042)→ RplW (COG0089)PRK14548 (PRK14548)→ |
| env-274 | JGI24707J26582_10001803 | + | 2105 | 2183 | **RNA**→ RplC (COG0087)rpl3p (PRK04231)→ RplD (COG0088)rpl4lp (PRK04042)→ RplW (COG0089)PRK14548 (PRK14548)→ RplB (COG0090)rpl2p (PRK09612)→ |
| env-275 | JGI24712J26585_10006974 | - | 5713 | 5635 | **RNA**→ RplC (COG0087)rpl3p (PRK04231)→ RplD (COG0088)rpl4lp (PRK04042)→ RplW (COG0089)PRK14548 (PRK14548)→ RplB (COG0090)rpl2p (PRK09612)→ RpsS (COG0185)rps19p (PRK04038)→ rpl22p (PRK04223)RplV (COG0091)→ RpsC (COG0092)rps3p (PRK04191)→ PRK14549 (PRK14549)Ribosomal_L29 (pfam00831)→ PRK03879 (PRK03879)UPF0086 (pfam01868)→ RpsQ (COG0186)rps17p (PRK08572)→ rpl14p (PRK08571)Ribosomal_L14 (pfam00238)→ |
| env-276 | JGI24709J26583_10619039 | + | 43 | 121 | **RNA**→ |
| env-277 | JGI24499J29688_1000702 | + | 823 | 901 | **RNA**→ RplC (COG0087)rpl3p (PRK04231)→ RplD (COG0088)rpl4lp (PRK04042)→ RplW (COG0089)PRK14548 (PRK14548)→ RplB (COG0090)rpl2p (PRK09612)→ RpsS (COG0185)rps19p (PRK04038)→ rpl22p (PRK04223)RplV (COG0091)→ RpsC (COG0092)rps3p (PRK04191)→ PRK14549 (PRK14549)Ribosomal_L29 (pfam00831)→ PRK03879 (PRK03879)UPF0086 (pfam01868)→ RpsQ (COG0186)rps17p (PRK08572)→ rpl14p (PRK08571)Ribosomal_L14 (pfam00238)→ rpl24p (PRK01191)KOW_RPL26 (cd06089)→ PRK04313 (PRK04313)KOW_RPS4 (cd06087)Ribosomal_S4e (pfam00900)→ RplE (COG0094)rpl5p (PRK04219)→ RpsN (COG0199)rps14P (PRK05766)→ rps8p (PRK04034)RpsH (COG0096)→ RplF (COG0097)rpl6p (PRK05518)→ rpl32e (PRK08562)Ribosomal_L32e (pfam01655)→ rpl19e (PRK08570)Ribosomal_L19e (pfam01280)→ RplR (COG0256)rpl18p (PRK08569)→ RpsE (COG0098)rps5p (PRK04044)→ Ribosomal_L7_archeal_euk (cd01657)rpl30p (PRK06049)→ rpl15p (PRK06419)Ribosomal_L27A (pfam00828)→ SecY (COG0201)PRK08568 (PRK08568)→ |
| env-278 | JGI24707J26582_10000621 | + | 14235 | 14314 | **RNA**→ ←hypo |
| env-279 | JGI24708J26588_10001024 | + | 13186 | 13265 | **RNA**→ ←hypo |
| env-280 | JGI24713J26584_10000622 | + | 13651 | 13730 | **RNA**→ ←hypo |
| env-281 | JGI24714J26587_10000570 | + | 13186 | 13265 | **RNA**→ ←hypo |
| env-282 | JGI24712J26585_10002902 | + | 13100 | 13179 | **RNA**→ ←hypo |
| env-283 | JGI24711J26586_10000985 | - | 9166 | 9087 | **RNA**→ RplC (COG0087)rpl3p (PRK04231)→ RplD (COG0088)rpl4lp (PRK04042)→ RplW (COG0089)PRK14548 (PRK14548)→ RplB (COG0090)rpl2p (PRK09612)→ RpsS (COG0185)rps19p (PRK04038)→ rpl22p (PRK04223)RplV (COG0091)→ RpsC (COG0092)rps3p (PRK04191)→ PRK14549 (PRK14549)Ribosomal_L29 (pfam00831)→ PRK03879 (PRK03879)UPF0086 (pfam01868)→ RpsQ (COG0186)rps17p (PRK08572)→ rpl14p (PRK08571)Ribosomal_L14 (pfam00238)→ RplX (COG0198)rpl24p (PRK01191)→ PRK04313 (PRK04313)KOW_RPS4 (cd06087)Ribosomal_S4e (pfam00900)→ RplE (COG0094)rpl5p (PRK04219)→ RpsN (COG0199)rps14P (PRK05766)→ rps8p (PRK04034)RpsH (COG0096)→ RplF (COG0097)rpl6p (PRK05518)→ rpl32e (PRK08562)Ribosomal_L32e (pfam01655)→ rpl19e (PRK08570)Ribosomal_L19e (pfam01280)→ |

| | | | | | |
|---|---|---|---|---|---|
| env-284 | JGI24709J26583_10001164 | - | 9166 | 9087 | **RNA**→ RplC (COG0087)rpl3p (PRK04231)→ RplD (COG0088)rpl4lp (PRK04042)→<br>RplW (COG0089)PRK14548 (PRK14548)→ RplB (COG0090)rpl2p (PRK09612)→<br>RpsS (COG0185)rps19p (PRK04038)→ rpl22p (PRK04223)RplV (COG0091)→<br>RpsC (COG0092)rps3p (PRK04191)→ PRK14549 (PRK14549)Ribosomal_L29 (pfam00831)→<br>PRK03879 (PRK03879)UPF0086 (pfam01868)→ RpsQ (COG0186)rps17p (PRK08572)→<br>rpl14p (PRK08571)Ribosomal_L14 (pfam00238)→ RplX (COG0198)rpl24p (PRK01191)→<br>PRK04313 (PRK04313)KOW_RPS4 (cd06087)Ribosomal_S4e (pfam00900)→ RplE (COG0094)rpl5p (PRK04219)→<br>RpsN (COG0199)rps14P (PRK05766)→ rps8p (PRK04034)RpsH (COG0096)→<br>RplF (COG0097)rpl6p (PRK05518)→ rpl32e (PRK08562)Ribosomal_L32e (pfam01655)→<br>rpl19e (PRK08570)Ribosomal_L19e (pfam01280)→ |
| env-285 | JGI24710J26742_10000974 | + | 13183 | 13262 | **RNA**→ ←hypo |
| env-286 | JGI24505J29691_1015564 | + | 211 | 290 | **RNA**→ ←hypo |
| env-287 | JGI24500J29687_10001710 | + | 1224 | 1303 | **RNA**→ RplC (COG0087)rpl3p (PRK04231)→ RplD (COG0088)rpl4lp (PRK04042)→<br>RplW (COG0089)PRK14548 (PRK14548)→ RplB (COG0090)rpl2p (PRK09612)→<br>RpsS (COG0185)rps19p (PRK04038)→ rpl22p (PRK04223)RplV (COG0091)→<br>RpsC (COG0092)rps3p (PRK04191)→ PRK14549 (PRK14549)Ribosomal_L29 (pfam00831)→<br>PRK03879 (PRK03879)UPF0086 (pfam01868)→ RpsQ (COG0186)rps17p (PRK08572)→<br>rpl14p (PRK08571)Ribosomal_L14 (pfam00238)→ RplX (COG0198)rpl24p (PRK01191)→<br>PRK04313 (PRK04313)KOW_RPS4 (cd06087)Ribosomal_S4e (pfam00900)→ RplE (COG0094)rpl5p (PRK04219)→<br>RpsN (COG0199)rps14P (PRK05766)→ rps8p (PRK04034)RpsH (COG0096)→<br>RplF (COG0097)rpl6p (PRK05518)→ rpl32e (PRK08562)Ribosomal_L32e (pfam01655)→<br>rpl19e (PRK08570)Ribosomal_L19e (pfam01280)→ RplR (COG0256)rpl18p (PRK08569)→<br>RpsE (COG0098)rps5p (PRK04044)→ Ribosomal_L7_archeal_euk (cd01657)rpl30p (PRK06049)→<br>rpl15p (PRK06419)Ribosomal_L27A (pfam00828)→ SecY (COG0201)PRK08568 (PRK08568)→ |
| env-288 | JGI24502J29692_10012380 | + | 876 | 955 | **RNA**→ RplC (COG0087)rpl3p (PRK04231)→ RplD (COG0088)rpl4lp (PRK04042)→<br>RplW (COG0089)PRK14548 (PRK14548)→ RplB (COG0090)rpl2p (PRK09612)→<br>RpsS (COG0185)rps19p (PRK04038)→ rpl22p (PRK04223)RplV (COG0091)→<br>RpsC (COG0092)rps3p (PRK04191)→ PRK14549 (PRK14549)Ribosomal_L29 (pfam00831)→<br>PRK03879 (PRK03879)UPF0086 (pfam01868)→ RpsQ (COG0186)rps17p (PRK08572)→<br>rpl14p (PRK08571)Ribosomal_L14 (pfam00238)→ RplX (COG0198)rpl24p (PRK01191)→<br>PRK04313 (PRK04313)KOW_RPS4 (cd06087)Ribosomal_S4e (pfam00900)→ RplE (COG0094)rpl5p (PRK04219)→<br>RpsN (COG0199)rps14P (PRK05766)→ rps8p (PRK04034)RpsH (COG0096)→<br>RplF (COG0097)rpl6p (PRK05518)→ rpl32e (PRK08562)Ribosomal_L32e (pfam01655)→<br>rpl19e (PRK08570)Ribosomal_L19e (pfam01280)→ RplR (COG0256)rpl18p (PRK08569)→<br>RpsE (COG0098)rps5p (PRK04044)→ Ribosomal_L7_archeal_euk (cd01657)rpl30p (PRK06049)→<br>rpl15p (PRK06419)Ribosomal_L27A (pfam00828)→ SecY (COG0201)PRK08568 (PRK08568)→ |
| env-289 | JGI24501J29690_1003097 | + | 211 | 290 | **RNA**→ RplC (COG0087)rpl3p (PRK04231)→ RplD (COG0088)rpl4lp (PRK04042)→<br>RplW (COG0089)PRK14548 (PRK14548)→ RplB (COG0090)rpl2p (PRK09612)→<br>RpsS (COG0185)rps19p (PRK04038)→ rpl22p (PRK04223)RplV (COG0091)→<br>RpsC (COG0092)rps3p (PRK04191)→ PRK14549 (PRK14549)Ribosomal_L29 (pfam00831)→<br>PRK03879 (PRK03879)UPF0086 (pfam01868)→ RpsQ (COG0186)rps17p (PRK08572)→<br>rpl14p (PRK08571)Ribosomal_L14 (pfam00238)→ RplX (COG0198)rpl24p (PRK01191)→<br>PRK04313 (PRK04313)KOW_RPS4 (cd06087)Ribosomal_S4e (pfam00900)→ RplE (COG0094)rpl5p (PRK04219)→<br>RpsN (COG0199)rps14P (PRK05766)→ rps8p (PRK04034)RpsH (COG0096)→<br>RplF (COG0097)rpl6p (PRK05518)→ rpl32e (PRK08562)Ribosomal_L32e (pfam01655)→<br>rpl19e (PRK08570)Ribosomal_L19e (pfam01280)→ RplR (COG0256)rpl18p (PRK08569)→<br>RpsE (COG0098)rps5p (PRK04044)→ Ribosomal_L7_archeal_euk (cd01657)rpl30p (PRK06049)→<br>rpl15p (PRK06419)Ribosomal_L27A (pfam00828)→ SecY (COG0201)PRK08568 (PRK08568)→ |

| | | | | | |
|---|---|---|---|---|---|
| env-290 | JGI24503J29689_10009320 | + | 876 | 955 | **RNA**→ RplC (COG0087)rpl3p (PRK04231)→ RplD (COG0088)rpl4lp (PRK04042)→<br>RplW (COG0089)PRK14548 (PRK14548)→ RplB (COG0090)rpl2p (PRK09612)→<br>RpsS (COG0185)rps19p (PRK04038)→ rpl22p (PRK04223)RplV (COG0091)→<br>RpsC (COG0092)rps3p (PRK04191)→ PRK14549 (PRK14549)Ribosomal_L29 (pfam00831)→<br>PRK03879 (PRK03879)UPF0086 (pfam01868)→ RpsQ (COG0186)rps17p (PRK08572)→<br>rpl14p (PRK08571)Ribosomal_L14 (pfam00238)→ RplX (COG0198)rpl24p (PRK01191)→<br>PRK04313 (PRK04313)KOW_RPS4 (cd06087)Ribosomal_S4e (pfam00900)→ RplE (COG0094)rpl5p (PRK04219)→<br>RpsN (COG0199)rps14P (PRK05766)→ rps8p (PRK04034)RpsH (COG0096)→<br>RplF (COG0097)rpl6p (PRK05518)→ rpl32e (PRK08562)Ribosomal_L32e (pfam01655)→<br>rpl19e (PRK08570)Ribosomal_L19e (pfam01280)→ RplR (COG0256)rpl18p (PRK08569)→<br>RpsE (COG0098)rps5p (PRK04044)→ Ribosomal_L7_archeal_euk (cd01657)rpl30p (PRK06049)→<br>rpl15p (PRK06419)Ribosomal_L27A (pfam00828)→ SecY (COG0201)PRK08568 (PRK08568)→ |
| env-291 | JGI24499J29688_1000521 | + | 876 | 955 | **RNA**→ RplC (COG0087)rpl3p (PRK04231)→ RplD (COG0088)rpl4lp (PRK04042)→<br>RplW (COG0089)PRK14548 (PRK14548)→ RplB (COG0090)rpl2p (PRK09612)→<br>RpsS (COG0185)rps19p (PRK04038)→ rpl22p (PRK04223)RplV (COG0091)→<br>RpsC (COG0092)rps3p (PRK04191)→ PRK14549 (PRK14549)Ribosomal_L29 (pfam00831)→<br>PRK03879 (PRK03879)UPF0086 (pfam01868)→ RpsQ (COG0186)rps17p (PRK08572)→<br>rpl14p (PRK08571)Ribosomal_L14 (pfam00238)→ RplX (COG0198)rpl24p (PRK01191)→<br>PRK04313 (PRK04313)KOW_RPS4 (cd06087)Ribosomal_S4e (pfam00900)→ RplE (COG0094)rpl5p (PRK04219)→<br>RpsN (COG0199)rps14P (PRK05766)→ rps8p (PRK04034)RpsH (COG0096)→<br>RplF (COG0097)rpl6p (PRK05518)→ rpl32e (PRK08562)Ribosomal_L32e (pfam01655)→<br>rpl19e (PRK08570)Ribosomal_L19e (pfam01280)→ RplR (COG0256)rpl18p (PRK08569)→<br>RpsE (COG0098)rps5p (PRK04044)→ Ribosomal_L7_archeal_euk (cd01657)rpl30p (PRK06049)→<br>rpl15p (PRK06419)Ribosomal_L27A (pfam00828)→ SecY (COG0201)PRK08568 (PRK08568)→ |
| env-292 | JGI24712J26585_10029187 | + | 2386 | 2464 | **RNA**→ hypo→ |
| env-293 | JGI24711J26586_10244524 | - | 365 | 287 | **RNA**→ ←hypo |
| env-294 | JGI24710J26742_10309618 | - | 365 | 287 | **RNA**→ ←hypo |
| env-295 | JGI24500J29687_10002376 | + | 168 | 246 | **RNA**→ ←hypo |
| env-296 | JGI24503J29689_10006492 | + | 104 | 182 | **RNA**→ rpl3p (PRK04231)Ribosomal_L3 (pfam00297)→ |
| Mbo-1-1 | NC_018227.2 | + | 1712758 | 1712836 | **RNA**→ RplC (COG0087)rpl3p (PRK04231)→ RplD (COG0088)rpl4lp (PRK04042)→<br>RplW (COG0089)PRK14548 (PRK14548)→ RplB (COG0090)rpl2p (PRK09612)→<br>RpsS (COG0185)rps19p (PRK04038)→ rpl22p (PRK04223)RplV (COG0091)→<br>RpsC (COG0092)rps3p (PRK04191)→ PRK14549 (PRK14549)Ribosomal_L29 (pfam00831)→<br>PRK03879 (PRK03879)UPF0086 (pfam01868)→ RpsQ (COG0186)rps17p (PRK08572)→<br>rpl14p (PRK08571)Ribosomal_L14 (pfam00238)→ RplX (COG0198)rpl24p (PRK01191)→<br>PRK04313 (PRK04313)KOW_RPS4 (cd06087)Ribosomal_S4e (pfam00900)→ RplE (COG0094)rpl5p (PRK04219)→<br>RpsN (COG0199)rps14P (PRK05766)→ rps8p (PRK04034)RpsH (COG0096)→<br>RplF (COG0097)rpl6p (PRK05518)→ rpl32e (PRK08562)Ribosomal_L32e (pfam01655)→<br>rpl19e (PRK08570)Ribosomal_L19e (pfam01280)→ RplR (COG0256)rpl18p (PRK08569)→<br>RpsE (COG0098)rps5p (PRK04044)→ |
| env-297 | JGI24501J29690_1006097 | + | 749 | 827 | **RNA**→ rpl3p (PRK04231)Ribosomal_L3 (pfam00297)→ |
| env-298 | JGI24707J26582_10054179 | + | 722 | 800 | **RNA**→ rpl3p (PRK04231)Ribosomal_L3 (pfam00297)→ |
| env-299 | JGI24708J26588_10030031 | + | 1058 | 1136 | **RNA**→ ←hypo |
| env-300 | JGI24709J26583_10416662 | - | 277 | 199 | **RNA**→ hypo→ |
| env-301 | JGI24502J29692_10003709 | + | 657 | 735 | **RNA**→ rpl3p (PRK04231)Ribosomal_L3 (pfam00297)→ |
| env-302 | JGI24503J29689_10003031 | + | 41 | 119 | **RNA**→ ←hypo |
| env-303 | JGI24710J26742_10068176 | + | 372 | 450 | **RNA**→ rpl3p (PRK04231)Ribosomal_L3 (pfam00297)→ |
| env-304 | JGI24709J26583_10014213 | + | 553 | 631 | **RNA**→ hypo→ RplC (COG0087)rpl3p (PRK04231)→ RplD (COG0088)rpl4lp (PRK04042)→<br>RplW (COG0089)PRK14548 (PRK14548)→ |
| env-305 | JGI24502J29692_10002344 | + | 84 | 165 | **RNA**→ RplC (COG0087)rpl3p (PRK04231)→ RplD (COG0088)rpl4lp (PRK04042)→ |
| env-306 | JGI24712J26585_10639760 | + | 51 | 132 | **RNA**→ ←hypo |
| env-307 | JGI24711J26586_10552659 | + | 51 | 132 | **RNA**→ ←hypo |
| env-308 | JGI24709J26583_10614246 | + | 51 | 132 | **RNA**→ ←hypo |
| env-309 | JGI24710J26742_10664497 | + | 51 | 132 | **RNA**→ ←hypo |

| | | | | | |
|---|---|---|---|---|---|
| Mma-1-1 | NC_009051.1 | + | 549081 | 549159 | **RNA**→ RplC (COG0087)rpl3p (PRK04231)→ RplD (COG0088)rpl4lp (PRK04042)→<br>RplW (COG0089)PRK14548 (PRK14548)→ RplB (COG0090)rpl2p (PRK09612)→<br>RpsS (COG0185)rps19p (PRK04038)→ rpl22p (PRK04223)RplV (COG0091)→<br>RpsC (COG0092)rps3p (PRK04191)→ PRK14549 (PRK14549)Ribosomal_L29 (pfam00831)→<br>PRK03879 (PRK03879)UPF0086 (pfam01868)→ RpsQ (COG0186)rps17p (PRK08572)→<br>rpl14p (PRK08571)Ribosomal_L14 (pfam00238)→ RplX (COG0198)rpl24p (PRK01191)→<br>PRK04313 (PRK04313)KOW_RPS4 (cd06087)Ribosomal_S4e (pfam00900)→ RplE (COG0094)rpl5p (PRK04219)→<br>RpsN (COG0199)rps14P (PRK05766)→ rps8p (PRK04034)RpsH (COG0096)→<br>RplF (COG0097)rpl6p (PRK05518)→ rpl32e (PRK08562)Ribosomal_L32_L32e (cd00513)→<br>rpl19e (PRK08570)Ribosomal_L19e (pfam01280)→ RplR (COG0256)rpl18p (PRK08569)→<br>RpsE (COG0098)rps5p (PRK04044)→ |
| Mse-1-1 | NZ_JXOJ01000001.1 | + | 373962 | 374040 | **RNA**→ RplC (COG0087)rpl3p (PRK04231)→ RplD (COG0088)rpl4lp (PRK04042)→<br>RplW (COG0089)PRK14548 (PRK14548)→ rpl2p (PRK09612)Ribosomal_L2_C (pfam03947)→<br>RpsS (COG0185)rps19p (PRK04038)→ rpl22p (PRK04223)RplV (COG0091)→<br>RpsC (COG0092)rps3p (PRK04191)→ PRK14549 (PRK14549)Ribosomal_L29 (pfam00831)→<br>PRK03879 (PRK03879)UPF0086 (pfam01868)→ RpsQ (COG0186)rps17p (PRK08572)→<br>rpl14p (PRK08571)Ribosomal_L14 (pfam00238)→ RplX (COG0198)rpl24p (PRK01191)→<br>PRK04313 (PRK04313)KOW_RPS4 (cd06087)Ribosomal_S4e (pfam00900)→ RplE (COG0094)rpl5p (PRK04219)→<br>RpsN (COG0199)rps14P (PRK05766)→ rps8p (PRK04034)RpsH (COG0096)→<br>RplF (COG0097)rpl6p (PRK05518)→ rpl32e (PRK08562)Ribosomal_L32e (pfam01655)→<br>rpl19e (PRK08570)Ribosomal_L19e (pfam01280)→ RplR (COG0256)rpl18p (PRK08569)→<br>RpsE (COG0098)rps5p (PRK04044)→ |
| Msp-1-1 | NZ_JMIO01000016.1 | + | 890 | 968 | **RNA**→ RplC (COG0087)rpl3p (PRK04231)→ RplD (COG0088)rpl4lp (PRK04042)→<br>RplW (COG0089)PRK14548 (PRK14548)→ RplB (COG0090)rpl2p (PRK09612)→<br>RpsS (COG0185)rps19p (PRK04038)→ rpl22p (PRK04223)RplV (COG0091)→<br>RpsC (COG0092)rps3p (PRK04191)→ PRK14549 (PRK14549)Ribosomal_L29 (pfam00831)→<br>PRK03879 (PRK03879)UPF0086 (pfam01868)→ RpsQ (COG0186)rps17p (PRK08572)→<br>rpl14p (PRK08571)Ribosomal_L14 (pfam00238)→ RplX (COG0198)rpl24p (PRK01191)→<br>PRK04313 (PRK04313)S4 (cd00165)KOW_RPS4 (cd06087)Ribosomal_S4e (pfam00900)→<br>RplE (COG0094)rpl5p (PRK04219)→ RpsN (COG0199)rps14P (PRK05766)→<br>rps8p (PRK04034)RpsH (COG0096)→ RplF (COG0097)rpl6p (PRK05518)→<br>rpl32e (PRK08562)Ribosomal_L32e (pfam01655)→ rpl19e (PRK08570)Ribosomal_L19e (pfam01280)→<br>RplR (COG0256)rpl18p (PRK08569)→ RpsE (COG0098)rps5p (PRK04044)→<br>Ribosomal_L7_archeal_euk (cd01657)rpl30p (PRK06049)→ rpl15p (PRK06419)Ribosomal_L27A (pfam00828)→<br>SecY (COG0201)PRK08568 (PRK08568)→ hypo→ |
| env-310 | JGI24712J26585_10057261 | - | 366 | 297 | **RNA**→ ←hypo |
| env-311 | JGI24710J26742_10004413 | - | 10050 | 9981 | **RNA**→ RplC (COG0087)rpl3p (PRK04231)→ RplD (COG0088)rpl4lp (PRK04042)→<br>RplW (COG0089)PRK14548 (PRK14548)→ RplB (COG0090)rpl2p (PRK09612)→<br>RpsS (COG0185)rps19p (PRK04038)→ rpl22p (PRK04223)RplV (COG0091)→<br>RpsC (COG0092)rps3p (PRK04191)→ PRK14549 (PRK14549)Ribosomal_L29 (pfam00831)→<br>PRK03879 (PRK03879)UPF0086 (pfam01868)→ RpsQ (COG0186)rps17p (PRK08572)→<br>rpl14p (PRK08571)Ribosomal_L14 (pfam00238)→ rpl24p (PRK01191)KOW_RPL26 (cd06089)→<br>PRK04313 (PRK04313)KOW_RPS4 (cd06087)Ribosomal_S4e (pfam00900)→ RplE (COG0094)rpl5p (PRK04219)→<br>RpsN (COG0199)rps14P (PRK05766)→ rps8p (PRK04034)RpsH (COG0096)→<br>RplF (COG0097)rpl6p (PRK05518)→ rpl32e (PRK08562)Ribosomal_L32e (pfam01655)→<br>rpl19e (PRK08570)Ribosomal_L19e (pfam01280)→ RplR (COG0256)rpl18p (PRK08569)→ |
| env-312 | JGI24500J29687_10026414 | + | 210 | 279 | **RNA**→ RplC (COG0087)rpl3p (PRK04231)→ RplD (COG0088)rpl4lp (PRK04042)→<br>RplW (COG0089)PRK14548 (PRK14548)→ RplB (COG0090)rpl2p (PRK09612)→<br>RpsS (COG0185)rps19p (PRK04038)→ rpl22p (PRK04223)RplV (COG0091)→<br>RpsC (COG0092)rps3p (PRK04191)→ PRK14549 (PRK14549)Ribosomal_L29 (pfam00831)→<br>PRK03879 (PRK03879)UPF0086 (pfam01868)→ RpsQ (COG0186)rps17p (PRK08572)→<br>rpl14p (PRK08571)Ribosomal_L14 (pfam00238)→ rpl24p (PRK01191)KOW_RPL26 (cd06089)→<br>PRK04313 (PRK04313)KOW_RPS4 (cd06087)Ribosomal_S4e (pfam00900)→ RplE (COG0094)rpl5p (PRK04219)→<br>RpsN (COG0199)rps14P (PRK05766)→ |
| env-313 | JGI24501J29690_1062315 | + | 210 | 279 | **RNA**→ ←hypo |

| | | | | | |
|---|---|---|---|---|---|
| env-314 | JGI24499J29688_1102173 | + | 190 | 259 | RNA→ ←hypo |
| env-315 | JGI24709J26583_10005356 | - | 7831 | 7762 | RNA→ RplC (COG0087)rpl3p (PRK04231)→ RplD (COG0088)rpl4lp (PRK04042)→ RplW (COG0089)PRK14548 (PRK14548)→ RplB (COG0090)rpl2p (PRK09612)→ RpsS (COG0185)rps19p (PRK04038)→ rpl22p (PRK04223)RplV (COG0091)→ RpsC (COG0092)rps3p (PRK04191)→ PRK14549 (PRK14549)Ribosomal_L29 (pfam00831)→ PRK03879 (PRK03879)UPF0086 (pfam01868)→ RpsQ (COG0186)rps17p (PRK08572)→ rpl14p (PRK08571)Ribosomal_L14 (pfam00238)→ rpl24p (PRK01191)KOW_RPL26 (cd06089)→ PRK04313 (PRK04313)KOW_RPS4 (cd06087)Ribosomal_S4e (pfam00900)→ RplE (COG0094)rpl5p (PRK04219)→ RpsN (COG0199)rps14P (PRK05766)→ rps8p (PRK04034)Ribosomal_S8 (pfam00410)→ |
| env-316 | JGI24711J26586_10160586 | + | 77 | 146 | RNA→ ←hypo |
| env-317 | JGI24500J29687_10008301 | + | 42 | 123 | RNA→ RplC (COG0087)rpl3p (PRK04231)→ RplD (COG0088)rpl4lp (PRK04042)→ RplW (COG0089)PRK14548 (PRK14548)→ RplB (COG0090)rpl2p (PRK09612)→ RpsS (COG0185)rps19p (PRK04038)→ rpl22p (PRK04223)RplV (COG0091)→ RpsC (COG0092)rps3p (PRK04191)→ PRK14549 (PRK14549)Ribosomal_L29 (pfam00831)→ PRK03879 (PRK03879)UPF0086 (pfam01868)→ RpsQ (COG0186)rps17p (PRK08572)→ rpl14p (PRK08571)Ribosomal_L14 (pfam00238)→ rpl24p (PRK01191)KOW_RPL26 (cd06089)→ PRK04313 (PRK04313)KOW_RPS4 (cd06087)Ribosomal_S4e (pfam00900)→ RplE (COG0094)rpl5p (PRK04219)→ RpsN (COG0199)rps14P (PRK05766)→ rps8p (PRK04034)RpsH (COG0096)→ RplF (COG0097)rpl6p (PRK05518)→ rpl32e (PRK08562)Ribosomal_L32e (pfam01655)→ rpl19e (PRK08570)Ribosomal_L19e (pfam01280)→ RplR (COG0256)rpl18p (PRK08569)→ |
| env-318 | JGI24502J29692_10009873 | + | 777 | 846 | RNA→ rpl3p (PRK04231)Ribosomal_L3 (pfam00297)→ |
| env-319 | JGI24503J29689_10009607 | + | 112 | 181 | RNA→ rpl3p (PRK04231)Ribosomal_L3 (pfam00297)→ |
| env-320 | JGI24712J26585_10076884 | - | 494 | 425 | RNA→ rpl3p (PRK04231)Ribosomal_L3 (pfam00297)→ |
| env-321 | JGI24500J29687_10085663 | + | 72 | 141 | RNA→ rpl3p (PRK04231)Ribosomal_L3 (pfam00297)→ |
| env-322 | JGI24711J26586_10010708 | - | 494 | 425 | RNA→ rpl3p (PRK04231)Ribosomal_L3 (pfam00297)→ |
| Mpa-1-1 | NC_011832.1 | + | 459047 | 459117 | RNA→ RplC (COG0087)rpl3p (PRK04231)→ RplD (COG0088)rpl4lp (PRK04042)→ RplW (COG0089)PRK14548 (PRK14548)→ rpl2p (PRK09612)Ribosomal_L2_C (pfam03947)→ RpsS (COG0185)rps19p (PRK04038)→ rpl22p (PRK04223)RplV (COG0091)→ RpsC (COG0092)rps3p (PRK04191)→ PRK14549 (PRK14549)Ribosomal_L29 (pfam00831)→ PRK03879 (PRK03879)UPF0086 (pfam01868)→ RpsQ (COG0186)rps17p (PRK08572)→ rpl14p (PRK08571)Ribosomal_L14 (pfam00238)→ rpl24p (PRK01191)KOW_RPL26 (cd06089)→ PRK04313 (PRK04313)S4 (cd00165)KOW_RPS4 (cd06087)Ribosomal_S4e (pfam00900)→ RplE (COG0094)rpl5p (PRK04219)→ RpsN (COG0199)rps14P (PRK05766)→ rps8p (PRK04034)RpsH (COG0096)→ RplF (COG0097)rpl6p (PRK05518)→ rpl32e (PRK08562)Ribosomal_L32e (pfam01655)→ rpl19e (PRK08570)Ribosomal_L19e (pfam01280)→ RplR (COG0256)rpl18p (PRK08569)→ RpsE (COG0098)rps5p (PRK04044)→ |
| env-323 | DHTCA2_GJDNVXK01BZYAY | - | 259 | 189 | RNA→ hypo→ |
| env-324 | 3300001580_10004417 | + | 2215 | 2285 | RNA→ hypo→ RplC (COG0087)rpl3p (PRK04231)→ RplD (COG0088)rpl4lp (PRK04042)→ RplW (COG0089)PRK14548 (PRK14548)→ RplB (COG0090)rpl2p (PRK09612)→ rps19p (PRK04038)Ribosomal_S19 (pfam00203)→ rpl22p (PRK04223)RplV (COG0091)→ RpsC (COG0092)rps3p (PRK04191)→ PRK14549 (PRK14549)Ribosomal_L29 (pfam00831)→ PRK03879 (PRK03879)UPF0086 (pfam01868)→ RpsQ (COG0186)rps17p (PRK08572)→ rpl14p (PRK08571)Ribosomal_L14 (pfam00238)→ rpl24p (PRK01191)KOW_RPL26 (cd06089)→ PRK04313 (PRK04313)S4 (cd00165)KOW_RPS4 (cd06087)Ribosomal_S4e (pfam00900)→ RplE (COG0094)rpl5p (PRK04219)→ RpsN (COG0199)rps14P (PRK05766)→ rps8p (PRK04034)RpsH (COG0096)→ RplF (COG0097)rpl6p (PRK05518)→ rpl32e (PRK08562)Ribosomal_L32e (pfam01655)→ rpl19e (PRK08570)Ribosomal_L19e (pfam01280)→ RplR (COG0256)rpl18p (PRK08569)→ RpsE (COG0098)rps5p (PRK04044)→ Ribosomal_L7_archeal_euk (cd01657)rpl30p (PRK06049)→ |
| env-325 | 3300001580_10080578 | - | 953 | 870 | RNA→ ←hypo |
| env-326 | 2232135516 | + | 86 | 169 | RNA→ |
| env-327 | 3300000558_11733700 | - | 1392 | 1309 | RNA→ ←hypo |
| env-328 | 3300000558_11835925 | - | 418 | 335 | RNA→ |

| | | | | | |
|---|---|---|---|---|---|
| env-329 | 3300000568_10049962 | - | 21413 | 21330 | **RNA**→ hypo→ RplC (COG0087)rpl3p (PRK04231)→ RplD (COG0088)rpl4lp (PRK04042)→<br>PRK14548 (PRK14548)Ribosomal_L23 (pfam00276)→ RplB (COG0090)rpl2p (PRK09612)→<br>rps19p (PRK04038)Ribosomal_S19 (pfam00203)→ rpl22p (PRK04223)RplV (COG0091)→<br>RpsC (COG0092)rps3p (PRK04191)→ PRK14549 (PRK14549)Ribosomal_L29 (pfam00831)→<br>PRK03879 (PRK03879)UPF0086 (pfam01868)→ RpsQ (COG0186)rps17p (PRK08572)→<br>rpl14p (PRK08571)Ribosomal_L14 (pfam00238)→ rpl24p (PRK01191)KOW_RPL26 (cd06089)→<br>PRK04313 (PRK04313)KOW_RPS4 (cd06087)Ribosomal_S4e (pfam00900)→<br>rpl5p (PRK04219)Ribosomal_L5 (pfam00281)Ribosomal_L5_C (pfam00673)→<br>RpsN (COG0199)rps14P (PRK05766)→ rps8p (PRK04034)RpsH (COG0096)→<br>RplF (COG0097)rpl6p (PRK05518)→ rpl32e (PRK08562)Ribosomal_L32e (pfam01655)→<br>rpl19e (PRK08570)Ribosomal_L19e (pfam01280)→ RplR (COG0256)rpl18p (PRK08569)→ |
| env-330 | 3300001605_10004100 | - | 1325 | 1242 | **RNA**→ ←hypo |
| env-331 | 3300000032_0004919 | + | 16062 | 16145 | **RNA**→ RplC (COG0087)rpl3p (PRK04231)→ RplD (COG0088)rpl4lp (PRK04042)→<br>PRK14548 (PRK14548)Ribosomal_L23 (pfam00276)→ RplB (COG0090)rpl2p (PRK09612)→<br>rps19p (PRK04038)Ribosomal_S19 (pfam00203)→ rpl22p (PRK04223)RplV (COG0091)→<br>RpsC (COG0092)rps3p (PRK04191)→ PRK14549 (PRK14549)Ribosomal_L29 (pfam00831)→<br>PRK03879 (PRK03879)UPF0086 (pfam01868)→ RpsQ (COG0186)rps17p (PRK08572)→<br>rpl14p (PRK08571)Ribosomal_L14 (pfam00238)→ rpl24p (PRK01191)KOW_RPL26 (cd06089)→<br>PRK04313 (PRK04313)RPS4A (COG1471)→ KOW (smart00739)KOW_RPS4 (cd06087)→<br>rpl5p (PRK04219)Ribosomal_L5 (pfam00281)Ribosomal_L5_C (pfam00673)→<br>RpsN (COG0199)rps14P (PRK05766)→ rps8p (PRK04034)RpsH (COG0096)→<br>RplF (COG0097)rpl6p (PRK05518)→ rpl32e (PRK08562)Ribosomal_L32e (pfam01655)→<br>rpl19e (PRK08570)Ribosomal_L19e (pfam01280)→ RplR (COG0256)rpl18p (PRK08569)→ |
| env-332 | 3300000185_104355 | - | 418 | 335 | **RNA**→ |
| Mfo-1-1 | NC_019943.1 | - | 2583107 | 2583023 | **RNA**→ RplC (COG0087)rpl3p (PRK04231)→ RplD (COG0088)rpl4lp (PRK04042)→<br>RplW (COG0089)PRK14548 (PRK14548)→ RplB (COG0090)rpl2p (PRK09612)→<br>rps19p (PRK04038)Ribosomal_S19 (pfam00203)→ rpl22p (PRK04223)RplV (COG0091)→<br>RpsC (COG0092)rps3p (PRK04191)→ PRK14549 (PRK14549)Ribosomal_L29 (pfam00831)→<br>PRK03879 (PRK03879)UPF0086 (pfam01868)→ RpsQ (COG0186)rps17p (PRK08572)→<br>rpl14p (PRK08571)Ribosomal_L14 (pfam00238)→ rpl24p (PRK01191)KOW_RPL26 (cd06089)→<br>PRK04313 (PRK04313)S4 (cd00165)KOW_RPS4 (cd06087)Ribosomal_S4e (pfam00900)→<br>rpl5p (PRK04219)Ribosomal_L5 (pfam00281)Ribosomal_L5_C (pfam00673)→<br>RpsN (COG0199)rps14P (PRK05766)→ rps8p (PRK04034)RpsH (COG0096)→<br>RplF (COG0097)rpl6p (PRK05518)→ rpl32e (PRK08562)Ribosomal_L32e (pfam01655)→<br>rpl19e (PRK08570)Ribosomal_L19e (pfam01280)→ RplR (COG0256)rpl18p (PRK08569)→ |
| env-333 | JGI20225J20221_1000316 | - | 54770 | 54691 | **RNA**→ RplC (COG0087)rpl3p (PRK04231)→ RplD (COG0088)rpl4lp (PRK04042)→<br>RplW (COG0089)PRK14548 (PRK14548)→ RplB (COG0090)rpl2p (PRK09612)→<br>rps19p (PRK04038)Ribosomal_S19 (pfam00203)→ rpl22p (PRK04223)RplV (COG0091)→<br>RpsC (COG0092)rps3p (PRK04191)→ PRK14549 (PRK14549)Ribosomal_L29 (pfam00831)→<br>PRK03879 (PRK03879)UPF0086 (pfam01868)→ RpsQ (COG0186)rps17p (PRK08572)→<br>rpl14p (PRK08571)Ribosomal_L14 (pfam00238)→ rpl24p (PRK01191)KOW_RPL26 (cd06089)→<br>PRK04313 (PRK04313)S4 (cd00165)KOW_RPS4 (cd06087)Ribosomal_S4e (pfam00900)→<br>RplE (COG0094)rpl5p (PRK04219)→ RpsN (COG0199)rps14P (PRK05766)→<br>rps8p (PRK04034)RpsH (COG0096)→ RplF (COG0097)rpl6p (PRK05518)→<br>rpl32e (PRK08562)Ribosomal_L32e (pfam01655)→ rpl19e (PRK08570)Ribosomal_L19e (pfam01280)→<br>RplR (COG0256)rpl18p (PRK08569)→ |
| env-334 | BMHB3a_c145790 | - | 1545 | 1466 | **RNA**→ RplC (COG0087)rpl3p (PRK04231)→ RplD (COG0088)rpl4lp (PRK04042)→ |
| env-335 | JGI24730J26740_1000194 | - | 6399 | 6320 | **RNA**→ RplC (COG0087)rpl3p (PRK04231)→ RplD (COG0088)rpl4lp (PRK04042)→<br>RplW (COG0089)PRK14548 (PRK14548)→ RplB (COG0090)rpl2p (PRK09612)→<br>rps19p (PRK04038)Ribosomal_S19 (pfam00203)→ rpl22p (PRK04223)RplV (COG0091)→<br>RpsC (COG0092)rps3p (PRK04191)→ PRK14549 (PRK14549)Ribosomal_L29 (pfam00831)→<br>PRK03879 (PRK03879)UPF0086 (pfam01868)→ RpsQ (COG0186)rps17p (PRK08572)→<br>rpl14p (PRK08571)Ribosomal_L14 (pfam00238)→ rpl24p (PRK01191)KOW_RPL26 (cd06089)→<br>PRK04313 (PRK04313)RPS4A (COG1471)→ |

| | | | | | |
|---|---|---|---|---|---|
| env-336 | JGI24730J26740_1000273 | - | 21541 | 21462 | **RNA**→ RplC (COG0087)rpl3p (PRK04231)→ RplD (COG0088)rpl4lp (PRK04042)→ RplW (COG0089)PRK14548 (PRK14548)→ RplB (COG0090)rpl2p (PRK09612)→ rps19p (PRK04038)Ribosomal_S19 (pfam00203)→ rpl22p (PRK04223)RplV (COG0091)→ RpsC (COG0092)rps3p (PRK04191)→ PRK14549 (PRK14549)Ribosomal_L29 (pfam00831)→ PRK03879 (PRK03879)UPF0086 (pfam01868)→ RpsQ (COG0186)rps17p (PRK08572)→ rpl14p (PRK08571)Ribosomal_L14 (pfam00238)→ rpl24p (PRK01191)KOW_RPL26 (cd06089)→ PRK04313 (PRK04313)S4 (cd00165)KOW_RPS4 (cd06087)Ribosomal_S4e (pfam00900)→ RplE (COG0094)rpl5p (PRK04219)→ RpsN (COG0199)rps14P (PRK05766)→ rps8p (PRK04034)RpsH (COG0096)→ RplF (COG0097)rpl6p (PRK05518)→ rpl32e (PRK08562)Ribosomal_L32e (pfam01655)→ rpl19e (PRK08570)Ribosomal_L19e (pfam01280)→ RplR (COG0256)rpl18p (PRK08569)→ |
| env-337 | 2013874846 | + | 4629 | 4707 | **RNA**→ RplC (COG0087)rpl3p (PRK04231)→ RplD (COG0088)rpl4lp (PRK04042)→ RplW (COG0089)PRK14548 (PRK14548)→ |
| env-338 | PRSSGFe2_Sequence0000003268 | + | 541 | 620 | **RNA**→ hypo→ |
| env-339 | WSSedL1Cam_c0014 | - | 25929 | 25851 | **RNA**→ RplC (COG0087)rpl3p (PRK04231)→ RplD (COG0088)rpl4lp (PRK04042)→ RplW (COG0089)PRK14548 (PRK14548)→ RplB (COG0090)rpl2p (PRK09612)→ rps19p (PRK04038)Ribosomal_S19 (pfam00203)→ rpl22p (PRK04223)RplV (COG0091)→ RpsC (COG0092)rps3p (PRK04191)→ PRK14549 (PRK14549)Ribosomal_L29 (pfam00831)→ PRK03879 (PRK03879)UPF0086 (pfam01868)→ RpsQ (COG0186)rps17p (PRK08572)→ rpl14p (PRK08571)Ribosomal_L14 (pfam00238)→ rpl24p (PRK01191)KOW_RPL26 (cd06089)→ PRK04313 (PRK04313)S4 (cd00165)KOW_RPS4 (cd06087)Ribosomal_S4e (pfam00900)→ RplE (COG0094)rpl5p (PRK04219)→ RpsN (COG0199)rps14P (PRK05766)→ rps8p (PRK04034)RpsH (COG0096)→ RplF (COG0097)rpl6p (PRK05518)→ rpl32e (PRK08562)Ribosomal_L32_L32e (cd00513)→ rpl19e (PRK08570)Ribosomal_L19e (pfam01280)→ RplR (COG0256)rpl18p (PRK08569)→ |
| CMe-1-1 | NC_009712.1 | + | 517727 | 517802 | **RNA**→ RplC (COG0087)rpl3p (PRK04231)→ RplD (COG0088)rpl4lp (PRK04042)→ RplW (COG0089)PRK14548 (PRK14548)→ RplB (COG0090)rpl2p (PRK09612)→ rps19p (PRK04038)Ribosomal_S19 (pfam00203)→ rpl22p (PRK04223)RplV (COG0091)→ RpsC (COG0092)rps3p (PRK04191)→ PRK14549 (PRK14549)Ribosomal_L29 (pfam00831)→ PRK03879 (PRK03879)UPF0086 (pfam01868)→ RpsQ (COG0186)rps17p (PRK08572)→ rpl14p (PRK08571)Ribosomal_L14 (pfam00238)→ rpl24p (PRK01191)KOW_RPL26 (cd06089)→ PRK04313 (PRK04313)S4 (cd00165)KOW_RPS4 (cd06087)Ribosomal_S4e (pfam00900)→ RplE (COG0094)rpl5p (PRK04219)→ RpsN (COG0199)rps14P (PRK05766)→ rps8p (PRK04034)RpsH (COG0096)→ RplF (COG0097)rpl6p (PRK05518)→ rpl32e (PRK08562)Ribosomal_L32_L32e (cd00513)→ rpl19e (PRK08570)Ribosomal_L19e (pfam01280)→ RplR (COG0256)rpl18p (PRK08569)→ |
| env-340 | 3300001567_10001587 | + | 11347 | 11431 | **RNA**→ rpl3p (PRK04231)Ribosomal_L3 (pfam00297)→ |
| env-341 | WSSedA2C_c003915 | - | 773 | 689 | **RNA**→ rpl3p (PRK04231)Ribosomal_L3 (pfam00297)→ |
| env-342 | WSSedA1TJGI_c001789 | - | 1725 | 1641 | **RNA**→ RplC (COG0087)rpl3p (PRK04231)→ RplD (COG0088)rpl4lp (PRK04042)→ |
| env-343 | WSSedB1B2_c003929 | - | 407 | 323 | **RNA**→ |
| env-344 | WSSedA1Ba_1001899 | - | 3122 | 3038 | **RNA**→ ←hypo |
| env-345 | WSSedA1Ba2_1001418 | - | 2992 | 2908 | **RNA**→ ←hypo |
| env-346 | WSSedA1Ba3_1004880 | - | 2426 | 2342 | **RNA**→ ←hypo |
| env-347 | WSSedA2Ca3_1003915 | - | 773 | 689 | **RNA**→ ←hypo |
| env-348 | JGI20214J14112_1009109 | + | 1012 | 1096 | **RNA**→ rpl3p (PRK04231)Ribosomal_L3 (pfam00297)→ |
| env-349 | JGI20213J14113_1038457 | - | 635 | 551 | **RNA**→ ←hypo |
| env-350 | B3_all_c_Cons72656 | + | 183 | 267 | **RNA**→ RplC (COG0087)rpl3p (PRK04231)→ RplD (COG0088)rpl4lp (PRK04042)→ |
| env-351 | 3300000568_10104459 | - | 1600 | 1514 | **RNA**→ RplC (COG0087)rpl3p (PRK04231)→ RplD (COG0088)rpl4lp (PRK04042)→ |
| env-352 | SakLake120mDRAFT_c0023947 | - | 758 | 684 | **RNA**→ rpl3p (PRK04231)Ribosomal_L3 (pfam00297)→ |
| env-353 | 3300000032_0022845 | - | 6964 | 6878 | **RNA**→ RplC (COG0087)rpl3p (PRK04231)→ RplD (COG0088)rpl4lp (PRK04042)→ |
| env-354 | 3300000568_10039005 | - | 399 | 313 | **RNA**→ ←hypo |
| env-355 | 3300001567_10000694 | + | 33701 | 33787 | **RNA**→ ←hypo |
| env-356 | 3300000558_11950893 | - | 214 | 128 | **RNA**→ ←hypo |
| env-357 | JGI26437J51864_10016855 | + | 1418 | 1504 | **RNA**→ |

| | | | | | |
|---|---|---|---|---|---|
| env-358 | WSSedA1Ba_1000820 | - | 2896 | 2810 | **RNA**→ RplC (COG0087)rpl3p (PRK04231)→ RplD (COG0088)rpl4lp (PRK04042)→ RplW (COG0089)PRK14548 (PRK14548)→ RplB (COG0090)rpl2p (PRK09612)→ |
| env-359 | JGI20214J14112_1000181 | - | 5712 | 5626 | **RNA**→ RplC (COG0087)rpl3p (PRK04231)→ RplD (COG0088)rpl4lp (PRK04042)→ RplW (COG0089)PRK14548 (PRK14548)→ RplB (COG0090)rpl2p (PRK09612)→ rps19p (PRK04038)Ribosomal_S19 (pfam00203)→ rpl22p (PRK04223)RplV (COG0091)→ RpsC (COG0092)rps3p (PRK04191)→ PRK14549 (PRK14549)Ribosomal_L29 (pfam00831)→ PRK03879 (PRK03879)UPF0086 (pfam01868)→ RpsQ (COG0186)rps17p (PRK08572)→ rpl14p (PRK08571)Ribosomal_L14 (pfam00238)→ |
| env-360 | WSSedB2Ca_1001724 | - | 887 | 801 | **RNA**→ rpl3p (PRK04231)Ribosomal_L3 (pfam00297)→ |
| env-361 | WSSedA2C_c001213 | + | 2849 | 2935 | **RNA**→ rpl3p (PRK04231)Ribosomal_L3 (pfam00297)→ |
| env-362 | WSSedA1TJGI_c002576 | + | 889 | 975 | **RNA**→ RplC (COG0087)rpl3p (PRK04231)→ RplD (COG0088)rpl4lp (PRK04042)→ RplW (COG0089)PRK14548 (PRK14548)→ RplB (COG0090)rpl2p (PRK09612)→ |
| env-363 | WSSedB1B2_c000733 | - | 1648 | 1562 | **RNA**→ rpl3p (PRK04231)Ribosomal_L3 (pfam00297)→ |
| env-364 | WSSedB2T_c003532 | + | 434 | 520 | **RNA**→ rpl3p (PRK04231)Ribosomal_L3 (pfam00297)→ |
| env-365 | WSSedA1Ba2_1000822 | - | 2885 | 2799 | **RNA**→ RplC (COG0087)rpl3p (PRK04231)→ RplD (COG0088)rpl4lp (PRK04042)→ RplW (COG0089)PRK14548 (PRK14548)→ RplB (COG0090)rpl2p (PRK09612)→ |
| env-366 | WSSedA1Ba3_1000489 | + | 1543 | 1629 | **RNA**→ RplC (COG0087)rpl3p (PRK04231)→ RplD (COG0088)rpl4lp (PRK04042)→ RplW (COG0089)PRK14548 (PRK14548)→ RplB (COG0090)rpl2p (PRK09612)→ rps19p (PRK04038)Ribosomal_S19 (pfam00203)→ rpl22p (PRK04223)RplV (COG0091)→ RpsC (COG0092)rps3p (PRK04191)→ PRK14549 (PRK14549)Ribosomal_L29 (pfam00831)→ PRK03879 (PRK03879)UPF0086 (pfam01868)→ RpsQ (COG0186)rps17p (PRK08572)→ rpl14p (PRK08571)Ribosomal_L14 (pfam00238)→ rpl24p (PRK01191)KOW_RPL26 (cd06089)→ PRK04313 (PRK04313)KOW_RPS4 (cd06087)Ribosomal_S4e (pfam00900)→ rpl5p (PRK04219)Ribosomal_L5 (pfam00281)Ribosomal_L5_C (pfam00673)→ RpsN (COG0199)rps14P (PRK05766)→ rps8p (PRK04034)RpsH (COG0096)→ RplF (COG0097)rpl6p (PRK05518)→ rpl32e (PRK08562)Ribosomal_L32e (pfam01655)→ rpl19e (PRK08570)Ribosomal_L19e (pfam01280)→ rpl18p (PRK08569)→ |
| env-367 | WSSedB2Ba_1001051 | - | 1502 | 1416 | **RNA**→ RplC (COG0087)rpl3p (PRK04231)→ RplD (COG0088)rpl4lp (PRK04042)→ |
| env-368 | WSSedA2Ca3_1001213 | + | 2849 | 2935 | **RNA**→ rpl3p (PRK04231)Ribosomal_L3 (pfam00297)→ |
| env-369 | JGI20213J14113_1000441 | - | 1013 | 927 | **RNA**→ rpl3p (PRK04231)Ribosomal_L3 (pfam00297)→ |
| env-370 | JGI26437J51864_10613048 | + | 135 | 220 | **RNA**→ hypo→ |
| env-371 | 3300000568_10163768 | + | 37 | 123 | **RNA**→ hypo→ |
| env-372 | JGI20213J14113_1352303 | + | 18 | 105 | **RNA**→ ←hypo |
| env-373 | WSSedB1B2_c001061 | - | 813 | 728 | **RNA**→ hypo→ |
| env-374 | JGI26437J51864_10024878 | + | 905 | 990 | **RNA**→ |
| env-375 | LWFCAn_GLQU6AQ01CDIKX | - | 257 | 171 | **RNA**→ hypo→ |
| env-376 | JGI24732J26686_1167705 | - | 325 | 240 | **RNA**→ hypo→ |
| env-377 | JGI24732J26686_1058151 | - | 388 | 303 | **RNA**→ hypo→ |
| env-378 | GYO3UBL02HBW2K | - | 172 | 86 | **RNA**→ hypo→ |
| env-379 | LWFCAn_GLO1YSU01C0WMX | + | 42 | 125 | **RNA**→ ←hypo |
| env-380 | 3300000558_10014112 | + | 936 | 1015 | **RNA**→ RplC (COG0087)rpl3p (PRK04231)→ RplD (COG0088)rpl4lp (PRK04042)→ RplW (COG0089)PRK14548 (PRK14548)→ RplB (COG0090)rpl2p (PRK09612)→ rps19p (PRK04038)Ribosomal_S19 (pfam00203)→ rpl22p (PRK04223)RplV (COG0091)→ RpsC (COG0092)rps3p (PRK04191)→ PRK14549 (PRK14549)Ribosomal_L29 (pfam00831)→ PRK03879 (PRK03879)UPF0086 (pfam01868)→ RpsQ (COG0186)rps17p (PRK08572)→ rpl14p (PRK08571)Ribosomal_L14 (pfam00238)→ rpl24p (PRK01191)KOW_RPL26 (cd06089)→ PRK04313 (PRK04313)KOW_RPS4 (cd06087)Ribosomal_S4e (pfam00900)→ rpl5p (PRK04219)Ribosomal_L5 (pfam00281)Ribosomal_L5_C (pfam00673)→ RpsN (COG0199)rps14P (PRK05766)→ rps8p (PRK04034)RpsH (COG0096)→ rpl6p (PRK05518)Ribosomal_L6 (pfam00347)→ |
| env-381 | 3300001580_10840377 | - | 216 | 137 | **RNA**→ ←hypo |
| env-382 | JGI20214J14112_1015362 | + | 514 | 593 | **RNA**→ rpl3p (PRK04231)Ribosomal_L3 (pfam00297)→ |
| env-383 | 3300001605_10335424 | - | 366 | 287 | **RNA**→ |
| env-384 | JGI26437J51864_10189827 | + | 91 | 170 | **RNA**→ ←hypo |
| env-385 | JGI26437J51864_10656079 | - | 220 | 141 | **RNA**→ ←hypo |

| | | | | | |
|---|---|---|---|---|---|
| env-386 | 3300001580_10004519 | - | 9169 | 9090 | **RNA**→ RplC (COG0087)rpl3p (PRK04231)→ RplD (COG0088)rpl4lp (PRK04042)→ RplW (COG0089)PRK14548 (PRK14548)→ RplB (COG0090)rpl2p (PRK09612)→ rps19p (PRK04038)Ribosomal_S19 (pfam00203)→ rpl22p (PRK04223)RplV (COG0091)→ RpsC (COG0092)rps3p (PRK04191)→ PRK14549 (PRK14549)Ribosomal_L29 (pfam00831)→ PRK03879 (PRK03879)UPF0086 (pfam01868)→ RpsQ (COG0186)rps17p (PRK08572)→ rpl14p (PRK08571)Ribosomal_L14 (pfam00238)→ rpl24p (PRK01191)KOW_RPL26 (cd06089)→ PRK04313 (PRK04313)KOW_RPS4 (cd06087)Ribosomal_S4e (pfam00900)→ rpl5p (PRK04219)Ribosomal_L5 (pfam00281)Ribosomal_L5_C (pfam00673)→ RpsN (COG0199)rps14P (PRK05766)→ rps8p (PRK04034)RpsH (COG0096)→ RplF (COG0097)rpl6p (PRK05518)→ rpl32e (PRK08562)Ribosomal_L32e (pfam01655)→ |
| env-387 | 3300000032_0388072 | - | 284 | 205 | **RNA**→ |
| env-388 | 3300001605_10952026 | + | 300 | 379 | **RNA**→ ←hypo |
| env-389 | B3_all_c_Cons98067 | + | 267 | 345 | **RNA**→ rpl3p (PRK04231)Ribosomal_L3 (pfam00297)→ |
| env-390 | LWAnNiSIP_F624WLL02IC2DN | - | 276 | 191 | **RNA**→ hypo→ |
| env-391 | LWAnN_F624WLL02IC2DN | - | 276 | 191 | **RNA**→ hypo→ |
| env-392 | JGI24728J21555_1000024 | + | 55297 | 55372 | **RNA**→ RplC (COG0087)rpl3p (PRK04231)→ RplD (COG0088)rpl4lp (PRK04042)→ RplW (COG0089)PRK14548 (PRK14548)→ RplB (COG0090)rpl2p (PRK09612)→ rps19p (PRK04038)Ribosomal_S19 (pfam00203)→ rpl22p (PRK04223)RplV (COG0091)→ RpsC (COG0092)rps3p (PRK04191)→ PRK14549 (PRK14549)Ribosomal_L29 (pfam00831)→ PRK03879 (PRK03879)UPF0086 (pfam01868)→ RpsQ (COG0186)rps17p (PRK08572)→ rpl14p (PRK08571)Ribosomal_L14 (pfam00238)→ rpl24p (PRK01191)KOW_RPL26 (cd06089)→ PRK04313 (PRK04313)KOW_RPS4 (cd06087)Ribosomal_S4e (pfam00900)→ rpl5p (PRK04219)Ribosomal_L5 (pfam00281)Ribosomal_L5_C (pfam00673)→ RpsN (COG0199)rps14P (PRK05766)→ rps8p (PRK04034)RpsH (COG0096)→ RplF (COG0097)rpl6p (PRK05518)→ rpl32e (PRK08562)Ribosomal_L32_L32e (cd00513)→ hypo→ hypo→ rpl19e (PRK08570)Ribosomal_L19e (pfam01280)→ RplR (COG0256)rpl18p (PRK08569)→ |
| env-393 | JGI24731J21663_1041451 | + | 763 | 838 | **RNA**→ |
| env-394 | JGI24732J26686_1000141 | + | 22138 | 22213 | **RNA**→ RplC (COG0087)rpl3p (PRK04231)→ RplD (COG0088)rpl4lp (PRK04042)→ RplW (COG0089)PRK14548 (PRK14548)→ RplB (COG0090)rpl2p (PRK09612)→ rps19p (PRK04038)Ribosomal_S19 (pfam00203)→ rpl22p (PRK04223)RplV (COG0091)→ RpsC (COG0092)rps3p (PRK04191)→ PRK14549 (PRK14549)Ribosomal_L29 (pfam00831)→ PRK03879 (PRK03879)UPF0086 (pfam01868)→ RpsQ (COG0186)rps17p (PRK08572)→ rpl14p (PRK08571)Ribosomal_L14 (pfam00238)→ rpl24p (PRK01191)KOW_RPL26 (cd06089)→ PRK04313 (PRK04313)KOW_RPS4 (cd06087)Ribosomal_S4e (pfam00900)→ rpl5p (PRK04219)Ribosomal_L5 (pfam00281)Ribosomal_L5_C (pfam00673)→ RpsN (COG0199)rps14P (PRK05766)→ rps8p (PRK04034)RpsH (COG0096)→ RplF (COG0097)rpl6p (PRK05518)→ rpl32e (PRK08562)Ribosomal_L32_L32e (cd00513)→ hypo→ hypo→ rpl19e (PRK08570)Ribosomal_L19e (pfam01280)→ RplR (COG0256)rpl18p (PRK08569)→ |
| env-395 | JGI24729J20445_1000265 | - | 14288 | 14211 | **RNA**→ RplC (COG0087)rpl3p (PRK04231)→ RplD (COG0088)rpl4lp (PRK04042)→ RplW (COG0089)PRK14548 (PRK14548)→ RplB (COG0090)rpl2p (PRK09612)→ rps19p (PRK04038)Ribosomal_S19 (pfam00203)→ rpl22p (PRK04223)RplV (COG0091)→ RpsC (COG0092)rps3p (PRK04191)→ PRK14549 (PRK14549)Ribosomal_L29 (pfam00831)→ PRK03879 (PRK03879)UPF0086 (pfam01868)→ RpsQ (COG0186)rps17p (PRK08572)→ rpl14p (PRK08571)Ribosomal_L14 (pfam00238)→ rpl24p (PRK01191)KOW_RPL26 (cd06089)→ PRK04313 (PRK04313)S4 (cd00165)KOW_RPS4 (cd06087)Ribosomal_S4e (pfam00900)→ RplE (COG0094)rpl5p (PRK04219)→ RpsN (COG0199)rps14P (PRK05766)→ rps8p (PRK04034)RpsH (COG0096)→ RplF (COG0097)rpl6p (PRK05518)→ rpl32e (PRK08562)Ribosomal_L32e (pfam01655)→ rpl19e (PRK08570)Ribosomal_L19e (pfam01280)→ RplR (COG0256)rpl18p (PRK08569)→ |

| | | | | | |
|---|---|---|---|---|---|
| env-396 | JGI24729J20445_1000376 | - | 22541 | 22464 | **RNA**→ RplC (COG0087)rpl3p (PRK04231)→ RplD (COG0088)rpl4lp (PRK04042)→ RplW (COG0089)PRK14548 (PRK14548)→ RplB (COG0090)rpl2p (PRK09612)→ rps19p (PRK04038)Ribosomal_S19 (pfam00203)→ rpl22p (PRK04223)RplV (COG0091)→ RpsC (COG0092)rps3p (PRK04191)→ PRK14549 (PRK14549)Ribosomal_L29 (pfam00831)→ PRK03879 (PRK03879)UPF0086 (pfam01868)→ RpsQ (COG0186)rps17p (PRK08572)→ rpl14p (PRK08571)Ribosomal_L14 (pfam00238)→ rpl24p (PRK01191)KOW_RPL26 (cd06089)→ PRK04313 (PRK04313)S4 (cd00165)KOW_RPS4 (cd06087)Ribosomal_S4e (pfam00900)→ RplE (COG0094)rpl5p (PRK04219)→ RpsN (COG0199)rps14P (PRK05766)→ rps8p (PRK04034)RpsH (COG0096)→ RplF (COG0097)rpl6p (PRK05518)→ rpl32e (PRK08562)Ribosomal_L32e (pfam01655)→ rpl19e (PRK08570)Ribosomal_L19e (pfam01280)→ RplR (COG0256)rpl18p (PRK08569)→ |
| env-397 | JGI24732J26686_1000166 | - | 22911 | 22834 | **RNA**→ RplC (COG0087)rpl3p (PRK04231)→ RplD (COG0088)rpl4lp (PRK04042)→ RplW (COG0089)PRK14548 (PRK14548)→ RplB (COG0090)rpl2p (PRK09612)→ rps19p (PRK04038)Ribosomal_S19 (pfam00203)→ rpl22p (PRK04223)RplV (COG0091)→ RpsC (COG0092)rps3p (PRK04191)→ PRK14549 (PRK14549)Ribosomal_L29 (pfam00831)→ PRK03879 (PRK03879)UPF0086 (pfam01868)→ RpsQ (COG0186)rps17p (PRK08572)→ rpl14p (PRK08571)Ribosomal_L14 (pfam00238)→ rpl24p (PRK01191)KOW_RPL26 (cd06089)→ PRK04313 (PRK04313)S4 (cd00165)KOW_RPS4 (cd06087)Ribosomal_S4e (pfam00900)→ RplE (COG0094)rpl5p (PRK04219)→ RpsN (COG0199)rps14P (PRK05766)→ rps8p (PRK04034)RpsH (COG0096)→ RplF (COG0097)rpl6p (PRK05518)→ rpl32e (PRK08562)Ribosomal_L32e (pfam01655)→ rpl19e (PRK08570)Ribosomal_L19e (pfam01280)→ RplR (COG0256)rpl18p (PRK08569)→ |
| env-398 | JGI24730J26740_1278979 | + | 62 | 139 | **RNA**→ ←hypo |
| env-399 | JGI20214J14112_1005972 | + | 608 | 694 | **RNA**→ hypo→ RplC (COG0087)rpl3p (PRK04231)→ |
| env-400 | 3300000558_10001908 | - | 3593 | 3508 | **RNA**→ hypo→ |
| env-401 | 3300001605_12127170 | + | 26 | 111 | **RNA**→ hypo→ |
| env-402 | JGI24732J26686_1168257 | - | 324 | 239 | **RNA**→ hypo→ |
| env-403 | JGI24729J20445_1001367 | - | 9125 | 9040 | **RNA**→ RplC (COG0087)rpl3p (PRK04231)→ RplD (COG0088)rpl4lp (PRK04042)→ RplW (COG0089)PRK14548 (PRK14548)→ RplB (COG0090)rpl2p (PRK09612)→ rps19p (PRK04038)Ribosomal_S19 (pfam00203)→ rpl22p (PRK04223)RplV (COG0091)→ RpsC (COG0092)rps3p (PRK04191)→ PRK14549 (PRK14549)Ribosomal_L29 (pfam00831)→ PRK03879 (PRK03879)UPF0086 (pfam01868)→ RpsQ (COG0186)rps17p (PRK08572)→ rpl14p (PRK08571)Ribosomal_L14 (pfam00238)→ rpl24p (PRK01191)KOW_RPL26 (cd06089)→ PRK04313 (PRK04313)S4 (cd00165)KOW_RPS4 (cd06087)Ribosomal_S4e (pfam00900)→ rpl5p (PRK04219)Ribosomal_L5 (pfam00281)Ribosomal_L5_C (pfam00673)→ RpsN (COG0199)rps14P (PRK05766)→ rps8p (PRK04034)RpsH (COG0096)→ RplF (COG0097)rpl6p (PRK05518)→ rpl32e (PRK08562)Ribosomal_L32e (pfam01655)→ |
| env-404 | JGI24730J26740_1168855 | + | 233 | 318 | **RNA**→ hypo→ |
| env-405 | JGI24728J21555_1218516 | - | 187 | 102 | **RNA**→ ←hypo |
| env-406 | JGI24729J20445_1004575 | + | 4026 | 4111 | **RNA**→ |
| env-407 | WSSedA1Ba3_1333478 | - | 172 | 89 | **RNA**→ hypo→ |

| | | | | | |
|---|---|---|---|---|---|
| Mli-2-1 | NZ_CM001436.1 | - | 2884644 | 2884571 | **RNA**→ RplC (COG0087)rpl3p (PRK04231)→ RplC (COG0087)rpl3p (PRK04231)→<br>RplD (COG0088)rpl4lp (PRK04042)→ RplD (COG0088)rpl4lp (PRK04042)→<br>RplW (COG0089)PRK14548 (PRK14548)→ RplW (COG0089)PRK14548 (PRK14548)→<br>RplB (COG0090)rpl2p (PRK09612)→ RplB (COG0090)rpl2p (PRK09612)→<br>rps19p (PRK04038)Ribosomal_S19 (pfam00203)→ rps19p (PRK04038)Ribosomal_S19 (pfam00203)→<br>rpl22p (PRK04223)RplV (COG0091)→ rpl22p (PRK04223)RplV (COG0091)→<br>RpsC (COG0092)rps3p (PRK04191)→ RpsC (COG0092)rps3p (PRK04191)→<br>PRK14549 (PRK14549)Ribosomal_L29 (pfam00831)→ PRK14549 (PRK14549)Ribosomal_L29 (pfam00831)→<br>PRK03879 (PRK03879)UPF0086 (pfam01868)→ PRK03879 (PRK03879)UPF0086 (pfam01868)→<br>RpsQ (COG0186)rps17p (PRK08572)→ RpsQ (COG0186)rps17p (PRK08572)→<br>rpl14p (PRK08571)Ribosomal_L14 (pfam00238)→ rpl14p (PRK08571)Ribosomal_L14 (pfam00238)→<br>rpl24p (PRK01191)KOW_RPL26 (cd06089)→ rpl24p (PRK01191)KOW_RPL26 (cd06089)→<br>PRK04313 (PRK04313)KOW_RPS4 (cd06087)Ribosomal_S4e (pfam00900)→<br>PRK04313 (PRK04313)KOW_RPS4 (cd06087)Ribosomal_S4e (pfam00900)→ RplE (COG0094)rpl5p (PRK04219)→<br>RplE (COG0094)rpl5p (PRK04219)→ RpsN (COG0199)rps14P (PRK05766)→<br>RpsN (COG0199)rps14P (PRK05766)→ rps8p (PRK04034)RpsH (COG0096)→<br>rps8p (PRK04034)RpsH (COG0096)→ RplF (COG0097)uL6_arch (TIGR03653)→<br>RplF (COG0097)uL6_arch (TIGR03653)→ rpl32e (PRK08562)Ribosomal_L32e (pfam01655)→<br>rpl32e (PRK08562)Ribosomal_L32e (pfam01655)→ rpl19e (PRK08570)Ribosomal_L19e (pfam01280)→<br>rpl19e (PRK08570)Ribosomal_L19e (pfam01280)→ RplR (COG0256)rpl18p (PRK08569)→<br>RplR (COG0256)rpl18p (PRK08569)→ RpsE (COG0098)rps5p (PRK04044)→<br>RpsE (COG0098)rps5p (PRK04044)→ |
| Mpe-1-1 | NC_014507.1 | + | 2206565 | 2206637 | **RNA**→ RplC (COG0087)rpl3p (PRK04231)→ RplD (COG0088)rpl4lp (PRK04042)→<br>RplW (COG0089)PRK14548 (PRK14548)→ RplB (COG0090)rpl2p (PRK09612)→<br>RpsS (COG0185)rps19p (PRK04038)→ rpl22p (PRK04223)RplV (COG0091)→<br>RpsC (COG0092)rps3p (PRK04191)→ PRK14549 (PRK14549)Ribosomal_L29 (pfam00831)→<br>PRK03879 (PRK03879)UPF0086 (pfam01868)→ RpsQ (COG0186)rps17p (PRK08572)→<br>rpl14p (PRK08571)Ribosomal_L14 (pfam00238)→ rpl24p (PRK01191)KOW_RPL26 (cd06089)→<br>PRK04313 (PRK04313)KOW_RPS4 (cd06087)Ribosomal_S4e (pfam00900)→ RplE (COG0094)rpl5p (PRK04219)→<br>RpsN (COG0199)rps14P (PRK05766)→ rps8p (PRK04034)RpsH (COG0096)→<br>RplF (COG0097)rpl6p (PRK05518)→ hypo→ rpl32e (PRK08562)Ribosomal_L32e (pfam01655)→<br>rpl19e (PRK08570)Ribosomal_L19e (pfam01280)→ RplR (COG0256)rpl18p (PRK08569)→<br>RpsE (COG0098)rps5p (PRK04044)→ |
| Mpa-2-1 | NZ_KN360930.1 | - | 82807 | 82734 | **RNA**→ ←hypo |
| env-408 | M940CN_1020281 | + | 3372 | 3442 | **RNA**→ rpl3p (PRK04231)Ribosomal_L3 (pfam00297)→ |
| Mmo-1-1 | NZ_JOMF01000007.1 | + | 9542 | 9636 | **RNA**→ RplC (COG0087)rpl3p (PRK04231)→ RplD (COG0088)rpl4lp (PRK04042)→<br>RplW (COG0089)PRK14548 (PRK14548)→ RplB (COG0090)rpl2p (PRK09612)→<br>RpsS (COG0185)rps19p (PRK04038)→ rpl22p (PRK04223)RplV (COG0091)→<br>RpsC (COG0092)rps3p (PRK04191)→ PRK14549 (PRK14549)Ribosomal_L29 (pfam00831)→<br>POP4 (COG1588)PRK03879 (PRK03879)→ RpsQ (COG0186)rps17p (PRK08572)→<br>rpl14p (PRK08571)Ribosomal_L14 (pfam00238)→ RplX (COG0198)rpl24p (PRK01191)→<br>PRK04313 (PRK04313)KOW_RPS4 (cd06087)Ribosomal_S4e (pfam00900)→ RplE (COG0094)rpl5p (PRK04219)→<br>RpsN (COG0199)rps14P (PRK05766)→ rps8p (PRK04034)RpsH (COG0096)→<br>RplF (COG0097)rpl6p (PRK05518)→ rpl32e (PRK08562)Ribosomal_L32e (pfam01655)→<br>rpl19e (PRK08570)Ribosomal_L19e (pfam01280)→ RplR (COG0256)rpl18p (PRK08569)→<br>RpsE (COG0098)rps5p (PRK04044)→ |
| env-409 | RUMENNODE_4185760_38501 | - | 73103 | 73009 | **RNA**→ RplC (COG0087)rpl3p (PRK04231)→ RplD (COG0088)rpl4lp (PRK04042)→<br>RplW (COG0089)PRK14548 (PRK14548)→ RplB (COG0090)rpl2p (PRK09612)→ rpl2p (PRK09612)→<br>RpsS (COG0185)rps19p (PRK04038)→ rpl22p (PRK04223)RplV (COG0091)→<br>KH (smart00322)archeal_30S_S3_KH (cd02411)→ rps3p (PRK04191)Ribosomal_S3_C (pfam00189)→<br>PRK14549 (PRK14549)Ribosomal_L29 (pfam00831)→ POP4 (COG1588)PRK03879 (PRK03879)→<br>RpsQ (COG0186)rps17p (PRK08572)→ rpl14p (PRK08571)Ribosomal_L14 (pfam00238)→<br>RplX (COG0198)rpl24p (PRK01191)→ PRK04313 (PRK04313)Ribosomal_S4e (pfam00900)→<br>PRK04313 (PRK04313)KOW_RPS4 (cd06087)→ RplE (COG0094)rpl5p (PRK04219)→<br>RpsN (COG0199)rps14P (PRK05766)→ rps8p (PRK04034)RpsH (COG0096)→<br>RplF (COG0097)rpl6p (PRK05518)→ |

| | | | | | |
|---|---|---|---|---|---|
| env-410 | SL_5KL_010_SED_10008304 | + | 2725 | 2796 | **RNA**→ RplC (COG0087)rpl3p (PRK04231)→ RplD (COG0088)rpl4lp (PRK04042)→ |
| env-411 | LCWY01003686.1 | + | 978 | 1049 | **RNA**→ hypo→ |
| env-412 | LCWZ01000078.1 | + | 773 | 844 | **RNA**→ RplC (COG0087)rpl3p (PRK04231)→ RplD (COG0088)rpl4lp (PRK04042)→ |
| | | | | | RplW (COG0089)PRK14548 (PRK14548)→ RplB (COG0090)rpl2p (PRK09612)→ |
| | | | | | RpsS (COG0185)rps19p (PRK04038)→ rpl22p (PRK04223)RplV (COG0091)→ |
| | | | | | RpsC (COG0092)rps3p (PRK04191)→ PRK14549 (PRK14549)Ribosomal_L29 (pfam00831)→ |
| | | | | | PRK03879 (PRK03879)UPF0086 (pfam01868)→ RpsQ (COG0186)rps17p (PRK08572)→ |
| | | | | | rpl14p (PRK08571)Ribosomal_L14 (pfam00238)→ rpl24p (PRK01191)KOW_RPL26 (cd06089)→ |
| | | | | | PRK04313 (PRK04313)KOW_RPS4 (cd06087)S4 (pfam01479)Ribosomal_S4e (pfam00900)→ |
| | | | | | rpl5p (PRK04219)Ribosomal_L5 (pfam00281)Ribosomal_L5_C (pfam00673)→ |
| | | | | | RpsN (COG0199)rps14P (PRK05766)→ rps8p (PRK04034)→ rps8p (PRK04034)Ribosomal_S8 (pfam00410)→ |
| | | | | | RplF (COG0097)rpl6p (PRK05518)→ hypo→ Rpl32e (COG1717)rpl32e (PRK08562)→ |
| | | | | | rpl19e (PRK08570)Ribosomal_L19e (pfam01280)→ RplR (COG0256)rpl18p (PRK08569)→ |
| | | | | | RpsE (COG0098)rps5p (PRK04044)→ Ribosomal_L30_like (cd00355)rpl30p (PRK06049)→ |
| | | | | | Ribosomal_L7_archeal_euk (cd01657)rpl30p (PRK06049)→ RplO (COG0200)rpl15p (PRK06419)→ |
| Mco-1-1 | NC_017034.1 | + | 1276328 | 1276418 | **RNA**→ MTH1 (COG2106)→ |
| env-413 | JGI24024J18818_10045249 | + | 1393 | 1489 | **RNA**→ Methyltrn_RNA_3 (pfam02598)→ |
| env-414 | LGVD01025758.1 | - | 1753 | 1657 | **RNA**→ Methyltrn_RNA_3 (pfam02598)→ |
| env-415 | ASHM260b_GM9OZ8V02GZRNU | + | 167 | 263 | **RNA**→ hypo→ |
| env-416 | LGVE01111852.1 | - | 441 | 345 | **RNA**→ Methyltrn_RNA_3 (pfam02598)→ |
| env-417 | LGVF01232254.1 | - | 138 | 42 | **RNA**→ Methyltrn_RNA_3 (pfam02598)→ |
| env-418 | LGOV01013195.1 | + | 3349 | 3445 | **RNA**→ Methyltrn_RNA_3 (pfam02598)→ RplC (COG0087)rpl3p (PRK04231)→ |
| | | | | | RplD (COG0088)rpl4lp (PRK04042)→ RplW (COG0089)PRK14548 (PRK14548)→ |
| | | | | | RplB (COG0090)rpl2p (PRK09612)→ RpsS (COG0185)rps19p (PRK04038)→ |
| | | | | | rpl22p (PRK04223)Ribosomal_L22 (pfam00237)→ rps3p (PRK04191)Ribosomal_S3_C (pfam00189)→ |
| | | | | | PRK14549 (PRK14549)Ribosomal_L29 (pfam00831)→ PRK03879 (PRK03879)UPF0086 (pfam01868)→ |
| | | | | | RpsQ (COG0186)rps17p (PRK08572)→ rpl14p (PRK08571)Ribosomal_L14 (pfam00238)→ |
| | | | | | rpl24p (PRK01191)KOW_RPL26 (cd06089)→ |
| | | | | | PRK04313 (PRK04313)KOW_RPS4 (cd06087)Ribosomal_S4e (pfam00900)→ |
| | | | | | rpl5p (PRK04219)Ribosomal_L5_C (pfam00673)→ RpsN (COG0199)rps14P (PRK05766)→ |
| | | | | | rps8p (PRK04034)RpsH (COG0096)→ RplF (COG0097)rpl6p (PRK05518)→ |
| | | | | | Rpl32e (COG1717)rpl32e (PRK08562)→ |
| env-419 | LGVC01002321.1 | + | 1612 | 1708 | **RNA**→ Methyltrn_RNA_3 (pfam02598)→ RplC (COG0087)rpl3p (PRK04231)→ |
| | | | | | RplD (COG0088)rpl4lp (PRK04042)→ RplW (COG0089)PRK14548 (PRK14548)→ |
| | | | | | RplB (COG0090)rpl2p (PRK09612)→ RpsS (COG0185)rps19p (PRK04038)→ |
| | | | | | rpl22p (PRK04223)RplV (COG0091)→ RpsC (COG0092)rps3p (PRK04191)→ |
| | | | | | PRK14549 (PRK14549)Ribosomal_L29 (pfam00831)→ PRK03879 (PRK03879)UPF0086 (pfam01868)→ |
| | | | | | RpsQ (COG0186)rps17p (PRK08572)→ rpl14p (PRK08571)Ribosomal_L14 (pfam00238)→ |
| | | | | | rpl24p (PRK01191)KOW_RPL26 (cd06089)→ |
| | | | | | PRK04313 (PRK04313)KOW_RPS4 (cd06087)Ribosomal_S4e (pfam00900)→ RplE (COG0094)rpl5p (PRK04219)→ |
| | | | | | RpsN (COG0199)rps14P (PRK05766)→ rps8p (PRK04034)RpsH (COG0096)→ |
| | | | | | RplF (COG0097)rpl6p (PRK05518)→ rpl32e (PRK08562)Ribosomal_L32_L32e (cd00513)→ |
| | | | | | rpl19e (PRK08570)Ribosomal_L19e (pfam01280)→ RplR (COG0256)rpl18p (PRK08569)→ |
| | | | | | RpsE (COG0098)rps5p (PRK04044)→ Ribosomal_L7_archeal_euk (cd01657)rpl30p (PRK06049)→ |
| | | | | | RplO (COG0200)rpl15p (PRK06419)→ SecY (COG0201)PRK08568 (PRK08568)→ |
| | | | | | adk (PRK00279)ADK (cd01428)→ COG1422 (COG1422)spore_lon_C (TIGR02903)→ |
| | | | | | CmkB (COG1102)PRK04182 (PRK04182)→ |
| | | | | | PRK04270 (PRK04270)PseudoU_synth_hDyskerin (cd02572)PUA (pfam01472)→ hypo→ hypo→ hypo→ |

| | | | | | |
|---|---|---|---|---|---|
| env-420 | LGVD01004949.1 | + | 1392 | 1488 | **RNA**→ Methyltrn_RNA_3 (pfam02598)→ RplC (COG0087)rpl3p (PRK04231)→<br>rpl4lp (PRK04042)Ribosomal_L4 (pfam00573)→ RplW (COG0089)PRK14548 (PRK14548)→<br>rpl2p (PRK09612)Ribosomal_L2_C (pfam03947)→ RpsS (COG0185)rps19p (PRK04038)→<br>rpl22p (PRK04223)RplV (COG0091)→ rps3p (PRK04191)Ribosomal_S3_C (pfam00189)→<br>PRK14549 (PRK14549)Ribosomal_L29 (pfam00831)→ PRK03879 (PRK03879)UPF0086 (pfam01868)→<br>RpsQ (COG0186)rps17p (PRK08572)→ rpl14p (PRK08571)Ribosomal_L14 (pfam00238)→ rpl24p (PRK01191)→<br>PRK04313 (PRK04313)KOW_RPS4 (cd06087)Ribosomal_S4e (pfam00900)→ RplE (COG0094)rpl5p (PRK04219)→<br>RpsN (COG0199)rps14P (PRK05766)→ rps8p (PRK04034)RpsH (COG0096)→<br>RplF (COG0097)rpl6p (PRK05518)→ rpl32e (PRK08562)Ribosomal_L32_L32e (cd00513)→<br>rpl19e (PRK08570)Ribosomal_L19e (pfam01280)→ RplR (COG0256)rpl18p (PRK08569)→<br>RpsE (COG0098)rps5p (PRK04044)→ RpmD (COG1841)rpl30p (PRK06049)→ |
| env-421 | LGVE01002492.1 | - | 11696 | 11600 | **RNA**→ Methyltrn_RNA_3 (pfam02598)→ RplC (COG0087)rpl3p (PRK04231)→<br>RplD (COG0088)rpl4lp (PRK04042)→ RplW (COG0089)PRK14548 (PRK14548)→<br>RplB (COG0090)rpl2p (PRK09612)→ rpl2p (PRK09612)Ribosomal_L2_C (pfam03947)→<br>RpsS (COG0185)rps19p (PRK04038)→ rpl22p (PRK04223)RplV (COG0091)→<br>rps3p (PRK04191)archeal_30S_S3_KH (cd02411)→ RpsC (COG0092)rps3p (PRK04191)→<br>RpmC (COG0255)PRK14549 (PRK14549)→ PRK03879 (PRK03879)UPF0086 (pfam01868)→<br>rps17p (PRK08572)→ rpl14p (PRK08571)Ribosomal_L14 (pfam00238)→<br>rpl24p (PRK01191)KOW_RPL26 (cd06089)→<br>PRK04313 (PRK04313)rpl5p (PRK04219)KOW_RPS4 (cd06087)Ribosomal_L5_C (pfam00673)Ribosomal_S4e (pfam00900)→<br>rps8p (PRK04034)RpsH (COG0096)→ RplF (COG0097)rpl6p (PRK05518)→<br>RplF (COG0097)rpl6p (PRK05518)→ Rpl32e (COG1717)rpl32e (PRK08562)→ |
| env-422 | LGVF01002618.1 | + | 131 | 227 | **RNA**→ Methyltrn_RNA_3 (pfam02598)→ rpl3p (PRK04231)Ribosomal_L3 (pfam00297)→<br>RplC (COG0087)rpl3p (PRK04231)→ |
| env-423 | LGVE01240744.1 | - | 496 | 400 | **RNA**→ Methyltrn_RNA_3 (pfam02598)→ |
| env-424 | LGVF01231129.1 | - | 567 | 471 | **RNA**→ Methyltrn_RNA_3 (pfam02598)→ |
| env-425 | LGOV01087460.1 | + | 959 | 1055 | **RNA**→ Methyltrn_RNA_3 (pfam02598)→ |
| env-426 | LGVE01063670.1 | + | 1228 | 1324 | **RNA**→ Methyltrn_RNA_3 (pfam02598)→ RplC (COG0087)rpl3p (PRK04231)→ |
| env-427 | LGVF01061963.1 | + | 239 | 335 | **RNA**→ Methyltrn_RNA_3 (pfam02598)→ rpl3p (PRK04231)Ribosomal_L3 (pfam00297)→ |
| env-428 | WSSedA1Ba3_1023920 | + | 778 | 871 | **RNA**→ Methyltrn_RNA_3 (pfam02598)→ |
| env-429 | WSSedA1Ba_1038552 | + | 407 | 500 | **RNA**→ MTH1 (COG2106)→ |
| Msh-1-1 | NZ_JONQ01000008.1 | - | 165643 | 165550 | **RNA**→ Methyltrn_RNA_3 (pfam02598)→ RplC (COG0087)rpl3p (PRK04231)→<br>RplD (COG0088)rpl4lp (PRK04042)→ RplW (COG0089)PRK14548 (PRK14548)→<br>RplB (COG0090)rpl2p (PRK09612)→ RpsS (COG0185)rps19p (PRK04038)→<br>rpl22p (PRK04223)RplV (COG0091)→ RpsC (COG0092)rps3p (PRK04191)→<br>PRK14549 (PRK14549)Ribosomal_L29_HIP (cd00427)→ PRK03879 (PRK03879)UPF0086 (pfam01868)→<br>RpsQ (COG0186)rps17p (PRK08572)→ rpl14p (PRK08571)Ribosomal_L14 (pfam00238)→<br>RplX (COG0198)rpl24p (PRK01191)→ PRK04313 (PRK04313)S4 (cd00165)Ribosomal_S4e (pfam00900)→<br>RplE (COG0094)rpl5p (PRK04219)→ RpsN (COG0199)rps14P (PRK05766)→<br>rps8p (PRK04034)RpsH (COG0096)→ RplF (COG0097)rpl6p (PRK05518)→<br>rpl32e (PRK08562)Ribosomal_L32e (pfam01655)→ rpl19e (PRK08570)Ribosomal_L19e (pfam01280)→<br>RplR (COG0256)rpl18p (PRK08569)→ |
| env-430 | taComm3_C1244 | - | 4081 | 3988 | **RNA**→ Methyltrn_RNA_3 (pfam02598)→ RplC (COG0087)rpl3p (PRK04231)→<br>RplD (COG0088)rpl4lp (PRK04042)→ RplW (COG0089)PRK14548 (PRK14548)→<br>RplB (COG0090)rpl2p (PRK09612)→ RpsS (COG0185)rps19p (PRK04038)→<br>rpl22p (PRK04223)RplV (COG0091)→ |
| env-431 | JGI12104J13512_1002612 | - | 3904 | 3811 | **RNA**→ Methyltrn_RNA_3 (pfam02598)→ RplC (COG0087)rpl3p (PRK04231)→<br>RplD (COG0088)rpl4lp (PRK04042)→ RplW (COG0089)PRK14548 (PRK14548)→<br>RplB (COG0090)rpl2p (PRK09612)→ RpsS (COG0185)rps19p (PRK04038)→<br>rpl22p (PRK04223)Ribosomal_L22 (pfam00237)→ |

| | | | | | |
|---|---|---|---|---|---|
| env-432 | JGI11944J13513_1000082 | - | 41802 | 41709 | **RNA**→ Methyltrn_RNA_3 (pfam02598)→ RplC (COG0087)rpl3p (PRK04231)→<br>RplD (COG0088)rpl4lp (PRK04042)→ RplW (COG0089)PRK14548 (PRK14548)→<br>RplB (COG0090)rpl2p (PRK09612)→ RpsS (COG0185)rps19p (PRK04038)→<br>rpl22p (PRK04223)RplV (COG0091)→ RpsC (COG0092)rps3p (PRK04191)→<br>PRK14549 (PRK14549)Ribosomal_L29 (pfam00831)→ PRK03879 (PRK03879)UPF0086 (pfam01868)→<br>RpsQ (COG0186)rps17p (PRK08572)→ rpl14p (PRK08571)Ribosomal_L14 (pfam00238)→<br>rpl24p (PRK01191)KOW_RPL26 (cd06089)→<br>PRK04313 (PRK04313)KOW_RPS4 (cd06087)S4 (pfam01479)Ribosomal_S4e (pfam00900)→<br>rpl5p (PRK04219)Ribosomal_L5 (pfam00281)Ribosomal_L5_C (pfam00673)→<br>RpsN (COG0199)rps14P (PRK05766)→ rps8p (PRK04034)RpsH (COG0096)→<br>RplF (COG0097)rpl6p (PRK05518)→ rpl32e (PRK08562)Ribosomal_L32e (pfam01655)→<br>rpl19e (PRK08570)Ribosomal_L19e (pfam01280)→ RplR (COG0256)rpl18p (PRK08569)→ |
| env-433 | JGI20179J14886_1000343 | - | 1356 | 1263 | **RNA**→ Methyltrn_RNA_3 (pfam02598)→ RplC (COG0087)rpl3p (PRK04231)→ |
| env-434 | 3300001580_10791145 | - | 337 | 244 | **RNA**→ MTH1 (COG2106)→ |
| env-435 | 3300000568_10023551 | - | 5356 | 5263 | **RNA**→ Methyltrn_RNA_3 (pfam02598)→ RplC (COG0087)rpl3p (PRK04231)→<br>RplD (COG0088)rpl4lp (PRK04042)→ RplW (COG0089)PRK14548 (PRK14548)→<br>RplB (COG0090)rpl2p (PRK09612)→ RpsS (COG0185)rps19p (PRK04038)→<br>rpl22p (PRK04223)RplV (COG0091)→ RpsC (COG0092)rps3p (PRK04191)→<br>PRK14549 (PRK14549)Ribosomal_L29 (pfam00831)→ PRK03879 (PRK03879)UPF0086 (pfam01868)→ |
| env-436 | 3300001592_10171977 | + | 123 | 216 | **RNA**→ MTH1 (COG2106)PRK10611 (PRK10611)→ rpl3p (PRK04231)Ribosomal_L3 (pfam00297)→ |
| env-437 | 3300000032_0536775 | - | 94 | 1 | **RNA**→ |
| env-438 | 3300001605_11573450 | - | 95 | 2 | **RNA**→ MTH1 (COG2106)→ |
| env-439 | 3300001567_10002499 | - | 7097 | 7004 | **RNA**→ Methyltrn_RNA_3 (pfam02598)→ RplC (COG0087)rpl3p (PRK04231)→<br>RplD (COG0088)rpl4lp (PRK04042)→ RplW (COG0089)PRK14548 (PRK14548)→<br>RplB (COG0090)rpl2p (PRK09612)→ RpsS (COG0185)rps19p (PRK04038)→<br>rpl22p (PRK04223)RplV (COG0091)→ RpsC (COG0092)rps3p (PRK04191)→<br>PRK14549 (PRK14549)Ribosomal_L29 (pfam00831)→ PRK03879 (PRK03879)UPF0086 (pfam01868)→<br>RpsQ (COG0186)rps17p (PRK08572)→ rpl14p (PRK08571)Ribosomal_L14 (pfam00238)→<br>KOW (smart00739)KOW_RPL26 (cd06089)→ PRK04313 (PRK04313)Ribosomal_S4e (pfam00900)→ |
| env-440 | 3300000558_10031507 | - | 264 | 171 | **RNA**→ MTH1 (COG2106)→ |
| env-441 | PCEOT_contig04628 | + | 1388 | 1481 | **RNA**→ Methyltrn_RNA_3 (pfam02598)→ RplC (COG0087)rpl3p (PRK04231)→<br>RplD (COG0088)rpl4lp (PRK04042)→ RplW (COG0089)PRK14548 (PRK14548)→<br>RplB (COG0090)rpl2p (PRK09612)→ RpsS (COG0185)rps19p (PRK04038)→<br>rpl22p (PRK04223)RplV (COG0091)→ RpsC (COG0092)rps3p (PRK04191)→<br>PRK14549 (PRK14549)Ribosomal_L29 (pfam00831)→ PRK03879 (PRK03879)UPF0086 (pfam01868)→ |
| env-442 | JGI12104J13512_1100715 | + | 77 | 170 | **RNA**→ MTH1 (COG2106)→ |
| Mco-2-1 | NC_015416.1 | - | 2306426 | 2306333 | **RNA**→ Methyltrn_RNA_3 (pfam02598)→ RplC (COG0087)rpl3p (PRK04231)→<br>RplD (COG0088)rpl4lp (PRK04042)→ RplW (COG0089)PRK14548 (PRK14548)→<br>RplB (COG0090)rpl2p (PRK09612)→ RpsS (COG0185)rps19p (PRK04038)→<br>rpl22p (PRK04223)RplV (COG0091)→ RpsC (COG0092)rps3p (PRK04191)→<br>PRK14549 (PRK14549)Ribosomal_L29 (pfam00831)→ PRK03879 (PRK03879)UPF0086 (pfam01868)→<br>RpsQ (COG0186)rps17p (PRK08572)→ rpl14p (PRK08571)Ribosomal_L14 (pfam00238)→<br>rpl24p (PRK01191)KOW_RPL26 (cd06089)→ rpl24p (PRK01191)KOW_RPL26 (cd06089)→<br>PRK04313 (PRK04313)KOW_RPS4 (cd06087)Ribosomal_S4e (pfam00900)→ RplE (COG0094)rpl5p (PRK04219)→<br>RpsN (COG0199)rps14P (PRK05766)→ rps8p (PRK04034)RpsH (COG0096)→<br>RplF (COG0097)rpl6p (PRK05518)→ rpl32e (PRK08562)Ribosomal_L32e (pfam01655)→<br>rpl19e (PRK08570)Ribosomal_L19e (pfam01280)→ RplR (COG0256)rpl18p (PRK08569)→ |
| env-443 | JGI24730J26740_1002326 | - | 3281 | 3188 | **RNA**→ Methyltrn_RNA_3 (pfam02598)→ RplC (COG0087)rpl3p (PRK04231)→<br>RplD (COG0088)rpl4lp (PRK04042)→ RplW (COG0089)PRK14548 (PRK14548)→<br>RplB (COG0090)rpl2p (PRK09612)→ |

| | | | | | |
|---|---|---|---|---|---|
| env-444 | JGI24732J26686_1000856 | + | 2537 | 2630 | **RNA**→ Methyltrn_RNA_3 (pfam02598)→ rpl3p (PRK04231)Ribosomal_L3 (pfam00297)→ rpl3p (PRK04231)Ribosomal_L3 (pfam00297)→ RplD (COG0088)rpl4lp (PRK04042)→ RplW (COG0089)PRK14548 (PRK14548)→ RplB (COG0090)rpl2p (PRK09612)→ RpsS (COG0185)rps19p (PRK04038)→ rpl22p (PRK04223)RplV (COG0091)→ RpsC (COG0092)rps3p (PRK04191)→ RpmC (COG0255)PRK14549 (PRK14549)→ POP4 (COG1588)PRK03879 (PRK03879)→ RpsQ (COG0186)rps17p (PRK08572)→ rpl14p (PRK08571)Ribosomal_L14 (pfam00238)→ rpl24p (PRK01191)KOW_RPL26 (cd06089)→ PRK04313 (PRK04313)KOW_RPS4 (cd06087)S4 (pfam01479)Ribosomal_S4e (pfam00900)→ rpl5p (PRK04219)Ribosomal_L5 (pfam00281)Ribosomal_L5_C (pfam00673)→ RpsN (COG0199)rps14P (PRK05766)→ rps8p (PRK04034)RpsH (COG0096)→ RplF (COG0097)rpl6p (PRK05518)→ rpl32e (PRK08562)Ribosomal_L32_L32e (cd00513)→ rpl19e (PRK08570)Ribosomal_L19e (pfam01280)→ RplR (COG0256)rpl18p (PRK08569)→ RpsE (COG0098)rps5p (PRK04044)→ Ribosomal_L7_archeal_euk (cd01657)rpl30p (PRK06049)→ rpl15p (PRK06419)→ |
| env-445 | JGI24730J26740_1077952 | - | 136 | 43 | **RNA**→ MTH1 (COG2106)→ |
| env-446 | JGI24729J20445_1000277 | + | 15893 | 15986 | **RNA**→ Methyltrn_RNA_3 (pfam02598)→ RplC (COG0087)rpl3p (PRK04231)→ RplD (COG0088)rpl4lp (PRK04042)→ RplW (COG0089)PRK14548 (PRK14548)→ RplB (COG0090)rpl2p (PRK09612)→ RpsS (COG0185)rps19p (PRK04038)→ rpl22p (PRK04223)RplV (COG0091)→ RpsC (COG0092)rps3p (PRK04191)→ PRK14549 (PRK14549)Ribosomal_L29 (pfam00831)→ PRK03879 (PRK03879)UPF0086 (pfam01868)→ RpsQ (COG0186)rps17p (PRK08572)→ rpl14p (PRK08571)Ribosomal_L14 (pfam00238)→ rpl24p (PRK01191)KOW_RPL26 (cd06089)→ PRK04313 (PRK04313)S4 (cd00165)KOW_RPS4 (cd06087)→ RplE (COG0094)rpl5p (PRK04219)→ RpsN (COG0199)rps14P (PRK05766)→ rps8p (PRK04034)RpsH (COG0096)→ RplF (COG0097)rpl6p (PRK05518)→ rpl32e (PRK08562)Ribosomal_L32e (pfam01655)→ rpl19e (PRK08570)Ribosomal_L19e (pfam01280)→ RplR (COG0256)rpl18p (PRK08569)→ |
| env-447 | JGI24732J26686_1234070 | + | 153 | 246 | **RNA**→ MTH1 (COG2106)→ |
| env-448 | SakLake120mDRAFT_c0020500 | - | 1276 | 1183 | **RNA**→ Methyltrn_RNA_3 (pfam02598)→ |
| env-449 | JGI24730J26740_1097811 | - | 201 | 108 | **RNA**→ Methyltrn_RNA_3 (pfam02598)→ |
| env-450 | JGI20198J14950_1052355 | + | 64 | 157 | **RNA**→ MTH1 (COG2106)→ |
| env-451 | JGI24729J20445_1001989 | - | 4749 | 4656 | **RNA**→ Methyltrn_RNA_3 (pfam02598)→ RplC (COG0087)rpl3p (PRK04231)→ RplD (COG0088)rpl4lp (PRK04042)→ RplW (COG0089)PRK14548 (PRK14548)→ RplB (COG0090)rpl2p (PRK09612)→ RpsS (COG0185)rps19p (PRK04038)→ rpl22p (PRK04223)RplV (COG0091)→ RpsC (COG0092)rps3p (PRK04191)→ |
| env-452 | JGI20214J14112_1145042 | + | 137 | 230 | **RNA**→ MTH1 (COG2106)→ |
| env-453 | JGI24732J26686_1001287 | - | 3712 | 3619 | **RNA**→ Methyltrn_RNA_3 (pfam02598)→ RplC (COG0087)rpl3p (PRK04231)→ RplD (COG0088)rpl4lp (PRK04042)→ RplW (COG0089)PRK14548 (PRK14548)→ rpl2p (PRK09612)Ribosomal_L2 (pfam00181)→ rpl2p (PRK09612)Ribosomal_L2_C (pfam03947)→ RpsS (COG0185)rps19p (PRK04038)→ |
| env-454 | JGI24730J26740_1003612 | - | 4510 | 4417 | **RNA**→ Methyltrn_RNA_3 (pfam02598)→ RplC (COG0087)rpl3p (PRK04231)→ RplD (COG0088)rpl4lp (PRK04042)→ RplW (COG0089)PRK14548 (PRK14548)→ RplB (COG0090)rpl2p (PRK09612)→ RpsS (COG0185)rps19p (PRK04038)→ rpl22p (PRK04223)RplV (COG0091)→ KH (smart00322)archeal_30S_S3_KH (cd02411)→ |
| env-455 | JGI24732J26686_1002014 | + | 4931 | 5024 | **RNA**→ Methyltrn_RNA_3 (pfam02598)→ RplC (COG0087)rpl3p (PRK04231)→ RplD (COG0088)rpl4lp (PRK04042)→ RplW (COG0089)PRK14548 (PRK14548)→ |
| env-456 | taComm3_C551 | - | 847 | 754 | **RNA**→ Methyltrn_RNA_3 (pfam02598)→ rpl3p (PRK04231)Ribosomal_L3 (pfam00297)→ |

| | | | | | |
|---|---|---|---|---|---|
| env-457 | JGI12104J13512_1000496 | - | 35195 | 35102 | **RNA**→ MTH1 (COG2106)→ RplC (COG0087)rpl3p (PRK04231)→ RplD (COG0088)rpl4lp (PRK04042)→ RplW (COG0089)PRK14548 (PRK14548)→ RplB (COG0090)rpl2p (PRK09612)→ RpsS (COG0185)rps19p (PRK04038)→ rpl22p (PRK04223)RplV (COG0091)→ RpsC (COG0092)rps3p (PRK04191)→ PRK14549 (PRK14549)Ribosomal_L29 (pfam00831)→ PRK03879 (PRK03879)UPF0086 (pfam01868)→ RpsQ (COG0186)rps17p (PRK08572)→ rpl14p (PRK08571)Ribosomal_L14 (pfam00238)→ rpl24p (PRK01191)KOW_RPL26 (cd06089)→ PRK04313 (PRK04313)KOW_RPS4 (cd06087)S4 (pfam01479)→ RplE (COG0094)rpl5p (PRK04219)→ RpsN (COG0199)rps14P (PRK05766)→ rps8p (PRK04034)RpsH (COG0096)→ RplF (COG0097)rpl6p (PRK05518)→ rpl32e (PRK08562)Ribosomal_L32e (pfam01655)→ rpl19e (PRK08570)Ribosomal_L19e (pfam01280)→ RplR (COG0256)rpl18p (PRK08569)→ RpsE (COG0098)rps5p (PRK04044)→ |
| env-458 | JGI12104J13512_1006964 | - | 3599 | 3506 | **RNA**→ Methyltrn_RNA_3 (pfam02598)→ RplC (COG0087)rpl3p (PRK04231)→ RplD (COG0088)rpl4lp (PRK04042)→ RplW (COG0089)PRK14548 (PRK14548)→ RplB (COG0090)rpl2p (PRK09612)→ rps19p (PRK04038)Ribosomal_S19 (pfam00203)→ |
| env-459 | JGI11944J13513_1000282 | + | 18417 | 18510 | **RNA**→ Methyltrn_RNA_3 (pfam02598)→ RplC (COG0087)rpl3p (PRK04231)→ RplD (COG0088)rpl4lp (PRK04042)→ RplW (COG0089)PRK14548 (PRK14548)→ RplB (COG0090)rpl2p (PRK09612)→ RpsS (COG0185)rps19p (PRK04038)→ rpl22p (PRK04223)RplV (COG0091)→ RpsC (COG0092)rps3p (PRK04191)→ PRK14549 (PRK14549)Ribosomal_L29 (pfam00831)→ PRK03879 (PRK03879)UPF0086 (pfam01868)→ RpsQ (COG0186)rps17p (PRK08572)→ rpl14p (PRK08571)Ribosomal_L14 (pfam00238)→ rpl24p (PRK01191)KOW_RPL26 (cd06089)→ PRK04313 (PRK04313)KOW_RPS4 (cd06087)S4 (pfam01479)→ RplE (COG0094)rpl5p (PRK04219)→ RpsN (COG0199)rps14P (PRK05766)→ rps8p (PRK04034)RpsH (COG0096)→ RplF (COG0097)rpl6p (PRK05518)→ rpl32e (PRK08562)Ribosomal_L32e (pfam01655)→ rpl19e (PRK08570)Ribosomal_L19e (pfam01280)→ RplR (COG0256)rpl18p (PRK08569)→ RpsE (COG0098)rps5p (PRK04044)→ |
| env-460 | WSSedA2C_c029104 | + | 89 | 182 | **RNA**→ MTH1 (COG2106)→ rpl3p (PRK04231)Ribosomal_L3 (pfam00297)→ |
| env-461 | WSSedA2Ca3_1029104 | + | 89 | 182 | **RNA**→ MTH1 (COG2106)→ rpl3p (PRK04231)Ribosomal_L3 (pfam00297)→ |
| env-462 | WSSedA1TJGI_c020031 | - | 764 | 671 | **RNA**→ MTH1 (COG2106)→ rpl3p (PRK04231)Ribosomal_L3 (pfam00297)→ |
| env-463 | WSSedA1Ba3_1007453 | + | 836 | 929 | **RNA**→ Methyltrn_RNA_3 (pfam02598)→ RplC (COG0087)rpl3p (PRK04231)→ rpl4lp (PRK04042)Ribosomal_L4 (pfam00573)→ |
| env-464 | JGI20213J14113_1028243 | - | 729 | 636 | **RNA**→ MTH1 (COG2106)→ rpl3p (PRK04231)Ribosomal_L3 (pfam00297)→ |
| env-465 | JGI20214J14112_1082845 | - | 335 | 242 | **RNA**→ MTH1 (COG2106)→ |
| env-466 | JGI20214J14112_1065024 | - | 197 | 104 | **RNA**→ MTH1 (COG2106)→ |
| env-467 | 3300000558_11706232 | - | 175 | 82 | **RNA**→ Methyltrn_RNA_3 (pfam02598)→ |
| env-468 | 3300001605_10001574 | + | 12383 | 12476 | **RNA**→ Methyltrn_RNA_3 (pfam02598)→ rpl3p (PRK04231)Ribosomal_L3 (pfam00297)→ rpl3p (PRK04231)Ribosomal_L3 (pfam00297)→ RplD (COG0088)rpl4lp (PRK04042)→ RplW (COG0089)PRK14548 (PRK14548)→ RplB (COG0090)rpl2p (PRK09612)→ RpsS (COG0185)rps19p (PRK04038)→ rpl22p (PRK04223)RplV (COG0091)→ RpsC (COG0092)rps3p (PRK04191)→ PRK14549 (PRK14549)Ribosomal_L29 (pfam00831)→ PRK03879 (PRK03879)UPF0086 (pfam01868)→ RpsQ (COG0186)rps17p (PRK08572)→ rpl14p (PRK08571)Ribosomal_L14 (pfam00238)→ rpl24p (PRK01191)KOW_RPL26 (cd06089)→ PRK04313 (PRK04313)KOW_RPS4 (cd06087)S4 (pfam01479)Ribosomal_S4e (pfam00900)→ rpl5p (PRK04219)Ribosomal_L5 (pfam00281)Ribosomal_L5_C (pfam00673)→ RpsN (COG0199)rps14P (PRK05766)→ rps8p (PRK04034)RpsH (COG0096)→ RplF (COG0097)rpl6p (PRK05518)→ rpl32e (PRK08562)Ribosomal_L32e (pfam01655)→ rpl19e (PRK08570)Ribosomal_L19e (pfam01280)→ RplR (COG0256)rpl18p (PRK08569)→ |
| env-469 | 2232004817 | + | 121 | 214 | **RNA**→ MTH1 (COG2106)→ |
| env-470 | 3300000558_10046922 | - | 175 | 82 | **RNA**→ Methyltrn_RNA_3 (pfam02598)→ |
| env-471 | 3300000032_0025728 | - | 2226 | 2133 | **RNA**→ Methyltrn_RNA_3 (pfam02598)→ RplC (COG0087)rpl3p (PRK04231)→ RplD (COG0088)rpl4lp (PRK04042)→ |
| env-472 | 3300000568_10127450 | - | 4713 | 4620 | **RNA**→ Methyltrn_RNA_3 (pfam02598)→ RplC (COG0087)rpl3p (PRK04231)→ RplD (COG0088)rpl4lp (PRK04042)→ RplW (COG0089)PRK14548 (PRK14548)→ RplB (COG0090)rpl2p (PRK09612)→ RpsS (COG0185)rps19p (PRK04038)→ rpl22p (PRK04223)RplV (COG0091)→ RpsC (COG0092)rps3p (PRK04191)→ |

| | | | | | |
|---|---|---|---|---|---|
| env-473 | 3300001580_10000904 | + | 13878 | 13971 | **RNA**→ Methyltrn_RNA_3 (pfam02598)→ RplC (COG0087)rpl3p (PRK04231)→ RplD (COG0088)rpl4lp (PRK04042)→ RplW (COG0089)PRK14548 (PRK14548)→ RplB (COG0090)rpl2p (PRK09612)→ RpsS (COG0185)rps19p (PRK04038)→ rpl22p (PRK04223)RplV (COG0091)→ RpsC (COG0092)rps3p (PRK04191)→ PRK14549 (PRK14549)Ribosomal_L29 (pfam00831)→ PRK03879 (PRK03879)UPF0086 (pfam01868)→ RpsQ (COG0186)rps17p (PRK08572)→ rpl14p (PRK08571)Ribosomal_L14 (pfam00238)→ rpl24p (PRK01191)KOW_RPL26 (cd06089)→ PRK04313 (PRK04313)KOW_RPS4 (cd06087)S4 (pfam01479)Ribosomal_S4e (pfam00900)→ rpl5p (PRK04219)Ribosomal_L5 (pfam00281)Ribosomal_L5_C (pfam00673)→ RpsN (COG0199)rps14P (PRK05766)→ rps8p (PRK04034)RpsH (COG0096)→ RplF (COG0097)rpl6p (PRK05518)→ rpl32e (PRK08562)Ribosomal_L32e (pfam01655)→ rpl19e (PRK08570)Ribosomal_L19e (pfam01280)→ RplR (COG0256)rpl18p (PRK08569)→ |
| env-474 | 3300001592_10239960 | + | 303 | 396 | **RNA**→ MTH1 (COG2106)→ |
| env-475 | 3300001580_10151383 | - | 980 | 887 | **RNA**→ MTH1 (COG2106)→ |
| Mha-1-1 | NC_017527.1 | - | 484979 | 484886 | **RNA**→ Methyltrn_RNA_3 (pfam02598)→ |
| env-476 | JGI12104J13512_1010054 | + | 871 | 964 | **RNA**→ Methyltrn_RNA_3 (pfam02598)→ RplC (COG0087)rpl3p (PRK04231)→ RplD (COG0088)rpl4lp (PRK04042)→ |
| env-477 | SakLake120mDRAFT_c0004292 | + | 2569 | 2662 | **RNA**→ Methyltrn_RNA_3 (pfam02598)→ RplC (COG0087)rpl3p (PRK04231)→ RplD (COG0088)rpl4lp (PRK04042)→ PRK14548 (PRK14548)→ RplB (COG0090)rpl2p (PRK09612)→ RpsS (COG0185)rps19p (PRK04038)→ rpl22p (PRK04223)RplV (COG0091)→ RpsC (COG0092)rps3p (PRK04191)→ |
| env-478 | JXWW01104647.1 | - | 233 | 140 | **RNA**→ Methyltrn_RNA_3 (pfam02598)→ |
| env-479 | JGI24025J20009_10001050 | - | 20258 | 20165 | **RNA**→ Methyltrn_RNA_3 (pfam02598)→ RplC (COG0087)rpl3p (PRK04231)→ RplD (COG0088)rpl4lp (PRK04042)→ RplW (COG0089)PRK14548 (PRK14548)→ RplB (COG0090)rpl2p (PRK09612)→ RpsS (COG0185)rps19p (PRK04038)→ rpl22p (PRK04223)RplV (COG0091)→ RpsC (COG0092)rps3p (PRK04191)rne (PRK10811)→ PRK14549 (PRK14549)Ribosomal_L29 (pfam00831)→ PRK03879 (PRK03879)UPF0086 (pfam01868)→ RpsQ (COG0186)rps17p (PRK08572)RING-CH-C4HC3_FANCL (cd16490)→ rpl14p (PRK08571)Ribosomal_L14 (pfam00238)→ RplX (COG0198)rpl24p (PRK01191)→ PRK04313 (PRK04313)KOW_RPS4 (cd06087)S4 (pfam01479)Ribosomal_S4e (pfam00900)→ RplE (COG0094)rpl5p (PRK04219)→ |
| env-480 | ASA129_contig07056 | - | 867 | 774 | **RNA**→ MTH1 (COG2106)→ |
| env-481 | JGI24020J35080_1000033 | + | 3315 | 3408 | **RNA**→ MTH1 (COG2106)→ RplC (COG0087)rpl3p (PRK04231)→ RplD (COG0088)rpl4lp (PRK04042)→ RplW (COG0089)PRK14548 (PRK14548)→ RplB (COG0090)rpl2p (PRK09612)→ RpsS (COG0185)rps19p (PRK04038)→ rpl22p (PRK04223)RplV (COG0091)→ RpsC (COG0092)rps3p (PRK04191)→ PRK14549 (PRK14549)Ribosomal_L29 (pfam00831)→ POP4 (COG1588)PRK03879 (PRK03879)→ RpsQ (COG0186)rps17p (PRK08572)→ rpl14p (PRK08571)Ribosomal_L14 (pfam00238)→ rpl24p (PRK01191)KOW_RPL26 (cd06089)→ PRK04313 (PRK04313)KOW_RPS4 (cd06087)S4 (pfam01479)Ribosomal_S4e (pfam00900)→ RplE (COG0094)rpl5p (PRK04219)→ hypo→ rps8p (PRK04034)RpsH (COG0096)→ RplF (COG0097)rpl6p (PRK05518)→ rpl32e (PRK08562)Ribosomal_L32e (pfam01655)→ rpl19e (PRK08570)Ribosomal_L19e (pfam01280)→ RplR (COG0256)rpl18p (PRK08569)→ RpsE (COG0098)rps5p (PRK04044)→ Ribosomal_L7_archeal_euk (cd01657)rpl30p (PRK06049)→ rpl15p (PRK06419)Ribosomal_L27A (pfam00828)→ SecY (COG0201)PRK08568 (PRK08568)→ adk (PRK00279)ADK (cd01428)→ COG1422 (COG1422)PRK02201 (PRK02201)→ CmkB (COG1102)PRK04182 (PRK04182)→ PRK04270 (PRK04270)PseudoU_synth_hDyskerin (cd02572)PUA (pfam01472)→ hypo→ hypo→ AAA (smart00382)CooC (cd02034)GTPase_EngA (TIGR03594)→ |

| | | | | | |
|---|---|---|---|---|---|
| env-482 | JGI24020J35080_1000258 | + | 14429 | 14513 | **RNA**→ MTH1 (COG2106)Raftlin (pfam15250)→ RplC (COG0087)rpl3p (PRK04231)→ RplD (COG0088)rpl4lp (PRK04042)→ RplW (COG0089)PRK14548 (PRK14548)→ RplB (COG0090)rpl2p (PRK09612)→ RpsS (COG0185)rps19p (PRK04038)→ rpl22p (PRK04223)RplV (COG0091)→ RpsC (COG0092)rps3p (PRK04191)→ PRK14549 (PRK14549)Ribosomal_L29 (pfam00831)→ POP4 (COG1588)PRK03879 (PRK03879)→ RpsQ (COG0186)rps17p (PRK08572)→ rpl14p (PRK08571)Ribosomal_L14 (pfam00238)→ rpl24p (PRK01191)KOW_RPL26 (cd06089)→ PRK04313 (PRK04313)KOW_RPS4 (cd06087)S4 (pfam01479)Ribosomal_S4e (pfam00900)→ rpl5p (PRK04219)Ribosomal_L5 (pfam00281)Ribosomal_L5_C (pfam00673)→ RpsN (COG0199)rps14P (PRK05766)→ rps8p (PRK04034)RpsH (COG0096)→ RplF (COG0097)rpl6p (PRK05518)→ Rpl32e (COG1717)rpl32e (PRK08562)HHH (pfam00633)HHH_5 (pfam14520)→ rpl19e (PRK08570)Ribosomal_L19e (pfam01280)→ RplR (COG0256)rpl18p (PRK08569)→ |
| env-483 | JGI20177J14857_1074829 | - | 299 | 212 | **RNA**→ MTH1 (COG2106)→ |
| env-484 | JGI20193J14888_1132815 | - | 101 | 14 | **RNA**→ Methyltrn_RNA_3 (pfam02598)→ |
| env-485 | JGI20191J14862_1083948 | + | 194 | 281 | **RNA**→ MTH1 (COG2106)→ |
| env-486 | JGI20195J14853_1011539 | - | 972 | 885 | **RNA**→ MTH1 (COG2106)→ rpl3p (PRK04231)Ribosomal_L3 (pfam00297)→ |
| env-487 | JGI20179J14886_1000011 | - | 26349 | 26262 | **RNA**→ MTH1 (COG2106)→ RplC (COG0087)rpl3p (PRK04231)→ RplD (COG0088)rpl4lp (PRK04042)→ RplW (COG0089)PRK14548 (PRK14548)→ RplB (COG0090)rpl2p (PRK09612)→ RpsS (COG0185)rps19p (PRK04038)→ rpl22p (PRK04223)RplV (COG0091)→ RpsC (COG0092)rps3p (PRK04191)→ RpmC (COG0255)PRK14549 (PRK14549)→ POP4 (COG1588)PRK03879 (PRK03879)→ RpsQ (COG0186)rps17p (PRK08572)→ rpl14p (PRK08571)Ribosomal_L14 (pfam00238)→ rpl24p (PRK01191)KOW_RPL26 (cd06089)→ PRK04313 (PRK04313)KOW_RPS4 (cd06087)S4 (pfam01479)Ribosomal_S4e (pfam00900)→ rpl5p (PRK04219)Ribosomal_L5 (pfam00281)Ribosomal_L5_C (pfam00673)→ |
| env-488 | JGI20199J14953_1000018 | + | 6725 | 6812 | **RNA**→ Methyltrn_RNA_3 (pfam02598)→ RplC (COG0087)rpl3p (PRK04231)→ RplD (COG0088)rpl4lp (PRK04042)→ RplW (COG0089)PRK14548 (PRK14548)→ RplB (COG0090)rpl2p (PRK09612)→ RpsS (COG0185)rps19p (PRK04038)→ rpl22p (PRK04223)RplV (COG0091)→ RpsC (COG0092)rps3p (PRK04191)→ RpmC (COG0255)PRK14549 (PRK14549)→ POP4 (COG1588)PRK03879 (PRK03879)→ RpsQ (COG0186)rps17p (PRK08572)→ |
| env-489 | JGI20201J14949_1002905 | - | 1578 | 1491 | **RNA**→ MTH1 (COG2106)→ RplC (COG0087)rpl3p (PRK04231)→ rpl4lp (PRK04042)→ |
| env-490 | JGI20197J15001_1000199 | - | 8178 | 8091 | **RNA**→ MTH1 (COG2106)→ RplC (COG0087)rpl3p (PRK04231)→ RplD (COG0088)rpl4lp (PRK04042)→ RplW (COG0089)PRK14548 (PRK14548)→ RplB (COG0090)rpl2p (PRK09612)→ RpsS (COG0185)rps19p (PRK04038)→ rpl22p (PRK04223)RplV (COG0091)→ RpsC (COG0092)rps3p (PRK04191)→ RpmC (COG0255)PRK14549 (PRK14549)→ POP4 (COG1588)PRK03879 (PRK03879)→ RpsQ (COG0186)rps17p (PRK08572)→ rpl14p (PRK08571)Ribosomal_L14 (pfam00238)→ rpl24p (PRK01191)KOW_RPL26 (cd06089)→ PRK04313 (PRK04313)KOW_RPS4 (cd06087)S4 (pfam01479)Ribosomal_S4e (pfam00900)→ rpl5p (PRK04219)Ribosomal_L5 (pfam00281)Ribosomal_L5_C (pfam00673)→ |
| env-491 | JGI20197J15136_1000199 | - | 8178 | 8091 | **RNA**→ MTH1 (COG2106)→ RplC (COG0087)rpl3p (PRK04231)→ RplD (COG0088)rpl4lp (PRK04042)→ RplW (COG0089)PRK14548 (PRK14548)→ RplB (COG0090)rpl2p (PRK09612)→ RpsS (COG0185)rps19p (PRK04038)→ rpl22p (PRK04223)RplV (COG0091)→ RpsC (COG0092)rps3p (PRK04191)→ RpmC (COG0255)PRK14549 (PRK14549)→ POP4 (COG1588)PRK03879 (PRK03879)→ RpsQ (COG0186)rps17p (PRK08572)→ rpl14p (PRK08571)Ribosomal_L14 (pfam00238)→ rpl24p (PRK01191)KOW_RPL26 (cd06089)→ PRK04313 (PRK04313)KOW_RPS4 (cd06087)S4 (pfam01479)Ribosomal_S4e (pfam00900)→ rpl5p (PRK04219)Ribosomal_L5 (pfam00281)Ribosomal_L5_C (pfam00673)→ |
| env-492 | AGCH01000237.1 | - | 413 | 326 | **RNA**→ MTH1 (COG2106)→ |
| env-493 | PermFrostAlaska_NODE_9622 | - | 413 | 326 | **RNA**→ MTH1 (COG2106)→ |

| | | | | | |
|---|---|---|---|---|---|
| env-494 | B3_all_c_Cons150562 | - | 21306 | 21219 | **RNA**→ MTH1 (COG2106)→ rpl3p (PRK04231)Ribosomal_L3 (pfam00297)→ RplC (COG0087)rpl3p (PRK04231)→ RplD (COG0088)rpl4lp (PRK04042)→ RplW (COG0089)PRK14548 (PRK14548)→ RplB (COG0090)rpl2p (PRK09612)→ RpsS (COG0185)rps19p (PRK04038)→ rpl22p (PRK04223)RplV (COG0091)→ RpsC (COG0092)rps3p (PRK04191)→ RpmC (COG0255)PRK14549 (PRK14549)→ POP4 (COG1588)PRK03879 (PRK03879)→ RpsQ (COG0186)rps17p (PRK08572)→ rpl14p (PRK08571)Ribosomal_L14 (pfam00238)→ rpl24p (PRK01191)KOW_RPL26 (cd06089)→ PRK04313 (PRK04313)KOW_RPS4 (cd06087)S4 (pfam01479)Ribosomal_S4e (pfam00900)→ rpl5p (PRK04219)Ribosomal_L5 (pfam00281)Ribosomal_L5_C (pfam00673)→ |
| env-495 | JGI20199J14953_1000162 | + | 190 | 277 | **RNA**→ MTH1 (COG2106)→ RplC (COG0087)rpl3p (PRK04231)→ RplD (COG0088)rpl4lp (PRK04042)→ RplW (COG0089)PRK14548 (PRK14548)→ RplB (COG0090)rpl2p (PRK09612)→ RpsS (COG0185)rps19p (PRK04038)→ rpl22p (PRK04223)RplV (COG0091)→ RpsC (COG0092)rps3p (PRK04191)→ RpmC (COG0255)PRK14549 (PRK14549)→ POP4 (COG1588)PRK03879 (PRK03879)→ RpsQ (COG0186)rps17p (PRK08572)→ rpl14p (PRK08571)Ribosomal_L14 (pfam00238)→ rpl24p (PRK01191)→ |
| env-496 | JGI20198J14950_1000001 | + | 50351 | 50438 | **RNA**→ Methyltrn_RNA_3 (pfam02598)→ RplC (COG0087)rpl3p (PRK04231)→ RplD (COG0088)rpl4lp (PRK04042)→ RplW (COG0089)PRK14548 (PRK14548)→ RplB (COG0090)rpl2p (PRK09612)→ RpsS (COG0185)rps19p (PRK04038)→ rpl22p (PRK04223)RplV (COG0091)→ RpsC (COG0092)rps3p (PRK04191)→ RpmC (COG0255)PRK14549 (PRK14549)→ POP4 (COG1588)PRK03879 (PRK03879)→ RpsQ (COG0186)rps17p (PRK08572)→ rpl14p (PRK08571)Ribosomal_L14 (pfam00238)→ RplX (COG0198)rpl24p (PRK01191)→ PRK04313 (PRK04313)KOW_RPS4 (cd06087)Ribosomal_S4e (pfam00900)→ rpl5p (PRK04219)Ribosomal_L5 (pfam00281)Ribosomal_L5_C (pfam00673)→ |
| env-497 | JGI20200J14955_1011738 | + | 613 | 700 | **RNA**→ MTH1 (COG2106)→ |
| env-498 | JGI20201J14949_1007869 | + | 187 | 274 | **RNA**→ MTH1 (COG2106)→ rpl3p (PRK04231)Ribosomal_L3 (pfam00297)→ |
| env-499 | JGI20197J15001_1000002 | - | 33762 | 33675 | **RNA**→ MTH1 (COG2106)→ RplC (COG0087)rpl3p (PRK04231)→ RplD (COG0088)rpl4lp (PRK04042)→ RplW (COG0089)PRK14548 (PRK14548)→ RplB (COG0090)rpl2p (PRK09612)→ RpsS (COG0185)rps19p (PRK04038)→ rpl22p (PRK04223)RplV (COG0091)→ RpsC (COG0092)rps3p (PRK04191)→ RpmC (COG0255)PRK14549 (PRK14549)→ POP4 (COG1588)PRK03879 (PRK03879)→ RpsQ (COG0186)rps17p (PRK08572)→ rpl14p (PRK08571)Ribosomal_L14 (pfam00238)→ RplX (COG0198)rpl24p (PRK01191)→ PRK04313 (PRK04313)KOW_RPS4 (cd06087)Ribosomal_S4e (pfam00900)→ rpl5p (PRK04219)Ribosomal_L5 (pfam00281)Ribosomal_L5_C (pfam00673)→ |
| env-500 | JGI20197J15136_1000002 | - | 33762 | 33675 | **RNA**→ MTH1 (COG2106)→ RplC (COG0087)rpl3p (PRK04231)→ RplD (COG0088)rpl4lp (PRK04042)→ RplW (COG0089)PRK14548 (PRK14548)→ RplB (COG0090)rpl2p (PRK09612)→ RpsS (COG0185)rps19p (PRK04038)→ rpl22p (PRK04223)RplV (COG0091)→ RpsC (COG0092)rps3p (PRK04191)→ RpmC (COG0255)PRK14549 (PRK14549)→ POP4 (COG1588)PRK03879 (PRK03879)→ RpsQ (COG0186)rps17p (PRK08572)→ rpl14p (PRK08571)Ribosomal_L14 (pfam00238)→ RplX (COG0198)rpl24p (PRK01191)→ PRK04313 (PRK04313)KOW_RPS4 (cd06087)Ribosomal_S4e (pfam00900)→ rpl5p (PRK04219)Ribosomal_L5 (pfam00281)Ribosomal_L5_C (pfam00673)→ |
| env-501 | AGCH01000005.1 | - | 11078 | 10991 | **RNA**→ Methyltrn_RNA_3 (pfam02598)→ rpl3p (PRK04231)Ribosomal_L3 (pfam00297)→ RplC (COG0087)rpl3p (PRK04231)→ RplD (COG0088)rpl4lp (PRK04042)→ RplW (COG0089)PRK14548 (PRK14548)→ RplB (COG0090)rpl2p (PRK09612)→ RpsS (COG0185)rps19p (PRK04038)→ rpl22p (PRK04223)RplV (COG0091)→ RpsC (COG0092)rps3p (PRK04191)→ RpmC (COG0255)PRK14549 (PRK14549)→ POP4 (COG1588)PRK03879 (PRK03879)→ RpsQ (COG0186)rps17p (PRK08572)→ rpl14p (PRK08571)Ribosomal_L14 (pfam00238)→ rpl24p (PRK01191)KOW_RPL26 (cd06089)→ PRK04313 (PRK04313)KOW_RPS4 (cd06087)S4 (pfam01479)Ribosomal_S4e (pfam00900)→ |

| | | | | | |
|---|---|---|---|---|---|
| env-502 | PermFrostAlaska_NODE_6124 | - | 23466 | 23379 | **RNA**→ Methyltrn_RNA_3 (pfam02598)→ rpl3p (PRK04231)Ribosomal_L3 (pfam00297)→ RplC (COG0087)rpl3p (PRK04231)→ RplD (COG0088)rpl4lp (PRK04042)→ RplW (COG0089)PRK14548 (PRK14548)→ RplB (COG0090)rpl2p (PRK09612)→ RpsS (COG0185)rps19p (PRK04038)→ rpl22p (PRK04223)RplV (COG0091)→ RpsC (COG0092)rps3p (PRK04191)→ RpmC (COG0255)PRK14549 (PRK14549)→ POP4 (COG1588)PRK03879 (PRK03879)→ RpsQ (COG0186)rps17p (PRK08572)→ rpl14p (PRK08571)Ribosomal_L14 (pfam00238)→ rpl24p (PRK01191)KOW_RPL26 (cd06089)→ PRK04313 (PRK04313)KOW_RPS4 (cd06087)S4 (pfam01479)Ribosomal_S4e (pfam00900)→ |
| env-503 | B3_all_c_Cons68376 | - | 6993 | 6906 | **RNA**→ Methyltrn_RNA_3 (pfam02598)→ RplC (COG0087)rpl3p (PRK04231)→ RplD (COG0088)rpl4lp (PRK04042)→ RplW (COG0089)PRK14548 (PRK14548)→ RplB (COG0090)rpl2p (PRK09612)→ RpsS (COG0185)rps19p (PRK04038)→ rpl22p (PRK04223)RplV (COG0091)→ RpsC (COG0092)rps3p (PRK04191)→ RpmC (COG0255)PRK14549 (PRK14549)→ POP4 (COG1588)PRK03879 (PRK03879)→ |
| env-504 | JGI20179J14886_1002439 | - | 264 | 177 | **RNA**→ Methyltrn_RNA_3 (pfam02598)→ |
| env-505 | JGI24728J21555_1000539 | + | 10047 | 10116 | **RNA**→ RplC (COG0087)rpl3p (PRK04231)→ RplD (COG0088)rpl4lp (PRK04042)→ RplW (COG0089)PRK14548 (PRK14548)→ RplB (COG0090)rpl2p (PRK09612)→ RpsS (COG0185)rps19p (PRK04038)→ rpl22p (PRK04223)RplV (COG0091)→ RpsC (COG0092)rps3p (PRK04191)→ PRK14549 (PRK14549)Ribosomal_L29 (pfam00831)→ PRK03879 (PRK03879)UPF0086 (pfam01868)→ RpsQ (COG0186)rps17p (PRK08572)→ rpl14p (PRK08571)Ribosomal_L14 (pfam00238)→ rpl24p (PRK01191)KOW_RPL26 (cd06089)→ PRK04313 (PRK04313)KOW_RPS4 (cd06087)Ribosomal_S4e (pfam00900)→ rpl5p (PRK04219)Ribosomal_L5 (pfam00281)Ribosomal_L5_C (pfam00673)→ RpsN (COG0199)rps14P (PRK05766)→ rps8p (PRK04034)RpsH (COG0096)→ RplF (COG0097)rpl6p (PRK05518)→ rpl32e (PRK08562)Ribosomal_L32e (pfam01655)→ |
| env-506 | JGI24728J21555_1000541 | - | 9030 | 8961 | **RNA**→ RplC (COG0087)rpl3p (PRK04231)→ RplD (COG0088)rpl4lp (PRK04042)→ RplW (COG0089)PRK14548 (PRK14548)→ RplB (COG0090)rpl2p (PRK09612)→ RpsS (COG0185)rps19p (PRK04038)→ rpl22p (PRK04223)RplV (COG0091)→ RpsC (COG0092)rps3p (PRK04191)→ PRK14549 (PRK14549)Ribosomal_L29 (pfam00831)→ PRK03879 (PRK03879)UPF0086 (pfam01868)→ RpsQ (COG0186)rps17p (PRK08572)→ rpl14p (PRK08571)Ribosomal_L14 (pfam00238)→ rpl24p (PRK01191)KOW_RPL26 (cd06089)→ PRK04313 (PRK04313)KOW_RPS4 (cd06087)Ribosomal_S4e (pfam00900)→ rpl5p (PRK04219)Ribosomal_L5 (pfam00281)Ribosomal_L5_C (pfam00673)→ RpsN (COG0199)rps14P (PRK05766)→ rps8p (PRK04034)RpsH (COG0096)→ RplF (COG0097)rpl6p (PRK05518)→ rpl32e (PRK08562)Ribosomal_L32e (pfam01655)→ |
| env-507 | JGI24728J21555_1000542 | - | 9049 | 8980 | **RNA**→ RplC (COG0087)rpl3p (PRK04231)→ RplD (COG0088)rpl4lp (PRK04042)→ RplW (COG0089)PRK14548 (PRK14548)→ RplB (COG0090)rpl2p (PRK09612)→ RpsS (COG0185)rps19p (PRK04038)→ rpl22p (PRK04223)RplV (COG0091)→ RpsC (COG0092)rps3p (PRK04191)→ PRK14549 (PRK14549)Ribosomal_L29 (pfam00831)→ PRK03879 (PRK03879)UPF0086 (pfam01868)→ RpsQ (COG0186)rps17p (PRK08572)→ rpl14p (PRK08571)Ribosomal_L14 (pfam00238)→ rpl24p (PRK01191)KOW_RPL26 (cd06089)→ PRK04313 (PRK04313)KOW_RPS4 (cd06087)Ribosomal_S4e (pfam00900)→ rpl5p (PRK04219)Ribosomal_L5 (pfam00281)Ribosomal_L5_C (pfam00673)→ RpsN (COG0199)rps14P (PRK05766)→ rps8p (PRK04034)RpsH (COG0096)→ RplF (COG0097)rpl6p (PRK05518)→ rpl32e (PRK08562)Ribosomal_L32e (pfam01655)→ |
| env-508 | taComm3_C12988 | + | 619 | 703 | **RNA**→ rpl3p (PRK04231)Ribosomal_L3 (pfam00297)→ |

| | | | | | |
|---|---|---|---|---|---|
| env-509 | JGI12104J13512_1000732 | + | 3737 | 3822 | **RNA**→ RplC (COG0087)rpl3p (PRK04231)→ RplD (COG0088)rpl4lp (PRK04042)→<br>RplW (COG0089)PRK14548 (PRK14548)→ RplB (COG0090)rpl2p (PRK09612)→<br>RpsS (COG0185)rps19p (PRK04038)→ rpl22p (PRK04223)RplV (COG0091)→<br>RpsC (COG0092)rps3p (PRK04191)→ PRK14549 (PRK14549)Ribosomal_L29 (pfam00831)→<br>PRK03879 (PRK03879)UPF0086 (pfam01868)→ RpsQ (COG0186)rps17p (PRK08572)→<br>rpl14p (PRK08571)Ribosomal_L14 (pfam00238)→ rpl24p (PRK01191)KOW_RPL26 (cd06089)→<br>PRK04313 (PRK04313)S4 (cd00165)KOW_RPS4 (cd06087)Ribosomal_S4e (pfam00900)→<br>RplE (COG0094)rpl5p (PRK04219)→ RpsN (COG0199)rps14P (PRK05766)→<br>rps8p (PRK04034)RpsH (COG0096)→ RplF (COG0097)rpl6p (PRK05518)→<br>rpl32e (PRK08562)Ribosomal_L32e (pfam01655)→ rpl19e (PRK08570)Ribosomal_L19e (pfam01280)→<br>RplR (COG0256)rpl18p (PRK08569)→ RpsE (COG0098)rps5p (PRK04044)→<br>Ribosomal_L7_archeal_euk (cd01657)rpl30p (PRK06049)→ rpl15p (PRK06419)Ribosomal_L27A (pfam00828)→<br>SecY (COG0201)PRK08568 (PRK08568)→ AdkA (COG2019)PRK04040 (PRK04040)→ DUF106 (pfam01956)→<br>CmkB (COG1102)PRK04182 (PRK04182)→ Lactamase_B (smart00849)DHPS-like_MBL-fold (cd07713)→ |
| env-510 | JGI11944J13513_1000175 | + | 47191 | 47276 | **RNA**→ RplC (COG0087)rpl3p (PRK04231)→ RplD (COG0088)rpl4lp (PRK04042)→<br>RplW (COG0089)PRK14548 (PRK14548)→ RplB (COG0090)rpl2p (PRK09612)→<br>RpsS (COG0185)rps19p (PRK04038)→ rpl22p (PRK04223)RplV (COG0091)→<br>RpsC (COG0092)rps3p (PRK04191)→ PRK14549 (PRK14549)Ribosomal_L29 (pfam00831)→<br>PRK03879 (PRK03879)UPF0086 (pfam01868)→ RpsQ (COG0186)rps17p (PRK08572)→<br>rpl14p (PRK08571)Ribosomal_L14 (pfam00238)→ rpl24p (PRK01191)KOW_RPL26 (cd06089)→<br>PRK04313 (PRK04313)S4 (cd00165)KOW_RPS4 (cd06087)Ribosomal_S4e (pfam00900)→<br>RplE (COG0094)rpl5p (PRK04219)→ RpsN (COG0199)rps14P (PRK05766)→<br>rps8p (PRK04034)RpsH (COG0096)→ RplF (COG0097)rpl6p (PRK05518)→<br>rpl32e (PRK08562)Ribosomal_L32e (pfam01655)→ rpl19e (PRK08570)Ribosomal_L19e (pfam01280)→<br>RplR (COG0256)rpl18p (PRK08569)→ RpsE (COG0098)rps5p (PRK04044)→ |
| Mta-1-1 | NZ_AGIY02000001.1 | - | 1752945 | 1752858 | **RNA**→ hypo→ RplC (COG0087)rpl3p (PRK04231)→ RplD (COG0088)rpl4lp (PRK04042)→<br>RplW (COG0089)PRK14548 (PRK14548)→ RplB (COG0090)rpl2p (PRK09612)→<br>RpsS (COG0185)rps19p (PRK04038)→ rpl22p (PRK04223)RplV (COG0091)→<br>RpsC (COG0092)rps3p (PRK04191)→ PRK14549 (PRK14549)Ribosomal_L29 (pfam00831)→<br>PRK03879 (PRK03879)UPF0086 (pfam01868)→ RpsQ (COG0186)rps17p (PRK08572)→<br>rpl14p (PRK08571)Ribosomal_L14 (pfam00238)→ rpl24p (PRK01191)KOW_RPL26 (cd06089)→<br>PRK04313 (PRK04313)S4 (cd00165)KOW_RPS4 (cd06087)Ribosomal_S4e (pfam00900)→<br>RplE (COG0094)rpl5p (PRK04219)→ RpsN (COG0199)rps14P (PRK05766)→<br>rps8p (PRK04034)RpsH (COG0096)→ RplF (COG0097)rpl6p (PRK05518)→<br>rpl32e (PRK08562)Ribosomal_L32e (pfam01655)→ rpl19e (PRK08570)Ribosomal_L19e (pfam01280)→<br>RplR (COG0256)rpl18p (PRK08569)→ RpsE (COG0098)rps5p (PRK04044)→ |
| env-511 | taComm3_C1003 | - | 71292 | 71205 | **RNA**→ hypo→ RplC (COG0087)rpl3p (PRK04231)→ RplD (COG0088)rpl4lp (PRK04042)→<br>RplW (COG0089)PRK14548 (PRK14548)→ RplB (COG0090)rpl2p (PRK09612)→<br>RpsS (COG0185)rps19p (PRK04038)→ rpl22p (PRK04223)RplV (COG0091)→<br>RpsC (COG0092)rps3p (PRK04191)→ PRK14549 (PRK14549)Ribosomal_L29 (pfam00831)→<br>PRK03879 (PRK03879)UPF0086 (pfam01868)→ RpsQ (COG0186)rps17p (PRK08572)→<br>rpl14p (PRK08571)Ribosomal_L14 (pfam00238)→ rpl24p (PRK01191)KOW_RPL26 (cd06089)→<br>PRK04313 (PRK04313)S4 (cd00165)KOW_RPS4 (cd06087)Ribosomal_S4e (pfam00900)→<br>RplE (COG0094)rpl5p (PRK04219)→ RpsN (COG0199)rps14P (PRK05766)→<br>rps8p (PRK04034)RpsH (COG0096)→ RplF (COG0097)rpl6p (PRK05518)→<br>rpl32e (PRK08562)Ribosomal_L32e (pfam01655)→ rpl19e (PRK08570)Ribosomal_L19e (pfam01280)→<br>RplR (COG0256)rpl18p (PRK08569)→ RpsE (COG0098)rps5p (PRK04044)→ |

| | | | | | |
|---|---|---|---|---|---|
| env-512 | JGI12104J13512_1000711 | - | 24857 | 24770 | **RNA**→ hypo→ RplC (COG0087)rpl3p (PRK04231)→ RplD (COG0088)rpl4lp (PRK04042)→ RplW (COG0089)PRK14548 (PRK14548)→ RplB (COG0090)rpl2p (PRK09612)→ RpsS (COG0185)rps19p (PRK04038)→ rpl22p (PRK04223)RplV (COG0091)→ RpsC (COG0092)rps3p (PRK04191)→ PRK14549 (PRK14549)Ribosomal_L29 (pfam00831)→ PRK03879 (PRK03879)UPF0086 (pfam01868)→ RpsQ (COG0186)rps17p (PRK08572)→ rpl14p (PRK08571)Ribosomal_L14 (pfam00238)→ rpl24p (PRK01191)KOW_RPL26 (cd06089)→ PRK04313 (PRK04313)S4 (cd00165)KOW_RPS4 (cd06087)Ribosomal_S4e (pfam00900)→ RplE (COG0094)rpl5p (PRK04219)→ RpsN (COG0199)rps14P (PRK05766)→ rps8p (PRK04034)RpsH (COG0096)→ RplF (COG0097)rpl6p (PRK05518)→ rpl32e (PRK08562)Ribosomal_L32e (pfam01655)→ rpl19e (PRK08570)Ribosomal_L19e (pfam01280)→ RplR (COG0256)rpl18p (PRK08569)→ RpsE (COG0098)rps5p (PRK04044)→ |
| env-513 | JGI11944J13513_1000170 | - | 24861 | 24774 | **RNA**→ hypo→ RplC (COG0087)rpl3p (PRK04231)→ RplD (COG0088)rpl4lp (PRK04042)→ RplW (COG0089)PRK14548 (PRK14548)→ RplB (COG0090)rpl2p (PRK09612)→ RpsS (COG0185)rps19p (PRK04038)→ rpl22p (PRK04223)RplV (COG0091)→ RpsC (COG0092)rps3p (PRK04191)→ PRK14549 (PRK14549)Ribosomal_L29 (pfam00831)→ PRK03879 (PRK03879)UPF0086 (pfam01868)→ RpsQ (COG0186)rps17p (PRK08572)→ rpl14p (PRK08571)Ribosomal_L14 (pfam00238)→ rpl24p (PRK01191)KOW_RPL26 (cd06089)→ PRK04313 (PRK04313)S4 (cd00165)KOW_RPS4 (cd06087)Ribosomal_S4e (pfam00900)→ RplE (COG0094)rpl5p (PRK04219)→ RpsN (COG0199)rps14P (PRK05766)→ rps8p (PRK04034)RpsH (COG0096)→ RplF (COG0097)rpl6p (PRK05518)→ rpl32e (PRK08562)Ribosomal_L32e (pfam01655)→ rpl19e (PRK08570)Ribosomal_L19e (pfam01280)→ RplR (COG0256)rpl18p (PRK08569)→ RpsE (COG0098)rps5p (PRK04044)→ |
| Hbo-1-1 | NC_014729.1 | + | 955608 | 955707 | **RNA**→ Methyltrn_RNA_3 (pfam02598)→ rpl3p (PRK04231)Ribosomal_L3 (pfam00297)→ RplD (COG0088)rpl4lp (PRK04042)→ RplW (COG0089)PRK14548 (PRK14548)→ RplB (COG0090)rpl2p (PRK09612)→ RpsS (COG0185)rps19p (PRK04038)→ rpl22p (PRK04223)RplV (COG0091)→ RpsC (COG0092)rps3p (PRK04191)→ PRK14549 (PRK14549)Ribosomal_L29 (pfam00831)→ PRK03879 (PRK03879)UPF0086 (pfam01868)→ RpsQ (COG0186)rps17p (PRK08572)→ rpl14p (PRK08571)Ribosomal_L14 (pfam00238)→ RplX (COG0198)rpl24p (PRK01191)→ PRK04313 (PRK04313)KOW_RPS4 (cd06087)Ribosomal_S4e (pfam00900)→ RplE (COG0094)rpl5p (PRK04219)→ RpsN (COG0199)rps14P (PRK05766)→ rps8p (PRK04034)RpsH (COG0096)→ RplF (COG0097)rpl6p (PRK05518)→ PRK12766 (PRK12766)RecA (COG0468)Rpl32e (COG1717)→ rpl19e (PRK08570)Ribosomal_L19e (pfam01280)→ |
| Ngr-1-1 | NC_019792.1 | - | 316138 | 316039 | **RNA**→ Methyltrn_RNA_3 (pfam02598)→ rpl3p (PRK04231)Ribosomal_L3 (pfam00297)→ RplD (COG0088)rpl4lp (PRK04042)→ RplW (COG0089)PRK14548 (PRK14548)→ RplB (COG0090)rpl2p (PRK09612)→ RpsS (COG0185)rps19p (PRK04038)→ rpl22p (PRK04223)RplV (COG0091)→ RpsC (COG0092)rps3p (PRK04191)→ RpmC (COG0255)PRK14549 (PRK14549)→ PRK03879 (PRK03879)UPF0086 (pfam01868)→ RpsQ (COG0186)rps17p (PRK08572)→ rpl14p (PRK08571)Ribosomal_L14 (pfam00238)→ rpl24p (PRK01191)KOW_RPL26 (cd06089)→ PRK04313 (PRK04313)S4 (cd00165)KOW_RPS4 (cd06087)Ribosomal_S4e (pfam00900)→ RplE (COG0094)rpl5p (PRK04219)→ RpsN (COG0199)rps14P (PRK05766)→ rps8p (PRK04034)RpsH (COG0096)→ RplF (COG0097)rpl6p (PRK05518)→ PRK12766 (PRK12766)RecA (COG0468)Rpl32e (COG1717)→ rpl19e (PRK08570)Ribosomal_L19e (pfam01280)→ |
| Hje-1-1 | NC_014297.1 | + | 971638 | 971737 | **RNA**→ Methyltrn_RNA_3 (pfam02598)→ RplC (COG0087)rpl3p (PRK04231)→ RplD (COG0088)rpl4lp (PRK04042)→ RplW (COG0089)PRK14548 (PRK14548)→ RplB (COG0090)rpl2p (PRK09612)→ RpsS (COG0185)rps19p (PRK04038)→ rpl22p (PRK04223)RplV (COG0091)→ RpsC (COG0092)rps3p (PRK04191)→ PRK14549 (PRK14549)Ribosomal_L29 (pfam00831)→ PRK03879 (PRK03879)UPF0086 (pfam01868)→ RpsQ (COG0186)rps17p (PRK08572)→ rpl14p (PRK08571)Ribosomal_L14 (pfam00238)→ rpl24p (PRK01191)KOW_RPL26 (cd06089)→ PRK04313 (PRK04313)KOW_RPS4 (cd06087)Ribosomal_S4e (pfam00900)→ RplE (COG0094)rpl5p (PRK04219)→ RpsN (COG0199)rps14P (PRK05766)→ rps8p (PRK04034)RpsH (COG0096)→ RplF (COG0097)rpl6p (PRK05518)→ PRK12766 (PRK12766)RecA (COG0468)Rpl32e (COG1717)→ rpl19e (PRK08570)Ribosomal_L19e (pfam01280)→ |

| | | | | | |
|---|---|---|---|---|---|
| Nma-1-1 | NC_013922.1 | + | 222997 | 223096 | **RNA**→ Methyltrn_RNA_3 (pfam02598)→ rpl3p (PRK04231)Ribosomal_L3 (pfam00297)→<br>RplD (COG0088)rpl4lp (PRK04042)→ RplW (COG0089)PRK14548 (PRK14548)→<br>RplB (COG0090)rpl2p (PRK09612)→ RpsS (COG0185)rps19p (PRK04038)→<br>rpl22p (PRK04223)RplV (COG0091)→ RpsC (COG0092)rps3p (PRK04191)→<br>RpmC (COG0255)PRK14549 (PRK14549)→ PRK03879 (PRK03879)UPF0086 (pfam01868)→<br>RpsQ (COG0186)rps17p (PRK08572)→ rpl14p (PRK08571)Ribosomal_L14 (pfam00238)→<br>rpl24p (PRK01191)KOW_RPL26 (cd06089)→<br>PRK04313 (PRK04313)S4 (cd00165)KOW_RPS4 (cd06087)Ribosomal_S4e (pfam00900)→<br>RplE (COG0094)rpl5p (PRK04219)→ RpsN (COG0199)rps14P (PRK05766)→<br>rps8p (PRK04034)RpsH (COG0096)→ RplF (COG0097)rpl6p (PRK05518)→<br>PRK12766 (PRK12766)RecA (COG0468)Rpl32e (COG1717)→ |
| Hal-1-1 | NZ_AOLL01000031.1 | - | 14961 | 14862 | **RNA**→ Methyltrn_RNA_3 (pfam02598)→ rpl3p (PRK04231)Ribosomal_L3 (pfam00297)→<br>RplD (COG0088)rpl4lp (PRK04042)→ RplW (COG0089)PRK14548 (PRK14548)→<br>RplB (COG0090)rpl2p (PRK09612)→ RpsS (COG0185)rps19p (PRK04038)→<br>rpl22p (PRK04223)RplV (COG0091)→ RpsC (COG0092)rps3p (PRK04191)→<br>PRK14549 (PRK14549)Ribosomal_L29 (pfam00831)→ PRK03879 (PRK03879)UPF0086 (pfam01868)→<br>RpsQ (COG0186)rps17p (PRK08572)→ rpl14p (PRK08571)Ribosomal_L14 (pfam00238)→<br>RplX (COG0198)rpl24p (PRK01191)→<br>PRK04313 (PRK04313)KOW_RPS4 (cd06087)Ribosomal_S4e (pfam00900)→ RplE (COG0094)rpl5p (PRK04219)→<br>RpsN (COG0199)rps14P (PRK05766)→ rps8p (PRK04034)RpsH (COG0096)→<br>RplF (COG0097)rpl6p (PRK05518)→ PRK12766 (PRK12766)RecA (COG0468)Rpl32e (COG1717)→<br>rpl19e (PRK08570)Ribosomal_L19e (pfam01280)→ |
| Hvo-1-1 | NC_013967.1 | - | 2419949 | 2419850 | **RNA**→ Methyltrn_RNA_3 (pfam02598)→ rpl3p (PRK04231)Ribosomal_L3 (pfam00297)→<br>RplD (COG0088)rpl4lp (PRK04042)→ RplW (COG0089)PRK14548 (PRK14548)→<br>RplB (COG0090)rpl2p (PRK09612)→ RpsS (COG0185)rps19p (PRK04038)→<br>rpl22p (PRK04223)RplV (COG0091)→ RpsC (COG0092)rps3p (PRK04191)→<br>PRK14549 (PRK14549)Ribosomal_L29 (pfam00831)→ PRK03879 (PRK03879)UPF0086 (pfam01868)→<br>RpsQ (COG0186)rps17p (PRK08572)→ rpl14p (PRK08571)Ribosomal_L14 (pfam00238)→<br>RplX (COG0198)rpl24p (PRK01191)→<br>PRK04313 (PRK04313)KOW_RPS4 (cd06087)Ribosomal_S4e (pfam00900)→ RplE (COG0094)rpl5p (PRK04219)→<br>RpsN (COG0199)rps14P (PRK05766)→ rps8p (PRK04034)RpsH (COG0096)→<br>RplF (COG0097)rpl6p (PRK05518)→ PRK12766 (PRK12766)RecA (COG0468)Rpl32e (COG1717)→<br>rpl19e (PRK08570)Ribosomal_L19e (pfam01280)→ |
| Nti-1-1 | NZ_KB913017.1 | - | 3677377 | 3677278 | **RNA**→ Methyltrn_RNA_3 (pfam02598)→ rpl3p (PRK04231)Ribosomal_L3 (pfam00297)→<br>RplC (COG0087)rpl3p (PRK04231)→ rpl3p (PRK04231)Ribosomal_L3 (pfam00297)→<br>RplD (COG0088)rpl4lp (PRK04042)→ RplW (COG0089)PRK14548 (PRK14548)→<br>RplB (COG0090)rpl2p (PRK09612)→ RpsS (COG0185)rps19p (PRK04038)→<br>rpl22p (PRK04223)RplV (COG0091)→ RpsC (COG0092)rps3p (PRK04191)→<br>RpmC (COG0255)PRK14549 (PRK14549)→ PRK03879 (PRK03879)UPF0086 (pfam01868)→<br>RpsQ (COG0186)rps17p (PRK08572)→ rpl14p (PRK08571)Ribosomal_L14 (pfam00238)→<br>rpl24p (PRK01191)KOW_RPL26 (cd06089)→<br>PRK04313 (PRK04313)KOW_RPS4 (cd06087)S4 (pfam01479)Ribosomal_S4e (pfam00900)→<br>RplE (COG0094)rpl5p (PRK04219)→ RpsN (COG0199)rps14P (PRK05766)→<br>rps8p (PRK04034)RpsH (COG0096)→ RplF (COG0097)rpl6p (PRK05518)→<br>PRK12766 (PRK12766)RecA (COG0468)Rpl32e (COG1717)→ rpl19e (PRK08570)Ribosomal_L19e (pfam01280)→ |

| | | | | | |
|---|---|---|---|---|---|
| Npe-1-1 | NC_019962.1 | + | 854999 | 855098 | **RNA**→ Methyltrn_RNA_3 (pfam02598)→ RplC (COG0087)rpl3p (PRK04231)→ RplD (COG0088)rpl4lp (PRK04042)→ RplW (COG0089)PRK14548 (PRK14548)→ RplB (COG0090)rpl2p (PRK09612)→ RpsS (COG0185)rps19p (PRK04038)→ rpl22p (PRK04223)RplV (COG0091)→ RpsC (COG0092)rps3p (PRK04191)→ RpmC (COG0255)PRK14549 (PRK14549)→ PRK03879 (PRK03879)UPF0086 (pfam01868)→ RpsQ (COG0186)rps17p (PRK08572)→ rpl14p (PRK08571)Ribosomal_L14 (pfam00238)→ rpl24p (PRK01191)KOW_RPL26 (cd06089)→ PRK04313 (PRK04313)KOW_RPS4 (cd06087)Ribosomal_S4e (pfam00900)→ RplE (COG0094)rpl5p (PRK04219)→ RpsN (COG0199)rps14P (PRK05766)→ rps8p (PRK04034)RpsH (COG0096)→ RplF (COG0097)rpl6p (PRK05518)→ UvrC (COG0322)Rpl32e (COG1717)rpl32e (PRK08562)HHH_5 (pfam14520)→ rpl19e (PRK08570)Ribosomal_L19e (pfam01280)→ |
| Hti-1-1 | NC_021921.1 | - | 1979794 | 1979695 | **RNA**→ Methyltrn_RNA_3 (pfam02598)→ RplC (COG0087)rpl3p (PRK04231)→ RplD (COG0088)rpl4lp (PRK04042)→ RplW (COG0089)PRK14548 (PRK14548)→ RplB (COG0090)rpl2p (PRK09612)→ RpsS (COG0185)rps19p (PRK04038)→ rpl22p (PRK04223)RplV (COG0091)→ RpsC (COG0092)rps3p (PRK04191)→ PRK14549 (PRK14549)Ribosomal_L29 (pfam00831)→ PRK03879 (PRK03879)UPF0086 (pfam01868)→ RpsQ (COG0186)rps17p (PRK08572)→ rpl14p (PRK08571)Ribosomal_L14 (pfam00238)→ RplX (COG0198)rpl24p (PRK01191)→ PRK04313 (PRK04313)KOW_RPS4 (cd06087)Ribosomal_S4e (pfam00900)→ RplE (COG0094)rpl5p (PRK04219)→ RpsN (COG0199)rps14P (PRK05766)→ rps8p (PRK04034)RpsH (COG0096)→ RplF (COG0097)rpl6p (PRK05518)→ |

#### 2.3 Conserved domains

Conserved domains found in protein-coding genes listed in Section 2.2 are shown below, with the first sentence in their description from the Conserved Domain Database (if any). Conserved domains associated with more than one L4-Archaeoglobi RNA are assigned a color, while others are shown in gray. The number in parentheses after the colored domain name is the number of occurrences in Section 2.2.

cd00141 (6) Nucleotidyltransferase (NT) domain of family X DNA Polymerases. [D/E]
cd00165 (46) S4/Hsp/ tRNA synthetase RNA-binding domain; The domain surface is populated by conserved, charged residues that define a likely RNA-binding site; Found in stress proteins, ribosomal proteins and tRNA synthetases; This may imply a hitherto unrecognized functional similarity between these three protein classes.
cd00355 (1) Ribosomal protein L30, which is found in eukaryotes and prokaryotes but not in archaea, is one of the smallest ribosomal proteins with a molecular mass of about 7kDa.
cd00427 (22) Ribosomal L29 protein/HIP.
cd00513 (12) Ribosomal_L32_L32e: L32 is a protein from the large subunit that contains a surface-exposed globular domain and a finger-like projection that extends into the RNA core to stabilize the tertiary structure.
cd01428 (2) Adenylate kinase (ADK) catalyzes the reversible phosphoryl transfer from adenosine triphosphates (ATP) to adenosine monophosphates (AMP) and to yield adenosine diphosphates (ADP).
cd01513 (1) Domain III of Elongation factor (EF) Tu (EF-TU) and related proteins.
cd01657 (30) Ribosomal protein L7, which is found in archaea and eukaryotes but not in prokaryotes, binds domain II of the 23S rRNA as well as the 5S rRNA and is one of five ribosomal proteins that mediate the interactions 5S rRNA makes with the ribosome.
cd01781 (1) Ubiquitin domain of AT-6, second repeat.
cd02034 (1) The accessory protein CooC, which contains a nucleotide-binding domain (P-loop) near the N-terminus, participates in the maturation of the nickel center of carbon monoxide dehydrogenase (CODH).
cd02411 (7) K homology RNA-binding domain (KH) of the archaeal 30S small ribosomal subunit S3 protein.
cd02572 (2) Pseudouridine synthase, human dyskerin like.
cd03586 (1) DNA Polymerase IV/Kappa.
cd03694 (1) Domain II of the GTPBP family of GTP binding proteins.
cd06087 (219) KOW motif of Ribosomal Protein S4 (RPS4).
cd06089 (159) KOW motif of Ribosomal Protein L26.
cd07713 (1) Methanocaldococcus jannaschii dihydropteroate synthase, Thermoanaerobacter tengcongensis Tflp, and related proteins; MBL-fold metallo hydrolase domain.
cd12188 (4) Saccharopine Dehydrogenase NAD-binding and catalytic domains.
cd16490 (1) RING-CH finger, H2 subclass (C4HC3-type), found in Fanconi anemia group L protein (FANCL) and similar proteins.
COG0087 (255) Ribosomal protein L3 [Translation, ribosomal structure and biogenesis]
COG0088 (321) Ribosomal protein L4 [Translation, ribosomal structure and biogenesis]
COG0089 (310) Ribosomal protein L23 [Translation, ribosomal structure and biogenesis]
COG0090 (299) Ribosomal protein L2 [Translation, ribosomal structure and biogenesis]
COG0091 (269) Ribosomal protein L22 [Translation, ribosomal structure and biogenesis]
COG0092 (256) Ribosomal protein S3 [Translation, ribosomal structure and biogenesis]
COG0094 (188) Ribosomal protein L5 [Translation, ribosomal structure and biogenesis]

COG0096 (210) Ribosomal protein S8 [Translation, ribosomal structure and biogenesis]
COG0097 (208) Ribosomal protein L6P/L9E [Translation, ribosomal structure and biogenesis]
COG0098 (59) Ribosomal protein S5 [Translation, ribosomal structure and biogenesis]
COG0185 (256) Ribosomal protein S19 [Translation, ribosomal structure and biogenesis]
COG0186 (242) Ribosomal protein S17 [Translation, ribosomal structure and biogenesis]
COG0198 (75) Ribosomal protein L24 [Translation, ribosomal structure and biogenesis]
COG0199 (213) Ribosomal protein S14 [Translation, ribosomal structure and biogenesis]
COG0200 (2) Ribosomal protein L15 [Translation, ribosomal structure and biogenesis]
COG0201 (25) Preprotein translocase subunit SecY [Intracellular trafficking, secretion, and vesicular transport]
COG0255 (60) Ribosomal protein L29 [Translation, ribosomal structure and biogenesis]
COG0256 (85) Ribosomal protein L18 [Translation, ribosomal structure and biogenesis]
COG0272 (4) NAD-dependent DNA ligase [Replication, recombination and repair]
COG0322 (25) Excinuclease UvrABC, nuclease subunit [Replication, recombination and repair]
COG0468 (99) RecA/RadA recombinase [Replication, recombination and repair]
COG1102 (4) Cytidylate kinase [Nucleotide transport and metabolism]
COG1422 (3) Uncharacterized archaeal membrane protein, DUF106 family, distantly related to YidC/Oxa1 [Function unknown]
COG1471 (2) Ribosomal protein S4E [Translation, ribosomal structure and biogenesis]
COG1555 (1) DNA uptake protein ComE and related DNA-binding proteins [Replication, recombination and repair]
COG1588 (34) RNase P/RNase MRP subunit p29 [Translation, ribosomal structure and biogenesis]
COG1717 (131) Ribosomal protein L32E [Translation, ribosomal structure and biogenesis]
COG1841 (1) Ribosomal protein L30/L7E [Translation, ribosomal structure and biogenesis]
COG2019 (2) Archaeal adenylate kinase [Nucleotide transport and metabolism]
COG2106 (89) Predicted RNA methylase MTH1, SPOUT superfamily [General function prediction only]
COG2251 (4) Predicted nuclease, RecB family [General function prediction only]
COG5271 (1) Midasin, AAA ATPase with vWA domain, involved in ribosome maturation [Translation, ribosomal structure and biogenesis]
pfam00181 (5) Ribosomal Proteins L2, RNA binding domain.
pfam00189 (3) Ribosomal protein S3, C-terminal domain.
pfam00203 (23) Ribosomal protein S19.
pfam00237 (3) Ribosomal protein L22p/L17e.
pfam00238 (239) Ribosomal protein L14p/L23e.
pfam00276 (4) Ribosomal protein L23.
pfam00281 (36) Ribosomal protein L5.
pfam00297 (180) Ribosomal protein L3.
pfam00347 (4) Ribosomal protein L6.
pfam00366 (3) Ribosomal protein S17.
pfam00410 (2) Ribosomal protein S8.
pfam00573 (16) Ribosomal protein L4/L1 family.
pfam00633 (2) Helix-hairpin-helix motif.
pfam00673 (35) ribosomal L5P family C-terminus.
pfam00828 (25) Ribosomal proteins 50S-L15, 50S-L18e, 60S-L27A.
pfam00831 (172) Ribosomal L29 protein.
pfam00900 (227) Ribosomal family S4e.
pfam01280 (169) Ribosomal protein L19e.
pfam01472 (2) PUA domain.
pfam01479 (47) S4 domain.
pfam01655 (63) Ribosomal protein L32.
pfam01868 (219) Domain of unknown function UPF0086.
pfam01956 (2) Integral membrane protein DUF106.
pfam02598 (347) Putative RNA methyltransferase.
pfam03947 (12) Ribosomal Proteins L2, C-terminal domain.
pfam04147 (1) Nop14-like family.
pfam09547 (1) Stage IV sporulation protein A (spore_IV_A).
pfam10559 (1) Plug domain of Sec61p.
pfam12226 (1) Turkey astrovirus capsid protein.
pfam14520 (61) Helix-hairpin-helix domain.
pfam15250 (1) Raftlin.
PHA03252 (1) DNA packaging tegument protein UL25; Provisional
PRK00279 (2) adenylate kinase; Reviewed
PRK01191 (235) 50S ribosomal protein L24P; Validated
PRK02201 (1) putative inner membrane protein translocase component YidC; Provisional
PRK03879 (253) ribonuclease P protein component 1; Validated
PRK04034 (214) 30S ribosomal protein S8P; Reviewed
PRK04038 (284) 30S ribosomal protein S19P; Provisional
PRK04040 (2) adenylate kinase; Provisional
PRK04042 (341) 50S ribosomal protein L4P; Provisional
PRK04044 (59) 30S ribosomal protein S5P; Reviewed
PRK04182 (4) cytidylate kinase; Provisional
PRK04191 (260) 30S ribosomal protein S3P; Reviewed
PRK04219 (226) 50S ribosomal protein L5P; Reviewed
PRK04223 (272) 50S ribosomal protein L22P; Reviewed
PRK04231 (435) 50S ribosomal protein L3P; Reviewed
PRK04270 (2) H/ACA RNA-protein complex component Cbf5p; Reviewed
PRK04313 (234) 30S ribosomal protein S4e; Validated
PRK05518 (209) 50S ribosomal protein L6P; Reviewed
PRK05766 (213) 30S ribosomal protein S14P; Reviewed
PRK05883 (2) acyl carrier protein; Validated
PRK06049 (32) 50S ribosomal protein L30P; Reviewed
PRK06419 (28) 50S ribosomal protein L15P; Reviewed
PRK07233 (1) hypothetical protein; Provisional
PRK08562 (116) 50S ribosomal protein L32e; Validated
PRK08568 (25) preprotein translocase subunit SecY; Reviewed
PRK08569 (86) 50S ribosomal protein L18P; Reviewed
PRK08570 (169) 50S ribosomal protein L19e; Reviewed
PRK08571 (240) 50S ribosomal protein L14P; Reviewed
PRK08572 (246) 30S ribosomal protein S17P; Reviewed
PRK09612 (316) 50S ribosomal protein L2P; Validated
PRK10575 (3) iron-hydroxamate transporter ATP-binding subunit; Provisional
PRK10611 (1) chemotaxis methyltransferase CheR; Provisional
PRK10811 (1) ribonuclease E; Reviewed
PRK12317 (1) elongation factor 1-alpha; Reviewed
PRK12442 (1) translation initiation factor IF-1; Reviewed
PRK12766 (86) 50S ribosomal protein L32e; Provisional
PRK13493 (1) chemoreceptor glutamine deamidase CheD; Provisional
PRK14548 (313) 50S ribosomal protein L23P; Provisional
PRK14549 (254) 50S ribosomal protein L29P; Provisional
smart00322 (6) K homology RNA-binding domain.
smart00382 (1) ATPases associated with a variety of cellular activities.

smart00739 (2) KOW (Kyprides, Ouzounis, Woese) motif.
smart00849 (1) Metallo-beta-lactamase superfamily.
TIGR01393 (1) elongation factor 4. [Unknown function, General]
TIGR02903 (1) ATP-dependent protease, Lon family. [Protein fate, Degradation of proteins, peptides, and glycopeptides, Cellular processes, Sporulation and germination]
TIGR03594 (1) ribosome-associated GTPase EngA. [Protein synthesis, Other]
TIGR03636 (2) ribosomal protein uL23, archaeal form. [Protein synthesis, Ribosomal proteins: synthesis and modification]
TIGR03653 (3) ribosomal protein uL6, archaeal form. [Protein synthesis, Ribosomal proteins: synthesis and modification]

#### 2.4 Multiple-sequence alignment

Each L4-Archaeoglobi RNA is denoted by an abbreviation (like "Eco-1-1") that refers to a taxonomy given in Section 2.1. The alignment may include sequences containing the RNA motif, as well as flanking sequence. The L4-Archaeoglobi RNA itself is denoted by the line underneath marked 5′ and 3′ on either end. Nucleotides in flanking sequences (i.e., not part of the motif) are written in gray letters. Stems of predicted rho-independent transcription terminators, if any, are shaded yellow. (Note: terminator predictions have not been analyzed manually, and many are likely to be false positives. Terminator predictions are those of the RNIE software.) Nucleotides predicted to function (as DNA) as transcription-factor binding sites, if any, are shaded in green. (But note: these predictions are manually annotated, so they might be under-predicted.) Annotated start codons, if any, are shaded green. (Note: start codons are frequently misannotated, especially in environmental samples.) Nucleotides proposed to basepair as part of the consensus structure are shaded in color when they comprise Watson-Crick or G-U pairs. Otherwise they are shaded gray. Conserved stems are also indicated at the bottom of the alignment by angle brackets, where matching < and > denote base-paired columns. Below these angle brackets, the symbol "2" denotes base pairs exhibiting covariation according to the statistically well-founded R-scape method. "1" denotes base pairs exhibiting covariation according to R2R's simplistic method. "0" denotes base pairs that are not observed to mutate and "?" denotes base pairs that have a significant frequency of non-canonical nucleotides for Watson-Crick or G-U pairs (> 5%). Below these base pair annotation is the consensus sequence: "R" = "A" or "G", "Y" = "C" or "U", red nucleotides: nucleotide identity conserved more than 97% of the time, black nucleotides: 90%, gray nucleotides: 75%, red circle (•): nucleotide is present 97% of the time, black circle (•): 90%, gray circle (•): 75%, white circle (◦): 50%. All percentages of sequences just described (e.g. 97% conserved) assume that sequences have been weighted by the GSC algorithm implemented by the Infernal software package.

Duplicate sequences: the following putative homologs are not shown in the alignment because their sequences are identical to a homolog already shown: Hdi-2-1, Hgi-2-1, Hha-1-2, Hhi-1-1, Hje-2-1, Hla-1-2, Hla-3-1, Hma-1-1, Hme-1-2, Hpa-1-2, Hsa-3-1, Hsi-1-1, Hsp-1-1, Hsp-4-1, Hsp-5-1, Hte-2-1, Hvo-1-1, Hwa-2-1, Mco-2-1, Mhu-1-1, Nal-1-1, Nsp-1-1, env-100, env-111, env-112, env-12, env-121, env-124, env-125, env-132, env-133, env-141, env-142, env-143, env-144, env-16, env-162, env-163, env-168, env-171, env-177, env-18, env-19, env-192, env-194, env-195, env-20, env-200, env-202, env-205, env-208, env-21, env-211, env-212, env-213, env-22, env-229, env-230, env-234, env-235, env-236, env-239, env-26, env-262, env-263, env-264, env-267, env-273, env-276, env-279, env-280, env-281, env-282, env-283, env-284, env-285, env-286, env-287, env-288, env-289, env-290, env-291, env-294, env-299, env-300, env-301, env-302, env-307, env-308, env-309, env-311, env-312, env-313, env-314, env-319, env-327, env-328, env-329, env-330, env-336, env-342, env-343, env-344, env-345, env-346, env-347, env-348, env-362, env-363, env-364, env-365, env-366, env-367, env-368, env-369, env-388, env-391, env-393, env-394, env-396, env-397, env-398, env-401, env-404, env-411, env-412, env-417, env-419, env-42, env-420, env-421, env-422, env-424, env-426, env-427, env-431, env-432, env-438, env-447, env-453, env-454, env-457, env-458, env-459, env-46, env-461, env-463, env-47, env-470, env-473, env-48, env-487, env-488, env-489, env-490, env-491, env-493, env-494, env-496, env-497, env-498, env-499, env-500, env-501, env-502, env-503, env-506, env-507, env-51, env-510, env-513, env-52, env-53, env-60, env-61, env-66, env-67, env-68, env-69, env-7, env-73, env-75, env-85, env-9, env-97, har-1-1

alignment positions 1 · · · 79

```
Mli-1-1  GUGAUGCCCCGGCUGCCCCUUAUGGGGGCACAACCGAGAAGGACUCUAAUCCAGUCCCAAGGAGCUGACAGACGCUGGA
env-1    UUCCCAGAUCCGUUCUCAUUCGUAAUACGAAAUGAGACCAGAAAGGAGUUGUACAAACAACAGGGCUGAAUUAGUGGGA
env-2    CAAUGUCACGAAGGAUAUAUCGGUUGGAGAAUAUAGAGAGGGAUUAGUACUCGAAAGGUCUAGCGAAGAGAAAGUAGGU
env-3    UAAUCUCGAUAAAACAACGGAAGUCGGUGAAAUAAGAGAGGGGUUUGUGUUAGAGAAGUUUGGUGAUGAGAGAGUAGGU
env-4    AUCGGGGACGGAUCGCCGUGCCCGGUCAUGUUAUGGGACGAAUGCCGUGCUUCUCGCACGGUGGCUGAUAGACGCUGGG
env-5    AUCGGGGGCGGAUCGCCGUGCCCGGUCAUGUUAUGGGACGAAUGCCGUGCUUCUCGCACGGUGGCUGAUAGACGCUGGG
env-6    GAAUGUUCCAUUUAGAGGCCAUUUAGGCUUUGAAAUGAGCAAUAAUUGCAUAAUAUAGCAAUGGGCUGAAAGAGUGGGA
env-8    UAAUUGAACCUAAUCUGACUUUUUUCGGUCAAGACGCGGUCAUAAAUUGCUUUAAUUGCAAUGGGCUGAAAGAGUGGGA

         ...............................................................................
         ...............................................................................
         •••••••••••••••••••••••••••••••••••••RR•••RRR•••••••••••••RRR•••GY•R••RRRGY•GGA
```

```
env-10    UGAAUGUACUGAUUCUGGCUUCUUUUAGCUAUGAAGCGGUCAAAAAUUGCUUUAAUUGCAAUGGGCUGAUAGAGUGGGA
env-11    UGAAUGUACUGAUUCUGGCUUCUUUUAGCUAUGAAGCGGUCAAAAAUUGCUUUAAUUGCAAUGGGCUGAUAGAGUGGGA
env-13    CUGAGGUCACGGCCCUCUCAUCUGACAGGGCCGUGGUUAUGGGAGUUGUGAGGAGAUCACAACCGGCUGACAUGGUGGA
env-14    GAAUGGUGUGAAGAUCCCGAUCCCGGGAUCAUAAACCGAUAGGAGUUGUGCCAGGGCACAACGGGCUGACAGAGCUGGA
env-15    CCUCCGCGUGUCGUCUACGACCGGCUCUGAGUCAACUGAUUCAGGGUCGUUAACACAGGGAAUCGUGACUGAGGUCGGA
Hwa-1-1   CCUCCGCGUGUCAUCUACGACCGGCUCUGAGUCAACUGAUUCAGGGUCGUUAACACAGGGAAUCGUGACCGAGGUCGGA
env-17    CCUCCGCGUGUCAUCUACGACCGGCUCUGAGUCAACUGAUUCAGGGUCGUUAACACAGGGAAUCGUGACCGAGGUCGGA
env-23    ACUCCGCGUCCCGUCGCAGACCGGCUCGACUCCAGACGGAGCCGAGUCGUCAACACAGGGAAUCGUGACCAAGGUAGGA
env-24    ...................................AGUCCGAACCCGAGUACAGAGAAGGCGUGGUGACGAAAGUCGGA
env-25    GCUCCGCACGCGUUCACAGACCGGCUCCGGAUCAGAUGGUUCGGGGUCGUUAAGACAGGGAAUCGUGACCGAGGUCGGA
env-27    GAUGCUGCCGACGGCCGAGGCCGGCGCCGGAUCGGAGGGUCCGGGGUCGUUAAUAGACGGAAUCGUGACCGAGGUCGGA
env-28    GCUUCGCGCCACGACACAGACCGGCUCCGAAUCAGACGGUUCGGGGUCGUCAAAACACGGAAUCGUGACCGAGGUCGGA
env-29    GCUUCGCGCCACGACACAGACCGGCUCCGAAUCAGACGGUUCGGGGUCGUCAAAACACGGAAUCGUGACCGAGGUCGGA
env-30    GCUUCGCGCCACGACACAGACCGGCUCCGAAUCAGACGGUUCGGGGUCGUCAAAACACGGAAUCGUGACCGAGGUCGGA
env-31    GCUCCGCGUGCGUUCACAGACCGGCUCCGGAUCCGAGGGUUCGGGGUCGUUAAGACAGGGAAUCGUGACCGAGGUCGGA
Hut-1-1   GCUCCGCGUUGCGACCCAGACCGGCUCCGGAUCGGACGAUUCGGGGUCGUUAAGACAGGGAAUCGUGACCGAGGUCGGA
env-32    GAUGCAGCCGUCGGCCGAGGCCGGCUCCGGAUCGCAGGGUCCGGGGUCGUUAAUAGACGGAAUCGUGACCGAGGUCGGA
env-33    GAUGCAGCCGUCGGCCGAGGCCGGCUCCGGAUCGCAGGGUCCGGGGUCGUUAAUAGACGGAAUCGUGACCGAGGUCGGA
env-34    GAUGCAGCCAUCGGCCGAGGCCGGCUCCGGAUCGCAGGGUCCGGGGUCGUUAAUAGACGGAAUCGUGACCGAGGUCGGA
env-35    GAUGCAGCCGUCGGCCGAGGCCGGCUCCGGAUCGCAGGGUCCGGGGUCGUUAAUAGACGGAAUCGUGACCGAGGUCGGA
env-36    GCUCCGCGUGGUGACACAGACCGGCUCCGGAUCCGAGGGUUCGGGGUCGUUAAGACAGGGAAUCGUGACCGAGGUCGGA
env-37    GCUCCGCGUCACAACACGGACCGGCUCCGGAUCGGAGGGUUCGGGGUCGUUAAGACAGGGAAUCGUGACCGAUGUCGGU
env-38    .............................................GUCGUUAAGACAGGGAAUCGUGACCGAUGUCGGU
env-39    GCUCCGCGUCACAACACGGACCGGCUCCGGAUCGGAGGGUUCGGGGUCGUUAAGACAGGGAAUCGUGACCGAUGUCGGU
env-40    GCUUCGGCUGCCGACAUGGACCGGCUCCGAAUCAAAGGAUUCGGGGUCGAUAAGAGAAGGAAUCGUGACCGAGGUCGGA
env-41    CCUCCGUAUCCCGUCUAUGACCGGCUCUGAAUCAGAUGAUUCAGAGUCGUUAACACAAGGAAUCGUGACCGAGGUCGGA
env-43    GCUCCGCGUGCGUUCACAGACCGGCUCCGGAUCCGAGGGUUCGGGGUCGUUAAGACAGGGAAUCGUGACCGAGGUCGGC
env-44    GCUCCGCGCCACGCCACAGACCGGCUCCGGAUCGAACGGUCCGGGGUCGUCAAGACGAGGGAUCGUGACCGAGGUCGGA
env-45    ACCGCUCCGCGCAUCAUCACGGACAUCGACCGACCCAGACGGGCCGGUCGAACAAACGGGAAUCGUGACCGAGGUCGGA
env-49    CCCGUUGCGGUGCUCGUCGCAGACCACAGCCGAUUCGUCCGAGUCGGCUGUCAACCAGGGAAUCGUGACCGAGGUCGGA
env-50    GCCGCUCCGGGUUUCGUCACAGACCGUAGCCGACCCGAACGGGCCGGCUACACACCAAGGAAUCGUGACCGAGGUCGGA
env-54    CCCGUUGGGCGUCUCGUCGCAGACCACAGCCGAUUCGUCCGAGUCGGCUGUCAACCACGGAAUCGUGACCGAGGUCGGA
env-55    UCCGCUCCGCGUUUCGUCACGGACGGUAACCGAUCCAAACGGAUCGGUUACGCACCAAGGAAUCGUGACCAAGGUCGGA
env-56    ...........................................CCGGCUACACACCAAGGAAUCGUGACCAAGGUCGGA
env-57    CCGGGUUUCGUCACAGACCGUAGCCGGUCCAGACGACGAUGGACCGGCUACACACCAAGGAAUCGUGACCAAGGUCGGA
env-58    ACCGCUCCGCGUUUCGUCACAGACCGUAGCCAACCCGGAUGGGCCGGCUACACACCAGGGAAUCGUGACCAAGGUCGGA
env-59    UCCGCUCCGGGUUUCGUCACAGACCGUAGCCGAUCCAGCCGGAUCGGUUACACAUCAAGGAAUCGUGACCAAGGUCGGA
env-62    UCCGCUCCGGGUUUCGUCACAGACCGUAGCCGAUCCAGCCGGAUCGGUUACACAUCAAGGAAUCGUGACCAAGGUCGGA
env-63    UCCGCUCCGGGUUUCGUCACAGACCGUAGCCGAUCCAGCCGGAUCGGUUACACAUCAAGGAAUCGUGACCAAGGUCGGA
env-64    ...................ACAGACCGUAGCCGAUCCAGCCGGAUCGGUUACACAUCAAGGAAUCGUGACCAGGUCGGA
env-65    UCCGCUCCGGGUUUCGUCACAGACCGUAGCCGAUCCAGCCGGAUCGGUUACACAUCAAGGAAUCGUGACCAAGGUCGGA
env-70    .....................AGACCGUAGCCGAUCCAGCCGGAUCGGUUACACAUCAAGGAAUCGUGACCAAGGUCGGA
env-71    .................CACAGACCGUAGCCGAUCCAGCCGGAUCGGUUACACAUCAAGGAAUCGUGACCAAGGUCGGA
env-72    GCUCCGUGCCACGUCACAGGCCGGCUCCGGAUCGAACGGUCCGGGGUCGAGAAGACAAGGGAUCGUGACCGAGGUCGGA
env-74    GCUCCGUGCCACGUCACAGACCGGCUCCGGAUCGAACGGUCCGGGGUCGAGAAGACAAGGGAUCGUGACCGAGGUCGGA
env-76    GCUCCACCUGGGUGCACAGACCGGCUCCGGAUCGAACGGUUCGGGGUCGUCAAGACAGGGCAUCGUGACCGAGGUCGGA
env-77    GCUCCACCUGGGUGCACGGACCGGCUCCGGAUCCGACGGUUCGGGGUCGUCAAGACAGGGCAUCGUGACCGAGGUCGGA
env-78    CGCUCCACCUGGGUGCACGGACCGGCUCCGGAUCCGACGUUCGGGGUCGUCAAGACAGGGCAUCGUGACCGAGGUCGGA
Hsa-1-1   GCUCCACGUGACACCACGGACCGGCUCCGGAUCGAACGAUUCGGGGUCGUUGAGACAGGGACUCGUGACCGAGGUCGGA
Hgi-1-1   CGUUCUCGUGGCGUCUACGACCGCGGACGACUCUGACGAGUCGCCCGCGUUGCGAGAAGGAAUCGUGACCGAGGUCGGA
Hsu-2-1   CGUUCUCGUCGCGUCUACGACCGCGGACGACUCCGACGAGUCGCCCGCGUUGCGAGAAGGAAUCGUGACCGAGGUCGGA
Hpr-1-1   CGUUCUCGUGGCGUCUACGACCGCGGACGACUCUGACGAGUCGCCCGCGUUGCGAGAAGGAAUCGUGACCGAGGUCGGA
Hsp-8-1   CGUUCUCGUCGCGUCUACGACCGCGGACGACUCCGACGAGUCGCCCGCGUUGCGAGAAGGAAUCGUGACCGAGGUCGGA
Hde-1-1   CGUUCUCGUCGCGUCUACGACCGCGGACGACUCCGACGAGUCGCCCGCGUUGCGAGAAGGAAUCGUGACCGAGGUCGGA
Hlu-1-1   CGUUCUCGUCGCGUCUACGACCGCGGACGACUCCGACGAGUCGCCCGCGUUGCGAGAAGGAAUCGUGACCGAGGUCGGA
Hal-2-1   CGUUCUCGUGGCGUCUACGACCGCGGACGGCUCUGACGAGUCGCCCGCGUUGCGAGAAGGAAUCGUGACCGAGGUCGGA
Hsp-6-1   CGUUCUCGUGGCGUCUACGACCGCGGACGACUCCGACGAGUCGCCCGCGUUGCGAGAAGGAAUCGUGACCGAGGUCGGA
Hme-1-1   GGUCCUGGUAUCGUCUACGACCGCGGGCGACUCUGACGAGACGCCCGCGUUGCGAGAAGGAAUCGUGACCGAGGUCGGA
Hla-2-1   CGUUCUCGUCUCGUCUACGACCGCGGACGACUCUGACGAGUUGCCCGCGUUGCGAGAAGGAAUCGUGACCGAGGUCGGA
Hel-1-1   CGUUCUCGUCUCGUCUACGACCGCGGACGACUCUGACGAGUUGCCCGCGUUGCGAGAAGGAAUCGUGACCGAGGUCGGA
Hmu-1-1   GGUCCUGGUAUCGUCUACGACCGCGGGCGACUCUGACGAGACGCCCGCGUUGCGAGAAGGAAUCGUGACCGAGGUCGGA
env-79    GCUCCGCGUCACAACACGGACCGGCUCCGGAUCGGAGGGUUCGGGGUCGUUAAGACAGGGAAUCGUGACCGAUGUCGGU
env-80    GCUCCGCGCGCGUUCACAGACCGGCUCCGGAUCGGAGGGUUCGGGGUCGUUAAGACAGGGAAUCGUGACGAAGGCCGGA
env-81    GCUCCGCGCGCGUUCACAGACCGGCUCCGGAUCGGAGGGUUCGGGGUCGUUAAGACAGGGAAUCGUGACGAAGGCCGGA
env-82    GCUCCGCGCGUUGUCACAGACCGGCUCCGGAUCAGACGGUUCGGGGUCGUUAAGACAGGGAAUCGUGACGAAGGCCGGA
env-83    GCUCCGCGCGUGUUCACAGACCGGCUCCGGAUCGGACGGUUCGGGGUCGUUAAGACAGGGAAUCGUGACGAAGGCCGGA
Hla-4-1   GCUCCGUGCCACGUCACAGACCGGCUCCGAAUCCGACGAUUCGGGGUCGAAAAGACAAGGAAUCGUGACCGAGGUCGGA
Ssp-1-1   GCUCCACGCCUCUUCACGGACCGGCUCCGGAUCGAGCGGUCCGGGGUCGUUGAGACAGGGUAUCGUGACCGAGGUCGGA
env-84    GCUCCGUCUCUCGGCAUGGACCGGCUCCGAAUCGAGCGGUUCGGGGUCGAUACGACAGGGAGUAGUGACCCAGGUCGGA
env-86    GCUCCGUGCCACCUCACAGACCGGCUCCGAAUCGGAGGGUUCGGGGUCGUCAAGACAAGGGAUCGUGACCGAGGUCGGA
Hsp-11-1  GCUCCGCGCACCGUCACAGACCGGCUCCGAAUCCGACGAUUCGGGGUCGAAAAGAUACGGAAUCGUGACCGAGGUCGGA
env-87    ........................................UUCGGGGCGUUAACACAGGGAAUCGUGACCAAGGUCGGA
env-88    ACCGCUCCGCGUUUCGUCACAGACCGUAGCCAACCCGGAUGGGCCGGCUACACACCAGGGAAUCGUGACCAAGGUCGGA
env-89    GCUCCGCAUCGCAACACAGACCGGCUCCGGAUCGGAGGGUUCGGGGUCGUUAAGACAGGGAAUCGUGACCGAGGUCGGA
Hka-1-1   GCUCCGCGUCCGUUCACAGACCGGCUCCGGAUCUGAGGGUUCGGGGUCGUUAAGACAGGGAAUCGUGACCGAGGUCGGA
Hmu-2-1   GCUCCGCGUCCGUUCACAGACCGGCUCCGGAUCUGAGGGUUCGGGGUCGUUAAGACAGGGAAUCGUGACCGAGGUCGGA
Hca-3-1   GCUCCGGGUCCGCUCAUGGACCGGCUCCGGUUCGGACGAAUCGGGGUCGUUAAGACAGGGAAUCGUGACCGAGGUCGGU
Hsa-5-1   GCUUCGCGCCGUCUCACGGACCGGCUCCGAAUCGCAAGGUUCGGGGUCGUCAAGACAGGGAAUCGUGACCGAGGUCGGC
env-90    GCUUCGCACCGUUUCACGGACCGGCUCCGGAUCACAGAAUCCGGGGUCGUUAAGACAGGGAAUCGUGACCGAGGUCGGC
Hag-1-1   GCUUCGCGCCGUCUCACGGACCGGCUCCGAAUCCCAAGGUUCGGGGUCGUCAAGACAGGGAAUCGUGACCGAGGUCGGC
Hsa-4-1   GCUUCGCGCCGUCUCACGGACCGGCUCCGAAUCCCAAGGUUCGGGGUCGUCAAGACAGGGAAUCGUGACCGAGGUCGGC
env-91    GCUCCGCGUGCGUUCACAGACCGGCUCCGGACCGAAGCGUUCGGGGUCGUCAAGACAGGGAAUCGUGACCGAGGUCGGA
env-92    CCCGUUGCGGCUCUCGUCGCAGACCACAGCCGAUUCGUCCGAGUCGGCUGUCAACCAGGGAAUCGUGACCGAGGUCGGA
env-93    CCCGUUGCGGCUCUCGUCGCAGACCACAGCCGAUUCGUCCGAGUCGGCUGUCAACCAGGGAAUCGUGACCGAGGUCGGA

          ...............................................................................
          ...............................................................................
          •••••••••••••••••••••••••••••••••••RR•••RRR••••••••••••••••RRR•••GY•R••RRGY•GGA
```

```
env-94     CCCGUUGCGGCUCUCGUCGCAGACCACAGCCGAUUCGUCCGAGUCGGCUGUCAACCAGGGAAUCGUGACCGAGGUCGGA
env-95     CCCGUUGCGGCUCUCGUCGCAGACCACAGCCGAUUCGUCCGAGUCGGCUGUCAACCAGGGAAUCGUGACCGAGGUCGGA
env-96     GCCCUUGCGGCUCUCGUCGCAGACCACAGCCGAUUCGUCCGAGUCGGCUGUCAACCAGGGAAUCGUGACCGAGGUCGGA
env-98     GCCCUUGCGGCUCUCGUCGCAGACCACAGCCGAUUCGUCCGAGUCGGCUGUCAACCAGGGAAUCGUGACCGAGGUCGGA
env-99     CCCAUUGCGGUGCUCGUCGCAGACCACAGCCGAUUCGUCCGAGUCGGCUGUCAACCAGGGAAUCGUGACCGAGGUCGGA
env-101    GCCGCUCCGCGUUUCGUCACAGACGGUAGCCGACCCAGACGGGACGGCUACACACCAAGGAAUCGUGACCAAGGUCGGA
env-102    ...........................................CCCGCUACACACCAAGGAAUCGUGACCAAGGUCGGA
env-103    .............................................GGUUACGCACCAAGGAAUCGUGACCAAGGUCGGA
env-104    GCUCCGGGUUUCGUCACAGACCGUAGCCGAUCCGGACGACGGACCGGCUACACACCAAGGAAUCGUGACCAAGGUCGGA
env-105    CCGCGUCUCGUCGCAGACCGUAGCCGAUCCAGACGACGACGGACCGGCUACACACCAAGGAAUCGUGACCAAGGUCGGA
env-106    ....................................................CACCAAGGAAUCGUGACCAAGGUCGGA
env-107    .........................UCCGGACCUGAGCGUCCGGGGUCGUUACAAGAGGGAAUCGUGACCGAGGUCGGA
env-108    ....AUGCCGCCGCUGCUCCCGUCGCGGACGGGUUCCGAGUCAUCGGAACUGCGAGAAGGAAUCGUGACCGAGGUCGGA
env-109    GCUCCGCGUCUCGUCGCAGACCGGCUCCGGAUCGGACGGUUCGGGGUCGUAAAGACAGGGAAUCGUGACCGAGGUCGGA
CHa-1-1    CGUGCUCCCGCCGAUGCGGCCCGACUCAUCCGGCCCCGACGGGUCGGAGCGACGAGCCGGAAUCGUGACCGAGGUCGGA
Hpa-2-1    CCUCCUCGUCUCGUCUACGACCGGCUCCGAAUCGAACGAUUCGGGGUCGUUACAACAGGGAAUCGUGACCGAGGUCGGA
env-110    GCUUCGCGCCGCGUCACAGACCGGCUCCGAAUCGAACGGUUCGGGGUCGUCAAGACAAGGGAUCGUGACCGAGGUCGGA
Nhu-1-1    GCUCCGCGCCACGUCACAGACCGGCUCCGAAUCAGACGGUUCGGGGUCGUUAAGACAAGGGAUCGUGACCGAGGUCGGA
env-113    GCUCCGCGUGCCGUGGCGGACCGGCUCGACCCCUAGCGGGGAAGAGUCGAAAACACAGGGACUCGUGACCGAGGUCGGA
env-114    CCCGUUGCGGUGCUCGUCGCAGACCACAGCCGAUUCGUCCGAGUCGGCUGUCAACCAGGGAAUCGUGACCGAGGUCGGA
env-115    CCCGUUGCGGUGCUCGUCGCAGACCACAGCCGAUUCGUCCGAGUCGGCUGUUAACCAGGGAAUCGUGACCGAGGUCGGA
env-116    ................................CGGACGACGGACCGGCUACACACCAAGGAAUCGUGACCAAGGUCGGA
env-117    ......................GCCGAUCCAGGCGAUGACGGACCGGCUACACACCAAGGAAUCGUGACCAAGGUCGGA
env-118    .....................................GACGGACCGGCUACACACCAAGGAAUCGUGACCAAGGUCGGA
env-119    GCCGCUCCGCGUUUCGUCACAGACCGUAGCCGACCCAGACGGGGCGGCUACACACCAAGGAAUCGUGACCAAGGUCGGA
env-120    UGUGCUCCCGCCGUUGUUGCCAGACACCUCCGGCCCAGACGGGGCGGAGCUACGAGACGGAAUCGUGACCGAGGUCGGA
env-122    CGUGCUCCCGCCGCUGUUGCCAGACACCUCCGGCCCAGACGGGGCGGAGCUGCGAGACGGAAUCGUGACCGAGGUCGGA
env-123    CGUGCUCCCGCCGUUGUUGCCAGACACCUCCGGCCCAGACGGGGCGGAGCUCCGAGACGGAAUCGUGACCGAGGUCGGA
env-126    ..........CCGGCUACGACCGGCUCCGAAUCCAACGGUUCGGGGUCGUUAACACAGGGAAUCGUGACCGAGGUCGGA
env-127    CCUCCGCGUCCCGUCUACGACCGGCUCCGAAUCCAACGGUUCGGGGUCGUUAACACAGGGAAUCGUGACCGAGGUCGGA
env-128    CCUCCGCGUCCCGUCUACGACCGGCUCCGAAUCCAACGGUUCGGGGUCGUUAACACAGGGAAUCGUGACCGAGGUCGGA
env-129    GCUUCGCGCGCUCUCACGGACCGGCUCCGGAUCCACCGAUCCGGGGUCGUUAAGACAGGGAAUCGUGACCGAGGUCGGA
Hna-1-1    CCUUCGCGUCUCGUCACGGACCGGCUCCGGACCGGAGCGUCCGGGGUCGUUACAAGAGGGAAUCGUGACCGAGGUCGGA
env-130    CCUUCGCGUCUCGUCACGGACCGGCUCCGGACCGGAGCGUCCGGGGUCGUUACAAGAGGGAAUCGUGACCGAGGUCGGA
env-131    CCCGCUCCGCGUCUCCCACGGACCGGCUCGGCGUCUGAACGGCCCGAGUCGAGAGAAGGAAUCGUGACCGAGGUCGGA
env-134    GCUCCGCGUCCGUUCACAGACCGGCUCCGGAUCGGACGGUUCGGGGUCGUUAAGACAGGGAAUCGUGACCGAGGUCGGA
env-135    ..................GACCGGCUCCGAAUCUACCGGUUCGGGGUCGUCAAGACAAGGAAUCGUGACCGAGGUCGGA
Nph-1-1    GCUCCGCGUGCGUUCACAGACCGGCUCCGGAUCGGACGGUUCGGGGUCGUUAAGACAGGGAAUCGUGACCGAGGUCGGA
env-136    GCUCCUCGUCCGUUCGCAGACCGGCUCCGGAUCGGACGGUUCGGGGUCGUUAAGACAGGGAAUCGUGACCGAGGUCGGA
env-137    GCUCCGCGUUCGUUCACAGACCGGCUCCGGAUCGGACGGUUCGGGGUCGUUAAGACAGGGAAUCGUGACCGAGGUCGGA
env-138    GCUCCGCGUACCAUGGCGGACCGGCUCGCCCCCAGACGGGAGCGAGUCGACAACACAGGGACUCGUGACCGAGGUCGGA
env-139    .................GGACCGGCUCGCCCCCCGACGGGGAGGAGUCGACAACACAGGGACUCGUGACCGAGGUCGGA
env-140    GCUUCGCGUGUCUUGGCGGACCGGCUCGCCCCCCGACGGGGGCGAGUCGACAACACAGGGACUCGUGACCGAGGUCGGA
env-145    GCUCCGCGUGUCGUGGCGGACCGGCUCCCCCCCCGACGGGGCGGAGUCGAAAACACAGGGACUCGUGACCGAGGUCGGA
env-146    .................................................AAAACACAGGGACUCGUGACCGAGGUCGGA
env-147    GCUCCGCGUGUCGUGGCGGACCGGCUCGUCCCUCGACGGGGAGGAGUCGACAACACAGGGACUCGUGACCGAGGUCGGA
env-148    GCUCCGCGUGCCGUGGCGGACCGGCUCGUCCCCCGACGGGGAGGAGUCGAUAACACAGGGACUCGUGACCGAGGUCGGA
Hsp-9-1    GCUCCGCGUGCCGUGGCGGACCGGCUCGACCCCUAGCGGGGGAGAGUCGAAAACACAGGGACUCGUGACCGAGGUCGGA
Hez-1-1    GCUCCGCGUGCCGUGGCGGACCGGCUCGACCCCUAGCGGGGAAGAGUCGAAAACACAGGGACUCGUGACCGAGGUCGGA
env-149    .........................................AAGAGUCGAAAACACAGGGACUCGUGACCGAGGUCGGA
env-150    .........................................AAGAGUCGAAAACACAGGGACUCGUGACCGAGGUCGGA
env-151    .....................................................CACAGGGACUCGUGACCGAGGUCGGA
Hco-1-1    GCUCCGCGUGCCGUGGCGGACCGGCUCGACCCCUAGCGGGGAAGAGUCGAAAACACAGGGACUCGUGACCGAGGUCGGA
env-152    GCUCCGCGUGCCGUGGCGGACCGGCUCGACCCCUAGCGGGGAAGAGUCGAAAACACAGGGACUCGUGACCGAGGUCGGA
env-153    GCUCCGCGUGCCGUGGCAGACCGGCUCGGCCCCCGACGGGGCGGAGUCGAAAACACAGGGACUCGUGACCGAGGUCGGA
env-154    ..............................................................UCGUGACCGAGGUCGGA
env-155    ....................................................ACACAGGGACUCGUGACCGAGGUCGGA
env-156    ...........................................GAGUCGAAAACACAGGGACUCGUGACCGAGGUCGGA
Hsp-10-1   GCUCCGCGUGCCGUGGCGGACCGGCUCGACCCCGAGCGGGGAAGAGUCGAAAACACAGGGACUCGUGACCGAGGUCGGA
Hli-1-1    GCUCCGCGUGCCGUGGCGGACCGGCUCGACCCCUAGCGGGGAAGAGUCGAAAACACAGGGACUCGUGACCGAGGUCGGA
Hte-1-1    GCUCCGCGUGCCGUGGCGGACCGGCUCGACCCCUAGCGGGGGAGAGUCGAAAACACAGGGACUCGUGACCGAGGUCGGA
Hko-1-1    GCUCCGCGUGCCGUGGCGGACCGGCUCGACCCCUAGCGGGGGAGAGUCGAAAACACAGGGACUCGUGACCGAGGUCGGA
Hli-2-1    GCUCCGCGUGCCGUGGCGGACCGGCUCGACCCCUAGCGGGGGAGAGUCGAAAACACAGGGACUCGUGACCGAGGUCGGA
Hsa-7-1    GCUCCGCGUGCCGUGGCGGACCGGCUCGACCCCUAGCGGGGAAGAGUCGAAAACACAGGGACUCGUGACCGAGGUCGGA
Hca-2-1    GCUCCGCGUGCCGUGGCGGACCGGCUCGACCCCUAACGGGGAAGAGUCGAAAACACAGGGACUCGUGACCGAGGUCGGA
Hsp-7-1    GCUCCGCGUGCCGUGGCGGACCGGCUCGACCCCUAGCGGGGGAGAGUCGAAAACACAGGGACUCGUGACCGAGGUCGGA
Hha-2-1    GCUCCGCGUGCCGUGGCGGACCGGCUCGACCCCUAGCGGGGAAGAGUCGAAAACACAGGGACUCGUGACCGAGGUCGGA
Hdi-1-1    GCUCCGCGUGCCGUGGCGGACCGGCUCGACCCCUAGCGGGGGAGAGUCGAAAACACAGGGACUCGUGACCGAGGUCGGA
Hsa-9-1    CGCUCCGCGUGCCGUGGCGGACCGGCUCGACCCUAGCGGGGAAGAGUCGAAAACACAGGGACUCGUGACCGAGGUCGGA
Har-2-1    GCUCCGCGUGCCGUGGCGGACCGGCUCGACCCCUAGCGGGGGAGAGUCGAAAACACAGGGACUCGUGACCGAGGUCGGA
env-157    .............................CCCCUAGCGGGGNAGAGUCGAAAACACAGGGACUCGUGACCGAGGUCGGA
env-158    .............................................................CUCGUGACCGAGGUCGGA
env-159    GCUCCGCGUGCCGUGGCGGACCGGCUCGACUCCUAGCGGAGAAGAGUCGAAAACACAGGGACUCGUGACCGAGGUCGGA
env-160    GCUCCGCGUGUCAUGGCGGACCGGCUCGUCCCCCGACGGGGAGGAGUCGACAACACAGGGACUCGUGACCGAGGUCGGA
h3A-1-1    GCUUCGUGUCCCGUCACGGACCGGCUCGCCCCCCGACGGGGACGAGUCGUCAACACAGGGACUCGUGACCGAGGUCGGA
Hai-1-1    GCUCCGCGUGCCGUGGCGGACCGGCUCGACCCCUAGCGGGGAAGAGUCGAAAACACAGGGACUCGUGACCGAGGUCGGA
env-161    GCUCCGCGUGCCGUGGCGGACCGGCUCGACCCCUAGCGGGGAAGAGUCGAAAACACAGGGACUCGUGACCGAGGUCGGA
Hho-1-1    GCUCCGCGUGCUGUGGCGGACCGGCUCGACCCCUAGCGGGGGAGAGUCGAAAACACAGGGACUCGUGACCGAGGUCGGA
env-164    .............................................GUCGAAAACACAGGGACUCGUGACCGAGGUCGGA
env-165    ........................CUCGCCCCCAGACGGGGGCGAGUCGACAACACAGGGACUCGUGACCGAGGUCGGA
env-166    GCUCCGCGUACCAUGGCGGACCGGCUCGCCCCCAGACGGGAGCGAGUCGACAACACAGGGACUCGUGACCGAGGUCGGA
env-167    ..............GGCGGACCGGCUCGCCCCCAGACGGGAGCGAGUCGACAACACAGGGACUCGUGACCGAGGUCGGA
env-169    .........GCCGUGGCGGACCGGCUCGACCCCUAGCGGGGAAGAGUCGAAAACACAGGGACUCGUGACCGAGGUCGGA
env-170    ................................................................GUGACCGAGGUCGGA

           ...............................................................................
           ...............................................................................
           •••••••••••••••••••••••••••••••••••••RR•••RRR•••••••••••••RRR•••GY•R••RRRGY•GGA
```

```
env-172   GCUCCGCGUACCAUGGCGGACCGGCUCGCCCCCAGACGGGAGCGAGUCGACAACACAGGGACUCGUGACCGAGGUCGGA
env-173   GCUUCACGCCUUGUCGCAGACCGGCUCUGGAUCAAGUGAUCCGGGGUCGUUAAGACAGGGAAUCGUGACCGAGGUCGGA
Hru-1-1   CCUCCGCGUCCCGUCUAUGACCGGCUCCGAAUCAGACGGUUCGGGGUCGUUAACACAGGGAAUCGUGACCGAGGUCGGA
env-174   CCUCCGCGUCCCGUCUAUGACCGGCUCCGAAUCAGACGGUUCGGGGUCGUUAACACAGGGAAUCGUGACCGAGGUCGGA
Hru-2-1   CCCCAUGCGUGUCUCCCCACGGACCGGCUCCGAGGGUGAAACCCCGGAGUCGAGAGAAGGAAUCGUGACCGAGGUCGGA
env-175   GCUUCGUGUCCGUGCACAGACCGGCUCCGGAUCAGAUGGUUCGGGGUCGUUAAGACAGGGAAUCGUGACCGAGGUCGGA
env-176   GCUCCGGGUUCGUUCACAGACCGGCUCCGGAUCGGACGGUUCGGGGUCGUUAAGACAGGGAAUCGUGACCGAGGUCGGA
env-178   GCUCCGAGUUCGUUCACAGACCGGCUCCGGAUCGGACGGUUCGGGGUCGUUAAGACAGGGAAUCGUGACCGAGGUCGGA
env-179   GCUUCGUGUCCGUGCACAGACCGGCUCCGGAUCGGACGGUUCGGGGUCGUUAAGACAGGGAAUCGUGACCGAGGUCGGA
env-180   GCUCCGCGUCCGUACACAGACCGGCUCCGGAUCGGAUGGUUCGGGGUCGUUAAGACAGGGAAUCGUGACCGAGGUCGGA
Nmo-1-1   GCUUCGUGUCCGUACACAGACCGGCUCCGGAUCAGACGGUUCGGGGUCGUUAAGACAGGGAAUCGUGACCGAGGUCGGA
Hha-1-1   GCUUCGCCCCGUCCCACGGACCGGCUCCGGACCACAGAGUUCGGGGUCGUUAAGACAGGGAAUCGUGACCGAGGUCGGC
env-181   GCUCCGUGCCACGUCACAGACCGGCUCCGAAUCUGACGGUUCGGGGUCGUUAAAACAAGGGAUCGUGACCGAGGUCGGA
Hpa-1-1   GCUCCGCGCCACAUCACAGACCGGCUCCGAAUCGGAGGGUUCGGGGUCGUUAAGACAGGGAAUCGUGACCGAGGUCGGA
Hci-1-1   GCUCCGCGCACCAUCGCAGACCGGCUCCGAAUCCGACGAUUCGGGGUCGUUAAGACAGGGAAUCGUGACCGAGGUCGGA
env-182   GCUCCGCGCACCGUCGCAGACCGGCUCCGAAUCUGACGAUUCGGGGUCGACAAGACAGGGAAUCGUGACCGAGGUCGGA
env-183   CGCUUCGCGCGGCCGCUCGGACCGGCUCCGAAUCCACAGUUCGGGGUCGUCAAGACAGGGAAUCGUGACCGAGGUCGGC
env-184   GCUUCGGCAGACGCCACAGACCGGCUCCGGAUCCAGCGAUUCGGGGUCGUCAAGACAGGGAAUCGUGACCGAGGUCGGA
env-185   GCUUCGCGUCCCAUCACAGACCGGCUCCGAAUCGAACGGUUCGGGGUCGAUAAGACGAGGAAUCGUGACCGAGGUCGGA
Hdj-1-1   GCUCCGCGCUGUGUCACGGACCGGCUCCGAAUCCGACGGUUCGGGGUCGUUAAGACAAGGGAUCGUGACCGAGGUCGGA
Hsa-10-1  GCUCCGCGCCAUGUCACAGACCGGCUCCGAAUCUACCGGUUCGGGGUCGUCAAGACAAGGGAUCGUGACCGAGGUCGGA
Nsu-1-1   GCUCCGCGCCAUGUCACAGACCGGCUCCGAAUCUACCGGUUCGGGGUCGUCAAGACAAGGAAUCGUGACCGAGGUCGGA
Nba-1-1   GCUCCGCGCCAUGUCACAGACCGGCUCCGAAUCUACCGGUUCGGGGUCGUCAAGACAAGGAAUCGUGACCGAGGUCGGA
Nin-1-1   GCUCCGCGCGGUGUCACAGACCGGCUCCGAAUCGAACGGUUCGGGGUCGUUAAGACAAGGGAUCGUGACCGAGGUCGGA
env-186   GCUCCGCGCACCGUCACAGACCGGCUCCGAAUCAGACGAUUCGGGGUCGACAAGACAGGGAAUCGUGACCGAGGUCGGA
env-187   GCUCCGCGCACCGUCGCAGACCGGCUCCGAAUCUGACGAUUCGGGGUCGACAAGACAGGGAAUCGUGACCGAGGUCGGA
env-188   GCUCCGCGCACCGUCACAGACCGGCUCCGAAUCGAACGAUUCGGGGUCGUCAAGACAGGGAAUCGUGACCGAGGUCGGA
env-189   ...CCGCGCACCGUCACAGACCGGCUCCGAAUCGAACGGUUCGGGGUCGACAAGACAGGGAAUCGUGACCGAGGUCGGA
env-190   GCUCCGUGCCCCGUCACAGACCGGCUCCGAAUCGAACGGUUCGGGGUCGUCAAAACAAGGAAUCGUGACCGAGGUCGGA
env-191   GCUCCGUGCCCCGUCACAGACCGGCUCCGAAUCGAACGGUUCGGGGUCGUCAAAACAAGGAAUCGUGACCGAGGUCGGA
Hla-1-1   GCUCCGCGCCGUGUCACGGACCGGCUCCGAAUCGAACGAUUCGGGGUCGUUAAGACAAGGGAUCGUGACCGAGGUCGGA
env-193   GCUCCGUGCCCCGUCACAGACCGGCUCCGAAUCGGACGGUUCGGGGUCGUCAAGACAAGGAAUCGUGACCGAGGUCGGA
Hac-1-1   GCUCCGCCUCUCGUCAUGGACCGGCUCCGAAUCGAGCGGUUCGGGGUCGCAUCGAGAAGGGAUCGUGACCCAGGUCGGA
Hac-1-2   GCUCCGCCUCUCGUCAUGGACCGGCUCCGAAUCGAGCGGUUCGGGGUCGCAUCGAGAAGGGAUCGUGACCCAGGUCGGA
env-196   GCUCCGUCUCUCGUCAUGGACCGGCUCAGACUCGAGCGGGUCUGGGUCGUUACGACAGGGAAUCGUGACCCAGGUCGGA
env-197   ...............................................................CGUGACCCAGGUCGGA
env-198   .................................................................UGACCCAGGUCGGA
Hsp-2-1   GCUCCGUCUCUCGUCAUGGACCGGCUCAGACUCGAGCGGGUCUGGGUCGUUACGACAGGGAAUCGUGACCCAGGUCGGA
env-199   GCUGCGCGCCGUGUCACGGACCGGCUCCGGACCACAGAGUUCGGGGUCGUUGAGACAGGGAAUCGUGACCGAGGUCGGC
Hsu-1-1   GCUUCGCGUUUCAGCAUGGACCGGCUCCGAAUCGAGCGGUUCGGGGUCGUUACGACAGGGAAUAGUGACCCAGGUCGGA
env-201   GCUCCGCGUGCGUUCACAGACCGGCUCCGGAUCUGAGGGUUCGGGGUCGUUAAGACAGGGAAUCGUGACCGAGGUCGGA
env-203   GCUCCGCGUGCGUUCACAGACCGGCUCCGGAUCUGAGGGUUCGGGGUCGUUAAGACAGGGAAUCGUGACCGAGGUCGGA
Hca-1-1   GCUCCGCGUGCGUUCACAGACCGGCUCCGGAUCUGAGGGUUCGGGGUCGUUAAGACAGGGAAUCGUGACCGAGGUCGGA
Ham-1-1   GCUCCGCGUGCGUUCACAGACCGGCUCCGGAUCUGAGGGUUCGGGGUCGUUAAGACAGGGAAUCGUGACCGAGGUCGGA
Hhi-2-1   GCUCCGCGUGCGUUCACAGACCGGCUCCGGAUCUGAGGGUUCGGGGUCGUUAAGACAGGGAAUCGUGACCGAGGUCGGA
Hsa-2-1   GCUCCGCGUGCGUUCACAGACCGGCUCCGGAUCUGAGGGUUCGGGGUCGUUAAGACAGGGAAUCGUGACCGAGGUCGGA
Har-1-1   GCUCCGCGUGCGUUCACAGACCGGCUCCGGAUCUGAGGGUUCGGGGUCGUUAAGACAGGGAAUCGUGACCGAGGUCGGA
Hva-1-1   GCUCCGCGUGCGUUCACAGACCGGCUCCGGAUCUGAGGGUUCGGGGUCGUUAAGACAGGGAAUCGUGACCGAGGUCGGA
Hja-1-1   GCUCCGCGUGCGUUCACAGACCGGCUCCGGAUCUGAGGGUUCGGGGUCGUUAAGACAGGGAAUCGUGACCGAGGUCGGA
env-204   CCCAUUGGGCGUCUCGUCGCAGACCACGGCCGGUUCGUCCGAAUCGGCCGUCCACCACGGAAUCGUGACCGAGGUCGGA
Hse-1-1   GCUUCGCGCGGCCGCUCGGACCGGCUCCGGAUCGAACGGUUCGGGGUCGUCAAGACAGGGAAUCGUGACCGAGGUCGGC
Hth-1-1   GCUUCGCGCGGCCGCUCGGACCGGCUCCGAAUCGAACGGUUCGGGGUCGUUAAGACAGGGAAUCGUGACCGAGGUCGGC
Hmo-1-1   GCUUCGCGCGGCCGCUCGGACCGGCUCCGAAUCGAACGGUUCGGGGUCGUCAAGACAGGGAAUCGUGACCGAGGUCGGC
env-206   GCUGCGUGCGACGUCAUGGACCGGCUCCGGAUCCAACGGUUCGGGGUCGUUAAGACAGGGAAUCGUGACCGAGGUCGGA
env-207   CGUGCUCCCGCCCGUGCUUCCGGAUCGGUCCGGCCCGGAUGGGUCGGAACUCCGAGACGGAAUCGUGACCGAGGUCGGA
env-209   CCUCCGAAUCGUGUCUACGACCGGCUCCGACUCGGACGAGUCGGGGUCGUUAACACAGGGAAUCGUGACUGAGGUCGGA
Nta-1-1   GCUCCGCGCCAUGUCACAGACCGGCUCCGAAUCUACCGGUUCGGGGUCGUUAAGACAAGGGAUCGUGACCGAGGUCGGA
Nas-1-1   GCUCCGCGCCGUGUCACAGACCGGCUCCGAAUCUACCGGUUCGGGGUCGUUAAGACAAGGGAUCGUGACCGAGGUCGGA
Noc-1-1   GCUCCGUGCCGUGUCACGGACCGGCUCCGAAUCGAACGAUUCGGGGUCGUCAAGACAAGGAAUCGUGACCGAGGUCGGA
Nje-1-1   GCUCCGCGCCGUGUCACAGACCGGCUCCGAAUCGGACGAUUCGGGGUCGUCAAGACAAGGAAUCGUGACCGAGGUCGGA
Nam-1-1   GCUCCGCGCCGUGUCACAGACCGGCUCCGAAUCGAACGGUUCGGGGUCGUCAAGACAAGGAAUCGUGACCGAGGUCGGA
env-210   GCUCCGCGCCAUGUCACAGACCGGCUCCGAAUCAGACGGUUCGGGGUCGUCAAGACAAGGAAUCGUGACCGAGGUCGGA
Hsp-3-1   GCUCCGUCUCUCGUCAUGGACCGGCUCAGACUCGAGCGGGUCUGGGUCGUUACGACAGGGAAUCGUGACCCAGGUCGGA
Nch-1-1   GCUCCGCGCCACGUCACAGACCGGCUCCGAAUCAGACGGUUCGGGGUCGUUAAGACAAGGGAUCGUGACCGAGGUCGGA
Nae-1-1   GCUCCGCGCCAUGUCACAGACCGGCUCCGAAUCUACCGGUUCGGGGUCGUUAAGACAAGGGAUCGUGACCGAGGUCGGA
env-214   GCUCCGCGCCGUGUCACAGACCGGCUCCGAAUCGAACGGUUCGGGGUCGUCAAGACAAGGAAUCGUGACCGAGGUCGGA
env-215   ............................GAAUCGAACGGUUCGGGGUCGUCAAGACAAGGAAUCGUGACCGAGGUCGGA
env-216   GCUGCUCCCCGCGUCACGGACCGGCUCCGAAUCGAACGAUUCGGGGUCGUUAAGACAGGGAAUCGUGACCGAGGUCGGA
Hni-1-1   GCUGCGUGCGGCGUCACGGACCGGCUCCGAAUCGAACGAUUCGGGGUCGUUAAGACAAGGGAUCGUGACCGAGGUCGGA
Has-1-1   GCUCCGCGCUUCAUCACGGACCGGCUCCGAAUCGAACGGUUCGGGGUCGUUAAGACGAGGGAUCGUGACCGAGGUCGGA
env-217   GCUCCGCGCCAUGUCACAGACCGGCUCCGAAUCUACCGGUUCGGGGUCGUCAAGACAAGGAAUCGUGACCGAGGUCGGA
Nal-2-1   GCUCCGCGCCAUGUCACAGACCGGCUCCGAAUCUACCGGUUCGGGGUCGUCAAGACAAGGAAUCGUGACCGAGGUCGGA
Npa-1-1   GCUCCGCGCCAUGUCACAGACCGGCUCCGAAUCUACCGGUUCGGGGUCGUCAAGACAAGGAAUCGUGACCGAGGUCGGA
Nga-1-1   GCUCCGCGCCAUGUCACAGACCGGCUCCGAAUCUACCGGUUCGGGGUCGUCAAGACAAGGAAUCGUGACCGAGGUCGGA
Nsp-2-1   GCUCCGCGCCAUGUCACAGACCGGCUCCGAAUCUACCGGUUCGGGGUCGUCAAGACAAGGAAUCGUGACCGAGGUCGGA
Htu-1-1   GCUCCGCGCCAUGUCACAGACCGGCUCCGAAUCUACCGGUUCGGGGUCGUCAAGACAAGGGAUCGUGACCGAGGUCGGA
Hsa-8-1   GCUCCGCGCCAUGUCACAGACCGGCUCCGAAUCUACCGGUUCGGGGUCGUCAAGACAAGGGAUCGUGACCGAGGUCGGA
Hth-2-1   GCUCCGCGCGGUGUCACAGACCGGCUCCGAAUCGAACGGUUCGGGGUCGUCAAGACAAGGGAUCGUGACCGAGGUCGGA
Nve-1-1   GCUCCGCGCCAUGUCACAGACCGGCUCCGAAUCUACCGGUUCGGGGUCGUCAAGACAAGGAAUCGUGACCGAGGUCGGA
Hli-3-1   GCUCCGCGCGGUGUCACAGACCGGCUCCGAAUCGAACGGUUCGGGGUCGUCAAGACAAGGGAUCGUGACCGAGGUCGGA
Hxa-1-1   GCUGCGCGCGGUGUCACAGACCGGCUCCGAAUCGAACGAUUCGGGGUCGUUAAGACAAGGGAUCGUGACCGAGGUCGGA
Hgo-1-1   GCUCCGCGCCGUGUCACGGACCGGCUCCGAAUCGAACGAUUCGGGGUCGUUAAGACAAGGGAUCGUGACCGAGGUCGGA
env-218   GCUCCGCGCCACGUCACAGACCGGCUCCGAAUCGAACGGUUCGGGGUCGUCAAGACAAGGGAUCGUGACCGAGGUCGGA
env-219   ...........GUCACAGACCGGCUCCGAAUCUGACGGUUCGGGGUCGUUAAGACAAGGGAUCGUGACCGAGGUCGGA

          ...............................................................................
          ...............................................................................
          •••••••••••••••••••••••••••••••••••••RR•••RRR•••••••••••••RRR•••GY•R••RRRGY•GGA
```

```
env-220    GCUUCGGCAGACGCCACAGACCGGCUCCGGAUCCAGCGAUUCGGGGUCGUCAACACAGGGAAUCGUGACCGAGGUCGGA
env-221    GCUCCGCGCCAUGUCACAGACCGGCUCCGAAUCUGACGGUUCGGGGUCGUCAAGACAAGGGAUCGUGACCGAGGUCGGA
Hsa-6-1    CCUGCGCGUCUCCCCACAGACCGGCUCCGAAUCAGACGGUUCGGGGUCGUUAACACAGGGAAUCGUGACCGAGGUCGGA
env-222    GCUCCUCGUCCGUGCACAGACCGGCUCCGGAUCGAACGGUUCGGGGUCGUUAAGACAGGGAAUCGUGACCGAGGUCGGA
env-223    GCUCCGCGUGCGUUCACAGACCGGCUCCGGAUCGAACGGUUCGGGGUCGUUAAGACAGGGAAUCGUGACCGAGGUCGGA
env-224    ..............................................UCGUUAAGACAGGGAAUCGUGACCGAGGUCGGA
env-225    UCUUAGAAUCUUACCACAGGCCGGCUCUGGAUCAGAAGAUUCAGAGUCGUUAAGACAAGGAAUAGUGACCCAGGUCGGA
env-226    UCUUAGAAUUUUACCACAGGCCGGCUCUGGAUCAGAAGAUUCAGAGUCGUUAAGACAAGGAAUAGUGACCCAGGUCGGA
env-227    UCUUAGAAUUUUACCACAGGCCGGCUCUGGAUCAGAAGAUUCCGAGUCGUUAAGACAAGGAAUAGUGACCCAGGUCGGA
env-228    CCUCCGCGUGUCGUCUACGACCGGCUCUGAGUCAACUGAUUCGGGGUCGUUAACACAGGGAAUCGUGACCGAGGUCGGA
env-231    GUCGGGGGUAUGGACAUGAAGCGUUCAUACCUUUGCAUGAGUGCGUGGUGAGCAAUCAUCACCAGCUGACGUGGUUGGA
env-232    UCGGGGGUAUGGACAUGAAGCGUUCAUACCUUUGCAUGAGUGUCGUGGUGAGCAAUCAUCACCAGCUGACGUGGUUGGA
env-233    GUCGGGGGUAUGGACAUGAAGCGUUCAUACCUUUGCAUGAGUGCGUGGUGAGCAAUCAUCACCAGCUGACGUGGUUGGA
env-237    CUCAUAGGUGAGUGACUCCCGGAGGGGAGUUACCGAGAAGGAGUUGUGGACUAAUUCACAACGGGCUGACAUAGCUGGA
env-238    CCCUCCACUCAGAACUUCUCAUCAUCCGCUGAAAAGCGAGUGGAGGGAGUUUCGAGAGGGAGUAGUCGUAAAGGUCGCA
env-240    CAUCCCACCGUUGCGCAUACCCUCUCAUACAGUUUCGAAAGAAUCGGAAUACCGCGAAGGUAUUGUGACCAAUGUCGGG
env-241    AUCAUCUGACCUCACUGAUGGAGAAUUCCGCGUUGGGGUCGUGGUCGCAAGACCAAAGAAGAGCGAGACAAAAGUCGGA
env-242    CAUGCCUGGUGUUUCCUGCAGCCUGCAGGGAAAACUCACAUGGAGUUGUGAUCCGUCACAACGGGAAGACAGAGUUGGA
env-243    CAUGCCUGGUGUUUCCUGCAGCCUGCAGGGAAAACUCACAUGGAGUUGUGAUCCGUCACAACGGGAAGACAGAGUUGGA
env-244    AUGGGUGGGCGUGCGCCCCCGUGUACGGGGCGCAGCCGAGAAGGAGUUGUGAUCAUUCACAACGGGCUGACACGGUGGA
env-245    AUAUGUGGCGGGUGCCGUCCAUGCGGGCGGCACAGCCGAGAAGGAGUUGUGUUCAUUCACAACGGGCUGACACGGUGGA
env-246    AUAUGUGGCGGGUGCCGUCCAUGCGGGCGGCACAGCCGAGAAGGAGUUGUGUUCAUUCACAACGGGCUGACACGGUGGA
env-247    AUAUGUGGCGGGUGCCGUCCAUGCGGGCGGCACAGCCGAGAAGGAGUUGUGUUCAUUCACAACGGGCUGACACGGUGGA
env-248    AUAUGUGGCGGGUGCCGUCCAUGCGGGCGGCACAGCCGAGAAGGAGUUGUGUUCAUUCACAACGGGCUGACACGGUGGA
env-249    CAGGUAGUGUCAUCCGGAGAUCCCGGGUGGCGUAACCGAGACGGAGUUGUGAUCACUCACAACGGGCUGACUUAGUAGA
env-250    CAGGUAGCGCCAUCCGGAGAUUCCGGGUGGCUUCACCGAGACGGAGUUGUGAUCAUUCACAACGGGCUGACACAGUAGA
env-251    AAGAGAGUAUCAUUCUGCUGCCACGUGUGGCGCAGGCUGAUGAGUUCUGCGAUAUCGCAGACGAGAUGACCAUUGCAGG
env-252    AAGAGAGUAUCACCCUGCUACCGAUUGUGGUGCAGGCUGGUGAGUUCUGCGAUAUCGCAGACGAGAUGACCAUUGCAGG
env-253    AAGAUAGCAUCAACCUGUAACCAAUCGCGGUGCAGGAUGGUGAGUUCUGCGAUAACGCAGACGAGAUGACCAUUGCAGG
env-254    CAUCAUAGGGUCAUACCCUCAGGAGGGAAAACAAGAAGGAGUUGUGGUCAGACACCACAACGGGCUGACAGAGGUAGGU
env-255    UUGGUGGAUCCCUGCUCAUCAUGAGCGGGGGUAUCCCGAGAAGGAGUUGUGAUCCGUCACAACAGGCUGACAGAGUGGG
env-256    GAAGGUCGAACCCUCCACAUGGAGGGCCGAGUGUCGAUAUGGAGUUGUGCUCGUCACGAACGGGAUGACAGGAGCUGGA
env-257    GCGAAGGUCGAACCCUCCACAUGGAGGGCCGAUGUCGAUAUGGAGUUGUGCUCGUCACAACGGGAUGACAGGAGCUGGA
env-258    GCGAAGGUCGAACCCUCCACAUGGAGGGCCGAUGUCGAUAUGGAGUUGUGCUCGUCACAACGGGAUGACAGGAGCUGGA
env-259    GCGAAGGUCGAACCCUCCACAUGGAGGGCCGAUGUCGAUAUGGAGUUGUGCUCGUCACAACGGGAUGACAGGAGCUGGA
env-260    GCGAAGGUCGAACCCUCCACAUGGAGGGCCGAUGUCGAUAUGGAGUUGUGCUCGUCACAACGGGAUGACAGGAGCUGGA
env-261    GCGAAGGUCGAACCCUCCUCCCGGAGGGCCGAUGUCGAUACGGAGUUGUGCUCGUCACAACGGGAUGACAGGAGCUGGA
env-265    CACAAGGUCGAACCCUCCUCACGGAGGGCCGAUGUCGAUAUGGAGUUGUGCUCGUCACAACGGGAUGACAGGAGCUGGA
env-266    GUGGAGAUUGAACCCUCCUUACGGAGGGCAGGUGUCGAUAUGGAGUUGUGCUUGGCACAACGGGACGACAGGAGCUGGA
env-268    UGGAGAUUGAACCCUCCUUACGGAGGGCAGGUAGUCGAUAUGGAGUUGUGCUUGGCACAACGGGACGACAGGAGCUGGA
env-269    GUGGAGAUUGAACCCUCCUUACGGAGGGCAGGUGUCGAUAUGGAGUUGUGCUUGGCACAACGGGACGACAGGAGCUGGA
env-270    UGGAGAUUGAACCCUCCUUACGGAGGGACAGGUGUCGAUAUGGAGUUGUGCUUGGCACAACGGGACGACAGGAGCUGGA
env-271    UCGAAGGUCGAACCCUCCAGAUGGAGGGCGAUAUCGAUUGAGGAGUUGUGCUCGUCACAACGGGAUGACAGGAGCUGGA
env-272    UCGAAGGUCGAACCCUCCAGAUGGAGGGCGAUAUCGAUUGAGGAGUUGUGCUCGUCACAACGGGAUGACAGGAGCUGGA
env-274    UCGAAGGUCGAACCCUCCAGAUGGAGGGCGAUAUCGAUUGAGGAGUUGUGCUCGUCACAACGGGAUGACAGGAGCUGGA
env-275    ......................................UGAGGAGUUGUGCUCGUCACAACGGGAUGACAGGAGCUGGA
env-277    UCGAAGGUCGAACCCUCCAGAUGGAGGGCGAUAUCGAUUGAGGAGUUGUGCUCGUCACAACGGGAUGACAGGAGCUGGA
env-278    GCGAAGGUUGAACCCUCCAGAUGGAGGGCGAUGUCGAUUAAGGAGUUGUGCUCGUCACAACGGGAUGACAGGAGCUGGA
env-292    CAAAGGUCGAACCCUCCACAUGGAGGGCCGAUGUCGAUUGAGGAGUUGUGCUCGUCACAACGGGAUGACAGGAGCUGGA
env-293    ...............................GAUGUCGAUAGGAGUUGUGCUCGUCACAACGGGAUGACAGGAGCUGGA
env-295    CACAAAGGUCGAACCCUCCACAUGGAGGGCCGAUGUCGAUAGGAGUUGUGCUCGUCACAACGGGAUGACAGGAGCUGGA
env-296    CACAAAGGUCGAACCCUCCACAUGGAGGGCCGAUGUCGAUAGGAGUUGUGCUCGUCACAACGGGAUGACAGGAGCUGGA
Mbo-1-1    CGCAAAGGUCGAACCCUCCACAUGGAGGGCCGAUGUCGAUAGGAGUUGUGCUCGUCACAACGGGAUGACAGGAGCUGGA
env-297    CACAAAGGUCGAACCCUCCACAUGGAGGGCCGAUGUCGAUAGGAGUUGUGCUCGUCACAACGGGAUGACAGGAGCUGGA
env-298    ........................................AGGAGUUGUGCUCGUCACAACGGGAUGACAGGAGCUGGA
env-303    CAAAGGUCGAACCCUCCACAUGGAGGGCCGAUGUCGAUUGAGGAGUUGUGCUCGUCACAACGGGAUGACAGGAGCUGGA
env-304    CGCAAAGGUCGAACCCUCCACAUGGAGGGCCGAUGUCGAUAGGAGUUGUGCUCGUCACAACGGGAUGACAGGAGCUGGA
env-305    UCGAAGGUCGAACCCUCCAGAUGGAGGGCGAUAUCGAUUGAGGAGUUGUGCUCGUCACAACGGGAUGACAGGAGCUGGA
env-306    ..............................AUAUCGAUUGAGGAGUUGUGCUCGUCACAACGGGAUGACAGGAGCUGGA
Mma-1-1    GACAAGGUCGAACCCUCCUCACGGAGGGCCGAUGUCGAUAUGGAGUUGUGCUCGUCACAACGGGAUGACAGGAGCUGGA
Mse-1-1    CACAAGGUCGAACCCUCCUCACGGAGGGCCGAUGUCGAUAUGGAGUUGUGCUCGUCACAACGGGAUGACAGGAGCUGGA
Msp-1-1    GACAAGGUCGAACCCUCCUCACGGAGGGCCGAUGUCGAUAUGGAGUUGUGCUCGUCACAACGGGAUGACAGGAGCUGGA
env-310    AGCAGGUCGAACCCUCCUUGCGGAGGGGCGACGUCGAUUGAGGAGUUGUGCUCGUCACAACGGGAUGACAGGAGCUGGA
env-315    AGCAGGUCGAACCCUCCUUGCGGAGGGGCGACGUCGAUUGAGGAGUUGUGCUCGUCACAACGGGAUGACAGGAGCUGGA
env-316    ....GGUCGAACCCUCCUUGCGGAGGGGCGACGUCGAUUGAGGAGUUGUGCUCGUCACAACGGGAUGACAGGAGCUGGA
env-317    .......................................GAGGAGUUGUGCUCGUCACAACGGGAUGACAGGAGCUGGA
env-318    AGCAGGUCGAACCCUCCUUGCGGAGGGGCGACGUCGAUUGAGGAGUUGUGCUCGUCACAACGGGAUGACAGGAGCUGGA
env-320    AGCAGGUCGAACCCUCCUUGCGGAGGGGCGACGUCGAUUGAGGAGUUGUGCUCGUCACAACGGGAUGACAGGAGCUGGA
env-321    .........AACCCUCCUUGCGGAGGGGCGACGUCGAUUGAGGAGUUGUGCUCGUCACAACGGGAUGACAGGAGCUGGA
env-322    AGCAGGUCGAACCCUCCUUGCGGAGGGGCGACGUCGAUUGAGGAGUUGUGCUCGUCACAACGGGAUGACAGGAGCUGGA
Mpa-1-1    AUACGGUCGCACCCGACCCUGAUGGUCGGGAAUGCCGAGAGGAGUUGUGAGCGAUCACAACGGGCUGACAGAAGCUGGA
env-323    ..............................AAUGCCGAGAGGAGUUGUGAGCGAUCACAACGGGCUGACAGAAGCUGGA
env-324    UUGAUUGGUGAACUCAUCAGUAACCUGAUGAGAUCCCGAGAAGGAGUUGUGAUUCGUCACAACAGGCUGACAGAGUGGG
env-325    CUCGAUUGGUGAACCCUCCUUCAGGAGGGGUAUACCGAGAAGGAGUUGUGAUCCGUCACAACAGGCUGACAGAGUAGGA
env-326    CUCGAUUGGUGAACCCUCCUUCAGGAGGGGUAUACCGAGAAGGAGUUGUGAUCCGUCACAACAGGCUGACAGAGUAGGA
env-331    CUCGAUUGGUGAACCCUCCUUCAGGAGGGGUAUACCGAGAAGGAGUUGUGAUCCGUCACAACAGGCUGACAGAGUAGGA
env-332    CUCGAUUGGUGAACCCUCCUUCAGGAGGGGUAUACCGAGAAGGAGUUGUGAUCCGUCACAACAGGCUGACAGAGUAGGA
Mfo-1-1    CGAUUGGUGGAUCCCCGUAUAACAACGGGGGUAUCCCGAGAAGGAGUUGUGAUUCGUCACAACAGGCUGACAGAGUGGG
env-333    CGAUUGGUGACCCCCGCAGACAGUGCGGGGGAUCCCGAGAAGGAGUUGUGAUCCGUCACAACGGGCUGACAGAGUGGGA
env-334    GAUUGGUGACCCCUGCAGAAAUAUGCGGGGGAUCCCGAGAAGGAGUUGUGAUCCGUCACAACGGGCUGACAAAGUGGGA
env-335    GAUUGGUGACCCUCGCAGGAAUCUGCGGGGGAUCCCGAGAAGGAGUUGUGAUCCGUCACAACGGGCUGACAAAGUGGGA
env-337    GAUUGGUGACCCCCGCAGAAAUAUGCGGGGGAUCCCGAGAAGGAGUUGUGAUCCGUCACAACGGGCUGACAAAGUGGGA
env-338    CGAUUGGUGACCCCCGCAGACAUUGCGGGGGAUCCCGAGAAGGAGUUGUGAUCCGUCACAACGGGCUGACAGAGUGGGA
env-339    CGAUUGGUGACCCCCGCACACAAUGCGGGGGAUCCCGAGAAGGAGUUGUGAUCCGUCACAACGGGCUGACAGAGUGGGA

           ...............................................................................
           ...............................................................................
           ●●●●●●●●●●●●●●●●●●●●●●●●●●●●●●●●●●●●●RR●●●RRR●●●●●●●●●●●●●RRR●●●GY●R●●RRRGY●GGA
```

```
CMe-1-1   CGAUUGGUGACCUCCUCAUUUCGUGAGGAGGAUCCCGAGAAGGAGUUGUGAUCCGUCACAACGGGCUGACAGAGUUGGA
env-340   GAUUGGUGAACCUCGCAGAAAUGCGGGGGGCAACCCGAGAAGGAGUUGUGAUUCGUCACAACAGGCUGACAGAGUAGGA
env-341   GAUUGGUGAACCCCGCAUACAGGUGGGGGUCAACCCGAGAAGGAGUUGUGUAUCGUCACAACAGGCUGACUGAGUGGGA
env-349   .......................................................................GAGUGGGA
env-350   CGAUUGGUGAACUCCGCGUUAUUGCGGGGGAAUCCCGAGUAGGAGUUGUGAUUAAUCACAACAGGCUGACUGAGUAGGA
env-351   GAUUGGUGAACCUCGCAGCAAUGCGGGGGGCAACCCGAGAAGGAGUUGUGAUUCGUCACAACAGGCUGACAGAGUAGGA
env-352   CCUUGAUUGGUGAACUCAUCCUAACGGAUGGGAUCCCGAGAAGGAGUUGUGAUUCGUCACAACAGGCUGACAGAGUGGG
env-353   GAUUGGUGAACUCCGCUUAAUUGCGGGGGGAAACCCGAGAAGGAGUUGUGACUCGUCACAACAGGUUGACUGAGUGGGA
env-354   GAUUGGUGAACUCCGCUUAAUUGCGGGGGGAAACCCGAGAAGGAGUUGUGACUCGUCACAACAGGUUGACUGAGUGGGA
env-355   GAUUGGUGAACUCCGUUUAAUUGCGGGGGGAAACCCGAGAAGGAGUUGUGACUCGUCACAACAGGUUGACUGAGUGGGA
env-356   CGAUUGGUGGACUCCGCUUAAUUGCGGGGGAAACCCGAGAAGGAGUUGUGACUCGUCACAACAGGUUGACUGAGUGGGA
env-357   GAUUGGUGAACUCCGCUUCAUNGCGGGNGGAAACCCGAGAAGGAGUUGUGACUCGUCACAACAGGCUGACUAAGUGGGA
env-358   CGAUUGGUGAACUCCGCAUCAGAGCGGAGGAAACCCGAGAAGGAGUUGUGACUCGUCACAACAGGCUGACUAAGUGGGA
env-359   CGAUUGGUGAACUCCGCAUCAGAGCGGAGGAAACCCGAGAAGGAGUUGUGACUCGUCACAACAGGCUGACUAAGUGGGA
env-360   CGAUUGGUGAACUCCGCAUCAGAGCGGAGGAAACCCGAGAAGGAGUUGUGACUCGUCACAACAGGCUGACUAAGUGGGA
env-361   CGAUUGGUGAACUCCGCAUCAGAGCGGAGGAAACCCGAGAAGGAGUUGUGACUCGUCACAACAGGCUGACUAAGUGGGA
env-370   GAUUGGUGAACUCCGCUUCAUUGCGGGGGGAAACCCGAGAAGGAGUUGUGACUCGUCACAACAGGCUGACUAAGUGGGA
env-371   ............................................GUUGUGACUCGUCACAACAGGCUGACUAAGUGGGA
env-372   ...............................................................GGCUGACUGAGUGGGA
env-373   GAUUGGUGAACCCCGCAUACAAGCGGGGGUCAACCCGAGAAGGAGUUGUGUAUCGUCACAACAGGCUGACUGAGUGGGA
env-374   CGAUUGGUGAACCCCGCCUCAUUGCGGGGGAAACCCGAGAAGGAGUUGUGAUUCGUCACAACAGGCUGACUACGUGGGA
env-375   ....UCAAUUGGUGCCCUCUCCGAGAAGGUGAGGACCGAGAAGGAGUUGUGAUUCGUCACAACAGGCUGACAGAGUGGG
env-376   ........................................................UCACAACGGGCUGACAGAGUGGG
env-377   CCUCAAUUGGUGCCCUCUCCGCAAGGUGAGGUAUACCGAGAAGGAGUUGUGAUUCGUCACAACAGGCUGACAGAGUGGG
env-378   UCGAUUGGUGCCCCUCUUAAAAAUGGGAGGGUAAACCGAGAAGGAGUUGUGAUUCGUCACAACAGGCUGACAGAGUGGG
env-379   .......................................GAAGGAGUUGUGAUUCGUCACAACAGGCUGACUGAGUGGG
env-380   CUCGAUUGGUGACCCCUCCUUAUGGAGGGCUGAACCGAGAAGGAGUUGUGAUUCGUCACAACAGGCUGACAGAGUAGGA
env-381   CUCGAUUGGUGACCCCUCCUUAUGGAGGGCUGAACCGAGAAGGAGUUGUGAUUCGUCACAACAGGCUGACAGAGUAGGA
env-382   CUCGAUUGGUGACCCCUCCUUAUGGAGGGCUGAACCGAGAAGGAGUUGUGAUUCGUCACAACAGGCUGACAGAGUAGGA
env-383   CUCGAUUGGUGACCCCUCCUUAUGGGGGGCUGAACCGAGAAGGAGUUGUGAUNCGUCACAACAGGCUGACAGAGUAGGA
env-384   CUCGAUUGGUGACCCCUCCUUAUGGAGGGCUGAACCGAGAAGGAGUUGUGAUUCGUCACAACAGGCUGACAGAGUAGGA
env-385   ..................CUUACGGAGGGCUAAACCGAGAAGGAGUUGUGAUCCGUCACAACGGGCUGACAGAGUAGGA
env-386   CUCGAUUGGUGAACCCUCCUUACGGAGGGCUAAACCGAGAAGGAGUUGUGAUCCGUCACAACGGGCUGACAGAGUAGGA
env-387   CUCGAUUGGUGAACCCUCCUUACGGAGGGCUAAACCGAGAAGGAGUUGUGAUCCGUCACAACGGGCUGACAGAGUAGGA
env-389   CUCGAUUGGUGACCCCUCCUUACGGAGGGUUAAACCGAGAAGGAGUUGUGAUUCGUCACAACAGGCUGACAGAGUAGGA
env-390   CGAUUGGUGAACCCCGUGUAAACACGGGGUCAACCCGAGAAGGAGUUGUGAUUCGUCACAACAGGCUGACUGAGUGGGA
env-392   CCCGAUUGGUGGAUCCCCCGUAUGGGGGGGUAUCCCGAGAAGGAGUUGUGAUUCGUCACAACAGGCUGACAGAGUGGG
env-395   ...................AAAAAGGAUGGGUAUCCCGAGAAGGAGUUGUGAUUCGUCACAACAGGCUGACCGAGUGGG
env-399   CUCGAUUGGUGAACCUUCCUUUAGGAAGGGUAUACCGAGAAAGGAGUUGUGAUUCGUCACAACGGGCUGACAGAGUGGG
env-400   .......................................................UCACAACAGGCUGACAGAGUAGGA
env-402   ........................................................UCACAACGGGCUGACAGAGUGGG
env-403   CUCGAUUGGUGCCUCUCCCCACAAGGAGGGGUAAACCGAGAAGGAGUUGUGAUUCGUCACAACGGGCUGACAGAGUGGG
env-405   ..............CCCGCCUCACCGCGGGUAUACCGAGAAGGAGUUGUGCAUCGUCACAACAGGCUGACAGAGUGGGA
env-406   CUUCGAUUGGUGAACCCGCCUCACCGCGGGUAUACCGAGAAGGAGUUGUGCAUCGUCACAACAGGCUGACAGAGUGGGA
env-407   GAUUGGUGAACUCCGCAGGAAUACGGAGGGCAACCCGAGAAGGAGUUGUGAUUCGUCACAACAGGCUGACAGAGUUGGA
Mli-2-1   AUGUGGUUGCCCCGGGUAUUUAUGCCGGGAUUUGCCGAGAUUAAUGUCCGGUUACGCCGGACGGGCUGACGAUGUAGAA
Mpe-1-1   CAACGUAAUCGCUGUCCGGAGAUUGCCGAGAAAGGACAAAAAGCCUAUGAAGGCUUUCUGUUCGGCUGACAAAGUAGGA
Mpa-2-1   CAACGUAAUCGCUGUCCGGAGAUUGCCGAGAAAGGACAAAAAGCCUAUGAAGGCUUUCUGUUCGGCUGACAAAGUAGGA
env-408   ACAACAUUUUUGCUGUCCGGAGAUUGCCGAGAAUGGACUGAAAGCCGCAGAGGCUGCAAGUCCGGCUGACAAAGUAGGA
Mmo-1-1   UUUCCGGUGAUUGCCGAGAUAGUAGGUCACACAAAAUUCCGACGGGAAUUUACAAGUGUGACGGGCUGACAACGUAGGA
env-409   UUUCCGGUGAUUGCCGAGAUAGUAGGUCACACAAAAUUCCGACGGGAAUUUACAAGUGUGACGGGCUGACAACGUAGGA
env-410   GAUCAACCGGCUCUGCUACGUAAGGUAGUAUAGCGGGUGAAUGCUUGCGACAAUUCGCAGGCCGACGACCACUAGCAGG
Mco-1-1   CGGAGUCAUCCCGCCACUGAGGAUACCAUCUCAUACUAGCGAAGAGGAAUACCGAGAAGGUAUUGUGACCAAAGUCGGC
env-413   GAACCUCAACGAAGGAGAAUUCCGGGUUGGGGUAGUGGUCCCCGGACCGCAAAAAAAGAAACGCGAGAAUAAAGUCGGA
env-414   GAACCUCAACGAAGGAGAAUUCCGGGUUGGGGUAGUGGUCCCGGGACCGCAAAAAAAGAAAAGCGAGAAAAAAAGUCGGA
env-415   GAACCUCAACGAAGGAGAAUUCCGGGUUGGGGUAGUGGUCCCAGGACCGCAAAAAAAGAAAAGCGAGAAAAAAAGUCGGA
env-416   GAACCUCAACGAAGGAGAAUUCCGGGUUGGGGUAGUGGUCCCGGGACCACACAAAAAGAAAAGCGAGAAAAAAAGUCGGA
env-418   ACACAUCAACGUAGGAGAAUACCGCGUUGGAGUAGUGGUUUCACAACCAAAAAAAAAAAUUAGCGAGAAAAAAGUAGGA
env-423   ACACAUCAACGUUGGAGAAUACCGCGUUGGAGUAGUGGUUCCACAACCAAAAAGAAAAAACAGCGAGAAAGAAGUAGGA
env-425   ACACAUCAACGUAGGAGAAUACCGCGUUGGAGUAGUGGUUAGACAACCAGGAAAAAAAAACAGCGAGAAAAAAAGUAGGA
env-428   UCAUCAUCCCAUAAACAAAAUGACAUCAGCGCUCAAGAUCGGCGAGUACCGAGUCGGAGCAGUCGUCGAAAGCGUCGGA
env-429   UCAUCAUCCCAUAAACAAAAUGACAUCAGCGCUCAAGAUCGGCGAGUACCGAGUCGGAGCAGUCGUCGAAAGCGUCGGA
Msh-1-1   GCCACACCAUCCGACGACAUCGGACACAUCGAAAACGAACAUCGGAGACAUCCGAACAGGUGUAGUGACCAAGGUAGGA
env-430   UCAUCAUCCCAUCAACUCAAAAUCAGAAUCGCUCAAGAUUGGCGAGUUUCGCGUCGGAGCAGUCGUUGAAAGCGUCGGA
env-433   ACAUCAUCCCAUAAACUCAAAAUCAGAAUCGCUCAAGAUUGGCGAGUUUCGCGUCGGAGCAGUCGUUGAAAGCGUCGGA
env-434   .........................................................AGCAGUCGUUGAAAGCGUCGGA
env-435   UCAUCAUCCCAUCAACUCAAAAUCAGAAUCGCUCAAGAUUGGCGAGUUUCGCGUCGGAGCAGUCGUUGAAAGCGUCGGA
env-436   CAUCAUCCCAUCAACUCAAAAAUCAGAAUCGCUCAAGAUUGGCGAGUUUCGCGUCGGAGCAGUCGUUGAAAGCGUCGGA
env-437   UCAUCAUCCCAUCAACUCAAAAUCAGAAUCGCUCAAGAUUGGCGAGUUUCGCGUCGGAGCAGUCGUUGAAAGCGUCGGA
env-439   UCAUCAUCCCAUCAACUCAAAAUCAGAAUCGCUCAAGAUUGGCGAGUUUCGCGUCGGAGCAGUCGUUGAAAGCGUCGGA
env-440   NCAUCAUCCCAUNAACUCAAAAUCAGAAUCGCUCAAGAUUGGCGAGUUUCGCGUCGGAGCAGUCGUUGAAAGCGUCGGA
env-441   UCAUCAUCCCAUCAACUCAAAAUCAGAAUCGCUCAAGAUUGGCGAGUUUCGCGUCGGAGCAGUCGUUGAAAGCGUCGGA
env-442   ....CAUCCCAUCAACUCAAAAUCAGAAUCGCUCAAGAUUGGCGAGUUUCGCGUCGGAGCAGUCGUUGAAAGCGUCGGA
env-443   UCAUCAUCCCACAAACUCAAAAUCAGAAUCGCUCAAGAUUGGCGAGUUUCGCGUCGGAGUUGUCGUUGAAAGCGUCGGA
env-444   UAAUCAUCCGUUGAACUCAAAAACAUCAGCGCUCAAAAUUGGCGAGUACCGAGUCGGAGCAGUCGUCGAAAGCGUCGGA
env-445   UCAUCAUCCCAUAAACUCAAGAUCAGAAUCGCUCAAGAUUGGCGAGUUUCGCGUCGGAGCAGUCGUUGAAAGCGUCGGA
env-446   UCAUCAUCCAUUAAACUCAAGAUCAGAAUCGCUCAAGAUUGGCGAGUUUCGCGUCGGAGCAGUCGUUGAAAGCGUCGGA
env-448   CCAUCAUCCUGUGAACUCAGAGAGCACAACGCUCAGAGUUGGCGAGUUCAGAGUCGGAGCAGUCGUCGAAAGCGUCGGA
env-449   UCAUCAUCCAAUCAACUCAAAGACGUCAGCGCUCAAGAUCGGCGAGUACCGAGUCGGAGCAGUCGUCGAAAGCGUCGGA
env-450   .................CAAAAUCAGAAUCGCUCAAGAUUGGCGAGUUUCGCGUCGGAGCAGUCGUUGAAAGCGUCGGA
env-451   UCAUCAUCCAAUAAACUCAAAAUCGGAAUCGCUCAAGAUUGGCGAGUUUCGCGUCGGAGCAGUCGUUGAAAGCGUCGGA
env-452   UCAUCAUCCAAUAAACUCAAAAUCGGAAUCGCUCAAGAUUGGCGAGUUUCGCGUCGGAGCAGUCGUUGAAAGCGUCGGA
env-455   UCAUCAUCCCAUAAACUCAAAAUCGGAAUCGCUCAAGAUUGGCGAGUUUCGCGUCGGAGCAGUCGUUGAAAGCGUCGGA
env-456   ...............CUCAAAAUCAGAAUCGCUCAAGAUUGGCGAGUUUCGCGUCGGAGCAGUCGUUGAAAGCGUCGGA

          ...............................................................................
          ...............................................................................
          •••••••••••••••••••••••••••••••••••••RR•••RRR•••••••••••••RRR•••GY•R••RRGY•GGA
```

```
env-460   UCAUCAUCCCAUCAACUCAAAAUCAGAAGCUCUCAAGAUCGGCGAGUUUCGCGUCGGAGCAGUCGUUGAAAGCGUCGGA
env-462   UCAUCAUCCCAUCAACUCAAAAUCAGAAGCUCUCAAGAUCGGCGAGUUUCGCGUCGGAGCAGUCGUUGAAAGCGUCGGA
env-464   .................................................UCGCGUCGGAGCAGUCGUUGAAAGCGUCGGA
env-465   UCAUCAUCCCAUAAACUCAAAAGCAGAAGCUCUCAAGAUCGGCGAGUUUCGCGUCGGAGCAGUCGUUGAAAGCGUCGGA
env-466   UCAUCAUCCAUUAAACUCAAAAUCAGAAUCGCUCAAGAUUGGCGAGUUUCGCGUCGGAGCAGUCGUUGAAAGCGUCGGA
env-467   UCAUCAUCCCACAAGCUCAAAGGUUUCAACGCUCAAGGUCGGCGAGUUUCGAGUAGGUUUGGUAGUCGAUGAAGUGGGA
env-468   UCAUCAUCCCACAAGCUCAAAGGUUUCAACGCUCAAGGUCGGCGAGUUUCGAGUAGGUUUGGUAGUCGACGAAGUGGGA
env-469   UCAUCAUCCCACAAGCUCAAAGGUUUCAACGCUCAAGGUCGGCGAGUUUCGAGUAGGUUUGGUAGUCGACGAAGUGGGA
env-471   UCAUCAUCCCACAAGCUCAAAGGUUUCAACGCUCAAGGUCGGCGAGUUUCGAGUAGGUUUGGUAGUCGACGAAGUGGGA
env-472   UCAUCAUCCCACAAGCUCAAAGGUUUCAACGCUCAAGGUCGGCGAGUUUCGAGUAGGUUUGGUAGUCGACGAAGUGGGA
env-474   UCAUCAUCCCACAAGCUCAAAGGUUUCAACGCUCAAGGUCGGCGAGUAUCGAGUAGGUUUAGUAGUCGAUGAAGUGGGA
env-475   UCAUCAUCCCAAAAGCUCAAAGGUUUCAACGCUCAAGGUCGGCGAGUAUCGAGUAGGUUUAGUAGUCGACGAAGUGGGA
Mha-1-1   UCAUCAUCCCAAAAGCUCAAAGGUUUCAACGCUCAAGGUCGGCGAGUAUCGAGUAGGUUUAGUAGUCGACGAAGUGGGA
env-476   UCAUCAUCCCACAAGCUCAAAGGUUUCAACGCUCAAGGUCGGCGAGUAUCGAGUAGGUUUAGUAGUCGACGAAGUGGGA
env-477   UCAUCAUCCCACAAGCUCAAAGGUUUCAACGCUCAAGGUCGGCGAGUUCAGAGUAGGUUUAGUAGUCGAUGAAGUGGGA
env-478   UCAUCAUCCGCUGAACUCAAAGUCGUCAGCGCUCAAGGUCGGCGAGUUCAGAGUCGGAGCUGUAGUCGAAAGCGUCGGA
env-479   ACGCACAGCACAUCAUCCCAAGGUUUCAACGCUCGAAGUCGGCGAGUUCAGAGUAGGAUUGGUUGUUGACGACGUCGGA
env-480   ....................GAGGUCACAACGCUCGAAGUCGGCGAGUUCAGAGUAGGAUUGGUUGUUGACGACGUCGGA
env-481   CCCCUUACGAACGCCACAUCAUCCAACAAGCUCAUCCGGGGUCGGAGAAAUCCGUCCCGGCGUGGUGACAAAAGUGGAG
env-482   CCCUCACCAUCCGGUAGGUUCCAGUGUAGAAAGCCUUUCCACUGGUGAAAUCAGGGAAGGCGUGGUAAGAAAGGUAGGG
env-483   .........................................CAAGAUAACGUCCGUGAUGGAAUUGUGAUAGACGUCGGA
env-484   GGUUGUCCCCCCUUUAAAAACCCGUCAUCAUGUGCACGAGCAAGAUAACGUCCGUGAUGGAAUUGUGAUAGACGUCGGA
env-485   GGUUGUCCCCCCUUUAAAAACCCGUCAUCAUGUGCACGAGCAAGAUAACGUCCGUGAUGGAAUUGUGAUAGACGUCGGA
env-486   GGUUGUCCCCCCUUUAAAAACCCGUCAUCAUGUGCACGAGCAAGAUAACGUCCGUGAUGGAAUUGUGAUAGACGUCGGA
env-492   GGUUGUCCCCCCUUUAAAAACCCGUCAUCAUGUGCACGAGCAAGAUAACGUCCGUGAUGGAAUUGUGAUAGACGUCGGA
env-495   GGUUGUCCCCCCGUUGAAGACCCGCCAUCAUGUGCACGAGUAUGAUAACGUUCGUGAUGGAAUUGUGAUAGACGUCGGA
env-504   GGUGGUUCCCCCGUUGAAGACCCCUCAUCACGCGCGGCCGCGUGAUGACAUUCGAGAAGGAAUUGUGAUAGACGUCGGA
env-505   UGACGGUCAAAUCGCAGAAAAAUAAAAUUUCUGCGGAGUGACAGGAUUGCGUUCAUCGCAAUGGGCUGAUAUAGUGGGA
env-508   GUCGGGGGCACGGGUCCCGGAUGGCCCGUGCCUUUGUAUGAGUAUGUGGUGAGCAAUCACCACCGGACGACACGGUGGA
env-509   GUCAGGGACAUGCGUUCCAGGCAACGCGUGUCUUUGCGUGAGAAUGUGGUGCAUGGUCACCACCGGCUGACACGGUGUA
Mta-1-1   GGUCAGGGACACGGGUUCCAGGAACGCGUGUCUUUGCGUGAGAAUGUGGUGAAUGGUCACCACCGGCUGACAGGGUGUC
env-511   GGUCAGGGACACGGGUUCCAGGAACGCGUGUCUUUGCGUGAGAAUGUGGUGAAUGGUCACCACCGGCUGACAGGGUGUC
env-512   GGUCAGGGACACGGGUUCCAGGAACGCGUGUCUUUGCGUGAGAAUGUGGUGAAUGGUCACCACCGGCUGACAGGGUGUC
Hbo-1-1   CCUGCUCGUUUCGUCUACGACCGGCUCCGAAUCAGACGAUUCGGGGUCGUUACAACAGGGAAUCGUGACCGAGGUCGGA
Ngr-1-1   GCUCCGCGCCACGUCACAGACCGGCUCCGAAUCGAACGGUUCGGGGUCGACAAGACAAGGGAUCGUGACCGAGGUCGGA
Hje-1-1   GCUUCGCGCCGUGUCACGGACCGGCUCCGGAUCGAACGGUUCGGGGUCGUUAAGACAGGGACUCGUGACCGAGGUCGGA
Nma-1-1   ACUCCGCGCCACGUCACAGACCGGCUCCGAAUCAGACGGUUCGGGGUCGUUAAGACAAGGGAUCGUGACCGAGGUCGGA
Hal-1-1   CGUUCUCGUCGCGUCUACGACCGCGGACGACUCCGACGAGUCGCCCGCGUUGCGAGAAGGAAUCGUGACCGAGGUCGGA
Nti-1-1   GCUCCGCGCCAUGUCACAGACCGGCUCCGAAUCUGACGGUUCGGGGUCGUCAAGACAAGGAAUCGUGACCGAGGUCGGA
Npe-1-1   GCUCCGCGCGGUGUCACAGACCGGCUCCGAAUCGAACGGUUCGGGGUCGUCAAGACAAGGGAUCGUGACCGAGGUCGGA
Hti-1-1   GCUCCGCGUUGCGACCCAGACCGGCUCCGGAUCGGACGAUUCGGGGUCGUUAAGACAGGGAAUCGUGACCGAGGUCGGA
          ................................................................................
          ................................................................................
          ●●●●●●●●●●●●●●●●●●●●●●●●●●●●●●●●●●●●●●RR●●●RRR●●●●●●●●●●●●●RRR●●●GY●R●●RRRGY●GGA
```

alignment positions 80 · · · 259

```
Mli-1-1   UCUGAU...CCGUGU.AAU..GGGCCUCC...GCC......CGUGCA.........GGC.GAGCCCGG....CCGGC.........................CCCGU.....................................G.CCGGCAU...GCACAACG.......AUCAGGA
env-1     UGUGAU...GAUAGC.AA...GGGUUGAA..AGCCA.....UCUG..........UGGCUUUGCCCGUG...UCUGCA.....UGAAGG.............AAAUA.....................CCU.CU............G.UAGACCU...GCUGAAAC.......GUCACGC
env-2     UCUGAC.UACUGUGCUA....GGGUGGAU...AUUG.....GCCU..........UGAU.GAGCCAGC....CCUCC......UCGUUGAAAAU........GAAGAA..............GUU...AGCGA............G.GAAGAAC...GCAUAACU.......GUCAGAA
env-3     UCUGAU.UACUGCGC.AA...GGGUGUAUAUAGGUC.....UUGACC........GACC.A.GCCCUU....CUCGC......UGAUU..............GUGAAG.....................GAUUA...........AG.CGAGAAACA.GCGCAUAACU.....GUCAGGA
env-4     GCUGAG.GCAGCAGC.A....GGGUGGAC...GUC......AUG...........GAC.G.GCCCU.....GCCCG.........................CACGA......................................C.GGGCGCU...GCUGCA.........UUCAGGA
env-5     GCUGAG.GCAGCAGC.A....GGGUGGAC...GUC......AUG...........GAC.G.GCCCU.....GCCCG.........................CACGA......................................C.GGGCGCU...GCUGCA.........UUCAGGA
env-6     UGUGAU..GACGGC..AA...GGGUGGAC...G........AUG.............C.G.GCCCGA....AUCC..........................UUAACC........................................GGAUAA.....GCCGGAGA......AUCACGC
env-8     UGUGAU...GUCGGC.AA...GGGUGGAC...G........AUG.............C.G.GCCCGA....GUCCU.........................GGUAAC....................................A.GGAUAA....GCCGAAAG.......AUCACGC
env-10    UGUGAU...GUCGGC.AA...GGGUGGAC...G........AUG.............C.G.GCCCGG....AUCCU.....G...................GUUU..............................C.............A.GGAUAA....GCCGAAGU.......AUCACGC
env-11    UGUGAU...GUCGGC.AA...GGGUGGAC...G........AUG.............C.G.GCCCGG....AUCCU.....G...................GUUU..............................C.............A.GGAUAA....GCCGAAGU.......AUCACGC
env-13    UCUGAA.AACGGCAC.AAC..GGGUGGAC...GUAAUCU..UUGA........AGA.UAC.GAGCCCUC....AUCCGG.....CGGA...............UCAA........................UCCG..........GC.GGAUAU....GUGCCCGG.......UUCAGGA
env-14    UCUGAC.GAAUCCGC.G....GGGUGGAC...GCACG....AAC..........CGUGC.GAGCCCAG....UUCUG......GUUAU..............AUCGG.......................AUA.C.............C.GGAA.....GCGGAGAUAGUAACCGUCAGGA
env-15    UCUGAU.GACCGCGUCC....GGGUCAAU...UGUG.....AAC...........UGCA.ACACCCAA....UUUCU......CUCCAUGUACCAGAC....UCGAU...........GUCUG.UGC...GAG...........AA.GGGAC.....ACGCGUUGCC.....AUCAGAA
Hwa-1-1   CCUGAU.GACCGCGUUC....GGGUCAAU...UGUG.....AAU...........UGCA.ACACCCGG....UCUCU......CUCC.AU..GUGCC.....AGAAUC...........GGUAUCUGUACGAG...........AG.GGGAC.....ACGCGUUGCU.....AUCAGGA
env-17    CCUGAU.GACCGCGUUC....GGGUCAAU...UGUG.....AAU...........UGCA.ACACCCGG....UCUCU......CUC...CAUGUGCC.....AGAAUC...........GGUAUCUGUACGAG...........AG.GGGAC.....ACGCGUUGCU.....AUCAGGA
env-23    CCUGAU.GAUCGCGUCC....GGGUCAAU...UGCGC....CCUC.........GCGCA.ACACCCGA....UCUCU......UUGCUCGUGCCCGACG..GCC............UGUCGG.....UCCGA.........AAGG.GAGA......GCGCGUCACA.....CUCAGGG
env-24    ACUGAC.GGACGCGUCU....GGGUCAAU...UGCG.....GAA...........CGCA.ACACCCGG....UCGCCC.....UUCGC..GUCCCC......GAUGAG...........GGGGAUACGGGAG............G.GGGA......ACGCGUCACC.....CUCAGAG
env-25    GCUGAU.GGGCGCGUCC....GGGUCAAU...UGCG.....GUC...........UGCA.ACACCCAA....UCUCG......CUA.CCGG.UGCCGG....AUGGA...........CUGGCAGCCGGCGAG............C.GUGA......GCGCGUCACC.....GUCAGGG
env-27    UCUGAA.GGCCGCGUCU....GGGUCAAU...UGCG.....ACA...........UGCA.ACACCCAC....UCGCC......CUCAUC......GCCCCACCUGAG........GU.GGGCACACUGGAUGAG............G.GCGA......CCGCGUCACC.....AUCAGGA
env-28    CCUGAC.AACCGCGUCC....GGGUCAAU...UGCG.....GCU...........UGCA.ACACCCAA....UCUCG......CUCCCCGUUGGACCA....GAAA.............UGG.CCAUAAAGGAG............G.GAGA......GCGCGUAACA.....GUCAGGA
env-29    CCUGAC.AACCGCGUCC....GGGUCAAU...UGCG.....GCU...........UGCA.ACACCCAC....UCUCG......CUCCCCGUUGGGCCA....GAAA.............UGGCCC.UCACAGAG............G.GAGA......ACGCGUGACA.....GUCAGGA
env-30    CCUGAC.AACCGCGUCC....GGGUCAAU...UGCG.....GCU...........UGCA.ACACCCAA....UCUCG......CUCCCCGUUGGACCA....GAAA.............UGGCCC.UCGAGGAG............G.GAGA......GCGCGUAACA.....GUCAGGA
env-31    GCUGAU.GGGCGCGUCC....GGGUCAAU...UGCG.....GUA...........UGCA.ACACCCGA....UCUCU......CU...CCACGUUCCAGA..CAG............UCUGGA.CGUGGCUCAG............G.GAGA......GCGCGUGACC.....GUCAGGG
Hut-1-1   UCUGAU.GGGCGCGUCC....GGGUCAAU...UGCG.....GCA...........UGCA.ACACCCGAU.....CUCC......CUC....GCCGUUCCGG..GCGA...........CCGGA.CGAUAAAGGAG............G.GGGA......ACGCGUCACC.....GUCAGGA
env-32    UCUGAA.GGCCGCGUCU....GGGUCAAU...UGCG.....AAC...........UGCA.ACACCCAC....UCGCC......CUCAUC..GC...CCCACCCGA.........GGUGGGACCGCUGGAUGAG............G.GCGA......CCGCGUCACC.....AUCAGGA
env-33    UCUGAA.GGCCGCGUCU....GGGUCAAU...UGCG.....AUC...........UGCA.ACACCCAC....UCGCC......CUCAUC.GC....UCCACCCGA.........GGUGGGACCGGUGGAUGAG............G.GUGA......CCGCGUCACC.....AUCAGGA
env-34    UCUGAA.GGCCGCGUCU....GGGUCAAU...UGCG.....AGC...........UGCA.ACACCCAC....UCGCC......CUCAUC..GC...CCCACCUGA.........GGUGGGACCGCUGGAUGAG............G.GCGA......CCGCGUCACC.....AUCAGGA
env-35    UCUGAA.GGCCGCGUCU....GGGUCAAU...UGCG.....AUC...........UGCA.ACACCCAC....UCGCC......CUCAUC.GC....CCCACCCGA.........GGUGGGACCGGUGGAUGAG............G.GUGA......CCGCGUCACC.....AUCAGGA
env-36    GCUGAU.GGCCGCGUAC....GGGUCAAU...UGCG.....GAC...........UGCA.ACACCCGG....UCUCC......CU..GCCGGUCCCGUCCG.GAA...........UGGACG.AACCGGUCGAG............G.GGGA......ACGCGUCUCC.....GUCAGGG
env-37    GCUGAU.GGGCGCGUCU....GGGUCAAU...UGCG.....GAC...........UGCA.ACACCCAC....UCUCU......CUCCC.......CGUCCC.GAAC..........GGGAUGGAACUCACGGAG............G.GAGA......CCGCGUUUCC.....GUCAGGG
env-38    GCUGAU.GGGCGCGUCU....GGGUCAAU...UGCG.....GAC...........UGCA.ACACCCAC....UCUCU......CUCCC.......CGUCCC.GGAU..........GGGAUGGAACUCACGGAG............G.GAGA......ACGCGUUUCC.....GUCAGGG
env-39    GCUGAU.GGGCGCGUCU....GGGUCAAU...UGCG.....GAC...........UGCA.ACACCCAC....UCUCU......CUCCC.......CGUCCC.AGAU..........GGGAUGGAACUCACGGAG............G.GGGA......ACGCGUUUCC.....GUCAGGG
env-40    GCUGAU.GGGCGCGUCC....GGGUCAAU...UGCG.....GCU...........UGCA.ACACCCGA....UCUCC......CUU..GUGGUGG.......CGGAUGGCGAU.........CCGCCGGCAAAG............G.GAGA......ACGCGUGACC.....GUCAGAA

          .<<<<<....<<<<<<.....<<<<...<...<<<<<<<<<.........>>>>>>>>>.>.>>>>......<<<<<......<<<<<<<<<<<<<<<<<<<.............>>>>>>>>>>>>>>>>>>>............>.>>>>.....>>>>>>..........>>>>>..
          .20.....22222.....001...2...22221110..........01112222.2.100......22222......222222..2.2.1111110.............0111111.2.2..222222............2.2222.....22222..........02..
          YCUGAY-○R●●RCGY○Y----GGGU●RAY---●●○○------●●●○----------○R●●-R○RCCCR●----UYYC●------●●○○○-------------●●●○--------------------○○○○○●-----------R-GRRR------GCGY●●●●○○-----RUCAGGR
          5'———————————————————————————————————————————————————————————————————————————————————————————3'
```

```
env-41    UCUGAA.GGCCGCGUUC....GGGUCAAU...UGUG.....AAC..........UACA.ACACCCAA....UCUCG......CUCG.UGCAAC..CGUCA.ACAA..........UGACG....CUCACUGAG............G.GAGA......ACGCGUUGCC.....AUCAGAA
env-43    ACUGAU.GGACGCGUUC....GGGUCAAU...UGCG.....GAU..........UGCA.ACACCCGA....UCUCC......CUCC.CC.GUUCCCG....AUGGC............UGGGAACUGGCCGAG............G.GGGA......GCGCGUCUCC.....GUCAGGG
env-44    CCUGAA.GGCCGCGUCC....GGGUCAAU...UGCG.....GAC..........UGCA.ACACCCGA....UCUCC......CUCCCCGUACCGGCA....GAAA..............UGUCGGU.CGAGGAG............G.GAGA......GCGCGUGACC.....GUCAGGA
env-45    CCUGAA.GGCCGCGUCC....GGGUCAAU...UGCG.....GAC..........UGCA.ACACCCGA....UCUCC......CU.GCUUG.UGCCACCA.ACGA...........UGGAGGAUCCAAGUAAG............G.GAGA......ACGCGUCACC.....GUCAGGA
env-49    UCUGAA.GGCCGCGUCC....GGGUCACU...UGCG.....GAC..........UGCA.ACACCCGA....UCUCC......CUCGC..CGUGCCGCCA.UCGA...........UGGCGACACCGGCCGAG............G.GGGA......GCGCGUCACC.....GUCAGGA
env-50    UCUGAA.GAGCGCGUUC....GGGUCACU...UGCG.....GAC..........UGCA.ACACCCGA....UCUCU......CUCCU..GGUGCCGCCG.UCGA...........UGGCGACACCCGCGGAG............G.GAGA......GCGCGUCACC.....AUCAGGA
env-54    CCUGAA.GGCCGCGUCC....GGGUCACU...UGCG.....GAC..........UGCA.ACACCCGA....UCUCC......CUCAU..GGUGCCGCCG.GCAA...........UGGCGACACCGACCGAG............G.GGGA......GCGCGUCACC.....GUCAGGA
env-55    UCUGAA.GGCCGCGUUC....GGGUCACU...UGCG.....GAC..........UGCA.ACACCCGA....UCUCC......CUCGG..CGUGCCGCCG.GAGA...........UGGCGACACCCACUGAG............G.GAGA......GCGCGUCACC.....GUCAGGA
env-56    UCUGAA.GGCCGCGUUC....GGGUCACU...UGCG.....GAC..........UGCA.ACACCCGA....UCUCC......CUCAU..GGUGCCGCCG.GAUA...........UGGCGACACCCACUGAG............G.GAGA......GCGCGUCACC.....GUCAGGA
env-57    UCUGAA.GGCCGCGUUC....GGGUCACU...UGCG.....GAC..........UGCA.ACACCCGA....UCUCC......CUUAU..GGUGCCGCCG.GAUA...........UGGCGACACCCACUGAG............G.GGGA......GCGCGUCACC.....GUCAGGA
env-58    UCUGAA.GGCCGCGUCU....GGGUCAAU...UGCG.....GAC..........UGCA.ACACCCAA....UCUCG......CUCAU..GGUGCCGCCG.AAUA...........UGGCGACACCGCAGGAG............G.GAGA......GCGCGUCACC.....GUCAGGA
env-59    UCUGAA.GGCCGCGUUC....GGGUCAAU...UGCG.....GAC..........UGCA.ACACCCGA....UCUCG......CUCAU..GGUGCCGCCG.AAUA...........UGGCGACACCCGCUGAG............G.GGGA......GCGCGUCACC.....GUCAGGA
env-62    UCUGAA.GGCCGCGUUC....GGGUCAAU...UGCG.....GAC..........UGCA.ACACCCGA....UCUCG......CUCAU..GGUGCCGCCG.AAUA...........UGGCGACACCCGCUGAG............G.GGGA......GCGCGUCACC.....GUCAGGA
env-63    UCUGAA.GGCCGCGUUC....GGGUCAAU...UGCG.....GAC..........UGCA.ACACCCGA....UCUCG......CUCAU..GGUGCCGCCG.AAUA...........UGGCGACACCCGCUGAG............G.GGGA......GCGCGUCACC.....GUCAGGA
env-64    UCUGAA.GGCCGCGUUC....GGGUCAAU...UGCG.....GAC..........UGCA.ACACCCGA....UCUCG......CUCAU..GGUGCCGCCG.AAUA...........UGGCGACACCCGCUGAG............G.GGGA......GCGCGUCACC.....GUCAGGA
env-65    UCUGAA.GGCCGCGUUC....GGGUCAAU...UGCG.....GAC..........UGCA.ACACCCGA....UCUCG......CUCAU..GGUGCCGCCG.AAUA...........UGGCGACACCCGCUGAG............G.GGGA......GCGCGUCACC.....GUCAGGA
env-70    UCUGAA.GGCCGCGUUC....GGGUCAAU...UGCG.....GAC..........UGCA.ACACCCGA....UCUCG......CUCAU..GGUGCCGCCG.AAUA...........UGGCGACACCCGCUGAG............G.GGGA......GCGCGUCACC.....GUCAGGA
env-71    UCUGAA.GGCCGCGUUC....GGGUCAAU...UGCG.....GAC..........UGCA.ACACCCGA....UCUCG......CUCAU..GGUGCCGCCG.AAUA...........UGGCGACACCCGCUGAG............G.GGGA......GCGCGUCACC.....GUCAGGA
env-72    CCUGAU.CGGCGCGUCC....GGGUCAAU...UGCG.....GAC..........UGCA.ACACCCGA....UCUCC......CUCGCCGUUCCUCCC....GACA.............GGGAGGUACG.CGAG............G.GAGA......GCGCGUGACA.....AUCAGGA
env-74    CCUGAU.CGGCGCGUCC....GGGUCAAU...UGCG.....GAC..........UGCA.ACACCCGA....UCUCC......CUCGCCGUCCCUCCC....GACA.............GGGAGGUCA.GCGAG............G.GAGA......GCGCGUGACA.....AUCAGGA
env-76    CCUGAG.GAGCGCGUCC....GGGUCAAU...UGCG.....GCC..........UGCA.ACACCCGA....UCUCC......CUGC..CGGUGCCUCCA.GGAA...........UGGAGGUGCCGCGCGAG............G.GGGA......ACGCGUGACU.....AUCAGGG
env-77    CCUGAG.GAGCGCGUCC....GGGUCAAU...UGCG.....GCC..........UGCA.ACACCCGA....UCUCC......CUCC..CGGUGCCGCCA.GGAA...........UGGCGGCCCCGAGCGAG............G.GGGA......ACGCGUAACC.....GUCAGGG
env-78    CCUGAG.GAGCGCGUCC....GGGUCAAU...UGCG.....GCC..........UGCA.ACACCCGA....UCUCC......CUCC..CGGUGCCGCCA.GGAA...........UGGCGGCCCCGAGCGAG............G.GGGA......ACGCGUAACC.....GUCAGGG
Hsa-1-1   UCUGAU.GGGCGCGUUC....GGGUCAAU...UGCG.....GCA..........UGCA.ACACCCGA....UCUCU......CUCGGUGUGCCUCC.....UUCCAU.............GGAGGU.CGCCGAG............G.GGGA......GCGCGUUGCC.....GUCAGGA
Hgi-1-1   CCUGAC.GACCGCGUUC....GGGUCAAU...UGCG.....GAA..........UGCA.ACACCCGA....UCUCCC....UGUUCGUCCC.........UCCC...................GGGAUGGACC....UCACAGAGG.GAGA......GCGCGUCGCU.....GUCAGGA
Hsu-2-1   CCUGAC.GACCGCGUUC....GGGUCAAU...UGCG.....GAA..........UGCA.ACACCCGA....UCUCCC....UGUUCGUCCC.........UCCC...................GGGAUGGACC....UCACAGAGG.GAGA......GCGCGUCGCU.....GUCAGGA
Hpr-1-1   CCUGAC.GACCGCGUUC....GGGUCAAU...UGCG.....GAA..........UGCA.ACACCCGA....UCUCCC....UGUUCGUCCC.........UCCC...................GGGAUGGACC....UCACAGAGG.GAGA......GCGCGUCGCU.....GUCAGGA
Hsp-8-1   CCUGAC.GACCGCGUUC....GGGUCAAU...UGCG.....GAA..........UGCA.ACACCCGA....UCUCCC....UGUUCGUCCC.........UCCC...................GGGAUGGACC....UCACAGAGG.GAGA......GCGCGUCGCU.....GUCAGGA
Hde-1-1   CCUGAC.GACCGCGUUC....GGGUCAAU...UGCG.....GAA..........UGCA.ACACCCGA....UCUCCC....UGUUCGUCCC.........UCCC...................GGGAUGGACC....UCACAGAGG.GAGA......GCGCGUCGCU.....GUCAGGA
Hlu-1-1   CCUGAC.GACCGCGUUC....GGGUCAAU...UGCG.....GAA..........UGCA.ACACCCGA....UCUCCC....UGUUCGUCCC.........UCCC...................GGGAUGGACC....UCACAGAGG.GAGA......GCGCGUCGCU.....GUCAGGA
Hal-2-1   CCUGAC.GACCGCGUUC....GGGUCAAU...UGCG.....GAA..........UGCA.ACACCCGA....UCUCCC....UGUUCGUCCC.........UCCC...................GGGAUGGACC....UCACAGAGG.GAGA......GCGCGUCGCU.....GUCAGGA
Hsp-6-1   CCUGAC.GACCGCGUUC....GGGUCAAU...UGCG.....GAA..........UGCA.ACACCCGA....UCUCCC....UGUUCGUCCC.........UCCC...................GGGAUGGACC....UCACAGAGG.GAGA......GCGCGUCGCU.....GUCAGGA
Hme-1-1   CCUGAC.GACCGCGUUC....GGGUCAAU...UGCG.....GAA..........UGCA.ACACCCGA....UCUCCC....UGUUCAUCCC.........UCCG...................GGGAUGGAUA....UUGCAAAGG.GGGA......GCGCGUCGCU.....GUCAGGA
Hla-2-1   CCUGAC.GACCGCGUUC....GGGUCAAU...UGCG.....GAA..........UGCA.ACACCCGA....UCUCCC....UGUUCGUCCC.........UCCC...................GGGAUGGACG.....CCACAGAGGGAGA......GCGCGUCGCU.....GUCAGGA
Hel-1-1   CCUGAC.GACCGCGUUC....GGGUCAAU...UGCG.....GAA..........UGCA.ACACCCGA....UCUCCC....UGUUCGUCCC.........UCCC...................GGGAUGGACG.....CCACAGAGGGAGA......GCGCGUCGCU.....GUCAGGA
Hmu-1-1   CCUGAC.GACCGCGUUC....GGGUCAAU...UGCG.....GAA..........UGCA.ACACCCGA....UCUCC.....CUGUUCACCCC........UCCG...................GGGAUGAAC.G....UUUCAAAGG.GGGA......GCGCGUCGCU.....GUCAGGA
env-79    GCUGAU.GGGCGCGUCU....GGGUCAAU...UGCG.....GAC..........UGCA.ACACCCACU....CUCU......CUCC........CCGUCCCGAAC.........GGGAUGG.AACUCACGGAG............G.GGGG......CGCGUUUCC.....GUCAGGG
env-80    ACUGAU.GGGCGCGUCC....GGGUCAAU...UGCG.....GAA..........UGCA.ACACCCGA....UCUCCC.....UUC.........CCGUCCCGAAU.........GGGAUGG.AUAUCACAGAG............G.GGGA......GCGUGUCACC.....GUCAGAG
env-81    ACUGAU.GGGCGCGUCC....GGGUCAAU...UGCG.....GAA..........UGCA.ACACCCGA....UCUCC......CUUCC.......CGUCC.CACU..........GGGAUGCAAGUACGGAA............G.GAGA......GCGUGUCGCC.....GUCAGAG
env-82    ACUGAU.GGGCACGUCC....GGGUCAAU...UGCG.....GAC..........UGCA.ACACCCGA....UCUCC......CUUCC.......CGUCC.GGAU..........GGGAUGGAUCUCACAGAG............G.GGGA......GCGUGUCACC.....GUCAGAG
env-83    ACUGAU.GGGCACGUCC....GGGUCAAU...UGCG.....GAC..........UGCA.ACACCCGA....UCUCC......CUUCC.......CGUCC.GGAU..........GGGAUAGAUCUCACAGAG............G.GGGA......GCGUGUCACC.....GUCAGAG
Hla-4-1   CCUGAU.CAACGCGUCC....GGGUCAAU...UGCG.....GAC..........UGCA.ACACCCGA....UCUCU......CUCAACG..UGCCAUC....GGA..............GAUGGCACUCGCGGAG............G.GAGA......GCGCGUGACA.....GUCAGGA
Ssp-1-1   CCUGAU.GGGCGCGUAC....GGGUCAAU...UGCG.....GCG..........UGCA.ACACCCGA....UCUCC......CU.GCCGGUUCCCC.....ACGAAC............GGGACAUCGGCGAG............G.GGGA......GCGUGUGACC.....GUCAGGG
env-84    UCUGAC.GGGCGCGUUC....GGGUCAAU...UGCG.....GAA..........UGCA.ACACCCGA....UCUCC......CUCC.ACACG.CCAC....CCGAC.............GUGGCCGUGUCGGAG............G.GGGA......GCGCGUGACC.....AUCAGGG
env-86    CCUGAA.GGGCGCGUCC....GGGUCAAU...UGCG.....GCU..........UGCA.ACACCCGA....UCUCU......CUCAA..UGUGCCG.....CCAUCGA.............UGGCACUCGAAGAG............G.GAGA......GCGCGUGACC.....GUCAGGA
Hsp-11-1  CCUGAU.CAACGCGUCC....GGGUCAAU...UGCG.....GAC..........UGCA.ACACCCGA....UCUCU......CUGAA..CGUGCCGUC....GGA..............GAUGGCACUCGACGAG............G.GAGA......GCGCGUGACA.....GUCAGGA
env-87    CCUGAG.GACCGCGUUC....GGGUCACU...UGCG.....GAA..........CGCA.ACACCCGA....UCUCC......CUUC..UCGCGCCC.....GGAC..............GGGCGCUGACGGAG............G.GGGA......GCGCGUCACC.....AUCAGGG
env-88    UCUGAA.GGCCGCGUCU....GGGUCAAU...UGCG.....GAC..........UGCA.ACACCCAA....UCUCG......CUCAU..GGUGCCGCCG.AAUA...........UGGCGACACCCAAGAG............G.GAGA......GCGCGUCACC.....GUCAGGA
env-89    GCUGAU.GGGCGCGCUC....GGGUCAAU...UGCG.....GUC..........UGCA.ACACCCAA....UCUCC......CU..GCCGGUCCCGUCCG.GAA...........UGGAUGGGCUGGUUGAG............G.GGGA......ACGCGUCUCC.....GUCAGGG
Hka-1-1   GCUGAU.GGGCGCGUUC....GGGUCAAU...UGCG.....GAC..........UGCA.ACACCCGA....UCUCU......CUC.CCGGUACCUUCCG..CGA.............UGGAAGUGCCGGCGAG............G.GGGA......GCGCGUCACC.....GUCAGGG
Hmu-2-1   GCUGAU.GGGCGCGUUC....GGGUCAAU...UGCG.....GAC..........UGCA.ACACCCGA....UCUCU......CUC.CCGGUACCUUCCG..CGA.............UGGAAGUGCCGGCGAG............G.GGGA......GCGCGUCACC.....GUCAGGG
Hca-3-1   GCUGAU.GAACGCGUAC....GGGUCAAU...UGCG.....GCC..........UGCA.ACACCCAG....UCUCG......CUCCCCGUGCCCUCCG...GAA.............UGGAGGUCGGGG..AG............G.GGGA......GCGCGUCUCC.....GUCAGGG
Hsa-5-1   CCUGAU.GGACGCGUUC....GGGUCAAU...UGCG.....GAC..........UGCA.ACACCCGA....UCUCA......CUGGUC.GAUCCCAC....CGAC..............GUGGGACUCGACGAG............G.GGGA......GCGUGUGACC.....GUCAGGA
env-90    CCUGAU.GGACGCGUUC....GGGUCAAU...UGCG.....GCU..........UGCA.ACACCCGA....UCUCG......CUGGUC.GAUCCCAC....CGAC..............GUGGGACUCGACGAG............G.GAGA......GCGUGUGACC.....GUCAGGA
Hag-1-1   CCUGAU.GGACGCGUUC....GGGUCAAU...UGCG.....GAC..........UGCA.ACACCCGA....UCUCG......CUGGUC.GAUCCCAC....CGAC..............GUGGGACUCGACGAG............G.GAGA......GCGUGUGACC.....GUCAGGA
Hsa-4-1   CCUGAU.GGACGCGUUC....GGGUCAAU...UGCG.....GAC..........UGCA.ACACCCGA....UCUCG......CUGGUC.GAUCCCAC....CGAC..............GUGGGACUCGACGAG............G.GGGA......GCGUGUGACC.....GUCAGGA
env-91    GCUGAU.GGCCGCGUUC....GGGUCAAU...UGCG.....GAC..........UGCA.ACACCCGG....UCUCA......CUCC..CGGUGUCGCC....CGCGAU..........GGCGACGCCGUCGGAG............G.GGGA......GCGUGUCACC.....GUCAGCA
env-92    UCUGAA.GGCCGCGUCC....GGGUCACU...UGCG.....GAC..........UGCA.ACACCCGA....UCUCU......CUCA..CGGUGCCGCCG.UCGA...........UGGCGACACCGGCUGAG............G.GGGA......GCGCGUCACC.....GUCAGGA
env-93    UCUGAA.GGCCGCGUCC....GGGUCACU...UGCG.....GAC..........UGCA.ACACCCGA....UCUCU......CUCA..CGGUGCCGCCG.UCGA...........UGGCGACACCGGCUGAG............G.GGGA......GCGCGUCACC.....GUCAGGA
env-94    UCUGAA.GGCCGCGUCC....GGGUCACU...UGCG.....GAC..........UGCA.ACACCCGA....UCUCU......CUCA..CGGUGCCGCCG.UCGA...........UGGCGACACCGGCUGAG............G.GGGA......GCGCGUCACC.....GUCAGGA
env-95    UCUGAA.GGCCGCGUCC....GGGUCACU...UGCG.....GAC..........UGCA.ACACCCGA....UCUCU......CUCA..CGGUGCCGCCG.UCGA...........UGGCGACACCGGCUGAG............G.GGGA......GCGCGUCACC.....GUCAGGA
env-96    UCUGAA.GGCCGCGUCC....GGGUCACU...UGCG.....GAC..........UGCA.ACACCCGA....UCUCU......CUCA..CGGUGCCGCCG.UCGA...........UGGCGACACCGGCUGAG............G.GGGA......GCGCGUCACC.....GUCAGGA
env-98    UCUGAA.GGCCGCGUCC....GGGUCACU...UGCG.....GAC..........UGCA.ACACCCGA....UCUCC......CUCA..CGGUGCCGCCG.UCGA...........UGGCGACACCGGCCGAG............G.GGGA......GCGCGUCACC.....GUCAGGA
env-99    UCUGAA.GGCCGCGUCC....GGGUCACU...UGCG.....GAC..........UGCA.ACACCCGA....UCUCU......CUCA..CGGUGCCGCCG.UCGA...........UGGCGACACCGGCCGAG............G.GGGA......GCGCGUCACC.....GUCAGAA
env-101   UCUGAA.GGCCGCGUUC....GGGUCACU...UGCG.....GAC..........UGCA.ACACCCGA....UCUCC......CUCA..UGGUGCCGCCG.GAGA...........UGGCAGCGCCGAGCGAG............G.GGGA......GCGCGUCACC.....GUCAGGA
env-102   UCUGAA.GGCCGCGUUC....GGGUCACU...UGCG.....GAC..........UGCA.ACACCCGA....UCUCC......CUCA..UGGUGCCGCCG.AAUA...........UGGCGACGCCAGCCGAG............G.GGGA......GCGCGUCACC.....GUCAGGA
env-103   UCUGAA.GGCCGCGUUC....GGGUCACU...UGCG.....GAC..........UGCA.ACACCCGA....UCUCC......CUCA..UGGUGCCGCCG.AAUA...........UGGCGACGCCAGCCGAG............G.GGGA......GCGCGUCACC.....GUCAGGA
env-104   UCUGAA.GGCCGCGUUC....GGGUCACU...UGCG.....GAC..........UGCA.ACACCCGA....UCUCC......CUCA..UGGUGCCGCCG.GAUA...........UGGCGACGCCAGCCGAG............G.GGGA......GCGCGUCACC.....GUCAGGA
env-105   UCUGAA.GGCCGCGUUC....GGGUCACU...UGCG.....GAC..........UGCA.ACACCCGA....UCUCC......CUCA..UGGUGCCGCCG.GAUA...........UGGCGACGCCAGCCGAG............G.GGGA......GCGCGUCACC.....GUCAGGA
env-106   UCUGAA.GGCCGCGUUC....GGGUCACU...UGCG.....GAC..........UGCA.ACACCCGA....UCUCC......CUCA..UGGUGCCGCCG.GAUA...........UGGCGACGCCAGCCGAG............G.GGGA......GCGCGUCACC.....GUCAGGA
env-107   UCUGAC.GGCCGCGUCC....GGGUCAAU...UGCG.....GAC..........UGCA.ACACCCGA....UCUCC......CUCCUCUCCCC........GAC...................GGAUGGAGGUG......UCCGAAG.GAGA......GCGCGUCGCC.....AUCAGGA
env-108   UCUGAG.GGACGCGUUC....GGGUCAAU...UGCG.....GAC..........UGCA.ACACCCGA....UCUCU......CUCCACACGCCUCCC....GGAC..............GGGAGCUGU.CGGAG............G.GGGA......GCGCGCAACC.....AUCAGGA
env-109   UCUGAC.GGGCGCGUCC....GGGUCAAU...UGCG.....GCA..........UGCA.ACACCCGA....UCUCCCU.....CCUC..GCUCCC.......GACC.................GGGAGUUCGAGG..........UCG.GAGA......GCGCGUUACC.....GUCAGGG
CHa-1-1   CCUGAA.GGCCGCGUUC....GGGUCAAU...UGCG.....GAC..........UGCA.ACACCCGA....UCUCCCUC....CACCUG..CCUCCC.....GGAC..............GGGAGGCGCAGGAG............G.GGGA......GCGCGUCACC.....GUCAGGG
Hpa-2-1   CCUGAA.GGUCGCGUUC....GGGUCAAU...UGCG.....GAC..........UGCA.ACACCCGG....UCUCC......CUCUA..CGCUCCGU.....CCGGU............ACGGAGUUGGAGGAG............G.GAGA......GCGCGUCGCC.....AUCAGGA
env-110   CCUGAU.CGGCGCGUCC....GGGUCAAU...UGCG.....GCU..........UGCA.ACACCCGA....UCUCG......CUCAACGUGCCUCCA....CGAA.............UGGAGGUU.ACGGAG............G.GAGA......GCGCGUGACA.....GUCAGGA
Nhu-1-1   CCUGAU.CAACGCGUGC....GGGUCAAU...UGCG.....GCU..........UGCA.ACACCCGA....UCUCC......CUCAACG..UACCGCCG..AAAA...........UGGCGGUCUCCG...AG............G.GAGA......GCGCGUGACU.....AUCAGGA
env-113   CCUGAA.GGCCGCGUCC....GGGUCAAU...UCCG.....CGCUGC.......GGGA.ACACCCGA....UCUCC......CUGCUCGUCCCCUCCG...GAA.............UGGAGGCCGAGC.CGG.............GGGA......ACGCGUCACC.....AUCAGGG
env-114   UCUGAA.GGCCGCGUCC....GGGUCACU...UGCG.....GAC..........UGCA.ACACCCGA....UCUCC......CUCGCCGUGC..CGCCG.UCGA...........UGGCGACACCGGCCGAG............G.GGGA......GCGCGUCACC.....GUCAGGA
env-115   UCUGAA.GGCCGCGUCC....GGGUCACU...UGCG.....GAC..........UGCA.ACACCCGA....UCUCC......CUCGCCGUGC..CGCCG.UCGA...........UGGCGACACCGGCCGAG............G.GGGA......GCGCGUCACC.....GUCAGGA
env-116   UCUGAA.GGCCGCGUUC....GGGUCACU...UGCG.....GAC..........UGCA.ACACCCGA....UCUCC......CUUGUGGUGC..CGCCG.GAGA...........UGGCGACACCCACUGAG............G.GGGA......GCGCGUCACC.....GUCAGGA
env-117   UCUGAA.GGCCGCGUUC....GGGUCACU...UGCG.....GAC..........UGCA.ACACCCGA....UCUCC......CUCAUGGUGC..CGCCG.GAUA...........UGGCGACGCCCACUGAG............G.GGGA......GCGCGUCACC.....GUCAGGA
env-118   UCUGAA.GGCCGCGUUC....GGGUCACU...UGCG.....GAC..........UGCA.ACACCCGA....UCUCU......CUCAUGGUGC..CGCCG.GAUA...........UGGCGACACCCACUGAG............G.GAGA......GCGCGUCACC.....GUCAGGA

          .<<<<<....<<<<<<.....<<<<...<...<<<<<<<<<..........>>>>>>>>>.>.>>>>......<<<<<......<<<<<<<<<<<<<<<<<<<<.............>>>>>>>>>>>>>>>>>>>>............>.>>>>......>>>>>>.........>>>>>..
          .20.......22222.....001...2...22221110..........01112222.2.100......22222......222222..2.2.1111110.............0111111.2.2..222222............2.2222......22222..........02..
          YCUGAY-•R••RCGY•Y----GGGU•RAY---••••-----•••◦---------◦R••-R•RCCCR•----UYYC•-------•••◦◦---------------••••-----------------------◦◦◦◦◦•-----------R-GRRR-----GCGY•••••◦◦-----RUCAGGR
          5'                                                                                                                                                                              3'
```

```
env-119    UCUGAA.GGCCGCGUUC....GGGUCACU...UGCG.....GAC..........UGCA.ACACCCGA....UCUCC......CUUGUGUGC..CGCCG..AAUA..........UGGCGACACCCGCUGCG............G.GAGA......GCGCGUCACC.....GUCAGGA
env-120    CCUGAG.GGCCGCGUUC....GGGUCAAU...UGCG.....GAC..........UGCA.ACACCCGA....UCUCCCU....CCGGGCGCCCC.......CAGAC...............GGGGUGCCCGA.......CGAGG.GGGA.....GCGUGUCACC.....GUCAGGG
env-122    CCUGAA.GGCCGCGUUC....GGGUCAAU...UGCG.....GAC..........UGCA.ACACCCGA....UCUCCCU....CCGGGCACCCC.......CAGAC...............GAGGUGCCCGC.......CGAGG.GGGA.....GCGCGUCACC.....GUCAGGG
env-123    CCUGAA.GGCCGCGUCC....GGGUCAAU...UGCG.....GAC..........UGCA.ACACCCGA....UCUCCCU....CCGGGCACCCC.......CAGAC...............GGGGUGCCCGC.......CGAGG.GGGA.....GCGCGUCACC.....GUCAGGG
env-126    CCUGAC.GGCCGCGUCC....GGGUCAAU...UGCG.....AAC..........UGCA.ACACCCAACC..GCCCU......CCGUGCGCCUCCG......UCGA............UGGAGG..UGCGG.........GAG.GGGCA....ACGCGUCGCC.....AUCAGGA
env-127    CCUGAC.GGCCGCGUCC....GGGUCAAU...UGCG.....AAC..........UGCA.ACACCCAACCA.CCCUU......CGUGCGCCUCCG.......UCGA..............UGGAGGUGCGGG.............A.GGGGCA....ACGCGUCGCC.....AUCAGGA
env-128    CCUGAC.GGCCGCGUCC....GGGUCAAU...UGCG.....AAC..........UGCA.ACACCCAACCA.CCCUU......CGUGCGCCUCCG.......UCGA..............UGGAGGUGCGGG.............A.GGGGCA....ACGCGUCGCC.....AUCAGGA
env-129    CCUGAU.GACCGCGUUC....GGGUCACU...UGCG.....GAC..........UGCA.ACACCCGA....UCUCA......CUCGG.CGUG.CCG.....AAAGAAA..........UGGAGGUCGCCGAG............G.GGGA......GCGCGUCACC.....GUCAGGA
Hna-1-1    CCUGAC.GGCCGCGUUC....GGGUCAAU...UGCG.....GAC..........UGCA.ACACCCGA....UCUCC......CUCCU.CAC.UCCGG....ACGGA...........CUGGAAGUCAAGGAG............G.GGGA......GCGCGUCGCC.....AUCAGGA
env-130    CCUGAC.GGCCGCGUCC....GGGUCAAU...UGCG.....GAC..........UGCA.ACACCCAA....UCUCC......CUCCU...CCUCC......CGGAUGGGAU.........GGAGGUGUCGGAC............G.GAGA......GCGCGUCGCC.....AUCAGGA
env-131    CCUGAC.GACCGCGUCC....GGGUCAAU...UGCG.....GCU..........UGCA.ACACCCGAUC..UCCCU......CCACGUCCGUCG......UCGA..............UGACGGU.CGUGG..........AAG.GGGA......GCGCGUCACU.....GUCAGGG
env-134    CCUGAC.GGGCGCGUCC....GGGUCAAU...UGCG.....GAC..........UGCA.ACACCCGA....UCUCG......CUCCCCGU..GCCUCC...GUC.............GGAGGAGCUCGAGGAG............G.GAGA......ACGCGUCACC.....AUCAGGG
env-135    CCUGAA.GGGCGCGUCC....GGGUCAAU...UGCG.....GAC..........UGCA.ACACCCCGA...UCUCC......CUCAACAUACCGCCG....AACA............UGGCGG..UCGAGGG...........AG.GGGGA.....GCGCGUGACC.....GUCAGGA
Nph-1-1    CCUGAC.GGGCGCGUCC....GGGUCAAU...UGCG.....GAA..........UGCA.ACACCCGA....UCUCU......CUCCCCGUCCCUCC.....CUCGAU.............GGAGG.UCGGGAAG............G.GAGA......GCGCGUCACC.....AUCAGGG
env-136    UCUGAC.GAGCGCGUCC....GGGUCAAU...UGCG.....GAC..........UGCA.ACACCCGA....UCUCC......CUCCCCGUUCCUCCA....UCGA............UGGA.GGUCGAGGAA............G.GGGA......ACGCGUCACC.....AUCAGGG
env-137    UCUGAC.AAGCGCGUCC....GGGUCAAU...UGCG.....GAC..........UGCA.ACACCCGA....UCUCC......CUCCCCGUUCCUCCA....UCGA............UGGA.GGUCGAGGAA............G.GGGA......ACGCGUCACC.....AUCAGGG
env-138    CCUGAA.GGCCGCGUCC....GGGUCAAU...UCCC.....CGCUGC........GGGA.ACACCCGA....UCUCCCU....GCUCGUCCCUCCG......AGA..............UGGAGGUG.CGAGC..........GG.GAGA......GCGCGUCACC.....AUCAGGG
env-139    CCUGAA.GGCCGCGUCC....GGGUCAAU...UCCC.....CGCUGC........GGGA.ACACCCGA....UCUCCCU....GCUCGUUCCCUCCG.....AGA..............UGGGGGU..CGAGU.........CGG.GGGA......GCGCGUUACC.....AUCAGGG
env-140    CCUGAA.GGCCGCGUCC....GGGUCAAU...UCCC.....CGCUGC........GGGA.ACACCCGA....UCUCCCUGC..UCGUCC..CCGACG.....AUA..............UGUCGGUCGAGCCG............G.GGGA......GCGCGUCACC.....AUCAGGG
env-145    CCUGAA.GGCCGCGUUC....GGGUCAAU...UCCCC....GCUG.........GGGGA.ACACCCGA....UCUCCCUGC..UCGUCC..CCUCCG.....AAA..............UGGAGGUCGAGCGG............G.GGGA......ACGCGUCACC.....AUCAGGG
env-146    CCUGAA.GGCCGCGUCC....GGGUCAAU...UCCC.....CGCUGC........GGGA.ACACCCGA....UCUCCCUGC..UCGUCC..CCUCCG.....GAA..............UGGAGGUUGAGCAA............G.GAGA......GCGCGUCACC.....AUCAGGG
env-147    CCUGAA.GGCCGCGUUC....GGGUCAAU...UCCC.....CGCUGC........GGGA.ACACCCGA....UCUCCCUGC..UCGUCC..CCUCCG.....GAA..............UGGAGGUCGAGCAG............G.GGGA......GCGCGUCACC.....AUCAGGG
env-148    CCUGAA.GGCCGCGUUC....GGGUCAAU...UCCCC....GCUG.........GGGGA.ACACCCGA....UCUCCCUGC..UCGUUC..CCUCCG.....GAC..............UGGAGGUCGAGCAG............G.GGGA......GCGCGUCACC.....AUCAGGG
Hsp-9-1    CCUGAA.GGCCGCGUCC....GGGUCAAU...UCCC.....CGCUGC........GGGA.ACACCCGA....UCUCCCUGC..UCGUUC..CCUCCG.....ACA..............UGGAGGUCGAGCAG............G.GGGA......GCGCGUCACC.....AUCAGGG
Hez-1-1    CCUGAA.GGCCGCGUCC....GGGUCAAU...UCCC.....CGCUGC........GGGA.ACACCCGA....UCUCCCUGC..UCGUCC..CCUCCG.....AAA..............UGGAGGUCGAGCGG............G.GGGA......GCGCGUCACC.....AUCAGGG
env-149    CCUGAA.GGCCGCGUCC....GGGUCAAU...UCCC.....CGCUGC........GGGA.ACACCCGA....UCUCCCUGC..UCGUGC..CCUCCG.....GAA..............UGGAGGUCGAGCGG............G.GGGA......GCGCGUCACC.....AUCAGGG
env-150    CCUGAA.GGCCGCGUCC....GGGUCAAU...UCCC.....CGCUGC........GGGA.ACACCCGA....UCUCCCUGC..UCGUGC..CCUCCG.....GAA..............UGGAGGUCGAGCGG............G.GGGA......GCGCGUCACC.....AUCAGGG
env-151    CCUGAA.GGCCGCGUCC....GGGUCAAU...UCCC.....CGCUGC........GGGA.ACACCCGA....UCUCCCUGC..UCGUCC..CCUCCG.....GAA..............UGGAGGUCGAGCGG............G.GGGA......ACGCGUCACC.....AUCAGGG
Hco-1-1    CCUGAA.GGCCGCGUCC....GGGUCAAU...UCCC.....CGCUGC........GGGA.ACACCCGA....UCUCCCUGC..UCGUCC..CCUCCG.....GAA..............UGGAGGUCGUGCGG............G.GGGA......GCGCGUCACC.....AUCAGGG
env-152    CCUGAA.GGCCGCGUCC....GGGUCAAU...UCCC.....CGCUGC........GGGA.ACACCCGA....UCUCCCUGC..UCGUCC..CCUCCG.....GAA..............UGGAGGUCGAGCGG............G.GGGA......GCGCGUCACC.....AUCAGGG
env-153    CCUGAA.GGCCGCGUCC....GGGUCAAU...UCCC.....CGCUGC........GGGA.ACACCCGA....UCUCCCUGC..UCGUCC..CCUCCG.....GAA..............UGGAGGUCGAGCGG............G.GGGA......GCGCGUCACC.....AUCAGGG
env-154    CCUGAA.GGCCGCGUCC....GGGUCAAU...UCCC.....CGCUGC........GGGA.ACACCCGA....UCUCCCUGC..UCGUCC..CCUCCG.....GAA..............UGGAGGUCGAGCGG............G.GGGA......GCGCGUCACC.....AUCAGGG
env-155    CCUGAA.GGCCGCGUCC....GGGUCAAU...UCCC.....CGCUGC........GGGA.ACACCCGA....UCUCCCUGC..UCGUCC..CCUCCG.....GAA..............UGGAGGUCGAGCGG............G.GGGA......GCGCGUCACC.....AUCAGGG
env-156    CCUGAA.GGCCGCGUCC....GGGUCAAU...UCCC.....CGCUGC........GGGA.ACACCCGA....UCUCCCUGC..UCGUCC..CCUCCG.....GAA..............UGGAGGUCGAGCGG............G.GGGA......GCGCGUCACC.....AUCAGGG
Hsp-10-1   CCUGAA.GGCCGCGUCC....GGGUCAAU...UCCC.....CGCUGC........GGGA.ACACCCGA....UCUCCCUGC..UCGUCC..CCUCCG.....GAA..............UGGAGGUCGAGCGG............G.GGGA......GCGCGUCACC.....AUCAGGG
Hli-1-1    CCUGAA.GGCCGCGUCC....GGGUCAAU...UCCC.....CGCUGC........GGGA.ACACCCGA....UCUCCCUGC..UCGUUC..CCUCCG.....GAA..............UGGAGGUCGAGCAA............G.GGGA......GCGCGUCACC.....AUCAGGG
Hte-1-1    CCUGAA.GGCCGCGUCC....GGGUCAAU...UCCCC....GCUG.........GGGGA.ACACCCGA....UCUCCCUGC..UCGUUC..CCUCCG.....GAA..............UGGAGGUCGAGCAG............G.GGGA......GCGCGUCACC.....AUCAGGG
Hko-1-1    CCUGAA.GGCCGCGUCC....GGGUCAAU...UCCC.....CGCUGC........GGGA.ACACCCGA....UCUCCCUGC..UCGUUC..CCUCCG.....GAA..............UGGAGGUCGAGCAG............G.GGGA......GCGCGUCACC.....AUCAGGG
Hli-2-1    CCUGAA.GGCCGCGUCC....GGGUCAAU...UCCC.....CGCUGC........GGGA.ACACCCGA....UCUCCCUGC..UCGUUC..CCUCCG.....GAA..............UGGAGGUCGAGCAG............G.GGGA......GCGCGUCACC.....AUCAGGG
Hsa-7-1    CCUGAA.GGCCGCGUCC....GGGUCAAU...UCCC.....CGCUGC........GGGA.ACACCCGA....UCUCCCUGC..UCGUUC..CCUCCG.....GAA..............UGGAGGUCGAGCAG............G.GGGA......GCGCGUCACC.....AUCAGGG
Hca-2-1    CCUGAA.GGCCGCGUCC....GGGUCAAU...UCCC.....CGCUGC........GGGA.ACACCCGA....UCUCCCUGC..UCGUUC..CCUCCG.....GAA..............UGGAGGUCGAGCAG............G.GGGA......GCGCGUCACC.....AUCAGGG
Hsp-7-1    CCUGAA.GGCCGCGUCC....GGGUCAAU...UCCC.....CGCUGC........GGGA.ACACCCGA....UCUCCCUGC..UCGUUC..CCUCCG.....GAA..............UGGAGGUCGAGCAG............G.GGGA......GCGCGUCACC.....AUCAGGG
Hha-2-1    CCUGAA.GGCCGCGUCC....GGGUCAAU...UCCC.....CGCUGC........GGGA.ACACCCGA....UCUCCCUGC..UCGUUC..CCUCCG.....GAA..............UGGAGGUCGAGCAG............G.GGGA......GCGCGUCACC.....AUCAGGG
Hdi-1-1    CCUGAA.GGCCGCGUCC....GGGUCAAU...UCCC.....CGCUGC........GGGA.ACACCCGA....UCUCCCUGC..UCGUUC..CCUCCG.....GAA..............UGGAGGUCGAGCAG............G.GGGA......GCGCGUCACC.....AUCAGGG
Hsa-9-1    CCUGAA.GGCCGCGUCC....GGGUCAAU...UCCC.....CGCUGC........GGGA.ACACCCGA....UCUCCCUGC..UCGUUC..CCUCCG.....GAA..............UGGAGGUCGAGCAG............G.GGGA......GCGCGUCACC.....AUCAGGG
Har-2-1    CCUGAA.GGCCGCGUCC....GGGUCAAU...UCCC.....CGCUGC........GGGA.ACACCCGA....UCUCCCUGC..UCGUUC..CCUCCG.....GAA..............UGGAGGUCGAGCAG............G.GGGA......GCGCGUCACC.....AUCAGGG
env-157    CCUGAA.GGCCGCGUCC....GGGUCAAU...UCCC.....CGCUGC........GGGA.ACACCCGA....UCUCCCUGC..UCGUUC..CCUCCG.....GAA..............UGGAGGUCGAGCAG............G.GGGA......GCGCGUCACC.....AUCAGGG
env-158    CCUGAA.GGCCGCGUCC....GGGUCAAU...UCCC.....CGCUGC........GGGA.ACACCCGA....UCUCCCUGC..UCGUUC..CCUCCG.....GAA..............UGGAGGUCGAGCGU............G.GAGA......GCGCGUCACC.....AUCAGGG
env-159    CCUGAA.GGCCGCGUCC....GGGUCAAU...UCCC.....CGCUGC........GGGA.ACACCCGA....UCUCCCUGC..UCGUUC..CCUCCG.....GAA..............UGGAGGUCGAGCGU............G.GAGA......GCGCGUCACC.....AUCAGGG
env-160    CCUGAA.GGCCGCGUUC....GGGUCAAU...UCCC.....CGCUGC........GGGA.ACACCCAA....UCUCCCUGC..UCGUCC..CCUCCG.....GAA..............UGGAGGUUGAGCGG............G.GGGA......GCGCGUCACC.....AUCAGGG
h3A-1-1    CCUGAC.GGCCGCGUUC....GGGUCAAU...UCCCC....GCUG.........GGGGA.ACACCCGA....UCUCCCUGC..UCGUUC..CCGACG.....GAC..............UGUCGGUCGAGCGG............G.GGGA......GCGCGUCACC.....AUCAGGA
Hai-1-1    CCUGAA.GGCCGCGUCC....GGGUCAAU...UCCC.....CGCUGC........GGGA.ACACCCGA....UCUCC......CUGCUCGUGCCCUCCG...GAA............UGGAGG..UCGAGCAG............G.GGGA......GCGCGUCACC.....AUCAGGG
env-161    CCUGAA.GGCCGCGUCC....GGGUCAAU...UCCC.....CGCUGC........GGGA.ACACCCGA....UCUCC......CUGCUCGUCCCUCCG...GAA............UGGAGG..UCGAGCAG............G.GGGA......GCGCGUCACC.....AUCAGGG
Hho-1-1    CCUGAA.GGCCGCGUCC....GGGUCAAU...UCCC.....CGCUGC........GGGA.ACACCCGA....UCUCC......CUGCUCGUCCCUCCG...ACA............UGGAGG..UCGAGCAG............G.GGGA......GCGCGUCACC.....AUCAGGG
env-164    CCUGAA.GGCCGCGUCC....GGGUCAAU...UCCC.....CGCUGC........GGGA.ACACCCGA....UCUCCCUG...CUCGUC.CCCGCCG.....ACA..............UGGCGGUCGAGCCG............G.GGGA......GCGCGUCACC.....AUCAGGG
env-165    CCUGAA.GGCCGCGUCC....GGGUCAAU...UCCC.....CGCUGC........GGGA.ACACCCAA....UCUCC......CUGCUCGUCCCUCCG...AGG............UGGAGG.UGCGGGC.G............G.GAGA......GCGCGUCACC.....AUCAGGG
env-166    CCUGAA.GGCCGCGUCC....GGGUCAAU...UCCC.....CGCUGC........GGGA.ACACCCAA....UCUCC......CUGCUCGUCCCUCCG...AGG............UGGAGG.UGCGGGC.G............G.GAGA......GCGCGUCACC.....AUCAGGG
env-167    CCUGAA.GGCCGCGUCC....GGGUCAAU...UCCC.....CGCUGC........GGGA.ACACCCGA....UCUCCCUGCU.CGUCC...CCUCCA.....ACA..............UGGAGGUGCGGGCG............G.GAGA......GCGCGUCACC.....AUCAGGG
env-169    CCUGAA.GGCCGCGUCC....GGGUCAAU...UCCC.....CGCUGC........GGGA.ACACCCGA....UCUCCCUGCU.CGUCC...CCUCCA.....ACA..............UGGAGGUGCGGGCG............G.GAGA......GCGCGUCACC.....AUCAGGG
env-170    CCUGAA.GGCCGCGUCC....GGGUCAAU...UCCC.....CGCUGC........GGGA.ACACCCGA....UCUCCCUGCU.CGUCC...CCUCCA.....ACA..............UGGAGGUGCGGGCG............G.GAGA......GCGCGUCACC.....AUCAGGG
env-172    CCUGAA.GGCCGCGUCC....GGGUCAAU...UCCC.....CGCUGC........GGGA.ACACCCGA....UCUCCCUGCU.CGUCC...CCUCCA.....ACA..............UGGAGGUGCGGGCG............G.GAGA......GCGCGUCACC.....AUCAGGG
env-173    CCUGAU.GGGCGCGUUC....GGGUCAAU...UGCG.....GCC..........UGCA.ACACCCGA....UCUCGCUC...CCCGUGCCGUCA.......AAAA...............UGUCGGUACAGG..........AGG.GAGA......GCGCGUAACC.....GUCAGGG
Hru-1-1    CCUGAC.GGCCGCGUUC....GGGUCAAU...UGCG.....GAC..........UGCA.ACACCCGG....UCUCU......CUGCUCGCGCCUCCG....UCGG............UGGAGGUC.CAGCAG............G.GGGA......GCGCGUCGCC.....AUCAGGA
env-174    CCUGAC.GGCCGCGUUC....GGGUCAAU...UGCG.....AAC..........UGCA.ACACCCGG....UCUC.GCUG..CUCGU..GCCUCCG.....UCGA..............UGGAGGUCGAGGAG..............GGGAC.....GCGCGUCGCC.....AUCAGGA
Hru-2-1    CCUGAC.GACCGCGUCC....GGGUCAAU...UGCG.....GAC..........UGCA.ACACCCGAUC..UCCCUG.....CUCGU..CCCGCCG.....UCGA..............UGGCGGUCGAGGAG............G.GGGA......GCGCGUCACA.....GUCAGGG
env-175    UCUGAC.GGGCGCGUCC....GGGUCGAA...UUGCG....GAC.........UGCAA..CACCCGAUC..UCCCUG.....CCGAUACCGCCA.......UCGA................UGGCGGUAUCGG..........CAG.GGGA......ACGCGUCACC.....AUCAGGG
env-176    UCUGAC.GGGCGCGUCC....GGGUCAAU...UGCG.....GAC..........UGCA.ACACCCGAUC..UCCCUA.....CCGGUGCCGC.........AGACGUUG.............GCGGUGUCGG..........AAG.GGGA......ACGCGUCACC.....AUCAGGG
env-178    UCUGAC.GGGCGCGUCC....GGGUCAAU...UGCG.....GAC..........UGCA.ACACCCGAUC..UCCCUG.....CCGGUACCGCCG.......ACGA................UGGCGGUGUCGG..........AAG.GGGA......ACGCGUCACC.....AUCAGGG
env-179    UCUGAC.GGGCGCGUCC....GGGUCAAU...UGCG.....GAC..........UGCA.ACACCCGAUC..UCCCUG.....CCGGUACCGCCG.......UCGA................UGGCGGUCUCGG..........CCG.GGGA......ACGCGUCACC.....AUCAGGG
env-180    UCUGAC.GGACGCGUCC....GGGUCAAU...UGCG.....GAC..........UGCA.ACACCCGAUC..UCCCU......UCCGGUGCCGCCG......UCGA................UGGCGGUUUCGGA...........GG.GGGA......GCGCGUCACC.....AUCAGGG
Nmo-1-1    UCUGAC.GGGCGCGUCC....GGGUCAAU...UGCG.....GAC..........UGCA.ACACCCGAUC..UCCCU......GCCGGUGCCGCCG......UCGA................UGGCGGUCUCGGC...........CG.GGGA......ACGCGUCACC.....AUCAGGG
Hha-1-1    CCUGAU.GGGCGCGUUC....GGGUCAAU...UGCG.....GAC..........UGCA.ACACCCGG....UCUCG......CUCGUCGUGCCUCC.....CGGAGU..........GGAGGUC.GGCGAG............G.GGGA......ACGCGUGACC.....GUCAGGA
env-181    CCUGAU.CGGCGCGUCC....GGGUCAAU...UGCG.....GCG..........UGCA.ACACCCGA....UCUCCCUCAA.CU..G..UCCGUCG.....GAAA..............UGACGGUCACCGAG............G.GGGA......GCGCGUGACG.....AUCAGGA
Hpa-1-1    CCUGAA.GGGCGCGUUC....GGGUCAAU...UGCG.....GAC..........UGCA.ACACCCGA....UCUCACUCG..UCGU..UCCGUCG......CAAA................UGACGGUCGGCGA..........GG.GAGA......ACGCGUUACC.....GUCAGGA
Hci-1-1    CCUGAA.GGGCGCGUCC....GGGUCAAU...UGCG.....GAU..........UGCA.ACACCCGAUU..UCGCUC.....GUCGU..CCCGUCG......CAAA..............UGACGGUCGGCGAG............G.GGGA......GCGCGUUACC.....GUCAGGA
env-182    CCUGAC.GGGCGCGUAC....GGGUCAAU...UG.CG....GCU.........UGACA.ACACCCGA....UCUCGCUUG..UCGUC.CCUCCG.......AAAA................UGGAGUCGACGA...........GG.GGGA......GCGCGUAACA.....GUCAGGA
env-183    CCUGAU.GGGCGCGUUC....GGGUCAAU...UGCG.....GCU..........UGCA.ACACCCGA....UCUCGCUCG..UCGU.GCCCUCCG......ACA.................UGGAGGUCGACGA............G.GGGA......GCGCGUGACC.....GUCAGGA
env-184    UCUGAC.GGGCGUGUCC....GGGUCAAU...UGCG.....GAC..........UGCA.ACACCCGA....UCUCCCUCGU.CGUU..CCCUCCG......GAA.................UGGAGGUCGAG.G..........AGG.GGGA......GCGCGUGACC.....GUCAGUA
env-185    CCUGAU.CAACGCGUGC....GGGUCAAU...UGCG.....GCU..........UGCA.ACACCCGG....UCUCCCUCAA.CACG....CCUCC.......CGGUAU...............GGAGGUCGCCGAG............G.GGGA......ACGCGUGACC.....GUCAGGA
Hdj-1-1    CCUGAA.GGGCGCGUCC....GGGUCAAU...UGCG.....GAC..........UGCA.ACACCCGA....UCUCCCUC...AACGUC..CCUCCG......AAAA..............UGGAGGUCGACGAG............G.GGGA......GCGCGUGACC.....GUCAGGA
Hsa-10-1   CCUGAA.GGGCGCGUCC....GGGUCAAU...UGCG.....GAC..........UGCA.ACACCCGA....UCUCCCUC...AAUGUC..CCUCCG......AAAA..............UGGAGGUCGACGAG............G.GGGA......GCGCGUGACC.....GUCAGGA
Nsu-1-1    CCUGAA.GGGCGCGUCC....GGGUCAAU...UGCG.....GCU..........UGCA.ACACCCGA....UCUCCCUC...AACGUC..CCUCCG......AAAA..............UGGAGGUCGACGAG............G.GGGA......GCGCGUGACC.....GUCAGGA
Nba-1-1    CCUGAA.GGGCGCGUCC....GGGUCAAU...UGCG.....GCU..........UGCA.ACACCCGA....UCUCCCUCA..ACGUU..CCUCCG......AAAA................UGGAGGUCGACGA...........GG.GGGA......GCGCGUGACC.....GUCAGGA

           .<<<<<....<<<<<<......<<<<...<...<<<<<<<<<...........>>>>>>>>>.>.>>>>......<<<<<......<<<<<<<<<<<<<<<<<<<............>>>>>>>>>>>>>>>>>>.............>.>>>>.....>>>>>>..........>>>>>..
           .20.......22222.......001...2...22221110..........01112222.2.100.......22222......222222..2.2.1111110.............0111111.2.2..222222.............2.2222.....22222..........02..
           YCUGAY-oR••RCGYoY----GGGU•RAY---•••o-----•••o----------oR••-RoRCCCR•----UYYC•------•••oo---------------•••o--------------------oooooo•------------R-GRRR------GCGY••••oo-----RUCAGGR
           5'                                                                                                                                                                                      3'
```

```
Nin-1-1   CCUGAA.GGGCGCGUCC....GGGUCAAU...UGCG.....GCU...........UGCA.ACACCCGA....UCUCCCUCAA.CACG....CCUCCG.....GAUA.............UGGAGGUCUCCGAG............G.GGGA......GCGCGUGACC.....GUCAGGA
env-186   CCUGAC.GGGCGCGUAC....GGGUCAAU...UGCG.....GCU...........UGCA.ACACCCGA....UCUCGCUC...GUCGUC..CCUCCG.....CAAA.............UGGAGGUCGACGAG............G.GGGA......GCGCGUAACA.....GUCAGGA
env-187   CCUGAC.GGGCGCGUAC....GGGUCAAU...UGCG.....GCU...........UGCA.ACACCCGA....UCUCGCUU...GUCGUC..CCUCCG.....AAAAA............UGGAGGUCGACGAG............G.GGGA......GCGCGUAACA.....GUCAGGA
env-188   CCUGAC.GGGCGCGUAC....GGGUCAAU...UGCG.....GCU...........UGCA.ACACCCGA....UCUCGCUG...GUCGUC..CCUCCG.....AAAA.............UGGAGGUCGACGAG............G.GAGA......GCGCGUAACA.....GUCAGGA
env-189   CCUGAC.GGGCGCGUAC....GGGUCAAU...UGCG.....GCU...........UGCA.ACACCCGA....UCUCGCUG...GUCGUC..CCUCCG.....AAAA.............UGGAGGUCGACGAG............G.GAGA......GCGCGUAACA.....GUCAGGA
env-190   CCUGAA.GGGCGCGUCC....GGGUCAAU...UGCG.....GAC...........UGCA.ACACCCGA....UCUCCCU....CAACA.CGCCUCC......CGGAAU............GGAGGUCGACGAG............G.GAGA......GCGCGUGACC.....GUCAGGA
env-191   CCUGAA.GGGCGCGUCC....GGGUCAAU...UGCG.....GAC...........UGCA.ACACCCGA....UCUCCCU....CAACA.CGCCUCC......CGGAAU............GGAGGUCGACGAG............G.GAGA......GCGCGUGACC.....GUCAGGA
Hla-1-1   CCUGAA.GGGCGCGUCC....GGGUCAAU...UGCG.....GAA...........UGCA.ACACCCGA....UCUCCCU....CAACAC.CCCUCC......UGACAU............GGAGGUCGAAGAG............G.GGGA......GCGCGUGACC.....GUCAGGA
env-193   CCUGAA.GGGCGCGUCC....GGGUCAAU...UGCG.....GCU...........UGCA.ACACCCAA....UCGCUCUCAA.CACA....CCUCC......CGGAAU............GGAGGUCGCUGAG............G.GCGA......GCGCGUGACC.....GUCAGGA
Hac-1-1   UCUGAA.GGGCGCGUUC....GGGUCAAU...UGCG.....GCA...........UGCA.ACACCCGA....UCUCC......CUCC...ACCUCC......CCGGCGAUAU........GGAGGUUUCGGAG............G.GGGA......GCGCGUGACC.....AUCAGGG
Hac-1-2   UCUGAA.GGGCGCGUUC....GGGUCAAU...UGCG.....GCA...........UGCA.ACACCCGA....UCUCC......CUCC...ACCUCC......CCGGCGAUAU........GGAGGUUUCGGAG............G.GGGA......GCGCGUGACC.....AUCAGGG
env-196   UCUGAA.GGGCGCGUUC....GGGUCAAU...UGCG.....GCA...........UGCA.ACACCCGA....UCUCCCUC...CACCA..GCCUCC......CGGUAU............GGAGGUUGUGGAG............G.GGGA......GCGCGUGACC.....AUCAGGG
env-197   UCUGAA.GGGCGCGUUC....GGGUCAAU...UGCG.....GCA...........UGCA.ACACCCGA....UCUCCCUC...CACCA..GCCUCC......CGGUAU............GGAGGUUGUGGAG............G.GGGA......GCGCGUGACC.....AUCAGGG
env-198   UCUGAA.GGGCGCGUUC....GGGUCAAU...UGCG.....GCA...........UGCA.ACACCCGA....UCUCCCUC...CACCA..GCCUCC......CGGUAU............GGAGGUUGUGGAG............G.GGGA......GCGCGUGACC.....AUCAGGG
Hsp-2-1   UCUGAA.GGGCGCGUUC....GGGUCAAU...UGCG.....GCA...........UGCA.ACACCCGA....UCUCCCUC...CACCA..GCCUCC......CGGUAU............GGAGGUUGUGGAG............G.GGGA......GCGCGUGACC.....AUCAGGG
env-199   CCUGAU.GGGCGCGUUC....GGGUCAAU...UGCG.....GCC...........UGCA.ACACCCGA....UCUCA......CUCGUGACCACC.......GACGA..............GGUGG..ACGAG............G.GGGA......GCGCGUGACC.....GUCAGGA
Hsu-1-1   UCUGAC.GGGCGCGUUC....GGGUCAAU...UGCG.....GAU...........UGCA.ACACCCAA....UCUCCCUC...CAC.GUC.CCGCC......CUCGAU............GGCGGUCGCGGAG............G.GGGA......ACGCGUGACC.....AUCAGGG
env-201   GCUGAU.GGGCGCGUCC....GGGUCAAU...UGCG.....GUA...........UGCA.ACACCCGA....UCUCC......CU..UCC.CGUCC......CAGCGGAUAU........GGACGUGGACCAG............G.GGGA......GCGCGUGACC.....GUCAGGG
env-203   GCUGAU.GGGCGCGUCC....GGGUCAAU...UGCG.....GUA...........UGCA.ACACCCGA....UCUCC......CU..UCC.CGUCC......CAGCGGAUAU........GGACGUGGACCAG............G.GGGA......GCGCGUGACC.....GUCAGGG
Hca-1-1   GCUGAU.GGGCGCGUCC....GGGUCAAU...UGCG.....GCA...........UGCA.ACACCCGA....UCUCC......CU..UCC.CGUCC......CAGCGGAUAU........GGACGUGGAACAG............G.GAGA......GCGCGUGACC.....GUCAGGG
Ham-1-1   GCUGAU.GGGCGCGUCC....GGGUCAAU...UGCG.....GCA...........UGCA.ACACCCGA....UCUCC......CU..UCC.CGUCC......CAGCGGAUAU........GGAUGUGGAACAG............G.GAGA......GCGCGUGACC.....GUCAGGG
Hhi-2-1   GCUGAU.GGGCGCGUCC....GGGUCAAU...UGCG.....GCA...........UGCA.ACACCCGA....UCUCU......CU..UCC.CGUCC......CAGCGGAUAU........GGACGUGGAACAG............G.GGGA......GCGCGUGACC.....GUCAGGG
Hsa-2-1   GCUGAU.GGGCGCGUCC....GGGUCAAU...UGCG.....GCA...........UGCA.ACACCCGA....UCUCC......CU..UCC.CGUCC......CAGCGGAUAU........GGACGUGGAACAG............G.GGGA......GCGCGUGACC.....GUCAGGG
Har-1-1   GCUGAU.GGGCGCGUCC....GGGUCAAU...UGCG.....GCA...........UGCA.ACACCCGA....UCUCC......CU..UCC.CGUCC......CAGCGGAUAU........GGACGUGGAACAG............G.GGGA......GCGCGUGACC.....GUCAGGG
Hva-1-1   GCUGAU.GGGCGCGUCC....GGGUCAAU...UGCG.....GCA...........UGCA.ACACCCGA....UCUCC......CU..UCC.CGUCC......CAGCGGAUAU........GGACGUGGAACAG............G.GGGA......GCGCGUGACC.....GUCAGGG
Hja-1-1   GCUGAU.GGGCGCGUCC....GGGUCAAU...UGCG.....GCA...........UGCA.ACACCCGA....UCUCC......CU..UCC.CGUCC......CAGCGGCUAU........GGACGUGGAACAG............G.GGGA......GCGCGUGACC.....GUCAGGG
env-204   CCUGAA.GGCCGCGUUC....GGGUCACU...UGCG.....GAC...........UGCA.ACACCCGA....UCUCC......CUCGGCGU..GCCGCCG..GAGA..........UGGCGACACCCGCCGAG............G.GGGA......GCGCGUCACC.....GUCAGGA
Hse-1-1   CCUGAU.GGGCGCGUUC....GGGUCAAU...UGCG.....GAC...........UGCA.ACACCCGA....UCUCG......CUCGUCGUGCCUCC.....CGAGAU...........GGAGGU.CGACGAG............G.GGGA......GCGCGUGACC.....GUCAGGA
Hth-1-1   CCUGAU.GGGCGCGUUC....GGGUCAAU...UGCG.....GAC...........UGCA.ACACCCGA....UCUCG......CUCGUCGCACCGCCG....GAGA............UGGCGGU.CGACGAG............G.GGGA......GCGCGUGACC.....GUCAGGA
Hmo-1-1   CCUGAU.GGGCGCGUUC....GGGUCAAU...UGCG.....GAC...........UGCA.ACACCCGA....UCUCG......CUCGUCGCACCGCCG....GAGA............UGGCGGU.CGACGAG............G.GGGA......GCGCGUGACC.....GUCAGGA
env-206   CCUGAC.CGGCGCGUCC....GGGUCAAU...UGCG.....GCA...........UGCA.ACACCCGAUC..UCCCU......CGUCGCCCGCC........CGAAAC.............GGCGAU.CGACG..........AGG.GGGA......GCGCGUGACA.....GUCAGGA
env-207   CCUGAC.GGCCGCGUUC....GGGUCACU...UGCG.....GAC...........UGCA.ACACCCGA....UCUCG......CUCCGUACCACG.......CGCGA..............CGUG..GUGGAG............G.GGGA......GCGCGUCACC.....GUCAGUG
env-209   CCUGAA.GGCCGCGUUC....GGGUCACU...UGCG.....GAC...........UGCA.ACACCCGA....UCUCG......CUCCUCGUCCCCCCCG...AGA............UGGCGGUC..GAGGAG............G.GAGA......GCGCGUCACC.....AUCAGGA
Nta-1-1   CCUGAU.CAACGCGUGC....GGGUCAAU...UGCG.....GCU...........UGCA.ACACCCGA....UCUCC......CUCAACGUGCCGCCG....AAAA............UGGCG.CUCGACGAG............G.GCGA......GCGCGUGACU.....AUCAGGA
Nas-1-1   CCUGAU.CAACGCGUGC....GGGUCAAU...UGCG.....GCU...........UGCA.ACACCCGA....UCUCC......CUCAACGUGCCGCCG....AAAA............UGGCG.CUCGACGAG............G.GCGA......GCGCGUGACU.....AUCAGGA
Noc-1-1   CCUGAU.CAACGCGUGC....GGGUCAAU...UGCG.....GCU...........UGCA.ACACCCGA....UCUCC......CUCAACGUGCCUCC.....CUCGAU...........GGAG.CUCGAAGAG............G.GGGA......GCGCGUGACC.....GUCAGGA
Nje-1-1   CCUGAU.CAACGCGUGC....GGGUCAAU...UGCG.....GCU...........UGCA.ACACCCGA....UCUCC......CUCAACGUGCCUCC.....CGGUAU...........GGAG.CUCGAGGAG............G.GGGA......GCGCGUAACC.....GUCAGGG
Nam-1-1   CCUGAU.CAACGCGUGC....GGGUCAAU...UGCG.....GCU...........UGCA.ACACCCGA....UCUCC......CUCAACGUGCCUCC.....CGGUAU...........GGAG.CUCGAGGAG............G.GGGA......GCGCGUGACC.....GUCAGGA
env-210   CCUGAU.CAACGCGUCC....GGGUCAAU...UGCG.....GCU...........UGCA.ACACCCGA....UCUCC......CUCAACGUUCCGCCA....GGAA............UGGCG.GUCGACGAG............G.GAGA......GCGCGUGACC.....GUCAGGA
Hsp-3-1   UCUGAA.GGGCGCGUUC....GGGUCAAU...UGCG.....GCA...........UGCA.ACACCCGA....UCUCC......CUCCACGAACCACCA....GGAA............UGGCGG.UCUCGGAA............G.GAGA......GCGCGUGACC.....AUCAGGG
Nch-1-1   CCUGAU.CAACGCGUGC....GGGUCAAU...UGCG.....GCU...........UGCA.ACACCCGA....UCUCC......CUCAACGUACCGCCG....AAAA............UGGCGGU.CUCCGUG............G.GAGA......GCGCGUGACU.....AUCAGGA
Nae-1-1   CCUGAU.CAACGCGUGC....GGGUCAAU...UGCG.....GCU...........UGCA.ACACCCGA....UCUCC......CUCAACGUGCCGCCG....AAAA............UGGCGGU.CGACGAG............G.GCGA......GCGCGUGACU.....AUCAGGA
env-214   CCUGAU.CAACGCGUGC....GGGUCAAU...UGCG.....GCU...........UGCA.ACACCCGA....UCUCC......CUCAACGUGCCUCC.....CGGUAU...........GGAGGU.CGACGAG............G.GGGA......GCGCGUGACU.....GUCAGGA
env-215   CCUGAU.CAACGCGUGC....GGGUCAAU...UGCG.....GCU...........UGCA.ACACCCGA....UCUCC......CUCAACGUGCCUCC.....CGGUAU...........GGAGGU.CGACGAG............G.GGGA......GCGCGUGACU.....GUCAGGA
env-216   CCUGAA.GGGCGCGUCC....GGGUCAAU...UGCG.....GAC...........UGCA.ACACCCGA....UCUCC......CUCAAUACGCCUCC.....CGGAAU...........GGAGGU.GGCUGAG............G.GAGA......GCGCGUGACC.....GUCAGGG
Hni-1-1   CCUGAA.GGGCGCGUCC....GGGUCAAU...UGCG.....GCA...........UGCA.ACACCCGA....UCUCC......CUCAAUACACCUCC.....CGACAU...........GGAGGU.CUCGGAG............G.GGGA......GCGCGUGACC.....GUCAGGA
Has-1-1   CCUGAA.GGCCGCGUCC....GGGUCAAU...UGCG.....GAC...........UGCA.ACACCCGA....UCUCC......CUCAACGUGCCACCG....AAAA............UGGUGGU.CGAUGAG............G.GGGA......GCGCGUGACC.....GUCAGGA
env-217   CCUGAA.GGGCGCGUCC....GGGUCAAU...UGCG.....GAC...........UGCA.ACACCCGA....UCUCC......CUCAACAUACCGCCG....AACA............UGGCGGU.CGAAGAG............G.GGGA......GCGCGUGACC.....GUCAGGA
Nal-2-1   CCUGAA.GGGCGCGUCC....GGGUCAAU...UGCG.....GAC...........UGCA.ACACCCGA....UCUCC......CUCAACGUACCGCCG....AAAA............UGGCGGU.CGAAGAG............G.GGGA......GCGCGUGACC.....GUCAGGA
Npa-1-1   CCUGAA.GGGCGCGUCC....GGGUCAAU...UGCG.....GAC...........UGCA.ACACCCGA....UCUCC......CUCAACGUACCGCCG....AAAA............UGGCGGU.CGAAGAG............G.GGGA......GCGCGUGACC.....GUCAGGA
Nga-1-1   CCUGAA.GGGCGCGUCC....GGGUCAAU...UGCG.....GAC...........UGCA.ACACCCGA....UCUCC......CUCAACGUACCGCCG....AAAA............UGGCGGU.CGAAGAG............G.GGGA......GCGCGUGACC.....GUCAGGA
Nsp-2-1   CCUGAA.GGGCGCGUCC....GGGUCAAU...UGCG.....GAC...........UGCA.ACACCCGA....UCUCC......CUCAACGUACCGCCG....AAAA............UGGCGGU.CGAAGAG............G.GGGA......GCGCGUGACC.....GUCAGGA
Htu-1-1   CCUGAA.GGGCGCGUCC....GGGUCAAU...UGCG.....GAU...........UGCA.ACACCCGA....UCUCC......CUCAACGUACCGCCG....AAAA............UGGCGGU.CGAGGAG............G.GGGA......GCGCGUGACC.....GUCAGGA
Hsa-8-1   CCUGAA.GGGCGCGUCC....GGGUCAAU...UGCG.....GAC...........UGCA.ACACCCGA....UCUCC......CUCAACGUACCGCCG....AAAA............UGGCGGU.CGACGAG............G.GGGA......GCGCGUGACC.....GUCAGGA
Hth-2-1   CCUGAA.GGGCGCGUCC....GGGUCAAU...UGCG.....GAC...........UGCA.ACACCCGA....UCUCC......CUCAACGUACCGCCG....AAAA............UGGCGGU.CGAGGAG............G.GGGA......GCGCGUGACC.....GUCAGGA
Nve-1-1   CCUGAA.GGGCGCGUCC....GGGUCAAU...UGCG.....GAC...........UGCA.ACACCCGA....UCUCC......CUCAACGUACCGCCG....AAAA............UGGCGGU.CGAGGAG............G.GGGA......GCGCGUGACC.....GUCAGGA
Hli-3-1   CCUGAA.GGGCGCGUCC....GGGUCAAU...UGCG.....GAC...........UGCA.ACACCCGA....UCUCC......CUCAACACACCUCC.....UGACAU...........GGAGGU.CGACGAG............G.GAGA......GCGCGUGACC.....GUCAGGA
Hxa-1-1   CCUGAA.GGGCGCGUCC....GGGUCAAU...UGCG.....GAC...........UGCA.ACACCCGA....UCUCC......CUCAACACGCCUCCG....GAUA............UGGAGGU.CGACGAG............G.GGGA......GCGCGUGACC.....GUCAGGA
Hgo-1-1   CCUGAU.CAUCGCGUCC....GGGUCAAU...UGCG.....GCU...........UGCA.ACACCCGA....UCUCG......CUCAACGUACCGCC.....CGAAAU...........GGCGGU.CGCUGAG............G.GAGA......ACGCGUGACA.....GUCAGGA
env-218   CCUGAU.CAUCGCGUCC....GGGUCAAU...UGCG.....GCU...........UGCA.ACACCCGA....UCUCC......CUCAACGUACCACCA....GAAA............UGGCGGU.CGAGGAG............G.GAGA......ACGCGUGACA.....GUCAGGA
env-219   CCUGAU.CGGCGCGUCC....GGGUCAAU...UGCG.....GAU...........UGCA.ACACCCGA....UCUCC......CUCACUGUGCCGUCA....GCAC............UGACGGU.CGCCGAG............G.GUGA......GCGCGUGACG.....AUCAGGA
env-220   UCUGAC.GGGCGUGUCC....GGGUCAAU...UGCG.....GAC...........UGCA.ACACCCGA....UCUCC......CUCGUUGUUCCCUCCG...GAA.............UGGAGGU....CGAG.........GCGG.GAGA......ACGCGUGACC.....GUCAGUA
env-221   CCUGAA.GGGCGCGUCC....GGGUCAAU...UGCG.....GCU...........UGCA.ACACCCGA....UCUCC......CUCAACGUUCCGCCG....GGAA............UGGCGGUCG.AAGAG............G.GGGA......GCGCGUGACC.....GUCAGGA
Hsa-6-1   CCUGAA.GGCCGCGUCC....GGGUCAAU...UGCG.....GAA...........UGCA.ACACCCGA....UCUCCCUCA..UCACCC.CCGACG......GAA...............UGUCGGUGGGCGA...........GG.GAGA......GCGCGUCACC.....GUCAGGA
env-222   UCUGAC.CGACGCGUCC....GGGUCAAU...UGCG.....GAC...........UGCA.ACACCCUAU...CUCCC......UUCCCGUCCCGCCG.....UCGA.............UGGU..GGUCGAGA...........AG.GGCGA.....ACGCGUCACU.....GUCAGGG
env-223   UCUGAU.GGGCGCGUCC....GGGUCAAU...UGCG.....GCU...........UGCA.ACACCCGA....UCUUGC.....UGCCGGU.GCCA.......GACGACA............UGGCCGCCGGCA..........AGC.GGGA......GCGCGUGACC.....GUCAGGG
env-224   GCUGAC.GGGCGCGUCC....GGGUCAAU...UGCG.....GCC...........UGCA.ACACCCAA....UCUCGC.....UGCCGG.UGCCGG......AUGGA.............UUGGCAGUCGGCG..........AGC.GCGA......GCGCGUCACC.....GUCAGGG
env-225   CCUGAU.AAUCGCGUUU....GGGUAAUU...UGCG.....GAC...........UGCA.AAACCCAAUU..UCCCU......AACAAUCCCCA........CUAAUA..............UGGAUAUUGUA...........GA.GGGACA....ACGCGUUUCU.....GUCAGAA
env-226   CCUGAU.AAUCGCGUUU....GGGUAAUU...UGUG.....GAC...........UGCA.AAACCCAAUU..UCCCU......AACAAUUCCCA........CUAAUA..............UGGAUAUUGUA...........GA.GGGACA....ACGCGUUUCU.....GUCAGAA
env-227   CCUGAU.AAUCGCGUUU....GGGUAAUU...UGUG.....GAC...........UACA.AAGCCCAA....UUUCUC.....UC.ACAAUACCUCC.....UAACAU...........GGA..UGUUGUCGA...........UG.GACA......ACGCGUUUCU.....GUCAGAG
env-228   CCUGAU.AACCGCGUUC....GGGUCAAU...UGUG.....AAU...........UGCA.ACACCCAGUCUCCCUUCAUGU..GCCAGA.............AUCAAUA..................UCUGUC.........CGAG.AAGGAAC...ACGCGUUGCU.....AUCAGGA
env-231   UCUGAA.CGUGGCGUAAA...GGGUGGAC...GUAAACCC.AGAU......GGG.UUAU.GUGCCCAU....UCCCU......GCCAU..............CCCA......................AUGGC..........CAG.GUGA......ACGCCCAC.......AUCAGGA
env-232   UCUGAA.CGUGGCGUAAA...GGGU.GAC...GUAAACCC.AGAU......GGG.UUAU.GUGCCCAU....UCCCU......GCCAU..............CCCA......................AUGGC..........CAG.GUGA......ACGCCCAC.......AUCAGGA
env-233   UCUGAA.CGUGGCGUAAA...GGGUGGAC...GUAAACCC.CAGAU.....GGG.UUAU.GUGCCCAU....UCCCU......GCCAU..............CCCA......................AUGGC..........CAG.GUGA......ACGCCCAC.......AUCAGGA
env-237   UCUGAU.GAGUGUGUCAU...GGGUGGAC...G........AUACCAAAA........C.G.GCCCG.....CUCUG......AAAAG..............ACAAUUGC..................UCUUU...........AC.AGGG.....GACACAUAAGU.....AUCAGGA
env-238   UCUGAC.GACCGCGUAU....GGGUCGAU...GUCG.....GUCU..........CGAC.A.GCCCAU....CACUC......GCAAAAG............GUAAAG..................CAAAGGU............G.GGUGAAC...GCGUGACA.......GUCAGGU
env-240   UCUGAC.CGGAGCGUUC....GGGUCGAU...GUCG.....GUAU..........CGAC.A.GCCCAAU...CUCCCUGG...AGACGCCU...........GAAA...................GGGCG.CU......GAAGAAG.GGAGA.....GCGCAUAUCA.....GUCAGAA
env-241   UCUGAC.ACAGUCACAU....GGGUCGAU...AUU......GGUGUA.........GAU.A.GCCCGG....UUCCCCUCAA.ACCUUCC............UCAA....................GGAAGGU......GCAAAAG.GGCGAGC...GUGUCAAU.......GUCAGGA
env-242   ACUGAU.GUGUGCGCUCAU..GGGCCAAC...GUU......GAAG...........AAC.GUGCCCGG....UUCAC......CGU................GAAA........................ACG...........GG.AGAA.....AGCGCCCCA.......GUCAGAA
env-243   ACUGAU.GUGUGCGCUCAU..GGGCCAAC...GUU......GAAG...........AAC.GUGCCCGG....UUCAC......CGU................GAAA........................ACG...........GG.AGAA.....AGCGCCCCA.......GUCAGAA
env-244   UCUGAU...AAGCGCAC....GGGUGGAC...GUG......GUCUCUA........CAC.GAGCCCGC....AUGUG......CCGC...............AGG........................GCGG............U.ACAUAU....GCGCGAGA.......CUCAGUA
env-245   ACUGAU...CAGCGC.AG...GGGUGGAC...GUGC.....UCU...........GCAC.GAGCCCGC....GUGUA......UCAGCA.............UCA......................UACUGG............U.ACAUCC....GCGCGAUA.......GUCAGGU
env-246   ACUGAU...CAGCGC.AG...GGGUGGAC...GUGC.....UCU...........GCAC.GAGCCCGC....GUGUA......UCAGCA.............UCA......................UACUGG............U.ACAUCC....GCGCGAUA.......GUCAGGU

          .<<<<<....<<<<<<.....<<<<...<...<<<<<<<<<.........>>>>>>>>>.>.>>>>......<<<<<......<<<<<<<<<<<<<<<<<<<............>>>>>>>>>>>>>>>>>>>............>.>>>>.....>>>>>>..........>>>>>..
          .20.......22222......001...2...22221110..........01112222.2.100.......22222......222222..2.2.1111110............0111111.2.2..222222............2.2222.....22222..........02..
          YCUGAY-•R••RCGY•Y----GGGU•RAY---••••-----•••○---------○R••-R•RCCCR•----UYYC•------•••○○-------------••••--------------------○○○○○•-----------R-GRRR------GCGY••••○○-----RUCAGGR
          5′                                                                                                                                                                               3′
```

```
env-247    ACUGAU...CAGCGC.AG...GGGUGGAC...GUGC.....UCU..........GCAC.GAGCCCGC....GUGUA......UCAGCA............UCA.....................UACUGG...........U.ACAUCC....GCGCGAUA.......GUCAGGU
env-248    ACUGAU...CAGCGC.AG...GGGUGGAC...GUGC.....UCU..........GCAC.GAGCCCGC....GUGUA......UCAGCA............UCA.....................UACUGG...........U.ACAUCC....GCGCGAUA.......GUCAGGU
env-249    UCUGAU..AUCACGCAAA...GGGUGGAC...GUCUU....CCC.........GGAAC.GAGCCCGU....UUCUG......CCA...............UGC.........................UGG...........C.AGAAAAC...GCGUGAAA.......GUCAGGA
env-250    UCUGAU..AUCACGCAAA...GGGUGGAC...GUCUU....CCC.........GGAAC.GAGCCCGU....UUUUG......CC................AUACA........................GG...........C.AGAAAAC...GCGUGAAC.......AUCAGGA
env-251    UCUGAUUGAACGCGC.C....GGGCGAAC...GU.......GAAA............AC.GAGCCCAC....UUUCU......GC................AUA..........................GC..........CG.GAAA.....GCGCGUCCGG.....GUCAGGA
env-252    UCUGAUUGAACGCGC.C....GGGCGAAC...GU.......GAAA............AC.GAGCCCAC....UUUCU......G.................AAUACA........................C...........A.GAAA.....GCGCGUCCGG.....GUCAGGA
env-253    UCUGAUUGAACGCGC.C....GGGCGAAC...GU.......GAAA............AC.GAGCCCAC....UUUCU......GA................ACUA.........................UC...........A.GAAA.....GCGCGUCAGG.....GUCAGGA
env-254    ACUGAU.CAAGCCGCCU....GGGUCGAC...GUUG.....GUG..........CAAC.GAGCCCUG....UUCCU......AUGGCCG...........AUGUAA..................UGCCCAU...........A.GGAAG....GUGGAUCACG.....GUCAGAA
env-255    UCUGAU..CAGCGCGUAUU..GGGCGAAC...GUU......CGUGC...........AAC.GAGCCCUU....CUUCA......UACAGC............CAUUAAU..................GAUGUG...........AU.GAAGAU...GCGCCAUU.......GUCAGGA
env-256    UCUGAU.CGAGACGA.U....GGGCCAAC...GC.......CAGUGCA..........GC.GCGCCCGG....UUCCG......GGAA..............CACUC......................UUUC...........C.GGAA.....ACGUCGGAGA.....AUCAGGA
env-257    UCUGAU.CGAGACGA.U....GGGCCAAC...GC.......CAGUGCA..........GC.GCGCCCGG....UUCCG......GGAA..............CACUC......................UUUC...........C.GGAA.....ACGUCGGAGA.....AUCAGGA
env-258    UCUGAU.CGAGACGA.U....GGGCCAAC...GC.......CAGUGCA..........GC.GCGCCCGG....UUCCG......GGAA..............CACUC......................UUUC...........C.GGAA.....ACGUCGGAGA.....AUCAGGA
env-259    UCUGAU.CGAGACGA.U....GGGCCAAC...GC.......CAGUGCA..........GC.GCGCCCGG....UUCCG......GGAA..............CACUC......................UUUC...........C.GGAA.....ACGUCGGAGA.....AUCAGGA
env-260    UCUGAU.CGAGACGA.U....GGGCCAAC...GC.......CAGUGCA..........GC.GCGCCCGG....UUCCG......GGAA..............CACUC......................UUUC...........C.GGAA.....ACGUCGGAGA.....AUCAGGA
env-261    UCUGAU.CGAGACGC.U....GGGCCAAC...GC.......CUGUGCA..........GC.GUGCCCGA....UUCCA......GGAGA.............ACG.......................UUGCC...........U.GGGA.....GCGUCGAAGA.....AUCAGGA
env-265    UCUGAU.CGAGACGC.U....GGGCCAAC...GC.......CAGUGCA..........GC.GCGCCCGG....UUCC.......GGGGA.............GAAUG.....................UCCCC.............GGAA.....GCGUCGAAGU.....GUCAGGA
env-266    UCUGAU.CGAGACGC.U....GGGCCAAC...GC.......CAGUGCA..........GC.GCGCCCGG....UUUC.......CGGGA.............UAAAUC....................UUCCC.............GGGA.....GCGUCAGAGA.....GUCAGGA
env-268    UCUGAU.CGAGACGC.U....GGGCCAAC...GC.......CAGUGCA..........GC.GCGCCCGG....UUUC.......CGGGA.............UAAAUC....................UUCCC.............GGGA.....GCGUCAGAGA.....GUCAGGA
env-269    UCUGAU.CGAGACGC.U....GGGCCAAC...GC.......CAGUGCA..........GC.GCGCCCGG....UUUC.......CGGGA.............UAAAUC....................UUCCC.............GGGA.....GCGUCAGAGA.....GUCAGGA
env-270    UCUGAU.CGAGACGC.U....GGGCCAAC...GC.......CAGUGCA..........GC.GCGCCCGG....UUUC.......CGGGA.............UAAAUC....................UUCCC.............GGGA.....GCGUCAGAGA.....GUCAGGA
env-271    UCUGAU.CGAGACGU.U....GGGCCAAC...GC.......CAGUGCA..........GC.GCGCCCGG....UUCC.......AGAGA.............ACA.......................UCUCU..........CC..GGAA.....ACGUCAAAGA.....GUCAGGA
env-272    UCUGAU.CGAGACGU.U....GGGCCAAC...GC.......CAGUGCA..........GC.GCGCCCGG....UUCC.......AGAGA.............ACA.......................UCUCU..........CC..GGAA.....ACGUCAAAGA.....GUCAGGA
env-274    UCUGAU.CGAGACGU.U....GGGCCAAC...GU.......CAGUGCA..........GC.GCGCCCGG....UUCC.......AGAGA.............ACA.......................UCUCU..........CC..GGAA.....ACGUCAAAGA.....GUCAGGA
env-275    UCUGAU.CGAGACGU.U....GGGCCAAC...GC.......CAGUGCA..........GC.GCGCCCGG....UUCC.......AGAGA.............ACA.......................UCUCU..........CC..GGAA.....ACGCCAAAGA.....GUCAGGA
env-277    UCUGAU.CGAGACGU.U....GGGCCAAC...GC.......CAGUGCA..........GC.GCGCCCGG....UUCC.......AGAGA.............ACA.......................UCUCU..........CC..GGAA.....ACGCCAAAGA.....GUCAGGA
env-278    UCUGAU.CGAGACGU.U....GGGCCAAC...GC.......CGGUGCA..........GC.GCGCCCGG....UUCC.......GGGAA.............CAGGCU....................UUCCC..........CC..GGAA.....ACGUUGAAGA.....AUCAGGA
env-292    UCUGAU.CGAGACGU.U....GGGCCAAC...GC.......CGGUGCA..........GC.GAGCCCGU....UUCCG......GGAA..............CAAGU......................UUCC...........C.GAAA.....GCGUCGAAGA.....GUCAGGA
env-293    UCUGAU.CGAGACGU.U....GGGCCAAC...GC.......CGGUGCA..........GC.GAGCCCGU....UUCCG......GGAA..............CAAGU......................UUCC...........C.GAAA.....GCGUCGAAGA.....GUCAGGA
env-295    UCUGAU.CGAGACGU.U....GGGCCAAC...GC.......CGGUGCA..........GC.GAGCCCGU....UUCCG......GGAA..............CAAGU......................UUCC...........C.GAAA.....GCGUCGAAGA.....GUCAGGA
env-296    UCUGAU.CGAGACGU.U....GGGCCAAC...GC.......CGGUGCA..........GC.GAGCCCGU....UUCCG......GGAA..............CAAGU......................UUCC...........C.GAAA.....GCGUCGAAGA.....GUCAGGA
Mbo-1-1    UCUGAU.CGAGACGU.U....GGGCCAAC...GC.......CGGUGCA..........GC.GAGCCCGU....UUCCG......GGAA..............CAAGU......................UUCC...........C.GAAA.....GCGUCGAAGA.....GUCAGGA
env-297    UCUGAU.CGAGACGU.U....GGGCCAAC...GC.......CGGUGCA..........GC.GAGCCCGU....UUCC.......GGGAA.............CAGGU.....................UUCCC.............GAAA.....GCGUCGAAGA.....GUCAGGA
env-298    UCUGAU.CGAGACGU.U....GGGCCAAC...GC.......CGGUGCA..........GC.GAGCCCGU....UUCC.......GGGAA.............CAGGU.....................UUCCC.............GAAA.....GCGUCGAAGA.....GUCAGGA
env-303    UCUGAU.CGAGACGU.U....GGGCCAAC...GC.......CGGUGCA..........GC.GAGCCCGU....UUCC.......GGGAA.............CAGGU.....................UUCCC.............GAAA.....GCGUCGAAGA.....GUCAGGA
env-304    UCUGAU.CGAGACGU.U....GGGCCAAC...GC.......CGGUGCA..........GC.GAGCCCGU....UUCC.......GGGAA.............CANGU.....................UUCCC.............GAAA.....GCGUCGAAGA.....GUCAGGA
env-305    UCUGAU.CGAGACGU.U....GGGCCAAC...GC.......CAGUGCA..........GC.GCGCCCGG....UUCCA......GAGA..............AGAAUAUC...................UCUC...........C.GGAA.....ACGUCGAAGA.....GUCAGGA
env-306    UCUGAU.CGAGACGU.U....GGGCCAAC...GC.......CAGUGCA..........GC.GCGCCCGG....UUCCA......GAGA..............AGAAUAUC...................UCUC...........C.GGAA.....ACGUCGAAGA.....GUCAGGA
Mma-1-1    UCUGAU.CGAGACGU.U....GGGCCAAC...GC.......CAGUGCA..........GC.GCGCCCGG....UUCCG......GGAA..............GAAUG......................UUCC...........C.GGGA.....GCGUCGAAGA.....GUCAGGA
Mse-1-1    UCUGAU.CGAGACGC.U....GGGCCAAC...GC.......CAGUGCA..........GC.GCGCCCGG....UUCCG......GGAA..............GAGUG......................UUCC...........C.GGGA.....GCGUCGAAGA.....GUCAGGA
Msp-1-1    UCUGAU.CGAGACGC.U....GGGCCAAC...GC.......CAGUGCA..........GC.GCGCCCGG....UUCCG......GGAA..............GAGUG......................UUCC...........C.GGGA.....GCGUCGAAGA.....GUCAGGA
env-310    UCUGAU.CGAGACGU.C....GGGCCAAC...GC.......CAGUGCA..........GC.GAGCCCGU....UUCCU.......................GAG.........................................A.GGAGA....GCGUCGAAGA.....GUCAGGA
env-315    UCUGAU.CGAGACGU.C....GGGCCAAC...GC.......CAGUGCA..........GC.GAGCCCGU....UUCCU.......................GAG.........................................A.GGAGA....GCGUCGAAGA.....GUCAGGA
env-316    UCUGAU.CGAGACGU.C....GGGCCAAC...GC.......CAGUGCA..........GC.GAGCCCGU....UUCCU.......................GAG.........................................A.GGAGA....GCGUCGAAGA.....GUCAGGA
env-317    UCUGAU.CGAGACGU.U....GGGCCAAC...GC.......CAGUGCA..........GC.GCGCCCGG....UUCC.......AGAGA.............ACAAUA....................UCUCU..........CC..GGAA.....ACGUCAAAGA.....GUCAGGA
env-318    UCUGAU.CGAGGCGC.A....GGGCCAAC...GC.......CAGUGCA..........GC.GAGCCCGU....UUUCU.......................GAAA........................................G.GAGA.....GCGUCGAAGA.....GUCAGGA
env-320    UCUGAU.CGAGGCGC.A....GGGCCAAC...GC.......CAGUGCA..........GC.GAGCCCGU....UUUCU.......................GAAA........................................G.GAGA.....GCGUCGAAGA.....GUCAGGA
env-321    UCUGAU.CGAGGCGC.A....GGGCCAAC...GC.......CAGUGCA..........GC.GAGCCCGU....UUUCU.......................GAAA........................................G.GAGA.....GCGUCGAAGA.....GUCAGGA
env-322    UCUGAU.CGAGGCGC.A....GGGCCAAC...GC.......CAGUGCA..........GC.GAGCCCGU....UUUCU.......................GAAA........................................G.GAGA.....GCGUCGAAGA.....GUCAGGA
Mpa-1-1    UCUGAA....GGACAGU....GGGUGGAC...GUU......CGUGC...........AAC.GAGCCCGU....UCCUG......CA................CAC..........................UG...........C.AGGAGAU...UGUCCUG........UUCAGGA
env-323    UCUGAA...GGACAG.U....GGGUGGAC...GUU......CGUGC...........AAC.GAGCCCGU....UCCUG......CA................CUC..........................UG...........C.AGGAGA....UGUGCCUG.......UUCAGGA
env-324    UCUGAU....ACCGGGCAU..GGGUGGAC...GUU......CGUGC...........AAC.GAGCCCGU....UCCUG......A.................ACAU..........................U...........C.AGGAU.....GCGUUCAU.......UUCAGGA
env-325    UCUGAC.CAGGACGCAU....GGGUCGAC...GUU......CGUGC...........AAC.GAGCCCGU....UCCUUCU....GCUGCA............UCAG....................UGUAGC...........A.GGGAU.....GCGCCAUU.......GUCAGGA
env-326    UCUGAC.CAGGACGCAU....GGGUCGAC...GUU......CGUGC...........AAC.GAGCCCGU....UCCUUCU....GCUGCA............UCAG....................UGUAGC...........A.GGGAU.....GCGCCAUU.......GUCAGGA
env-331    UCUGAC.CAGGACGCAU....GGGUCGAC...GUU......CGUGC...........AAC.GAGCCCGU....UCCUUCU....GCUGCA............UCAG....................UGUAGC...........A.GGGAU.....GCGCCAUU.......GUCAGGA
env-332    UCUGAC.CAGGACGCAU....GGGUCGAC...GUU......CGUGC...........AAC.GAGCCCGU....UCCUUCU....GCUGCA............UCAG....................UGUAGC...........A.GGGAU.....GCGCCAUU.......GUCAGGA
Mfo-1-1    UCUGAU..CAGGCGCAU....GGGCGAAC...GUU......CGUGC...........AAC.GAGCCCCU....UCUUCGU....GCGGAC............UAAU....................GUCCGC.........GUG.AAGAU.....GCGCCAUU.......GUCAGGA
env-333    UCUGAC.CGUGACGCAU....GGGUGGAC...GUUU.....CCAUAAA........AAAC.GAGCCUGU....UCCCC......GG................AAUUAC.......................CC...........G.GGGAU.....GCGUCAAU.......GUCAGGG
env-334    UCUGAC.CGUGACGCAU....GGGUGGAC...GUUUU....CAUAAA.......AAAAC.GAGCCUGU....UCCCC......GG................AAUAC........................CC...........G.GGGAU.....GCGUCAAU.......GUCAGGG
env-335    UCUGAC.CGUGACGCAU....GGGUGGAC...GUUUU....CAUAAA.......AAAAC.GAGCCUGU....UCCCC......GG................AAUAC........................CC...........G.GGGAU.....GCGUCAAU.......GUCAGGG
env-337    UCUGAC.CGUGACGCAU....GGGUGGAC...GUUUU....CAUAA........AAAAC.GAGCCUGU....UCCCC......GG................AAUAC........................CC...........G.GGGAU.....GCGUCAAU.......GUCAGGG
env-338    UCUGAC.CGUGACGCAC....GGGUGGAC...GUUUC....CAAGA........GAAAC.GAGCCUGU....UCCCC......GG................AAUUAC.......................CC...........G.GGGAU.....GCGUCAAU.......GUCAGGG
env-339    UCUGAC.CGUGACGCAC....GGGUGGAC...GUUUC....CAAGA........GAAAC.GAGCCUGU....UCCCU......GG................AAUAC........................CC...........G.GGGAU.....GCGUCAAU.......GUCAGGG
CMe-1-1    UCUGAC.CGCGACGCAU....GGGUGGAC...GUUU.....AAA............AAAC.GAGCCUGG....UUCCC......UGA...............AUCA........................UCA...........G.GGAAU.....GCGUCAGU.......GUCAGGG
env-340    ACUGAC.AGGUACGCAU....GGGUCGAC...GUU......CGUGC...........AAC.GAGCCCGU....UCCC.GC....UGCUU.............CAAUAAU....................GGGCA..........GA..AGGAU.....GCGUUAUU.......GUCAGGA
env-341    UCUGAC.CGGUACGCAU....GGGUCGAC...GUUU.....GAAC...........AAAC.GAGCCCAU....UCCACA.....UGCUCU............AAAA.....................AGUGCA..........CG.UGGAU.....GCGUUAUU.......GUCAGGA
env-349    UCUGAC.CGGUACGCAU....GGGUCGAC...GUUU.....GAAC...........AAAC.GAGCCCAU....UCCACA.....UGCUCU............AAAA.....................AGUGCA..........CG.UGGAU.....GCGUUAUU.......GUCAGGA
env-350    UCUGAC.CGAUACGCA.....GGGUCGAC...GUU......CGUGC...........AAC.GAGCCCGC....UCCCG......CUGCUU............CCCUCAAA..................GAGUAG...........A.AGGAU.....GCGUUAUU.......GUCAGGA
env-351    UCUGAC.CGGUACGCAU....GGGUCGAC...GUU......CGUGC...........AAC.GAGCCCGU....UCCCG......CUGUUU............CAAUAAUUU.................GGGCAG...........A.AGGAU.....GCGUUAUU.......GUCAGGA
env-352    UCUGAU.UACCGCGCAU....GGGUCGAC...GUU......CGUGC...........AAC.GAGCCCGU....UCCCC......C.................UGAAUAC.......................G...........G.GGGAU.....GCGUCAUU.......AUCAGGG
env-353    UCUGAU.AAGCACGCAU....GGGUCGAC...GUUU.....GAAC...........AAAC.GAGCCCGU....UCCAA......UUGCUC............AAAUCAUA..................GGGUGA...........A.UGGAU.....GCGUUAUU.......GUCAGGA
env-354    UCUGAU.AAGCACGCAU....GGGUCGAC...GUUU.....GAAC...........AAAC.GAGCCCGU....UCCAA......UUGCUC............AAAUUAUA..................GGGCGG...........A.UGGAU.....GCGUUAUU.......GUCAGGA
env-355    UCUGAU.AAGCACGCAU....GGGUCGAC...GUUU.....GAAC...........AAAC.GAGCCCGU....UCCAA......UUGCUC............AAAUCAUA..................GGGCGG...........A.UGGAU.....GCGUUAUU.......GUCAGGA
env-356    UCUGAU.AAGCACGCAU....GGGUCGAC...GUUU.....GAAC...........AAAC.GAGCCCGU....UCCAA......UUGCUC............AAAUCAUA..................GGGCGG...........A.UGGAU.....GCGUUAUU.......GUCAGGA
env-357    UCUGAC.CAGCACGCAU....GGGUCGAC...GUUU.....AAAC...........AAAC.GAGCCCGU....UCCAC......CUGUUCU...........CAUCAU..................GGAGCAG...........A.UGGAU.....GCGUUAUU.......GUCAGGA
env-358    UCUGAC.CAGCACGCAU....GGGUCGAC...GUUU.....GAAC...........AAAC.GAGCCCGU....UCCAC......CUGCUCU...........CCUCAU..................GGAGAAG...........A.UGGAU.....GCGUUAUU.......GUCAGGA
env-359    UCUGAC.CAGCACGCAU....GGGUCGAC...GUUU.....GAAC...........AAAC.GAGCCCGU....UCCAC......CUGCUCU...........CCUCAU..................GGAGAAG...........A.UGGAU.....GCGUUAUU.......GUCAGGA
env-360    UCUGAC.CAGCACGCAU....GGGUCGAC...GUUU.....GAAC...........AAAC.GAGCCCGU....UCCAC......CUGCUCU...........CCUCAU..................GGAGAAG...........A.UGGAU.....GCGUUAUU.......GUCAGGA
env-361    UCUGAC.CAGCACGCAU....GGGUCGAC...GUUU.....GAAC...........AAAC.GAGCCCGU....UCCAC......CUGCUCU...........CCUCAU..................GGAGAAG...........A.UGGAU.....GCGUUAUU.......GUCAGGA
env-370    UCUGAC.CAGCACGCAU....GGGUCGAC...GUUU.....GAAC...........AAAC.GAGCCCGU....UCCAA......UUGCCCCA..........UCA.....................UGGGGCAG...........A.UGGAU.....GCGUUAUU.......GUCAGGA
env-371    UCUGAC.CAGCACGCAU....GGGUCGAC...GUUU.....GAAC...........AAAC.GAGCCCGU....UCCAA......UUGCUCUG..........AUCA....................CGGAGCAG...........A.UGGAU.....GCGUUAUU.......GUCAGGA
env-372    UCUGAC.CGGUACGCAU....GGGUCGAC...GUUU.....GAAC...........AAAC.GAGCCCGU....UCCAA......AUGCUC............UAAUAAAAU.................GAGCCU...........C.UGGAU.....GCGUUAUU.......GUCAGGA
env-373    UCUGAC.CGGUACGCAU....GGGUCGAC...GUUU.....GAAU...........AGAC.GAGCCCGU....UCCAA......AUGCUCUA..........AAA....................UAGAGCAU...........C.UGGAU.....GCGUUAUU.......GUCAGGA
env-374    UCUGAC.CAGCACGUAU....GGGUCGAC...GUU......AAAAA...........AAC.GAGCCCGU....UCCAU......CCGCUC............UAAAAAAU..................GUGCAG...........A.UGGAU.....GCGUUAUU.......GUCAGGA

           .<<<<<....<<<<<<.....<<<<...<...<<<<<<<<<.........>>>>>>>>>.>.>>>>.....<<<<<......<<<<<<<<<<<<<<<<<<............>>>>>>>>>>>>>>>>>>............>.>>>>.....>>>>>>..........>>>>>..
           .20.......22222......001...2...22221110..........01112222.2.100......22222......222222..2.2.1111110............0111111.2.2..222222............2.2222.....22222...........02..
           YCUGAY-•R••RCGY•Y----GGGU•RAY---•••○------•••○----------○R••-R•RCCCR•----UYYC•-------•••○○--------------•••○--------------------○○○○○•------------R-GRRR------GCGY•••••○○----RUCAGGR
           5'                                                                                                                                                                                        3'
```

```
env-375   UCUGAU.UAGGACGCAU....GGGUCGAC...GUU.....CGUGC.........AAC.GAGCCCGU....UCCCU......CUGCUGU...........AAUAAUA.................GCAGCAG...........G.AGGAU.....GCACCAAU.......AUCAGGA
env-376   UCUGAU.UAGGACGCAU....GGGUCGAC...GUU.....CGUGC.........AAC.GAGCCCUU....UCCCU......AUGCUGC...........AACAGA..................GCAGCAG...........G.UGGAU.....GCGCCAAU.......AUCAGGA
env-377   UCUGAU.UAGGACGCAU....GGGUCGAC...GUU.....CGUGC.........AAC.GAGCCCGU....UCCCU......AUGCUGC...........AACAAA..................GCAGCAG...........G.UGGAU.....GCGCCAAU.......AUCAGGA
env-378   UCUGAC.UAAGACGCAU....GGGUCGAC...GUU.....CGUGC.........AAC.GAGCCCUU....UCCCU......CUGCUGC...........AACAUGA.................GCAGCAG...........G.UGGAU.....GCGCCAUU.......GUCAGGA
env-379   UCUGAC.CAGGACGCAU....GGGUCGAC...GUU.....CGUGC.........AAC.GAGCCCUU....UCCCU......CUGCUGC...........AAAA....................GCAGCAG...........G.CGGAU.....GCGCCAUU.......GUCAGGA
env-380   UCUGAC.CAGGACGCAU....GGGUCGAC...GUU.....CGUGC.........AAC.GAGCCCCU....UCCUC......CUGC..............UGAU.......................GCAG...........G.UGGAU.....GCGACCAAAA.....GUCAGGA
env-381   UCUGAC.CAGGACGCAU....GGGUCGAC...GUU.....CGUGC.........AAC.GAGCCCCU....UCCUC......CUGC..............UGAU.......................GCAG...........G.UGGAU.....GCGACCAAAA.....GUCAGGA
env-382   UCUGAC.CGGGACGCAU....GGGUCGAC...GUU.....CGUGC.........AAC.GAGCCCCU....UCUGC......CUGC..............UAAC.......................GCAG...........G.UGGAU.....GCGACCAAAA.....GUCAGGA
env-383   UCUGAC.CAGGACGCAU....GGGUCGAC...GUU.....CGUGC.........AAC.GAGCCCCU....UCCGC......CUGC..............UUAU.......................GCAG...........G.UGGAU.....GCGACCAAAA.....GUCAGGA
env-384   UCUGAC.CAGGACGCAU....GGGUCGAC...GUU.....CGUGC.........AAC.GAGCCCCC....UUCAC......CUGC..............UGAC.......................GCAG...........G.AGGAU.....GCGACCAGAA.....GUCAGGA
env-385   UCUGAC.CAGGACGCAA....GGGUCGAC...GUU.....CGUGC.........AAC.GAGCCCAU....UCCAC......CUGC..............AUUAA......................GCAG...........G.UGGAU.....GCCCCAAAA......GUCAGGA
env-386   UCUGAC.CAGGACGCAU....GGGUCGAC...GUU.....CGUGC.........AAC.GAGCCCAU....UCCAC......CUGCU.............AUA.......................AGCAG...........G.UGGAU.....GCCCCAAAA......GUCAGGA
env-387   UCUGAC.CAGGACGCAU....GGGUCGAC...GUU.....CGUGC.........AAC.GAGCCCAU....UCCAC......CUGCU.............AUA.......................AGCAG...........G.UGGAU.....GCCCCAAAA......GUCAGGA
env-389   UCUGAC.CAGGAUGCAU....GGGUCGAC...GUU.....CGUGC.........AAC.GAGCCCUC....UCCGC......CUG...............CUAAGC......................CAG...........G.CGGAU.....GCGCCAAAA......GUCAGGA
env-390   UCUGAC.CAGCACGCAU....GGGUCGAC...GUU.....CAUGC.........AAC.GAGCCCGU....UUCAU......CUGCUUCA..........AACA....................UGAGGCAG...........A.UGGAU.....GCGUUAUU.......GUCAGGA
env-392   UCUGAU..CAGGCGCAU....GGGCCAAC...GUU.....CGUGC.........AAC.GAGCCCCC....UUCAC......GUG...............AAAAC.......................CAC...........G.UGAAU.....GCGCCAUU.......AUCAGGA
env-395   UCUGAU.UACGACGCAU....GGGCCGAC...GUU.....CGUGC.........AAC.GAGCCCUC....UCCAC......CUGC..............UAAC.......................GCAG...........G.UGGAU.....GCGCCAUU.......AUCAGGA
env-399   UCUGAU.CUGGCCGCAU....GGGUCGAC...GUU.....CGUGC.........AAC.GAGCCCGU....UCCGCGAU...GUUAC.............CUUUGA....................GUAAC.........AAA.CGGAU.....GCGCCAAU.......GUCAGGA
env-400   UCUGAC.CAGGACGCAU....GGGUCGAC...GUU.....CGUGC.........AAC.GAGCCCC.....UCUCCU.....UACAACC...........UGAC....................GGUAGUA..........AG.GAGAU.....GCGACCAAAA.....GUCAGGA
env-402   UCUGAU.UAGGACGCAU....GGGUCGAC...GUU.....CGUGC.........AAC.GAGCCCUU....UCCCUAU....GCUGC.............AACAGA......................GCAGC.........AGG.UGGAU.....GCGCUAAU.......AUCAGGA
env-403   UCUGAU.UAGGACGCAU....GGGUCGAC...GUU.....CGUGC.........AAC.GAGCCCUU....UCCCUAU....GCUGC.............AACAGA......................GCAGC.........AGG.UGGAU.....GCGCUAAU.......AUCAGGA
env-405   UCUGAU.UAGGACGCAU....GGGUCGAC...GUU.....CGUGC.........AAC.GAGCCCUU....UCCCCCG....UCACA.............GGAUA......................UGUGA........GUUG.GGGAU.....GCGCCAUU.......AUCAGGA
env-406   UCUGAU.UAGGACGCAU....GGGUCGAC...GUU.....CGUGC.........AAC.GAGCCCUU....UCCCCCG....UCACA.............GGAUA......................UGUGA........GUUG.GGGAU.....GCGCCAUU.......AUCAGGA
env-407   UCUGAU.UGGGACGCAU....GGGUCGAC...GUU.....CGUGC.........AAC.GAGCCCGU....UCCGC......UGCC..............UCAUUGAAU...................GGCA...........GAAGGAU.....GCGUUAUU.......GUCAGGA
Mli-2-1   UUUGAU.GUAUGCGCAUAU..GGGUGGAC...GAA.....AUUU..........UUC.G.GCCCUU....ACUC...............................UUAAAU.................................GAGUAGU...GCAUAUUAAG.....GUCAGAA
Mpe-1-1   UCUGAU.AAAUGCGCCUCAA.GGGUGGAC...GAA.....AUU...........UUC.G.GCCCUU....UAUCC..............................GACG...........................................G.GAUA.....GCGCACAAAUAG...GUCAGAA
Mpa-2-1   UCUGAU.AAAUGCGCUUUUGAGGGUGGAC...GAA.....AUU...........UUC.G.GCCCUU....UAUCC..............................GACG...........................................G.GAUA.....GCGCACAAAUAG...GUCAGAA
env-408   UCUGAU.UAAUGCGCUCA...GGGUGGAC...GAA.....AUAU..........UUC.G.GCCCUU....UAUCC..............................GAAG...........................................G.GAUA.....GCGCAUAAUAG....GUCAGAA
Mmo-1-1   UCUGAU.UAAUGCGCCCAA..GGGUGGAC..AGAUGCG....GAUG.......UGCAUC.GGGCCCUU....UCUU.A.....AAGGCA............CAAG....................UGUCUU.........AA..AAGA.....GCGCAAAUUACUG..GUCAGAA
env-409   UCUGAU.UAAUGCGCCCAA..GGGUGGAC..AGAUGCG....GAUG.......UGCAUC.GGGCCCUU....UCUU.A.....AAGGCA............CAAA....................UGUCUU.........AA..AAGA.....GCGCAAAUUACUG..GUCAGAA
env-410   UCUGAU.UGACGCGC.C....GGGCCAAC...G.......CACCGU...........C.GGCCCAU.....UUCCC......GG................CAGU.........................CC...........G.GGAA.....GCGCGCUUAGG....GUCAGGA
Mco-1-1   ACUGAC.GGAAACGCAU....GGGUCGAU...GUCG....GAAG.........CGAC.A.GCCCGGCAAUGCUCCC......CGAC..............GCAAAA......................GUCG.......AGAAG.GGGCAGC...GCGUGACA.......GUCAGGA
env-413   UCUGAC.AAAAAUGCAU....GGGUCGAU...AUCG....GAAU.........CGAU.A.GCCCGGUC..CGCCUCGAA..GCACCUU...........CAAGA...................AAAGUGC......GAACU.GGCGAGC...GCGUUAAU.......GUCAGGA
env-414   UCUGAC.AAAAGCGCAU....GGGUCGAU...AUC.....GGAAUA.........GAU.A.GCCCGGUC..CGCC.UCGAA.GCACCUUC..........AAG....................GAAAGUGC......GAACU.GGCGAGC...GCGUUAAU.......GUCAGAA
env-415   UCUGAC.AAAAGCGCAU....GGGUCGAU...AUCG....GAAU.........CGAU.A.GCCCGGUC..CGCCUCGAA..GCACCUUC..........AAG....................GAAAGUGC......GAACU.GGCGA....GCGCGUUAAN.....GUCAGAA
env-416   UCUGAC.ACAGGCGCAU....GGGUUGAU...AUCG....GAAU.........CGAU.A.GCCCGGUC..CGCCUCGAA..GCACCUUC..........AAG....................GAAAGUGC........GAAAUGGCGA...GCGCGUUAAU.....GUCAGAA
env-418   UCUGAU.AAAUAUGCAU....GGGUCGAU...AUC.....GGAAUA.........GAU.A.GCCCGG....UUCUCUUAAAGUACUUC...........AGCAG....................GAAGUAU......CAAUA.GGAGA....GCGCGUUAAU.....GUCAGAA
env-423   UCUGAC.AAAUUCGUAU....GGGUCGAU...AUC.....GGAAUA.........GAU.A.GCCCGGU....UCGCCUUGAAGUACUUUC..........GAA...................GGAAGUAU.....CAAUAGG.AGAAC....GCGUUAAU.......GUCAGGA
env-425   UCUGAC.AAAUUCGUAU....GGGUCGAU...AUC.....GGAAUA.........GAU.A.GCCCGGUU..CGCCU......UGGAGUACUUC.......CGAAA...............GAAGUAU..CA...........AUA.GGAGAAC...GCGUUAAU.......GUCAGGA
env-428   UCUGAU.GGCGGCGCAU....GGGUGGAA...AUCG....GUGU.........CGAU..CGCCCAA....UUCCC......CUUCGAAC..........UGAUGAGAU...............GUUCGAAG..........CU.GGAAA....ACGCCUAAAU.....GUCAGAA
env-429   UCUGAU.GGCGGCGCAU....GGGUGGAA...AUCG....GUGU.........CGAU..CGCCCAA....UUCCC......CUUCGAAC..........UGAUGAGAU...............GUUCGAAG..........CU.GGAAA....ACGCCUAAAU.....GUCAGAA
Msh-1-1   UCUGAC.GGCAGCGCAU....GGGUCGA...ACUCG....GACU.........CGAG..AGCCCA.....GCUCC......ACU..CAGAAACCC.....GAAA.............GG...UGUGAAGGU...........G.GGGCA....GCGCAUAGAC.....GUCAGGA
env-430   UCUGAU.GGCGGCGUCU....GGGUGGAA...AUCG....GAAU.........CAA..UCGCCCAAUUC.CCCUUCGCAC.CGGAGAA...........AAG......................UUC.CCG...........GUAGGGCA....GCGCCUAAGU.....GUCAGAA
env-433   UCUGAU.GGCGGCGUCU....GGGUGGAA...AUCG....GAAU.........CAA..UCGCCCUAU...UCCCCUUCG..CACCGG............AGAAAAGUAC................UCGGUG...........G.GGCA.....GCGCCUAAGU.....GUCAGAA
env-434   UCUGAU.GGCGGCGUAU....GGGUGGAA...AUCG....GAAU.........CAAU..CGCCCAAU...UCCCCUUCGC.ACCGGA............GAAAAUUA..................UCCGGU.........AG.GGCA.....ACGCCUAAAC.....GUCAGAA
env-435   UCUGAU.GGCGGCGUAU....GGGUGGAA...AUCG....GAAU.........CAAU..CGCCCAAU...UCCCCUUCGC.ACCGGA............GAAAAGUA..................UCCGGU.........AG.GGCA.....ACGCCUAAAC.....GUCAGA.
env-436   UCUGAU.GGCGGCGUAU....GGGUGGAA...AUCG....GAAU.........CAAU..CGCCCAAU...UCCCCUUCGC.ACCGGA............GAAAAGUA..................UCCGGU.........AG.GGCA.....ACGCCUAAAC.....GUCAGAA
env-437   UCUGAU.GGCGGCGUAU....GGGUGGAA...AUCG....GAAU.........CAAU..CGCCCAAU...UCCCCUUCGC.ACCGGA............GAAAAGUA..................UCCGGU.........AG.GGCA.....ACGCCUAAAC.....GUCAG..
env-439   UCUGAU.GGCGGCGUAU....GGGUGGAA...AUCG....GAAU.........CAAU..CGCCCAAU...UCCCCUUCGC.ACCGGA............GAAAAGUA..................UCCGGU.........AG.GGCA.....ACGCCUAAAC.....GUCAGAA
env-440   UCUGAU.GGCGGCGUAU....GGGUGGAA...AUCG....GAAU.........CAAU..CGCCCAAU...UCCCCUUCGC.ACCGGA............GAAAAGUA..................UCCGGU.........AG.GGCA.....ACGCCUAAAC.....GUCAGAA
env-441   UCUGAU.GGCGGCGUAU....GGGUGGAA...AUCG....GAAU.........CAAU..CGCCCAAU...UCCCCUUCGC.ACCGGA............GAAAAGUA..................UCCGGU.........AG.GGCA.....ACGCCUAAAC.....GUCAGAA
env-442   UCUGAU.GGCGGCGUAU....GGGUGGAA...AUCG....GAAU.........CAAU..CGCCCAAU...UCCCCUUCGC.ACCGGA............GAAAAGUA..................UCCGGU.........AG.GGCA.....ACGCCUAAAC.....GUCAGAA
env-443   UCUGAU.AGCGGCGUCU....GGGUGGAA...AUCG....GAAU.........CAA..UCGCCCAAU...UCCCCUUCGC.ACCGA.............AAGUAGGUAC.................UCGGU.........AG.GGGA.....GCGCCUNAAU.....GUCAGAA
env-444   UCUGAU.AAAGGCGUCU....GGGUGGAA...AUCG....GAGU.........UGA..UCGCCCAAUA..CCCCUA.....GUGUC.............GAAAGA......................GAUAU.....ACAAGCC.GGGGCA....ACGCCUAAAU.....GUCAGAA
env-445   UCUGAU.GGCGGCGUUU....GGGUGGAA...AUCG....GAAU.........CGAU..CGCCCAAUU...CCCCU......CCG..CACC..........GGAGGCA..................GGUACUCGG............U..GGGGCA....GCGCCUUAAC.....GUCAGAA
env-446   UCUGAU.GGCGGCGUUU....GGGUGGAA...AUCG....GAAU.........CGAU..CGCCCAAUU...CCCCU......CCG..CACC..........GGAGGCA..................GGUACUCGG............U..GGGGCA....GCGCCUUAAC.....GUCAGAA
env-448   UCUGAU.GGCGGUGCAU....GGGUGGAA...AUCG....GUAU.........CGAU..CGCCCAGU....GCCUCUG....CGAACC............GAGAAUGA...................GGUUCG........ACCG.GGGCA....GCGCCUCAAC.....GUCAGAA
env-449   UCUGAU.GGUGGCGCAU....GGGUGGAA...AUCG....GUGU.........CGAU..CGCCCAAUAC..CCCUC......CGCA..............CUGAGGAGAAAUA..................UGCG...........GUAGGGAG....ACGCCUGAAC.....GUCAGAA
env-450   UCUGAU.GGCGGCGUCU....GGGUGGAA...AUCG....GAAU.........CAAU..CGCCCUAUU...CCCCU......UCG..UA....CC.......GGU...............GGUAAUACUCGG............U..GGGGCA....GCGCCUAAAU.....GUCAGAA
env-451   UCUGAU.GGCGGCGUAU....GGGUGGAA...AUCG....GAAU.........CGAU..CGCCCAAUU...CCCCU......CCGC..ACC..........GUGGGUA..................GGUACUCGG............U..GGGGCA....GCGCCUAAGU.....GUCAGAA
env-452   UCUGAU.GGCGGCGUAU....GGGUGGAA...AUCG....GAAU.........CGAU..CGCCCAAUU...CCCCU......CCGC..ACC..........GUGGGUA..................GGUACUCGG............U..GGGGCA....GCGCCUAAGU.....GUCAGAA
env-455   UCUGAU.GGCGGCGUCU....GGGUGGAA...AUCG....GAAU.........CGAU..CGCCCAAUU...CCCCU......CCGC..ACC..........GUGGGUA..................GGUACUCGG............U..GGGGCA....GCGCCUAAAU.....GUCAGAA
env-456   UCUGAU.GGCGGCGUCU....GGGUGGAA...AUCG....GAAU.........CGAU..CGCCCAAUU...CCCCU......CCGC..ACC..........GCGGGUA..................GGUACUCGG............U..GGGGCA....ACGCCUAAAU.....GUCAGAA
env-460   UCUGAU.GACGGCGUAU....GGGUGGAA...AUCG....GAAU.........CGAU..CGCCCAAUU...CCCCUUC....GCACC.............GAUAA......................GGUGU.....ACCAGUUG.GGGGA....GCGCCUUGAC.....GUCAGAA
env-462   UCUGAU.GACGGCGUAU....GGGUGGAA...AUCG....GAAU.........CGAU..CGCCCAAUU...CCCCUCC....GCACC.............GAUAA......................GGUGU.....ACCAGUUG.GGGGA....GCGCCUUGAC.....GUCAGAA
env-464   UCUGAU.GACGGCGUAU....GGGUGGAA...AUCG....GAAU.........CGAU..CGCCCAA.....UUCCCCUCC..GCACC.............GAUAA......................GGUGU.....ACCAGGCG.GGGGA....GCGCCUUGAC.....GUCAGAA
env-465   UCUGAU.GACGGCGUAU....GGGUGGAA...AUCG....GAAU.........CGAU..CGCCCAAUU...CCCCUC.....CGCACC............GAUAA.....................GGUGUG......CCAGGUG.GGGGA....GCGCCUUGAC.....GUCAGAA
env-466   UCUGAU.GGCGGCGUCU....GGGUGGAA...AUCG....GAGU.........CGAU..CGCCCAAUU...CCCCUCC....GCACC.............AAUGG......................GGUGU......UCGAGAUGGGGCA....GCGCCUAAGU.....GUCAGAA
env-467   UCUGAC.GACGGCGCAU....GGGUGGAG...AUAG....GUGU.........UGAG.C.GCCCAU....UGCCCCUCAG.GACCGAAA...........AGCGG..................UUUCGGUU............GG.GGCA.....GCGCCUGAAC.....GUCAGAG
env-468   UCUGAC.GACGGCGCAU....GGGUGGAG...AUAG....GUGU.........UGAG.C.GCCCAU....UGCCCCUCAG.GACCGAAA...........AGCGG..................UUUCGGUU............GG.GGCA.....GCGCCUGAAC.....GUCAGAG
env-469   UCUGAC.GACGGCGCAU....GGGUGGAG...AUAG....GUGU.........UGAG.C.GCCCAU....UGCCCCUCAG.GACCGAAA...........AGCGG..................UUUCGGUU............GG.GGCA.....GCGCCUGAAC.....GUCAGAA
env-471   UCUGAC.GACGGCGCAU....GGGUGGAG...AUAG....GUGU.........UGAG.C.GCCCAU....UGCCCCUCAG.GACCGAAA...........AGCGG..................UUUCGGUU............GG.GGCA.....GCGCCUGAAC.....GUCAGAN
env-472   UCUGAC.GACGGCGCAU....GGGUGGAG...AUAG....GUGU.........UGAG.C.GCCCAU....UGCCCCUCAG.GACCGAAA...........AGCGG..................UUUCGGUU............GG.GGCA.....GCGCCUGAAC.....GUCAGAA
env-474   UCUGAC.AACGGCGCAU....GGGUGGAG...AUAG....GUGU.........UGAG.C.GCCCAU....UGCCC......CUCAAGACCGC.......AUC..................GCGGUACAGAG.........CAG.GGCA.....GCGCCUGAAC.....GUCAGAG
env-475   UCUGAC.GACGGCGCAU....GGGUGGAG...AUAG....GUGU.........UGAG.C.GCCCUU....UGCCC......CUCAGAACCGC.......GAA..................GCGGUUUUCUG.........CUG.GGCA.....GCGUCUGAAC.....GUCAGAG
Mha-1-1   UCUGAC.GACGGCGCAU....GGGUGGAG...AUAG....GUGU.........UGAG.C.GCCCUU....UGCCC......CUCAGAACCGC.......GAA..................GCGGUUUUCUG.........CUG.GGCA.....GCGUCUGAAC.....GUCAGAG
env-476   UCUGAC.GACGGCGCAU....GGGUGGAG...AUAG....GUGU.........UGAG.C.GCCCUU....UGCCC......CUGAGAAC..........CGCUCUGC.................GGUUUCAG.......ACUG.GGCA.....GCGUCUGAAC.....GUCAGAA
env-477   UCUGAC.AACGGCGCAU....GGGUGGAC...AUAG....GUG..........UUGA.GUGCCCAU....UGCCCCUCAAGACCGC..............UCG..........................GCGGU....ACGAAGUGG.GGCA.....GCGCCUGAAC.....GUCAGAG
env-478   UCUGAU.AAAGGCGUCU....GGGUGGAA...AUCG....GAGU.........CGAU..CGCCCUG.....UGCCC......CUCGUC............ACA...........................GACGAGAAAUACAAGUUGG.GGCA.....GCGCCUGAAC.....GUCAGGA
env-479   UCUGAU.GGUGGCGCAA....GGGUGGAA...CUCG....GAGU.........CGAG.U.GCCCGG....UGCCCCUCC..CUCACC............AAGAAAAC...................GGUGAG.......ACAG.GGCA.....GCGCCUGAAC.....GUCAGAA
env-480   UCUGAU.GGUGGCGCAA....GGGUAGAA...CUCG....GAGU.........UGAG.U.GCCCGG....UGCCCCUCCA.CACC..............AAGAGAAC.....................GGUG.....AGACGAG.GGCA.....GCGCCUGAAC.....GUCAGAA
env-481   CCUGAC.GGAAGCGCAU....GGGUAGAG...CUCG....GGUU.........UGAG.C.GCCCAGCGUACUUC.......CUGAGCCU..........AAAG.....................GGGCAAAG...........G.AAGGAGA...GCGCGUGAAC.....GUCAGGA
env-482   CCUGAC.GGUGGCGUUU....GGGUAGAU...AUAG....GUG..........UUGG.ACGCCUUG....CCCUC......CUCAA.............AAAUGG......................UAAAG.........CUG.GGCA.....ACGUGUGACC.....GUCAGGG

          .<<<<....<<<<<.....<<<<...<...<<<<<<<<<..........>>>>>>>>>.>.>>>>......<<<<......<<<<<<<<<<<<<<<<<<.............>>>>>>>>>>>>>>>>>>...........>.>>>>.....>>>>>...........>>>>>..
          .20......22222......001...2...22221110..........01112222.2.100.......22222......222222..2.2.1111110.............0111111.2.2..222222...........2.2222.....22222...........02..
          YCUGAY-•R••RCGY•Y---GGGU•RAY---••••-----•••○---------○R••-R•RCCCR•----UYYC•-------••○○○-------------••••○---------------------○○○○○•-----------R-GRRR-----GCGY•••••○○-----RUCAGGR
          5'                                                                                                                                                                              3'
```

```
env-483    ACUGGC....UGCGCUU....GGGUGGAA...AUC......GGUUUG.........GAU.U.GCCCAG....CCUUU......UUGUGCA.CG.........AAUA..................CGCUGCGCAA............G.GGGGA.....GCGCGUGACA.....GUUAAGA
env-484    ACUGGC....UGCGCUU....GGGUGGAA...AUC......GGUUUG.........GAU.U.GCCCAG....CCUUU......UUGUGCA.CG.........AAUA..................CGCUGCGCAA............G.GGGGA.....GCGCGUGACA.....GUUAAGA
env-485    ACUGGC....UGCGCUU....GGGUGGAA...AUC......GGUUUG.........GAU.U.GCCCAG....CCUUU......UUGUGCA.CG.........AAUA..................CGCUGCGCAA............G.GGGGA.....GCGCGUGACA.....GUUAAGA
env-486    ACUGGC....UGCGCUU....GGGUGGAA...AUC......GGUUUG.........GAU.U.GCCCAG....CCUUU......UUGUGCA.CG.........AAUA..................CGCUGCGCAA............G.GGGGA.....GCGCGUGACA.....GUUAAGA
env-492    ACUGGC....UGCGCUU....GGGUGGAA...AUC......GGUUUG.........GAU.U.GCCCAG....CCUUU......UUGUGCA.CG.........AAUA..................CGCUGCGCAA............G.GGGGA.....GCGCGUGACA.....GUUAAGA
env-495    ACUGAC....UGCGCUU....GGGUUGAA...AUC......GGUUUG.........GAU.U.GCCCAG....CCUUU......UUGUGCA.CG.........AAGA..................CGCUGCGCAA............A.GGGGA.....GCGCGUGACA.....GUCAAGA
env-504    ACUGAC....UGCGCUU....GGGUUGAA...AUC......GGUUUG.........GAU.U.GCCCGG....CCCUU......UUGCGAACCG.........CGA...................CGUUGCGCAA............A.GGGGA.....GCGCGUGACA.....GUUAAGA
env-505    UGUGAU.UUCGGCAA......GGGUGGAC...G........AUG..............C.G.GCCCAU....UACCU......GUGA...............UUGA........................UCAC............G.GGUU......AAGCCUGAAU.....UUCACGC
env-508    UCUGAA.AGCGGCGAGAA...GGGCGGAC...GUAUCU...UCUG........GGAUAU.GAGCCCGG....UACC.......AGCCA..............GCAGG......................UGGCA..............GGUAC.....ACGCCCGU.......UUCAGGA
env-509    ACUGAAAACAGGCAUGCG...GGGUGGAC...GUAUCC...CCUG........GGAUAU.GAGCCCAU....AAUCU......GCCUU..............GAA........................ACGGC............A.GGUUC.....AUGCUGGA.......UUCAGGA
Mta-1-1    GCUGAA.ACAGGCAUGCA...GGGUGGAC...GAAUCC...CCUG........GGAUAC.GGGCCCGU....CAUCU......GCC................CAUAAAAAGA...................GGC............A.GGUGC.....AUGCCGGA.......UUCAGGA
env-511    GCUGAA.ACAGGCAUGCA...GGGUGGAC...GAAUCC...CCUG........GGAUAC.GGGCCCGU....CAUCU......GCC................CAUAAAAAGA...................GGC............A.GGUGC.....AUGCCGGA.......UUCAGGA
env-512    GCUGAA.ACAGGCAUGCA...GGGUGGAC...GAAUCC...CCUG........GGAUAC.GGGCCCGU....CAUCU......GCC................CAUAAAAAGA...................GGC............A.GGUGC.....AUGCCGGA.......UUCAGGA
Hbo-1-1    CCUGAC.GGCCGCGUUC....GGGUCAAU...UGCG.....GAA...........UGCA.ACACCCGA....UCUCC......CUCUACACGCCAUCG...GAUA.............UGAUGGUGAA.AGAG............G.GGGA......GCGCGUCGCC.....AUCAGGA
Ngr-1-1    CCUGAA.GGGCGCGUCC....GGGUCAAU...UGCG.....GCU...........UGCA.ACACCCGA....UCUCC......CUCAACACACCUCC.....CGACGU............GGAGGU.CGCCGAG............G.GGGA......GCGCGUGACC.....AUCAGGA
Hje-1-1    CCUGAA.GGGCGCGUAC....GGGUCAAU...UGCG.....GAC...........UGCA.ACACCCAC....UCUCC......CUCAUAACGCCUCC.....CGGAAU............GGAGGU.CCAGCAG............G.GGGA......ACGCGUGACC.....GUCAGGA
Nma-1-1    CCUGAU.CAACGCGUGC....GGGUCAAU...UGCG.....GCU...........UGCA.ACACCCGA....UCUCC......CUCAAUGUACCGCCG...AAAA.............UGGCGGU.CUCCGUG............G.GAGA......GCGCGUGACU.....AUCAGGA
Hal-1-1    CCUGAC.GACCGCGUUC....GGGUCAAU...UGCG.....GAA...........UGCA.ACACCCGA....UCUCCC.....UGUUCGUCCC.........UCCC..................GGGAUGGACC....UCACAGAGG.GAGA......GCGCGUCGCU.....GUCAGGA
Nti-1-1    CCUGAA.GGGCGCGUCC....GGGUCAAU...UGCG.....GCU...........UGCA.ACACCCGA....UCUCC......CUCAACGUUCCUCCG...AAAA.............UGGAGGU.CGAAGAG............G.GGGA......GCGCGUGACC.....GUCAGGA
Npe-1-1    CCUGAA.GGGCGCGUCC....GGGUCAAU...UGCG.....GAC...........UGCA.ACACCCGA....UCUCC......CUCAACGUACCGCCG...AAAA.............UGGCGGU.CGAGGAG............G.GAGA......GCGCGUGACC.....GUCAGGA
Hti-1-1    UCUGAU.GGGCGCGUCC....GGGUCAAU...UGCG.....GUA...........UGCA.ACACCCGA....UCUCC......CUC....GCCGUUCCGG..GCGA...........CCGGA.CGAUACGUGAG............G.GGGA......ACGCGUCACC.....GUCAGGA
           .<<<<<....<<<<<<.....<<<<...<...<<<<<<<<<.........>>>>>>>>>.>.>>>>......<<<<<......<<<<<<<<<<<<<<<<<<<.............>>>>>>>>>>>>>>>>>>>............>.>>>>.....>>>>>>..........>>>>>..
           .20.......22222......001...2...22221110..........01112222.2.100.......22222......222222..2.2.1111110..............0111111.2.2..222222............2.2222......22222..........02..
           YCUGAY-∘R●●RCGY∘Y----GGGU●RAY---●●∘○-----●●●○----------○R●●-R∘RCCCR●----UYYC●------●●∘∘○--------------●●●∘----------------------○○∘∘∘●------------R-GRRR------GCGY●●●●○○-----RUCAGGR
           5′                                                                                                                                                                                3′
```

alignment positions 260 · · · 439

```
Mli-1-1    UCUCUUCGCGAGACGAGGUAUUCCGGUGCACACCGGAAUGAUCUCCAUGACAUGGAGACCGCGACCCACGGCACAUCACUCCCGACACCACCGCAGCAGGACUCCAGAUGAUCUGCUGCGCGGAAGAAGCGUGCCCGUUCGAUCUGUCCGGACCCCGGUAACGGGGCGAGGAGUGGGCCG
env-1      AUCUCUUGCUUGAGACCGGUCGUUAUAUGAGAAUUGCAGAGAAGCAUUCUAACCGCAUCACGGGGAAAGGAUUUCUGGCAGGAUAUUCCUGCAUGAAGACCAUUGCGGUGCCAGUGUGAUGUUUUUGGAGAAUGCUUCGUAGUCACUCUCAUCUCUAUCAUACCUGCCUGAUCAGCAGAU
env-2      CUAUUUCCUGCGAGAGCCCUGUUAAGGUACGUAAGGUUAACAAAUCUGAUGUUCCCUAUUACUGGGGAUACGAUACUACCGUUUGCAAAAAUCUUGAGAAAAAGAUUUCAGAUCUUCGGGAUGAGGAUGGAGCAUAAUUUCCACCUCCCGAUAUGGUGAAAAAGUUAGUACAGAAAAUG
env-3      CUAUUUCCUGCGAUGACAAAAUUGUGGUUGAAAAGGUCCCGCCCAAUGAUGUACCCUGGUAUUGGGGAUUUGACGUAGGUAUCUGUGACCUAGAUACUGAGCUUAAGAACCUAAAACAAGGGGAUUGGAGUGUAAUAUCCACUUCUAGAAAAGGAGAGGUGGUUUCAAAUAGCUCCAUUG
env-4      UCUCUUUGCGAGACAGGUUGGAGUCGGCGAUUCACGCCGACAGGUGCUCCUCCAU.............................................................................................................................
env-5      UCUCUUUGCGAGACAGGUUGGAGUCGGCGAUUCACGCCGACAGGUGCUCCUUCAUACCGGUCACGGAGUCUGCAGACGGGAUGCUGCAGAACGGGGCCGGGCAUGCGGAGAGUUACAACUGGCACUCUCUUUGACGAUCCGCGGCCUUUUCAGUCGACGGCAUUCUCCUGCCGGGUCAGC
env-6      AUCUUACGAGAGACCGGUAGUCGGACAUUCCGGCUAAAGAUCUGUUUACACGCGAGAGAAUUUGUUCUGGC.............................................................................................................
env-8      AUCUUACGAGAGACCGGUAGCCGCAAGAUAAAGGCAAGAUCUGUCUACACGCGAGAGUGUUCAUUCUAGCAGGAUUCCCUGCAUAUCCGAAAAGGGACUCCGUGUUGUAGGGAGAGAUCUGGUCAUGUCUUCUGGCAAGUAGCAGUUAUGGCGAAAGCGCAUAUACAGCUCAUGUCCCGU
env-10     AUCUUACGAGAGACCGGUAGCUGGACGAUACAAAUCCGAUCUGUCUACACAUGUGAGGAACAUUCUGGCAGUUUAUUCUGCACAUCCGAAACGGAACUCCAUGAGGUAGGAAGAGAUCUGAUCGUAGGUCCGGCAUGCAGCAGUUAUGGCGAAAGCGCAUAUACAGCUCAUGUCCCGUGU
env-11     AUCUUACGAGAGACCGGUCGCUGGAUGAUACAAAUCCGAUCUGUCUACACAUGUGAGGAUCAUUCUGGCAGUAUUCUGCACAUCCGAAACGGAACUCCAUGAGGUAGGAAGAGAUCUGAUCGUAUGUCCGGCAUGCAGCAGUUAUGGCGAAAGCGCAUAUACAGCUCAUGUCCCGUGUCU
env-13     UCUAUUCGCGAGACUGGUUCAUCCGGAUGACGGAGCGGACAUGACCGACUCCACCGGAGAGAUUCCGAAAGAGGGAAGGAUCCUCUGCAGGUAUCGGAACGCCACAGUACCCCCGCGGUAAUGGACGAAGAGAUUUCGUUCUCGUCCUGUACCGCAUGAUCAGUGUCGGAAGCGUCACUC
env-14     UCUCUUCGCGAAACGAAUGCUCCCGAGGGAAGAACCGGACAUCGCCGGAUCAUCCUUUUAGGAAUAUCGGGCGAGGGAGAAGAACGCUUUGAGUGAGCGGUAUUCUCCUUCUCCACGAUCAGCACACGGCAGCACAGCCAUUUCCUGCGGCACGAACACUCGAAUAGAUGUAAGUGAGUU
env-15     UUUCUUCGCGAGAACCGGUCCGAGCACGGAUCGUCGAUACUCCACUUUCGGGGUUUGAUGUAUCCUGCUUGGAUCUUGUAGAUGCACUCGACCACGCCACUGCAGGGAUCACCGUUGCGACAUCCAGACAUGGUGAGAUGAUAACCACCGGGCGAUUGCCAGAACUUACCGCCCGUAUCG
Hwa-1-1    UCUCUUCGCGAGAACCGGUCCGGGCACGGAUCGUCGAUACUCCACUUUCGGGGUUUGAUGUCUCCUGCCUGGACCUUGUGGAUGCACUUGAGCACGCUGCUGCAGGUAUCACCGUUGCGACGUCCAGACACGGUGACACAGUAACCACAGGGCGAUUGCCAGAACUUGCCGCACGCGUCA
env-17     UCUCUUCGCGAGAACCGGUCCGGGCACGGAUCGUCGAUACUCCACUUUCGGGGUUUGAUGUCUCCUGCCUGGACCUUGUGGAUGCACUUGAGCACGCUGCUGCAGGUAUCACCGUUGCGACGUCCAGACACGGUGACACAGUAACCACAGGGCGAUUGCCAGAACUUGCCGCACGCGUCA
env-23     UAUCUUCGCGAGAUCCACUCCGCGCCCGCAUCACCGGCUAUCCCGAGGCUGGAUUCCAGGUGACCGGCGCGGAGCUCACGGAAGUCAUUGCCGAGGCUGGACUGACAAUUGCCGCCUCACGACACGGACAGGAGUUAUCCGUGUCCCGGCUAGGCGAGGUAGUCCAGACGGUGUCGGAGG
env-24     UCUCUUCGAGAGAGCCUUUGCGUGCCGAGAUAGCCGAGGACGCAGAUUCGCCCUACGGCGGAUACGAGGUCGAACGCGCACCGCUCGACGACUGGCUCGACUCCUACGAAGGGGUCGCCGUCUGUACGUCGAGACGGGGCAAGGAGGUCGAACUGUCUGACCUCCGUGUCGACGACGAAA
env-25     UCUCUUCGCGGGAGCCGGUCCGGGCAAAACUUGUCGACCAGUCUCCACCGGGAUAUGUCGUCGAGCGCGCGGACCUCGACACAGUGCUGGCGCGGGCGGACGGCCUAACCGUCGCAGCAUCGCGACACGGCCGGCCGCUCACGCCCGAUCGGGUAACCCGGUUGGGCGAGCGCGCCGACG
env-27     UCUCUUCGAGAGACCCACUCCAAGCACGGAUCGUGCAGCGCGCUCCGGCGACGUUCGCCGUCGAGACGGCGACGCUCCCCGAGCUGCUCCAGCGACCCGACGCCGGGGUAUGCAUCGCGACGUCCCGCCACGGGACGCCGCUGACGCAGGCACACCUGGCGACGCUCACCACGGCGAUCG
env-28     UCUCUUCGCGACGACCGGUCCGGGCAAAGAUCGUCGACACGGACCCUUCGGGGCUCGUUGUCGACCGCACGGACCUUCAGGCAGCACUCGGCCGUGAGGACGCUGGCGUUCGGAUCGCGUCGUCCCGCUACGGUGAACAUCUCACCGUGGGGCGACUCGAGACGCUGGCCGGAGAUGUCG
env-29     UCUCUUCGCGACGACCGGUCCGGGCAAAAAUCGUCGACACGGACCCUUCGGGGCUCGUUGUCGACCGCACGGGCCUUCAGGCAGCACUCAGCCGUGAGGAUGCUGGCGUUUGCAUCGCGUCAUCCCGCUACGGUGAACAUCUCACUGUCGGACGACUCGAGACGCUGGCCGGACAAGUCG
env-30     UCUCUUCGCGACGACCGGUCCGGGCAAAGAUCGUCGACACGGACCCUUCGGGGCUCGUUGUCGACCGCGCGGACCUUCAGGCAGCACUCGGCCGUGAGGACGCUGGCGUUCGGAUCGCGUCGUCCCGCUACGGUGAACAUCUCACCGUGGGGCGACUCGAGACGCUGGCCGGAGAUGUCG
env-31     UCUCUUCGCGACGGCCGGUUCGGGCGAAACUCGUCGACGCCCCACUCUCGGGGUUCGACGUCGUUUCCGCGCGGGCGAGGGGGAGCGCGUGACCGUCAGGGUCUCUUCGCGACGGCCGGUCCGGGCGAAACUCGUCGACGCCCCCAUAUCGGGGUUCGACGUCGUUUCCGCGGACCUCGA
Hut-1-1    UCUCUUCGAGACGACCGGUUCGCGCGAAGAUCGUCGAUCAGCCCCCACCGGGCUAUCAGGUCGACCGUGCGAACCUUGACGCUGCGCUCUCGCGGCCCGACGCCGGCGUACGGAUCGCCGCGUCCCGACACGGCCAGUCGCUCACGACCGAGCGACUCGGGCGACUCGCCGGGACGAUCG
env-32     UCUCUUCGAGAGACCCACUCCAAGCACGGAUCGUCGAGCGCGCUCCGGCGACGUUCGCCGUCGAGGCGGCGGCGCUGGAUGCGCUGCUCGACCGACCCGAUGCCGGGGUGCCAUCGCGACGUCCCGCCACGGGACGCCGCUGACGCAGGCACACCUGGCUGCGCUCACCACGGCGAUCG
env-33     UCUCUUCGAGAGACCCACUCCAAGCACGGAUCGUGGAGCGCGCACCCGCGACGUUCGCCGUCGAGACGGCGGCGCUGCCCGAGCUGCUCGGGCGGCCCGACGCGGGGGUGUCCAUCGCGACGUCCCGCCAUGGGACGCCGCUGACACAGGCACACCUGGCGACGCUCACUGCCGCGAUCG
env-34     UCUCUUCGAGAGACCCACUCCAAGCACGGAUCGUCGAGCGCGCACCCGCGACGUUCGCCGUCGAGACGGCGACGUUGGCGGAGCUGCUCGGGCGGCCCGACGCCGGGGUGUGCAUCGCGACGUCCCGACACGGGACGGCGCUGACGCAGGCACACCUGGCUGCGCUCACCGCGGCGAUCG
env-35     UCUCUUCGAGAGACCCACUCCAAGCCCGGAUCGUCGAGCACGCACCGGCGACGUUCGCCGUCGAGACGGCGACGCUCACCGAGCAGCUCCAGCGACCCGACGCCGGGGUGCGCAUCGCGACGUCCCGCCACGGGACGCCGCUGACGCAGGCACACCUGGCGACGCUCACCACGGCGAUCG
env-36     UCUCUUCGCGAGAGCCGGUUCGGGCGAAACUCGUCGACGAGCCACAGCCUGGCUACGUCGUCGACACCGCGACACUCGACAGCGCACUCGCCCGUGCAGACGCCGGGUUCACUAUCGCCUCGUCGAGAUACGGCGAGCGCCUGCGCACAGGCCGUCUCGACGAACUUGCCGCGAGAACCA
env-37     UCUCUUCGCGAGAGCCGGUUCGGGCGAAACUAACUGAUGAGCCACUUCCUGGCUACGCAGUGGACGCCGCGAACCUCGAUUCGGCAGUUACACGCGACGACGCUGGGUUCACCAUCGCGACUUCGCGGUUUGGUGAGCCAGUGACCGCGUCGACACUCUCCACGCUCGUCCCCAGCACAC
env-38     UCUCUUCGCGAGAGCCGGUUCGGGCGAAACUGACCGAAGAGCCAGUCGCUGGUUACUCGGUAGACGCCGCGAACCUCGAUUCGGCAGUUAGCCGCGACGACGCUGGGUUCACUAUCGCAACUUCGCGAUUUGGUGAGCCAGUUACCACGUCGGCGCUCGAAACGCUCGUUCCCAGUACGG
env-39     UCUCUUCGCGAGAGCCGGUUCGGGCGAAACUGACCGAAGAGCCAGUCGCUGGUUACUCGGUAGACGCCGCGAACCUCGAUUCGGCAGUUAGCCGCGACGACGCUGGGUUCACUAUCGCAACUUCGCGAUUUGGUGAGCCAGUUACCACGUCGGCGCUCGAAACGCUCGUUCCCAGUACGG
env-40     UCUCUUCGCGAGAACCGGUUCGGGCAAAACUCCUCGAUGAGGAGCCACCCGGACCACGUGUCUCCGAACGAUCGCUAUCGUCGGCGCUCAACGCCGAUGAUGCAGGGCUCCGAAUCGCCACCUCACGGCUGGGCCACCCGCUCA....................................
env-41     UCUCUUCGCGAGAACCAGUCCGGGCACGGCUCAUUGAUGAGCCACUACCGGGCUUCCAGGUCUCUCGUAUGGACCUGUCAGACGCACUCAAUCGAGAUGAAGCAGGGGUUCGAAUCGCAACCUCUCGACAUGGCACUGCUAUUACCCCUUCACUGAUGUCCACGCUGUCUGCACGGAUAG
env-43     UCUCUUCGCGACGGCCGGUUCGGGCGAAACUCGUCGACGAGCCCACUCCAGGGUUCGCCGUCGACGCUUCGAACCUGCAGACGGCACUCGCGCGUGACGACGUCGGACUCGCCAUCGCCGCUUCGCGGCUCGGCGAGGAUCUGACC..................................
env-44     UCUCUUCGCGACGACC....................................................................................................................................................................
env-45     UCUCUUCGCGAGAACCGGUCCGUGCUCGCCUUGUCAACGAGGCCCCCCGGGCUUCGAUGUGAAGGCCAUGGACCUGGCGGAAGCACUCACCACCGAAUCCGGAUUACGGAUUGCCACGUCUCGAUAUGGAGAACCACUCUCCAUUUCGCGGCUUGGAGAACUCGCCCCGCAGACGCGGG
env-49     UCUCUUCGAGAGAACCGGUUCGCGCGCGCCUCGUCGAUAAGCCCACACCGGGCUUCGACGUGGCGACUGUGAACCUGUCGGCCGCCCUUGCCGACGCCGCCGGACGACGGAUCGCCACCUCCCGUCAUGGACAGGAGUUGUCCAUCGGGCGACUGGCGGAGUUCCGCCCCGACUACCGCG
env-50     UCUCUUCGAGAGAACCGGUUCGUGCCCGCCUCGUCGACGAGGCCCCCCGGGCUUCGUCGUCGACCGCAUGGACCUGUCGGAAGCGCUUGCCGACGAACCGGGGCUGACGAUCGCCACAUCCGCGUAUGGACAGACGCUGUCGCAUGCGCGACUCGGCACCCUCCUCGCCGACGUUCGCG
env-54     UCUCUUCGAGAGACCCG...................................................................................................................................................................
env-55     UCUCUUCGAGAGAACCGGUCCGUGCACGACUCGUCGACGAGCCCAUAUCGGGGCUUGUUGUCGAAUGUGCGGACCUCUCGGCUGCCCUCGACGACGAAUCAGGCGUAACGGUCGCCGCCUCCGAGUAUGGGGAACUACUCUCCCAUACUCGACUCGGCGACCUCCUUCCGGAUCUCCGGG
env-56     UCUCUUCGAGAGAACCGGUCCGUGCACGACUCGUCGACGAGCCCAUCUCGGGGCUCGUUGUCGAGUGUAUGGACCUCUCGGCUGCCCUCAGCGACGAAUCAGGCGUGCGGAUCGCUGCCUCCGAGUACGGGGAGGAACUCUCCCACACUCGGCUCGGCGAACUCCUUCCGGAUCUCCGGG
env-57     UCUCUUCGAGAGAACCGGUCCGUGCACGACUCGUCGACGAACCCAUAUCGGGGCUCGUUGUCGAAUGUAUGGACCUCUCGGCUGCCCUCAGCGACGAAUCGGGCCUAACGGUUGCCGCCUCCGAGUACGGGGAAACACUCUCCCAUACUCGACUCGGCGAACUCCUUCCGGAACUCCGGG
env-58     UCUCUUCGCGAGAACCGGUCCGUGCCCGCCUCGUCGACGAACCCGCGCCGGGCUUCGUCGUCGACAGCAUGGACCUGUCGGCAGCCCUCAACGGAGAAUCGGGGCUACGGAUCGCCACGUCCGCUUAUGGACAGCCACUGUCCAUCGAGCGGCUUGGCAAACUCCGUCCGCAACUGGCGG
env-59     UCUC................................................................................................................................................................................
env-62     UCUCUUCGAGAGAACCGGUCCGUGCACGGCUCGUCGACGAGGCCCACCGGGCUUCGACGUCGAAAGCACGGACCUCUCGGCAGCCCUCGACGACGAAUCGGGGCUACGGAUCGCCACGUCAGCCUAUGGGGAAGAACUGUCCCAUGGUCGACUCGGUGAUCUCCUUCCGCAGAUCGCGG
env-63     UCUCUUCGAGAGAACCGGUCCGUGCACGGCUCGUCGACGAGGCCCACCGGGCUUCGACGUCGAAUGCACGGACCUCUCGGACGCCCUCGCUGAAGAAUCGGGGCUACGGAUCGCCACGUCAGCCUAUGGAGAAGAACUCUCCCAUGGUCGACUCGGUGAUCUCCUUCCGCAGCUCGCGG
env-64     UCUCUUCGAGAGAACCGGUCCGUGCACGGCUCGUCGACGAGGCCCACCGGGCUUCGACGUCGAAUGCACGGACCUCUCGGACGCCCUCGCUGAAGAAUCGGGGCUACGGAUCGCCACGUCAGCCUAUGGAGAAGAACUCUCCCAUGGUCGACUCGGUGAUCUCCUUCCGCAGCUCGCGG
env-65     UCUCUUCGAGAGAACCGGUCCGUGCACGGCUCGUCGACGAGGCCCACCGGGCUUCGACGUCGAAUGCACGGACCUCUCGGACGCCCUCGCUGAAGAAUCGGGGCUACGGAUCGCCACGUCAGCCUAUGGAGAAGAACUCUCCCAUGGUCGACUCGGUGAUCUCCUUCCGCAGCUCGCGG
env-70     UCUCUUCGAGAGAACCGGUCCGUGCACGGCUCGUCGACGAGGCCCACCGGGCUUCGACGUCGAAUGCACGGACCUCUCGGACGCCCUGCUGAAGAAUCGGGCUACGGAUCGCCACGUCAGCCUAUGGAGAAGAACUCUCCCAUAGGUCGACUCGGUGAUCUCCUUCCGCAGCUCGC...
env-71     UCUCUUCGAGAGAACCGGUCCGUGCACGGCUCGUCGACGAGGCCCACCGGGCUUCGACGUCGAAUGCACGGACCUCUCGGACGCCCCUCGCUGAAGAAUCGGGGCUACGGAUCGCCACGUCAGCCUAUGGAGAAGAACUCUCCCAUGGUCGACUCGGUGAUCUCCUUCCGCAGCUCGCG
env-72     UCUCUUCGCGACGACCGGUCCGGGCGAAGCUCGUCGACGACCCCCUCCCGGGGCUCGCCGUCGAGUGGACGGACCUAUCGGCAGCGCUCGGCCGUGAGGACGCCGGCGUUCGGAUCGCGGCCUCCAGACACGGUGAACCGCUCACCGUCGGAGGCCUCGAGACGCUGGUCGGACGCGUCG
env-74     UCUCUUCGCGACGACCGGUCCGGGCGAAGCUCGUCGACGACCCCCUCCCGGGGCUCGCCGUCGAGUGGACGGACCUCUCGGCAGCACUCGGCCGUGAGGACGCCGGCGUUCGGAUCGCGGCUUCCAGACACGGUGAACCGCUCACCGUCGGAGGCCUCGAGACGCUGGUCGGGCACGUCG

           ....................................................................................................................................................................................
           ....................................................................................................................................................................................
           UCUYUUYR●GRRRR●YR●●●●●●●Y●●●R●●●RY●RR●R●●●●●●●●●●●●●●●●●●●●●●●●●●●●●●●●●●●●●●●●●●R●●●●●●●●●●●●●●●●●●●●●●●●●●●●●●●●●●●●●●●●●●●●●●●●●●●●●●●●●●●●●●●●●●●●●●●●●●●●●●●●●●●●●●●●●●●●●●●●●
```

| | |
|---|---|
| env-76 | UUGCCUCGCGCGAACCGGUCCGCGCCAGGCUCGUCGACGAACCCCAGCCGGGGUUCG........................................................................................................................... |
| env-77 | UUUCAUCGCGCGAACCGGUCCGCGCCAAGCUCGUCGACGAACCCCAGCCGGGGUUCGACGUCGAACGCGCGGGCCUCGCUGCAACGCUCGCCCGAGCCGACGCCGGCGUCCGCAUCGCCGCGUCCCGGUUCGGGGCCGAGCUCUCCGUCGCC............................ |
| env-78 | UUUCAUCGCGCGAACCGGUCCGCGCCAAGCUCGUCGACGAACCCCAGCCGGGGUUCGACGUCGAACGCGCGGGCCUCGCUGCAACGCUCGCCCGAGCCGACGCCGGCGUCCGCAUCG............................................................... |
| Hsa-1-1 | UCUAUUCGCGAAGACCGGUCCGCGCGAAGCUCGUCGACGAACCCCUUCCGGGGUACGCCGUCGAUCGGGCGGACCUGGACGAAGCGCUCUCGCGUGCCGACGCUGGCGUUCGGAUCGCCUCCUCGCGACACGGUGUCGAAGCGACGACUGGCCGUCUCGGCCAGUUGAUCGAGACGACAC |
| Hgi-1-1 | UCUCUUCGCGAGAACCGGUCCGUGCGCGGAUCGUCGACGAGCCCCUCCCGGGCUUCGACGUAUCCCGCAUGGACCUCACGGAAGCACUCGGCAGGCCAGAUGCGGGCGUGCGAAUCGCCACGUCUCGCUAUGGCGAACCGCUGUCGGUCCCCACAUUGGGCGACUUGACGGCCCGCAUCA |
| Hsu-2-1 | UCUCUUCGCGAGAACCGGUCCGUGCGCGGAUCGUCGACGAGCCCCGUCCGGGCUUCGACGUAUCCCGCAUGGACCUCACGGAAGCACUCGGCAGGCCGGAUGCGGGCGUGCGAAUCGCCACGUCUCGCUAUGGCGAACCGCUGUCGGUCCCCACAUUGGGUGACUUGACGGCCCGCAUCA |
| Hpr-1-1 | UCUCUUCGCGAGAACCGGUCCGUGCGCGGAUCGUCGACGAGCCCCUCCCGGGCUUCGACGUAUCCCGCAUGGACCUCACGGAAGCACUCGGCAGGCCAGAUGCGGGCGUGCGAAUCGCCACGUCUCGCUAUGGCGAACCGCUGUCGGUCCCCACAUUGGGUGACUUGACGGCCCGCAUCA |
| Hsp-8-1 | UCUCUUCGCGAGAACCGGUCCGUGCGCGGAUCGUCGACGAGCCCCUCCCGGGCUUCGACGUAUCCCGCAUGGACCUCACGGAAGCACUCGGCAGGCCGGAUGCGGGCGUGCGAAUCGCCACGUCUCGCUAUGGCGAACCGCUGUCGGUCCCCAGAUUGGGCGACCUGACGGCCCGCAUCA |
| Hde-1-1 | UCUCUUCGCGAGAACCGGUCCGUGCGCGGAUCGUCGACGAGCCCCUUCCGGGCUUCGACGUAUCCCGCAUGGACCUCACGGAAGCACUCGGCAGGCCGGAUGCGGGCGUGCGAAUCGCCACGUCUCGCUAUGGCGAACCGCUGUCGGUCCCCACAUUGGGUGACCUGACGGCCCGCAUCA |
| Hlu-1-1 | UCUCUUCGCGAGAACCGGUCCGUGCGCGGAUCGUCGACGAGCCCCUCCCGGGCUUCGACGUAUCCCGCAUGGACCUCACGGAAGCACUCGGCAGGCCGGAUGCGGGCGUGCGAAUCGCCACGUCUCGCUAUGGCGAACCGCUGUCGGUCCCCACAUUGGGCGACUUGACGGCCCGCAUCA |
| Hal-2-1 | UCUCUUCGCGAGAACCGGUCCGUGCGCGGAUCGUCGACGAGCCCCUCCCGGGCUUCGACGUAUCCCGCAUGGACCUCACGGAAGCACUCGGCAGGCCAGAUGCGGGCGUGCGAAUCGCCACGUCUCGCUAUGGCGAACCGCUGUCGGUCCCCACAUUGGGUGACUUGACGGCCCGCAUCA |
| Hsp-6-1 | UCUCUUCGCGAGAACCGGUCCGUGCGCGGAUCGUCGACGAGCCCCUCCCGGGCUUCGACGUAUCCCGCAUGGACCUCACGGAAGCACUCGGCAGGCCGGAUGCGGGCGUGCGAAUCGCCACGUCUCGCUAUGGCGAACCGCUGUCGGUCCCCACAUUGGGCGACCUGACGGCCCGCAUCA |
| Hme-1-1 | UCUCUUCGCGAGAACCGGUCCGUGCGAGGAUCGUCGACGAGCCCCUUCCGGGCUUCGACGUAUCCCGCAUGGACCUCACGGAAGCACUCGGCAGGCCAGAUGCGGGCGUGCGAAUCGCCACGUCUCGCUAUGGCGAGUCGCUGUCGGUCCCCCGACUGGGUGACCUGACGGCCCGCAUCA |
| Hla-2-1 | UCUCUUCGCGAGAACCGGUCCGUGCGAGGAUCGUCGACGAGCCCCUUCCGGGGUUCGACGUAUCCCGCAUGGACCUCACGGAAGCUCUCGGCAGACCGGAUGCGGGCGUGCGAAUCGCCACGUCUCGCUAUGGUGAGCCGCUUUCGGUCCCCCGACUGGGGAACUGACAGCCCGUAUCG |
| Hel-1-1 | UCUCUUCGCGAGAACCGGUCCGUGCGAGGAUCGUCGACGAGCCCCUUCCGGGGUUCGACGUAUCCCGCAUGGACCUCACGGAAGCUCUCGGCAGACCGGAUGCGGGCGUGCGAAUCGCCACGUCUCGCUAUGGUGAGCCGCUUUCGGUCCCCCGACUGGGGAACUGACAGCCCGUAUCG |
| Hmu-1-1 | UCUCUUCGCGAGAACCGGUCCGUGCGAGGAUCGUCGAUGAACCCCUUCCGGGGUUCGACGUAUCCCGCAUGGACCUCACGGAAGCACUCGGCAGGCCAGAUGCGGGCGUGCGAAUCGCCACGUCUCGCUAUGGCGAGUCGCUGUCGGUCCCCCGACUGGGCGACCUGACGGCCCGCAUCA |
| env-79 | UCUCUUCGCGAGAGCCGGUUCGGGCGAAACUGACCGAGGAGCCAGUCGCUGGUUACUCGGUAGACGCCGCGAGCCUCGAUUCGGCAGUUAGCCGCGACGACGCUGGUUUCACCAUCGCUACUUCGCGAUUUGGUGAGCCAGUUACCACGUCGACACUCAACACACUCGUUCCCAGCACGG |
| env-80 | UCUCUUCGCGAGAGCCGGUCCGAGCAAAGUUAGUCGACGUCGACCCAUCUGGGUUCGUUGUCGACCGCUCGGACCUCGAUACGGCAGUGUCGCGCGCGGAUGCAGGUCUCACUAUCGCCGCUUCGCGGUACGGUGAGCAACUCACGCCGCAGCGCCUCGACGAGAUCGCCCCCAGUGUC |
| env-81 | UCUCUUCGCGAGAGCCAAUCCGAGCAAAACUCGUCGACACCUCACUCACGGGGUAUCACGUCGACCGCUCGGAGAUCGAUACGGCAGUCGACCGCUCGAAUGCAGGACUCACUAUUGCCGCCUCGCGGUAUGGUAAGUCGCUUGCGCCGGGUCGCCUCGACGACAUCGCACGCCAGUGUC |
| env-82 | UCUCUUCGCGAGAGCCGGUCCGAGCAAAGCUCGUCGACGUCGACCCAUCUGGGUUCGUCGUCGACCGCUCGGACCUCGAUACGGCAGUGUCGCGCUCGGAUGCAGGACUCGCUAUCGCCGCUUCGCGGUACGGCGAGGAACUCACACCGAAGCGCCUCGACGAGAUCGCCCCACAGUGUC |
| env-83 | UCUCUUCGCGAGAGCCGGUCCGAGCAAAGCUCGUCGACGUCGACCCAUCUGGGUUCGUCGUCGACCGCUCGGACCUCGAUACGGCAGUGUCGCGCUCGGAUGCAGGACUCGCUAUCGCCGCUUCGCGGUACGGCGAGGAACUCACACCGAAGCGCCUCGACGAGAUCGCCCCACAGUGUC |
| Hla-4-1 | UCUCUUCGCGACGACCGGUCCGUGCGAAAAUCGUCGACAAGCCCACACAGGGGCUAUCGAUCGAGCAAACGGACCUUUCGGCGGCUCUCAGCCGUGAGGACGCCGGCGUCCGAAUCGCAGCUUCCCGGUUCGGUGAACCGCUCACCGUGGGGAAACUCGAGACGCUGGCCGGACGACUCG |
| Ssp-1-1 | UCUCUUCGCGACGACCGGUCCGCGCCAAGAUCGUCGACGAGCCCCUCCCGGGCUUCGAGGUUGAUCGCGCGGACCUGCCGACGGCCAUCGAGGACUCCCGGAACACGUGUCGCAUCGCCGCGUCCCGCCACGGGACGCCGCUGACCGCGACCCGACUGACGGACCUCGUCGGACAUCUCC |
| env-84 | UCUCUUCGCGACGACCGGUUCGCGCGAAACUCGUCGAUGAGCCCCGUCCGGGCUUUAUCGUCGAACGCGUGGACCUCCGGGACGCUCUCGACCGGCCCGACG.............................................................................. |
| env-86 | UCUCUUCGCGACGACCGGUCCGUGCCAAGCUCGUCGACGAGCCCCCCUCGGGGCUGUCGGUCGAGCGCGCGGACCUGCGGGCAGCUCUCGGCCGUGAGGAUGCUGGCGUCUGCAUCGCGUCGUCCCGGUUCGGUGAGGAACUCACCGUCGGGCGACUCGGGACGCUGGCCGGACGCAUCG |
| Hsp-11-1 | UCUCUUCGCGACGACCGGUCCGUGCGAAAAUCGUCGGCAAGCCCACACAGGGGUUGUCGAUCGAGCAAACGGACCUUUCGGCGGCUCUCAGCCGUGAGGAUGCCGGCGUUCGAAUCGCAGCUUCCCGGUUCGGUGAAGCGCUCACCGUGGGGAAACUCGAGACGCUGGCCGGACGAAUCG |
| env-87 | UCUCUUCCGUAGAACCGGUCCGCGCCCGCAUCGUCGACGAGCCCCCUCCGGGGCCGACGGUGCUGGAACAGGAUCUGUCGACAACGCUCGGAGCGGCGGACGCGGGCUGUGUAUCGCGGCCUCGCGCCACGGCUCGCCGCUCACGGUCGGCCGUCUGGCCGACCUGCCGGGCUCCGGCG |
| env-88 | UCUCUUCGCGAGAACCGGUCCGUGCCCGCCUCGUCGACGAACCCGCGCCGGGCUUCGUCGUCGACAGCAUGGACCUGUCGGCAGCCCUUUCGGUCGAAUCGGGGCUACGGAUCGCCACAUCCGCUUAUGGACAGCCACUGUCCAUCGAGCGGCUUGGCCAACUCCGUCCGCAACUGGCGG |
| env-89 | UCUCUU.............................................................................................................................................................................. |
| Hka-1-1 | UCUCUUCGCGACGGCCAGUCCGCGCAAAGCUCGUCGACGAGCCCCUCACCGGGUACGUCGUCGACACCGCGGACGUGGAUACGGCACUCGAACGGUCCGACGCCGGGUUCGCCAUCGCCGCAUCGCGGUAUGGUGAGGAACUCGGCGUCGACCGGCUCGGCCAACUGGUCGAGCGCCGGG |
| Hmu-2-1 | UCUCUUCGCGACGGCCAGUCCGCGCAAAGCUCGUCGACGAGCCCCUCACCGGGUACGUCGUCGACACCGCGGACGUGGAUACGGCACUCGAACGGUCCGACGCCGGGUUCGCCAUCGCCGCAUCGCGGCAUGGUGAGGAACUCGGCGUCGACCGGCUCGGCCAACUGGUCGAGCGCCGGG |
| Hca-3-1 | UCUCUUCGCGACGACCGGUACGGGCGAAGCUCGUCGACGAGCCCGCGGCAGGGUUCGUCGUCGACUCGACCGAUAUCGACGACGCACUCGCGCGUCCGGACGCCGGGCUGACCAUCGCGGCCUCUCGCUACGGCGAGGAGCUGACCGUCACGAAGCUCGACCAGCUGGUCGAACGGCUUC |
| Hsa-5-1 | UCUCUUCGCGAGAGCCGGUCCGCGCGCGGAUCGUCGACGAGCCCGCACCCGGGUUCGCCGUCGAGCGCGCGGACCUCUCGGCAGCGCUCGGCCGUGAGGACGCCGGGCUCCGGAUCGCCACGUCGCGCCACGGUGAGGCGCUCACGACGGAUCGACUCGGAGUGCUGGCCGGACGGUCG |
| env-90 | UCUCUUCGCGAGAGCCGGUUCGCGCACGGAUCGUCGACGAG........................................................................................................................................... |
| Hag-1-1 | UCUCUUCGCGAGAGCCGGUUCGCGCGCGGAUCGUCGACGAGCCCGCACCCGGGUUCGCCGUCGAGCGUGCGGACCUCUCGGCAGCGCUCGGCCGUGAGGACGCCGGGCUCCGGAUCGCCACGUCGCGCCACGGUGAGGCGCUCACGACGGAGCGACUCGGGGUGCUGGCCGGACGGUCG |
| Hsa-4-1 | UCUCUUCGCGAGAGCCGGUCCGCGCGCGGAUCGUCGACGAGCCCCUUCCGGGGUUCGUCGUCGAGCGCGCGGACCUCUCGGCAGCGCUCGGCCGUGAGGACGCCGGGCUCCGGAUCGCCACGUCGCGCCACGGUCAGGCGCUCACGACGGAGCGACUCGGAGUGCUGGCCGGACGGUCG |
| env-91 | UCUCUUCGCGAGAGCCGGUCCGGGCGAAGCUUAUCGACCAGCCCCCGUCGGGCUACGCUGUCGACGCCGCGGACCUCGAUACAGCGCUCGCGCGCGAGGACGCAGGACUGAGAAUCGCGUCCUCGCGGCACGGCGAGCGCCUGCGGCCGGGUCGACUCGACAACCUCGCGGCGGCGUGCC |
| env-92 | UCUCUUCGAGAGAACCGGUUCGCGCGCGCCUCGUCGANGAGCCCACACCGGGCUUCGACGUGGCGACUGUGAACCUGUCGGCCGCCCUUGCCGACGCCGCCGGACGACGGAUCGCCACCUCCCGUCAUGGACAGGAGCUGUCCAUCACGCGACUGGCGGAGUUCCGUCCCGACUACCGGG |
| env-93 | UCUCUUCGAGAGAACCGGUUCGCGCGCGCCUCGUCGANGANCCCACACCGGGCUUCGACGUGGCGACUGUGAACCUGUCGGCCGCCCUUGCCGACGCCGCCGGACGACGGAUCGCCACC............................................................. |
| env-94 | UCUCUUCGAGAGAACCGGUUCGCGCGCGCCUCGUCGACGAGCCCACACCGGGCUUCGACGUGGCGACUGUGAACCUGUCGGCCGCCCUUGCCGACGCCGCCGGACGACGGAUCGCCACCUCCCGUCAUGGACAGGAGCUGUCCAUCA................................. |
| env-95 | UCUCUUCGAGAGAACCGGUUCGCGCGCGCCUCGUCGACGAGCCCACACCGGGCUUCGACGUGGCGACUGUGAACCUGUCGGCCGCCCUUGCCGACGCCGCCGGACGACGGAUCGCCACCUCCCGUCAUGGACAGGAGCUGUCCAUCNCGCGACUGGCGGAGUUCCGUCCCGACUACCGGG |
| env-96 | UCUCUUCGAGAGAACCGGUUCGCGCGCGCCUCGUCGACGAGCCCACACCGGGCUUCGACGUGGCGACUGUGAACCUGUCGGCCGCCCUUGCCGACGCCGCCGGACGACGGAUCGCCACGUCCCGUCAUGGACAGAUGCUGUCCAUCGCGCGACUGGCUGAGUCCCGCCCGACUAACCGGG |
| env-98 | UCUCUUCGAGAGAACCGGUUCGCGCGCGCCUCGUCGACGAGCCCACACCGGGCUUCGACGUGGCGACUGUGAACCUGUCGGCCGCCCUUGCCGACGCCGCCGGACGACGGAUCGCCACGUCCCGUCAUGGACAGCAGCUGUCCAUCGCGCGACUGGCGGAGUUCCGCCCCGACUACUGCG |
| env-99 | UCUCUUCGAGAGAACCGGUUCGCGCGCGCCUCGUCGACAAGCCCACACCGGGCUUCGAUGUGGCGACUGUGAACCUGUCGGCCGCCCUUGCCGACGCCGCCGGACGACGGAUCGCCACCUCCCGUCAUGGACAGCAGCUGUCCAUCGGGCGACUGGCCGAGUUCCGUCCCGCACGGACG |
| env-101 | UCUCUUCGAGAGAACCGGUCCGUGCACGACUCGUCGACGAGCCCAUAUCGGGGUUCGACGUCGAAUGCAUGGGCCUCUCGGCUGCCCUCAGCGACGAAUCGGGGCUCCGGAUCGCCGCCUCCGAGUACGGGAAAGAGCUGUCCCACACUCGGCUCGGCGACCUCCUGCCGCAGUUGGCAG |
| env-102 | UCUCUUCGAGAGAACCGGUUCGUGCACGACUCGUCGACGAACCCACAACGGGGUUCGACGUCGAAUGUACGGGCCUCUCGGCUGCCCUCAGCGACGAAUCGGGCCUCACAAUCGCCACCUCGGCCUACGGAGACAAACUCUCUUCUGGCCGACUUGGCGAACUCCUUCCGGAACUCCGGG |
| env-103 | UCUCUUCGAGAGAACCGGUUCGUGCACGACUCGUCGACGAACCCACAACGGGGUUCGACGUCGAAUGUACGGGCCUCUCGGCUGCCCUCAGCGACGAAUCGGGCCUCACAAUCGCCACCUCGGCCUACGGAGACAAACUCUCUUCUGGCCGGCUUGGCGAACUCCUUUCGGAACUCCGGG |
| env-104 | UCUCUUCGAGAGAACCGGUCCGUGCACGCCUCGUCGACGAACCCAUAUCGGGGCUCGUUGUCGAACGUGCGGACCUCUCGGCUGCCCUCAGCGACGAAUCGGGCGCAACGAUCGCCACCUCGGCCUACGGGGAGACGCUCUCCCACGGUCGACUCGGCGAACUCCUUC............ |
| env-105 | UCUCUUCGAGAGAACCGGUCCGUGCACGACUCGUCGACGAACCCAUAUCGGGGUUCGACGUCGAAUGUACGGGCCUCUCGGCUGCCCUCAGCGACGAAUCGGGCCUCACAAUCGCCACCUCGGCCUACGGAGACAAACUCUCUUCUGGCCGGCUUGGCGAACUCCUUUCGGAACUCCGGG |
| env-106 | UCUCUUCGAGAGAACCGGUCCGUGCACGACUCGUCGACGAGCCCAUAUCGGGGCUCGUUGUCGAAUGUAUGGGCCUCUCGGCUGCCCUCAGCGACGAAUCAGGCCUCACGGUUGCCGCCUCCGAGUACGGGGAGAAACUCACCCAUACUCGACUCGGCGGACUCCUUCCGGAACUCCGGG |
| env-107 | UCUCUUCGAGAGAGCCGGUCCGCGCACGGAUCGUCGACGAACCCUUGCCUGGGUUCGAGAUCCAGCGUGCGGACCUGCCGGAAGUGCUGGCCCGGCCCGACGCCGGGGCUC..................................................................... |
| env-108 | UCUCUUCGCGAGAGCCGGUCCGAGCACGGAUCGUGGACGAGCCCGCACCGGGCUACACGAUCCUGAC................................................................................................................. |
| env-109 | UCUCUUCGCGACGGCCGGUCCGCGCCAAGAUCGUCGACG............................................................................................................................................. |
| CHa-1-1 | UCUCUUCGAGAGAACCCGUCCGCGCCCGCAUCGUGGACGACCCCCACCGGGGUUCCGCGUCGCGGGCGAGGACGUCGCGGACGCCGCCGCCGCCGCCGAACUGGCGGUCGCCACCUCGCGGCACGGACGCCCGCUGUCCGUGUCGGGACUGAGCGAUCUCCACGCGCGACUGGACGACA |
| Hpa-2-1 | UCUCUUCGCGAGAACCGGUCCGUGCGCGGAUCGUGGACGAGCCCGUUCCGGGGUUCUCCAUCGCCCGCACGGACCUCGAGGAAGCACUCGGCCGCUCCGACGCGGGCGUCACUAUCGCGACGUCCCGCCACGGAGAGUCGCUGACGGUGCCGAAGCUAGCGGAACUCUCCCCGCGAAUCG |
| env-110 | UCUCUUCGCGACGACCGGUCCGAGCGAAACUCGAAGACAGCCCCCUCCCGGGGCUUGCUAUCGAGCGGACGGACCUACAGGCAGCACUCAGCCGUGAGGACGCCGGCGUCUGCAUCGCGUCGUCUCGCUACGGUGAACCGCUCACCGUCGGGCGACUCGAGACGCUGGCCGGACGGAUCG |
| Nhu-1-1 | UCUCUUCGCGACGACCGGUCCGGGCGAAACUCGUCGACGAGCCCCUCCCGGGGCUUACCGUCGAGCGGACGGACCUUCAGGCGGCUCUCAGCCGUGAGGACGCCGGCGUUCGAAUUGCGGCUUCCCGAUUCGGUGCAGAACUCACCGUCGGGCGGCUCGGAACGCUGGCCGGACACACAG |
| env-113 | UCUCUUCGAGAGAACCGGUCCGCGCCCGUAUCACCGGGAAGCCCGAGGCCGGCUUCCAGGUCGCGGCCGCGGACCUGUCGGAAGCGCUCGCCGGCGACGGACUGGCCGUCGCAACGUCGCGACACGGAGCGGCACUCUCCGUCUCGCGGCUCGGCGACCUCGUUUCGGACGCGCGGAUG |
| env-114 | UCUCUUCGAGAGAACCGGUUCGCGCGCGCCUCGUCGAUAAGCCCACACCGGGCUUCGACGUGGCGACUGUGAACCUGUCGGCCGCCCUUGCCGACGCCGCCGGACGACGGAUCGCCACCUCCCGUCAUGGACAGGAGUUGUCCAUCGGGCGGCUGGCGGAGUUCCGCCCCGACUACCGCG |
| env-115 | UCUCUUCGAGAGAACCGGUUCGCGCGCGCCUCGUCGAUAAGC.......................................................................................................................................... |
| env-116 | UCUCUUCGAGAGAACCGGUCCGUGCACGACUCGUCGACGAGCCCACAACGGGGUUCGACGUCGAAUGUACGGGCCUCUCGGCUGCCCUCAGCGAUGAAUCAGGCAUAACGGUCGCCGCCUCCGAGUAUGGCGAGAUACUCUCGCAUGCGCGGCUCGGCGAACUCCUUCCGGAUCUCCGGG |
| env-117 | UCUCUUCGAGAGAACCGGUCCGUGCACGNCUCGUCGACGAACCCAUAUCGGGGCUCGUUGUCGAACGUGCGGACCUCUCGGCUGCCCUCGACGACGAAUCGGGCGUAACGGUUGCCGCCUCCGAGUAUGGGGAGAAACUCUCCCACACUCGACUCGGCGACCUCCUUCCGGGUCUCCGGG |
| env-118 | UCUCUUCGAGAGAACCGGUCCGUGCACGACUCGUCGACGAACCCAUAUCGGGGCUCGUUGUCGAACGUGCGGACCUCUCGGCUGCCCUCAGCGAUGAAUCAGGCAUAACGGUCGCCGCCUCCGAGUAUGGCGAGAUACUCUCGCAUGCGCGGCUCGGCGAACUCCUUCCGGAUCUCCGGG |
| env-119 | UCUCUUCGAGAGAACCGGUCCGUGCACGACUCGUCGACGAACCC........................................................................................................................................ |
| env-120 | UCUCUUCGCGAGAGCCCGUUCGCGCCAAGCUCGUCGACGACACGGUGUUGUCCGUCGAGCGCGCGGACGUGGGGAGCAUCACGGCCGACGCCGACGUGUCGAUCGCCGCCUCCCGCCACGGGGAACGUCUCUCCGUGGCGCGGUUGGAGACGGUGACGGCCCGACUACGCGACGCAGACC |
| env-122 | UCUCUUCGCGAGAGCCCGUUCGCGCCAAGCUCGUCGACGACACCGCGUUGUCGGUCGAGCGCGCGGACGUGGGGAGCGUCACGGCCGACGCCGACGUGUCGAUCGCCGCCUCCCGUCACGGGGACACACUCUCCGUGCCGGCGCUGGAGACGGUGACGACGCAGUUGGCGACGGCCGACC |
| env-123 | UCUCUUCGCGAGAGCCGGUUCGCGCCAAACUCGUCGACGACACCGUGUUGUCGGUCGAGCGCGCGGACCUGGGGAGCGUCACAGCCGACGCCGACGUGUCGAUCGCCGCCUCCCGCCACGGGGACACACUCUCCGUGCCGGCGUUGGAGGUGGUGACGACGGAGUUGGUGGCGGCAGACC |
| env-126 | UCUCUUCGCGAGAACCGGUCCGUGCGCGGAUCGUCGAUCGGCCCCUUCGGGGCUUCGACGUCUCCCGUAUGGACCUCGAGGAAGCACUCGGCCGCGCCGACGCGGGGGUCGUAAUCGCGACGUCCCGCCACGGCGAGGCGCUGACCACCACGCGGCUGCCGGAUCUCGCCGCCCGUAUCG |
| env-127 | UCUCU............................................................................................................................................................................... |
| env-128 | UCUCUUCGCGAGAACCGGUCCGUGCGCGGAUCGUCAAUCGGCCCCUUCCGGGCUUCGACGUGUCCCGUAUGGACCUCGAGGAAGCACUCGGCCGCGCCGACACGGGUGUCGCUAUCGCGACAUCCCGCCACGGCGAGGCGCUGACCACCACGCGGCUGCCGGAUCUCGCCGCCCGCAUCG |
| env-129 | UCUCUUCGCGACGACCGGUCCGCGCACGGCUGGUGAACGGGCCAACGGGGCUCACAGUCGAGCGUGUGGACCUAUCGGCGGCGCUCGGCCGUGAGGACGCCGGCGUCCGUAUCGCGACGUCGAGGCUCGGUGAACCGCUCACCGUCGCUCGACUGCGGACGCUGGCCGGACGGUCGAGG |
| Hna-1-1 | UCUCUUCGAGAGAGCCGGUCCGCGCACGGAUCGUCGACGAACCCCUGCCGGGGUUCGAGGUCCAACGCGCGGACCUGCCGGACGUGCUGGCCCGCCCCGACGCCGGGGUUCGGAUCGCCACCUCCGUCCACGGGACGGAACUGUCGGUCGGUCGCCUGACGGAACUCGUCGGGCGGAUCG |
| env-130 | UCUCUUCGAGAGAGCCGGUCCGCGCACGGAUCGUCGACGACCCCCUACCGGGGUUCGAGGUCCAACGCGCGGACCUGCCGGAAGUGCUGGCCCGCCCCGAUGCGGGAGNUCGGAUCGCCACGUCCGUUCACGGAACGGAACUGUCGGUCGGUCGCCUGACCGGGUUGGUCGGGCGCAUCG |
| env-131 | UCUCUUCGCGAGAACCCGUCCGCGCCCGCCUCGUCGACGAGCCCACCGCGGGCUACGUUGUCGAGGCCGCGGACAUCCAGGAAACCCUCGCCCGUCAGAACGCGGGCGUCCGGAUCGGCACCUCACGCCACGGUGAUGACCUCACCGUGUCGCGACUCGCCGAACUGGGCUCGCAGUGUC |
| env-134 | UCUCUUCGCGACGGCCGGUUCGCGCGCGGAUCGUCGACGAACCCCCACCGGGGUUCGCCGUCGAGCGCGCGACACUCGAUGACGCCCUCGCCAGGGAGGACGCAGGCGUG...................................................................... |
| env-135 | UCUCUUCGCGACGACCGGUCCGGGCGAAGCUCGUCGAU.............................................................................................................................................. |
| Nph-1-1 | UCUCUUCGCGAGAGCCGGUUCGCGCAAAGCUCGUCGACGAAUCCCCACCGGGAUUCGCCGUCGACUGUGCGACCAUCGAAGCCAGUCUCGCCCGCCCGGACGCCGGCGUCCGUAUCGCCGCGUCCCGGCAUGGGGAGACGCUUUCGGUCGACCGUCUCGACGGUAUCGUCGGUCGGACGA |
| env-136 | UCUCUUCGCGAGAGCCGGUUCGCGCAAAGCUCGUGGCCGAGCCCCCACCGGGGUUCGCCGUCGAGCGCGCGACCAUCGCAGACUGCCUCACCCGUCCGGACGCUGGCGUCCGUAUCGCCGCGUCCAGACACGGCGAGGUGUUGACGGUCGAUCUGCUCGGCGACAUCGUCGAGCGAUCGA |
| env-137 | UCUCUUCGCGAGAGCCGGUUCGCGCAAAGCUCGUGGCCGAGCCCCCACCGGGGUUCGCCGUCGAGCGCGCGACCAUCGCAGACUGCCUCACCCGUCCGGACGCUGGCGUCCGUAUCGCCGCGUCCAGACACGGCGAGGUGUUGACGGUCGAUCUGCUCGGCGACAUCGUCGAGCGAUCGA |
| env-138 | UCUCUUCGAGAGAACCGGUCCGCGCCCGGAUCACCGGGAAGCCGUCUGCUGGCUUCCAGGUGGUGGGUGCGGACCUGUCGGAAGCGCUCGUCGAUGGCGGACUCGCCGUCGCAACGUCGCGACACGGAGCGAACCUCUCCGUCUCGCGGCUCGGCGAACUCGUCGCCGACGCGUCGAAGG |
| env-139 | UCUCUUCGAGAGAACCGAUCCGCGCCCGUAUCACUGGAAAGCCGGCGGUCGGCUUCCAGGUCGUGGACGCGGACCUGUCGGAAGCACUCGACGCCGACGGACUCGCCGUCGCCACGUCGCGACACGGGGAGCAGCUCUCCGUUUCGCGGCUCGGUGAGAUCGUCACGGACGCACGGACGG |
| env-140 | UCUCUUCGAGAGAACCGGUCCGCGCCCGCAUCACCGGAAAGCCGGCGGACGGCUUUCAGGUCGUUAGCGCGGACCUGUCGGCAGCACUCGAUGACGGACUCGCCAUCGCAACGUCGCGACACGGGGAAUCGCUCUCCGUCUCGCGGCUCGGCGAGAUCGUCGCGGACGCACGGGAGGCUG |
| env-145 | UCUCUUCGAGAGAACCGGUCCGCGCCCGCAUCACCGGAAAGCCGGAGGACGGCUUUCAGGUCGUGGCCGCGGACCUGUCGGAAGCGCUCACGGGCGACGAACUCGCCAUCGCGACGUCGCGGCACGGGGAACCGCUCUCCGUCCCGCGGCUCGGCGAACUCGUCGCCGGCGCACGGAGG |
| env-146 | UCUCUUCGAGAGAACCAGUCCGCGCCCGCAUCACCGGGAAGCCGGAGGUCGGUUUCCAGGUCGUGGGCGCGAGGCUGUCGGAAGCGCUCGCCGGCGACGGAGUGGCCGUCGCGACGUCGCGACACGGAGAAGAACUCUCCGUCUCGCGGCUCGGCGACCUCGUUUCGGACGCGCGGAAGG |
| | .................................................................................................................................................................................... |
| | .................................................................................................................................................................................... |
| | UCUYUUYR•GRRRR•YR•••••••Y•••R•••RY•RR•R•••••••••••••••••••••••••••••••••••••••••R••••••••••••••••••••••••••••••••••••••••••••••••••••••••••••••••••••••••••••••••••••••••••••••••••••••• |

```
env-147    UCUCUUCGAGAGAACCGGUCCGCGCCCGCAUCACCGGAACGCCGGCGGACGGCUUUCAGGUCGUGGGAGCGGACCUUUCGGAAGCGCUCGGCGGCGACGGACUCGCCGUCGCGACAUCGCGGUACGGGGAGGAGCUCUCCGUCCCGCGACUGGGCGAUAUCGUCUCGGACGCACGCGAAG
env-148    UCUCUUCGAGAGAACCGGUCCGCGCCCGCAUCACCGGGAAGCCGGAGGACGGUUUCCAGGUCGUGGGUGCGGACCUUUCGGAAGCGCUCGACGGCGACGGACUCGCCGUCGCAACGUCACGACACGGGGAAGAGCUCUCCGUCUCGCGGCUGGGCGACAUCGUCUCGGACGCACGGGAGG
Hsp-9-1    UCUCUUCGAGAGAACCGGUCCGCGCCCGCAUCGCCGGGAAGCCCGAGGAGGGUUUCCAGGUCGUGGCCGCGGACCUGUCGGAAGCGCUCGCCGGCGACGGACUGGCCGUCGCGACGUCGCGACACGGAGCAGAACUCUCCGUCUCGCGGCUCGGCGACCUCGUCUCGGACGCGCGGAAGG
Hez-1-1    UCUCUUCGAGAGAACCGGUCCGCGCCCGCAUCACCGGGAAGCCGGAGGUCGGCUUCCAGGUCGCGGCCGCGGACCUGUCGGAAGCGCUCGCCGGCGACGGACUGGCCGUCGCGACGUCGCGACACGGAGCGGAACUCUCCGUCUCGCGGCUCGGCGACCUCGUCUCGGACGCCCGCGAGG
env-149    UCUCUUCGAGAGAACCGGUCCGCGCCCGCAUCACCGGGAAGCCGGAGGACGGUUUCCAGGUCGUGGACGCGGACCUGUCGGAAGCGCUCGCCGGCGACGGACUGGCCGUCGCGACGUCGCGACACGGGGAGGAACUCUCCGUCUCGCGGCUCGGCG........................
env-150    UCUCUUCGAGAGAACCGGUCCGCGCCCGCAUCACCGGGAAGCCGGAGGACGGUUUCCAGGUCGUGGACGCGGACCUGUCGGAAGCGCUCGCCGGCGACGGCUUGGCCGUCGCGACGUCGCGACACGGGGAGGAACUCUCCGUCUCGCGGCUC............................
env-151    UCUCUUCGAGAGAACCGGUCCGCGCCCGCAUCACCGGGAAGCCGGAGGACGGUUUCCAGGUCGCGGCCGCGGACCUGUCGGAAGCGCUCGCCGGCGACGGACUGGCCGUCGCGACGUCGCGACACGGAGCGGAACUCUCCGUCUCGCGGCUCGGCGACCUCGUCUCGGACGCCCGCGAGG
Hco-1-1    UCUCUUCGAGAGAACCGGUCCGCGCCCGCAUCACCGGGAAGCCGGAGGUCGGUUUCCAGGUCGCGGCCGCGGACCUGUCGGAAGCGCUCGCCGGCGACGGACUGGCCGUCGCGACGUCGCGACACGGAGGGGAACUCUCCGUCUCGCGGCUCGGCGACCUCGUCUCGGACGCCCGCGAGG
env-152    UCUCUUCGAGAGAACCGGUCCGCGCCCGCAUCACCGGGAAGCCGGAGGACGGUUUCCAGGUCGUGGACGCGGACCUGUCGGAAGCGCUCGCCGGCGACGGACUGGCCGUCGCGACGUCGCGACACGGGGAGGAACUCUCCGUCUCGCGGCUCGGCGGCAUCGUCUCGGACGCGCGGAAGG
env-153    UCUCUUCGAGAGAACCGGUCCGCGCCCGCAUCACCGGGAAGCCGGAGGUCGGCUUCCAGGUCGCGGCCGCGGACCUGUCGGAAGCGCUCGCCGGCGACGGACUGGCCGUCGCGACGUCGCGACACGGAGCGGAACUCUCCGUCUCGCGGCUCGGCGACCUCGUCUCGGACGCCCGCGAGG
env-154    UCUCUUCGAGAGAACCGGUCCGCGCCCGCAUCACCGGGAAGCCGGAGGUCGGCUUCCAGGUCGCGGCCGCGGACCUGUCGGAAGCGCUCGCCGGCGACGGACUGGCCGUCGCGACGUCGCGACACGGAGCGGAACUCUCCGUCUCGCGGCUCGGCGACCUCGUUUCGGACGCCCGCGAGG
env-155    UCUCUUCGAGAGAACCGGUCCGCGCCCGCAUCACCGGGAAGCCGGAGGCCGGCUUCCAGGUCGCGGCCGCGGACCUGUCGGAAGCGCUCGCCGGCGACGGACUGGCCGUCGCAACGUCGCGACACGGAGCGGAACUCUCCGUCUCGCGGCUCGGCGACCUCGUUUCGGACGCCCGCGAGG
env-156    UCUCUUCGAGAGAACCGGUCCGCGCCCGCAUCACCGGGAAGCCGGAGGUCGGCUUCCAGGUCGCGGCCGCGGACCUGUCGGAAGCGCUCGCCGGCGACGGACUGGCCGUCGCGACGUCGCGACACGGAGCGGAACUCUCCGUCUCGCGGCUCGGCGACCUCGUCUCGGACGCCCGCGAGG
Hsp-10-1   UCUCUUCGAGAGAACCGGUCCGCGCCCGCAUCACCGGGAAGCCGGAGGACGGUUUCCAGGUCGUGGCCGCGGACCUAUCGGAAGCGCUCGCCGGCGACGGACUGACCGUCGCCACGUCGCGACACGGGGAGGAACUCUCCGUCUCGCGGCUCGGCGACGUCGUCUCGGACGCGCGGAAGG
Hli-1-1    UCUCUUCGAGAGAACCGGUCCGCGCCCGCAUCACCGGGAAGCCCGAGGAGGGUUUCCAGGUCGUGGGAGCGGACCUGUCGGAAGCGCUCGCCGGCGACGGCUUGGCCGUCGCGACGUCGCGACACGGGGAGGAACUCUCCGUCUCGCGGCUCGGCGGCAUCGUCUCGGACGCGCGGAAGG
Hte-1-1    UCUCUUCGAGAGAACCGGUCCGCGCCCGCAUCACCGGGAAGCCCGAGGCGGGCUUCCAGGUCGUGGCCGCGGACCUGUCGGAAGCGCUCGCCGGCGACGGACUGGCCGUCGCGACGUCGCGACACGGGGAGGAACUCUCCGUCUCGCGGCUCGGCGACCUCGUCUCGGACGCCCGCGAGG
Hko-1-1    UCUCUUCGAGAGAACCGGUCCGCGCCCGCAUCACCGGGAAGCCGGAGGACGGCUUCCAGGUCGUGGAAGCGGACCUGUCGGAAGCGCUCGCCGGCGACGGUUUGGCCGUCGCGACGUCGCGACACGGGGAGGAACUCUCCGUCUCGCGGCUCGGCACCAUCGUCUCGGACGCGCGGAGG
Hli-2-1    UCUCUUCGAGAGAACCGGUCCGCGCCCGCAUCGCCGGGAAGCCCGAGGAGGGGUUCCAGGUCGUGGCCGCGGACCUGUCGGAAGCGCUCGCCGGCGACGGAGUGGCCGUCGCGACGUCGCGACACGGAGAAUCGCUCUCCGUCUCGCGGCUCGGCGACCUCGUCUCGGACGCGCGGAAGG
Hsa-7-1    UCUCUUCGAGAGAACCGGUCCGCGCCCGCAUCACCGGGAAGCCGGAGGACGGUUUCCAGGUCGUGGAAGCGGACCUGUCGGAAGCGCUCGCCGGCGACGGCUUGGCCGUCGCGACGUCGCGACACGGGGAGGAACUCUCCGUCUCGCGGCUUGGCACCAUCGUCUCGGACGCGCGGAGG
Hca-2-1    UCUCUUCGAGAGAACCGGUCCGCGCCCGCAUCGCCGGGAAGCCCGAGGAGGGGUUCCAGGUCGUGGCCGCGGACCUGUCGGAAGCGCUCGCCGGCGACGGACUGGCCGUCGCGACGUCGCGACACGGAGAAUCGCUCUCCGUCUCGCGGCUCGGCGACCUCGUCUCGGACGCCCGCGAGG
Hsp-7-1    UCUCUUCGAGAGAACCGGUCCGCGCCCGCAUCGCCGGGAAGCCCGAGGAGGGCUUCCAGGUCGUGGCCGCGGACCUGUCGGAAGCGCUCGCCGGCGACGGAGUGGCCGUCGCGACGUCGCGACACGGAGAAGCACUCUCCGUCUCGCGGCUCGGCGACCUCGUCUCGGACGCCCGCGAGG
Hha-2-1    UCUCUUCGAGAGAACCGGUCCGCGCCCGCAUCACCGGGAAGCCCGAGGAGGGGUUCCAAGUCGUGGGGGCGGACCUGUCGGAAGCGCUCGCCGGCGACGGCUUGGCCGUCGCGACGUCGCGACACGGAGAAGAGCUCUCCGUCUCGCGGCUCGGCACCAUCGUCUCGGACGCGCGGAAGG
Hdi-1-1    UCUCUUCGAGAGAACCGGUCCGCGCCCGCAUCGCCGGGAAGCCCGAGGAGGGGUUCCAGGUCGUGGCCGCGGACCUGUCGGAAGCGCUCGCCGGCGACGGAGUGGCCGUCGCGACGUCGCGACACGGAGAAUCGCUCUCCGUCUCGCGGCUCGGCGACCUCGUCUCGGACGCGCGGAAGG
Hsa-9-1    UCUCUUCGAGAGAACCGGUCCGCGCCCGCAUCACCGGGAAGCCCGAGGAGGGCUUCCAGGUCGUGGACGCGGACCUGUCGGAAGCGCUCGCCGGCGACGGUUUGGCCGUCGCGACGUCGCGACACGGGGAGGAACUCUCCGUCUCGCGGCUCGGCACCAUCGUCUCGGACGCGCGGAAGG
Har-2-1    UCUCUUCGAGAGAACCGGUCCGCGCCCGCAUCGCCGGGAAGCCCGAGGAGGGGUUCCAGGUCGUGGCCGCGGACCUGUCGGAAGCGCUCGCCGGCGACGGAGUGGCCGUCGCGACGUCGCGACACGGAGAAUCGCUCUCCGUCUCGCGGCUCGGCGACCUCGUCUCGGACGCGCGGAAGG
env-157    UCUCUUCGAGAGAACCGGUCCGCGCCCGCAUCACCGGGAAGCCGGAGGUCGGCUUCCAGGUCGUGGCCGCGGACCUGUCGGAAGCGCUCGCCGGCGACGGACUGGCCGUCGCGACGUCGCGACACGGAGCGGAACUCUCCGUCUCGCGGCUCGGCGACCU...................
env-158    UCUCUUCGAGAGAACCGGUCCGUGCCCGCAUCACCGGGAAGCCGGAGGUCGGUUUCCAGGUCGUGGCGCGGAUCUGUCGGAAGCGCUUGCCGGCGACGGAGUGGCCGUCGCGACGUCGCGACACGGAGAAGCGCUCUCCGUGUCGCGUCUCGGCGACCUCGUUUCGGACGCGCGAGAGG
env-159    UCUCUUCGAGAGAACCGGUCCGUGCCCGCAUCACCGGGAAGCCGGAGGUCGGUUUCCAGGUCGUGGGCGCGGAUCUGUCGGAAGCGCUUGCCGGCGACGGAGUGGCCGUCGCGACGUCGCGACACGGAGAAGCGCUCUCCGUGUCGCGUCUCGGCGACCUCGUUUCGGACGCGCGAGAGG
env-160    UCUCUUCGAGAGAACCGGUCCGCGCCCGCAUCACCGGAAAACCGGCGAACGGUUUUCAGGUCGUGGGAGCGGACCUUUCGGAAGCGCUCGGCGGCGACGGACUCGCCGUCGCGACGUCGCGGUACGGGGAGGAGCUCUCCGUCUCGCGACUGGGCGAUAUCGUCUCGGACGCGCGAAG
h3A-1-1    UCUCUUCGAGAGAACCGGUCCGCGCCCGCAUCACCGGGAAGCCGGAGGACGGUUUCCAGGUCGUUGGCGCGGACCUUCCGGAGGUGCUCGCCGACGGCGGGCUCGCCAUCGCGACGUCACGACACGGGGAGCCGCUCUCCGUGUCGCGGCUCGCCGAACUCGCCCCGCGGGUGCAGGAGG
Hai-1-1    UCUCUUCGAGAGAACCGGUCCGCGCCCGCAUCACCGGGAAGCCCGAGGAGGGUUUCCAGGUCGUGGGUGCGGACCUGUCGGAAGCGCUCGCCGGCGACGGCUUGGCCGUCGCGACGUCGCGGCACGGAGAAGAGCUCUCCGUCUCGCGGCUCGGCGACCUCGUCUCGGACGCACGGAGG
env-161    UCUCUUCGAGAGAACCGGUCCGCGCCCGCAUCACCGGGAAGCCCGAGGAGGGUUUCCAGGUCGUGGAAGCGGACCUGUCGGAAGCGCUCGCCGGCGACGGCUUGGCCGUCGCGACGUCGCGACACGGAGAGGAACUCUCCGUCUCGCGGCUCGGCGACAUCGUCUCGGACGCGCGGAAGG
Hho-1-1    UCUCUUCGAGAGAACCGGUCCGCGCCCGCAUCGCCGGGAAGCCCGAGGCAGGCUUCCAGGUCGUGGCCGCGGACCUGUCGGAAGCGCUCGCCGGCGACGGACUGGCCGUCGCGACGUCGCGACACGGGGAGGAACUCUCCGUCUCGCGGCUCGGCGACCUCGUCUCGGACGCCCGCGAGG
env-164    UCUCUUCGAGAGAACCGGUCCGCGCCCGCAUCACCGGGAAGCCGUCGGAGGGCUUCCAGGUCGUGACCGCGGACCUGUCGGCAGCGCUCGCCGACGACGGACUCGCCGUCGCCACGUCACGUUCGGAGGAGGAGCUCUCCGUCUCGCGGCUCGGCGGAUCGUCACGGACGCACGGGGG
env-165    UCUCUUCGAGAGAACCGGUCCGCGCCCGGAUCACCGGGAAGCCGUCUGCUGGCUUCCAAGUGGUGGGUGCGGACCUGUCGGAAGCGCUCGCCGAUGGCGGGCUCGCCGUCGCAACGUCGCGACACGGAGAAUCGCUCUCCGUCUCGCGGCUCGGCGAACU..................
env-166    UCUCUUCGAGAGAACCGGUCCGCGCCCGGAUCACCGGGAAGCCGUCUGCUGGCUUCCAAGUGGUGGGUGCGGACCUGUCGGAAGCGCUCGCCGAUGGCGGGCUCGCCGUCGCAACGUCGCGACACGGAGAAUCGCUCC....................................
env-167    UCUCUUCGAGAGAACCGGUCCGCGCCCGGAUCACCGGGAAGCCAUCCGCUGGCUUCCAGGUGGUGGGUGCGGACCUGUCGGAAGCGCUCGCCGAUGGCGGGCUCGCCGUCGCAACGUCGCGACACGGAGAAUCGCUCUCCGUCUCGCGGCUCGGCGAACUCGUCGCCGACGCGGCAAAGG
env-169    UCUCUUCGAGAGAACCGGUCCGCGCCCGGAUCACCGGGAAGCCAUCCGCUGGCUUCCAGGUGGUGGGUGCGGACCUGUCGGAAGCGCUCGCCGAUGGCGGGCUCGCCGUCGCAACGUCGCGACACGGAGAAUCGCUCUCCGUCUCGCGGCUCGGCGAACUCGUCGCCGACGCGGCAAAGG
env-170    UCUCUUCGAGAGAACCGGUCCGCGCCCGGAUCACCGGGAAGCCAUCCGCUGGCUUCCAGGUGGUGGGUGCGGACCUGUCGGAAGCGCUCGCCGAUGGCGGGCUCGCCGUCGCAACGUCGCGACACGGAGAAUCGCUCC....................................
env-172    UCUCUUCGAGAGAACCGGUCCGCGCCCGGAUCACCGGGAAGCCAUCCGCUGGCUUCCAGGUGGUGGGUGCGGACCUGUCGGAAGCGCUCGCCGAUGGCGGGCUCGCCGUCGCAACGUCGCGACACGGAGAAUCGCUCUCCGUCUCGCGGCUCGGCGAGCU..................
env-173    UCUCUUCGCGACGACCCGUCCGCGCGAAACUUGCUGACGAGCCCGUUUCGGGCUUUCAAGUUUAUCGUGCGGACCUCCCGGAAGCUCUCAGCUGGUCCGAUACGGGACUCUGCAUUGCAGCGUCUCGGUAUGGCCAGGAUUUGACCGUUGAUACUCUCGACGAGUUAACGUCGGAGAUCA
Hru-1-1    UCUCUUCGCGAGAACCGGUCCGUGCGCGGAUCGUCGACGAGCCCGUUCGGGCUUCGACGUGUCCCGUAUGGACCUCGAGGAAGCACUCGGCCACGCCGACGCGGGCGUCGCGAUUGCGACGUCCCGCUUCGGCGAGGCGCUGACCACGAGUCGGCUGCCGCAGCUCGCCUCGCGCGUCG
env-174    UCUCUUCGCGAGAACCGGUCCGUGCGCGGAUCG......................................................................................................................................................
Hru-2-1    UCUCUUCGCGAGAACCCGUCCGCGCACGGCUGGUCGACGAGGACCCACCCGGGUUUACCGUCGACGCUCGGGGACAUCCAGGAAGCCCUGGCCCGCGACGACGCGGGCCUCUGCAUCGGCACGUCGCGCCACGGGGACGAACUCACCCGUUCGCGGCUGACCGAACUCGCCCCGCGCU
env-175    UCUCUUCGCGAGAACCGGUUCGCGCGAAAUCGUGGCCGAACCCCCACCGGGGUUCGCCGUUGAGCGCGCGGCCAUCGGCAACUGCCUUGCCCGGCCGGACGCUGGCAUUCGAAUUGCCGCGUCCAGGCACGGGGA...........................................
env-176    UCUCUUCGCGAGAGCCGGUUCGCGCGAAACUCGUGGCGGACGCCCCACCGGGCUUCGCCGUCGAGCGCGCGACCGUCGACGAGUGCCUCUCCCGGUCGGACGCCGGCGUCCGUAUCGCCGCGUCCAGACACGGGGAGGUGCUGACGGUCGCCCGACUCGGGGAACUCUCCGAGCGGACGA
env-178    UCUCUUCGCGAGAGCCGGUUCGCGCGAAACUCGUGGCGGACGCCCCACCGGGCUUCGCCGUCGAGCGCGCGACCGUCGACGAGUGCCUCUCCCGGUCGGACGCCGGCGUCCGUAUCGCCGCGUCCAGACACGGGGAGGUGCUGACGGUCGCUCGGCUCGGAGAACUCUCCGAGCGGACGA
env-179    UCUCUUCGCGAGAGCCGGUCCGCGCGAAACUCGUGGCCGAGCCCCCACCGGGGUUCGCCGUCGAGCGCGCGGCCAUCGACGACUGCCUCGCCCGUCCGGACGCCGGCGUCCGAAUCGCCGCGUCCAGACACGGGGAGGUACUGACGGUCGAUCGGCUCGGGGAACUCUCCGGGCGGACGA
env-180    UCUCUUCGCGAGAGCCGGUC...................................................................................................................................................................
Nmo-1-1    UCUCUUCGCGAGAGCCGGUUCGCGCGAAACUCGUGGCGGACCCCUCACCGGGGUUCGCCGUCGAGCGUGCGACCAUCGACGACUGCCUCGCCCGUCCGGACGCCGGCGUCCGAAUCGCCGCGUCCAGACACGGGGAGGUACUGACGGUCGAUCGGCUCGGGGAACUCUCCGGGCGGACGG
Hha-1-1    UAUCUUCGCGAGAACCGGUCCGCGCACGGAUCGUCGACGAACCCGCACCCGGGUUCGCCGUCGACCGCACGGACCUCCCGGCGGCGCUCGGCCGUGAGGAUGCCGGGCUCCGGGUCGCCACGUCGCGCCACGGCGAGGCGCUGACGGCGGACGACCUCGGAGCACUGGCCGGCGGGCCCG
env-181    UCUCUUCGCGACGACCGGUCCGGGCGAAACUCGUCGACGACCCCCUCCCGGGGCUCGCCGUCGAGCGGACGGACCUCUCCGCAGCACUCGGCCGUGAGGACGCCGGCGUCACCAUUGCUGCCUCUCGAUUCGGUGAGAUUCUCACCGUGGGGCGGCUGGACACGC..............
Hpa-1-1    UCUCUUCGAGAAGACCGGUCCGUGCGAAGCUCGUGGACGCCCCCGUUCCGGGGCUCACAGUGACGCGCAUGGACCUCUCGGCGGCCCUCGACCGCGACGAUGCUGGCGUCAGAAUCGCCACAUCGCGUUACGGAGAACCGCUCUCCGUCGGACGGCGGCUUGAAGAAGUCGUCACGCGAACGA
Hci-1-1    UCUCUUCGAGAAGACCGGUUCGUGCGAAGCUCGUGGACGCCCCGCAGUCGGGCUACACAGUGAUGCGCAUGAACCUCUCGGCGGCCCUCGACCGCGACGAUGCUGGCGUUCGAAUCGCCACGUCUCGCUUCGGAGAAUCGCUCUCCGUGGCGCGACUCGACGAAAUCGUUGCGCAAACCG
env-182    UCUCUUCGCG.............................................................................................................................................................................
env-183    UAUCUUCGAGAGAACCGGUCCGCGCGAAUCACGGACGAAGCCCACCGGGUUUCGCCGUCGAAC.........................................................................................................................
env-184    UCUCUUCGCGAGAUCCGGUCCGGGCCCGACUCACCGAGGAGCCCCCGCCGGGCUACUCGGUCGGGCCGUCGGACCUUUCG......................................................................................................
env-185    UCUCUUCGCGACGACCGGUCCGGGCGAAGCUCGUAGACGAGCCCCUCCCGGGGCUCGCCGUCGAGUCGACGGACCUAUCGGCGGCUCUC..............................................................................................
Hdj-1-1    UCUCUUCGCGACGACCGGUCCGGGCGAAGCUCGUCGACGGCCGGCCCCUCCCGGGGCUGUCGGUCGAGCGGACGGACCUUUCGGCAGCACUCGGCCGUGAGGACGCCGGCGUCUGCAUCGCCGCCUCCCGCUACGGUGAACGACUCACCGUCGGGCGGCUCGAGACGCUGGCCGGGACGAUCG
Hsa-10-1   UCUCUUCGCGACGACCGGUCCGGGCGAAGCUCGUCGAUCGGCCCCUUCCGGGGCUAUCGAUUGAGCAGACGGACCUUUCGGCAGCACUCGGCCGUGAGGACGCCGGCGUCCGUAUCGCCGCCUCCCGUUACGGUGAAGAACUCACCGUGGGGCGGCUCGAGACGCUGGCCGGGCGCGUCG
Nsu-1-1    UCUCUUCGCGACGACCGGUCCGGGCGAAACUCGAGAACAAGCCCCUCCCGGGGCUUGCAGUCGAGCGGACGGACCUUUCCGCAGCACUCGGCCGUGAGGACGCUGGCGUCUGCAUCGCUGCCUCCCGAUUCGGUGAAGAACUCACCGUCGGGCGGCUCGAGACGUUGGCCGGACGCAUCG
Nba-1-1    UCUCUUCGCGACGACCGGUCCGAGCGAAACUCGUCGAGGACAAACCCCUCCCGGGGCUGUCAGUCGAGCGGACGGACCUUUCCGCAGCACUCGGCCGUGAGGACGCUGGCGUCUGCAUCGCCGCGUUCCCGAUUCGGUGAAGAGCUCACCGUCGGGCGGCUUGGACGCUGGCCGGACGCAUCG
Nin-1-1    UCUCUUCGCGACGACCGGUCCGGGCGAAACUCGUCGACGAACCCCUUCCGGGGCUCGCAGUCGAGCGGACGGACCUUUCGACGGCACUCGGCCGUGAGGACGCCGGCGUCUGUAUUGCGUCGUCUCGCUACGGUAUCGAACUUACCGUCGGACGGCUCGAGACGCUGGCCGGACGCGUCG
env-186    UCUCUUCGCGAGAGCCGGUCCGUGCGAGGCUGGUCGACGAGCCCGUUUCGGGGCUCUCGAUCCAUCGCAUGGACCUAUCGGAAGCGCUCGGCCGUGAGGACGCCGGUGUUCGGAUCGCCGACGUCGAGGUUCGGCGAGAACCUUAGCGUCUCGCGACUCGGAACGCUGGCCGGACGGAUCG
env-187    UCUCUUCGCGAGAGCCGGU....................................................................................................................................................................
env-188    UCUCUUCGCGAGAGCCGGUCCGUGCGAAGCUGGUCGACGAGCCCAUUUCGGGGCUCUCGAUCCAUCGCAUGGACCUAUCGGAAGCGCUCGGCCGUGAGGACGCCGGUGUCCGAAUCGCGACGUCGAGGUUCGGCGAGAACCUUAGCGUCUCGCGACUCGGGACACUGGCCGGACGAAUCG
env-189    UCUCUUCGCGAGAGCCGGUCCGUGCGAAGCUGGUCGACGAGCCCAUUUCGGGGCUCUCGAUCCAUCGCAUGGACCUAUCGGAAGCGCUCGGCCGUGAGGACGCCGGUGUCCGAAUCGCGACGUCGAGGUUCGGCGAGAACCUUAGCGUCUCGCGACUCGGGACACUGGCCGG........
env-190    UCUCUUCGCGACGACCGGUCCGGGCGAAACUCGUCGACGAAUCCCCGCCGGGG..................................................................................................................................
env-191    UCUCUUCGCGACGACCGGUCCGGGCGAAACUCGUCGACGAAUCCCCACCGGGAUUCGCCGUCGAGCGGACGGACCUCCCGGCAGCACUCGGCCGUGAGGACGCCGGCGUUCGUAUCGCCGCCUCCCGAUUCGGUGAACCGCUCACCAUCGGACGGCUCGAGACGCUGGCCGGGGACAUCG
Hla-1-1    UCUCUUCGCGACGACCGGUCCGGGCGAAGCUCGUCGAUCGCCCCCUUCCGGGGCUUUCAGUCGAGCGGACGGACCUAUCGGCAGCACUCGGCCGUGAGGACGCCGGCGUCCGCAUCGCCGCCUCCCGACACGGUGAGGAACUCACCGUGGGGCGGCUCGAGACGCUGGCCGGACGCAUCG
env-193    UCUCUUCGCGACGACCGGUCCGGGCGAAACUCGUCGACGAAUCCCCACCGGGAUUUACCGUCGAGCGGACGGACCUCUCGGCAGCACUCGGCCGUGAGGACGCUGGCGUUCGUAUCGCCGCGUUCUCGCUACGGUGAGCAACUCACCGUGGAACGGCUC.....................
Hac-1-1    UAUCUUCGAGACGACCGGUCCGGGCGAAGAUCAUCGAGGAGGCCCCACCGGGCUUUCAG............................................................................................................................
Hac-1-2    UAUCUUCGAGACGACCGGUCCGGGCGAAGAUCAUCGAGGAGGCCCCACCGGGCUUUCAGGUCUUCACCCGGCCCAUCGGGGACGAACUCGACCGUCCGGACGCGGGGUUCCGGGUCGCCACAUCACGCUUCGGGGUGCCAGUCGGCGCCGCGAACGUGGGCGACCUCACCGCCGCGUCC
env-196    UCUCUUCGCGACGACCGGU....................................................................................................................................................................
env-197    UCUCUUCGCGACGACCGGUCCGUGCGAAACUGGUCGACGAACCCCUCCCGGGGUUCUCGGUCGAACGCACGGCCCUCGGUGACGCUCUCGA.........................................................................................
env-198    UCUCUUCGCGACGACCGGUCCGUGCGAAACUGGUCGACGAACCCCUCCCGGGGUUCUCGGUCGAACGCACGACCAUCGGGGACGCCCUCGACCGCCCCGACGCGGGCGUUCGCAUCGCUACCUCCCGCCACGGGACGCCGCUUUCGGCGGCGUCCCUCGGGGGGUACGUCGAACGCAUCG
Hsp-2-1    UCUCUUCGCGACGACCGGUCCGUGCGAAACUGGUCGACGAACCCCUCCCGGGGUUCUCGGUCGAACGCACGACCAUCGGGGACGCCCUCGACCGCCCCGACGCGGGCGUUCGCAUCGCUACCUCCCGCCACGGGACGCCGCUUUCGGCGGCGUCCCUCGGGGGGUACGUCGAACGCAUCG
env-199    UCUCUUCGAGAGACCGGUCCGGGCGAAGAUCGUCGACGAUCGAUCGGUUGGUCGGUCGACCGUGCGGACCUGCCGGCGCACUCGGCCGUGAGGACGCCGGACUCCGGAUCGCCGCCUCACGCCACGGCGACGCGUGGUGAGAGCGGUGACAUCGGGAGUCUGGCCGGUCGCGACGAGG
Hsu-1-1    UCUCUUCGCGACGACCGGUUCGUGCGAAGCUGGUCGACGAGCCCCUUCCGGGGUUCGUCGUCACUCGGACGAACCUCCGGGAUGCCCUCGAUCGCCCCGACGCAGGCUAUCGCGUUGCCACCUCCCGACACGGGACACCGGUUACGGUGUCCGGACUGUCGGGAAUCAAAGAACGCGUCG
env-201    UCUCUUCGCGACGGCCGGUCCGGGCGAAACUCGUCGACGCCCCCACAACGGGAUUCGACGUUGUCGCCGCGGACCUCGAUGCGGCACUCUCGCGCGACGAUGCCGGGCUCACUAUCGCGUCCUCGCGAUACGGUGAGCCGGUAACGUCGACCCGUCUC......................
env-203    UCUCUUCGCGACGGCCGGUCCGGGCGAAACUCGUCGACGCCCCCACAACGGGAUUCGACGUUGUCGCCGCGGACCUCGAUGCGGCACUCUCGCGCGACGAUGCCGGGCUCACUAUCGCGUCCUCGCGAUACGGUGAGCCGGUAACGUCGACCCGUCUCGGCCAGCUGGUC..........
Hca-1-1    UCUCUUCGCGACGGCCGGUCCGGGCGAAACUCGUCGACGCCCCCACAACGGGAUUCGACGUUGUCGCCGCGGACCUCGAUGCGGCACUCUCGCGCGACGAUGCCGGGCUCACUAUCGCGUCCUCGCGAUACGGCGAGCCGGUAACGUCGACCCGUCUCGGCCAGUUGGCCGAGCGGCGCG
Ham-1-1    UCUCUUCGCGACGGCCGGUCCGGGCGAAACUCGUCGACGCCCCCACAACGGGAUUCGACGUUGUCGCCGCGGACCUCGAUGCGGCACUCUCGCGCGACGAUGCCGGGCUCACUAUCGCGUCCUCGCGAUACGGCGAGCCGGUAACGUCGACCCGUCUCGGCCAGUUGGCCGAGCGGCGCG
Hhi-2-1    UCUCUUCGCGACGGCCGGUCCGGGCGAAACUCGUCGACGCCCCCACAACGGGAUUCGACGUUGUCGCCGCGGACCUCGAUGCGGCACUCUCGCGCGACGAUGCCGGGCUCACUAUCGCGUCCUCGCGAUACGGCGAGCCGGUAACGUCGACCCGUCUCGGCCAGUUGGCCGAGCGGCGCG

           ...........................................................................................................................................................................................
           ...........................................................................................................................................................................................
           UCUYUUYR•GRRRR•YR•••••••Y•••R•••RY•RR•R••••••••••••••••••••••••••••••••••••••••••R•••••••••••••••••••••••••••••••••••••••••••••••••••••••••••••••••••••••••••••••••••••••••••••••••••••••••
```

| | |
|---|---|
| Hsa-2-1 | [illegible] |
| Har-1-1 | [illegible] |
| Hva-1-1 | [illegible] |
| Hja-1-1 | [illegible] |
| env-204 | [illegible] |
| Hse-1-1 | [illegible] |
| Hth-1-1 | [illegible] |
| Hmo-1-1 | [illegible] |
| env-206 | [illegible] |
| env-207 | [illegible] |
| env-209 | [illegible] |
| Nta-1-1 | [illegible] |
| Nas-1-1 | [illegible] |
| Noc-1-1 | [illegible] |
| Nje-1-1 | [illegible] |
| Nam-1-1 | [illegible] |
| env-210 | [illegible] |
| Hsp-3-1 | [illegible] |
| Nch-1-1 | [illegible] |
| Nae-1-1 | [illegible] |
| env-214 | [illegible] |
| env-215 | [illegible] |
| env-216 | [illegible] |
| Hni-1-1 | [illegible] |
| Has-1-1 | [illegible] |
| env-217 | [illegible] |
| Nal-2-1 | [illegible] |
| Npa-1-1 | [illegible] |
| Nga-1-1 | [illegible] |
| Nsp-2-1 | [illegible] |
| Htu-1-1 | [illegible] |
| Hsa-8-1 | [illegible] |
| Hth-2-1 | [illegible] |
| Nve-1-1 | [illegible] |
| Hli-3-1 | [illegible] |
| Hxa-1-1 | [illegible] |
| Hgo-1-1 | [illegible] |
| env-218 | [illegible] |
| env-219 | [illegible] |
| env-220 | [illegible] |
| env-221 | [illegible] |
| Hsa-6-1 | [illegible] |
| env-222 | [illegible] |
| env-223 | [illegible] |
| env-224 | [illegible] |
| env-225 | [illegible] |
| env-226 | [illegible] |
| env-227 | [illegible] |
| env-228 | [illegible] |
| env-231 | [illegible] |
| env-232 | [illegible] |
| env-233 | [illegible] |
| env-237 | [illegible] |
| env-238 | [illegible] |
| env-240 | [illegible] |
| env-241 | [illegible] |
| env-242 | [illegible] |
| env-243 | [illegible] |
| env-244 | [illegible] |
| env-245 | [illegible] |
| env-246 | [illegible] |
| env-247 | [illegible] |
| env-248 | [illegible] |
| env-249 | [illegible] |
| env-250 | [illegible] |
| env-251 | [illegible] |
| env-252 | [illegible] |
| env-253 | [illegible] |
| env-254 | [illegible] |
| env-255 | [illegible] |
| env-256 | [illegible] |
| env-257 | [illegible] |
| env-258 | [illegible] |
| env-259 | [illegible] |
| env-260 | [illegible] |
| env-261 | [illegible] |
| env-265 | [illegible] |
| env-266 | [illegible] |
| env-268 | [illegible] |
| env-269 | [illegible] |
| env-270 | [illegible] |
| env-271 | [illegible] |
| env-272 | [illegible] |
| env-274 | [illegible] |

```
env-275   UCUAUUAGCGACGACUGGACACGGACCCGGUGCACGCCGGGGCGGUGGCGGGAGCGAUAGGAGAGGAUCGUCUCUCCCCCGUCGCCUC..........................................................................................................
env-277   UCUAUUAGCGACGACUGGACACGGACCCGGUGCACGCCGGGGCGGUGGCGGGAGCGAUAGGAGAGGAUCGUCUCUCCCCCGUCGCCUCCAUCCGACUGUUUCCCUUCACGCCGUGCCAUGAUGGCACGACGUUCGAUGUCAUGCCCAAGUGGGUUCACCACAAUUUGCCCGAUCUGCGAC
env-278   UCUAUUAGCGACGACUGGACGUACCCGGCGCACACCGGGCCGGUGGCGGGGAAGAUAGGAGGAGAUGGGCGUCUCCCCCGCCGCCUCCAUCCAACUGUAUCCCUUCACGCCGCGGCAAAACGGCGUGGCGUUCGAUGUCAUGCCCGAUACGGGUUCGCCGUAAUGCCCGACCUGCGGCGG
env-292   UCUAUUAGCGACGACUGGACGUACCCGGUGCACACCGGGCCGGUGGCGGGAGCGGUAGGAGACGAUCGUCUCCCCCGUGCCUGCCGCCUCCAUCCAACUGUUUCCCUUCACGCCGCGCCAUUCUGGCGCGUCGUUCGAUGUCAUGCCCGAUACGGGUUCGCCGUAAUGCCCGACCGUGCC
env-293   UCUAUUAGCGACGACUGGACGUACCCGGUGCACACCGGGCCGGUGGCGGGAGCGGUAGGAGACGAUCGUCUCCCCCGUGCCUGCCGCCUCCAUCCAACUGUUUCCCUUCACGCCGCGCCAUUCUGGCGCGUCGUUCGAUGUCAUGCCCGAUACGGGUUCGCCGUAAUGCCCGACCGUGCC
env-295   UCUAUUAGCGACGACUGGACGUACCCGGUGCACACCGGGCCGGUGGCGGGAGCGGUAGGAGACGAUCGUCUCCCCCGUGCCUGCCGCCUCCAUCCAACUGUUUCCCUUCACGCCGCGCCAUUCUGGCGCGUCGUUCGAUGUCAUGCCCGAUACGGGUUCGCCGUAAUGCCCGACCGUGCC
env-296   UCUAUUAGCGACGACUGGACGUACCCGGUGCACACCGGGCCGGUGGCGGGAGCGGUAGGAGACGAUCGUCUCCCCCGUGCCUGCCGCCUCCAUCCAACUGUUUCCCUUCACGCCGCGCCAGUUCUGGCGCGUCGUUCGAUGUCAUGCCCGAUACGGGUUCGCCGUAAUGCCCGACCGUGCC
Mbo-1-1   UCUAUUAGCGACGACUGGACGUACCCGGUGCACACCGGGCUGGUGGCGGGAGCGAUAGGAGACGAUCGUCUCCCCCGUGCCUGCCGCCUCCAUCCAACUGUUUCCCUUCACGCCGCGCCGUUCUGGCGCGUCGUUCGAUGUCAUGCCCGAUACGGGUUCGCCGUAAUGCCCGACCGUGCC
env-297   UCUAUUAGCGACGACUGGACGUACCCGGUGCACACCGGGCCGGUGGCGGGAGCGAUAGGAGACGAGUGUCUCCCCGACCCCGCGGCCUCCAUCCAACUGUUUCCCUUCACGACGCGCUAGAACUGCGCGCCGUUCGAUGUCAUGCCCGAUACGGGUUCGCCGUAAUGCCCGACCGUGC
env-298   UCUAUUAGCGACGACUGGACGUACCCGGUGCACACCGGGCCGGUGGCGGGAGCGAUAGGAGACGAGUGUCUCCCCGACCCCGCGGCCUCCAUCCAACUGUUUCCCUUCACGACGCGCUAGAACUGCGCGCCGUUCGAUGUCAUGCCCGAUACGGGUUCGCCGUAAUGCCCGACCGUGCCG
env-303   UCUAUUAGCGACGACUGGACGUACCCGGUGCACACCGGGCCGGUGGCGGGAGCGAUAGGAGACGAGUGUCUCCCCGACCCCGCGGCCUCCAUCCAACUGUUUCCCUUCACGACGCGCUAGAACUGCGCGCCGUUCGAUGUCAUGCCCGAUACGGGUUCGCCGUAAUGCCCGACCGUGCCG
env-304   UCUAUUAGCGACGACUGGACGUACCCGGUGCACACCGGGCCGGUGGCGGGAGCGNUAGGAGACGAUCGUCUCCCCCGUGCCUGCCGCCUCCAUCCAACUGUUUCCCUUCACGCCGCGCCAUUCUGGCGCGNCGUUCGAUGUCAUGCCCGAUACGGGUUCGCCGUAAUGCCCGACCGUGCC
env-305   UCUAUUAGCGACGACUGGACACGGACCCGGUGCACGCCGGGGCGGUGGCGGGAGCGAUAGGAGAGGAUCGUCUCUCCCCCGUCGCCUCCAUCCGACUGUUUCCCUUCACGCCGUGCCAUGAUGGCACGACGUUCGAUGUCAUGCCCAAGUGGGUUCACCACAAUUUGCCCGAUCUGCGAC
env-306   UCUAUUAGCGACGACUGGACACGGACCCGGUGCACGCCGGGGCGGUGGCGGGAGCGAUAGGAGAGGAUCGUCUCUCCCCCGUCGCCUCCA.......................................................................................................
Mma-1-1   UCUAUUGGCGACGACUGGAAUGCACCCGGUGCACACCGGGACGGCAGCAGAGAGAACGGUCGAGGCGCACGGAGACGAACGUCUCUGUGACUUCGCGUUCCGCUGCCUCCAUCAACUGUUUCCCUUACGCUACGGCAAAAUGGCGUGGCGUUCGAUAUCCCGCCCCAUCCGGGUUCGCCA
Mse-1-1   UCUAUUGGCGACGACUGGAAUGCACCCGGUGCACACCGGGACGGCAGCAGAGAGAGCGGUCGAGGCGCACGGAGACGAAUGUCUUUGUGACUUCACGUUCCGCUGCCUCCAUCAACUGUUUCCCUUACGCUACGGCAAAACGGCGUGGCGUUCGAUAUCCCGCCCCAUCCGGGUUCGCCA
Msp-1-1   UCUAUUGGCGACGACUGGAAUGCACCCGGUGCACACCGGGACGGCAGCAGAGAGAGCGGUCGAGGCGCACAGAGACGAAUGUCUCUGUGACUUCACGUUCCGCUGCCUCCAUCAACUGUAUCCCUUACGCUACGGCAACAUGGCGUGGCGUUCGAUGUCCCGCCCCAUCCGGGUUCGCCA
env-310   UCUAUUAGCGAC..................................................................................................................................................................................
env-315   UCUAUUAGCGACGACUGGACGCACCCGGUGCACGCCGGGCCGGUGGCGGGAGCGAUCAGGAGACAGGCGUCUCUCUGACCCCCGCUGCCACCAUCCAACUGUUUCCCAUCACGUCGCGUCAGAACGGCGCGUCGUUCGAUGUCAUGCCCGAUACGGGUUCGCCGUAAUGUGCCUGAUCUG
env-316   UCUAUUAGCGACGACUGGACGCACCCGGUGCACGCCGGGCCGGUGGCGGGAGCGAUCAGGAGACAGGCGUCUCUCUGACCCCGCUGCCACCAUCCAACUGUUUCCCAUCACGUCGCGUCAGAACGGCGCGUCGUUCGAUGUCAUGCCCGAUACGGGUUCGCCGUAAUGUGCCUGAUCUG
env-317   UCUAUUAGCGACGACUGGACACGGACCCGGUGCACGCCGGGGCGGUGGCGGGAGCGAUAGGAGAGGAUCGUCUCUCCCCCGUCGCCUCCAUCCGACUGUUUCCCUUCACGCCGUGCCAUGAUGGCACGACGUUCGAUGUCAUGCCCAAGUGGGUUCACCACAAUUUGCCCGAUCUGCGAC
env-318   UCUAUUAGCGACGACUGGGCAUACCCGGUGCACGCCGGGCCGGUGGCGGGAGCGAUUCAGGAGACAGGCGUCUCUCCUGACCCCCGCUAUCACCAUCCAACUGUUUCCCAUCACGCCGCGCCGGAACGGCACGUCGUUCGAUGUCACACCCGAUACGGGUUCGCCGCAAUGUGCCUGACC
env-320   UCUAUUAGCGACGACUGGGCAUACCCGGUGCACACCGGGCCGGUGGCGGGAGCGAUGCAGGAGACAGGCGUCUCCUGACCCCCCGCUAUCACCAUCCAACUGUUUCCCAUCACGCCGCGCCGGAACGGCGCGUCGUUCGAUGUCACACCCGAUACGGGUUCGCCGCAAUGUGCCUGACC
env-321   UCUAUUAGCGACGACUGGGCAUACCCGGUGCACGCCGGGCCGGUGGCGGGAGCGAUUCAGGAGACAGACGUCUCCCUGACCCCCGCUAUCACCAUCCAACUGUUUCCCAUCACGCCGCGCCGGAACGGCGCGNCGUUCGAUGUCACACCCGAUACGGGUUCGCCGCAAUGUGCCUGACC
env-322   UCUAUUAGCGACGACUGGGCAUACCCGGUGCACNCCGGGCCGGUGGCGGGAGCGAUNCAGGAGACAGGCGUCUCCUGACCCCCCGCUAUCACCAUCCAACUGUUUCCCAUCACGCCGCGCCGGAACGGCNCGUCGUUCGAUGUCACACCCGAUACGGGUUCGCCGCAAUGUGCCUGACC
Mpa-1-1   UCUAUUCGCGAGACAGGUACAUUCUGGCGCACUCCAGAAGAUGCUCACCAGGAAGGAACGAUUCCGCUCUCUGUGAAAGGGGGGCUGAUCGCUCUGGACUGGCAUAAUCCUCAAGCGAAGUGCACUGCUCAAUCUCGGGUUUACGAUGCGGCAUCGUCCCGUCGAUAAACAUGAACAAAC
env-323   UCUAUUCGCGAGACAGGUACAUUCUGGCGCACUCCAGAAGAUGCUCACCAUGAAAGGACGGUCCUGCCCCUGCAGAAGGUGGCAGAUCGCCCUGGACUGGCAUAAUCCUCAAGCGAAGUGCACUGCUCAAUCUCGGGUUUACGAUGGGGCAUCGUCCCGUCGGUAAGCAUCAACAAACGC
env-324   UCUUCUUUCGCGAGACAGGAAAUACCCGGUGCACACCGGAUCAUGCCGAAGAGACAAUAUCUUCUGCAUCCUCGUAACCGGACGAACCGACAGAGCAGGGAUAACCAGGAUUAUCUGCCACGAACUGGCGGAUAUCCCAAAAAAUUCCCCGAUCUUCGGCGUGCCGCGUGUUGGCUUAAG
env-325   ACUUCUUUCGCGAGACAGGAAUGCUGGUGCACACCAGUAGGGCAAUAACACAAGCUCAUUCGUGCAUCUCCAGAAACGGGUGAAUGGUAACCAUGACCCGCUGCCUUGAAAAACCACGUGGUUUUUCAUGGCAUGCCGUGGUCGGGCAUAAAAACCACCGUUCCCGCGUGUUGGCUUAAC
env-326   ACUUCUUUCGCGAGACAGGAAUGCUGGUGCACACCAGUAGGGCAAUAACACAAGCUCAUUCGUGCAUCUCCAGAAACGGGUGAAUGGUAACCAUGACCCGCUGCCUUGAAAAACCACGUGGUUUUUCAUGGCAUGCCGUGGUCGGGCAUAAAAACCACCGUUCCCGCGUGUUGGCUUAAC
env-331   ACUUCUUUCGCGAGACAGGAAUGCUGGUGCACACCAGUAGGGCAAUAACACAAGCUCAUUCGUGCAUCUCCAGAAACGGGUGAAUGGUAACCAUGACCCGCUGCCUUAGAAAAACCACGUGGUUUUUCAUGGCAUGCCGUGGUCGGGCAUAAAAACCACCGUUCCCGCGUGUUGGCUUAA
env-332   ACUUCUUUCGCGAGACAGGAAUGCUGGUGCACACCAGUAGGGCAAUAACACAAGCUCAUUCGUGCAUCUCCAGAAACGGGUGAAUGGUAACCAUGACCCGCUGCCUUGAAAAACCACGUGGUUUUUCAUGGCAUGCCGUGGUCGGGCAUAAAAACCACCGUUCCCGCGUGUUGGCUUAAC
Mfo-1-1   UCUUCUUUCGCGAGACAGGAAUACCGGUGCACACCGGUCCAGCGGUGGGAGUAAUCUCAUACGCACAUCCCCGAAACGAUGAACCAGUCUGCGGUCGGCAGUGUAAUGUAUCCCUACCGGAUCCUCUGCACUGCCUGUCGCGACUGCCUUUCAUCUCCGUCGCGUGUUGGCAUCUGUAAC
env-333   UCUUCUUUCGCGAACAAGAAAAAAACACCAAAACCGGGGAGCCGGGCAGUAAUAUGCCCCGGCCUCCCGGGAUAACGGUGUGCCGGCAAAGGGAAAACAACCGGUUUCAGGAGGUUUUCCCGCAUUUUGCCGGUCGCGUGUUGGCUUACCAGACAGGCACGUGAUAUCACUUCAGUCUCG
env-334   UCUUCUUUCGCGAAAAAAGCAAUACCGAAAAUAUCCGGGAACGAGGCACACAAAUGUCUCGUCCCAGAUAUAACGGUGUACCGACAAAGGGACAAUUACCGGUUCCGGUAGUUUUCCCUCAUGGCGGUCGCGUGUUGGCUUACCAAGACAGGCACGUGAUAUCACUUCAGUCUCGGAAUC
env-335   UCUUCUUUCGCGAAAAAGGAAACACUGAAUGUAUCAGGGAACGGGACACCCCGUGUCCCCUUCUGAUAUAACGGUGUGCCGGCAAAAGGAAAAUUACCGGUUCCGGUGGUUUUCCGGUAUGCUGGUCGCGUGUUGGCUUACCAAGACAGGCACGUGAUAUCACUUCAGUCUCGGAAUCA
env-337   UCUUCUUUCGCGAAAAAAGCAAUACCGAAAAUAUCCGGGAACGAGGCACACAAAUGUCUCGUCCCAGAUAUAACGGUGUACCGACAAAGGGACAAUUACCGGUUCCCGGUAGUUUUCCCUCAUGGCGGUCGCGUGUUGGCUUACCAAGACAGGCACGUGAUAUCACUUCAGUCUCGGAAU
env-338   UCUUCUUUCGCGAAAAAGAAAAGCACCCGAACCGGGGAUACGGGCAGGAAUGUGCCCGGCUUUCCGGUUAACGGUGUGCCGGCAAAGGAAAAACAACCGGUUUUCCGGUGGUUUUUCCCGUAUCUUGCCGGUCGCGUGUUGGCUUACCAGACAGGCACGUGAUAUCACUUCAGUCUCGGA
env-339   UCUUCUUUCGCGAAAACGAAAAAAAACACCCGAACCGGGAAUCCGGGCAGUAAUGUGCCCGGCUUUUCGGUGAAGGUGUGCCGGCAAAGGAAAAACCACCGGUUUCCGGUGGUUUUUUUAUAUUUUGCCGGUCGCGUGUUGGCUUACCAGACAGGCACGUGAUAUCACUUCAGUCUCGGA
CMe-1-1   UCUUCUUUCGCGAGGAAAAAAAGCGGAGAUCCCUUGAGGGUGGGAGAAAGCUCUCUCCCUCUGCAGGUCUCCAGGCUAACCGGUUCACGGGCAGGGUGAGUAUGCUCCCGCGCAUCACGUGCGGGAAAAAUCGUCCUGCAAGAUGACACCGGAUCGCGUGUUGGCUUACCGGACAGGCAAG
env-340   UCUUCUUUCGCGAGACAGGAAUGCUGGUGCACAUCAGCUCAUGGAUGAAACAUCAUCCACAUGUCUGUAAGCUACAGAAAACAGGGAUAUUCAGUACUACAUGAAAUCCGCUUGCCGCGGAUCCUCGUUGUACUUGAGUUACCUCGUUUUUUCGAUGGCGUCGCGUGUUGGCUCAUUAAA
env-341   UCUUCUUUCGCGAGACAGGAAUGCUGGUGAAUACCAGCUCAUGGAUGAGGAAAUUAUCAUCCAUAUCUCCUUAAGCAGAGGAAAACCGGACGUUCAAUAAUACAAAAAAUCCGAAUAUAUGGAUCAUUGUAUUUUUUUGAAUUCGCCGGUUCUCUUCUCUGGCGUCGCGUGUUGGCAUCA
env-349   UCUUCUUUCGCGAGACAGGAAUGCUGGUGAAUACCAGCUCAUGGAUGAGGAAAUUAUCAUCCAUAUCUCCUUAAGCAGAGGAAAACCGGACGUUCAAUAAUACAAAAAAUCCGAAUAUAUGGAUCAUUGUAUGUAUUUUGNAUUCNCCGUUUUUCUUCUCUGGCGUCGCGUGUUGGCAA
env-350   UCUUGUUUCGCGAGACUUAGGCAAGCUGGUGCACAUCAGCUCAUGGAUGAGGAAAAAAACAUCCCUAUUGUCCGUAACCGGAAAAGAAUGUCAAGAUCUUCAAUACAACCUUCAAAUCCGUGUUAUCGGAUCGGUGUAUAAUUGAACGAUCCUGGUUCACCGGCAUCGCGUGUUGGCUUCAG
env-351   UCUUCUUUCGCGAGACAGGAAUGCUGGUGCACAUCAGCUCAUGGGUGAAAAAUCUCAUCCAAAUGUCCGUAACCCAAUGAAAAACCGGAUGUUCGAUACUCCAUGAAAUCCGCUUAACGCGGAUCAAGGAAGUCCCGGAGACAUCCUGUUAUUCAGAUGGCGUCGCGUGUUGGCUAUAUC
env-352   UCUUCUUUCGCGAGACAGGAAUGCCGGUGCACACCGGCGCAGGCGUUUGAGUUCAUCUCUUCCGCAUCUCCGUAACCGGGCAAACUGACAGAACGAGCGCAUACGGAUAAGGGCCCACAUAAAGGAUCCCUCGUAUAGCUCACUAUUGUCGGCAUGUCCGUGUCGCGUGUUGGCUUUAU
env-353   UCUUCUUUCGCGAGACACAGAAUGCGGUGCAUACCAGCUCAUGGAUUGGGACAGUCUCAUCCAUAUGUCCGUAGCCGGAUGAAAAACCGGAGUUCGAUAAUGCAUAAAAAAUCCGCAACCGCGGAUUAGUGUAUUACCCGAGGUAUCCGGUUAACCGACUGGCGUCGCGUGUUGGCAAC
env-354   UCUUCUUUCGCGAGACACAGAAUGCUGGUGCAUACCAGCUCAUGGAUUGGGACAGUCUCAUCCAUAUGUCCGUAGCCGGAUGAAAAAACCGGAGUUCGAUAAUGCAUAAAAAAUCCGCAACCGCGGAUUAGUGUAUUACCCGAGGUAUCCGGUUAACCGACUGGCGUCGCGUGUUGGCAA
env-355   UCUUCUUUCGCGAGACACAGAAUGCUGGUGCAGACCAGCUCAUGGAUUGGGACAGUCUCAUCCAUAUGUCCGUAGCCGGAUGAAAAAACCGGAGUUCGAUAAUACAUAAAAAUCCGCAACCGCGGAUUAGUGUAUUACCCGAGGUAUCCGGUUAACCGACUGGCGUCGCGUGUUGGCAAC
env-356   UCUUCUUUCGCGAGACACAGAAUGCUGGUGCAUACCAGCUCAUGGAUUGGGACAGUCUCAUCCAUAUGUCCGUAGCCGGAUGAAAAAACCGGAGUUCGAUAAUGCAUAAAAAAUCCGCAACCGCGG................................................
env-357   UCUUCUUUCGCGAGACAGGAAAGCUGGUGUAUAUCAGCUCUGGGAUGAGAAAAAUAUCUCCCUAUGUCCGUUGG.........................................................................................................
env-358   UCUUCUUUCGCGAGACAGAAAAGCUGGUGUAAUUCAGCUUUUGGAUGAGAAAAAUAUCAUCCGUAUGUCCGUAGCCACACGAAAAAUCCGGACGUUCAUGAAUACCGAAAAAUCCCGCAACAGUGGGAUAGCGUAUUACCUGAGUUGUCCGGAUUUCUCACCAGGCUUCGCGUGUUGGCA
env-359   UCUUCUUUCGCGAGACAGAAAAGCUGGNGUAAUUCAGCUUUUGGAUGAGAAAAAUAUCAUCNGUAUGUCCGUAGCCACACGAANAAUCCGGACGUUCAUGAAUACCCGAAAAAUCCCGCAACNGUGGGAUAGCGUAUUACCUGAGUUGUCCGNAUUUCUCACCAGGCUUCGCGUGUGGC
env-360   UCUUCUUUCGCGAGACAGAAAAGCUGGUGUAAUUCAGCUUUUGGAUGAGAAAAAUAUCAUCCGUAUGUCCGUAGCCACACGAAAAAUCCGGACGUUCAUGAAUACCCGAAAAAUCCCGCAACAGUGGGAUAGCGUAUUACCUGAGUUGUCCGGAUUUCUCACCAGGCUUCGCGUGUUGGC
env-361   UCUUCUUUCGCGAGACAGAAAAGCUGGUGUAAUUCAGCUUUUGGAUGAGAAAAAUAUCAUCUGUAUGUCCGUAGCCACAUGAAAAAUCCGGACGUUCAUGAAUACCCGAAAAAUCCCGCAACAGUGGGAUAGCGUAUUACCUGAGUUGUCCGGAUUUCUCACCAGGCUUCGCGUGUUGGC
env-370   UCUUCUUUCGCGAGACAGGAAAGCUGGUAUACAUCAGCUCUCGGAUGAGCAAAAUAUCAUCCCUAUGUC.............................................................................................................
env-371   UCUUCUUUCGCGAGACAGGAUUGCUGAUCAAGAAUCAGCUCAUGGAUGAGUAUAGUAUCAUCCAUAUGUCCAUUGCCCGUUGCAUAACCGGAUGUUCAAUCAUAUAUAAAAUCCACAAAAGUGGAUUCGUGUAUAACCUGAAUUCUCGGUUCUCUCAACUGGCGUCGCGUGUUGGCAUC
env-372   UCUUCUUUCGCGAGACAGGAACGCUGGUGCAUACCAGCUCAUGGAUGAGGAAAAUUUCAUCCAUAUCUCCUUAGGCAGAGGAAAAGCGGUCGUUCAAUAAUACAAAAAAAAUCCGUAUAUGCGGAUUAUUGUAUUUUUGUAUU..................................
env-373   UCUUCUUUCGCGAGACAGGAAUGCUGGUGCAUACCAGCUCAUGGAUGAUGAAAUUAUCAUCCAUAUCUCCUUAAGCAGAGGAAAAGCGGACGUUCAAUAAUACAAAAAAAUAAUACAAAAAAUCCGAAUAUGCGGAUCAUUGUAUUUUUUGUAUUCGCCGUUUUUCUUCUCUGGAGUCGC
env-374   UCUUCUUUCGCGAGACAGGAUUGCUGAUAAAAAAUCAGCGUAAGGAUGAGGCAUUAUCAUCCUAAUGUCUAUACCCCGAUAAGAAAACCGGAUAUCCGAUAAUACCUGAAAAACCCAUAAUCUGGGAACGUGUAUUACCUGGACUUUCCGGUUUCCUAAACCGGCGUCGCGUGUUGGCAUC
env-375   UCUUCUUUCGCGAGACAGGAAUGCCGGUGCACACUGGCACAGUGAUGCAACUUCUGUUGUAUCCCCUAUAAUUACGGACGAAAAUACCAGUAUCAGUUAUCAGAGCCAGUUAUCCGUUACUACGGAAUGACCGCGCUGAUGGUGUUGGUGUAAGUUGCCCGCUA...........
env-376   UCUUCUUUCGCGAGACAGGAAUACCGGUGCACACUGGUACAGCGAUGCAACUUCUGUUGCAUCCCCUUAAACUACGGACGAAAAUACCAAUAUCGUCACAGGCUGUAAUCCGUUUAACGGAUGAAUUCCCCGACAAUCUUGGUGUGUUAUUGUCCGUGUCGCGUGUUAGUUUCUGAAACAAU
env-377   UCUUCUUUCGCGAGACAGGAAUGCCGGUGCACACUGGCACAGCGAUGCAACUUCUGUUGCAUCCCCUUAAUUACGGACGAAAACCAGCAUCAGUCAUAAGAGCCGGACAUCCAUAACUUUGGAUUGACUUCGCUGAUGGCAGGUGUCGGUGUUAUUGUCCGUGUCGCGUGUUGGCUUCUGA
env-378   UCUUCUUUCGCGAGACAGGAAUGCCGGUGCACACUGGCAAAGCGAUGCAACUCUCGUUGCAUCCCCUAACUAUGGACGAAUAU.................................................................................................
env-379   UCUUCUUUCGCGAGACAGGAAUGCCGGUGCACACUGGCAUAAGCGAUGCAAUUAUCCAUUGCAUCCCCUUACUACGGACAAAUAUUUUCUCAUCGUCAUCAGAGCCGGUAAUCCAUAAAUAAGGAUUUGUGCCCUGAUGGCUGCUGAUAUACAAGUCCGUGGCCGCGUGUUAGUUUCUGG
env-380   ACUUCUUUCGCGAGACAGGUAAGCUGGUGCACACCAGUAGGGCAAUAAAAUAACGCUCAAUCGUGCAUCCCCAGAAACGGGUGGAUAUCGAUCAUGAUCCGCUACCCUCAAACCCCCCGGGUUAAAAGGGCAGACUGUCAUCCCCGGUACACACCACCCGUUCCCGCGUGUUGGCUUAAC
env-381   ACUUCUUUCGCGAGACAGGUAAGCUGGUGCACACCAGUAGGGCAAUACAAUAACGCUCAAUCGUGCAUCUCCAGAAACGGGUGGAUAUCGAUCAUGAUCCGCUACCCUCAAACCCUUCGGGUUAAAAGGGCAGA...........................................
env-382   ACUUCUUUCGCGAGACAGGAAAGCUGGUGCACACCAGUAGGGCAAUAAAAUAACGCUCAAUCGUGCAUCUCCAGAAACGGGUGGAUAUCGAUCAUGAUCCGCUACCCCCAAACCCCCCGGGGUUUUCAAGGGGCAGACGGUCAUAACGGUAUAAACCAUCCGUUCCCGCGUGUUGGCUU
env-383   ACUUCUUUCGCGAGACAGGAAAGCUGGUGCACACCAGUAGGGCAAUAAAAUAUCGCUCAAUCGUGCAUCCCCAGAAACGGGUGGAUAUCGAUCAUGAUCUGCUACCCUCAAACCCCCCGGGGUUAAAAGGGUCGACUGUCAUCAAAGGUACACACCACCCGUUCCCGCGUGUUGGCUUAA
env-384   ACUUCUUUCGCGAGACAGGAAUGCUGGUGCACACCAGUAGGGCAAAAAAAGCUCAAUUGUGCAUCUCCAGAAACGGGUGGAUAUUUACGAUGAUCCACUGCUGAUAACCCCCGCGGGGGUUUUCCAGGGCAGAACAUGAUCAUGGAUAUACACCACCCGUUCCCGCGUGUUGGCUUACCA
env-385   ACUUCUUUCGCGAGACAGGAAUGCUGGUGCACACCAGUAGGGCAAUACAUAACGCUCAAUCGUGCAUCUCCAGAAACGGGUGGAUAUCUGCCACGAUCCACUGCCAAAAUACUCUUGGCAGACCGUGAACAGAGGAGU.........................................
env-386   ACUUCUUUCGCGAGACAGGAAUGCUGGUGCACACCAGUAGGGCAAUACGUAACGCUCAAUCGUGCAUCUCCAGAAACGGGUGGAUAUCUGCCAUGACGCUGCCAACGAACUCUUGGCAGGCCGUGAACAGAGGAGUUAACCCCCCGUUCCCGCGUGUUGGCUUAACCGGAAGCGAAGCUA
env-387   ACUUCUUUCGCGAG................................................................................................................................................................................
env-389   ACUUCUUUCGCGAGACAGGAAUGCUGGUGCACACCAGUAGAGCGAUAUAAAAAAAGCUCAAUCGUGCAUCUCCAGAAACGGGUGAAUAUCCAAAAACGAUCCGCUGCAAGAAAAUCCCCCCGGGGUUUUUCCGGACAGACCGUGAUCACAGGUAUAAACCAUCCGUUCCCGCGUGUUGGC
env-390   UCUUCUUUCGCGAGACAGGAAUGCUGGUAAAUACCAGACCCUGGAUGAGAAAAAUAUCAUCCAUAAUGUCCGUAGCCAAAAAAAAUCGGAUAUCUCAAUAAUACGAAAAAAUCUGCAACUGCGGACUAGUGUAUUAUUUGUACUCUCCGAUUUCUCCCUCUGGCGUCCGCGUGUUGGCAUC
env-392   UCUUCUUUCGCGAGACAGGAACGCCGGUGCACACUGGCAUCGCCAUGAGAUCACAUCUAUGUGCAGC...............................................................................................................
env-395   UCUUCUUUCGCGAGACAGGUAUGCCGGUGCACAACCGGCUCAAGCGACGAGUAUUCCUUGUCGCAUCUCCGUAACCGGGCGAACAAUCCCCAGAGCGCAUACGGGUAAUCCACCCUGACGUGGUUUCCCGUG...........................................
env-399   UCUUCUUUCGCGAGACAGGAAUACUGGUGCACACCGGUACAGCGAUGCAAUAACCAUUGCAUCCCCUCAUCGGAUGAAUACAUCAGCUUCAGUCACCGGGCAAGGCAACCAAUACUGGUUUCAUGUCCCGAGAUAUGUCUGAUCUGUUGUCCGUCCCCGCGAGUUGGCUUCUGGAAUUA
env-400   ACUUCUUUCGCGAGACAGGAAUGCUGGUGCACACCAGUAGGGCAAUAGAAUAACGCUCAAUUGUGCAUCUCCAGAAACGGCUGGAUAUCGAUCAUGAUCCGCUGCC.......................................................................
env-402   UCUUCUUUCGCGAGACAGGAAUACCGGUGCACACUGGUACAGCGAUGCAACUUCUGUUGCAUCCCCUUAAACUACGGACGAAAAUACCAAUAUCGUUACAGGCUGUAAUCCGUUUAACGGAUGAAUUCGCCGAUGGUCUUGGUGUUAUUGUCCGUGUCGCGUGUUAGUUUCUGAAACAAUA
env-403   UCUUCUUUCGCGAGACAGGAAUACCGGUGCACACUGGUACAG......................................................................................................................................
env-405   UCUUCUUUCGCGAGACAGGCUUGCCGGUGCACUCCGGCAAUACGAUGAGUAUAUUCUCAUCCGAAUCACCAUAACCGGACGAAAUGUGUUCCAAUAUCG..............................................................................
env-406   UCUUCUUUCGCGAGACAGGCUU........................................................................................................................................................
env-407   UCUUCUUUCGCGAGACAGGAAAAGGCUGAUGCACAUCAGCUCAUGGAUGAAAAAUAUCAUCCCCAUGUCCGUAACCUCAUGAAAAC..............................................................................................

          ..............................................................................................................................................................................................
          ..............................................................................................................................................................................................
          UCUYUUYR•GRRRR•YR••••••Y•••R•••RY•RR•R••••••••••••••••••••••••••••••••••••••••••R•••••••••••••••••••••••••••••••••••••••••••••••••••••••••••••••••••••••••••••••••••••••••••••••••
```

```
Mli-2-1   UCUGUUUGAGAGUAGUAAGUUGCCCUUUAUAACGUAUAGGUGCACUGUCGGAUGUCUGUGAAGAUGCAGUUUAUCCCCGGCCGGGUUUAAAGCCUGACUGCUGUACAUUCGUUUCUGUUCCGAUUCAGUCGUUUACUGUCCUUUGCAGGAUUUAUUUCUGCUUCUCGGAAGAAUAUUGUG
Mpe-1-1   UCUUUAGAGAGAGAUUUCCGAUAACAGGAAAGACAAGGCAGAGGAAAAACGUCCUGUAUAACAGGCAGGUUCCCGGCCGAAUGCAGACAGCAUUACAGUCUGCCGCAGUUAUGGUUAUUCUCAGAUUCAGCAGACCGAAAAUCCUUUUUGGUCCGGCCGAAAUCCGUAAACCCGGAUUUC
Mpa-2-1   UCUUUAGAGAGAGAUUUCCGAUAACAGGAAAGACAAGGCAGAGGAAAAACGUCCUGUAACAACAGGCAGGUUCCCGGCCGAAUGCAGACAGCAUUACAGUCUGCCGCAGUUAUGGUUAUUCUCAGAUUCAACAGACCGAAAAUCCUUUUUGGUCCGGCCGAAAUCCGUAAACACGGAUUU
env-408   UCUUUAGAGAGAGAUUUCCGAUAACAGGAAAGACAAGGCAUGGGAAGAGCACCAUAUUUAAGGCAUGCUUCCGGCCGAAUGCAGACAGCAUUACCGUCCACCGCAGUUAUGGUUAUUCUCAGAUUCAGCAGACCGAAAUCCUUUUUGGUCCGGCCGAAAUCCGUUAAAACGGAUUUCUCG
Mmo-1-1   UCUGUUUGAGAGACGUGAUACAAAGAUCGGUCUGCAAAGUUAACAUGCAGACGGGUCUUGGGAAUAUUAAUUCCAAGGCAAUGCGAGGCAAAGUCAGUUUGGUCUGAAUACCAAGUAUUGUGCCCUGAAAUUACACGGAUAUUUCAGGUUCGAAAAAAGACCGACGAAGACCGUCUGUUA
env-409   UCUGUUUGAGAGACGUGAUACAAAGAUCGGUCUGCAAAGUUAACAUGCAGACGGGUCUUGGGAAUAUUAAUUCCAAGGCAAUGCGAGGCAAAAUCAGUUUGGUCUGAAUACCAAGUAUUGUGCCCUGAAAUUACACGGAUAUUUCAGGUUCGAAAAAAGACCGACGAAGACCGUCUGUUA
env-410   ACUCUUUGAGAGACCGGGAAACCCGUUAUCGGGAAAGAAAUAUCAAUUCAUGAAAGGAGACGCUUCAGGCAUCCUGAUGUAUCAGGCAGAGCCCUUUCGUCCAGCAUUCAUGAGCCAGAACGAUUUCACAGGUCCUCUAUUGCUCCGGCCAACGACUCCCGCCUACCAGCCGUUGCGGCU
Mco-1-1   UCUAUUCGCGAAGGCCGCUAACGGUCGAGCUCGUGAAGAGGAGCGACGUGCCCCUGUACUGGGGCUACGAAGUGCGUAUCGCCGAUACGCUCCACGAUGCGCUGGAAACCGAUGGCUUAAGGAUAGCGACGUCGAGGCUGGGGCAUCCCCUGGCCUGCGAGAUGCUAUCCGAGAUAAAAA
env-413   UCUUUUCGCGAAGACCAAUACAAGCAAAGCUGGUAACAAAGGACGAGAUCCCCAUGUACUGGGGAUACCGUACAGCAGUAGAAGACAGCCUGUCAGGGGCGCUGGAAAAGAUUUCAGAAGAAGGUGCCCUGAUAAUCGCCACAUCCCGCCAAGG..........................
env-414   UCUUUUCGCGAAGACCAAUACAAGCAAAGCUGGUAACAAAGGACGAGAUCCCCAUGUACUGGGGAUACCGUACAGCAGUAGAAGACAGCCUGUCAGGGGCGCUGGAAAAGAUUUCAGAAGAAAGUGCCCUGGUAAUCGCCACAUCCCGCCAGGGUAAGUUGCUCGACAACACACUCCUGG
env-415   UCUUUUCGCGAAGACCAAUACAAGUAAAGCUGGUAACAAGGACGAGAUCCCCAUGUACUGGGGAUACCGUACAGCAGUAGAAGACAGCCUGUCAGGGGCGCUGGAAAAGAUUUCAGAAGAAGGUGCCCUGAUAAU...........................................
env-416   UCUUUUCGCGAAGACCAAUACAAGCAAAGCUGGUAACAA.....................................................................................................................................................
env-418   UCUCUUUAAGACGACCAUUACAAGUAAAACUUGUAGAAAAAGAUGAGAUUCCACUAUAUUGGGGAUACUCAAUUAUAAUAGCAAAUAGCCUUUCUGAAGCUCUCAAAGAGCUUUCAAAACAAGGUGCACUGAUCAUAGUAACCGGACGACAGGGCCAGGUAUUAAAUACAAGUGUUCUAA
env-423   UCUCUUUAAGACGACCAAUACAAGCAGAACUUGUAGAAAAAAAUGAGAUUCCACUAUAUUGGGGAUACUCUACUAUAAUGGCAGAUAGCCUUUCUAAAGCUCUCAAAAAGCUUUCAAGGCAAAAUGCACUGAUCAUAGUAACUGCACGUCAGGGCCAGGUAUUAAAUACCAGCGUUCUAA
env-425   UCUCUUUAAGACGACCAAUACAAGCAAAACUUGUAGAAAAAGAUGAGAUUCCACUAUAUUGGGGAUACUCUACUAUAAUAGCAGAUAGUCUUUCUAAAGCUCUUAAAAAGCUUUCAAGGCAAAAUGCACAGAUCAUAGUAACUGCACGUCAGGGCCAGGUACUAGAUACCAGCGUUCUAA
env-428   UCUUUUCGCUAGACCCUCUCGCCGCUGAGCCUGUCGGUCGAAAAGAGAUCCCUCAAUACUGGGGGUAUGAAACAGAAGUAGUAGGUAGCCUGGAGAAAUGCCUAGAGGAUUAUCUGGGCUCAACUGAGGCUUUAGUGCUUCUAUCCUCCCGGAAGGGAAGAACUCUCACGACCCAACUCC
env-429   UCUUUUCGCUAGACCCCCUCGCCGCUGAGCCUGUCGGUCGAAAAGAGAUCCCUCAAUACUGGGGGUACGAGACAGAAGUAGUAGGUAGCCUGGAGAAAUACCUAGAGGAUUAUCUGGGCUCAAUUGAGGCUUUAGUGCUUCUGACCUCCAGGAAGGGCAGACCUCUUACGGCCGAACUCC
Msh-1-1   UCUUUUCGAAGACGCCACUUACAGUCGAGUAUAUAGCGAGGGAGCACAUACCCACAUACUGGGGCUACAGAACCCAUGUGUGCGGACCCCUCCACUCCACGCUCAGCGCCCUGCGCUCUCGGGGCUGGUGGGUGCUGGGCACAUCGAGGGAGGGAAGACCGCUGGACGCCAAUGCCCUCA
env-430   UCUUUUCGCAGGAACCGCUCGCCGCUGAGCCUGUGGACCAGAAGGAUAUACCCCAUUACUGGGGCUACGAGACCGAGAUAGUGGAUAGCCUGGAGAGCUAUCUGGCGUCAUUAGGAGAUGCCCGAAUCGUUCUCGCCUCCAGAAAUGGCCGGGCAGUCACGCCCCGGGUACUCCAGGAGA
env-433   UCUUUUCGCAUGAACCGCUCGCCGCUGAGCCUGUGGACCCGAAGGAGAUCCCUCAAUACUGGGGGUUCGAGACCGAGAUAGUGGAGAGCCUGGAAGGCUACUUAGCUGCAAAGCAUGAGCAGGGUAGCCAUGUGGUUCUCACCUCCCGGAAUGGAAAGGCCCUAACGCCUGAACUCCUCU
env-434   UCUUUUCGCAGGAACCGCUCGCCGCUGAGCCUGUGGACCAGAAGGACAUACCCCAUUACUGGGGGUACGAGACCGAGGUAGUGGAUAGCCUGGAGGGCUAUCUGGAAUCAAUGAGAGAUACCCGGAUCGUUCUCGCCUCCAGAAGCGGUCGGGCUAUAACGCCCCAGGUACUCCAGGAGU
env-435   ............................................................................................................................................................................................
env-436   UCUUUUCGCAGGAACCGCUCGCCGCUGAGCCUGUGGACCAGAAGGAUAUACCCCAUUACUGGGGGUACGAGACCGAGGUAGUGGAUAGCCUGGAGAGCUAUCUGGAAUCAAUGAGAGAUACCCGGAUCGUUCUUGCCUCCAGAAGCGGUCGGGCUAUAACGCCCCANAUACUCCAGGAGU
env-437   ............................................................................................................................................................................................
env-439   UCUUUUCGCAGGAACCGCUCGCCGCUGAGCCUGUUGACCAAAAGGAUAUACCCCAUUACUGGGGGUACGAGACCGAGGUAGUGGAUAGCCUGGAGNGCUAUCUGGAAUCAAUGAGAGAUACCCGGAUCGUUCUUGCCUCCAGAAGCGGUCGGGCUAUAACGCCCCAGGCACUUCUGGAGU
env-440   UCUUUUCGCAGGAACCGCUCGCCGCUGAGCCUGUNGACCANAAGGAUAUACCCCAUUACUGGGGGUACGAGACCGAGGUAGUGGAUAGCCUGGAGAGCUAUCUGGAAUCAAUGAGAGAUACCCGGAUCGUUCUUGCCUCCAGAAGCGGUCGGGCUAUAACGCCCCAGG............
env-441   UCUUUUCGCAGGAACCGCUCGCCGCUGAGCCUGUUGACCAAAAGGAUAUACCCCAUUACUGGGGGUACGAGACCGAGGUAGUGGAUAGCCUGGAGAGCUAUCUGGAAUCAAUGAGAGAUACCCGGAUCGUUCUUGCCUCCAGAAGCGGUCGGGCUAUAACGCCCCAGGCACUUCUGGAGU
env-442   UCUUUUCGCAGGAACCGCUCGCCGCUGAGCCUGUUGACCAAAAGGAUAUACCCCAUUACUGGGGGUACGAGACCGAGGUAGUGGAUAGCCUGGAGAGCUAUCUGGAAUCAAUGAGAGAUACCCGGAUCGUUCUUGCCUCCAGAAGCGGUCGGGCUAUAACGCCCCAGGCACUUCUGGAGU
env-443   UCUUUUCGCAGCAGCCGCUCGCCGCUGAGCCUGUGGACCAGAAGGAUAUCCCCCAUUACUGGGGGUACGAGACCGUUGUAGUGGAGAGCUUAGAGAGCUACCUUGCUGCAAAGCAUGAGCAGGGUGGCUACGUGGUUCUUGCCUCGCGGUACGGAAAAGGCAUAACGCCUGAAAUCCUCU
env-444   UCUUUUCGCUGAACCCGCUCGCCGCUGAGCCUGUGGAUAUGUUGGAGAUCCCAGAGUACUGGGGGUAUGAGACCGAGGUAGUCGGCAGCCUGGAGAAAUAUCUCCUGGCCAAGGAAGCCUUUGUCAUACUUACUUCCAAGAAGGGCAGGGUCAUAGAUCCAAAACUCCUUCAGAGCAUAG
env-445   UCUUUUCGCAGGAACCGCUCGCCGCUGAGCCUGUGGACCA....................................................................................................................................................
env-446   UCUUUUCGCAGGAACCGCUCGCCGCUGAGCCUGUGGACCA....................................................................................................................................................
env-448   UCUUUUCGCUGAGCCCGCUCGCCACUGAGCAAGUGGAUCGGAGUAAGAUUCCCGGCUAUUGGGGCUACCAGACGGUAUCGGUGGAGAGCGCUGAAGAGUACCUGCGCUCCUGGAACCCACGGGUGGUGGCUACAUCCCGAAAGGGACGGGUGGUGACUCCGGAUCUGCUGGAGCAGAUAG
env-449   UCUUUUCGCGAAACCCGCUCGCCGCUGAGCCUGUAGACAGGAAGGAAAUUCCCCAGUACUGGGGGUAUGAGACAGAGGUAGUAGAGAGCCUUGAGAAAUAUCUGG..........................................................................................
env-450   UCUUUUCGCAUGAACCGCUCGCCGCUGAGCCUGUGGACCCGAAGGAUAUCCCUCAUUACUGGGGAUUCGAGACCGAGAUAGUGGAGAGCCUGGAAGGCUACUUA...........................................................................................
env-451   UCUUUUCGCAUAACCCGCUCGCCGCUGAGCCAGUGGACCAGAAGGAGAUCCCUCAUUACUGGGGGUUUGAAACCGAAGUAGUGGAUAGCCUUGAAGGCUACCUGGCGUCGAAAGAUGCCGCCUCCGUGCUUCUCACCUCCAUAAAAGGAAAGGCGAUAACGCCAGAAAUCCUCCAGCAAA
env-452   UCUUUUCGCAUAACCCGCUCGCCGCUGAGCCAGUGGACCAGAAGGAGAUCCCUCAUUACUGGGGGUUUGAAACCGAAGUAGUGGAUAGCCUUGAAGGCUACCUGGCGUCGAAAGAUGCCGCCUCCGUGCUUCUCACCUCCAUAAAAGGAAAGGCGAUAACGCCAGAAAU...........
env-455   UCUUUUCGCAUAACCCGCUCGCCGCUGAGCCAGUGGACCAGAAGGAGAUCCCUCAUUACUGGGGGUACGAGACCGAAGUAGUGGACAGCCUUGAAGGCUACCUGGCAUCGAAAGAUGCCGCCUCCGUGCUUCUUACCUCCAUAAAAGGAAAGGCGAUAACGCCAGAAAUCCUCCAGCAAA
env-456   UCUUUUCGCGAAAACCGCUCGCCGCUGAGCCAGUAGACCGAAAGGAGAUCCCUCACUACUGGGGGUAUGAGAUAGAAGUAGUCGAUAGCCUUGAAGGCUACCUGGCGUCGAAGGACGACGCCUUUGUGCUUCUCACCUCCAGAAAAGGAGAGGCCAUAACGCCAGAAACCCUCCAGAAGA
env-460   UCUUUUCGCAUAACCCGCUCGCCGCUGAGCCUGUGGACCAAAAGGAGAUCCCUCAUUACUGGGGUUACGGGACCGAAGUAGUGGAAAGCCUUGAAGGCUACCUGGCAUCAAGAGACGAUGCCCAAGUGGUUCUAGCCUCCAGGAAUGGAAAGGCUAUCACGCCCGAAAUCCUCCAGAAGU
env-462   UCUUUUCGCAUAACCCGCUCGCCGCUGAGCCUGUGGACCAAAAGGAGAUCCCUCAUUACUGGGGUUACGGGACCGAAGUAGUGGAAAGCCUUGAAGGCUACCUGGCAUCAAGAGACGAUGCCCAAGUGGUUCUAGCCUCCAGGAAUGGAAAGGCUAUCACGCCCGAAAUCCUCCAGAAGU
env-464   UCUUUUCGCAUAACCCGCUCGCCGCUGAGCCUGUGGACCAAAAGGAGAUCCCUCAUUACUGGGGUUACGGGACCGAAGUAGUGGAAAGCCUUGAAGGCUACCUGGCAUCAAGAGACGAUGCCCAAGUGGUUCUAGCCUCCAGGAAUGGAAAGGCUAUCACGCCCGAAAUCCUCCAGAAGU
env-465   UCUUUUCGCAUAACCCGCUCGCCGCUGAGCCUGUGGACCAAAAGGAGAUCCCUCAUUACUGGGGUUACGGGACCGAAGUAGUGGAAAGCCUUGAAGGCUACCUGGCAUCAAGAGACGAUGCCCAAGUGGUUCUAGCCUCCAGGAAUGGAAAGGCUAUCACGCCCGAAAUCCUCCAAAGCU
env-466   UCUUUUCGCAUGAACCGCUCGCCGCUGAGCCUGUGGACCAAAAGGACAUCCCCAUCUACUGGGGGUAUGAGAUAGAGGCAGCACAGAGCCUGGAGGCCGCA...............................................................................................
env-467   UCUUUUCGCUGAAACCCCUCGCCGCGGAGCCGGUAGACCGAUCGGAGAUACCUCACUACUGGGGCUACGAGACAGAGGU.....................................................................................................................
env-468   UCUUUUCGCUGAAACCCCUCGCCGCGGAGCCGGUAGACCGAUCGGAGAUACCUCACUACUGGGGCUACGAGACAGAGGUCAUCCCGAACCUGGACGACCACGUCCGCUCGACAGAGAGGCUAGUCGUGGCCACGUCCAGGAGGGGCGCGCCAGCCACCCCAGAACUCCUGAGCCAAAUGG
env-469   UCUUUUCGCUGAAACCCCUCGCCGCGGAGCCGGUAGACCGAUCGGAGAUACCUCACUACUGGGGCUACGAGACAGAGGU.....................................................................................................................
env-471   UCUUUUCGCUGAAACCCCUCGCCGCGGAGCCGGUAGACCGAUCGGAGAUACCUCACUACUGGGGCUACGAGACAGAGGUCAUCCCGAACCUGGACGACCACGUCCGCUCGACAGAGAGGCUAGUCGUGGCCACGUCCAGGAGNGGCGCNCCAGCCACCCCAGAACUCCUGAGCCAAAUGG
env-472   UCUUUUCGCUGAAACCCCUCGCCGCGGAGCCGGUAGACCGAUCGGAGAUACCUCACUACUGGGGCUACGAGACAGAGGUCAUCCCGAACCUGGACGACCACGUCCGCUCGACAGAGAGGCUAGUCGUGGCCACGUCCAGGAGGGGCGCGCCAGCCACCCCAGAACUCCUGAGCCAAAUGG
env-474   UCUUUUCGCUGAAACCCCUCGCCGCGGAGCCGGUAGACCGAUCGGAGAUACCUCACUACUGGGGCUACGAGACAGAGGUCAUCCCGAACCUGGACGAACACGUCCGCUCUAGCGACGGGCUAGUGGUGGUCACGUCCAGGCGUGGUAAGCCAGCAACCCUAGACCUUCUGGCCCAGACGG
env-475   UCUUUUCGCUGAAACCCCUCGCCGCGGAGCCUGUAGACCGAUCGGAGAUACCCCAUUACUGGGGCUACGAGACCGAGGUCAUCCCAGACCUGGACGGGCACGUCCGCUCGAGCGAGCGGCUGGUGGUGGCCACGUCCAGAUUGGGGACGCCAGCAACCCCAGACCUCCUCGCGCGGGUGG
Mha-1-1   UCUUUUCGCUGAAACCCCUCGCCGCGGAGCCUGUAGACCGAUCGGAGAUACCCCAUUACUGGGGCUACGAGACCGAGGUCAUCCCGGACCUGGACGAGCACGUCCGCUCGAGCGGGCGGCUGGUGGUGGCCACGUCCAGAUUGGGGACGCCAGCAACCCCAGACCUCCUCGCGCGGGUGG
env-476   UCUUUUCGCUGAAACCCCUCGCCGCGGAGCCGGUAGACCGAUCGGAGAUACCCCACUACUGGGGCUACGAGACCGAGGUCAUCCCGAACCUGGACGAGCACGUCCGCUCGAGCGAGCGGCUGGUCGUGGCCACGUCCAGAUUGGGGACGUCAGCAACCCCAGACCUCCUCGCGCGGGUGG
env-477   UCUUUUCGCUGAAACCCCUCGCCGCGGAGCCGGUAGACCGAUCGGAGAUACCUCACUACUGGGGCUACGAAACAGAGGUCAUCCCGAACCUGGAAGAACACGUUCGCUCUAGCGAUAGGCUAGUCGUGGCCACGUCCAGGCAGGGCACGCUGGCGACCCCAGACCUCCUGAGGCAGACGG
env-478   UCUUUUCGCUAGAACCCCUCGCCGCUGAGCCCGUGGAUAAAAGAACAUCCCCCUGUACUGGGGGUACGAUACGAAGGUAGUGGGAAGCCUAGAGGAUUAUCUCAGGGCGACCAAGGCCUUCGUAGUGCUCACCUCU...........................................
env-479   UCUUCUCGCUGGACCCGCUCGCCGCGGAGCCCGUAGAUCAAAACGAGAUACCCCACUACUGGGGCUACGAGACUAUAGUCGUGGACGACCUGGAGGAGUACCUCCUGUCCACCGACGCCCGCCUCGUAGUCGCAUCGAGAAAAGGCAGACCGGCGACCCCAGAUCUCCUGGAGAAUAUGG
env-480   UCUUCUCGCUGGACCCGCUCGCCGCGGAGCCCGUAGACCAAAACGAGAUACCCCACUACUGGGGUUACGAGACGAUAGUCGUGGAUGACCUGGAGGAAUACCUCCAGUCUAUCGACGCCCUGCUCGUGGUCGCAUCGAGAAAAGGCAGACCGGCGACCCUAGAUCUCUUAGAGAAUAUGG
env-481   UCUUUUCGUUGACGCCCUUGGUGGUAGAGCGUGUCCAAAAAGACGAAAUCCCCAAAUAUUGGGGAUACACCGUAAGGAUAGAGGGAAACCUUACCUCUACACUCAAAAAAAUGAGGAAAGAAGGUAUGUACAUAAUAGGUACUUCAAAGUAUGGAGAUAGUGUGAGUAUGGAAAAACUUG
env-482   UCUGUUCGAAGAAACCGUUGGAAGUGGAACCGGUUGAAAAACCGGAAGUGUGGGGAUACCGGGUAAAAGAGGUCAAGCUGAGAGAGCUUGUCCAGAAGGAUCUGCUGAUAACCUCCAGAAAAUGUCCCUAUGCGGAUCCGCAGGGGAUAAAGAGGUAUAUCCAGUCGAGAGAUGAGAUAU
env-483   UCUUUUCGCGAGAUCCGCUCGAAGUUGAAGCGAUUCAAAAGCCACCUGAAUAUUGGGGAUACUCGGUUCGGGUGACAGCUAGCAUCAAAGAGGCACUGGAAGGGUUUGUGGUGACCGCCGCGAAGCGCGGACAGGUGAUAAACCCUGAAAUGUGCGAUGAAAUUAAAAGGAGAGGCACCG
env-484   UCUUUUCGCGA.........................................................................................................................................................................................
env-485   UCUUUUCGCGAGAUCCGCUCGAAGUUGAAGCGAUUCAAAAGCCACCUGAAUAUUGGGGAUACUCGGUUCGGGUGACAGCUAGCAUCA.............................................................................................................
env-486   UCUUUUCGCGAGAUCCGCUCGAAGUUGAAGCGAUUCAAAAGCCACCUGAAUAUUGGGGAUACUCGGUUCGGGUGACAGCUAGCAUCAAAGAGGCACUGGAAGGGUUUGUGGUGACCGCCGCGAAGCGCGGACAGGUGAUAAACCCUGAAAUGUGCGAUGAAAUUAAAAGGAGAGGCACCG
env-492   UCUUUUCGCGAGAUCCGCUCGAAGUUGAAGCGAUUCAAAAGCCACCUGAAUAUUGGGGAUACUCGGUUCGGGUGACCGCUAGCAUCAAAGAGGCACUGGAAGGGUUUGUGGUGACCGCCGCGAAGCGCGGACAGGUGAUAAACCCUGAAAUGUGCGAUGAAAUUAAAAGGAGAGGCACCG
env-495   UCUUUUCGCGAGAUCCGCUCGAAGUUGAAGCGAUUCAAAAGCCGCCUGGAUAUUGGGGAUAUUCGGUUCGGGUGGUAGCUAGCAUCAAAGAAGCACUGGAGGGGCUGGUGGUGACCACCGCGAAGCGCGGACAGGUAAUAAGCCCUAUAAUGCGAGAUGACAUUAAAAGGAGAGGCACCG
env-504   UCUUUUCGCGAGAUCCGCUUGAAGUUGGACUGACUCAGAAGCCGCCGCGUUAUUGGGGGUAUUCGGUGCGGGUGGUCGAUAGCAUCGCAGACGCGUUGGAGGGGUUGGUCAUAGCCACGUCGAGGCACGGACAGCCAAUAAGUCCCGCUAUACACAAUGAAAUAAGAAGGAGAA......
env-505   AUCUUACGAGAGACCGGUAUCUGAACGCAUUACAUCUGACUGCCUGACUACACGUCAAGGAUUUCGUUCUGGCACCCUGAGUAAUAAAGGGCGCAUAAGACUUCUGGUACCUUGACGCUGUGGGGAGAGGCAGGUCUGAACGUUCGUUCAGCUGAAGUUACGCUUCAUGCAACUUCUGCU
env-508   UCUCUUUCGCGAGACAGGUCACAGGGGAGACGGCCAGGAGCCAGAGGAAAGGCCGCCUUUCCGGGGUACUGCACCAGAAGGGGGGAUCUCCCUCUUCGCACCGCUGCGGCGAUGCGCCGGGCGCCUGGCUAGAGUCCGGGCGUCCUGCAGGCAUAUGCGUGCCAGCGGCCCGGAAACAUC
env-509   UCUCUUUGCGAGACAGGUCACAGGGAGGCAGUCGCUGCCAGGAGAAAGACUGCCUUUCCGGGGUACUGCACCGGGAGCGGGUAUCUUCCCCUCCAACCCUGCGGCGAAGAAUGGAUCGGGUGCCUGUGCAUGCAAUGCGGGGCAUCCUCCUAUCUUCCGCACGUCAGGAGCCCGGACACA
Mta-1-1   UCUCUUUGCGAGACAGGUCACAGGGAGACGGUCACUAACCAGGAACAGGGCCGUCUUUUUGGAGUACUGCACCUGGAGGGGGAAAACCCACUCCAUACUGCUGACGGAGGGAUAGACGGGUGCCUGGACAUCUAUCCAUCGCCCUUUCCCCCUCCGCGCGACAGGAGCCCGGACACGUCA
env-511   UCUCUUUGCGAGACAGGUCACAGGGAGACGGUCACUAACCAGGAACAGGGCCGUCUUUUUGGAGUACUGCACCUGGAGGGGGAAAACCCACUCCAUACUGCUGACGGAGGGAUAGACGGGUGCCUGGACAUCUAUCCAUCGCCCUUUCCCCCUCCGCGCGACAGGAGCCCGGACACGUCA
env-512   UCUCUUUGCGAGACAGGUCACAGGGAGACGGUCACUAACCAGGAACAGGGCCGUCUUUUUGGAGUACUGCACCUGGAGGGGGAAAACCCACUCCAUACUGCUGACGGAGGGAUAGACGGGUGCCUGGACAUCUAUCCAUCGCCCUUUCCCCCUCCGCGCGACAGGAGCCCGGACACGUCA
Hbo-1-1   UCUCUUCGCGAGAACCGGUCCGUGCGCGGAUUGUGGACGAGCCCGUUCCAGGGUUCGACGUAUCCCGCAUGGACCUCGAGGAAGCACUCGGCCGCGCCGAUGCGGGCAUCACCAUCGCUACGUCCCGCUUCGGUGAGUCGCUGACGGUGCCGAAACUGGCGGAUCUCUCCCCGCGAAUUG
Ngr-1-1   UCUCUUCGCGACGACCGGUCCGGGCGAAACUCGUCGAUUGCCCCCUUCCGGGGCUUUCGAUCGAGCGGACGGACCUAUCGGCAGCACUCGGCCGUGAGGAUGCUGGCGUCCGUAUUGCCGCCUCUCGAUUCGGUGAGCACCUCACCGUCGGGCGGCUCGAGACGCUGGCCGGACGCAUCG
Hje-1-1   UAUCUUCGCGACGACCGGUCCGUGCAAAGCUCACCGAGGAGCCUGCGGGGCUCGAAGUCGAGCGCACGGACCUUCCGGCAGCACUCGGCCGCCCCGACGCGGGCGUCCGGAUCGCGACUUCUCGCUUCGGCGAGGAGCUCACGACCGGACGUCUGCCGGGGCUUUCGGCGCGCAUCGAAG
Nma-1-1   UCUCUUCGCGACGACCGGUCCGGGCGAAACUCGUCGACGAGCCCCUCCCGGGGCUUACCGUCGAGCGGACGGACCUUCAGGCGGCUCUCAGCCGUGAGGACGCCGGCGUUCGAAUUGCGGCUUCCCGAUUCGGUGCAGAACUCACCGUCGGGCGGCUCGGAACGCUGGCCGGACACACAG
Hal-1-1   UCUCUUCGCGAGAACCGGUCCGUGCGCGGAUCGUCGACGAGCCCCUUCCGGGCUUCGACGUAUCCCGCAUGGACCUCACGGAAGCACUCGGCAGGCCGGAUGCGGGCGUGCGAAUCGCCACGUCUCGCUAUGGCGAACCGCUGUCGGUCCCCACAUUGGGCGACUUGACGGCCCGCAUCA
Nti-1-1   UCUCUUCGCGACGACCGGUCCGGGCGAAACUCGUCGACGAUCCCCUUCCGGGAUUGUCGGUCGAGCGGACGGACCUCUCCGCAGCACUCGGCCGUGAGGACGCCGGCGUCCGUAUCGCAGCUUCCCGGUUCGGUGAGAAUCUCACCGUCGGGCGGCUCGAGACGUUGGCCGGACGCGUCG
Npe-1-1   UCUCUUCGCGACGACCGGUCCGGGCGAAACUCGAAGACGUUCCCCUUCCGGGGCUCGCCGUCGAGCGGACGGACCUUUCGACAGCACUCGGCCGUGAGGACGCCGGCGUCUGCAUCGCCGCCUCCCGAUUCGGUGAAGAACUCACCGUCGGGCGGCUCGAGACGCUGGCCGGACGCAUCG
Hti-1-1   UCUCUUCGAGACGACCGGUUCGCGCGAAGAUCGUCGAUCAGCCCCCACCGGGCUAUCAGGUCGACCGUGCGAACCUUGACGCUGCGCUCUCGCGGCCCGACGCCGGCGUGCGGAUCGCCGCGUCCCGACACGGCCAGUCGCUCACGACCGAGCGACUCGGUCGACUCGUCGGAACGAUCG
          ............................................................................................................................................................................................
          ............................................................................................................................................................................................
          UCUYUUYR•GRRRR•YR•••••••Y•••R•••RY•RR•R••••••••••••••••••••••••••••••••••••••••••R••••••••••••••••••••••••••••••••••••••••••••••••••••••••••••••••••••••••••••••••••••••••••••••••••
```

alignment positions 440 · · · 557

```
Mli-1-1   GUCCCCGUUCGCAAGUGUUUGCGCAAUCCUGAGCAGGAAACCUGGGAUUCUCGGAGAGAACCAAAGCAGAGGAUUGAUAGGAACAUGGCGAAUAUACACAGGCCACGCAGGGGUUCCC
env-1     AUCAUAUCAUAAACUAUGAUAUGCCGGUUAGAAAAUAUCCAGCGGCAUCUUUCAUGAGGAUCAUGAACUAUGCCGGUUUAAGGAUUGUAAUAAUUAGGACUUACAUACAAUGCCACGA
env-2     UCGAUACAUAUCUUUCAGAUAAGGUUGCCUUGAUAUUUGGAUCCCCAGAUAAAGGAGUUGAUGCUUUCAUUGAUCAAGAUCAGCUUGUCGAUGAUUCUAUUAACACAGUUGUCAAGCA
env-3     ACCAGUAUCUUUCAGAUCGAACCGCAGUUAUAUUUGGAUCUCCAGACAGGGGAGUCAGCCAGAUAACUGGUAGGAUGGACCUUGUGGACCAGGAAAUAAACACAAUUCCAAACCAAGG
env-4     ......................................................................................................................
env-5     ACGACCACGCCGUCUCCCGUGCGGCAUCACAACCAGAUAGGACUGAUCGGACAAUGCCAAAAAUUAACAGGCCGCGGCGGGGAUCGCUUGCGUACAGCCCGAGAAAGCGGGCAAAGAG
env-6     ......................................................................................................................
env-8     GUCUCUUUCCUGCAUCGAAGCACUACCGGAAAUCGGGGUGUUCCCAGAUACAGUAAAGAAGGACUGAUUGAUACAUGCCAACUAUUCACAGACCUCGCAUGGGUUCCCUGGCAUAUAG
env-10    CUCUUUCCUGCAUCGAAGCAUACCGGAAUUCCGGUGUGUCCCCAGAUACAGAAAAGAAGGACUGAUUGAUACAUGCCAACAAUUCACAGACCUCGCAUGGGUUCUCUGGCAUACAGCC
env-11    CUUUCCUGCAUCGAAGCAUACCGGAAUUCUGGUGUGUCCCCAGAUACAGAAAAGAAGGACUGAUUGAUACAUGCCAACAAUUCACAGACCUCGCAUGGGUUCUCUGGCAUACAGCCCG
env-13    AUCCCCGGCUCAUAUGGCCGAUAUGGGCGAUUUACAUUUCCGGGAAACCGGAUUUGCGGUCGAAUAAGACCCGAAGAUUGAGUCAUGUAGAGGAUUGAUUGAACAAUGCCCAAGAUCA
env-14    GAACAGACCAUGCCGAAGAUCAACAGGCCACGCAGGGGUUCCCUUGCCUACAGUCCGCGAAAACGGGCCAAGAGUCCGAUACCACGGUACCAUUCCUGGGCGGACACUCGGGGAGAAC
env-15    ACCGGGCGGGAGCGACAAUCGUCUUUGGUGCGCCUGGUCGUGGGCUCCUAGCCAUCCUUGGCGAUGACAACCCACCAAAUACUCAAACGCAGAUGGGUGAGCACGCGAGUUCACACAU
Hwa-1-1   AUCGCGCGGGAGCGACAAUCGCUUUUGGUGCGCCUGGUCGUGGGCUUCCAGUCAUCCUUGGCGAUGAUAAUGCGCCGGAUAGCCAAACACAAACGGAUGAGAAAACGGUCUCACCCAC
env-17    AUCGCGCGGGAGCGACAAUCGCUUUUGGUGCGCCUGGUCGUGGGCUUCCAGUCAUCCUUGGCGAUGAUAAUGCGCCGGAUAGCCAAACACAAACGGAUGAGAAAACGGUCUCACCCGC
env-23    CCGACGGAACGGCUGUCGUCUUCGGUGCGCCUCAGCGGGGACUUCCGGCGAUGUUGGGGGUUGCGCCCGAGCAGGUGACCGGUGAGCGGGCAUCUGCCGGGCGUGACCAGUCCCAGUC
env-24    CAGACCUCGCCUUAGUCUUCGGCUCUCCCGAGCGCGGCGUUCACGAGAUUCUGAGGGAGCGUGACGCUUCCGUCGAUUUCGAUUACGUACUAAACACCGUGCCCGACCAGGGCGUAGA
env-25    GUCGGUUCAUGAUUGCGUUUGGCGCCCCCGAGCGUGGCCUCCCGGCGAUACUUGGGAUUGCUCCGGACGCGGUCCGCGGAGACGACCAGCGCGAGACUGGCCGGUUUGAUCUCUGGCU
env-27    CCGAGGAUGGGAUGACGGUCGUCUUCGGCGCCCCCGGACGGGGGCUGCUGGCGAUGGCCGACGCAGAUCCAGAUGCACUCGCUGCGGGGAUGUCCCCGCUGGGGGUGGAUCACUGGCU
env-28    AAGACGGCAUGACAGUCGCCUUCGGCUCGCCCGAGAGAGGGUU...........................................................................
env-29    AAGACGGCAUGACCGUCGCCUUCGGCUCGCCCGAGAGAGGGUUGCCCGAGAUACUCGACAUCGACGCGACCGCCGUUGAGGCGUCCACCGGGGACGACGGAGACAUCGAACCUGCCGA
env-30    AAGACGGCAUGACAGUCGCCUUCGGCUCGCCCGAGAGAGGGUUGCCCGAGAUACUCGGAAUCGACGCGACCGCCGUUGAGUCGUCCAGCAGGGACGACGGAGACGUCGAACCCGCGGC
env-31    UACGGCACUCUCGCGUGACGACGCCGGACUCACCAUCGCGUCAUCGCGAUACGGUGAGCUGGCGACGACUCGUCGACUCGGCCAACUGGCCGACCGACGAGACCGCG...........
Hut-1-1   CGUCGGCUGGCGACAUGACAGUCGCCUUUGGCGCGCCGGAGCGCGGCUUGCCGGCUAUCCUCGGCGUCGAACCAGGUGCUGUCGACCCGUCCACUGUGGACGAGUUGGCGGCAGCCGU
env-32    CCGAGGAUGGGAUGACGGUCGUCUUCGGCGCCCCCGGACGGGGGCUGCUGGCGAUGGCCGACGCAGAUCCGGAUGCACUCGCGGCGGGGAUGUCCCCGCUGGGGGUGGAUCGCUGGCU
env-33    CCGAGGAUGGGAUGACGGUCGUCUUCGGCGCCCCCGGACGGGGGCUGCUGGCGAUGGCCGACGCAGAUCCGGAUGCACUGUCGGCGGGGAUGUCCCCGCUGGGGGUGGAUCACUGGCU
env-34    CCGAGGAUGGGAUGACGGUCGUCUUCGGCGCCCCCGGACGGGGGCUGCUGGCGAUGGCCAACGCAGAUCCGGAUGCGCUGUCGGCGGGGAUGUCCCCGCUGGCGGUGGAUCACUGGCU
env-35    CCGAGGAUGGGAUGACGGUCGUCUUCGGCGCCCCCGGACGGGGGCUGCUGGCGAUGGCCGACGCAGAUCCAGAUGCACUCACCGCGGGGAUGUCCCCGCUGGGGGUGGAUCACUGGCU
env-36    GCACGGAUGCUGGCGAUGGCGAGGAUGGCGAACGGACCUACACAGUCGCGUUCGGUGCACCCGAGCGGGGCAUUCCGGCGGUUCUGGGCAUCGAGCCUGGAGCGGUUGGCGAAGACGA
env-37    ACGCUGGCAGCGAGGGUUCCGGUGUCGACUGUACCGUCGCCUUUGGCGCGCCUGAGCGCGGUCUGCCGGAGAUAUUGGGCAUUUCGCCCGAGUCGAUACGCGAAGCGGACUCCACCGA
env-38    AUGCUGGCAGCGAGGGUGUGAGUGUCGACCUGACCGUCGCCUUUGGCGCACCCGAGCGCGGCCUGCCGGAGAUUCUGGGCAUUUCGCCCGAGUCGAUACGCGAAGCUGACUCCACCGA
env-39    AUGCUGGCAGCGAGGGUGCGGGUGUCGACCUGACCGUCGCCUUUGGCGCACCCGAGCGCGGCC.......................................................
env-40    ......................................................................................................................
env-41    CGAAUCAAGGGGCAACAGUGGUGUUUGGUUCGCCUGGCCGAGGGCUUCCAGCUAUCCUUGGGGUCGACGUAGCAAGCGACUCUGAGACCACUGUCGAGCCCUCCGACGGUGUCGGGUU
env-43    ......................................................................................................................
env-44    ......................................................................................................................
env-45    CAGGACUUACCGUUGUCUUUGGAUCACCCGGUCGAGGGCUUCCAGAAAUCCUUGGGGUUACCCCCGAGGCCAUUACGGCCGCCGAAGGCGGUCCAGUCGAACCCGAUCCGGGGUUCGA
env-49    AGGGGCUGACCGUCGCGUUCGGCUCGCCGGGACGAGGGCUGCCGGACAUACUCGGGAUCCCACCCGAGGCCAUCUCGGCUGAGCAAUCGGCCGGGGCGUCCGAGCCAGACGGCUUGGA
env-50    AGGGGAGUAUGACCGUCGCCUUCGGUUCGCCCGGUAGGGGCCUUCCGCAAAUGCUCGGGAUCUCACCCGAGGCGGUUUCGGCCGCCAUCGCGGCCGACACUCACGUCGACACCGAUCC
env-54    ......................................................................................................................
env-55    AGGGGAUGACCGUCGCCUUUGGUUCGCCCGGACGAGGGUUGCCGGAAAUACUCGGGAUCGAACCCGAGGCAGUCUCGGCCGCAGUCGCGGCUGGAGACCAUGUUGAAUCCGACCCCG.
env-56    AGGGGAUGACCGUCGCCUUCGGUUCGCCCGGACGAGGGUUGCCGGAAAUACUCGGGAUCGAACCCGAGGC................................................
env-57    AGGGCGCAACUGUUGCGUUCGGCUCGCCUGGCAGAGGGUUGCCGGAAAUACUCGGGAUCGAACCCGAGGCAGU.............................................
env-58    AUGGAAUGACCGUCGCCUUCGGCUCGCCGGGACGGGGACUGCCGGAAAUACUCGGGAUCACUCCCGAGGCAGUCUCGGCCGCAAUCGCGGCUGGAGACCAUGUUGAAUCCGAUCCGGG
env-59    ......................................................................................................................
env-62    AUGGAGCAACCGUCGCCUUCGGCUCGCCCGGCAGAGGGCUUCCGGAAAUACUCGGGAUUGAACCCGAGGCAGUCUCGGCCGCAGUCGCGGCUGGAGACCAUGUCGACUCCGAUCCGGG
env-63    AUGGAGCAACCGUCGCCUUCGGUUCGCCCGGCAGAGGGCUUCCGGAAAUACUCGGGAUUGAACCCGAGGCAGUCUCGGCCGCAGUCGCGGCUGGAGACCACGUUGAAUCCGUUCCUGG
env-64    AUGGAGCAACCGUCGCCUUCGGUUCGCCCGGCAGAGGGCUUCCGGAAAUACUCGGGAUUGAACCCGAGGCAGUCUCGGCCGCAGUCGCGGCU..........................
env-65    AUGGAGCAACCGUCGCCUUCG.................................................................................................
env-70    ......................................................................................................................
env-71    GAUGGAGCAACCGUCGCCUUCGGUUCNCCCGGCAGAGGGCUUCCGGAAAUACUCGGGAUUGAACCCGAGGCAGUCUCGGCCGCAGUCGCGGCUGGAGACCACGUUGAAUC........
env-72    AGCGCGACGGGAUGACCGUCGCGUUCGGUGCGCCCGGGAGAGGGCUGCCGGCCAUCCUCGGCAUCGAGGAAUCUGCCGUCCCGUCGUCGACGAUAGACGACUGGGCCGGCCGCGACGA
env-74    AACGCGACGGUUAUACCGUCGCGUUCGGGGCGCCCGGGAGAGGGCUGCCGGCCAUCCUCGGCAUCGAGGAAUCGGCCGUCCCGUCGUCGUCGGUAGACGACUGGGCCGGCCGCGACGA
env-76    ......................................................................................................................
env-77    ......................................................................................................................
env-78    ......................................................................................................................
Hsa-1-1   AGUCGCGUGGCGAUCUGACCGUCGCGUUCGGUUCACCCGAGCGUGGGUUGCCCGCGAUCAUGGGGAUCGACCCCGAUGACGUCGGGCCGACUGAAAUCGAAGCGGUCACCGGAUCCGA
Hgi-1-1   CCGAUGCCGGCGGCAUGACCGUCGUCUUCGGGUCGCCGGGCCGUGGGCUUCCGGACAUGCUCGGGAUGUCUCCCGAGGAUGUUGCAGACGUCGAACCUUCCGUUGGUCCGGGGUUCGA
Hsu-2-1   CCGAUGCCGGCGGCAUGACCGUCGUCUUCGGGUCGCCGGGCCGUGGGCUUCCGGACAUGCUCGGGAUGCCUCCCGAGGAUGUUGCAGACGUCGAACCUUCCGUUGGUCCGGGGUUCGA
Hpr-1-1   CCGAUGUCGGCGGCAUGACCGUCGUCUUCGGGUCGCCGGGCCGUGGGCUUCCGGACAUGCUCGGGAUGUCUCCCGAGGAUGUUGCAGACGUCGAACCUUCCGUUGGUCCGGGGUUCGA
Hsp-8-1   CCGAUGCCGGCGGCAUGACCGUCGUCUUCGGGUCGCCGGGCCGUGGGCUUCCGGACAUGCUCGGGAUGCCUCCCGAGGAUGUUGCAGACGUCGAACCUUCCGUUGGUCCGGGGUUCGA
Hde-1-1   CCGACGCCGGCGGUAUGACCGUCGUCUUCGGGUCGCCGGGCCGUGGGCUUCCGGACAUGCUCGGGAUGUCUCCCGAGGAUGUUGCAGACGUCGAACCUUCCGUUGGUCCGGGGUUCGA
Hlu-1-1   CCGAUGCCGGCGGCAUGACCGUCGUCUUCGGGUCGCCGGGCCGUGGGCUUCCGGACAUGCUCGGGAUGCCUCCCGAGGAUGUUGCAGACGUCGAACCUUCCGUUGGUCCGGGGUUCGA
Hal-2-1   CCGAUGUCGGCGGCAUGACCGUCGUCUUCGGGUCGCCGGGCCGUGGGCUUCCGGACAUGCUCGGGAUGUCUCCCGAGGAUGUUGCAGACGUCGAACCUUCCGUUGGUCCGGGGUUCGA
Hsp-6-1   CCGAGGCCGGCGGCAUGACCGUCGUCUUCGGGUCGCCGGGCCGUGGGCUUCCGGACAUGCUCGGGAUGUCUCCCGAGGAUGUUGCAGACGUCGAACCUUCCGUUGGUCCGGGGUUCGA
Hme-1-1   CCGAUGCCGGCGGUAUGACCGUCGCCUUCGGUUCACCAGGCCGUGGGCUUCCGGACAUGCUUGGGAUGUCUCCCGAGGAUGUUGCAGACGUCGAACCUUCCGUUGGUCCGGGGUUCGA
Hla-2-1   CCGACGCCGACGGUAUGACCGUCGUGUUCGGUUCGCCGGGCCGUGGGCUUCCGGACAUGCUGGGGAUGACUCCCGAGGAUGUCGCAGACGUCGAACCCUCCGUUGGUCCGGGGUUCGA
Hel-1-1   CCGACGCCGACGGCAUGACCGUCGUGUUCGGUUCGCCGGGCCGUGGGCUUCCGGACAUGCUGGGGAUGACUCCCGAGGAUGUCGCAGACGUCGAACCCUCCGUUGGUCCGGGGUUCGA
Hmu-1-1   CCGAUGCCGGCGGUAUGACCGUCGUCUUCGGUUCUCCCGGCCGUGGGCUUCCGGACAUGCUCGGGAUGUCUCCCGAGGAUGUUGCAGACGUCGAACCUUCCGUUGGUCCGGGGUUCGA
env-79    AUGCUGGCAGCGAGGGUUCCGGUGCCGACCUGACCGUCGCCUUUGGCGCACCCGAGCGCGGCCUGCCGGAGAUUUUGGGCAUUUCGCCCGAGUCGAUACGCGAAGCUGACUCCACCGA
env-80    GCCGAGAGUCGACGACAGGCGGUGAUCUGGCCGAUUCCGCGUUCACAGUCGCCUUUG.............................................................
env-81    ACACAGCGGGUGUGAACACAGUCGAUGGCGAGGCUGACUCGGAGACCGAGCACGCCUAUACAGUCGCGUUCGGUGCACCCGAGCGUGGAUUGCCAGAAAUCCUGGGCACAACGCCCGA
env-82    GCCGUGAGUCGGCGACAGGCGGUGAUCUAGCCGAUUCCGCGUUCACAGUCGCCUUUGGCGCACCAGAGCGCGGUUUGCCGGAGAUUCUCGGAUUCACACCCGAGAAGAUUCGCGAAGA
env-83    GCCGUGAGUCGGCGACAGGCGGUGAUCUAGCCGAUUCCGCGUUCACAGUCGCCUUUGGCGCACCAGAGCGCGGUUUGCCGGAGAUUCUCGGAUUCACACCCGAGAAGAUUCGCGAAGA
Hla-4-1   AGCGGGACGGUCUCACCGUCGCGUUCGGUGCGCCCGAGAGAGGGCUGCCGGCAAUCCUCGGCAUCGAUGCCGACGCCGUACAGAAAGACGGCGUCGCGCGACUCGAUGCCGCGGACGA
Ssp-1-1   AAACCGAGGGGCUCACCCUCGCGCUGGGGGCGCCCGAGCGGGGGCUUCCGGAGAUCCUCGACGUCGACGCCGAUUCGGUCGACGACUCACCGAGUGCGCUUCGUCGCGCGAUCGAGUC
env-84    ......................................................................................................................
env-86    ACGACGACGGGAUGACCGUCGCCUUCGGUGCGCCCGAGAGAGGGCUGCCGGCGAUCCUCGAUAUCGAGGAUGCUGGCGAGGCCCUAUCCGGGUCUCGUUUCGGCGAAGCGGGCGACGG
Hsp-11-1  AGCGGGACGGUCUCACCGUCGCGUUCGGCGCGCCCGAGAGAGGGCUGCCGGCAAUCCUCGGCAUCGACGCCGACACCGUGCGAACGGAUGGUGUCGCGCGACUCGAUGCCGCGGACGA
env-87    GCGACCUCACGGUCGCCUUCGGCGCGCCCGAACGAGGGCUGCCGGCGAUCCUCGGCCUCGCGGUCGAGGACGUGAGCGCGGGCGAGUCCACGGCUGGAUUCGACCGCUGGCUCAACAC
env-88    AUGGAAUGACCGUUGCCUUCGGCUCGCCGGGACGGGGACUGCCGGAAAUACUCGGGAUCACUCCCGAGGCAGUCUCGGCCGCAACCGCGGCUGGAGACCAUGUUGAAUCCGAUCCGGG

          ......................................................................................................................
          ......................................................................................................................
          ••••••••••••••••••••••••••••••••••••••••••••••••••••••••••••••••••••••••••••••••••••••••••••••••••••••••••••••••••••••
```

```
env-89     ........................................................................................................................
Hka-1-1    ACGACGCGGGCCACAUGACAGUGGCCUUCGGCGCUCCGGAGCGCGGGUUGCCAGCGAUAUUCGGAUUGGACCCGGACGACGCCGUCCCAGACGGCACAGGUGACGACGACGCCGGGUU
Hmu-2-1    ACGACGCGGGCCACAUGACAGUGGCCUUCGGCGCUCCGGAGCGCGGGUUGCCAGCGAUAUUCGGAUUGGACCCGGACGACGCCGUCCCAGACGGCACAGGUGACGACGACGCCGGGUU
Hca-3-1    GUGGCGGAGACGAUGGGAACGCAUCCGGUGUCGAGACGGGCGGAUCCGACGAAUCUGACGCAUCCGACGACGAGGCGGCGCCCCUGGGUGACCUGACAGUCGCCUUCGGUGCGCCCGA
Hsa-5-1    AAGACGGCAUGACCGUCGCCUUCGGCGCGCCCGAGAGAGGGCUGCCGGCGAUGCUUGGCAUCGACACCGGGUCGGUGGCGAGUGGCAGCGAGACCGACGAGCCUGCGGGGUUCGAUCG
env-90     ........................................................................................................................
Hag-1-1    AAGACGGCAUGACCGUCGCCUUCGGCGCGCCCGAGCGAGGGCUGCCGGCGAUGCUCGGCAUCGACGAGGCGUCGGUGGCGAGUGCGGACGAGACCGGGUCGGACCCUGCGGGGUUCGA
Hsa-4-1    AAGACGGCAUGACUGUCGCCUUCGGCGCGCCCGAGCGAGGGCUGCCGGCGAUGCUCGGCAUCGACACGGCGUCGGUGGCGAGUGCGGACGAGACCGGGUCGGACCCUGGAGGGUUCGA
env-91     GCGAGACUGUCGACAGCGACGGCCGACCCCCGUACACGGUCGCGUUCGGAGCGCCCGAACGUGGCCUGCCGGCGAUCCGCGGACUCGACCCGGAGGCCGUCCGGGCGGGCUCGCGCGG
env-92     AUGGGCUGACCGUUGCUUUCGGCUCGCCGGGACGAGGGCUGCCGGAGAUACUCGGGGUUGACCCCGAGACAA................................................
env-93     ........................................................................................................................
env-94     ........................................................................................................................
env-95     AUGGGCUGACCGUUGCUUUCGGCUCG..............................................................................................
env-96     AUGGGCUGACGUCGCUUCGGCGCGCGGGCCGCGGGCUGCCGGAGAUACUCGGGAUCCCACCCGAAGGCUAUCUCGGCUGCUGACGCCG................................
env-98     AGGGGCUGACCGUCGCCUUCGGCGCGCCGGGCCGCGGGCUGCCGGAGAUACUCGGGAUCCCACCCGAGGCAAUCUCGGC.........................................
env-99     GGCUGACCGUCGCCUUCGGCGCGCCGGAACGAGGGCUGCCGGAGAUACUCGGGAUUCCACCCGAGGCUAUCUCGGCUGCUGACGCAGCCGGGGCGUCCAAGCCGUAUGGAUUGGACGA
env-101    GGGGGAUGUCCGUCGCCUUCGGUUCGCCCGGCAGAGGGCUGCCGGAAAUACUCGGGAUCGAACCCGAGGCAGUCU.............................................
env-102    AGGGCGCAACCGUUGCGUUCGGCUCGCCAGGCAGAGGGUUGCCGGAAAUACUCGGGAUCAAACCCGAGGCAGUCUCAGCCGCAC....................................
env-103    AGGGCGCAACCGUUGCGUUCGGCUCGCCAGGCAGAGGGUUGCCGGAAAUACUCGGGAUCAAACCCGAGGCAGUCUCAGCCGCACUCGCGGCUGGAGACCAUGUUGAAUCCGACCCCGG
env-104    ........................................................................................................................
env-105    AGGGCGCAACCGUUGCGUUCGGCUCGCCAGGCAGAGGGUUGCCGGAAAUACUCGGGAUCAAACCCGAGGCAGUCUCAGCCGCACUCGCGGCUGGAGACCAUGUUGAAUCCGACCCCGG
env-106    AGGGGAUGACCGUUGGCUUUGGUUCGCCCGGACGAGGGUUGCCGGAAAUACUCGGGAUCGAACCCGAGGCAGUCUCGGCCGCAGUCGCGGCUGGGCAGCACGUUGAAUCCACGCCGGG
env-107    ........................................................................................................................
env-108    ........................................................................................................................
env-109    ........................................................................................................................
CHa-1-1    CGCACGUCGCCGUCGCGUUCGGUUCGCCGGGGCGGGGGCUUCCGGCCAUCCUCGGGAUGUCCCCCGAGGAGGUGCCGGUCGAACCCGGAGACGCAGACGACUCGGGGUUCGACCUCUG
Hpa-2-1    ACCGGGAGGGAGCCACCGUCGUCUUCGGUUCGCCAGGCCGCGGGCUUCCGGACAUUCUCGGCAUCAACGCCGAGGAAGUCACAGUCGAACCCUCCGACGGUCCGGGGUUCGACCUCUG
env-110    AACGUGACGGAAUGACCGUCGCCUUCGGCUCGCCCGAAAGAGGGCUGCCAGAUAUCCUCGGUAUCGAGGUGUCGGCCAUCGAGGCGUCGAACGAAGACGCCGGUGGCUCGCCUGCGAC
Nhu-1-1    AACGCGAUACUACCCGCGACGGAUACACAAUCGCGUUCGGUGCGCCCGAGAGAGGGCUGCCUGCCAUCCUCGAACUCGAGGAUACCACCGUCGAGUCGUUGUCGGCAUGGCCAUCGUG
env-113    CCGGCGGCUACACCGUCGCCUUCGGCGCGCCCGAACGGGGGCUUCCGGAGAUGCUCGGGCUCUCGCCCGACGACGUUCGGGCGGCCGUCGCCGACGGUCGCCCGGUCGAACCCGAUCC
env-114    AGGGGCUGACCGUCGCGUUCGGCUCGCCGGGACGAGGGCUGCCGGACAUACUCGGGAUCGCCCCCGAGACAAUCUCGGCUGAGCAAUCGGCCGGGGCGUCCGAGCCAGACGGCUUGGA
env-115    ........................................................................................................................
env-116    AGGGGAUGACCGUCGCCUUCGGUUCGCCCGGACGAGGGUUGCCGGAAAUACUCGGGAUCGAACCCGAGGC..................................................
env-117    AGGGGAUGACCGUCGCCUUUGGUUCACCCGGACGAGGGUUGCCGGAAAUACUCGGGAUCGAACCCGAGGCAGUCUCGGCCGCAGUCGCG...............................
env-118    AGGGGAUGACCGUCGCCUUCGGUUCGCCCGGACGAGGGUUGCCGGAAAUACUCGGGAU..............................................................
env-119    ........................................................................................................................
env-120    GCGUCGCAGUCGCGUUCGGCGCCCCGGCGCGUGGGCUCCCCGCCAUCUUCGGGAUCGAGGGUGGACCGGUCGACGCCGAUACGCGGUUCGACCGGUGGCUGAACACGGUUCCAGACCA
env-122    GCGUCGCCGUCGCGUUCGGCGCGCCGGCGCGGGGGCUCCCCGCCAUCCUCGGGUACGAGGAAGGCCCGGUCGACCCCGCCGACACGCGGUUCGACCGGUGGCUGAACACGGUUCCGGA
env-123    GCGUCGCCGUCGCGUUCGGCGCGCCGGCGCGGGGGCUCCCCGCCAUCCUCGGGUACGAGGACGGGCCGGUCGACCCCGCGGACACGCGGUUCGACCGGUGGCUGAACACGGUUCCGGA
env-126    AGCGGGACGGCGCGACAGUCGUGUUCGGGUCGCCCGGCCGCGGGCUUCCGGGCCAUUCUCGGCGACGGCGUCGAUGUCGACGCCCGAUCCGAGGUGGUGGU..................
env-127    ........................................................................................................................
env-128    AGCGGGAUGGCGCGACGGUCGCGUUCGGGUCGCC......................................................................................
env-129    GCGACGGCACCACCGUCGUCUUCGGCUCGCCCGCGAGAGGGCUGCCGGAGAUCCUCGGGGUGGACCCCGAGGUGGUCGAGCGGUCGGGAGACGACGUCGAACCCGCUGGGGGGUUCGA
Hna-1-1    GCGACGGUGAUCUGACGGUCGCCUUCGGGGCGCCCGGGCGCGGCCUUCCGGCCAUCCUCGACGUGGCUGUCGAGGAAGUAGCCGCCGGCGGCGGCGAGGGCGUCGAACCCGAUCCGGG
env-130    ACGACGCGGGCGAUCUGACGGUCGCCUUCGGGGCGCCCGGGCGCGGCCUUCCGGCCAUCCUCGACGUGGCUGUCGAGGACGUAGCCGCACGCGGCGGCGAGGGCGUCGAACCCGGUGA
env-131    GGGACGAUGGGGCGACCAUCGUUUUCGGCUCGCCGGGGCGAGGGAUUCCGGAGAUUCUCGGGGUGUCCCCCGAUGAUCUCCCCGUCGAACCCGACUCACCCGGGUUCGACCUCUGGCU
env-134    ........................................................................................................................
env-135    ........................................................................................................................
Nph-1-1    CCGCCAACGGAUUCACGGUCGCCUUCGGCGCCCCCGAACGGGGGCUGCCGGAGAUACUCGGCUGUGAGCCUCGCGAGGAGGCUAUCGGCGAUGAGCCCGAAGCGCGGUUCGACCUUUG
env-136    CCGACGACGGACUCACGGUCGUCUUCGGAGCCCCCGAACGGGGACUGCCGGAGAUACUCGACUGUGAGCCUCGCGCGGAGACUAUCGGCGACGAACCGAAUGCGCGGUUCGACCUUUG
env-137    CCGACGACGGACUCACGGUCGUCUUCGGAGCCCCCGAACGGGGACUGCCGGAGAUACUCGACUGUGAGCCUCGCGCGGAGACUAUCGGCGACGAACCGAAUGCGCGGUUCGACCUUUG
env-138    CUGGCGGCUACACCGUUGCCUUC.................................................................................................
env-139    CCGGCGGCUACGUCGUCGCCGUCGGUGCGCCCGAACGAGGGCUUCCGGACGUUCUCGGGCUCACACCCGAGGACAUCCGCGCGGCCGUCGCCGACGGUCGCACGGUCGAACCCGAUCC
env-140    GCGGAUACACCGUCGCCUUCGGUGCGCCCGGUCGAGGGCUGCCGGAAAUGCUCGGGCUGUCGCCCGAGGAGCUCCGCGCGGCCGUCGCCGACGGUCGCCCGGUCGAACCCGAUCCGGG
env-145    CCGGC...................................................................................................................
env-146    CCGGCGGCUACACCGUCGCUUUCGGCUCACCCGAACGCGGGCUUCCAGAAAUGCUCGGGCUAUCGCCCGACGACAUUCGAGCGGCCGUCGCCGACGGUCGCUCGGUCGAACCCGAUCC
env-147    CCGGCGGGUACACCGUCGCCUUCGGCGCGCCCGAACGAGGGCUUCCGGAGAUGCUCGGGGUAUCACCCGAGACAGUCCAGGCGGCCGUCGCCGACGGCCGCCGU...............
env-148    CCGGCGGAUACACCGUCGCCUUCGGUGCGCCCGAACGAGGGCUUCCGGAGAUGCUCGGGCUGUCGCCCGAGGACAUCCGCACGGCCGUCUCAGACGGCCGCCGAGUCGAACCCGAUCC
Hsp-9-1    CCGGCGGGUACACCGUCGCCUUCGGCGCGCCCGAGCGGGGGCUUCCGGAGAUGCUCGGGCUUUCGCCCGACCGCAUUCGGGCGGCCGUCGCCGACGGCCGCUCGGUCGAACCCGAUCC
Hez-1-1    CCGGCGGUUACACCGUCGCCUUCGGCGCGCCCGAGCGGGGGCUUCCGGAGAUGCUCGGGCUUCCGCCCGACCGCAUUCGGGCGGCCGUCGCCGACGGCCGCUCGGUCGAACCCGAUCC
env-149    ........................................................................................................................
env-150    ........................................................................................................................
env-151    CCGGCGGUUACACCGUCGCCUUCGGCGCACCCGAGCGGGGGCUUCCGGAGAUGCUCGGGCUUUCGCCCGACCGCAUUCGGGCGGCCGUCGCCGACGGCCGCUCGGUCGAACCCGAUCC
Hco-1-1    CCGGCGGUUACACCGUCGCCUUCGGCGCGCCCGAGCGGGGGCUUCCGGAGAUGCUCGGGCUUUCGCCCGACCGCAUUCGGUCGGCCGUCGCCGACGGCCGUCCGGUCGAACCCGAUCC
env-152    CCGGCGGCUAC.............................................................................................................
env-153    CCGGCGGUUACACCGUCGCCUUCGGCGCGCCCGAGCGGGGGCUUCCGGAGAUGCUCGGGCUUCCGCCCGACCGCAUUCGGGCGG....................................
env-154    CCGGCGGUUACACCGUCGCCUUCGGCGCGCCCGAGCGGGGGCUUCCGGAGAUGCUCGGGCUUUCGCCCGACCGCAUUCGGGCGGCCGUCGCCGACGGCCGCUCGGUCGAACCCGAUCC
env-155    CCGGCGGCUACACCGUCGCCUUCGGCGCGCCCGAACGGGGGCUUCCGGAGAUGCUCGGGCUCUCGCCCGACCGCAUUCGGGCGGCCGUCGCCGACGGCCGCCCGGUCGAACCCGAUCC
env-156    CCGGCGGUUACACCGUCGCCUUCGGCGCGCCCGAGCGGGGGCUUCCGGAGAUGCUCGGGCUUCCGCCCGACCGCAUUCGGGCGGCCGUCGCCGACGGCCGCUCGGUCGAACCCGAUCC
Hsp-10-1   CCGGCGGCUACACCGUCGCCUUCGGCGCGCCCGAGCGGGGGCUUCCGGAGAUGCUCGGGCUGUCGCCCGACGACAUUCGGACGGCCGUUGCCGACGGUCGCCCGGUCGAACCCGAUCC
Hli-1-1    CCGGCGGGUACACCGUCGCCUUCGGCUCGCCCGAACGAGGGCUUCCGGAGAUGCUCGGGCUUUCGCCCGACGGCAUUCGGGCGGCCGUUGCCGACGGCCGCUCGGUCGAACCCGAUCC
Hte-1-1    CAGGCGGUUACACCGUCGCCUUCGGCGCGCCCGAGCGGGGGCUUCCGGAGAUGCUCGGGCUCUCGCCCGACCGCAUUCGAGCGGCCGUCGCCGACGGCCGCUCGGUCGAACCCGAUCC
Hko-1-1    CCGGCGGGUACACCGUCGCCUUCGGCUCGCCCGAGCGAGGGCUUCCGGAGAUGCUCGGGCUUUCGCCCGACGGCAUUCGGGCGGCCGUUGCCGACGGCCGCUCGGUCGAACCCGAUCC
Hli-2-1    CCGGCAGUUACACCGUCGCCUUCGGCGCGCCCGAGCGGGGGCUUCCGGAGAUGCUCGGGCUUUCGCCCGACCGCAUUCGGGCGGCCGUCGCCGACGGCCGCUCGGUCGAACCCGAUCC
Hsa-7-1    CCGGCGGGUACACCGUCGCCUUCGGCUCGCCCGAGCGAGGGCUUCCGGAGAUGCUCGGGCUCUCGCCCGACGGCAUUCGGGCGGCUGUCGCCGACGGCCGCUCGGUCGAACCCGAUCC
Hca-2-1    CCGGCGGUUACACCGUCGCCUUCGGGGCGCCCGAGCGGGGGCUUCCGGAGAUGCUCGGGCUUUCGCCCGACCGCAUUCGGGCGGCCGUCGCCGACGGCCGCUCGGUCGAACCCGAUCC
Hsp-7-1    CCGGCGGUUACACCGUCGCCUUCGGGGCGCCCGAGCGGGGGCUUCCGGAGAUGCUCGGGCUUUCGCCCGACCGCAUUCGGGCGGCCGUCGCCGACGGCCGCUCGGUCGAACCCGAUCC
Hha-2-1    CCGGCGGCUACACCGUCGCCUUCGGCUCGCCCGAGCGAGGGCUUCCGGAGAUGCUCGGGCUUUCGCCCGACAGCAUUCGGGCGGCCGCCGCCGACGGCCGCUCGGUCGAACCCGAUCC
Hdi-1-1    CCGGCGGUUACACCGUCGUCUUCGGCGCGCCCGAGCGGGGGCUUCCGGAGAUGCUCGGGCUUUCGCCCGACCGCAUUCGGGCGGCCGUCGCCGACGGCCGCUCGGUCGAACCCGAUCC
Hsa-9-1    CCGGCGGCUACACCGUCGCCUUCGGCUCGCCCGAGCGAGGGCUUCCGGAGAUGCUCGGGCUUUCGCCCGACGGCAUUCGGGCGGCCGUCGCCGACGGCCGCUCGGUCGAACCCGAUCC
Har-2-1    CCGGCGGUUACACCGUCGCCUUCGGCGCGCCCGAGCGGGGGCUUCCGGAGAUGCUCGGGCUUUCGCCCGACCGCAUUCGGGCGGCCGUCGCCGACGGCCGCUCGGUCGAACCCGAUCC
env-157    ........................................................................................................................
env-158    CGGGCGGUUACACCGUCGCCUUUGGCUCGCCCGAACGAGGGCUUCCGGAGAUGCUCGGGCUGUCGCCCGACAGCAUUCGGGCGGCCGUCGCCGACGGCCGCCCGGUCGAACCCGAUCC
env-159    CGGGCGGUUACACCGUCGCCUUUGGCUCGCCCGAACGAGGGCUUCCGGAGAUGCUCGGGCUGUCGCCCGACAGCAUUCGGGCGGCCGUCGCCGACGGCCGCCCGGUCGAACCCGAUCC

           ........................................................................................................................
           ........................................................................................................................
           ........................................................................................................................
```

```
env-160    CCGGCGGGUACACCGUCGCUUUCGGCGCGCCCGAACGGGGGCUUCCGGAGAUGCUCGGGGUAUCACCCGAGACAGUCCAGGCGGCCGUCGCCGACGGCCGCC................
h3A-1-1    CGGACGAUCCGGUCGUCGCCUUCGGCGCGCCCGGCAGAGGGCUUCCGGAGAUCCUCGGGGUCCGCCCCGAGGACGUCACGGCCGCAGCCGGCGGCCCCGUCGAACCCGAUCCGGGGUU
Hai-1-1    CCGGUGGCUACGUCAUCGCCUUCGGUGCGCCCGAACGAGGGCUUCCGGAGAUGCUCGGGCUUUCGCCCGACGGCAUUCGGGCGGCCGUCGCCGACGGCCGUCCGGUCGAACCCGAUCC
env-161    CCGGCGGCUACACCGUCGCCUUCGGCUCGCCCGAACGAGGGCUUCCGGAGAUGCUCGGGCUUUCGCCCGACGGCAUUCGGGCGGCCGUUGCCGACGGCCGCUCGGUCGAACCCGAUCC
Hho-1-1    CCGGCGGUUACACCGUCGCCUUCGGCGCGCCCGAGCGGGGGCUUCCGGAGAUGCUCGGGCUCUCGCCCGACCGCAUUCGGGCGGCCGUCGCCGACGGCCGCUCGGUCGAACCCGAUCC
env-164    CCGGCGGCUACGUCGUCGCCUUCGGUGCGCCCGGUCGAGGGCUGCCGGACAUGCUCGGGCUGUCGCCCGAGGAUCUCCGCGCGGCCGUCGCCGACGGCCGCCCGGUCG..........
env-165    ..............................................................................................................................
env-166    ..............................................................................................................................
env-167    CUGGCGGCUACACCGUCGCCUUUGGCGCGCCGGAGCGGGGGCUUCCGGAGAUGCUCGGGUGUCAACCCGAGGACGUUCACACGGCCGUCGAAGACGGUCGUCGAGUCGAACCCGAUCC
env-169    CUGGCGGCUACACCGUCGCCUUUGGCGCGCCGGAGCGGGGGCUUCCGGAGAUGCUCGGG............................................................
env-170    ..............................................................................................................................
env-172    ..............................................................................................................................
env-173    GCCAACGAGGUCUCACAAUCGCGUUUGGCGCUCCACAGCGGGGACUUCCGGAC..................................................................
Hru-1-1    AGCGCGAGGGCGCGACAGUCGUCUUCGGGUCGCCCGGUCGUGGGCUUCCGGCCAUCCUCGGCGUCGACGUCGAGGAUGCGGUCGAACCCGCAGACGGUCCGGGGUUCGAUCUCUGGCU
env-174    ..............................................................................................................................
Hru-2-1    ACCGGGACGGCGCCACCGUCGUCUUCGGCUCGCCGGGCCGCGGGCUUCCGGAGAUCUUGGGGGUAUCCCCCGACGAUCUCCCCGUCGAACCCGACUCGCCCGGGUUCGACCUCUGGCU
env-175    ..............................................................................................................................
env-176    CAGACGAUGGGUUCACGGUCGCCUUCGGCGCCCCCGAACGGGGGUUGCCGGAGAUACUCGACUGCGAACCGAGCGCGGAGACCAUCGGCGACGAACCGAACGUGCGGUUCGACCUUUG
env-178    CCGACGACGGGUUCACGGUCGCUUUCGGCGCCCCCGAACGGGGGUUGCCGGAGAUACUCGACUGUGAGCCGGGCGCGGAGGCCAUCGGCGACGAACCGAACGUGCGGUUCGACCUUUG
env-179    CCGAGGACG..............................................................................................................
env-180    ..............................................................................................................................
Nmo-1-1    CCGACGGACUCACGGUCGCCUUCGGCGCCCCCGAACGGGGGUUGCCGGAGAUACUCGACUGUGAGCCACGCGCGGAGGCCAUCGGCGACGAACCGAACGCGCGGUUCGACCUUUGGCU
Hha-1-1    GCGGCGACCUGACAGUCGCCUUCGGGGCCCCCGAACGAGGGCUGCCGGCGAUGCUCGGCAUCGACGUCGAGUCGGUGACGAGCGCCGAACCGAACGGCGGGUUCGACCGCUGGCUCAA
env-181    ..............................................................................................................................
Hpa-1-1    ACCGCGACGGAAUGACCGUCGCGUUCGGUUCGCCGGGUCGAGGACUGCCGGAGAUAUUCGGGGUCCCCCCCGAAGGUAUCUCCGCGGUCGAAUCCGGCGUACCAGCCCGGUUCGACCU
Hci-1-1    AACGCGACGGCAUGACCGUCGUGUUCGGCUCGCCCGGUCGGGGACUGCCGGAGAUACUCGGGGUUGAACCCGAGCGUAUCUCCGCGGUCGAAUCCGGCGUACCAGCCCGGUUCGACCU
env-182    ..............................................................................................................................
env-183    ..............................................................................................................................
env-184    ..............................................................................................................................
env-185    ..............................................................................................................................
Hdj-1-1    AACGGGACGGGAUGACCGUCGCGUUCGGUGCACCCGAGCGAGGGUUGCCGGAUAUCCUCGGAAUCGAGGCGUCAGCCGUCCCCUCGUCCGACGAGGGUGACGCUGACGACGGAGUCGA
Hsa-10-1   AGCGUGACGGUAUGACCGUCGCGUUCGGGGCGCCCGAGAGAGGGCUGCCGGACAUCCUCGGAAUCGAGGCAUCGGCCGUCACCGCGUCGGACGACUCCGACGACGGGGUGGACGAUGA
Nsu-1-1    AUCGUGACGGGCUGACCGUCGCAUUUGGUGCGCCCGAGAGAGGGCUGCCGGAUAUCCUCGGAAUCGAGGCAUCGGCUGUCAACGCGUCGUCGGGACAAGGCGACGCGGCCGAAUCGUC
Nba-1-1    AUCGUGACGGGCUGACUGUCGCGUUCGGUGCGCCCGAGAGAGGGCUGCCGGAUAUCCUCGGAAUCGAGGCAUCAGCUGUCAACGCGUCGUCGGGACAAGGCGACGCGGCCGAAUCGUC
Nin-1-1    GCGACGGACUGACCGUCGCCUUUGGUGCGCCCGAGAGAGGGCUGCCGGAUAUCCUCGGAAUCGAGGCAUCCGCCAUCGACGCUGACCCAUCGGGGCAGCUGGACACCGCGGCGGAUGA
env-186    AAGCCGACGGGAUGACCGUCGCCUUCGGCUCGCCCGAGAGAGGGCUUCCGGAGAUGCUCGACGGGGUAACUGUCGAGCAACGAGCGGACGGGGACCGAGUCCCCGGGAACGGAGACGA
env-187    ..............................................................................................................................
env-188    AAGCCGACGGGAUGACCGUCGCCUUCGGCUCGCCCGAGAGAGGGCUUCCGGAGAUGCUCGACGGGGUAACUGUCGAGCAACGAGCGGACGGGGACGCAGUCCCCGGGAACGGAGACGA
env-189    ..............................................................................................................................
env-190    ..............................................................................................................................
env-191    AACGGGACGGGCUCACCGUCGCCUUCGGUGCCCCCGAACGAGGGUUGCCGGACAUCCUCGACAUCGACGAGGCGGCUGUCGAAGCCGCCGUCGAUGGCGAGGGCGACGCAAACGUCGA
Hla-1-1    AGGACGACGGUCUGACCGUCGCCUUCGGCGCGCCCGAGAGAGGGCUGCCGGAUAUCCUCGGAAUCGAGGCGUCCGCCGUCGCCGUCGAUCCGGAUCCAUCGGAUCGAGGGGAUUCCGA
env-193    ..............................................................................................................................
Hac-1-1    ..............................................................................................................................
Hac-1-2    ACGAGGACGGACUGACCGUGGCGUUCGGCUCGCCGGAACGCGGGCUGCCGCCGAUGCUCGGCCUCACCGCCGAGGCGGUCAGGGAGUUCGACUCCGCCCAGUCCACCCGCGCACCCGG
env-196    ..............................................................................................................................
env-197    ..............................................................................................................................
env-198    CCCGGGACGGACUGACAGUCGCCUUCGGUGCACCCGAACGUGGGCUUCCGCCCAUGCUCGGACUCACCGAGGACGAGGUCAGCGAGGCCGUCGCCGACCCCAACUCCGACGCACCCGC
Hsp-2-1    CCCGGGACGGACUGACAGUCGCCUUCGGUGCACCCGAACGUGGGCUUCCGCCCAUGCUCGGACUCACCGAGGACGAGGUCAGCGAGGCCGUCGCCGACCCCAACUCCGACGCACCCGC
env-199    ACGGCGUCACCGUUGCCUUCGGCGCGCCCGAGCGAGGGCUGCCGGCGAUACUCGGUAUCGACCCAGCGCGGGUCGGUGCCGACUCCCCGCCAGGGGACUUCGACCGCUGGCUGAACAC
Hsu-1-1    CCGCCGACGGACUGACCGUCGCCUUCGGAGCGCCGGGUCGUGGCCUCCCCGACAUCUUGGACGUCGACGUCCAGGAAAUCGGGGCCGCGACCGGCUCCACGACGGUCGGCGAGGGUGC
env-201    ..............................................................................................................................
env-203    ..............................................................................................................................
Hca-1-1    ACGCCGAGGGCGGCAUGACCGUCGCCUUCGGGGCACCCGAGCGCGGGUUACCGUCGAUACUCGACGUUGCCCCGGACGCCGUCGGGGGAGACCAGACGAGCGACGAACCAGAAGGGUU
Ham-1-1    ACGCCGAGGGCGGCAUGACCGUCGCCUUCGGGGCACCCGAGCGCGGGUUACCGUCGAUACUCGACGUUGCCCCGGACGCCGUCGGGGGAGACCAGACGAGCGACGAACCAGAAGGGUU
Hhi-2-1    ACGCCGAGGGCGGCAUGACCGUCGCCUUCGGGGCACCCGAGCGCGGGUUACCGUCGAUACUCGACGUUGCCCCGGACGCCGUCGGGGGAGACCAGACGAGCGACGAACCAGAAGGGUU
Hsa-2-1    ACGCCGAGGGCGGCAUGACCGUCGCCUUCGGGGCACCCGAGCGCGGGUUACCGUCGAUACUCGACGUUGCCCGGACGCGUCGGGGGAGACCAGACGAGCGACGAACCAGAAGGGUUCG
Har-1-1    ACGCCGAGGGCGGCAUGACAGUCGCCUUCGGGGCACCCGAGCGCGGGUUACCGUCGAUACUCGACGUUGCCCCGGACGCCGUCGGGGGAGACCAGACGAGCGACGAACCAGAAGGGUU
Hva-1-1    ACGCCGAGGGCGGCAUGACCGUCGCCUUCGGGGCACCCGAGCGCGGGUUACCGUCGAUACUCGACGUUGCCCCGGACGCCGUCGGGGGAGACCAGACGAGCGACGAACCAGAAGGGUU
Hja-1-1    ACGCCGAGGGCGGCAUGACUGUCGCCUUCGGGGCACCCGAGCGCGGGUUACCGUCGAUACUCGACGUUGCCCCGGACGCCGUCGGGGGAGACCAGACGAGCGACGAACCAGAAGGGUU
env-204    ACGGACUCACCGUCGCCUUCGGCGCGCCACAACGGGGGCUGCCGGCGAUUCUCGGGAUCGAACCCGAGGUCAUUUCGGCCGAACCAUCGGCCGACACGUCCGAGCCAGCCGGGUCGGA
Hse-1-1    ACGACGGACUGACCGUCGCCUUCGGCGCGCCCGAGCGAGGGCUGCCGGCCAUGCUCGGCAUCGAGCACGAAUCGGUGUCGAGCGCCGAAUCGAACCACGCGGGGUUCGACUGCUGGCU
Hth-1-1    ACGACGGACUGACCGUCGCCUUCGGCGCGCCCGAGCGAGGGCUGCCGGCGAUGCUCGACAUCGACGCGGCGUCGGUGGCGAGCGACUCACCCCACGAGGGGUUCGACUGCUGGCUCAA
Hmo-1-1    ACGACGGGCUGACCGUCGCCUUCGGCGCGCCCGAGCGAGGGCUGCCGGCGAUGCUCGACAUCGACGCGGCGUCGGUGGCGAGCGAUACACCCCACGCGGGGUUCGACUGCUGGCUCAA
env-206    AAGCCGACGGGAUGACCGUCGCCUUCGGCGCGCCCGAGAGAGGGCUGCCGGAGAUCCUCGAUCUGGCGAUCGAGGCUGUCGGCG..................................
env-207    AACGAGUCGCCGUGGCUUUCGGCGCGCCGGCGCGAGGCUUGCCCGCCAUCCUCGACGGGGUGGCGGACGACGAGCGGAUCGACGCGGCGGAGACGCCGUUCGACCGCUGGUUGAACAC
env-209    CAGGAGGCG..............................................................................................................
Nta-1-1    ACCGCGACGGGAUGACCGUCGCCUUCGGCGCGCCCGAGAGAGGGCUGCCGGACAUCCUCGAGUUCGAAUCCGAGGACGCCGUCGACACGUUGUCGUCCUGGCCGCCGCGGUCAGACGA
Nas-1-1    ACCGCGACGGGAUGACCGUCGCCUUCGGCGCGCCCGAGAGAGGGCUGCCGGACAUCCUCGAGUUCGAAUCCGAGGACGCCGUCGACACACUGUCGUCCUGGCCACCGCGGCCGGACGA
Noc-1-1    AGCGCGACGGGAUGACCGUCGCCUUCGGGGCACCCGAGAGAGGGCUGCCGGCCAUCCUCGGAAUGGACGAGGAGACCGUCGACGCGCUGUCCCGCUGGGACGGCGUCGACGGUGGCGG
Nje-1-1    AGCGCGACGGGAUGACCGUCGCCUUCGGGGCACCCGAGAGAGGGCUGCCGGCAAUCCUCGGAAUGGACGAGGAGGCCGUCGACGCGCUGUCCCGCCGGGACGGCGUCGACGGCGACGG
Nam-1-1    AGCGCGACGGGAUGACCGUCGUCUUCGGGGCACCCGAGAGAGGGCUGCCGGACAUCCUCGGAAUGGACGAGGAGGCCGUCGACGCGCUGUCCAACUGGGACGGCGUCGACGGUGACGG
env-210    AAGACGACGGGAUGACCGUCGCGUUCGGUGCGCCCGAGAGAGGGCUGCCGGCUAUCCUCGGAAUCGAGGCAUCGGCUAUCGAGCCGUCCGGCGAGGACGAGCCGGCCGAUGACGGAGU
Hsp-3-1    CCAGGGACGGCGUGACCGUUGCGUUCGGAGCACCGGAACGUGGGCUGCCGCCCAUGCUUGGUGUUUCGGCGGACGCGGUCAACGAGUCGGUCACCGACUCAUCCGCAGACGCCCCCGC
Nch-1-1    AACGCGAUACUACCCGCGACGGAUACACAAUCGCGUUCGGUGCGCCCGAAAGAGGGCUGCCUGCCAUCCUCGAACUCGAGGAUACCACCGUCGAGUCGUUGUCGGCAUGGCCAUCGUG
Nae-1-1    AGCGCGACGGGAUGACCGUCGCCUUCGGCGCGCCCGAGAGAGGGCUGCCGGACAUCCUCGAGUUCGAAUCCGAGGACGCCGUCGACACGUUGUCGUCCUGGCCGCCGCGGUCAGACGA
env-214    ..............................................................................................................................
env-215    AGCGCGACGGGAUGACCGUCGCCUUCGGGGCACCCGAGAGAGGGCUGCCGGACAUCCUCGGAAUGGCCGAGGAGACC..........................................
env-216    AGGACGAGGGA............................................................................................................
Hni-1-1    AAGCCGACGGGAUGACCGUCGCUUUCGGUGCGCCCGAGAGAGGGCUGCCGGACAUCCUCGGAAUCGAGGCAUCCGACGUUGCGUCGGCCCACGCAGGCGACGGGCCGGAGAGCGCGGA
Has-1-1    AGCGCGACGGCAUGACCGUUGCGUUCGGUGCGCCCGAGAGAGGGCUGCCGGACAUCCUCGGAAUCGAGGAGUCACGUGCCGGCGCGUCGUCCAGCGGGGACGGCGGGGCGGCAGACGG
env-217    ..............................................................................................................................
Nal-2-1    AGCGCGACGGGAUGACCGUCGCCUUCGGCGCGCCCGAGAGAGGGCUGCCGGCUAUCCUCGGAAUCGAGGAAUCGGCCAUCGAACCGUCGUCCGGACAAGACGACGCGAUCGAACCGUC
Npa-1-1    AGCGCGACGGGAUGACCGUCGCCUUCGGCGCGCCCGAGAGAGGGCUGCCGGCUAUCCUCGGAAUCGAGGAAUCGGCCAUCGAAGCGUCGUCCGGACAAGACGACGCGAUCGAACCGUC

           ..............................................................................................................................
           ..............................................................................................................................
           ..............................................................................................................................
```

```
Nga-1-1   AGCGCGACGGGAUGACCGUCGCCUUCGGCGCGCCCGAGAGAGGGCUGCCGGCUAUCCUCGGAAUCGAGGAAUCGGCCAUCGACGCGUCGUCCGGACGAGACGACGCGGUCGACCCGUC
Nsp-2-1   AGCGCGACGGGAUGACCGUCGCCUUCGGCGCGCCCGAGAGAGGGCUGCCGGCUAUCCUCGGAAUCGAGGAAUCGGCCAUCGACGCGUCGUCCGGACGAGACGACGCGGUCGACCCGUC
Htu-1-1   AGGGCGACGGGAUGACCGUCGCCUUCGGCGCGCCCGAGAGAGGGCUGCCUGCUAUCCUCGGAAUCGAGGCAUCGGCCAUCGAAGCGUCGUCGGGACAAGGCGACGCGAUCGAAUCGUC
Hsa-8-1   AGGGUGACGGGAUGACCGUCGCCUUCGGCGCGCCCGAGAGAGGGCUGCCUGCUAUCCUCGGAAUCGAGGCAUCGGCCAUCGAAGCGUCGUCGGGACAAGGCGACGUGAUCGAACCGUC
Hth-2-1   AGGGCGAUGGGAUGACCGUCGCGUUCGGUGCGCCCGAACGAGGGCUGCCGGAUAUCCUCGAAAUCGAGGCAUCCGCCGUCGGGGCCCCGAAGGACGCGGCCGCCGGCGAGGAUGACGG
Nve-1-1   AGCGCGACGGGAUGACCGUUGCCUUCGGCGCGCCCGAGAGAGGGCUGCCGGCUAUCCUCGGAAUCGAGGAAUCGGCCAUCGAAGCGUCGUCCGGACAAGGCGACGCGGCCGAUAACGG
Hli-3-1   AGGACGACGGGAUGACCGUCGCCUUUGGCGCGCCCGAGCGAGGGUUGCCGGAUAUCCUCGGAAUCGAGGCAUCCGCCGUCGGGGCUGUCCAGGAGACAGCCACGGACGGGGAUGACGG
Hxa-1-1   AUCGCGACGGUUUGACCGUCGCCUUCGGAGCGCCCGAGAGAGGGCUGCCGGCCAUCCUCGGAAUCGAGGCGUCCGACGUCGCUGCCGCAUACGGCGGCGACGCGGACGACGGCAACGU
Hgo-1-1   AGGCCGACGGGAUGACCGUCGCCUUCGGCGCGCCCGAGAGAGGGCUGCCGAGCAUCCUCGGAAUCGAGGCGUCGGCCGUCGAGUCGUCGUCCACCGGGGACGACGCCGGACGAUCGUC
env-218   ........................................................................................................................
env-219   AGCAGGACGGAUACACCGUCACGUUCGGAGCGCCCGAUAGAGGGCUGCCGGAUAUCCUCGAUAUCGAGGCGUCGGCCGUCGCGUCGUCAGCGGACACCAGCGGCGGUGGCGACGACGU
env-220   AGGGACUCACAGUCGCCUUUGGCGCGCCCGAACGCGGUCUACCGGCCAUUCUUGGCGUCGAGGCCGGCGAUCCCGAAGCGGGAUUCGACCUGUGGCUGAACACGGUUCCGAACCAGGG
env-221   AACGAGACGGGAUGACCGUCGCGUUCGGUGCGCCCGAGAGAGGGCUGCCGGCUAUCCUCGGAAUCGAGGAAUCGGCUGUCGAGUCGUCCAGUGAGGACGGGGCAGCCGUCGACGGAGU
Hsa-6-1   ACGACGGAAUGACCGUCGUCUUCGGAUCCCCCGGACGAGGGCUUCCGGACAUCCUCGACGUCGCCGUCGAGGACCUCCCCACCAGUACCGGUGGGGCGAGCGUCGGACCCGACAGCGG
env-222   ........................................................................................................................
env-223   ........................................................................................................................
env-224   ........................................................................................................................
env-225   AUCGGGAGGGAAUGACAGUUGCAUUUGGGGCCCCCGGCAGGGGUUUGCCAACGAUUCUUGGGUUAGACCCCAUAGAAAUACUUGAGCCAGAUAUCCCUGGCCCCAAGUUCGAUUGCUG
env-226   ........................................................................................................................
env-227   ........................................................................................................................
env-228   AUCGGGCGGGAGCGACAAUUGCCUUCGGUGCGCCUGGUCGUGGGCUCCCAGUCAUCCUUGGCGAUGACAAUGCGCCGGAUAGCCGAACGCAAACGGGGAACCAAACAGCCUCACCCGU
env-231   CCUCACUCAUCUCCGGUGUGUAUCGGCCGAUAUAGGCGAAUUCAACCGGUAUACCGGUAUGUCGUGGUCAAAUACCACAAAUCCAGGAAAUUAGUGCAGGAUUGAUAGAACCAUGCCA
env-232   ........................................................................................................................
env-233   CCUCACUCAUCUCCGGUGUGUAUCGGCCGAUAUAGGCGAAUUCAACCGGUAUACCGGUAUGUCGUGGUCAAAUACCACAAAUCCAGGAAAUUAGUGCA.....................
env-237   GGACCCGGUGGAAAUCCCCGGAAAAUCCAGAAGAAAGGACACCUGUUACCAGGUAGUUCC............................................................
env-238   AGAAUGGAUACACUCUAAUCUUUGGCUCACCUCUGGGUGGAAUUCCCAGAGAUCACUGGGAGAAAUUUCACAUGGUUGUGAAUUUCAUUCCAGAUCAGGAGACCGAAACUGUGAGGAC
env-240   CAAAUGCGAAGCAAAGGGAUAAGGUUUCCGUGGCGUUCGGCAGCCCUCGAAAGGGCUGGAUGCUAUCUUACUCGAUGAAGGGCACAAGCUUGAGGAUCAUUCCAGUUACGUGGUGAA
env-241   AGGAAUUGGGCAAGCAAGCGAGGUUAAAGUCAUAUACUGUUGUGGUAUUCGGUUCACCCAAACAGGGUGUGGAGUCCAUAUUAGCAAGAGAGGGUACAGAACUCUCAAAAUAUCAGUG
env-242   GAAAUUCCGGCAUGCACGAUGGAACGAAGGAAUGAUUGAAAAAUGCCGAAAAUAAACAGGCCACGCAGAGGAUCUCUUGCAUUCAGCCCCCGGAAGAGGGCAAAGAGCCCGGUGCCCA
env-243   ........................................................................................................................
env-244   GUAUGGUGAAGCCGCGGUCCGGAAUACUGCACGGAAAGUUCAGAUCUCAUACGAGGAUUGAUAAAAAAUGCCGAAGAUCAACACACCGCGAAAGGGAUCCCUCGCAUUCAGCCCCAG
env-245   CCGUACACANAUUACCCCGGUUUUCCGCGGGAGAGGUUGCGGCAGGUGCAGCACUGAAUUGUCAGAUUAGAGAAGGAUUGAUGAAAAAAUGCCGAAGAUCAACACACCGCGAAAAGGA
env-246   CCGUACACACAUUACCCCGGUUUUCCGCGGGAGAGGUUGCGGCAGGUGCAGCACUGAAUUGUCAGAUUAGAGAAGGAUUGAUGAAAAAAUGCCGAAGAUCAACACACCGCGAAAAGGA
env-247   CGUACACAAAUUACCCCGGUUUUCCGCGGGAGAGGUUGCGGCAGGUGCAGCACUGAAUUGUCAGAUUAGAGAAGGAUUGAUGAAAAAAUGCCGAAGAUCAACACACCGCGAAAAGGAU
env-248   GCCGUACACAACUUACCCCGGUUUUCCGCGGGAGAGGUUGCGGCAGGUGCAGCACUGAAUUGUCAGAUUAGAGAAGGAUUGAUGAAAAAAUGCCGAAGAUCAACACACCGCGAAAAGG
env-249   AUCUCGGAUCCCGGACCG......................................................................................................
env-250   CUCGGAUCCAAAGACCGGUGGAUUAUUUUACCGGUAAAUGACAGAGCCUGGAGGUUCCCCGGUAAAAAUUACCGGGCCCUCCGCAUGUACGGGCACAAUUGUCCCUGCAUCAAAAGAC
env-251   CGGUGCAGAGCGGGCAUACGCCCUUGAUAGAAAGAUUGAAGGAUUGAAGAAUACAUGCCGAAGAUUAACAGGCCACGUUAUGGUUCACUCGCAUUCAGCCCACGGAAACGUGCACGAA
env-252   CACGGUGCAGAAUGGGUAUACGCCCUUGAUAGAAAGACUUGAAGGAUUGAAGAAUACAUGCCGAAGAUUCACAGGCCACGUUCAGGUUCACUCGCAUUCAGCCCGCGGACACGUGCUC
env-253   UUACCCGGUGCAGAAUGGGCGUACGCCCUUGUUAGAAAGACUUGAAGGAUUGAAGAAUACAUGCCGAAGAUUCACAGGCCACGUUCAGGUUCACUCGCAUUCAGCCCACGGAAGCGUG
env-254   ........................................................................................................................
env-255   CCUGCAACAGGUCUCACAGAGUCUCGGAUGAAUCUCUCAAACGUGUCGCAAUGAGAAUAACCUACACGCAAAGGUUAGGAUGGAUUAACACAUGCCAAAAAUCAACAGACCACGCCGU
env-256   ........................................................................................................................
env-257   AGGCCGAUCGUGCCGGCAGAUCCGGCGCGAGGUCCGGCGCACGCAUCGCUCUGCAGUAUACCUGCAGUCGAUGCACCACGCUACAAUGCACGAAGGAUUGAUUGAAACAUGCCGAAGA
env-258   AAGGCCGAUCGUGCCGGCAGAUCCGGCGCGAGGUCCGGCGCACGCAUCGCUCUGCAGUAUACCUGCAGUCGAUGCACCACGCUACAAUGCACGAAGGAUUGAUUGAAACAUGCCGAAG
env-259   AGGCCGAUCGUGCCGGCAGAUCCGGCGCGAGGUCCGGCGCACGCAUCGCUCUGCAGUAUACCUGCAGUCGAUGCACCACGCUACAAUGCACGAAGGAUUGAUUGAAACAU........
env-260   AGACCGAUCGUGCCGGCAGAUCCGGCGCGAGGUCCGGCGCACGCAUCGCUCUGCAGUAUACCUGCAGUCGAUGCACCACGCUACAAUGCACGAAGGAUUGAUUGAAACAUGCCGAAGA
env-261   CACCAGAAGAAUCCAUCGUGUCGGGUACCCGGCGCACGGGGUUCGGCGCAGCAUCUCACUACAACGCACGAAGGGUUGAUUGAAACAUGCCGAAUAUACACAGACCACGCAGAGGAUC
env-265   AAAGGGCUUUCGUGCCGGAACAUCCGGCACGGAGCCCGGCGAGUGCAUCGCUCUGCAGCAGGCCUGCGGCCGAUGCAUCACGCUACAACGCACGAAGGGUUGAUUGAAACAUGCCGAA
env-266   CGGAAGCCCGGCGCAGGCAUCGCUCUGCAACAGAGUGGUGCGUCACGCUAUAACCACGAAGGGUUGAUUGAAACAUGCCCAAGAUUAACAGACCACGCAGAGGAUCCCUCGCGUAUAG
env-268   ........................................................................................................................
env-269   CGGAAGCCCGGCGCAGGCAUCGCUCUGCAACAGAGUGGUGCGUCACGCUAUAACCACGAAGGGUUGAUUGAAACAUGCCCAAGAUUAACAGACCACGCAGAGG...............
env-270   ACGGAAGCCCGGCGCAGGCAUCGCUCUGCAACAGAGUGGUGCGUCACGCUAUAACCACGAAGGGUUGAUUGAAACAUGCCCAAGAUUAACAGACCACGCAGAGGNUCCCUCGCGUAUA
env-271   UGCCUCUCCGGCAGGGAUCAUGCAGUCGGUGCAUCAGACUAGUUGCACGAAGGGUUGAUUGAAACAUGCCGAAGUAUAACAGGCCACGCAGAGGAUCCCUUGCAUACAGUCCAAGGAA
env-272   CUGCCUCUCCGGCAGGGAUCAUGCAGUCGGUGCAUCAGACUAGUUGCACGAAGGGUUGAUUGAAACAUGCCGAAGUAUAACAGGCCACGCAGAGGAUCCCUUGCAUACAGUCCAAGGA
env-274   CUGCCUCUCCGGCAGGGAUCAUGCAGUCGGUGCAUCAGACUAGUUGCACGAAGGGUUGAUUGAAACAUGCCGAAGUAUAACAGGCCACGCAGAGGAUCCCUUGCAUACAGUCCAAGGA
env-275   ........................................................................................................................
env-277   CUGCCUCUCCGGCGGGGAUCAUGCAGUCGGUGCAUCAGACUAGUUGCACGAAGGGUUGAUUGAAACAUGCCGAAGUAUAACAGGCCACGCAGAGGAUCCCUUGCAUACAGUCCAAGGA
env-278   CCUGUCCGGCAGCGAUUUGGUGCGUUCUGCAGUCCAUGCAGCAGUACGAAUCGCUACGCAGUCGGUGCAUCUCAGACUACAGUUGCACGAAGGGUUGAUUAAAAACAUGCCGAAGUAU
env-292   GGCCCGUCCGGCAGAGGAUUCGGUGCAUCACACUACAGUUGCACGAAGGGUUGAUUGAAACAUGCCGAAGUAUAACAGGCCACGCAGGGGAUCCCUCGCAUACA.............
env-293   GGCCCGUCCGGCAGAGGAUUCGGUGCAUCACACUACAGUUGCACGAAGGGUUGAUUGAAACAUGCCGAAGUAUAACAGGCCACGCAGGGGAUCCCUCGCAUACA.............
env-295   GGCCCGUCCGGCAGAGGAUUCGGUGCAUCACACUACAGUUGCACGAAGGGUUGAUUGAAACAUGCCGAAGUAUAACAGGCCACGCAGGGGAUCCCUCGCAUACAGCCCGAGAAAACGG
env-296   CGGCCCGUCCGGCAGAGGAUUCGGUGCAUCACACUACAGUUGCACGAAGGGUUGAUUGAAACAUGCCGAAGUAUAACAGGCCACGCAGGGGAUCCCUCGCAUACAGCCCGAGAAAACG
Mbo-1-1   GGCCCGUCCGGCAGAGGAUUCGGUGCAUCACACUACAGUUGCACGAAGGGUUGAUUGAAACAUGCCGAAGUAUAACAGGCCACGCAGGGGAUCCCUCGCAUACAGCCCGAGAAAACGG
env-297   CGGUCCGUCCGGCAGAGGAUUCGGUGCAUCACACUACAGUUGCACGAAGGGUUGAUUGAAACAUGCCGAAGUAUAACAGGCCACGCAGGGGAUCCCUCGCAUACAGCCCGAGAAAACG
env-298   GUCCGUCCGGCAGAGG........................................................................................................
env-303   GUCCGUCCGGCAGAGGAUUCGGUGCAUCACACUACAGUUGCACGAAGGGUUGAUUGAAACAUGCCGAAGUAUAACAGGCCACGCAGGGGAUCCCUCGCAUACAGCCCGAGAAAACGGG
env-304   GGNCCGUCCGGCAGAGGAUUCGGUGCAUCACACUACAGUUGCACGAAGGGUUGAUUGAAACAUGCCGAAGUAUAACAGGCCACGCAGGGGAUCCCUCGCAUACAGCCCGAGAAAACGG
env-305   CUGCCUCUCCGGCAGGGAUCAUGCAGUCGGUGCAUCAGACUAGUUGCACGAAGGGUUGAUUGAAACAUGCCGAAGUAUAACAGGCCACGCAGAGGAUCCCUUGCAUACAGUCCAAGGA
env-306   ........................................................................................................................
Mma-1-1   GAGGGCUUACCGUGCCGGUGCGUCCGGUACGGGGCCCGGCGCACGCAUCGCGCUGCAGCAAGCCUGCAGUCGAUGCCUCACGCUACAACGCACGAAGGGUUGAUUGAAAAAUGCCGAA
Mse-1-1   GAGGGCUUACCAUGCCGGUGCGUCCGGCACGGGGCCCGGCGCACGCAUCGCACUGCAGCGAGCCUGCAGUCGAUGCCUCACGCUAUAACGCACGAAGGGUUGAUUGAAAAAUGCCGAA
Msp-1-1   GAGGCUUAUCGUGCCGGUGCGUCCGGCACGGAGCCCGGCGCACGCAUCGCACUGCAGCGAGCCUGCAGUCGAUGCCUCACGCUACAACGCACGAAGGGUUGAUUGAAAAAUGCCGAAG
env-310   ........................................................................................................................
env-315   GCCGGCUCGCCCGGCAGAGGAUCUGGCAUAUCAACACUAGUUGCACGAAGGGUUGAUUGAAACAUGCCGAAAUAUAACAGGCCACGCAGAGGAUCCCUCGCGUACAGCCCGAGGAAAC
env-316   GCCGGCUCGCCCGGCAGAGGAUCUGGCAUAUCAACACUAGUUGCACGAAGGGUUGAUUGAAACAUGCCGAAAUAUAACAGGCCACGCAGAGGAUCCCUCGCGUACAGCCCGAGGAAAC
env-317   CUGCCUCUCCGGCAGGGAUCAUGCAGUCGGUGCAUCAGACUAGUUGCACGAAGGGUUGAUUGAAACAUGCCGAAGUAUAACAGGCCACGCAGAGGAUCCCUUGCAUACAGUCCAAGGA
env-318   UAGCCGGCCCGUCCGGCAGAGGAUAUGGCACAUCAACACUAGUUGCACGAAGGGUUGAUUGAAACAUGCCGAAAUAUAACAGGCCACGCAGAGGAUCCCUCGCGUACAGCCCGAGGAA
env-320   UGGCCGGCCCGUCCGGCAGAGGAUCUGGCACAUCAACACUAGUUGCACGAAGGGUUGAUUGAAACAUGCCGAAAUAUAACAGGCCACGCAGAGGAUCCCUCGCGUACAGCCCGAGGAA
env-321   UAGCCGGCCCGUCCGGCAGAGGAUAUGGCACAUCAACACUAGUUGCACGAAGGGUUGAUUGAAACAUGCCGAAAUAUAACAGGCCACGCAGAGGAUCCCUCGCGUACAGCCCGAGGAA
env-322   UNGCCGGCCCGUCCGGCAGAGGAUNUGGCACAUCAACACUAGUUGCACGAAGGGUUGAUUGAAACAUGCCGAAAUAUAACAGGCCACGCAGAGGAUCCCUCGCGUACAGCCCGAGGAA
Mpa-1-1   GCAGAGAAGGAUUGAUUAAGACAUGGCAAAUAUACACAGGCCACGUAGAGGAUCACUUGCAUACAGUCCUCGCAAGCGUGCAAAGAGCCAGGUUCCAAAGUAUCACUCUUGGCCUGCC
env-323   AGAGAA..................................................................................................................
env-324   AGCGAAAAAUAUCACACAGUCUCGGAUUUCAUAAAAAAUUCGUGUCGCAUUUGAGAAUAACCUACACGCAAAAAGGAUUGAUUACCACAUGCCAAAAAUCAACAGACCACGCAGAGGC

          ........................................................................................................................
          ........................................................................................................................
          ........................................................................................................................
```

```
env-325   CGGAAGCGAAGCUAUCACACAAUCUCGGAUUCAUCUCACAAUCGUGUCGCAAUGAGAAUAACCUACACGCAAAAAGGAUUGAUUCACACAUGCCAAAAAUCAACAGACCACGCCGCGG
env-326   CGGAAGCGAAGCUAUCACACAAUCUCGGAUUCAUCUCACAAUCGUGUCGCAAUGAGAAUAACCUA.......................................................
env-331   CCGGAAGCGAAGCUAUCACACAAUCUCGGAUUCAUCUCACAAUCGUGUCGCAAUGAGAAUAACCUACACGCAAAAAGGAUUGAUUCACACAUGCCAAAAAUCAACAGACCACGUCGCG
env-332   CGGAAGCGAAGCUAUCACACAAUCUCGGAUUCAUCUCACAAUCGUGUCGCAAUGAGAAUAACCUACACGCAAAAAGGAUUGAUUCACACAUGCCAAAAAUCAACAGACCACGUCGCGG
Mfo-1-1   CUGCAAAGGUUUCAGAAGAGUCUCGGAUGAAUCUCUCAAACGUGUCGCAAUGAGAACAACCUACACGCAAAAGUCAGGAUGGAUUAAUACAUGCCAAAAAUUAACAGACCACGCCGUG
env-333   GAAUCAUCUUUGAAAAGUUCGUGUCACAAAAAAACAAAAAAACACGAAUGAAAAGGAUUGAUUGACACAUGCCAAAGAUAAACAGACCACGUCGAGGCUCGCUGGCAUUCAGCCCACG
env-334   AUCUUUGAAAAGUUCGUGUCACGCAAAAUUUCGAAAAAAAACACGAAUGAAAAGGAUUGAUUGACACAUGCCAAAGAUAAACAGACCACGCCGAGGCUCGCUGGCAUUCAGCCCACGG
env-335   UCUUUGAAAAGUUCGUGUCACACAAAAUUUACAAAAAAACACGAAUGAAAAGGAUUGAUUGACACAUGCCAAAGAUAAACAGACCACGCCGAGGCUCGCUGGCAUUCAGCCCACGGAA
env-337   CAUCUUUGAAAAGUUCGUGUCACGCAAAAUUUCGAAAAAAAACACGAAUGAAAAGGAUUGAUUGACACAUGCCAAAGAUAAACAGACCACGCCGAGGCUCGCUGGCAUUCAGCCCACG
env-338   AUCAUCUUUGAAAAGUUCGUGUCAUAAAAAAAAUCAAAAAAAACACGAAUGAAAAGGAUUGAUUGACACAUGCCAAAGAUAAACAGACCACGUCGAGGCUCGCUGGCAUUCAGCCCAC
env-339   AUCAUCUUUGAAAAAUUCGUGUCACCAAAAAAAAUCAAAAAAACACGAAUGAAAAGGAUUGAUUGACACAUGCCAAAGAUAAACAGACCACGUCGAGGCUCGCUGGCAUUCAGCCCAC
CMe-1-1   UGAGAGCACUUCCGUCUCGGAUUCAUCUCUCUAGUUCGUGUCAAAAAGAUCCAAAACACGAAUGUACAGGAAUGAUUGACACAUGCCAAAGAUAAACAGACCACGCCGAGGCUCGCUG
env-340   ACUAACAGGUAUCACACGGUCUCGGAUUAAUCGUAAUUCGUGUCGAAUUGGGAAUAACCUACACGCAGAAAAAGGAUUGAUUAACACAUGCCAAAAAUAAACAGACCACGCCGUGGUU
env-341   CAAACACAGUUAUCACGCUGUCUCGGAUUAAUCGUAAUUCGUGUCACAAUGGGAAUCACCCACACGCACAAAAAGGAUUGAUUAACACAUGCCAAAAAUAAACAGACCACGCCGUGGU
env-349   CAAAAACACAGUUAUCACACUGUCUCGGAUUAAUCGUAAUUCGUGUCACAAUGGGAAUCACCCACACGCACAAAAAGGAUUGAUUAACACAUGCCAAAAAUAAACAGACCACGCCGUG
env-350   CAACAAACAACUGAUUAUAAAUCUCGGAUUAAUCUAAAAUUCGUGUCGCAAUGGGAACAACCUACACGCAGAAAAAGGAUGGAUUAACACAUGCCAAAAAUAAACCGACCACGCCGUG
env-351   AACAGACGAAGGCAUCGUCUCGGAUUAAUCGUAAUUCGUGUCAAAAUGGGAAUAACCUACACGCAGAAAAAGGAUUGAUUAACACAUGCCAAAAAUAAACAGACCACGCCGUGGUUCU
env-352   CACGAAGUUAUCACACAUCUCGGAUUUAAAAAAAAAAUUCGUGUCGCACUUGAGAAUAACCUACACGCAAAAAGGAUUGAUAAACACAUGCCAAAAAUCAACAGACCACGCCGCGGCU
env-353   UUAACCAGACAAAGUUAUCACACAGUCUCGGAUUAAUCAAAAAUUCGUGUCGCAUUAGGAAUAACCUACACGCAGAAAAAGGAUUGAUUAACACAUGCCAAAAAUUAACAGACCACGC
env-354   CUUAACCAGACAAAGUUAUCACACAGUCUCGGAUUAAUCAAAAAUUCGUGUCGCAUUAGGAAUAACCUACACGCAGAAAAAGGAUUGAUUAACACAUGCCAAAAAUAAACAGACCACG
env-355   UUAACCAGACAAAGUUAUCACACAGUCUCGGAUUAAUCAAAAAUUCGUGUCGCAUAAGGAAUAACCUACACGCAGAAAAAGGAUUGAUUAACACAUGCCAAAAAUUAACAGACCACGC
env-356   ........................................................................................................................
env-357   ........................................................................................................................
env-358   UCUUAACCAGACAACGUUAUCACACGGUCUCGGAUUAAUCAAAAAUUCGUGUCGCAAUGGGAAUAACCUACACGCAGAAAAAGGAUUGAUUAACACAUGCCAAAAAUAAACAGACCAC
env-359   AUCUUAACCAGACAACGUUAUCACACGGUCUCGGAUUAAUCAAAAAUUCGUGUCGCAAUGGGAAUAACCUACACGCAGAAAAAGGAUUGAUUAACACAUGCCAAAAAUAAACAGACCA
env-360   AUCUUAACCAGACAACGUUAUCACACGGUCUCGGAUUAAUCAAAAAUUCGUGUCGCAAUGGGAAUAACCUACACGCAGAAAAAGGAUUGAUUAACACAUGCCAAAAAUAAACAGACCA
env-361   AUCUUAACCAGACAACGUUAUCACACGGUCUCGGAUUAAUCAAAAAUUCGUGUCGCAAUGGGAAUAACCUACACGCAGAAAAAGGAUUGAUUAACACAUGCCAAAAAUAAACAGACCA
env-370   ........................................................................................................................
env-371   UUAACCAGAUACGGUUAUCACACCGUCUCGGAUUAAUCGUAAUUCGUGUCGCAAUGGGAAUAACCUACACGCAGAAAAAGGAUUGAUUAACACAUGCCAAAAAUAAACAGACCAC...
env-372   ........................................................................................................................
env-373   GUGUUGGCAUCAUAAACACAGUUAUCACACUGUCUCGGAUUAAUCGUAAUUCGUGUCAUAAUGGGAAUCACCCACACGCACAAAAAGGAUUGAUUAACACAUGCCAAAAAUAAACAGA
env-374   UUAACCAGACAAAGUUAUCACACGGUCUCGGAUUAAUCGUAUUUCGUGUCGAAAUGGGAAUAACCUACACGCAGAAAAAGGAUUGAUUAACACAUGCCAAAAAUAAACA.........
env-375   ........................................................................................................................
env-376   AUUUUGUUUCAAAAGCUCGGAUUAAUCUCAAAUUCGUGUCGCAAUCAGAAUAACCUA...............................................................
env-377   AACAAUAUUUUGUUUCAAAAGCUCGGAUUAAUCUCAAAUUCGUGUCGCAAUCAGAAUAACCUACACGCAAAUAAAGGAUUGAUUAACACAUGCCAAAAAUUAACAGACCUCGCCGUGG
env-378   ........................................................................................................................
env-379   AACAAUACGUUUCAAAAGCUCGGAUUAAUCUUAAAAAUCGUGUCGCAAUCAGAAUAACCUACACGCUAAAAAAGGAUUGAUUAACACAUGCCAAAAAUUAACAGACCCCGCCGUGGUU
env-380   CGAAAGCGAAGCUAUCACACAAUCUCGGAUUAAUCUCACAAACGUGUCGCAAUGAGAAUAACCUACACGCAAAAAGGAUUGAUUCACACAUGCCAAAAAUCAACAGACCACGCCGCGG
env-381   ........................................................................................................................
env-382   AACCGGAAGCGAAGCUAUCACACAAUCUCGGAUUAAUCUCACAAAACGUGUCGCAAUGAGAAUAACCUACACGCAAAAAGGAUUGAUUCACACAUGCCAAAAAUUAACAGACCACGCC
env-383   CCGAAAGCGAAGCUAUCACACAAUCUCGGAUUAAUCUCACAAACGUGUCGCAAUGAGAAUAACCUACACGCAAAAAGGAUUGAUUCACACAUGCCAAAAAUCAA..............
env-384   GAAGCGAAGCUAUCACACAAUCUCGGAUUAAUCUCAAAAAUCGUGUCGCAAUGAGAAUAACCUACACGCAAAAAGGAUUGAUUCACACAUGCCAAAAAUUAACAGACCACGCCGUGGC
env-385   ........................................................................................................................
env-386   UCACACAAUCUCGGAUUCAUCUCACAAAACGUGUCGCAAUGAGAAUAACCUACACGCAAAAAGGAUUGAUUCACACAUGCCAAAAAUCAACAGACCACGCCGCGGCUCUCUUGCAUUC
env-387   ........................................................................................................................
env-389   UUAACGGAAGCGAAGCUAUCACACAUCUCGGAUUAAUCUUAUCAAAACGUGUCGCAAUGAGAAUAACCUACACGCAAAAAGGAUUGAUUCACACAUGCCAAAAAUCAACAGACCACGC
env-390   AUAAACAA................................................................................................................
env-392   ........................................................................................................................
env-395   ........................................................................................................................
env-399   UCAUUGUUCCAAAAGUCUCGGAUUAAUCUCACAUACGUGUCGCUAUCAGAAUAACCUACACGCAGAAGAGGAUUGAUUUACACAUGCCAAAAAUUAACAGACCACGCCGUGGAUCUCU
env-400   ........................................................................................................................
env-402   UUUUGUUUCAAAAGCUCGGAUUAAUCUCAAAUUCGUGUCGCAAUCAGAAUAACCUA................................................................
env-403   ........................................................................................................................
env-405   ........................................................................................................................
env-406   ........................................................................................................................
env-407   ........................................................................................................................
Mli-2-1   AAAAUUUAGUUUGUAGGACUGAUCGAAAAAUGCCCCAAACAACCAGACCACGCAGGGGAUCCCUUGCAUACAGCCCCCGCAAGCGUGCAAAAAGCCAGGUACCAAAGUACCAGUCAUG
Mpe-1-1   UCGGAAAAAAGUUCAAAGAAUAAACAGGACUGAUUGAUAAAUGCCUGGAACAACCAGACCACGCAGAGGAUCACUUGCAUACAGCCCGCGCAAAAGGGCGAAGAGUCAGGUACCGAAG
Mpa-2-1   CUCGGAAAAAAGUUCAAAGAAUAAACAGGACUGAUUGAUAAAUGCCUGGAACAACCAGACCACGCAGAGGAUCACUUGCAUACAGCCCGCGCAAAAGGGCGAAGAGUCAGGUACCGAA
env-408   GGAAAAAAGUUCAAAGAACAAAUAGGACUGAUUGAUAAAUGCCUGGAAAAACCAGACCACGCAGAGGAUCACUUGCAUACAGCCCGCGCAAAAGGGCAAAGAGUCAGGUACCUAAGUA
Mmo-1-1   CACCCAAAAAUGUCCGAAUUAUCUUUCGAUACCGAAAUUCGCAUAAAUACGGCAUGACGCCCUCUGUGAUGCGAAAAACAACGGAAAAACGAAACAGGAUAUUCGGAAUCUCGGAAAA
env-409   CACCCAAAAAUGUCCGAAGUAUCUUUCGAUACCGAAAUUCGCAUAAAUACGGCAUGACGCCCUCUGUGAUGCGAAAAACAACGGAAAAACGAAACAGGAUAUUCGGAAUCUCGGAAAA
env-410   GGGUGCAGCGC.............................................................................................................
Mco-1-1   GCAAAAUCCGGGAUAAGGUAUCCGUCGCAUUUGGAAGCCCCUCGAAGGGGCUGGAGCAGCUCCUGCACGAUGAAGGCCACAAGCUUGAGGACCAUUCCGAUUGCGUGGUGAACUCCAU
env-413   ........................................................................................................................
env-414   CACAGAUUGGUGAGAGGACAAAGCAGUCCAAUCAUACUGCUGUGGUAUUUGGCUCACCCAGGCAAGGUUUAGAGACCAUACUGGACAAUGAGGGUUUCAAGCUCUCAGACUUCCAUUG
env-415   ........................................................................................................................
env-416   ........................................................................................................................
env-418   CAAACAUCGGUAGUAUGAUCAGACAAUCAUCAUUAACUGCUAUUGUAUUCGGAUCACCCAAACAAGGUGUGGAAUCGAUAUUGGCAAAUGAGGGAACACAACUCUCACAUAUCCAGUG
env-423   AACACGUCAGUGGUUUGAUCAGACAUUCAUCAUCAACUGCUAUUGUAUUCGGGUCACCCAAACAAGGUGUGGAAUCGAUAUUGGCAAAUGAGGGCACACAACUCUCACAAAUUCAGUG
env-425   AACAUGUCAGUGGUUUGAUCAGACAUUCAUCAUUAACUGCUAUUGUAUUCGGGUCACCCAAACAAGGUGUGGAAUCGAUAUUGGCAAAUGAGGGUACUCAACUCUCACAAAUUCAGUG
env-428   UGCAAAAGAUAGGAAAUGCAGGCCAAAGCGACCUUGCGGUGGCGUUCGGCUCCCCUGCACGAGGCGUUGAUGCAUUCUUGAGCCAGAGGCUCAUGGAAAGAUGCUGCAUGAUAAAUAC
env-429   UGCAAAGAUUAGGAAAAGCAGGCCAAAGCGACCUUGCGGUGGCAUUCGGCUCCCCUGCACGAGGCGUUGAUGCAUU............................................
Msh-1-1   UCGCUCUCAAAGGGCACAUGAGCGGCAGGGUCUGUGUGGUGUUCGGCUCCCAGAAGAGAGGCAUCAGAGAGAUGCUAAGUGCCCAUGUGGGCGAUGGAUGGAGGCAGCACUUCGAUGA
env-430   UGGGCCAAUCAGGCAAAAAUCGCGACCUUGCGGUGGCCUUCGGCUCACCCGCAAGGGGUCUGGAUGCGUUCUUGAGUGAAGAAUCAAUGAAGAGGUAUGAAAUGAUCAACACCAUUCC
env-433   CAAAAUUGGGCCACUCAGUUCAGAAGCGCGAUCUUGCGGUGGCCUUCGGCUCACCCGCACGAGGCGUGGAUGCCUUCUUGAGCAAAGAAUCGAUGGACAGGUAUGAAAUGAUCAACAC
env-434   UGGUCCAAUCGGGCUGCAAAUGCGACCUUGCGGUGGCCUUCGGCUCACCCGCACGAGGGCU...........................................................
env-435   ........................................................................................................................
env-436   UGGUCCAAUCGGGCUGCAAAUGCGACCUUGCGGUGGCCUUUGGCUCACCCGCACGAGGGCUAGAUGCGUUCUUGAGUAAAGAAUCAAUGAAGAGGUAUGAAAUGAUCAACACCAUUCC
env-437   ........................................................................................................................
env-439   UGGUCCAAUCGGGCUGCAAAUGCGACCUUGCGGUGGCCUUUGGCUCACCCGCACGAGGGCUAGAUGCGUUCUUGAGUAAAGAAUCAAUGAAGAGGUAUGAAAUGAUCAACACCAUUCC
env-440   ........................................................................................................................
env-441   UGGUCCAAUCGGGCAACAAACGCGACCUUGCGGUGGCCUUUGGCUCACCCGCACGAGGGCUAGAUGCGUUCUUGAGUAAAGAAUCAAUGAAGAGGUAUGAAAUGAUCAACACCAUUCC
env-442   UGGUCCAAUCGGGCAACAAA....................................................................................................

          ........................................................................................................................
          ........................................................................................................................
          ........................................................................................................................
```

| | |
|---|---|
| env-443 | CCAACAUGGGACACACCGGCUAUCAGCGCGAUCUUGCGGUCGUCUUCGGCUCUCCCUCACGGGGCGUAGAUGCAUUCUUGAGCAAAGAGUCAAUGGAAAGAUAUGAAAUGAUCAACAC |
| env-444 | GCAGAUCCAGUCAGAAGAGCGAUCUUGCAGUGGUAUUCGGCUCCCCUUCGCAAGGCGUUGAUGCGUUCUUGAGCGAUGAAGUCAAGGACAGGUAUUGCAUGAUUAACACCAUUCCGCA |
| env-445 | ...................................................................................................................... |
| env-446 | ...................................................................................................................... |
| env-448 | AGCGAUCCAGCAAGCGAGGCCUUGCGGUUGUCUUUGGCUCCCCAUCAAGAGGCGUUGAUGCGUUCUUGAGUCGAGAUAUGUUGGAGAGAUGUUGUAUGAUCAACACCAUUCCGCAUCA |
| env-449 | ...................................................................................................................... |
| env-450 | ...................................................................................................................... |
| env-451 | UGGGUCAAUUGGGCCAGAAGCGCGAUCUUGUGGUGGUAUUCGGCUCACCCGCACGAGGCGUAGAUGCGUUCUUGAGUUGUGAAGCAAUUGGCAGAUAUGACAUGAUUAACACCAUUCC |
| env-452 | ...................................................................................................................... |
| env-455 | UGGGUCAAUUGGACCAGAAGCGCGAUCUUGCGGUGGUAUUCGGCUCACCAGCACGAGGCGUAGAUGCGUUCUUGAGUAAGGAAGCUAUUGGCAGGUAUGACAUGAUUAACACCAUUCC |
| env-456 | UAGGUCGAUUGGGCCAGAAGCGCGAUCUUGCGGUGGUAUUCGGCUCACCCGCACGAGGCGUAGAUGCGUUCUUGAGUUGUGAAGCAAUGAACAGGUACGACAUGAUCAACACCAUUCC |
| env-460 | UGGGUCAACUAGGCCAGAAGCGUGAUCUUGCGGUGGUAUUCGGCUCACCCUCACGAGGCGUAGAUGCGUUCUUGAGCAAGGAAACAAUGGACAGGUAUGACAUGAUUAACACCAUUCC |
| env-462 | UGGGUCAACUAGGCCAGAAGCGUGAUCUUGCGGUGGUAUUCGGCUCACCCUCACGAGGCGUAGAUGCGUUCUUGAGCAAGGAAACAAUGGACAGGUAUGACAUGAUUAACACCAUUCC |
| env-464 | UGGGUCAACUAGGCCAGAAGCGUGAUCUUGCGGUGGUAUUCGGCUCACCCUCACGAGGCGUAGAUGCGUUCUUGAGCAAGGAAACAAUGGACAGGUAUGACAUGAUUAACACCAUUCC |
| env-465 | UAGGUCAAUUAGGCCAGAAGCGUGAUCUUGCGGUGGUAUUCGGCUCACCCUCACGAGGC........................................................... |
| env-466 | ...................................................................................................................... |
| env-467 | ...................................................................................................................... |
| env-468 | UCCAUGAGGGCCGAAGGAGAAGGGGUCUUGAGGUGGUCUUCGGCUCGCCCGCACGCGGCGUUGAUGCGUUCUUGAGCAGCGAAACGAUGGAAGGAUGUUGCAUGAUUAACACCAUACC |
| env-469 | ...................................................................................................................... |
| env-471 | UCCAUGAGGGCCGANGGAGANGGGGUCUUGAGGUGGUCUUCGGCUCGCCCGCACGCGGCGUUGAUGCGUUCUUGAGCAGCGAAACGAUGGAAGGAUGUUGCAUGAUUAACACCAUACC |
| env-472 | UCCAUGAGGGCCGAAGGAGAAGGGGUCUUGAGGUGGUCUUCGGCUCGCCCGCACGCGGCGUUGAUGCGUUCUUGAGCAGCGAAACGAUGGAAGGAUGUUGCAUGAUUAACACCAUACC |
| env-474 | UGCGUCAGGGCCAACGGCGUGGUCUGGAGGUGGUCUUCGGCUCGCCCGCACGCGGCGUUGAUGCGUUCUUGAGCAGCGAAACGAUGGAAGGAUGUUGCGUGAUUAACAC......... |
| env-475 | UCCGCCAGGGCCAAGGGCGAGGUCUGGAACUGGUCUACGGAUCGCCCGCUCGCGGCGUUGAUGCGUUCUUGAGCAGCGAAACGAUGGAAGGAUGUUGCGUGAUUAACACCAUACCGCA |
| Mha-1-1 | UCCGCCAGGGCCAAGGGCGAGGUCUGGAACUGGUCUACGGAUCGCCCGCUCGCGGCGUUGAUGCGUUCUUGAGCAGCGAAACGAUGGAAGGAUGUUGCGUGAUUAACACCAUACCGCA |
| env-476 | UCCGCCAGGGCCAAAGGAGAGGUCUGGAACUGGUCUACGGCUCGCCCGCUCGCGGCGUUGAUGCGUUCUUGAGCAGCGAAACGAUGGAAGGAUGUUGCGUGAUUAACACCAUUCCGCA |
| env-477 | UACGUCAAGGCCGAAAGCAUGGUCUGGAGGUGGUCUUCGGCUCGCCCGCACGCGGCGUUGAUGCAUUCUUGAGCAGCGAAACGAUGGAAGGAUGUUGCGUGAUUAACACCAUACCGCA |
| env-478 | ...................................................................................................................... |
| env-479 | CCGUACACUGCAACCUGGUGGCGAUAUUCGGUUCACCAGCUCGGGGCGUCGACCGGUUCUUGAGCGACGAAAUGAUGGAUAGAUGCUGCGUAAUCAACACAAUUCCGCAUCAGGGAAC |
| env-480 | CCGUACACCGCAACCUUGUGGCGAUAUUCGGUUCACCAGCUCGGGGCGUCGACCGGUUCUUGAGCGACGAAAUGAUGGAUAGAUGCUGCGUAAUCAACACAAUUCCGCAUCAGGGAAC |
| env-481 | UGUCCCUUAAGGACAGGGGAGACCUGUGUAUAGUGUUCGGUUCACCAAGGAAAGGACUACUGGAAAUGGUGGAGAGAAAUAUGUUCGACGCAGUGCUGAACACAAUACCAAAUCAAGG |
| env-482 | GUGUGAUAUUUGGCUCUGCCAGAAAAGGAGUAUUUGAUAUGGCAAAGCAGGAAAAAGUCGAACUGAAUGGAACAUGUUGGAAUUUCGUUCCUGAACAGGGGACAGAAAGUGUGAGAAC |
| env-483 | GCUUGACAACCUUCGUCUUUGGCGCUCCA......................................................................................... |
| env-484 | ...................................................................................................................... |
| env-485 | ...................................................................................................................... |
| env-486 | GCUUGACAACCUUCGUCUUUGGCGCUCCAAACCGCGGAGUAGAAGCGCUUUUAAACGAAGCCGGGCUAUCGAUCGAGGAAGCGUCAGACUUUGUCGUUAAUACGAUUCCUGACCAAGG |
| env-492 | GCUUGACAACCUUUGUCUUUGGCGCUCCAAACCGCGGAGUAGAAGCGCUUUUAAACGAAGCCGGGCUAUCGAUCGAGGAAGCGUCAGACUUUGUCGUUAAUACGAUUCCUGACCAAGG |
| env-495 | GCUUGA................................................................................................................ |
| env-504 | ...................................................................................................................... |
| env-505 | AUAAGGUAUACUUCGUGUCUCUGAUCCUGCAUCGACUAUCAUUCUGGCAGAACUGCCUGAAUGAUCUGAUACAGACAGAAGGACUGAUUGAAAAAUGCCAAACGUACACAGACCCCGC |
| env-508 | ACUCAUAUCCGGUGUAACAAUCGGCCGAUAUUAGGCGAGUUCAACCGAUUAGUUUCGGUUCCGGCCAAAAAAAUGGCCAACGUUCCAGUCAUUAGUGCAGGAUUGAUUGAACCAUGCC |
| env-509 | UCACUAUCUCCGGUAUACUCUGGCCAAAGAGUAGGCGAAUUGACCCGGAACUCCGGGAAAGGCCACCGCAGGUGGCUCCAGUGAACCGCACGUGAAGGAUUGAUUGAAACAUGCCCAA |
| Mta-1-1 | CACAUCUCCGGUAUGCUCUGGCCGAAGAUAGGCGAAAAAACCGGAAAUACCGGUAAAGGCCACAAGGAGUGGCCUGCGACACAAUGCAAGUGCAGGAUUGAUUGAAACAUGCCCAAGA |
| env-511 | CACAUCUCCGGUAUGCUCUGGCCGAAGAUAGGCGAAAAAACCGGAAAUACCGGUAAAGGCCACAAGGAGUGGCCUGCGACACAAUGCAAGUGCAGGAUUGAUUGAAACAUGCCCAAGA |
| env-512 | CACAUCUCCGGCAUGCUCUGGCCGAAGAUAGGCGAAAAAACCGGAAAUACCGGUAAAGGCCACAAGGAGUGGCCUGCGACACAAUGCAAGUGCAGGAUUGAUUGAAACAUGCCCAAGA |
| Hbo-1-1 | AGCGGGAAGGGGCGACCGUCGUCUUCGGUUCACCCGGUCGUGGGCUUCCAGACAUCCUCGGCAUCGACGCCGAGGAAGUCACAGUCGAACCCUCCGACGGUCCGGGGUUCGACCUCUG |
| Ngr-1-1 | AGGGCGACGGGAUGACCGUCGCCUUCGGCGCGCCCGAGAGAGGGCUGCCGGACAUCCUCGGAAUCGAGGCGUCUGCCAUCUCCUCGAGCGACGAGGGGGGCGACGACGGAGUCGAACC |
| Hje-1-1 | CCGACGGAACGACCGUCGCCUUCGGUUCGCCGGGCCGCGGCCUGCCGGACAUACUGGACGUGAACGCCGAGGCCGUCGGCGACCAGCCGGCGGACGGUGUCGAACCCGGUUCGGACGG |
| Nma-1-1 | AACGCGAUACUACCCGCGACGGAUUCACAAUCGCGUUCGGUGCGCCCGAGAGAGGGCUGCCUGCCAUCCUCGAACUCGAGGAUACCACCGUCGAGUCGUUGUCGGCAUGGCCAUCGUG |
| Hal-1-1 | CCGAUGCCGGCGGCAUGACCGUCGUCUUCGGGUCGCCGGGCCGUGGGCUUCCGGACAUGCUCGGGAUGCCUCCCGAGGAUGUUGCAGACGUCGAACCUUCCGUUGGUCCGGGGUUCGA |
| Nti-1-1 | AACGGGACGGGAUGACCGUCGCGUUCGGGGCGCCCGAACGAGGGCUGCCGGACAUCCUCGACAUCGAGGCUUCCGCUGUCGCCCCCGCGGAUGGAACGGGCGACGCGGACGACGGAGU |
| Npe-1-1 | AGGGCGAUGGGAUGACCGUCGCGUUCGGUGCGCCCGAACGAGGGCUGCCGGAUAUCCUCGAAAUCGAGGCAUCCGCCGUCGGGGCCCCGAAGGACGCGGCCGCCGGCGAGGAUGACGG |
| Hti-1-1 | CGUCGGCUGGCGACAUGACAGUCGCCUUUGGCGCGCCGGAGCGCGGCUUGCCGGCUAUCCUCGGCGUCGAACCGGGUGCUGUCGACCCGUGCACCGUGGACGAGUUGGCGGCAGCCGU |
| | ...................................................................................................................... |
| | ...................................................................................................................... |
| | ...................................................................................................................... |

### 3 L13-Bacteroidia

#### 3.1 Taxa

The taxonomy of each organism containing a putative L13-Bacteroidia RNA is listed, with abbreviations identifying each hit (e.g., "Eco-1-1" and "Eco-1-2" might hypothetically represent two distinct RNAs in *E. coli*). The abbreviations will be used to identify each individual L13-Bacteroidia RNA in Sections 3.2 and 3.4:.

| **abbrev. of hits** | **taxonomy of species** |
|---|---|
| Bsp-1-1 | Bacteria Bacteroidetes Bacteroidia Bacteroidales Bacteroidaceae *Bacteroides sp.* Ga6A1 |
| Bsp-2-1 | Bacteria Bacteroidetes Bacteroidia Bacteroidales Bacteroidaceae *Bacteroides sp.* Ga6A2 |
| Psp-1-1 | Bacteria Bacteroidetes Bacteroidia Bacteroidales Prevotellaceae Alloprevotella *Prevotella sp. oral taxon* 473 str. F0040 |
| Pcl-1-1 | Bacteria Bacteroidetes Bacteroidia Bacteroidales Prevotellaceae *Paraprevotella clara* YIT 11840 |
| Pxy-1-1 | Bacteria Bacteroidetes Bacteroidia Bacteroidales Prevotellaceae *Paraprevotella xylaniphila* YIT 11841 |
| Hse-1-1 | Bacteria Bacteroidetes Bacteroidia Bacteroidales Prevotellaceae Prevotella *Hallella seregens* ATCC 51272 |

| | |
|---|---|
| Pal-1-1 to Pal-1-2 | Bacteria Bacteroidetes Bacteroidia Bacteroidales Prevotellaceae *Prevotella albensis* DSM 11370 = JCM 12258 |
| Pam-1-1 | Bacteria Bacteroidetes Bacteroidia Bacteroidales Prevotellaceae *Prevotella amnii* CRIS 21A-A |
| Pam-2-1 | Bacteria Bacteroidetes Bacteroidia Bacteroidales Prevotellaceae *Prevotella amnii* DNF00058 |
| Pam-3-1 | Bacteria Bacteroidetes Bacteroidia Bacteroidales Prevotellaceae *Prevotella amnii* DSM 23384 |
| Pam-4-1 | Bacteria Bacteroidetes Bacteroidia Bacteroidales Prevotellaceae *Prevotella amnii* DSM 23384 = JCM 14753 |
| Pau-1-1 | Bacteria Bacteroidetes Bacteroidia Bacteroidales Prevotellaceae *Prevotella aurantiaca* JCM 15754 |
| Pba-1-1 to Pba-1-2 | Bacteria Bacteroidetes Bacteroidia Bacteroidales Prevotellaceae *Prevotella baroniae* DSM 16972 = JCM 13447 |
| Pba-2-1 | Bacteria Bacteroidetes Bacteroidia Bacteroidales Prevotellaceae *Prevotella baroniae* F0067 |
| Pbe-1-1 | Bacteria Bacteroidetes Bacteroidia Bacteroidales Prevotellaceae *Prevotella bergensis* DSM 17361 |
| Pbi-1-1 | Bacteria Bacteroidetes Bacteroidia Bacteroidales Prevotellaceae *Prevotella bivia* |
| Pbi-2-1 | Bacteria Bacteroidetes Bacteroidia Bacteroidales Prevotellaceae *Prevotella bivia* DNF00188 |
| Pbi-3-1 | Bacteria Bacteroidetes Bacteroidia Bacteroidales Prevotellaceae *Prevotella bivia* DNF00320 |
| Pbi-4-1 | Bacteria Bacteroidetes Bacteroidia Bacteroidales Prevotellaceae *Prevotella bivia* DNF00650 |
| Pbi-5-1 | Bacteria Bacteroidetes Bacteroidia Bacteroidales Prevotellaceae *Prevotella bivia* DSM 20514 |
| Pbi-6-1 | Bacteria Bacteroidetes Bacteroidia Bacteroidales Prevotellaceae *Prevotella bivia* JCVIHMP010 |
| Pbr-1-1 | Bacteria Bacteroidetes Bacteroidia Bacteroidales Prevotellaceae *Prevotella brevis* ATCC 19188 |
| Pbr-2-1 | Bacteria Bacteroidetes Bacteroidia Bacteroidales Prevotellaceae *Prevotella brevis* P6B11 |
| Pbr-3-1 | Bacteria Bacteroidetes Bacteroidia Bacteroidales Prevotellaceae *Prevotella bryantii* B14 |
| Pbr-4-1 | Bacteria Bacteroidetes Bacteroidia Bacteroidales Prevotellaceae *Prevotella bryantii* C21a |
| Pbu-1-1 | Bacteria Bacteroidetes Bacteroidia Bacteroidales Prevotellaceae *Prevotella buccae* |
| Pbu-2-1 | Bacteria Bacteroidetes Bacteroidia Bacteroidales Prevotellaceae *Prevotella buccae* ATCC 33574 |
| Pbu-3-1 | Bacteria Bacteroidetes Bacteroidia Bacteroidales Prevotellaceae *Prevotella buccae* D17 |
| Pbu-4-1 | Bacteria Bacteroidetes Bacteroidia Bacteroidales Prevotellaceae *Prevotella buccalis* ATCC 35310 |
| Pbu-5-1 | Bacteria Bacteroidetes Bacteroidia Bacteroidales Prevotellaceae *Prevotella buccalis* DNF00853 |
| Pbu-6-1 | Bacteria Bacteroidetes Bacteroidia Bacteroidales Prevotellaceae *Prevotella buccalis* DNF00985 |
| Pco-1-1 | Bacteria Bacteroidetes Bacteroidia Bacteroidales Prevotellaceae *Prevotella conceptionensis* 9403948 |
| Pco-2-1 | Bacteria Bacteroidetes Bacteroidia Bacteroidales Prevotellaceae *Prevotella copri* DSM 18205 |
| Pco-3-1 to Pco-3-2 | Bacteria Bacteroidetes Bacteroidia Bacteroidales Prevotellaceae *Prevotella corporis* DSM 18810 = JCM 8529 |
| Pde-1-1 | Bacteria Bacteroidetes Bacteroidia Bacteroidales Prevotellaceae *Prevotella dentalis* DSM 3688 |
| Pde-2-1 | Bacteria Bacteroidetes Bacteroidia Bacteroidales Prevotellaceae *Prevotella dentasini* JCM 15908 |
| Pde-3-1 | Bacteria Bacteroidetes Bacteroidia Bacteroidales Prevotellaceae *Prevotella denticola* CRIS 18C-A |
| Pde-4-1 | Bacteria Bacteroidetes Bacteroidia Bacteroidales Prevotellaceae *Prevotella denticola* DNF00960 |
| Pde-5-1 to Pde-5-2 | Bacteria Bacteroidetes Bacteroidia Bacteroidales Prevotellaceae *Prevotella denticola* DSM 20614 = JCM 13449 |
| Pde-6-1 | Bacteria Bacteroidetes Bacteroidia Bacteroidales Prevotellaceae *Prevotella denticola* F0289 |
| Pdi-1-1 | Bacteria Bacteroidetes Bacteroidia Bacteroidales Prevotellaceae *Prevotella disiens* DNF00882 |
| Pdi-2-1 | Bacteria Bacteroidetes Bacteroidia Bacteroidales Prevotellaceae *Prevotella disiens* FB035-09AN |
| Pdi-3-1 | Bacteria Bacteroidetes Bacteroidia Bacteroidales Prevotellaceae *Prevotella disiens* JCM 6334 = ATCC 29426 |
| Pen-1-1 | Bacteria Bacteroidetes Bacteroidia Bacteroidales Prevotellaceae *Prevotella enoeca* JCM 12259 |
| Pfa-1-1 | Bacteria Bacteroidetes Bacteroidia Bacteroidales Prevotellaceae *Prevotella falsenii* DSM 22864 = JCM 15124 |
| Pfu-1-1 | Bacteria Bacteroidetes Bacteroidia Bacteroidales Prevotellaceae *Prevotella fusca* JCM 17724 |
| Phi-1-1 | Bacteria Bacteroidetes Bacteroidia Bacteroidales Prevotellaceae *Prevotella histicola* F0411 |
| Phi-2-1 to Phi-2-2 | Bacteria Bacteroidetes Bacteroidia Bacteroidales Prevotellaceae *Prevotella histicola* JCM 15637 = DNF00424 |
| Pin-1-1 | Bacteria Bacteroidetes Bacteroidia Bacteroidales Prevotellaceae *Prevotella intermedia* 17 |
| Pin-2-1 | Bacteria Bacteroidetes Bacteroidia Bacteroidales Prevotellaceae *Prevotella intermedia* ATCC 25611 = DSM 20706 |
| Pin-3-1 | Bacteria Bacteroidetes Bacteroidia Bacteroidales Prevotellaceae *Prevotella intermedia* ZT |
| Plo-1-1 | Bacteria Bacteroidetes Bacteroidia Bacteroidales Prevotellaceae *Prevotella loescheii* DSM 19665 = JCM 12249 = ATCC 15930 |
| Pma-1-1 | Bacteria Bacteroidetes Bacteroidia Bacteroidales Prevotellaceae *Prevotella maculosa* DSM 19339 |
| Pma-2-1 | Bacteria Bacteroidetes Bacteroidia Bacteroidales Prevotellaceae *Prevotella maculosa* DSM 19339 = JCM 15638 |
| Pma-3-1 | Bacteria Bacteroidetes Bacteroidia Bacteroidales Prevotellaceae *Prevotella maculosa* OT 289 |
| Pma-4-1 | Bacteria Bacteroidetes Bacteroidia Bacteroidales Prevotellaceae *Prevotella marshii* DSM 16973 |
| Pma-5-1 | Bacteria Bacteroidetes Bacteroidia Bacteroidales Prevotellaceae *Prevotella marshii* DSM 16973 = JCM 13450 |
| Pme-1-1 | Bacteria Bacteroidetes Bacteroidia Bacteroidales Prevotellaceae *Prevotella melaninogenica* ATCC 25845 |
| Pme-2-1 | Bacteria Bacteroidetes Bacteroidia Bacteroidales Prevotellaceae *Prevotella melaninogenica* D18 |
| Pme-3-1 | Bacteria Bacteroidetes Bacteroidia Bacteroidales Prevotellaceae *Prevotella melaninogenica* DNF00666 |
| Pmi-1-1 | Bacteria Bacteroidetes Bacteroidia Bacteroidales Prevotellaceae *Prevotella micans* DSM 21469 |
| Pmi-2-1 | Bacteria Bacteroidetes Bacteroidia Bacteroidales Prevotellaceae *Prevotella micans* DSM 21469 = JCM 16134 |
| Pmi-3-1 | Bacteria Bacteroidetes Bacteroidia Bacteroidales Prevotellaceae *Prevotella micans* F0438 |
| Pmu-1-1 | Bacteria Bacteroidetes Bacteroidia Bacteroidales Prevotellaceae *Prevotella multiformis* DSM 16608 |
| Pmu-2-1 | Bacteria Bacteroidetes Bacteroidia Bacteroidales Prevotellaceae *Prevotella multisaccharivorax* DSM 17128 |
| Pna-1-1 | Bacteria Bacteroidetes Bacteroidia Bacteroidales Prevotellaceae *Prevotella nanceiensis* DSM 19126 |
| Pna-2-1 | Bacteria Bacteroidetes Bacteroidia Bacteroidales Prevotellaceae *Prevotella nanceiensis* DSM 19126 = JCM 15639 |
| Pni-1-1 | Bacteria Bacteroidetes Bacteroidia Bacteroidales Prevotellaceae *Prevotella nigrescens* ATCC 33563 |
| Pni-2-1 | Bacteria Bacteroidetes Bacteroidia Bacteroidales Prevotellaceae *Prevotella nigrescens* CC14M |
| Pni-3-1 | Bacteria Bacteroidetes Bacteroidia Bacteroidales Prevotellaceae *Prevotella nigrescens* F0103 |
| Por-1-1 | Bacteria Bacteroidetes Bacteroidia Bacteroidales Prevotellaceae *Prevotella oralis* ATCC 33269 |

| | |
|---|---|
| Por-2-1 | Bacteria Bacteroidetes Bacteroidia Bacteroidales Prevotellaceae *Prevotella oralis* CC98A |
| Por-3-1 | Bacteria Bacteroidetes Bacteroidia Bacteroidales Prevotellaceae *Prevotella oralis* HGA0225 |
| Por-4-1 | Bacteria Bacteroidetes Bacteroidia Bacteroidales Prevotellaceae *Prevotella oris* C735 |
| Por-5-1 | Bacteria Bacteroidetes Bacteroidia Bacteroidales Prevotellaceae *Prevotella oris* DSM 18711 |
| Por-6-1 | Bacteria Bacteroidetes Bacteroidia Bacteroidales Prevotellaceae *Prevotella oris* DSM 18711 = JCM 12252 |
| Por-7-1 | Bacteria Bacteroidetes Bacteroidia Bacteroidales Prevotellaceae *Prevotella oris* F0302 |
| Por-8-1 | Bacteria Bacteroidetes Bacteroidia Bacteroidales Prevotellaceae *Prevotella oryzae* DSM 17970 |
| Pou-1-1 | Bacteria Bacteroidetes Bacteroidia Bacteroidales Prevotellaceae *Prevotella oulorum* F0390 |
| Pou-2-1 | Bacteria Bacteroidetes Bacteroidia Bacteroidales Prevotellaceae *Prevotella oulorum* JCM 14966 |
| Ppa-1-1 | Bacteria Bacteroidetes Bacteroidia Bacteroidales Prevotellaceae *Prevotella pallens* ATCC 700821 |
| Ppa-2-1 | Bacteria Bacteroidetes Bacteroidia Bacteroidales Prevotellaceae *Prevotella paludivivens* DSM 17968 |
| Ppa-3-1 | Bacteria Bacteroidetes Bacteroidia Bacteroidales Prevotellaceae *Prevotella paludivivens* DSM 17968 = JCM 13650 |
| Ppl-1-1 | Bacteria Bacteroidetes Bacteroidia Bacteroidales Prevotellaceae *Prevotella pleuritidis* F0068 |
| Ppl-2-1 | Bacteria Bacteroidetes Bacteroidia Bacteroidales Prevotellaceae *Prevotella pleuritidis* JCM 14110 |
| Pru-1-1 | Bacteria Bacteroidetes Bacteroidia Bacteroidales Prevotellaceae *Prevotella ruminicola* 23 |
| Pru-2-1 | Bacteria Bacteroidetes Bacteroidia Bacteroidales Prevotellaceae *Prevotella ruminicola* Ga6B6 |
| Psa-1-1 | Bacteria Bacteroidetes Bacteroidia Bacteroidales Prevotellaceae *Prevotella saccharolytica* F0055 |
| Psa-2-1 | Bacteria Bacteroidetes Bacteroidia Bacteroidales Prevotellaceae *Prevotella salivae* DSM 15606 |
| Psa-3-1 | Bacteria Bacteroidetes Bacteroidia Bacteroidales Prevotellaceae *Prevotella salivae* F0493 |
| Psc-1-1 | Bacteria Bacteroidetes Bacteroidia Bacteroidales Prevotellaceae *Prevotella scopos* JCM 17725 |
| Psh-1-1 | Bacteria Bacteroidetes Bacteroidia Bacteroidales Prevotellaceae *Prevotella shahii* DSM 15611 = JCM 12083 |
| Psp-2-1 | Bacteria Bacteroidetes Bacteroidia Bacteroidales Prevotellaceae *Prevotella sp.* 109 |
| Psp-3-1 | Bacteria Bacteroidetes Bacteroidia Bacteroidales Prevotellaceae *Prevotella sp.* AGR2160 |
| Psp-4-1 | Bacteria Bacteroidetes Bacteroidia Bacteroidales Prevotellaceae *Prevotella sp.* BV3P1 |
| Psp-5-1 | Bacteria Bacteroidetes Bacteroidia Bacteroidales Prevotellaceae *Prevotella sp.* C561 |
| Psp-6-1 | Bacteria Bacteroidetes Bacteroidia Bacteroidales Prevotellaceae *Prevotella sp.* F0091 |
| Psp-7-1 | Bacteria Bacteroidetes Bacteroidia Bacteroidales Prevotellaceae *Prevotella sp.* FD3004 |
| Psp-8-1 | Bacteria Bacteroidetes Bacteroidia Bacteroidales Prevotellaceae *Prevotella sp.* HJM029 |
| Psp-9-1 | Bacteria Bacteroidetes Bacteroidia Bacteroidales Prevotellaceae *Prevotella sp.* HUN102 |
| Psp-10-1 | Bacteria Bacteroidetes Bacteroidia Bacteroidales Prevotellaceae *Prevotella sp.* ICM33 |
| Psp-11-1 | Bacteria Bacteroidetes Bacteroidia Bacteroidales Prevotellaceae *Prevotella sp.* MA2016 |
| Psp-12-1 | Bacteria Bacteroidetes Bacteroidia Bacteroidales Prevotellaceae *Prevotella sp.* MSX73 |
| Psp-23-1 | Bacteria Bacteroidetes Bacteroidia Bacteroidales Prevotellaceae *Prevotella sp. oral taxon* 299 str. F0039 |
| Psp-24-1 | Bacteria Bacteroidetes Bacteroidia Bacteroidales Prevotellaceae *Prevotella sp. oral taxon* 306 str. F0472 |
| Psp-25-1 | Bacteria Bacteroidetes Bacteroidia Bacteroidales Prevotellaceae *Prevotella sp. oral taxon* 317 str. F0108 |
| Psp-26-1 | Bacteria Bacteroidetes Bacteroidia Bacteroidales Prevotellaceae *Prevotella sp. oral taxon* 472 str. F0295 |
| Psp-13-1 | Bacteria Bacteroidetes Bacteroidia Bacteroidales Prevotellaceae *Prevotella sp.* P4-65 |
| Psp-14-1 | Bacteria Bacteroidetes Bacteroidia Bacteroidales Prevotellaceae *Prevotella sp.* P4-76 |
| Psp-15-1 | Bacteria Bacteroidetes Bacteroidia Bacteroidales Prevotellaceae *Prevotella sp.* P5-119 |
| Psp-16-1 | Bacteria Bacteroidetes Bacteroidia Bacteroidales Prevotellaceae *Prevotella sp.* P5-125 |
| Psp-17-1 | Bacteria Bacteroidetes Bacteroidia Bacteroidales Prevotellaceae *Prevotella sp.* P5-60 |
| Psp-18-1 | Bacteria Bacteroidetes Bacteroidia Bacteroidales Prevotellaceae *Prevotella sp.* P6B1 |
| Psp-19-1 | Bacteria Bacteroidetes Bacteroidia Bacteroidales Prevotellaceae *Prevotella sp.* P6B4 |
| Psp-20-1 | Bacteria Bacteroidetes Bacteroidia Bacteroidales Prevotellaceae *Prevotella sp.* RM4 |
| Psp-21-1 | Bacteria Bacteroidetes Bacteroidia Bacteroidales Prevotellaceae *Prevotella sp.* S7 MS 2 |
| Psp-22-1 | Bacteria Bacteroidetes Bacteroidia Bacteroidales Prevotellaceae *Prevotella sp.* S7-1-8 |
| Pst-1-1 | Bacteria Bacteroidetes Bacteroidia Bacteroidales Prevotellaceae *Prevotella stercorea* DSM 18206 |
| Pti-1-1 | Bacteria Bacteroidetes Bacteroidia Bacteroidales Prevotellaceae *Prevotella timonensis* 4401737 = DSM 22865 = JCM 15640 |
| Pti-2-1 | Bacteria Bacteroidetes Bacteroidia Bacteroidales Prevotellaceae *Prevotella timonensis* CRIS 5C-B1 |
| Pti-3-1 | Bacteria Bacteroidetes Bacteroidia Bacteroidales Prevotellaceae *Prevotella timonensis* S9-PR14 |
| Pve-1-1 | Bacteria Bacteroidetes Bacteroidia Bacteroidales Prevotellaceae *Prevotella veroralis* DSM 19559 |
| Pve-2-1 | Bacteria Bacteroidetes Bacteroidia Bacteroidales Prevotellaceae *Prevotella veroralis* DSM 19559 = JCM 6290 |
| Pve-3-1 | Bacteria Bacteroidetes Bacteroidia Bacteroidales Prevotellaceae *Prevotella veroralis* F0319 |
| env-1 to env-1259 | environmental samples |

#### 3.2 Gene contexts

Each L13-Bacteroidia RNA (indicated by "RNA→") is listed. For each hit, the downstream genes predicted to reside in a regulated operon are listed. If the nearest downstream gene is encoding in the opposite strand (and therefore presumed to not be a part of a regulated operon), then that gene is still depicted. Some environmental sequences and some RefSeq entries lack gene annotations, and so no genes are available for such sequences. The direction of each gene is indicated with an arrow (→), and each predicted conserved domain in the gene is named. Conserved domains associated with more than one L13-Bacteroidia RNA are assigned a color; other domains are gray. Information about these conserved domains is given in Section 3.3. The accession of the sequence containing each L13-Bacteroidia RNA is given in the column named "Seq. accession". Accessions beginning with "NC_", "NS_", "NW_" or "NZ_" are contained in RefSeq. Other accession refer to environmental samples. Nucleotide coordinates are given for the 5′ and 3′ boundaries of each L13-Bacteroidia RNA. If the 5′ coordinate is greater than the 3′ coordinate, the RNA is present on the reverse-complement strand of the containing genomic DNA sequence. Each hit is denoted by an abbreviation (like "Eco-1-1") that refers to a taxonomy given in Section 3.1.

| abbrev. | Seq. accession | | 5′ at | 3′ at | genes |
|---|---|---|---|---|---|
| env-1 | AMCI01008330.1 | - | 1066 | 991 | RNA→ rplM (PRK09216)Ribosomal_L13 (pfam00572)→ rpsI (PRK00132)Ribosomal_S9 (pfam00380)→ |
| env-2 | AUXO012860029.1 | - | 1571 | 1495 | RNA→ rplM (PRK09216)Ribosomal_L13 (pfam00572)→ rpsI (PRK00132)Ribosomal_S9 (pfam00380)→ |
| env-3 | SRS011529_WUGC_scaffold_27015 | - | 1315 | 1235 | RNA→ rplM (PRK09216)Ribosomal_L13 (pfam00572)→ rpsI (PRK00132)Ribosomal_S9 (pfam00380)→ |
| env-4 | RUMENNODE_4362630_1 | + | 14588 | 14659 | RNA→ rplM (PRK09216)Ribosomal_L13 (pfam00572)→ rplM (PRK09216)Ribosomal_L13 (pfam00572)→ rpsI (PRK00132)Ribosomal_S9 (pfam00380)→ |
| env-5 | scaffold15734_5_MH0046 | + | 243 | 321 | RNA→ rplM (PRK09216)Ribosomal_L13 (pfam00572)→ rpsI (PRK00132)Ribosomal_S9 (pfam00380)→ |
| env-6 | SRMUA_GNVGR9Q01D7VOE | - | 237 | 168 | RNA→ rplM (PRK09216)RplM (COG0102)→ |
| env-7 | AUXO010978888.1 | + | 545 | 616 | RNA→ rplM (PRK09216)Ribosomal_L13 (pfam00572)→ rpsI (PRK00132)Ribosomal_S9 (pfam00380)→ |
| env-8 | SRS048164_C2255498 | + | 534 | 617 | RNA→ |
| env-9 | scaffold6072_39_MH0053 | + | 450 | 533 | RNA→ |
| env-10 | SRS019910_WUGC_scaffold_74502 | + | 208 | 291 | RNA→ rplM (PRK09216)Ribosomal_L13 (pfam00572)→ rpsI (PRK00132)Ribosomal_S9 (pfam00380)→ |
| env-11 | scaffold204458_1_MH0006 | - | 1372 | 1304 | RNA→ rplM (PRK09216)Ribosomal_L13 (pfam00572)→ rpsI (PRK00132)Ribosomal_S9 (pfam00380)→ |
| env-12 | UnmappedStool_Broad_scaffold_711238 | - | 140 | 72 | RNA→ hypo→ |
| env-13 | 4491686.3.479386 | + | 127 | 205 | RNA→ |
| env-14 | scaffold15346_2_O2.UC-24 | + | 302 | 371 | RNA→ rplM (PRK09216)Ribosomal_L13 (pfam00572)→ |
| env-15 | HCF12C_103092 | - | 189 | 116 | RNA→ |
| env-16 | AUXO015828536.1 | + | 340 | 412 | RNA→ rplM (PRK09216)Ribosomal_L13 (pfam00572)→ |
| env-17 | DOM022_scaffold34730_2 | - | 1753 | 1684 | RNA→ rplM (PRK09216)Ribosomal_L13 (pfam00572)→ rpsI (PRK00132)Ribosomal_S9 (pfam00380)→ |
| env-18 | scaffold19504_1_MH0036 | - | 69802 | 69732 | RNA→ rplM (PRK09216)Ribosomal_L13 (pfam00572)→ rpsI (PRK00132)Ribosomal_S9 (pfam00380)→ |
| env-19 | scaffold4731_12_MH0057 | - | 597 | 527 | RNA→ rplM (PRK09216)Ribosomal_L13 (pfam00572)→ |
| env-20 | DLM014_scaffold3130_5 | - | 5881 | 5811 | RNA→ rplM (PRK09216)Ribosomal_L13 (pfam00572)→ rpsI (PRK00132)Ribosomal_S9 (pfam00380)→ |
| Pxy-1-1 | NZ_GL883827.1 | - | 169388 | 169318 | RNA→ rplM (PRK09216)Ribosomal_L13 (pfam00572)→ rpsI (PRK00132)Ribosomal_S9 (pfam00380)→ |
| env-21 | SRS048164_WUGC_scaffold_29212 | + | 2023 | 2093 | RNA→ rplM (PRK09216)Ribosomal_L13 (pfam00572)→ |
| env-22 | scaffold5875_2_MH0020 | + | 21461 | 21531 | RNA→ rplM (PRK09216)Ribosomal_L13 (pfam00572)→ rpsI (PRK00132)Ribosomal_S9 (pfam00380)→ |
| env-23 | scaffold18015_8_MH0044 | + | 3289 | 3359 | RNA→ rplM (PRK09216)Ribosomal_L13 (pfam00572)→ rpsI (PRK00132)Ribosomal_S9 (pfam00380)→ |
| env-24 | scaffold39182_1_MH0047 | - | 395 | 325 | RNA→ rplM (PRK09216)Ribosomal_L13 (pfam00572)→ |
| env-25 | scaffold33768_3_MH0073 | + | 2463 | 2533 | RNA→ rplM (PRK09216)Ribosomal_L13 (pfam00572)→ |
| env-26 | scaffold6160_1_V1.UC-19 | + | 273 | 343 | RNA→ rplM (PRK09216)Ribosomal_L13 (pfam00572)→ |
| env-27 | scaffold2340_6_MH0035 | + | 477 | 547 | RNA→ rplM (PRK09216)Ribosomal_L13 (pfam00572)→ |
| env-28 | scaffold54231_2_MH0050 | + | 1169 | 1239 | RNA→ rplM (PRK09216)Ribosomal_L13 (pfam00572)→ |
| env-29 | scaffold58113_4_MH0056 | - | 1209 | 1139 | RNA→ rplM (PRK09216)Ribosomal_L13 (pfam00572)→ rpsI (PRK00132)Ribosomal_S9 (pfam00380)→ |
| env-30 | scaffold2091_1_MH0058 | - | 156 | 86 | RNA→ |
| env-31 | scaffold16985_10_MH0059 | - | 522 | 452 | RNA→ rplM (PRK09216)Ribosomal_L13 (pfam00572)→ |
| env-32 | scaffold41222_2_MH0070 | - | 522 | 452 | RNA→ rplM (PRK09216)Ribosomal_L13 (pfam00572)→ |
| env-33 | scaffold1773_9_MH0082 | + | 1494 | 1564 | RNA→ rplM (PRK09216)Ribosomal_L13 (pfam00572)→ rpsI (PRK00132)Ribosomal_S9 (pfam00380)→ rpsB (PRK05299)Ribosomal_S2 (pfam00318)→ |
| env-34 | scaffold14401_2_O2.UC-12 | + | 160 | 230 | RNA→ rplM (PRK09216)Ribosomal_L13 (pfam00572)→ rpsI (PRK00132)Ribosomal_S9 (pfam00380)→ |
| env-35 | 4491401.3_NODE_125 | + | 6382 | 6452 | RNA→ rplM (PRK09216)Ribosomal_L13 (pfam00572)→ rpsI (PRK00132)Ribosomal_S9 (pfam00380)→ |
| env-36 | 4491414.3_NODE_4995 | + | 3654 | 3724 | RNA→ rplM (PRK09216)Ribosomal_L13 (pfam00572)→ rpsI (PRK00132)Ribosomal_S9 (pfam00380)→ |
| env-37 | 4491415.3_NODE_16304 | - | 35739 | 35669 | RNA→ rplM (PRK09216)Ribosomal_L13 (pfam00572)→ rpsI (PRK00132)Ribosomal_S9 (pfam00380)→ |
| env-38 | 4491417.3_NODE_72813 | + | 28708 | 28778 | RNA→ rplM (PRK09216)Ribosomal_L13 (pfam00572)→ rpsI (PRK00132)Ribosomal_S9 (pfam00380)→ |
| env-39 | 4491419.3_NODE_42377 | - | 198572 | 198502 | RNA→ rplM (PRK09216)Ribosomal_L13 (pfam00572)→ rpsI (PRK00132)Ribosomal_S9 (pfam00380)→ |
| env-40 | 4491477.3_NODE_70256 | - | 8117 | 8047 | RNA→ rplM (PRK09216)Ribosomal_L13 (pfam00572)→ |

| | | | | | |
|---|---|---|---|---|---|
| env-41 | DLM006_scaffold552_72 | - | 772 | 702 | **RNA**→ rplM (PRK09216)Ribosomal_L13 (pfam00572)→ rpsI (PRK00132)Ribosomal_S9 (pfam00380)→ |
| env-42 | DLM028_scaffold52693_6 | + | 193 | 263 | **RNA**→ rplM (PRK09216)Ribosomal_L13 (pfam00572)→ rpsI (PRK00132)Ribosomal_S9 (pfam00380)→ |
| env-43 | DOM010_scaffold9237_5 | + | 12401 | 12471 | **RNA**→ rplM (PRK09216)Ribosomal_L13 (pfam00572)→ rpsI (PRK00132)Ribosomal_S9 (pfam00380)→ |
| env-44 | NLM008_scaffold4989_13 | + | 788 | 858 | **RNA**→ rplM (PRK09216)Ribosomal_L13 (pfam00572)→ |
| env-45 | SRS011529_WUGC_scaffold_21913 | + | 1369 | 1439 | **RNA**→ rplM (PRK09216)Ribosomal_L13 (pfam00572)→ rpsI (PRK00132)Ribosomal_S9 (pfam00380)→ |
| env-46 | SRS013521_LANL_scaffold_6003 | - | 8813 | 8743 | **RNA**→ rplM (PRK09216)Ribosomal_L13 (pfam00572)→ rpsI (PRK00132)Ribosomal_S9 (pfam00380)→ |
| env-47 | SRS013687_C2637478 | - | 685 | 615 | **RNA**→ rplM (PRK09216)Ribosomal_L13 (pfam00572)→ |
| env-48 | SRS013800_Baylor_scaffold_15627 | + | 1409 | 1479 | **RNA**→ rplM (PRK09216)Ribosomal_L13 (pfam00572)→ |
| env-49 | SRS015133_WUGC_scaffold_36455 | - | 685 | 615 | **RNA**→ rplM (PRK09216)Ribosomal_L13 (pfam00572)→ |
| env-50 | SRS015782_WUGC_scaffold_38346 | + | 1363 | 1433 | **RNA**→ rplM (PRK09216)Ribosomal_L13 (pfam00572)→ |
| env-51 | SRS017307_Baylor_scaffold_15330 | - | 1112 | 1042 | **RNA**→ rplM (PRK09216)Ribosomal_L13 (pfam00572)→ rpsI (PRK00132)Ribosomal_S9 (pfam00380)→ |
| env-52 | SRS017433_Baylor_scaffold_19937 | - | 547 | 477 | **RNA**→ rplM (PRK09216)Ribosomal_L13 (pfam00572)→ |
| env-53 | SRS017701_Baylor_scaffold_13140 | - | 970 | 900 | **RNA**→ rplM (PRK09216)Ribosomal_L13 (pfam00572)→ rpsI (PRK00132)Ribosomal_S9 (pfam00380)→ |
| env-54 | SRS023914_C2116633 | - | 685 | 615 | **RNA**→ rplM (PRK09216)Ribosomal_L13 (pfam00572)→ |
| env-55 | SRS024435_LANL_scaffold_1171 | - | 1412 | 1342 | **RNA**→ rplM (PRK09216)Ribosomal_L13 (pfam00572)→ rpsI (PRK00132)Ribosomal_S9 (pfam00380)→ |
| env-56 | SRS045004_LANL_scaffold_14319 | + | 1359 | 1429 | **RNA**→ rplM (PRK09216)Ribosomal_L13 (pfam00572)→ rpsI (PRK00132)Ribosomal_S9 (pfam00380)→ |
| env-57 | SRS045645_C2019336 | + | 871 | 941 | **RNA**→ |
| env-58 | SRS047044_C1551280 | - | 547 | 477 | **RNA**→ rplM (PRK09216)Ribosomal_L13 (pfam00572)→ |
| env-59 | SRS049712_C15572285 | + | 896 | 966 | **RNA**→ rplM (PRK09216)Ribosomal_L13 (pfam00572)→ |
| env-60 | SRS049995_LANL_scaffold_2946 | - | 33940 | 33870 | **RNA**→ rplM (PRK09216)Ribosomal_L13 (pfam00572)→ rpsI (PRK00132)Ribosomal_S9 (pfam00380)→ |
| env-61 | SRS063985_LANL_scaffold_39061 | - | 685 | 615 | **RNA**→ rplM (PRK09216)Ribosomal_L13 (pfam00572)→ |
| env-62 | SRS078176_C2299272 | - | 333 | 263 | **RNA**→ rplM (PRK09216)Ribosomal_L13 (pfam00572)→ |
| Pcl-1-1 | NZ_JH376625.1 | + | 66716 | 66786 | **RNA**→ rplM (PRK09216)Ribosomal_L13 (pfam00572)→ rpsI (PRK00132)Ribosomal_S9 (pfam00380)→ |
| env-63 | scaffold5721_2_MH0077 | - | 792 | 722 | **RNA**→ rplM (PRK09216)Ribosomal_L13 (pfam00572)→ rpsI (PRK00132)Ribosomal_S9 (pfam00380)→ |
| env-64 | scaffold32997_1_MH0002 | - | 109 | 39 | **RNA**→ |
| env-65 | scaffold77706_1_MH0069 | - | 138 | 68 | **RNA**→ |
| env-66 | NLM006_scaffold34944_6 | + | 1503 | 1573 | **RNA**→ rplM (PRK09216)Ribosomal_L13 (pfam00572)→ rpsI (PRK00132)Ribosomal_S9 (pfam00380)→ |
| env-67 | NOF008_scaffold67812_2 | - | 140 | 70 | **RNA**→ |
| env-68 | NOF010_scaffold32157_6 | + | 458 | 528 | **RNA**→ rplM (PRK09216)Ribosomal_L13 (pfam00572)→ rpsI (PRK00132)Ribosomal_S9 (pfam00380)→ |
| env-69 | SRS017821_Baylor_scaffold_12585 | + | 5448 | 5518 | **RNA**→ rplM (PRK09216)Ribosomal_L13 (pfam00572)→ rpsI (PRK00132)Ribosomal_S9 (pfam00380)→ |
| env-70 | AUXO011504091.1 | - | 895 | 820 | **RNA**→ rplM (PRK09216)Ribosomal_L13 (pfam00572)→ |
| env-71 | AUXO012576065.1 | + | 558 | 627 | **RNA**→ rplM (PRK09216)Ribosomal_L13 (pfam00572)→ rpsI (PRK00132)Ribosomal_S9 (pfam00380)→ |
| env-72 | AUXO017805936.1 | + | 175 | 245 | **RNA**→ rplM (PRK09216)Ribosomal_L13 (pfam00572)→ |
| env-73 | AUXO015702136.1 | - | 1049 | 974 | **RNA**→ rplM (PRK09216)Ribosomal_L13 (pfam00572)→ rpsI (PRK00132)Ribosomal_S9 (pfam00380)→ |
| env-74 | AUXO010123457.1 | + | 71 | 143 | **RNA**→ rplM (PRK09216)Ribosomal_L13 (pfam00572)→ rpsI (PRK00132)Ribosomal_S9 (pfam00380)→ |
| env-75 | CAM_READ_0099979911 | + | 161 | 230 | **RNA**→ rplM (PRK09216)Ribosomal_L13 (pfam00572)→ |
| env-76 | NOM008_scaffold29131_6 | - | 3871 | 3801 | **RNA**→ rplM (PRK09216)Ribosomal_L13 (pfam00572)→ rpsI (PRK00132)Ribosomal_S9 (pfam00380)→ |
| env-77 | AUXO013980254.1 | - | 161 | 92 | **RNA**→ hypo→ |
| env-78 | AUXO014577836.1 | - | 989 | 916 | **RNA**→ rplM (PRK09216)Ribosomal_L13 (pfam00572)→ rpsI (PRK00132)Ribosomal_S9 (pfam00380)→ |
| env-79 | 2013414608 | - | 600 | 529 | **RNA**→ rplM (PRK09216)Ribosomal_L13 (pfam00572)→ rplM (PRK09216)Ribosomal_L13 (pfam00572)→ |
| env-80 | pre-rum113-01766 | + | 19308 | 19380 | **RNA**→ rplM (PRK09216)Ribosomal_L13 (pfam00572)→ rpsI (PRK00132)Ribosomal_S9 (pfam00380)→ |
| Pbr-3-1 | NZ_ADWO01000013.1 | - | 1784 | 1707 | **RNA**→ rplM (PRK09216)Ribosomal_L13 (pfam00572)→ rplM (PRK09216)Ribosomal_L13 (pfam00572)→ rpsI (PRK00132)Ribosomal_S9 (pfam00380)→ rpsI (PRK00132)Ribosomal_S9 (pfam00380)→ |
| Pbr-4-1 | NZ_AUKF01000007.1 | - | 102040 | 101963 | **RNA**→ rplM (PRK09216)Ribosomal_L13 (pfam00572)→ rpsI (PRK00132)Ribosomal_S9 (pfam00380)→ |
| env-81 | AUXO013781889.1 | + | 110 | 187 | **RNA**→ rplM (PRK09216)Ribosomal_L13 (pfam00572)→ rpsI (PRK00132)Ribosomal_S9 (pfam00380)→ |
| env-82 | AUXO016535493.1 | + | 268 | 342 | **RNA**→ rplM (PRK09216)Ribosomal_L13 (pfam00572)→ rpsI (PRK00132)Ribosomal_S9 (pfam00380)→ |
| env-83 | AUXO012398056.1 | + | 1683 | 1756 | **RNA**→ rplM (PRK09216)Ribosomal_L13 (pfam00572)→ rpsI (PRK00132)Ribosomal_S9 (pfam00380)→ |
| env-84 | scaffold11907_5_MH0040 | - | 2845 | 2774 | **RNA**→ rplM (PRK09216)Ribosomal_L13 (pfam00572)→ rpsI (PRK00132)Ribosomal_S9 (pfam00380)→ |
| env-85 | 4491405.3_NODE_26186 | + | 3884 | 3955 | **RNA**→ rplM (PRK09216)Ribosomal_L13 (pfam00572)→ rpsI (PRK00132)Ribosomal_S9 (pfam00380)→ |
| env-86 | scaffold25515_12_MH0059 | + | 284 | 355 | **RNA**→ rplM (PRK09216)Ribosomal_L13 (pfam00572)→ rpsI (PRK00132)Ribosomal_S9 (pfam00380)→ |
| env-87 | AUXO015466288.1 | + | 1512 | 1584 | **RNA**→ rplM (PRK09216)Ribosomal_L13 (pfam00572)→ rpsI (PRK00132)Ribosomal_S9 (pfam00380)→ |
| Pal-1-1 | NZ_AUFP01000001.1 | - | 13559 | 13481 | **RNA**→ rplM (PRK09216)Ribosomal_L13 (pfam00572)→ rpsI (PRK00132)Ribosomal_S9 (pfam00380)→ |

| | | | | | |
|---|---|---|---|---|---|
| Pal-1-2 | NZ_BAJD01000011.1 | + | 60623 | 60701 | **RNA**→ rplM (PRK09216)Ribosomal_L13 (pfam00572)→ rpsI (PRK00132)Ribosomal_S9 (pfam00380)→ |
| env-88 | SRS062878_C3036997 | - | 256 | 183 | **RNA**→ hypo→ rplM (PRK09216)RplM (COG0102)→ |
| env-89 | Tongue_dor_WUGI_scaffold_10412 | + | 328 | 401 | **RNA**→ rplM (PRK09216)Ribosomal_L13 (pfam00572)→ rpsI (PRK00132)Ribosomal_S9 (pfam00380)→ |
| env-90 | SRS014470_WUGC_scaffold_46671 | - | 1515 | 1442 | **RNA**→ rplM (PRK09216)Ribosomal_L13 (pfam00572)→ rpsI (PRK00132)Ribosomal_S9 (pfam00380)→ |
| env-91 | SRS018439_C2372430 | + | 460 | 533 | **RNA**→ rplM (PRK09216)Ribosomal_L13 (pfam00572)→ |
| env-92 | SRS019607_WUGC_scaffold_53821 | - | 2535 | 2462 | **RNA**→ rplM (PRK09216)Ribosomal_L13 (pfam00572)→ rpsI (PRK00132)Ribosomal_S9 (pfam00380)→ |
| env-93 | SRS019894_C3844330 | - | 310 | 237 | **RNA**→ rplM (PRK09216)Ribosomal_L13 (cd00392)→ |
| env-94 | SRS047219_C2319273 | + | 489 | 562 | **RNA**→ rplM (PRK09216)Ribosomal_L13 (pfam00572)→ rpsI (PRK00132)Ribosomal_S9 (pfam00380)→ |
| env-95 | SRS057539_LANL_scaffold_61402 | - | 1835 | 1762 | **RNA**→ rplM (PRK09216)Ribosomal_L13 (pfam00572)→ rpsI (PRK00132)Ribosomal_S9 (pfam00380)→ |
| env-96 | SRS057791_C3471145 | + | 135 | 208 | **RNA**→ |
| env-97 | SRS062544_LANL_scaffold_41679 | - | 1006 | 933 | **RNA**→ rplM (PRK09216)Ribosomal_L13 (pfam00572)→ rpsI (PRK00132)Ribosomal_S9 (pfam00380)→ |
| env-98 | Throat_LANL_C5730482 | + | 257 | 330 | **RNA**→ |
| env-99 | SRS011306_Baylor_scaffold_83447 | + | 6688 | 6761 | **RNA**→ rplM (PRK09216)Ribosomal_L13 (pfam00572)→ rpsI (PRK00132)Ribosomal_S9 (pfam00380)→ |
| env-100 | SRS015038_C1882965 | - | 403 | 330 | **RNA**→ rplM (PRK09216)Ribosomal_L13 (pfam00572)→ |
| env-101 | SRS015762_C3563104 | - | 158 | 85 | **RNA**→ |
| env-102 | SRS015893_WUGC_scaffold_41335 | - | 2217 | 2144 | **RNA**→ rplM (PRK09216)Ribosomal_L13 (pfam00572)→ rpsI (PRK00132)Ribosomal_S9 (pfam00380)→ |
| env-103 | SRS015941_C3475421 | + | 479 | 552 | **RNA**→ rplM (PRK09216)→ |
| env-104 | SRS016002_WUGC_scaffold_14080 | - | 451 | 378 | **RNA**→ rplM (PRK09216)Ribosomal_L13 (pfam00572)→ |
| env-105 | SRS018357_Baylor_scaffold_26234 | + | 880 | 953 | **RNA**→ hypo→ |
| env-106 | SRS018591_WUGC_scaffold_33810 | - | 202 | 129 | **RNA**→ hypo→ |
| env-107 | SRS018739_C3556239 | + | 325 | 398 | **RNA**→ |
| env-108 | SRS019022_C2654036 | - | 1005 | 932 | **RNA**→ rplM (PRK09216)Ribosomal_L13 (pfam00572)→ rpsI (PRK00132)Ribosomal_S9 (pfam00380)→ |
| env-109 | SRS019026_C1997533 | - | 401 | 328 | **RNA**→ rplM (PRK09216)Ribosomal_L13 (pfam00572)→ |
| env-110 | SRS019027_WUGC_scaffold_18825 | + | 340 | 413 | **RNA**→ rplM (PRK09216)Ribosomal_L13 (pfam00572)→ |
| env-111 | SRS019045_WUGC_scaffold_59681 | - | 2127 | 2054 | **RNA**→ rplM (PRK09216)Ribosomal_L13 (pfam00572)→ rpsI (PRK00132)Ribosomal_S9 (pfam00380)→ |
| env-112 | SRS019974_Baylor_scaffold_58098 | + | 480 | 553 | **RNA**→ rplM (PRK09216)Ribosomal_L13 (pfam00572)→ rpsI (PRK00132)Ribosomal_S9 (pfam00380)→ |
| env-113 | SRS020856_C2589008 | + | 484 | 557 | **RNA**→ |
| env-114 | SRS022621_C4094963 | - | 158 | 85 | **RNA**→ hypo→ |
| env-115 | SRS023352_C4653467 | - | 740 | 667 | **RNA**→ rplM (PRK09216)Ribosomal_L13 (pfam00572)→ rpsI (PRK00132)Ribosomal_S9 (pfam00380)→ |
| env-116 | SRS024138_C2697359 | + | 474 | 547 | **RNA**→ |
| env-117 | SRS024277_LANL_scaffold_45644 | + | 325 | 398 | **RNA**→ rplM (PRK09216)Ribosomal_L13 (pfam00572)→ rpsI (PRK00132)Ribosomal_S9 (pfam00380)→ |
| env-118 | SRS024580_C3091241 | - | 952 | 879 | **RNA**→ rplM (PRK09216)Ribosomal_L13 (pfam00572)→ rpsI (PRK00132)Ribosomal_S9 (pfam00380)→ |
| env-119 | SRS047824_WUGC_scaffold_40341 | - | 1220 | 1147 | **RNA**→ rplM (PRK09216)Ribosomal_L13 (pfam00572)→ rpsI (PRK00132)Ribosomal_S9 (pfam00380)→ |
| env-120 | SRS048411_C6958491 | - | 407 | 334 | **RNA**→ rplM (PRK09216)Ribosomal_L13 (pfam00572)→ |
| env-121 | SRS048791_C3141610 | + | 191 | 264 | **RNA**→ rplM (PRK09216)RplM (COG0102)→ |
| env-122 | SRS053603_LANL_scaffold_55573 | + | 1162 | 1235 | **RNA**→ rplM (PRK09216)Ribosomal_L13 (pfam00572)→ rpsI (PRK00132)Ribosomal_S9 (pfam00380)→ |
| env-123 | SRS058336_C2151893 | + | 935 | 1008 | **RNA**→ |
| env-124 | SRS064423_LANL_scaffold_69482 | - | 3254 | 3181 | **RNA**→ rplM (PRK09216)Ribosomal_L13 (pfam00572)→ rpsI (PRK00132)Ribosomal_S9 (pfam00380)→ |
| env-125 | SRS065278_C2565433 | - | 3238 | 3165 | **RNA**→ rplM (PRK09216)Ribosomal_L13 (pfam00572)→ rpsI (PRK00132)Ribosomal_S9 (pfam00380)→ |
| env-126 | SRS016740_C2737901 | - | 1286 | 1213 | **RNA**→ hypo→ rplM (PRK09216)Ribosomal_L13 (pfam00572)→ rpsI (PRK00132)Ribosomal_S9 (pfam00380)→ |
| env-127 | SRS047210_C3305333 | - | 1003 | 930 | **RNA**→ rplM (PRK09216)Ribosomal_L13 (pfam00572)→ rpsI (PRK00132)Ribosomal_S9 (pfam00380)→ |
| env-128 | SRS064774_C2715190 | + | 51 | 124 | **RNA**→ hypo→ rplM (PRK09216)Ribosomal_L13 (pfam00572)→ |
| env-129 | SRS014692_C1518495 | - | 385 | 312 | **RNA**→ rplM (PRK09216)Ribosomal_L13 (pfam00572)→ |
| env-130 | SRS014684_WUGC_scaffold_43074 | + | 2486 | 2559 | **RNA**→ rplM (PRK09216)Ribosomal_L13 (pfam00572)→ rpsI (PRK00132)Ribosomal_S9 (pfam00380)→ |
| env-131 | SRS014689_WUGC_scaffold_12190 | + | 2627 | 2700 | **RNA**→ rplM (PRK09216)Ribosomal_L13 (pfam00572)→ rpsI (PRK00132)Ribosomal_S9 (pfam00380)→ |
| env-132 | Supragingival_plaque_LANL_scaffold_252697 | - | 158 | 85 | **RNA**→ |
| env-133 | SRS013705_Baylor_scaffold_2470 | - | 1400 | 1327 | **RNA**→ rplM (PRK09216)Ribosomal_L13 (pfam00572)→ rpsI (PRK00132)Ribosomal_S9 (pfam00380)→ |
| env-134 | SRS014271_C2964309 | + | 4142 | 4215 | **RNA**→ |
| env-135 | SRS015215_WUGC_scaffold_38570 | + | 157 | 230 | **RNA**→ rplM (PRK09216)Ribosomal_L13 (pfam00572)→ |
| env-136 | SRS016342_C2855557 | - | 402 | 329 | **RNA**→ rplM (PRK09216)Ribosomal_L13 (pfam00572)→ |
| env-137 | SRS017808_Baylor_scaffold_55041 | - | 1325 | 1252 | **RNA**→ rplM (PRK09216)Ribosomal_L13 (pfam00572)→ rpsI (PRK00132)Ribosomal_S9 (pfam00380)→ |
| env-138 | SRS018791_C2310859 | + | 489 | 562 | **RNA**→ rplM (PRK09216)Ribosomal_L13 (pfam00572)→ rpsI (PRK00132)Ribosomal_S9 (pfam00380)→ |

| | | | | | |
|---|---|---|---|---|---|
| env-139 | SRS020220_Baylor_scaffold_52325 | - | 173 | 100 | **RNA**→ |
| env-140 | SRS021954_Baylor_scaffold_48638 | + | 2555 | 2628 | **RNA**→ rplM (PRK09216)Ribosomal_L13 (pfam00572)→ rpsI (PRK00132)Ribosomal_S9 (pfam00380)→ |
| env-141 | SRS023617_Baylor_scaffold_62383 | + | 2343 | 2416 | **RNA**→ rplM (PRK09216)Ribosomal_L13 (pfam00572)→ rpsI (PRK00132)Ribosomal_S9 (pfam00380)→ |
| env-142 | SRS023926_Baylor_scaffold_35364 | + | 6413 | 6486 | **RNA**→ rplM (PRK09216)Ribosomal_L13 (pfam00572)→ rpsI (PRK00132)Ribosomal_S9 (pfam00380)→ |
| env-143 | SRS024081_C2596496 | - | 158 | 85 | **RNA**→ |
| env-144 | SRS044373_C2896605 | + | 144 | 217 | **RNA**→ rplM (PRK09216)Ribosomal_L13 (pfam00572)→ rpsI (PRK00132)Ribosomal_S9 (pfam00380)→ |
| env-145 | SRS044486_C2263445 | - | 551 | 478 | **RNA**→ rplM (PRK09216)Ribosomal_L13 (pfam00572)→ |
| env-146 | SRS050244_C4466451 | - | 293 | 220 | **RNA**→ rplM (PRK09216)Ribosomal_L13 (pfam00572)→ |
| env-147 | SRS056892_C1498053 | - | 740 | 667 | **RNA**→ rplM (PRK09216)Ribosomal_L13 (pfam00572)→ rpsI (PRK00132)Ribosomal_S9 (pfam00380)→ |
| env-148 | SRS015797_C2641570 | + | 936 | 1009 | **RNA**→ |
| env-149 | SRS019219_C2933673 | + | 150 | 223 | **RNA**→ rplM (PRK09216)Ribosomal_L13 (pfam00572)→ rpsI (PRK00132)Ribosomal_S9 (pfam00380)→ |
| env-150 | SRS024318_LANL_scaffold_22081 | - | 1688 | 1615 | **RNA**→ rplM (PRK09216)Ribosomal_L13 (pfam00572)→ rpsI (PRK00132)Ribosomal_S9 (pfam00380)→ |
| env-151 | SRS042643_WUGC_scaffold_65245 | - | 159 | 86 | **RNA**→ |
| env-152 | SRS043663_LANL_scaffold_40074 | - | 6366 | 6293 | **RNA**→ rplM (PRK09216)Ribosomal_L13 (pfam00572)→ rpsI (PRK00132)Ribosomal_S9 (pfam00380)→ |
| env-153 | SRS053854_LANL_scaffold_53869 | + | 18601 | 18674 | **RNA**→ rplM (PRK09216)Ribosomal_L13 (pfam00572)→ rpsI (PRK00132)Ribosomal_S9 (pfam00380)→ |
| env-154 | SRS062761_LANL_scaffold_7973 | + | 6020 | 6093 | **RNA**→ rplM (PRK09216)Ribosomal_L13 (pfam00572)→ rpsI (PRK00132)Ribosomal_S9 (pfam00380)→ |
| env-155 | SRS075404_C3410168 | + | 150 | 223 | **RNA**→ |
| env-156 | SRS023835_Baylor_scaffold_45883 | - | 172 | 99 | **RNA**→ |
| env-157 | SRS049147_LANL_scaffold_19223 | - | 1397 | 1324 | **RNA**→ rplM (PRK09216)Ribosomal_L13 (pfam00572)→ rpsI (PRK00132)Ribosomal_S9 (pfam00380)→ |
| env-158 | SRS020856_Baylor_scaffold_43228 | + | 1090 | 1165 | **RNA**→ |
| env-159 | SRS023557_C1740022 | + | 539 | 614 | **RNA**→ |
| env-160 | SRS023930_Baylor_scaffold_1355 | + | 765 | 840 | **RNA**→ rplM (PRK09216)Ribosomal_L13 (pfam00572)→ rpsI (PRK00132)Ribosomal_S9 (pfam00380)→ |
| env-161 | SRS013946_WUGC_scaffold_1088 | + | 79348 | 79423 | **RNA**→ rplM (PRK09216)Ribosomal_L13 (pfam00572)→ rpsI (PRK00132)Ribosomal_S9 (pfam00380)→ |
| env-162 | SRS019124_C1404027 | - | 844 | 769 | **RNA**→ rplM (PRK09216)Ribosomal_L13 (pfam00572)→ rpsI (PRK00132)Ribosomal_S9 (pfam00380)→ |
| Psp-23-1 | NC_022111.1 | - | 1426745 | 1426670 | **RNA**→ rplM (PRK09216)Ribosomal_L13 (pfam00572)→ rpsI (PRK00132)Ribosomal_S9 (pfam00380)→ |
| env-163 | Keratinized_gingiva_LANL_scaffold_26598 | - | 469 | 394 | **RNA**→ rplM (PRK09216)Ribosomal_L13 (pfam00572)→ |
| env-164 | SRS045127_LANL_scaffold_9552 | - | 310 | 235 | **RNA**→ rplM (PRK09216)Ribosomal_L13 (pfam00572)→ |
| env-165 | SRS057539_LANL_scaffold_47289 | + | 1537 | 1612 | **RNA**→ rplM (PRK09216)Ribosomal_L13 (pfam00572)→ rpsI (PRK00132)Ribosomal_S9 (pfam00380)→ |
| env-166 | SRS064774_LANL_scaffold_66778 | - | 205 | 130 | **RNA**→ rplM (PRK09216)→ |
| env-167 | SRS024375_LANL_scaffold_43091 | - | 2175 | 2100 | **RNA**→ rplM (PRK09216)Ribosomal_L13 (pfam00572)→ rpsI (PRK00132)Ribosomal_S9 (pfam00380)→ |
| env-168 | SRS045715_LANL_scaffold_76474 | + | 674 | 749 | **RNA**→ rplM (PRK09216)Ribosomal_L13 (pfam00572)→ rpsI (PRK00132)Ribosomal_S9 (pfam00380)→ |
| env-169 | SRS013818_Baylor_scaffold_58539 | - | 3474 | 3399 | **RNA**→ rplM (PRK09216)Ribosomal_L13 (pfam00572)→ rpsI (PRK00132)Ribosomal_S9 (pfam00380)→ |
| env-170 | SRS018439_Baylor_scaffold_27830 | + | 14827 | 14902 | **RNA**→ rplM (PRK09216)Ribosomal_L13 (pfam00572)→ rpsI (PRK00132)Ribosomal_S9 (pfam00380)→ |
| env-171 | Palatine_Tonsils_LANL_C6199093 | - | 206 | 131 | **RNA**→ rplM (PRK09216)→ |
| env-172 | SRS016740_Baylor_scaffold_45058 | - | 3466 | 3391 | **RNA**→ rplM (PRK09216)Ribosomal_L13 (pfam00572)→ rpsI (PRK00132)Ribosomal_S9 (pfam00380)→ |
| env-173 | SRS011306_Baylor_scaffold_30322 | - | 1178 | 1103 | **RNA**→ rplM (PRK09216)Ribosomal_L13 (pfam00572)→ rpsI (PRK00132)Ribosomal_S9 (pfam00380)→ |
| env-174 | SRS013164_Baylor_scaffold_48360 | - | 1235 | 1160 | **RNA**→ rplM (PRK09216)Ribosomal_L13 (pfam00572)→ rpsI (PRK00132)Ribosomal_S9 (pfam00380)→ |
| env-175 | SRS013942_C1226381 | + | 56 | 131 | **RNA**→ rplM (PRK09216)Ribosomal_L13 (pfam00572)→ |
| env-176 | SRS013947_WUGC_scaffold_26275 | + | 3554 | 3629 | **RNA**→ rplM (PRK09216)Ribosomal_L13 (pfam00572)→ rpsI (PRK00132)Ribosomal_S9 (pfam00380)→ |
| env-177 | SRS013948_C1540062 | - | 1341 | 1266 | **RNA**→ rplM (PRK09216)Ribosomal_L13 (pfam00572)→ rpsI (PRK00132)Ribosomal_S9 (pfam00380)→ |
| env-178 | SRS014888_C2952726 | - | 278 | 203 | **RNA**→ rplM (PRK09216)RplM (COG0102)→ |
| env-179 | SRS015762_WUGC_scaffold_24890 | - | 155 | 80 | **RNA**→ |
| env-180 | SRS016002_WUGC_scaffold_61027 | + | 308 | 383 | **RNA**→ rplM (PRK09216)Ribosomal_L13 (pfam00572)→ rpsI (PRK00132)Ribosomal_S9 (pfam00380)→ |
| env-181 | SRS016086_WUGC_scaffold_26845 | - | 1575 | 1500 | **RNA**→ rplM (PRK09216)Ribosomal_L13 (pfam00572)→ rpsI (PRK00132)Ribosomal_S9 (pfam00380)→ |
| env-182 | SRS016225_C2845363 | + | 174 | 249 | **RNA**→ |
| env-183 | SRS016569_Baylor_scaffold_56398 | - | 14475 | 14400 | **RNA**→ rplM (PRK09216)Ribosomal_L13 (pfam00572)→ rpsI (PRK00132)Ribosomal_S9 (pfam00380)→ |
| env-184 | SRS017808_C3912739 | + | 987 | 1062 | **RNA**→ rplM (PRK09216)Ribosomal_L13 (pfam00572)→ rpsI (PRK00132)Ribosomal_S9 (pfam00380)→ |
| env-185 | SRS018145_Baylor_scaffold_43542 | - | 203 | 128 | **RNA**→ rplM (PRK09216)→ |
| env-186 | SRS019022_WUGC_scaffold_45455 | + | 194 | 269 | **RNA**→ rplM (PRK09216)Ribosomal_L13 (pfam00572)→ rpsI (PRK00132)Ribosomal_S9 (pfam00380)→ |
| env-187 | SRS019045_C2719629 | + | 63 | 138 | **RNA**→ rplM (PRK09216)Ribosomal_L13 (pfam00572)→ rpsI (PRK00132)Ribosomal_S9 (pfam00380)→ |
| env-188 | SRS019126_WUGC_scaffold_13188 | + | 379 | 454 | **RNA**→ rplM (PRK09216)Ribosomal_L13 (pfam00572)→ rpsI (PRK00132)Ribosomal_S9 (pfam00380)→ |
| env-189 | SRS019894_C3913439 | - | 664 | 589 | **RNA**→ rplM (PRK09216)Ribosomal_L13 (pfam00572)→ rpsI (PRK00132)rps9 (CHL00079)→ |

| | | | | | |
|---|---|---|---|---|---|
| env-190 | SRS019974_Baylor_scaffold_2117 | + | 182 | 257 | **RNA**→ rplM (PRK09216)Ribosomal_L13 (pfam00572)→ rpsI (PRK00132)Ribosomal_S9 (pfam00380)→ |
| env-191 | SRS021954_C3044878 | + | 70 | 145 | **RNA**→ rplM (PRK09216)Ribosomal_L13 (pfam00572)→ rpsI (PRK00132)Ribosomal_S9 (pfam00380)→ |
| env-192 | SRS023958_C1974288 | + | 58 | 133 | **RNA**→ rplM (PRK09216)Ribosomal_L13 (pfam00572)→ |
| env-193 | SRS024015_Baylor_scaffold_54415 | + | 4601 | 4676 | **RNA**→ rplM (PRK09216)Ribosomal_L13 (pfam00572)→ rpsI (PRK00132)Ribosomal_S9 (pfam00380)→ |
| env-194 | SRS024277_C2835398 | + | 56 | 131 | **RNA**→ rplM (PRK09216)Ribosomal_L13 (pfam00572)→ rpsI (PRK00132)Ribosomal_S9 (pfam00380)→ |
| env-195 | SRS048411_C6973885 | + | 102 | 177 | **RNA**→ rplM (PRK09216)Ribosomal_L13 (pfam00572)→ rpsI (PRK00132)Ribosomal_S9 (pfam00380)→ |
| env-196 | SRS048791_C3265861 | - | 1162 | 1087 | **RNA**→ rplM (PRK09216)Ribosomal_L13 (pfam00572)→ rpsI (PRK00132)Ribosomal_S9 (pfam00380)→ |
| env-197 | SRS052227_C2916121 | - | 1045 | 970 | **RNA**→ rplM (PRK09216)Ribosomal_L13 (pfam00572)→ rpsI (PRK00132)Ribosomal_S9 (pfam00380)→ |
| env-198 | SRS053603_LANL_scaffold_51146 | - | 1182 | 1107 | **RNA**→ rplM (PRK09216)Ribosomal_L13 (pfam00572)→ rpsI (PRK00132)Ribosomal_S9 (pfam00380)→ |
| env-199 | SRS054687_C3493368 | + | 1536 | 1611 | **RNA**→ rplM (PRK09216)Ribosomal_L13 (pfam00572)→ |
| env-200 | SRS056323_C2562669 | - | 459 | 384 | **RNA**→ rplM (PRK09216)Ribosomal_L13 (pfam00572)→ |
| env-201 | SRS057355_C2358048 | - | 1102 | 1027 | **RNA**→ rplM (PRK09216)Ribosomal_L13 (pfam00572)→ rpsI (PRK00132)Ribosomal_S9 (pfam00380)→ |
| env-202 | SRS063193_LANL_scaffold_66856 | - | 3099 | 3024 | **RNA**→ rplM (PRK09216)Ribosomal_L13 (pfam00572)→ rpsI (PRK00132)Ribosomal_S9 (pfam00380)→ |
| env-203 | SRS064423_LANL_scaffold_47969 | - | 3499 | 3424 | **RNA**→ rplM (PRK09216)Ribosomal_L13 (pfam00572)→ rpsI (PRK00132)Ribosomal_S9 (pfam00380)→ |
| env-204 | SRS077736_LANL_scaffold_38236 | + | 42461 | 42536 | **RNA**→ rplM (PRK09216)Ribosomal_L13 (pfam00572)→ rpsI (PRK00132)Ribosomal_S9 (pfam00380)→ |
| env-205 | Tongue_dor_WUGI_scaffold_8501 | + | 269 | 344 | **RNA**→ rplM (PRK09216)Ribosomal_L13 (pfam00572)→ rpsI (PRK00132)Ribosomal_S9 (pfam00380)→ |
| env-206 | SRS014470_C3713445 | + | 547 | 622 | **RNA**→ rplM (PRK09216)RplM (COG0102)→ |
| env-207 | SRS018300_C3341863 | - | 280 | 205 | **RNA**→ rplM (PRK09216)RplM (COG0102)→ |
| env-208 | SRS024441_LANL_scaffold_48252 | + | 856 | 931 | **RNA**→ |
| env-209 | SRS075404_C3567588 | + | 297 | 372 | **RNA**→ rplM (PRK09216)RplM (COG0102)→ |
| env-210 | SRS011086_WUGC_scaffold_15953 | - | 4147 | 4072 | **RNA**→ rplM (PRK09216)Ribosomal_L13 (pfam00572)→ rpsI (PRK00132)Ribosomal_S9 (pfam00380)→ |
| env-211 | SRS013879_WUGC_scaffold_7871 | + | 156 | 231 | **RNA**→ rplM (PRK09216)Ribosomal_L13 (pfam00572)→ rpsI (PRK00132)Ribosomal_S9 (pfam00380)→ |
| env-212 | SRS011243_Baylor_scaffold_40702 | - | 1370 | 1295 | **RNA**→ rplM (PRK09216)Ribosomal_L13 (pfam00572)→ rpsI (PRK00132)Ribosomal_S9 (pfam00380)→ |
| env-213 | SRS014573_WUGC_scaffold_49775 | - | 908 | 833 | **RNA**→ rplM (PRK09216)Ribosomal_L13 (pfam00572)→ rpsI (PRK00132)Ribosomal_S9 (pfam00380)→ |
| env-214 | SRS015797_WUGC_scaffold_1453 | - | 1656 | 1581 | **RNA**→ rplM (PRK09216)Ribosomal_L13 (pfam00572)→ rpsI (PRK00132)Ribosomal_S9 (pfam00380)→ |
| env-215 | SRS016037_WUGC_scaffold_35398 | + | 102 | 177 | **RNA**→ rplM (PRK09216)Ribosomal_L13 (pfam00572)→ |
| env-216 | SRS021496_Baylor_scaffold_24398 | - | 5055 | 4980 | **RNA**→ rplM (PRK09216)Ribosomal_L13 (pfam00572)→ rpsI (PRK00132)Ribosomal_S9 (pfam00380)→ |
| env-217 | SRS024138_Baylor_scaffold_896 | + | 1270 | 1345 | **RNA**→ rplM (PRK09216)Ribosomal_L13 (pfam00572)→ rpsI (PRK00132)Ribosomal_S9 (pfam00380)→ |
| env-218 | SRS042643_WUGC_scaffold_46473 | + | 11248 | 11323 | **RNA**→ rplM (PRK09216)Ribosomal_L13 (pfam00572)→ rpsI (PRK00132)Ribosomal_S9 (pfam00380)→ |
| env-219 | SRS047219_C2146198 | + | 105 | 180 | **RNA**→ rplM (PRK09216)RplM (COG0102)→ |
| env-220 | SRS049147_LANL_scaffold_18610 | + | 946 | 1021 | **RNA**→ rplM (PRK09216)Ribosomal_L13 (pfam00572)→ rpsI (PRK00132)Ribosomal_S9 (pfam00380)→ |
| env-221 | SRS055426_LANL_scaffold_19831 | - | 3585 | 3510 | **RNA**→ rplM (PRK09216)Ribosomal_L13 (pfam00572)→ rpsI (PRK00132)Ribosomal_S9 (pfam00380)→ hypo→ |
| env-222 | Buccal_mucosa_LANL_scaffold_78460 | - | 760 | 685 | **RNA**→ rplM (PRK09216)Ribosomal_L13 (pfam00572)→ rpsI (PRK00132)Ribosomal_S9 (pfam00380)→ |
| env-223 | SRS014271_WUGC_scaffold_629 | + | 43 | 118 | **RNA**→ rplM (PRK09216)Ribosomal_L13 (pfam00572)→ rpsI (PRK00132)Ribosomal_S9 (pfam00380)→ |
| env-224 | SRS062544_LANL_scaffold_4602 | - | 1246 | 1171 | **RNA**→ rplM (PRK09216)Ribosomal_L13 (pfam00572)→ rpsI (PRK00132)Ribosomal_S9 (pfam00380)→ |
| env-225 | SRS065278_LANL_scaffold_63970 | - | 3725 | 3650 | **RNA**→ rplM (PRK09216)Ribosomal_L13 (pfam00572)→ rpsI (PRK00132)Ribosomal_S9 (pfam00380)→ |
| env-226 | SRS024081_LANL_scaffold_32043 | + | 406 | 481 | **RNA**→ rplM (PRK09216)Ribosomal_L13 (pfam00572)→ rpsI (PRK00132)Ribosomal_S9 (pfam00380)→ |
| env-227 | SRS022719_LANL_scaffold_53966 | + | 5202 | 5277 | **RNA**→ rplM (PRK09216)Ribosomal_L13 (pfam00572)→ rpsI (PRK00132)Ribosomal_S9 (pfam00380)→ |
| env-228 | SRS044662_LANL_scaffold_14148 | - | 15885 | 15810 | **RNA**→ rplM (PRK09216)Ribosomal_L13 (pfam00572)→ rpsI (PRK00132)Ribosomal_S9 (pfam00380)→ |
| env-229 | SRS050669_LANL_scaffold_25595 | + | 601 | 676 | **RNA**→ rplM (PRK09216)Ribosomal_L13 (pfam00572)→ rpsI (PRK00132)Ribosomal_S9 (pfam00380)→ |
| env-230 | SRS058336_LANL_scaffold_14781 | - | 102 | 27 | **RNA**→ hypo→ |
| env-231 | SRS016342_Baylor_scaffold_48981 | + | 82 | 157 | **RNA**→ rplM (PRK09216)Ribosomal_L13 (pfam00572)→ rpsI (PRK00132)Ribosomal_S9 (pfam00380)→ |
| env-232 | SRS018739_WUGC_scaffold_60144 | - | 1366 | 1291 | **RNA**→ rplM (PRK09216)Ribosomal_L13 (pfam00572)→ rpsI (PRK00132)Ribosomal_S9 (pfam00380)→ |
| env-233 | SRS019126_C2043960 | + | 254 | 329 | **RNA**→ |
| Pna-2-1 | NZ_BAKD01000002.1 | - | 398642 | 398567 | **RNA**→ rplM (PRK09216)Ribosomal_L13 (pfam00572)→ rpsI (PRK00132)Ribosomal_S9 (pfam00380)→ |
| Pna-1-1 | NZ_KB904328.1 | + | 132132 | 132207 | **RNA**→ rplM (PRK09216)Ribosomal_L13 (pfam00572)→ rpsI (PRK00132)Ribosomal_S9 (pfam00380)→ |
| env-234 | SRS011140_Baylor_scaffold_53572 | - | 2158 | 2083 | **RNA**→ rplM (PRK09216)Ribosomal_L13 (pfam00572)→ rpsI (PRK00132)Ribosomal_S9 (pfam00380)→ |
| env-235 | SRS015434_C5898829 | + | 238 | 313 | **RNA**→ rplM (PRK09216)Ribosomal_L13 (pfam00572)→ |
| env-236 | SRS015644_WUGC_scaffold_48566 | + | 1536 | 1611 | **RNA**→ rplM (PRK09216)Ribosomal_L13 (pfam00572)→ rpsI (PRK00132)Ribosomal_S9 (pfam00380)→ |
| env-237 | SRS019077_WUGC_scaffold_47444 | - | 372 | 297 | **RNA**→ rplM (PRK09216)Ribosomal_L13 (pfam00572)→ |
| env-238 | SRS019127_C1659470 | + | 56 | 131 | **RNA**→ rplM (PRK09216)Ribosomal_L13 (pfam00572)→ rpsI (PRK00132)Ribosomal_S9 (pfam00380)→ |
| env-239 | SRS020856_Baylor_scaffold_12760 | - | 1113 | 1038 | **RNA**→ rplM (PRK09216)Ribosomal_L13 (pfam00572)→ rpsI (PRK00132)Ribosomal_S9 (pfam00380)→ |

| | | | | | |
|---|---|---|---|---|---|
| env-240 | SRS022077_Baylor_scaffold_47573 | + | 56 | 131 | **RNA**→ rplM (PRK09216)Ribosomal_L13 (pfam00572)→ rpsI (PRK00132)Ribosomal_S9 (pfam00380)→ |
| env-241 | SRS022621_Baylor_scaffold_3806 | - | 26534 | 26459 | **RNA**→ rplM (PRK09216)Ribosomal_L13 (pfam00572)→ rpsI (PRK00132)Ribosomal_S9 (pfam00380)→ |
| env-242 | SRS023617_Baylor_scaffold_52938 | - | 36529 | 36454 | **RNA**→ rplM (PRK09216)Ribosomal_L13 (pfam00572)→ rpsI (PRK00132)Ribosomal_S9 (pfam00380)→ |
| env-243 | SRS024580_C3074526 | - | 620 | 545 | **RNA**→ rplM (PRK09216)Ribosomal_L13 (pfam00572)→ |
| env-244 | SRS042131_WUGC_scaffold_8821 | - | 37335 | 37260 | **RNA**→ rplM (PRK09216)Ribosomal_L13 (pfam00572)→ rpsI (PRK00132)Ribosomal_S9 (pfam00380)→ |
| env-245 | SRS050244_LANL_scaffold_95985 | + | 1714 | 1789 | **RNA**→ rplM (PRK09216)Ribosomal_L13 (pfam00572)→ rpsI (PRK00132)Ribosomal_S9 (pfam00380)→ |
| env-246 | SRS019607_C2757825 | - | 1575 | 1500 | **RNA**→ rplM (PRK09216)Ribosomal_L13 (pfam00572)→ rpsI (PRK00132)Ribosomal_S9 (pfam00380)→ |
| env-247 | SRS023835_Baylor_scaffold_30646 | - | 382 | 307 | **RNA**→ rplM (PRK09216)Ribosomal_L13 (pfam00572)→ |
| env-248 | SRS057205_LANL_scaffold_32691 | + | 1114 | 1189 | **RNA**→ |
| env-249 | SRS012279_C2758674 | + | 3094 | 3169 | **RNA**→ rplM (PRK09216)Ribosomal_L13 (pfam00572)→ rpsI (PRK00132)Ribosomal_S9 (pfam00380)→ |
| env-250 | SRS014692_C1492733 | + | 63 | 138 | **RNA**→ rplM (PRK09216)Ribosomal_L13 (pfam00572)→ |
| env-251 | SRS019026_C1981803 | + | 132 | 207 | **RNA**→ rplM (PRK09216)Ribosomal_L13 (pfam00572)→ rpsI (PRK00132)Ribosomal_S9 (pfam00380)→ |
| env-252 | SRS019027_C1968276 | + | 63 | 138 | **RNA**→ rplM (PRK09216)Ribosomal_L13 (pfam00572)→ rpsI (PRK00132)Ribosomal_S9 (pfam00380)→ |
| env-253 | SRS019327_WUGC_scaffold_24766 | - | 2945 | 2870 | **RNA**→ rplM (PRK09216)Ribosomal_L13 (pfam00572)→ rpsI (PRK00132)Ribosomal_S9 (pfam00380)→ |
| env-254 | SRS019389_C2252888 | + | 2434 | 2509 | **RNA**→ |
| env-255 | SRS023352_C4687326 | - | 1370 | 1295 | **RNA**→ rplM (PRK09216)Ribosomal_L13 (pfam00572)→ rpsI (PRK00132)Ribosomal_S9 (pfam00380)→ |
| env-256 | SRS047210_C3287465 | - | 670 | 595 | **RNA**→ rplM (PRK09216)Ribosomal_L13 (pfam00572)→ rpsI (PRK00132)rps9 (CHL00079)→ |
| env-257 | SRS057791_LANL_scaffold_82241 | + | 477 | 552 | **RNA**→ rplM (PRK09216)Ribosomal_L13 (pfam00572)→ rpsI (PRK00132)Ribosomal_S9 (pfam00380)→ |
| env-258 | SRS014684_WUGC_scaffold_44011 | + | 3280 | 3355 | **RNA**→ rplM (PRK09216)Ribosomal_L13 (pfam00572)→ rpsI (PRK00132)Ribosomal_S9 (pfam00380)→ |
| env-259 | SRS017533_Baylor_scaffold_27038 | - | 80 | 5 | **RNA**→ |
| env-260 | SRS043663_LANL_scaffold_17693 | + | 210 | 285 | **RNA**→ rplM (PRK09216)Ribosomal_L13 (pfam00572)→ |
| env-261 | SRS062761_C4207862 | - | 374 | 299 | **RNA**→ rplM (PRK09216)Ribosomal_L13 (pfam00572)→ |
| env-262 | SRS013946_WUGC_scaffold_14040 | + | 788 | 863 | **RNA**→ rplM (PRK09216)Ribosomal_L13 (cd00392)→ |
| env-263 | SRS019125_WUGC_scaffold_9864 | - | 274 | 199 | **RNA**→ rplM (PRK09216)RplM (COG0102)→ |
| env-264 | SRS013506_Baylor_scaffold_8126 | - | 301 | 226 | **RNA**→ rplM (PRK09216)Ribosomal_L13 (cd00392)→ |
| env-265 | SRS014473_C1126965 | + | 422 | 497 | **RNA**→ rplM (PRK09216)Ribosomal_L13 (pfam00572)→ rpsI (PRK00132)Ribosomal_S9 (pfam00380)→ |
| env-266 | AUXO014368639.1 | - | 674 | 600 | **RNA**→ rplM (PRK09216)Ribosomal_L13 (pfam00572)→ |
| env-267 | AUXO015562770.1 | + | 235 | 303 | **RNA**→ rplM (PRK09216)Ribosomal_L13 (pfam00572)→ rpsI (PRK00132)Ribosomal_S9 (pfam00380)→ |
| Bsp-2-1 | NZ_JNKX01000045.1 | - | 3997 | 3929 | **RNA**→ rplM (PRK09216)Ribosomal_L13 (pfam00572)→ rpsI (PRK00132)Ribosomal_S9 (pfam00380)→ |
| Bsp-1-1 | NZ_JNLA01000048.1 | + | 27849 | 27917 | **RNA**→ rplM (PRK09216)Ribosomal_L13 (pfam00572)→ rpsI (PRK00132)Ribosomal_S9 (pfam00380)→ |
| env-268 | SRS023352_C4496944 | - | 123 | 46 | **RNA**→ hypo→ |
| env-269 | SRS062544_C2920745 | + | 463 | 540 | **RNA**→ |
| env-270 | SRS019219_WUGC_scaffold_426 | - | 214 | 137 | **RNA**→ hypo→ |
| env-271 | SRS051791_LANL_scaffold_54765 | - | 217 | 140 | **RNA**→ |
| env-272 | SRS015057_WUGC_scaffold_61382 | + | 542 | 619 | **RNA**→ |
| env-273 | SRS048411_C6924690 | + | 505 | 582 | **RNA**→ |
| env-274 | SRS057791_C3531782 | + | 268 | 345 | **RNA**→ |
| env-275 | SRS045715_LANL_scaffold_76924 | + | 360 | 437 | **RNA**→ |
| env-276 | SRS015272_C3500034 | - | 214 | 137 | **RNA**→ |
| env-277 | SRS018791_C2278485 | + | 773 | 850 | **RNA**→ |
| env-278 | SRS011255_C2362793 | - | 117 | 40 | **RNA**→ |
| env-279 | SRS013950_WUGC_scaffold_2138 | - | 220 | 143 | **RNA**→ |
| env-280 | SRS013836_C4630228 | + | 244 | 321 | **RNA**→ |
| env-281 | SRS023841_C1489733 | - | 221 | 144 | **RNA**→ hypo→ |
| env-282 | Supragingival_plaque_LANL_scaffold_194841 | - | 474 | 397 | **RNA**→ rplM (PRK09216)Ribosomal_L13 (pfam00572)→ |
| env-283 | SRS017691_C5410429 | + | 2077 | 2154 | **RNA**→ |
| env-284 | SRS024561_C2727733 | + | 380 | 457 | **RNA**→ |
| env-285 | SRS024277_C2727216 | - | 173 | 96 | **RNA**→ hypo→ |
| env-286 | SRS016319_C3957492 | - | 123 | 46 | **RNA**→ hypo→ |
| env-287 | SRS042131_C2217398 | + | 340 | 417 | **RNA**→ |
| env-288 | SRS043663_C4080713 | + | 808 | 885 | **RNA**→ |
| env-289 | SRS062761_C4201374 | + | 277 | 354 | **RNA**→ |

| | | | | | |
|---|---|---|---|---|---|
| env-290 | SRS015797_C2619071 | - | 219 | 142 | **RNA**→ |
| env-291 | SRS019607_C2703769 | - | 184 | 107 | **RNA**→ ←hypo |
| env-292 | SRS021496_Baylor_scaffold_55246 | + | 49 | 126 | **RNA**→ rplM (PRK09216)Ribosomal_L13 (pfam00572)→ |
| env-293 | SRS024081_C2601354 | + | 658 | 735 | **RNA**→ |
| env-294 | SRS049389_C4506064 | + | 176 | 253 | **RNA**→ |
| env-295 | SRS075404_C3622013 | + | 648 | 725 | **RNA**→ |
| env-296 | SRS018739_C3603893 | + | 489 | 566 | **RNA**→ |
| env-297 | SRS020220_C3576851 | - | 115 | 38 | **RNA**→ hypo→ |
| env-298 | SRS011306_C3387646 | + | 94 | 171 | **RNA**→ |
| env-299 | SRS019071_C5065527 | - | 138 | 61 | **RNA**→ hypo→ |
| env-300 | SRS019327_WUGC_scaffold_20822 | + | 376 | 453 | **RNA**→ |
| env-301 | Supragingival_plaque_LANL_C182575899 | - | 117 | 40 | **RNA**→ hypo→ |
| env-302 | SRS016002_C2496321 | - | 179 | 102 | **RNA**→ hypo→ |
| env-303 | SRS019126_C2138744 | + | 361 | 438 | **RNA**→ |
| env-304 | SRS019127_C1661984 | + | 1415 | 1492 | **RNA**→ |
| env-305 | SRS017209_Baylor_scaffold_64433 | - | 214 | 137 | **RNA**→ |
| env-306 | SRS024318_LANL_scaffold_131 | - | 214 | 137 | **RNA**→ |
| env-307 | SRS057205_LANL_scaffold_46232 | - | 410 | 333 | **RNA**→ rplM (PRK09216)Ribosomal_L13 (pfam00572)→ |
| env-308 | SRS018357_C3783001 | - | 179 | 102 | **RNA**→ hypo→ |
| env-309 | SRS020856_C2544654 | + | 239 | 316 | **RNA**→ |
| env-310 | SRS044373_WUGC_scaffold_63614 | - | 123 | 46 | **RNA**→ |
| env-311 | SRS014271_C2955858 | + | 2330 | 2407 | **RNA**→ |
| env-312 | SRS050244_C4442009 | + | 418 | 495 | **RNA**→ |
| env-313 | SRS062544_C3030084 | + | 2332 | 2409 | **RNA**→ |
| env-314 | SRS013164_C2312325 | - | 119 | 42 | **RNA**→ hypo→ |
| env-315 | SRS016225_C2929058 | + | 310 | 388 | **RNA**→ |
| env-316 | 4491686.3.521687 | + | 140 | 220 | **RNA**→ ←hypo |
| env-317 | scaffold7222_1_MH0052 | - | 220 | 140 | **RNA**→ |
| env-318 | 4491686.3.475604 | - | 103 | 23 | **RNA**→ ←hypo |
| env-319 | 2013397958 | - | 676 | 600 | **RNA**→ rplM (PRK09216)Ribosomal_L13 (pfam00572)→ rplM (PRK09216)Ribosomal_L13 (pfam00572)→ |
| Psp-3-1 | NZ_AUJK01000003.1 | + | 63931 | 64008 | **RNA**→ rplM (PRK09216)Ribosomal_L13 (pfam00572)→ rpsI (PRK00132)Ribosomal_S9 (pfam00380)→ |
| env-320 | scaffold26454_4_MH0084 | + | 5212 | 5288 | **RNA**→ rplM (PRK09216)Ribosomal_L13 (pfam00572)→ rpsI (PRK00132)Ribosomal_S9 (pfam00380)→ |
| env-321 | scaffold66021_2_MH0081 | + | 3270 | 3346 | **RNA**→ rplM (PRK09216)Ribosomal_L13 (pfam00572)→ rpsI (PRK00132)Ribosomal_S9 (pfam00380)→ |
| env-322 | scaffold25402_2_MH0054 | + | 2416 | 2492 | **RNA**→ rplM (PRK09216)Ribosomal_L13 (pfam00572)→ rpsI (PRK00132)Ribosomal_S9 (pfam00380)→ |
| env-323 | scaffold8724_1_O2.UC-16 | - | 106 | 30 | **RNA**→ |
| env-324 | SRS015782_WUGC_scaffold_20008 | - | 24364 | 24288 | **RNA**→ rplM (PRK09216)Ribosomal_L13 (pfam00572)→ rpsI (PRK00132)Ribosomal_S9 (pfam00380)→ |
| env-325 | SRS047044_WUGC_scaffold_15995 | - | 1654 | 1578 | **RNA**→ rplM (PRK09216)Ribosomal_L13 (pfam00572)→ rpsI (PRK00132)Ribosomal_S9 (pfam00380)→ |
| env-326 | scaffold34413_1_V1.CD-13 | + | 111 | 187 | **RNA**→ rplM (PRK09216)Ribosomal_L13 (pfam00572)→ rpsI (PRK00132)Ribosomal_S9 (pfam00380)→ |
| env-327 | scaffold69522_1_MH0044 | + | 6237 | 6313 | **RNA**→ rplM (PRK09216)Ribosomal_L13 (pfam00572)→ rpsI (PRK00132)Ribosomal_S9 (pfam00380)→ |
| env-328 | scaffold59624_1_MH0052 | + | 3663 | 3739 | **RNA**→ ←hypo |
| env-329 | scaffold15726_3_MH0032 | - | 12257 | 12181 | **RNA**→ rplM (PRK09216)Ribosomal_L13 (pfam00572)→ rpsI (PRK00132)Ribosomal_S9 (pfam00380)→ |
| env-330 | scaffold58986_1_MH0033 | - | 3162 | 3086 | **RNA**→ rplM (PRK09216)Ribosomal_L13 (pfam00572)→ rpsI (PRK00132)Ribosomal_S9 (pfam00380)→ |
| env-331 | scaffold11675_1_MH0026 | + | 12207 | 12283 | **RNA**→ rplM (PRK09216)Ribosomal_L13 (pfam00572)→ rpsI (PRK00132)Ribosomal_S9 (pfam00380)→ |
| env-332 | NLM025_scaffold6375_35 | - | 965 | 889 | **RNA**→ rplM (PRK09216)Ribosomal_L13 (pfam00572)→ rpsI (PRK00132)Ribosomal_S9 (pfam00380)→ |
| env-333 | SRS013951_WUGC_scaffold_10163 | - | 35684 | 35608 | **RNA**→ rplM (PRK09216)Ribosomal_L13 (pfam00572)→ rpsI (PRK00132)Ribosomal_S9 (pfam00380)→ |
| env-334 | SRS019161_WUGC_scaffold_63747 | + | 921 | 997 | **RNA**→ rplM (PRK09216)Ribosomal_L13 (pfam00572)→ rpsI (PRK00132)Ribosomal_S9 (pfam00380)→ |
| env-335 | SRS050422_LANL_scaffold_39679 | - | 16737 | 16661 | **RNA**→ rplM (PRK09216)Ribosomal_L13 (pfam00572)→ rpsI (PRK00132)Ribosomal_S9 (pfam00380)→ |
| env-336 | scaffold3922_2_MH0001 | - | 532 | 456 | **RNA**→ rplM (PRK09216)Ribosomal_L13 (pfam00572)→ |
| env-337 | scaffold12670_1_MH0041 | + | 6023 | 6099 | **RNA**→ rplM (PRK09216)Ribosomal_L13 (pfam00572)→ rpsI (PRK00132)Ribosomal_S9 (pfam00380)→ |
| env-338 | 4491401.3_NODE_59631 | + | 440 | 516 | **RNA**→ rplM (PRK09216)Ribosomal_L13 (pfam00572)→ |
| env-339 | scaffold24499_3_MH0018 | - | 199 | 123 | **RNA**→ |
| env-340 | SRS048164_WUGC_scaffold_7139 | - | 11028 | 10952 | **RNA**→ rplM (PRK09216)Ribosomal_L13 (pfam00572)→ rpsI (PRK00132)Ribosomal_S9 (pfam00380)→ |

| | | | | | |
|---|---|---|---|---|---|
| env-341 | scaffold13081_6_MH0009 | - | 3032 | 2956 | **RNA**→ rplM (PRK09216)Ribosomal_L13 (pfam00572)→ rpsI (PRK00132)Ribosomal_S9 (pfam00380)→ |
| env-342 | scaffold29722_3_MH0034 | + | 2377 | 2453 | **RNA**→ rplM (PRK09216)Ribosomal_L13 (pfam00572)→ rpsI (PRK00132)Ribosomal_S9 (pfam00380)→ |
| env-343 | scaffold19923_2_O2.UC-12 | - | 430 | 354 | **RNA**→ rplM (PRK09216)Ribosomal_L13 (pfam00572)→ |
| env-344 | SRS015794_Baylor_scaffold_29113 | - | 896 | 820 | **RNA**→ rplM (PRK09216)Ribosomal_L13 (pfam00572)→ rpsI (PRK00132)Ribosomal_S9 (pfam00380)→ |
| env-345 | scaffold988_4_MH0046 | - | 10774 | 10698 | **RNA**→ rplM (PRK09216)Ribosomal_L13 (pfam00572)→ rpsI (PRK00132)Ribosomal_S9 (pfam00380)→ |
| env-346 | scaffold60646_3_MH0076 | - | 3034 | 2958 | **RNA**→ rplM (PRK09216)Ribosomal_L13 (pfam00572)→ rpsI (PRK00132)Ribosomal_S9 (pfam00380)→ |
| env-347 | scaffold60041_3_MH0056 | - | 523 | 447 | **RNA**→ rplM (PRK09216)Ribosomal_L13 (pfam00572)→ |
| env-348 | scaffold62900_7_O2.UC-19 | + | 1505 | 1581 | **RNA**→ rplM (PRK09216)Ribosomal_L13 (pfam00572)→ rpsI (PRK00132)Ribosomal_S9 (pfam00380)→ |
| env-349 | Subgingival_plaque_LANL_C11191810 | + | 401 | 479 | **RNA**→ |
| Pma-3-1 | NZ_JH594503.1 | - | 275251 | 275173 | **RNA**→ rplM (PRK09216)Ribosomal_L13 (pfam00572)→ rpsI (PRK00132)Ribosomal_S9 (pfam00380)→ |
| env-350 | SRS024087_C3572528 | - | 219 | 141 | **RNA**→ |
| env-351 | SRS024289_LANL_scaffold_69854 | - | 837 | 759 | **RNA**→ rplM (PRK09216)Ribosomal_L13 (pfam00572)→ rpsI (PRK00132)Ribosomal_S9 (pfam00380)→ |
| env-352 | SRS047634_LANL_scaffold_34142 | - | 1677 | 1599 | **RNA**→ rplM (PRK09216)Ribosomal_L13 (pfam00572)→ rpsI (PRK00132)Ribosomal_S9 (pfam00380)→ |
| Pma-1-1 | NZ_KB908319.1 | + | 316856 | 316934 | **RNA**→ rplM (PRK09216)Ribosomal_L13 (pfam00572)→ rpsI (PRK00132)Ribosomal_S9 (pfam00380)→ |
| Pma-2-1 | NZ_BAJZ01000003.1 | + | 9704 | 9782 | **RNA**→ rplM (PRK09216)Ribosomal_L13 (pfam00572)→ rpsI (PRK00132)Ribosomal_S9 (pfam00380)→ |
| env-353 | SRS018443_Baylor_scaffold_42450 | + | 2749 | 2827 | **RNA**→ |
| env-354 | SRS019980_C3029122 | + | 888 | 966 | **RNA**→ |
| env-355 | SRS017227_Baylor_scaffold_86068 | - | 121 | 43 | **RNA**→ |
| env-356 | SRS049268_LANL_scaffold_27761 | + | 405 | 483 | **RNA**→ |
| env-357 | SRS017120_Baylor_scaffold_16174 | + | 172 | 250 | **RNA**→ rplM (PRK09216)Ribosomal_L13 (pfam00572)→ rpsI (PRK00132)Ribosomal_S9 (pfam00380)→ |
| env-358 | SRS024318_LANL_scaffold_13466 | - | 227 | 149 | **RNA**→ hypo→ |
| env-359 | SRS018791_WUGC_scaffold_24827 | + | 398 | 476 | **RNA**→ rplM (PRK09216)Ribosomal_L13 (pfam00572)→ rpsI (PRK00132)Ribosomal_S9 (pfam00380)→ |
| env-360 | SRS075404_C3518551 | + | 334 | 412 | **RNA**→ |
| env-361 | SRS015395_C2316301 | + | 133 | 211 | **RNA**→ |
| env-362 | SRS019122_C5835162 | + | 454 | 532 | **RNA**→ |
| env-363 | SRS011086_WUGC_scaffold_41294 | + | 17651 | 17729 | **RNA**→ rplM (PRK09216)Ribosomal_L13 (pfam00572)→ rpsI (PRK00132)Ribosomal_S9 (pfam00380)→ |
| env-364 | SRS020334_Baylor_scaffold_26466 | - | 656 | 578 | **RNA**→ rplM (PRK09216)Ribosomal_L13 (pfam00572)→ |
| env-365 | SRS023617_Baylor_scaffold_72410 | + | 1232 | 1310 | **RNA**→ |
| env-366 | SRS047210_C3259279 | - | 125 | 47 | **RNA**→ |
| env-367 | SRS050244_LANL_scaffold_47339 | - | 253 | 175 | **RNA**→ |
| env-368 | SRS014689_C1337541 | - | 253 | 175 | **RNA**→ |
| env-369 | SRS016225_C2877068 | + | 184 | 262 | **RNA**→ |
| env-370 | SRS018591_WUGC_scaffold_20233 | + | 10037 | 10115 | **RNA**→ rplM (PRK09216)Ribosomal_L13 (pfam00572)→ rpsI (PRK00132)Ribosomal_S9 (pfam00380)→ |
| env-371 | SRS013705_C3658225 | + | 116 | 194 | **RNA**→ |
| env-372 | SRS019607_WUGC_scaffold_9276 | - | 440 | 362 | **RNA**→ rplM (PRK09216)Ribosomal_L13 (pfam00572)→ |
| env-373 | SRS014684_C3202863 | + | 288 | 366 | **RNA**→ |
| env-374 | SRS047824_WUGC_scaffold_43430 | + | 1021 | 1099 | **RNA**→ rplM (PRK09216)Ribosomal_L13 (pfam00572)→ rpsI (PRK00132)Ribosomal_S9 (pfam00380)→ |
| env-375 | SRS014573_WUGC_scaffold_51555 | - | 550 | 472 | **RNA**→ rplM (PRK09216)Ribosomal_L13 (pfam00572)→ |
| env-376 | SRS015209_WUGC_scaffold_7145 | - | 1222 | 1144 | **RNA**→ rplM (PRK09216)Ribosomal_L13 (pfam00572)→ rpsI (PRK00132)Ribosomal_S9 (pfam00380)→ |
| env-377 | SRS021496_Baylor_scaffold_21245 | + | 106 | 184 | **RNA**→ rplM (PRK09216)Ribosomal_L13 (pfam00572)→ rpsI (PRK00132)Ribosomal_S9 (pfam00380)→ |
| env-378 | SRS023835_Baylor_scaffold_21120 | - | 440 | 362 | **RNA**→ rplM (PRK09216)Ribosomal_L13 (pfam00572)→ |
| env-379 | SRS047219_WUGC_scaffold_3870 | + | 3309 | 3387 | **RNA**→ rplM (PRK09216)Ribosomal_L13 (pfam00572)→ rpsI (PRK00132)Ribosomal_S9 (pfam00380)→ |
| env-380 | SRS048791_LANL_scaffold_13616 | + | 156 | 234 | **RNA**→ |
| env-381 | SRS051791_LANL_scaffold_6037 | + | 5220 | 5298 | **RNA**→ rplM (PRK09216)Ribosomal_L13 (pfam00572)→ rpsI (PRK00132)Ribosomal_S9 (pfam00380)→ |
| env-382 | SRS062544_LANL_scaffold_33552 | + | 2985 | 3063 | **RNA**→ rplM (PRK09216)Ribosomal_L13 (pfam00572)→ rpsI (PRK00132)Ribosomal_S9 (pfam00380)→ |
| env-383 | SRS063193_LANL_scaffold_17717 | - | 550 | 472 | **RNA**→ rplM (PRK09216)Ribosomal_L13 (pfam00572)→ |
| env-384 | SRS023926_Baylor_scaffold_61302 | - | 446 | 368 | **RNA**→ rplM (PRK09216)Ribosomal_L13 (pfam00572)→ |
| env-385 | SRS019126_C2050939 | + | 106 | 184 | **RNA**→ |
| env-386 | SRS014692_C1429530 | + | 228 | 306 | **RNA**→ |
| env-387 | SRS023352_LANL_scaffold_12203 | + | 574 | 652 | **RNA**→ rplM (PRK09216)Ribosomal_L13 (pfam00572)→ rpsI (PRK00132)Ribosomal_S9 (pfam00380)→ |
| env-388 | SRS053603_LANL_scaffold_56678 | - | 125 | 47 | **RNA**→ hypo→ |
| env-389 | SRS075404_LANL_scaffold_32190 | + | 334 | 412 | **RNA**→ rplM (PRK09216)Ribosomal_L13 (pfam00572)→ |

| | | | | | |
|---|---|---|---|---|---|
| env-390 | SRS019026_C1988327 | + | 1161 | 1239 | **RNA**→ |
| env-391 | SRS042131_WUGC_scaffold_38955 | - | 396 | 318 | **RNA**→ rplM (PRK09216)Ribosomal_L13 (pfam00572)→ |
| env-392 | SRS015038_WUGC_scaffold_15341 | - | 253 | 175 | **RNA**→ |
| env-393 | SRS016002_WUGC_scaffold_10716 | + | 1413 | 1491 | **RNA**→ rplM (PRK09216)Ribosomal_L13 (pfam00572)→ rplM (PRK09216)Ribosomal_L13 (pfam00572)→ rpsI (PRK00132)Ribosomal_S9 (pfam00380)→ |
| env-394 | SRS056323_C2617078 | - | 253 | 175 | **RNA**→ |
| env-395 | SRS065278_C2566571 | - | 253 | 175 | **RNA**→ |
| env-396 | SRS019974_C3363040 | + | 862 | 940 | **RNA**→ |
| Psa-2-1 | NZ_GL629647.1 | + | 2253698 | 2253776 | **RNA**→ rplM (PRK09216)Ribosomal_L13 (pfam00572)→ rpsI (PRK00132)Ribosomal_S9 (pfam00380)→ |
| env-397 | SRS016086_C2389321 | - | 101 | 23 | **RNA**→ hypo→ |
| env-398 | SRS020856_Baylor_scaffold_13408 | + | 234 | 312 | **RNA**→ rplM (PRK09216)Ribosomal_L13 (pfam00572)→ rpsI (PRK00132)Ribosomal_S9 (pfam00380)→ |
| env-399 | SRS014271_WUGC_scaffold_18759 | + | 466 | 544 | **RNA**→ rplM (PRK09216)Ribosomal_L13 (pfam00572)→ |
| env-400 | SRS015057_WUGC_scaffold_34905 | + | 1097 | 1175 | **RNA**→ rplM (PRK09216)Ribosomal_L13 (pfam00572)→ |
| Psa-3-1 | NZ_AWGW01000017.1 | + | 92405 | 92483 | **RNA**→ rplM (PRK09216)Ribosomal_L13 (pfam00572)→ rpsI (PRK00132)Ribosomal_S9 (pfam00380)→ |
| env-401 | SRS012279_C2722110 | + | 914 | 992 | **RNA**→ |
| env-402 | SRS024015_Baylor_scaffold_12438 | + | 2443 | 2521 | **RNA**→ rplM (PRK09216)Ribosomal_L13 (pfam00572)→ rpsI (PRK00132)Ribosomal_S9 (pfam00380)→ |
| env-403 | SRS013234_C2993348 | - | 253 | 175 | **RNA**→ |
| env-404 | SRS024081_LANL_scaffold_75049 | + | 9042 | 9120 | **RNA**→ |
| env-405 | SRS057205_C3268011 | + | 63 | 141 | **RNA**→ |
| env-406 | Tongue_dor_WUGI_scaffold_48438 | - | 373 | 295 | **RNA**→ rplM (PRK09216)Ribosomal_L13 (pfam00572)→ |
| env-407 | SRS022143_WUGC_scaffold_32861 | + | 47 | 125 | **RNA**→ rplM (PRK09216)Ribosomal_L13 (pfam00572)→ |
| env-408 | SRS016541_Baylor_scaffold_9219 | - | 350 | 274 | **RNA**→ rplM (PRK09216)Ribosomal_L13 (pfam00572)→ |
| env-409 | SRS016575_C3620180 | + | 304 | 380 | **RNA**→ |
| env-410 | SRS017691_Baylor_scaffold_1218 | - | 223 | 147 | **RNA**→ |
| env-411 | SRS049318_C6476930 | + | 709 | 785 | **RNA**→ |
| env-412 | SRS016569_Baylor_scaffold_25639 | - | 252 | 175 | **RNA**→ |
| Por-6-1 | NZ_BAJC01000029.1 | + | 9652 | 9728 | **RNA**→ hypo→ rplM (PRK09216)Ribosomal_L13 (pfam00572)→ rpsI (PRK00132)Ribosomal_S9 (pfam00380)→ |
| Por-5-1 | NZ_KB898581.1 | - | 166045 | 165968 | **RNA**→ rplM (PRK09216)Ribosomal_L13 (pfam00572)→ rpsI (PRK00132)Ribosomal_S9 (pfam00380)→ |
| env-413 | SRS013723_C3447116 | - | 123 | 46 | **RNA**→ |
| env-414 | SRS013170_Baylor_scaffold_49654 | - | 224 | 147 | **RNA**→ hypo→ |
| env-415 | SRS015055_C1639257 | - | 224 | 147 | **RNA**→ hypo→ |
| env-416 | SRS015062_C160810 | - | 234 | 157 | **RNA**→ hypo→ |
| env-417 | SRS023964_C2594004 | - | 224 | 147 | **RNA**→ |
| Por-4-1 | NZ_GL349571.1 | + | 7135 | 7212 | **RNA**→ rplM (PRK09216)Ribosomal_L13 (pfam00572)→ rplM (PRK09216)Ribosomal_L13 (pfam00572)→ rpsI (PRK00132)Ribosomal_S9 (pfam00380)→ rpsI (PRK00132)Ribosomal_S9 (pfam00380)→ |
| Por-7-1 | NZ_GG703883.1 | + | 239161 | 239238 | **RNA**→ rplM (PRK09216)Ribosomal_L13 (pfam00572)→ rpsI (PRK00132)Ribosomal_S9 (pfam00380)→ |
| env-418 | Buccal_mucosa_LANL_C65077335 | + | 342 | 419 | **RNA**→ hypo→ |
| env-419 | SRS019980_C3064054 | - | 118 | 41 | **RNA**→ hypo→ |
| env-420 | SRS019029_C3217778 | + | 42 | 119 | **RNA**→ hypo→ rplM (PRK09216)Ribosomal_L13 (pfam00572)→ |
| env-421 | SRS024087_C3239609 | + | 108 | 185 | **RNA**→ |
| env-422 | SRS014476_C3433381 | + | 868 | 945 | **RNA**→ |
| env-423 | Subgingival_plaque_LANL_C11098963 | - | 118 | 41 | **RNA**→ hypo→ |
| env-424 | SRS024649_LANL_scaffold_18754 | + | 580 | 657 | **RNA**→ |
| Psp-8-1 | NZ_AZHT01000177.1 | + | 6805 | 6883 | **RNA**→ rplM (PRK09216)Ribosomal_L13 (pfam00572)→ rpsI (PRK00132)Ribosomal_S9 (pfam00380)→ |
| env-425 | SRS019219_WUGC_scaffold_64631 | - | 18808 | 18730 | **RNA**→ rplM (PRK09216)Ribosomal_L13 (pfam00572)→ rpsI (PRK00132)Ribosomal_S9 (pfam00380)→ |
| env-426 | SRS014271_C2786805 | - | 121 | 43 | **RNA**→ hypo→ |
| env-427 | SRS024318_LANL_scaffold_76469 | + | 3580 | 3658 | **RNA**→ |
| env-428 | SRS062544_C2827308 | - | 121 | 43 | **RNA**→ hypo→ |
| env-429 | SRS017120_Baylor_scaffold_66317 | - | 655 | 577 | **RNA**→ rplM (PRK09216)Ribosomal_L13 (pfam00572)→ |
| env-430 | SRS042643_WUGC_scaffold_1740 | - | 1427 | 1349 | **RNA**→ rplM (PRK09216)Ribosomal_L13 (pfam00572)→ rpsI (PRK00132)Ribosomal_S9 (pfam00380)→ |
| env-431 | SRS018739_WUGC_scaffold_55643 | + | 229 | 307 | **RNA**→ rplM (PRK09216)Ribosomal_L13 (pfam00572)→ |
| env-432 | SRS050244_LANL_scaffold_95751 | + | 228 | 306 | **RNA**→ rplM (PRK09216)Ribosomal_L13 (pfam00572)→ rpsI (PRK00132)Ribosomal_S9 (pfam00380)→ |
| env-433 | SRS019026_C1859658 | + | 229 | 307 | **RNA**→ |

| | | | | | |
|---|---|---|---|---|---|
| env-434 | SRS044373_C2740995 | - | 121 | 43 | **RNA**→ hypo→ |
| env-435 | SRS021496_Baylor_scaffold_66287 | + | 228 | 306 | **RNA**→ rplM (PRK09216)Ribosomal_L13 (pfam00572)→ rpsI (PRK00132)Ribosomal_S9 (pfam00380)→ |
| env-436 | SRS023352_LANL_scaffold_79269 | - | 463 | 385 | **RNA**→ rplM (PRK09216)Ribosomal_L13 (pfam00572)→ |
| env-437 | SRS022143_WUGC_scaffold_73957 | + | 77 | 155 | **RNA**→ rplM (PRK09216)Ribosomal_L13 (pfam00572)→ rpsI (PRK00132)Ribosomal_S9 (pfam00380)→ |
| env-438 | SRS015893_C2357989 | - | 655 | 577 | **RNA**→ rplM (PRK09216)Ribosomal_L13 (pfam00572)→ |
| env-439 | SRS015057_WUGC_scaffold_71704 | + | 230 | 308 | **RNA**→ rplM (PRK09216)Ribosomal_L13 (pfam00572)→ |
| env-440 | SRS024355_LANL_scaffold_106705 | - | 2853 | 2775 | **RNA**→ rplM (PRK09216)Ribosomal_L13 (pfam00572)→ rpsI (PRK00132)Ribosomal_S9 (pfam00380)→ |
| env-441 | SRS058053_C5276896 | - | 119 | 41 | **RNA**→ |
| env-442 | SRS023595_C6791388 | - | 141 | 63 | **RNA**→ hypo→ |
| env-443 | SRS047219_C2204013 | + | 361 | 439 | **RNA**→ |
| Pou-1-1 | NZ_JH114217.1 | + | 161819 | 161897 | **RNA**→ rplM (PRK09216)Ribosomal_L13 (pfam00572)→ rpsI (PRK00132)Ribosomal_S9 (pfam00380)→ |
| env-444 | SRS019980_C2940732 | + | 377 | 455 | **RNA**→ |
| env-445 | SRS022536_LANL_scaffold_97402 | - | 1401 | 1323 | **RNA**→ rplM (PRK09216)Ribosomal_L13 (pfam00572)→ rpsI (PRK00132)Ribosomal_S9 (pfam00380)→ |
| env-446 | SRS018591_C1244473 | - | 270 | 192 | **RNA**→ rplM (PRK09216)→ |
| env-447 | SRS063999_LANL_scaffold_89325 | + | 324 | 402 | **RNA**→ rplM (PRK09216)Ribosomal_L13 (pfam00572)→ rpsI (PRK00132)Ribosomal_S9 (pfam00380)→ |
| Pou-2-1 | NZ_BAJQ01000079.1 | - | 3607 | 3529 | **RNA**→ rplM (PRK09216)Ribosomal_L13 (pfam00572)→ rpsI (PRK00132)Ribosomal_S9 (pfam00380)→ |
| env-448 | AUXO010375897.1 | + | 341 | 415 | **RNA**→ rplM (PRK09216)Ribosomal_L13 (pfam00572)→ rpsI (PRK00132)Ribosomal_S9 (pfam00380)→ |
| env-449 | SRS063603_C4098342 | + | 290 | 366 | **RNA**→ |
| Pba-2-1 | NZ_AWEY01000020.1 | - | 92342 | 92266 | **RNA**→ rplM (PRK09216)Ribosomal_L13 (pfam00572)→ rpsI (PRK00132)Ribosomal_S9 (pfam00380)→ |
| Pba-1-1 | NZ_AUFQ01000002.1 | - | 90788 | 90712 | **RNA**→ rplM (PRK09216)Ribosomal_L13 (pfam00572)→ rpsI (PRK00132)Ribosomal_S9 (pfam00380)→ |
| Pba-1-2 | NZ_BAJF01000002.1 | + | 155272 | 155348 | **RNA**→ rplM (PRK09216)Ribosomal_L13 (pfam00572)→ rpsI (PRK00132)Ribosomal_S9 (pfam00380)→ |
| env-450 | SRS049389_C4455900 | + | 102 | 178 | **RNA**→ |
| Pde-2-1 | NZ_BAKG01000016.1 | + | 37012 | 37087 | **RNA**→ rplM (PRK09216)Ribosomal_L13 (pfam00572)→ rpsI (PRK00132)Ribosomal_S9 (pfam00380)→ |
| Pbe-1-1 | NZ_GG704782.1 | + | 371273 | 371346 | **RNA**→ rplM (PRK09216)Ribosomal_L13 (pfam00572)→ rpsI (PRK00132)Ribosomal_S9 (pfam00380)→ |
| env-451 | AUXO016035411.1 | - | 430 | 357 | **RNA**→ rplM (PRK09216)Ribosomal_L13 (pfam00572)→ |
| env-452 | AUXO017177372.1 | + | 741 | 814 | **RNA**→ rplM (PRK09216)Ribosomal_L13 (pfam00572)→ rpsI (PRK00132)Ribosomal_S9 (pfam00380)→ |
| env-453 | AUXO011185619.1 | + | 157 | 231 | **RNA**→ rplM (PRK09216)Ribosomal_L13 (pfam00572)→ rpsI (PRK00132)Ribosomal_S9 (pfam00380)→ |
| env-454 | SRS022725_LANL_scaffold_110518 | - | 1243 | 1167 | **RNA**→ rplM (PRK09216)Ribosomal_L13 (pfam00572)→ rpsI (PRK00132)Ribosomal_S9 (pfam00380)→ |
| env-455 | SRS047634_LANL_scaffold_138961 | - | 948 | 872 | **RNA**→ rplM (PRK09216)Ribosomal_L13 (pfam00572)→ rpsI (PRK00132)Ribosomal_S9 (pfam00380)→ |
| env-456 | SRS063603_LANL_scaffold_20726 | + | 2347 | 2423 | **RNA**→ rplM (PRK09216)Ribosomal_L13 (pfam00572)→ rpsI (PRK00132)Ribosomal_S9 (pfam00380)→ |
| env-457 | SRS058808_LANL_scaffold_70845 | - | 2078 | 2005 | **RNA**→ rplM (PRK09216)Ribosomal_L13 (pfam00572)→ rpsI (PRK00132)Ribosomal_S9 (pfam00380)→ |
| env-458 | SRS022602_Baylor_scaffold_3137 | + | 97 | 171 | **RNA**→ rplM (PRK09216)Ribosomal_L13 (pfam00572)→ |
| env-459 | SRS013252_Baylor_scaffold_54444 | - | 7007 | 6933 | **RNA**→ rplM (PRK09216)Ribosomal_L13 (pfam00572)→ rpsI (PRK00132)Ribosomal_S9 (pfam00380)→ |
| env-460 | SRS017511_C3369784 | - | 87 | 13 | **RNA**→ hypo→ |
| env-461 | SRS063603_LANL_scaffold_102327 | + | 763 | 837 | **RNA**→ |
| Psa-1-1 | NZ_KB290984.1 | - | 62870 | 62796 | **RNA**→ rplM (PRK09216)Ribosomal_L13 (pfam00572)→ rpsI (PRK00132)Ribosomal_S9 (pfam00380)→ |
| env-462 | SRS049268_C4307889 | - | 590 | 516 | **RNA**→ rplM (PRK09216)Ribosomal_L13 (pfam00572)→ |
| env-463 | SRS017227_C5254572 | - | 533 | 459 | **RNA**→ rplM (PRK09216)Ribosomal_L13 (pfam00572)→ |
| env-464 | SRS018665_C3785311 | + | 95 | 169 | **RNA**→ rplM (PRK09216)Ribosomal_L13 (pfam00572)→ rpsI (PRK00132)Ribosomal_S9 (pfam00380)→ |
| env-465 | SRS051941_C3920117 | - | 311 | 237 | **RNA**→ rplM (PRK09216)Ribosomal_L13 (pfam00572)→ |
| env-466 | HCF12C_28379 | - | 230 | 155 | **RNA**→ hypo→ |
| env-467 | 08_07_GOS_DNA_C8001741_1 | + | 143 | 218 | **RNA**→ rplM (PRK09216)Ribosomal_L13 (pfam00572)→ |
| env-468 | AUXO013741021.1 | + | 250 | 324 | **RNA**→ hypo→ |
| env-469 | AUXO016321942.1 | + | 109 | 183 | **RNA**→ rplM (PRK09216)Ribosomal_L13 (pfam00572)→ rpsI (PRK00132)Ribosomal_S9 (pfam00380)→ |
| env-470 | AUXO016352636.1 | + | 21 | 94 | **RNA**→ rplM (PRK09216)Ribosomal_L13 (pfam00572)→ rpsI (PRK00132)Ribosomal_S9 (pfam00380)→ |
| Ppa-3-1 | NZ_BAJH01000014.1 | + | 85099 | 85176 | **RNA**→ rplM (PRK09216)Ribosomal_L13 (pfam00572)→ rpsI (PRK00132)Ribosomal_S9 (pfam00380)→ |
| Ppa-2-1 | NZ_KB890632.1 | + | 85379 | 85456 | **RNA**→ rplM (PRK09216)Ribosomal_L13 (pfam00572)→ rpsI (PRK00132)Ribosomal_S9 (pfam00380)→ |
| env-471 | 2205195008 | + | 38 | 115 | **RNA**→ |
| env-472 | 2157459302 | + | 38 | 115 | **RNA**→ |
| env-473 | JGI20225J20221_1004075 | + | 804 | 881 | **RNA**→ rplM (PRK09216)Ribosomal_L13 (pfam00572)→ rpsI (PRK00132)Ribosomal_S9 (pfam00380)→ |
| env-474 | BMHB3a_c149604 | - | 2595 | 2518 | **RNA**→ rplM (PRK09216)Ribosomal_L13 (pfam00572)→ rpsI (PRK00132)Ribosomal_S9 (pfam00380)→ |
| env-475 | JGI20225J20221_1000003 | - | 388531 | 388454 | **RNA**→ rplM (PRK09216)Ribosomal_L13 (pfam00572)→ rpsI (PRK00132)Ribosomal_S9 (pfam00380)→ |

| | | | | | |
|---|---|---|---|---|---|
| env-476 | BMHBC_68715 | + | 149 | 226 | **RNA**→ rplM (PRK09216)Ribosomal_L13 (pfam00572)→ rpsI (PRK00132)Ribosomal_S9 (pfam00380)→ |
| env-477 | BMHBC_2_1802482 | + | 57 | 134 | **RNA**→ |
| env-478 | BMHBC_2_3001399 | + | 57 | 134 | **RNA**→ |
| env-479 | BMHBC_2_2370598 | + | 57 | 134 | **RNA**→ |
| env-480 | BMHBC_2_1178038 | + | 57 | 134 | **RNA**→ |
| env-481 | BMHBC_2_612469 | + | 57 | 134 | **RNA**→ |
| env-482 | BMHBC_2_3578336 | + | 57 | 134 | **RNA**→ |
| env-483 | HCE12Call500_c0056283 | + | 177 | 251 | **RNA**→ rplM (PRK09216)Ribosomal_L13 (pfam00572)→ rpsI (PRK00132)Ribosomal_S9 (pfam00380)→ |
| env-484 | HCF12C_221933 | - | 28484 | 28410 | **RNA**→ rplM (PRK09216)Ribosomal_L13 (pfam00572)→ rpsI (PRK00132)Ribosomal_S9 (pfam00380)→ |
| env-485 | HCF14C_11080 | - | 2194 | 2120 | **RNA**→ rplM (PRK09216)Ribosomal_L13 (pfam00572)→ rpsI (PRK00132)Ribosomal_S9 (pfam00380)→ |
| env-486 | AUXO012434654.1 | - | 366 | 290 | **RNA**→ rplM (PRK09216)RplM (COG0102)Ribosomal_L13 (cd00392)→ |
| env-487 | AUXO011866134.1 | + | 7 | 83 | **RNA**→ ←hypo |
| env-488 | AUXO017757733.1 | - | 1325 | 1250 | **RNA**→ rplM (PRK09216)Ribosomal_L13 (pfam00572)→ rpsI (PRK00132)Ribosomal_S9 (pfam00380)→ |
| env-489 | AUXO011774349.1 | - | 771 | 700 | **RNA**→ rplM (PRK09216)Ribosomal_L13 (pfam00572)→ rpsI (PRK00132)Ribosomal_S9 (pfam00380)→ |
| env-490 | SRS013542_LANL_scaffold_1766 | - | 69128 | 69055 | **RNA**→ rplM (PRK09216)Ribosomal_L13 (pfam00572)→ rpsI (PRK00132)Ribosomal_S9 (pfam00380)→ |
| env-491 | SRS023604_Baylor_scaffold_50 | + | 320 | 393 | **RNA**→ rplM (PRK09216)Ribosomal_L13 (pfam00572)→ rpsI (PRK00132)Ribosomal_S9 (pfam00380)→ |
| env-492 | SRS017497_Baylor_scaffold_193 | - | 91994 | 91921 | **RNA**→ rplM (PRK09216)Ribosomal_L13 (pfam00572)→ rpsI (PRK00132)Ribosomal_S9 (pfam00380)→ |
| env-493 | SRS023468_LANL_scaffold_8569 | + | 176 | 249 | **RNA**→ rplM (PRK09216)Ribosomal_L13 (pfam00572)→ rpsI (PRK00132)Ribosomal_S9 (pfam00380)→ |
| Pam-1-1 | NZ_ADFQ01000027.1 | + | 26891 | 26964 | **RNA**→ rplM (PRK09216)Ribosomal_L13 (pfam00572)→ rplM (PRK09216)Ribosomal_L13 (pfam00572)→ rpsI (PRK00132)Ribosomal_S9 (pfam00380)→ rpsI (PRK00132)Ribosomal_S9 (pfam00380)→ |
| Pam-2-1 | NZ_JRNU01000014.1 | - | 45509 | 45436 | **RNA**→ rplM (PRK09216)Ribosomal_L13 (pfam00572)→ rpsI (PRK00132)Ribosomal_S9 (pfam00380)→ |
| Pam-3-1 | NZ_KB905270.1 | + | 27167 | 27240 | **RNA**→ rplM (PRK09216)Ribosomal_L13 (pfam00572)→ rpsI (PRK00132)Ribosomal_S9 (pfam00380)→ |
| Pam-4-1 | NZ_BAJP01000004.1 | - | 90359 | 90286 | **RNA**→ rplM (PRK09216)Ribosomal_L13 (pfam00572)→ rpsI (PRK00132)Ribosomal_S9 (pfam00380)→ |
| Pbi-1-1 | NZ_JUWN01000307.1 | + | 2136 | 2209 | **RNA**→ rplM (PRK09216)Ribosomal_L13 (pfam00572)→ rpsI (PRK00132)Ribosomal_S9 (pfam00380)→ |
| Pbi-6-1 | NZ_ADFO01000036.1 | - | 4358 | 4285 | **RNA**→ rplM (PRK09216)Ribosomal_L13 (pfam00572)→ rplM (PRK09216)Ribosomal_L13 (pfam00572)→ rpsI (PRK00132)Ribosomal_S9 (pfam00380)→ rpsI (PRK00132)Ribosomal_S9 (pfam00380)→ |
| Pbi-5-1 | NZ_JH660660.1 | + | 893597 | 893670 | **RNA**→ rplM (PRK09216)Ribosomal_L13 (pfam00572)→ rpsI (PRK00132)Ribosomal_S9 (pfam00380)→ |
| Pbi-2-1 | NZ_JRNF01000004.1 | + | 30225 | 30298 | **RNA**→ rplM (PRK09216)Ribosomal_L13 (pfam00572)→ rpsI (PRK00132)Ribosomal_S9 (pfam00380)→ |
| Pbi-4-1 | NZ_JRNM01000029.1 | + | 14173 | 14246 | **RNA**→ rplM (PRK09216)Ribosomal_L13 (pfam00572)→ rpsI (PRK00132)Ribosomal_S9 (pfam00380)→ |
| Pbi-3-1 | NZ_JRNQ01000001.1 | - | 38223 | 38150 | **RNA**→ rplM (PRK09216)Ribosomal_L13 (pfam00572)→ rpsI (PRK00132)Ribosomal_S9 (pfam00380)→ |
| Pve-1-1 | NZ_KB898330.1 | + | 50120 | 50193 | **RNA**→ rplM (PRK09216)Ribosomal_L13 (pfam00572)→ rpsI (PRK00132)Ribosomal_S9 (pfam00380)→ |
| Pve-2-1 | NZ_BAIP01000005.1 | + | 49830 | 49903 | **RNA**→ rplM (PRK09216)Ribosomal_L13 (pfam00572)→ rpsI (PRK00132)Ribosomal_S9 (pfam00380)→ |
| Pve-3-1 | NZ_GG698713.1 | - | 155542 | 155469 | **RNA**→ rplM (PRK09216)Ribosomal_L13 (pfam00572)→ rpsI (PRK00132)Ribosomal_S9 (pfam00380)→ |
| env-494 | SRS013836_C4770317 | - | 112 | 39 | **RNA**→ hypo→ |
| env-495 | SRS014689_WUGC_scaffold_10524 | - | 158 | 84 | **RNA**→ hypo→ |
| env-496 | SRS019045_WUGC_scaffold_55373 | + | 7079 | 7153 | **RNA**→ |
| env-497 | SRS064329_LANL_scaffold_12626 | + | 135 | 209 | **RNA**→ rplM (PRK09216)Ribosomal_L13 (pfam00572)→ |
| Psc-1-1 | NZ_BAKP01000004.1 | + | 58256 | 58330 | **RNA**→ rplM (PRK09216)Ribosomal_L13 (pfam00572)→ rpsI (PRK00132)Ribosomal_S9 (pfam00380)→ |
| env-498 | SRS017808_Baylor_scaffold_59118 | + | 835 | 909 | **RNA**→ |
| env-499 | SRS044373_WUGC_scaffold_55808 | + | 931 | 1005 | **RNA**→ |
| env-500 | SRS053603_LANL_scaffold_86116 | + | 1113 | 1187 | **RNA**→ |
| Psp-6-1 | NZ_KI259601.1 | + | 30175 | 30249 | **RNA**→ rplM (PRK09216)Ribosomal_L13 (pfam00572)→ rpsI (PRK00132)Ribosomal_S9 (pfam00380)→ |
| Pme-3-1 | NZ_JRNS01000294.1 | + | 180 | 255 | **RNA**→ rplM (PRK09216)Ribosomal_L13 (pfam00572)→ rpsI (PRK00132)Ribosomal_S9 (pfam00380)→ |
| env-501 | SRS024318_LANL_scaffold_11533 | - | 620 | 547 | **RNA**→ rplM (PRK09216)Ribosomal_L13 (pfam00572)→ rpsI (PRK00132)Ribosomal_S9 (pfam00380)→ |
| env-502 | SRS017439_Baylor_scaffold_50412 | - | 108 | 36 | **RNA**→ |
| env-503 | SRS011086_WUGC_scaffold_40282 | - | 108 | 36 | **RNA**→ hypo→ |
| env-504 | SRS014124_WUGC_scaffold_14407 | - | 802 | 730 | **RNA**→ rplM (PRK09216)Ribosomal_L13 (pfam00572)→ rplM (PRK09216)Ribosomal_L13 (pfam00572)→ rpsI (PRK00132)Ribosomal_S9 (pfam00380)→ |
| env-505 | SRS017533_Baylor_scaffold_7923 | + | 285 | 357 | **RNA**→ rplM (PRK09216)Ribosomal_L13 (pfam00572)→ |
| env-506 | SRS049147_LANL_scaffold_62005 | + | 1217 | 1289 | **RNA**→ |
| Phi-1-1 | NZ_JH376764.1 | + | 326001 | 326073 | **RNA**→ hypo→ rplM (PRK09216)Ribosomal_L13 (pfam00572)→ rpsI (PRK00132)Ribosomal_S9 (pfam00380)→ |
| env-507 | SRS017076_C2635852 | + | 287 | 359 | **RNA**→ hypo→ |
| Phi-2-1 | NZ_BAJX01000005.1 | - | 144658 | 144586 | **RNA**→ rplM (PRK09216)Ribosomal_L13 (pfam00572)→ rpsI (PRK00132)Ribosomal_S9 (pfam00380)→ |

| | | | | | |
|---|---|---|---|---|---|
| env-508 | SRS013234_Baylor_scaffold_28796 | - | 148 | 76 | **RNA**→ |
| env-509 | SRS024637_LANL_scaffold_306 | + | 443 | 515 | **RNA**→ rplM (PRK09216)Ribosomal_L13 (pfam00572)→ rpsI (PRK00132)Ribosomal_S9 (pfam00380)→ |
| Phi-2-2 | NZ_JRNJ01000036.1 | - | 45722 | 45650 | **RNA**→ rplM (PRK09216)Ribosomal_L13 (pfam00572)→ rpsI (PRK00132)Ribosomal_S9 (pfam00380)→ |
| env-510 | SRS014573_WUGC_scaffold_34708 | - | 1134 | 1060 | **RNA**→ rplM (PRK09216)Ribosomal_L13 (pfam00572)→ rpsI (PRK00132)Ribosomal_S9 (pfam00380)→ |
| env-511 | SRS021496_Baylor_scaffold_22917 | + | 96 | 170 | **RNA**→ rplM (PRK09216)Ribosomal_L13 (pfam00572)→ |
| env-512 | SRS015644_WUGC_scaffold_31522 | - | 876 | 802 | **RNA**→ rplM (PRK09216)Ribosomal_L13 (pfam00572)→ rpsI (PRK00132)Ribosomal_S9 (pfam00380)→ |
| env-513 | SRS042131_C2183549 | + | 266 | 340 | **RNA**→ |
| env-514 | SRS022719_LANL_scaffold_14486 | + | 176535 | 176609 | **RNA**→ rplM (PRK09216)Ribosomal_L13 (pfam00572)→ rpsI (PRK00132)Ribosomal_S9 (pfam00380)→ |
| env-515 | SRS044662_LANL_scaffold_9271 | + | 199524 | 199598 | **RNA**→ rplM (PRK09216)Ribosomal_L13 (pfam00572)→ rpsI (PRK00132)Ribosomal_S9 (pfam00380)→ |
| env-516 | SRS013879_C2814589 | - | 110 | 36 | **RNA**→ |
| env-517 | SRS018145_Baylor_scaffold_16176 | - | 11602 | 11528 | **RNA**→ rplM (PRK09216)Ribosomal_L13 (pfam00572)→ rpsI (PRK00132)Ribosomal_S9 (pfam00380)→ |
| env-518 | SRS057692_LANL_scaffold_51493 | + | 229 | 303 | **RNA**→ |
| env-519 | SRS016002_WUGC_scaffold_30040 | + | 6147 | 6221 | **RNA**→ |
| Psp-5-1 | NZ_JH114141.1 | + | 226443 | 226517 | **RNA**→ rplM (PRK09216)Ribosomal_L13 (pfam00572)→ rpsI (PRK00132)Ribosomal_S9 (pfam00380)→ |
| env-520 | SRS017713_Baylor_scaffold_12488 | - | 158 | 84 | **RNA**→ hypo→ |
| env-521 | SRS056622_LANL_scaffold_17923 | - | 158 | 84 | **RNA**→ hypo→ |
| env-522 | SRS022625_Baylor_scaffold_4609 | - | 158 | 84 | **RNA**→ hypo→ |
| env-523 | SRS016037_WUGC_scaffold_34392 | + | 1842 | 1916 | **RNA**→ rplM (PRK09216)Ribosomal_L13 (pfam00572)→ rpsI (PRK00132)Ribosomal_S9 (pfam00380)→ |
| env-524 | SRS054687_LANL_scaffold_40466 | - | 149 | 75 | **RNA**→ hypo→ |
| env-525 | SRS016569_Baylor_scaffold_12365 | + | 198 | 272 | **RNA**→ rplM (PRK09216)Ribosomal_L13 (pfam00572)→ rplM (PRK09216)Ribosomal_L13 (pfam00572)→ rpsI (PRK00132)Ribosomal_S9 (pfam00380)→ |
| env-526 | SRS043663_LANL_scaffold_8456 | + | 196 | 270 | **RNA**→ rplM (PRK09216)Ribosomal_L13 (pfam00572)→ |
| env-527 | SRS011086_WUGC_scaffold_9083 | - | 2222 | 2148 | **RNA**→ rplM (PRK09216)Ribosomal_L13 (pfam00572)→ rpsI (PRK00132)Ribosomal_S9 (pfam00380)→ |
| env-528 | SRS024441_LANL_scaffold_8252 | - | 965 | 891 | **RNA**→ rplM (PRK09216)Ribosomal_L13 (pfam00572)→ rpsI (PRK00132)Ribosomal_S9 (pfam00380)→ |
| env-529 | SRS045127_LANL_scaffold_25864 | + | 4 | 78 | **RNA**→ rplM (PRK09216)Ribosomal_L13 (pfam00572)→ |
| env-530 | SRS052227_C2913848 | - | 999 | 925 | **RNA**→ rplM (PRK09216)Ribosomal_L13 (pfam00572)→ rpsI (PRK00132)Ribosomal_S9 (pfam00380)→ |
| Psp-10-1 | NZ_JACD01000007.1 | + | 195948 | 196022 | **RNA**→ rplM (PRK09216)Ribosomal_L13 (pfam00572)→ rpsI (PRK00132)Ribosomal_S9 (pfam00380)→ |
| env-531 | SRS018591_WUGC_scaffold_24424 | + | 1589 | 1663 | **RNA**→ |
| env-532 | SRS013947_C1601091 | - | 178 | 104 | **RNA**→ hypo→ |
| env-533 | SRS019120_C1333688 | + | 439 | 513 | **RNA**→ |
| env-534 | SRS017209_Baylor_scaffold_33086 | + | 1225 | 1299 | **RNA**→ |
| env-535 | SRS018394_C4052969 | + | 130 | 204 | **RNA**→ rplM (PRK09216)Ribosomal_L13 (pfam00572)→ rpsI (PRK00132)Ribosomal_S9 (pfam00380)→ |
| env-536 | SRS055378_LANL_scaffold_77236 | - | 397 | 323 | **RNA**→ rplM (PRK09216)Ribosomal_L13 (pfam00572)→ |
| env-537 | SRS013164_Baylor_scaffold_24536 | - | 1111 | 1037 | **RNA**→ rplM (PRK09216)Ribosomal_L13 (pfam00572)→ rpsI (PRK00132)Ribosomal_S9 (pfam00380)→ |
| env-538 | SRS011243_Baylor_scaffold_14041 | + | 7051 | 7125 | **RNA**→ rplM (PRK09216)Ribosomal_L13 (pfam00572)→ rpsI (PRK00132)Ribosomal_S9 (pfam00380)→ |
| env-539 | SRS011247_C1360241 | + | 440 | 514 | **RNA**→ rplM (PRK09216)Ribosomal_L13 (pfam00572)→ |
| env-540 | SRS013502_Baylor_scaffold_7651 | - | 3320 | 3246 | **RNA**→ rplM (PRK09216)Ribosomal_L13 (pfam00572)→ rpsI (PRK00132)Ribosomal_S9 (pfam00380)→ |
| env-541 | SRS013879_WUGC_scaffold_9024 | - | 172 | 98 | **RNA**→ |
| env-542 | SRS014124_WUGC_scaffold_11710 | - | 948 | 874 | **RNA**→ rplM (PRK09216)Ribosomal_L13 (pfam00572)→ rplM (PRK09216)Ribosomal_L13 (pfam00572)→ rpsI (PRK00132)Ribosomal_S9 (pfam00380)→ rpsI (PRK00132)Ribosomal_S9 (pfam00380)→ |
| env-543 | SRS014689_WUGC_scaffold_10523 | - | 158 | 84 | **RNA**→ |
| env-544 | SRS016501_C1543222 | + | 266 | 340 | **RNA**→ rplM (PRK09216)RplM (COG0102)→ |
| env-545 | SRS019022_WUGC_scaffold_10171 | - | 190 | 116 | **RNA**→ |
| env-546 | SRS019045_WUGC_scaffold_18044 | + | 6417 | 6491 | **RNA**→ |
| env-547 | SRS023557_Baylor_scaffold_343 | - | 20477 | 20403 | **RNA**→ rplM (PRK09216)Ribosomal_L13 (pfam00572)→ rpsI (PRK00132)Ribosomal_S9 (pfam00380)→ |
| env-548 | SRS024580_LANL_scaffold_5027 | + | 1876 | 1950 | **RNA**→ rplM (PRK09216)Ribosomal_L13 (pfam00572)→ rpsI (PRK00132)Ribosomal_S9 (pfam00380)→ |
| env-549 | SRS057355_LANL_scaffold_16577 | - | 735 | 661 | **RNA**→ rplM (PRK09216)Ribosomal_L13 (pfam00572)→ rpsI (PRK00132)Ribosomal_S9 (pfam00380)→ |
| env-550 | SRS063288_LANL_scaffold_61854 | + | 267 | 341 | **RNA**→ rplM (PRK09216)Ribosomal_L13 (pfam00572)→ rpsI (PRK00132)Ribosomal_S9 (pfam00380)→ |
| env-551 | SRS077736_LANL_scaffold_11170 | - | 5468 | 5394 | **RNA**→ rplM (PRK09216)Ribosomal_L13 (pfam00572)→ rpsI (PRK00132)Ribosomal_S9 (pfam00380)→ |
| Pme-2-1 | NZ_GG740010.1 | + | 211066 | 211140 | **RNA**→ rplM (PRK09216)Ribosomal_L13 (pfam00572)→ rplM (PRK09216)Ribosomal_L13 (pfam00572)→ rpsI (PRK00132)Ribosomal_S9 (pfam00380)→ rpsI (PRK00132)Ribosomal_S9 (pfam00380)→ |
| Pme-1-1 | NC_014370.1 | - | 173052 | 172978 | **RNA**→ rplM (PRK09216)Ribosomal_L13 (pfam00572)→ rpsI (PRK00132)Ribosomal_S9 (pfam00380)→ |
| env-552 | SRS024447_LANL_scaffold_5458 | - | 189 | 115 | **RNA**→ hypo→ |

| | | | | | |
|---|---|---|---|---|---|
| env-553 | SRS018739_WUGC_scaffold_1229 | - | 110 | 36 | **RNA**→ hypo→ |
| env-554 | SRS021954_Baylor_scaffold_773 | + | 402 | 476 | **RNA**→ rplM (PRK09216)Ribosomal_L13 (pfam00572)→ |
| env-555 | SRS020858_C742461 | + | 146 | 220 | **RNA**→ |
| env-556 | SRS055426_LANL_scaffold_45579 | - | 373 | 299 | **RNA**→ rplM (PRK09216)Ribosomal_L13 (pfam00572)→ |
| env-557 | SRS013818_Baylor_scaffold_15170 | + | 1030 | 1104 | **RNA**→ rplM (PRK09216)Ribosomal_L13 (pfam00572)→ |
| env-558 | SRS022530_LANL_scaffold_8517 | - | 2577 | 2503 | **RNA**→ rplM (PRK09216)Ribosomal_L13 (pfam00572)→ rpsI (PRK00132)Ribosomal_S9 (pfam00380)→ |
| env-559 | SRS018969_WUGC_scaffold_48352 | - | 1834 | 1760 | **RNA**→ rplM (PRK09216)Ribosomal_L13 (pfam00572)→ rpsI (PRK00132)Ribosomal_S9 (pfam00380)→ |
| env-560 | SRS019029_C3094948 | + | 207 | 281 | **RNA**→ |
| env-561 | SRS057539_LANL_scaffold_11258 | + | 1634 | 1708 | **RNA**→ rplM (PRK09216)Ribosomal_L13 (pfam00572)→ |
| Pfu-1-1 | NZ_CP012074.1 | - | 335584 | 335511 | **RNA**→ rplM (PRK09216)Ribosomal_L13 (pfam00572)→ rpsI (PRK00132)Ribosomal_S9 (pfam00380)→ |
| Psp-24-1 | NZ_AJIN01000058.1 | + | 75456 | 75530 | **RNA**→ rplM (PRK09216)Ribosomal_L13 (pfam00572)→ rplM (PRK09216)Ribosomal_L13 (pfam00572)→ rpsI (PRK00132)Ribosomal_S9 (pfam00380)→ rpsI (PRK00132)Ribosomal_S9 (pfam00380)→ |
| env-562 | SRS018357_Baylor_scaffold_53049 | + | 659 | 733 | **RNA**→ |
| env-563 | SRS044486_WUGC_scaffold_29347 | + | 207 | 281 | **RNA**→ |
| env-564 | SRS042643_WUGC_scaffold_65829 | + | 2043 | 2117 | **RNA**→ |
| Pde-3-1 | NZ_AEXO01000095.1 | + | 287046 | 287120 | **RNA**→ rplM (PRK09216)Ribosomal_L13 (pfam00572)→ rplM (PRK09216)Ribosomal_L13 (pfam00572)→ rpsI (PRK00132)Ribosomal_S9 (pfam00380)→ rpsI (PRK00132)Ribosomal_S9 (pfam00380)→ |
| env-565 | SRS017691_C5408377 | + | 1907 | 1981 | **RNA**→ |
| Pde-4-1 | NZ_JRNO01000001.1 | - | 3358 | 3284 | **RNA**→ rplM (PRK09216)Ribosomal_L13 (pfam00572)→ rpsI (PRK00132)Ribosomal_S9 (pfam00380)→ |
| Pde-5-1 | NZ_ATWK01000001.1 | + | 151366 | 151440 | **RNA**→ rplM (PRK09216)Ribosomal_L13 (pfam00572)→ rpsI (PRK00132)Ribosomal_S9 (pfam00380)→ |
| Pde-5-2 | NZ_BAJG01000001.1 | + | 303181 | 303255 | **RNA**→ rplM (PRK09216)Ribosomal_L13 (pfam00572)→ rpsI (PRK00132)Ribosomal_S9 (pfam00380)→ |
| env-566 | SRS013836_Baylor_scaffold_11239 | + | 579 | 653 | **RNA**→ |
| env-567 | SRS011255_Baylor_scaffold_46592 | + | 1416 | 1490 | **RNA**→ |
| Pde-6-1 | NC_015311.1 | + | 832827 | 832901 | **RNA**→ rplM (PRK09216)Ribosomal_L13 (pfam00572)→ rpsI (PRK00132)Ribosomal_S9 (pfam00380)→ |
| env-568 | SRS013836_C4811161 | - | 112 | 38 | **RNA**→ |
| env-569 | SRS019029_WUGC_scaffold_26302 | + | 226 | 300 | **RNA**→ |
| Pmu-1-1 | NZ_GL872283.1 | + | 37707 | 37781 | **RNA**→ rplM (PRK09216)Ribosomal_L13 (pfam00572)→ rpsI (PRK00132)Ribosomal_S9 (pfam00380)→ |
| env-570 | AUXO010614209.1 | + | 217 | 289 | **RNA**→ |
| env-571 | Supragingival_plaque_LANL_C181911197 | - | 124 | 49 | **RNA**→ hypo→ |
| env-572 | SRS018739_C3456198 | - | 221 | 146 | **RNA**→ hypo→ |
| env-573 | Supragingival_plaque_LANL_C182733701 | + | 533 | 608 | **RNA**→ |
| env-574 | MA40A_contig28992 | - | 33539 | 33466 | **RNA**→ rplM (PRK09216)Ribosomal_L13 (pfam00572)→ rpsI (PRK00132)Ribosomal_S9 (pfam00380)→ |
| env-575 | SRS023604_C759437 | + | 77 | 149 | **RNA**→ rplM (PRK09216)Ribosomal_L13 (pfam00572)→ |
| env-576 | SRS023468_LANL_scaffold_9905 | + | 61199 | 61271 | **RNA**→ rplM (PRK09216)Ribosomal_L13 (pfam00572)→ rpsI (PRK00132)Ribosomal_S9 (pfam00380)→ |
| Psp-22-1 | NZ_JRNC01000011.1 | + | 61278 | 61350 | **RNA**→ rplM (PRK09216)Ribosomal_L13 (pfam00572)→ rpsI (PRK00132)Ribosomal_S9 (pfam00380)→ |
| env-577 | RUMENNODE_143372_1 | - | 3480 | 3407 | **RNA**→ rplM (PRK09216)Ribosomal_L13 (pfam00572)→ rpsI (PRK00132)Ribosomal_S9 (pfam00380)→ |
| env-578 | SRS049318_LANL_scaffold_15105 | - | 3354 | 3278 | **RNA**→ rplM (PRK09216)Ribosomal_L13 (pfam00572)→ rpsI (PRK00132)Ribosomal_S9 (pfam00380)→ |
| env-579 | SRS017691_C5027809 | - | 118 | 42 | **RNA**→ hypo→ |
| Pmi-3-1 | NZ_JH594521.1 | - | 42001 | 41925 | **RNA**→ rplM (PRK09216)Ribosomal_L13 (pfam00572)→ rpsI (PRK00132)Ribosomal_S9 (pfam00380)→ |
| Pmi-1-1 | NZ_KB892256.1 | - | 41862 | 41786 | **RNA**→ rplM (PRK09216)Ribosomal_L13 (pfam00572)→ rpsI (PRK00132)Ribosomal_S9 (pfam00380)→ |
| Pmi-2-1 | NZ_BAKH01000001.1 | + | 67606 | 67682 | **RNA**→ rplM (PRK09216)Ribosomal_L13 (pfam00572)→ rpsI (PRK00132)Ribosomal_S9 (pfam00380)→ |
| env-580 | SRS047634_LANL_scaffold_139898 | + | 902 | 978 | **RNA**→ rplM (PRK09216)Ribosomal_L13 (pfam00572)→ rpsI (PRK00132)Ribosomal_S9 (pfam00380)→ |
| env-581 | SRS055378_C3881430 | - | 175 | 99 | **RNA**→ hypo→ |
| env-582 | SRS022602_Baylor_scaffold_67890 | - | 3320 | 3244 | **RNA**→ rplM (PRK09216)Ribosomal_L13 (pfam00572)→ rpsI (PRK00132)Ribosomal_S9 (pfam00380)→ |
| Pin-1-1 | NC_017861.1 | - | 192856 | 192781 | **RNA**→ rplM (PRK09216)Ribosomal_L13 (pfam00572)→ rpsI (PRK00132)Ribosomal_S9 (pfam00380)→ |
| env-583 | SRS014692_WUGC_scaffold_15999 | + | 4242 | 4316 | **RNA**→ rplM (PRK09216)Ribosomal_L13 (pfam00572)→ rpsI (PRK00132)Ribosomal_S9 (pfam00380)→ |
| env-584 | SRS014684_WUGC_scaffold_44181 | + | 3215 | 3289 | **RNA**→ |
| env-585 | SRS014689_WUGC_scaffold_31060 | + | 16578 | 16652 | **RNA**→ |
| env-586 | SRS019026_WUGC_scaffold_25915 | - | 1526 | 1452 | **RNA**→ rplM (PRK09216)Ribosomal_L13 (pfam00572)→ rpsI (PRK00132)Ribosomal_S9 (pfam00380)→ |
| env-587 | SRS019027_C1975919 | + | 1325 | 1399 | **RNA**→ |
| env-588 | SRS022530_LANL_scaffold_21668 | - | 23768 | 23694 | **RNA**→ rplM (PRK09216)Ribosomal_L13 (pfam00572)→ rpsI (PRK00132)Ribosomal_S9 (pfam00380)→ |
| env-589 | SRS043663_C4103193 | + | 3216 | 3290 | **RNA**→ |
| env-590 | SRS044373_WUGC_scaffold_63271 | + | 16814 | 16888 | **RNA**→ rplM (PRK09216)Ribosomal_L13 (pfam00572)→ rpsI (PRK00132)Ribosomal_S9 (pfam00380)→ |

| | | | | | |
|---|---|---|---|---|---|
| env-591 | SRS048791_LANL_scaffold_39630 | - | 1512 | 1438 | **RNA**→ rplM (PRK09216)Ribosomal_L13 (pfam00572)→ rpsI (PRK00132)Ribosomal_S9 (pfam00380)→ |
| env-592 | SRS047824_WUGC_scaffold_53533 | - | 614 | 540 | **RNA**→ rplM (PRK09216)Ribosomal_L13 (pfam00572)→ |
| env-593 | SRS057205_LANL_scaffold_58665 | + | 670 | 744 | **RNA**→ rplM (PRK09216)RplM (COG0102)→ |
| env-594 | SRS012279_Baylor_scaffold_56590 | - | 3134 | 3060 | **RNA**→ rplM (PRK09216)Ribosomal_L13 (pfam00572)→ rpsI (PRK00132)Ribosomal_S9 (pfam00380)→ |
| env-595 | SRS011306_Baylor_scaffold_89003 | - | 10148 | 10074 | **RNA**→ rplM (PRK09216)Ribosomal_L13 (pfam00572)→ rpsI (PRK00132)Ribosomal_S9 (pfam00380)→ |
| env-596 | SRS075404_LANL_scaffold_64891 | + | 238 | 312 | **RNA**→ |
| env-597 | SRS023352_LANL_scaffold_62464 | - | 47439 | 47365 | **RNA**→ rplM (PRK09216)Ribosomal_L13 (pfam00572)→ rpsI (PRK00132)Ribosomal_S9 (pfam00380)→ |
| env-598 | SRS063603_C4102126 | + | 332 | 406 | **RNA**→ |
| env-599 | SRS064329_C2725335 | + | 3215 | 3289 | **RNA**→ |
| env-600 | SRS062878_LANL_scaffold_70775 | - | 8062 | 7988 | **RNA**→ rplM (PRK09216)Ribosomal_L13 (pfam00572)→ rpsI (PRK00132)Ribosomal_S9 (pfam00380)→ |
| env-601 | Subgingival_plaque_LANL_C11028051 | + | 219 | 293 | **RNA**→ |
| env-602 | SRS014691_C2755366 | + | 2883 | 2957 | **RNA**→ |
| env-603 | SRS015272_C3490354 | + | 339 | 413 | **RNA**→ |
| env-604 | SRS015644_WUGC_scaffold_48472 | + | 933 | 1007 | **RNA**→ rplM (PRK09216)Ribosomal_L13 (pfam00572)→ rpsI (PRK00132)Ribosomal_S9 (pfam00380)→ |
| env-605 | SRS018357_Baylor_scaffold_17708 | - | 24023 | 23949 | **RNA**→ rplM (PRK09216)Ribosomal_L13 (pfam00572)→ rpsI (PRK00132)Ribosomal_S9 (pfam00380)→ |
| env-606 | SRS019974_Baylor_scaffold_54950 | + | 3215 | 3289 | **RNA**→ rplM (PRK09216)Ribosomal_L13 (pfam00572)→ rpsI (PRK00132)Ribosomal_S9 (pfam00380)→ |
| env-607 | SRS023617_Baylor_scaffold_30918 | + | 5850 | 5924 | **RNA**→ rplM (PRK09216)Ribosomal_L13 (pfam00572)→ rpsI (PRK00132)Ribosomal_S9 (pfam00380)→ |
| env-608 | SRS024375_C2325854 | + | 138 | 212 | **RNA**→ |
| env-609 | SRS042131_WUGC_scaffold_50435 | + | 3411 | 3485 | **RNA**→ |
| env-610 | SRS047210_WUGC_scaffold_57327 | + | 9864 | 9938 | **RNA**→ rplM (PRK09216)Ribosomal_L13 (pfam00572)→ rpsI (PRK00132)Ribosomal_S9 (pfam00380)→ |
| env-611 | SRS050244_LANL_scaffold_37949 | - | 1334 | 1260 | **RNA**→ rplM (PRK09216)Ribosomal_L13 (pfam00572)→ rpsI (PRK00132)Ribosomal_S9 (pfam00380)→ |
| env-612 | SRS053603_LANL_scaffold_49200 | + | 16206 | 16280 | **RNA**→ rplM (PRK09216)Ribosomal_L13 (pfam00572)→ rpsI (PRK00132)Ribosomal_S9 (pfam00380)→ |
| env-613 | SRS017076_C2680416 | + | 454 | 528 | **RNA**→ |
| env-614 | SRS013946_C747973 | - | 1186 | 1112 | **RNA**→ rplM (PRK09216)Ribosomal_L13 (pfam00572)→ rpsI (PRK00132)Ribosomal_S9 (pfam00380)→ |
| env-615 | SRS014470_WUGC_scaffold_48816 | - | 9526 | 9452 | **RNA**→ rplM (PRK09216)Ribosomal_L13 (pfam00572)→ rpsI (PRK00132)Ribosomal_S9 (pfam00380)→ |
| env-616 | SRS014573_C3755401 | + | 2926 | 3000 | **RNA**→ |
| env-617 | SRS015038_WUGC_scaffold_38040 | + | 3215 | 3289 | **RNA**→ |
| env-618 | SRS015057_WUGC_scaffold_8828 | + | 2155 | 2229 | **RNA**→ rplM (PRK09216)Ribosomal_L13 (pfam00572)→ rpsI (PRK00132)Ribosomal_S9 (pfam00380)→ |
| env-619 | SRS015209_WUGC_scaffold_49105 | + | 45106 | 45180 | **RNA**→ rplM (PRK09216)Ribosomal_L13 (pfam00572)→ rpsI (PRK00132)Ribosomal_S9 (pfam00380)→ |
| env-620 | SRS015893_WUGC_scaffold_32763 | + | 3119 | 3193 | **RNA**→ rplM (PRK09216)Ribosomal_L13 (pfam00572)→ rpsI (PRK00132)Ribosomal_S9 (pfam00380)→ |
| env-621 | SRS016002_WUGC_scaffold_17242 | + | 35903 | 35977 | **RNA**→ rplM (PRK09216)Ribosomal_L13 (pfam00572)→ rpsI (PRK00132)Ribosomal_S9 (pfam00380)→ |
| env-622 | SRS017120_Baylor_scaffold_72849 | + | 96 | 170 | **RNA**→ rplM (PRK09216)Ribosomal_L13 (pfam00572)→ rpsI (PRK00132)Ribosomal_S9 (pfam00380)→ |
| env-623 | SRS017439_Baylor_scaffold_52553 | - | 1177 | 1103 | **RNA**→ rplM (PRK09216)Ribosomal_L13 (pfam00572)→ rpsI (PRK00132)Ribosomal_S9 (pfam00380)→ |
| env-624 | SRS018300_Baylor_scaffold_51866 | + | 3216 | 3290 | **RNA**→ rplM (PRK09216)Ribosomal_L13 (pfam00572)→ rpsI (PRK00132)Ribosomal_S9 (pfam00380)→ |
| env-625 | SRS018439_Baylor_scaffold_51883 | - | 3196 | 3122 | **RNA**→ rplM (PRK09216)Ribosomal_L13 (pfam00572)→ rpsI (PRK00132)Ribosomal_S9 (pfam00380)→ |
| env-626 | SRS018739_WUGC_scaffold_60459 | - | 1177 | 1103 | **RNA**→ rplM (PRK09216)Ribosomal_L13 (pfam00572)→ rpsI (PRK00132)Ribosomal_S9 (pfam00380)→ |
| env-627 | SRS019126_WUGC_scaffold_41627 | - | 1177 | 1103 | **RNA**→ rplM (PRK09216)Ribosomal_L13 (pfam00572)→ rpsI (PRK00132)Ribosomal_S9 (pfam00380)→ |
| env-628 | SRS019127_C1673699 | + | 3216 | 3290 | **RNA**→ |
| env-629 | SRS019219_WUGC_scaffold_22098 | - | 3099 | 3025 | **RNA**→ rplM (PRK09216)Ribosomal_L13 (pfam00572)→ rpsI (PRK00132)Ribosomal_S9 (pfam00380)→ |
| env-630 | SRS020220_Baylor_scaffold_52465 | - | 1526 | 1452 | **RNA**→ rplM (PRK09216)Ribosomal_L13 (pfam00572)→ rpsI (PRK00132)Ribosomal_S9 (pfam00380)→ |
| env-631 | SRS020856_Baylor_scaffold_35690 | - | 3199 | 3125 | **RNA**→ rplM (PRK09216)Ribosomal_L13 (pfam00572)→ rpsI (PRK00132)Ribosomal_S9 (pfam00380)→ |
| env-632 | SRS022143_WUGC_scaffold_32909 | - | 9534 | 9460 | **RNA**→ rplM (PRK09216)Ribosomal_L13 (pfam00572)→ rpsI (PRK00132)Ribosomal_S9 (pfam00380)→ |
| env-633 | SRS022621_C4119557 | + | 2940 | 3014 | **RNA**→ |
| env-634 | SRS024318_LANL_scaffold_74074 | - | 11441 | 11367 | **RNA**→ rplM (PRK09216)Ribosomal_L13 (pfam00572)→ rpsI (PRK00132)Ribosomal_S9 (pfam00380)→ |
| env-635 | SRS024580_C3038766 | - | 171 | 97 | **RNA**→ |
| env-636 | SRS024637_LANL_scaffold_30251 | + | 233 | 307 | **RNA**→ |
| env-637 | SRS042910_C5625461 | + | 3216 | 3290 | **RNA**→ |
| env-638 | SRS045715_LANL_scaffold_89135 | + | 3431 | 3505 | **RNA**→ rplM (PRK09216)Ribosomal_L13 (pfam00572)→ |
| env-639 | SRS049147_LANL_scaffold_58601 | + | 95 | 169 | **RNA**→ rplM (PRK09216)Ribosomal_L13 (pfam00572)→ rpsI (PRK00132)Ribosomal_S9 (pfam00380)→ |
| env-640 | SRS053854_C1915023 | + | 140 | 214 | **RNA**→ |
| env-641 | SRS054687_C3415499 | - | 171 | 97 | **RNA**→ hypo→ |
| env-642 | SRS055426_LANL_scaffold_56878 | + | 12351 | 12425 | **RNA**→ rplM (PRK09216)Ribosomal_L13 (pfam00572)→ |

| | | | | | |
|---|---|---|---|---|---|
| env-643 | SRS056622_LANL_scaffold_38514 | - | 47437 | 47363 | **RNA**→ rplM (PRK09216)Ribosomal_L13 (pfam00572)→ rpsI (PRK00132)Ribosomal_S9 (pfam00380)→ |
| env-644 | SRS058336_C2178143 | + | 3215 | 3289 | **RNA**→ |
| env-645 | SRS063193_LANL_scaffold_53070 | + | 2926 | 3000 | **RNA**→ rplM (PRK09216)Ribosomal_L13 (pfam00572)→ rpsI (PRK00132)Ribosomal_S9 (pfam00380)→ |
| env-646 | SRS064423_LANL_scaffold_66525 | - | 11227 | 11153 | **RNA**→ rplM (PRK09216)Ribosomal_L13 (pfam00572)→ rpsI (PRK00132)Ribosomal_S9 (pfam00380)→ |
| env-647 | SRS065278_C2564619 | + | 3215 | 3289 | **RNA**→ |
| Ppa-1-1 | NZ_GL982513.1 | + | 723490 | 723564 | **RNA**→ rplM (PRK09216)Ribosomal_L13 (pfam00572)→ rpsI (PRK00132)Ribosomal_S9 (pfam00380)→ |
| env-648 | SRS018791_WUGC_scaffold_38475 | - | 1334 | 1260 | **RNA**→ rplM (PRK09216)Ribosomal_L13 (pfam00572)→ rpsI (PRK00132)Ribosomal_S9 (pfam00380)→ |
| env-649 | SRS022077_Baylor_scaffold_52460 | - | 1526 | 1452 | **RNA**→ rplM (PRK09216)Ribosomal_L13 (pfam00572)→ rpsI (PRK00132)Ribosomal_S9 (pfam00380)→ |
| env-650 | SRS024015_Baylor_scaffold_54468 | - | 5401 | 5327 | **RNA**→ rplM (PRK09216)Ribosomal_L13 (pfam00572)→ rpsI (PRK00132)Ribosomal_S9 (pfam00380)→ |
| env-651 | SRS016319_WUGC_scaffold_54116 | + | 3215 | 3289 | **RNA**→ rplM (PRK09216)Ribosomal_L13 (pfam00572)→ rpsI (PRK00132)Ribosomal_S9 (pfam00380)→ |
| env-652 | SRS047219_WUGC_scaffold_57943 | + | 16790 | 16864 | **RNA**→ rplM (PRK09216)Ribosomal_L13 (pfam00572)→ rpsI (PRK00132)Ribosomal_S9 (pfam00380)→ |
| env-653 | Buccal_mucosa_LANL_C64807355 | - | 110 | 36 | **RNA**→ hypo→ |
| env-654 | Contig7238817.1 | - | 110 | 36 | **RNA**→ |
| env-655 | SRS015434_C5938921 | - | 110 | 36 | **RNA**→ |
| env-656 | SRS017808_Baylor_scaffold_37180 | + | 3883 | 3957 | **RNA**→ |
| env-657 | SRS016740_C2751815 | - | 117 | 43 | **RNA**→ |
| env-658 | SRS023930_C2173199 | - | 90 | 16 | **RNA**→ |
| env-659 | SRS013705_Baylor_scaffold_102904 | + | 23297 | 23371 | **RNA**→ |
| env-660 | SRS013879_C2856030 | + | 1392 | 1466 | **RNA**→ |
| env-661 | SRS015762_WUGC_scaffold_41259 | + | 774 | 848 | **RNA**→ rplM (PRK09216)Ribosomal_L13 (pfam00572)→ rpsI (PRK00132)Ribosomal_S9 (pfam00380)→ |
| env-662 | SRS015797_WUGC_scaffold_34883 | - | 23901 | 23827 | **RNA**→ rplM (PRK09216)Ribosomal_L13 (pfam00572)→ rpsI (PRK00132)Ribosomal_S9 (pfam00380)→ |
| env-663 | SRS015941_WUGC_scaffold_78227 | + | 3216 | 3290 | **RNA**→ rplM (PRK09216)Ribosomal_L13 (pfam00572)→ rpsI (PRK00132)Ribosomal_S9 (pfam00380)→ |
| env-664 | SRS016086_WUGC_scaffold_26855 | - | 171 | 97 | **RNA**→ |
| env-665 | SRS016225_C2803421 | - | 171 | 97 | **RNA**→ hypo→ |
| env-666 | SRS016569_Baylor_scaffold_56485 | + | 16533 | 16607 | **RNA**→ rplM (PRK09216)Ribosomal_L13 (pfam00572)→ rpsI (PRK00132)Ribosomal_S9 (pfam00380)→ |
| env-667 | SRS021496_Baylor_scaffold_85386 | - | 12316 | 12242 | **RNA**→ rplM (PRK09216)Ribosomal_L13 (pfam00572)→ rpsI (PRK00132)Ribosomal_S9 (pfam00380)→ |
| env-668 | SRS021954_C2960878 | + | 284 | 358 | **RNA**→ |
| env-669 | SRS023926_C2818602 | + | 2890 | 2964 | **RNA**→ |
| env-670 | SRS024081_LANL_scaffold_39378 | + | 40034 | 40108 | **RNA**→ rplM (PRK09216)Ribosomal_L13 (pfam00572)→ rpsI (PRK00132)Ribosomal_S9 (pfam00380)→ |
| env-671 | SRS042643_WUGC_scaffold_26426 | - | 1334 | 1260 | **RNA**→ rplM (PRK09216)Ribosomal_L13 (pfam00572)→ rpsI (PRK00132)Ribosomal_S9 (pfam00380)→ |
| env-672 | SRS051791_LANL_scaffold_12138 | - | 47738 | 47664 | **RNA**→ rplM (PRK09216)Ribosomal_L13 (pfam00572)→ rpsI (PRK00132)Ribosomal_S9 (pfam00380)→ |
| env-673 | SRS057539_LANL_scaffold_35057 | + | 49731 | 49805 | **RNA**→ rplM (PRK09216)Ribosomal_L13 (pfam00572)→ rpsI (PRK00132)Ribosomal_S9 (pfam00380)→ |
| env-674 | SRS063288_C2542184 | + | 84 | 158 | **RNA**→ rplM (PRK09216)Ribosomal_L13 (pfam00572)→ |
| env-675 | SRS064774_LANL_scaffold_17002 | - | 16294 | 16220 | **RNA**→ rplM (PRK09216)Ribosomal_L13 (pfam00572)→ rpsI (PRK00132)Ribosomal_S9 (pfam00380)→ |
| env-676 | UnmappedStool_Broad_C252508257 | - | 155 | 79 | **RNA**→ hypo→ |
| Pco-3-1 | NZ_AUME01000007.1 | - | 36087 | 36011 | **RNA**→ rplM (PRK09216)Ribosomal_L13 (pfam00572)→ rpsI (PRK00132)Ribosomal_S9 (pfam00380)→ |
| Pco-3-2 | NZ_BAIT01000003.1 | + | 74722 | 74798 | **RNA**→ rplM (PRK09216)Ribosomal_L13 (pfam00572)→ rpsI (PRK00132)Ribosomal_S9 (pfam00380)→ |
| Pni-1-1 | NZ_GL982464.1 | + | 2020144 | 2020219 | **RNA**→ rplM (PRK09216)Ribosomal_L13 (pfam00572)→ rplM (PRK09216)Ribosomal_L13 (pfam00572)→ rpsI (PRK00132)Ribosomal_S9 (pfam00380)→ |
| env-677 | SRS012285_Baylor_scaffold_36585 | + | 18214 | 18289 | **RNA**→ rplM (PRK09216)Ribosomal_L13 (pfam00572)→ rpsI (PRK00132)Ribosomal_S9 (pfam00380)→ |
| env-678 | SRS022083_C1814115 | - | 190 | 115 | **RNA**→ hypo→ |
| env-679 | SRS024144_C1709965 | - | 224 | 149 | **RNA**→ rplM (PRK09216)RplM (COG0102)Ribosomal_L13 (cd00392)Ribosomal_L13 (pfam00572)→ |
| env-680 | SRS063932_LANL_scaffold_105725 | + | 15855 | 15930 | **RNA**→ |
| env-681 | Subgingival_plaque_LANL_C10982129 | - | 147 | 72 | **RNA**→ hypo→ |
| env-682 | SRS022725_LANL_scaffold_18998 | - | 20108 | 20033 | **RNA**→ rplM (PRK09216)Ribosomal_L13 (pfam00572)→ rpsI (PRK00132)Ribosomal_S9 (pfam00380)→ |
| env-683 | SRS063215_C2206217 | + | 314 | 389 | **RNA**→ hypo→ |
| env-684 | SRS045313_LANL_scaffold_12927 | + | 11427 | 11502 | **RNA**→ |
| env-685 | SRS058053_LANL_scaffold_46648 | + | 2496 | 2571 | **RNA**→ rplM (PRK09216)Ribosomal_L13 (pfam00572)→ rpsI (PRK00132)Ribosomal_S9 (pfam00380)→ |
| env-686 | SRS015215_WUGC_scaffold_60845 | + | 31443 | 31518 | **RNA**→ |
| env-687 | SRS016575_C3771021 | - | 171 | 96 | **RNA**→ hypo→ |
| env-688 | SRS017227_Baylor_scaffold_128981 | - | 4583 | 4508 | **RNA**→ rplM (PRK09216)Ribosomal_L13 (pfam00572)→ rpsI (PRK00132)Ribosomal_S9 (pfam00380)→ |
| env-689 | SRS018573_C2468474 | - | 171 | 96 | **RNA**→ hypo→ |

| | | | | | |
|---|---|---|---|---|---|
| env-690 | SRS045197_C3607732 | + | 2688 | 2763 | **RNA**→ |
| env-691 | SRS047113_C5438044 | + | 4885 | 4960 | **RNA**→ |
| env-692 | SRS049318_LANL_scaffold_180129 | + | 334 | 409 | **RNA**→ rplM (PRK09216)Ribosomal_L13 (pfam00572)→ |
| env-693 | SRS051930_C3291029 | + | 3901 | 3976 | **RNA**→ |
| env-694 | SRS057022_LANL_scaffold_21753 | + | 985 | 1060 | **RNA**→ rplM (PRK09216)Ribosomal_L13 (pfam00572)→ |
| env-695 | SRS064449_C4104329 | + | 731 | 806 | **RNA**→ |
| env-696 | SRS014691_C2755332 | - | 159 | 84 | **RNA**→ hypo→ |
| env-697 | SRS019029_WUGC_scaffold_30607 | - | 318 | 243 | **RNA**→ rplM (PRK09216)Ribosomal_L13 (pfam00572)→ |
| env-698 | SRS011255_Baylor_scaffold_75811 | - | 105 | 30 | **RNA**→ |
| env-699 | SRS023938_C2346941 | - | 137 | 62 | **RNA**→ |
| Pni-3-1 | NZ_ADGJ01000034.1 | + | 4034 | 4109 | **RNA**→ rplM (PRK09216)Ribosomal_L13 (pfam00572)→ rpsI (PRK00132)Ribosomal_S9 (pfam00380)→ |
| env-700 | SRS013533_PGA_scaffold_55845 | + | 31911 | 31986 | **RNA**→ ←hypo |
| env-701 | SRS013950_WUGC_scaffold_19463 | + | 6777 | 6852 | **RNA**→ rplM (PRK09216)Ribosomal_L13 (pfam00572)→ rpsI (PRK00132)Ribosomal_S9 (pfam00380)→ |
| env-702 | SRS015574_C3429056 | + | 765 | 840 | **RNA**→ |
| env-703 | SRS019980_Baylor_scaffold_52314 | - | 426 | 351 | **RNA**→ rplM (PRK09216)Ribosomal_L13 (pfam00572)→ |
| env-704 | SRS022536_LANL_scaffold_126332 | + | 32859 | 32934 | **RNA**→ rplM (PRK09216)Ribosomal_L13 (pfam00572)→ rpsI (PRK00132)Ribosomal_S9 (pfam00380)→ |
| env-705 | SRS023595_C7191239 | - | 113 | 38 | **RNA**→ |
| env-706 | SRS024087_LANL_scaffold_6797 | - | 49193 | 49118 | **RNA**→ rplM (PRK09216)Ribosomal_L13 (pfam00572)→ rpsI (PRK00132)Ribosomal_S9 (pfam00380)→ |
| env-707 | SRS058808_C2875695 | - | 162 | 87 | **RNA**→ |
| env-708 | SRS065099_LANL_scaffold_67594 | - | 171 | 96 | **RNA**→ hypo→ |
| Pni-2-1 | NZ_KI669452.1 | - | 66638 | 66563 | **RNA**→ rplM (PRK09216)Ribosomal_L13 (pfam00572)→ rpsI (PRK00132)Ribosomal_S9 (pfam00380)→ |
| env-709 | SRS057355_C2357028 | + | 1923 | 1998 | **RNA**→ |
| env-710 | SRS014573_C3689837 | + | 580 | 655 | **RNA**→ |
| env-711 | SRS013164_Baylor_scaffold_6044 | + | 24960 | 25035 | **RNA**→ rplM (PRK09216)Ribosomal_L13 (pfam00572)→ rpsI (PRK00132)Ribosomal_S9 (pfam00380)→ |
| env-712 | SRS015434_C5939871 | + | 1383 | 1458 | **RNA**→ hypo→ |
| env-713 | SRS016037_WUGC_scaffold_40495 | - | 1517 | 1442 | **RNA**→ rplM (PRK09216)Ribosomal_L13 (pfam00572)→ rpsI (PRK00132)Ribosomal_S9 (pfam00380)→ |
| env-714 | SRS017209_C2612132 | - | 171 | 96 | **RNA**→ hypo→ |
| env-715 | SRS018145_Baylor_scaffold_45351 | + | 15741 | 15816 | **RNA**→ rplM (PRK09216)Ribosomal_L13 (pfam00572)→ rpsI (PRK00132)Ribosomal_S9 (pfam00380)→ |
| env-716 | SRS019071_WUGC_scaffold_1653 | - | 393 | 318 | **RNA**→ rplM (PRK09216)Ribosomal_L13 (pfam00572)→ |
| env-717 | SRS019327_WUGC_scaffold_7031 | - | 9945 | 9870 | **RNA**→ rplM (PRK09216)Ribosomal_L13 (pfam00572)→ rpsI (PRK00132)Ribosomal_S9 (pfam00380)→ |
| env-718 | SRS022719_LANL_scaffold_29884 | - | 15205 | 15130 | **RNA**→ rplM (PRK09216)Ribosomal_L13 (pfam00572)→ rpsI (PRK00132)Ribosomal_S9 (pfam00380)→ |
| env-719 | SRS023958_Baylor_scaffold_35671 | + | 24960 | 25035 | **RNA**→ rplM (PRK09216)Ribosomal_L13 (pfam00572)→ rpsI (PRK00132)Ribosomal_S9 (pfam00380)→ |
| env-720 | SRS044662_LANL_scaffold_26917 | - | 15127 | 15052 | **RNA**→ rplM (PRK09216)Ribosomal_L13 (pfam00572)→ rpsI (PRK00132)Ribosomal_S9 (pfam00380)→ |
| env-721 | SRS049389_C4741864 | + | 3211 | 3286 | **RNA**→ hypo→ |
| env-722 | SRS050669_C2732573 | - | 171 | 96 | **RNA**→ hypo→ |
| env-723 | SRS057692_LANL_scaffold_59063 | + | 15741 | 15816 | **RNA**→ rplM (PRK09216)Ribosomal_L13 (pfam00572)→ rpsI (PRK00132)Ribosomal_S9 (pfam00380)→ |
| env-724 | SRS043422_C2606710 | - | 418 | 343 | **RNA**→ rplM (PRK09216)Ribosomal_L13 (pfam00572)→ |
| Pau-1-1 | NZ_BAKF01000067.1 | - | 163 | 88 | **RNA**→ |
| Psp-9-1 | NZ_JIAF01000004.1 | - | 988952 | 988877 | **RNA**→ rplM (PRK09216)Ribosomal_L13 (pfam00572)→ rpsI (PRK00132)Ribosomal_S9 (pfam00380)→ |
| env-725 | SRS055401_C2262988 | - | 171 | 96 | **RNA**→ hypo→ |
| env-726 | SRS016746_C3112314 | - | 171 | 96 | **RNA**→ hypo→ |
| env-727 | Contig7208859.1 | - | 113 | 38 | **RNA**→ |
| env-728 | SRS048411_C6994262 | + | 2390 | 2465 | **RNA**→ |
| env-729 | SRS013723_C3473767 | + | 936 | 1011 | **RNA**→ hypo→ |
| env-730 | SRS020226_C3504182 | + | 3062 | 3137 | **RNA**→ hypo→ |
| env-731 | SRS022149_C2844760 | - | 185 | 110 | **RNA**→ hypo→ |
| env-732 | SRS023595_Baylor_scaffold_12538 | - | 315 | 240 | **RNA**→ rplM (PRK09216)Ribosomal_L13 (pfam00572)→ |
| env-733 | SRS017511_Baylor_scaffold_37568 | - | 3386 | 3311 | **RNA**→ rplM (PRK09216)Ribosomal_L13 (pfam00572)→ rpsI (PRK00132)Ribosomal_S9 (pfam00380)→ |
| env-734 | SRS062761_C4185168 | - | 157 | 82 | **RNA**→ hypo→ |
| env-735 | SRS063603_LANL_scaffold_16033 | + | 3063 | 3138 | **RNA**→ rplM (PRK09216)Ribosomal_L13 (pfam00572)→ rpsI (PRK00132)Ribosomal_S9 (pfam00380)→ |
| env-736 | SRS049268_C4330408 | + | 785 | 860 | **RNA**→ hypo→ |
| Pin-3-1 | NZ_ATMK01000012.1 | - | 3560 | 3485 | **RNA**→ rplM (PRK09216)Ribosomal_L13 (pfam00572)→ rpsI (PRK00132)Ribosomal_S9 (pfam00380)→ |

| | | | | | |
|---|---|---|---|---|---|
| Pfa-1-1 | NZ_BAJY01000023.1 | + | 18148 | 18223 | **RNA**→ rplM (PRK09216)Ribosomal_L13 (pfam00572)→ rpsI (PRK00132)Ribosomal_S9 (pfam00380)→ |
| env-737 | SRS049318_LANL_scaffold_111644 | + | 3483 | 3558 | **RNA**→ rplM (PRK09216)Ribosomal_L13 (pfam00572)→ rpsI (PRK00132)Ribosomal_S9 (pfam00380)→ |
| Pin-2-1 | NZ_KE392224.1 | + | 52909 | 52984 | **RNA**→ rplM (PRK09216)Ribosomal_L13 (pfam00572)→ rpsI (PRK00132)Ribosomal_S9 (pfam00380)→ |
| env-738 | SRS014474_C725551 | + | 364 | 439 | **RNA**→ |
| env-739 | SRS014477_C1641189 | + | 1209 | 1284 | **RNA**→ |
| env-740 | SRS015061_C521155 | - | 3425 | 3350 | **RNA**→ rplM (PRK09216)Ribosomal_L13 (pfam00572)→ rpsI (PRK00132)Ribosomal_S9 (pfam00380)→ |
| env-741 | SRS015062_WUGC_scaffold_566 | - | 3425 | 3350 | **RNA**→ rplM (PRK09216)Ribosomal_L13 (pfam00572)→ rpsI (PRK00132)Ribosomal_S9 (pfam00380)→ |
| env-742 | SRS047634_C4505245 | + | 2722 | 2797 | **RNA**→ |
| Pdi-3-1 | NZ_KI259320.1 | - | 18576 | 18501 | **RNA**→ rplM (PRK09216)Ribosomal_L13 (pfam00572)→ rpsI (PRK00132)Ribosomal_S9 (pfam00380)→ |
| env-743 | SRS017521_Baylor_scaffold_47871 | - | 90 | 15 | **RNA**→ |
| Pdi-2-1 | NZ_AEDO01000037.1 | + | 25023 | 25098 | **RNA**→ rplM (PRK09216)Ribosomal_L13 (pfam00572)→ rplM (PRK09216)Ribosomal_L13 (pfam00572)→ rpsI (PRK00132)Ribosomal_S9 (pfam00380)→ rpsI (PRK00132)Ribosomal_S9 (pfam00380)→ |
| Pdi-1-1 | NZ_JRNR01000099.1 | - | 3529 | 3454 | **RNA**→ rplM (PRK09216)Ribosomal_L13 (pfam00572)→ rpsI (PRK00132)Ribosomal_S9 (pfam00380)→ |
| env-744 | NOM017_scaffold152_1 | + | 2821 | 2896 | **RNA**→ |
| env-745 | 4448812.3_Contig1872 | + | 367 | 442 | **RNA**→ rplM (PRK09216)Ribosomal_L13 (pfam00572)→ rpsI (PRK00132)Ribosomal_S9 (pfam00380)→ |
| env-746 | DLM003_scaffold159_2 | + | 2840 | 2915 | **RNA**→ rplM (PRK09216)Ribosomal_L13 (pfam00572)→ rpsI (PRK00132)Ribosomal_S9 (pfam00380)→ |
| env-747 | NOM022_scaffold34272_1 | + | 2699 | 2774 | **RNA**→ rplM (PRK09216)Ribosomal_L13 (pfam00572)→ rpsI (PRK00132)Ribosomal_S9 (pfam00380)→ |
| env-748 | contig104641 | + | 10740 | 10815 | **RNA**→ rplM (PRK09216)Ribosomal_L13 (pfam00572)→ |
| env-749 | DLM001_scaffold46771_1 | - | 555 | 480 | **RNA**→ rplM (PRK09216)Ribosomal_L13 (pfam00572)→ |
| env-750 | DOM003_scaffold20629_2 | + | 164 | 239 | **RNA**→ rplM (PRK09216)Ribosomal_L13 (pfam00572)→ |
| env-751 | NLM001_scaffold44488_1 | + | 2699 | 2774 | **RNA**→ rplM (PRK09216)Ribosomal_L13 (pfam00572)→ rpsI (PRK00132)Ribosomal_S9 (pfam00380)→ |
| env-752 | SRS011271_WUGC_scaffold_12531 | + | 603 | 678 | **RNA**→ |
| env-753 | 4448818.3_Contig13931 | - | 222 | 147 | **RNA**→ rplM (PRK09216)→ |
| env-754 | NLM006_scaffold2689_1 | - | 559 | 484 | **RNA**→ rplM (PRK09216)Ribosomal_L13 (pfam00572)→ |
| env-755 | SRS015794_Baylor_scaffold_8969 | + | 743 | 818 | **RNA**→ rplM (PRK09216)Ribosomal_L13 (pfam00572)→ |
| env-756 | DOF011_scaffold4575_1 | + | 2870 | 2945 | **RNA**→ rplM (PRK09216)Ribosomal_L13 (pfam00572)→ rpsI (PRK00132)Ribosomal_S9 (pfam00380)→ |
| env-757 | NLM004_scaffold39342_1 | - | 326 | 250 | **RNA**→ rplM (PRK09216)Ribosomal_L13 (pfam00572)→ |
| env-758 | DLF003_scaffold48930_1 | + | 25126 | 25201 | **RNA**→ rplM (PRK09216)Ribosomal_L13 (pfam00572)→ |
| env-759 | NLM004_scaffold4959_1 | - | 319 | 244 | **RNA**→ rplM (PRK09216)Ribosomal_L13 (pfam00572)→ |
| env-760 | NOM012_scaffold33926_2 | - | 272 | 197 | **RNA**→ rplM (PRK09216)Ribosomal_L13 (pfam00572)→ |
| env-761 | scaffold3211_3_V1.CD-2 | + | 538 | 613 | **RNA**→ |
| env-762 | SRS019910_WUGC_scaffold_54703 | + | 204 | 279 | **RNA**→ |
| env-763 | DLM001_scaffold46772_1 | - | 551 | 476 | **RNA**→ rplM (PRK09216)Ribosomal_L13 (pfam00572)→ |
| env-764 | DOM016_scaffold20994_2 | + | 58303 | 58378 | **RNA**→ rplM (PRK09216)Ribosomal_L13 (pfam00572)→ rpsI (PRK00132)Ribosomal_S9 (pfam00380)→ |
| env-765 | SRS013521_LANL_scaffold_8254 | - | 121 | 46 | **RNA**→ |
| env-766 | NLM016_scaffold16524_1 | - | 542 | 467 | **RNA**→ rplM (PRK09216)Ribosomal_L13 (pfam00572)→ |
| env-767 | 4491403.3_NODE_17602 | - | 322 | 247 | **RNA**→ rplM (PRK09216)Ribosomal_L13 (pfam00572)→ |
| env-768 | SRS049959_WUGC_scaffold_5722 | - | 414 | 338 | **RNA**→ rplM (PRK09216)Ribosomal_L13 (pfam00572)→ |
| env-769 | scaffold28598_1_V1.UC-17 | - | 126 | 50 | **RNA**→ |
| env-770 | scaffold2660_2_MH0049 | + | 139 | 215 | **RNA**→ rplM (PRK09216)Ribosomal_L13 (pfam00572)→ |
| env-771 | 4491403.3_NODE_60160 | - | 980 | 904 | **RNA**→ rplM (PRK09216)Ribosomal_L13 (pfam00572)→ rpsI (PRK00132)Ribosomal_S9 (pfam00380)→ |
| env-772 | SRS013521_LANL_scaffold_15414 | - | 158 | 82 | **RNA**→ hypo→ |
| env-773 | SRS049995_LANL_scaffold_36980 | + | 752 | 828 | **RNA**→ |
| env-774 | BAAX01002455.1 | + | 95 | 171 | **RNA**→ rplM (PRK09216)Ribosomal_L13 (pfam00572)→ rpsI (PRK00132)Ribosomal_S9 (pfam00380)→ rpsI (PRK00132)Ribosomal_S9 (pfam00380)→ |
| env-775 | SRS015794_C2240237 | + | 338 | 414 | **RNA**→ |
| env-776 | DLM006_scaffold20724_1 | + | 4613 | 4689 | **RNA**→ rplM (PRK09216)Ribosomal_L13 (pfam00572)→ rpsI (PRK00132)Ribosomal_S9 (pfam00380)→ |
| env-777 | NLF006_scaffold9001_1 | - | 11920 | 11844 | **RNA**→ rplM (PRK09216)Ribosomal_L13 (pfam00572)→ rpsI (PRK00132)Ribosomal_S9 (pfam00380)→ |
| env-778 | NLM002_scaffold28362_1 | + | 827 | 903 | **RNA**→ rplM (PRK09216)Ribosomal_L13 (pfam00572)→ rpsI (PRK00132)Ribosomal_S9 (pfam00380)→ |
| env-779 | NLM004_scaffold34280_1 | + | 818 | 894 | **RNA**→ rplM (PRK09216)Ribosomal_L13 (pfam00572)→ |
| env-780 | NLM006_scaffold7359_1 | + | 197 | 273 | **RNA**→ rplM (PRK09216)Ribosomal_L13 (pfam00572)→ |
| env-781 | DLM002_scaffold24298_1 | + | 12011 | 12086 | **RNA**→ rplM (PRK09216)Ribosomal_L13 (pfam00572)→ rpsI (PRK00132)Ribosomal_S9 (pfam00380)→ |
| env-782 | NOF010_scaffold54121_1 | + | 59215 | 59290 | **RNA**→ rplM (PRK09216)Ribosomal_L13 (pfam00572)→ rpsI (PRK00132)Ribosomal_S9 (pfam00380)→ |

| | | | | | |
|---|---|---|---|---|---|
| env-783 | NOM009_scaffold53764_1 | + | 59215 | 59290 | **RNA**→ rplM (PRK09216)Ribosomal_L13 (pfam00572)→ rpsI (PRK00132)Ribosomal_S9 (pfam00380)→ |
| env-784 | NOM017_scaffold40285_1 | + | 1439 | 1514 | **RNA**→ rplM (PRK09216)Ribosomal_L13 (pfam00572)→ |
| env-785 | NLM004_scaffold29467_1 | + | 19562 | 19637 | **RNA**→ rplM (PRK09216)Ribosomal_L13 (pfam00572)→ |
| env-786 | SRS022609_Baylor_scaffold_13243 | + | 21067 | 21142 | **RNA**→ rplM (PRK09216)Ribosomal_L13 (pfam00572)→ rpsI (PRK00132)Ribosomal_S9 (pfam00380)→ |
| env-787 | 4491403.3_NODE_11716 | + | 384 | 459 | **RNA**→ |
| env-788 | DLM027_scaffold23115_4 | + | 11864 | 11939 | **RNA**→ rplM (PRK09216)Ribosomal_L13 (pfam00572)→ rpsI (PRK00132)Ribosomal_S9 (pfam00380)→ |
| env-789 | 4448819.3_Contig631 | - | 7314 | 7239 | **RNA**→ rplM (PRK09216)Ribosomal_L13 (pfam00572)→ rpsI (PRK00132)Ribosomal_S9 (pfam00380)→ |
| env-790 | 4448812.3_Contig639 | + | 2967 | 3042 | **RNA**→ rplM (PRK09216)Ribosomal_L13 (pfam00572)→ rpsI (PRK00132)Ribosomal_S9 (pfam00380)→ |
| env-791 | scaffold18566_1_MH0009 | - | 109 | 34 | **RNA**→ |
| env-792 | NLM016_scaffold16376_1 | - | 143 | 68 | **RNA**→ |
| env-793 | scaffold58558_1_MH0065 | + | 213 | 288 | **RNA**→ rplM (PRK09216)Ribosomal_L13 (pfam00572)→ rpsI (PRK00132)Ribosomal_S9 (pfam00380)→ |
| env-794 | 4491477.3_NODE_2169 | - | 8743 | 8668 | **RNA**→ rplM (PRK09216)Ribosomal_L13 (pfam00572)→ rpsI (PRK00132)Ribosomal_S9 (pfam00380)→ |
| env-795 | SRS015782_WUGC_scaffold_26114 | - | 11843 | 11768 | **RNA**→ rplM (PRK09216)Ribosomal_L13 (pfam00572)→ rpsI (PRK00132)Ribosomal_S9 (pfam00380)→ |
| env-796 | SRS024132_Baylor_scaffold_32360 | - | 11900 | 11825 | **RNA**→ rplM (PRK09216)Ribosomal_L13 (pfam00572)→ rpsI (PRK00132)Ribosomal_S9 (pfam00380)→ |
| env-797 | scaffold43386_1_MH0050 | + | 966 | 1041 | **RNA**→ |
| env-798 | 4448813.3_Contig493 | - | 5921 | 5846 | **RNA**→ rplM (PRK09216)Ribosomal_L13 (pfam00572)→ rpsI (PRK00132)Ribosomal_S9 (pfam00380)→ |
| env-799 | NOF001_scaffold377_4 | - | 5305 | 5230 | **RNA**→ rplM (PRK09216)Ribosomal_L13 (pfam00572)→ rpsI (PRK00132)Ribosomal_S9 (pfam00380)→ |
| env-800 | CAM_READ_0099651871 | - | 384 | 309 | **RNA**→ rplM (PRK09216)Ribosomal_L13 (pfam00572)→ |
| env-801 | SRS047044_WUGC_scaffold_16328 | + | 257 | 332 | **RNA**→ rplM (PRK09216)Ribosomal_L13 (pfam00572)→ |
| env-802 | 4444165.3.02693 | + | 265 | 340 | **RNA**→ rplM (PRK09216)Ribosomal_L13 (pfam00572)→ |
| env-803 | 4448807.3_Contig1253 | - | 1893 | 1817 | **RNA**→ rplM (PRK09216)Ribosomal_L13 (pfam00572)→ rpsI (PRK00132)Ribosomal_S9 (pfam00380)→ |
| env-804 | NLM008_scaffold29501_1 | - | 12251 | 12176 | **RNA**→ rplM (PRK09216)Ribosomal_L13 (pfam00572)→ rpsI (PRK00132)Ribosomal_S9 (pfam00380)→ |
| env-805 | NLM024_scaffold18343_2 | + | 16425 | 16500 | **RNA**→ rplM (PRK09216)Ribosomal_L13 (pfam00572)→ rpsI (PRK00132)Ribosomal_S9 (pfam00380)→ |
| env-806 | NOM013_scaffold55258_1 | - | 10498 | 10423 | **RNA**→ rplM (PRK09216)Ribosomal_L13 (pfam00572)→ rpsI (PRK00132)Ribosomal_S9 (pfam00380)→ |
| env-807 | SRS017849_C4402944 | - | 468 | 393 | **RNA**→ rplM (PRK09216)Ribosomal_L13 (pfam00572)→ |
| env-808 | C4444188_1_MH0077 | + | 224 | 299 | **RNA**→ rplM (PRK09216)Ribosomal_L13 (pfam00572)→ |
| env-809 | 4448817.3_Contig838 | - | 764 | 689 | **RNA**→ rplM (PRK09216)Ribosomal_L13 (pfam00572)→ rpsI (PRK00132)Ribosomal_S9 (pfam00380)→ |
| env-810 | 4491404.3_NODE_16218 | + | 95 | 170 | **RNA**→ rplM (PRK09216)Ribosomal_L13 (pfam00572)→ rpsI (PRK00132)Ribosomal_S9 (pfam00380)→ |
| env-811 | 4491414.3_NODE_1809 | + | 43317 | 43392 | **RNA**→ rplM (PRK09216)Ribosomal_L13 (pfam00572)→ rpsI (PRK00132)Ribosomal_S9 (pfam00380)→ |
| env-812 | 4491421.3_NODE_2987 | + | 40037 | 40112 | **RNA**→ rplM (PRK09216)Ribosomal_L13 (pfam00572)→ rpsI (PRK00132)Ribosomal_S9 (pfam00380)→ |
| env-813 | DLF006_scaffold45333_1 | + | 259 | 334 | **RNA**→ rplM (PRK09216)Ribosomal_L13 (pfam00572)→ |
| env-814 | DLM005_scaffold13338_1 | + | 240 | 315 | **RNA**→ rplM (PRK09216)Ribosomal_L13 (pfam00572)→ rpsI (PRK00132)Ribosomal_S9 (pfam00380)→ |
| env-815 | DLM017_scaffold38149_2 | - | 11964 | 11889 | **RNA**→ rplM (PRK09216)Ribosomal_L13 (pfam00572)→ rpsI (PRK00132)Ribosomal_S9 (pfam00380)→ |
| env-816 | DOM001_scaffold19626_3 | + | 109863 | 109938 | **RNA**→ rplM (PRK09216)Ribosomal_L13 (pfam00572)→ rpsI (PRK00132)Ribosomal_S9 (pfam00380)→ |
| env-817 | DOM003_scaffold20232_1 | + | 1899 | 1974 | **RNA**→ rplM (PRK09216)Ribosomal_L13 (pfam00572)→ |
| env-818 | NOM019_scaffold53163_1 | + | 23837 | 23912 | **RNA**→ rplM (PRK09216)Ribosomal_L13 (pfam00572)→ rpsI (PRK00132)Ribosomal_S9 (pfam00380)→ |
| env-819 | SRS019910_WUGC_scaffold_10772 | + | 132 | 207 | **RNA**→ rplM (PRK09216)Ribosomal_L13 (pfam00572)→ |
| env-820 | SRS053398_LANL_scaffold_37336 | + | 287 | 362 | **RNA**→ rplM (PRK09216)Ribosomal_L13 (pfam00572)→ rpsI (PRK00132)Ribosomal_S9 (pfam00380)→ |
| env-821 | 4448820.3_Contig_1858 | - | 2732 | 2657 | **RNA**→ rplM (PRK09216)Ribosomal_L13 (pfam00572)→ rpsI (PRK00132)Ribosomal_S9 (pfam00380)→ |
| env-822 | DLF003_scaffold47495_1 | + | 445 | 520 | **RNA**→ rplM (PRK09216)Ribosomal_L13 (pfam00572)→ |
| env-823 | DOM023_scaffold26204_1 | - | 12481 | 12406 | **RNA**→ rplM (PRK09216)Ribosomal_L13 (pfam00572)→ rpsI (PRK00132)Ribosomal_S9 (pfam00380)→ |
| env-824 | 4448818.3_Contig459 | - | 19229 | 19154 | **RNA**→ rplM (PRK09216)Ribosomal_L13 (pfam00572)→ rpsI (PRK00132)Ribosomal_S9 (pfam00380)→ |
| env-825 | 4448811.3_Contig642 | - | 2634 | 2559 | **RNA**→ rplM (PRK09216)RplM (COG0102)→ rplM (PRK09216)Ribosomal_L13 (pfam00572)→ rpsI (PRK00132)Ribosomal_S9 (pfam00380)→ |
| env-826 | 4448814.3_Contig2119 | + | 771 | 846 | **RNA**→ rplM (PRK09216)Ribosomal_L13 (pfam00572)→ |
| env-827 | scaffold12317_1_MH0026 | - | 272 | 197 | **RNA**→ rplM (PRK09216)Ribosomal_L13 (pfam00572)→ |
| env-828 | scaffold30210_1_MH0049 | - | 536 | 461 | **RNA**→ rplM (PRK09216)Ribosomal_L13 (pfam00572)→ |
| env-829 | 4448810.3.47000 | - | 13461 | 13386 | **RNA**→ rplM (PRK09216)Ribosomal_L13 (pfam00572)→ rpsI (PRK00132)Ribosomal_S9 (pfam00380)→ |
| env-830 | 4448816.3_Contig1564 | - | 700 | 625 | **RNA**→ rplM (PRK09216)Ribosomal_L13 (pfam00572)→ rplM (PRK09216)Ribosomal_L13 (pfam00572)→ rpsI (PRK00132)Ribosomal_S9 (pfam00380)→ |
| env-831 | NOM001_scaffold36808_1 | - | 12342 | 12267 | **RNA**→ rplM (PRK09216)Ribosomal_L13 (pfam00572)→ rpsI (PRK00132)Ribosomal_S9 (pfam00380)→ |
| env-832 | CAM_READ_0099565819 | + | 27 | 102 | **RNA**→ rplM (PRK09216)Ribosomal_L13 (pfam00572)→ |
| env-833 | 4448808.3_Contig33004 | - | 288 | 214 | **RNA**→ rplM (PRK09216)RplM (COG0102)→ |

| | | | | | |
|---|---|---|---|---|---|
| env-834 | NLM016_scaffold13879_1 | - | 134 | 59 | **RNA**→ hypo→ |
| env-835 | scaffold9468_1_O2.UC-12 | + | 3921 | 3996 | **RNA**→ rplM (PRK09216)Ribosomal_L13 (pfam00572)→ |
| env-836 | NLM006_scaffold9242_1 | + | 439 | 514 | **RNA**→ rplM (PRK09216)RplM (COG0102)→ |
| env-837 | NLM009_scaffold10030_1 | + | 15733 | 15808 | **RNA**→ rplM (PRK09216)Ribosomal_L13 (pfam00572)→ rpsI (PRK00132)Ribosomal_S9 (pfam00380)→ |
| Pco-2-1 | NZ_GG703855.1 | - | 96347 | 96272 | **RNA**→ rplM (PRK09216)Ribosomal_L13 (pfam00572)→ rpsI (PRK00132)Ribosomal_S9 (pfam00380)→ |
| env-838 | contig13883 | - | 85 | 10 | **RNA**→ hypo→ |
| env-839 | DLF004_scaffold32113_1 | - | 12265 | 12190 | **RNA**→ rplM (PRK09216)Ribosomal_L13 (pfam00572)→ rpsI (PRK00132)Ribosomal_S9 (pfam00380)→ |
| env-840 | SRS013687_Baylor_scaffold_46077 | - | 545 | 470 | **RNA**→ rplM (PRK09216)Ribosomal_L13 (pfam00572)→ |
| env-841 | SRS023914_Baylor_scaffold_16462 | + | 15313 | 15388 | **RNA**→ rplM (PRK09216)Ribosomal_L13 (pfam00572)→ rpsI (PRK00132)Ribosomal_S9 (pfam00380)→ |
| env-842 | SRS049959_WUGC_scaffold_10211 | - | 449 | 374 | **RNA**→ rplM (PRK09216)Ribosomal_L13 (pfam00572)→ |
| env-843 | scaffold2008_3_V1.UC-21 | + | 224 | 299 | **RNA**→ rplM (PRK09216)Ribosomal_L13 (pfam00572)→ rpsI (PRK00132)Ribosomal_S9 (pfam00380)→ |
| env-844 | NOM020_scaffold12283_2 | - | 11931 | 11856 | **RNA**→ rplM (PRK09216)Ribosomal_L13 (pfam00572)→ rpsI (PRK00132)Ribosomal_S9 (pfam00380)→ |
| Por-8-1 | NZ_KK073873.1 | + | 3030252 | 3030326 | **RNA**→ rplM (PRK09216)Ribosomal_L13 (pfam00572)→ rpsI (PRK00132)Ribosomal_S9 (pfam00380)→ |
| Psp-21-1 | NZ_JRPT01000012.1 | - | 33492 | 33419 | **RNA**→ rplM (PRK09216)Ribosomal_L13 (pfam00572)→ rpsI (PRK00132)Ribosomal_S9 (pfam00380)→ |
| env-845 | RUMENNODE_2697187_1 | - | 3085 | 3011 | **RNA**→ rplM (PRK09216)Ribosomal_L13 (pfam00572)→ rpsI (PRK00132)Ribosomal_S9 (pfam00380)→ |
| env-846 | AUXO015280558.1 | - | 453 | 379 | **RNA**→ rplM (PRK09216)Ribosomal_L13 (pfam00572)→ |
| env-847 | AUXO011297008.1 | - | 1734 | 1660 | **RNA**→ rplM (PRK09216)Ribosomal_L13 (pfam00572)→ rpsI (PRK00132)Ribosomal_S9 (pfam00380)→ |
| env-848 | SRS013723_Baylor_scaffold_19570 | + | 4339 | 4413 | **RNA**→ rplM (PRK09216)Ribosomal_L13 (pfam00572)→ rpsI (PRK00132)Ribosomal_S9 (pfam00380)→ |
| env-849 | SRS012285_C2809459 | - | 149 | 75 | **RNA**→ hypo→ |
| env-850 | SRS019980_C2790887 | - | 135 | 61 | **RNA**→ hypo→ |
| env-851 | SRS016043_WUGC_scaffold_31124 | + | 1754 | 1828 | **RNA**→ rplM (PRK09216)Ribosomal_L13 (pfam00572)→ |
| env-852 | SRS011152_Baylor_scaffold_57690 | - | 154 | 80 | **RNA**→ |
| env-853 | SRS013836_C4801934 | + | 838 | 912 | **RNA**→ |
| env-854 | SRS045313_C1296217 | - | 154 | 80 | **RNA**→ hypo→ |
| env-855 | SRS022725_LANL_scaffold_20670 | - | 412 | 338 | **RNA**→ rplM (PRK09216)Ribosomal_L13 (pfam00572)→ |
| env-856 | SRS022149_C2846288 | + | 4393 | 4467 | **RNA**→ |
| env-857 | SRS051930_C3273277 | + | 1215 | 1289 | **RNA**→ |
| env-858 | SRS042984_LANL_scaffold_16758 | + | 9998 | 10072 | **RNA**→ rplM (PRK09216)Ribosomal_L13 (pfam00572)→ rpsI (PRK00132)Ribosomal_S9 (pfam00380)→ |
| env-859 | SRS052876_LANL_scaffold_54535 | + | 954 | 1028 | **RNA**→ |
| Pco-1-1 | NZ_HE999443.1 | - | 281229 | 281155 | **RNA**→ rplM (PRK09216)Ribosomal_L13 (pfam00572)→ rpsI (PRK00132)Ribosomal_S9 (pfam00380)→ |
| env-860 | SRS014578_WUGC_scaffold_18370 | - | 7328 | 7254 | **RNA**→ rplM (PRK09216)Ribosomal_L13 (pfam00572)→ rpsI (PRK00132)Ribosomal_S9 (pfam00380)→ |
| env-861 | SRS013170_Baylor_scaffold_112597 | + | 2727 | 2801 | **RNA**→ |
| env-862 | SRS043239_C1976103 | - | 186 | 112 | **RNA**→ hypo→ |
| env-863 | SRS063999_LANL_scaffold_2422 | - | 7865 | 7791 | **RNA**→ rplM (PRK09216)Ribosomal_L13 (pfam00572)→ rpsI (PRK00132)Ribosomal_S9 (pfam00380)→ |
| env-864 | SRS016360_Baylor_scaffold_11199 | - | 720 | 646 | **RNA**→ rplM (PRK09216)Ribosomal_L13 (pfam00572)→ rpsI (PRK00132)Ribosomal_S9 (pfam00380)→ |
| env-865 | SRS019128_C4357049 | - | 303 | 229 | **RNA**→ rplM (PRK09216)Ribosomal_L13 (pfam00572)→ |
| env-866 | SRS024087_LANL_scaffold_47659 | - | 135 | 61 | **RNA**→ |
| env-867 | SRS063932_LANL_scaffold_92613 | + | 4336 | 4410 | **RNA**→ |
| env-868 | SRS024289_LANL_scaffold_70016 | - | 135 | 61 | **RNA**→ |
| env-869 | SRS018443_Baylor_scaffold_65820 | + | 8016 | 8090 | **RNA**→ rplM (PRK09216)Ribosomal_L13 (pfam00572)→ rpsI (PRK00132)Ribosomal_S9 (pfam00380)→ |
| env-870 | SRS018337_C2365684 | + | 512 | 586 | **RNA**→ |
| env-871 | SRS063215_LANL_scaffold_21409 | - | 945 | 871 | **RNA**→ rplM (PRK09216)Ribosomal_L13 (pfam00572)→ rpsI (PRK00132)Ribosomal_S9 (pfam00380)→ |
| env-872 | SRS014107_WUGC_scaffold_50706 | - | 134 | 60 | **RNA**→ |
| env-873 | SRS015044_WUGC_scaffold_67840 | - | 135 | 61 | **RNA**→ |
| env-874 | SRS016331_C4131247 | - | 156 | 82 | **RNA**→ |
| env-875 | SRS017139_C3804675 | - | 135 | 61 | **RNA**→ |
| env-876 | SRS024355_LANL_scaffold_42842 | - | 1441 | 1367 | **RNA**→ rplM (PRK09216)Ribosomal_L13 (pfam00572)→ rpsI (PRK00132)Ribosomal_S9 (pfam00380)→ |
| Psp-25-1 | NZ_GG740073.1 | - | 461425 | 461351 | **RNA**→ rplM (PRK09216)Ribosomal_L13 (pfam00572)→ rplM (PRK09216)Ribosomal_L13 (pfam00572)→ rpsI (PRK00132)Ribosomal_S9 (pfam00380)→ rpsI (PRK00132)Ribosomal_S9 (pfam00380)→ |
| env-877 | SRS011343_C2242542 | + | 1051 | 1125 | **RNA**→ |
| env-878 | SRS014691_C2503925 | - | 145 | 71 | **RNA**→ hypo→ |
| env-879 | SRS019028_WUGC_scaffold_45064 | - | 145 | 71 | **RNA**→ hypo→ |

| | | | | | |
|---|---|---|---|---|---|
| env-880 | SRS015574_WUGC_scaffold_63161 | + | 871 | 945 | **RNA**→ rplM (PRK09216)Ribosomal_L13 (pfam00572)→ |
| env-881 | SRS016746_Baylor_scaffold_73191 | + | 1882 | 1956 | **RNA**→ |
| env-882 | SRS017511_Baylor_scaffold_92834 | - | 135 | 61 | **RNA**→ |
| env-883 | SRS023358_C3462583 | + | 482 | 556 | **RNA**→ |
| env-884 | SRS049318_LANL_scaffold_32648 | + | 9208 | 9282 | **RNA**→ |
| env-885 | SRS050628_LANL_scaffold_20687 | - | 978 | 904 | **RNA**→ rplM (PRK09216)Ribosomal_L13 (pfam00572)→ rpsI (PRK00132)Ribosomal_S9 (pfam00380)→ |
| env-886 | SRS022719_LANL_scaffold_45285 | - | 804 | 729 | **RNA**→ rplM (PRK09216)Ribosomal_L13 (pfam00572)→ rpsI (PRK00132)Ribosomal_S9 (pfam00380)→ |
| env-887 | SRS019129_WUGC_scaffold_43257 | - | 338 | 263 | **RNA**→ rplM (PRK09216)Ribosomal_L13 (pfam00572)→ |
| env-888 | SRS016043_C2115867 | - | 218 | 143 | **RNA**→ rplM (PRK09216)RplM (COG0102)→ |
| env-889 | SRS013533_PGA_scaffold_117699 | + | 3103 | 3178 | **RNA**→ |
| env-890 | SRS017511_Baylor_scaffold_92627 | + | 951 | 1026 | **RNA**→ |
| env-891 | SRS021477_C2588523 | - | 323 | 248 | **RNA**→ rplM (PRK09216)Ribosomal_L13 (pfam00572)→ |
| env-892 | SRS013170_Baylor_scaffold_67347 | - | 307 | 232 | **RNA**→ rplM (PRK09216)Ribosomal_L13 (pfam00572)→ |
| env-893 | SRS064449_C4126558 | - | 118 | 43 | **RNA**→ |
| env-894 | SRS053917_LANL_scaffold_88433 | + | 607 | 682 | **RNA**→ rplM (PRK09216)Ribosomal_L13 (pfam00572)→ |
| env-895 | SRS013252_C3525919 | - | 153 | 78 | **RNA**→ hypo→ |
| env-896 | SRS015803_C2675896 | + | 3433 | 3508 | **RNA**→ rplM (PRK09216)Ribosomal_L13 (pfam00572)→ rpsI (PRK00132)Ribosomal_S9 (pfam00380)→ |
| env-897 | SRS018394_C4057209 | - | 982 | 907 | **RNA**→ rplM (PRK09216)Ribosomal_L13 (pfam00572)→ rpsI (PRK00132)Ribosomal_S9 (pfam00380)→ |
| env-898 | SRS024649_LANL_scaffold_51754 | - | 635 | 560 | **RNA**→ rplM (PRK09216)Ribosomal_L13 (pfam00572)→ |
| env-899 | SRS054430_LANL_scaffold_20956 | - | 988 | 913 | **RNA**→ rplM (PRK09216)Ribosomal_L13 (pfam00572)→ rpsI (PRK00132)Ribosomal_S9 (pfam00380)→ |
| Plo-1-1 | NZ_KB899213.1 | + | 249357 | 249432 | **RNA**→ rplM (PRK09216)Ribosomal_L13 (pfam00572)→ rpsI (PRK00132)Ribosomal_S9 (pfam00380)→ |
| env-900 | SRS017445_Baylor_scaffold_83464 | + | 1528 | 1603 | **RNA**→ rplM (PRK09216)Ribosomal_L13 (pfam00572)→ |
| env-901 | SRS018573_C2480980 | - | 139 | 64 | **RNA**→ |
| env-902 | SRS045197_WUGC_scaffold_65903 | + | 7998 | 8073 | **RNA**→ rplM (PRK09216)Ribosomal_L13 (pfam00572)→ rpsI (PRK00132)Ribosomal_S9 (pfam00380)→ |
| Psp-26-1 | NZ_GG704825.1 | - | 516881 | 516806 | **RNA**→ rplM (PRK09216)Ribosomal_L13 (pfam00572)→ rpsI (PRK00132)Ribosomal_S9 (pfam00380)→ hypo→ rpsB (PRK05299)Ribosomal_S2 (pfam00318)→ tsf (PRK09377)UBA_EF-Ts (cd14275)EF_TS (pfam00889)→ |
| env-903 | SRS023841_Baylor_scaffold_21872 | - | 947 | 872 | **RNA**→ rplM (PRK09216)Ribosomal_L13 (pfam00572)→ rpsI (PRK00132)Ribosomal_S9 (pfam00380)→ |
| env-904 | SRS011098_C1871947 | - | 560 | 485 | **RNA**→ rplM (PRK09216)Ribosomal_L13 (pfam00572)→ |
| env-905 | SRS011255_C2493253 | - | 151 | 76 | **RNA**→ |
| env-906 | SRS016746_C2995254 | + | 311 | 386 | **RNA**→ |
| env-907 | SRS017227_C5088817 | + | 278 | 353 | **RNA**→ |
| env-908 | SRS018157_Baylor_scaffold_33504 | + | 7670 | 7745 | **RNA**→ rplM (PRK09216)Ribosomal_L13 (pfam00572)→ rpsI (PRK00132)Ribosomal_S9 (pfam00380)→ |
| env-909 | SRS018665_C3707955 | + | 500 | 575 | **RNA**→ |
| env-910 | SRS019906_C2276695 | + | 7839 | 7914 | **RNA**→ |
| env-911 | SRS023595_C6887425 | - | 139 | 64 | **RNA**→ hypo→ |
| env-912 | SRS023938_C2241954 | - | 119 | 44 | **RNA**→ hypo→ |
| env-913 | SRS024087_LANL_scaffold_42583 | - | 139 | 64 | **RNA**→ |
| env-914 | SRS051378_C1873917 | - | 1295 | 1220 | **RNA**→ rplM (PRK09216)Ribosomal_L13 (pfam00572)→ rpsI (PRK00132)Ribosomal_S9 (pfam00380)→ |
| env-915 | SRS051941_LANL_scaffold_77484 | - | 153 | 78 | **RNA**→ |
| env-916 | SRS052604_LANL_scaffold_13013 | + | 539 | 614 | **RNA**→ rplM (PRK09216)Ribosomal_L13 (pfam00572)→ rpsI (PRK00132)Ribosomal_S9 (pfam00380)→ |
| env-917 | SRS055401_C2209469 | + | 195 | 270 | **RNA**→ |
| env-918 | SRS058053_LANL_scaffold_60299 | + | 304 | 379 | **RNA**→ |
| env-919 | SRS063603_C4249750 | - | 118 | 43 | **RNA**→ |
| env-920 | SRS063932_LANL_scaffold_98774 | - | 139 | 64 | **RNA**→ |
| env-921 | SRS017808_C3874880 | - | 146 | 71 | **RNA**→ |
| env-922 | SRS017439_Baylor_scaffold_51404 | - | 632 | 557 | **RNA**→ rplM (PRK09216)Ribosomal_L13 (pfam00572)→ |
| env-923 | SRS075404_LANL_scaffold_60611 | + | 3391 | 3466 | **RNA**→ |
| env-924 | SRS012279_Baylor_scaffold_46820 | + | 3560 | 3635 | **RNA**→ rplM (PRK09216)Ribosomal_L13 (pfam00572)→ rpsI (PRK00132)Ribosomal_S9 (pfam00380)→ |
| env-925 | SRS013705_Baylor_scaffold_63054 | + | 3114 | 3189 | **RNA**→ rplM (PRK09216)Ribosomal_L13 (pfam00572)→ rpsI (PRK00132)Ribosomal_S9 (pfam00380)→ |
| env-926 | SRS013879_C2765042 | + | 277 | 352 | **RNA**→ |
| env-927 | SRS017209_Baylor_scaffold_40939 | + | 72651 | 72726 | **RNA**→ rplM (PRK09216)Ribosomal_L13 (pfam00572)→ rpsI (PRK00132)Ribosomal_S9 (pfam00380)→ |
| env-928 | SRS018357_Baylor_scaffold_46794 | + | 98 | 173 | **RNA**→ rplM (PRK09216)Ribosomal_L13 (pfam00572)→ |

| | | | | | |
|---|---|---|---|---|---|
| env-929 | SRS022077_Baylor_scaffold_41018 | + | 3560 | 3635 | **RNA**→ rplM (PRK09216)Ribosomal_L13 (pfam00572)→ rpsI (PRK00132)Ribosomal_S9 (pfam00380)→ |
| env-930 | SRS023926_Baylor_scaffold_55027 | + | 3644 | 3719 | **RNA**→ rplM (PRK09216)Ribosomal_L13 (pfam00572)→ rpsI (PRK00132)Ribosomal_S9 (pfam00380)→ |
| env-931 | SRS050669_LANL_scaffold_50571 | + | 3562 | 3637 | **RNA**→ rplM (PRK09216)Ribosomal_L13 (pfam00572)→ rpsI (PRK00132)Ribosomal_S9 (pfam00380)→ |
| env-932 | SRS064329_LANL_scaffold_50907 | - | 985 | 910 | **RNA**→ rplM (PRK09216)Ribosomal_L13 (pfam00572)→ rpsI (PRK00132)Ribosomal_S9 (pfam00380)→ |
| env-933 | Tongue_dor_WUGI_scaffold_12918 | + | 1043 | 1118 | **RNA**→ |
| env-934 | SRS011306_Baylor_scaffold_62841 | - | 35842 | 35767 | **RNA**→ rplM (PRK09216)Ribosomal_L13 (pfam00572)→ rpsI (PRK00132)Ribosomal_S9 (pfam00380)→ |
| env-935 | SRS014573_WUGC_scaffold_21848 | + | 3270 | 3345 | **RNA**→ rplM (PRK09216)Ribosomal_L13 (pfam00572)→ |
| env-936 | SRS015762_C3664821 | + | 2929 | 3004 | **RNA**→ |
| env-937 | SRS016002_C2493893 | - | 135 | 60 | **RNA**→ hypo→ |
| env-938 | SRS018300_Baylor_scaffold_10118 | + | 3538 | 3613 | **RNA**→ rplM (PRK09216)Ribosomal_L13 (pfam00572)→ rpsI (PRK00132)Ribosomal_S9 (pfam00380)→ |
| env-939 | SRS018739_WUGC_scaffold_27638 | + | 3156 | 3231 | **RNA**→ rplM (PRK09216)Ribosomal_L13 (pfam00572)→ rpsI (PRK00132)Ribosomal_S9 (pfam00380)→ |
| env-940 | SRS019389_C2124919 | - | 146 | 71 | **RNA**→ hypo→ |
| env-941 | SRS024081_LANL_scaffold_17499 | + | 2653 | 2728 | **RNA**→ rplM (PRK09216)Ribosomal_L13 (pfam00572)→ rpsI (PRK00132)Ribosomal_S9 (pfam00380)→ |
| env-942 | SRS024441_LANL_scaffold_33890 | + | 6153 | 6228 | **RNA**→ rplM (PRK09216)Ribosomal_L13 (pfam00572)→ rpsI (PRK00132)Ribosomal_S9 (pfam00380)→ |
| env-943 | SRS049389_WUGC_scaffold_44708 | - | 146 | 71 | **RNA**→ |
| env-944 | SRS053603_LANL_scaffold_56769 | + | 3538 | 3613 | **RNA**→ |
| env-945 | SRS057205_LANL_scaffold_36133 | - | 1438 | 1363 | **RNA**→ rplM (PRK09216)Ribosomal_L13 (pfam00572)→ rpsI (PRK00132)Ribosomal_S9 (pfam00380)→ |
| env-946 | SRS063193_LANL_scaffold_48783 | + | 3570 | 3645 | **RNA**→ rplM (PRK09216)Ribosomal_L13 (pfam00572)→ |
| env-947 | SRS064774_C2783916 | - | 146 | 71 | **RNA**→ hypo→ |
| Psh-1-1 | NZ_BAIZ01000030.1 | - | 35483 | 35408 | **RNA**→ rplM (PRK09216)Ribosomal_L13 (pfam00572)→ rpsI (PRK00132)Ribosomal_S9 (pfam00380)→ |
| env-948 | SRS022143_WUGC_scaffold_91746 | - | 632 | 557 | **RNA**→ rplM (PRK09216)Ribosomal_L13 (pfam00572)→ |
| env-949 | SRS019327_WUGC_scaffold_70870 | - | 10558 | 10483 | **RNA**→ rplM (PRK09216)Ribosomal_L13 (pfam00572)→ rpsI (PRK00132)Ribosomal_S9 (pfam00380)→ |
| env-950 | SRS015434_C5949372 | - | 146 | 71 | **RNA**→ |
| env-951 | SRS018439_Baylor_scaffold_22146 | - | 17449 | 17374 | **RNA**→ rplM (PRK09216)Ribosomal_L13 (pfam00572)→ rpsI (PRK00132)Ribosomal_S9 (pfam00380)→ |
| env-952 | SRS019071_C5217890 | + | 1981 | 2056 | **RNA**→ |
| env-953 | SRS013164_Baylor_scaffold_59130 | - | 979 | 904 | **RNA**→ rplM (PRK09216)Ribosomal_L13 (pfam00572)→ rpsI (PRK00132)Ribosomal_S9 (pfam00380)→ |
| env-954 | SRS014888_C3104655 | - | 146 | 71 | **RNA**→ |
| env-955 | SRS023958_Baylor_scaffold_31813 | + | 3560 | 3635 | **RNA**→ rplM (PRK09216)Ribosomal_L13 (pfam00572)→ |
| env-956 | SRS057355_C2359830 | - | 146 | 71 | **RNA**→ |
| env-957 | SRS021496_C2846806 | - | 146 | 71 | **RNA**→ |
| env-958 | SRS022530_LANL_scaffold_60444 | - | 7595 | 7520 | **RNA**→ rplM (PRK09216)Ribosomal_L13 (pfam00572)→ rpsI (PRK00132)Ribosomal_S9 (pfam00380)→ |
| env-959 | SRS063288_C2558656 | + | 371 | 446 | **RNA**→ |
| env-960 | scaffold15263_2_MH0055 | - | 943 | 868 | **RNA**→ rplM (PRK09216)Ribosomal_L13 (pfam00572)→ rpsI (PRK00132)Ribosomal_S9 (pfam00380)→ |
| env-961 | scaffold18695_5_MH0014 | + | 2193 | 2268 | **RNA**→ rplM (PRK09216)Ribosomal_L13 (pfam00572)→ rpsI (PRK00132)Ribosomal_S9 (pfam00380)→ |
| env-962 | scaffold58703_4_MH0042 | + | 509 | 584 | **RNA**→ rplM (PRK09216)Ribosomal_L13 (pfam00572)→ |
| env-963 | scaffold410_3_MH0063 | - | 5281 | 5206 | **RNA**→ rplM (PRK09216)Ribosomal_L13 (pfam00572)→ rpsI (PRK00132)Ribosomal_S9 (pfam00380)→ |
| env-964 | scaffold1879_1_O2.UC-24 | + | 1299 | 1374 | **RNA**→ rplM (PRK09216)Ribosomal_L13 (pfam00572)→ rpsI (PRK00132)Ribosomal_S9 (pfam00380)→ |
| env-965 | scaffold6667_6_V1.UC-14 | - | 958 | 883 | **RNA**→ rplM (PRK09216)Ribosomal_L13 (pfam00572)→ rpsI (PRK00132)Ribosomal_S9 (pfam00380)→ |
| env-966 | 4491482.3_NODE_74973 | - | 6204 | 6129 | **RNA**→ rplM (PRK09216)Ribosomal_L13 (pfam00572)→ rpsI (PRK00132)Ribosomal_S9 (pfam00380)→ |
| env-967 | BAAX01036178.1 | - | 165 | 88 | **RNA**→ |
| env-968 | 4491403.3_NODE_60304 | - | 979 | 902 | **RNA**→ rplM (PRK09216)Ribosomal_L13 (pfam00572)→ rplM (PRK09216)Ribosomal_L13 (pfam00572)→ rpsI (PRK00132)Ribosomal_S9 (pfam00380)→ rpsI (PRK00132)Ribosomal_S9 (pfam00380)→ |
| env-969 | DLF003_scaffold24849_4 | - | 8212 | 8135 | **RNA**→ rplM (PRK09216)Ribosomal_L13 (pfam00572)→ rpsI (PRK00132)Ribosomal_S9 (pfam00380)→ |
| env-970 | NLM016_scaffold8572_3 | - | 1028 | 951 | **RNA**→ rplM (PRK09216)Ribosomal_L13 (pfam00572)→ rpsI (PRK00132)Ribosomal_S9 (pfam00380)→ |
| env-971 | 4491404.3_NODE_3637 | + | 6552 | 6629 | **RNA**→ rplM (PRK09216)Ribosomal_L13 (pfam00572)→ |
| env-972 | DLM006_scaffold334_9 | + | 10545 | 10622 | **RNA**→ rplM (PRK09216)Ribosomal_L13 (pfam00572)→ rpsI (PRK00132)Ribosomal_S9 (pfam00380)→ |
| env-973 | scaffold78050_1_O2.UC-19 | + | 162 | 239 | **RNA**→ rplM (PRK09216)Ribosomal_L13 (pfam00572)→ rpsI (PRK00132)Ribosomal_S9 (pfam00380)→ |
| env-974 | DOM016_scaffold1189_4 | + | 9750 | 9827 | **RNA**→ rplM (PRK09216)Ribosomal_L13 (pfam00572)→ rpsI (PRK00132)Ribosomal_S9 (pfam00380)→ |
| env-975 | DOM022_scaffold19179_2 | + | 11637 | 11714 | **RNA**→ rplM (PRK09216)Ribosomal_L13 (pfam00572)→ rpsI (PRK00132)Ribosomal_S9 (pfam00380)→ |
| env-976 | SRS011529_WUGC_scaffold_9481 | + | 35 | 112 | **RNA**→ rplM (PRK09216)Ribosomal_L13 (pfam00572)→ rpsI (PRK00132)Ribosomal_S9 (pfam00380)→ |
| Pst-1-1 | NZ_JH379462.1 | - | 10330 | 10253 | **RNA**→ rplM (PRK09216)Ribosomal_L13 (pfam00572)→ rpsI (PRK00132)Ribosomal_S9 (pfam00380)→ |
| env-977 | NLF006_scaffold10643_1 | - | 1262 | 1185 | **RNA**→ rplM (PRK09216)Ribosomal_L13 (pfam00572)→ rpsI (PRK00132)Ribosomal_S9 (pfam00380)→ |

| | | | | | |
|---|---|---|---|---|---|
| env-978 | NLM029_scaffold46472_1 | - | 9492 | 9415 | **RNA**→ rplM (PRK09216)Ribosomal_L13 (pfam00572)→ rpsI (PRK00132)Ribosomal_S9 (pfam00380)→ |
| env-979 | NLF005_scaffold222_5 | + | 9773 | 9850 | **RNA**→ rplM (PRK09216)Ribosomal_L13 (pfam00572)→ rpsI (PRK00132)Ribosomal_S9 (pfam00380)→ |
| env-980 | NOM007_scaffold112_4 | + | 15977 | 16054 | **RNA**→ rplM (PRK09216)Ribosomal_L13 (pfam00572)→ rpsI (PRK00132)Ribosomal_S9 (pfam00380)→ |
| env-981 | NLM025_scaffold10432_10 | - | 8972 | 8895 | **RNA**→ rplM (PRK09216)Ribosomal_L13 (pfam00572)→ rpsI (PRK00132)Ribosomal_S9 (pfam00380)→ |
| env-982 | DLM002_scaffold10936_3 | + | 5488 | 5565 | **RNA**→ rplM (PRK09216)Ribosomal_L13 (pfam00572)→ rpsI (PRK00132)Ribosomal_S9 (pfam00380)→ |
| env-983 | NLM009_scaffold4804_2 | + | 26579 | 26656 | **RNA**→ rplM (PRK09216)Ribosomal_L13 (pfam00572)→ rpsI (PRK00132)Ribosomal_S9 (pfam00380)→ |
| env-984 | NLM002_scaffold27297_2 | - | 9435 | 9358 | **RNA**→ rplM (PRK09216)Ribosomal_L13 (pfam00572)→ rpsI (PRK00132)Ribosomal_S9 (pfam00380)→ |
| env-985 | UnmappedStool_Broad_scaffold_727222 | - | 162 | 82 | **RNA**→ hypo→ |
| env-986 | 4491404.3_NODE_8492 | + | 545 | 625 | **RNA**→ rplM (PRK09216)Ribosomal_L13 (pfam00572)→ |
| env-987 | SRS016541_C1524265 | + | 279 | 359 | **RNA**→ |
| env-988 | SRS013687_Baylor_scaffold_57705 | + | 279 | 359 | **RNA**→ rplM (PRK09216)Ribosomal_L13 (pfam00572)→ |
| env-989 | SRS023914_Baylor_scaffold_14686 | + | 279 | 359 | **RNA**→ rplM (PRK09216)Ribosomal_L13 (pfam00572)→ rpsI (PRK00132)Ribosomal_S9 (pfam00380)→ |
| env-990 | BAAX01006449.1 | - | 273 | 193 | **RNA**→ rplM (PRK09216)Ribosomal_L13 (pfam00572)→ |
| env-991 | NLF006_scaffold165_1 | - | 402 | 322 | **RNA**→ rplM (PRK09216)Ribosomal_L13 (pfam00572)→ |
| env-992 | 4491403.3_NODE_60304 | - | 780 | 700 | **RNA**→ rplM (PRK09216)Ribosomal_L13 (pfam00572)→ rplM (PRK09216)Ribosomal_L13 (pfam00572)→ rpsI (PRK00132)Ribosomal_S9 (pfam00380)→ rpsI (PRK00132)Ribosomal_S9 (pfam00380)→ |
| env-993 | SRS049959_C3758114 | - | 124 | 44 | **RNA**→ hypo→ |
| env-994 | scaffold29644_4_O2.UC-13 | - | 6421 | 6346 | **RNA**→ rplM (PRK09216)Ribosomal_L13 (pfam00572)→ rpsI (PRK00132)Ribosomal_S9 (pfam00380)→ |
| Pbu-3-1 | NZ_GG739928.1 | - | 179846 | 179770 | **RNA**→ rplM (PRK09216)Ribosomal_L13 (pfam00572)→ rplM (PRK09216)Ribosomal_L13 (pfam00572)→ rpsI (PRK00132)Ribosomal_S9 (pfam00380)→ rpsI (PRK00132)Ribosomal_S9 (pfam00380)→ |
| Pbu-1-1 | NZ_JVYX01000008.1 | - | 5815 | 5739 | **RNA**→ rplM (PRK09216)Ribosomal_L13 (pfam00572)→ rpsI (PRK00132)Ribosomal_S9 (pfam00380)→ |
| Pbu-2-1 | NZ_GL586311.1 | + | 1240091 | 1240167 | **RNA**→ rplM (PRK09216)Ribosomal_L13 (pfam00572)→ rpsI (PRK00132)Ribosomal_S9 (pfam00380)→ |
| Psp-12-1 | NZ_ALJQ01000010.1 | - | 159165 | 159089 | **RNA**→ rplM (PRK09216)Ribosomal_L13 (pfam00572)→ rplM (PRK09216)Ribosomal_L13 (pfam00572)→ rpsI (PRK00132)Ribosomal_S9 (pfam00380)→ rpsI (PRK00132)Ribosomal_S9 (pfam00380)→ |
| env-995 | AUXO015968780.1 | + | 216 | 290 | **RNA**→ |
| env-996 | AUXO017534585.1 | - | 968 | 894 | **RNA**→ rplM (PRK09216)Ribosomal_L13 (pfam00572)→ rpsI (PRK00132)Ribosomal_S9 (pfam00380)→ |
| env-997 | pre-rum113-00662 | - | 3481 | 3406 | **RNA**→ rplM (PRK09216)Ribosomal_L13 (pfam00572)→ rpsI (PRK00132)Ribosomal_S9 (pfam00380)→ |
| env-998 | pre-rum114-03875 | + | 2977 | 3052 | **RNA**→ rplM (PRK09216)Ribosomal_L13 (pfam00572)→ rpsI (PRK00132)Ribosomal_S9 (pfam00380)→ |
| env-999 | HCE12Call500_c0260910 | + | 4902 | 4978 | **RNA**→ rplM (PRK09216)Ribosomal_L13 (pfam00572)→ |
| env-1000 | HCF12C_329458 | - | 179 | 103 | **RNA**→ ←hypo |
| Por-1-1 | NZ_GL833119.1 | - | 125193 | 125119 | **RNA**→ rplM (PRK09216)Ribosomal_L13 (pfam00572)→ rpsI (PRK00132)Ribosomal_S9 (pfam00380)→ |
| Por-3-1 | NZ_KE340302.1 | - | 130474 | 130400 | **RNA**→ rplM (PRK09216)Ribosomal_L13 (pfam00572)→ rpsI (PRK00132)Ribosomal_S9 (pfam00380)→ |
| env-1001 | SRS013836_C4829201 | - | 117 | 43 | **RNA**→ |
| Por-2-1 | NZ_KI669542.1 | - | 130434 | 130360 | **RNA**→ rplM (PRK09216)Ribosomal_L13 (pfam00572)→ rpsI (PRK00132)Ribosomal_S9 (pfam00380)→ |
| env-1002 | SRS023604_Baylor_scaffold_3034 | + | 34418 | 34492 | **RNA**→ rplM (PRK09216)Ribosomal_L13 (pfam00572)→ rpsI (PRK00132)Ribosomal_S9 (pfam00380)→ |
| env-1003 | SRS013542_C165903 | - | 485 | 411 | **RNA**→ rplM (PRK09216)Ribosomal_L13 (pfam00572)→ |
| Pti-3-1 | NZ_JRPQ01000080.1 | + | 19409 | 19483 | **RNA**→ rplM (PRK09216)Ribosomal_L13 (pfam00572)→ rpsI (PRK00132)Ribosomal_S9 (pfam00380)→ |
| env-1004 | SRS011111_Baylor_scaffold_4265 | + | 5684 | 5758 | **RNA**→ rplM (PRK09216)Ribosomal_L13 (pfam00572)→ |
| Pti-1-1 | NZ_HG417093.1 | - | 162971 | 162897 | **RNA**→ rplM (PRK09216)Ribosomal_L13 (pfam00572)→ rpsI (PRK00132)Ribosomal_S9 (pfam00380)→ |
| Pti-2-1 | NZ_ADEF01000048.1 | + | 153828 | 153902 | **RNA**→ rplM (PRK09216)Ribosomal_L13 (pfam00572)→ rplM (PRK09216)Ribosomal_L13 (pfam00572)→ rpsI (PRK00132)Ribosomal_S9 (pfam00380)→ rpsI (PRK00132)Ribosomal_S9 (pfam00380)→ |
| env-1005 | SRS023468_LANL_scaffold_1144 | - | 3752 | 3678 | **RNA**→ rplM (PRK09216)Ribosomal_L13 (pfam00572)→ rpsI (PRK00132)Ribosomal_S9 (pfam00380)→ |
| env-1006 | SRS023604_Baylor_scaffold_2736 | - | 116 | 42 | **RNA**→ hypo→ |
| Psp-4-1 | NZ_AWXC01000037.1 | - | 65697 | 65623 | **RNA**→ rplM (PRK09216)Ribosomal_L13 (pfam00572)→ rpsI (PRK00132)Ribosomal_S9 (pfam00380)→ |
| Pbu-6-1 | NZ_JRNP01000028.1 | + | 52030 | 52104 | **RNA**→ rplM (PRK09216)Ribosomal_L13 (pfam00572)→ rpsI (PRK00132)Ribosomal_S9 (pfam00380)→ |
| env-1007 | SRS011111_Baylor_scaffold_12781 | - | 1177 | 1103 | **RNA**→ rplM (PRK09216)Ribosomal_L13 (pfam00572)→ rpsI (PRK00132)Ribosomal_S9 (pfam00380)→ |
| Pbu-5-1 | NZ_JRNN01000034.1 | + | 33581 | 33655 | **RNA**→ rplM (PRK09216)Ribosomal_L13 (pfam00572)→ rpsI (PRK00132)Ribosomal_S9 (pfam00380)→ |
| Pbu-4-1 | NZ_ADEG01000023.1 | - | 35564 | 35490 | **RNA**→ rplM (PRK09216)Ribosomal_L13 (pfam00572)→ rplM (PRK09216)Ribosomal_L13 (pfam00572)→ rpsI (PRK00132)Ribosomal_S9 (pfam00380)→ rpsI (PRK00132)Ribosomal_S9 (pfam00380)→ |
| env-1008 | SRS017511_Baylor_scaffold_48340 | - | 333 | 258 | **RNA**→ hypo→ rplM (PRK09216)Ribosomal_L13 (pfam00572)→ |
| Pen-1-1 | NZ_BAIX01000001.1 | + | 113639 | 113714 | **RNA**→ rplM (PRK09216)Ribosomal_L13 (pfam00572)→ rpsI (PRK00132)Ribosomal_S9 (pfam00380)→ |
| env-1009 | SRS013950_C2181444 | - | 1427 | 1352 | **RNA**→ rplM (PRK09216)Ribosomal_L13 (pfam00572)→ rpsI (PRK00132)Ribosomal_S9 (pfam00380)→ |
| env-1010 | SRS014107_WUGC_scaffold_54517 | - | 223 | 148 | **RNA**→ rplM (PRK09216)RplM (COG0102)→ |

| | | | | | |
|---|---|---|---|---|---|
| env-1011 | SRS014476_C3414245 | + | 514 | 589 | **RNA**→ rplM (PRK09216)RplM (COG0102)→ |
| env-1012 | SRS014691_C2727318 | + | 851 | 926 | **RNA**→ |
| env-1013 | SRS016746_Baylor_scaffold_78063 | + | 1074 | 1149 | **RNA**→ rplM (PRK09216)Ribosomal_L13 (pfam00572)→ |
| env-1014 | SRS017227_Baylor_scaffold_162432 | - | 14350 | 14275 | **RNA**→ rplM (PRK09216)Ribosomal_L13 (pfam00572)→ rpsI (PRK00132)Ribosomal_S9 (pfam00380)→ |
| env-1015 | SRS019029_WUGC_scaffold_53097 | + | 1214 | 1289 | **RNA**→ rplM (PRK09216)Ribosomal_L13 (pfam00572)→ |
| env-1016 | SRS047634_LANL_scaffold_137848 | + | 8745 | 8820 | **RNA**→ |
| Ppl-1-1 | NZ_AWET01000036.1 | + | 123358 | 123433 | **RNA**→ rplM (PRK09216)Ribosomal_L13 (pfam00572)→ rpsI (PRK00132)Ribosomal_S9 (pfam00380)→ |
| env-1017 | Buccal_mucosa_LANL_C64865474 | + | 28 | 103 | **RNA**→ rplM (PRK09216)Ribosomal_L13 (pfam00572)→ |
| env-1018 | SRS011255_C2371241 | - | 446 | 371 | **RNA**→ rplM (PRK09216)Ribosomal_L13 (pfam00572)→ |
| env-1019 | SRS047113_LANL_scaffold_81006 | + | 28 | 103 | **RNA**→ hypo→ rplM (PRK09216)Ribosomal_L13 (pfam00572)→ rpsI (PRK00132)Ribosomal_S9 (pfam00380)→ |
| env-1020 | SRS058808_C2744514 | + | 90 | 165 | **RNA**→ hypo→ rplM (PRK09216)Ribosomal_L13 (pfam00572)→ |
| Ppl-2-1 | NZ_BAJN01000002.1 | - | 253813 | 253738 | **RNA**→ rplM (PRK09216)Ribosomal_L13 (pfam00572)→ rpsI (PRK00132)Ribosomal_S9 (pfam00380)→ |
| env-1021 | scaffold29515_4_MH0046 | - | 478 | 404 | **RNA**→ rplM (PRK09216)Ribosomal_L13 (pfam00572)→ |
| env-1022 | scaffold6118_1_MH0049 | + | 1778 | 1852 | **RNA**→ rplM (PRK09216)Ribosomal_L13 (pfam00572)→ rpsI (PRK00132)Ribosomal_S9 (pfam00380)→ |
| env-1023 | scaffold121376_1_MH0056 | - | 1807 | 1733 | **RNA**→ rplM (PRK09216)Ribosomal_L13 (pfam00572)→ rpsI (PRK00132)Ribosomal_S9 (pfam00380)→ |
| env-1024 | 4448812.3_Contig33336 | + | 223 | 300 | **RNA**→ |
| env-1025 | DLF003_scaffold46021_1 | - | 4697 | 4623 | **RNA**→ rplM (PRK09216)Ribosomal_L13 (pfam00572)→ rpsI (PRK00132)Ribosomal_S9 (pfam00380)→ |
| env-1026 | NOF001_scaffold14758_1 | - | 4682 | 4608 | **RNA**→ rplM (PRK09216)Ribosomal_L13 (pfam00572)→ rpsI (PRK00132)Ribosomal_S9 (pfam00380)→ |
| env-1027 | SRS017227_C4992648 | - | 155 | 81 | **RNA**→ hypo→ |
| Pma-4-1 | NZ_GL397214.1 | - | 1714297 | 1714223 | **RNA**→ rplM (PRK09216)Ribosomal_L13 (pfam00572)→ rpsI (PRK00132)Ribosomal_S9 (pfam00380)→ |
| env-1028 | SRS023595_Baylor_scaffold_66299 | + | 21 | 95 | **RNA**→ rplM (PRK09216)Ribosomal_L13 (pfam00572)→ rpsI (PRK00132)Ribosomal_S9 (pfam00380)→ |
| Pma-5-1 | NZ_BAJI01000002.1 | - | 152336 | 152262 | **RNA**→ rplM (PRK09216)Ribosomal_L13 (pfam00572)→ hypo→ rpsI (PRK00132)Ribosomal_S9 (pfam00380)→ |
| env-1029 | AUXO014575475.1 | + | 800 | 874 | **RNA**→ rplM (PRK09216)Ribosomal_L13 (pfam00572)→ rpsI (PRK00132)Ribosomal_S9 (pfam00380)→ |
| env-1030 | RUMENNODE_3847963_1 | + | 42999 | 43073 | **RNA**→ rplM (PRK09216)Ribosomal_L13 (pfam00572)→ rplM (PRK09216)Ribosomal_L13 (pfam00572)→ |
| env-1031 | AUXO014586155.1 | + | 7411 | 7485 | **RNA**→ rplM (PRK09216)Ribosomal_L13 (pfam00572)→ rpsI (PRK00132)Ribosomal_S9 (pfam00380)→ |
| env-1032 | HCF12C_125474 | - | 35705 | 35631 | **RNA**→ rplM (PRK09216)Ribosomal_L13 (pfam00572)→ rpsI (PRK00132)Ribosomal_S9 (pfam00380)→ |
| env-1033 | HCE12Call500_c0173985 | + | 472 | 546 | **RNA**→ rplM (PRK09216)Ribosomal_L13 (pfam00572)→ rpsI (PRK00132)Ribosomal_S9 (pfam00380)→ |
| env-1034 | NLF006_scaffold31027_2 | - | 1626 | 1552 | **RNA**→ rplM (PRK09216)Ribosomal_L13 (pfam00572)→ rpsI (PRK00132)Ribosomal_S9 (pfam00380)→ |
| env-1035 | AUXO013075926.1 | + | 551 | 625 | **RNA**→ rplM (PRK09216)Ribosomal_L13 (pfam00572)→ rpsI (PRK00132)Ribosomal_S9 (pfam00380)→ |
| env-1036 | RUMENNODE_4167426_1 | + | 15701 | 15775 | **RNA**→ rplM (PRK09216)Ribosomal_L13 (pfam00572)→ rpsI (PRK00132)Ribosomal_S9 (pfam00380)→ |
| env-1037 | AUXO014456099.1 | - | 1496 | 1422 | **RNA**→ rplM (PRK09216)Ribosomal_L13 (pfam00572)→ rpsI (PRK00132)Ribosomal_S9 (pfam00380)→ |
| env-1038 | scaffold2367_2_V1.UC-14 | - | 1330 | 1257 | **RNA**→ rplM (PRK09216)Ribosomal_L13 (pfam00572)→ rpsI (PRK00132)Ribosomal_S9 (pfam00380)→ |
| env-1039 | 4491414.3_NODE_1526 | - | 5046 | 4973 | **RNA**→ rplM (PRK09216)Ribosomal_L13 (pfam00572)→ rpsI (PRK00132)Ribosomal_S9 (pfam00380)→ |
| env-1040 | DLF003_scaffold16038_8 | - | 1088 | 1015 | **RNA**→ rplM (PRK09216)Ribosomal_L13 (pfam00572)→ rpsI (PRK00132)Ribosomal_S9 (pfam00380)→ |
| env-1041 | DLF005_scaffold3173_4 | - | 4285 | 4212 | **RNA**→ rplM (PRK09216)Ribosomal_L13 (pfam00572)→ rpsI (PRK00132)Ribosomal_S9 (pfam00380)→ |
| env-1042 | NOM012_scaffold23711_5 | + | 2538 | 2611 | **RNA**→ rplM (PRK09216)Ribosomal_L13 (pfam00572)→ rpsI (PRK00132)Ribosomal_S9 (pfam00380)→ |
| env-1043 | scaffold1280_4_V1.UC-13 | - | 11294 | 11221 | **RNA**→ rplM (PRK09216)Ribosomal_L13 (pfam00572)→ rpsI (PRK00132)Ribosomal_S9 (pfam00380)→ |
| env-1044 | SRS015663_WUGC_scaffold_11548 | + | 5547 | 5620 | **RNA**→ rplM (PRK09216)Ribosomal_L13 (pfam00572)→ rpsI (PRK00132)Ribosomal_S9 (pfam00380)→ |
| env-1045 | SRS051031_LANL_scaffold_24636 | - | 16175 | 16102 | **RNA**→ rplM (PRK09216)Ribosomal_L13 (pfam00572)→ rpsI (PRK00132)Ribosomal_S9 (pfam00380)→ |
| env-1046 | scaffold8699_1_MH0021 | + | 371 | 444 | **RNA**→ rplM (PRK09216)Ribosomal_L13 (pfam00572)→ |
| env-1047 | C1729250_1_MH0022 | - | 754 | 681 | **RNA**→ rplM (PRK09216)Ribosomal_L13 (pfam00572)→ rpsI (PRK00132)Ribosomal_S9 (pfam00380)→ |
| env-1048 | scaffold49_1_MH0057 | + | 5529 | 5602 | **RNA**→ rplM (PRK09216)Ribosomal_L13 (pfam00572)→ rpsI (PRK00132)Ribosomal_S9 (pfam00380)→ |
| env-1049 | scaffold98913_1_O2.UC-19 | - | 593 | 520 | **RNA**→ rplM (PRK09216)Ribosomal_L13 (pfam00572)→ |
| env-1050 | scaffold80421_1_O2.UC-1 | + | 180 | 253 | **RNA**→ rplM (PRK09216)Ribosomal_L13 (pfam00572)→ |
| env-1051 | DLM018_scaffold38383_1 | + | 191 | 264 | **RNA**→ rplM (PRK09216)Ribosomal_L13 (pfam00572)→ rpsI (PRK00132)Ribosomal_S9 (pfam00380)→ |
| env-1052 | 4491401.3_NODE_2070 | - | 1171 | 1098 | **RNA**→ rplM (PRK09216)Ribosomal_L13 (pfam00572)→ rpsI (PRK00132)Ribosomal_S9 (pfam00380)→ |
| env-1053 | 4491477.3_NODE_12413 | + | 2389 | 2462 | **RNA**→ rplM (PRK09216)Ribosomal_L13 (pfam00572)→ rpsI (PRK00132)Ribosomal_S9 (pfam00380)→ |
| Psp-2-1 | NZ_LFQU01000025.1 | + | 6048 | 6121 | **RNA**→ rplM (PRK09216)Ribosomal_L13 (pfam00572)→ rpsI (PRK00132)Ribosomal_S9 (pfam00380)→ |
| env-1054 | 4491405.3_NODE_34345 | + | 86865 | 86938 | **RNA**→ rplM (PRK09216)Ribosomal_L13 (pfam00572)→ rpsI (PRK00132)Ribosomal_S9 (pfam00380)→ |
| env-1055 | scaffold4912_4_MH0034 | - | 2299 | 2226 | **RNA**→ rplM (PRK09216)Ribosomal_L13 (pfam00572)→ rpsI (PRK00132)Ribosomal_S9 (pfam00380)→ |
| env-1056 | C1986582_1_MH0079 | + | 198 | 271 | **RNA**→ rplM (PRK09216)Ribosomal_L13 (pfam00572)→ |
| env-1057 | scaffold47598_1_O2.UC-11 | + | 2719 | 2792 | **RNA**→ rplM (PRK09216)Ribosomal_L13 (pfam00572)→ rpsI (PRK00132)Ribosomal_S9 (pfam00380)→ |

| | | | | | |
|---|---|---|---|---|---|
| env-1058 | AUXO010723907.1 | - | 1439 | 1366 | **RNA**→ rplM (PRK09216)Ribosomal_L13 (pfam00572)→ rpsI (PRK00132)Ribosomal_S9 (pfam00380)→ |
| env-1059 | SRM_contig03116 | + | 29 | 103 | **RNA**→ rplM (PRK09216)Ribosomal_L13 (pfam00572)→ |
| env-1060 | AUXO013680712.1 | + | 1792 | 1865 | **RNA**→ rplM (PRK09216)Ribosomal_L13 (pfam00572)→ rpsI (PRK00132)Ribosomal_S9 (pfam00380)→ |
| env-1061 | SRS049959_WUGC_scaffold_56943 | - | 909 | 836 | **RNA**→ rplM (PRK09216)Ribosomal_L13 (pfam00572)→ rpsI (PRK00132)Ribosomal_S9 (pfam00380)→ |
| Psp-13-1 | NZ_JXQH01000086.1 | + | 71385 | 71458 | **RNA**→ rplM (PRK09216)Ribosomal_L13 (pfam00572)→ rpsI (PRK00132)Ribosomal_S9 (pfam00380)→ |
| Psp-14-1 | NZ_JXQI01000060.1 | + | 39882 | 39955 | **RNA**→ rplM (PRK09216)Ribosomal_L13 (pfam00572)→ rpsI (PRK00132)Ribosomal_S9 (pfam00380)→ |
| env-1062 | scaffold253617_1_MH0006 | + | 404 | 477 | **RNA**→ |
| Psp-17-1 | NZ_JXQJ01000163.1 | + | 35397 | 35470 | **RNA**→ rplM (PRK09216)Ribosomal_L13 (pfam00572)→ rpsI (PRK00132)Ribosomal_S9 (pfam00380)→ |
| env-1063 | NOM022_scaffold35257_3 | + | 8385 | 8458 | **RNA**→ rplM (PRK09216)Ribosomal_L13 (pfam00572)→ |
| Psp-15-1 | NZ_JXQK01000088.1 | + | 79922 | 79995 | **RNA**→ rplM (PRK09216)Ribosomal_L13 (pfam00572)→ rpsI (PRK00132)Ribosomal_S9 (pfam00380)→ |
| Psp-16-1 | NZ_JXQL01000086.1 | - | 112978 | 112905 | **RNA**→ rplM (PRK09216)Ribosomal_L13 (pfam00572)→ rpsI (PRK00132)Ribosomal_S9 (pfam00380)→ |
| env-1064 | BAAX01026741.1 | - | 768 | 695 | **RNA**→ rplM (PRK09216)Ribosomal_L13 (pfam00572)→ |
| env-1065 | scaffold15697_4_MH0030 | - | 142 | 69 | **RNA**→ |
| env-1066 | NOM022_scaffold26603_2 | - | 448 | 375 | **RNA**→ rplM (PRK09216)Ribosomal_L13 (pfam00572)→ |
| env-1067 | AUXO015787294.1 | - | 1451 | 1378 | **RNA**→ rplM (PRK09216)Ribosomal_L13 (pfam00572)→ rpsI (PRK00132)Ribosomal_S9 (pfam00380)→ |
| Hse-1-1 | NZ_JADU01000007.1 | + | 92405 | 92478 | **RNA**→ rplM (PRK09216)Ribosomal_L13 (pfam00572)→ rpsI (PRK00132)Ribosomal_S9 (pfam00380)→ |
| env-1068 | scaffold4605_11_MH0050 | - | 3079 | 3003 | **RNA**→ rplM (PRK09216)Ribosomal_L13 (pfam00572)→ rpsI (PRK00132)Ribosomal_S9 (pfam00380)→ |
| env-1069 | HCE12Call500_c0256376 | - | 434 | 359 | **RNA**→ rplM (PRK09216)Ribosomal_L13 (pfam00572)→ |
| env-1070 | HCF14C_2_1_1_newblercontig23395 | + | 452 | 527 | **RNA**→ rplM (PRK09216)Ribosomal_L13 (pfam00572)→ |
| env-1071 | AUXO017008710.1 | + | 176 | 252 | **RNA**→ hypo→ |
| env-1072 | HCF14C_3223 | - | 3241 | 3167 | **RNA**→ rplM (PRK09216)Ribosomal_L13 (pfam00572)→ rpsI (PRK00132)Ribosomal_S9 (pfam00380)→ |
| env-1073 | HCF12C_138759 | - | 454 | 380 | **RNA**→ rplM (PRK09216)Ribosomal_L13 (pfam00572)→ |
| env-1074 | HCF14C_3378 | + | 16846 | 16920 | **RNA**→ |
| env-1075 | HCF12C_432612 | + | 209 | 283 | **RNA**→ rplM (PRK09216)Ribosomal_L13 (pfam00572)→ rpsI (PRK00132)Ribosomal_S9 (pfam00380)→ |
| env-1076 | AUXO011672201.1 | - | 118 | 44 | **RNA**→ |
| env-1077 | AUXO010615907.1 | - | 963 | 889 | **RNA**→ rplM (PRK09216)Ribosomal_L13 (pfam00572)→ rpsI (PRK00132)Ribosomal_S9 (pfam00380)→ |
| env-1078 | RUMENNODE_3874741_74000 | + | 4185 | 4261 | **RNA**→ rplM (PRK09216)Ribosomal_L13 (pfam00572)→ |
| env-1079 | AUXO013969103.1 | + | 2307 | 2382 | **RNA**→ |
| env-1080 | AUXO015357845.1 | + | 1768 | 1843 | **RNA**→ rplM (PRK09216)Ribosomal_L13 (pfam00572)→ rpsI (PRK00132)Ribosomal_S9 (pfam00380)→ |
| env-1081 | SRM_contig10604 | + | 931 | 1006 | **RNA**→ |
| env-1082 | SRMUA_GNVGR9Q01BHOFT | - | 77 | 2 | **RNA**→ |
| env-1083 | SRMUA_GNVGR9Q01BGO0O | - | 154 | 79 | **RNA**→ |
| env-1084 | AUXO010497420.1 | + | 284 | 358 | **RNA**→ rplM (PRK09216)Ribosomal_L13 (pfam00572)→ rpsI (PRK00132)Ribosomal_S9 (pfam00380)→ |
| env-1085 | AUXO018287067.1 | - | 792 | 718 | **RNA**→ rplM (PRK09216)Ribosomal_L13 (pfam00572)→ rpsI (PRK00132)Ribosomal_S9 (pfam00380)→ |
| env-1086 | AUXO015587525.1 | + | 974 | 1048 | **RNA**→ |
| env-1087 | AUXO011220652.1 | + | 3323 | 3399 | **RNA**→ |
| env-1088 | AUXO012321124.1 | - | 85 | 11 | **RNA**→ |
| env-1089 | AUXO014609123.1 | - | 80 | 6 | **RNA**→ |
| env-1090 | AUXO018327041.1 | + | 2387 | 2463 | **RNA**→ |
| env-1091 | AUXO018475375.1 | - | 90 | 14 | **RNA**→ ←hypo |
| env-1092 | SRMUA_GMSDS1U03GX2P3 | + | 122 | 197 | **RNA**→ rplM (PRK09216)Ribosomal_L13 (pfam00572)→ |
| env-1093 | AUXO015617850.1 | + | 121 | 196 | **RNA**→ rplM (PRK09216)Ribosomal_L13 (pfam00572)→ rpsI (PRK00132)Ribosomal_S9 (pfam00380)→ |
| env-1094 | AUXO010064957.1 | - | 160 | 84 | **RNA**→ hypo→ |
| env-1095 | AUXO012890930.1 | - | 277 | 202 | **RNA**→ |
| env-1096 | RUMENNODE_1622158_1 | + | 23986 | 24061 | **RNA**→ rplM (PRK09216)Ribosomal_L13 (pfam00572)→ rpsI (PRK00132)Ribosomal_S9 (pfam00380)→ |
| env-1097 | AUXO012504799.1 | + | 349 | 424 | **RNA**→ |
| env-1098 | RUMENNODE_4144373_1 | - | 20587 | 20512 | **RNA**→ rplM (PRK09216)Ribosomal_L13 (pfam00572)→ rplM (PRK09216)Ribosomal_L13 (pfam00572)→ rpsI (PRK00132)Ribosomal_S9 (pfam00380)→ |
| Psp-7-1 | NZ_KN050820.1 | - | 266447 | 266372 | **RNA**→ rplM (PRK09216)Ribosomal_L13 (pfam00572)→ rpsI (PRK00132)Ribosomal_S9 (pfam00380)→ |
| env-1099 | 4491686.3.305921 | - | 489 | 414 | **RNA**→ rplM (PRK09216)Ribosomal_L13 (pfam00572)→ |
| env-1100 | 4491686.3.312252 | + | 219 | 294 | **RNA**→ rplM (PRK09216)Ribosomal_L13 (pfam00572)→ |
| Psp-11-1 | NZ_JHUW01000006.1 | + | 275505 | 275580 | **RNA**→ rplM (PRK09216)Ribosomal_L13 (pfam00572)→ rpsI (PRK00132)Ribosomal_S9 (pfam00380)→ |

| | | | | | |
|---|---|---|---|---|---|
| Psp-19-1 | NZ_JHVE01000005.1 | + | 85620 | 85695 | **RNA**→ rplM (PRK09216)Ribosomal_L13 (pfam00572)→ rpsI (PRK00132)Ribosomal_S9 (pfam00380)→ |
| Psp-18-1 | NZ_JNKF01000010.1 | + | 588125 | 588200 | **RNA**→ rplM (PRK09216)Ribosomal_L13 (pfam00572)→ rpsI (PRK00132)Ribosomal_S9 (pfam00380)→ |
| env-1101 | 4491686.3.93290 | - | 116 | 41 | **RNA**→ |
| Pru-1-1 | NC_014033.1 | + | 3193901 | 3193976 | **RNA**→ rplM (PRK09216)Ribosomal_L13 (pfam00572)→ rpsI (PRK00132)Ribosomal_S9 (pfam00380)→ |
| Pru-2-1 | NZ_JHXD01000002.1 | + | 95378 | 95453 | **RNA**→ rplM (PRK09216)Ribosomal_L13 (pfam00572)→ rpsI (PRK00132)Ribosomal_S9 (pfam00380)→ |
| Psp-20-1 | NZ_JNKG01000008.1 | + | 135579 | 135654 | **RNA**→ rplM (PRK09216)Ribosomal_L13 (pfam00572)→ rpsI (PRK00132)Ribosomal_S9 (pfam00380)→ |
| env-1102 | AUXO018400480.1 | - | 139 | 64 | **RNA**→ |
| env-1103 | AUXO015757529.1 | - | 664 | 588 | **RNA**→ rplM (PRK09216)Ribosomal_L13 (pfam00572)→ |
| env-1104 | AUXO015472756.1 | - | 170 | 95 | **RNA**→ |
| Pbr-1-1 | NZ_JHXM01000009.1 | - | 128416 | 128341 | **RNA**→ rplM (PRK09216)Ribosomal_L13 (pfam00572)→ rpsI (PRK00132)Ribosomal_S9 (pfam00380)→ |
| Pbr-2-1 | NZ_KK211353.1 | - | 323686 | 323611 | **RNA**→ rplM (PRK09216)Ribosomal_L13 (pfam00572)→ rpsI (PRK00132)Ribosomal_S9 (pfam00380)→ |
| env-1105 | AUXO010774619.1 | - | 135 | 60 | **RNA**→ |
| env-1106 | AUXO016205324.1 | + | 977 | 1053 | **RNA**→ |
| env-1107 | AUXO018699643.1 | - | 84 | 9 | **RNA**→ |
| env-1108 | AUXO010272107.1 | + | 282 | 357 | **RNA**→ ←hypo |
| env-1109 | AUXO014911040.1 | + | 6911 | 6987 | **RNA**→ rplM (PRK09216)Ribosomal_L13 (pfam00572)→ rpsI (PRK00132)Ribosomal_S9 (pfam00380)→ |
| env-1110 | AUXO018573799.1 | - | 2207 | 2131 | **RNA**→ rplM (PRK09216)Ribosomal_L13 (pfam00572)→ rpsI (PRK00132)Ribosomal_S9 (pfam00380)→ |
| env-1111 | AUXO016034740.1 | - | 861 | 785 | **RNA**→ rplM (PRK09216)Ribosomal_L13 (pfam00572)→ rpsI (PRK00132)Ribosomal_S9 (pfam00380)→ |
| env-1112 | AUXO013985143.1 | + | 1048 | 1123 | **RNA**→ rplM (PRK09216)Ribosomal_L13 (pfam00572)→ |
| env-1113 | AUXO012106665.1 | + | 1076 | 1151 | **RNA**→ rplM (PRK09216)Ribosomal_L13 (pfam00572)→ rpsI (PRK00132)Ribosomal_S9 (pfam00380)→ |
| env-1114 | AUXO010611192.1 | + | 2306 | 2381 | **RNA**→ rplM (PRK09216)Ribosomal_L13 (pfam00572)→ rpsI (PRK00132)Ribosomal_S9 (pfam00380)→ |
| env-1115 | AUXO017780793.1 | + | 2412 | 2486 | **RNA**→ rplM (PRK09216)Ribosomal_L13 (pfam00572)→ |
| env-1116 | AUXO016964339.1 | - | 2517 | 2443 | **RNA**→ ←hypo |
| env-1117 | DLM014_scaffold29163_3 | - | 749 | 677 | **RNA**→ rplM (PRK09216)Ribosomal_L13 (pfam00572)→ rpsI (PRK00132)Ribosomal_S9 (pfam00380)→ |
| env-1118 | AUXO012501382.1 | + | 1300 | 1372 | **RNA**→ rplM (PRK09216)Ribosomal_L13 (pfam00572)→ |
| env-1119 | RUMENNODE_4186824_1 | - | 20452 | 20380 | **RNA**→ rplM (PRK09216)Ribosomal_L13 (pfam00572)→ rpsI (PRK00132)Ribosomal_S9 (pfam00380)→ |
| env-1120 | HCE12Call500_c0163880 | - | 1019 | 945 | **RNA**→ rplM (PRK09216)Ribosomal_L13 (pfam00572)→ rpsI (PRK00132)Ribosomal_S9 (pfam00380)→ |
| env-1121 | HCF14C_7730 | + | 2022 | 2096 | **RNA**→ rplM (PRK09216)Ribosomal_L13 (pfam00572)→ rpsI (PRK00132)Ribosomal_S9 (pfam00380)→ |
| env-1122 | HCF12C_211171 | + | 370 | 445 | **RNA**→ rplM (PRK09216)Ribosomal_L13 (pfam00572)→ |
| env-1123 | 4491686.3.18377 | - | 3185 | 3110 | **RNA**→ rplM (PRK09216)Ribosomal_L13 (pfam00572)→ rpsI (PRK00132)Ribosomal_S9 (pfam00380)→ |
| env-1124 | 4491686.3.998042 | + | 30719 | 30794 | **RNA**→ rplM (PRK09216)Ribosomal_L13 (pfam00572)→ rpsI (PRK00132)Ribosomal_S9 (pfam00380)→ |
| env-1125 | HCF12C_268453 | + | 254 | 328 | **RNA**→ |
| env-1126 | HCF12C_148665 | + | 852 | 926 | **RNA**→ hypo→ |
| env-1127 | HCE12Call500_c0204626 | - | 679 | 605 | **RNA**→ rplM (PRK09216)Ribosomal_L13 (pfam00572)→ rpsI (PRK00132)rps9 (CHL00079)→ |
| env-1128 | HCF12C_185472 | + | 268 | 342 | **RNA**→ rplM (PRK09216)RplM (COG0102)→ |
| env-1129 | 4491421.3_NODE_9264 | + | 154 | 230 | **RNA**→ rplM (PRK09216)Ribosomal_L13 (pfam00572)→ |
| env-1130 | AUXO015981657.1 | - | 1275 | 1203 | **RNA**→ rplM (PRK09216)Ribosomal_L13 (pfam00572)→ rpsI (PRK00132)Ribosomal_S9 (pfam00380)→ |
| env-1131 | AUXO010201767.1 | + | 271 | 345 | **RNA**→ rplM (PRK09216)Ribosomal_L13 (pfam00572)→ |
| env-1132 | AUXO014390899.1 | - | 563 | 489 | **RNA**→ rplM (PRK09216)Ribosomal_L13 (pfam00572)→ |
| env-1133 | AUXO015967901.1 | + | 641 | 712 | **RNA**→ rplM (PRK09216)Ribosomal_L13 (pfam00572)→ rpsI (PRK00132)Ribosomal_S9 (pfam00380)→ |
| env-1134 | AUXO015633410.1 | + | 206 | 279 | **RNA**→ rplM (PRK09216)Ribosomal_L13 (pfam00572)→ rpsI (PRK00132)Ribosomal_S9 (pfam00380)→ |
| env-1135 | AUXO017839357.1 | + | 144 | 211 | **RNA**→ rplM (PRK09216)Ribosomal_L13 (pfam00572)→ |
| env-1136 | AUXO016134445.1 | + | 192 | 259 | **RNA**→ rplM (PRK09216)Ribosomal_L13 (pfam00572)→ rpsI (PRK00132)Ribosomal_S9 (pfam00380)→ |
| env-1137 | SRS064449_C3977517 | + | 160 | 232 | **RNA**→ rplM (PRK09216)RplM (COG0102)→ |
| env-1138 | SRS013170_C4245439 | - | 179 | 107 | **RNA**→ |
| env-1139 | SRS019071_C4979592 | + | 177 | 249 | **RNA**→ |
| env-1140 | SRS063999_C3760459 | + | 1331 | 1403 | **RNA**→ rplM (PRK09216)Ribosomal_L13 (pfam00572)→ rpsI (PRK00132)Ribosomal_S9 (pfam00380)→ |
| env-1141 | SRS053917_LANL_scaffold_97956 | + | 384 | 456 | **RNA**→ rplM (PRK09216)Ribosomal_L13 (pfam00572)→ |
| env-1142 | SRS049318_LANL_scaffold_147506 | + | 27 | 99 | **RNA**→ rplM (PRK09216)Ribosomal_L13 (pfam00572)→ |
| env-1143 | SRS047634_C4471531 | - | 1001 | 929 | **RNA**→ rplM (PRK09216)Ribosomal_L13 (pfam00572)→ rpsI (PRK00132)Ribosomal_S9 (pfam00380)→ |
| env-1144 | SRS022602_Baylor_scaffold_89900 | - | 465 | 393 | **RNA**→ rplM (PRK09216)Ribosomal_L13 (pfam00572)→ |
| env-1145 | SRS019028_C2913254 | - | 170 | 98 | **RNA**→ |

| | | | | | |
|---|---|---|---|---|---|
| env-1146 | SRS063603_LANL_scaffold_35578 | + | 6849 | 6921 | **RNA**→ rplM (PRK09216)Ribosomal_L13 (pfam00572)→ rpsI (PRK00132)Ribosomal_S9 (pfam00380)→ |
| env-1147 | SRS016225_WUGC_scaffold_29763 | + | 2155 | 2226 | **RNA**→ |
| env-1148 | SRS022621_Baylor_scaffold_78299 | + | 1192 | 1263 | **RNA**→ rplM (PRK09216)Ribosomal_L13 (pfam00572)→ |
| env-1149 | SRS022719_LANL_scaffold_24264 | - | 1070 | 999 | **RNA**→ rplM (PRK09216)Ribosomal_L13 (pfam00572)→ rpsI (PRK00132)Ribosomal_S9 (pfam00380)→ rpsI (PRK00132)Ribosomal_S9 (pfam00380)→ |
| env-1150 | SRS023926_C2746934 | + | 199 | 270 | **RNA**→ rplM (PRK09216)Ribosomal_L13 (pfam00572)→ |
| env-1151 | SRS014888_C3094079 | - | 1005 | 933 | **RNA**→ rplM (PRK09216)Ribosomal_L13 (pfam00572)→ rpsI (PRK00132)Ribosomal_S9 (pfam00380)→ |
| env-1152 | SRS057355_LANL_scaffold_38534 | + | 74 | 146 | **RNA**→ rplM (PRK09216)Ribosomal_L13 (pfam00572)→ |
| env-1153 | SRS018739_C3681903 | - | 158 | 86 | **RNA**→ |
| env-1154 | SRS013164_Baylor_scaffold_53891 | - | 1272 | 1200 | **RNA**→ rplM (PRK09216)Ribosomal_L13 (pfam00572)→ rpsI (PRK00132)Ribosomal_S9 (pfam00380)→ |
| env-1155 | SRS015797_WUGC_scaffold_34484 | - | 665 | 593 | **RNA**→ rplM (PRK09216)Ribosomal_L13 (pfam00572)→ rpsI (PRK00132)Ribosomal_S9 (pfam00380)→ |
| env-1156 | SRS042131_WUGC_scaffold_59323 | - | 801 | 729 | **RNA**→ rplM (PRK09216)Ribosomal_L13 (pfam00572)→ rpsI (PRK00132)Ribosomal_S9 (pfam00380)→ |
| env-1157 | SRS024441_LANL_scaffold_55878 | - | 1004 | 932 | **RNA**→ rplM (PRK09216)Ribosomal_L13 (pfam00572)→ rpsI (PRK00132)Ribosomal_S9 (pfam00380)→ |
| env-1158 | SRS016037_WUGC_scaffold_44051 | - | 27077 | 27005 | **RNA**→ rplM (PRK09216)Ribosomal_L13 (pfam00572)→ rpsI (PRK00132)Ribosomal_S9 (pfam00380)→ |
| env-1159 | SRS018357_Baylor_scaffold_49795 | + | 563 | 635 | **RNA**→ |
| env-1160 | SRS044662_LANL_scaffold_9532 | + | 25254 | 25326 | **RNA**→ rplM (PRK09216)Ribosomal_L13 (pfam00572)→ rpsI (PRK00132)Ribosomal_S9 (pfam00380)→ |
| env-1161 | SRS045127_LANL_scaffold_15711 | - | 3179 | 3107 | **RNA**→ rplM (PRK09216)Ribosomal_L13 (pfam00572)→ rpsI (PRK00132)Ribosomal_S9 (pfam00380)→ |
| env-1162 | SRS049147_C4127564 | + | 678 | 750 | **RNA**→ |
| env-1163 | Palatine_Tonsils_LANL_C6209475 | + | 331 | 403 | **RNA**→ rplM (PRK09216)Ribosomal_L13 (pfam00572)→ |
| env-1164 | SRS015434_C5847974 | - | 383 | 311 | **RNA**→ rplM (PRK09216)Ribosomal_L13 (pfam00572)→ |
| env-1165 | SRS017533_Baylor_scaffold_51088 | + | 1318 | 1390 | **RNA**→ |
| env-1166 | SRS019327_C2789739 | + | 1129 | 1201 | **RNA**→ rplM (PRK09216)Ribosomal_L13 (pfam00572)→ |
| env-1167 | Buccal_mucosa_LANL_scaffold_123568 | - | 308 | 236 | **RNA**→ rplM (PRK09216)Ribosomal_L13 (cd00392)→ |
| env-1168 | SRS015985_C1563967 | + | 138 | 210 | **RNA**→ rplM (PRK09216)Ribosomal_L13 (pfam00572)→ |
| env-1169 | SRS062878_LANL_scaffold_81556 | + | 2146 | 2218 | **RNA**→ rplM (PRK09216)RplM (COG0102)→ |
| env-1170 | SRS075404_C3564214 | - | 158 | 86 | **RNA**→ hypo→ |
| env-1171 | Throat_LANL_C5768501 | + | 142 | 214 | **RNA**→ rplM (PRK09216)Ribosomal_L13 (pfam00572)→ |
| env-1172 | SRS011306_Baylor_scaffold_57099 | + | 475 | 547 | **RNA**→ rplM (PRK09216)RplM (COG0102)→ |
| env-1173 | SRS013879_WUGC_scaffold_12081 | + | 2721 | 2793 | **RNA**→ rplM (PRK09216)Ribosomal_L13 (pfam00572)→ rpsI (PRK00132)Ribosomal_S9 (pfam00380)→ |
| env-1174 | SRS013947_C1628089 | - | 350 | 278 | **RNA**→ rplM (PRK09216)Ribosomal_L13 (pfam00572)→ |
| env-1175 | SRS013948_WUGC_scaffold_13434 | + | 25402 | 25474 | **RNA**→ rplM (PRK09216)Ribosomal_L13 (pfam00572)→ |
| env-1176 | SRS014271_C2902322 | - | 158 | 86 | **RNA**→ |
| env-1177 | SRS015762_WUGC_scaffold_30677 | - | 1005 | 933 | **RNA**→ rplM (PRK09216)Ribosomal_L13 (pfam00572)→ rpsI (PRK00132)Ribosomal_S9 (pfam00380)→ |
| env-1178 | SRS016002_C2530304 | + | 27 | 99 | **RNA**→ rplM (PRK09216)Ribosomal_L13 (pfam00572)→ |
| env-1179 | SRS016569_C2383578 | + | 27 | 99 | **RNA**→ rplM (PRK09216)Ribosomal_L13 (pfam00572)→ |
| env-1180 | SRS017209_Baylor_scaffold_3818 | + | 25398 | 25470 | **RNA**→ rplM (PRK09216)Ribosomal_L13 (pfam00572)→ rpsI (PRK00132)Ribosomal_S9 (pfam00380)→ |
| env-1181 | SRS018145_Baylor_scaffold_64 | + | 25195 | 25267 | **RNA**→ rplM (PRK09216)Ribosomal_L13 (pfam00572)→ rpsI (PRK00132)Ribosomal_S9 (pfam00380)→ |
| env-1182 | SRS018439_Baylor_scaffold_41344 | - | 879 | 807 | **RNA**→ rplM (PRK09216)Ribosomal_L13 (pfam00572)→ rpsI (PRK00132)Ribosomal_S9 (pfam00380)→ |
| env-1183 | SRS019071_C4950970 | - | 163 | 91 | **RNA**→ hypo→ |
| env-1184 | SRS019120_C1301558 | + | 33 | 105 | **RNA**→ rplM (PRK09216)Ribosomal_L13 (pfam00572)→ |
| env-1185 | SRS019126_WUGC_scaffold_35353 | + | 161 | 233 | **RNA**→ rplM (PRK09216)Ribosomal_L13 (pfam00572)→ rpsI (PRK00132)Ribosomal_S9 (pfam00380)→ |
| env-1186 | SRS023958_Baylor_scaffold_34017 | + | 4806 | 4878 | **RNA**→ rplM (PRK09216)Ribosomal_L13 (pfam00572)→ rpsI (PRK00132)Ribosomal_S9 (pfam00380)→ |
| env-1187 | SRS024081_C2561546 | + | 437 | 509 | **RNA**→ |
| env-1188 | SRS050669_LANL_scaffold_65278 | - | 28287 | 28215 | **RNA**→ rplM (PRK09216)Ribosomal_L13 (pfam00572)→ rpsI (PRK00132)Ribosomal_S9 (pfam00380)→ |
| env-1189 | SRS057205_LANL_scaffold_2683 | - | 1005 | 933 | **RNA**→ rplM (PRK09216)Ribosomal_L13 (pfam00572)→ rpsI (PRK00132)Ribosomal_S9 (pfam00380)→ |
| env-1190 | SRS057692_C2560902 | + | 134 | 206 | **RNA**→ |
| env-1191 | SRS058336_LANL_scaffold_19337 | - | 5666 | 5594 | **RNA**→ rplM (PRK09216)Ribosomal_L13 (pfam00572)→ rpsI (PRK00132)Ribosomal_S9 (pfam00380)→ |
| env-1192 | SRS015057_WUGC_scaffold_40957 | - | 277 | 205 | **RNA**→ rplM (PRK09216)Ribosomal_L13 (pfam00572)→ |
| env-1193 | SRS063603_LANL_scaffold_3918 | - | 1001 | 931 | **RNA**→ rplM (PRK09216)Ribosomal_L13 (pfam00572)→ rpsI (PRK00132)Ribosomal_S9 (pfam00380)→ |
| env-1194 | SRS016746_Baylor_scaffold_3854 | + | 350 | 421 | **RNA**→ |
| Psp-1-1 | NZ_KB290717.1 | - | 518225 | 518153 | **RNA**→ rplM (PRK09216)Ribosomal_L13 (pfam00572)→ rpsI (PRK00132)Ribosomal_S9 (pfam00380)→ |
| env-1195 | SRS013949_C3355882 | - | 134 | 64 | **RNA**→ hypo→ |

| | | | | | |
|---|---|---|---|---|---|
| env-1196 | SRS024087_C3225267 | - | 91 | 21 | RNA→ hypo→ |
| env-1197 | SRS013950_C2065648 | - | 258 | 188 | RNA→ rplM (PRK09216)RplM (COG0102)→ |
| env-1198 | SRS018157_C4805273 | - | 576 | 506 | RNA→ rplM (PRK09216)Ribosomal_L13 (pfam00572)→ |
| env-1199 | SRS019128_C4141267 | + | 154 | 224 | RNA→ |
| env-1200 | SRS047634_LANL_scaffold_18390 | + | 479 | 549 | RNA→ rplM (PRK09216)Ribosomal_L13 (pfam00572)→ rpsI (PRK00132)Ribosomal_S9 (pfam00380)→ |
| env-1201 | SRS055401_C2292367 | - | 364 | 294 | RNA→ rplM (PRK09216)Ribosomal_L13 (pfam00572)→ |
| env-1202 | SRS063932_LANL_scaffold_79042 | + | 232 | 302 | RNA→ rplM (PRK09216)Ribosomal_L13 (pfam00572)→ rpsI (PRK00132)Ribosomal_S9 (pfam00380)→ |
| env-1203 | SRS015803_WUGC_scaffold_2190 | + | 350 | 420 | RNA→ rplM (PRK09216)Ribosomal_L13 (pfam00572)→ rpsI (PRK00132)Ribosomal_S9 (pfam00380)→ |
| env-1204 | SRS024289_LANL_scaffold_56240 | + | 232 | 302 | RNA→ rplM (PRK09216)Ribosomal_L13 (pfam00572)→ rpsI (PRK00132)Ribosomal_S9 (pfam00380)→ |
| env-1205 | SRS023938_C2238754 | - | 301 | 231 | RNA→ rplM (PRK09216)Ribosomal_L13 (pfam00572)→ |
| env-1206 | SRS012285_C2904644 | - | 382 | 310 | RNA→ rplM (PRK09216)Ribosomal_L13 (pfam00572)→ |
| env-1207 | SRS017227_Baylor_scaffold_121114 | + | 26 | 98 | RNA→ rplM (PRK09216)Ribosomal_L13 (pfam00572)→ rpsI (PRK00132)Ribosomal_S9 (pfam00380)→ |
| env-1208 | SRS023595_Baylor_scaffold_24496 | + | 230 | 302 | RNA→ rplM (PRK09216)Ribosomal_L13 (pfam00572)→ rpsI (PRK00132)Ribosomal_S9 (pfam00380)→ |
| env-1209 | SRS047113_C5260445 | - | 272 | 200 | RNA→ rplM (PRK09216)RplM (COG0102)→ |
| env-1210 | SRS050628_C1880365 | + | 114 | 186 | RNA→ rplM (PRK09216)Ribosomal_L13 (pfam00572)→ |
| env-1211 | SRS019028_C3034176 | - | 899 | 827 | RNA→ rplM (PRK09216)Ribosomal_L13 (pfam00572)→ rpsI (PRK00132)Ribosomal_S9 (pfam00380)→ |
| env-1212 | SRS019029_C3106614 | + | 230 | 302 | RNA→ |
| env-1213 | SRS016037_C2055413 | - | 3265 | 3192 | RNA→ rplM (PRK09216)Ribosomal_L13 (pfam00572)→ rpsI (PRK00132)Ribosomal_S9 (pfam00380)→ |
| env-1214 | SRS022719_LANL_scaffold_71174 | - | 5214 | 5141 | RNA→ rplM (PRK09216)Ribosomal_L13 (pfam00572)→ rpsI (PRK00132)Ribosomal_S9 (pfam00380)→ |
| env-1215 | SRS044662_LANL_scaffold_46735 | - | 2254 | 2181 | RNA→ rplM (PRK09216)Ribosomal_L13 (pfam00572)→ rpsI (PRK00132)Ribosomal_S9 (pfam00380)→ |
| env-1216 | SRS013164_Baylor_scaffold_32078 | + | 335 | 407 | RNA→ rplM (PRK09216)Ribosomal_L13 (pfam00572)→ rpsI (PRK00132)Ribosomal_S9 (pfam00380)→ |
| env-1217 | SRS013879_WUGC_scaffold_46467 | - | 1318 | 1246 | RNA→ rplM (PRK09216)Ribosomal_L13 (pfam00572)→ rpsI (PRK00132)Ribosomal_S9 (pfam00380)→ |
| env-1218 | SRS015434_C5929714 | - | 1009 | 937 | RNA→ rplM (PRK09216)Ribosomal_L13 (pfam00572)→ rpsI (PRK00132)Ribosomal_S9 (pfam00380)→ |
| env-1219 | SRS019071_C5025454 | - | 172 | 100 | RNA→ hypo→ |
| env-1220 | SRS023958_Baylor_scaffold_23657 | + | 335 | 407 | RNA→ rplM (PRK09216)Ribosomal_L13 (pfam00572)→ rpsI (PRK00132)Ribosomal_S9 (pfam00380)→ |
| env-1221 | SRS049389_WUGC_scaffold_42978 | - | 343 | 271 | RNA→ rplM (PRK09216)Ribosomal_L13 (pfam00572)→ |
| env-1222 | SRS052227_LANL_scaffold_7065 | - | 172 | 100 | RNA→ |
| env-1223 | SRS011140_C5077184 | + | 62 | 134 | RNA→ rplM (PRK09216)Ribosomal_L13 (pfam00572)→ |
| env-1224 | SRS016225_WUGC_scaffold_16679 | - | 161 | 89 | RNA→ |
| env-1225 | SRS017439_C3816727 | + | 159 | 231 | RNA→ rplM (PRK09216)Ribosomal_L13 (pfam00572)→ rpsI (PRK00132)Ribosomal_S9 (pfam00380)→ |
| env-1226 | SRS018357_Baylor_scaffold_50190 | - | 1318 | 1246 | RNA→ rplM (PRK09216)Ribosomal_L13 (pfam00572)→ rpsI (PRK00132)Ribosomal_S9 (pfam00380)→ |
| env-1227 | SRS019327_WUGC_scaffold_31 | + | 300 | 372 | RNA→ rplM (PRK09216)Ribosomal_L13 (pfam00572)→ rpsI (PRK00132)Ribosomal_S9 (pfam00380)→ |
| env-1228 | SRS013239_Baylor_scaffold_8550 | + | 223 | 295 | RNA→ rplM (PRK09216)Ribosomal_L13 (pfam00572)→ rpsI (PRK00132)Ribosomal_S9 (pfam00380)→ |
| env-1229 | SRS013252_C3407129 | + | 146 | 218 | RNA→ |
| env-1230 | SRS019329_C822464 | - | 1070 | 998 | RNA→ rplM (PRK09216)Ribosomal_L13 (pfam00572)→ rpsI (PRK00132)Ribosomal_S9 (pfam00380)→ |
| env-1231 | SRS045049_C1487558 | - | 103 | 31 | RNA→ |
| env-1232 | SRS046686_C923094 | + | 169 | 241 | RNA→ rplM (PRK09216)Ribosomal_L13 (pfam00572)→ rpsI (PRK00132)Ribosomal_S9 (pfam00380)→ |
| env-1233 | SRS012279_C2693900 | - | 583 | 511 | RNA→ rplM (PRK09216)Ribosomal_L13 (pfam00572)→ |
| env-1234 | SRS014573_WUGC_scaffold_5954 | + | 353 | 425 | RNA→ rplM (PRK09216)Ribosomal_L13 (pfam00572)→ rpsI (PRK00132)Ribosomal_S9 (pfam00380)→ |
| env-1235 | SRS014888_C3088705 | - | 1002 | 930 | RNA→ rplM (PRK09216)Ribosomal_L13 (pfam00572)→ rpsI (PRK00132)Ribosomal_S9 (pfam00380)→ |
| env-1236 | SRS018145_Baylor_scaffold_49105 | - | 1318 | 1246 | RNA→ rplM (PRK09216)Ribosomal_L13 (pfam00572)→ rpsI (PRK00132)Ribosomal_S9 (pfam00380)→ |
| env-1237 | SRS019122_C5893890 | + | 223 | 295 | RNA→ rplM (PRK09216)Ribosomal_L13 (pfam00572)→ rpsI (PRK00132)Ribosomal_S9 (pfam00380)→ |
| env-1238 | SRS019126_C2101879 | - | 343 | 271 | RNA→ rplM (PRK09216)Ribosomal_L13 (pfam00572)→ |
| env-1239 | SRS021496_Baylor_scaffold_82682 | + | 222 | 294 | RNA→ rplM (PRK09216)Ribosomal_L13 (pfam00572)→ rpsI (PRK00132)Ribosomal_S9 (pfam00380)→ |
| env-1240 | SRS054569_C1464969 | - | 199 | 127 | RNA→ hypo→ |
| env-1241 | SRS057205_LANL_scaffold_57883 | - | 1002 | 930 | RNA→ rplM (PRK09216)Ribosomal_L13 (pfam00572)→ rpsI (PRK00132)Ribosomal_S9 (pfam00380)→ |
| env-1242 | SRS057692_LANL_scaffold_10341 | + | 335 | 407 | RNA→ rplM (PRK09216)Ribosomal_L13 (pfam00572)→ rpsI (PRK00132)Ribosomal_S9 (pfam00380)→ |
| env-1243 | SRS018739_C3498499 | - | 172 | 100 | RNA→ hypo→ |
| env-1244 | SRS013946_WUGC_scaffold_9812 | + | 797 | 869 | RNA→ |
| env-1245 | SRS017127_Baylor_scaffold_10764 | - | 3240 | 3168 | RNA→ rplM (PRK09216)Ribosomal_L13 (pfam00572)→ rpsI (PRK00132)Ribosomal_S9 (pfam00380)→ |
| env-1246 | SRS019120_C1331431 | + | 223 | 295 | RNA→ rplM (PRK09216)Ribosomal_L13 (pfam00572)→ |
| env-1247 | SRS019124_WUGC_scaffold_15768 | + | 336 | 408 | RNA→ rplM (PRK09216)Ribosomal_L13 (pfam00572)→ rpsI (PRK00132)Ribosomal_S9 (pfam00380)→ |

| | | | | | |
|---|---|---|---|---|---|
| env-1248 | SRS019125_WUGC_scaffold_5306 | + | 336 | 408 | RNA→ rplM (PRK09216)Ribosomal_L13 (pfam00572)→ rpsI (PRK00132)Ribosomal_S9 (pfam00380)→ |
| env-1249 | SRS019128_C4205376 | + | 199 | 271 | RNA→ |
| env-1250 | SRS047113_C5255867 | - | 272 | 200 | RNA→ rplM (PRK09216)RplM (COG0102)→ |
| env-1251 | SRS014473_WUGC_scaffold_265 | - | 32420 | 32348 | RNA→ rplM (PRK09216)Ribosomal_L13 (pfam00572)→ rpsI (PRK00132)Ribosomal_S9 (pfam00380)→ |
| env-1252 | SRS015060_WUGC_scaffold_459 | + | 456 | 528 | RNA→ rplM (PRK09216)Ribosomal_L13 (pfam00572)→ rpsI (PRK00132)Ribosomal_S9 (pfam00380)→ |
| env-1253 | SRS063272_LANL_scaffold_10757 | - | 7611 | 7539 | RNA→ rplM (PRK09216)Ribosomal_L13 (pfam00572)→ rpsI (PRK00132)Ribosomal_S9 (pfam00380)→ |
| env-1254 | SRS017215_C2622978 | + | 37 | 109 | RNA→ rplM (PRK09216)Ribosomal_L13 (pfam00572)→ rpsI (PRK00132)Ribosomal_S9 (pfam00380)→ |
| env-1255 | SRS049318_C6258073 | - | 173 | 101 | RNA→ hypo→ |
| env-1256 | SRS016225_WUGC_scaffold_16862 | - | 163 | 90 | RNA→ hypo→ |
| env-1257 | SRS018300_C3300755 | - | 410 | 337 | RNA→ rplM (PRK09216)Ribosomal_L13 (pfam00572)→ |
| env-1258 | SRS013506_Baylor_scaffold_6896 | + | 328 | 400 | RNA→ rplM (PRK09216)Ribosomal_L13 (pfam00572)→ rpsI (PRK00132)Ribosomal_S9 (pfam00380)→ |
| env-1259 | SRS049147_C4035837 | - | 173 | 101 | RNA→ hypo→ |
| Pmu-2-1 | NZ_GL945017.1 | + | 791406 | 791482 | RNA→ rplM (PRK09216)Ribosomal_L13 (pfam00572)→ rplM (PRK09216)Ribosomal_L13 (pfam00572)→ rpsI (PRK00132)Ribosomal_S9 (pfam00380)→ rpsI (PRK00132)Ribosomal_S9 (pfam00380)→ |
| Pde-1-1 | NC_019960.1 | - | 1444430 | 1444357 | RNA→ rplM (PRK09216)Ribosomal_L13 (pfam00572)→ rpsI (PRK00132)Ribosomal_S9 (pfam00380)→ |

#### 3.3 Conserved domains

Conserved domains found in protein-coding genes listed in Section 3.2 are shown below, with the first sentence in their description from the Conserved Domain Database (if any). Conserved domains associated with more than one L13-Bacteroidia RNA are assigned a color, while others are shown in gray. The number in parentheses after the colored domain name is the number of occurrences in Section 3.2.

cd00392 (6) Ribosomal protein L13.
cd14275 (1) UBA domain found in elongation factor Ts (EF-Ts) from bacteria, chloroplasts and mitochondria of eukaryotes.
CHL00079 (3) ribosomal protein S9
COG0102 (26) Ribosomal protein L13 [Translation, ribosomal structure and biogenesis]
pfam00318 (2) Ribosomal protein S2.
pfam00380 (712) Ribosomal protein S9/S16.
pfam00572 (955) Ribosomal protein L13.
pfam00889 (1) Elongation factor TS.
PRK00132 (715) 30S ribosomal protein S9; Reviewed
PRK05299 (2) 30S ribosomal protein S2; Provisional
PRK09216 (990) 50S ribosomal protein L13; Reviewed
PRK09377 (1) elongation factor Ts; Provisional

#### 3.4 Multiple-sequence alignment

Each L13-Bacteroidia RNA is denoted by an abbreviation (like "Eco-1-1") that refers to a taxonomy given in Section 3.1. The alignment may include sequences containing the RNA motif, as well as flanking sequence. The L13-Bacteroidia RNA itself is denoted by the line underneath marked 5′ and 3′ on either end. Nucleotides in flanking sequences (i.e., not part of the motif) are written in gray letters. Stems of predicted rho-independent transcription terminators, if any, are shaded yellow. (Note: terminator predictions have not been analyzed manually, and many are likely to be false positives. Terminator predictions are those of the RNIE software.) Nucleotides predicted to function (as DNA) as transcription-factor binding sites, if any, are shaded in green. (But note: these predictions are manually annotated, so they might be under-predicted.) Annotated start codons, if any, are shaded green. (Note: start codons are frequently misannotated, especially in environmental samples.) Nucleotides proposed to basepair as part of the consensus structure are shaded in color when they comprise Watson-Crick or G-U pairs. Otherwise they are shaded gray. Conserved stems are also indicated at the bottom of the alignment by angle brackets, where matching $<$ and $>$ denote base-paired columns. Below these angle brackets, the symbol "2" denotes base pairs exhibiting covariation according to the statistically well-founded R-scape method. "1" denotes base pairs exhibiting covariation according to R2R's simplistic method. "0" denotes base pairs that are not observed to mutate and "?" denotes base pairs that have a significant frequency of non-canonical nucleotides for Watson-Crick or G-U pairs ($> 5\%$). Below these base pair annotation is the consensus sequence: "R" = "A" or "G", "Y" = "C" or "U", red nucleotides: nucleotide identity conserved more than 97% of the time, black nucleotides: 90%, gray nucleotides: 75%, red circle (•): nucleotide is present 97% of the time, black circle (•): 90%, gray circle (•): 75%, white circle (◦): 50%. All percentages of sequences just described (e.g. 97% conserved) assume that sequences have been weighted by the GSC algorithm implemented by the Infernal software package.

Duplicate sequences: the following putative homologs are not shown in the alignment because their sequences are identical to a homolog already shown: Bsp-1-1, Pal-1-2, Pam-2-1, Pam-4-1, Pba-1-2, Pbi-2-1, Pbi-3-1, Pbi-4-1, Pbi-5-1, Pbi-6-1, Pbr-4-1, Pbu-6-1, Pcl-1-1, Pco-1-1, Pco-3-2, Pde-5-2, Phi-1-1, Pin-2-1, Plo-1-1, Pma-2-1, Pma-5-1, Pme-1-1, Pme-2-1, Pmi-1-1, Pmi-2-1, Pmi-3-1, Pna-1-1, Pna-2-1, Pni-2-1, Por-3-1, Por-4-1, Por-7-1, Ppa-1-1, Ppa-2-1, Ppl-1-1, Pru-2-1, Psa-1-1, Psa-2-1, Psa-3-1, Psh-1-1, Psp-10-1, Psp-14-1, Psp-18-1, Psp-20-1, Psp-22-1, Psp-23-1, Pst-1-1, Pve-2-1, Pxy-1-1, env-100, env-1000, env-101, env-1010, env-1011, env-1012, env-1013, env-1014, env-1015, env-1016, env-1018, env-1019, env-102, env-1026, env-103, env-1030, env-1036, env-104, env-1041, env-1042, env-1045, env-1047, env-1048, env-1049, env-105, env-1050, env-1051, env-1057, env-106, env-107, env-1070, env-1073, env-108, env-1082, env-109, env-110, env-111, env-112, env-1121, env-1124, env-1128, env-113, env-114, env-1142, env-1146, env-115, env-1150, env-1152, env-116, env-1160, env-1161, env-1162, env-1168, env-1169, env-117, env-1170, env-1171, env-1172, env-1173, env-1174, env-1175, env-1176, env-1177, env-1178, env-1179, env-118, env-1180, env-1181, env-1182, env-1183, env-1184, env-1185, env-1186, env-1187, env-1188, env-1189, env-119, env-1190, env-1191, env-1196, env-1198, env-1199, env-120, env-1200, env-1201, env-1202, env-1204, env-1207, env-1208, env-1209, env-121, env-1210, env-1212, env-1214, env-1215, env-1217, env-1218, env-1219, env-122, env-1220, env-1221, env-1222, env-1225, env-1226, env-1227, env-1229, env-123, env-1230, env-1231, env-1232, env-1234, env-1235, env-1236, env-1237, env-1238, env-1239, env-124, env-1240, env-1241, env-1242, env-1246, env-1247, env-1248, env-1249, env-125, env-1252, env-1253, env-1255, env-1257, env-127, env-128, env-130, env-131, env-133, env-134, env-135, env-136, env-137, env-138, env-139, env-140, env-141, env-142, env-143, env-144, env-145, env-146, env-147, env-149, env-150, env-151, env-152, env-153, env-154, env-156, env-161, env-162, env-172, env-173, env-174, env-175, env-176, env-177, env-178, env-179, env-180, env-181, env-182, env-183, env-184, env-185, env-186, env-187, env-188, env-189, env-19, env-190, env-191, env-192, env-193, env-194, env-195, env-196, env-197, env-198, env-199, env-20, env-200, env-201, env-202, env-203, env-204, env-214, env-215, env-216, env-217, env-218, env-219, env-220, env-221, env-223, env-224, env-225, env-228, env-23, env-235, env-236, env-237, env-238, env-239, env-24, env-240, env-241, env-242, env-243, env-244, env-245, env-247, env-25, env-251, env-252, env-253, env-254, env-255, env-256, env-261, env-263, env-271, env-274, env-277, env-28, env-282, env-283, env-284, env-289, env-29, env-294, env-296, env-30, env-300, env-303, env-304, env-306, env-31, env-312, env-313, env-32, env-325, env-33, env-332, env-333, env-334, env-335, env-34, env-340, env-342, env-343, env-344, env-346, env-35, env-352, env-356, env-358, env-36, env-364, env-365, env-366, env-367, env-37, env-372, env-374, env-376, env-377, env-378, env-379, env-38, env-380, env-381, env-382, env-383, env-39, env-391, env-393, env-394, env-395, env-40, env-404, env-405, env-407, env-409, env-41, env-415, env-416, env-417, env-42, env-424, env-427, env-428, env-43, env-430, env-44, env-45, env-456, env-459, env-46, env-460, env-461, env-465, env-47, env-472, env-475, env-478, env-479, env-48, env-480, env-481, env-482, env-484, env-49, env-496, env-499, env-50, env-500, env-505, env-506, env-51, env-513, env-515, env-518, env-52, env-524, env-528, env-529, env-53, env-530, env-533, env-536, env-539, env-54, env-540, env-541, env-542, env-543, env-544, env-545, env-546, env-547, env-548, env-549, env-55, env-550, env-551, env-556, env-56, env-57, env-58, env-581, env-584, env-585, env-586, env-587, env-588, env-59, env-590, env-591, env-60, env-601, env-602, env-603, env-604, env-605, env-606, env-607, env-608, env-609, env-61, env-610, env-611, env-612, env-614, env-615, env-616, env-617, env-618, env-619, env-62, env-620, env-621, env-622, env-623, env-624, env-625, env-626, env-627, env-628, env-629, env-630, env-631, env-632, env-633, env-634, env-635, env-636, env-637, env-638, env-639, env-640, env-641, env-642, env-643, env-644, env-645, env-646, env-647, env-649, env-65, env-650, env-654, env-655, env-656, env-658, env-659, env-66, env-660, env-661, env-662, env-663, env-664, env-665, env-666, env-667, env-668, env-669, env-67, env-670, env-671, env-672, env-673, env-674, env-675, env-678, env-68, env-685, env-687, env-688, env-689, env-69, env-690, env-691, env-692, env-693, env-694, env-695, env-697, env-701, env-702, env-703, env-704, env-705, env-706, env-707, env-708, env-712, env-713, env-714, env-715, env-716, env-717, env-718, env-719, env-720, env-721, env-722, env-723, env-728, env-730, env-731, env-732, env-739, env-740, env-741, env-750, env-764, env-773, env-777, env-778, env-779, env-782, env-783, env-784, env-786, env-792, env-794, env-795, env-796, env-805, env-806, env-807, env-809, env-810, env-811, env-812, env-813, env-814, env-815, env-816, env-817, env-818, env-819, env-820, env-823, env-841, env-844, env-85, env-850, env-853, env-854, env-863, env-867, env-871, env-879, env-884, env-89, env-895, env-896, env-897, env-898, env-899, env-90, env-904, env-905, env-906, env-907, env-908, env-909, env-91, env-910, env-911, env-912, env-913, env-914, env-915, env-916, env-917, env-918, env-919, env-92, env-920, env-925, env-926, env-927, env-928, env-929, env-93, env-930, env-931, env-933, env-934, env-935, env-936, env-937, env-938, env-939, env-94, env-940, env-941, env-942, env-943, env-944, env-945, env-946, env-947, env-95, env-951, env-952, env-954, env-955, env-956, env-959, env-96, env-962, env-963, env-964, env-965, env-966, env-969, env-97, env-974, env-975, env-976, env-980, env-987, env-988, env-989, env-99, env-991, env-998

alignment positions 1 · · · 79

```
env-1   CACCGGUUUCAACAGGAAACUUACAGCGAAGACCUUGCCAAUAAAUAAAAUAGUUGUACUUUUGCGCGCCGAUUAUUAU
env-2   CGUUCAGAUACACAAAAAGUUGCAAAAAAGUUUUGUCGGUUCACAGAAAGUUCGUACCUUUGCACGCCGAUUAUUAUCA
env-3   ACGCCGCAAAAUCACGAAAAAAAGACAAGUAGAUAUUUGCCAUGAAUAAAAAAAAUAGUACUUUUGCACGCCGAUUAUU
env-4   CCAAAGAAGUAGCAAAAAUAAGUAAAAAAGUUUUGUCAGUUCGUGAAAAAGCCGUACCUUUGCACGCCGAUUAUUAUCA
env-5   AUAAUAUAUAAACAAACAUAACAACACGCCUCACAGCAUAAGAAUACUCAUUAUAUAUAUGUAACUUUGCAAACUGGAU
env-6   AAUGAAAGUGCGAAAAAAACUUAAAAAAAUCUUUGUUGGAUGGGGGAAAAGUGCUAACUUUGCAAGCCGAUUAUUAUCAA

        ...............................................................................
        ...............................................................................
        ●●●●●●●●●●●●●●●●●●●●●●●●●●●●●●●●●●●●●●●●●●●●●●●●●R●●●●●●●●●●Y●Y●●●●●●●●●●●●●Y●●
```

```
env-7      AGUAUGAAAGUGCGAAAAAAACUUAAAAAAUCUUGGUCAGAUGGUGGAAAAGUGCUAACUUUGCAAGCCGAUUAUUAUC
env-8      AACGAAUUACGGCAAAAAAUCAUAAAGAAAUCAUCGCCACAUAAAAAAUAAAAUUGUAAUUUUGCACACCGAUUAUUAUC
env-9      AACGAAUUACGGCAAAAAAUCAUAAAGAAAUCAUCGCCACAUAAAAAAUAAAAUCGUAAUUUUGCGCACCGAUUAUUAUC
env-10     AACGAAUUACGGCAAAAAAUCAUAAAGAAAUCAUCGCCACAUAAAAAAUAAAAUUGUAAUUUUGCGCACCGAUUAUUAUC
env-11     AAAAACCAGCAAAAAAGUCGCGGCAGGGGUUUGCUGUUCUGUGAAAAAGUCAUACCUUUGCACGCCGAUUAUUAUCAAG
env-12     CCUUAAAUAAAAAUAAUUGGCUGUAAACCAUUGUGGUUUGGCAUAAAAAAUCGUAACUUUGCAAGCCGAUUAUUAUCAAA
env-13     GAAAUCUGACAAAAAAAAACGCUCAGCUUUGCUUGCUGAAUAAAAAGAAAGUCGUACUUUUGCACGCCGAUUAUUAUCA
env-14     CAGGGAAGAAAAGAAAAACAACGGGGGGAACCUUGCCAUUCCAAUGAAUUUUAGUACUUUUGCACGCCGAUUAUUAUCA
env-15     AAAACAUAAAAAAUGCUAUCGUUGCAUAAAUUUUCAUCUUUUUUAUAAGGAAAAAGUAUUAACUUUGCACCCCGAAAUCG
env-16     GGGGGAAAAGCAAAAAAAAAGCAAAAAAUGCUUGCCAAUUCAGGAAAAAGCAGUACCUUUGCAAGCCGAUUAUUAUCAA
env-17     UAUAAUACUGAUAAAAAAAGUUCUUUUUUUGUUGGCUGUUUGGUAAUAAAAUAGUAAUUUUCCAAGCCGAUUAUUAUCAA
env-18     AAGAAAAGGUGAAAAAGUUUGUGGAUUUUGUUGGACGUUUGAUUGAAAAGUAGUAAUUUUGCAAGCCGAUUAUUAUCAA
env-21     AAGAAAGGGAGAAAAAGUUUGUGGAUUUUGUUGGCCGUUUGAUUGAAAAGUAGUAAUUUUGCAAGCCGAUUAUUAUCAA
env-22     AAGAAAGGGAGAAAAAGUUUGUGGAUUUUGUUGGCCGUUUGAUUGAAAAGUAGUAAUUUUGCAAGCCGAUUAUUAUCAA
env-26     AAGAAAGGGAGAAAAAGUUUGUGGAUUUUGUUGGCCGUUUGAUUGAAAAGUAGUAAUUUUGCAAGCCGAUUAUUAUCAA
env-27     AAGAAAGGGAGAAAAAGUUUGUGGAUUUUGUUGGCCGUUUGAUUGAAAAGUAGUAAUUUUGCAAGCCGAUUAUUAUCAA
env-63     AAGAAAGGGAGAAAAAGUUUGUGGAUUUUGUUGGCCGUUUGAUUGAAAAGUAGUAAUUGUAGAAGCCGAUUAUUAUCAA
env-64     AAGAAAGGGAGAAAAAGUUUGUGGAUUUUGUUGGCCGUUUGAUUGAAAAGUAGUAAUUUUGCAAGCCGAUUAUUAUCAA
env-70     AAUGUGCUCCAAACAAAAAAAAUCGGCAAAAAAACUUGUCAAUUCACGAAAAAGCAGUACCUUUGCAAGCCGAUUAUUAU
env-71     GUGAGAAGAAAAAAUUUUUGCAAAAAUACUUGUGGAUUCACGAAAAAGCAGUACCUUUGCAAGCCGAUUAUUAUCAGAG
env-72     UUUCACAUAAAAAAAGUUACGCAAAACGCUUGCUGAAUGGCAUAAAAGUCGUACCUUUGCACGCCGAUUAUUAUCAAGG
env-73     ACUAACUUUUUCUCAAAAAAAAAUACCUAAAUCGCUUGUCAAUUCACGAAAAAGCAGUACCUUUGCAAGCCGAUUAUUAU
env-74     ..........AUUAAAAAAAAACGAGAAAUUCCUUUGCCAUGUCAGCGGGAAACCGUACCUUUGCAAGCCGAUUAUUAUC
env-75     AAUUUUUUGUGAAAUAAUCGGUUAGAAAUGCUUGUUAUGUGACUAUAAAUUACUAUCUUUGCAAGCCGAUUAUUAUCAA
env-76     AUUUUUUUAUAAAAAUAAUCGGUGCAUUUUGCUUGUUAUGUCCCUAUAAAUUACUAUCUUUGCAAGCCGAUUAUUAUCA
env-77     CUAUUUUAUGAUUCAUUUUCAUAAAAAAAUGCUUGCCAGUUUCAGAAAAAGUGGUACCUUUGCACGCCGAUUAUUAUCAA
env-78     GCACCUAAGGAAGAAAAAAACUGCCCGCAACGCUUGCAGAUUAGGGAAAAAGUUGUACCUUUGCACGCCGAUUAUUAUCA
env-79     GUAGCGAGCCCUUCAAAAAAAGCACGAAAUGCUUUGUUAUGUAACAGAAAAUGACUACCUUUGCAAGCCGAUUAUUAUC
env-80     GACUGUGUCAUGGGAAAAAUAUCGGCCAAAAACUUUGUGGUUCAGGGAAAAAUGACUACCUUUGCAAGCCGAUUAUUAU
Pbr-3-1    AGUAUUUAAAACGUCUUUUUCUAAAGAAUGUGUAGAAGGGUUAUUUCAAAUAAAAUAUGUACUUUUGCACUCUGAAUUG
env-81     AGUAUUUAAAACGUCUUUUUCUAAAGAAUGUGUAGAAGGGUUAUUUCAAAUAAAAUAUGUACUUUUGCACUCUGAAUUG
env-82     AGUAUAAAUAUUAAUAUGCUAAAAGAAUAUGCUUUCUUGAAUAUCAAUUAAUAGUUGUACCUUUGCAGUCCGAUUAUUA
env-83     UUUGAAAAUAAACAAAAAACACGAAAAAACUUUUGUCAGAAUAUAGGAAAGUCGUAUCUUUGCACGCCGAUUAUUAUCG
env-84     GCAAAACCUCUGAAAAUAUUUAUGAUAAACAUGCGACGGAAUGUAAAUAAAUUCGUACUUUUGCACGCCGAUUAUUAUC
env-86     GCAAAACCUCUGAAAAUAUUUAUGAUAAACAUGCGACGGAAUGUAAAUAAAUUCGUACUUUUGCACGCCGAUUAUUAUC
env-87     GCCUCAAGAAGAAAAAAAGUAUCACCCGGGCUUGCAGAAUAAGGAAAAAGUUGUACCUUUGCAAGCCGAUUAUUAUCAA
Pal-1-1    AAUUUAUAAAAUGACAUACUUGCAAGACAAGUGUUGAUGCUCGUUUUAAAUAAAAUAUGUACCUUUGCACACCGAAUUG
env-88     CCAUGAAAAGAAGAUUUUCAAGUUCACAAGCUUGGCUAUUUCGCAGAAUAUAAGUACAUUUGCAUGCCGAUUAUUAUCA
env-98     CCAUGAAAAGAAGAUUUUCAAGUUCACAAGCUUGGCUAUUUCGCAGAAUAUAAGUACAUUUGCAUGCCGAUUAUUAUCA
env-126    ..............................CUUGGCUAUUUCGCAGAAUAUAAGUACAUUUGCAUGCCGAUUAUUAUCA
env-129    CCAUGAAAAGAAGAUUUUCAAGUUCACAAGCUUGGCUAUUUCGCAGAAUAUAAGUACAUUUGCAUGCCGAUUAUUAUCA
env-132    CCAUGAAAAGAAGAUUUUCAAGUUCACAAGCUUGGCUAUUUCGCAGAAUAUAAGUACAUUUGCAUGCCGAUUAUUAUCA
env-148    CCAUGAAAAGAAGAUUUUCAAGUUCACAAGCUUGGCUAUUUCGCAGAAUAUAAGUACAUUUGCAUGCCGAUUAUUAUCA
env-155    CCAUGAAAAGAAGAUUUUCAAGUUCACAAGCUUGGCUAUUUCGCAGAAUAUAAGUACAUUUGCAUGCCGAUUAUUAUCA
env-157    CCAUGAAAAGAAGAUUUUCAAGUUCACAAGCUUGGCUAUUUCGCAGAAUAUAAGUACAUUUGCAUGCCGAUUAUUAUCA
env-158    UAAAUAAAUAAUCACAUUCAUAGACAAUCACUUAUCCUUAUAAUAAAAAGAAAGUAAUACCUUUGUAGUCCGAUUAUUA
env-159    UAAAUAAAUAAUCACAUUCAUAGACAAUCACUUAUCCUUAUAAUAAAAAGAAAGUAAUACCUUUGUAGUCCGAUUAUUA
env-160    UAAAUAAAUAUACACAAGCAAACACAACCAUAUCUCCUUAUAAUAAAAAGAAAGUAAUACCUUUGUAGUCCGAUUAUUA
env-163    .................................CUCCUUAUAAUAAAAAGAAAGUAAUACCUUUGUAGUCCGAUUAUUA
env-164    NNNNNNNNNNNNNNNNNNNNNNNNNNNNCACUUAUCCUUAUAAUAAAAAGAAAGUAAUACCUUUGUAGUCCGAUUAUUA
env-165    UAAAUAAAUAAUCACAUUCAUAGACAAUCACUUAUCCUUAUAAUAAAAAGAAAGUAAUACCUUUGUAGUCCGAUUAUUA
env-166    UAAAUAAAUAAUCACAUCCACACACAAUCACUUAUCCUUAUAAUAAAAAGAAAGUAAUACCUUUGUAGUCCGAUUAUUA
env-167    UAAAUAAAUAAUCACACCCACAGACAAUCACUUAUCCUUAUAAUAAAAAGAAAGUAAUACCUUUGUAGUCCGAUUAUUA
env-168    UAAAUAAAUAAUUACAUCCAUAGACAAUCACUUAUCCUUAUAAUAAAAAGAAAGUAAUACCUUUGUAGUCCGAUUAUUA
env-169    .......AUAAUCAUAUCCACACACAAUCACUUAUCCUUAUAAUAAAAAGAAAGUAAUACCUUUGUAGUCCGAUUAUUA
env-170    UAAAUAAAUAACCACAUCCACACACAAUCACUUAUCCUUAUAAUAAAAAGAAAGUAAUACCUUUGUAGUCCGAUUAUUA
env-171    ..........................AUCACUUAUCCUUAUAAUAAAAAGAAAGUAAUACCUUUGUAGUCCGAUUAUUA
env-205    UAAAUAAAUAAUCGCAUCCAUAGAUAAUCACUUAUCCUUAUAAUAAAAAGAAAGUAAUACCUUUGUAGUCCGAUUAUUA
env-206    UAAAUAAAUAAUCACAUCCGCGCACAAUCACUUAUCCUUAUAAUAAAAAGAAAGUAAUACCUUUGUAGUCCGAUUAUUA
env-207    UAAAUAAAUAAUCACAUCUAUAGACAAUCACUUAUCCUUAUAAUAAAAAGAAAGUAAUACCUUUGUAGUCCGAUUAUUA
env-208    UAAAUAAAUAAUCACAGUCAUAGACAAUCACUUAUCCUUAUAAUAAAAAGAAAGUAAUACCUUUGUAGUCCGAUUAUUA
env-209    UAAAUAAAUAAUCACAUCCACGAACAACCACUUAUCCUUAUAAUAAAAAGAAAGUAAUACCUUUGUAGUCCGAUUAUUA
env-210    .................CCAUAGACAAUCACUUAUCCUUAUAAUAAAAAGAAAGUAAUACCUUUGUAGUCCGAUUAUUA
env-211    AAUCGCUCCUUUUACACAUAUAUUAUAUAUAAGGGNNNNNNNNNNNNNNNAAAGUAAUACCUUUGUAGUCCGAUUAUUA
env-212    UAAAUAAAUAAUCACAUCCAUAGACAAUCACUUAUCCUUAUAAUAAAAAGAAAGUAAUACCUUUGUAGUCCGAUUAUUA
env-213    ...........UCGCAUCCACAAACAACCACUUAUCCGUAUAAUAAAAAGAAAGUAAUACCUUUGUAGUCCGAUUAUUA
env-222    ............................................................CUUUGUAGUCCGAUUAUUA
env-226    UAAAUAAAUAAUCACAUCCAUAGACAACCACUUAUCCUUAUAAUAAAAAGAAAGUAAUACCUUUGUAGUCCGAUUAUUA
env-227    UAAAUAAAUAAUCACAUCCACAGAUAAUCACUUAUCCUUAUAAUAAAAAGAAAGUAAUACCUUUGUAGUCCGAUUAUUA
env-229    UAAAUAAAUAAUCACAUCCACAGAUAAUCACUUAUCCUUAUAAUAAAAAGAAAGUAAUACCUUUGUAGUCCGAUUAUUA
env-230    UAAAUAAAUAAUCGCAUCCAUAGAUAAUCACUUAUCCUUAUAAUAAAAAGAAAGUAAUACCUUUGUAGUCCGAUUAUUA
env-231    UAAAUAAAUAAUCACAUCCAUAGAUAAUCACUUAUCCUUAUAAUAAAAAGAAAGUAAUACCUUUGUAGUCCGAUUAUUA
env-232    UAAAUAAAUAAUCAUGUCCACAAACAACCACUUAUCCUUAUAAUAAAAAGAAAGUAAUACCUUUGUAGUCCGAUUAUUA
env-233    UAAAUAAAUAAUCACAUCCACAGAUAAUCACUUAUCCUUAUAAUAAAAAGAAAGUAAUACCUUUGUAGUCCGAUUAUUA
env-234    .........................AAUCACUUAUCCUUAUAAUAAAAAGAAAGUAAUACCUUUGUAGUCCGAUUAUUA
env-246    UAAAUAAAUAAUCACAUCCACGAACAACCACUUAUCCUUAUAAUAAAAAGAAAGUAAUACCUUUGUAGUCCGAUUAUUA
env-248    UAAAUAAAUAAUUACAUCCACAGAUAAUCACUUAUCCUUAUAAUAAAAAGAAAGUAAUACCUUUGUAGUCCGAUUAUUA
env-249    UAAAUAAAUAAUCGCAUCCAUAGAUAAUCACUUAUCCUUAUAAUAAAAAGAAAGUAAUACCUUUGUAGUCCGAUUAUUA
env-250    ..................CAUAGACAAUCACUUAUCCUUAUAAUAAAAAGAAAGUAAUACCUUUGUAGUCCGAUUAUUA
env-257    AAAAAUGUUAAUAAUAAAUAAAUAAUCACACCCAUAGACAAUCANNNNNNNNNNNNNNNNNNNNNNNNNNNNAUUAUUA
env-258    UAAAUAAAUAAUCACAUUCAUAGACAAUCACUUAUCCUUAUAAUAAAAAGAAAGUAAUACCUUUGUAGUCCGAUUAUUA
env-259    UAAAUAAAUAAUCACAUCCAUAGAUAAUCACUUAUCCUUAUAAUAAAAAGAAAGUAAUACCUUUGUAGUCCGAUUAUUA
env-260    UAAAUAAAUAAUCACAUCCACACACAAUCACUUAUCCUUAUAAUAAAAAGAAAGUAAUACCUUUGUAGUCCGAUUAUUA
env-262    GAUUGUAAAAUAAGGAAAUCAUAAAAAGUCUAUUGCCUCUUUAAGAAAAGAAAUUGCUAUCUUUGUAGUCCGAUUAUUA

           ...............................................................................
           ...............................................................................
           ●●●●●●●●●●●●●●●●●●●●●●●●●●●●●●●●●●●●●●●●●●●●●●●●●R●●●●●●●●●●Y●Y●●●●●●●●●●●●●Y●●
```

| | |
|---|---|
| env-264 | AAUUGCAAAAUAAGAAAGUUGUAAAAAGCCUAUUACCUCUUUAAGAAAAGAAAUUGCUAUCUUUGUAGUCCGAUUAUUA |
| env-265 | AACCGUAAAAUAAGGAAAUCAUAAAAAGCCUAUUACCUCUUUAAGAAAAGAAAUUGCUAUCUUUGUAGUCCGAUUAUUA |
| env-266 | AUGUGCCCGCGCGCGGAAAUACAUAAAAUUAUUUGGUAGUUAGGGCGCUAUUUCGUACCUUUGCAAGCCGAUUAUUAUC |
| env-267 | UUGAAAACUUCAGAAAAUUGUUGAAAAACUUUGCCGGUUCGUCAAAAAUUAGUACUUUUGCAAGCCGAUUAUUAUCAAG |
| Bsp-2-1 | UUGAAAACUUCAGAAAAUUGUUGAAAAACUUUGCCAGUUCGUCAAAAAUUAGUACUUUUGCAAGCCGAUUAUUAUCAAG |
| env-268 | AUAAAUAUAAACAAUAUCAUUGUGGAGCCUUGAUAGCCUCUCGGCCUGAAUAAAAUAUGUAACUUUGCAUAUCGAAUUG |
| env-269 | AUAAAUAUAAACAAUAUCAUUGUGGAGUCUUGAUUGCCUCUCGACCUGAAUAAAAUAUGUAACUUUGCACAUCGAAUUG |
| env-270 | AUAAAUAUAAACAAUGGCAUUGCAGAGCCAUGAUAGCCUCUCGGCCUGAAUAAAAUAUGUAACUUUGCAUAUCGAAUUG |
| env-272 | AUAAAUAUAAACAAUGGCAUUGCAGAGCCAUGAUAGCCUCUCGGCCUGAAUAAAAUAUGUAACUUUGCACAUCGAAUUG |
| env-273 | AUAAAUAUAAACAAUGGCAUUGCGGAGCCAUGAUAGCCUCUUGGCCUGAAUAAAAUAUGUAACUUUGCACAUCGAAUUG |
| env-275 | AUAAAUAUAAACAAUAUCAUUGUGGAGCCUUGAUAGCCUCUCGGCCUGAAUAAAAUAUGUAACUUUGCACAUCGAAUUG |
| env-276 | AUAAAUAUAAACAAUGGCAUUGCAGAGCCAUGAUAGCCUCUCGGCCUGAAUAAAAUAUGUAACUUUGCACAUCGAAUUG |
| env-278 | AUAAUCGUAAAUAUUCGGUUUCGCCGCUUUUCAGGCAGGAGUCUGCUUAAAUAAAUACGUAUCUUUGCAAUCCGAAUUG |
| env-279 | AUAAUCGUAAAUAUUCAGUUUCGCCGCUUUUCAGGCAGGAGUCUGCCUAAAUAAAUAUGUAUCUUUGCAAUUCGAAUUG |
| env-280 | AUAAUCGUAAAUAUUCAGUUUCGUCGCUUUUCAGGCAGGAGUCUGCCUAAAUAAAUAUGUAUCUUUGCAAUCCGAAUUG |
| env-281 | AUAAUCGUAAAUAUUCAGUUUCGCCGCUUUUCAGGCAGGAGUCUGCCUAAAUAAAUAUGUAUCUUUGCAAUCCGAAUUG |
| env-285 | UAAUCGUAAAUCUUCAGUUUUGCGACGUUUCCUGCAAGAGGUAAGCUAAAUAAAAUAUGUAACUUUGCAUUCCGAAUUG |
| env-286 | UAAUCGUAAAUCUUCAGUUUUGCGACGUUUCCUGCAAGAGGUAAGCUAAAUAAAAUAUGUAACUUUGCAUUCCGAAUUG |
| env-287 | UAAUCGUAAAUCUUCGGUUUUGUGACGUUUCCUGCAAGAGGUAAGCUAAAUAAAAUAUGUAACUUUGCAUUCCGAAUUG |
| env-288 | UAAUCGUAAAUCUUUAGUUUUGCGACGUUUCGUGCAAGAGGUAAGCUAAAUAAAAUAUGUAACUUUGCAUUCCGAAUUG |
| env-290 | UAAUCGUAAAUCUUCAGUUUUGCGACGUUUCCUGCAAGAGGUAAGCUAAAUAAAAUAUGUAACUUUGCAUUCCGAAUUG |
| env-291 | UAAUCGUAAAUCUUCAGUUUUGCGACGUUUCCUGCAAGAGGUAAGCUAAAUAAAAUAUGUAACUUUGCAUUCCGAAUUG |
| env-292 | .................................UGCAAGAGGUAAGCUAAAUAAAAUAUGUAACUUUGCAUUCCGAAUUG |
| env-293 | UAAUCGUAAAUCUUCAGUUUUGCGACGUUUUGUGCAAGAGGUAAGCUAAAUAAAAUAUGUAACUUUGCAUUCCGAAUUG |
| env-295 | UAAUCGUAAAUCUUUAGUUUUGCGACGUUUCGUGCAAGAGGUAAGCUAAAUAAAAUAUGUAACUUUGCAUUCCGAAUUG |
| env-297 | UAAUCGUAAAUCUUCAGUUUUGCGACGUUUCGUGCAAGAGGUAAGCUAAAUAAAAUAUGUAACUUUGCAUUCCGAAUUG |
| env-298 | UAAUCGUAAAUCUUCAGUUUUGCGACGUUUCCUGCAAGAGGUAAGCUAAAUAAAAUAUGUAACUUUGCAUUCCGAAUUG |
| env-299 | UAAUCGUAAAUCUUCGGUUUUGCGACGUUUCCUGCAAGAGGUAAGCUAAAUAAAAUAUGUAACUUUGCAUUCCGAAUUG |
| env-301 | UAAUCGUAAAUCUUCAGUUUUGUGACGUUUCCUGCAAGAGGUAAGCUAAAUAAAAUAUGUAACUUUGCAUUCCGAAUUG |
| env-302 | UAAUCGUAAAUCUUCAGUUUUACGACGUUUCCUGCAAGAGGUAAGCUAAAUAAAAUAUGUAACUUUGCAUUCCGAAUUG |
| env-305 | UAAUCGUAAAUCUUCAGUUUUGCGACGUUUCCUGCAAGAGGUAAGCUAAAUAAAAUAUGUAACUUUGCAUUCCGAAUUG |
| env-307 | UAAUCGUAAAUCUUCAGUUUUGCGACGUUUCGUGCAAGAGGUAAGCUAAAUAAAAUAUGUAACUUUGCAUUCCGAAUUG |
| env-308 | UAAUCGUAAAUCUUCAGUUUUGCGACGUUUCGUGCAAGAGGUAAGCUAAAUAAAAUAUGUAACUUUGCAUUCCGAAUUG |
| env-309 | UAAUCGUAAAUCUUCAGUUUUGCGACGUUUUAUGCAAGAGGUAAGCUAAAUAAAAUAUGUAACUUUGCAUUCCGAAUUG |
| env-310 | UAAUCGUAAAUCUUCAGUUUUGCGACGUUUUGUGCAAGAGGUAAGCUAAAUAAAAUAUGUAACUUUGCAUUCCGAAUUG |
| env-311 | UAAUUGUAAAUAUAUGGCUUGACAACGUUUAAGACAAGGGCUAACCUAAAUAAAAUAUGUAACUUUGCAUUCCGAAUUG |
| env-314 | UAAUCGUAAAUCUUCGGUUUUGCGACGUUUCCUGCAAGAGGUAAGCUAAAUAAAAUAUGUAACUUUGCAUUCCGAAUUG |
| env-315 | AUAAUCGUAAAUCUUUAGUUUUGCGACGUUUUGUGCAAGAUGUAAGCUAAAUAAAAUAUGUAACUUUGCAUUCCGAAUU |
| env-316 | UAAUCAAUAAACAGAAGCAGAAAAGAACGCUCCACAGACAAGCGUCUCAAAUAAAAGUGGUAACUUUGCAAACCGAUUG |
| env-317 | UAAUCAAUAAACAAAGGCAGAAAAGACCAUUCCACCGGCGAGUGUCUCAAAUAAAAGUGGUAACUUUGCAAACCGAUUG |
| env-318 | UAAUCAAUAAACAAAGGCNGAAAAAACCAUUCCACCGGCGAGUGUCUCAAAUAAAAGUGGUAACUUUGCAAACCGAUUG |
| env-319 | AAUAAUCAUAAAUCAACCUUAUCACAAGACUUGCAGAAAGCUCAUUUACAAUAAAAUAUGUACCUUUGCACCCCGAUUG |
| Psp-3-1 | AUUGAGUUAAUAAUUCAUAAAUAUAGUAUCUUCUCCUCGCGCGAUCUUAAAUAAAAGUUGUACCUUUGCACACCGAUUG |
| env-320 | AAUAUUCGUAAAAAUGCAGGUCACACUCAUCCGCCCCGUGGCACUGCCAUAUAAAAUAUGUAACUUUGCACCCCGACUU |
| env-321 | AAUAUUCGUAAAAAUGCAGGUCGCGUCCAUCCGCCCCGUGGCACUGCCAUAUAAAAUGUGUAACUUUGCACCCCGACUU |
| env-322 | AAUAUUCGUAAAAAUGCAGGUCGCGUCCAUCCGCCCCGUGGCACUGCCAUAUAAAAUAUGUAACUUUGCACCCCGACUU |
| env-323 | AAUAUUCGUAAAAAUGCAGGUCGCGUCCAUCCGCCCCGUGGCACUGCCAUAUAAAAUAUGUAACUUUGCACCCCGACUU |
| env-324 | AAUAUUCGUAAAAAUGCAGGUCGCGUCCAUCCGCCCCGUGGCACUGCCAUAUAAAAUAUGUAACUUUGCACCCCGACUU |
| env-326 | AAUAUUCGUAAAAAUGCAGGUCGCGUCCAUCCACCCCGUGGCACUGCCAUAUAAAAUAUGUAACUUUGCACCCCGACUU |
| env-327 | AAUAUUCGUAAAAAUGCAGGUCGCGUCCAUCCGCCCCGUGGCACUGCAAUAUAAAAUAUGUAACUUUGCACCCCGACUU |
| env-328 | AAUAUUCGUAAAAAUGCAGGUCGCAUCCAUUCGCCCCGAGGCACUGCCAUAUAAAAUAUGUAACUUUGCACCCCGACUU |
| env-329 | AAUAUUCGUAAAAAUGCAGGCCACACCCAUCCGCCCCGUGGCACUGCCAUAUAAAAUAUGUAACUUUGCACCCCGACUU |
| env-330 | AAUAUUCGUAAAAAUGCAGGUCACAUCCACUCGCCCCGUGGCACUGCCAUAUAAAAUAUGUAACUUUGCACCCCGACUU |
| env-331 | AAUAUUCGUAAAAAUGCAGGUCGCGUCCAUCCGCCCCGUGGCACUGCCAUAUAAAAUAUGUAACUUUGCACCCCGACUU |
| env-336 | AAUAUUCGUAAAAAUGCAGGUCGCGUCCAUUCGCCCCGUGGCACUGCCAUAUAAAAUAUGUAACUUUGCACCCCGACUU |
| env-337 | AAUAUUUGUAAAAAUGCAGGUCGCGUCCAUCCGCCCCGUGGCACUGCCAUAUAAAAUAUGUAACUUUGCACCCCGACUU |
| env-338 | AAUAUUCGUAAAAAUGCAGGCCACACCCAUCCGCCCCGUGGCGCUGCCAUAUAAAAUAUGUAACUUUGCACCCCGACUU |
| env-339 | AAUAUUCGUAAAAAUGCAGGCCACGCCCAUCCGCCCCGUGGCACUGCCAUAUAAAAUAUGUAACUUUGCACCCCGACUU |
| env-341 | AAUAUUCGUAAAAAUGCAGGUCGCGUCCAUUCGCCCCGUGGCACUGCCAUAUAAAAUAUGUAACUUUGCACCCCGACUU |
| env-345 | AAUAUUCGUAAAAAUGCAGGCCACGCCCACCCGCCCCGUGGCACUGCCAUAUAAAAUAUGUAACUUUGCACCCCGACUU |
| env-347 | AAUAUUCGUAAAAAUGCAGGCCACACCCAUCCGCCCCGUGGCACUGCCAUAUAAAAUAUGUAACUUUGCACCCCGACUU |
| env-348 | AAUAUUCGUAAAAAUGCAGGCCACAUCCAUUCGCCCCGUGGCACUGCCAUAUAAAAUAUGUAACUUUGCACCCCGACUU |
| env-349 | AAGUUUUUAAUUCAGCCGUUAUUGAUGGAUAGAGACGCGCGUUUCUAUCCAUAAAAUAUGUAACUUUGCAAACCGAUUG |
| Pma-3-1 | AAGUUUUUAAUUCAGCCGUUAUUGAUGGAUAGAGACGCGCGUUUCUAUCCAUAAAAUAUGUAACUUUGCAAACCGAUUG |
| env-350 | AAGUUUCUAAUUCGGCCAUUAUUGAUGGAUAGAGACGCGCGUUUCUAUCCAUAAAAUAUGUAACUUUGCAAACCGAUUG |
| env-351 | AAGUUUUUAAUUCAGCCGUUAUUGAUGGAUAGAGACGCGCGUUUCUAUCCAUAAAAUAUGUAACUUUGCAAACCGAUUG |
| Pma-1-1 | AAGUUUCUAAUUCAGCCGUUAUUGAUGGAUAGAGACGCGCGUUUCUAUCCAUAAAAUAUGUAACUUUGCAAACCGAUUG |
| env-353 | AAGUUUUUAAUUCGGCCGUUAUUGAUGGAUAGAGACGCGCGUUUCUAUCCAUAAAAUAUGUAACUUUGCAAACCGAUUG |
| env-354 | AAGUUUUUAAUUCGACCGUUAUUGAUGGAUAGAGACGCGCGUUUCUAUCCAUAAAAUAUGUAACUUUGCAAACCGAUUG |
| env-355 | AAGUUUUUAAUUCAGCCGUUAUUGAUGGAUAGAGACGCGCGUUUCUAUCCAUAAAAUAUGUAACUUUGCAAACCGAUUG |
| env-357 | AUAUUCGUAAUCUGAAUGCUUUUGCCUGCUAAUACUUUUCGAUUCUUCCUAUAAAAUAUGUAUCUUUGCAAACCGAUUA |
| env-359 | AUAUUCGUAACCAGGAUGCUUUUGUCUGCUAAUGCUUUUCGAUUCUUCCUAUAAAAUAUGUAUCUUUGCAAACCGAUUA |
| env-360 | AUAUUCGUAAUCAGAAUGCUUUUGUCUGCUAAUACUUUUCGGUUCUUCCUAUAAAAUAUGUAUCUUUGCAAACCGAUUA |
| env-361 | AUAUUCGUAAUCAGAAUGCUUUUGUCUGCUAAUACUUUUCGGUUCUUCCUAUAAAAUAUGUAUCUUUGCAAACCGAUUA |
| env-362 | AUAUUCGUAAUCAGAAUGCUUUUGCCUGCUAAUACUUUUCGGUUCUUCCUAUAAAAUAUGUAUCUUUGCAAACCGAUUA |
| env-363 | AUAUUCGUAACCAGAAUGCUUUUGUCUGCUAAUACUUUUCGGUUCUUCCUAUAAAAUAUGUAUCUUUGCAAACCGAUUA |
| env-368 | AUAUUCGUAAUCAGGAUGCUUUUGUCUGCUAAUACUUUUCGGUUCUUCCUAUAAAAUAUGUAUCUUUGCAAACCGAUUA |
| env-369 | AUAUUCGUAACCGGGAUGCUUUUGUCUGCUAAUGCUUUUCGGUUCUUCCUAUAAAAUAUGUAUCUUUGCAAACCGAUUA |
| env-370 | AUAUUCGUAACCAGGAUGCUUUUGUCUGCUAAUGCUUUUCGGUUCUUCCUAUAAAAUAUGUAUCUUUGCAAACCGAUUA |
| env-371 | AUAUUCGUAAUCAGAAUGCUUUUGUCUGCUAAUACUUUUCGGUUCUUUCUAUAAAAUAUGUAUCUUUGCAAACCGAUUA |
| env-373 | AUAUUCGUAAUCAGGAUGCUUUUGUCUGCUAAUACUUUUCGGUUCUUCCUAUAAAAUAUGUAUCUUUGCAAACCGAUUA |
| env-375 | ..........................................GUUCUUCCUAUAAAAUAUGUAUCUUUGCAAACCGAUUA |
| env-384 | AUAUCCGUAAUCAGAAUACUUUUGUCUGCUAAUACUUUUCGGUUCUUCCUAUAAAAUAUGUAUCUUUGCAAACCGAUUA |
| env-385 | AUAUUCGUAAUCAGAAUGCUUUUGCCUGCUAAUACUUUUCGGUUCUUCCUAUAAAAUAUGUAUCUUUGCAAACCGAUUA |
| | ................................................................................ |
| | ................................................................................ |
| | ●●●●●●●●●●●●●●●●●●●●●●●●●●●●●●●●●●●●●●●●●●●●●●●●●●R●●●●●●●●●●Y●Y●●●●●●●●●●●●●Y●● |

```
env-386  AUAUUCGUAAUCAGGAUGCUUUUGUCUGCUAAUACUUUUCGGUUCUUCCUAUAAAAUAUGUAUCUUUGCAAACCGAUUA
env-387  AUAUUCAUAAUUGGAAUGCUUUUGCCUGCUAAUACUUUUCGGUUCUUCCUAUAAAAUAUGUAUCUUUGCAAACCGAUUA
env-388  AUAUUUGUAAUUGGGAUGCUUUUAUCUGCUCAUACUUUUCGGUUCUUCCUAUAAAAUAUGUAUCUUUGCAAACCGAUUA
env-389  AUAUUCGUAACCAGAAUGCUUUUGUCUGCUAAUGCUUUUCGGUUCUUCCUAUAAAAUAUGUAUCUUUGCAAACCGAUUA
env-390  ...........................................................................AUUA
env-392  AUAUUCGUAACCAGGAUGCUUUUGUCUGCUAAUGCUUUUCGAUUCUUCCUAUAAAAUAUGUAUCUUUGCAAACCGAUUA
env-396  AUAUUCGUAACCAGAAUGCUUUUGUCUGCUAAUGCUUUUCGGUUCUUCCUAUAAAAUAUGUAUCUUUGCAAACCGAUUA
env-397  AUAUUUGUAAUUGGAAUGCUUUUAUCUGCUCAUACUUUUCGGUUCUUCCUAUAAAAUAUGUAUCUUUGCAAACCGAUUA
env-398  AUAUUCGUAACCGGGAUGCUUUUGUCUGCUCAUACUUUUCGGUUCUUCCUAUAAAAUAUGUAUCUUUGCAAACCGAUUA
env-399  UUGUGUAUCUUGGGUUGUAGGGAUUGAUUUCAUGAAAUGAAUAAUAUUANNNNNNNNNNGUAUCUUUGCAAACCGAUUA
env-400  AUAUUCGUAACCAGAAUGCUUUUGUCUGCUAAUGCUUUUCGAUUCUUCCUAUAAAAUAUGUAUCUUUGCAAACCGAUUA
env-401  AUAUUCGUAAUCUGAAUGCUUUUGCCUGCUAAUACUUUUCGAUUCUUCCUAUAAAAUAUGUAUCUUUGCAAACCGAUUA
env-402  AUAUUUGUAAUUGGAAUGCUUUUAUCUGCUCAUACUUUUCGGUUCUUCCUAUAAAAUAUGUAUCUUUGCAAACCGAUUA
env-403  ..................CUUUUGUCUGCUAAUACUUUUCGGUUCUUCCUAUAAAAUAUGUAUCUUUGCAAACCGAUUA
env-406  ....................................CUUUUCGGUUCUUCCUAUAAAAUAUGUAUCUUUGCAAACCGAUUA
env-408  UUAUGUAAUUAGAAUAGAUUUACAGUCUAUUACAUUAUGCUUCUUUCUUUAAAAUAUGUACCUUUGCAAACCGAUUGCU
env-410  UUGUGUAAUUGGGAUAGAUUUACAGUCUAUUACAUUAUGCUUCUUUCUUUAAAAUAUGUACCUUUGCAAACCGAUUGCU
env-411  UUGUGUAAUUAGAAUAGAUUUACAGUCUAUUACAUUAUGCUUCUUUCUUUAAAAUAUGUACCUUUGCAAACCGAUUGCU
env-412  UAUUCGUAACCAGAAUACUUUUGUCUGCCAAUGCUUUUCGAUUCUUCCUAUAAAAUAUGUAUCUUUGCAAACCGAUUAC
Por-6-1  GUUAUGUAAUUAGAAUGGAUUUGCAGUCUAUUACAUUAUGCUUCUUUCUUUAAAAUAUGUACCUUUGCAAACCGAUUGC
Por-5-1  GUUAUGUAAUUAGAAUGGAUUUGCAGUCUAUUACAUUAUGCUUCUUUCUUUAAAAUAUGUACCUUUGCAAACCGAUUGC
env-413  GUUAUGUAAUUAGAAUAGAUUUACAGUCUAUUACAUUAUGCUUCUUUCUUUAAAAUAUGUACCUUUGCAAACCGAUUGC
env-414  GUUAUGUAAUUAGAAUGGAUUUGCAGUCUAUUACAUUAUGCUUCUUUCUUUAAAAUAUGUACCUUUGCAAACCGAUUGC
env-418  GUUAUGUAAUUAGAAUGGAUUUGCAGUCUAUUACAUUAUGCUUCUUUCUUUAAAAUAUGUACCUUUGCAAACCGAUUGC
env-419  GUUGUGUAAUUAGAAUGGAUUUACAGUCUAUUACAUUAUGCUUCUUUCUUUAAAAUAUGUACCUUUGCAAACCGAUUGC
env-420  .......................................GCUUCUUUCUUUAAAAUAUGUACCUUUGCAAACCGAUUGC
env-421  GUUAUGUAAUUAGAAUAGAUUUACAGUCUAUUACAUUAUGCUUCUUUCUUUAAAAUAUGUACCUUUGCAAACCGAUUGC
env-422  GUUGUGUAAUUAGAAUAGAUUUGCAGUCUAUUACAUUAUGCUUCUUUCUUUAAAAUAUGUACCUUUGCAAACCGAUUGC
env-423  GUUAUAUAAUUAGAAUGGAUUUGCAGUCUAUUACAUUAUGCUUCUUUCUUUAAAAUAUGUACCUUUGCAAACCGAUUGC
Psp-8-1  AUAUAUAUAAGAUGAGGUGUUUUAGGGGGCACUGUGAAUACAUCCCUCUCAUAAAAUAUGUAACUUUGCACACCGAUUG
env-425  AUAUAUAUAAGACGAGGCGUUUUAGGGGGCGCUGUGAAUACAUCCCUCUCAUAAAAUAUGUAACUUUGCACACCGAUUG
env-426  AUAUAUAUAAGACGAGGCGUUUUAGGGGGCGCUGUGAAUACAUCCCUCUCAUAAAAUAUGUAACUUUGCAUACCGAUUG
env-429  AUAUAUAUAAGACGAGGCGUUUUAGGGGGCGCUGUGAAUACAUCCCUCUCAUAAAAUAUGUAACUUUGCAUACCGAUUG
env-431  AUAUAUAUAAGAUGAGGUGUUUUAGGGGGCGCUGUGAAUACAUCCCUCUCAUAAAAUAUGUAACUUUGCACACCGAUUG
env-432  AUAUAUAUAAGAUGAGGUGUUUUAGGGGGCGCUGUGAAUACAUCCCUCUCAUAAAAUAUGUAACUUUGCAUACCGAUUG
env-433  AUAUAUGUAAGAUGAGGUGUUUUAGGGGGCACUAUGAAUACAUCCCUCUCAUAAAAUAUGUAACUUUGCAAACCGAUUG
env-434  AUAUAUAUAAGAUGAGGUGUUUUAGGGGGCGCUGUGAAUACAUCCCUCUCAUAAAAUAUGUAACUUUGCAUACCGAUUG
env-435  AUAUAUAUAAGAUGAGGUGUUUUAGGGGGUACUGUGAAUACAUCCCUCUCAUAAAAUAUGUAACUUUGCAUACCGAUUG
env-436  AUAUAUGUAAUAUAUGGUACUUUGGAGGGCGUUCCGAGUGCAUCCCUGUUAUAAAAUAUGUAACUUUGCACACCGAUUG
env-437  ....AUGUAAUAUAUGGUACUUUGGAGGGCGUUCCGAGUGCAUCCCUGUUAUAAAAUAUGUAACUUUGCACACCGAUUG
env-438  AUAUAUAUAAGAUGAGGUGUUUUAGGGGGCACUGUGAAUACAUCCCUCUCAUAAAAUAUGUAACUUUGCACACCGAUUG
env-439  AUAUAUAUAAGAUGAGGCGUUUUAGGGGGUACUGUGAAUACAUCCCUCUCAUAAAAUAUGUAACUUUGCACACCGAUUG
env-440  AUAUAUGUAAUAUGUAGUGCUUUGGAGGGCGUUCCGGGUGCAUCCCUGUUAUAAAAUAUGUAUCUUUGCACGCCGAUUG
env-441  AUAUAUGUAAUAUAUGGUGCUUUGAAGGGUGUUCCGGGUGCAUCCCUGUUAUAAAAUAUGUAUCUUUGCACGCCGAUUG
env-442  AUAUAUGUAAUAUAUGGUGCUUUGGAGGGCGUUCCGAGUGCAUCCCUGUUAUAAAAUAUGUAUCUUUGCACGCCGAUUG
env-443  AUAUAUGUAAUAUAUGGCGCUUUGGAGGGUGUUCCGAGUGCAUCCCUGUUAUAAAAUAUGUAUCUUUGCACGCCGAUUG
Pou-1-1  AUAUAUGUAAUAUAUGGUGCUUUGGAGGGCGUCCCGAGUGCAUCCCUGUUAUAAAAUAUGUAUCUUUGCACGCCGAUUG
env-444  AUAUAUGUAAUAUAUAGUGCUUUGGAGAGCGUUCCGGGUGCAUCCCUGUUAUAAAAUAUGUAUCUUUGCACGCCGAUUG
env-445  AUAUAUGUAAUAUAUAGUGCUUUGGAGGGUGUUCCGAGUGCAUCCCUGUUAUAAAAUAUGUAUCUUUGCACGCCGAUUG
env-446  AUAUAUGUAAUAUAUGGUGCUUUGGAGGGUGCUCAGGGUGCAUCCCUGUUAUAAAAUAUGUAUCUUUGCACGCCGAUUG
env-447  AUAUAUGUAAUAUAUGGUGCUUUGGAGGGCGUUCCGAGUGCAUCCCUGUUAUAAAAUAUGUAUCUUUGCACGCCGAUUG
Pou-2-1  AUAUAUGUAAUAUAUGGUACUUUGGAGGGCGUUCCGAGUGCAUCUCUGUUAUAAAAUAUGUAUCUUUGCACGCCGAUUG
env-448  AAUCAUAAUUCAUACAUUUUUUCUGUAUAAAUACCUUGUAAUUAAAAUAAAAUACAUACUUUUGCAGACCGAUUAUUAU
env-449  UGUUUUGUAAAAUAAAAGCCGCGUUCUCCAUGCCGUGGCCCCAUCUCGUCAUAAAAUAUGUAACUUUGCAGUCCGAUUG
Pba-2-1  AUGUUUUGUAAAAUAAAGCCGCGUUCUCCAUGCCGUGGCCCCAUCUCCUCAUAAAAUAUGUAACUUUGCAGUCCGAUUG
Pba-1-1  UGUUUUGUAAAAUAAAAGCCGCGUUCUCCAUGCCGUGGCCCCAUCUCGUCAUAAAAUAUGUAACUUUGCAGUCCGAUUG
env-450  UGUUUUGUAAUUUGUGUAGUGUGAAAUAUGUCCCGUGAAUCAAACUCGUAAUAAAAUAUGUAACUUUGCAGUCCGAUUG
Pde-2-1  UCAUAAAAGGAAAGUCGUGGCAUGCCCGCGGUGCUGCUACUCUAAUUAAAAUGUGUACUUUUGCAGACCGAUUAUUAAA
Pbe-1-1  CUUGUUAAAACAAAGCUCUGUGGGAGUGUUUUGUUAGUAUGCACGAAAUAAAAUAGGUAAUUUUGCAGACCGAUUGUUU
env-451  UUUCAUAGUCGGAAAUUGAUAGGUGAGGCUUUUACUAUAUAAAAUGUGUACCUUUGCAGUCCGAUUAUUAAAAUUGUUG
env-452  ACAUAAAACAUUGCUUCCGUAGGCUCCUUCGGAGGCUCAUCACCAUAUUAAAUACCUACCUUUGCAGUCCGAUUAUUAU
env-453  AGGUAGAAAACGCAAAUUUUUAUCGAAAAAAGUUGUUGAUUAAUAAAAAAUGCAUACUUUUGCAGUCCGAUUAUUAUUU
env-454  AUUCUUAACGCUUGAUCUCUCCUCUUCUCCUCGGGUUCGUUUUCCCAUAGAAAUACGUACCUUUGCAGUCCGAUUAUUA
env-455  AUUCUUAACGCUUGAUCUCUCCUCUUCUCCUCGGGUUCGUUUUCCCAUAGAAAUACGUACCUUUGCAGUCCGAUUAUUA
env-457  UUCGUAAAAUCUACUUUUCUUUCGCAGUAGGUUGGCUUGUUGCUCUAUAUAAAUAUGUAACUUUGCAGUCCGAUUAUUA
env-458  UUCGUAAAAUCUACUUUUCUUUCACAGUGGGUUGGCUUGUUGCUCUAUAUAAAUAUGUAACUUUGCAGUCCGAUUAUUA
env-462  UUCGUAAAAUCUACUUUUCUUUCGCAGUGGGUUGGCUUGUUGCUCUAUAUAAAUAUGUAACUUUGCAGUCCGAUUAUUA
env-463  UUCGUAAAAUGCACUUUUCUUUCGCAGUGGGUUGGUUUGUUGCUCUAUAUAAAUAUGUAACUUUGCAGUCCGAUUAUUA
env-464  UUCGUAAAAUACACUUUUCUUUCGCAGUGGGUUGGUUUGUUGCUCUAUAUAAAUAUGUAACUUUGCAGUCCGAUUAUUA
env-466  UAUAAAAAUCGCAUUUUAGUAAGCGAAAACGAGCAUUUAAUCACCGGAAAAUGCGUACUUUUGCAGACCGAUUUUAUCC
env-467  UACAUAAAAACAACAUAUUUAACAUCCCGUCAUAUAUUAAAUAAUGAGAAAAAGCAGUACCUUUGCAAUCCGAUUAUUU
env-468  AUAUUAAAUUGUGUGGUUUCUGUGACCAUCAGGAUGCGAUUAUGAAUAAAAAAUAUGUAACUUUGCAGUCCGAUUAUUA
env-469  AUGUUAAAUGUGAUGUCUUGGAAUCUUGGAGAGUGCGUUAAAAGGAUAAAAAUGAGUACCUUUGCAGUCCGAUUAUUAC
env-470  ............................................................UUUGCACACCGAUUAUUUU
Ppa-3-1  UAACUGUAAAUUUAAGGAUUGCGUCAUGCUCUUGCAUCAUGAAGUAAGAAUAAAAUAUGUACCUUUGCACACCGAUUAC
env-471  ...........................................GUAAGAAUAAAAUAUGUACCUUUGCACACCGAUUAC
env-473  UAACUAUAAAUUCAAGGAUUGAGCCAUGCUCUUGCAUCAUGAAGUAAGAAUAAAAUAUGUACCUUUGCACACCGAUUAC
env-474  UAACUGUAAAUUUAAGGAUUGCGUCAUGCUCUUGCAUCAUGAAGUAAGAAUAAAAUAUGUACCUUUGCACACCGAUUAC
env-476  UAACUGUAAAUUUAAGGAUUGCAUCAUGCUCUUGCAUCAUGAAGUAAGAAUAAAAUAUGUACCUUUGCACACCGAUUAC
env-477  ........................CAUGCUCUUGCAUCAUGAAGUAAGAAUAAAAUAUGUACCUUUGCACACCGAUUAC
env-483  AAACUAUAAAAAUCGCUUAUAAAAGCGAAUCGGCGCUUGCGGUAUGAAUAAAAAGUUGUACCUUUGCACCCCGAAACGG
env-485  AAACUAUAAAAAUCGCUUAUAAAAGCGAAUCGGUGUUUGUGGUAUGAAUAAAAAGUUGUACCUUUGCACCCCGAAACGG
env-486  CAUAAUUAUUCGUUUUCAAGGAUUGGUGAUGGCGGGACUCUGAAAGAAAAGCAGUACCUUUGCAGUCCGAUUAUUACCA
env-487  ..........................................................................GAUUU

         ...............................................................................
         ...............................................................................
         ●●●●●●●●●●●●●●●●●●●●●●●●●●●●●●●●●●●●●●●●●●●●●●●●●R●●●●●●●●●●Y●Y●●●●●●●●●●●●●Y●●
```

```
env-488    AAUAAGUAUAAAUCUAUGCAAAUAAUCCGCUUUUACGCUCCACACCUUAUUUAAAAUGUGUACCUUUGCACCUCGAUUU
env-489    CCUUAAUUUGAAACCAUCUAUUCCUCUUUACAUUUCCUCAUAAUAAAUAAAGUGAGUACCUUUGCAGUCCGAUUAUUAU
env-490    UUGUAAAAAACAGGAGGAUUGGUGCUAUAGUUGUGUUUUCACAUCAUUUAAAUUAGUACCUUUGUAGUCCGAUUAUUAU
env-491    UUAUAAAAAAUGGAAGGAUUGGUGCUGUAGUUGUGUUUUCACAUCAUUUAAAUUAGUACCUUUGUAGUCCGAUUAUUAU
env-492    UUAUAAAAAAUAGGAGGAUAGGUGCUGUAGUUGUGUUUUCACAUCAUUUAAAUUAGUACCUUUGUAGUCCGAUUAUUAU
env-493    UCAUAAAAAAUGGAAGGGUUGGUGCUGUAGUUGUGUUUUCACAUCAUUUAAAUUAGUACCUUUGUAGUCCGAUUAUUAU
Pam-1-1    UUAUAAAAAAUGGAAGGAUUGGUGCUGUAGUUGUGUUUUCACAUCAUUUAAAUUAGUACCUUUGUAGUCCGAUUAUUAU
Pam-3-1    UUAUAAAAAAUGGAGGGAUUGGUGCUGUAGUUGUGUUUUCACAUCAUUUAAAUUAGUACCUUUGUAGUCCGAUUAUUAU
Pbi-1-1    AUUUAUAAAAAUACUAUCUCUCAAAAUGCUAUCUCUCUUUUCACAUCAUUAAUUAGUACCUUUGCAGUCCGAUUAUUAU
Pve-1-1    UCGUAAAUAAUACAGAAUGAUUAUAGGAAAAGUAUAUAUAUAAGGUAGGAAAUAAUUACCUUUGCAGUCCGAUUAUUAU
Pve-3-1    UCGUAAAUAACAUAGAAUGAUUAUAGGAAAAGUAUAUAUAUAAGGUAGAAAACAAUUACCUUUGCAGUCCGAUUAUUAU
env-494    UCGUAAAUAACGCACAAUGAUUAUAGGAAAAGUAUAUAUAUAAGGUAGAAAACAAUUACCUUUGCAGUCCGAUUAUUAU
env-495    UCAUAAACAAUACAUAUCUAUGCUAUUCAAACCUUUUAUAUAAGGUUGAAAAUAAGUACCUUUGCAGUCCGAUUAUUAA
env-497    UCAUAAACAACACAUAUCUAUCCUAUUCAAACCUUUUAUAUAAGGUUGAAAAUAAGUACCUUUGCAGUCCGAUUAUUAA
Psc-1-1    AUAUAAACAUUACAUACUUACACGAGGUAACAUUAGUGUAUAAGGUUGAAAAUAAGUACCUUUGCAGUCCGAUUAUUAA
env-498    AUAUAAACAUUACAUACUUACACGAGGUAACAUUAGUGUAUAAGGUUGAAAAUAAGUACCUUUGCAGUCCGAUUAUUAA
Psp-6-1    ACCUAAAAAUCAUACACUUAAACAAGCUAACAUUAUUAUAUAAGGUUGAAAAUAAGUACCUUUGCAGUCCGAUUAUUAA
Pme-3-1    AAUCAUAAACUACAUAUAAUCUUACUAUCCAAAAAAUUAUAUAAGGUUGAAAAAAAGUACCUUUGCAGUCCGAUUAUUA
env-501    UGUAUAAAUUAUUUAUAACGAUGAUUGCCAUUCUUAUAUAUAAGGUUGGAAAUGACUACCUUUGCAGUCCGAUUAUUAU
env-502    UAUAUAAAUUAUUUAUAACGAUGAUUGCCAUUCUUAUAUAUAAGGUUGGAAAUGACUACCUUUGCAGUCCGAUUAUUAU
env-503    UAUAUAAAUUAUUUACAACGGUGAUUGCCAUUCUUAUAUAUAAGGUUGGAAAUGACUACCUUUGCAGUCCGAUUAUUAU
env-504    UAUAUAAAUUAUUUAUAACGAUGAUUGCCAUUCUUAUAUAUAAGGUUGGAAAUGACUACCUUUGCAGUCCGAUUAUUAU
env-507    UAUAUAAAUUAUUUAUAACGAUGAUUGCCAUUCUUAUAUAUAAGGUUGGAAAUGACUACCUUUGCAGUCCGAUUAUUAU
Phi-2-1    UAUAUAAAUUAUUUAUAACGAUGAUUGCCAUUCUUAUAUAUAAGGUUGGAAAUGACUAUCUUUGCAGUCCGAUUAUUAU
env-508    UAUAUAAAUUAUUUAUAACGAUGAUUGCCAUUCUUAUAUAUAAGGUUGGAAAUGACUACCUUUGCAGUCCGAUUAUUAU
env-509    UAUAUAAAUUAUUUAUAACGAUGAUUGCCAUUCUUAUAUAUAAGGUUGGAAAUGACUACCUUUGCAGUCCGAUUAUUAU
Phi-2-2    UAUAUAAAUUAUUUAUAACGAUGAUUGCCAUUCUUAUAUAUAAGGUUGGAAAUGACUACCUUUGCAGUCCGAUUAUUAU
env-510    UCAUAAACUACAUAUAAUCUUACUAUCCAAAACAAUUAUAUAAGGUUGAAAAUAAGUACCUUUGCAGUCCGAUUAUUAA
env-511    UCAUAAACUACAUAUAAUCUUACUAUCCAAAACAAUUAUAUAAGGUUGAAAAUAAGUACCUUUGCAGUCCGAUUAUUAA
env-512    UCAUAAACUACAUAUAAUCUUACUAUCCAAAACAAUUAUAUAAGGUUGAAAAUAAGUACCUUUGCAGUCCGAUUAUUAA
env-514    CAUAAACUACAUACAAUCCUACCAUUCAAAACAAUUAUAUAAGGUUGAAAAUAAGUACCUUUGCAGUCCGAUUAUUAAA
env-516    CAUAAACUACAUACAAUCCUACCAUUCAAAACAAUUAUAUAAGGUUGAAAAUAAGUACCUUUGCAGUCCGAUUAUUAAA
env-517    CCUAAACUACAUACAAUCCUACCAUUCAAAACAAUUAUAUAAGGUUGAAAAUAAGUACCUUUGCAGUCCGAUUAUUAAA
env-519    CAUAAACAACACAUAUCUAUCCUAUUCAAACCUUUUAUAUAAGGUUGAAAAUAAGUACCUUUGCAGUCCGAUUAUUAAA
Psp-5-1    UCAUAAAAAAUACGUAGCUAUGCUAUUCAAACCUUUUAUAUAAGGUUGAAAAUAAGUACCUUUGCAGUCCGAUUAUUAA
env-520    UCAUAAACAAUACGUAGCUAUGCUAUUCAAACCUUUUAUAUAAGGUUGAAAAUAAGUAUCUUUGCAGCCCGAUUAUUAA
env-521    UCAUAAACAAUACAUAUCUAUGCUAUUCAAACCUUUUAUAUAAGGUUGAAAAUAAGUACCUUUGCAGUCCGAUUAUUAA
env-522    UCAUAAACAACACAUAUCUAUGCUAUUCAAACCUUUUAUAUAAGGUUGAAAAUAAGUACCUUUGCAGUCCGAUUAUUAA
env-523    UCAUAAACUACAUAUAAUCUUACUAUCCAAAACAAUUAUAUAAGGUUGAAAAUAAGUACCUUUGCAGUCCGAUUAUUAA
env-525    UCAUAAACUACAUAUAAUCUUACUAUCCAAAACAAUUAUAUAAGGUUGAAAAUAAGUACCUUUGCAGUCCGAUUAUUAA
env-526    NNNNNNNNNNCACAUAAUCCUACUAUCCAAAACAAUUAUAUAAGGUUGAAAAUAAGUACCUUUGCAGUCCGAUUAUUAA
env-527    .............................................................................AA
env-531    UCAUAAACUACACAUAAUCCUACUAUCCAAAACAAUUAUAUAAGGUUGAAAAUAAGUACCUUUGCAGUCCGAUUAUUAA
env-532    UCAUAAACUACAUAUAAUCCUACUAUCCAAAACAAUUAUAUAAGGUUGAAAAUAAGUACCUUUGCAGUCCGAUUAUUAA
env-534    UCAUAAACUACAUAUAAUCUUACUAUCCAAAACAAUUAUAUAAGGUUGAAAAUAAGUACCUUUGCAGUCCGAUUAUUAA
env-535    UCAUAAACUACAUAUAAUCCUACUAUCCAAAACAAUUAUAUAAAGUUGAAAAUAAGUACCUUUGCAGUCCGAUUAUUAA
env-537    UCAUAAACUACAUAUAAUCCUACUAUCCAAAACAAUUAUAUAAGGUUGAAAAUAAGUACCUUUGCAGUCCGAUUAUUAA
env-538    UCAUAAACUACAUAUAAUCUUACUAUCCAAAACAAUUAUAUAAGGUUGAAAAUAAGUACCUUUGCAGUCCGAUUAUUAA
env-552    UCAUAAACUACAUAUAAUCUUACUAUCCAAAACAAUUAUAUAAGGUUGAAAAUAAGUACCUUUGCAGUCCGAUUAUUAA
env-553    UCAUAAACAACACAUAUCUAUCCUAUUCAAACCUUUUAUAUAAGGUUGAAAAUAAGUACCUUUGCAGUCCGAUUAUUAA
env-554    CAUAAACUACAUAUAAUCCUACUAUCCAAAACAAUUAUAUAACGUUGAAAAUAAGUACCUUUGCAGUCCGAUUAUUAAA
env-555    UCAUAAACUACAUAUAAUCUUACUAUCCAAAACAAUUAUAUAAGGUUGAAAAUAAGUACCUUUGCAGUCCGAUUAUUAA
env-557    UCAUAAACUACAUAUAAUCCUACUAUCCAAAACAAUUAUAUAAGGUUAAAAAUAAGUACCUUUGCAGUCCGAUUAUUAA
env-558    UCAUAAACUACAUAUAAUCUUACUAUCCAAAACAAUUAUAUAAGGUUGAAAAUAAGUACCUUUGCAGUCCGAUUAUUAA
env-559    UCAUAAACUACAUAUAAUCUUACUAUCCAAAACAAUUAUAUAAGGUUGAAAAUAAGUACCUUUGCAGUCCGAUUAUUAA
env-560    UCAUAAACUAUAUAUAAUCCUACUACCCAAAACAAUUAUAUAAGGUUGAAAAUAAGUACCUUUGCAGUCCGAUUAUUAA
env-561    UCAUAAACUACAUAUAAUCCUACUAUCCAAAACAAUUAUAUAAGGUUGAAAAUAAGUACCUUUGCAGUCCGAUUAUUAA
Pfu-1-1    AUCAUAAACAACACGCCAUCAAACAUUCAAAUAUAAUUAUAAAUGUAGAAAAUAAGUACCUUUGCAGUCCGAUUAUUAU
Psp-24-1   AAUCGUAAAUAACACAGAAUGGUUAUAGGAAAUCAUAUAUAUAAGGUAGAAAACAAUUACCUUUGCAGUCCGAUUAUUA
env-562    AUCGUAAAUAACACAGAAUGAUUAUAGGAAAAGUAUAUAUAUAAGGUAGAAAACAAUUACCUUUGCAGUCCGAUUAUUA
env-563    AAUCGUAAAUAACACAGAAUGAUUAUAGGAAAUCAUAUAUAUAAGGUAGAAAACAAUUACCUUUGCAGUCCGAUUAUUA
env-564    AAUCGUAAAUAACACAGAAUGAUUAUAGGAAAUCAUAUAUAUAAGGUAGAAAACAAUUACCUUUGCAGUCCGAUUAUUA
Pde-3-1    CGUAAAAAGCAAACGUCGGUUCCUUGCUCCUUAUAGAAUGAGGUUGGAAAUGAGUACCUUUGCAGUCCGAUUAUUAUGG
env-565    CGUAAAAAGCAAACGUCGGUCCCUUGCUCCUUAUAGAAUGAGGUUGGAAAUGAGUACCUUUGCAGUCCGAUUAUUAUGG
Pde-4-1    CGUAAAAAGCAAGCGUCGGUCCCUUGCUCCUUAUAGAAUGAGGUUGGAAAUGAGUACCUUUGCAGUCCGAUUAUUAUGG
Pde-5-1    CGUAAAAAGCAAACGUCGGUCCCUUGCUCCUUAUAGAAUGAGGUUGGAAAUGAGUACCUUUGCAGUCCGAUUAUUAUGG
env-566    CGUAAAAAGCAAACGCCGGUCCCUUGCUCCUUAUAGAAUGAGGUUGGAAAUGAGUACCUUUGCAGUCCGAUUAUUAUGG
env-567    CGUAAAAAGCAAACGCCGGUCCCUUGCUCCUUAUAGAAUGAGGUUGGAAAUGAGUACCUUUGCAGUCCGAUUAUUAUGG
Pde-6-1    CGUAAAAAGCAAACGCCGGUCCCUUGCUCCUUAUAGAAUGAGGUUGGAAAUGAGUACCUUUGCAGUCCGAUUAUUAUGG
env-568    AUCGUAAAUAAUGGGUGCGGAAAACAUCACCCUUAUAAUAUAAGGUUGGAAAUAAGUACCUUUGCAGUCCGAUUAUUAU
env-569    AUCGUAAAUAAUGGGUGCGGAAAACAUCACCCUUAUAAUAUAAGGUUGGAAAUAAGUACCUUUGCAGUCCGAUUAUUAU
Pmu-1-1    AUCGUAAAUAAUGGGUGCGGAAAACAUCACCCUUAUAAUAUAAGGUUGGAAAUAAGUACCUUUGCAGUCCGAUUAUUAU
env-570    UCAUAAACUUAUAAGUUUGGAGAUGGUAUAGAGCAAUCUCUUAUACAUAAAAUGUAUAUCUUUGCAGUCCGAUUAUUUC
env-571    AAUUUAUAAAUCCGUUCGUUUUGUCUCCUGCUGCCCUUGUCUGAUUUAUAAAUAAUGAGUAAUUUUGCAAUCCGAUUGA
env-572    AAUAUAUAAAACGAUUUGUCUUGCAUCCCGUCGUUCUUGUCGGAUUUAUAAAUAAUAGGUAAUUUUGCAAUCCGAUUGA
env-573    AAUAUAUAAAACGAUUUGUCUUGCAUCCCGUCGUUCUUGUCGGAUUUAUAAAUAAUAGGUAAUUUUGCAAUCCGAUUGA
env-574    GUUUAAAACUGUAACACUUGCUUUGUCGGAUACUUCUCUAUGCCGAAUAAAAUGAGUAAUUUUGCAGACCGAUUAUUUC
env-575    ....UAAAUAUAGAUUUUUUGGUUGUGUUUGAUAGUUCAGUCAUCUAUAAAAUAAGUACUUUUGCAUACCGAUUAUUUU
env-576    UUUUUAAAUAUUGGCUUUUUGGUUGUUUUUGAUCGUUUAGUCAUCUGUAAAAUAAGUACUUUUGCAUACCGAUUAUUUU
env-577    ................CCGAGCAGACGCAACAUGCAAGUAUUGUAAGAAAUAUGUGUACCUUUGCAGUCCGAUUAUUUC
env-578    UACGUAAAAGACUAAUUAGCUUUUAAGGUAGUAUUUCCAAAAGUUUCCUAAAUGUGUACUUUUGCGCACCGAUUAUUAU
env-579    UACGUAAAAGACUAAUUAGCUUUUAAGGUAGUAUUUCCAAAAGUUUCCUAAAUGUGUACUUUUGCGCACCGAUUAUUAU
env-580    UACGUAAAAGACUAAUUAGCUUUUAAGGUAGUAUUUCCAAAAGUUUCCUAAAUGUGUACUUUUGCGCACCGAUUAUUAU
env-582    UACGUAAAAGACUAAUUAGCUUUUAAGGUAGUAUUUCCAAAAGUUUCCUAAAUGUGUACUUUUGCGCACCGAUUAUUAU

           ...............................................................................
           ...............................................................................
           ●●●●●●●●●●●●●●●●●●●●●●●●●●●●●●●●●●●●●●●●●●●●●●●●●R●●●●●●●●●●Y●Y●●●●●●●●●●●●●Y●●
```

```
Pin-1-1   AUAAUUAAUAUCAAUGCUUUCACGCUGUUAUGCGUUUCGUACCUAAUAAAAAAUGAGUAAUUUUGCAGACCGAUUAUUU
env-583   AUGAUUAAGAUAUAAGCAUCUACGGAUUUAUCCACUUUGUAAAUAAUAAAAAAUGAGUAAUUUUGCGAACCGAUUAUUU
env-589   AUGAUUAAGAUAUAAGCAUCUACGGAUUUAUCCACUUUGUAAAUAAUAAAAAAUGAGUAAUUUUGCGAACCGAUUAUUU
env-592   AUAAUUAAGAUAUAAGCGUCUACGGAUUUAUCCACUUUGUAAAUAAUAAAAAAUGAGUAAUUUUGCGAACCGAUUAUUU
env-593   AUAAUUAAGAUAUAAGCGUCUACGGAUUUAUCCACUUUGUAAAUAAUAAAAAAUGAGUAAUUUUGCGAACCGAUUAUUU
env-594   AUAAUUAAGAUAUAAGCGUCUGUGGAUUUAUCCACUUUGUAAAUAAUAAAAAAUGAGUAAUUUUGCGCACCGAUUAUUU
env-595   AUAAUUAAGAUAUAAGCGUCUACGGAUUUAUCCACUUUGUAAAUAAUAAAAAAUGAGUAAUUUUGCGAACCGAUUAUUU
env-596   AUGAUUAAGAUAUAAGCAUCUACGGAUUUAUCCACUUUGUAAAUAAUAAAAAAUGAGUAAUUUUGCGAACCGAUUAUUU
env-597   AUAAUUAAGAUAUAAGCAUCUACGGAUUUAUCCACUUUGUAAAUAAUAAAAAAUGAGUAAUUUUGCGAACCGAUUAUUU
env-598   AUAAUUAAGAUAUAAGCAUCUACGGAUUUAUCCACUUUGUAAAUAAUAAAAAAUGAGUAAUUUUGCGAACCGAUUAUUU
env-599   AUAAUUAAGAUAUAAGCGUCUACGGAUUUAUCCACUUUGUAAAUAAUAAAAAAUGAGUAAUUUUGCGAACCGAUUAUUU
env-600   AUAAUUAAGAUAUAAGCAUCUGCGGAUUUAUCCACUUUGUAAAUAAUAAAAAAUGAGUAAUUUUGCGAACCGAUUAUUU
env-613   AUAAUUAAGAUAUAAGCAUCUACGGAUUUAUCCACUUUGUAAAUAAUAAAAAAUGAGUAAUUUUGCGAACCGAUUAUUU
env-648   AUAAUUAAGAUAUAAGCGUCUGUGGAUUUAUCCACUUUGUAAAUAAUAAAAAAUGAGUAAUUUUGCGCACCGAUUAUUU
env-651   AUGAUUAAGAUAUAAGCAUCUGCGGAUUUAUCCACUUUGUAAAUAAUAAAAAAUGAGUAAUUUUGCGAACCGAUUAUUU
env-652   AUAAUUAAGAUAUAAGCAUCUGCGGAUUUAUCCACUUUGUAAAUAAUAAAAAAUGAGUAAUUUUGCGAACCGAUUAUUU
env-653   AUAAUUAAGAUAUAAGCAUCUACGGAUUUAUCCACUUUGUAAAUAAUAAAAAAUGAGUAAUUUUGCGAACCGAUUAUUU
env-657   AUAAUUAAGAUAUAAGCGUCUACGGAUUUAUCCACUUUGUAAAUAAUAAAAAAUGAGUAAUUUUGCGAACCGAUUAUUU
env-676   AUAAUAAAUGAACGGAAUUAAUCCCUGACGAUAGUGAUUCUGGUAUAUAAAAUGAGUAAUUUUGCAGACCGAUUAUUUU
Pco-3-1   AUAAUAAAUGAACGGAAUUAAUCCCUGACGAUAGUGAUUCUGGUAUAUAAAAUGAGUAAUUUUGCAGACCGAUUAUUUU
Pni-1-1   AUAAUUAAUAAGCAGACUUUCGCAGAGUUAUACACUUUGUAACCAAUAAAAAAUGAGUAAUUUUGCAGACCGAUUAUUC
env-677   AUGAUUAAUAAGCAAACUUUCGCAGAGUUAUACACUUUGUAACCAAUAAAAAAUGAGUAAUUUUGCAGACCGAUUAUUC
env-679   AUGAUUAAUAAGUAAACUUUCGCAGAGUUAUACACUUUGUAACCAAUAAAAAAUGAGUAAUUUUGCAGACCGAUUAUUC
env-680   AUGAUUAAUAAGUAAACUUUCGCAGAGUUAUACACUUUGUAACCAAUAAAAAAUGAGUAAUUUUGCAGACCGAUUAUUC
env-681   AUAAUUAAUAAGCAAACUUUCGCAGAGUUAUACACUUUGUAACCAAUAAAAAAUGAGUAAUUUUGCAGACCGAUUAUUC
env-682   AUGAUUAAUAAGUAGAUUUUCGCAGAGUUAUACACUUUGUAACCAAUAAAAAAUGAGUAAUUUUGCAGACCGAUUAUUC
env-683   AUGAUUAAUAAGUAGACUUUCGCAGAGUUAUACACUUUGUAACCAAUAAAAAAUGAGUAAUUUUGCAGACCGAUUAUUC
env-684   AUGAUUAAUAAGCAAACUUUCGCAGAGUUAUACACUUUGUAACCAAUAAAAAAUGAGUAAUUUUGCAGACCGAUUAUUC
env-686   AUAAUUAAUAAGCAGACUUUCGCAGAGUUAUACACUUUGUAACCAAUAAAAAAUGAGUAAUUUUGCAGACCGAUUAUUC
env-696   AUAAUUAAUAAGCAAACUUUCGCAGAGUUAUACACUUUGUAACCAAUAAAAAAUGAGUAAUUUUGCAGACCGAUUAUUC
env-698   AUGAUUAAUAAGCAAACUUUCGCAGAGUUAUACACUUUGUAACCAAUAAAAAAUGAGUAAUUUUGCAGACCGAUUAUUC
env-699   AUGAUUAAUAAGCAAACUUUCGCAGAGUUAUACACUUUGUAACCAAUAAAAAAUGAGUAAUUUUGCAGACCGAUUAUUC
Pni-3-1   AUGAUUAAUAAGCAGACUUUCGCAGAGUUAUACACUUUGUAACCAAUAAAAAAUGAGUAAUUUUGCAGACCGAUUAUUC
env-700   AUGAUUAAUAAGUAAACUUUCGCAGAGUUAUACACUUUGUAACCAAUAAAAAAUGAGUAAUUUUGCAGACCGAUUAUUC
env-709   AUAUUUAAUACAAAUACUUUCACGCGGCUACACAUUUCGUAGCUGAUAAAAAAUGAGUAAUUUUGCAGACCGAUUAUUU
env-710   UAUUUAAUACAAAUACUUUCAUGCGGCUACACAUUUCGUAGCUGAUAAAAAAAUGAGUAAUUUUGCAGACCGAUUAUUU
env-711   AUAUUUAAUACAAAUACUUUCAUGCGGCUACACAUUUCGUAGCUGAUAAAAAAUGAGUAAUUUUGCAGACCGAUUAUUU
env-724   ................................................AAAAAAUGAGUAAUUUUGCAGACCGAUUAUUU
Pau-1-1   AUAUUUAAUACAAAUACUUUCACGCGGCUACACAUUUCGUAGCUGAUAAAAAAUGAGUAAUUUUGCAGACCGAUUAUUU
Psp-9-1   UAUCUAAAAACGGCAAAUUCGGAUAUAAGGGAGAAUGGCAACAUAAAUAAAAAAUGAGUAAUUUUGCAGACCGAUUAUUU
env-725   AUAAUUAAUACCAACGCUUUCACGCUGUUAUGCGUUUCGUACCUAAUAAAAAAUGAGUAAUUUUGCAGACCGAUUAUUU
env-726   AUAAUUAAUACCAACGCUUUCACGCUGUUAUGCGUUUCGUACCUAAUAAAAAAUGAGUAAUUUUGCAGACCGAUUAUUU
env-727   AUAAUUAAUAUCAAUACUUUCACACUGUUAUGCGUUUCGUACCUAAUAAAAAAUGAGUAAUUUUGCAGACCGAUUAUUU
env-729   AUAAUUAAUACCAACGCUUUCACGCUGUUAUGCGUUUCGUACCUAAUAAAAAAUGAGUAAUUUUGCAGACCGAUUAUUU
env-733   AUAAUUAAUAUCAACGCUUUCACGCUGUUAUGCGUUUCGUACCUAAUAAAAAAUGAGUAAUUUUGCAGACCGAUUAUUU
env-734   AUAAUUAAUACCAACGCUUUCACGCUGUUAUGCGUUUCGUACCUAAUAAAAAAUGAGUAAUUUUGCAGACCGAUUAUUU
env-735   AUGAUUAAUACCAACGCUUUCACGCUGUUAUGCGUUUCGUACCUAAUAAAAAAUGAGUAAUUUUGCAGACCGAUUAUUU
env-736   AUAAUUAAUACCAACACUUUCACGCUGUUAUGCGUUUCAUACCUAAUAAAAAAUGAGUAAUUUUGCAGACCGAUUAUUU
Pin-3-1   AUAAUUAAUACCAAUGCUUUCGCGCUGUUAUGCGUUUCGUACCUAAUAAAAAAUGAGUAAUUUUGCAGACCGAUUAUUU
Pfa-1-1   AUAAUUAAUACUAACACUUUCGCACUGUUAUGCGUUUUGUAGCUAAUAAAAAAUGAGUAAUUUUGCAGACCGAUUAUUU
env-737   AUAAUUAAUACCAACGCUUUCACGCUGUUAUGCGUUUCGUACCUAAUAAAAAAUGAGUAAUUUUGCAGACCGAUUAUUU
env-738   UAAUUAAUACCAACGCUUUCACGCUGUUAUGCGUUUCGUACCUAAAUAAAAAAUGAGUAAUUUUGCAGACCGAUUAUUU
env-742   AUAAUUAAUACCAACGCUUUCACGCUGUUAUGCGUUUCGUACCUAAUAAAAAAUGAGUAAUUUUGCAGACCGAUUAUUU
Pdi-3-1   AUUUGUAAAUAAUUCCCUUUGGCGGCUGCAUCGCAACUCUACCUAAUAAAAAAUGAGUAAUUUUGCAGACCGAUUAUUU
env-743   AUUUGUAAAUAAUUCCCUUUGGCGGCUGCAUCGCAACUCUACCUAAUAAAAAAUGAGUAAUUUUGCAGACCGAUUAUUU
Pdi-2-1   AUUUGUAAAUAAUUACCUUUGGCGGCUGCAUCGCAACUCUACCUAAUAAAAAAUGAGUAAUUUUGCAGACCGAUUAUUU
Pdi-1-1   AUUUGUAAAUAAUUCCCUUUGGCGGCUGCAUCGCAACUCUACCUAAUAAAAAAUGAGUAAUUUUGCAGACCGAUUAUUU
env-744   AUUAUAAAAGAUAGGGGGCUGCGGCUCGUAACUUACACAUAAUCAAAUAAAAAUGUGUAAUUUUGCAGUCCGAUUAUUU
env-745   AAUUAUAAAAGAUAAGGACUACGGAGCGUAACUUACACAUAAUCAAAUAAAAAUGUGUAAUUUUGCAGUCCGAUUAUUU
env-746   AAUUAUAAAAGAUAAGGACUGCGGCUCGUAACUUGCACAUAAUCAAAUAAAAAUGUGUAAUUUUGCAGUCCGAUUAUUU
env-747   UAUUAUAAAAGAUAGGGGCUGCGGCUCGUAACUUACACAUAAUCAAAUAAAAAUGUGUAAUUUUGCAGUCCGAUUAUUU
env-748   AAUUAUAAAAGAUAGGGGCUGCGACUCGUAACUUACACAUAAUCAAAUAAAAAUGUGUAAUUUUGCAGUCCGAUUAUUU
env-749   AAUUAUAAAAGAUAGGGGCUGCGGUUCGUAACUUACACAUAAUCAAAUAAAAAUGUGUAAUUUUGCAGUCCGAUUAUUU
env-751   UAUUAUAAAAGAUAGGGACUGCGGCUCGUAACUUACACAUAAUCAAAUAAAAAUGUGUAAUUUUGCAGUCCGAUUAUUU
env-752   UAUUAUAAAAGAUAGGGGCUGCGGCUCGUAACUUACACAUAAUCAAAUAAAAAUGUGUAAUUUUGCAGUCCGAUUAUUU
env-753   AAUUAUAAAAGAUAAGGACUACGGAGCGUAACUUACACAUAAUCAAAUAAAAAUGUGUAAUUUUGCAGUCCGAUUAUUU
env-754   AAUUAUAAAAGAUAAGGACUGCGGCUCGUAACUUACACAUAAUCAAAUAAAAAUGUGUAAUUUUGCAGUCCGAUUAUUU
env-755   AAUCAUAAAAGAUGGGGGCUGUGACUCGUAACUUACACAUAAUCAAAUAAAAAUGUGUAAUUUUGCAGUCCGAUUAUUU
env-756   UAUUAUAAAAGAUAGGGACUGCGGCUCGUAACUUACACAUAAUCAAAUAAAAAUGUGUAAUUUUGCAGUCCGAUUAUUU
env-757   AACUAUAAAAGAUAAGGAGGUGGAAUCGUAACCUACACAUAAUCAAAUAAAAAUGUGUAAUUUUGCAGUCCGAUUAUUU
env-758   AUUCAUAAAAGAUGGGGUGCAGGACUCGUAACAUUCACAUAAUCAAAUAAAAAUGUGUAAUUUUGCAGUCCGAUUAUUU
env-759   UUCAUAAAAGUUUGGGGGUUAAAACUCGUAACCUACACAUAAUCAAAUAAAAAUGUGUAAUUUUGCAGUCCGAUUAUUU
env-760   AAUUAUAAAAGAUAGGGGCUGCGGCUCGUAACUUACACAUAAUCAAAUAAAAAUGUGUAAUUUUGCAGUCCGAUUAUUU
env-761   AUUUAUAAAAAUUUUGGUUGAAAACUCGUAACUUACACAUAAUUAAAUUUAAAUGUGUACUUUUGCAGUCCGAUUAUAU
env-762   AUUUAUAAAAGUUUUGAUUGAAAACUCGUAACUUACACAUAAUCAAAUAAAAAUGUGUAAUUUUGCAGUCCGAUUAUUU
env-763   AUUCAUAAAAGUUUGGGGUUAAAACUCGUAACUUACACAUAAUUAAAUAAAAAUGUGUAAUUUUGCAGUCCGAUUAUUU
env-765   GCAGCUGAUAGGAUAUGUGCAGUAGCCUUGCUAUGUGGCGUGAAAAGAUGUNNNNNNNNNNNNUGCAGUCCGAUUAUUU
env-766   AUUUAUAAAAGUUUUGAUUGAAAACUCGUAACCUACACAUAAUCAAAUAAAAAUGUGUAAUUUUGCAGUCCGAUUAUUU
env-767   AUUCAUAAAAGUUUGGGGUUAAAACUCGUAACAUACACAUAAUUAAAUAAAAAUGUGUAAUUUUGCAGUCCGAUUAUUU
env-768   UCUAUAAAAGUAAAACACUUGAGUUUGUAACUUACACAUAAUUAAAUAAAAAUGUGUACUUUUGCAGUCCGAUUAUUUA
env-769   UCCGUAAAAAUAAUACUUGAGGACUGCAGAGUUACACAUAAUUAAAUAAAAAUGUGUACUUUUGCAGUCCGAUUAUUUG
env-770   UCUAUAAAAGUAAAGCACUUGAGUUUGUAACUUACACAUAAUUAAAUAAAAAUGUGUACUUUUGCAGUCCGAUUAUUUA
env-771   UCUAUAAAAGUAAAACACUUGAGUUUGUAACUUACACAUAAUUAAAUAAAAAUGUGUACUUUUGCAGUCCAAUUAUUUA
env-772   UCUAUAAAAGUAAAACACUUGAGUUUGUAACUUACACAUAAUUAAAUAAAAAUGUGUACUUUUGCAGUCCGAUUAUUUG

          ................................................................................
          ................................................................................
          ●●●●●●●●●●●●●●●●●●●●●●●●●●●●●●●●●●●●●●●●●●●●●●●●●●R●●●●●●●●●●Y●Y●●●●●●●●●●●●●Y●●
```

```
env-774   AAUUAUAAAAAUUUUGGUUGAAACUUCGUAACUUACACAUAAUGAAAUAAAAAGGGUACUGUUGCAGUCCGAUUAUUUG
env-775   UCUAUAAAAGUAAAACGCUUGAGUUUGUAACUUACACAUAAUUAAAUAAAAAUGUGUACUUUUGCAGUCCGAUUAUUUG
env-776   UCUAUAAAAGUAAAACGCUUGAGUUUGUAACUUACACAUAAUUAAAUAAAAAUGUGUACUUUUGCAGUCCGAUUAUUUG
env-780   UUUAUAAAAAUUUUGGUUGAAAACUCGUAACUUACACAUAAUUAAAUAAAAAUGUGUACUUUUGCAGUCCGAUUAUUUG
env-781   AUUUAUAAAAGUUUGGGGUUAAAACUCGUAACCUACACAUAAUCAAAUAAAAAUGUGUAAUUUUGCAGUCCGAUUAUUU
env-785   AUUCAUAAAAGUUUGGGGUUAAAACUCGUAACCUACACAUAAUCAAAUAAAAAUGUGUAAUUUUGCAGUCCGAUUAUUU
env-787   AUUCAUAAAAGUUUGGGAUUAAAACUCGUAACCUACACAUAAUCAAAUAAAAAUGUGUAAUUUUGCAGUCCGAUUAUUU
env-788   AUUUAUAAAAGUUUGGGGUUAAAACUCCUAACCUACACAUAAUCAAAUAAAAAUGUGUAAUUUUGCAGUCCGAUUAUUU
env-789   AUUCAUAAAAGUUURGGGUUAAAACUCGUAACCUACACAUAAUCAAAUAAAAAUGUGUAAUUUUGCAGUCCGAUUAUUU
env-790   AUUYAUAAAAGUUUAGGGUUAAAACUCGUAACCUACACAUAAUCAAAUAAAAAUGUGUAAUUUUGCAGUCCGAUUAUUU
env-791   AUUCAUAAAAGUUUGGGGUUAAAACUCGUAACCUACACAUAAUCAAAUAAAAAUGUGUAAUUUUGCAGUCCGAUUAUUU
env-793   AUUUAUAAAAGUUUAGGGUUAAAACUCGUAACCUACACAUAAUCAAAUAAAAAUGUGUAAUUUUGCAGUCCGAUUAUUU
env-797   AUUUAUAAAAGUUUAGGGUUAAAACUCGUAACCUACACAUAAUCAAAUAAAAAUGUGUAAUUUUGCAGUCCGAUUAUUU
env-798   AUUUAUAAAAGUUUAGGGUUAAAACUCGUAACCUACACAUAAUCAAAUAAAAAUGUGUAAUUUUGCAGUCCGAUUAUUU
env-799   AUUCAUAAAAGAUGGGGUGCAAGACUCGUAACCUACACAUAAUCAAAUAAAAAUGUGUAAUUUUGCAGUCCGAUUAUUU
env-800   .CAUAAAAAGUUUGGGGUUAAAAACUCGUAACCUACACAUAAUCAAAUAAAAAUGUGUAAUUUUGCAGUCCGAUUAUUU
env-801   NNNNNNNNNNNUUUAGGGUUAAAACUCGUAACCUACACAUAAUCAAAUAAAAAUGUGUAAUUUUGCAGUCCGAUUAUUU
env-802   UUUAUAAAAAGUUUAGGGUUAAAACUCGUAACCUACACAUAAUCAAAUAAAAAUGUGUAAUUUUGCAGUCCGAUUAUUU
env-803   AUUUAUAAAAGUUUAGGGUUAAAACUCGUAACCUACACAUAAUCAAAUAAAAAUGUGUAAUUUUGCAGUCCGAUUAUUU
env-804   AUUCAUAAAAGUUUGGGAUUAAAACUCGUAACCUACACAUAAUCAAAUAAAAAUGUGUAAUUUUGCAGUCCGAUUAUUU
env-808   AUUCAUAAAAGUUUGGGGUUAAAACUCGUAACCUACACAUAAUCAAAUAAAAAUGUGUAAUUUUGCAGUCCGAUUAUUU
env-821   UUYAUAAAAGUUURGGGUUAAAAACUCGUAACCUACACAUAAUCAAAUAAAAAUGUGUAAUUUUGCARUCCGAUUAUUU
env-822   AUUUAUAAAAAUUCAGGGUUAAAACUCGUAACCUACACAUAAUCAAAUAAAAAUGUGUAAUUUUGCAGUCCGAUUAUUU
env-824   AUUUAUAAAAGUUUAGGGUUAAAACUCGUAACCUACACAUAAUCAAAUAAAAAUGUGUAAUUUUGCAGUCCGAUUAUUU
env-825   AUUUAUAAAAGUUUAGGGUUAAAACUCGUAACCUACACAUAAUCAAAUAAAAAUGUGUAAUUUUGCAGUCCGAUUAUUU
env-826   AUUCAUAAAAGUUURGGGUUAAAACUCGUAACCUACACAUAAUCAAAUAAAAAUGUGUAAUUUUGCAGUCCGAUUAUUU
env-827   AUUCAUAAAAGUUUGGGGUUAAAAAUCGUAACCUACACAUAAUCAAAGAAAAAUGUGUAAUUUUGCAGUCCGAUUAUUU
env-828   ....................................ACAUAAUCAAAUAAAAAUGUGUAAUUUUGCAGUCCGAUUAUUU
env-829   AUUCAUAAAAGUUUGGGGUUAAAACUCGUAACCUACACAUAAUCAAAUAAAAAUGUGUAAUUUUGCAGUCCGAUUAUUU
env-830   UCAUAAAAGUUUGGGGUUAAAACUCGUAACCUACACAUAAUCAAAUAAAAAAUGUGUAAUUUUGCARUCCGAUUAUUUC
env-831   AUUCAUAAAAGUUUGAGAUUAAAACUCGUAACCUACACAUAAUCAAAUAAAAAUGUGUAAUUUUGCAGUCCGAUUAUUU
env-832   ......................................................GUGUAAUUUUGCAGUCCGAUUAUUU
env-833   AAUAUUCAUAAGUUUAGGGUUAAAACUCGUAACCUACACAUAAUCAAAUAAAAUGUGUAAUUUUGCAGUCCGAUUAUUU
env-834   AUUUAUAAAAUACCUGCUUUUAAGUUUGUAACCUACACAAAAUCAAAUAAAAAUGUGUAAUUUUGCAGUCCGAUUAUUU
env-835   AUUCAUAAAAGUUUGGGGUUAAAACUCGUAACCUACACAUAAUCAAAAAAAUAUGUGUAAUUUUGCAGUCCGAUUAUUU
env-836   UUUCAUAAAAGUUUGGGUUUAAAACUCGUAACCUACACAUAAUCAAAUAAAAAUGUGUAAUUUUGCAGUCCGAUUAUUU
env-837   UUUUAUAAAAGUUUGGGUUUAAAACUCGUAACCUACACAUAAUCAAAUAAAAAUGUGUAAUUUUGCAGUCCGAUUAUUU
Pco-2-1   UUUUAUAAAAGUUUGGGUUUGAAACUCGUAACCUACACAUAAUCAAAUAAAAAUGUGUAACUUUGCAGUCCGAUUAUUU
env-838   UUUAUAAAAGUUUGGGGUUAAGACUCGUAACCUACACAUAAUCAAAUAAAAAAUGUGUAAUUUUGCAGUCCGAUUAUUU
env-839   AUUUAUAAAAGUUUGGGAUUAAAACUCGUAACCUACACAUAAUCAAAUAAAAAUGUGUAAUUUUGCAGUCCGAUUAUUU
env-840   AUUCAUAAAAGUUUGGGGUUAAGACUCGUAACCUACACAUAAUCAAAUAAAAAUGUGUAAUUUUGCAGUCCGAUUAUUU
env-842   AUUCAUAAAAGUUUGGGGUUAAAACUCGUAACCUACACAUAAUCAAAAAAAUAUGUGUAAUUUUGCAGUCCGAUUAUUU
env-843   AUUUAUAAAAGUUUGGGGUUAAAACUCGUAACCUACACAUAAUCAAAUAAAAAUGUGUAAUUUUGCAGUCCGAUUAUUU
Por-8-1   AUGAUUAAAUAGAUAAUUUUAAUCAUUUCUUAUUUUCCAACUUACGAUUAUACUUCGUACCUUUGCAGUCCGAUUAUUU
Psp-21-1  AAUAAUAAAAUAGGAGAUUUGUAAUAGCUCUGAUUAUACCAUGUAAAUAAAACGUCGUAACUUUGCAGUCCGAUUAUUU
env-845   NNNNNNNNNNNNNNNNNNNNUGCUCCCAUGCUGUCUUCGCCAUUCGGCAGAAUCUUCGUACCUUUGCAGUCCGAUUAUU
env-846   AAUAUUAAAUCAACACCUUUUACCCCCAUGCUAUCUUCGCCAUUCGGCAGAAUCUUCGUACCUUUGCAGUCCGAUUAUU
env-847   GUUAAAUGUUUAGCCAAAUUACGCAUUACGCAUUACGCUUUACGCAUUAAAAUCGUACCUUUGCAGUCCGAUUAUUACA
env-848   ACGUAACAACAAAGUUAAAUAGCUGUCGCACUUAUUAGUUAAAGCAUAUUAAUAUGUACCUUUGCAGUCCGAUUAUUAU
env-849   ACGUAACAACAAAGUUAAAUAGCUGUCGCACUUAUUAGUUAAAGCAUAUUAAUAUGUACCUUUGCAGUCCGAUUAUUAU
env-851   GCGUAACGACAAAGUUAAAUAGCUGUCGCACUUAUUAGUUAAAGCAUAUUAAUAUGUACCUUUGCAGUCCGAUUAUUAU
env-852   ACGUAACGACAAAGUUAAAUAGCCAUCGUACUUAUUAGUUAAAGCAUAUUAAUAUGUACCUUUGCAGUCCGAUUAUUAU
env-855   ACGUAACGGUAAAGUUAAAUAGCUGUCGCACUUAUUAGUUAAAGCAUAUUAAUAUGUACCUUUGCAGUCCGAUUAUUAU
env-856   ACAUAACGAUAAAGUUAAAUAGCUGUCGCACUUAUUAGUUAAAGCAUAUUAAUACGUACCUUUGCAGUCCGAUUAUUAU
env-857   ACAUAACGAUAAAGUUAAAUAGCUGUCGCACUUAUUAGUUAAAGCAUAUUAAUAUGUACCUUUGCAGUCCGAUUAUUAU
env-858   ACAUAACGAUAAAGUUAAAUAGCUGUCGCACUUAUUAGUUAAAGCAUAUUAAUACGUACCUUUGCAGUCCGAUUAUUAU
env-859   ACGUAACGACAAAGUUAAAUAGCUGUCGCACUUAUUAGUUAAAGCAUAUUAAUAUGUACCUUUGCAGUCCGAUUAUUAU
env-860   ACAUAACGACAAGGUUAAAUAUGCGCCGCAUCUAUUAGUUAAAGCAUAUUAAUAUGUACCUUUGCAGUCCGAUUAUUAU
env-861   GCGUAACGACAAAGUUAAAUAGCUGUCGCACUUAUUAGUUAAAGCAUAUUAAUAUGUACCUUUGCAGUCCGAUUAUUAU
env-862   ACAUAACGGCAAAGUUAAAUAGCAAUCGCACUUAUUAGUUAAAGCAUAUUAAUAUGUACCUUUGCAGUCCGAUUAUUAU
env-864   ACGUAACGGCAAAGUUAAAUAUGCGCCGCAUUUAUUAGUUAAAGCAUAUUAAUAUGUACCUUUGCAGUCCGAUUAUUAU
env-865   ACGUAAUGACAAAGUUAAAUAUGCGCCGCAUUUAUUAGUUAAAGCAUAUUAAUAUGUACCUUUGCAGUCCGAUUAUUAU
env-866   ACAUAACGAUAAGGUUAAAUAGCUGUCGCACUUAUUAGUUAAAGCAUAUUAAUAUGUACCUUUGCAGUCCGAUUAUUAU
env-868   ACAUAACGAUAAGGUUAAAUAGCUGUCGCUCUUAUUAGUUAAAGCAUAUUAAUAUGUACCUUUGCAGUCCGAUUAUUAU
env-869   ACGUAACGACAAGGUUAAAUAUGCGCCGCAUUUAUUAGUUAAAGCAUAUUAAUAUGUACCUUUGCAGUCCGAUUAUUAU
env-870   ACAUAACGGCAAAGUUAAAUAUGCGCCGCAUUUAUUAGUUAAAGCAUAUUAAUAUGUACCUUUGCAGUCCGAUUAUUAU
env-872   AUUUAUCGUGNNNNNNNNNNNNNNNNNNNNNNNNNNNNNNNNNNNNNNNNNAAUACGUACCUUUGCAGUCCGAUUAUUAU
env-873   ACAUAACGGUAAAGUUAAAUAGCCAUCGCACUUAUUAGUUAAAGCAUAUUAAUACGUACCUUUGCAGUCCGAUUAUUAU
env-874   AUAUAACGACAAAGUUAAAUAGCUGUCGCACUUAUUAGUUAAAGCAUAUUAAUAUGUACCUUUGCAGUCCGAUUAUUAU
env-875   ACAUAACGGCAAGGUUAAAUAGCUGUCGCACUUAUUAGUUAAAGCAUAUUAAUAUGUACCUUUGCAGUCCGAUUAUUAU
env-876   ACAUAACGGCAAGGUUAAAUAGCUGUCGCACUUAUUAGUUAAAGCAUAUUAAUAUGUACCUUUGCAGUCCGAUUAUUAU
Psp-25-1  ACGUAACGAUAAAGUUAAAUAGCUGUCGCACUUAUUAGUUAAAGCAUAUUAAUACGUACCUUUGCAGUCCGAUUAUUAU
env-877   ACAUAACGACAAAGUUAAAUAGCUGUCGCACUUAUUAGUUAAAGCAUAUUAAUAUGUACCUUUGCAGUCCGAUUAUUAU
env-878   ACAUAAUGGCAAAGUUAAAUAGCAAUCGCACUUAUUAGUUAAAGCAUAUUAAUAUGUACCUUUGCAGUCCGAUUAUUAU
env-880   ACAUAACGACAAAGUUAAAUAGCCAUCGUACUUAUUAGUUAAAGCAUAUUAAUAUGUACCUUUGCAGUCCGAUUAUUAU
env-881   ACGUAACGAUAAAGUUAAAUAGCUGUCGCACUUAUUAGUUAAAGCAUAUUAAUACGUACCUUUGCAGUCCGAUUAUUAU
env-882   ACAUAACGGCAAAGUUAAAUAGCUGUCGCACUUAUUAGUUAAAGCAUAUUAAUAUGUACCUUUGCAGUCCGAUUAUUAU
env-883   ACGUAACGAUAAAGUUAAAUAGCUGUCGCACUUAUUAGUUAAAGCAUAUUAAUAUGUACCUUUGCAGUCCGAUUAUUAU
env-885   ACAUAACGGUAAAGUUAAAUAGCCAUCGCACUUAUUAGUUAAAGCAUAUUAAUACGUACCUUUGCAGUCCGAUUAUUAU
env-886   AGCGUAACGACAAAGUUAAAUAGCUGUCGCACUUAUUAGUUAAAGCACAUUAAUUCGUACCUUUGCAGUCCGAUUAUUA
env-887   UAUAUAACGCUAAGGUUAAAUAUGCGAUGCGCUUAUUAGUUAAAGCAUAUUAAUGCGUACCUUUGCAGUCCGAUUAUUA
env-888   UAUAUAACGCUAAGGUUAAAUAUGCGAUGCACUUAUUAGUUAAAGCAUAUUAAUGCGUACCUUUGCAGUCCGAUUAUUA
env-889   UAUAUAACUCUAAGGUUAAAUAUGCGAUGCGCUUAUUAGUUAAAGCAUAUUAAUGCGUACCUUUGCAGUCCGAUUAUUA
env-890   UAUAUAACGCUAAGGUUAAAUAUGCGAUGCGCUUAUUAGUUAAAGCAUAUUAAUGCGUACCUUUGCAGUCCGAUUAUUA

          ...............................................................................
          ...............................................................................
          ●●●●●●●●●●●●●●●●●●●●●●●●●●●●●●●●●●●●●●●●●●●●●●●●●R●●●●●●●●●●Y●Y●●●●●●●●●●●●●Y●●
```

```
env-891    UAUAUAACUCUAAGGUUAAAUAUGCGAUGCGUUUAUUAGUUAAAGCAUAUUAAUGCGUACCUUUGCAGUCCGAUUAUUA
env-892    UAUAUAAUGCUAAGGUUAAAUAUGCGAUGCGUUUAUUAGUUAAAGCAUAUUAAUGCGUACCUUUGCAGUCCGAUUAUUA
env-893    UAUAUAACGCUAAGGUUAAAUAUGCGAUAUGCUUAUUAGUUAAAGCAUAUUAAUGCGUACCUUUGCAGUCCGAUUAUUA
env-894    UAUAUAACGCUAAGGUUAAAUAUGCGAUGCGUUUAUUAGUUAAAGCAUAUUAAUGCGUACCUUUGCAGUCCGAUUAUUA
env-900    UAUAUAACGCUAAGGUUAAAUAUGCGAUGCGUUUAUUCGUUAAAGCAUAUUAAUGCGUACCUUUGCAGUCCGAUUAUUA
env-901    UAUAUAACGCUAAGGUUAAAUAUGCGAUGCGUUUAUUCGUUAAAGCAUAUUAAUGCGUACCUUUGCAGUCCGAUUAUUA
env-902    UAUAUAACGCUAAGGUUAAAUAUGCGAUGCGUUUAUUCGUUAAAGCAUAUUAAUGCGUACCUUUGCAGUCCGAUUAUUA
Psp-26-1   UAUAUAACGCUAAGGUUAAAUAUGCGAUGCGUUUAUUAGUUAAAGCAUAUUAAUGUGUACCUUUGCAGUCCGAUUAUUA
env-903    UAUAUAACGCUAAGGUUAAAUAUGCGAUGCGCUUAUUAGUUAAAGCAUAUUAAUGCGUACCUUUGCAGUCCGAUUAUUA
env-921    AACGUAACGACAAAGUUAAAUAGCUGUCGUACUUAUUAGUUAAAGCACAUUAAUUCGUACCUUUGCAGUCCGAUUAUUA
env-922    AGCGUAACGACAAAGUUAAAUAACUGUCGCACUUAUUAGUUAAAGCACAUUAAUUCGUACCUUUGCAGUCCGAUUAUUA
env-923    AACGUAACAACAAAGUUAAAUAGCUGUCGCACUUAUUAGUUAAAGCACAUUAAUUCGUACCUUUGCAGUCCGAUUAUUA
env-924    AGCGUAACGACAAAGUUAAAUAGCUGUCGCACUUAUUAGUUAAAGCACAUUAAUUCGUACCUUUGCAGUCCGAUUAUUA
env-932    AACGUAACGACAAAGUUAAAUAGCUGUCGCACUUAUUAGUUAAAGCACAUUAAUUCGUACCUUUGCAGUCCGAUUAUUA
env-948    ACGUAACGACAAAGUUAAAUAGCUGUCGUACUUAUUAGUUAAAGCACAUUAAUUCGUACCUUUGCAGUCCGAUUAUUAU
env-949    AACGUAACAACAAAGUUAAAUAGCUGUCGCACUUAUUAGUUAAAGCACAUUAAUUCGUACCUUUGCAGUCCGAUUAUUA
env-950    AGCGUAACGACAAAGUUAAAUAACUGUCGCACUUAUUAGUUAAAGCACAUUAAUUCGUACCUUUGCAGUCCGAUUAUUA
env-953    AACGUAACGACAAAGUUAAAUAGCUGUCGCACUUAUUAGUUAAAGCACAUUAAUUCGUACCUUUGCAGUCCGAUUAUUA
env-957    AACGUAACGGCAAAGUUAAAUAGCUGUCGCACUUAUUAGUUAAAGCACAUUAAUUCGUACCUUUGCAGUCCGAUUAUUA
env-958    AACGUAACGGCAAAGUUAAAUAGCUGUCGCACUUAUUAGUUAAAGCACAUUAAUUCGUACCUUUGCAGUCCGAUUAUUA
env-960    UUCAUUAAUAUUACUUUCCCCUCGCCGUUCCCGUUUGCUCCCGCUGAAUAAAUUAAGUAAUUUUGCAGUCCGAUUAUUU
env-961    UUCAUUAAUAUUACUUUCCCCGCGCCGUUCCCGUUUGCUCCCGCUGAAUAAAUUAAGUAAUUUUGCAGUCCGAUUAUUU
env-967    UAAUUCUAAAUAUUCGCCGAUGCUUGCUGUAUGUCUCUUUUAUAUAAUAAAAAAUGUGUAAUUUUGCAGACCGAUUAUU
env-968    UAAUUCUAAAUAUUCGCCGAUGCUUGCUGUAUGUCUCUUUUAUAUAAUAAAAAAUGUGUAAUUUUGCAGACCGAUUAUU
env-970    UAAUUCUAAAUAUUCGCCAAUGCUUGCUGUAUGUCUCUUUUAUAUAAUAAAAAAUGUGUAAUUUUGCAGACCGAUUAUU
env-971    UAAUUCUAAAUAUUCGCCAAUGCGUGCUGUAUGUCUCUUUUAUAUAAUAAAAAAUGUGUAAUUUUGCAGACCGAUUAUU
env-972    UAAUUCUAAAUAUUCGUUGCUGCUUAUUGUAUAUCACUUCUAUAUAAUAAAAAAUGUGUAAUUUUGCAGACCGAUUAUU
env-973    ..............................................AUAAAAAAUGUGUAAUUUUGCAGACCGAUUAUU
env-977    UAAUUCUAAAUAUUCGUUGCCGCUUAUUGUAUAUCACUUCUAUAUAAUAAAAAAUGUGUAAUUUUGCAGACCGAUUAUU
env-978    UAAUUCUAAAUAUUCGUUGCUGCUUAUUGCAUAUUACUUCUAUAUAAUAAAAAAUGUGUAAUUUUGCAGACCGAUUAUU
env-979    UAAUUCUAAAUAUUCGUUGCUGCUUAUCGUAUCUCACUUCUAUAUAAUAAAAAAUGUGUAAUUUUGCAGACCGAUUAUU
env-981    UAAUUCUAAAUAUUCGUUGCUGCUUAUUGUAUAUCACUUCUAUAUAAUAAAAAAUGUGUAAUUUUGCAGACCGAUUAUU
env-982    UAAUUCUAAAUAUUCGUUGCUGCUUAUUGCAUGUUACUUCUAUAUAAUAAAAAAUGUGUAAUUUUGCAGACCGAUUAUU
env-983    UAAUUCUAAAUAUUCGUUGCCGCUUAUUGCAUAUCACUUCUAUAUAAUAAAAAAUGUGUAAUUUUGCAGACCGAUUAUU
env-984    UAAUUCUAAAUAUUCGUUGCUGCUUAUUGCAUAUUACUUCUAUACAAUAAAAAAUGUGUAAUUUUGCAGACCGAUUAUU
env-985    UAACUUUAAAUAUUACACUUUGUGAGCUGUCUUGUUUUUGUAUAUAAUAAAUAAUGAGUAAUUUUGCGGACCGAUUAUU
env-986    UAAUUUUAAAUAUUACGCUUUGUGAGCUGUCUUGUUUUUGUAUAUAAUAAAUAAUGAGUAAUUUUGCGGACCGAUUAUU
env-990    UAAAUGUAAAUAAGCGGGCUGCUUAAAAGCAUGAUUUCUUUAUAUAAUAAAAAAUGAGUAAUUUUGCGGACCGAUUAUU
env-992    UUAAUUAAAAAAAUGAACACUUUAAGUUACAAGACUAUUUCCGUAAAUAAGGAAACAGCUAANNNNNNNNNNGAUUAUU
env-993    UAAACGUAAAUAUACAGUAUGCUUAAAAGCAUCAUUUCUCUAUAUAAUAAAAAAUGAGUAAUUUUGCGGACCGAUUAUU
env-994    UUUAUUAAUAUUAUAUUCUUCCCCGAUUCCGUUCUUGCUUUCAAAAAAUAAAAUAUGUAAUUUUGCAGUCCGAUUAUUU
Pbu-3-1    UAGUAUUUAAACGAUUGUGUUCUUGAUGCGAUAAUGGUGCGUUGUUUUUAAUAAAAUAUGUAACUUUGCACUCCGAUUG
Pbu-1-1    UAGUGUUUAAACGAUUGUAUUCUUGAUGCGAUAAUGGUGCGUUGUUUUUAAUAAAAUAUGUAACUUUGCACUCCGAUUG
Pbu-2-1    UAGUGUUUAAACGAUUGUGUUCUUGAUGUGAUAAUGGUGCGUUGUUUUUAAUAAAAUAUGUAACUUUGCACUCCGAUUG
Psp-12-1   UAGUGUUUAAACGAUUGUGUUCUUGAUGCGAUAAUGGUGCGUUGUUUUUAAUAAAAUAUGUAACUUUGCACUCCGAUUG
env-995    UCAUCCAAAUAGGAUAAAAAAGUCAUUUCUUGUUUGCGAAACACAUAAAAUGCGUACCUUUGCAGUCCGAUUAUUACAU
env-996    AUAAAUCUUACGAUAAAUUGUGAUGCUUUUUGUUUGCGAUAUAUAUAAAAUGCGUACCUUUGCAGUCCGAUUAUUACAA
env-997    UCAAAAAAUAGUUCUCCCUUACACAUCUCCCGUUAAAUAUAAGUCUGAAAUGUUUUGUAUCUUUGCAGUCCGAUUAUUU
env-999    UACAUAAAAUACACACAUUUGACCUUCAGAAGUAAAUAACAAAAGAAAAAGUAUAAGUAACUUUGCAGACCGAUUAUUU
Por-1-1    UUAUAAAAUUGGUUUUAAGCGUAUCUUAUACAGUACUCUAAUCUGAAAAAUAUACGUACUUUUGCAGACCGAUUAUUUU
env-1001   UUAUAAAAUUGGUUUUAAGCGUAUCUUAUACAGUACUCUAAUCUGAAAAAUAUACGUACUUUUGCAGACCGAUUAUUUU
Por-2-1    UUAUAAAAUUGGUUUUAAGCGUAUCUUAUACAGUACUCUAAUCUGAAAAAUAUACGUACUUUUGCAGACCGAUUAUUUU
env-1002   AGAUAUAAUUAAACGGGCUGGAUUGUACCUUUUCUUUGCACAUUACUAUUAAAUAUGUAUUUUUGCAGACCGAUUAUUU
env-1003   AGAUAUAAUUAAAAGGGCUGGAUAGUACCUUUUCUUUGCACUUUAAUGUUAAAUAUGUAUUUUUGCAGACCGAUUAUUU
Pti-3-1    AGAUAUAAUUAAACAGGCUGGUUUGUACCUUUUCUUUGCACUUUACAGUUAAAUAUGUAUUUUUGCAGACCGAUUAUUU
env-1004   AGAUAUAAUUAAACGGGCUGGAUUGUACCUUUUCUUUGCACAUUACUAUUAAAUAUGUAUUUUUGCAGACCGAUUAUUU
Pti-1-1    AGAUAUAAUUAAGCGUGCGGGGUUGUAUCUUUUCUUUGCACUUUACUAUUAAAUAUGUACUUUUGCAGACCGAUUAUUU
Pti-2-1    AGAUAUAAUUAAGCGUGCGGGGUUGUAUCUUUUCUUUGCACUUUACUAUUAAAUAUGUACUUUUGCAGACCGAUUAUUU
env-1005   AGAUAUAAUUAAACGGGCUGGAUUGUACCUUUUCUUUGCACAUUACUAUUAAAUAUGUACUUUUGCAGACCGAUUAUUU
env-1006   AUUAUAAAAUAAUGGCUAUUGAGAGAGUCUAAUGUUGGCUAUGUUUAUAUAUAUGUGUAUUUUUGCAGACCGAUUAUUU
Psp-4-1    AUUAUAAAAUAAUAGCUAUUGAGAGAGUCUAAUGUUGGCUAUGUUUAUAGAUAUGUGUAUUUUUGCAGACCGAUUAUUU
env-1007   AUUAUAAAAUAGCAGCUAUUAAGAGAGCCUAAUGUUGGCUAUGUUAAUAGAUAUGUGUAUUUUUGCAGACCGAUUAUUU
Pbu-5-1    AUUAUAAAAUAAUAGCUAUUAAGAGAGUCUAAUGUUGGCUAUGUUUAUAGAUAUGUGUAUUUUUGCAGACCGAUUAUUU
Pbu-4-1    AUUAUAAAAUAACGGCUAUUAAGAGAGCCUAAUGUUGGCUAUGUUUAUAGAUAUGUGUAUUUUUGCAGACCGAUUAUUU
env-1008   UACAUAAUUCACAAGAUUUCAUCCUGGGUCUUACCUGUGUUUUAUUUAUAAAAUAUGUAACUUUGCAGUCCGAUUAUUU
Pen-1-1    UUCGUAAUUUAUAGCAUCUCGUUUUUUGCCCGACUAGCGUUUCUCUUAUAAAAUAUGUAACUUUGCAGUCCGAUUAUUU
env-1009   UACAUAAUUCACAAGAUUUCAUCCUGGGUCUUACCUGUGUUUUAUUUAUAAAAUAUGUAACUUUGCAGUCCGAUUAUUU
env-1017   ...............................................................UGCAGUCCGAUUAUUU
env-1020   UACAUAAUUCACAAAAUUUCAUCCUGGGUCUUACCUGUGUUUUAUUUAUAAAAUAUGUAACUUUGCAGUCCGAUUAUUU
Ppl-2-1    UGCAUAAUUCACAAGAUUUCAUCCUGGGUCUUACCUGUGUUUUAUUUAUAAAAUAUGUAACUUUGCAGUCCGAUUAUUU
env-1021   UUCAUAAAUCUUCAUCAUUGGGAGCAUCCCGAUAAUGGCUGAUAUAAAAAUAAUGUGUAAUUUUGCACGCCGAUUAUUU
env-1022   UUCAUAAAUAUUCAUCAUUGGAAGCGCCUCGAUAAUGGCUGAUAUAAAAAUAAUGUGUAAUUUUGCACGCCGAUUAUUU
env-1023   UUCAUAAAUCUUCAUCAUUGGGAGCGUCUCGAUAAUGGCUGAUAUAAAAAUAAUGUGUAAUUUUGCACGCCGAUUAUUU
env-1024   GAUUUUAAAUCAAAUAGCUUCAUGCGCUCAUAACUUGCUGAUAUAAGAUAUAAUGUGUAAUUUUGCAGACCGAUUAUUU
env-1025   GUUUUUAAAUCAAAUGGUCUCAUGCGCUCAUAACUUGCUGAUAUAAGAUAUAAUGUGUAAUUUUGCAGACCGAUUAUUU
env-1027   AUGUUAAAAAACCGUAACCAACAAGCGAAAAUGAGUAUAUAUUCAAACAUUACUUCGUACCUUUGCAGUCCGAUUAUUU
Pma-4-1    GUGUUAAAAAACCAUAACUAAUAAGCAAAAAUGAGUAUGUAUUCAAACAUUACUUCGUACCUUUGCAGUCCGAUUAUUU
env-1028   ............................................................CUUUGCAGUCCGAUUAUUU
env-1029   CACAAAUUUAAAAUUUAGAGGGCAUCUGUUUUUCAUUACGAUAUGAAAAAUGAGUACCUUUGCAGUCCGAUUAUUAUUU
env-1031   CUCAAUUUGAAAAUAUUUGAUGGUGUAUGUUUUUCAAUACGCGAUGAAAAAUGUGUACCUUUGCAGUCCGAUUAUUAUU
env-1032   UUGUUUAAAUUUCGUCUUUUUCGCGUUUUAUUACGCGCGUAGAAGGGUUGGAAUGUGUACUUUUGCAGACCGAUUAUUU
env-1033   UUGUUUAAAUUUCGUCUUUUUCGCGUUUUAUUACGCGCGUAGAAGCGUUGGAAUGUGUACUUUUGCAGACCGAUUAUUU
env-1034   AUCCUAAUUAUCAGCCCCUUACCCACCUCCAUUCUUGGCCCAAUAAAAGAUAAUGUGUAAUUUUGCAGACCGAUUAUUU
env-1035   UCUAUAAAUAUGAUUGUCCGAUGGCAGAAUGAUAACAUGCAGGUAAAAUAAAAUAAGUACUUUUGCAGACCGAUUAUUU

           ...............................................................................
           ...............................................................................
           ●●●●●●●●●●●●●●●●●●●●●●●●●●●●●●●●●●●●●●●●●●●●●●●●●R●●●●●●●●●●Y●Y●●●●●●●●●●●●●Y●●
```

```
env-1037   ..............AGCUUUAAGAAGAGCUAUCGAUGUGAAAAAUAAAUAAAAUGUGUACCUUUGCAGUCCGAUUAUUA
env-1038   AAUAUAAUACAACACAUUCAACGCCCUUCCCUGCGAACGAGUUAUAAUAAAAAUAUGUACCUUUGCAGUCCGAUUAUUA
env-1039   AAUAUAAUACAACACAUUCCGCGCCCUUCCCUGCGAACGAGUUAUAAUAAAAAUAUGUACCUUUACAGUCCGAUUAUUA
env-1040   AAUAUAAUACAACACAUUCAACGCCCUUCCCAGCGAACGAGUUAUAAUAAAAAUAUGUACCUUUGCAGUCCGAUUAUUA
env-1043   AAUAUAAUACAGCACAUUCAACGCCCUUCCCUGCGAACGAGUUAUAAUAAAAAUAUGUACCUUUGCAGUCCGAUUAUUA
env-1044   AAUAUAAUACAGCACAUUCAGCGCCCUUCCCUGCGAACGAGUUAUAAUAAAAAUAUGUACCUUUGCAGUCCGAUUAUUA
env-1046   AAUAUAAUACAACACAUUCAACGCCCUUCCCUGCGAACGAGUUAUAAUAAAAAUAUGUACCUUUGCAGUCCGAUUAUUA
env-1052   AAUAUAACUAAAAGGUAUUAAAAGCAUACAGCUAUUCAUUCUUAUUAAUAAAAUAUGUACCUUUGCAGUCCGAUUAUUA
env-1053   AAUAUAAUACAACACAUUCAACGCCCUUCCCUGCGAACGAGUUAUAAUAAAAAUGUGUACCUUUGCAGUCCGAUUAUUA
Psp-2-1    AAUAUAAUACAACACGUUCAACGCCCUUCCCUGCGAACGAGUUAUAAUAAAAAUAUGUACCUUUGCAGUCCGAUUAUUA
env-1054   AAUAUAAUACAACACGUUCAGCGCCCUUCCCUGCGAACGAGUUAUAAUAAAAAUAUGUACCUUUGCAGUCCGAUUAUUA
env-1055   AAUAUAAUACAACACAUUCCGCGCCCUUCCCUGCGAACGAGUUAUAAUAAAAAUAUGUACCUUUGCAGUCCGAUUAUUA
env-1056   AAUAUAAUACAACACAUUCAGCGCCCUUCCCUGCGAACGAGUUAUAAUAAAAAUAUGUACCUUUGCAGUCCGAUUAUUA
env-1058   AAUGUUAAUGUUUUAUAAUUUGUGGUUUUGAGCAUCUCGUAAUCCUUAUAAAAUUAGUACCUUUGCAGUCCGAUUAUUA
env-1059   ....................................................AGUAGUACCUUUGCAGUCCGAUUAUUA
env-1060   CCGCUAAAAAUUAUUAAUUUCGUUUUCUCCCUAACCUCACAUAAUAAAUAAAAGUAGUACCUUUGCAGUCCGAUUAUUA
env-1061   UCAUAAAAAUAAAAUUCCAACAAGUCUUGUAUUAUAAAUAUAAAGGAAAAAUUCAUACCUUUGCAAUCCGAUUAUUAAA
Psp-13-1   UCAUAAAAAUAAAAUCCUAACGAGUCUUGUAUUAUAAAUAUAAAGGAAAAAUUCAUACCUUUGCAAUCCGAUUAUUAAA
env-1062   UCAUAAAAAUAAAAUUCUAACGAGUCUUGUAUUAUAAAUAUAAAGGAAAAAUUCAUACCUUUGCAAUCCGAUUAUUAAA
Psp-17-1   UCAUAAAAAUAAAAUCCUAACAAGUCUUGUAUUAUAAAUAUAAAGGAAAAAUUCAUACCUUUGCAAUCCGAUUAUUAAA
env-1063   UCAUAAAAAUAAAACUCUAACGAGUCUUGUAUUAUAAAUAUAAAGGAAAAAUUCAUACCUUUGCAAUCCGAUUAUUAAA
Psp-15-1   UCAUAAAAAUAAAAUUCUAACGAGUCUUGUAUUAUAAAUAUAAAGGAAAAAUUCAUACCUUUGCAAUCCGAUUAUUAAA
Psp-16-1   UCAUAAAAAUAAAAUUCUAACGAGUCUUGUAUUAUAAAUAUAAAGGAAAAAUUCAUACCUUUGCAAUCCGAUUAUUAAA
env-1064   AUAUAAAAACAAAGCUUACAUUCUUCCAUAUAUAAUGUACAUAUAAAAAUUAUUCAUACCUUUGCAAUCCGAUAAUUUA
env-1065   AUAUAAAAUCACAGUUUGCAUUCUUUCAUAUAUAAUGUACAUACAAAAAUUAUUCAUACCUUUGCAAUCCGAUUAUUUA
env-1066   AUAUAAAAACAAAGCUUACAUUCUUCCAUAUAUAAUGUACAUAUAAAAAUUAUUCAUACCUUUGCAAUCCGAUUAUUUA
env-1067   .................UUUCUUGGUGCACUGAAUUCAGCGGACCAAUAAAAAUGUGUACCUUUGCAGUCCGAUUAUUA
Hse-1-1    CUUAAUCCAUGGGCGUUUUCACCGUUUUGUCGUGCGUUGUACGUCAUAAAAUGUGUAAUUUUGCAGAACCGAUUAUUUC
env-1068   AAUAUAAAACAAAGGUCGCACCGGCGUCCAAUCACAUUUUAUUAUGGAGAAAAAUACUACCUUUGCAGACCGAUUAUUG
env-1069   GCAUUUUCUUCACGCGUGAGGAGAUGCAUUUGCGAGAAAAUAAGUAAUUUUGCAGUCCGAUUAAUUCAAUCCGAUUAAA
env-1071   AAUAAUCACAAAAACGCAAUAUCAAGACCUACAUUAUGCUGCUAUGAAAAGAAAAAGCUGUACCUUUGCAGUCCGAUUA
env-1072   AACCUUAAAAGAUUGGUUCUUAAGCAGAAUGACGGACAGGUUUUCACCUGAAAACCACUACUUUUGCAGUCCAAACAAA
env-1074   ACCUUAAAAGAUUGAUUCUUAAGCAGAAUGACGGACAGGUUUUCAUCUGAAAACCACUACUUUUUGCAGUCCAAACAAA
env-1075   AACCUUAAAAGAUUGAUUCUUAAGCAGAAUGACGGACAGGUUUUCAUCUGAAAACCACUACUUUUGCAGUCCAAACAAA
env-1076   UUCAUAAAAAAACGGCGUCGAUGGUAGGAGAUUUAUUGCCGAUAGACAAUUAAUUAUUACCUUUGCAGUCCGAUUAUUAU
env-1077   ........................................................UACCUUUGCAGUCCGAUUAUUAU
env-1078   AAUUAUCAUAAAAGUUUGGUUUCGGAAGGGGAAAGGGCUUCGCAUAUUAACAAAUAUGUGUACCUUUGCAGUCCGAUUA
env-1079   AAUAUCUAUAAAAUACCUAUCCUGCGUGGGUGGUGCUUUCUUAGAAAAGAAAAUUCUUCGUACCUUUGCAGUCCGAUUU
env-1080   AUAAUCAUAAAUCUUUCAUCGCUUAUCGCUUAUCACUUGUUGCUUGUCAUUUAAAAUAUGUACCUUUGCAGUCCGAUUU
env-1081   AAUAAUCAUAAUCCGAGGCAGAUUCUUCACGCUUCACUCUUCACUCUUCACUUAAAAUAUGUACCUUUGCAGUCCGAUU
env-1083   AAUAAUCAUAAUCCGAGGCAGAUUCUUCACGCUUCACUCUUCACUCUUCACUUAAAAUAUGUACCUUUGCAGUCCGAUU
env-1084   AAAAACAUAAAAGAGACGUUUUUCGUUGUUACGCGUACAUUUAAAAGAAGAAAAAGCCGUACCUUUGCAGUCCGAUUAC
env-1085   ..........................................................GUACCUUUGCAGUCCGAUUAU
env-1086   AUAAUAUUAAAUCUUUAACUUAUCCAGUUUCUUAUUUUCCAGAUUCAAGAGAAAAGCUGUACCUUUGCAGUCCGAUUAU
env-1087   AAUUACAGCUGAUUGGCUCUAUUUCACUCUUUUCACUUUCCACUUUUCACUUUUCUUCGUACCUUUGCAGUCCGAUUAU
env-1088   AUAAUAUUAAAUCUUUAACUUAUCCAGUUCUUCGUUUCCUAGAUUCAAGAGAAAAACCGUACCUUUGCAGUCCGAUUAU
env-1089   AUAAUAUUAAAUCUUUAACUUAUCCAGUUUCUUGUUUUCAUGAUUCAAGAGAAAAGCCGUACCUUUGCAGUCCGAUUAU
env-1090   AUAAUCCUAAAAAUACACCUAUUUGCUUUCCUGACGUACAUUAUUAAGGAGAAAAGCCGUACCUUUGCAGUCCGAUUAU
env-1091   AAUAAUCAUAAAAAUUCUUUUCGUCGGACUGCAAAGGCGCGCACGUAUAAUAAAAUAUGUGUACCUUUGCAGUCCGAUUA
env-1092   AAAAUAUAAAUCUUUACUUUCAAUGGCUUUCUCGUGCCGUGAGAAAGUAGAAAAAGUAGUACCUUUGCAGUCCGAUUAU
env-1093   AAGUACGCCUUUUGCCCUCUUUUUUCACUUUUCACUUUUCACUUUUCAUUUUUUCGUCGUACCUUUGCAGUCCGAUUAU
env-1094   AAUAACACCAUUUUUACUUCCUUUAAGGCUCCUACUCUACUCAUUUCUCGUUUUUUAGCAGUACCUUUGCAGUCCGAUU
env-1095   AUUCAUAAAUCUUUGACUUCUGACAGGUUUCCUCGCGCGAUAUUAGAAAGAAAUAGCGUACCUUUGCAGUCCGAUUAUU
env-1096   AUAAUCAUAAAUCUUCCGUACGCCAGACUAAAAUAGCCUCGCGUACGGAAAAAUGUUCGUACCUUUGCAGUCCGAUUAU
env-1097   AAAUGCAAGCUUGAAGCCUAUUUAACAUUCCGCACCACACUUUUUUCAAUGAAAAACCGUACCUUUGCAGUCCGAUUAU
env-1098   AUAAUCAUAAAAGUUCGUGAAUAAAGAGGGUAAAGACACUCUAAAAUACAAAAUAUGUGUACCUUUGCACGCCGAUUAU
Psp-7-1    AAAAAUAUAAAUAUCUAUUUCUUGGACUGCAAAGGUGUGCAUGAAUAAAUAAAUAUGUGUACCUUUGCAGUCCGAUUAU
env-1099   UUUGUCUAAAUUUGACGCUUUUUUCUUUUGCGCGUACCUUAUAUAAUAAAAAUAGUUAGUACCUUUGCACACCGAUUUA
env-1100   AUUUGUCUAAAUUUGACGCUUUUUUCUUUUGCGCGUACCUUAUAUAAUAAAAAUAGUUAGUACCUUUGCACACCGAUUU
Psp-11-1   AUUUGUCUAAAUUUGACGCUUUUUUCUUUUGCGCGUACAUUAUAUAAUAAAAAUAGUUAGUACCUUUGCACACCGAUUU
Psp-19-1   AUUUCUCUAAAUUUGACGCUUUUUUCUUUUGCGCGUACCUUAUAUAAUAAAAAUAGUUAGUACCUUUGCACACCGAUUU
env-1101   AUUUCUCUAAAUUUAGCGCUUUUUUCUUUUGCGCGUACAUUAUAUAAUAAAAAUAGUUAGUACCUUUGCACACCGAUUU
Pru-1-1    AUUUGUCUAAAUUUGACGCUUUUUUCUUUUGCGCGUACCUUAUAUAAUAAAAAUAGUUAGUACCUUUGCACACCGAUUU
env-1102   AUAAUCAUAAAUAGUAAUAUCCAUAGCUGUCGCAGGCACGUAAAUUAAGAAAUAGUUCGUACCUUUGCAGUCCGAUUAU
env-1103   AAAUAUAAAUAACAGGGCGCUAAAAGGCAAAUAUAUCGUGUAAAACGCGAAAAAGCUGUACCUUUGCAGUCCGAUUAUU
env-1104   UAAUCAUAAAUACAGACUAUAAAUCCCAAUUCUCAAUUCUCAAUUCUUAAUUUAUUAGUACCUUUGCAGUCCGAUUAUU
Pbr-1-1    AACUAUAAUUUUAUGGUUACUGCAUGGAUGUAUGCUUCUUAAUUUCAAUUAAUGUGAGUACCUUUGCAGUCCGAUUAUU
Pbr-2-1    AACUAUAAUUUUAUGGUUGCUGCAUGGAUGUAUGCUUCUUAAUUUCAAUUAAUGUGAGUACCUUUGCAGUCCGAUUAUU
env-1105   AAAAUAAUUGCCGAGGUAAAAACGCGGCAAACCCUCUUACAGUCUCAAUAAAAAUAAUUACCUUUGCAGUCCGAUUAUU
env-1106   AAAACGUAAAUGUUAAGGUAGUGAGAGAUUCCAUCGCGCGAUAUAUGACGAAAAAGCAGUACCUUUGCAGUCCGAUUAU
env-1107   AAUCGUAAAAGUUGGGUCCUGGCUGGCUGCUGAUGGUCGAUGAAUGGCAGAAAAGUAGUACCUUUGCAGUCCGAUUAUU
env-1108   AAGCAUAAAUGUUUAGCUUCAAAAAUAUUAUAGGAGCCUGUAAAAGAAGAAAAAGUAGUACCUUUGCAGUCCGAUUUAG
env-1109   CUUUUAUAAAAAUACACUUUCCCGCUCGCUUCCGUAAGCGGAAUCAUCAUUUAAUUUGUACCUUUGCAGUCCGAUUUAA
env-1110   AAUCAUAAAUCUUAUGCCGUUUGCUGUUCUCGGUUUCUCGUUUUCCAUUUAAACUGCGUACCUUUGCAGUCCGAUUUUG
env-1111   AAUCAUAAUUUGAUGGUCUUUUCUCCUUUCUCAUUACUUCUUUCUCAUAUUAUCUCACUACCUUUGCAGUCCGAUUUUU
env-1112   AUCAUAAAAGUUGCGUCCUAAUAGCUGCGUUAUACCUCAUAUAAUGAGAAAAAGCAGUACCUUUGCAGUCCGAUUAUUG
env-1113   AAGUGCAGUUUUGCGCUCGCGCGAUACCCCUUAAAUGGAAAAAUAUGUGUAAUUUUGCACUCCGAUUUCCGAUUAUCGC
env-1114   UAUGCGGUUGCUUAGCUGGCAUGAUAGCCUCUAAGUAAAAAAAUAUGUGUAAUUUUGCACUCCGAUUUCCGAUUAUCGC
env-1115   ACUGCACUAAUGAAGGUCAUAGCAUAGCUACUAUGUCAAUAAAUAUGUGUAAUUUUGCACUCCGAUUUCCGAUUAUCGC
env-1116   AUAAAUUAGACAGCAAACACAGCAACGGAAACAACAUAUUCCAUAUAAAAUAUGUACUUUUGCAGUCCGAUUAUUAAAA
env-1117   AUACGUAAAAGAUGAAACAAAAUGUGUCGCGCGUAUAUUAAAUAAUGAGAAAAGCCGUAACUUUGCAGUCCGAUUAUUU
env-1118   UUUCUUAAAAAUACAGACUGCAAGGAAACUCAUUGCACAGAAGUGCGGAAAAAUGAGUACCUUUGCAGUCCGAUUAUUA
env-1119   UUUCUUAAAAAUACGGACUGCAAGGAAACUCAUUGCACAGAAGUGCGGAAAAAUGAGUACCUUUGCAGUCCGAUUAUUA
env-1120   AAAACGUAAAUCUGAUGAUUUCAUAAAAGUUCACGCGCGAUAUUUAAAAUAAAAUGCUUACCUUUGCAGUCCGAAAUCA

           ...............................................................................
           ...............................................................................
           ●●●●●●●●●●●●●●●●●●●●●●●●●●●●●●●●●●●●●●●●●●●●●●●●●R●●●●●●●●●●Y●Y●●●●●●●●●●●●●Y●●
```

```
env-1122  AAAACGUAAUCUGAUGAUUUCAUAAAAGUUCACGCGCGAAUAUUUAAAAAUAAAUGCUUACCUUUGCAGUCCGAAAUCA
env-1123  CAGCCAAGCAAACAGUCAGAAAAUCAUCAAAUUGUUUUCUUGUUUCGAAAAAGUAUCUACCUUUGCAGUCCGAUUAUUA
env-1125  AAAACAUAAAAGUUGCGCUUUCGCUCGCCGUGUCAAUCGCAAUGCGAAGAAAAAUCAGUACCUUUGCAGUCCGAUUAUU
env-1126  AAAACGUAAAUCUUAAUUUGUUUGCCCCUCUUUGGAUUGGUAAACGAAGAAAAAGCCGUACCUUUGCAGUCCGAUUAUU
env-1127  AAAACGUAAAUCUCACACUGUUUGCCCCUCUUUGGGUGGUUAAACGGAGAAAAAUGAGUACCUUUGCAGUCCGAUUAUU
env-1129  UGCAUAAAUAUUAUAGCAGAAUAGGGUGCAUGUUACGCAUAUUAUAAAAAUAAUGUGUAAUUUUGCAGUCCGAUUAUUU
env-1130  UUUGCUGAGGGAAUGAAAAAUUUUAAAUAUAUUUGGUGUAUUCGGUGAUUUUUCGUACCUUUGCAAGCCGAUUAUUAUC
env-1131  GUAUACUCGCGGUUGAAAAUUUGGAAAAGUUUUGUUAGUUACGCUUUUUUUCGUACCUUUGCACGCCGAUUAUUAUCA
env-1132  UGUAUGCAUGCGCGAAAAAAACAAGGAAAGUUUUGCCAGUUAAAAGAAUUUUCGUACCUUUGCACGCCGAUUAUUAUCA
env-1133  AUGCCAAGAAGAAAAAAAGUCUUGCCUGUGCUUGCAGAAUUGGGAAAAAGUUGUACCUUUGCACGCCGAUUAUUAUCAA
env-1134  GUGGUUCGGAGACAAAAAAGAAAGGAAACUUUUGGUCAGUUAAUAUAAAAAUCGUACCUUUGCAAGCCGAUUAUUAUCA
env-1135  CCUGAAAAUUAUUUUUUUCAGUGGAAAUGCUUGCCGUUUCCGGAAAAAGUCGUACCUUUGCACGCCGAUUAUUGUCUAC
env-1136  CAAUAAUUAAUAAAAAAAGCAAGGAGAUGCUUGUCGGUUCAGAAAAAAGUUGUACCUUUGCACGCCGAUUAUUGUCUAA
env-1137  UCAACGAUUCCAUAUGAAAAAUAGGAGGAAGUUUGGCUGUUUCAGUAGAAAUGCGUAAUUUUGCAUGCCGAUUAUUAUC
env-1138  UCGACGAUUCUAUAUGAAAAAUAGGAGAGAGUUUGGCUGUUUCAGUAGAAAUGCGUAAUUUUGCAUGCCGAUUAUUAUC
env-1139  UCAACGAUUCCAUAUGAAAAAUAGGAGGAAGUUUGGCUGUUUCAGUAGAAAUGCGUAAUUUUGCAUGCCGAUUAUUAUC
env-1140  UCGACGAUUCUAUAUGAAAAAUAGGAGGAAGUUUGGCUGUUUCAGUAGAAAUGCGUAAUUUUGCAUGCCGAUUAUUAUC
env-1141  ......................................................GUAAUUUUGCAUGCCGAUUAUUAUC
env-1143  UCGACGAUUCUAUAUGAAAAAUAGGAGGAAGUUUGGCUGUUUCAGUAGAAAUGCGUAAUUUUGCAUGCCGAUUAUUAUC
env-1144  UCGACGAUUCUAUAUGAAAAAUAGGAGGAAGUUUGGCUGUUUCAGUAGAAAUGCGUAAUUUUGCAUGCCGAUUAUUAUC
env-1145  UCGACGAUUCUAUAUGAAAAAUAGGAGGAAGUUUGGCUGUUUCAGUAGAAAUGCGUAAUUUUGCAUGCCGAUUAUUAUC
env-1147  CGCCGACAUGCGCUUUUUGCCCUCACCUGCUUGGCUAUUUCGCAGAAUAUAAGUACAUUUGCAUGCCGAUUAUUAUCAA
env-1148  CGCCGACAUGCGCUUUUCGCCCUCACCUGCUUGGCUAUUUCGCAGAAUAUAAGUACAUUUGCAUGCCGAUUAUUAUCAA
env-1149  CGCCGACAUGCGCUUUUCGCCCUCACCUGCUUGGCUAUUUCGCAGAAUAUAAGUACAUUUGCAUGCCGAUUAUUAUCAA
env-1151  .......CUGCGCUUUUCGCCCUCACCUUCUUGGCUAUUUCGCAGAAUAUAAGUACAUUUGCAUGCCGAUUAUUAUCAA
env-1153  CGUCGACCUUCGCUUUUCGCCCUCACCGUCUUGGCUAUUUCGCAGAAUAUAAGUACAUUUGCAUGCCGAUUAUUAUCAA
env-1154  CGUCGACCUGCGCUUUUCGCCCUCACGUUCUUGGCUAUUUCGCAGAAUAUAAGUACAUUUGCAUGCCGAUUAUUAUCAA
env-1155  CGUCGACCUGCGCUUUUCGCCCUCACCUUCUUGGCUAUUUCGCAGAAUAUAAGUACAUUUGCAUGCCGAUUAUUAUCAA
env-1156  CAUGAAAAAAAGAUUUUCAAGUUCACAAGCUUGGCUAUUUCGCAGAAUAUAAGUACAUUUGCAUGCCGAUUAUUAUCAA
env-1157  CGUCGACCUGCGCUUUUCGCCCUCACCUUCUUGGCUAUUUCGCAGAAUAUAAGUACAUUUGCAUGCCGAUUAUUAUCAA
env-1158  CGUCGACCUGCGCUUUUCGCCCUCACCUUCUUGGCUAUUUCGCAGAAUAUAAGUACAUUUGCAUGCCGAUUAUUAUCAA
env-1159  CGCCGACAUGCGCUUUUCGCCCUCACCUGCUUGGCUAUUUCGCAGAAUAUAAGUACAUUUGCAUGCCGAUUAUUAUCAA
env-1163  CGCCGACCUGCGCUUUUCGCCCUCACCUUCUUGGCUAUUUCGCAGAAAAUAAGUACAUUUGCAUGCCGAUUAUUAUCAA
env-1164  CAUGAAAAGAAGAUUUUCAAGUUCACAAGCUUGGCUAUUUCGCAGAAUAUAAGUACAUUUGCAUGCCGAUUAUUAUCAA
env-1165  CGUCGACCAGCGCUUUUCGCCCUCACCUUCUUGGCUAUUUCGCAGAAUAUAAGUACAUUUGCAUGCCGAUUAUUAUCAA
env-1166  CGUCGACCUGCGCUUUUCGUCCUCACCUUCUUGGCUAUUUCGCAGAAUAUAAGUACAUUUGCAUGCCGAUUAUUAUCAA
env-1167  ......................................................ACAUUUGCAUGCCGAUUAUUAUCAA
env-1192  CAUGAAAAGAAGAUUUUCAAGUUCACAAGCUUGGCUAUUUCGCAGAAUAUAAGUACAUUUGCAUGCCGAUUAUUAUCAA
env-1193  UAAGUUUUGCAUUUAUAGAAGGAAAUGUUUGGUAGAUUCGUGGAAAAUAAGUAAUUUUGCAUGCCGAUUAUUAUCAAGU
env-1194  AAGUUUUGCAUUUAUAGAGGGAAAUGUUUGGUAGAUUCGUGGAAAAUAAGUAAUUUUGCAUGCCGAUUAUUAUCAAGUU
Psp-1-1   AAGUUUUGCAUUUAUAGAGGGAAAUGUUUGGUAGAUUCGUGGAAAAUAAGUAAUUUUGCAUGCCGAUUAUUAUCAAGUU
env-1195  AAGUUUUGCAUUUAUAGAGGGAAAUGUUUGGUAGAUUCGUGGAAAAUAAGUAAUUUUGCAUGCCGAUUAUUAUCAAGUU
env-1197  AAGUUUUGCAUUUAUAGAGGGAAAUGUUUGGUAGAUUCGUGGAAAAUAAGUAAUUUUGCAUGCCGAUUAUUAUCAAGUU
env-1203  AAGUUUUGCAUUUAUAGAAGGAAAUGUUUGGUAGAUUCGUGGAAAAUAAGUAAUUUUGCAUGCCGAUUAUUAUCAAGUU
env-1205  AAGUUUUGCAUUUAUGGAAGGAAAUGUUUGGUAGAUUCGUGGAAAAUAAGUAAUUUUGCAUGCCGAUUAUUAUCAAGUU
env-1206  .......................................................UUUUGCAUGCCGAUUAUUAUCAAG
env-1211  CUAAGUUUUGCAUUUAUAGAAGGAAAUGUUUGGUAGAUUCGUGGAAAAUAAGUAAUUUUGCAUGCCGAUUAUUAUCAAG
env-1213  GGGGGAGAAGUUGCUUUUCUCCGAGAAAUGUUUGGUGGUAUCACGAGAAAUAAGUAAUUUUGCAUGCCGAUUAUUAUCA
env-1216  GGCGUUUGCAGUGCUUUUCUCCGAGAAAUGUUUGGUGGAAUCAUGAGAAAUGAGUAAUUUUGCGUGCCGAUUAUUAUCA
env-1223  ...................UCUGAGAAAUGUUUGGUGGAAUCAUGAGAAAUGAGUAAUUUUGCGUGCCGAUUAUUAUCA
env-1224  GGCGUUUGCAGUGCUUUUCUCUGAGAAAUGUUUGGUGGAAUCAUGAGAAAUGAGUAAUUUUGCGUGCCGAUUAUUAUCA
env-1228  GUGUGAUAGUGUGCUUUUCUCUGAGAAAUGUUUGGUGGUAUCAUGAGAAAUGAGUAAUUUUGCGUGCCGAUUAUUAUCA
env-1233  GGCGUUUGCAGUGCUUUUCUCUGAGAAAUGUUUGGUGGAAUCAUGAGAAAUGAGUAAUUUUGCGUGCCGAUUAUUAUCA
env-1243  GGCGUUUGCAGUGCUUUUCUCCGAGAAAUGUUUGGUGGAAUCAUGAGAAAUGAGUAAUUUUGCGUGCCGAUUAUUAUCA
env-1244  GGGUGUUAUAGUGCUUUUCUCUGAGAAAUGUUUGGUGGAAUCAUGAGAAAUGAGUAAUUUUGCAUGCCGAUUAUUAUCA
env-1245  GGGUGUUAUAGUGCUUUUCUCUGAGAAAUGUUUGGUGGAAUCAUGAGAAAUGAGUAAUUUUGCAUGCCGAUUAUUAUCA
env-1250  GUGUGAUAGUGUGCUUUUCUCUGAGAAAUGUUUGGUGGAAUCAUGAGAAAUGAGUAAUUUUGCGUGCCGAUUAUUAUCA
env-1251  GUGUGAUAGUGUGCUUUUCUCUGAGAAAUGUUUGGUGGUAUCAUGAGAAAUGAGUAAUUUUGCGUGCCGAUUAUUAUCA
env-1254  ............................................GAGAAAUGAGUAAUUUUGCGUGCCGAUUAUUAUCA
env-1256  GGCGUUUGCAGUGCUUUUCUCCGAGAAAUGUUUGGUGGUGUCACGAGAAAUAAGUAAUUUUGCAUGCCGAUUAUUAUCA
env-1258  GGGUAUUCUAGUGCUUUUCUAUGAGAAAAGUUUGGCAGAAUCAGGAGAAAUGAGUAAUUUUGCGUGCCGAUUAUUAUCA
env-1259  GGGUGUUAUAGUGCUUUUCUCUGAGAAAAGUUUGGCAGAAUCAGGAGAAAUGAGUAAUUUUGCGUGCCGAUUAUUAUCA
Pmu-2-1   UUAUUUAUAAUUAAACCAAAUAAUAAUAAUUCAUUCUUUUCCGCAUCAACUGAUAUGUGUACCUUUGCACUGCUGAUUG
Pde-1-1   CUUAAUCCAUGGGCGUUUUCACCGUUUUGCCGUGCGUUGUACGUCAUAAAAUAUGUAAUUUUGCAGAACCGAUUAUUUC
          ...............................................................................
          ...............................................................................
          ●●●●●●●●●●●●●●●●●●●●●●●●●●●●●●●●●●●●●●●●●●●●●●●●●R●●●●●●●●●●Y●Y●●●●●●●●●●●●Y●●
```

alignment positions 80 · · · 259

```
env-1   CAAGGGU.......................ACCCUUAAUGCCCAACCAAGAGUUAAUAGUU..AC..GUCC.....UAUUUUGGCCUGGACG.UAAUGUUGGUAAGCUCUUGGAAAGAUAAACAUUAGUAGUUACAAUUUAAUUCAAACAAAAGUGGACACUUUAAGUUACAAGACCA
env-2   AGGGAGGCGC..AGAU...........GAUCCUCCC....AAAGUGGGCGUGUAGAUAGCG..CGU.UGUU.........UUGGCC.GACAC.GUGUGGUCUGUG.AAUCGCUCAGAGAAAAAACUACAUCUCCAUCGUGGCCGCUCUCCUUUGCGAGGUGGUGUACGAUAGUGGAGUG
env-3   AUCUGGC.....CGGCCAAAA.........GCCAGC.....CACGCCGCGGAUUAAUAGUU..GC..AUCC....GUUUUAUUGGCCAGGAUG.UAAUGGUUAGGU.AAUAUUCGUGGCAAAAGAUAAACAUUAGUAGUUACAAUUUAAAACAAACAAAAGUGGACACUUUAAGUUACAA
env-4   GAGAGAGUC...AUUA............G.UUCUCU....CAAAAGGGUGUGUAGAUAGCGU..G..AACU.........UUGGCC.GGUUC..UGUGGUCUGUG.AAUCACUCAAAUAAACUAAUCUCCGCCGCUAAUCCCGAAAGGGAGAACGGUAAGGGAGUGAUUUUUCGUGAUG
env-5   UGGGGCGACACUCUU..........AGAAUCGCUCC....AGCAUGAGCGUGUUAAUAGUG..UC..GCGU.........AAGGCC.AUGCG.UCAUGGUUAGGG.AAUCGCCCAUGCGUUCAGGGUUAAAGCCAUGGAUGGUCUCUUUUAGACUGGAGGCACCGCGAGGCCCCUGUUC
env-6   GGGAUGC.....GAAAA.............GCGUCC....UCUACAAGUGUGUAGAUAGCGU.CC..AUA..........UUGGCC..UAUG.GCGUGGUCUGUG.AAUCACUUGUGAAAAUGAAGAAUUGAACAAUUAAUUAGUAGUAACAAAAUCAAGAAUUUAAAGAUGAACACUU
env-7   AGGGGACGC...GAAA............GUG.CCCC....CAUAUAAGUGUGUAGAUAGCGU.CC..AUA..........UUGGCC..UGUG.GCGUGGUCUGUG.AAUCACUUGUAAAAAAACAAGAAUUGAACAAUUAAUUAGUAGUAACAAAAAUUUUAGAGAUGAACACUUUAAG
env-8   AACAGGC.....AACAAAAA..........GCCUGU....UAAUCCCACGUGUUAAUAGCU..GU..AUCUAUUAAAGAAUUUGGCC.UGAUG.CAGUGGUUAGUG.AAUCGUGGUGCGUCGUCAUGCGGCGCAAAAAAACUUUAUGGC................................
env-9   AACAGGC.....AACAAAAA..........GCCUGU....UAAUCCCACGUGUUAAUAGCU..GU..AUCUAUUAAAGAAUUUGGCC.UGAUG.CAGUGGUUAGUG.AAUCGUGGUGCGUCAUCAUGCGGCGCAAAAAAAAACUUUAUGGCAAACCCGAAGAAAAACUUAGAACGGAAGG
env-10  AACAGGC.....AACAAAAA..........GCCUGU....UAAUCCCACGUGUUAAUAGCU..GU..AUCUAUUAAAGAAUUUGGCC.UGAUG.CAGUGGUUAGUG.AAUCGUGGUGCGUCGUCAUGCGGCGCAAAAAAACUUUAUGGCAAACCCGAAGAAAAACUUAGAACGGAAGGCAA

        .<<<<<<<<<<<............>>>>>>>>>>>................<<<<<.<<<..<<..<<<<<..............>>>>>>.>>>>.>>>>>.........................................................................
        .222222.1111............1111.222222................02200.122..22..2222...............22222.2221.00220.........................................................................
        ●GGGG●○-----R●●●○○○○----------○●CCCC----○○●●●●●●YGUGU●AAUAGY●--●Y--RYYU---------UUGGCC-GRRY●-●●RYGGUU●GUG-AAUCR●●●●●●●RR●●R●●●●●●●●●●●●●●●●●●●●●●●●●●●●●●●●●●●●●●●●●●●●●●●●●●●●●●●
         5′                                                                                                      3′
```

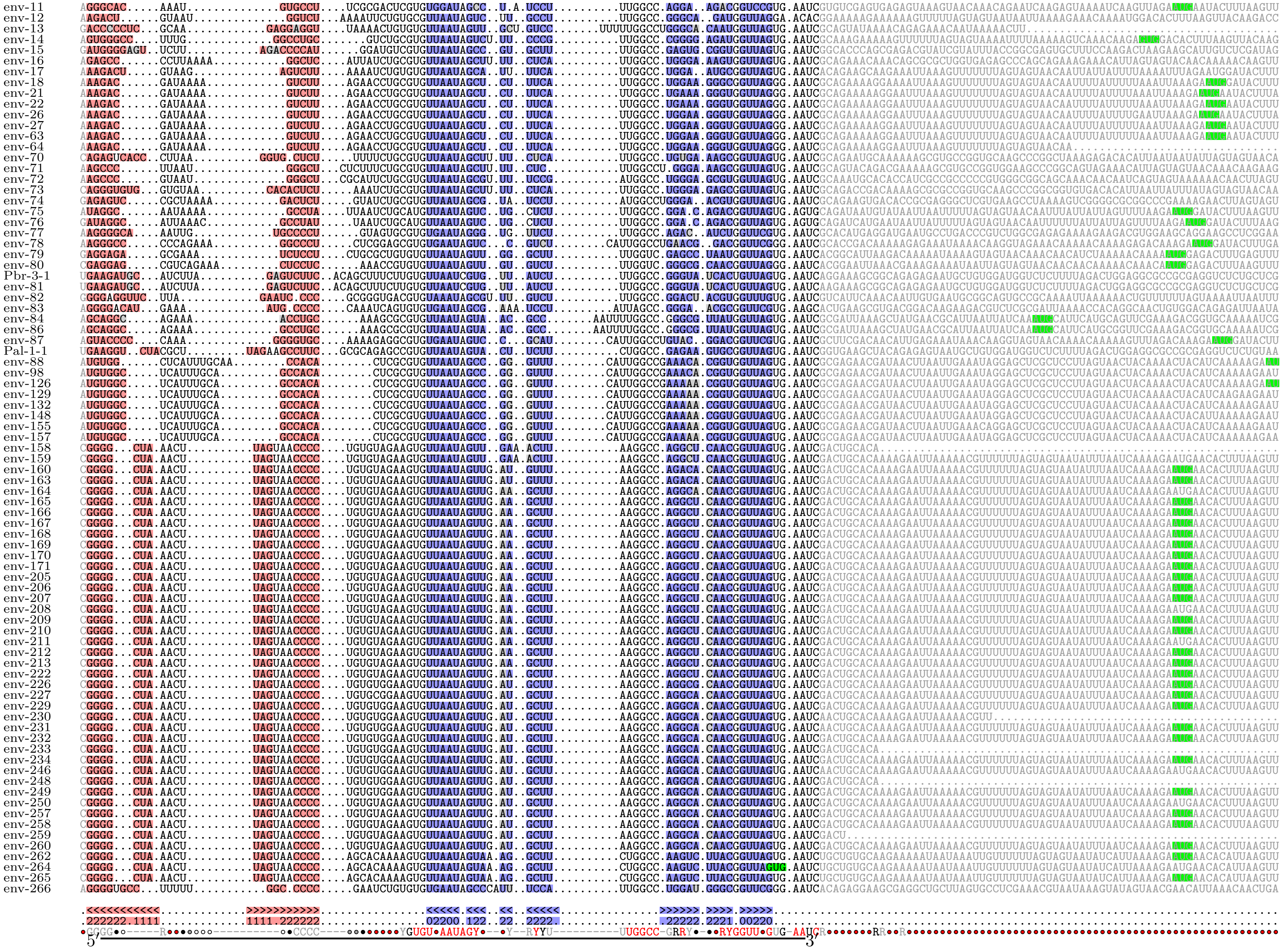

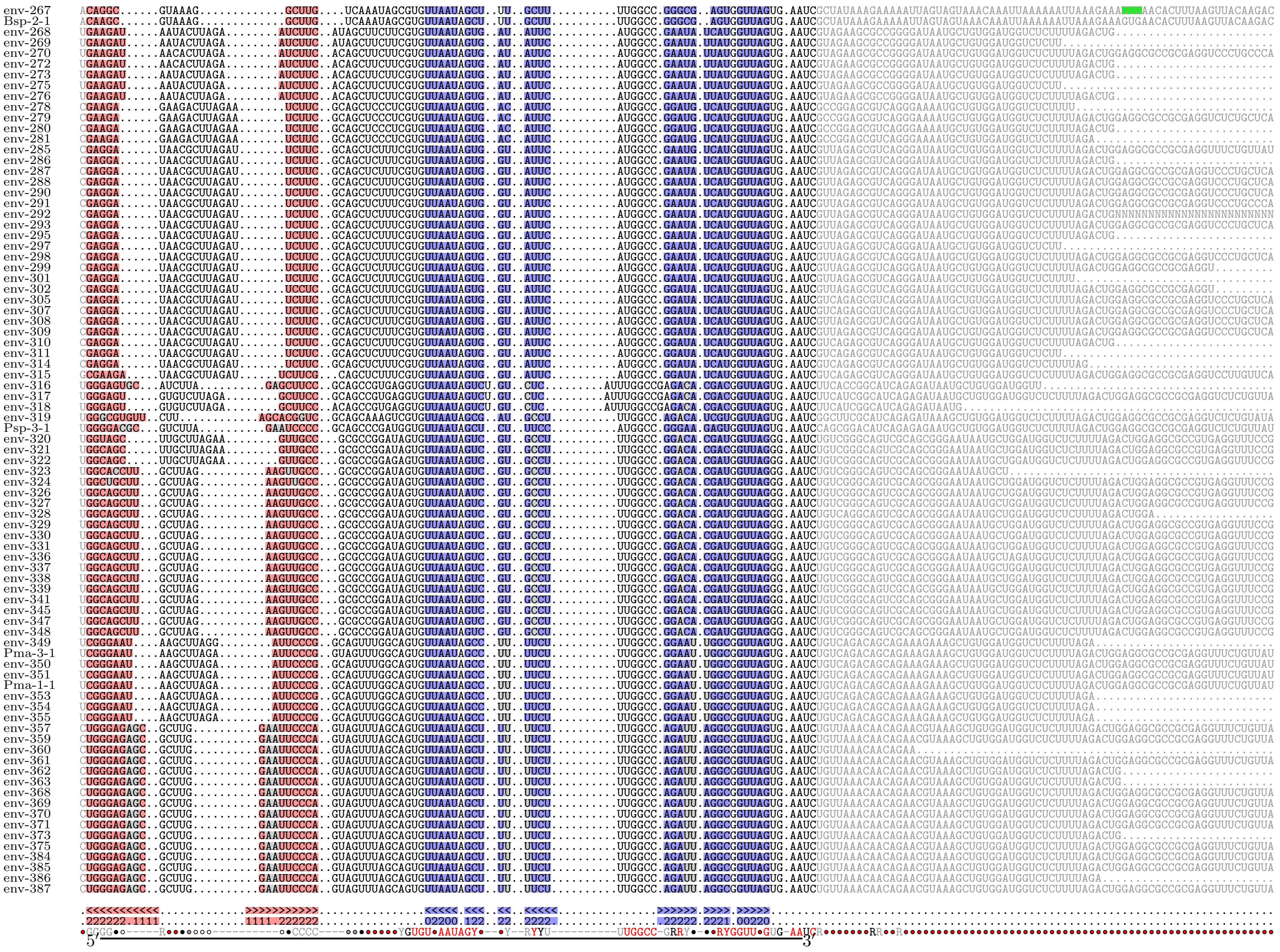

```
env-388  CUGGGAGAGC..GCUUG.........GAAUUCCCA..GUAGUUUAGCAGUGUUAAUAGCU..UU..UUCU.........UUGGCC.AGAUU.AGGCGGUUAGUG.AAUCUGUUAAACAACAGAACGUAAAGCUGUGGAUGGUCUCUUUUAGACUG.......................
env-389  CUGGGAGAGC..GCUUG.........GAAUUCCCA..GUAGUUUAGCAGUGUUAAUAGCC..UU..UUCU.........UUGGCC.AGAUU.AGGCGUUUAGUG.AAUCUGUUAAACAACAGAACGUAAAGCUGUGGAUGGUCUCUUUUAGACUGGAGGCGCCGCGAGGUUUCUGUUA
env-390  CUGGGAGAGC..GCUUG.........GAAUUCCCA..GUAGUUUAGCAGUGUUAAUAGCC..UU..UUCU.........UUGGCC.AGAUU.AGGUGGUUAGUG.AAUCUGUUAAACAACAGAACGUAAAGCUGUGGAUGGUCUCUUUUAGACUGGAGGCGCCGCGAGGUUUCUGUUA
env-392  CUGGGAGAGC..GCUUG.........GAAUUCCCA..GUAGUUUAGCAGUGUUAAUAGCC..UU..UUCU.........UUGGCC.AGAUU.AGGCGGUUAGUG.AAUCUGUUAAACAACAGAACGUAAAGCUGUGGAUGGUCUCUUUUAGACUGGAGGCGCCGCGAGGUUUCUGUUA
env-396  CUGGGAGAGC..GCUUG.........GAAUUCCCA..GUAGUUUAGCAGUGUUAAUAGCC..UU..UUCU.........UUGGCC.AGAUU.AGGCGGUUAGUG.AAUCUGUUAAACAACAGAACGUAAAGCUGUGGAUGGUCUCUUUUAGACUGGAGGCGCCGCGAGGUUUCUGUUA
env-397  CUGGGAGAGC..GCUUG.........GAAUUCCCA..GUAGUUUAGCAGUGUUAAUAGCC..UU..UUCU.........UUGGCC.AGAUU.AGGCGGUUAGUG.AAUCUGUUAAACAACAGAACGUAAAG.............................................
env-398  CUGGGAGAGC..GCUUG.........GAAUUCCCA..GUAGUUUAGCAGUGUUAAUAGCC..UU..UUCU.........UUGGCC.AGAUU.AGGCGGUUAGUG.AAUCUGUUAAACAACAGAACGUAAAGCUGUGGAUGGUCUCUUUUAGACUGGAGGCGCCGCGAGGUUUCUGUUA
env-399  CUGGGAGAGC..GCUUG.........GAAUUCCCA..GUAGUUUAGCAGUGUUAAUAGCC..UU..UUCU.........UUGGCC.AGAUU.AGGCGGUUAGUG.AAUCUGUUAAACAACAGAACGUAAAGCUGUGGAUGGUCUCUUUUAGACUGGAGGCGCCGCGAGGUUUCUGUUA
env-400  CUGGGAGAGC..GCUUG.........GAAUUCCCA..GUAGUUUAGCAGUGUUAAUAGCC..UU..UUCU.........UUGGCC.AGAUU.AGGCGGUUAGUG.AAUCUGUUAAACAACAGAACGUAAAGCUGUGGAUGGUCUCUUUUAGACUGGAGGCGCCGCGAGGUUUCUGUUA
env-401  CUGGGAGAGC..GCUUG.........GAAUUCCCA..GUAGUUUAGCAGUGUUAAUAGCC..UU..UUCU.........UUGGCC.AGAUU.AGGCGGUUAGUG.AAUCUGUUAAACAACAGAACGUAAAGCUGUGGAUGGUCUCUUUUAGACUGGAGGCGCCGCGAGGUUUCUGUUA
env-402  CUGGGAGAGC..GCUUG.........GAAUUCCCA..GUAGUUUAGCAGUGUUAAUAGCC..UU..UUCU.........UUGGCC.AGAUU.AGGCGGUUAGUG.AAUCUGUUAAACAACAGAACGUAAAGCUGUGGAUGGUCUCUUUUAGACUGGAGGCGCCGCGAGGUUUCUGUUA
env-403  CUGGGAGAGC..GCUUG.........GAAUUCCCA..GUAGUUUAGCAGUGUUAAUAGCC..UU..UUCU.........UUGGCC.AGAUU.AGGCGGUUAGUG.AAUCUGUUAAACAACAGAACGUAAAGCUGUGGAUGGUCUCUUUUAGACUGGAGGCGCCGCGAGGUUUCUGUUA
env-406  CUGGGAGAGC..GCUUG.........GAAUUCCCA..GUAGUUUAGCAGUGUUAAUAGCC..UU..UUCU.........UUGGCC.AGAUU.AGGCGGUUAGUG.AAUCUGUUAAACAACAGAACGUAAAGCUGUGGAUGGUCUCUUUUAGACUGGAGGCGCCGCGAGGUUUCUGUUA
env-408  GGGAAA.....GCACUUGGAA.........UUUCC.GGCAGUCUGGCAGUGUUAAUAGCC..UU..UUCU.........UUGGCC.GGAUA.AGGCGGUUAGUG.AAUCUGUCGGACAACAGAACAAAAGCUGUGGAUGGUCUCUUUUAGACUGGAGGCGCCGCGAGGUUUCUGUUAU
env-410  GGGAAA.....GCACUUGGAA.........UUUCC.GGCAGUCUGGCAGUGUUAAUAGCC..UU..UUCU.........UUGGCC.GGAUA.AGGCGGUUAGUG.AAUCUGUCGGACAACAGAACAAAAGCUGUGGAUGGUCUCUUUUAGACUGGAGGCGCCGCGAGGUUUCUGUUAU
env-411  GGGAAA.....GCACUUGGAA.........UUUCC.GGCAGUCUGGCAGUGUUAAUAGCC..UU..UUCU.........UUGGCC.GGAUA.AGGCGGUUAGUG.AAUCUGUCGGGCAACAGAACAAAAGCUGUGGAUGGUCUCUUUUAGACUGGAGGCGCCGCGAGGUUUCUGUUAU
env-412  UGGGAGAGC...GCUUG..........GAAUUCCC.UGUAGUUUAGCAGUGUUAAUAGCC..UU..UUCU.........UUGGCC.AGAUU.AGGCGGUUAGUG.AAUCUGUUAAACAACAGAACGUAAAGCUGUGGAUGGUCUCUUUUAGACUGGAGGCGCCGCGAGGUUUCUGUUA
Por-6-1  UGGGAAAGC...GCUUG..........GAAUUUCC.GGCAGUCUGGCAGUGUUAAUAGCC..UU..UUCU.........UUGGC..GGAUA.AGGCGGUUAGUG.AAUUUGUCGGACAACAGGACAAAAGCUGUGGAUGGUCUCUUUUAGACUGGAGGCGCCGCGAGGUUUCUGUUAU
Por-5-1  UGGGAAAGC...GCUUG..........GAAUUUCC.GGCAGUCUGGCAGUGUUAAUAGCC..UU..UUCU.........UUGGCC.GGAUA.AGGCGGUUAGUG.AAUUUGUCGGACAACAGGACAAAAGCUGUGGAUGGUCUCUUUUAGACUGGAGGCGCCGCGAGGUUUCUGUUAU
env-413  UGGGAAAGC...GCUUG..........GAAUUUCC.GGCAGUCUGGCAGUGUUAAUAGCC..UU..UUCU.........UUGGCC.GGAUA.AGGCGGUUAGUG.AAUCUGUCGGACAACAGAACAAAAGCUGUGGAUGGUCUCUUUUAGACUG.......................
env-414  UGGGAAAGC...GCUUG..........GAAUUUCC.GGCAGUCUGGCAGUGUUAAUAGCC..UU..UUCU.........UUGGCC.GGAUA.AGGCGGUUAGUG.AAUCUGUCGGACAACAGAACAAAAGCUGUGGAUGGUCUCUUUUAGACUGGAGGCGCCGCGAGGUUUCUGUUAU
env-418  UGGGAAAGC...GCUUG..........GAAUUUCC.GGCAGUCUGGCAGUGUUAAUAGCC..UU..UUCU.........UUGGCC.GGAUA.AGGCGGUUAGUG.AAUCUGUCGGACAACAGAACAAAAGCUGUGGAUGGUCUCUUUUA............................
env-419  UGGGAAAGC...GCUUG..........GAAUUUCC.GGCAGUCUGGCAGUGUUAAUAGCC..UU..UUCU.........UUGGCC.GGAUA.AGGCGGUUAGUG.AAUCUGUCGGACAACAGAACAAAAGCUGUGGAUGGUCUCUUUUA............................
env-420  UGGGAAAGC...GCUUG..........GAAUUUCC.GGCAGUCUGGCAGUGUUAAUAGCC..UU..UUCU.........UUGGCC.GGAUA.AGGCGGUUAGUG.AAUCUGUCGGACAACAGAACAAAAGCUGUGGAUGGUCUCUUUUAGACUGGAGGCGCCGCGAGGUUUCUUUGUUAU
env-421  UGGGAAAGC...GCUUG..........GAAUUUCC.GGCAGUCUGGCAGUGUUAAUAGCC..UU..UUCU.........UUGGCC.GGAUA.AGGCGGUUAGUG.AAUCUGUCGGACAACAGAACAAAAGCUGUGGAUGGUCUCUUUUAGACUGGAGGCGCCGCGAGGUUUCUGUUAU
env-422  UGGGAAAGC...GCUUG..........GAAUUUCC.GGCAGUCUGGCAGUGUUAAUAGCC..UU..UUCU.........UUGGCC.GGAUA.AGGCGGUUAGUG.AAUCUGUCGGACA..........................................................
env-423  UGGGAAAGC...GCUUG..........GAAUUUCC.GGCAGUCUGGCAGUGUUAAUAGCC..UU..UUCU.........UUGGCC.GGAUA.AGGCGGUUAGUG.AAUCUGUCGGACAACAGAACAAAAGCUGUGGAUGGUCUCUUUUA............................
Psp-8-1  UUGAGAAAGC..GCUUA.........GAAUCUCA..GCAGUUCAGUGGUGUUAAUAGCG..UU..CUC.........UCUGGCC.UGAUG.ACGCGGUUAGUG.AAUCUGCUGAUCAGCAAAGCAAAAGCUGUGGAUGGUCUCUUUUAGACUGGAGGCGCCGCGAGGUCUUUGUUAU
env-425  UUGAGAAAGC..GCUUA.........GAAUUCUCA..GCAGUUCAGUGGUGUUAAUAGCG..UU..CUCU.........CCGGCC.AGAUG.ACGCGGUUAGUG.AAUCUGCUGAUCAGCAAAGCAAAAGCUGUGGAUGGUCUCUUUUAGACUGGAGGCGCCGCGAGGUCUUUGUUAU
env-426  UUGAGAAAGC..GCUUA.........GAAAUCUCA..GCAGUUCAGUGGUGUUAAUAGCG..UU..CUCU.........CUGGCC.AGAUG.ACGCGGUUAGUG.AAUCUGCUGAUCAGCAAAGCAAAAGCUGUGGAUGGUCUCUUUUAGA..........................
env-429  UUGAGAAAGC..GCUUA.........GAAAUCUCA..GCAGUUCAGUGGUGUUAAUAGCG..UU..CUCU.........CUGGCC.AGAUG.ACGCGGUUAGUG.AAUCUGCUGAUCAGCAAAGCAAAAGCUGUGGAUGGUCUCUUUUAGACUGGAGGCGCCGCGAGGUCUUUGUUAU
env-431  UUGAGAAAGC..GCUUA.........GAAAUCUCA..GCAGUUCAGUGGUGUUAAUAGCG..UU..CUCU.........CUGGCC.AGAUG.ACGCGGUUAGUG.AAUCUGCUGAUCAGCAAAGCAAAAGCUGUGGAUGGUCUCUUUUAGACUGGAGGCGCCGCGAGGUCUUUGUUAU
env-432  UUGAGAAAGC..GCUUA.........GAAAUCUCA..GCAGUUCAGUGGUGUUAAUAGCG..UU..CUCU.........CUGGCC.AGAUG.ACGCGGUUAGUG.AAUCUGCUGAUCAGCAAAGCAAAAGCUGUGGAUGGUCUCUUUUAGACUGGAGGCGCCGCGAGGUCUUUGUUAU
env-433  UUGAGAAAGC..GCUUA.........GAAAUCUCA..GCAGUUCAGUGGUGUUAAUAGCG..UU..CUCU.........CUGGCC.AGAUG.ACGCGGUUAGUG.AAUCUGCUGAUCAGCAAAGCAAAAGCUGUGGAUGGUCUCUUUUAGACUGGAGGCGCCGCGAGGUC.......
env-434  UUGAGAAAGC..GCUUA.........GAAAUCUCA..GCAGUUCAGUGGUGUUAAUAGCG..UU..CUCU.........CUGGCC.AGAUG.ACGCGGUUAGUG.AAUCUGCUGAUCAGCAAAGCAAAAGCUGUGGAUGGUCUCUUUUAGA..........................
env-435  UUGAGAAAGC..GCUUA.........GAAAUCUCA..GCAGUUCAGUGGUGUUAAUAGCG..UU..CUCU.........CUGGCC.AGAUG.ACGCGGUUAGUG.AAUCUACUGAUCAGCAAAGCAAAAGCUGUGGAUGGUCUCUUUUAGACUGGAGGCGCCGCGAGGUCUUUGUUAU
env-436  UUGAGAAAGC..GCUUA.........GAAUUCUCA..GCGGUUCAGUGGUGUUAAUAGCG..UU..CUCU.........CUGGCC.AGAUG.ACGCGGUUAGUG.AAUCUGCUGAUCAGCAAAGCAAAAGCUGUGGAUGGUCUCUUUUAGACUGGAGGCGCCGCGAGGUCUUUGUUAU
env-437  UUGAGAAAGC..GCUUA.........GAAUUCUCA..GCGGUUCAGUGGUGUUAAUAGCG..UU..CUCU.........CUGGCC.AGAUG.ACGCGGUUAGUG.AAUCUGCUGAUCAGCAAAGCAAAAGCUGUGGAUGGUCUCUUUUAGACUGGAGGCGCCGCGAGGUCUUUGUUAU
env-438  UUGAGAAAGC..GCUUA.........GAAAUCUCA..GCAGUUCAGUGGUGUUAAUAGCG..UU..CUCU.........CUGGCC.AGAUG.ACGCGGUUAGUG.AAUCUGCUGAUCAGCAAAGCAAAAGCUGUGGAUGGUCUCUUUUAGACUGGAGGCGCCGCGAGGUCUUUGUUAU
env-439  UUGAGAAAGC..GCUUA.........GAAUUCUCA..GCAGUUCAGUGGUGUUAAUAGCG..UU..CUCU.........CUGGCC.AGAUG.ACGCGGUUAGUG.AAUCUGCUGAUCAGCAAAGCAAAAGCUGUGGAUGGUCUCUUUUAGACUGGAGGCGCCGCGAGGUCUUUGUUAU
env-440  UUGAGAA.....AGUGCUUAGAA......UUCUCA..GCAGUUCAGUGGUGUUAAUAGCG..UU..CUCU.........CUGGCC.AGAUG.ACGCGGUUAGUG.AAUCUGCUGAUCAGCAAAGCAAAAGCUGUGGAUGGUCUCUUUUAGACUGGAGGCGCCGCGAGGUCUUUGUUAU
env-441  UUGAGAA.....AGUGCUUAGAA......UUCUCA..GCAGUUCAGUGGUGUUAAUAGCG..UU..CUCU.........CUGGCC.AGAUG.ACGCGGUUAGUG.AAUCUGCUGAUCAGCAAAGCAAAAGCUGUGGAUGGUCUCUUUUA............................
env-442  UUGAGAA.....AGUGCUUAGAA......UUCUCA..GCAGUUCAGUGGUGUUAAUAGCG..UU..CUCU.........CUGGCC.AGAUG.ACGCGGUUAGUG.AAUCUGCUGAUCAGCAAAGCAAAAGCUGUGGAUGGUCUCUUUUAGACUGGAGGCGCCGCGAGGUCU.......
env-443  UUGAGAA.....AGUGCUUAGAA......UUCUCA..GCAGUUCAGUGGUGUUAAUAGCG..UU..CUCU.........CUGGCC.AGAUG.ACGCGGUUAGUG.AAUCUGCUGAUCAGCAAAGCAAAAGCUGUGGAUGGUCUCUUUUAGA..........................
Pou-1-1  UUGAGAA.....AGUGCUUAGAA......UUCUCA..GCAGUUCAGUGGUGUUAAUAGCG..UU..CUCU.........CUGGCC.AGAUG.ACGCGGUUAGUG.AAUCUGCUGAUCAGCAAAGCAAAAGCUGUGGAUGGUCUCUUUUAGACUGGAGGCGCCGCGAGGUCUUUGUUAU
env-444  UUGAGAA.....AGUGCUUAGAA......UUCUCA..GCAGUUCAGUGGUGUUAAUAGCG..UU..CUCU.........CUGGCC.AGAUG.ACGCGGUUAGUG.AAUCUGCUGAUCAGCAAAGCAAAAGCUGUGGAUGGUCUCUUUUA............................
env-445  UUGAGAA.....AGUGCUUAGAA......UUCUCA..GCAGUUCAGUGGUGUUAAUAGCG..UU..CUCU.........CUGGCC.AGAUG.ACGCGGUUAGUG.AAUCUGCUGAUCAGCAAAGCAAAAGCUGUGGAUGGUCUCUUUUAGACUGGAGGCGCCGCGAGGUCUUUGUUAU
env-446  UUGAGAA.....AGUGCUUAGAA......UUCUCA..GCAGUUCAGUGGUGUUAAUAGCG..UU..CUCU.........CUGGCC.AGAUG.ACGCGGUUAGUG.AAUCUGCUGAUCAACAAAGCAAAAGCUGUGGAUGGUCUCUUUUAGACUGGAGGCGCCGCGAGGUCUUUGUUAU
env-447  UUGAGAA.....AGUGCUUAGAA......UUCUCA..GCAGUUCAGUGGUGUUAAUAGCG..UU..CUCU.........CUGGCC.AGAUG.ACGCGGUUAGUG.AAUCUGCUGAUCAGCAAAGCAAAAGCUGUGGAUGGUCUCUUUUAGACUGGAGGCGCCGCGAGGUCUUUGUUAU
Pou-2-1  UUGAGAA.....AGUGCUUAGAA......UUCUCA..GCAGUUCAGUGGUGUUAAUAGCG..UU..CUCU.........CUGGCC.AGAUG.ACGCGGUUAGUG.AAUCUGCUGAUCAGCAAAGCAAAAGCUGUGGAUGGUCUCUUUUAGACUGGAGGCGCCGCGAGGUCUUUGUUAU
env-448  UUGGGG......AACACUUAGAA.......CCCCA....CGUACAGAAGUGUUAAUAGCG..GU..GCUU.........UUGGCC.GAACA.GCUGGUUAGUG.AAUCGUCUGUACAAGCUUGAUAAAAAAAACGUUUUUUAGUAGUAAAUAUUUUCAAAAUUAGAUUUAACACAUUAA
env-449  CGGGAAUAC...GCUUAG.........G.AUUCUC..AUAGCGUGGUCGUGUUAAUAGUU..CU..UCUU.........UUGGCC.GAGGA.GAGCGGUUAGUG.AAUCGGCCGUCGCAUCAGAGAUAAUGCUGUGGAUGGUCUCUUUUAGACUGGAGG..................
Pba-2-1  UGGGAAUAC...GCUUAG.........G.AUUCUC..AUAGCGUGGUCGUGUUAAUAGUU..CU..UCUU.........UUGGCC.GAGGA.GAGCGGUUAGUG.AAUCGGCCGUCGCAUCAGAGAUAAUGCUGUGGAUGGUCUCUUUUAGACUGGAGGCGCCGCGAGGUCUCUGUUA
Pba-1-1  UGGGAAUAC...GCUUAG.........G.AUUCUC..AUAGCGUGGUCGUGUUAAUAGUU..CU..UCUU.........UUGGCC.GAGGA.GAGCGGUUAGUG.AAUCGGCCGUCGCAUCAGAGAUAAUGCUGUGGAUGGUCUCUUUUAGACUGGAGGCGCCGCGAGGUCUCUGUUA
env-450  UGGGAAUAC...GCUUAG.........G.AUUCUC..AUAGCGCGGUCGUGUUAAUAGUGU..UU..UCUU.........UUGGCC.GAGAA.GAGCGGUUAGGG.AAUCGGCUGUCGCGUCAGAGAUAAUGCUGUGGAUGGUCUCUUUUAGACUGGAGGCGCCGCGAGGUCUGUUA
Pde-2-1  UGGAGGUU....GAUACUUAG.........AGCCUCC.....GGCACGUGCGUGUUAAUAGUGU.CA..GCUU.........UUGGCC.GAGCU.GUACGGUUAGUG.AAUCUCACGGGCAACAAAAGAAAACGUUUUUUAGUAGUAAAUAUUAUUUUUUUAGAUUUGACACUUUAAGU
Pbe-1-1  CAGGGGC.....ACGCUUAGA.........GCCCCU.....CGCACCUUGUGUUAAUAGUG..AU..GCCU.........UUGGCC.GGGCA.UCACGGUUAGUG.AAUCAAGGGUGCAGCAGAGCGAAAGCCAUGGAUGGUCUCUUUUAGACUGGAGGCGCCGUGAGGUCUCUGUAUA
env-451  AAGGGG......AAACCUUAGAA........CCCCU.....UUGCGAGAGUGUUAAUAGUCA.AU..AGCU..........UGGCC.GGCUU.GAUCGGUUAGUG.AAUCGCUCGCAAGAAAAAUUAAGUUUUUUAGUAGUAAAAUUAAAAUUUUAAUUUGACACUUUAAGUUACAAG
env-452  UGGGGG......AAGGCUUAGAG.......CCCCC.....AGCCCGGGAGUGUUAAUAGUG..CU..GCA..........UUGGCC.AUGCA.CCACGGUUAGUG.AAUCGCCCGUGCAAAGUUCAAGAGCCUGAAGCGAGAAGUUUCAGGUGUGCGUUCUUGAGCGUCAGCCCGAGUC
env-453  GGGGGU......GGCUUAGA..........GCCCCC...GCUUGCGGAAGUGUUAAUAGUA..GU..GUUU.........UUGGCC.GGACA.UUACGGUUAGUG.AAUCGACCGCAAAUAAAGAGAAAAAACGUUUUUUAGUAGUAAUUACAAACAUUAAAUUUAACACUUUAAGUUA
env-454  UGGAGGUU....AACUUAG...........GACCUCC..GCGUACAGUUCGUGUUAAUAGUC..AC..AUCU.........UUGGCC.AGGUG.UGACGGUUAGUG.AAUCGGCUGUCGAAGAAAAACGUUUUUUAGUAGUAUUACAAUUUUAAAAUUGAAAUUUGACACUUUAAGUUAC
env-455  UGGAGGUU....AACUUAG...........GACCUCC..GCGUACAGUUCGUGUUAAUAGUC..AU..AUCU.........UUGGCC.AGGUG.UGACGGUUAGUG.AAUCGGCUGUCGAAGAAAAACGUUUUUUAGUAGUAUUACAAUUUUAAAAUUGAAAAUUGGACACUUUAAGUUAC
env-457  CGGGGGU.....AACUUAGG..........GCCUCC....AUACACAGUUCGUGUUAAUAGUU..GU..ACUU.........UUGGCC.GAGUG.CAGCGGUUAGUG.AAUCGGCUGUAAAAAUAAGAUAUAAACGUUUUUUAGUAGUAUUACAAUUUUAAUUUAAAUUUGACACUUUAAG
env-458  CGGGGGU.....AACUUAGG..........GCCUCC....ACUACAGUUCGUGUUAAUAGUU..GU..ACUU.........UUGGCC.GAGUG.CAACGGUUAGUG.AAUCGGCUGUAAAAAC.............................................................
env-462  CGGGGGU.....AACUUAGG..........GCCUCC....ACUACAGUUCGUGUUAAUAGUU..GU..ACUU.........UUGGCC.GAGUG.CAACGGUUAGUG.AAUCGGCUGUAAAACAAGAUAUAAACGUUUUUUAGUAGUAUUACAAUUUUAAUUUAAAUUUGACACUUUAAGU
env-463  CGGGGGU.....AACUUAGG..........GCCUCC....ACUACAGUUCGUGUUAAUAGUU..GU..ACUU.........UUGGCC.GAGUG.CAACGGUUAGUG.AAUCGGCUGUAAAAACAAGAUAUAAACGUUUUUUAGUAGUAUUACAAUUUUAAUUUAAAUUUGACACUUUAAGU
env-464  CGGGGGU.....AACUUAGG..........GCCUCC....ACUACAGUUCGUGUUAAUAGUU..GU..ACUU.........UUGGCC.GAGUG.CAACGGUUAGUG.AAUCGGCUGUAAAAACAAGAUAUAAACGUUUUUUAGUAGUAUUACAAUUUUAAUUUAAAUUUGACACUUUAAG
env-466  UGAGGGCAC...GCUUA.........GAGCCCUC.....GUCGGGCGUGUGUUAAUAGCGA.GU..GUC..........AUUUGGCCUGGACA.GCGUGGUUAGAG.AAUCGCUCGAAGACAUUCGAACACCGCAGGGGAACGCCGUUUCUCAUGCAGGUUAUUUUUUGUCAUCACCAA
env-467  CUGGGG......AACACUUAGAGC.......CCCCA.....CGGACGUAGUGUUAAUAGUCG.CU..GUCU.........UUGGCC.GGGCA.CGACGGUUAGUG.AAUCGACGUCCAAACGGCAGAAAGCACACAUGCCAUAUAUAUUAAUGUAUGACAAAGUCUGUAAAAAGGCGUG
env-468  CGAGGGAGU...ACUUA...........GGUCCCUC....AGAAGCGGUGUGUUAAUAGUG..AU..GCUU.........UUGGCC.AGGCA.UGACGGUUAGUG.AAUCACGGCUUCACGACACAGAAAAUUGUUUUUUAGUAGUAAAUUUAAAAUUUGAAUGAACACUUUAAG...
env-469  CGGGGAGU....ACUUAG..........GC.CCCCC....AGGAGCGACGUGUUAAUAGUU..GU..GCUU.........UUGGCC.AGGCA.UAGUGGUUAGUG.AAUCGAAGCCCCAACGAAGAAAAAACGUUUUUUAGUAGUAAACCAAAAGAAAUUUAACACUUUAAGUUACAA
env-470  UUGGGUAC....ACUUA...........GAGCCCA...GUUUGGAGGCAGUGUUAAUAGUC..GA..GCGU.........UUGGCC.GCGC..CGAUGGUUAGUG.AAUCUGCGUUCAGCAAAAAUGUUUUUUAGUAGUAAUAAAAUUUUCUAAAAGUCAAGAUUUGACACUUUAAGU
Ppa-3-1  AGGGG.......CGACUCUUAGAAA.......CCCC.GGUAGUGCAAUGGUGUUAAUAGCU..UU..ACUU.........UUGGCC.GAGUA.UCGCAGUUAGUG.AAUCUAUUGCACGGCAGAGAAAAAGCUAUGGAUGGUCUCUUUUAGACUGGAGGCGCCGCGAGGUCUCUGUUAU
env-471  AGGGG.......CGACUCUUAGAAA.......CCCC.AGUAGUGCAAUGGUGUUAAUAGCC..UU..ACUU.........UUGGCC.GAGUA.UCGCAGUUAGUG.AAUCUAUUGCACGGCAGAGAAAAAGCUAUGGAUGGUCUCUUUUAGACUGGAGGCGCCGUGAGGUCUCUGUUAU
env-473  AGGGG.......CGACUCUUAGAAA.......CCCC.AGUAGUGCAAUGGUGUUAAUAGCU..UU..ACUU.........UUGGCC.GAGUA.UCGCAGUUAGUG.AAUCUAUUGCACGGCAGAGAAAAAGCUUUGGAUGGUCUCUUUUAGACUGGAGGCGCCGAGAGGUCUCUGUUAU
env-474  AGGGG.......CGACUCUUAGAAA.......CCCC.AGUAGUGCAAUGGUGUUAAUAGCU..UU..ACUU.........UUGGCC.GAGUA.UCGCAGUUAGUG.AAUCUAUUGCACGGCAGAGAAAAAGCUUUGGAUGGUCUCUUUUAGACUGGAGGCGCCGAGAGGUCUCUGUUAU
env-476  AGGGGCGACU..CUU...........AGGAACCCC.AGUAGUGCAAUGGUGUUAAUAACU..UU..ACUU.........UUGGCC.GAGUA.UCGCAGUUAGUG.AAUCUAUUGCACGGCAGAGCAAAAGCUUUGGAUGGUCUCUUUUAGACUGGAGGCGCCGAGAGGUCUCUGUUAU
env-477  AGGGGCGACU..CUU...........AGGAACCCC.AGUAGUGCAAUGGUGUUAAUAACU..UU..ACUU.........UUGGCC.GAGUA.UCGCAGUUAGUG.AAUCUAUUGCACGGCAGAGCAAAAGCUUUGGAUGGUCUCUUUUAGACUGGAGGCGCCGAGAGGUCUCUGUUAU
env-483  AUGGGG......GAAGUCUUAGAA.......CCCCA.....UGGAUGCCGUGUUAAUAGCG..AU..GCCU.........UUGGCC.GGGCA.UGGCGGUUAGUG.AAUCGGCAUUCACGAGACGCAUCGCAUAAAGACCGGCGAGUGCCUUCCAAGACUAAAAGGCAUAGUCUCGAUA
env-485  AUGGGG......GAAGUCUUAGAA.......CCCCA.....UGGAUGCCGUGUUAAUAGCU..GU..GCCU.........UUGGCC.GGGCA.UGGCGGUUAGUG.AAUCGGCAUUCACGAGACGCAUCGCAUAAAGACCGGCGAGUGCCUUCCAAGACUAAAAGGCAUAGUCUCGAUA
env-486  AGGGGGU.....AAAUCCUUAGA.......GCCCCC....AGCGUGGCCGUGUUAAUAGUG..GU..GUCU.........UUGGCC.GGGCA.UCAUGGUUAGUG.AAUCGGCUGCGCAGUCAUCGAGGUUUUGUAAGCGGGAGCACCCAGUGGCUCGCUUGCUUGUUUGUGUGCGCUG
env-487  AGGGGA......AAACACUUAGAA........UCCCC...GUGGACUAGAGUGUGAAUAGCA..AC..GUCU.........UUGGCC.GGACG.UCGUGGUUCGGG.AAUCGCUGGUUCAACCAUCCGCAACACGCCUGUUUACAGGCCCCGCGGAGCGCACGUUAGACUAACAACCGGG
env-488  AGGGGAAACU..CUU...........AAGAUCCCC...GUGAACAGGGGUGUGAAUAGCG..GC..GUCU.........UUGGCC.GGGCG.AAGCGGUUAGUG.AAUCGUCUGUUCAAGCACAAGUUAGACUAACAGCCGGGAGUUAGACCUCGGAUGGGAGUCAAUGUGCGUACAU

         .<<<<<<<<<<<............>>>>>>>>>>>...............<<<<<.<<<..<<..<<<<<...............>>>>>>.>>>>.>>>>>.................................................................
         .222222.1111............1111.222222...............02200.122..22..2222...............22222.2221.00220................................................................
         •GGGG••-----R•••oooo----------o•CCCC----oo•••••••YGUGU•AAUAGY•--•Y--RYYU---------UUGGCC-GRRY•-••RYGGUU•GUG-AAUCR•••••••RR••R•••••••••••••••••••••••••••••••••••••••••••
          5'                                                                                                              3'
```

```
env-489    UGGGGUACU...CUU.............AGAACCCC.....GGAGAAGCGUGUUAAUAGUC..GU..GCUU.........UUGGCC.GGGCA.CGAUGGUUAGGG.AAUCGCCCUCCAAAGUAUGAAGAAAUUGUUUUUUAGUAGUAAAAUUUUUAAAUCGAAAUGAACACAUUAAGUU
env-490    GGGGAUUUU...CUU.............AGAAUCCC....GUACGUAACAUGUUAAUAGCA..UG..GUUU.........UUGGCC.GAGCU.GUGUGGUUAGGUAAAGCAUGUGCGUGAGUUAAUAAAAACGUUUUUUAGUAGUAAAUUUAAUUAAAAAAAUGAACACUUUAAGUUAC
env-491    GGGGAUUUU...CUU.............AGAAUCCC....GUACGUAACAUGUUAAUAGCA..UG..GUUU.........UUGGCC.GAGCU.GUGUGGUUAGGUAAAGCAUGUGCGUGAAUUAAUAAAAACGUUUUUUAGUAGUAAAUUUAAUUAAAAAAAUGAACACUUUAAGUUAU
env-492    GGGGAUUUU...CUU.............AGAAUCCC....GUACGUAACAUGUUAAUAGCA..UG..GUUU.........UUGGCC.GAGCU.GUGUGGUUAGGUAAAGCAUGUGCGUGAGUUAAUAAAAACGUUUUUUAGUAGUAAAUUUAAUUAAAAAAAUGAACACUUUAAGUUAC
env-493    GGGGAUUUU...CUU.............AGAAUCCC....GUACGUAACAUGUUAAUAGCA..UG..GUUU.........UUGGCC.GAGCU.GUGUGGUUAGGUAAAGCAUGUGCGUGAAUUAAUAAAAACGUUUUUUAGUAGUAAAUUUAAUUAAAAAAAUGAACACUUUAAGUUAC
Pam-1-1    GGGGAUUUU...CUU.............AGAAUCCC....GUACGUAACAUGUUAAUAGCA..UG..GUUU.........UUGGCC.GAGCU.GUGUGGUUAGGUAAAGCAUGUGCGUGAAUUAAUAAAAACGUUUUUUAGUAGUAAAUUUAAUUAAAAAAAUGAACACUUUAAGUUAC
Pam-3-1    GGGGAUUUU...CUU.............AGAAUCCC....GUACGUAACAUGUUAAUAGCA..UG..GUUU.........UUGGCC.GAGCU.GUGUGGUUAGGUAAAGCAUGUGCGUGAAUUAAUAAAAACGUUUUUUAGUAGUAAAUUUAAUUAAAAAAAUGAACACUUUAAGUUAC
Pbi-1-1    GGGGAUUUU...CUU.............AGAAUCCC....AUACGUUACGUGUUAAUAACA..UA..GUCU.........UUGGCC.GGGCU.AUGUGGUUAGGAAAAGCAUGUGCGUGAAUUAAUAAAAACGUUUUUUAGUAGUAAAAUUUAAUUUUUAAAAUGGACACUUUAAGUUA
Pve-1-1    AGGGGUACU...CUU.............AGAGCCCC....AUACGAUGCAUGUUAAUAGUG..UU..GUCU.........UUGGCC.GGGCG.ACAUGGUUAGGUAAGGCAUGUUCGUGAAUAGAAAACGUUUUUUAGUAGUAAAAUUAUUAAUUUUUUAGAAUGGACACUUUAAGUUA
Pve-3-1    AGGGGUACU...CUU.............AGAGCCCC....AUACGAUACAUGUUAAUAGUG..UU..GUCU.........UUGGCC.GGGCG.ACAUGGUUAGGAAAAGCAUGAUCGUGAAAUAGAAAACGUUUUUUAGUAGUAAAAUUAUUAAUUUUUUAGAAUGGACACUUUAAGUU
env-494    AGGGGUACU...CUU.............AGAGCCCC....AUACGAUGCAUGUUAAUAGUG..UU..GUCU.........UUGGCC.GGGCG.ACAUGGUUAGGGAAAGCAUGAUCGUGAAAUAGAAAACGUUUUUUAGUAGUAAAAU................................
env-495    CGGGG.......AAACACUUAGAA........CCCC....AUACGAUACGUGUUAAUAGUG..UU..GUCU.........UUGGCC.GGACA.GCAUGGUUAGGUCAAGCACAGUCGUAGAAAUAGAAAACGUUUUUUAGUAGUAAAAUUAUUAAUUUUUUAGAAUGAACACUUUAAGU
env-497    UGGGG.......AUACACUUAGAA........CCCC....AUACGAUACGUGUUAAUAGUG..UU..GUCU.........UUGGCC.GGGCA.GCAUGGUUAGGUCAAGCACAGUCGUAAAAAUAGAAAACGUUUUUUAGUAGUAAAAUUAUUAAUUUUUUAGAAUGGACACUUUAAGU
Psc-1-1    AGGGG.......GAAUUCUUAGAA........CCCC....AUACGAUACGUGUUAAUAGUG..UU..GUCU.........UUGGCC.GGGCA.AUAUGGUUAGGAAAAGCACAAUCGUAGAAAUAGAAAACGUUUUUUAGUAGUAAAAUUAUUAAUUUUUUAGAAUGGACACUUUAAGUU
env-498    AGGGG.......GAAUUCUUAGAA........CCCC....AUACGAUACGUGUUAAUAGUG..UU..GUCU.........UUGGCC.GGGCA.AUAUGGUUAGGAAAAGCACAAUCGUAGAAAUAGAAAACGUUUUUUA.......................................
Psp-6-1    AGGGG.......GAAUUCUUAGAA........CCCC....AUACGAUGCGUGUUAAUAGUG..UU..GUCU.........UUGGCC.GGGCA.ACAUGGUUAGGAAAAGCACAGUCGUAGAAACAGAAAACGUUUUUUAGUAGUAAAAUUAUUAAUUUUUUGAAUGGACACUUUAAGU
Pme-3-1    AGGGG.......AAUUUUCUUAGAA.......CCCC....AUACGAUACGUGUUAAUAGUG..UU..GUCU.........UUGGCC.GGGCA.GCAUGGUUAGGUCAAGCAUUGUCGUAAAAAUAGAAAACGUUUUUUAGUAGUAAAAUUAUUAAUUUUUUAGAAUGGACACUUUAAGU
env-501    AGGGGU......AAUCUUAGA..........GCCCC....AUACGAUUCAUGUUAAUAGUG..UU..GUCU.........UUGGCC.GGGCA.GCAUGGUUAGGUCAAGCACAGUCGUAGAAAUAGAAAACGUUUUUUAGUAGUAAAAUUAUUAAUUUUUUAGAAUGGACACUUUAAGU
env-502    AGGGGU......AAUCUUAGA..........GCCCC....AUACGAUUCAUGUUAAUAGUG..UU..GUCU.........UUGGCC.GGGCA.GCAUGGUUAGGU.AAACACAUGAUCGUAAAAUAGAAAACGUUUUUUAGUAGU..................................
env-503    AGGGGU......AAUCUUAGA..........GCCCC....AUACGAUUCAUGUUAAUAGUG..UU..GUCU.........UUGGCC.GGGCG.GCAUGGUUAGGU.AAACACAUGAUCGUAAAAUAGAAAACGUUUUUUAGUAGU..................................
env-504    AGGGGU......AAUCUUAGA..........GCCCC....AUACGAUUCAUGUUAAUAGUG..UU..GUCU.........UUGGCC.GGGCG.GCAUGGUUAGGU.AAACACAUGGUCGUAAAAUAGAAAACGUUUUUUAGUAGUAAAAUUAUUAAUUUUUUAGAAUGGACACUUUAAG
env-507    AGGGGU......AAUCUUAGA..........GCCCC....AUACGAUUCAUGUUAAUAGUG..UU..GUCU.........UUGGCC.GGGCG.GCAUGGUUAGGU.AAACACAUGGUCGUAAAAUAGAAAACGUUUUUUAGUAGU..................................
Phi-2-1    AGGGGU......AAUCUUAGA..........GCCCC....AUACGAUUCAUGUUAAUAGUG..UU..GUCU.........UUGGCC.GGGCG.GCAUGGUUAGGU.AAACACAUGGUCGUAAAAUAGAAAACGUUUUUUAGUAGUAAAAUUAUUAAUUUUUUAGAAUGGACACUUUAAG
env-508    AGGGGU......AAUCUUAGA..........GCCCC....AUACGAUUCAUGUUAAUAGUG..UU..GUCU.........UUGGCC.GGGCG.GCAUGGUUAGGU.AAACACAUGGUCGUAAAAUAGAAAACGUUUUUUAGUAGUAAAAUUAUUAAUUUUUUAGAAUGGACACUUUAAG
env-509    AGGGGU......AAUCUUAGA..........GCCCC....AUACGAUUCAUGUUAAUAGUG..UU..GUCU.........UUGGCC.GGGCG.GCAUGGUUAGGU.AAACACAUGGUCGUAAAAUAGAAAACGUUUUUUAGUAGUAAAAUUAUUAAUUUUUUAUGANNNNNNNNNAGCC
Phi-2-2    AGGGGU......AAUCUUAGA..........GCCCC....AUACGAUUCAUGUUAAUAGUG..UU..GUCU.........UUGGCC.GGGCG.GCAUGGUUAGGU.AAACACAUGGUCGUUAAAUAGAAAACGUUUUUUAGUAGUAAAAUUAUUAAUUUUUUAGAAUGGACACUUUAAG
env-510    GGGGGAUUU...UCUU............AGAACCCC....AUAUGAUACGUGUUAAUAGUG..UU..GUCU.........UUGGCC.GGGCA.GCAUGGUUAGGUCAAGCACAGUCGUAAAAAUAGAAAACGUUUUUUAGUAGUAAAAUUAUUAAUUUUUUAGAAUGGACACUUUAAGU
env-511    GGGGGAUUU...UCUU............AGAGCCCC....AUACGAUACGUGUUAAUAGUG..UU..GUCU.........UUGGCC.GGGCA.GCAUGGUUAGGUCAAGCACAGUCGUAAAAACAGAAAACGUUUUUUAGUAGUAAAAUUAUUAAUUUUUUAGAAUGGACACUUUAAGU
env-512    GGGGGAUUU...UCUU............AGAACCCC....AUACGAUACGUGUUAAUAGUG..UU..GUCU.........UUGGCC.AGGCA.GCAUGGUUAGGUCAAGCACAGUCGUAAAAAUAGAAAACGUUUUUUAGUAGUAAAAUUAUUAAUUUUUUAGAAUGGACACUUUAAGU
env-514    GGGGGAUUU...UCUU............AGAACCCC....AUACGAUACGUGUUAAUAGUG..UU..GUCU.........UUGGCC.GGGCA.GCAUGGUUAGGACAAGCACCGUCGUAGAAAUAGAAAACGUUUUUUAGUAGUAAAAUUAUUAAUUUUUUAGAAUGGACACUUUAAGU
env-516    GGGGGAUUU...UCUU............AGAACCCC....AUACGAUUCGUGUUAAUAGUG..UU..GUCU.........UUGGCC.GGGCA.GCAUGGUUAGGUCAAGCACAGUCGUAGAAAUAGAAAACGUUUUUUAGUAGUA................................
env-517    GGGGGAUUU...UCUU............AGAACCCC....AUACGAUUCGUGUUAAUAGUG..UU..GUCU.........UUGGCC.GGGCA.GCAUGGUUAGGACAAGCACAGUCGUAGAAAUAGAAAACGUUUUUUAGUAGUAAAAUUAUU.........................
env-519    GGGGGAUUU...UCUU............AGAACCCC....AUACGAUACGUGUUAAUAGUG..UU..GUCU.........UUGGCC.GGACA.GCAUGGUUAGGUAAAGCACAGUCGUAGAAAUAGAAAACGUUUUUUAGUAGUAAAAUUAUUAAUUUUUUAGAAUGGACACUUUAAGU
Psp-5-1    CGGGGAU.....ACACUUAG..........AACCCC....AUACGAUACGUGUUAAUAGUG..UU..GUCU.........UUGGCC.GGACA.GCAUGGUUAGGUCAAGCACAGUCGUAGAAAUAGAAAACGUUUUUUAGUAGUAAAAUUAUUAAUUUUUUAGAAUGGACACUUUAAGU
env-520    CGGGGAU.....ACACUUAG..........AACCCC....AUACGAUACGUGUUAAUAGUG..UU..GUCU.........UUGGCC.GGACA.GCAUGGUUAGGUCAAGCACAGUCGUAGAAAUAGAAAACGUUUUUUAGUAGUAAAAUUAUUAAUUUUUUAGAAUGAACACUUUAAGU
env-521    CGGGGAU.....ACACUUAG..........AACCCC....AUACGAUACGUGUUAAUAGUG..UU..GUCU.........UUGGCC.GGACA.GCAUGGUUAGGUCAAGCACAGUCGUAGAAAUAGAAAACGUUUUUUAGUAGUAAAAUUAUUAAUUUUUUAGAAUGAACACUUUAAGU
env-522    CGGGGAU.....ACACUUAG..........AACCCC....AUACGAUACGUGUUAAUAGUG..UU..GUCU.........UUGGCC.GGACA.GCAUGGUUAGGUCAAGCACAGUCGUAGAAAUAGAAAACGUUUUUUAGUAGUAAAAUUAUUAAUUUUUUAGAAUGAACACUUUAAGU
env-523    CGGGGAU.....UUUCUUAG..........AACCCC....AUACGAUACGUGUUAAUAGUG..CU..GUCU.........UUGGCC.GGGCA.GCAUGGUUAGGUCAAGCACAGUCGUAGAAAUAGAAAACGUUUUUUAGUAGUAAAAUUAUUAAUUUUUUAGAAUGGACACUUUAAGU
env-525    GGGGGAU.....UUUCUUAG..........AACCCC....AUACGAUACGUGUUAAUAGUG..CU..GUCU.........UUGGCC.GGGCA.GCAUGGUUAGGUCAAGCACAGUCGUAGAAAUAGAAAACGUUUUUUAGUAGUNNNNNNNNNNNNNNNNNNNNNNNNNNNNNNNNNNNN
env-526    CGGGGAU.....UUUCUUAG..........AACCCC....AUACGAUACGUGUUAAUAGUG..UU..GUCU.........UUGGCC.GGGCA.GCAUGGUUAGGUCAAGCACAGUCGUAAAAAUAGAAAACGUUUUUUAGUAGUAAAAUUAUUAAUUUUUUAGAAUGGACACUUUAAGUU
env-527    GGGGGAU.....UUUCUUAG..........AACCCC....AUACGAUACGUGUUAAUAGUG..UU..GUCU.........UUGGCC.GGGCA.GCAUGGUUAGGUCAAGCACAGUCGUAAAAAUAGAAAACGUUUUUUAGUAGUAAAAUUAUUAAUUUUUUAGAAUGGACACUUUAAGU
env-531    CGGGGAU.....UUUCUUAG..........AACCCC....AUACGAUACGUGUUAAUAGUG..UU..GUCU.........UUGGCC.GGGCA.GCAUGGUUAGGUCAAGCACAGUCGUAAAAAUAGAAAACGUUUUUUAGUAGUAAAAUUAUUAAUUUUUUAGAAUGGACACUUUAAGU
env-532    CGGGGAU.....UUUCUUAG..........AACCCC....AUACGAUACGUGUUAAUAGUG..UU..GUCU.........UUGGCC.GGGCA.GCAUGGUUAGGUCAAGCACAGUCGUAAAAAUAGAAAACGUUUUUUAGUAGUAAAAUUAUUAAUUUUUUAGAAUGGACACUUUAAGUU
env-534    GGGGGAU.....UUUCUUAG..........AACCCC....AUACGAUACGUGUUAAUAGUG..UU..GUCU.........UUGGCC.GGGCA.GCAUGGUUAGGUCAAGCACAGUCGUAAAAAUAGAAAACGUUUUUUAGUAGUAAAAUUAUUAAUUUUUUAAAAUGGACACUUUAAGU
env-535    CGGGGAU.....UUUCUUAG..........AACCCC....AUACGAUACGUGUUAAUAGUG..UU..GUCU.........UUGGCC.GGGCA.GCAUGGUUAGGUCAAGCACAGUCGUAAAAAUAGAAAACGUUUUUUAGUAGUAAAAUUAUUAAUUUUUUAGAAUGGACACUUUAAGU
env-537    CGGGGAU.....UUUCUUAG..........AACCCC....AUACGAUACGUGUUAAUAGUG..UU..GUCU.........UUGGCC.GGGCA.GCAUGGUUAGGUCAAGCACAGUCGUAAAAAUAGAAAACGUUUUUUAGUAGUAAAAUUAUUAAUUUUUUAGAAUGGACACUUUAAGU
env-538    CGGGGAU.....UUUCUUAG..........AACCCC....AUACGAUACGUGUUAAUAGUG..UU..GUCU.........UUGGCC.GGGCA.GCAUGGUUAGGUCAAGCACAGUCGUAAAAAUAGAAAACGUUUUUUAGUAGUAAAAUUAUUAAUUUUUUAGAAUGGACACUUUAAGU
env-552    CGGGGAU.....UUUCUUAG..........AACCCC....AUACGAUACGUGUUAAUAGUG..UU..GUCU.........UUGGCC.GGGCA.GCAUGGUUAGGUCAAGCACAGUCGUAAAAAUAGAAAACGUUUUUUAGUAGUAAAAUUAUUAAUUUUUUAGAAUGGACACUUUAAGUU
env-553    CGGGGAU.....UUUCUUAG..........AACCCC....AUACGAUACGUGUUAAUAGUG..UU..GUCU.........UUGGCC.GGGCA.GCAUGGUUAGGUCAAGCACAGUCGUAAAAAUAGAAAACGUUUUUUAGUAGUA................................
env-554    GGGGGAU.....UUUCUUAG..........AACCCC....AUACGAUACGUGUUAAUAGUG..UU..GUCU.........UUGGCC.GGGCA.GCAUGGUUAGGUCAAGCACAGUCGUAAAAAUAGAAAACGUUUUUUAGUAGUAAAAUUAUUAAUUUUUUAGAAUGGACACUUUAAGU
env-555    GGGGGAU.....UUUCUUAG..........AACCCC....AUACGAUACGUGUUAAUAGUG..UU..GUCU.........UUGGCC.GGGCA.GCAUGGUUAGGUCAAGCACAGUCGUAGAAAUAGAAAACGUUUUUUAGUAGUAAAAUUAUUAAUUUUUUAGAAUGGACACUUUAAGU
env-557    CGGGGAU.....UUUCUUAG..........AACCCC....AUACGAUACGUGUUAAUAGUG..UU..GUCU.........UUGGCC.GGGCA.GCAUGGUUAGGUCAAGCACAGUCGUAAAAAUAGAAAACGUUUUUUAGUAGUAAAAUUAUUAAUUUUUUAGAAUGGACACUUUAAGU
env-558    GGGGGAU.....UUUCUUAG..........AACCCC....AUACGAUACGUGUUAAUAGUG..UU..GUCU.........UUGGCC.GGGCA.GCAUGGUUAGGUCAAGCACAGUCGUAAAAAUAGAAAACGUUUUUUAGUAGUAAAAUUAUUAAUUUUUUAGAAUGGACANNNNNNNNN
env-559    CGGGGAU.....UUUCUUAG..........AACCCC....AUACGAUACGUGUUAAUAGUG..UU..GUCU.........UUGGCC.GGGCA.GCAUGGUUAGGUCAAGCACAGUCGUAAAAAUAGAAAACGUUUUUUAGUAGUAAAAUUAUUAAUUUUUUAGAAUGGACACUUUAAGU
env-560    CGGGGAU.....UUUCUUAG..........AACCCC....AUACGAUACGUGUUAAUAGUG..UU..GUCU.........UUGGCC.GGGCA.GCAUGGUUAGGUCAAGCACAGUCGUAAAAAUAGAAAACGUUUUUUAGUAG..................................
env-561    CGGGGAU.....UUUCUUAG..........AACCCC....AUACGAUACGUGUUAAUAGUG..UU..GUCU.........UUGGCC.GGGCA.GCAUGGUUAGGUCAAGCACAGUCGUAAAAAUAGAAAACGUUUUUUAGUAGUAAAAUUAUUAAUUUUUUAAAAUGGACACUUUAAGU
Pfu-1-1    AGGGGU......AGUCUUAGA..........ACCCC....GUACGAUGCGUGUUAAUAGUG..UU..GUCU.........UUGGCC.GGGCA.GCAUGGUUAGGAAAGGCACAGUCGUAAAACAGAAAACGUUUUUUAGUAGUAAAAUUAUUAAUUUUUUAGAAUGGACACUUUAAGUU
Psp-24-1   UAGGGGU.....ACUCUUAGA.........GCCCCA.....UACGAAUCAUGUUAAUAGUG..UU..GUCU.........UUGGCC.GGGCG.ACAUGGUUAGGCAAAGCAUGUUCGUGAAAUAGAAAACGUUUUUUAGUAGUAAAAUUAUUAAUUUUUUAAAAUGGACACUUUAAGUU
env-562    UAGGGGU.....ACUCUUAGA.........GCCCCA.....UACGAAUCAUGUUAAUAGUG..UU..GUCU.........UUGGCC.GGGCG.ACAUGGUUAGGCAAAGCAUGUUCGUGAAAUAGAAAACGUUUUUUAGUAGU..................................
env-563    UAGGGGU.....ACUCUUAGA.........GCCCCA.....UACGAAUCAUGUUAAUAGUG..UU..GUCU.........UUGGCC.GGGCG.ACAUGGUUAGGCAAAGCAUGUUCGUGAAAUAGAAAACGUUUUUUAGUAGU..................................
env-564    UAGGGGUACU..CUU............AGAGCCCCU.....UACGAAUCAUGUUAAUAGUG..UU..GUCU.........UUGGCC.GGGCG.ACAUGGUUAGGCAAAGCAUAUUCGUGAAAUAGAAAACGUUUUUUAGUAGUAAAAUUAUUAAUUUUUUAAAAUGGACACUUUAAGUU
Pde-3-1    UGGGGUAC....ACUUA...........GAACCCC...CGUACGGUCUGUGUUAAUAGUG..CU..GUCU.........UUGGCC.GGGCA.GCAUGGUUAGGUUAAGCGCAGCCGUAAAAAGAAAACGUUUUUUAGUAGUAAAAUUAUUAAUUUUCAAGAAUGGACACUUUAAGUU
env-565    UGGGGUAC....ACUUA...........GAACCCC...CGUACGGUCUGUGUUAAUAGUG..CU..GUCU.........UUGGCC.GGGCA.GCAUGGUUAGGUCAAGCGCAGCCGUAAAAAGAAAACGUUUUUUAGUAGUAAAAU.................................
Pde-4-1    UGGGGUAC....ACUUA...........GAACCCC...CGUACGGUCUGUGUUAAUAGUG..CU..GUCU.........UUGGCC.GGGCA.GCAUGGUUAGGUCAAGCGCAGCCGUAAAAAGAAAACGUUUUUUAGUAGUAAAAUUAUUAAUUUUUUAAGAAUGGACACUUUAAGUU
Pde-5-1    UGGGGUAC....ACUUA...........GAACCCC...CGUACGGUCUGUGUUAAUAGUG..CU..GUCU.........UUGGCC.GGGCA.GCAUGGUUAGGUCAAGCGCAGCCGUAAAAAGAAAACGUUUUUUAGUAGUAAAAUUAUUAAUUUUUUAAGAAUGGACACUUUAAGUUA
env-566    UGGGGUAC....ACUUA...........GAACCCC...CGUACGGUCUGUGUUAAUAGUG..CU..GUCU.........UUGGCC.GGGCA.GCAUGGUUAGGUCAAGCGCAGCCGUAAAAAGAAAACGUUUUUUAGUAGUAAAAUUAUUAAUUUUUUAAGAAUGGACACUUUAAGUUA
env-567    UGGGGUAC....ACUUA...........GAACCCC...UGUACGGUCUGUGUUAAUAGUG..CU..GUCU.........UUGGCC.GGGCA.GCAUGGUUAGGUCAAGCGCAGCCGUAAAAAAGAAAACGUUUUUUAGUAGUAAAAUUAUUAAUUUUUUAAGAAUGGACACUUUAAGUU
Pde-6-1    UGGGGUAC....ACUUA...........GAACCCC...CGUACGGUCUGUGUUAAUAGUG..CU..GUCU.........UUGGCC.GGGCA.GCAUGGUUAGGUCAAGUGCAGCCGUAAAAAGAAAACGUUUUUUAGUAGUAAAAUUAUUAAUUUUUUAAGAAUGGACACUUUAAGUUA
env-568    UGUGGGUAC...ACUUA..........GAGCCC.C....GUACGGCUCGUGUUAAUAGUG..CU..GUCU.........UUGGCC.GGGCA.GCAUGGUUAGGACAAGCACGGUCGUAGAAAGAAAACGUUUUUUAGUAGUAAAAU...............................
env-569    UGUGGGUAC...ACUUA..........GAGCCC.C....GUACGGCUCGUGUUAAUAGUG..CU..GUCU.........UUGGCC.GGGCA.GCAUGGUUAGGACAAGCACGGUCGUAGAAAGAAAACGUUUUUUAGUAGUAAAAUUAUUAAUUUUUUAGAAUGGACACUUUAAGUUA
Pmu-1-1    UGUGGGUAC...ACUUA..........GAGCCC.C....GUACGGCUCGUGUUAAUAGUG..CU..GUCU.........UUGGCC.GGGCA.GCAUGGUUAGGACAAGCACGGUCGUAGAAAGAAAACGUUUUUUAGUAGUAAAAUUAUUAAUUUUUUAGAAUGGACACUUUAAGUUA
env-570    AGGGGCACU...CUU.............AGAACCCC....UGUGUCGUUGUGUUAAUAGCGC.UU..GCUU..........UGGCC.GAGCA.UCGUGGUUAGUG.AAUCAAUGGCA............................................................
env-571    UUGGGGCAC...GCUUA...........GAGCCCCA...AGACAUGGUUGUGUUAAUAGCG..UC..GCCU.........UUGGCC.GGACG.GCGCGGUUAGUG.AAUCAGCCGUGCGGCAGAGCGAAAGCCAUGGAUGGUCUCUUUUAGACUGGAG.....................
env-572    UUGGGGCAC...GCUUA...........GAGCCCCA...AGGCAUGGUUGUGUUAAUAGCG..UC..GCCU.........UUGGCC.GGACG.GCGUGGUUAGUG.AAUCAAUCGUGUAGCAGAGCGAAAGCUAUGGAUGGUCUCUUUUAGACUGGAGGCGCCGUGAGGUCUCUGCUGG
env-573    UUGGGGCAC...GCUUA...........GAGCCCCA...AGGCAUGGUUGUGUUAAUAGCG..UC..GCCU.........UUGGCC.GGACG.GCGUGGUUAGUG.AAUCAAUCGUGUAGCAGAGCGAAAGCUAUGGAUGGUCUCUUUUAGA.............................
env-574    CGGGGCAC....GCUUA............GAGCCCC...ACGCACAGCAGUGUUAAUAGCG..GC..GCCU.........UUGGCC.GGACG.UCGUGGUUAGUG.AAUCGGCCGUGCAGCAGAGCUAACGCCAUGGAUGGUCUCUUUUAGACUGGAGGCGCCGUGAGGUCUCUGUCAA
env-575    UGGGGCAC....GCUUA............GAGUCCC...UUGCGCAUUGUGUUAAUAGUG..GU..GCCU.........UUGGCC.GGACA.UCGUGGUUAGUG.AAUCAAUGGCGCGACAGAGCGAAAGCCAUGGAUGGUCUCUUUUAGACUGGAGGCGCCGUGAGGUCUCUGUUUU
env-576    UGGGGCAC....GCUUA............GAGUCCC...UUGCGCAUUGUGUUAAUAGCG..AU..GCCU.........UUGGCC.GGACA.UCGUGGUUAGUG.AAUCAAUGGCGCGACAGAGCGAAAGCCAUGGAUGGUCUCUUUUAGACUGGAGGCGCCGUGAGGUCUCUGUUUU
env-577    UGGGGU......GAACUUAGA..........GCCCC...GUGGAUGUGCGUGUUAAUAGCC..GU..GCCU.........UUGGCC.GGACA.UGGUGGUUAGUG.AAUCGUCGUCCGCAGAGAAAAUGUUAAUUAACAAAAGGUUUUUUAGUAGUUUUUAUUUAGAAAUGAACACU
env-578    UGAGGGC.....AGUCUUAGA.........GCCCUU....GGCACAUACGUGUUAAUAGUGUCAU..GCCU.........UUGGCC.GAGCA.ACAUGGUUAGUG.AAUCGUAUGAGCAAAGAAUUAGAAAACGUUUUUUAGUAGUAAAAUUUAAUUUCACAGAAUGGACACUUUAAG
env-579    UGAGGGC.....AGUCUUAGA.........GCCCUU....GGCACAUACGUGUUAAUAGUGUCAU..GCUU.........UUGGCC.GAGCA.ACAUGGUUAGUG.AAUCGUAUGAGCAAAGAAUUAGAAAACGUUUUUUAGUAGUAAAA..............................
env-580    UGAGGGC.....AGUCUUAGA.........GCCCUU....GGCACAUGCGUGUUAAUAGUGUCAU..GCCU.........UUGGCC.GAGCA.ACAUGGUUAGUG.AAUCGUAUGAGCAAAGAAUUAGAAAACGUUUUUUAGUAGUAAAAUUUAAUUUCACAGAAUGGACACUUUAAG
env-582    UGAGGGC.....AGUCUUAGA.........GCCCUU....AGCACAUGCGUGUUAAUAGUGUCAU..GCCU.........UUGGCC.GAGCA.ACAUGGUUAGUG.AAUCGUAUGAGCAAAGAAUUAGAAAACGUUUUUUAGUAGUAAAAUUUAAUUUCACAGAAUGGACACUUUAAG

           .<<<<<<<<<<<............>>>>>>>>>>>...............<<<<<.<<<..<<..<<<<<..............>>>>>>.>>>>.>>>>>.............................................................................
           .222222.1111............1111.222222...............02200.122..22..2222...............22222.2221.00220.............................................................................
           •GGGG•∘-----R••••∘∘∘∘---------∘•CCCC---∘∘•••••YGUGU•AAUAGY•--•Y--RYYU---------UUGGCC-GRRY•-••RYGGUU•GUG-AAUCR•••••••RR••R••••••••••••••••••••••••••••••••••••••••••••••••••••
            5′                                                                                                               3′
```

```
Pin-1-1  CGAGGG......UGAGCUUAGAU........CCCUC...AGCACGUUGUGUGUUAAUAGUG..UU..GCUU..........UUGGCC.GAGCA.GCAUGGUUAGUG.AAUCCGACGGGCAAGAAUUAAAAACGUUUUUUAGUAGUAAAAUUUAAAAUUAAAAAUGAACACUUUAAGUUA
env-583  CGAGAGU.....GAACUUAGA.........ACUCUC...AGCACAUUGUGUGUUAAUAGCG..UU..ACUU..........UUGGCC.GGGUU.ACGUGGUUAGAG.AAU.GCAAUGAGCAAGUAUUAAAAACGUUUUUUAGUAGUAAAAUUUAAAAUUUAAAAAUGGACACUUUAAGUU
env-589  CGAGAGU.....GAACUUAGA.........ACUCUC...AGCACAUUGUGUGUUAAUAGCG..UU..GCUU..........UUGGCC.GGGUU.ACGUGGUUAGAG.AAU.GCAAUGAGCAAGUAUUAAAAACGUUUUUUAGUAGUAAAAUUUAAAAUUUAAAAAUGGACACUUUAAGUU
env-592  CGAGAGU.....GAACUUAGA.........ACUCUC...AGCACAUUGUGUGUUAAUAGCG..UU..GCUU..........UUGGCC.GGGUU.ACGUGGUUAGAG.AAU.GCAAUGAGCAAGUAUUAAAAACGUUUUUUAGUAGUAAAAUUUAAAANAGGAAACAGCUAAGAAAGAGUG
env-593  CGAGAGU.....GAACUUAGA.........ACUCUC...AGCACAUUGUGUGUUAAUAGCG..UU..GCUU..........UUGGCC.GGGUU.ACGUGGUUAGAG.AAU.GCAAUGAGCAAGUAUUAAAAACGUUUUUUAGUAGUAAAAUUUAAAAUUUAAAAAUGGACACUUUAAGUU
env-594  CGAGAGU.....GAACUUAGA.........ACUCUC...AGCACAUUGUAUGUUAAUAGCG..UU..ACUU..........UUGGCC.GGGUU.ACGUGGUUAGAG.AAU.GCAAUGAGCAAGUAUNNNNNNNNNNNNNNNAGGAAACAGCUAAGAAAGAGUGGGUAGUAGUAGAUGCUAC
env-595  CGAGAGU.....GAACUUAGA.........ACUCUC...AGCACAUUGUAUGUUAAUAGCG..UU..ACUU..........UUGGCC.GGGUU.ACGUGGUUAGAG.AAU.GCAAUGAGCAAGUAUUAAAAACGUUUUUUAGUAGUAAAAUUUAAAAUUUAAAAAUGGACACUUUAAGUU
env-596  CGAGAGU.....GAACUUAGA.........ACUCUC...AGCACAUUGUAUGUUAAUAGCG..UU..ACUU..........UUGGCC.GGGUU.ACGUGGUUAGAG.AAU.GCAAUGAGCAAGUAUUAAAAACGUUUUUUAGUAGUAAAAUUUAAAAUUUAAAAAUGGACACUUUAAGUU
env-597  CGAGAGU.....GAACUUAGA.........ACUCUC...AGCACAUUGUAUGUUAAUAGCG..UU..ACUU..........UUGGCC.GGGUU.ACGUGGUUAGAG.AAU.GCAAUGAGCAAGUAUNAGGAAACAGCUAAGAAAGAGUGGGUAGUAGUAGAUGCUACAGACCAGAUAGUU
env-598  AGAGAGU.....GAACUUAGA.........ACUCUC...AGCACAUUGUAUGUUAAUAGCG..UU..ACUU..........UUGGCC.GGGUU.ACGUGGUUAGAG.AAU.GCAAUGAGCAAGUA.......................................................
env-599  CGAGAGU.....GAACUUAGA.........ACUCUC...AGCACAUUGUAUGUUAAUAGCG..UU..ACUU..........UUGGCC.GGGUU.ACGUGGUUAGAG.AAU.GCAAUGAGCAAGUAUUAAAAACGUUUUUUAGUAGUAAAAUUUAAAAUUUAAAAAUGGACACUUUAAGUU
env-600  CGAGAGU.....GAACUUAGA.........ACUCUC...AGCACAUUGUAUGUUAAUAGCG..UU..ACUU..........UUGGCC.GGGUU.ACGUGGUUAGAG.AAU.GCAAUGAGCAAGUAUUAAAAACGUUUUUUAGUAGUAAAAUUUAAAAUUUAAAAAUGGACACUUUAAGUU
env-613  CGAGAGU.....GAACUUAGA.........ACUCUC...AGCACAUUGUAUGUUAAUAGCG..UU..ACUU..........UUGGCC.GGGUU.ACGUGGUUAGAG.AAU.GCAAUGAGCAAGUAUUAAAAACGUUUUUUAGUAGUAAAAUUUAAAAUUUAAAAAUGGACACUUUAAGUU
env-648  CGAGAGU.....GAACUUAGA.........ACUCUC...AGCACAUUGUAUGUUAAUAGCG..UU..ACUU..........UUGGCC.GGGUU.ACGUGGUUAGAG.AAU.GCAAUGAGCAAGUAUUAAAAACGUUUUUUAGUAGUAAAAUUUAAAAUUUAAAAAUGGACACUUUAAGUU
env-651  CGAGAGU.....GAACUUAGA.........ACUCUC...AGCACAUUGUAUGUUAAUAGCG..UU..ACUU..........UUGGCC.GGGUU.ACGUGGUUAGAG.AAU.GCAAUGAGCAAGUAUUAAAAACGUUUUUUAGUAGUAAAAUUUAAAAUUUAAAAAUGGACACUUUAAGUU
env-652  CGAGAGU.....GAACUUAGA.........ACUCUC...AGCACAUUGUAUGUUAAUAGCG..UU..ACUU..........UUGGCC.GGGUU.ACGUGGUUAGAG.AAU.GCAAUGAGCAAGUAUUAAAAACGUUUUUUAGUAGUNAGUAGUAGAUGCUACAGACCAGAUAGUUGGUCG
env-653  CGAGAGU.....GAACUUAGA.........ACUCUC...AGCACAUUGUAUGUUAAUAGCG..UU..ACUU..........UUGGCC.GGGUU.ACGUGGUUAGAG.AAU.GCAAUGAGCAAGUAUUAAAAACGUUUUUUAGUAGU..................................
env-657  CGAGAGU.....GAACUUAGA.........ACUCUC...AGCACAUUGUAUGUUAAUAGCG..UU..ACUU..........UUGGCC.GGGUU.ACGUGGUUAGAG.AAU.GCAAUGAGCAAGUAU......................................................
env-676  UGAGGGU.....GUACUUAGA.........GCCCUC...UGCACGAUGUGUGUUAAUAGUAG.UG..GCUU..........UUGGCC.GAGCC.GCACGGUUAGAG.AAUCACACGGGCAAAAUUAGAAAACGUUUUUUAGUAGUAAAAUUAUUAAUUUUUUAGAAUGGACACUUUAAGU
Pco-3-1  CGAGGGU.....GUACUUAGA.........GCCCUC...UGCACGAUGUGUGUUAAUAGUAG.UG..GCUU..........UUGGCC.GAGCC.GCACGGUUAGAG.AAUCACACGGGCAAAAUUAGAAAACGUUUUUUAGUAGUAAAAUUAUUAAUUUUUUAGAAUGGACACUUUAAGU
Pni-1-1  CGAGGGU.....GAACUUAGA.........GCCCUC...AGCACGUUGUGUGUUAAUAGCG..CU..GCUU..........UUGGCC.AGGCU.GCGUGGUUAGUG.AAUCCGACGGGCAAGCAUUAAAAACGUUUUUUAGUAGUAAAAUUUAAAAUUAAAAAUGAUACUUUAAGUUA
env-677  CGAGGGU.....GAACUUAGA.........ACCCUC...AGCACGUUGUGUGUUAAUAGCG..CU..GCUU..........UUGGCC.GGGCU.GCGUGGUUAGUG.AAUCCGACGGGCAAGCAUUAAAAACGUUUUUUAGUAGUAAAAUUUAAAAUUAAAAAUGGAUACUUUAAGUUA
env-679  CGAGGGU.....GAACUUAGA.........GCCCUC...AGCACGUUGUGUGUUAAUAGCG..CU..GCUU..........UUGGCC.GGGCU.GUGUGGUUAGUG.AAUCCGACGGGCAAGCAUUAAAAACGUUUUUUAGUAGUAAAAUUUAAAAUUAAAAAUGGAUACUUUAAGUUA
env-680  CGAGGGU.....GAACUUAGA.........GCCCUC...AGCACGUUGUGUGUUAAUAGCG..CU..GCUU..........UUGGCC.GGGCU.GCGUGGUUAGUG.AAUCCGACGGGCAAGCAUUAAAAACGUUUUUUAGUAGUAAAAUUUAAAAUUAAAAAUGGACACUUUAAGUUA
env-681  CGAGGGU.....GAACUUAGA.........GCCCUC...AGCACGUUGUGUGUUAAUAGCG..CU..GCUU..........UUGGCC.GGGCU.GCGUGGUUAGUG.AAUCCGACGGGCAAGCAUUAAAAACGUUUUUUAGUAGUAAAAUUUAAAAUUAAAAAUGGACACUUUAAGUUA
env-682  CGAGGGU.....GAACUUAGA.........GCCCUC...AGCACGUUGUGUGUUAAUAGCG..CU..GCUU..........UUGGCC.GGGCU.GCGUGGUUAGUG.AAUCCGACGGGCAAGCAUUAAAAACGUUUUUUAGUAGUAAAAUUUAAAAUUAAAAAUGGAUACUUUAAGUUA
env-683  CGAGGGU.....GAACUUAGA.........GCCCUC...AGCACGUUGUGUGUUAAUAGCG..CU..GCUU..........UUGGCC.GGGCU.GCGUGGUUAGUG.AAUCCGACGGGCAAGCAUUAAAAACGUUUUUUAGUAGUAAAAUUUAAAAUUAAAAAUGGAUACUUUAAGUUA
env-684  CGAGGGU.....GAACUUAGA.........GCCCUC...AGCACGUUGUGUGUUAAUAGCG..CU..GCUU..........UUGGCC.GGGCU.GCGUGGUUAGUG.AAUCCGACGGGCAAGCAUUAAAAACGUUUUUUAGUAGUAAAAUUUAAAAUUAAAAAUGGAUACUUUAAGUUA
env-686  CGAGGGU.....GAACUUAGA.........GCCCUC...AGCACGUUGUGUGUUAAUAGCG..CU..GCUU..........UUGGCC.GGGCU.GCGUGGUUAGUG.AAUCCGACGGGCAAGCAUUAAAAACGUUUUUUAGUAGUAAAAUUUAAAAUUAAAAAUGGAUACUUUAAGUUA
env-696  CGAGGGU.....GAACUUAGA.........GCCCUC...AGCACGUUGUGUGUUAAUAGCG..CU..GCUU..........UUGGCC.GGGCU.GCGUGGUUAGUG.AAUCCGACGGGCAAGCAUUAAAAACGUUUUUUAGUAGUAAAAUUUAAAAUUAAAAAUGGAUACUUUAAGUUA
env-698  CGAGGGU.....GAACUUAGA.........GCCCUC...AGCACGUUGUGUGUUAAUAGCG..CU..GCUU..........UUGGCC.GGGCU.GCGUGGUUAGUG.AAUCCGACGGGCAAGCAUUAAAAACGUUUUUUA.......................................
env-699  CGAGGGU.....GAACUUAGA.........GCCCUC...AGCACGUUGUGUGUUAAUAGCG..CU..GCUU..........UUGGCC.GGGCU.GCGUGGUUAGUG.AAUCCGACGGGCAAGCAUUAAAAACGUUUUUUAGUAGUAAAAUUUAAAAUUAAAAAUGGAUACU........
Pni-3-1  CGAGGGU.....GAACUUAGA.........GCCCUC...AGCACGUUGUGUGUUAAUAGCG..CU..GCUU..........UUGGCC.GGGCU.GCGUGGUUAGUG.AAUCCGACGGGCAAGCAUUAAAAACGUUUUUUAGUAGUAAAAUUUAAAAUUAAAAAUGGAUACUUUAAGUUA
env-700  CGAGGGU.....GAACUUAGA.........GCCCUC...AGCACGUUGUGUGUUAAUAGCG..CU..GCUU..........UUGGCC.GGGCU.GCGUGGUUAGUG.AAUCCGACGGGCAAGCAUUAAAAACGUUUUUUAGUAGUAAA...............................
env-709  CGAGGGU.....GAACUUAGA.........GCCCUC...AGCACGUUGUGUGUUAAUAGUG..CC..GCUU..........UUGGCC.GGGCU.GCAUGGUUAGUG.AAUCCAACGAGCAAGAAUUAAAAACGUUUUUUAGUAGUAAA...............................
env-710  CGAGGGU.....GAACUUAGA.........GCUCUC...AGCACGUUGUGUGUUAAUAGUG..CC..GCUU..........UUGGCC.GGGCU.GCAUGGUUAGUG.AAUCCAACGAGCAAGAAUUAAAAACGUUUUUUAGUAGUAAAAUUUAAAAUUAAAAAUGAACACUUUAAGUUA
env-711  CGAGGGU.....GAACUUAGA.........GCUCUC...AGCACGUUGUGUGUUAAUAGUG..CC..GCUU..........UUGGCC.GGGCU.GCAUGGUUAGUG.AAUCCAACGAGCAAGAAUUAAAAACGUUUUUUAGUAGUAAAAUUUAAAAUUAAAAAUGAACACUUUAAGUUA
env-724  CGAGGGU.....GAACUUAGA.........GCUCUC...AGCACGUUGUGUGUUAAUAGUG..CC..GCUU..........UUGGCC.GGGCU.GCAUGGUUAGUG.AAUCCAACGAGCAAGAAUUAAAAACGUUUUUUAGUAGUAAAAUUUAAAAUUAAAAAUGAACACUUUAAGUUA
Pau-1-1  CGAGGGU.....GAACUUAGA.........GCUCUC...AGCACGUUGUGUGUUAAUAGUG..CC..GCUU..........UUGGCC.GGGCU.GCAUGGUUAGUG.AAUCCAACGAGCAAGAAUUAAAAACGUUUUUUAGUAGUAAAAUUUAAAAAUUAAAAAAUUGAAACACAUUUA
Psp-9-1  CGAGGGU.....GUACUUAGA.........GCCCUC...AGCACGUGGUGUGUUAAUAGUG..CU..GCUU..........UUGGCC.GAGCU.GCACGGUUAGUG.AAUCACACGGGCAAAGAACAAAAACGUUUUUUAGUAGUAAAAUUAAAAUUUAAAAUGACACUUUAAGUUAC
env-725  CGAGGGU.....GAACUUAGA.........GCCCUC...AGCACGUUGUGUGUUAAUAGUG..UU..GCUU..........UUGGCC.GAGCA.GUAUGGUUAGUG.AAUCCGACGGGCAAGAAUUAAAAACGUUUUUUAGUAGUAAAAUUUAAAAUUAAAAAUGAACACUUUAAGUUA
env-726  CGAGGGU.....GAACUUAGA.........GCCCUC...AGCACGUUGUGUGUUAAUAGUG..UU..GCUU..........UUGGCC.GAGCA.GUAUGGUUAGUG.AAUCCGACGGGCAAGAAUUAAAAACGUUUUUUAGUAGUAAAAUUUAAAAUUAAAAAUGGAUACUUUAAGUUA
env-727  CGAGGGU.....AAGCUUAGA.........GCCCUC...AGCACGUUGUGUGUUAAUAGUG..UU..GCUU..........UUGGCC.GAGCU.GCAUGGUUAGUG.AAUCCGACGGGCAAGAAUUAAAAACGUUUUUUAGUAGUAAA...............................
env-729  CGAGGGU.....GAGCUUAGA.........GCCCUC...GGCACGUUGUGUGUUAAUAGUG..UU..GCUU..........UUGGCC.GAGCU.GCAUGGUUAGUG.AAUCCGACGGGCAAGAAUUAAAAACGUUUUUUAGUAGUAAAAUUUAAAAUUAAAAAUGAACACUUUAAGUUA
env-733  CGAGGGU.....GAGCUUAGA.........GCCCUC...AGCACGUUGUGUGUUAAUAGUG..UU..GCUU..........UUGGCC.GAGCU.GCAUGGUUAGUG.AAUCCGACGGGCAAGAAUUAAAAACGUUUUUUAGUAGUAAAAUUUAAAAUUAAAAAUGAACACUUUAAGUUA
env-734  CGAGGGU.....GAGCUUAGA.........GCCCUC...AGCACGUUGUGUGUUAAUAGUG..UU..GCUU..........UUGGCC.GAGCU.GCAUGGUUAGUG.AAUCCGACGGGCAAGAAUUAAAAACGUUUUUUAGUAGUAAAAUUUAAAAUUAAAAAUGAACACUUUAAGUUA
env-735  CGAGGGU.....GAGCUUAGA.........GCCCUC...AGCACGUUGUGUGUUAAUAGUG..UU..GCUU..........UUGGCC.GAGCU.GCAUGGUUAGUG.AAUCCGACGGGCAAGAAUUAAAAACGUUUUUUAGUAGUAAAAUUUAAAAUUAAAAAUGAACACUUUAAGUUA
env-736  CGAGGGU.....GAGCUUAGA.........ACCCUC...AGCACGUUGUGUGUUAAUAGUG..UU..GCUU..........UUGGCC.GAGCU.GCAUGGUUAGUG.AAUCCGACGGGCAAGAAUUAAAAACGUUUUUUAGUAGUAAAAUUUAAAAUUAAAAAUGAACACUUUAAGUUA
Pin-3-1  CGAGGGU.....GAACUUAGA.........GCCCUC...AGCACGUUGUGUGUUAAUAGUG..UU..GCUU..........UUGGCC.GAGCU.GCAUGGUUAGUG.AAUCCGACGGGCAAGAAUUAAAAACGUUUUUUAGUAGUAAAAUUUAAAAUUAAAAAUGAAUACUUUAAGUUA
Pfa-1-1  CGAGGGU.....GAACUUAGA.........GCCCUC...AGCACGUUGUGUGUUAAUAGUG..UU..GCUU..........UUGGCC.GAGCU.GCAUGGUUAGUG.AAUCCGACGGGCAAGAAUUAAAAACGUUUUUUAGUAGUAAAAUUUAAAAUUAAAAUGAACACUUUAAGUUAU
env-737  CGAGGGU.....GAACUUAGA.........GCCCUC...AGCACGUUGUGUGUUAAUAGUG..UU..GCUU..........UUGGCC.GAGCU.GCAUGGUUAGUG.AAUCCGACGGGCAAGAAUUAAAAACGUUUUUUAGUAGUAAAAUUUAAAAUUAAAAAUGAACACUUUAAGUUA
env-738  CGAGGGU.....GAACUUAGA.........GCCCUC...AGCACGUUGUGUGUUAAUAGUG..UU..GCUU..........UUGGCC.GAGCU.GCACGGUUAGUG.AAUCCGACGGGCAAGAAUUAAAAACGUUUUUUAGUAGUAAAAUUUAAAAUUAAAAAUGAACACUUUAAGUUA
env-742  CGAGGGU.....GAACUUAGA.........GCCCUC...AGCACGUUGUGUGUUAAUAGUG..UU..GCUU..........UUGGCC.GAGCU.GCACGGUUAGUG.AAUCCGACGGGCAAGAAUUAAAAACGUUUUUUAGUAGUAAAAUUUAAAAUUAAAAAUGGAUACUUUAAGUUA
Pdi-3-1  CGAGGGU.....GAACUUAGA.........ACCCUC...AGCUCGUUGUGUGUUAAUAGUG..UA..ACUU..........UUGGCC.GAGUC.GCAUGGUUAGUG.AAUCCAAUGAGCAAGAAUUAAAAACGUUUUUUAGUAGUAAAAUUUAAAAUUUAAAAUGAACACUUUAAGUUAC
env-743  CGAGGGU.....GAACUUAGA.........ACCCUC...AGCUCGUUGUGUGUUAAUAGUG..UA..ACUU..........UUGGCC.GAGUC.GCAUGGUUAGUG.AAUCCAAUGAGCAAGAAU......................................................
Pdi-2-1  CGAGGGU.....GAACUUAGA.........ACCCUC...AGCUUGUUGUGUGUUAAUAGUG..UA..ACUU..........UUGGCC.GAGUC.GCAUGGUUAGUG.AAUCCAAUGAGCAAGAAUUAAAAACGUUUUUUAGUAGUAAAAUUUAAAAUUUAAAAUGAACACUUUAAGUUAC
Pdi-1-1  CGAGGGU.....GAACUUAGA.........ACCCUC...AGCUUGUUGUGUGUUAAUAGUG..UA..ACUU..........UUGGCC.GAGUC.GCAUGGUUAGUG.AAUCCAAUGAGCAAGAAUUAAAAACGUUUUUUAGUAGUAAAAUUUAAAAUUUAAAAUGAACACUUUAAGUUAC
env-744  CGGGGAU.....ACACUUAGA.........GUCCCC...UGGGUAUGGUGUGUUAAUAGCA..CA..UCCU..........GUGGCC.GGGAU.UUGCGGUUAGUG.AAUCACCUAUCCGAAAUUAAGUGAGUAAGAAAAGAUUAAAACGUUUUUUAGUAGUAAAUUUAAAUUUUAAAAU
env-745  CGGGGAU.....ACACUUAGA.........GUCCCC...UGGGUAUGGUGUGUUAAUAGCA..CA..UCCU..........GUGGCC.GGGAU.UUGCGGUUAGUG.AAUCACCUAUCCGAAAUUAAGUGAGUAAAAAAGAUUAAAACGUUUUUUAGUAGUAAAUUUAAAUUUUAAAAUG
env-746  CGGGGAU.....ACACUUAGA.........GUCCCC...UGGGUAUGGUGUGUUAAUAGCA..CA..UCCU..........GUGGCC.GGGAU.UUGCGGUUAGUG.AAUCACCUAUCCGAAAAUUAAGUGAGUAAGAAAAGAUUAAAACGUUUUUUAGUAGUAAAUUUAAAUUUUAAAA
env-747  CGGGGAU.....ACACUUAGA.........GUCCCC...UGGGUAUGGUGUGUUAAUAGCA..CA..UCCU..........GUGGCC.GGGAU.UUGCGGUUAGUG.AAUCACCUAUCCGAAAAUUAAGUGAGUAAGAAAAGAUUAAAACGUUUUUUAGUAGUAAAUUUAAAUUUUAAAA
env-748  CGGGGAU.....ACACUUAGA.........GUCCCC...UGGGUAUGGUGUGUUAAUAGCA..CA..UCCU..........GUGGCC.GGGAU.UUGCGGUUAGUG.AAUCACCUAUCCGAAAUUAAGUGAGUAAGAAAAGAUUAAAACGUUUUUUAGUAGUAAAUUUAAAUUUUAAAAU
env-749  CGGGGAU.....ACACUUAGA.........GUCCCC...UGGGUAUGGUGUGUUAAUAGCA..CA..UCCU..........GUGGCC.GGGAU.UUGCGGUUAGUG.AAUCACCUAUCCGAAAUUAAGUGAGUAAGAAAAGAUUAAAACGUUUUUUAGUAGUAAAUUUAAAUUUUAAAAU
env-751  CGGGGAU.....ACACUUAGA.........GUCCCC...UGGGUAUGGUGUGUUAAUAGCA..CA..UCCU..........GUGGCC.GGGAU.UUGCGGUUAGUG.AAUCACCUAUCCGAAAUUAAGUGAGUAAGAAAAGAUUAAAACGUUUUUUAGUAGUAAAUUUAAAUUUUAAAAU
env-752  CGGGGAU.....ACACUUAGA.........GUCCCC...UGGGUAUGGUGUGUUAAUAGCA..CA..UCCU..........GUGGCC.GGGAU.UUGCGGUUAGUG.AAUCACCUAUCCGAAAUUAAGUGAGUAAGAAAAGAUUAAAACGUUUUUUAGUAGUAAAUUUA..........
env-753  CGGGGAU.....ACACUUAGA.........GUCCCC...UGGGUAUGGUGUGUUAAUAGCA..CA..UCCU..........GUGGCC.GGGAU.UUGCGGUUAGUG.AAUCACCUAUCCGAAAUUAAGUGAGUAAAAAAGAUUAAAACGUUUUUUAGUAGUAAAUUUAAAUUUUAAAAUGGA
env-754  CGGGGAU.....ACACUUAGA.........GUCCCC...UGGGUAUGGUGUGUUAAUAGCA..CA..UCCU..........GUGGCC.GGGAU.UUGCGGUUAGUG.AAUCACCUAUCCGAAAUUAAGUGAGUAAGAAAAGAUUAAAACGUUUUUUAGUAGUAAAUUUAAAUUUUAAAAU
env-755  CGGGGAU.....ACACUUAGA.........GUCCCC...UGGGUAUGGUGUGUUAAUAGCA..CA..UCCU..........GUGGCC.GGGAU.UUGCGGUUAGUG.AAUCACCUAUCCGAAAUUAAGUGAGUAAGAAAAGAUUAAAACGUUUUUUAGUAGUAAAUUUANNNNNNNNNUU
env-756  CGGGGAU.....ACACUUAGA.........GUCCCC...UGGGUAUGGUGUGUUAAUAGCA..CA..UCCU..........GUGGCC.GGGAU.UUGCGGUUAGUG.AAUCACCUAUCCGAAAAUUAAGUGAGUAAGAAAAGAUUAAAACGUUUUUUAGUAGUAAAUUUAAAUUUUAAAA
env-757  CGGGGAUAC...ACUUA...........GAGUCCCC...UGGAUAGCGUGUGUUAAUAGCAA.CG..UCCU..........GUGGCC.GGGAU.UUGCGGUUAGUG.AAUCACCUAUCCAAAUAGUGAGUAAAAAGAUUAAAAACGUUUUUUAGUAGUAAUUUAAAUUUUAAAAUGGACA
env-758  CGGGGAUAC...ACUUA...........GAGUCCCC...UGGGUAUGGUGUGUUAAUAGCA..CG..UCCU..........GUGGCC.GGGAU.UUGCGGUUAGAG.AAUCACCUAUCCGAAACAUAAGUGAGUAAGAAAAAGAUUAAAAACGUUUUUUAGUAGUAAAUUUAAAUUUUAA
env-759  CGGGGAUAC...ACUUA...........GAGUCCCC...UGGAUAUGGUGUGUUAAUAGCA..CU..UCCU..........GUGGCC.GGGAU.UUGCGGUUAGUG.AAUCACCUAUCCGAAACAUAAGUGAAUUAAGAAAAAGAUUAAAAACGUUUUUUAGUAGUAAAUUUAAAUUUUA
env-760  CGGGGAUAC...GCUUA...........GAGUCCCC...UGGGUAUGGUGUGUUAAUAGCA..CA..UCCU..........GUGGCC.GGGAU.UUGCGGUUAGUG.AAUCACCUAUCCGAAAUUAAGUGAGUAAGAAAAGAUUAAAACGUUUUUUAGUAGUAAAUUUAAAUUUUAAAAU
env-761  CGGGGAUAC...ACUUA...........GAGUCCCC....UGGACUGGUGUGUUAAUAGCAA.UC..UCCU..........GUGGCC.GGGAA.UUGCGGUUAGUG.AAUCGCCUGUCUAAAAGUGAGUAAAAAGAUUUAAAAACGUUUUUUAGUAGU......................
env-762  CGGGGAUAC...ACUUA...........GAGUCCCC....UGGAUUGGUGUGUUAAUAGCAA.UG..UCCU..........GUGGCC.GGGAU.UUGCGGUUAGAG.AAUCACUUGUCCGAAAUAGUGAGUAAAAAGAUUAAAAACGUUUUUUAGUAGUAAUUUAAAUUUUAAAAUGGAC
env-763  CGGGGAUAC...ACUUA...........GAGUCCCC....UGGACUGGUGUGUUAAUAGCAA.UG..UCCU..........GUGGCC.GGGAU.UUGCGGUUAGAG.AAUCACCUGUCCGAAAUAGUGAGUAAAAAGAUUAAAAACGUUUUUUAGUAGUAAUUUAAAUUUUAAAAUGGAC
env-765  CGGGGAUAC...ACUUA...........GAGUCCCC....UGGACUGGUGUGUUAAUAGCAA.UG..UCCU..........GUGGCC.GGGAU.UUGCGGUUAGAG.AAUCACCUGUCCGAAAUAGUGAGUAAAAAGAUUAAAAACGUUUUUUAGU........................
env-766  CGGGGAUAC...ACUUA...........GAGUCCCC....UGGACUGGUGUGUUAAUAGCAA.UG..UCCU..........GUGGCC.GGGAU.UUGCGGUUAGAG.AAUCACCUGUCCGAAAUAGUGAGUAAAAAGAUUAAAAACGUUUUUUAGUAGUAAUUUAAAUUUUAAAAUGGAC
env-767  CGGGGAUAC...ACUUA...........GAGUCCCC....UGGACUGGUGUGUUAAUAGCAA.UG..UCCU..........GUGGCC.GGGAU.UUGCGGUUAGAG.AAUCACCUGUCCGAAAUAGUGAGUAAAAAGAUUAAAAACGUUUUUUAGUAGUAAUUUAAAUUUUAAAAUGGAC
env-768  CGGGGAUAC...GCUUA...........GAGUCCCC...UGGGCAGUGUGUGUUAAUAGCAA.UC..UCCU..........GUGGCC.GGGAA.UUGCGGUUAGUG.AAUCGCCUGUCUAAAAGUGAGUAAAAAGAUUUAAAAACGUUUUUUAGUAGUAAUUUAAAAUUUAAAAUGGACA
env-769  CGGGGAUAC...GCUUA...........GAGUCCCC...UGGGCAGUGUGUGUUAAUAGCAA.UC..UCCU..........GUGGCC.GGGAA.UUGCGGUUAGUG.AAUCGCCUGUCUAAAAGUGAGUAAAAAGAUUUAAAAACGUUUUUUAGUAGUAA....................
env-770  CGGGGAUAC...GCUUA...........GAGUCCCC...UGGGCAGUGUGUGUUAAUAGCAA.UC..UCCU..........GUGGCC.GGGAA.UUGCGGUUAGUG.AAUCGCCUGUCUAAAAGUGAGUAAAAAGAUUUAAAAACGUUUUUUAGUAGUAAUUUAAAAUUUAAAAUGGACA
env-771  CGGGGAUAC...GCUUA...........GAGUCCCC...UGGGCAGUGUGUGUUAAUAGCAA.UC..UCCU..........GUGGCC.GGGAA.UUGCGGUUAGUG.AAUCGCCUGUCUAAAAGUGAGUAAAAAGAUUUAAAAACGUUUUUUAGUAGUAAUUUAAAAUUUAAAAUGGACA

         .<<<<<<<<<<<.............>>>>>>>>>>>................<<<<<.<<<..<<..<<<<<................>>>>>>.>>>>.>>>>>.............................................................
         .222222.1111.............1111.222222................02200.122..22..2222.................22222.2221.00220.............................................................
         •GGGG•∘-----R••••∘∘∘----------∘•CCCC----••••••••YGUGU•AAUAGY•--•Y--RYYU----------UUGGCC-GRRY•-••RYGGUU•GUG-AAUCR•••••••RR••R••••••••••••••••••••••••••••••••••••••••••••••••••••••••
          5′                                                                                                            3′
```

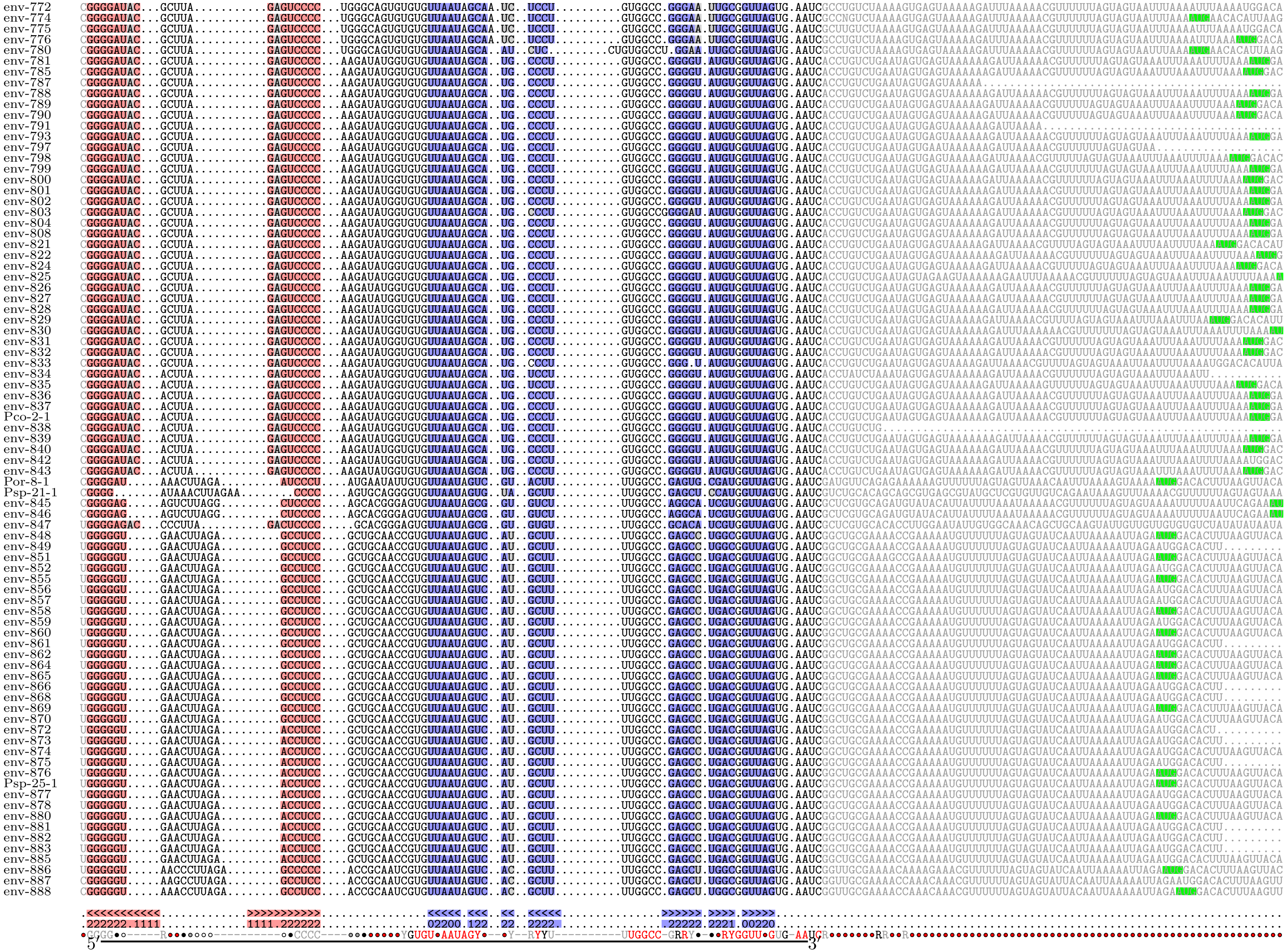
env-772
env-774
env-775
env-776
env-780
env-781
env-785
env-787
env-788
env-789
env-790
env-791
env-793
env-797
env-798
env-799
env-800
env-801
env-802
env-803
env-804
env-808
env-821
env-822
env-824
env-825
env-826
env-827
env-828
env-829
env-830
env-831
env-832
env-833
env-834
env-835
env-836
env-837
Pco-2-1
env-838
env-839
env-840
env-842
env-843
Por-8-1
Psp-21-1
env-845
env-846
env-847
env-848
env-849
env-851
env-852
env-855
env-856
env-857
env-858
env-859
env-860
env-861
env-862
env-864
env-865
env-866
env-868
env-869
env-870
env-872
env-873
env-874
env-875
env-876
Psp-25-1
env-877
env-878
env-880
env-881
env-882
env-883
env-885
env-886
env-887
env-888
5′
3′

```
env-889    CGGGGGU.....AAACCUUAGA........GCCUCC....ACCGCAAUCGUGUUAAUAGUC..AC..GCUU..........UUGGCC.GAGCU.UGGCGGUUAGUG.AAUCGGUUGCGAAAACCAAACAAACGUUUUUUAGUAGUAUUACAAU...........................
env-890    CGGGGGU.....AAACCUUAGA........GCCUCC....ACCGCAAUCGUGUUAAUAGUC..AC..GCUU..........UUGGCC.GAGCU.UGGCGGUUAGUG.AAUCGGUUGCGAAAACCAAACAAAUGUUUUUUAGUAGUAUUACAAUUAAAAAUUAGAAUGGACACUU......
env-891    CGGGGGU.....AAACCUUAGA........GCCUCC....ACCGCAAUCGUGUUAAUAGUC..AC..GCUU..........UUGGCC.GAGCU.UGGCGGUUAGUG.AAUCGGUUGCGAAAACCAAACAAACGUUUUUUAGUAGUAUUACAAUUAAAAAUUAGAAUGGACACUUUAAGUU
env-892    CGGGGGU.....AAACCUUAGA........GCCUCC....ACCGCAAUCGUGUUAAUAGUC..AC..GCUU..........UUGGCC.GAGCU.UGGCGGUUAGUG.AAUCGGUUGCGAAAACCAAACAAACGUUUUUUAGUAGUAUUACAAUUAAAAAUUAGAAUGGACACUUUAAGUU
env-893    CGGGGGU.....AAACCUUAGA........GCCUCC....ACCGCAAUCGUGUUAAUAGUC..AC..GCUU..........UUGGCC.GAGCU.UGGCGGUUAGUG.AAUCGGUUGCGAAAACCAAACAAACGUUUUUUAGUAGUAUUACAAU...........................
env-894    CGGGGGU.....AAACCUUAGA........GCCUCC....ACCGCAAUCGUGUUAAUAGUC..AC..GCUU..........UUGGCC.GAGCU.UGGCGGUUAGUG.AAUCGGUUGCGAAAACCAAACAAACGUUUUUUAGUAGUAUUACAAUUAAAAAUUAGAAUGGACACUUUAAGUU
env-900    CGGGGGU.....AAACCUUAGA........GCCUCC....ACCGCAAUCGUGUUAAUAGUC..AC..GCUU..........UUGGCC.GAGCU.UGGCGGUUAGUG.AAUCGGUUGCGAAAACCAAACAAACGUUUUUUAGUAGUAUUACAAUUAAAAAUUAGAAUGGACACUUUAAGUU
env-901    CGGGGGU.....AAACCUUAGA........GCCUCC....ACCGCAAUCGUGUUAAUAGUC..AC..GCUU..........UUGGCC.GAGCU.UGGCGGUUAGUG.AAUCGGUUGCGAAAACCAAACAAACGCUUUUUAGUAGUAUUACAAUUAAAAAUUAGAAUGGACACUU......
env-902    CGGGGGU.....AAACCUUAGA........GCCUCC....ACCGCAAUCGUGUUAAUAGUC..AC..GCUU..........UUGGCC.GAGCU.UGGCGGUUAGUG.AAUCGGUUGCGAAAACCAAACAAACGCUUUUUAGUAGUAUUACAAUUAAAAAUUAGAAUGGACACUUUAAGUU
Psp-26-1   CGGGGGU.....AAACCUUAGA........GCCUCC....ACCGCAAUCGUGUUAAUAGUC..AC..GCUU..........UUGGCC.GAGCU.UGGCGGUUAGUG.AAUCGGUUGCGAAAACCAAACAAACGUUUUUUAGUAGUAUUACAAUUAAAAAUUAGAAUGGACACUUUAAGUU
env-903    CGGGGGU.....AAACCUUAGA........GCCUCC....ACCGCAAUCGUGUUAAUAGUC..AC..GCUU..........UUGGCC.GAGCU.UGGCGGUUAGUG.AAUCGGUUGCGAAAACCAAACAAACGUUUUUUAGUAGUAUUACAAU...........................
env-921    UGGGGGU.....AACCCUUAGA........GCCUCC....GCUGCAACCGUGUUAAUAGUC..AU..GCUU..........UUGGCC.GAGCC.UGACGGUUAGUG.AAUCGGCUGCGAAAACCGAAAAAUGUUUUUUAGUAGUAUCAAUUAAAAAUUAGAAUGGACACUUUAAGUUACA
env-922    UGGGGGU.....AACCCUUAGA........GCCUCC....GCUGCAACCGUGUUAAUAGUC..AU..GCUU..........UUGGCC.GAGCC.UGACGGUUAGUG.AAUCGGCUGCGAAAACCGAAAAAUGUUUUUUAGUAGUAUCAAUUAAAAAUUAGAAUGGACACUUUAAGUUACA
env-923    UGGGGGU.....AACCCUUAGA........GCCUCC....GCUGCAACCGUGUUAAUAGUC..AU..GCUU..........UUGGCC.GAGCC.UGACGGUUAGUG.AAUCGGCUGCGAAAACCGAAAAAUGUUUUUUAGUAGUAUCAAUUAAAAAUUAGAAUGGACACUUUAAGUUACA
env-924    UGGGGGU.....AACCCUUAGA........GCCUCC....GCUGCAACCGUGUUAAUAGUC..AU..GCUU..........UUGGCC.GAGCC.UGACGGUUAGUG.AAUCGGCUGCGAAAACCGAAAAAUGUUUUUUAGUAGUAUCAAUUAAAAAUUAGAAUGGACACUUUAAGUUACA
env-932    UGGGGGU.....AACCCUUAGA........GCCUCC....GCUGCAACCGUGUUAAUAGUC..AU..GCUU..........UUGGCC.GAGCC.UGACGGUUAGUG.AAUCGGCUGCGAAAACCGAAAAAUGUUUUUUAGUAGUAUCAAUUAAAAAUUAGAAUGGACACU..........
env-948    GGGGGGU.....AACCCUUAGA........GCCUCC....GCUGCAACCGUGUUAAUAGUC..AU..GCUU..........UUGGCC.GAGCC.UGACGGUUAGUG.AAUCGGCUGCGAAAACCGAAAAAUGUUUUUUAGUAGUAUCAAUUAAAAAUUAGAAUGGACACUUUAAGUUACA
env-949    UGGGGGU.....AACCCUUAGA........GCCUCC....GCUGCAACCGUGUUAAUAGUC..AU..GCUU..........UUGGCC.GAGCC.UGACGGUUAGUG.AAUCGGCUGCGAAAACCGAAAAAUGUUUUUUAGUAGUAUCAAUUAAAAAUUAGAAUGGACACUUUAAGUUACA
env-950    UGGGGGU.....AACCCUUAGA........GCCUCC....GCUGCAACCGUGUUAAUAGUC..AU..GCUU..........UUGGCC.GAGCC.UGACGGUUAGUG.AAUCGGCUGCGAAAACCGAAAAAUGUUUUUUAGUAGUAUCAAUUAAAAAUUAGAAUGGACACUUUAAGUUACA
env-953    UGGGGGU.....AACCCUUAGA........GCCUCC....ACUGCAACCGUGUUAAUAGUC..AU..GCUU..........UUGGCC.GAGCC.UGACGGUUCGUG.AAUCGGCUGCGAAAACCGAAAAAUGUUUUUUAGUAGUAUCAAUUAAAAAUUAGAAUGGACACUUUAAGUUACA
env-957    UGGGGGU.....AACCCUUAGA........GCCUCC....ACUGCAACCGUGUUAAUAGUC..AU..GCUU..........UUGGCC.GAGCC.UGACGGUUAGUG.AAUCGGCUGCGAAAACCGAAAAAUGUUUUUUAGUAGUAUCAAUUAAAAAUUAGAAUGGACACUUUAAGUUACA
env-958    UGGGGGU.....AACCCUUAGA........GCCUCC....ACUGCAACCGUGUUAAUAGUC..AU..GCUU..........UUGGCC.GAGCC.UGACGGUUAGUG.AAUCGGCUGCGAAAACCGAAAAAUGUUUUUUAGUAGUAUCAAUUAAAAAUUAGAAUGGACACUUUAAGUUACA
env-960    CGGGGAU.....ACACUUAGA.........GUCCCC....AGCGUGAGAGUGUUAAUAGUGG.AG..GUCU..........UUGGCC.GGACC.UCACGGUUAGUG.AAUCGCUCGCGCAAAGUUAUAAUGUUUUUUAGUAGUAAUAUUUAAAUUAAACAAUGAACACUUUAAGUUACAA
env-961    CGGGGAU.....ACACUUAGA.........GUCCCC....AGCGUGAGAGUGUUAAUAGUGG.AG..GUCU..........UUGGCC.GGACC.UCACGGUUAGUG.AAUCGCUCGCGCAAAGUUAUAAUGUUUUUUAGUAGUAAUAUUUAAAUUAAACAAUGAACACUUUAAGUUACAA
env-967    CGGGGAUAC...ACUUA...........GAGUCCCC....UGCAUGAGGGUGUUAAUAGCGC.UU..GCC........GAAAAGGCCA.GGCG.GCGUGGUUAGUG.AAUCGCUCGUGCAAGAAAAAAUGAAUGUUUUUUAGUAGUAAUAUUUUUAAUUACAACAAUGAACGACAACUUC
env-968    CGGGGAUAC...ACUUA...........GAGUCCCC....UGCAUGAGGGUGUUAAUAGCGC.UU..GCC.....G...AAAAGGCCA.GGCG.GCGUGGUUAGUG.AAUCGCUCGUGCAAGAAAAAUAAAUGUUUUUUAGUAGUAAUAUUUUUAAUUAAAAAAAUGAACACUUUAAGUU
env-970    CGGGGAUAC...ACUUA...........GAGUCCCC....UGCAUGAGGGUGUUAAUAGCGC.UU..GCC.....G...AAAAGGCCA.GGCG.GCGUGGUUAGUG.AAUCGCUCGUGCAAGAAAAAUAAAUGUUUUUUAGUAGUAAUAUUUUUAAUUAAAAAAAUGAACACUUUAAGUU
env-971    CGGGGAUAC...ACUUA...........GAGUCCCC....UGCAUGAGGGUGUUAAUAGCGC.UU..GCC.....G...AAAAGGCCA.GGCG.GCGUGGUUAGUG.AAUCGCUCGUGCAAGAAAAAUAAAUGUUUUUUAGUAGUAAUAUUUUUAAUUAAAAAAAUGAACACUUUAAGUU
env-972    UGGGGAUAC...ACUUA...........GAGUCCCC....UGCAUGAGGGUGUUAAUAGCGC.UU..GUC.....G...AAAAGGCCA.GACG.GCGUGGUUAGUG.AAUCGCUCGUGCAAGAAAGAUAAAUGUUUUUUAGUAGUAAUAUUUUAAUUAAAAAAAUGAACACUUUAAGUUA
env-973    UGGGGAUAC...ACUUA...........GAGUCCCC....UGCAUGAGGGUGUUAAUAGCGU.UU..GUC.....G...AAAAGGCCA.GACG.GCGUGGUUAGUG.AAUCGCUCGUGCAAGAAAAAUAAAUGUUUUUUAGUAGUAAUAUUUUAAUUAAAAAAAAUGAACACUUUAAGUUA
env-977    UGGGGAUAC...ACUUA...........GAGUCCCC....UGCAUGAGGGUGUUAAUAGCGU.UU..GUC.....G...AAAAGGCCA.GACG.GCGUGGUUAGUG.AAUCGCUCGUGCAAGAAAAAUAAAUGUUUUUUAGUAGUAAUAUUUUAAUUAAAAAAAAUGAACACUUUAAGUUA
env-978    UGGGGAUAC...ACUUA...........GAGUCCCC....UGCAUGAGGGUGUUAAUAGCGU.UU..GUC.....G...AAAAGGCCA.GACG.GCGUGGUUAGUG.AAUCGCUCGUGCAAGAAAAAUAAAUGUUUUUUAGUAGUAAUAUUUUAAUUAAAAAAAUGAACACUUUAAGUUA
env-979    GGGGGAUAC...ACUUA...........GAGUCCCC....UGCAUGAGGGUGUUAAUAGCGU.UU..GUC.....G...AAAAGGCCA.GACG.GCGUGGUUAGUG.AAUCGCUCGUGCAAGAAAAAUAAAUGUUUUUUAGUAGUAAUAUUUUAAUUAAAAAAAUGAACACUUUAAGUUA
env-981    UGGGGAUAC...ACUUA...........GAGUCCCC....UGCAUGAGGGUGUUAAUAGCGU.UU..GUC.....G...AAAAGGCCA.GACG.GCGUGGUUAGUG.AAUCGCUCGUGCAAGAAAAAUAAAUGUUUUUUAGUAGUAAUAUUUUAAUUAAAAAAAUGAACACUUUAAGUUA
env-982    UGGGGAUAC...ACUUA...........GAGUCCCC....UGCAUGAGGGUGUUAAUAGCGU.UU..GUC.....G...AAAAGGCCA.GACG.GCGUGGUUAGUG.AAUCGCUCGUGCAAGAAAAAUAAAUGUUUUUUAGUAGUAAUAUUUUAAUUAAAAAAAUGAACACUUUAAGUUA
env-983    UGGGGAUAC...ACUUA...........GAGUCCCC....UGCAUGAGGGUGUUAAUAGCGU.UU..GUC.....G...AAAAGGCCA.GACG.GCGUGGUUAGUG.AAUCGCUCGUGCAAGAAAAAUAAAUGUUUUUUAGUAGUAAUAUUUUAAUUAAAAAAAAUGAACACUUUAAGUUA
env-984    UGGGGAUAC...ACUUA...........GAGUCCCC....UGCAUGAGGGUGUUAAUAGCGU.UU..GUC.....G...AAAAGGCCA.GACG.GCGUGGUUAGUG.AAUCGCUCGUGCAAGAAAAAUAAAUGUUUUUUAGUAGUAAUAUUUUAAUUAAAAAAAUGAACACUUUAAGUUA
env-985    CGGGGAUAC...ACUUA...........GAGUCCCC....UGCAUGAGGGUGUUAAUAGCGU.UC..GCU.....AAAGAAUAGGCCA.AGCG.GCGUGGUUAGUG.AAUCGCUCGUGCAAAAAAGAAGAAACGUUUUUUAGUAGUAAUUUUUAAAUUAAACAAUGAACACUUUAAGUUA
env-986    CGGGGAUAC...ACUUA...........GAGUCCCC....UGCAUGAGGGUGUUAAUAGCGU.UC..GCU.....AAAGAAUAGGCCA.AGCG.GCGUGGUUAGUG.AAUCGCUCGUGCAAAAAAGAAGAAACGUUUUUUAGUAGUAAUUUUUAAAUUAAACAAUGAACACUUUAAGUUA
env-990    UGGGGAUAC...ACUUA...........GAGUCCCC....UGCAUGAGGGUGUUAAUAGCGC.UC..GUUAU.....AGAAAAGGCCA.AACG.GCGUGGUUAGUG.AAUCGCUCGUGCAACAAAAAUAAAACGUUUUUUAGUAGUAAUUUUUAAAUUAAACAAUGAACACUUUAAGUUA
env-992    UGGGGAUAC...ACUUA...........GAGUCCCC....UGCAUGAGGGUGUUAAUAGCGC.UC..GUUAU.....AGAAAAGGCCA.AACG.GCGUGGUUAGUG.AAUCGCUCGUGCAACAAAAAUAAAACGUUUUUUAGUAGUAAUUUUUAAAUUAAACAAUGAACACUUUAAGUUA
env-993    UGGGGAUAC...ACUUA...........GAGUCCCC....UGCAUGAGGGUGUUAAUAGCGC.UC..GUUAU.....AGAAAAGGCCA.AACG.GCGUGGUUAGUG.AAUCGCUCGUGCAACAAAAAUAAAACGUUUUUUAGUAGUAAUUUUUA.........................
env-994    CGGGGAUAC...ACUUA...........GAGUCCCC....CGCGUGAGAGUGUUAAUAGUGU.UG..GUCU..........UUGGCC.GGACC.ACGCGGUUAGUG.AAUCGCUCGCGCAAAGUUAUAAUGUUUUUUAGUAGUAAUAUUUAAAUUAAACAAUGAACACUUUAAGUUACAA
Pbu-3-1    UAGGGAU.....UGACUUAGA.........GUCCCU..UCUGCACAGUCGUGUUAAUAGUG..AU..ACUU..........UUGGCC.GAGUA.UCAUGGUUAGUG.AAUCGGUUGUGCGACAGAGAUAAAGCUAUGGAUGGUCUCUUUUAGACUGGAGGCGCCGUGAGGUCUCUGUGAC
Pbu-1-1    UAGGGAUU....GACUUAG..........AGUCCCU..UCUGCGCAGUCGUGUUAAUAGUG..AU..ACUU..........UUGGCC.GAGUA.UCAUGGUUAGUG.AAUCGGUUGUGCGACAGAGAUAAAGCUAUGGAUGGUCUCUUUUAGACUGGAGGCGCCGUGAGGUCUCUGUGAC
Pbu-2-1    UAGGGAUU....GACUUAG..........AGUCCCU..UCUGCGCAGUCGUGUUAAUAGUG..AU..ACUU..........UUGGCC.GAGUA.UCAUGGUUAGUG.AAUCGGUUGUGCGACAGAGAUAAAGCUAUGGAUGGUCUCUUUUAGACUGGAGGCGCCGUGAGGUCUCUGUGAC
Psp-12-1   UAGGGAUU....GACUUAG..........AGUCCCU..UCUGCGCAGUCGUGUUAAUAGUG..AU..ACUU..........UUGGCC.GAGUA.UCAUGGUUAGUG.AAUCGGUUGUGCGACAGAGAUAAAGCUAUGGAUGGUCUCUUUUAGACUGGAGGCGCCGUGAGGUCUCUGUGAC
env-995    UGGGGAU.....GAACUUAGA.........GUCCCU....GGUACGGGAGUGUUAAUAGUG..AA..GCUU..........UUGGCC.GAGUU.UCAUGGUUAGUG.AAUCGCUCGUACAACAUUCCGAUAAGCGGGAGCACAGCAUGCUCGCUUGUGUUUUU.................
env-996    UGGGGAU.....GAACUUAGA.........GUCCCC....GGUACGGGAGUGUUAAUAGUG..AG..GCUU..........UUGGCC.GGGCU.UCAUGGUUAGUG.AAUCGCUCGUACGAUGAUUUCGAUAAGCGGGAGCGCAGCAUGCUCGCUUGUGUUUUUGCGUGUCGGGGUCAGA
env-997    CGGGGAU.....GAACUUAGA.........GUCCCC...AAGUACGGAAGUGUUAAUAGUG..GC..ACUU..........UUGGCC.GGGUG.UCAUGGUUAGUG.AAUCGACCGUACGAAGACAAGUCUUCAGGUGUUGCAAAGGCAGUAGAUGUCUGUGCGUCUUUCCGUUUGGGCU
env-999    CGGGGAU.....GAACUUAGA.........GUCCCC..AACGUAUGGGAGUGUUAAUAGUC..GC..ACUU..........UUGGCC.AGGUU.CGACGGUUAGUG.AAUCGCUCAUACAAACAAAAAAGCAGAGUCAAAAAGUUGAUUUUUGAACUCUGUGUGUUGUGUUGCGCUACAU
Por-1-1    CGGGGAU.....GAACUUAGA.........GUCCCC....UGUGCGGACGUGUUAAUAGCA..GU..ACUU..........UUGGCC.GAGUG.UUGCGGUUAGGG.AAUCGCUCGUACAAGUAUAAGAUUAAAAACGUUUUUUAGUAGUAAAUUUAAAUUUUAAAAUGAACACUUUAAG
env-1001   CGGGGAU.....GAACUUAGA.........GUCCCC....UGUGCGGACGUGUUAAUAGCA..GC..ACUU..........UUGGCC.GAGUG.UUGCGGUUAGGG.AAUCGCUCGUACAAGUAUAAGAUUAAAAACGUUUUUUAGUAGUAAA...........................
Por-2-1    CGGGGAU.....GAACUUAGA.........GUCCCC....UGUGCGGACGUGUUAAUAGCA..GC..ACUU..........UUGGCC.GAGUG.UUGCGGUUAGGG.AAUCGCUCGUACAAGUAUAAGAUUAAAAACGUUUUUUAGUAGUAAAUUUAAAUUUUAAAAUGAACACUUUAAG
env-1002   CGGGGAU.....GAACUUAGA.........GUCCCC....UGUGCGGUCGUGUUAAUAGUG..AU..GCUU..........UUGGCC.GAGUG.UUACGGUUAGUG.AAUCGCCCGUGCAAGUGAGAAAAAAAAAUGUUUUUUAGUAGUAAAAUUUAAAAGUUUAAUGGACACUUUAAGUU
env-1003   CGGGGAU.....GAACUUAGA.........GUCCCC....UGUGCGGUCGUGUUAAUAGUA..AU..GCUU..........UUGGCC.GAGUG.UUACGGUUAGUG.AAUCGCCCGUGCAAGUGAGAAAAAAAAAUGUUUUUUAGUAGUAAAAUUUAAAAGUUUAAUGGACACUUUAAGUU
Pti-3-1    CGGGGAU.....GAACUUAGA.........GUCCCC....UGUGCGGUCGUGUUAAUAGUA..AU..GCUU..........UUGGCC.GAGUG.UUACGGUUAGUG.AAUCGCCCGUGCAAGUGAGAAAAAAAAAUGUUUUUUAGUAGUAAAAUUUAAAAGUUUAAUGGACACUUUAAGUU
env-1004   CGGGGAU.....GAACUUAGA.........GUCCCC....UGUGCGGUCGUGUUAAUAGUA..AU..GCUU..........UUGGCC.GAGUG.UUACGGUUAGUG.AAUCGCCCGUGCAAGUGAGAAAAAAAUAAUGUUUUUUAGUAGUAAAAUUUAAAAGUUUAAUGGACACUUUAAGU
Pti-1-1    CGGGGAU.....GAACUUAGA.........GUCCCC....UGUGCGGUCGUGUUAAUAGUA..AU..GCUU..........UUGGCC.GAGUG.UUACGGUUAGUG.AAUCGCCCGUGCAAAUGAGAAAAAAAAAUGUUUUUUAGUAGUAAAAUUUAAAAGUUUAAUGGACACUUUAAGUU
Pti-2-1    CGGGGAU.....GAACUUAGA.........GUCCCC....UGUGCGGUCGUGUUAAUAGUA..AU..GCUU..........UUGGCC.GAGUG.UUACGGUUAGUG.AAUCGCCCGUGCAAGUGAGAAAAAAAAAUGUUUUUUAGUAGUAAAAUUUAAAAGUUUAAUGGACACUUUAAGUU
env-1005   CGGGGAU.....GAACUUAGA.........GUCCCC....UGUGCGGUCGUGUUAAUAGUA..AU..GCUU..........UUGGCC.GAGUG.UUACGGUUAGUG.AAUCGCCCGUGCAAGUGAGAAAAAAAAAUGUUUUUUAGUAGUAAAAUUUAAAAGUUUAAUGGACACUUUAAGUU
env-1006   CGGGGAU.....GAACUUAGA.........GUCCCC....UGUGCGGAUGUGUUAAUAGUG..GU..GCUU..........UUGGCC.GAGCA.UCACGGUUAGUG.AAUCGUUCGUACAUGAGUGAAAUAAAAACGUUUUUUAGUAGUAAA...........................
Psp-4-1    CGGGGAU.....GAACUUAGA.........GUCCCC....UGUGCGGAUGUGUUAAUAGUG..GU..GCUU..........UUGGCC.GAGCA.UCACGGUUAGUG.AAUCGUUCGUACAUGAGUGAAAUAAAAACGUUUUUUAGUAGUAAAAUUUAAAUUUUUAAUGGACACAUUAAGU
env-1007   CGGGGAU.....GAACUUAGA.........GUCCCC....UGUGCGGAUGUGUUAAUAGUG..GU..GCUU..........UUGGCC.GAGCA.UCACGGUUAGUG.AAUCGUUCGCACAUGAGUGAAAUAAAAACGUUUUUUAGUAGUAAAAUUUAAUUUUUUAAUGGACACAUUAAGU
Pbu-5-1    CGGGGAU.....GAACUUAGA.........GUCCCC....UGUGCGGAUGUGUUAAUAGUG..GU..GCUU..........UUGGCC.GAGCA.UCACGGUUAGUG.AAUCGUUCGUACAUGAGUGAAAUAAAAACGUUUUUUAGUAGUAAAAUUUAAAUUUUUAAUGGACACAUUAAGU
Pbu-4-1    CGGGGAU.....GAACUUAGA.........GUCCCC....UGUGCGGAUGUGUUAAUAGUG..GU..GCUU..........UUGGCC.GAGCA.UCACGGUUAGUG.AAUCGUUCGUACAUGAGUGAAAUAAAAACGUUUUUUAGUAGUAAAAUUUAAAUUUUUAAUGGACACAUUAAGU
env-1008   CGGGGAU.....GAACUUAGA.........GUCCCC...ACUUUCGGGCGUGUUAAUAGUC..GU..GCUU..........UUGGCC.GAGCG.CGACGGUUAGUG.AAUCGUCCGGAGCCAACGAAUGAAAUGUUUUUUAGUAGUAAAAUUGAAAAGAAGAAAUGGACACUUUAAGUUA
Pen-1-1    CGGGGAU.....GAACUUAGA.........GUCCCC...ACUUUCGGGCGUGUUAAUAGUC..GU..GCUU..........UUGGCC.GAGCG.CGACGGUUAGUG.AAUCGUCCGGAGCCAACGAAUGAAAUGUUUUUUAGUAGUAAAAUUGAAAAGAAGAAAUGGACACUUUAAGUUA
env-1009   CGGGGAU.....GAACUUAGA.........GUCCCC...ACUUUCGGGCGUGUUAAUAGUC..GU..GCUU..........UUGGCC.GAGCG.CGACGGUUAGUG.AAUCGUCCGGAGCCAACGAAUGAAAUGUUUUUUAGUAGUAAAAUUGAAAAGAAGAAAUGGACACUUUAAGUUA
env-1017   CGGGGAU.....GAACUUAGA.........GUCCCC...ACUUUCGGGCGUGUUAAUAGUC..GU..GCUU..........UUGGCC.GAGCG.CGACGGUUAGUG.AAUCGUCCGGAGCCAACGAAUGAAAUGUUUUUUAGUAGUAAAAUUGAAAAGAAGAAAUGGACACUUUAAGUUA
env-1020   CGGGGAU.....GAACUUAGA.........GUCCCC...ACUUUCGGGCGUGUUAAUAGUC..GU..GCUU..........UUGGCC.GAGCG.CGACGGUUAGUG.AAUCGUCCGGAGCCAACGAAUGAAAUGUUUUUUAGUAGUAAAAUUGAAAAGAAGAAAUGGACACUUUAAGUUA
Ppl-2-1    CGGGGAU.....GAACUUAGA.........GUCCCC...ACUUUCGGGCGUGUUAAUAGUC..GU..GCUU..........UUGGCC.GAGCG.CGACGGUUAGUG.AAUCGUCCGGAGCCAACGAAUGAAAUGUUUUUUAGUAGUAAAAUUGAAAAGAAGAAAUGGACACUUUAAGUUA
env-1021   CGGGGAC.....ACACUUAGA.........GUCCCC....CGUGUGAGCGUGUUAAUAGUG..GC..GCUU..........UUGGCC.GAGCG.UCAUGGUUAGUG.AAUCGCCUGCACAUCAAGAGAAAUAAAAACGUUUUUUAGUAGUAAAAAUUAAAAUUAAAAAUGAACACUUUAA
env-1022   CGGGGAC.....ACACUUAGA.........GUCCCC....CGUGUGAGCGUGUUAAUAGUG..GC..GCUU..........UUGGCC.GAGCG.UCAUGGUUAGUG.AAUCGCCUGCACAUCAAGAGAAAUAAAAACGUUUUUUAGUAGUAAAAAUUAAAAUUAAAAAUGAACACUUUAA
env-1023   CGGGGAC.....ACACUUAGA.........GUCCCC....CGUGUGAGCGUGUUAAUAGUG..GC..GCUU..........UUGGCC.GAGCG.UCAUGGUUAGUG.AAUCGCCUGCACAUCAAGAGAAAUAAAAACGUUUUUUAGUAGUAAAAAUUAAAAUUAAAAAUGAACACUUUAA
env-1024   CGGGGG.C....ACACUUAGAA.......GACCCCC....UGUGCUGGUGUGUUAAUAGUG..AC..UCUU.........UUGGUCC.GAGAG.CCAUGGUUAGUG.AAUCGCUUGCACAAAGUAAAUAAUAAUGUUUUUUAGUAGUAAAAUUUAAUUUUAAAAU..............
env-1025   CGGGGGC.....ACACUUAGA.........GCCCCC....UGUGCUGGUGUGUUAAUAGUG..AC..UCUU..........UUGGCC.GAGAG.CCAUGGUUAGUG.AAUCGCUUGCACAAAGUAAAAUAAUAAUGUUUUUUAGUAGUAAAAUUUAAAUUUUAAAAUGAACACUUUAAGU
env-1027   CGGGGAUAC...GCUUA...........GAGUCCCC....AAGGGUAUCGUGUUAAUAGUU..GU..GUCU..........UUGGCC.GGGUA.CGACGGUUAGUG.AAUCGGUAUUCUAAACGAAGUAAAUAAGUUUUUUAGUAGUACAAACAAUUCAAUUGAAGAUGAACACUUUAAG
Pma-4-1    CGGGGAUAC...GCUUA...........GAGUCCCC....AAGGGUAUCGUGUUAAUAGUU..GU..GUCU..........UUGGCC.GGGCA.CGACGGUUAGUG.AAUCGGUAUUCUAAACGAAGUAAAUAAGUUUUUUAGUAGUACAAACAAUUCAAUUGAAGAUGAACACUUUAAG
env-1028   CGGGGAUAC...GCUUA...........GAGUCCCC....AAGGGUAUCGUGUUAAUAGUU..GU..GUCU..........UUGGCC.GGGCA.CGACGGUUAGUG.AAUCGGUAUUCUAAACGAAGUAAAUAAGUUUUUUAGUAGUACAAACAAUUCAAUUGAAGAUGAACACUUUAAG
env-1029   GGGGGAUAC...ACUUA...........GAGUCCCC....AGUACAGGAGUGUUAAUAGCG..GU..GUCU..........UUGGCC.GGGCA.CGGUGGUUAGUG.AAUCGCUUGUACAACAGUGAUAAAAAGGAAAAAACGUUUUUUAGUAGUAAAAUUUAAAUUUUAAAAUGGACAC
env-1031   GGGGGAUAC...GCUUA...........GAGUCCCC....AGUACAGGAGUGUUAAUAGCU..GU..GUCU..........UUGGCC.GGGCA.UGGUGGUUAGUG.AAUCGCUUGUACAACGUGAUAAAAAGGAAAAAAACGUUUUUUAGUAGUAAAAUUUAAAAUUUAAAAUGGACAC
env-1032   CGGGGAUAC...ACUUA...........GAGUCCCC....CGUACGAGCGUGUAAAUAGUG..GU..CCUU..........UUGGCC.GAGUC.CCACGGUUUGUG.AAUCGCUCGCACACAGCAAGGUAAGGACGUUCCGCAAGGAGUGUUGUUCCUUAUCGUGUUGUUAUGCGGUGUC

           .<<<<<<<<<<<.............>>>>>>>>>>>................<<<<<.<<<..<<..<<<<<...............>>>>>>.>>>>.>>>>>...........................................................................
           .222222.1111.............1111.222222................02200.122..22..2222...............22222.2221.00220...........................................................................
           •GGGG•○-----R••••○○○----------○•CCCC----••••••••YGUGU•AAUAGY•--•Y--RYYU---------UUGGCC-GRRY•-••RYGGUU•GUG-AAUCR•••••••RR••R••••••••••••••••••••••••••••••••••••••••••••••••••••••
            5′                                                                                                  3′
```

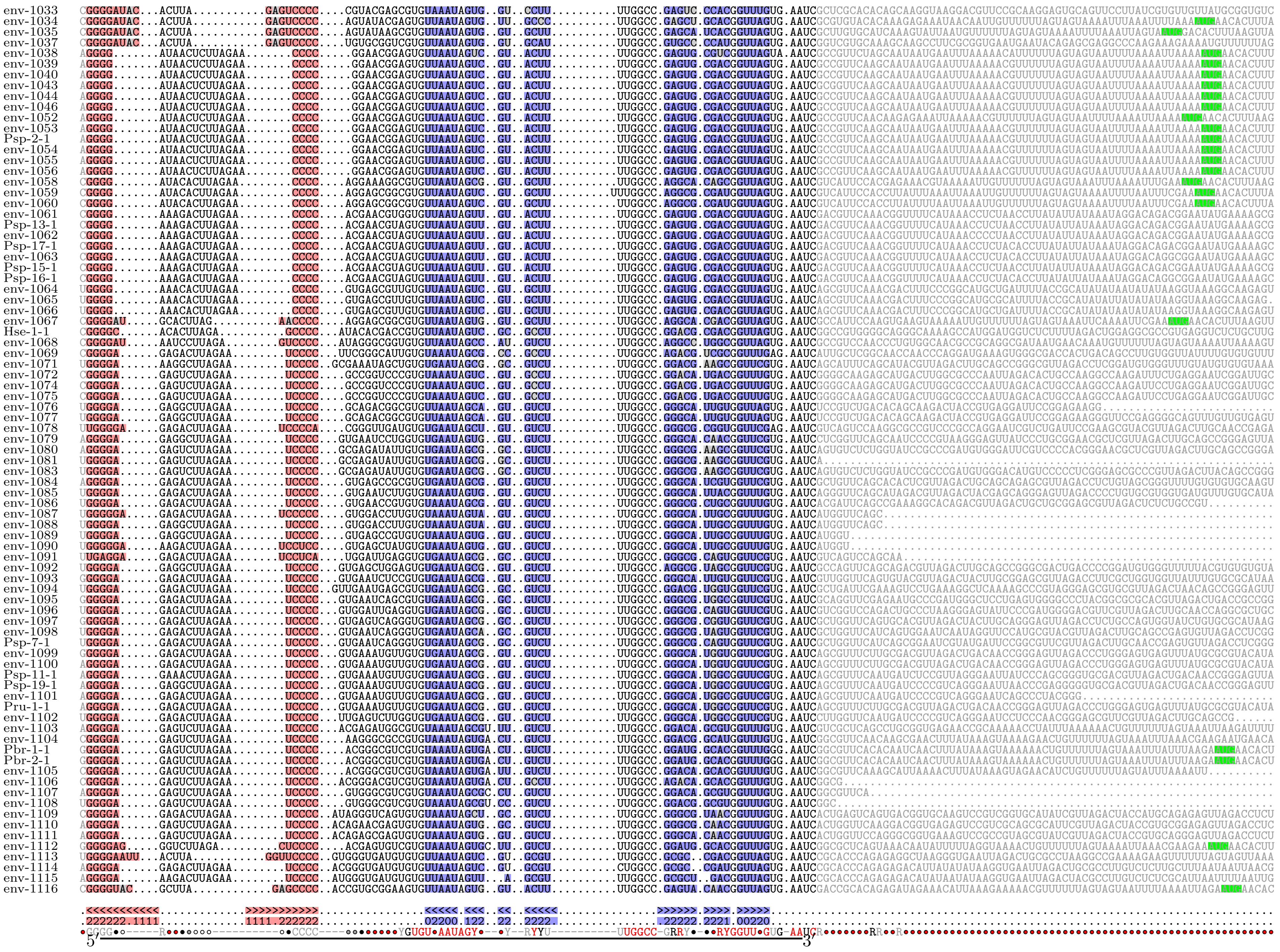

```
env-1117  CGGGGAAC....ACUUA............GAUCCCC....AUGGCCGUAGUGUUAAUAGUC..AU..GUCU..........CUGGCC.GGGCA.UGGCGGUUAGUG.AAUCUGCGUCCAGACGGCAGACAGCCGGAGAGCGACGGUUCUUCCGGCGGUUGGUCGUGAGAAAGAAUUGAAA
env-1118  UGGGGCAC....ACUUA............GAACCCC...GCGAGCGAUGGUGUUAAUAGUC..AU..GCGU..........UUGGCC.GCGCA..GACGGUUAGUG.AAUCCGCGUUCGAAACAAAUGUUUUUUAGUAGUAUUUUUAAAAAGAUUUUUAAAGUUAAAUGAACACUUUAAG
env-1119  UGGGGCAC....ACUUA............GAACCCC...GCGAGCGAUGGUGUUAAUAGUC..AU..GCGU..........UUGGCC.GCGCA..GACGGUUAGUG.AAUCCGCGUUCGAAACAAUUGUUUUUUAGUAGUAUUUUUAAAAAGAUUUUUAAAGUUAAAUGAACACUUUAAG
env-1120  CGGGGG......AAGUCUUAG..........CCCCC..CGCCGGGCAAAGUGUUAAUAGCG..CC..GCUUG..........UGGC.CGAGCG.UCGCGGUUAGUG.AAUCUGCGCCCGACCGAGAGUCUCCCGAGACUAAGAGCCUCUCGCAAAGAAUGUUUUUUAGUAGUAUAAAUUU
env-1122  CGGGGG......AAGUCUUACG.........CCCCC..CGCCGGGCAAAGUGUUAAUAGCG..CC..GCUUG..........UGGC.CGAGCG.UCGCGGUUAGUG.AAUCUGCGCCCGACCGAGAGUCUCCCGAGACUAAGAGCCUCUCGCAAAGAAUGUUUUUAGUAGUAUAAAUUUU
env-1123  UGGAGCG.....AGUCUUAGA.........CGCUCC....GGUGCUGACGUGUUAAUAGUG..GC..AUCU..........UUGGCC.GGGUGACCAUGGUUAGUG.AAUCGAAAGCACAGCGACACAUAAGGUGAGCAGCACAGCGUUCGCUCUGUUUGUUGUGCGCCUGCGUGUCGAA
env-1125  UGGGGGAU....AGCUUAG..........GUCCUCC....GCUCUUGGCGUGUUAAUAGCG..UU..GCUU..........UUGGCC.GAGCA.UCGUGGUUAGUG.AAUCGUCAGGAAAGAGACAACAGAAAAGAAAACGUUUUUUAGUAGUAAUUUUUUAACUUAGAAAUGAACACAU
env-1126  UGAAGGAU....GUCUUAG..........GUCCUUC....GGUCUCGUCGUGUUAAUAGCU..GU..ACUU..........UUGGCC.GAGUG.CGGUGGUUAGUG.AAUCGUCGGGGCAAGCAAAGAGUUUUUUAGUAGUAAAGAAAGAAUUAGAGUUUAGAAAUGAACACGUUAAGUU
env-1127  UGAAGGAU....GUCUUAG..........GUCCUUC....GGUCUCGUCGUGUUAAUAGCG..GU..ACUU..........UUGGCC.GAGUG.CAGUGGUUAGUG.AAUCGUCGGGGCAAGCAAAAAGUUUUUUAGUAGUAAAAAGUAUUUUUAGAGUUAGAAAUGAACACGUUAAGUU
env-1129  CGGGGAU.....GUGCUUAGA.........GUCCCC...AGGAUGUGGCGUGUUAAUAGAG..GUA.UUCU..........GUGGCC.GGAAG.CCUUGGUUAGUG.AAUCGCCUGUCCAAAGAAAAGUGAGAUUAUAAAAACGUUUUUUAGUAGUAAAAUUUAAAAGUUAAAAAUGGAC
env-1130  AGGGCAC.....CCCAAAAA..........GUGCCC......AAACUGCGUGUGAAUAGCC..CGU.UUCA..........UUGGCC.UGGAC.GGACGGUUCGUG.AAUCGCAGAGAAUCAAGAAGUUAAACGUAGAAAACGAACAUUAAAAGAAAUGGACACACUGAGUCAGAAGACC
env-1131  GGGGU..ACC..CUCU...........GGUAUACCC.....AAAUCUGCGUGUAAAUAGUC..CGUUUCCA..........UUGGCC.UGGAC.GGGCGGUUUGUG.AAUCGCAGAACAAGUAGUAAUGAACAAUAAAGAAUUAAAAAACUAAAAUGGACACACUUAGUCAGAAGACCAC
env-1132  GGGGUACCC...CUCUGA..........GCGUACCC.......AUCUGCGUGUGAAUAGUC..CGUUUCCA..........UUGGCC.UGGAC.GGGUGGUUCGUG.AAUCGCAGAAGAGAACAAAAAACGUAGUAACGAACAAAAGAAACUAAAAAACAAAAAAUGGACACACUGAGUC
env-1133  AGGACCC.....CUAAA.............GGGCCC....UCUCGAAGCGUGUGAAUAGUC..CGC.AUCA..........UUGGCC.UGAAC.GGACGGUUCGUG.AAUCGCUCGACAAAAUGAGAAAUAAACUAGGUAGAAACAAACAAAAGUAAGACAAAGAAUGGAUACUUUGAGU
env-1134  AGAGGGG.....CUAAAAUA..........CCCCUC......AAACUGCGUGUAAAUAGUU..CGUUUCCA..........UUGGCC.UGGAC.GGAUGGUUUGUG.AAUCGCAGAAAACAAGAAAAAUUAAACGUAGUAACGAACAAAACAAAAGAAAUGGACACACUGAGUCAGAAGA
env-1135  AGGGGC......GAAUUAU............GCCCC......AAAGUGCGUGUGAAUAGUU..CG..CUCU..........UUGGCC.AGA.C.GGACGGUUCGUG.AAUCGCGCUGAAAAUGAACUUAGUAGAAAACAAAAAGAAAAAACAAAAUGGACACAUUGAGUUACAAGACCAC
env-1136  CGGGGC......AAAAACA............GCCCC......AAAGUGCGUGUGAAUAGUG..U...GCUU.........AUUGGCC.AGGU..ACACGGUUCGUG.AAUCGCGCUGAAGAAGAGUUAAAUUUAGUAGAGAACAAAAAGAAAAGAACAAAGAAAUGGACACAUUGAGUUA
env-1137  AAGUGGCCUC..UUUU...........GAGG.CACU.......AUCUGCGUGUUAAUAGCU..AC..GUUC.........UUUGGCC.GAACG.UGGUGGUUAGUG.AAUCGCAGAGAAAAGUGAUAAUUUAUUAGAUUUAGGGAUUAAUUCCCCUUAGUAACUACAAACAAUUAACAAC
env-1138  AAGUGGCCUC..UUUU...........GAGG.CACU.......AUCUGUGUGUUAAUAGCU..GC..GUUC.........UUUGGCC.GAACG.UGGUGGUUAGUG.AAUCGCAGAGAAAAGUGAUAAUUUAUUAGAUUUAGGGAUUAAUUCCCCUUAGUAACUACAAACAAUUAACAAC
env-1139  AAGUGGCCUC..UUUU...........GAGG.CACU.......AUCUGUGUGUUAAUAGCU..GC..GUUC.........UUUGGCC.GAACG.UGGUGGUUAGUG.AAUCGCAGAGAAAAAUGAUAAUUUAUUAGAUUUAGGGAAUCAAUUCCCCUUAGUAACUACAAACAAUUAACAA
env-1140  AAGUGGCCUC..UUUU...........GAGG.CGCU.......AUCUGUGUGUUAAUAGCU..GC..GUUC.........UUUGGCC.GAACG.UGGUGGUUAGUG.AAUCGCAGAGAAAAAUGAUAAUUUAUUAGAUUUAGGGAAUCAAUUCCCCUUAGUAACUACAAACAAUUAACAA
env-1141  AAGUGGCCUC..UUUU...........GAGG.CGCU.......AUCUGUGUGUUAAUAGCU..AC..GUUC.........UUUGGCC.GAACG.UGGUGGUUAGUG.AAUCGCAGAGAAAAAUGAUAAUUUAUUAGAAUUAGGGAUUCAAUUCCCCUUAGUAACUACAAACAAUUAACAA
env-1143  AAGUGGCCUC..UUUU...........GAGG.CGCU.......AUCUGUGUGUUAAUAGCU..AC..GUUC.........UUUGGCC.GAACG.UGGUGGUUAGUG.AAUCGCAGAGAAAAAUGAUAAUUUAUUAGAUUUAGGGAUUCAAUUCCCCUUAGUAACUACAAACAAUUAACAA
env-1144  AAGUGGCCUC..UUUU...........GAGG.CGCU.......AUCUGUGUGUUAAUAGCU..AC..GUUC.........UUUGGCC.GAACG.UGGUGGUUAGUG.AAUCGCAGAGAAAAAUGAUAAUUUAUUAGAUUUAGGGAAUCAAUUCCCCUUAGUAACUACAAACAAUUAACAA
env-1145  AAGUGGCCUC..UUUU...........GAGG.CGCU.......AUCUGUGUGUUAAUAGCU..AC..GUUC.........UUUGGCC.GAACG.UGGUGGUUAGUG.AAUCGCAGAGAAAAAUGAUAAUUUAUUAGAAUUAGGGAUUCAAUUCCCCUUAGUAACUACAAACAAUUAACAA
env-1147  AGUGCCUC....AAAA.............GAGUCAC.....G..GCCGCGUGUUAAUAGCC..GA..GUUU........CAUUGGCCGAAAAU.CGGUGGUUAGUG.AAUCGCGGAAACGAUAACUUAAUUGAAUUAGGAGUACGCUCCUUAGUAACUACAAAAAUCUCAACAAAAAGAA
env-1148  AGUGCCUC....AAAA.............GAGUCAC.....G..GCCGCGUGUUAAUAGCC..GA..GUUU........CAUUGGCCGAAAAU.CGGUGGUUAGUG.AAUCGCGGAAACGAUAACUUAAUUGAAUUAGGAGUACGCUCCUUAGUAACUACAAAAUCUCAACAAAAAGAAU
env-1149  AGUGCCUC....AAAA.............GAGUCAC.....G..GCCGCGUGUUAAUAGCC..GA..GUUU........CAUUGGCCGAAAAU.CGGUGGUUAGUG.AAUCGCGGAAACGAUAACUUAAUUGAAUUAGGAGUACGCUCCUUAGUAACUACAAAAAUCUCAACAAAAAGAA
env-1151  AGUGC.CUC...AUAA............GAGAGCAC.......AGCCGCGUGUUAAUAGCC..GA..GUUUC........AUUGGCCGAAAAU.CGGUGGUUAGUG.AAUCGCGGAAACGAUAACUUAAUUGAAAUAGGAGUACGCUCCUUAGUAACUACAAAAAUCUCAACAAAAAGAA
env-1153  AGUGC.CUC...AAAA............GAGUGCAC.......GGCCGCGUGUUAAUAGCC..GA..GUUUC........AUUGGCCGAAAAU.CGGUGGUUAGUG.AAUCGCGGAAACGAUAACUUAAUUGAAUUAGGAGUACGCUCCUUAGUAACUACAAAAAUCUCAACAAAAAGAA
env-1154  AGUGC.CUC...AAAA............GAGUGCAC.......GGCCGCGUGUUAAUAGCC..GA..GUUUC........AUUGGCCGAAAAU.CGGUGGUUAGUG.AAUCGCGGAAACGAUAACUUAAUUGAAAUAGGAGUACGCUCCUUAGUAACUACAAAAUCUCAACAAAAAGAAU
env-1155  AGUGC.CUC...AAAA............GAGUGCAC.......GGCCGCGUGUUAAUAGCC..GA..GUUUC........AUUGGCCGAAAAU.CGGUGGUUAGUG.AAUCGCGGAAACGNNNNNNNNAAGCUACAAGACCAUUUCUGCCAACAAGGAAACCGCACAGAAAGAGUGGGUC
env-1156  AGUGC.CUC...AAAA............GAGUGCAC.......GGCCGCGUGUUAAUAGCC..GA..GUUUC........AUUGGCCGAAAAU.CGGUGGUUAGUG.AAUCGCGGAAACGAUAACUUAAUUGAAUUAGGAGUACGCUCCUUAGUAACUACAAAAAUCUCAACAAAAAGAA
env-1157  AGUGC.CUC...AAAA............GAGUGCAC.......GGCCGCGUGUUAAUAGCC..GA..GUUUC........AUUGGCCGAAAAU.CGGUGGUUAGUG.AAUCGCGGAAACGAUAACUUAAUUGAAAUAGGAGUACGCUCCUUAGUAACUACAAAAUCUCAACAAAAAGAAU
env-1158  AGUGC.CUC...AAAA............GAGUGCAC.......GGCCGCGUGUUAAUAGCC..GA..GUUUC........AUUGGCCGAAAAU.CGGUGGUUAGUG.AAUCGCGGAAACGAUAACUUAAUUGAAAUAGGAGUACGCUCCUUAGUAACUACAAAAAUCUCAACAAAAAGAA
env-1159  AGUGC.CUC...AAAA............GAGUGCAC.......GGCCGCGUGUUAAUAGCC..GA..GUUUC........AUUGGCCGAAAAU.CGGUGGUUAGUG.AAUCGCGGAAACGAUAACUUAAUUGAAUUAGGAGUACGCUCCUUAGUAACUACAAAAAUCUCAACAAAAAGAA
env-1163  AGUGC.CUC...AAAA............GAGUGCAC.......GGCCGCGUGUUAAUAGCC..GA..GUUUC........AUUGGCCGAAAAU.CGGUGGUUAGUG.AAUCGCGGAAACGAUAACUUAAUUGAAAUAGGAGUACGCUCCUUAGUAACUACAAAAAUCUCAACAAAAAGAA
env-1164  AGUGC.CUC...AAAA............GAGUGCAC.......GGCCGCGUGUUAAUAGCC..GA..GUUUC........AUUGGCCGAAAAU.CGGUGGUUAGUG.AAUCGCGGAAACGAUAACUUAAUUGAAAUAGGAGUACGCUCCUUAGUAACUACAAAAAUCUCAACAAAAAGAA
env-1165  AGUGC.CUC...AAAA............GAGUGCAC.......GGCCGCGUGUUAAUAGCC..GA..GUUUC........AUUGGCCGAAAAU.CGGUGGUUAGUG.AAUCGCGGAAACGAUAACUUAAUUGAAAUAGGAGUACGCUCCUUAGUAACUACAAAAAUCUCAACAAAAAGAA
env-1166  AGUGC.CUC...AAAA............GAGUGCAC.......GGCCGCGUGUUAAUAGCC..GA..GUUUC........AUUGGCCGAAAAU.CGGUGGUUAGUG.AAUCGCGGAAACGAUAACUUAAUUGAAUCUAGGAGUACGCUCCUUAGUAACUACAAAAAUCUCAACAAAAAGA
env-1167  AGUGC.CUC...AAAA............GAGUGCAC.......GGCCGCGUGUUAAUAGCC..GA..GUUUC........AUUGGCCGAAAAU.CGGUGGUUAGUG.AAUCGCGGAAACGAUAACUUAAUUGAAAUAGGAGUACGCUCCUUAGUAACUACAAAAAUCUCAACAAAAAGAA
env-1192  UGUGGCU.C...AUUU............GCAGCCAC.......ACUCGCGUAUUAAUAGCC..GG..GUUU........CAUUGGCC.GAAAAACGGUGGUUAGUG.AAUCGCGAGAACGAUAACUUAAUUGAAAUAGGAGCUCGCUCCUUAGUAANNNNNNNNNNNNNNAAGCUACAAGA
env-1193  UGCCCCA.....AUUG..............UGGGGU...UACACUCACGUGUUAAUAGCC..UA..GUUC.........UUUGGCC.GAAUU.UGGUGGUUAGUG.AAUCGUGAGAAAAAGAUAAUUCAUUGAUUUAUGGGCAUUUGCCCACUUAGUAACUACAAAUUAAUCACUUAGA
env-1194  GCCCCAA.....UUU...............UUGGGG..GUUACACUCACGUGUUAAUAGCC..UA..GUUC.........UUUGGCC.GAACU.UGGUGGUUAGUG.AAUCGUGAGAAAAAGAUAAUUCAUUGAUUUAUGGGCAUUUGCCCACUUAGUAACUACAAAUUAAUCACUUAGA
Psp-1-1   GCCCCA......AUUUUG.............UGGGG..GUUACACUCACGUGUUAAUAGCC..UA..GUUC.........UUUGGCC.GAACU.UGGUGGUUAGUG.AAUCGUGAGAAAAAGAUAAUUCAUUGAUUUAUGGGCAUUUGCCCACUUAGUAACUACAAAUUAAUCACUUAGA
env-1195  GCCCCA......AUUG...............UGGGG..GUUACACUCACGUGUUAAUAGCC..UA..GUUC.........UUUGGCC.GAACU.UGGUGGUUAGUG.AAUCGUGAGAAAAAGAUAAUUCAU.................................................
env-1197  GCCCCA......AUUG...............UGGGG..GUUACACUCACGUGUUAAUAGCC..UA..GUUC.........UUUGGCC.GAACU.UGGUGGUUAGUG.AAUCGUGAGAAAAAGAUAAUUCAUUGAUUUAUGGGCAUUUGCCCACUUAGUAACUACAAAUUAAUCACUUAGA
env-1203  GCCCCA......AUUG...............UGGGG..GUUACACUCACGUGUUAAUAGCC..UA..GUUC.........UUUGGCC.GAACU.UGGUGGUUAGUG.AAUCGUGAGAAAAAGAUAAUUCAUUGAUUUAUGGGCAUUUGCCCACUUAGUAACUACAAAUUAAUCACUUAGA
env-1205  GCCCCA......AUUG...............UGGGG..GUUACACUCACGUGUUAAUAGCC..UA..GUUC.........UUUGGCC.GAACU.UGGUGGUUAGUG.AAUCGUGAGAAAAAGAUAAUUCAUUGAUUUAUGGGCAUUUGCCCACUUAGUAACUACAAAUUAAUCACUUAGA
env-1206  UUGCCCC.....AAUUGUGG..........GGGGUA......CACUCACGUGUUAAUAGCC..UA..GUUC.........UUUGGCC.GAACU.UGGUGGUUAGUG.AAUCGUGAGAAAAAGAUAAUUCAUUGAUUUAUGGGCAUUUGCCCACUUAGUAACUACAAAUUAAUCACUUAGA
env-1211  UUGCCCC.....AAUUGUGG..........GGGGUA......CACUCACGUGUUAAUAGCC..UA..GUUC.........UUUGGCC.GAACU.UGGUGGUUAGUG.AAUCGUGAGAAAAAGAUAAUUCAUUG...............................................
env-1213  AGUUGCCCCA..AUUG...........UGGGGUUAC.......ACUCACGUGUUAAUAGCC..UU..GUUC.........UUUGGCC.GAACU.UGGUGGUUAGUG.AAUCGUGAGAAAAAGAUAAUUCAUUGAUUUAUGGGCAUUCGCCCACUUAGUAACUACAAAUUAAUCACUUAGA
env-1216  AGUUACCUC...GUUUU...........GAGGUUGC.......ACUUGCGUGUUAAUAGCC..AA..GUUC.........UUUGGCC.GAAUU.UGGUGGUUAGAG.AAUCGCAAGAAAAAGAUAAUUUAUUGAUUUAUGGGCGAAAGCCCACUUAGUAACUACAAAUUAAUCAUUAGAA
env-1223  AGUUACCUC...GUUUU...........GAGGUUGC.......ACUUGCGUGUUAAUAGCC..AA..GUUC.........UUUGGCC.GAAUU.UGGUGGUUAGAG.AAUCGCAAGAAAAAGAUAAUUUAUUGAUUUAUGGGCGAAAGCCCACUUAGUAACUACAAAUUAAUCAUUAGAA
env-1224  AGUUACCUC...GUUUU...........GAGGUUGC.......ACUUGCGUGUUAAUAGCC..AA..GUUC.........UUUGGCC.GAAUU.UGGUGGUUAGAG.AAUCGCAAGAAAAAGAUAAUUUAUUGAUUUAUGGGCGAAAGCCCACUUAGUAACUACAAAUUAAUCAUUAGAA
env-1228  AGUUACCUCA..UUU............UGAGGUUGC.......ACUUGCGUGUUAAUAGCC..AA..GUUC.........UUUGGCC.GAAUU.UGGUGGUUAGAG.AAUCGCAAGAAAAAGAUAAUUUAUUGAUUUAUGG.......................................
env-1233  AGUUACCUCA..UUU............UGAGGUUGC.......ACUUGCGUGUUAAUAGCC..AA..GUUC.........UUUGGCC.GAAUU.UGGUGGUUAGAG.AAUCGCAAGAAAAAGAUAAUUUAUUGAUUUAUGGGCGAAAGCCCACUUAGUAACUACAAAUUAAUCAUUAGAA
env-1243  AGUUACCUCA..UUU............UGAGGUUGC.......ACUUGCGUGUUAAUAGCC..AA..GUUC.........UUUGGCC.GAAUU.UGGUGGUUAGAG.AAUCGCAAGAAAAAGAUAAUUUAUUGAUUUAUGGGCGAAAGCCCACUUAGUAACUACAAAUUAAUCAUUAGAA
env-1244  AGUUGCCUCA..UUU............UGAGGUUGC.......ACUUGCGUGUUAAUAGCC..AA..GUUC.........UUUGGCC.GAAUU.UGGUGGUUAGAG.AAUCGCAAGAAAAAGAUAAUUUAUUGAUUUA..........................................
env-1245  AGUUGCCUCA..UUU............UGAGGUUGC.......ACUUGCGUGUUAAUAGCC..AA..GUUC.........UUUGGCC.GAAUU.UGGUGGUUAGAG.AAUCGCAAGAAAAAGAUAAUUUAUUGAUUUAUGGGCGAAAGCCCACUUAGUAACUACAAAUUAAUCAUUUAGA
env-1250  AGUUGCCUCA..UUU............UGAGGUUGC.......ACUUGCGUGUUAAUAGCC..AA..GUUC.........UUUGGCC.GAAUU.UGGUGGUUAGAG.AAUCGCAAGAAAAAGAUAAUUUAUUGAUUUAUGGGCGAAUGCCCACUUAGUAACUACAAAUUAAUCAUUUAGA
env-1251  AGUUGCCUCA..UUU............UGAGGUUGC.......ACUUGCGUGUUAAUAGCC..AA..GUUC.........UUUGGCC.GAAUU.UGGUGGUUAGAG.AAUCGCAAGAAAAAGAUAAUUUAUUGAUUUAUGGGCGAAAGCCCACUUAGUAACUACAAAUUAAUCAUUUAGA
env-1254  AGUUGCCUCA..UUU............UGAGGUUGC.......ACUUGCGUGUUAAUAGCC..AA..GUUC.........UUUGGCC.GAAUU.UGGUGGUUAGAG.AAUCGCAAGAAAAAGAUAAUUUAUUGAUUUAUGGGCGAAUGCCCACUUAGUAACUACAAAUUAAUCAUUUAGA
env-1256  AGUUGCCUCA..UUUU...........UGAGGUUGC.......ACUUGCGUGUUAAUAGCC..AA..GCUC.........UUUGGCC.GAACU.UGGUGGUUAGAG.AAUCGCAAGAAAAAGAUAAUUUAUUGAUUUAUGGGCAUUCGCCCACUUAGUAACUACAAAUUAAUCACUUAGA
env-1258  AGUUGCCUC...UUUUU...........GAGGUCGC.......ACUUGCGUGUUAAUAGCC..AA..GCUC.........UUUGGCC.GAACU.UGGUGGUUAGAG.AAUCGCAAGAAAAAGAUAAUUUAUUGAUUUAUGGGCGAAAGCCCACUUAGUAACUACAAAUUAAUCAUUUAGA
env-1259  AGUUGCCUC...UUUUU...........GAGGUCGC.......ACUUGCGUGUUAAUAGCC..AA..GCUC.........UUUGGCC.GAACU.UGGUGGUUAGAG.AAUCGCAAGAAAAAGAUAAUUUAUUGAUUUAUGGGCGAAAGCCCACUUAGUAACUACAAAUUAAUCAUUUAGA
Pmu-2-1   GGGCAGCU....GUCUUAG..........AGCUGCC..ACGGCAUUGGCGUGUUAAUAGGG..CU..GUGC..........AUGGCC.GUACA.GUCUGGUUAGAG.AAUCGCCCGUGCGCUCAGGGUUAAAGCUGUGGAUGGUCUCUUUUAGACUGGAGGCACCGCGAGGCCCCUGUUC
Pde-1-1   CGGGGC......ACACUUAGA..........GCCCC...AUACACGACCGUGUUAAUAGUC..GC..GCCU..........UUGGCC.GGACG.CGACGGUUAGUG.AAUCGGCCGUGGGGCAGGGCAAAAGCCAUGGAUGGUCUCUUUUAGACUGGAGGCGCCGUGAGGUCUCUGCCUG
          .<<<<<<<<<<<.............>>>>>>>>>>>................<<<<<.<<<..<<..<<<<<...............>>>>>>.>>>>.>>>>>............................................................................
          .222222.1111.............1111.222222................02200.122..22..2222.................22222.2221.00220............................................................................
          •GGGG•∘-----R•••∘∘∘∘----------∘•CCCC----∘∘•••••••YGUGU•AAUAGY•--•Y--RYYU----------UUGGCC-GRRY•-••RYGGUU•GUG-AAUCR•••••••RR••R••••••••••••••••••••••••••••••••••••••••••••••••••••••••
           5′                                                                                                    3′
```

alignment positions 260 · · · 267

```
env-1     UUUCUGCU
env-2     UGUUUUUA
env-3     GACCAUUU
env-4     UUAAAUGA
env-5     AUCUGAAG
env-6     UAAGUUUC

          ........
          ........
          ••••R•••
```

| | |
|---|---|
| env-7 | UUUCAAGA |
| env-8 | ........ |
| env-9 | CAAGCCUA |
| env-10 | GCCUAAUG |
| env-11 | ACAAGACC |
| env-12 | AU...... |
| env-13 | ........ |
| env-14 | ACCAUUUC |
| env-15 | UAAACGAU |
| env-16 | UAACAAAA |
| env-17 | AAGUUACA |
| env-18 | AAGUUACA |
| env-21 | AGUUACAA |
| env-22 | AAGUUACA |
| env-26 | AGUUACAA |
| env-27 | AAGUUACA |
| env-63 | AAGUUACA |
| env-64 | ........ |
| env-70 | AACAGAAG |
| env-71 | UAUUAACA |
| env-72 | UAAGAAAC |
| env-73 | ACAGAAGU |
| env-74 | AAAACAAA |
| env-75 | ACAAGACC |
| env-76 | UUACAAGA |
| env-77 | AGAUAAAA |
| env-78 | GCUACAAG |
| env-79 | CAAGACCA |
| env-80 | CAAGACCA |
| Pbr-3-1 | UUCAAUAG |
| env-81 | UUCAAUAG |
| env-82 | UAAGAAUG |
| env-83 | AUUAAAAG |
| env-84 | UCAUGAAU |
| env-86 | UCAUGAAU |
| env-87 | UGAGCUAC |
| Pal-1-1 | GAUUCGAC |
| env-88 | GGACACUU |
| env-98 | GGACACUU |
| env-126 | GGACACUU |
| env-129 | GGACACUU |
| env-132 | GGACACUU |
| env-148 | GGACACUU |
| env-155 | GGACACUU |
| env-157 | UGGACACU |
| env-158 | ........ |
| env-159 | ACAAAACC |
| env-160 | ACAAAACC |
| env-163 | ACAAAACC |
| env-164 | ACAAAACC |
| env-165 | ACAAAACC |
| env-166 | ACAAAACC |
| env-167 | ACAAAACC |
| env-168 | ACAAAACC |
| env-169 | ACAAAACC |
| env-170 | ACAAAACC |
| env-171 | ACAAAACC |
| env-205 | ACAAAACC |
| env-206 | ACAAAACC |
| env-207 | ACAAAACC |
| env-208 | ACAAAACC |
| env-209 | ACAAAACC |
| env-210 | ACAAAACC |
| env-211 | ACAAAACC |
| env-212 | ACAAAACC |
| env-213 | ACAAAACC |
| env-222 | ACAAAACC |
| env-226 | ACAAAACC |
| env-227 | ACAAAACC |
| env-229 | ACAAAACC |
| env-230 | ........ |
| env-231 | ACAAAACC |
| env-232 | ACAAAACC |
| env-233 | ........ |
| env-234 | ACAAAACC |
| env-246 | ACAAAACC |
| env-248 | ........ |
| env-249 | ACAAAACC |
| env-250 | ACAAAACC |
| env-257 | ACAAAACC |
| env-258 | ACAAAACC |
| env-259 | ........ |
| env-260 | ACAAAACC |
| env-262 | ACAAAACC |
| | ........ |
| | ........ |
| | ••••R••• |

| | |
|---|---|
| env-264 | ACAAAACC |
| env-265 | ACAAAACC |
| env-266 | AAUGGACA |
| env-267 | CAUUUCCG |
| Bsp-2-1 | CAUUUCCG |
| env-268 | ........ |
| env-269 | ........ |
| env-270 | UUCAAACC |
| env-272 | ........ |
| env-273 | ........ |
| env-275 | ........ |
| env-276 | ........ |
| env-278 | ........ |
| env-279 | UUCAGACA |
| env-280 | ........ |
| env-281 | ........ |
| env-285 | CAUUGAAU |
| env-286 | ........ |
| env-287 | UUCAGGUA |
| env-288 | UUCAGGUA |
| env-290 | UUCAAGUA |
| env-291 | UUCAAACC |
| env-292 | NNNNNNNN |
| env-293 | UUCAGGUA |
| env-295 | ........ |
| env-297 | ........ |
| env-298 | UUCAGGUA |
| env-299 | ........ |
| env-301 | ........ |
| env-302 | ........ |
| env-305 | UUCAGGUA |
| env-307 | UUCAGGUA |
| env-308 | UUCAGGUA |
| env-309 | UUCAGGUA |
| env-310 | ........ |
| env-311 | ........ |
| env-314 | ........ |
| env-315 | UUCAAGUA |
| env-316 | ........ |
| env-317 | UCACAAAG |
| env-318 | ........ |
| env-319 | UUCAAUAC |
| Psp-3-1 | CAUGUGUU |
| env-320 | CUAUAUCA |
| env-321 | CUAUAUCA |
| env-322 | CUAUAUCA |
| env-323 | ........ |
| env-324 | CUAUAUCA |
| env-326 | CUAUAUCA |
| env-327 | CUAUAUCA |
| env-328 | ........ |
| env-329 | CUAUAUCA |
| env-330 | CUAUAUCA |
| env-331 | CUAUAUCA |
| env-336 | CUAUAUCA |
| env-337 | CUAUAUCA |
| env-338 | CUAUAUCA |
| env-339 | CUAUAUCA |
| env-341 | CUAUAUCA |
| env-345 | CUAUAUCA |
| env-347 | CUAUAACA |
| env-348 | CUAUAUCA |
| env-349 | ........ |
| Pma-3-1 | CAAAGCAU |
| env-350 | CAAAGCAU |
| env-351 | CAAAGCAU |
| Pma-1-1 | CAAAGCAU |
| env-353 | ........ |
| env-354 | ........ |
| env-355 | ........ |
| env-357 | UCAUUGAU |
| env-359 | UCAUUGAU |
| env-360 | ........ |
| env-361 | UCAUUGAU |
| env-362 | ........ |
| env-363 | ........ |
| env-368 | UCAUUGAU |
| env-369 | UCAUUGAU |
| env-370 | UCAUUGAU |
| env-371 | UCAUUGAU |
| env-373 | ........ |
| env-375 | UCAUUGAU |
| env-384 | UCAUUGAU |
| env-385 | UCAUUGAU |
| | ........ |
| | ........ |
| | ●●●●R●●● |

| | |
|---|---|
| env-386 | ........ |
| env-387 | UCAUUGAU |
| env-388 | ........ |
| env-389 | UCAUUGAU |
| env-390 | UCAUUGAU |
| env-392 | UCAUUGAU |
| env-396 | UCAUUGAU |
| env-397 | ........ |
| env-398 | UCAUUGAU |
| env-399 | UCAUUGAU |
| env-400 | UCAUUGAU |
| env-401 | UCAUUGAU |
| env-402 | UCAUUGAU |
| env-403 | UCAUUGAU |
| env-406 | UCAUUGAU |
| env-408 | CGAAAAGU |
| env-410 | CGAAAAGU |
| env-411 | CGAAAAGU |
| env-412 | UCAUUGAU |
| Por-6-1 | CGAAAAGU |
| Por-5-1 | CGAAAAGU |
| env-413 | ........ |
| env-414 | CGAAAAGU |
| env-418 | ........ |
| env-419 | ........ |
| env-420 | CGAAGGUA |
| env-421 | CGAAAAGU |
| env-422 | ........ |
| env-423 | ........ |
| Psp-8-1 | CGAAGGUA |
| env-425 | CGAAGGUA |
| env-426 | ........ |
| env-429 | CGAAGGUA |
| env-431 | CGAAGGUA |
| env-432 | CGAAGGUA |
| env-433 | ........ |
| env-434 | ........ |
| env-435 | CGAAGGUA |
| env-436 | CGAAGGUA |
| env-437 | CGAAGGUA |
| env-438 | CGAAGGUA |
| env-439 | CGAAGGUA |
| env-440 | CGAAGGUA |
| env-441 | ........ |
| env-442 | ........ |
| env-443 | ........ |
| Pou-1-1 | CGAAGGUA |
| env-444 | ........ |
| env-445 | CGAAGGUA |
| env-446 | CGAAGGUA |
| env-447 | CGAAGGUA |
| Pou-2-1 | CGAAGGUA |
| env-448 | GUUACAAG |
| env-449 | ........ |
| Pba-2-1 | UCGAAUAC |
| Pba-1-1 | UCGAAUAC |
| env-450 | UCGAAUGU |
| Pde-2-1 | UACAAGAC |
| Pbe-1-1 | AAUAAGGU |
| env-451 | ACUAUUUC |
| env-452 | GCACAAGC |
| env-453 | CAAGACUG |
| env-454 | AAGACUAU |
| env-455 | AAGACUAU |
| env-457 | UUAUAAAA |
| env-458 | ........ |
| env-462 | UAUAAAAC |
| env-463 | UAUAAAAC |
| env-464 | UUAUAAAA |
| env-466 | AAUGAGUU |
| env-467 | UGGCGGUA |
| env-468 | ........ |
| env-469 | GACAAUUU |
| env-470 | ACAAGACA |
| Ppa-3-1 | AACAAUGU |
| env-471 | AAUAAUGU |
| env-473 | AAUAAUGU |
| env-474 | AAUAAUGU |
| env-476 | AAUAAUGU |
| env-477 | AAUAAUGU |
| env-483 | UAAGGAAC |
| env-485 | UAAUAGGA |
| env-486 | AGCAAUAC |
| env-487 | AGUUAGAC |
| | ........ |
| | ........ |
| | ●●●●R●●● |

| | |
|---|---|
| env-488 | AACAAAUU |
| env-489 | ACAAGACU |
| env-490 | AAGACUAU |
| env-491 | AAGACUAU |
| env-492 | AAGACUAU |
| env-493 | AAGACUAU |
| Pam-1-1 | AAGACUAU |
| Pam-3-1 | AAGACUAU |
| Pbi-1-1 | CAAGACUA |
| Pve-1-1 | CAAGACUA |
| Pve-3-1 | ACAAGACU |
| env-494 | ........ |
| env-495 | UACAAGAC |
| env-497 | UACAAGAC |
| Psc-1-1 | ACAAGACU |
| env-498 | ........ |
| Psp-6-1 | UACAAGAC |
| Pme-3-1 | UACAAGAC |
| env-501 | UACAAGAC |
| env-502 | ........ |
| env-503 | ........ |
| env-504 | UUACAAGA |
| env-507 | ........ |
| Phi-2-1 | UUACAAGA |
| env-508 | UUACAA.. |
| env-509 | AACUUUCA |
| Phi-2-2 | UUACAAGA |
| env-510 | UACAAGAC |
| env-511 | UACAANNN |
| env-512 | UACAAGAC |
| env-514 | UACAAGAC |
| env-516 | ........ |
| env-517 | ........ |
| env-519 | UACAAGAC |
| Psp-5-1 | UACAAGAC |
| env-520 | UACAAGAC |
| env-521 | UACAAGAC |
| env-522 | UACAAGAC |
| env-523 | UACAA... |
| env-525 | CUUUGCUC |
| env-526 | ACAAGACU |
| env-527 | UACAAGAC |
| env-531 | UACAAGAC |
| env-532 | ACAAGACU |
| env-534 | UACA.... |
| env-535 | UACAAGAC |
| env-537 | UACAAGAC |
| env-538 | UACAAGAC |
| env-552 | ACAAGACU |
| env-553 | ........ |
| env-554 | UACAAGAC |
| env-555 | UACAAGAC |
| env-557 | UACAAGAC |
| env-558 | NNNNNNNN |
| env-559 | UACAAGAC |
| env-560 | ........ |
| env-561 | UACAAGAC |
| Pfu-1-1 | ACAAGACU |
| Psp-24-1 | ACAAGACU |
| env-562 | ........ |
| env-563 | ........ |
| env-564 | ACA..... |
| Pde-3-1 | ACAAGACU |
| env-565 | ........ |
| Pde-4-1 | ACAAGACU |
| Pde-5-1 | CAAGACUA |
| env-566 | CAAGA... |
| env-567 | ACAAGA.. |
| Pde-6-1 | CAAGACUA |
| env-568 | ........ |
| env-569 | CAA..... |
| Pmu-1-1 | CAAGACUA |
| env-570 | ........ |
| env-571 | ........ |
| env-572 | AAUUAAUA |
| env-573 | ........ |
| env-574 | UAACAGGU |
| env-575 | AUCAGGUA |
| env-576 | AUCAGGUA |
| env-577 | UAAGUUAC |
| env-578 | UUAUAAGA |
| env-579 | ........ |
| env-580 | UUAUAAGA |
| env-582 | UUAUAAGA |
| | ........ |
| | ........ |
| | ••••R••• |

```
Pin-1-1    UAAGACUA
env-583    AU......
env-589    AUAAGACU
env-592    GGUAGUAG
env-593    AUAAGACU
env-594    AGACCAGA
env-595    AUAAGACU
env-596    AU......
env-597    GGUCGCCU
env-598    ........
env-599    AUAAGAC.
env-600    AUAAGACU
env-613    AUAAGACU
env-648    AUAAGACU
env-651    AUAAGACU
env-652    CCUCUGCU
env-653    ........
env-657    ........
env-676    UACAAGAC
Pco-3-1    UACAAGAC
Pni-1-1    UAAGACUA
env-677    UAAGACUA
env-679    UAAGACUA
env-680    CAAGACU.
env-681    CA......
env-682    UAAGACUA
env-683    UAAGACUA
env-684    UAAGACUA
env-686    UAAGACUA
env-696    UAAGACUA
env-698    ........
env-699    ........
Pni-3-1    UAAGACUA
env-700    ........
env-709    ........
env-710    UAAGACUA
env-711    UAAGACUA
env-724    UAAGACUA
Pau-1-1    AGUUAUUA
Psp-9-1    AAGACUAU
env-725    UAAGACUA
env-726    UAAGACUA
env-727    ........
env-729    UAAGACUA
env-733    UAAGACUA
env-734    UAAGACUA
env-735    UAAGACUA
env-736    UAAGACUA
Pin-3-1    UAAGACUA
Pfa-1-1    AAGACUAU
env-737    UAAGACUA
env-738    UAAGACUA
env-742    UAAGACUA
Pdi-3-1    AAGACUAU
env-743    ........
Pdi-2-1    AAGACUAU
Pdi-1-1    AAGACUAU
env-744    GGACACA.
env-745    GACACAUU
env-746    UGGACACA
env-747    UGGACACA
env-748    GGACACAU
env-749    GGACACAU
env-751    GGACACAU
env-752    ........
env-753    CACAUUAA
env-754    GGACACAU
env-755    GUAAUCGA
env-756    UGGACACA
env-757    CAUUAAGU
env-758    AAUGGACA
env-759    AAAUGGAC
env-760    GGACACAU
env-761    ........
env-762    ACAUUAAG
env-763    ACAUUAAG
env-765    ........
env-766    ACAUUAAG
env-767    ACAUUAAG
env-768    CAUUAAGU
env-769    ........
env-770    CAUUAAGU
env-771    CAUUAAGU
env-772    CAUUAAGU

           ........
           ........
           ••••R•••
```

| | |
|---|---|
| env-774 | UUUCAAGA |
| env-775 | CAUUAAGU |
| env-776 | CAUUAAGU |
| env-780 | UUUCAAGA |
| env-781 | CACAUUAA |
| env-785 | ACAUUAAG |
| env-787 | ........ |
| env-788 | CACAUUAA |
| env-789 | CAUUAAGU |
| env-790 | CAUUAAGU |
| env-791 | ........ |
| env-793 | CACAUUAA |
| env-797 | ........ |
| env-798 | AUUAAGUU |
| env-799 | CACAUUAA |
| env-800 | ACAUUAAG |
| env-801 | CACAUUAA |
| env-802 | CACAUUAA |
| env-803 | ACAUUAAG |
| env-804 | CACAUUAA |
| env-808 | CACAUUAA |
| env-821 | UAAGUUAC |
| env-822 | ACACAUUA |
| env-824 | CAUUAAGU |
| env-825 | UGGACACA |
| env-826 | CACAUUAA |
| env-827 | CACAUUAA |
| env-828 | CACAUUAA |
| env-829 | AAGUUACA |
| env-830 | GGACACAU |
| env-831 | ACAUUAAG |
| env-832 | ACAUUAAG |
| env-833 | AGUUACAA |
| env-834 | ........ |
| env-835 | CAUUAAGU |
| env-836 | CACAUUAA |
| env-837 | CACAUUAA |
| Pco-2-1 | CACAUUAA |
| env-838 | ........ |
| env-839 | CACAUUAA |
| env-840 | ACAUUAAG |
| env-842 | ACAUUAAG |
| env-843 | ACAUUAAG |
| Por-8-1 | AGACCAUU |
| Psp-21-1 | AUCAAAUC |
| env-845 | GGAUACUU |
| env-846 | GGAUACUU |
| env-847 | AGGUGGAA |
| env-848 | AGACUAUU |
| env-849 | ........ |
| env-851 | AGACUAUU |
| env-852 | A....... |
| env-855 | AGACUAUU |
| env-856 | AGACUGUU |
| env-857 | A....... |
| env-858 | AGACUAUU |
| env-859 | AGACUAUU |
| env-860 | AGACUAUU |
| env-861 | ........ |
| env-862 | AGACUAUU |
| env-864 | AGACUAUU |
| env-865 | AGACUAUU |
| env-866 | ........ |
| env-868 | ........ |
| env-869 | AGACUAUU |
| env-870 | AGACUAUU |
| env-872 | ........ |
| env-873 | ........ |
| env-874 | AGACUAUU |
| env-875 | ........ |
| env-876 | AGACUAUU |
| Psp-25-1 | AGACUAUU |
| env-877 | AGACUAUU |
| env-878 | A....... |
| env-880 | AGACUAUU |
| env-881 | ........ |
| env-882 | ........ |
| env-883 | ........ |
| env-885 | AGACUAUU |
| env-886 | AAGACUAU |
| env-887 | ACAAGACU |
| env-888 | ACAAGACU |
| env-889 | ........ |
| env-890 | ........ |
| | ........ |
| | ........ |
| | ....R... |

| | |
|---|---|
| env-891 | ACAAGACU |
| env-892 | ACAAGACU |
| env-893 | ........ |
| env-894 | ACAAGACU |
| env-900 | ACAAGACU |
| env-901 | ........ |
| env-902 | ACAAGACU |
| Psp-26-1 | ACAAGACU |
| env-903 | ........ |
| env-921 | A....... |
| env-922 | AGACUAUU |
| env-923 | A....... |
| env-924 | AGACUAUU |
| env-932 | ........ |
| env-948 | AGACUAUU |
| env-949 | AGACUAUU |
| env-950 | A....... |
| env-953 | A....... |
| env-957 | A....... |
| env-958 | AGACUAUU |
| env-960 | GACUGUUU |
| env-961 | GACUGUUU |
| env-967 | AGCCUAGA |
| env-968 | ACAAGACU |
| env-970 | ACAAGACU |
| env-971 | ACAAGACU |
| env-972 | CAAGACUA |
| env-973 | CAAGACUA |
| env-977 | CAAGACUA |
| env-978 | CAAGACUA |
| env-979 | CAAGACUA |
| env-981 | CAAGACUA |
| env-982 | CAAGACUA |
| env-983 | CAAGACUA |
| env-984 | CAAGACUA |
| env-985 | CAAGACUA |
| env-986 | CAAGACUA |
| env-990 | CAAGACUA |
| env-992 | CAAGACUA |
| env-993 | ........ |
| env-994 | GACUGUUU |
| Pbu-3-1 | UGUAAUUA |
| Pbu-1-1 | UGUAAUUA |
| Pbu-2-1 | UGUAAUUA |
| Psp-12-1 | UGUAAUUA |
| env-995 | ........ |
| env-996 | GAAAGUAA |
| env-997 | UGUCGAAA |
| env-999 | UAUAAUAU |
| Por-1-1 | UUUCAAAA |
| env-1001 | ........ |
| Por-2-1 | UUUCAAAA |
| env-1002 | AUAAGACU |
| env-1003 | AUAAGACU |
| Pti-3-1 | AUAAGACU |
| env-1004 | UAUAAGAC |
| Pti-1-1 | AUAAGACU |
| Pti-2-1 | AUAAGACU |
| env-1005 | AUAAGACU |
| env-1006 | ........ |
| Psp-4-1 | UACAAGAC |
| env-1007 | UACAAGAC |
| Pbu-5-1 | UACAAGAC |
| Pbu-4-1 | UACAAGAC |
| env-1008 | NNNNNNNN |
| Pen-1-1 | CAAAACUA |
| env-1009 | CAAAACUA |
| env-1017 | CAAAACUA |
| env-1020 | CAAAACUA |
| Ppl-2-1 | CAAAACUA |
| env-1021 | GUUACAAG |
| env-1022 | GUUACAAG |
| env-1023 | GUUACAAG |
| env-1024 | ........ |
| env-1025 | UACAAGAC |
| env-1027 | UUACAAGA |
| Pma-4-1 | UUACAAGA |
| env-1028 | UUACAAGA |
| env-1029 | UUUAAGUU |
| env-1031 | UUUAAGUU |
| env-1032 | UAUUGAAG |
| env-1033 | UAUUGAAG |
| env-1034 | AGUUACAA |
| env-1035 | CAAGACUA |
| | ........ |
| | ........ |
| | ●●●●R●●● |

| | |
|---|---|
| env-1037 | UAGUAAAU |
| env-1038 | AAGUUACA |
| env-1039 | AAGUUACA |
| env-1040 | AAGUUACA |
| env-1043 | AAGUUACA |
| env-1044 | AAGUUACA |
| env-1046 | AAGUUACA |
| env-1052 | UUACAAAA |
| env-1053 | AAGUUACA |
| Psp-2-1 | AAGUUACA |
| env-1054 | AAGUUACA |
| env-1055 | AAGUUACA |
| env-1056 | AAGUUACA |
| env-1058 | UUACAAGA |
| env-1059 | AGUUACAA |
| env-1060 | AGUUACAA |
| env-1061 | UGAAGAAA |
| Psp-13-1 | UGAAGAAA |
| env-1062 | UGAAGAAA |
| Psp-17-1 | GUGAAGAA |
| env-1063 | GUGAAGAA |
| Psp-15-1 | UGAAGAAA |
| Psp-16-1 | GUGAAGAA |
| env-1064 | UGGAAAGA |
| env-1065 | ........ |
| env-1066 | UGGAAAGA |
| env-1067 | ACAAGACU |
| Hse-1-1 | CAUCAAGG |
| env-1068 | UAAAAUGA |
| env-1069 | AACAAUUC |
| env-1071 | AUAAUUAA |
| env-1072 | UCGGAUGC |
| env-1074 | UCGGAUGC |
| env-1075 | UCGGAUGC |
| env-1076 | ........ |
| env-1077 | GAUUAUUU |
| env-1078 | GUUAGACC |
| env-1079 | GACCUCGG |
| env-1080 | GUUAGACC |
| env-1081 | ........ |
| env-1083 | AGUUAGAC |
| env-1084 | AAGAGAUA |
| env-1085 | AGAGAGUA |
| env-1086 | ........ |
| env-1087 | ........ |
| env-1088 | ........ |
| env-1089 | ........ |
| env-1090 | ........ |
| env-1091 | ........ |
| env-1092 | AUAUAAAU |
| env-1093 | GAGAGAAA |
| env-1094 | AGACCUCG |
| env-1095 | GAGUUAGA |
| env-1096 | GCCUGGGA |
| env-1097 | AGAACAAA |
| env-1098 | AUGUGGGU |
| Psp-7-1 | UGUGAGUU |
| env-1099 | UUUAAUUA |
| env-1100 | UUUAAUUA |
| Psp-11-1 | GACCCUGG |
| Psp-19-1 | AGACCCUG |
| env-1101 | ........ |
| Pru-1-1 | UUUAAUUA |
| env-1102 | ........ |
| env-1103 | UUGAAAUG |
| env-1104 | CUUUAAGU |
| Pbr-1-1 | UUAAGUUA |
| Pbr-2-1 | UUAAGUUA |
| env-1105 | ........ |
| env-1106 | ........ |
| env-1107 | ........ |
| env-1108 | ........ |
| env-1109 | GUGGGUGG |
| env-1110 | UGCGGGUG |
| env-1111 | GCGGGUGG |
| env-1112 | UAAGUUAC |
| env-1113 | AAUUAUCG |
| env-1114 | UGAUAAAA |
| env-1115 | UGAUAAAA |
| env-1116 | UUUAAGUU |
| env-1117 | AUGUUUUU |
| env-1118 | UUACAAGA |
| env-1119 | UUACAAGA |
| env-1120 | UAAGAAUU |
| | ........ |
| | ........ |
| | ••••R••• |

| | |
|---|---|
| env-1122 | AAGAAUUA |
| env-1123 | CGAAAUAA |
| env-1125 | UAAGUUAC |
| env-1126 | ACAAGACU |
| env-1127 | ACAAGACU |
| env-1129 | ACAUUAAG |
| env-1130 | ACUUAUGU |
| env-1131 | UAAUGUGA |
| env-1132 | AGAAGACC |
| env-1133 | UACAAGAC |
| env-1134 | CCACAAAC |
| env-1135 | AAAUGUGA |
| env-1136 | CAAGACCA |
| env-1137 | UUAGAAUG |
| env-1138 | UUAGAAUG |
| env-1139 | CUUAGAAU |
| env-1140 | CUUAGAAU |
| env-1141 | CUUAGAAU |
| env-1143 | CUUAGAAU |
| env-1144 | CUUAGAAU |
| env-1145 | CUUAGAAU |
| env-1147 | UGGACACU |
| env-1148 | GGACACUU |
| env-1149 | UGGACACU |
| env-1151 | UGGACACU |
| env-1153 | UGGACACU |
| env-1154 | GGACACUU |
| env-1155 | AUCGUUGA |
| env-1156 | UGGACACU |
| env-1157 | GGACACUU |
| env-1158 | UGGACACU |
| env-1159 | UGGACACU |
| env-1163 | UGGACACU |
| env-1164 | UGGACACU |
| env-1165 | UGGACACU |
| env-1166 | AUGGACAC |
| env-1167 | UGGACACU |
| env-1192 | CCAUUUCU |
| env-1193 | AUGGAUAC |
| env-1194 | AUGGAUAC |
| Psp-1-1 | AUGGAUAC |
| env-1195 | ........ |
| env-1197 | AUGGAUAC |
| env-1203 | AUGGAUAC |
| env-1205 | AUGGAUAC |
| env-1206 | AUGGAUAC |
| env-1211 | ........ |
| env-1213 | AUGGAUAC |
| env-1216 | UGGAUACU |
| env-1223 | UGGAUACU |
| env-1224 | UGGAUACU |
| env-1228 | ........ |
| env-1233 | UGGAUACU |
| env-1243 | UGGAUACU |
| env-1244 | ........ |
| env-1245 | AUGGAUAC |
| env-1250 | AUGGAUAC |
| env-1251 | AUGGAUAC |
| env-1254 | AUGGAUAC |
| env-1256 | AUGGAUAC |
| env-1258 | AUGGAUAC |
| env-1259 | AUGGAUAC |
| Pmu-2-1 | GUCAACAG |
| Pde-1-1 | GUAUCAAG |
| | ........ |
| | ........ |
| | ●●●●R●●● |

### 4 eL15-Euryarchaeota

#### 4.1 Taxa

The taxonomy of each organism containing a putative eL15-Euryarchaeota RNA is listed, with abbreviations identifying each hit (e.g., "Eco-1-1" and "Eco-1-2" might hypothetically represent two distinct RNAs in *E. coli*). The abbreviations will be used to identify each individual eL15-Euryarchaeota RNA in Sections 4.2 and 4.4:.

| abbrev. of hits | taxonomy of species |
|---|---|
| har-1-1 | halophilic archaeon DL31 Archaea |
| ume-1-1 | Archaea Euryarchaeota environmental samples uncultured methanogenic archaeon RC-I |
| h3A-1-1 | Archaea Euryarchaeota Halobacteria Halobacteriales haloarchaeon 3A1_DGR |
| Hsa-1-1 | Archaea Euryarchaeota Halobacteria Halobacteriales Haloarculaceae *Halapricum salinum* |
| Hsp-1-1 | Archaea Euryarchaeota Halobacteria Halobacteriales Haloarculaceae *Haloarcula sp.* CBA1115 |
| CHa-1-1 | Archaea Euryarchaeota Halobacteria Halobacteriales Halobacteriaceae *Candidatus* Halobonum tyrrellensis G22 |
| Hci-1-1 | Archaea Euryarchaeota Halobacteria Halobacteriales Halobacteriaceae *Haladaptatus cibarius* D43 |
| Hpa-1-1 to Hpa-1-2 | Archaea Euryarchaeota Halobacteria Halobacteriales Halobacteriaceae *Haladaptatus paucihalophilus* DX253 |
| Hje-1-1 | Archaea Euryarchaeota Halobacteria Halobacteriales Halobacteriaceae *Halalkalicoccus jeotgali* B3 |
| Hsu-1-1 | Archaea Euryarchaeota Halobacteria Halobacteriales Halobacteriaceae *Halanaeroarchaeum sulfurireducens* |
| Hac-1-1 to Hac-1-2 | Archaea Euryarchaeota Halobacteria Halobacteriales Halobacteriaceae *Halarchaeum acidiphilum* MH1-52-1 |
| Ham-1-1 | Archaea Euryarchaeota Halobacteria Halobacteriales Halobacteriaceae *Haloarcula amylolytica* JCM 13557 |
| Har-1-1 | Archaea Euryarchaeota Halobacteria Halobacteriales Halobacteriaceae *Haloarcula argentinensis* DSM 12282 |
| Hca-1-1 | Archaea Euryarchaeota Halobacteria Halobacteriales Halobacteriaceae *Haloarcula californiae* ATCC 33799 |
| Hhi-1-1 | Archaea Euryarchaeota Halobacteria Halobacteriales Halobacteriaceae *Haloarcula hispanica* ATCC 33960 |
| Hhi-2-1 | Archaea Euryarchaeota Halobacteria Halobacteriales Halobacteriaceae *Haloarcula hispanica* N601 |
| Hja-1-1 | Archaea Euryarchaeota Halobacteria Halobacteriales Halobacteriaceae *Haloarcula japonica* DSM 6131 |
| Hma-1-1 | Archaea Euryarchaeota Halobacteria Halobacteriales Halobacteriaceae *Haloarcula marismortui* ATCC 43049 |
| Hsa-2-1 | Archaea Euryarchaeota Halobacteria Halobacteriales Halobacteriaceae *Haloarcula salaria* |
| Hsi-1-1 | Archaea Euryarchaeota Halobacteria Halobacteriales Halobacteriaceae *Haloarcula sinaiiensis* ATCC 33800 |
| Hva-1-1 | Archaea Euryarchaeota Halobacteria Halobacteriales Halobacteriaceae *Haloarcula vallismortis* ATCC 29715 |
| Hsa-3-1 | Archaea Euryarchaeota Halobacteria Halobacteriales Halobacteriaceae *Halobacterium salinarum* R1 |
| Hsp-2-1 | Archaea Euryarchaeota Halobacteria Halobacteriales Halobacteriaceae *Halobacterium sp.* DL1 |
| Hsp-3-1 | Archaea Euryarchaeota Halobacteria Halobacteriales Halobacteriaceae *Halobacterium sp.* NRC-1 |
| Hla-1-1 to Hla-1-2 | Archaea Euryarchaeota Halobacteria Halobacteriales Halobacteriaceae *Halobiforma lacisalsi* AJ5 |
| Hni-1-1 | Archaea Euryarchaeota Halobacteria Halobacteriales Halobacteriaceae *Halobiforma nitratireducens* JCM 10879 |
| Hha-1-1 to Hha-1-2 | Archaea Euryarchaeota Halobacteria Halobacteriales Halobacteriaceae *Halococcus hamelinensis* 100A6 |
| Hmo-1-1 | Archaea Euryarchaeota Halobacteria Halobacteriales Halobacteriaceae *Halococcus morrhuae* DSM 1307 |
| Hsa-4-1 | Archaea Euryarchaeota Halobacteria Halobacteriales Halobacteriaceae *Halococcus saccharolyticus* DSM 5350 |
| Hsa-5-1 | Archaea Euryarchaeota Halobacteria Halobacteriales Halobacteriaceae *Halococcus salifodinae* DSM 8989 |
| Hth-1-1 | Archaea Euryarchaeota Halobacteria Halobacteriales Halobacteriaceae *Halococcus thailandensis* JCM 13552 |
| Hal-1-1 | Archaea Euryarchaeota Halobacteria Halobacteriales Halobacteriaceae *Haloferax alexandrinus* JCM 10717 |
| Hde-1-1 | Archaea Euryarchaeota Halobacteria Halobacteriales Halobacteriaceae *Haloferax denitrificans* ATCC 35960 |
| Hel-1-1 | Archaea Euryarchaeota Halobacteria Halobacteriales Halobacteriaceae *Haloferax elongans* ATCC BAA-1513 |
| Hgi-1-1 | Archaea Euryarchaeota Halobacteria Halobacteriales Halobacteriaceae *Haloferax gibbonsii* ATCC 33959 |
| Hla-2-1 | Archaea Euryarchaeota Halobacteria Halobacteriales Halobacteriaceae *Haloferax larsenii* JCM 13917 |
| Hlu-1-1 | Archaea Euryarchaeota Halobacteria Halobacteriales Halobacteriaceae *Haloferax lucentense* DSM 14919 |
| Hme-1-1 to Hme-1-2 | Archaea Euryarchaeota Halobacteria Halobacteriales Halobacteriaceae *Haloferax mediterranei* ATCC 33500 |
| Hmu-1-1 | Archaea Euryarchaeota Halobacteria Halobacteriales Halobacteriaceae *Haloferax mucosum* ATCC BAA-1512 |
| Hpr-1-1 | Archaea Euryarchaeota Halobacteria Halobacteriales Halobacteriaceae *Haloferax prahovense* DSM 18310 |
| Hsp-4-1 | Archaea Euryarchaeota Halobacteria Halobacteriales Halobacteriaceae *Haloferax sp.* ATCC BAA-644 |
| Hsp-5-1 | Archaea Euryarchaeota Halobacteria Halobacteriales Halobacteriaceae *Haloferax sp.* ATCC BAA-645 |
| Hsp-6-1 | Archaea Euryarchaeota Halobacteria Halobacteriales Halobacteriaceae *Haloferax sp.* ATCC BAA-646 |
| Hsp-7-1 | Archaea Euryarchaeota Halobacteria Halobacteriales Halobacteriaceae *Haloferax sp.* BAB2207 |
| Hsu-2-1 | Archaea Euryarchaeota Halobacteria Halobacteriales Halobacteriaceae *Haloferax sulfurifontis* ATCC BAA-897 |
| Hvo-1-1 | Archaea Euryarchaeota Halobacteria Halobacteriales Halobacteriaceae *Haloferax volcanii* DS2 |
| Hbo-1-1 | Archaea Euryarchaeota Halobacteria Halobacteriales Halobacteriaceae *Halogeometricum borinquense* DSM 11551 |
| Hsa-6-1 | Archaea Euryarchaeota Halobacteria Halobacteriales Halobacteriaceae *Halogranum salarium* B-1 |
| Hka-1-1 | Archaea Euryarchaeota Halobacteria Halobacteriales Halobacteriaceae *Halomicrobium katesii* DSM 19301 |
| Hmu-2-1 | Archaea Euryarchaeota Halobacteria Halobacteriales Halobacteriaceae *Halomicrobium mukohataei* DSM 12286 |
| Hxa-1-1 | Archaea Euryarchaeota Halobacteria Halobacteriales Halobacteriaceae *Halopiger xanaduensis* SH-6 |
| Hwa-1-1 | Archaea Euryarchaeota Halobacteria Halobacteriales Halobacteriaceae *Haloquadratum walsbyi* C23 |
| Hwa-2-1 | Archaea Euryarchaeota Halobacteria Halobacteriales Halobacteriaceae *Haloquadratum walsbyi* DSM 16790 |
| Hti-1-1 | Archaea Euryarchaeota Halobacteria Halobacteriales Halobacteriaceae *Halorhabdus tiamatea* SARL4B |
| Hut-1-1 | Archaea Euryarchaeota Halobacteria Halobacteriales Halobacteriaceae *Halorhabdus utahensis* DSM 12940 |
| Hai-1-1 | Archaea Euryarchaeota Halobacteria Halobacteriales Halobacteriaceae *Halorubrum aidingense* JCM 13560 |
| Har-2-1 | Archaea Euryarchaeota Halobacteria Halobacteriales Halobacteriaceae *Halorubrum arcis* JCM 13916 |
| Hca-2-1 | Archaea Euryarchaeota Halobacteria Halobacteriales Halobacteriaceae *Halorubrum californiensis* DSM 19288 |
| Hco-1-1 | Archaea Euryarchaeota Halobacteria Halobacteriales Halobacteriaceae *Halorubrum coriense* DSM 10284 |
| Hdi-1-1 | Archaea Euryarchaeota Halobacteria Halobacteriales Halobacteriaceae *Halorubrum distributum* JCM 10118 |
| Hdi-2-1 | Archaea Euryarchaeota Halobacteria Halobacteriales Halobacteriaceae *Halorubrum distributum* JCM 9100 |
| Hho-1-1 | Archaea Euryarchaeota Halobacteria Halobacteriales Halobacteriaceae *Halorubrum hochstenium* ATCC 700873 |
| Hko-1-1 | Archaea Euryarchaeota Halobacteria Halobacteriales Halobacteriaceae *Halorubrum kocurii* JCM 14978 |
| Hla-3-1 | Archaea Euryarchaeota Halobacteria Halobacteriales Halobacteriaceae *Halorubrum lacusprofundi* ATCC 49239 |

| | |
|---|---|
| Hli-1-1 | Archaea Euryarchaeota Halobacteria Halobacteriales Halobacteriaceae *Halorubrum lipolyticum* DSM 21995 |
| Hli-2-1 | Archaea Euryarchaeota Halobacteria Halobacteriales Halobacteriaceae *Halorubrum litoreum* JCM 13561 |
| Hsa-7-1 | Archaea Euryarchaeota Halobacteria Halobacteriales Halobacteriaceae *Halorubrum saccharovorum* DSM 1137 |
| Hsp-8-1 | Archaea Euryarchaeota Halobacteria Halobacteriales Halobacteriaceae *Halorubrum sp.* T3 |
| Hte-1-1 | Archaea Euryarchaeota Halobacteria Halobacteriales Halobacteriaceae *Halorubrum tebenquichense* DSM 14210 |
| Hte-2-1 | Archaea Euryarchaeota Halobacteria Halobacteriales Halobacteriaceae *Halorubrum terrestre* JCM 10247 |
| Hpa-2-1 | Archaea Euryarchaeota Halobacteria Halobacteriales Halobacteriaceae *Halosarcina pallida* JCM 14848 |
| Hca-3-1 | Archaea Euryarchaeota Halobacteria Halobacteriales Halobacteriaceae *Halosimplex carlsbadense* 2-9-1 |
| Hli-3-1 | Archaea Euryarchaeota Halobacteria Halobacteriales Halobacteriaceae *Haloterrigena limicola* JCM 13563 |
| Hsa-8-1 | Archaea Euryarchaeota Halobacteria Halobacteriales Halobacteriaceae *Haloterrigena salina* JCM 13891 |
| Hth-2-1 | Archaea Euryarchaeota Halobacteria Halobacteriales Halobacteriaceae *Haloterrigena thermotolerans* DSM 11522 |
| Htu-1-1 | Archaea Euryarchaeota Halobacteria Halobacteriales Halobacteriaceae *Haloterrigena turkmenica* DSM 5511 |
| Has-1-1 | Archaea Euryarchaeota Halobacteria Halobacteriales Halobacteriaceae *Halovivax asiaticus* JCM 14624 |
| Nae-1-1 | Archaea Euryarchaeota Halobacteria Halobacteriales Halobacteriaceae *Natrialba aegyptia* DSM 13077 |
| Nas-1-1 | Archaea Euryarchaeota Halobacteria Halobacteriales Halobacteriaceae *Natrialba asiatica* DSM 12278 |
| Nch-1-1 | Archaea Euryarchaeota Halobacteria Halobacteriales Halobacteriaceae *Natrialba chahannaoensis* JCM 10990 |
| Nhu-1-1 | Archaea Euryarchaeota Halobacteria Halobacteriales Halobacteriaceae *Natrialba hulunbeirensis* JCM 10989 |
| Nma-1-1 | Archaea Euryarchaeota Halobacteria Halobacteriales Halobacteriaceae *Natrialba magadii* ATCC 43099 |
| Nta-1-1 | Archaea Euryarchaeota Halobacteria Halobacteriales Halobacteriaceae *Natrialba taiwanensis* DSM 12281 |
| Nal-1-1 | Archaea Euryarchaeota Halobacteria Halobacteriales Halobacteriaceae *Natrinema altunense* JCM 12890 |
| Nga-1-1 | Archaea Euryarchaeota Halobacteria Halobacteriales Halobacteriaceae *Natrinema gari* JCM 14663 |
| Npa-1-1 | Archaea Euryarchaeota Halobacteria Halobacteriales Halobacteriaceae *Natrinema pallidum* DSM 3751 |
| Npe-1-1 | Archaea Euryarchaeota Halobacteria Halobacteriales Halobacteriaceae *Natrinema pellirubrum* DSM 15624 |
| Nsp-1-1 | Archaea Euryarchaeota Halobacteria Halobacteriales Halobacteriaceae *Natrinema sp.* J7-2 |
| Nve-1-1 | Archaea Euryarchaeota Halobacteria Halobacteriales Halobacteriaceae *Natrinema versiforme* JCM 10478 |
| Ngr-1-1 | Archaea Euryarchaeota Halobacteria Halobacteriales Halobacteriaceae *Natronobacterium gregoryi* SP2 |
| Nam-1-1 | Archaea Euryarchaeota Halobacteria Halobacteriales Halobacteriaceae *Natronococcus amylolyticus* DSM 10524 |
| Nje-1-1 | Archaea Euryarchaeota Halobacteria Halobacteriales Halobacteriaceae *Natronococcus jeotgali* DSM 18795 |
| Noc-1-1 | Archaea Euryarchaeota Halobacteria Halobacteriales Halobacteriaceae *Natronococcus occultus* SP4 |
| Nin-1-1 | Archaea Euryarchaeota Halobacteria Halobacteriales Halobacteriaceae *Natronolimnobius innermongolicus* JCM 12255 |
| Nmo-1-1 | Archaea Euryarchaeota Halobacteria Halobacteriales Halobacteriaceae *Natronomonas moolapensis* 8.8.11 |
| Nph-1-1 | Archaea Euryarchaeota Halobacteria Halobacteriales Halobacteriaceae *Natronomonas pharaonis* DSM 2160 |
| Nba-1-1 | Archaea Euryarchaeota Halobacteria Halobacteriales Halobacteriaceae *Natronorubrum bangense* JCM 10635 |
| Nsu-1-1 | Archaea Euryarchaeota Halobacteria Halobacteriales Halobacteriaceae *Natronorubrum sulfidifaciens* JCM 14089 |
| Nti-1-1 | Archaea Euryarchaeota Halobacteria Halobacteriales Halobacteriaceae *Natronorubrum tibetense* GA33 |
| Ssp-1-1 | Archaea Euryarchaeota Halobacteria Halobacteriales Halobacteriaceae *Salinarchaeum sp.* Harcht-Bsk1 |
| Hag-1-1 | Archaea Euryarchaeota Halobacteria Halobacteriales Halococcaceae *Halococcus agarilyticus* |
| Hse-1-1 | Archaea Euryarchaeota Halobacteria Halobacteriales Halococcaceae *Halococcus sediminicola* |
| Hru-1-1 | Archaea Euryarchaeota Halobacteria Haloferacales Haloferacaceae *Halobellus rufus* |
| Hal-2-1 | Archaea Euryarchaeota Halobacteria Haloferacales Haloferacaceae *Haloferax alexandrinus* |
| Hgi-2-1 | Archaea Euryarchaeota Halobacteria Haloferacales Haloferacaceae *Haloferax gibbonsii* |
| Hme-1-1 to Hme-1-2 | Archaea Euryarchaeota Halobacteria Haloferacales Haloferacaceae *Haloferax mediterranei* ATCC 33500 |
| Hsp-9-1 | Archaea Euryarchaeota Halobacteria Haloferacales Haloferacaceae *Haloferax sp.* ATB1 |
| Hna-1-1 | Archaea Euryarchaeota Halobacteria Haloferacales Haloferacaceae *Haloplanus natans* DSM 17983 |
| Hru-2-1 | Archaea Euryarchaeota Halobacteria Haloferacales Halorubraceae *Halolamina rubra* |
| Hez-1-1 | Archaea Euryarchaeota Halobacteria Haloferacales Halorubraceae *Halorubrum ezzemoulense* DSM 17463 |
| Hha-2-1 | Archaea Euryarchaeota Halobacteria Haloferacales Halorubraceae *Halorubrum halophilum* |
| Hsa-9-1 | Archaea Euryarchaeota Halobacteria Haloferacales Halorubraceae *Halorubrum saccharovorum* |
| Hsp-10-1 | Archaea Euryarchaeota Halobacteria Haloferacales Halorubraceae *Halorubrum sp.* BV1 |
| Hdj-1-1 | Archaea Euryarchaeota Halobacteria Natrialbales Natrialbaceae *Halopiger djelfimassiliensis* |
| Hgo-1-1 | Archaea Euryarchaeota Halobacteria Natrialbales Natrialbaceae *Halopiger goleimassiliensis* |
| Hsa-10-1 | Archaea Euryarchaeota Halobacteria Natrialbales Natrialbaceae *Halopiger salifodinae* |
| Hla-4-1 | Archaea Euryarchaeota Halobacteria Natrialbales Natrialbaceae *Halostagnicola larsenii* XH-48 |
| Hsp-11-1 | Archaea Euryarchaeota Halobacteria Natrialbales Natrialbaceae *Halostagnicola sp.* A56 |
| Hje-2-1 | Archaea Euryarchaeota Halobacteria Natrialbales Natrialbaceae *Haloterrigena jeotgali* A29 |
| Nal-2-1 | Archaea Euryarchaeota Halobacteria Natrialbales Natrialbaceae *Natrinema altunense* |
| Nsp-2-1 | Archaea Euryarchaeota Halobacteria Natrialbales Natrialbaceae *Natrinema sp.* J7-1 |
| Mar-1-1 | Archaea Euryarchaeota Methanobacteria Methanobacteriales Methanobacteriaceae *Methanobacterium arcticum* |
| Mfo-1-1 | Archaea Euryarchaeota Methanobacteria Methanobacteriales Methanobacteriaceae *Methanobacterium formicicum* |
| Mfo-2-1 | Archaea Euryarchaeota Methanobacteria Methanobacteriales Methanobacteriaceae *Methanobacterium formicicum* DSM 3637 |
| Msp-1-1 | Archaea Euryarchaeota Methanobacteria Methanobacteriales Methanobacteriaceae *Methanobacterium sp.* AL-21 |
| Msp-2-1 | Archaea Euryarchaeota Methanobacteria Methanobacteriales Methanobacteriaceae *Methanobacterium sp.* MB1 |
| Msp-3-1 | Archaea Euryarchaeota Methanobacteria Methanobacteriales Methanobacteriaceae *Methanobacterium sp.* SMA-27 |
| Msp-4-1 | Archaea Euryarchaeota Methanobacteria Methanobacteriales Methanobacteriaceae *Methanobacterium sp.* SWAN-1 |
| Mve-1-1 | Archaea Euryarchaeota Methanobacteria Methanobacteriales Methanobacteriaceae *Methanobacterium veterum* |

| | |
|---|---|
| Mor-1-1 | Archaea Euryarchaeota Methanobacteria Methanobacteriales Methanobacteriaceae *Methanobrevibacter oralis* JMR01 |
| Msm-1-1 | Archaea Euryarchaeota Methanobacteria Methanobacteriales Methanobacteriaceae *Methanobrevibacter smithii* |
| Msm-2-1 | Archaea Euryarchaeota Methanobacteria Methanobacteriales Methanobacteriaceae *Methanobrevibacter smithii* ATCC 35061 |
| Msm-3-1 | Archaea Euryarchaeota Methanobacteria Methanobacteriales Methanobacteriaceae *Methanobrevibacter smithii* DSM 2374 |
| Msm-4-1 | Archaea Euryarchaeota Methanobacteria Methanobacteriales Methanobacteriaceae *Methanobrevibacter smithii* DSM 2375 |
| Msm-5-1 | Archaea Euryarchaeota Methanobacteria Methanobacteriales Methanobacteriaceae *Methanobrevibacter smithii* TS145A |
| Msm-6-1 | Archaea Euryarchaeota Methanobacteria Methanobacteriales Methanobacteriaceae *Methanobrevibacter smithii* TS145B |
| Msm-7-1 | Archaea Euryarchaeota Methanobacteria Methanobacteriales Methanobacteriaceae *Methanobrevibacter smithii* TS146A |
| Msm-8-1 | Archaea Euryarchaeota Methanobacteria Methanobacteriales Methanobacteriaceae *Methanobrevibacter smithii* TS146B |
| Msm-9-1 | Archaea Euryarchaeota Methanobacteria Methanobacteriales Methanobacteriaceae *Methanobrevibacter smithii* TS146C |
| Msm-10-1 | Archaea Euryarchaeota Methanobacteria Methanobacteriales Methanobacteriaceae *Methanobrevibacter smithii* TS146E |
| Msm-11-1 | Archaea Euryarchaeota Methanobacteria Methanobacteriales Methanobacteriaceae *Methanobrevibacter smithii* TS147A |
| Msm-12-1 | Archaea Euryarchaeota Methanobacteria Methanobacteriales Methanobacteriaceae *Methanobrevibacter smithii* TS147B |
| Msm-13-1 | Archaea Euryarchaeota Methanobacteria Methanobacteriales Methanobacteriaceae *Methanobrevibacter smithii* TS147C |
| Msm-14-1 | Archaea Euryarchaeota Methanobacteria Methanobacteriales Methanobacteriaceae *Methanobrevibacter smithii* TS94A |
| Msm-15-1 | Archaea Euryarchaeota Methanobacteria Methanobacteriales Methanobacteriaceae *Methanobrevibacter smithii* TS94B |
| Msm-16-1 | Archaea Euryarchaeota Methanobacteria Methanobacteriales Methanobacteriaceae *Methanobrevibacter smithii* TS94C |
| Msm-17-1 | Archaea Euryarchaeota Methanobacteria Methanobacteriales Methanobacteriaceae *Methanobrevibacter smithii* TS95A |
| Msm-18-1 | Archaea Euryarchaeota Methanobacteria Methanobacteriales Methanobacteriaceae *Methanobrevibacter smithii* TS95B |
| Msm-19-1 | Archaea Euryarchaeota Methanobacteria Methanobacteriales Methanobacteriaceae *Methanobrevibacter smithii* TS95C |
| Msm-20-1 | Archaea Euryarchaeota Methanobacteria Methanobacteriales Methanobacteriaceae *Methanobrevibacter smithii* TS95D |
| Msm-21-1 | Archaea Euryarchaeota Methanobacteria Methanobacteriales Methanobacteriaceae *Methanobrevibacter smithii* TS96A |
| Msm-22-1 | Archaea Euryarchaeota Methanobacteria Methanobacteriales Methanobacteriaceae *Methanobrevibacter smithii* TS96B |
| Msm-23-1 | Archaea Euryarchaeota Methanobacteria Methanobacteriales Methanobacteriaceae *Methanobrevibacter smithii* TS96C |
| Mae-1-1 | Archaea Euryarchaeota Methanococci Methanococcales Methanococcaceae *Methanococcus aeolicus* Nankai-3 |
| Mvo-1-1 | Archaea Euryarchaeota Methanococci Methanococcales Methanococcaceae *Methanococcus voltae* A3 |
| Mco-1-1 | Archaea Euryarchaeota Methanomicrobia Methanocellales Methanocellaceae *Methanocella conradii* HZ254 |
| CMe-1-1 | Archaea Euryarchaeota Methanomicrobia Methanomicrobiales Generaincertae sedis *Candidatus* Methanoregula boonei 6A8 |
| Mpa-1-1 | Archaea Euryarchaeota Methanomicrobia Methanomicrobiales Generaincertae sedis *Methanosphaerula palustris* E1-9c |
| Mma-1-1 | Archaea Euryarchaeota Methanomicrobia Methanomicrobiales Methanomicrobiaceae *Methanoculleus marisnigri* JR1 |
| Mse-1-1 | Archaea Euryarchaeota Methanomicrobia Methanomicrobiales Methanomicrobiaceae *Methanoculleus sediminis* |
| Msp-5-1 | Archaea Euryarchaeota Methanomicrobia Methanomicrobiales Methanomicrobiaceae *Methanoculleus sp.* MH98A |
| Mli-1-1 | Archaea Euryarchaeota Methanomicrobia Methanomicrobiales Methanomicrobiaceae *Methanofollis liminatans* DSM 4140 |
| Mta-1-1 | Archaea Euryarchaeota Methanomicrobia Methanomicrobiales Methanoregulaceae *Methanolinea tarda* NOBI-1 |
| Mhu-1-1 | Archaea Euryarchaeota Methanomicrobia Methanomicrobiales Methanospirillaceae *Methanospirillum hungatei* JF-1 |
| CMe-2-1 | Archaea Euryarchaeota Methanomicrobia Methanosarcinales Candidatus Methanoperedenaceae *Candidatus* Methanoperedens nitroreducens |
| Mbu-1-1 | Archaea Euryarchaeota Methanomicrobia Methanosarcinales Methanosarcinaceae *Methanococcoides burtonii* DSM 6242 |
| Mme-1-1 | Archaea Euryarchaeota Methanomicrobia Methanosarcinales Methanosarcinaceae *Methanococcoides methylutens* |
| Mme-2-1 | Archaea Euryarchaeota Methanomicrobia Methanosarcinales Methanosarcinaceae *Methanococcoides methylutens* MM1 |
| Mev-1-1 | Archaea Euryarchaeota Methanomicrobia Methanosarcinales Methanosarcinaceae *Methanohalobium evestigatum* Z-7303 |
| Mma-2-1 | Archaea Euryarchaeota Methanomicrobia Methanosarcinales Methanosarcinaceae *Methanohalophilus mahii* DSM 5219 |
| Mps-1-1 | Archaea Euryarchaeota Methanomicrobia Methanosarcinales Methanosarcinaceae *Methanolobus psychrophilus* R15 |
| Mti-1-1 | Archaea Euryarchaeota Methanomicrobia Methanosarcinales Methanosarcinaceae *Methanolobus tindarius* DSM 2278 |
| Mho-1-1 | Archaea Euryarchaeota Methanomicrobia Methanosarcinales Methanosarcinaceae *Methanomethylovorans hollandica* DSM 15978 |
| Mzh-1-1 | Archaea Euryarchaeota Methanomicrobia Methanosarcinales Methanosarcinaceae *Methanosalsum zhilinae* DSM 4017 |
| Mac-1-1 | Archaea Euryarchaeota Methanomicrobia Methanosarcinales Methanosarcinaceae *Methanosarcina acetivorans* C2A |
| Mba-1-1 | Archaea Euryarchaeota Methanomicrobia Methanosarcinales Methanosarcinaceae *Methanosarcina barkeri* 227 |
| Mba-2-1 | Archaea Euryarchaeota Methanomicrobia Methanosarcinales Methanosarcinaceae *Methanosarcina barkeri* 3 |
| Mba-3-1 | Archaea Euryarchaeota Methanomicrobia Methanosarcinales Methanosarcinaceae *Methanosarcina barkeri* CM1 |
| Mba-4-1 | Archaea Euryarchaeota Methanomicrobia Methanosarcinales Methanosarcinaceae *Methanosarcina barkeri* MS |
| Mba-5-1 | Archaea Euryarchaeota Methanomicrobia Methanosarcinales Methanosarcinaceae *Methanosarcina barkeri str.* Fusaro |
| Mba-6-1 | Archaea Euryarchaeota Methanomicrobia Methanosarcinales Methanosarcinaceae *Methanosarcina barkeri str.* Wiesmoor |
| Mho-2-1 | Archaea Euryarchaeota Methanomicrobia Methanosarcinales Methanosarcinaceae *Methanosarcina horonobensis* HB-1 = JCM 15518 |
| Mla-1-1 | Archaea Euryarchaeota Methanomicrobia Methanosarcinales Methanosarcinaceae *Methanosarcina lacustris* Z-7289 |
| Mma-3-1 to Mma-3-56 | Archaea Euryarchaeota Methanomicrobia Methanosarcinales Methanosarcinaceae *Methanosarcina mazei* |
| Mma-4-1 | Archaea Euryarchaeota Methanomicrobia Methanosarcinales Methanosarcinaceae *Methanosarcina mazei* C16 |
| Mma-5-1 | Archaea Euryarchaeota Methanomicrobia Methanosarcinales Methanosarcinaceae *Methanosarcina mazei* Go1 |
| Mma-6-1 | Archaea Euryarchaeota Methanomicrobia Methanosarcinales Methanosarcinaceae *Methanosarcina mazei* LYC |
| Mma-7-1 | Archaea Euryarchaeota Methanomicrobia Methanosarcinales Methanosarcinaceae *Methanosarcina mazei* S-6 |
| Mma-8-1 | Archaea Euryarchaeota Methanomicrobia Methanosarcinales Methanosarcinaceae *Methanosarcina mazei* SarPi |
| Mma-9-1 | Archaea Euryarchaeota Methanomicrobia Methanosarcinales Methanosarcinaceae *Methanosarcina mazei* WWM610 |
| Msi-1-1 | Archaea Euryarchaeota Methanomicrobia Methanosarcinales Methanosarcinaceae *Methanosarcina siciliae* C2J |
| Msi-2-1 | Archaea Euryarchaeota Methanomicrobia Methanosarcinales Methanosarcinaceae *Methanosarcina siciliae* T4/M |
| Mso-1-1 | Archaea Euryarchaeota Methanomicrobia Methanosarcinales Methanosarcinaceae *Methanosarcina soligelidi* |
| Msp-6-1 | Archaea Euryarchaeota Methanomicrobia Methanosarcinales Methanosarcinaceae *Methanosarcina sp.* 1.H.A.2.2 |

| | |
|---|---|
| Msp-7-1 | Archaea Euryarchaeota Methanomicrobia Methanosarcinales Methanosarcinaceae *Methanosarcina sp.* 1.H.T.1A.1 |
| Msp-8-1 | Archaea Euryarchaeota Methanomicrobia Methanosarcinales Methanosarcinaceae *Methanosarcina sp.* 2.H.A.1B.4 |
| Msp-9-1 | Archaea Euryarchaeota Methanomicrobia Methanosarcinales Methanosarcinaceae *Methanosarcina sp.* 2.H.T.1A.15 |
| Msp-10-1 | Archaea Euryarchaeota Methanomicrobia Methanosarcinales Methanosarcinaceae *Methanosarcina sp.* 2.H.T.1A.3 |
| Msp-11-1 | Archaea Euryarchaeota Methanomicrobia Methanosarcinales Methanosarcinaceae *Methanosarcina sp.* 2.H.T.1A.6 |
| Msp-12-1 | Archaea Euryarchaeota Methanomicrobia Methanosarcinales Methanosarcinaceae *Methanosarcina sp.* 2.H.T.1A.8 |
| Msp-13-1 | Archaea Euryarchaeota Methanomicrobia Methanosarcinales Methanosarcinaceae *Methanosarcina sp.* Kolksee |
| Msp-14-1 | Archaea Euryarchaeota Methanomicrobia Methanosarcinales Methanosarcinaceae *Methanosarcina sp.* MTP4 |
| Msp-15-1 | Archaea Euryarchaeota Methanomicrobia Methanosarcinales Methanosarcinaceae *Methanosarcina sp.* WH1 |
| Msp-16-1 | Archaea Euryarchaeota Methanomicrobia Methanosarcinales Methanosarcinaceae *Methanosarcina sp.* WWM596 |
| Mth-1-1 | Archaea Euryarchaeota Methanomicrobia Methanosarcinales Methanosarcinaceae *Methanosarcina thermophila* TM-1 |
| Mva-1-1 | Archaea Euryarchaeota Methanomicrobia Methanosarcinales Methanosarcinaceae *Methanosarcina vacuolata* Z-761 |
| Msh-1-1 | Archaea Euryarchaeota Methanomicrobia Methanosarcinales Methermicoccaceae *Methermicoccus shengliensis* DSM 18856 |
| Pfe-1-1 | Archaea Euryarchaeota Thermococci Thermococcales Thermococcaceae *Palaeococcus ferrophilus* DSM 13482 |
| Ppa-1-1 | Archaea Euryarchaeota Thermococci Thermococcales Thermococcaceae *Palaeococcus pacificus* DY20341 |
| Pab-1-1 | Archaea Euryarchaeota Thermococci Thermococcales Thermococcaceae *Pyrococcus abyssi* GE5 |
| Pfu-1-1 | Archaea Euryarchaeota Thermococci Thermococcales Thermococcaceae *Pyrococcus furiosus* COM1 |
| Pfu-2-1 | Archaea Euryarchaeota Thermococci Thermococcales Thermococcaceae *Pyrococcus furiosus* DSM 3638 |
| Pho-1-1 | Archaea Euryarchaeota Thermococci Thermococcales Thermococcaceae *Pyrococcus horikoshii* OT3 |
| Psp-1-1 | Archaea Euryarchaeota Thermococci Thermococcales Thermococcaceae *Pyrococcus sp.* NA2 |
| Psp-2-1 | Archaea Euryarchaeota Thermococci Thermococcales Thermococcaceae *Pyrococcus sp.* ST04 |
| Pya-1-1 | Archaea Euryarchaeota Thermococci Thermococcales Thermococcaceae *Pyrococcus yayanosii* CH1 |
| Tba-1-1 | Archaea Euryarchaeota Thermococci Thermococcales Thermococcaceae *Thermococcus barophilus* MP |
| Teu-1-1 | Archaea Euryarchaeota Thermococci Thermococcales Thermococcaceae *Thermococcus eurythermalis* |
| Tga-1-1 | Archaea Euryarchaeota Thermococci Thermococcales Thermococcaceae *Thermococcus gammatolerans* EJ3 |
| Tko-1-1 | Archaea Euryarchaeota Thermococci Thermococcales Thermococcaceae *Thermococcus kodakarensis* KOD1 |
| Tli-1-1 | Archaea Euryarchaeota Thermococci Thermococcales Thermococcaceae *Thermococcus litoralis* DSM 5473 |
| Tna-1-1 | Archaea Euryarchaeota Thermococci Thermococcales Thermococcaceae *Thermococcus nautili* |
| Ton-1-1 | Archaea Euryarchaeota Thermococci Thermococcales Thermococcaceae *Thermococcus onnurineus* NA1 |
| Tpa-1-1 | Archaea Euryarchaeota Thermococci Thermococcales Thermococcaceae *Thermococcus paralvinellae* |
| Tsi-1-1 | Archaea Euryarchaeota Thermococci Thermococcales Thermococcaceae *Thermococcus sibiricus* MM 739 |
| Tsp-1-1 | Archaea Euryarchaeota Thermococci Thermococcales Thermococcaceae *Thermococcus sp.* 4557 |
| Tsp-2-1 | Archaea Euryarchaeota Thermococci Thermococcales Thermococcaceae *Thermococcus sp.* AM4 |
| Tsp-3-1 | Archaea Euryarchaeota Thermococci Thermococcales Thermococcaceae *Thermococcus sp.* CL1 |
| Tsp-4-1 | Archaea Euryarchaeota Thermococci Thermococcales Thermococcaceae *Thermococcus sp.* PK |
| Tzi-1-1 | Archaea Euryarchaeota Thermococci Thermococcales Thermococcaceae *Thermococcus zilligii* AN1 |
| env-1 to env-713 | environmental samples |

#### 4.2 Gene contexts

Each eL15-Euryarchaeota RNA (indicated by "RNA→") is listed. For each hit, the downstream genes predicted to reside in a regulated operon are listed. If the nearest downstream gene is encoding in the opposite strand (and therefore presumed to not be a part of a regulated operon), then that gene is still depicted. Some environmental sequences and some RefSeq entries lack gene annotations, and so no genes are available for such sequences. The direction of each gene is indicated with an arrow (→), and each predicted conserved domain in the gene is named. Conserved domains associated with more than one eL15-Euryarchaeota RNA are assigned a color; other domains are gray. Information about these conserved domains is given in Section 4.3. The accession of the sequence containing each eL15-Euryarchaeota RNA is given in the column named "Seq. accession". Accessions beginning with "NC_", "NS_", "NW_" or "NZ_" are contained in RefSeq. Other accession refer to environmental samples. Nucleotide coordinates are given for the 5′ and 3′ boundaries of each eL15-Euryarchaeota RNA. If the 5′ coordinate is greater than the 3′ coordinate, the RNA is present on the reverse-complement strand of the containing genomic DNA sequence. Each hit is denoted by an abbreviation (like "Eco-1-1") that refers to a taxonomy given in Section 4.1.

| abbrev. | Seq. accession | 5′ at | 3′ at | genes |
|---|---|---|---|---|

| | | | | | |
|---|---|---|---|---|---|
| env-1 | JGI24721J44947_10004045 | - | 13735 | 13690 | **RNA**→ PRK04243 (PRK04243)Ribosomal_L15e (pfam00827)→ |
| | | | | | PRK14555 (PRK14555)RNA_binding (pfam01877)→ RPP1 (COG1603)PRK00912 (PRK00912)→ |
| | | | | | PRK03717 (PRK03717)POP5 (COG1369)→ PRK03996 (PRK03996)proteasome_alpha_archeal (cd03756)→ |
| | | | | | Sdo1 (COG1500)PRK13760 (PRK13760)→ Rrp4 (COG1097)PRK04163 (PRK04163)→ |
| | | | | | RNase_PH_archRRP41 (cd11366)PRK03983 (PRK03983)→ |
| | | | | | RNase_PH_archRRP42 (cd11365)PRK04282 (PRK04282)→ RPL43A (COG1997)rpl37ae (PRK03976)→ |
| | | | | | RPC10 (COG1996)rpoP (PRK00398)→ Brix (smart00879)IMP4 (COG2136)→ |
| | | | | | Pcc1 (COG2892)Pcc1 (pfam09341)→ gimC_beta (TIGR02338)GimC (COG1382)→ |
| env-2 | JGI24721J26819_10075606 | - | 1121 | 1075 | **RNA**→ PRK04243 (PRK04243)Ribosomal_L15e (pfam00827)→ |
| | | | | | PRK14555 (PRK14555)RNA_binding (pfam01877)→ PRK00912 (PRK00912)→ |
| Mma-2-1 | NC_014002.1 | + | 819315 | 819361 | **RNA**→ PRK04243 (PRK04243)Ribosomal_L15e (pfam00827)→ |
| | | | | | PRK14555 (PRK14555)RNA_binding (pfam01877)→ PRK00912 (PRK00912)RNase_P_p30 (pfam01876)→ |
| | | | | | PRK03717 (PRK03717)RNase_P_Rpp14 (pfam01900)→ |
| | | | | | PRK03996 (PRK03996)proteasome_alpha_archeal (cd03756)→ Sdo1 (COG1500)PRK13760 (PRK13760)→ |
| | | | | | Rrp4 (COG1097)PRK04163 (PRK04163)→ RNase_PH_archRRP41 (cd11366)PRK03983 (PRK03983)→ |
| | | | | | RNase_PH_archRRP42 (cd11365)PRK04282 (PRK04282)→ rpl37ae (PRK03976)Ribosomal_L37ae (pfam01780) |
| | | | | | RPC10 (COG1996)rpoP (PRK00398)→ IMP4 (COG2136)PRK00933 (PRK00933)→ |
| | | | | | Pcc1 (COG2892)Pcc1 (pfam09341)→ hypo→ |
| env-3 | APHM01012746.1 | - | 3591 | 3541 | **RNA**→ PRK04243 (PRK04243)Ribosomal_L15e (pfam00827)→ |
| env-4 | LTJ07As871_J07ABscf098854 | + | 859526 | 859576 | **RNA**→ PRK04243 (PRK04243)Ribosomal_L15e (pfam00827)→ |
| Hha-1-1 | NZ_AJRK01000373.1 | + | 10855 | 10927 | **RNA**→ PRK04243 (PRK04243)Ribosomal_L15e (pfam00827)→ |
| | | | | | PRK04243 (PRK04243)Ribosomal_L15e (pfam00827)→ FrnE (COG2761)Thioredoxin_4 (pfam13462)→ |
| | | | | | FrnE (COG2761)Thioredoxin_4 (pfam13462)→ |
| Hha-1-2 | NZ_AOMB01000014.1 | + | 37016 | 37088 | **RNA**→ PRK04243 (PRK04243)Ribosomal_L15e (pfam00827)→ FrnE (COG2761)Thioredoxin_4 (pfam13462)– |
| env-5 | SS_3KL_010_SOIL_10297861 | - | 306 | 257 | **RNA**→ ←hypo |
| env-6 | SS_3KL_010_SOIL_10032413 | - | 1596 | 1549 | **RNA**→ PRK04243 (PRK04243)Ribosomal_L15e (pfam00827)→ |
| env-7 | APHM01002430.1 | + | 679 | 749 | **RNA**→ PRK04243 (PRK04243)Ribosomal_L15e (pfam00827)→ |
| | | | | | PRK10089 (PRK10089)tRNA_bindingDomain (cd02153)→ |
| env-8 | LTJ07As871_J07ABscf098776 | - | 39554 | 39484 | **RNA**→ PRK04243 (PRK04243)Ribosomal_L15e (pfam00827)→ |
| | | | | | PRK10089 (PRK10089)tRNA_bindingDomain (cd02153)→ |
| env-9 | GSLNARP_GFPJP1N02HFTYD | - | 319 | 187 | **RNA**→ PRK04243 (PRK04243)Ribosomal_L15e (pfam00827)→ |
| env-10 | GSLNARP_GFPJP1N02IOKU2 | + | 246 | 309 | **RNA**→ PRK04243 (PRK04243)Ribosomal_L15e (pfam00827)→ |
| Hsa-6-1 | NZ_ALJD01000003.1 | - | 716758 | 716671 | **RNA**→ PRK04243 (PRK04243)Ribosomal_L15e (pfam00827)→ |
| | | | | | PRK04243 (PRK04243)Ribosomal_L15e (pfam00827)→ |
| h3A-1-1 | NZ_KK033114.1 | + | 399056 | 399132 | **RNA**→ PRK04243 (PRK04243)Ribosomal_L15e (pfam00827)→ |
| env-11 | P_1C_Sed_2_UnCty_1134943 | + | 65 | 158 | **RNA**→ RPL15A (COG1632)PRK04243 (PRK04243)→ |
| env-12 | P_1C_Liq_2_UnCty_1030158 | - | 327 | 255 | **RNA**→ RPL15A (COG1632)PRK04243 (PRK04243)→ |
| env-13 | P_A23_Liq_2_Fmt_1019611 | - | 352 | 280 | **RNA**→ RPL15A (COG1632)PRK04243 (PRK04243)→ |
| env-14 | P_1C_Sed_2_UnCty_1034182 | + | 193 | 285 | **RNA**→ PRK04243 (PRK04243)Ribosomal_L15e (pfam00827)→ |
| env-15 | ADL5mRS1u_contig05278 | - | 96 | 32 | **RNA**→ |
| har-1-1 | NC_015954.1 | + | 2102369 | 2102433 | **RNA**→ PRK04243 (PRK04243)Ribosomal_L15e (pfam00827)→ |
| env-16 | ADL20m3uS_contig11659 | - | 8862 | 8798 | **RNA**→ PRK04243 (PRK04243)Ribosomal_L15e (pfam00827)→ |
| env-17 | 2140918017-4741 | - | 11103 | 11039 | **RNA**→ PRK04243 (PRK04243)Ribosomal_L15e (pfam00827)→ |
| env-18 | 2140918027-00084 | - | 11554 | 11490 | **RNA**→ PRK04243 (PRK04243)Ribosomal_L15e (pfam00827)→ |
| env-19 | P_A23_Liq_1_Fmt_1000005 | + | 6223 | 6278 | **RNA**→ PRK04243 (PRK04243)Ribosomal_L15e (pfam00827)→ |
| | | | | | ThiF_MoeB_HesA_family (cd00757)adenyl_thiF (TIGR02356)→ |
| env-20 | P_A23_Liq_3_Fmt_1000005 | + | 8005 | 8060 | **RNA**→ PRK04243 (PRK04243)Ribosomal_L15e (pfam00827)→ |
| | | | | | ThiF_MoeB_HesA_family (cd00757)adenyl_thiF (TIGR02356)→ |
| env-21 | P_A23_Liq_2_Fmt_1000009 | - | 43137 | 43082 | **RNA**→ PRK04243 (PRK04243)Ribosomal_L15e (pfam00827)→ |
| | | | | | ThiF_MoeB_HesA_family (cd00757)adenyl_thiF (TIGR02356)→ |
| Hwa-2-1 | NC_008212.1 | + | 526505 | 526560 | **RNA**→ PRK04243 (PRK04243)Ribosomal_L15e (pfam00827)→ |
| | | | | | ThiF_MoeB_HesA_family (cd00757)adenyl_thiF (TIGR02356)→ |
| | | | | | PRK08762 (PRK08762)ThiF_MoeB_HesA_family (cd00757)→ |
| env-22 | GSLNARP_contig12467 | - | 1205 | 1150 | **RNA**→ PRK04243 (PRK04243)Ribosomal_L15e (pfam00827)→ |
| | | | | | PRK08762 (PRK08762)ThiF_MoeB_HesA_family (cd00757)→ |

| | | | | | |
|---|---|---|---|---|---|
| Hwa-1-1 | NC_017459.1 | + | 520849 | 520904 | **RNA**→ PRK04243 (PRK04243)Ribosomal_L15e (pfam00827)→ |
| | | | | | ThiF_MoeB_HesA_family (cd00757)adenyl_thiF (TIGR02356)→ |
| | | | | | PRK08762 (PRK08762)ThiF_MoeB_HesA_family (cd00757)→ |
| env-23 | APHM01021410.1 | + | 37065 | 37122 | **RNA**→ PRK04243 (PRK04243)Ribosomal_L15e (pfam00827)→ |
| | | | | | PRK08762 (PRK08762)ThiF_MoeB_HesA_family (cd00757)→ |
| env-24 | LTJ07As871_J07ABscf098855 | + | 941476 | 941533 | **RNA**→ PRK04243 (PRK04243)Ribosomal_L15e (pfam00827)→ |
| env-25 | APHM01021802.1 | + | 241797 | 241852 | **RNA**→ PRK04243 (PRK04243)Ribosomal_L15e (pfam00827)→ |
| | | | | | ThiF_MoeB_HesA_family (cd00757)adenyl_thiF (TIGR02356)→ |
| env-26 | LTJ07As871_J07ABscf098874 | + | 856037 | 856092 | **RNA**→ PRK04243 (PRK04243)Ribosomal_L15e (pfam00827)→ |
| | | | | | ThiF_MoeB_HesA_family (cd00757)adenyl_thiF (TIGR02356)→ |
| env-27 | JXWS01023088.1 | - | 1497 | 1442 | **RNA**→ PRK04243 (PRK04243)Ribosomal_L15e (pfam00827)→ |
| | | | | | ThiF_MoeB_HesA_family (cd00757)adenyl_thiF (TIGR02356)→ |
| env-28 | P_A23_Liq_2_Fmt_1049687 | + | 77 | 131 | **RNA**→ PRK04243 (PRK04243)Ribosomal_L15e (pfam00827)→ |
| env-29 | P_A23_Liq_1_Fmt_1040915 | - | 222 | 168 | **RNA**→ PRK04243 (PRK04243)Ribosomal_L15e (pfam00827)→ |
| env-30 | P_A23_Liq_3_Fmt_1034622 | + | 56 | 110 | **RNA**→ RPL15A (COG1632)PRK04243 (PRK04243)→ |
| env-31 | GSLNARP_GFPJP1N02J0WUN | + | 62 | 117 | **RNA**→ RPL15A (COG1632)PRK04243 (PRK04243)→ |
| Hru-1-1 | NZ_BBJO01000020.1 | + | 43857 | 43911 | **RNA**→ PRK04243 (PRK04243)Ribosomal_L15e (pfam00827)→ |
| env-32 | JXWS01113690.1 | + | 469 | 523 | **RNA**→ RPL15A (COG1632)PRK04243 (PRK04243)→ |
| env-33 | GSLNARP_GGGT6729_b1 | + | 216 | 270 | **RNA**→ RPL15A (COG1632)PRK04243 (PRK04243)→ |
| Hpa-2-1 | NZ_AOIV01000027.1 | - | 154749 | 154689 | **RNA**→ PRK04243 (PRK04243)Ribosomal_L15e (pfam00827)→ |
| env-34 | GSLNAS_contig09763 | + | 194 | 284 | **RNA**→ RPL15A (COG1632)PRK04243 (PRK04243)→ |
| env-35 | SL_5KL_010_BRINE_10000873 | + | 7980 | 8030 | **RNA**→ PRK04243 (PRK04243)Ribosomal_L15e (pfam00827)→ |
| env-36 | SL_6KL_011_BRINE_1000559 | + | 8034 | 8084 | **RNA**→ PRK04243 (PRK04243)Ribosomal_L15e (pfam00827)→ |
| env-37 | SL_4KL_010_BRINE_10001420 | - | 8559 | 8509 | **RNA**→ PRK04243 (PRK04243)Ribosomal_L15e (pfam00827)→ |
| env-38 | SL_7KL_010_BRINE_10000923 | - | 9479 | 9429 | **RNA**→ PRK04243 (PRK04243)Ribosomal_L15e (pfam00827)→ |
| env-39 | GSLNARP_contig11045 | + | 688 | 814 | **RNA**→ PRK04243 (PRK04243)Ribosomal_L15e (pfam00827)→ |
| env-40 | APHM01004060.1 | - | 1754 | 1627 | **RNA**→ PRK04243 (PRK04243)Ribosomal_L15e (pfam00827)→ |
| env-41 | LTJ07As871_J07ABscf087768 | + | 8333 | 8460 | **RNA**→ PRK04243 (PRK04243)Ribosomal_L15e (pfam00827)→ |
| Hac-1-1 | NZ_BATA01000010.1 | - | 40529 | 40466 | **RNA**→ PRK04243 (PRK04243)Ribosomal_L15e (pfam00827)→ |
| Hac-1-2 | NZ_BANO01000201.1 | + | 3030 | 3093 | **RNA**→ PRK04243 (PRK04243)Ribosomal_L15e (pfam00827)→ |
| env-42 | JXWS01007612.1 | + | 354 | 427 | **RNA**→ PRK04243 (PRK04243)Ribosomal_L15e (pfam00827)→ |
| env-43 | P_A23_Liq_1_Fmt_1012496 | + | 495 | 568 | **RNA**→ PRK04243 (PRK04243)Ribosomal_L15e (pfam00827)→ |
| env-44 | P_A23_Liq_3_Fmt_1032483 | - | 283 | 210 | **RNA**→ RPL15A (COG1632)PRK04243 (PRK04243)→ |
| env-45 | P_1C_Liq_3_UnCty_1034822 | + | 91 | 164 | **RNA**→ RPL15A (COG1632)PRK04243 (PRK04243)→ |
| env-46 | P_1C_Liq_1_UnCty_1087024 | - | 228 | 155 | **RNA**→ PRK04243 (PRK04243)Ribosomal_L15e (pfam00827)→ |
| Hna-1-1 | NZ_KE386573.1 | + | 519442 | 519497 | **RNA**→ PRK04243 (PRK04243)Ribosomal_L15e (pfam00827)→ |
| env-47 | P_1C_Sed_2_UnCty_1011141 | + | 185 | 240 | **RNA**→ PRK04243 (PRK04243)Ribosomal_L15e (pfam00827)→ |
| env-48 | GSLSAAI_contig05270 | + | 171 | 218 | **RNA**→ PRK04243 (PRK04243)Ribosomal_L15e (pfam00827)→ |
| | | | | | PRK08762 (PRK08762)ThiF_MoeB_HesA_family (cd00757)→ |
| env-49 | GSLSAAI_contig13583 | - | 272 | 225 | **RNA**→ PRK04243 (PRK04243)Ribosomal_L15e (pfam00827)→ |
| env-50 | GSLSAAI_contig05339 | - | 621 | 574 | **RNA**→ PRK04243 (PRK04243)Ribosomal_L15e (pfam00827)→ |
| env-51 | GSLSAAI_contig11105 | + | 1628 | 1675 | **RNA**→ PRK04243 (PRK04243)Ribosomal_L15e (pfam00827)→ |
| | | | | | PRK08762 (PRK08762)ThiF_MoeB_HesA_family (cd00757)→ |
| env-52 | GSLSAAI_contig11811 | - | 21562 | 21515 | **RNA**→ PRK04243 (PRK04243)Ribosomal_L15e (pfam00827)→ |
| | | | | | PRK08762 (PRK08762)ThiF_MoeB_HesA_family (cd00757)→ PRK05399 (PRK05399)MutS_I (pfam01624)→ |
| | | | | | MUTSd (smart00533)MutS (COG0249)→ MUTSac (smart00534)MutS (COG0249)→ |
| | | | | | HATPase_c (smart00387)HATPase_c (cd00075)→ DNA_mis_repair (pfam01119)MutL_C (pfam08676)→ |
| env-53 | PR_CR_10_Liq_1_inCR_1003279 | - | 5806 | 5759 | **RNA**→ PRK04243 (PRK04243)Ribosomal_L15e (pfam00827)→ |
| | | | | | PRK08762 (PRK08762)ThiF_MoeB_HesA_family (cd00757)→ |
| | | | | | MUTSac (smart00534)MUTSd (smart00533)MutS (COG0249)→ MutL (COG0323)mutL (PRK00095)→ |
| env-54 | PR_CR_10_Liq_3_inCR_1104247 | - | 209 | 162 | **RNA**→ RPL15A (COG1632)PRK04243 (PRK04243)→ |
| env-55 | SL_6KL_011_BRINE_1017367 | + | 1682 | 1774 | **RNA**→ PRK04243 (PRK04243)Ribosomal_L15e (pfam00827)→ |
| env-56 | SL_4KL_010_BRINE_10011168 | - | 1073 | 981 | **RNA**→ PRK04243 (PRK04243)Ribosomal_L15e (pfam00827)→ |
| env-57 | SL_6KL_011_BRINE_1001181 | - | 10407 | 10313 | **RNA**→ PRK04243 (PRK04243)Ribosomal_L15e (pfam00827)→ |

| | | | | | |
|---|---|---|---|---|---|
| env-58 | SL_5KL_010_SED_10002058 | - | 10404 | 10310 | **RNA**→ PRK04243 (PRK04243)Ribosomal_L15e (pfam00827)→ |
| env-59 | SL_8KL_010_SED_10019608 | + | 2296 | 2390 | **RNA**→ PRK04243 (PRK04243)Ribosomal_L15e (pfam00827)→ |
| env-60 | SL_4KL_010_BRINE_10010798 | - | 3106 | 3012 | **RNA**→ PRK04243 (PRK04243)Ribosomal_L15e (pfam00827)→ |
| env-61 | SL_6KL_011_BRINE_1000334 | + | 4498 | 4594 | **RNA**→ PRK04243 (PRK04243)Ribosomal_L15e (pfam00827)→ |
| env-62 | SL_8KL_010_SED_10044433 | - | 1782 | 1686 | **RNA**→ PRK04243 (PRK04243)Ribosomal_L15e (pfam00827)→ |
| env-63 | SL_6KL_011_BRINE_1254331 | + | 116 | 210 | **RNA**→ |
| env-64 | SL_5KL_010_BRINE_10001910 | - | 7669 | 7575 | **RNA**→ PRK04243 (PRK04243)Ribosomal_L15e (pfam00827)→ |
| env-65 | SL_4KL_010_BRINE_10067600 | - | 209 | 106 | **RNA**→ RPL15A (COG1632)PRK04243 (PRK04243)→ |
| env-66 | SL_6KL_011_BRINE_1036440 | - | 1105 | 1005 | **RNA**→ PRK04243 (PRK04243)Ribosomal_L15e (pfam00827)→ |
| Hai-1-1 | NZ_AOJI01000011.1 | + | 21092 | 21169 | **RNA**→ PRK04243 (PRK04243)Ribosomal_L15e (pfam00827)→ |
| Hco-1-1 | NZ_AOJL01000012.1 | + | 32123 | 32204 | **RNA**→ PRK04243 (PRK04243)Ribosomal_L15e (pfam00827)→ |
| Hsp-8-1 | NZ_JH815329.1 | + | 137323 | 137405 | **RNA**→ PRK04243 (PRK04243)Ribosomal_L15e (pfam00827)→ |
| Hte-1-1 | NZ_AOJD01000051.1 | + | 33749 | 33831 | **RNA**→ PRK04243 (PRK04243)Ribosomal_L15e (pfam00827)→ |
| Hho-1-1 | NZ_AOJO01000019.1 | - | 62900 | 62818 | **RNA**→ PRK04243 (PRK04243)Ribosomal_L15e (pfam00827)→ |
| env-67 | P_1C_Liq_3_UnCty_1013660 | - | 413 | 331 | **RNA**→ RPL15A (COG1632)PRK04243 (PRK04243)→ |
| env-68 | P_1C_Liq_1_UnCty_1013722 | - | 796 | 714 | **RNA**→ PRK04243 (PRK04243)Ribosomal_L15e (pfam00827)→ |
| env-69 | P_1C_Liq_2_UnCty_1006954 | - | 774 | 692 | **RNA**→ PRK04243 (PRK04243)Ribosomal_L15e (pfam00827)→ |
| Hca-2-1 | NZ_AOJK01000041.1 | + | 102530 | 102612 | **RNA**→ PRK04243 (PRK04243)Ribosomal_L15e (pfam00827)→ |
| env-70 | P_A23_Liq_1_Fmt_1003600 | - | 1435 | 1353 | **RNA**→ PRK04243 (PRK04243)Ribosomal_L15e (pfam00827)→ |
| env-71 | P_A23_Liq_3_Fmt_1003343 | + | 1237 | 1319 | **RNA**→ PRK04243 (PRK04243)Ribosomal_L15e (pfam00827)→ |
| env-72 | P_A23_Liq_2_Fmt_1003882 | - | 1692 | 1610 | **RNA**→ PRK04243 (PRK04243)Ribosomal_L15e (pfam00827)→ |
| Hez-1-1 | NZ_ATXS01000005.1 | + | 87176 | 87258 | **RNA**→ PRK04243 (PRK04243)Ribosomal_L15e (pfam00827)→ |
| Hli-2-1 | NZ_AOJF01000031.1 | - | 113040 | 112960 | **RNA**→ PRK04243 (PRK04243)Ribosomal_L15e (pfam00827)→ |
| Har-2-1 | NZ_AOJJ01000054.1 | - | 111488 | 111408 | **RNA**→ PRK04243 (PRK04243)Ribosomal_L15e (pfam00827)→ |
| Hdi-1-1 | NZ_AOJN01000039.1 | - | 113054 | 112974 | **RNA**→ PRK04243 (PRK04243)Ribosomal_L15e (pfam00827)→ |
| Hte-2-1 | NZ_AOIW01000042.1 | - | 36897 | 36817 | **RNA**→ PRK04243 (PRK04243)Ribosomal_L15e (pfam00827)→ |
| Hdi-2-1 | NZ_AOJM01000058.1 | - | 27117 | 27037 | **RNA**→ PRK04243 (PRK04243)Ribosomal_L15e (pfam00827)→ |
| env-73 | P_1C_Liq_1_UnCty_1078077 | - | 117 | 36 | **RNA**→ hypo→ |
| Hli-1-1 | NZ_AOJG01000017.1 | - | 108989 | 108896 | **RNA**→ PRK04243 (PRK04243)Ribosomal_L15e (pfam00827)→ |
| Hha-2-1 | NZ_BBJP01000018.1 | + | 13726 | 13819 | **RNA**→ PRK04243 (PRK04243)Ribosomal_L15e (pfam00827)→ |
| Hsa-9-1 | NZ_JNFH02000023.1 | + | 21503 | 21596 | **RNA**→ PRK04243 (PRK04243)Ribosomal_L15e (pfam00827)→ |
| env-74 | ADL20m3uS_contig07384 | + | 1833 | 1926 | **RNA**→ PRK04243 (PRK04243)Ribosomal_L15e (pfam00827)→ |
| env-75 | 2140918017-2486 | + | 2582 | 2675 | **RNA**→ PRK04243 (PRK04243)Ribosomal_L15e (pfam00827)→ |
| env-76 | 2140918027-05980 | - | 1029 | 936 | **RNA**→ PRK04243 (PRK04243)Ribosomal_L15e (pfam00827)→ |
| Hla-3-1 | NC_012029.1 | + | 432908 | 433001 | **RNA**→ PRK04243 (PRK04243)Ribosomal_L15e (pfam00827)→ |
| Hko-1-1 | NZ_AOJH01000069.1 | + | 3726 | 3819 | **RNA**→ PRK04243 (PRK04243)Ribosomal_L15e (pfam00827)→ |
| env-77 | GQKUMWE01AOE6H | - | 136 | 43 | **RNA**→ hypo→ |
| env-78 | GQKUMWE02FUCCT | + | 205 | 298 | **RNA**→ hypo→ |
| Hsa-7-1 | NZ_AOJE01000060.1 | + | 111633 | 111732 | **RNA**→ PRK04243 (PRK04243)Ribosomal_L15e (pfam00827)→ |
| env-79 | P_1C_Liq_2_UnCty_1069180 | - | 260 | 182 | **RNA**→ PRK04243 (PRK04243)Ribosomal_L15e (pfam00827)→ |
| env-80 | P_A23_Liq_1_Fmt_1002719 | - | 2388 | 2327 | **RNA**→ PRK04243 (PRK04243)Ribosomal_L15e (pfam00827)→ |
| env-81 | P_A23_Liq_3_Fmt_1003378 | - | 2423 | 2362 | **RNA**→ PRK04243 (PRK04243)Ribosomal_L15e (pfam00827)→ |
| env-82 | P_A23_Liq_2_Fmt_1009559 | - | 879 | 818 | **RNA**→ PRK04243 (PRK04243)Ribosomal_L15e (pfam00827)→ |
| env-83 | P_1C_Sed_2_UnCty_1017312 | + | 398 | 479 | **RNA**→ PRK04243 (PRK04243)Ribosomal_L15e (pfam00827)→ |
| env-84 | P_1C_Liq_1_UnCty_1034993 | + | 194 | 275 | **RNA**→ RPL15A (COG1632)PRK04243 (PRK04243)→ |
| env-85 | P_1C_Liq_2_UnCty_1023529 | - | 401 | 320 | **RNA**→ RPL15A (COG1632)PRK04243 (PRK04243)→ |
| env-86 | P_1C_Liq_3_UnCty_1021498 | - | 346 | 265 | **RNA**→ RPL15A (COG1632)PRK04243 (PRK04243)→ |
| env-87 | P_1C_Sed_2_UnCty_1004595 | + | 1788 | 1869 | **RNA**→ PRK04243 (PRK04243)Ribosomal_L15e (pfam00827)→ |
| env-88 | P_1C_Sed_2_UnCty_1006549 | + | 274 | 355 | **RNA**→ PRK04243 (PRK04243)Ribosomal_L15e (pfam00827)→ |
| env-89 | SL_6KL_011_BRINE_1064466 | - | 575 | 484 | **RNA**→ PRK04243 (PRK04243)Ribosomal_L15e (pfam00827)→ |
| env-90 | SL_6KL_011_BRINE_1008323 | - | 510 | 419 | **RNA**→ RPL15A (COG1632)PRK04243 (PRK04243)→ |
| env-91 | SL_6KL_011_BRINE_1063445 | + | 299 | 393 | **RNA**→ RPL15A (COG1632)PRK04243 (PRK04243)→ |

| | | | | | |
|---|---|---|---|---|---|
| env-92 | SL_6KL_011_BRINE_1001542 | - | 7921 | 7827 | **RNA**→ PRK04243 (PRK04243)Ribosomal_L15e (pfam00827)→ |
| env-93 | SL_6KL_011_BRINE_1006605 | + | 4992 | 5093 | **RNA**→ RPL15A (COG1632)PRK04243 (PRK04243)→ |
| env-94 | SL_6KL_011_BRINE_1010435 | - | 890 | 821 | **RNA**→ PRK04243 (PRK04243)Ribosomal_L15e (pfam00827)→ |
| Hmu-1-1 | NZ_AOLN01000018.1 | + | 760151 | 760214 | **RNA**→ PRK04243 (PRK04243)Ribosomal_L15e (pfam00827)→ |
| Hel-1-1 | NZ_AOLK01000011.1 | - | 499306 | 499243 | **RNA**→ PRK04243 (PRK04243)Ribosomal_L15e (pfam00827)→ |
| Hla-2-1 | NZ_AOLI01000030.1 | - | 9206 | 9143 | **RNA**→ PRK04243 (PRK04243)Ribosomal_L15e (pfam00827)→ |
| Hde-1-1 | NZ_AOLP01000017.1 | + | 194706 | 194766 | **RNA**→ PRK04243 (PRK04243)Ribosomal_L15e (pfam00827)→ |
| Hal-2-1 | NZ_LK053000.1 | - | 1797714 | 1797654 | **RNA**→ PRK04243 (PRK04243)Ribosomal_L15e (pfam00827)→ |
| Hpr-1-1 | NZ_AOLG01000009.1 | - | 95328 | 95268 | **RNA**→ PRK04243 (PRK04243)Ribosomal_L15e (pfam00827)→ |
| Hgi-1-1 | NZ_AOLJ01000011.1 | + | 406353 | 406413 | **RNA**→ PRK04243 (PRK04243)Ribosomal_L15e (pfam00827)→ |
| Hgi-2-1 | NZ_CP011947.1 | - | 470623 | 470563 | **RNA**→ PRK04243 (PRK04243)Ribosomal_L15e (pfam00827)→ |
| Hsp-9-1 | NZ_JPES01000007.1 | - | 10173 | 10113 | **RNA**→ PRK04243 (PRK04243)Ribosomal_L15e (pfam00827)→ |
| Hsu-2-1 | NZ_AOLM01000018.1 | + | 49470 | 49530 | **RNA**→ PRK04243 (PRK04243)Ribosomal_L15e (pfam00827)→ |
| Hsp-6-1 | NZ_AOLD01000031.1 | + | 8987 | 9047 | **RNA**→ PRK04243 (PRK04243)Ribosomal_L15e (pfam00827)→ |
| Hsp-4-1 | NZ_AOLF01000036.1 | - | 67323 | 67263 | **RNA**→ PRK04243 (PRK04243)Ribosomal_L15e (pfam00827)→ |
| Hsp-5-1 | NZ_AOLE01000023.1 | + | 9123 | 9183 | **RNA**→ PRK04243 (PRK04243)Ribosomal_L15e (pfam00827)→ |
| Hme-1-1 | NZ_AOLO01000007.1 | + | 387185 | 387264 | **RNA**→ PRK04243 (PRK04243)Ribosomal_L15e (pfam00827)→ |
| Hme-1-2 | NC_017941.2 | - | 494773 | 494694 | **RNA**→ PRK04243 (PRK04243)Ribosomal_L15e (pfam00827)→ |
| Hru-2-1 | NZ_BBJN01000010.1 | - | 27162 | 27075 | **RNA**→ PRK04243 (PRK04243)Ribosomal_L15e (pfam00827)→ |
| env-95 | P_A23_Liq_1_Fmt_1000360 | + | 2899 | 2970 | **RNA**→ PRK04243 (PRK04243)Ribosomal_L15e (pfam00827)→ |
| env-96 | P_A23_Liq_3_Fmt_1000054 | + | 17691 | 17762 | **RNA**→ PRK04243 (PRK04243)Ribosomal_L15e (pfam00827)→ |
| env-97 | P_A23_Liq_2_Fmt_1000005 | + | 59053 | 59124 | **RNA**→ PRK04243 (PRK04243)Ribosomal_L15e (pfam00827)→ |
| env-98 | JXWS01130832.1 | - | 481 | 414 | **RNA**→ RPL15A (COG1632)PRK04243 (PRK04243)→ |
| env-99 | APHM01022051.1 | + | 3638 | 3699 | **RNA**→ PRK04243 (PRK04243)Ribosomal_L15e (pfam00827)→ |
| env-100 | LTJ07As871_J07ABscf098878 | + | 1502510 | 1502571 | **RNA**→ PRK04243 (PRK04243)Ribosomal_L15e (pfam00827)→ |
| CHa-1-1 | NZ_ASGZ01000052.1 | + | 24839 | 24927 | **RNA**→ PRK04243 (PRK04243)Ribosomal_L15e (pfam00827)→ GldE (TIGR03520)DUF21 (pfam01595)→ |
| env-101 | SL_4KL_010_BRINE_10493865 | - | 158 | 111 | **RNA**→ RPL15A (COG1632)PRK04243 (PRK04243)→ |
| env-102 | SL_6KL_011_BRINE_1000129 | + | 15997 | 16046 | **RNA**→ PRK04243 (PRK04243)Ribosomal_L15e (pfam00827)→ |
| | | | | | PRK08762 (PRK08762)ThiF_MoeB_HesA_family (cd00757)→ NikR (COG0864)NikR_C (pfam08753)→ |
| env-103 | SL_4KL_010_BRINE_10001303 | + | 2439 | 2488 | **RNA**→ PRK04243 (PRK04243)Ribosomal_L15e (pfam00827)→ |
| | | | | | PRK08762 (PRK08762)ThiF_MoeB_HesA_family (cd00757)→ NikR (COG0864)NikR_C (pfam08753)→ |
| env-104 | SL_4KL_010_BRINE_10001201 | + | 2902 | 2951 | **RNA**→ PRK04243 (PRK04243)Ribosomal_L15e (pfam00827)→ |
| | | | | | ThiF_MoeB_HesA_family (cd00757)adenyl_thiF (TIGR02356)→ NikR (COG0864)NikR_C (pfam08753)→ |
| env-105 | SL_5KL_010_BRINE_10000041 | + | 10173 | 10222 | **RNA**→ PRK04243 (PRK04243)Ribosomal_L15e (pfam00827)→ |
| env-106 | SL_6KL_011_BRINE_1002627 | - | 6475 | 6426 | **RNA**→ PRK04243 (PRK04243)Ribosomal_L15e (pfam00827)→ |
| env-107 | SL_5KL_010_SED_10011446 | + | 130 | 179 | **RNA**→ PRK04243 (PRK04243)Ribosomal_L15e (pfam00827)→ |
| env-108 | SL_8KL_010_SED_10489198 | - | 248 | 199 | **RNA**→ PRK04243 (PRK04243)Ribosomal_L15e (pfam00827)→ |
| env-109 | SL_4KL_010_BRINE_10000898 | + | 3047 | 3096 | **RNA**→ PRK04243 (PRK04243)Ribosomal_L15e (pfam00827)→ |
| env-110 | SL_6KL_011_BRINE_1001896 | - | 92 | 43 | **RNA**→ ←hypo |
| env-111 | SL_4KL_010_BRINE_10030276 | + | 2918 | 2967 | **RNA**→ hypo→ |
| env-112 | P_1C_Liq_3_UnCty_1038130 | - | 250 | 201 | **RNA**→ PRK04243 (PRK04243)Ribosomal_L15e (pfam00827)→ |
| env-113 | P_1C_Liq_2_UnCty_1042794 | + | 154 | 203 | **RNA**→ PRK04243 (PRK04243)Ribosomal_L15e (pfam00827)→ |
| env-114 | P_1C_Liq_1_UnCty_1078870 | + | 144 | 193 | **RNA**→ PRK04243 (PRK04243)Ribosomal_L15e (pfam00827)→ |
| env-115 | P_A23_Liq_3_Fmt_1006374 | - | 68 | 19 | **RNA**→ |
| env-116 | JXWS01040251.1 | + | 738 | 787 | **RNA**→ |
| env-117 | P_A23_Liq_3_Fmt_1005856 | + | 1663 | 1712 | **RNA**→ |
| env-118 | P_A23_Liq_2_Fmt_1003095 | - | 2041 | 1992 | **RNA**→ PRK04243 (PRK04243)Ribosomal_L15e (pfam00827)→ |
| env-119 | APHM01020319.1 | - | 3958 | 3909 | **RNA**→ PRK04243 (PRK04243)Ribosomal_L15e (pfam00827)→ |
| env-120 | LTJ07As871_J07ABscf098871 | + | 227603 | 227652 | **RNA**→ PRK04243 (PRK04243)Ribosomal_L15e (pfam00827)→ |
| env-121 | P_A23_Liq_1_Fmt_1002579 | - | 586 | 537 | **RNA**→ PRK04243 (PRK04243)Ribosomal_L15e (pfam00827)→ |
| env-122 | P_A23_Liq_1_Fmt_1006552 | - | 756 | 707 | **RNA**→ PRK04243 (PRK04243)Ribosomal_L15e (pfam00827)→ |
| env-123 | P_A23_Liq_2_Fmt_1001062 | + | 420 | 469 | **RNA**→ PRK04243 (PRK04243)Ribosomal_L15e (pfam00827)→ |
| env-124 | GSLNARP_contig28426 | + | 697 | 746 | **RNA**→ PRK04243 (PRK04243)Ribosomal_L15e (pfam00827)→ |

| | | | | | |
|---|---|---|---|---|---|
| env-125 | P_A23_Liq_3_Fmt_1000673 | - | 2025 | 1976 | **RNA**→ PRK04243 (PRK04243)Ribosomal_L15e (pfam00827)→ |
| env-126 | ADL20m3uS_contig11627 | + | 9912 | 9962 | **RNA**→ PRK04243 (PRK04243)Ribosomal_L15e (pfam00827)→ |
| env-127 | ADL5mRS1u_contig00042 | - | 12742 | 12692 | **RNA**→ PRK04243 (PRK04243)Ribosomal_L15e (pfam00827)→ |
| env-128 | 2140918027-00046 | - | 29012 | 28962 | **RNA**→ PRK04243 (PRK04243)Ribosomal_L15e (pfam00827)→ |
| env-129 | P_1C_Sed_2_UnCty_1108461 | + | 263 | 312 | **RNA**→ PRK04243 (PRK04243)Ribosomal_L15e (pfam00827)→ |
| env-130 | ADL20m3uS_contig09914 | + | 12790 | 12839 | **RNA**→ PRK04243 (PRK04243)Ribosomal_L15e (pfam00827)→ |
| env-131 | ADL5mRS1u_contig00058 | - | 5253 | 5204 | **RNA**→ PRK04243 (PRK04243)Ribosomal_L15e (pfam00827)→ |
| env-132 | 2140918017-271 | - | 51697 | 51648 | **RNA**→ PRK04243 (PRK04243)Ribosomal_L15e (pfam00827)→ |
| env-133 | 2140918017-8071 | - | 5288 | 5239 | **RNA**→ PRK04243 (PRK04243)Ribosomal_L15e (pfam00827)→ |
| env-134 | 2140918027-00195 | - | 5210 | 5161 | **RNA**→ PRK04243 (PRK04243)Ribosomal_L15e (pfam00827)→ |
| env-135 | SL_4KL_010_BRINE_10074326 | - | 160 | 113 | **RNA**→ PRK04243 (PRK04243)Ribosomal_L15e (pfam00827)→ |
| env-136 | SL_6KL_011_BRINE_1020994 | - | 164 | 117 | **RNA**→ PRK04243 (PRK04243)Ribosomal_L15e (pfam00827)→ |
| env-137 | SL_6KL_011_BRINE_1123850 | + | 243 | 291 | **RNA**→ PRK04243 (PRK04243)Ribosomal_L15e (pfam00827)→ |
| env-138 | SL_8KL_010_SED_10020664 | - | 519 | 471 | **RNA**→ PRK04243 (PRK04243)Ribosomal_L15e (pfam00827)→ |
| env-139 | SL_8KL_010_SED_10132433 | - | 343 | 294 | **RNA**→ RPL15A (COG1632)PRK04243 (PRK04243)→ |
| env-140 | SL_5KL_010_SED_10120051 | + | 387 | 435 | **RNA**→ |
| env-141 | SL_5KL_010_BRINE_10075440 | - | 64 | 16 | **RNA**→ hypo→ |
| env-142 | SL_5KL_010_SED_10001678 | - | 13040 | 12992 | **RNA**→ PRK04243 (PRK04243)Ribosomal_L15e (pfam00827)→ |
| | | | | | MCP_signal (cd11386)MA (smart00283)PRK10935 (PRK10935)HAMP (pfam00672)→ |
| | | | | | ThiF_MoeB_HesA_family (cd00757)adenyl_thiF (TIGR02356)→ PRK04460 (PRK04460)ParD (COG3609)→ |
| | | | | | TIGR00697 (TIGR00697)Vut_1 (pfam02592)→ |
| env-143 | SL_7KL_010_BRINE_10383585 | + | 15 | 63 | **RNA**→ RPL15A (COG1632)PRK04243 (PRK04243)→ |
| env-144 | SL_5KL_010_SED_10002997 | - | 7066 | 7018 | **RNA**→ PRK04243 (PRK04243)Ribosomal_L15e (pfam00827)→ |
| | | | | | MA (smart00283)Tar (COG0840)PRK10935 (PRK10935)HAMP (pfam00672)→ |
| | | | | | ThiF_MoeB_HesA_family (cd00757)adenyl_thiF (TIGR02356)→ PRK04460 (PRK04460)ParD (COG3609)→ |
| | | | | | TIGR00697 (TIGR00697)Vut_1 (pfam02592)→ YhaK (COG1741)→ |
| env-145 | SL_5KL_010_BRINE_10088461 | + | 207 | 254 | **RNA**→ PRK04243 (PRK04243)Ribosomal_L15e (pfam00827)→ |
| env-146 | SL_6KL_011_BRINE_1001921 | - | 5758 | 5709 | **RNA**→ PRK04243 (PRK04243)Ribosomal_L15e (pfam00827)→ |
| | | | | | MA (smart00283)Tar (COG0840)HAMP (pfam00672)→ |
| | | | | | ThiF_MoeB_HesA_family (cd00757)adenyl_thiF (TIGR02356)→ PRK04460 (PRK04460)ParD (COG3609)→ |
| | | | | | TIGR00697 (TIGR00697)Vut_1 (pfam02592)→ YhaK (COG1741)→ |
| env-147 | SL_6KL_011_BRINE_1002375 | - | 67 | 20 | **RNA**→ |
| env-148 | SL_4KL_010_BRINE_10002475 | + | 10303 | 10350 | **RNA**→ RPL15A (COG1632)PRK04243 (PRK04243)→ |
| env-149 | SL_5KL_010_BRINE_10000147 | - | 15060 | 15012 | **RNA**→ PRK04243 (PRK04243)Ribosomal_L15e (pfam00827)→ |
| | | | | | MA (smart00283)Tar (COG0840)HAMP (pfam00672)→ |
| | | | | | ThiF_MoeB_HesA_family (cd00757)adenyl_thiF (TIGR02356)→ PRK04460 (PRK04460)ParD (COG3609)→ |
| | | | | | TIGR00697 (TIGR00697)Vut_1 (pfam02592)→ |
| env-150 | SL_6KL_011_BRINE_1000125 | + | 13899 | 13947 | **RNA**→ PRK04243 (PRK04243)Ribosomal_L15e (pfam00827)→ |
| | | | | | MA (smart00283)Tar (COG0840)HAMP (pfam00672)→ |
| | | | | | ThiF_MoeB_HesA_family (cd00757)adenyl_thiF (TIGR02356)→ PRK04460 (PRK04460)ParD (COG3609)→ |
| | | | | | TIGR00697 (TIGR00697)Vut_1 (pfam02592)→ |
| env-151 | SL_4KL_010_BRINE_10000557 | + | 10772 | 10820 | **RNA**→ PRK04243 (PRK04243)Ribosomal_L15e (pfam00827)→ |
| | | | | | MA (smart00283)Tar (COG0840)HAMP (pfam00672)→ |
| | | | | | ThiF_MoeB_HesA_family (cd00757)adenyl_thiF (TIGR02356)→ PRK04460 (PRK04460)ParD (COG3609)→ |
| | | | | | TIGR00697 (TIGR00697)Vut_1 (pfam02592)→ |
| env-152 | SL_4KL_010_BRINE_10273491 | - | 277 | 229 | **RNA**→ ←hypo |
| env-153 | P_A23_Liq_1_Fmt_1052201 | - | 109 | 60 | **RNA**→ hypo→ |
| env-154 | P_A23_Liq_3_Fmt_1050467 | + | 211 | 260 | **RNA**→ hypo→ |
| env-155 | P_A23_Liq_2_Fmt_1033762 | + | 198 | 247 | **RNA**→ PRK04243 (PRK04243)Ribosomal_L15e (pfam00827)→ |
| env-156 | APHM01021601.1 | - | 2022 | 1973 | **RNA**→ PRK04243 (PRK04243)Ribosomal_L15e (pfam00827)→ |
| env-157 | LTJ07As871_J07ABscf098869 | - | 39692 | 39643 | **RNA**→ PRK04243 (PRK04243)Ribosomal_L15e (pfam00827)→ |
| env-158 | JXWU01132649.1 | - | 361 | 312 | **RNA**→ RPL15A (COG1632)PRK04243 (PRK04243)→ |
| env-159 | SL_4KL_010_BRINE_10652442 | - | 116 | 67 | **RNA**→ |
| Hsp-10-1 | NZ_KN050825.1 | + | 204974 | 205055 | **RNA**→ PRK04243 (PRK04243)Ribosomal_L15e (pfam00827)→ |

| | | | | | |
|---|---|---|---|---|---|
| Hse-1-1 | NZ_BBMP01000007.1 | - | 359177 | 359090 | **RNA**→ PRK04243 (PRK04243)Ribosomal_L15e (pfam00827)→ |
| | | | | | DsbA_FrnE (cd03024)Thioredoxin_4 (pfam13462)→ |
| Hmo-1-1 | NZ_AOMC01000142.1 | - | 76777 | 76718 | **RNA**→ PRK04243 (PRK04243)Ribosomal_L15e (pfam00827)→ |
| | | | | | DsbA_FrnE (cd03024)Thioredoxin_4 (pfam13462)→ |
| Hth-1-1 | NZ_AOMF01000174.1 | - | 199637 | 199578 | **RNA**→ PRK04243 (PRK04243)Ribosomal_L15e (pfam00827)→ |
| Nae-1-1 | NZ_AOIP01000031.1 | + | 430123 | 430196 | **RNA**→ PRK04243 (PRK04243)Ribosomal_L15e (pfam00827)→ |
| Nta-1-1 | NZ_AOIL01000013.1 | + | 203216 | 203289 | **RNA**→ PRK04243 (PRK04243)Ribosomal_L15e (pfam00827)→ |
| Nas-1-1 | NZ_AOIO01000020.1 | - | 6198 | 6125 | **RNA**→ PRK04243 (PRK04243)Ribosomal_L15e (pfam00827)→ |
| env-160 | GSLNAS_F7I09SK02JROTM | - | 174 | 92 | **RNA**→ ←hypo |
| env-161 | GSLNARP_GFPJP1N02H548U | + | 202 | 278 | **RNA**→ RPL15A (COG1632)PRK04243 (PRK04243)→ |
| env-162 | SL_8KL_010_SED_10010026 | + | 6182 | 6238 | **RNA**→ RPL15A (COG1632)PRK04243 (PRK04243)→ |
| env-163 | SL_4KL_010_BRINE_10000926 | + | 6176 | 6232 | **RNA**→ PRK04243 (PRK04243)Ribosomal_L15e (pfam00827)→ |
| env-164 | SL_5KL_010_BRINE_10000029 | + | 1081 | 1137 | **RNA**→ PRK04243 (PRK04243)Ribosomal_L15e (pfam00827)→ |
| env-165 | SL_6KL_011_BRINE_1183487 | + | 36 | 92 | **RNA**→ RPL15A (COG1632)PRK04243 (PRK04243)→ |
| env-166 | SL_5KL_010_SED_10001187 | - | 8518 | 8462 | **RNA**→ PRK04243 (PRK04243)Ribosomal_L15e (pfam00827)→ |
| env-167 | SL_4KL_010_BRINE_10007163 | + | 5927 | 5983 | **RNA**→ PRK04243 (PRK04243)Ribosomal_L15e (pfam00827)→ |
| env-168 | SL_4KL_010_BRINE_10050687 | - | 1683 | 1627 | **RNA**→ PRK04243 (PRK04243)Ribosomal_L15e (pfam00827)→ |
| env-169 | SL_8KL_010_SED_10059739 | - | 1479 | 1423 | **RNA**→ PRK04243 (PRK04243)Ribosomal_L15e (pfam00827)→ |
| env-170 | SL_5KL_010_BRINE_10000179 | + | 3935 | 3995 | **RNA**→ PRK04243 (PRK04243)Ribosomal_L15e (pfam00827)→ |
| env-171 | SL_6KL_011_BRINE_1206958 | - | 208 | 148 | **RNA**→ RPL15A (COG1632)PRK04243 (PRK04243)→ |
| Hla-4-1 | NZ_CP007055.1 | - | 515819 | 515738 | **RNA**→ PRK04243 (PRK04243)Ribosomal_L15e (pfam00827)→ |
| Hsp-11-1 | NZ_JMIP02000020.1 | + | 43217 | 43284 | **RNA**→ PRK04243 (PRK04243)Ribosomal_L15e (pfam00827)→ |
| Hsp-3-1 | NC_002607.1 | - | 153435 | 153378 | **RNA**→ PRK04243 (PRK04243)Ribosomal_L15e (pfam00827)→ |
| Hsa-3-1 | NC_010364.1 | - | 153597 | 153540 | **RNA**→ PRK04243 (PRK04243)Ribosomal_L15e (pfam00827)→ |
| env-172 | SL_4KL_010_BRINE_10049878 | + | 1656 | 1711 | **RNA**→ RPL15A (COG1632)PRK04243 (PRK04243)→ |
| Hsa-1-1 | NZ_BBMO01000001.1 | - | 1407520 | 1407456 | **RNA**→ PRK04243 (PRK04243)Ribosomal_L15e (pfam00827)→ |
| env-173 | ADL20m3uS_GLR497S01AM9VW | - | 353 | 303 | **RNA**→ RPL15A (COG1632)PRK04243 (PRK04243)→ |
| env-174 | 2140918017-22757 | - | 213 | 163 | **RNA**→ PRK04243 (PRK04243)Ribosomal_L15e (pfam00827)→ |
| env-175 | GSLNAS_F7I09SK02IM9H7 | + | 19 | 70 | **RNA**→ PRK04243 (PRK04243)Ribosomal_L15e (pfam00827)→ |
| env-176 | P_1C_Sed_2_UnCty_1039923 | - | 750 | 700 | **RNA**→ PRK04243 (PRK04243)Ribosomal_L15e (pfam00827)→ |
| Hpa-1-1 | NZ_AEMG01000025.1 | - | 56166 | 56103 | **RNA**→ PRK04243 (PRK04243)Ribosomal_L15e (pfam00827)→ |
| | | | | | PRK04243 (PRK04243)Ribosomal_L15e (pfam00827)→ DUF1211 (pfam06736)→ DUF1211 (pfam06736)→ |
| Hpa-1-2 | NZ_AQXI01000001.1 | - | 2637650 | 2637587 | **RNA**→ PRK04243 (PRK04243)Ribosomal_L15e (pfam00827)→ DUF1211 (pfam06736)→ |
| env-177 | SL_4KL_010_BRINE_10364003 | - | 267 | 151 | **RNA**→ RPL15A (COG1632)PRK04243 (PRK04243)→ |
| env-178 | SL_4KL_010_BRINE_10015962 | - | 1055 | 974 | **RNA**→ PRK04243 (PRK04243)Ribosomal_L15e (pfam00827)→ |
| env-179 | SL_4KL_010_BRINE_10475926 | + | 13 | 62 | **RNA**→ RPL15A (COG1632)PRK04243 (PRK04243)→ |
| env-180 | SL_4KL_010_BRINE_10013350 | - | 4649 | 4598 | **RNA**→ PRK04243 (PRK04243)Ribosomal_L15e (pfam00827)→ |
| env-181 | SL_4KL_010_BRINE_10006798 | - | 2037 | 1873 | **RNA**→ PRK04243 (PRK04243)Ribosomal_L15e (pfam00827)→ |
| env-182 | SL_4KL_010_BRINE_10145615 | + | 98 | 148 | **RNA**→ PRK04243 (PRK04243)Ribosomal_L15e (pfam00827)→ |
| Ssp-1-1 | NC_021313.1 | + | 780000 | 780118 | **RNA**→ PRK04243 (PRK04243)Ribosomal_L15e (pfam00827)→ |
| env-183 | JXWU01070236.1 | - | 388 | 249 | **RNA**→ PRK04243 (PRK04243)Ribosomal_L15e (pfam00827)→ |
| Nph-1-1 | NC_007426.1 | - | 125427 | 125378 | **RNA**→ PRK04243 (PRK04243)Ribosomal_L15e (pfam00827)→ |
| env-184 | SL_4KL_010_BRINE_10509292 | + | 104 | 196 | **RNA**→ RPL15A (COG1632)PRK04243 (PRK04243)→ |
| Hgo-1-1 | NZ_HG315690.1 | - | 2398586 | 2398507 | **RNA**→ PRK04243 (PRK04243)Ribosomal_L15e (pfam00827)→ PRK15005 (PRK15005)USP_Like (cd00293)→ |
| | | | | | ars_reduc_gluta (TIGR02689)LMWPc (pfam01451)→ |
| Hxa-1-1 | NC_015666.1 | + | 1583029 | 1583107 | **RNA**→ PRK04243 (PRK04243)Ribosomal_L15e (pfam00827)→ |
| env-185 | SL_4KL_010_BRINE_10004067 | + | 1089 | 1146 | **RNA**→ PRK04243 (PRK04243)Ribosomal_L15e (pfam00827)→ PRK15005 (PRK15005)USP_Like (cd00293)→ |
| env-186 | SL_5KL_010_BRINE_10021750 | + | 302 | 359 | **RNA**→ PRK04243 (PRK04243)Ribosomal_L15e (pfam00827)→ PRK15005 (PRK15005)USP_Like (cd00293)→ |
| env-187 | SL_8KL_010_SED_10621895 | - | 164 | 104 | **RNA**→ RPL15A (COG1632)PRK04243 (PRK04243)→ |
| Nin-1-1 | NZ_AOHZ01000075.1 | - | 121628 | 121567 | **RNA**→ PRK04243 (PRK04243)Ribosomal_L15e (pfam00827)→ |
| env-188 | SL_5KL_010_BRINE_10005738 | - | 1188 | 1131 | **RNA**→ PRK04243 (PRK04243)Ribosomal_L15e (pfam00827)→ |
| env-189 | SL_6KL_011_BRINE_1053602 | + | 271 | 328 | **RNA**→ PRK04243 (PRK04243)Ribosomal_L15e (pfam00827)→ |
| env-190 | SL_8KL_010_SED_10028257 | - | 828 | 771 | **RNA**→ PRK04243 (PRK04243)Ribosomal_L15e (pfam00827)→ |

| | | | | | |
|---|---|---|---|---|---|
| env-191 | SL_4KL_010_BRINE_10066144 | - | 810 | 753 | **RNA**→ PRK04243 (PRK04243)Ribosomal_L15e (pfam00827)→ |
| env-192 | SS_3KL_010_SOIL_10028172 | - | 1924 | 1854 | **RNA**→ ←hypo |
| env-193 | SL_3KL_010_SED_10125154 | + | 137 | 207 | **RNA**→ PRK04243 (PRK04243)Ribosomal_L15e (pfam00827)→ |
| Nje-1-1 | NZ_AOIA01000114.1 | + | 146675 | 146744 | **RNA**→ PRK04243 (PRK04243)Ribosomal_L15e (pfam00827)→ |
| Nam-1-1 | NZ_AOIB01000026.1 | + | 119391 | 119460 | **RNA**→ PRK04243 (PRK04243)Ribosomal_L15e (pfam00827)→ |
| Noc-1-1 | NC_019974.1 | - | 1797311 | 1797242 | **RNA**→ PRK04243 (PRK04243)Ribosomal_L15e (pfam00827)→ |
| env-194 | GSLNARP_GFPJP1N02GQ822 | + | 449 | 495 | **RNA**→ |
| env-195 | SL_4KL_010_BRINE_10350018 | - | 232 | 148 | **RNA**→ RPL15A (COG1632)PRK04243 (PRK04243)→ |
| env-196 | SL_4KL_010_BRINE_10097165 | - | 841 | 785 | **RNA**→ PRK04243 (PRK04243)Ribosomal_L15e (pfam00827)→ |
| env-197 | SL_5KL_010_BRINE_10001159 | - | 6520 | 6463 | **RNA**→ PRK04243 (PRK04243)Ribosomal_L15e (pfam00827)→ |
| env-198 | SL_5KL_010_SED_10028357 | + | 365 | 422 | **RNA**→ PRK04243 (PRK04243)Ribosomal_L15e (pfam00827)→ |
| env-199 | SL_6KL_011_BRINE_1084724 | - | 464 | 407 | **RNA**→ RPL15A (COG1632)PRK04243 (PRK04243)→ |
| env-200 | SL_8KL_010_SED_10260913 | + | 164 | 221 | **RNA**→ RPL15A (COG1632)PRK04243 (PRK04243)→ |
| env-201 | SL_4KL_010_BRINE_10003303 | + | 1476 | 1533 | **RNA**→ PRK04243 (PRK04243)Ribosomal_L15e (pfam00827)→ |
| env-202 | GSLNAS_F7I09SK02F8ECV | - | 271 | 209 | **RNA**→ RPL15A (COG1632)PRK04243 (PRK04243)→ |
| Hla-1-1 | NZ_AGFZ01000059.1 | - | 22715 | 22654 | **RNA**→ hypo→ hypo→ PRK04243 (PRK04243)Ribosomal_L15e (pfam00827)→ PRK04243 (PRK04243)Ribosomal_L15e (pfam00827)→ |
| Hla-1-2 | NZ_AOLZ01000073.1 | - | 22804 | 22743 | **RNA**→ hypo→ PRK04243 (PRK04243)Ribosomal_L15e (pfam00827)→ |
| Hva-1-1 | NZ_AOLQ01000004.1 | - | 73870 | 73819 | **RNA**→ PRK04243 (PRK04243)Ribosomal_L15e (pfam00827)→ |
| Hsi-1-1 | NZ_AOLR01000005.1 | + | 181787 | 181840 | **RNA**→ PRK04243 (PRK04243)Ribosomal_L15e (pfam00827)→ |
| Hma-1-1 | NC_006396.1 | + | 1849657 | 1849710 | **RNA**→ PRK04243 (PRK04243)Ribosomal_L15e (pfam00827)→ |
| Ham-1-1 | NZ_AOLW01000010.1 | - | 93451 | 93398 | **RNA**→ PRK04243 (PRK04243)Ribosomal_L15e (pfam00827)→ |
| env-203 | P_A23_Liq_1_Fmt_1025305 | + | 182 | 235 | **RNA**→ PRK04243 (PRK04243)Ribosomal_L15e (pfam00827)→ |
| env-204 | P_A23_Liq_3_Fmt_1015036 | - | 204 | 151 | **RNA**→ PRK04243 (PRK04243)Ribosomal_L15e (pfam00827)→ |
| env-205 | P_A23_Liq_2_Fmt_1015236 | + | 488 | 541 | **RNA**→ PRK04243 (PRK04243)Ribosomal_L15e (pfam00827)→ |
| Hca-1-1 | NZ_AOLS01000038.1 | - | 57729 | 57676 | **RNA**→ PRK04243 (PRK04243)Ribosomal_L15e (pfam00827)→ |
| Hsa-2-1 | NZ_JXAO01000004.1 | + | 115750 | 115803 | **RNA**→ hypo→ |
| Hhi-2-1 | NC_023013.1 | + | 2490914 | 2490967 | **RNA**→ PRK04243 (PRK04243)Ribosomal_L15e (pfam00827)→ |
| Hhi-1-1 | NC_015948.1 | + | 2479481 | 2479534 | **RNA**→ PRK04243 (PRK04243)Ribosomal_L15e (pfam00827)→ |
| Hsp-1-1 | NZ_CP010529.1 | + | 1092738 | 1092791 | **RNA**→ PRK04243 (PRK04243)Ribosomal_L15e (pfam00827)→ |
| Hja-1-1 | NZ_AOLY01000007.1 | - | 129303 | 129250 | **RNA**→ PRK04243 (PRK04243)Ribosomal_L15e (pfam00827)→ |
| Har-1-1 | NZ_AOLX01000025.1 | - | 30531 | 30480 | **RNA**→ PRK04243 (PRK04243)Ribosomal_L15e (pfam00827)→ |
| env-206 | GSLNAS_F7I09SK02I6FWL | + | 298 | 351 | **RNA**→ PRK04243 (PRK04243)Ribosomal_L15e (pfam00827)→ |
| Hmu-2-1 | NC_013202.1 | - | 272589 | 272518 | **RNA**→ PRK04243 (PRK04243)Ribosomal_L15e (pfam00827)→ |
| Hka-1-1 | NZ_KB905378.1 | + | 614167 | 614238 | **RNA**→ PRK04243 (PRK04243)Ribosomal_L15e (pfam00827)→ |
| env-207 | SL_6KL_011_BRINE_1037538 | - | 695 | 635 | **RNA**→ ←hypo |
| env-208 | SL_6KL_011_BRINE_1012586 | - | 1128 | 1038 | **RNA**→ PRK04243 (PRK04243)Ribosomal_L15e (pfam00827)→ |
| env-209 | SL_4KL_010_BRINE_10018197 | + | 2238 | 2328 | **RNA**→ PRK04243 (PRK04243)Ribosomal_L15e (pfam00827)→ |
| env-210 | SL_6KL_011_BRINE_1090145 | + | 342 | 402 | **RNA**→ PRK04243 (PRK04243)Ribosomal_L15e (pfam00827)→ |
| env-211 | SL_4KL_010_BRINE_10055587 | - | 1221 | 1161 | **RNA**→ PRK04243 (PRK04243)Ribosomal_L15e (pfam00827)→ |
| env-212 | SL_6KL_011_BRINE_1018015 | - | 1651 | 1591 | **RNA**→ PRK04243 (PRK04243)Ribosomal_L15e (pfam00827)→ |
| env-213 | SL_4KL_010_BRINE_10006569 | + | 5039 | 5099 | **RNA**→ PRK04243 (PRK04243)Ribosomal_L15e (pfam00827)→ |
| env-214 | SL_6KL_011_BRINE_1094071 | - | 230 | 150 | **RNA**→ RPL15A (COG1632)PRK04243 (PRK04243)→ |
| env-215 | GSLSAAI_GFP0ZZ301EUXSI | + | 466 | 528 | **RNA**→ ←hypo |
| env-216 | SL_4KL_010_BRINE_10104860 | + | 909 | 969 | **RNA**→ RPL15A (COG1632)PRK04243 (PRK04243)→ |
| env-217 | GSLNARP_contig03416 | - | 112 | 53 | **RNA**→ |
| env-218 | GSLNAS_F7I09SK02HZKWC | - | 504 | 445 | **RNA**→ RPL15A (COG1632)PRK04243 (PRK04243)→ |
| env-219 | GSLNAS_F7I09SK02GL76P | - | 421 | 362 | **RNA**→ RPL15A (COG1632)PRK04243 (PRK04243)→ |
| env-220 | P_A23_Liq_1_Fmt_1037775 | + | 87 | 146 | **RNA**→ PRK04243 (PRK04243)Ribosomal_L15e (pfam00827)→ |
| Nmo-1-1 | NC_020388.1 | + | 780497 | 780546 | **RNA**→ PRK04243 (PRK04243)Ribosomal_L15e (pfam00827)→ |
| env-221 | APHM01007304.1 | + | 2549 | 2595 | **RNA**→ PRK04243 (PRK04243)Ribosomal_L15e (pfam00827)→ PRK08671 (PRK08671)MetAP2 (cd01088)→ |
| env-222 | LTJ07As871_J07ABscf089578 | - | 54536 | 54490 | **RNA**→ PRK04243 (PRK04243)Ribosomal_L15e (pfam00827)→ |

| | | | | | |
|---|---|---|---|---|---|
| env-223 | GSLNARP_contig28880 | + | 3300 | 3346 | **RNA**→ PRK04243 (PRK04243)Ribosomal_L15e (pfam00827)→ PRK08671 (PRK08671)APP_MetAP (cd01066 |
| env-224 | APHM01013436.1 | + | 115 | 161 | **RNA**→ PRK04243 (PRK04243)Ribosomal_L15e (pfam00827)→ |
| env-225 | LTJ07As871_J07ABscf098875 | + | 1708553 | 1708599 | **RNA**→ PRK04243 (PRK04243)Ribosomal_L15e (pfam00827)→ |
| env-226 | SL_4KL_010_BRINE_10204549 | + | 36 | 97 | **RNA**→ RPL15A (COG1632)PRK04243 (PRK04243)→ |
| env-227 | JXWU01037046.1 | - | 538 | 484 | **RNA**→ RPL15A (COG1632)PRK04243 (PRK04243)→ |
| env-228 | SS_3KL_010_SOIL_10003299 | + | 3588 | 3637 | **RNA**→ PRK04243 (PRK04243)Ribosomal_L15e (pfam00827)→ |
| env-229 | SS_3KL_010_SOIL_10003445 | + | 3378 | 3426 | **RNA**→ PRK04243 (PRK04243)Ribosomal_L15e (pfam00827)→ |
| Hci-1-1 | NZ_JDTH01000002.1 | - | 1455055 | 1454992 | **RNA**→ PRK04243 (PRK04243)Ribosomal_L15e (pfam00827)→ |
| Hut-1-1 | NC_013158.1 | - | 2429098 | 2429049 | **RNA**→ PRK04243 (PRK04243)Ribosomal_L15e (pfam00827)→ |
| env-230 | GSLNARP_GFPJP1N02FZ9S5 | + | 70 | 153 | **RNA**→ PRK04243 (PRK04243)Ribosomal_L15e (pfam00827)→ |
| env-231 | SL_8KL_010_SED_10278449 | - | 167 | 118 | **RNA**→ RPL15A (COG1632)PRK04243 (PRK04243)→ |
| env-232 | SL_4KL_010_BRINE_10001615 | - | 5266 | 5217 | **RNA**→ PRK04243 (PRK04243)Ribosomal_L15e (pfam00827)→ |
| env-233 | SL_6KL_011_BRINE_1000077 | - | 3468 | 3419 | **RNA**→ PRK04243 (PRK04243)Ribosomal_L15e (pfam00827)→ hypo→ hypo→ |
| env-234 | SL_6KL_011_BRINE_1009755 | - | 3271 | 3210 | **RNA**→ PRK04243 (PRK04243)Ribosomal_L15e (pfam00827)→ |
| env-235 | SL_8KL_010_SED_10678060 | - | 174 | 125 | **RNA**→ RPL15A (COG1632)PRK04243 (PRK04243)→ |
| env-236 | SL_4KL_010_BRINE_10654803 | + | 125 | 172 | **RNA**→ RPL15A (COG1632)PRK04243 (PRK04243)→ |
| Nba-1-1 | NZ_AOHY01000007.1 | + | 209062 | 209133 | **RNA**→ PRK04243 (PRK04243)Ribosomal_L15e (pfam00827)→ |
| env-237 | SL_3KL_010_SED_11113557 | - | 168 | 97 | **RNA**→ RPL15A (COG1632)PRK04243 (PRK04243)→ |
| Nsu-1-1 | NZ_AOHX01000026.1 | + | 190456 | 190532 | **RNA**→ PRK04243 (PRK04243)Ribosomal_L15e (pfam00827)→ |
| Npa-1-1 | NZ_AOII01000068.1 | + | 9874 | 9946 | **RNA**→ PRK04243 (PRK04243)Ribosomal_L15e (pfam00827)→ |
| Nal-2-1 | NZ_JNCS01000001.1 | + | 188067 | 188139 | **RNA**→ PRK04243 (PRK04243)Ribosomal_L15e (pfam00827)→ |
| Nal-1-1 | NZ_AOIK01000025.1 | - | 13704 | 13632 | **RNA**→ PRK04243 (PRK04243)Ribosomal_L15e (pfam00827)→ |
| Nga-1-1 | NZ_AOIJ01000043.1 | + | 26494 | 26566 | **RNA**→ PRK04243 (PRK04243)Ribosomal_L15e (pfam00827)→ |
| Nsp-2-1 | NZ_AJVG01000013.1 | + | 23678 | 23750 | **RNA**→ PRK04243 (PRK04243)Ribosomal_L15e (pfam00827)→ |
| Nsp-1-1 | NC_018224.1 | + | 1880458 | 1880530 | **RNA**→ PRK04243 (PRK04243)Ribosomal_L15e (pfam00827)→ |
| Hdj-1-1 | NZ_HG315684.1 | - | 683477 | 683408 | **RNA**→ PRK04243 (PRK04243)Ribosomal_L15e (pfam00827)→ |
| Hli-3-1 | NZ_AOIT01000029.1 | - | 228185 | 228113 | **RNA**→ PRK04243 (PRK04243)Ribosomal_L15e (pfam00827)→ |
| Hje-2-1 | NZ_JDTG01000014.1 | - | 528274 | 528203 | **RNA**→ PRK04243 (PRK04243)Ribosomal_L15e (pfam00827)→ |
| Hth-2-1 | NZ_AOIR01000063.1 | - | 74675 | 74604 | **RNA**→ PRK04243 (PRK04243)Ribosomal_L15e (pfam00827)→ |
| Hsa-10-1 | NZ_JROF01000002.1 | + | 114940 | 115012 | **RNA**→ PRK04243 (PRK04243)Ribosomal_L15e (pfam00827)→ |
| env-238 | SL_5KL_010_BRINE_10014771 | + | 1979 | 2040 | **RNA**→ PRK04243 (PRK04243)Ribosomal_L15e (pfam00827)→ |
| env-239 | SL_5KL_010_BRINE_10003441 | + | 1496 | 1557 | **RNA**→ PRK04243 (PRK04243)Ribosomal_L15e (pfam00827)→ |
| env-240 | SL_6KL_011_BRINE_1002462 | - | 7480 | 7419 | **RNA**→ PRK04243 (PRK04243)Ribosomal_L15e (pfam00827)→ |
| env-241 | SL_8KL_010_SED_10108063 | - | 214 | 153 | **RNA**→ PRK04243 (PRK04243)Ribosomal_L15e (pfam00827)→ |
| env-242 | SL_4KL_010_BRINE_10000366 | + | 1598 | 1659 | **RNA**→ PRK04243 (PRK04243)Ribosomal_L15e (pfam00827)→ |
| Hni-1-1 | NZ_AOMA01000119.1 | + | 3513 | 3588 | **RNA**→ PRK04243 (PRK04243)Ribosomal_L15e (pfam00827)→ |
| Nch-1-1 | NZ_AOIN01000080.1 | + | 5529 | 5592 | **RNA**→ PRK04243 (PRK04243)Ribosomal_L15e (pfam00827)→ |
| Nhu-1-1 | NZ_AOIM01000013.1 | + | 217184 | 217247 | **RNA**→ PRK04243 (PRK04243)Ribosomal_L15e (pfam00827)→ |
| Nve-1-1 | NZ_AOID01000026.1 | - | 208079 | 208007 | **RNA**→ PRK04243 (PRK04243)Ribosomal_L15e (pfam00827)→ |
| Hsa-8-1 | NZ_AOIS01000035.1 | + | 13267 | 13339 | **RNA**→ PRK04243 (PRK04243)Ribosomal_L15e (pfam00827)→ |
| Htu-1-1 | NC_013743.1 | - | 2869593 | 2869521 | **RNA**→ PRK04243 (PRK04243)Ribosomal_L15e (pfam00827)→ |
| env-243 | SL_8KL_010_SED_10556862 | - | 191 | 117 | **RNA**→ RPL15A (COG1632)PRK04243 (PRK04243)→ |
| Hca-3-1 | NZ_AOIU01000004.1 | + | 46196 | 46248 | **RNA**→ PRK04243 (PRK04243)Ribosomal_L15e (pfam00827)→ |
| env-244 | JXWS01111537.1 | - | 180 | 72 | **RNA**→ ←Form_Nir_trans (pfam01226) |
| env-245 | SL_4KL_010_BRINE_10017818 | - | 3866 | 3781 | **RNA**→ PRK04243 (PRK04243)Ribosomal_L15e (pfam00827)→ PRK15005 (PRK15005)USP_Like (cd00293)→ |
| env-246 | GSLNAS_F7I09SK02G2VWW | + | 218 | 297 | **RNA**→ PRK04243 (PRK04243)Ribosomal_L15e (pfam00827)→ |
| env-247 | SL_5KL_010_BRINE_10003121 | + | 300 | 385 | **RNA**→ PRK04243 (PRK04243)Ribosomal_L15e (pfam00827)→ |
| env-248 | SL_5KL_010_BRINE_10044475 | - | 707 | 612 | **RNA**→ PRK04243 (PRK04243)Ribosomal_L15e (pfam00827)→ |
| env-249 | SL_4KL_010_BRINE_10064659 | + | 812 | 918 | **RNA**→ RPL15A (COG1632)PRK04243 (PRK04243)→ RPL15A (COG1632)PRK04243 (PRK04243)→ |
| env-250 | GSLNARP_GFPJP1N02IRMJO | + | 320 | 372 | **RNA**→ PRK04243 (PRK04243)Ribosomal_L15e (pfam00827)→ |
| env-251 | APHM01020421.1 | + | 442 | 499 | **RNA**→ PRK04243 (PRK04243)Ribosomal_L15e (pfam00827)→ |
| env-252 | LTJ07As871_J07ABscf098852 | - | 1211893 | 1211836 | **RNA**→ PRK04243 (PRK04243)Ribosomal_L15e (pfam00827)→ |

| | | | | | |
|---|---|---|---|---|---|
| env-253 | GSLNARP_GFPJP1N02H78XW | + | 40 | 86 | **RNA**→ RPL15A (COG1632)PRK04243 (PRK04243)→ |
| Hsu-1-1 | NZ_CP008874.1 | - | 516965 | 516913 | **RNA**→ PRK04243 (PRK04243)Ribosomal_L15e (pfam00827)→ |
| Hsp-2-1 | NZ_CP007060.1 | - | 692145 | 692080 | **RNA**→ PRK04243 (PRK04243)Ribosomal_L15e (pfam00827)→ |
| Has-1-1 | NZ_AOIQ01000019.1 | + | 72263 | 72320 | **RNA**→ PRK04243 (PRK04243)Ribosomal_L15e (pfam00827)→ |
| env-254 | SS_3KL_010_SOIL_10043763 | - | 169 | 95 | **RNA**→ RPL15A (COG1632)PRK04243 (PRK04243)→ |
| Hag-1-1 | NZ_BAFM01000013.1 | + | 96977 | 97051 | **RNA**→ PRK04243 (PRK04243)Ribosomal_L15e (pfam00827)→ FrnE (COG2761)Thioredoxin_4 (pfam13462)– |
| Hsa-4-1 | NZ_AOMD01000030.1 | + | 286601 | 286667 | **RNA**→ PRK04243 (PRK04243)Ribosomal_L15e (pfam00827)→ hypo→ hypo→ |
| | | | | | FrnE (COG2761)Thioredoxin_4 (pfam13462)→ |
| Hsa-5-1 | NZ_AOME01000076.1 | + | 93813 | 93880 | **RNA**→ PRK04243 (PRK04243)Ribosomal_L15e (pfam00827)→ FrnE (COG2761)Thioredoxin_4 (pfam13462)– |
| env-255 | CERQ01166877.1 | - | 487 | 436 | **RNA**→ RPL15A (COG1632)PRK04243 (PRK04243)→ |
| env-256 | CETH01173410.1 | + | 12810 | 12860 | **RNA**→ PRK04243 (PRK04243)Ribosomal_L15e (pfam00827)→ PreA (COG1146)Fer4_7 (pfam12838)→ |
| env-257 | CETM01021463.1 | - | 36859 | 36809 | **RNA**→ PRK04243 (PRK04243)Ribosomal_L15e (pfam00827)→ PreA (COG1146)Fer4_7 (pfam12838)→ |
| env-258 | CEUQ01031760.1 | + | 6382 | 6432 | **RNA**→ PRK04243 (PRK04243)Ribosomal_L15e (pfam00827)→ PreA (COG1146)Fer4_7 (pfam12838)→ |
| env-259 | CEUS01376465.1 | + | 263 | 313 | **RNA**→ RPL15A (COG1632)PRK04243 (PRK04243)→ |
| env-260 | CEUY01138208.1 | - | 851 | 801 | **RNA**→ PRK04243 (PRK04243)Ribosomal_L15e (pfam00827)→ hypo→ |
| env-261 | CEUZ01171605.1 | + | 1156 | 1206 | **RNA**→ RPL15A (COG1632)PRK04243 (PRK04243)→ |
| env-262 | JCVI_SCAF_1096628214146 | - | 161 | 111 | **RNA**→ |
| env-263 | CENY01171985.1 | - | 10152 | 10102 | **RNA**→ PRK04243 (PRK04243)Ribosomal_L15e (pfam00827)→ PreA (COG1146)Fer4_13 (pfam13370)→ |
| env-264 | CEOQ01117861.1 | + | 720 | 770 | **RNA**→ RPL15A (COG1632)PRK04243 (PRK04243)→ |
| env-265 | CEUL01353856.1 | + | 7 | 57 | **RNA**→ PRK04243 (PRK04243)Ribosomal_L15e (pfam00827)→ PreA (COG1146)Fer4_7 (pfam12838)→ |
| env-266 | CENY01152858.1 | + | 14934 | 14985 | **RNA**→ PRK04243 (PRK04243)Ribosomal_L15e (pfam00827)→ PreA (COG1146)Fer4_9 (pfam13187)→ |
| Mae-1-1 | NC_009635.1 | - | 1562691 | 1562646 | **RNA**→ PRK04243 (PRK04243)Ribosomal_L15e (pfam00827)→ |
| env-267 | PBDCA2_FIDWTPW02QZKOZ | + | 178 | 232 | **RNA**→ hypo→ RPL15A (COG1632)PRK04243 (PRK04243)→ |
| env-268 | JGI24422J19971_10000787 | - | 11928 | 11884 | **RNA**→ PRK04243 (PRK04243)Ribosomal_L15e (pfam00827)→ |
| env-269 | JGI2172J19969_10313737 | - | 395 | 351 | **RNA**→ RPL15A (COG1632)PRK04243 (PRK04243)→ |
| env-270 | LGVF01069618.1 | - | 525 | 473 | **RNA**→ PRK04243 (PRK04243)Ribosomal_L15e (pfam00827)→ |
| env-271 | LGVE01065530.1 | - | 193 | 142 | **RNA**→ PRK04243 (PRK04243)Ribosomal_L15e (pfam00827)→ |
| env-272 | LGVF01108952.1 | + | 1406 | 1457 | **RNA**→ PRK04243 (PRK04243)Ribosomal_L15e (pfam00827)→ |
| env-273 | LGVF01076430.1 | - | 3041 | 2990 | **RNA**→ PRK04243 (PRK04243)Ribosomal_L15e (pfam00827)→ |
| env-274 | JGI24721J26819_10009006 | + | 6561 | 6611 | **RNA**→ PRK04243 (PRK04243)Ribosomal_L15e (pfam00827)→ PRK12277 (PRK12277)RPL13 (COG4352)→ |
| | | | | | PRK14555 (PRK14555)RNA_binding (pfam01877)→ RPP1 (COG1603)PRK00912 (PRK00912)→ |
| | | | | | PRK03717 (PRK03717)RNase_P_Rpp14 (pfam01900)→ |
| env-275 | JGI24721J44947_10030901 | - | 78 | 28 | **RNA**→ |
| env-276 | JGI24721J44947_10028419 | + | 536 | 586 | **RNA**→ PRK04243 (PRK04243)Ribosomal_L15e (pfam00827)→ PRK12277 (PRK12277)RPL13 (COG4352)→ |
| | | | | | PRK14555 (PRK14555)RNA_binding (pfam01877)→ RPP1 (COG1603)PRK00912 (PRK00912)→ |
| | | | | | PRK03717 (PRK03717)RNase_P_Rpp14 (pfam01900)→ |
| env-277 | JGI24721J44947_10022851 | + | 2001 | 2047 | **RNA**→ PRK04243 (PRK04243)Ribosomal_L15e (pfam00827)→ PRK12277 (PRK12277)RPL13 (COG4352)→ |
| | | | | | PRK14555 (PRK14555)RNA_binding (pfam01877)→ hypo→ RPP1 (COG1603)PRK00912 (PRK00912)→ |
| | | | | | PRK03717 (PRK03717)RNase_P_Rpp14 (pfam01900)→ |
| env-278 | JGI20130J15003_1001487 | + | 10993 | 11039 | **RNA**→ PRK04243 (PRK04243)Ribosomal_L15e (pfam00827)→ PRK12277 (PRK12277)RPL13 (COG4352)→ |
| | | | | | PRK14555 (PRK14555)RNA_binding (pfam01877)→ RPP1 (COG1603)PRK00912 (PRK00912)→ |
| | | | | | PRK03717 (PRK03717)RNase_P_Rpp14 (pfam01900)→ |
| env-279 | JGI20130J15133_1001487 | + | 10993 | 11039 | **RNA**→ PRK04243 (PRK04243)Ribosomal_L15e (pfam00827)→ PRK12277 (PRK12277)RPL13 (COG4352)→ |
| | | | | | PRK14555 (PRK14555)RNA_binding (pfam01877)→ RPP1 (COG1603)PRK00912 (PRK00912)→ |
| | | | | | PRK03717 (PRK03717)RNase_P_Rpp14 (pfam01900)→ |
| env-280 | JGI24721J26819_10027382 | - | 1728 | 1682 | **RNA**→ PRK04243 (PRK04243)Ribosomal_L15e (pfam00827)→ PRK12277 (PRK12277)RPL13 (COG4352)→ |
| | | | | | PRK14555 (PRK14555)RNA_binding (pfam01877)→ PRK00912 (PRK00912)RNase_P_p30 (pfam01876)→ |
| env-281 | JGI2172J19969_10022280 | + | 1906 | 1952 | **RNA**→ RPL15A (COG1632)PRK04243 (PRK04243)→ |
| env-282 | SMTZ1_10021852 | + | 5904 | 5950 | **RNA**→ RPL15A (COG1632)PRK04243 (PRK04243)→ |
| env-283 | JGI20214J14112_1187723 | + | 274 | 322 | **RNA**→ |
| env-284 | WSSedA2C_c189965 | + | 176 | 224 | **RNA**→ RPL15A (COG1632)PRK04243 (PRK04243)→ |
| env-285 | WSSedA2Ca3_1189965 | + | 176 | 224 | **RNA**→ RPL15A (COG1632)PRK04243 (PRK04243)→ |
| env-286 | WSSedB1B2_c027137 | - | 242 | 194 | **RNA**→ PRK04243 (PRK04243)Ribosomal_L15e (pfam00827)→ |
| env-287 | JGI24721J26819_10037939 | + | 1348 | 1398 | **RNA**→ PRK04243 (PRK04243)Ribosomal_L15e (pfam00827)→ PRK12277 (PRK12277)RPL13 (COG4352)→ |

| | | | | | |
|---|---|---|---|---|---|
| env-288 | JGI24721J44947_10000270 | - | 47013 | 46963 | **RNA**→ PRK04243 (PRK04243)Ribosomal_L15e (pfam00827)→ PRK12277 (PRK12277)RPL13 (COG4352)→ |
| env-289 | JGI2160J19893_10105365 | - | 280 | 230 | **RNA**→ RPL15A (COG1632)PRK04243 (PRK04243)→ |
| env-290 | JGI2171J19970_10224488 | + | 281 | 331 | **RNA**→ RPL15A (COG1632)PRK04243 (PRK04243)→ |
| env-291 | JGI2173J19968_10055779 | + | 876 | 926 | **RNA**→ RPL15A (COG1632)PRK04243 (PRK04243)→ |
| env-292 | SMTZ1_10000331 | - | 33118 | 33068 | **RNA**→ PRK04243 (PRK04243)Ribosomal_L15e (pfam00827)→ PRK12277 (PRK12277)RPL13 (COG4352)→ PRK14555 (PRK14555)RNA_binding (pfam01877)→ RPP1 (COG1603)PRK00912 (PRK00912)→ PRK03717 (PRK03717)RNase_P_Rpp14 (pfam01900)→ |
| env-293 | JGI2171J19970_10312620 | + | 332 | 382 | **RNA**→ RPL15A (COG1632)PRK04243 (PRK04243)→ |
| env-294 | SMTZ1_10014562 | - | 4684 | 4634 | **RNA**→ PRK04243 (PRK04243)Ribosomal_L15e (pfam00827)→ PRK12277 (PRK12277)RPL13 (COG4352)→ PRK14555 (PRK14555)RNA_binding (pfam01877)→ RPP1 (COG1603)PRK00912 (PRK00912)→ PRK03717 (PRK03717)RNase_P_Rpp14 (pfam01900)→ |
| env-295 | WSSedA1TJGI_c051973 | + | 58 | 106 | **RNA**→ RPL15A (COG1632)PRK04243 (PRK04243)→ |
| env-296 | WSSedA1Ba3_1047821 | - | 762 | 714 | **RNA**→ PRK04243 (PRK04243)Ribosomal_L15e (pfam00827)→ hypo→ |
| env-297 | JGI20171J14444_1002001 | + | 1726 | 1771 | **RNA**→ PRK04243 (PRK04243)Ribosomal_L15e (pfam00827)→ |
| env-298 | JGI20165J15291_1047466 | - | 228 | 183 | **RNA**→ PRK04243 (PRK04243)Ribosomal_L15e (pfam00827)→ |
| env-299 | JGI20164J15296_1001389 | + | 654 | 699 | **RNA**→ PRK04243 (PRK04243)Ribosomal_L15e (pfam00827)→ |
| env-300 | JGI20163J15578_10000609 | - | 5418 | 5373 | **RNA**→ PRK04243 (PRK04243)Ribosomal_L15e (pfam00827)→ |
| env-301 | JGI20164J15296_1023034 | - | 338 | 292 | **RNA**→ RPL15A (COG1632)PRK04243 (PRK04243)→ |
| env-302 | JGI20164J15296_1001061 | + | 1474 | 1520 | **RNA**→ PRK04243 (PRK04243)Ribosomal_L15e (pfam00827)→ |
| env-303 | JGI20164J15296_1003383 | - | 130 | 84 | **RNA**→ |
| env-304 | WSSedA2C_c120722 | - | 217 | 169 | **RNA**→ PRK04243 (PRK04243)Ribosomal_L15e (pfam00827)→ |
| env-305 | WSSedA1Ba2_1101229 | - | 82 | 34 | **RNA**→ hypo→ |
| env-306 | WSSedA1Ba3_1039983 | - | 339 | 291 | **RNA**→ RPL15A (COG1632)PRK04243 (PRK04243)→ |
| env-307 | WSSedA2Ca3_1120722 | - | 217 | 169 | **RNA**→ PRK04243 (PRK04243)Ribosomal_L15e (pfam00827)→ |
| env-308 | JREQ01002797.1 | + | 594 | 639 | **RNA**→ RPL15A (COG1632)PRK04243 (PRK04243)→ |
| env-309 | JREQ01001068.1 | + | 594 | 639 | **RNA**→ RPL15A (COG1632)PRK04243 (PRK04243)→ RPL15A (COG1632)PRK04243 (PRK04243)→ |
| env-310 | JREQ01004635.1 | - | 265 | 221 | **RNA**→ PRK04243 (PRK04243)Ribosomal_L15e (pfam00827)→ |
| env-311 | JGI24422J19971_10442158 | + | 33 | 79 | **RNA**→ RPL15A (COG1632)PRK04243 (PRK04243)→ |
| env-312 | JGI2172J19969_10165153 | + | 155 | 200 | **RNA**→ RPL15A (COG1632)PRK04243 (PRK04243)→ |
| env-313 | JGI2173J19968_10019714 | - | 204 | 159 | **RNA**→ PRK04243 (PRK04243)Ribosomal_L15e (pfam00827)→ |
| env-314 | JGI2171J19970_10593475 | + | 159 | 204 | **RNA**→ PRK04243 (PRK04243)Ribosomal_L15e (pfam00827)→ |
| env-315 | JGI24422J19971_10429142 | - | 262 | 217 | **RNA**→ PRK04243 (PRK04243)Ribosomal_L15e (pfam00827)→ |
| env-316 | SMTZ1_10013484 | - | 9454 | 9409 | **RNA**→ PRK04243 (PRK04243)Ribosomal_L15e (pfam00827)→ PRK14555 (PRK14555)RNA_binding (pfam01877)→ RPP1 (COG1603)PRK00912 (PRK00912)→ PRK03717 (PRK03717)RNase_P_Rpp14 (pfam01900)→ |
| env-317 | JGI2172J19969_10374243 | - | 221 | 175 | **RNA**→ PRK04243 (PRK04243)Ribosomal_L15e (pfam00827)→ |
| env-318 | JGI2173J19968_10119450 | - | 690 | 644 | **RNA**→ hypo→ PRK04243 (PRK04243)Ribosomal_L15e (pfam00827)→ |
| env-319 | JGI2171J19970_10117997 | - | 822 | 776 | **RNA**→ hypo→ PRK04243 (PRK04243)Ribosomal_L15e (pfam00827)→ RNA_binding (pfam01877)→ |
| env-320 | SMTZ1_10010538 | - | 5263 | 5217 | **RNA**→ PRK04243 (PRK04243)Ribosomal_L15e (pfam00827)→ PRK14555 (PRK14555)RNA_binding (pfam01877)→ RPP1 (COG1603)PRK00912 (PRK00912)→ PRK03717 (PRK03717)RNase_P_Rpp14 (pfam01900)→ |
| env-321 | TB_LI09_3_1376570 | + | 4 | 50 | **RNA**→ PRK04243 (PRK04243)Ribosomal_L15e (pfam00827)→ |
| env-322 | LGVD01087542.1 | - | 817 | 771 | **RNA**→ RPL15A (COG1632)PRK04243 (PRK04243)→ RPL15A (COG1632)PRK04243 (PRK04243)→ RNA_binding (pfam01877)→ |
| env-323 | JGI24024J18818_10296176 | - | 207 | 161 | **RNA**→ hypo→ |
| env-324 | SMTZ1_10000031 | + | 24101 | 24147 | **RNA**→ PRK04243 (PRK04243)Ribosomal_L15e (pfam00827)→ PRK14555 (PRK14555)RNA_binding (pfam01877)→ PRK00912 (PRK00912)RNase_P_p30 (pfam01876)→ PRK03717 (PRK03717)RNase_P_Rpp14 (pfam01900)→ |
| env-325 | JGI2173J19968_10468069 | - | 221 | 175 | **RNA**→ PRK04243 (PRK04243)Ribosomal_L15e (pfam00827)→ |
| env-326 | LGVF01282102.1 | - | 355 | 309 | **RNA**→ RPL15A (COG1632)PRK04243 (PRK04243)→ |
| env-327 | LGVE01222292.1 | - | 389 | 343 | **RNA**→ RPL15A (COG1632)PRK04243 (PRK04243)→ |
| env-328 | LGOV01067113.1 | + | 3689 | 3735 | **RNA**→ PRK04243 (PRK04243)Ribosomal_L15e (pfam00827)→ PRK14555 (PRK14555)RNA_binding (pfam01877)→ PRK00912 (PRK00912)RNase_P_p30 (pfam01876)→ POP5 (COG1369)→ |

| | | | | | |
|---|---|---|---|---|---|
| Pab-1-1 | NC_000868.1 | + | 818641 | 818684 | **RNA**→ PRK04243 (PRK04243)Ribosomal_L15e (pfam00827)→ hydrog_prot (TIGR00072)H2MP_MemB-H2evol (cd06067)→ |
| Psp-2-1 | NC_017946.1 | - | 888644 | 888600 | **RNA**→ PRK04243 (PRK04243)Ribosomal_L15e (pfam00827)→ |
| Tga-1-1 | NC_012804.1 | + | 1735891 | 1735934 | **RNA**→ PRK04243 (PRK04243)Ribosomal_L15e (pfam00827)→ |
| Tsp-3-1 | NC_018015.1 | - | 513605 | 513561 | **RNA**→ PRK04243 (PRK04243)Ribosomal_L15e (pfam00827)→ |
| Tsp-2-1 | NC_016051.1 | - | 1259118 | 1259075 | **RNA**→ PRK04243 (PRK04243)Ribosomal_L15e (pfam00827)→ |
| Tna-1-1 | NZ_CP007264.1 | + | 1944164 | 1944207 | **RNA**→ PRK04243 (PRK04243)Ribosomal_L15e (pfam00827)→ |
| Teu-1-1 | NZ_CP008887.1 | + | 1002663 | 1002706 | **RNA**→ PRK04243 (PRK04243)Ribosomal_L15e (pfam00827)→ |
| env-329 | 3300001592_11448690 | + | 60 | 103 | **RNA**→ PRK04243 (PRK04243)Ribosomal_L15e (pfam00827)→ |
| Tsp-4-1 | NZ_KE387149.1 | + | 630216 | 630259 | **RNA**→ PRK04243 (PRK04243)Ribosomal_L15e (pfam00827)→ |
| Tsi-1-1 | NC_012883.1 | + | 1077704 | 1077747 | **RNA**→ PRK04243 (PRK04243)Ribosomal_L15e (pfam00827)→ |
| Tli-1-1 | NC_022084.1 | + | 51943 | 51986 | **RNA**→ PRK04243 (PRK04243)Ribosomal_L15e (pfam00827)→ |
| env-330 | JGI24019J35510_1000020 | - | 119249 | 119205 | **RNA**→ PRK04243 (PRK04243)Ribosomal_L15e (pfam00827)→ |
| Tba-1-1 | NC_014804.1 | + | 122047 | 122091 | **RNA**→ PRK04243 (PRK04243)Ribosomal_L15e (pfam00827)→ |
| Tpa-1-1 | NZ_CP006965.1 | + | 229409 | 229452 | **RNA**→ PRK04243 (PRK04243)Ribosomal_L15e (pfam00827)→ |
| Ppa-1-1 | NZ_CP006019.1 | + | 356037 | 356081 | **RNA**→ PRK04243 (PRK04243)Ribosomal_L15e (pfam00827)→ COG1325 (COG1325)PRK14555 (PRK14555) PRK03892 (PRK03892)RPP1 (COG1603)→ PRK03717 (PRK03717)POP5 (COG1369)→ TIGR00341 (TIGR00341)DUF389 (pfam04087)→ |
| Pfu-2-1 | NC_003413.1 | - | 849639 | 849596 | **RNA**→ PRK04243 (PRK04243)Ribosomal_L15e (pfam00827)→ |
| Pfu-1-1 | NC_018092.1 | - | 651734 | 651691 | **RNA**→ PRK04243 (PRK04243)Ribosomal_L15e (pfam00827)→ |
| Pho-1-1 | NC_000961.1 | + | 891553 | 891596 | **RNA**→ PRK04243 (PRK04243)Ribosomal_L15e (pfam00827)→ PRK04243 (PRK04243)Ribosomal_L15e (pfam00827)→ hydrog_prot (TIGR00072)H2MP_MemB-H2evol (cd06067)→ |
| Psp-1-1 | NC_015474.1 | + | 1545307 | 1545351 | **RNA**→ PRK04243 (PRK04243)Ribosomal_L15e (pfam00827)→ hydrog_prot (TIGR00072)H2MP_MemB-H2evol (cd06067)→ |
| Pya-1-1 | NC_015680.1 | + | 224726 | 224769 | **RNA**→ PRK04243 (PRK04243)Ribosomal_L15e (pfam00827)→ |
| Tko-1-1 | NC_006624.1 | - | 1280545 | 1280502 | **RNA**→ PRK04243 (PRK04243)Ribosomal_L15e (pfam00827)→ |
| Tsp-1-1 | NC_015865.1 | - | 869933 | 869890 | **RNA**→ PRK04243 (PRK04243)Ribosomal_L15e (pfam00827)→ |
| Pfe-1-1 | NZ_LANF01000010.1 | - | 104355 | 104312 | **RNA**→ PRK04243 (PRK04243)Ribosomal_L15e (pfam00827)→ |
| Ton-1-1 | NC_011529.1 | + | 120097 | 120140 | **RNA**→ PRK04243 (PRK04243)Ribosomal_L15e (pfam00827)→ COG1325 (COG1325)PRK14555 (PRK14555) PRK03892 (PRK03892)RPP1 (COG1603)→ |
| Tzi-1-1 | NZ_AJLF01000001.1 | - | 197475 | 197432 | **RNA**→ PRK04243 (PRK04243)Ribosomal_L15e (pfam00827)→ PRK04243 (PRK04243)Ribosomal_L15e (pfam00827)→ |
| env-331 | JGI24721J44947_10092924 | + | 379 | 432 | **RNA**→ PRK04243 (PRK04243)Ribosomal_L15e (pfam00827)→ PRK12277 (PRK12277)RPL13 (COG4352)→ PRK14555 (PRK14555)RNA_binding (pfam01877)→ PRK00912 (PRK00912)RNase_P_p30 (pfam01876)→ |
| env-332 | JGI24721J26819_10019109 | + | 3371 | 3424 | **RNA**→ PRK04243 (PRK04243)Ribosomal_L15e (pfam00827)→ PRK12277 (PRK12277)RPL13 (COG4352)→ PRK14555 (PRK14555)RNA_binding (pfam01877)→ PRK00912 (PRK00912)RNase_P_p30 (pfam01876)→ |
| env-333 | JGI24422J19971_10125801 | - | 282 | 234 | **RNA**→ RPL15A (COG1632)PRK04243 (PRK04243)→ |
| env-334 | JGI24721J26819_10947299 | + | 173 | 218 | **RNA**→ |
| env-335 | JGI24721J44947_10473589 | - | 321 | 276 | **RNA**→ PRK04243 (PRK04243)Ribosomal_L15e (pfam00827)→ |
| env-336 | JGI24721J26819_10001491 | + | 21482 | 21529 | **RNA**→ PRK04243 (PRK04243)Ribosomal_L15e (pfam00827)→ PRK12277 (PRK12277)RPL13 (COG4352)→ PRK14555 (PRK14555)RNA_binding (pfam01877)→ RPP1 (COG1603)PRK00912 (PRK00912)→ PRK03717 (PRK03717)RNase_P_Rpp14 (pfam01900)→ |
| env-337 | JGI24721J44947_10032959 | - | 4920 | 4873 | **RNA**→ PRK04243 (PRK04243)Ribosomal_L15e (pfam00827)→ PRK12277 (PRK12277)RPL13 (COG4352)→ PRK14555 (PRK14555)RNA_binding (pfam01877)→ RPP1 (COG1603)PRK00912 (PRK00912)→ PRK03717 (PRK03717)RNase_P_Rpp14 (pfam01900)→ |
| env-338 | TB_LI09_3_1044718 | + | 801 | 846 | **RNA**→ PRK04243 (PRK04243)Ribosomal_L15e (pfam00827)→ |
| env-339 | JGI24721J26819_10004799 | - | 3984 | 3939 | **RNA**→ PRK04243 (PRK04243)Ribosomal_L15e (pfam00827)→ PRK12277 (PRK12277)RPL13 (COG4352)→ PRK14555 (PRK14555)RNA_binding (pfam01877)→ RPP1 (COG1603)PRK00912 (PRK00912)→ PRK03717 (PRK03717)RNase_P_Rpp14 (pfam01900)→ |
| env-340 | JGI24721J44947_10004827 | - | 9632 | 9587 | **RNA**→ PRK04243 (PRK04243)Ribosomal_L15e (pfam00827)→ PRK12277 (PRK12277)RPL13 (COG4352)→ PRK14555 (PRK14555)RNA_binding (pfam01877)→ RPP1 (COG1603)PRK00912 (PRK00912)→ PRK03717 (PRK03717)RNase_P_Rpp14 (pfam01900)→ |
| env-341 | JGI20130J15003_1091000 | + | 354 | 398 | **RNA**→ PRK04243 (PRK04243)Ribosomal_L15e (pfam00827)→ |
| env-342 | JGI20130J15133_1091000 | + | 354 | 398 | **RNA**→ PRK04243 (PRK04243)Ribosomal_L15e (pfam00827)→ |

| | | | | | |
|---|---|---|---|---|---|
| env-343 | JGI24721J44947_10014353 | - | 2950 | 2906 | **RNA**→ PRK04243 (PRK04243)Ribosomal_L15e (pfam00827)→ PRK12277 (PRK12277)RPL13 (COG4352)→ PRK14555 (PRK14555)RNA_binding (pfam01877)→ RPP1 (COG1603)PRK00912 (PRK00912)→ PRK03717 (PRK03717)RNase_P_Rpp14 (pfam01900)→ |
| env-344 | JGI20130J15003_1005034 | + | 5585 | 5630 | **RNA**→ PRK04243 (PRK04243)Ribosomal_L15e (pfam00827)→ PRK12277 (PRK12277)RPL13 (COG4352)→ PRK14555 (PRK14555)RNA_binding (pfam01877)→ PRK00912 (PRK00912)RNase_P_p30 (pfam01876)→ RPP1 (COG1603)PRK00912 (PRK00912)→ PRK03717 (PRK03717)RNase_P_Rpp14 (pfam01900)→ |
| env-345 | JGI20130J15133_1005034 | + | 5585 | 5630 | **RNA**→ PRK04243 (PRK04243)Ribosomal_L15e (pfam00827)→ PRK12277 (PRK12277)RPL13 (COG4352)→ PRK14555 (PRK14555)RNA_binding (pfam01877)→ PRK00912 (PRK00912)RNase_P_p30 (pfam01876)→ RPP1 (COG1603)PRK00912 (PRK00912)→ PRK03717 (PRK03717)RNase_P_Rpp14 (pfam01900)→ |
| env-346 | JGI24721J26819_10047889 | - | 1496 | 1451 | **RNA**→ PRK04243 (PRK04243)Ribosomal_L15e (pfam00827)→ PRK12277 (PRK12277)RPL13 (COG4352)→ PRK14555 (PRK14555)RNA_binding (pfam01877)→ Acyl-CoA_dh_N (pfam02771)→ |
| env-347 | JGI24721J44947_10203480 | + | 445 | 490 | **RNA**→ PRK04243 (PRK04243)Ribosomal_L15e (pfam00827)→ |
| env-348 | JREQ01006527.1 | - | 305 | 258 | **RNA**→ PRK04243 (PRK04243)Ribosomal_L15e (pfam00827)→ |
| env-349 | JGI2172J19969_10000042 | - | 4265 | 4219 | **RNA**→ RPL15A (COG1632)PRK04243 (PRK04243)→ PRK12277 (PRK12277)RPL13 (COG4352)→ PRK14555 (PRK14555)RNA_binding (pfam01877)→ RPP1 (COG1603)PRK00912 (PRK00912)→ PRK03717 (PRK03717)RNase_P_Rpp14 (pfam01900)→ |
| env-350 | JGI2173J19968_10000091 | + | 26877 | 26923 | **RNA**→ RPL15A (COG1632)PRK04243 (PRK04243)→ PRK12277 (PRK12277)RPL13 (COG4352)→ PRK14555 (PRK14555)RNA_binding (pfam01877)→ PRK00912 (PRK00912)RNase_P_p30 (pfam01876)→ |
| env-351 | JGI2171J19970_10013243 | - | 1906 | 1860 | **RNA**→ RPL15A (COG1632)PRK04243 (PRK04243)→ PRK12277 (PRK12277)RPL13 (COG4352)→ PRK14555 (PRK14555)RNA_binding (pfam01877)→ PRK00912 (PRK00912)RNase_P_p30 (pfam01876)→ |
| env-352 | JGI24422J19971_10240577 | + | 294 | 340 | **RNA**→ RPL15A (COG1632)PRK04243 (PRK04243)→ |
| env-353 | JGI24422J19971_10004830 | + | 6226 | 6272 | **RNA**→ PRK04243 (PRK04243)Ribosomal_L15e (pfam00827)→ PRK12277 (PRK12277)RPL13 (COG4352)→ PRK14555 (PRK14555)RNA_binding (pfam01877)→ PRK00912 (PRK00912)RNase_P_p30 (pfam01876)→ PRK03717 (PRK03717)POP5 (COG1369)→ |
| env-354 | JGI2172J19969_10181609 | - | 256 | 210 | **RNA**→ PRK04243 (PRK04243)Ribosomal_L15e (pfam00827)→ |
| env-355 | JGI24422J19971_10009670 | - | 3736 | 3690 | **RNA**→ PRK04243 (PRK04243)Ribosomal_L15e (pfam00827)→ PRK12277 (PRK12277)RPL13 (COG4352)→ PRK14555 (PRK14555)RNA_binding (pfam01877)→ RPP1 (COG1603)PRK00912 (PRK00912)→ PRK03717 (PRK03717)RNase_P_Rpp14 (pfam01900)→ |
| env-356 | JGI2172J19969_10028492 | + | 394 | 440 | **RNA**→ PRK04243 (PRK04243)Ribosomal_L15e (pfam00827)→ PRK12277 (PRK12277)RPL13 (COG4352)→ PRK14555 (PRK14555)RNA_binding (pfam01877)→ PRK00912 (PRK00912)→ |
| env-357 | JGI24422J19971_10492128 | - | 124 | 78 | **RNA**→ ←hypo |
| env-358 | JGI24422J19971_10371442 | + | 38 | 84 | **RNA**→ RPL15A (COG1632)PRK04243 (PRK04243)→ |
| env-359 | JGI24019J35510_1000149 | + | 57719 | 57764 | **RNA**→ PRK04243 (PRK04243)Ribosomal_L15e (pfam00827)→ PRK12277 (PRK12277)RPL13 (COG4352)→ PRK14555 (PRK14555)RNA_binding (pfam01877)→ PRK00912 (PRK00912)RNase_P_p30 (pfam01876)→ |
| env-360 | JGI24020J35080_1000786 | + | 20256 | 20301 | **RNA**→ |
| env-361 | JGI24020J35080_1002222 | - | 8791 | 8746 | **RNA**→ PRK04243 (PRK04243)Ribosomal_L15e (pfam00827)→ PRK12277 (PRK12277)RPL13 (COG4352)→ PRK14555 (PRK14555)RNA_binding (pfam01877)→ PRK00912 (PRK00912)RNase_P_p30 (pfam01876)→ PRK03717 (PRK03717)RNase_P_Rpp14 (pfam01900)→ |
| env-362 | JGI24422J19971_10000253 | - | 1868 | 1822 | **RNA**→ PRK04243 (PRK04243)Ribosomal_L15e (pfam00827)→ PRK12277 (PRK12277)RPL13 (COG4352)→ PRK14555 (PRK14555)RNA_binding (pfam01877)→ PRK00912 (PRK00912)RNase_P_p30 (pfam01876)→ |
| env-363 | JGI26339J46600_10000361 | + | 3851 | 3897 | **RNA**→ PRK04243 (PRK04243)Ribosomal_L15e (pfam00827)→ PRK12277 (PRK12277)RPL13 (COG4352)→ PRK14555 (PRK14555)RNA_binding (pfam01877)→ RPP1 (COG1603)PRK00912 (PRK00912)→ PRK03717 (PRK03717)RNase_P_Rpp14 (pfam01900)→ |
| env-364 | JGI24025J20009_10035112 | - | 1739 | 1692 | **RNA**→ PRK04243 (PRK04243)Ribosomal_L15e (pfam00827)→ |
| env-365 | JGI24422J19971_10447265 | - | 312 | 266 | **RNA**→ RPL15A (COG1632)PRK04243 (PRK04243)→ |
| env-366 | JGI24422J19971_10005604 | - | 2467 | 2422 | **RNA**→ hypo→ PRK04243 (PRK04243)Ribosomal_L15e (pfam00827)→ PRK12277 (PRK12277)RPL13 (COG4352)→ PRK14555 (PRK14555)RNA_binding (pfam01877)→ RPP1 (COG1603)PRK00912 (PRK00912)→ PRK03717 (PRK03717)RNase_P_Rpp14 (pfam01900)→ |
| env-367 | JGI2171J19970_10311510 | + | 423 | 468 | **RNA**→ hypo→ |
| env-368 | JGI24422J19971_10296037 | - | 379 | 334 | **RNA**→ hypo→ RPL15A (COG1632)PRK04243 (PRK04243)→ |
| env-369 | JGI24422J19971_10083467 | + | 1365 | 1411 | **RNA**→ RPL15A (COG1632)PRK04243 (PRK04243)→ |
| env-370 | JGI24422J19971_10004103 | - | 1823 | 1778 | **RNA**→ hypo→ PRK04243 (PRK04243)Ribosomal_L15e (pfam00827)→ PRK12277 (PRK12277)RPL13 (COG4352)→ PRK14555 (PRK14555)RNA_binding (pfam01877)→ PRK00912 (PRK00912)RNase_P_p30 (pfam01876)→ |
| env-371 | JGI2172J19969_10042350 | + | 1081 | 1126 | **RNA**→ hypo→ RPL15A (COG1632)PRK04243 (PRK04243)→ |
| env-372 | JGI2173J19968_10455679 | - | 124 | 79 | **RNA**→ hypo→ |

| | | | | | |
|---|---|---|---|---|---|
| env-373 | JGI2171J19970_10583661 | + | 269 | 314 | **RNA**→ hypo→ |
| env-374 | SMTZ1_10006093 | + | 9757 | 9802 | **RNA**→ hypo→ PRK04243 (PRK04243)Ribosomal_L15e (pfam00827)→ PRK12277 (PRK12277)RPL13 (COG4352)→ PRK14555 (PRK14555)RNA_binding (pfam01877)→ PRK00912 (PRK00912)RNase_P_p30 (pfam01876)→ PRK03717 (PRK03717)RNase_P_Rpp14 (pfam01900)→ |
| env-375 | JGI24025J20009_10133169 | - | 100 | 54 | **RNA**→ ←hypo |
| env-376 | JGI24422J19971_10019282 | - | 678 | 632 | **RNA**→ PRK04243 (PRK04243)Ribosomal_L15e (pfam00827)→ |
| env-377 | JGI24025J20009_10005934 | - | 75 | 29 | **RNA**→ |
| env-378 | JGI24025J20009_10025887 | + | 2637 | 2683 | **RNA**→ |
| env-379 | JGI24025J20009_10011070 | + | 4857 | 4903 | **RNA**→ |
| env-380 | JGI24025J20009_10053632 | + | 1424 | 1470 | **RNA**→ ←hypo |
| env-381 | JGI24422J19971_10774031 | - | 188 | 142 | **RNA**→ PRK04243 (PRK04243)Ribosomal_L15e (pfam00827)→ |
| env-382 | JGI2171J19970_10721007 | - | 71 | 25 | **RNA**→ |
| env-383 | JGI24025J20009_10067810 | - | 1101 | 1055 | **RNA**→ PRK04243 (PRK04243)Ribosomal_L15e (pfam00827)→ PRK12277 (PRK12277)RPL13 (COG4352)→ COG1325 (COG1325)PRK14555 (PRK14555)→ |
| env-384 | JGI24025J20009_10468419 | - | 121 | 75 | **RNA**→ |
| env-385 | JGI2172J19969_10020584 | - | 1314 | 1266 | **RNA**→ hypo→ PRK04243 (PRK04243)Ribosomal_L15e (pfam00827)→ PRK12277 (PRK12277)RPL13 (COG4352)→ PRK14555 (PRK14555)RNA_binding (pfam01877)→ |
| env-386 | JGI24422J19971_10090327 | - | 1116 | 1068 | **RNA**→ PRK04243 (PRK04243)Ribosomal_L15e (pfam00827)→ PRK12277 (PRK12277)RPL13 (COG4352)→ COG1325 (COG1325)PRK14555 (PRK14555)→ |
| env-387 | JGI24422J19971_10132992 | - | 55 | 7 | **RNA**→ hypo→ |
| env-388 | JGI24721J26819_10042702 | + | 773 | 820 | **RNA**→ PRK04243 (PRK04243)Ribosomal_L15e (pfam00827)→ PRK12277 (PRK12277)RPL13 (COG4352)→ PRK14555 (PRK14555)RNA_binding (pfam01877)→ RPP1 (COG1603)PRK00912 (PRK00912)→ |
| env-389 | JGI24721J44947_10035793 | + | 3282 | 3329 | **RNA**→ PRK04243 (PRK04243)Ribosomal_L15e (pfam00827)→ PRK12277 (PRK12277)RPL13 (COG4352)→ PRK14555 (PRK14555)RNA_binding (pfam01877)→ |
| env-390 | JGI24721J26819_10782576 | - | 189 | 142 | **RNA**→ PRK04243 (PRK04243)Ribosomal_L15e (pfam00827)→ |
| env-391 | JGI24721J44947_11017790 | - | 189 | 142 | **RNA**→ PRK04243 (PRK04243)Ribosomal_L15e (pfam00827)→ |
| env-392 | GBSCES77__Contig_41123 | - | 1948 | 1904 | **RNA**→ PRK04243 (PRK04243)Ribosomal_L15e (pfam00827)→ PRK12277 (PRK12277)RPL13 (COG4352)→ PRK14555 (PRK14555)RNA_binding (pfam01877)→ RPP1 (COG1603)PRK00912 (PRK00912)→ |
| env-393 | GBSCES77__Contig_16200 | - | 2430 | 2386 | **RNA**→ PRK04243 (PRK04243)Ribosomal_L15e (pfam00827)→ PRK12277 (PRK12277)RPL13 (COG4352)→ PRK14555 (PRK14555)RNA_binding (pfam01877)→ RPP1 (COG1603)PRK00912 (PRK00912)→ PRK03717 (PRK03717)RNase_P_Rpp14 (pfam01900)→ |
| env-394 | GBSCES77__Contig_3030 | + | 791 | 835 | **RNA**→ PRK04243 (PRK04243)Ribosomal_L15e (pfam00827)→ PRK12277 (PRK12277)RPL13 (COG4352)→ PRK14555 (PRK14555)RNA_binding (pfam01877)→ RPP1 (COG1603)PRK00912 (PRK00912)→ PRK03717 (PRK03717)RNase_P_Rpp14 (pfam01900)→ |
| env-395 | GBSCES77__Contig_34313 | + | 671 | 715 | **RNA**→ PRK04243 (PRK04243)Ribosomal_L15e (pfam00827)→ PRK12277 (PRK12277)RPL13 (COG4352)→ PRK14555 (PRK14555)RNA_binding (pfam01877)→ RPP1 (COG1603)PRK00912 (PRK00912)→ PRK03717 (PRK03717)RNase_P_Rpp14 (pfam01900)→ |
| env-396 | GBSCES77__Contig_24520 | - | 2097 | 2053 | **RNA**→ PRK04243 (PRK04243)Ribosomal_L15e (pfam00827)→ PRK12277 (PRK12277)RPL13 (COG4352)→ PRK14555 (PRK14555)RNA_binding (pfam01877)→ RPP1 (COG1603)PRK00912 (PRK00912)→ |
| env-397 | GBSCES77__Contig_45658 | + | 559 | 603 | **RNA**→ PRK04243 (PRK04243)Ribosomal_L15e (pfam00827)→ PRK12277 (PRK12277)RPL13 (COG4352)→ PRK14555 (PRK14555)RNA_binding (pfam01877)→ RPP1 (COG1603)PRK00912 (PRK00912)→ PRK03717 (PRK03717)RNase_P_Rpp14 (pfam01900)→ |
| env-398 | GBSCES77__Contig_11719 | - | 2430 | 2386 | **RNA**→ PRK04243 (PRK04243)Ribosomal_L15e (pfam00827)→ PRK12277 (PRK12277)RPL13 (COG4352)→ PRK14555 (PRK14555)RNA_binding (pfam01877)→ RPP1 (COG1603)PRK00912 (PRK00912)→ PRK03717 (PRK03717)RNase_P_Rpp14 (pfam01900)→ |
| env-399 | GBSCES77__Contig_6779 | + | 671 | 715 | **RNA**→ PRK04243 (PRK04243)Ribosomal_L15e (pfam00827)→ PRK12277 (PRK12277)RPL13 (COG4352)→ PRK14555 (PRK14555)RNA_binding (pfam01877)→ RPP1 (COG1603)PRK00912 (PRK00912)→ |
| env-400 | GBSCES77__Contig_20802 | - | 2430 | 2386 | **RNA**→ PRK04243 (PRK04243)Ribosomal_L15e (pfam00827)→ PRK12277 (PRK12277)RPL13 (COG4352)→ PRK14555 (PRK14555)RNA_binding (pfam01877)→ RPP1 (COG1603)PRK00912 (PRK00912)→ PRK03717 (PRK03717)RNase_P_Rpp14 (pfam01900)→ |
| env-401 | GBSCES77__Contig_29831 | - | 2430 | 2386 | **RNA**→ PRK04243 (PRK04243)Ribosomal_L15e (pfam00827)→ PRK12277 (PRK12277)RPL13 (COG4352)→ PRK14555 (PRK14555)RNA_binding (pfam01877)→ RPP1 (COG1603)PRK00912 (PRK00912)→ PRK03717 (PRK03717)RNase_P_Rpp14 (pfam01900)→ |
| env-402 | JGI20130J15003_1003712 | - | 4413 | 4369 | **RNA**→ PRK04243 (PRK04243)Ribosomal_L15e (pfam00827)→ PRK12277 (PRK12277)RPL13 (COG4352)→ PRK14555 (PRK14555)RNA_binding (pfam01877)→ RPP1 (COG1603)PRK00912 (PRK00912)→ PRK03717 (PRK03717)RNase_P_Rpp14 (pfam01900)→ |

| | | | | | |
|---|---|---|---|---|---|
| env-403 | JGI20130J15133_1003712 | - | 4413 | 4369 | **RNA**→ PRK04243 (PRK04243)Ribosomal_L15e (pfam00827)→ PRK12277 (PRK12277)RPL13 (COG4352)→ PRK14555 (PRK14555)RNA_binding (pfam01877)→ RPP1 (COG1603)PRK00912 (PRK00912)→ PRK03717 (PRK03717)RNase_P_Rpp14 (pfam01900)→ |
| env-404 | JGI24721J26819_10021412 | - | 955 | 911 | **RNA**→ PRK04243 (PRK04243)Ribosomal_L15e (pfam00827)→ PRK12277 (PRK12277)RPL13 (COG4352)→ |
| env-405 | JGI24721J44947_10036157 | + | 3704 | 3748 | **RNA**→ PRK04243 (PRK04243)Ribosomal_L15e (pfam00827)→ PRK12277 (PRK12277)RPL13 (COG4352)→ |
| env-406 | JGI20130J15003_1001502 | + | 15316 | 15361 | **RNA**→ PRK04243 (PRK04243)Ribosomal_L15e (pfam00827)→ PRK12277 (PRK12277)RPL13 (COG4352)→ PRK14555 (PRK14555)RNA_binding (pfam01877)→ RPP1 (COG1603)PRK00912 (PRK00912)→ PRK03717 (PRK03717)RNase_P_Rpp14 (pfam01900)→ |
| env-407 | JGI20130J15133_1001502 | + | 15316 | 15361 | **RNA**→ PRK04243 (PRK04243)Ribosomal_L15e (pfam00827)→ PRK12277 (PRK12277)RPL13 (COG4352)→ PRK14555 (PRK14555)RNA_binding (pfam01877)→ RPP1 (COG1603)PRK00912 (PRK00912)→ PRK03717 (PRK03717)RNase_P_Rpp14 (pfam01900)→ |
| env-408 | JGI24721J26819_10470799 | + | 270 | 315 | **RNA**→ |
| env-409 | JGI24721J44947_10040429 | + | 409 | 454 | **RNA**→ PRK04243 (PRK04243)Ribosomal_L15e (pfam00827)→ PRK12277 (PRK12277)RPL13 (COG4352)→ PRK14555 (PRK14555)RNA_binding (pfam01877)→ RPP1 (COG1603)PRK00912 (PRK00912)→ PRK03717 (PRK03717)RNase_P_Rpp14 (pfam01900)→ |
| env-410 | JGI20130J15003_1014452 | - | 2679 | 2634 | **RNA**→ PRK04243 (PRK04243)Ribosomal_L15e (pfam00827)→ PRK12277 (PRK12277)RPL13 (COG4352)→ PRK14555 (PRK14555)RNA_binding (pfam01877)→ PRK00912 (PRK00912)RNase_P_p30 (pfam01876)→ PRK03717 (PRK03717)RNase_P_Rpp14 (pfam01900)→ |
| env-411 | JGI20130J15133_1014452 | - | 2679 | 2634 | **RNA**→ PRK04243 (PRK04243)Ribosomal_L15e (pfam00827)→ PRK12277 (PRK12277)RPL13 (COG4352)→ PRK14555 (PRK14555)RNA_binding (pfam01877)→ PRK00912 (PRK00912)RNase_P_p30 (pfam01876)→ PRK03717 (PRK03717)RNase_P_Rpp14 (pfam01900)→ |
| env-412 | JGI24721J26819_10066975 | - | 1900 | 1855 | **RNA**→ PRK04243 (PRK04243)Ribosomal_L15e (pfam00827)→ PRK12277 (PRK12277)RPL13 (COG4352)→ PRK14555 (PRK14555)RNA_binding (pfam01877)→ PRK00912 (PRK00912)RNase_P_p30 (pfam01876)→ |
| env-413 | JGI24721J44947_10053920 | - | 3312 | 3267 | **RNA**→ RPL15A (COG1632)PRK04243 (PRK04243)→ RPL15A (COG1632)PRK04243 (PRK04243)→ PRK12277 (PRK12277)RPL13 (COG4352)→ PRK14555 (PRK14555)RNA_binding (pfam01877)→ PRK00912 (PRK00912)RNase_P_p30 (pfam01876)→ PRK03717 (PRK03717)RNase_P_Rpp14 (pfam01900)→ |
| env-414 | JGI24721J26819_10007452 | + | 1045 | 1090 | **RNA**→ PRK04243 (PRK04243)Ribosomal_L15e (pfam00827)→ PRK12277 (PRK12277)RPL13 (COG4352)→ PRK14555 (PRK14555)RNA_binding (pfam01877)→ RPP1 (COG1603)PRK00912 (PRK00912)→ PRK03717 (PRK03717)RNase_P_Rpp14 (pfam01900)→ |
| env-415 | JGI24721J44947_10012689 | + | 1045 | 1091 | **RNA**→ PRK04243 (PRK04243)Ribosomal_L15e (pfam00827)→ PRK12277 (PRK12277)RPL13 (COG4352)→ PRK14555 (PRK14555)RNA_binding (pfam01877)→ RPP1 (COG1603)PRK00912 (PRK00912)→ PRK03717 (PRK03717)RNase_P_Rpp14 (pfam01900)→ |
| env-416 | JGI24721J26819_10000566 | + | 1047 | 1092 | **RNA**→ PRK04243 (PRK04243)Ribosomal_L15e (pfam00827)→ PRK12277 (PRK12277)RPL13 (COG4352)→ PRK14555 (PRK14555)RNA_binding (pfam01877)→ RPP1 (COG1603)PRK00912 (PRK00912)→ PRK03717 (PRK03717)RNase_P_Rpp14 (pfam01900)→ |
| env-417 | JGI24721J44947_10000929 | - | 39531 | 39486 | **RNA**→ PRK04243 (PRK04243)Ribosomal_L15e (pfam00827)→ PRK12277 (PRK12277)RPL13 (COG4352)→ PRK14555 (PRK14555)RNA_binding (pfam01877)→ RPP1 (COG1603)PRK00912 (PRK00912)→ PRK03717 (PRK03717)RNase_P_Rpp14 (pfam01900)→ |
| env-418 | JGI20130J15003_1040014 | + | 320 | 364 | **RNA**→ PRK04243 (PRK04243)Ribosomal_L15e (pfam00827)→ PRK12277 (PRK12277)RPL13 (COG4352)→ |
| env-419 | JGI20130J15133_1040014 | + | 320 | 364 | **RNA**→ PRK04243 (PRK04243)Ribosomal_L15e (pfam00827)→ PRK12277 (PRK12277)RPL13 (COG4352)→ |
| env-420 | JGI24721J26819_10008710 | - | 5175 | 5131 | **RNA**→ PRK04243 (PRK04243)Ribosomal_L15e (pfam00827)→ PRK12277 (PRK12277)RPL13 (COG4352)→ PRK14555 (PRK14555)RNA_binding (pfam01877)→ RPP1 (COG1603)PRK00912 (PRK00912)→ PRK03717 (PRK03717)RNase_P_Rpp14 (pfam01900)→ |
| env-421 | JGI24721J44947_10014925 | - | 5175 | 5131 | **RNA**→ PRK04243 (PRK04243)Ribosomal_L15e (pfam00827)→ PRK12277 (PRK12277)RPL13 (COG4352)→ PRK14555 (PRK14555)RNA_binding (pfam01877)→ RPP1 (COG1603)PRK00912 (PRK00912)→ PRK03717 (PRK03717)RNase_P_Rpp14 (pfam01900)→ |
| env-422 | JGI20130J15003_1006708 | + | 320 | 364 | **RNA**→ PRK04243 (PRK04243)Ribosomal_L15e (pfam00827)→ PRK12277 (PRK12277)RPL13 (COG4352)→ PRK14555 (PRK14555)RNA_binding (pfam01877)→ RPP1 (COG1603)PRK00912 (PRK00912)→ PRK03717 (PRK03717)RNase_P_Rpp14 (pfam01900)→ |
| env-423 | JGI20130J15133_1006708 | + | 320 | 364 | **RNA**→ PRK04243 (PRK04243)Ribosomal_L15e (pfam00827)→ PRK12277 (PRK12277)RPL13 (COG4352)→ PRK14555 (PRK14555)RNA_binding (pfam01877)→ RPP1 (COG1603)PRK00912 (PRK00912)→ PRK03717 (PRK03717)RNase_P_Rpp14 (pfam01900)→ |
| env-424 | JGI24422J19971_10000261 | - | 3582 | 3531 | **RNA**→ PRK04243 (PRK04243)Ribosomal_L15e (pfam00827)→ PRK12277 (PRK12277)RPL13 (COG4352)→ PRK14555 (PRK14555)RNA_binding (pfam01877)→ RPP1 (COG1603)PRK00912 (PRK00912)→ PRK03717 (PRK03717)RNase_P_Rpp14 (pfam01900)→ |
| env-425 | JGI2173J19968_10016998 | + | 1834 | 1885 | **RNA**→ PRK04243 (PRK04243)Ribosomal_L15e (pfam00827)→ PRK12277 (PRK12277)RPL13 (COG4352)→ |

| | | | | | |
|---|---|---|---|---|---|
| env-426 | SMTZ1_10000482 | + | 8104 | 8155 | RNA→ PRK04243 (PRK04243)Ribosomal_L15e (pfam00827)→ PRK12277 (PRK12277)RPL13 (COG4352)→ PRK14555 (PRK14555)RNA_binding (pfam01877)→ RPP1 (COG1603)PRK00912 (PRK00912)→ PRK03717 (PRK03717)RNase_P_Rpp14 (pfam01900)→ |
| env-427 | JGI2171J19970_10271144 | - | 353 | 302 | RNA→ RPL15A (COG1632)PRK04243 (PRK04243)→ |
| env-428 | SMTZ1_10032630 | - | 6487 | 6436 | RNA→ ←hypo |
| env-429 | JGI2172J19969_10033071 | + | 406 | 457 | RNA→ ←hypo |
| env-430 | JGI2171J19970_10083154 | - | 519 | 468 | RNA→ ←hypo |
| env-431 | JGI24422J19971_10005257 | + | 6290 | 6341 | RNA→ PRK04243 (PRK04243)Ribosomal_L15e (pfam00827)→ PRK12277 (PRK12277)RPL13 (COG4352)→ PRK14555 (PRK14555)RNA_binding (pfam01877)→ PRK00912 (PRK00912)RNase_P_p30 (pfam01876)→ |
| env-432 | JGI24422J19971_10022574 | + | 1413 | 1464 | RNA→ PRK04243 (PRK04243)Ribosomal_L15e (pfam00827)→ PRK12277 (PRK12277)Ribosomal_L13e (pfam01294)→ PRK14555 (PRK14555)RNA_binding (pfam01877)→ RPP1 (COG1603)PRK00912 (PRK00912)→ PRK03717 (PRK03717)RNase_P_Rpp14 (pfam01900)→ |
| env-433 | JGI2172J19969_10002202 | - | 186 | 135 | RNA→ PRK04243 (PRK04243)Ribosomal_L15e (pfam00827)→ |
| env-434 | JGI2172J19969_10005325 | + | 99 | 150 | RNA→ PRK04243 (PRK04243)Ribosomal_L15e (pfam00827)→ PRK12277 (PRK12277)RPL13 (COG4352)→ PRK14555 (PRK14555)RNA_binding (pfam01877)→ RPP1 (COG1603)PRK00912 (PRK00912)→ PRK03717 (PRK03717)RNase_P_Rpp14 (pfam01900)→ |
| env-435 | JGI2173J19968_10000262 | + | 3449 | 3500 | RNA→ PRK04243 (PRK04243)Ribosomal_L15e (pfam00827)→ PRK12277 (PRK12277)RPL13 (COG4352)→ PRK14555 (PRK14555)RNA_binding (pfam01877)→ RPP1 (COG1603)PRK00912 (PRK00912)→ PRK03717 (PRK03717)RNase_P_Rpp14 (pfam01900)→ |
| env-436 | JGI2171J19970_10002036 | + | 2023 | 2074 | RNA→ PRK04243 (PRK04243)Ribosomal_L15e (pfam00827)→ PRK12277 (PRK12277)RPL13 (COG4352)→ PRK14555 (PRK14555)RNA_binding (pfam01877)→ RPP1 (COG1603)PRK00912 (PRK00912)→ PRK03717 (PRK03717)RNase_P_Rpp14 (pfam01900)→ |
| env-437 | JGI24422J19971_10243708 | + | 627 | 678 | RNA→ |
| env-438 | JGI24721J44947_10007540 | - | 2578 | 2525 | RNA→ PRK04243 (PRK04243)Ribosomal_L15e (pfam00827)→ PRK12277 (PRK12277)RPL13 (COG4352)→ PRK14555 (PRK14555)RNA_binding (pfam01877)→ RPP1 (COG1603)PRK00912 (PRK00912)→ PRK03717 (PRK03717)RNase_P_Rpp14 (pfam01900)→ |
| env-439 | JGI24721J26819_10005385 | + | 10350 | 10403 | RNA→ PRK04243 (PRK04243)Ribosomal_L15e (pfam00827)→ PRK12277 (PRK12277)RPL13 (COG4352)→ PRK14555 (PRK14555)RNA_binding (pfam01877)→ RPP1 (COG1603)PRK00912 (PRK00912)→ PRK03717 (PRK03717)RNase_P_Rpp14 (pfam01900)→ |
| env-440 | JGI2172J19969_10330441 | + | 133 | 180 | RNA→ RPL15A (COG1632)PRK04243 (PRK04243)→ |
| env-441 | JGI24422J19971_10230678 | - | 541 | 494 | RNA→ PRK04243 (PRK04243)Ribosomal_L15e (pfam00827)→ |
| env-442 | JGI24422J19971_10330695 | - | 518 | 473 | RNA→ PRK04243 (PRK04243)Ribosomal_L15e (pfam00827)→ |
| env-443 | JGI2171J19970_10787211 | - | 272 | 227 | RNA→ RPL15A (COG1632)PRK04243 (PRK04243)→ |
| env-444 | JGI2172J19969_10018448 | + | 977 | 1022 | RNA→ PRK04243 (PRK04243)Ribosomal_L15e (pfam00827)→ |
| env-445 | SMTZ1_10005895 | - | 12232 | 12187 | RNA→ hypo→ PRK04243 (PRK04243)Ribosomal_L15e (pfam00827)→ |
| env-446 | JGI2171J19970_10523375 | - | 233 | 185 | RNA→ PRK04243 (PRK04243)Ribosomal_L15e (pfam00827)→ |
| env-447 | JGI24721J26819_10011755 | + | 847 | 896 | RNA→ PRK04243 (PRK04243)Ribosomal_L15e (pfam00827)→ PRK12277 (PRK12277)RPL13 (COG4352)→ PRK14555 (PRK14555)RNA_binding (pfam01877)→ RPP1 (COG1603)PRK00912 (PRK00912)→ PRK03717 (PRK03717)RNase_P_Rpp14 (pfam01900)→ |
| env-448 | JGI24721J44947_10000482 | - | 52170 | 52121 | RNA→ PRK04243 (PRK04243)Ribosomal_L15e (pfam00827)→ PRK12277 (PRK12277)RPL13 (COG4352)→ PRK14555 (PRK14555)RNA_binding (pfam01877)→ RPP1 (COG1603)PRK00912 (PRK00912)→ PRK03717 (PRK03717)RNase_P_Rpp14 (pfam01900)→ |
| env-449 | JGI2173J19968_10000733 | + | 4783 | 4829 | RNA→ PRK04243 (PRK04243)Ribosomal_L15e (pfam00827)→ |
| env-450 | JGI2172J19969_10000762 | - | 8526 | 8480 | RNA→ PRK04243 (PRK04243)Ribosomal_L15e (pfam00827)→ |
| env-451 | JGI2171J19970_10187156 | - | 317 | 271 | RNA→ RPL15A (COG1632)PRK04243 (PRK04243)→ |
| env-452 | JGI24422J19971_10000606 | + | 10914 | 10960 | RNA→ PRK04243 (PRK04243)Ribosomal_L15e (pfam00827)→ |
| env-453 | NCBI_PBSM_READ_68102390 | + | 1020 | 1063 | RNA→ |
| Mvo-1-1 | NC_014222.1 | - | 1392579 | 1392536 | RNA→ PRK04243 (PRK04243)Ribosomal_L15e (pfam00827)→ |
| env-454 | SakLake120mDRAFT_c0000339 | + | 19224 | 19273 | RNA→ RPL15A (COG1632)PRK04243 (PRK04243)→ RPP1 (COG1603)PRK00912 (PRK00912)→ RNase_P_Rpp14 (pfam01900)→ PRK03996 (PRK03996)proteasome_alpha_archeal (cd03756)→ Sdo1 (COG1500)PRK13760 (PRK13760)→ rpl37ae (PRK03976)Ribosomal_L37ae (pfam01780)→ |

| | | | | | |
|---|---|---|---|---|---|
| ume-1-1 | NC_009464.1 | - | 994905 | 994839 | **RNA**→ PRK04243 (PRK04243)Ribosomal_L15e (pfam00827)→ COG1325 (COG1325)PRK14555 (PRK14555)<br>PRK00912 (PRK00912)RNase_P_p30 (pfam01876)→ PRK03717 (PRK03717)RNase_P_Rpp14 (pfam01900)→<br>PRK03996 (PRK03996)proteasome_alpha_archeal (cd03756)→ Sdo1 (COG1500)PRK13760 (PRK13760)→<br>Rrp4 (COG1097)PRK04163 (PRK04163)→ RNase_PH_archRRP41 (cd11366)PRK03983 (PRK03983)→<br>RNase_PH_archRRP42 (cd11365)PRK04282 (PRK04282)→ rpl37ae (PRK03976)Ribosomal_L37ae (pfam01780)<br>RPC10 (COG1996)rpoP (PRK00398)→ Pcc1 (COG2892)Pcc1 (pfam09341)→<br>gimC_beta (TIGR02338)Prefoldin_2 (pfam01920)→ hypo→ |
| CMe-2-1 | NZ_JMIY01000003.1 | + | 337389 | 337437 | **RNA**→ PRK04243 (PRK04243)Ribosomal_L15e (pfam00827)→<br>PRK14555 (PRK14555)RNA_binding (pfam01877)→ |
| Mev-1-1 | NC_014253.1 | - | 592639 | 592588 | **RNA**→ PRK04243 (PRK04243)Ribosomal_L15e (pfam00827)→<br>PRK14555 (PRK14555)RNA_binding (pfam01877)→ PRK00912 (PRK00912)RNase_P_p30 (pfam01876)→<br>PRK03717 (PRK03717)POP5 (COG1369)→ PRK03996 (PRK03996)proteasome_alpha_archeal (cd03756)→<br>Sdo1 (COG1500)PRK13760 (PRK13760)→ hypo→ hypo→ hypo→ |
| env-455 | WSSedA2C_c207490 | + | 72 | 126 | **RNA**→ PRK04243 (PRK04243)Ribosomal_L15e (pfam00827)→ |
| env-456 | WSSedA2Ca3_1207490 | + | 72 | 126 | **RNA**→ PRK04243 (PRK04243)Ribosomal_L15e (pfam00827)→ |
| env-457 | WSSedA1TJGI_c024695 | + | 624 | 678 | **RNA**→ PRK04243 (PRK04243)Ribosomal_L15e (pfam00827)→ |
| env-458 | WSSedA1Ba2_1000622 | + | 1087 | 1141 | **RNA**→ PRK04243 (PRK04243)Ribosomal_L15e (pfam00827)→ RPP1 (COG1603)PRK00912 (PRK00912)→<br>PRK03717 (PRK03717)RNase_P_Rpp14 (pfam01900)→<br>PRK03996 (PRK03996)proteasome_alpha_archeal (cd03756)→ Sdo1 (COG1500)PRK13760 (PRK13760)→<br>rpl37ae (PRK03976)Ribosomal_L37ae (pfam01780)→ |
| env-459 | WSSedA1Ba3_1003811 | + | 2343 | 2397 | **RNA**→ PRK04243 (PRK04243)Ribosomal_L15e (pfam00827)→ RPP1 (COG1603)PRK00912 (PRK00912)→<br>PRK03717 (PRK03717)RNase_P_Rpp14 (pfam01900)→ |
| env-460 | JGI26437J51864_10646690 | + | 117 | 172 | **RNA**→ |
| env-461 | WSSedB1B2_c010830 | + | 452 | 507 | **RNA**→ RPL15A (COG1632)PRK04243 (PRK04243)→ |
| env-462 | JGI20214J14112_1483376 | + | 5 | 54 | **RNA**→ PRK04243 (PRK04243)Ribosomal_L15e (pfam00827)→ |
| Mzh-1-1 | NC_015676.1 | - | 1281814 | 1281760 | **RNA**→ PRK04243 (PRK04243)Ribosomal_L15e (pfam00827)→<br>PRK14555 (PRK14555)RNA_binding (pfam01877)→ RPP1 (COG1603)PRK00912 (PRK00912)→<br>PRK03717 (PRK03717)RNase_P_Rpp14 (pfam01900)→<br>PRK03996 (PRK03996)proteasome_alpha_archeal (cd03756)→ Sdo1 (COG1500)PRK13760 (PRK13760)→<br>Rrp4 (COG1097)PRK04163 (PRK04163)→ RNase_PH_archRRP41 (cd11366)PRK03983 (PRK03983)→<br>RNase_PH_archRRP42 (cd11365)PRK04282 (PRK04282)→ rpl37ae (PRK03976)Ribosomal_L37ae (pfam01780)<br>RPC10 (COG1996)rpoP (PRK00398)→ Brix (smart00879)IMP4 (COG2136)→<br>Pcc1 (COG2892)Pcc1 (pfam09341)→ gimC_beta (TIGR02338)Prefoldin_2 (pfam01920)→<br>PRK14538 (PRK14538)DHH (pfam01368)→ PaaK (COG1541)AMP-binding_C_2 (pfam14535)→ |
| env-463 | JGI24020J35080_1007792 | + | 2787 | 2832 | **RNA**→ RPL15A (COG1632)PRK04243 (PRK04243)→ |
| env-464 | JGI24019J35510_1000013 | + | 160353 | 160398 | **RNA**→ PRK04243 (PRK04243)Ribosomal_L15e (pfam00827)→<br>PRK14555 (PRK14555)RNA_binding (pfam01877)→ RPP1 (COG1603)PRK00912 (PRK00912)→<br>PRK03717 (PRK03717)RNase_P_Rpp14 (pfam01900)→<br>PRK03996 (PRK03996)proteasome_alpha_archeal (cd03756)→ Sdo1 (COG1500)PRK13760 (PRK13760)→<br>Rrp4 (COG1097)PRK04163 (PRK04163)→ RNase_PH_archRRP41 (cd11366)PRK03983 (PRK03983)→<br>RNase_PH_archRRP42 (cd11365)PRK04282 (PRK04282)→ rpl37ae (PRK03976)Ribosomal_L37ae (pfam01780)<br>Pcc1 (COG2892)Pcc1 (pfam09341)→ gimC_beta (TIGR02338)GimC (COG1382)→ |
| env-465 | JGI24724J26744_10037895 | - | 1732 | 1679 | **RNA**→ PRK04243 (PRK04243)Ribosomal_L15e (pfam00827)→<br>ABC_ATPase (cd00267)AAA (smart00382)NapF (COG1145)Fer4_7 (pfam12838)→ |
| env-466 | LWFCAn_GLQU6AQ01BL2KP | - | 265 | 213 | **RNA**→ PRK04243 (PRK04243)Ribosomal_L15e (pfam00827)→ |
| env-467 | JGI24730J26740_1001090 | + | 9520 | 9567 | **RNA**→ PRK04243 (PRK04243)Ribosomal_L15e (pfam00827)→<br>PRK14555 (PRK14555)RNA_binding (pfam01877)→ |
| env-468 | JGI24730J26740_1002557 | - | 5350 | 5303 | **RNA**→ PRK04243 (PRK04243)Ribosomal_L15e (pfam00827)→<br>PRK14555 (PRK14555)RNA_binding (pfam01877)→ |
| env-469 | taComm3_C6796 | + | 6853 | 6906 | **RNA**→ PRK04243 (PRK04243)Ribosomal_L15e (pfam00827)→<br>PRK00912 (PRK00912)RNase_P_p30 (pfam01876)→ PRK03717 (PRK03717)RNase_P_Rpp14 (pfam01900)→<br>PRK03996 (PRK03996)proteasome_alpha_archeal (cd03756)→ Sdo1 (COG1500)PRK13760 (PRK13760)→<br>rpl37ae (PRK03976)Ribosomal_L37ae (pfam01780)→ RPC10 (COG1996)rpoP (PRK00398)→ |

| | | | | | |
|---|---|---|---|---|---|
| env-470 | JGI12104J13512_1001091 | + | 6473 | 6526 | **RNA**→ PRK04243 (PRK04243)Ribosomal_L15e (pfam00827)→ PRK00912 (PRK00912)RNase_P_p30 (pfam01876)→ PRK03717 (PRK03717)RNase_P_Rpp14 (pfam01900)→ PRK03996 (PRK03996)proteasome_alpha_archeal (cd03756)→ Sdo1 (COG1500)PRK13760 (PRK13760)→ rpl37ae (PRK03976)Ribosomal_L37ae (pfam01780)→ RPC10 (COG1996)rpoP (PRK00398)→ |
| env-471 | JGI11944J13513_1001226 | + | 6477 | 6530 | **RNA**→ PRK04243 (PRK04243)Ribosomal_L15e (pfam00827)→ PRK00912 (PRK00912)RNase_P_p30 (pfam01876)→ PRK03717 (PRK03717)RNase_P_Rpp14 (pfam01900)→ PRK03996 (PRK03996)proteasome_alpha_archeal (cd03756)→ Sdo1 (COG1500)PRK13760 (PRK13760)→ rpl37ae (PRK03976)Ribosomal_L37ae (pfam01780)→ RPC10 (COG1996)rpoP (PRK00398)→ |
| Mta-1-1 | NZ_AGIY02000001.1 | + | 329402 | 329455 | **RNA**→ PRK04243 (PRK04243)Ribosomal_L15e (pfam00827)→ PRK00912 (PRK00912)RNase_P_p30 (pfam01876)→ PRK00912 (PRK00912)RNase_P_p30 (pfam01876)→ PRK03717 (PRK03717)RNase_P_Rpp14 (pfam01900)→ PRK03996 (PRK03996)proteasome_alpha_archeal (cd03756)→ Sdo1 (COG1500)PRK13760 (PRK13760)→ rpl37ae (PRK03976)Ribosomal_L37ae (pfam01780)→ RPC10 (COG1996)rpoP (PRK00398)→ |
| env-472 | TB_LI09_4_10122878 | + | 413 | 459 | **RNA**→ RPL15A (COG1632)PRK04243 (PRK04243)→ |
| env-473 | TB_LI09_4_10800703 | + | 157 | 203 | **RNA**→ hypo→ |
| env-474 | BDMC2_contig02027 | - | 540 | 492 | **RNA**→ RPL15A (COG1632)PRK04243 (PRK04243)→ PRK04243 (PRK04243)Ribosomal_L15e (pfam00827)→ |
| Mpa-1-1 | NC_011832.1 | - | 2661123 | 2661072 | **RNA**→ PRK04243 (PRK04243)Ribosomal_L15e (pfam00827)→ RPP1 (COG1603)PRK00912 (PRK00912)→ PRK03717 (PRK03717)RNase_P_Rpp14 (pfam01900)→ PRK03996 (PRK03996)proteasome_alpha_archeal (cd03756)→ Sdo1 (COG1500)PRK13760 (PRK13760)→ rpl37ae (PRK03976)Ribosomal_L37ae (pfam01780)→ RPC10 (COG1996)rpoP (PRK00398)→ IMP4 (COG2136)PRK00933 (PRK00933)→ Pcc1 (COG2892)PRK14887 (PRK14887)→ yhbS (COG3153)Acetyltransf_7 (pfam13508)→ |
| env-475 | JGI20225J20221_1000037 | + | 100793 | 100845 | **RNA**→ PRK04243 (PRK04243)Ribosomal_L15e (pfam00827)→ PRK00912 (PRK00912)RNase_P_p30 (pfam01876)→ PRK03717 (PRK03717)RNase_P_Rpp14 (pfam01900)→ PRK03996 (PRK03996)proteasome_alpha_archeal (cd03756)→ Sdo1 (COG1500)PRK13760 (PRK13760)→ rpl37ae (PRK03976)Ribosomal_L37ae (pfam01780)→ RPC10 (COG1996)rpoP (PRK00398)→ Pcc1 (COG2892)Pcc1 (pfam09341)→ gimC_beta (TIGR02338)Prefoldin_2 (pfam01920)→ |
| env-476 | WSSedL1Cam_c0003 | - | 103534 | 103482 | **RNA**→ PRK04243 (PRK04243)Ribosomal_L15e (pfam00827)→ PRK00912 (PRK00912)RNase_P_p30 (pfam01876)→ PRK03717 (PRK03717)RNase_P_Rpp14 (pfam01900)→ PRK03996 (PRK03996)proteasome_alpha_archeal (cd03756)→ Sdo1 (COG1500)PRK13760 (PRK13760)→ rpl37ae (PRK03976)Ribosomal_L37ae (pfam01780)→ RPC10 (COG1996)rpoP (PRK00398)→ Pcc1 (COG2892)Pcc1 (pfam09341)→ gimC_beta (TIGR02338)Prefoldin_2 (pfam01920)→ |
| env-477 | JGI24729J20445_1007040 | + | 2798 | 2848 | **RNA**→ ←hypo |
| env-478 | JGI24730J26740_1000148 | - | 22730 | 22680 | **RNA**→ PRK04243 (PRK04243)Ribosomal_L15e (pfam00827)→ RPP1 (COG1603)PRK00912 (PRK00912)→ PRK03717 (PRK03717)RNase_P_Rpp14 (pfam01900)→ PRK03996 (PRK03996)proteasome_alpha_archeal (cd03756)→ Sdo1 (COG1500)PRK13760 (PRK13760)→ rpl37ae (PRK03976)Ribosomal_L37ae (pfam01780)→ RPC10 (COG1996)rpoP (PRK00398)→ PRK00933 (PRK00933)→ Pcc1 (COG2892)Pcc1 (pfam09341)→ |
| env-479 | 3300000558_10037240 | - | 1302 | 1247 | **RNA**→ PRK04243 (PRK04243)Ribosomal_L15e (pfam00827)→ PRK00912 (PRK00912)RNase_P_p30 (pfam01876)→ |
| env-480 | 3300000568_10759524 | - | 62 | 6 | **RNA**→ |
| env-481 | 3300000032_0445971 | - | 908 | 852 | **RNA**→ PRK04243 (PRK04243)Ribosomal_L15e (pfam00827)→ PRK00912 (PRK00912)→ |
| env-482 | 3300000568_10762646 | - | 251 | 195 | **RNA**→ PRK04243 (PRK04243)Ribosomal_L15e (pfam00827)→ |
| env-483 | 3300000568_10812672 | + | 106 | 162 | **RNA**→ PRK04243 (PRK04243)Ribosomal_L15e (pfam00827)→ |
| env-484 | 3300000568_10718528 | + | 26 | 82 | **RNA**→ PRK04243 (PRK04243)Ribosomal_L15e (pfam00827)→ |
| env-485 | 3300001567_10000590 | + | 50837 | 50893 | **RNA**→ PRK04243 (PRK04243)Ribosomal_L15e (pfam00827)→ PRK00912 (PRK00912)RNase_P_p30 (pfam01876)→ PRK03717 (PRK03717)RNase_P_Rpp14 (pfam01900)→ PRK03996 (PRK03996)proteasome_alpha_archeal (cd03756)→ Sdo1 (COG1500)PRK13760 (PRK13760)→ rpl37ae (PRK03976)Ribosomal_L37ae (pfam01780)→ RPC10 (COG1996)rpoP (PRK00398)→ Pcc1 (COG2892)Pcc1 (pfam09341)→ |
| env-486 | 3300000568_10082522 | + | 368 | 421 | **RNA**→ PRK04243 (PRK04243)Ribosomal_L15e (pfam00827)→ |
| env-487 | 3300001567_10000995 | - | 27743 | 27691 | **RNA**→ PRK04243 (PRK04243)Ribosomal_L15e (pfam00827)→ PRK00912 (PRK00912)RNase_P_p30 (pfam01876)→ PRK03717 (PRK03717)RNase_P_Rpp14 (pfam01900)→ PRK03996 (PRK03996)proteasome_alpha_archeal (cd03756)→ Sdo1 (COG1500)PRK13760 (PRK13760)→ rpl37ae (PRK03976)Ribosomal_L37ae (pfam01780)→ RPC10 (COG1996)rpoP (PRK00398)→ Pcc1 (COG2892)Pcc1 (pfam09341)→ |

| | | | | | |
|---|---|---|---|---|---|
| env-488 | JGI26437J51864_10005597 | - | 1881 | 1829 | **RNA**→ PRK04243 (PRK04243)Ribosomal_L15e (pfam00827)→ RPP1 (COG1603)PRK00912 (PRK00912)→ PRK03717 (PRK03717)RNase_P_Rpp14 (pfam01900)→ |
| env-489 | lwComb_BCIB16813_x1 | + | 697 | 749 | **RNA**→ |
| env-490 | WSSedA1Ba3_1028801 | + | 186 | 238 | **RNA**→ PRK04243 (PRK04243)Ribosomal_L15e (pfam00827)→ PRK00912 (PRK00912)RNase_P_p30 (pfam01876)→ |
| env-491 | WSSedA1Ba3_1008109 | + | 557 | 609 | **RNA**→ PRK04243 (PRK04243)Ribosomal_L15e (pfam00827)→ PRK00912 (PRK00912)RNase_P_p30 (pfam01876)→ PRK03717 (PRK03717)RNase_P_Rpp14 (pfam01900)→ Proteasome_A_N (smart00948)PRE1 (COG0638)→ |
| env-492 | WSSedA1TJGI_c023358 | - | 396 | 344 | **RNA**→ RPL15A (COG1632)PRK04243 (PRK04243)→ |
| env-493 | WSSedA1Ba_1000153 | - | 7580 | 7528 | **RNA**→ PRK04243 (PRK04243)Ribosomal_L15e (pfam00827)→ PRK00912 (PRK00912)RNase_P_p30 (pfam01876)→ PRK03717 (PRK03717)RNase_P_Rpp14 (pfam01900)→ PRK03996 (PRK03996)proteasome_alpha_archeal (cd03756)→ Sdo1 (COG1500)PRK13760 (PRK13760)→ rpl37ae (PRK03976)Ribosomal_L37ae (pfam01780)→ RPC10 (COG1996)rpoP (PRK00398)→ Pcc1 (COG2892)Pcc1 (pfam09341)→ |
| env-494 | WSSedA1Ba2_1001361 | - | 527 | 475 | **RNA**→ RPL15A (COG1632)PRK04243 (PRK04243)→ |
| env-495 | JGI26437J51864_10016078 | - | 877 | 823 | **RNA**→ PRK04243 (PRK04243)Ribosomal_L15e (pfam00827)→ PRK00912 (PRK00912)→ |
| env-496 | LWFCAn_GLQU6AQ01EP314 | - | 386 | 333 | **RNA**→ RPL15A (COG1632)PRK04243 (PRK04243)→ |
| env-497 | JGI24729J20445_1011542 | + | 712 | 765 | **RNA**→ PRK04243 (PRK04243)Ribosomal_L15e (pfam00827)→ PRK00912 (PRK00912)RNase_P_p30 (pfam01876)→ |
| env-498 | JGI24728J21555_1031680 | - | 431 | 378 | **RNA**→ RPL15A (COG1632)PRK04243 (PRK04243)→ |
| env-499 | 3300001580_10004398 | - | 7500 | 7446 | **RNA**→ PRK04243 (PRK04243)Ribosomal_L15e (pfam00827)→ RPP1 (COG1603)PRK00912 (PRK00912)→ PRK03717 (PRK03717)RNase_P_Rpp14 (pfam01900)→ PRK03996 (PRK03996)proteasome_alpha_archeal (cd03756)→ Sdo1 (COG1500)PRK13760 (PRK13760)→ rpl37ae (PRK03976)Ribosomal_L37ae (pfam01780)→ RPC10 (COG1996)rpoP (PRK00398)→ Pcc1 (COG2892)Pcc1 (pfam09341)→ gimC_beta (TIGR02338)Prefoldin_2 (pfam01920)→ spoT_relA (TIGR00691)SpoT (COG0317)→ |
| env-500 | JGI24729J20445_1004823 | - | 1184 | 1131 | **RNA**→ PRK04243 (PRK04243)Ribosomal_L15e (pfam00827)→ RPP1 (COG1603)PRK00912 (PRK00912)→ |
| env-501 | JGI24732J26686_1002952 | - | 3151 | 3098 | **RNA**→ PRK04243 (PRK04243)Ribosomal_L15e (pfam00827)→ RPP1 (COG1603)PRK00912 (PRK00912)→ PRK03717 (PRK03717)RNase_P_Rpp14 (pfam01900)→ PRK03996 (PRK03996)proteasome_alpha_archeal (cd03756)→ Sdo1 (COG1500)PRK13760 (PRK13760)→ |
| env-502 | JGI24732J26686_1094881 | - | 48 | 1 | **RNA**→ |
| env-503 | JGI24729J20445_1001141 | + | 9710 | 9763 | **RNA**→ PRK04243 (PRK04243)Ribosomal_L15e (pfam00827)→ RPP1 (COG1603)PRK00912 (PRK00912)→ |
| env-504 | JGI24732J26686_1139212 | - | 359 | 306 | **RNA**→ RPL15A (COG1632)PRK04243 (PRK04243)→ |
| env-505 | TB_GS10_10_10036752 | + | 1072 | 1125 | **RNA**→ PRK04243 (PRK04243)Ribosomal_L15e (pfam00827)→ RPP1 (COG1603)PRK00912 (PRK00912)→ |
| env-506 | JGI24730J26740_1018719 | + | 53 | 106 | **RNA**→ PRK04243 (PRK04243)Ribosomal_L15e (pfam00827)→ RPP1 (COG1603)PRK00912 (PRK00912)→ RNase_P_Rpp14 (pfam01900)→ |
| env-507 | SakLake120mDRAFT_c0042489 | - | 330 | 276 | **RNA**→ ←hypo |
| env-508 | 3300001605_10380848 | + | 193 | 245 | **RNA**→ PRK04243 (PRK04243)Ribosomal_L15e (pfam00827)→ |
| env-509 | 3300000032_0318192 | - | 212 | 159 | **RNA**→ PRK04243 (PRK04243)Ribosomal_L15e (pfam00827)→ |
| env-510 | 3300001580_10019781 | - | 923 | 870 | **RNA**→ PRK04243 (PRK04243)Ribosomal_L15e (pfam00827)→ hypo→ |
| env-511 | 3300000558_10012475 | + | 6222 | 6275 | **RNA**→ RPL15A (COG1632)PRK04243 (PRK04243)→ |
| env-512 | 3300001567_10108382 | + | 569 | 622 | **RNA**→ PRK04243 (PRK04243)Ribosomal_L15e (pfam00827)→ |
| env-513 | 3300001580_10224100 | + | 558 | 611 | **RNA**→ RPL15A (COG1632)PRK04243 (PRK04243)→ |
| env-514 | JGI26437J51864_10657425 | - | 97 | 44 | **RNA**→ hypo→ |
| env-515 | 3300001580_10162759 | + | 554 | 608 | **RNA**→ RPL15A (COG1632)PRK04243 (PRK04243)→ |
| env-516 | 3300001567_10107871 | + | 571 | 625 | **RNA**→ PRK04243 (PRK04243)Ribosomal_L15e (pfam00827)→ |
| env-517 | JGI24728J21555_1000054 | - | 38560 | 38510 | **RNA**→ PRK04243 (PRK04243)Ribosomal_L15e (pfam00827)→ RPP1 (COG1603)PRK00912 (PRK00912)→ PRK03717 (PRK03717)RNase_P_Rpp14 (pfam01900)→ PRK03996 (PRK03996)proteasome_alpha_archeal (cd03756)→ Sdo1 (COG1500)PRK13760 (PRK13760)→ rpl37ae (PRK03976)Ribosomal_L37ae (pfam01780)→ |
| env-518 | JGI24728J21555_1000137 | - | 38579 | 38529 | **RNA**→ PRK04243 (PRK04243)Ribosomal_L15e (pfam00827)→ RPP1 (COG1603)PRK00912 (PRK00912)→ PRK03717 (PRK03717)RNase_P_Rpp14 (pfam01900)→ PRK03996 (PRK03996)proteasome_alpha_archeal (cd03756)→ Sdo1 (COG1500)PRK13760 (PRK13760)→ rpl37ae (PRK03976)Ribosomal_L37ae (pfam01780)→ |

| | | | | | |
|---|---|---|---|---|---|
| env-519 | JGI24728J21555_1000138 | + | 5955 | 6005 | **RNA**→ PRK04243 (PRK04243)Ribosomal_L15e (pfam00827)→ RPP1 (COG1603)PRK00912 (PRK00912)→ PRK03717 (PRK03717)RNase_P_Rpp14 (pfam01900)→ PRK03996 (PRK03996)proteasome_alpha_archeal (cd03756)→ Sdo1 (COG1500)PRK13760 (PRK13760)→ rpl37ae (PRK03976)Ribosomal_L37ae (pfam01780)→ |
| env-520 | JGI24728J21555_1000139 | + | 5947 | 5997 | **RNA**→ PRK04243 (PRK04243)Ribosomal_L15e (pfam00827)→ RPP1 (COG1603)PRK00912 (PRK00912)→ PRK03717 (PRK03717)RNase_P_Rpp14 (pfam01900)→ PRK03996 (PRK03996)proteasome_alpha_archeal (cd03756)→ Sdo1 (COG1500)PRK13760 (PRK13760)→ rpl37ae (PRK03976)Ribosomal_L37ae (pfam01780)→ |
| env-521 | JGI24728J21555_1000140 | - | 38568 | 38518 | **RNA**→ PRK04243 (PRK04243)Ribosomal_L15e (pfam00827)→ RPP1 (COG1603)PRK00912 (PRK00912)→ PRK03717 (PRK03717)RNase_P_Rpp14 (pfam01900)→ PRK03996 (PRK03996)proteasome_alpha_archeal (cd03756)→ Sdo1 (COG1500)PRK13760 (PRK13760)→ rpl37ae (PRK03976)Ribosomal_L37ae (pfam01780)→ |
| env-522 | SL_5KL_010_SED_10012660 | - | 440 | 389 | **RNA**→ RPL15A (COG1632)PRK04243 (PRK04243)→ |
| env-523 | LCWY01000735.1 | - | 1783 | 1732 | **RNA**→ PRK04243 (PRK04243)Ribosomal_L15e (pfam00827)→ PRK00912 (PRK00912)RNase_P_p30 (pfam01876)→ PRK03717 (PRK03717)RNase_P_Rpp14 (pfam01900)→ |
| env-524 | LCWZ01000388.1 | - | 3340 | 3289 | **RNA**→ PRK04243 (PRK04243)Ribosomal_L15e (pfam00827)→ PRK00912 (PRK00912)RNase_P_p30 (pfam01876)→ PRK03717 (PRK03717)RNase_P_Rpp14 (pfam01900)→ PRK03996 (PRK03996)proteasome_alpha_archeal (cd03756)→ PRK13760 (PRK13760)SBDS (pfam01172)→ Sdo1 (COG1500)SBDS_C (pfam09377)→ hypo→ |
| env-525 | SL_5KL_010_SED_10011794 | + | 2661 | 2714 | **RNA**→ PRK04243 (PRK04243)Ribosomal_L15e (pfam00827)→ |
| env-526 | SL_3KL_010_SED_10302459 | + | 187 | 240 | **RNA**→ RPL15A (COG1632)PRK04243 (PRK04243)→ |
| env-527 | LCWZ01004261.1 | - | 1068 | 1015 | **RNA**→ RPL15A (COG1632)PRK04243 (PRK04243)→ PRK00912 (PRK00912)RNase_P_p30 (pfam01876)→ |
| env-528 | JGI2065J20421_1127642 | + | 79 | 128 | **RNA**→ PRK04243 (PRK04243)Ribosomal_L15e (pfam00827)→ |
| env-529 | 2013877062 | + | 961 | 1010 | **RNA**→ PRK04243 (PRK04243)Ribosomal_L15e (pfam00827)→ PRK00912 (PRK00912)RNase_P_p30 (pfam01876)→ |
| env-530 | BMHB3a_c117030 | + | 389 | 438 | **RNA**→ PRK04243 (PRK04243)Ribosomal_L15e (pfam00827)→ |
| env-531 | BMHB3a_c11385 | - | 71 | 22 | **RNA**→ hypo→ |
| env-532 | JGI24730J26740_1000056 | + | 41101 | 41154 | **RNA**→ PRK04243 (PRK04243)Ribosomal_L15e (pfam00827)→ PRK00912 (PRK00912)RNase_P_p30 (pfam01876)→ PRK03717 (PRK03717)RNase_P_Rpp14 (pfam01900)→ PRK03996 (PRK03996)proteasome_alpha_archeal (cd03756)→ Sdo1 (COG1500)PRK13760 (PRK13760)→ rpl37ae (PRK03976)Ribosomal_L37ae (pfam01780)→ RPC10 (COG1996)rpoP (PRK00398)→ Pcc1 (COG2892)Pcc1 (pfam09341)→ gimC_beta (TIGR02338)GimC (COG1382)→ |
| env-533 | JGI24730J26740_1000097 | - | 49515 | 49462 | **RNA**→ PRK04243 (PRK04243)Ribosomal_L15e (pfam00827)→ PRK00912 (PRK00912)RNase_P_p30 (pfam01876)→ PRK03717 (PRK03717)RNase_P_Rpp14 (pfam01900)→ PRK03996 (PRK03996)proteasome_alpha_archeal (cd03756)→ Sdo1 (COG1500)PRK13760 (PRK13760)→ rpl37ae (PRK03976)Ribosomal_L37ae (pfam01780)→ RPC10 (COG1996)rpoP (PRK00398)→ Pcc1 (COG2892)Pcc1 (pfam09341)→ gimC_beta (TIGR02338)GimC (COG1382)→ |
| CMe-1-1 | NC_009712.1 | - | 1906137 | 1906085 | **RNA**→ PRK04243 (PRK04243)Ribosomal_L15e (pfam00827)→ PRK00912 (PRK00912)RNase_P_p30 (pfam01876)→ PRK03717 (PRK03717)RNase_P_Rpp14 (pfam01900)→ PRK03996 (PRK03996)proteasome_alpha_archeal (cd03756)→ Sdo1 (COG1500)PRK13760 (PRK13760)→ rpl37ae (PRK03976)Ribosomal_L37ae (pfam01780)→ RPC10 (COG1996)rpoP (PRK00398)→ Pcc1 (COG2892)Pcc1 (pfam09341)→ PRK09343 (PRK09343)GimC (COG1382)→ |
| env-534 | PRSSGFe2_Sequence0000007570 | - | 3311 | 3258 | **RNA**→ PRK04243 (PRK04243)Ribosomal_L15e (pfam00827)→ PRK00912 (PRK00912)RNase_P_p30 (pfam01876)→ PRK03717 (PRK03717)RNase_P_Rpp14 (pfam01900)→ PRK03996 (PRK03996)proteasome_alpha_archeal (cd03756)→ Sdo1 (COG1500)PRK13760 (PRK13760)→ |
| env-535 | JGI26339J46600_10050974 | + | 104 | 156 | **RNA**→ RPL15A (COG1632)PRK04243 (PRK04243)→ PRK00912 (PRK00912)→ |
| env-536 | SL_3KL_010_SED_10003966 | - | 9606 | 9558 | **RNA**→ PRK04243 (PRK04243)Ribosomal_L15e (pfam00827)→ RPP1 (COG1603)PRK00912 (PRK00912)→ PRK03717 (PRK03717)RNase_P_Rpp14 (pfam01900)→ PRK03996 (PRK03996)proteasome_alpha_archeal (cd03756)→ Sdo1 (COG1500)PRK13760 (PRK13760)→ rpl37ae (PRK03976)Ribosomal_L37ae (pfam01780)→ RPC10 (COG1996)rpoP (PRK00398)→ IMP4 (COG2136)PRK00933 (PRK00933)→ Pcc1 (COG2892)Pcc1 (pfam09341)→ PRK09343 (PRK09343)GimC (COG1382)→ |
| env-537 | SL_3KL_010_SED_10034542 | + | 3130 | 3178 | **RNA**→ RPL15A (COG1632)PRK04243 (PRK04243)→ |

| | | | | | |
|---|---|---|---|---|---|
| env-538 | JGI12104J13512_1003036 | - | 8013 | 7965 | **RNA**→ PRK04243 (PRK04243)Ribosomal_L15e (pfam00827)→<br>PRK00912 (PRK00912)RNase_P_p30 (pfam01876)→ PRK03717 (PRK03717)RNase_P_Rpp14 (pfam01900)→<br>PRK03996 (PRK03996)proteasome_alpha_archeal (cd03756)→ Sdo1 (COG1500)PRK13760 (PRK13760)→<br>rpl37ae (PRK03976)Ribosomal_L37ae (pfam01780)→ |
| env-539 | JGI11944J13513_1000334 | + | 17500 | 17548 | **RNA**→ PRK04243 (PRK04243)Ribosomal_L15e (pfam00827)→<br>PRK00912 (PRK00912)RNase_P_p30 (pfam01876)→ PRK03717 (PRK03717)RNase_P_Rpp14 (pfam01900)→<br>PRK03996 (PRK03996)proteasome_alpha_archeal (cd03756)→ Sdo1 (COG1500)PRK13760 (PRK13760)→<br>rpl37ae (PRK03976)Ribosomal_L37ae (pfam01780)→ |
| env-540 | PCEOT_contig21575 | - | 446 | 398 | **RNA**→ RPL15A (COG1632)PRK04243 (PRK04243)→ |
| Mhu-1-1 | NC_007796.1 | + | 2543565 | 2543613 | **RNA**→ PRK04243 (PRK04243)Ribosomal_L15e (pfam00827)→ RPP1 (COG1603)PRK00912 (PRK00912)→<br>PRK03717 (PRK03717)RNase_P_Rpp14 (pfam01900)→<br>PRK03996 (PRK03996)proteasome_alpha_archeal (cd03756)→ Sdo1 (COG1500)PRK13760 (PRK13760)→<br>rpl37ae (PRK03976)Ribosomal_L37ae (pfam01780)→ RPC10 (COG1996)rpoP (PRK00398)→<br>PRK00933 (PRK00933)→ Pcc1 (COG2892)Pcc1 (pfam09341)→ PRK09343 (PRK09343)Prefoldin_2 (pfam0192<br>hypo→ hypo→ hypo→<br>PRK14960 (PRK14960)CARDB (pfam07705)NosD (pfam05048)Beta_helix (pfam13229)DUF4832 (pfam16116)-<br>PRK14960 (PRK14960)CARDB (pfam07705)NosD (pfam05048)Beta_helix (pfam13229)DUF4832 (pfam16116)-<br>PRK14960 (PRK14960)CARDB (pfam07705)NosD (pfam05048)Beta_helix (pfam13229)DUF4832 (pfam16116)- |
| env-541 | JGI24730J26740_1152247 | - | 310 | 262 | **RNA**→ PRK04243 (PRK04243)Ribosomal_L15e (pfam00827)→ |
| env-542 | 3300001580_10002056 | + | 12087 | 12136 | **RNA**→ PRK04243 (PRK04243)Ribosomal_L15e (pfam00827)→<br>PRK00912 (PRK00912)RNase_P_p30 (pfam01876)→ PRK03717 (PRK03717)RNase_P_Rpp14 (pfam01900)→<br>PRK03996 (PRK03996)proteasome_alpha_archeal (cd03756)→ Sdo1 (COG1500)PRK13760 (PRK13760)→<br>rpl37ae (PRK03976)Ribosomal_L37ae (pfam01780)→ |
| env-543 | JGI26437J51864_10233459 | - | 229 | 178 | **RNA**→ PRK04243 (PRK04243)Ribosomal_L15e (pfam00827)→ |
| env-544 | JGI24730J26740_1087991 | - | 503 | 453 | **RNA**→ RPL15A (COG1632)PRK04243 (PRK04243)→ |
| env-545 | JGI24731J21663_1016982 | - | 1473 | 1423 | **RNA**→ PRK04243 (PRK04243)Ribosomal_L15e (pfam00827)→ RPP1 (COG1603)PRK00912 (PRK00912)→<br>hypo→ |
| env-546 | JGI24732J26686_1102613 | - | 402 | 352 | **RNA**→ RPL15A (COG1632)PRK04243 (PRK04243)→ |
| Mli-1-1 | NZ_CM001555.1 | + | 718118 | 718169 | **RNA**→ PRK04243 (PRK04243)Ribosomal_L15e (pfam00827)→<br>PRK04243 (PRK04243)Ribosomal_L15e (pfam00827)→ PRK00912 (PRK00912)RNase_P_p30 (pfam01876)→<br>PRK00912 (PRK00912)RNase_P_p30 (pfam01876)→ PRK03717 (PRK03717)RNase_P_Rpp14 (pfam01900)→<br>PRK03717 (PRK03717)RNase_P_Rpp14 (pfam01900)→<br>PRK03996 (PRK03996)proteasome_alpha_archeal (cd03756)→<br>PRK03996 (PRK03996)proteasome_alpha_archeal (cd03756)→ Sdo1 (COG1500)PRK13760 (PRK13760)→<br>Sdo1 (COG1500)PRK13760 (PRK13760)→ rpl37ae (PRK03976)Ribosomal_L37ae (pfam01780)→<br>rpl37ae (PRK03976)Ribosomal_L37ae (pfam01780)→ RPC10 (COG1996)rpoP (PRK00398)→<br>RPC10 (COG1996)rpoP (PRK00398)→ IMP4 (COG2136)PRK00933 (PRK00933)→<br>IMP4 (COG2136)PRK00933 (PRK00933)→ Pcc1 (COG2892)Pcc1 (pfam09341)→<br>Pcc1 (COG2892)Pcc1 (pfam09341)→ gimC_beta (TIGR02338)GimC (COG1382)→<br>gimC_beta (TIGR02338)GimC (COG1382)→ |
| env-547 | JGI24712J26585_10534437 | + | 165 | 211 | **RNA**→ |
| env-548 | JGI24711J26586_10063235 | + | 193 | 239 | **RNA**→ PRK04243 (PRK04243)Ribosomal_L15e (pfam00827)→<br>PRK00912 (PRK00912)RNase_P_p30 (pfam01876)→ |
| env-549 | JGI24709J26583_10002362 | + | 11095 | 11141 | **RNA**→ RPL15A (COG1632)PRK04243 (PRK04243)→ PRK00912 (PRK00912)RNase_P_p30 (pfam01876)→<br>PRK03717 (PRK03717)RNase_P_Rpp14 (pfam01900)→<br>PRK03996 (PRK03996)proteasome_alpha_archeal (cd03756)→ Sdo1 (COG1500)PRK13760 (PRK13760)→<br>rpl37ae (PRK03976)Ribosomal_L37ae (pfam01780)→ |
| env-550 | JGI24710J26742_10002715 | - | 4756 | 4710 | **RNA**→ RPL15A (COG1632)PRK04243 (PRK04243)→ PRK00912 (PRK00912)RNase_P_p30 (pfam01876)→<br>PRK03717 (PRK03717)RNase_P_Rpp14 (pfam01900)→<br>PRK03996 (PRK03996)proteasome_alpha_archeal (cd03756)→ Sdo1 (COG1500)PRK13760 (PRK13760)→<br>rpl37ae (PRK03976)Ribosomal_L37ae (pfam01780)→ |
| env-551 | JGI24500J29687_10032170 | + | 390 | 436 | **RNA**→ RPL15A (COG1632)PRK04243 (PRK04243)→ PRK00912 (PRK00912)RNase_P_p30 (pfam01876)→<br>PRK03717 (PRK03717)RNase_P_Rpp14 (pfam01900)→<br>PRK03996 (PRK03996)proteasome_alpha_archeal (cd03756)→ Sdo1 (COG1500)PRK13760 (PRK13760)→ |

| | | | | | |
|---|---|---|---|---|---|
| Mse-1-1 | NZ_JXOJ01000004.1 | + | 7876 | 7922 | **RNA**→ hypo→ PRK04243 (PRK04243)Ribosomal_L15e (pfam00827)→<br>PRK00912 (PRK00912)RNase_P_p30 (pfam01876)→ PRK03717 (PRK03717)RNase_P_Rpp14 (pfam01900)→<br>PRK03996 (PRK03996)proteasome_alpha_archeal (cd03756)→ Sdo1 (COG1500)PRK13760 (PRK13760)→<br>rpl37ae (PRK03976)Ribosomal_L37ae (pfam01780)→ RPC10 (COG1996)rpoP (PRK00398)→<br>IMP4 (COG2136)PRK00933 (PRK00933)→ Pcc1 (COG2892)Pcc1 (pfam09341)→ |
| Mma-1-1 | NC_009051.1 | - | 1680920 | 1680874 | **RNA**→ PRK04243 (PRK04243)Ribosomal_L15e (pfam00827)→<br>PRK00912 (PRK00912)RNase_P_p30 (pfam01876)→ PRK03717 (PRK03717)RNase_P_Rpp14 (pfam01900)→<br>PRK03996 (PRK03996)proteasome_alpha_archeal (cd03756)→ Sdo1 (COG1500)PRK13760 (PRK13760)→<br>rpl37ae (PRK03976)Ribosomal_L37ae (pfam01780)→ RPC10 (COG1996)rpoP (PRK00398)→<br>IMP4 (COG2136)PRK00933 (PRK00933)→ Pcc1 (COG2892)Pcc1 (pfam09341)→ |
| Msp-5-1 | NZ_JMIO01000049.1 | + | 18527 | 18573 | **RNA**→ PRK04243 (PRK04243)Ribosomal_L15e (pfam00827)→ RPP1 (COG1603)PRK00912 (PRK00912)→<br>PRK03717 (PRK03717)RNase_P_Rpp14 (pfam01900)→<br>PRK03996 (PRK03996)proteasome_alpha_archeal (cd03756)→ Sdo1 (COG1500)PRK13760 (PRK13760)→<br>rpl37ae (PRK03976)Ribosomal_L37ae (pfam01780)→ RPC10 (COG1996)rpoP (PRK00398)→<br>IMP4 (COG2136)PRK00933 (PRK00933)→ hypo→ Pcc1 (COG2892)Pcc1 (pfam09341)→ |
| env-552 | JGI24712J26585_10228661 | + | 308 | 354 | **RNA**→ PRK04243 (PRK04243)Ribosomal_L15e (pfam00827)→ |
| env-553 | JGI24707J26582_10055176 | - | 84 | 39 | **RNA**→ ←hypo |
| env-554 | JGI24708J26588_10022103 | - | 1842 | 1797 | **RNA**→ PRK04243 (PRK04243)Ribosomal_L15e (pfam00827)→ RPP1 (COG1603)PRK00912 (PRK00912)→<br>PRK03717 (PRK03717)RNase_P_Rpp14 (pfam01900)→ |
| env-555 | JGI24712J26585_10005651 | + | 1497 | 1542 | **RNA**→ PRK04243 (PRK04243)Ribosomal_L15e (pfam00827)→ RPP1 (COG1603)PRK00912 (PRK00912)→<br>PRK03717 (PRK03717)RNase_P_Rpp14 (pfam01900)→<br>PRK03996 (PRK03996)proteasome_alpha_archeal (cd03756)→ Sdo1 (COG1500)PRK13760 (PRK13760)→<br>Sdo1 (COG1500)PRK13760 (PRK13760)→ rpl37ae (PRK03976)Ribosomal_L37ae (pfam01780)→ |
| env-556 | JGI24710J26742_10007141 | + | 2776 | 2821 | **RNA**→ PRK04243 (PRK04243)Ribosomal_L15e (pfam00827)→ RPP1 (COG1603)PRK00912 (PRK00912)→<br>PRK03717 (PRK03717)RNase_P_Rpp14 (pfam01900)→<br>PRK03996 (PRK03996)proteasome_alpha_archeal (cd03756)→ Sdo1 (COG1500)PRK13760 (PRK13760)→<br>rpl37ae (PRK03976)Ribosomal_L37ae (pfam01780)→ |
| env-557 | JGI24502J29692_10004908 | + | 2312 | 2357 | **RNA**→ PRK04243 (PRK04243)Ribosomal_L15e (pfam00827)→ RPP1 (COG1603)PRK00912 (PRK00912)→<br>PRK03717 (PRK03717)RNase_P_Rpp14 (pfam01900)→<br>PRK03996 (PRK03996)proteasome_alpha_archeal (cd03756)→ Sdo1 (COG1500)PRK13760 (PRK13760)→<br>rpl37ae (PRK03976)Ribosomal_L37ae (pfam01780)→ |
| env-558 | JGI24503J29689_10004405 | + | 2321 | 2366 | **RNA**→ PRK04243 (PRK04243)Ribosomal_L15e (pfam00827)→ RPP1 (COG1603)PRK00912 (PRK00912)→<br>PRK03717 (PRK03717)RNase_P_Rpp14 (pfam01900)→<br>PRK03996 (PRK03996)proteasome_alpha_archeal (cd03756)→ Sdo1 (COG1500)PRK13760 (PRK13760)→<br>rpl37ae (PRK03976)Ribosomal_L37ae (pfam01780)→ |
| env-559 | JGI24707J26582_10007021 | + | 73 | 118 | **RNA**→ PRK04243 (PRK04243)Ribosomal_L15e (pfam00827)→ RPP1 (COG1603)PRK00912 (PRK00912)→<br>PRK03717 (PRK03717)RNase_P_Rpp14 (pfam01900)→<br>PRK03996 (PRK03996)proteasome_alpha_archeal (cd03756)→ Sdo1 (COG1500)PRK13760 (PRK13760)→<br>rpl37ae (PRK03976)Ribosomal_L37ae (pfam01780)→ |
| env-560 | JGI24708J26588_10445994 | - | 46 | 1 | **RNA**→ |
| env-561 | JGI24711J26586_10009933 | - | 1842 | 1797 | **RNA**→ PRK04243 (PRK04243)Ribosomal_L15e (pfam00827)→ RPP1 (COG1603)PRK00912 (PRK00912)→<br>PRK03717 (PRK03717)RNase_P_Rpp14 (pfam01900)→ |
| env-562 | JGI24709J26583_10016669 | + | 2203 | 2248 | **RNA**→ PRK04243 (PRK04243)Ribosomal_L15e (pfam00827)→<br>PRK00912 (PRK00912)RNase_P_p30 (pfam01876)→ RPP1 (COG1603)PRK00912 (PRK00912)→<br>PRK03717 (PRK03717)RNase_P_Rpp14 (pfam01900)→ |
| env-563 | JGI24500J29687_10006466 | + | 556 | 601 | **RNA**→ PRK04243 (PRK04243)Ribosomal_L15e (pfam00827)→ RPP1 (COG1603)PRK00912 (PRK00912)→<br>PRK03717 (PRK03717)RNase_P_Rpp14 (pfam01900)→ Proteasome_A_N (smart00948)PRE1 (COG0638)→ |
| env-564 | 3300000568_10010387 | + | 293 | 339 | **RNA**→ PRK04243 (PRK04243)Ribosomal_L15e (pfam00827)→ |
| env-565 | 3300000032_0492979 | + | 132 | 178 | **RNA**→ |
| env-566 | 3300000558_10049236 | + | 193 | 239 | **RNA**→ PRK04243 (PRK04243)Ribosomal_L15e (pfam00827)→<br>PRK00912 (PRK00912)RNase_P_p30 (pfam01876)→ PRK03717 (PRK03717)RNase_P_Rpp14 (pfam01900)→<br>PRK03996 (PRK03996)proteasome_alpha_archeal (cd03756)→ Sdo1 (COG1500)PRK13760 (PRK13760)→<br>rpl37ae (PRK03976)Ribosomal_L37ae (pfam01780)→ |
| env-567 | 3300000032_0511128 | + | 210 | 256 | **RNA**→ |
| env-568 | 3300000032_0037457 | - | 630 | 584 | **RNA**→ PRK04243 (PRK04243)Ribosomal_L15e (pfam00827)→ |
| env-569 | 3300001592_10229257 | + | 50 | 96 | **RNA**→ PRK04243 (PRK04243)Ribosomal_L15e (pfam00827)→ |

| | | | | | |
|---|---|---|---|---|---|
| env-570 | 3300000507_100942 | + | 577 | 624 | **RNA**→ PRK04243 (PRK04243)Ribosomal_L15e (pfam00827)→ PRK00912 (PRK00912)RNase_P_p30 (pfam01876)→ PRK03717 (PRK03717)RNase_P_Rpp14 (pfam01900)→ PRK03996 (PRK03996)proteasome_alpha_archeal (cd03756)→ Sdo1 (COG1500)PRK13760 (PRK13760)→ rpl37ae (PRK03976)Ribosomal_L37ae (pfam01780)→ |
| env-571 | JGI24713J26584_10001369 | + | 2226 | 2272 | **RNA**→ RPL15A (COG1632)PRK04243 (PRK04243)→ RPP1 (COG1603)PRK00912 (PRK00912)→ PRK03717 (PRK03717)RNase_P_Rpp14 (pfam01900)→ PRK03996 (PRK03996)proteasome_alpha_archeal (cd03756)→ Sdo1 (COG1500)PRK13760 (PRK13760)→ rpl37ae (PRK03976)Ribosomal_L37ae (pfam01780)→ |
| env-572 | JGI24714J26587_10001520 | - | 13960 | 13913 | **RNA**→ RPL15A (COG1632)PRK04243 (PRK04243)→ RPP1 (COG1603)PRK00912 (PRK00912)→ PRK03717 (PRK03717)RNase_P_Rpp14 (pfam01900)→ PRK03996 (PRK03996)proteasome_alpha_archeal (cd03756)→ Sdo1 (COG1500)PRK13760 (PRK13760)→ rpl37ae (PRK03976)Ribosomal_L37ae (pfam01780)→ |
| env-573 | JGI24504J29685_1025823 | + | 107 | 154 | **RNA**→ RPL15A (COG1632)PRK04243 (PRK04243)→ |
| env-574 | 3300001592_10015540 | - | 752 | 702 | **RNA**→ PRK04243 (PRK04243)Ribosomal_L15e (pfam00827)→ |
| env-575 | 3300000032_0000829 | + | 1108 | 1158 | **RNA**→ PRK04243 (PRK04243)Ribosomal_L15e (pfam00827)→ PRK00912 (PRK00912)RNase_P_p30 (pfam01876)→ PRK03717 (PRK03717)RNase_P_Rpp14 (pfam01900)→ PRK03996 (PRK03996)proteasome_alpha_archeal (cd03756)→ Sdo1 (COG1500)PRK13760 (PRK13760)→ rpl37ae (PRK03976)Ribosomal_L37ae (pfam01780)→ RPC10 (COG1996)rpoP (PRK00398)→ PRK00933 (PRK00933)→ Pcc1 (COG2892)PRK14887 (PRK14887)→ gimC_beta (TIGR02338)GimC (COG13[illegible] |
| env-576 | 3300000558_12043292 | + | 53 | 104 | **RNA**→ PRK04243 (PRK04243)Ribosomal_L15e (pfam00827)→ |
| env-577 | 3300000568_10821221 | - | 73 | 23 | **RNA**→ hypo→ |
| env-578 | 3300001567_10009047 | + | 577 | 629 | **RNA**→ PRK04243 (PRK04243)Ribosomal_L15e (pfam00827)→ RPP1 (COG1603)PRK00912 (PRK00912)→ PRK03717 (PRK03717)RNase_P_Rpp14 (pfam01900)→ PRK03996 (PRK03996)proteasome_alpha_archeal (cd03756)→ Sdo1 (COG1500)PRK13760 (PRK13760)→ rpl37ae (PRK03976)Ribosomal_L37ae (pfam01780)→ |
| env-579 | 3300001592_10165112 | + | 124 | 175 | **RNA**→ PRK04243 (PRK04243)Ribosomal_L15e (pfam00827)→ PRK00912 (PRK00912)RNase_P_p30 (pfam01876)→ |
| env-580 | 3300001567_10003826 | + | 5679 | 5730 | **RNA**→ PRK04243 (PRK04243)Ribosomal_L15e (pfam00827)→ RPP1 (COG1603)PRK00912 (PRK00912)→ PRK03717 (PRK03717)RNase_P_Rpp14 (pfam01900)→ PRK03996 (PRK03996)proteasome_alpha_archeal (cd03756)→ Sdo1 (COG1500)PRK13760 (PRK13760)→ rpl37ae (PRK03976)Ribosomal_L37ae (pfam01780)→ |
| env-581 | 3300000032_0010281 | - | 14322 | 14272 | **RNA**→ PRK04243 (PRK04243)Ribosomal_L15e (pfam00827)→ RPP1 (COG1603)PRK00912 (PRK00912)→ PRK03717 (PRK03717)RNase_P_Rpp14 (pfam01900)→ PRK03996 (PRK03996)proteasome_alpha_archeal (cd03756)→ Sdo1 (COG1500)PRK13760 (PRK13760)→ rpl37ae (PRK03976)Ribosomal_L37ae (pfam01780)→ RPC10 (COG1996)rpoP (PRK00398)→ |
| env-582 | 3300000558_11443494 | - | 245 | 195 | **RNA**→ PRK04243 (PRK04243)Ribosomal_L15e (pfam00827)→ |
| env-583 | 3300000568_10668294 | - | 19377 | 19327 | **RNA**→ PRK04243 (PRK04243)Ribosomal_L15e (pfam00827)→ RPP1 (COG1603)PRK00912 (PRK00912)→ PRK03717 (PRK03717)RNase_P_Rpp14 (pfam01900)→ PRK03996 (PRK03996)proteasome_alpha_archeal (cd03756)→ Sdo1 (COG1500)PRK13760 (PRK13760)→ rpl37ae (PRK03976)Ribosomal_L37ae (pfam01780)→ RPC10 (COG1996)rpoP (PRK00398)→ |
| env-584 | 3300001580_10001349 | - | 19377 | 19327 | **RNA**→ PRK04243 (PRK04243)Ribosomal_L15e (pfam00827)→ RPP1 (COG1603)PRK00912 (PRK00912)→ PRK03717 (PRK03717)RNase_P_Rpp14 (pfam01900)→ PRK03996 (PRK03996)proteasome_alpha_archeal (cd03756)→ Sdo1 (COG1500)PRK13760 (PRK13760)→ rpl37ae (PRK03976)Ribosomal_L37ae (pfam01780)→ RPC10 (COG1996)rpoP (PRK00398)→ |
| env-585 | JGI12104J13512_1001025 | + | 6838 | 6890 | **RNA**→ RPL15A (COG1632)PRK04243 (PRK04243)→ PRK00912 (PRK00912)RNase_P_p30 (pfam01876)→ PRK03717 (PRK03717)RNase_P_Rpp14 (pfam01900)→ PRK03996 (PRK03996)proteasome_alpha_archeal (cd03756)→ Sdo1 (COG1500)PRK13760 (PRK13760)→ rpl37ae (PRK03976)Ribosomal_L37ae (pfam01780)→ RPC10 (COG1996)rpoP (PRK00398)→ |
| env-586 | JGI11944J13513_1001244 | + | 5853 | 5905 | **RNA**→ RPL15A (COG1632)PRK04243 (PRK04243)→ PRK00912 (PRK00912)RNase_P_p30 (pfam01876)→ PRK03717 (PRK03717)RNase_P_Rpp14 (pfam01900)→ PRK03996 (PRK03996)proteasome_alpha_archeal (cd03756)→ Sdo1 (COG1500)PRK13760 (PRK13760)→ rpl37ae (PRK03976)Ribosomal_L37ae (pfam01780)→ RPC10 (COG1996)rpoP (PRK00398)→ |
| env-587 | taComm3_C10759 | - | 580 | 529 | **RNA**→ PRK04243 (PRK04243)Ribosomal_L15e (pfam00827)→ |
| env-588 | 3300001592_10896941 | - | 243 | 192 | **RNA**→ PRK04243 (PRK04243)Ribosomal_L15e (pfam00827)→ |
| env-589 | 2227094145 | - | 323 | 269 | **RNA**→ ←hypo |
| env-590 | 2227164185 | + | 1201 | 1252 | **RNA**→ ←hypo |

| | | | | | |
|---|---|---|---|---|---|
| env-591 | 2226969280 | + | 1286 | 1337 | **RNA**→ RPL15A (COG1632)PRK04243 (PRK04243)→ |
| env-592 | 2227071043 | + | 381 | 432 | **RNA**→ PRK04243 (PRK04243)Ribosomal_L15e (pfam00827)→ |
| env-593 | IMNBL1_c0001816 | - | 461 | 410 | **RNA**→ RPL15A (COG1632)PRK04243 (PRK04243)→ |
| env-594 | JGI20163J15578_10187233 | - | 825 | 772 | **RNA**→ PRK04243 (PRK04243)Ribosomal_L15e (pfam00827)→ |
| env-595 | JGI996J12353_1000527 | - | 5644 | 5593 | **RNA**→ PRK04243 (PRK04243)Ribosomal_L15e (pfam00827)→ PRK14555 (PRK14555)RNA_binding (pfam01877)→ RPP1 (COG1603)PRK00912 (PRK00912)→ |
| env-596 | 2227035939 | + | 2727 | 2778 | **RNA**→ RPL15A (COG1632)PRK04243 (PRK04243)→ |
| env-597 | 2227538537 | - | 5296 | 5245 | **RNA**→ PRK04243 (PRK04243)Ribosomal_L15e (pfam00827)→ PRK14555 (PRK14555)RNA_binding (pfam01877)→ RPP1 (COG1603)PRK00912 (PRK00912)→ |
| env-598 | IMNBL1_c0047771 | - | 55 | 4 | **RNA**→ |
| env-599 | 2226957925 | - | 288 | 237 | **RNA**→ RPL15A (COG1632)PRK04243 (PRK04243)→ |
| env-600 | IMNBL1_c0018460 | - | 1848 | 1797 | **RNA**→ PRK04243 (PRK04243)Ribosomal_L15e (pfam00827)→ PRK14555 (PRK14555)RNA_binding (pfam01877)→ RPP1 (COG1603)PRK00912 (PRK00912)→ |
| env-601 | JGI994J12354_10008059 | + | 6285 | 6336 | **RNA**→ PRK04243 (PRK04243)Ribosomal_L15e (pfam00827)→ PRK14555 (PRK14555)RNA_binding (pfam01877)→ RPP1 (COG1603)PRK00912 (PRK00912)→ |
| env-602 | 2227477430 | - | 52 | 1 | **RNA**→ |
| env-603 | 2227211080 | - | 136 | 82 | **RNA**→ |
| env-604 | 2227517699 | + | 3271 | 3325 | **RNA**→ hypo→ |
| env-605 | IMNBL1_c0279633 | + | 244 | 298 | **RNA**→ PRK04243 (PRK04243)Ribosomal_L15e (pfam00827)→ |
| env-606 | JGI994J12354_10081286 | - | 57 | 3 | **RNA**→ ←hypo |
| env-607 | 2227618780 | - | 56 | 2 | **RNA**→ |
| env-608 | IMNBL1_c0051091 | + | 1242 | 1296 | **RNA**→ hypo→ |
| env-609 | JGI994J12354_10078750 | - | 57 | 3 | **RNA**→ |
| env-610 | IMNBL1_c0049435 | + | 1175 | 1227 | **RNA**→ PRK04243 (PRK04243)Ribosomal_L15e (pfam00827)→ |
| env-611 | 2227139717 | - | 365 | 311 | **RNA**→ ←hypo |
| env-612 | JGI994J12354_10115912 | - | 1015 | 961 | **RNA**→ PRK04243 (PRK04243)Ribosomal_L15e (pfam00827)→ PRK14555 (PRK14555)RNA_binding (pfam01877)→ |
| env-613 | IMNBL1_c0291723 | + | 33 | 87 | **RNA**→ RPL15A (COG1632)PRK04243 (PRK04243)→ |
| env-614 | IMNBL1_c0045419 | + | 1265 | 1319 | **RNA**→ PRK04243 (PRK04243)Ribosomal_L15e (pfam00827)→ |
| env-615 | JGI994J12354_10079817 | - | 170 | 116 | **RNA**→ PRK04243 (PRK04243)Ribosomal_L15e (pfam00827)→ |
| env-616 | 3300000594_10398 | + | 31 | 76 | **RNA**→ PRK04243 (PRK04243)Ribosomal_L15e (pfam00827)→ PRK14555 (PRK14555)RNA_binding (pfam01877)→ PRK00912 (PRK00912)RNase_P_p30 (pfam01876)→ |
| Mco-1-1 | NC_017034.1 | + | 356660 | 356705 | **RNA**→ PRK04243 (PRK04243)Ribosomal_L15e (pfam00827)→ PRK14555 (PRK14555)RNA_binding (pfam01877)→ RPP1 (COG1603)PRK00912 (PRK00912)→ BcsA (COG1215)glyco_like_mftF (TIGR04283)→ |
| env-617 | SL_5KL_010_SED_10351708 | + | 176 | 228 | **RNA**→ |
| env-618 | SL_8KL_010_SED_10426921 | + | 67 | 119 | **RNA**→ PRK04243 (PRK04243)Ribosomal_L15e (pfam00827)→ |
| Mps-1-1 | NC_018876.1 | + | 2608031 | 2608076 | **RNA**→ PRK04243 (PRK04243)Ribosomal_L15e (pfam00827)→ COG1325 (COG1325)PRK14555 (PRK14555) RPP1 (COG1603)PRK00912 (PRK00912)→ PRK03717 (PRK03717)RNase_P_Rpp14 (pfam01900)→ PRK03996 (PRK03996)proteasome_alpha_archeal (cd03756)→ Sdo1 (COG1500)PRK13760 (PRK13760)→ Rrp4 (COG1097)PRK04163 (PRK04163)Pup (pfam05639)→ RNase_PH_archRRP41 (cd11366)PRK03983 (PRK03983)→ RNase_PH_archRRP42 (cd11365)PRK04282 (PRK04282)→ rpl37ae (PRK03976)Ribosomal_L37ae (pfam01780) RPC10 (COG1996)rpoP (PRK00398)→ |
| env-619 | JGI25321J50212_10071092 | - | 517 | 472 | **RNA**→ RPL15A (COG1632)PRK04243 (PRK04243)→ |
| env-620 | 3300001567_10059738 | + | 145 | 190 | **RNA**→ PRK04243 (PRK04243)Ribosomal_L15e (pfam00827)→ COG1325 (COG1325)PRK14555 (PRK14555) |
| Mti-1-1 | NZ_AZAJ01000001.1 | - | 571920 | 571858 | **RNA**→ PRK04243 (PRK04243)Ribosomal_L15e (pfam00827)→ PRK14555 (PRK14555)RNA_binding (pfam01877)→ PRK00912 (PRK00912)RNase_P_p30 (pfam01876)→ PRK03717 (PRK03717)POP5 (COG1369)→ PRK03996 (PRK03996)proteasome_alpha_archeal (cd03756)→ Sdo1 (COG1500)PRK13760 (PRK13760)→ Rrp4 (COG1097)PRK04163 (PRK04163)→ RNase_PH_archRRP41 (cd11366)PRK03983 (PRK03983)→ RNase_PH_archRRP42 (cd11365)PRK04282 (PRK04282)→ rpl37ae (PRK03976)Ribosomal_L37ae (pfam01780) RPC10 (COG1996)rpoP (PRK00398)→ |
| env-621 | 2231757710 | + | 31 | 79 | **RNA**→ PRK04243 (PRK04243)Ribosomal_L15e (pfam00827)→ |

| | | | | | |
|---|---|---|---|---|---|
| Mba-2-1 | NZ_CP009517.1 | - | 1228703 | 1228652 | RNA→ PRK04243 (PRK04243)Ribosomal_L15e (pfam00827)→ COG1325 (COG1325)PRK14555 (PRK14555) RPP1 (COG1603)PRK00912 (PRK00912)→ PRK03717 (PRK03717)POP5 (COG1369)→ |
| env-622 | B4_c_Cons7318 | + | 381 | 432 | RNA→ PRK04243 (PRK04243)Ribosomal_L15e (pfam00827)→ |
| env-623 | JGI20225J20221_1000854 | - | 92739 | 92688 | RNA→ PRK04243 (PRK04243)Ribosomal_L15e (pfam00827)→ COG1325 (COG1325)PRK14555 (PRK14555) RPP1 (COG1603)PRK00912 (PRK00912)→ PRK03717 (PRK03717)POP5 (COG1369)→ |
| env-624 | 3300000507_111820 | - | 641 | 590 | RNA→ PRK04243 (PRK04243)Ribosomal_L15e (pfam00827)→ |
| env-625 | JGI24714J26587_10012885 | - | 2658 | 2606 | RNA→ PRK04243 (PRK04243)Ribosomal_L15e (pfam00827)→ COG1325 (COG1325)PRK14555 (PRK14555) RPP1 (COG1603)PRK00912 (PRK00912)→ PRK03717 (PRK03717)POP5 (COG1369)→ |
| env-626 | JGI24712J26585_10034019 | - | 1360 | 1308 | RNA→ PRK04243 (PRK04243)Ribosomal_L15e (pfam00827)→ COG1325 (COG1325)PRK14555 (PRK14555) RPP1 (COG1603)PRK00912 (PRK00912)→ |
| env-627 | JGI24711J26586_10062120 | + | 435 | 487 | RNA→ PRK04243 (PRK04243)Ribosomal_L15e (pfam00827)→ |
| Mho-2-1 | NZ_CP009516.1 | - | 3291833 | 3291782 | RNA→ PRK04243 (PRK04243)Ribosomal_L15e (pfam00827)→ COG1325 (COG1325)PRK14555 (PRK14555) RPP1 (COG1603)PRK00912 (PRK00912)→ PRK03717 (PRK03717)POP5 (COG1369)→ |
| env-628 | JGI24731J21663_1002020 | - | 12930 | 12879 | RNA→ PRK04243 (PRK04243)Ribosomal_L15e (pfam00827)→ COG1325 (COG1325)PRK14555 (PRK14555) RPP1 (COG1603)PRK00912 (PRK00912)→ PRK03717 (PRK03717)POP5 (COG1369)→ |
| env-629 | JGI24731J21663_1031455 | - | 306 | 255 | RNA→ RPL15A (COG1632)PRK04243 (PRK04243)→ |
| env-630 | JGI12419J13241_1124316 | + | 56 | 108 | RNA→ RPL15A (COG1632)PRK04243 (PRK04243)→ |
| Mth-1-1 | NZ_CP009501.1 | - | 381745 | 381693 | RNA→ PRK04243 (PRK04243)Ribosomal_L15e (pfam00827)→ COG1325 (COG1325)PRK14555 (PRK14555) RPP1 (COG1603)PRK00912 (PRK00912)→ PRK03717 (PRK03717)POP5 (COG1369)→ |
| Msp-13-1 | NZ_CP009524.1 | - | 1406720 | 1406669 | RNA→ PRK04243 (PRK04243)Ribosomal_L15e (pfam00827)→ COG1325 (COG1325)PRK14555 (PRK14555) RPP1 (COG1603)PRK00912 (PRK00912)→ PRK03717 (PRK03717)POP5 (COG1369)→ |
| Mva-1-1 | NZ_CP009520.1 | - | 1438747 | 1438696 | RNA→ COG4585 (COG4585)→ PRK04243 (PRK04243)Ribosomal_L15e (pfam00827)→ COG1325 (COG1325)PRK14555 (PRK14555)→ RPP1 (COG1603)PRK00912 (PRK00912)→ PRK03717 (PRK03717)POP5 (COG1369)→ |
| Mba-4-1 | NZ_CP009528.1 | - | 2670750 | 2670699 | RNA→ PRK04243 (PRK04243)Ribosomal_L15e (pfam00827)→ COG1325 (COG1325)PRK14555 (PRK14555) RPP1 (COG1603)PRK00912 (PRK00912)→ PRK03717 (PRK03717)POP5 (COG1369)→ |
| Mba-1-1 | NZ_CP009530.1 | + | 476717 | 476768 | RNA→ PRK04243 (PRK04243)Ribosomal_L15e (pfam00827)→ COG1325 (COG1325)PRK14555 (PRK14555) RPP1 (COG1603)PRK00912 (PRK00912)→ PRK03717 (PRK03717)POP5 (COG1369)→ |
| Mba-3-1 | NZ_CP008746.1 | + | 468339 | 468390 | RNA→ PRK04243 (PRK04243)Ribosomal_L15e (pfam00827)→ COG1325 (COG1325)PRK14555 (PRK14555) RPP1 (COG1603)PRK00912 (PRK00912)→ PRK03717 (PRK03717)POP5 (COG1369)→ |
| env-631 | BMHB3a_c34562 | + | 135 | 186 | RNA→ |
| env-632 | JGI2065J20421_1016166 | + | 872 | 923 | RNA→ PRK04243 (PRK04243)Ribosomal_L15e (pfam00827)→ PRK14555 (PRK14555)RNA_binding (pfam01877)→ |
| env-633 | B3_all_c_Cons125890 | + | 461 | 512 | RNA→ PRK04243 (PRK04243)Ribosomal_L15e (pfam00827)→ |
| Mla-1-1 | NZ_CP009515.1 | + | 2312125 | 2312176 | RNA→ PRK04243 (PRK04243)Ribosomal_L15e (pfam00827)→ COG1325 (COG1325)PRK14555 (PRK14555) RPP1 (COG1603)PRK00912 (PRK00912)→ PRK03717 (PRK03717)POP5 (COG1369)→ |
| Mba-5-1 | NC_007355.1 | + | 3156160 | 3156211 | RNA→ PRK04243 (PRK04243)Ribosomal_L15e (pfam00827)→ COG1325 (COG1325)PRK14555 (PRK14555) RPP1 (COG1603)PRK00912 (PRK00912)→ PRK03717 (PRK03717)POP5 (COG1369)→ |
| Mba-6-1 | NZ_CP009526.1 | - | 2909575 | 2909524 | RNA→ PRK04243 (PRK04243)Ribosomal_L15e (pfam00827)→ COG1325 (COG1325)PRK14555 (PRK14555) RPP1 (COG1603)PRK00912 (PRK00912)→ PRK03717 (PRK03717)POP5 (COG1369)→ |
| Msp-7-1 | NZ_JJQY01000019.1 | - | 28722 | 28671 | RNA→ PRK04243 (PRK04243)Ribosomal_L15e (pfam00827)→ COG1325 (COG1325)PRK14555 (PRK14555) RPP1 (COG1603)PRK00912 (PRK00912)→ PRK03717 (PRK03717)POP5 (COG1369)→ |
| Msp-8-1 | NZ_JJOV01000053.1 | - | 61527 | 61476 | RNA→ PRK04243 (PRK04243)Ribosomal_L15e (pfam00827)→ COG1325 (COG1325)PRK14555 (PRK14555) RPP1 (COG1603)PRK00912 (PRK00912)→ PRK03717 (PRK03717)POP5 (COG1369)→ |
| Msp-10-1 | NZ_JJOX01000057.1 | - | 29319 | 29268 | RNA→ PRK04243 (PRK04243)Ribosomal_L15e (pfam00827)→ COG1325 (COG1325)PRK14555 (PRK14555) RPP1 (COG1603)PRK00912 (PRK00912)→ PRK03717 (PRK03717)POP5 (COG1369)→ |
| Msp-11-1 | NZ_JJOY01000041.1 | - | 29252 | 29201 | RNA→ PRK04243 (PRK04243)Ribosomal_L15e (pfam00827)→ COG1325 (COG1325)PRK14555 (PRK14555) RPP1 (COG1603)PRK00912 (PRK00912)→ PRK03717 (PRK03717)POP5 (COG1369)→ |
| Msp-6-1 | NZ_JJQL01000038.1 | - | 47137 | 47086 | RNA→ PRK04243 (PRK04243)Ribosomal_L15e (pfam00827)→ COG1325 (COG1325)PRK14555 (PRK14555) RPP1 (COG1603)PRK00912 (PRK00912)→ PRK03717 (PRK03717)POP5 (COG1369)→ |
| Msp-12-1 | NZ_JJOZ01000046.1 | - | 28637 | 28586 | RNA→ PRK04243 (PRK04243)Ribosomal_L15e (pfam00827)→ COG1325 (COG1325)PRK14555 (PRK14555) RPP1 (COG1603)PRK00912 (PRK00912)→ PRK03717 (PRK03717)POP5 (COG1369)→ |
| Msp-9-1 | NZ_JJOW01000294.1 | - | 20540 | 20489 | RNA→ PRK04243 (PRK04243)Ribosomal_L15e (pfam00827)→ COG1325 (COG1325)PRK14555 (PRK14555) RPP1 (COG1603)PRK00912 (PRK00912)→ PRK03717 (PRK03717)POP5 (COG1369)→ |
| Msp-16-1 | NZ_CP009503.1 | + | 3735833 | 3735884 | RNA→ PRK04243 (PRK04243)Ribosomal_L15e (pfam00827)→ COG1325 (COG1325)PRK14555 (PRK14555) RPP1 (COG1603)PRK00912 (PRK00912)→ PRK03717 (PRK03717)POP5 (COG1369)→ |
| Msp-15-1 | NZ_CP009504.1 | - | 2456027 | 2455976 | RNA→ PRK04243 (PRK04243)Ribosomal_L15e (pfam00827)→ COG1325 (COG1325)PRK14555 (PRK14555) RPP1 (COG1603)PRK00912 (PRK00912)→ PRK03717 (PRK03717)POP5 (COG1369)→ |

| | | | | | |
|---|---|---|---|---|---|
| env-634 | JGI24731J21663_1041287 | + | 802 | 854 | **RNA**→ ←hypo |
| Mso-1-1 | NZ_JQLR01000001.1 | + | 2442416 | 2442467 | **RNA**→ PRK04243 (PRK04243)Ribosomal_L15e (pfam00827)→ COG1325 (COG1325)PRK14555 (PRK14555)<br>RPP1 (COG1603)PRK00912 (PRK00912)→ PRK03717 (PRK03717)POP5 (COG1369)→ |
| Mma-5-1 | NC_003901.1 | + | 3119311 | 3119362 | **RNA**→ PRK04243 (PRK04243)Ribosomal_L15e (pfam00827)→ COG1325 (COG1325)PRK14555 (PRK14555)<br>RPP1 (COG1603)PRK00912 (PRK00912)→ PRK03717 (PRK03717)POP5 (COG1369)→ |
| Mma-6-1 | NZ_CP009513.1 | + | 3747266 | 3747317 | **RNA**→ PRK04243 (PRK04243)Ribosomal_L15e (pfam00827)→ COG1325 (COG1325)PRK14555 (PRK14555)<br>RPP1 (COG1603)PRK00912 (PRK00912)→ PRK03717 (PRK03717)POP5 (COG1369)→ |
| Mma-8-1 | NZ_CP009511.1 | - | 2504533 | 2504482 | **RNA**→ PRK04243 (PRK04243)Ribosomal_L15e (pfam00827)→ COG1325 (COG1325)PRK14555 (PRK14555)<br>RPP1 (COG1603)PRK00912 (PRK00912)→ PRK03717 (PRK03717)POP5 (COG1369)→ |
| Mma-7-1 | NZ_CP009512.1 | - | 2625933 | 2625882 | **RNA**→ PRK04243 (PRK04243)Ribosomal_L15e (pfam00827)→ COG1325 (COG1325)PRK14555 (PRK14555)<br>RPP1 (COG1603)PRK00912 (PRK00912)→ PRK03717 (PRK03717)POP5 (COG1369)→ |
| Mma-3-1 | NZ_JJPB01000050.1 | + | 11936 | 11987 | **RNA**→ PRK04243 (PRK04243)Ribosomal_L15e (pfam00827)→ COG1325 (COG1325)PRK14555 (PRK14555)<br>RPP1 (COG1603)PRK00912 (PRK00912)→ PRK03717 (PRK03717)POP5 (COG1369)→ |
| Mma-3-2 | NZ_JJQI01000092.1 | - | 16421 | 16370 | **RNA**→ PRK04243 (PRK04243)Ribosomal_L15e (pfam00827)→ COG1325 (COG1325)PRK14555 (PRK14555)<br>RPP1 (COG1603)PRK00912 (PRK00912)→ PRK03717 (PRK03717)POP5 (COG1369)→ |
| Mma-3-3 | NZ_JJQC01000141.1 | - | 42369 | 42318 | **RNA**→ PRK04243 (PRK04243)Ribosomal_L15e (pfam00827)→ COG1325 (COG1325)PRK14555 (PRK14555)<br>RPP1 (COG1603)PRK00912 (PRK00912)→ PRK03717 (PRK03717)POP5 (COG1369)→ |
| Mma-3-4 | NZ_JJPX01000042.1 | + | 11441 | 11492 | **RNA**→ PRK04243 (PRK04243)Ribosomal_L15e (pfam00827)→ COG1325 (COG1325)PRK14555 (PRK14555)<br>RPP1 (COG1603)PRK00912 (PRK00912)→ PRK03717 (PRK03717)POP5 (COG1369)→ |
| Mma-3-5 | NZ_JJQA01000093.1 | - | 42369 | 42318 | **RNA**→ PRK04243 (PRK04243)Ribosomal_L15e (pfam00827)→ COG1325 (COG1325)PRK14555 (PRK14555)<br>RPP1 (COG1603)PRK00912 (PRK00912)→ PRK03717 (PRK03717)POP5 (COG1369)→ |
| Mma-3-6 | NZ_JJPN01000151.1 | + | 11194 | 11245 | **RNA**→ PRK04243 (PRK04243)Ribosomal_L15e (pfam00827)→ COG1325 (COG1325)PRK14555 (PRK14555)<br>RPP1 (COG1603)PRK00912 (PRK00912)→ PRK03717 (PRK03717)POP5 (COG1369)→ |
| Mma-3-7 | NZ_JJOS01000070.1 | + | 20296 | 20347 | **RNA**→ PRK04243 (PRK04243)Ribosomal_L15e (pfam00827)→ COG1325 (COG1325)PRK14555 (PRK14555)<br>RPP1 (COG1603)PRK00912 (PRK00912)→ PRK03717 (PRK03717)POP5 (COG1369)→ |
| Mma-3-8 | NZ_JJPG01000087.1 | + | 11936 | 11987 | **RNA**→ PRK04243 (PRK04243)Ribosomal_L15e (pfam00827)→ COG1325 (COG1325)PRK14555 (PRK14555)<br>RPP1 (COG1603)PRK00912 (PRK00912)→ PRK03717 (PRK03717)POP5 (COG1369)→ |
| Mma-3-9 | NZ_JJPP01000085.1 | + | 31236 | 31287 | **RNA**→ PRK04243 (PRK04243)Ribosomal_L15e (pfam00827)→ COG1325 (COG1325)PRK14555 (PRK14555)<br>RPP1 (COG1603)PRK00912 (PRK00912)→ PRK03717 (PRK03717)POP5 (COG1369)→ |
| Mma-3-10 | NZ_JJQB01000132.1 | - | 42369 | 42318 | **RNA**→ PRK04243 (PRK04243)Ribosomal_L15e (pfam00827)→ COG1325 (COG1325)PRK14555 (PRK14555)<br>RPP1 (COG1603)PRK00912 (PRK00912)→ PRK03717 (PRK03717)POP5 (COG1369)→ |
| Mma-3-11 | NZ_JJQG01000125.1 | + | 110287 | 110338 | **RNA**→ PRK04243 (PRK04243)Ribosomal_L15e (pfam00827)→ COG1325 (COG1325)PRK14555 (PRK14555)<br>RPP1 (COG1603)PRK00912 (PRK00912)→ PRK03717 (PRK03717)POP5 (COG1369)→ |
| Mma-3-12 | NZ_JJPF01000029.1 | - | 32163 | 32112 | **RNA**→ PRK04243 (PRK04243)Ribosomal_L15e (pfam00827)→ COG1325 (COG1325)PRK14555 (PRK14555)<br>RPP1 (COG1603)PRK00912 (PRK00912)→ PRK03717 (PRK03717)POP5 (COG1369)→ |
| Mma-3-13 | NZ_JJQR01000115.1 | + | 47927 | 47978 | **RNA**→ PRK04243 (PRK04243)Ribosomal_L15e (pfam00827)→ COG1325 (COG1325)PRK14555 (PRK14555)<br>RPP1 (COG1603)PRK00912 (PRK00912)→ PRK03717 (PRK03717)POP5 (COG1369)→ |
| Mma-3-14 | NZ_JJQW01000009.1 | - | 43611 | 43560 | **RNA**→ PRK04243 (PRK04243)Ribosomal_L15e (pfam00827)→ COG1325 (COG1325)PRK14555 (PRK14555)<br>RPP1 (COG1603)PRK00912 (PRK00912)→ PRK03717 (PRK03717)POP5 (COG1369)→ |
| Mma-3-15 | NZ_JJPO01000048.1 | + | 11937 | 11988 | **RNA**→ PRK04243 (PRK04243)Ribosomal_L15e (pfam00827)→ COG1325 (COG1325)PRK14555 (PRK14555)<br>RPP1 (COG1603)PRK00912 (PRK00912)→ PRK03717 (PRK03717)POP5 (COG1369)→ |
| Mma-3-16 | NZ_JJQX01000051.1 | + | 8135 | 8186 | **RNA**→ PRK04243 (PRK04243)Ribosomal_L15e (pfam00827)→ COG1325 (COG1325)PRK14555 (PRK14555)<br>RPP1 (COG1603)PRK00912 (PRK00912)→ PRK03717 (PRK03717)POP5 (COG1369)→ |
| Mma-3-17 | NZ_JJQE01000025.1 | - | 33596 | 33545 | **RNA**→ PRK04243 (PRK04243)Ribosomal_L15e (pfam00827)→ COG1325 (COG1325)PRK14555 (PRK14555)<br>RPP1 (COG1603)PRK00912 (PRK00912)→ PRK03717 (PRK03717)POP5 (COG1369)→ |
| Mma-3-18 | NZ_JJRA01000090.1 | - | 35290 | 35239 | **RNA**→ PRK04243 (PRK04243)Ribosomal_L15e (pfam00827)→ COG1325 (COG1325)PRK14555 (PRK14555)<br>RPP1 (COG1603)PRK00912 (PRK00912)→ PRK03717 (PRK03717)POP5 (COG1369)→ |
| Mma-3-19 | NZ_JJRB01000023.1 | - | 35592 | 35541 | **RNA**→ PRK04243 (PRK04243)Ribosomal_L15e (pfam00827)→ COG1325 (COG1325)PRK14555 (PRK14555)<br>RPP1 (COG1603)PRK00912 (PRK00912)→ PRK03717 (PRK03717)POP5 (COG1369)→ |
| Mma-3-20 | NZ_JJOT01000028.1 | + | 142723 | 142774 | **RNA**→ PRK04243 (PRK04243)Ribosomal_L15e (pfam00827)→ COG1325 (COG1325)PRK14555 (PRK14555)<br>RPP1 (COG1603)PRK00912 (PRK00912)→ PRK03717 (PRK03717)POP5 (COG1369)→ |
| Mma-3-21 | NZ_JJQS01000135.1 | - | 34925 | 34874 | **RNA**→ PRK04243 (PRK04243)Ribosomal_L15e (pfam00827)→ COG1325 (COG1325)PRK14555 (PRK14555)<br>RPP1 (COG1603)PRK00912 (PRK00912)→ PRK03717 (PRK03717)POP5 (COG1369)→ |
| Mma-3-22 | NZ_JJPC01000138.1 | + | 16225 | 16276 | **RNA**→ PRK04243 (PRK04243)Ribosomal_L15e (pfam00827)→ COG1325 (COG1325)PRK14555 (PRK14555)<br>RPP1 (COG1603)PRK00912 (PRK00912)→ PRK03717 (PRK03717)POP5 (COG1369)→ |
| Mma-3-23 | NZ_JJOU01000016.1 | - | 43691 | 43640 | **RNA**→ PRK04243 (PRK04243)Ribosomal_L15e (pfam00827)→ COG1325 (COG1325)PRK14555 (PRK14555)<br>RPP1 (COG1603)PRK00912 (PRK00912)→ PRK03717 (PRK03717)POP5 (COG1369)→ |

| | | | | | |
|---|---|---|---|---|---|
| Mma-3-24 | NZ_JJPS01000107.1 | - | 6231 | 6180 | **RNA**→ PRK04243 (PRK04243)Ribosomal_L15e (pfam00827)→ COG1325 (COG1325)PRK14555 (PRK14555)<br>RPP1 (COG1603)PRK00912 (PRK00912)→ PRK03717 (PRK03717)POP5 (COG1369)→ |
| Mma-3-25 | NZ_JJPY01000103.1 | - | 42289 | 42238 | **RNA**→ PRK04243 (PRK04243)Ribosomal_L15e (pfam00827)→ COG1325 (COG1325)PRK14555 (PRK14555)<br>RPP1 (COG1603)PRK00912 (PRK00912)→ PRK03717 (PRK03717)POP5 (COG1369)→ |
| Mma-3-26 | NZ_JJOR01000097.1 | + | 48050 | 48101 | **RNA**→ PRK04243 (PRK04243)Ribosomal_L15e (pfam00827)→ COG1325 (COG1325)PRK14555 (PRK14555)<br>RPP1 (COG1603)PRK00912 (PRK00912)→ PRK03717 (PRK03717)POP5 (COG1369)→ |
| Mma-3-27 | NZ_JJQD01000017.1 | + | 37333 | 37384 | **RNA**→ PRK04243 (PRK04243)Ribosomal_L15e (pfam00827)→ COG1325 (COG1325)PRK14555 (PRK14555)<br>RPP1 (COG1603)PRK00912 (PRK00912)→ PRK03717 (PRK03717)POP5 (COG1369)→ |
| Mma-3-28 | NZ_JJQM01000066.1 | + | 47345 | 47396 | **RNA**→ PRK04243 (PRK04243)Ribosomal_L15e (pfam00827)→ COG1325 (COG1325)PRK14555 (PRK14555)<br>RPP1 (COG1603)PRK00912 (PRK00912)→ PRK03717 (PRK03717)POP5 (COG1369)→ |
| Mma-3-29 | NZ_JJPA01000100.1 | - | 34395 | 34344 | **RNA**→ PRK04243 (PRK04243)Ribosomal_L15e (pfam00827)→ COG1325 (COG1325)PRK14555 (PRK14555)<br>RPP1 (COG1603)PRK00912 (PRK00912)→ PRK03717 (PRK03717)POP5 (COG1369)→ |
| Mma-3-30 | NZ_JJPR01000056.1 | - | 42079 | 42028 | **RNA**→ PRK04243 (PRK04243)Ribosomal_L15e (pfam00827)→ COG1325 (COG1325)PRK14555 (PRK14555)<br>RPP1 (COG1603)PRK00912 (PRK00912)→ PRK03717 (PRK03717)POP5 (COG1369)→ |
| Mma-3-31 | NZ_JJPH01000044.1 | - | 32163 | 32112 | **RNA**→ PRK04243 (PRK04243)Ribosomal_L15e (pfam00827)→ COG1325 (COG1325)PRK14555 (PRK14555)<br>RPP1 (COG1603)PRK00912 (PRK00912)→ PRK03717 (PRK03717)POP5 (COG1369)→ |
| Mma-3-32 | NZ_JJPT01000015.1 | - | 33938 | 33887 | **RNA**→ PRK04243 (PRK04243)Ribosomal_L15e (pfam00827)→ COG1325 (COG1325)PRK14555 (PRK14555)<br>RPP1 (COG1603)PRK00912 (PRK00912)→ PRK03717 (PRK03717)POP5 (COG1369)→ |
| Mma-3-33 | NZ_JJPW01000129.1 | - | 35393 | 35342 | **RNA**→ PRK04243 (PRK04243)Ribosomal_L15e (pfam00827)→ COG1325 (COG1325)PRK14555 (PRK14555)<br>RPP1 (COG1603)PRK00912 (PRK00912)→ PRK03717 (PRK03717)POP5 (COG1369)→ |
| Mma-3-34 | NZ_JJQQ01000052.1 | - | 35254 | 35203 | **RNA**→ PRK04243 (PRK04243)Ribosomal_L15e (pfam00827)→ COG1325 (COG1325)PRK14555 (PRK14555)<br>RPP1 (COG1603)PRK00912 (PRK00912)→ PRK03717 (PRK03717)POP5 (COG1369)→ |
| Mma-3-35 | NZ_JJQZ01000044.1 | - | 45155 | 45104 | **RNA**→ PRK04243 (PRK04243)Ribosomal_L15e (pfam00827)→ COG1325 (COG1325)PRK14555 (PRK14555)<br>RPP1 (COG1603)PRK00912 (PRK00912)→ PRK03717 (PRK03717)POP5 (COG1369)→ |
| Mma-3-36 | NZ_JJPI01000027.1 | + | 21733 | 21784 | **RNA**→ PRK04243 (PRK04243)Ribosomal_L15e (pfam00827)→ COG1325 (COG1325)PRK14555 (PRK14555)<br>RPP1 (COG1603)PRK00912 (PRK00912)→ PRK03717 (PRK03717)POP5 (COG1369)→ |
| Mma-3-37 | NZ_JJPQ01000027.1 | + | 11208 | 11259 | **RNA**→ PRK04243 (PRK04243)Ribosomal_L15e (pfam00827)→ COG1325 (COG1325)PRK14555 (PRK14555)<br>RPP1 (COG1603)PRK00912 (PRK00912)→ PRK03717 (PRK03717)POP5 (COG1369)→ |
| Mma-3-38 | NZ_JJQH01000164.1 | - | 45155 | 45104 | **RNA**→ PRK04243 (PRK04243)Ribosomal_L15e (pfam00827)→ COG1325 (COG1325)PRK14555 (PRK14555)<br>RPP1 (COG1603)PRK00912 (PRK00912)→ PRK03717 (PRK03717)POP5 (COG1369)→ |
| Mma-3-39 | NZ_JJPD01000048.1 | + | 11958 | 12009 | **RNA**→ PRK04243 (PRK04243)Ribosomal_L15e (pfam00827)→ COG1325 (COG1325)PRK14555 (PRK14555)<br>RPP1 (COG1603)PRK00912 (PRK00912)→ PRK03717 (PRK03717)POP5 (COG1369)→ |
| Mma-3-40 | NZ_JJPJ01000102.1 | - | 35123 | 35072 | **RNA**→ PRK04243 (PRK04243)Ribosomal_L15e (pfam00827)→ COG1325 (COG1325)PRK14555 (PRK14555)<br>RPP1 (COG1603)PRK00912 (PRK00912)→ PRK03717 (PRK03717)POP5 (COG1369)→ |
| Mma-3-41 | NZ_JJPK01000008.1 | + | 160849 | 160900 | **RNA**→ PRK04243 (PRK04243)Ribosomal_L15e (pfam00827)→ COG1325 (COG1325)PRK14555 (PRK14555)<br>RPP1 (COG1603)PRK00912 (PRK00912)→ PRK03717 (PRK03717)POP5 (COG1369)→ |
| Mma-3-42 | NZ_JJPZ01000008.1 | + | 11958 | 12009 | **RNA**→ PRK04243 (PRK04243)Ribosomal_L15e (pfam00827)→ COG1325 (COG1325)PRK14555 (PRK14555)<br>RPP1 (COG1603)PRK00912 (PRK00912)→ PRK03717 (PRK03717)POP5 (COG1369)→ |
| Mma-3-43 | NZ_JJQF01000099.1 | - | 13178 | 13127 | **RNA**→ PRK04243 (PRK04243)Ribosomal_L15e (pfam00827)→ COG1325 (COG1325)PRK14555 (PRK14555)<br>RPP1 (COG1603)PRK00912 (PRK00912)→ PRK03717 (PRK03717)POP5 (COG1369)→ |
| Mma-3-44 | NZ_JJPL01000149.1 | + | 76874 | 76925 | **RNA**→ PRK04243 (PRK04243)Ribosomal_L15e (pfam00827)→ COG1325 (COG1325)PRK14555 (PRK14555)<br>RPP1 (COG1603)PRK00912 (PRK00912)→ PRK03717 (PRK03717)POP5 (COG1369)→ |
| Mma-3-45 | NZ_JJPV01000067.1 | - | 142282 | 142231 | **RNA**→ PRK04243 (PRK04243)Ribosomal_L15e (pfam00827)→ COG1325 (COG1325)PRK14555 (PRK14555)<br>RPP1 (COG1603)PRK00912 (PRK00912)→ PRK03717 (PRK03717)POP5 (COG1369)→ |
| Mma-3-46 | NZ_JJPM01000067.1 | + | 8095 | 8146 | **RNA**→ PRK04243 (PRK04243)Ribosomal_L15e (pfam00827)→ COG1325 (COG1325)PRK14555 (PRK14555)<br>RPP1 (COG1603)PRK00912 (PRK00912)→ PRK03717 (PRK03717)POP5 (COG1369)→ |
| Mma-3-47 | NZ_JJPU01000129.1 | - | 54102 | 54051 | **RNA**→ PRK04243 (PRK04243)Ribosomal_L15e (pfam00827)→ COG1325 (COG1325)PRK14555 (PRK14555)<br>RPP1 (COG1603)PRK00912 (PRK00912)→ PRK03717 (PRK03717)POP5 (COG1369)→ |
| Mma-3-48 | NZ_JJQU01000233.1 | + | 21011 | 21062 | **RNA**→ PRK04243 (PRK04243)Ribosomal_L15e (pfam00827)→ COG1325 (COG1325)PRK14555 (PRK14555)<br>RPP1 (COG1603)PRK00912 (PRK00912)→ PRK03717 (PRK03717)POP5 (COG1369)→ |
| Mma-3-49 | NZ_JJPE01000037.1 | - | 32163 | 32112 | **RNA**→ PRK04243 (PRK04243)Ribosomal_L15e (pfam00827)→ COG1325 (COG1325)PRK14555 (PRK14555)<br>RPP1 (COG1603)PRK00912 (PRK00912)→ PRK03717 (PRK03717)POP5 (COG1369)→ |
| Mma-3-50 | NZ_JJQK01000170.1 | + | 11911 | 11962 | **RNA**→ PRK04243 (PRK04243)Ribosomal_L15e (pfam00827)→ COG1325 (COG1325)PRK14555 (PRK14555)<br>RPP1 (COG1603)PRK00912 (PRK00912)→ PRK03717 (PRK03717)POP5 (COG1369)→ |
| Mma-3-51 | NZ_JJQO01000228.1 | + | 47525 | 47576 | **RNA**→ PRK04243 (PRK04243)Ribosomal_L15e (pfam00827)→ COG1325 (COG1325)PRK14555 (PRK14555)<br>RPP1 (COG1603)PRK00912 (PRK00912)→ PRK03717 (PRK03717)POP5 (COG1369)→ |
| Mma-3-52 | NZ_JJQJ01000089.1 | - | 22779 | 22728 | **RNA**→ PRK04243 (PRK04243)Ribosomal_L15e (pfam00827)→ COG1325 (COG1325)PRK14555 (PRK14555)<br>RPP1 (COG1603)PRK00912 (PRK00912)→ PRK03717 (PRK03717)POP5 (COG1369)→ |

| | | | | | |
|---|---|---|---|---|---|
| Mma-3-53 | NZ_JJQP01000172.1 | + | 39888 | 39939 | **RNA**→ PRK04243 (PRK04243)Ribosomal_L15e (pfam00827)→ COG1325 (COG1325)PRK14555 (PRK14555)<br>RPP1 (COG1603)PRK00912 (PRK00912)→ PRK03717 (PRK03717)POP5 (COG1369)→ |
| Mma-3-54 | NZ_JJQN01000109.1 | + | 11937 | 11988 | **RNA**→ PRK04243 (PRK04243)Ribosomal_L15e (pfam00827)→ COG1325 (COG1325)PRK14555 (PRK14555)<br>RPP1 (COG1603)PRK00912 (PRK00912)→ PRK03717 (PRK03717)POP5 (COG1369)→ |
| Mma-3-55 | NZ_JJQT01000136.1 | - | 6188 | 6137 | **RNA**→ PRK04243 (PRK04243)Ribosomal_L15e (pfam00827)→ COG1325 (COG1325)PRK14555 (PRK14555)<br>RPP1 (COG1603)PRK00912 (PRK00912)→ PRK03717 (PRK03717)POP5 (COG1369)→ |
| Mma-3-56 | NZ_JJQV01000102.1 | - | 35394 | 35343 | **RNA**→ PRK04243 (PRK04243)Ribosomal_L15e (pfam00827)→ COG1325 (COG1325)PRK14555 (PRK14555)<br>RPP1 (COG1603)PRK00912 (PRK00912)→ PRK03717 (PRK03717)POP5 (COG1369)→ |
| Mma-9-1 | NZ_CP009509.1 | + | 2909485 | 2909536 | **RNA**→ PRK04243 (PRK04243)Ribosomal_L15e (pfam00827)→ COG1325 (COG1325)PRK14555 (PRK14555)<br>RPP1 (COG1603)PRK00912 (PRK00912)→ PRK03717 (PRK03717)POP5 (COG1369)→ |
| Mma-4-1 | NZ_CP009514.1 | + | 1235377 | 1235428 | **RNA**→ PRK04243 (PRK04243)Ribosomal_L15e (pfam00827)→ COG1325 (COG1325)PRK14555 (PRK14555)<br>RPP1 (COG1603)PRK00912 (PRK00912)→ PRK03717 (PRK03717)POP5 (COG1369)→ |
| Mac-1-1 | NC_003552.1 | - | 2189802 | 2189751 | **RNA**→ PRK04243 (PRK04243)Ribosomal_L15e (pfam00827)→ COG1325 (COG1325)PRK14555 (PRK14555)<br>RPP1 (COG1603)PRK00912 (PRK00912)→ PRK03717 (PRK03717)POP5 (COG1369)→ |
| Msi-2-1 | NZ_CP009506.1 | - | 3147028 | 3146977 | **RNA**→ PRK04243 (PRK04243)Ribosomal_L15e (pfam00827)→ COG1325 (COG1325)PRK14555 (PRK14555)<br>RPP1 (COG1603)PRK00912 (PRK00912)→ PRK03717 (PRK03717)POP5 (COG1369)→ |
| env-635 | WSSedB2T_c104918 | + | 74 | 125 | **RNA**→ RPL15A (COG1632)PRK04243 (PRK04243)→ |
| Msi-1-1 | NZ_CP009508.1 | - | 2149794 | 2149743 | **RNA**→ PRK04243 (PRK04243)Ribosomal_L15e (pfam00827)→ COG1325 (COG1325)PRK14555 (PRK14555)<br>RPP1 (COG1603)PRK00912 (PRK00912)→ PRK03717 (PRK03717)POP5 (COG1369)→ |
| env-636 | M940CN_1003840 | - | 2106 | 2055 | **RNA**→ PRK04243 (PRK04243)Ribosomal_L15e (pfam00827)→ COG1325 (COG1325)PRK14555 (PRK14555)<br>RPP1 (COG1603)PRK00912 (PRK00912)→ PRK03717 (PRK03717)RNase_P_Rpp14 (pfam01900)→ |
| Msp-14-1 | NZ_CP009505.1 | - | 2602304 | 2602254 | **RNA**→ ←hypo |
| env-637 | 3300001567_10002926 | - | 2776 | 2730 | **RNA**→ PRK04243 (PRK04243)Ribosomal_L15e (pfam00827)→ COG1325 (COG1325)PRK14555 (PRK14555)<br>RPP1 (COG1603)PRK00912 (PRK00912)→ PRK03717 (PRK03717)POP5 (COG1369)→<br>PRK03996 (PRK03996)proteasome_alpha (cd01911)→ |
| env-638 | DHTCA2_contig15787 | + | 370 | 416 | **RNA**→ |
| env-639 | AMWB01080572.1 | + | 1326 | 1372 | **RNA**→ PRK04243 (PRK04243)Ribosomal_L15e (pfam00827)→ |
| env-640 | 3300001580_10062933 | + | 412 | 459 | **RNA**→ PRK04243 (PRK04243)Ribosomal_L15e (pfam00827)→<br>PRK14555 (PRK14555)RNA_binding (pfam01877)→ RPP1 (COG1603)PRK00912 (PRK00912)→ |
| Mho-1-1 | NC_019977.1 | + | 1869779 | 1869826 | **RNA**→ PRK04243 (PRK04243)Ribosomal_L15e (pfam00827)→<br>PRK14555 (PRK14555)RNA_binding (pfam01877)→ RPP1 (COG1603)PRK00912 (PRK00912)→<br>PRK03717 (PRK03717)POP5 (COG1369)→ PRK03996 (PRK03996)proteasome_alpha_archeal (cd03756)→<br>Sdo1 (COG1500)PRK13760 (PRK13760)→ Rrp4 (COG1097)PRK04163 (PRK04163)→<br>RNase_PH_archRRP41 (cd11366)PRK03983 (PRK03983)PRK05658 (PRK05658)→<br>RNase_PH_archRRP42 (cd11365)PRK04282 (PRK04282)→ rpl37ae (PRK03976)Ribosomal_L37ae (pfam01780)→<br>RPC10 (COG1996)rpoP (PRK00398)→ Brix (smart00879)IMP4 (COG2136)→<br>Pcc1 (COG2892)Pcc1 (pfam09341)→ gimC_beta (TIGR02338)GimC (COG1382)→<br>PRK05427 (PRK05427)DHH (pfam01368)→ PaaK (COG1541)AMP-binding_C_2 (pfam14535)→ |
| env-641 | AMWB01126899.1 | + | 489 | 536 | **RNA**→ PRK04243 (PRK04243)Ribosomal_L15e (pfam00827)→<br>PRK14555 (PRK14555)RNA_binding (pfam01877)→ PRK00912 (PRK00912)RNase_P_p30 (pfam01876)→<br>PRK03717 (PRK03717)POP5 (COG1369)→ PRK03996 (PRK03996)proteasome_alpha_archeal (cd03756)→ |
| Mbu-1-1 | NC_007955.1 | + | 176502 | 176552 | **RNA**→ PRK04243 (PRK04243)Ribosomal_L15e (pfam00827)→<br>PRK14555 (PRK14555)RNA_binding (pfam01877)→ PRK00912 (PRK00912)RNase_P_p30 (pfam01876)→<br>PRK03717 (PRK03717)RNase_P_Rpp14 (pfam01900)→ |
| env-642 | AZIB01003834.1 | + | 91 | 141 | **RNA**→ PRK04243 (PRK04243)Ribosomal_L15e (pfam00827)→<br>PRK14555 (PRK14555)RNA_binding (pfam01877)→ |
| Mme-2-1 | NZ_CP009518.1 | + | 794999 | 795052 | **RNA**→ PRK04243 (PRK04243)Ribosomal_L15e (pfam00827)→ COG1325 (COG1325)PRK14555 (PRK14555)→<br>RPP1 (COG1603)PRK00912 (PRK00912)→ PRK03717 (PRK03717)RNase_P_Rpp14 (pfam01900)→ |
| Mme-1-1 | NZ_JRHO01000010.1 | + | 86627 | 86680 | **RNA**→ PRK04243 (PRK04243)Ribosomal_L15e (pfam00827)→ COG1325 (COG1325)PRK14555 (PRK14555)→<br>PRK00912 (PRK00912)RNase_P_p30 (pfam01876)→ PRK03717 (PRK03717)POP5 (COG1369)→ |
| Msh-1-1 | NZ_JONQ01000008.1 | + | 45804 | 45849 | **RNA**→ RPL15A (COG1632)PRK04243 (PRK04243)→ PRK14555 (PRK14555)RNA_binding (pfam01877)→<br>RPP1 (COG1603)PRK00912 (PRK00912)→ |
| env-643 | HCF12C_96986 | - | 646 | 605 | **RNA**→ PRK04243 (PRK04243)Ribosomal_L15e (pfam00827)→ |
| env-644 | AUXO016695324.1 | - | 1274 | 1233 | **RNA**→ PRK04243 (PRK04243)Ribosomal_L15e (pfam00827)→ |
| env-645 | JGI12104J13512_1011404 | - | 2541 | 2501 | **RNA**→ PRK04243 (PRK04243)Ribosomal_L15e (pfam00827)→<br>PRK14555 (PRK14555)RNA_binding (pfam01877)→ RPP1 (COG1603)PRK00912 (PRK00912)→<br>PRK03717 (PRK03717)POP5 (COG1369)→ |

| | | | | | |
|---|---|---|---|---|---|
| env-646 | 3300001580_10676249 | + | 109 | 149 | **RNA**→ PRK04243 (PRK04243)Ribosomal_L15e (pfam00827)→ |
| env-647 | 08_07_CONTROL_DNA_scaffold24236_7 | - | 6049 | 6007 | **RNA**→ PRK04243 (PRK04243)Ribosomal_L15e (pfam00827)→ PRK14555 (PRK14555)RNA_binding (pfam01877)→ |
| env-648 | 08_07_XOS_DNA_scaffold878_1 | + | 2078 | 2120 | **RNA**→ PRK04243 (PRK04243)Ribosomal_L15e (pfam00827)→ |
| env-649 | O8_19_POLYDEXT_DNA_scaffold34267_3 | - | 167 | 125 | **RNA**→ PRK04243 (PRK04243)Ribosomal_L15e (pfam00827)→ |
| env-650 | JGI994J12354_10354549 | - | 293 | 251 | **RNA**→ PRK04243 (PRK04243)Ribosomal_L15e (pfam00827)→ |
| env-651 | JGI24713J26584_10000181 | - | 19302 | 19259 | **RNA**→ PRK04243 (PRK04243)Ribosomal_L15e (pfam00827)→ COG1325 (COG1325)PRK14555 (PRK14555)→ RPP1 (COG1603)PRK00912 (PRK00912)→ PRK03717 (PRK03717)POP5 (COG1369)→ PRK03996 (PRK03996)proteasome_alpha_archeal (cd03756)→ Sdo1 (COG1500)PRK13760 (PRK13760)→ Rrp4 (COG1097)PRK04163 (PRK04163)→ RNase_PH_archRRP41 (cd11366)PRK03983 (PRK03983)→ RNase_PH_archRRP42 (cd11365)PRK04282 (PRK04282)→ |
| env-652 | JGI24714J26587_10000218 | + | 19819 | 19862 | **RNA**→ PRK04243 (PRK04243)Ribosomal_L15e (pfam00827)→ COG1325 (COG1325)PRK14555 (PRK14555)→ RPP1 (COG1603)PRK00912 (PRK00912)→ PRK03717 (PRK03717)POP5 (COG1369)→ PRK03996 (PRK03996)proteasome_alpha_archeal (cd03756)→ Sdo1 (COG1500)PRK13760 (PRK13760)→ Rrp4 (COG1097)PRK04163 (PRK04163)→ RNase_PH_archRRP41 (cd11366)PRK03983 (PRK03983)→ RNase_PH_archRRP42 (cd11365)PRK04282 (PRK04282)→ |
| env-653 | JGI24504J29685_1000272 | - | 114106 | 114063 | **RNA**→ PRK04243 (PRK04243)Ribosomal_L15e (pfam00827)→ COG1325 (COG1325)PRK14555 (PRK14555)→ RPP1 (COG1603)PRK00912 (PRK00912)→ PRK03717 (PRK03717)POP5 (COG1369)→ PRK03996 (PRK03996)proteasome_alpha_archeal (cd03756)→ Sdo1 (COG1500)PRK13760 (PRK13760)→ Rrp4 (COG1097)PRK04163 (PRK04163)→ RNase_PH_archRRP41 (cd11366)PRK03983 (PRK03983)→ RNase_PH_archRRP42 (cd11365)PRK04282 (PRK04282)→ |
| env-654 | JGI24504J29685_1000273 | + | 20634 | 20677 | **RNA**→ PRK04243 (PRK04243)Ribosomal_L15e (pfam00827)→ COG1325 (COG1325)PRK14555 (PRK14555)→ RPP1 (COG1603)PRK00912 (PRK00912)→ PRK03717 (PRK03717)POP5 (COG1369)→ PRK03996 (PRK03996)proteasome_alpha_archeal (cd03756)→ Sdo1 (COG1500)PRK13760 (PRK13760)→ Rrp4 (COG1097)PRK04163 (PRK04163)→ RNase_PH_archRRP41 (cd11366)PRK03983 (PRK03983)→ RNase_PH_archRRP42 (cd11365)PRK04282 (PRK04282)→ |
| env-655 | JGI24505J29691_1005987 | + | 2006 | 2049 | **RNA**→ PRK04243 (PRK04243)Ribosomal_L15e (pfam00827)→ COG1325 (COG1325)PRK14555 (PRK14555)→ RPP1 (COG1603)PRK00912 (PRK00912)→ PRK03717 (PRK03717)POP5 (COG1369)→ PRK03996 (PRK03996)proteasome_alpha_archeal (cd03756)→ Sdo1 (COG1500)PRK13760 (PRK13760)→ Rrp4 (COG1097)PRK04163 (PRK04163)→ RNase_PH_archRRP41 (cd11366)PRK03983 (PRK03983)→ RNase_PH_archRRP42 (cd11365)PRK04282 (PRK04282)→ |
| env-656 | RUMENNODE_3862964_1 | + | 19057 | 19104 | **RNA**→ hypo→ PRK04243 (PRK04243)Ribosomal_L15e (pfam00827)→ PRK14555 (PRK14555)RNA_binding (pfam01877)→ hypo→ |
| env-657 | AUXO010246320.1 | + | 2947 | 2993 | **RNA**→ |
| env-658 | RUMENNODE_3957055_1 | + | 1849 | 1895 | **RNA**→ PRK04243 (PRK04243)Ribosomal_L15e (pfam00827)→ PRK14555 (PRK14555)RNA_binding (pfam01877)→ |
| Mve-1-1 | NZ_JQJK01000008.1 | + | 40819 | 40860 | **RNA**→ PRK04243 (PRK04243)Ribosomal_L15e (pfam00827)→ PRK14555 (PRK14555)RNA_binding (pfam01877)→ RPP1 (COG1603)PRK00912 (PRK00912)→ PRK03717 (PRK03717)RNase_P_Rpp14 (pfam01900)→ |
| Mar-1-1 | NZ_JQKN01000001.1 | + | 40819 | 40860 | **RNA**→ PRK04243 (PRK04243)Ribosomal_L15e (pfam00827)→ PRK14555 (PRK14555)RNA_binding (pfam01877)→ RPP1 (COG1603)PRK00912 (PRK00912)→ PRK03717 (PRK03717)RNase_P_Rpp14 (pfam01900)→ |
| env-659 | JGI24731J21663_1049964 | - | 360 | 319 | **RNA**→ RPL15A (COG1632)PRK04243 (PRK04243)→ |
| env-660 | B3_all_c_Cons195859 | + | 487 | 528 | **RNA**→ RPL15A (COG1632)PRK04243 (PRK04243)→ |
| Msp-4-1 | NC_015574.1 | + | 503676 | 503718 | **RNA**→ PRK04243 (PRK04243)Ribosomal_L15e (pfam00827)→ PRK14555 (PRK14555)RNA_binding (pfam01877)→ RPP1 (COG1603)PRK00912 (PRK00912)→ |
| env-661 | 2014734625 | + | 11972 | 12018 | **RNA**→ PRK04243 (PRK04243)Ribosomal_L15e (pfam00827)→ PRK14555 (PRK14555)RNA_binding (pfam01877)→ RPP1 (COG1603)PRK00912 (PRK00912)→ |
| env-662 | ANASMECb_4085297_C2853 | + | 11972 | 12018 | **RNA**→ PRK04243 (PRK04243)Ribosomal_L15e (pfam00827)→ PRK14555 (PRK14555)RNA_binding (pfam01877)→ RPP1 (COG1603)PRK00912 (PRK00912)→ |
| env-663 | 3300001580_10131017 | - | 51 | 5 | **RNA**→ |
| env-664 | 3300001592_10138106 | - | 985 | 939 | **RNA**→ PRK04243 (PRK04243)Ribosomal_L15e (pfam00827)→ PRK14555 (PRK14555)RNA_binding (pfam01877)→ |
| env-665 | JGI24712J26585_10348213 | - | 47 | 4 | **RNA**→ hypo→ |

| | | | | | |
|---|---|---|---|---|---|
| env-666 | JGI24709J26583_10002022 | - | 8087 | 8044 | **RNA**→ PRK04243 (PRK04243)Ribosomal_L15e (pfam00827)→<br>PRK14555 (PRK14555)RNA_binding (pfam01877)→ RPP1 (COG1603)PRK00912 (PRK00912)→<br>PRK03717 (PRK03717)RNase_P_Rpp14 (pfam01900)→<br>PRK03996 (PRK03996)proteasome_alpha_archeal (cd03756)→ Sdo1 (COG1500)PRK13760 (PRK13760)→<br>Rrp4 (COG1097)PRK04163 (PRK04163)→ RNase_PH_archRRP41 (cd11366)PRK03983 (PRK03983)→<br>RNase_PH_archRRP42 (cd11365)PRK04282 (PRK04282)→ |
| env-667 | JGI24710J26742_10000204 | + | 18022 | 18065 | **RNA**→ PRK04243 (PRK04243)Ribosomal_L15e (pfam00827)→<br>PRK14555 (PRK14555)RNA_binding (pfam01877)→ RPP1 (COG1603)PRK00912 (PRK00912)→<br>PRK03717 (PRK03717)RNase_P_Rpp14 (pfam01900)→<br>PRK03996 (PRK03996)proteasome_alpha_archeal (cd03756)→ Sdo1 (COG1500)PRK13760 (PRK13760)→<br>Rrp4 (COG1097)PRK04163 (PRK04163)→ RNase_PH_archRRP41 (cd11366)PRK03983 (PRK03983)→<br>RNase_PH_archRRP42 (cd11365)PRK04282 (PRK04282)→ |
| env-668 | JGI24500J29687_10014623 | + | 1789 | 1832 | **RNA**→ hypo→ PRK04243 (PRK04243)Ribosomal_L15e (pfam00827)→<br>PRK14555 (PRK14555)RNA_binding (pfam01877)→ RPP1 (COG1603)PRK00912 (PRK00912)→<br>PRK03717 (PRK03717)RNase_P_Rpp14 (pfam01900)→<br>PRK03996 (PRK03996)proteasome_alpha_archeal (cd03756)→ Sdo1 (COG1500)PRK13760 (PRK13760)→<br>Rrp4 (COG1097)PRK04163 (PRK04163)→ RNase_PH_archRRP41 (cd11366)PRK03983 (PRK03983)→<br>RNase_PH_archRRP42 (cd11365)PRK04282 (PRK04282)→ |
| env-669 | JGI24501J29690_1010023 | + | 224 | 267 | **RNA**→ hypo→ PRK04243 (PRK04243)Ribosomal_L15e (pfam00827)→<br>PRK14555 (PRK14555)RNA_binding (pfam01877)→ RPP1 (COG1603)PRK00912 (PRK00912)→<br>PRK03717 (PRK03717)RNase_P_Rpp14 (pfam01900)→<br>PRK03996 (PRK03996)proteasome_alpha_archeal (cd03756)→ Sdo1 (COG1500)PRK13760 (PRK13760)→<br>Rrp4 (COG1097)PRK04163 (PRK04163)→ RNase_PH_archRRP41 (cd11366)PRK03983 (PRK03983)→<br>RNase_PH_archRRP42 (cd11365)PRK04282 (PRK04282)→ |
| env-670 | JGI24711J26586_10150374 | - | 270 | 227 | **RNA**→ PRK04243 (PRK04243)Ribosomal_L15e (pfam00827)→ |
| env-671 | JGI24731J21663_1027378 | - | 1115 | 1069 | **RNA**→ PRK04243 (PRK04243)Ribosomal_L15e (pfam00827)→<br>PRK14555 (PRK14555)RNA_binding (pfam01877)→ |
| env-672 | JGI24729J20445_1009801 | + | 789 | 835 | **RNA**→ PRK04243 (PRK04243)Ribosomal_L15e (pfam00827)→<br>PRK14555 (PRK14555)RNA_binding (pfam01877)→ RPP1 (COG1603)PRK00912 (PRK00912)→ |
| Mfo-2-1 | NZ_AMPO01000006.1 | + | 4040 | 4086 | **RNA**→ PRK04243 (PRK04243)Ribosomal_L15e (pfam00827)→<br>PRK04243 (PRK04243)Ribosomal_L15e (pfam00827)→ PRK14555 (PRK14555)RNA_binding (pfam01877)→<br>PRK14555 (PRK14555)RNA_binding (pfam01877)→ RPP1 (COG1603)PRK00912 (PRK00912)→<br>RPP1 (COG1603)PRK00912 (PRK00912)→ |
| env-673 | JGI20214J14112_1380804 | - | 244 | 198 | **RNA**→ RPL15A (COG1632)PRK04243 (PRK04243)→ |
| env-674 | JGI24731J21663_1005210 | + | 4307 | 4353 | **RNA**→ PRK04243 (PRK04243)Ribosomal_L15e (pfam00827)→<br>PRK14555 (PRK14555)RNA_binding (pfam01877)→ RPP1 (COG1603)PRK00912 (PRK00912)→ |
| Msp-2-1 | NC_023044.1 | + | 529281 | 529327 | **RNA**→ PRK04243 (PRK04243)Ribosomal_L15e (pfam00827)→<br>PRK14555 (PRK14555)RNA_binding (pfam01877)→ RPP1 (COG1603)PRK00912 (PRK00912)→<br>PRK03717 (PRK03717)RNase_P_Rpp14 (pfam01900)→<br>PRK03996 (PRK03996)proteasome_alpha_archeal (cd03756)→ Sdo1 (COG1500)PRK13760 (PRK13760)→<br>Rrp4 (COG1097)PRK04163 (PRK04163)→ RNase_PH_archRRP41 (cd11366)PRK03983 (PRK03983)→<br>RNase_PH_archRRP42 (cd11365)PRK04282 (PRK04282)→ |
| Mfo-1-1 | NZ_CP006933.1 | - | 1785014 | 1784968 | **RNA**→ PRK04243 (PRK04243)Ribosomal_L15e (pfam00827)→<br>PRK14555 (PRK14555)RNA_binding (pfam01877)→ RPP1 (COG1603)PRK00912 (PRK00912)→<br>PRK03717 (PRK03717)POP5 (COG1369)→ PRK03996 (PRK03996)proteasome_alpha_archeal (cd03756)→<br>Sdo1 (COG1500)PRK13760 (PRK13760)→ Rrp4 (COG1097)PRK04163 (PRK04163)→<br>RNase_PH_archRRP41 (cd11366)PRK03983 (PRK03983)→<br>RNase_PH_archRRP42 (cd11365)PRK04282 (PRK04282)→ |
| env-675 | PRSSGFe2_Sequence0000019669 | + | 951 | 992 | **RNA**→ PRK04243 (PRK04243)Ribosomal_L15e (pfam00827)→<br>PRK14555 (PRK14555)RNA_binding (pfam01877)→ |
| env-676 | JGI12104J13512_1023983 | - | 1071 | 1030 | **RNA**→ PRK04243 (PRK04243)Ribosomal_L15e (pfam00827)→<br>PRK14555 (PRK14555)RNA_binding (pfam01877)→ |
| env-677 | JGI11944J13513_1001397 | - | 15451 | 15410 | **RNA**→ PRK04243 (PRK04243)Ribosomal_L15e (pfam00827)→<br>PRK14555 (PRK14555)RNA_binding (pfam01877)→ PRK00912 (PRK00912)RNase_P_p30 (pfam01876)→<br>PRK03717 (PRK03717)POP5 (COG1369)→ |
| env-678 | 3300001592_10385652 | - | 169 | 128 | **RNA**→ PRK04243 (PRK04243)Ribosomal_L15e (pfam00827)→ |

| | | | | | |
|---|---|---|---|---|---|
| Msp-1-1 | NC_015216.1 | - | 2225955 | 2225907 | **RNA**→ PRK04243 (PRK04243)Ribosomal_L15e (pfam00827)→<br>PRK14555 (PRK14555)RNA_binding (pfam01877)→ RPP1 (COG1603)PRK00912 (PRK00912)→<br>RPP1 (COG1603)PRK00912 (PRK00912)→ PRK03717 (PRK03717)RNase_P_Rpp14 (pfam01900)→ |
| env-679 | JGI2065J20421_1000005 | + | 287784 | 287832 | **RNA**→ PRK04243 (PRK04243)Ribosomal_L15e (pfam00827)→<br>PRK14555 (PRK14555)RNA_binding (pfam01877)→ RPP1 (COG1603)PRK00912 (PRK00912)→ |
| env-680 | WSSedB2Ca_1123116 | - | 192 | 151 | **RNA**→ PRK04243 (PRK04243)Ribosomal_L15e (pfam00827)→ |
| Msp-3-1 | NZ_JQLY01000001.1 | + | 314798 | 314839 | **RNA**→ PRK04243 (PRK04243)Ribosomal_L15e (pfam00827)→<br>PRK14555 (PRK14555)RNA_binding (pfam01877)→ RPP1 (COG1603)PRK00912 (PRK00912)→<br>PRK03717 (PRK03717)POP5 (COG1369)→ |
| env-681 | B3_all_c_Cons30296 | + | 231 | 272 | **RNA**→ PRK04243 (PRK04243)Ribosomal_L15e (pfam00827)→ |
| env-682 | B3_all_c_Cons42794 | + | 295 | 336 | **RNA**→ RPL15A (COG1632)PRK04243 (PRK04243)→ |
| env-683 | SL_5KL_010_SED_10126917 | - | 228 | 187 | **RNA**→ PRK04243 (PRK04243)Ribosomal_L15e (pfam00827)→ |
| Mor-1-1 | NZ_HG796200.1 | - | 398447 | 398403 | **RNA**→ PRK04243 (PRK04243)Ribosomal_L15e (pfam00827)→<br>PRK14555 (PRK14555)RNA_binding (pfam01877)→ |
| Msm-14-1 | NZ_AELU01000003.1 | - | 71106 | 71058 | **RNA**→ PRK04243 (PRK04243)Ribosomal_L15e (pfam00827)→<br>PRK14555 (PRK14555)RNA_binding (pfam01877)→ YurZ (COG0599)→ |
| Msm-4-1 | NZ_DS996911.1 | + | 39506 | 39554 | **RNA**→ PRK04243 (PRK04243)Ribosomal_L15e (pfam00827)→<br>PRK14555 (PRK14555)RNA_binding (pfam01877)→ |
| env-684 | scaffold116291_3_V1.CD-8 | - | 666 | 618 | **RNA**→ PRK04243 (PRK04243)Ribosomal_L15e (pfam00827)→ |
| env-685 | 4491416.3_NODE_1031 | + | 40283 | 40331 | **RNA**→ PRK04243 (PRK04243)Ribosomal_L15e (pfam00827)→<br>PRK14555 (PRK14555)RNA_binding (pfam01877)→ YurZ (COG0599)→ |
| env-686 | scaffold101749_2_MH0011 | + | 6770 | 6818 | **RNA**→ PRK04243 (PRK04243)Ribosomal_L15e (pfam00827)→<br>PRK14555 (PRK14555)RNA_binding (pfam01877)→ YurZ (COG0599)→ |
| env-687 | scaffold45427_5_O2.UC-12 | + | 989 | 1037 | **RNA**→ PRK04243 (PRK04243)Ribosomal_L15e (pfam00827)→ |
| env-688 | scaffold45612_1_V1.CD-9 | - | 1221 | 1173 | **RNA**→ PRK04243 (PRK04243)Ribosomal_L15e (pfam00827)→<br>PRK14555 (PRK14555)RNA_binding (pfam01877)→ |
| env-689 | scaffold99220_1_V1.UC-8 | - | 16733 | 16685 | **RNA**→ PRK04243 (PRK04243)Ribosomal_L15e (pfam00827)→<br>PRK14555 (PRK14555)RNA_binding (pfam01877)→ |
| env-690 | UnmappedStool_Broad_scaffold_1133206 | + | 660 | 708 | **RNA**→ PRK04243 (PRK04243)Ribosomal_L15e (pfam00827)→<br>PRK14555 (PRK14555)RNA_binding (pfam01877)→ |
| env-691 | DLM022_scaffold90350_1 | + | 654 | 702 | **RNA**→ PRK04243 (PRK04243)Ribosomal_L15e (pfam00827)→<br>PRK14555 (PRK14555)RNA_binding (pfam01877)→ YurZ (COG0599)→ |
| env-692 | ADJS01014849.1 | - | 6089 | 6041 | **RNA**→ PRK04243 (PRK04243)Ribosomal_L15e (pfam00827)→<br>PRK14555 (PRK14555)RNA_binding (pfam01877)→ |
| env-693 | SRS014235_C3325073 | + | 168 | 216 | **RNA**→ PRK04243 (PRK04243)Ribosomal_L15e (pfam00827)→ |
| env-694 | SRS014979_C3300304 | - | 2323 | 2275 | **RNA**→ PRK04243 (PRK04243)Ribosomal_L15e (pfam00827)→<br>PRK14555 (PRK14555)RNA_binding (pfam01877)→ |
| Msm-1-1 | NZ_CCXV01000062.1 | - | 3684 | 3636 | **RNA**→ PRK04243 (PRK04243)Ribosomal_L15e (pfam00827)→<br>PRK14555 (PRK14555)RNA_binding (pfam01877)→ |
| Msm-19-1 | NZ_AELZ01000004.1 | - | 93410 | 93362 | **RNA**→ PRK04243 (PRK04243)Ribosomal_L15e (pfam00827)→<br>PRK14555 (PRK14555)RNA_binding (pfam01877)→ |
| Msm-11-1 | NZ_AELR01000006.1 | - | 74375 | 74327 | **RNA**→ PRK04243 (PRK04243)Ribosomal_L15e (pfam00827)→<br>PRK14555 (PRK14555)RNA_binding (pfam01877)→ |
| Msm-23-1 | NZ_AEMD01000013.1 | - | 30363 | 30315 | **RNA**→ PRK04243 (PRK04243)Ribosomal_L15e (pfam00827)→<br>PRK14555 (PRK14555)RNA_binding (pfam01877)→ |
| Msm-12-1 | NZ_AELS01000005.1 | - | 55986 | 55938 | **RNA**→ PRK04243 (PRK04243)Ribosomal_L15e (pfam00827)→<br>PRK14555 (PRK14555)RNA_binding (pfam01877)→ |
| Msm-13-1 | NZ_AELT01000005.1 | - | 60811 | 60763 | **RNA**→ PRK04243 (PRK04243)Ribosomal_L15e (pfam00827)→<br>PRK14555 (PRK14555)RNA_binding (pfam01877)→ |
| Msm-22-1 | NZ_AEMC01000004.1 | - | 93153 | 93105 | **RNA**→ PRK04243 (PRK04243)Ribosomal_L15e (pfam00827)→<br>PRK14555 (PRK14555)RNA_binding (pfam01877)→ |
| Msm-21-1 | NZ_AEMB01000003.1 | - | 93149 | 93101 | **RNA**→ PRK04243 (PRK04243)Ribosomal_L15e (pfam00827)→<br>PRK14555 (PRK14555)RNA_binding (pfam01877)→ |
| Msm-20-1 | NZ_AEMA01000007.1 | - | 93143 | 93095 | **RNA**→ PRK04243 (PRK04243)Ribosomal_L15e (pfam00827)→<br>PRK14555 (PRK14555)RNA_binding (pfam01877)→ |
| Msm-10-1 | NZ_AELQ01000008.1 | - | 55971 | 55923 | **RNA**→ PRK04243 (PRK04243)Ribosomal_L15e (pfam00827)→<br>PRK14555 (PRK14555)RNA_binding (pfam01877)→ |

| | | | | | |
|---|---|---|---|---|---|
| Msm-18-1 | NZ_AELY01000006.1 | - | 93416 | 93368 | **RNA**→ PRK04243 (PRK04243)Ribosomal_L15e (pfam00827)→ |
| | | | | | PRK14555 (PRK14555)RNA_binding (pfam01877)→ |
| Msm-9-1 | NZ_AELO01000021.1 | - | 3502 | 3454 | **RNA**→ PRK04243 (PRK04243)Ribosomal_L15e (pfam00827)→ |
| | | | | | PRK14555 (PRK14555)RNA_binding (pfam01877)→ |
| Msm-5-1 | NZ_AEKU01000009.1 | - | 30411 | 30363 | **RNA**→ PRK04243 (PRK04243)Ribosomal_L15e (pfam00827)→ |
| | | | | | PRK14555 (PRK14555)RNA_binding (pfam01877)→ |
| Msm-17-1 | NZ_AELX01000007.1 | - | 93439 | 93391 | **RNA**→ RPL15A (COG1632)PRK04243 (PRK04243)→ hypo→ hypo→ |
| | | | | | PRK14555 (PRK14555)RNA_binding (pfam01877)→ |
| Msm-8-1 | NZ_AELN01000005.1 | - | 30375 | 30327 | **RNA**→ PRK04243 (PRK04243)Ribosomal_L15e (pfam00827)→ |
| | | | | | PRK14555 (PRK14555)RNA_binding (pfam01877)→ |
| Msm-2-1 | NC_009515.1 | - | 288824 | 288776 | **RNA**→ PRK04243 (PRK04243)Ribosomal_L15e (pfam00827)→ |
| | | | | | PRK14555 (PRK14555)RNA_binding (pfam01877)→ YurZ (COG0599)→ |
| Msm-3-1 | NZ_GG704759.1 | - | 1679032 | 1678984 | **RNA**→ PRK04243 (PRK04243)Ribosomal_L15e (pfam00827)→ |
| | | | | | PRK14555 (PRK14555)RNA_binding (pfam01877)→ |
| env-695 | scaffold43629_1_V1.CD-14 | + | 589 | 637 | **RNA**→ RPL15A (COG1632)PRK04243 (PRK04243)→ |
| Msm-16-1 | NZ_AELW01000005.1 | - | 70811 | 70763 | **RNA**→ PRK04243 (PRK04243)Ribosomal_L15e (pfam00827)→ |
| | | | | | PRK14555 (PRK14555)RNA_binding (pfam01877)→ YurZ (COG0599)→ |
| Msm-15-1 | NZ_AELV01000008.1 | - | 55039 | 54991 | **RNA**→ PRK04243 (PRK04243)Ribosomal_L15e (pfam00827)→ |
| | | | | | PRK14555 (PRK14555)RNA_binding (pfam01877)→ YurZ (COG0599)→ |
| Msm-6-1 | NZ_AELL01000010.1 | - | 30405 | 30357 | **RNA**→ PRK04243 (PRK04243)Ribosomal_L15e (pfam00827)→ |
| | | | | | PRK14555 (PRK14555)RNA_binding (pfam01877)→ |
| Msm-7-1 | NZ_AELM01000013.1 | - | 30412 | 30364 | **RNA**→ PRK04243 (PRK04243)Ribosomal_L15e (pfam00827)→ |
| | | | | | PRK14555 (PRK14555)RNA_binding (pfam01877)→ |
| env-696 | JGI26339J46600_10088026 | - | 219 | 169 | **RNA**→ PRK04243 (PRK04243)Ribosomal_L15e (pfam00827)→ |
| env-697 | JGI24730J26740_1120805 | + | 4 | 51 | **RNA**→ RPL15A (COG1632)PRK04243 (PRK04243)→ |
| env-698 | LGVF01127568.1 | + | 648 | 697 | **RNA**→ |
| env-699 | LGOV01033634.1 | - | 1312 | 1261 | **RNA**→ PRK04243 (PRK04243)Ribosomal_L15e (pfam00827)→ |
| | | | | | PRK14555 (PRK14555)RNA_binding (pfam01877)→ PRK00912 (PRK00912)RNase_P_p30 (pfam01876)→ |
| env-700 | LGVE01048678.1 | - | 619 | 570 | **RNA**→ PRK04243 (PRK04243)Ribosomal_L15e (pfam00827)→ |
| env-701 | LGVF01407700.1 | + | 445 | 496 | **RNA**→ RPL15A (COG1632)PRK04243 (PRK04243)→ |
| env-702 | LGVE01104961.1 | - | 973 | 923 | **RNA**→ PRK04243 (PRK04243)Ribosomal_L15e (pfam00827)→ |
| env-703 | LGVF01373635.1 | - | 2320 | 2270 | **RNA**→ PRK04243 (PRK04243)Ribosomal_L15e (pfam00827)→ |
| env-704 | LGVE01091035.1 | + | 101 | 150 | **RNA**→ PRK04243 (PRK04243)Ribosomal_L15e (pfam00827)→ |
| env-705 | LGVF01174437.1 | - | 990 | 941 | **RNA**→ PRK04243 (PRK04243)Ribosomal_L15e (pfam00827)→ |
| env-706 | LGVE01094124.1 | + | 632 | 681 | **RNA**→ PRK04243 (PRK04243)Ribosomal_L15e (pfam00827)→ |
| env-707 | JGI24024J18818_10368636 | + | 225 | 274 | **RNA**→ |
| env-708 | LGVF01194351.1 | - | 1341 | 1292 | **RNA**→ PRK04243 (PRK04243)Ribosomal_L15e (pfam00827)→ |
| env-709 | PBDCA2_contig12847 | - | 760 | 713 | **RNA**→ PRK04243 (PRK04243)Ribosomal_L15e (pfam00827)→ |
| | | | | | PRK14555 (PRK14555)RNA_binding (pfam01877)→ |
| env-710 | JGI2161J19892_1008470 | - | 767 | 717 | **RNA**→ RPL15A (COG1632)PRK04243 (PRK04243)→ |
| env-711 | JGI2172J19969_10002521 | + | 546 | 596 | **RNA**→ RPL15A (COG1632)PRK04243 (PRK04243)→ |
| env-712 | JGI24422J19971_10970220 | + | 95 | 145 | **RNA**→ RPL15A (COG1632)PRK04243 (PRK04243)→ |
| env-713 | JGI2160J19893_10000003 | - | 51272 | 51222 | **RNA**→ RPL15A (COG1632)PRK04243 (PRK04243)→ |
| Ngr-1-1 | NC_019792.1 | - | 2417457 | 2417388 | **RNA**→ PRK04243 (PRK04243)Ribosomal_L15e (pfam00827)→ |
| Hti-1-1 | NC_021921.1 | - | 2128296 | 2128249 | **RNA**→ PRK04243 (PRK04243)Ribosomal_L15e (pfam00827)→ |
| Npe-1-1 | NC_019962.1 | - | 782478 | 782407 | **RNA**→ PRK04243 (PRK04243)Ribosomal_L15e (pfam00827)→ |
| Hsp-7-1 | NZ_ANPG01000166.1 | - | 1829 | 1769 | **RNA**→ PRK04243 (PRK04243)Ribosomal_L15e (pfam00827)→ |
| Hlu-1-1 | NZ_AOLH01000025.1 | + | 31699 | 31759 | **RNA**→ PRK04243 (PRK04243)Ribosomal_L15e (pfam00827)→ |
| Hal-1-1 | NZ_AOLL01000011.1 | + | 546240 | 546300 | **RNA**→ PRK04243 (PRK04243)Ribosomal_L15e (pfam00827)→ |
| Hvo-1-1 | NC_013967.1 | - | 495895 | 495835 | **RNA**→ PRK04243 (PRK04243)Ribosomal_L15e (pfam00827)→ |
| Hje-1-1 | NC_014297.1 | + | 2507160 | 2507224 | **RNA**→ PRK04243 (PRK04243)Ribosomal_L15e (pfam00827)→ |
| | | | | | ThiF_MoeB_HesA_family (cd00757)adenyl_thiF (TIGR02356)→ |
| Hbo-1-1 | NC_014729.1 | + | 2535983 | 2536036 | **RNA**→ PRK04243 (PRK04243)Ribosomal_L15e (pfam00827)→ |
| Nma-1-1 | NC_013922.1 | - | 988031 | 987966 | **RNA**→ PRK04243 (PRK04243)Ribosomal_L15e (pfam00827)→ |
| Nti-1-1 | NZ_KB913017.1 | + | 2799470 | 2799545 | **RNA**→ PRK04243 (PRK04243)Ribosomal_L15e (pfam00827)→ |

#### 4.3 Conserved domains

Conserved domains found in protein-coding genes listed in Section 4.2 are shown below, with the first sentence in their description from the Conserved Domain Database (if any). Conserved domains associated with more than one eL15-Euryarchaeota RNA are assigned a color, while others are shown in gray. The number in parentheses after the colored domain name is the number of occurrences in Section 4.2.

cd00075 (1) Histidine kinase-like ATPases; This family includes several ATP-binding proteins for example: histidine kinase, DNA gyrase B, topoisomerases, heat shock protein HSP90, phytochrome-like ATPases and DNA mismatch repair proteins
cd00267 (1) ATP-binding cassette transporter nucleotide-binding domain.
cd00293 (4) Usp: Universal stress protein family.
cd00757 (26) ThiF_MoeB_HesA.
cd01066 (1) A family including aminopeptidase P, aminopeptidase M, and prolidase.
cd01088 (1) Methionine Aminopeptidase 2.
cd01911 (1) proteasome alpha subunit.
cd02153 (2) The tRNA binding domain is also known as the Myf domain in literature.
cd03024 (2) DsbA family, FrnE subfamily; FrnE is a DsbA-like protein containing a CXXC motif.
cd03756 (76) proteasome_alpha_archeal.
cd06067 (3) Endopeptidases belonging to membrane-bound hydrogen evolving hydrogenase group. [NiFe]
cd11365 (19) RRP42 subunit of archaeal exosome.
cd11366 (19) RRP41 subunit of archaeal exosome.
cd11386 (1) Methyl-accepting chemotaxis protein (MCP), signaling domain.
COG0249 (3) DNA mismatch repair ATPase MutS [Replication, recombination and repair]
COG0317 (1) (p)ppGpp synthase/hydrolase, HD superfamily [Signal transduction mechanisms, Transcription]
COG0323 (1) DNA mismatch repair ATPase MutL [Replication, recombination and repair]
COG0599 (7) Uncharacterized conserved protein YurZ, alkylhydroperoxidase/carboxymuconolactone decarboxylase family [General function prediction only]
COG0638 (2) 20S proteasome, alpha and beta subunits [Posttranslational modification, protein turnover, chaperones]
COG0840 (5) Methyl-accepting chemotaxis protein [Cell motility, Signal transduction mechanisms]
COG0864 (3) Metal-responsive transcriptional regulator, contains CopG/Arc/MetJ DNA-binding domain [Transcription]
COG1097 (19) Exosome complex RNA-binding protein Rrp4, contains S1 and KH domains [Translation, ribosomal structure and biogenesis]
COG1145 (1) Ferredoxin [Energy production and conversion]
COG1146 (6) NAD-dependent dihydropyrimidine dehydrogenase, PreA subunit [Nucleotide transport and metabolism]
COG1215 (1) Glycosyltransferase, catalytic subunit of cellulose synthase and poly-beta-1,6-N-acetylglucosamine synthase [Cell motility]
COG1325 (106) Exosome subunit, RNA binding protein with dsRBD fold [Translation, ribosomal structure and biogenesis]
COG1369 (108) RNase P/RNase MRP subunit POP5 [Translation, ribosomal structure and biogenesis]
COG1382 (10) Prefoldin, chaperonin cofactor [Posttranslational modification, protein turnover, chaperones]
COG1500 (76) Ribosome maturation protein Sdo1 [Translation, ribosomal structure and biogenesis]
COG1541 (2) Phenylacetate-coenzyme A ligase PaaK, adenylate-forming domain family [Coenzyme transport and metabolism]
COG1603 (223) RNase P/RNase MRP subunit p30 [Translation, ribosomal structure and biogenesis]
COG1632 (147) Ribosomal protein L15E [Translation, ribosomal structure and biogenesis]
COG1741 (2) Redox-sensitive bicupin YhaK, pirin superfamily [General function prediction only]
COG1996 (35) DNA-directed RNA polymerase, subunit RPC12/RpoP, contains C4-type Zn-finger [Transcription]
COG1997 (1) Ribosomal protein L37AE/L43A [Translation, ribosomal structure and biogenesis]
COG2136 (11) rRNA maturation protein Rpf1, contains Brix/IMP4 (anticodon-binding) domain [Translation, ribosomal structure and biogenesis]
COG2761 (6) Predicted dithiol-disulfide isomerase, DsbA family [Posttranslational modification, protein turnover, chaperones]
COG2892 (25) tRNA threonylcarbamoyladenosine modification (KEOPS) complex, Pcc1 subunit [Translation, ribosomal structure and biogenesis]
COG3153 (1) Predicted N-acetyltransferase YhbS [General function prediction only]
COG3609 (6) Transcriptional regulator, contains Arc/MetJ-type RHH (ribbon-helix-helix) DNA-binding domain [Transcription]
COG4352 (80) Ribosomal protein L13E [Translation, ribosomal structure and biogenesis]
COG4585 (1) Signal transduction histidine kinase [Signal transduction mechanisms]
pfam00672 (6) HAMP domain.
pfam00827 (780) Ribosomal L15.
pfam01119 (1) DNA mismatch repair protein, C-terminal domain.
pfam01172 (1) Shwachman-Bodian-Diamond syndrome (SBDS) protein.
pfam01226 (1) Formate/nitrite transporter.
pfam01294 (1) Ribosomal protein L13e.
pfam01368 (2) DHH family.
pfam01451 (1) Low molecular weight phosphotyrosine protein phosphatase.
pfam01595 (1) Domain of unknown function DUF21.
pfam01624 (1) MutS domain I.
pfam01780 (58) Ribosomal L37ae protein family.
pfam01876 (70) RNase P subunit p30.
pfam01877 (163) RNA binding.
pfam01900 (136) Rpp14/Pop5 family.
pfam01920 (6) Prefoldin subunit.
pfam02592 (6) Putative vitamin uptake transporter.
pfam02771 (1) Acyl-CoA dehydrogenase, N-terminal domain.
pfam04087 (1) Domain of unknown function (DUF389).
pfam05048 (3) Periplasmic copper-binding protein (NosD).

pfam05639 (1) Pup-like protein. [EQ]
pfam06736 (3) Protein of unknown function (DUF1211).
pfam07705 (3) CARDB.
pfam08676 (1) MutL C terminal dimerisation domain.
pfam08753 (3) NikR C terminal nickel binding domain.
pfam09341 (23) Transcription factor Pcc1.
pfam09377 (1) SBDS protein C-terminal domain.
pfam12838 (5) 4Fe-4S dicluster domain.
pfam13187 (1) 4Fe-4S dicluster domain.
pfam13229 (3) Right handed beta helix region.
pfam13370 (1) 4Fe-4S single cluster domain of Ferredoxin I. [4Fe-4S]
pfam13462 (8) Thioredoxin.
pfam13508 (1) Acetyltransferase (GNAT) domain.
pfam14535 (2) AMP-binding enzyme C-terminal domain.
pfam16116 (3) Domain of unknown function (DUF4832).
PRK00095 (1) DNA mismatch repair protein; Reviewed
PRK00398 (35) DNA-directed RNA polymerase subunit P; Provisional
PRK00912 (296) ribonuclease P protein component 3; Provisional
PRK00933 (11) ribosomal biogenesis protein; Validated
PRK03717 (241) ribonuclease P protein component 2; Provisional
PRK03892 (2) ribonuclease P protein component 3; Provisional
PRK03976 (59) 50S ribosomal protein L37Ae; Reviewed
PRK03983 (19) exosome complex exonuclease Rrp41; Provisional
PRK03996 (77) proteasome subunit alpha; Provisional
PRK04163 (19) exosome complex RNA-binding protein Rrp4; Provisional
PRK04243 (927) 50S ribosomal protein L15e; Validated
PRK04282 (19) exosome complex RNA-binding protein Rrp42; Provisional
PRK04460 (6) nickel responsive regulator; Provisional
PRK05399 (1) DNA mismatch repair protein MutS; Provisional
PRK05427 (1) putative manganese-dependent inorganic pyrophosphatase; Provisional
PRK05658 (1) RNA polymerase sigma factor RpoD; Validated
PRK08671 (2) methionine aminopeptidase; Provisional
PRK08762 (10) molybdopterin biosynthesis protein MoeB; Validated
PRK09343 (3) prefoldin subunit beta; Provisional
PRK10089 (2) tRNA-binding protein; Provisional
PRK10935 (2) nitrate/nitrite sensor protein NarQ; Provisional
PRK12277 (81) 50S ribosomal protein L13e; Provisional
PRK13760 (76) putative RNA-associated protein; Provisional
PRK14538 (1) putative bifunctional signaling protein/50S ribosomal protein L9; Provisional
PRK14555 (267) hypothetical protein; Provisional
PRK14887 (2) KEOPS complex Pcc1-like subunit; Provisional
PRK14960 (3) DNA polymerase III subunits gamma and tau; Provisional
PRK15005 (4) universal stress protein F; Provisional
smart00283 (6) Methyl-accepting chemotaxis-like domains (chemotaxis sensory transducer).
smart00382 (1) ATPases associated with a variety of cellular activities.
smart00387 (1) Histidine kinase-like ATPases.
smart00533 (2) DNA-binding domain of DNA mismatch repair MUTS family.
smart00534 (2) ATPase domain of DNA mismatch repair MUTS family.
smart00879 (3) The Brix domain is found in a number of eukaryotic proteins.
smart00948 (2) Proteasome subunit A N-terminal signature Add an annotation.
TIGR00072 (3) hydrogenase maturation protease.
TIGR00341 (1) TIGR00341 family protein. [Hypothetical proteins, Conserved]
TIGR00691 (1) (p)ppGpp synthetase, RelA/SpoT family. [Cellular processes, Adaptations to atypical conditions]
TIGR00697 (6) conserved hypothetical integral membrane protein. [Hypothetical proteins, Conserved]
TIGR02338 (13) prefoldin, beta subunit, archaeal.
TIGR02356 (16) thiazole biosynthesis adenylyltransferase ThiF, E. [Biosynthesis of cofactors, prosthetic groups, and carriers, Thiamine]
TIGR02689 (1) arsenate reductase, glutathione/glutaredoxin type. [Cellular processes, Detoxification]
TIGR03520 (1) gliding motility-associated protein GldE.
TIGR04283 (1) transferase 2, rSAM/selenodomain-associated. [Unknown function, Enzymes of unknown specificity]

#### 4.4 Multiple-sequence alignment

Each eL15-Euryarchaeota RNA is denoted by an abbreviation (like "Eco-1-1") that refers to a taxonomy given in Section 4.1. The alignment may include sequences containing the RNA motif, as well as flanking sequence. The eL15-Euryarchaeota RNA itself is denoted by the line underneath marked 5′ and 3′ on either end. Nucleotides in flanking sequences (i.e., not part of the motif) are written in gray letters. Stems of predicted rho-independent transcription terminators, if any, are shaded yellow. (Note: terminator predictions have not been analyzed manually, and many are likely to be false positives. Terminator predictions are those of the RNIE software.) Nucleotides predicted to function (as DNA) as transcription-factor binding sites, if any, are shaded in green. (But note: these predictions are manually annotated, so they might be under-predicted.) Annotated start codons, if any, are shaded green. (Note: start codons are frequently misannotated, especially in environmental samples.) Nucleotides proposed to basepair as part of the consensus structure are shaded in color when they comprise Watson-Crick or G-U pairs. Otherwise they are shaded gray. Conserved stems are also indicated at the bottom of the alignment by angle brackets, where matching $<$ and $>$ denote base-paired columns. Below these angle brackets, the symbol "2" denotes base pairs exhibiting covariation according to the statistically well-founded R-scape method. "1" denotes base pairs exhibiting covariation according to R2R's simplistic method. "0" denotes base pairs that are not observed to mutate and "?" denotes base pairs that have a significant frequency of non-canonical nucleotides for Watson-Crick or G-U pairs ($> 5\%$). Below these base pair annotation is the consensus sequence: "R" = "A" or "G", "Y" = "C" or "U", red nucleotides: nucleotide identity conserved more than 97% of the time, black nucleotides: 90%, gray nucleotides: 75%, red circle (•): nucleotide is present 97% of the time, black circle (•): 90%, gray circle (•): 75%, white circle (◦): 50%. All percentages of sequences just described (e.g. 97% conserved) assume that sequences have been weighted by the GSC algorithm implemented by the Infernal software package.

Duplicate sequences: the following putative homologs are not shown in the alignment because their sequences are identical to a homolog already shown: Hac-1-2, Hal-1-1, Hdi-1-1, Hdi-2-1, Hez-1-1, Hha-1-2, Hhi-1-1, Hhi-2-1, Hla-1-2, Hla-3-1, Hlu-1-1, Hme-1-2, Hpa-1-2, Hsa-3-1, Hsp-1-1, Hsp-4-1, Hsp-5-1, Hte-2-1, Hvo-1-1, Hwa-2-1, Mar-1-1, Mba-1-1, Mba-3-1, Mba-6-1, Mho-1-1, Mhu-1-1, Mma-3-10, Mma-3-11, Mma-3-12, Mma-3-13, Mma-3-14, Mma-3-15, Mma-3-16, Mma-3-17, Mma-3-18, Mma-3-19, Mma-3-2, Mma-3-20, Mma-3-21, Mma-3-22, Mma-3-23, Mma-3-24, Mma-3-25, Mma-3-26, Mma-3-27, Mma-3-28, Mma-3-29, Mma-3-3, Mma-3-30, Mma-3-31, Mma-3-32, Mma-3-33, Mma-3-34, Mma-3-35, Mma-3-36, Mma-3-37, Mma-3-38, Mma-3-39, Mma-3-4, Mma-3-40, Mma-3-41, Mma-3-42, Mma-3-43, Mma-3-44, Mma-3-45, Mma-3-46, Mma-3-47, Mma-3-48, Mma-3-49, Mma-3-5, Mma-3-50, Mma-3-51, Mma-3-52, Mma-3-53, Mma-3-54, Mma-3-55, Mma-3-56, Mma-3-6, Mma-3-7, Mma-3-8, Mma-3-9, Mma-4-1, Mma-5-1, Mma-6-1, Mma-7-1, Mma-8-1, Mma-9-1, Msm-1-1, Msm-10-1, Msm-11-1, Msm-12-1, Msm-13-1, Msm-15-1, Msm-17-1, Msm-18-1, Msm-19-1, Msm-2-1, Msm-20-1, Msm-21-1, Msm-22-1, Msm-23-1, Msm-3-1, Msm-5-1, Msm-7-1, Msm-8-1, Msm-9-1, Msp-10-1, Msp-11-1, Msp-12-1, Msp-15-1, Msp-6-1, Msp-8-1, Msp-9-1, Nal-1-1, Nsp-1-1, Nsp-2-1, Pfu-1-1, env-100, env-103, env-106, env-107, env-108, env-109, env-111, env-113, env-118, env-120, env-123, env-124, env-127, env-128, env-13, env-131, env-132, env-133, env-134, env-143, env-150, env-151, env-154, env-155, env-157, env-163, env-165, env-166, env-167, env-168, env-17, env-171, env-18, env-190, env-191, env-198, env-20, env-200, env-204, env-205, env-209, env-21, env-211, env-222, env-225, env-24, env-240, env-241, env-242, env-252, env-257, env-258, env-259, env-26, env-260, env-261, env-263, env-264, env-265, env-272, env-275, env-279, env-280, env-282, env-285, env-288, env-292, env-294, env-300, env-303, env-305, env-306, env-307, env-313, env-314, env-315, env-316, env-320, env-335, env-337, env-340, env-342, env-343, env-345, env-346, env-347, env-350, env-351, env-352, env-355, env-357, env-36, env-361, env-368, env-37, env-372, env-373, env-374, env-38, env-389, env-391, env-393, env-394, env-395, env-396, env-397, env-398, env-399, env-4, env-400, env-401, env-403, env-404, env-405, env-407, env-409, env-41, env-411, env-415, env-417, env-419, env-421, env-423, env-426, env-430, env-431, env-434, env-435, env-436, env-437, env-44, env-445, env-448, env-451, env-452, env-456, env-458, env-459, env-46, env-464, env-468, env-470, env-471, env-478, env-482, env-483, env-493, env-494, env-498, env-501, env-504, env-510, env-518, env-519, env-520, env-521, env-523, env-524, env-526, env-527, env-533, env-54, env-548, env-549, env-550, env-551, env-554, env-555, env-556, env-557, env-558, env-56, env-562, env-566, env-569, env-572, env-573, env-58, env-582, env-583, env-584, env-586, env-59, env-592, env-593, env-60, env-600, env-601, env-614, env-615, env-618, env-62, env-626, env-627, env-632, env-648, env-649, env-652, env-653, env-654, env-655, env-662, env-666, env-667, env-668, env-669, env-677, env-678, env-68, env-687, env-688, env-689, env-69, env-690, env-691, env-692, env-693, env-694, env-703, env-705, env-71, env-711, env-712, env-713, env-72, env-75, env-76, env-8, env-81, env-82, env-85, env-88, env-96, env-97, har-1-1

alignment positions 1 · · · 180

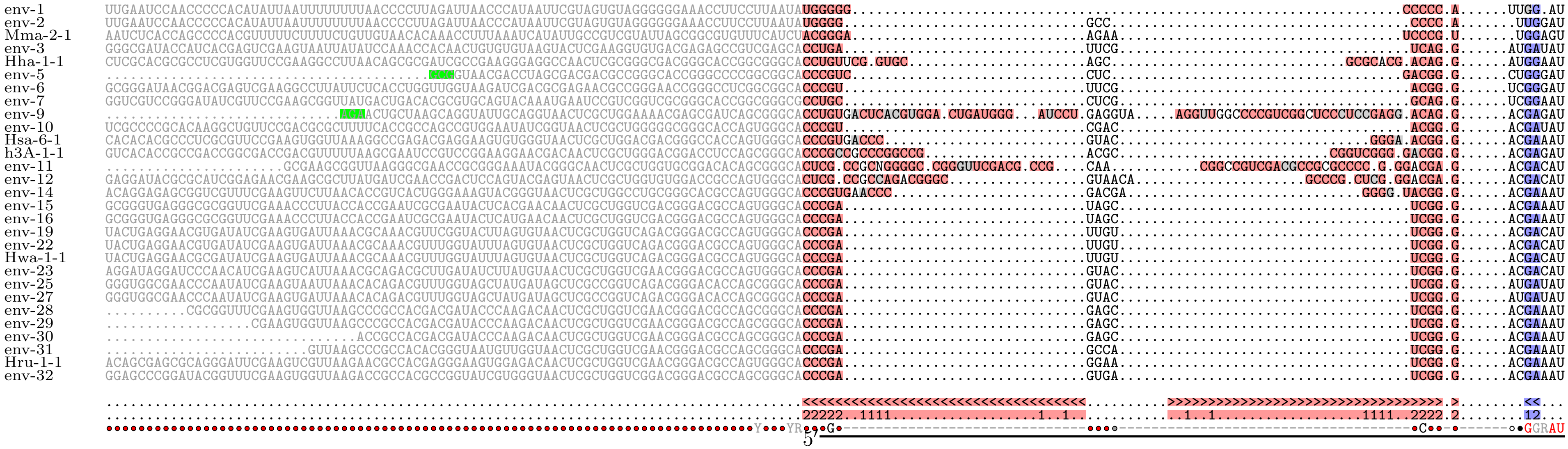

```
env-33   CGGCGGACGCUGCGGUUUCGAAGUCGUUAAGACUGCCACGGCAGUAACAAAUAUAACUCGCUGGUCGGACGGGACGCCAGUGGGCACCCGA...............................GAGA.....................................UCGG.G......ACGAAAU
Hpa-2-1  GCACGGCGUCGACCGAUUCGAAGUCGUUAAAGGCGAACCGACGGUAGAGUCGGCAACUCGCUGGUCAGACGGGACGCCAGCGGGCACCCGA...............................GCAC.....................................UCGG.G......ACGACAU
env-34   GCACGCCGCUCGGGCCGUUCCGACGCGCUUUUAGCCCGGAGCGGCGGAUCGAGAGGUAACUCGCUGGGGGCGGGCACCAGCGGGCACCCGU...............................ACAC.....................................ACGG.G......ACGACAU
env-35   ACCCACAGAAGGGGUGUUCCGAAGCCCUUAAUGUACCCACACGGGUAGAACACAUAACUCGCUGGCAGACGGGCUCCCAGCGGGCACCCGAAC.............................ACC....................................GUUCGG.G......ACGAAAU
env-39   CGGACCGGUCGACCAGGAACGAAGCCUUUAGAAGCAGUACAGGCGGAUAUACUGUAACUCGCUGGCUGACGGGCGAUCAGCGGGCACCCGUGACCCACAGGGCAGCA.....GGCGA...CUC.........UCGCCACUAGUUGCUGCUC.GUGGGA.ACGG.G......ACGAAAU
env-40   CGGACCGGUCGACCAGGAACGAAGCCUUUAGGAGCAGUACAGGCGGAUAUACUGUAACUCGCUGGCUGACGGGCGAUCAGCGGGCACCCGUGACCCACAGGGCAGCA.....GGCGA...CUC.........UCGCCACUCGUUGCUGCCC.GUGGGA.ACGG.G......ACGAAAU
Hac-1-1  CGGGACCGUGGCCGCGCGCGUUCCGAAGCCGUUAAAGCCGCCACGACGGUAGAUGGGAUAACUCGUCGGUGCGGGCCCGACGGGCACCCGCCGCGACG........................AUCC..............................CGUCGUGACGG.G......ACGACAU
env-42   GGCUGGGCGUGCAUCGAAGAACGAAGCGCUUAUGAUCGAACCGGCACCAGUACGAGUAACUCGCUGGUGUGGACCGCCAGUGGGCACUCGCCGCCGGCCGAGCGU.................GAC........................ACGCUCGCUCGG.GACGA.G......AUGACAU
env-43   GGCUGAGCGUGCAUCGAAGAACGAAGCGCUUAUGAUCGAACCCGCACCAGUACGAGUAACUCGCUGGUGUGGACCGCCAGUGGGCACUCGCCGCCGGACGGGCG..................CGACA.......................CGCUCGCUCGG.GACGA.G......ACGACAU
env-45   GAGUAUACGCGCAUCGGAGAACGAAGCGCUUAUGAUCGAACCGGUUCCAGUACGAGUAACUCGCUGGUGUGGACCGCCAGUGGGCACUCGCCGCCGGACGGGCG..................CGACA.......................CGCCCGCGCGG.GACGA.G......ACGACAU
Hna-1-1  AGACGGUGCGCGCGGCGAGUCGAAGUUCUUAAACGGAGCCGGGCGGUAGACCGACCAACUCGCUGGCUGACGGGCCCCAGCGGGCACCCGA...............................CUCG.....................................UCGG.G......ACGACAU
env-47   AGACGGUGCGUGCGGCGAGUCGAAGUUCUUAAACCGAAACAGGGGGUAGAACCGCCAACUCGCUGGUCGACGGGCCCCAGCGGGCACCCGA...............................CUCG.....................................UCGG.G......ACGACAU
env-48   GCGGUGACCCAUGGGCAUUCCGAAGCGGUUUUACUCGCGACUCGGGUGAUUGCAGCACUGACUGGCUCGCGGGACACCAGCAGGCACCCGA...............................UCCGU....................................UCGG.G......AUGACAU
env-49   GCGGCGACUCAUGGGCAUUCCGAAGCGGUUUUACUCGCGACUCGGGUGAUUGCAGCACUGACUGGCUCGCGGGACACCAGCAGGCACCCGA...............................UCCGU....................................UCGG.G......AUGACAU
env-50   AUCACAGCCCAUGGGCAUUCCGAAGCGGUUUUACUCGCCACUUGGAUGGUGGGUACACUGACUGGCUCGCGGGGCACCAGCAGGCACCCGA...............................UCUGU....................................UCGG.G......AUGACAU
env-51   GCAGCGACCCAUCACCGUUCCGAAGCGGUUUUACUCGCGACUCGGAUGGUUGAGGUACUGACUGGCUCGCGGGACACCAGCAGGCACCCGA...............................UCCGU....................................UCGG.G......AUGACAU
env-52   GAUUUGCUCCAUCGGCAAUCCGAAGCGGUUUUACUCGCAUCUCGGGUCUUUUGGCUACUGACUGGCUCGCGGGACACCAGCAGGCACCCGA...............................UCCGU....................................UCGG.G......AUGACAU
env-53   GAUUUGCUCCAUCGGCAAUCCGAAGCGGUUUUACUCGCAUCUCGGGUCUUUUGGCUACUGACUGGCUCGCGGGACACCAGCAGGCACCCGA...............................UCAGU....................................UCGG.G......AUGACAU
env-55   CACCACCCACGCGAGCCCCGUAGCGAAGCGGUUAAGGGCGAACCGCGGGAAGUACGGACAACUCGCUGGUGCGGACACAGCGGGCACUCGCCGCCGGAGCCGGGUCGCGACGCGGAG...GUCG.........CUCCGC.UCGCU.CCC.GC.CCGG.GACGA.G......ACGAAAU
env-57   CACUCAUGGGCUGCGACGGAGCGAAGCCCUUACAACCGAACGCGCGGUAUGGCCGAUAACUCGCUGGGACGGACGUCCAGCGGGCACUCGCCGCCGGGAAGUGGCG..UGGGUCGAC....ACGGG.........GUCGAUCCGUCCGCC....CCCGG.GACGA.G......ACGAAAU
env-61   UACGCAGGGGCGGCGACGGAGCGAAGCCCUUACAACCGAACGGACGGUACCACCAGUAACUCGCUGGGACGGACGUCCAGCGGGCACUCGCCGCCGGACCGAAGGU.CGAGUCGACG....UCA..........CGUCGAC.CGCACCGCCGGACCGG.GACGA.G......ACGAAAU
env-63   CAUACACGGGCCACGACGGAGCGAAGCCCUUACAACCGAACGCGCGGUAUGGCCGAUAACUCGCUGGGACGGACGUCCAGCGGGCACUCGCCGCCGGGGAGUGGCGUGGGUCGAC......ACGAG..........GUCGAUCCGU.CCGC.CCCCGG.GACGA.G......ACGAAAU
env-64   CACUCAUGGGCUGCGACGGAGCGAAGCCCUUACAACCGAACGCGCGGUAUGGCCGAUAACUCGCUGGGACGGACGUCCAGCGGGCACUCGCCGCCGGGGAGUGGCGUGGGUCGAC......ACGGG..........GUCGAUCCGU.CCGC.CCCCGG.GACGA.G......ACGAAAU
env-65   GGCGCAGGGACGGCGACGGAGCGAAGCCCUUACAACCGAACGCGCGGUACGACCGGUAACUCGCUGGGACGGACGUCCAGCGGGCACUCGCCGCCGGACC.GAGGCGUC....GGGUCGACGCGGAG....UCAACCCCGUCGACGCCGAGGGUCCGG.GACGA.G......ACGAAAU
env-66   CCGACGCAUGCCCGUAUCAAGGCGAAUCGCUUAAGAACGACGGCGGGGAAACUUUGAUAACUCGCUGGUUGGACGUCCAGCGGGCACUCGCCGCCGGAUCGCGG.CGCCGGG.GCGAC...CGCGCA......GUCGCUUCCGUCGACACCGAUCCGG.GACGA.G......ACGAAAU
Hai-1-1  CCACACGCACGCGAACCCUGGAGCGAAGCGCUUAACAGCGAACCGACGGAAAUACGGACAACUCGCUGGUGCGGACACAGCGGGCACUCGCCACCGGCGC.CGGGGA..............CGCGC...................UCCCUGUGC.CCGG.GACGA.G......ACGACAU
Hco-1-1  GUCUCCCCACGCGACCCGCGGAGCGAAGCGGUUAAGGGCGAACCACGGGAAUUCUCGGUAACUCGCUGGUGCGGACACAGCGGGCACUCGCCGCCGGGGC.CGCGGCCGG..........ACCGA.................CC.GCCGCGCGCCCGGGA..CGA.G......ACGACAU
Hsp-8-1  GGUUCCCCACGCGACCCGCGGAGCGAAGCGGUUAAGGGCGGACCGCGGGUAUUUCUGCCAACUCGCUGGUGCGGACACAGCGGGCACUCGCCGCCGGGGC.CGCGGCCGG..........ACCGA.................CC.GCCGCACGCCCCGG.GACGA.G......ACGACAU
Hte-1-1  GCGCUCGCACGCGAUCCGGGGAACGAAGCCGUUAAGGGCGAACCGCGGGUACUGCCGAUAACUCGCUGGUGCGGACACAGCGGGCACUCGCCGCCGGGAC.CGCCGUCGG..........AUCGA.................CCGAC.GCGCGUCCCGG.GACGA.G......ACGACAU
Hho-1-1  GCGCUCGCACGCGAUCCGGGGAACGAAGCCGUUAAGGGCGAACCGCGGGUACUGCCGACAACUCGCUGGUGCGGACACAGCGGGCACUCGCCGCUGGGAC.CGCCGUCGG..........AUCGA.................CCGAC.GCGCGUCCCGG.GACGA.G......ACGACAU
env-67   GGCCACCCACGCGACCCGCGGAGCGAAGCGGUUAAGGGCGAACCGCGGGAAUUCACGCUAACUCGCUGGUGCGGACACAGCGGGCACUCGCCGCCGGGAC.CGCGGUCGG..........ACCGA.................CCG.CCGCGCGCCCCGG.GACGA.G......ACGACAU
Hca-2-1  CGUCCCCCACGCGACCCGCGGAGCGAAGCGGUUAAGAGUGAACCGCGGAUAAUUUCGCCAACUCGCUGGUGCGGACACAGCGGGCACUCGCCGCCGAGGC.CGCGGUCGG..........ACCGA.................CCG.CCGCGCGCCCCGG.GACGA.G......ACGACAU
env-70   CGCAACCCACGCGACCCGCGGAGCGAAGCGGUUAAGAGUGAACCGCGGAUAUUUCCGUCAACUCGCUGGUGCGGACACAGCGGGCACUCGCCGCCGGGGC.CGCGGUCGG..........ACCGA.................CCG.CCGCGCGCCCCGG.GACGA.G......ACGACAU
Hli-2-1  CACUCCCCACGCGACCCGCGGAGCGAAGCGGUUAAGGGCGGACCGCGGGUAUUUCCGCCAACUCGCUGGUGCGGACACAGCGGGCACUCGCCGCCGAGGCGUGCGGUCG...........GAC...................CGACCGCACGCCCCGG.GACGA.G......ACGACAU
Har-2-1  CUCUCCCCACGCGACCCGCGGAGCGAAGCGGUUAAGGGCGGACCGCGGGUAUUUCCGCCAACUCGCUGGUGCGGACACAGCGGGCACUCGCCGCCGAGGCGUGCGGUCG...........GAC...................CGACCGCACGCCCCGG.GACGA.G......ACGACAU
env-73   GCACACGCACACGAACGCCGUAGCGAAGCGGUUAAGGGCGAACCACAGGUAAAUGAGAUAACUCGCUGGUGCGGACACAGCGGGCACUCGCCGCCGAGG.CCGAGGUCA...........CCGU..................UGAUCUCGUAUCUCGG.GACGA.G......ACGACAU
Hli-1-1  CUAUCCCCACGCGAACGCCGUAGCGAAGCGGUUAAGGGCGAACCGCGGGAAAUACGGGCAACUCGCUGGUGCGGACACAGCGGGCACUCGCCGCCGGGGC.CGGCCUCGACG.CCG.....ACG...........CGAUCGUCGACGUCGCGCCCCGG.GACGA.G......ACGACAU
Hha-2-1  UCACCCGCACGCGAACGCCGUAGCGAAGCGGUUAAGGGCGAACCACGGGAAAUACGGACAACUCGCUGGUGCGGACACAGCGGGCACUCGCCGCCGGAGC.CGGCCUUGACG.CC.....GCGAU..........GGUCGUCAACGCUGCGCCCCGG.GACGA.G......ACGACAU
Hsa-9-1  UCACCCGCACGCGAACGCCGUAGCGAAGCGGUUAAGGGCGAACCACGGGAAAUACAGACAACUCGCUGGUGCGGACACAGCGGGCACUCGCCGCCGGAGC.CGGCCUUGACG.CC.....GCGAU..........GGUCGUCAACGCUGCGCCCCGG.GACGA.G......ACGACAU
env-74   CAUUCCCCACGCGACCGCCGUUGCGAAGCGGUUAAGGGCGAACCGCGGGAAAUACGGGUAACUCGCUGGUGCGGACACAGCGGGCACUCGCCGCCGGAGC.CGGCUUCGACG.CC.....GCGAU..........GGCCGUCGACGCCGCGCCCCGG.GACGA.G......ACGACAU
Hko-1-1  CGACCCCCACGCGAACGCCGUAGCGAAGCGGUUAAGGGCGAACCGCGGGAACUACGGACAACUCGCUGGUGCGGACACAGCGGGCACUCGCCGCCGGGGC.CGGGUUCGACG.CC.....GCGAU..........GGCCGUCGACGUCGCGCCCCGG.GACGA.G......ACGACAU
env-77   CAUCCCCCACGCGACCGCCGUUGCGAAGCGGUUAAGGGCGAACCGCGGGAAAUACGGGCAACUCGCUGCUGCGGACACAGCGGGCACUCGCCGCCGGGGC.CGACAUCGACG.CC.....GCGAU..........GGCCGUCGACGCCGCGCUCCGG.GACGA.G......ACGACAU
env-78   CAUCCCCCACGCGACCGCCGUUGCGAAGCGGUUAAGGGCGAACCGCGGGAAAUACGGGCAACUCGCUGCUGCGGACACAGCGGGCACUCGCCGCCGGGGC.CGACAUCGACG.CC.....GCGAU..........GGCCGUCGACGCCGCGCUCCGG.GACGA.G......ACGACAU
Hsa-7-1  UUAUUUGCACGCGAACGCCGUAGCGAAGCGGUUAAGGGCGAACCGCGGGAAGUACGGGCAACUCGCUGGUGCGGACACAGCGGGCACUCGCCGCCGGGAC.CGGCUUCGACG.CCGACG..CCAC........CGU.GGCCGUCGACGCCGAGUCCCGG.GACGA.G......ACGACAU
env-79   ......................GCGAAGCCGUUAAGAGCGAACCGCAGGUAUUCAAGAUAACUCGCUGGUGCGGACACAGCGGGCACUCGCCGCCGGG..AACGUCGG............UUCGG...................UCGACGCACGCCCGG.GACGA.G......ACGACAU
env-80   GGGCUCGCACGCGGUCCGCGGAACGAAGCGAUUAAACCCCGACCGCGGGUACACGUGAUAACUCGCUGUCGCGGACACAGCGGGCGCUCGCCGCCGAGC...................CGGAC..........................GCUCGG.GACGA.G......ACGACAU
env-83   GACUCCGCACGCGACCCGCGGAGCGAAGCGGUUAAGGGCGAACCACGGGAAAUUUCGACAACUCGCUGGUGCGGACACAGCGGGCACUCGCCGCCGGGGCCGCGGCCGG.............ACGGA...................CCGCCGCGCGCCCGG.GACGA.G......ACGACAU
env-84   CCGCACCCACGCGAACGCCGUAGCGAAGCGGUUAAGGGCGAACCGCGGGAAAUACGGGCAACUCGCUGGUGCGGACACAGCGGGCACUCGCCGCCGGGCG.UCGGGACC............GCAGG..................GGUCUCGCUC.CCCGG.GACGA.G......ACGACAU
env-86   CCGCACCCACGCGAACGCCGUAGCGAAGCGGUUAAGGGCGAACCGCGGGAAAUACGGGCAACUCGCUGGUGCGGACACAGCGGGCACUCGCCGCCGGGCG.UCGGGACC............GCAGG..................GGUCUCGCUC.CCCGG.GACGA.G......ACGACAU
env-87   CCGCACCCACGCGAACGCCGUAGCGAAGCGGUUAAGGGCGAACCGCGGAAAAUACGGGCAACUCGCUGGUGCGGACACAGCGGGCACUCGCCGCCGGGCG.UCGGGACC............GCAGG..................GGUCUCGCUC.CCCGG.GACGA.G......ACGACAU
env-89   UGUGACGUUCAAAUCGCGGAGCGAAGCGCUUAAACUCGGUGGCUGAAGAUAUGUAGUAACUCGCUGGGACGGACGUCCAGCGGGCACUCGCCACCGGAUCGCAAACGGAGCCC.......CGUCGU..........GGUCUCCAUCU.CGACCCGG.GACGA.G......ACGAAAU
env-90   GCUGUCGCGGGAACGGCGGAGCGAAGCCCUUAACGUCGACGACGGGGAAUGUCGGACAACUCGCUGGAACGGACGUUCAGCGGGCACUCGCCGCCGAGUCG.AGUCC.CGGUCCA.....UCG..........UGGGCCGUGGUCGAUGACUCGG.GACGA.G......ACGAGAU
env-91   GCUUACGCGGGAACGCCGGAGCGAAGCGCUUAAACUCCACGCCGGGGAAACGACAGUAACUCGCUGGUACGGACGUCCAGCGGGCACUCGCCGCCGGAUCG.CGGGCAACAGUCCG....UUCG.........CGGACCGCC.UCCGACGAUCCGG.GACGA.G......ACGAAAU
env-92   GCUGUCGCGGGAACCCCGGAACGAAGCGCUUAAACUCCACGCCGGGGAAACGAGAGUAACUCGCUGGUACGGACGUCCAGCGGGCACUCGCCGCCGGAUCG.CGGGCAACAGUCCG....UUCG.........CGGACCGCC.CCCGACGACCCGG.GACGA.G......ACGAAAU
env-93   GGCGCAGGGACGGCGACGGAGCGAAGCCCUUACAACCGAACGCGCGGUACGACCGGUAACUCGCUGGGACGGACGUCCAGCGGGCACUCGCCGCCGGACCGAGGCGU...CGGGUCGAC..CCAUC.......GUCGAUCCGGUCACGCCGCGUUCCGG.GACGA.G......ACGAAAU
env-94   CUGACGCACGCUCGGCGCGGAACGAAGCCCUUACAACCGAACGGUGGGUACGUUCGCCAACUCGCUGGAACGGACGUCAGCGGGCACUCGCCGCCGCGCCGGG...................GGA.......................CCCGACGCGG.GACGA.G......ACGAAAU
Hmu-1-1  GCACGGGUGCUCUGAAUCCGAAGUUCUUAACCCUCCCCUGUUGGUAGGUCAGCCAACUCGCUGGCAUGCGGGCACGCCAGAGGGCACCCGU...............................CACGA....................................ACGG.G......ACGAAAU
Hel-1-1  GCACGCCAGCUCUCAAUCCGAAGUUCUUAACCCUCCCCUGUUGGUAGUUCCGCCAACUCGCUGGCAUGCGGGCACGCCAGAGGGCACCCGU...............................CACGU....................................ACGG.G......ACGAAAU
Hla-2-1  ACACGCCAGCUCUCAAUCCGAAGUUCUUAACCCUCCCCUGUUGGUAGUUCCGCCAACUCGCUGGCAUGCGGGCACGCCAGAGGGCACCCGU...............................CACGU....................................ACGG.G......ACGAAAU
Hde-1-1  CGCGCGCCGCGAGAAUCCGAAGUCCUUAACCCUCCCCUGCCGGUAAGUUCGCUAACUCGCUGGCAAUGCGGGCACGCCAGAGGGCACCCGU...............................CACGC....................................ACGG.G......ACGAAAU
Hal-2-1  CAUGCCAGUUCCCAAUCCGAAGUCCUUAACCCUCCCCUGUUGGUAGUUCCGCCAACUCGCUGGCAAUGCGGGCACGCCAGAGGGCACCCGU...............................CACGA....................................ACGG.G......ACGAAAU
Hpr-1-1  CAUGCCAGUUCCCAAUCCGAAGUCCUUAACCCUCCCCUGUUGGUAGUUCCGCCAACUCGCUGGCAAUGCGGGCACGCCAGAGGGCACCCGU...............................CACGA....................................ACGG.G......ACGAAAU
Hgi-1-1  CGCGCGUCGCUCGAAUCCGAAGUCCUUAACCCUCCCCUGUUGGUAGUUCCGCCAACUCGCUGGCAAUGCGGGCACGCCAGAGGGCACCCGU...............................CACGA....................................ACGG.G......ACGAAAU
Hgi-2-1  GCAUGCUGUUUCCAAUCCGAAGUCCUUAACCCUCCCCUGUUGGUAGUUCCGCCAACUCGCUGGCAAUGCGGGCACGCCAGAGGGCACCCGU...............................CACGA....................................ACGG.G......ACGAAAU
Hsp-9-1  CACACCCGCUGUUAAUCCGAAGUUCUUAACCCUCCCCAUAGGGAAAAUCCGCCAACUCGCUGGCAACGCGGGCACGCCAGAGGGCACCCGU...............................CACGC....................................ACGG.G......ACGAAAU
Hsu-2-1  CACGCGUCGCGAGAAUCCGAAGUCCUUAACCCUCCCCUGUUGGUAGUUCCGCCAACUCGCUGGCAAUGCGGGCACGCCAGAGGGCACCCGU...............................CACGC....................................ACGG.G......ACGAAAU
Hsp-6-1  CGCGCGUCGCUCGAAUUCGAAGUCCUUAACCCUCCCCUGUUGGUAGUUCCGCUAACUCGCUGGCAACGCGGGCACGCCAGAGGGCACCCGU...............................CACGC....................................ACGG.G......ACGAAAU
Hme-1-1  ACACAGUCGCUGUGAAUCCGAAGUUCUUAACCCUCCCCUAUUGGUAGGUUCGGCAACUCGCUGGCAUGCGGGCACGCCAGAGGGCACCCGU...............................CACGU....................................ACGG.G......ACGAAAU
Hru-2-1  CGGUGAGUCGGGGCGCUUCGAAACCCUUACCACCGAAUCCCGAGUAUCCACGCCCAACUCGCUGGUCGACGGGACGCCAGUGGGCACCCGA...............................UAGC.....................................UCGG.G......ACGAAAU
env-95   GGGGGGCCCCGUGGUGCUUCGAAGUGUUCAAGACCGACGACAGCGAACGACGGGUAACUCGUCGGCCAGUGGGCCGCCGGCGGGCACCCGG...............................UUCG.....................................CCGG.G......AGGAGAU
env-98   CCCGGCCGUCGUGGCGCUUCGAAGUGUUCAAGAGCGGCCACGGCGAACAACCGGUAACUCGCCGGCCAGUGGGCCGCCGGUGGGCACCCGG...............................UUCG.....................................CCGG.G......ACGAGAU
env-99   CCCACCGGUCACGGCGCUUCGAAGUGUUGAAACCCGACACCACCGAACCGUGAGCAACUCGCCGGCCAGUGGGCCGCCGGCGGGCACCCGG...............................UUCG.....................................CCGG.G......ACGAGAU
CHa-1-1  CGCACGGUCGAGCGGCGUUCGAAGUGCUUAACAGCGCCGCACCGGUACGUCCGCCAACUCGCUGGCACGUGGGCCGCCAGCGGGCACCCGG...............................UCCGC....................................CCGG.G......ACGACAU
env-101  ACCCACAGGGAGGUUGGUCCGAAGCCAUUAAACGCCUCUCGGAAGUAGAACCCCUAACUCGCUGGCAAACGGGCCACCAGUGGGCACCCGG...............................UUCG.....................................CCGG.G......ACGAAAU
env-102  CUGACGCAGGAGAAACGGUUCGAAGCUAUUAAACGGCUCUCAGCAGUAGAUCCGGCAACUCACUGAGGACGGGCCACCAGUGGGCACCCGG...............................UUCG.....................................CCGG.G......ACGAAAU
env-104  CUGACGCGGGAAGGACGGUUCGAAGCUAUUAAACGGCUCUCAGCAGUAGGUCCGCUAACUCACUGAGGACGGGCCACCAGUGGGCACCCGG...............................UUCG.....................................CCGG.G......ACGAAAU
env-105  ..................................ACACCUCUCAGCAGUAGAGUCGCCAACUCACUGUGACGGGCCUCAGUGGGCACCCGG...............................UUCG.....................................CCGG.G......ACGAAAU
env-110  ACCACGCACCAGCAGAAACCCGAAGCUGUUAAACGCCUCUCAGCAGUAGGUUCGCCAACUCACUGCGGACGGGCCACCAGUGGGCACCCGG...............................UUCG.....................................CCGG.G......ACGAAAU
env-112  CCCGCGGUGCGGCGAGUCCGAAGCGGUUAAACGCGCCUCGGAGAUAGAACAGGCAACUCACUGCGGUGACGGGCUACCAGUGGGCACCCGG...............................AUUU.....................................CCGG.G......ACGAAAU
env-114  CUCGCGGUGCGGCGAGUCCGAAGCGGUUAAACGCGCCUCGGAGAUAGAACAGGCAACUCACUGCGGUGACGGGCUACCAGUGGGCACCCGG...............................AUUU.....................................CCGG.G......ACGAAAU
env-115  CUCGUGGUGCAGCCAGUCCGAAGCGGUUAAACGCAUCUCGGAGGUACCACAGGUAACUCACUGCGGUGACGGGCUACCAGUGGGCACCCGG...............................UUCG.....................................CCGG.G......ACGAAAU
env-116  CUCGCGGUACAGCAAGUCCGAAGCCGUUAAACAGGUCUCGGGGGUAGGAAAGCCAACUCACUGCGGUGACGGGCUACCAGUGGGCACCCGG...............................UUCG.....................................CCGG.G......ACGAAAU
env-117  CUCUCGGUGCAGCGAGUCCGAAGCGGUUAAACGGGCUUCGGAGGUAGAACAGGCAACUCACUGCGGUGACGGGCUACCAGUGGGCACCCGG...............................UUUG.....................................CCGG.G......ACGAAAU

         ......................................................................................<<<<<<<<<<<<<<<<<<<<<<<<<<<<<<<<<<<<..........>>>>>>>>>>>>>>>>>>>>>>>>>>>>>>>>>>>.>........<<...
         ......................................................................................22222..1111..................1..1............1..1.................1111..2222.2........12...
         ••••••••••••••••••••••••••••••••••••••••••••••••••••••••••••••••••••••••••••••••••Y•••YR•••G•---------------------------•••○-------------------------------•C••-•------○GGRAU
                                                                                               5′
```

```
env-119   CCCGCGGUACAGCAAGUCCGAAGCGGUUAAACGCGCUUCGAGGGUAGAAUAGACAACUCACUGCGGUGACGGGCUACCAGUGGGCACCCGG..............................UUUG....................................CCGG.G......ACGAAAU
env-121   CUCUCGGUGCAGCGAGUCCGAAGCGGUUAAACGGGCUUCGGAGGUAGAACAGGCAACUCACUGCGGUGACGGGCUACCAGUGGGCACCCGG..............................UUUG....................................CCGG.G......ACGAAAU
env-122   CACGCGGUACAGCAAGUCCGAAGCGGUUAAACGGGUCUCGGAGGUAGAGACGCCAACUCACUGCGGUGACGGGCUACCAGUGGGCACCCGG..............................UUUG....................................CCGG.G......ACGAAAU
env-125   CUCGUGGUGCGGCAAGUCCGAAGCGAUUAAACGAGUCUCGGCGGUAACACAGGUAACUCACUGCGGUGACGGGCUACCAGUGGGCACCCGG..............................UUUG....................................CCGG.G......ACGAAAU
env-126   AUCUCGCACCACUACGAACCCGAAGCAAUUAAACAGGUCUCGGCAGUAGAAUGCCUAACUCGCUGGAGACGGGCCACCAGUGGGCACCCGG..............................UUCG....................................CCGG.G......ACGAAAU
env-129   UACACACAACACCGGCAACCCGAAGCCAUUAAACGGGUCUCGGCAGUAGAACGCCUAACUCGCUGGCAGCGGGCCACCAGUGGGCACCCGG..............................UUCG....................................CCGG.G......ACGAAAU
env-130   AUCUCGCACCACCACCAACCCGAAGCCAUUAAACAGGUCUCGGCAGUAGACACGCCAACUCGCUGGAGACGGGCCACCAGUGGGCACCCGG..............................UUCG....................................CCGG.G......ACGAAAU
env-135   GCCCACAGAGCUACCAGUCCGAAGCCAUUAAACGCCUCUCGACAGUAGAACACCCAACUCGCUGGCAAACGGGCCACCAGUGGGCACCCGG..............................UUCG....................................CCGG.G......ACGAAAU
env-136   CCACGUACUACCAGCAACCCGAAGCCAUUAAACGGCCUUCGACAGUAGAACGCCUAACUCGCUGGCAAACGGGCCACCAGUGGGCACCCGG..............................UUCG....................................CCGG.G......ACGAAAU
env-137   UCUCGUGGCGACAGCGGUCCGAAGCCAUUAAACGCCUCUCGGAAGUAGAAGGGCUAACUCGCUGGCAAACGGGCCACCAGUGGGCACCCGG..............................UUCG....................................CCGG.G......ACGAAAU
env-138   UCUCGUGGCGACAGCGGUCCGAAGCCAUUAAACGCCUCUCGGGAGUAGAACGGCUAACUCGCUGGCAAACGGGCCACCAGUGGGCACCCGG..............................UUCG....................................CCGG.G......ACGAAAU
env-139   UGCCUUGCCGAGAGCGGUCCGAAGCCAUUAAACGGCUCUCCGAAGUAGAACGCCUAACUCGCUGGCAAACGGGCCACCAGUGGGCACCCGG..............................UUCG....................................CCGG.G......ACGACAU
env-140   CCACGCACAGCUAGCAACCCGAAGCCGUUAAACGACUCUCCGAAGUAUUCACACCAACUCGCUGGCAAACGGGCCACCAGUGGGCACCCGG..............................UUCG....................................CCGG.G......ACGAAAU
env-141   CCCCACACCAGCAGAAACCCGAAGCCAUUAAACGCCUCUCGACAGUAGAACGCCUAACUCGCUGGCAAACGGGCCACCAGUGGGCACCCGG..............................UUCG....................................CCGG.G......ACGAAAU
env-142   ........................................................................GCCACCAGUGGGCACCCGG..............................UUCG....................................CCGG.G......ACGAAAU
env-144   CCCCACACCAGCAGAAACCCGAAGCCAUUAAACGCCUCUCGACAGUAGAACGCCUAACUCGCUGGCAAACGGGCCACCAGUGGGCACCCGG..............................UUCG....................................CCGG.G......ACGAAAU
env-145   UCUCGUGGCGGCAGCGGUCCGAAGCCAUUAAACGCCUCUCGGAAGUAGAAGGGCUAACUCGCUGGCAAACGGGCCACCAGUGGGCACCCGG..............................UUCG....................................CCGG.G......ACGAAAU
env-146   ACCCACAGAGCGGUUGGUCCGAAGCCAUUAAACGCCUCUCGGAAGUAGAACGCCUAACUCGCUGGCAAACGGGCCACCAGUGGGCACCCGG..............................UUCG....................................CCGG.G......ACGAAAU
env-147   ACUCACAGAGCGACUGGUCCGAAGCUAUUAAACGCCCUUCGACAGUAGAACGCCUAACUCGCUGGAAAACGGGCCACCAGUGGGCACCCGG..............................UUCG....................................CCGG.G......ACGAGAU
env-148   ACUCACAGAGCGACUGGUCCGAAGCUAUUAAACGCCCUUCGACAGUAGAACGCCUAACUCGCUGGAAAACGGGCCACCAGUGGGCACCCGG..............................UUCG....................................CCGG.G......ACGAAAU
env-149   CCACGCACAGCUAGCAACCCGAAGCCGUUAAACGACUCUCCGAAGUAUUCACACCAACUCGCUGGCAAACGGGCCACCAGUGGGCACCCGG..............................UUCG....................................CCGG.G......ACGAAAU
env-152   CUCACAGAGCGACUGGUCCGAAGCCAUUAAACGCCUCUCGGGAGUAGAGACGCCUAACUCGCUGGCAAACGGGCCACCAGUGGGCACCCGG..............................UUCG....................................CCGG.G......ACGAAAU
env-153   CGCGUGUCGCGGCGAGUCCGAAGCGGUUAAACCGGUUUCGGCGGUAGAACAGCCAACUCACUGUGGAACGGGCCACCAGUGGGCACCCGG..............................UACG....................................CCGG.G......ACGAAAU
env-156   CGCUCGGCACAGCGAGUCCGAAGCGGUUAAACCGGUUUCGACGGUAUAGCAGGCAACUCACUGUGGAACGGGCCACCAGUGGGCACCCGG..............................UACG....................................CCGG.G......ACGAAAU
env-158   CCCGUCGGCCCGCCCGGCUUUCGAAGUCCUUAUGCGCCACAGACGGGUAUGGCCGCCAACUCGCUGGUCGCGGGCACCAGCGGGCACCCGC..............................UUCG....................................GCGG.G......ACGACAU
env-159   CUGUUGUCACGCGUCGUUUCGAAGCCCUUAACCGCCGACCCGAAGUAGCAUCGAUAACUCGCUGGCUGGUGGGCCGCCAGUGGGCACCCGG..............................UACG....................................CCGG.G......ACGAAAU
Hsp-10-1  GGUUCGCACACACGGCGCGUAGCGAAGCCGUUAAAACCUAUCCCGGGGAAGCUACGGGUAACUCGCUGGUGCGGACACAGCGGGCACUCGCCGCCGGCCGCGUCGACGG............CUCG..................CCGCCGACGC.GCCGG.GACGA.G......ACGACAU
Hse-1-1   GGCGACCGGACGAACGGUUUCGAAGCGCUUAACAGCGAACCGCCGAAACGGACGACAACUCGCUGGGGACCGGGCACCGGCGGGCACCUGCUCGUGUG.......................AACGC............................CACAC.AGCGG.G......CCGGUAU
Hmo-1-1   GGCGAGCCCGGCGGUGGGUUUCGAAGGCCUUAACAGCGAUUGCGCGGAAGGAUCGGUAACUCGCUGGUACCGGGCACCGGCGGGCACCUGCUCGCGUG.......................AACC.............................CACGC.AGCGG.G......UCGGUAU
Hth-1-1   GGCGGUCCUGGCGGCGGGUUUCGAAGGCCUUAACAGCGAUUGCCCGGAAGGAUCGGUAACUCGCUGGUAUCGGGCACCGGCGGGCACCUGCUCGCGUG.......................AACC.............................CACGC.AGCGG.G......UCGGUAU
Nae-1-1   UGUUAUACUCGAAACGCUUCGAAGCCCUUAUACUCGGGAGUUGGAUAGAGUAUCGUAACUCGCUGGGCCGCGGGCACCAGCGGGCACCUGU.CCGGC........................ACGAC.............................GCCGCCACAG.G......ACGGAAU
Nta-1-1   UGUCACACUCGAAACGCUUCGAAGCCCUUAUACUCGGGAAUUGGAUAGAGUAUCGUAACUCGCUGGGCCGCGGGCACCAGCGGGCACCUGU.CCGGC........................ACGAC.............................GCCGCCACAG.G......ACGGAAU
Nas-1-1   CGUUACACUCGAAACGCUUCGAAGCCCUUAUACUCGGGAAUUGGAUAGAGUACCAUAACUCGCUGGGCCGCGGGCACCAGCGGGCACCUGU.CCGGC........................ACGAC.............................GCCGCCACAG.G......ACGGAAU
env-160   CGAGAGCACGGGACGGAGUUCGAAGCGCUUUUGUCCGCCAGCCGGAAAUCUGUGGUAUCUCGCUGGGACGCGGGCGCCAGCGGGCACCUGC..............................CUGU....................................GCAG.G......UCGGGAU
env-161   CCGGCGCCCCGGGCACGGUCCGAAGCCCUUUUGCCCGAAAGCCGGGAACAGACGGUAACUCGCUGGGACGCGGGCGCCAGCGGGCACCUGU..............................CUGU....................................GCAG.G......UCGGGAU
env-162   CCCUGCCCUCGCCUGCGCUUCGAAGCCCUUAUGCCCGCCACAGGGAUAUCACGAGGUAACUCGUCGAGAGCGGGCUUCGGCGGGCACCUGUUCGGC.........................AGCC...............................GCC.ACAGG.G......AUGGAAU
env-164   ...................................................CGGGGUAACUCGUCGAGAGCGGGCUUCGGCGGGCACCUGUUCGGC.........................ACAC...............................GCC..ACAG.G.....CAUGGAAU
env-169   CGGAUUCGAACACGCCGCUUCGAAGGGCUUAUACCCGCCACGAGGGAAUCUCGAGGUAACUCGUCGAGAGCGGGCUUCGGCGGGCACCUGUUCGGC.........................ACAC...............................GCC..ACAG.G.....CAUGGAAU
env-170   ...........GGUGGGCUUCGAAGCGCUUAUACUCGACAGUUGGGUACGUUCGGUAACUCGCUGGUCGCGGGCGUCAGCGGGCACCUGUUCGACG........................UCU...............................CGUC..ACAG.G.....UACGGAAU
Hla-4-1   GACUCGAGGCGUGACGCUUCGAAGCCCUUAUACUCGCAAGUUGGAUAGUUGCAGACGACUCGCUGGUCCGCGGGCGACAGCGGGCACCUGUUCGACACACGUC..................ACAG........................GUCGGAAGUC..ACAG.G......UCGGAAU
Hsp-11-1  GGCCCGAGGCGCGACGCUUCGAAGCCCUUAUACUCGCAAGUUGGAUAGUUGCAGACGACUCGCUGGUCCGCGGGCGACAGCGGGCACCUGUUCGAC.........................GCAC...............................GUC..ACAG.G......UCGGGAU
Hsp-3-1   GCACGGACGUGGCGUCGUUCCGAAGUUCUUAACCCCGCCGGCGGAGUACCGAACGGUAACUCGCUGGUGGCGGGCUCCAGCGGGCGCCUGAC.............................GACGC..................................GUCGG.G......CCGGGAU
env-172   AACCAUGCAUUCGGCGCUUCGAAGCGCUUAUACGACCCUCUCGACUAUCUGAACCUGACUCGGUGGCACGCGGGCGCCAGCGGGCACCUGAACGCA.........................CAC................................UGC..UCAG.G......UCGGGAU
Hsa-1-1   CGAACGACCUGUAACUCAGUUCGAAGCCCUUAAACUCGCCAACGCAGAAGACUCGAUAACUCGCUGGUGGCGGGCUCCAGCGGGCACCUGCACGGAGCG......................UGAU............................CGCUCCGCUCAG.G......CUGGAAU
env-173   ...........GGCGGGUUCCGAAACCCUUUACCUGACGACGCGGGAAGUUCUGGUAACUCGCUGGACGACGGGCACCAGUGGGCACCUGC..............................CAGAUA..................................GCAG.G...UCAUCGGAAU
env-174   UCUCGUGCGCUGGCGGGUUCCGAAACCCUUUACCUGACGACGCGGGAAGUUCUGGUAACUCGCUGGACGACGGGCACCAGUGGGCACCUGC..............................CAGAUA..................................GCAG.G...UCAUCGGAAU
env-175   ....................................................................GACGGGCACCAGUGGGCACCUGU..............................CAUCG...................................ACAG.G...UCAUCGGAAU
env-176   ....GGACGGCGGCGGGUUCCGAAACCCUUUACCUGCCGACGGGGGAAUUUCGACCAACUCGCUGGUCGACGGGCACCAGCGGGCACCUGUC.............................GUC....................................GACAG.G...UCAUCGGAAU
Hpa-1-1   CUGGUUGUGACGAGCGAAGACGAAGCUCUUAAACCCGCCAACCGGGAACGUUCGAUAACUCGCUGGGAAACGGGCACCAGUGGGCACCUGC..............................AACU....................................GCAG.G......ACGGAAU
env-177   .GCCGUAACGAUUCGUCUUCGAAGCCCUUUUGUCCUCAACUGGAAAAGAACAGGAUAGCUAACUGGACGACGGGCACCAGUGGGCGCCUGU..............................UUCAGC..................................ACAG.G......UCGGGAU
env-178   CGCCCCUGACGAACCGCGUUUCGAAGCCCUUAACAGCCGCCCUGGAUAAUUUCGGGUAACUCGCUGGAGACGGGCAACAGCGGGCACCUGU..............................ACAC....................................ACAG.G......UCGGGAU
env-179   ..........................................................................CGCCAGCGGGCACCUGA..............................GAUAC...................................UCAG.G......UCGGGAU
env-180   AACACACCAGCUGUGGUUCGAAGUCCUUUUAUCCGCAAGUGCGGUAGAACCGUCAACUCGCUGGGCGACACGGGCACCAGCGGGCACCUGU..............................UGCAU...................................ACAG.G......CUGGGAU
env-181   AAUACUGCAACAUGAGUUCGAAGUCCUUUUAUUUACAACCGCGAUAUUUUCACCAACUCGCUGGGUGACACGGGCACCAGCGGGCACCUGU..............................CAACU...................................ACAG.G......CUGGGAU
env-182   CGAGUAGCCAGCCGAGUUCGACAUCCUUUUAUCCGUAACCGGGAUAGAUCCGGCAACUCGCUGGCGAACACGGGCACCAGCGGGCACCUGU..............................ACUC....................................ACAG.G......ACGGGAU
Ssp-1-1   ACCGGUCGUCGACCGCCAUUCGAAGCGUUUAACCACCGAACGGGCGGAGAAUGGACAACUCGCUGGCAGUCGGGCUCCAGCGGGCACCCGUC.............................UCGC...................................GACGG.G......CCGGGAU
env-183   ACCUCCGUCGAGCGCCAUUCGAAGCGUUUAACCACCGAACGGGCGGAUGAACGGACAACUCGCUGGCAGUCGGGCUCCAGCGGGCACCCGUC.............................UCGC...................................GACGG.G......CCGGGAU
Nph-1-1   GUUACAGCCGCUACCGUUCCGAAGCGCUUAACCACGCCAGCGGCCACGGUUGAAACGACUCGCUGGAUGACGGGCACCAGCGGGCACCUGU..............................CGAC....................................ACAG.G......CUGGAAU
env-184   CCCAGGGAGACGACUGCUUCGAAGCGCUUAUACUCCGGACUUGGAUAGAUACGAUAGACUCGCUGGACCGCGGGCGACAGCGGGCACCUGUUCGGC.........................AACU...............................GCC..ACAG.G......UCGGGAU
Hgo-1-1   CACGCCCCCUUCGACGGUUCGAAGCGCUUAUACGGGAACGCUGGGUACGUUCGGGCAACUCGCUGGGUCGCGGGCUCCAGCGGGCACCUGUUCACGCGGCGGUGC................AGUCGAC...................GCACCGUUGCC...ACAG.G......UAGGGAU
Hxa-1-1   ACAUCCCGUUUCGACGGUUCGAAGCGCUUAUACGGGAACGCUGGGUACGUUCGGGCAACUCGCUGGGUCGCGGGCGCCAGCGGGCACCUGUUCACGCGGCGCUGCA...............GUGA.....................UGCGGCGCCGCC...ACAG.G......UCGGGAU
env-185   ACGGGCCGUUUCGGCGGUUCGAAGGCCUUAUCAACGGCUGCUUUGUACGUUUGGGUGACUCGCCGAGUCGCGGGCGUCGGCGGGCACCUGUCCGGC.........................AUCU...............................GCCC.ACAG.G......UCGGGAU
env-186   ACGGGCCGUUUCGGCGGUUCGAAGGCCUUAUGAACGGCUGCUUUGUAGGUUCGGGUGACUCGCCGAGUCGCGGGCGUCGGCGGGCACCUGUCCGGC.........................AUCU...............................GCCC.ACAG.G......UCGGGAU
env-187   GAUCGAUUCCGCCGCUUCGAAGCGCUUAUAGGCGAACGCUGGAUACGUCCUGUCAGACUCGCUGGGCACGCGGACGCCAGCGGGCACCUGUCCGGC.........................UCAC...............................GCCC.ACAG.G......UCGGGAU
Nin-1-1   GCCUUCGCCCGAAGAGUUCGAAGGCCUUAUACUCGAGAGUUGGCUAUGCUGCGGUAGACUCGCUGAGUCGCGGGCAUCAGCGGGCACCUGUUCGGC.........................ACAC...............................GCCCCACAG.G......UCGGGAU
env-188   CGCGAGCGCGUGAGCGGUUCGAAGGCCUUAUGAAAGACAGCCGGCUAGCCGGAGGCAACCUGUCGGUCGGCGGGCACCGGCGGGCACCCGCACGGC.........................ACGC...............................GCCA..CGG.G......UCGGGAU
env-189   CGAGCGCGUGAGAGCGGUUCGAAGCCCUUAUCACCGACAGCCGGCAACCCUGAGGUAACCUGUCGGUCGGCGGGCACCGACGGGCACCCGUACGGC.........................ACGC...............................GCC..ACGG.G......UCGGGAU
env-192   .......................GCGUUUAUACUCGGCAGUCCGAUAGAUCAGGACGACUCGCUGGGUCGCGGGCGCCGGCGGGCACCUGUUCGGC.........................AUUCGAC............................GCCC.ACAG.G......UCGGGAU
env-193   GGGCGACGUCGCCACGGUUCGAAGCGUUUAUACUCGGCAGUCCGAUAGAUCAGGACGACUCGCUGGGUCGCGGGCGCCGGCGGGCACCUGUUCGGC.........................AUUCGAC............................GCCC.ACAG.G......UCGGGAU
Nje-1-1   CCGCGCCGUCGACGCGGUUCGAAGCGUUUAUACUCGGGAGCUGGGUAGAUCGGGACGACUCGCUGGGUCGCGGGCACCGGCGGGCACCUGUUCGGCA........................GUGA..............................UGUCC.ACAG.G......UCGGGAU
Nam-1-1   GGGCGACCUCGUCGCGGUUCGAAGCGUUUAUACUCGGCAGUCCGAUAGAUCAGGAUAACUCGCUGGGUCGCGGGCGCCGGCGGGCACCUGUUCGGU.........................AUCGAC.............................GCCC.ACAG.G......UCGGGAU
Noc-1-1   CGUCAGCGUCGUCGCGGUUCGAAGCGUUUAUACUCGGUAGCUGGGUAGAUAGGGACGACUCGCUGGGUCGCGGGCGCCGGCGGGCACCUGUACGGC.........................AUCGAC.............................GCCC.ACAG.G......UCGGGAU
env-194   GGUAUCGCGUUCCGGCGGGCUUCGAAGCGCUUUUGAUAGGAGACAGCGAAGAUUGGACAACUCGCUGGGACGGGCACCAGCGGGCACCUGU..............................CUGU....................................ACAG.G......UCGGGAU
env-195   GCCGCGGAUAGGUCAGCUUCGAAGCCCUUUUGUCCUCAAAUCAGAAAGGAUAGCGUAGCUAACUGGACGACGGGCACCAGUGGGCACCUGU..............................CACAGA..................................ACAG.G......UCGGGAU
env-196   GCACAGCGAGCAGUGAUAUCGACGCCCUUUUGUUNGAAACCACGAAAUCUCGUGUAACUCGCUGGGUAAACGGGCACCAGUGGGCACCUGU..............................UCCAU...................................ACAG.G......UUGGGAU
env-197   ACACACCAAGCAGUGAUAUCGACGCCCUUUUGUUUGAAACCACGAAAUUCCCUGUAACUCGCUGGGAGAACGGGCACCAGCGGGCACCUGU..............................UCCAU...................................ACAG.G......UCGGGAU
env-199   ACACAGCCAGCAGUGAUAUCGACGCCCUUUUGUUCGAAACCACGAAAUCUCGUGUAACUCGCUGGGUAAACGGGCACCAGUGGGCACCUGU..............................UCCAU...................................ACAG.G......UUGGGAU
env-201   ACAUAGCCAGCAAGAAUAUCGACGCCCUUUUGUUUGAAACCACGAAAUUUUCUGUAACUCGCUGGGUAAACGGGCACCAGUGGGCACCUGU..............................UCCAU...................................ACAG.G......UCGGGAU
env-202   ..CCACGACCGGCGGAGUUCGAAGUCCUUUUUGUCCGAUAGCAGGGUAGAAUCGGUAACUCGCUGGCGGACGGGCACCAGCGGGCACCUGU..............................CACAU...................................ACAG.G......UCGGGAU
Hla-1-1   AGACAGGGUCAGGGCGGUUCGAAGCGCUUAUACACCGCCGCUGAGUACGUUUCGACGACUCGCUGGGUCGCGGGCGCCAGCGGGCACCUGU.UCGGC........................ACCGAC............................GCCGCCACAG.G......UCGGGAU
Hva-1-1   UCACACCCGUGAGUCUGGUCCGAACCGCUUAAGAGUCCGAACGGCAAAUUGCGGAUAACUCGCUGGACGACGGGCACCAGCGGGCACCUGC..............................GUGU....................................GCAG.G......UCGGGAU
Hsi-1-1   CUCAUCCACAAGUCUGGUCCGAACCGUUUAAGAGUCCGAGCGGCAAACUCCGGGGUAACUCGCUGGACAACGGGCACCAGCGGGCACCUGC..............................GUGU....................................GCAG.G......UCGGGAU
Hma-1-1   CUCAUCCACAAGUCUGGUCCGAACCGUUUAAGACUCCGAGCGGCAAACUCCGGGGUAACUCGCUGGACAACGGGCACCAGCGGGCACCUGC..............................GUGU....................................GCAG.G......UCGGGAU

          ......................................................................................<<<<<<<<<<<<<<<<<<<<<<<<<<<<<<<<<<<..........>>>>>>>>>>>>>>>>>>>>>>>>>>>>>>>>>>.>........<<...
          ......................................................................................22222..1111..................1..1............1..1..................1111..2222.2........12...
          ●●●●●●●●●●●●●●●●●●●●●●●●●●●●●●●●●●●●●●●●●●●●●●●●●●●●●●●●●●●●●●●●●●●●●●●●●●●●●●●●●●Y●●●YR●●●G●------------------------------●●●○-----------------------------------●C●●-●------○●GGRAU
                                                                                                5′
```

```
Ham-1-1    UCACAUCCGCGAGUCUGGUCCGAACCGUUUAAGGCUCCGAACGGCAAACUCCGGGUAACUCGCUGGACGACGGGCACCAGCGGGCACCUGC..............................GUGU...................................GCAG.G......UCGGGAU
env-203    UCACACCCGCGAGUCGGGUCCGAACCGCUUAAGAGUCCGAACGGCAAACUCCGGGUAACUCGCUGGACGACGGGCACCAGCGGGCACCUGC..............................GUGU...................................GCAG.G......CCGGGAU
Hca-1-1    CUCAUCCACAAGUCUGGUCCGAACCGUUUAAGAGUCCGAGCGGCAAACUCCGGGGUAACUCGCUGGACGACGGGCACCAGCGGGCACCUGC..............................GUGU...................................GCAG.G......UCGGGAU
Hsa-2-1    UCACAUCCGCGAGUCUGGUCCGAACCGUUUAAGACUCCGAACGGCAAACUCCGGGUAACUCGCUGGACGACGGGCACCAGCGGGCACCUGC..............................GUGU...................................GCAG.G......UCGGGAU
Hja-1-1    CCCAUCCCUAAGUCCGGUCCGAACCGCUUAAGAGUCCGAACGGCAAACUCCGGAGUAACUCGCUGGACGACGGGCACCAGCGGGCACCUGC..............................GUGU...................................GCAG.G......UCGGGAU
Har-1-1    CGCAUCCACGAGUCUGGUCCGAACCGCUUAAGAGUCCGAACGGCAAACUCCGGAGUAACUCGCUGGGCGACGGGCACCAGCGGGCACCUGC..............................GUGU...................................GCAG.G......UCGGGAU
env-206    UGAGCGGCUCCCGGCGGAGUUCGAAGGCCUUUUGGCCGCCGGCGGGGUACCGCCUGUAACUCGCUGGAGACGGGCACCAGCGGGCACCUGC..............................CUGU...................................GCAG.G......ACGGGAU
Hmu-2-1    AGCCCGUCUCGAUCAGCUUCGAAUGGCUUAAGGGCGCAAGCCGGGAAGCAGUGAGUAACUCGCUGGACGACGGGCACCAGCGGGCACCUGCAUGGACCG......................AUCGA..........................CGGGUCGUACAG.G......UCGGGAU
Hka-1-1    AGACCGUCUCGAUCAGCUUCGAAUGGCUUAAGGGCGCAAGCCGGGAAGCAGUGGGUAACUCGCUGGACGACGGGCACCAGCGGGCACCUGCAUGGACCG......................AUCGA..........................CGGGUCGUACAG.G......UCGGGAU
env-207    CUCAACGGACGGCGGGGUUUCGAAGCCCUUAACAGCGGUUGCGGUAAAGUUGGAGCAACUCGCCGGACGACGGGCACCGGUGGGCACCUGU..............................CUAC...................................GCAG.G......UCGGGAU
env-208    ACCAGUUGUCGACCGCAUUUCGAAGCCCUUAACCCCUGCUGUGAUAAGUUUCGAACAACUCGCCGGACGACGGGCAUCGGCGGGCACCUGU..............................CUAC...................................GCAG.G......UCGGGAU
env-210    CGCGGAAACCCGUUCGAUUUCGAAGUGUUUAACCCCGUCAGUCGUAAAAAUCAGGCAACUCGCUGGACGACGGGCACCAGCGGGCGCCUGU..............................CCAC...................................ACAG.G......UCGGGAU
env-212    CGCAAGGCCCCGAUCGGUUUCGAAGUGUUUAACCCCGGCAGUCGUGAAUCUCCCGUAACUCGCUGGACGACGGGCACCAGCGGGCGCCUGU..............................CUAC...................................GCAG.G......UCGGGAU
env-213    CGUGAGAUCCCGUUCGGUUUCGAAGUGUUUAACCCCGACAGUCGUGAAUCUCCCGUAACUCGCUGGACGACGGGCACCAGCGGGCGCCUGU..............................CUAC...................................GCAG.G......UCGGGAU
env-214    CGACGUUUUGAGGAUACUUUCGAAGCCCUUAACCCCACUAGCGAAGAAUCUUAGUCAACGCGCCGGACGACGGGCACCGGUGGGCACCUGU..............................CUAC...................................ACAG.G......UCGGAAU
env-215    AUCCGGCGCGUAAGACGUUUCGAAGCCCUUAACCGCGAUUAUUGUGAAGAGACGGUAACUCGCUGGACGACGGGCACCAGUGGGCACCUGU..............................CUAC...................................GCAG.G......UCGGGAU
env-216    CGGCCAGUUCGAGCGCUUUCGAAGCGCUUAACCCCGACAAUACAGAAAAUCUGGGUAACUCGCUGGCGGACGGGCACCAGCGGGCACCUGU..............................CUAC...................................GCAG.G......UCGGGAU
env-217    CUUGACCGUCACGGGCUUUCGAAGCCCUUAACCGACACAGCAGCGAAUUCCACGGUAACUCGCCGGAGGACGGGCACCGGUGGGCACCUGU..............................CUAC...................................ACAG.G......UCGGGAU
env-218    ........................GCUUAACCGACACAGCGGUGAAUUUAAUCGUAACUCGCCGGACGACGGGCGACCGGUGGGCACCUGU..............................CUAC...................................GCAG.G......UCGGGAU
env-219    CGUGGGUGCCACAGGCUUUCGAAGCGCUUAACCGACACAGCGGUGAAUUUAAUCGUAACUCGCCGGACGACGGGCACCGGUGGGCACCUGU..............................CUAC...................................GCAG.G......UCGGGAU
env-220    CGUGGGUGCCGCAGGCUUUCGAAGCGCUUAACCGACACAGCGGUGAAUUUAAGCGUAACUCGCCGGACGACGGGCACCGGUGGGCACCUGU..............................CUAC...................................GCAG.G......UCGGGAU
Nmo-1-1    CGUCGACCGCCACGGGCUUUCGAAGCGCUUAACCGACCCAGCGGUGAAUUCCGGGUAACGCGCCGGACGACGGGCACCGGUGGGCACCUGU..............................CUAC...................................ACAG.G......UCGGGAU
env-221    GGACUCGGGCUGUACGGGGUUCGAAGCGUUUUUCCACGACGGCGAUGAACUGUGGGUAACUCGCUGGAGACGGGCACCAGCGGGCGCCUGU..............................CUGU...................................ACAG.G......UCGGGAU
env-223    GUAUCGCCCUCCGGUGGGCUUCGAAGCGUUUUUUAUCCGAGACGCGAAAGUUCAGGUAACUCGCUGGAGACGGGCACCAGCGGGCACCUGU..............................CUGU...................................ACAG.G......UCGGGAU
env-224    CACCACCGGUGUUCGGACCGCUUCGAAGUAUUUUAACUUAAAACUGAGAAGUAUAUACAACUCGCUGGGACGGGCACCAGCGGGCACCUGU..............................CUGU...................................ACAG.G......UCGGGAU
env-226    ...................................................AGCCUAACUCGCUGGACGACGGGCACCAGUGGGCACCUGU..............................CCAC...................................ACAG.G......UAGGGAU
env-227    .......................................AAUACGGAACUAUCGAUAACUCGCUGGACGACGGGCACCAGUGGGCACCUGC..............................GUGU...................................GCAG.G......UCGGGAU
env-228    CGAAUCGAGCGGCGCGGUUCGAAGCUCUUAAACGCCCAGUGCGACGUAUCUCGGGUAGACUCGCUGGAGACGGGCACCAGCGGGCACCCGUC.............................GCG...................................GACGG.G......CCGGGAU
env-229    CGAGGCGAGCGGCGCGGUUCGAAGCUCUUAACCCUCCCGUAGGACGUAGCUCUGGUAGACUCGCUGGGGACGGGCACCAGCGGGCACCCGUC.............................GCGC..................................GACGG.G......CCGGGAU
Hci-1-1    AUGUACGACCGUGCGAAAGACGAAGGCCUUAAACCCCCGAACGGGAUAUCUUGGGUAACUCGCUGGUCAACGGGCACCAGUGGGCACCUGC..............................AAUC...................................GCAG.G......ACGGAAU
Hut-1-1    AUCUCACUCACGGAAGUUCGAAUUCCUUAAGAGGACGACCGAAGUAUGAGUGAGUAACUCGCUGGUCGGCGGGCAGCCAGCGGGCACCUGU..............................UUAG...................................GCAG.G......UCGGAAU
env-230    .................UUCCGAAGCCCUUAGAACCCCGAGCGAAGAAGAGGCGUAUAACUCGUCGGUCGCGGGCACCGGCGGGCACCUGC..............................UACG...................................GCAG.G......ACGGAAU
env-231    GGAGUCUCGCAGGUGCUUUCGAAGGUCUUAAACUCCGAACAGCGGUAUUCCAGAGUAACUCGCUGGACGACGGGCACCAGCGGGCACCUGUC.............................AUC...................................GACAG.G......ACGGAAU
env-232    GAUGGAUCGACGCUGCUUUCGAAGGCCUUAAACUCCAAACGGCGGUACUCACGGGUAACUCGCUGGACGACGGGCACCGGCGGGCACCUGUC.............................AUC...................................GACAG.G......ACGGAAU
env-233    GACGUAUCGAGGGGUCUUUCGAAGGUCUUAAACUCCGAACGACGGUAUCCAACUGUAACUCGCUGGACGACGGGCACCAGCGGGCACCUGUC.............................AUC...................................GACAG.G......ACGGAAU
env-234    AACAGCCGCAGCCUCACUUCGAAGCCCUUUUGUCCACAAAUCGGAAAGGAGAGCACACCACGCUGGACGACGGGCACCAGCGGGCACCUGUC.............................ACAG..................................GACAG.G......ACGGGAU
env-235    ...........CCCUGUAUCGAAGUGCUUAAGCGAAUCUCAUGGGUAAAUUAUAGUAUCUCGCUGGAAGACGGGCACCAGCGGGCACCUGUC.............................UCU...................................GACAG.G......UCGGGAU
env-236    CUGACGCCCGGGGGUUUCGAAGCGCUUAAACGAAGCACAGGGGAAUAUAUCAUAAGUUCUCGCUGGCGGACGGGCACCAGCGGGCACCUGUC.............................AUC...................................GACAG.G......CCGGGAU
Nba-1-1    ACAGCCACUUCCGGAGUUCGAAGCCCUUAUACUCGAGAGUUGGAUAGACUGUCGUAGACUCGCUGGGUCGCGGGCGUCAGCGGGCACCUGU.GCA..........................GCAUUG.............................UGCUACAG.G......UCGGGAU
env-237    ACUGCCGUUCCCGGAGUUCGAAGCCCUUAUACUCGAGAGUUGGAUAGAUUGUCGUAGACUCGCUGGGUCGCGGGCGUCAGCGGGCACCUGU.GCA..........................GCAUUG.............................UGCUACAG.G......UCGGGAU
Nsu-1-1    CCGUCUGGUCCGGGAGUUCGAAGGCCUUAUACUCGAGAGUUGGGUAGAUGGUCGUAGACUCGCUGGGUCGCGGGCGUCAGCGGGCACCUGCGCAGCAC.......................GUCUCU..........................GUGCUA..CAG.G......UCGGGAU
Npa-1-1    CACGGGCGUUCUCGGAGUUCGAAGCCCUUAUACUCGAGAGUUGGGAAGUGUGAGUAGACUCGCUGGUUCGCGGGCAUCAGCGGGCACCUGU..CCGA........................CGG..............................UCGGUCACAG.G......UCGGGAU
Nal-2-1    CCACGGGCGUCUCGGAGUUCGAAGCCCUUAUACUCGAGAGUUGGGAAGUGUGAGUAGACUCGCUGGUUCGCGGGCAUCAGCGGGCACCUGU..CCGA........................CGG..............................UCGGUCACAG.G......UCGGGAU
Nga-1-1    CAUGGCUGUUCUCGGAGUUCGAAGCCCUUAUACUCGAGAGUUGGGAAGUGUGAGUAGACUCGCUGGUUCGCGGGCAUCAGCGGGCACCUGU..CCGA........................CGG..............................UCGGUCACAG.G......UCGGGAU
Hdj-1-1    ACGCAGGGUGUGGCCGGUUCGAAGCGCUUAUACUCGAGAGCUAGAUAGUUACGGUAGACUCGCUGGUUCGCGGGCAUCAGCGGGCACCUGCACGGC.........................UUU...............................GCC..GCAG.G......UCGGGAU
Hli-3-1    GUAGCGCGCUCGGGGAGUUCGAAGCCCUUAUACUCGAGAGUUGGGUAGAUCCAGUAGACUCGCUGGUUCGCGGGCAUCAGCGGGCACCUGCACGGU.........................ACCGG.............................GCC..GCAG.G......UCGGGAU
Hje-2-1    ACGCCGCGUUCCCGGAGUUCGAAGCGCUUAUACUCGAGAGCCGGGAAGAGAGGGUAGACUCGCUGGUUCGCGGGCAUCAGCGGGCACCUGCACGACG........................CCG..............................CGUCA..CAG.G......UCGGGAU
Hth-2-1    ACGCCGCGUUCCCGGAGUUCGAAGCGCUUAUACUCGAGAGCCGGGAAGAGUGGGUAGACUCGCUGGUUCGCGGGCAUCAGCGGGCACCUG...CACG........................ACGC.............................CGUGUCACAG.G......UCGGGAU
Hsa-10-1   CGUUACCGUCGGCGUCGUUCGAAGCCCUUAUACUCGAGAGUUGGGUAGGUCCGGUAGACUCGCUGGUUCGCGGGCAUCAGCGGGCACCUGUACGACG........................CCG..............................CGUC..ACAG.G......UCGGGAU
env-238    GCCGUUUGUUUCGACGCUUCGAAGCGCUUAUACACGAACGCUGGAUAGAACCGACAGACUCGCUGGGCUGCGGGCACCAGCGGGCACCUGU.UCGGC........................UCGAC............................GCCGCCACAG.G......ACGGGAU
env-239    GCCGUUUGUUUCGACGCUUCGAAGCGCUUAUACACGAACGCUGGAUAGAACCGACAGACUCGCUGGGCUGCGGGCACCAGUGGGCACCUGU.UCGGC........................UCGAC............................GCCGCCACAG.G......ACGGGAU
Hni-1-1    CACGCCACUUCCGAUGGUUCGAAGCGCUUAUACACGGCGGCUGGGUACGUUUCGACGACUCGCUGGGUCGCGGGCACCAGCGGGCACCUGU.UCGGC........................AACCGAC..........................GCCGCCACAG.G......CCGGGAU
Nch-1-1    CAACACGCUCCGCCUGCUUCGAAGCCCUUAUACUCGGGAGUUGAAUAGGUGAAACUGACUCGCUGGAUCGCGGGCACCAGCGGGCACCUGU.UCGGC........................ACGCGAC..........................GCCGCCACAG.G......ACGGGAU
Nhu-1-1    CGACACGGUCCGCACGCUUCGAAGCCCUUAUACUCGGGAGUUGAAUAGGUGAAACCGACUCGCUGGAUCGCGGGCACCAGCGGGCACCUGU.UCGGC........................ACACGAC..........................GCCGCCACAG.G......ACGGGAU
Nve-1-1    CACGGCGUCUCACGGAGUUCGAAGCCCUUAUACUCGAGAGUUGGGAAGGGUGAGUAGACUCGCUGGUUCGCGGGCAUCAGCGGGCACCUGU.ACGGC........................ACC..............................GCCGCCACAG.G......UCGGGAU
Hsa-8-1    GCGGCGUGGCCGCGCAGUUCGAAGCCCUUAUACUCGAGAGUUGGGAAGAGAGAGUAGACUCGCUGGGUCGCGGGCGCCAGCGGGCACCUGU.ACGGC........................ACC..............................GCCGCCACAG.G......UCGGGAU
Htu-1-1    GCUCGCCGUUCCCGCAGUUCGAAGCCCUUAUACUCGAGAGUUGGGAAGAGAAGGUAGACUCGCUGGGUCGCGGGCGCCAGCGGGCACCUGU.ACGGC........................AUU..............................GCCGCCACAG.G......UCGGGAU
env-243    GUGACCGGCCACGGAGUUCGAAGGCCUUAUACUCGAGAGUUGGCUAUCACGACGUAGACUCGCUGAGUCGCGGGCAUCAGCGGGCACCUGU.UCGGCG.......................CGG.............................CGCCACCACAG.G......UCGGGAU
Hca-3-1    GGUGCGCCGUCGCGAGUUCGGAUCGCUUAAGGGCGCGAGCGGGGAACUGGCGAGUAACUCGCUGGGCGGUCGGGCACCAGCGGGCGCCUGC..............................CUGU...................................GCAG.G......UCGGGAU
env-244    AACCCACGCCAGCGGAGUUCGAAGUGUUUAACCCCGGGACCGGGGAAGGACGGGGUAACUCGCUGGUCGACGGGCACCAGCGGGCACCUGCCGCGUCGCGUUUCGAGGGG...........ACG.................CCCCUCG...CGCGAUCCA.CAG.G......ACGGGAU
env-245    AAUACGGCGAGCACGUGGUUCGAAGCGCUUAUAUUUGCGACGCUGAUAGCUGCUGUUGACUUACUGGCGGCGGGCACCAGUGGGCACCUGCACGG..........................GUUCGA.............................CC..GCAG.G......UCGGGAU
env-246    CGAGCGGGGCGGCGCCACUCCGAAGCUCUUUUGCCCGAAAGCGGGGAAGAGAGGAUAACUCGCUGGGACGCGGGCGCCAGCGGGCACCUGC..............................CUGU...................................GCAG.G......UCGGGAU
env-247    UGUGGCUAAACCGGCGGUUCGAAGCCCUUAUACUCACCACUCCGGUAGCUGUGAGUGACUGACUGGACGACGGGCACCAGUGGGCACCUGCACGG..........................CUAGA..............................CC..GCAG.G......UCGGGAU
env-248    AAAUCGGAUACUGACGCUUCGAAGCGCUUAUACUCGCCACUCCGGUAGCUCUGGGUGACUGACUGGACGGCGGGCACCAGUGGGCACCUGCACGG..........................CUAGA..............................CC..GCAG.G......UCGGGAU
env-249    AAAGCGGAUACAGACGCUUCGAAGCGCUUAUACUCGCCACUCCGGUAGCUCUGGGUGACUGACUGGACGGCGGGCACCAGUGGGCACCUGCACGG..........................CUAGA..............................CC..GCAG.G......UCGGGAU
env-250    GUUCCGGGACCGGCCGAGUUCGAAGCGUUUUAGCCCGGCACUCGAGUAGGUUGUGUAUCUCGCUGGCGGACGGGCACCAGCGGGCGCCUGC..............................GUGU...................................GCAG.G......UCGGGAU
env-251    UCCGGGACCGGCCGACGAGUUCGAAGCGUUUUAGCCUGGCACUCGCAUAACUGUCCAUCUCGCUGGCGAACGGGCACCAGCGGGCGCCUGC..............................GUGU...................................GCAG.G......UCGGAAU
env-253    ...............................................AUUUCUGAUAACUCGCUGGCGGACGGGCACCAGCGGGCACCUGC..............................CUGU...................................GCAG.G......CCGGGAU
Hsu-1-1    AACCGGCCUGACCGCGAUUCGAAGUCCUUUUAGUCGAACCGCGCCGAUCAUACGAUAUCUCGCUGGUGAGCGGGCUCCAGCGGGCUCCCGA..............................UAGCC..................................UCGG.G......ACGGGAU
Hsp-2-1    CCUUCGACGUCGAGUAGUUCCGAAGUUCUUAACCCCGACCGAUUCGAACUCAGGGUAACUCGCUGGCGAGCGGGCUCCAGCGGGCUCCUGAC.............................CACA..................................GUCAG.G......ACGGGAU
Has-1-1    ACUGCGGACGACAGCGCUUCGAAGCCCUUAUGAAACGACGUUCGGUAGUAUGGGUAGCUACGCUGGGCGGCGGGCACCAGCGGGCACCCGUACGG..........................GUUAU..............................CC..ACGG.G......UUGGAAU
env-254    CCGUACACCGCCGCUCGCUUCGAAGCCCUUAUGAAACCGGACGGGAAACCAUCGGUAGACUCGUCGGCAGCGGGCACCGACGGGCACCCGUACGGC.........................AACC..............................GCCC.ACGG.G......CCGGGAU
Hag-1-1    GACCGGGUCGAUGGAGGAUGUUUCGAAGCCCUUAACAGCGCCACUGGCGGAGUUCCGGUGACUCGCCGGGCGGGCACCGGCGGGCACCUGCG.UGCGA.......................UGU.............................UCGCAUUGCAG.G......UCGGAAU
Hsa-4-1    UCGGCCAAGGACCGUGGCGUUCCGAAGCGCUUAACAGCGCCACUGGCGGAGAUCCAGUGACUCGCUGGGACGGGCACCGGCGGGCACCUGCG.............................UGAUAC................................UGCAG.G......UCGGAAU
Hsa-5-1    CCGGACGAGGACCGCGGCGUUCCGAAGCGCUUAACAGCGCCACUGGCGGAGAUCCAGUGACUCGCUGGGACGGGCACCGGCGGGCACCUGUGG............................CGAUA................................CUGCAG.G......UCGGAAU
env-255    UCACUGAUGUUUGUUUCGAAGUGCUUAAAAGGGUCGACAGCUGCAUGAGGAGUACAGAUUCGCCGGGAUGCGGAACCGGCGGGGCGCCUGU..............................AUAAC..................................UCAG.G.UUGGUGCGGGAU
env-256    CGAUUGACGUCUGUUUCGAAGUGCUUAAAAGGAUCGAAAGCUGCAUGAGGAGUACAGAUUCGCCGGAAUGCGGAACCGGUGGGGCGCCUGU..............................AUAAC..................................UCAG.G.UUGAUGCGGGAU
env-262    ................................................................................GGGGCGCCUGU..............................AUAAC..................................UCAG.G.UUGAUGCGGGAU
env-266    CUAAGGACAUUUUUUUCGAAGUGCUUAAAAGGGUCGGAAGCUGCAUGAUGAAUACAGAUUCGUCGGGUAGCGGAACCGGCGGGGCGCCUGU..............................AUAGA..................................UCAG.G.UUGACGCGGGAU
Mae-1-1    AACCGUUAAAUUAAUAUACUAAAUACCAUACAAGAUAAUGGGCAUAUUAUUACAUAUAGGAUAGAAAUAAAAACAACAGACAAUAACGGGA..............................GUCAAA.................................UCCC.G.....UCGGGCAU
env-267    GGGCGAAGGCUAUCGUUGCGGGGCCGGUUUAUUCUGCGAAAUAUUUAUAUUAUAGUACCUGUUUAUUGUACCCACAGUUUCGUUCGCGGGA..............................UCCG...................................UCCC.G......CCGGGAU
env-268    UGAAAGAAGCUAGCCUAGUGGAUAAGCCAUUGAGAACUUGAAAAGGUUUAUUAACAAUCCACUGGCUUCAUACCUACGAGCUAUCUACGGA..............................UGUCGU.................................UCCG.U.......UGGGAU
env-269    .....................................................ACAAUCCACUGGCUUCAUACCUACGAGCUAUCUACGGA..............................UGUCGU.................................UCCG.U.......UGGGAU

           ......................................................................................<<<<<<<<<<<<<<<<<<<<<<<<<<<<<<<<<<<.........>>>>>>>>>>>>>>>>>>>>>>>>>>>>>>>>>>.>........<<...
           ......................................................................................22222..1111..................1..1...........1..1....................1111.2222.2........12...
           ●●●●●●●●●●●●●●●●●●●●●●●●●●●●●●●●●●●●●●●●●●●●●●●●●●●●●●●●●●●●●●●●●●●●●●●●●●●●●●●●Y●●●YR●●●G●------------------------------●●●○-----------------------------------●C●●-●------○●GGRAU
                                                                                                 5′
```

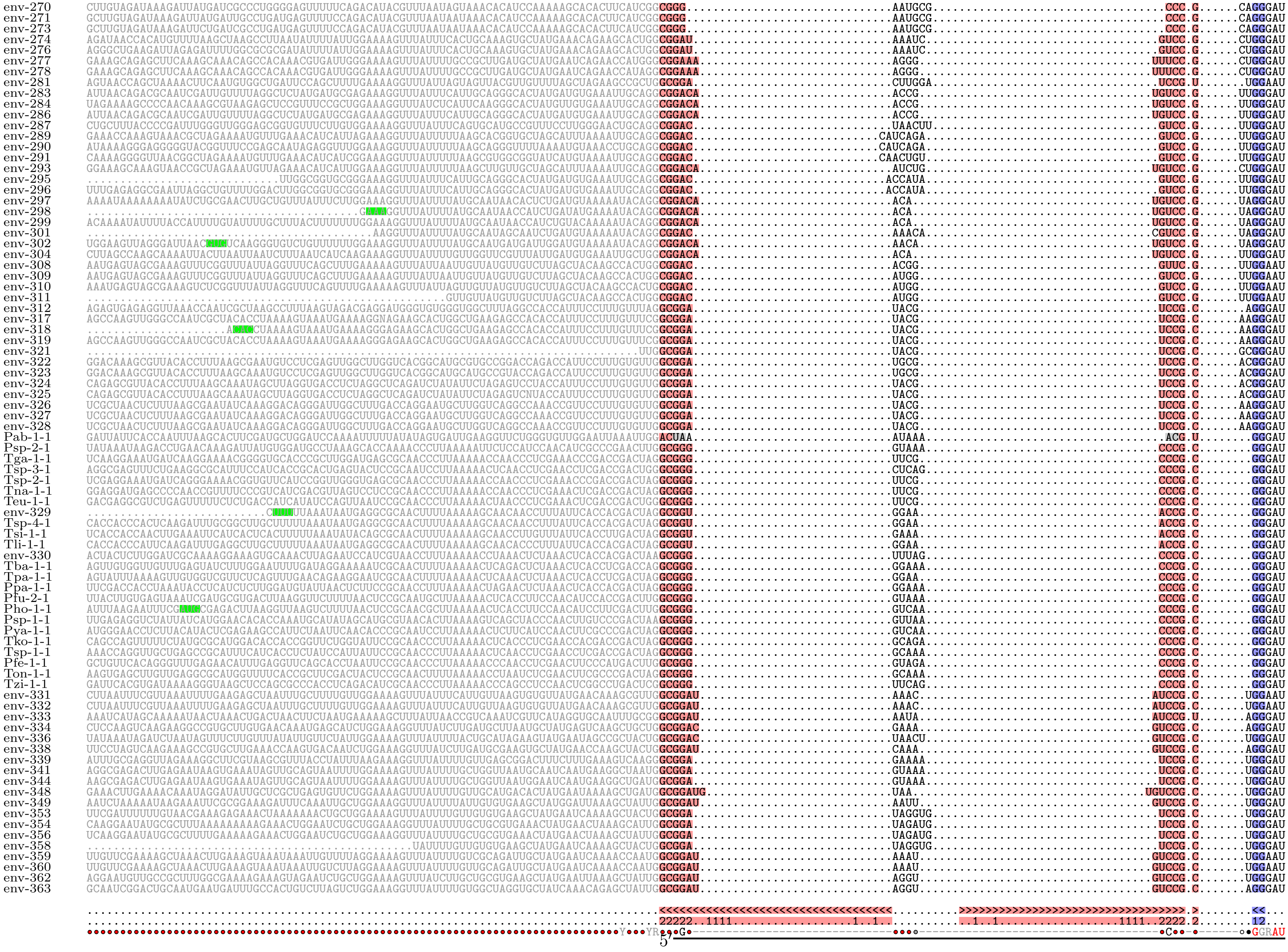
env-270
env-271
env-273
env-274
env-276
env-277
env-278
env-281
env-283
env-284
env-286
env-287
env-289
env-290
env-291
env-293
env-295
env-296
env-297
env-298
env-299
env-301
env-302
env-304
env-308
env-309
env-310
env-311
env-312
env-317
env-318
env-319
env-321
env-322
env-323
env-324
env-325
env-326
env-327
env-328
Pab-1-1
Psp-2-1
Tga-1-1
Tsp-3-1
Tsp-2-1
Tna-1-1
Teu-1-1
env-329
Tsp-4-1
Tsi-1-1
Tli-1-1
env-330
Tba-1-1
Tpa-1-1
Ppa-1-1
Pfu-2-1
Pho-1-1
Psp-1-1
Pya-1-1
Tko-1-1
Tsp-1-1
Pfe-1-1
Ton-1-1
Tzi-1-1
env-331
env-332
env-333
env-334
env-336
env-338
env-339
env-341
env-344
env-348
env-349
env-353
env-354
env-356
env-358
env-359
env-360
env-362
env-363
5'

```
env-364 CUGAGGCAAGGCUUGAAAAUAAACCAGAGUGUGACUGAAGCCGGAAAAGUUUAUUAUGUUGCGUGAAGCUAUGAAUCAAGCUAUCGGCGGAUG..........................AAAG..................................GUCCG.C......UGGGAU
env-365 AGGAAUAUCGUCGAUUUGGAAAAAAGAAACUGGAAUCUUCUGGAAAAGUUUAUUUUGCUGCAUGAAACUAUGAAUCAAAGCUAUUGGCGGAU...........................AGAU..................................GUCCG.C......UGGGAU
env-366 CAAAGCAAAGCUUGAAAACAAUUUGAAUGUGCCUGAGAUUCUGGAAAGGUUUAUUUUGAUGCGUGGAACUAUGAAUCAAAGCUACUGGCGGAC...........................GGGU..................................GUCCG.C......UGGGAU
env-367 AAAAGCGGGGCUUGAAAACAAUUUGAAUGUGCCUGAGAUUCUGGAAAAGUUUAUUUUGAUGCGUGAAACUAUGAAUCAAAGCUACUGGCGGAU...........................AGGU..................................GUCCG.C......UGGGAU
env-369 AGCAAAACUUGAAAACAAAUGAACAUUGCUAGCUGAGUACUGGAAAAGUUUAUUUUGUUGCAUGAAGCUAUGAAUUAAAGCUACUGGCGGAU...........................AGGU..................................GUCCG.C......UGGGAU
env-370 CAAAGCGGAGCUUGAAAACAAUUUGAAUGUGCCUGAGAUUCUGGAAAGGUUUAUUUUGAUGCGUGAAACUAUGAAUCAAAGCUACUGGCGGAU...........................GGGU..................................GUCCG.C......UGGGAU
env-371 CAAAGCGGAGCUUGAAAACAAUUUGAAUGUACCUGAGAUUCUGGAAAAGUUUAUUUUGCUGCGUGAAACUAUGAAUCAAAGCUACUGGCGGAU...........................GGGU..................................GUCCG.C......UGGGAU
env-375 ACCACGCUAUUAAAUUUUUCAAAGAACANCCCCAUUUAACUGGAAAAGUUUAUUUCACUGUAAGAUGCUAUGAAUNAAAACAAUUGGCGGAU...........................AAAA..................................GUCCG.C......UGGGAU
env-376 UCUAUCACGCUAUUAAAUUUUUCAAACAGUCCCCUUUAGCUGGAAAAGUUUAUUUCAUUGUAAGAUGCUAUGAAUCAAUCAAAACAAUUGGCGGAU...........................AGAA..................................GUCCG.C......UGGGAU
env-377 GUUACCACGCUAUUAAAUUUUUCAAACAGCCCCCUUUAUCUGGAAAAGUUUAUUUCACUGUAAGAUGCUAUCAAUCAAUCAAAACAAUUGGCGGA............................UGAAAG..................................UCCG.C......UGGGAU
env-378 ACCAUACUAUUAAAUUUUUCAAAGAACAGCCCCAUUCAACUGGAAAAGUUUAUUUCACUGUAAGAUGCUAUGAAUCAAAACAAUUGGCGGA............................UGAAAG..................................UCCG.C......UGGGAU
env-379 AACUUUAGAAAGCGAAAACAGAUAGAGAAUGAAACCUAUUGGAAAAGUUUAUUUCAUUGUAAGAUGCUAUCAAUCAAAACAAUUGGCGGA............................UGAAAG..................................UCCG.C......UGGGAU
env-380 UCUACCACGCUAUUAAAUUUUUCAAAUAGUCCCUUUUAGCUGGAAAAGUUUAUUUCAUUGUAAGAUGUUAUGAAUCAAAACAAUUGGCGGA............................UGAAAG..................................UCCG.C......UGGGAU
env-381 UCUACCACGCUAUUAAAUUUUUCAAACAGUCCUCUUUAGCUGGAAAAGUUUAUUUCAUUGUAAGAUGCUAUGAAUCAAAACAAUUGGCGGA............................UAAAAG..................................UCCG.C......UGGGAU
env-382 ACCGCACUAUUAAAUUUUUCAAAGAACAGCCCCAUUUAGCUGGAAAAGUUUAUUUCAUUGUAAGAUGCUAUCAAUCAAAACAAUUGGCGGA............................UAAAAG..................................UCCG.C......UGGGAU
env-383 UCUACCACGCUAUUAAAUUUUUCAAACAGUCCCUUUUAGCUGGAAAAGUUUAUUUCAUUGUAAGAUGCUAUGAAUUAAAACAAUUGGCGGA............................UAAAAG..................................UCCG.C......UGGGAU
env-384 UCUACCACGCUAUUAAAUUUUUCAAACAGUCCCUUUUAGCUGGAAAAGUUUAUUUCAUUGUAAGAUGCUAUGAAUAAAACAAUUGGCGGA............................UAAAAA..................................UCCG.C......UGGGAU
env-385 GGCGAAGCUUGAAAACAAUUUAACAUUGCCAGUUUAGCGCUGGAAAAGUUUAUUUUGCCGCGUGAAGCUAUGAAUCAAAGCUACUGGCGGAU...........................AAGU..................................GUCCG.U......UGGAAU
env-386 GGCGAAGCUUGAAGACAAAUGAACAUUGCCAGUUUAGCGCUGGAAAAGUUUAUUUUGCCGUGAAGGUAUGAAUCAAAGCUACUGGCGGAU...........................AAGU..................................GUCCG.U......UGGAAU
env-387 GGCGAAGCUUGAAAACAAUUUAACAUUGCCAGUUUAGCGUUGGAAAAGUUUAUUUUGCCGCGUGAAGCUAUGAAUCAAAGCUACUGGCGGAU...........................AAGU..................................GUCCG.U......UGGAAU
env-388 GUAAAAUUAACUGAAGAAAAUGAAUAAGUGACUGUUUUCUGGAAAAGUUUAUUUUACUGCAUGAUGCUAUGAAUCAAAGCUAUUGGCGGA............................UUAGAUG................................UCCG.C......UGGGAU
env-390 GUAAAAUUAACUAAAAGAAAAUGAAUAAGUGACUGUUUUCUGGAAAAGUUUAUUUUACUGCAUGAUGCUAUGAAUCAAAGCUAUUGGCGGA............................UUAGACA................................UCCG.C......UGGGAU
env-392 GGCCGUGCUUGAGGAUAAGCCGAAAUAGGCUGUUCGCCUUUGGAAAAGUUUAUUUUGUCGGCUUGUGAUUUAUUGAGAGCCGAGCGGCGGA............................CUUCA..................................UCCG.C......UGGGAU
env-402 GGCCGUGCUUGAGGAUAAGCCGAAAUAGGCUGUUCGCCUUUGGAAAAGUUUAUUUUGUCGGCUUGUGAAUGAUUGAGAGCCGAGCGGCGGA............................CUUCA..................................UCCG.C......UGGGAU
env-406 GUUAAAAAAGCUGUGCUUGAGGCUAAAAAUGGUUAACUUGCUGGAAAAGUUUAUUUUUGUGGUUGGGCAAUCAUUGAGAGCCAGUGGCGGAU...........................AGAA..................................AUCCG.C......UGGGAU
env-408 GUUAAAAAAGCUGUGCUUGAGACUAAAAAUGGUUAACUUGCUGGAAAAGUUUAUUUUUGUGGUUGGGCAAUCAUUGAGAGCCAGUGGCGGAU...........................GGAA..................................AUCCG.C......UGGGAU
env-410 UUAAAAAGGCCGUGCUUGAGGCUAAGACCUAACAGCCUUGUUGGAAAAGUUUAUUUUGUUGGUUGGGGAAUCAUUGAGAGCCAGUGGCGGAU...........................GAGA..................................AUCCG.C......UGGGAU
env-412 ...................................................................AAUCAUUGAGAGCCAGUGGCGGAU...........................GAGA..................................AUCCG.C......UGGGAU
env-413 .................................................................GGGAAUCAUUGAGAGCCAGUGGCGGAU...........................GAGA..................................AUCCG.C......UGGGAU
env-414 AGAGCUUGAAGACAAGACAAGAUAGGCGCCUCAAGCUUUUGGAAAAGUUUAUUUUGCUGGUUGGCGAAUCAUUGAAAGCCGAGCGGCGGAC...........................UUU...................................GUCCG.C......UGGGAU
env-416 GAGCUUGCAGACAAGGCAAAAUAGGCACUUCAAGCCCUUUUGGAAAAGUUUAUUUUGCUGGUUGGCGAAUCAUUGAAAGCCGAGCGGCGGAC...........................UUU...................................GUCCG.C......UGGGAU
env-418 GCUGAACUUGAGGACAAGCCGAAAUAGGCAGUUUGCCUCUUGGAAAAGUUUAUUUUGCUGGCUUGUGAAUGAUUGAGAGCCGAGCGGCGGAC...........................AAG...................................GUCCG.C.....UAGG.AU
env-420 AGCUGAACUUGAGGACAAGCCAAAAUAGGCAGUUUGCCUUUGGAAAAGUUUAUUUUGCUGGCUUGUGAAUGAUUGAGAGCCGAGCGGCGGAC...........................AAG...................................GUCCG.C......UGGGAU
env-422 GCUGAACUUGAGGACAAGCCAAAAUAGGCGGUUUGCCUUUUGGAAAAGUUUAUUUUGCUGGCUUGUGAAUGAUUGAGAGCCGAGCGGCGGAC...........................AAG...................................GUCCG.C......UGGGAU
env-424 UCAAGAAAUCUUGAAGGCUAGAAAAUGUUUGAAACAUCAUAGAAAAGGUUUAUUUUAAGCACUGCGCUAACAUUCAAAAUUGCAGGCGGACU..........................ACCAGA................................GUCCG.C......UGGGAU
env-425 UUUAUAUAAGAGAGAGUAGUCUAAUACAGUAAUAGCGUUUUGGAAAGGUUUAUUUUAGGCAUGGUGCUAAGAUGUAAAAUUGCAGGCGGAC...........................CAGCAGA...............................GUCCG.C......UGGGAU
env-427 UUUAUAUAAGAGAGAGUAGUCUAAUACAGCAAUAGCGUUUUGGAAAGGUUUAUUUUAGGCAUGGUGCUAAGAUGUAAAAUUGCAGGCGGAC...........................CAGCAGA...............................GUCCG.C......UGGGAU
env-428 GCUUAAGAACCUUUCGUCACACGCACAGCGCUAGCGUUCUAGAAAAGGUUUAUUUUUAACCACGGCGCUAACAUUUAAAAUUGCAGGCGGAC...........................CAACCGU...............................GUCCG.C......UGGGAU
env-429 GUAUAGGAGGCUAUAGUCUCCGCAGAGCACUAGGGUUCUGGAAAAGUUUAUUUUUAACCACGGCGCUAACAUUCAAAAUUGCAGGCGGAC...........................CAGCUGU...............................GUCCG.C......UGGGAU
env-432 AUAUAGAGGGGCUAUAGUCUCCGCAGAGCACUAGUGUUCUGGAAAGGUUUAUUUUUAACCACGGCGCUAACAUUGCAGAAUUGCAGGCGGAC...........................CAACCGU...............................GUCCG.C......UGGGAU
env-433 AUAUAGAGGGGCUAUAGUCACCGCAGAGCACCAGAGUUCUGGAAAAGUUUAUUUUUAACCACGGCGCUAACAUUCAAAAUUGCAGGCGGAC...........................CAACCGU...............................GUCCG.C......UGGGAU
env-438 UCAUUUUAAUUCGCACUUCUGAAGAGUGAUUUGACUUUUGUUGGAAAAGUUUAUUUCAUAGGUAAGUGUUAUGAAACAAAACAUUGGCGGAU...........................AACA..................................GUCCG.C......UGGAAU
env-439 UCAUUUUAAUUCGCACUUCUGAAGAGUGAUUUGACUUUUGUGGAAAAGUUUAUUUCAUAGGUAACUGUUAUGAAACAAAACAUUGGCGGAU...........................AACA..................................GUCCG.C......UGGAAU
env-440 GGUUGUAGCUAAUAAAUUCAUUGUAGAAGGCUGACUGCGCGCGCAAAAGAUUUAUUAACAUUCGUCAUGUGUUAACUGGCUGCUGGCGGAU...........................GCCAA.................................AUCCG.U......UGGAAU
env-441 UUGAACUGUGGCUAAUAAAUUCAUUGCUGGCUGUAUGUUUAGAAAAAGCUUUAUUAGGAUUCUCAUUGUGUUAGCUACAAGCUGUUGGCGGAU...........................GCUCG.................................GUCCG.C......UGGAAU
env-442 ......GCUGUGGUUAAUAAAUUCAUUGCUUGUGCCUGUUGAGAAAAGCUUUAUUAGGAUUCUUGGCUUGUUACCUAANGCUGUUGGCGGAU...........................AUU...................................GUCCG.C......UGGAAU
env-443 ................................................UUUUUUAGGAUUCUUGGCUUGUUACCUAAAGCUGUUGGCGGAU...........................AUU...................................GUCCG.C......UGGAAU
env-444 CCUAGUCUGUGGCUAAUAAAUUCACUGCUGGUCCAAGCUUAGAAAAAGCUUUAUUAGGACUCUUCAUUUGUUACCUAAAAUUGUUAGCGGA............................CACGA..................................UCCG.C......UGGAAU
env-446 CCUCAGCUGCGCACNCANUCGCUUGGACCCACGAAGCAUCAGAAAAAACUUUAUUAGGAUUCCUUGGGUGUAAGCUAGGACUACAGGCGGAUC..........................CUUC.................................GAUCCG.U......UGGGAU
env-447 CUACGGAUUUAUCCGUGGGUAGCUGACGUUAUGCUCUGUGCUGGAAAAGGUUUAUUUUUAACCGUCUGAUUAGCAUGUGCGUUGCUUGCGGACG..........................GUGA.................................UGUCCG.C.....UUGGGAU
env-449 CUCAGUGUAGGUAGUAAGCNCGCGUCGGCGGCGACGAAGCCGAAAAAGCUUUAAUAGACGUCUGUUUGUUAGCUACAAACUAUUGGCGGAUG..........................GGC..................................CAUCCG.C......UGGAAU
env-450 CUCAGUGUAGGUAGUAAGCCGCGUCGGCGGCGACGAAGCCGAAAAAGCUUUAAUAGACGUCUGUUUGUUAGCUACAAACUAUUGGCGGAUG..........................GGC..................................CAUCCG.C......UGGAAU
env-453 ACGGAAACCAAGGGGCCAGGGGCCGGGGGGGCCCAAAAAAUCCCAACAGGUCGGAUUUUUGGGGAAACCCCCAAAACUUUGGGGGGGGG.............................CAGAA..................................CCCG.C.........GGGAA
Mvo-1-1 UUUUUAACAAUAUAUUUUUUGAACGUUAACUUAAUAUACUUGAGUGUCUUAAUUAGGUAAGAUAACCUAAAAGGUUCUCACAUUAACGGA.............................GUAA..................................UCCG.G.....UAGGGAAU
env-454 UUUUUGACACUGAGAAAGAACAGAAGCACGUGAAUCGGUGACUUCGAUAGGUAUGUAGCAAUAACUGCAAACCAUACUGGAAUCACGGGA.............................UCCG..................................UCCCG.G.....AGAGGAAU
ume-1-1 CAUGUGCCAGCAGGCUGCUUAUAGCGCCCGGACAAUACGAAAAUAUUUAUAUUAUAGGACCUGUUUAUUGUUCCAACGGUUUCAGUCGCGGA.............................UACGU..................................UCCG.G......CCGGGAU
CMe-2-1 UGCUGAAGGAAUUGAAAUAAGAACCGAAGGUGGACGAAGAUUCAAGAAAUCUUUAACUACAAUGAAAUAGCUUAAAGUCCCUGGUAGCGGGA............................GAAAA.................................UCCCG.C.......AGGAAU
Mev-1-1 UUGUAUUUUUGUAUGAUAAUAUAACUCAGUAUAGUUGCAUAAAAAACUUUAAAUCAUAUUAAGUUUGUAGUGCACUUGUUUCAUUUACGGGA............................UGUUA.................................UCCCG.U.......CGGAAU
env-455 ..............GCUCUNAUGGGUCCNGNAACUGCUUCUGCCGGAGUACUUUAAUAGACCUCCGGUUAAAGUAUGUGGGAACAUGGGA.............................UCCG..................................UCCG.A..UUGAAAGGAAU
env-457 ACACGGGUGCGAGUUGCUCUCAUGGGUCCCGCAACUGCUUCUGCCGGAGUACUUUAAUAGACCUCCGGUUAAAGUAUGUGGGAACAUGGGA.............................UCCG..................................UCCG.A..UUGAAAGGAAU
env-460 GCGCAAAGCUCGUGAUGCUGAUGGUCCUCGUCAACUGCCCGUGCCGGAGUACUUUAAUAGACCUCCGGUUAAAGUAUGUGGGAACAUGGGA.............................UCCG..................................UCCG.A..UUGAAAGGAAU
env-461 AACAAAGUGCUCGUAGUUCGUAUGGUUCCGGCACCUGCUUCUUCCGGAGUACUUUAAUAGACCUCCGGUUAAAGUAUGUGGGAACAUGGGA.............................UCCG..................................UCCG.A..UUGAAAGGAAU
env-462 ............................................................................................GUCGCGGGA.............................GUGA..................................UCCG.G......CCGGAAU
Mzh-1-1 ACAAAUGCUACUGCAUUACAUAGAAACUGAACAAAACACAAAAACUUAAAUGCAAUCAGGUAUGUAGUAGCGGAUGUUUCAUUAUCGGGA.............................UAUAU.................................UCCG.G......UAGGGAU
env-463 UCAAAGGAAUAAAAAACUUUGGACAUUCCCUAGAACGUCAAUAGUAACAUUUAAAUUCGAGAAUUACAUUUGAAACACUGAAGUGGCGGGA.............................GAGA..................................UCCG.G......CUGGGAU
env-465 AUUUACUACUAUCCUUGCAGAUGAGAAGUACCUUUUCAUUUAGAUAGCUUUAAUACUGAACGGACUGCACUUCAUAUACCCUAACCGUGGGA.............................GAGA..................................UCCG.G......CUGGAAU
env-466 AGGGCGUAUAAUUCCUCUGGCAGCAAUGGAUUGACACGAUAAUGCCGGUGUACUUUAUUACCCUGCCAGUCCAAUUUUAUGGGAAUCGCGG..............................AUUCGUA................................CCCG.U...UGAAAGGAAU
env-467 AAGCUUUAAAUAUGGGGUAAAAAUUCUGAUUGUACCUGGGAUAUCUUUAAUACAAUGAAAUAGUUUAAAGCUUCUCAAGCGGCGGGA.............................GGAGA.................................UCCG.G......CAGGGAU
env-469 CAUCGGGGAAGAGUGACCGGAAAGGGCACUCUCCAUUCCUGUAACCGCGCACUUUAAGUGCGUCCCUGCCGAUUCAUAUAGGUCUUACGGGA............................UGCG..................................UCCCG.U.....CAAGGAAU
Mta-1-1 CAUCGGGGAAGAGUGACCGGAAAGGGCACUCUCCAUUCCUGCAACCGCGCACUUUAAGUGCGUCCCUGCCGAUUCAUAUAGGUCUUACGGGA............................UGCG..................................UCCCG.U.....CAAGGAAU
env-472 CCUCACGCUCUCCGCGGGACACGCAAAACCGCCCGGUUCAGGGCAAAGCAUAUAAGCCGCGCAUCCCAACGGGGCGCCCCUAGUGCCGGGG.............................CCCG..................................CCCG.G......GUGGGAU
env-473 AGCCAGAAAGGCACAAAAAGCACGUAAAACCGCCCGCUUCGGGACAAAGCAUAUAAACAGCCCUCACAAUGGGGCGCCCCUAGUGCCGGGG.............................UACG..................................CCCG.G......GUGGGAU
env-474 ACCGUCUUCCCCCAUCUGUGAAUUUCUCGCCCAAAGAUACCGAAGCCCAUUGUUUUAAUACCCCAUCCAUCGAAAUAUACAGACCACGGGAU...........................AGU...................................AUCCG.G.....AAAGGAAU
Mpa-1-1 GCGAUAUCCGGGUGAUCGAGAAGAUCAAGGGCUGACCAGGAUCACCGCUGUUAUUAAUAACCGGGUGGUCGACAUAUAAAGGAAUACGGGA.............................UCCG..................................UCCG.G.....UAAAGGAAU
env-475 GGCAGUGAACUCCCCCUGCCAAUUAUGGUCUCUCUCGGCACCUGCCGAGUACUUUAAUACCCUGCCUCUGUCCAAUUCUAUAGAACCGCGGGA............................UUCG..................................UCCCG.U...UAAAAGGAAU
env-476 GUCAUUAUUGCCGGUUAUAAGUGUCCAUGAAUGGAAUGAAUCCGUCCCUCCCUUUAUUACCUCUCCUCCAUCCAAUUCUAUAGAACCGCGGGA............................UUCG..................................UCCCG.U...UAAAAGGAAU
env-477 CGAUAAGGGUGCAGGCAGGAACCCCGGGAUACCAGGUUUCCGGCACCAUUCUUUUAAUAGGGCCCCGGUUAAUGUAUUAAGGAACACGGGA.............................UCCG..................................UCCG.G....UUAAGGAAU
env-479 ACGGCGAAAAAGGGCGCGUGUUAAUAUUCCGGUCUUUCUCCAGCCCCCACUCUUUAUUACCCCCAUAUCGUAGUAUAGGAAUGCGGGA..............................UACG..................................UCCCG.UGUUGAAAAGGAAU
env-480 GCGCAAGAGUACCUGAUAGUAUAAUGAUCCCAGCUGAUGCAGGAGCCGGUGUAUUUAAUAUGGCUCCGGUUAAAGUAUAUAGGAAUGCGGGA............................UACG..................................UCCCG.U...UGAAAGGAAU
env-481 GCGCAAGAGUACCUGAUAGUAUAAUGAUCCCCGGAGAUGCAGGAGCCGGUGUAUUUAAUACGGCUCCGGUUAAAGUAUAUGGGAAUGCGGGA............................UACG..................................UCCCG.U...UGAAAGGAAU
env-484 ..................................................................GGCUCCGGUUAAAGUAUAUAGGAAUGCGGGA............................UACG..................................UCCCG.U...UGAAAGGAAU
env-485 GCGCAAGAGUACUUGAUAGUAUAAUGAUCCCCGGAGAUGCAGGAGCCGGUGUAUUUAAUACGGCUCCGGUUAAAGUAUAUGGGAAUACGGGA............................UACG..................................UCCCG.U...UGAAAGGAAU
env-486 CACAAAGGUCCCUGACACCAGGAUCACCCGGAAGGGAGGAGGAGCCGGAGUACUUUAAUAGCCCUCCGGUUAAAGUAUGUGGGAAUACGGGA............................UCUG..................................UCCCG.U...UGAAAGGAAU

.........................................................................................<<<<<<<<<<<<<<<<<<<<<<<<<<<<<<<<<<<..........>>>>>>>>>>>>>>>>>>>>>>>>>>>>>>>>>.>........<<...
.........................................................................................22222..1111.................1..1............1..1.................1111..2222.2........12...
......................................................................................Y...YR...G..........................................................C........GGRAU
5'
```

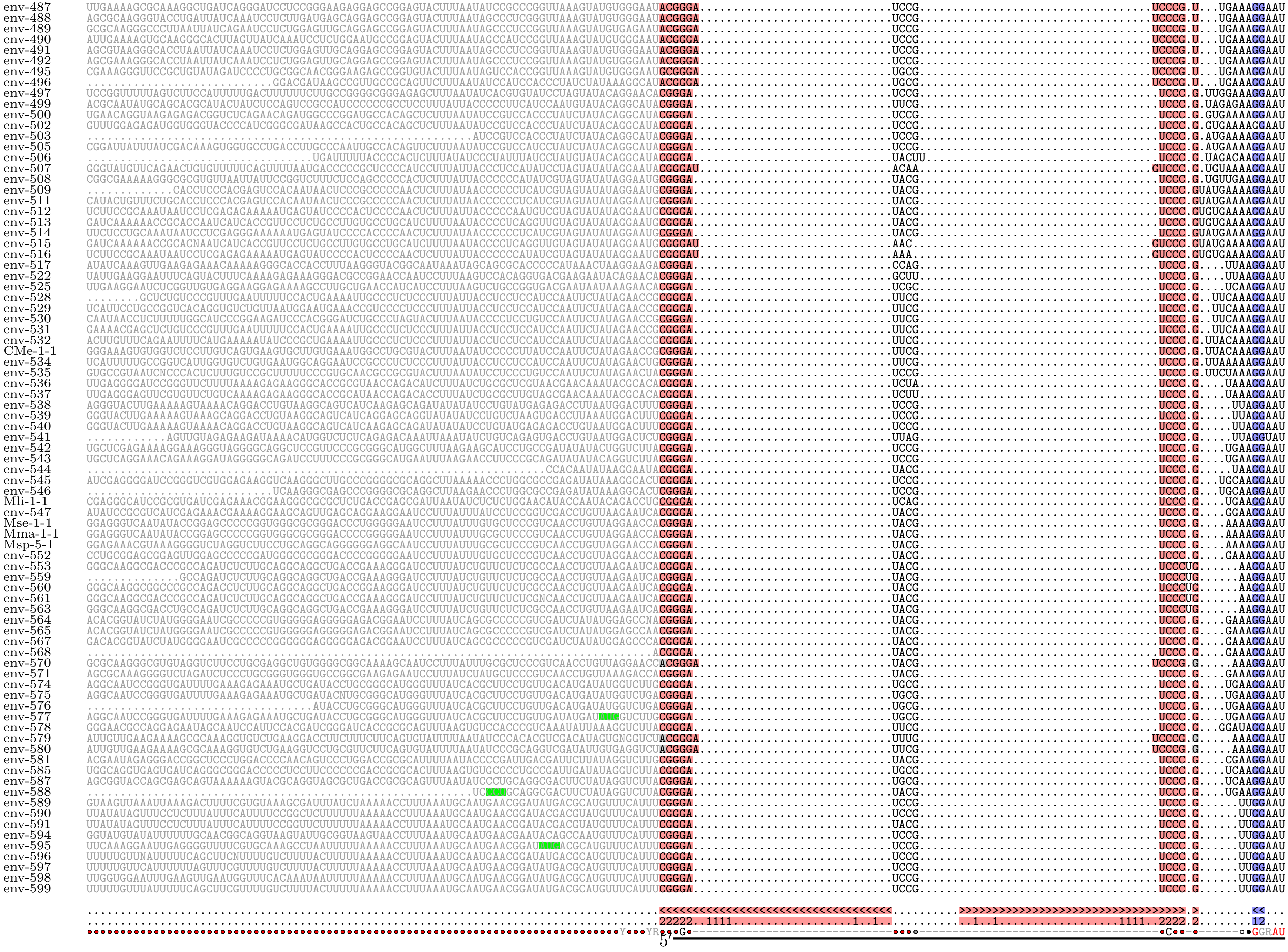

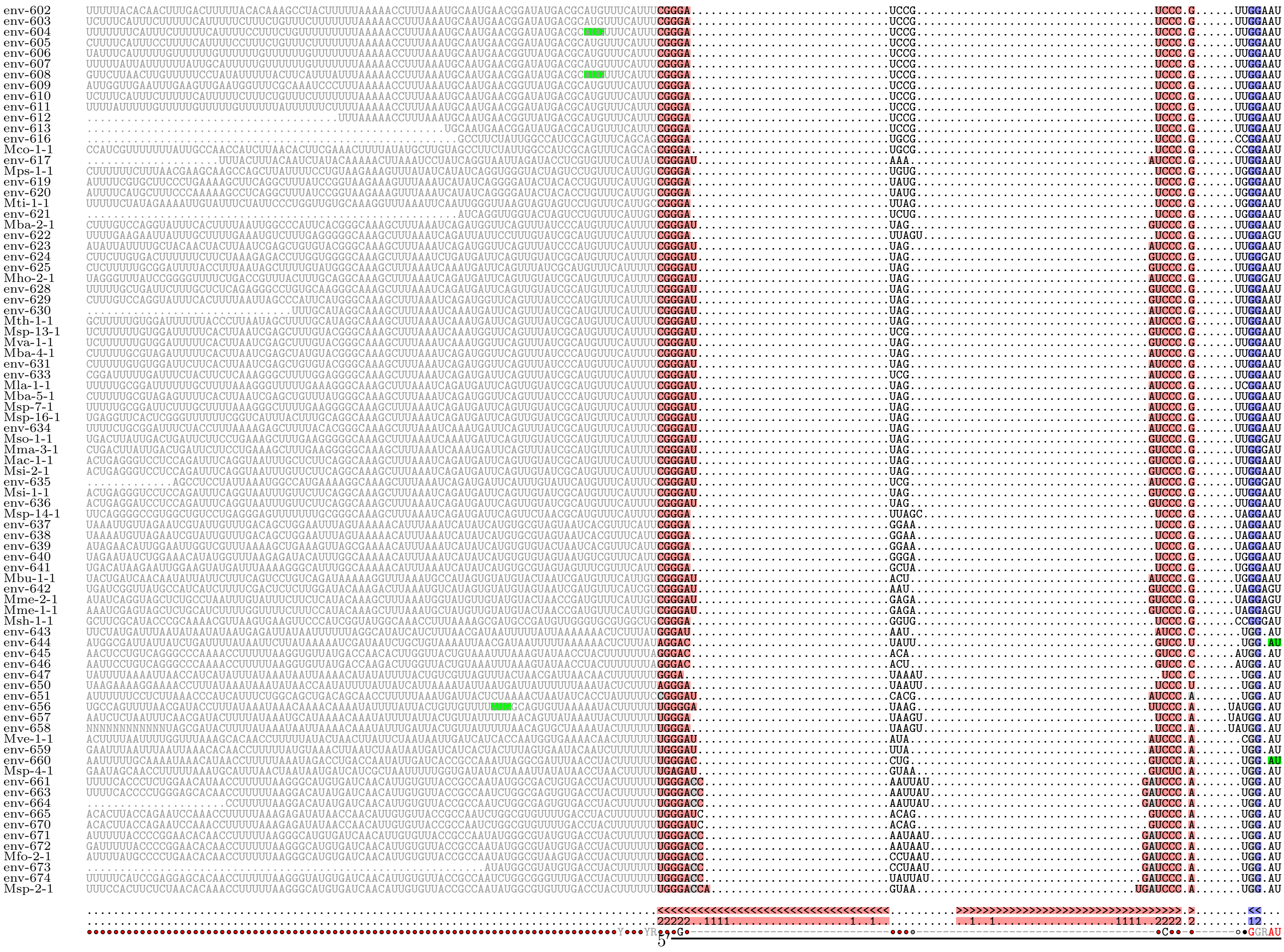
env-602
env-603
env-604
env-605
env-606
env-607
env-608
env-609
env-610
env-611
env-612
env-613
env-616
Mco-1-1
env-617
Mps-1-1
env-619
env-620
Mti-1-1
env-621
Mba-2-1
env-622
env-623
env-624
env-625
Mho-2-1
env-628
env-629
env-630
Mth-1-1
Msp-13-1
Mva-1-1
Mba-4-1
env-631
env-633
Mla-1-1
Mba-5-1
Msp-7-1
Msp-16-1
env-634
Mso-1-1
Mma-3-1
Mac-1-1
Msi-2-1
env-635
Msi-1-1
env-636
Msp-14-1
env-637
env-638
env-639
env-640
env-641
Mbu-1-1
env-642
Mme-2-1
Mme-1-1
Msh-1-1
env-643
env-644
env-645
env-646
env-647
env-650
env-651
env-656
env-657
env-658
Mve-1-1
env-659
env-660
Msp-4-1
env-661
env-663
env-664
env-665
env-670
env-671
env-672
Mfo-2-1
env-673
env-674
Msp-2-1
5'

```
Mfo-1-1    UUCCCUUAUCCCCAACACAAACCUUUUUAAGGGCAUGUGAUCAACAUUGUGUUACCGCCAAUCUGGCGUGUUUGACCUACUUUUUUUGGGACCA...........................GUUA..................................UGAUCCC.A......UGG.AU
env-675    AAUGUCAAAUUUUUUAGAGCACAACCUUUUUAUACUUACUUAUUCUAAUAGUUGAUCAUCACCAACUGGUGACAAACAACUUUUUUUGGGA..............................AACAA....................................UCCC.A......UGG.AU
env-676    AAUGUUUUACAUUUAUCAUAACCUUUUUAAGUAGUGGUGAUCAAUAUUUAUCACCGCCAAUUUGGCGAUAAAAAACAACUUUUUUUGGGA..............................UAAAU....................................UCCC.A......UGG.AU
Msp-1-1    AUCAAUAGUGAUCACCGCCAGUUUUGGCGAUUAAAUAAUUGACUAAAACUUGAUUGCUUACAUUGAAGUUAUAUAAAACUUUUUUUGGGAU.............................AAGUAAAACC..............................AUCCC.A......UGG.AU
env-679    UCAAUAUUGAUCACCGCCAGUUUUGGCGAUUAAAUCAAUUGAAUAAAACUUGAUUGCUUACAUUGAAGUUAUAUAAAACUUUUUUUGGGAU.............................AAGUAAAACC..............................AUCCC.A......UGG.AU
env-680    AUCAAUAUUUAUCACCGCCAAUUAUGGCGAAUAAAUCUAGUAAAUAGAACUUGAUUGCUUACAUUGAAGUUAUAUACAACUUUUUUUGGGAU.............................UUG.....................................AUCCC.A......UGG.AU
Msp-3-1    AUCAAUAUUGAUCACCGCCAAUUCUGGCGAUUAAAUCUAGUAAAUAAAACUUGAUUGCUUACAUUGAAGUUAUAUACAACUUUUUUUGGGAU.............................AGU.....................................AUCCC.A......UGG.AU
env-681    CAACAUUGAUCACCGCCAAUUCUGGCUAUUUAAAUUUAGUAGUUAAAACUUGAUUGCUUUACAUAGAAGUUAUAUACAACUUUUUUUGGGAU.............................AUU.....................................AUCCC.A......UGG.AU
env-682    CAACAUUGAUCACCGCCAAUUUUGGCGAUUUAAAUUUAGAAGUUAAAACUUGAUUGCUUUACAUUGAAGUUAUUUACAACUUUUUUUGGGAU.............................AAU.....................................AUCCC.A......UGG.AU
env-683    CGGUCUGUAUUUUUUAGCACAAACUUUUUAAGUAUUCAUGAUCAACAUUGAUCACCGCCAAUUGGUGGAUAAUAAAUACUCUUUUUAGGGAU.............................CAA.....................................GUCCC.U......UGG.AU
Mor-1-1    UUUUAAUAAAGUCAAAUAAACGAUAUCUUUAUAAACAUAUAAUUGCUAAUAUUUAAUUACUGCCAAUUUGGUAGAAUAACUUUUUUUGGGA..............................UAUUU.....................................UCCC.A.....UUGG.AU
Msm-14-1   CUUUAUAUGUUUAACCAUAACUUUAUAAAUAAACAAAACAAAUAUUUUAUUAUCGUUGAUUUAUUGACGAUUACAUAACUUUUUUCGAGAC.............................AAU.....................................GUCUC.A......UGG.AU
Msm-4-1    UCUUUAUAUGUUUAACCAUAACUUUAUAAAUAAACAAAACAAAUAUUUUAUUAUCGUUGAUUUAUUGACGAUUGCAUAACUUUUUUCGAGAC.............................AAU.....................................GUCUC.A......UGG.AU
env-684    UCUUUAUAUGUUUAACCAUAACUUUAUAAAUAAACAAAACAAAUAUUUUAUUAUCGUUGAUUUAUUGACGAUUAUAUAACUUUUUUCGAGAC.............................AAU.....................................GUCUC.A......UGG.AU
env-685    UCUUUAUAAAUUUGACCAAAACUUUAUAAAUAAAUAAAACAAAUAUUUUAUUAUCGUUAAUUUAUUGACGAUUACAUAACUUUUUUCGAGAC.............................AAU.....................................GUCUC.A......UGG.AU
env-686    UCUUUAUAUGUUUAACCAUAACUUUAUAAAUAAACAAAACAAAUAUUUUAUUAUCGUUGAUUUAUUGACGAUUACAUAACUUUUUUCGAGAC.............................AAU.....................................GUCUC.A......UGG.AU
env-695    UCUUUAUAUGUUUAACCAUAACUUUAUAAAUAAACAAAACAAAUAUUUUAUUAUCGUUAAUUUAUUGACGAUUACAUAACUUUUUUCGAGAC.............................AAU.....................................GUCUC.A......UGG.AU
Msm-16-1   UCUUUAUAUGUUUAACCAUAACUUUAUAAAUAAACAAAACAAAUAUUUUAUUAUCGUUGAUUUAUUGACGAUUACAUAACUUUUUUCGAGAC.............................AAU.....................................GUCUC.A......UGG.AU
Msm-6-1    CUUUAUAUGUUUAACCAUAACUUUAUAAAUAAACAAAACAAAUAUUUUAUUAUCGUUGAUUUAUUGACGAUUACAUAACUUUUUUCGAGAC.............................AAU.....................................GUCUC.A......UGG.AU
env-696    UGAAUUCCGAAUGUAGGGCAACAAUCUUUUUAUUAUUCGAAACGUUUAUAUGCUAAUACAAGUUAUUCUGCAAGGCAGUUUCAAUCGCGGGA.............................UACG....................................UCCCG.C......CGGAAU
env-697    ...................................................................................UGGCGGGA..............................GAAAA.....................................UCCC.G.....CAGGGAU
env-698    CGCUUACAUAUGAUCAAUGAUCUCUUACCCAACCUUUAUAACUCAUAAAAGUAUCUAUACCUUUUUACAACAAGCCAGUUUCAUCGCGGGAU.............................AAAU....................................AUCCC.G.....CAGGGAU
env-699    CACUUACAUAUGAUCAAUGAUCUCUUACCCAACCUUUAUAACCCAUAAAAGAAUUUAUAACCUUUUACAACAAGCCAGUUUCAUUGGCGGGA.............................UAAAAUC.................................UCCCG.C......AGGGAU
env-700    AGCAGUGUGUAGUACCAAAUUCCAUCCCCAACCUUUAUUACCCAUUAAUGUAUCUAUAAUCCUCUCACAACAUGCCAGUUUCAUAGGCGGGA.............................UACG....................................UCCCG.C......AGGAAU
env-701    UCCAAAUUUCAUACCGAUGCAUCAUCCCCAACCUUUAUUACCCAUUAAUGUAUCUAUAAUCCACUCAAAACAUGCCAGUUUCAUAGGCGGGA.............................UACG....................................UCCCG.C......AGGGAU
env-702    UCCAAAUUUCAUACCGAUGCAUCAUCCCCAACCUUUAUUACCCAUUAACGUAUCUAUAAUCCACUCACAGCAUGCCAGUUUCAUAGGCGGGA.............................UACG....................................UCCCG.C......AGGGAU
env-704    ...........................................AUUAACGUAUCUAUAAUCCCCUUACAACAUGCCAGUUUCAUAGGCGGGA.............................UACG....................................UCCCG.C......AGGGAU
env-706    ACCAAAUUCCAUCCCGAUGCAUCAUCCCCAACAUUUAUUACCCAUUAACGUAUCUAUAAUCCCCUCACAAGAUGCCAGUUUCAUAGGCGGGA.............................UACG....................................UCCCG.C......AGGGAU
env-707    AUUAAAUUUCAUACCAAUACAUCAUCCCCAACAUUUAUUACCCAUUAAUGUAUCUAUAAUCCUCUCAGAACAUGCCAGUUUCAUAGGCGGGA.............................UACG....................................UCCCG.C......AGGGAU
env-708    NNNNNNNNNCAUCCCGAUGCAUCAUCCCCAACAUUUAUUACCCAUUAACGUAUCUAUAAUCCCCUCACAAGAUGCCAGUUUCAUAGGCGGGA.............................UACG....................................UCCCG.C......AGGGAU
env-709    UUUGCGCAGGAAGCCAUAAACCAGUUCAACAGUUUUACGAAAUAUUUAUAUCAUAGUACCUGAUUAUUCUGCCUACGGUUUCGUUCGCGGGA.............................UCCG....................................UCCCG.C......CGGAAU
env-710    GGAAUAACAAUCUUUUAAAUGGUUUUUUUUCUGAUAAUUCAUAGAAAAGGUUUAUUUUUAAGCAAUGUUGUGAUUAAAAAUUGAAAACGGAC.............................CUGUAGA.................................GUCCG.U......AGGGAU
Ngr-1-1    GCCGUGUCGGCGACGGUUCGAAGCGCUUAUACUCGAGAGUUGAGUAAAGACCGAUAGACUCGCUGUAUCGCGGGCGACAGCGGGCACCUGUACGGC.........................UUU.................................GCC..ACAG.G.....UCGGGAU
Hti-1-1    UCCCGGCCUGGCGGGAGUUCGAAUCGCUUAAGGGUGCCAGCACGGUAGUACCGAUAACUCGCUGGUCGGCGGGCAGCCAGCGGGCACCUGU..............................UUAG.....................................GCAG.G.....ACGGAAU
Npe-1-1    UCGCUGCGUUCCCGGAGUUCGAAGCGCUUAUACCCGAGAGCCGGGAAGGGAGGGUAGACUCGCUGGUUCGCGGGCAUCAGCGGGCACCUG...CACG........................ACGC...............................CGUGUCACAG.G.....UCGGGAU
Hsp-7-1    CGCGCGCCGUGAGAAUCCGAAGUCCUUAACCCUCCCCUGUUGGUAGUUCCGCCAACUCGCUGGCAACGCGGGCACGCCAGAGGGCACCCGU..............................CACGC....................................ACGG.G.....ACGAAAU
Hje-1-1    ACCGGCGAAACGACGCGGGACCGAAGGGCUUAACCGUGAUUCGGGGGUAGAUCCUCCACUCGCUGGGAAACGGGCACCAGCAGGCACCUGCUACC..........................GUCG.................................GGUCGCAG.G.....CAGGAAU
Hbo-1-1    GACACGUGCCGGCAGAUUCGAAGUCGUUAAAGGCGAACCGAGGGAAGAGUCGGUAACUCGCUGGCCGGACGGGACGCCAGCGGGCACCCGA..............................GAAC.....................................UCGG.G.....ACGAAAU
Nma-1-1    CAACAUGGUCCGCCCGCUUCGAAGCCCUUAUACUCGGGAGUUGAAUAGGUGAAACUGACUCGCUGGAUCGCGGGCACCAGCGGGCACCUGU.UCGGC........................ACAUUCGAC..........................GCCGCCACAG.G.....ACGGGAU
Nti-1-1    AAGCACGCUCCCGGAGUUCGAAGCCCUUAUACUCGAGAGUUGGGUAGACUGUCGUAGACUCGCUGAGUCGCGGGCAUCAGCGGGCACCUGU.UCGGCG.......................UCGA..............................CGCCACCACAG.G.....UCGGGAU
           ......................................................................................<<<<<<<<<<<<<<<<<<<<<<<<<<<<<<<<<<<...........>>>>>>>>>>>>>>>>>>>>>>>>>>>>>>>>>>.>......<<....
           ......................................................................................22222..1111..................1..1.............1..1..................1111..2222.2......12....
           ●●●●●●●●●●●●●●●●●●●●●●●●●●●●●●●●●●●●●●●●●●●●●●●●●●●●●●●●●●●●●●●●●●●●●●●●●●●●●●●Y●●●YR●●●G●----------------------------●●●○------------------------------------------●C●●-●------○●GGRAU
                                                                                              5′
```

alignment positions 181 · · · 360

```
env-1      GU.GGCC.............................................................................................................................................................................
env-2      GU.GGCC.............................................................................................................................................................................
Mma-2-1    GU.GGCC.............................................................................................................................................................................
env-3      GU.GGGUGA...........................................................................................................................................................................
Hha-1-1    GU.GGCCUGUGCGGU.....................................................................................................................................................................
env-5      GU.GACCGG...........................................................................................................................................................................
env-6      GU.GACCGU...........................................................................................................................................................................
env-7      GU.GGCC..ACCCUCGUGGU................................................................................................................................................................
env-9      GU.GCCCAGA.CUAGC.GUC.CC.............................................................................................................................................................
env-10     GU.GGCUCAAGCGGUCC...................................................................................................................................................................
Hsa-6-1    GU.GGUUGCGC.GGCGAGGACGAC............................................................................................................................................................
h3A-1-1    GU.GGUUCCGCGGC......................................................................................................................................................................
env-11     GU.GGCCGC...........................................................................................................................................................................
env-12     GU.GGCCGG...........................................................................................................................................................................
env-14     GU.GACG.CGGGAGACGUGACGUCUC.........................................................................................................................................................
env-15     GU.GGUCGGC.CGGGCGC..................................................................................................................................................................
env-16     GU.GGUCGGC.CGGGCGC..................................................................................................................................................................
env-19     GU.GGGAAUACGU.......................................................................................................................................................................
env-22     GU.GGGAAUACGU.......................................................................................................................................................................
Hwa-1-1    GU.GGGAAUACGU.......................................................................................................................................................................
env-23     GU.GGGAAGACGUC......................................................................................................................................................................
env-25     GU.GGGAAUGCGUC......................................................................................................................................................................
env-27     GU.GGGAAUGCGUC......................................................................................................................................................................
env-28     GU.GGGCCGGCG........................................................................................................................................................................
env-29     GU.GGGCCGGCG........................................................................................................................................................................
env-30     GU.GGGCCGGCG........................................................................................................................................................................
env-31     GU.GGGUCGGCGU.......................................................................................................................................................................
Hru-1-1    GU.GGGUCGGCGU.......................................................................................................................................................................
env-32     GU.GGGUCGGCGU.......................................................................................................................................................................
env-33     GU.GGGUCGACGU.......................................................................................................................................................................
Hpa-2-1    GU.GGAGUGCGGUC......................................................................................................................................................................
env-34     GU.GGCUC.................................................................GGGUGGUCC.......ACC..................GGACCACC.GCAA......CCGA..........GCGGC................GGA.........
env-35     GU.GGCGG............................................................................................................................................................................

           ...<<<<<<<<<<<<<<<<<<<<<<<<<<<<<<<<<<<<<<<<<<<<<<<<<<<<<<<<<<<<<<<<<<<...<<<<<<<<<<<<<<<<............>>>>>>>>>>>>>>>>.....<<<<<<<...>>>>>>>.<<<<<<<<<<<<<<<<<<<<<<<<<<.......>>>>>
           ...22222..................1.1..000000000.0000000...000.000...00000.......1111111...100000.............000001...1111111.....000000....000000.1011..1.11....111.10000000.......00000
           GU-GRYY○------------------------------------------------------------------------------------------------------------------------------------------------------------------------------
```

```
env-39   GU.GGCCUGUCUGAG..CCGGG.CUC..........................................................................................................................................................
env-40   GU.GGCCUGUCUGAG..CCGGG..CUC.........................................................................................................................................................
Hac-1-1  GU.GGUGCGC..........................................................................................................................................................................
env-42   GU.GGCCGG...........................................................................................................................................................................
env-43   GU.GGCCGG...........................................................................................................................................................................
env-45   GU.GGCCGG...........................................................................................................................................................................
Hna-1-1  GU.AGC.CCGCCG.......................................................................................................................................................................
env-47   GU.GGC.UCGCCG.......................................................................................................................................................................
env-48   GU.GGCGC............................................................................................................................................................................
env-49   GU.GGCGC............................................................................................................................................................................
env-50   GU.GGCGC............................................................................................................................................................................
env-51   GU.GGCGC............................................................................................................................................................................
env-52   GU.GGCGC............................................................................................................................................................................
env-53   GU.GGCGC............................................................................................................................................................................
env-55   GU.GGCCC............................................................................................................................................................................
env-57   GU.GGCCGC...........................................................................................................................................................................
env-61   GU.GGCCG............................................................................................................................................................................
env-63   GU.GGCCGC...........................................................................................................................................................................
env-64   GU.GGCCGC...........................................................................................................................................................................
env-65   GU.GGCCGC...........................................................................................................................................................................
env-66   GU.GGCCGC...........................................................................................................................................................................
Hai-1-1  GU.GGCCGC...........................................................................................................................................................................
Hco-1-1  GU.GGCC.............................................................................................................................................................................
Hsp-8-1  GU.GGCC.............................................................................................................................................................................
Hte-1-1  GU.GGCC.............................................................................................................................................................................
Hho-1-1  GU.GGCC.............................................................................................................................................................................
env-67   GU.GGCC.............................................................................................................................................................................
Hca-2-1  GU.GGCC.............................................................................................................................................................................
env-70   GU.GGCC.............................................................................................................................................................................
Hli-2-1  GU.GGCC.............................................................................................................................................................................
Har-2-1  GU.GGCC.............................................................................................................................................................................
env-73   GU.GGCCGC...........................................................................................................................................................................
Hli-1-1  GU.GGCCGC...........................................................................................................................................................................
Hha-2-1  GU.GGCCGC...........................................................................................................................................................................
Hsa-9-1  GU.GGCCGC...........................................................................................................................................................................
env-74   GU.GGCCGC...........................................................................................................................................................................
Hko-1-1  GU.GGCCGC...........................................................................................................................................................................
env-77   GU.GGCCGC...........................................................................................................................................................................
env-78   GU.GGCCGC...........................................................................................................................................................................
Hsa-7-1  GU.GGCCGC...........................................................................................................................................................................
env-79   GU.GGCC.............................................................................................................................................................................
env-80   GU.GGCC.............................................................................................................................................................................
env-83   GU.GGCC.............................................................................................................................................................................
env-84   GU.GGCCGC...........................................................................................................................................................................
env-86   GU.GGCCGC...........................................................................................................................................................................
env-87   GU.GGCCGC...........................................................................................................................................................................
env-89   GU.GGCCGC...........................................................................................................................................................................
env-90   GU.GGCCGC...........................................................................................................................................................................
env-91   GU.GGCCGC...........................................................................................................................................................................
env-92   GU.GGCCGC...........................................................................................................................................................................
env-93   GU.GGCCGC...........................................................................................................................................................................
env-94   GU.GGCCGC...........................................................................................................................................................................
Hmu-1-1  GU.GGUC.AUCGAGGGA...................................................................................................................................................................
Hel-1-1  GU.GGUC.AUCGAGGGA...................................................................................................................................................................
Hla-2-1  GU.GGUC.AUCGAGGGA...................................................................................................................................................................
Hde-1-1  GU.GGUCU.CAGGGG.....................................................................................................................................................................
Hal-2-1  GU.GGUCU.CAGGGG.....................................................................................................................................................................
Hpr-1-1  GU.GGUCU.CAGGGG.....................................................................................................................................................................
Hgi-1-1  GU.GGUCU.CAGGGG.....................................................................................................................................................................
Hgi-2-1  GU.GGUCU.CAGGGG.....................................................................................................................................................................
Hsp-9-1  GU.GGUCU.CAGGGG.....................................................................................................................................................................
Hsu-2-1  GU.GGUCU.CAGGGG.....................................................................................................................................................................
Hsp-6-1  GU.GGUCU.CAGGG......................................................................................................................................................................
Hme-1-1  GU.GGCCGA.C.GGGGAAUUCCCCGUCUG.......................................................................................................................................................
Hru-2-1  GU.GGUCGAAAGAGGCG..CGACG.CGCCU......................................................................................................................................................
env-95   GU.GGUCUGGGUGGCGCGGAC...............................................................................................................................................................
env-98   GU.GGUCUGGGUGACGCGG.................................................................................................................................................................
env-99   GU.GGUCUGGGUGAC.....................................................................................................................................................................
CHa-1-1  GU.GGUCGGCGGGCG......UCGUCCG........................................................................................................................................................
env-101  GU.GGUCG............................................................................................................................................................................
env-102  GU.GGUCGG...........................................................................................................................................................................
env-104  GU.GGUCGG...........................................................................................................................................................................
env-105  GU.GGUCGG...........................................................................................................................................................................
env-110  GU.GGUCGG...........................................................................................................................................................................
env-112  GU.GGUCGG...........................................................................................................................................................................
env-114  GU.GGUCGG...........................................................................................................................................................................
env-115  GU.GGUCGG...........................................................................................................................................................................
env-116  GU.GGUCGG...........................................................................................................................................................................
env-117  GU.GGUCGG...........................................................................................................................................................................
env-119  GU.GGUCGG...........................................................................................................................................................................
env-121  GU.GGUCGG...........................................................................................................................................................................
env-122  GU.GGUCGG...........................................................................................................................................................................
env-125  GU.GGUCGG...........................................................................................................................................................................

         ...<<<<<<<<<<<<<<<<<<<<<<<<<<<<<<<<<<<<<<<<<<<<<<<<<<<<<<<<<<<<<<<<<<<...<<<<<<<<<<<<<<<<..............>>>>>>>>>>>>>>>>.....<<<<<<<...>>>>>>>.<<<<<<<<<<<<<<<<<<<<<<<<<<.......>>>>>
         ...22222..................1.1..000000000.0000000...000.000...00000.......1111111...100000..............000001...1111111.....000000....000000.1011..1.11....111.10000000.......00000
         GU-GRYYo------------------------------------------------------------------------------------------------------------------------------------------------------------------------------------
```

```
env-126    GU.GGUCGGG..........................................................................................................................................................................
env-129    GU.GGUCGGA..........................................................................................................................................................................
env-130    GU.GGUCGG...........................................................................................................................................................................
env-135    GU.GGUCGA...........................................................................................................................................................................
env-136    GU.GGUCGA...........................................................................................................................................................................
env-137    GU.GGUCGA...........................................................................................................................................................................
env-138    GU.GGUCGA...........................................................................................................................................................................
env-139    GU.GGUCGA...........................................................................................................................................................................
env-140    GU.GGUCGA...........................................................................................................................................................................
env-141    GU.GGUCGA...........................................................................................................................................................................
env-142    GU.GGUCGA...........................................................................................................................................................................
env-144    GU.GGUCGA...........................................................................................................................................................................
env-145    GU.GGUCGA...........................................................................................................................................................................
env-146    GU.GGUCGA...........................................................................................................................................................................
env-147    GU.GGUCGA...........................................................................................................................................................................
env-148    GU.GGUCGA...........................................................................................................................................................................
env-149    GU.GGUCGA...........................................................................................................................................................................
env-152    GU.GGUCGA...........................................................................................................................................................................
env-153    GU.GGUCGG...........................................................................................................................................................................
env-156    GU.GGUCGG...........................................................................................................................................................................
env-158    GU.GGCCGGU..........................................................................................................................................................................
env-159    GU.GGCCGCC..........................................................................................................................................................................
Hsp-10-1   GU.AGCCGC...........................................................................................................................................................................
Hse-1-1    GU.GGCCU.GUGGCG.CGACCGU.............................................................................................................................................................
Hmo-1-1    GU.GGCCC............................................................................................................................................................................
Hth-1-1    GU.GGCCC............................................................................................................................................................................
Nae-1-1    GU.GAUCCG.CGAGG.....................................................................................................................................................................
Nta-1-1    GU.GAUCCG.CGGGGG....................................................................................................................................................................
Nas-1-1    GU.GAUCCG.CGGGGA....................................................................................................................................................................
env-160    GU.GGCU...............................................................GG.CGGGGUC.........UCU....................GACCCCG.GGCA..................GGCGG.....................CACCG.......
env-161    GU.GACCAGC...............................................................GGGGUC..........GAA.....................GACCCC.AAAA..................CGGCG.....................UCGC........
env-162    GU.GGGCC............................................................................................................................................................................
env-164    GU.GGGCC............................................................................................................................................................................
env-169    GU.GGGCC............................................................................................................................................................................
env-170    GU.GGACCGC..........................................................................................................................................................................
Hla-4-1    GU.GGAACGCGAGAG.....................................................................................................................................................................
Hsp-11-1   GU.GGAACGCGGGAG.....................................................................................................................................................................
Hsp-3-1    GU.GAUUCUCGG........................................................................................................................................................................
env-172    GU.GAUCCU...........................................................................................................................................................................
Hsa-1-1    GU.GGUCGG...........................................................................................................................................................................
env-173    GU.GGUCU............................................................................................................................................................................
env-174    GU.GGUCU............................................................................................................................................................................
env-175    GU.GGUCCC...........................................................................................................................................................................
env-176    GU.GGUCC............................................................................................................................................................................
Hpa-1-1    GU.GGCGUGG..UCGAUU..................................................................................................................................................................
env-177    GU.GAACCG................................................................GGAGGGU...GAGAUCGAA...........GAUCUCGCAGCCCUCC....ACGAGAUUUCCAAUCUCGCCGGGG.....................CCUUCGA.....
env-178    GU.GACCUGCG..CGGCUUCGGCCGC..........................................................................................................................................................
env-179    GU.GAUCUC...........................................................................................................................................................................
env-180    GU.GAACUCC..........................................................................................................................................................................
env-181    GU.GAACCGACGGCGGGUUUC.CCGCCUC..AUACUGGUGACUAUAAGAGCGUG.GAAAAUUCCGACCCU..............................................................................................................
env-182    GU.GAACCUG..........................................................................................................................................................................
Ssp-1-1    GU.GACCC..............................................................UGGCCGACGGACGC.....CGAC...............GCGUUCGUCGG.GAGA..................GGCG.ACGCGC..GUGCCG.......GCA.........
env-183    GU.GACCC...UGGC..........................................................CGGCCGACGCC.....GAC................GGCGUCGGUCGGGGGA..................GGCGACGGUCGAGGCCGAUCGCACCGAUCCG..CGGUG
Nph-1-1    GU.GACCGGC..........................................................................................................................................................................
env-184    GU.GAUUGACAC.ACCACUCGAGUC...........................................................................................................................................................
Hgo-1-1    GU.GAUCCCGA.........................................................................................................................................................................
Hxa-1-1    GU.GAUCUCGA.........................................................................................................................................................................
env-185    GU.GAUCUC...........................................................................................................................................................................
env-186    GU.GAUCUC...........................................................................................................................................................................
env-187    GU.GAUCUCGC.........................................................................................................................................................................
Nin-1-1    GU.GAUCCCG..........................................................................................................................................................................
env-188    GU.GAUCCGG..........................................................................................................................................................................
env-189    GU.GAUCCGG..........................................................................................................................................................................
env-192    GU.GAUCCG.CGGG......................................................................................................................................................................
env-193    GU.GAUCCG.CGGG......................................................................................................................................................................
Nje-1-1    GU.GAUCC.GCGGG......................................................................................................................................................................
Nam-1-1    GU.GAUCC.GCGGG......................................................................................................................................................................
Noc-1-1    GU.GAUCC.GCGGG......................................................................................................................................................................
env-194    GU.GGCCC............................................................................................................................................................................
env-195    GU.GAACC...GCGAGGUCGAAGAUCUC........................................................................................................................................................
env-196    GU.GAACUAUCGC.......................................................................................................................................................................
env-197    GU.GAACUAUCGC.......................................................................................................................................................................
env-199    GU.GAACUAUCGC.......................................................................................................................................................................
env-201    GU.GAACUAUCGC.......................................................................................................................................................................
env-202    GU.GAACCGC.CGAGG....................................................................................................................................................................
Hla-1-1    GU.GAUCCC...........................................................................................................................................................................
Hva-1-1    GU.GAGUCGC..........................................................................................................................................................................
Hsi-1-1    GU.GAGUCGCGC........................................................................................................................................................................
Hma-1-1    GU.GAGUCGCGC........................................................................................................................................................................
Ham-1-1    GU.GAGUCGCGU........................................................................................................................................................................
env-203    GU.GAGUCGCGC........................................................................................................................................................................
Hca-1-1    GU.GAGUCGCGC........................................................................................................................................................................
Hsa-2-1    GU.GAGUCGCGC........................................................................................................................................................................

           ...<<<<<<<<<<<<<<<<<<<<<<<<<<<<<<<<<<<<<<<<<<<<<<<<<<<<<<<<<<<<<<<<<<<...<<<<<<<<<<<<<<<<..............>>>>>>>>>>>>>>>>.....<<<<<<<...>>>>>>>.<<<<<<<<<<<<<<<<<<<<<<<<<<.......>>>>>
           ...22222..................1.1..000000000.0000000...000.000...00000.......1111111...100000..............000001...1111111.....000000....000000.1011..1.11....111.10000000.......00000
           GU-GRYY∘------------------------------------------------------------------------------------------------------------------------------------------------------------------------
```

```
Hja-1-1    GU.GAGUCGCGC........................................................................................................................................................................
Har-1-1    GU.GAGUCGCG.........................................................................................................................................................................
env-206    GU.GAGUCGCG.........................................................................................................................................................................
Hmu-2-1    GU.GACUCCGGA........................................................................................................................................................................
Hka-1-1    GU.GACUCCGGA........................................................................................................................................................................
env-207    GU.GACCCCCGCGG......................................................................................................................................................................
env-208    GU.GAUUCGCGUUCGUGGCCGACG..GCCGC.....................................................................................................................................................
env-210    GU.GAUCC.CCAGCC.....................................................................................................................................................................
env-212    GU.GAUUCCC.AGC......................................................................................................................................................................
env-213    GU.GAUUCCC.AGCC.....................................................................................................................................................................
env-214    GU.GACUCCCG.CGGGGC...GGCCGU.........................................................................................................................................................
env-215    GU.GAUUCCCACCGCC....................................................................................................................................................................
env-216    GU.GAUUCCC..CGCC....................................................................................................................................................................
env-217    GU.GACUCCGCGGGC.....................................................................................................................................................................
env-218    GU.GACUCCGCGGGC.....................................................................................................................................................................
env-219    GU.GACUCCGCGGGC.....................................................................................................................................................................
env-220    GU.GACUCCGCGGGC.....................................................................................................................................................................
Nmo-1-1    GU.GACUCCG..........................................................................................................................................................................
env-221    GU.GGUCC............................................................................................................................................................................
env-223    GU.GGCCC............................................................................................................................................................................
env-224    GU.GGCCU............................................................................................................................................................................
env-226    GU.GACUCUCGGCGUC....................................................................................................................................................................
env-227    GU.GAGGCGGCG........................................................................................................................................................................
env-228    GU.GGCCGC...........................................................................................................................................................................
env-229    GU.GGCCG............................................................................................................................................................................
Hci-1-1    GU.GGCAGGCGGGAUUC...................................................................................................................................................................
Hut-1-1    GU.GGCCGGC..........................................................................................................................................................................
env-230    GU.GGCCGGCGGGGG..CGAAAGCCCCC........................................................................................................................................................
env-231    GU.GGCCGG...........................................................................................................................................................................
env-232    GU.GGCCGG...........................................................................................................................................................................
env-233    GU.GGCCGG...........................................................................................................................................................................
env-234    GU.GAACC..CCGAG.....................................................................................................................................................................
env-235    GU.GACUUC...........................................................................................................................................................................
env-236    GU.GACCC............................................................................................................................................................................
Nba-1-1    GU.GAUCC...GCGGGGC..................................................................................................................................................................
env-237    GU.GAUCC...GCGGGGC..................................................................................................................................................................
Nsu-1-1    GU.GAUCC...GCGGGGC..................................................................................................................................................................
Npa-1-1    GU.GAUCC...GCGGGGC..................................................................................................................................................................
Nal-2-1    GU.GAUCC...GCGGGGC..................................................................................................................................................................
Nga-1-1    GU.GAUCC...GCGGGGC..................................................................................................................................................................
Hdj-1-1    GU.GAUCC...GCGGGGC..................................................................................................................................................................
Hli-3-1    GU.GAUCC...GCGGGGC..................................................................................................................................................................
Hje-2-1    GU.GAUCC...GCGGGGC..................................................................................................................................................................
Hth-2-1    GU.GAUCC...GCGGGGC..................................................................................................................................................................
Hsa-10-1   GU.GAUCC...GCGGGGC..................................................................................................................................................................
env-238    GU.GAUCCC...........................................................................................................................................................................
env-239    GU.GAUCCC...........................................................................................................................................................................
Hni-1-1    GU.GAUCC..GCGGGGC...................................................................................................................................................................
Nch-1-1    GU.GAUCCGA..........................................................................................................................................................................
Nhu-1-1    GU.GAUCCGA..........................................................................................................................................................................
Nve-1-1    GU.GAUCC...GCGGGGC..................................................................................................................................................................
Hsa-8-1    GU.GAUCC..GC.GGGG...................................................................................................................................................................
Htu-1-1    GU.GAUCCG...CGGGG...................................................................................................................................................................
env-243    GU.GAUCC...GCGGGG...................................................................................................................................................................
Hca-3-1    GU.GAUCGCCG.........................................................................................................................................................................
env-244    GU.GAACCGCGCUCUCGGUUCGGAGGUC........................................................................................................................................................
env-245    GU.GAUC.C.ACGGGAGCGGUCUCG...........................................................................................................................................................
env-246    GU.GACC.GGCGAGACCUUCGGUCUCGU........................................................................................................................................................
env-247    GU.GAUC...............................................................CG.CGAGUA..........CGG.....................UACUCGAGUGA..................GGG.......................ACCGC.......
env-248    GU.GAUCCG................................................................CGAGCG..........CGAUA...................CGCUCGAGAGA..................GGUGACAG..................AUG.........
env-249    GU.GAUC...............................................................CG.CGAGCGU.........GCAUGCGAUACGAG.........GCGCUCGAGAGA..................GGUGACAG..................CGG.........
env-250    GU.GACCC.CG.........................................................................................................................................................................
env-251    GU.GACCCUU..........................................................................................................................................................................
env-253    GU.GACCC............................................................................................................................................................................
Hsu-1-1    GU.GAUCCGC..........................................................................................................................................................................
Hsp-2-1    GU.GAUCCGUGAGGCGC...................................................................................................................................................................
Has-1-1    GU.GGGCCGG..........................................................................................................................................................................
env-254    GU.GGACCUGAUUCGAGG..................................................................................................................................................................
Hag-1-1    GU.GGCCAGCGUGGUCC...................................................................................................................................................................
Hsa-4-1    GU.GGCCAGCGUGGUCC...................................................................................................................................................................
Hsa-5-1    GU.GGCCAGCGUGGUCC...................................................................................................................................................................
env-255    GU.GGCU.............................................................................................................................................................................
env-256    GU.GGCU.............................................................................................................................................................................
env-262    GU.GGCU.............................................................................................................................................................................
env-266    GU.GGCU.............................................................................................................................................................................
Mae-1-1    GU.GGC..............................................................................................................................................................................
env-267    GU.GGACGUUCC........................................................................................................................................................................
env-268    GU.AAC..............................................................................................................................................................................
env-269    GU.AAC..............................................................................................................................................................................
env-270    GU.GACUGC...........................................................................................................................................................................
env-271    GU.GACUGC...........................................................................................................................................................................
env-273    GU.GACUGC...........................................................................................................................................................................
env-274    GU.AGCCUU...........................................................................................................................................................................

           ...<<<<<<<<<<<<<<<<<<<<<<<<<<<<<<<<<<<<<<<<<<<<<<<<<<<<<<<<<<<<<<<<<<<...<<<<<<<<<<<<<<<<..............>>>>>>>>>>>>>>>>.....<<<<<<<...>>>>>>>.<<<<<<<<<<<<<<<<<<<<<<<<<<.......>>>>>
           ...22222..................1.1..000000000.0000000...000.000...00000.......1111111...100000..............000001...1111111.....000000....000000.1011..1.11....111.10000000.......00000
           GU-GRYY∘------------------------------------------------------------------------------------------------------------------------------------------------------------------------
```

```
env-276  GU.AGCCUU...................................................................................................................................................................................
env-277  GU.AGCC.....................................................................................................................................................................................
env-278  GU.AGCC.....................................................................................................................................................................................
env-281  GU.GACC.....................................................................................................................................................................................
env-283  GU.GGUCC....................................................................................................................................................................................
env-284  GU.GGUCC....................................................................................................................................................................................
env-286  GU.GGUCC....................................................................................................................................................................................
env-287  GU.GGUCCC...................................................................................................................................................................................
env-289  GU.GGCCC....................................................................................................................................................................................
env-290  GU.GGCCCA...................................................................................................................................................................................
env-291  GU.GGCCC....................................................................................................................................................................................
env-293  GU.GGCCCU...................................................................................................................................................................................
env-295  GU.GGUCC....................................................................................................................................................................................
env-296  GU.GGUCC....................................................................................................................................................................................
env-297  GU.AGCC.....................................................................................................................................................................................
env-298  GU.GGCC.....................................................................................................................................................................................
env-299  GU.GGCC.....................................................................................................................................................................................
env-301  GU.GGCC.....................................................................................................................................................................................
env-302  GU.GGCC.....................................................................................................................................................................................
env-304  GU.GGCCC....................................................................................................................................................................................
env-308  GU.AACCU....................................................................................................................................................................................
env-309  GU.AACCU....................................................................................................................................................................................
env-310  GU.AACC.....................................................................................................................................................................................
env-311  GU.AACC.....................................................................................................................................................................................
env-312  GU.GGCCC....................................................................................................................................................................................
env-317  GU.GGCCC....................................................................................................................................................................................
env-318  GU.GGCCC....................................................................................................................................................................................
env-319  GU.GGCCC....................................................................................................................................................................................
env-321  GU.GGCC.....................................................................................................................................................................................
env-322  GU.GGCCC....................................................................................................................................................................................
env-323  GU.GGCCC....................................................................................................................................................................................
env-324  GU.GGCC.....................................................................................................................................................................................
env-325  GU.GGCC.....................................................................................................................................................................................
env-326  GU.GGCCC....................................................................................................................................................................................
env-327  GU.GGCCC....................................................................................................................................................................................
env-328  GU.GGCCC....................................................................................................................................................................................
Pab-1-1  GU.CCCC.....................................................................................................................................................................................
Psp-2-1  GU.UCCC.....................................................................................................................................................................................
Tga-1-1  GU.UCCG.....................................................................................................................................................................................
Tsp-3-1  GU.UCCG.....................................................................................................................................................................................
Tsp-2-1  GU.UCCG.....................................................................................................................................................................................
Tna-1-1  GU.UCCG.....................................................................................................................................................................................
Teu-1-1  GU.UCCG.....................................................................................................................................................................................
env-329  GU.UCCG.....................................................................................................................................................................................
Tsp-4-1  GU.UCCG.....................................................................................................................................................................................
Tsi-1-1  GU.UCCG.....................................................................................................................................................................................
Tli-1-1  GU.UCCG.....................................................................................................................................................................................
env-330  GU.UCCG.....................................................................................................................................................................................
Tba-1-1  GU.UCCG.....................................................................................................................................................................................
Tpa-1-1  GU.UCCG.....................................................................................................................................................................................
Ppa-1-1  GU.UCCG.....................................................................................................................................................................................
Pfu-2-1  GU.CCCC.....................................................................................................................................................................................
Pho-1-1  GU.CCCC.....................................................................................................................................................................................
Psp-1-1  GU.CCCC.....................................................................................................................................................................................
Pya-1-1  GU.UCCC.....................................................................................................................................................................................
Tko-1-1  GU.UCCC.....................................................................................................................................................................................
Tsp-1-1  GU.UCCC.....................................................................................................................................................................................
Pfe-1-1  GU.UCCC.....................................................................................................................................................................................
Ton-1-1  GU.UCCC.....................................................................................................................................................................................
Tzi-1-1  GU.UCCC.....................................................................................................................................................................................
env-331  GU.AACCCU...................................................................................................................................................................................
env-332  GU.AACCCU...................................................................................................................................................................................
env-333  GU.AACCC....................................................................................................................................................................................
env-334  GU.GGCC.....................................................................................................................................................................................
env-336  GU.AGCC.....................................................................................................................................................................................
env-338  GU.GGCC.....................................................................................................................................................................................
env-339  GU.AACC.....................................................................................................................................................................................
env-341  GU.AACC.....................................................................................................................................................................................
env-344  GU.AACC.....................................................................................................................................................................................
env-348  GU.AGCC.....................................................................................................................................................................................
env-349  GU.AGCC.....................................................................................................................................................................................
env-353  GU.AGCC.....................................................................................................................................................................................
env-354  GU.AGCC.....................................................................................................................................................................................
env-356  GU.AGCC.....................................................................................................................................................................................
env-358  GU.AGCC.....................................................................................................................................................................................
env-359  GU.GGCC.....................................................................................................................................................................................
env-360  GU.GGCC.....................................................................................................................................................................................
env-362  GU.GGCC.....................................................................................................................................................................................
env-363  GU.GGCC.....................................................................................................................................................................................
env-364  GU.AGCC.....................................................................................................................................................................................
env-365  GU.AGCC.....................................................................................................................................................................................
env-366  GU.AGCC.....................................................................................................................................................................................
env-367  GU.AGCC.....................................................................................................................................................................................

         ...<<<<<<<<<<<<<<<<<<<<<<<<<<<<<<<<<<<<<<<<<<<<<<<<<<<<<<<<<<<<<<<<<<<...<<<<<<<<<<<<<<<<..............>>>>>>>>>>>>>>>>.....<<<<<<<...>>>>>>>.<<<<<<<<<<<<<<<<<<<<<<<<<<.......>>>>>
         ...22222..................1.1..000000000.0000000...000.000...00000.......1111111...100000..............000001...1111111.....000000....000000.1011..1.11....111.10000000.......00000
         GU-GRYYo------------------------------------------------------------------------------------------------------------------------------------------------------------------------------------
```

```
env-369    GU.AGCC.............................................................................................................................................................................
env-370    GU.GGCC.............................................................................................................................................................................
env-371    GU.GGCC.............................................................................................................................................................................
env-375    GU.AGCC.............................................................................................................................................................................
env-376    GU.AGCC.............................................................................................................................................................................
env-377    GU.AGCC.............................................................................................................................................................................
env-378    GU.AGCC.............................................................................................................................................................................
env-379    GU.AGCC.............................................................................................................................................................................
env-380    GU.AGCC.............................................................................................................................................................................
env-381    GU.AGCC.............................................................................................................................................................................
env-382    GU.AGCC.............................................................................................................................................................................
env-383    GU.AGCC.............................................................................................................................................................................
env-384    GU.AGCC.............................................................................................................................................................................
env-385    GU.AGCCUA...........................................................................................................................................................................
env-386    GU.AGCCUA...........................................................................................................................................................................
env-387    GU.AGCCUA...........................................................................................................................................................................
env-388    GU.AGCC.............................................................................................................................................................................
env-390    GU.AGCC.............................................................................................................................................................................
env-392    GU.AACC.............................................................................................................................................................................
env-402    GU.AACC.............................................................................................................................................................................
env-406    GU.AACC.............................................................................................................................................................................
env-408    GU.AACC.............................................................................................................................................................................
env-410    GU.AACC.............................................................................................................................................................................
env-412    GU.AACC.............................................................................................................................................................................
env-413    GU.AACC.............................................................................................................................................................................
env-414    GU.AACCA............................................................................................................................................................................
env-416    GU.AACCA............................................................................................................................................................................
env-418    GU.AACC.............................................................................................................................................................................
env-420    GU.AACC.............................................................................................................................................................................
env-422    GU.AACC.............................................................................................................................................................................
env-424    GU.AGCCC............................................................................................................................................................................
env-425    GU.AGCCC............................................................................................................................................................................
env-427    GU.AGCCC............................................................................................................................................................................
env-428    GU.GGCCCA...........................................................................................................................................................................
env-429    GU.GGCCC............................................................................................................................................................................
env-432    GU.GGCCC............................................................................................................................................................................
env-433    GU.GGCCC............................................................................................................................................................................
env-438    GU.AACCCU...........................................................................................................................................................................
env-439    GU.AACCCU...........................................................................................................................................................................
env-440    GU.AACCU............................................................................................................................................................................
env-441    GU.AACCU............................................................................................................................................................................
env-442    GU.AACCU............................................................................................................................................................................
env-443    GU.AACCU............................................................................................................................................................................
env-444    GU.AACCU............................................................................................................................................................................
env-446    GU.AACCU............................................................................................................................................................................
env-447    GU.AGCCU............................................................................................................................................................................
env-449    GU.AACC.............................................................................................................................................................................
env-450    GU.AACC.............................................................................................................................................................................
env-453    AU.UCCC.............................................................................................................................................................................
Mvo-1-1    GUAGC...............................................................................................................................................................................
env-454    GU.GGUCC............................................................................................................................................................................
ume-1-1    GU.GGCCUAUCC........................................................................................................................................................................
CMe-2-1    GU.GGCCC............................................................................................................................................................................
Mev-1-1    GU.GACUCA...........................................................................................................................................................................
env-455    GUAGCUUUU...........................................................................................................................................................................
env-457    GUAGCUUUU...........................................................................................................................................................................
env-460    GUAGCUUUUU..........................................................................................................................................................................
env-461    GUAGCUUUUU..........................................................................................................................................................................
env-462    GU.GGCUUA...........................................................................................................................................................................
Mzh-1-1    GU.GAUUCGUU.........................................................................................................................................................................
env-463    GU.GGCCA............................................................................................................................................................................
env-465    GU.GGCCCUGA.........................................................................................................................................................................
env-466    GU.AGCCC............................................................................................................................................................................
env-467    GU.GGCUC............................................................................................................................................................................
env-469    GU.GGCCCGG..........................................................................................................................................................................
Mta-1-1    GU.GGCCCGG..........................................................................................................................................................................
env-472    GU.GGCCC............................................................................................................................................................................
env-473    GU.GGCCC............................................................................................................................................................................
env-474    GU.AGCUU............................................................................................................................................................................
Mpa-1-1    GU.AGCUCAA..........................................................................................................................................................................
env-475    GU.AGCCUG...........................................................................................................................................................................
env-476    GUAGCCUG............................................................................................................................................................................
env-477    GUAGCACG............................................................................................................................................................................
env-479    GUAGCCC.............................................................................................................................................................................
env-480    GUAGCUCU............................................................................................................................................................................
env-481    GUAGCUCU............................................................................................................................................................................
env-484    GUAGCUCU............................................................................................................................................................................
env-485    GUAGCUCU............................................................................................................................................................................
env-486    GUAGCUC.............................................................................................................................................................................
env-487    GUAGCUC.............................................................................................................................................................................
env-488    GUAGCUC.............................................................................................................................................................................
env-489    GUAGCUC.............................................................................................................................................................................
env-490    GUAGCUC.............................................................................................................................................................................

           ...<<<<<<<<<<<<<<<<<<<<<<<<<<<<<<<<<<<<<<<<<<<<<<<<<<<<<<<<<<<<<<<<<<<...<<<<<<<<<<<<<<<<..............>>>>>>>>>>>>>>>>.....<<<<<<<...>>>>>>>.<<<<<<<<<<<<<<<<<<<<<<<<<<.......>>>>>
           ...22222..................1.1..000000000.0000000...000.000...00000.......1111111...100000..............000001...1111111.....000000....000000.1011..1.11....111.10000000.......00000
           GU-GRYY∘------------------------------------------------------------------------------------------------------------------------------------------------------------------------
```

```
env-491    GUAGCUC.........................................................................................................................................................................
env-492    GUAGCUC.........................................................................................................................................................................
env-495    GUAGCUCUG.......................................................................................................................................................................
env-496    GUAGCUCA........................................................................................................................................................................
env-497    GUAGCUCA........................................................................................................................................................................
env-499    GUAGCUCA........................................................................................................................................................................
env-500    GUAGCUCA........................................................................................................................................................................
env-502    GUAGCUCA........................................................................................................................................................................
env-503    GUAGCUCA........................................................................................................................................................................
env-505    GUAGCUCA........................................................................................................................................................................
env-506    GUAGCCC.........................................................................................................................................................................
env-507    GUAGCCC.........................................................................................................................................................................
env-508    GUAGCCC.........................................................................................................................................................................
env-509    GUAGCCC.........................................................................................................................................................................
env-511    GUAGCCC.........................................................................................................................................................................
env-512    GUAGCCC.........................................................................................................................................................................
env-513    GUAGCCC.........................................................................................................................................................................
env-514    GUAGCCC.........................................................................................................................................................................
env-515    GUAGCCC.........................................................................................................................................................................
env-516    GUAGCCC.........................................................................................................................................................................
env-517    GU.GGCCUG.......................................................................................................................................................................
env-522    GU.GGCUU.C......................................................................................................................................................................
env-525    GU.GGCUU.C......................................................................................................................................................................
env-528    GU.AGCC.........................................................................................................................................................................
env-529    GU.AGCC.........................................................................................................................................................................
env-530    GU.AGCC.........................................................................................................................................................................
env-531    GU.AGCC.........................................................................................................................................................................
env-532    GUAGCCC.........................................................................................................................................................................
CMe-1-1    GUAGCCC.........................................................................................................................................................................
env-534    GUAGCCC.........................................................................................................................................................................
env-535    GUAGCCC.........................................................................................................................................................................
env-536    GU.GGCUC........................................................................................................................................................................
env-537    GU.GGCUC........................................................................................................................................................................
env-538    GU.AGCCUG.......................................................................................................................................................................
env-539    GU.AGCCUG.......................................................................................................................................................................
env-540    GU.AGCCUG.......................................................................................................................................................................
env-541    GU.GGCCU........................................................................................................................................................................
env-542    GU.GGCGUC.......................................................................................................................................................................
env-543    GU.GGCGUUU......................................................................................................................................................................
env-544    GU.AGCCUG.......................................................................................................................................................................
env-545    GU.GGCGG........................................................................................................................................................................
env-546    GU.GGCGG........................................................................................................................................................................
Mli-1-1    GU.GGCGC........................................................................................................................................................................
env-547    GU.GGCG.........................................................................................................................................................................
Mse-1-1    GU.GGCG.........................................................................................................................................................................
Mma-1-1    GU.GGCG.........................................................................................................................................................................
Msp-5-1    GU.GGCG.........................................................................................................................................................................
env-552    GU.GGCG.........................................................................................................................................................................
env-553    GU.GGCG.........................................................................................................................................................................
env-559    GU.GGCG.........................................................................................................................................................................
env-560    GU.GGCG.........................................................................................................................................................................
env-561    GU.GGCG.........................................................................................................................................................................
env-563    GU.GGCG.........................................................................................................................................................................
env-564    GU.GGCG.........................................................................................................................................................................
env-565    GU.GGCG.........................................................................................................................................................................
env-567    GU.GGCG.........................................................................................................................................................................
env-568    GU.GGCG.........................................................................................................................................................................
env-570    GU.GGCG.........................................................................................................................................................................
env-571    GU.GGCG.........................................................................................................................................................................
env-574    GU.GGCGUG.......................................................................................................................................................................
env-575    GU.GGCGUG.......................................................................................................................................................................
env-576    GU.GGCGUG.......................................................................................................................................................................
env-577    GU.GGCGUG.......................................................................................................................................................................
env-578    GU.GGCUCA.......................................................................................................................................................................
env-579    GU.GGCUC........................................................................................................................................................................
env-580    GU.GGCUCA.......................................................................................................................................................................
env-581    GU.GGCCCG.......................................................................................................................................................................
env-585    GU.GGCCCG.......................................................................................................................................................................
env-587    GU.GGCUCG.......................................................................................................................................................................
env-588    GU.GGCUCG.......................................................................................................................................................................
env-589    G..UAGCCGC......................................................................................................................................................................
env-590    GU.GGCUG........................................................................................................................................................................
env-591    GU.GGCUG........................................................................................................................................................................
env-594    GU.GGCUGC.......................................................................................................................................................................
env-595    GU.GGCUGC.......................................................................................................................................................................
env-596    GU.GGCUGC.......................................................................................................................................................................
env-597    GU.GGCUGC.......................................................................................................................................................................
env-598    GU.GGCUGC.......................................................................................................................................................................
env-599    GU.GGCUGC.......................................................................................................................................................................
env-602    GU.GGCUGC.......................................................................................................................................................................
env-603    GU.GGCCGC.......................................................................................................................................................................
env-604    GU.GGCCGC.......................................................................................................................................................................
env-605    GU.GGCCGC.......................................................................................................................................................................

           ...<<<<<<<<<<<<<<<<<<<<<<<<<<<<<<<<<<<<<<<<<<<<<<<<<<<<<<<<<<<<<<<<<<<...<<<<<<<<<<<<<<<<..............>>>>>>>>>>>>>>>>.....<<<<<<<...>>>>>>>.<<<<<<<<<<<<<<<<<<<<<<<<<<.......>>>>>
           ...22222..................1.1..000000000.0000000...000.000...00000.......1111111...100000..............000001...1111111.....000000....000000.1011..1.11....111.10000000.......00000
           GU-GRYY∘------------------------------------------------------------------------------------------------------------------------------------------------------------------------
```

```
env-606   GU.GGCCGC..........................................................................................................................................................................
env-607   GU.GGCCGC..........................................................................................................................................................................
env-608   GU.GGCCGC..........................................................................................................................................................................
env-609   GU.GGCCGC..........................................................................................................................................................................
env-610   GU.GGCCGC..........................................................................................................................................................................
env-611   GU.GGCCGC..........................................................................................................................................................................
env-612   GU.GGCCGC..........................................................................................................................................................................
env-613   GU.GGCCGC..........................................................................................................................................................................
env-616   GU.GGCUA...........................................................................................................................................................................
Mco-1-1   GU.GGCUA...........................................................................................................................................................................
env-617   GU.GAUCUUU.........................................................................................................................................................................
Mps-1-1   GU.GGCUU...........................................................................................................................................................................
env-619   GU.GGCUU...........................................................................................................................................................................
env-620   GU.GGCUU...........................................................................................................................................................................
Mti-1-1   G..UAGCUUUUG.......................................................................................................................................................................
env-621   G..UAGCUU..........................................................................................................................................................................
Mba-2-1   G..UAGUCCUG........................................................................................................................................................................
env-622   GU.GACCCUG.........................................................................................................................................................................
env-623   GU.GGUCCUG.........................................................................................................................................................................
env-624   GU.GGUCCUG.........................................................................................................................................................................
env-625   GU.GGUCCUGU........................................................................................................................................................................
Mho-2-1   GU.GGUCCUG.........................................................................................................................................................................
env-628   GU.GGUCCUG.........................................................................................................................................................................
env-629   GU.GGUCCUG.........................................................................................................................................................................
env-630   GU.GGUCCUGU........................................................................................................................................................................
Mth-1-1   GU.GGUCCUGU........................................................................................................................................................................
Msp-13-1  GU.GGUCCUG.........................................................................................................................................................................
Mva-1-1   GU.GGUCCUG.........................................................................................................................................................................
Mba-4-1   GU.GGUCCUG.........................................................................................................................................................................
env-631   GU.GGUCCUG.........................................................................................................................................................................
env-633   GU.GGUCCUG.........................................................................................................................................................................
Mla-1-1   GU.GGUCCUG.........................................................................................................................................................................
Mba-5-1   GU.GGUCCUG.........................................................................................................................................................................
Msp-7-1   GU.GGUCCUG.........................................................................................................................................................................
Msp-16-1  GU.GGUCCUG.........................................................................................................................................................................
env-634   GU.GGUCCUGC........................................................................................................................................................................
Mso-1-1   G.UGGUCCUG.........................................................................................................................................................................
Mma-3-1   G.UGGUCCUG.........................................................................................................................................................................
Mac-1-1   G.UGGCCCUG.........................................................................................................................................................................
Msi-2-1   G.UGGCCCUG.........................................................................................................................................................................
env-635   GU.GGCCUUG.........................................................................................................................................................................
Msi-1-1   GUGGCCCUG..........................................................................................................................................................................
env-636   GUGGCCCUG..........................................................................................................................................................................
Msp-14-1  GU.GGCUCG..........................................................................................................................................................................
env-637   GU.GGUCC...........................................................................................................................................................................
env-638   GU.GGUCC...........................................................................................................................................................................
env-639   G..UAGUCC..........................................................................................................................................................................
env-640   G..UAGCC...........................................................................................................................................................................
env-641   GUGGCCCA...........................................................................................................................................................................
Mbu-1-1   G.UGGCUGAA.........................................................................................................................................................................
env-642   GU.GGCUGAA.........................................................................................................................................................................
Mme-2-1   GU.GGCCGAAG........................................................................................................................................................................
Mme-1-1   GU.GGCCGAAG........................................................................................................................................................................
Msh-1-1   GU.GGCUC...........................................................................................................................................................................
env-643   GU.GGCU............................................................................................................................................................................
env-644   GU.GGC.............................................................................................................................................................................
env-645   GU.GGC.............................................................................................................................................................................
env-646   GU.GGC.............................................................................................................................................................................
env-647   GU.GGCC............................................................................................................................................................................
env-650   GU.GGCU............................................................................................................................................................................
env-651   GU.GGC.............................................................................................................................................................................
env-656   GU.GGCC............................................................................................................................................................................
env-657   GU.GGCC............................................................................................................................................................................
env-658   GU.GGCC............................................................................................................................................................................
Mve-1-1   GU.GGC.............................................................................................................................................................................
env-659   GU.GGC.............................................................................................................................................................................
env-660   GU.GGC.............................................................................................................................................................................
Msp-4-1   GU.GGC.............................................................................................................................................................................
env-661   GU.GGC.............................................................................................................................................................................
env-663   GU.GGC.............................................................................................................................................................................
env-664   GU.GGC.............................................................................................................................................................................
env-665   GU.GGC.............................................................................................................................................................................
env-670   GU.GGC.............................................................................................................................................................................
env-671   GU.GGC.............................................................................................................................................................................
env-672   GU.GGC.............................................................................................................................................................................
Mfo-2-1   GU.GGC.............................................................................................................................................................................
env-673   GU.GGC.............................................................................................................................................................................
env-674   GU.GGC.............................................................................................................................................................................
Msp-2-1   GU.GGC.............................................................................................................................................................................
Mfo-1-1   GU.GGC.............................................................................................................................................................................
env-675   GU.GGC.............................................................................................................................................................................
env-676   GU.GGC.............................................................................................................................................................................
Msp-1-1   GU.GGC.............................................................................................................................................................................

          ...<<<<<<<<<<<<<<<<<<<<<<<<<<<<<<<<<<<<<<<<<<<<<<<<<<<<<<<<<<<<<<<<<<<...<<<<<<<<<<<<<<<<..............>>>>>>>>>>>>>>>>.....<<<<<<<...>>>>>>>.<<<<<<<<<<<<<<<<<<<<<<<<<<.......>>>>>
          ...22222.................1.1..000000000.0000000...000.000...00000.......1111111...100000..............000001...1111111.....000000....000000.1011..1.11....111.10000000.......00000
          GU-GRYY∘-------------------------------------------------------------------------------------------------------------------------------------------------------------------------------
```

```
env-679    GU.GGC...........................................................................................
env-680    GU.GGC...........................................................................................
Msp-3-1    GU.GGC...........................................................................................
env-681    GU.GGC...........................................................................................
env-682    GU.GGC...........................................................................................
env-683    GU.GGC...........................................................................................
Mor-1-1    GU.GGCU..........................................................................................
Msm-14-1   GU.GGCUUUU.......................................................................................
Msm-4-1    GU.GGCUUUU.......................................................................................
env-684    GU.GGCUUUU.......................................................................................
env-685    GU.GGCUUUU.......................................................................................
env-686    GU.GGCUUUU.......................................................................................
env-695    GU.GGCUUUU.......................................................................................
Msm-16-1   GU.GGCUUUU.......................................................................................
Msm-6-1    GU.GGCUUUU.......................................................................................
env-696    GU.GGCUUA........................................................................................
env-697    GU.GGCCU.........................................................................................
env-698    GU.GGCCU.........................................................................................
env-699    GU.GGCCU.........................................................................................
env-700    GU.GGCC..........................................................................................
env-701    GU.GGCCC.........................................................................................
env-702    GU.GGCCC.........................................................................................
env-704    GU.GGCCC.........................................................................................
env-706    GU.GGCCC.........................................................................................
env-707    GU.GGCCC.........................................................................................
env-708    GU.GGCCC.........................................................................................
env-709    GU.GGCCC.........................................................................................
env-710    GU.GGCUU.........................................................................................
Ngr-1-1    GU.GAUC.CG.CGGGGC................................................................................
Hti-1-1    GU.GGCCGC........................................................................................
Npe-1-1    GU.GAUCC...GCGGGGC...............................................................................
Hsp-7-1    GU.GGUCU.CAGGGG..................................................................................
Hje-1-1    GU.GGUCCGCGC.....................................................................................
Hbo-1-1    GU.GGAGGGCGU.....................................................................................
Nma-1-1    GU.GAUCCGA.......................................................................................
Nti-1-1    GU.GAUCC...GCGGGG................................................................................
           ...<<<<<<<<<<<<<<<<<<<<<<<<<<<<<<<<<<<<<<<<<<<<<<<<<<<<<<<<<<<<<<<<<<<<<...<<<<<<<<<<<<<<<<..............>>>>>>>>>>>>>>>>.....<<<<<<<...>>>>>>>.<<<<<<<<<<<<<<<<<<<<<<<<<<.......>>>>>
           ...22222.................1.1..000000000.0000000...000.000...00000.......1111111...100000..............000001...1111111.....000000....000000.1011..1.11....111.10000000.......00000
           GU-GRYY∘-------------------------------------------------------------------------------------------
```

alignment positions 361 · · · 512

```
env-1      .....................CAUUUGGA.................................................GGCU.GAACAAAUCAUUAGGGUGAAAUAUC........................AUG
env-2      .....................CAUUUGGA.................................................GGCU.GAACAAAUCAUUAGGGUGAAAUAUC........................AUG
Mma-2-1    .....................UUUAU....................................................GGCU.GAACCGUCAAUGAAUGAAACAAGGUGAAAUC..................AUG
env-3      .....................AUACAG.................................................UCACUC.GAAUCGAAACCGCUCGUCAGAGACAACA.....................AUG
Hha-1-1    .....................CGGG.............................................ACCGCAC.GGCU.GAACCACCAACGCCCGGAGGCACAUC.......................AUG
env-5      .....................UUGA...................................................CCGGUU.GAACCACUAACGCCCCGGGAACCGAAAGACC..................AUG
env-6      .....................UUC....................................................ACGGUU.GAACCACCAACGCCCCGGGAAGGUACACCAA..................AUG
env-7      .....................GAAGGG...................................ACCGCCCCGGAACGGCU.GAACCAUCGAGCCCGCGGUGACAACA.......................AUG
env-9      .....................GACUCGAA...........................GGAGGCUGCUGGAUCUGGGU.GAAUCGCAAGCGCUCGUGGAAUACACA.....................AUG
env-10     .....................GAC.................................................GGACCACGAGGGCC.GAAUCGCAAGCGCCCGCGGUAGACCA......................AUG
Hsa-6-1    .....................GACGU.................................GUCGUCUUCGUUGGCGGAACU.GAAUCGCAAGCGUCCGUGGUGAUUACA.....................AUG
h3A-1-1    .....................GUC.......................................................GUCGCGGGACU.GAAUCGCAAGCGUCCGUGGACAACCC....................AUG
env-11     .....................UUC....................................................GCGGCU.GAAUCGCAACCGUCCGCGGACAACAU......................AUG
env-12     .....................AGA....................................................CCGGCC.GAAUCGCAAGCGUCCACAGGCAACC.......................AUG
env-14     .....................CGG...................................GAGGCGGCCCCACCGCCGACGUU.GAAUCGCAAGCGCCCGCGGUGAAACAU.....................AUG
env-15     .....................AAC...............................................GCGCCCCUGCCGACU.GAAUCGCAAGCGCCCGCAAUCUAACC......................AUG
env-16     .....................AAC...............................................GCGCCCCUGCCGACU.GAAUCGCAAGCGCCCGCAAUCUAACC......................AUG
env-19     .....................CAUUG...................................................ACGU..UCCC.GAAUCACAAGCGCCCGUGGUGAAAACAG...................AUG
env-22     .....................CAUUG...................................................ACGU..UCCC.GAAUCACAAGCGCCCGUGGUGAAAACAG...................AUG
Hwa-1-1    .....................CAUUG...................................................ACGU..UCCC.GAAUCACAAGCGCCCGUGGUGAAAACAG...................AUG
env-23     .....................AAU.....................................................GACGUCUUCCC.GAAUCGCAAGCGCCCGUGGUGAAAACAG..................AUG
env-25     .....................AUU.....................................................GACGUC..CCC.GAAUCGCAAGCGCCCGUGGUGAAAAUAC..................AUG
env-27     .....................AUU.....................................................GACGUC..CCC.GAAUCGCAAGCGCCCGUGGUGAAAAUAC..................AUG
env-28     .....................UCGAC.....................................................CGUC.GCCC.GAAUCGCAAGCGCCCGUGGUGACACCA...................AUG
env-29     .....................UCGAC.....................................................CGUC.GCCC.GAAUCGCAAGCGCCCGUGGUGACACCA...................AUG
env-30     .....................UCGAC.....................................................CGUC.GCCC.GAAUCGCAAGCGCCCGUGGUGACACCA...................AUG
env-31     .....................CGA.......................................................ACGCCCGCCC.GAAUCGCAAGCGCCCGUGGUGACACCA..................AUG
Hru-1-1    .....................CGA.......................................................ACGCCG.CCC.GAAUCGAAAGCGCCCGUGGUGAAACCA..................AUG
env-32     .....................CGA.......................................................ACGUCG.CCC.GAAUCGCAAGCGCCCGUGGUGAAACCA..................AUG
env-33     .....................CGA.......................................................ACGUCG.CCC.GAAUCGCAAGCGCCCGUGGUGAAACCA..................AUG
Hpa-2-1    .....................GACGGUA..................................................GAGCGC.CUCC.GAAUCGCAAGCGCCCGUGGUGAAACU..................AUG
env-34     ............GCCGC....GAGGGGA.......................................................GAGCU.GAAUCGCAAGCGCCCGCGGUACACCA....................AUG
env-35     .....................UAUAU.........................................................CCGCU.GAAUCGCAAGCGCCCGUGGUGAUACAU...................AUG
env-39     .....................CGCCGU..................................GAACCCCGACUCUCAAUCAGGCU.GAAUCGCAAGCGUCCGUGGUACACAUC.....................AUG
env-40     .....................CACGGU.................................GAACCCCCGACUCUCAAUCAGGCU.GAAUCGCAAGCGUCCGUGGAACACAUC.....................AUG
Hac-1-1    .....................GUC.........................................................GCGCACU.GAAUCGCAAGCGCCCGCGGAACACACU...................AUG
env-42     .....................AGA...........................................................CCGGCC.GAAUCGCAAGCGUCCGCAGGCACCC.....................AUG

           >>>>>>>>>>>>>>>>>>>>>..........>>>>>>>>>>>>>>>>>>>>>>>>>>>>>>>>>>>>>>>>>>>>>>>>>>>>>>>>>>>>>>....>>.................................................
           001.111....11.1..1101.............00000...000.000...0000000.000000000..1.1...................22222....21.................................................
           ---------------------••••∘--------------------------------------------------------------∘RRYU-GAACCR•RRR••••••••R•••••••••∘∘-------------------AUG
                                                                                                              3'
```

```
env-43    .....................GGA..........................................................CCGGCC.GAAUCGCAAGCGUCCGCAGGCAACC.........................AUG
env-45    .....................GGA..........................................................CCGGCC.GAAUCGCAAGCGUCCGCAGGCAACC.........................AUG
Hna-1-1   .....................ACUU.....................................................CGGCGGCGCC.GAAUCGCAAGCGCCCGUGGGUAACUC........................AUG
env-47    .....................ACUN.....................................................CGGCGACGCU.GAAUCGCAAGCGCCCGUGGUGAAACCC.......................AUG
env-48    .....................AGGU..........................................................GCGCU.GAAUCGCAACGGCCCGCGGUGAAACC........................AUG
env-49    .....................AGGU..........................................................GCGCU.GAAUCGCAACGGCCCGCGGUGAAACC........................AUG
env-50    .....................AGAU..........................................................GCGCU.GAAUCGCAACGGCCCGCGGUGAAACC........................AUG
env-51    .....................AGAU..........................................................GCGCU.GAAUCGCAACGGCCCGCGGUGAAACC........................AUG
env-52    .....................AGAC..........................................................GCGCU.GAAUCGCAAGCGCCCGCGGUGAAACC........................AUG
env-53    .....................AGAC..........................................................GCGCU.GAAUCGCAAGCGCCCGCGGUGAAACC........................AUG
env-55    .....................UUC...........................................................GGGCU.GAAUCGCAACCGUCCGCGGACACACCA.......................AUG
env-57    .....................AUGC.........................................................GCGGCU.GAAUCGCAAACGUCCGUGGACAACACCC......................AUG
env-61    .....................CAUGCA........................................................CGGCU.GAAUCGCAAACGUCCGUGGACACGACC.......................AUG
env-63    .....................AUGC.........................................................GCGGCU.GAAUCGCAAACGUCCGUGGACAACACCC......................AUG
env-64    .....................AUGC.........................................................GCGGCU.GAAUCGCAAACGUCCGUGGACAACACCC......................AUG
env-65    .....................AUGC.........................................................GCGGCU.GAAUCGCAACCGUCCGUGGACACGAACC......................AUG
env-66    .....................CUC..........................................................GCGGCU.GAAUCGCAACCGUCCGUGGACAACCA........................AUG
Hai-1-1   .....................CUC..........................................................GCGGCU.GAAUCGCAACCGUCCGCGGAUACACCA.......................AUG
Hco-1-1   .....................CCUCGC.........................................................GGCU.GAAUCGCAACCGUCCGCAGACACACA........................AUG
Hsp-8-1   .....................CCUCGC.........................................................GGCU.GAAUCGCAACCGUCCGCGGACACACA........................AUG
Hte-1-1   .....................CCUCGC.........................................................GGCU.GAAUCGCAACCGUCCGCGGACACACA........................AUG
Hho-1-1   .....................CCUCGC.........................................................GGCU.GAAUCGCAACCGUCCGCGGACACACA........................AUG
env-67    .....................CCUCGC.........................................................GGCU.GAAUCGCAACCGUCCGCGGACACACA........................AUG
Hca-2-1   .....................CCUCGC.........................................................GGCU.GAAUCGCAACCGUCCGCGCAACACACA.......................AUG
env-70    .....................CCUCGC.........................................................GGCU.GAAUCGCAACCGUCCGCGGACAACA.........................AUG
Hli-2-1   .....................CCUCGC.........................................................GGCU.GAAUCGCAACCGUCCGCGCAACACACA.......................AUG
Har-2-1   .....................CCUCGC.........................................................GGCU.GAAUCGCAACCGUCCGCGCAACACACA.......................AUG
env-73    .....................CUC..........................................................GCGGCU.GAAUCGCAACCGUCCGCGGACAACAU........................AUG
Hli-1-1   .....................UUC..........................................................GCGGCU.GAAUCGCAACCGUCCGCGGACAACAU........................AUG
Hha-2-1   .....................UUC..........................................................GCGGCU.GAAUCGCAACCGUCCGCGGACAACAU........................AUG
Hsa-9-1   .....................UUC..........................................................GCGGCU.GAAUCGCAACCGUCCGCGGACAACAU........................AUG
env-74    .....................UUC..........................................................GCGGCU.GAAUCGCAACCGUCCGCGGACAACAU........................AUG
Hko-1-1   .....................UUC..........................................................GCGGCU.GAAUCGCAACCGUCCGCGGACAACAU........................AUG
env-77    .....................UUC..........................................................GCGGCU.GAAUCGCAACCGUCCGCGGACAACAU........................AUG
env-78    .....................CUC..........................................................GCGGCU.GAAUCGCAACCGUCCGCGGACAACAU........................AUG
Hsa-7-1   .....................CUC..........................................................GCGGCU.GAAUCGCAACCGUCCGCGGACAACAU........................AUG
env-79    .....................CCUCGC.........................................................GGCU.GAAUCGCAACCGUCCGCGGACAACAU........................AUG
env-80    .....................CUCGC..........................................................GGCU.GAAUCGCAACCGUCCGCACACACACA........................AUG
env-83    .....................CCUCGC.........................................................GGCU.GAAUCGCAACCGUCCGCAGACACACA........................AUG
env-84    .....................UUC..........................................................GCGGCU.GAAUCGCAACCGUCCGCGGACACACCA.......................AUG
env-86    .....................UUC..........................................................GCGGCU.GAAUCGCAACCGUCCGCGGACACACAU.......................AUG
env-87    .....................UUC..........................................................GCGGCU.GAAUCGCAACCGUCCGCGGACACACAU.......................AUG
env-89    .....................CUC..........................................................GCGGCU.GAAUCGCAAUCGUCCGUGGACGAUACA.......................AUG
env-90    .....................CUC..........................................................GCGGCU.GAAUCGCAAGCGUCCGUGGACAACACA.......................AUG
env-91    .....................CUC..........................................................GCGGCU.GAAUCGCAAGCGUCCGUGGACAACACA.......................AUG
env-92    .....................CUC..........................................................GCGGCU.GAAUCGCAAGCGUCCGUGGACGAUACA.......................AUG
env-93    .....................AUGC.........................................................GCGGCU.GAAUCGCAAACGUCCGUGGACACGAACC......................AUG
env-94    .....................UUC..........................................................GCGGCU.GAAUCGCAAGCGUCCGUGGACACGACA.......................AUG
Hmu-1-1   .....................GUG..................................................UCCCUCCACCGACU.GAAUCGCAAGCGCCCGCGGUGAUAUACA......................AUG
Hel-1-1   .....................UGG..................................................UCCCUCAACCGACU.GAAUCGCAAGCGCCCGCGGUGAUAUACA......................AUG
Hla-2-1   .....................UGG..................................................UCCCUCAUUCGACU.GAAUCGCAAGCGCCCGCGGUGAUAUACA......................AUG
Hde-1-1   .....................GAUU....................................................CCCCGACGGACU.GAAUCGCAAGCGCCCGCGGUGAUACACA......................AUG
Hal-2-1   .....................AGUA....................................................CCCCGACGGACU.GAAUCGCAAGCGCCCGCGGUGAUAUACA......................AUG
Hpr-1-1   .....................AGUU....................................................CCCCGACGGACU.GAAUCGCAAGCGCCCGCGGUGAUAUACA......................AUG
Hgi-1-1   .....................AGUU....................................................CCCCGACGGACU.GAAUCGCAAGCGCCCGCGGUGAUAUACA......................AUG
Hgi-2-1   .....................AGUU....................................................CCCCGACGGACU.GAAUCGCAAGCGCCCGCGGUGAUAUACA......................AUG
Hsp-9-1   .....................AGUU....................................................CCCCGACGGACU.GAAUCGCAAGCGCCCGCGGUGAUACACA......................AUG
Hsu-2-1   .....................AGUU....................................................CCCCGACGGACU.GAAUCGCAAGCGCCCGCGGUGAUACACA......................AUG
Hsp-6-1   .....................AAGUUC...................................................CCCGACGGACU.GAAUCGCAAGCGCCCGCGGUGAUACACA......................AUG
Hme-1-1   .....................GGG......................................CGGACG.......CCCCAGAAUGGCU.GAAUCGCAAGCGCCCGCGGUGAUAUACA......................AUG
Hru-2-1   .....................CCCG....................................AGGCGGCGGAGCCGCCGACAUUCGACU.GAAUCGCAAGCGCCCGCACAGAACC.........................AUG
env-95    .....................GGC................................................GUUCGCGCCGCCACGACU.GAAUCGCAAGCGCCCGCAGGAGACACU.....................AUG
env-98    .....................ACU..................................................CCGCGUCGUCGCGACU.GAAUCGCAAGCGCCCGCAGGAGACACU.....................AUG
env-99    .....................GGACG....................................................GUCGCCACGACU.GAAUCGCAAGCGCCCGCAGGAACACU.......................AUG
CHa-1-1   .....................ACGCCCGAGGG...........................CGGGGGACCCGCCCGACCGCCGACU.GAAUCGCAAGCGCCCGCGGAGCAACCC.....................AUG
env-101   .....................UCGUU..........................................................CGACU.GAAUCGCAAGCGCCCGAGGAUACCA.........................AUG
env-102   .....................GUGAU.........................................................CCGACU.GAAUCGCAAGCGCCCGAGGAUAACCU........................AUG
env-104   .....................GUGAU.........................................................CCGACU.GAAUCGCAAGCGCCCGAGGAUAACCA........................AUG
env-105   .....................AUGAA.........................................................CCGACU.GAAUCGCAAGCGCCCGAGGACAAUCU........................AUG
env-110   .....................AUGAA.........................................................CCGACU.GAAUCGCAAGCGCCCGAGGACAAUCU........................AUG
env-112   .....................CCUGA.........................................................CCGACU.GAAUCGCAAGCGCCCGAGGACAACU.........................AUG
env-114   .....................CCUGA.........................................................CCGACU.GAAUCGCAAGCGCCCGAGGACAACU.........................AUG
env-115   .....................CCUGA.........................................................CCGACU.GAAUCGCAAGCGCCCGAGGACAACU.........................AUG
env-116   .....................CCUGA.........................................................CCGACU.GAAUCGCAAGCGCCCGAGGACAACU.........................AUG
env-117   .....................CUUGA.........................................................CCGACU.GAAUCGCAAGCGCCCGAGGACAACU.........................AUG
env-119   .....................CCUGA.........................................................CCGACU.GAAUCGCAAGCGCCCGAGGACAACU.........................AUG
env-121   .....................CCUGA.........................................................CCGACU.GAAUCGCAAGCGCCCGAGGACAACU.........................AUG
env-122   .....................CCUGA.........................................................CCGACU.GAAUCGCAAGCGCCCGAGGACAACU.........................AUG
env-125   .....................CCUGA.........................................................CCGACU.GAAUCGCAAGCGCCCGAGGACAACU.........................AUG
env-126   .....................AUUU.........................................................UCCGACU.GAAUCGCAAGCGCCCGAGGAUAACA.........................AUG
env-129   .....................AUU..........................................................UCCGACU.GAAUCGCAAGCGCCCGAGGACAACU.........................AUG
env-130   .....................GAUUU.........................................................CCGACU.GAAUCGCAAGCGCCCGAGGACAACA.........................AUG
env-135   .....................GUU...........................................................UCGACU.GAAUCGCAAGCGCCCGAGGACAACA.........................AUG

          >>>>>>>>>>>>>>>>>>>>>...........>>>>>>>>>>>>>>>>>>>>>>>>>>>>>>>>>>>>>>>>>>>>>>>>>>>>>>>>....>>................................................
          001.111....11.1..1101...............00000...000.000...0000000.000000000..1.1..................22222....21................................................
          ---------------------●●●○-----------------------------------------------------------○RRYU-GAACCR●RRR●●●●●●●●R●●●●●●●○○--------------------AUG
                                                                                                         3′
```

```
env-136   ....................AUU..........UCGACU.GAAUCGCAAGCGCCCGAGGACAAAUA..........AUG
env-137   ....................CUGU.........UCGACU.GAAUCGCAAGCGCCCGAGGACAACCA..........AUG
env-138   ....................CUCU.........UCGACU.GAAUCGCAAGCGCCCGAGGACAAACA..........AUG
env-139   ....................CUCCU........UCGACU.GAAUCGCAAGCGCCCGAGGACAAACA..........AUG
env-140   ....................CUCC.........UCGACU.GAAUCGCAAGCGCCCGAGGACAAACA..........AUG
env-141   ....................CUCC.........UCGACU.GAAUCGCAAGCGCCCGAGGACAAAC............
env-142   ....................CUCC.........UCGACU.GAAUCGCAAGCGCCCGAGGACAAACA..........AUG
env-144   ....................CUCC.........UCGACU.GAAUCGCAAGCGCCCGAGGACAAACA..........AUG
env-145   ....................CUU..........UCGACU.GAAUCGCAAGCGCCCGAGGACAAACA..........AUG
env-146   ....................CUUGU........UCGACU.GAAUCGCAAGCGCCCGAGGACAACCA..........AUG
env-147   ....................CGU..........UCGACU.GAAUCGCAAGCGCCCGAGGACAACCA..........AUG
env-148   ....................CGU..........UCGACU.GAAUCGCAAGCGCCCGAGGACAACCA..........AUG
env-149   ....................CUCC.........UCGACU.GAAUCGCAAGCGCCCGAGGAUAACA...........AUG
env-152   ....................CUCU.........UCGACU.GAAUCGCAAGCGCCCGAGGAAAACA...........AUG
env-153   ....................CUCGA........CCGACU.GAAUCGCAAGCGCCCGACGAACU.............AUG
env-156   ....................CUUGA........CCGACU.GAAUCGCAAGCGCCCGAGACAACU............AUG
env-158   ....................UCG.........ACCGGCU.GAAUCGCAAGCGCCCGCGGUGAAACG..........AUG
env-159   ....................UCC.........GGCGGCU.GAAUCGCAAGCGCCCGCGGUGAACCAC.........AUG
Hsp-10-1  ....................CUC..........GCGGCU.GAAUCGCAACCGUCCGCGAACACACA..........AUG
Hse-1-1   ....................GCG.....GCGGUCGUCGCCCGAGGGCU.GAACCACCAACGCCCGCAGAUAACC...AUG
Hmo-1-1   ....................UCAC..........GGGCU.GAACCCUCAACGCCCGCAGACAACC............AUG
Hth-1-1   ....................UCAC..........GGGCU.GAACCCGCAACGCCCGCAGACAACC............AUG
Nae-1-1   ....................CAGCGC.....CCUCGCCGGGUU.GAACCAUCACCGCCCGCGGUGAAUACA......AUG
Nta-1-1   ....................AUCG......UCCUCGCCGGGUU.GAACCAUCACCGCCCGCGGUGAAUACA......AUG
Nas-1-1   ....................AUCG......UCCUCGCCGGGUU.GAACCAUCACCGCCCGCGGUGAAUACA......AUG
env-160   ...............CCGCCGGAG...........AGCU.GAACCACACAAGGCCCGCACGCCAACC.........AUG
env-161   ...............CGCCG...............GGUU.GAACCACGCAAGGCCCGCACGCCAACA.........AUG
env-162   ....................CUCGA..........GGCCU.GAACCAACGACGCCCGCGGAUACACU.........AUG
env-164   ....................CUCGA..........GGCCU.GAACCAACGACGCCCGCGGACAACCU.........AUG
env-169   ....................CUCGA..........GGCCU.GAACCAACGACGCCCGCGGACAACCU.........AUG
env-170   ....................UCUC.........GUGGUCU.GAACCGGGAAACGCCCGCAGGUAAAACG.......AUG
Hla-4-1   ....................AUC.......CUCUCGAGUUCU.GAACCAGGAAACGCCCGCAGCAAAACACA....AUG
Hsp-11-1  ....................ACU........CUCUCGAGUUCU.GAACCAGGAAACGCCCGCAGGUAUACA......AUG
Hsp-3-1   ....................UGCG.........CCGGGAAUU.GAACCACACAACGCCCGCGGACACAGAACA....AUG
env-172   ....................CUCG...........AGGAUU.GAACCAGGAACUGCCCGCAGCACAACU.......AUG
Hsa-1-1   ....................GCGA...........CCGACU.GAACCACUCGACGCCCGCGGUGAAAUC.......AUG
env-173   ....................UCG.............GGACU.GAACCGGCCAACGCCCGUGGUGAUACACG.....AUG
env-174   ....................UCG.............GGACU.GAACCGGCCAACGCCCGUGGUGAUACACG.....AUG
env-175   ....................UCC............GGGACU.GAACCGGCAAACGCCCGUGGUGAUAGACC.....AUG
env-176   ....................CUCG............GGACU.GAACCGGCAAACGCCCGUGGUGAUAGACC.....AUG
Hpa-1-1   ....................GUG......AAUCGAUUCCACGCU.GAACCACUCAACGCCCGUGGUGAUAGACC..AUG
env-177   ...............CCCCGCA.............UGGUUU.GAACCACGCAACGCCCGUGGUGAUAGACAU....AUG
env-178   ....................GGGGAG....GCGGAUC.GGCCGCCGAACGGUU.GAACCACACGACGCCCGUGGUGAACACACU..AUG
env-179   ....................UCAC...........GGGAUU.GAACCAGGAAACGCCCGCAAGAAAACC.......AUG
env-180   ....................GCUC..........GGAGUUU.GAACCGUACAACGCCCGUGGGUGAAACCACA...AUG
env-181   ....................ACGGG....GGUCGGAUA.UUCACACAAAUUUAUAGCCACCAGUAUCAGUGGUGGCGAAGCCACCAG.GGUUU.GAACCACACAACGCCCGUGGUGAUACCAC..AUG
env-182   ....................AAUC..........CAGGUUU.GAACCACUCAACGCCCGUGGGUGAAACCACA...AUG
Ssp-1-1   ..CGGCACCACCGCG.ACGCCGAUC............GGGUU.GAACCACCAAACGCCCGAGGAGAAUACA......AUG
env-183   CGACCGGGCCCGACCGCUGCCGAUC............GGGUU.GAACCACCAAACGCCCGAGGAGAACACA......AUG
Nph-1-1   ....................UCC............GCCGGUU.GAACCACGAAACGCCCGUGGUGAACAUC.......AUG
env-184   ....................CCGAGAACU....GACUCGAGUGGGCGUGAAAAUU.GAACCAGAAAACGCCCGCAGAGAAUACCA..AUG
Hgo-1-1   ....................AAC..........UCGGGAUU.GAACCAGGAAACGCCCGCGGGUGAAACC......AUG
Hxa-1-1   ....................AGU..........UCGGGAUU.GAACCAGGAAACGCCCGCAGGUGAUAAUC......AUG
env-185   ....................ACUC............GAGAUU.GAACCAGGAAACGCCCGCAGAAACCU........AUG
env-186   ....................ACUC............GAGAUU.GAACCAGGAAACGCCCGCAGAAACCU........AUG
env-187   ....................GAC............GCGGGAUU.GAACCAGGAAACGUCCGCGGGUGAAACU.....AUG
Nin-1-1   ....................GAGGU...........CGGGAUU.GAACCAGGAAACGCCCGCGGUGAUUACC.....AUG
env-188   ....................AGG.............CCGGAUU.GAACCAGGAAACGCCCGCGGUGAACACG.....AUG
env-189   ....................AGG.............CCGGAUU.GAACCAGGAAACGCCCGCGGUGAACACG.....AUG
env-192   ....................GACGA........CCCGCCGGGUU.GAACCAGGAAACGUCCGCAGUGAUCACG....AUG
env-193   ....................GACGA........CCCGCCGGGUU.GAACCAGGAAACGUCCGCAGUGAUCACG....AUG
Nje-1-1   ....................GUCGA........CCCGCUGGGUU.GAACCAGGAAACGCCCGCAGUGAUCACG....AUG
Nam-1-1   ....................GCCGG........CCCGCUGGGUU.GAACCAGGAAACGCCCGCAGUGAUCACG....AUG
Noc-1-1   ....................ACCGA........CCCGCUGGGUU.GAACCAGGAAACGCCCGCAGUGAUCACG....AUG
env-194   ....................GCGA..............GGGCU.GAACCAGGCAACG.....................
env-195   ....................GCC......GAGAUUGGAAAUCUCGCUACGGUUU.GAACCACGCAACGCCCGUGGUGAUAGACAU..AUG
env-196   ....................CCC..........GCGAUGGUUU.GAACCACGCAACGCCCGUGGUGAUACCA.....AUG
env-197   ....................ACCC.........GCGAUGGUUU.GAACCACGCAACGCCCGUGGUGAUAGCA.....AUG
env-199   ....................ACCC.........GCGAUGGUUU.GAACCACGCAACGCCCGUGGUGAUAACA.....AUG
env-201   ....................ACCC.........GCGAUGGUUU.GAACCACGCAACGCCCGUGGUGAUAACA.....AUG
env-202   ....................ACGU........CCCCGUGCGGUUU.GAACCACGCAACGCCCGUGGUGACAAUC...AUG
Hla-1-1   ....................CUUC...........GGGAUU.GAACCAGGAAACGCCCGCGGGUGAUAACC......AUG
Hva-1-1   ....................AGACC..........GCGACUU.GAACCACGCAACGCCCGUGGUGAAUACAU.....AUG
Hsi-1-1   ....................CGC..........GCGCGACUU.GAACCACGCAACGCCCGUGGUGAAUACAU.....AUG
Hma-1-1   ....................CGC..........GCGCGACUU.GAACCACGCAACGCCCGUGGUGAAUACAU.....AUG
Ham-1-1   ....................CGA..........GCGCGACUU.GAACCACGCAACGCCCGUGGUGAAUACAU.....AUG
env-203   ....................CGA..........GCGCGACUU.GAACCACGCAACGCCCGUGGUGAAUACAU.....AUG
Hca-1-1   ....................CGA..........GCGCGACUU.GAACCACGCAACGCCCGUGGUGAAUACAU.....AUG
Hsa-2-1   ....................CGA..........GCGCGACUU.GAACCACGCAACGCCCGUGGUGAAUACAU.....AUG
Hja-1-1   ....................CGA..........GCGCGACUU.GAACCACGCAACGCCCGUGGUGAAUACAU.....AUG
Har-1-1   ....................AAA...........CGCGACUU.GAACCACGCAACGCCCGUGGUGAAUACAU.....AUG
env-206   ....................GACCG.........CGCGACUU.GAACCACGCAACGCCCGAGGUGAAACCC......AUG
Hmu-2-1   ....................CGAU.........UCCGGCGUU.GAACCACGCAACGCCCGUGGUGAUACACAC....AUG

          >>>>>>>>>>>>>>>>>>>>...........>>>>>>>>>>>>>>>>>>>>>>>>>>>>>>>>>>>>>>>>>>>>>>>>>>>....>>..........
          001.111....11.1..1101...........00000...000.000...0000000.000000000..1.1..................22222....21..........
          --------------------●●●○------------------------○RRYU-GAACCR●RRR●●●●●●●●R●●●●●●●●○○----------AUG
                                                                      5'
```

```
Hka-1-1    ....................GGAU..................................................................UCCGGCGUU.GAACCACGCAACGCCCGUGGUGAUACACAC....................AUG
env-207    ....................GCGGUU..............................................................CCGCGGGGGUU.GAACCACGCAACGCCCGUGGUGAAACC.......................AUG
env-208    ....................GAGGAG..............................................GCGGCACCGCCG.CC.GAAACGGGAAUU.GAACCACACAACGCCCGUGGUGAAACC.......................AUG
env-210    ....................GUCGG...............................................................GGCUCGAGGAUU.GAACCAGCAAACGCCCGUGGUGAAACC.......................AUG
env-212    ....................CGUCGAA..............................................................GCUCGGGAAUU.GAACCACACAACGCCCGUGGUGAAACC.......................AUG
env-213    ....................GUCGG...............................................................GGCUCGGGAAUU.GAACCACACAACGCCCGUGGUGAAACC.......................AUG
env-214    ....................GAGA...................................................ACGGCCACAGCGCCGAUGGGGGUU.GAACCACACAACGCCCGUGGUGACACAU......................AUG
env-215    ....................GCAC..............................................................GGAUCCGGGAAUU.GAACCACACAACGCCCGUGGUGAAACU.......................AUG
env-216    ....................GGAU..............................................................GGCGUCGGGAAUU.GAACCACACGACGCCCGUGGUGAAACC.......................AUG
env-217    ....................GGC.................................................................GCUCGCGGGGUU.GAACCAUACAACGCCCGUGGUGAAACC.......................AUG
env-218    ....................GGC.................................................................GCUCGCGGGGUU.GAACCAUACAACGCCCGUGGUGAAACC.......................AUG
env-219    ....................GGC.................................................................GCUCGCGGGGUU.GAACCAUACAACGCCCGUGGUGAAACC.......................AUG
env-220    ....................GGC.................................................................GCUCGCGGGGUU.GAACCAUACAACGCCCGUGGUGAAACC.......................AUG
Nmo-1-1    ....................CGG......................................................................CGGAGUU.GAACCACACAACGCCCGUGGUGAAACU.......................AUG
env-221    ....................GAAG......................................................................GGGCU.GAACCAGCAAACGCCCGUGGUGAAACAG......................AUG
env-223    ....................GCGA......................................................................GGGCU.GAACCAGACAACGCCCGUGGUGAGACAG......................AUG
env-224    ....................ACGU......................................................................GGGCU.GAACCAACAAACGCCCGUGGUGGUGAGACCA...................AUG
env-226    ....................UUU...............................................................GGCGUCGAGAGUU.GAACCAUGCAACGCCCGUGGUGAAACC.......................AUG
env-227    ....................ACCA..................................................................CGCCGCCUU.GAACCACACAACGCCCGUGGUGAUACC.......................AUG
env-228    ....................AGAC.....................................................................GCGGCU.GAACCACGAAACGCCCGUGGUGAAACACG.....................AUG
env-229    ....................CACA......................................................................CGGCU.GAACCACCAAACGCCCGUGGUGAAACACG.....................AUG
Hci-1-1    ....................GUA..............................................................GAAUCCUGCCCGCU.GAACCACUCAACGCCCGUGGUGAUAGACC.....................AUG
Hut-1-1    ....................GGA.....................................................................GCCGGCU.GAACCACCCAACGCCCGCGGUGAAAGAU......................AUG
env-230    ....................GGAGA....................................................GGCGGC..ACGACCGCCGACCGGCU.GAACCACGCAACGCCCGCGGUGACAAC.......................AUG
env-231    ....................GCUC.....................................................................CCGGCU.GAACCACCCAACGCCCGUGGUGAAACAU......................AUG
env-232    ....................ACUC.....................................................................CCGGCU.GAACCACGCAACGCCCGUGGUGAAACAU......................AUG
env-233    ....................ACUC.....................................................................CCGGCU.GAACCACCCAACGCCCGUGGUGAAACAU......................AUG
env-234    ....................ACCGA..............................................................CUCGCACGGUUU.GAACCACUCGACGCCCGUGGUGAGACAAC.....................AUG
env-235    ....................ACAC.....................................................................GGAGUU.GAACCACUUGACGCCCGUGGUGAUAAACU.....................AUG
env-236    ....................ACGC......................................................................GGGUU.GAACCACCUGACGCCCGUGGUGAUAUAC......................AUG
Nba-1-1    ....................AUG.............................................................GCCCCGGUCAGGAUU.GAACCAGGAAACGCCCGCGGGAACAUACA.....................AUG
env-237    ....................AUG.............................................................GCCCCGGUCAGGAUU.GAACCAGGAAACGCCCGCGGGAACAUACA.....................AUG
Nsu-1-1    ....................AUG.............................................................GCCCCGGUCGGGAUU.GAACCAGAAAACGCCCGCGGGAACAUACA.....................AUG
Npa-1-1    ....................ACGC............................................................GCUCCGGUCGGGAUU.GAACCACGAAACGCCCGCGGGUGAAUACAG....................AUG
Nal-2-1    ....................GCGC............................................................GCCCCGGUCGGGAUU.GAACCACGAAACGCCCGCGGGUGAAUACAG....................AUG
Nga-1-1    ....................ACAC............................................................GCUCCGGUCGGGAUU.GAACCACGAAACGCCCGCGGGUGAAUACAG....................AUG
Hdj-1-1    ....................AUG.............................................................GCCCCGGUCGGGAUU.GAACCAGGAAACGCCCGCGGGUGAACAUAC....................AUG
Hli-3-1    ....................AUGU............................................................GCCCCGGUCGGGAUU.GAACCAGGAAACGCCCGCGGGUGAAUACG.....................AUG
Hje-2-1    ....................UCG.............................................................GCCCCGGUCGGGAUU.GAACCAGCAAACGCCCGCGGGUGAAUACA.....................AUG
Hth-2-1    ....................UUG.............................................................GCCCCGGUCGGGAUU.GAACCAGCAAACGCCCGCGGGUGAAUACA.....................AUG
Hsa-10-1   ....................GCCG............................................................GCCCCGGUCAGGAUU.GAACCAGGAAACGCCCGCGAGUGAUAAUAC....................AUG
env-238    ....................UGGUU....................................................................GGGAUU.GAACCACGAAACGCCCGCGGUGAUUACG......................AUG
env-239    ....................UGGUU....................................................................GGGAUU.GAACCACGAAACGCCCGCGGUGAUUACG......................AUG
Hni-1-1    ....................CCC...............................................................GCCCCGUUGGGAUU.GAACCAGGAAACGCCCGCGGUGAUACAU......................AUG
Nch-1-1    ....................CAC.....................................................................UCGGAUU.GAACCAGGAAACGCCCGCGGUGAUAAUCA.....................AUG
Nhu-1-1    ....................CGC.....................................................................UCGGAUU.GAACCAGGAAACGCCCGCGGUGACAAUCA.....................AUG
Nve-1-1    ....................UCG.............................................................GCCCCGGUCGGGAUU.GAACCAGCAAACGCCCGCGGAGAAUACA......................AUG
Hsa-8-1    ....................ACGGG.............................................................CCCCGGCCAGGAUU.GAACCAGGAAACGCCCGCGGUGAUACAC......................AUG
Htu-1-1    ....................ACCGC.............................................................CCCCGGUCGGGAUU.GAACCAGGAAACGCCCGCGGUGAUACAC......................AUG
env-243    ....................ACCGA.............................................................CCCCGGUCGGGAUU.GAACCAGGCAACGCCCGCGGUGAUAGCU......................AUG
Hca-3-1    ....................GCGA..................................................................CGGCGGUU.GAACCACGCAACGCCCGGCGGUGAACACCC....................AUG
env-244    ....................GGA.....................................................GACCUCC..........CGCGGUUU.GAACCACUCGACGCCCGUGGUGAUAGCCC.....................AUG
env-245    ....................GAGA.......................................................CGAG.CCGUCACCGGCGUGAUU.GAACCACUCAUUGCCCGCGGUGAUACCA......................AUG
env-246    ....................GGAG...................................................GCGGCACCG......CCGCCGGGUU.GAACCACGCAAGGCCCGCACGCCAACC.......................AUG
env-247    .................CCCGAGGC.......................................................................GAUU.GAACCAAGAACGCCCGUAGGUACACU........................AUG
env-248    .............CUGUCACCGA.....................................................................CGC.GAUU.GAACCAGGAAAGCCCGCAGGCAAACU........................AUG
env-249    .............UUGUCGCCGAGGC......................................................................GAUU.GAACCAGGAAAGCCCGCAGGCAAACU........................AUG
env-250    ....................UUCCG...................................................................UGAGGGUU.GAACCACACACGCCCGUGGUGACAGAUC......................AUG
env-251    ....................UCUCAGCCUAC..............................................................GAGGGUU.GAACCACUCACGCCCGUGGUGACAGAUU......................AUG
env-253    ....................GCGA......................................................................GGGUU.GAACCACGAACGCCCGUGGUGACAGAC.......................AUG
Hsu-1-1    ....................CUCGA...................................................................GCGGAUU.GAACCACCAAACGCCCGCGAGACAAUC.......................AUG
Hsp-2-1    ....................UGC...............................................................GUGCCUCGCGGAUU.GAACCACGCAACGCCCGCGGACAACAC.......................AUG
Has-1-1    ....................CUGA....................................................................CCGGUCU.GAACCAGGAACGCCCGCGGUGAUACAC.......................AUG
env-254    ....................ACGU............................................................CCUCGAUGC.GGUCU.GAACCACGAACGCCCGCGGUGAAACG........................AUG
Hag-1-1    ....................AUC..............................................................GGGCCACGC.GGCU.GAACCACCAACGCCCGCAGACAACAC........................AUG
Hsa-4-1    ....................CCC..............................................................GGACCACGC.GGCU.GAACCAUCAACGCCCGCAAACAACAC........................AUG
Hsa-5-1    ....................GGU..............................................................GGACCACGC.GGCU.GAACCAUCAACGCCCGCAGCAACC..........................AUG
env-255    ....................UUCAC.....................................................................GGCU.GAACCGUUAAGAAUCCGCGAGGUUAGCAA.....................AUG
env-256    ....................UUAC......................................................................GGCU.GAACCGUUAAGAAUCCGCGAGGUUAGCAA.....................AUG
env-262    ....................UUAC......................................................................GGCU.GAACCGUUAAGAAUCCGCGAGGUUAGCAA.....................AUG
env-266    ....................UUUGU.....................................................................GGCU.GAACCGUUAAGAAUCCGCGAGGUUAAGAA.....................AUG
Mae-1-1    ....................AAGA.......................................................................GCU.GAACCCAUAAACAACAAUUACAAUAACUUCCAACAACCAAUUAUAUUAUAUG
env-267    ....................UGAU..................................................................GGAACGUUU.GAACCACGAAACGCGAAUGCACAGGAGACACGAAACUA...........AUG
env-268    ....................AGAUG.....................................................................GUU.GAACCACAAAUGGAGGGGAGAUUCCACUAG....................AUG
env-269    ....................AGAUG.....................................................................GUU.GAACCACAAAUGGAGGGGAGAUUCCACUAG....................AUG
env-270    ....................GAAAAAAA...............................................................GCAGUU.GAACCAUAAAAUAGGUGAAUUUGUAA........................AUG
env-271    ....................GAAAAAG................................................................GCAGUU.GAACCAUAAAAUAGGUGAAUUUGUAA........................AUG
env-273    ....................GAAAAAG................................................................GCAGUU.GAACCAUAAAAUAGGUGAAUUUGUAA........................AUG
env-274    ....................ACAAA..................................................................AAGGCU.GAACCAUAAACUAAUUGAGAGGUGAAAAU.....................AUG
env-276    ....................ACAAA..................................................................AAGGCU.GAACCAUAAACUAAUUGAGAGGUGAAAAU.....................AUG
env-277    ....................GCAG......................................................................GGCU.GAACCGUAAACAAGUGUGAGGUGAAAA.......................AUG
env-278    ....................GCAG......................................................................GGCU.GAACCGUAAACAAGUGUGAGGUGAAAA.......................AUG
env-281    ....................UUAAU.....................................................................GGUU.GAACCGUAGACUUGAAGGAGAGUUGAGA......................AUG

           >>>>>>>>>>>>>>>>>>>>...........>>>>>>>>>>>>>>>>>>>>>>>>>>>>>>>>>>>>>>>>>>>>>>>>>>>>>>>>>>>>....>>.............................................
           001.111....11.1..1101...............00000...000.000...0000000.000000000..1.1..................22222....21.............................................
           --------------------●●●○------------------------------------------------------------------○RRYU-GAACCR●RRR●●●●●●●●R●●●●●●●●○○------------------AUG
                                                                                                                   3'
```

```
env-283  ....................UUCG..........................GGACU.GAACCAUAAGCAAACAAUACAGAUAGGAUGAAAAAUAU...........AUG
env-284  ....................UUCG..........................GGACU.GAACCAUAAGCAAACAAUACAGAUAGGAUGAAAAAUAU...........AUG
env-286  ....................UUCG..........................GGACU.GAACCAUAAGCAAACAAUACAGAUAGGAUGAAAAACAU...........AUG
env-287  ....................UUUG.........................GGGACU.GAACCGCAAACAGAAAAAAACAACAAAAAGUGAUUAAAC...........AUG
env-289  ....................AGUAC.........................GGGCU.GAACCGCAAACAUAAGAUUAGAGGUGAAAAAAAGAAAG............AUG
env-290  ....................UAU..........................UGGGCU.GAACCGUAAACUUAAGAUUAGAGGUGAAAAAAAGAAAG............AUG
env-291  ....................AAACA.........................GGGCU.GAACCGAAAACAUAAACUGAGAGGUGAAAAAGGAAAG............AUG
env-293  ....................AAC..........................GGGGCU.GAACCGUAAACUUAAGAUGUGAGGUGAAAGAAGAAAG............AUG
env-295  ....................UUCG..........................GGACU.GAACCGUAAGCAACAAAACACAUAGGAGAAAAAAAUUUAA..........AUG
env-296  ....................UUCG..........................GGACU.GAACCGUAAGCAACAAAACACAUAGGAGAAAAAAAUUUAA..........AUG
env-297  ....................GAAA...........................GGCU.GAACCGUAAACAAAUAAUACACAUAAUAAACAUAG..............GUG
env-298  ....................GAAA...........................GGCU.GAACCGUAAACAAUUAACACACAGGUGAAGAAACAAA............AUG
env-299  ....................GAAA...........................GGCU.GAACCGUAAACAAUUAACACACAGGUGAAGAAACAAA............AUG
env-301  ....................GAAA...........................GGCU.GAACCAUAAACAAAUAAUACUUAG.........................GUG
env-302  ....................GAGA...........................GGCU.GAACCAUAAACAAGUAAUACUUAGGUGAAAAAC................AUG
env-304  ....................CACGU.........................GGGCU.GAACCGUAAGCAUAAAACACAUUGGAGGUAAAAAAU.............AUG
env-308  ....................UAA...........................AGGUU.GAACCGUAAACGUAAAGGAGAAUUAGACA....................AUG
env-309  ....................UAA...........................AGGUU.GAACCGUAAACGUAAAGGAGGAUUUAGAAACGGCUUACAAGUAU.....UUG
env-310  ....................UAAA...........................GGUU.GAACCGUAAACGUAAAGGAGGAU..........................UUG
env-311  ....................CUUAGA.........................GGUU.GAACCGUAAGAAUAGGAGAAUUAGAGA......................AUG
env-312  ....................CAGA..........................GGGCU.GAACCACAAACAAAGCGAAUAGAUCACAACAGAGG..............AUG
env-317  ....................GCAG..........................GGGCU.GAACCACAAACAAAGCGAAUGAAUUACAGCAAAGG..............AUG
env-318  ....................GCAG..........................GGGCU.GAACCACAAACAAAGCGAAUGAAUUACAGCAAAGG..............AUG
env-319  ....................GCAG..........................GGGCU.GAACCACAAACAAAGCGAAUGAAUUACAGCAAAGG..............AUG
env-321  ....................CUCUGA.........................GGCU.GAACCACAAACAAAGUGAAUCUAUUACGACAAAGG..............AUG
env-322  ....................UUAG..........................GGGCU.GAACCACAAACAAAGUGAAUAUAUUACGACAAAGG..............AUG
env-323  ....................UUAG..........................GGGCU.GAACCACAAACAAAGCGAAUAUAUUACGACAAAGG..............AUG
env-324  ....................CAUAGA.........................GGCU.GAACCACAAACAAAGUGAAUAUAGUUACAGCAAAGG.............AUG
env-325  ....................CAUAGA.........................GGCU.GAACCACAAACAAAGUGAAUAUAGUUACAGCAAAGG.............AUG
env-326  ....................AUUA..........................GGGCU.GAACCACAAACAAAGCGAAUUUAAUCACUGCAAAGGAAGGCAA......AUG
env-327  ....................AUUA..........................GGGCU.GAACCACAAACAAAGCGAAUUAAAUCACGGCAAAGGAAGGCAA......UUG
env-328  ....................AUUA..........................GGGCU.GAACCACAAACAAAGCGAAUUUAAUCACGGCAAAGGAAGGCAA......UUG
Pab-1-1  ....................GUAAU..........................GGGG.GAACCACGAGGGAAGGAGGGAAAGAA.......................AUG
Psp-2-1  ....................GAGUU..........................GGGA.GAACCACAAACAGGAGGGAAAAGGG........................AUG
Tga-1-1  ....................CUCUG..........................CGGA.GAACCACAAACAAGGAAGGAGGGUGCAAG....................AUG
Tsp-3-1  ....................CACUG..........................CGGA.GAACCACAAACGUGGAAGGAGGGUGCAAG....................AUG
Tsp-2-1  ....................CUCUG..........................CGGA.GAACCACAAACGUGGGAGGGAAUUGAA......................AUG
Tna-1-1  ....................CUCUG..........................CGGA.GAACCACAAACGUGGGAGGGAAUUGAA......................AUG
Teu-1-1  ....................CUCUG..........................CGGA.GAACCACAAACAGGGAAGAGGUGAAAGAG....................AUG
env-329  ....................CUCUG..........................CGGA.GAACCACAAACAGCGAAAGAUGAGGUGGAAAAA................AUG
Tsp-4-1  ....................CUCUG..........................CGGA.GAACCACAAACAGCGAAAGAUGAGGUGGAAAAA................AUG
Tsi-1-1  ....................CUCUA..........................CGGA.GAACCAUAAACAGUGAAAGACGAGGUGAAAAAA................AUG
Tli-1-1  ....................CUCUG..........................CGGA.GAACCACAAACAGUGAAGAACGAGGUGAAAAAG................AUG
env-330  ....................CUCUG..........................CGGA.GAACCACAAACAGUGAAAGCGAGGUGAAGAAG..................AUG
Tba-1-1  ....................CUCUG..........................CGGA.GAACCACAAACAGCGAAAGCGAGGUGGAAGGA..................AUG
Tpa-1-1  ....................CUCUG..........................CGGA.GAACCACAAACAGCGAAAGCGAGGUGGAAAGA..................AUG
Ppa-1-1  ....................CUAAG..........................CGGA.GAACCAUAAACAGCAAAAACGAGGUGAAAAAG..................AUG
Pfu-2-1  ....................GGAU...........................GGGG.GAACCACAAACUAGGGAGGGAGAGCA........................AUG
Pho-1-1  ....................GGAU...........................GGGG.GAACCACAAACCAGGGAGGGAGAGCA........................AUG
Psp-1-1  ....................GAGCC..........................GGGG.GAACCACAAACUAGGGAGGGAGCAAG........................AUG
Pya-1-1  ....................GGAC...........................GGGA.GAACCACAAACGAGGAAGGGGUGAAAGGA....................AUG
Tko-1-1  ....................GGAA...........................GGGA.GAACCACAAACAGGGAAGAGGUGAACGAG....................AUG
Tsp-1-1  ....................GUAA...........................GGGA.GAACCACAAACAUGGAAGAGGUGAACGAG....................AUG
Pfe-1-1  ....................GUAA...........................GGGA.GAACCACAAACAUGGAAGGAGGGAACGAG....................AUG
Ton-1-1  ....................GUAA...........................GGGA.GAACCACAAACGAGGAAGGAGGGAGCAAG....................AUG
Tzi-1-1  ....................GGAA...........................GGGA.GAACCACAAACGGGGAAGAGGUGUGAAAG....................AUG
env-331  ....................ACAAGCAA.....................AGGGUU.GAACCAUAAACUAAGAAGGAGGAAUGCUUUAAA................AUG
env-332  ....................AUAAGCAA.....................AGGGUU.GAACCAUAAACUAAGAAGGAGGAAUGCUUUAAA................AUG
env-333  ....................AAACG.........................GGGUU.GAACCGCAAACAAAGGAGGAACAAUGAAU....................AUG
env-334  ....................GCAA...........................GGCU.GAACCAUAAACAACCGAACACACGUGAACUGAA................AUG
env-336  ....................AUAUA..........................GGCU.GAACCGUAAACUGUUGAGGAGUGAAACUAGAA..................AUG
env-338  ....................GCAA...........................GGCU.GAACCAUAAACAAGUGGCACAAGUGAACAAAG..................AUG
env-339  ....................GUUUA..........................GGUU.GAACCAUAAACUAGUGGAGGUGAGA........................AUG
env-341  ....................GAAA...........................GGUU.GAACCGUAAACUAGAAGCGAGGUGAAAAAGGAAA...............AUG
env-344  ....................GAAAA..........................GGUU.GAACCGUAAACUAGAAGCGAGGUGAAAAAGGAAA...............AUG
env-348  ....................GUAUA..........................GGCU.GAACCAUAAACCAAGAAAGCGAGGUGAAAAAA..................AUG
env-349  ....................GCAUA..........................GGCU.GAACCAUAAACCGAAAAGGCGAAGUGAAAAAA..................AUG
env-353  ....................GUAUA..........................GGCU.GAACCAUAAACUAAGAACGAGGUGAAAAA....................AUG
env-354  ....................GUAUA..........................GGCU.GAACCGAAAACUAGAAGCGAGGUGGAAAA....................AUG
env-356  ....................GUAUA..........................GGCU.GAACCAUAAACUAAAAGCGAGGUGGAAAA....................AUG
env-358  ....................GUAUA..........................GGCU.GAACCAUAAACCAAGAGAGCGAGGUGAAAAA..................AUG
env-359  ....................UUAA...........................GGCU.GAACCAUAAACUAAAAACGAGGUGAAAAA....................AUG
env-360  ....................UUAA...........................GGCU.GAACCAUAAACUAAAAACGAGGUGAAAAA....................AUG
env-362  ....................GCGUA..........................GGCU.GAACCAUAAACUAAAAGCGAGGUGGAAAA....................AUG
env-363  ....................GAGAA..........................GGCU.GAACCAUAAACUAACGUUGAGGUUGAAAGAAA..................UUG
env-364  ....................GUGCA..........................GGCU.GAACCGUAAACCAAAAGCGAGGUGAAAAUGA..................AUG
env-365  ....................GUACA..........................GGCU.GAACCGAAAACUAAAAGCGAGGUGGAAAA....................AUG
env-366  ....................UAUG...........................GGCU.GAACCAUAAACUAAAAGGCGAGGUAAAAAAA..................AUG
env-367  ....................UUUA...........................GGCU.GAACCAUAAACUGAAUGGCGAGAUGAAAAA...................AUG
env-369  ....................GUAUU..........................GGCU.GAACCAUAAACUAAAAGGCGAGGUGAAAAA...................AUG
env-370  ....................UAUA...........................GGCU.GAACCAUAAACUAAAUGGCGAGGUGAAAAA...................AUG
env-371  ....................UAUA...........................GGCU.GAACCAUAAACUAAAUGGCGAGGUGAAAAA...................AUG
env-375  ....................GCAUA..........................GGCU.GAACCAUAAACUAAAGAGUGAGGUGAAAAA...................AUG

         >>>>>>>>>>>>>>>>>>>>..........>>>>>>>>>>>>>>>>>>>>>>>>>>>>>>>>>>>>>>>>>>>>>>>>>>>....>>.......................................
         001.111....11.1..1101..........00000..000.000...0000000.000000000..1.1..................22222....21.......................................
         --------------------●●●○---------------------------------------------------○RRYU-GAACCR●RRR●●●●●●●●R●●●●●●●○○------------------AUG
                                                                                              3′
```

```
env-376   .....................GCGUA.....................................................................GGCU.GAACCAUAAACUAAAGAGUGAGGUGAAAAA...................AUG
env-377   .....................GCAUA.....................................................................GGCU.GAACCAUAAACUAAAGAGUGAGGUGAAAAA...................AUG
env-378   .....................GCAUA.....................................................................GGCU.GAACCAUAAACUAAAGAGUGAGGUGAAAAA...................AUG
env-379   .....................GCGUA.....................................................................GGCU.GAACCAUAAACUAAAGAGUGAGGUGAAAAA...................AUG
env-380   .....................GCGUA.....................................................................GGCU.GAACCAUAAACUAAAGAGUGAGGUGAAAAA...................AUG
env-381   .....................GCAGA.....................................................................GGCU.GAACCAUAAACUAAAGAGUGAGGUGAAAAA...................AUG
env-382   .....................GCAUA.....................................................................GGCU.GAACCAUAAACUAAAGAGUGAGGUGAAAAA...................AUG
env-383   .....................GCAUA.....................................................................GGCU.GAACCAUAAAGUAAAGAGUGAGGUGAAAAA...................AUG
env-384   .....................GCAUA.....................................................................GGCU.GAACCAUAAACUAAAGAGUGAGGUGAAAAA...................AUG
env-385   .....................UAA.....................................................................UAGGCU.GAACCAUAAACUAAGAGGCGAGGUGAGAGA...................AUG
env-386   .....................UAA.....................................................................UAGGCU.GAACCAUAAACUAAGAGGCGAGGUGAGAGA...................AUG
env-387   .....................UAA.....................................................................UAGGCU.GAACCAUAAACUAAGA....................................
env-388   .....................GUAAA.....................................................................GGCU.GAACCAUAAACUAGUGUGAGGUGAAUUAAA...................AUG
env-390   .....................GUAAA.....................................................................GGCU.GAACCAUAAACUAGUGUGAGGUGAAUAAA....................AUG
env-392   .....................GCAU......................................................................GGUU.GAACCAUAAACUAGUGCGAGGUGAAAAG.....................AUG
env-402   .....................GCAU......................................................................GGUU.GAACCAUAAACCAGUGCGAGGUGAAAAG.....................AUG
env-406   .....................GUUU......................................................................GGUU.GAACCAUAAACAAUAAUGAGGUGAAAAACAA..................AUG
env-408   .....................GUUU......................................................................GGUU.GAACCAUAAACAAUAAUGAGGUGAAAAACAA..................AUG
env-410   .....................AUUU......................................................................GGUU.GAACCAUAAACCGUGGGCGAGGUGAAAAAG...................UUG
env-412   .....................AUUU......................................................................GGUU.GAACCAUAAACUGUGGCGAGGUGAAAAAGG...................UUG
env-413   .....................AUUU......................................................................GGUU.GAACCAUAAACUGUGGCGAGGUGAAAAAGG...................UUG
env-414   .....................UCA......................................................................UGGUU.GAACCGUAAACUAAGCGAGGUGAAAAA......................AUG
env-416   .....................UCA......................................................................UGGUU.GAACCGUAAACUGAAGCGAGGUGAAAAA.....................AUG
env-418   .....................GUAA......................................................................GGUU.GAACUAUAAACUGGUGCGAGGUGAAAAG.....................GUG
env-420   .....................GUAA......................................................................GGUU.GAACCAUAAACUAGUGUGAGGUGAAAAA.....................GUG
env-422   .....................GUAA......................................................................GGUU.GAACCAUAAACUAGUGUGAGGUGAAAAA.....................GUG
env-424   .....................AAACC....................................................................GGGCU.GAACCGCAAACAUAAGAUGAGAGGUGAAAAAAGAAAG............AUG
env-425   .....................AGCAC....................................................................GGGCU.GAACCGUAAACGUAAGAUGAGAGGUGAAAAAAGAAAG............AUG
env-427   .....................AGCAC....................................................................GGGCU.GAACCGUAAACGUAAGAUGAGAGGUGAAAAAAGAAAG............AUG
env-428   .....................AAC.....................................................................UGGGCU.GAACCGUAAGCAUAAAAUGUGAGGUGAAAAAAGAAG.............AUG
env-429   .....................AGCAC....................................................................GGGCU.GAACCGUAAACAUUAGAUGUGAGGUGAAAAAAGAGAG............AUG
env-432   .....................AACAC....................................................................GGGCU.GAACCGCAAACAUAAGAUGUGAGGUGAAAAAAGAAAG............AUG
env-433   .....................AAACC....................................................................GGGCU.GAACCGCAAACAUAAGAUGAGAGGUGAAAAAAGAAAG............AUG
env-438   .....................AAUCAAAA................................................................AGGGUU.GAACCAUAAACUNAAAAGUGAUUAAAA......................AUG
env-439   .....................AAUCAAAA................................................................AGGGUU.GAACCAUAAACUGAAAAGUGAUUAAAA......................AUG
env-440   .....................AUA......................................................................AGGUU.GAACCAUAAACAGGAGAGCAGAUAGG.......................AUG
env-441   .....................UUA......................................................................AGGUU.GAACCACAAACUGGAGAAUGCAUAA........................AUG
env-442   .....................UUA......................................................................AGGUU.GAACCGUAAACUGGAGAAUACGUAAG.......................AUG
env-443   .....................UUA......................................................................AGGUU.GAACCGUAAACUGGAGAAUACGUAAG.......................AUG
env-444   .....................UAA......................................................................AGGUU.GAACCACAAACUGAGGAACCAUAA.........................AUG
env-446   .....................UUA......................................................................AGGUU.GAACCUCAAACUGAGGAAUUGGA..........................AUG
env-447   .....................UGA......................................................................AGGCU.GAACCGUUAAACCGUUGAGGAGGAAACAUA...................GUG
env-449   .....................GUAA......................................................................GGUU.GAACCGUUAAGCUGGAGAUCACUUAG.......................AUG
env-450   .....................GUAA......................................................................GGUU.GAACCGUUAAGCUGGAGAUCACUUAG.......................AUG
env-453   .....................GCAAU.....................................................................UGGG.GAAC.AAAAACGGGGAGGAGGGGAGAAAAAGGA...................
Mvo-1-1   .....................GAAU........................................................................GCCGAACCCGGAAACAAAAUAUUGACUACAGUGAGUGAUAAA..........AUG
env-454   .....................UUCU.....................................................................GGACU.GAACCUAGGUGAAAAUAAUGGUACGAUCAAUGU................AUG
ume-1-1   .....................AUAUUCAUCAUGACA......................................................GGAUAGGUU.GAACCAGGAAACACUAUUCAAAGGAGACACGAAACUA............AUG
CMe-2-1   .....................AUAA.....................................................................GGGCU.GAACCGUAAAACUUAGGUGAAUACAAUGAAAUCA...............AUG
Mev-1-1   .....................AAAUA...................................................................UGAGUU.GAACCAUGAAACUAAAAUGAAACCAGGUGAGAAAA..............UUG
env-455   .....................UAAA....................................................................AAAAGCCGAACCUUAGGUGAAUAUC...............................AUG
env-457   .....................UAAA....................................................................AAAAGCCGAACCUUAGGUGAAUAUC...............................AUG
env-460   .....................AAA....................................................................AAAAAGCCGAACCUCAGGUGAAUAUC...............................AUG
env-461   .....................AAA....................................................................AAAAAGCCGAACCUUAGGUGAAUAUC...............................AUG
env-462   .....................AGACC...................................................................UAAGCU.GAACCCUUGAAACUUUUAUCGGAGACGAAACUUA...............AUG
Mzh-1-1   .....................AAACC.................................................................GAUGAAUU.GAACCAUGAAACUAAGAUGAAACAAAGGUGACCAAA.............UUG
env-463   .....................UUU......................................................................UGGCU.GAACCGUAAAACUGCGGAUGGUGAGUGGG....................GUG
env-465   .....................AAAAAG.................................................................UCGGGCU.GAACCGUGAAAGGUGAUUAAAAG..........................AUG
env-466   .....................GUAA.....................................................................GGGCU.GAACCUUGAGGUGACACUAA.............................AUG
env-467   .....................AACA.....................................................................GAGCU.GAACCGUAAAACUUUAGGUGAAUACA.......................AUG
env-469   .....................UCUU...................................................................CCGGGCU.GAACCUGAAAGGUGUAAAAAUGGCACGAUCA..................AUG
Mta-1-1   .....................UCUU...................................................................CCGGGCU.GAACCUGAAAGGUGUAAAA..............................AUG
env-472   .....................GCGA.....................................................................GGGCU.GAACCGUAAAACCAUGGAGCAACGCC.......................AUG
env-473   .....................GAAA.....................................................................GGGCU.GAACCGUAAAACCAUGGAGCAAUGCC.......................AUG
env-474   .....................AUCC.....................................................................AAGCU.GAACCUUGAGGUGAACCGAUUCAAUGGCAAAAUCA..............AUG
Mpa-1-1   .....................AAU....................................................................UUGAGCU.GAACCUUGGUGAAGAUCAUGUCAAAAUCUAUGU................AUG
env-475   .....................UCA.....................................................................CAGGCU.GAACCUUGAGGUGUUAUAACAC...........................AUG
env-476   .....................UUA......................................................................CAGGCCGAACCUUGAGGUGUUAUAACAC...........................AUG
env-477   .....................GAAA.....................................................................CGUGCCGAACCUGAGGUGAAAUCCAUAUGGCAAAAUCA.................AUG
env-479   .....................UCUUU.....................................................................GGGCCGAACCUNAGGUGAUUACC...............................AUG
env-480   .....................CAAAAAA..................................................................GGGGCCGAACCUUAGGUGAUU.....................................
env-481   .....................CAAAAAA..................................................................GGGGCCGAACCUUAGGUGAUUACC...............................AUG
env-484   .....................CAAAAAA..................................................................GGGGCCGAACCUUAGGUGAUUACC...............................AUG
env-485   .....................CAAAAAA..................................................................GGGGCCGAACCUUAGGUGAUUACC...............................AUG
env-486   .....................GUUAAA....................................................................GAGCCGAACCUUAGGUGAUUAUC...............................AUG
env-487   .....................GUAAA.....................................................................GAGCCGAACCUUAGGUGAUUAUC...............................AUG
env-488   .....................CACGU.....................................................................GAGCCGAACCUUAGGAGAUCAAC...............................AUG
env-489   .....................CACGU.....................................................................GAGCCGAACCUUAGGAGAUCAAC...............................AUG
env-490   .....................UACGU.....................................................................GAGCCGAACCUUAGGUGAUUAUC...............................AUG
env-491   .....................CACGU.....................................................................GAGCCGAACCUUAGGUGAGUAUC...............................AUG
env-492   .....................CACGU.....................................................................GAGCCGAACCUUAGGUGAGUAUC...............................AUG
env-495   .....................UAA.....................................................................CAGAGCCGAACCUUAGGUGANUAUC...............................AUG
env-496   .....................GCAA.....................................................................UGAGCCGAACCUUAGGUGAUCACC...............................AUG

          >>>>>>>>>>>>>>>>>>>>>...........>>>>>>>>>>>>>>>>>>>>>>>>>>>>>>>>>>>>>>>>>>>>>>>>>>>>>>>>>>>>>>>>>>>>....>>...............................................
          001.111....11.1..1101...............00000..000.000...0000000.000000000..1.1...................22222....21...............................................
          ---------------------●●●○---------------------------------------------------------------------○RRYU-GAACCR●RRR●●●●●●●●R●●●●●●●○○---------------------AUG
                                                                                                               3′
```

```
env-497   ....................CCAA...........................UGAGCCGAACCUUACAGGUGAUCAGACUAUGGUUAAAUCG..............AUG
env-499   ....................GGAAA..........................UGAGCCGAACCUUAGGUGAUUACCAUGGUAAAAUCA..................AUG
env-500   ....................GCAA...........................UGAGCCGAACCUUAGGUGAUUACCAUGGUAAAAUCA..................AUG
env-502   ....................GCAA...........................UGAGCCGAAC.............................................
env-503   ....................GCAA...........................UGAGCCGAACCUUAGGUGAUUACCAUGGUAAAAUCG..................AUG
env-505   ....................GCAA...........................UGAGCCGAACCUUAGGUGAUUACCAUGGUAAAAUCG..................AUG
env-506   ....................UCUUU...........................GGGCCGAACCUUAGGUGAUUACC...............................AUG
env-507   ....................UCUUU...........................GGGCCGAACCUUAGGUGAUUACCAUGGUAAAAUCG..................AUG
env-508   ....................UCUUU...........................GGGCCGAACCUCAGGUGAUUACCAUGGUAAAAUCA..................AUG
env-509   ....................UCUUU...........................GGGCCGAACCUUAGGUGAUCACCAUGGUAAAAUCG..................AUG
env-511   ....................UCUUU...........................GGGCCGAACCUUAGGUGAUCACCAUGGUAAAAUCG..................AUG
env-512   ....................UCUUU...........................GGGCCGAACCUUAGGUGAUCACCAUGGUAAAAUCG..................AUG
env-513   ....................UCUUU...........................GGGCCGAACCUUAGGUGAUCACCAUGGUAAAAUCG..................AUG
env-514   ....................UCUUU...........................GGGCCGAACCUUAGGUGAUUACCAUGGUAAAAUCA..................AUG
env-515   ....................UCUUU...........................GGGCCGAACCUUAGGUGAUCACCAUGGUAAAAUCG..................AUG
env-516   ....................UCUUU...........................GGGCCGAACCUUAGGUGAUCACC...............................AUG
env-517   ....................AUAU..........................CAGGCU.GAACCUUUGGAGAGAUC................................AUG
env-522   ....................AUUG.........................GCAAGCC.GAACCUGGAGCAUAAU.................................AUG
env-525   ....................AAUUCA.......................GUAAGCC.GAACCUGGAGAAUAAC.................................AUG
env-528   ....................UUACC...........................GGCU.GAACCUUGAGGUGUCAAUAACACAUGGUAAAAUCA.............AUG
env-529   ....................UUACC...........................GGCU.GAACCUUGAGGUGUCAAUAACACAUGGUAAAAUCA.............AUG
env-530   ....................UUACC...........................GGCU.GAACCUUGAGGUGUCAAUAACACAUGGUAAAAUCA.............AUG
env-531   ....................UUACC...........................GGCU.GAACCUUGAGGUGUCAAUAACACAUGGUAAA...................
env-532   ....................UUACUU..........................GGGCCGAACCUUGAGGUGUUAUAACACAUGGUAAAAUCG..............AUG
CMe-1-1   ....................GUCAU...........................GGGCCGAACCUUGAGGUGUUAUCGCACAUGGUAAAAUCG..............AUG
env-534   ....................UUACGU..........................GGGCCGAACCUUGAGGUGUUAUAACACAUGGUAAAAUCG..............AUG
env-535   ....................GCGAU...........................GGGCCGAACCUUGAGGUGUUAUAACAGAUGGUAAAAUCG..............AUG
env-536   ....................AAAA...........................GAGCU.GAACCUGGAGCAACUAUGUCACGAUCU......................AUG
env-537   ....................AUAC...........................GAGCU.GAACCUGGAGCAACUAUGUCACGAUCA......................AUG
env-538   ....................AUU..........................CAGGCU.GAACCUGGUGAACCGC..................................AUG
env-539   ....................AUU..........................CAGGCU.GAACCUGGUGAACCGC..................................AUG
env-540   ....................AUU..........................CAGGCU.GAACCUGGUGAACCGC..................................AUG
env-541   ....................GAAAA...........................AGGCU.GAACCUGGAGAAUUUAUGUCAAAAUCGAUGUAUUCGU..........AUG
env-542   ....................UCU..........................GACGCU.GAACCUGAUGGUGAUCUACCAUGGCGAAAUCGAUGU............AUG
env-543   ....................UUC.........................AGACGCU.GAACCUGAGGCUGAUUGUCUAUGGCGAAAUCGAUGU............AUG
env-544   ....................AAAAA........................UAGGCU.GAACCUUGGAGAUAUC..................................AUG
env-545   ....................UUCAU...........................CCGCU.GAACCUCAGGUGAUCGCA................................AUG
env-546   ....................UUCAU...........................CCGCU.GAACCUCAGGUGAUCGCA................................AUG
Mli-1-1   ....................GAAACCA.........................GCGCU.GAACCUGAAGGUGAUUAUCAAUGGCAAAGUCAAUGU............AUG
env-547   ....................GAAA.............................CGCU.GAACCUGAGGUGAUUUGAAAAAUGGCAAAAUCGAUGU...........AUG
Mse-1-1   ....................GAAA.............................CGCU.GAACCUGAGGUGAUUCUAAACAUGGCAAAAUCGAUGU...........AUG
Mma-1-1   ....................GAAA.............................CGCU.GAACCUGAGGUGAUUCUAAACAUGGCAAAAUCGAUGU...........AUG
Msp-5-1   ....................GAAA.............................CGCU.GAACCUGAGGUGAUUCUAAACAUGGCAAAAUCGAUGU...........AUG
env-552   ....................GAAA.............................CGCU.GAACCUGAGGUGAUUCUAAACAUGGCAAAAUCGAUGU...........AUG
env-553   ....................GAAA.............................CGCU.GAACCUGAGGUGAUUUGUGAAUGACAAAAUCG.................AUG
env-559   ....................GAAA.............................CGCU.GAACCUGAGGUGAUUUGUGAAUGACAAAAUCG.................AUG
env-560   ....................GAAA.............................CGCU.GAACCUGAGG.........................................
env-561   ....................GAAA.............................CGCU.GAACCUGAGGUGAUUUGUGAAUGACAAAAUCG.................AUG
env-563   ....................GAAA.............................CGCU.GAACCUGAGGUGAUUUGUGAAUGACAAAAUCG.................AUG
env-564   ....................GAAA.............................CGCU.GAACCUGAGGUGAUUGAGAGCAUGGCAAAAUCGAUGU...........AUG
env-565   ....................GAAA.............................CGCU.GAACCUGAGGUGAUUGAGAGCAUGGCAAAAUCGAUGU...........AUG
env-567   ....................GAAA.............................CGCU.GAACCUGAGGUGAUUGAGAGCAUGGCAAAAUCGAUGU...........AUG
env-568   ....................GAAA.............................CGCU.GAACCUGAGGUGAUUGAGAGCAUGGCAAAAUCGAUGU...........AUG
env-570   ....................GCAA.............................CGCU.GAACCUGAGGUGAUUCUAAACAUGGCAAAAUCGAUGU...........AUG
env-571   ....................GAAA.............................CGCU.GAACCUGAGGUGAUAAAAGAAUGGCAAAGUCAAUGU............AUG
env-574   ....................GGAA.........................CACGCU.GAACCUGGAUGGUGACAGAGCAUGNCGAAAUCGAUGU............AUG
env-575   ....................GCAA.........................CACGCU.GAACCUGGAUGGUGACAGAGUAUGUCGAAAUCGAUGU............AUG
env-576   ....................GCAA.........................CACGCUUGAACCUGGAUGGUGACAGAGUAUGUCGAAAUCGAUGU............AUG
env-577   ....................GGAA.........................CACGCU.GAACCUGGAUGUGACAGAGC..............................AUG
env-578   ....................UCUAC........................UGAGCU.GAACCUGAAGGUGAAGAAAU..............................AUG
env-579   ....................UCCAUU...........................GAGCU.GAACCUGAGGUGAUUUGUUAUGGCAAAAUCGAUGU............AUG
env-580   ....................CCAU.........................UGAGCU.GAACCUGAGGUGAUUUGUAAAUGGCAAAAUCG..................AUG
env-581   ....................UCUU.........................CGGGCU.GAACCUGUAGGUGCAAGAAUC.............................AUG
env-585   ....................UUCCAU.......................CGGGCU.GAACCUGAAAGGUGUGAGAA..............................AUG
env-587   ....................UCUUU........................CGAGCU.GAACCUGAAAGGUGCAAGAACC.............................AUG
env-588   ....................UCUUU........................CGAGCU.GAACCUGAAAGGUGCAAGAACC.............................AUG
env-589   ....................GGAAAAAAAG..................GCGGCUG..AACCGUGAAACUCUCUUAUUAAAG........................GUG
env-590   ....................UAAUAACGU.......................CAGCU.GAACCGUGAAACUGUAUAAAAUUGGUGAAACAACA.............AUG
env-591   ....................UAAUAACGU.......................CAGCU.GAACCGUGAAACUGUAUAAAAUUGGUGAAACAACA.............AUG
env-594   ....................UAAAAGCAU.....................GCGGCU.GAACCGUGAAACAAUCAUAAAUUGGUGAAACACA...............AUG
env-595   ....................AAAAACA.......................GUAGCU.GAACCGUGAAACUAUAAACUAUUGGUGAAACAAA...............AUG
env-596   ....................AAAAACA.......................GUAGCU.GAACCGUGAAACUAUAAACUAUUGGUGAAACAAA...............AUG
env-597   ....................AAAAACA.......................GUAGCU.GAACCGUGAAACUAUAAACUAUUGGUGAAACAAA...............AUG
env-598   ....................AAAAACA.......................GUAGCU.GAACCGUGAAACU.......................................
env-599   ....................AAAAACA.......................GUAGCU.GAACCGUGAAACUAUAAACUAUUGGUGAAACAAA...............AUG
env-602   ....................AAAAACA.......................GUAGCU.GAACCGUGAA..........................................
env-603   ....................GAAAAAAAAG....................GCGGCU.GAACCGUGAAACUCUCUUAUUAAAGGUGAAACAAA...............AUG
env-604   ....................GGAAAAAAAG....................GCGGCU.GAACCGUGAAACUCUCUUAUUAAAGGUGAAACAAA...............AUG
env-605   ....................GGAAAAAAAG....................GCGGCU.GAACCGUGAAACUCUCUUAUUAAAGGUGAAACAAA...............AUG
env-606   ....................AGAAAAAAAG....................GCGGCU.GAACCGUGAAAC........................................
env-607   ....................AGAAAAAAAG....................GCGGCU.GAACCGUGAAA.........................................
env-608   ....................AGAAAAAAAG....................GCGGCU.GAACCGUGAAACUCUCUU..................................
env-609   ....................AGAAAAAAAG....................GCGGCU.GAACCGUGAAAC........................................

          >>>>>>>>>>>>>>>>>>>>..........>>>>>>>>>>>>>>>>>>>>>>>>>>>>>>>>>>>>>>>>>>>>>>>>>>>....>>.................
          001.111....11.1..1101..........00000...000.000...0000000.000000000..1.1.................22222....21..........
          --------------------●●●○------------------------------------○RRYU-GAACCR●RRR●●●●●●●●R●●●●●●●●○○-----------AUG
                                                                                           3'
```

```
env-610   .....................GAUAAAAG.......................GCGGCU.GAACCGUGAAACUCUAUUAUUAUAGGUGAAACAAA..............AUG
env-611   .....................AGAAAAAAA.....................GGCGGCU.GAACCGUGAAACUCUCUUACUAAAGGUGAAACAAA..............AUG
env-612   .....................AGAAAAAAAA....................GGCGGCU.GAACCGUGAAACUCUCUUAUUUAAAGGUGAAACAAA.............AUG
env-613   .....................AGAAAAAAAA....................GGCGGCU.GAACCGUGAAACUCUCUUAUUUAAAGGUGAAACAAA.............AUG
env-616   .....................ACA............................UAGCU.GAACCGUGAAACCAUGAUGGAGAAUCGAAAC..................AUG
Mco-1-1   .....................ACA............................UAGCU.GAACCGUGAAACCAUGAUGGAGAAUCGAAAC..................AUG
env-617   .....................AAAGA........................AAAGAUU.GAACCAUGAAACUAAAAAUGGAACAGGUGAAUAAUU.............UUG
Mps-1-1   .....................UUG............................AAGCU.GAACCGUGAAACAAAGAAAUGAAACAAGGUGAACGUAU...........UUG
env-619   .....................UUG............................AAGCU.GAACCGUGAAACACAGAAAUGAAACAAGGUGAAUAUAU...........UUG
env-620   .....................UUG............................AAGCU.GAACCGUGAAACACAGAAAUGAAACAAGGUGA.................AUG
Mti-1-1   .....................UAAUAAUUAGAAGC...............CAAAAGCUG..AACCGUGAAACUAGAAAUGAAACUUAG....................GUG
env-621   .....................UGAUAU........................AAGCUG..AACCGUGAAACAAGAAGAUGAAACAAG....................GUG
Mba-2-1   .....................CAGG.........................CAGGGCUG..AACCGUGAAACAUAAGUGAAACUUG......................GUG
env-622   .....................CAGA.........................CAGGGUU.GAACCGUGAAACAUAAAUGAAACUUGGUGUUUUUAA..............UUG
env-623   .....................CAGG.........................CAGGGCU.GAACCGUGAAACAAAGUGAAACUUG........................GUG
env-624   .....................CAGG.........................CAGGACU.GAACCGUGAAACGUAAGUGAAACUUG.......................GUG
env-625   .....................AGG.........................GCAGGGCU.GAACCAUGAAACAAAAGUGAAACUUG.......................GUG
Mho-2-1   .....................CUGG.........................CAGGGCU.GAACCGUGAAACAUAAGUGAAACUUG.......................GUG
env-628   .....................CUGG.........................CAGGGCU.GAACCGUGAAACAUAAGUGAAACCUG.......................GUG
env-629   .....................CAAG.........................CAGGGCU.GAACCGUGAAACAUAAGUGAAACUUG.......................GUG
env-630   .....................AGG.........................GCAGGGCU.GAACCGUGAAACAAAAGUGAAACUUG.......................GUG
Mth-1-1   .....................AGG.........................GCAGGGCU.GAACCGUGAAACAAAAGUGAAACUUG.......................GUG
Msp-13-1  .....................CAGG.........................CAGGGCU.GAACCGUGAAACAAAAGUGAAACCUG.......................GUG
Mva-1-1   .....................CAGG.........................CAGGGCU.GAACCGUGAAACAAAAGUGAAACCUG.......................GUG
Mba-4-1   .....................CUGG.........................CAGGGCU.GAACCGUGAAACAAAAGUGAAACCUG.......................GUG
env-631   .....................CAGG.........................CAGGGCU.GAACCGUGAAACAAAAGUGAAACCUG.......................GUG
env-633   .....................CAGG.........................CAGGGCU.GAACCGUGAAACAUAAGUGAAACUUG.......................GUG
Mla-1-1   .....................CAGG.........................CAGGGCU.GAACCAUGAAACAUAAGUGAAACUUG.......................GUG
Mba-5-1   .....................CAGG.........................CAGGGCU.GAACCGUGAAACAAAAGUGAAACUUG.......................GUG
Msp-7-1   .....................CAGG.........................CAGGGCU.GAACCAUGAAACAUAAGUGAAACUUG.......................GUG
Msp-16-1  .....................CAGG.........................CAGGGCU.GAACCGUGAAACAUAAGUGAAACUUG.......................GUG
env-634   .....................AAG.........................GCAGGGCU.GAACCGUGAAACAAAAGUGAAACUUG.......................GUG
Mso-1-1   .....................CUGG.........................CAGGGCUG.AACCGUGAAACAUAAGUGAAACUUG.......................GUG
Mma-3-1   .....................CUGG.........................CAGGGCUG.AACCGUGAAACAUAAGUGAAACUUG.......................GUG
Mac-1-1   .....................CAGG.........................CAGGGCUG.AACCGUGAAACAAAAGUGAAACUUG.......................GUG
Msi-2-1   .....................CAGG.........................CAGGGCUG.AACCGUGAAACAAAAGUGAAACUUG.......................GUG
env-635   .....................CAGG.........................CAGGGCU.GAACCGUGAAGCAAAAAGUGAAACUUG......................GUG
Msi-1-1   .....................CAGG..........................CAGGGUUGAACCGUGAAACAAAAGUGAAACUUGGUGUUUAAA..............UUG
env-636   .....................CAGG..........................CAGGGUUGAACCGUGAAACAAAAGUGAAACUUGGUGUUUAAA..............UUG
Msp-14-1  .....................AGACC.........................CGGGCU.GAACCGUGAAACAAGAAGUGAAACUUGGUGUUAUAAAA...........UUG
env-637   .....................GACA...........................GGACU.GAACCGUGAAACUGAAAAUGAAACAG.......................GUG
env-638   .....................GACA...........................GGACU.GAACCGUGAAACUGAAAAUGAAACAG.......................GUG
env-639   .....................GAUA..........................GGACUG..AACCGUGAAACUUAAAAUGAAACAG.......................GUG
env-640   .....................GGAAAAA........................GGCUG..AACCGUGAAACUAAAAAUGAAACAAG......................GUG
env-641   .....................AAA............................UGGGUUGAACCGUGAAACUGAAAAUGAAACAAGGUGAACAUA............UUG
Mbu-1-1   .....................AAC.........................UUCAGCUG.AACCGUGAAACUAACAUGAAACAAGGUGAAUAAA..............UUG
env-642   .....................AAC.........................UUCAGCU.GAACCGUGAAACUAACAUGAAACAAG.......................GUG
Mme-2-1   .....................UAA........................CUUUGGCU.GAACCGUGAAACUAACAUGAAACUAAGGUGAAAAAU............UUG
Mme-1-1   .....................UAA........................CUUUGGCU.GAACCGUGAAACUAACGUGAAACAAGGUGAAUAAA.............UUG
Msh-1-1   .....................ACU............................GAGCU.GAACCAUAAAGCAGAGAUGGAGGCACUGU...................AUG
env-643   .....................UUAA............................AGCU.GAACGAGAGGUUAUAAAAUG............................AUG
env-644   .....................GUAAA............................GCU.GAACGAGAGGUUAUUAAAAUG...........................AUG
env-645   .....................GAAA.............................GCU.GAACGAAAAGAGGUGUUUUAA...........................UUG
env-646   .....................GAAG.............................GCU.GAACAAAAAGAGGUGUUUUAA...........................UUG
env-647   .....................GAGAA...........................GGCU.GAACAAAAAGGUGUUUAUU.............................AUG
env-650   .....................UUA.............................AGCU.GAACAAGAGGUGUUAAU...............................UUG
env-651   .....................GUAAA............................GCU.GAACAGAUAGAGGUGUAAUUA...........................AUG
env-656   .....................CUAGU...........................GGCU.GAACAAAAAGGUGUUAUUU.............................AUG
env-657   .....................CUCGU...........................GGCU.GAACAAAAAGGUGUUAUUU.............................AUG
env-658   .....................CUCGU...........................GGCU.GAACAAAAGGUGUUAUU...............................AUG
Mve-1-1   .....................GCAA.............................GCU.GAACAAAAGAGGUGUUUUA.............................AUG
env-659   .....................GCAA.............................GCU.GAACAAAAGAGGUGUUUUA.............................AUG
env-660   .....................GCAA.............................GCU.GAACAAAAGAGGUGUUUUA.............................AUG
Msp-4-1   .....................GAAA.............................GCU.GAACAAAAGAGGUGUUUUA.............................AUG
env-661   .....................GAAA.............................GCU.GAACAAAAGAGGUGUAUAUUAA..........................AUG
env-663   .....................GAAA.............................GCU.GAACAAAAGAGGUG..................................
env-664   .....................GAAA.............................GCU.GAACAAAAGAGGUGUAUAUUAA..........................AUG
env-665   .....................GAAA.............................GCU.GAACAAAAAGAGG...................................
env-670   .....................GAAA.............................GCU.GAACAAAAAGAGGUGUAUUA............................AUG
env-671   .....................GAAA.............................GCU.GAACAAAAAGAGGUGUAUUUA...........................AUG
env-672   .....................GAAA.............................GCU.GAACAAAAAGAGGUGUAUUUA...........................AUG
Mfo-2-1   .....................GAAA.............................GCU.GAACAAAAAGAGGUGUAUUUA...........................AUG
env-673   .....................GAAA.............................GCU.GAACAAAAAGAGGUGUAUUUA...........................AUG
env-674   .....................GAAA.............................GCU.GAACAAAAAGAGGUGUAUAUUAA.........................AUG
Msp-2-1   .....................GAAA.............................GCU.GAACAAAAAGAGGUGUAUAUUAA.........................AUG
Mfo-1-1   .....................GAAA.............................GCU.GAACAAAAAGAGGUGUAUAUUAA.........................AUG
env-675   .....................GCAA.............................GCU.GAACAAAGAGGUGUUUUUA.............................AUG
env-676   .....................GAAA.............................GCU.GAACAAAGAGGUGUUUAA..............................AUG
Msp-1-1   .....................GCAA.............................GCU.GAACAAAGAGGUGUAAUUUA............................AUG
env-679   .....................GCAA.............................GCU.GAACAAAGAGGUGUAAUUUA............................AUG
env-680   .....................GCAA.............................GCU.GAACAAAGAGGUGUUAAUUA............................AUG
Msp-3-1   .....................GCAA.............................GCU.GAACAAAGAGGUGUUAAGUA............................AUG
env-681   .....................GCAA.............................GCU.GAACAAAGAGGUGUUAAUUA............................AUG

          >>>>>>>>>>>>>>>>>>>>>..........>>>>>>>>>>>>>>>>>>>>>>>>>>>>>>>>>>>>>>>>>>>>>>>>>>>>>>>....>>..........................
          001.111....11.1..1101..........(...00000..000.000...0000000.000000000..1.1.................22222....21..........................
          --------------------●●●○---------------------------------------------------○RRYU-GAACCR●RRR●●●●●●●●R●○○○○○○○○----------AUG
                                                                                                     3′
```

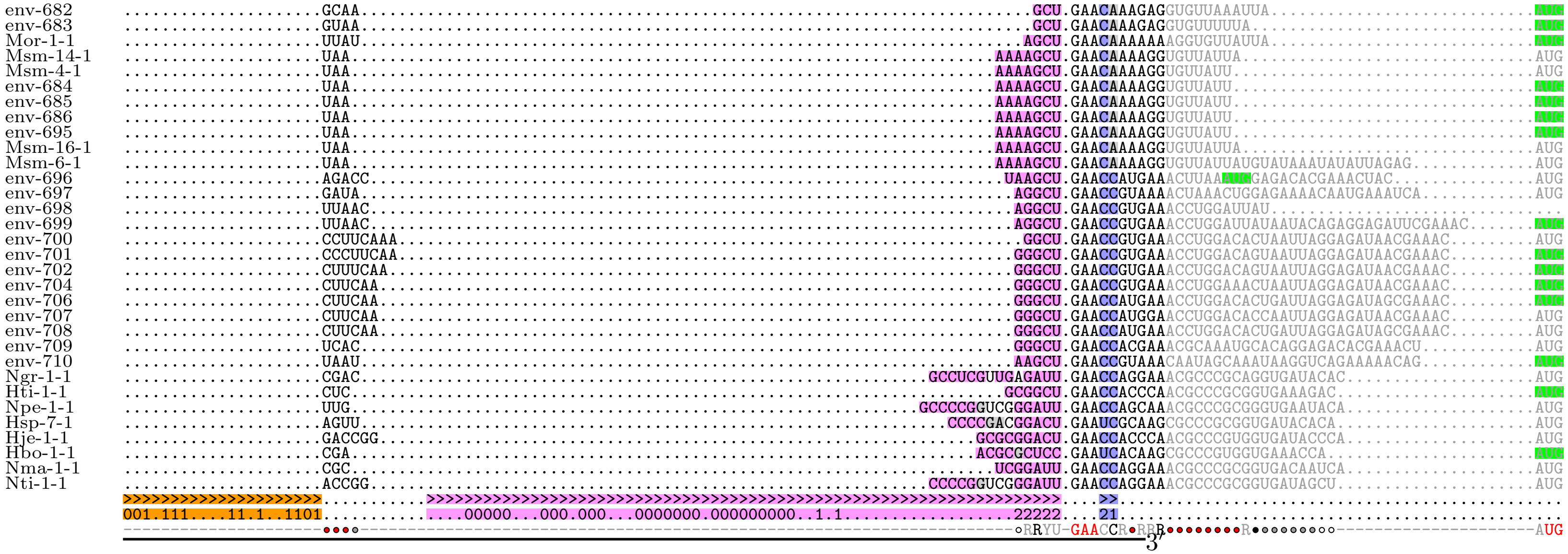

### 5 L17-Actino-Proteobacteria

#### 5.1 Taxa

The taxonomy of each organism containing a putative L17-Actino-Proteobacteria RNA is listed, with abbreviations identifying each hit (e.g., "Eco-1-1" and "Eco-1-2" might hypothetically represent two distinct RNAs in *E. coli*). The abbreviations will be used to identify each individual L17-Actino-Proteobacteria RNA in Sections 5.3 and 5.5:.

| **abbrev. of hits** | **taxonomy of species** |
|---|---|
| Aal-1-1 | Bacteria Actinobacteria Actinobacteridae Actinomycetales Propionibacterineae Nocardioidaceae *Actinopolymorpha alba* DSM 45243 |
| Tbi-1-1 | Bacteria Actinobacteria Actinobacteridae Actinomycetales Pseudonocardineae Pseudonocardiaceae *Thermobispora bispora* DSM 43833 |
| Sac-1-1 | Bacteria Actinobacteria Actinobacteridae Actinomycetales Streptomycineae Streptomycetaceae *Streptomyces acidiscabies* 84-104 |
| Saf-1-1 | Bacteria Actinobacteria Actinobacteridae Actinomycetales Streptomycineae Streptomycetaceae *Streptomyces afghaniensis* 772 |
| Sal-1-1 | Bacteria Actinobacteria Actinobacteridae Actinomycetales Streptomycineae Streptomycetaceae *Streptomyces albulus* CCRC 11814 |
| Sal-2-1 | Bacteria Actinobacteria Actinobacteridae Actinomycetales Streptomycineae Streptomycetaceae *Streptomyces albus* J1074 |
| Sau-1-1 | Bacteria Actinobacteria Actinobacteridae Actinomycetales Streptomycineae Streptomycetaceae *Streptomyces aurantiacus* JA 4570 |
| Sau-2-1 | Bacteria Actinobacteria Actinobacteridae Actinomycetales Streptomycineae Streptomycetaceae *Streptomyces auratus* AGR0001 |
| Sav-1-1 | Bacteria Actinobacteria Actinobacteridae Actinomycetales Streptomycineae Streptomycetaceae *Streptomyces avermitilis* MA-4680 |
| Sbi-1-1 | Bacteria Actinobacteria Actinobacteridae Actinomycetales Streptomycineae Streptomycetaceae *Streptomyces bingchenggensis* BCW-1 |
| Sbo-1-1 to Sbo-1-2 | Bacteria Actinobacteria Actinobacteridae Actinomycetales Streptomycineae Streptomycetaceae *Streptomyces bottropensis* ATCC 25435 |
| Sca-1-1 | Bacteria Actinobacteria Actinobacteridae Actinomycetales Streptomycineae Streptomycetaceae *Streptomyces canus* 299MFChir4.1 |
| Sca-2-1 to Sca-2-2 | Bacteria Actinobacteria Actinobacteridae Actinomycetales Streptomycineae Streptomycetaceae *Streptomyces cattleya* NRRL 8057 = DSM 46488 |
| Sch-1-1 | Bacteria Actinobacteria Actinobacteridae Actinomycetales Streptomycineae Streptomycetaceae *Streptomyces chartreusis* NRRL 12338 |
| Scl-1-1 to Scl-1-2 | Bacteria Actinobacteria Actinobacteridae Actinomycetales Streptomycineae Streptomycetaceae *Streptomyces clavuligerus* ATCC 27064 |
| Sco-1-1 | Bacteria Actinobacteria Actinobacteridae Actinomycetales Streptomycineae Streptomycetaceae *Streptomyces coelicoflavus* ZG0656 |
| Sco-2-1 | Bacteria Actinobacteria Actinobacteridae Actinomycetales Streptomycineae Streptomycetaceae *Streptomyces coelicolor* A3(2) |
| Sco-3-1 | Bacteria Actinobacteria Actinobacteridae Actinomycetales Streptomycineae Streptomycetaceae *Streptomyces collinus* Tu 365 |
| Sda-1-1 | Bacteria Actinobacteria Actinobacteridae Actinomycetales Streptomycineae Streptomycetaceae *Streptomyces davawensis* JCM 4913 |
| Sfl-1-1 | Bacteria Actinobacteria Actinobacteridae Actinomycetales Streptomycineae Streptomycetaceae *Streptomyces flavogriseus* ATCC 33331 |
| Sfu-1-1 | Bacteria Actinobacteria Actinobacteridae Actinomycetales Streptomycineae Streptomycetaceae *Streptomyces fulvissimus* DSM 40593 |

| | |
|---|---|
| Sga-1-1 | Bacteria Actinobacteria Actinobacteridae Actinomycetales Streptomycineae Streptomycetaceae *Streptomyces gancidicus* BKS 13-15 |
| Sgh-1-1 | Bacteria Actinobacteria Actinobacteridae Actinomycetales Streptomycineae Streptomycetaceae *Streptomyces ghanaensis* ATCC 14672 |
| Sgl-1-1 | Bacteria Actinobacteria Actinobacteridae Actinomycetales Streptomycineae Streptomycetaceae *Streptomyces globisporus* C-1027 |
| Sgr-1-1 | Bacteria Actinobacteria Actinobacteridae Actinomycetales Streptomycineae Streptomycetaceae *Streptomyces griseoaurantiacus* M045 |
| Sgr-2-1 | Bacteria Actinobacteria Actinobacteridae Actinomycetales Streptomycineae Streptomycetaceae *Streptomyces griseoflavus* Tu4000 |
| Sgr-4-1 | Bacteria Actinobacteria Actinobacteridae Actinomycetales Streptomycineae Streptomycetaceae *Streptomyces griseus subsp. griseus* NBRC 13350 |
| Sgr-3-1 | Bacteria Actinobacteria Actinobacteridae Actinomycetales Streptomycineae Streptomycetaceae *Streptomyces griseus* XylebKG-1 |
| Shy-1-1 | Bacteria Actinobacteria Actinobacteridae Actinomycetales Streptomycineae Streptomycetaceae *Streptomyces hygroscopicus* ATCC 53653 |
| Shy-2-1 | Bacteria Actinobacteria Actinobacteridae Actinomycetales Streptomycineae Streptomycetaceae *Streptomyces hygroscopicus subsp. jinggangensis* 5008 |
| Shy-3-1 | Bacteria Actinobacteria Actinobacteridae Actinomycetales Streptomycineae Streptomycetaceae *Streptomyces hygroscopicus subsp. jinggangensis* TL01 |
| Sip-1-1 | Bacteria Actinobacteria Actinobacteridae Actinomycetales Streptomycineae Streptomycetaceae *Streptomyces ipomoeae* 91-03 |
| Sli-1-1 | Bacteria Actinobacteria Actinobacteridae Actinomycetales Streptomycineae Streptomycetaceae *Streptomyces lividans* 1326 |
| Sli-2-1 to Sli-2-2 | Bacteria Actinobacteria Actinobacteridae Actinomycetales Streptomycineae Streptomycetaceae *Streptomyces lividans* TK24 |
| Smo-1-1 | Bacteria Actinobacteria Actinobacteridae Actinomycetales Streptomycineae Streptomycetaceae *Streptomyces mobaraensis* NBRC 13819 = DSM 40847 |
| Sni-1-1 | Bacteria Actinobacteria Actinobacteridae Actinomycetales Streptomycineae Streptomycetaceae *Streptomyces niveus* NCIMB 11891 |
| Spr-1-1 | Bacteria Actinobacteria Actinobacteridae Actinomycetales Streptomycineae Streptomycetaceae *Streptomyces pristinaespiralis* ATCC 25486 |
| Spr-2-1 | Bacteria Actinobacteria Actinobacteridae Actinomycetales Streptomycineae Streptomycetaceae *Streptomyces prunicolor* NBRC 13075 |
| Spu-1-1 | Bacteria Actinobacteria Actinobacteridae Actinomycetales Streptomycineae Streptomycetaceae *Streptomyces purpureus* KA281 |
| Sri-1-1 | Bacteria Actinobacteria Actinobacteridae Actinomycetales Streptomycineae Streptomycetaceae *Streptomyces rimosus subsp. rimosus* ATCC 10970 |
| Sro-1-1 | Bacteria Actinobacteria Actinobacteridae Actinomycetales Streptomycineae Streptomycetaceae *Streptomyces roseochromogenes subsp. oscitans* DS 12.976 |
| Sro-2-1 | Bacteria Actinobacteria Actinobacteridae Actinomycetales Streptomycineae Streptomycetaceae *Streptomyces roseosporus* NRRL 15998 |
| Ssc-1-1 | Bacteria Actinobacteria Actinobacteridae Actinomycetales Streptomycineae Streptomycetaceae *Streptomyces scabiei* 87.22 |
| Ssc-2-1 | Bacteria Actinobacteria Actinobacteridae Actinomycetales Streptomycineae Streptomycetaceae *Streptomyces scabrisporus* DSM 41855 |
| Sso-1-1 | Bacteria Actinobacteria Actinobacteridae Actinomycetales Streptomycineae Streptomycetaceae *Streptomyces somaliensis* DSM 40738 |
| Ssp-1-1 | Bacteria Actinobacteria Actinobacteridae Actinomycetales Streptomycineae Streptomycetaceae *Streptomyces sp.* 303MFCol5.2 |
| Ssp-2-1 | Bacteria Actinobacteria Actinobacteridae Actinomycetales Streptomycineae Streptomycetaceae *Streptomyces sp.* 351MFTsu5.1 |
| Ssp-3-1 | Bacteria Actinobacteria Actinobacteridae Actinomycetales Streptomycineae Streptomycetaceae *Streptomyces sp.* AA0539 |
| Ssp-4-1 | Bacteria Actinobacteria Actinobacteridae Actinomycetales Streptomycineae Streptomycetaceae *Streptomyces sp.* AA1529 |
| Ssp-6-1 | Bacteria Actinobacteria Actinobacteridae Actinomycetales Streptomycineae Streptomycetaceae *Streptomyces sp.* Amel2xE9 |
| Ssp-5-1 | Bacteria Actinobacteria Actinobacteridae Actinomycetales Streptomycineae Streptomycetaceae *Streptomyces sp.* ATexAB-D23 |
| Ssp-7-1 | Bacteria Actinobacteria Actinobacteridae Actinomycetales Streptomycineae Streptomycetaceae *Streptomyces sp.* BoleA5 |
| Ssp-8-1 | Bacteria Actinobacteria Actinobacteridae Actinomycetales Streptomycineae Streptomycetaceae *Streptomyces sp.* C |
| Ssp-17-1 | Bacteria Actinobacteria Actinobacteridae Actinomycetales Streptomycineae Streptomycetaceae *Streptomyces sp.* CcalMP-8W |
| Ssp-9-1 | Bacteria Actinobacteria Actinobacteridae Actinomycetales Streptomycineae Streptomycetaceae *Streptomyces sp.* CNB091 |
| Ssp-10-1 | Bacteria Actinobacteria Actinobacteridae Actinomycetales Streptomycineae Streptomycetaceae *Streptomyces sp.* CNQ766 |
| Ssp-11-1 | Bacteria Actinobacteria Actinobacteridae Actinomycetales Streptomycineae Streptomycetaceae *Streptomyces sp.* CNS335 |
| Ssp-12-1 | Bacteria Actinobacteria Actinobacteridae Actinomycetales Streptomycineae Streptomycetaceae *Streptomyces sp.* CNS615 |
| Ssp-13-1 | Bacteria Actinobacteria Actinobacteridae Actinomycetales Streptomycineae Streptomycetaceae *Streptomyces sp.* CNT302 |
| Ssp-14-1 | Bacteria Actinobacteria Actinobacteridae Actinomycetales Streptomycineae Streptomycetaceae *Streptomyces sp.* CNT372 |
| Ssp-15-1 | Bacteria Actinobacteria Actinobacteridae Actinomycetales Streptomycineae Streptomycetaceae *Streptomyces sp.* CNY228 |
| Ssp-16-1 | Bacteria Actinobacteria Actinobacteridae Actinomycetales Streptomycineae Streptomycetaceae *Streptomyces sp.* CNY243 |
| Ssp-18-1 | Bacteria Actinobacteria Actinobacteridae Actinomycetales Streptomycineae Streptomycetaceae *Streptomyces sp.* DvalAA-83 |
| Ssp-49-1 | Bacteria Actinobacteria Actinobacteridae Actinomycetales Streptomycineae Streptomycetaceae *Streptomyces sp. e14* |
| Ssp-19-1 | Bacteria Actinobacteria Actinobacteridae Actinomycetales Streptomycineae Streptomycetaceae *Streptomyces sp.* FxanaC1 |
| Ssp-20-1 | Bacteria Actinobacteria Actinobacteridae Actinomycetales Streptomycineae Streptomycetaceae *Streptomyces sp.* FxanaD5 |
| Ssp-21-1 | Bacteria Actinobacteria Actinobacteridae Actinomycetales Streptomycineae Streptomycetaceae *Streptomyces sp.* GBA 94-10 |
| Ssp-22-1 | Bacteria Actinobacteria Actinobacteridae Actinomycetales Streptomycineae Streptomycetaceae *Streptomyces sp.* HCCB10043 |
| Ssp-23-1 | Bacteria Actinobacteria Actinobacteridae Actinomycetales Streptomycineae Streptomycetaceae *Streptomyces sp.* HGB0020 |
| Ssp-25-1 | Bacteria Actinobacteria Actinobacteridae Actinomycetales Streptomycineae Streptomycetaceae *Streptomyces sp.* HmicA12 |
| Ssp-24-1 | Bacteria Actinobacteria Actinobacteridae Actinomycetales Streptomycineae Streptomycetaceae *Streptomyces sp.* HPH0547 |
| Ssp-26-1 | Bacteria Actinobacteria Actinobacteridae Actinomycetales Streptomycineae Streptomycetaceae *Streptomyces sp.* KhCrAH-244 |
| Ssp-27-1 | Bacteria Actinobacteria Actinobacteridae Actinomycetales Streptomycineae Streptomycetaceae *Streptomyces sp.* KhCrAH-340 |
| Ssp-28-1 | Bacteria Actinobacteria Actinobacteridae Actinomycetales Streptomycineae Streptomycetaceae *Streptomyces sp.* LaPpAH-108 |
| Ssp-29-1 | Bacteria Actinobacteria Actinobacteridae Actinomycetales Streptomycineae Streptomycetaceae *Streptomyces sp.* LaPpAH-165 |
| Ssp-30-1 | Bacteria Actinobacteria Actinobacteridae Actinomycetales Streptomycineae Streptomycetaceae *Streptomyces sp.* LaPpAH-202 |
| Ssp-31-1 | Bacteria Actinobacteria Actinobacteridae Actinomycetales Streptomycineae Streptomycetaceae *Streptomyces sp.* LaPpAH-95 |
| Ssp-32-1 to Ssp-32-2 | Bacteria Actinobacteria Actinobacteridae Actinomycetales Streptomycineae Streptomycetaceae *Streptomyces sp.* Mg1 |
| Ssp-33-1 | Bacteria Actinobacteria Actinobacteridae Actinomycetales Streptomycineae Streptomycetaceae *Streptomyces sp.* MspMP-M5 |
| Ssp-34-1 | Bacteria Actinobacteria Actinobacteridae Actinomycetales Streptomycineae Streptomycetaceae *Streptomyces sp.* PAMC26508 |
| Ssp-36-1 | Bacteria Actinobacteria Actinobacteridae Actinomycetales Streptomycineae Streptomycetaceae *Streptomyces sp.* PsTaAH-124 |
| Ssp-35-1 | Bacteria Actinobacteria Actinobacteridae Actinomycetales Streptomycineae Streptomycetaceae *Streptomyces sp.* PVA 94-07 |
| Ssp-37-1 | Bacteria Actinobacteria Actinobacteridae Actinomycetales Streptomycineae Streptomycetaceae *Streptomyces sp.* R1-NS-10 |
| Ssp-38-1 | Bacteria Actinobacteria Actinobacteridae Actinomycetales Streptomycineae Streptomycetaceae *Streptomyces sp.* S4 |
| Ssp-39-1 | Bacteria Actinobacteria Actinobacteridae Actinomycetales Streptomycineae Streptomycetaceae *Streptomyces sp.* SA3_actF |
| Ssp-40-1 | Bacteria Actinobacteria Actinobacteridae Actinomycetales Streptomycineae Streptomycetaceae *Streptomyces sp.* SA3_actG |

| | |
|---|---|
| Ssp-44-1 | Bacteria Actinobacteria Actinobacteridae Actinomycetales Streptomycineae Streptomycetaceae *Streptomyces sp.* ScaeMP-e10 |
| Ssp-45-1 | Bacteria Actinobacteria Actinobacteridae Actinomycetales Streptomycineae Streptomycetaceae *Streptomyces sp.* SirexAA-E |
| Ssp-41-1 | Bacteria Actinobacteria Actinobacteridae Actinomycetales Streptomycineae Streptomycetaceae *Streptomyces sp.* SPB74 |
| Ssp-42-1 | Bacteria Actinobacteria Actinobacteridae Actinomycetales Streptomycineae Streptomycetaceae *Streptomyces sp.* SPB78 |
| Ssp-43-1 | Bacteria Actinobacteria Actinobacteridae Actinomycetales Streptomycineae Streptomycetaceae *Streptomyces sp.* SS |
| Ssp-46-1 | Bacteria Actinobacteria Actinobacteridae Actinomycetales Streptomycineae Streptomycetaceae *Streptomyces sp.* TOR3209 |
| Ssp-47-1 | Bacteria Actinobacteria Actinobacteridae Actinomycetales Streptomycineae Streptomycetaceae *Streptomyces sp.* Tu6071 |
| Ssp-48-1 | Bacteria Actinobacteria Actinobacteridae Actinomycetales Streptomycineae Streptomycetaceae *Streptomyces sp.* W007 |
| Ssu-1-1 | Bacteria Actinobacteria Actinobacteridae Actinomycetales Streptomycineae Streptomycetaceae *Streptomyces sulphureus* DSM 40104 |
| Ssu-2-1 | Bacteria Actinobacteria Actinobacteridae Actinomycetales Streptomycineae Streptomycetaceae *Streptomyces sulphureus* L180 |
| Ssv-1-1 | Bacteria Actinobacteria Actinobacteridae Actinomycetales Streptomycineae Streptomycetaceae *Streptomyces sviceus* ATCC 29083 |
| Sts-1-1 | Bacteria Actinobacteria Actinobacteridae Actinomycetales Streptomycineae Streptomycetaceae *Streptomyces tsukubaensis* NRRL18488 |
| Stu-1-1 | Bacteria Actinobacteria Actinobacteridae Actinomycetales Streptomycineae Streptomycetaceae *Streptomyces turgidiscabies* Car8 |
| Sve-1-1 | Bacteria Actinobacteria Actinobacteridae Actinomycetales Streptomycineae Streptomycetaceae *Streptomyces venezuelae* ATCC 10712 |
| Svi-1-1 | Bacteria Actinobacteria Actinobacteridae Actinomycetales Streptomycineae Streptomycetaceae *Streptomyces violaceusniger* SPC6 |
| Svi-2-1 | Bacteria Actinobacteria Actinobacteridae Actinomycetales Streptomycineae Streptomycetaceae *Streptomyces violaceusniger* Tu 4113 |
| Svi-3-1 | Bacteria Actinobacteria Actinobacteridae Actinomycetales Streptomycineae Streptomycetaceae *Streptomyces viridochromogenes* DSM 40736 |
| Svi-4-1 | Bacteria Actinobacteria Actinobacteridae Actinomycetales Streptomycineae Streptomycetaceae *Streptomyces viridochromogenes* Tue57 |
| Svi-5-1 | Bacteria Actinobacteria Actinobacteridae Actinomycetales Streptomycineae Streptomycetaceae *Streptomyces viridosporus* T7A |
| Svi-6-1 | Bacteria Actinobacteria Actinobacteridae Actinomycetales Streptomycineae Streptomycetaceae *Streptomyces vitaminophilus* DSM 41686 |
| Szi-1-1 | Bacteria Actinobacteria Actinobacteridae Actinomycetales Streptomycineae Streptomycetaceae *Streptomyces zinciresistens* K42 |
| Sba-1-1 | Bacteria Actinobacteria Actinobacteridae Actinomycetales Streptomycineae *Streptomycetaceae bacterium* MP113-05 |
| Nco-1-1 | Bacteria Actinobacteria Actinobacteridae Actinomycetales Streptosporangineae Streptosporangiaceae *Nonomuraea coxensis* DSM 45129 |
| Sro-3-1 | Bacteria Actinobacteria Actinobacteridae Actinomycetales Streptosporangineae Streptosporangiaceae *Streptosporangium roseum* DSM 43021 |
| Aat-1-1 | Bacteria Actinobacteria Actinobacteridae Actinomycetales Streptosporangineae Thermomonosporaceae *Actinomadura atramentaria* DSM 43919 |
| Afl-1-1 | Bacteria Actinobacteria Actinobacteridae Actinomycetales Streptosporangineae Thermomonosporaceae *Actinomadura flavalba* DSM 45200 |
| Aac-1-1 | Bacteria Actinobacteria Catenulisporales Actinospicaceae *Actinospica acidiphila* |
| Nal-1-1 | Bacteria Actinobacteria Propionibacteriales Nocardioidaceae *Nocardioides alkalitolerans* DSM 16699 |
| Nha-1-1 | Bacteria Actinobacteria Propionibacteriales Nocardioidaceae *Nocardioides halotolerans* DSM 19273 |
| Nin-1-1 | Bacteria Actinobacteria Propionibacteriales Nocardioidaceae *Nocardioides insulae* DSM 17944 |
| Nlu-1-1 | Bacteria Actinobacteria Propionibacteriales Nocardioidaceae *Nocardioides luteus* |
| Nsp-1-1 | Bacteria Actinobacteria Propionibacteriales Nocardioidaceae *Nocardioides sp.* J54 |
| Nsp-2-1 | Bacteria Actinobacteria Propionibacteriales Nocardioidaceae *Nocardioides sp.* URHA0020 |
| Nsp-3-1 | Bacteria Actinobacteria Propionibacteriales Nocardioidaceae *Nocardioides sp.* URHA0032 |
| Psi-1-1 | Bacteria Actinobacteria Propionibacteriales Nocardioidaceae *Pimelobacter simplex* |
| Ssp-50-1 | Bacteria Actinobacteria Pseudonocardiales Pseudonocardiaceae *Saccharothrix sp.* ST-888 |
| Kau-1-1 to Kau-1-2 | Bacteria Actinobacteria Streptomycetales Streptomycetaceae *Kitasatospora aureofaciens* |
| Kaz-1-1 | Bacteria Actinobacteria Streptomycetales Streptomycetaceae *Kitasatospora azatica* KCTC 9699 |
| Kme-1-1 | Bacteria Actinobacteria Streptomycetales Streptomycetaceae *Kitasatospora mediocidica* KCTC 9733 |
| Kph-1-1 to Kph-1-2 | Bacteria Actinobacteria Streptomycetales Streptomycetaceae *Kitasatospora phosalacinea* |
| Kpu-1-1 | Bacteria Actinobacteria Streptomycetales Streptomycetaceae *Kitasatospora purpeofusca* |
| Ksp-1-1 | Bacteria Actinobacteria Streptomycetales Streptomycetaceae *Kitasatospora sp.* MBT63 |
| Ksp-2-1 | Bacteria Actinobacteria Streptomycetales Streptomycetaceae *Kitasatospora sp.* MBT66 |
| Ksp-3-1 | Bacteria Actinobacteria Streptomycetales Streptomycetaceae *Kitasatospora sp.* MY 5-36 |
| Ksp-4-1 | Bacteria Actinobacteria Streptomycetales Streptomycetaceae *Kitasatospora sp.* NRRL B-11411 |
| Sal-3-1 | Bacteria Actinobacteria Streptomycetales Streptomycetaceae *Streptacidiphilus albus* |
| Sal-4-1 | Bacteria Actinobacteria Streptomycetales Streptomycetaceae *Streptacidiphilus albus* JL83 |
| Sca-3-1 | Bacteria Actinobacteria Streptomycetales Streptomycetaceae *Streptacidiphilus carbonis* |
| Sje-1-1 | Bacteria Actinobacteria Streptomycetales Streptomycetaceae *Streptacidiphilus jeojiense* |
| Sor-1-1 | Bacteria Actinobacteria Streptomycetales Streptomycetaceae *Streptacidiphilus oryzae* TH49 |
| [pa-1-1 | Bacteria Actinobacteria Streptomycetales Streptomycetaceae Streptomyces [Kitasatospora] papulosa |
| Asp-1-1 | Bacteria Actinobacteria Streptomycetales Streptomycetaceae Streptomyces *Actinosporangium sp.* NRRL B-3428 |
| Sac-2-1 | Bacteria Actinobacteria Streptomycetales Streptomycetaceae *Streptomyces achromogenes subsp. achromogenes* |
| Sac-3-1 | Bacteria Actinobacteria Streptomycetales Streptomycetaceae *Streptomyces acidiscabies* |
| Sah-1-1 | Bacteria Actinobacteria Streptomycetales Streptomycetaceae *Streptomyces ahygroscopicus subsp. wuyiensis* |
| Sal-5-1 | Bacteria Actinobacteria Streptomycetales Streptomycetaceae Streptomyces Streptomyces albidoflavus group *Streptomyces albidoflavus* |
| Sal-6-1 | Bacteria Actinobacteria Streptomycetales Streptomycetaceae *Streptomyces alboflavus* |
| Sal-7-1 | Bacteria Actinobacteria Streptomycetales Streptomycetaceae *Streptomyces alboviridis* |
| Sal-8-1 | Bacteria Actinobacteria Streptomycetales Streptomycetaceae *Streptomyces albulus* |
| Sal-9-1 | Bacteria Actinobacteria Streptomycetales Streptomycetaceae *Streptomyces albulus* PD-1 |
| Sal-10-1 | Bacteria Actinobacteria Streptomycetales Streptomycetaceae *Streptomyces albulus* ZPM |
| Sal-11-1 | Bacteria Actinobacteria Streptomycetales Streptomycetaceae *Streptomyces albus* |
| Sal-12-1 to Sal-12-5 | Bacteria Actinobacteria Streptomycetales Streptomycetaceae *Streptomyces albus subsp. albus* |
| Sam-1-1 | Bacteria Actinobacteria Streptomycetales Streptomycetaceae *Streptomyces ambofaciens* ATCC 23877 |
| San-1-1 | Bacteria Actinobacteria Streptomycetales Streptomycetaceae *Streptomyces antibioticus* |

| | |
|---|---|
| San-2-1 to San-2-2 | Bacteria Actinobacteria Streptomycetales Streptomycetaceae *Streptomyces anulatus* |
| Sat-1-1 | Bacteria Actinobacteria Streptomycetales Streptomycetaceae *Streptomyces atratus* |
| Sat-2-1 | Bacteria Actinobacteria Streptomycetales Streptomycetaceae *Streptomyces atroolivaceus* |
| Sau-3-1 to Sau-3-2 | Bacteria Actinobacteria Streptomycetales Streptomycetaceae *Streptomyces aureocirculatus* |
| Sau-4-1 | Bacteria Actinobacteria Streptomycetales Streptomycetaceae *Streptomyces aureus* |
| Sav-2-1 to Sav-2-2 | Bacteria Actinobacteria Streptomycetales Streptomycetaceae *Streptomyces avermitilis* MA-4680 = NBRC 14893 |
| Sav-3-1 | Bacteria Actinobacteria Streptomycetales Streptomycetaceae *Streptomyces avicenniae* |
| Sba-2-1 | Bacteria Actinobacteria Streptomycetales Streptomycetaceae *Streptomyces baarnensis* |
| Sbi-2-1 to Sbi-2-2 | Bacteria Actinobacteria Streptomycetales Streptomycetaceae *Streptomyces bicolor* |
| Sbi-3-1 to Sbi-3-3 | Bacteria Actinobacteria Streptomycetales Streptomycetaceae *Streptomyces bikiniensis* |
| Sbr-1-1 | Bacteria Actinobacteria Streptomycetales Streptomycetaceae *Streptomyces brasiliensis* |
| Sca-4-1 | Bacteria Actinobacteria Streptomycetales Streptomycetaceae *Streptomyces caatingaensis* |
| Sca-5-1 to Sca-5-2 | Bacteria Actinobacteria Streptomycetales Streptomycetaceae *Streptomyces californicus* |
| Sca-6-1 to Sca-6-2 | Bacteria Actinobacteria Streptomycetales Streptomycetaceae *Streptomyces capuensis* |
| Sca-7-1 | Bacteria Actinobacteria Streptomycetales Streptomycetaceae *Streptomyces catenulae* |
| Sce-1-1 | Bacteria Actinobacteria Streptomycetales Streptomycetaceae *Streptomyces celluloflavus* |
| Sce-2-1 | Bacteria Actinobacteria Streptomycetales Streptomycetaceae *Streptomyces cellulosae* |
| Scy-1-1 | Bacteria Actinobacteria Streptomycetales Streptomycetaceae *Streptomyces cyaneofuscatus* |
| Scy-2-1 | Bacteria Actinobacteria Streptomycetales Streptomycetaceae *Streptomyces cyaneogriseus subsp. noncyanogenus* |
| Sde-1-1 | Bacteria Actinobacteria Streptomycetales Streptomycetaceae *Streptomyces decoyicus* |
| Sdu-1-1 | Bacteria Actinobacteria Streptomycetales Streptomycetaceae *Streptomyces durhamensis* |
| Ser-1-1 | Bacteria Actinobacteria Streptomycetales Streptomycetaceae *Streptomyces erythrochromogenes* |
| Seu-1-1 | Bacteria Actinobacteria Streptomycetales Streptomycetaceae *Streptomyces europaeiscabiei* |
| Sex-1-1 | Bacteria Actinobacteria Streptomycetales Streptomycetaceae *Streptomyces exfoliatus* |
| Sex-2-1 | Bacteria Actinobacteria Streptomycetales Streptomycetaceae *Streptomyces exfoliatus* DSM 41693 |
| Sfl-2-1 | Bacteria Actinobacteria Streptomycetales Streptomycetaceae *Streptomyces flaveolus* |
| Sfl-3-1 | Bacteria Actinobacteria Streptomycetales Streptomycetaceae *Streptomyces flaveus* |
| Sfl-4-1 | Bacteria Actinobacteria Streptomycetales Streptomycetaceae *Streptomyces flavidovirens* DSM 40150 |
| Sfl-5-1 | Bacteria Actinobacteria Streptomycetales Streptomycetaceae *Streptomyces flavochromogenes* |
| Sfl-6-1 | Bacteria Actinobacteria Streptomycetales Streptomycetaceae *Streptomyces flavotricini* |
| Sfl-7-1 | Bacteria Actinobacteria Streptomycetales Streptomycetaceae *Streptomyces flavovariabilis* |
| Sfl-8-1 | Bacteria Actinobacteria Streptomycetales Streptomycetaceae *Streptomyces flavovirens* |
| Sfl-9-1 | Bacteria Actinobacteria Streptomycetales Streptomycetaceae *Streptomyces floridae* |
| Sfr-1-1 to Sfr-1-2 | Bacteria Actinobacteria Streptomycetales Streptomycetaceae *Streptomyces fradiae* |
| Sfu-2-1 | Bacteria Actinobacteria Streptomycetales Streptomycetaceae *Streptomyces fulvoviolaceus* |
| Sfu-3-1 | Bacteria Actinobacteria Streptomycetales Streptomycetaceae *Streptomyces fulvoviridis* |
| Sga-2-1 | Bacteria Actinobacteria Streptomycetales Streptomycetaceae *Streptomyces galbus* |
| Sgl-2-1 | Bacteria Actinobacteria Streptomycetales Streptomycetaceae *Streptomyces glaucescens* |
| Sgl-3-1 to Sgl-3-2 | Bacteria Actinobacteria Streptomycetales Streptomycetaceae *Streptomyces globisporus subsp. globisporus* |
| Sgr-5-1 | Bacteria Actinobacteria Streptomycetales Streptomycetaceae *Streptomyces graminis* |
| Sgr-6-1 | Bacteria Actinobacteria Streptomycetales Streptomycetaceae *Streptomyces griseoflavus* |
| Sgr-7-1 | Bacteria Actinobacteria Streptomycetales Streptomycetaceae *Streptomyces griseofuscus* |
| Sgr-8-1 | Bacteria Actinobacteria Streptomycetales Streptomycetaceae *Streptomyces griseolus* |
| Sgr-9-1 | Bacteria Actinobacteria Streptomycetales Streptomycetaceae *Streptomyces griseoluteus* |
| Sgr-10-1 | Bacteria Actinobacteria Streptomycetales Streptomycetaceae *Streptomyces griseorubens* |
| Sgr-11-1 | Bacteria Actinobacteria Streptomycetales Streptomycetaceae *Streptomyces griseus* |
| Sgr-12-1 to Sgr-12-9 | Bacteria Actinobacteria Streptomycetales Streptomycetaceae *Streptomyces griseus subsp. griseus* |
| Sgr-13-1 to Sgr-13-5 | Bacteria Actinobacteria Streptomycetales Streptomycetaceae *Streptomyces griseus subsp. rhodochrous* |
| Sha-1-1 | Bacteria Actinobacteria Streptomycetales Streptomycetaceae *Streptomyces halstedii* |
| Shy-4-1 | Bacteria Actinobacteria Streptomycetales Streptomycetaceae *Streptomyces hygroscopicus subsp. hygroscopicus* |
| Sia-1-1 | Bacteria Actinobacteria Streptomycetales Streptomycetaceae *Streptomyces iakyrus* |
| Sir-1-1 | Bacteria Actinobacteria Streptomycetales Streptomycetaceae *Streptomyces iranensis* |
| Ska-1-1 to Ska-1-2 | Bacteria Actinobacteria Streptomycetales Streptomycetaceae *Streptomyces katrae* |
| Sla-1-1 | Bacteria Actinobacteria Streptomycetales Streptomycetaceae *Streptomyces lavendulae* |
| Sla-2-1 to Sla-2-4 | Bacteria Actinobacteria Streptomycetales Streptomycetaceae *Streptomyces lavendulae subsp. lavendulae* |
| Sla-3-1 | Bacteria Actinobacteria Streptomycetales Streptomycetaceae *Streptomyces lavenduligriseus* |
| Sle-1-1 | Bacteria Actinobacteria Streptomycetales Streptomycetaceae *Streptomyces leeuwenhoekii* |
| Sli-2-1 to Sli-2-2 | Bacteria Actinobacteria Streptomycetales Streptomycetaceae *Streptomyces lividans* TK24 |
| Slu-1-1 | Bacteria Actinobacteria Streptomycetales Streptomycetaceae *Streptomyces luteus* |
| Sly-1-1 | Bacteria Actinobacteria Streptomycetales Streptomycetaceae *Streptomyces lydicus* |
| Sly-2-1 | Bacteria Actinobacteria Streptomycetales Streptomycetaceae *Streptomyces lydicus* A02 |
| Sme-1-1 | Bacteria Actinobacteria Streptomycetales Streptomycetaceae *Streptomyces mediolani* |
| Sme-2-1 | Bacteria Actinobacteria Streptomycetales Streptomycetaceae *Streptomyces megasporus* |
| Smi-1-1 | Bacteria Actinobacteria Streptomycetales Streptomycetaceae *Streptomyces mirabilis* |
| Smo-2-1 | Bacteria Actinobacteria Streptomycetales Streptomycetaceae *Streptomyces monomycini* |

| | |
|---|---|
| Sna-1-1 | Bacteria Actinobacteria Streptomycetales Streptomycetaceae *Streptomyces natalensis* |
| Sna-2-1 | Bacteria Actinobacteria Streptomycetales Streptomycetaceae *Streptomyces natalensis* ATCC 27448 |
| Sni-2-1 | Bacteria Actinobacteria Streptomycetales Streptomycetaceae *Streptomyces niger* |
| Sno-1-1 | Bacteria Actinobacteria Streptomycetales Streptomycetaceae *Streptomyces nodosus* |
| Sno-2-1 | Bacteria Actinobacteria Streptomycetales Streptomycetaceae *Streptomyces novaecaesareae* |
| Soc-1-1 | Bacteria Actinobacteria Streptomycetales Streptomycetaceae *Streptomyces ochraceiscleroticus* |
| Sod-1-1 | Bacteria Actinobacteria Streptomycetales Streptomycetaceae *Streptomyces odonnellii* |
| Sol-1-1 | Bacteria Actinobacteria Streptomycetales Streptomycetaceae *Streptomyces olindensis* |
| Sol-2-1 to Sol-2-2 | Bacteria Actinobacteria Streptomycetales Streptomycetaceae *Streptomyces olivaceus* |
| Spe-1-1 | Bacteria Actinobacteria Streptomycetales Streptomycetaceae *Streptomyces peruviensis* |
| Spe-2-1 | Bacteria Actinobacteria Streptomycetales Streptomycetaceae *Streptomyces peucetius* |
| Spl-1-1 to Spl-1-2 | Bacteria Actinobacteria Streptomycetales Streptomycetaceae *Streptomyces pluripotens* |
| Spu-2-1 to Spu-2-2 | Bacteria Actinobacteria Streptomycetales Streptomycetaceae *Streptomyces puniceus* |
| Spu-3-1 to Spu-3-2 | Bacteria Actinobacteria Streptomycetales Streptomycetaceae *Streptomyces purpeochromogenes* |
| Spy-1-1 | Bacteria Actinobacteria Streptomycetales Streptomycetaceae *Streptomyces pyridomyceticus* |
| Sre-1-1 | Bacteria Actinobacteria Streptomycetales Streptomycetaceae *Streptomyces resistomycificus* |
| Sri-2-1 to Sri-2-2 | Bacteria Actinobacteria Streptomycetales Streptomycetaceae *Streptomyces rimosus* |
| Sri-3-1 | Bacteria Actinobacteria Streptomycetales Streptomycetaceae *Streptomyces rimosus* R6-500 |
| Sri-4-1 to Sri-4-20 | Bacteria Actinobacteria Streptomycetales Streptomycetaceae *Streptomyces rimosus subsp. rimosus* |
| Sro-4-1 | Bacteria Actinobacteria Streptomycetales Streptomycetaceae *Streptomyces roseosporus* NRRL 11379 |
| Sro-5-1 | Bacteria Actinobacteria Streptomycetales Streptomycetaceae *Streptomyces roseoverticillatus* |
| Sro-6-1 | Bacteria Actinobacteria Streptomycetales Streptomycetaceae *Streptomyces roseus* |
| Sru-1-1 to Sru-1-2 | Bacteria Actinobacteria Streptomycetales Streptomycetaceae *Streptomyces ruber* |
| Ssc-3-1 to Ssc-3-2 | Bacteria Actinobacteria Streptomycetales Streptomycetaceae *Streptomyces scabiei* |
| Ssc-4-1 to Ssc-4-2 | Bacteria Actinobacteria Streptomycetales Streptomycetaceae *Streptomyces sclerotialus* |
| Ssc-5-1 | Bacteria Actinobacteria Streptomycetales Streptomycetaceae *Streptomyces scopuliridis* RB72 |
| Sse-1-1 | Bacteria Actinobacteria Streptomycetales Streptomycetaceae *Streptomyces seoulensis* |
| Ssp-51-1 | Bacteria Actinobacteria Streptomycetales Streptomycetaceae *Streptomyces sp.* 142MFCol3.1 |
| Ssp-52-1 | Bacteria Actinobacteria Streptomycetales Streptomycetaceae *Streptomyces sp.* 150FB |
| Ssp-53-1 | Bacteria Actinobacteria Streptomycetales Streptomycetaceae *Streptomyces sp.* 769 |
| Ssp-55-1 | Bacteria Actinobacteria Streptomycetales Streptomycetaceae *Streptomyces sp.* AcH 505 |
| Ssp-54-1 | Bacteria Actinobacteria Streptomycetales Streptomycetaceae *Streptomyces sp.* AW19M42 |
| Ssp-56-1 | Bacteria Actinobacteria Streptomycetales Streptomycetaceae *Streptomyces sp.* CNH099 |
| Ssp-57-1 | Bacteria Actinobacteria Streptomycetales Streptomycetaceae *Streptomyces sp.* CNH189 |
| Ssp-58-1 | Bacteria Actinobacteria Streptomycetales Streptomycetaceae *Streptomyces sp.* CNH287 |
| Ssp-59-1 | Bacteria Actinobacteria Streptomycetales Streptomycetaceae *Streptomyces sp.* CNQ-509 |
| Ssp-60-1 | Bacteria Actinobacteria Streptomycetales Streptomycetaceae *Streptomyces sp.* CNQ-525 |
| Ssp-61-1 | Bacteria Actinobacteria Streptomycetales Streptomycetaceae *Streptomyces sp.* CNQ329 |
| Ssp-62-1 | Bacteria Actinobacteria Streptomycetales Streptomycetaceae *Streptomyces sp.* CNQ431 |
| Ssp-63-1 | Bacteria Actinobacteria Streptomycetales Streptomycetaceae *Streptomyces sp.* CNQ865 |
| Ssp-64-1 | Bacteria Actinobacteria Streptomycetales Streptomycetaceae *Streptomyces sp.* CNR698 |
| Ssp-65-1 | Bacteria Actinobacteria Streptomycetales Streptomycetaceae *Streptomyces sp.* CNS606 |
| Ssp-66-1 | Bacteria Actinobacteria Streptomycetales Streptomycetaceae *Streptomyces sp.* CNS654 |
| Ssp-67-1 | Bacteria Actinobacteria Streptomycetales Streptomycetaceae *Streptomyces sp.* CNT318 |
| Ssp-68-1 | Bacteria Actinobacteria Streptomycetales Streptomycetaceae *Streptomyces sp.* CNT360 |
| Ssp-69-1 | Bacteria Actinobacteria Streptomycetales Streptomycetaceae *Streptomyces sp.* CNT371 |
| Ssp-70-1 | Bacteria Actinobacteria Streptomycetales Streptomycetaceae *Streptomyces sp.* CT34 |
| Ssp-71-1 | Bacteria Actinobacteria Streptomycetales Streptomycetaceae *Streptomyces sp.* DpondAA-B6 |
| Ssp-73-1 | Bacteria Actinobacteria Streptomycetales Streptomycetaceae *Streptomyces sp.* FxanaA7 |
| Ssp-72-1 | Bacteria Actinobacteria Streptomycetales Streptomycetaceae *Streptomyces sp.* FXJ7.023 |
| Ssp-74-1 | Bacteria Actinobacteria Streptomycetales Streptomycetaceae *Streptomyces sp.* GXT6 |
| Ssp-75-1 | Bacteria Actinobacteria Streptomycetales Streptomycetaceae *Streptomyces sp.* HNS054 |
| Ssp-76-1 | Bacteria Actinobacteria Streptomycetales Streptomycetaceae *Streptomyces sp.* JS01 |
| Ssp-77-1 | Bacteria Actinobacteria Streptomycetales Streptomycetaceae *Streptomyces sp.* KE1 |
| Ssp-78-1 | Bacteria Actinobacteria Streptomycetales Streptomycetaceae *Streptomyces sp.* KhCrAH-337 |
| Ssp-79-1 | Bacteria Actinobacteria Streptomycetales Streptomycetaceae *Streptomyces sp.* KhCrAH-40 |
| Ssp-80-1 | Bacteria Actinobacteria Streptomycetales Streptomycetaceae *Streptomyces sp.* M10 |
| Ssp-81-1 | Bacteria Actinobacteria Streptomycetales Streptomycetaceae *Streptomyces sp.* MBT28 |
| Ssp-32-1 to Ssp-32-2 | Bacteria Actinobacteria Streptomycetales Streptomycetaceae *Streptomyces sp.* Mg1 |
| Ssp-82-1 | Bacteria Actinobacteria Streptomycetales Streptomycetaceae *Streptomyces sp.* MNU77 |
| Ssp-83-1 | Bacteria Actinobacteria Streptomycetales Streptomycetaceae *Streptomyces sp.* MUSC 125 |
| Ssp-84-1 | Bacteria Actinobacteria Streptomycetales Streptomycetaceae *Streptomyces sp.* MUSC119T |
| Ssp-85-1 | Bacteria Actinobacteria Streptomycetales Streptomycetaceae *Streptomyces sp.* MUSC136T |
| Ssp-86-1 | Bacteria Actinobacteria Streptomycetales Streptomycetaceae *Streptomyces sp.* MUSC149T |
| Ssp-87-1 | Bacteria Actinobacteria Streptomycetales Streptomycetaceae *Streptomyces sp.* MUSC164 |

| | |
|---|---|
| Ssp-88-1 | Bacteria Actinobacteria Streptomycetales Streptomycetaceae *Streptomyces sp.* NBRC 110027 |
| Ssp-89-1 | Bacteria Actinobacteria Streptomycetales Streptomycetaceae *Streptomyces sp.* NBRC 110035 |
| Ssp-90-1 | Bacteria Actinobacteria Streptomycetales Streptomycetaceae *Streptomyces sp.* NRRL B-11253 |
| Ssp-91-1 | Bacteria Actinobacteria Streptomycetales Streptomycetaceae *Streptomyces sp.* NRRL B-12105 |
| Ssp-92-1 | Bacteria Actinobacteria Streptomycetales Streptomycetaceae *Streptomyces sp.* NRRL B-1322 |
| Ssp-93-1 | Bacteria Actinobacteria Streptomycetales Streptomycetaceae *Streptomyces sp.* NRRL B-1347 |
| Ssp-94-1 | Bacteria Actinobacteria Streptomycetales Streptomycetaceae *Streptomyces sp.* NRRL B-1381 |
| Ssp-95-1 | Bacteria Actinobacteria Streptomycetales Streptomycetaceae *Streptomyces sp.* NRRL B-24051 |
| Ssp-96-1 | Bacteria Actinobacteria Streptomycetales Streptomycetaceae *Streptomyces sp.* NRRL B-24484 |
| Ssp-97-1 | Bacteria Actinobacteria Streptomycetales Streptomycetaceae *Streptomyces sp.* NRRL B-24720 |
| Ssp-98-1 | Bacteria Actinobacteria Streptomycetales Streptomycetaceae *Streptomyces sp.* NRRL B-3229 |
| Ssp-99-1 | Bacteria Actinobacteria Streptomycetales Streptomycetaceae *Streptomyces sp.* NRRL B-3253 |
| Ssp-100-1 | Bacteria Actinobacteria Streptomycetales Streptomycetaceae *Streptomyces sp.* NRRL F-2202 |
| Ssp-101-1 | Bacteria Actinobacteria Streptomycetales Streptomycetaceae *Streptomyces sp.* NRRL F-2305 |
| Ssp-102-1 | Bacteria Actinobacteria Streptomycetales Streptomycetaceae *Streptomyces sp.* NRRL F-2580 |
| Ssp-103-1 | Bacteria Actinobacteria Streptomycetales Streptomycetaceae *Streptomyces sp.* NRRL F-2664 |
| Ssp-104-1 | Bacteria Actinobacteria Streptomycetales Streptomycetaceae *Streptomyces sp.* NRRL F-2747 |
| Ssp-105-1 | Bacteria Actinobacteria Streptomycetales Streptomycetaceae *Streptomyces sp.* NRRL F-2799 |
| Ssp-106-1 | Bacteria Actinobacteria Streptomycetales Streptomycetaceae *Streptomyces sp.* NRRL F-2890 |
| Ssp-107-1 | Bacteria Actinobacteria Streptomycetales Streptomycetaceae *Streptomyces sp.* NRRL F-3218 |
| Ssp-108-1 | Bacteria Actinobacteria Streptomycetales Streptomycetaceae *Streptomyces sp.* NRRL F-3273 |
| Ssp-109-1 | Bacteria Actinobacteria Streptomycetales Streptomycetaceae *Streptomyces sp.* NRRL F-3307 |
| Ssp-110-1 | Bacteria Actinobacteria Streptomycetales Streptomycetaceae *Streptomyces sp.* NRRL F-4335 |
| Ssp-111-1 | Bacteria Actinobacteria Streptomycetales Streptomycetaceae *Streptomyces sp.* NRRL F-4428 |
| Ssp-112-1 | Bacteria Actinobacteria Streptomycetales Streptomycetaceae *Streptomyces sp.* NRRL F-4474 |
| Ssp-113-1 | Bacteria Actinobacteria Streptomycetales Streptomycetaceae *Streptomyces sp.* NRRL F-4835 |
| Ssp-114-1 | Bacteria Actinobacteria Streptomycetales Streptomycetaceae *Streptomyces sp.* NRRL F-5008 |
| Ssp-115-1 | Bacteria Actinobacteria Streptomycetales Streptomycetaceae *Streptomyces sp.* NRRL F-5053 |
| Ssp-116-1 | Bacteria Actinobacteria Streptomycetales Streptomycetaceae *Streptomyces sp.* NRRL F-5065 |
| Ssp-117-1 | Bacteria Actinobacteria Streptomycetales Streptomycetaceae *Streptomyces sp.* NRRL F-5123 |
| Ssp-118-1 | Bacteria Actinobacteria Streptomycetales Streptomycetaceae *Streptomyces sp.* NRRL F-5126 |
| Ssp-119-1 | Bacteria Actinobacteria Streptomycetales Streptomycetaceae *Streptomyces sp.* NRRL F-5135 |
| Ssp-120-1 | Bacteria Actinobacteria Streptomycetales Streptomycetaceae *Streptomyces sp.* NRRL F-5140 |
| Ssp-121-1 | Bacteria Actinobacteria Streptomycetales Streptomycetaceae *Streptomyces sp.* NRRL F-5193 |
| Ssp-122-1 | Bacteria Actinobacteria Streptomycetales Streptomycetaceae *Streptomyces sp.* NRRL F-525 |
| Ssp-123-1 | Bacteria Actinobacteria Streptomycetales Streptomycetaceae *Streptomyces sp.* NRRL F-5527 |
| Ssp-124-1 | Bacteria Actinobacteria Streptomycetales Streptomycetaceae *Streptomyces sp.* NRRL F-5555 |
| Ssp-125-1 | Bacteria Actinobacteria Streptomycetales Streptomycetaceae *Streptomyces sp.* NRRL F-5630 |
| Ssp-126-1 | Bacteria Actinobacteria Streptomycetales Streptomycetaceae *Streptomyces sp.* NRRL F-5635 |
| Ssp-127-1 | Bacteria Actinobacteria Streptomycetales Streptomycetaceae *Streptomyces sp.* NRRL F-5639 |
| Ssp-128-1 | Bacteria Actinobacteria Streptomycetales Streptomycetaceae *Streptomyces sp.* NRRL F-5650 |
| Ssp-129-1 | Bacteria Actinobacteria Streptomycetales Streptomycetaceae *Streptomyces sp.* NRRL F-5681 |
| Ssp-130-1 | Bacteria Actinobacteria Streptomycetales Streptomycetaceae *Streptomyces sp.* NRRL F-5702 |
| Ssp-131-1 | Bacteria Actinobacteria Streptomycetales Streptomycetaceae *Streptomyces sp.* NRRL F-5727 |
| Ssp-132-1 | Bacteria Actinobacteria Streptomycetales Streptomycetaceae *Streptomyces sp.* NRRL F-5917 |
| Ssp-133-1 | Bacteria Actinobacteria Streptomycetales Streptomycetaceae *Streptomyces sp.* NRRL F-6131 |
| Ssp-134-1 | Bacteria Actinobacteria Streptomycetales Streptomycetaceae *Streptomyces sp.* NRRL F-6628 |
| Ssp-135-1 | Bacteria Actinobacteria Streptomycetales Streptomycetaceae *Streptomyces sp.* NRRL F-6674 |
| Ssp-136-1 | Bacteria Actinobacteria Streptomycetales Streptomycetaceae *Streptomyces sp.* NRRL F-6676 |
| Ssp-137-1 | Bacteria Actinobacteria Streptomycetales Streptomycetaceae *Streptomyces sp.* NRRL F-6677 |
| Ssp-138-1 | Bacteria Actinobacteria Streptomycetales Streptomycetaceae *Streptomyces sp.* NRRL S-1022 |
| Ssp-139-1 | Bacteria Actinobacteria Streptomycetales Streptomycetaceae *Streptomyces sp.* NRRL S-104 |
| Ssp-140-1 | Bacteria Actinobacteria Streptomycetales Streptomycetaceae *Streptomyces sp.* NRRL S-118 |
| Ssp-141-1 | Bacteria Actinobacteria Streptomycetales Streptomycetaceae *Streptomyces sp.* NRRL S-1314 |
| Ssp-142-1 | Bacteria Actinobacteria Streptomycetales Streptomycetaceae *Streptomyces sp.* NRRL S-1448 |
| Ssp-143-1 | Bacteria Actinobacteria Streptomycetales Streptomycetaceae *Streptomyces sp.* NRRL S-146 |
| Ssp-144-1 | Bacteria Actinobacteria Streptomycetales Streptomycetaceae *Streptomyces sp.* NRRL S-15 |
| Ssp-145-1 | Bacteria Actinobacteria Streptomycetales Streptomycetaceae *Streptomyces sp.* NRRL S-1813 |
| Ssp-146-1 | Bacteria Actinobacteria Streptomycetales Streptomycetaceae *Streptomyces sp.* NRRL S-1824 |
| Ssp-147-1 | Bacteria Actinobacteria Streptomycetales Streptomycetaceae *Streptomyces sp.* NRRL S-1831 |
| Ssp-148-1 | Bacteria Actinobacteria Streptomycetales Streptomycetaceae *Streptomyces sp.* NRRL S-1868 |
| Ssp-149-1 | Bacteria Actinobacteria Streptomycetales Streptomycetaceae *Streptomyces sp.* NRRL S-1896 |
| Ssp-150-1 | Bacteria Actinobacteria Streptomycetales Streptomycetaceae *Streptomyces sp.* NRRL S-237 |
| Ssp-151-1 | Bacteria Actinobacteria Streptomycetales Streptomycetaceae *Streptomyces sp.* NRRL S-241 |
| Ssp-152-1 | Bacteria Actinobacteria Streptomycetales Streptomycetaceae *Streptomyces sp.* NRRL S-244 |

| | |
|---|---|
| Ssp-153-1 | Bacteria Actinobacteria Streptomycetales Streptomycetaceae *Streptomyces sp.* NRRL S-31 |
| Ssp-154-1 | Bacteria Actinobacteria Streptomycetales Streptomycetaceae *Streptomyces sp.* NRRL S-325 |
| Ssp-155-1 | Bacteria Actinobacteria Streptomycetales Streptomycetaceae *Streptomyces sp.* NRRL S-337 |
| Ssp-156-1 | Bacteria Actinobacteria Streptomycetales Streptomycetaceae *Streptomyces sp.* NRRL S-340 |
| Ssp-157-1 | Bacteria Actinobacteria Streptomycetales Streptomycetaceae *Streptomyces sp.* NRRL S-350 |
| Ssp-158-1 | Bacteria Actinobacteria Streptomycetales Streptomycetaceae *Streptomyces sp.* NRRL S-37 |
| Ssp-159-1 | Bacteria Actinobacteria Streptomycetales Streptomycetaceae *Streptomyces sp.* NRRL S-378 |
| Ssp-160-1 | Bacteria Actinobacteria Streptomycetales Streptomycetaceae *Streptomyces sp.* NRRL S-384 |
| Ssp-161-1 | Bacteria Actinobacteria Streptomycetales Streptomycetaceae *Streptomyces sp.* NRRL S-455 |
| Ssp-162-1 | Bacteria Actinobacteria Streptomycetales Streptomycetaceae *Streptomyces sp.* NRRL S-474 |
| Ssp-163-1 | Bacteria Actinobacteria Streptomycetales Streptomycetaceae *Streptomyces sp.* NRRL S-475 |
| Ssp-164-1 | Bacteria Actinobacteria Streptomycetales Streptomycetaceae *Streptomyces sp.* NRRL S-481 |
| Ssp-165-1 | Bacteria Actinobacteria Streptomycetales Streptomycetaceae *Streptomyces sp.* NRRL S-495 |
| Ssp-166-1 | Bacteria Actinobacteria Streptomycetales Streptomycetaceae *Streptomyces sp.* NRRL S-515 |
| Ssp-167-1 | Bacteria Actinobacteria Streptomycetales Streptomycetaceae *Streptomyces sp.* NRRL S-575 |
| Ssp-168-1 | Bacteria Actinobacteria Streptomycetales Streptomycetaceae *Streptomyces sp.* NRRL S-623 |
| Ssp-169-1 | Bacteria Actinobacteria Streptomycetales Streptomycetaceae *Streptomyces sp.* NRRL S-646 |
| Ssp-170-1 | Bacteria Actinobacteria Streptomycetales Streptomycetaceae *Streptomyces sp.* NRRL S-813 |
| Ssp-171-1 | Bacteria Actinobacteria Streptomycetales Streptomycetaceae *Streptomyces sp.* NRRL S-87 |
| Ssp-172-1 | Bacteria Actinobacteria Streptomycetales Streptomycetaceae *Streptomyces sp.* NRRL S-920 |
| Ssp-173-1 | Bacteria Actinobacteria Streptomycetales Streptomycetaceae *Streptomyces sp.* NRRL S-98 |
| Ssp-174-1 | Bacteria Actinobacteria Streptomycetales Streptomycetaceae *Streptomyces sp.* NRRL WC-3540 |
| Ssp-175-1 | Bacteria Actinobacteria Streptomycetales Streptomycetaceae *Streptomyces sp.* NRRL WC-3549 |
| Ssp-176-1 | Bacteria Actinobacteria Streptomycetales Streptomycetaceae *Streptomyces sp.* NRRL WC-3626 |
| Ssp-177-1 | Bacteria Actinobacteria Streptomycetales Streptomycetaceae *Streptomyces sp.* NRRL WC-3641 |
| Ssp-178-1 | Bacteria Actinobacteria Streptomycetales Streptomycetaceae *Streptomyces sp.* NRRL WC-3683 |
| Ssp-179-1 | Bacteria Actinobacteria Streptomycetales Streptomycetaceae *Streptomyces sp.* NRRL WC-3702 |
| Ssp-180-1 | Bacteria Actinobacteria Streptomycetales Streptomycetaceae *Streptomyces sp.* NRRL WC-3703 |
| Ssp-181-1 | Bacteria Actinobacteria Streptomycetales Streptomycetaceae *Streptomyces sp.* NRRL WC-3719 |
| Ssp-182-1 | Bacteria Actinobacteria Streptomycetales Streptomycetaceae *Streptomyces sp.* NRRL WC-3725 |
| Ssp-183-1 | Bacteria Actinobacteria Streptomycetales Streptomycetaceae *Streptomyces sp.* NRRL WC-3742 |
| Ssp-184-1 to Ssp-184-2 | Bacteria Actinobacteria Streptomycetales Streptomycetaceae *Streptomyces sp.* NRRL WC-3744 |
| Ssp-185-1 | Bacteria Actinobacteria Streptomycetales Streptomycetaceae *Streptomyces sp.* NRRL WC-3773 |
| Ssp-186-1 | Bacteria Actinobacteria Streptomycetales Streptomycetaceae *Streptomyces sp.* NRRL WC-3774 |
| Ssp-187-1 | Bacteria Actinobacteria Streptomycetales Streptomycetaceae *Streptomyces sp.* NRRL WC-3795 |
| Ssp-188-1 | Bacteria Actinobacteria Streptomycetales Streptomycetaceae *Streptomyces sp.* NTK 937 |
| Ssp-189-1 | Bacteria Actinobacteria Streptomycetales Streptomycetaceae *Streptomyces sp.* PCS3-D2 |
| Ssp-190-1 | Bacteria Actinobacteria Streptomycetales Streptomycetaceae *Streptomyces sp.* PRh5 |
| Ssp-191-1 | Bacteria Actinobacteria Streptomycetales Streptomycetaceae *Streptomyces sp.* RSD-27 |
| Ssp-192-1 | Bacteria Actinobacteria Streptomycetales Streptomycetaceae *Streptomyces sp.* SBT349 |
| Ssp-193-1 | Bacteria Actinobacteria Streptomycetales Streptomycetaceae *Streptomyces sp.* SolWspMP-sol2th |
| Ssp-194-1 | Bacteria Actinobacteria Streptomycetales Streptomycetaceae *Streptomyces sp.* TAA040 |
| Ssp-195-1 | Bacteria Actinobacteria Streptomycetales Streptomycetaceae *Streptomyces sp.* TAA204 |
| Ssp-196-1 | Bacteria Actinobacteria Streptomycetales Streptomycetaceae *Streptomyces sp.* Tu 6176 |
| Ssp-197-1 | Bacteria Actinobacteria Streptomycetales Streptomycetaceae *Streptomyces sp.* UNC401CLCol |
| Ssp-198-1 | Bacteria Actinobacteria Streptomycetales Streptomycetaceae *Streptomyces sp.* URHA0041 |
| Ssp-199-1 | Bacteria Actinobacteria Streptomycetales Streptomycetaceae *Streptomyces sp.* WM6386 |
| Ssp-200-1 | Bacteria Actinobacteria Streptomycetales Streptomycetaceae *Streptomyces sp.* WM6391 |
| Ssp-201-1 | Bacteria Actinobacteria Streptomycetales Streptomycetaceae *Streptomyces sp.* WMMB 714 |
| Sst-1-1 | Bacteria Actinobacteria Streptomycetales Streptomycetaceae *Streptomyces stelliscabiei* |
| Sto-1-1 | Bacteria Actinobacteria Streptomycetales Streptomycetaceae *Streptomyces toyocaensis* |
| Sva-1-1 | Bacteria Actinobacteria Streptomycetales Streptomycetaceae *Streptomyces variegatus* |
| Sva-2-1 to Sva-2-2 | Bacteria Actinobacteria Streptomycetales Streptomycetaceae *Streptomyces varsoviensis* |
| Svi-7-1 | Bacteria Actinobacteria Streptomycetales Streptomycetaceae *Streptomyces vietnamensis* |
| Svi-8-1 | Bacteria Actinobacteria Streptomycetales Streptomycetaceae *Streptomyces vinaceus* |
| Svi-9-1 to Svi-9-2 | Bacteria Actinobacteria Streptomycetales Streptomycetaceae *Streptomyces violaceoruber* |
| Svi-10-1 | Bacteria Actinobacteria Streptomycetales Streptomycetaceae *Streptomyces violaceorubidus* |
| Svi-11-1 | Bacteria Actinobacteria Streptomycetales Streptomycetaceae *Streptomyces violens* |
| Svi-12-1 to Svi-12-3 | Bacteria Actinobacteria Streptomycetales Streptomycetaceae *Streptomyces virginiae* |
| Svi-13-1 to Svi-13-4 | Bacteria Actinobacteria Streptomycetales Streptomycetaceae *Streptomyces viridochromogenes* |
| Swa-1-1 | Bacteria Actinobacteria Streptomycetales Streptomycetaceae *Streptomyces wadayamensis* |
| Swe-1-1 | Bacteria Actinobacteria Streptomycetales Streptomycetaceae *Streptomyces wedmorensis* |
| Sxa-1-1 to Sxa-1-2 | Bacteria Actinobacteria Streptomycetales Streptomycetaceae *Streptomyces xanthophaeus* |
| Sxi-1-1 | Bacteria Actinobacteria Streptomycetales Streptomycetaceae *Streptomyces xiamenensis* |
| Sxi-2-1 | Bacteria Actinobacteria Streptomycetales Streptomycetaceae *Streptomyces xinghaiensis* S187 |

| | |
|---|---|
| Sxy-1-1 | Bacteria Actinobacteria Streptomycetales Streptomycetaceae *Streptomyces xylophagus* |
| Sya-1-1 | Bacteria Actinobacteria Streptomycetales Streptomycetaceae *Streptomyces yangpuensis* |
| Sye-1-1 | Bacteria Actinobacteria Streptomycetales Streptomycetaceae *Streptomyces yeochonensis* CN732 |
| Sye-2-1 | Bacteria Actinobacteria Streptomycetales Streptomycetaceae *Streptomyces yerevanensis* |
| Hcr-1-1 | Bacteria Actinobacteria Streptosporangiales Streptosporangiaceae *Herbidospora cretacea* |
| Mro-1-1 | Bacteria Actinobacteria Streptosporangiales Streptosporangiaceae *Microbispora rosea subsp. nonnitritogenes* |
| Msp-1-1 | Bacteria Actinobacteria Streptosporangiales Streptosporangiaceae *Microbispora sp.* ATCC PTA-5024 |
| Mgl-1-1 | Bacteria Actinobacteria Streptosporangiales Streptosporangiaceae *Microtetraspora glauca* |
| Nca-1-1 | Bacteria Actinobacteria Streptosporangiales Streptosporangiaceae *Nonomuraea candida* |
| Nsp-4-1 | Bacteria Actinobacteria Streptosporangiales Streptosporangiaceae *Nonomuraea sp.* SBT364 |
| Sam-2-1 | Bacteria Actinobacteria Streptosporangiales Streptosporangiaceae *Streptosporangium amethystogenes* |
| Sro-7-1 | Bacteria Actinobacteria Streptosporangiales Streptosporangiaceae *Streptosporangium roseum* |
| Ama-1-1 | Bacteria Actinobacteria Streptosporangiales Thermomonosporaceae *Actinomadura madurae* LIID-AJ290 |
| Ari-1-1 | Bacteria Actinobacteria Streptosporangiales Thermomonosporaceae *Actinomadura rifamycini* DSM 43936 |
| Sal-13-1 | Bacteria Actinobacteria Streptosporangiales Thermomonosporaceae *Spirillospora albida* |
| [br-1-1 | Bacteria Proteobacteria Betaproteobacteria Burkholderiales [Polyangium] brachysporum |
| Rge-1-1 | Bacteria Proteobacteria Betaproteobacteria Burkholderiales *Rubrivivax gelatinosus* CBS |
| Age-1-1 | Bacteria Proteobacteria Deltaproteobacteria Myxococcales Cystobacterineae Archangiaceae *Archangium gephyra* |
| Avi-1-1 | Bacteria Proteobacteria Deltaproteobacteria Myxococcales Cystobacterineae Archangiaceae *Archangium violaceum* Cb vi76 |
| Hmi-1-1 | Bacteria Proteobacteria Deltaproteobacteria Myxococcales Cystobacterineae Archangiaceae *Hyalangium minutum* |
| Cfu-1-1 | Bacteria Proteobacteria Deltaproteobacteria Myxococcales Cystobacterineae Cystobacteraceae *Cystobacter fuscus* DSM 2262 |
| Sau-5-1 | Bacteria Proteobacteria Deltaproteobacteria Myxococcales Cystobacterineae Cystobacteraceae *Stigmatella aurantiaca* DW4/3-1 |
| Cco-1-1 | Bacteria Proteobacteria Deltaproteobacteria Myxococcales Cystobacterineae Myxococcaceae *Corallococcus coralloides* DSM 2259 |
| Mfu-1-1 | Bacteria Proteobacteria Deltaproteobacteria Myxococcales Cystobacterineae Myxococcaceae *Myxococcus fulvus* 124B02 |
| Mfu-2-1 | Bacteria Proteobacteria Deltaproteobacteria Myxococcales Cystobacterineae Myxococcaceae *Myxococcus fulvus* HW-1 |
| Mha-1-1 | Bacteria Proteobacteria Deltaproteobacteria Myxococcales Cystobacterineae Myxococcaceae *Myxococcus hansupus* |
| Msp-2-1 | Bacteria Proteobacteria Deltaproteobacteria Myxococcales Cystobacterineae Myxococcaceae *Myxococcus sp. (contaminant ex* DSM 436) |
| Mst-1-1 | Bacteria Proteobacteria Deltaproteobacteria Myxococcales Cystobacterineae Myxococcaceae *Myxococcus stipitatus* DSM 14675 |
| Mxa-1-1 | Bacteria Proteobacteria Deltaproteobacteria Myxococcales Cystobacterineae Myxococcaceae *Myxococcus xanthus* DK 1622 |
| Mxa-2-1 | Bacteria Proteobacteria Deltaproteobacteria Myxococcales Cystobacterineae Myxococcaceae *Myxococcus xanthus* DZ2 |
| Mxa-3-1 | Bacteria Proteobacteria Deltaproteobacteria Myxococcales Cystobacterineae Myxococcaceae *Myxococcus xanthus* DZF1 |
| env-1 to env-41 | environmental samples |

#### 5.2 Notes

Superscript numbers are used to annotate any L17-Actino-Proteobacteria RNA that has special characteristics (described below). These numeric annotations will be used in Sections 5.3 and 5.5:. The meaning of the numbers are as follows.

[1] Left hairpin does not conform well to the characteristics of those in most other sequences.

#### 5.3 Gene contexts

Each L17-Actino-Proteobacteria RNA (indicated by "RNA→") is listed. For each hit, the downstream genes predicted to reside in a regulated operon are listed. If the nearest downstream gene is encoding in the opposite strand (and therefore presumed to not be a part of a regulated operon), then that gene is still depicted. Some environmental sequences and some RefSeq entries lack gene annotations, and so no genes are available for such sequences. The direction of each gene is indicated with an arrow (→), and each predicted conserved domain in the gene is named. Conserved domains associated with more than one L17-Actino-Proteobacteria RNA are assigned a color; other domains are gray. Information about these conserved domains is given in Section 5.4. The accession of the sequence containing each L17-Actino-Proteobacteria RNA is given in the column named "Seq. accession". Accessions beginning with "NC_", "NS_", "NW_" or "NZ_" are contained in RefSeq. Other accession refer to environmental samples. Nucleotide coordinates are given for the 5′ and 3′ boundaries of each L17-Actino-Proteobacteria RNA. If the 5′ coordinate is greater than the 3′ coordinate, the RNA is present on the reverse-complement strand of the containing genomic DNA sequence. Each hit is denoted by an abbreviation (like "Eco-1-1") that refers to a taxonomy given in Section 5.1. (Superscript numbers refer to annotations listed in Section 5.2)

| abbrev. | Seq. accession | | 5′ at | 3′ at | genes |
|---|---|---|---|---|---|
| env-1 | C687J26621_10007310 | - | 562 | 514 | **RNA**→ RplQ (COG0203)rplQ (PRK05591)→ hypo→ |
| env-2 | C687J26621_10071554 | - | 1140 | 1092 | **RNA**→ hypo→ RplQ (COG0203)rplQ (PRK05591)→ hypo→ hypo→ |
| [br-1-1 | NZ_CP011371.1 | - | 5726771 | 5726711 | **RNA**→ RplQ (COG0203)rplQ (PRK05591)→ |
| env-3 | soil_Contig5330 | + | 734 | 782 | **RNA**→ RplQ (COG0203)rplQ (PRK05591)→ hypo→ hypo→ hypo→ |
| Nal-1-1 | NZ_AUFN01000013.1 | - | 290198 | 290141 | **RNA**→ RplQ (COG0203)rplQ (PRK05591)→ hypo→ |
| Nsp-1-1 | NZ_JAER01000028.1 | + | 47081 | 47133 | **RNA**→ RplQ (COG0203)rplQ (PRK05591)→ |
| Mxa-1-1 | NC_008095.1 | + | 3870176 | 3870231 | **RNA**→ RplQ (COG0203)rplQ (PRK05591)→ |
| Mxa-2-1 | NZ_AKYI02000032.1 | + | 15862 | 15917 | **RNA**→ RplQ (COG0203)rplQ (PRK05591)→ |
| Mxa-3-1 | NZ_AOBT01000010.1 | - | 325703 | 325648 | **RNA**→ RplQ (COG0203)rplQ (PRK05591)→ |
| env-4 | G312J29652_10012153 | - | 190 | 133 | **RNA**→ RplQ (COG0203)rplQ (PRK05591)→ |
| Cfu-1-1 | NZ_ANAH02000033.1 | - | 26146 | 26089 | **RNA**→ RplQ (COG0203)rplQ (PRK05591)→ |
| Avi-1-1 | NZ_JPMI01000295.1 | + | 16333 | 16389 | **RNA**→ RplQ (COG0203)rplQ (PRK05591)→ |
| Age-1-1 | NZ_CP011509.1 | + | 5915327 | 5915383 | **RNA**→ RplQ (COG0203)rplQ (PRK05591)→ |
| Hmi-1-1 | NZ_JMCB01000003.1 | + | 16437 | 16494 | **RNA**→ RplQ (COG0203)rplQ (PRK05591)→ |
| env-5 | JGI12467J12023_1013060 | + | 231 | 287 | **RNA**→ RplQ (COG0203)rplQ (PRK05591)→ |
| Sau-5-1 | NC_014623.1 | + | 4510803 | 4510859 | **RNA**→ RplQ (COG0203)rplQ (PRK05591)→ |
| Mfu-1-1 | NZ_CP006003.1 | - | 6488472 | 6488416 | **RNA**→ RplQ (COG0203)rplQ (PRK05591)→ |
| Mst-1-1 | NC_020126.1 | - | 5952427 | 5952371 | **RNA**→ RplQ (COG0203)rplQ (PRK05591)→ |
| Msp-2-1 | NZ_AKXT02000011.1 | - | 416054 | 415998 | **RNA**→ RplQ (COG0203)rplQ (PRK05591)→ |
| Mha-1-1 | NZ_CP012109.1 | - | 4586241 | 4586185 | **RNA**→ RplQ (COG0203)rplQ (PRK05591)→ |
| Mfu-2-1 | NC_015711.1 | + | 5782024 | 5782080 | **RNA**→ RplQ (COG0203)rplQ (PRK05591)→ |
| env-6 | JGI12467J12023_1015101 | + | 1120 | 1177 | **RNA**→ RplQ (COG0203)rplQ (PRK05591)→ |
| Cco-1-1 | NC_017030.1 | + | 3764344 | 3764400 | **RNA**→ RplQ (COG0203)rplQ (PRK05591)→ |
| env-7 | SRB_FXPBX6M01ETLHE | - | 521 | 467 | **RNA**→ RplQ (COG0203)rplQ (PRK05591)→ |
| env-8 | LWAnN_GIDYKCY01AF5CK | - | 393 | 338 | **RNA**→ RplQ (COG0203)rplQ (PRK05591)→ |
| Ama-1-1 | NZ_AWOO02000003.1 | - | 173008 | 172954 | **RNA**→ rplQ (PRK05591)Ribosomal_L17 (pfam01196)→ |
| env-9 | topACOD_F95O9CU02FUYVO | + | 189 | 240 | **RNA**→ rplQ (PRK05591)Ribosomal_L17 (pfam01196)→ |
| env-10 | botACOD_contig00616 | + | 47 | 98 | **RNA**→ rplQ (PRK05591)Ribosomal_L17 (pfam01196)→ PseudoU_synth_EcTruA (cd02570)truA (PRK00021)→ |
| | | | | | AdoMet_MTases (cd02440)Methyltransf_25 (pfam13649)→ |
| | | | | | AAA (smart00382)Uup (COG0488)ABC_tran_Xtn (pfam12848)→ |
| Ssp-68-1 | NZ_AUKO01000030.1 | + | 121030 | 121087 | **RNA**→ rplQ (PRK05591)Ribosomal_L17 (pfam01196)→ |
| [1]env-11 | topACOD_F95O9CU01C589N | - | 233 | 171 | **RNA**→ rplQ (PRK05591)Ribosomal_L17 (pfam01196)→ |
| env-12 | DCrCPGB_c521527 | + | 223 | 278 | **RNA**→ RplQ (COG0203)rplQ (PRK05591)→ |
| Ssp-96-1 | NZ_JOAI01000002.1 | - | 176853 | 176802 | **RNA**→ rplQ (PRK05591)Ribosomal_L17 (pfam01196)→ PseudoU_synth_EcTruA (cd02570)truA (PRK00021)→ |
| Ksp-1-1 | NZ_JAIZ01000648.1 | - | 13562 | 13510 | **RNA**→ rplQ (PRK05591)Ribosomal_L17 (pfam01196)→ PseudoU_synth_EcTruA (cd02570)truA (PRK00021)→ |
| | | | | | AAA (smart00382)Uup (COG0488)→ |
| Kme-1-1 | NZ_JQLN01000005.1 | + | 2502927 | 2502979 | **RNA**→ rplQ (PRK05591)Ribosomal_L17 (pfam01196)→ PseudoU_synth_EcTruA (cd02570)truA (PRK00021)→ |
| | | | | | AAA (smart00382)Uup (COG0488)AAA_21 (pfam13304)→ |
| Rge-1-1 | NZ_AJFF01000004.1 | + | 16232 | 16281 | **RNA**→ RplQ (COG0203)rplQ (PRK05591)→ |
| Sca-3-1 | NZ_BBPM01000018.1 | - | 109868 | 109813 | **RNA**→ rplQ (PRK05591)Ribosomal_L17 (pfam01196)→ PseudoU_synth_EcTruA (cd02570)truA (PRK00021)→ |
| Sal-3-1 | NZ_BBPL01000005.1 | + | 86334 | 86389 | **RNA**→ rplQ (PRK05591)Ribosomal_L17 (pfam01196)→ |
| Sal-4-1 | NZ_JQML01000001.1 | + | 5666634 | 5666689 | **RNA**→ rplQ (PRK05591)Ribosomal_L17 (pfam01196)→ |
| env-13 | JGI24770J26754_10005505 | - | 4344 | 4287 | **RNA**→ RplQ (COG0203)rplQ (PRK05591)PRK11855 (PRK11855)→ |
| | | | | | PRK09434 (PRK09434)bac_FRK (cd01167)→ CarboxypepD_reg (pfam13620)→ |
| Sav-3-1 | NZ_JOEK01000005.1 | - | 58397 | 58339 | **RNA**→ RplQ (COG0203)rplQ (PRK05591)→ |
| [1]Ssp-195-1 | NZ_AUKW01000004.1 | - | 554930 | 554861 | **RNA**→ PRK13428 (PRK13428)rplQ (PRK05591)Ribosomal_L17 (pfam01196)→ |
| | | | | | PseudoU_synth_EcTruA (cd02570)truA (PRK00021)→ |
| Nsp-4-1 | NZ_LAVL01000256.1 | - | 2722 | 2668 | **RNA**→ rplQ (PRK05591)Ribosomal_L17 (pfam01196)→ PseudoU_synth_EcTruA (cd02570)truA (PRK00021)→ |
| env-14 | botACOD_F64RS5002G1I32 | + | 29 | 84 | **RNA**→ rplQ (PRK05591)Ribosomal_L17 (pfam01196)→ |
| Nco-1-1 | NZ_KB903951.1 | - | 119743 | 119681 | **RNA**→ rplQ (PRK05591)Ribosomal_L17 (pfam01196)→ PseudoU_synth_EcTruA (cd02570)truA (PRK00021)→ |
| Sje-1-1 | NZ_JOEH01000005.1 | - | 70730 | 70674 | **RNA**→ rplQ (PRK05591)Ribosomal_L17 (pfam01196)→ PseudoU_synth_EcTruA (cd02570)truA (PRK00021)→ |
| Kaz-1-1 | NZ_JQMO01000003.1 | + | 3548080 | 3548135 | **RNA**→ rplQ (PRK05591)Ribosomal_L17 (pfam01196)→ |

| | | | | | |
|---|---|---|---|---|---|
| Kph-1-1 | NZ_JNWZ01000014.1 | - | 38553 | 38499 | RNA→ rplQ (PRK05591)Ribosomal_L17 (pfam01196)→ PseudoU_synth_EcTruA (cd02570)truA (PRK00021)→ AAA (smart00382)Uup (COG0488)→ |
| Kph-1-2 | NZ_JNYE01000006.1 | + | 139183 | 139237 | RNA→ rplQ (PRK05591)Ribosomal_L17 (pfam01196)→ |
| Sno-2-1 | NZ_JNWQ01000041.1 | - | 6189 | 6134 | RNA→ rplQ (PRK05591)Ribosomal_L17 (pfam01196)→ |
| Ksp-3-1 | NZ_LFVW01000009.1 | + | 42129 | 42184 | RNA→ rplQ (PRK05591)Ribosomal_L17 (pfam01196)→ PseudoU_synth_EcTruA (cd02570)truA (PRK00021)→ AAA (smart00382)Uup (COG0488)AAA_21 (pfam13304)→ |
| Spy-1-1 | NZ_KL573373.1 | + | 195480 | 195535 | RNA→ rplQ (PRK05591)Ribosomal_L17 (pfam01196)→ |
| Ksp-4-1 | NZ_JOGH01000001.1 | - | 191862 | 191808 | RNA→ rplQ (PRK05591)Ribosomal_L17 (pfam01196)→ PseudoU_synth_EcTruA (cd02570)truA (PRK00021)→ AAA (smart00382)Uup (COG0488)→ |
| Ssp-165-1 | NZ_JZWY01000017.1 | - | 18645 | 18591 | RNA→ rplQ (PRK05591)Ribosomal_L17 (pfam01196)→ |
| Ssp-133-1 | NZ_JOHN01000002.1 | + | 49938 | 49992 | RNA→ rplQ (PRK05591)Ribosomal_L17 (pfam01196)→ PseudoU_synth_EcTruA (cd02570)truA (PRK00021)→ AAA (smart00382)Uup (COG0488)→ |
| Ssp-50-1 | NZ_JYJF01000115.1 | - | 10192 | 10138 | RNA→ rplQ (PRK05591)Ribosomal_L17 (pfam01196)→ PseudoU_synth_EcTruA (cd02570)truA (PRK00021)→ |
| Ssp-157-1 | NZ_JOHO01000003.1 | + | 268650 | 268704 | RNA→ rplQ (PRK05591)Ribosomal_L17 (pfam01196)→ PseudoU_synth_EcTruA (cd02570)truA (PRK00021)→ AAA (smart00382)Uup (COG0488)AAA_21 (pfam13304)→ |
| Ssp-160-1 | NZ_JOJA01000001.1 | + | 115341 | 115395 | RNA→ rplQ (PRK05591)Ribosomal_L17 (pfam01196)→ PseudoU_synth_EcTruA (cd02570)truA (PRK00021)→ |
| Ksp-2-1 | NZ_JAIY01000008.1 | - | 87931 | 87877 | RNA→ rplQ (PRK05591)Ribosomal_L17 (pfam01196)→ PseudoU_synth_EcTruA (cd02570)truA (PRK00021)→ AAA (smart00382)Uup (COG0488)→ |
| Kpu-1-1 | NZ_JODS01000006.1 | + | 94156 | 94210 | RNA→ rplQ (PRK05591)Ribosomal_L17 (pfam01196)→ |
| Ssp-183-1 | NZ_JOCF01000001.1 | - | 67177 | 67123 | RNA→ rplQ (PRK05591)Ribosomal_L17 (pfam01196)→ PseudoU_synth_EcTruA (cd02570)truA (PRK00021)→ AAA (smart00382)Uup (COG0488)AAA_21 (pfam13304)→ |
| Sam-2-1 | NZ_JOEQ01000010.1 | - | 266003 | 265948 | RNA→ rplQ (PRK05591)Ribosomal_L17 (pfam01196)→ PseudoU_synth_EcTruA (cd02570)truA (PRK00021)→ |
| Sro-7-1 | NZ_JOEP01000065.1 | + | 28073 | 28126 | RNA→ rplQ (PRK05591)Ribosomal_L17 (pfam01196)→ PseudoU_synth_EcTruA (cd02570)truA (PRK00021)→ |
| Ssp-87-1 | NZ_LAKD01000081.1 | + | 62702 | 62751 | RNA→ rplQ (PRK05591)Ribosomal_L17 (pfam01196)→ PseudoU_synth_EcTruA (cd02570)truA (PRK00021)→ |
| Ssp-190-1 | NZ_JABQ01000020.1 | + | 41705 | 41754 | RNA→ rplQ (PRK05591)Ribosomal_L17 (pfam01196)→ PseudoU_synth_EcTruA (cd02570)truA (PRK00021)→ |
| Sir-1-1 | NZ_LK022848.1 | - | 5397417 | 5397368 | RNA→ rplQ (PRK05591)Ribosomal_L17 (pfam01196)→ PseudoU_synth_EcTruA (cd02570)truA (PRK00021)→ |
| Shy-4-1 | NZ_JOIK01000025.1 | + | 178719 | 178768 | RNA→ rplQ (PRK05591)Ribosomal_L17 (pfam01196)→ PseudoU_synth_EcTruA (cd02570)truA (PRK00021)→ |
| Sro-3-1 | NC_013595.1 | + | 1131580 | 1131634 | RNA→ rplQ (PRK05591)Ribosomal_L17 (pfam01196)→ PseudoU_synth_EcTruA (cd02570)truA (PRK00021)→ |
| Nin-1-1 | NZ_KE383928.1 | + | 830235 | 830308 | RNA→ rplQ (PRK05591)Ribosomal_L17 (pfam01196)→ |
| Nca-1-1 | NZ_JOAG01000016.1 | + | 29392 | 29446 | RNA→ rplQ (PRK05591)Ribosomal_L17 (pfam01196)→ |
| [1]Ssp-192-1 | NZ_LAVK01000035.1 | + | 24842 | 24902 | RNA→ rplQ (PRK05591)Ribosomal_L17 (pfam01196)→ PseudoU_synth_EcTruA (cd02570)truA (PRK00021)→ |
| Ssp-172-1 | NZ_JODF01000016.1 | - | 82667 | 82612 | RNA→ rplQ (PRK05591)Ribosomal_L17 (pfam01196)→ PseudoU_synth_EcTruA (cd02570)truA (PRK00021)→ |
| Ssp-93-1 | NZ_JOJM01000060.1 | - | 2808 | 2751 | RNA→ rplQ (PRK05591)Ribosomal_L17 (pfam01196)→ |
| Sau-3-1 | NZ_JOAP01000029.1 | + | 78987 | 79043 | RNA→ rplQ (PRK05591)Ribosomal_L17 (pfam01196)→ |
| Sau-3-2 | NZ_JOAR01000028.1 | + | 78983 | 79039 | RNA→ rplQ (PRK05591)Ribosomal_L17 (pfam01196)→ |
| Sal-6-1 | NZ_JNXT01000010.1 | - | 3945 | 3889 | RNA→ rplQ (PRK05591)Ribosomal_L17 (pfam01196)→ PseudoU_synth_EcTruA (cd02570)truA (PRK00021)→ |
| Sru-1-1 | NZ_JODN01000016.1 | - | 4008 | 3952 | RNA→ rplQ (PRK05591)Ribosomal_L17 (pfam01196)→ PseudoU_synth_EcTruA (cd02570)truA (PRK00021)→ |
| Sru-1-2 | NZ_JOAQ01000016.1 | - | 4052 | 3996 | RNA→ rplQ (PRK05591)Ribosomal_L17 (pfam01196)→ PseudoU_synth_EcTruA (cd02570)truA (PRK00021)→ |
| Sau-1-1 | NZ_AOPZ01000478.1 | - | 3734 | 3676 | RNA→ ←hypo |
| Ssp-106-1 | NZ_JOIG01000009.1 | - | 261110 | 261058 | RNA→ rplQ (PRK05591)Ribosomal_L17 (pfam01196)→ PseudoU_synth_EcTruA (cd02570)truA (PRK00021)→ |
| Sxi-1-1 | NZ_CP009922.2 | - | 2234636 | 2234584 | RNA→ rplQ (PRK05591)Ribosomal_L17 (pfam01196)→ PseudoU_synth_EcTruA (cd02570)truA (PRK00021)→ |
| Ssp-3-1 | NZ_ALNP01000008.1 | + | 75229 | 75281 | RNA→ rplQ (PRK05591)Ribosomal_L17 (pfam01196)→ PseudoU_synth_EcTruA (cd02570)truA (PRK00021)→ |
| env-15 | JGI12419J13241_1048700 | + | 807 | 860 | RNA→ |
| env-16 | JGI12341J12878_1001749 | + | 4790 | 4843 | RNA→ rplQ (PRK05591)Ribosomal_L17 (pfam01196)→ |
| env-17 | JGI12298J13902_1000008 | - | 41137 | 41084 | RNA→ rplQ (PRK05591)Ribosomal_L17 (pfam01196)→ |
| Tbi-1-1 | NC_014165.1 | + | 644331 | 644384 | RNA→ rplQ (PRK05591)Ribosomal_L17 (pfam01196)→ |
| env-18 | JGI12419J13241_1045219 | - | 901 | 848 | RNA→ rplQ (PRK05591)Ribosomal_L17 (pfam01196)→ |
| Ssp-118-1 | NZ_JOFZ01000006.1 | - | 529759 | 529704 | RNA→ rplQ (PRK05591)Ribosomal_L17 (pfam01196)→ |
| env-19 | 2200133452 | + | 1231 | 1291 | RNA→ rplQ (PRK05591)Ribosomal_L17 (pfam01196)→ |
| env-20 | 2200130945 | - | 4098 | 4038 | RNA→ rplQ (PRK05591)Ribosomal_L17 (pfam01196)→ |
| env-21 | JGI12419J13241_1059560 | - | 305 | 245 | RNA→ rplQ (PRK05591)Ribosomal_L17 (pfam01196)→ |
| Sor-1-1 | NZ_JQMQ01000005.1 | + | 4289657 | 4289704 | RNA→ rplQ (PRK05591)Ribosomal_L17 (pfam01196)→ PseudoU_synth_EcTruA (cd02570)truA (PRK00021)→ |
| Ssp-64-1 | NZ_AZXC01000006.1 | - | 241043 | 240986 | RNA→ rplQ (PRK05591)rpsP (PRK14521)Ribosomal_L17 (pfam01196)→ PseudoU_synth_EcTruA (cd02570)truA (PRK00021)→ |

| | | | | | |
|---|---|---|---|---|---|
| Ssp-12-1 | NZ_KB846744.1 | - | 49103 | 49046 | RNA→ rplQ (PRK05591)Ribosomal_L17 (pfam01196)→ PseudoU_synth_EcTruA (cd02570)truA (PRK00021)→ |
| Ssp-13-1 | NZ_KB898271.1 | - | 505841 | 505784 | RNA→ rplQ (PRK05591)Ribosomal_L17 (pfam01196)→ PseudoU_synth_EcTruA (cd02570)truA (PRK00021)→ |
| Ssp-61-1 | NZ_AXVU01000018.1 | - | 122599 | 122542 | RNA→ rplQ (PRK05591)Ribosomal_L17 (pfam01196)→ PseudoU_synth_EcTruA (cd02570)truA (PRK00021)→ |
| Ssp-59-1 | NZ_CP011492.1 | + | 4931209 | 4931266 | RNA→ rplQ (PRK05591)Ribosomal_L17 (pfam01196)→ PseudoU_synth_EcTruA (cd02570)truA (PRK00021)→ |
| Ssp-16-1 | NZ_KB897717.1 | - | 411625 | 411568 | RNA→ rplQ (PRK05591)Ribosomal_L17 (pfam01196)→ PseudoU_synth_EcTruA (cd02570)truA (PRK00021)→ |
| Ssp-10-1 | NZ_KB897364.1 | - | 373832 | 373775 | RNA→ rplQ (PRK05591)Ribosomal_L17 (pfam01196)→ PseudoU_synth_EcTruA (cd02570)truA (PRK00021)→ |
| Ssp-11-1 | NZ_KB897623.1 | + | 230842 | 230899 | RNA→ rplQ (PRK05591)Ribosomal_L17 (pfam01196)→ PseudoU_synth_EcTruA (cd02570)truA (PRK00021)→ |
| Ssp-63-1 | NZ_AUKP01000008.1 | + | 18089 | 18146 | RNA→ rplQ (PRK05591)Ribosomal_L17 (pfam01196)→ PseudoU_synth_EcTruA (cd02570)truA (PRK00021)→ |
| Ssp-60-1 | NZ_JNID01000001.1 | + | 69239 | 69296 | RNA→ rplQ (PRK05591)Ribosomal_L17 (pfam01196)→ PseudoU_synth_EcTruA (cd02570)truA (PRK00021)→ |
| Ssp-69-1 | NZ_AZWZ01000009.1 | - | 266579 | 266522 | RNA→ rplQ (PRK05591)Ribosomal_L17 (pfam01196)→ PseudoU_synth_EcTruA (cd02570)truA (PRK00021)→ hypo→ |
| Ssp-56-1 | NZ_AZWL01000015.1 | - | 17420 | 17364 | RNA→ rplQ (PRK05591)Ribosomal_L17 (pfam01196)→ PseudoU_synth_EcTruA (cd02570)truA (PRK00021)→ |
| Hcr-1-1 | NZ_JODQ01000002.1 | - | 560025 | 559979 | RNA→ rplQ (PRK05591)Ribosomal_L17 (pfam01196)→ PseudoU_synth_EcTruA (cd02570)truA (PRK00021)→ |
| Msp-1-1 | NZ_KI866524.1 | - | 954571 | 954521 | RNA→ rplQ (PRK05591)Ribosomal_L17 (pfam01196)→ PseudoU_synth_EcTruA (cd02570)truA (PRK00021)→ |
| Mro-1-1 | NZ_JNZQ01000013.1 | + | 26247 | 26297 | RNA→ rplQ (PRK05591)Ribosomal_L17 (pfam01196)→ PseudoU_synth_EcTruA (cd02570)truA (PRK00021)→ |
| Ssp-171-1 | NZ_JOGB01000027.1 | - | 2040 | 1986 | RNA→ rplQ (PRK05591)Ribosomal_L17 (pfam01196)→ PseudoU_synth_EcTruA (cd02570)truA (PRK00021)→ |
| Ssp-43-1 | NZ_JH941042.1 | + | 296021 | 296076 | RNA→ rplQ (PRK05591)Ribosomal_L17 (pfam01196)→ PseudoU_synth_EcTruA (cd02570)truA (PRK00021)→ |
| Svi-13-1 | NZ_LGUP01000370.1 | - | 89952 | 89897 | RNA→ rplQ (PRK05591)Ribosomal_L17 (pfam01196)→ PseudoU_synth_EcTruA (cd02570)truA (PRK00021)→ |
| Swe-1-1 | NZ_JNWK01000006.1 | - | 89924 | 89869 | RNA→ rplQ (PRK05591)Ribosomal_L17 (pfam01196)→ PseudoU_synth_EcTruA (cd02570)truA (PRK00021)→ |
| Ssp-73-1 | NZ_LACL01000017.1 | + | 268699 | 268754 | RNA→ rplQ (PRK05591)Ribosomal_L17 (pfam01196)→ |
| Ssc-3-1 | NZ_KL997422.1 | + | 626325 | 626380 | RNA→ rplQ (PRK05591)Ribosomal_L17 (pfam01196)→ |
| Kau-1-1 | NZ_JODU01000011.1 | - | 303136 | 303081 | RNA→ rplQ (PRK05591)Ribosomal_L17 (pfam01196)→ |
| Sgr-1-1 | NZ_AEYX01000036.1 | - | 159801 | 159746 | RNA→ rplQ (PRK05591)Ribosomal_L17 (pfam01196)→ rplQ (PRK05591)Ribosomal_L17 (pfam01196)→ PseudoU_synth_EcTruA (cd02570)truA (PRK00021)→ PseudoU_synth_EcTruA (cd02570)truA (PRK00021)→ |
| Sto-1-1 | NZ_JFCB01000003.1 | + | 275848 | 275903 | RNA→ rplQ (PRK05591)Ribosomal_L17 (pfam01196)→ |
| Ssp-114-1 | NZ_JOHW01000009.1 | - | 153796 | 153741 | RNA→ rplQ (PRK05591)Ribosomal_L17 (pfam01196)→ PseudoU_synth_EcTruA (cd02570)truA (PRK00021)→ |
| Ssc-4-1 | NZ_JODX01000002.1 | + | 54572 | 54627 | RNA→ rplQ (PRK05591)Ribosomal_L17 (pfam01196)→ PseudoU_synth_EcTruA (cd02570)truA (PRK00021)→ |
| Ssp-1-1 | NZ_KB911732.1 | - | 106280 | 106225 | RNA→ rplQ (PRK05591)Ribosomal_L17 (pfam01196)→ |
| Sau-4-1 | NZ_KL647039.1 | - | 228013 | 227958 | RNA→ rplQ (PRK05591)Ribosomal_L17 (pfam01196)→ PseudoU_synth_EcTruA (cd02570)truA (PRK00021)→ |
| Spr-1-1 | NZ_CM000950.1 | - | 3302692 | 3302637 | RNA→ hypo→ rplQ (PRK05591)Ribosomal_L17 (pfam01196)→ hypo→ PseudoU_synth_EcTruA (cd02570)truA (PRK00021)→ |
| Ssp-189-1 | NZ_JDUZ01000002.1 | + | 233594 | 233648 | RNA→ rplQ (PRK05591)Ribosomal_L17 (pfam01196)→ PseudoU_synth_EcTruA (cd02570)truA (PRK00021)→ AAA (smart00382)Uup (COG0488)AAA_21 (pfam13304)→ DUF2029 (pfam09594)→ |
| Sfl-6-1 | NZ_JNXV01000034.1 | + | 16887 | 16941 | RNA→ rplQ (PRK05591)Ribosomal_L17 (pfam01196)→ |
| Ssp-103-1 | NZ_JOFX01000039.1 | + | 152601 | 152655 | RNA→ rplQ (PRK05591)Ribosomal_L17 (pfam01196)→ |
| Ssp-111-1 | NZ_JYJI01000223.1 | + | 17460 | 17514 | RNA→ rplQ (PRK05591)Ribosomal_L17 (pfam01196)→ |
| Ssp-159-1 | NZ_JOBX01000009.1 | + | 138190 | 138244 | RNA→ rplQ (PRK05591)Ribosomal_L17 (pfam01196)→ |
| Ser-1-1 | NZ_JNZG01000024.1 | + | 137960 | 138014 | RNA→ rplQ (PRK05591)Ribosomal_L17 (pfam01196)→ PseudoU_synth_EcTruA (cd02570)truA (PRK00021)→ AAA (smart00382)Uup (COG0488)AAA_21 (pfam13304)→ DUF2029 (pfam09594)→ |
| Sya-1-1 | NZ_LBMK01000005.1 | - | 318921 | 318867 | RNA→ rplQ (PRK05591)Ribosomal_L17 (pfam01196)→ |
| Svi-12-1 | NZ_LGUV01000155.1 | - | 24360 | 24306 | RNA→ rplQ (PRK05591)Ribosomal_L17 (pfam01196)→ PseudoU_synth_EcTruA (cd02570)truA (PRK00021)→ AAA (smart00382)Uup (COG0488)AAA_21 (pfam13304)→ DUF2029 (pfam09594)→ |
| Ssp-151-1 | NZ_JOCY01000032.1 | - | 24441 | 24387 | RNA→ rplQ (PRK05591)Ribosomal_L17 (pfam01196)→ PseudoU_synth_EcTruA (cd02570)truA (PRK00021)→ AAA (smart00382)Uup (COG0488)AAA_21 (pfam13304)→ DUF2029 (pfam09594)→ |
| Ssp-110-1 | NZ_JOFY01000047.1 | - | 27927 | 27873 | RNA→ rplQ (PRK05591)Ribosomal_L17 (pfam01196)→ PseudoU_synth_EcTruA (cd02570)truA (PRK00021)→ AAA (smart00382)Uup (COG0488)AAA_21 (pfam13304)→ DUF2029 (pfam09594)→ |
| Ssp-112-1 | NZ_JOIB01000007.1 | + | 127472 | 127526 | RNA→ rplQ (PRK05591)Ribosomal_L17 (pfam01196)→ PseudoU_synth_EcTruA (cd02570)truA (PRK00021)→ AAA (smart00382)Uup (COG0488)AAA_21 (pfam13304)→ DUF2029 (pfam09594)→ |
| Ssp-102-1 | NZ_JOIR01000010.1 | + | 206080 | 206134 | RNA→ rplQ (PRK05591)Ribosomal_L17 (pfam01196)→ PseudoU_synth_EcTruA (cd02570)truA (PRK00021)→ AAA (smart00382)Uup (COG0488)AAA_21 (pfam13304)→ DUF2029 (pfam09594)→ |
| Sla-2-1 | NZ_JOBU01000029.1 | + | 79242 | 79296 | RNA→ rplQ (PRK05591)Ribosomal_L17 (pfam01196)→ |
| Ssp-92-1 | NZ_JOHF01000013.1 | + | 131540 | 131594 | RNA→ rplQ (PRK05591)Ribosomal_L17 (pfam01196)→ |
| Ssp-139-1 | NZ_JZWW01000181.1 | - | 23883 | 23829 | RNA→ rplQ (PRK05591)Ribosomal_L17 (pfam01196)→ |
| Ssp-173-1 | NZ_JOJD01000018.1 | - | 24958 | 24904 | RNA→ rplQ (PRK05591)Ribosomal_L17 (pfam01196)→ |

| | | | | | |
|---|---|---|---|---|---|
| Svi-12-2 | NZ_JOAK01000021.1 | - | 24986 | 24932 | **RNA**→ rplQ (PRK05591)Ribosomal_L17 (pfam01196)→ |
| Ssp-8-1 | NZ_GG657750.1 | + | 4667597 | 4667651 | **RNA**→ rplQ (PRK05591)Ribosomal_L17 (pfam01196)→ rplQ (PRK05591)Ribosomal_L17 (pfam01196)→ |
| Svi-7-1 | NZ_CP010407.1 | + | 5084066 | 5084122 | **RNA**→ rplQ (PRK05591)Ribosomal_L17 (pfam01196)→ PseudoU_synth_EcTruA (cd02570)truA (PRK00021)→ |
| Sgr-12-1 | NZ_JNZT01000074.1 | - | 4616 | 4560 | **RNA**→ rplQ (PRK05591)Ribosomal_L17 (pfam01196)→ PseudoU_synth_EcTruA (cd02570)truA (PRK00021)→ |
| Sgr-12-2 | NZ_JOEN01000002.1 | - | 181210 | 181154 | **RNA**→ rplQ (PRK05591)Ribosomal_L17 (pfam01196)→ PseudoU_synth_EcTruA (cd02570)truA (PRK00021)→ |
| Sfl-5-1 | NZ_JNZO01000012.1 | - | 190895 | 190839 | **RNA**→ rplQ (PRK05591)Ribosomal_L17 (pfam01196)→ PseudoU_synth_EcTruA (cd02570)truA (PRK00021)→ |
| Sbi-3-1 | NZ_JNWL01000008.1 | + | 233987 | 234043 | **RNA**→ rplQ (PRK05591)Ribosomal_L17 (pfam01196)→ PseudoU_synth_EcTruA (cd02570)truA (PRK00021)→ |
| Sbi-3-2 | NZ_JOAU01000005.1 | + | 321503 | 321559 | **RNA**→ rplQ (PRK05591)Ribosomal_L17 (pfam01196)→ PseudoU_synth_EcTruA (cd02570)truA (PRK00021)→ |
| Sbi-3-3 | NZ_JNXJ01000008.1 | + | 234102 | 234158 | **RNA**→ rplQ (PRK05591)Ribosomal_L17 (pfam01196)→ PseudoU_synth_EcTruA (cd02570)truA (PRK00021)→ |
| Sex-1-1 | NZ_KL647056.1 | + | 327108 | 327164 | **RNA**→ rplQ (PRK05591)Ribosomal_L17 (pfam01196)→ PseudoU_synth_EcTruA (cd02570)truA (PRK00021)→ |
| Sgl-3-1 | NZ_JODW01000006.1 | - | 235779 | 235723 | **RNA**→ rplQ (PRK05591)Ribosomal_L17 (pfam01196)→ PseudoU_synth_EcTruA (cd02570)truA (PRK00021)→ |
| Sgr-9-1 | NZ_JOBE01000026.1 | + | 78758 | 78814 | **RNA**→ rplQ (PRK05591)Ribosomal_L17 (pfam01196)→ PseudoU_synth_EcTruA (cd02570)truA (PRK00021)→ |
| Ssp-121-1 | NZ_JOHZ01000078.1 | + | 27510 | 27566 | **RNA**→ rplQ (PRK05591)Ribosomal_L17 (pfam01196)→ PseudoU_synth_EcTruA (cd02570)truA (PRK00021)→ |
| Sve-1-1 | NC_018750.1 | + | 4750596 | 4750652 | **RNA**→ rplQ (PRK05591)Ribosomal_L17 (pfam01196)→ PseudoU_synth_EcTruA (cd02570)truA (PRK00021)→ |
| Sav-2-1 | NZ_JZJK01000077.1 | - | 121007 | 120951 | **RNA**→ rplQ (PRK05591)Ribosomal_L17 (pfam01196)→ PseudoU_synth_EcTruA (cd02570)truA (PRK00021)→ |
| Sav-1-1 | NC_003155.4 | + | 6003039 | 6003095 | **RNA**→ rplQ (PRK05591)Ribosomal_L17 (pfam01196)→ PseudoU_synth_EcTruA (cd02570)truA (PRK00021)→ |
| Sav-2-2 | NZ_BAVY01000021.1 | - | 121478 | 121422 | **RNA**→ rplQ (PRK05591)Ribosomal_L17 (pfam01196)→ PseudoU_synth_EcTruA (cd02570)truA (PRK00021)→ |
| Mgl-1-1 | NZ_JOFO01000009.1 | + | 79442 | 79498 | **RNA**→ rplQ (PRK05591)Ribosomal_L17 (pfam01196)→ PseudoU_synth_EcTruA (cd02570)truA (PRK00021)→ |
| Asp-1-1 | NZ_JOGW01000014.1 | - | 37012 | 36956 | **RNA**→ rplQ (PRK05591)Ribosomal_L17 (pfam01196)→ |
| Stu-1-1 | NZ_AEJB01000201.1 | - | 20150 | 20094 | **RNA**→ rplQ (PRK05591)Ribosomal_L17 (pfam01196)→ PseudoU_synth_EcTruA (cd02570)truA (PRK00021)→ |
| Sye-2-1 | NZ_JNWJ01000097.1 | + | 561 | 617 | **RNA**→ rplQ (PRK05591)Ribosomal_L17 (pfam01196)→ PseudoU_synth_EcTruA (cd02570)truA (PRK00021)→ |
| Sdu-1-1 | NZ_JNXR01000027.1 | + | 434 | 490 | **RNA**→ rplQ (PRK05591)Ribosomal_L17 (pfam01196)→ PseudoU_synth_EcTruA (cd02570)truA (PRK00021)→ |
| Sgh-1-1 | NZ_DS999641.1 | - | 3242980 | 3242924 | **RNA**→ rplQ (PRK05591)Ribosomal_L17 (pfam01196)→ rplQ (PRK05591)Ribosomal_L17 (pfam01196)→ PseudoU_synth_EcTruA (cd02570)truA (PRK00021)→ PseudoU_synth_EcTruA (cd02570)truA (PRK00021)→ |
| Svi-5-1 | NZ_JH993790.1 | - | 1541255 | 1541199 | **RNA**→ rplQ (PRK05591)Ribosomal_L17 (pfam01196)→ PseudoU_synth_EcTruA (cd02570)truA (PRK00021)→ |
| Ssp-153-1 | NZ_JOCB01000058.1 | + | 23941 | 23997 | **RNA**→ rplQ (PRK05591)Ribosomal_L17 (pfam01196)→ PseudoU_synth_EcTruA (cd02570)truA (PRK00021)→ |
| Ssp-86-1 | NZ_LAVA01000018.1 | + | 78943 | 78999 | **RNA**→ rplQ (PRK05591)Ribosomal_L17 (pfam01196)→ PseudoU_synth_EcTruA (cd02570)truA (PRK00021)→ |
| Sro-1-1 | NZ_CM002285.1 | - | 3697722 | 3697666 | **RNA**→ rplQ (PRK05591)Ribosomal_L17 (pfam01196)→ PseudoU_synth_EcTruA (cd02570)truA (PRK00021)→ |
| Sgr-7-1 | NZ_JOFU01000024.1 | + | 46860 | 46916 | **RNA**→ rplQ (PRK05591)Ribosomal_L17 (pfam01196)→ |
| Ssp-28-1 | NZ_KB891987.1 | + | 241284 | 241340 | **RNA**→ rplQ (PRK05591)Ribosomal_L17 (pfam01196)→ PseudoU_synth_EcTruA (cd02570)truA (PRK00021)→ |
| Ssp-176-1 | NZ_JOCA01000010.1 | + | 82614 | 82670 | **RNA**→ rplQ (PRK05591)Ribosomal_L17 (pfam01196)→ |
| Ssp-14-1 | NZ_KB897780.1 | - | 288156 | 288100 | **RNA**→ rplQ (PRK05591)Ribosomal_L17 (pfam01196)→ |
| Sac-2-1 | NZ_JODT01000001.1 | - | 203810 | 203754 | **RNA**→ rplQ (PRK05591)Ribosomal_L17 (pfam01196)→ PseudoU_synth_EcTruA (cd02570)truA (PRK00021)→ |
| Sbo-1-1 | NZ_KB405078.1 | - | 1073332 | 1073276 | **RNA**→ rplQ (PRK05591)Ribosomal_L17 (pfam01196)→ |
| Sbo-1-2 | NZ_KB911581.1 | + | 1052469 | 1052525 | **RNA**→ rplQ (PRK05591)Ribosomal_L17 (pfam01196)→ |
| Ssp-2-1 | NZ_KB911779.1 | - | 228049 | 227993 | **RNA**→ rplQ (PRK05591)Ribosomal_L17 (pfam01196)→ PseudoU_synth_EcTruA (cd02570)truA (PRK00021)→ |
| Ssp-98-1 | NZ_JOGP01000006.1 | - | 287417 | 287361 | **RNA**→ rplQ (PRK05591)Ribosomal_L17 (pfam01196)→ |
| Sac-3-1 | NZ_KQ257828.1 | + | 111389 | 111445 | **RNA**→ rplQ (PRK05591)Ribosomal_L17 (pfam01196)→ PseudoU_synth_EcTruA (cd02570)truA (PRK00021)→ |
| Sac-1-1 | NZ_AHBF01000045.1 | - | 10486 | 10430 | **RNA**→ rplQ (PRK05591)Ribosomal_L17 (pfam01196)→ rplQ (PRK05591)Ribosomal_L17 (pfam01196)→ PseudoU_synth_EcTruA (cd02570)truA (PRK00021)→ PseudoU_synth_EcTruA (cd02570)truA (PRK00021)→ |
| Sca-7-1 | NZ_JODY01000013.1 | - | 8155 | 8099 | **RNA**→ rplQ (PRK05591)Ribosomal_L17 (pfam01196)→ PseudoU_synth_EcTruA (cd02570)truA (PRK00021)→ |
| Sgr-2-1 | NZ_GG657758.1 | - | 3229434 | 3229378 | **RNA**→ rplQ (PRK05591)Ribosomal_L17 (pfam01196)→ rplQ (PRK05591)Ribosomal_L17 (pfam01196)→ |
| Ssp-105-1 | NZ_JOIF01000015.1 | + | 61537 | 61593 | **RNA**→ rplQ (PRK05591)Ribosomal_L17 (pfam01196)→ PseudoU_synth_EcTruA (cd02570)truA (PRK00021)→ |
| Ssp-85-1 | NZ_LBDA01000016.1 | + | 39978 | 40034 | **RNA**→ rplQ (PRK05591)Ribosomal_L17 (pfam01196)→ PseudoU_synth_EcTruA (cd02570)truA (PRK00021)→ |
| Sco-3-1 | NC_021985.1 | + | 5098374 | 5098430 | **RNA**→ rplQ (PRK05591)Ribosomal_L17 (pfam01196)→ PseudoU_synth_EcTruA (cd02570)truA (PRK00021)→ |
| Sse-1-1 | NZ_JNXP01000008.1 | - | 210569 | 210513 | **RNA**→ rplQ (PRK05591)Ribosomal_L17 (pfam01196)→ PseudoU_synth_EcTruA (cd02570)truA (PRK00021)→ |
| Sre-1-1 | NZ_KL575664.1 | - | 76522 | 76466 | **RNA**→ RplQ (COG0203)rplQ (PRK05591)→ |
| Spe-1-1 | NZ_KL575016.1 | + | 244282 | 244340 | **RNA**→ rplQ (PRK05591)Ribosomal_L17 (pfam01196)→ |
| Sbi-2-1 | NZ_JOFS01000017.1 | - | 10150 | 10092 | **RNA**→ rplQ (PRK05591)Ribosomal_L17 (pfam01196)→ |
| Sbi-2-2 | NZ_JOFR01000006.1 | - | 10415 | 10357 | **RNA**→ rplQ (PRK05591)Ribosomal_L17 (pfam01196)→ |
| Ssp-83-1 | NZ_JUIG01000050.1 | - | 40198 | 40142 | **RNA**→ rplQ (PRK05591)Ribosomal_L17 (pfam01196)→ PseudoU_synth_EcTruA (cd02570)truA (PRK00021)→ |
| Spl-1-1 | NZ_JTDH01000075.1 | - | 3887 | 3831 | **RNA**→ rplQ (PRK05591)Ribosomal_L17 (pfam01196)→ PseudoU_synth_EcTruA (cd02570)truA (PRK00021)→ |
| Spl-1-2 | NZ_JUIF01000014.1 | - | 27221 | 27165 | **RNA**→ rplQ (PRK05591)Ribosomal_L17 (pfam01196)→ PseudoU_synth_EcTruA (cd02570)truA (PRK00021)→ |

| | | | | | |
|---|---|---|---|---|---|
| Sno-1-1 | NZ_CP009313.1 | + | 4387573 | 4387629 | **RNA**→ rplQ (PRK05591)Ribosomal_L17 (pfam01196)→ |
| San-1-1 | NZ_LGUX01000327.1 | + | 17957 | 18013 | **RNA**→ rplQ (PRK05591)Ribosomal_L17 (pfam01196)→ PseudoU_synth_EcTruA (cd02570)truA (PRK00021)→ |
| Sfl-3-1 | NZ_JOCU01000004.1 | + | 188839 | 188895 | **RNA**→ rplQ (PRK05591)Ribosomal_L17 (pfam01196)→ PseudoU_synth_EcTruA (cd02570)truA (PRK00021)→ |
| Ssp-109-1 | NZ_JOIC01000006.1 | + | 63381 | 63437 | **RNA**→ rplQ (PRK05591)Ribosomal_L17 (pfam01196)→ PseudoU_synth_EcTruA (cd02570)truA (PRK00021)→ |
| Sfl-2-1 | NZ_JNWV01000004.1 | + | 188944 | 189000 | **RNA**→ rplQ (PRK05591)Ribosomal_L17 (pfam01196)→ PseudoU_synth_EcTruA (cd02570)truA (PRK00021)→ |
| Sla-3-1 | NZ_JOBD01000005.1 | + | 187241 | 187297 | **RNA**→ rplQ (PRK05591)Ribosomal_L17 (pfam01196)→ PseudoU_synth_EcTruA (cd02570)truA (PRK00021)→ |
| Ssp-74-1 | NZ_KI601353.1 | - | 672230 | 672174 | **RNA**→ rplQ (PRK05591)Ribosomal_L17 (pfam01196)→ PseudoU_synth_EcTruA (cd02570)truA (PRK00021)→ |
| Ssp-162-1 | NZ_KL585947.1 | + | 251029 | 251085 | **RNA**→ rplQ (PRK05591)Ribosomal_L17 (pfam01196)→ |
| Ssp-164-1 | NZ_KL585169.1 | - | 174791 | 174735 | **RNA**→ rplQ (PRK05591)Ribosomal_L17 (pfam01196)→ |
| Ssp-161-1 | NZ_JOCT01000004.1 | - | 557487 | 557431 | **RNA**→ rplQ (PRK05591)Ribosomal_L17 (pfam01196)→ |
| Sgl-2-1 | NZ_CP009438.1 | + | 4600285 | 4600341 | **RNA**→ rplQ (PRK05591)Ribosomal_L17 (pfam01196)→ |
| Ssp-143-1 | NZ_JOAW01000083.1 | + | 21714 | 21772 | **RNA**→ rplQ (PRK05591)Ribosomal_L17 (pfam01196)→ |
| Ssp-36-1 | NZ_KB892158.1 | + | 80443 | 80499 | **RNA**→ rplQ (PRK05591)Ribosomal_L17 (pfam01196)→ |
| Ssp-196-1 | NZ_KK106988.1 | + | 235730 | 235786 | **RNA**→ rplQ (PRK05591)Ribosomal_L17 (pfam01196)→ |
| Ssp-200-1 | NZ_JXWX01000073.1 | - | 12207 | 12151 | **RNA**→ rplQ (PRK05591)Ribosomal_L17 (pfam01196)→ |
| Sli-2-1 | NZ_GG657756.1 | - | 3314931 | 3314875 | **RNA**→ rplQ (PRK05591)Ribosomal_L17 (pfam01196)→ rplQ (PRK05591)Ribosomal_L17 (pfam01196)→ |
| Sli-1-1 | NZ_CM001889.1 | + | 5192600 | 5192656 | **RNA**→ rplQ (PRK05591)Ribosomal_L17 (pfam01196)→ |
| Sli-2-2 | NZ_CP009124.1 | - | 3347250 | 3347194 | **RNA**→ rplQ (PRK05591)Ribosomal_L17 (pfam01196)→ |
| Sco-2-1 | NC_003888.3 | + | 5143469 | 5143525 | **RNA**→ rplQ (PRK05591)Ribosomal_L17 (pfam01196)→ |
| Svi-9-1 | NZ_JOCE01000019.1 | - | 83006 | 82950 | **RNA**→ rplQ (PRK05591)Ribosomal_L17 (pfam01196)→ |
| Sco-1-1 | NZ_AHGS01001102.1 | + | 34290 | 34346 | **RNA**→ rplQ (PRK05591)Ribosomal_L17 (pfam01196)→ |
| Ssp-128-1 | NZ_JOGV01000002.1 | + | 114246 | 114302 | **RNA**→ rplQ (PRK05591)Ribosomal_L17 (pfam01196)→ |
| Sol-2-1 | NZ_JOFH01000023.1 | - | 178524 | 178468 | **RNA**→ rplQ (PRK05591)Ribosomal_L17 (pfam01196)→ PseudoU_synth_EcTruA (cd02570)truA (PRK00021)→ |
| Ssp-72-1 | NZ_APIV01000001.1 | - | 622171 | 622115 | **RNA**→ rplQ (PRK05591)Ribosomal_L17 (pfam01196)→ PseudoU_synth_EcTruA (cd02570)truA (PRK00021)→ |
| Ssp-124-1 | NZ_JOHJ01000006.1 | + | 93024 | 93080 | **RNA**→ rplQ (PRK05591)Ribosomal_L17 (pfam01196)→ |
| Ssp-147-1 | NZ_JOGT01000009.1 | - | 100768 | 100712 | **RNA**→ rplQ (PRK05591)Ribosomal_L17 (pfam01196)→ |
| Ssp-126-1 | NZ_JOGJ01000017.1 | + | 292846 | 292902 | **RNA**→ rplQ (PRK05591)Ribosomal_L17 (pfam01196)→ |
| Sce-2-1 | NZ_JOEV01000018.1 | - | 142923 | 142867 | **RNA**→ rplQ (PRK05591)Ribosomal_L17 (pfam01196)→ PseudoU_synth_EcTruA (cd02570)truA (PRK00021)→ |
| Smo-1-1 | NZ_AORZ01000051.1 | - | 35230 | 35174 | **RNA**→ rplQ (PRK05591)Ribosomal_L17 (pfam01196)→ PseudoU_synth_EcTruA (cd02570)truA (PRK00021)→ |
| Svi-9-2 | NZ_JODI01000022.1 | + | 100216 | 100272 | **RNA**→ rplQ (PRK05591)Ribosomal_L17 (pfam01196)→ PseudoU_synth_EcTruA (cd02570)truA (PRK00021)→ |
| Sso-1-1 | NZ_AJJM01000095.1 | - | 4025 | 3969 | **RNA**→ rplQ (PRK05591)Ribosomal_L17 (pfam01196)→ rplQ (PRK05591)Ribosomal_L17 (pfam01196)→ |
| Soc-1-1 | NZ_JOAX01000002.1 | + | 178274 | 178330 | **RNA**→ rplQ (PRK05591)Ribosomal_L17 (pfam01196)→ PseudoU_synth_EcTruA (cd02570)truA (PRK00021)→ |
| Sni-2-1 | NZ_JOFQ01000004.1 | - | 303573 | 303517 | **RNA**→ rplQ (PRK05591)Ribosomal_L17 (pfam01196)→ PseudoU_synth_EcTruA (cd02570)truA (PRK00021)→ |
| Svi-11-1 | NZ_JOBH01000014.1 | + | 179111 | 179167 | **RNA**→ rplQ (PRK05591)Ribosomal_L17 (pfam01196)→ PseudoU_synth_EcTruA (cd02570)truA (PRK00021)→ |
| Sex-2-1 | NZ_AZSS01000011.1 | + | 62972 | 63028 | **RNA**→ rplQ (PRK05591)Ribosomal_L17 (pfam01196)→ PseudoU_synth_EcTruA (cd02570)truA (PRK00021)→ |
| Ssp-6-1 | NZ_KB912947.1 | - | 14396 | 14340 | **RNA**→ rplQ (PRK05591)Ribosomal_L17 (pfam01196)→ PseudoU_synth_EcTruA (cd02570)truA (PRK00021)→ |
| Ssp-101-1 | NZ_JOFW01000029.1 | - | 52272 | 52216 | **RNA**→ rplQ (PRK05591)Ribosomal_L17 (pfam01196)→ |
| Ssp-49-1 | NZ_GG753626.1 | + | 3590861 | 3590917 | **RNA**→ rplQ (PRK05591)Ribosomal_L17 (pfam01196)→ rplQ (PRK05591)Ribosomal_L17 (pfam01196)→<br>PseudoU_synth_EcTruA (cd02570)truA (PRK00021)→ PseudoU_synth_EcTruA (cd02570)truA (PRK00021)→ |
| Sal-11-1 | NZ_CP010519.1 | + | 5393200 | 5393256 | **RNA**→ rplQ (PRK05591)Ribosomal_L17 (pfam01196)→ |
| Shy-2-1 | NC_017765.1 | + | 6630461 | 6630517 | **RNA**→ rplQ (PRK05591)Ribosomal_L17 (pfam01196)→ PseudoU_synth_EcTruA (cd02570)truA (PRK00021)→ |
| Shy-3-1 | NC_020895.1 | + | 6324153 | 6324209 | **RNA**→ rplQ (PRK05591)Ribosomal_L17 (pfam01196)→ PseudoU_synth_EcTruA (cd02570)truA (PRK00021)→ |
| Ssp-46-1 | NZ_AGNH01000472.1 | - | 11085 | 11029 | **RNA**→ rplQ (PRK05591)Ribosomal_L17 (pfam01196)→ |
| Ssp-187-1 | NZ_JOCG01000011.1 | + | 26447 | 26503 | **RNA**→ rplQ (PRK05591)Ribosomal_L17 (pfam01196)→ |
| Ssp-156-1 | NZ_JOIY01000026.1 | + | 57550 | 57606 | **RNA**→ rplQ (PRK05591)Ribosomal_L17 (pfam01196)→ |
| Sgr-10-1 | NZ_KL503830.1 | + | 4506022 | 4506078 | **RNA**→ rplQ (PRK05591)Ribosomal_L17 (pfam01196)→ |
| Aac-1-1 | NZ_JNYX01000001.1 | - | 551094 | 551038 | **RNA**→ rplQ (PRK05591)Ribosomal_L17 (pfam01196)→ |
| Ssp-197-1 | NZ_JMLN01000037.1 | + | 22969 | 23025 | **RNA**→ rplQ (PRK05591)Ribosomal_L17 (pfam01196)→ |
| Ssp-20-1 | NZ_KB891532.1 | + | 21043 | 21099 | **RNA**→ rplQ (PRK05591)Ribosomal_L17 (pfam01196)→ |
| Ssp-123-1 | NZ_JOHL01000062.1 | - | 14420 | 14364 | **RNA**→ rplQ (PRK05591)Ribosomal_L17 (pfam01196)→ |
| Ssp-141-1 | NZ_JOHU01000049.1 | - | 16428 | 16372 | **RNA**→ rplQ (PRK05591)Ribosomal_L17 (pfam01196)→ |
| Sga-1-1 | NZ_AOHP01000041.1 | - | 49677 | 49621 | **RNA**→ rplQ (PRK05591)Ribosomal_L17 (pfam01196)→ |

| | | | | | | | |
|---|---|---|---|---|---|---|---|
| Ssp-81-1 | NZ_LARV01000087.1 | + | 36996 | 37052 | **RNA**→ | rplQ (PRK05591)Ribosomal_L17 (pfam01196)→ | |
| Ssp-113-1 | NZ_KL591910.1 | + | 68579 | 68635 | **RNA**→ | rplQ (PRK05591)Ribosomal_L17 (pfam01196)→ | |
| Ssp-57-1 | NZ_AZUS01000002.1 | + | 290899 | 290955 | **RNA**→ | rplQ (PRK05591)Ribosomal_L17 (pfam01196)→ | |
| Ssp-149-1 | NZ_JOIV01000008.1 | + | 53324 | 53380 | **RNA**→ | rplQ (PRK05591)Ribosomal_L17 (pfam01196)→ | PseudoU_synth_EcTruA (cd02570)truA (PRK00021)→ |
| Ssp-178-1 | NZ_JOJG01000009.1 | - | 357423 | 357367 | **RNA**→ | rplQ (PRK05591)Ribosomal_L17 (pfam01196)→ | PseudoU_synth_EcTruA (cd02570)truA (PRK00021)→ |
| Sah-1-1 | NZ_JXYI02000085.1 | + | 175249 | 175305 | **RNA**→ | rplQ (PRK05591)Ribosomal_L17 (pfam01196)→ | PseudoU_synth_EcTruA (cd02570)truA (PRK00021)→ |
| Sal-8-1 | NZ_CP007574.1 | - | 4322438 | 4322382 | **RNA**→ | rplQ (PRK05591)Ribosomal_L17 (pfam01196)→ | PseudoU_synth_EcTruA (cd02570)truA (PRK00021)→ |
| Sal-10-1 | NZ_CP006871.1 | - | 4620900 | 4620844 | **RNA**→ | rplQ (PRK05591)Ribosomal_L17 (pfam01196)→ | PseudoU_synth_EcTruA (cd02570)truA (PRK00021)→ |
| Sal-1-1 | NZ_AROY02000047.1 | + | 56170 | 56226 | **RNA**→ | rplQ (PRK05591)Ribosomal_L17 (pfam01196)→ | PseudoU_synth_EcTruA (cd02570)truA (PRK00021)→ |
| Sal-9-1 | NZ_AXDB02000042.1 | - | 10629 | 10573 | **RNA**→ | rplQ (PRK05591)Ribosomal_L17 (pfam01196)→ | PseudoU_synth_EcTruA (cd02570)truA (PRK00021)→ |
| Svi-10-1 | NZ_JODM01000001.1 | + | 289559 | 289615 | **RNA**→ | rplQ (PRK05591)Ribosomal_L17 (pfam01196)→ | |
| Sch-1-1 | NZ_JH164842.1 | - | 142738 | 142680 | **RNA**→ | rplQ (PRK05591)Ribosomal_L17 (pfam01196)→ | |
| Sia-1-1 | NZ_JNXI01000009.1 | - | 19967 | 19909 | **RNA**→ | rplQ (PRK05591)Ribosomal_L17 (pfam01196)→ | |
| Saf-1-1 | NZ_KE354180.1 | - | 5829 | 5771 | **RNA**→ | rplQ (PRK05591)Ribosomal_L17 (pfam01196)→ | |
| Ssp-89-1 | NZ_BBNN01000017.1 | + | 134818 | 134874 | **RNA**→ | rplQ (PRK05591)Ribosomal_L17 (pfam01196)→ | |
| Sva-1-1 | NZ_JYJH01000002.1 | + | 426 | 482 | **RNA**→ | rplQ (PRK05591)Ribosomal_L17 (pfam01196)→ | |
| Sfl-7-1 | NZ_JNXD01000002.1 | - | 531073 | 531017 | **RNA**→ | rplQ (PRK05591)Ribosomal_L17 (pfam01196)→ | |
| Ssp-158-1 | NZ_JOIZ01000007.1 | + | 222840 | 222896 | **RNA**→ | rplQ (PRK05591)Ribosomal_L17 (pfam01196)→ | |
| Sro-5-1 | NZ_JOFL01000019.1 | - | 115762 | 115706 | **RNA**→ | rplQ (PRK05591)Ribosomal_L17 (pfam01196)→ | |
| Ssp-116-1 | NZ_JOHV01000022.1 | - | 72383 | 72327 | **RNA**→ | rplQ (PRK05591)Ribosomal_L17 (pfam01196)→ | PseudoU_synth_EcTruA (cd02570)truA (PRK00021)→ |
| env-22 | JGI25153J46596_10079674 | + | 416 | 472 | **RNA**→ | rplQ (PRK05591)Ribosomal_L17 (pfam01196)→ | |
| Sam-1-1 | NZ_CP012382.1 | + | 4777985 | 4778041 | **RNA**→ | rplQ (PRK05591)Ribosomal_L17 (pfam01196)→ | PseudoU_synth_EcTruA (cd02570)truA (PRK00021)→ |
| Ssp-51-1 | NZ_AUKV01000055.1 | + | 421 | 477 | **RNA**→ | rplQ (PRK05591)Ribosomal_L17 (pfam01196)→ | PseudoU_synth_EcTruA (cd02570)truA (PRK00021)→ |
| Slu-1-1 | NZ_KN039947.1 | - | 2174179 | 2174123 | **RNA**→ | rplQ (PRK05591)Ribosomal_L17 (pfam01196)→ | PseudoU_synth_EcTruA (cd02570)truA (PRK00021)→ |
| Sna-2-1 | NZ_JRKI01000063.1 | + | 196029 | 196085 | **RNA**→ | rplQ (PRK05591)Ribosomal_L17 (pfam01196)→ | |
| Sna-1-1 | NZ_KL570007.1 | - | 55047 | 54991 | **RNA**→ | rplQ (PRK05591)Ribosomal_L17 (pfam01196)→ | |
| Ssp-75-1 | NZ_LDZX01000014.1 | - | 12861 | 12805 | **RNA**→ | rplQ (PRK05591)Ribosomal_L17 (pfam01196)→ | |
| env-23 | JGI12419J13241_1019540 | - | 599 | 543 | **RNA**→ | rplQ (PRK05591)Ribosomal_L17 (pfam01196)→ | |
| Ssp-182-1 | NZ_JOJH01000005.1 | - | 219931 | 219875 | **RNA**→ | rplQ (PRK05591)Ribosomal_L17 (pfam01196)→ | PseudoU_synth_EcTruA (cd02570)truA (PRK00021)→ |
| Scy-2-1 | NZ_CP010849.1 | + | 4633351 | 4633407 | **RNA**→ | rplQ (PRK05591)Ribosomal_L17 (pfam01196)→ | PseudoU_synth_EcTruA (cd02570)truA (PRK00021)→ |
| Ssp-184-1 | NZ_JNXK01000001.1 | + | 184828 | 184884 | **RNA**→ | rplQ (PRK05591)Ribosomal_L17 (pfam01196)→ | PseudoU_synth_EcTruA (cd02570)truA (PRK00021)→ |
| Ssp-184-2 | NZ_JOCJ01000007.1 | + | 53530 | 53586 | **RNA**→ | rplQ (PRK05591)Ribosomal_L17 (pfam01196)→ | PseudoU_synth_EcTruA (cd02570)truA (PRK00021)→ |
| Svi-4-1 | NZ_AMLP01000111.1 | - | 27781 | 27725 | **RNA**→ | rplQ (PRK05591)Ribosomal_L17 (pfam01196)→ | |
| Ssp-163-1 | NZ_JOJB01000037.1 | + | 425 | 481 | **RNA**→ | rplQ (PRK05591)Ribosomal_L17 (pfam01196)→ | PseudoU_synth_EcTruA (cd02570)truA (PRK00021)→ |
| Ssp-177-1 | NZ_KL579031.1 | - | 16656 | 16600 | **RNA**→ | rplQ (PRK05591)Ribosomal_L17 (pfam01196)→ | |
| Sol-1-1 | NZ_JJOH01000017.1 | + | 251041 | 251097 | **RNA**→ | rplQ (PRK05591)Ribosomal_L17 (pfam01196)→ | |
| Svi-3-1 | NZ_GG657757.1 | + | 5367559 | 5367615 | **RNA**→ | rplQ (PRK05591)Ribosomal_L17 (pfam01196)→ | rplQ (PRK05591)Ribosomal_L17 (pfam01196)→ |
| Szi-1-1 | NZ_AGBF01000029.1 | - | 28163 | 28107 | **RNA**→ | rplQ (PRK05591)Ribosomal_L17 (pfam01196)→ | rplQ (PRK05591)Ribosomal_L17 (pfam01196)→ |
| Svi-13-2 | NZ_LFNT01000154.1 | + | 521 | 577 | **RNA**→ | rplQ (PRK05591)Ribosomal_L17 (pfam01196)→ | |
| Svi-13-3 | NZ_LGUR01000037.1 | + | 521 | 577 | **RNA**→ | rplQ (PRK05591)Ribosomal_L17 (pfam01196)→ | |
| Svi-13-4 | NZ_LGUQ01000054.1 | - | 6858 | 6802 | **RNA**→ | rplQ (PRK05591)Ribosomal_L17 (pfam01196)→ | |
| Ssp-120-1 | NZ_JOIA01001474.1 | + | 156 | 212 | **RNA**→ | rplQ (PRK05591)Ribosomal_L17 (pfam01196)→ | |
| Ssp-131-1 | NZ_JOGX01000044.1 | + | 26042 | 26098 | **RNA**→ | rplQ (PRK05591)Ribosomal_L17 (pfam01196)→ | PseudoU_synth_EcTruA (cd02570)truA (PRK00021)→ |
| Sca-4-1 | NZ_LFXA01000017.1 | + | 733261 | 733317 | **RNA**→ | rplQ (PRK05591)Ribosomal_L17 (pfam01196)→ | PseudoU_synth_EcTruA (cd02570)truA (PRK00021)→ |
| Sba-1-1 | NZ_KI547048.1 | + | 192269 | 192328 | **RNA**→ | rplQ (PRK05591)Ribosomal_L17 (pfam01196)→ | |
| Ssp-65-1 | NZ_AUFD01000010.1 | - | 185036 | 184977 | **RNA**→ | rplQ (PRK05591)Ribosomal_L17 (pfam01196)→ | |
| Sme-2-1 | NZ_JODL01000029.1 | - | 20518 | 20462 | **RNA**→ | rplQ (PRK05591)Ribosomal_L17 (pfam01196)→ | PseudoU_synth_EcTruA (cd02570)truA (PRK00021)→ |
| Sgr-12-3 | NZ_JNZI01000001.1 | - | 441683 | 441627 | **RNA**→ | rplQ (PRK05591)Ribosomal_L17 (pfam01196)→ | PseudoU_synth_EcTruA (cd02570)truA (PRK00021)→ |
| Sgr-12-4 | NZ_JOGN01000053.1 | - | 20537 | 20481 | **RNA**→ | rplQ (PRK05591)Ribosomal_L17 (pfam01196)→ | PseudoU_synth_EcTruA (cd02570)truA (PRK00021)→ |
| Ssp-15-1 | NZ_KB898225.1 | + | 40192 | 40248 | **RNA**→ | rplQ (PRK05591)Ribosomal_L17 (pfam01196)→ | PseudoU_synth_EcTruA (cd02570)truA (PRK00021)→ |
| Ssp-38-1 | NZ_FR873697.1 | - | 1509259 | 1509203 | **RNA**→ | rplQ (PRK05591)Ribosomal_L17 (pfam01196)→ | PseudoU_synth_EcTruA (cd02570)truA (PRK00021)→ |
| Sal-5-1 | NZ_JOII01000001.1 | - | 312744 | 312688 | **RNA**→ | rplQ (PRK05591)Ribosomal_L17 (pfam01196)→ | PseudoU_synth_EcTruA (cd02570)truA (PRK00021)→ |

| | | | | | |
|---|---|---|---|---|---|
| Ssp-99-1 | NZ_JOGQ01000003.1 | + | 125525 | 125581 | **RNA**→ rplQ (PRK05591)Ribosomal_L17 (pfam01196)→ PseudoU_synth_EcTruA (cd02570)truA (PRK00021)→ |
| Ssp-77-1 | NZ_LAYX01000002.1 | - | 159453 | 159397 | **RNA**→ rplQ (PRK05591)Ribosomal_L17 (pfam01196)→ PseudoU_synth_EcTruA (cd02570)truA (PRK00021)→ |
| Swa-1-1 | NZ_JHDU01000025.1 | - | 87726 | 87670 | **RNA**→ rplQ (PRK05591)Ribosomal_L17 (pfam01196)→ PseudoU_synth_EcTruA (cd02570)truA (PRK00021)→ |
| Ssp-30-1 | NZ_KB890704.1 | - | 981330 | 981274 | **RNA**→ rplQ (PRK05591)Ribosomal_L17 (pfam01196)→ PseudoU_synth_EcTruA (cd02570)truA (PRK00021)→ |
| Ssp-134-1 | NZ_JOHX01000007.1 | - | 127009 | 126953 | **RNA**→ rplQ (PRK05591)Ribosomal_L17 (pfam01196)→ PseudoU_synth_EcTruA (cd02570)truA (PRK00021)→ |
| Ssp-62-1 | NZ_JTCK01000085.1 | + | 32179 | 32235 | **RNA**→ rplQ (PRK05591)Ribosomal_L17 (pfam01196)→ PseudoU_synth_EcTruA (cd02570)truA (PRK00021)→ |
| Ssp-21-1 | NZ_CM002271.1 | + | 4349395 | 4349451 | **RNA**→ rplQ (PRK05591)Ribosomal_L17 (pfam01196)→ PseudoU_synth_EcTruA (cd02570)truA (PRK00021)→ |
| Ssp-35-1 | NZ_CM002273.1 | + | 4354309 | 4354365 | **RNA**→ rplQ (PRK05591)Ribosomal_L17 (pfam01196)→ PseudoU_synth_EcTruA (cd02570)truA (PRK00021)→ |
| Ssp-80-1 | NZ_KN549146.1 | + | 250775 | 250831 | **RNA**→ rplQ (PRK05591)Ribosomal_L17 (pfam01196)→ PseudoU_synth_EcTruA (cd02570)truA (PRK00021)→ |
| Sgr-12-5 | NZ_JOGU01000007.1 | - | 148467 | 148411 | **RNA**→ rplQ (PRK05591)Ribosomal_L17 (pfam01196)→ PseudoU_synth_EcTruA (cd02570)truA (PRK00021)→ |
| Sva-2-1 | NZ_JOBF01000008.1 | - | 20052 | 19996 | **RNA**→ rplQ (PRK05591)Ribosomal_L17 (pfam01196)→ |
| Sva-2-2 | NZ_JOFN01000040.1 | - | 18282 | 18226 | **RNA**→ rplQ (PRK05591)Ribosomal_L17 (pfam01196)→ |
| Ssp-58-1 | NZ_AUKQ01000015.1 | + | 52320 | 52377 | **RNA**→ rplQ (PRK05591)Ribosomal_L17 (pfam01196)→ |
| Sal-12-1 | NZ_JOID01000003.1 | + | 68800 | 68856 | **RNA**→ rplQ (PRK05591)Ribosomal_L17 (pfam01196)→ |
| Sfr-1-1 | NZ_LGSP01000027.1 | - | 10382 | 10326 | **RNA**→ rplQ (PRK05591)Ribosomal_L17 (pfam01196)→ |
| Sfr-1-2 | NZ_JNAD01000026.1 | - | 10369 | 10313 | **RNA**→ rplQ (PRK05591)Ribosomal_L17 (pfam01196)→ |
| Sxi-2-1 | NZ_AFRP01001524.1 | + | 40341 | 40397 | **RNA**→ rplQ (PRK05591)Ribosomal_L17 (pfam01196)→ |
| Ssp-119-1 | NZ_JOHR01000002.1 | + | 230095 | 230151 | **RNA**→ rplQ (PRK05591)Ribosomal_L17 (pfam01196)→ |
| Ssp-33-1 | NZ_KB891885.1 | - | 239596 | 239541 | **RNA**→ rplQ (PRK05591)Ribosomal_L17 (pfam01196)→ PseudoU_synth_EcTruA (cd02570)truA (PRK00021)→ |
| Ssp-37-1 | NZ_BARG01000058.1 | - | 624 | 569 | **RNA**→ rplQ (PRK05591)Ribosomal_L17 (pfam01196)→ |
| Ssp-170-1 | NZ_JODG01000026.1 | + | 329 | 385 | **RNA**→ rplQ (PRK05591)Ribosomal_L17 (pfam01196)→ PseudoU_synth_EcTruA (cd02570)truA (PRK00021)→ Ugd (COG1004)NDP-sugDHase (TIGR03026)→ |
| Smi-1-1 | NZ_KN050740.1 | - | 288662 | 288606 | **RNA**→ rplQ (PRK05591)Ribosomal_L17 (pfam01196)→ PseudoU_synth_EcTruA (cd02570)truA (PRK00021)→ |
| Ssp-138-1 | NZ_JOIU01000016.1 | + | 88730 | 88786 | **RNA**→ rplQ (PRK05591)Ribosomal_L17 (pfam01196)→ PseudoU_synth_EcTruA (cd02570)truA (PRK00021)→ |
| Sga-2-1 | NZ_JRHJ01000054.1 | + | 156382 | 156438 | **RNA**→ rplQ (PRK05591)Ribosomal_L17 (pfam01196)→ |
| env-24 | JGI24740J21852_10018017 | - | 980 | 924 | **RNA**→ rplQ (PRK05591)Ribosomal_L17 (pfam01196)→ |
| env-25 | JGI24739J22299_10011386 | - | 2321 | 2265 | **RNA**→ rplQ (PRK05591)Ribosomal_L17 (pfam01196)→ |
| env-26 | JGI24737J22298_10030274 | - | 711 | 655 | **RNA**→ rplQ (PRK05591)Ribosomal_L17 (pfam01196)→ hypo→ |
| env-27 | JGI24735J21928_10022680 | + | 1260 | 1316 | **RNA**→ rplQ (PRK05591)Ribosomal_L17 (pfam01196)→ |
| env-28 | JGI24738J21930_10016389 | + | 901 | 957 | **RNA**→ rplQ (PRK05591)Ribosomal_L17 (pfam01196)→ |
| env-29 | JGI24738J21930_10017841 | - | 666 | 610 | **RNA**→ rplQ (PRK05591)Ribosomal_L17 (pfam01196)→ |
| Sca-1-1 | NZ_KB911586.1 | + | 111863 | 111919 | **RNA**→ rplQ (PRK05591)Ribosomal_L17 (pfam01196)→ PseudoU_synth_EcTruA (cd02570)truA (PRK00021)→ |
| Ssp-23-1 | NZ_KE150428.1 | + | 48833 | 48889 | **RNA**→ rplQ (PRK05591)Ribosomal_L17 (pfam01196)→ PseudoU_synth_EcTruA (cd02570)truA (PRK00021)→ |
| Ssp-135-1 | NZ_JNXF01000033.1 | - | 14094 | 14038 | **RNA**→ rplQ (PRK05591)Ribosomal_L17 (pfam01196)→ |
| Ssp-137-1 | NZ_JOHP01000033.1 | - | 14040 | 13984 | **RNA**→ rplQ (PRK05591)Ribosomal_L17 (pfam01196)→ |
| Ssp-91-1 | NZ_JNZW01000034.1 | - | 14086 | 14030 | **RNA**→ rplQ (PRK05591)Ribosomal_L17 (pfam01196)→ |
| Ssp-136-1 | NZ_JOHS01000031.1 | - | 14033 | 13977 | **RNA**→ rplQ (PRK05591)Ribosomal_L17 (pfam01196)→ |
| Ssp-169-1 | NZ_JODC01000003.1 | - | 760731 | 760675 | **RNA**→ rplQ (PRK05591)Ribosomal_L17 (pfam01196)→ |
| env-30 | JGI24736J21556_1066001 | + | 409 | 465 | **RNA**→ |
| Ssv-1-1 | NZ_CM000951.1 | + | 5617327 | 5617383 | **RNA**→ rplQ (PRK05591)Ribosomal_L17 (pfam01196)→ hypo→ PseudoU_synth_EcTruA (cd02570)truA (PRK00021)→ hypo→ |
| Ssp-186-1 | NZ_JOBZ01000048.1 | + | 11344 | 11400 | **RNA**→ rplQ (PRK05591)Ribosomal_L17 (pfam01196)→ |
| Sda-1-1 | NC_020504.1 | - | 4173417 | 4173361 | **RNA**→ rplQ (PRK05591)Ribosomal_L17 (pfam01196)→ PseudoU_synth_EcTruA (cd02570)truA (PRK00021)→ |
| Sgr-12-6 | NZ_JOJE01000032.1 | - | 42148 | 42092 | **RNA**→ rplQ (PRK05591)Ribosomal_L17 (pfam01196)→ |
| Ssp-199-1 | NZ_JXTE01000103.1 | - | 19424 | 19368 | **RNA**→ rplQ (PRK05591)Ribosomal_L17 (pfam01196)→ |
| env-31 | JGI25160J50197_1027159 | - | 1290 | 1235 | **RNA**→ rplQ (PRK05591)Ribosomal_L17 (pfam01196)→ |
| Ssp-88-1 | NZ_BBNO01000005.1 | + | 169561 | 169616 | **RNA**→ rplQ (PRK05591)Ribosomal_L17 (pfam01196)→ PseudoU_synth_EcTruA (cd02570)truA (PRK00021)→ |
| Ssp-19-1 | NZ_AQWO01000012.1 | - | 42118 | 42063 | **RNA**→ rplQ (PRK05591)Ribosomal_L17 (pfam01196)→ PseudoU_synth_EcTruA (cd02570)truA (PRK00021)→ |
| Sfu-2-1 | NZ_JOEY01000005.1 | - | 283078 | 283023 | **RNA**→ rplQ (PRK05591)Ribosomal_L17 (pfam01196)→ PseudoU_synth_EcTruA (cd02570)truA (PRK00021)→ Glyco_hydro_26 (pfam02156)→ |
| Sau-2-1 | NZ_JH725387.1 | + | 3734051 | 3734106 | **RNA**→ rplQ (PRK05591)Ribosomal_L17 (pfam01196)→ |
| Sxy-1-1 | NZ_JNWO01000013.1 | + | 27275 | 27330 | **RNA**→ rplQ (PRK05591)Ribosomal_L17 (pfam01196)→ |
| Ssp-122-1 | NZ_JNXE01000005.1 | + | 80283 | 80338 | **RNA**→ rplQ (PRK05591)Ribosomal_L17 (pfam01196)→ |

| | | | | | |
|---|---|---|---|---|---|
| Spr-2-1 | NZ_BARF01000005.1 | + | 85303 | 85358 | **RNA**→ rplQ (PRK05591)Ribosomal_L17 (pfam01196)→ |
| Ssp-84-1 | NZ_LBMU01000061.1 | - | 29957 | 29901 | **RNA**→ rplQ (PRK05591)Ribosomal_L17 (pfam01196)→ |
| Sip-1-1 | NZ_AEJC01000092.1 | + | 9205 | 9260 | **RNA**→ rplQ (PRK05591)Ribosomal_L17 (pfam01196)→ PseudoU_synth_EcTruA (cd02570)truA (PRK00021)→ |
| Sly-2-1 | NZ_CP007699.1 | + | 4988677 | 4988732 | **RNA**→ rplQ (PRK05591)Ribosomal_L17 (pfam01196)→ PseudoU_synth_EcTruA (cd02570)truA (PRK00021)→ |
| Ssc-2-1 | NZ_KB889574.1 | - | 55636 | 55581 | **RNA**→ rplQ (PRK05591)Ribosomal_L17 (pfam01196)→ |
| Sye-1-1 | NZ_JQNR01000005.1 | + | 1956765 | 1956820 | **RNA**→ rplQ (PRK05591)Ribosomal_L17 (pfam01196)→ |
| Svi-6-1 | NZ_KB904651.1 | - | 121812 | 121756 | **RNA**→ rplQ (PRK05591)Ribosomal_L17 (pfam01196)→ |
| Sni-1-1 | NZ_CM002280.1 | - | 3417483 | 3417429 | **RNA**→ rplQ (PRK05591)Ribosomal_L17 (pfam01196)→ PseudoU_synth_EcTruA (cd02570)truA (PRK00021)→ |
| Ssu-2-1 | NZ_AJTQ01000307.1 | + | 13549 | 13604 | **RNA**→ RplQ (COG0203)rplQ (PRK05591)→ |
| Ssu-1-1 | NZ_KB905817.1 | - | 189202 | 189147 | **RNA**→ RplQ (COG0203)rplQ (PRK05591)→ PseudoU_synth_EcTruA (cd02570)truA (PRK00021)→ |
| Svi-2-1 | NC_015957.1 | + | 705115 | 705170 | **RNA**→ rplQ (PRK05591)Ribosomal_L17 (pfam01196)→ PseudoU_synth_EcTruA (cd02570)truA (PRK00021)→ |
| Sfl-4-1 | NZ_AUBE01000006.1 | - | 238624 | 238569 | **RNA**→ rplQ (PRK05591)Ribosomal_L17 (pfam01196)→ PseudoU_synth_EcTruA (cd02570)truA (PRK00021)→ |
| Spu-1-1 | NZ_KB913030.1 | + | 4401513 | 4401568 | **RNA**→ rplQ (PRK05591)Ribosomal_L17 (pfam01196)→ PseudoU_synth_EcTruA (cd02570)truA (PRK00021)→ |
| Ssc-4-2 | NZ_JOBC01000019.1 | - | 77220 | 77165 | **RNA**→ rplQ (PRK05591)Ribosomal_L17 (pfam01196)→ PseudoU_synth_EcTruA (cd02570)truA (PRK00021)→ |
| Shy-1-1 | NZ_GG657754.1 | - | 4571299 | 4571244 | **RNA**→ rplQ (PRK05591)Ribosomal_L17 (pfam01196)→ rplQ (PRK05591)Ribosomal_L17 (pfam01196)→ |
| env-32 | JGI25160J50197_1040719 | - | 1000 | 945 | **RNA**→ rplQ (PRK05591)Ribosomal_L17 (pfam01196)→ truA (PRK00021)PseudoU_synth_1 (pfam01416)→ |
| Sbi-1-1 | NC_016582.1 | + | 7517266 | 7517321 | **RNA**→ rplQ (PRK05591)Ribosomal_L17 (pfam01196)→ |
| Afl-1-1 | NZ_KB894099.1 | - | 208768 | 208711 | **RNA**→ rplQ (PRK05591)Ribosomal_L17 (pfam01196)→ |
| Ssp-55-1 | NZ_JTIY01000001.1 | - | 1514095 | 1514041 | **RNA**→ rplQ (PRK05591)Ribosomal_L17 (pfam01196)→ |
| Ssp-152-1 | NZ_JOCX01000005.1 | + | 52944 | 52998 | **RNA**→ rplQ (PRK05591)Ribosomal_L17 (pfam01196)→ |
| Ssp-104-1 | NZ_JOIS01000020.1 | - | 101942 | 101888 | **RNA**→ rplQ (PRK05591)Ribosomal_L17 (pfam01196)→ |
| Ssp-166-1 | NZ_JODE01000005.1 | + | 46633 | 46687 | **RNA**→ rplQ (PRK05591)Ribosomal_L17 (pfam01196)→ PseudoU_synth_EcTruA (cd02570)truA (PRK00021)→ |
| | | | | | AAA (smart00382)Uup (COG0488)AAA_21 (pfam13304)→ DUF2029 (pfam09594)→ DUF2029 (pfam09594)→ |
| Ssp-167-1 | NZ_JODJ01000029.1 | + | 46652 | 46706 | **RNA**→ rplQ (PRK05591)Ribosomal_L17 (pfam01196)→ PseudoU_synth_EcTruA (cd02570)truA (PRK00021)→ |
| | | | | | AAA (smart00382)Uup (COG0488)AAA_21 (pfam13304)→ DUF2029 (pfam09594)→ DUF2029 (pfam09594)→ |
| Ssp-191-1 | NZ_JWZS01000804.1 | + | 24884 | 24938 | **RNA**→ rplQ (PRK05591)Ribosomal_L17 (pfam01196)→ PseudoU_synth_EcTruA (cd02570)truA (PRK00021)→ |
| | | | | | AAA (smart00382)Uup (COG0488)AAA_21 (pfam13304)→ hypo→ |
| Sxa-1-1 | NZ_JNZH01000037.1 | - | 123110 | 123056 | **RNA**→ rplQ (PRK05591)Ribosomal_L17 (pfam01196)→ PseudoU_synth_EcTruA (cd02570)truA (PRK00021)→ |
| | | | | | AAA (smart00382)Uup (COG0488)→ DUF2029 (pfam09594)→ |
| Sxa-1-2 | NZ_JOFT01000047.1 | + | 32685 | 32739 | **RNA**→ rplQ (PRK05591)Ribosomal_L17 (pfam01196)→ PseudoU_synth_EcTruA (cd02570)truA (PRK00021)→ |
| | | | | | AAA (smart00382)Uup (COG0488)→ DUF2029 (pfam09594)→ |
| Sla-1-1 | NZ_JNXL01000004.1 | + | 16844 | 16898 | **RNA**→ rplQ (PRK05591)Ribosomal_L17 (pfam01196)→ |
| Svi-12-3 | NZ_JNYC01000038.1 | + | 47873 | 47927 | **RNA**→ rplQ (PRK05591)Ribosomal_L17 (pfam01196)→ PseudoU_synth_EcTruA (cd02570)truA (PRK00021)→ |
| | | | | | AAA (smart00382)Uup (COG0488)AAA_21 (pfam13304)→ |
| Sla-2-2 | NZ_JOEW01000015.1 | + | 48662 | 48716 | **RNA**→ rplQ (PRK05591)Ribosomal_L17 (pfam01196)→ PseudoU_synth_EcTruA (cd02570)truA (PRK00021)→ |
| | | | | | AAA (smart00382)Uup (COG0488)AAA_21 (pfam13304)→ DUF2029 (pfam09594)→ |
| Ssp-150-1 | NZ_JODA01000027.1 | - | 24120 | 24066 | **RNA**→ rplQ (PRK05591)Ribosomal_L17 (pfam01196)→ PseudoU_synth_EcTruA (cd02570)truA (PRK00021)→ |
| | | | | | AAA (smart00382)Uup (COG0488)AAA_21 (pfam13304)→ DUF2029 (pfam09594)→ |
| Ssp-32-1 | NZ_DS570396.1 | - | 78156 | 78102 | **RNA**→ rplQ (PRK05591)Ribosomal_L17 (pfam01196)→ rplQ (PRK05591)Ribosomal_L17 (pfam01196)→ |
| | | | | | PseudoU_synth_EcTruA (cd02570)truA (PRK00021)→ PseudoU_synth_EcTruA (cd02570)truA (PRK00021)→ |
| | | | | | AAA (smart00382)Uup (COG0488)AAA_21 (pfam13304)→ |
| | | | | | AAA (smart00382)Uup (COG0488)AAA_21 (pfam13304)→ DUF2029 (pfam09594)→ DUF2029 (pfam09594)→ |
| Ssp-32-2 | NZ_CP011664.1 | + | 4787937 | 4787991 | **RNA**→ rplQ (PRK05591)Ribosomal_L17 (pfam01196)→ PseudoU_synth_EcTruA (cd02570)truA (PRK00021)→ |
| | | | | | AAA (smart00382)Uup (COG0488)AAA_21 (pfam13304)→ DUF2029 (pfam09594)→ |
| Ska-1-1 | NZ_JNZY01000017.1 | + | 138776 | 138830 | **RNA**→ rplQ (PRK05591)Ribosomal_L17 (pfam01196)→ |
| Sro-6-1 | NZ_LFML01000045.1 | + | 17342 | 17396 | **RNA**→ rplQ (PRK05591)Ribosomal_L17 (pfam01196)→ |
| Ssp-194-1 | NZ_AUKR01000012.1 | + | 193841 | 193895 | **RNA**→ rplQ (PRK05591)Ribosomal_L17 (pfam01196)→ PseudoU_synth_EcTruA (cd02570)truA (PRK00021)→ |
| env-33 | LWSO_NODE_580 | + | 2805 | 2860 | **RNA**→ rplQ (PRK05591)Ribosomal_L17 (pfam01196)→ PseudoU_synth_EcTruA (cd02570)truA (PRK00021)→ |
| Ssp-181-1 | NZ_JOCD01000044.1 | - | 26202 | 26147 | **RNA**→ rplQ (PRK05591)Ribosomal_L17 (pfam01196)→ |
| Ssp-7-1 | NZ_KB892128.1 | + | 655 | 710 | **RNA**→ rplQ (PRK05591)Ribosomal_L17 (pfam01196)→ PseudoU_synth_EcTruA (cd02570)truA (PRK00021)→ |
| Ssp-140-1 | NZ_KL591014.1 | + | 86235 | 86290 | **RNA**→ rplQ (PRK05591)Ribosomal_L17 (pfam01196)→ PseudoU_synth_EcTruA (cd02570)truA (PRK00021)→ |
| Svi-1-1 | NZ_ASHX01000001.1 | - | 1241638 | 1241583 | **RNA**→ rplQ (PRK05591)Ribosomal_L17 (pfam01196)→ |
| Sca-2-1 | NC_016111.1 | + | 3889184 | 3889239 | **RNA**→ rplQ (PRK05591)Ribosomal_L17 (pfam01196)→ PseudoU_synth_EcTruA (cd02570)truA (PRK00021)→ |
| Sca-2-2 | NC_017586.1 | + | 3888995 | 3889050 | **RNA**→ rplQ (PRK05591)Ribosomal_L17 (pfam01196)→ PseudoU_synth_EcTruA (cd02570)truA (PRK00021)→ |

| | | | | | |
|---|---|---|---|---|---|
| Sts-1-1 | NZ_AJSZ01000399.1 | - | 24785 | 24730 | **RNA**→ rplQ (PRK05591)Ribosomal_L17 (pfam01196)→ rplQ (PRK05591)Ribosomal_L17 (pfam01196)→ |
| Ssp-25-1 | NZ_KB891797.1 | + | 527129 | 527185 | **RNA**→ rplQ (PRK05591)Ribosomal_L17 (pfam01196)→ PseudoU_synth_EcTruA (cd02570)truA (PRK00021)→ |
| Ska-1-2 | NZ_JZWV01000003.1 | + | 52624 | 52679 | **RNA**→ rplQ (PRK05591)Ribosomal_L17 (pfam01196)→ PseudoU_synth_EcTruA (cd02570)truA (PRK00021)→ |
| | | | | | AAA (smart00382)Uup (COG0488)AAA_21 (pfam13304)→ DUF2029 (pfam09594)→ |
| Smo-2-1 | NZ_JNYL01000032.1 | - | 28345 | 28290 | **RNA**→ rplQ (PRK05591)Ribosomal_L17 (pfam01196)→ |
| Ssp-70-1 | NZ_KN708638.1 | - | 3669055 | 3669000 | **RNA**→ rplQ (PRK05591)Ribosomal_L17 (pfam01196)→ |
| Sba-2-1 | NZ_KL573554.1 | + | 64146 | 64201 | **RNA**→ rplQ (PRK05591)Ribosomal_L17 (pfam01196)→ PseudoU_synth_EcTruA (cd02570)truA (PRK00021)→ |
| Sme-1-1 | NZ_JOJK01000001.1 | - | 178778 | 178723 | **RNA**→ rplQ (PRK05591)Ribosomal_L17 (pfam01196)→ PseudoU_synth_EcTruA (cd02570)truA (PRK00021)→ |
| Sgr-11-1 | NZ_JYBE01000004.1 | - | 64041 | 63986 | **RNA**→ rplQ (PRK05591)Ribosomal_L17 (pfam01196)→ PseudoU_synth_EcTruA (cd02570)truA (PRK00021)→ |
| Sgl-1-1 | NZ_JH689267.1 | - | 288602 | 288547 | **RNA**→ rplQ (PRK05591)Ribosomal_L17 (pfam01196)→ PseudoU_synth_EcTruA (cd02570)truA (PRK00021)→ |
| Ssp-76-1 | NZ_JPWW01000002.1 | + | 1014580 | 1014635 | **RNA**→ rplQ (PRK05591)Ribosomal_L17 (pfam01196)→ PseudoU_synth_EcTruA (cd02570)truA (PRK00021)→ |
| Sgr-12-7 | NZ_JOGA01000004.1 | - | 151218 | 151163 | **RNA**→ rplQ (PRK05591)Ribosomal_L17 (pfam01196)→ PseudoU_synth_EcTruA (cd02570)truA (PRK00021)→ |
| Sgl-3-2 | NZ_JNZK01000003.1 | - | 116076 | 116021 | **RNA**→ rplQ (PRK05591)Ribosomal_L17 (pfam01196)→ PseudoU_synth_EcTruA (cd02570)truA (PRK00021)→ |
| Sgr-4-1 | NC_010572.1 | - | 3318808 | 3318753 | **RNA**→ rplQ (PRK05591)Ribosomal_L17 (pfam01196)→ PseudoU_synth_EcTruA (cd02570)truA (PRK00021)→ |
| San-2-1 | NZ_JNXS01000008.1 | + | 26146 | 26201 | **RNA**→ rplQ (PRK05591)Ribosomal_L17 (pfam01196)→ PseudoU_synth_EcTruA (cd02570)truA (PRK00021)→ |
| San-2-2 | NZ_JOEZ01000006.1 | - | 175251 | 175196 | **RNA**→ rplQ (PRK05591)Ribosomal_L17 (pfam01196)→ PseudoU_synth_EcTruA (cd02570)truA (PRK00021)→ |
| Sgr-3-1 | NZ_GL877172.1 | - | 3511493 | 3511438 | **RNA**→ rplQ (PRK05591)Ribosomal_L17 (pfam01196)→ rplQ (PRK05591)Ribosomal_L17 (pfam01196)→ |
| | | | | | PseudoU_synth_EcTruA (cd02570)truA (PRK00021)→ PseudoU_synth_EcTruA (cd02570)truA (PRK00021)→ |
| Sgr-12-8 | NZ_JOBR01000006.1 | + | 349597 | 349652 | **RNA**→ rplQ (PRK05591)Ribosomal_L17 (pfam01196)→ PseudoU_synth_EcTruA (cd02570)truA (PRK00021)→ |
| Ssp-9-1 | NZ_KB899005.1 | + | 126861 | 126916 | **RNA**→ rplQ (PRK05591)Ribosomal_L17 (pfam01196)→ PseudoU_synth_EcTruA (cd02570)truA (PRK00021)→ |
| Seu-1-1 | NZ_KQ257782.1 | - | 24925 | 24870 | **RNA**→ rplQ (PRK05591)Ribosomal_L17 (pfam01196)→ PseudoU_synth_EcTruA (cd02570)truA (PRK00021)→ |
| Ssp-82-1 | NZ_JXUQ01000001.1 | + | 4992228 | 4992283 | **RNA**→ rplQ (PRK05591)Ribosomal_L17 (pfam01196)→ PseudoU_synth_EcTruA (cd02570)truA (PRK00021)→ |
| Sal-12-2 | NZ_JOED01000012.1 | + | 368027 | 368082 | **RNA**→ rplQ (PRK05591)Ribosomal_L17 (pfam01196)→ PseudoU_synth_EcTruA (cd02570)truA (PRK00021)→ |
| Ssp-48-1 | NZ_AGSW01000026.1 | - | 4221 | 4166 | **RNA**→ rplQ (PRK05591)Ribosomal_L17 (pfam01196)→ rplQ (PRK05591)Ribosomal_L17 (pfam01196)→ |
| | | | | | PseudoU_synth_EcTruA (cd02570)truA (PRK00021)→ PseudoU_synth_EcTruA (cd02570)truA (PRK00021)→ |
| Sha-1-1 | NZ_JOAZ01000028.1 | - | 8990 | 8935 | **RNA**→ rplQ (PRK05591)Ribosomal_L17 (pfam01196)→ PseudoU_synth_EcTruA (cd02570)truA (PRK00021)→ |
| Sgr-8-1 | NZ_JOFC01000040.1 | - | 9004 | 8949 | **RNA**→ rplQ (PRK05591)Ribosomal_L17 (pfam01196)→ PseudoU_synth_EcTruA (cd02570)truA (PRK00021)→ |
| Ssp-45-1 | NC_015953.1 | + | 4421046 | 4421101 | **RNA**→ rplQ (PRK05591)Ribosomal_L17 (pfam01196)→ PseudoU_synth_EcTruA (cd02570)truA (PRK00021)→ |
| Ssp-18-1 | NZ_ARPI01000009.1 | + | 69587 | 69642 | **RNA**→ rplQ (PRK05591)Ribosomal_L17 (pfam01196)→ PseudoU_synth_EcTruA (cd02570)truA (PRK00021)→ |
| Ssp-71-1 | NZ_AZVE01000001.1 | - | 707671 | 707616 | **RNA**→ rplQ (PRK05591)Ribosomal_L17 (pfam01196)→ PseudoU_synth_EcTruA (cd02570)truA (PRK00021)→ |
| Ssp-175-1 | NZ_KL585382.1 | + | 76745 | 76800 | **RNA**→ rplQ (PRK05591)Ribosomal_L17 (pfam01196)→ PseudoU_synth_EcTruA (cd02570)truA (PRK00021)→ |
| Ssp-188-1 | NZ_JJOB01000001.1 | + | 3882743 | 3882798 | **RNA**→ rplQ (PRK05591)Ribosomal_L17 (pfam01196)→ PseudoU_synth_EcTruA (cd02570)truA (PRK00021)→ |
| Ssp-52-1 | NZ_JTHL01000001.1 | + | 5725538 | 5725593 | **RNA**→ rplQ (PRK05591)Ribosomal_L17 (pfam01196)→ |
| Sri-4-1 | NZ_JOCQ01000089.1 | - | 23248 | 23193 | **RNA**→ rplQ (PRK05591)Ribosomal_L17 (pfam01196)→ |
| Ssp-145-1 | NZ_JOHB01000009.1 | + | 199860 | 199915 | **RNA**→ rplQ (PRK05591)Ribosomal_L17 (pfam01196)→ |
| env-34 | JGI25160J50197_1029655 | - | 174 | 119 | **RNA**→ |
| Sly-1-1 | NZ_JNZA01000027.1 | - | 104738 | 104683 | **RNA**→ rplQ (PRK05591)Ribosomal_L17 (pfam01196)→ |
| Ssp-142-1 | NZ_JOGE01000029.1 | + | 58699 | 58754 | **RNA**→ rplQ (PRK05591)Ribosomal_L17 (pfam01196)→ |
| Ssp-155-1 | NZ_JOIX01000020.1 | + | 81701 | 81756 | **RNA**→ rplQ (PRK05591)Ribosomal_L17 (pfam01196)→ PseudoU_synth_EcTruA (cd02570)truA (PRK00021)→ |
| Ssp-53-1 | NZ_CP003987.1 | + | 5490097 | 5490152 | **RNA**→ rplQ (PRK05591)Ribosomal_L17 (pfam01196)→ PseudoU_synth_EcTruA (cd02570)truA (PRK00021)→ |
| Sat-2-1 | NZ_JNXG01000002.1 | + | 464448 | 464503 | **RNA**→ rplQ (PRK05591)Ribosomal_L17 (pfam01196)→ PseudoU_synth_EcTruA (cd02570)truA (PRK00021)→ |
| Ssp-95-1 | NZ_JOAE01000005.1 | - | 277533 | 277478 | **RNA**→ rplQ (PRK05591)Ribosomal_L17 (pfam01196)→ PseudoU_synth_EcTruA (cd02570)truA (PRK00021)→ |
| Sfl-1-1 | NC_016114.1 | - | 2965273 | 2965218 | **RNA**→ rplQ (PRK05591)Ribosomal_L17 (pfam01196)→ PseudoU_synth_EcTruA (cd02570)truA (PRK00021)→ |
| Ssp-34-1 | NC_021055.1 | + | 4462927 | 4462982 | **RNA**→ rplQ (PRK05591)Ribosomal_L17 (pfam01196)→ PseudoU_synth_EcTruA (cd02570)truA (PRK00021)→ |
| [pa-1-1 | NZ_JNYQ01000004.1 | - | 851086 | 851031 | **RNA**→ rplQ (PRK05591)Ribosomal_L17 (pfam01196)→ PseudoU_synth_EcTruA (cd02570)truA (PRK00021)→ |
| Sol-2-2 | NZ_JNWM01000003.1 | + | 440846 | 440901 | **RNA**→ rplQ (PRK05591)Ribosomal_L17 (pfam01196)→ PseudoU_synth_EcTruA (cd02570)truA (PRK00021)→ |
| Sfu-3-1 | NZ_JNXH01000004.1 | - | 821119 | 821064 | **RNA**→ rplQ (PRK05591)Ribosomal_L17 (pfam01196)→ PseudoU_synth_EcTruA (cd02570)truA (PRK00021)→ |
| Sfl-8-1 | NZ_JOAB01000003.1 | - | 795932 | 795877 | **RNA**→ rplQ (PRK05591)Ribosomal_L17 (pfam01196)→ PseudoU_synth_EcTruA (cd02570)truA (PRK00021)→ |
| Ssp-154-1 | NZ_JOIW01000004.1 | - | 790730 | 790675 | **RNA**→ rplQ (PRK05591)Ribosomal_L17 (pfam01196)→ PseudoU_synth_EcTruA (cd02570)truA (PRK00021)→ |
| Sce-1-1 | NZ_JOEL01000031.1 | + | 46103 | 46158 | **RNA**→ rplQ (PRK05591)Ribosomal_L17 (pfam01196)→ |
| Ssc-5-1 | NZ_JOEI01000008.1 | + | 148665 | 148720 | **RNA**→ rplQ (PRK05591)Ribosomal_L17 (pfam01196)→ PseudoU_synth_EcTruA (cd02570)truA (PRK00021)→ |
| Ssp-144-1 | NZ_KL589850.1 | + | 71321 | 71376 | **RNA**→ rplQ (PRK05591)Ribosomal_L17 (pfam01196)→ PseudoU_synth_EcTruA (cd02570)truA (PRK00021)→ |
| Ssp-54-1 | NZ_CBRG010000181.1 | + | 9589 | 9644 | **RNA**→ rplQ (PRK05591)Ribosomal_L17 (pfam01196)→ PseudoU_synth_EcTruA (cd02570)truA (PRK00021)→ |

| | | | | | |
|---|---|---|---|---|---|
| Sgr-13-1 | NZ_JOFF01000044.1 | - | 4277 | 4222 | **RNA**→ rplQ (PRK05591)Ribosomal_L17 (pfam01196)→ PseudoU_synth_EcTruA (cd02570)truA (PRK00021)→ |
| Sfl-9-1 | NZ_JOAC01000001.1 | + | 308726 | 308781 | **RNA**→ rplQ (PRK05591)Ribosomal_L17 (pfam01196)→ PseudoU_synth_EcTruA (cd02570)truA (PRK00021)→ |
| Sro-4-1 | NZ_ABYX02000001.1 | + | 4766062 | 4766117 | **RNA**→ rplQ (PRK05591)Ribosomal_L17 (pfam01196)→ PseudoU_synth_EcTruA (cd02570)truA (PRK00021)→ |
| Ssp-22-1 | NZ_AWOQ01000620.1 | + | 11739 | 11794 | **RNA**→ rplQ (PRK05591)Ribosomal_L17 (pfam01196)→ |
| Sro-2-1 | NZ_DS999644.1 | + | 4713512 | 4713567 | **RNA**→ rplQ (PRK05591)Ribosomal_L17 (pfam01196)→ rplQ (PRK05591)Ribosomal_L17 (pfam01196)→ PseudoU_synth_EcTruA (cd02570)truA (PRK00021)→ PseudoU_synth_EcTruA (cd02570)truA (PRK00021)→ |
| Sal-12-3 | NZ_JNWW01000015.1 | - | 121666 | 121611 | **RNA**→ rplQ (PRK05591)Ribosomal_L17 (pfam01196)→ PseudoU_synth_EcTruA (cd02570)truA (PRK00021)→ |
| Ssp-174-1 | NZ_JOCW01000027.1 | + | 95259 | 95314 | **RNA**→ rplQ (PRK05591)Ribosomal_L17 (pfam01196)→ PseudoU_synth_EcTruA (cd02570)truA (PRK00021)→ |
| Ssp-107-1 | NZ_JOIP01000001.1 | + | 298835 | 298890 | **RNA**→ rplQ (PRK05591)Ribosomal_L17 (pfam01196)→ PseudoU_synth_EcTruA (cd02570)truA (PRK00021)→ |
| Ssp-108-1 | NZ_JOIO01000005.1 | + | 89666 | 89721 | **RNA**→ rplQ (PRK05591)Ribosomal_L17 (pfam01196)→ PseudoU_synth_EcTruA (cd02570)truA (PRK00021)→ |
| Sgr-13-2 | NZ_LGUZ01000023.1 | - | 4252 | 4197 | **RNA**→ rplQ (PRK05591)Ribosomal_L17 (pfam01196)→ PseudoU_synth_EcTruA (cd02570)truA (PRK00021)→ |
| Spu-3-1 | NZ_JOBG01000005.1 | - | 224288 | 224233 | **RNA**→ rplQ (PRK05591)Ribosomal_L17 (pfam01196)→ PseudoU_synth_EcTruA (cd02570)truA (PRK00021)→ |
| Sgr-13-3 | NZ_JOFE01000006.1 | + | 95269 | 95324 | **RNA**→ rplQ (PRK05591)Ribosomal_L17 (pfam01196)→ PseudoU_synth_EcTruA (cd02570)truA (PRK00021)→ |
| Sgr-13-4 | NZ_JOFD01000016.1 | - | 125262 | 125207 | **RNA**→ rplQ (PRK05591)Ribosomal_L17 (pfam01196)→ PseudoU_synth_EcTruA (cd02570)truA (PRK00021)→ |
| Spu-3-2 | NZ_JODK01000012.1 | + | 27097 | 27152 | **RNA**→ rplQ (PRK05591)Ribosomal_L17 (pfam01196)→ PseudoU_synth_EcTruA (cd02570)truA (PRK00021)→ |
| Sbr-1-1 | NZ_JNXB01000005.1 | + | 80235 | 80290 | **RNA**→ rplQ (PRK05591)Ribosomal_L17 (pfam01196)→ PseudoU_synth_EcTruA (cd02570)truA (PRK00021)→ |
| Sca-5-1 | NZ_KL573598.1 | - | 234310 | 234255 | **RNA**→ rplQ (PRK05591)Ribosomal_L17 (pfam01196)→ PseudoU_synth_EcTruA (cd02570)truA (PRK00021)→ |
| Sca-5-2 | NZ_KL569913.1 | + | 82305 | 82360 | **RNA**→ rplQ (PRK05591)Ribosomal_L17 (pfam01196)→ PseudoU_synth_EcTruA (cd02570)truA (PRK00021)→ |
| Sgr-13-5 | NZ_JODO01000006.1 | + | 95248 | 95303 | **RNA**→ rplQ (PRK05591)Ribosomal_L17 (pfam01196)→ PseudoU_synth_EcTruA (cd02570)truA (PRK00021)→ |
| Ssp-129-1 | NZ_JOHA01000009.1 | + | 79533 | 79588 | **RNA**→ rplQ (PRK05591)Ribosomal_L17 (pfam01196)→ PseudoU_synth_EcTruA (cd02570)truA (PRK00021)→ |
| Ssp-130-1 | NZ_JOHD01000004.1 | + | 34514 | 34569 | **RNA**→ rplQ (PRK05591)Ribosomal_L17 (pfam01196)→ PseudoU_synth_EcTruA (cd02570)truA (PRK00021)→ |
| Ssp-100-1 | NZ_JOIH01000008.1 | - | 104878 | 104823 | **RNA**→ rplQ (PRK05591)Ribosomal_L17 (pfam01196)→ PseudoU_synth_EcTruA (cd02570)truA (PRK00021)→ |
| Spu-2-1 | NZ_JOFA01000009.1 | + | 27215 | 27270 | **RNA**→ rplQ (PRK05591)Ribosomal_L17 (pfam01196)→ PseudoU_synth_EcTruA (cd02570)truA (PRK00021)→ |
| Sgr-12-9 | NZ_JOIT01000002.1 | + | 49057 | 49112 | **RNA**→ rplQ (PRK05591)Ribosomal_L17 (pfam01196)→ PseudoU_synth_EcTruA (cd02570)truA (PRK00021)→ |
| Spu-2-2 | NZ_JOBQ01000009.1 | + | 27228 | 27283 | **RNA**→ rplQ (PRK05591)Ribosomal_L17 (pfam01196)→ PseudoU_synth_EcTruA (cd02570)truA (PRK00021)→ |
| Svi-8-1 | NZ_JNYP01000002.1 | + | 382717 | 382772 | **RNA**→ rplQ (PRK05591)Ribosomal_L17 (pfam01196)→ |
| Ssp-66-1 | NZ_JNLT01000004.1 | - | 111420 | 111365 | **RNA**→ rplQ (PRK05591)Ribosomal_L17 (pfam01196)→ PseudoU_synth_EcTruA (cd02570)truA (PRK00021)→ |
| Ssp-94-1 | NZ_JOHG01000019.1 | - | 96015 | 95960 | **RNA**→ rplQ (PRK05591)Ribosomal_L17 (pfam01196)→ PseudoU_synth_EcTruA (cd02570)truA (PRK00021)→ |
| Ssp-17-1 | NZ_KB890938.1 | + | 398321 | 398376 | **RNA**→ rplQ (PRK05591)Ribosomal_L17 (pfam01196)→ PseudoU_synth_EcTruA (cd02570)truA (PRK00021)→ |
| Ssp-193-1 | NZ_ARPF01000004.1 | - | 200548 | 200493 | **RNA**→ rplQ (PRK05591)Ribosomal_L17 (pfam01196)→ PseudoU_synth_EcTruA (cd02570)truA (PRK00021)→ |
| Sat-1-1 | NZ_JQJU01000001.1 | + | 70024 | 70079 | **RNA**→ rplQ (PRK05591)Ribosomal_L17 (pfam01196)→ PseudoU_synth_EcTruA (cd02570)truA (PRK00021)→ |
| Ssp-44-1 | NZ_KB891296.1 | - | 1243298 | 1243243 | **RNA**→ rplQ (PRK05591)Ribosomal_L17 (pfam01196)→ PseudoU_synth_EcTruA (cd02570)truA (PRK00021)→ |
| Scy-1-1 | NZ_JOEM01000001.1 | - | 651664 | 651609 | **RNA**→ rplQ (PRK05591)Ribosomal_L17 (pfam01196)→ PseudoU_synth_EcTruA (cd02570)truA (PRK00021)→ |
| Sfu-1-1 | NC_021177.1 | + | 4972092 | 4972147 | **RNA**→ rplQ (PRK05591)Ribosomal_L17 (pfam01196)→ PseudoU_synth_EcTruA (cd02570)truA (PRK00021)→ |
| Sal-7-1 | NZ_JNWU01000006.1 | - | 157973 | 157918 | **RNA**→ rplQ (PRK05591)Ribosomal_L17 (pfam01196)→ PseudoU_synth_EcTruA (cd02570)truA (PRK00021)→ |
| Ssp-168-1 | NZ_KL647092.1 | - | 163335 | 163280 | **RNA**→ rplQ (PRK05591)Ribosomal_L17 (pfam01196)→ PseudoU_synth_EcTruA (cd02570)truA (PRK00021)→ |
| Ssp-97-1 | NZ_JOJO01000001.1 | + | 71929 | 71984 | **RNA**→ rplQ (PRK05591)Ribosomal_L17 (pfam01196)→ PseudoU_synth_EcTruA (cd02570)truA (PRK00021)→ |
| Ssp-146-1 | NZ_JOGZ01000044.1 | - | 52593 | 52538 | **RNA**→ rplQ (PRK05591)Ribosomal_L17 (pfam01196)→ PseudoU_synth_EcTruA (cd02570)truA (PRK00021)→ |
| Sod-1-1 | NZ_LATD01000297.1 | + | 18268 | 18323 | **RNA**→ rplQ (PRK05591)Ribosomal_L17 (pfam01196)→ PseudoU_synth_EcTruA (cd02570)truA (PRK00021)→ |
| Sri-4-2 | NZ_JOBO01000012.1 | + | 88122 | 88177 | **RNA**→ rplQ (PRK05591)Ribosomal_L17 (pfam01196)→ PseudoU_synth_EcTruA (cd02570)truA (PRK00021)→ |
| Sgr-6-1 | NZ_LGUW01000024.1 | + | 68205 | 68259 | **RNA**→ rplQ (PRK05591)Ribosomal_L17 (pfam01196)→ PseudoU_synth_EcTruA (cd02570)truA (PRK00021)→ |
| Sri-4-3 | NZ_JOBV01000009.1 | + | 106610 | 106664 | **RNA**→ rplQ (PRK05591)Ribosomal_L17 (pfam01196)→ PseudoU_synth_EcTruA (cd02570)truA (PRK00021)→ |
| Sri-4-4 | NZ_JOES01000007.1 | + | 106606 | 106660 | **RNA**→ rplQ (PRK05591)Ribosomal_L17 (pfam01196)→ PseudoU_synth_EcTruA (cd02570)truA (PRK00021)→ |
| Sri-4-5 | NZ_JNWX01000006.1 | + | 69237 | 69291 | **RNA**→ rplQ (PRK05591)Ribosomal_L17 (pfam01196)→ PseudoU_synth_EcTruA (cd02570)truA (PRK00021)→ |
| Sri-2-1 | NZ_JMGX01000035.1 | + | 49554 | 49609 | **RNA**→ rplQ (PRK05591)Ribosomal_L17 (pfam01196)→ PseudoU_synth_EcTruA (cd02570)truA (PRK00021)→ |
| Kau-1-2 | NZ_LGUY01000192.1 | + | 63444 | 63499 | **RNA**→ rplQ (PRK05591)Ribosomal_L17 (pfam01196)→ PseudoU_synth_EcTruA (cd02570)truA (PRK00021)→ |
| Ssp-179-1 | NZ_JOCC01000033.1 | + | 69292 | 69347 | **RNA**→ rplQ (PRK05591)Ribosomal_L17 (pfam01196)→ PseudoU_synth_EcTruA (cd02570)truA (PRK00021)→ |
| Sri-4-6 | NZ_JOEO01000033.1 | - | 28104 | 28049 | **RNA**→ rplQ (PRK05591)Ribosomal_L17 (pfam01196)→ PseudoU_synth_EcTruA (cd02570)truA (PRK00021)→ |
| Sri-4-7 | NZ_JNYK01000036.1 | - | 28104 | 28049 | **RNA**→ rplQ (PRK05591)Ribosomal_L17 (pfam01196)→ PseudoU_synth_EcTruA (cd02570)truA (PRK00021)→ |
| Sri-4-8 | NZ_JOBN01000012.1 | + | 209341 | 209396 | **RNA**→ rplQ (PRK05591)Ribosomal_L17 (pfam01196)→ PseudoU_synth_EcTruA (cd02570)truA (PRK00021)→ |
| Sri-4-9 | NZ_JOCL01000017.1 | + | 107763 | 107818 | **RNA**→ rplQ (PRK05591)Ribosomal_L17 (pfam01196)→ PseudoU_synth_EcTruA (cd02570)truA (PRK00021)→ |
| Sri-4-10 | NZ_JOCR01000028.1 | + | 69299 | 69354 | **RNA**→ rplQ (PRK05591)Ribosomal_L17 (pfam01196)→ PseudoU_synth_EcTruA (cd02570)truA (PRK00021)→ |

| | | | | | |
|---|---|---|---|---|---|
| Sri-4-11 | NZ_JOCO01000017.1 | + | 107815 | 107870 | RNA→ rplQ (PRK05591)Ribosomal_L17 (pfam01196)→ PseudoU_synth_EcTruA (cd02570)truA (PRK00021)→ |
| Sri-4-12 | NZ_JOBS01000018.1 | + | 107803 | 107858 | RNA→ rplQ (PRK05591)Ribosomal_L17 (pfam01196)→ PseudoU_synth_EcTruA (cd02570)truA (PRK00021)→ |
| Ssp-180-1 | NZ_JOCH01000032.1 | + | 69299 | 69354 | RNA→ rplQ (PRK05591)Ribosomal_L17 (pfam01196)→ PseudoU_synth_EcTruA (cd02570)truA (PRK00021)→ |
| Sca-6-1 | NZ_JNWP01000055.1 | + | 21186 | 21241 | RNA→ rplQ (PRK05591)Ribosomal_L17 (pfam01196)→ PseudoU_synth_EcTruA (cd02570)truA (PRK00021)→ |
| Sca-6-2 | NZ_JOFM01000031.1 | + | 69322 | 69377 | RNA→ rplQ (PRK05591)Ribosomal_L17 (pfam01196)→ PseudoU_synth_EcTruA (cd02570)truA (PRK00021)→ |
| Sri-4-13 | NZ_JOBT01000154.1 | - | 8051 | 7996 | RNA→ rplQ (PRK05591)Ribosomal_L17 (pfam01196)→ PseudoU_synth_EcTruA (cd02570)truA (PRK00021)→ |
| Sri-4-14 | NZ_JOCM01000018.1 | + | 107761 | 107816 | RNA→ rplQ (PRK05591)Ribosomal_L17 (pfam01196)→ PseudoU_synth_EcTruA (cd02570)truA (PRK00021)→ |
| Sri-4-15 | NZ_JOCP01000018.1 | - | 28099 | 28044 | RNA→ rplQ (PRK05591)Ribosomal_L17 (pfam01196)→ PseudoU_synth_EcTruA (cd02570)truA (PRK00021)→ |
| Sri-4-16 | NZ_JNYR01000006.1 | + | 209435 | 209490 | RNA→ rplQ (PRK05591)Ribosomal_L17 (pfam01196)→ PseudoU_synth_EcTruA (cd02570)truA (PRK00021)→ |
| Sla-2-3 | NZ_JOEX01000032.1 | + | 107616 | 107671 | RNA→ rplQ (PRK05591)Ribosomal_L17 (pfam01196)→ PseudoU_synth_EcTruA (cd02570)truA (PRK00021)→ |
| Sla-2-4 | NZ_JOCN01000020.1 | + | 107890 | 107945 | RNA→ rplQ (PRK05591)Ribosomal_L17 (pfam01196)→ PseudoU_synth_EcTruA (cd02570)truA (PRK00021)→ |
| Sri-1-1 | NZ_ANSJ01000031.1 | + | 47278 | 47333 | RNA→ rplQ (PRK05591)Ribosomal_L17 (pfam01196)→ PseudoU_synth_EcTruA (cd02570)truA (PRK00021)→ |
| Sri-3-1 | NZ_JJNO01000013.1 | + | 209385 | 209440 | RNA→ rplQ (PRK05591)Ribosomal_L17 (pfam01196)→ PseudoU_synth_EcTruA (cd02570)truA (PRK00021)→ |
| Sri-2-2 | NZ_JMGY01000011.1 | + | 209215 | 209270 | RNA→ rplQ (PRK05591)Ribosomal_L17 (pfam01196)→ PseudoU_synth_EcTruA (cd02570)truA (PRK00021)→ |
| Sri-4-17 | NZ_JOET01000019.1 | + | 107682 | 107737 | RNA→ rplQ (PRK05591)Ribosomal_L17 (pfam01196)→ PseudoU_synth_EcTruA (cd02570)truA (PRK00021)→ |
| Ssp-90-1 | NZ_JNWN01000026.1 | - | 28254 | 28199 | RNA→ rplQ (PRK05591)Ribosomal_L17 (pfam01196)→ |
| Spe-2-1 | NZ_JOCK01000016.1 | + | 107552 | 107607 | RNA→ rplQ (PRK05591)Ribosomal_L17 (pfam01196)→ |
| Sri-4-18 | NZ_JOBW01000066.1 | + | 21254 | 21309 | RNA→ rplQ (PRK05591)Ribosomal_L17 (pfam01196)→ PseudoU_synth_EcTruA (cd02570)truA (PRK00021)→ |
| Ssp-185-1 | NZ_JOJI01000014.1 | + | 111307 | 111361 | RNA→ rplQ (PRK05591)Ribosomal_L17 (pfam01196)→ PseudoU_synth_EcTruA (cd02570)truA (PRK00021)→ |
| Sri-4-19 | NZ_JOBP01000017.1 | - | 140212 | 140157 | RNA→ rplQ (PRK05591)Ribosomal_L17 (pfam01196)→ PseudoU_synth_EcTruA (cd02570)truA (PRK00021)→ |
| Sri-4-20 | NZ_JOJJ01000026.1 | + | 70453 | 70508 | RNA→ rplQ (PRK05591)Ribosomal_L17 (pfam01196)→ PseudoU_synth_EcTruA (cd02570)truA (PRK00021)→ |
| Ssp-5-1 | NZ_KB892002.1 | - | 454475 | 454420 | RNA→ rplQ (PRK05591)Ribosomal_L17 (pfam01196)→ PseudoU_synth_EcTruA (cd02570)truA (PRK00021)→ |
| Ssp-29-1 | NZ_KB891785.1 | - | 425407 | 425352 | RNA→ rplQ (PRK05591)Ribosomal_L17 (pfam01196)→ PseudoU_synth_EcTruA (cd02570)truA (PRK00021)→ |
| Ssp-31-1 | NZ_AQWQ01000002.1 | - | 425407 | 425352 | RNA→ rplQ (PRK05591)Ribosomal_L17 (pfam01196)→ PseudoU_synth_EcTruA (cd02570)truA (PRK00021)→ |
| Ssp-26-1 | NZ_KB891594.1 | - | 382830 | 382775 | RNA→ rplQ (PRK05591)Ribosomal_L17 (pfam01196)→ PseudoU_synth_EcTruA (cd02570)truA (PRK00021)→ |
| Ssp-27-1 | NZ_KB891406.1 | - | 1044742 | 1044687 | RNA→ rplQ (PRK05591)Ribosomal_L17 (pfam01196)→ PseudoU_synth_EcTruA (cd02570)truA (PRK00021)→ |
| Ssp-79-1 | NZ_KI911349.1 | + | 472182 | 472237 | RNA→ rplQ (PRK05591)Ribosomal_L17 (pfam01196)→ PseudoU_synth_EcTruA (cd02570)truA (PRK00021)→ |
| Ssp-78-1 | NZ_AZVF01000001.1 | - | 1050889 | 1050834 | RNA→ rplQ (PRK05591)Ribosomal_L17 (pfam01196)→ PseudoU_synth_EcTruA (cd02570)truA (PRK00021)→ |
| Aat-1-1 | NZ_KB907214.1 | - | 233058 | 233000 | RNA→ rplQ (PRK05591)Ribosomal_L17 (pfam01196)→ |
| Ssp-117-1 | NZ_JOHY01000071.1 | - | 9294 | 9239 | RNA→ rplQ (PRK05591)Ribosomal_L17 (pfam01196)→ PseudoU_synth_EcTruA (cd02570)truA (PRK00021)→ |
| Ssp-198-1 | NZ_KL370770.1 | + | 196586 | 196640 | RNA→ rplQ (PRK05591)Ribosomal_L17 (pfam01196)→ PseudoU_synth_EcTruA (cd02570)truA (PRK00021)→ |
| Ssp-115-1 | NZ_JOHT01000007.1 | + | 90429 | 90475 | RNA→ rplQ (PRK05591)Ribosomal_L17 (pfam01196)→ PseudoU_synth_EcTruA (cd02570)truA (PRK00021)→ |
| Ssp-148-1 | NZ_JOGD01000029.1 | + | 57793 | 57839 | RNA→ rplQ (PRK05591)Ribosomal_L17 (pfam01196)→ PseudoU_synth_EcTruA (cd02570)truA (PRK00021)→ |
| Psi-1-1 | NZ_CP009896.1 | - | 4283040 | 4282985 | RNA→ rplQ (PRK05591)Ribosomal_L17 (pfam01196)→ IspH (COG0761)ispH (PRK01045)→ PRK11630 (PRK11630)SUA5 (COG0009)→ |
| Ssp-4-1 | NZ_ALAP01000018.1 | - | 152336 | 152283 | RNA→ rplQ (PRK05591)Ribosomal_L17 (pfam01196)→ PseudoU_synth_EcTruA (cd02570)truA (PRK00021)→ |
| Ssp-67-1 | NZ_AUKN01000012.1 | + | 38678 | 38731 | RNA→ rplQ (PRK05591)Ribosomal_L17 (pfam01196)→ PseudoU_synth_EcTruA (cd02570)truA (PRK00021)→ |
| Ssp-201-1 | NZ_LAND01000020.1 | + | 47240 | 47292 | RNA→ rplQ (PRK05591)Ribosomal_L17 (pfam01196)→ PseudoU_synth_EcTruA (cd02570)truA (PRK00021)→ |
| Aal-1-1 | NZ_KB892746.1 | + | 281585 | 281636 | RNA→ rplQ (PRK05591)Ribosomal_L17 (pfam01196)→ rplQ (PRK05591)Ribosomal_L17 (pfam01196)→ PseudoU_synth_EcTruA (cd02570)truA (PRK00021)→ AdoMet_MTases (cd02440)Methyltransf_25 (pfam13649)→ hypo→ hypo→ |
| Sal-13-1 | NZ_JOFJ01000024.1 | - | 50986 | 50914 | RNA→ RplQ (COG0203)rplQ (PRK05591)→ |
| Ari-1-1 | NZ_AULB01000026.1 | + | 70386 | 70458 | RNA→ rplQ (PRK05591)Ribosomal_L17 (pfam01196)→ |
| Ssp-40-1 | NZ_ADXA01000020.1 | - | 61314 | 61258 | RNA→ rplQ (PRK05591)Ribosomal_L17 (pfam01196)→ rplQ (PRK05591)Ribosomal_L17 (pfam01196)→ PseudoU_synth_EcTruA (cd02570)truA (PRK00021)→ PseudoU_synth_EcTruA (cd02570)truA (PRK00021)→ |
| Ssp-39-1 | NZ_ADXB01001219.1 | + | 56 | 112 | RNA→ rplQ (PRK05591)Ribosomal_L17 (pfam01196)→ rplQ (PRK05591)Ribosomal_L17 (pfam01196)→ |
| Ssp-42-1 | NZ_GG657742.1 | - | 2775010 | 2774954 | RNA→ rplQ (PRK05591)Ribosomal_L17 (pfam01196)→ rplQ (PRK05591)Ribosomal_L17 (pfam01196)→ PseudoU_synth_EcTruA (cd02570)truA (PRK00021)→ PseudoU_synth_EcTruA (cd02570)truA (PRK00021)→ |
| Ssp-47-1 | NZ_CM001165.1 | + | 4663630 | 4663686 | RNA→ ←hypo |
| Ssp-41-1 | NZ_GG770539.1 | + | 4279054 | 4279112 | RNA→ hypo→ rplQ (PRK05591)Ribosomal_L17 (pfam01196)→ PseudoU_synth_EcTruA (cd02570)truA (PRK00021)→ hypo→ |
| Ssp-125-1 | NZ_JOGO01000002.1 | - | 47335 | 47279 | RNA→ rplQ (PRK05591)Ribosomal_L17 (pfam01196)→ PseudoU_synth_EcTruA (cd02570)truA (PRK00021)→ |
| Nsp-3-1 | NZ_JIAV01000015.1 | - | 114208 | 114159 | RNA→ RplQ (COG0203)rplQ (PRK05591)rne (PRK10811)→ |

| | | | | | |
|---|---|---|---|---|---|
| Nsp-2-1 | NZ_KK211169.1 | + | 269460 | 269509 | **RNA**→ RplQ (COG0203)rplQ (PRK05591)→ |
| env-35 | JGI12075J13256_1000799 | - | 577 | 527 | **RNA**→ rplQ (PRK05591)Ribosomal_L17 (pfam01196)→ |
| env-36 | JGI12112J13074_1005592 | - | 506 | 456 | **RNA**→ RplQ (COG0203)rplQ (PRK05591)→ |
| env-37 | JGI12271J12027_1176655 | + | 141 | 191 | **RNA**→ |
| env-38 | JGI12396J12026_1276148 | + | 100 | 150 | **RNA**→ hypo→ |
| env-39 | JGI12397J12024_1184993 | + | 151 | 201 | **RNA**→ |
| env-40 | JGI12271J12027_1236889 | + | 33 | 83 | **RNA**→ rplQ (PRK05591)Ribosomal_L17 (pfam01196)→ |
| Nlu-1-1 | NZ_JZDQ01000118.1 | + | 6491 | 6537 | **RNA**→ rplQ (PRK05591)Ribosomal_L17 (pfam01196)→ |
| env-41 | JXWU01108663.1 | + | 303 | 347 | **RNA**→ rplQ (PRK05591)Ribosomal_L17 (pfam01196)→ |
| Nha-1-1 | NZ_AUGT01000009.1 | + | 577134 | 577185 | **RNA**→ RplQ (COG0203)rplQ (PRK05591)→ |
| Sst-1-1 | NZ_KQ257875.1 | + | 79579 | 79635 | **RNA**→ rplQ (PRK05591)Ribosomal_L17 (pfam01196)→ |
| Sle-1-1 | NZ_LN831790.1 | - | 3535110 | 3535054 | **RNA**→ rplQ (PRK05591)Ribosomal_L17 (pfam01196)→ PseudoU_synth_EcTruA (cd02570)truA (PRK00021)→ |
| Sal-2-1 | NC_020990.1 | + | 4279609 | 4279665 | **RNA**→ rplQ (PRK05591)Ribosomal_L17 (pfam01196)→ PseudoU_synth_EcTruA (cd02570)truA (PRK00021)→ |
| Ssp-24-1 | NZ_KE150476.1 | - | 126365 | 126312 | **RNA**→ ←hypo |
| Ssp-132-1 | NZ_JOHQ01000016.1 | + | 22026 | 22079 | **RNA**→ ←hypo |
| Ssp-127-1 | NZ_JOGK01000040.1 | + | 21913 | 21966 | **RNA**→ ←hypo |
| Sgr-5-1 | NZ_JNWT01000058.1 | + | 21899 | 21952 | **RNA**→ ←hypo |
| Sal-12-4 | NZ_JODZ01000010.1 | + | 169503 | 169556 | **RNA**→ ←hypo |
| Sal-12-5 | NZ_JODR01000016.1 | + | 224011 | 224064 | **RNA**→ ←hypo |
| Scl-1-1 | NZ_CM001015.1 | + | 4251346 | 4251401 | **RNA**→ rplQ (PRK05591)Ribosomal_L17 (pfam01196)→ |
| Scl-1-2 | NZ_CM000913.1 | + | 4257377 | 4257432 | **RNA**→ rplQ (PRK05591)Ribosomal_L17 (pfam01196)→ rplQ (PRK05591)Ribosomal_L17 (pfam01196)→ |
| Ssc-1-1 | NC_013929.1 | - | 4144210 | 4144152 | **RNA**→ rplQ (PRK05591)Ribosomal_L17 (pfam01196)→ |
| Ssc-3-2 | NZ_KL997514.1 | + | 21465 | 21523 | **RNA**→ rplQ (PRK05591)Ribosomal_L17 (pfam01196)→ |
| Sde-1-1 | NZ_KL570245.1 | + | 195754 | 195809 | **RNA**→ rplQ (PRK05591)Ribosomal_L17 (pfam01196)→ |

#### 5.4 Conserved domains

Conserved domains found in protein-coding genes listed in Section 5.3 are shown below, with the first sentence in their description from the Conserved Domain Database (if any). Conserved domains associated with more than one L17-Actino-Proteobacteria RNA are assigned a color, while others are shown in gray. The symbols 'd' and 'u' (if any) to the left of the colored domain name indicate whether the domain occurs upstream or downstream of the RNA motif. Domains marked with lower-case 'd' occur (at least once) present downstream of some L17-Actino-Proteobacteria RNA in the opposite orientation. Domains marked with capital 'D' occur downstream and in the same orientation (i.e., a potential *cis*-regulatory arrangement). Domains marked with a 'u' occur upstream of the RNA in either orientation. If the 'd'/'u' symbols are missing, then the domain is downstream and in the same strand. (This mode is used for motifs predicted as *cis*-regulatory.) The number in parentheses after the colored domain name is the number of occurrences in Section 5.3.

cd01167 (1) Fructokinases (FRKs) mainly from bacteria and plants are enzymes with high specificity for fructose, as are all FRKs, but they catalyzes the conversion of fructose to fructose-6-phosphate, which is an entry point into glycolysis via conversion into glucose-6-phosphate. [or ketohexokinases (KHKs)]
cd02440 (2) S-adenosylmethionine-dependent methyltransferases (SAM or AdoMet-MTase), class I; AdoMet-MTases are enzymes that use S-adenosyl-L-methionine (SAM or AdoMet) as a substrate for methyltransfer, creating the product S-adenosyl-L-homocysteine (AdoHcy).
cd02570 (329) Eukaryotic and bacterial pseudouridine synthases similar to E.
COG0009 (1) tRNA A37 threonylcarbamoyladenosine synthetase subunit TsaC/SUA5/YrdC [Translation, ribosomal structure and biogenesis]
COG0203 (37) Ribosomal protein L17 [Translation, ribosomal structure and biogenesis]
COG0488 (29) ATPase components of ABC transporters with duplicated ATPase domains [General function prediction only]
COG0761 (1) 4-Hydroxy-3-methylbut-2-enyl diphosphate reductase IspH [Lipid transport and metabolism]
COG1004 (1) UDP-glucose 6-dehydrogenase [Cell wall/membrane/envelope biogenesis]
pfam01196 (524) Ribosomal protein L17.
pfam01416 (1) tRNA pseudouridine synthase.
pfam02156 (1) Glycosyl hydrolase family 26.
pfam09594 (19) Protein of unknown function (DUF2029).
pfam12848 (1) ABC transporter.
pfam13304 (21) AAA domain, putative AbiEii toxin, Type IV TA system.
pfam13620 (1) Carboxypeptidase regulatory-like domain.
pfam13649 (2) Methyltransferase domain.
PRK00021 (330) tRNA pseudouridine synthase A; Validated

PRK01045 (1) 4-hydroxy-3-methylbut-2-enyl diphosphate reductase; Reviewed
PRK05591 (561) 50S ribosomal protein L17; Validated
PRK09434 (1) aminoimidazole riboside kinase; Provisional
PRK10811 (1) ribonuclease E; Reviewed
PRK11630 (1) hypothetical protein; Provisional
PRK11855 (1) dihydrolipoamide acetyltransferase; Reviewed
PRK13428 (1) F0F1 ATP synthase subunit delta; Provisional
PRK14521 (1) 30S ribosomal protein S16; Provisional
smart00382 (29) ATPases associated with a variety of cellular activities.
TIGR03026 (1) nucleotide sugar dehydrogenase.

#### 5.5 Multiple-sequence alignment

Each L17-Actino-Proteobacteria RNA is denoted by an abbreviation (like "Eco-1-1") that refers to a taxonomy given in Section 5.1. (Superscript numbers refer to annotations listed in Section 5.2) The alignment may include sequences containing the RNA motif, as well as flanking sequence. The L17-Actino-Proteobacteria RNA itself is denoted by the line underneath marked 5′ and 3′ on either end. Nucleotides in flanking sequences (i.e., not part of the motif) are written in gray letters. Stems of predicted rho-independent transcription terminators, if any, are shaded yellow. (Note: terminator predictions have not been analyzed manually, and many are likely to be false positives. Terminator predictions are those of the RNIE software.) Nucleotides predicted to function (as DNA) as transcription-factor binding sites, if any, are shaded in green. (But note: these predictions are manually annotated, so they might be under-predicted.) Annotated start codons, if any, are shaded green. (Note: start codons are frequently misannotated, especially in environmental samples.) Nucleotides proposed to basepair as part of the consensus structure are shaded in color when they comprise Watson-Crick or G-U pairs. Otherwise they are shaded gray. Conserved stems are also indicated at the bottom of the alignment by angle brackets, where matching < and > denote base-paired columns. Below these angle brackets, the symbol "2" denotes base pairs exhibiting covariation according to the statistically well-founded R-scape method. "1" denotes base pairs exhibiting covariation according to R2R's simplistic method. "0" denotes base pairs that are not observed to mutate and "?" denotes base pairs that have a significant frequency of non-canonical nucleotides for Watson-Crick or G-U pairs (> 5%). Below these base pair annotation is the consensus sequence: "R" = "A" or "G", "Y" = "C" or "U", red nucleotides: nucleotide identity conserved more than 97% of the time, black nucleotides: 90%, gray nucleotides: 75%, red circle (•): nucleotide is present 97% of the time, black circle (•): 90%, gray circle (•): 75%, white circle (◦): 50%. All percentages of sequences just described (e.g. 97% conserved) assume that sequences have been weighted by the GSC algorithm implemented by the Infernal software package.

Duplicate sequences: the following putative homologs are not shown in the alignment because their sequences are identical to a homolog already shown: Kau-1-2, Kpu-1-1, Mha-1-1, Mxa-2-1, Mxa-3-1, Sac-1-1, Sal-1-1, Sal-10-1, Sal-12-4, Sal-12-5, Sal-4-1, Sal-8-1, Sal-9-1, San-2-2, Sau-3-2, Sav-1-1, Sav-2-2, Sbi-2-2, Sbi-3-2, Sbi-3-3, Sbo-1-2, Sbr-1-1, Sca-2-2, Sca-5-1, Sca-5-2, Sca-6-1, Sca-6-2, Scl-1-2, Sfl-2-1, Sfl-3-1, Sfl-7-1, Sfr-1-2, Sfu-3-1, Sga-1-1, Sgr-12-2, Sgr-12-8, Sgr-12-9, Sgr-13-2, Sgr-13-3, Sgr-13-4, Sgr-13-5, Sgr-5-1, Sgr-8-1, Shy-3-1, Sla-2-3, Sla-2-4, Sli-1-1, Sli-2-1, Sli-2-2, Sna-1-1, Sol-2-2, Spe-2-1, Spl-1-2, Spu-2-2, Spu-3-1, Spu-3-2, Sri-1-1, Sri-2-2, Sri-3-1, Sri-4-10, Sri-4-11, Sri-4-12, Sri-4-13, Sri-4-14, Sri-4-15, Sri-4-16, Sri-4-17, Sri-4-20, Sri-4-3, Sri-4-4, Sri-4-6, Sri-4-7, Sri-4-8, Sri-4-9, Sro-2-1, Sru-1-1, Sru-1-2, Ssc-3-2, Ssp-10-1, Ssp-100-1, Ssp-107-1, Ssp-108-1, Ssp-109-1, Ssp-11-1, Ssp-126-1, Ssp-127-1, Ssp-130-1, Ssp-132-1, Ssp-136-1, Ssp-137-1, Ssp-147-1, Ssp-148-1, Ssp-167-1, Ssp-173-1, Ssp-178-1, Ssp-179-1, Ssp-18-1, Ssp-180-1, Ssp-181-1, Ssp-184-2, Ssp-196-1, Ssp-22-1, Ssp-27-1, Ssp-31-1, Ssp-32-2, Ssp-34-1, Ssp-35-1, Ssp-60-1, Ssp-63-1, Ssp-67-1, Ssp-78-1, Ssp-79-1, Ssp-88-1, Ssp-91-1, Ssp-92-1, Ssp-99-1, Sva-2-2, Svi-12-2, Svi-13-3, Svi-13-4, Swe-1-1, Sxa-1-2, Sxi-1-1, Tbi-1-1, env-17, env-25, env-26, env-27, env-28, env-29, env-36

alignment positions 1 · · · 79

```
env-1     AGUGCUCGCUUCGCGCGGCCUGACCUUGGGUAUGAAGCUCGAAAACUGGCCGCCGGCCGGUCUCGACAAACACUGAACU
env-2     ..............................................UGGCCGCCGGCCGGUCUCGACAAACACUGAACU
[br-1-1   CGUGCGGCUCGAGAACUGGCCGCCGUCCGGCCUCGACAAGCGUUAAAAAAUCGGGGUGUCGGCUGCAGGCCGGCACCCC
env-3     GCGCGGCCUGACCUUGGGCAUGAAACUGGAAUCGUGGCCGCCCGCCGGCCUGGACAAACGUUAAUUCAACCCGGUGAAA
Nal-1-1   CGCAGGCCGCCCUCGCGGCGUACGGCGACGACGACGACGACGCGUUCGUCGAGGACGAGCAGUACUGAUCCAGCCCCAC
Nsp-1-1   CCUCCGCCUGGCGGCAAGACACGAGAGACGCAGCCGUCGGGUGGGGCCUGGCCCCGCCUGGCGGCAAGACAAGUGAAGA
Mxa-1-1   GCCUCCCCACGUGGCCGCGGUACGGCGGUCAGCGGCGCGAGGGGAGGGGGCUGGGACGUCUUCCGGUCCGAAUGACAGU
env-4     UGCUUUAACCCGCGCGUCCGGGGCGGCACGGCCGCAUGCCCCGGAUGGACGGGGACCCUCACCGGUCCAAUACAUUGA
Cfu-1-1   GCGAGUACCUGCCCACCCGGGGCGGCAUGGUUUCAUGCCCCGGGACGAGCAGGGGGCCCUCACCGGCCCAACCCCGCAG
Avi-1-1   UCGCAGUACCUCGUCCCUCGGGGCGGCAUGGCCGUAUACCCCGGGGGAGGAGGGGGCCUUCCCAGGCCCGAUACAGCAA
Age-1-1   CCGCAGUCCCUCGUCCCUCGGGGCGGCAUGGCCGCAUACCCCGGGGGAGGAGGGGGCCCUCCCAGGCCCAACACAGCAA
Hmi-1-1   UCGGCCCGCGCCGUGACACGGCUUCACGCGGCGCGGGAAGAGGGGGCCCCUCCCCAUAAAAGUGGGGGCCUAUUGGCAA

          ...............................................................................
          ...............................................................................
          •••••••••••••••••••••••••••••••••••••••••••••••••••••••••••••••••••••••••••••••
```

```
  env-5       UCUGAAGCACCUGCUCUCGCGCCGCGACACAGCCGCAGGCGGCGCGAGGGGAGGGGGCCCGUAAGGCCCAACAAGGCAA
  Sau-5-1     UGUCGUACCUCACGCCUCGCCGCGACACGGCUUCACACGGCGGGGAGUGAGGGGGGCCCCUGAAAAGGCUCCUUGGCAA
  Mfu-1-1     CCCCGCGCGCCCGCGACAUGGCGGCAAGCGGCGCGCGGAGGGAAGGAGCCCGGUGAAGUCCUCCGGGUUCGACGAGCAG
  Mst-1-1     UUUCCUCCGUGCCCGCGACAUGGUUGCAGGCGGCGCGGAGGGAAGGGACCUGGUGAGACCCACCUGGUCCAAUCAGCAG
  Msp-2-1     ACCUCCUCACGCGUCCGCGGGCACGGCGGUCAGCGGCGCGAGGGGAGGGGGCUGGGACGUCUUCCGGCCCGAAUGACAG
  Mfu-2-1     GCCUCCCCACGCGGCCGCGGGCACGGCGGUCAGCGGCGCGAGGGGAGGGGGCCGGGACGUCUUCCAGCCCGAAUGACAG
  env-6       CUCGGUCCACGCCGUGACACGGCUUCACGCGGCGCGGGCAGAGGGGGCCUCCCUCCAUAAACGGGAGGCCCAAUGGCAA
  Cco-1-1     CACCUGCCUUGGCUGCCCACCGACACGGCUUCACGGUGGCGCGAAGGCAGGGGGCCCUGGUACACAGGGCCCGAUUCAA
  env-7       ..........................................................GCCGCGAGGCACGGUCGACGAA
  env-8       AGAACUGGCCGCCGCAGAACCUGGACAAGCGCUAGAGCUCUGACAAUACGAAAGGCGGCCCAAGUGCCGUCCCAGCGUG
  Ama-1-1     GGACAACUACGGCGACGACGACCAGAGCUACGCCGAGACCGAGCAGUACUAGACACCCGAUUCCGGCGGCCGGGGGCAC
  env-9       CGGAGUGCGACCAUGAGUACCGAGCACGAAGCGAGCGCGACCGUCAGUGCAGAGUCCGAGCGGCGCGCGGAUCCCAGAC
  env-10      ...................................GCGCGACCGUCAGUGCUGAGUCCGAGCGGCGCGCCGAUCCCAGAC
1 Ssp-68-1    CUGCGGGCCGGCUGCGGCCUGCCGCUCCCACGCGGCGAAGCCGCAUGGGAGCACAGCCCCGCGCCCUUCGGGGGCGCA
  env-11      AGACCGAGCAGUACUGACCGGACCGCGGCGGCCGGGGCCGCGUUCCCCGGCCGUCCGGGCAGCACAGGCGCGCGGGCGU
  env-12      GCAGUUGGCGCGGUCACAACUACAGAUUGAUUGGCGGCUGCGGGCCGAAACUCUUGGUUUCGGGUUCGCUUCGCCACAC
  Ssp-96-1    CUACGCGGAGACCGAGCAGUACUAAGAAUUCGGGACGGGGGCGGCCCUUCGCGGGCCGCCCCCGCCACCGUGCAAGUCG
  Ksp-1-1     GCGGAGACCGAGCAGUACUAAGAGUUUGGGACGGGGGCGGCCCCUCGCGGACCGCCCCCGCUUCAAUGCACACAAGCCG
  Kme-1-1     UUACGCGGAGACCGAGCAGUACUGAGUUUGGUCCGGGACGGCUCCCACCGGAGCCGUCCCGGUUCAUGAAGGAAAGUCG
  Rge-1-1     AGCGUUGAGAUCGCGGGCCGCUCGCGGCUCGUCGUCGUAAGAACCGCUGCGGCUCCCGGACGGGAGCCGCGGUGUCGAU
  Sca-3-1     UGAUCCAGCACUGAUCACCGUUCAGGUGGUACGGGACCGCUCCGGCGGUCCCGUACUCUUGAUGGCUGUAUGACCAAGC
  Sal-3-1     AGACCGAGCAGUACUGAGAAACACCUGGGUACGGGACCGCCUCGGCGGUCCCGUACUCUUGAAGGCUGUGCAACCAAGU
  env-13      GACGACGACGACGACGCGUUCGUCGAGGACGAGCAGUACUGAUCCGUCCGGGAGGCUCCGCCCUCCCGGACACCCCCGA
1 Sav-3-1     GACGCGUUCGGCGCGGAGGACGACGGCGACGCGGGCUUCGUCGAGACCGAGCAGUACUGACCCCUCGGGGUCGCCAACC
  Ssp-195-1   AGACCGAGCAGUACUGAUCCGACGGGGCGUGCGACCGGCCGCGGCCGGCCCCACGCCCCGGUCCCCGUCCGGCGAUCCG
  Nsp-4-1     ACCGCGGUGGCCGGCGGCGGCUACGACGACGACGACAGCGCGUACGUCGAGACCGAGCAGUACUGAUUUUCCCGAUAGC
  env-14      ......................................................GGAGAGGGGCCCGCGCGGUGAACGUA
  Nco-1-1     CGCGGUGGCCGGCGGCGGCUACGACGACGACGACAGCGCGUACGUCGAGACCGAGCAGUACUGAUCACACACUGAUAGC
  Sje-1-1     GACCGAGCAGUACUGAGCCUGAUCGAGUGGUACGGGACCGCCCCGGCGGUCCCGUACCCUUGAAGGCUGUGCAACCAAG
  Kaz-1-1     GGACGGGGGCGGCCCUCGGGUCGCCCCUGUUCGAUGCACACAGGGCUCUGCGGGGGACAACCCCCAGACCCCCACUAAG
  Kph-1-1     GGCUACGCGGAGACCGAGCAGUACUGAGUUUCGACACCGGGGGCGGUCCUCGCCGGACCGCCCCCGGUGGCGUGAGAAG
  Kph-1-2     GCUACGCGGAGACCGAGCAGUACUGAGUUUCGACACCGGGGGCAGCCCUUCGCCGGGCCGCCCCCGGUGGCGUGAGAAG
  Sno-2-1     GGUUACGCCGAGACCGAGCAGUACUAAGAGAUUUGGCCGCGGGGGCGGUUCCACGGAGCCGCCCCCGUCGUCACGGAAG
  Ksp-3-1     GCACUUCGCCGAGACCGAGCAGUACUGAGAGAAUUGGCCGCGGGGGCGGUUCCACGAACCGCCCCCGUCGUCGUGAAAG
  Spy-1-1     GGUUACGCCGAGACCGAGCAGUACUAAGAGAUUUGGCCGCGGGGGCGGUUCCACGGAGCCGCCCCCGUCGUCACGGAAG
  Ksp-4-1     AGACCGAGCAGUACUGAGUUCCCUGAAGUCUUGACGCCGGGGGCGGUUCCCAGUGGAUCGCCCCCGGGGCCAUGAAAAG
  Ssp-165-1   GGUUACGCGGAGACCGAGCAGUACUGAGAGACUUGACCGCGGGGGCGGUUCCCCGGAACCGCCCCCGUCGUCGUGAAAG
  Ssp-133-1   CGGGUUACGCGGAGACCGAGCAGUACUGAGAGACUUGACCGCGGGGCGGUUCCCCGGAACCGCCCCGCUGUCGUGAAAG
  Ssp-50-1    AGGGCUACGCGGAGACCGAGCAGUACUAAGAGUUUGAGUCGGGGGCGACUCGCCACGGGUCGCCCCCGAUCCAUGCAAG
  Ssp-157-1   GGCUACGCGGAGACCGAGCAGUACUAAGAGACUUGGCCGCGGGGGCGGUUCCCCGGAACCGCCCCCGUCGUCCUGGAAG
  Ssp-160-1   GCUACGCGGAGACCGAGCAGUACUGAGUUUCGACACCGGGGGCGGUCCUUCGCCGGACCGCCUCCGGUGGCGUGAGAAG
  Ksp-2-1     GGUUACGCGGAGACCGAGCAGUACUGAGAGACUUGACCGCGGGGGCGGUUCCCAGGAACCGCCUCCGUUGUCGUGAAAG
  Ssp-183-1   CGGGUUACGCGGAGACCGAGCAGUACUGAGAGACUUGACCGCGGGGCGGUUCCCCGGAACCGCCCCGCUGUCGUGAAAG
  Sam-2-1     CCGAGCGCGGUGGCCGGUGGCGGCUACGACGACGACGACAGCGCCUACGUCGAGACCGAGCAGUACUGAUUUUUCAAUA
  Sro-7-1     CCCAGCGCGGUGGCCGGCGGCGGCUACGACGACGACGACAGCGCGUACGUCGAGACCGAGCAGUACUGAUUUUUCUGUA
  Ssp-87-1    CGGACGCCGGUUUCGUCGAGACCGAGCAGUACUAGAACCUCCCGGUCCGCUGUGGCUGAGGUCUUGUCCUCAAUCGCCG
  Ssp-190-1   GCGGACGCCGGUUUCGUCGAGACCGAGCAGUACUAGAACCUGCUGGUCCCUGUGGUUGAGGUCUUGUCCUCAAUCGCCG
  Sir-1-1     CGGACGCUGGAUUCGUCGAGACCGAGCAGUACUAGAACCUGCUGGUCCGCUGUGGUUGAGGUCUUGUCCUCAACCGCCG
  Shy-4-1     CGGACGCCGGUUUCGUGGAGACCGAGCAGUACUAGAACCCGCUGGUCCGCUGUGGCUGAGGCCGCGUCCUCAGCCACCG
  Sro-3-1     CCCCAGCGCGGUGGCCGGCGGCGGCUACGACGACGACGACAGCGCGUACGUCGAGACCGAGCAGUACUGAUUUUUCUGU
  Nin-1-1     GUUCGUCGAGCCCGGCCGGUGGAUGCAAGCGGCACUGUCCACCGGCUGCCGCGCGACAACUGAAGAUCCGGUGGCCGAG
1 Nca-1-1     ACGACAGCGCGUACGUCGAGACCGAGCAGUACUGAUUCACCGUCAGCCCCCGCGUCUGAUCGCGGGGGCUCAAUAACGG
  Ssp-192-1   UCGCGCCGCCACCGGUACCUGAUACGGCCGGUGCGGCUCGCCUCCGGUGGGGAAGCCGUGUGCCGCCCCCGCCGGGGCC
  Ssp-172-1   GACGACGCCGACGCCGGGUUCGUCGAGACCGAGCAGUACUAAAUUCUCUGCGGACCGGCUCGGGCCGGUCGCGCAGUUC
  Ssp-93-1    CGAGACCGAGCAGUACUAAGAGCUCGGGACCGGCCGCUUGAUUUUGGCUGCGCGUGCGUUGUGGUUGUUCGCGCAGUUC
  Sau-3-1     CGAGACCGAGCAGUACUGAGAGCUCGGGCCUCGCGGUUCGAUUGUGGCUGCGGACCGGCUUCGGCCGGUCGCGCAGCUC
  Sal-6-1     CGAGACCGAGCAGUACUGAGAGCUCGGGACUGGCGGAUUGAUUUUGGCUGCGCGUGCGUCUUGAUUGCGCGCGCAGUUC
  Sau-1-1     UCGAGACCGAGCAGUACUAAGAGCUCGGGACUGACGGUUCGUUUUGGCUGCGGGGCCGUUGUGGCUGGUCGCGCAGUUC
  Ssp-106-1   AGGACGACGGGGACACGGGCUUCGCGGAGACCGAGCAGUACUGACGAAGCGGUACUGAGCGGCCCCCGCGGCUGACAGU
  Ssp-3-1     GACGCCUUCGGGGCCGAGGACGACGGGGACACGGGCUUCGCGGAGACCGAGCAGUACUGACGGUCCUGCGUCUCACAGU
  env-15      GCGGCCCUCGGCACGGUGACCGCCCGGCCCGGCCGGGGCGGCACCGCCGCCAGCGGUCUGCCUGGCAAGCCAUGAAGGU
  env-16      GCGGCCCUCGGCACGGUGACCGCCCGGCCCGGCCGGGGCGGCACCGCCGCCAGCGGUCUGCCUGGCAAGCCAUGAAGGU
  env-18      ...................................................................AAGCCAUGAAGGU
  Ssp-118-1   GCCGCGCGGCACCGCCGUGCUCGCGGCGUCACGGGGCGGCGUGCUGAACGCCGCACGCCGUACGCCGCACACACGAAGC
  env-19      CACCGGGCGCCGGACUGCUCGGCGGUAGUGAGCCCUCCGGAUCGCGGCCACCGGCCCGUUCCGGGAGAUUGUCAAAGAG
  env-20      CACCGGGCGCCGGACUGCUCGGCGGUAGUGAGCCCUCCGGAUCGCGGCCACCGGCCCGUUCCGGGAGAUUGUCAAAGAG
  env-21      ACCGGACGCCCGGACUGUUCGGCGGCAGUGAGCCCUCCGGAUCGCGGUGACCGCCCGAUCCGGCAGACAUGUCAAAGAG
  Sor-1-1     ACCGAGCAGUACUGAGAACACAUCGUCCCGUCGGCGUCCCCGCCGGCGUGAUGCCGUAAGUCGUACGGCGUGCGCGGAC
  Ssp-64-1    GUGGCUGCCUCCGGGGCCACGGCCCCGGGCUCCGGGCGCGAGAGCCCGGCACAGAUGUCCCCGGUACCGCCGGGGGGUG
  Ssp-12-1    GGGUGGCUGCCUCCGGGGCCACGGCCCCGGGCUCCGGGCGCGAGCCCGGCACAGAUGUCCCCGGCACCGCCGGGGGGUG
  Ssp-13-1    GGGUGGCUGCCUCCGGGGCCACGGCCCCGGGCUCCGGGCGCGAGCCCGGCACAGAUGUCCCCGGCACCGCCGGGGGGUG
  Ssp-61-1    CCCGACCUCCGCCGCCGACACCUUCGGCGCCGACGACGACGCCGACGCGGGCUUCGUCGAGACCGAGCAGUACUGAAGU
  Ssp-59-1    CCCCACCUCGGCCGCCGACACCUUCGGCGCCGACGACGACGCCGACGCCGGCUUCGUCGAGACCGAGCAGUACUGAUGU
  Ssp-16-1    CCCCACCUCGGCCGCCGACACCUUCGGCGCCGACGACGACGCCGACGCCGGCUUCGUCGAGACCGAGCAGUACUGAUGU
  Ssp-69-1    CCCCACCUCGGCCGCCGACACCUUCGGCGCCGACGACGACGCCGACGCCGGCUUCGUCGAGACCGAGCAGUACUGAUGU
  Ssp-56-1    GCCGCCGACACCUUCGGCGCCGAGGACGACGCCGACGCGGGCUUCGUCGAGACCGAGCAGUACUGAGCAGCACUGAUGU
  Hcr-1-1     CGCUGUCGCCGGUGGCGGCUACGACGACGAGGACGGCGGCUUCAUCGAGACCGAGCAGUACUGAGUUUUCGCAGAUAGA
  Msp-1-1     CGUCACCGAUCGCUGCCGGUAGGCACCACCGCAUCACCGGCCGCCCCGCGUCGUACGGCGUGGGCGGCGAUGACAGCC
  Mro-1-1     CUGAUCGCCCGUCUCCGAGCAGCGGCACCACCGUCUCGUGCCGCCCGUCCAGGGCGGCAUGAGGCGGCGAUGACAAGUC
  Ssp-171-1   CCGACGCCGGCUUCGUCGAGACCGAGCAGUACUGAUCCACGCUUGCGGGAGGCCCGGGCACUGCCCGGGCCCCCCGUAC
  Ssp-43-1    GAGCAGUACUAGUAGAUCGUCUGCGGGCCGGCUUCGGCCGGUCGCGCAGUUCCCCGCGCCCCUUACGGGGCGCGGGUCU
  Svi-13-1    GAGCAGUACUAGUAGAUCGUCUGCGGGCCGGCUUCGGCCGGUCGCGCAGUUCCCCGCGCCCCUUACGGGGCGCGGGUCU
  Ssp-73-1    CGUGGCUGGUCGCGCAGUUCCCCGCGCCCCUUACGGGGCGCGGUCCUCCGAGGCGCUUCGAAAGAGGAACCCCGGAUCU
  Ssc-3-1     GGUGCGGGUGCGCUGUGGCUUGUCGCGCAGUUCCCCGUGCCCCUUCCGGGAGCGUCCCUUCGGGACUGCUCCCGGAUCU

              ................................................................................
              ................................................................................
              ●●●●●●●●●●●●●●●●●●●●●●●●●●●●●●●●●●●●●●●●●●●●●●●●●●●●●●●●●●●●●●●●●●●●●●●●●●●●●●●●
```

| | |
|---|---|
| Kau-1-1 | GGUUCGACGGUUCGUUUUGGCUGCCGGCCCGCUGUGGCUGGUCGCGCAGUUCCCCGCGCCCCUUUGGGGCGCGGGGUCU |
| Sgr-1-1 | GCGCGGGGCGGCUGUGGCUUGACGCGCAGUUCCCCGUGCCCCUUCCGGGCACGGCCCCUCGGGGCCCGUGUCCGGAUCU |
| Sto-1-1 | GGUGCGGGUGCGCUGUGGCUUGUCGCGCAGUUCCCCGCGCCCCUUCCGGGAGCGCCCCUUCGGGGCUGCUUCCGGAUCU |
| Ssp-114-1 | GGCUGUGGCUGGUCGCGCAGUUCCCCGCGCCCCUUUGGGGCGCGGACCACUCCGGGCGUCCGAAAAGGCGCCCGGAUCU |
| Ssc-4-1 | GGUGCGGGUGCACUGUGGCUGAGCGCGCAGUUCCCCGCGCCCCUUCCGGGAGCGCCCCUGCGGGGCGACUCCCGGAUCU |
| Ssp-1-1 | CCGGGUGCGGGUGCGCUGUGGCUUGUCGCGCAGUUCCCCGCGCCCCUUCCGACUCGCCCCUUCGGGGCUCGUCGGAUCU |
| Sau-4-1 | GACCGGCUGUGGCUGAUCGCGCAGUUCCCCGCGCCCCUUACGGGGCUCGGUACCUCCGGCGCUGAAAAGUGCCGGAUCU |
| Spr-1-1 | CCGACGACGACGCGGACGCGGGCUUCGUCGAGACCGAGCAGUACUGAGAAGGCUUGCGGGGAGCUCCGGCUCCCCGCAC |
| Ssp-189-1 | CGACGACGACGCGGACGCCGGUUUCGUCGAGACCGAGCAGUACUGAGAAAGCCUGCGGGGGGCCUCGCCCCCCGCACCC |
| Sfl-6-1 | GACGACGACGCGGACGCCGGUUUCGUCGAGACCGAGCAGUACUGAGAAAGCUUGCGGGGGUCUCGCGCCCCCCGCACCC |
| Ssp-103-1 | CGACGACGACGCGGACGCCGGUUUCGUCGAGACCGAGCAGUACUGAGAAAGCUUGCGGGGGGCCCCGCCCCCCGCACCC |
| Ssp-111-1 | GACGACGACGCGGACGCCGGUUUCGUCGAGACCGAGCAGUACUGAGAAAGCUUGCGGGGGGCUCGCGCCCCCCGCACCC |
| Ssp-159-1 | CGACGACGACGCGGACGCCGGUUUCGUCGAGACCGAGCAGUACUGAGAAAGCUUGCGGGGGGCUCGCGCCCCCCGCACC |
| Ser-1-1 | GACGACGACGCGGACGCCGGUUUCGUCGAGACCGAGCAGUACUGAGAAAGCUUGCGGGGGGCUCACGCCCCCCGCACCC |
| Sya-1-1 | GACGACGACGCGGACGCCGGUUUCGUCGAGACCGAGCAGUACUGAGAAAGCUUGCGGGGGGCUCGCGCCCCCCGUACCC |
| Svi-12-1 | CCGACGACGACGCGGACGCCGGUUUCGUCGAGACCGAGCAGUACUAGAAAAGCUUGUGGGGGGCUCACGCCCCCCACAC |
| Ssp-151-1 | CCGACGACGACGCGGACGCCGGUUUCGUCGAGACCGAGCAGUACUAGAAAAGCUUGUGGGGGGCUCGCGCCCCCCACAC |
| Ssp-110-1 | GCCGACGACGACGCGGACGCCGGUUUCGUCGAGACCGAGCAGUACUAGAAAGCUUGUGGGGGGCUCGCGCCCCCCACAC |
| Ssp-112-1 | GACGACGACGCGGACGCCGGUUUCGUCGAGACCGAGCAGUACUGAGAAAGCUUGCGGGGGGCUCACGCCCCCCGCACCC |
| Ssp-102-1 | CCGACGACGACGCGGACGCCGGUUUCGUCGAGACCGAGCAGUACUAGAAAACUUGUGGGGGGCUCACGCCCCCCGCACC |
| Sla-2-1 | CGACGACGACGCGGACGCCGGUUUCGUCGAGACCGAGCAGUACUGAGAAAGCUUGUGGGGGGCUCACGCCCCCCACACA |
| Ssp-139-1 | CCGACGACGACGCGGACGCCGGUUUCGUCGAGACCGAGCAGUACUGAGAAAGCUUGUGGGGGGCUCGCGCCCCCCACAC |
| Ssp-8-1 | CCGACGACGACGCGGACGCCGGUUUCGUCGAGACCGAGCAGUACUGAGAAAGCUUGCGGGGGGCUCGCCCCCCCGCACCC |
| Svi-7-1 | GACCGACGGUCCAGAUUUGGCUGCGGACCGGCUUCGGCUGGUCGCGCAGUUCCCCGCGCCCCUAACGGGGCGCGGGGUC |
| Sgr-12-1 | UACUGACGGCCGCAGUUCCGCUGCCGGUCCGCUCCGGCUGACCGCGCAGUUCCCCGCGCCCCUCACGGGGCGCGGGGUC |
| Sfl-5-1 | CCCGGGUGCGGGAGCGCUGUGCUUGAUCGCGCAGUUCCCCGCGCCCCUUCCGGGCGCCCCUUCGGGGGCUGCCCGGAUC |
| Sbi-3-1 | GAUCGCGCUCACACGGCGGAGCCGUACAUCGGCUCCGCCCCGCGCCCCUUCCGGGCGCCUCCACGAGGCCUUCCGGAUC |
| Sex-1-1 | AGCAGUACUAGUAGAUCGUCUGCGGGCCGGUUUCGGCUGGUCGCGCAGUUCCCCGCGCCCCUUAACAGGGGCGCGGGUC |
| Sgl-3-1 | ACUGAGGUCGUAGUUCUUGCUGCGGACCGGCCUCGGCCGGUCACGCAGUUCCCCGCGCCCCUUCCGGGGGCGCGGGGUC |
| Sgr-9-1 | GGACCGACGGUCCAGAUUCGCCUGCGGGCCGGCUGUGGCUGGUCGCGCAGUUCCCCGCGCCCCUGAGGGGGCGCGGGUC |
| Ssp-121-1 | UCGGGUCCGUACUCGGGUGCGGGUGGACUGUCGCUGGUCGCGCAGUUCCCCGCGCCCCUUCCGGGGCCCUGCCCCGGUC |
| Sve-1-1 | AGUACUAGUCGCAGUCUUGCUGCGGGCCGGCUUCGGCUGGUCGCGCAGUUCCCCGCGCCCCUGACCGGGGCGCGGGGUC |
| Sav-2-1 | GGGUGCGGGUGCGCUGUGGCUUGUCGCGCAGUUCCCCGCGCCCCUUCCGGGGGCGCCUCUACGAGGCUGCCCCCGGAUC |
| Mgl-1-1 | GGGUCCGUACUCGGGUGCGGGUGGACUGUGGCUGUUCGCGCAGUUCCCCGCGCCCCUUUCCGGGGCGUACGCCCCGGUC |
| Asp-1-1 | GCGGACGCGGGUUUCGUGGAGACCGAGCAGUACUGAUCGGUCCCCCUUCCGGGAGCCGCCUUCGGGAGGCUCCCGGAUC |
| Stu-1-1 | GGGUGCGGGUGCGCUGUGCUUGAUCGCGCAGUUCCUCGCGCCCCUUCCGGGAUCGCCCCUUCGUGGCUGCUCCCGGAUC |
| Sye-2-1 | GCGGAGCCGCAAAUUGACACAGCCCGCGCCCCUGAAGGGGCGCGGGUGCUCCGGGCGCCUUGAAAAGGGUGCCCGGAUC |
| Sdu-1-1 | CCGGCUUCGGCUGGUCGCGCAGUUCCCCGCGCCCCUUCGGGGGCGCGGGUCUUCCGGUGCUCGAAGGAGCGCCCGGAUC |
| Sgh-1-1 | GAUCGGAUCCUAGGAAGCGGGCGCGCCAUGCGCGCCCGCUCCCCUUCCGGGAGCUCCCCCACGGGGUGGCUCCCGGAUC |
| Svi-5-1 | GAUCGGAUCCUAGGAAGCGGGCGCGCCGUGCGCGCCCGCUCCCCUUCCGGGAGCUCCCCCACGGGGUGGCUCCCGGAUC |
| Ssp-153-1 | CCGGGUUCGUCGAGACCGAGCAGUACUAGGAGCUCGGGACCGACGGUCCUCCGGGCGCCCCCGCGGGGACGCCCGGAUC |
| Ssp-86-1 | CGAGGAUCGGCAGCACUGCCACUGCGGGUCGGCUGUGGCUGGUCGCGCAGUUCCCCGCGCCCCUCUGGGGCGCGGGGUC |
| Sro-1-1 | ACGCGGACGCGGGUUUCGUGGAGACCGAGCAGUACUGAUCUCCACAGCCUCCGGGGUUCGCGUCAGCGGACCCCGGAUC |
| Sgr-7-1 | GGGCCUCAGGGCUCGAUUUCGUCUGCCGGCCGGCUGCGGCUGGUCGCGCAGUUCCCCGCGCCCCUUCGGGGCGCGGGUC |
| Ssp-28-1 | GGUGCGGGAGCGCUGUGCUUGAUCGCGCAGUUCCCCGCGCCCCUUUCCGAGCAGGCCCCCUUCGGGGCCCGCUCGGAUC |
| Ssp-176-1 | CGUUUUUCGGUUGCGGGUCGGCUGUGGCUGGUCGCGCAGUUCCCCGCGUCCCUUCCAGGCCCCUUCGGGGGCCUGGAUC |
| Ssp-14-1 | GCAGCCGCACAGGCAGCACCGCCCCGCGCCCCUGACGGUGCGCCCUUCCGGGGGCGCACCCUGACGGGGGGCGCGGGUC |
| Sac-2-1 | CGACGCGGACGCGGGUUUCGUGGAGACCGAGCAGUACUGAGUCUCUGCUUCCGGGGUUCGCGUCAGCGGACCCCGGAUC |
| Sbo-1-1 | CGGGCGCGGGUGCGCUGUGCUUGAUCGCGCAGUUCCCCGCGCCGCUUCCGGGCAGGCCCUUCGGGGCCUCGCCCGGAUC |
| Ssp-2-1 | ACGCGGACGCCGGGUUCGUGGAGACCGAGCAGUACUAAGCGCUCCGCACCCCUUCCAGGCCCCUUCGGGGGCCUGGAUC |
| Ssp-98-1 | CGGGUGCGCUGUGGCUUGUCGCGCAGUUCCCCGCGCCCCUUCCGGACGAUGCGCCCCUUCGGGGCUUGUCGUCCGGAUC |
| Sac-3-1 | GCGCUGUGGCUUGUCGCGCAGUUCCCCGCGCCCCUAAGGGCGAUGCCCCUUUCCGGAGUUUCGUAAGAAGCCUCGGAUC |
| Sca-7-1 | GCGGUUCCGUUCUUGCUGCGGGUCGGCUGUGGCCGGUCGCGCAGUUCCCCGCGCCCCUCUUUGCGAGGGGCGCGGGGUC |
| Sgr-2-1 | CGUACUGCCCGUGCGGGCCAGCUGUGGCUGGUCGCGCAGUUCCCCGCGCCCCUUCCAGGCCCCUUCGGGGGCCUGGAUC |
| Ssp-105-1 | GCGUCUGCGGCCGGCAGUGGCCGGUCGCCCAGUUCCCCGCGCCCCUUCCGAGCGGGCCCCUUCGGGGCCGGCUCGGAUC |
| Ssp-85-1 | CGGCUGCGGCCGGCCGCGCGGUUCCCCGCGCCCCUUCGGGAGCGCGGGCCCUCCGGGAGUCCGCGAGGACUCCCGGAUC |
| Sco-3-1 | CCGGCUGUGGCCGGUCGCGCAGUUCCCCGCGCCCCUUCGGGGGCGCGGACCCUCCGGGAUCGAACGUGAACCCCGGAUC |
| Sse-1-1 | CGUCUGCGGGCCGGCUGUGACUGGUUCGCGCAGUUCCCCGCACCCCUUCCGAGCGGGCCCUUCAGGGCCUGCUCGGAUC |
| Sre-1-1 | NNNGGCUGGUCGCGCAGUUCCCCGCGCCCCUUACGGGGCGCGGAUCCUCCGGGGUGCUUCGAAAGAGGCGCCCCGGAUC |
| Spe-1-1 | AGCAGUACUGAUCGGUUUCGGCUGCGGGCCCGCUGUGACUGGUCACGCAGUUCCCCGCGCCCCUAUGGGGCGCGGGGUC |
| Sbi-2-1 | GGACCGACUGCCGGUUCUUGGCUGCGGGCCCGCUGUGGCUGGUCGCGCAGUUCCCCGCGCCCCUUUGGGGCGCGGGGUC |
| Ssp-83-1 | GACACGGACGCGGGUUUCGUGGAGACCGAGCAGUACUGAUCGGUACCCCUUCCGGGGGCCUCACCGAGGCCCCCGGAUC |
| Spl-1-1 | GACACGGACGCGGGUUUCGUGGAGACCGAGCAGUACUGAUCGGUACCCCUUCCGGGGGCCUCACCGAGGCCCCCGGAUC |
| Sno-1-1 | GUCCGUUGUGGCCGGUCGCGCAGUUCCCCGUGCCCCUUUGGGGCGCGGGGCCUCCGGGAGUCGUCAGGACGACCGGAUC |
| San-1-1 | GUUUCUGCGGGCCGGUUGUGGCUGGCCGCGCAGUUCCCCGCGCCCCUUCCGGGGCGCCCCUUCGGGGACGCCCCGGAUC |
| Sla-3-1 | CGUACUGCGUCUGCGGGCCGGUUGUGGCUGGUCGCGCAGUUCCCCGCGCCCCUUCCAGGCCCCUUCGGGGGCCUGGAUC |
| Ssp-74-1 | GACGCGGACUCGGGUUUCGUGGAGACCGAGCAGUACUAGGUCUCCGUUCUCCGGGGCCUGCGACAGUGGGCCCCGGAUC |
| Ssp-162-1 | NNNNNNNNNNNNNNNNNNNNNNNNGGCUGGUCGCGCAGUUCCCCGCGCCCCUUCCAGGCCCCUUCGGGGGCCUGGAUC |
| Ssp-164-1 | CGUACUGCGUCUGCGGGCCGGUCGUGGCUGGUCGCGCAGUUCCCCGCGCCCCUUCCAGGCCCCUUCGGGGGCCUGGAUC |
| Ssp-161-1 | CGUACUGCGUCUGCGGGCCGGUCGUGGCUGGUCGCGCAGUUCCCCGCGCCCCUUCCAGGCCCCUUCGGGGGCCUGGAUC |
| Sgl-2-1 | UGAGGGUCGCCGCGCCGUGUCGCGCGGUUGCCCGCACCCCGCUCCGAGCGCGGCCCCUCGCGGGGUCAGCGCCCGGAUC |
| Ssp-143-1 | CUGGUCGCGCCCACGCGACGGAGUCGCAUAUCGACACCGCCCCGCGCCCCUUCCAGGCCCCCUACGGGGGGCCUGGAUC |
| Ssp-36-1 | GUCCGAUGCCGGGUGCGGGGGCGCUGUGCCUGAUCGCGCAGUUCCCCGCGCCCCUUCCGGGGCUGGAAGGCCCCGGAUC |
| Ssp-200-1 | GCGAAGCCGCACAGGGGUACAGCCCCGCGCCCCUUGGGGGUGCGGACCCUCCGGGGCGCUUGCGAAGGCGCCCCGGAUC |
| Sco-2-1 | GCGAAGCCGCACAGGGGUACAGCCCCGCGCCCCUUUGGGGUGCGGACCCUCCGGGGCGCUUGCGAAGGCGCCCCGGAUC |
| Svi-9-1 | GCGAAGCCGCACAGGGGUACAGCCCCGCGCCCCUUUGGGGUGCGGGCCCUCCGGGGCGCUUGCGAAGGCGCCCCGGAUC |
| Sco-1-1 | GACGCGGACGCGGGCUUCGUGGAGACCGAGCAGUACUGAUCGGCACCACUUCCGGGGGCCUCGUGAAGGCCCCCGGAUC |
| Ssp-128-1 | GGUUCGACGGUUCGUUUUCGACUGCCGGCCGGUUGUGGCCGGUCGCGCAGUUCCCCGUGCCCCAUGGGGGCGCGGGGUC |
| Sol-2-1 | GGCUGCGGCUGGUCGCGCAGUUCUCCGAGCCCCUUCGGGGCGCGGAUCCUCCGGGGCGUCUGCGAAGACGCCCCGGAUC |
| Ssp-72-1 | GGCUGUGGCUGGUCGCGCAGUUCCGCGUACCCCUUCGGGGCGCGGAUCCUCCGGGGCGUCUGCGAAGAGGCCCCGGAUC |
| Ssp-124-1 | CGGCUGUGGCCGGUCGCGCAGUUCCCCGCGCCCCUUCGGGCGGGGACCCUCCGGGGCGUGUGAAAAGACGCCCCGGAUC |
| Sce-2-1 | GACGCGGACGCGGGUUUCGUGGAGACCGAGCAGUACUGAUCGGUGCCACCUCCGGGGGUCUCGCGCAGACCCCCGGAUC |
| Smo-1-1 | GGCCCGCUGUGGCUGGUCGCGCAGUUCCCCGCGCCCCUCGGGGCGCGGGCCCUUCCGGGCCCCGAAAGAGGCUCGGAUC |
| Svi-9-2 | GACGCGGACGCGGGUUUCGUGGAGACCGAGCAGUACUGAUCGGUGCCACCUCCGGGGGUCUCGCGCAGACCCCCGGAUC |
| | ................................................................................ |
| | ................................................................................ |
| | ●●●●●●●●●●●●●●●●●●●●●●●●●●●●●●●●●●●●●●●●●●●●●●●●●●●●●●●●●●●●●●●●●●●●●●●●●●●●●●●● |

```
Sso-1-1      UCCCACGAGGUCCGACCGCUCCCGCGGACCGGCUCCGGCCGGCCGCGGAGCUCCCCGCGCCCCCCUGGGGGCGCGGGUC
Soc-1-1      GGCCUAACGGCCUGAUUUUGACUGCCGGCCCGCUGUGGCUGGUCGCGCAGUUCCCCGCGCCCCUUCGGGGGCGCGGGUC
Sni-2-1      GGCCUGACGGUCUGAUUUUGCUGCGGACCGGCUUCGGCUGGUCGCGCAGUUCCCCGCGCCCCUUACGGGGCGCGGGGUC
Svi-11-1     CGGCCCGCUGUGGCUGGUCGCGCCCACGCGGCAGAGCCGCAGAUCAACACCUUCCCGCGCCCCUUCGGGGGCGCGGGUC
Sex-2-1      UCCUGAGGAUCGAUUGACCCCUGCCGGCCGGUUUCGGCUGGUCGCGCAGUUCCCCGCGCCCCUUGUUGGGCGCGGGGUC
Ssp-6-1      GGGCGCGGGUGCGCUGUGCUUGAUCGCGCAGCUCCUCGCGUCCCUUCCGGGAGCGCCCCUUCGGGGCUGCUCCCGGAUC
Ssp-101-1    UCGGCUGUGGCCGGUCGCGCAGUUCCCCGCGCCCCUCUGGGGCGCGGACCCUCCGGGCGUCAGAAAUGGCGCCCGGAUC
Ssp-49-1     UAUCGACACCGCCCCGCGCCCCUGUCGGGAGCGCCCCUGCGGGGCGGCUCCCGACUCCUCCGGGGCGCGUCCCCGGAUC
Sal-11-1     CGGGUGCGGGUGCGCUGCGGUACUCACGCAGCUCCCCGCGCCCCUGUCCGGACCGCCCCUUCGGGGGCGGAUCCGGAUC
Shy-2-1      GGCCCGCCGGACCGUUUUCGGCUGCCGGCCCGCUGUGGCUGGUCGCGCAGUUCCCCGCGCCCCUUGCGGGCGCGGGGUC
Ssp-46-1     GUCCGUUGCGGCCGGUCGCGCAGUUCCCCGCGCGCUUCGGGGCGCGGAUCCUCCGGGGUGUCGAAAGGCGCCCCGGAUC
Ssp-187-1    GUCCGUCGUGGCCGGUCGCGCAGUUCCCCGCGCCCCUCGGGGCGCGGACCCUCCGGGGCGUCGAAAGGCGCCCCGGAUC
Ssp-156-1    UCGCGCAGUUCCCCGCGCCCCUGUCGGGACGCCCUUCGGGCUGCCCCGACACCCUCCGGAGCUGAAGAAGCUCCGGAUC
Sgr-10-1     GUUUCGGCUGGUCGCGCAGUUCCCCGCGCCCCUUCGGGGGCGCGGGUCCUCCAGGACGUCUGUGAAGGCGCUCUGGAUC
Aac-1-1      GUUUCGGCUGGUCGCGCAGUUCCCCGCGCCCCUUCGGGGGCGCGGGUCCUCCAGGACGUCUGUGAAGGCGCCCUGGAUC
Ssp-197-1    CGUACGCGGGUGCGGGUGCGCUGUGCUUGACCGCGCAGUUCCCCGCGCCCCUUUCCGGGCCCCUUCGGGGGUCCGGAUC
Ssp-20-1     CGUACGCGGGUGCGGGUGCGCUGUGCUUGACCGCGCAGUUCCCCGCGCCCCUUUCCGGGCCCCUUCGAGGGUCCGGAUC
Ssp-123-1    CCGGCUGUGGCUGGUCGCGCAGUUCCCCGCGCCCCUUCGGGGCGCGGACCCUCCGGGGCGUCGCAAGGCGCCCCGGAUC
Ssp-141-1    CCGGCUGUGGCUGGUCGCGCAGUUCCCCGCGCCCCUUCGGGGCGCGGACCCUCCGGGGCGUCGCAAGGCGCCCCGGAUC
Ssp-81-1     GUCCGUCGUGGCCGGUCGCGCAGUUCCCCGCGCCCUUCGGGGCGCGGAUCCUCCGGGGCGUCGAAAGGCGCCCCGGAUC
Ssp-113-1    GUCCGUUGUGGCCGGUCGCGCAGUUCCCCGCGCCCUUCGGGGCGCGGAUCCUCCGGGGCGUCGAAAGGCGCCCCGGAUC
Ssp-57-1     GUUUCGGCUGGUCGCGCGGUCCCCCGCGCCCCUUCGGGGGCGCGGGUCCUCCAGGACGUCGGUGAAGGCGUCCUGGAUC
Ssp-149-1    CGGAGCCGCAUAUGGGCACAGCCCCGCGACCCUUCGGGGCCGCGGACCCUCCGGGGCGCCUGACAAGGCGCCCCGGAUC
Sah-1-1      GGUCCUCGACCGGCCACGCGGUUCCCCGCGCCCCUUCUGGGGGCGCGGGUCCUCCGGGCCCCGAGCAGGGGUCCGGAUC
Svi-10-1     GUGCGGGUGCGCUGUGGCUUGUCGCGCAGUUCCCCGCGCCCCUUCCGAGCACGGUCCCUCCGGGACCUUUGCUCGGAUC
Sch-1-1      CGACGCGGACGCCGGGUUCGUGGAGACCGAGCAGUACUGAGUCUCUGCUUCCGGGGUCCGCGUCAGCGGACCCCGGAUC
Sia-1-1      CUCGGGUGCGGGUGCGCUGUGGCUUGUCGCGCAGUUCCCCGCGCCCCUUCCGAGCAGCCCCUUCGGGGCUGCUCGGAUC
Saf-1-1      CGACGCGGACGCGGGCUUCGUGGAGACCGAGCAGUACUGAGUCUCUGCUUCCGGGGUUCGCGUCAGCGGACCCCGGAUC
Ssp-89-1     GGUUCCGUUUCGGCUGGUCGCGCCCACGCGACGGAGCCGCACAUCGGCACCUCCCCGCGCCCCUGUGGGGCGCGGGGUC
Sva-1-1      GGUGCGGGAGCGCUGUGCUUGAUCGCGCAGUUUCCCGCGCCCCUUCCGGACGGGCCCCUGCGGGGGCCUCGUCCGGAUC
Ssp-158-1    CGCCCACGCGACGGAGUCGCAUAUCGAUACCGCCCGCGCCCCUUCCGAGCACGGUCCCUUCGGACCUGUGCUCGGAUC
Sro-5-1      UGCGGCCCGGCUUCGGCUGGGCGCGCAGUUCCCCGCGCCCCUUACGGGGCGCGGACCCUCCGGGGCGCAGCCCCGGAUC
Ssp-116-1    UACUGAUUGACCCGGCCCGUCCGGCUCGAUCGGCGCUGACCGGUCCCGCCUCCGGGGGCCCCGCGCGGGCCCCCGGAUC
env-22       UGUGGAGACCGAGCAGUACUGAUCGGUCUCCGGCUGCGGCUGGUCGCGCAGCUCCCCGCGCCCCCCGGGGCGCGGGGUC
Sam-1-1      GGGUGCGGGUGGGCUGUGGCUGGUCGCGCAGUUCCCCGCGCCCCUUCCGGACGGGCCCCUUCGGGGCCCCGUCCGGAUC
Ssp-51-1     CGUACCCGGGCGCGGGUGCGCUGUGGCUGAUCGCGCAGUUCCCCGCGCCCCUUUCCAGGCCCCUUCGGGAGCCUGGAUC
Slu-1-1      GCGGAGUCGUAUACCGACACCGCCCCGCGCUCCUUCGGGGGCGCGGAUCCUCCGGGGCGCCCGAAAGGCGCCCCGGAUC
Sna-2-1      GGCCGGCUGUGGCUGGUCGCGCAGUUCCCCGCGCCCCUUACGGGGCGCGGCCCCUCCGGGCCCCACUGGGGCCCGGAUC
Ssp-75-1     CCGUACGCGGGUGCGGGUGCGCUGUGCUUGAUCGCGCAGUUCCCCGCGCCCCUCUCCGGGCCCCUGCGGGGCCCGGAUC
env-23       UGCGGGCGCGCCGCGCUUGAUCGCACGGCUCCCCGCACCUCUUCCGGGCGCGGUCCCCUCGCGGGCCCGUGCCCGGAUC
Ssp-182-1    CGGAGCCGCAUAUGGGCACAGCCCCGCGACCCUUCGGGGCCGCGGACCCUCCGGGGCGCCUGACAAGGCGCCCCGGAUC
Scy-2-1      GGGUGCGGGUGCGCUGUGCUUGAUCGCGCAGUUCCCCGCGCCCCCUCCGGGAGCGCCCCUCGGGGACGGCUCCCGGAUC
Ssp-184-1    CCGUUGUGGCUGGUCGCAGUUCCCCGCGACCCUUCGGGGCCGCGGACCCUCCGGGGUGCCUGACAAGGCGCCCCGGAUC
Svi-4-1      GCGGGUGCGCUGUGCCUGAUCGCGCAGUUCCCCGCGCCCCUUCCGGACGACAGCCCCUUCGGGGCUUGUCGCCCGGAUC
Ssp-163-1    GGGUGCGGGUGCGCUGUGCUUGAUCGCGCAGUUCCCCGCGCCCCUUCCGGGAGCGUCCCUUCGGGGCGGCUCCCGGAUC
Ssp-177-1    UCCGUUGUGGCUGGUCGCACAGUUCCCCGCGCCCCUUACGGGGCGCGGUCCCUUCCGGAGUUGAAAAGAACCCCGGAUC
Sol-1-1      UCCGUUGUGGCUGGUCGCACAGUUCCCCGCGCCCCUUACGGGGCGCGGUCCCUUCCGGAGUUGAAAAGAACCCCGGAUC
Svi-3-1      CGAUGUGGCUGAUCGCGCAGUUCCCCGCGCCCCUUACGGGGCGCGGAUCCUUCCGGGCGUCUGAAAGGACGCCCGGAUC
Szi-1-1      GCCCCCCGGGGCCGGUCGCGCGGGUUCCGCACCCCUCGGGGCGCGGACCCUCCGGGCGUCCGACAGGGCGCCCGGAUC
Svi-13-2     UUCUGCGGGCCGGUUUCGGCUGGUCGCGCAGUUCCCCGCGCCCCUUCCGGGAGCGCCCCUUCGGGGCUGCUCCCGGAUC
Ssp-120-1    CUUCUGCGGGCCGGUUUCGGCUGGUCGCGCAGUUCCCCGCGCCCCUUCCGGGAGCGCCCUUCGGGGCCGCUCCCGGAUC
Ssp-131-1    UACUGACGGUCGUAGUUCUGCUGCGGGUCCGCUUCGGCUGAUCGCGCAGUUCCCCGCGCCCCUUACGGGGCACGGGGUC
Sca-4-1      GUACUAAGAGCUCGGGUCCGACCCACGGUUCUUGGGCGCGGGUGCGCUGUGGCUUGUCGCGCAGUUCCCCGCGCCCCUU
Sba-1-1      GCGGGCGGAACCGACGACGUCGGGUCCGCCCCGCCCGUGGGGCCUCCGGGACGACGUUCGGCCCGAACGUCGUCCCGGG
Ssp-65-1     GACUUCCGGCCUCCGGGCCGGGCGACCGGCCGGUUCACACCGGCCGGCUCCGGGGCGGCGGUACGCCGACCGUCCCGGG
Sme-2-1      GGCACUCCGACCGGGCGCGGUGCCGACCGCGCCCGGUCGGCCCACGCUUCCGGGGCCGCCGCUGCCGCGCGCUCCGGGU
Sgr-12-3     GGGUCCGUCGGACAGACUGAAGUCGGUCUGGGGCGGUUGAGGGGUCCGGGUGGCUCUUCGGAGUCGUUCGGGUCGAGAC
Sgr-12-4     UGGAAAAGAUGUCCUCAAUCGCCGGACGGGCUUUAAGUGGCUCGCUGGGCAGGCUUCUGGCCGACGGUUGAGGUGAGAC
Ssp-15-1     GGGUCCGUUGGACAGGCUGAAGUCGGUCUGGGGCGGUUGAGGGGUCCGGGUGGCUCUUCGGAGUCGUUCGGGUCGAGAC
Ssp-38-1     GGGUCUGUCGGACAGGCUGAAGUCGGUCUGGGGCGGUUGAGGGGUCCGGGUGGCUCUUCGGAGUCGUUCGGGUCGAGAC
Sal-5-1      GGGUCCGUCGGACAGGCUGCAGUCGGUCUGGGGCGGUUGAGGGGUCCGGGUGGCUCUUUGGAGUCGUUCGGGUCGAGAC
Ssp-77-1     AUGUCCUCAAUCGCCGGACGGGCUUAGUGGGGCUCGUCGGACGGGCUCAGUCGGGCCCGUCUCGCGGUUGAGGUGAGAC
Swa-1-1      GGGUCCGUCGGACAGGCUGAAGUCGGUCUGGGGCGGUUGAGGGGUCCGGGUGGCUCUUCGGAGUCGUUCGGGUCGAGAC
Ssp-30-1     GGGUCCGUCGGACGGGCUGAAGUUGGUCUGGGGCGGUUGAGGGGUCCGGGUGGCUCUUCGGAGUUGUUCGGGUCGGGAC
Ssp-134-1    GGGUUCGUCGGACAGGCUGAAGUUGGUCUGGGGCGGUUGAGGGGUCCGGGUGGCUCUUCGGAGUCGUUCGGGUCGAGAC
Ssp-62-1     UCGGGUGCGUGGCCGGUUGAAAAGAUGUCCUCAAUCGCCGGACGGGCUUUAUCUGGCACGUUUCGCGGUUGAGGGGGAC
Ssp-21-1     UUGAAAAGAUGUCCUCAAUCGCCGGACGGGCUUUAUGUGGCUCGCCGGACAGGCUUUUGGCCGACGGUUGAGGUGAGAC
Ssp-80-1     UUGAAAAGAUGUCCUCAAUCGCCGGACGGGCUUUAGGUGGCUCGCCGGGCAGGCUUUCGGCCGACGGUUGAGGUGAGAC
Sgr-12-5     UUGAGAAGAUGUCCUCAAUCGCCGGACGGGCUUUAGGUGGCUCGCCGGGCAGGCUUUCGGCCGACGGUUGAGGUGAGAC
Sva-2-1      UCCUCAGCUACGGCUGAGGCUUUGUCCUCAAGUGCCGGACGGGCUUGGUUUGGCCUGGGCCGUUCGGCGAACGCACGCU
Ssp-58-1     GUCGGGUCCCGGCCCGCACCCCCGCCCGCGGACCACGCACGGUCCGCGGGCAUUCCGGGGCGGCGCGUGCCGCCUCGGG
Sal-12-1     CUCACGGCUGAGGUGAGGGCGGAGCCGCCCGGCGCUCCGAGCCCGGACGCAAGCCUCCGGGAACCGCCGGUUCCCGGGU
Sfr-1-1      CGGCACCGGGUGCCGGGGGUGCCGCCCCGCCGGGCGCCCGGCCCGGACGCAAGCCUCCGGGAACCGCCGGUUCCCGGGU
Sxi-2-1      CCGGCACCGGGUGCCGGGGUGCCGCCCCGCCGGGCGCGCGGCCCGGACGCAAGCCUCCGGGAACCGCCGGUUCCCGGGU
Ssp-119-1    CGGCGCGGACGACGACGCGGACGCGGGCUUCGUGGAGACCGAGCAGUACUGAAACACCUCCGGAGCGUCAGCUCCGGGU
Ssp-33-1     UCGAGACCGAGCAGUACUGACGUCUCGGGGGCUCUGCCCCCGGACCCCCGCUCCUCAAACGCCGGAGGGGCUGGGUAUU
Ssp-37-1     GGCCGCUGCCGACGCCUUCGGCGCCGACGACGACGUGGACGCGGGUUUUGUGGAGACCGAGCAGUACUGAUCGGUUGUU
Ssp-170-1    GGCCGCUGCCGACGCCUUCGGCGCCGACGACGACGCGGACGCGGGUUUCGUGGAGACCGAGCAGUACUGAUCGGUCGUU
Smi-1-1      GCCGCUGCGGACGCGUUCGGCGCGGACGACGACGCGGACGCGGGUUUCGUGGAGACCGAGCAGUACUGAUCGGUCGUUU
Ssp-138-1    GGCCGCUGCCGACGCCUUCGGCGCCGACGACGACGCGGACGCGGGUUUCGUGGAGACCGAGCAGUACUGAUCGGUCGUU
Sga-2-1      CGACGCCUUCGGCGCCGACGACGACGCGGACGCGGGUUUCGUGGAGACCGAGCAGUACUGACCGUACUGACCGGUCGUU
env-24       CGCCGCCGCCGACGCCUUCGGCGCCGACGACGACGCGGACGCGGGUUUCGUGGAGACCGAGCAGUACUGACCGGUCUUU
Sca-1-1      GGCCGCCGCCGACGCCUUCGGCGCCGACGACGACGCGGACGCGGGUUUCGUGGAGACCGAGCAGUACUGAUCGGUCGUU
Ssp-23-1     GCGCCGACGACGACGCGGACGCGGGUUUCGUGGAGACCGAGCAGUACUGAGCCCUACGGGUCUGUGCUGAACGGUCGUU
```

```
Ssp-135-1   GGCCGCUGCCGACGCCUUCGGCGCCGACGACGACGCGGACGCGGGUUUCGUGGAGACCGAGCAGUACUGAUCGGUCGAU
Ssp-169-1   CGCCGCCGCCGACGCCUUCGGCGCCGACGACGACGCGGACGCGGGUUUCGUGGAGACCGAGCAGUACUGAUCGGUCUUU
env-30      CGCCGCCGCCGACGCCUUCGGCGCCGACGACGACGCGGACGCGGGUUUCGUGGAGACCGAGCAGUACUGACCGGUCUUU
Ssv-1-1     GCCGCUGCCGACGCCUUCGGCGCCGACGACGACGCGGACGCGGGUUUCGUGGAGACCGAGCAGUACUGAUCGGUCGUUU
Ssp-186-1   GCCGCUGCCGACGCCUUCGGCGCCGACGACGACGCGGACGCGGGUUUCGUGGAGACCGAGCAGUACUGAUCGGUCGUUU
Sda-1-1     ACGCCUUCGGCGCCGACGACGACGCGGACGCGGGUUUCGUGGAGACCGAGCAGUACUGAUCAGUACUUGAUCGGUCGAU
Sgr-12-6    CGCCGCCGCCGACGCCUUCGGCGCCGACGACGACGCGGACGCGGGUUUCGUGGAGACCGAGCAGUACUGAUCGGUUGUU
Ssp-199-1   CGCCGCCGCCGACGCCUUCGGUGCCGACGACGACGCGGACGCGGGUUUUGUGGAGACCGAGCAGUACUGAUCGGUCGUU
env-31      UCGCUGGUCGCGCAGUUCCCCGCGCCCCUUACAGGGCGCGGGGGUCCCGCCCCUCUCCUCGCGGGAGGGGCUGGUUGUU
Ssp-19-1    UGGCCGGUCGCGCAGUUCCCCGCGCCCUUUACAGGGCGCGGGGGUCCCGUCCCUCUCCUCGCAGGAGGGGCUGGUUGUU
Sfu-2-1     CGCCGCCGCCGACGCCUUCGGCGCCGACGACGACGCGGACGCGGGUUUCGUGGAGACCGAGCAGUACUGAUCGGUUGUU
Sau-2-1     AGACGGCCGGAUUGUGACUGCGCGGCCGCUGUGGCUGGUCGCGCAGUUCCCCGCGCCCCUUACGGGGGCGCUGGUUGUU
Sxy-1-1     CGCUGCCGCCGACGCCUUUGGCGCCGACGACGACGUGGACGCGGGUUUCGUGGAGACCGAGCAGUACUGAUCGGUUGUU
Ssp-122-1   CGCCGCCGCCGAUGCCUUUGGCGCCGACGACGAUGUGGACGCGGGUUUCGUGGAGACCGAGCAGUACUGAUCGGUUGUU
Spr-2-1     CGCUGCCGCCGACGCCUUUGGCGCCGACGACGACGUGGACGCGGGUUUCGUGGAGACCGAGCAGUACUGAUCGGUUGUU
Ssp-84-1    ACGCCUUCGGCGCCGACGACGACGCGGACGCGGGUUUCGUGGAGACCGAGCAGUACUGAUCAGUACCUGACCGGUCGUU
Sip-1-1     UCGGCGGUUCGUUUUAGGCUGCGCGACCGCUGUGGCUGGUCGCGCAGUUCCCCGCGCCCCUUACGGGGCGCUGGUUGUU
Sly-2-1     CUGACGGUCUGAUUGUGGCUGCGGGCCGGCUUCGGCCGGUCGCGCAGUUCCCCGCGCCCCUUUCGGGGCGCUGGUUGUU
Ssc-2-1     CGGACAGCUUCGGCGACGAGGACGACCACGGCUUCGCGGAGACCGAGCAGUACUGAGCUUCGCCGCGAGGCGUAGCCAC
Sye-1-1     CCCUUUCCCGGGUACACACCCGCGUCCUUCCGGACGUACCGCGCCCUGCGGGGCGCACUCCGCGCCCUCGGGCGCGUUG
Svi-6-1     GCGGACGCCUUCGGCACCGAGGACGACGCGGACGCCGGGUUCGUCGAGACCGAGCAGUACUGAGGCGACCCGCUGAAUA
Sni-1-1     CCGCCGCCGCCGACGCCUUCGGCGCGGACGACGACGCGGACGCCGGGUUCGUGGAGACCGAGCAGUACUGAAAAGCCUU
Ssu-2-1     GAGCCGUUCUCGGCCCGCGCACCCGGCGUCCGCCCGGUGCGCGGGCCGAGCGCCGUAGGCGCCGGUGGCGCGUACGGAU
Ssu-1-1     GAGCCGUUCUCGGCCCGCGCACCCGGCGUCCGCCCGGUGCGCGGGCCGAGCGCCGUAGGCGCCCGUGGCGCGUACGGAC
Svi-2-1     CUGAGGCGCUGUCCUCAAUCGCCGGACGGGCUUGAUUUUGCUGAGCUUGAUCGAGUCCGUGCGGCGGGCGACGAAAAUU
Sfl-4-1     CCGACCGCCGCCGCCGACGCCUUCGGCGCCGACGACGACGCGGACGCGGGUUUCGUCGAGACCGAGCAGUACUAGAACU
Spu-1-1     GGGCUUCGUCGAGACCGAGCAGUACUAAGCAGUACCCGGUGGUCCGGGGCCCGCGUGGCCCCGGACCACCGUAUGGACU
Ssc-4-2     CGCCUUCGGCGCCGACGACGACGCGGACGCCGGGUUCGUCGAGACCGAGCAGUACUGAGCCGACCGGCUCAGCGACACU
Shy-1-1     CCGACCGCCGCCGCGGACGCCUUCGGCGCGGACGAUGACGCGGAUGCCGGUUUCGUGGAGACCGAGCAGUACUGAUACC
env-32      ......GCCGCCGCCGACGCCUUCGGCGCGGACGACGACGCGGAUGCCGGUUUCGUGGAGACCGAGCAGUACUGAUACC
Sbi-1-1     GUUUCGUCGAGACCGAGCAGUACUGAGCCCUGGGGCCCCGAGCUUCGCGCUCGGGGCCCUGAGCUGAAGCCUGAAGACU
Afl-1-1     UUUCGGCAGUGCUGCUGCCGGGGGUCUGGGGGUUGUCCCCCCGAAGUGACACGCACAGUACUGACCCUCGGAGCACGAC
Ssp-55-1    CGCCGACGCCUUCGGCGCCGACGACGACGCGGACGCGGGUUUCGUGGAGACCGAGCAGUACUGAGCAGUCCCUCGAAUU
Ssp-152-1   CGACGCCUUCGGCGCCGACGACGACGCGGACGCCGGUUUCGUCGAGACCGAGCAGUACUGAGCAGCACCCGUAGAACUU
Ssp-104-1   CGACGCCUUCGGCGCCGACGACGACGCGGACGCCGGUUUCGUGGAGACCGAGCAGUACUGAGCAGCACCCGUUGAACUU
Ssp-166-1   CCGAGCAGUACUGAUUCAUCCCGCCGCCGGCGGCCGAGGCUUUCGCCCAGGUCGCCGGCCGGGCUUGAACCGAACACUU
Ssp-191-1   GCCGCCGCCGACGCCUUCGGCGCCGACGACGACGCGGACGCCGGUUUCGUCGAGACCGAGCAGUACUGAGAAACGACUU
Sxa-1-1     UUCGGCGCCGACGACGACGCGGACGCCGGUUUCGUCGAGACCGAGCAGUACUAGUCCCACCCCGCACCAACGAAGACUU
Sla-1-1     CGAGCAGUACUGAUCCAUCCCGCCGCCGGCGGCCGGGGCCUUCGGCCCGGGUCGCCGGCCGGGCUUGGACCCAAGACUU
Svi-12-3    GAGCAGUACUGAUCCGUCCCGCCGCCGGCGGCCGAGGUCUUCGGACCGCGGUCGCCGGCCGGGCUCGGAACCAAGACUU
Sla-2-2     CGACGCCUUCGGCGCCGACGACGACGCGGACGCCGGUUUCGUCGAGACCGAGCAGUACUAAGCAGCACCCGUAGAUCUU
Ssp-150-1   GAGCAGUACUGAUUCAUCCCGCCGCCGGCGGCCGAGGUCCGCGGACCGUGGUCGCCGGCCGGGCUUGGAACCAAGACUU
Ssp-32-1    CGACGCCUUCGGCGCCGACGACGACGCGGACGCCGGUUUCGUCGAGACCGAGCAGUACUAAGCACCACCCGUAGAACUU
Ska-1-1     GCCGCUGCCGACGCCUUCGGCGCCGACGACGACGCGGACGCCGGUUUCGUCGAGACCGAGCAGUACUAAGAAAAGACUU
Sro-6-1     GCCGCCGCCGACGCCUUCGGCGCCGACGACGACGCGGACGCCGGUUUCGUCGAGACCGAGCAGUACUAAGAGAAGACUU
Ssp-194-1   CGGACGGGGCCCGGCCCCGUCCGGAACCCCCUCGCGGGCGCUCCCGGACACGGAAGCACCCGCGAGGGGCACACGCACU
env-33      CCUUCGGCGCCGACGACGACGCGGACGCCGGCUUCGUCGAGACCGAGCAGUACUGAGCCGCACGGCUCGGGUGACAACU
Ssp-7-1     ACCUCCGCCGCCGACACCUUCGGUGCGGACGACGACGCCGACGCCGGCUUCGUCGAGACCGAGCAGUACUGAUCAGACU
Ssp-140-1   CGCGGGCUUCGUCGAGACCGAGCAGUACUGAUCCGCCGCUGAUCCCCUCACCGGGAUCAGCGCGGUUCAUGCGACAUCU
Svi-1-1     CGACCGCCGCCGCCGACGCCUUCGGCGCCGACGACGACGCGGACGCGGGCUUCGUCGAGACCGAGCAGUACUGAUCACU
Sca-2-1     CGACGCGUUCGGCGCGGACGACGACGACGCCGGCUUCGUCGAGACCGAGCAGUACUGAUCCGCGAGCGGUACUGAGACU
Sts-1-1     CCAGCGCCCCUAUCGGGGCACACCCUCCUGCGCCCGUGACGGGCGCAGGAACCCCCGCUUCUCGCGGGGCACACACACU
Ssp-25-1    ACCGCCGCCGCCGACGCCUUCGGCGCCGACGACGACGCGGACGCCGGUUUCGUGGAGACCGAGCAGUACUGAUGUGACU
Ska-1-2     CGCCGCCGCCGACGCCUUCGGCGCGGACGACGACGCCGACGCCGGCUUCGUCGAGACCGAGCAGUACUAAGAAAUGACU
Smo-2-1     UUCGGCCGGGCUGGAUGAAUGAGCCUCGCCGAGACUGGAAGCGUCGGGGUAGCCCGGCCGCUUCCAGUGCGAACGAUCU
Ssp-70-1    GUUUCGUGGAGACCGAGCAGUACUGAGCGUGACCGACUGGGGUCCGGGCGGUGUCCCGGGCCCCAGCCGUGUGAAAUCU
Sba-2-1     NNNNNNNNNNNNNNNNNNNNNNNNNNNNNNNNNNNNNNCGUCGGGCAGGCUGAAUUUGGCUGAUGUCCGGCAGCUAUGACU
Sme-1-1     AUCGCCGGACGGGCUUGAUUUUGGCUGAUGUCCGGCGUGAGCGUCGGGCGGGCUUGAAUGCCGUCCGGCACGUAUGACU
Sgr-11-1    AUCGCCGGACGGGCUUGAUUUCGGCUGAUGUCCGGCGUGAGCGUCGGGCGGGCUUGAAUGCCGUCCGGCACGUAUGACU
Sgl-1-1     UUGAGGGUGCCCCCGUGCCGGGGCUGAGGCUUUGUCCUCAAUCGCCGGACGGGUUUGAUUUUGUCCGGCAGCUAUGACU
Ssp-76-1    GCCGGACGGGCUUGAUUUUGGCUGAUGUCCGGCGUGAGCGUCGGGCGGGCUUGAAUGCGGCCGUCCGGCACGUAUGACU
Sgr-12-7    AAUCGCCGGACGGGCUGAAUUUGGCUGAUGUCCGGCGCGAACGUCGGGCGGGCUUGAACGCCGUCCGGCAGCUAUGACU
Sgl-3-2     CUCGAUCGCCGGACGGGCUUGUUUUCGGCUGAUGUCCGGCGAACGUCGGGCGGGCUGAUUCUGGCUCGCGCCGAAGACU
Sgr-4-1     UCGCGGGUGCCCCGGGCCGGGGCUGAGGCUCUGUCCUCAAACGCCGGGCGGGCUUGAUCGCCGCCCGGCAGCCAUGACU
San-2-1     CCGGACGGGCUUGAUUUCGGCUGAUGCCCGGCGAAAGUCGGGCGGGCUGGAAACAUCGAGCCCGACCGCACGUAUGACU
Sgr-3-1     UCGCGGGUGCCCCGGGCCGGGGCUGAGGCUUUGUCCUCAAACGCCGGGCGGGCUUGACCGCCGCCCGGCAGCCAUGACU
Ssp-9-1     UUGGCUUAUGUCCGGCGAGCGUCGGGCGGGCUUGAGUGUGGCUGAUGCCGGACUCCGAGCCCGAACGGCAGCUAUGACU
Seu-1-1     CCGGACAGGCUUGAUUUUGGCUGAUGUCCGGCGAACGUCGGGCGGGCUUGAAAGCCGUGCCCGACCGGCACGUAUGACU
Ssp-82-1    GGUUUUGUGGGUGCCCCCGGGCCGGGCUGAGGCUUUGUCCUCAAUCGCCGGACGGGCUUGCCGUCCGGCAGCUAUGACU
Sal-12-2    CGGACGGGCUUGAAUUUGGCUGAUGUCCGGCGUGAACGUCGGGCAGGCUGAAUUUGGCUGAUGUCCGGCAGCUAUGACU
Ssp-48-1    UCAAUCGCCGGACGGGCUUGAUUUUGGCUGAUGUCCGGCGAACGUCGGGCGGGCUGAUUCCGGCUCACGUCCGAAGACU
Sha-1-1     GAGGCCGUGUCCUCAGGCGCCGGACGGCCCGGUCUCGUGGCCCCCUGGGCCCGAUGACGGCCGUGCGGCACGCACGACU
Ssp-45-1    AGACCGAGCAGUACUGAGCCGUUCCCCGGCCGGGGCGCCCUGUGAGGGGCGUCCCCGUGCCGGGUGGGGCCGCACGACU
Ssp-71-1    GAGGCCGUGGCCUCAGGCGCCGGACGGACCGGUCUCGUGGCCCCCUGGGCCCGAUGACGGCCGUGCGGCACGCACGACU
Ssp-175-1   GACCGAGCAGUACUGACCGUGUUCCGGCAGGGGCGCCCGUUCUGACGGGUGCCCCCGUGCCGGAUCCGGCAGCAGGACU
Ssp-188-1   CGAGGCCGUGGCCUCAGGCGCCGGACGGACCGGUCUCGUGGCCCCCUGGGCCCGAUGACGGCCGUGCGGCACGCACGACU
Ssp-52-1    UUCGGCGCCGACGACGACGUGGACGCCGGGUUCGUGGAGACCGAGCAGUACUGAGCCGGUCGCACGAUCGCACGAAUCU
Sri-4-1     CGCCUUCGGCGCCGACGACGACGCGGACGCCGGUUUCGUGGAGACCGAGCAGUACUGAGCCCAAGGGCUCGCGCAAUCU
Ssp-145-1   GAGACCGAGCAGUACUGAGCCUGCGGGUUCGUUGCGGCUGAGGCUUUGUCCUCAAUCGCCGGACGGGCUGAUUUGAUCU
env-34      GCUUCGUGGAGACCGAGCAGUACUGAGUUUGUCCGGCGGAGGUCCGGGGUGUCCCCCCGGGCCUCCGCCGUGUGAGUCU
Sly-1-1     GUUUCGUGGAGACCGAGCAGUACUGAGUUUGUCCGGCUGGGGUCCGGGGCGCUGCCCCGGACCCCAGUCGUGUGAAUCU
Ssp-142-1   GGUUCGUGGAGACCGAGCAGUACUGAGUUCGACCGGCAGGGGUCCGGGAGCCUUUCCCGGGCCCUUGCCGUGUGCAUCU
Ssp-155-1   GGGUUCGUGGAGACCGAGCAGUACUGAGUUCGACCGGCAGGGGUCCGGGAGCCUUCCCGGGCCCCUGCCGUGUGCAUCU
Ssp-53-1    CUCGAACGCCGAACGGGCUGAUGCAUGAACCCCCUGCUGGGGGCCCGGGGGUCGUCCCCCGGGCGAAUGCAGUGUGACU
```

```
Sat-2-1     AGACCGAGCAGUACUGAGCCGCAUCCCGGACGGGGGUGCCUGAUCUUCCAGGUGCCCCCGUGCCGGGCCAGUAGGGACU
Ssp-95-1    GAGCAGUACUGAUCCGCUUCCCGGCCAGGGGCGCCCGCUGACGCGGAGCCCCUGGGCCGGUCGCGGACCGUUCGAGACU
Sfl-1-1     GAGCAGUACUGAUCCGCUUCCCGGCCAGGGGCGCCCGCUCACGCGGAGCCCCUGGGCCGGUCGCGGACCGUUCAAGACU
Jpa-1-1     GAGCAGUACUGAUCCGCUUCCCGGCCAGGGGCGCCCGCUGACGCGGAGCCCCUGGGCCGGUCGCGGACCGUUCAAGACU
Sfl-8-1     GAGCAGUACUGAUCCGCUUCCCGGCCAGGGGCGCCCGCUCACGCGGAGCCCCUGGGCCGGUCGCGGACCGUUCAAGACU
Ssp-154-1   GAGCAGUACUGAUCCGCUUCCCGGCCAGGGGCGCCCGCUCACGCGGAGCCCCUGCGCCGGCCGCGGACCGUUCAAGACU
Sce-1-1     GACGCCUUCGGCGCCGACGACGACGCGGACGCCGGGUUCGUGGAGACCGAGCAGUACUGAGCCGCGGGCUCGUGAAUCU
Ssc-5-1     CGCGGGUUUCGUGGAGACCGAGCAGUACUAAGGGUUCCGGGUCCGGGGCUCGGACAGCCCGGACCGGCCUGAUUUCGCU
Ssp-144-1   UCCGGGGCGCCCCCGCCGGGGCCGAGGGCUUCGUCCUCGAUCGCCGGCCGGGCUCGGACGAGCCCGGCCGGCAAGAACU
Ssp-54-1    CGAGGGCUCUGUCCUCAGGCGCCGGACGGGCUUGAUAUUCGGGCUGAUGUCCGAUGACAGCCGUAAGGCUCGUAAGACU
Sgr-13-1    GCGCCCCGGGCCGGGGCUGAGGCUUGGUCCUCAAUCGCCGGACGGGCUUGAAUUCGGCCGCUGUCCGGCACGUACGACU
Sfl-9-1     GCGCCCCGGUCCGGGGCUGAGGCUUUGUCCUCAUUCGCCGGACGGGCUUGAAUUCGGCCGCUGUCCGGCACGUACGACU
Sro-4-1     GUGCCCCGUGCCGGGCUGAGGCUUUGUCCUCAAUCGCCGUUCGGGCUUGAUAUAUCAAGCCCGAACGGCAGCUAUGACU
Sal-12-3    AUCGCCGGACGGGCUUGAUUUUGGCUGAUGUCCGGCGUGGACGUCGGGCGGGCUUGAACAUCGUCCGGCAGCUAUGACU
Ssp-174-1   GCGCCCCGGGCCGGGGCUGAGGCUUUGUCCUCAAUCGCCGGACAGGCUUGAAUUCGGCCGCUGUCCGGCACGUACGACU
Ssp-129-1   GCGCCCCGGGCCGGGGCUGAGGCUUUGUCCUCAAUCGCCGGACAGGCUUGAAUUCGGCCGCUGUCCGGCACGUACGACU
Spu-2-1     GCGCCCCGGGCCGGGGCUGAGGCUUUGUCCUCAAUCGCCGGACAGGCUUGAAUUCGGCCGCUGUCCGGCACGUACGACU
Svi-8-1     GCGCCCCGGGCCGGGGCUGAGGCUUUGUCCUCAUUCGCCGGACAGGCUUGCAUUCGGCCGCUGUCCGGCACGUACGACU
Ssp-66-1    GUGCCCCGUGCCGGGCUGAGGCUUUGUCCUCAAUCGCCGGACGGGCUUGAUUUAUCAAGCCCGAACGGCAGCUAUGACU
Ssp-94-1    GGCGCCCCGGGCCGGGCUGAGGCUUUGUCCUCAAUCGCCGGACAGGCUUGAAUUCGGCCGCUGUCCGGCACGUACGACU
Ssp-17-1    CGAUGCCUUCGGUGCCGACGACGACGCGGACGCCGGUUUCGUGGAGACCGAGCAGUACUGAGCACCACCCCGUACGACU
Ssp-193-1   CGACGCCUUCGGUGCCGACGACGACGCGGACGCCGGUUUCGUGGAGACCGAGCAGUACUGAGCACCACCCCGUACGACU
Sat-1-1     CCUCAACAGCCGGACGGGCUUGAUUUCGGCCGAUGACUGAUCUCAUCGGCCGAGAUGGUGUCCGGAUGGCCCAUGGACU
Ssp-44-1    UUCGAGGGUGCCCCGGGCCGGGGCUGAGGCUUUGUCCUCAAUCGCCGGACGGGCCUGAUGUCGUCCGGCAGCCAUGACU
Scy-1-1     GGUGCCCCGGGCCGGGGCUGAGGCUUUGUCCUCAAUCGCCGGAUGGGCUGAAUGUGGCUGAUGUCCGGCAGCUAUGACU
Sfu-1-1     GGUUUCGAGGGUGCCCCGGGCCGGGGCUGAGGCUUUGUCCCCAAUCGCCGGACGGGCUUGAUGUCCGGCAGCUAUGACU
Sal-7-1     GUGCCCCCGUGCCGGGAUGAGGCUUUGUCCUCAAUCGCCGGACGGGCUUGAAUUUGGCUGAUGUCCGGCAGCUAUGACU
Ssp-168-1   GUGCCCCCGUGCCGGGAUGAGGCUUUGUCCUCAAUCGCCGGACGGGCUUGAAUUCGGCUGAUGUCCGGCAGCUAUGACU
Ssp-97-1    UAUCCGGGCGCCCGGACCGGGGUGAGGCUCCGGCCUCGAUAGCCGGACGGGCUCGAUGAGCCCCGACGGCCCAUGAUCU
Ssp-146-1   UUUCCGGGCGCCCCGACCGGGGUGAGGCUCCGGCCUCGAUCGCCGGACGGGCUCGAUGAGCCCCGACGGCCCAUGAUCU
Sod-1-1     CAGUACUGAGCGGCCCCGGGCCCGGGGCUGGGGCUUUGUCCUCAAUCGCCGGACGGGCUUGAUUUCGUCGCUUGAUUUU
Sri-4-2     GGCCCUGGGGCCUCGUCCUCAAUCUCCCCCAGCUGCCGCUGGGAGGUGCCCCCAGGACGGGCUUGAAAUGAAAGAGACU
Sgr-6-1     CGGCCAGGUUUGAUGGAUGAAGCCGGCCGAAGCUGGAAGCGGCGGGGGCGACCCGGCCGCUUCCAGCGCGAUAAGAACU
Sri-4-5     CCGCCGUGUUCGGGUGCAGGCUUGAUCGGCCCUGGGGCCUCGUCCUCAAACGCCGGACGGGCUUGAAAUGAAAAAGACU
Sri-2-1     CAAACGCCGGACGGGCUGGAUAGGGAUGAGCUGCGCUCAGCCACGACAGCCCAGCGGGCCCCCGGCCCGCGUGCAAUCU
Ssp-90-1    GCCCGCCGUGUUCGGGUGCAGGCUUAUCGGCCUGCGGCCUCGUCCUCAAACGCCGGACGGGCUUGAAAUGAAAAAGACU
Sri-4-18    CGGCCAGGUUUGAUGGAUGAAGCCGGCCGAAGCUGGAAGCGGCGGGGGCGACCCGGCCGCUUCCAGCGCGAAAAGAACU
Ssp-185-1   CGGCCCUGAGGCCUCGUCCUCAAACUCCCCCAGCUACCGCUGGGAGGUGCCCCCAGGACGGGCUUGAAUCGAAAGAUCU
Sri-4-19    CGGCCAGGUUUGAUGGAUGAAGCCGGCCGAAGCUGGAAGCGGCGGGGGCGACCCGGCCGCUUCCAGCGCGAAAAGAACU
Ssp-5-1     UUGAUCUUCGGGCUGAGGCCCGAGAUUUCGGGUGCGGAUGCCGACUCGUGAGUGAGUCGGUCCGGCGACAGCACACACU
Ssp-29-1    UGUGGGCGCCCCGGCCCGGGUGAGGGCUUUUGUCCUCAAUCGCCGGACGGGCUUGAUCGGUCCGGCGCGAUCACACACU
Ssp-26-1    GCCGGACGGGCUUGAUUUUCGCCCGAUGCCGCGUGCGGAUCCGACUCGCGAGAGUCGGUCCGGCGACAGCACACACACU
Aat-1-1     GCAACGACCGCCGCGUCGUUUCUCAGUACCGCGGGGCUGUGGGGGUCGUCCCCCACAGGUGACACGCCACACGCAACAC
Ssp-117-1   GUGUAGUUGCCCCUCGGGGCAGAAUGGAAGCCUCCCAGGGCGGCAGCCCUGGGGACUCCGGGGCUCGGCCCCGGGACAC
Ssp-198-1   GGCGCCGACGACGACGUGGACGCGGGCUUCGUCGAGACCGAGCAGUACUGAGCCCCCCGAGGCGCAGCCUCGGGACCCC
Ssp-115-1   UGGUCGCGCAGUUCCCCGCGCCCCUCUCGGGGCGCAUCCCCGUCGGGCGUUCCGUCCGGCGCGGUUCGUGAGCCCGGU
Psi-1-1     CUCCCGACACGCUGCCGGCCUCGCGGUCGAGGUCCGCUGAGCGAGAGAUCGCUCGGCGGCCAGACCACCCACCCAAUGG
Ssp-4-1     CGCCGGGUUCGUCGAGACCGAGCAGUACUGAGCGACUGCCGACCGGCGGGGGCUCCGCCCGGAGCCCCCGCGAACCCCU
Ssp-201-1   CCUUCGGGCCCGCUCCCUGGCCGGGAAUCCGCCCGGUCCGGGAUUCCCCUCGGCCCUCGCGGACGAGGGGGCACCCCCU
Aal-1-1     CUCCUUCGACGAAGAGGACGAGAGCUUCGCGGAGACCGAGCAGUACUAACCCUCAACACAGUCCGCAACAACCGCCCCA
Sal-13-1    CGACCCGAGCGCGGCGGCCGACGCCUACAGCGACGACGACCAGAGCUACGCCGAGACCGAGCAGUACUAGAAACCCGAG
Ari-1-1     CGACCCGAGCGCGGCGGCCGACAACUACGGCGACGACGACCAGAGCUACGCCGAGACCGAGCAGUACUAGAACCCAGAU
Ssp-40-1    CGCUUCGCGCUCGUCCUCAAGCGCCGGACGGGCUGGAUUCUUUUCCGCCCGUACCGGAUUCCGGACUUCGGGCCGGAUC
Ssp-39-1    .........................GGACGGGCUGGAUUCUUUUCCGCCCGUACCGGAUUCCGGACUUCGGGCCGGAUC
Ssp-42-1    CGCUUCGCGCUCGUCCUCAAGCGCCGGACGGGCUGGAUUCUUUUCCGCCCGUACCGGAUUCCGGACUUCGGGCCGGAUC
Ssp-47-1    CGCUUGGCGCUCGUCCUCAAGCGCCGGACGGGCUGGAUUCUUUUCCGCCCGUACCGGAUUCCGGACUUCGGGCCGGAUC
Ssp-41-1    CGCUUCGCGCUCGUUCUCAAGCGCCGGACGGGCUGGAAUUUUUCCCGCCCGUGCCGGAUUCCGGACUUCGGGCCGGAUC
Ssp-125-1   GCGCUUCGCGCUCGUCCUCAAGCGCCGGACGGGCUGGAUUCUUCCCGCCCGCACCGGAUUCCGGACUUCGGGCCGGAUC
Nsp-3-1     GACCCGCACGCCGCGCUCGCGGCGUACGGCGAUGACGACGACGACGCCUUCGUCGAGGACGAGCAGUACUGAUCCCAAA
Nsp-2-1     ACGACGCGUUCGUCGAGGACGAGCAGUACUGAUCCGCUCCUGAGGCAGGCGCUCGUGCGCCUGCCUCAGACCAGCCAAA
env-35      GACGCGUUCGUCGAGGACGAGCAGUACUGAGCACCACCUGAUCGAUCCCGCCGGGGCCGCAGCCGCGGCCCCGGCCAAA
env-37      GACCCGCAGGCUGCGCUCGCGGCGUACGGCGACGACGACGACGACGCGUUUGUCGAGGACGAGCAGUACUGAUCCCCAA
env-38      GACCCGCAGGCUGCGCUCGCGGCGUACGGCGACGACGACGACGACGCGUUUGUCGAGGACGAGCAGUACUGAUCCCCAA
env-39      GACCCGCAGGCUGCGCUCGCGGCGUACGGCGACGACGACGACGACGCGUUUGUCGAGGACGAGCAGUACUGAUCCCCAA
env-40      ................................................GCGUCGAGGACGAGCAGUACUGAGAACGAAA
Nlu-1-1     UCCGCCGCCCUCGCCGCCUACGGUGACGAUGACGACGACGCGUUCGUCGAGGACGAGCAGUACUGAGUUUCCACCAAAA
env-41      CGUCGAGGACGAGCAGUACUGAGUUGUAGCGCCGGGCCAGGCGGCUUCGCCGCGGCCCGGCGUUAACCCAUAAAGAAAA
Nha-1-1     CGAGCAGCUCUGACUGGAUUGCGCGAGCGUGCCGGCCGAGCGCUGUGGUCGGCACCUCGCACCAGUCACACAAGUGAAA
Sst-1-1     CGGGUGCGGGUGCGCUGUGCUUGAUCGCGCGGUUCCCCGCACCCCUUCCGGGCAGGCCCUUCGGGGCCCUGCCCGGAUC
Sle-1-1     GGGUGCGGGUGCGCUGUGCUCGACCGCGCAGUUCCCCGCGCCCCCUCCGGGAGCGCCCCUCGGGGACGGCUCCCGGAUC
Sal-2-1     GGGUCCGUCGGACAGGCUGAAGUUGGUCUGGGGCGGUUGAGGGGUCCGGGUGGCUCUUUGGAGUCGUUCGGGUCGAGAC
Ssp-24-1    GCGGGCCGGCUGCGGCCGGCCGUGCGGCUCCCCGCGCCCCUUUCGGGGCACCCCGCGCCGCACGGGCGCGGGGCCGACU
Scl-1-1     GCGGGGACGGUGCGCCGCCGUUGUGGCUGAGCGCGCGGGUCCCUGCGCCCCUGGCGGGGCCUUCGGGCCCCGGCACACU
Ssc-1-1     GGACCGACUGCCGGUUCUUGGCUGCCGGCCCGCUGUGGCUGGUCGCGCAGUUCCCCGCGCCCCUAUGGGGCGCGGGGUC
Sde-1-1     GGUUCGUGGAGACCGAGCAGUACUGAGUUUGUCCGGCUGGGGUCCGGGGCCGCGUGCCCCGGGCCUCAGCCGUGAGCCU
            ...............................................................................
            ...............................................................................
            ●●●●●●●●●●●●●●●●●●●●●●●●●●●●●●●●●●●●●●●●●●●●●●●●●●●●●●●●●●●●●●●●●●●●●●●●●●●●●●●
```

alignment positions 80 · · · 259

```
.<<<<<<<<<<<<<<<<.......................>>>>>>>>>>>>>>>>.......<<<..<<<<<<<..........>>>>>>>..>>>.......................................................................................
.2222222221110000.......................0000111222222222.......122...221100..........001122...221.......................................................................................
YC●●●●●R●●-------R●YY--------------------------●●Y●●●●●G●●●----CUGA-C●CCGGUACCUGAYACGGCCGG●G--CAGR●●●●-------------------------------●AGGAGARR●●●●-----AUGCC●●R●CCCRCCAAGGGYGCCCGYCU
 5'                                                                                           3'
```

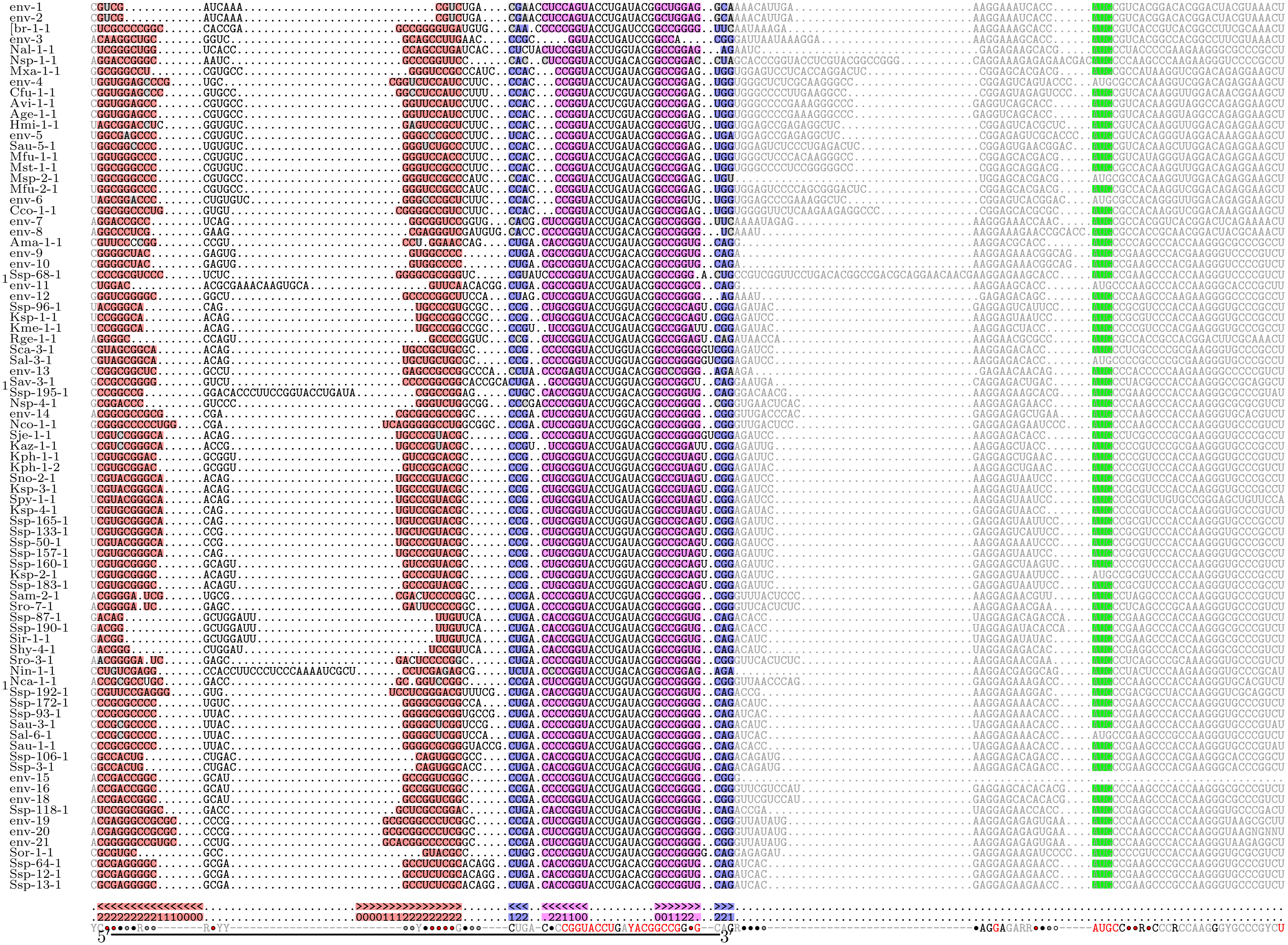

env-1
env-2
[br-1-1
env-3
Nal-1-1
Nsp-1-1
Mxa-1-1
env-4
Cfu-1-1
Avi-1-1
Age-1-1
Hmi-1-1
env-5
Sau-5-1
Mfu-1-1
Mst-1-1
Msp-2-1
Mfu-2-1
env-6
Cco-1-1
env-7
env-8
Ama-1-1
env-9
env-10
Ssp-68-1
env-11
env-12
Ssp-96-1
Ksp-1-1
Kme-1-1
Rge-1-1
Sca-3-1
Sal-3-1
env-13
Sav-3-1
Ssp-195-1
Nsp-4-1
env-14
Nco-1-1
Sje-1-1
Kaz-1-1
Kph-1-1
Kph-1-2
Sno-2-1
Ksp-3-1
Spy-1-1
Ksp-4-1
Ssp-165-1
Ssp-133-1
Ssp-50-1
Ssp-157-1
Ssp-160-1
Ksp-2-1
Ssp-183-1
Sam-2-1
Sro-7-1
Ssp-87-1
Ssp-190-1
Sir-1-1
Shy-4-1
Sro-3-1
Nin-1-1
Nca-1-1
Ssp-192-1
Ssp-172-1
Ssp-93-1
Sau-3-1
Sal-6-1
Sau-1-1
Ssp-106-1
Ssp-3-1
env-15
env-16
env-18
Ssp-118-1
env-19
env-20
env-21
Sor-1-1
Ssp-64-1
Ssp-12-1
Ssp-13-1
5'
3'

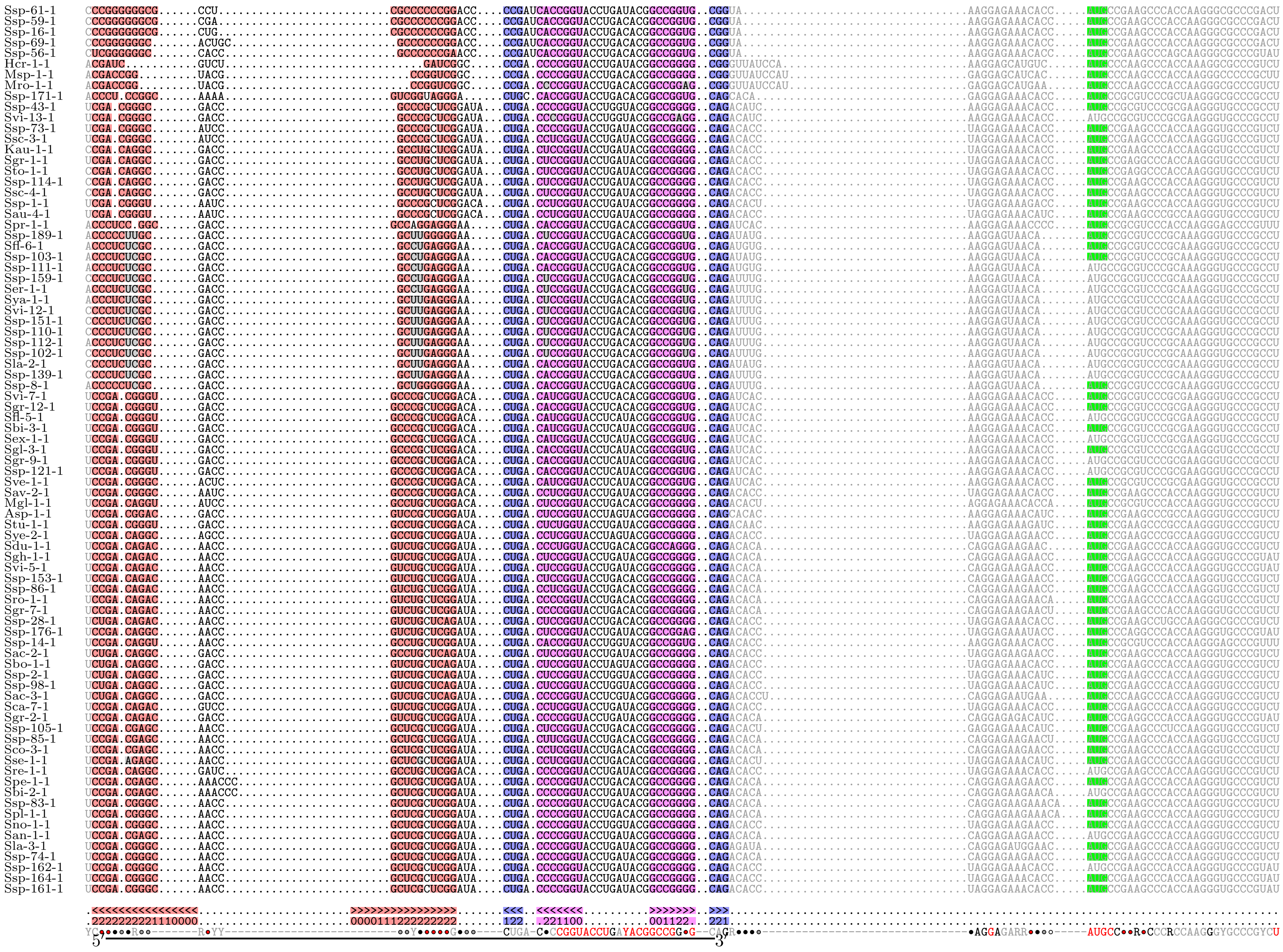

```
Sgl-2-1     UCCGA.CGGGC......AACC.........................GCUCGCUCGGAUA....CUGA.CCCCGGUACCUGAUACGGCCGGGG..CAGACACA...............................CAGGAGAAGAACC.....AUGCCGAAGCCCACCAAGGGUGCCCGUCU
Ssp-143-1   UCCGA.CGGGU......AAACCC.......................GCUCGCUCGGAUA....CUGA.CCCCGGUACCUGAUACGGCCGGGG..CAGACACC...............................UAGGAGAAGUUCC.....AUGCCGAAGCCCACCAAGGGUGCCCGUAU
Ssp-36-1    UCCGA.CAGGC......GACC.........................GCCUGCUCGGAUA....CUGA.CUCCGGUACCUCGUACGGCCGGGG..CAGACACC...............................UAGGAGAAACAUC.....AUGCCGAGGCCCACCAAGGGUGCCCGUCU
Ssp-200-1   UCCGA.CAGGC......GGCC.........................GCCUGCUCGGAUA....CUGA.CUUCGGUACCUGAUACGGCCGGGG..CAGACACC...............................UAGGAGAAACAUC.....AUGCCGAAGCCCAGCAAGGGUGCCCGUCU
Sco-2-1     UCCGA.CAGGC......GGCC.........................GCCUGCUCGGAUA....CUGA.CUUCGGUACCUGAUACGGCCGGGG..CAGACACC...............................UAGGAGAAACAUC.....AUGCCGAAGCCCAGCAAGGGUGCCCGUCU
Svi-9-1     UCCGA.CAGGC......GGCC.........................GCCUGCUCGGAUA....CUGA.CUUCGGUACCUGAUACGGCCGGGG..CAGACACC...............................UAGGAGAAACAUC.....AUGCCGAAGCCCAGCAAGGGUGCCCGUCU
Sco-1-1     UCCGA.CAGGC......GGCC.........................GCCUGCUCGGAUA....CUGA.CUUCGGUACCUGAUACGGCCGGGG..CAGACACC...............................UAGGAGAAACACC.....AUGCCGAAGCCCACCAAGGGUGCCCGUCU
Ssp-128-1   UCCGA.CAGGC......GGCC.........................GCCUGCUCGGAUA....CUGA.CCCCGGUACCUGAUACGGCCGGGG..CAGACACC...............................UAGGAGAAACACC.....AUGCCGAAGCCCACCAAGGGUGCCCGUCU
Sol-2-1     UCCGA.CAGGC......GGCC.........................GCCUGCUCGGAUA....CUGA.CCUCGGUACCUGAUACGGCCGGGG..CAGACACC...............................UAGGAGAAACAUC.....AUGCCGAAGCCCACCAAGGGUGCCCGUCU
Ssp-72-1    UCCGA.CAGGC......GGCC.........................GCCUGCUCGGAUA....CUGA.CCUCGGUACCUGAUACGGCCGGGG..CAGACACC...............................UAGGAGAAACAUC.....AUGCCGAAGCCCACCAAGGGUGCCCGUCU
Ssp-124-1   UCCGA.CAGGC......GGCC.........................GCCUGCUCGGAUA....CUGA.CCUCGGUACCUGAUACGGCCGGGG..CAGACACC...............................UAGGAGAAACACC.....AUGCCGAAGCCCACCAAGGGUGCCCGUCU
Sce-2-1     UCCGA.CGGGC......GACC.........................GCCCGCUCGGAUA....CUGA.CUCCGGUACCUGAUACGGCCGGGG..CAGACCCC...............................UAGGAGAAAGACC.....AUGCCGAAGCCCACCAAGGGUGCCCGUCU
Smo-1-1     UCCGA.CGGGC......GACC.........................GCCCGCUCGGAUA....CUGA.CUCCGGUACCUGAUACGGCCGGGG..CAGACACC...............................UAGGAGAAACACC.....AUGCCGCGUCCCACCAAGGGUGCCCGUCU
Svi-9-2     UCCGA.CGGGC......GACC.........................GCCCGCUCGGAUA....CUGA.CUCCGGUACCUGAUACGGCCGGGG..CAGACACC...............................UAGGAGAAAGACC.....AUGCCGAAGCCCACCAAGGGUGCCCGUCU
Sso-1-1     UCCGA.CGGGC......GACC.........................GCCCGCUCGGAUA....CUGA.CCCCGGUACCUGGUACGGCCGGGG..CAGACACC...............................AAGGAGAAACACC.....AUGCCGCGUCCCGCGAAGGGUGCCCGCCU
Soc-1-1     UCCGA.CGGGC......GACC.........................GCCCGCUCGGAUA....CUGA.CCCCGGUACCUGAUACGGCUGGGG..CAGACACA...............................GAGGAGAAACACC.....AUGCCGCGUCCCGCCAAGGGUGCCCGUCU
Sni-2-1     UCCGA.CGGGC......GACC.........................GCCCGCUCGGAUA....CUGA.CCCCGGUACCUGAUACGGCUGGGG..CAGACACA...............................GAGGAGAAACACC.....AUGCCGCGUCCCGCCAAGGGUGCCCGUCU
Svi-11-1    UCCGA.CGGGC......GACC.........................GCCCGCUCGGAUA....CUGA.CCCCGGUACCUGAUACGGCUGGGG..CAGACACA...............................GAGGAGAAACACC.....AUGCCGCGUCCCGCCAAGGGUGCCCGUCU
Sex-2-1     UCCGA.CGGGU......GACC.........................GCCCGCUCGGAUA....CUGA.CCCCGGUACCUGAUACGGCCGGGG..CAGACACC...............................AAGGAGAAACACC.....AUGCCGCGUCCCGCGAAGGGUGCCCGCCU
Ssp-6-1     UCCGA.CGGGC......GACC.........................GCCCGCUCGGAUA....CUGA.CCCCGGUACCUGAUACGGCCGGGG..CAGACACC...............................UAGGAGAAACACC.....AUGCCGAAGCCCACCAAGGGUGCCCGUCU
Ssp-101-1   UCCGA.CGGGC......GACC.........................GCCCGCUCGGAUA....CUGA.CCCCGGUACCUGAUACGGCCGGGG..CAGACACC...............................UAGGAGAGACACC.....AUGCCGAGGCCCACCAAGGGCGCCCGUAU
Ssp-49-1    UCCGA.CGGGC......GACC.........................GCCCGCUCGGAUA....CUGA.CCCCGGUACCUGAUACGGCCGGGG..CAGACACA...............................CAGGAGAAGAACC.....AUGCCGAAGCCCACCAAGGGUGCCCGUCU
Sal-11-1    UCCGA.CGGGC......GACC.........................GCCUGCUCGGACA....CUGA.CCCCGGUACCUGAUACGGCCGGGG..CAGACAUG...............................GAGGAGCAACACC.....AUGCCGAAGCCCACCAAGGGUGCCCGUUU
Shy-2-1     UCCGA.CAGGC......GACC.........................GCCUGCUCGGAUA....CUGA.CUCCGGUACCUGACACGGCCGGGG..CAGACACA...............................CAGGAGAAAGAAC.....AUGCCGAAGCCCACCAAGGGUGCCCGUCU
Ssp-46-1    UCCGA.CAGGC......GACC.........................GCCUGCUCGGAUA....CUGA.CUCCGGUACCUGAUACGGCCGGGG..CAGACACC...............................UAGGAGAAUCACC.....AUGCCGAAGCCCACCAAGGGUGCCCGUCU
Ssp-187-1   UCCGA.CAGGC......GACC.........................GCCUGCUCGGAUA....CUGA.CUCCGGUACCUGAUACGGCCGGGG..CAGACACC...............................UAGGAGAAUCACC.....AUGCCGAAGCCCACCAAGGGUGCCCGUCU
Ssp-156-1   UCCGA.CAGGC......GACC.........................GCCUGCUCGGAUA....CUGA.CUCCGGUACCUGAUACGGCCGGGG..CAGACACC...............................UAGGAGAAACAUC.....AUGCCGAGGCCCACCAAGGGUGCCCGUCU
Sgr-10-1    UCCGA.CAGGC......GACC.........................GCCUGCUCGGAUA....CUGA.CUCCGGUACCUGAUACGGCCGGGG..CAGACACA...............................CAGGAGAGUCACU.....AUGCCGAGGCCCACCAAGGGUGCCCGUAU
Aac-1-1     UCCGA.CAGGC......GACC.........................GCCUGCUCGGAUA....CUGA.CUCCGGUACCUGAUACGGCCGGGG..CAGACACA...............................CAGGAGAGUCACU.....AUGCCGAGGCCCACCAAGGGUGCCCGUAU
Ssp-197-1   UCCGA.CAGGC......GACC.........................GCCUGCUCGGAUA....CUGA.CUCCGGUACCUGAUACGGCCGGGG..CAGACACA...............................CAGGAGAGUCACU.....AUGCCGAGGCCCACCAAGGGUGCCCGUCU
Ssp-20-1    UCCGA.CAGGC......GACC.........................GCCUGCUCGGAUA....CUGA.CUCCGGUACCUGAUACGGCCGGGG..CAGACACA...............................CAGGAGAGUCACU.....AUGCCGAGGCCCACCAAGGGUGCCCGUCU
Ssp-123-1   UCCGA.CAGGC......GACC.........................GCCUGCUCGGAUA....CUGA.CUCCGGUACCUGAUACGGCCGGGG..CAGACACC...............................UAGGAGAUACACC.....AUGCCGAGGCCCACCAAGGGUGCCCGUCU
Ssp-141-1   UCCGA.CAGGC......GACC.........................GCCUGCUCGGAUA....CUGA.CUCCGGUACCUGAUACGGCCGGGG..CAGACACC...............................UAGGAGAUACACC.....AUGCCGAGGCCCACCAAGGGUGCCCGUCU
Ssp-81-1    UCCGA.CAGGC......GACC.........................GCCUGCUCGGAUA....CUGA.CUCCGGUACCUGAUACGGCCGGGG..CAGACACC...............................UAGGAGAAUCACC.....AUGCCGAAGCCCACCAAGGGUGCCCGUCU
Ssp-113-1   UCCGA.CAGGC......GACC.........................GCCUGCUCGGAUA....CUGA.CUCCGGUACCUGAUACGGCCGGGG..CAGACACC...............................UAGGAGAAUCACC.....AUGCCGAAGCCCACCAAGGGUGCCCGUCU
Ssp-57-1    UCCGA.CAGGC......GACC.........................GCCUGCUCGGAUA....CUGA.CUCCGGUACCUGAUACGGCCGGGG..CAGACACA...............................CAGGAGAGUCAUC.....AUGCCGAGGCCCACCAAGGGUGCCCGUAU
Ssp-149-1   UCCGA.CAGGC......GACC.........................GCCUGCUCGGAUA....CUGA.CUCCGGUACCUGAUACGGCCGGGG..CAGACACA...............................CAGGAGAAGAACC.....AUGCCGAAGCCCACCAAGGGUGCCCGUCU
Sah-1-1     UCCGA.CAGGC......AACC.........................GCCUGCUCGGAUA....CUGA.CUCCGGUACCUGAUACGGCCGGGG..CAGACGUUU..............................GAGGAGAAACACC.....AUGCCGAAGCCCGCCAAGGGUGCCCGUCU
Svi-10-1    UCCGA.CAGGC......AACC.........................GCCUGCUCGGAUA....CUGA.CUCCGGUACCUGAUACGGCCGGGG..CAGACACC...............................UAGGAGAAACACC.....AUGCCGAAGCCCACCAAGGGUGCCCGUUU
Sch-1-1     UCCGA.CAGGC......AAACCC.......................GUCUGCUCGGAUA....CUGA.CCCCGGUACCUGAUACGGCCGGGG..CAGACACC...............................UAGGAGAAACACC.....AUGCCGAAGCCCACCAAGGGUGCCCGUCU
Sia-1-1     UCCGA.CAGGC......AAACCC.......................GCCUGCUCGGAUA....CUGA.CCCCGGUACCUGAUACGGCCGGGG..CAGACACC...............................UAGGAGAAGCACC.....AUGCCGAAGCCCACCAAGGGUGCCCGUAU
Saf-1-1     UCCGA.CAGGC......AAACCC.......................GCCUGCUCGGAUA....CUGA.CCCCGGUACCUGAUACGGCCGGGG..CAGACACC...............................UAGGAGAAACACC.....AUGCCGAAGCCCACCAAGGGUGCCCGUAU
Ssp-89-1    UCCGA.CAGGC......AACC.........................GCCUGCUCGGAUA....CUGA.CCCCGGUACCUGAUACGGCCGGGG..CAGACACC...............................UAGGAGAGACAGC.....AUGCCGAAGCCCACCAAGGGUGCCCGUAU
Sva-1-1     UCCGA.CGGGC......AACC.........................GCCUGCUCGGAUA....CUGA.CCCCGGUACCUGAUACGGCCGGGG..CAGACACC...............................UAGGAGAAACACC.....AUGCCGAAGCCCACCAAGGGUGCCCGUAU
Ssp-158-1   UCCGA.CGGGC......GACC.........................GCCUGCUCGGAUA....CUGA.CCCCGGUACCUGAUACGGCCGGGG..CAGACACC...............................UAGGAGAAACACC.....AUGCCGAGGCCCACCAAGGGUGCCCGUAU
Sro-5-1     UCCGA.CAGGC......AACC.........................GCCUGCUCGGAUA....CUGA.CCUCGGUACCUGAUACGGCCGGGG..CAGACUA................................AAGGAGAAACACC.....AUGCCGCGUCCCACCAAGGGUGCCCGUCU
Ssp-116-1   UCCGA.CAGGC......GACC.........................GCCUGCUCGGAUA....CUGA.CCCCGGUACCUGGUACGGCCGGGG..CAGACACC...............................UAGGAGAAUCACC.....AUGCCGAAGCCCGCCAAGGGUGCCCGUCU
env-22      UCCGA.CAGGC......GCCC.........................GCCUGCUCGGAUA....CUGA.CCUCGGUACCUGAUACGGCCGGGG..CAGACACA...............................CAGGAGAAGAACU.....AUGCCGAAGCCCACCAAGGGUGCCCGUCU
Sam-1-1     UCCGA.CAGGC......GACC.........................GCCUGCUCGGAUA....CUGA.CCUCGGUACCUGAUACGGCCGGGG..CAGACACC...............................UAGGAGAAACACC.....AUGCCGAAGCCCAGCAAGGGUGCCCGUAU
Ssp-51-1    UCCGA.CAGGC......GACC.........................GCCUGCUCGGAUA....CUGA.CCUCGGUACCUGAUACGGCCGGGG..CAGACACA...............................CAGGAGAAGAACA.....AUGCCGAAGCCCACCAAGGGUGCCCGUCU
Slu-1-1     UCCGA.CAGGC......GACC.........................GCCUGCUCGGAUA....CUGA.CCUCGGUACCUGAUACGGCCGGGG..CAGACACC...............................UAGGAGAAUCACC.....AUGCCGAAGCCCACCAAGGGUGCCCGUCU
Sna-2-1     UCCGA.CAGGC......GACC.........................GCCUGCUCGGAUA....CUGA.CCCCGGUACCUGAUACGGCCGGGG..CAGACACC...............................AAGGAGAAACACC.....AUGCCGAAGCCUUCCAAGGGUGCCCGUCU
Ssp-75-1    UCCGA.CAGGC......GACC.........................GCCUGCUCGGAUA....CUGA.CCCCGGUACCUGAUACGGCCGGGG..CAGACACA...............................CAGGAGAGUCACU.....AUGCCGAGGCCCACCAAGGGUGCCCGUCU
env-23      UCCGA.CAGGC......GACC.........................GCCUGCUCGGAUA....CUGA.CCCCGGUACCUGAUACGGCCGGGG..CAGACACC...............................UAGGAGAAACACG.....AUGCCGAAGCCCACCAAGGGUGCCCGUCU
Ssp-182-1   UCCGA.CAGGC......GACC.........................GCCUGCUCGGAUA....CUGA.CCCCGGUACCUGAUACGGCCGGGG..CAGACACA...............................CAGGAGAAGAACC.....AUGCCGAAGCCCACCAAGGGUGCCCGUCU
Scy-2-1     UCCGA.CAGGC......GACC.........................GCCUGCUCGGAUA....CUGA.CCCCGGUACCUGAUACGGCCGGGG..CAGACACC...............................UAGGAGAAACACC.....AUGCCGAAGCCCACCAAGGGUGCCCGUCU
Ssp-184-1   UCCGA.CAGGC......GACC.........................GCCUGCUCGGAUA....CUGA.CCCCGGUACCUGAUACGGCCGGGG..CAGACACA...............................CAGGAGAAGAACC.....AUGCCGAAGCCCACCAAGGGUGCCCGUCU
Svi-4-1     UCUGA.CAGGC......AACC.........................GCCUGCUCAGAUA....CUGA.CCCCGGUACCUGAUACGGCCGGGG..CAGACACA...............................CAGGAGAAGAACA.....AUGCCGAAGCCCACCAAGGGUGCCCGUCU
Ssp-163-1   UCUGA.CAGAC......AACC.........................GUCUGCUCAGAUA....CUGA.CCCCGGUACCUGAUACGGCCGGGG..CAGACACC...............................UAGGAGAAGUUCC.....AUGCCGAAGCCCACCAAGGGUGCCCGUAU
Ssp-177-1   UCUGA.CAGAC......AACC.........................GUCUGCUCAGAUA....CUGA.CCCCGGUACCUGAUACGGCCGGGG..CAGACACC...............................UAGGAGAAACACC.....AUGCCGAAGCCCACCAAGGGUGCCCGUAU
Sol-1-1     UCUGA.CAGAC......AACC.........................GUCUGCUCAGAUA....CUGA.CCCCGGUACCUGAUACGGCCGGGG..CAGACACC...............................UAGGAGAAACACC.....AUGCCGAAGCCCACCAAGGGUGCCCGUAU
Svi-3-1     UCUGA.CAGAC......AACC.........................GUCUGCUCAGAUA....CUGA.CCUCGGUACCUGAUACGGCCGGGG..CAGACACC...............................UAGGAGAAACACC.....AUGCCGAAGCCCACCAAGGGUGCCCGUCU
Szi-1-1     UCCGA.CAGGU......AACC.........................GCCUGCUCGGAUA....CUGA.CCCCGGUACCUCAUACGGCCGGGG..CAGACACA...............................CAGGAGAUUGAC......AUGCCGAAGCCCACCAAGGGUGCCCGUCU
Svi-13-2    UCCGA.CAGGU......AACC.........................GCCUGCUCGGAUA....CUGA.CCUCGGUACCUCAUACGGCCGGGG..CAGACAUA...............................CAGGAGAUUGAC......AUGCCGAAGCCCACCAAGGGUGCCCGUCU
Ssp-120-1   UCCGA.CAGGU......AACC.........................GCCUGCUCGGAUA....CUGA.CCUCGGUACCUCAUACGGCCGGGG..CAGACAUA...............................CAGGAGAUUGAC......AUGCCGAAGCCCACCAAGGGUGCCCGUCU
Ssp-131-1   UCCGA.CAGGU......AACC.........................GCCUGCUCGGAUA....CUGA.CACCGGUACCUCAUACGGCCGGUG..CAGAUCAC...............................AAGGAGAAACACC.....AUGCCGCGUCCCGCGAAGGGUGCCCGCCU
Sca-4-1     UCCG.GCGGGC......AACC.........................GUCCGCUCGGAUA....CUGA.CUCCGGUACCUGAUACGGCCGGGG..CAGACAUC...............................AAGGAGAAACACC.....AUGCCGCGUCCCACCAAGGGUGCCCGUCU
Sba-1-1     CCCGUGCGGAC......GACCU........................GUCCGUACGGACCC...CUGA.CACCGGUACCUGACACGGCCGGUG..CAGUAACC...............................AAGGAGAAGCACC.....AUGCCGAAGCCCACCAAGGGUGCCCGUCU
Ssp-65-1    CCCGUGCGGACG.....GCC.........................CGUCCGUACGGACCC...CUGA.CACCGGUACCUGACACGGCCGGUG..CAGUAACC...............................AAGGAGAAGCACC.....AUGCCGAAGCCCACCAAGGGUGCCCGUCU
Sme-2-1     UCCG.GCGGAC......ACAA.........................GUCCGUACGGACA....CUGA.CAUCGGUACCUGAUACGGCCGGUG..CAGACACGA..............................GAGGAACGGAUCA.....AUGCCGAAGCCCCCCAAGGGCGCCCGCCU
Sgr-12-3    UCCG.GCGGAC......ACAA.........................GUUCGUGCGGACA....CUGU.CAUCGGUACCUGAUACGGCCGGUG..CAGAUCAC...............................AAGGAGAAAGACC.....AUGCCGAAGCCCACCAAGGGUGCCCGUCU
Sgr-12-4    UCCG.GCGGAC......ACAA.........................GUUCGUGCGGACA....CUGU.CAUCGGUACCUGAUACGGCCGGUG..CAGAUCAC...............................AAGGAGAAAGACC.....AUGCCGAAGCCCACCAAGGGUGCCCGUCU
Ssp-15-1    UCCG.GCGGAC......ACAA.........................GUUCGUGCGGACA....CUGU.CAUCGGUACCUGAUACGGCCGGUG..CAGAUCAC...............................AAGGAGAAAGACC.....AUGCCGAAGCCCACCAAGGGUGCCCGUCU
Ssp-38-1    UCCG.GCGGAC......ACAA.........................GUUCGUGCGGACA....CUGU.CAUCGGUACCUGAUACGGCCGGUG..CAGAUCAC...............................AAGGAGAAAGACC.....AUGCCGAAGCCCACCAAGGGUGCCCGUCU
Sal-5-1     UCCG.GCGGAC......ACAA.........................GUUCGUGCGGACA....CUGU.CAUCGGUACCUGAUACGGCCGGUG..CAGAUCAC...............................AAGGAGAAAGACC.....AUGCCGAAGCCCACCAAGGGUGCCCGUCU
Ssp-77-1    UCCG.GCGGAC......ACAA.........................GUUCGUGCGGACA....CUGU.CAUCGGUACCUGAUACGGCCGGUG..CAGAUCAC...............................AAGGAGAAAGACC.....AUGCCGAAGCCCACCAAGGGUGCCCGUCU
Swa-1-1     UCCG.GCGGAC......ACAA.........................GUUCGUGCGGACA....CUGU.CAUCGGUACCUGAUACGGCCGGUG..CAGAUCAC...............................AAGGAGAAAGACC.....AUGCCGAAGCCCACCAAGGGUGCCCGUCU
Ssp-30-1    UCCG.GCGGAC......ACAA.........................GUUCGUGCGGACA....CUGU.CAUCGGUACCUGAUACGGCCGGUG..CAGAUCAC...............................AAGGAGAAAGACC.....AUGCCGAAGCCCACCAAGGGUGCCCGUCU
Ssp-134-1   UCCG.GCGGAC......ACAA.........................GUUCGUGCGGACA....CUGU.CAUCGGUACCUGAUACGGCCGGUG..CAGAUCAC...............................AAGGAGAAAGACC.....AUGCCGAAGCCCACCAAGGGUGCCCGUCU
Ssp-62-1    UCCG.GCGGAC......ACAA.........................GUUCGUGCGGACA....CUGU.CAUCGGUACCUGGUACGGCCGGUG..CAGAUCAC...............................AAGGAGAAAGACC.....AUGCCGAAGCCCACCAAGGGUGCCCGUCU
Ssp-21-1    UCCG.GCGGAC......ACAA.........................GUUCGUGCGGACA....CUGU.CAUCGGUACCUGGUACGGCCGGUG..CAGAUCAC...............................AAGGAGAAAGACC.....AUGCCGAAGCCCACCAAGGGUGCCCGUCU
Ssp-80-1    UCCG.GCGGAC......ACAA.........................GUUCGUGCGGACA....CUGU.CAUCGGUACCUGGUACGGCCGGUG..CAGAUCAC...............................AAGGAGAAAGACC.....AUGCCGAAGCCCACCAAGGGUGCCCGUCU
Sgr-12-5    UCCG.GCGGAC......ACAA.........................GUUCGUGCGGACA....CUGU.CAUCGGUACCUGGUACGGCCGGUG..CAGAUCAC...............................AAGGAGAAAGACC.....AUGCCGAAGCCCACCAAGGGUGCCCGUCU

            .<<<<<<<<<<<<<<<<.......................>>>>>>>>>>>>>>>>.......<<<..<<<<<<<..........>>>>>>>..>>>...................................................................................
            .2222222221110000.......................0000111222222222.......122...221100..........001122...221...................................................................................
            YC●●●○●R○○-------R●YY--------------------------○○Y●●●●●G●○○----CUGA-C●CCGGUACCUGAYACGGCCGG●G--CAGR●●●○-------------------------------●AGGAGARR●●○○-----AUGCC●●R●CCCRCCAAGGGYGCCCGYCU
             5′————————————————————————————————————————————————————————————————————————————————3′
```

```
Sva-2-1    UCCG.GCGGGC......AAAC.........................GCUCGUACGGACA....CUGA.CACCGGUACCUGAUACGGCCGGUG..CAGAAAUG...............................AAGGAGAAAGACC.....AUGCCGAAGCCCGCCAAGGGUGCCCGUCU
Ssp-58-1   CCCGUACGGGC......AGCA.........................GCCCGUGCGGACC....CUGA.CACCGGUACCUGAUACGGCCGGUG..CAGAGACCC..............................AAGGAGAAAGAC......AUGCCGAAGCCCACCAAGGGUGCCCGUCU
Sal-12-1   UCCG.GCGGGC......AUCA.........................GCCCGUACGGACA....CUGA.CACCGGUACCUGAUACGGCCGGUG..CAGAGACU...............................GAGGAGAAACAC......AUGCCGAAGCCCGCCAAGGGUGCCCGUCU
Sfr-1-1    UCCG.GCGGGC......ACGA.........................GCCCGUACGGACA....CUGA.CACCGGUACCUGAUACGGCCGGUG..CAGAGACU...............................GAGGAGACACAC......AUGCCGAAGCCCGCCAAGGGUGCCCGUCU
Sxi-2-1    UCCG.GCGGGC......ACGA.........................GCCCGUACGGACA....CUGA.CACCGGUACCUGAUACGGCCGGUG..CAGAGACU...............................GAGGAGACACAC......AUGCCGAAGCCCGCCAAGGGUGCCCGUCU
Ssp-119-1  UCCGG.CGGGU......AAUC.........................GCCCGUCCGGGAA....CUGA.CACCGGUACCUGAUACGGCCGGUG..CAGAUACG...............................AAGGAGUCACACCC....AUGCCGAAGCCCACCAAGGGUGCCCGUCU
Ssp-33-1   GCCGGUGAGC.......GCUC..........................GCUCAUCGGGUC....CUGA.CACCGGUACCUGAUACGGCCGGUG..CAGCAAUG...............................AAGGAGAUACACC.....AUGCCGAAGCCCACCAAGGGCGCCCGUCU
Ssp-37-1   GCCGGUGAGC.......GUCC..........................GCUCACCGGGUC....CUGA.CCCCGGUACCUCAUACGGCCGGGG..CAGAUACC...............................AAGGAGAAAGAA......AUGCCGAAGCCCACCAAGGGUGCCCGUCU
Ssp-170-1  GCCGGUGAGC.......AUCC..........................GCUCAUCGGGUUC...CUGA.CCCCGGUACCUGACACGGCCGGGG..CAGACACA...............................CAGGAGAAGAACA.....AUGCCGAAGCCCACCAAGGGUGCCCGUCU
Smi-1-1    GCCGGUGAGC.......ACCC..........................GCUCAUCGGGUUC...CUGA.CCCCGGUACCUGAUACGGCCGGGG..CAGACACA...............................CAGGAGAUAGAAC.....AUGCCGAAGCCCACCAAGGGUGCCCGUCU
Ssp-138-1  GCCGGUGAGC.......AUCC..........................GCUCAUCGGGUUC...CUGA.CCCCGGUACCUGAUACGGCCGGGG..CAGACACA...............................CAGGAGAAAGAAC.....AUGCCGAAGCCCACCAAGGGUGCCCGUCU
Sga-2-1    GCCGGUGAGC.......AUCC..........................GCUCAUCGGGUUC...CUGA.CCCCGGUACCUGAUACGGCCGGGG..CAGACACA...............................CAGGAGAAGAACA.....AUGCCGAAGCCCACCAAGGGUGCCCGUCU
env-24     GCCGGUGAGC.......AUCC..........................GCUCAUCGGGUUC...CUGA.CCCCGGUACCUGAUACGGCCGGGG..CAGACACA...............................CAGGAGAAGAACC.....AUGCCGAAGCCCACCAAGGGUGCCCGUCU
Sca-1-1    GCCGGUGAGC.......AUCC..........................GCUCAUCGGGUUC...CUGA.CCCCGGUACCUGAUACGGCCGGGG..CAGACACA...............................CAGGAGAUAGAAA.....AUGCCGAAGCCCACCAAGGGUGCCCGUAU
Ssp-23-1   GCCGGUGAGC.......AUCC..........................GCUCAUCGGGUUC...CUGA.CCCCGGUACCUGAUACGGCCGGGG..CAGACACA...............................CAGGAGAAAGAA......AUGCCGAAGCCCACCAAGGGUGCCCGUCU
Ssp-135-1  GCCGGUGAGC.......AUCC..........................GCUCAUCGGGUUC...CUGA.CCCCGGUACCUGAUACGGCCGGGG..CAGACACA...............................CAGGAGAAAGAAC.....AUGCCGAAGCCCACCAAGGGUGCCCGUCU
Ssp-169-1  GCCGGUGAGC.......AUCC..........................GCUCAUCGGGUUC...CUGA.CCCCGGUACCUGAUACGGCCGGGG..CAGACACA...............................CAGGAGAAGAACA.....AUGCCGAAGCCCACCAAGGGUGCCCGUCU
env-30     GCCGGUGAGC.......AUCC..........................GCUCAUCGGGUUC...CUGA.CCCCGGUACCUGAUACGGCCGGGG..CAGACACA...............................CAGGAGAAGAACC.....AUGCCGAAGCCCACCAAGGGUGCCCGUCU
Ssv-1-1    GCCGGUGAGC.......AUCC..........................GCUCAUCGGGUUC...CUGA.CCCCGGUACCUGAUACGGCCGGGG..CAGACACA...............................CAGGAGAUAGAAA.....AUGCCGAAGCCCACCAAGGGUGCCCGUAU
Ssp-186-1  GCCGGUGAGC.......AUCC..........................GCUCAUCGGGUUC...CUGA.CCCCGGUACCUGAUACGGCCGGGG..CAGACACA...............................CAGGAGAUAGAAA.....AUGCCGAAGCCCACCAAGGGUGCCCGUAU
Sda-1-1    GCCGGUGAGC.......AUCC..........................GCUCAUCGGGUUC...CUGA.CCCCGGUACCUGAUACGGCCGGGG..CAGACACA...............................CAGGAGAUUGAA.....AAUGCCGAAGCCCACCAAGGGUGCCCGUC
Sgr-12-6   GCCGGUGAGC.......AUCC..........................GCUCACCGGGUUC...CUGA.CCCCGGUACCUGAUACGGCCGGGG..CAGAUACC...............................AAGGAGAAAGAA......AUGCCGAAGCCCGCCAAGGGUGCCCGUCU
Ssp-199-1  GCCGGUGAGC.......AUCC..........................GCUCACCGGGUUC...CUGA.CCCCGGUACCUGAUACGGCCGGGG..CAGACC.................................AAGGAGAAAGAA......AUGCCGAAGCCCACCAAGGGUGCCCGUCU
env-31     GCCGGUGAGC.......AACC..........................GCUCGCCGGGUC....CUGA.CCUCGGUACCUGAUACGGCCGGGG..CAGAUACC...............................AAGGAGAAAGAAC.....AUGCCGAAGCCCGCCAAGGGUGCCCGUCU
Ssp-19-1   GCCGGUGAGC.......AACC..........................GCUCACCGGGUC....CUGA.CCUCGGUACCUGAUACGGCCGGGG..CAGAUACC...............................AAGGAGAAAGAAA.....AUGCCGAAGCCCGCCAAGGGUGCCCGUCU
Sfu-2-1    GCCGGUGAGC.......AUCC..........................GCUCACCGGGUC....CUGA.CCCCGGUACCUGAUACGGCCGGGG..CAGAUGCC...............................AAGGAGAAAGAA......AUGCCGAAGCCCACCAAGGGUGCCCGUCU
Sau-2-1    GCCGGUGAGC.......AACC..........................GCUCACCGGGUC....CUGA.CCCCGGUACCUGAUACGGCCGGGG..CAGAUACC...............................AAGGAGAAAGAA......AUGCCGAAGCCCGCCAAGGGUGCCCGUCU
Sxy-1-1    GCCGGUGAGC.......AUCC..........................GCUCGCCGGGUC....CUGA.CCCCGGUACCUGAUACGGCCGGGG..CAGAUACC...............................AAGGAGAAAGAA......AUGCCGAAGCCCUCCAAGGGUGCCCGUCU
Ssp-122-1  GCCGGUGAGC.......GUCC..........................GCUCGCCGGGUC....CUGA.CCCCGGUACCUGAUACGGCCGGGG..CAGAUACC...............................AAGGAGAAAGAA......AUGCCGAAGCCCACCAAGGGUGCCCGUCU
Spr-2-1    GCCGGUGAGC.......GUCC..........................GCUCGCCGGGUC....CUGA.CCCCGGUACCUGAUACGGCCGGGG..CAGAUACC...............................AAGGAGAAAGAA......AUGCCGAAGCCCACCAAGGGUGCCCGUCU
Ssp-84-1   GCCGGUGGGC.......AUCC..........................GCUCACCGGGUUC...CUGA.CCCCGGUACCUGACACGGCCGGGG..CAGACACA...............................CAGGAGAAGAACC.....AUGCCGAAGCCCACCAAGGGUGCCCGUCU
Sip-1-1    GCCGGUGAGC.......AUCC..........................GCUCAUCGGGCC....CUGA.CCUCGGUACCUCAUACGGCCGGGG..CAGAUACC...............................AAGGAGAAAGAAA.....AUGCCGAAGCCCACCAAGGGUGCCCGUCU
Sly-2-1    GCCGGUGAGC.......GACC..........................GCUCAUCGGGUC....CUGA.CCCCGGUACCUAGUACGGCCGGGG..CAGAUACC...............................AAGGAGAAACACC.....AUGCCGAAGCCCUCCAAGGGUGCCCGUAU
Ssc-2-1    CACCCGGGCC.......GACC..........................GGCCCGGGUGGA....CUGA.CGCCGGUACCUGACACGGCCGGCG..CAGGAAUAUC.............................UAGGAGAGAAC.......AUGCCCAGGCCCACCAAGGGUCCGCGCCU
Sye-1-1    ACGGGUCGGC.......GGCC..........................GCCGACCCGCCA....CUGA.CACCGGUACCUGACACGGCCGGUG..CAGCGAAC...............................AAGGAGAAUGAC......AUGCCCCAGCCCGCGAAGGGUGCCCGCCU
Svi-6-1    UCGGCGGGGC.......AACCCC........................GCCCCGCCGAA.....CUGA.CACCGGUACCUGACACGGCCGGUG..CAGACCUG...............................GAGGAGAGACACAACC..AUGCCGAAGCCCACCAAGGGUCCCCGUCU
Sni-1-1    UCCGCGGAC........GGCC...........................GUUCGCGGGGAA...CUGA.CACCGGUACCUGAUACGGCCGGUG..CAGAUAUG...............................AAGGAGUAACACC.....AUGCCGAAGCCCGCCAAGGGUGCCCGUCU
Ssu-2-1    ACCGGCGGGGC......GCCC.........................GCCUCGUCGGC......CUGA.CACCGGUACCUGACACGGCCGGUG..CAGAGUUC...............................AAGGAGAGAGAAG.....AUGCCGAAGCCCACCAAGGGUCCCCGUCU
Ssu-1-1    UCCGGCGGGGC......GCCC.........................GCCUCGUCGGC......CUGA.CACCGGUACCUGACACGGCCGGUG..CAGAGUUC...............................AAGGAGAGAGAAG.....AUGCCGAAGCCCACCAAGGGUCCCCGUCU
Svi-2-1    UCCGGCGGGC.......GGCC..........................GCUCGCCGGUAA....CUGA.CACUGGUACCUGAUACGGCCGGUG..CAGACACC...............................UAGGAGAUACACCA....AUGCCGAAGCCCGCCAAGGGCGCCCGUCU
Sfl-4-1    UCCGGCGGGC.......AGCA..........................GCUCGCCGGUAA....CUGA.CACCGGUACCUGAUACGGCCGGUG..CAGACAC................................AAGGAGAAACACC.....AUGCCGCGUCCCGCAAAGGGUGCCCGUCU
Spu-1-1    UCCGGCGGGC.......GGUC..........................GCCCGUCGGUAA....CUGA.CACCGGUACCUGACACGGCCGGUG..CAGAGAC................................GAGGAGAAACACC.....AUGCCGCGUCCCACCAAGGGAGCCCGUCU
Ssc-4-2    UCCGGCGGGC.......GUCC..........................GCUCGUCGGUAA....CUGA.CACCGGUACCUGACACGGCCGGUG..CAGACAG................................AAGGAGAAACACC.....AUGCCGCGUCCCGCCAAGGGAGCCCGUCU
Shy-1-1    UCCGGCGGGC.......ACAC..........................GCUCGUCGGUAA....CUGA.CACCGGUACCUGAUACGGCCGGUG..CAGACAUC...............................UAGGAGAUACACC.....AUGCCGAAGCCCACCAAGGGCGCCCGUCU
env-32     UCCGGCGGGC.......ACAC..........................GCUCGUCGGUAA....CUGA.CACCGGUACCUGAUACGGCCGGUG..CAGACAUC...............................UAGGAGAUACACC.....AUGCCGAAGCCCGCCAAGGGCGCCCGUCU
Sbi-1-1    UCCGGCGGGC.......ACUC..........................GCCCGUCGGUAA....CUGA.CACCGGUACCUCAUACGGCCGGUG..CAGACCA................................AAGGAGAAACA.......AUGCCGAGGCCCACCAAGGGUGCCCGUCU
Afl-1-1    CCCCGGGGCC.......GCCC..........................GGCCCCGGGGCAAG..CUGA.CGUCGGUACCUGAUACGGCCGGCG..CAGG...................................AAGGACGAGAC.......AUGCCUCAGCCCACCAAGGGUGCCCGCCU
Ssp-55-1   UCCCGGGGC........GGCC...........................GCUCCGGGGGAA...CUGA.CAUCGGUACCUGAUACGGCCGGUG..CAGAUAUG...............................AAGGAGUAACACC.....AUGCCUCAGCCCGCCAAGGGUGCCCGUCU
Ssp-152-1  UCCCGCGGC........AGCC...........................GCCGCGGGGGAA...CUGA.CACCGGUACCUGACACGGCCGGUG..CAGAUAUG...............................AAGGAGUAACACC.....AUGCCGCGUCCCGCAAAGGGUGCCCGUCU
Ssp-104-1  UCCCGCGGC........AGCC...........................GCCGCGGGGGAA...CUGA.CACCGGUACCUGACACGGCCGGUG..CAGAAAUG...............................AAGGAGUAACACC.....AUGCCGCGUCCCGCAAAGGGUGCCCGUCU
Ssp-166-1  UCCCGCGGC........AUCC...........................GCCGCGGGGGAA...CUGA.CACCGGUACCUGACACGGCCGGUG..CAGAUAUG...............................AAGGAGUAACACC.....AUGCCGCGUCCCGCAAAGGGUGCCCGUCU
Ssp-191-1  UCCCGGGGC........GUCC...........................GCCCCGGGGGAA...CUGA.CACCGGUACCUGACACGGCCGGUG..CAGAUAUG...............................AAGGAGUAACACC.....AUGCCGCGUCCCGCUAAGGGUGCCCGUCU
Sxa-1-1    UCCCGCGGC........AUCC...........................GCCCCGGGGGAA...CUGA.CACCGGUACCUGACACGGCCGGUG..CAGAUGUG...............................AAGGAGUAACACC.....AUGCCGCGUCCCGCAAAGGGUGCCCGUCU
Sla-1-1    UCCCGCGGC........AUCC...........................GCCUCGGGGGAA...CUGA.CACCGGUACCUGACACGGCCGGUG..CAGAUAUG...............................AAGGAGUAACACC.....AUGCCGCGUCCCGCAAAGGGUGCCCGCCU
Svi-12-3   UCCCGCGGC........AUCC...........................GCCUCGGGGGAA...CUGA.CACCGGUACCUGACACGGCCGGUG..CAGAUAUG...............................AAGGAGUAACACC.....AUGCCGCGUCCCGCAAAGGGUGCCCGUCU
Sla-2-2    UCCCGCGGC........AUCC...........................GCCUCGGGGGAA...CUGA.CACCGGUACCUGACACGGCCGGUG..CAGAUAUG...............................AAGGAGUAACACC.....AUGCCGCGUCCCGCAAAGGGUGCCCGCCU
Ssp-150-1  UCCCGCGGC........AUCC...........................GCCUCGGGGGAA...CUGA.CACCGGUACCUGACACGGCCGGUG..CAGAUAUG...............................AAGGAGUAACACC.....AUGCCGCGUCCCGCAAAGGGUGCCCGUCU
Ssp-32-1   UCCCGCGGC........GUCC...........................GCCCCGGGGGAA...CUGA.CACCGGUACCUGACACGGCCGGUG..CAGAUAUG...............................AAGGAGUAUCACC.....AUGCCGCGUCCCGCAAAGGGUGCCCGUCU
Ska-1-1    UCCCGGGGC........AGCC...........................GCCCCGGGGGAA...CUGA.CACCGGUACCUGACACGGCCGGUG..CAGAUAUG...............................AAGGAGUAACACC.....AUGCCGCGUCCCGCAAAGGGUGCCCGUCU
Sro-6-1    UCCCGGGGC........AGCC...........................GCCCCGGGGGAA...CUGA.CACCGGUACCUGACACGGCCGGUG..CAGAUAUG...............................AAGGAGUAACACC.....AUGCCGCGUCCCGCAAAGGGUGCCCGUCU
Ssp-194-1  CCGUCCGGGC.......GUCC..........................GCCCGGACGGC.....CUGA.CACCGGUACCUGAUACGGCCGGUG..CAGAGAAA...............................GAGGAGAAGCAAC.....AUGCCGAAGCCCACCAAGGGCGCCCGUAU
env-33     GCCCGCAGGC.......GUCC..........................GCCUGCGGGGCC....CUGA.CCCCGGUACCUGACACGGCCGGGG..CAGAUCAC...............................AAGGAGAAACACC.....AUGCCGAAGCCCGCCAAGGGUGCCCGUCU
Ssp-7-1    GCCCCUGGGC.......GUCC..........................GCCCGGGGGGCC....CUGA.CACCGGUACCUGACACGGCCGGUG..CAGAUCA................................GAGGAGAAACACC.....AUGCCGAAGCCCGCCAAGGGUGCCCGUCU
Ssp-140-1  GCCCGCAGGC.......GUCC..........................GCCUGCGGGGUC....CUGA.CACCGGUACCUGAUACGGCCGGUG..CAGAUUUC...............................AAGGAGAAACACC.....AUGCCGAAGCCCGCCAAGGGUGCCCGUCU
Svi-1-1    GCCCGCGGGC.......GUCC..........................GCCCGCGGGGCC....CUGA.CACCGGUACCUGAUACGGCCGGUG..CAGAUCAC...............................AAGGAGAAACACC.....AUGCCGAAGCCCGCCAAGGGUGCCCGUCU
Sca-2-1    GCCCGCAGGC.......GUCC..........................GCCCGCGGGGCC....CUGA.CACCGGUACCUGAUACGGCCGGUG..CAGAUAG................................AAGGAGAAACCC......AUGCCGAAGCCCACCAAGGGCGCCCGUCU
Sts-1-1    GCCCGCAGGC.......GGCC..........................GCCUGUGGGGUC....CUGA.CAUCGGUACCUGAUACGGCCGGUG..CAGAUUG................................AAGGAGAAACACC.....AUGCCGAAGCCCGCCAAGGGCGCCCGUCU
Ssp-25-1   GCCCGCAGGC.......AAUCC.........................GCCUGCGGGGCC....CUGA.CCCCGGUACCUGAUACGGCCGGGG..CAGACACC...............................UAGGAGAAACACC.....AUGCCGAAGCCCACCAAGGGUGCCCGUCU
Ska-1-2    UCCCCGGGGC.......AUCC..........................GCCCCGGGGGAA....CUGA.CACCGGUACCUGACACGGCCGGUG..CAGAUAUG...............................AAGGAGUAACA.......AUGCCGCGUCCCGCAAAGGGUGCCCGCCU
Smo-2-1    UCUACGUGGC.......GGCC..........................GCCACGUGGGAA....CUGA.CACCGGUACCUGAUACGGCCGGUG..CAGCAAUG...............................AAGGAGAAAUACC.....AUGCCGAAGCCCGCCAAGGGUGCCCGUCU
Ssp-70-1   UCCCCGGGGC.......GGCC..........................GCCUCGGGGGAA....CUGA.CACCGGUACCUGAUACGGCCGGUG..CAGCAAUG...............................AAGGAGAUACACC.....AUGCCGAAGCCCGCCAAGGGCGCCCGUCU
Sba-2-1    UCCGUGAGGC.......GUCC..........................GCCUUGCGGGAA....CUGA.CAUCGGUACCUGAUACGGCCGGUG..CAGCACAC...............................AAGGAGAAUCACC.....AUGCCGCGUCCCGCCAAGGGUGCCCGUCU
Sme-1-1    UCCGUGAGGC.......GACC..........................GCCUUGCGGGAA....CUGA.CACCGGUACCUGAUACGGCCGGUG..CAGCACAC...............................AAGGAGAAUCACC.....AUGCCGCGUCCCGCCAAGGGUGCCCGUCU
Sgr-11-1   UCCGUGAGGC.......GACC..........................GCCUUGCGGGAA....CUGA.CACCGGUACCUGAUACGGCCGGUG..CAGCACAC...............................AAGGAGAAUCACC.....AUGCCGCGUCCCGCCAAGGGUGCCCGUCU
Sgl-1-1    UCCGUGAGGC.......GACC..........................GCCUCGCGGGAA....CUGA.CACCGGUACCUGAUACGGCCGGUG..CAGCACAC...............................AAGGAGAAUCACC.....AUGCCGCGUCCCGCCAAGGGUGCCCGUCU
Ssp-76-1   UCCGUGAGGC.......GACC..........................GCCUCGCGGGAA....CUGA.CACCGGUACCUGAUACGGCCGGUG..CAGCACAC...............................AAGGAGAAUCACC.....AUGCCGCGUCCCGCCAAGGGUGCCCGUCU
Sgr-12-7   UCCGUGAGGC.......GACC..........................GCCUCGCGGGAA....CUGA.CACCGGUACCUGAUACGGCCGGUG..CAGCACAC...............................AAGGAGAAUCACC.....AUGCCGCGUCCCGCCAAGGGUGCCCGUCU
Sgl-3-2    UCCGUGAGGC.......GACC..........................GCCUCGCGGGAA....CUGA.CACCGGUACCUGAUACGGCCGGUG..CAGCACAC...............................AAGGAGAAUCACC.....AUGCCGCGUCCCGCCAAGGGUGCCCGUCU
Sgr-4-1    UCCGGGAGGC.......GUCC..........................GCCUCGCGGGAA....CUGA.CAUCGGUACCUGAUACGGCCGGUG..CAGCACAC...............................AAGGAGAAUCACC.....AUGCCGCGUCCCGCAAAGGGUGCCCGUCU
San-2-1    UCCGGGAGGC.......GUCC..........................GCCUCGCGGGAA....CUGA.CAUCGGUACCUGAUACGGCCGGUG..CAGCACAC...............................AAGGAGAAUCACC.....AUGCCGCGUCCCGCAAAGGGUGCCCGUCU
Sgr-3-1    UCCGGGAGGC.......GUCC..........................GCCUCGCGGGAA....CUGA.CAUCGGUACCUGAUACGGCCGGUG..CAGCACAC...............................AAGGAGAAUCACC.....AUGCCGCGUCCCGCAAAGGGUGCCCGUCU
Ssp-9-1    UCCGGGAGGC.......GUCC..........................GCCUCGCGGGAA....CUGA.CACCGGUACCUGAUACGGCCGGUG..CAGCACAC...............................AAGGAGAAUCACC.....AUGCCGCGUCCCGCAAAGGGUGCCCGUCU
Seu-1-1    UCCGGGAGGC.......GUCC..........................GCCUCGCGGGAA....CUGA.CACCGGUACCUGAUACGGCCGGUG..CAGCACAC...............................AAGGAGAAUCACC.....AUGCCGCGUCCCGCAAAGGGUGCCCGUCU

           .<<<<<<<<<<<<<<<<.......................>>>>>>>>>>>>>>>>.......<<<..<<<<<<<..........>>>>>>>..>>>...................................................................................
           .2222222221110000.......................0000111222222222.......122...221100..........001122...221...................................................................................
           YC●●●○●R○○-------R●YY--------------------------○○Y●●●●●G●○○----CUGA-C●CCGGUACCUGAYACGGCCGG●G--CAGR●●●○-------------------------------●AGGAGARR●●○○-----AUGCC●●R●CCCRCCAAGGGYGCCCGYCU
            5′                                                                                  3′
```

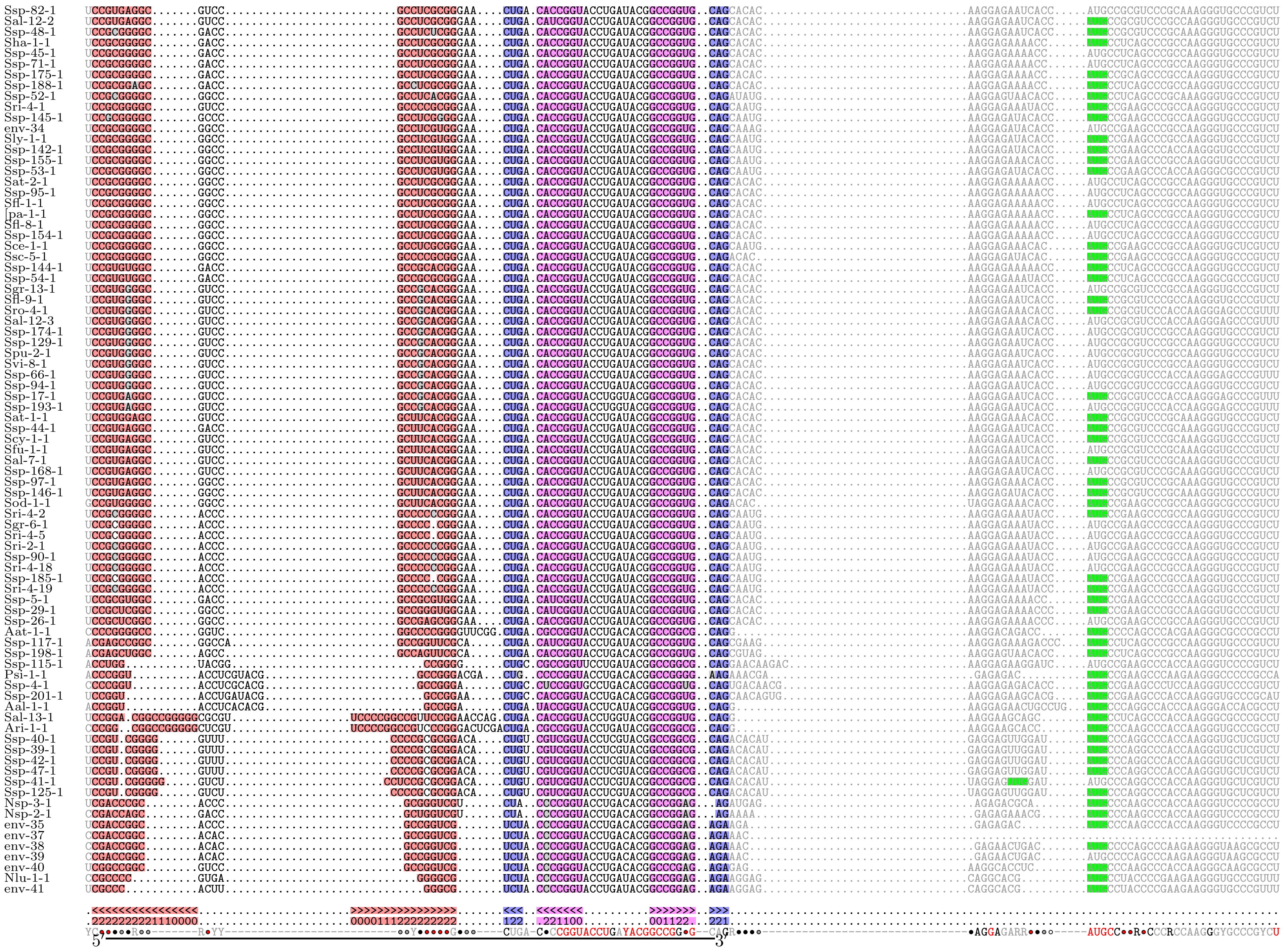
Ssp-82-1
Sal-12-2
Ssp-48-1
Sha-1-1
Ssp-45-1
Ssp-71-1
Ssp-175-1
Ssp-188-1
Ssp-52-1
Sri-4-1
Ssp-145-1
env-34
Sly-1-1
Ssp-142-1
Ssp-155-1
Ssp-53-1
Sat-2-1
Ssp-95-1
Sfl-1-1
Jpa-1-1
Sfl-8-1
Ssp-154-1
Sce-1-1
Ssc-5-1
Ssp-144-1
Ssp-54-1
Sgr-13-1
Sfl-9-1
Sro-4-1
Sal-12-3
Ssp-174-1
Ssp-129-1
Spu-2-1
Svi-8-1
Ssp-66-1
Ssp-94-1
Ssp-17-1
Ssp-193-1
Sat-1-1
Ssp-44-1
Scy-1-1
Sfu-1-1
Sal-7-1
Ssp-168-1
Ssp-97-1
Ssp-146-1
Sod-1-1
Sri-4-2
Sgr-6-1
Sri-4-5
Sri-2-1
Ssp-90-1
Sri-4-18
Ssp-185-1
Sri-4-19
Ssp-5-1
Ssp-29-1
Ssp-26-1
Aat-1-1
Ssp-117-1
Ssp-198-1
Ssp-115-1
Psi-1-1
Ssp-4-1
Ssp-201-1
Aal-1-1
Sal-13-1
Ari-1-1
Ssp-40-1
Ssp-39-1
Ssp-42-1
Ssp-47-1
Ssp-41-1
Ssp-125-1
Nsp-3-1
Nsp-2-1
env-35
env-37
env-38
env-39
env-40
Nlu-1-1
env-41
5'
3'

```
Nha-1-1   UCGGCUGACC.......GCAC.........................GGUCGGCCGU......CUA..CCCCGGUACCUGACACGGCCGGAG...AGAUGAGAG.............................AAGGACG...........AUGCCCACUCCCAAGAAGGGUGCCCGCCU
Sst-1-1   UCUGA.CAGGC......GACC........................GUCUGCUCAGAUA....CUGA.CUCCGGUACCUAGUACGGCCGGGG..CAGACACC.............................UAGGAGAAACACC.....AUGCCGAAGCCCACCAAGGGUGCCCGUCU
Sle-1-1   UCCGA.CAGGC......GACC........................GCCUGCUCGGAUA....CUGA.CCCCGGUACCUGAUACGGCCGGGG..CAGACACC.............................UAGGAGAAACACC.....AUGCCGAAGCCCACCAAGGGUGCCCGUCU
Sal-2-1   UCCG.GCGGAC......ACAA........................GUUCGUGCGGACA....CUGU.CAUCGGUACCUGAUACGGCCGGUG..CAGAUCAC.............................AAGGAGAAAGACC.....AUGCCGAAGCCCACCAAGGGUGCCCGUCU
Ssp-24-1  CCCCGGU..........ACCUGAUACG......................GCCGGGA....CUGC.CGCCGGUGCCUGAUACGGCCGGCG..CAGGGACAACG..........................AAGGAGAAGCACC.....AUGCCGAAGCCCACCAAGGGUCCCCGUCU
Scl-1-1   GCCCGUGGGC.......AUCC.........................GCCCGCGGGGCC....CUGA.CACCGGUACCUGACACGGCCGGUG..CAGCACAC.............................AAGGAGAAACACC.....AUGCCGAAGCCCGCCAAGGGUGCCCGUCU
Ssc-1-1   UCCGA.CGAGC......AAACCC......................GCUCGCUCGGAUA....CUGA.CCCCGGUACCUGAUACGGCCGGGG..CAGACACC.............................UAGGAGAAACACC.....AUGCCGAAGCCCACCAAGGGUGCCCGUCU
Sde-1-1   UCCGCGGGGC.......GCCC.........................GCCUCGUGGGAA....CUGA.CACCGGUACCUGAUACGGCCGGUG..CAGCAAUG.............................AAGGAGAUACACC.....AUGCCGAAGCCCGCCAAGGGUGCCCGUCU
          .<<<<<<<<<<<<<<<<.....................>>>>>>>>>>>>>>>>......<<<..<<<<<<<..........>>>>>>>..>>>.......................................................................
          .2222222221110000.....................0000111222222222......122...221100..........001122...221.......................................................................
          YC•••••R••-------R•YY-------------------------••Y•••••G•••----CUGA-C•CCGGUACCUGAYACGGCCGG•G--CAGR••••-----------------------------•AGGAGARR••••-----AUGCC••R•CCCRCCAAGGGYGCCCGYCU
          5'____________________________________________________________________________________________3'
```

alignment positions 260 · · · 439

```
env-1      CAAUCGCACCAGCGAGCACCGCCUCGCCAUGCUGCGCAACAUGAUGAACUCGCUGCUCACGCACGAAGUCAUCAAGACCACGCUGCCCAAGGCCAAAGAGCUGCGCCGCGUCGUGGAACCCAUGAUCACCCUGGCCAAGGUCGCGACCGUGGCAAACCGUCGUCUGGCUUUGACCGCCU
env-2      CAAUCGCACCAGCGAGCACCGCCUCGCCAUGCUGCGCAACAUGAUGAACUCGCUGCUCACGCACGAAGUCAUCAAGACCACGCUGCCCAAGGCCAAAGAGCUGCGCCGCGUCGUGGAACCCAUGAUCACCCUGGCCAAGGUCGCGACCGUGGCAAACCGUCGUCUGGCUUUGACCGCCU
[br-1-1    CAACCGCACCUCCGAGCACCGCCUGGCGAUGCUCCGCAACAUGGCCAACUCGCUGAUCGAGCACGAGGCCAUCAAGACCACGGUGCCCAAGGCCAAGGAACUGCGUCGCGUCGUCGAGCCGCUGAUCACCCUCGGCAAGGAGCCGACCGUCGCCAACAAGCGCCUGGCGUUCUCGCGCCU
env-3      CAACCGCACCAGCGAGCACCGCCUCGCCAUGCUGCGCAACAUGAUGAACUCGCUGCUCGAGCACGAGGUCAUCAAGACCACCGUCCCCAAAGCCAAGGAAUUGCGCCGCGUGGUCGAGCCCAUGAUCACGCUUGCCAAGGAACCGACCGUCGCCAACAAGCGUCUGGCGUUCGAUCGCCU
Nal-1-1    CGGCGGCAGCCCCGCGCACCAGCGCCUCAUCCUCGCGAACCUGGCCACGGACCUGUUCCGCUACGGCCGGAUCACGACGACCGAGUCGAAGGCGCGCACGCUGCGUCCGCACGCCGAGAAGCUCAUCACCAAGGCGAAGAAGGCCCACCUGGGCGAGAACCCGCUGGCCCAGCGCCGCGA
Nsp-1-1    CGGCGGCAGCCCGGCGCACCAGCGCCUCAUCCUCGCGAACCUCGCCACCUCGCUCUUCGAGCACGGCCGGAUCACGACGACCGAGGCCAAGGCCCGCACCCUGCGGCCCUACGCCGAGAAGCUGAUCACCAAGGCCAAGAAGGCCCACGCCGGCGAGAACCCGCUGCACCAGCGCCGCGA
Mxa-1-1    UCACCGCAGCACGAGCCACCGGCUCGCGAUGCUCAACAACAUGGUGACCUCGCUGCUGGAGCACCAGGCCAUCCGCACCACGCUUCCCAAGGCCAAGGAGGCCCGGAAGAUCGCGGAGCGGAUCAUCACCCUGGGCAAGCGUGGCGGCCUCGCCAACGUUCGCCUGGCGGCUCGUACGGU
env-4      CCACCGCACCACGAGCCACCGGCUCGCGAUGCUCCACAACAUGGUCACCUCGCUGCUCGAGCACGAGGCCA.............................................................................................................
Cfu-1-1    CCACCGCACCACGAGCCACCGGCUCGCGAUGCUGCACAACAUGGUCACCUCGCUGCUCGAGCACGAGGCCAUCCGCACCACGCUGCCCAAGGCCAAGGAGGCCCGGGCCCUCGCCGAGCGCAUCAUCACCCUCGGCAAGCGCGGCGGUCUGUCCAAUGUGCGCCUCGCGGAGCGCACCGU
Avi-1-1    CCACCGCACCACGAGCCACCGGCUCGCGAUGCUCCACAACAUGGUCACCUCGCUGCUCGAGCACGAGGCCAUCCGCACCACGCUGCCCAAGGCCAAGGAGGCCCGGGCGAUCGCCGAGCGCAUCAUCACCCUCGGCAAGCGCGGUGGCCUGGCCAACGUGCGCCUCGCGGAGCGGACGGU
Age-1-1    CCACCGCACCACGAGCCACCGGCUCGCGAUGCUCCACAACAUGGUCACCUCGCUGCUCGAGCACGAGGCCAUCCGCACCACGCUGCCCAAGGCCAAGGAGGCCCGGGCGAUCGCCGAGCGCAUCAUCACCCUCGGCAAGCGCGGUGGCCUGGCCAACGUGCGCCUCGCGGAGCGGACAGU
Hmi-1-1    UCACCGCACCACGCCGCACCGGCUCGCGAUGCUCAACAACAUGGUCACCUCGCUGCUCGAGCACGAGGCGAUCCGCACCACCGUCCCCAAGGCCAAGGAGGCCCGGAAGAUGGCGGAGCGGAUCAUCACGCUCGGCAAGCGCGGCGGGCUGGCCAAUGUUCGUCUGGCUGCCCGAAUCGU
env-5      GCACCGCACCACGAGCCACCGGCUCGCGAUGCUCAACAACAUGGUCACCUCGCUGCUCGAGCACGAGGCCAUUCGCACCACCGUCCCCAAGGCCAAGGAGGCCCGGAAGAUGGCGGAGCGCAUCAUCACCCUCGGCAAGCGCGGUGGGCUGGCCAAUGUUCGCCUCGCGGCGCGCAUCGU
Sau-5-1    GCACCGCACCACGAGCCACCGGCUCGCGAUGCUCAACAACAUGGUCACCUCGCUGCUCGAGCACGAGGCCAUCCGCACCACGCUGCCCAAGGCCAAGGAGGCCCGGAAGCUGGCGGAGCGCAUCAUCACGCUCGGCAAGCGCGGUGGUCUGGCCAAUGUCCGUCUCGCGUCCCGGACCGU
Mfu-1-1    GCACCGCACCACGAGCCACCGUCUCGCGAUGCUGAACAACAUGGUCACCUCGCUGCUCGAGCACCAGGCCAUCCGCACCACGCUGCCCAAGGCCAAGGAGGCCCGGAAGCUGGCGGAGCGCAUCAUCACGCUGGGUAAGCGUGGUGGGCUCUCCAACGUGCGCCUGGCGGCCCGGACCGU
Mst-1-1    GCACCGCACCACGAGCCACCGGCUCGCGAUGUUGAACAACAUGGUCACCUCGCUGCUCGAGCACCAGGCCAUCCGCACCACGCUGCCCAAGGCCAAGGAGGCCCGGAAGCUGGCGGAGCGCAUCAUCACGCUGGGUAAGCGUGGUGGUCUCUCCAACGUGCGCCUGGCGGCCCGGACGGU
Msp-2-1    GCACCGCACCACGAGCCACCGUCUCGCGAUGCUGAACAACAUGGUGACUUCGCUGCUGGAGCACCAGGCCAUCCGCACCACGCUUCCCAAGGCCAAGGAGGCCCGGAAGAUCGCGGAGCGGAUCAUCACCCUGGGCAAGCGUGGCGGCCUCGCCAACGUUCGCCUGGCGGCCCGGACGGU
Mfu-2-1    UCACCGCACCACGAGCCACCGGCUCGCGAUGCUCAACAACAUGGUGACCUCGCUGCUGGAGCACCAGGCCAUCCGCACCACGCUUCCCAAGGCCAAGGAGGCCCGGAAGAUCGCGGAGCGGAUCAUCACCCUGGGCAAGCGUGGCGGUCUCGCCAACGUUCGCCUGGCGGCCCGGACGGU
env-6      UCACCGCACCACGCCGCACCGGCUCGCGAUGCUCAACAACAUGGUCACCUCGCUGCUCGAGCACGAGGCGAUCCGCACCACCGUCCCCAAGGCCAAGGAGGCCCGGAAGAUGGCGGAGCGGAUCAUCACGCUCGGCAAGCGCGGCGGGCUGGCCAAUGUCCGUCUGGCUUCCCGGAUCGU
Cco-1-1    GCACCGCACCACGAGCCACCGUCUCGCGAUGCUGAACAACAUGGUCACGUCCCUGCUGGAGCACGGGGCCAUCCGCACCACCGUUCCCAAGGCGAAGGAAGUCCGUCCGCUGGCCGAGCGCAUCAUCACGCUCGCCAAGCGCGGUGGCCUGUCCAACGUGCGCCUUGCUGCCCGCACCGU
env-7      CAACCGCACGUCGGAGCACCGCCUGGCGAUGUUCCGCAACAUGUGCAAUUCGUUGAUUCAACACGAGGCGAUCAAGACGACGGUGCCGAAGGCGAAGGAACUGCGCCGGCUGGUCGAGCCGCUGAUCACGCUCGGCAAGGAACCGACGCUCGCGAACCGGCGGCUGGCUUUCAACCGCAC
env-8      CAACCGCACCAGUGAGCAUCGCCAGGCGAUGUUGCGCAACAUGUGCAAUUCGUUGCUGCAGCACGAAGCCAUCAAGACCACGCUGCCCAAGGCCAAGGAAUUGCGCCGCGUGGUCGAGCCGCUCAUCACGCUGGCCAAGGAAGCGACACUGGCGAACCGCCGUCUGGCUUUCGAUCGCAC
Ama-1-1    CGGCGGUAGCGCGGCGCACCAGAAGCUGAUCCUGGCGAACCUGGCCACGGCCCUGUUCGAGCACGGCCGCAUCACCACCACCGAGGCCAAGGCCCGCCGGGUCCGCCCGCUGGCGGAGAAGCUGAUCACCAAGGCCAAGAAGGGCGACCUGCACAACCGGCGCCUGGUGGCCCGCACGAU
env-9      CGGCGGGAGCCCGGCACACGAACGGUUGAUCCUGGCGAACCUCGCGACCGCGUUGUUCGAGCACGGAUCGAUCCGGACGACCGAGGCGAAGGCGCGGCGGGUGCGGCCACUGGCCGAGCGGCUGAUCACCAAGGCCAAGAAGGGUGAUCUGCACAACCGCCGUCAGGUGCACGCGGUG..
env-10     CGGCGGGAGCCCGGCACACGAACGGUUGAUCCUGGCGAACCUCGCGACCGCGUUGUUCGAGCACGGAUCGAUCCGGACGACCGAGGCGAAGGCGCGGCGGGUGCGGCCACUGGCCGAGCGGCUGAUCACCAAGGCCAAGAAGGGUGAUCUGCACAACCGCCGUCAGGUGCACGCGGUGAU
1 Ssp-68-1 GGGCGGCAGUGCCGCGCACCAGAAGGCGAUGCUGGGCAACCUGGCGACCGCGCUGUUCGAGCACGGCCGCAUCACGACGACCGAGGCGAAGGCCCGCCGCCUGCGUCCGGUCGCCGAGCGUCUGAUCACCAAGGCGAAGAAGGGCGACCUGCACAACCGUCGGCAGGUCAUGCGCACCGU
env-11     CGGCGGCAGCCCCGCUCACCAGCGGCGGAUGCUGGCGAACCUGGCGACGGCGCUGUUCGAGCACGGCCGCAUCACCACCACCGAGGCCAAGGCCCGGCGGGUGCGGCCGCUCGCGGAGAAGCUGAUCAC..................................................
env-12     CGGCGGCAGCCCGGCCCACCAGCGGCUCAUCCUCCGCAACCUGGCCACCUCGCUCUUCGAGCACGGCCGGAUCACCACCACGGAGGCCAAGGCGCA...................................................................................
Ssp-96-1   CGGCGGCGGCCCGCACCACGAGCCCCUGCUGCUCGCCGGUCUGGCCCGUGAGCUGUUCCAGUACGGCCGCAUCACCACCACCGAGGCCAAGGCCCGCCGCCUGCGCCCGCUGGCGGAGAAGCUGAUCACCAAGGCGAAGAAGGGCGACAUCCACAACCGUCGCCUGGUGCGCAAGACCAU
Ksp-1-1    CGGCGGCGGCCCGCACCACGAGCCGCUGCUGCUCGCCGGUCUGUGCCGGGAGCUGUUCCAGUACGGCCGCAUCACCACGACCGAGGCCAAGGCCCGUCGGAUGCGCCCGCUGGCGGAGAAGCUGAUCACCAAGGCGAAGAAGGGCGACAUCCACAACCGUCGCCUGGUGCGCAAGACCAU
Kme-1-1    CGGCGGUGGCCCGCACCACGAGCCGCUGCUGCUGGCCGGCCUGUGCCGCGAGCUGUUCACCCACGGCCGCAUCACCACGACCGAGGCCAAGGCCCGUCGGAUGCGCCCGCUGGCCGAGCGCCUGAUCACCAAGGCGAAGGUCGGUGACAUCCACAACCGUCGUAUCGUCCGCAAGACGAU
Rge-1-1    GAACCGCACCAGCAGCCAUCGCCUCGCGAUGCUGCGCAACAUGGCCAACUCGCUGAUCCGCCACGAGGCGAUCAAGACGACGGUGCCGAAGGCCAAGGAACUGCGCGCCGUCGUCGAGCCGCUGAUCACGCUGGGCAAGACCCCGACGCUGGCCAACCGCCGCCUGGCUUUCGACCGCAC
Sca-3-1    CGGUGGCGGUCCCUCCCACGAGCGUCUGCUGCUGGCCGGUCUGGCCCGCGAGCUGUUCCAGUACGGCCGGAUCACCACCACCGAGGCCAAGGCUCGCCGCCUGCGCCCGGUGGCGGAGAAGCUGAUCACCAAGGCCAAGAAGGGUGACAUCCACAACCGCCGUGAGGUCCGCAAGACCAU
Sal-3-1    CGGCGGCGGCCCUGCCCACGAGCGUCUGCUGCUCGCCGGUCUGGCCCGUGAGCUGUUCCAGUACGGCCGCAUCACCACCACCGAGGCCAAGGCUCGCCGCCUGCGCCCGGUGGCGGAGAAGCUGAUCACCAAGGCCAAGAAGGGUGACAUCCACAACCGCCGUGAGGUCCGCAAGACCAU
env-13     GGGGGGCAGCGCCGCGCACCAGCGCCUGAUCCUGCGCAACCUGGCCACGUCGCUGUUCGAGCACGGCGCAGUGACCACGACCGAGGCGAAGGCCCGCAUGCUGCGGCCCUACGCCGAGAAGCUGAUCACCAAGGCCAAGAAGGGCGACCUGCACAACCGUCGCCUCGUGCUGCGCGAGCU
1 Sav-3-1  CGGCGGCAGUGCCGCUCACGAGCGCAUGCUGCUGGCGAACCUCGCCACCUCGCUGUUCGAGCACGGCCGCAUCACCACCACCGUGGCCAAGGCCAGGCGGCUGCGUCCGGUCGCCGAGCGGCUCAUCACCAAGGCGAAGAAGGGCGACCUGCACAACCGUCGCCAGGUGAUGCAGACCAU
Ssp-195-1  GGGCGGCAGCGCCUCGCACCAGAAGGCCAUGCUGGCGAACCUCGCCACGGACCUCUUCGAGUACGGGCGCAUCACCACCACCGAGACGCGGGCCCGCCGCCUGCGUCCCGUGGCGGAGCGGCUGAUCACCAAGGCGAAGAACGGCGACCUUCACAACCGCCGUCAGGUCAUGCGUACGGU
Nsp-4-1    UGGCGGCAGUCCGGCGCACGAGCGGCUGAUCCUGGCCAACCUUGCCACGGACCUGUUCCGCCACGGCAAGAUCCGCACCACGGUCACCAAGGCCAAGCGCCUUCGUCCGGUGGCGGAGCGCCUGAUCACCAAGGCGAAGAAGGGCGACAUCCACAACCGUCGCCAGGUGCUCACCGUCGU
env-14     UGGCGGCAGCCCGGCGCACGAGCGGUUGAUCCUGGCGAACCUGGCCACCGAUCUGUUCCGCCACGGCAAGAUCCGCACGACGGUCGCCAAGGCCAAGCGCCUGCGCCCGCUGGCGGAGCGCCUGAUCACCAAGGCGAAGAAGGGCGACAUCCACAACCGUCGCCAGGUCCUGACCGUCGU
Nco-1-1    UGGCGGCAGCCCGGCGCACGAGCGGCUGAUCCUGGCCAACCUGGCGACCGACCUGUUCCGGCACGGCAAGAUCCGCACGACGGUCGCGAAGGCCAAGCGCCUGCGUCCGGUCGCGGAGCGCCUGAUCACCAAGGCGAAGAAGGGCGACAUCCACAACCGUCGCCAGGUCCUGACCGUCGU
Sje-1-1    CGGUGGCGGUCCCUCCCACGAGCGUCUGCUGCUGGCCGGUCUGGCCCGCGAGCUCUUCCAGUACGGCCGGAUCACCACCACCGAGGCCAAGGCUCGCCGCCUGCGCCCGGUGGCGGAGAAGCUGAUCACCAAGGCCAAGAAGGGUGACAUCCACAACCGCCGUGAGGUCCGCAAGACCAU
Kaz-1-1    CGGCGGUGGCCCGCACCACGAGCCGCUGCUGCUCGCCGGCCUCUGCCGGGAGCUCUUCCAGUACGGCCGCAUCACCACGACCGAGGCCAAGGCCCGUCGGAUGCGCCCGCUGGCGGAGAAGCUGAUCACCAAGGCCAAGGCCGGUGACAUCCACAACCGUCGCAUCGUCCGCAAGACGAU
Kph-1-1    CGGCGGCGGCCCGUCCCACGAGCCGCUGCUGCUGGCCGGCCUGGCCCGCGAGCUGUUCCAGUACGGCCGCAUCACCACCACCGAGGCCAAGGCCCGCCGCCUGCGCCCGCUGGCGGAGAAGCUGAUCACCAAGGCGAAGAAGGGCGACAUCCACAACCGUCGCCUGGUGCGCAAGACCAU
Kph-1-2    CGGCGGCGGCCCGUCCCACGAGCCGCUGCUGCUGGCCGGCCUGGCCCGCGAGCUGUUCCAGUACGGCCGCAUCACCACCACCGAGGCCAAGGCCCGCCGCCUGCGCCCGCUGGCGGAGAAGCUGAUCACCAAGGCGAAGAAGGGCGACAUCCACAACCGUCGCCUGGUGCGCAAGACCAU
Sno-2-1    CGGCGGCGGCCCGCACCACGAGCCGCUGCUGCUCGCCGGUCUGUGCCGGGAGCUGUUCCAGUACGGCCGCAUCACCACGACCGAGGCCAAGGCCCGUCGGAUGCGCCCGCUGGCGGAGAAGCUGAUCACCAAGGCGAAGAAGGGCGACAUCCACAACCGUCGCCUGGUGCGCAAGACCAU
Ksp-3-1    CGGCGGCGGCCCGCACCACGAGCCGCUGCUGCUCGCCGGUCUGUGCCGGGAGCUGUUCCAGUACGGCCGCAUCACCACGACCGAGGCCAAGGCCCGUCGGAUGCGCCCGCUGGCGGAGAAGCUGAUCACCAAGGCGAAGAAGGGCGACAUCCACAACCGUCGCCUGGUGCGCAAGACCAU
Spy-1-1    GUACGGCCGCAUCACCACGACCGAGGCCAAGGCCCGUCGGAUGCGCCCGCUGGCGGAGAAGCUGAUCACCAAGGCGAAGAAGGGCGACAUCCACAACCGUCGCCUGGUGCGCAAGACCAUCACCGACAUCUCGGUGCUGCACACCCUCUUCACCGAGAUCGCGCCGCGCUACGAGAACCG
Ksp-4-1    CGGCGGCGGCCCGUCCCACGAGCCGCUCCUGCUGGCCGGCCUGGCCCGCGAGCUGUUCCAGUACGGCCGCAUCACCACCACCGAGGCCAAGGCCCGUCGCCUGCGCCCGCUGGCGGAGAAGCUGAUCACCAAGGCGAAGAAGGGCGACAUCCACAACCGUCGCCUGGUGCGCAAGACCAU
Ssp-165-1  CGGCGGCGGCCCGCACCACGAGCCGCUGCUGCUCGCCGGUCUGUGCCGUGAGCUGUUCCAGUACGGCCGCAUCACCACGACCGAGGCCAAGGCCCGUCGGAUGCGCCCGCUGGCGGAGAAGCUGAUCACCAAGGCGAAGAAGGGCGACAUCCACAACCGUCGCCUGGUGCGCAAGACCAU
Ssp-133-1  CGGCGGCGGCCCGCACCACGAGCCGCUGCUGCUCGCCGGUCUGUGCCGGGAGCUGUUCCAGUACGGCCGCAUCACCACGACCGAGGCCAAGGCCCGUCGGAUGCGCCCGCUGGCGGAGAAGCUGAUCACCAAGGCGAAGAAGGGCGACAUCCACAACCGUCGUCUGGUUCGCAAGACCAU
Ssp-50-1   CGGCGGCGGCCCGCACCACGAGCCGCUGCUGCUCGCCGGUCUGUGCCGGGAGCUGUUCCAGUACGGCCGCAUCACCACGACCGAGGCCAAGGCCCGUCGGAUGCGCCCGCUGGCGGAGAAGCUGAUCACCAAGGCGAAGAAGGGCGACAUCCACAACCGUCGCCUGGUUCGCAAGACCAU
Ssp-157-1  CGGCGGCGGCCCGCACCACGAGCCGCUGCUGCUCGCCGGUCUGUGCCGGGAGCUGUUCCAGUACGGCCGCAUCACCACGACCGAGGCGAAGGCCCGUCGGAUGCGCCCGCUGGCGGAGAAGCUGAUCACCAAGGCGAAGAAGGGCGACAUCCACAACCGUCGCCUGGUGCGCAAGACGAU
Ssp-160-1  CGGCGGCGGCCCGUCCCACGAGCCGCUGCUGCUGGCCGGCCUGGCCCGCGAGCUGUUCCAGUACGGCCGCAUCACCACCACCGAGGCCAAGGCCCGCCGCCUGCGCCCGCUGGCGGAGAAGCUGAUCACCAAGGCGAAGAAGGGCGACAUCCACAACCGUCGCCUGGUGCGCAAGACCAU
Ksp-2-1    CGGCGGCGGCCCGCACCACGAGCCGCUGCUGCUCGCCGGUCUGUGCCGGGAGCUGUUCCAGUACGGCCGCAUCACCACGACCGAGGCCAAGGCCCGUCGGAUGCGCCCGCUGGCGGAGAAGCUGAUCACCAAGGCGAAGAAGGGCGACAUCCACAACCGUCGCCUGGUGCGCAAGACCAU
Ssp-183-1  CGGCGGCGGCCCGCACCACGAGCCGCUGCUGCUCGCCGGUCUGUGCCGGGAGCUCUUCCAGCACGGCCGCAUCACCACGACCGAGGCCAAGGCCCGUCGGAUGCGCCCGCUGGCCGAGAAGCUGAUCACCAAGGCGAAGAAGGGCGACCUGCACAACCGUCGCCUGGUGCGUAAGACCAU
Sam-2-1    CGGCGGCAGCCCGGCCCACGAGCGGCUGAUCCUGGCGAACCUCGCCACCCAGCUGUUCCAGCACGGCCGGAUCAAGACCACCGAGGCCAAGGCGAAGCGUCUGCGGCCGCUGGCGGAGAAGCUGAUCACCAAGGCGAAGAAGGGCGACAUGCACAACCGUCGCCAGGUGCUGACGGUCGU
Sro-7-1    CGGCGGAAGCCCGGCUCACGAGCGGCUGAUCCUGGCGAACCUCGCCACCCAGCUGUUCCAGCACGGCCGGAUCAAGACCACGGUGGCCAAGGCCAAGCGUCUGCGCCCGCUGGCGGAGCGGCUGAUCACCAAGGCGAAGAAGGGCGACAUGCACAACCGUCGCCAGGUGCUGACCGUCGU
Ssp-87-1   GGGCGGCAGCGCCGCGCACGAGCGGCUCCUGCUGGCGAACCUCGCCACCGCGCUGUUCGAGCACGGCCGGAUCACCACCACCGAGGCGAAGGCCCGCCGCCUGCGCCCGGUCGCGGAGAAGCUGGUCACCAAGGCGAAGAAGGGCGACCUUCACAACCGCCGUCUGGUGCUCCAGACCAU
Ssp-190-1  GGGCGGCAGCGCUGCGCACGAGCGGCUCCUGCUGGCGAACCUCGCCACCGCGCUGUUCGAGCACGGCCGGAUCACCACCACCGAGGCAAAGGCCCGCCGCCUGCGCCCGGUCGCGGAGAAGCUGGUCACCAAGGCGAAGAAGGGCGACCUUCACAACCGCCGUCUGGUGCUCCAGACCAU
Sir-1-1    GGGCGGCAGCGCUGCGCACGAGCGGCUCCUGCUGGCGAACCUCGCCACCGCGCUGUUCGAGCACGGCCGGAUCACCACCACCGAGGCGAAGGCCCGCCGCCUGCGCCCGGUCGCGGAGAAGCUGGUCACCAAGGCGAAGAAGGGCGACCUUCACAACCGCCGUCUGGUGCUCCAGACCAU
Shy-4-1    GGGCGGCAGCGCUGCGCACGAGCGGCUCCUGCUGGCGAACCUCGCCACCGCGCUGUUCGAGCACGGCCGGAUCACCACCACCGAGGCGAAGGCCCGCCGCCUGCGCCCGGUCGCGGAGAAGCUGGUCACCAAGGCGAAGAAGGGCGACCUUCACAACCGCCGUCUGGUGCUCCAGACCAU
Sro-3-1    CGGCGGAAGCCCGGCUCACGAGCGGCUGAUCCUGGCGAACCUCGCCACCCAGCUGUUCCAGCACGGCCGGAUCAAGACCACGGUGGCCAAGGCCAAGCGUCUGCGCCCGCUGGCGGAGCGGCUGAUCACCAAGGCGAAGAAGGGCGACAUGCACAACCGUCGCCAGGUGCUGACCGUCGU
Nin-1-1    CGGCGGUAGCCCGGCCCACCAGAAGCUCAUCAUCGCCAACCUGGCGACCGCGCUCUUCGAGCACGGCCGGAUCACCACCACCGAGGCGAAGGCGCGACUGCUGCGUCCGCACGCGGAGAAGCUGAUCACCAAGGCGAGGAAGGGCGACCUGCACAACCGCCGCGAGGUCCUCAAGACCAU
1 Nca-1-1  UGGCGGCAGCCCGGCGCACGAGCGGCUGAUCCUGGCCAACCUGGCCACGGACCUGUUCCGCCACGGCAAGAUCCGCACGACGGUCGCGAAGGCCAAGCGCCUGCGUCCGCUGGCCGAGCGCCUGAUCACCAAGGCGAAGAAGGGCGACAUCCACAACCGGCGUCAGGUCCUGACCGUCGU
Ssp-192-1  GGGUGGCAGCCCCGCCCACGAGCGGAUGCUGGUGGCGAACCUGGCCACCGCGCUCUUCGAGCACGGCCGCAUCACCACCACCGUGGCCAAGGCCCGGCGGCUGCGUCCGGUGGCCGAGCGCCUGAUCACCAAGGCGAAGAAGGGCGACCUGCACAACCGCCGCCAGGUGAUGCAGACCAU
Ssp-172-1  GGGCGGCAGCGCCGCGCACGAGAAGCUUCUGCUGGCGAACCUCGCCAAGCAGCUCUUCGAGUACGGCCGCAUCACCACCACCGAGGCGAAGGCCCGUCGCCUGCGCCCGUACGCGGAGCGUCUGGUCACCAAGGCGAAGAAGGGUGACCUUCACAACCGCCGCCAGGUGCUGUCCGUCAU

           ....................................................................................................................................................................................
           ....................................................................................................................................................................................
           •GGCGGCAGCRC•GCGCAC•AG•R•CU••U•CU•RY•AACCU•GC•A••••GCU•UUCGAGCACGGCCG•AUCAC•AC•ACCGAGGC•AAGGCCGYCG•CUGCGYCCG••CGC•GAGCR•CUGRUCACCAAGGCGAAGAAGGGCGAC•UYCACAACCGYCGYC•GGU•YU•YRGRY•AU
```

| | |
|---|---|
| Ssp-93-1 | GGGCGGCAGCGCCGCGCACGAGAAGCUGAUGCUGGCGAACCUCGCCAAGGAUCUCUUCGAGCACGGCCGCAUCACCACGACCGAGGCGAAGGCCCGUCGCCUGCGUCCGUUCGCGGAGCGUCUGGUCACCAAGGCGAAGAAGGGCGACCUUCACAACCGCCGUCAGGUGCUCCAGGUCAU |
| Sau-3-1 | GGGCGGCAGUGCCGCGCACCAGAAGCUGAUGCUGGCGAACCUCGCCAAGGAUCUCUUCGAGCACGGCCGCAUCACCACCACCGAGGCCAAGGCCCGUCGUCUGCGUCCCUACGCGGAGCGUCUGGUCACCAAGGCGAAGAAGGGCGACCUUCACAACCGCCGCCAGGUCCUCCAGGUCAU |
| Sal-6-1 | GGGCGGCAGCGCCGCGCACGAGAAGCUUCUGCUGGCGAACCUCGCCAAGCAGCUCUUCGAGCACGGCCGCAUCACCACCACCGAGGCCAAGGCCCGUCGUCUGCGUCCCUACGCGGAGCGUCUGGUCACCAAGGCGAAGAAGGGCGACCUUCACAACCGCCGCCAGGUCCUCCAGGUCAU |
| Sau-1-1 | GGGCGGCAGUGCCGCGCACCAGAAGCUGAUGCUGGCGAACCUCGCCAAGGAUCUCUUCGAGCACGGCCGCAUCACCACCACCGAGGCCAAGGCCCGUCGUCUGCGUCCCUACGCGGAGCGUCUGGUCACCAAGGCGAAGAAGGGCGACCUUCACAACCGCCGCCAGGUCCUCCAGGUCAU |
| Ssp-106-1 | CGGUGGCAGCCCCGCCCACGAGCGGCUCAUGCUGGCGAACCUCGCCACCGCGCUGUUCGAGCACGGCCGGAUCACCACCACGGUCGCCAAGGCCCGCGGCUGCGUCCGGCUGCGCCGAGCGGCUGAUCACCAAGGCGAAGAAGGGCGACCUGCACAACCGCCGUCAGGUGAUGCGCACCGU |
| Ssp-3-1 | CGGUGGCGGCCCGGCCCACGAGCGGCUCAUGCUGGCGAACCUCGCCACGGCGCUGUUCGAGCACGGCCGGAUCACCACCACGGUGGCGAAGGCGCGCCGGCUGCGUCCGGUGGCCGAGCGGCUGAUCACCAAGGCGAAGAAGGGCGACCUGCACAACCGCCGUCAGGUGAUGCGCACCAU |
| env-15 | .............................................................................................................................................................................. |
| env-16 | CGGCGGCAGCCCCGGCCCACGAGCGGCUGAUCCUGGCGAACCUCGCCACGCAGCUGUUCCAGCACGGCCGGAUCCGCACCACGGAGGCGAAGGCCAAGCGUCUGCGCCCGUUCGCUGAGCGGUUGAUCACCAAGGCAAAGAAGGGCGACCUGCACAACCGCCGCCAGGUGCUGGCGGUCAU |
| env-18 | CGGCGGCAGCCCCGGCCCACGAGCGGCUGAUCCUGGCGAACCUCGCCACGCAGCUGUUCCAGCACGGCCGGAUCCGCACCACGGAGGCGAAGGCCAAGCGUCUGCGCCCGUUCGCUGAGCGGUUGAUCACCAAGGCAAAGAAGGGCGACCUGCACAACCGCCGCCAGGUGCUGGCGGUCAU |
| Ssp-118-1 | GGGCGGCAGCGCCGCGCACGAGAAGCUGCUCCUGCUCGCGAACCUCGCGAAGUCGCUCUUCGAGCACGGCCGCAUCACCACGACCGAGGCGAAGGCCCGCCGCCUGCGUCCGGUCGCGGAGCGCCUGAUCACCAAGGCGAAGAAGGGCGACAUCCACAACCGUCGCCUGGUGCUGCGCACGAU |
| env-19 | CGGCGGCAGCCCCGGCGCACGAGCGGCUGAUCCUGGCGAACCUCGCCACCCAGCUGUUCCAGCACGGCCGCAUCCGCACCACCGAGGCGAAGGCCAAGCGUCUGCGCCCGCUGGCGGAGAGGCUGAUCACCAAGGCGAAGAAGGGCGACAUGCACAACCGUCGCCAGGUGCUCACCGUCGU |
| env-20 | CGGCGGCAGCCCGGCGCACGAGCGGCUNAUCCUGGCGAACCUCGCCACNCAGCUNUUCCAGCACGGCCGCAUCCGNACCACCGAGGCGAAGGCCAAGCGUCUGCGCCCGCUGGCGGAGNGGCUGGUCAUCACCAAGGCGAAGAAGGGCGACAUGCACAACCGUCGCCAGGUGCUCACCNUCGU |
| env-21 | CGGCGGCAGCCCGGCGCACGAGCGGCUCAUCCUGGCGAACCUCGCCACGCAGCUCUUCCAGCACGGCCGCAUCCGUACCACCGAGGCGAAGGCCAAGCGUCUGCGCCCGCUGGCGGAGCGGCUGAUCACCAAGGCGAAGAAGGGCGACAUGCACAACCGUCGCCAGGUGCUCACCAUCGU |
| Sor-1-1 | CGGCGGUAGCCCGGCGCACGAGCGGCUGCUGCUCGCGAACCUGGCCACCGCGCUCUUCGAGCACGGCCGCAUCACCACCACCGAGGCCAAGGCCCGCCGCCUGCGCCCGCUGGCGGAGAAGCUGAUCACCAAGGCGAAGAAGGGCGACCUCCACAACCGCCGCCUGGUGCGCAAGACCGU |
| Ssp-64-1 | GGGCGGCAGCGCCGCGCACGAGAAGCUGCUUCUCGCGAACCUCGCGAAGUCGCUCUUCGAGCACGGCCGCAUCACCACGACCGAGGCCAAGGCCCGCCGCCUGCGUCCGGUCGCCGAGCGCCUGAUCACCAAGGCGAAGAAGGGCGACAUCCACAACCGUCGCCUGGUGCUGCAGACGAU |
| Ssp-12-1 | GGGCGGCAGCGCCGCGCACGAGAAGCUGCUUCUCGCGAACCUCGCGAAGUCGCUCUUCGAGCACGGCCGCAUCACCACGACCGAGGCCAAGGCCCGCCGCCUGCGUCCGGUCGCCGAGCGCCUGAUCACCAAGGCGAAGAAGGGCGACAUCCACAACCGUCGCCUGGUGCUGCAGACGAU |
| Ssp-13-1 | GGGCGGCAGCGCCGCGCACGAGAAGCUGCUUCUCGCGAACCUCGCGAAGUCGCUCUUCGAGCACGGUCGCAUCACCACGACCGAGGCCAAGGCCCGCCGCCUGCGUCCGGUCGCCGAGCGCCUGAUCACCAAGGCGAAGAAGGGCGACAUCCACAACCGUCGCCUGGUGCUGCAGACGAU |
| Ssp-61-1 | GGGCGGCGGCGCGGCGCACGAGCGCGCUAUGCUGCGGAACCUGGCCACCGCUCUCUUCGAGCACGGUCGCAUCACGACGACCGAGGCCAAGGCCCGCAGGCUGCGUCCGUACGCCGAGCGGCUGGUGACCAAGGCGAAGAAGGGCGACCUUCACAACCGCCGGCAGGUCAUGCAGCUCAU |
| Ssp-59-1 | GGGCGGCGGCGCGGCGCACGAGCGCGCCAUGCUGCGGAACCUGGCCACCGCUCUCUUCGAGCACGGUCGCAUCACCACGACCGAGGCCAAGGCCCGCAGGCUGCGUCCGUACGCCGAGCGGCUGGUGACCAAGGCGAAGAAGGGUGACCUUCACAACCGCCGGCAGGUCAUGCAGCUCAU |
| Ssp-16-1 | GGGCGGCGGCGCGGCGCACGAGCGCGCCAUGCUGCGGAACCUGGCCACCGCUCUCUUCGAGCACGGUCGCAUCACGACGACCGAGGCCAAGGCCCGCAGGCUGCGUCCGUACGCCGAGCGGCUGGUGACCAAGGCGAAGAAGGGCGACCUUCACAACCGCCGGCAGGUCAUGCAGCUCAU |
| Ssp-69-1 | GGGCGGCGGCGCGGCGCACGAGCGCGCCAUGCUGCGGAACCUGGCCACCGCUCUCUUCGAGCACGGUCGCAUCACGACGACCGAGGCCAAGGCCCGCAGGCUGCGUCCGUACGCCGAGCGGCUGGUGACCAAGGCGAAGAAGGGCGACCUUCACAACCGCCGGCAGGUCAUGCAGCUCAU |
| Ssp-56-1 | GGGCGGCGGCGCGGCGCACGAGCGCGCCAUGCUGCGGAACCUGGCCACCGCUCUCUUCGAGCACGGUCGCAUCACCACCACCGAGGCCAAGGCCCGCAGGCUGCGUCCGUACGCCGAGCGACUGGUGACCAAGGCGAAGAAGGGCGACCUGCACAACCGCCGCCAGGUCAUGCAGCUGAU |
| Hcr-1-1 | CGGUGGUUCCCCGGCGCACGAGAAGCUGAUCCUGUCGAACCUCGCGACGCAGCUCUUCCAGCACGGGAAGAUCAAGACCACCGAGGCCAAGGCCCAAGCGCCUGCGCCCGCUGGCUGAGCGGCUGAUCACCAAGGCGAAGAAGGGCGACAUCCACAACCGUCGCCAGGUGCUGACCGUCGU |
| Msp-1-1 | CGGCGGCAGCCCGGCCCACGAGCGGCUGAUCCUGGCGAACCUCGCCACCCAGCUGUUCCAGAACGGCCGGAUCCGCACCACCGAGGCCAAGGCGAAGCGUCUGCGCCCGCUCGCGGAGAAGCUGAUCACCAAGGCGAAGAAGGGCGACAUCCACAACCGUCGCCAGGUGCUGACCGUCGU |
| Mro-1-1 | CGGUGGCAGCCCGUCCCACGAGCGGCUGAUCCUGGCGAACCUCGCCACGGCGCUGUUCCAGAACGGCCGGAUCACCACCACCGAGGCCAAGGCCCAGGCGUCUCCGCCCGCUCGCGGAGAAGCUGAUCACCAAGGCGAAGAAGGGCGACAUCCACAACCGUCGCCAGGUGCUGACGGUCGU |
| Ssp-171-1 | CGGCGGCUCCGCCGCGCACGAGAAGCUGCUGCUCGCCAACCUGGCGAAGUCGCUCUUCGAGCACGGCCGCAUCACCACGACCGAGGCCAAGGCCCGCCGCCUGCGUCCGGUCGCCGAGCGCCUGAUCACCAAGGCGAAGAAGGGCGACAUCCACAACCGUCGCCUGGUGCUGCAGACGAU |
| Ssp-43-1 | CGGCGGUUCCGCCGCGCACGAGAAGCUGCUGCUCGCCAACCUGGCGAAGUCGCUCUUCGAGCACGGCCGCAUCACCACGACCGAGGCCAAGGCCCGUCGCCUGCGUCCGGUCGCCGAGCGCCUGAUCACCAAGGCGAAGAAGGGCGACAUCCACAACCGUCGCCUGGUGCUGCAGUCGAU |
| Svi-13-1 | CGGCGGUUCCGCCGCGCACGAGAAGCUGCUGCUCGCCAACCUGGCGAAGUCGCUCUUCGAGCACGGCCGCAUCACCACGACCGAGGCCAAGGCCCGUCGCCUGCGUCCGGUCGCCGAGCGCCUGAUCACCAAGGCGAAGAAGGGCGACAUCCACAACCGUCGCCUGGUGCUGCAGUCGAU |
| Ssp-73-1 | GGGCGGCAGCGCCGCGCACGAGAAGCUGCUCUCGCGAACCUCGCGAAGUCGCUCUUCGAGCACGGCCGGAUCACCACCACCGAGGCGAAGGCCCGCCGCCUGCGUCCGUACGCCGAGCGCCUGAUCACCAAGGCGAAGAAGGGCGACCUUCACAACCGCCGCCAGGUGCUCCAGGUCAU |
| Ssc-3-1 | GGGCGGCGGCGCCGCGCACGAGAAGCUGCUCUCGCGAACCUCGCGAAGGCGCUCUUCGAGCACGGCCGCAUCACCACCACCGAGGCGAAGGCCCGCCGUCUGCGGCCGUACGCCGAGCGUCUGGUCACCAAGGCGAAGAAGGGCGACCUUCACAACCGCCGUCAGGUGCUCCAGGUCAU |
| Kau-1-1 | GGGCGGCAGUGCCGCGCACGAGAAGCUGCUCUCGCCAACCUGGCGAAGAGCCUCUUCGAGCACGGCCGGAUCACCACCACCGAGGCGAAGGCGCGCAAGCUGCGUCCGUACGCCGAGCGUCUGGUCACCAAGGCGAAGAAGGGCGACCUUCACAACCGCCGUCAGGUGCUCCAGGUGAU |
| Sgr-1-1 | GGGCGGCAGCGCCGCGCACGAGAAGCUCCUCUCGCGAACCUCGCGAAGGCGCUCUUCGAGCACGGCCGCAUCACCACCACCGAGGCGAAGGCGCGCAAGCUGCGUCCGUACGCCGAGCGUCUGGUCACCAAGGCGAAGAAGGGCGACCUUCACAACCGCCGUCAGGUGCUCCAGGUGAU |
| Sto-1-1 | GGGCGGCAGCGCCGCGCACGAGAAGCUGAUGCUCGCGAACCUCGCGAAGGCGCUUUUCGAGCACGGCCGCAUCACCACCACCGAGGCGAAGGCGCGCAAGCUGCGUCCGUACGCCGAGCGUCUGGUCACCAAGGCGAAGAAGGGCGACCUUCACAACCGCCGUCAGGUGCUCCAGGUGAU |
| Ssp-114-1 | GGGCGGCAGCGCCGCGCACGAGAAGCUGCUCUCGCGAACCUCGCGAAGGCGCUCUUCGAGCACGGCCGCAUCACCACCACCGAGGCGAAGGCCCGCCGCCUGCGCCCGUACGCCGAGCGACUGGUCACCAAGGCGAAGAAGGGCGACCUUCACAACCGCCGUCAGGUGCUCCAGGUGAU |
| Ssc-4-1 | GGGCGGCAGCGCCGCGCACGAGAAGCUGCUCUCGCGAACCUCGCGAAGUCGCUGUUCGAGCACGGCCGUAUCACCACCACCGAGGCGAAGGCCCGCCGUCUGCGGCCGUACGCCGAGCGUCUGGUCACCAAGGCGAAGAAGGGCGACCUUCACAACCGCCGUCAGGUGCUCCAGGUCAU |
| Ssp-1-1 | GGGCGGCAGCGCCGCGCACGAGAAGCUGCUCUCGCGAACCUCGCGAAGUCGCUGUUCGAGCACGGCCGCAUCACCACCACCGAGGCGAAGGCGCGCAAGCUGCGUCCGUACGCCGAGCGUCUGGUCACCAAGGCGAAGAAGGGCGACCUUCACAACCGCCGUCAGGUGCUCCAGGUGAU |
| Sau-4-1 | GGGCGGCAGCGCCGCGCACGAGAAGCUGCUCUCGCGAACCUCGCGAAGUCGCUCUUCGAGCACGGCCGCAUCACCACCACCGAGGCGAAGGCCCGCCGCCUGCGUCCCUACGCGGAGCGUCUGGUCACCAAGGCGAAGAAGGGUGACCUUCACAACCGCCGUCAGGUGCUCUCGGUGAU |
| Spr-1-1 | CGGUGGCAGCGCCGCUCACGAGAAGCUUCUUCUCGCGAACCUCGCCGAAGUCGCUGUUCGAGCACGGCCGCAUCACCACGACCGAGGCCAAGGCCCGCCGCCUGCGUCCGGUCGCCGAGCGCCUGAUCACCAAGGCGAAGAAGGGCGACAUCCACAACCGUCGCCUGGUGCUGCAGACGAU |
| Ssp-189-1 | GGGCGGCUCCGCCGCGCACGAGAAGCACCUCCUCGCGAACCUCGCGAAGGCGCUCUUCGAGCACGGCCGCAUCACCACCACCGAGGCCAAGGCCCGCCGCCUGCGUCCCUACGCCGAGCGUCUGGUCACCAAGGCCAAGAAGGGCGACAUCCACAACCGUCGCCUGGUGCUGCAGACGAU |
| Sfl-6-1 | GGGCGGCUCCGCCGCGCACACGAGAAGCACCUGCUCGCGAACCUCGCGAAGGCGCUCUUCGAGCACGGCCGCAUCACCACCACCGAGGCCAAGGCCCGCCGCCUGCGUCCCUACGCCGAGCGUCUGGUCACCAAGGCCAAGAAGGGCGACAUCCACAACCGUCGCCUGGUGCUGCAGACGAU |
| Ssp-103-1 | GGGCGGCUCCGCCGCGCACGAGAAGCACCUCCUCGCGAACCUCGCGAAGGCGCUCUUCGAGCACGGCCGCAUCACCACCACCGAGGCCAAGGCCCGCCGCCUGCGUCCCUACGCCGAGCGUCUGGUCACCAAGGCCAAGAAGGGCGACAUCCACAACCGUCGCCUGGUGCUGCAGACGAU |
| Ssp-111-1 | GGGCGGCUCCGCCGCGCACACGAGAAGCACCUGCUCGCGAACCUCGCGAAGGCGCUCUUCGAGCACGGCCGCAUCACCACCACCGAGGCCAAGGCCCGCCGCCUGCGUCCCUACGCCGAGCGUCUGGUCACCAAGGCCAAGAAGGGCGACAUCCACAACCGUCGCCUGGUGCUGCAGACGAU |
| Ssp-159-1 | GGGCGGCUCCGCCGCGCACGAGAAGCACCUGCUCGCGAACCUCGCGAAGGCGCUCUUCGAGCACGGCCGCAUCACCACCACCGAGGCCAAGGCCCGCCGCCUGCGUCCCUACGCCGAGCGUCUGGUCACCAAGGCCAAGAAGGGCGACAUCCACAACCGUCGCCUGGUGCUGCAGACGAU |
| Ser-1-1 | GGGCGGCUCCGCCGCGCACGAGAAGCACCUCCUCGCGAACCUCGCGAAGGCGCUCUUCGAGCACGGCCGCAUCACCACCACCGAGGCCAAGGCCCGCCGCCUGCGUCCCUACGCCGAGCGUCUGGUCACCAAGGCCAAGAAGGGCGACAUCCACAACCGUCGCCUGGUGCUGCAGACGAU |
| Sya-1-1 | GGGCGGCUCCGCCGCGCACGAGAAGCACCUGCUCGCGAACCUCGCGAAGGCGCUCUUCGAGCACGGCCGCAUCACCACCACCGAGGCCAAGGCCCGCCGCCUGCGUCCCUACGCCGAGCGUCUGGUCACCAAGGCCAAGAAGGGCGACAUCCACAACCGUCGCCUGGUGCUGCAGACGAU |
| Svi-12-1 | GGGCGGCUCCGCCGCGCACGAGAAGCACCUCCUCGCGAACCUCGCGAAGGCGCUCUUCGAGCACGGCCGCAUCACCACCACCGAGGCCAAGGCCCGCCGCCUGCGUCCCUACGCCGAGCGUCUGGUCACCAAGGCCAAGAAGGGCGACAUCCACAACCGUCGCCUGGUGCUGCAGACGAU |
| Ssp-151-1 | GGGCGGCUCCGCCGCGCACGAGAAGCACCUCCUCGCGAACCUCGCGAAGGCGCUCUUCGAGCACGGCCGCAUCACCACCACCGAGGCCAAGGCCCGUCGCCUGCGUCCCUACGCCGAGCGUCUGGUCACCAAGGCCAAGAAGGGCGACAUCCACAACCGUCGCCUGGUGCUGCAGACGAU |
| Ssp-110-1 | GGGCGGCUCCGCCGCGCACGAGAAGCACCUCCUCGCGAACCUCGCGAAGGCGCUCUUCGAGCACGGCCGGAUCACCACCACCGAGGCCAAGGCCCGUCGCCUGCGUCCCUACGCCGAGCGUCUGGUCACCAAGGCCAAGAAGGGCGACAUCCACAACCGUCGCCUGGUGCUGCAGACGAU |
| Ssp-112-1 | GGGCGGCUCCGCCGCGCACGAGAAGCACCUCCUCGCGAACCUCGCGAAGGCGCUCUUCGAGCACGGCCGCAUCACCACCACCGAGGCCAAGGCCCGCCGCCUGCGUCCCUACGCCGAGCGUCUGGUCACCAAGGCCAAGAAGGGCGACAUCCACAACCGUCGCCUGGUGCUGCAGACGAU |
| Ssp-102-1 | GGGCGGCUCCGCCGCGCACGAGAAGCACCUCCUCGCGAACCUCGCGAAGGCGCUCUUCGAGCACGGCCGCAUCACCACCACCGAGGCCAAGGCCCGUCGCCUGCGUCCCUACGCCGAGCGUCUGGUCACCAAGGCCAAGAAGGGCGACAUCCACAACCGUCGCCUGGUGCUGCAGACGAU |
| Sla-2-1 | GGGCGGCUCCGCCGCGCACACGAGAAGCACCUGCUCGCGAACCUCGCGAAGGCGCUCUUCGAGCACGGCCGCAUCACCACCACCGAGGCCAAGGCCCGCCGCCUGCGUCCCUACGCCGAGCGUCUGGUCACCAAGGCCAAGAAGGGCGACAUCCACAACCGUCGCCUGGUGCUGCAGACGAU |
| Ssp-139-1 | GGGCGGCUCCGCCGCACACGAGAAGCACCUGCUCGCGAACCUCGCGAAGGCGCUCUUCGAGCACGGCCGCAUCACCACCACCGAGGCCAAGGCCCGCCGCCUGCGUCCCUACGCCGAGCGUCUGGUCACCAAGGCCAAGAAGGGCGACAUCCACAACCGUCGCCUGGUGCUGCAGACGAU |
| Ssp-8-1 | GGGCGGCUCCGCCGCGCACGAGAAGCACCUCCUCGCGAACCUCGCGAAGGCCCUCUUCGAGCACGGCCGCAUCACCACCACCGAGGCCAAGGCCCGCCGCCUGCGUCCCUACGCCGAGCGUCUGGUCACCAAGGCGAAGAAGGGCGACAUCCACAACCGUCGCCUGGUGCUGCAGACGAU |
| Svi-7-1 | CGGCGGUUCCGCCGCGCACGAGAAGCUGCUCCUCGCCAACCUGGCGAAGUCGCUCUUCGAGCACGGCCGCAUCACCACGACCGAGGCCAAGGCCCGUCGCCUGCGUCCGGUCGCCGAGCGCCUGAUCACCAAGGCGAAGAAGGGCGACAUCCACAACCGUCGCCUGGUGCUGCAGACGAU |
| Sgr-12-1 | CGGCGGUUCCGCCGCUCACGAGAAGCUGCUCCUCGCCAACCUGGCGAAGUCGCUGUUCGAGCACGGCCGCAUCACGACGACCGAGGCCAAGGCCCGCCGUCUGCGUCCGGUCGCCGAGCGCCUGAUCACCAAGGCGAAGAAGGGCGACAUCCACAACCGUCGCCUGGUGCUGCAGACGAU |
| Sfl-5-1 | CGGCGGUUCCGCCGCGCACGAGAAGCUGCUCCUCGCCAACCUGGCGAAGUCGCUCUUCGAGCACGGCCGCAUCACCACGACCGAGGCCAAGGCCCGUCGCCUGCGUCCGGUCGCCGAGCGCCUGAUCACCAAGGCGAAGAAGGGCGACAUCCACAACCGUCGCCUGGUGCUGCAGACGAU |
| Sbi-3-1 | CGGCGGUUCCGCCGCGCACGAGAAGCUGCUCCUCGCCAACCUGGCGAAGUCGCUGUUCGAGCACGGCCGCAUCACCACGACCGAGGCCAAGGCCCGUCGCCUGCGUCCGGUCGCCGAGCGCCUGAUCACCAAGGCGAAGAAGGGCGACAUCCACAACCGUCGCCUGGUGCUGCAGACGAU |
| Sex-1-1 | CGGCGGUUCCGCCGCGCACGAGAAGCUGCUCCUCGCCAACCUGGCGAAGUCGCUCUUCGAGCACGGCCGCAUCACCACGACCGAGGCCAAGGCCCGUCGCCUGCGUCCGGUCGCCGAGCGCCUGAUCACCAAGGCGAAGAAGGGCGACAUCCACAACCGUCGCCUGGUGCUGCAGACGAU |
| Sgl-3-1 | CGGCGGUUCCGCCGCGCACGAGAAGCUGCUCCUCGCCAACCUGGCGAAGUCGCUCUUCGAGCACGGCCGCAUCACCACGACCGAGGCCAAGGCCCGUCGCCUGCGUCCGGUCGCCGAGCGCCUGAUCACCAAGGCGAAGAAGGGCGACAUCCACAACCGUCGCCUGGUGCUGCAGUCGAU |
| Sgr-9-1 | CGGCGGUUCCGCCGCGCACGAGAAGCUGCUCCUCGCCAACCUGGCGAAGUCGCUCUUCGAGCACGGCCGCAUCACCACGACCGAGGCCAAGGCCCGUCGCCUGCGUCCGGUCGCCGAGCGCCUGAUCACCAAGGCGAAGAAGGGCGACAUCCACAACCGUCGCCUCGUGCUGCAGACGAU |
| Ssp-121-1 | CGGCGGUUCCGCCGCUCACGAGAAGCUGCUGCUCGCCAACCUGGCGAAGUCGCUGUUCGAGCACGGCCGCAUCACGACGACCGAGGCCAAGGCCCGCCGCCUGCGUCCGCUCGCCGAGCGCCUGAUCACCAAGGCGAAGAAGGGCGACAUCCACAACCGUCGCCUGGUGCUGCAGACGAU |
| Sve-1-1 | CGGCGGUUCCGCCGCGCACGAGAAGCUGCUCCUCGCCAACCUGGCGAAGUCGCUCUUCGAGCACGGCCGCAUCACCACGACCGAGGCCAAGGCCCGUCGCCUGCGUCCGGUCGCCGAGCGCCUGAUCACCAAGGCGAAGAAGGGCGACAUCCACAACCGUCGCCUGGUGCUGCAGACGAU |
| Sav-2-1 | GGGCGGCAGCGCCGCGCACGAGAAGCUGCUCCUCGCGAACCUCGCGAAGUCGCUCUUCGAGCACGGCCGCAUCACCACCACCGAGGCGAAGGCCCGCCGCCUGCGCCCGUACGCCGAGCGUCUGGUCACCAAGGCGAAGAAGGGUGACCUUCACAACCGCCGCCAGGUGCUCCAGGUCAU |
| Mgl-1-1 | CGGCGGUUCCGCCGCGCACGAGAAGCUGCUCCUCGCCAACCUGGCGAAGUCGCUCUUCGAGCACGGCCGCAUCACGACGACCGAGGCCAAGGCCCGCCGCCUGCGUCCGGUCGCCGAGCGCCUGAUCACCAAGGCGAAGAAGGGCGACAUCCACAACCGUCGCCUGGUGCUGCAGACGAU |
| Asp-1-1 | GGGCGGCAGCGCUGCGCACGAGCGUCACAUGCUGGCGAACCUCGCCAAGUCGCUGUUCGAGCACGGCCGUAUCACGACGACCGAGGCCAAGGCCCGCCGCCUGCGUCCGGUCGCCGAGCGCCUGAUCACCAAGGCGAAGAAGGGCGACAUCCACAACCGCCGCCAGGUGCUGCAGACGAU |
| Stu-1-1 | GGGCGGCAGUGCCGCGCACGAGAAGCUGCUUCUUGCGAACCUCGCGAAGUCGCUCUUCGAGCACGGCAAGAUCACCACCACCGAGGCGAAGGCGCGCCGCCUGCGUCCGUACGCCGAGCGUCUGGUCACCAAGGCGAAGAAGGGUGACCUUCACAACCGCCGUCAGGUGCUCCAGGUCAU |
| Sye-2-1 | GGGCGGCGGUGCCGCGCACGAGAAGCUGCUUCUUGCGAACCUUGCGAAGGCGCUGUUCGAGCACGGCCGUAUCACCACGACCGAGGCGAAGGCCCGCCGGCUGCGGCCGUACGCCGAGCGUCUGGUCACCAAGGCGAAGAAGGGCGACCUUCACAACCGCCGUCAGGUGCUUCAGGUCAU |
| Sdu-1-1 | GGGCGGCAGCGCCGCGCACGAGAAGCUGCUCCUCGCGAACCUCGCGAAGAGCCUCUUCGAGCACGGCCGUAUCACCACCACCGAGGCGAAGGCCCGUCGCCUGCGCCCGUACGCCGAGCGUCUGAUCACCAAGGCGAAGAAGGGUGACCUUCACAACCGCCGCCAGGUGCUCCAGGUCAU |
| Sgh-1-1 | GGGCGGCAGCGCCGCGCACGAGAAGCUGAUGCUCGCGAACCUCGCGAAGUCGCUCUUCGAGCACGGCCGCAUCACCACCACCGAGGCGAAGGCGCGCAAGCUGCGCCCGUACGCCGAGCGUCUGAUCACCAAGGCGAAGAAGGGCGACCUUCACAACCGCCGUCAGGUGCUCCAGGUCAU |
| Svi-5-1 | GGGCGGCAGCGCCGCGCACGAGAAGCUGAUGCUCGCGAACCUCGCGAAGUCGCUCUUCGAGCACGGCCGCAUCACCACCACCGAGGCGAAGGCGCGCAAGCUGCGCCCGUACGCCGAGCGUCUGAUCACCAAGGCGAAGAAGGGCGACCUUCACAACCGCCGUCAGGUGCUCCAGGUCAU |
| Ssp-153-1 | GGGCGGCAGCGCCGCGCACGAGAAGCUCCUCCUCGCGAACCUCGCGAAGAGCCUCUUCGAGCACGGCCGUAUCACCACCACCGAGGCGAAGGCCCGCCGCCUGCGCCCGUACGCCGAGCGUCUGAUCACCAAGGCGAAGAAGGGCGACCUUCACAACCGCCGCCAGGUGCUCCAGGUCAU |
| Ssp-86-1 | GGGCGGCAGCGCCGCGCACGAGAAGCUGCUCCUCGCGAACCUCGCGAAGGCGCUGUUCGAGCACGGCCGUAUCACCACCACCGAGGCGAAGGCCCGCCGCCUGCGCCCGUACGCCGAGCGUCUGGUCACCAAGGCGAAGAAGGGCGACCUUCACAACCGCCGUCAGGUGCUCCAGGUCAU |
| Sro-1-1 | GGGCGGCAGCGCCGCGCACGAGAAGCUGCUCCUCGCGAACCUCGCGAAGGCGCUCUUCGAGCACGGCCGUAUCACCACCACCGAGGCGAAGGCCCGCCGCCUGCGCCCGUACGCCGAGCGUCUGAUCACCAAGGCGAAGAAGGGCGACCUUCACAACCGCCGUCAGGUGCUCCAGGUCAU |
| Sgr-7-1 | GGGCGGCAGCGCCGCGCACGAGAAGCUGCUCCUCGCGAACCUCGCGAAGAGCCUCUUCGAGCACGGCCGCAUCACCACGACCGAGGCGAAGGCCCGCCGCCUGCGCCCGUACGCCGAGCGUCUGGUCACCAAGGCGAAGAAGGGCGACCUUCACAACCGCCGUCAGGUGCUCCAGGUCAU |
| Ssp-28-1 | GGGCGGCAGUGCCGCGCACGAGAAGCUGCUGCUCGCGAACCUCGCGAAGAGCCUCUUCGAGCACGGCCGUAUCACCACCACCGAGGCGAAGGCCCGCCGCCUGCGCCCGUACGCCGAGCGUCUGGUCACCAAGGCGAAGAAGGGUGACCUUCACAACCGCCGCCAGGUGCUCCAGGUCAU |
| Ssp-176-1 | GGGCGGCAGCGCCGCGCACGAGAAGCUCCUCCUCGCGAACCUCGCGAAGGCGCUGUUCGAGCACGGUCGCAUCACCACCACCGAGGCGAAGGCGCGCAAGCUGCGUCCGUACGCCGAGCGUCUGGUCACCAAGGCGAAGAAGGGCGACCUUCACAACCGCCGUCAGGUGCUCCAGGUGAU |
| Ssp-14-1 | CGGUGGCAGCGCCGCUCACGAGAAGCUGCUUCUCGCGAACCUGGCGAAGUCGCUGUUCGAGCACGGCCGCAUCACCACGACCGAGGCGAAGGCCCGCCGCCUGCGUCCGGUCGCGGAGCGCCUGAUCACCAAGGCGAAGAAGGGCGACAUCCACAACCGUCGCCUGGUGCUGCAGAGCAU |
| Sac-2-1 | GGGCGGCAGCGCCGCGCACGAGAAGCUGCUCCUCGCGAACCUCGCGAAGAGCCUCUUCGAGCACGGCAAGAUCACCACCACCGAGGCGAAGGCCCGCCGCCUGCGCCCGUACGCCGAGCGUCUGAUCACCAAGGCGAAGAAGGGUGACCUUCACAACCGCCGCCAGGUGCUCCAGGUGAU |
| Sbo-1-1 | GGGCGGCAGCGCCGCGCACGAGAAGCUGCUCCUCGCGAACCUCGCGAAGAGCCUCUUCGAGCACGGCCGCAUCACCACCACCGAGGCGAAGGCCCGCCGUCUGCGGCCGUACGCCGAGCGUCUGGUCACCAAGGCGAAGAAGGGCGACCUUCACAACCGCCGUCAGGUGCUCCAGGUCAU |
| Ssp-2-1 | GGGCGGCAGCGCCGCGCACGAGAAGCUGCUGCUCGCGAACCUCGCGAAGAGCCUCUUCGAGCACGGCCGUAUCACCACCACCGAGGCGAAGGCGCGCAAGCUGCGUCCGUACGCCGAGCGUCUGGUCACCAAGGCGAAGAAGGGCGACCUUCACAACCGCCGUCAGGUGCUUCAGGUGAU |
| Ssp-98-1 | GGGCGGCAGUGCCGCGCACGAGAAGCUGCUGCUCGCGAACCUCGCGAAGAGCCUCUUCGAGCACGGCCGUAUCACCACCACCGAGGCGAAGGCCCGUCGGCUGCGGCCGUACGCCGAGCGUCUGGUCACCAAGGCGAAGAAGGGCGACCUUCACAACCGCCGUCAGGUGCUCCAGGUGAU |
| Sac-3-1 | CGGUGGUAGUGCUUCUCACGAGAAGCUGCUUCUGGCGAACCUUGCCAAGUCGCUCUUCGAGCACGGUCGGAUCACCACGACCGAGGCGAAGGCGCGUCGGCUGCGGCCGUACGCCGAGCGGCUGGUGACCAAGGCGAAGAAGGGUGACCUUCACAACCGUCGCCAGGUGCUUCAGGUCAU |
| Sca-7-1 | GGGCGGUAGCGCCGCGCACGAGCGUCUCAUGCUGCGCAACCUGGCCACCGCGCUCUUCGAGCACGGCCGCAUCACGACGACCGAGGCCAAGGCCCGCCGCCUGCGUCCGUACGCCGAGCGGCUGGUGACCAAGGCGAAGAAGGGUGACCUUCACAACCGUCGCCAGGUCAUGCAGUUGAU |
| Sgr-2-1 | GGGCGGCAGUGCCGCGCACGAGAAGCUGCUUCUCGCGAACCUCGCGAAGGCGCUGUUCGAGCACGGCCGCAUCACCACCACCGAGGCGAAGGCGCGCAAGCUGCGUCCGUACGCCGAGCGUCUGGUCACCAAGGCGAAGAAGGGCGACCUUCACAACCGCCGUCAGGUUCUCCAGGUCAU |

..........................................................................................................................................................................

..........................................................................................................................................................................

•GGCGGCAGCRC•GCGCAC•AG•R•CU••U•CU•RY•AACCU•GC•A••••GCU•UUCGAGCACGGCCG•AUCAC•AC•ACCGAGGC•AAGGCCCGYCG•CUGCGYCCG••CGC•GAGCR•CUGRUCACCAAGGCGAAGAAGGGCGAC•UYCACAACCGYCGYC•GGU•YU•YRGRY•AU

| | |
|---|---|
| Ssp-105-1 | GGGCGGCAGUGCCGCGCACGAGAAGCUGCUCCUCGCGAACCUCGCGAAGAGCCUCUUCGAGCACGGCCGCAUCACCACCACCGAGGCGAAGGCGCGCCGUCUGCGCCCGUACGCCGAGCGUCUGGUCACCAAGGCGAAGAAGGGCGACCUUCACAACCGCCGUCAGGUGCUCCAGGUGAU |
| Ssp-85-1 | GGGCGGCAGCGCCGCGCACGAGAAGCUGCUCCUCGCGAACCUCGCGAAGAGCCUCUUCGAGCACGGCCGCAUCACCACCACCGAGGCGAAGGCCCGCCGUCUGCGCCCGUACGCCGAGCGUCUGGUCACCAAGGCGAAGAAGGGCGACCUUCACAACCGCCGUCAGGUGCUCCAGGUAU |
| Sco-3-1 | GGGCGGCAGCGCCGCGCACGAGAAGCUGCUCCUCGCGAACCUCGCGAAGUCGCUCUUCGAGCACGGCCGCAUCACCACCACCGAGGCGAAGGCCCGCCGCCUGCGCCCGUACGCCGAGCGUCUGGUCACCAAGGCGAAGAAGGGCGACCUUCACAACCGCCGUCAGGUGCUCCAGGUCAU |
| Sse-1-1 | GGGCGGCAGCGCCGCGCACGAGAAGCUGCUUCUCGCGAACCUCGCGAAGAGCCUCUUCGAGCACGGCCGCAUCACCACCACCGAGGCGAAGGCGCGCCGUCUGCGCCCGUACGCCGAGCGUCUGGUCACCAAGGCGAAGAAGGGCGACCUUCACAACCGCCGUCAGGUGCUCCAGGUGAU |
| Sre-1-1 | GGGCGGCAGCGCCGCGCACGAGAAGCUGCUUGUCGCGAACCUCGCGAAGGCGCUGUUCGAGCACGGCCGUAUCACCACCACCGAGGCGAAGGCGCGCAAGCUGCGCCCGUACGCCGAGCGUCUGAUCACCAAGGCGAAGAAGGGUGACCUUCACAACCGCCGCCAGGUGCUCCAGGUGAU |
| Spe-1-1 | GGGCGGCAGUGCCGCGCACGAGAAGCUGCUCCUCGCGAACCUCGCGAAGGCGCUCUUCGAGCACGGUCGUAUCACCACCACCGAGGCGAAGGCCCGCCGUCUGCGGCCGUACGCCGAGCGUCUGGUCACCAAGGCGAAGAAGGGCGACCUUCACAACCGCCGUCAGGUGCUCCAGGUCAU |
| Sbi-2-1 | GGGCGGCAGUGCCGCGCACGAGAAGCUGCUCCUCGCGAACCUGGCGAAGAGCCUCUUCGAGCACGGCCGGAUCACCACCACCGAGGCGAAGGCCCGCCGUCUGCGGCCGUACGCCGAGCGUCUGAUCACCAAGGCGAAGAAGGGUGACCUUCACAACCGCCGUCAGGUGCUCCAGGUGAU |
| Ssp-83-1 | GGGCGGCAGCGCCGCGCACGAGAAGCUGCUCCUCGCGAACCUCGCGAAGGCGCUCUUCGAGCACGGCCGCAUCACCACCACCGAGGCGAAGGCCCGCCGCCUGCGCCCGUACGCCGAGCGUCUGGUCACCAAGGCGAAGAAGGGCGACCUUCACAACCGCCGUCAGGUGCUCCAGGUCAU |
| Spl-1-1 | GGGCGGCAGCGCCGCGCACGAGAAGCUGCUCCUCGCGAACCUCGCGAAGGCGCUCUUCGAGCACGGCCGCAUCACCACCACCGAGGCGAAGGCCCGCCGCCUACGCCCGUACGCCGAGCGUCUGGUCACCAAGGCGAAGAAGGGCGACCUUCACAACCGCCGUCAGGUGCUCCAGGUCAU |
| Sno-1-1 | GGGCGGCAGCGCCGCGCACGAGAAGCUGCUCCUCGCGAACCUCGCGAAGGCGCUGUUCGAGCACGGCCGUAUCACCACCACCGAGGCGAAGGCGCGCCGUCUGCGCCCGUACGCCGAGCGUCUGGUCACCAAGGCGAAGAAGGGUGACCUUCACAACCGCCGCCAGGUGCUCCAGGUGAU |
| San-1-1 | GGGCGGCAGCGCCGCGCACGAGAAGCUGCUCCUCGCGAACCUGGCGAAGAGCCUCUUCGAGCACGGCCGUAUCACCACCACCGAGGCGAAGGCGCGCCGUCUGCGUCCGUACGCCGAGCGUCUGAUCACCAAGGCGAAGAAGGGCGACCUUCACAACCGCCGCCAGGUGCUUCAGGUCAU |
| Sla-3-1 | GGGCGGCAGCGCCGCGCACGAGAAGCUGCUCCUCGCGAACCUCGCGAAGAGCCUCUUCGAGCACGGCAAGAUCACCACCACCGAGGCGAAGGCGCGCCGUCUGCGCCCGUACGCCGAGCGUCUGAUCACCAAGGCGAAGAAGGGUGACCUUCACAACCGCCGCCAGGUGCUUCAGGUGAU |
| Ssp-74-1 | GGGCGGCAGCGCCGCGCACGAGAAGCUGCUCCUCGCGAACCUCGCGAAGUCGCUGUUCGAGCACGGCCGCAUCACCACCACCGAGGCGAAGGCCCGCCGCCUGCGCCCGUACGCCGAGCGUCUGAUCACCAAGGCGAAGAAGGGCGACCUUCACAACCGCCGUCAGGUGCUCCAGGUCAU |
| Ssp-162-1 | GGGCGGCAGCGCCGCGCACGAGAAGCUGAUGCUCGCGAACCUCGCGAAGUCGCUGUUCGAGCACGGCCGUAUCACCACCACCGAGGCGAAGGCCCGCCGCCUGCGUCCGUACGCCGAGCGUCUGGUCACCAAGGCGAAGAAGGGCGACCUUCACAACCGCCGUCAGGUGCUCCAGGUCAU |
| Ssp-164-1 | GGGCGGCAGCGCCGCGCACGAGAAGCUGAUGCUCGCGAACCUCGCGAAGUCGCUGUUCGAGCACGGUCGUAUCACCACCACCGAGGCGAAGGCCCGCCGCCUGCGUCCGUACGCCGAGCGUCUGGUCACCAAGGCGAAGAAGGGCGACCUUCACAACCGCCGUCAGGUGCUCCAGGUCAU |
| Ssp-161-1 | GGGCGGCAGCGCCGCGCACGAGAAGCUGAUGCUCGCGAACCUCGCGAAGUCGCUGUUCGAGCACGGUCGUAUCACCACCACCGAGGCGAAGGCCCGCCGCCUGCGUCCGUACGCCGAGCGUCUGGUCACCAAGGCGAAGAAGGGCGACCUUCACAACCGCCGUCAGGUGCUCCAGGUCAU |
| Sgl-2-1 | GGGCGGCAGUGCCGCGCACGAGAAGCUGCUGCUCGCGAACCUCGCGAAGUCGCUCUUCGAGCACGGCAAGAUCACGACCACCGGAGGCGAAGGCCCGCCGCCUGCGCCCGUACGCCGAGCGUCUGAUCACCAAGGCGAAGAAGGGCGACCUUCACAACCGCCGCCAGGUGCUCCAGGUCAU |
| Ssp-143-1 | GGGCGGCAGCGCCGCGCACGAGAAGCUGCUCCUCGCGAACCUCGCGAAGGCGCUCUUCGAGCACGGCCGCAUCACCACCACCGAGGCGAAGGCGCGCAAGCUGCGUCCGUACGCCGAGCGUCUGGUCACCAAGGCGAAGAAGGGCGACCUUCACAACCGCCGUCAGGUGCUCCAGGUCAU |
| Ssp-36-1 | GGGCGGCAGCGCCGCGCACGAGAAGCUCCUCCUCGCGAACCUCGCGAAGGCGCUGUUCGAGCACGGCCGUAUCACCACCACCGAGGCGAAGGCCCGCCGUCUGCGCCCGUACGCCGAGCGUCUGGUCACCAAGGCGAAGAAGGGCGACCUUCACAACCGCCGUCAGGUGCUCCAGGUCAU |
| Ssp-200-1 | GGGCGGCAGUGCCGCGCACGAGAAGCUGCUCCUCGCGAACCUGGCGAAGAGCCUCUUCGAGCACGGCCGGAUCACCACCACCGAGGCGAAGGCGCGCAAGCUGCGUCCGUACGCCGAGCGUCUGGUCACCAAGGCGAAGAAGGGCGACCUUCACAACCGCCGUCAGGUGCUCCAGGUGAU |
| Sco-2-1 | GGGCGGCAGUGCCGCGCACGAGAAGCUGCUCCUCGCGAACCUGGCGAAGAGCCUCUUCGAGCACGGCCGGAUCACCACCACCGAGGCGAAGGCGCGCAAGCUGCGCCCGUACGCCGAGCGUCUGGUCACCAAGGCGAAGAAGGGCGACCUUCACAACCGCCGUCAGGUGCUCCAGGUGAU |
| Svi-9-1 | GGGCGGCAGUGCCGCGCACGAGAAGCUGCUCCUCGCGAACCUGGCGAAGAGCCUCUUCGAGCACGGCCGCAUCACCACCACCGAGGCGAAGGCGCGCAAGCUGCGUCCGUACGCCGAGCGUCUGGUCACCAAGGCGAAGAAGGGCGACCUUCACAACCGCCGUCAGGUGCUCCAGGUGAU |
| Sco-1-1 | GGGCGGCAGCGCCGCGCACGAGAAGCUGCUCCUCGCGAACCUGGCGAAGAGCCUCUUCGAGCACGGCAAGAUCACCACCACCGAGGCGAAGGCGCGCAAGCUGCGUCCGUACGCCGAGCGUCUGGUCACCAAGGCGAAGAAGGGCGACCUUCACAACCGCCGUCAGGUGCUCCAGGUGAU |
| Ssp-128-1 | GGGCGGCAGCGCCGCGCACGAGAAGCUCCUCCUCGCCAACCUGGCGAAGAGCCUCUUCGAGCACGGCCGCAUCACCACCACCGAGGCGAAGGCGCGCAAGCUGCGUCCGUACGCCGAGCGUCUGGUCACCAAGGCGAAGAAGGGCGACCUUCACAACCGCCGUCAGGUGCUCCAGGUGAU |
| Sol-2-1 | GGGCGGCAGUGCCGCGCACGAGAAGCUGCUCAUCGCGAACCUCGCGAAGUCGCUGUUCGAGCACGGCCGCAUCACCACCACCGAGGCGAAGGCGCGCAAGCUGCGUCCGUACGCCGAGCGUCUGGUCACCAAGGCGAAGAAGGGCGACCUUCACAACCGCCGUCAGGUGCUCCAGGUGAU |
| Ssp-72-1 | GGGCGGCAGUGCCGCGCACGAGAAGCUGCUCAUCGCGAACCUCGCGAAGUCGCUGUUCGAGCACGGCCGCAUCACCACCACCGAGGCGAAGGCGCGCAAGCUGCGUCCGUACGCCGAGCGUCUGGUCACCAAGGCGAAGAAGGGCGACCUUCACAACCGCCGUCAGGUGCUCCAGGUGAU |
| Ssp-124-1 | GGGCGGCAGCGCCGCGCACGAGAAGCUGCUCCUCGCCAACCUGGCGAAGAGCCUCUUCGAGCACGGCCGCAUCACCACCACCGAGGCGAAGGCGCGCAAGCUGCGUCCGUACGCCGAGCGUCUGGUCACCAAGGCGAAGAAGGGCGACCUUCACAACCGCCGUCAGGUGCUCCAGGUGAU |
| Sce-2-1 | GGGCGGCAGUGCCGCGCACGAGAAGCUGCUCCUCGCGAACCUCGCGAAGUCGCUCUUCGAGCACGGCCGCAUCACCACCACCGAGGCGAAGGCGCGCAGGCUGCGCCCGUACGCCGAGCGUCUGGUCACCAAGGCGAAGAAGGGCGACCUUCACAACCGCCGUCAGGUGCUCCAGGUGAU |
| Smo-1-1 | GGGCGGCGGUGCCGCGCACGAGAAGCUGAUGCUGGCGAACCUCGCCCGCUCCUCUUCGAGCACGGCCGCAUCACCACCACCGAGGCCAAGGCCCGCGUCUGCGUCCGGUGGCGGAGCGCCUGAUCACCAAGGCGAAGAAGGGCGACAUCCACAACCGUCGCCAGGUGCUGCAGACGAU |
| Svi-9-2 | GGGCGGCAGUGCCGCGCACGAGAAGCUGCUCCUCGCGAACCUCGCGAAGUCGCUCUUCGAGCACGGCCGCAUCACCACCACCGAGGCGAAGGCGCGCAGGCUGCGUCCGUACGCCGAGCGUCUGGUCACCAAGGCGAAGAAGGGCGACCUUCACAACCGCCGUCAGGUGCUCCAGGUGAU |
| Sso-1-1 | CGGCGGUUCCGCCGCGCACGAGAAGCUGCUCCUCGCGAACCUCGCGAAGUCGCUCUUCGAGCACGGCCGCAUCACCACGACCGAGGCCAAGGCCCGCCGCCUGCGCCCGUCGCGGAGCGCCUGAUCACCAAGGCGAAGAAGGGCGACAUCCACAACCGUCGCCUGGUGCUGCAGACGAU |
| Soc-1-1 | GGGCGGCAGCGCCGCGCACGAGCGUCUGCUGCUCGCGAACCUCGCGAAGUCGCUGUUCGAGCACGGCCGCAUCACCACGACCGAGGCCAAGGCCCGCCGCCUGCGCCCGUCGCGGAGAAGCUGAUCACCAAGGCGAAGAAGGGCGACAUCCACAACCGUCGCCUGGUGCUGGCCACGAU |
| Sni-2-1 | GGGCGGCAGCGCCGCGCACGAGCGUCUGCUGCUCGCGAACCUCGCGAAGUCGCUGUUCGAGCACGGCCGCAUCACCACGACCGAGGCCAAGGCCCGCCGCCUGCGCCCGUCGCGGAGAAGCUGAUCACCAAGGCGAAGAAGGGCGACAUCCACAACCGUCGCCUGGUGCUGGCCACGAU |
| Svi-11-1 | GGGCGGCAGCGCCGCGCACGAGCGUCUGCUGCUCGCGAACCUCGCGAAGUCGCUGUUCGAGCACGGCCGCAUCACCACGACCGAGGCCAAGGCCCGCCGCCUGCGCCCGUCGCGGAGAAGCUGAUCACCAAGGCGAAGAAGGGCGACAUCCACAACCGUCGCCUGGUGCUGGCCACGAU |
| Sex-2-1 | CGGCGGUUCCGCCGCGCACGAGCGUCUGCUGCUCGCCAACCUGGCGAAGUCCCUCUUCGAGCACGGCCGCAUCACGACGACCGAGGCCAAGGCCCGUCGCCUGCGUCCGUCGCGGAGCGCCUGAUCUCCAAGGCGAAGAAGGGCGACAUCCACAACCGUCGCCUGGUGCUGCAGACGAU |
| Ssp-6-1 | GGGCGGCAGUGCCGCGCACGAGAAGCUGCUCCUCGCGAACCUCGCGAAGGCGCUCUUCGAGCACGGCCGCAUCACCACCACCGAGGCGAAGGCGCGCAAGCUGCGUCCGUACGCCGAGCGUCUGAUCACCAAGGCGAAGAAGGGCGACCUUCACAACCGCCGCCAGGUGCUCCAGGUCAU |
| Ssp-101-1 | GGGCGGCAGCGCCGCGCACGAGAAGCUGCUCCUCGCGAACCUCGCGAAGGCGCUGUUCGAGCACGGCCGUAUCACCACCACCGAGGCGAAGGCGCGCAAGCUGCGUCCGUACGCCGAGCGUCUGGUCACCAAGGCGAAGAAGGGCGACCUUCACAACCGCCGUCAGGUGCUCCAGGUCAU |
| Ssp-49-1 | GGGCGGCAGUGCCGCGCACGAGAAGCUGCUCCUCGCGAACCUCGCGAAGGCGCUCUUCGAGCACGGCCGCAUCACCACCACCGAGGCGAAGGCGCGCCGUCUGCGUCCCUACGCCGAGCGUCUGAUCACCAAGGCGAAGAAGGGCGACCUUCACAACCGCCGCCAGGUGCUCCAGGUCAU |
| Sal-11-1 | UGGCGGCAGUGCCGCGCACGAGAAGCUGCUCCUCGCGAACCUCGCGAAGUCGCUCUUCGAGCACGGCCGCAUCACCACCACCGAGGCGAAGGCGCGCCGUCUGCGGCCCUACGCCGAGCGUCUGGUCACCAAGGCGAAGAAGGGCGACCUUCACAACCGCCGCCAGGUGCUCCAGGUCAU |
| Shy-2-1 | GGGCGGCAGCGCCGCGCACGAGAAGCUCCUCCUCGCGAACCUGGCGAAGAGCCUCUUCGAGCACGGUCGCAUCACCACCACCGAGGCGAAGGCGCGCCGUCUGCGGCCCGUACGCCGAGCGUCUGAUCACCAAGGCGAAGAAGGGCGACCUUCACAACCGCCGUCAGGUGCUCCAGGUCAU |
| Ssp-46-1 | GGGCGGCAGCGCCGCGCACGAGAAGCUGCUCCUCGCGAACCUCGCGAAGAGCCUCUUCGAGCACGGCCGGAUCACCACCACCGAGGCGAAGGCGCGCAAGCUGCGUCCGUACGCCGAGCGUCUGGUCACCAAGGCGAAGAAGGGCGACCUUCACAACCGCCGUCAGGUGCUCCAGGUGAU |
| Ssp-187-1 | GGGCGGCAGCGCCGCGCACGAGAAGCUGCUCCUCGCGAACCUCGCGAAGAGCCUCUUCGAGCACGGCCGGAUCACCACCACCGAGGCGAAGGCGCGCAAGCUGCGUCCGUACGCCGAGCGUCUGGUCACCAAGGCGAAGAAGGGCGACCUUCACAACCGCCGUCAGGUGCUCCAGGUGAU |
| Ssp-156-1 | GGGCGGCAGCGCCGCGCACGAGAAGCUCCUCCUCGCGAACCUCGCGAAGAGCCUCUUCGAGCACGGCCGCAUCACCACCACCGAGGCGAAGGCCCGCCGCCUGCGUCCGUACGCCGAGCGUCUGGUCACCAAGGCGAAGAAGGGCGACCUUCACAACCGCCGUCAGGUGCUCCAGGUCAU |
| Sgr-10-1 | GGGCGGCAGCGCCGCGCACGAGAAGCUCCUCCUCGCGAACCUCGCGAAGGCGCUCUUCGAGCACGGCCGCAUCACCACCACCGAGGCGAAGGCGCGCAAGCUGCGUCCGUACGCCGAGCGUCUGGUCACCAAGGCGAAGAAGGGCGACCUUCACAACCGCCGCCAGGUGCUCCAGGUCAU |
| Aac-1-1 | GGGCGGCAGCGCCGCGCACGAGAAGCUCCUCCUCGCGAACCUCGCGAAGGCGCUCUUCGAGCACGGCCGCAUCACCACCACCGAGGCGAAGGCGCGCAAGCUGCGUCCCUACGCCGAGCGUCUGGUCACCAAGGCGAAGAAGGGCGACCUUCACAACCGCCGCCAGGUGCUCCAGGUGAU |
| Ssp-197-1 | GGGCGGCAGCGCCGCGCACGAGAAGCUUCUCCUCGCGAACCUCGCGAAGGCGCUUUUCGAGCACGGCCGCAUCACCACCACCGAGGCGAAGGCGCGCAAGCUGCGUCCCUACGCCGAGCGUCUGGUCACCAAGGCGAAGAAGGGCGACCUUCACAACCGCCGUCAGGUGCUCCAGGUGAU |
| Ssp-20-1 | GGGCGGCAGCGCCGCGCACGAGAAGCUUCUCCUCGCGAACCUCGCGAAGGCGCUUUUCGAGCACGGCCGCAUCACCACCACCGAGGCGAAGGCGCGCAAGCUGCGUCCCUACGCCGAGCGUCUGGUCACCAAGGCGAAGAAGGGCGACCUUCACAACCGCCGUCAGGUGCUCCAGGUGAU |
| Ssp-123-1 | GGGCGGCAGCGCCGCGCACGAGAAGCUCAUGCUCGCGAACCUCGCGAAGGCGCUUUUCGAGCACGGCCGCAUCACCACCACCGAGGCGAAGGCGCGCAAGCUGCGUCCGUACGCCGAGCGUCUGGUCACCAAGGCGAAGAAGGGCGACCUUCACAACCGCCGUCAGGUGCUCCAGGUCAU |
| Ssp-141-1 | GGGCGGCAGCGCCGCGCACGAGAAGCUCAUGCUCGCGAACCUCGCGAAGGCGCUUUUCGAGCACGGCCGCAUCACCACCACCGAGGCGAAGGCGCGCAAGCUGCGUCCGUACGCCGAGCGUCUGGUCACCAAGGCGAAGAAGGGCGACCUUCACAACCGCCGUCAGGUGCUCCAGGUCAU |
| Ssp-81-1 | GGGCGGCAGCGCCGCGCACGAGAAGCUGCUCCUCGCGAACCUCGCGAAGAGCCUCUUCGAGCACGGCCGGAUCACCACCACCGAGGCGAAGGCGCGCAAGCUGCGUCCGUACGCCGAGCGUCUGGUCACCAAGGCGAAGAAGGGCGACCUUCACAACCGCCGUCAGGUGCUCCAGGUGAU |
| Ssp-113-1 | GGGCGGCAGCGCCGCGCACGAGAAGCUGCUCCUCGCGAACCUCGCGAAGAGCCUCUUCGAGCACGGCCGGAUCACCACCACCGAGGCGAAGGCGCGCAAGCUGCGUCCGUACGCCGAGCGUCUGGUCACCAAGGCGAAGAAGGGCGACCUUCACAACCGCCGUCAGGUGCUCCAGGUGAU |
| Ssp-57-1 | GGGCGGCAGUGCCGCGCACGAGAAGCUGCUUCUCGCGAACCUCGCCAAGGCGCUGUUCGAGCACGGCCGCAUCACCACCACCGAGGCGAAGGCGCGCAAGCUGCGUCCGUACGCCGAGCGUCUGGUCACCAAGGCGAAGAAGGGCGACCUUCACAACCGCCGUCAGGUGCUCCAGGUGAU |
| Ssp-149-1 | GGGCGGCAGCGCCGCGCACGAGAAGCUGCUCCUCGCGAACCUGGCGAAGAGCCUCUUCGAGCACGGCCGUAUCACCACCACCGAGGCGAAGGCGCGCCGUCUGCGUCCGUACGCCGAGCGUCUGAUCACCAAGGCGAAGAAGGGCGACCUUCACAACCGCCGCCAGGUGCUUCAGGUCAU |
| Sah-1-1 | GGGCGGCAGCGCUUCCCACGAGAAGGCGCUCCUGCGCAACCUGUCGACCAGCCUCUUCGAGCACGGCCGCAUCACGACGACCGAGGCCAAGGCCCGCCGUCUGCGUCCGUUCGCGGAGCGCCUGGUGACCAAGGCGAAGAAGGGCGACCUUCACAACCGCCGCCAGGUCAUGGAGCUGAU |
| Svi-10-1 | GGGCGGCAGCGCCGCGCACGAGAAGCUGCUCCUCGCGAACCUGGCGAAGAGCCUCUUCGAGCACGGCCGGAUCACCACCACCGAGGCGAAGGCGCGCAAGCUGCGUCCGUACGCCGAGCGUCUGGUCACCAAGGCGAAGAAGGGCGACCUUCACAACCGCCGCCAGGUGCUCCAGGUGAU |
| Sch-1-1 | GGGCGGCAGCGCCGCGCACGAGAAGCUGAUGCUCGCGAACCUCGCGAAGUCGCUGUUCGAGCACGGCCGGAUCACCACCACCGAGGCGAAGGCGCGCAAGCUGCGUCCGUACGCCGAGCGUCUGGUCACCAAGGCGAAGAAGGGCGACCUUCACAACCGCCGUCAGGUGCUCCAGGUCAU |
| Sia-1-1 | GGGCGGCAGCGCCGCGCACGAGAAGCUGAUGCUCGCGAACCUCGCGAAGUCGCUCUUCGAGCACGGCCGCAUCACCACCACCGAGGCCAAGGCCCGCAAGCUGCGGCCGUACGCCGAGCGUCUGGUCACCAAGGCGAAGAAGGGCGACCUUCACAACCGCCGUCAGGUGCUCCAGGUCAU |
| Saf-1-1 | GGGCGGCAGCGCCGCGCACGAGAAGCUGCUCCUCGCGAACCUCGCGAAGGCGCUGUUCGAGCACGGCCGCAUCACCACCACCGAGGCGAAGGCGCGCAAGCUGCGUCCGUACGCCGAGCGUCUGAUCACCAAGGCGAAGAAGGGCGACCUUCACAACCGCCGCCAGGUGCUCCAGGUCAU |
| Ssp-89-1 | GGGCGGCAGCGCCGCGCACGAGAAGCUGCUCCUCGCGAACCUCGCGAAGUCGCUCUUCGAGCACGGCCGGAUCACCACCACCGAGGCGAAGGCGCGCAGGCUGCGUCCGUACGCCGAGCGCCUGGUCACCAAGGCGAAGAAGGGCGACCUUCACAACCGCCGCCAGGUGCUCCAGGUCAU |
| Sva-1-1 | GGGCGGCGGCGCCGCGCACGAGAAGCUGCUCCUCGCGAACCUCGCGAAGGCGCUGUUCGAGCACGGUCGCAUCACCACCACCGAGGCGAAGGCCCGCAGGCUGCGGCCGUACGCCGAGCGUCUGGUCACCAAGGCGAAGAAGGGCGACCUUCACAACCGCCGCCAGGUGCUCCAGGUCAU |
| Ssp-158-1 | GGGCGGCAGCGCCGCGCACGAGAAGCUCCUCCUCGCGAACCUCGCGAAGGCGCUGUUCGAGCACGGCCGCAUCACCACCACCGAGGCGAAGGCGCGCAAGCUGCGUCCGUACGCCGAGCGUCUGGUCACCAAGGCGAAGAAGGGCGACCUUCACAACCGCCGUCAGGUGCUCCAGGUGAU |
| Sro-5-1 | GGGCGGCGGUGCCGCGCACGAGAAGCUGAUGCUGGCGAACCUCGCCAAGUCGCUCUUCGAGCACGGCCGCAUCACGACGACCGAGGCCAAGGCCCGCCGCCUGCGUCCGGUCGCGGAGCGCCUGAUCACCAAGGCGAAGAAGGGCGACAUCCACAACCGUCGCCAGGUGCUGCAGACGAU |
| Ssp-116-1 | GGGCGGCAGCGCCGCGCACGAGAAGCUGCUCCUCGCGAACCUCGCGAAGUCGCUCUUCGAGCACGGCCGCAUCAAGACGACCGAGUCGAAGGCCCGCCGUCUGCGGCCCGUACGCCGAGCGUCUGAUCACCAAGGCGAAGAAGGGCGACCUUCACAACCGCCGCCAGGUGCUCCAGGUCAU |
| env-22 | GGGCGGCAGUGCCGCGCACGAGAAGCUGCUUCUCGCGAACCUCGCGAAGUCGCUCUUCGAGCACGGCCGUAUCACCACCACCGAGGCGAAGGCCCGCCGCCUGCGGCCGUACGCCGAGCGUCUGGUCACCAAGGCGAAGAAGGGCGACCUUCACAACCGCCGUCAGGUGCUCCAGGUCAU |
| Sam-1-1 | GGGCGGCAGCGCCGCGCACGAGAAGCUGCUCCUCGCGAACCUCGCGAAGUCGCUCUUCGAGCACGGCAAGAUCAAGACCACCGAGGCGAAGGCGCGCAAGCUGCGUCCGUACGCCGAGCGUCUGGUCACCAAGGCGAAGAAGGGCGACCUUCACAACCGCCGUCAGGUGCUCCAGGUGAU |
| Ssp-51-1 | GGGCGGCAGUGCCGCGCACGAGAAGCUGCUCCUCGCGAACCUCGCGAAGUCGCUCUUCGAGCACGGCCGCAUCACCACCACCGAGGCGAAGGCCCGUCGCCUGCGCCCGUACGCCGAGCGUCUGGUCACCAAGGCGAAGAAGGGUGACCUUCACAACCGCCGCCAGGUGCUGUCGGUGAU |
| Slu-1-1 | GGGCGGCAGCGCCGCGCACGAGAAGCUGCUCCUCGCGAACCUCGCGAAGGCGCUGUUCGAGCACGGCCGGAUCACCACCACCGAGGCGAAGGCGCGCAAGCUGCGUCCGUACGCCGAGCGUCUGGUCACCAAGGCGAAGAAGGGCGACCUUCACAACCGCCGUCAGGUGCUCCAGGUGAU |
| Sna-2-1 | GGGCGGCAGCGCGUCGCACGAGAAGCGAUGCUGCGCAACCUGGCGACCAGCCUCUUCGAGCACGGCCGCAUCACGACGACCGAGGCCAAGGCCCGUCGUCUGCGUCCGUACGCCGAGCGUCUGGUGACCAAGGCGAAGAAGGGCGACCUUCACAACCGCCGUCAGGUCAUGCAGCUGAU |
| Ssp-75-1 | GGGCGGCAGCGCCGCGCACGAGAAGCUCAUGCUCGCGAACCUCGCGAAGGCGCUUUUCGAGCACGGCCGCAUCACCACCACCGAGGCGAAGGCGCGCAAGCUGCGUCCCUACGCCGAGCGUCUGGUCACCAAGGCGAAGAAGGGCGACCUUCACAACCGCCGUCAGGUGCUCCAGGUGAU |
| env-23 | GGGCGGCAGUGCCGCGCACGAGAAGCUGAUGCUCGCGAACCUCGCGAAGUCGCUGUUCGAGCACGGCCGCAUCACCACCACCGAGGCGAAGGCCCGCCGUCUGCGGCCGUACGCCGAGCGUCUGAUCACCAAGGCGAAGAAGGGCGACCUUCACAACCGCCGCCAGGUGCUCCAGGUCAU |
| Ssp-182-1 | GGGCGGCAGCGCCGCGCACGAGAAGCUGCUCCUCGCGAACCUGGCGAAGAGCCUCUUCGAGCACGGCCGUAUCACCACCACCGAGGCGAAGGCGCGCCGUCUGCGUCCGUACGCCGAGCGUCUGAUCACCAAGGCGAAGAAGGGCGACCUUCACAACCGCCGCCAGGUGCUUCAGGUCAU |
| Scy-2-1 | GGGCGGCAGCGCCGCGCACGAGAAGCUGAUGCUCGCGAACCUCGCGAAGUCGCUCUUCGAGCACGGCCGUAUCACCACGACCGAGGCGAAGGCCCGCCGUCUGCGGCCGUACGCCGAGCGUCUGAUCACCAAGGCGAAGAAGGGCGACCUUCACAACCGCCGUCAGGUGCUCCAGGUCAU |
| Ssp-184-1 | GGGCGGCAGCGCCGCGCACGAGAAGUUGCUCCUCGCGAACCUGGCGAAGAGCCUCUUCGAGCACGGCCGUAUCACCACCACCGAGGCGAAGGCGCGCCGUCUGCGUCCGUACGCCGAGCGUCUGAUCACCAAGGCGAAGAAGGGCGACCUUCACAACCGCCGCCAGGUGCUUCAGGUCAU |
| Svi-4-1 | GGGCGGCAGCGCCGCGCACGAGAAGCUCCUCCUCGCGAACCUCGCGAAGAGCCUCUUCGAGCACGGCCGUAUCACCACCACCGAGGCGAAGGCCCGCCGUCUGCGGCCGUACGCCGAGCGUCUGAUCACCAAGGCGAAGAAGGGCGACCUUCACAACCGCCGUCAGGUGCUCCAGGUCAU |
| Ssp-163-1 | GGGCGGCAGCGCCGCGCACGAGAAGCUGCUCCUCGCGAACCUCGCGAAGGCGCUCUUCGAGCACGGCCGCAUCACCACCACCGAGGCGAAGGCGCGCAAGCUGCGUCCGUACGCCGAGCGUCUGGUCACCAAGGCGAAGAAGGGCGACCUUCACAACCGCCGUCAGGUGCUCCAGGUCAU |
| Ssp-177-1 | GGGCGGCAGCGCCGCGCACGAGAAGCUGAUGCUCGCGAACCUCGCGAAGUCGCUGUUCGAGCACGGUCGUAUCACCACCACCGAGGCCAAGGCGCGCAAGCUGCGGCCGUACGCCGAGCGUCUGGUCACCAAGGCGAAGAAGGGCGACCUUCACAACCGCCGUCAGGUGCUCCAGGUCAU |
| Sol-1-1 | GGGCGGCAGCGCCGCGCACGAGAAGCUGAUGCUCGCGAACCUCGCGAAGUCGCUGUUCGAGCACGGUCGUAUCACCACCACCGAGGCCAAGGCGCGCAAGCUGCGGCCGUACGCCGAGCGUCUGGUCACCAAGGCGAAGAAGGGCGACCUUCACAACCGCCGUCAGGUGCUCCAGGUCAU |
| Svi-3-1 | GGGCGGCAGCGCCGCGCACGAGAAGCUGAUGCUCGCGAACCUCGCGAAGUCGCUCUUCGAGCACGGCCGCAUCACCACCACCGAGGCGAAGGCUCGCCGCCUGCGUCCCUACGCCGAGCGUCUGGUCACCAAGGCGAAGAAGGGCGACCUUCACAACCGCCGUCAGGUGCUCCAGGUCAU |
| Szi-1-1 | GGGCGGCAGCGCCGCGCACGAGAAGCUCCUCCUCGCGAACCUCGCGAAGGCGCUCUUCGAGCACGGCCGCAUCACCACCACGGAGGCGAAGGCCCGCCGCCUGCGCCCGUACGCCGAGCGUCUGGUCACCAAGGCGAAGAAGGGCGACCUUCACAACCGCCGUCAGGUGCUCCAGGUGAU |
| Svi-13-2 | GGGCGGCAGCGCCGCGCACGAGAAGCUGCUCCUCGCGAACCUCGCGAAGGCGCUCUUCGAGCACGGCCGUAUCACCACCACCGAGGCGAAGGCCCGCCGUCUGCGGCCGUACGCCGAGCGUCUGGUCACCAAGGCGAAGAAGGGCGACCUUCACAACCGCCGUCAGGUGCUCCAGGUCAU |
| Ssp-120-1 | GGGCGGCAGUGCCGCGCACGAGAAGCUGCUUCUCGCGAACCUCGCGAAGGCGCUCUUCGAGCACGGCCGUAUCACCACCACCGAGGCGAAGGCCCGCCGUCUGCGGCCCUACGCCGAGCGUCUGGUGACCAAGGCGAAGAAGGGCGACCUUCACAACCGCCGUCAGGUGCUCCAGGUCAU |
| Ssp-131-1 | CGGCGGUUCCGCCGCGCACGAGAAGCUGCUCCUCGCCAACCUGGCGAAGUCGCUGUUCGAGCACGGCCGCAUCACGACGACCGAGGCCAAGGCCCGCCGUCUGCGUCCCGUCGCGGAGCGCCUGAUCACCAAGGCGAAGAAGGGCGACAUCCACAACCGUCGCCUGGUGCUGCAGACGAU |
| Sca-4-1 | GGGCGGCAGCGCCGCGCACGAGAAGCUGAUGCUGGCGAACCUCGCCCGCUCGCUCUUCGAGAACGGCCGCAUCACCACCACCGAGGCCAAGGCCCGCCGCCUGCGCCCGGUCGCGGAGCGCCUGAUCACCAAGGCGAAGAAGGGCGACAUCCACAACCGUCGCCAGGUGCUGCAGACGAU |
| Sba-1-1 | GGGCGGCAGCGCCGCGCACCAGAAGGCCAUGCUGGCGAACCUCGCCACGAACCUUUUUGAGCACGGCCGGAUCACCACCACCGAGGCCAAGGCCCGUCGCCUCCGCCCGGUGGCGGAGCGUCUGAUCACCAAGGCGAAGAAGGGUGACCUUCACAACCGCCGCCAGGUCAUGCGCACGGU |

.........................................................................................................................................................................

.........................................................................................................................................................................

•GGCGGCAGCRC•GCGCAC•AG•R•CU••U•CU•RY•AACCU•GC•A••••GCU•UUCGAGCACGGCCG•AUCAC•AC•ACCGAGGC•AAGGCCCGYCG•CUGCGYCCG••CGC•GAGCR•CUGRUCACCAAGGCGAAGAAGGGCGAC•UYCACAACCGYCGYC•GGU•YU•YRGRY•AU

```
Ssp-65-1   GGGCGGCAGCGCCGCGCACCAGAAGGCCAUGCUGGCGAACCUCGCCACCGACCUUUUCGAGCACGGCCGCAUCACCACCACCGAGGCGAAGGCCCGCCGCCUGCGCCCGGUCGCGGAGCGUCUGAUCACCAAGGCGAAGAAGGGCGACCUUCACAACCGCCGCCAGGUCAUGCGCACGGU
Sme-2-1    CGGUGGCGGCCCCGCGCACGAGAAGGCCAUGCUGGCGAACCUCGCCACCAACCUCUUCGAGCACGGCCGCAUCAAGACCACCGAGGCCAAGGCCCGUCGGCUGCGCCCGGUGGCGGAGCGUCUGAUCACCAAGGCGAAGAAGGGCGACCUUCACAACCGCCGCCAGGUGAUGCGGACGAU
Sgr-12-3   GGGCGGCAGCGCCGCGCACGAGCGGCUCAUGCUCGCGAACCUCGCGCGCUCGCUGUUCGAGCACGGCCGGAUCAAGACGACCGAGGCGAAGGCGCGCCGUCUGCGUCCCUACGCCGAGCGUCUGGUCACCAAGGCGAAGAAGGGCGACCUUCACAACCGCCGCCAGGUUCUCCAGGUCAU
Sgr-12-4   GGGCGGCAGCGCCGCGCACGAGCGGCUCAUGCUCGCGAACCUCGCGCGCUCGCUGUUCGAGCACGGCCGGAUCAAGACGACCGAGGCGAAGGCGCGCCGUCUGCGUCCCUACGCCGAGCGUCUGGUCACCAAGGCGAAGAAGGGCGACCUUCACAACCGCCGCCAGGUUCUCCAGGUCAU
Ssp-15-1   GGGCGGCAGCGCCGCGCACGAGCGGCUCAUGCUCGCGAACCUCGCGCGCUCGCUGUUCGAGCACGGCCGGAUCAAGACGACCGAGGCGAAGGCGCGCCGUCUGCGUCCCUACGCCGAGCGUCUGGUCACCAAGGCGAAGAAGGGCGACCUUCACAACCGCCGCCAGGUUCUCCAGGUCAU
Ssp-38-1   GGGCGGCAGCGCCGCGCACGAGCGGCUCAUGCUCGCGAACCUCGCGCGCUCGCUGUUCGAGCACGGCCGGAUCAAGACGACCGAGGCGAAGGCGCGCCGUCUGCGUCCCUACGCCGAGCGUCUGGUCACCAAGGCGAAGAAGGGCGACCUUCACAACCGCCGCCAGGUUCUCCAGGUCAU
Sal-5-1    GGGCGGCAGCGCCGCGCACGAGCGGCUCAUGCUCGCGAACCUCGCGCGCUCGCUGUUCGAGCACGGCCGGAUCAAGACGACCGAGGCGAAGGCGCGCCGUCUGCGUCCCUACGCCGAGCGUCUGGUCACCAAGGCGAAGAAGGGCGACCUUCACAACCGCCGCCAGGUUCUCCAGGUCAU
Ssp-77-1   GGGCGGCAGCGCCGCGCACGAGCGGCUCAUGCUCGCGAACCUCGCGCGCUCGCUGUUCGAGCACGGCCGGAUCAAGACGACCGAGGCGAAGGCGCGCCGCCUGCGUCCGUACGCCGAGCGUCUGGUCACCAAGGCGAAGAAGGGCGACCUUCACAACCGCCGUCAGGUUCUCCAGGUCAU
Swa-1-1    GGGCGGCAGCGCCGCGCACGAGCGGCUCAUGCUCGCGAACCUCGCGCGCUCGCUGUUCGAGCACGGCCGGAUCAAGACGACCGAGGCGAAGGCGCGCCGUCUGCGUCCCUACGCCGAGCGUCUGGUCACCAAGGCGAAGAAGGGCGACCUUCACAACCGCCGCCAGGUUCUCCAGGUCAU
Ssp-30-1   GGGCGGCAGCGCCGCGCACGAGCGGCUCAUGCUCGCGAACCUCGCGCGCUCGCUGUUCGAGCACGGCCGGAUCAAGACGACCGAGGCGAAGGCGCGCCGUCUGCGUCCCUACGCCGAGCGUCUGGUCACCAAGGCGAAGAAGGGCGACCUUCACAACCGCCGCCAGGUUCUCCAGGUCAU
Ssp-134-1  GGGCGGCAGCGCCGCGCACGAGCGGCUCAUGCUCGCGAACCUCGCGCGCUCGCUGUUCGAGCACGGCCGGAUCAAGACGACCGAGGCGAAGGCGCGCCGUCUGCGUCCCUACGCCGAGCGUCUGGUCACCAAGGCGAAGAAGGGCGACCUUCACAACCGCCGCCAGGUUCUCCAGGUCAU
Ssp-62-1   GGGCGGCAGCGCCGCGCACGAGCGGCUCAUGCUCGCGAACCUCGCGCGCUCGCUGUUCGAGCACGGCCGGAUCAAGACGACCGAGGCGAAGGCGCGCCGUCUGCGUCCGUACGCCGAGCGUCUGGUCACCAAGGCGAAGAAGGGCGACCUUCACAACCGUCGCCAGGUUCUCCAGGUCAU
Ssp-21-1   GGGCGGCAGCGCCGCGCACGAGCGGCUCAUGCUCGCGAACCUCGCGCGCUCGCUGUUCGAGCACGGCCGGAUCAAGACGACCGAGGCGAAGGCGCGCCGUCUGCGUCCGUACGCCGAGCGUCUGGUCACCAAGGCGAAGAAGGGCGACCUUCACAACCGUCGCCAGGUUCUCCAGGUCAU
Ssp-80-1   GGGCGGCAGCGCCGCGCACGAGCGGCUCAUGCUCGCGAACCUCGCGCGCUCGCUGUUCGAGCACGGCCGGAUCAAGACGACCGAGGCGAAGGCGCGCCGUCUGCGUCCGUACGCCGAGCGUCUGGUCACCAAGGCGAAGAAGGGCGACCUUCACAACCGCCGUCAGGUUCUCCAGGUCAU
Sgr-12-5   GGGCGGCAGCGCCGCGCACGAGCGGCUCAUGCUCGCGAACCUCGCGCGCUCGCUGUUCGAGCACGGCCGGAUCAAGACGACCGAGGCGAAGGCGCGCCGUCUGCGUCCGUACGCCGAGCGUCUGGUCACCAAGGCGAAGAAGGGCGACCUUCACAACCGCCGUCAGGUUCUCCAGGUCAU
Sva-2-1    GGGCGGCAGCGCCGCGCACGAGCGUCACCUGCUCGCGAACCUCGCGAAGUCGCUGUUCGAGCACGGCCGCAUCACCACCACCGAGGCCAAGGCGCGCCGGCUCCGUCCGGUCGCCGAGCGUCUGGUGACCAAGGCGAAGAAGGGCGACAUCCACAACCGUCGCCAGGUUCUUCAGACGAU
Ssp-58-1   GGGCGGCAGUGCCGCGCACCAGAAGGCGAUGCUGGGGAACCUCGCGACCAGCCUCUUCGAGCACGGCCGCAUCACCACGACCGAGGCGAAGGCACGCCGCCUGCGCCCGUUCGCGGAGCAGCUGAUCACCAAGGCGAAGAAGGGCGACCUUCACAACCGGCGGCUGGUCAUGCGUACGGU
Sal-12-1   GGGCGGCAGCGCCGCGCACGAGCGGCUGAUGCUGGCGAACCUCGCCACCGCGCUCUUCGAGCACGGCCGCAUCACCACCACCGAGGCCAAGGCCCGCCGGCUGCGCCCGGUGGCCGAGCGUCUGAUCACCAAGGCGAAGAAGGGUGACCUUCACAACCGCCGCCAGGUGCUCCAGACGGU
Sfr-1-1    GGGCGGCAGCGCCGCGCACGAGCGGCUGAUGCUGGCGAACCUCGCCACCGCGCUCUUCGAGCACGGCCGCAUCACCACCACCGAGGCCAAGGCCCGCCGGCUGCGCCCGGUGGCCGAGCGUCUGAUCACCAAGGCGAAGAAGGGCGACCUUCACAACCGCCGCCAGGUGCUCCAGACGGU
Sxi-2-1    GGGCGGCAGCGCCGCGCACGAGCGGCUGAUGCUGGCGAACCUCGCCACCGCGCUCUUCGAGCACGGCCGCAUCACCACCACCGAGGCCAAGGCCCGCCGGCUGCGCCCGGUGGCCGAGCGUCUGAUCACCAAGGCGAAGAAGGGCGACCUUCACAACCGCCGCCAGGUGCUCCAGACGGU
Ssp-119-1  GGGCGGCAGCGCCGCGCACGAGAAGCUGCUCCUCGCGAACCUCGCGAAGGCGCUCUUCGAGCACGGCCGCAUCACCACGACCGAGGCCAAGGCCCGCCGCCUGCGCCCCGUCGCGGAGCGCUUCAUCACCAAGGCGAAGAAGGGCGACAUCCACAACCGUCGCCAGGUGCUGCGGUCGAU
Ssp-33-1   GGGCGGCAGCGCUGCGCACGAGAAGGCGCUCCUGCGCAACCUGGCGACCAACCUCUUCGAGCACGGCCGCAUCCGGACGACCGAGGCCAAGGCCCGUCGUCUGCGUCCGUUCGCGGAGCGUCUGGUGACCAAGGCGAAGAAGGGCGACCUUCACAACCGCCGCCAGGUCAUGGAGCUGAU
Ssp-37-1   GGGCGGCAGCGCCGCGCACGAGAAGCUGCUCAUCGCGAACCUCGCGAAGAGCCUCUUCGAGCACGGCCGUAUCACCACCACCGAGGCGAAGGCCCGCAAGCUGCGGCCGUACGCCGAGCGUCUGGUCACCAAGGCGAAGAAGGGCGACCUUCACAACCGCCGUCAGGUGCUCCAGGUGAU
Ssp-170-1  GGGCGGCAGUGCCGCGCACGAGAAGCUGCUCCUCGCGAACCUCGCGAAGGCGCUGUUCGAGCACGGCCGUAUCACCACCACCGAGGCGAAGGCCCGCCGCCUGCGCCCGUACGCCGAGCGUCUGGUCACCAAGGCGAAGAAGGGCGACCUUCACAACCGCCGUCAGGUGCUCCAGGUCAU
Smi-1-1    GGGCGGCAGCGCCGCGCACGAGAAGCUGCUCCUCGCGAACCUCGCGAAGAGCCUCUUCGAGCACGGCCGUAUCACCACCACCGAGGCGAAGGCGCGCAAGCUGCGCCCGUACGCCGAGCGUCUGGUCACCAAGGCGAAGAAGGGCGACCUUCACAACCGCCGUCAGGUGCUCCAGGUGAU
Ssp-138-1  GGGCGGCAGCGCCGCGCACGAGAAGCUCCUCCUCGCCAACCUGGCGAAGAGCCUCUUCGAGCACGGCCGUAUCACCACCACCGAGGCGAAGGCCCGCCGUCUGCGGCCGUACGCCGAGCGUCUGGUCACCAAGGCGAAGAAGGGCGACCUUCACAACCGCCGUCAGGUGCUCCAGGUCAU
Sga-2-1    GGGCGGCAGUGCCGCGCACGAGAAGCUGCUGCUCGCGAACCUCGCGAAGUCGCUCUUCGAGCACGGCCGGAUCACCACCACCGAGGCGAAGGCCCGCCGUCUGCGGCCCUACGCCGAGCGUCUGGUCACCAAGGCGAAGAAGGGCGACCUUCACAACCGCCGCCAGGUGCUCCAGGUCAU
env-24     GGGCGGCAGCGCCGCGCACGAGAAGCUGCUCCUCGCGAACCUCGCGAAGGCGCUCUUCGAGCACGGCCGUAUCACCACCACCGAGGCGAAGGCCCGCCGUCUGCGGCCGUACGCCGAGCGUCUGGUGACCAAGGCGAAGAAGGGCGACCUUCACAACCGCCGCCAGGUGCUGCAGGUCAU
Sca-1-1    GGGCGGCAGCGCCGCGCACGAGAAGCUGCUCCUCGCGAACCUGGCGAAGAGCCUCUUCGAGCACGGCCGUAUCACCACCACCGAGGCGAAGGCGCGCAAGCUGCGUCCGUACGCCGAGCGUCUGGUCACCAAGGCGAAGAAGGGCGACCUUCACAACCGCCGUCAGGUGCUCCAGGUGAU
Ssp-23-1   GGGCGGUAGUGCCGCGCACGAGAAGCUGCUCCUCGCGAACCUCGCGAAGGCGCUCUUCGAGCACGGCCGCAUCACCACCACCGAGGCGAAGGCCCGCCGUCUGCGGCCGUACGCCGAGCGUCUGGUGACCAAGGCGAAGAAGGGCGACCUUCACAACCGCCGCCAGGUGCUGCAGGUCAU
Ssp-135-1  GGGCGGCAGUGCCGCGCACGAGAAGCUGCUCCUCGCGAACCUCGCGAAGGCGCUCUUCGAGCACGGCCGCAUCACCACCACCGAGGCCAAGGCGCGCCGGCUGCGGCCGUACGCCGAGCGUCUGGUGACCAAGGCGAAGAAGGGCGACCUUCACAACCGCCGUCAGGUGUUGCAGGUCAU
Ssp-169-1  GGGCGGCAGCGCCGCGCACGAGAAGCUGCUCCUCGCGAACCUCGCGAAGUCGCUGUUCGAGCACGGCCGUAUCACCACCACCGAGGCGAAGGCCCGCCGUCUGCGGCCGUACGCCGAGCGUCUGGUGACCAAGGCGAAGAAGGGCGACCUUCACAACCGCCGCCAGGUGCUGCAGGUCAU
env-30     GGGCGGCAGCGCCGCGCACGAGAAGCUGCUCCUCGCGAACCUCGCGAAGG..................................................................................................................................
Ssv-1-1    GGGCGGCAGCGCCGCGCACGAGAAGCUGCUCCUCGCGAACCUUGCGAAGAGCCUCUUCGAGCACGGCCGUAUCACCACCACCGAGGCGAAGGCGCGCAAGCUGCGUCCGUACGCCGAGCGUCUGGUCACCAAGGCGAAGAAGGGCGACCUUCACAACCGCCGUCAGGUGCUCCAGGUGAU
Ssp-186-1  GGGCGGCAGCGCCGCGCACGAGAAGCUGCUCCUCGCGAACCUCGCGAAGAGCCUCUUCGAGCACGGCCGUAUCACCACCACCGAGGCGAAGGCGCGCAAGCUGCGUCCCUACGCCGAGCGUCUGGUCACCAAGGCGAAGAAGGGCGACCUUCACAACCGCCGUCAGGUGCUCCAGGUGAU
Sda-1-1    UGGGCGGCAGCGCCGCGCACGAGAAGCUGCUCCUCGCGAACCUCGCGAAGGCGCUCUUCGAGCACGGCCGCAUCACCACCACCGAGGCGAAGGCCCGCCGCCUGCGGCCGUACGCCGAGCGUCUGGUCACCAAGGCGAAGAAGGGCGACCUUCACAACCGCCGUCAGGUGCUCCAGGUGA
Sgr-12-6   GGGCGGCAGUGCCGCGCACGAGAAGCUGCUGCUCGCGAACCUCGCGAAGAGCCUCUUCGAGCACGGCCGCAUCACCACCACCGAGGCGAAGGCCCGCCGCCUGCGCCCGUACGCCGAGCGUCUGGUCACCAAGGCGAAGAAGGGCGACCUUCACAACCGCCGUCAGGUGCUCCAGGUGAU
Ssp-199-1  GGGCGGCAGCGCCGCGCACGAGAAGCUCCUCCUCGCGAACCUCGCGAAGAGCCUCUUCGAGCACGGCCGUAUCACCACCACCGAGGCGAAGGCGCGCAAGCUGCGCCCGUACGCCGAGCGUCUGGUCACCAAGGCGAAGAAGGGCGACCUUCACAACCGCCGUCAGGUGCUCCAGGUGAU
env-31     GGGCGGCAGCGCUGCGCACGAGCGCCUCAUGCUGCGCAACCUGGCCACCGCGCUCUUCGAGCACGGCCGCAUCACGACGACCGAGGCCAAGGCCCGUCGUCUGCGUCCGUACGCGGAGCGCCUGGUGACCAAGGCGAAGAAGGGCGACCUUCACAACCGCCGUCAGGUCAUGCAGCUGAU
Ssp-19-1   GGGCGGCAGCGCUGCGCACGAGCGCCUCAUGCUGCGCAACCUGGCCACCGCGCUCUUCGAGCACGGCCGCAUCACGACGACCGAGGCCAAGGCCCGUCGUCUGCGUCCGUACGCGGAGCGUCUGGUGACCAAGGCGAAGAAGGGCGACCUUCACAACCGCCGUCAGGUCAUGCAGCUGAU
Sfu-2-1    GGGCGGCAGCGCCGCGCACGAGAAGCUGCUCCUCGCGAACCUCGCGAAGAGCCUCUUCGAGCACGGCCGUAUCACCACCACCGAGGCGAAGGCGCGCAAGCUGCGUCCGUACGCCGAGCGUCUGGUCACCAAGGCGAAGAAGGGCGACCUUCACAACCGCCGUCAGGUGCUCCAGACCAU
Sau-2-1    GGGCGGCAGCGCCGCGCACGAGCGUCUGAUGCUGCGCAACCUGGCCACCGCUCUCUUCGAGCACGGCCGCAUCACGACGACCGAGGCCAAGGCCCGUCGUCUGCGUCCGUACGCGGAGCGCCUGGUGACCAAGGCGAAGAAGGCGGACCUGCACAACCGCCGCCAGGUCAUGCAGCUGAU
Sxy-1-1    GGGCGGCAGUGCCGCGCACGAGAAGCUGCUUCUCGCGAACCUCGCGAAGUCGCUCUUCGAGCACGGCCGUAUCACCACCACCGAGGCGAAGGCCCGCAAGCUGCGGCCGUACGCCGAGCGUCUGAUCACCAAGGGCAAGAAGGGCGACCUUCACAACCGCCGUCAGGUGCUUCAGGUGAU
Ssp-122-1  GGGCGGCAGUGCCGCGCACGAGAAGCUGCUGCUCGCGAACCUUGCGAAGAGCCUCUUCGAGCACGGCCGUAUCACCACCACCGAGGCGAAGGCCCGCAAGCUGCGGCCGUACGCCGAGCGUCUGAUCACCAAGGGCAAGAAGGGCGACCUUCACAACCGCCGUCAGGUGCUUCAGGUGAU
Spr-2-1    GGGCGGCAGUGCCUCGCACGAGAAGCUGCUCCUCGCGAACCUCGCGAAGUCGCUUUUCGAGCACGGCCGUAUCACCACCACCGAGGCGAAGGCCCGCAAGCUGCGGCCGUACGCCGAGCGUCUGAUCACCAAGGGCAAGAAGGGCGACCUUCACAACCGCCGUCAGGUGCUUCAGGUGAU
Ssp-84-1   GGGCGGCAGCGCCGCGCACGAGAAGCUGCUCCUCGCGAACCUCGCGAAGUCUCUCUUCGAGCACGGCCGUAUCACCACCACCGAGGCGAAGGCCCGCCGCCUGCGCCCGUACGCCGAGCGUCUGGUCACCAAGGCGAAGAAGGGCGACCUUCACAACCGCCGUCAGGUGCUCCAGGUCAU
Sip-1-1    GGGCGGCAGCGCCGCGCACGAGAAGCUGCUUCUUGCGAACCUCGCCAAGAGCCUUUUCGAGCACGGUCGUAUCACCACCACCGAGGCGAAGGCCCGUCGGCUGCGGCCGUACGCCGAGCGCCUGAUCACCAAGGCGAAGAAGGGUGACCUUCACAACCGCCGUCAGGUGCUCUCGGUCAU
Sly-2-1    GGGCGGCAGCGCGUCGCACGAGAAGGCGAUGCUGCGCAACCUGGCGACCAGCCUCUUCGAGCACGGCCGCAUCACGACGACCGAGGCCAAGGCCCGUCGUCUGCGUCCGUACGCCGAGCGUCUGGUGACCAAGGCGAAGAAGGGCGACCUUCACAACCGCCGUCAGGUCAUGCAGCUGAU
Ssc-2-1    CGGCGGCGGCCCGUCCCACGAGCGGCUGAUCCUGGCCGCUCUCGCGACGGCGCUGUUCGAGCACGGUCGGAUCACCACGACCGAGGCCAAGGCGCGUCGGCUGCGUCCCGUCGCCGAGCGACUGAUCACCAAGGCGAAGAAGGGCGACCUGCACAACCGUCGCCAGGUGCUCACGAUCGU
Sye-1-1    CGGCGGCAGCGCCUCGCACGAGAAGCUGCUGCUCGCGAACCUCGCGAAGUCGCUCUUCGAGCACGGCCGGAUCACCACCACCGAGGCCAAGGCCCGGCGGCUGCGUCCGGUGGCCGAGCGGCUGAUCACCAAGGCGAAGAAGGGCGACAUCCACAACCGUCGCCAGGUGCUCCAGACGAU
Svi-6-1    GGGCGGCAGCCCGGCGCACGAGCGGCUGAUGCUGGCCAACCUGGCCACGUCGCUGUUCGAGCACGGCCGCAUCACCACCACCGAGGCGAAGGCCCGUCGGCUGCGCCCGGUGGCCGAGCGCCUGAUCACCAAGGCCAGGCGCGGCGACCUGCACGCCCGUCGCCAGGUGCUCGCGGUCAU
Sni-1-1    GGGCGGCAGCGCCGCGCACGAGCGGCUGCUUCUCGCGAACCUCGCGAAGGCGCUGUUCGAGCACGGCAAGAUCACGACGACCGAGGCCAAGGCCCGCCGCCUGCGUCCGGUCGCCGAGCGUCUGGUCACCAAGGCGAAGAAGGGCGACAUCCACAACCGUCGCCUCGUUCUCCAGACGAU
Ssu-2-1    CGGUGGCAGUGCCGCGCACCAGAAGGCCAUGCUGGCGAACCUCGCCACGGACCUCUUCGAGUACGGCCGUAUCACCACCACGGAGUCGCGGGCCCGGAAGCUGCGGCCGGUCGCCGAGCGGCUGAUCACCAAGGCGAAGAAGGGCGACCUGCACAACCGUCGCCAGGUCAUGCGCACCAU
Ssu-1-1    CGGUGGCAGUGCCGCGCACCAGAAGGCCAUGCUGGCGAACCUCGCCACGGACCUCUUCGAGUACGGCCGUAUCACCACCACUGAGUCGCGGGCCCGGAAGCUGCGGCCCGUCGCCGAGCGGCUGAUCACCAAGGCGAAGAAGGGCGACCUGCACAACCGUCGCCAGGUGAUGCGCACCAU
Svi-2-1    GGGCGGCAGCGCUGCGCACGAGCGGCUCCUGCUGGCGAACCUCGCCACCGCGCUGUUCGAGCACGGCCGGAUCACCACCACCGAGGCGAAGGCCCGCCGCCUGCGCCCGGUCGCGGAGAAGCUGGUCACCAAGGCGAAGAAGGGCGACCUUCACAACCGCCGUCUGGUGCUCCAGACCGU
Sfl-4-1    GGGCGGCAGCGCCGCGCACGAGCGUCUGCUCCUCGCCAACCUGGCGAAGUCGCUGUUCGAGCACGGCCGCAUCACGACGACCGAGGCCAAGGCCCGCCGCCUGCGCCCCGUCGCCGAGCGUCUGGUCACCAAGGCGAAGAAGGGCGACAUCCACAACCGUCGCCUGGUGCUGCAGACGAU
Spu-1-1    GGGCGGCAGCGCCGCGCACGAGAAGCUGCUUCUCGCCAACCUGGCGAAGUCGCUCUUCGAGCACGGCCGCAUCACCACGACCGAGGCCAAGGCCCGCCGCCUGCGUCCGGUCGCCGAGCGCCUGAUCACCAAGGCGAAGAAGGGCGACAUCCACAACCGUCGCCUGGUGCUGCAGACCAU
Ssc-4-2    GGGCGGCAGCGCUUCGCACGAGCGUCUGAUGCUGGCGAACCUCGCGACCGCGCUCUUCGAGCACGGCCGCAUCACCACGACCGAGGCCAAGGCCCGCCGCCUGCGCCCGGUCGCGGAGAAGCUGGUCACCAAGGCGAAGAAGGGCGACCUUCACAACCGCCGUCUGGUGCUCCAGACCAU
Shy-1-1    GGGCGGCAGCGCUGCGCACGAGCGGCUCCUGCUGGCGAACCUCGCCACCGCGCUGUUCGAGCACGGCCGGAUCACCACCACCGAGGCGAAGGCCCGCCGCCUGCGCCCGGUCGCGGAGAAGCUGGUCACCAAGGCGAAGAAGGGCGACCUUCACAACCGCCGUCUGGUGCUCCAGACCGU
env-32     GGGCGGCAGCGCUGCGCACGAGCGGCUCCUGCUGGCGAACCUCGCCACCGCGCUGUUCGAGCACGGCCGGAUCACCACCACCGAGGCGAAGGCCCGCCGCCUGCGCCCGGUCGCGGAGAAGCUGGUCACCAAGGCGAAGAAGGGCGACCUUCACAACCGCCGUCUGGUGCUCCAGACCGU
Sbi-1-1    GGGCGGCAGCGCGGCGCACGAGCGGCUGAUGCUGGCGAACCUCGCCACCUCGCUGUUCGAGCACGGCCGCAUCAAGACCACCGAGGCGAAGGCCCGCCGCCUGCGCCCGGUCGCGGAGAAGCUGAUCACCAAGGCGAAGAAGGGCGACCUUCACAACCGCCGUCUGGUGCUCCAGACCAU
Afl-1-1    CGGCGGUAGCGCCGCGCACCAGCGUCUGAUCCUCGCGAACCUGGCGACGUCGCUGUUCGAGCACGGCCGCAUCACCACCACCGAGGCCAAGGCCCGCCGGGUCCGCCCGCUGGCGGAGAAGCUGAUCACCAAGGCCAAGAAGGGCGACCUGCACAACCGUCGCCAGGUCGCGCAGGUCAU
Ssp-55-1   GGGCGGCAGCGGCGCGCACGAGAAGCUUCUCCUCGCGAACCUCGCGAAGUCGCUCUUCGAGCACGGCCGUAUCACCACCACCGAGGCGAAGGCCCGCCGUGUGCGGCCCGUCGCCGAGCGUCUGAUCACCAAGGCGAAGCGGGCGACAUCCACAACCGUCGCCAGGUGCUGCAGUCGAU
Ssp-152-1  GGGCGGCAGCGCCGCGCACGAGAAGCACCUUCUCGCGAACCUCGCGAAGGCGCUCUUCGAGCACGGCCGCAUCACCACCACCGAGGCCAAGGCCCGCCGCCUGCGUCCCUACGCCGAGCGUCUGGUCACCAAGGCCAAGAAGGGCGACAUCCACAACCGUCGCCUGGUGCUGCAGACGAU
Ssp-104-1  GGGCGGCUCCGCCGCGCACGAGAAGCACCUUCUCGCGAACCUCGCGAAGGCGCUCUUCGAGCACGGCCGCAUCACCACCACCGAGGCCAAGGCCCGCCGCCUGCGUCCCUACGCCGAGCGUCUGGUCACCAAGGCCAAGAAGGGCGACAUCCACAACCGUCGCCUGGUGCUGCAGACGAU
Ssp-166-1  GGGCGGCAGCGCCGCGCACGAGAAGCACCUCCUCGCGAACCUCGCGAAGGCGCUCUUCGAGCACGGCCGCAUCACCACCACCGAGGCCAAGGCCCGUCGCCUGCGUCCCUACGCCGAGCGUCUGGUCACCAAGGCCAAGAAGGGCGACAUCCACAACCGUCGCCUGGUGCUGCAGACGAU
Ssp-191-1  GGGCGGCUCCGCCGCGCACGAGAAGCACCUUCUCGCGAACCUCGCGAAGGCGCUCUUCGAGCACGGCCGCAUCACCACCACCGAGGCCAAGGCCCGCCGCCUGCGUCCCUACGCCGAGCGUCUGGUCACCAAGGCCAAGAAGGGCGACAUCCACAACCGUCGCCUGGUGCUGCAGACGAU
Sxa-1-1    GGGCGGCAGCGCUGCGCACGAGAAGCACCUCCUCGCGAACCUCGCGAAGUCGCUCUUCGAGCACGGCCGCAUCACCACCACCGAGGCCAAGGCCCGCCGCCUGCGUCCCUACGCCGAGCGUCUGGUCACCAAGGCCAAGAAGGGCGACAUCCACAACCGUCGCCUGGUGCUGCAGACGAU
Sla-1-1    GGGCGGUUCCGCCGCGCACGAGAAGCACCUCCUCGCGAACCUCGCGAAGUCGCUCUUCGAGCACGGCCGCAUCACCACCACCGAGGCCAAGGCCCGCCGCCUGCGUCCCUACGCCGAGCGUCUGGUCACCAAGGCCAAGAAGGGCGACAUCCACAACCGUCGCCUGGUGCUGCAGACGAU
Svi-12-3   GGGCGGCAGCGCCGCGCACGAGAAGCACCUCCUCGCGAACCUCGCGAAGGCGCUCUUCGAGCACGGCCGCAUCACCACCACCGAGGCCAAGGCCCGCCGCCUGCGUCCCUACGCCGAGCGUCUGGUCACCAAGGCCAAGAAGGGCGACAUCCACAACCGUCGCCUGGUGCUGCAGACGAU
Sla-2-2    GGGCGGUUCCGCCGCGCACGAGAAGCACCUCCUCGCGAACCUCGCGAAGGCGCUCUUCGAGCACGGCCGCAUCACCACCACCGAGGCCAAGGCCCGUCGCCUGCGUCCCUACGCCGAGCGUCUGGUCACCAAGGCCAAGAAGGGCGACAUCCACAACCGUCGCCUGGUGCUGCAGACGAU
Ssp-150-1  GGGCGGUUCCGCCGCGCACGAGAAGCACCUCCUCGCGAACCUCGCGAAGGCGCUCUUCGAGCACGGCCGCAUCACCACCACCGAGGCCAAGGCCCGCCGCCUGCGUCCCUACGCCGAGCGUCUGGUCACCAAGGCCAAGAAGGGCGACAUCCACAACCGUCGCCUGGUGCUGCAGACGAU
Ssp-32-1   GGGCGGCAGCGCCGCGCACGAGAAGCACCUCCUCGCGAACCUCGCGAAGUCGCUCUUCGAGCACGGCCGCAUCACGACGACCGAGGCCAAGGCCCGCCGCCUGCGUCCGGUCGCCGAGCGUCUGAUCACCAAGGCGAAGAAGGGCGACAUCCACAACCGUCGCCUCGUUCUGCAGACGAU
Ska-1-1    GGGCGGCUCCGCCGCGCACGAGAAGCACCUUCUCGCGAACCUCGCGAAGGCGCUCUUCGAGCACGGCCGCAUCACCACCACCGAGGCCAAGGCCCGCCGCCUGCGUCCCUACGCCGAGCGUCUGGUCACCAAGGCCAAGAAGGGCGACAUCCACAACCGUCGCCUGGUGCUGCAGACGAU
Sro-6-1    GGGCGGCUCCGCCGCGCACGAGAAGCACCUUCUCGCGAACCUCGCGAAGGCGCUCUUCGAGCACGGCCGCAUCACCACCACCGAGGCCAAGGCCCGCCGCCUGCGUCCCUACGCCGAGCGUCUGGUCACCAAGGCCAAGAAGGGCGACAUCCACAACCGUCGCCUGGUGCUGCAGACGAU
Ssp-194-1  GGGCGGCGGCGCCGCGCACCAGAAGGCGAUGCUGGGCAACCUUGCCACCAGCCUUUUCGAGCACGGCCGCAUCACCACCACCGAGGCGAAGGCACGCCGCCUGCGUCCGGUGGCGGAGCGUCUGAUCACCAAGGCGAAGAAGGGCGACCUUCACAACCGCCGCCAGGUCAUGCGCACGGU
env-33     GGGCGGCAGCGCCGCGCACGAGAAGCUGCUUCUCGCGAACCUCGCGAAGUCGCUGUUCGAGCACGGCCGCAUCACCACGACCGAGGCCAAGGCCCGCCGCCUGCGUCCGGUCGCCGAGCGCCUGAUCACCAAGGCGAAGAAGGGCGACAUCCACAACCGUCGCCUGGUGCUGCAGACGAU
Ssp-7-1    GGGCGGCAGCGCCGCGCACGAGCGUCUGCUUCUCGCGAACCUCGCGAAGUCGCUGUUCGAGCACGGCCGCAUCACCACCACCGAGGCCAAGGCGCGCCGUCUGCGUCCGGUCGCGGAGCGCCUGAUCACCAAGGCGAAGAAGGGCGACAUCCACAACCGUCGCCUGGUGCUGCAGACGAU
Ssp-140-1  GGGCGGCAGCGCCGCGCACGAGAAGCUGCUUCUCGCGAACCUCGCGAAGUCGCUGUUCGAGCACGGCCGCAUCACCACGACCGAGGCCAAGGCCCGCCGCCUGCGUCCGGUCGCCGAGCGCCUGAUCACCAAGGCGAAGAAGGGCGACAUCCACAACCGUCGCCUGGUGCUGCAGACGAU
Svi-1-1    GGGCGGCAGCGCCGCGCACGAGAAGCUGCUCCUCGCGAACCUCGCGAAGUCGCUCUUCGAGCACGGCCGCAUCACCACGACCGAGGCCAAGGCCCGCCGCCUGCGUCCGGUCGCCGAGCGCCUGAUCACCAAGGCGAAGAAGGGCGACAUCCACAACCGUCGCCUGGUUCUCCAGACGAU
Sca-2-1    GGGCGGCAGCGCGGCCCACGAGAAGCUGCUGCUGGCGAACCUCGCCAAGUCGCUCUUCGAGCACGGCCGGAUCACCACCACCGAGGCCAAGGCGCGCCGGCUGCGCCCGGUCGCGGAGAAGCUGAUCACCAAGGCGAAGAAGGGCGACCUGCACAACCGUCGCCUGGUGCUGGAGGUCAU
Sts-1-1    GGGCGGCAGCGCUGCGCACGAGCGGCUGCUCCUCGCGAACCUCGCGAAGUCGCUGUUCGAGCACGGCCGGAUCACCACGACCGAGGCGAAGGCCCGCCGCCUGCGUCCGGUCGCCGAGCGGCUGAUCACCAAGGCGAAGAAGGGCGACAUCCACAACCGUCGUCUCGUUCUUCAGACGAU
Ssp-25-1   GGGCGGCAGCGCCGCGCACGAGAAGCUGCUCCUCGCGAACCUCGCGAAGUCGCUCUUCGAGCACGGCAAGAUCACGACGACCGAGGCCAAGGCCCGUCGUCUGCGCCCGUACGCGGAGCGUCUGGUCACCAAGGCGAAGAAGGGCGACCUUCACAACCGCCGUCAGGUGCUCACGGUCAU
Ska-1-2    GGGCGGCUCCGCCGCGCACGAGAAGCACCUCCUCGCGAACCUCGCGAAGGCCCUCUUCGAGCACGGCCGCAUCACCACCACCGAGGCCAAGGCCCGCCGCCUGCGUCCCUACGCCGAGCGUCUGGUCACCAAGGCCAAGAAGGGCGACAUCCACAACCGUCGCCUGGUGCUGCAGACGAU

           ....................................................................................................................................................................................
           ....................................................................................................................................................................................
           •GGCGGCAGCRC•GCGCAC•AG•R•CU••U•CU•RY•AACCU•GC•A••••GCU•UUCGAGCACGGCCG•AUCAC•AC•ACCGAGGC•AAGGCCCGYCG•CUGCGYCCG••CGC•GAGCR•CUGRUCACCAAGGCGAAGAAGGGCGAC•UYCACAACCGYCGYC•GGU•YU•YRGRY•AU
```

| | |
|---|---|
| Smo-2-1 | GGGCGGCAGCGCAGCGCACGAGCGUCUCAUGCUGCGCAACCUGGCCACCGCGCUGUUCGAGCACGGCCGCAUCACGACGACCGAGGCCAAGGCCCGUCGUCUGCGUCCGUACGCCGAGCGUCUGGUGACCAAGGCGAAGAAGGGUGACCUUCACAACCGUCGCCAGGUCAUGCAGCUGAU |
| Ssp-70-1 | GGGCGGCAGCGCUGCGCACGAGAAGGCGCUCCUGCGCAACCUGGCCACCAACCUCUUCGAGCACGGCCGCAUCACGACGACCGAGGCCAAGGCCCGUCGUCUGCGUCCGUUCGCGGAGCGUCUGGUGACCAAGGCGAAGAAGGGUGACCUGCACAACCGCCGUCAGGUCAUGGAGCUGAU |
| Sba-2-1 | GGGCGGCAGCGCCGCGCACGAGAAGCUGCUCCUCGCCAACCUGGCGAAGUCGCUGUUCGAGCACGGCCGCAUCACGACGACCGAGGCCAAGGCCCGCCGCCUGCGCCCCGUCGCCGAGCGCCUCGUCACCAAGGCGAAGAAGGGCGACAUCCACAACCGUCGCCUGGUGCUCCAGACGAU |
| Sme-1-1 | GGGCGGCAGCGCCGCGCACGAGCGUCUGCUCCUCGCCAACCUGGCGAAGUCGCUGUUCGAGCACGGCCGCAUCACGACGACCGAGGCCAAGGCUCGCCGCCUGCGCCCCGUCGCCGAGCGCCUCGUCACCAAGGCGAAGAAGGGCGACAUCCACAACCGUCGCCUGGUGCUCCAGACGAU |
| Sgr-11-1 | GGGCGGCAGCGCCGCGCACGAGCGUCUGCUCCUCGCCAACCUGGCGAAGUCGCUGUUCGAGCACGGCCGCAUCACGACGACCGAGGCCAAGGCUCGCCGCCUGCGCCCCGUCGCCGAGCGCCUCGUCACCAAGGCGAAGAAGGGCGACAUCCACAACCGUCGCCUGGUGCUCCAGACGAU |
| Sgl-1-1 | GGGCGGCAGCGCCGCGCACGAGCGUCUGCUCCUCGCCAACCUGGCGAAGUCGCUGUUCGAGCACGGCCGCAUCACGACGACCGAGGCCAAGGCUCGCCGCCUGCGCCCCGUCGCCGAGCGCCUCGUCACCAAGGCGAAGAAGGGCGACAUCCACAACCGUCGCCUGGUGCUCCAGACGAU |
| Ssp-76-1 | GGGCGGCAGCGCCGCGCACGAGCGUCUGCUCCUCGCCAACCUGGCGAAGUCGCUGUUCGAGCACGGCCGCAUCACGACGACCGAGGCCAAGGCUCGCCGCCUGCGCCCCGUCGCCGAGCGCCUCGUCACCAAGGCGAAGAAGGGCGACAUCCACAACCGUCGCCUGGUGCUCCAGACGAU |
| Sgr-12-7 | GGGCGGCAGCGCCGCGCACGAGCGUCUGCUCCUCGCCAACCUGGCGAAGUCGCUGUUCGAGCACGGCCGCAUCACGACGACCGAGGCCAAGGCUCGCCGCCUGCGCCCCGUCGCCGAGCGCCUCGUCACCAAGGCGAAGAAGGGCGACAUCCACAACCGUCGCCUGGUGCUCCAGACGAU |
| Sgl-3-2 | GGGCGGCAGCGCCGCGCACGAGCGUCUGCUCCUCGCCAACCUGGCGAAGUCGCUGUUCGAGCACGGCCGCAUCACGACGACCGAGGCCAAGGCCCGCCGCCUGCGCCCCGUCGCCGAGCGCCUCGUCACCAAGGCGAAGAAGGGCGACAUCCACAACCGUCGCCUGGUGCUCCAGACGAU |
| Sgr-4-1 | GGGCGGCAGCGCCGCGCACGAGAAGCUGCUCCUCGCCAACCUGGCGAAGUCGCUGUUCGAGCACGGCCGCAUCACGACGACCGAGGCCAAGGCCCGCCGCCUGCGUCCGGUCGCCGAGCGCCUCGUCACCAAGGCGAAGAAGGGCGACAUCCACAACCGUCGCCUGGUGCUCCAGACGAU |
| San-2-1 | GGGCGGCAGCGCCGCGCACGAGCGUCUGCUCCUCGCCAACCUGGCGAAGUCGCUGUUCGAGCACGGCCGCAUCACGACGACCGAGGCCAAGGCCCGCCGCCUGCGUCCGGUCGCCGAGCGCCUCGUCACCAAGGCGAAGAAGGGCGACAUCCACAACCGUCGCCUGGUGCUCCAGACGAU |
| Sgr-3-1 | GGGCGGCAGCGCCGCGCACGAGAAGCUGCUCCUCGCCAACCUGGCGAAGUCGCUGUUCGAGCACGGCCGCAUCACGACGACCGAGGCCAAGGCCCGCCGCCUGCGUCCGGUCGCCGAGCGCCUCGUCACCAAGGCGAAGAAGGGCGACAUCCACAACCGUCGCCUGGUGCUCCAGACGAU |
| Ssp-9-1 | GGGCGGCAGCGCCGCGCACGAGCGUCUGCUCCUCGCCAACCUGGCGAAGUCGCUGUUCGAGCACGGCCGCAUCACGACGACCGAGGCGAAGGCCCGUCGCCUGCGUCCGGUCGCCGAGCGCCUCGUCACCAAGGCGAAGAAGGGCGACAUCCACAACCGUCGCCUGGUGCUCCAGACGAU |
| Seu-1-1 | GGGCGGCAGCGCCGCGCACGAGCGUCUGCUUCUCGCCAACCUGGCGAAGUCGCUGUUCGAGCACGGCCGCAUCACGACGACCGAGGCCAAGGCCCGUCGCCUGCGUCCGGUCGCCGAGCGCCUCGUCACCAAGGCGAAGAAGGGCGACAUCCACAACCGUCGCCUGGUGCUCCAGACGAU |
| Ssp-82-1 | GGGCGGCAGCGCCGCGCACGAGCGUCUGCUUCUCGCCAACCUGGCGAAGUCGCUGUUCGAGCACGGCCGCAUCACGACGACCGAGGCCAAGGCCCGUCGCCUGCGCCCCGUCGCCGAGCGCCUCGUCACCAAGGCGAAGAAGGGCGACAUCCACAACCGUCGCCUGGUGCUCCAGACGAU |
| Sal-12-2 | GGGCGGCAGCGCCGCGCACGAGAAGCUGCUCCUCGCCAACCUGGCGAAGUCGCUGUUCGAGCACGGCCGCAUCACGACGACCGAGGCCAAGGCCCGCCGCCUGCGCCCCGUCGCCGAGCGCCUCGUCACCAAGGCGAAGAAGGGCGACAUCCACAACCGUCGCCUGGUGCUCCAGACGAU |
| Ssp-48-1 | GGGCGGCAGCGCCGCGCACGAGCGUCUGCUUCUCGCCAACCUGGCGAAGUCGCUGUUCGAGCACGGCCGCAUCACGACGACCGAGGCCAAGGCCCGUCGCCUGCGUCCGGUCGCCGAGCGCCUCGUCACCAAGGCGAAGAAGGGCGACAUCCACAACCGUCGCCUGGUGCUCCAGACGAU |
| Sha-1-1 | GGGCGGCAGCGCCGCGCACGAGAAGCUUCUUCUCGUCAACCUCGCGAAGGCGCUGUUCGAGCACGGCCGCAUCACCACGACCGAGGCCAAGGCCCGCCGCCUGCGUCCGGUCGCCGAGCGCUUCAUCACCAAGGCGAAGAAGGGCGACAUCCACAACCGUCGCCUGGUGCUGCAGUCGAU |
| Ssp-45-1 | GGGCGGCAGCGCCGCGCACGAGAAGCUUCUUCUCAUCAACCUCGCGAAGUCGCUGUUCGAGCACGGCCGCAUCACCACGACCGAGGCCAAGGCCCGCCGCCUGCGUCCGGUCGCCGAGCGUUUCAUCACCAAGGCGAAGAAGGGCGACAUCCACAACCGUCGCCUGGUGCUGCAGUCGAU |
| Ssp-71-1 | GGGCGGCAGCGCCGCGCACGAGAAGCUUCUUCUCGUCAACCUCGCGAAGGCGCUGUUCGAGCACGGCCGCAUCACCACGACCGAGGCCAAGGCCCGCCGCCUGCGUCCGGUCGCCGAGCGCUUCAUCACCAAGGCGAAGAAGGGCGACAUCCACAACCGUCGCCUGGUGCUGCAGUCGAU |
| Ssp-175-1 | GGGCGGCAGCGCCGCGCACGAGAAGCUUCUUCUCAUCAACCUCGCGAAGUCGCUGUUCGAGCACGGCCGCAUCACCACGACCGAGGCCAAGGCCCGCCGCCUGCGUCCGGUCGCCGAGCGUUUCAUCACCAAGGCGAAGAAGGGCGACAUCCACAACCGUCGCCUGGUGCUGCAGUCGAU |
| Ssp-188-1 | GGGCGGCAGCGCCGCGCACGAGAAGCUUCUUCUCGUCAACCUCGCGAAGGCGCUGUUCGAGCACGGCCGCAUCACCACGACCGAGGCCAAGGCCCGCCGCCUGCGGCCGGUCGCCGAGCGCUUCAUCACCAAGGCGAAGAAGGGCGACAUCCACAACCGUCGCCUGGUGCUGCAGUCGAU |
| Ssp-52-1 | GGGCGGCAGCGGCGCGCACGAGAAGCUGCUCCUCGCGAACCUCGCGAAGGCGCUCUUCGAGCACGGCCGCAUCACGACGACCGAGGCCAAGGCCCGCCGCCUGCGUCCGGUCGCCGAGCGUCUCGUCACCAAGGCGAAGAAGGGCGACAUGCACAACCGUCGCCUGGUGCUUCAGACGAU |
| Sri-4-1 | GGGCGGCAGCGCAGCGCACGAGCGUCUCAUGCUGCGCAACCUGGCCACCGCGCUGUUCGAGCACGGCCGCAUCACGACGACCGAGGCCAAGGCCCGUCGUCUGCGUCCGUACGCCGAGCGUCUGGUGACCAAGGCGAAGAAGGGUGACCUUCACAACCGCCGCCAGGUCAUGCAGCUGAU |
| Ssp-145-1 | GGGCGGCAGCGCUGCGCACGAGCGCCUCAUGCUGCGCAACCUGGCCACCUCGCUGUUCGAGCACGGCCGCAUCACGACGACCGAGGCCAAGGCCCGUCGUCUGCGUCCGUACGCGGAGCGCCUGGUGACCAAGGCGAAGAAGGGCGACCUUCACAACCGCCGUCAGGUCAUGCAGCUGAU |
| env-34 | GGGCGGCAGCGCUGCGCACGAGCGCCUCAUGCUGCGCAACCUGGCCACCGCGCUCUUCGAGCACGGCCGCA............................................................................................................. |
| Sly-1-1 | GGGCGGCAGCGCCGCGCACGAGCGUCUGAUGCUGCGCAACCUGGCCACCGCUCUCUUCGAGCACGGCCGCAUCACGACGACCGAGGCCAAGGCCCGUCGUCUGCGUCCGUACGCGGAGCGCCUGGUGACCAAGGCGAAGAAGGCGGACCUGCACAACCGCCGCCAGGUCAUGCAGCUGAU |
| Ssp-142-1 | GGGCGGCAGCGCUGCGCACGAGCGUGCCAUGCUGCGCAACCUGGCGACCAACCUCUUCGAGCACGGCCGCAUCACGACGACCGAGGCCAAGGCCCGCCGUCUGCGUCCGUACGCGGAGCGUCUGGUGACCAAGGCGAAGAAGGGCGACCUUCACAACCGCCGUCAGGUCAUGCAGCUGAU |
| Ssp-155-1 | GGGCGGCAGCGCGGCGCACGAGCGUCUCAUGCUGCGCAACCUGGCCACCGCGCUCUUCGAGCACGGCCGCAUCACGACGACCGAGGCCAAGGCCCGUCGUCUGCGUCCGUACGCGGAGCGUCUGGUGACCAAGGCGAAGAAGGGCGACCUUCACAACCGCCGUCAGGUCAUGCAGCUGAU |
| Ssp-53-1 | GGGCGGCAGCGCUGCGCACGAGAAGGCGCUCCUGCGCAACCUGGCGACCAACCUCUUCGAGCACGGCCGCAUCACGACGACCGAGGCCAAGGCCCGUCGUCUGCGUCCGUUCGCGGAGCGUCUGGUGACCAAGGCGAAGAAGGGCGACCUUCACAACCGCCGCCAGGUCAUGGAGCUGAU |
| Sat-2-1 | GGGCGGCAGCGCUGCGCACGAGAAGCUUCUUCUCAUCAACCUCGCGAAGUCGCUGUUCGAGCACGGCCGCAUCACCACGACCGAGGCCAAGGCCCGCCGCCUGCGUCCGGUCGCCGAGCGUUUCAUCACCAAGGCGAAGAAGGGCGACAUCCACAACCGUCGCCUGGUGCUGCAGUCGAU |
| Ssp-95-1 | GGGCGGCAGCGCUGCGCACGAGAAGCUUCUUCUCAUCAACCUCGCGAAGUCGCUGUUCGAGCACGGCCGCAUCACCACGACCGAGGCCAAGGCCCGCCGCCUGCGUCCGGUCGCCGAGCGUUUCAUCACCAAGGCCAAGAAGGGCGACAUCCACAACCGUCGCCUGGUGCUGCAGUCGAU |
| Sfl-1-1 | GGGCGGCAGCGCUGCGCACGAGAAGCUUCUUCUCAUCAACCUCGCGAAGUCGCUGUUCGAGCACGGCCGCAUCACCACGACCGAGGCCAAGGCCCGCCGCCUGCGUCCGGUCGCCGAGCGUUUCAUCACCAAGGCCAAGAAGGGCGACAUCCACAACCGUCGCCUGGUGCUGCAGUCGAU |
| [pa-1-1 | GGGCGGCAGCGCUGCGCACGAGAAGCUUCUUCUCAUCAACCUCGCGAAGUCGCUGUUCGAGCACGGCCGCAUCACCACGACCGAGGCCAAGGCCCGCCGCCUGCGUCCGGUCGCCGAGCGUUUCAUCACCAAGGCCAAGAAGGGCGACAUCCACAACCGUCGCCUGGUGCUGCAGUCGAU |
| Sfl-8-1 | GGGCGGCAGCGCUGCGCACGAGAAGCUUCUUCUCAUCAACCUCGCGAAGUCGCUGUUCGAGCACGGCCGCAUCACCACGACCGAGGCCAAGGCGCGCCGCCUGCGUCCGGUCGCCGAGCGUUUCAUCACCAAGGCCAAGAAGGGCGACAUCCACAACCGUCGCCUGGUGCUGCAGUCGAU |
| Ssp-154-1 | GGGCGGCAGCGCUGCGCACGAGAAGCUUCUUCUCAUCAACCUCGCGAAGUCGCUGUUCGAGCACGGCCGCAUCACCACGACCGAGGCCAAGGCCCGCCGCCUGCGUCCGGUCGCCGAGCGUUUCAUCACCAAGGCCAAGAAGGGCGACAUCCACAACCGUCGCCUGGUGCUGCAGUCGAU |
| Sce-1-1 | GGGUGGCGGCGCCGCCCACGAGCGGCUCAUGCUGCGCAACCUGGCCACCGCUCUCUUCGAGCACGGUCGCAUCAAGACGACCGAGGCCAAGGCGCGCCGGCUGCGCCCCUACGCCGAGCGUCUGGUGACCAAGGCGAAGAAGGGCGACCUGCACAACCGCCGUCAGGUCAUGCAGCUGAU |
| Ssc-5-1 | GGGCGGCAGCGCCGCGCACGAGAAGCUGCUCCUCGCGAACCUCGCGAAGUCGCUCUUCGAGCACGGCCGCAUCACCACGACCGAGGCCAAGGCCCGCCGCCUGCGCCCGAUCGCGGAGCGUUUCAUCACCAAGGCGAAGAAGGGCGACAUCCACAACCGUCGCCUGGUGCUGCAGUCGAU |
| Ssp-144-1 | GGGCGGCAGCGCUGCGCACGAGAAGCUUCUUCUCAUCAACCUCGCGAAGUCGCUGUUCGAGCACGGCCGCAUCACCACGACCGAGGCCAAGGCCCGUCGCCUGCGUCCGGUCGCCGAGCGUUUCAUCACCAAGGCGAAGAAGGGCGACAUCCACAACCGUCGCCUGGUGCUGCAGUCGAU |
| Ssp-54-1 | GGGCGGCAGCGCUGCGCACGAGAAGCUUCUUCUCAACAACCUCGCGAAGUCGCUGUUCGAGCACGGCCGCAUCACCACGACCGAGGCCAAGGCCCGCCGCCUGCGUCCGGUCGCCGAGCGUUUCAUCACCAAGGCGAAGAAGGGCGACAUCCACAACCGUCGCCUGGUGCUGCAGUCGAU |
| Sgr-13-1 | GGGCGGCAGCGCCGCGCACGAGAAGCUGCUCCUCGCCAACCUGGCGAAGUCGCUGUUCGAGCACGGCCGCAUCACGACGACCGAGGCCAAGGCCCGCCGCCUGCGCCCCGUCGCCGAGCGCCUCGUCACCAAGGCGAAGAAGGGCGACAUCCACAACCGUCGCCUGGUGCUCCAGACGAU |
| Sfl-9-1 | GGGCGGCAGCGCCGCGCACGAGAAGCUGCUCCUCGCCAACCUGGCGAAGUCGCUGUUCGAGCACGGCCGCAUCACGACGACCGAGGCCAAGGCCCGCCGCCUGCGCCCCGUCGCCGAGCGCCUCGUCACCAAGGCGAAGAAGGGCGACAUCCACAACCGUCGCCUGGUGCUCCAGACGAU |
| Sro-4-1 | CGGCGGUAGCGCCGCUCACGAGAAGCUGCUUCUCGCGAACCUGGCGAAGUCGCUGUUCGAGCACGGCCGCAUCACGACGACCGAGGCCAAGGCCCGCCGCCUGCGCCCCGUCGCCGAGCGCCUCGUCACCAAGGCGAAGAAGGGCGACAUCCACAACCGUCGCCUGGUGCUCCAGACGAU |
| Sal-12-3 | CGGCGGUAGCGCCGCUCACGAGAAGCUGCUUCUCGCGAACCUGGCGAAGUCGCUGUUCGAGCACGGCCGCAUCACGACGACCGAGGCCAAGGCCCGCCGCCUGCGCCCCGUCGCCGAGCGCCUCGUCACCAAGGCGAAGAAGGGCGACAUCCACAACCGUCGCCUGGUGCUCCAGACGAU |
| Ssp-174-1 | GGGCGGCAGCGCCGCGCACGAGAAGCUGCUCCUCGCCAACCUGGCGAAGUCGCUGUUCGAGCACGGCCGCAUCACGACGACCGAGGCCAAGGCCCGCCGCCUGCGCCCCGUCGCCGAGCGCCUCGUCACCAAGGCGAAGAAGGGCGACAUCCACAACCGUCGCCUGGUGCUCCAGACGAU |
| Ssp-129-1 | GGGCGGCAGCGCCGCGCACGAGAAGCUGCUCCUCGCCAACCUGGCGAAGUCGCUGUUCGAGCACGGCCGCAUCACGACGACCGAGGCCAAGGCCCGCCGCCUGCGCCCCGUCGCCGAGCGCCUCGUCACCAAGGCGAAGAAGGGCGACAUCCACAACCGUCGCCUGGUGCUCCAGACGAU |
| Spu-2-1 | GGGCGGCAGCGCCGCGCACGAGAAGCUGCUCCUCGCCAACCUGGCGAAGUCGCUGUUCGAGCACGGCCGCAUCACGACGACCGAGGCCAAGGCUCGCCGCCUGCGCCCCGUCGCCGAGCGCCUCGUCACCAAGGCGAAGAAGGGCGACAUCCACAACCGUCGCCUGGUGCUCCAGACGAU |
| Svi-8-1 | GGGCGGCAGCGCCGCGCACGAGAAGCUGCUCCUCGCCAACCUGGCGAAGUCGCUGUUCGAGCACGGCCGCAUCACGACGACCGAGGCCAAGGCCCGCCGCCUGCGCCCCGUCGCCGAGCGCCUCGUCACCAAGGCGAAGAAGGGCGACAUCCACAACCGUCGCCUGGUGCUCCAGACGAU |
| Ssp-66-1 | CGGCGGUAGCGCCGCUCACGAGAAGCUGCUUCUCGCGAACCUGGCGAAGUCGCUGUUCGAGCACGGCCGCAUCACGACGACCGAGGCCAAGGCCCGCCGCCUGCGCCCCGUCGCCGAGCGCCUCGUCACCAAGGCGAAGAAGGGCGACAUCCACAACCGUCGCCUGGUGCUCCAGACGAU |
| Ssp-94-1 | GGGCGGCAGCGCCGCGCACGAGAAGCUGCUCCUCGCCAACCUGGCGAAGUCGCUGUUCGAGCACGGCCGCAUCACGACGACCGAGGCCAAGGCCCGCCGCCUGCGCCCCGUCGCCGAGCGCCUCGUCACCAAGGCGAAGAAGGGCGACAUCCACAACCGUCGCCUGGUGCUCCAGACGAU |
| Ssp-17-1 | CGGCGGUAGCGCCGCUCACGAGAAGCUGCUUCUCGCGAACCUGGCGAAGUCGCUGUUCGAGCACGGCCGCAUCACGACGACCGAGGCCAAGGCCCGCCGCCUGCGCCCCGUCGCCGAGCGCCUCGUCACCAAGGCGAAGAAGGGCGACAUCCACAACCGUCGCCUGGUGCUCCAGACGAU |
| Ssp-193-1 | CGGCGGUAGCGCCGCUCACGAGAAGCUGCUUCUCGCGAACCUGGCGAAGUCGCUGUUCGAGCACGGCCGCAUCACGACGACCGAGGCCAAGGCCCGCCGCCUGCGCCCCGUCGCCGAGCGCCUCGUCACCAAGGCGAAGAAGGGCGACAUCCACAACCGUCGCCUGGUGCUCCAGACGAU |
| Sat-1-1 | GGGCGGCAGCGCCGCGCACGAGCGUCUGCUUCUCGCGAACCUCGCGAAGUCGCUGUUCGAGCACGGCCGCAUCACGACGACCGAGGCCAAGGCCCGCCGCCUGCGUCCGGUCGCCGAGCGUCUGAUCACCAAGGCGAAGAAGGGCGACAUCCACAACCGUCGCCUCGUUCUGCAGACGAU |
| Ssp-44-1 | GGGCGGCAGCGCCGCGCACGAGCGUCUGCUCCUCGCCAACCUGGCGAAGUCGCUGUUCGAGCACGGCCGCAUCACGACGACCGAGGCCAAGGCCCGCCGCCUGCGCCCCGUCGCCGAGCGCCUCGUCACCAAGGCGAAGAAGGGCGACAUCCACAACCGUCGCCUGGUGCUGCAGACGAU |
| Scy-1-1 | GGGCGGCAGCGCCGCGCACGAGCGUCUGCUCCUCGCCAACCUGGCGAAGUCGCUGUUCGAGCACGGCCGCAUCACGACGACCGAGGCCAAGGCCCGCCGCCUGCGCCCCGUCGCCGAGCGCCUCGUCACCAAGGCGAAGAAGGGCGACAUCCACAACCGUCGCCUGGUGCUGCAGACGAU |
| Sfu-1-1 | GGGCGGCAGCGCCGCGCACGAGCGUCUGCUCCUCGCCAACCUGGCGAAGUCGCUGUUCGAGCACGGCCGCAUCACGACGACCGAGGCCAAGGCCCGCCGCCUGCGUCCGGUCGCCGAGCGCCUCGUCACCAAGGCGAAGAAGGGCGACAUCCACAACCGUCGCCUGGUGCUGCAGACGAU |
| Sal-7-1 | GGGCGGCAGCGCCGCGCACGAGCGUCUGCUCCUCGCCAACCUGGCGAAGUCGCUGUUCGAGCACGGCCGCAUCACGACGACCGAGGCCAAGGCCCGCCGCCUGCGUCCGGUCGCCGAGCGCCUCGUCACCAAGGCGAAGAAGGGCGACAUCCACAACCGUCGCCUGGUGCUGCAGACGAU |
| Ssp-168-1 | GGGCGGCAGCGCCGCGCACGAGCGUCUGCUCCUCGCCAACCUGGCGAAGUCGCUGUUCGAGCACGGCCGCAUCACGACGACCGAGGCCAAGGCCCGCCGCCUGCGUCCGGUCGCCGAGCGCCUCGUCACCAAGGCGAAGAAGGGCGACAUCCACAACCGUCGCCUGGUGCUGCAGACGAU |
| Ssp-97-1 | GGGCGGCAGCGCCGCGCACGAGAAGCUUCUUCUCAACAACCUUGCGAAGUCGCUGUUCGAGCACGGCCGCAUCACGACGACCGAGGCCAAGGCCCGCCGCCUGCGUCCGGUCGCCGAGCGUUUCAUCACCAAGGCGAAGAAGGGCGACAUCCACAACCGUCGCCUCGUUCUGCAGUCGAU |
| Ssp-146-1 | GGGCGGCAGCGCCGCGCACGAGCGUCUGCUUCUCGCGAACCUCGCGAAGUCGCUGUUCGAGCACGGCCGCAUCACGACGACCGAGGCCAAGGCCCGCCGCCUGCGUCCGGUCGCCGAGCGUCUGAUCACCAAGGCGAAGAAGGGCGACAUCCACAACCGUCGCCUGGUUCUGCAGUCGAU |
| Sod-1-1 | GGGCGGCAGCGCCGCGCACGAGAAGCUGCUUCUCGCGAACCUCGCGAAGUCGCUCUUCGAGCACGGCCGCAUCACCACGACCGAGGCCAAGGCCCGCCGCCUGCGUCCGGUCGCGGAGCGCUUCAUCACCAAGGCGAAGAAGGGCGACAUCCACAACCGUCGCCUGGUGCUCCGUUCGAU |
| Sri-4-2 | GGGCGGCAGCGCUGCGCACGAGCGUCUUAUGCUGCGCAACCUGGCCACCAGCCUGUUCGAGCACGGCCGCAUCACGACGACCGAGGCCAAGGCCCGUCGUCUGCGUCCGUACGCCGAGCGUCUGGUGACCAAGGCGAAGAAGGGUGACCUUCACAACCGCCGUCAGGUCAUGCAGCUGAU |
| Sgr-6-1 | GGGCGGCAGCGCUGCGCACGAGCGUGCGAUGCUGCGCAACCUGGCCACCAACCUGUUCGAGCACGGCCGCAUCACGACGACCGAGGCCAAGGCCCGUCGGCUGCGUCCGUACGCCGAGCGUCUGGUGACCAAGGCGAAGAAGGGUGACCUUCACAACCGCCGCCAGGUCAUGCAGCUGAU |
| Sri-4-5 | GGGCGGCAGCGCUGCGCACGAGCGUCUUAUGCUGCGCAACCUGGCCACCAGCCUGUUCGAGCACGGCCGCAUCACGACGACCGAGGCCAAGGCCCGUCGUCUGCGUCCGUACGCCGAGCGUCUGGUGACCAAGGCGAAGAAGGGUGACCUUCACAACCGCCGUCAGGUCAUGCAGCUGAU |
| Sri-2-1 | GGGCGGCAGCGCUGCGCACGAGCGUCUUAUGCUGCGCAACCUGGCCACCAGCCUGUUCGAGCACGGCCGCAUCACGACGACCGAGGCCAAGGCCCGUCGUCUGCGUCCGUACGCCGAGCGUCUGGUGACCAAGGCGAAGAAGGGUGACCUUCACAACCGCCGUCAGGUCAUGCAGCUGAU |
| Ssp-90-1 | GGGCGGCAGCGCUGCGCACGAGCGUCUUAUGCUGCGCAACCUGGCCACCAGCCUGUUCGAGCACGGCCGCAUCACGACGACCGAGGCCAAGGCCCGUCGUCUGCGUCCGUACGCCGAGCGUCUGGUGACCAAGGCGAAGAAGGGUGACCUUCACAACCGCCGUCAGGUCAUGCAGCUGAU |
| Sri-4-18 | GGGCGGCAGCGCUGCGCACGAGCGUCUUAUGCUGCGCAACCUGGCCACCAGCCUGUUCGAGCACGGCCGCAUCACGACGACCGAGGCCAAGGCCCGUCGUCUGCGUCCGUACGCCGAGCGCCUGGUGACCAAGGCGAAGAAGGGUGACCUUCACAACCGCCGUCAGGUCAUGCAGCUGAU |
| Ssp-185-1 | GGGCGGCAGCGCCGCGCACGAGCGUCUCAUGCUGCGCAACCUGGCCACCAGCCUCUUCGAGCACGGCCGGAUCACGACGACCGAGGCCAAGGCCCGUCGUCUGCGUCCGUACGCCGAGCGUCUGGUGACCAAGGCGAAGAAGGGUGACCUUCACAACCGCCGCCAGGUCAUGCAGCUGAU |
| Sri-4-19 | GGGCGGCAGCGCUGCGCACGAGCGUCUGAUGCUGCGCAACCUGGCCACCAGCCUGUUCGAGCACGGCCGCAUCACGACGACCGAGGCCAAGGCCCGUCGUCUGCGUCCGUACGCCGAGCGUCUGGUGACCAAGGCGAAGAAGGGUGACCUUCACAACCGCCGCCAGGUCAUGCAGCUGAU |
| Ssp-5-1 | GGGCGGCAGCGCAGCGCACGAGAAGCUUCUGCUGAACAACCUCGCGAAGUCGCUGUUCGAGCACGGCCGCAUCACCACGACCGAGGCCAAGGCCCGCCGCCUGCGUCCGGUCGCGGAGCGCUUCAUCACCAAGGCGAAGAAGGGCGACAUCCACAACCGUCGCCUGGUGCUGCAGUCGAU |
| Ssp-29-1 | GGGCGGCAGCGCCGCGCACGAGAAGCUUCUUCUCAACAACCUCGCGAAGUCGCUGUUCGAGCACGGCCGCAUCACCACGACCGAGGCCAAGGCCCGCCGCCUGCGUCCGGUCGCGGAGCGUUUCAUCACCAAGGCGAAGAAGGGCGACAUCCACAACCGUCGCCUGGUGCUGCAGUCGAU |
| Ssp-26-1 | GGGCGGCAGCGCCGCGCACGAGAAGCUUCUUCUCAACAACCUCGCGAAGUCGCUGUUCGAGCACGGCCGCAUCACCACGACCGAGGCCAAGGCCCGCCGCCUGCGUCCGGUCGCGGAGCGUUUCAUCACCAAGGCGAAGAAGGGCGACAUCCACAACCGUCGCCUGGUGCUUCAGUCGAU |
| Aat-1-1 | CGGCGGCAGCGCCGCGCACCAGCGGCUCAUCCUCGCCAACCUGGCGACGGCGCUGUUCGAGCACGGCCGCAUCACCACGACCGAGGCCAAGGCCCGCCGGGUCCGCCCGCUCGCGGAGAAGCUGAUCACCAAGGCCAAGAAGGGCGACCUGCACAACCGUCGCCAGGUCGCGGCCGUGAU |
| Ssp-117-1 | GGGCGGCAGCGCCGCGCACGAGAAGCUGCUCCUCGCGAACCUCGCGAAGUCGCUCUUCGAGCACGGCCGCAUCACCACCACCGAGGCCAAGGCGCGCCGGCUGCGCCCGGUCGCGGAGCGUCUGAUCACCAAGGCGAAGAAGGGCGACAUCCACAACCGUCGCCAGGUCCUGCAGGUCAU |
| Ssp-198-1 | GGGCGGCAGCGCCGCGCACGAGAAGCUGCUCCUCGCGAACCUCGCGAAGUCGCUCUUCGAGCACGGCCGCAUCACCACCACCGAGGCCAAGGCGCGCCGGCUGCGCCCGGUCGCGGAGCGCCUGAUCACCAAGGCGAAGAAGGGCGACAUCCACAACCGUCGCCAGGUGCUGCAGACCAU |
| Ssp-115-1 | GGGCGGCAGCGCCUCGCACCAGAAGGCGAUGCUGGGCAACCUGGCCACCGCGCUGUUCGAGCACGGCCGCAUCACGACGACCGAGGCCAAGGCCCGUCGUCUGCGUCCGGUCGCGGAGCGGCUGAUCACCAAGGCGAAGAAGGGCGACCUGCACAACCGCCGGCAGGUCAUGCGCACGAU |
| Psi-1-1 | CGGCGGUAGCGCCGCGCACCAGCGCCUCAUCCUGUCGAACCUGGCCACCGCGCUCUUCGAGCACGGCCGGAUCACGACGACCGAGGCGAAGGCCCGCACGCUGCGUCCGUACGCGGAGAAGCUCAUCACCAAGGCGAAGAAGGCCCACGCCGGCGAGAACCCGCUGCACCAGCGCCGCGA |
| Ssp-4-1 | GGGCGGCAGCGCCGCACACCAGAAGGCGAUGCUGGGCAACCUGGCGACCGCGCUGUUCGAGCACGGCCGCAUCACGACGACCGAGGCGAAGGCCCGCCGCCUGCGUCCGGUCGCCGAGCGUCUGAUCACCAAGGCGAAGAAGGGCGACCUGCACAACCGUCGCCAGGUCAUGCGCACCGU |
| Ssp-201-1 | GGGCGGCAGCGCCUCGCACCAGAAGGCCAUGCUGGCGAACCUCGCCACGGACCUUUUCGAGUACGGGCGCAUCACCACCACCGAGGCGCGGGCCCGCCGUCUGCGUCCCGUGGCGGAGCGUCUGAUCACCAAGGCGAAGAAGGGCGACCUUCACAACCGCCGCCAGGUCAUGCGCACGGU |
| Aal-1-1 | CGGCGGAAGCCCAGCGCACGAGCGCCUGAUGCUCGCCAACCUCGCCACCUCGUUGUUCCGCCACGGGCGGAUCACGACGACGGAGACGAAGGCCCGGCGGCUGCGCCCGUACGCCGAGCGACUCAUCACCAAGGCCAAGCGUGGUGACCUGCACGCGCCGCCAGGUGCACUCCGUGAU |
| Sal-13-1 | CGGCGGCUCCGCGGCGCACCAGAAGCUGAUCCUGGCGAACCUGGCCACCGCGCUGUUCGAGCACGGCCGGAUCACCACCACCGAGGCGAAGGCCCGCCGGCUCCGUCCGCUGGCGGAGAAGCUGAUCACGAAGGCCAAGCGCGGCGACCUGCACAACCGUCGCCUCGUGGCCCAGACGAU |
| Ari-1-1 | CGGCGGUUCCGCCGCGCACCAGCGCCUCAUCCUGGCGAACCUGGCCACGGCGCUGUUCGAGCACGGCAGCAUCACCACCACCGAGGCCAAGGCCCGGCGGGUGCGUCCGCUGGCGGAGAAGCUGAUCACCAAGGCCAAGAAGGGCGACCUGCACAACCGUCGCCAGGUGGCCAAGACGAU |
| Ssp-40-1 | CGGCGGCGGUGCCGCGCACGAGAAGCUGCUGCUCGCGAACCUCGCGAAGGCGCUGUUCGAGCACGGCAAGAUCACCACGACCGAGGCGAAGGCGCGGCGGCUGCGGCCCUACGCCGAGCGUCUGGUGACCAAGGCGAAGAAGGGUGACCUUCACAACCGCCGCCAGGUCCUCCAGGUCAU |
| Ssp-39-1 | CGGCGGCGGUGCCGCGCACGAGAAGCUGCUGCUCGCGAACCUCGCGAAGGCGCUGUUCGAGCACGGCAAGAUCACCACGACCGAGGCGAAGGCGCGGCGGCUGCGGCCCUACGCCGAGCGUCUGGUGACCAAGGCGAAGAAGGGUGACCUUCACAACCGCCGCCAGGUCCUCCAGGUCAU |
| | .................................................................................................................................................................................... |
| | .................................................................................................................................................................................... |
| | •GGCGGCAGCRC•GCGCAC•AG•R•CU••U•CU•RY•AACCU•GC•A••••GCU•UUCGAGCACGGCCG•AUCAC•AC•ACCGAGGC•AAGGCCCGYCG•CUGCGYCCG••CGC•GAGCR•CUGRUCACCAAGGCGAAGAAGGGCGAC•UYCACAACCGYCGYC•GGU•YU•YRGRY•AU |

```
Ssp-42-1   CGGCGGCGGUGCCGCGCACGAGAAGCUGUUGCUCGCGAACCUCGCGAAGGCGCUGUUCGAGCACGGCAAGAUCACCACGACCGAGGCGAAGGCGCGGCGGCUGCGGCCCUACGCCGAGCGUCUGGUGACCAAGGCGAAGAAGGGUGACCUUCACAACCGCCGCCAGGUCCUCCAGGUCAU
Ssp-47-1   CGGCGGCGGUGCCGCGCACGAGAAGCUGCUGCUCGCGAACCUCGCGAAGGCGCUGUUCGAGCACGGCAAGAUCACCACGACCGAGGCGAAGGCGCGGCGGCUGCGGCCCUACGCCGAGCGUCUGGUGACCAAGGCGAAGAAGGGUGACCUUCACAACCGCCGUCAGGUCCUCCAGGUCAU
Ssp-41-1   CGGCGGCGGUGCCGCGCACGAGAAGCUGCUGCUCGCGAACCUCGCGAAGGCGCUGUUCGAGCACGGCAAGAUCACCACGACCGAGGCGAAGGCGCGGCGGCUGCGGCCCUACGCCGAGCGACUGGUGACCAAGGCGAAGAAGGGCGACCUUCACAACCGCCGCCAGGUCCUCCAGGUCAU
Ssp-125-1  CGGCGGCGGUGCCGCGCACGAGAAGCUGCUGCUCGCGAACCUCGCGAAGGCUCUCUUCGAGCACGGCAGGAUCACCACGACCGAGGCGAAGGCGCGGCGGCUGCGGCCCUACGCCGAGCGUCUGGUGACCAAGGCGAAGAAGGGCGACCUUCACAACCGCCGCCAGGUCCUCCAGGUCAU
Nsp-3-1    CGGCGGUAGCCCGUCCCACCAGCGCCUGAUCCUCGCGAACCUGGCCACCCAGCUCUUCGAGCACGGCCGGAUCACCACCACCGAGGCCAAGGCUCGCGCGCUGCGCCCGCACGCCGAGAAGCUGAUCACCAAGGCGAAGAAGGGCGACCUGCACAACCGUCGCGAGGUCCUCAAGACCAU
Nsp-2-1    CGGCGGUAGCGCCGCUCACCAGCGCCUGAUCCUGUCGAACCUCUCGACCCAGCUCUUCGAGCACGGCCGGAUCACGACGACCGAGGCCCGGGCCCGCGCCCUGCGUCCGGUCGCCGAGAAGCUGAUCACGAAGGCCAAGAAGGGCGACCUGCACAACCGUCGCGAGGUCCUCAAGACCAU
env-35     CGGCGGCAGCCCCUCGCACCAGCGCCUGAUCCUGUCGAACCUGGCCACCGCCCUGUUCGAGAACGGUCGCAUCACCACCACCGAGUCGCGCGCCCGUGCGCUGCGCCCGCUGGCGGAGAAGCUGAUCACCAAGGCCAAGAAGGGCGACCUGCACAACCGCCGCGAGGUCCUCAAGACCAU
env-37     ..............................................................................................................................................................................................
env-38     GGGCGGUAGCGCCGCCCACCAGCGCCUCAUCCUCG...........................................................................................................................................................
env-39     ..............................................................................................................................................................................................
env-40     CGGCGGCAGCGCGGCGCACCAGCGCCUCAUGCUGUCCAACCUGGCGACCGCCCUGUUCGAGCACGGCCGGAUCACCACCACCGAGACCCGGGCGCGCAUGCUGCGCCCGCUCGCGGAGAAGCUGAUCACC...........................................................
Nlu-1-1    CGGCGGCUCGCCCGCGCACCAGAAGCUGAUCCUCUCGAACCUCGCCACCGCGCUGUUCGAGAACGGCCGCAUCACCACCACCGAGGCCAAGGCCCGGCACCUCCGCCCCGUCGCGGAGAAGCUGAUCACCAAGGCCAAGAAGGGUGACCUGCACAACCGUCGCGAGGUCCUCAAGACCAU
env-41     CGGCGGUUCGCCCGCGCACCAGAAGCUGAUCCUCUCGAACCUCGCCACCGCGCUGUUCGAGAACGGCCGCAUCACCACCACCGAGGCCAAGGCCCGGCACCUCCGCCCCGUCGCGGAGAAGCUGAUCACCAAGGCCAAGAAGGGUGACCUGCACAACCGUCGCGAGGUCCUCAAGACCAU
Nha-1-1    CGGCGGCAGCCCCUCGCACCAGAGGCUGAUCGUCUCGAACCUCGCCACGGCGCUGUUCGAGCACGGCCGGAUCACGACCACCGAGGCGCGCGCCCGCGUGCUGCGGCCGGUCGCGGAGAAGCUGAUCACCAAGGCCAAGCGGGGCGACCUGCACAACCGGCGCGAGGUCCUCAAGACCAU
Sst-1-1    GGGCGGCAGCGCCGCGCACGAGAAGCUGCUCCUCGCGAACCUCGCGAAGAGCCUCUUCGAGCACGGCCGCAUCACCACCACCGAGGCGAAGGCCCGCCGUCUGCGGCCCUACGCCGAGCGUCUGGUCACCAAGGCGAAGAAGGGCGACCUUCACAACCGCCGCCAGGUGCUCCAGGUCAU
Sle-1-1    GGGCGGCAGCGCCGCGCACGAGAAGCUGAUGCUCGCGAACCUCGCGAAGUCGCUCUUCGAGCACGGUCGUAUCACCACCACCGAGGCGAAGGCCCGCCGUCUGCGGCCGUACGCCGAGCGUCUGAUCACCAAGGCGAAGAAGGGCGACCUUCACAACCGCCGCCAGGUGCUCCAGGUCAU
Sal-2-1    GGGCGGCAGCGCCGCGCACGAGCGGCUCAUGCUCGCGAACCUCGCGCGCUCGCUGUUCGAGCACGGCCGGAUCAAGACGACCGAGGCGAAGGCGCGCGCCGUCUGCGUCCCUACGCCGAGCGUCUGGUCACCAAGGCGAAGAAGGGCGACCUUCACAACCGCCGCCAGGUUCUCCAGGUCAU
Ssp-24-1   GGGCGGCAGCGCCUCGCACCAGAAGGCGAUGCUGGGCAACCUGGCGACCGCGCUGUUCGAGCACGGCCGCAUCACGACGACCGAGGCGAAGGCCCGCCGCCUGCGUCCGGUCGCCGAGCGCCUGAUCACCAAGGCGAAGAAGGGUGACCUGCACAACCGUCGUCAGGUCAUGCGCACCAU
Scl-1-1    GGGCGGCAGCGCCGCGCACGAGCGUCUGCUCCUCGCGAACCUCGCGAAGGCGCUGUUCGAGCACGGCCGGAUCACCACGACCGAGGCCAAGGCCCGCCGCCUGCGUCCGGUCGCCGAGCGCCUGAUCACCAAGGCGAAGAAGGGCGACAUCCACAACCGUCGCCUGGUGCUUCAGACGAU
Ssc-1-1    GGGCGGCAGUGCCGCGCACGAGAAGCUGCUCCUCGCGAACCUCGCGAAGAGCCUCUUCGAGCACGGCCGCAUCACCACCACCGAGGCGAAGGCCCGCCGUCUGCGGCCGUACGCCGAGCGUCUGAUCACCAAGGGCAAGAAGGGCGACCUUCACAACCGCCGUCAGGUGCUUCAGGUGAU
Sde-1-1    GGGCGGCAGCGCUGCGCACGAGCGCCUCAUGCUGCGCAACCUGGCCACCUCGCUGUUCGAGCACGGCCGCAUCACGACGACCGAGGCCAAGGCCCGUCGUCUGCGUCCGUACGCGGAGCGCCUGGUGACCAAGGCGAAGAAGGGCGACCUUCACAACCGCCGUCAGGUCAUGCAGCUGAU
           ..............................................................................................................................................................................................
           ..............................................................................................................................................................................................
           •GGCGGCAGCRC•GCGCAC•AG•R•CU••U•CU•RY•AACCU•GC•A••••GCU•UUCGAGCACGGCCG•AUCAC•AC•ACCGAGGC•AAGGCCCGYCG•CUGCGYCCG••CGC•GAGCR•CUGRUCACCAAGGCGAAGAAGGCGAC•UYCACAACCGYCGYC•GGU•YU•YRGRY•AU
```

alignment positions 440 · · · 533

```
env-1       GCGCGACCGCGAUGUGGUCGUCAAGCUCUUCAACGAGCUGGGCCCGCGCUUCGCAGCCCGCCCGGGUGGCUAUACCCGCAUCCUGAAGAUGGGC
env-2       GCGCGACCGCGAUGUGGUCGUCAAGCUCUUCAACGAGCUGGGCCCGCGCUUCGCAGCCCGCCCGGGUGGCUAUACCCGCAUCCUGAAGAUGGGC
[br-1-1     GCGCAACCGCGACAACGUCGUGAAGCUGUUCGAAGUCCUGGGCCCGCGUUUCAACGCUCGCCCGGGCGGCUACACGCGCAUCCUGAAGAUGGGU
env-3       GCGCGACCGCGACAUCGUCACCAAGCUGUUCGCCGAACUGGGCCCGCGCUACCAGACGCGUCCGGGCGGCUACACCCGCAUCCUGAAGAUGGGC
Nal-1-1     GGUCCUCAAGACCAUCCGUGACAAGGGCGUCGUGCACGUGCUCUUCACGGAGAUCGCGCCCACCUUCGCCGAGCGUCCGGGCGGGUACACCCGC
Nsp-1-1     GGUCCUCAAGGUCAUCCGGGACAAGGGUGUCGUGCACACCCUCUUCACGGAGAUCGCGCCGACCUUCGCCGACCGCCCGGGCGGCUACACCCGC
Mxa-1-1     CAAGGACCGCGACGUGCUGCAGAAGGUGUUCGGCGAGUACAAGGACCGGUACGCCAGCCGUCCCGGUGGCUACACCCGCAUCGUUCGGCUCGGC
env-4       ..............................................................................................
Cfu-1-1     CCGCAAUCGCGUCAUCCUGAAGAAGGUCUUCGGGGAGUACAAGGAGCGCUACGCCAGCCGUCCGGGUGGUUACACCCGCAUCGUCCGGCUCGGG
Avi-1-1     UCGCAACCGCGUCAUCCUCCAGAAGGUGUUCAGCGAGUACAAGGACCGGUACGCCAACCGCCCGGGCGGGUACACCCGCAUCGUGCGCCUCGGA
Age-1-1     UCGCAACCGCGUCAUCCUCCAGAAGGUGUUCAGCGAGUACAAGGAUCGUUACGCCAGCCGCCCGGGCGGGUACACCCGCAUCGUGCGCCUCGGC
Hmi-1-1     GAAGGACAAGGACGUCCUCCAGAAGGUCUUCAGCACCUACAAGGACCGCUACGCCAGCCGUAAGGGCGGCUACACCCGCAUCAUCAAGCUCGGC
env-5       GAAGGACAAGAACGUCCUUCAGAAGGUGUUUGGCCCGUACAAGGAGCGCUACGCCAACCGCAAUGGCGGCUACACCCGCAUCGUGAAGCUGGGC
Sau-5-1     GAAGGACAAGAACGUCCUCCAGAAGGUCUUCAGCGAGUACAAGGACCGCUACAGCACCCGCAAGGGUGGCUACACCCGCAUCAUCAAGCUCGGU
Mfu-1-1     GAAGGACCGUGACGUGCUGAAGAAGGUCUUCAGCGAGUACAAGGACCGCUACGCCAACCGUCCCGGCGGCUACACCCGCAUCGUUCGUCUCGGC
Mst-1-1     GAAGGACCGUGACGUGCUGAAGAAGGUCUUCAGCGAGUACAAGGACCGGUACGCCACGCGUCCCGGCGGCUACACCCGCAUCAUCCGCCUCGGC
Msp-2-1     GAAGGACCGCGAGGUGCUGCAGAAGGUGUUCAGCGAGUACAAGGACCGUUACGCCAGCCGUCCCGGUGGCUACACGCGCAUCAUCCGGCUCGGC
Mfu-2-1     CAAGGACCGCGACGUGCUGCAGAAGGUGUUCGGCGAGUACAAGGACCGGUACGCCAACCGUCCCGGCGGCUACACCCGCAUCGUCCGGCUCGGC
env-6       GAAGGACAAGAACGUCCUCCAGAAGGUCUUCAGCACCUACAAGGACCGCUACGCCACCCGCAAGGGCGGCUACACCCGCAUCAUCAAGCUCGGC
Cco-1-1     GAAGGACCGCACGGUCCUCCAGAAGGUGUUCAGCGAGUACAAGGAGCGUUACGCCUCCCGUCCGGGUGGCUACACCCGCAUCGUGCGCCUGGGC
env-7       CCGGGAUCGCGCCAUCGUCAGCAAGCUGUUCGACGAACUCGGCCCGCGCUACAAGGCCCGGCCCGGCGGCUACACACGGAUCCUGAAGAUGGGC
env-8       GCGCGACCGCGACAUCGUCACCAAGCUGUUUGGCGAGUUGGGACCACGCUACAAGGCACGCCCGGGCGGCUACACCCGGAUUCUCAAGAUGGGA
Ama-1-1     CCGCGACAAGGGCGUGCUGCACGAGCUCUUCACCGAGAUCGCGCCGCGCUUCGAGAACCGGCCCGGCGGCUACACCCGCAUCACCAAGAUCGGC
env-9       ..............................................................................................
env-10      CCGGAACAAGGACGUGGUGCAUCAGCUGUUCACGGCGAUCGGUCCCAGCCAGGCCGCGCGUCCGGGCGGGUACACCAGGAUCACCAAGGUCGGU
1 Ssp-68-1  CCUCGACAAGAGCGUCGUGCACACCCUCUUCACCGAGAUCGGUCCGCGGUACGAGAACCGUCCCGGUGGCUACACCCGCAUCACCAAGAUCGGC
env-11      ..............................................................................................
env-12      ..............................................................................................
Ssp-96-1    CACCGACAUCUCGGUGCUGCACACCCUCUUCACCGAGAUCGCGCCGCGCUUCGAGAACCGCCCCGGCGGUUACACCCGCAUCACCAAGAUCGGC
Ksp-1-1     CACCGACGUCUCGGUGCUGCACACCCUCUUCACCGAGAUCGCGCCGCGCUACGAGAACCGCCCGGGUGGCUACACCCGUAUCACCAAGAUCGGU
Kme-1-1     CACCGACGUCUCCGUGCUGCACACCCUCUUCACCGAGAUCGCGCCGCGCUACGAGAACCGCCCGGGUGGCUACACCCGUAUCACCAAGAUCGGU
Rge-1-1     CCGUGACCGCGAAGUCGUCACCAAGCUGUUCAACGAACUGGGCCCGCGCUACGCCGCGCGUCCGGGCGGCUACACCCGCAUCCUGAAGAUGGGC
Sca-3-1     CACCGACGUGGCCGUGCUGCACACCCUCUUCACCGAGAUCGCUCCUCGCUACGAGAACCGUCCCGGUGGCUACACCCGGAUCACCAAGGUCGGC
Sal-3-1     CACCGACGUGGCUGUGCUGCACACCCUCUUCACCGAGAUCGGUCCUCGCUACGAGAACCGCCCCGGUGGCUACACCCGCAUCACCAAGAUCGGC
env-13      CUACAAGAGCGUCGACCAGAAGAAGCUCAGCGAGGACGGCGUGCUGGUGGUCAACCAGUACCCCCAGCACCGGCUGUUCACCGAGAUCGCGCCG
1 Sav-3-1   CCGUGACAAGAGCGUGGUUCACACCCUCUUCACGGAGAUCGGCCCGAAGUUCGCCAACCGGCCGGGUGGCUACACCCGCAUCACCAAGCUCGGC
Ssp-195-1   CAAGGACAAGAGCGUGGUGCACACGCUCUUCACCGAGAUCGGCCCGCGCUACGAGAACCGGCCGGGCGGCUACACCCGCAUCACCAAGAUCGGU
Nsp-4-1     CAAGGACAAGGGCGUCGUUCACCACCUCUUCACCGAGAUCGCGCCGACGUUCGCCGAGCGUCCGGGCGGCUACACCCGCAUCACCAAGGUCGGG
env-14      UCGGGACAAGGGCGUCGUGCACCACCUGUUCACCGAGAUCGCGACGACGUUCGCCGAGCGU.................................
Nco-1-1     CAGGGACAAGGGCGUCGUUCACCACCUCUUCACCGAGAUCGCGACGACGUUCGCCGAGCGUCCGGGCGGCUACACCCGGAUCACCAAGAUCGGU
Sje-1-1     CACCGACGUGGCUGUGCUGCACACCCUCUUCACCGAGAUCGGUCCUCGCUACGAGAACCGUCCCGGUGGCUACACCCGCAUCACCAAGAUCGGU
Kaz-1-1     CACCGACGUCUCCGUGCUGCACACCCUCUUCACCGAGAUCGCGCCGCGCUACGAGAACCGUCCCGGUGGCUACACCCGCAUCACCAAGAUCGGU
Kph-1-1     CACCGACGUGUCGGUGCUGCACACCCUCUUCACCGAGAUCGCCCCGCGGUUCGAGAACCGCCCCGGCGGCUACACCCGCAUCACCAAGAUCGGC
Kph-1-2     CACCGACGUGUCGGUGCUGCACACCCUCUUCACCGAGAUCGCCCCGCGGUUCGAGAACCGCCCCGGCGGCUACACCCGCAUCACCAAGAUCGGC
Sno-2-1     CACCGACAUCUCGGUGCUGCACACCCUCUUCACCGAGAUCGCGCCGCGCUACGAGAACCGCCCGGGUGGCUACACCCGCAUCACCAAGAUCGGU
Ksp-3-1     CACCGACGUGUCGGUGCUGCACACCCUCUUCACCGAGAUCGCGCCGCGCUACGAGAACCGCCCGGGUGGUUACACCCGUAUCACCAAGAUCGGU
Spy-1-1     CCCGGGUGGCUACACCCGCAUCACCAAGAUCGGUCCCCGUCGUGGCGACAACGCCCCGAUGGCCGUGAUCGAGCUGGUCGAGGCGCUGACCGUC
Ksp-4-1     CACCGACGUGUCGGUGCUGCACACCCUCUUCACCGAGAUCGCCCCGCGGUUCGAGAACCGUCCCGGCGGCUACACCCGCAUCACCAAGAUCGGC
Ssp-165-1   CACCGACAUCUCGGUGCUGCACACCCUCUUCACCGAGAUCGCGCCGCGCUACGAGAACCGCCCGGGUGGCUACACCCGUAUCACCAAGAUCGGU
Ssp-133-1   CACCGACGUCUCGGUGCUGCACACCCUCUUCACCGAGAUCGCGCCGCGCUACGAGAACCGCCCGGGUGGCUACACCCGCAUCACCAAGAUCGGU
Ssp-50-1    CACCGACGUCUCGGUGCUGCACACCCUCUUCACCGAGAUCGCGCCGCGCUACGAGAACCGCCCGGGUGGUUACACCCGUAUCACCAAGAUCGGU
Ssp-157-1   CACCGACAUCGCCGUGCUGCACACCCUCUUCACCGAGAUCGCGCCGCGCUACGAGAACCGCCCGGGUGGCUACACCCGCAUCACCAAGAUCGGU
Ssp-160-1   CACCGACGUGUCGGUGCUGCACACCCUCUUCACCGAGAUCGCCCCGCGGUUCGAGAACCGCCCCGGCGGCUACACCCGCAUCACCAAGAUCGGC
Ksp-2-1     CACCGACAUCUCGGUGCUGCACACCCUCUUCACCGAGAUCGCGCCGCGCUACGAGAACCGCCCGGGUGGUUACACCCGUAUCACCAAGAUCGGU

            ..............................................................................................
            ..............................................................................................
            C•••GACAAGRGCRUCGU•CACAC•CUCUUCACCGAGAUCG••CCGCG•UACG•RRA•CGYCC•GGUGGYUACACCCGYAUCACCAAGAUCGGY
```

```
Ssp-183-1  CACCGACGUGUCGGUCCUGCACACCCUCUUCACCGAGAUCGCGCCGCGCUACGAGAACCGCCCGGGUGGUUACACCCGUAUCACCAAGAUCGGU
Sam-2-1    CAGGGACAAGGGUGUCGUCCACCACCUCUUCACCGAGAUCGCCUCGACCUUCGCCGAGCGUCCCGGUGGUUACACCCGGAUCACCAAGGUCGGC
Sro-7-1    CCGUGACAAGGGUGUCGUCCACCACCUGUUCACGGAGAUCGCGCCGACCUUCGCCGAGCGUCCGGGCGGCUACACCCGUAUCACGAAGGUCGGC
Ssp-87-1   CCGCGACAAGAGCGUCGUGCACACCCUCUUCACCGAGAUCGGCCCGCGCUACGAGAACCGGCCCGGUGGCUACACCCGCAUCACCAAGAUCGGU
Ssp-190-1  CCGUGACAAGAGCGUCGUGCACACUCUCUUCACCGAGAUCGGGCCGCGCUACGAGAACCGGCCGGGUGGCUACACCCGCAUCACCAAGAUCGGU
Sir-1-1    CCGCGACAAGAGCGUCGUGCACACUCUCUUCACCGAGAUCGGGCCGCGCUACGAGAACCGGCCGGGUGGCUACACCCGCAUCACCAAGAUCGGU
Shy-4-1    CCGCGACAAGAGCGUCGUGCACACCCUCUUCACCGAGAUCGCGCCGCGCUACGAGAACCGGCCGGGUGGCUACACCCGCAUCACCAAGAUCGGU
Sro-3-1    CCGUGACAAGGGUGUCGUCCACCACCUGUUCACGGAGAUCGCGCCGACCUUCGCCGAGCGUCCGGGCGGCUACACCCGCAUCACGAAGGUCGGC
Nin-1-1    CCGCGACAAGUCGAUCGUGCACGUGCUCUUCACCGACAUCGCCGAGCAGUUCGCGGACCGUCCGGGUGGCUACACCCGGAUCACCAAGAUCGGC
1 Nca-1-1  UCGGGACAAGGGCGUCGUGCACCACCUCUUCACCGAGCUGGCGACGACGUUCGCCGAGCGGCCGGUGGUUACACCCGGAUCACCAAGAUCGGU
Ssp-192-1  CCGCGACAAGAGCGUGGUCCACACGCUGUUCACGGAGAUCGGGCCCAAGUUCGCCAACCGGCCCGGCGGUUACACCCGCAUCACCAAGAUCGGC
Ssp-172-1  CACCGACAAGAGCAUCGUGCACACGCUCUUCACGGAGAUCGGCCCGCGCUACGAGAACCGCCCGGGUGGCUACACCCGUAUCACCAAGAUCGGU
Ssp-93-1   CACGGACAAGAGCGUCGUCCACACGCUCUUCGCCGAGAUCGGCCCGCGCUACGAGAACCGCCCGGGUGGCUACACCCGUAUCACCAAGAUCGGU
Sau-3-1    CUCGGACAAGAGCAUCGUCCACACGCUCUUCACCGAGAUCGGCCCGCGCUACGAGAACCGCCCGGGUGGCUACACCCGUAUCACCAAGAUCGGU
Sal-6-1    CACCGACAAGAGCAUCGUGCACACGCUCUUCACGGAGAUCGGCCCGCGCUACGAGAACCGCCCGGGUGGCUACACCCGCAUCACCAAGAUCGGU
Sau-1-1    CUCGGACAAGAGCGUCGUCCACACGCUCUUCACCGAGAUCGGCCCGCGCUACGAGAACCGCCCGGGUGGCUACACCCGCAUCACCAAGAUCGGU
Ssp-106-1  CACCGACAAGAGCGUCGUGCACACCCUCUUCACCGAGAUCGGUCCGCAGUUCGCGAACCGUCCGGGCGGCUACACCCGGAUCACCAAGAUCGGC
Ssp-3-1    CACCGACAAGAGCGUCGUGCACACCCUCUUCACCGAGAUCGGUCCGCAGUUCGCGAACCGGCCGGGCGGCUACACCCGCAUCACCAAGAUCGGU
env-15     ................................................................................................
env-16     CCGUGACAAGAGCGUCGUCCACCACCUCUUCACCGAGAUCGCGCCGACGUUCGCCGAGCGUCCCGGUGGCUACACCCGGAUCACCAAGAUCGGC
env-18     CCGUGACAAGAGCGUCGUCCACCACCUCUUCACCGAGAUCGCGCCGACGUUCGCCGAGCGUCCCGGUGGCUACACCCGGAUCACCAAGAUCGGC
Ssp-118-1  CACCGACAAGAGCGUCGUCCACACGCUCUUCGCCGAGAUCGGCCCGCGCUACGAGAACCGCCCGGGUGGCUACACCCGGAUCACCAAGAUCGGC
env-19     GCGGGACAAGGGCGUCGUCCACCACCUCUUCACCGAGAUCGCUCCGGCGUUCGCCCAGCGUCCGGGCGGGUACACCCGGAUCACCAAGAUCGGC
env-20     GCGGGACAAGGGCGUCGUCCACCACCUCUUCACCGAGAUCGCUCCGGCGUUCGCCCAGCGUCCGGGCGGNUACACCCGGAUCACCAAGAUCGGC
env-21     GCGGGACAAGGGC...................................................................................
Sor-1-1    GACCAACGUCGCGGUGCUGCACACCCUCUUCACCGAGAUCGCUCCGCGGUACGAGAACCGCCCGGGCGGCUACACCCGCAUCACCAAGAUCGGU
Ssp-64-1   CACGGACAAGAGCAUCGUCCACACGCUGUUCACCGAGAUCGCCCCGCGCUACGCCGAGCGUCCGGGUGGCUACACCCGGAUCACCAAGAUCGGC
Ssp-12-1   CACGGACAAGAGCAUCGUCCACACGCUGUUUACCGAGAUCGCCCCGCGCUACGCCGAGCGUCCGGGUGGCUACACCCGGAUCACCAAGAUCGGC
Ssp-13-1   CACGGACAAGAGCAUCGUCCACACGCUGUUCACCGAGAUCGCCCCGCGCUACGCCGAGCGUCCGGGUGGCUACACCCGGAUCACCAAGAUCGGC
Ssp-61-1   CUCGGACAAGAGCGUCGUGCACACGCUGUUCACCGAGAUCGGUCCGCGGUACGAGAACCGCCCGGGCGGCUACACCCGGAUCACCAAGAUCGGC
Ssp-59-1   CUCGGACAAGAGCGUGGUCCACACGCUCUUCACCGAGAUCGGCCCGCGGUACGACAACCGCCCGGGCGGCUACACCCGGAUCACCAAGAUCGGC
Ssp-16-1   CUCGGACAAGAGCGUGGUGCACACGCUGUUCACCGAGAUCGGUCCGCGGUACGACAACCGCCCGGGCGGCUACACCCGGAUCACCAAGAUCGGC
Ssp-69-1   CUCGGACAAGAGCGUGGUGCACACGCUGUUCACCGAGAUCGGCCCGCGGUACGACAACCGCCCGGGCGGCUACACCCGGAUCACCAAGAUCGGC
Ssp-56-1   CUCGGACAAGAGCGUCGUGCACACGCUCUUCACCGAGAUCGGCCCGCGGUACGAGAACCGCCCGGGUGGCUACACCCGGAUCACCAAGAUCGGU
Hcr-1-1    CCGUGACAAGGGCGUCGUCCACCACCUCUUCACCGAGAUCGCCCCCAGCUUCGCGGAGCGUCCCGGUGGCUACACCCGCAUCACCAAGAUCGGU
Msp-1-1    CCGCGACAAGGGCGUCGUCCACCACCUCUUCACCGAGAUCGCCCCCACCUUUGCCGAGCGCCCGGGCGGCUACACCCGGAUCACCAAGCUUGGC
Mro-1-1    CCGCGACAAGGGCGUCGUCCACCACCUCUUCACCGAGAUCGCCCCGACGUACGCCGAGCGUCCCGGUGGCUACACCCGGAUCACCAAGCUCGGC
Ssp-171-1  CACCGACAAGGGCGUCGUCCACACGCUCUUCACCGAGAUCGCGCCGCGCUACUCGGAGCGUCCGGGUGGUUACACCCGCAUCACCAAGAUCGGC
Ssp-43-1   CACCGACAAGGGGAUCGUCCACACCCUCUUCACCGAGAUCGCCCCGCGUUACUCGGAGCGUCCGGGUGGCUACACCCGCAUCACCAAGAUCGGC
Svi-13-1   CACCGACAAGGGGAUCGUCCACACCCUCUUCACCGAGAUCGCCCCGCGUUACUCGGAGCGUCCGGGUGGCUACACCCGCAUCACCAAGAUCGGC
Ssp-73-1   CACGGACAAGAGCGUCGUGCACACGCUCUUCACCGAGAUCGGCCCGCGGUACGAGAACCGUCCGGGUGGCUACACCCGGAUCACCAAGAUCGGU
Ssc-3-1    CACGGACAAGAGCAUCGUGCACACGCUCUUCACCGAGAUCGGCCCGCGGUACGAGAACCGUCCGGGUGGCUACACCCGUAUCACCAAGAUCGGU
Kau-1-1    CACGGACAAGAGCGUCGUCCACACGCUCUUCACCGAGAUCGGCCCGCGCUACGAGAACCGUCCCGGUGGCUACACGCGGAUCACCAAGAUCGGC
Sgr-1-1    CACGGACAAGAGCAUCGUCCACACGCUCUUCACCGAGAUCGGCCCGCGCUACGAGAACCGCCCGGGUGGCUACACCCGCAUCACCAAGAUCGGU
Sto-1-1    CACGGACAAGAGCAUCGUCCACACGCUCUUCACCGAGAUCGGCCCGCGGUACGAGAACCGUCCCGGUGGCUACACCCGCAUCACCAAGAUCGGC
Ssp-114-1  CACGGACAAGAGCAUCGUGCACACGCUCUUCACCGAGAUCGGCCCGCGCUACGAGAACCGCCCGGGUGGCUACACCCGUAUCACCAAGAUCGGU
Ssc-4-1    CACGGACAAGGGCAUCGUCCACACGCUCUUCACCGAGAUCGGCCCGCGGUACGAGAACCGUCCGGGUGGCUACACCCGCAUCACCAAGAUCGGU
Ssp-1-1    CACGGACAAGAGCAUCGUGCACACGCUCUUCACCGAGAUCGGCCCGCGUUACGAGAACCGCCCCGGUGGCUACACCCGCAUCACCAAGAUCGGU
Sau-4-1    CACGGACAAGAGCAUCGUGCACACGCUCUUCACCGAGAUCGGCCCGCGCUACGAGAACCGCCCGGGUGGUUACACCCGUAUCACCAAGAUCGGU
Spr-1-1    CACGGACAAGGGCAUCGUGCACACGCUCUUCACCGAGAUCGCUCCGCGUUACGCCGAGCGUCCGGGUGGCUACACCCGUAUCACCAAGAUCGGC
Ssp-189-1  CACGGACAAGAGCAUCGUGCACACGCUGUUCACCGAGAUCGCCCCGCGCUACGAGAACCGCCCCGGUGGUUACACGCGCAUCACCAAGAUCGGC
Sfl-6-1    CACGGACAAGAGCAUCGUGCACACGCUGUUCACCGAGAUCGCCCCGCGCUACGAGAACCGCCCCGGUGGUUACACGCGCAUCACCAAGAUCGGC
Ssp-103-1  CACGGACAAGAGCAUCGUGCACACGCUGUUCACCGAGAUCGCCCCGCGCUACGAGAACCGCCCCGGUGGUUACACGCGCAUCACCAAGAUCGGC
Ssp-111-1  CACGGACAAGAGCAUCGUGCACACGCUGUUCACCGAGAUCGCCCCGCGCUACGAGAACCGCCCCGGUGGUUACACGCGCAUCACCAAGAUCGGC
Ssp-159-1  CACGGACAAGAGCAUCGUGCACACGCUGUUCACCGAGAUCGCCCCGCGCUACGAGAACCGCCCCGGUGGUUACACGCGCAUCACCAAGAUCGGC
Ser-1-1    CACGGACAAGAGCAUCGUCCACACGCUCUUCACCGAGAUCGCCCCGCGCUACGAGAACCGCCCCGGUGGUUACACGCGCAUCACCAAGAUCGGC
Sya-1-1    CACGGACAAGAGCAUCGUGCACACGCUGUUCACCGAGAUCGCCCCGCGCUACGAGAACCGCCCCGGUGGUUACACGCGCAUCACCAAGAUCGGC
Svi-12-1   CACGGACAAGAGCAUCGUGCACACGCUGUUCACCGAGAUCGCCCCGCGCUACGAGAACCGCCCCGGUGGUUACACGCGCAUCACCAAGAUCGGC
Ssp-151-1  CACGGACAAGAGCAUCGUGCACACGCUGUUCACCGAGAUCGCCCCGCGCUACGAGAACCGCCCCGGUGGUUACACGCGCAUCACCAAGAUCGGC
Ssp-110-1  CACGGACAAGAGCAUCGUGCACACGCUGUUCACCGAGAUCGCCCCGCGCUACGAGAACCGCCCCGGUGGUUACACGCGCAUCACCAAGAUCGGC
Ssp-112-1  CACGGACAAGAGCAUCGUGCACACGCUGUUCACCGAGAUCGCCCCGCGCUACGAGAACCGCCCCGGUGGUUACACGCGCAUCACCAAGAUCGGC
Ssp-102-1  CACGGACAAGAGCAUCGUGCACACGCUGUUCACCGAGAUCGCCCCGCGCUACGAGAACCGCCCCGGUGGUUACACGCGCAUCACCAAGAUCGGC
Sla-2-1    CACGGACAAGAGCAUCGUGCACACGCUGUUCACCGAGAUCGCCCCGCGCUACGAGAACCGCCCCGGUGGUUACACGCGCAUCACCAAGAUCGGC
Ssp-139-1  CACGGACAAGAGCAUCGUGCACACGCUGUUCACCGAGAUCGCCCCGCGCUACGAGAACCGCCCCGGUGGUUACACGCGCAUCACCAAGAUCGGC
Ssp-8-1    CACGGACAAGAGCAUCGUGCACACGCUCUUCACCGAGAUCGCCCCGCGCUACGAGAACCGCCCCGGUGGUUACACGCGCAUCACCAAGAUCGGC
Svi-7-1    CACCGACAAGGGCAUCGUCCACACCCUCUUCACCGAGAUCGCCCCGCGUUACUCGGAGCGUCCGGGUGGCUACACCCGCAUCACCAAGAUCGGC
Sgr-12-1   CACCGACAAGGGCAUCGUCCACACCCUCUUCACCGAGAUCGCCCCGCGCUACUCGGAGCGUCCGGGUGGUUACACCCGCAUCACCAAGAUCGGC
Sfl-5-1    CACCGACAAGGGCAUCGUCCACACCCUCUUCACCGAGAUCGCCCCGCGCUACUCCGAGCGCCCGGGUGGCUACACCCGUAUCACCAAGAUCGGC
Sbi-3-1    CACCGACAAGGGCAUCGUCCACACCCUCUUCACCGAGAUCGCCCCGCGUUACUCGGAGCGUCCGGGUGGCUACACCCGCAUCACCAAGAUCGGC
Sex-1-1    CACCGACAAGGGCAUCGUCCACACCCUCUUCACCGAGAUCGCGCCGCGCUACUCCGAGCGCCCCGGUGGCUACACCCGGAUCACCAAGAUCGGC
Sgl-3-1    CACCGACAAGGGGAUCGUCCACACGCUCUUCACCGAGAUCGCCCCGCGUUACUCGGAGCGUCCGGGUGGCUACACCCGCAUCACCAAGAUCGGC
Sgr-9-1    CACCGACAAGGGCAUCGUCCACACCCUCUUCACCGAGAUCGCGCCGCGCUACUCCGAGCGCCCGGGUGGCUACACCCGUAUCACCAAGAUCGGC
Ssp-121-1  CACCGACAAGGGCAUCGUCCACACCCUCUUCACCGAGAUCGCCCCGCGUUACUCGGAGCGUCCGGGUGGUUACACCCGCAUCACGAAGAUCGGC
Sve-1-1    CACCGACAAGGGCAUCGUCCACACCCUCUUCACCGAGAUCGCCCCGCGUUACUCGGAGCGCCCGGGUGGCUACACCCGCAUCACCAAGAUCGGC
Sav-2-1    CACGGACAAGAGCGUCGUGCACACGCUCUUCACCGAGAUCGGCCCGCGCUACGAGAACCGUCCCGGUGGCUACACCCGCAUCACCAAGAUCGGU
Mgl-1-1    CACCGACAAGGGCAUCGUCCACACCCUCUUCACCGAGAUCGCCCCGCGCUACGCGGAGCGUCCGGGUGGUUACACCCGCAUCACGAAGAUCGGC
Asp-1-1    CACCGACAAGGGCAUCGUCCACACCCUCUUCACCGAGAUCGCGCCGCGGUACACCGAGCGUCCGGGUGGUUACACCCGUAUCACCAAGAUCGGC
Stu-1-1    CUCGGACAAGAGCAUCGUGCACACGCUCUUCACCGAGAUCGGCCCGCGCUACGAGAACCGCCCGGGUGGUUACACCCGUAUCACCAAGAUCGGU
Sye-2-1    CACGGACAAGAGCAUCGUGCACACGCUCUUCACCGAGAUCGGCCCGCGGUAUGAGAACCGUCCGGGUGGCUACACCCGUAUCACCAAGAUCGGU
Sdu-1-1    CACGGACAAGGGCGUCGUCCACACGCUCUUCACCGAGAUCGGCCCGCGUUACGAGAACCGUCCGGGUGGCUACACCCGCAUCACCAAGAUCGGU
Sgh-1-1    CACGGACAAGGGCAUCGUCCACACGCUCUUCACCGAGAUCGGCCCGCGGUACGAGAACCGUCCGGGCGGCUACACCCGCAUCACCAAGAUCGGU
Svi-5-1    CACGGACAAGGGCAUCGUCCACACGCUCUUCACCGAGAUCGGCCCGCGGUACGAGAACCGUCCGGGCGGCUACACCCGCAUCACCAAGAUCGGU
Ssp-153-1  CACGGACAAGGGCAUCGUCCACACGCUCUUCACCGAGAUCGGCCCGCGGUACGAGAACCGUCCCGGUGGUUACACCCGCAUCACCAAGAUCGGU
Ssp-86-1   CACGGACAAGAGCGUCGUCCACACGCUCUUCACCGAGAUCGGCCCGCGCUACGAGAACCGCCCCGGUGGCUACACCCGCAUCACCAAGAUCGGU

           ................................................................................................
           ................................................................................................
           C•••GACAAGRGCRUCGU•CACAC•CUCUUCACCGAGAUCG••CCGCG•UACG•RRA•CGYCC•GGUGGYUACACCCGYAUCACCAAGAUCGGY
```

| | |
|---|---|
| Sro-1-1 | CACGGACAAGGGCGUCGUCCACACGCUCUUCACCGAGAUCGGCCCGCGGUACGAGAACCGUCCGGGUGGCUACACCCGCAUCACCAAGAUCGGU |
| Sgr-7-1 | CACGGACAAGAGCAUCGUGCACACGCUCUUCACCGAGAUCGGCCCGCGGUACGAGAACCGUCCCGGUGGCUACACCCGCAUCACCAAGAUCGGU |
| Ssp-28-1 | CACGGACAAGGGCGUCGUCCACACGCUCUUCACCGAGAUCGCCCCGCGCUACGAGAACCGCCCCGGUGGUUACACCCGCAUCACCAAGAUCGGU |
| Ssp-176-1 | CACGGACAAGGGCAUCGUCCACACGCUCUUCACCGAGAUCGGCCCGCGUUACGAGAACCGUCCCGGUGGCUACACCCGCAUCACCAAGAUCGGU |
| Ssp-14-1 | CACGGACAAGGGUGUCGUGCACACCCUGUUCACCGAGAUCGCUCCGCGUUACGCCGAGCGUCCCGGCGGUUACACCCGCAUCACCAAGAUCGGC |
| Sac-2-1 | CACGGACAAGAGCGUCGUCCACACGCUCUUCACCGAGAUCGGCCCGCGCUACGAGAACCGUCCGGGUGGCUACACCCGCAUCACCAAGAUCGGU |
| Sbo-1-1 | CACGGACAAGAGCAUCGUGCACACGCUCUUCACCGAGAUCGGCCCGCGGUACGAGAACCGUCCCGGUGGCUACACCCGCAUCACCAAGAUCGGU |
| Ssp-2-1 | CACGGACAAGAGCGUCGUCCACACGCUCUUCACCGAGAUCGGCCCGCGCUACGAGAACCGUCCGGGUGGUUACACCCGUAUCACCAAGAUCGGU |
| Ssp-98-1 | CACGGACAAGAGCGUCGUCCACACGCUCUUCACCGAGAUCGGCCCGCGCUACGAGAACCGUCCGGGUGGUUACACCCGUAUCACCAAGAUCGGU |
| Sac-3-1 | CACGGACAAGAGCAUCGUCCACACGCUCUUCACCGAGAUCGGCCCGCGGUACGAGAACCGUCCCGGUGGUUACACCCGUAUCACCAAGAUCGGU |
| Sca-7-1 | CUCGGACAAGGGUGUCGUGCACACGCUGUUCACCGAGAUCGCCCCGCGGUUCGAGAACCGCCCGGGUGGUUACACCCGCAUCACCAAGAUCGGU |
| Sgr-2-1 | CACGGACAAGAGCAUCGUCCACACGCUCUUCACCGAGAUCGGCCCGCGCUACGAGAACCGUCCCGGUGGCUACACCCGCAUCACCAAGAUCGGU |
| Ssp-105-1 | CACGGACAAGAGCGUCGUCCACACGCUCUUCGCCGAGAUCGGCCCGCGCUACGAGAACCGUCCCGGUGGUUACACCCGCAUCACCAAGAUCGGU |
| Ssp-85-1 | CACGGACAAGAGCAUCGUGCACACGCUCUUCACCGAGAUCGGCCCGCGGUACGAGAACCGUCCCGGUGGCUACACCCGCAUCACCAAGAUCGGU |
| Sco-3-1 | CACGGACAAGAGCAUCGUCCACACGCUCUUCACCGAGAUCGGCCCGCGGUACGAGAACCGUCCCGGUGGCUACACCCGCAUCACCAAGAUCGGU |
| Sse-1-1 | CACGGACAAGAGCGUCGUCCACACGCUCUUCGCCGAGAUCGGCCCGCGCUACGAGAACCGCCCCGGUGGUUACACCCGCAUCACCAAGAUCGGU |
| Sre-1-1 | CACGGACAAGAGCAUCGUGCACACGCUCUUCACCGAGAUCGGCCCGCGCUACGAGAACCGUCCCGGUGGCUACACCCGCAUCACCAAGAUCGGU |
| Spe-1-1 | CACGGACAAGAGCGUCGUCCACACGCUCUUCACCGAGAUCGGCCCGCGGUACGAGAACCGUCCGGGUGGUUACACCCGUAUCACCAAGAUCGGU |
| Sbi-2-1 | CACGGACAAGGGCAUCGUCCACACGCUCUUCACCGAGAUCGGCCCGCGGUACGAGAACCGUCCGGGUGGCUACACCCGCAUCACCAAGAUCGGU |
| Ssp-83-1 | CACGGACAAGAGCAUCGUCCACACGCUCUUCACCGAGAUCGGCCCGCGUUACGAGAACCGUCCGGGUGGUUACACCCGCAUCACCAAGAUCGGU |
| Spl-1-1 | CACGGACAAGAGCAUCGUCCACACGCUCUUCACCGAGAUCGGCCCGCGUUACGAGAACCGUCCGGGUGGUUACACCCGCAUCACCAAGAUCGGU |
| Sno-1-1 | CACGGACAAGAGCAUCGUCCACACGCUCUUCACCGAGAUCGGCCCGCGCUACGAGAACCGUCCGGGUGGUUACACCCGGAUCACCAAGAUCGGU |
| San-1-1 | CACGGACAAGAGCGUCGUCCACACGCUCUUCACCGAGAUCGGCCCGCGGUACGAGAACCGUCCCGGUGGCUACACCCGCAUCACCAAGAUCGGU |
| Sla-3-1 | CACGGACAAGAGCGUCGUCCACACGCUCUUCACCGAGAUCGGCCCGCGCUACGAGAACCGUCCCGGUGGUUACACCCGCAUCACCAAGAUCGGC |
| Ssp-74-1 | CACGGACAAGAGCGUCGUCCACACGCUCUUCACCGAGAUCGGCCCGCGCUACGAGAACCGUCCGGGUGGCUACACCCGCAUCACCAAGAUCGGC |
| Ssp-162-1 | CACGGACAAGAGCGUCGUCCACACGCUCUUCACCGAGAUCGGCCCGCGCUACGAGAACCGUCCGGGUGGCUACACCCGCAUCACCAAGAUCGGU |
| Ssp-164-1 | CACGGACAAGAGCGUCGUCCACACGCUCUUCACCGAGAUCGGCCCGCGCUACGAGAACCGUCCGGGUGGCUACACCCGCAUCACCAAGAUCGGU |
| Ssp-161-1 | CACGGACAAGAGCGUCGUCCACACGCUCUUCACCGAGAUCGGCCCGCGCAACGAGAACCGUCCGGGUGGCUACACCCGCAUCACCAAGAUCGGU |
| Sgl-2-1 | CACGGACAAGAGCGUCGUCCACACGCUCUUCACCGAGAUCGGCCCGCGCUACGAGAACCGCCCGGGUGGCUACACCCGCAUCACCAAGAUCGGU |
| Ssp-143-1 | CACGGACAAGAGCAUCGUGCACACGCUCUUCACCGAGAUCGGUCCGCGCUACGAGAACCGUCCGGGUGGCUACACCCGCAUCACCAAGAUCGGU |
| Ssp-36-1 | CACGGACAAGAGCGUCGUCCACACGCUCUUCACCGAGAUCGGCCCGCGGUACGAGAACCGUCCCGGUGGCUACACCCGCAUCACCAAGAUCGGU |
| Ssp-200-1 | CACGGACAAGAGCAUCGUCCACACGCUCUUCACCGAGAUCGGCCCGCGCUACGAGAACCGUCCCGGUGGCUACACCCGGAUCACCAAGAUCGGU |
| Sco-2-1 | CACGGACAAGAGCAUCGUCCACACGCUCUUCACCGAGAUCGGCCCGCGCUACGAGAACCGUCCCGGUGGCUACACCCGGAUCACCAAGAUCGGU |
| Svi-9-1 | CACGGACAAGAGCAUCGUCCACACGCUCUUCACCGAGAUCGGCCCGCGCUACGAGAACCGUCCCGGUGGCUACACCCGGAUCACCAAGAUCGGU |
| Sco-1-1 | CACGGACAAGAGCAUCGUGCACACGCUCUUCACCGAGAUCGGCCCGCGCUACGAGAACCGUCCCGGUGGCUACACCCGGAUCACCAAGAUCGGU |
| Ssp-128-1 | CACGGACAAGAGCGUCGUCCACACGCUCUUCACCGAGAUCGGCCCGCGCUACGAGAACCGUCCCGGUGGCUACACCCGGAUCACCAAGAUCGGC |
| Sol-2-1 | CUCGGACAAGAGCGUCGUCCACACGCUCUUCACCGAGAUCGGCCCGCGCUACGAGAACCGUCCCGGUGGUUACACCCGGAUCACCAAGAUCGGU |
| Ssp-72-1 | CUCGGACAAGAGCGUCGUCCACACGCUCUUCACCGAGAUCGGCCCGCGCUACGAGAACCGUCCCGGUGGCUACACCCGGAUCACCAAGAUCGGU |
| Ssp-124-1 | CACGGACAAGAGCGUCGUCCACACGCUCUUCACCGAGAUCGGCCCGCGCUACGAGAACCGUCCCGGUGGCUACACCCGGAUCACCAAGAUCGGC |
| Sce-2-1 | CACGGACAAGAGCAUCGUCCACACGCUCUUCACCGAGAUCGGCCCGCGCUACGAGAACCGUCCCGGUGGCUACACCCGCAUCACCAAGAUCGGU |
| Smo-1-1 | CACCGACAAGGGCAUCGUCCACACCCUCUUCACCGAGAUCGCUCCGCGCUACGCCGAGCGUCCGGGUGGCUACACCCGUAUCACCAAGAUCGGU |
| Svi-9-2 | CACGGACAAGAGCAUCGUCCACACGCUCUUCACCGAGAUCGGCCCGCGCUACGAGAACCGUCCCGGUGGCUACACCCGCAUCACCAAGAUCGGU |
| Sso-1-1 | CACGGACAAGGGCAUCGUCCACACGCUCUUCACCGAGAUCGCCCCGCGCUACGCCGAGCGCCCGGGUGGCUACACCCGCAUCACCAAGAUCGGC |
| Soc-1-1 | CACGGACAAGAGCAUCGUGCACACGCUGUUCACCGAGAUCGCCCCGCGCUACGCCGAGCGUCCGGGUGGCUACACCCGUAUCACCAAGAUCGGC |
| Sni-2-1 | CACGGACAAGAGCAUCGUGCACACGCUGUUCACCGAGAUCGCGCCGCGCUACGCCGAGCGUCCGGGUGGCUACACCCGUAUCACCAAGAUCGGC |
| Svi-11-1 | CACGGACAAGAGCAUCGUGCACACGCUGUUCACCGAGAUCGCGCCGCGCUACGCCGAGCGUCCGGGUGGCUACACCCGUAUCACCAAGAUCGGC |
| Sex-2-1 | CACCGACAAGGGCAUCGUCCACACCCUCUUCACCGAGAUCGCCCCGCGUUACUCGGAGCGUCCGGGUGGUUACACCCGUAUCACCAAGAUCGGC |
| Ssp-6-1 | CACGGACAAGGGCAUCGUCCACACGCUCUUCACCGAGAUCGGCCCGCGGUACGAGAACCGUCCCGGUGGCUACACCCGCAUCACCAAGGUCGGC |
| Ssp-101-1 | CACGGACAAGAGCAUCGUCCACACGCUCUUCACCGAGAUCGGCCCGCGGUACGAGAACCGUCCCGGUGGCUACACCCGCAUCACCAAGAUCGGU |
| Ssp-49-1 | CACGGACAAGGGCAUCGUCCACACGCUCUUCACCGAGAUCGGCCCGCGGUACGAGAACCGUCCCGGUGGCUACACCCGCAUCACCAAGGUCGGC |
| Sal-11-1 | CACCGACAAGAGCGUCGUGCACACGCUCUUCACGGAGAUCGGCCCGCGGUACGAGAACCGCCCGGGUGGCUACACCCGUAUCACCAAGAUCGGU |
| Shy-2-1 | CACGGACAAGAGCAUCGUCCACACGCUCUUCACCGAGAUCGGCCCGCGGUACGAGAACCGUCCCGGUGGCUACACCCGCAUCACCAAGAUCGGU |
| Ssp-46-1 | CACGGACAAGAGCAUCGUCCACACGCUCUUCACCGAGAUCGGCCCGCGCUACGAGAACCGUCCCGGUGGCUACACCCGGAUCACCAAGAUCGGU |
| Ssp-187-1 | CACGGACAAGAGCAUCGUCCACACGCUCUUCACCGAGAUCGGCCCGCGCUACGAGAACCGUCCCGGUGGCUACACCCGGAUCACCAAGAUCGGU |
| Ssp-156-1 | CACGGACAAGAGCGUCGUCCACACCCUCUUCACCGAGAUCGGCCCGCGGUACGAGAACCGUCCCGGUGGCUACACCCGCAUCACCAAGAUCGGU |
| Sgr-10-1 | CACGGACAAGAGCAUCGUCCACACGCUCUUCACCGAGAUCGGCCCGCGGUACGAGAACCGUCCGGUGGCUACACCCGCAUCACCAAGAUCGGU |
| Aac-1-1 | CACGGACAAGAGCAUCGUCCACACGCUCUUCACCGAGAUCGGCCCGCGGUACGAGAACCGUCCCGGUGGCUACACCCGCAUCACCAAGAUCGGU |
| Ssp-197-1 | CACGGACAAGAGCAUCGUCCACACGCUCUUCACCGAGAUCGGCCCGCGGUACGAGAACCGUCCCGGUGGCUACACCCGCAUCACCAAGAUCGGU |
| Ssp-20-1 | CACGGACAAGAGCAUCGUCCACACGCUCUUCACCGAGAUCGGCCCGCGGUACGAGAACCGUCCCGGUGGCUACACCCGCAUCACCAAGAUCGGU |
| Ssp-123-1 | CACGGACAAGAGCAUCGUGCACACGCUCUUCACCGAGAUCGGCCCGCGGUACGAGAACCGUCCCGGUGGCUACACCCGCAUCACCAAGAUCGGU |
| Ssp-141-1 | CACGGACAAGAGCAUCGUCCACACGCUCUUCACCGAGAUCGGCCCGCGGUACGAGAACCGUCCCGGUGGCUACACCCGCAUCACCAAGAUCGGU |
| Ssp-81-1 | CACGGACAAGAGCAUCGUCCACACGCUCUUCACCGAGAUCGGCCCGCGCUACGAGAACCGUCCCGGUGGCUACACCCGGAUCACCAAGAUCGGU |
| Ssp-113-1 | CACGGACAAGAGCAUCGUCCACACGCUCUUCACCGAGAUCGGCCCGCGCUACGAGAACCGUCCCGGUGGCUACACCCGGAUCACCAAGAUCGGU |
| Ssp-57-1 | CACGGACAAGAGCAUCGUCCACACGCUCUUCACCGAGAUCGGCCCGCGGUACGAGAACCGUCCCGGUGGCUACACCCGCAUCACCAAGAUCGGU |
| Ssp-149-1 | CACGGACAAGGGCGUCGUCCACACGCUCUUCACCGAGAUCGGCCCGCGGUACGAGAACCGUCCCGGUGGCUACACCCGCAUCACCAAGAUCGGU |
| Sah-1-1 | CACGGACAAGAGCGUCGUCCACACGCUCUUCACCGAGAUCGCGCCCCGUUUCUCGGAGCGUCCGGGUGGUUACACCCGCAUCACCAAGAUCGGC |
| Svi-10-1 | CACGGACAAGAGCAUCGUCCACACGCUCUUCACCGAGAUCGGCCCGCGCUACGAGAACCGUCCCGGUGGCUACACCCGGAUCACCAAGAUCGGU |
| Sch-1-1 | CACGGACAAGAGCAUCGUCCACACGCUCUUCACCGAGAUCGGCCCGCGUUACGAGAACCGUCCGGGUGGCUACACCCGCAUCACCAAGAUCGGU |
| Sia-1-1 | CACGGACAAGAGCAUCGUGCACACGCUCUUCACCGAGAUCGGCCCGCGCUACGAGAACCGUCCGGGUGGUUACACCCGCAUCACCAAGAUCGGU |
| Saf-1-1 | CACGGACAAGAGCAUCGUCCACACGCUCUUCACCGAGAUCGGCCCGCGCUACGAGAACCGUCCGGGUGGCUACACCCGCAUCACCAAGAUCGGU |
| Ssp-89-1 | CACGGACAAGAGCAUCGUGCACACGCUCUUCACCGAGAUCGGCCCGCGGUACGAGAACCGGCCGGGUGGCUACACCCGCAUCACCAAGAUCGGU |
| Sva-1-1 | CACGGACAAGAGCAUCGUGCACACGCUCUUCACCGAGAUCGGCCCGCGCUACGAGAACCGUCCGGGUGGCUACACCCGCAUCACCAAGAUCGGU |
| Ssp-158-1 | CACGGACAAGAGCAUCGUCCACACGCUCUUCACCGAGAUCGGCCCGCGGUACGAGAACCGUCCCGGUGGCUACACCCGCAUCACCAAGAUCGGU |
| Sro-5-1 | CACGGACAAGGGCAUCGUGCACGUGCUCUUCACCGAGAUCGCUCCGCGCUUCGCCGAGCGUCCGGGUGGCUACACCCGCAUCACCAAGAUCGGC |
| Ssp-116-1 | CACGGACAAGGGCAUCGUCCACACGCUCUUCACCGAGAUCGGCCCGCGGUACGAGAACCGCCCGGGUGGCUACACCCGGAUCACCAAGGUGGGC |
| env-22 | CACGGACAAGAGCGUCGUCCACACGCUCUUCACCGAGAUCGGCCCGCGGUACGAGAACCGUCCGGGUGGCUACACGCGCAUCACCAAGAUCGGU |
| Sam-1-1 | CACGGACAAGAGCAUCGUGCACACGCUCUUCACCGAGAUCGGCCCGCGCUACGAGAACCGUCCCGGUGGCUACACCCGGAUCACCAAGGUCGGC |
| Ssp-51-1 | CACGGACAAGAGCGUCGUGCACACGCUCUUCACCGAGAUCGGCCCGCGCUACGAGAACCGCCCGGGUGGUUACACCCGUAUCACCAAGAUCGGU |
| Slu-1-1 | CACGGACAAGAGCAUCGUCCACACGCUCUUCACCGAGAUCGGCCCGCGCUACGAGAACCGUCCCGGUGGCUACACCCGGAUCACCAAGAUCGGU |
| Sna-2-1 | CUCGGACAAGAGCGUCGUCCACACGCUCUUCACCGAGAUCGCCCCGCGGUUCUCGGAGCGUCCGGGUGGCUACACCCGUAUCACCAAGAUCGGU |
| Ssp-75-1 | CACGGACAAGAGCAUCGUCCACACGCUCUUCACCGAGAUCGGCCCGCGGUACGAGAACCGUCCCGGUGGCUACACCCGCAUCACCAAGAUCGGU |
| env-23 | CACGGACAAGAGCGUCGUGCACACGCUCUUCACCGAGAUCGGCCCGCGGUACGAGAACCGUCCGGGUGGCUACACCCGCAUCACCAAGAUCGGU |
| Ssp-182-1 | CACGGACAAGAGCGUCGUCCACACGCUCUUCACCGAGAUCGGCCCGCGGUACGAGAACCGUCCCGGUGGCUACACCCGCAUCACCAAGAUCGGU |
| Scy-2-1 | CACGGAUAAGAGCAUCGUCCACACGCUCUUCACCGAGAUCGGCCCGCGGUACGAGAACCGCCCGGGCGGCUACACGCGUAUCACCAAGAUCGGC |
| | ................................................................................................ |
| | ................................................................................................ |
| | C•••GACAAGRGCRUCGU•CACAC•CUCUUCACCGAGAUCG••CCGCG•UACG•RRA•CGYCC•GGUGGYUACACCCGYAUCACCAAGAUCGGY |

```
Ssp-184-1   CACGGACAAGAGCGUCGUCCACACGCUCUUCACCGAGAUCGGCCCGCGGUACGAGAACCGUCCCGGUGGCUACACCCGCAUCACCAAGAUCGGU
Svi-4-1     CACGGACAAGGGCAUCGUCCACACGCUCUUCACCGAGAUCGGCCCGCGGUACGAGAACCGUCCGGGUGGUUACACCCGUAUCACCAAGAUCGGU
Ssp-163-1   CUCGGACAAGAGCAUCGUGCACACGCUCUUCACCGAGAUCGGCCCGCGCUACGAGAACCGUCCGGGUGGCUACACCCGCAUCACCAAGAUCGGU
Ssp-177-1   CACGGACAAGAGCGUUGUCCACACGCUCUUCACCGAGAUCGGCCCGCGCUACGAGAACCGUCCGGGUGGCUACACCCGCAUCACCAAGAUCGGC
Sol-1-1     CACGGACAAGAGCGUCGUCCACACGCUCUUCACCGAGAUCGGCCCGCGCUACGAGAACCGUCCGGGUGGCUACACCCGCAUCACCAAGAUCGGC
Svi-3-1     CACGGACAAGAGCAUCGUCCACACGCUCUUCACCGAGAUCGGCCCGCGCUACGAGAACCGUCCGGGUGGCUACACCCGCAUCACCAAGAUCGGU
Szi-1-1     CACGGACAAGAGCAUCGUCCACACGCUCUUCACCGAGAUCGGCCCGCGCUACGAGAACCGUCCCGGUGGCUACACCCGCAUCACCAAGGUCGGC
Svi-13-2    CACGGACAAGAGCGUCGUCCACACGCUCUUCACCGAGAUCGGUCCGCGGUACGAGAACCGUCCGGGUGGUUACACCCGUAUCACCAAGAUCGGU
Ssp-120-1   CACGGACAAGAGCGUCGUCCACACGCUCUUCACCGAGAUCGGCCCGCGCUACGAGAACCGUCCGGGUGGCUACACCCGCAUCACCAAGAUCGGU
Ssp-131-1   CACCGACAAGGGCAUCGUCCACACCCUCUUCACCGAGAUCGCCCCGCGCUACUCGGAGCGUCCGGGUGGUUACACCCGCAUCACGAAGAUCGGC
Sca-4-1     CACCGACAAGGGCGUCGUCCACACCCUCUUCACCGAGAUCGCUCCGCGCUACGCCGAGCGUCCGGGUGGCUACACCCGCAUCACCAAGAUCGGU
Sba-1-1     CCUGGACAAGAGCGUCGUCCACACGCUCUUCACCGAGAUCGGCCCGCGCUACGAGAACCGUCCGGGUGGCUACACCCGCAUCACCAAGGUCGGC
Ssp-65-1    CCUGGACAAGGGCGUCGUGCACACGCUCUUCACCGAGAUCGGUCCCCGCUACGAGAACCGUCCGGGUGGCUACACCCGCAUCACCAAGAUCGGC
Sme-2-1     CCUCGACAAGAGCGUCGUCCACACGCUCUUCACCGAGAUCGCCCCGCGCUACGAGAACCGUCCGGGUGGCUACACCCGCAUCACCAAGAUCGGU
Sgr-12-3    CACGGACAAGAGCGUCGUCCACACGCUCUUCACCGAGAUCGGCCCCCGUUACGAGAACCGCCCGGGUGGCUACACCCGGAUCACCAAGGUCGGU
Sgr-12-4    CACGGACAAGAGCGUCGUCCACACGCUCUUCACCGAGAUCGGCCCCCGUUACGAGAACCGCCCGGGUGGCUACACCCGGAUCACCAAGGUCGGU
Ssp-15-1    CACGGACAAGAGCGUCGUCCACACGCUCUUCACCGAGAUCGGCCCCCGUUACGAGAACCGCCCGGGUGGCUACACCCGGAUCACCAAGGUCGGU
Ssp-38-1    CACGGACAAGAGCGUCGUCCACACGCUCUUCACCGAGAUCGGCCCCCGUUACGAGAACCGCCCGGGUGGCUACACCCGGAUCACCAAGGUCGGU
Sal-5-1     CACGGACAAGAGCGUCGUCCACACGCUCUUCACCGAGAUCGGCCCCCGUUACGAGAACCGCCCGGGUGGCUACACCCGGAUCACCAAGGUCGGU
Ssp-77-1    CACGGACAAGAGCGUCGUCCACACGCUCUUCACCGAGAUCGGCCCCCGUUACGAGAACCGCCCGGGUGGCUACACCCGGAUCACCAAGGUCGGU
Swa-1-1     CACGGACAAGAGCGUCGUCCACACGCUCUUCACCGAGAUCGGCCCCCGUUACGAGAACCGCCCGGGUGGCUACACCCGGAUCACCAAGGUCGGU
Ssp-30-1    CACGGACAAGAGCGUCGUCCACACGCUCUUCACCGAGAUCGGCCCCCGUUACGAGAACCGCCCGGGUGGCUACACCCGGAUCACCAAGGUCGGU
Ssp-134-1   CACGGACAAGAGCGUCGUCCACACGCUCUUCACCGAGAUCGGCCCCCGUUACGAGAACCGCCCGGGUGGCUACACCCGGAUCACCAAGGUCGGU
Ssp-62-1    CACGGACAAGAGCGUCGUCCACACGCUCUUCACCGAGAUCGGCCCCCGUUACGAGAACCGCCCGGGUGGUUACACCCGGAUCACCAAGGUCGGU
Ssp-21-1    CACGGACAAGAGCGUCGUCCACACGCUCUUCACCGAGAUCGGCCCCCGUUACGAGAACCGCCCGGGUGGUUACACCCGGAUCACCAAGGUCGGU
Ssp-80-1    CACGGACAAGAGCGUCGUCCACACGCUCUUCACCGAGAUCGGCCCCCGUUACGAGAACCGCCCGGGUGGCUACACCCGGAUCACCAAGGUCGGU
Sgr-12-5    CACGGACAAGAGCGUCGUCCACACGCUCUUCACCGAGAUCGGCCCCCGUUACGAGAACCGCCCGGGUGGCUACACCCGGAUCACCAAGGUCGGU
Sva-2-1     CACCGACAAGAGCGUCGUCCACACGCUCUUCACGGAGAUCGCGCCGCGCUACGAGAACCGCCCCGGCGGGUACACGCGCAUCACCAAGAUCGGC
Ssp-58-1    CCUGGACAAGAGCGUGGUCCACACGCUGUUCACCGAGAUCGGCCCGCGCUACGAGAACCGGCCCGGUGGCUACACCCGUAUCACCAAGAUCGGC
Sal-12-1    CCGCGACAAGGGUGUCGUCCACACCCUGUUCACGGAGAUCGGCCCGCGGUACGAGAACCGUCCGGGUGGCUACACCCGGAUCACCAAGAUCGGC
Sfr-1-1     CCGCGACAAGGGUGUCGUCCACACCCUGUUCACGGAGAUCGGCCCGCGGUACGAGAACCGUCCGGGUGGCUACACCCGGAUCACCAAGAUCGGC
Sxi-2-1     CCGCGACAAGGGUGUCGUCCACACCCUGUUCACGGAGAUCGGCCCGCGGUACGAGAACCGUCCGGGUGGCUACACCCGGAUCACCAAGAUCGGC
Ssp-119-1   CACGGACAAGAGCGUCGUGCACACGCUCUUCACCGAGAUCGCCCCGCGCUACGAGAACCGCCCCGGUGGUUACACGCGCAUCACCAAGAUCGGC
Ssp-33-1    CACGGACAAGAGCGUCGUGCACACGCUCUUCACCGAGAUCGCGCCGCGGUUCGCGGAGCGUCCGGGUGGCUACACCCGUAUCACCAAGAUCGGC
Ssp-37-1    CACGGACAAGAGCGUCGUCCACACGCUCUUCACCGAGAUCGGCCCGCGCUACGAGAACCGUCCCGGUGGCUACACCCGUAUCACCAAGAUCGGU
Ssp-170-1   CACGGACAAGAGCGUCGUCCACACGCUCUUCACCGAGAUCGGCCCGCGUUACGAGAACCGUCCGGGUGGCUACACCCGCAUCACCAAGAUCGGU
Smi-1-1     CACGGACAAGAGCGUCGUGCACACGCUCUUCACCGAGAUCGGCCCGCGCUACGAGAACCGUCCCGGUGGCUACACCCGCAUCACCAAGAUCGGU
Ssp-138-1   CACGGACAAGAGCGUCGUCCACACGCUCUUCACCGAGAUCGGCCCGCGGUACGAGAACCGUCCGGGUGGUUACACCCGCAUCACCAAGAUCGGU
Sga-2-1     CACGGACAAGAGCGUCGUGCACACGCUCUUCACCGAGAUCGGCCCGCGGUACGAGAACCGUCCGGGUGGCUACACCCGGAUCACCAAGAUCGGU
env-24      CACGGACAAGAGCGUCGUCCACACGCUCUUCACCGAGAUCGGCCCGCGCUACGAGAACCGUCCGGGUGGCUACACCCGCAUCACCAAGAUCGGU
Sca-1-1     CACGGACAAGAGCAUCGUGCACACGCUCUUCACCGAGAUCGGCCCGCGCUACGAGAACCGUCCCGGUGGCUACACCCGCAUCACCAAGAUCGGU
Ssp-23-1    CACGGACAAGAGCGUCGUCCACACGCUCUUCACCGAGAUCGGCCCGCGGUACGAGAACCGUCCGGGUGGCUACACCCGUAUCACCAAGAUCGGU
Ssp-135-1   CACGGACAAGAGCAUCGUCCACACGCUCUUCACCGAGAUCGGCCCGCGCUACGAGAACCGUCCGGGUGGCUACACCCGCAUCACCAAGAUCGGU
Ssp-169-1   CACGGACAAGAGCGUCGUCCACACGCUCUUCACCGAGAUCGGCCCGCGCUACGAGAACCGUCCGGGUGGCUACACCCGCAUCACCAAGAUCGGU
env-30      ................................................................................................
Ssv-1-1     CACGGACAAGAGCAUCGUGCACACGCUCUUCACCGAGAUCGGCCCGCGCUACGAGAACCGUCCCGGUGGCUACACCCGCAUCACCAAGAUCGGU
Ssp-186-1   CACGGACAAGAGCAUCGUGCACACGCUCUUCACCGAGAUCGGCCCGCGCUACGAGAACCGUCCCGGUGGCUACACCCGCAUCACCAAGAUCGGU
Sda-1-1     UCACGGACAAGAGCGUCGUCCACACGCUCUUCACCGAGAUCGGCCCGCGCUACGAGAACCGUCCGGGUGGCUACACCCGUAUCACCAAGAUCGG
Sgr-12-6    CACGGACAAGAGCGUCGUCCACACGCUCUUCACCGAGAUCGGCCCGCGCUACGAGAACCGUCCGGGUGGCUACACCCGCAUCACCAAGAUCGGU
Ssp-199-1   CACGGACAAGAGCAUCGUCCACACGCUUUUCACCGAGAUCGGCCCGCGCUACGAGAACCGUCCCGGUGGCUACACCCGCAUCACCAAGAUCGGU
env-31      CUCGGACAAGAGCGUCGUGCACACGCUCUUCACGGAGAUCGCCCCGCGGUUCGAGAACCGCCCGGGUGGCUACACCCGUAUCACCAAGAUCGGU
Ssp-19-1    CUCGGACAAGAGCGUCGUGCACACGCUCUUCACGGAGAUCGCCCCGCGGUUCGAGAACCGCCCGGGUGGCUACACCCGUAUCACCAAGAUCGGU
Sfu-2-1     CACGGACAAGAGCGUCGUCCACACGCUCUUCACCGAGAUCGGCCCGCGCUACGAGAACCGCCCGGGUGGCUACACCCGCAUCACCAAGAUCGGU
Sau-2-1     CUCGGACAAGAGCGUCGUCCACACGCUGUUCACCGAGAUCGCCCCGCGGUUCGAGAACCGCCCGGGUGGCUACACCCGUAUCACCAAGAUCGGU
Sxy-1-1     CACGGACAAGAGCGUCGUCCACACGCUCUUCACCGAGAUCGGCCCGCGCUACGAGAACCGUCCCGGUGGCUACACCCGUAUCACCAAGAUCGGU
Ssp-122-1   CACGGACAAGAGCAUCGUCCACACGCUCUUCACCGAGAUCGGCCCGCGCUACGAGAACCGUCCCGGUGGCUACACCCGUAUCACCAAGAUCGGU
Spr-2-1     CACGGACAAGAGCAUCGUCCACACGCUCUUCACCGAGAUCGGCCCGCGCUACGAGAACCGUCCCGGUGGCUACACCCGUAUCACCAAGAUCGGU
Ssp-84-1    CACGGACAAGAGCAUCGUGCACACGCUCUUCACCGAGAUCGGCCCCCGCUACGAGAACCGUCCGGGUGGCUACACCCGCAUCACCAAGAUCGGU
Sip-1-1     CACGGACAAGGGCGUCGUGCACACGCUCUUCACCGAGAUCGGCCCGCGCUACGAGAACCGUCCGGGUGGCUACACCCGUAUCACCAAGAUCGGU
Sly-2-1     CUCGGACAAGAGCGUCGUCCACACGCUCUUCGUGGAGAUCGCGCCGCGGUUCGCGGAGCGUCCGGGUGGCUACACCCGCAUCACCAAGAUCGGC
Ssc-2-1     UCGUGACAAGAGCGUCGUGCACUCGCUGUUCGAGGAGAUCGCUCCCCGGUACGAGAACCGCCCGGGCGGUUACACGCGCAUCACCAAGAUCGGC
Sye-1-1     CACGGACAAGGGCAUCGUGCACACGCUCUUCACCGAGAUCGCCCCCCGGUACGAGAACCGUCCCGGUGGCUACACCCGGAUCACCAAGAUCGGC
Svi-6-1     CCGGGACAAGGGUGUGGUGCACACCCUGUUCACCGAGAUCGGUCCCCGGUUCGAGAACCGUCCGGGUGGCUACACGCGCAUCACCAAGAUCGGU
Sni-1-1     CACGGACAAGAGCAUCGUGCACACGCUGUUCACCGAGAUCGCUCCGCGUUACGAGAACCGCCCCGGUGGCUACACCCGCAUCACCAAGAUCGGC
Ssu-2-1     CACCGACAAGGGCGUGGUGCACACCCUCUUCACCGAGAUCGGCCCGCGGUACGAGAACCGCCCCGGCGGUUAUACCCGCAUCACCAAGAUCGGC
Ssu-1-1     CACCGACAAGGGCGUGGUGCACACCCUCUUCACCGAGAUCGGCCCGCGGUACGAGAACCGCCCCGGCGGUUACACCCGCAUCACCAAGAUCGGC
Svi-2-1     CCGCGACAAGAGCGUCGUGCACACCCUCUUCACCGAGAUCGGCCCGCGCUACGAGAACCGGCCGGGUGGCUACACCCGGAUCACCAAGAUCGGU
Sfl-4-1     CACGGACAAGAGCGUCGUCCACACGCUCUUCACCGAGAUCGCCCCGCGGUACGAGAACCGCCCCGGUGGUUACACCCGCAUCACGAAGAUCGGC
Spu-1-1     CACGGACAAGGGCGUCGUCCACACGCUCUUCACCGAGAUCGCUCCGCGUUACGCCGAGCGUCCGGGUGGCUACACCCGCAUCACCAAGGUCGGC
Ssc-4-2     CCGUGACAAGAGCGUCGUGCACACGCUCUUCACGGAGAUCGCGCCGCGCUUCGCGGAGCGUCCGGGUGGCUACACCCGCAUCACCAAGAUCGGC
Shy-1-1     CCGGGACAAGAGCGUCGUGCACACUCUCUUCACCGAGAUCGCGCCGCGCUACGAGAACCGGCCGGGUGGCUACACCCGCAUCACCAAGAUCGGC
env-32      CCGGGACAAGAGCGUCGUGCACACUCUCUUCACCGAGAUCGCGCCGCGCUACGAGAACCGGCCGGGUGGCUACACCCGCAUCACCAAGAUCGGC
Sbi-1-1     CCGUGACAAGAGCGUGGUGCACACUCUCUUCACGGAGAUCGGCCCGCGUUACGAGAACCGCCCGGGUGGCUACACCCGUAUCACCAAGAUCGGU
Afl-1-1     UCGCGACAAGAGCGUCGUGCACGAGCUGUUCACGGUGAUCGGCCCGCAGUUCGCGGCUCGUCCGGGUGGCUACACCCGCAUCACCAAGCUCGGC
Ssp-55-1    CACCGACAAGUCGGUCGUGCACACCCUCUUCACCGAGAUCGCGCCGCGGUACGAGAACCGCCCCGGUGGCUACACCCGCAUCACCAAGAUCGGC
Ssp-152-1   CACGGACAAGAGCAUCGUGCACACGCUGUUCACCGAGAUCGCCCCGCGCUACGAGAACCGCCCCGGUGGUUACACGCGCAUCACCAAGAUCGGC
Ssp-104-1   CACGGACAAGAGCAUCGUGCACACGCUGUUCACCGAGAUCGCCCCGCGCUACGAGAACCGCCCCGGUGGUUACACGCGCAUCACCAAGAUCGGC
Ssp-166-1   CACGGACAAGAGCAUCGUGCACACGCUGUUCACCGAGAUCGCCCCGCGCUACGAGAACCGCCCCGGUGGUUACACGCGCAUCACCAAGAUCGGC
Ssp-191-1   CACGGACAAGAGCAUCGUGCACACGCUGUUCACCGAGAUCGCCCCGCGCUACGAGAACCGCCCCGGUGGUUACACGCGCAUCACCAAGAUCGGC
Sxa-1-1     CACGGACAAGAGCAUCGUGCACACGCUGUUCACCGAGAUCGCCCCGCGCUACGAGAACCGCCCCGGUGGUUACACGCGCAUCACCAAGAUCGGC
Sla-1-1     CACGGACAAGAGCAUCGUGCACACGCUGUUCACCGAGAUCGCCCCGCGCUACGAGAACCGCCCCGGUGGUUACACGCGCAUCACCAAGAUCGGC
Svi-12-3    CACGGACAAGAGCAUCGUGCACACGCUGUUCACCGAGAUCGCCCCGCGCUACGAGAACCGCCCCGGUGGUUACACGCGCAUCACCAAGAUCGGC
Sla-2-2     CACGGACAAGAGCAUCGUGCACACGCUGUUCACCGAGAUCGCCCCGCGCUACGAGAACCGCCCCGGUGGUUACACGCGCAUCACCAAGAUCGGC
Ssp-150-1   CACGGACAAGAGCAUCGUGCACACGCUGUUCACCGAGAUCGCCCCGCGCUACGAGAACCGCCCCGGUGGUUACACGCGCAUCACCAAGAUCGGC

            ................................................................................................
            ................................................................................................
            C•••GACAAGRGCRUCGU•CACAC•CUCUUCACCGAGAUCG••CCGCG•UACG•RRA•CGYCC•GGUGGYUACACCCGYAUCACCAAGAUCGGY
```

```
Ssp-32-1    CACGGACAAGAGCAUCGUGCACACGCUGUUCACCGAGAUCGCCCCGCGUUACGAGAACCGCCCCGGUGGUUACACGCGCAUCACCAAGAUCGGC
Ska-1-1     CACGGACAAGAGCAUCGUGCACACGCUGUUCACCGAGAUCGCCCCGCGCUACGAGAACCGCCCCGGUGGUUACACGCGCAUCACCAAGAUCGGC
Sro-6-1     CACGGACAAGAGCAUCGUGCACACGCUGUUCACCGAGAUCGCCCCGCGCUACGAGAACCGCCCCGGUGGUUACACGCGCAUCACCAAGAUCGGC
Ssp-194-1   CCUGGACAAGGGCGUCGUCCACACGCUCUUCACCGAGAUCGGCCCGCGGUACGAGAACCGUCCGGGCGGCUACACCCGCAUCACCAAGAUCGGC
env-33      CACGGACAAGGGCAUCGUCCACACGCUCUUCACCGAGAUCGCCCCGCGCUACGCCGAGCGUCCGGGUGGCUACACCCGCAUCACCAAGAUCGGC
Ssp-7-1     CACGGACAAGAGCAUCGUCCACACGCUCUUCACCGAGAUCGCCCCGCGGUACGAGAACCGCCCGGGUGGUUACACCCGCAUCACCAAGAUCGGU
Ssp-140-1   CACGGACAAGGGCAUCGUCCACACGCUCUUCACCGAGAUCGCUCCGCGUUACGCCGAGCGCCCGGGUGGCUACACCCGCAUCACCAAGAUCGGC
Svi-1-1     CACGGACAAGAGCAUCGUCCACACGCUCUUCACCGAGAUCGCCCCGCGCUACGCCGAGCGCCCGGGUGGUUACACCCGCAUCACCAAGAUCGGC
Sca-2-1     CACCGACAAGAGCGUCGUCCACACGCUCUUCACGGAGAUCGCCCCCCGGUACGAGAACCGCCCGGGUGGUUACACCCGCAUCACCAAGAUCGGC
Sts-1-1     CACGGACAAGGGCAUCGUGCACGCGCUCUUCACCGAGAUCGCCCCGCGCUACGCCGAGCGCCCGGGUGGCUACACCCGGAUCACCAAGAUCGGC
Ssp-25-1    CACGGACAAGAGCGUCGUCCACACGCUCUUCACCGAGAUCGCCCCGCGCUACGAGAACCGUCCGGGUGGUUACACCCGUAUCACCAAGAUCGGU
Ska-1-2     CACGGACAAGAGCAUCGUGCACACGCUGUUCACCGAGAUCGCCCCGCGCUACGAGAACCGCCCCGGUGGUUACACGCGCAUCACCAAGAUCGGC
Smo-2-1     CUCGGACAAGAGCGUCGUGCACACGCUCUUCACGGAGAUCGCCCCGCGCUUCGAGGAGCGCCCGGGUGGCUACACCCGCAUCACCAAGAUCGGC
Ssp-70-1    CACGGACAAGAGCGUCGUGCACACGCUCUUCACCGAGAUCGCCCCGCGCUUCGCCGAGCGUCCGGGUGGCUACACCCGCAUCACCAAGAUCGGC
Sba-2-1     CACGGACAAGAGCGUCGUCCACACGCUCUUCACCGAGAUCGCUCCCCGGUACGAGAACCGCCCCGGUGGUUACACCCGUAUCACCAAGAUCGGC
Sme-1-1     CACGGACAAGAGCGUCGUCCACACGCUCUUCACCGAGAUCGCUCCCCGGUACGAGAACCGCCCCGGUGGUUACACCCGUAUCACCAAGAUCGGC
Sgr-11-1    CACGGACAAGAGCGUCGUCCACACGCUCUUCACCGAGAUCGCUCCCCGGUACGAGAACCGCCCCGGUGGUUACACCCGUAUCACCAAGAUCGGC
Sgl-1-1     CACGGACAAGAGCGUCGUCCACACGCUCUUCACCGAGAUCGCUCCCCGGUACGAGAACCGCCCCGGUGGUUACACCCGUAUCACCAAGAUCGGC
Ssp-76-1    CACGGACAAGAGCGUCGUCCACACGCUCUUCACCGAGAUCGCUCCCCGGUACGAGAACCGCCCCGGUGGUUACACCCGUAUCACCAAGAUCGGC
Sgr-12-7    CACGGACAAGAGCGUCGUCCACACGCUCUUCACCGAGAUCGCUCCCCGGUACGAGAACCGCCCCGGUGGUUACACCCGUAUCACCAAGAUCGGC
Sgl-3-2     CACGGACAAGAGCGUCGUCCACACGCUCUUCACCGAGAUCGCUCCCCGGUACGAGAACCGCCCCGGUGGUUACACCCGUAUCACCAAGAUCGGC
Sgr-4-1     CACGGACAAGAGCGUCGUCCACACGCUCUUCACCGAGAUCGCUCCCCGGUACGAGAACCGCCCCGGUGGUUACACCCGUAUCACCAAGAUCGGC
San-2-1     CACGGACAAGAGCAUCGUCCACACGCUCUUCACCGAGAUCGCUCCCCGAUACGAGAACCGCCCCGGUGGUUACACCCGUAUCACCAAGAUCGGC
Sgr-3-1     CACGGACAAGAGCGUCGUCCACACGCUCUUCACCGAGAUCGCUCCCCGGUACGAGAACCGCCCCGGUGGUUACACCCGUAUCACCAAGAUCGGC
Ssp-9-1     CACGGACAAGAGCAUCGUCCACACGCUCUUCACCGAGAUCGCUCCCCGAUACGAGAACCGCCCCGGUGGUUACACCCGUAUCACCAAGAUCGGC
Seu-1-1     CACGGACAAGAGCAUCGUCCACACGCUCUUCACCGAGAUCGCUCCCCGAUACGAGAACCGCCCCGGUGGUUACACCCGUAUCACCAAGAUCGGC
Ssp-82-1    CACGGACAAGAGCAUCGUCCACACGCUCUUCACCGAGAUCGCUCCCCGGUACGAGAACCGCCCCGGUGGUUACACCCGUAUCACCAAGAUCGGC
Sal-12-2    CACGGACAAGAGCGUCGUCCACACGCUCUUCACCGAGAUCGCUCCCCGGUACGAGAACCGCCCCGGUGGUUACACCCGUAUCACCAAGAUCGGC
Ssp-48-1    CACGGACAAGAGCAUCGUCCACACGCUCUUCACCGAGAUCGCUCCCCGAUACGAGAACCGCCCCGGUGGUUACACCCGUAUCACCAAGAUCGGC
Sha-1-1     CACGGACAAGAGCAUCGUGCACACGCUCUUCACCGAGAUCGCCCCGCGGUACGAGAACCGCCCCGGUGGUUACACCCGCAUCACCAAGAUCGGC
Ssp-45-1    CACGGACAAGAGCAUCGUGCACACGCUCUUCACCGAGAUCGCCCCGCGGUACGAGAACCGCCCCGGUGGUUACACCCGCAUCACCAAGAUCGGC
Ssp-71-1    CACGGACAAGAGCAUCGUGCACACGCUCUUCACCGAGAUCGCCCCGCGGUACGAGAACCGCCCCGGUGGUUACACCCGCAUCACCAAGAUCGGC
Ssp-175-1   CACGGACAAGAGCAUCGUGCACACGCUCUUCACCGAGAUCGCGCCGCGGUACGAGAACCGCCCCGGUGGUUACACCCGCAUCACCAAGAUCGGC
Ssp-188-1   CACGGACAAGAGCAUCGUGCACACGCUCUUCACCGAGAUCGCCCCGCGGUACGAGAACCGCCCCGGUGGUUACACCCGCAUCACCAAGAUCGGC
Ssp-52-1    CACGGACAAGAGCAUCGUGCACACGCUCUUCACCGAGAUCGCCCCGCGCUACGAGAACCGCCCCGGUGGCUACACGCGCAUCACCAAGAUCGGC
Sri-4-1     CUCGGACAAGAGCGUCGUGCACACGCUCUUCACGGAGAUCGCCCCGCGCUUCGAGGAGCGCCCGGGUGGCUACACCCGUAUCACCAAGAUCGGC
Ssp-145-1   CUCGGACAAGAGCGUCGUGCACACGCUCUUCACGGAGAUCGCCCCGCGGUUCGAGAACCGCCCGGGUGGCUACACCCGUAUCACCAAGAUCGGU
env-34      ..............................................................................................
Sly-1-1     CUCGGACAAGAGCGUCGUGCACACGCUCUUCACCGAGAUCGCCCCGCGGUUCGAGAACCGCCCGGGUGGCUACACCCGCAUCACCAAGAUCGGU
Ssp-142-1   CUCGGACAAGAGCGUCGUGCACACGCUCUUCACGGAGAUCGCGCCGCGGUUCGCGGAGCGUCCGGGUGGCUACACCCGUAUCACCAAGAUCGGC
Ssp-155-1   CUCGGACAAGAGCGUCGUGCACACGCUCUUCACGGAGAUCGCGCCGCGGUUCGCGGAGCGUCCGGGUGGCUACACCCGUAUCACCAAGAUCGGC
Ssp-53-1    CACGGACAAGAGCGUCGUGCACACGCUCUUCACCGAGAUCGCGCCGCGGUUCGCGGAGCGUCCGGGUGGCUACACCCGUAUCACCAAGAUCGGC
Sat-2-1     CACGGACAAGAGCAUCGUGCACACGCUCUUCACCGAGAUCGCCCCGCGGUACGAGAACCGCCCCGGUGGUUACACCCGCAUCACCAAGAUCGGC
Ssp-95-1    CACGGACAAGAGCAUCGUGCACACGCUCUUCACCGAGAUCGCCCCGCGGUACGAGAACCGCCCCGGUGGUUACACCCGCAUCACCAAGAUCGGC
Sfl-1-1     CACGGACAAGAGCAUCGUGCACACGCUCUUCACCGAGAUCGCCCCGCGGUACGAGAACCGCCCCGGUGGUUACACCCGCAUCACCAAGAUCGGC
[pa-1-1     CACGGACAAGAGCAUCGUGCACACGCUCUUCACCGAGAUCGCCCCGCGGUACGAGAACCGCCCCGGUGGUUACACCCGCAUCACCAAGAUCGGC
Sfl-8-1     CACGGACAAGAGCAUCGUGCACACGCUCUUCACCGAGAUCGCCCCGCGGUACGAGAACCGCCCCGGUGGUUACACCCGCAUCACCAAGAUCGGC
Ssp-154-1   CACGGACAAGAGCAUCGUGCACACGCUCUUCACCGAGAUCGCCCCGCGGUACGAGAACCGCCCCGGUGGUUACACCCGCAUCACCAAGAUCGGC
Sce-1-1     CUCGGACAAGAGCGUCGUGCACACGCUCUUCACCGAGAUCGGUCCGCGGUUCGAGAACCGCCCGGGUGGCUACACCCGGAUCACCAAGAUCGGU
Ssc-5-1     CACGGACAAGAGCGUCGUCCACACGCUCUUCACCGAGAUCGCCCCGCGGUACGAGAACCGCCCCGGUGGUUACACGCGUAUCACCAAGAUCGGC
Ssp-144-1   CACGGACAAGGGCAUCGUGCACACGCUCUUCACCGAGAUCGCCCCGCGGUACGAGAACCGCCCCGGUGGUUACACCCGCAUCACCAAGAUCGGC
Ssp-54-1    CACGGACAAGGGCGUCGUCCACACGCUCUUCACCGAGAUCGCCCCGCGGUACGAGAACCGCCCCGGUGGUUACACCCGUAUCACCAAGAUCGGC
Sgr-13-1    CACGGACAAGAGCGUCGUCCACACGCUCUUCACCGAGAUCGCCCCCCGGUACGAGAACCGCCCCGGUGGUUACACCCGUAUCACCAAGAUCGGC
Sfl-9-1     CACGGACAAGAGCGUCGUCCACACGCUCUUCACCGAGAUCGCCCCCCGGUACGAGAACCGCCCCGGUGGUUACACCCGUAUCACCAAGAUCGGC
Sro-4-1     CACGGACAAGAGCGUCGUCCACACGCUCUUCACCGAGAUCGCUCCCCGGUACGAGAACCGCCCCGGUGGUUACACCCGUAUCACCAAGAUCGGC
Sal-12-3    CACGGACAAGAGCGUCGUCCACACGCUCUUCACCGAGAUCGCUCCCCGGUACGAGAACCGCCCCGGUGGUUACACCCGUAUCACCAAGAUCGGC
Ssp-174-1   CACGGACAAGAGCGUCGUCCACACGCUCUUCACCGAGAUCGCUCCCCGGUACGAGAACCGCCCCGGUGGUUACACCCGUAUCACCAAGAUCGGC
Ssp-129-1   CACGGACAAGAGCGUCGUCCACACGCUCUUCACCGAGAUCGCCCCCCGGUACGAGAACCGCCCCGGUGGUUACACCCGUAUCACCAAGAUCGGC
Spu-2-1     CACGGACAAGAGCGUCGUCCACACGCUCUUCACCGAGAUCGCUCCCCGGUACGAGAACCGCCCCGGUGGUUACACCCGUAUCACCAAGAUCGGC
Svi-8-1     CACGGACAAGAGCGUCGUCCACACGCUCUUCACCGAGAUCGCCCCCCGGUACGAGAACCGCCCCGGUGGUUACACCCGUAUCACCAAGAUCGGC
Ssp-66-1    CACGGACAAGAGCGUCGUCCACACGCUCUUCACCGAGAUCGCCCCCCGGUACGAGAACCGCCCCGGUGGUUACACCCGUAUCACCAAGAUCGGC
Ssp-94-1    CACGGACAAGAGCGUCGUCCACACGCUCUUCACCGAGAUCGCUCCCCGGUACGAGAACCGCCCCGGUGGUUACACCCGUAUCACCAAGAUCGGC
Ssp-17-1    CACGGACAAGAGCGUCGUCCACACGCUCUUCACCGAGAUCGCGCCCCGGUACGAGAACCGCCCCGGUGGUUACACCCGUAUCACCAAGAUCGGC
Ssp-193-1   CACGGACAAGAGCGUCGUCCACACGCUCUUCACCGAGAUCGCGCCCCGGUACGAGAACCGCCCCGGUGGUUACACCCGUAUCACCAAGAUCGGC
Sat-1-1     CACGGACAAGAGCAUCGUGCACACGCUCUUCACCGAGAUCGCCCCGCGCUACGAGAACCGCCCCGGUGGUUACACGCGCAUCACCAAGAUCGGC
Ssp-44-1    CACGGACAAGAGCGUCGUCCACACGCUCUUCACCGAGAUCGCCCCCCGGUACGAGAACCGCCCCGGUGGUUACACGCGUAUCACCAAGAUCGGC
Scy-1-1     CACGGACAAGAGCAUCGUCCACACGCUCUUCACCGAGAUCGCCCCCCGUUACGAGAACCGCCCCGGUGGUUACACGCGUAUCACCAAGAUCGGC
Sfu-1-1     CACGGACAAGAGCAUCGUGCACACGCUCUUCACCGAGAUCGCCCCCCGAUACGAGAACCGCCCCGGUGGUUACACGCGUAUCACCAAGAUCGGC
Sal-7-1     CACGGACAAGAGCAUCGUGCACACGCUCUUCACCGAGAUCGCCCCCCGCUACGAGAACCGCCCCGGUGGUUACACGCGUAUCACCAAGAUCGGC
Ssp-168-1   CACGGACAAGAGCAUCGUGCACACGCUCUUCACCGAGAUCGCCCCCCGGUACGAGAACCGCCCCGGUGGUUACACGCGUAUCACCAAGAUCGGC
Ssp-97-1    CACGGACAAGGGCAUCGUGCACACGCUCUUCACCGAGAUCGCCCCGCGCUACGAGAACCGCCCUGGUGGUUACACGCGUAUCACCAAGAUCGGC
Ssp-146-1   CACGGACAAGGGCAUCGUGCACACGCUGUUCACCGAGAUCGCCCCGCGCUACGAGAACCGCCCCGGUGGUUACACGCGCAUCACCAAGAUUGGC
Sod-1-1     CACGGACAAGGGCAUCGUGCACACGCUCUUCACCGAGAUCGCGCCGCGGUACGAGAACCGCCCCGGUGGUUACACCCGUAUCACCAAGAUCGGC
Sri-4-2     CUCGGACAAGAGCGUCGUGCACACGCUCUUCACGGAGAUCGCGCCGCGCUUCGCGGAGCGUCCGGGUGGUUACACCCGUAUCACCAAGAUCGGC
Sgr-6-1     CUCGGACAAGAGCGUCGUGCACACGCUCUUCACGGAGAUCGCGCCGCGCUUCGCGGAGCGUCCGGGUGGCUACACCCGUAUCACCAAGAUCGGC
Sri-4-5     CUCGGACAAGAGCGUCGUGCACACGCUCUUCACGGAGAUCGCGCCGCGCUUCGCGGAGCGUCCGGGUGGUUACACCCGUAUCACCAAGAUCGGC
Sri-2-1     CUCGGACAAGAGCGUCGUGCACACGCUCUUCACGGAGAUCGCGCCGCGCUUCGCGGAGCGUCCGGGUGGUUACACCCGUAUCACCAAGAUCGGC
Ssp-90-1    CUCGGACAAGAGCGUCGUGCACACGCUCUUCACGGAGAUCGCGCCGCGCUUCGCGGAGCGUCCGGGUGGUUACACCCGUAUCACCAAGAUCGGC
Sri-4-18    CUCGGACAAGAGCGUCGUGCACACGCUCUUCACGGAGAUCGCGCCGCGCUUCGCGGAGCGUCCGGGUGGUUACACCCGUAUCACCAAGAUCGGC
Ssp-185-1   CUCGGACAAGAGCGUCGUGCACACGCUCUUCACGGAGAUCGCGCCGCGCUUCUCGGAGCGUCCGGGUGGUUACACCCGUAUCACCAAGAUCGGC
Sri-4-19    CUCGGACAAGAGCGUCGUGCACACGCUCUUCACGGAGAUCGCGCCGCGCUUCGCGGAGCGUCCGGGUGGUUACACCCGUAUCACCAAGAUCGGC
Ssp-5-1     CACGGACAAGGGCAUCGUUCACACGCUCUUCACCGAGAUCGCCCCGCGGUACGAGAACCGCCCCGGUGGUUACACCCGCAUCACCAAGAUCGGC
Ssp-29-1    CACGGACAAGGGCGUCGUCCACACGCUCUUCACCGAGAUCGCCCCGCGGUACGAGAACCGCCCCGGUGGUUACACCCGCAUCACCAAGAUCGGC
Ssp-26-1    CACGGACAAGGGCAUCGUCCACACGCUCUUCACCGAGAUCGCCCCGCGGUACGAGAACCGCCCCGGUGGUUACACCCGCAUCACCAAGAUCGGC

            ..............................................................................................
            ..............................................................................................
            C•••GACAAGRGCRUCGU•CACAC•CUCUUCACCGAGAUCG••CCGCG•UACG•RRA•CGYCC•GGUGGYUACACCCGYAUCACCAAGAUCGGY
```

```
Aat-1-1     CCGCGACAAGGGCGUGGUCCACGAGCUGUUCACCGAGAUCGCGCCGCGGUUCGCGAACCGUCCGGGCGGCUACACCCGGAUCACCAAGCUCGGG
Ssp-117-1   CACGGACAAGAGCAUCGUGCACACGCUCUUCACCGAGAUCGGCCCGCGGUACGAGAACCGCCCGGGUGGUUACACCCGCAUCACCAAGGUCGGC
Ssp-198-1   CACGGACAAGGGCGUCGUGCACACGCUCUUCACCGAGAUCGGCCCGCGGUACGAGAACCGCCCGGGUGGCUACACCCGCAUCACCAAGAUCGGU
Ssp-115-1   CCUCGACAAGAGCGUCGUGCACACCCUCUUCACCGAGAUCGGUCCGCGCUACGAGAACCGUCCCGGUGGCUACACCCGCAUCACCAAGAUCGGC
Psi-1-1     GGUCCUCAAGGUCAUCCGGGACAAGGGUGUCGUGCACACGCUCUUCACCGAGAUCGCGCCGACGUUCUCCGAGCGCCCGGGCGGCUACACCCGG
Ssp-4-1     CCUGGACAAGAGCGUCGUCCACACGCUCUUCACCGAGAUCGGUCCCCGGUACGAGAACCGUCCCGGUGGCUACACCCGCAUCACCAAGAUCGGC
Ssp-201-1   CAAGGACAAGAGCGUGGUCCACACGCUGUUCACCGAGAUCGGCCCGCGCUACGAGAACCGGCCGGGCGGCUACACCCGCAUCACCAAGAUCGGU
Aal-1-1     CCGCGACAAGGACAUCGUCCACACGCUGUUCGCGGAGAUCGGGCCGCUGUACGCCACUCGGCCCGGCGGCUACACCCGCAUCACGAAGGUCGGC
Sal-13-1    CCGCGACAAGGGCGUCGUCCACACGCUCUUCACCGAGAUCGCGCCGACCUUCGAGGCCCGGCCGGGCGGCUACACGCGCAUCACCAAGAUCGGC
Ari-1-1     CCGCGACAAGGGCGUCGUGCACGAGCUCUUCACGGAGAUCGCGCCGAAGUUCGAGAACCGUCCGGGCGGCUACACGCGCAUCACCAAGAUCGGG
Ssp-40-1    CACCGACAAGAGCGUCGUCCACACGCUCUUCACGGAGAUCGGCCCGCGUUACGAGAACCGCCCGGGUGGCUACACCCGUAUCACCAAGAUCGGC
Ssp-39-1    CACCGACAAGAGCGUCGUCCACACGCUCUUCACGGAGAUCGGCCCGCGUUACGAGAACCGCCCGGGUGGCUACACCCGUAUCACCAAGAUCGGC
Ssp-42-1    CACCGACAAGAGCGUCGUCCACACGCUCUUCACGGAGAUCGGCCCGCGUUACGAGAACCGCCCGGGUGGCUACACCCGUAUCACCAAGAUCGGC
Ssp-47-1    CACCGACAAGAGCGUCGUCCACACGCUCUUCACGGAGAUCGGCCCGCGUUACGAGAACCGCCCGGGUGGCUACACCCGUAUCACCAAGAUCGGC
Ssp-41-1    CACCGACAAGAGCGUCGUCCACACGCUCUUCACGGAGAUCGGCCCGCGUUACGAGAACCGUCCGGGUGGCUACACCCGUAUCACCAAGAUCGGC
Ssp-125-1   CACCGACAAGAGCGUCGUCCACACGCUCUUCACGGAGAUCGGCCCGCGUUACGAGAACCGCCCGGGUGGCUACACCCGGAUCACCAAGAUCGGC
Nsp-3-1     CCGCGACAAGUCGGUCGUGCACACGCUCUUCACGGACAUCGCGCCGACGUUCUCGGAGCGUCCCGGUGGCUACACCCGGAUCACCAAGAUCGGU
Nsp-2-1     CCGCGACAAGUCGGUCGUCCACGUGCUCUUCACCGAGAUCGCGGUCACCUUCGCCGAGCGUCCGGGUGGCUACACCCGGAUCACCAAGAUCGGU
env-35      CCGCGACAAGGGUGUCGUGCACGUCCUGUUCACCGAGAUCGCCCCGACCUUCUCCGAGCGUCCCGGUGGCUACACCCGCAUCACCAAGAUCGGC
env-37      ..............................................................................................
env-38      ..............................................................................................
env-39      ..............................................................................................
env-40      ..............................................................................................
Nlu-1-1     CCGCGACAAGUCCGUCGUGCACACCCUCUUCACCGAGAUCGCCGAGCAGUUCGCGGAGCGUCCGGGUGGCUACACCCGGAUCACCAAGAUCGGU
env-41      CCGCGACAAGUCCGUCGUGCACACCCUCUUCACCGAGAUCGCCGAGCAGUU...........................................
Nha-1-1     CCGCGACAAGUCGGUCGUGCACACGCUGUUCACCGAGAUCGCGCCCACCUUCGAGGAGCGUCCCGGGGGCUACACCCGGAUCACCAAGAUCGGU
Sst-1-1     CACGGACAAGAGCGUCGUGCACACGCUCUUCACCGAGAUCGGCCCGCGGUACGAGAACCGUCCCGGUGGCUACACCCGCAUCACCAAGAUCGGU
Sle-1-1     CACGGACAAGGGCAUCGUCCACACGCUCUUCACCGAGAUCGGCCCGCGGUACGAGAACCGCCCGGGUGGCUACACCCGGAUCACCAAGAUCGGC
Sal-2-1     CACGGACAAGAGCGUCGUCCACACGCUCUUCACCGAGAUCGGCCCCCGUUACGAGAACCGCCCGGGUGGCUACACCCGGAUCACCAAGGUCGGU
Ssp-24-1    CCUCGACAAGGGCGUCGUGCACACCCUCUUCACCGAGAUCGGCCCGCGGUACGAGAACCGCCCCGGCGGUUACACCCGCAUCACCAAGAUCGGU
Scl-1-1     CACGGACAAGGGCAUCGUCCACGCCCUGUUCACCGAGAUCGCUCCGCGGUACGCCGAGCGUCCGGGUGGUUACACCCGUAUCACCAAGAUCGGC
Ssc-1-1     CACGGACAAGAGCGUCGUCCACACGCUCUUCACCGAGAUCGGCCCGCGCUACGAGAACCGUCCGGGUGGUUACACCCGUAUCACCAAGAUCGGU
Sde-1-1     CUCGGACAAGAGCGUCGUGCACACGCUCUUCACGGAGAUCGCCCCGCGGUUCGAGAACCGCCCGGGUGGCUACACCCGUAUCACCAAGAUCGGU
            ..............................................................................................
            ..............................................................................................
            C•••GACAAGRGCRUCGU•CACAC•CUCUUCACCGAGAUCG••CCGCG•UACG•RRA•CGYCC•GGUGGYUACACCCGYAUCACCAAGAUCGGY
```

### 6 L19-Flavobacteria

#### 6.1 Taxa

The taxonomy of each organism containing a putative L19-Flavobacteria RNA is listed, with abbreviations identifying each hit (e.g., "Eco-1-1" and "Eco-1-2" might hypothetically represent two distinct RNAs in *E. coli*). The abbreviations will be used to identify each individual L19-Flavobacteria RNA in Sections 6.2 and 6.4:.

| abbrev. of hits | taxonomy of species |
|---|---|
| Fba-1-1 | Bacteria Bacteroidetes *Flavobacteria bacterium* BBFL7 |
| Fba-2-1 | Bacteria Bacteroidetes Flavobacteria Flavobacteriales *Flavobacteria bacterium* BAL38 |
| Cgi-1-1 | Bacteria Bacteroidetes Flavobacteria Flavobacteriales Flavobacteriaceae *Capnocytophaga gingivalis* ATCC 33624 |
| Coc-1-1 | Bacteria Bacteroidetes Flavobacteria Flavobacteriales Flavobacteriaceae *Capnocytophaga ochracea* DSM 7271 |
| Csp-1-1 | Bacteria Bacteroidetes Flavobacteria Flavobacteriales Flavobacteriaceae *Capnocytophaga sputigena* ATCC 33612 |
| Cat-1-1 | Bacteria Bacteroidetes Flavobacteria Flavobacteriales Flavobacteriaceae *Croceibacter atlanticus* HTCC2559 |
| Fba-3-1 | Bacteria Bacteroidetes Flavobacteria Flavobacteriales *Flavobacteriaceae bacterium* 3519-10 |
| Fjo-1-1 | Bacteria Bacteroidetes Flavobacteria Flavobacteriales Flavobacteriaceae *Flavobacterium johnsoniae* UW101 |
| Gfo-1-1 | Bacteria Bacteroidetes Flavobacteria Flavobacteriales Flavobacteriaceae *Gramella forsetii* KT0803 |
| Kal-1-1 | Bacteria Bacteroidetes Flavobacteria Flavobacteriales Flavobacteriaceae *Kordia algicida* OT-1 |
| Lbl-1-1 | Bacteria Bacteroidetes Flavobacteria Flavobacteriales Flavobacteriaceae *Leeuwenhoekiella blandensis* MED217 |
| Pir-1-1 | Bacteria Bacteroidetes Flavobacteria Flavobacteriales Flavobacteriaceae *Polaribacter irgensii* 23-P |
| Rbi-1-1 | Bacteria Bacteroidetes Flavobacteria Flavobacteriales Flavobacteriaceae *Robiginitalea biformata* HTCC2501 |
| Zpr-1-1 | Bacteria Bacteroidetes Flavobacteria Flavobacteriales Flavobacteriaceae *Zunongwangia profunda* SM-A87 |
| Fba-4-1 | Bacteria Bacteroidetes Flavobacteria *Flavobacteriales bacterium* ALC-1 |
| Fba-5-1 | Bacteria Bacteroidetes Flavobacteria *Flavobacteriales bacterium* HTCC2170 |
| Oho-1-1 | Bacteria Bacteroidetes Flavobacteriia Flavobacteriales Cryomorphaceae *Owenweeksia hongkongensis* DSM 17368 |
| Aca-1-1 | Bacteria Bacteroidetes Flavobacteriia Flavobacteriales Flavobacteriaceae *Aequorivita capsosiphonis* DSM 23843 |
| Ale-1-1 to Ale-1-2 | Bacteria Bacteroidetes Flavobacteriia Flavobacteriales Flavobacteriaceae *Algibacter lectus* |
| Aag-1-1 | Bacteria Bacteroidetes Flavobacteriia Flavobacteriales Flavobacteriaceae *Aquimarina agarilytica* ZC1 |
| Aat-1-1 | Bacteria Bacteroidetes Flavobacteriia Flavobacteriales Flavobacteriaceae *Aquimarina atlantica* |

| | |
|---|---|
| Ala-1-1 | Bacteria Bacteroidetes Flavobacteriia Flavobacteriales Flavobacteriaceae *Aquimarina latercula* DSM 2041 |
| Ama-1-1 | Bacteria Bacteroidetes Flavobacteriia Flavobacteriales Flavobacteriaceae *Aquimarina macrocephali* JAMB N27 |
| Ame-1-1 | Bacteria Bacteroidetes Flavobacteriia Flavobacteriales Flavobacteriaceae *Aquimarina megaterium* XH134 |
| Amu-1-1 | Bacteria Bacteroidetes Flavobacteriia Flavobacteriales Flavobacteriaceae *Aquimarina muelleri* DSM 19832 |
| Apa-1-1 | Bacteria Bacteroidetes Flavobacteriia Flavobacteriales Flavobacteriaceae *Aquimarina pacifica* |
| Aal-1-1 | Bacteria Bacteroidetes Flavobacteriia Flavobacteriales Flavobacteriaceae *Arenibacter algicola* |
| Ace-1-1 | Bacteria Bacteroidetes Flavobacteriia Flavobacteriales Flavobacteriaceae *Arenibacter certesii* DSM 19833 |
| Ala-2-1 | Bacteria Bacteroidetes Flavobacteriia Flavobacteriales Flavobacteriaceae *Arenibacter latericius* DSM 15913 |
| Bar-1-1 | Bacteria Bacteroidetes Flavobacteriia Flavobacteriales Flavobacteriaceae *Bizionia argentinensis* JUB59 |
| Cca-1-1 to Cca-1-3 | Bacteria Bacteroidetes Flavobacteriia Flavobacteriales Flavobacteriaceae *Capnocytophaga canis* |
| Cgr-1-1 | Bacteria Bacteroidetes Flavobacteriia Flavobacteriales Flavobacteriaceae *Capnocytophaga granulosa* ATCC 51502 |
| Coc-2-1 | Bacteria Bacteroidetes Flavobacteriia Flavobacteriales Flavobacteriaceae *Capnocytophaga ochracea* |
| Coc-3-1 | Bacteria Bacteroidetes Flavobacteriia Flavobacteriales Flavobacteriaceae *Capnocytophaga ochracea* F0287 |
| Coc-4-1 | Bacteria Bacteroidetes Flavobacteriia Flavobacteriales Flavobacteriaceae *Capnocytophaga ochracea str.* Holt 25 |
| Csp-2-1 | Bacteria Bacteroidetes Flavobacteriia Flavobacteriales Flavobacteriaceae *Capnocytophaga sp.* CM59 |
| Csp-3-1 | Bacteria Bacteroidetes Flavobacteriia Flavobacteriales Flavobacteriaceae *Capnocytophaga sp. oral taxon* 324 str. F0483 |
| Csp-4-1 | Bacteria Bacteroidetes Flavobacteriia Flavobacteriales Flavobacteriaceae *Capnocytophaga sp. oral taxon* 326 str. F0382 |
| Csp-5-1 | Bacteria Bacteroidetes Flavobacteriia Flavobacteriales Flavobacteriaceae *Capnocytophaga sp. oral taxon* 329 str. F0087 |
| Csp-6-1 | Bacteria Bacteroidetes Flavobacteriia Flavobacteriales Flavobacteriaceae *Capnocytophaga sp. oral taxon* 332 str. F0381 |
| Csp-7-1 | Bacteria Bacteroidetes Flavobacteriia Flavobacteriales Flavobacteriaceae *Capnocytophaga sp. oral taxon* 335 str. F0486 |
| Csp-8-1 | Bacteria Bacteroidetes Flavobacteriia Flavobacteriales Flavobacteriaceae *Capnocytophaga sp. oral taxon* 336 str. F0502 |
| Csp-9-1 | Bacteria Bacteroidetes Flavobacteriia Flavobacteriales Flavobacteriaceae *Capnocytophaga sp. oral taxon* 380 str. F0488 |
| Csp-10-1 | Bacteria Bacteroidetes Flavobacteriia Flavobacteriales Flavobacteriaceae *Capnocytophaga sp. oral taxon* 412 str. F0487 |
| Csp-11-1 | Bacteria Bacteroidetes Flavobacteriia Flavobacteriales Flavobacteriaceae *Capnocytophaga sp. oral taxon* 863 str. F0517 |
| Cal-1-1 | Bacteria Bacteroidetes Flavobacteriia Flavobacteriales Flavobacteriaceae *Cellulophaga algicola* DSM 14237 |
| Cba-1-1 | Bacteria Bacteroidetes Flavobacteriia Flavobacteriales Flavobacteriaceae *Cellulophaga baltica* 13 |
| Cba-2-1 | Bacteria Bacteroidetes Flavobacteriia Flavobacteriales Flavobacteriaceae *Cellulophaga baltica* 18 |
| Cba-3-1 | Bacteria Bacteroidetes Flavobacteriia Flavobacteriales Flavobacteriaceae *Cellulophaga baltica* 4 |
| Cba-4-1 | Bacteria Bacteroidetes Flavobacteriia Flavobacteriales Flavobacteriaceae *Cellulophaga baltica* NN016038 |
| Csp-12-1 | Bacteria Bacteroidetes Flavobacteriia Flavobacteriales Flavobacteriaceae *Cellulophaga sp.* E6(2014) |
| Csp-13-1 | Bacteria Bacteroidetes Flavobacteriia Flavobacteriales Flavobacteriaceae *Cellulophaga sp.* Hel_I_12 |
| Cca-2-1 | Bacteria Bacteroidetes Flavobacteriia Flavobacteriales Flavobacteriaceae *Chryseobacterium caeni* DSM 17710 |
| Cgr-2-1 | Bacteria Bacteroidetes Flavobacteriia Flavobacteriales Flavobacteriaceae *Chryseobacterium gregarium* DSM 19109 |
| Cha-1-1 | Bacteria Bacteroidetes Flavobacteriia Flavobacteriales Flavobacteriaceae *Chryseobacterium halperniae* |
| Chi-1-1 | Bacteria Bacteroidetes Flavobacteriia Flavobacteriales Flavobacteriaceae *Chryseobacterium hispalense* DSM 25574 |
| Cko-1-1 | Bacteria Bacteroidetes Flavobacteriia Flavobacteriales Flavobacteriaceae *Chryseobacterium koreense* CCUG 49689 |
| Cpa-1-1 | Bacteria Bacteroidetes Flavobacteriia Flavobacteriales Flavobacteriaceae *Chryseobacterium palustre* DSM 21579 |
| Csp-14-1 | Bacteria Bacteroidetes Flavobacteriia Flavobacteriales Flavobacteriaceae *Chryseobacterium sp.* FH1 |
| Csp-15-1 | Bacteria Bacteroidetes Flavobacteriia Flavobacteriales Flavobacteriaceae *Chryseobacterium sp.* FH2 |
| Csp-16-1 | Bacteria Bacteroidetes Flavobacteriia Flavobacteriales Flavobacteriaceae *Chryseobacterium sp.* OV259 |
| Csp-17-1 | Bacteria Bacteroidetes Flavobacteriia Flavobacteriales Flavobacteriaceae *Chryseobacterium sp.* YR005 |
| Csp-18-1 | Bacteria Bacteroidetes Flavobacteriia Flavobacteriales Flavobacteriaceae *Chryseobacterium sp.* YR460 |
| Cte-1-1 | Bacteria Bacteroidetes Flavobacteriia Flavobacteriales Flavobacteriaceae *Chryseobacterium tenax* DSM 16811 |
| Ddo-1-1 | Bacteria Bacteroidetes Flavobacteriia Flavobacteriales Flavobacteriaceae *Dokdonia donghaensis* DSW-1 |
| Dsp-1-1 | Bacteria Bacteroidetes Flavobacteriia Flavobacteriales Flavobacteriaceae *Dokdonia sp.* MED134 |
| Dsp-2-1 | Bacteria Bacteroidetes Flavobacteriia Flavobacteriales Flavobacteriaceae *Dokdonia sp.* PRO95 |
| Ean-1-1 to Ean-1-4 | Bacteria Bacteroidetes Flavobacteriia Flavobacteriales Flavobacteriaceae *Elizabethkingia anophelis* |
| Ean-2-1 | Bacteria Bacteroidetes Flavobacteriia Flavobacteriales Flavobacteriaceae *Elizabethkingia anophelis* Ag1 |
| Ean-3-1 | Bacteria Bacteroidetes Flavobacteriia Flavobacteriales Flavobacteriaceae *Elizabethkingia anophelis* FMS-007 |
| Ean-4-1 | Bacteria Bacteroidetes Flavobacteriia Flavobacteriales Flavobacteriaceae *Elizabethkingia anophelis* NUH1 |
| Ean-5-1 | Bacteria Bacteroidetes Flavobacteriia Flavobacteriales Flavobacteriaceae *Elizabethkingia anophelis* NUH11 |
| Ean-6-1 | Bacteria Bacteroidetes Flavobacteriia Flavobacteriales Flavobacteriaceae *Elizabethkingia anophelis* NUH4 |
| Ean-7-1 | Bacteria Bacteroidetes Flavobacteriia Flavobacteriales Flavobacteriaceae *Elizabethkingia anophelis* NUH6 |
| Ean-8-1 | Bacteria Bacteroidetes Flavobacteriia Flavobacteriales Flavobacteriaceae *Elizabethkingia anophelis* NUHP1 |
| Ean-9-1 | Bacteria Bacteroidetes Flavobacteriia Flavobacteriales Flavobacteriaceae *Elizabethkingia anophelis* NUHP2 |
| Ean-10-1 | Bacteria Bacteroidetes Flavobacteriia Flavobacteriales Flavobacteriaceae *Elizabethkingia anophelis* NUHP3 |
| Ean-11-1 | Bacteria Bacteroidetes Flavobacteriia Flavobacteriales Flavobacteriaceae *Elizabethkingia anophelis* PW2809 |
| Ean-12-1 | Bacteria Bacteroidetes Flavobacteriia Flavobacteriales Flavobacteriaceae *Elizabethkingia anophelis* R26 |
| Eme-1-1 | Bacteria Bacteroidetes Flavobacteriia Flavobacteriales Flavobacteriaceae *Elizabethkingia meningoseptica* 502 |
| Eme-2-1 to Eme-2-2 | Bacteria Bacteroidetes Flavobacteriia Flavobacteriales Flavobacteriaceae *Elizabethkingia meningoseptica* ATCC 13253 = NBRC 12535 |
| Emi-1-1 | Bacteria Bacteroidetes Flavobacteriia Flavobacteriales Flavobacteriaceae *Elizabethkingia miricola* |
| Ead-1-1 | Bacteria Bacteroidetes Flavobacteriia Flavobacteriales Flavobacteriaceae *Eudoraea adriatica* DSM 19308 |
| Fic-1-1 | Bacteria Bacteroidetes Flavobacteriia Flavobacteriales Flavobacteriaceae *Flaviramulus ichthyoenteri* Th78 |
| Fba-6-1 | Bacteria Bacteroidetes Flavobacteriia Flavobacteriales *Flavobacteriaceae bacterium* HQM9 |
| Fba-7-1 | Bacteria Bacteroidetes Flavobacteriia Flavobacteriales *Flavobacteriaceae bacterium* P7-3-5 |

| | |
|---|---|
| Fba-8-1 | Bacteria Bacteroidetes Flavobacteriia Flavobacteriales *Flavobacteriaceae bacterium* S85 |
| Fan-1-1 | Bacteria Bacteroidetes Flavobacteriia Flavobacteriales Flavobacteriaceae *Flavobacterium antarcticum* DSM 19726 |
| Faq-1-1 | Bacteria Bacteroidetes Flavobacteriia Flavobacteriales Flavobacteriaceae *Flavobacterium aquatile* LMG 4008 = ATCC 11947 |
| Fbr-1-1 | Bacteria Bacteroidetes Flavobacteriia Flavobacteriales Flavobacteriaceae *Flavobacterium branchiophilum* FL-15 |
| Fca-1-1 to Fca-1-2 | Bacteria Bacteroidetes Flavobacteriia Flavobacteriales Flavobacteriaceae *Flavobacterium cauense* R2A-7 |
| Fch-1-1 | Bacteria Bacteroidetes Flavobacteriia Flavobacteriales Flavobacteriaceae *Flavobacterium chungangense* LMG 26729 |
| Fco-1-1 | Bacteria Bacteroidetes Flavobacteriia Flavobacteriales Flavobacteriaceae *Flavobacterium columnare* ATCC 49512 |
| Fda-1-1 | Bacteria Bacteroidetes Flavobacteriia Flavobacteriales Flavobacteriaceae *Flavobacterium daejeonense* DSM 17708 |
| Fde-1-1 | Bacteria Bacteroidetes Flavobacteriia Flavobacteriales Flavobacteriaceae *Flavobacterium denitrificans* DSM 15936 |
| Fen-1-1 to Fen-1-2 | Bacteria Bacteroidetes Flavobacteriia Flavobacteriales Flavobacteriaceae *Flavobacterium enshiense* DK69 |
| Ffi-1-1 | Bacteria Bacteroidetes Flavobacteriia Flavobacteriales Flavobacteriaceae *Flavobacterium filum* DSM 17961 |
| Ffr-1-1 | Bacteria Bacteroidetes Flavobacteriia Flavobacteriales Flavobacteriaceae *Flavobacterium frigidarium* DSM 17623 |
| Ffr-2-1 | Bacteria Bacteroidetes Flavobacteriia Flavobacteriales Flavobacteriaceae *Flavobacterium frigoris* PS1 |
| Fge-1-1 | Bacteria Bacteroidetes Flavobacteriia Flavobacteriales Flavobacteriaceae *Flavobacterium gelidilacus* DSM 15343 |
| Fgi-1-1 | Bacteria Bacteroidetes Flavobacteriia Flavobacteriales Flavobacteriaceae *Flavobacterium gilvum* |
| Fhi-1-1 | Bacteria Bacteroidetes Flavobacteriia Flavobacteriales Flavobacteriaceae *Flavobacterium hibernum* |
| Fin-1-1 | Bacteria Bacteroidetes Flavobacteriia Flavobacteriales Flavobacteriaceae *Flavobacterium indicum* GPTSA100-9 |
| Fli-1-1 | Bacteria Bacteroidetes Flavobacteriia Flavobacteriales Flavobacteriaceae *Flavobacterium limnosediminis* JC2902 |
| Fps-1-1 to Fps-1-10 | Bacteria Bacteroidetes Flavobacteriia Flavobacteriales Flavobacteriaceae *Flavobacterium psychrophilum* |
| Fps-2-1 | Bacteria Bacteroidetes Flavobacteriia Flavobacteriales Flavobacteriaceae *Flavobacterium psychrophilum* FPG101 |
| Fps-3-1 | Bacteria Bacteroidetes Flavobacteriia Flavobacteriales Flavobacteriaceae *Flavobacterium psychrophilum* FPG3 |
| Fps-4-1 | Bacteria Bacteroidetes Flavobacteriia Flavobacteriales Flavobacteriaceae *Flavobacterium psychrophilum* JIP02/86 |
| Fre-1-1 | Bacteria Bacteroidetes Flavobacteriia Flavobacteriales Flavobacteriaceae *Flavobacterium reichenbachii* |
| Fsa-1-1 | Bacteria Bacteroidetes Flavobacteriia Flavobacteriales Flavobacteriaceae *Flavobacterium saliperosum* S13 |
| Fsa-2-1 | Bacteria Bacteroidetes Flavobacteriia Flavobacteriales Flavobacteriaceae *Flavobacterium sasangense* DSM 21067 |
| Fse-1-1 | Bacteria Bacteroidetes Flavobacteriia Flavobacteriales Flavobacteriaceae *Flavobacterium seoulense* |
| Fso-1-1 | Bacteria Bacteroidetes Flavobacteriia Flavobacteriales Flavobacteriaceae *Flavobacterium soli* DSM 19725 |
| Fsp-1-1 | Bacteria Bacteroidetes Flavobacteriia Flavobacteriales Flavobacteriaceae *Flavobacterium sp.* 83 |
| Fsp-2-1 | Bacteria Bacteroidetes Flavobacteriia Flavobacteriales Flavobacteriaceae *Flavobacterium sp.* ABG |
| Fsp-3-1 | Bacteria Bacteroidetes Flavobacteriia Flavobacteriales Flavobacteriaceae *Flavobacterium sp.* ACAM 123 |
| Fsp-4-1 | Bacteria Bacteroidetes Flavobacteriia Flavobacteriales Flavobacteriaceae *Flavobacterium sp.* AED |
| Fsp-5-1 | Bacteria Bacteroidetes Flavobacteriia Flavobacteriales Flavobacteriaceae *Flavobacterium sp.* B17 |
| Fsp-6-1 | Bacteria Bacteroidetes Flavobacteriia Flavobacteriales Flavobacteriaceae *Flavobacterium sp.* CF136 |
| Fsp-7-1 | Bacteria Bacteroidetes Flavobacteriia Flavobacteriales Flavobacteriaceae *Flavobacterium sp.* F52 |
| Fsp-8-1 | Bacteria Bacteroidetes Flavobacteriia Flavobacteriales Flavobacteriaceae *Flavobacterium sp.* Fl |
| Fsp-9-1 | Bacteria Bacteroidetes Flavobacteriia Flavobacteriales Flavobacteriaceae *Flavobacterium sp.* KJJ |
| Fsp-10-1 | Bacteria Bacteroidetes Flavobacteriia Flavobacteriales Flavobacteriaceae *Flavobacterium sp.* MEB061 |
| Fsp-11-1 | Bacteria Bacteroidetes Flavobacteriia Flavobacteriales Flavobacteriaceae *Flavobacterium sp.* URHB0058 |
| Fsp-12-1 | Bacteria Bacteroidetes Flavobacteriia Flavobacteriales Flavobacteriaceae *Flavobacterium sp.* WG21 |
| Fsu-1-1 | Bacteria Bacteroidetes Flavobacteriia Flavobacteriales Flavobacteriaceae *Flavobacterium succinicans* LMG 10402 |
| Fsu-2-1 to Fsu-2-2 | Bacteria Bacteroidetes Flavobacteriia Flavobacteriales Flavobacteriaceae *Flavobacterium suncheonense* GH29-5 = DSM 17707 |
| Fte-1-1 | Bacteria Bacteroidetes Flavobacteriia Flavobacteriales Flavobacteriaceae *Flavobacterium tegetincola* DSM 22377 |
| Fag-1-1 | Bacteria Bacteroidetes Flavobacteriia Flavobacteriales Flavobacteriaceae *Formosa agariphila* KMM 3901 |
| Fsp-13-1 | Bacteria Bacteroidetes Flavobacteriia Flavobacteriales Flavobacteriaceae *Formosa sp.* AK20 |
| Gsa-1-1 | Bacteria Bacteroidetes Flavobacteriia Flavobacteriales Flavobacteriaceae *Gaetbulibacter saemankumensis* DSM 17032 |
| Gsp-1-1 | Bacteria Bacteroidetes Flavobacteriia Flavobacteriales Flavobacteriaceae *Galbibacter sp. ck*-I2-15 |
| Gme-1-1 | Bacteria Bacteroidetes Flavobacteriia Flavobacteriales Flavobacteriaceae *Gelidibacter mesophilus* DSM 14095 |
| Gli-1-1 | Bacteria Bacteroidetes Flavobacteriia Flavobacteriales Flavobacteriaceae *Gillisia limnaea* DSM 15749 |
| Gsp-2-1 | Bacteria Bacteroidetes Flavobacteriia Flavobacteriales Flavobacteriaceae *Gillisia sp.* CAL575 |
| Gsp-3-1 | Bacteria Bacteroidetes Flavobacteriia Flavobacteriales Flavobacteriaceae *Gillisia sp.* CBA3202 |
| Gsp-4-1 | Bacteria Bacteroidetes Flavobacteriia Flavobacteriales Flavobacteriaceae *Gillisia sp.* Hel_I_29 |
| Gsp-5-1 | Bacteria Bacteroidetes Flavobacteriia Flavobacteriales Flavobacteriaceae *Gillisia sp.* JM1 |
| Gec-1-1 | Bacteria Bacteroidetes Flavobacteriia Flavobacteriales Flavobacteriaceae *Gramella echinicola* DSM 19838 |
| Gpo-1-1 | Bacteria Bacteroidetes Flavobacteriia Flavobacteriales Flavobacteriaceae *Gramella portivictoriae* DSM 23547 |
| Iha-1-1 | Bacteria Bacteroidetes Flavobacteriia Flavobacteriales Flavobacteriaceae *Imtechella halotolerans* K1 |
| Jpa-1-1 to Jpa-1-3 | Bacteria Bacteroidetes Flavobacteriia Flavobacteriales Flavobacteriaceae *Jejuia pallidilutea* |
| Jma-1-1 | Bacteria Bacteroidetes Flavobacteriia Flavobacteriales Flavobacteriaceae *Joostella marina* DSM 19592 |
| Kje-1-1 | Bacteria Bacteroidetes Flavobacteriia Flavobacteriales Flavobacteriaceae *Kordia jejudonensis* |
| Kzh-1-1 | Bacteria Bacteroidetes Flavobacteriia Flavobacteriales Flavobacteriaceae *Kordia zhangzhouensis* |
| Ksp-1-1 | Bacteria Bacteroidetes Flavobacteriia Flavobacteriales Flavobacteriaceae *Krokinobacter sp.* 4H-3-7-5 |
| Lja-1-1 | Bacteria Bacteroidetes Flavobacteriia Flavobacteriales Flavobacteriaceae *Lacinutrix jangbogonensis* |
| Lsp-1-1 | Bacteria Bacteroidetes Flavobacteriia Flavobacteriales Flavobacteriaceae *Lacinutrix sp.* 5H-3-7-4 |
| Lsp-2-1 | Bacteria Bacteroidetes Flavobacteriia Flavobacteriales Flavobacteriaceae *Lacinutrix sp.* Hel_I_90 |
| Lsp-3-1 | Bacteria Bacteroidetes Flavobacteriia Flavobacteriales Flavobacteriaceae *Leeuwenhoekiella sp.* Hel_I_48 |
| Lsp-4-1 | Bacteria Bacteroidetes Flavobacteriia Flavobacteriales Flavobacteriaceae *Leeuwenhoekiella sp.* MAR_2009_132 |

| | |
|---|---|
| Myu-1-1 | Bacteria Bacteroidetes Flavobacteriia Flavobacteriales Flavobacteriaceae *Mangrovimonas yunxiaonensis* |
| Man-1-1 | Bacteria Bacteroidetes Flavobacteriia Flavobacteriales Flavobacteriaceae *Maribacter antarcticus* DSM 21422 |
| Mfo-1-1 | Bacteria Bacteroidetes Flavobacteriia Flavobacteriales Flavobacteriaceae *Maribacter forsetii* DSM 18668 |
| Msp-1-1 | Bacteria Bacteroidetes Flavobacteriia Flavobacteriales Flavobacteriaceae *Maribacter sp.* Hel_I_7 |
| Mth-1-1 | Bacteria Bacteroidetes Flavobacteriia Flavobacteriales Flavobacteriaceae *Maribacter thermophilus* |
| Mze-1-1 | Bacteria Bacteroidetes Flavobacteriia Flavobacteriales Flavobacteriaceae *Mesoflavibacter zeaxanthinifaciens* DSM 18436 |
| Mze-2-1 | Bacteria Bacteroidetes Flavobacteriia Flavobacteriales Flavobacteriaceae *Mesoflavibacter zeaxanthinifaciens* S86 |
| Mmo-1-1 | Bacteria Bacteroidetes Flavobacteriia Flavobacteriales Flavobacteriaceae *Mesonia mobilis* DSM 19841 |
| Mlu-1-1 | Bacteria Bacteroidetes Flavobacteriia Flavobacteriales Flavobacteriaceae *Muricauda lutaonensis* |
| Msp-2-1 | Bacteria Bacteroidetes Flavobacteriia Flavobacteriales Flavobacteriaceae *Muricauda sp.* MAR_2010_75 |
| Min-1-1 | Bacteria Bacteroidetes Flavobacteriia Flavobacteriales Flavobacteriaceae *Myroides injenensis* M09-0166 |
| Ndo-1-1 | Bacteria Bacteroidetes Flavobacteriia Flavobacteriales Flavobacteriaceae *Nonlabens dokdonensis* DSW-6 |
| Nma-1-1 | Bacteria Bacteroidetes Flavobacteriia Flavobacteriales Flavobacteriaceae *Nonlabens marinus* S1-08 |
| Nse-1-1 | Bacteria Bacteroidetes Flavobacteriia Flavobacteriales Flavobacteriaceae *Nonlabens sediminis* |
| Nul-1-1 to Nul-1-6 | Bacteria Bacteroidetes Flavobacteriia Flavobacteriales Flavobacteriaceae *Nonlabens ulvanivorans* |
| Oma-1-1 | Bacteria Bacteroidetes Flavobacteriia Flavobacteriales Flavobacteriaceae *Olleya marilimosa* CAM030 |
| Osp-1-1 | Bacteria Bacteroidetes Flavobacteriia Flavobacteriales Flavobacteriaceae *Olleya sp.* VCSA23 |
| Osp-2-1 | Bacteria Bacteroidetes Flavobacteriia Flavobacteriales Flavobacteriaceae *Olleya sp.* VCSM12 |
| Orh-1-1 | Bacteria Bacteroidetes Flavobacteriia Flavobacteriales Flavobacteriaceae *Ornithobacterium rhinotracheale* DSM 15997 |
| Orh-2-1 | Bacteria Bacteroidetes Flavobacteriia Flavobacteriales Flavobacteriaceae *Ornithobacterium rhinotracheale* H06-030791 |
| Orh-3-1 | Bacteria Bacteroidetes Flavobacteriia Flavobacteriales Flavobacteriaceae *Ornithobacterium rhinotracheale* ORT-UMN 88 |
| Psp-1-1 | Bacteria Bacteroidetes Flavobacteriia Flavobacteriales Flavobacteriaceae *Polaribacter sp.* Hel1_33_49 |
| Psp-2-1 | Bacteria Bacteroidetes Flavobacteriia Flavobacteriales Flavobacteriaceae *Polaribacter sp.* Hel_I_88 |
| Psp-3-1 | Bacteria Bacteroidetes Flavobacteriia Flavobacteriales Flavobacteriaceae *Polaribacter sp.* MED152 |
| Pgo-1-1 | Bacteria Bacteroidetes Flavobacteriia Flavobacteriales Flavobacteriaceae *Psychroflexus gondwanensis* ACAM 44 |
| Pto-1-1 | Bacteria Bacteroidetes Flavobacteriia Flavobacteriales Flavobacteriaceae *Psychroflexus torquis* ATCC 700755 |
| Ptr-1-1 | Bacteria Bacteroidetes Flavobacteriia Flavobacteriales Flavobacteriaceae *Psychroflexus tropicus* DSM 15496 |
| Pbu-1-1 | Bacteria Bacteroidetes Flavobacteriia Flavobacteriales Flavobacteriaceae *Psychroserpens burtonensis* DSM 12212 |
| Pda-1-1 | Bacteria Bacteroidetes Flavobacteriia Flavobacteriales Flavobacteriaceae *Psychroserpens damuponensis* |
| Pja-1-1 | Bacteria Bacteroidetes Flavobacteriia Flavobacteriales Flavobacteriaceae *Psychroserpens jangbogonensis* |
| Pme-1-1 | Bacteria Bacteroidetes Flavobacteriia Flavobacteriales Flavobacteriaceae *Psychroserpens mesophilus* |
| Psp-4-1 | Bacteria Bacteroidetes Flavobacteriia Flavobacteriales Flavobacteriaceae *Psychroserpens sp.* Hel_I_66 |
| Ran-1-1 to Ran-1-2 | Bacteria Bacteroidetes Flavobacteriia Flavobacteriales Flavobacteriaceae *Riemerella anatipestifer* |
| Ran-2-1 to Ran-2-2 | Bacteria Bacteroidetes Flavobacteriia Flavobacteriales Flavobacteriaceae *Riemerella anatipestifer* ATCC 11845 = DSM 15868 |
| Ran-3-1 | Bacteria Bacteroidetes Flavobacteriia Flavobacteriales Flavobacteriaceae *Riemerella anatipestifer* CH3 |
| Ran-4-1 | Bacteria Bacteroidetes Flavobacteriia Flavobacteriales Flavobacteriaceae *Riemerella anatipestifer* RA-CH-1 |
| Ran-5-1 | Bacteria Bacteroidetes Flavobacteriia Flavobacteriales Flavobacteriaceae *Riemerella anatipestifer* RA-CH-2 |
| Ran-6-1 | Bacteria Bacteroidetes Flavobacteriia Flavobacteriales Flavobacteriaceae *Riemerella anatipestifer* RA-GD |
| Ran-7-1 | Bacteria Bacteroidetes Flavobacteriia Flavobacteriales Flavobacteriaceae *Riemerella anatipestifer* RA-SG |
| Ran-8-1 | Bacteria Bacteroidetes Flavobacteriia Flavobacteriales Flavobacteriaceae *Riemerella anatipestifer* RA-YM |
| Ran-9-1 | Bacteria Bacteroidetes Flavobacteriia Flavobacteriales Flavobacteriaceae *Riemerella anatipestifer* Yb2 |
| Rco-1-1 | Bacteria Bacteroidetes Flavobacteriia Flavobacteriales Flavobacteriaceae *Riemerella columbina* DSM 16469 |
| Ssp-1-1 | Bacteria Bacteroidetes Flavobacteriia Flavobacteriales Flavobacteriaceae *Salegentibacter sp.* Hel_I_6 |
| Sxi-1-1 | Bacteria Bacteroidetes Flavobacteriia Flavobacteriales Flavobacteriaceae *Salinimicrobium xinjiangense* DSM 19287 |
| Ssp-2-1 | Bacteria Bacteroidetes Flavobacteriia Flavobacteriales Flavobacteriaceae *Sediminibacter sp.* Hel_I_10 |
| Sze-1-1 | Bacteria Bacteroidetes Flavobacteriia Flavobacteriales Flavobacteriaceae *Siansivirga zeaxanthinifaciens* CC-SAMT-1 |
| Tna-1-1 | Bacteria Bacteroidetes Flavobacteriia Flavobacteriales Flavobacteriaceae *Tamlana nanhaiensis* |
| Tse-1-1 | Bacteria Bacteroidetes Flavobacteriia Flavobacteriales Flavobacteriaceae *Tamlana sedimentorum* |
| Tma-1-1 | Bacteria Bacteroidetes Flavobacteriia Flavobacteriales Flavobacteriaceae *Tenacibaculum maritimum* NCIMB 2154 |
| Tme-1-1 | Bacteria Bacteroidetes Flavobacteriia Flavobacteriales Flavobacteriaceae *Tenacibaculum mesophilum* |
| Tov-1-1 | Bacteria Bacteroidetes Flavobacteriia Flavobacteriales Flavobacteriaceae *Tenacibaculum ovolyticum* DSM 18103 |
| Wma-1-1 | Bacteria Bacteroidetes Flavobacteriia Flavobacteriales Flavobacteriaceae *Weeksella massiliensis* |
| Wvi-1-1 | Bacteria Bacteroidetes Flavobacteriia Flavobacteriales Flavobacteriaceae *Weeksella virosa* DSM 16922 |
| Wps-1-1 | Bacteria Bacteroidetes Flavobacteriia Flavobacteriales Flavobacteriaceae *Winogradskyella psychrotolerans* RS-3 |
| Wsp-1-1 | Bacteria Bacteroidetes Flavobacteriia Flavobacteriales Flavobacteriaceae *Winogradskyella sp.* PG-2 |
| Zam-1-1 | Bacteria Bacteroidetes Flavobacteriia Flavobacteriales Flavobacteriaceae *Zhouia amylolytica* AD3 |
| Zga-1-1 | Bacteria Bacteroidetes Flavobacteriia Flavobacteriales Flavobacteriaceae *Zobellia galactanivorans* |
| Zul-1-1 | Bacteria Bacteroidetes Flavobacteriia Flavobacteriales Flavobacteriaceae *Zobellia uliginosa* |
| env-1 to env-2123 | environmental samples |

#### 6.2 Gene contexts

Each L19-Flavobacteria RNA (indicated by “RNA→”) is listed. For each hit, the downstream genes predicted to reside in a regulated operon are listed. If the nearest downstream gene is encoding in the opposite strand (and therefore presumed to not be a part of a regulated operon), then that gene is still depicted. Some environmental sequences and some RefSeq entries lack gene annotations, and so no genes are available for such sequences. The direction of each gene is indicated with an arrow (→), and each predicted conserved domain in the gene is named. Conserved domains associated with more than one L19-Flavobacteria RNA are assigned a color; other domains are gray. Information about these conserved domains is given in Section 6.3. The accession of the sequence containing each L19-Flavobacteria RNA is given in the column named “Seq. accession”. Accessions beginning with “NC_”, “NS_”, “NW_” or “NZ_” are contained in RefSeq. Other accession refer to environmental samples. Nucleotide coordinates are given for the 5′ and 3′ boundaries of each L19-Flavobacteria RNA. If the 5′ coordinate is greater than the 3′ coordinate, the RNA is present on the reverse-complement strand of the containing genomic DNA sequence. Each hit is denoted by an abbreviation (like “Eco-1-1”) that refers to a taxonomy given in Section 6.1.

| abbrev. | Seq. accession | | 5′ at | 3′ at | genes |
|---|---|---|---|---|---|
| env-1 | DelMOWin2010_c10054634 | + | 824 | 851 | RNA→ rplS (PRK05338)Ribosomal_L19 (pfam01245)→ |
| env-2 | JCVI_SCAF_1101668113425 | - | 1241 | 1214 | RNA→ rplS (PRK05338)Ribosomal_L19 (pfam01245)→ |
| env-3 | scaffold96896_1_MH0080 | + | 1611 | 1645 | RNA→ rplS (PRK05338)Ribosomal_L19 (pfam01245)→ |
| env-4 | JCVI_SCAF_1101668324204 | + | 356 | 380 | RNA→ rplS (PRK05338)Ribosomal_L19 (pfam01245)→ PRK08299 (PRK08299)→ |
| env-5 | CEVX01024844.1 | + | 3189 | 3216 | RNA→ rplS (PRK05338)Ribosomal_L19 (pfam01245)→ |
| env-6 | JCVI_SCAF_1101668025146 | + | 75 | 102 | RNA→ rplS (PRK05338)Ribosomal_L19 (pfam01245)→ |
| env-7 | scaffold46008_4_MH0055 | + | 3799 | 3835 | RNA→ rplS (PRK05338)Ribosomal_L19 (pfam01245)→ |
| env-8 | 2236876009_110915 | - | 51 | 24 | RNA→ |
| env-9 | 2236876011_102929 | - | 302 | 275 | RNA→ rplS (PRK05338)Ribosomal_L19 (pfam01245)→ |
| env-10 | 2236876009_236522 | + | 137 | 164 | RNA→ rplS (PRK05338)Ribosomal_L19 (pfam01245)→ |
| env-11 | 2236876010_0006057 | - | 121 | 94 | RNA→ hypo→ |
| env-12 | SI47jul10_100m_c1059181 | + | 191 | 218 | RNA→ rplS (PRK05338)Ribosomal_L19 (pfam01245)→ |
| env-13 | SI39nov09_120m_c1230431 | - | 223 | 196 | RNA→ rplS (PRK05338)Ribosomal_L19 (pfam01245)→ |
| env-14 | SI36aug09_135m_c1030014 | + | 671 | 698 | RNA→ rplS (PRK05338)Ribosomal_L19 (pfam01245)→ |
| env-15 | SI53jan11_100m_c1051597 | + | 314 | 341 | RNA→ rplS (PRK05338)Ribosomal_L19 (pfam01245)→ |
| env-16 | SI60aug11_100m_c1037386 | - | 486 | 459 | RNA→ rplS (PRK05338)Ribosomal_L19 (pfam01245)→ |
| env-17 | SI47jul10_120m_c1000746 | + | 8693 | 8720 | RNA→ rplS (PRK05338)Ribosomal_L19 (pfam01245)→ |
| env-18 | SI53jan11_150m_c1118125 | - | 192 | 165 | RNA→ rplS (PRK05338)Ribosomal_L19 (pfam01245)→ |
| env-19 | SI34jun09_120m_1000436 | + | 12936 | 12963 | RNA→ rplS (PRK05338)Ribosomal_L19 (pfam01245)→ |
| env-20 | SI34jun09_135m_1001026 | + | 8736 | 8763 | RNA→ rplS (PRK05338)Ribosomal_L19 (pfam01245)→ |
| env-21 | SI36aug09_100m_1040962 | - | 274 | 247 | RNA→ rplS (PRK05338)Ribosomal_L19 (pfam01245)→ |
| env-22 | SI36aug09_120m_1000931 | - | 2702 | 2675 | RNA→ rplS (PRK05338)Ribosomal_L19 (pfam01245)→ |
| env-23 | SI34jun09_100m_1000067 | - | 31230 | 31203 | RNA→ rplS (PRK05338)Ribosomal_L19 (pfam01245)→ |
| env-24 | F10SI0310a_Sequence0000020675 | + | 300 | 327 | RNA→ rplS (PRK05338)Ribosomal_L19 (pfam01245)→ |
| env-25 | F10SI03100a_Sequence0000002003 | - | 449 | 422 | RNA→ rplS (PRK05338)Ribosomal_L19 (pfam01245)→ |
| env-26 | F10SI03120a_Sequence0000009017 | - | 632 | 605 | RNA→ rplS (PRK05338)Ribosomal_L19 (pfam01245)→ |
| env-27 | JCVI_SCAF_1096627066532 | - | 2055 | 2028 | RNA→ rplS (PRK05338)Ribosomal_L19 (pfam01245)→ |
| env-28 | CEWG01100961.1 | + | 836 | 863 | RNA→ rplS (PRK05338)Ribosomal_L19 (pfam01245)→ |
| env-29 | A09P0410a_Sequence0000049938 | - | 828 | 801 | RNA→ rplS (PRK05338)Ribosomal_L19 (pfam01245)→ |
| env-30 | JCVI_SCAF_1096627807607 | - | 1865 | 1838 | RNA→ rplS (PRK05338)Ribosomal_L19 (pfam01245)→ |
| env-31 | CENP01449906.1 | + | 6989 | 7016 | RNA→ rplS (PRK05338)Ribosomal_L19 (pfam01245)→ |
| env-32 | CEOK01161594.1 | - | 675 | 648 | RNA→ rplS (PRK05338)Ribosomal_L19 (pfam01245)→ |
| env-33 | CESO01045118.1 | - | 1107 | 1080 | RNA→ rplS (PRK05338)Ribosomal_L19 (pfam01245)→ |
| env-34 | LPaug08P2610m_c1001156 | + | 4888 | 4915 | RNA→ rplS (PRK05338)Ribosomal_L19 (pfam01245)→ |
| env-35 | CESF01045010.1 | + | 37 | 64 | RNA→ rplS (PRK05338)Ribosomal_L19 (pfam01245)→ |
| env-36 | CESI01106464.1 | - | 2060 | 2033 | RNA→ rplS (PRK05338)Ribosomal_L19 (pfam01245)→ |
| env-37 | CESU01171607.1 | - | 76 | 49 | RNA→ |
| env-38 | DelMOSum2011_c10436309 | - | 43 | 16 | RNA→ ←hypo |
| env-39 | CEWO01376889.1 | - | 537 | 510 | RNA→ rplS (PRK05338)Ribosomal_L19 (pfam01245)→ |

| | | | | | |
|---|---|---|---|---|---|
| env-40 | JGI20156J14371_10000063 | - | 46374 | 46347 | **RNA**→ rplS (PRK05338)Ribosomal_L19 (pfam01245)→ |
| env-41 | CENG01060425.1 | - | 1443 | 1416 | **RNA**→ hypo→ rplS (PRK05338)Ribosomal_L19 (pfam01245)→ |
| env-42 | CEOM01373684.1 | + | 885 | 912 | **RNA**→ rplS (PRK05338)Ribosomal_L19 (pfam01245)→ |
| env-43 | CENJ01127951.1 | + | 724 | 751 | **RNA**→ rplS (PRK05338)Ribosomal_L19 (pfam01245)→ |
| env-44 | JCVI_SCAF_1096627276236 | + | 31293 | 31320 | **RNA**→ rplS (PRK05338)Ribosomal_L19 (pfam01245)→ |
| env-45 | P_2C_Liq_1_UnCty_1015735 | - | 1527 | 1500 | **RNA**→ rplS (PRK05338)Ribosomal_L19 (pfam01245)→ |
| env-46 | SRS022609_C3248272 | - | 149 | 114 | **RNA**→ rplS (PRK05338)→ |
| env-47 | NLM026_scaffold33221_1 | - | 492 | 457 | **RNA**→ hypo→ rplS (PRK05338)Ribosomal_L19 (pfam01245)→ |
| env-48 | SRS013476_C3069736 | + | 704 | 739 | **RNA**→ rplS (PRK05338)Ribosomal_L19 (pfam01245)→ |
| env-49 | NOF004_scaffold34062_6 | + | 254 | 289 | **RNA**→ rplS (PRK05338)Ribosomal_L19 (pfam01245)→ |
| env-50 | SRS020233_C4251492 | - | 197 | 162 | **RNA**→ rplS (PRK05338)Ribosomal_L19 (pfam01245)→ |
| env-51 | JGI20154J14316_10000121 | + | 19996 | 20023 | **RNA**→ rplS (PRK05338)Ribosomal_L19 (pfam01245)→ DUF1015 (pfam06245)→ |
| | | | | | TIGR00044 (TIGR00044)PLPDE_III_YBL036c_like (cd00635)→ ValS (COG0525)valS (PRK05729)→ |
| env-52 | JGI20153J14295_10000586 | + | 20619 | 20646 | **RNA**→ rplS (PRK05338)Ribosomal_L19 (pfam01245)→ DUF1015 (pfam06245)→ |
| | | | | | TIGR00044 (TIGR00044)PLPDE_III_YBL036c_like (cd00635)→ ValS (COG0525)valS (PRK05729)→ |
| env-53 | JGI20153J14318_10000586 | + | 20619 | 20646 | **RNA**→ rplS (PRK05338)Ribosomal_L19 (pfam01245)→ DUF1015 (pfam06245)→ |
| | | | | | TIGR00044 (TIGR00044)PLPDE_III_YBL036c_like (cd00635)→ ValS (COG0525)valS (PRK05729)→ |
| env-54 | JGI20154J14316_10009206 | - | 959 | 931 | **RNA**→ rplS (PRK05338)Ribosomal_L19 (pfam01245)→ DUF1015 (pfam06245)→ |
| env-55 | DelMOSum2011_c10004732 | - | 2425 | 2397 | **RNA**→ rplS (PRK05338)Ribosomal_L19 (pfam01245)→ DUF1015 (pfam06245)→ |
| | | | | | SDR_c (cd05233)fabG (PRK05653)→ |
| env-56 | 2236876010_0079494 | - | 132 | 104 | **RNA**→ hypo→ |
| env-57 | CEWG01003263.1 | - | 1195 | 1167 | **RNA**→ rplS (PRK05338)Ribosomal_L19 (pfam01245)→ DUF1015 (pfam06245)→ |
| env-58 | 2236876009_196101 | - | 440 | 412 | **RNA**→ rplS (PRK05338)Ribosomal_L19 (pfam01245)→ |
| env-59 | JGI20154J14316_10002796 | - | 4419 | 4392 | **RNA**→ rplS (PRK05338)Ribosomal_L19 (pfam01245)→ DUF1015 (pfam06245)→ |
| | | | | | PRK07825 (PRK07825)adh_short (pfam00106)→ TIGR00044 (TIGR00044)PLPDE_III_YBL036c_like (cd00635)→ |
| env-60 | CGUI_GFCP3IQ02J1ARS | + | 207 | 234 | **RNA**→ rplS (PRK05338)Ribosomal_L19 (pfam01245)→ |
| env-61 | 2236876001_386712 | + | 198 | 225 | **RNA**→ rplS (PRK05338)Ribosomal_L19 (pfam01245)→ |
| env-62 | 2236876001_178730 | + | 31 | 58 | **RNA**→ rplS (PRK05338)Ribosomal_L19 (pfam01245)→ |
| env-63 | SI34jun09_10m_1002607 | - | 5144 | 5117 | **RNA**→ rplS (PRK05338)Ribosomal_L19 (pfam01245)→ |
| env-64 | JGI1221J11331_1002397 | - | 3466 | 3439 | **RNA**→ rplS (PRK05338)Ribosomal_L19 (pfam01245)→ |
| env-65 | BS_KBB_SWE26_205m_c1114174 | - | 142 | 115 | **RNA**→ rplS (PRK05338)Ribosomal_L19 (pfam01245)→ |
| env-66 | BS_KBA_SWE21_205m_10293102 | + | 80 | 107 | **RNA**→ ←hypo |
| env-67 | CEQG01051930.1 | + | 175 | 202 | **RNA**→ hypo→ rplS (PRK05338)Ribosomal_L19 (pfam01245)→ |
| Nul-1-1 | NZ_BBMK01000006.1 | + | 140006 | 140033 | **RNA**→ rplS (PRK05338)Ribosomal_L19 (pfam01245)→ |
| Fba-1-1 | NZ_CH672374.1 | + | 838999 | 839026 | **RNA**→ rplS (PRK05338)Ribosomal_L19 (pfam01245)→ |
| Nul-1-2 | NZ_BBLG01000002.1 | - | 286727 | 286700 | **RNA**→ rplS (PRK05338)Ribosomal_L19 (pfam01245)→ |
| Nul-1-3 | NZ_BBMJ01000007.1 | + | 150921 | 150948 | **RNA**→ rplS (PRK05338)Ribosomal_L19 (pfam01245)→ |
| Nul-1-4 | NZ_BBNT01000005.1 | - | 74527 | 74500 | **RNA**→ rplS (PRK05338)Ribosomal_L19 (pfam01245)→ |
| Nul-1-5 | NZ_JPJI01000032.1 | - | 1150068 | 1150041 | **RNA**→ rplS (PRK05338)Ribosomal_L19 (pfam01245)→ |
| Nul-1-6 | NZ_BBMM01000005.1 | + | 148362 | 148389 | **RNA**→ rplS (PRK05338)Ribosomal_L19 (pfam01245)→ |
| env-68 | TDF_OR_ARG04_113m_c1026478 | + | 123 | 150 | **RNA**→ rplS (PRK05338)Ribosomal_L19 (pfam01245)→ |
| Csp-18-1 | NZ_JUGG01000008.1 | - | 653525 | 653498 | **RNA**→ rplS (PRK05338)Ribosomal_L19 (pfam01245)→ |
| Csp-15-1 | NZ_LFNE01000002.1 | - | 475215 | 475188 | **RNA**→ rplS (PRK05338)Ribosomal_L19 (pfam01245)→ |
| Csp-16-1 | NZ_KN050755.1 | - | 362647 | 362620 | **RNA**→ rplS (PRK05338)Ribosomal_L19 (pfam01245)→ |
| Chi-1-1 | NZ_JARQ01000007.1 | - | 271126 | 271099 | **RNA**→ rplS (PRK05338)Ribosomal_L19 (pfam01245)→ |
| env-69 | APMI01120637.1 | - | 2839 | 2812 | **RNA**→ rplS (PRK05338)Ribosomal_L19 (pfam01245)→ |
| Cgr-2-1 | NZ_AUMU01000006.1 | - | 52910 | 52883 | **RNA**→ rplS (PRK05338)Ribosomal_L19 (pfam01245)→ |
| Csp-17-1 | NZ_JUGK01000011.1 | - | 424788 | 424761 | **RNA**→ rplS (PRK05338)Ribosomal_L19 (pfam01245)→ |
| Fsp-5-1 | NZ_BACY01000298.1 | - | 3551 | 3524 | **RNA**→ rplS (PRK05338)Ribosomal_L19 (pfam01245)→ |
| env-70 | 07_22_INULIN_DNA_scaffold76662_14 | - | 670 | 643 | **RNA**→ rplS (PRK05338)Ribosomal_L19 (pfam01245)→ |
| env-71 | JGI20153J14295_10392805 | + | 60 | 87 | **RNA**→ rplS (PRK05338)Ribosomal_L19 (pfam01245)→ |
| env-72 | JGI20153J14318_10392805 | + | 60 | 87 | **RNA**→ rplS (PRK05338)Ribosomal_L19 (pfam01245)→ |
| env-73 | JCVI_SCAF_1096627352169 | - | 5043 | 5016 | **RNA**→ rplS (PRK05338)Ribosomal_L19 (pfam01245)→ |

| | | | | | |
|---|---|---|---|---|---|
| env-74 | JCVI_SCAF_1096626620328 | - | 745 | 718 | **RNA**→ rplS (PRK05338)Ribosomal_L19 (pfam01245)→ |
| env-75 | CEWG01401482.1 | - | 206 | 179 | **RNA**→ rplS (PRK05338)Ribosomal_L19 (pfam01245)→ |
| env-76 | 84538774 | + | 290 | 317 | **RNA**→ rplS (PRK05338)Ribosomal_L19 (pfam01245)→ |
| env-77 | JCVI_READ_1106515117443 | + | 381 | 408 | **RNA**→ rplS (PRK05338)Ribosomal_L19 (pfam01245)→ |
| env-78 | 84539387 | - | 983 | 956 | **RNA**→ rplS (PRK05338)Ribosomal_L19 (pfam01245)→ |
| env-79 | JCVI_READ_1101854579158 | + | 714 | 741 | **RNA**→ |
| env-80 | JCVI_READ_1101754002299 | + | 718 | 745 | **RNA**→ |
| env-81 | CESE01025856.1 | + | 5377 | 5404 | **RNA**→ rplS (PRK05338)Ribosomal_L19 (pfam01245)→ |
| env-82 | CESI01208183.1 | + | 540 | 567 | **RNA**→ |
| env-83 | CESJ01033644.1 | - | 458 | 431 | **RNA**→ rplS (PRK05338)Ribosomal_L19 (pfam01245)→ |
| env-84 | CEST01017140.1 | + | 5896 | 5923 | **RNA**→ rplS (PRK05338)Ribosomal_L19 (pfam01245)→ |
| env-85 | CESU01147895.1 | - | 55 | 28 | **RNA**→ |
| env-86 | JCVI_SCAF_1096627480095 | - | 1803 | 1776 | **RNA**→ rplS (PRK05338)Ribosomal_L19 (pfam01245)→ |
| env-87 | DelMOSum2010_c10678701 | + | 177 | 204 | **RNA**→ |
| env-88 | CAM_READ_0235983637 | + | 371 | 398 | **RNA**→ |
| env-89 | JCVI_SCAF_1101668031618 | + | 670 | 697 | **RNA**→ rplS (PRK05338)Ribosomal_L19 (pfam01245)→ |
| env-90 | CERM01141528.1 | + | 522 | 549 | **RNA**→ |
| env-91 | JGI20157J14317_10131042 | + | 360 | 387 | **RNA**→ rplS (PRK05338)Ribosomal_L19 (pfam01245)→ |
| env-92 | 2236876002_127420 | + | 329 | 356 | **RNA**→ |
| env-93 | JCVI_SCAF_1096626613738 | - | 310 | 283 | **RNA**→ rplS (PRK05338)Ribosomal_L19 (pfam01245)→ |
| env-94 | 2236876006_283195 | - | 34 | 7 | **RNA**→ |
| env-95 | DelMOSum2010_c10002012 | + | 12245 | 12272 | **RNA**→ rplS (PRK05338)Ribosomal_L19 (pfam01245)→ |
| env-96 | DelMOSum2011_c10031706 | - | 525 | 498 | **RNA**→ rplS (PRK05338)Ribosomal_L19 (pfam01245)→ |
| env-97 | DelMOWin2010_c10001229 | + | 4276 | 4303 | **RNA**→ rplS (PRK05338)Ribosomal_L19 (pfam01245)→ |
| env-98 | JGI20152J14361_10056501 | - | 442 | 415 | **RNA**→ rplS (PRK05338)Ribosomal_L19 (pfam01245)→ |
| env-99 | JGI20156J14371_10026599 | + | 782 | 809 | **RNA**→ rplS (PRK05338)Ribosomal_L19 (pfam01245)→ |
| env-100 | JGI20154J14316_10083694 | + | 359 | 386 | **RNA**→ rplS (PRK05338)Ribosomal_L19 (pfam01245)→ |
| env-101 | JGI20160J14292_10012380 | - | 2489 | 2462 | **RNA**→ rplS (PRK05338)Ribosomal_L19 (pfam01245)→ |
| env-102 | JGI20159J14440_10010596 | + | 2893 | 2920 | **RNA**→ rplS (PRK05338)Ribosomal_L19 (pfam01245)→ |
| env-103 | JGI20155J14468_10002089 | - | 1100 | 1073 | **RNA**→ rplS (PRK05338)Ribosomal_L19 (pfam01245)→ |
| env-104 | JGI20158J14315_10029275 | + | 515 | 542 | **RNA**→ rplS (PRK05338)Ribosomal_L19 (pfam01245)→ |
| env-105 | A09P0410a_Sequence0000056231 | + | 784 | 811 | **RNA**→ |
| env-106 | F10SI0310a_Sequence0000007596 | - | 1376 | 1349 | **RNA**→ rplS (PRK05338)Ribosomal_L19 (pfam01245)→ |
| env-107 | JCVI_SCAF_1096626854527 | + | 2612 | 2639 | **RNA**→ rplS (PRK05338)Ribosomal_L19 (pfam01245)→ |
| env-108 | DelMOWin2010_c10025809 | + | 1404 | 1431 | **RNA**→ rplS (PRK05338)Ribosomal_L19 (pfam01245)→ |
| env-109 | JGI20152J14361_10004558 | + | 2419 | 2446 | **RNA**→ rplS (PRK05338)Ribosomal_L19 (pfam01245)→ |
| env-110 | JGI20151J14362_10005809 | + | 2820 | 2847 | **RNA**→ rplS (PRK05338)Ribosomal_L19 (pfam01245)→ |
| env-111 | JGI20153J14295_10006540 | - | 5010 | 4983 | **RNA**→ rplS (PRK05338)Ribosomal_L19 (pfam01245)→ |
| env-112 | JGI20153J14318_10006540 | - | 5010 | 4983 | **RNA**→ rplS (PRK05338)Ribosomal_L19 (pfam01245)→ |
| env-113 | JGI20159J14440_10023770 | - | 1237 | 1210 | **RNA**→ rplS (PRK05338)Ribosomal_L19 (pfam01245)→ |
| env-114 | JGI20155J14468_10003739 | + | 1801 | 1828 | **RNA**→ rplS (PRK05338)Ribosomal_L19 (pfam01245)→ |
| env-115 | JGI20158J14315_10006321 | + | 2255 | 2282 | **RNA**→ rplS (PRK05338)Ribosomal_L19 (pfam01245)→ |
| env-116 | CESS01145451.1 | - | 462 | 435 | **RNA**→ rplS (PRK05338)Ribosomal_L19 (pfam01245)→ |
| env-117 | JGI20156J14371_10212006 | + | 469 | 496 | **RNA**→ |
| env-118 | JGI20154J14316_10004022 | + | 6496 | 6523 | **RNA**→ rplS (PRK05338)Ribosomal_L19 (pfam01245)→ |
| env-119 | JGI20160J14292_10007636 | + | 2383 | 2410 | **RNA**→ rplS (PRK05338)Ribosomal_L19 (pfam01245)→ |
| env-120 | P_2C_Liq_3_UnCty_100387 | + | 12092 | 12119 | **RNA**→ rplS (PRK05338)Ribosomal_L19 (pfam01245)→ |
| env-121 | P_2C_Liq_1_UnCty_1000156 | + | 13091 | 13118 | **RNA**→ rplS (PRK05338)Ribosomal_L19 (pfam01245)→ |
| env-122 | P_2C_Liq_2_UnCty_1000768 | + | 4871 | 4898 | **RNA**→ rplS (PRK05338)Ribosomal_L19 (pfam01245)→ |
| env-123 | JGI20151J14362_10005693 | + | 5310 | 5337 | **RNA**→ rplS (PRK05338)Ribosomal_L19 (pfam01245)→ |
| env-124 | JGI20153J14295_10091054 | + | 386 | 413 | **RNA**→ rplS (PRK05338)Ribosomal_L19 (pfam01245)→ |
| env-125 | JGI20153J14318_10091054 | + | 386 | 413 | **RNA**→ rplS (PRK05338)Ribosomal_L19 (pfam01245)→ |

| | | | | | |
|---|---|---|---|---|---|
| env-126 | JGI11876J14442_10491868 | - | 201 | 174 | **RNA**→ rplS (PRK05338)Ribosomal_L19 (pfam01245)→ |
| env-127 | JGI11876J14442_10523508 | + | 238 | 265 | **RNA**→ rplS (PRK05338)Ribosomal_L19 (pfam01245)→ |
| env-128 | CEOS01057510.1 | + | 637 | 664 | **RNA**→ rplS (PRK05338)Ribosomal_L19 (pfam01245)→ |
| env-129 | CETR01239150.1 | + | 650 | 677 | **RNA**→ rplS (PRK05338)Ribosomal_L19 (pfam01245)→ |
| env-130 | CETS01121663.1 | - | 506 | 479 | **RNA**→ rplS (PRK05338)Ribosomal_L19 (pfam01245)→ |
| env-131 | CETZ01264369.1 | + | 15462 | 15489 | **RNA**→ rplS (PRK05338)Ribosomal_L19 (pfam01245)→ |
| env-132 | CEUA01417981.1 | - | 740 | 713 | **RNA**→ rplS (PRK05338)Ribosomal_L19 (pfam01245)→ |
| env-133 | CEVA01411844.1 | - | 180 | 153 | **RNA**→ rplS (PRK05338)Ribosomal_L19 (pfam01245)→ |
| Oho-1-1 | NC_016599.1 | - | 405639 | 405612 | **RNA**→ rplS (PRK05338)Ribosomal_L19 (pfam01245)→ hypo→ |
| env-134 | PR_CR_10_Liq_1_inCR_1003339 | + | 5067 | 5094 | **RNA**→ rplS (PRK05338)Ribosomal_L19 (pfam01245)→ hypo→ |
| env-135 | PR_CR_10_Liq_3_inCR_1015616 | + | 986 | 1013 | **RNA**→ rplS (PRK05338)Ribosomal_L19 (pfam01245)→ |
| env-136 | JCVI_SCAF_1101668700438 | + | 1308 | 1335 | **RNA**→ |
| env-137 | CETE01027368.1 | + | 993 | 1020 | **RNA**→ rplS (PRK05338)Ribosomal_L19 (pfam01245)→ |
| env-138 | CEWH01269799.1 | + | 175 | 202 | **RNA**→ rplS (PRK05338)Ribosomal_L19 (pfam01245)→ |
| env-139 | CEUH01076195.1 | - | 529 | 502 | **RNA**→ rplS (PRK05338)Ribosomal_L19 (pfam01245)→ |
| env-140 | CEVG01157966.1 | + | 2615 | 2642 | **RNA**→ rplS (PRK05338)Ribosomal_L19 (pfam01245)→ |
| env-141 | CEUU01157968.1 | - | 568 | 541 | **RNA**→ rplS (PRK05338)Ribosomal_L19 (pfam01245)→ |
| env-142 | CEVA01228795.1 | - | 1304 | 1277 | **RNA**→ rplS (PRK05338)Ribosomal_L19 (pfam01245)→ |
| env-143 | CEUW01090977.1 | - | 185 | 158 | **RNA**→ rplS (PRK05338)Ribosomal_L19 (pfam01245)→ |
| env-144 | CENG01005083.1 | + | 2607 | 2634 | **RNA**→ rplS (PRK05338)Ribosomal_L19 (pfam01245)→ |
| env-145 | CENU01397455.1 | - | 1003 | 976 | **RNA**→ rplS (PRK05338)Ribosomal_L19 (pfam01245)→ PRK08299 (PRK08299)→ |
| env-146 | CENJ01029369.1 | - | 130 | 103 | **RNA**→ rplS (PRK05338)Ribosomal_L19 (pfam01245)→ |
| Nse-1-1 | NZ_BBML01000004.1 | - | 136788 | 136761 | **RNA**→ rplS (PRK05338)Ribosomal_L19 (pfam01245)→ |
| env-147 | CENG01172060.1 | - | 288 | 261 | **RNA**→ rplS (PRK05338)Ribosomal_L19 (pfam01245)→ |
| env-148 | CENJ01036011.1 | - | 1206 | 1179 | **RNA**→ rplS (PRK05338)Ribosomal_L19 (pfam01245)→ PRK08299 (PRK08299)→ |
| env-149 | CENJ01107579.1 | - | 484 | 457 | **RNA**→ rplS (PRK05338)Ribosomal_L19 (pfam01245)→ |
| env-150 | JGI20151J14362_10016516 | - | 3534 | 3507 | **RNA**→ rplS (PRK05338)Ribosomal_L19 (pfam01245)→ PRK08299 (PRK08299)→ |
| env-151 | JGI20156J14371_10021029 | + | 1321 | 1348 | **RNA**→ rplS (PRK05338)Ribosomal_L19 (pfam01245)→ PRK08299 (PRK08299)→ |
| env-152 | JGI20160J14292_10169914 | + | 210 | 237 | **RNA**→ rplS (PRK05338)Ribosomal_L19 (pfam01245)→ |
| env-153 | JGI20159J14440_10065380 | + | 387 | 414 | **RNA**→ rplS (PRK05338)Ribosomal_L19 (pfam01245)→ PRK08299 (PRK08299)→ |
| env-154 | JGI20155J14468_10022096 | - | 399 | 372 | **RNA**→ rplS (PRK05338)Ribosomal_L19 (pfam01245)→ |
| env-155 | JGI20158J14315_10073378 | - | 1006 | 979 | **RNA**→ rplS (PRK05338)Ribosomal_L19 (pfam01245)→ PRK08299 (PRK08299)→ |
| env-156 | CEOS01040381.1 | - | 4293 | 4266 | **RNA**→ rplS (PRK05338)Ribosomal_L19 (pfam01245)→ |
| env-157 | CEOX01212680.1 | + | 1004 | 1031 | **RNA**→ |
| env-158 | CEPD01030902.1 | - | 4264 | 4237 | **RNA**→ rplS (PRK05338)Ribosomal_L19 (pfam01245)→ |
| env-159 | CERQ01076022.1 | - | 4315 | 4288 | **RNA**→ rplS (PRK05338)Ribosomal_L19 (pfam01245)→ |
| env-160 | CESL01149743.1 | - | 4963 | 4936 | **RNA**→ rplS (PRK05338)Ribosomal_L19 (pfam01245)→ |
| env-161 | CETA01027592.1 | - | 4529 | 4502 | **RNA**→ rplS (PRK05338)Ribosomal_L19 (pfam01245)→ |
| env-162 | CETB01172258.1 | + | 5225 | 5252 | **RNA**→ rplS (PRK05338)Ribosomal_L19 (pfam01245)→ |
| env-163 | CETG01164098.1 | - | 4952 | 4925 | **RNA**→ rplS (PRK05338)Ribosomal_L19 (pfam01245)→ |
| env-164 | CETM01103031.1 | - | 1792 | 1765 | **RNA**→ rplS (PRK05338)Ribosomal_L19 (pfam01245)→ |
| env-165 | CETO01013763.1 | + | 6548 | 6575 | **RNA**→ rplS (PRK05338)Ribosomal_L19 (pfam01245)→ |
| env-166 | CETR01123512.1 | - | 2833 | 2806 | **RNA**→ rplS (PRK05338)Ribosomal_L19 (pfam01245)→ |
| env-167 | CETS01046135.1 | + | 8383 | 8410 | **RNA**→ rplS (PRK05338)Ribosomal_L19 (pfam01245)→ |
| env-168 | CETT01303147.1 | + | 949 | 976 | **RNA**→ rplS (PRK05338)Ribosomal_L19 (pfam01245)→ |
| env-169 | CETU01309689.1 | - | 765 | 738 | **RNA**→ rplS (PRK05338)Ribosomal_L19 (pfam01245)→ |
| env-170 | CEUN01257627.1 | + | 429 | 456 | **RNA**→ rplS (PRK05338)Ribosomal_L19 (pfam01245)→ |
| env-171 | CEUX01208011.1 | - | 370 | 343 | **RNA**→ rplS (PRK05338)Ribosomal_L19 (pfam01245)→ |
| env-172 | CEUY01486419.1 | - | 3370 | 3343 | **RNA**→ rplS (PRK05338)Ribosomal_L19 (pfam01245)→ |
| env-173 | CEVI01034447.1 | - | 284 | 257 | **RNA**→ rplS (PRK05338)Ribosomal_L19 (pfam01245)→ |
| env-174 | CEVJ01148327.1 | - | 504 | 477 | **RNA**→ rplS (PRK05338)Ribosomal_L19 (pfam01245)→ |
| env-175 | CEVK01192335.1 | + | 325 | 352 | **RNA**→ rplS (PRK05338)Ribosomal_L19 (pfam01245)→ |

| | | | | | |
|---|---|---|---|---|---|
| env-176 | CEVM01083244.1 | - | 265 | 238 | **RNA**→ rplS (PRK05338)Ribosomal_L19 (pfam01245)→ |
| env-177 | CEWI01140434.1 | - | 2409 | 2382 | **RNA**→ rplS (PRK05338)Ribosomal_L19 (pfam01245)→ |
| Zpr-1-1 | NC_014041.1 | + | 3889655 | 3889682 | **RNA**→ rplS (PRK05338)Ribosomal_L19 (pfam01245)→ |
| env-178 | CEVA01498907.1 | + | 662 | 689 | **RNA**→ rplS (PRK05338)Ribosomal_L19 (pfam01245)→ |
| env-179 | Buccal_mucosa_LANL_scaffold_37880 | + | 293 | 320 | **RNA**→ rplS (PRK05338)Ribosomal_L19 (pfam01245)→ |
| env-180 | SRS018149_Baylor_scaffold_24394 | + | 14431 | 14458 | **RNA**→ rplS (PRK05338)Ribosomal_L19 (pfam01245)→ |
| env-181 | SRS054569_C1505048 | - | 440 | 413 | **RNA**→ rplS (PRK05338)Ribosomal_L19 (pfam01245)→ |
| env-182 | SRS024140_C1177031 | - | 285 | 258 | **RNA**→ rplS (PRK05338)Ribosomal_L19 (pfam01245)→ |
| env-183 | SRS052668_LANL_scaffold_1943 | - | 64691 | 64664 | **RNA**→ rplS (PRK05338)Ribosomal_L19 (pfam01245)→ |
| env-184 | SRS024347_C1548200 | - | 195 | 168 | **RNA**→ rplS (PRK05338)Ribosomal_L19 (pfam01245)→ |
| env-185 | SRS017127_Baylor_scaffold_3172 | - | 838 | 811 | **RNA**→ PLN02820 (PLN02820)MmdA (COG4799)→ rplS (PRK05338)Ribosomal_L19 (pfam01245)→ |
| env-186 | LGOV01048772.1 | + | 5860 | 5887 | **RNA**→ rplS (PRK05338)Ribosomal_L19 (pfam01245)→ |
| env-187 | LGVF01504154.1 | - | 463 | 436 | **RNA**→ rplS (PRK05338)Ribosomal_L19 (pfam01245)→ |
| env-188 | JCVI_SCAF_1096627098231 | + | 761 | 788 | **RNA**→ rplS (PRK05338)Ribosomal_L19 (pfam01245)→ monomer_idh (TIGR00178)IDH (pfam03971)→ |
| env-189 | CESM01261374.1 | - | 289 | 262 | **RNA**→ rplS (PRK05338)Ribosomal_L19 (pfam01245)→ |
| env-190 | CETE01135469.1 | - | 628 | 601 | **RNA**→ rplS (PRK05338)Ribosomal_L19 (pfam01245)→ monomer_idh (TIGR00178)IDH (pfam03971)→ |
| env-191 | CETQ01070376.1 | - | 697 | 670 | **RNA**→ rplS (PRK05338)Ribosomal_L19 (pfam01245)→ monomer_idh (TIGR00178)IDH (pfam03971)→ |
| env-192 | CEUH01016525.1 | + | 7232 | 7259 | **RNA**→ rplS (PRK05338)Ribosomal_L19 (pfam01245)→ monomer_idh (TIGR00178)IDH (pfam03971)→<br>PRK13484 (PRK13484)TonB_dep_Rec (pfam00593)Plug (pfam07715)CarbopepD_reg_2 (pfam13715)OMP_b-brl_3 (pfam1<br>DUF4249 (pfam14054)→ murQ (PRK05441)SIS_Etherase (cd05007)→ hypo→ |
| env-193 | CEWG01371477.1 | - | 849 | 822 | **RNA**→ rplS (PRK05338)Ribosomal_L19 (pfam01245)→ monomer_idh (TIGR00178)IDH (pfam03971)→ |
| env-194 | TahiMoana_1003731 | - | 598 | 571 | **RNA**→ rplS (PRK05338)Ribosomal_L19 (pfam01245)→ |
| env-195 | ElkS_mat_CD6A_1022331 | + | 121 | 148 | **RNA**→ rplS (PRK05338)Ribosomal_L19 (pfam01245)→ |
| env-196 | JGI11876J14442_10027587 | + | 312 | 339 | **RNA**→ rplS (PRK05338)Ribosomal_L19 (pfam01245)→ |
| env-197 | CEVK01107798.1 | - | 6100 | 6073 | **RNA**→ rplS (PRK05338)Ribosomal_L19 (pfam01245)→ |
| env-198 | CEVM01261662.1 | + | 6748 | 6775 | **RNA**→ rplS (PRK05338)Ribosomal_L19 (pfam01245)→ |
| env-199 | CEVO01248002.1 | - | 144 | 117 | **RNA**→ rplS (PRK05338)Ribosomal_L19 (pfam01245)→ |
| env-200 | LPaug08P2610m_c1119498 | + | 167 | 194 | **RNA**→ |
| env-201 | LPjun09P1210m_c1035922 | + | 137 | 164 | **RNA**→ rplS (PRK05338)Ribosomal_L19 (pfam01245)→ |
| env-202 | G312J29652_10010132 | + | 1022 | 1049 | **RNA**→ rplS (PRK05338)Ribosomal_L19 (pfam01245)→ |
| Rbi-1-1 | NC_013222.1 | + | 2791912 | 2791939 | **RNA**→ rplS (PRK05338)Ribosomal_L19 (pfam01245)→ |
| env-203 | JGI24024J18818_10269885 | - | 332 | 305 | **RNA**→ rplS (PRK05338)Ribosomal_L19 (pfam01245)→ |
| env-204 | P_1C_Liq_3_UnCty_1004041 | + | 444 | 471 | **RNA**→ rplS (PRK05338)Ribosomal_L19 (pfam01245)→ |
| env-205 | P_1C_Liq_1_UnCty_1002842 | - | 2900 | 2873 | **RNA**→ rplS (PRK05338)Ribosomal_L19 (pfam01245)→ |
| env-206 | P_2C_Liq_1_UnCty_1206961 | + | 13 | 40 | **RNA**→ rplS (PRK05338)Ribosomal_L19 (pfam01245)→ |
| env-207 | JGI20156J14371_10587087 | - | 192 | 165 | **RNA**→ rplS (PRK05338)Ribosomal_L19 (pfam01245)→ |
| env-208 | ElkS_mat_MD6A_1140673 | + | 80 | 107 | **RNA**→ rplS (PRK05338)Ribosomal_L19 (pfam01245)→ |
| env-209 | JGI11876J14442_10684723 | - | 250 | 223 | **RNA**→ rplS (PRK05338)Ribosomal_L19 (pfam01245)→ |
| env-210 | LM29OCT2010E_c89422 | + | 70 | 97 | **RNA**→ rplS (PRK05338)Ribosomal_L19 (pfam01245)→ |
| env-211 | JGI11876J14442_10078056 | - | 328 | 301 | **RNA**→ rplS (PRK05338)Ribosomal_L19 (pfam01245)→ |
| Gsp-1-1 | NZ_AMSG01000008.1 | + | 129752 | 129779 | **RNA**→ rplS (PRK05338)Ribosomal_L19 (pfam01245)→ rplS (PRK05338)Ribosomal_L19 (pfam01245)→ |
| env-212 | PR_CR_10_Liq_1_inCR_1041635 | - | 595 | 568 | **RNA**→ rplS (PRK05338)Ribosomal_L19 (pfam01245)→ |
| env-213 | PR_CR_10_Liq_3_inCR_1112054 | - | 41 | 14 | **RNA**→ |
| env-214 | ATED1026_F5B15RW02FGXY8 | - | 36 | 9 | **RNA**→ |
| env-215 | JGI24023J19991_10000222 | - | 18096 | 18069 | **RNA**→ rplS (PRK05338)Ribosomal_L19 (pfam01245)→ |
| env-216 | JGI24023J19991_10496689 | - | 276 | 249 | **RNA**→ hypo→ rplS (PRK05338)Ribosomal_L19 (pfam01245)→ |
| Man-1-1 | NZ_JHZC01000004.1 | - | 677641 | 677614 | **RNA**→ rplS (PRK05338)Ribosomal_L19 (pfam01245)→ |
| env-217 | NCBI_READ_1112493462061 | + | 858 | 885 | **RNA**→ |
| env-218 | JGI24723J26617_10000035 | - | 19920 | 19893 | **RNA**→ rplS (PRK05338)Ribosomal_L19 (pfam01245)→ |
| env-219 | UnmappedStool_Broad_C253267521 | + | 390 | 417 | **RNA**→ rplS (PRK05338)Ribosomal_L19 (pfam01245)→ |
| env-220 | SRS016095_C2602647 | + | 2397 | 2424 | **RNA**→ rplS (PRK05338)Ribosomal_L19 (pfam01245)→ hypo→ |
| env-221 | SRS019968_C2032833 | + | 279 | 306 | **RNA**→ rplS (PRK05338)Ribosomal_L19 (pfam01245)→ |
| env-222 | 4491417.3_NODE_24004 | + | 1215 | 1242 | **RNA**→ rplS (PRK05338)Ribosomal_L19 (pfam01245)→ |

| | | | | | |
|---|---|---|---|---|---|
| env-223 | UnmappedStool_Broad_C252323657 | + | 165 | 192 | **RNA**→ rplS (PRK05338)Ribosomal_L19 (pfam01245)→ |
| env-224 | SRS053917_LANL_scaffold_85389 | + | 54783 | 54810 | **RNA**→ rplS (PRK05338)Ribosomal_L19 (pfam01245)→ KAS_I_II (cd00834)Ketoacyl-synt_2 (pfam13723)→ PRK10925 (PRK10925)SodA (COG0605)→ |
| env-225 | SRS022725_LANL_scaffold_115907 | - | 16699 | 16672 | **RNA**→ rplS (PRK05338)Ribosomal_L19 (pfam01245)→ KAS_I_II (cd00834)Ketoacyl-synt_2 (pfam13723)→ PRK10925 (PRK10925)SodA (COG0605)→ |
| env-226 | SRS075410_LANL_scaffold_18149 | - | 1834 | 1807 | **RNA**→ rplS (PRK05338)Ribosomal_L19 (pfam01245)→ |
| env-227 | SRS023841_C1664739 | + | 283 | 310 | **RNA**→ rplS (PRK05338)Ribosomal_L19 (pfam01245)→ KAS_I_II (cd00834)fabF (TIGR03150)→ PRK10925 (PRK10925)SodA (COG0605)→ tonB_Cterm (TIGR01352)TonB_C (pfam03544)→ SCO (cd02968)→ DUF4325 (pfam14213)→ PIN (pfam01850)→ RsmA (COG0030)ksgA (PRK00274)→ |
| env-228 | Subgingival_plaque_LANL_C11225722 | - | 66 | 39 | **RNA**→ |
| env-229 | SRS011126_Baylor_scaffold_19796 | - | 15490 | 15463 | **RNA**→ rplS (PRK05338)Ribosomal_L19 (pfam01245)→ KAS_I_II (cd00834)fabF (TIGR03150)→ PRK10925 (PRK10925)SodA (COG0605)→ tonB_Cterm (TIGR01352)TonB_C (pfam03544)→ SCO (cd02968)→ DUF4325 (pfam14213)→ PIN (pfam01850)→ RsmA (COG0030)ksgA (PRK00274)→ |
| env-230 | SRS013949_C3491244 | + | 478 | 505 | **RNA**→ |
| env-231 | SRS015899_WUGC_scaffold_40977 | + | 29485 | 29512 | **RNA**→ rplS (PRK05338)Ribosomal_L19 (pfam01245)→ KAS_I_II (cd00834)fabF (TIGR03150)→ PRK10925 (PRK10925)SodA (COG0605)→ tonB_Cterm (TIGR01352)TonB_C (pfam03544)→ SCO (cd02968)→ DUF4325 (pfam14213)→ PIN (pfam01850)→ RsmA (COG0030)ksgA (PRK00274)→ |
| env-232 | SRS016297_C532705 | - | 142 | 115 | **RNA**→ |
| env-233 | SRS022083_C1794273 | + | 126 | 153 | **RNA**→ rplS (PRK05338)Ribosomal_L19 (pfam01245)→ |
| env-234 | SRS022536_LANL_scaffold_110861 | - | 15742 | 15715 | **RNA**→ rplS (PRK05338)Ribosomal_L19 (pfam01245)→ KAS_I_II (cd00834)fabF (TIGR03150)→ PRK10925 (PRK10925)SodA (COG0605)→ tonB_Cterm (TIGR01352)TonB_C (pfam03544)→ SCO (cd02968)→ DUF4325 (pfam14213)→ PIN (pfam01850)→ RsmA (COG0030)ksgA (PRK00274)→ |
| env-235 | SRS023938_C2099915 | + | 111 | 138 | **RNA**→ rplS (PRK05338)Ribosomal_L19 (pfam01245)→ |
| env-236 | SRS024381_C2283100 | + | 338 | 365 | **RNA**→ |
| env-237 | SRS042984_LANL_scaffold_49763 | - | 3257 | 3230 | **RNA**→ rplS (PRK05338)Ribosomal_L19 (pfam01245)→ KAS_I_II (cd00834)fabF (TIGR03150)→ PRK10925 (PRK10925)SodA (COG0605)→ tonB_Cterm (TIGR01352)TonB_C (pfam03544)→ hypo→ |
| env-238 | SRS049318_LANL_scaffold_97896 | + | 617 | 644 | **RNA**→ |
| env-239 | SRS065099_LANL_scaffold_36643 | - | 202 | 175 | **RNA**→ rplS (PRK05338)Ribosomal_L19 (pfam01245)→ |
| env-240 | SRS065310_C1333489 | + | 502 | 529 | **RNA**→ |
| env-241 | SRS075410_C3068469 | + | 2319 | 2346 | **RNA**→ |
| env-242 | SRS015803_C2605009 | + | 372 | 399 | **RNA**→ rplS (PRK05338)Ribosomal_L19 (pfam01245)→ |
| env-243 | SRS047634_LANL_scaffold_52163 | + | 1818 | 1845 | **RNA**→ |
| env-244 | SRS017691_Baylor_scaffold_41406 | + | 1771 | 1798 | **RNA**→ |
| env-245 | SRS055450_LANL_scaffold_77438 | + | 1416 | 1443 | **RNA**→ |
| Csp-2-1 | NZ_ALNN01000014.1 | - | 9694 | 9667 | **RNA**→ rplS (PRK05338)Ribosomal_L19 (pfam01245)→ rplS (PRK05338)Ribosomal_L19 (pfam01245)→ KAS_I_II (cd00834)Ketoacyl-synt_2 (pfam13723)→ KAS_I_II (cd00834)Ketoacyl-synt_2 (pfam13723)→ PRK10925 (PRK10925)SodA (COG0605)→ PRK10925 (PRK10925)SodA (COG0605)→ tonB_Cterm (TIGR01352)TonB_C (pfam03544)→ tonB_Cterm (TIGR01352)TonB_C (pfam03544)→ SCO (cd02968)→ SCO (cd02968)→ |
| env-246 | SRS050244_C4263116 | - | 66 | 39 | **RNA**→ |
| Cgr-1-1 | NZ_KE150260.1 | + | 63374 | 63401 | **RNA**→ rplS (PRK05338)Ribosomal_L19 (pfam01245)→ KAS_I_II (cd00834)Ketoacyl-synt_2 (pfam13723)→ PRK10925 (PRK10925)SodA (COG0605)→ tonB_Cterm (TIGR01352)TonB_C (pfam03544)→ SCO (cd02968)→ DUF4325 (pfam14213)→ PIN (pfam01850)→ RsmA (COG0030)ksgA (PRK00274)→ |
| env-247 | SRS019980_C3055592 | - | 63 | 36 | **RNA**→ |
| env-248 | SRS063603_LANL_scaffold_70638 | - | 72 | 45 | **RNA**→ |
| env-249 | SRS011255_C2494079 | - | 69 | 42 | **RNA**→ |
| env-250 | SRS014578_WUGC_scaffold_19186 | + | 5221 | 5248 | **RNA**→ rplS (PRK05338)Ribosomal_L19 (pfam01245)→ KAS_I_II (cd00834)fabF (TIGR03150)→ PRK10925 (PRK10925)SodA (COG0605)→ |
| env-251 | SRS017445_Baylor_scaffold_33849 | + | 184 | 211 | **RNA**→ rplS (PRK05338)Ribosomal_L19 (pfam01245)→ |
| env-252 | SRS017691_C5080407 | - | 78 | 51 | **RNA**→ hypo→ |
| env-253 | SRS018573_WUGC_scaffold_28515 | + | 3449 | 3476 | **RNA**→ rplS (PRK05338)Ribosomal_L19 (pfam01245)→ KAS_I_II (cd00834)fabF (TIGR03150)→ PRK10925 (PRK10925)Sod_Fe_N (pfam00081)→ |
| env-254 | SRS045197_WUGC_scaffold_27322 | - | 194 | 167 | **RNA**→ rplS (PRK05338)Ribosomal_L19 (pfam01245)→ |
| env-255 | SRS063932_LANL_scaffold_8547 | + | 316 | 343 | **RNA**→ rplS (PRK05338)Ribosomal_L19 (pfam01245)→ |
| env-256 | SRS022602_Baylor_scaffold_17361 | - | 665 | 638 | **RNA**→ rplS (PRK05338)Ribosomal_L19 (pfam01245)→ hypo→ |

| | | | | | |
|---|---|---|---|---|---|
| env-257 | SRS013252_C3528677 | + | 307 | 334 | RNA→ rplS (PRK05338)Ribosomal_L19 (pfam01245)→ |
| env-258 | SRS013705_C3926261 | + | 2029 | 2056 | RNA→ rplS (PRK05338)Ribosomal_L19 (pfam01245)→ |
| env-259 | SRS015434_C5837920 | + | 260 | 287 | RNA→ rplS (PRK05338)Ribosomal_L19 (pfam01245)→ |
| env-260 | SRS017139_Baylor_scaffold_77500 | + | 898 | 925 | RNA→ KAS_I_II (cd00834)fabF (TIGR03150)→ hypo→ PRK10925 (PRK10925)SodA (COG0605)→ |
| env-261 | SRS018157_Baylor_scaffold_33010 | + | 2985 | 3012 | RNA→ rplS (PRK05338)Ribosomal_L19 (pfam01245)→ |
| env-262 | SRS018300_Baylor_scaffold_56055 | + | 6669 | 6696 | RNA→ rplS (PRK05338)Ribosomal_L19 (pfam01245)→ |
| env-263 | SRS019028_WUGC_scaffold_24330 | + | 43055 | 43082 | RNA→ rplS (PRK05338)Ribosomal_L19 (pfam01245)→ |
| env-264 | SRS019029_WUGC_scaffold_43710 | + | 5639 | 5666 | RNA→ rplS (PRK05338)Ribosomal_L19 (pfam01245)→ |
| env-265 | SRS019071_C5218564 | + | 2113 | 2140 | RNA→ |
| env-266 | SRS023926_C2716754 | - | 232 | 205 | RNA→ rplS (PRK05338)Ribosomal_L19 (pfam01245)→ |
| env-267 | SRS024355_LANL_scaffold_16886 | - | 95 | 68 | RNA→ |
| env-268 | SRS024649_LANL_scaffold_7780 | - | 430 | 403 | RNA→ rplS (PRK05338)Ribosomal_L19 (pfam01245)→ |
| env-269 | SRS049318_LANL_scaffold_73392 | + | 507 | 534 | RNA→ |
| env-270 | SRS051941_LANL_scaffold_75018 | - | 1329 | 1302 | RNA→ rplS (PRK05338)Ribosomal_L19 (pfam01245)→ |
| env-271 | SRS052604_C1778119 | + | 204 | 231 | RNA→ rplS (PRK05338)Ribosomal_L19 (pfam01245)→ |
| env-272 | SRS055378_LANL_scaffold_12682 | + | 3851 | 3878 | RNA→ rplS (PRK05338)Ribosomal_L19 (pfam01245)→ |
| env-273 | SRS014477_C1574933 | + | 316 | 343 | RNA→ |
| env-274 | SRS017304_C1915201 | + | 220 | 247 | RNA→ rplS (PRK05338)Ribosomal_L19 (pfam01245)→ |
| env-275 | SRS045197_WUGC_scaffold_24835 | + | 1618 | 1645 | RNA→ |
| env-276 | SRS047113_C5057443 | - | 72 | 45 | RNA→ |
| Csp-11-1 | NZ_KE992168.1 | + | 41060 | 41087 | RNA→ rplS (PRK05338)Ribosomal_L19 (pfam01245)→ |
| env-277 | SRS015762_C3669651 | - | 1738 | 1711 | RNA→ rplS (PRK05338)Ribosomal_L19 (pfam01245)→ KAS_I_II (cd00834)Ketoacyl-synt_2 (pfam13723)→ PRK10925 (PRK10925)SodA (COG0605)→ |
| env-278 | SRS015989_C2177835 | - | 199 | 172 | RNA→ rplS (PRK05338)Ribosomal_L19 (pfam01245)→ |
| env-279 | SRS016746_Baylor_scaffold_79161 | - | 4503 | 4476 | RNA→ rplS (PRK05338)Ribosomal_L19 (pfam01245)→ |
| env-280 | SRS043772_WUGC_scaffold_6381 | + | 547 | 574 | RNA→ rplS (PRK05338)Ribosomal_L19 (pfam01245)→ |
| env-281 | SRS064449_LANL_scaffold_70561 | + | 3228 | 3255 | RNA→ |
| env-282 | SRS011098_C1795866 | + | 278 | 305 | RNA→ |
| env-283 | SRS012279_C2574426 | + | 101 | 128 | RNA→ rplS (PRK05338)Ribosomal_L19 (pfam01245)→ |
| env-284 | SRS013723_Baylor_scaffold_90380 | - | 302 | 275 | RNA→ rplS (PRK05338)Ribosomal_L19 (pfam01245)→ |
| env-285 | SRS013836_C4642922 | + | 91 | 118 | RNA→ rplS (PRK05338)Ribosomal_L19 (pfam01245)→ |
| env-286 | SRS014578_WUGC_scaffold_47009 | - | 8623 | 8596 | RNA→ rplS (PRK05338)Ribosomal_L19 (pfam01245)→ |
| env-287 | SRS014690_WUGC_scaffold_15252 | + | 1864 | 1891 | RNA→ rplS (PRK05338)Ribosomal_L19 (pfam01245)→ |
| env-288 | SRS014888_C3106699 | + | 96 | 123 | RNA→ rplS (PRK05338)Ribosomal_L19 (pfam01245)→ |
| env-289 | SRS015755_WUGC_scaffold_39561 | + | 2225 | 2252 | RNA→ |
| env-290 | SRS015762_WUGC_scaffold_50704 | + | 8795 | 8822 | RNA→ rplS (PRK05338)Ribosomal_L19 (pfam01245)→ |
| env-291 | SRS015797_WUGC_scaffold_33578 | + | 7661 | 7688 | RNA→ rplS (PRK05338)Ribosomal_L19 (pfam01245)→ |
| env-292 | SRS016319_C3872782 | + | 96 | 123 | RNA→ rplS (PRK05338)Ribosomal_L19 (pfam01245)→ |
| env-293 | SRS017511_Baylor_scaffold_29232 | + | 1941 | 1968 | RNA→ rplS (PRK05338)Ribosomal_L19 (pfam01245)→ |
| env-294 | SRS018300_Baylor_scaffold_56130 | + | 96 | 123 | RNA→ rplS (PRK05338)Ribosomal_L19 (pfam01245)→ |
| env-295 | SRS019028_WUGC_scaffold_30630 | - | 13299 | 13272 | RNA→ rplS (PRK05338)Ribosomal_L19 (pfam01245)→ |
| env-296 | SRS019029_WUGC_scaffold_51290 | - | 216 | 189 | RNA→ rplS (PRK05338)Ribosomal_L19 (pfam01245)→ |
| env-297 | SRS019122_C5922434 | + | 96 | 123 | RNA→ rplS (PRK05338)Ribosomal_L19 (pfam01245)→ |
| env-298 | SRS019127_WUGC_scaffold_19698 | - | 2353 | 2326 | RNA→ rplS (PRK05338)Ribosomal_L19 (pfam01245)→ KAS_I_II (cd00834)Ketoacyl-synt_2 (pfam13723)→ PRK10925 (PRK10925)SodA (COG0605)→ |
| env-299 | SRS019906_WUGC_scaffold_11203 | - | 334 | 307 | RNA→ |
| env-300 | SRS019980_Baylor_scaffold_24077 | + | 36052 | 36079 | RNA→ rplS (PRK05338)Ribosomal_L19 (pfam01245)→ |
| env-301 | SRS021496_Baylor_scaffold_80652 | - | 8962 | 8935 | RNA→ rplS (PRK05338)Ribosomal_L19 (pfam01245)→ KAS_I_II (cd00834)Ketoacyl-synt_2 (pfam13723)→ PRK10925 (PRK10925)SodA (COG0605)→ tonB_Cterm (TIGR01352)TonB_C (pfam03544)→ |
| env-302 | SRS022077_C2503359 | + | 101 | 128 | RNA→ rplS (PRK05338)Ribosomal_L19 (pfam01245)→ |
| env-303 | SRS022530_LANL_scaffold_60953 | + | 128871 | 128898 | RNA→ rplS (PRK05338)Ribosomal_L19 (pfam01245)→ KAS_I_II (cd00834)Ketoacyl-synt_2 (pfam13723)→ PRK10925 (PRK10925)SodA (COG0605)→ tonB_Cterm (TIGR01352)TonB_C (pfam03544)→ |
| env-304 | SRS023938_Baylor_scaffold_64946 | + | 867 | 894 | RNA→ rplS (PRK05338)Ribosomal_L19 (pfam01245)→ rplS (PRK05338)Ribosomal_L19 (pfam01245)→ |

| | | | | | |
|---|---|---|---|---|---|
| env-305 | SRS051378_LANL_scaffold_26894 | - | 581 | 554 | **RNA**→ rplS (PRK05338)Ribosomal_L19 (pfam01245)→ |
| env-306 | SRS052876_LANL_scaffold_30038 | + | 1896 | 1923 | **RNA**→ rplS (PRK05338)Ribosomal_L19 (pfam01245)→ |
| env-307 | SRS055378_LANL_scaffold_27817 | - | 196 | 169 | **RNA**→ rplS (PRK05338)Ribosomal_L19 (pfam01245)→ |
| env-308 | SRS063288_LANL_scaffold_21438 | - | 2311 | 2284 | **RNA**→ rplS (PRK05338)Ribosomal_L19 (pfam01245)→ KAS_I_II (cd00834)Ketoacyl-synt_2 (pfam13723)→ PRK10925 (PRK10925)SodA (COG0605)→ |
| env-309 | SRS063999_LANL_scaffold_36605 | + | 103 | 130 | **RNA**→ rplS (PRK05338)Ribosomal_L19 (pfam01245)→ |
| Cgi-1-1 | NZ_ACLQ01000031.1 | + | 74550 | 74577 | **RNA**→ rplS (PRK05338)Ribosomal_L19 (pfam01245)→ rplS (PRK05338)Ribosomal_L19 (pfam01245)→ KAS_I_II (cd00834)Ketoacyl-synt_2 (pfam13723)→ KAS_I_II (cd00834)Ketoacyl-synt_2 (pfam13723)→ PRK10925 (PRK10925)SodA (COG0605)→ PRK10925 (PRK10925)SodA (COG0605)→ tonB_Cterm (TIGR01352)TonB_C (pfam03544)→ tonB_Cterm (TIGR01352)TonB_C (pfam03544)→ |
| env-310 | SRS015158_WUGC_scaffold_7291 | - | 289 | 262 | **RNA**→ rplS (PRK05338)Ribosomal_L19 (pfam01245)→ |
| env-311 | SRS065310_C1342652 | - | 237 | 210 | **RNA**→ rplS (PRK05338)Ribosomal_L19 (pfam01245)→ |
| env-312 | SRS016200_WUGC_scaffold_37401 | + | 138 | 165 | **RNA**→ rplS (PRK05338)Ribosomal_L19 (pfam01245)→ |
| env-313 | SRS013705_C3910892 | - | 51 | 24 | **RNA**→ |
| env-314 | SRS017209_Baylor_scaffold_54331 | - | 82 | 55 | **RNA**→ |
| env-315 | SRS018439_C2415034 | + | 2567 | 2594 | **RNA**→ |
| env-316 | SRS018739_C3517180 | - | 170 | 143 | **RNA**→ rplS (PRK05338)Ribosomal_L19 (pfam01245)→ |
| env-317 | SRS019077_WUGC_scaffold_45325 | + | 94 | 121 | **RNA**→ rplS (PRK05338)Ribosomal_L19 (pfam01245)→ |
| env-318 | SRS023926_Baylor_scaffold_72969 | + | 206 | 233 | **RNA**→ rplS (PRK05338)Ribosomal_L19 (pfam01245)→ KAS_I_II (cd00834)Ketoacyl-synt_2 (pfam13723)→ PRK10925 (PRK10925)SodA (COG0605)→ hypo→ |
| env-319 | SRS024441_C2947315 | + | 96 | 123 | **RNA**→ rplS (PRK05338)Ribosomal_L19 (pfam01245)→ |
| env-320 | SRS050244_LANL_scaffold_54867 | + | 23174 | 23201 | **RNA**→ rplS (PRK05338)Ribosomal_L19 (pfam01245)→ |
| env-321 | SRS050669_LANL_scaffold_708 | + | 7741 | 7768 | **RNA**→ rplS (PRK05338)Ribosomal_L19 (pfam01245)→ KAS_I_II (cd00834)Ketoacyl-synt_2 (pfam13723)→ PRK10925 (PRK10925)SodA (COG0605)→ tonB_Cterm (TIGR01352)TonB_C (pfam03544)→ |
| env-322 | SRS062544_C3036364 | + | 1599 | 1626 | **RNA**→ rplS (PRK05338)Ribosomal_L19 (pfam01245)→ KAS_I_II (cd00834)Ketoacyl-synt_2 (pfam13723)→ PRK10925 (PRK10925)SodA (COG0605)→ |
| env-323 | SRS023964_Baylor_scaffold_63915 | - | 289 | 262 | **RNA**→ rplS (PRK05338)Ribosomal_L19 (pfam01245)→ |
| env-324 | SRS024355_LANL_scaffold_68198 | - | 289 | 262 | **RNA**→ rplS (PRK05338)Ribosomal_L19 (pfam01245)→ |
| env-325 | SRS042131_WUGC_scaffold_62340 | + | 1868 | 1895 | **RNA**→ rplS (PRK05338)Ribosomal_L19 (pfam01245)→ |
| env-326 | SRS075410_LANL_scaffold_45971 | - | 82 | 55 | **RNA**→ |
| env-327 | SRS022602_Baylor_scaffold_115301 | - | 611 | 584 | **RNA**→ rplS (PRK05338)Ribosomal_L19 (pfam01245)→ hypo→ |
| env-328 | SRS017445_Baylor_scaffold_77384 | + | 73 | 100 | **RNA**→ rplS (PRK05338)Ribosomal_L19 (pfam01245)→ |
| env-329 | SRS058336_C2180371 | - | 2178 | 2151 | **RNA**→ rplS (PRK05338)Ribosomal_L19 (pfam01245)→ KAS_I_II (cd00834)Ketoacyl-synt_2 (pfam13723)→ PRK10925 (PRK10925)SodA (COG0605)→ |
| env-330 | SRS016002_C2622796 | + | 98 | 125 | **RNA**→ rplS (PRK05338)Ribosomal_L19 (pfam01245)→ KAS_I_II (cd00834)Ketoacyl-synt_2 (pfam13723)→ |
| env-331 | SRS018394_Baylor_scaffold_70182 | + | 834 | 861 | **RNA**→ rplS (PRK05338)Ribosomal_L19 (pfam01245)→ |
| env-332 | SRS023841_Baylor_scaffold_23089 | - | 5503 | 5476 | **RNA**→ rplS (PRK05338)Ribosomal_L19 (pfam01245)→ KAS_I_II (cd00834)Ketoacyl-synt_2 (pfam13723)→ PRK10925 (PRK10925)SodA (COG0605)→ tonB_Cterm (TIGR01352)TonB_C (pfam03544)→ |
| env-333 | SRS013252_C3501854 | + | 222 | 249 | **RNA**→ rplS (PRK05338)Ribosomal_L19 (pfam01245)→ |
| env-334 | SRS013949_WUGC_scaffold_26347 | - | 289 | 262 | **RNA**→ rplS (PRK05338)Ribosomal_L19 (pfam01245)→ |
| env-335 | SRS013950_C2064126 | + | 253 | 280 | **RNA**→ rplS (PRK05338)Ribosomal_L19 (pfam01245)→ |
| env-336 | SRS015215_WUGC_scaffold_59667 | - | 1128 | 1101 | **RNA**→ rplS (PRK05338)Ribosomal_L19 (pfam01245)→ |
| env-337 | SRS015278_C1853906 | + | 1366 | 1393 | **RNA**→ rplS (PRK05338)Ribosomal_L19 (pfam01245)→ |
| env-338 | SRS015989_WUGC_scaffold_22267 | + | 1178 | 1205 | **RNA**→ rplS (PRK05338)Ribosomal_L19 (pfam01245)→ |
| env-339 | SRS016331_WUGC_scaffold_4458 | - | 4432 | 4405 | **RNA**→ rplS (PRK05338)Ribosomal_L19 (pfam01245)→ |
| env-340 | SRS016746_Baylor_scaffold_49712 | + | 3708 | 3735 | **RNA**→ rplS (PRK05338)Ribosomal_L19 (pfam01245)→ |
| env-341 | SRS018157_Baylor_scaffold_36967 | + | 16031 | 16058 | **RNA**→ rplS (PRK05338)Ribosomal_L19 (pfam01245)→ |
| env-342 | SRS018975_WUGC_scaffold_5756 | + | 23754 | 23781 | **RNA**→ rplS (PRK05338)Ribosomal_L19 (pfam01245)→ |
| env-343 | SRS019071_WUGC_scaffold_32884 | - | 289 | 262 | **RNA**→ rplS (PRK05338)Ribosomal_L19 (pfam01245)→ |
| env-344 | SRS020340_C2977594 | - | 289 | 262 | **RNA**→ rplS (PRK05338)Ribosomal_L19 (pfam01245)→ |
| env-345 | SRS024021_Baylor_scaffold_35803 | + | 10911 | 10938 | **RNA**→ rplS (PRK05338)Ribosomal_L19 (pfam01245)→ |
| env-346 | SRS024144_C1810569 | + | 1730 | 1757 | **RNA**→ rplS (PRK05338)Ribosomal_L19 (pfam01245)→ |
| env-347 | SRS024561_LANL_scaffold_19935 | + | 3850 | 3877 | **RNA**→ rplS (PRK05338)Ribosomal_L19 (pfam01245)→ |
| env-348 | SRS024649_C3635117 | + | 578 | 605 | **RNA**→ |

| | | | | | |
|---|---|---|---|---|---|
| env-349 | SRS042984_LANL_scaffold_17949 | - | 524 | 497 | **RNA**→ rplS (PRK05338)Ribosomal_L19 (pfam01245)→ |
| env-350 | SRS043755_C2384479 | - | 289 | 262 | **RNA**→ rplS (PRK05338)Ribosomal_L19 (pfam01245)→ |
| env-351 | SRS043772_WUGC_scaffold_25088 | + | 1877 | 1904 | **RNA**→ rplS (PRK05338)Ribosomal_L19 (pfam01245)→ |
| env-352 | SRS047265_WUGC_scaffold_21571 | + | 10737 | 10764 | **RNA**→ rplS (PRK05338)Ribosomal_L19 (pfam01245)→ KAS_I_II (cd00834)Ketoacyl-synt_2 (pfam13723)→ PRK10925 (PRK10925)SodA (COG0605)→ tonB_Cterm (TIGR01352)TonB_C (pfam03544)→ |
| env-353 | SRS047634_LANL_scaffold_65807 | - | 82 | 55 | **RNA**→ |
| env-354 | SRS051930_LANL_scaffold_29394 | - | 1258 | 1231 | **RNA**→ rplS (PRK05338)Ribosomal_L19 (pfam01245)→ |
| env-355 | SRS075406_C837804 | + | 4602 | 4629 | **RNA**→ rplS (PRK05338)Ribosomal_L19 (pfam01245)→ |
| env-356 | SRS016541_Baylor_scaffold_34066 | + | 23526 | 23553 | **RNA**→ rplS (PRK05338)Ribosomal_L19 (pfam01245)→ KAS_I_II (cd00834)Ketoacyl-synt_2 (pfam13723)→ PRK10925 (PRK10925)SodA (COG0605)→ tonB_Cterm (TIGR01352)TonB_C (pfam03544)→ |
| env-357 | SRS053630_LANL_scaffold_23956 | - | 14901 | 14874 | **RNA**→ rplS (PRK05338)Ribosomal_L19 (pfam01245)→ KAS_I_II (cd00834)Ketoacyl-synt_2 (pfam13723)→ PRK10925 (PRK10925)SodA (COG0605)→ tonB_Cterm (TIGR01352)TonB_C (pfam03544)→ |
| env-358 | SRS064329_C2710786 | - | 237 | 210 | **RNA**→ rplS (PRK05338)Ribosomal_L19 (pfam01245)→ |
| env-359 | SRS011126_Baylor_scaffold_64408 | + | 519 | 546 | **RNA**→ rplS (PRK05338)Ribosomal_L19 (pfam01245)→ |
| env-360 | SRS011343_Baylor_scaffold_21101 | + | 3877 | 3904 | **RNA**→ rplS (PRK05338)Ribosomal_L19 (pfam01245)→ |
| env-361 | SRS012285_Baylor_scaffold_35900 | - | 433 | 406 | **RNA**→ rplS (PRK05338)Ribosomal_L19 (pfam01245)→ |
| env-362 | SRS013533_C5373250 | + | 215 | 242 | **RNA**→ rplS (PRK05338)Ribosomal_L19 (pfam01245)→ |
| env-363 | SRS015899_WUGC_scaffold_32404 | + | 5473 | 5500 | **RNA**→ rplS (PRK05338)Ribosomal_L19 (pfam01245)→ |
| env-364 | SRS016043_WUGC_scaffold_15886 | - | 289 | 262 | **RNA**→ rplS (PRK05338)Ribosomal_L19 (pfam01245)→ |
| env-365 | SRS017227_Baylor_scaffold_5549 | + | 3292 | 3319 | **RNA**→ |
| env-366 | SRS018573_WUGC_scaffold_23933 | + | 4746 | 4773 | **RNA**→ rplS (PRK05338)Ribosomal_L19 (pfam01245)→ |
| env-367 | SRS019333_C1276128 | + | 649 | 676 | **RNA**→ |
| env-368 | SRS021477_Baylor_scaffold_72269 | + | 2598 | 2625 | **RNA**→ rplS (PRK05338)Ribosomal_L19 (pfam01245)→ |
| env-369 | SRS023538_Baylor_scaffold_41620 | + | 7749 | 7776 | **RNA**→ rplS (PRK05338)Ribosomal_L19 (pfam01245)→ KAS_I_II (cd00834)Ketoacyl-synt_2 (pfam13723)→ PRK10925 (PRK10925)SodA (COG0605)→ tonB_Cterm (TIGR01352)TonB_C (pfam03544)→ |
| env-370 | SRS024081_LANL_scaffold_74647 | - | 14542 | 14515 | **RNA**→ rplS (PRK05338)Ribosomal_L19 (pfam01245)→ KAS_I_II (cd00834)Ketoacyl-synt_2 (pfam13723)→ PRK10925 (PRK10925)SodA (COG0605)→ tonB_Cterm (TIGR01352)TonB_C (pfam03544)→ |
| env-371 | SRS024138_Baylor_scaffold_45959 | + | 4204 | 4231 | **RNA**→ rplS (PRK05338)Ribosomal_L19 (pfam01245)→ KAS_I_II (cd00834)Ketoacyl-synt_2 (pfam13723)→ PRK10925 (PRK10925)SodA (COG0605)→ tonB_Cterm (TIGR01352)TonB_C (pfam03544)→ |
| env-372 | SRS045197_WUGC_scaffold_31928 | - | 203 | 176 | **RNA**→ rplS (PRK05338)Ribosomal_L19 (pfam01245)→ |
| env-373 | SRS047113_LANL_scaffold_42767 | - | 289 | 262 | **RNA**→ rplS (PRK05338)Ribosomal_L19 (pfam01245)→ |
| env-374 | SRS049268_C4056582 | - | 82 | 55 | **RNA**→ hypo→ |
| env-375 | SRS049318_LANL_scaffold_162736 | - | 82 | 55 | **RNA**→ |
| env-376 | SRS053584_LANL_scaffold_34108 | + | 3382 | 3409 | **RNA**→ rplS (PRK05338)Ribosomal_L19 (pfam01245)→ |
| env-377 | SRS058053_LANL_scaffold_42593 | + | 127 | 154 | **RNA**→ rplS (PRK05338)Ribosomal_L19 (pfam01245)→ |
| env-378 | SRS063215_LANL_scaffold_9737 | - | 206 | 179 | **RNA**→ rplS (PRK05338)Ribosomal_L19 (pfam01245)→ |
| env-379 | SRS065099_LANL_scaffold_38497 | + | 1419 | 1446 | **RNA**→ rplS (PRK05338)Ribosomal_L19 (pfam01245)→ |
| env-380 | SRS015044_WUGC_scaffold_40958 | - | 82 | 55 | **RNA**→ |
| env-381 | SRS015470_WUGC_scaffold_48518 | + | 881 | 908 | **RNA**→ |
| env-382 | SRS015947_C1406781 | - | 1120 | 1093 | **RNA**→ rplS (PRK05338)Ribosomal_L19 (pfam01245)→ |
| env-383 | SRS052604_C1832355 | - | 290 | 263 | **RNA**→ rplS (PRK05338)Ribosomal_L19 (pfam01245)→ |
| env-384 | SRS055401_LANL_scaffold_4017 | + | 3753 | 3780 | **RNA**→ rplS (PRK05338)Ribosomal_L19 (pfam01245)→ KAS_I_II (cd00834)Ketoacyl-synt_2 (pfam13723)→ PRK10925 (PRK10925)SodA (COG0605)→ tonB_Cterm (TIGR01352)TonB_C (pfam03544)→ |
| env-385 | SRS016360_Baylor_scaffold_25143 | + | 778 | 805 | **RNA**→ rplS (PRK05338)Ribosomal_L19 (pfam01245)→ |
| env-386 | SRS017025_C1733308 | - | 289 | 262 | **RNA**→ rplS (PRK05338)Ribosomal_L19 (pfam01245)→ |
| env-387 | SRS019591_C1897347 | - | 576 | 549 | **RNA**→ rplS (PRK05338)Ribosomal_L19 (pfam01245)→ |
| env-388 | SRS050628_LANL_scaffold_19974 | + | 122 | 149 | **RNA**→ rplS (PRK05338)Ribosomal_L19 (pfam01245)→ KAS_I_II (cd00834)Ketoacyl-synt_2 (pfam13723)→ |
| env-389 | SRS058808_LANL_scaffold_17722 | - | 1470 | 1443 | **RNA**→ rplS (PRK05338)Ribosomal_L19 (pfam01245)→ |
| env-390 | SRS019225_WUGC_scaffold_3159 | - | 14652 | 14625 | **RNA**→ rplS (PRK05338)Ribosomal_L19 (pfam01245)→ KAS_I_II (cd00834)Ketoacyl-synt_2 (pfam13723)→ PRK10925 (PRK10925)SodA (COG0605)→ tonB_Cterm (TIGR01352)TonB_C (pfam03544)→ |
| env-391 | SRS014573_C3736643 | - | 387 | 360 | **RNA**→ rplS (PRK05338)Ribosomal_L19 (pfam01245)→ |
| env-392 | SRS045715_C4787013 | + | 279 | 306 | **RNA**→ rplS (PRK05338)Ribosomal_L19 (pfam01245)→ |
| env-393 | SRS015064_C1649023 | - | 315 | 288 | **RNA**→ rplS (PRK05338)Ribosomal_L19 (pfam01245)→ |
| env-394 | SRS013170_Baylor_scaffold_118831 | - | 45 | 18 | **RNA**→ |

| | | | | | |
|---|---|---|---|---|---|
| env-395 | SRS016575_Baylor_scaffold_24948 | + | 13 | 40 | **RNA**→ rplS (PRK05338)Ribosomal_L19 (pfam01245)→ |
| env-396 | SRS011098_Baylor_scaffold_7568 | + | 1143 | 1170 | **RNA**→ rplS (PRK05338)Ribosomal_L19 (pfam01245)→ |
| env-397 | SRS011255_Baylor_scaffold_19128 | + | 7753 | 7780 | **RNA**→ |
| env-398 | SRS016092_C1080935 | + | 130 | 157 | **RNA**→ rplS (PRK05338)Ribosomal_L19 (pfam01245)→ |
| env-399 | SRS063603_LANL_scaffold_85471 | + | 432 | 459 | **RNA**→ rplS (PRK05338)Ribosomal_L19 (pfam01245)→ |
| env-400 | JGI24723J26617_10169939 | + | 159 | 186 | **RNA**→ rplS (PRK05338)Ribosomal_L19 (pfam01245)→ |
| Aal-1-1 | NZ_JPOO01000003.1 | + | 1020008 | 1020035 | **RNA**→ rplS (PRK05338)Ribosomal_L19 (pfam01245)→ |
| env-401 | CENR01047393.1 | + | 64 | 91 | **RNA**→ rplS (PRK05338)Ribosomal_L19 (pfam01245)→ |
| env-402 | JGI24723J26617_10002127 | + | 8185 | 8212 | **RNA**→ rplS (PRK05338)Ribosomal_L19 (pfam01245)→ |
| env-403 | JGI24723J26617_10400028 | - | 167 | 140 | **RNA**→ rplS (PRK05338)Ribosomal_L19 (pfam01245)→ |
| env-404 | JGI11876J14442_10842250 | - | 169 | 142 | **RNA**→ rplS (PRK05338)Ribosomal_L19 (pfam01245)→ |
| env-405 | 3300001197_1009609 | + | 200 | 227 | **RNA**→ rplS (PRK05338)Ribosomal_L19 (pfam01245)→ |
| Cpa-1-1 | NZ_KE383813.1 | + | 351899 | 351926 | **RNA**→ rplS (PRK05338)Ribosomal_L19 (pfam01245)→ |
| env-406 | JGI20160J14292_10276556 | + | 252 | 281 | **RNA**→ rplS (PRK05338)Ribosomal_L19 (pfam01245)→ |
| env-407 | JGI26524J50256_1027228 | - | 1038 | 1011 | **RNA**→ rplS (PRK05338)Ribosomal_L19 (pfam01245)→ |
| env-408 | JGI26523J50269_1011208 | - | 1852 | 1825 | **RNA**→ rplS (PRK05338)Ribosomal_L19 (pfam01245)→ |
| env-409 | JGI26527J51213_1021506 | + | 117 | 144 | **RNA**→ rplS (PRK05338)Ribosomal_L19 (pfam01245)→ |
| env-410 | JGI26531J51214_10040747 | - | 1247 | 1220 | **RNA**→ rplS (PRK05338)Ribosomal_L19 (pfam01245)→ |
| env-411 | SwRhRL2b_contig_430118 | + | 1429 | 1456 | **RNA**→ rplS (PRK05338)Ribosomal_L19 (pfam01245)→ |
| Cte-1-1 | NZ_AUAA01000036.1 | + | 28922 | 28949 | **RNA**→ rplS (PRK05338)Ribosomal_L19 (pfam01245)→ |
| env-412 | APMI01002267.1 | - | 4164 | 4137 | **RNA**→ rplS (PRK05338)Ribosomal_L19 (pfam01245)→ |
| env-413 | JGI26530J50255_1090020 | + | 26 | 53 | **RNA**→ rplS (PRK05338)Ribosomal_L19 (pfam01245)→ |
| Cca-2-1 | NZ_AUFK01000009.1 | + | 927162 | 927189 | **RNA**→ rplS (PRK05338)Ribosomal_L19 (pfam01245)→ |
| env-414 | 2204596394 | - | 335 | 308 | **RNA**→ rplS (PRK05338)Ribosomal_L19 (pfam01245)→ |
| env-415 | 2158507912 | - | 335 | 308 | **RNA**→ rplS (PRK05338)Ribosomal_L19 (pfam01245)→ |
| env-416 | 3300001196_1012603 | - | 263 | 236 | **RNA**→ hypo→ rplS (PRK05338)Ribosomal_L19 (pfam01245)→ |
| env-417 | 3300001197_1004711 | - | 1007 | 980 | **RNA**→ rplS (PRK05338)Ribosomal_L19 (pfam01245)→ |
| env-418 | JGI26538J50259_1000010 | + | 108285 | 108312 | **RNA**→ hypo→ rplS (PRK05338)Ribosomal_L19 (pfam01245)→ |
| Fba-3-1 | NC_013062.1 | - | 907127 | 907100 | **RNA**→ rplS (PRK05338)Ribosomal_L19 (pfam01245)→ |
| Cha-1-1 | NZ_JPLY01000004.1 | - | 726335 | 726308 | **RNA**→ rplS (PRK05338)Ribosomal_L19 (pfam01245)→ |
| env-419 | 3300001592_10000850 | + | 8847 | 8874 | **RNA**→ hypo→ rplS (PRK05338)Ribosomal_L19 (pfam01245)→ |
| Cko-1-1 | NZ_LFNG01000001.1 | + | 195794 | 195821 | **RNA**→ rplS (PRK05338)Ribosomal_L19 (pfam01245)→ |
| Csp-14-1 | NZ_JPLZ01000003.1 | - | 97674 | 97647 | **RNA**→ rplS (PRK05338)Ribosomal_L19 (pfam01245)→ |
| env-420 | G312J29652_10118847 | + | 270 | 297 | **RNA**→ rplS (PRK05338)Ribosomal_L19 (pfam01245)→ |
| env-421 | KVRMV2_101465984 | + | 447 | 474 | **RNA**→ rplS (PRK05338)Ribosomal_L19 (pfam01245)→ |
| env-422 | JGI26530J50255_1091746 | - | 216 | 189 | **RNA**→ rplS (PRK05338)Ribosomal_L19 (pfam01245)→ |
| env-423 | JGI26529J51215_1001451 | + | 7912 | 7939 | **RNA**→ rplS (PRK05338)Ribosomal_L19 (pfam01245)→ |
| env-424 | CESG01229099.1 | + | 720 | 747 | **RNA**→ rplS (PRK05338)Ribosomal_L19 (pfam01245)→ |
| env-425 | CEQS01103232.1 | + | 83 | 110 | **RNA**→ rplS (PRK05338)Ribosomal_L19 (pfam01245)→ |
| env-426 | CEOF01031608.1 | - | 756 | 729 | **RNA**→ rplS (PRK05338)Ribosomal_L19 (pfam01245)→ PRK08299 (PRK08299)→ |
| env-427 | CENX01416069.1 | + | 1028 | 1055 | **RNA**→ rplS (PRK05338)Ribosomal_L19 (pfam01245)→ PRK08299 (PRK08299)→ |
| env-428 | CESI01090854.1 | + | 6360 | 6387 | **RNA**→ rplS (PRK05338)Ribosomal_L19 (pfam01245)→ |
| env-429 | CESU01215068.1 | - | 134 | 107 | **RNA**→ rplS (PRK05338)Ribosomal_L19 (pfam01245)→ |
| env-430 | CESI01241033.1 | - | 576 | 549 | **RNA**→ hypo→ rplS (PRK05338)Ribosomal_L19 (pfam01245)→ |
| Sze-1-1 | NZ_CP007202.1 | + | 3148679 | 3148706 | **RNA**→ rplS (PRK05338)Ribosomal_L19 (pfam01245)→ |
| env-431 | NCBI_READ_1112533034697 | + | 726 | 753 | **RNA**→ |
| env-432 | CEVX01297991.1 | - | 487 | 460 | **RNA**→ rplS (PRK05338)Ribosomal_L19 (pfam01245)→ |
| env-433 | CENF01074325.1 | - | 280 | 253 | **RNA**→ rplS (PRK05338)Ribosomal_L19 (pfam01245)→ |
| env-434 | CEVU01029800.1 | - | 121 | 94 | **RNA**→ hypo→ |
| env-435 | CESM01056441.1 | - | 321 | 294 | **RNA**→ rplS (PRK05338)Ribosomal_L19 (pfam01245)→ |
| env-436 | CEQN01349805.1 | - | 798 | 771 | **RNA**→ rplS (PRK05338)Ribosomal_L19 (pfam01245)→ PRK08299 (PRK08299)→ |
| env-437 | CEUA01051077.1 | + | 1051 | 1078 | **RNA**→ hypo→ |

| | | | | | |
|---|---|---|---|---|---|
| env-438 | CEOX01208336.1 | - | 960 | 933 | **RNA**→ rplS (PRK05338)Ribosomal_L19 (pfam01245)→ PRK08299 (PRK08299)→ |
| env-439 | CEPK01106220.1 | - | 708 | 681 | **RNA**→ rplS (PRK05338)Ribosomal_L19 (pfam01245)→ PRK08299 (PRK08299)→ |
| env-440 | CEQX01336274.1 | - | 961 | 934 | **RNA**→ rplS (PRK05338)Ribosomal_L19 (pfam01245)→ PRK08299 (PRK08299)→ |
| env-441 | CEVS01105966.1 | + | 832 | 859 | **RNA**→ rplS (PRK05338)Ribosomal_L19 (pfam01245)→ PRK08299 (PRK08299)→ |
| env-442 | CEOW01119178.1 | - | 642 | 615 | **RNA**→ rplS (PRK05338)Ribosomal_L19 (pfam01245)→ PRK08299 (PRK08299)→ |
| env-443 | CEPW01076239.1 | + | 1420 | 1447 | **RNA**→ rplS (PRK05338)Ribosomal_L19 (pfam01245)→ PRK08299 (PRK08299)→ |
| env-444 | CETU01284987.1 | + | 291 | 318 | **RNA**→ rplS (PRK05338)Ribosomal_L19 (pfam01245)→ |
| env-445 | CEVL01112834.1 | - | 1317 | 1290 | **RNA**→ rplS (PRK05338)Ribosomal_L19 (pfam01245)→ PRK08299 (PRK08299)→ |
| env-446 | CEVR01040825.1 | - | 1071 | 1044 | **RNA**→ rplS (PRK05338)Ribosomal_L19 (pfam01245)→ PRK08299 (PRK08299)→ |
| env-447 | CEVW01007019.1 | - | 234 | 207 | **RNA**→ rplS (PRK05338)Ribosomal_L19 (pfam01245)→ |
| env-448 | CEVY01168308.1 | + | 338 | 365 | **RNA**→ rplS (PRK05338)Ribosomal_L19 (pfam01245)→ |
| env-449 | CEVZ01408390.1 | - | 747 | 720 | **RNA**→ rplS (PRK05338)Ribosomal_L19 (pfam01245)→ PRK08299 (PRK08299)→ |
| env-450 | CEWO01314421.1 | - | 167 | 140 | **RNA**→ rplS (PRK05338)Ribosomal_L19 (pfam01245)→ |
| env-451 | CEVG01047150.1 | + | 617 | 644 | **RNA**→ |
| env-452 | CEWH01127616.1 | - | 502 | 475 | **RNA**→ rplS (PRK05338)Ribosomal_L19 (pfam01245)→ |
| env-453 | CETE01195508.1 | - | 433 | 406 | **RNA**→ rplS (PRK05338)Ribosomal_L19 (pfam01245)→ |
| env-454 | CESB01079027.1 | - | 492 | 465 | **RNA**→ rplS (PRK05338)Ribosomal_L19 (pfam01245)→ |
| env-455 | CEWH01141888.1 | + | 60 | 87 | **RNA**→ rplS (PRK05338)Ribosomal_L19 (pfam01245)→ |
| env-456 | JCVI_SCAF_1096627080784 | - | 1550 | 1523 | **RNA**→ rplS (PRK05338)Ribosomal_L19 (pfam01245)→ |
| env-457 | JCVI_SCAF_1096627154527 | - | 948 | 921 | **RNA**→ rplS (PRK05338)Ribosomal_L19 (pfam01245)→ |
| env-458 | 2236876001_368620 | + | 5 | 32 | **RNA**→ rplS (PRK05338)Ribosomal_L19 (pfam01245)→ |
| env-459 | CEVE01148832.1 | + | 48 | 75 | **RNA**→ rplS (PRK05338)Ribosomal_L19 (pfam01245)→ |
| env-460 | CEUF01225509.1 | - | 494 | 467 | **RNA**→ rplS (PRK05338)Ribosomal_L19 (pfam01245)→ |
| env-461 | CEVH01093382.1 | + | 145 | 172 | **RNA**→ rplS (PRK05338)Ribosomal_L19 (pfam01245)→ |
| env-462 | CENG01146222.1 | + | 169 | 196 | **RNA**→ rplS (PRK05338)Ribosomal_L19 (pfam01245)→ |
| env-463 | CEOM01318431.1 | - | 353 | 326 | **RNA**→ rplS (PRK05338)Ribosomal_L19 (pfam01245)→ |
| env-464 | CENO01076922.1 | - | 362 | 335 | **RNA**→ rplS (PRK05338)Ribosomal_L19 (pfam01245)→ |
| env-465 | CEOC01161351.1 | + | 852 | 879 | **RNA**→ rplS (PRK05338)Ribosomal_L19 (pfam01245)→ |
| env-466 | CEUQ01033343.1 | - | 386 | 359 | **RNA**→ rplS (PRK05338)Ribosomal_L19 (pfam01245)→ |
| env-467 | CEUV01047175.1 | + | 3626 | 3653 | **RNA**→ rplS (PRK05338)Ribosomal_L19 (pfam01245)→ |
| env-468 | CEVH01118179.1 | + | 204 | 231 | **RNA**→ rplS (PRK05338)Ribosomal_L19 (pfam01245)→ |
| env-469 | JCVI_READ_1108799335411 | - | 468 | 441 | **RNA**→ rplS (PRK05338)Ribosomal_L19 (pfam01245)→ |
| env-470 | JCVI_READ_1105333524717 | - | 686 | 659 | **RNA**→ rplS (PRK05338)Ribosomal_L19 (pfam01245)→ |
| env-471 | JCVI_SCAF_1096627165370 | - | 2607 | 2580 | **RNA**→ rplS (PRK05338)Ribosomal_L19 (pfam01245)→ |
| env-472 | CEOM01332252.1 | + | 48 | 75 | **RNA**→ rplS (PRK05338)Ribosomal_L19 (pfam01245)→ |
| env-473 | CEUR01223384.1 | - | 46 | 19 | **RNA**→ |
| env-474 | CEWK01258637.1 | - | 133 | 106 | **RNA**→ hypo→ |
| env-475 | JCVI_SCAF_1096627377686 | - | 4544 | 4517 | **RNA**→ rplS (PRK05338)Ribosomal_L19 (pfam01245)→ Icd (COG0538)PRK08299 (PRK08299)→ |
| env-476 | SI34jun09_10m_1057978 | - | 369 | 342 | **RNA**→ rplS (PRK05338)Ribosomal_L19 (pfam01245)→ |
| env-477 | CENG01034942.1 | - | 1486 | 1459 | **RNA**→ rplS (PRK05338)Ribosomal_L19 (pfam01245)→ |
| env-478 | CEOM01098376.1 | - | 235 | 208 | **RNA**→ rplS (PRK05338)Ribosomal_L19 (pfam01245)→ |
| env-479 | CEVS01030948.1 | + | 1746 | 1773 | **RNA**→ rplS (PRK05338)Ribosomal_L19 (pfam01245)→ |
| env-480 | CEWR01159377.1 | + | 50034 | 50061 | **RNA**→ rplS (PRK05338)Ribosomal_L19 (pfam01245)→ |
| env-481 | CESF01020515.1 | + | 988 | 1015 | **RNA**→ rplS (PRK05338)Ribosomal_L19 (pfam01245)→ |
| env-482 | CESO01050146.1 | + | 1060 | 1087 | **RNA**→ rplS (PRK05338)Ribosomal_L19 (pfam01245)→ |
| env-483 | KVRMV2_100000088 | - | 5600 | 5573 | **RNA**→ rplS (PRK05338)Ribosomal_L19 (pfam01245)→ |
| env-484 | KVRMV2_100034891 | - | 6394 | 6367 | **RNA**→ rplS (PRK05338)Ribosomal_L19 (pfam01245)→ |
| env-485 | CERW01158022.1 | + | 442 | 469 | **RNA**→ |
| env-486 | LPaug08P2610m_c1078019 | + | 6 | 33 | **RNA**→ rplS (PRK05338)Ribosomal_L19 (pfam01245)→ |
| env-487 | KGI_S1_ANT01_95m_c10010320 | + | 2206 | 2233 | **RNA**→ rplS (PRK05338)Ribosomal_L19 (pfam01245)→ |
| env-488 | KGI_S2_ANT04_2345m_c1008589 | - | 3087 | 3060 | **RNA**→ rplS (PRK05338)Ribosomal_L19 (pfam01245)→ |
| env-489 | KGI_S2_ANT05_2345m_c1137067 | - | 164 | 137 | **RNA**→ rplS (PRK05338)Ribosomal_L19 (pfam01245)→ |

| | | | | | |
|---|---|---|---|---|---|
| env-490 | KGI_S1_ANT03_95m_c1048383 | + | 153 | 180 | RNA→ rplS (PRK05338)Ribosomal_L19 (pfam01245)→ |
| env-491 | KGI_S1_ANT02_95m_c10003481 | - | 1313 | 1286 | RNA→ rplS (PRK05338)Ribosomal_L19 (pfam01245)→ |
| env-492 | KGI_S1_ANT02_95m_c10032222 | + | 261 | 288 | RNA→ rplS (PRK05338)Ribosomal_L19 (pfam01245)→ |
| env-493 | KGI_S1_ANT01_95m_c10010675 | + | 3983 | 4010 | RNA→ rplS (PRK05338)Ribosomal_L19 (pfam01245)→ |
| env-494 | KGI_S2_ANT05_2345m_c1144219 | + | 109 | 136 | RNA→ rplS (PRK05338)Ribosomal_L19 (pfam01245)→ |
| env-495 | KGI_S1_ANT02_95m_c10086448 | - | 710 | 683 | RNA→ rplS (PRK05338)Ribosomal_L19 (pfam01245)→ |
| env-496 | SI34jun09_10m_1080363 | + | 219 | 246 | RNA→ rplS (PRK05338)Ribosomal_L19 (pfam01245)→ |
| env-497 | JGI20157J14317_10015639 | + | 4166 | 4193 | RNA→ rplS (PRK05338)Ribosomal_L19 (pfam01245)→ |
| env-498 | CEPW01210536.1 | + | 127 | 154 | RNA→ rplS (PRK05338)Ribosomal_L19 (pfam01245)→ |
| env-499 | CEUA01108642.1 | + | 6252 | 6279 | RNA→ rplS (PRK05338)Ribosomal_L19 (pfam01245)→ PRK08299 (PRK08299)→ |
| env-500 | CEWA01174457.1 | + | 746 | 773 | RNA→ ←hypo |
| env-501 | JGI24023J19991_10003771 | - | 406 | 379 | RNA→ rplS (PRK05338)Ribosomal_L19 (pfam01245)→ |
| env-502 | JCVI_READ_1104230194226 | + | 528 | 555 | RNA→ rplS (PRK05338)Ribosomal_L19 (pfam01245)→ |
| env-503 | wf3_AHAI437_g1 | + | 799 | 826 | RNA→ |
| env-504 | 2236876001_449510 | + | 47 | 74 | RNA→ rplS (PRK05338)Ribosomal_L19 (pfam01245)→ |
| env-505 | TDF_MC_ARG01_113m_c1099489 | - | 124 | 97 | RNA→ hypo→ |
| env-506 | JGI20153J14295_10005629 | - | 2919 | 2892 | RNA→ rplS (PRK05338)Ribosomal_L19 (pfam01245)→ PRK08299 (PRK08299)→ PRK08299 (PRK08299)→ murQ (PRK05441)SIS_Etherase (cd05007)→ hypo→ |
| env-507 | JGI20153J14318_10005629 | - | 2919 | 2892 | RNA→ rplS (PRK05338)Ribosomal_L19 (pfam01245)→ PRK08299 (PRK08299)→ PRK08299 (PRK08299)→ murQ (PRK05441)SIS_Etherase (cd05007)→ hypo→ |
| env-508 | CEPT01227262.1 | - | 57 | 30 | RNA→ |
| env-509 | CESU01040866.1 | - | 1275 | 1248 | RNA→ rplS (PRK05338)Ribosomal_L19 (pfam01245)→ PRK08299 (PRK08299)→ |
| env-510 | JCVI_SCAF_1096627031829 | - | 1224 | 1197 | RNA→ rplS (PRK05338)Ribosomal_L19 (pfam01245)→ PRK08299 (PRK08299)→ |
| env-511 | CEVH01254860.1 | - | 217 | 190 | RNA→ rplS (PRK05338)Ribosomal_L19 (pfam01245)→ |
| env-512 | JGI24023J19991_10003593 | + | 2712 | 2739 | RNA→ rplS (PRK05338)Ribosomal_L19 (pfam01245)→ |
| env-513 | ASA120_GJFD58A02F7OV7 | + | 101 | 128 | RNA→ rplS (PRK05338)Ribosomal_L19 (pfam01245)→ |
| env-514 | 2236876001_074437 | + | 323 | 350 | RNA→ |
| env-515 | KGI_S1_ANT01_95m_c10456706 | - | 66 | 39 | RNA→ |
| env-516 | KGI_S1_ANT03_95m_c1008295 | - | 214 | 187 | RNA→ rplS (PRK05338)Ribosomal_L19 (pfam01245)→ |
| env-517 | SA_S1_NOR08_45m_c10345472 | - | 298 | 271 | RNA→ rplS (PRK05338)Ribosomal_L19 (pfam01245)→ |
| env-518 | KGI_S1_ANT03_95m_c1006880 | + | 307 | 334 | RNA→ rplS (PRK05338)Ribosomal_L19 (pfam01245)→ |
| env-519 | JGI20154J14316_10012696 | + | 4846 | 4873 | RNA→ rplS (PRK05338)Ribosomal_L19 (pfam01245)→ |
| env-520 | CEOO01015956.1 | + | 214 | 241 | RNA→ rplS (PRK05338)Ribosomal_L19 (pfam01245)→ PRK08299 (PRK08299)→ murQ (PRK05441)SIS_Etherase (cd05007)→ hypo→ |
| env-521 | CESR01281926.1 | + | 492 | 519 | RNA→ rplS (PRK05338)Ribosomal_L19 (pfam01245)→ |
| env-522 | CEVG01003056.1 | + | 390 | 417 | RNA→ hypo→ |
| env-523 | JGI24025J20009_10088571 | - | 715 | 688 | RNA→ rplS (PRK05338)Ribosomal_L19 (pfam01245)→ |
| env-524 | JCVI_READ_1108800202219 | - | 163 | 136 | RNA→ |
| Tma-1-1 | NZ_BAUG01000055.1 | + | 4233 | 4260 | RNA→ rplS (PRK05338)Ribosomal_L19 (pfam01245)→ |
| env-525 | 2236876001_552703 | - | 123 | 96 | RNA→ hypo→ |
| env-526 | KGI_S1_ANT01_95m_c10209358 | + | 393 | 420 | RNA→ |
| env-527 | P_2C_Liq_1_UnCty_1013314 | + | 1523 | 1550 | RNA→ rplS (PRK05338)Ribosomal_L19 (pfam01245)→ |
| env-528 | P_2C_Liq_1_UnCty_1013450 | - | 874 | 847 | RNA→ rplS (PRK05338)Ribosomal_L19 (pfam01245)→ |
| env-529 | JGI24024J18818_10000452 | - | 6510 | 6483 | RNA→ rplS (PRK05338)Ribosomal_L19 (pfam01245)→ |
| env-530 | CERI01232405.1 | - | 69 | 42 | RNA→ |
| env-531 | CEUB01095771.1 | - | 744 | 717 | RNA→ rplS (PRK05338)Ribosomal_L19 (pfam01245)→ PRK08299 (PRK08299)→ |
| env-532 | LGVF01272697.1 | + | 220 | 247 | RNA→ rplS (PRK05338)Ribosomal_L19 (pfam01245)→ |
| env-533 | JCVI_READ_1103242157104 | - | 160 | 133 | RNA→ |
| env-534 | JCVI_READ_1108800189128 | + | 253 | 280 | RNA→ rplS (PRK05338)Ribosomal_L19 (pfam01245)→ PRK08299 (PRK08299)→ |
| env-535 | 2236876001_291567 | - | 86 | 59 | RNA→ |
| env-536 | JGI20152J14361_10348201 | - | 184 | 157 | RNA→ rplS (PRK05338)Ribosomal_L19 (pfam01245)→ |
| env-537 | 2006938804 | + | 246 | 273 | RNA→ hypo→ |
| env-538 | JGI20156J14371_10009645 | + | 545 | 572 | RNA→ rplS (PRK05338)Ribosomal_L19 (pfam01245)→ |

| | | | | | |
|---|---|---|---|---|---|
| env-539 | JGI20157J14317_10000094 | + | 89205 | 89232 | **RNA**→ rplS (PRK05338)Ribosomal_L19 (pfam01245)→ |
| env-540 | DelMOSum2010_c10046964 | + | 738 | 765 | **RNA**→ rplS (PRK05338)Ribosomal_L19 (pfam01245)→ |
| env-541 | JGI20152J14361_10050080 | - | 464 | 437 | **RNA**→ rplS (PRK05338)Ribosomal_L19 (pfam01245)→ |
| env-542 | JGI20154J14316_10114600 | + | 178 | 205 | **RNA**→ rplS (PRK05338)Ribosomal_L19 (pfam01245)→ |
| env-543 | JGI20160J14292_10167608 | + | 546 | 573 | **RNA**→ |
| env-544 | JGI20159J14440_10045553 | - | 1068 | 1041 | **RNA**→ rplS (PRK05338)Ribosomal_L19 (pfam01245)→ |
| env-545 | JGI20155J14468_10052104 | - | 1080 | 1053 | **RNA**→ rplS (PRK05338)Ribosomal_L19 (pfam01245)→ |
| env-546 | JGI20158J14315_10097952 | - | 470 | 443 | **RNA**→ rplS (PRK05338)Ribosomal_L19 (pfam01245)→ |
| env-547 | F10SI0310a_Sequence0000013889 | - | 1032 | 1005 | **RNA**→ rplS (PRK05338)Ribosomal_L19 (pfam01245)→ |
| env-548 | DelMOSum2011_c10479874 | + | 149 | 176 | **RNA**→ rplS (PRK05338)Ribosomal_L19 (pfam01245)→ |
| env-549 | JGI20152J14361_10049778 | - | 470 | 443 | **RNA**→ rplS (PRK05338)Ribosomal_L19 (pfam01245)→ |
| env-550 | JGI20153J14295_10000026 | - | 1056 | 1029 | **RNA**→ rplS (PRK05338)Ribosomal_L19 (pfam01245)→ |
| env-551 | JGI20153J14318_10000026 | - | 1056 | 1029 | **RNA**→ rplS (PRK05338)Ribosomal_L19 (pfam01245)→ |
| env-552 | JGI20159J14440_10045424 | - | 1068 | 1041 | **RNA**→ rplS (PRK05338)Ribosomal_L19 (pfam01245)→ |
| env-553 | JGI20155J14468_10052697 | - | 1064 | 1037 | **RNA**→ rplS (PRK05338)Ribosomal_L19 (pfam01245)→ |
| env-554 | JGI20158J14315_10000044 | - | 470 | 443 | **RNA**→ rplS (PRK05338)Ribosomal_L19 (pfam01245)→ |
| env-555 | P_2C_Liq_1_UnCty_1031250 | - | 753 | 726 | **RNA**→ rplS (PRK05338)Ribosomal_L19 (pfam01245)→ |
| env-556 | P_2C_Liq_2_UnCty_1144299 | - | 190 | 163 | **RNA**→ rplS (PRK05338)Ribosomal_L19 (pfam01245)→ |
| env-557 | DelMOWin2010_c10191725 | + | 109 | 136 | **RNA**→ rplS (PRK05338)Ribosomal_L19 (pfam01245)→ |
| env-558 | PML_READ_00935591 | - | 131 | 104 | **RNA**→ ←hypo |
| env-559 | BS_KBA_SWE12_21m_c10032736 | + | 594 | 621 | **RNA**→ rplS (PRK05338)Ribosomal_L19 (pfam01245)→ |
| env-560 | BS_KBA_SWE21_205m_10115428 | - | 372 | 345 | **RNA**→ rplS (PRK05338)Ribosomal_L19 (pfam01245)→ |
| env-561 | BS_KBA_SWE02_21m_10146188 | + | 256 | 283 | **RNA**→ rplS (PRK05338)Ribosomal_L19 (pfam01245)→ |
| env-562 | SA_S1_NOR08_45m_c10057602 | - | 805 | 778 | **RNA**→ rplS (PRK05338)Ribosomal_L19 (pfam01245)→ |
| env-563 | SA_S2_NOR15_50m_c10029395 | + | 1634 | 1661 | **RNA**→ rplS (PRK05338)Ribosomal_L19 (pfam01245)→ |
| env-564 | SA_S1_NOR02_45m_c1073127 | - | 170 | 143 | **RNA**→ rplS (PRK05338)Ribosomal_L19 (pfam01245)→ |
| env-565 | SA_S2_NOR15_50m_c10237660 | - | 437 | 410 | **RNA**→ rplS (PRK05338)Ribosomal_L19 (pfam01245)→ |
| env-566 | SA_S1_NOR05_45m_c10015507 | + | 313 | 340 | **RNA**→ rplS (PRK05338)Ribosomal_L19 (pfam01245)→ |
| env-567 | SA_S1_NOR08_45m_c10002712 | + | 1145 | 1172 | **RNA**→ rplS (PRK05338)Ribosomal_L19 (pfam01245)→ |
| env-568 | SA_S2_NOR15_50m_c10347322 | - | 312 | 285 | **RNA**→ rplS (PRK05338)Ribosomal_L19 (pfam01245)→ |
| env-569 | SA_S1_NOR02_45m_c1174369 | + | 156 | 183 | **RNA**→ hypo→ |
| env-570 | KGI_S1_ANT01_95m_c10003353 | - | 6873 | 6846 | **RNA**→ rplS (PRK05338)Ribosomal_L19 (pfam01245)→ |
| env-571 | KGI_S1_ANT02_95m_c10361752 | - | 97 | 70 | **RNA**→ |
| env-572 | JGI1221J11331_1000054 | - | 29154 | 29127 | **RNA**→ rplS (PRK05338)Ribosomal_L19 (pfam01245)→ |
| env-573 | ASLM20b_GM71RDO02HMAJ4 | - | 59 | 32 | **RNA**→ hypo→ |
| Fba-5-1 | NC_014472.1 | + | 3298444 | 3298471 | **RNA**→ rplS (PRK05338)Ribosomal_L19 (pfam01245)→ |
| env-574 | KGI_S1_ANT01_95m_c10282318 | - | 277 | 250 | **RNA**→ hypo→ rplS (PRK05338)Ribosomal_L19 (pfam01245)→ |
| env-575 | SA_S1_NOR05_45m_c10063021 | + | 354 | 381 | **RNA**→ rplS (PRK05338)Ribosomal_L19 (pfam01245)→ |
| env-576 | SA_S1_NOR02_45m_c1006420 | + | 323 | 350 | **RNA**→ rplS (PRK05338)Ribosomal_L19 (pfam01245)→ |
| env-577 | KGI_S1_ANT02_95m_c10152699 | + | 34 | 61 | **RNA**→ hypo→ rplS (PRK05338)Ribosomal_L19 (pfam01245)→ |
| env-578 | CESI01056244.1 | + | 1354 | 1381 | **RNA**→ |
| Tme-1-1 | NZ_LDOD01000007.1 | - | 131191 | 131164 | **RNA**→ rplS (PRK05338)Ribosomal_L19 (pfam01245)→ |
| Zul-1-1 | NZ_JQMD01000002.1 | - | 686490 | 686463 | **RNA**→ rplS (PRK05338)Ribosomal_L19 (pfam01245)→ |
| env-579 | KGI_S1_ANT01_95m_c10220709 | - | 418 | 391 | **RNA**→ hypo→ rplS (PRK05338)Ribosomal_L19 (pfam01245)→ |
| env-580 | SA_S2_NOR13_50m_c1009519 | + | 390 | 417 | **RNA**→ rplS (PRK05338)Ribosomal_L19 (pfam01245)→ |
| env-581 | SA_S1_NOR05_45m_c10030739 | + | 81 | 108 | **RNA**→ rplS (PRK05338)Ribosomal_L19 (pfam01245)→ |
| env-582 | SA_S1_NOR08_45m_c10078726 | - | 177 | 150 | **RNA**→ rplS (PRK05338)Ribosomal_L19 (pfam01245)→ |
| env-583 | SA_S2_NOR15_50m_c10004324 | + | 2414 | 2441 | **RNA**→ rplS (PRK05338)Ribosomal_L19 (pfam01245)→ |
| env-584 | SA_S2_NOR18_50m_1017234 | + | 451 | 478 | **RNA**→ rplS (PRK05338)Ribosomal_L19 (pfam01245)→ |
| env-585 | SA_S1_NOR08_45m_c10100170 | + | 733 | 760 | **RNA**→ rplS (PRK05338)Ribosomal_L19 (pfam01245)→ |
| env-586 | SA_S2_NOR15_50m_c10049348 | + | 877 | 904 | **RNA**→ rplS (PRK05338)Ribosomal_L19 (pfam01245)→ |
| env-587 | TDF_MC_ARG02_113m_c1036547 | + | 146 | 173 | **RNA**→ rplS (PRK05338)Ribosomal_L19 (pfam01245)→ |

| | | | | | |
|---|---|---|---|---|---|
| env-588 | KGI_S1_ANT01_95m_c10011644 | - | 2952 | 2925 | **RNA**→ rplS (PRK05338)Ribosomal_L19 (pfam01245)→ |
| env-589 | KGI_S1_ANT03_95m_c1001414 | + | 244 | 271 | **RNA**→ rplS (PRK05338)Ribosomal_L19 (pfam01245)→ |
| env-590 | BS_KBA_SWE21_205m_10096012 | - | 541 | 514 | **RNA**→ rplS (PRK05338)Ribosomal_L19 (pfam01245)→ |
| env-591 | LGVF01642023.1 | + | 527 | 554 | **RNA**→ rplS (PRK05338)Ribosomal_L19 (pfam01245)→ |
| env-592 | DelMOSum2010_c10000064 | + | 13406 | 13433 | **RNA**→ rplS (PRK05338)Ribosomal_L19 (pfam01245)→ |
| env-593 | DelMOSum2011_c10042862 | + | 72 | 99 | **RNA**→ rplS (PRK05338)Ribosomal_L19 (pfam01245)→ |
| env-594 | SI34jun09_10m_1001399 | + | 1917 | 1944 | **RNA**→ rplS (PRK05338)Ribosomal_L19 (pfam01245)→ |
| env-595 | JGI20152J14361_10000209 | + | 40144 | 40171 | **RNA**→ rplS (PRK05338)Ribosomal_L19 (pfam01245)→ |
| env-596 | JGI20151J14362_10000555 | - | 1411 | 1384 | **RNA**→ rplS (PRK05338)Ribosomal_L19 (pfam01245)→ |
| env-597 | JGI20156J14371_10000042 | + | 40239 | 40266 | **RNA**→ rplS (PRK05338)Ribosomal_L19 (pfam01245)→ |
| env-598 | JGI20154J14316_10000040 | + | 40642 | 40669 | **RNA**→ rplS (PRK05338)Ribosomal_L19 (pfam01245)→ |
| env-599 | JGI20160J14292_10000074 | + | 5141 | 5168 | **RNA**→ rplS (PRK05338)Ribosomal_L19 (pfam01245)→ |
| env-600 | JGI20153J14295_10000199 | + | 26032 | 26059 | **RNA**→ rplS (PRK05338)Ribosomal_L19 (pfam01245)→ |
| env-601 | JGI20153J14318_10000199 | + | 26032 | 26059 | **RNA**→ rplS (PRK05338)Ribosomal_L19 (pfam01245)→ |
| env-602 | JGI20157J14317_10000069 | - | 62313 | 62286 | **RNA**→ rplS (PRK05338)Ribosomal_L19 (pfam01245)→ |
| env-603 | JGI20159J14440_10000007 | + | 39982 | 40009 | **RNA**→ rplS (PRK05338)Ribosomal_L19 (pfam01245)→ |
| env-604 | JGI20155J14468_10000017 | - | 82695 | 82668 | **RNA**→ rplS (PRK05338)Ribosomal_L19 (pfam01245)→ |
| env-605 | JGI20158J14315_10000602 | - | 6255 | 6228 | **RNA**→ rplS (PRK05338)Ribosomal_L19 (pfam01245)→ |
| env-606 | F10SI0310a_Sequence0000031405 | + | 45 | 72 | **RNA**→ rplS (PRK05338)Ribosomal_L19 (pfam01245)→ |
| Zga-1-1 | NC_015844.1 | + | 11699 | 11726 | **RNA**→ rplS (PRK05338)Ribosomal_L19 (pfam01245)→ |
| env-607 | TDF_OR_ARG04_113m_c1009311 | + | 312 | 339 | **RNA**→ rplS (PRK05338)Ribosomal_L19 (pfam01245)→ |
| env-608 | KGI_S2_ANT04_2345m_c1075036 | - | 569 | 542 | **RNA**→ rplS (PRK05338)Ribosomal_L19 (pfam01245)→ |
| env-609 | KGI_S1_ANT02_95m_c10135656 | - | 480 | 453 | **RNA**→ rplS (PRK05338)Ribosomal_L19 (pfam01245)→ |
| env-610 | KGI_S1_ANT02_95m_c10219286 | - | 147 | 120 | **RNA**→ rplS (PRK05338)Ribosomal_L19 (pfam01245)→ |
| env-611 | JGI24023J19991_10008759 | + | 3422 | 3449 | **RNA**→ rplS (PRK05338)Ribosomal_L19 (pfam01245)→ |
| Tov-1-1 | NZ_KE387163.1 | + | 593630 | 593657 | **RNA**→ rplS (PRK05338)Ribosomal_L19 (pfam01245)→ |
| env-612 | KGI_S1_ANT01_95m_c10030583 | + | 914 | 941 | **RNA**→ rplS (PRK05338)Ribosomal_L19 (pfam01245)→ |
| env-613 | KGI_S1_ANT02_95m_c10406104 | + | 132 | 159 | **RNA**→ rplS (PRK05338)Ribosomal_L19 (pfam01245)→ |
| env-614 | KGI_S1_ANT02_95m_c10445002 | + | 31 | 58 | **RNA**→ rplS (PRK05338)Ribosomal_L19 (pfam01245)→ |
| Ace-1-1 | NZ_KE386915.1 | - | 8797 | 8770 | **RNA**→ rplS (PRK05338)Ribosomal_L19 (pfam01245)→ |
| env-615 | DelMOSum2011_c10154912 | + | 8 | 35 | **RNA**→ rplS (PRK05338)Ribosomal_L19 (pfam01245)→ PRK08299 (PRK08299)→ |
| env-616 | CERS01121503.1 | + | 479 | 506 | **RNA**→ |
| env-617 | JGI20152J14361_10000298 | - | 36606 | 36579 | **RNA**→ rplS (PRK05338)Ribosomal_L19 (pfam01245)→ Icd (COG0538)PRK08299 (PRK08299)→<br>murQ (PRK05441)SIS_Etherase (cd05007)→ COX15-CtaA (pfam02628)→<br>TGS_ThrRS_N (cd01667)ThrS (COG0441)thrS (PRK00413)→ |
| env-618 | JGI20154J14316_10019853 | - | 413 | 386 | **RNA**→ rplS (PRK05338)Ribosomal_L19 (pfam01245)→ |
| env-619 | JGI20160J14292_10056708 | + | 610 | 637 | **RNA**→ rplS (PRK05338)Ribosomal_L19 (pfam01245)→ PRK08299 (PRK08299)→ |
| env-620 | JGI20153J14295_10004656 | + | 5844 | 5871 | **RNA**→ rplS (PRK05338)Ribosomal_L19 (pfam01245)→ Icd (COG0538)PRK08299 (PRK08299)→<br>murQ (PRK05441)SIS_Etherase (cd05007)→ COX15-CtaA (pfam02628)→ |
| env-621 | JGI20153J14318_10004656 | + | 5844 | 5871 | **RNA**→ rplS (PRK05338)Ribosomal_L19 (pfam01245)→ Icd (COG0538)PRK08299 (PRK08299)→<br>murQ (PRK05441)SIS_Etherase (cd05007)→ COX15-CtaA (pfam02628)→ |
| env-622 | JGI20157J14317_10171026 | - | 182 | 155 | **RNA**→ rplS (PRK05338)Ribosomal_L19 (pfam01245)→ |
| env-623 | JGI20153J14295_10241690 | + | 224 | 251 | **RNA**→ rplS (PRK05338)Ribosomal_L19 (pfam01245)→ |
| env-624 | JGI20153J14318_10241690 | + | 224 | 251 | **RNA**→ rplS (PRK05338)Ribosomal_L19 (pfam01245)→ |
| env-625 | JGI20156J14371_10014241 | - | 1035 | 1008 | **RNA**→ rplS (PRK05338)Ribosomal_L19 (pfam01245)→ PRK08299 (PRK08299)→ |
| env-626 | NCBI_READ_1112614356934 | + | 470 | 497 | **RNA**→ rplS (PRK05338)Ribosomal_L19 (pfam01245)→ |
| env-627 | JGI20152J14361_10011154 | - | 64 | 37 | **RNA**→ |
| env-628 | JGI20152J14361_10002150 | + | 8757 | 8784 | **RNA**→ rplS (PRK05338)Ribosomal_L19 (pfam01245)→ |
| env-629 | CESU01163506.1 | - | 117 | 90 | **RNA**→ hypo→ |
| env-630 | JGI20154J14316_10029844 | + | 675 | 702 | **RNA**→ rplS (PRK05338)Ribosomal_L19 (pfam01245)→ |
| env-631 | DelMOSum2011_c10007301 | - | 1729 | 1702 | **RNA**→ rplS (PRK05338)Ribosomal_L19 (pfam01245)→ |
| env-632 | JGI20156J14371_10057408 | + | 984 | 1011 | **RNA**→ rplS (PRK05338)Ribosomal_L19 (pfam01245)→ |
| env-633 | JGI20153J14295_10093405 | - | 123 | 96 | **RNA**→ |

| | | | | | |
|---|---|---|---|---|---|
| env-634 | JGI20153J14318_10093405 | - | 123 | 96 | **RNA**→ |
| env-635 | JGI20157J14317_10007733 | + | 5168 | 5195 | **RNA**→ rplS (PRK05338)Ribosomal_L19 (pfam01245)→ |
| env-636 | 2236876009_261411 | - | 77 | 50 | **RNA**→ hypo→ |
| env-637 | DelMOSum2010_c10004838 | - | 7637 | 7610 | **RNA**→ rplS (PRK05338)Ribosomal_L19 (pfam01245)→ |
| env-638 | JGI20152J14361_10046914 | + | 1115 | 1142 | **RNA**→ |
| env-639 | 2236876010_0651014 | + | 319 | 346 | **RNA**→ rplS (PRK05338)Ribosomal_L19 (pfam01245)→ |
| Fsp-12-1 | NZ_AMYW01000004.1 | + | 95479 | 95506 | **RNA**→ rplS (PRK05338)Ribosomal_L19 (pfam01245)→ |
| env-640 | JGI20152J14361_10056262 | + | 870 | 897 | **RNA**→ |
| env-641 | CESE01016993.1 | - | 115 | 88 | **RNA**→ |
| env-642 | CESI01086171.1 | + | 110 | 137 | **RNA**→ rplS (PRK05338)Ribosomal_L19 (pfam01245)→ |
| env-643 | CESJ01062511.1 | - | 5384 | 5357 | **RNA**→ rplS (PRK05338)Ribosomal_L19 (pfam01245)→ |
| env-644 | CESU01102979.1 | - | 419 | 392 | **RNA**→ rplS (PRK05338)Ribosomal_L19 (pfam01245)→ |
| env-645 | JGI20152J14361_10002317 | + | 11082 | 11109 | **RNA**→ rplS (PRK05338)Ribosomal_L19 (pfam01245)→ |
| env-646 | JGI20153J14295_10001978 | + | 15013 | 15040 | **RNA**→ |
| env-647 | JGI20153J14295_10015022 | + | 1046 | 1073 | **RNA**→ rplS (PRK05338)Ribosomal_L19 (pfam01245)→ |
| env-648 | JGI20153J14318_10001978 | + | 15013 | 15040 | **RNA**→ |
| env-649 | JGI20153J14318_10015022 | + | 1046 | 1073 | **RNA**→ rplS (PRK05338)Ribosomal_L19 (pfam01245)→ |
| env-650 | CERM01075739.1 | + | 5107 | 5134 | **RNA**→ rplS (PRK05338)Ribosomal_L19 (pfam01245)→ |
| env-651 | sludgePhrap_Contig8207 | - | 493 | 466 | **RNA**→ rplS (PRK05338)Ribosomal_L19 (pfam01245)→ |
| env-652 | 2156199154 | - | 535 | 508 | **RNA**→ rplS (PRK05338)Ribosomal_L19 (pfam01245)→ |
| env-653 | 2210285019 | + | 294 | 321 | **RNA**→ rplS (PRK05338)Ribosomal_L19 (pfam01245)→ |
| env-654 | JGI20160J14292_10046497 | + | 690 | 717 | **RNA**→ rplS (PRK05338)Ribosomal_L19 (pfam01245)→ |
| env-655 | CESE01253289.1 | - | 285 | 258 | **RNA**→ rplS (PRK05338)Ribosomal_L19 (pfam01245)→ |
| env-656 | CESJ01016245.1 | + | 3196 | 3223 | **RNA**→ Ribosomal_L19 (pfam01245)→ |
| env-657 | JGI20152J14361_10000482 | + | 17691 | 17718 | **RNA**→ rplS (PRK05338)Ribosomal_L19 (pfam01245)→ |
| env-658 | JGI20154J14316_10002680 | + | 12374 | 12401 | **RNA**→ rplS (PRK05338)Ribosomal_L19 (pfam01245)→ |
| env-659 | JGI20160J14292_10003989 | - | 962 | 935 | **RNA**→ rplS (PRK05338)Ribosomal_L19 (pfam01245)→ |
| env-660 | JGI20153J14295_10000826 | - | 7449 | 7422 | **RNA**→ rplS (PRK05338)Ribosomal_L19 (pfam01245)→ |
| env-661 | JGI20153J14318_10000826 | - | 7449 | 7422 | **RNA**→ rplS (PRK05338)Ribosomal_L19 (pfam01245)→ |
| env-662 | A09P04_1300_Sequence0000003336 | - | 2306 | 2279 | **RNA**→ rplS (PRK05338)Ribosomal_L19 (pfam01245)→ |
| env-663 | NCBI_BBAY_READ_1106073089325 | - | 898 | 871 | **RNA**→ rplS (PRK05338)Ribosomal_L19 (pfam01245)→ PRK08299 (PRK08299)→ |
| env-664 | P_1C_Liq_3_UnCty_1009162 | - | 198 | 171 | **RNA**→ rplS (PRK05338)Ribosomal_L19 (pfam01245)→ |
| env-665 | P_1C_Liq_1_UnCty_1001883 | - | 3248 | 3221 | **RNA**→ rplS (PRK05338)Ribosomal_L19 (pfam01245)→ |
| env-666 | P_1C_Liq_2_UnCty_1016458 | + | 549 | 576 | **RNA**→ |
| env-667 | NCBI_BBAY_READ_1106105074739 | + | 204 | 231 | **RNA**→ rplS (PRK05338)Ribosomal_L19 (pfam01245)→ PRK08299 (PRK08299)→ |
| env-668 | CENJ01092743.1 | - | 137 | 110 | **RNA**→ rplS (PRK05338)Ribosomal_L19 (pfam01245)→ |
| env-669 | NCBI_BBAY_READ_1106073037205 | + | 277 | 304 | **RNA**→ rplS (PRK05338)Ribosomal_L19 (pfam01245)→ PRK08299 (PRK08299)→ |
| env-670 | CERS01174496.1 | - | 38 | 11 | **RNA**→ |
| env-671 | CEVW01141574.1 | + | 1150 | 1177 | **RNA**→ rplS (PRK05338)Ribosomal_L19 (pfam01245)→ |
| env-672 | CEUA01411516.1 | + | 349 | 376 | **RNA**→ rplS (PRK05338)Ribosomal_L19 (pfam01245)→ |
| env-673 | CEQM01197603.1 | + | 392 | 419 | **RNA**→ rplS (PRK05338)Ribosomal_L19 (pfam01245)→ |
| env-674 | CEQN01023522.1 | + | 8459 | 8486 | **RNA**→ rplS (PRK05338)Ribosomal_L19 (pfam01245)→ |
| env-675 | CEQO01070212.1 | + | 1934 | 1961 | **RNA**→ rplS (PRK05338)Ribosomal_L19 (pfam01245)→ |
| env-676 | CEVS01116577.1 | - | 1094 | 1067 | **RNA**→ rplS (PRK05338)Ribosomal_L19 (pfam01245)→ |
| env-677 | CEVP01261667.1 | + | 2 | 29 | **RNA**→ rplS (PRK05338)Ribosomal_L19 (pfam01245)→ |
| env-678 | JCVI_READ_1104230344964 | - | 901 | 874 | **RNA**→ rplS (PRK05338)Ribosomal_L19 (pfam01245)→ |
| env-679 | CERW01179445.1 | + | 13 | 40 | **RNA**→ rplS (PRK05338)Ribosomal_L19 (pfam01245)→ |
| env-680 | DelMOWin2010_c10013017 | + | 4133 | 4160 | **RNA**→ rplS (PRK05338)Ribosomal_L19 (pfam01245)→ PRK08299 (PRK08299)→ |
| env-681 | JGI20157J14317_10033833 | + | 996 | 1023 | **RNA**→ rplS (PRK05338)Ribosomal_L19 (pfam01245)→ Icd (COG0538)PRK08299 (PRK08299)→ |
| env-682 | CEWH01233956.1 | - | 272 | 245 | **RNA**→ rplS (PRK05338)Ribosomal_L19 (pfam01245)→ |
| env-683 | JCVI_SCAF_1101668058877 | - | 1488 | 1461 | **RNA**→ rplS (PRK05338)Ribosomal_L19 (pfam01245)→ PRK08299 (PRK08299)→ |
| env-684 | CENT01032076.1 | - | 7307 | 7280 | **RNA**→ rplS (PRK05338)Ribosomal_L19 (pfam01245)→ |

| | | | | | |
|---|---|---|---|---|---|
| env-685 | CENG01052807.1 | + | 215 | 242 | **RNA**→ rplS (PRK05338)Ribosomal_L19 (pfam01245)→ |
| env-686 | CESE01258788.1 | + | 699 | 726 | **RNA**→ rplS (PRK05338)Ribosomal_L19 (pfam01245)→ |
| env-687 | CESJ01025343.1 | + | 3507 | 3534 | **RNA**→ rplS (PRK05338)Ribosomal_L19 (pfam01245)→ |
| Ean-11-1 | NZ_CBYE010000026.1 | - | 462809 | 462782 | **RNA**→ rplS (PRK05338)Ribosomal_L19 (pfam01245)→ PRK10546 (PRK10546)Nudix_Hydrolase_31 (cd04690)→ |
| Eme-1-1 | NZ_AVCQ01000015.1 | + | 91308 | 91335 | **RNA**→ rplS (PRK05338)Ribosomal_L19 (pfam01245)→ PRK08999 (PRK08999)Nudix_Hydrolase_31 (cd04690)→ |
| Ean-1-1 | NZ_JNCG01000035.1 | + | 11502 | 11529 | **RNA**→ rplS (PRK05338)Ribosomal_L19 (pfam01245)→ PRK08999 (PRK08999)Nudix_Hydrolase_31 (cd04690)→ |
| Ean-1-2 | NZ_LFKT01000011.1 | - | 207966 | 207939 | **RNA**→ rplS (PRK05338)Ribosomal_L19 (pfam01245)→ PRK08999 (PRK08999)Nudix_Hydrolase_31 (cd04690)→ |
| Ean-2-1 | NZ_AHHG01000005.1 | - | 82828 | 82801 | **RNA**→ rplS (PRK05338)Ribosomal_L19 (pfam01245)→ rplS (PRK05338)Ribosomal_L19 (pfam01245)→ PRK08999 (PRK08999)Nudix_Hydrolase_31 (cd04690)→ PRK08999 (PRK08999)Nudix_Hydrolase_31 (cd04690)→ |
| Ean-8-1 | NZ_CP007547.1 | + | 2704094 | 2704121 | **RNA**→ rplS (PRK05338)Ribosomal_L19 (pfam01245)→ PRK08999 (PRK08999)Nudix_Hydrolase_31 (cd04690)→ |
| Ean-3-1 | NZ_CP006576.1 | + | 2075235 | 2075262 | **RNA**→ rplS (PRK05338)Ribosomal_L19 (pfam01245)→ PRK08999 (PRK08999)Nudix_Hydrolase_31 (cd04690)→ |
| Ean-6-1 | NZ_ASYI01000012.1 | - | 167353 | 167326 | **RNA**→ rplS (PRK05338)Ribosomal_L19 (pfam01245)→ PRK08999 (PRK08999)Nudix_Hydrolase_31 (cd04690)→ |
| Ean-5-1 | NZ_ASYK01000009.1 | + | 92547 | 92574 | **RNA**→ rplS (PRK05338)Ribosomal_L19 (pfam01245)→ PRK08999 (PRK08999)Nudix_Hydrolase_31 (cd04690)→ |
| Ean-7-1 | NZ_ASYJ01000045.1 | - | 40079 | 40052 | **RNA**→ rplS (PRK05338)Ribosomal_L19 (pfam01245)→ PRK08999 (PRK08999)Nudix_Hydrolase_31 (cd04690)→ |
| Ean-10-1 | NZ_ASYG01000013.1 | - | 186964 | 186937 | **RNA**→ rplS (PRK05338)Ribosomal_L19 (pfam01245)→ PRK08999 (PRK08999)Nudix_Hydrolase_31 (cd04690)→ |
| Ean-9-1 | NZ_ASYF01000003.1 | + | 87715 | 87742 | **RNA**→ rplS (PRK05338)Ribosomal_L19 (pfam01245)→ PRK08999 (PRK08999)Nudix_Hydrolase_31 (cd04690)→ |
| Ean-4-1 | NZ_ASYH01000003.1 | + | 87833 | 87860 | **RNA**→ rplS (PRK05338)Ribosomal_L19 (pfam01245)→ PRK08999 (PRK08999)Nudix_Hydrolase_31 (cd04690)→ |
| Ean-1-3 | NZ_CCAC010000045.1 | + | 65940 | 65967 | **RNA**→ rplS (PRK05338)Ribosomal_L19 (pfam01245)→ PRK08999 (PRK08999)Nudix_Hydrolase_31 (cd04690)→ |
| Ean-1-4 | NZ_CCAB010000079.1 | + | 65942 | 65969 | **RNA**→ rplS (PRK05338)Ribosomal_L19 (pfam01245)→ PRK08999 (PRK08999)Nudix_Hydrolase_31 (cd04690)→ |
| Ean-12-1 | NZ_ANIW01000063.1 | - | 189045 | 189018 | **RNA**→ rplS (PRK05338)Ribosomal_L19 (pfam01245)→ PRK08999 (PRK08999)Nudix_Hydrolase_31 (cd04690)→ |
| Eme-2-1 | NZ_BARD01000013.1 | - | 28567 | 28540 | **RNA**→ rplS (PRK05338)Ribosomal_L19 (pfam01245)→ |
| Eme-2-2 | NZ_ASAN01000023.1 | - | 28545 | 28518 | **RNA**→ rplS (PRK05338)Ribosomal_L19 (pfam01245)→ |
| env-688 | LPaug09P1610m_c1098533 | + | 95 | 122 | **RNA**→ rplS (PRK05338)Ribosomal_L19 (pfam01245)→ |
| env-689 | JGI20157J14317_10000212 | - | 34865 | 34838 | **RNA**→ rplS (PRK05338)Ribosomal_L19 (pfam01245)→ Icd (COG0538)PRK08299 (PRK08299)→ |
| env-690 | LPaug08P2610m_c1056069 | + | 194 | 221 | **RNA**→ rplS (PRK05338)Ribosomal_L19 (pfam01245)→ |
| env-691 | CESU01106244.1 | - | 3668 | 3641 | **RNA**→ rplS (PRK05338)Ribosomal_L19 (pfam01245)→ Icd (COG0538)PRK08299 (PRK08299)→ |
| env-692 | wf3_Contig2666 | + | 306 | 333 | **RNA**→ rplS (PRK05338)Ribosomal_L19 (pfam01245)→ |
| env-693 | JGI11876J14442_10791786 | - | 254 | 227 | **RNA**→ rplS (PRK05338)Ribosomal_L19 (pfam01245)→ |
| env-694 | DelMOSum2011_c10050897 | - | 758 | 731 | **RNA**→ rplS (PRK05338)Ribosomal_L19 (pfam01245)→ |
| env-695 | JGI20153J14295_10078886 | + | 823 | 850 | **RNA**→ rplS (PRK05338)Ribosomal_L19 (pfam01245)→ |
| env-696 | JGI20153J14318_10078886 | + | 823 | 850 | **RNA**→ rplS (PRK05338)Ribosomal_L19 (pfam01245)→ |
| env-697 | JGI20153J14295_10079231 | - | 179 | 152 | **RNA**→ rplS (PRK05338)Ribosomal_L19 (pfam01245)→ |
| env-698 | JGI20153J14318_10079231 | - | 179 | 152 | **RNA**→ rplS (PRK05338)Ribosomal_L19 (pfam01245)→ |
| env-699 | SRS043422_C2637862 | - | 319 | 292 | **RNA**→ rplS (PRK05338)Ribosomal_L19 (pfam01245)→ |
| env-700 | SRS013506_C1070803 | - | 155 | 128 | **RNA**→ rplS (PRK05338)Ribosomal_L19 (pfam01245)→ |
| env-701 | SRS058105_C1803295 | + | 211 | 238 | **RNA**→ rplS (PRK05338)Ribosomal_L19 (pfam01245)→ |
| env-702 | CEQS01154894.1 | - | 496 | 469 | **RNA**→ rplS (PRK05338)Ribosomal_L19 (pfam01245)→ |
| env-703 | CETA01127273.1 | + | 1082 | 1109 | **RNA**→ rplS (PRK05338)Ribosomal_L19 (pfam01245)→ |
| env-704 | 2236876010_0239681 | + | 214 | 241 | **RNA**→ rplS (PRK05338)Ribosomal_L19 (pfam01245)→ |
| env-705 | 2236876010_0451387 | + | 219 | 246 | **RNA**→ rplS (PRK05338)Ribosomal_L19 (pfam01245)→ |
| env-706 | JGI20154J14316_10000655 | + | 31048 | 31075 | **RNA**→ rplS (PRK05338)Ribosomal_L19 (pfam01245)→ |
| env-707 | JGI20157J14317_10157594 | + | 564 | 591 | **RNA**→ |
| env-708 | CESI01155210.1 | - | 333 | 306 | **RNA**→ rplS (PRK05338)Ribosomal_L19 (pfam01245)→ |
| env-709 | CESU01106790.1 | - | 246 | 219 | **RNA**→ rplS (PRK05338)Ribosomal_L19 (pfam01245)→ |
| Gfo-1-1 | NC_008571.1 | - | 690781 | 690754 | **RNA**→ rplS (PRK05338)Ribosomal_L19 (pfam01245)→ |
| env-710 | DelMOSum2010_c10261886 | - | 392 | 365 | **RNA**→ rplS (PRK05338)Ribosomal_L19 (pfam01245)→ |
| env-711 | JGI20154J14316_10114945 | - | 324 | 297 | **RNA**→ rplS (PRK05338)Ribosomal_L19 (pfam01245)→ |
| env-712 | JGI24025J20009_10008959 | - | 4721 | 4694 | **RNA**→ rplS (PRK05338)Ribosomal_L19 (pfam01245)→ |
| env-713 | JGI20154J14316_10088205 | + | 421 | 448 | **RNA**→ rplS (PRK05338)Ribosomal_L19 (pfam01245)→ |
| env-714 | JGI20156J14371_10005271 | - | 8543 | 8516 | **RNA**→ rplS (PRK05338)Ribosomal_L19 (pfam01245)→ |
| env-715 | CETR01024806.1 | + | 447 | 474 | **RNA**→ rplS (PRK05338)Ribosomal_L19 (pfam01245)→ |
| env-716 | CXWF01011805.1 | - | 863 | 836 | **RNA**→ rplS (PRK05338)Ribosomal_L19 (pfam01245)→ hypo→ |

| | | | | | |
|---|---|---|---|---|---|
| Cat-1-1 | NC_014230.1 | - | 1512168 | 1512141 | **RNA**→ rplS (PRK05338)Ribosomal_L19 (pfam01245)→ |
| env-717 | KGI_S1_ANT02_95m_c10343606 | + | 218 | 245 | **RNA**→ rplS (PRK05338)Ribosomal_L19 (pfam01245)→ |
| env-718 | CEWR01292008.1 | - | 1347 | 1320 | **RNA**→ rplS (PRK05338)Ribosomal_L19 (pfam01245)→ |
| env-719 | JGI24723J26617_10003020 | + | 3244 | 3271 | **RNA**→ rplS (PRK05338)Ribosomal_L19 (pfam01245)→ |
| env-720 | SA_S1_NOR05_45m_c10349549 | + | 1 | 28 | **RNA**→ rplS (PRK05338)Ribosomal_L19 (pfam01245)→ |
| env-721 | SA_S2_NOR15_50m_c10137904 | + | 11 | 38 | **RNA**→ rplS (PRK05338)Ribosomal_L19 (pfam01245)→ |
| env-722 | SA_S1_NOR08_45m_c10597501 | - | 252 | 225 | **RNA**→ hypo→ rplS (PRK05338)Ribosomal_L19 (pfam01245)→ |
| env-723 | KGI_S2_ANT04_2345m_c1082035 | + | 110 | 137 | **RNA**→ rplS (PRK05338)Ribosomal_L19 (pfam01245)→ |
| env-724 | SA_S1_NOR08_45m_c10030363 | - | 789 | 762 | **RNA**→ rplS (PRK05338)Ribosomal_L19 (pfam01245)→ |
| env-725 | KGI_S1_ANT02_95m_c10228337 | - | 406 | 379 | **RNA**→ rplS (PRK05338)Ribosomal_L19 (pfam01245)→ |
| env-726 | JGI24024J18818_10002320 | + | 5869 | 5896 | **RNA**→ rplS (PRK05338)Ribosomal_L19 (pfam01245)→ |
| env-727 | KVRMV2_100472114 | - | 249 | 222 | **RNA**→ rplS (PRK05338)Ribosomal_L19 (pfam01245)→ |
| Tna-1-1 | NZ_JTDV01000005.1 | + | 417562 | 417589 | **RNA**→ rplS (PRK05338)Ribosomal_L19 (pfam01245)→ |
| Pbu-1-1 | NZ_AUDE01000014.1 | + | 155495 | 155522 | **RNA**→ rplS (PRK05338)Ribosomal_L19 (pfam01245)→ |
| env-728 | KGI_S1_ANT01_95m_c10008907 | - | 926 | 899 | **RNA**→ rplS (PRK05338)Ribosomal_L19 (pfam01245)→ |
| env-729 | KGI_S2_ANT06_2345m_c1016060 | - | 116 | 89 | **RNA**→ |
| env-730 | KGI_S2_ANT04_2345m_c1006248 | - | 1101 | 1074 | **RNA**→ rplS (PRK05338)Ribosomal_L19 (pfam01245)→ |
| env-731 | KGI_S1_ANT02_95m_c10014716 | - | 776 | 749 | **RNA**→ rplS (PRK05338)Ribosomal_L19 (pfam01245)→ |
| env-732 | SA_S1_NOR05_45m_c10011330 | + | 878 | 905 | **RNA**→ rplS (PRK05338)Ribosomal_L19 (pfam01245)→ |
| env-733 | SA_S2_NOR15_50m_c10030572 | + | 1601 | 1628 | **RNA**→ rplS (PRK05338)Ribosomal_L19 (pfam01245)→ |
| env-734 | SA_S1_NOR08_45m_c10005809 | + | 1419 | 1446 | **RNA**→ rplS (PRK05338)Ribosomal_L19 (pfam01245)→ |
| env-735 | KGI_S2_ANT04_2345m_c1312501 | - | 63 | 36 | **RNA**→ |
| env-736 | SA_S2_NOR18_50m_1150586 | - | 220 | 193 | **RNA**→ rplS (PRK05338)Ribosomal_L19 (pfam01245)→ |
| env-737 | KGI_S2_ANT04_2345m_c1135335 | + | 346 | 373 | **RNA**→ |
| env-738 | KGI_S1_ANT02_95m_c10003498 | - | 1873 | 1846 | **RNA**→ rplS (PRK05338)Ribosomal_L19 (pfam01245)→ |
| env-739 | TDF_MC_ARG02_113m_c1071523 | + | 280 | 307 | **RNA**→ ←hypo |
| env-740 | JGI24723J26617_10000135 | + | 62418 | 62445 | **RNA**→ rplS (PRK05338)Ribosomal_L19 (pfam01245)→ |
| Ale-1-1 | NZ_BBNQ01000002.1 | + | 52311 | 52338 | **RNA**→ rplS (PRK05338)Ribosomal_L19 (pfam01245)→ |
| Fic-1-1 | NZ_KN525716.1 | + | 641421 | 641448 | **RNA**→ rplS (PRK05338)Ribosomal_L19 (pfam01245)→ |
| env-741 | KGI_S1_ANT01_95m_c10145689 | + | 78 | 105 | **RNA**→ rplS (PRK05338)Ribosomal_L19 (pfam01245)→ |
| env-742 | KGI_S1_ANT01_95m_c10094256 | + | 98 | 125 | **RNA**→ rplS (PRK05338)Ribosomal_L19 (pfam01245)→ |
| env-743 | BS_KBA_SWE12_21m_c10430184 | - | 261 | 234 | **RNA**→ rplS (PRK05338)Ribosomal_L19 (pfam01245)→ |
| env-744 | SA_S1_NOR08_45m_c10000775 | + | 7919 | 7946 | **RNA**→ rplS (PRK05338)Ribosomal_L19 (pfam01245)→ |
| env-745 | SA_S2_NOR15_50m_c10050530 | + | 950 | 977 | **RNA**→ rplS (PRK05338)Ribosomal_L19 (pfam01245)→ |
| env-746 | SA_S1_NOR08_45m_c10816085 | + | 12 | 39 | **RNA**→ hypo→ rplS (PRK05338)Ribosomal_L19 (pfam01245)→ |
| env-747 | SA_S2_NOR15_50m_c10122310 | - | 573 | 546 | **RNA**→ rplS (PRK05338)Ribosomal_L19 (pfam01245)→ |
| env-748 | KGI_S1_ANT01_95m_c10075767 | + | 706 | 733 | **RNA**→ rplS (PRK05338)Ribosomal_L19 (pfam01245)→ |
| env-749 | KGI_S1_ANT02_95m_c10035975 | - | 1547 | 1520 | **RNA**→ rplS (PRK05338)Ribosomal_L19 (pfam01245)→ |
| env-750 | ElkS_mat_MD6A_1000445 | - | 13256 | 13229 | **RNA**→ rplS (PRK05338)Ribosomal_L19 (pfam01245)→ |
| env-751 | JGI11876J14442_10004783 | - | 7288 | 7261 | **RNA**→ rplS (PRK05338)Ribosomal_L19 (pfam01245)→ |
| env-752 | JGI24025J20009_10002662 | + | 3106 | 3133 | **RNA**→ rplS (PRK05338)Ribosomal_L19 (pfam01245)→ |
| env-753 | JGI24724J26744_10000817 | - | 7312 | 7285 | **RNA**→ rplS (PRK05338)Ribosomal_L19 (pfam01245)→ |
| env-754 | JGI24723J26617_10075658 | + | 845 | 872 | **RNA**→ rplS (PRK05338)Ribosomal_L19 (pfam01245)→ |
| Ale-1-2 | NZ_BBNU01000009.1 | - | 60878 | 60851 | **RNA**→ hypo→ hypo→ |
| Wsp-1-1 | NZ_AP014583.1 | + | 217643 | 217670 | **RNA**→ rplS (PRK05338)Ribosomal_L19 (pfam01245)→ |
| env-755 | DelMOSum2011_c10001390 | + | 13601 | 13628 | **RNA**→ rplS (PRK05338)Ribosomal_L19 (pfam01245)→ |
| env-756 | BS_KBA_SWE12_21m_c10000778 | - | 8562 | 8535 | **RNA**→ rplS (PRK05338)Ribosomal_L19 (pfam01245)→ |
| env-757 | BS_KBB_SWE26_205m_c1007554 | + | 1145 | 1172 | **RNA**→ rplS (PRK05338)Ribosomal_L19 (pfam01245)→ |
| env-758 | BS_KBA_SWE07_21m_c1035024 | - | 470 | 443 | **RNA**→ rplS (PRK05338)Ribosomal_L19 (pfam01245)→ |
| env-759 | BS_KBA_SWE21_205m_10000389 | - | 3010 | 2983 | **RNA**→ rplS (PRK05338)Ribosomal_L19 (pfam01245)→ |
| env-760 | BS_KBA_SWE02_21m_10003979 | + | 3505 | 3532 | **RNA**→ rplS (PRK05338)Ribosomal_L19 (pfam01245)→ |
| env-761 | SA_S1_NOR05_45m_c10404754 | - | 237 | 210 | **RNA**→ rplS (PRK05338)Ribosomal_L19 (pfam01245)→ |

| | | | | | |
|---|---|---|---|---|---|
| env-762 | SA_S1_NOR08_45m_c10035522 | + | 438 | 465 | **RNA**→ rplS (PRK05338)Ribosomal_L19 (pfam01245)→ |
| env-763 | SA_S2_NOR15_50m_c10123094 | + | 331 | 358 | **RNA**→ rplS (PRK05338)Ribosomal_L19 (pfam01245)→ |
| env-764 | SA_S2_NOR15_50m_c10622339 | - | 159 | 132 | **RNA**→ rplS (PRK05338)Ribosomal_L19 (pfam01245)→ |
| env-765 | BS_KBA_SWE02_21m_10429781 | + | 153 | 180 | **RNA**→ rplS (PRK05338)Ribosomal_L19 (pfam01245)→ |
| env-766 | JGI24723J26617_10219199 | + | 387 | 414 | **RNA**→ hypo→ |
| Pda-1-1 | NZ_JXJO01000016.1 | - | 240145 | 240118 | **RNA**→ rplS (PRK05338)Ribosomal_L19 (pfam01245)→ |
| Fba-7-1 | NZ_ALIH01000012.1 | + | 102806 | 102833 | **RNA**→ rplS (PRK05338)Ribosomal_L19 (pfam01245)→ |
| env-767 | SA_S2_NOR13_50m_c1039248 | + | 208 | 235 | **RNA**→ rplS (PRK05338)Ribosomal_L19 (pfam01245)→ |
| env-768 | SA_S1_NOR05_45m_c10021836 | - | 1574 | 1547 | **RNA**→ rplS (PRK05338)Ribosomal_L19 (pfam01245)→ |
| env-769 | SA_S1_NOR08_45m_c10009016 | - | 4028 | 4001 | **RNA**→ rplS (PRK05338)Ribosomal_L19 (pfam01245)→ |
| env-770 | SA_S2_NOR15_50m_c10078015 | - | 652 | 625 | **RNA**→ rplS (PRK05338)Ribosomal_L19 (pfam01245)→ |
| env-771 | TB_PC08_66_10107273 | + | 586 | 613 | **RNA**→ rplS (PRK05338)Ribosomal_L19 (pfam01245)→ |
| env-772 | BS_KBA_SWE12_21m_c10659546 | + | 51 | 78 | **RNA**→ hypo→ rplS (PRK05338)Ribosomal_L19 (pfam01245)→ |
| env-773 | TB_PC08_64_1001738 | + | 2654 | 2681 | **RNA**→ rplS (PRK05338)Ribosomal_L19 (pfam01245)→ |
| env-774 | TB_FS06_10DRAFT_1005018 | + | 3980 | 4007 | **RNA**→ rplS (PRK05338)Ribosomal_L19 (pfam01245)→ |
| env-775 | TB_PC08_66_10001095 | + | 2574 | 2601 | **RNA**→ rplS (PRK05338)Ribosomal_L19 (pfam01245)→ |
| env-776 | TB_GS10_10_10000237 | + | 23689 | 23716 | **RNA**→ rplS (PRK05338)Ribosomal_L19 (pfam01245)→ |
| env-777 | TB_GS10_10_10014454 | - | 4102 | 4075 | **RNA**→ rplS (PRK05338)Ribosomal_L19 (pfam01245)→ |
| env-778 | TB_GS09_5_10009762 | + | 4122 | 4149 | **RNA**→ rplS (PRK05338)Ribosomal_L19 (pfam01245)→ |
| env-779 | TB_PC08_3_1001100 | - | 12933 | 12906 | **RNA**→ rplS (PRK05338)Ribosomal_L19 (pfam01245)→ |
| env-780 | JGI24723J26617_10196616 | + | 45 | 72 | **RNA**→ rplS (PRK05338)Ribosomal_L19 (pfam01245)→ |
| env-781 | JGI20151J14362_10404994 | + | 205 | 232 | **RNA**→ |
| env-782 | JGI20155J14468_10372448 | + | 233 | 260 | **RNA**→ rplS (PRK05338)Ribosomal_L19 (pfam01245)→ |
| env-783 | 3300001592_10000620 | - | 7266 | 7239 | **RNA**→ rplS (PRK05338)Ribosomal_L19 (pfam01245)→ |
| env-784 | GBANfinal_contig20918 | + | 428 | 455 | **RNA**→ |
| Tse-1-1 | NZ_JTDW01000005.1 | + | 217195 | 217222 | **RNA**→ rplS (PRK05338)Ribosomal_L19 (pfam01245)→ |
| env-785 | SA_S1_NOR05_45m_c10008190 | - | 2600 | 2573 | **RNA**→ rplS (PRK05338)Ribosomal_L19 (pfam01245)→ |
| env-786 | SA_S1_NOR08_45m_c10005913 | - | 109 | 82 | **RNA**→ ←hypo |
| env-787 | SA_S2_NOR15_50m_c10028300 | - | 1363 | 1336 | **RNA**→ rplS (PRK05338)Ribosomal_L19 (pfam01245)→ |
| env-788 | CEQZ01051997.1 | - | 539 | 512 | **RNA**→ rplS (PRK05338)Ribosomal_L19 (pfam01245)→ |
| env-789 | CERU01034551.1 | - | 2620 | 2593 | **RNA**→ rplS (PRK05338)Ribosomal_L19 (pfam01245)→ |
| env-790 | JGI24025J20009_10046633 | + | 1505 | 1532 | **RNA**→ rplS (PRK05338)Ribosomal_L19 (pfam01245)→ |
| Psp-4-1 | NZ_JUGU01000001.1 | + | 1535967 | 1535994 | **RNA**→ rplS (PRK05338)Ribosomal_L19 (pfam01245)→ |
| Fsp-13-1 | NZ_ANLA01000019.1 | - | 4070 | 4043 | **RNA**→ rplS (PRK05338)Ribosomal_L19 (pfam01245)→ |
| env-791 | CEQG01218791.1 | + | 295 | 322 | **RNA**→ rplS (PRK05338)Ribosomal_L19 (pfam01245)→ |
| env-792 | CERD01073987.1 | + | 168 | 195 | **RNA**→ rplS (PRK05338)Ribosomal_L19 (pfam01245)→ |
| env-793 | JGI11876J14442_10006999 | + | 7283 | 7310 | **RNA**→ rplS (PRK05338)Ribosomal_L19 (pfam01245)→ |
| env-794 | JGI24723J26617_10011294 | + | 1041 | 1068 | **RNA**→ rplS (PRK05338)Ribosomal_L19 (pfam01245)→ |
| Jpa-1-1 | NZ_BBNS01000022.1 | + | 50201 | 50228 | **RNA**→ rplS (PRK05338)Ribosomal_L19 (pfam01245)→ |
| Pja-1-1 | NZ_JSWG01000016.1 | + | 19956 | 19983 | **RNA**→ rplS (PRK05338)Ribosomal_L19 (pfam01245)→ |
| env-795 | BS_KBA_SWE02_21m_10588502 | + | 82 | 109 | **RNA**→ rplS (PRK05338)Ribosomal_L19 (pfam01245)→ |
| env-796 | JGI11876J14442_10024983 | + | 2546 | 2573 | **RNA**→ rplS (PRK05338)Ribosomal_L19 (pfam01245)→ |
| env-797 | KVRMV2_101767380 | + | 513 | 540 | **RNA**→ |
| Pme-1-1 | NZ_JXJP01000027.1 | - | 9100 | 9073 | **RNA**→ rplS (PRK05338)Ribosomal_L19 (pfam01245)→ |
| env-798 | SA_S1_NOR08_45m_c10161364 | - | 92 | 65 | **RNA**→ ←hypo |
| env-799 | ElkS_mat_MD6A_1031343 | - | 868 | 841 | **RNA**→ rplS (PRK05338)Ribosomal_L19 (pfam01245)→ |
| Jpa-1-2 | NZ_BBNR01000023.1 | - | 12164 | 12137 | **RNA**→ rplS (PRK05338)Ribosomal_L19 (pfam01245)→ |
| Jpa-1-3 | NZ_BBNY01000007.1 | + | 50312 | 50339 | **RNA**→ rplS (PRK05338)Ribosomal_L19 (pfam01245)→ |
| env-800 | DelMOSum2010_c10067390 | - | 753 | 726 | **RNA**→ rplS (PRK05338)Ribosomal_L19 (pfam01245)→ |
| env-801 | JGI20154J14316_10000408 | + | 38586 | 38613 | **RNA**→ rplS (PRK05338)Ribosomal_L19 (pfam01245)→ |
| env-802 | JGI20160J14292_10000275 | + | 38583 | 38610 | **RNA**→ rplS (PRK05338)Ribosomal_L19 (pfam01245)→ |
| env-803 | JGI20157J14317_10000632 | + | 26198 | 26225 | **RNA**→ rplS (PRK05338)Ribosomal_L19 (pfam01245)→ |

| | | | | | |
|---|---|---|---|---|---|
| env-804 | JGI24724J26744_10000045 | - | 14321 | 14294 | **RNA**→ rplS (PRK05338)Ribosomal_L19 (pfam01245)→ |
| env-805 | JGI20157J14317_10002549 | + | 15190 | 15217 | **RNA**→ rplS (PRK05338)Ribosomal_L19 (pfam01245)→ |
| env-806 | JGI24724J26744_10018347 | - | 3457 | 3430 | **RNA**→ rplS (PRK05338)Ribosomal_L19 (pfam01245)→ |
| env-807 | JGI20153J14295_10046331 | - | 387 | 360 | **RNA**→ rplS (PRK05338)Ribosomal_L19 (pfam01245)→ |
| env-808 | JGI20153J14318_10046331 | - | 387 | 360 | **RNA**→ rplS (PRK05338)Ribosomal_L19 (pfam01245)→ |
| env-809 | SA_S1_NOR08_45m_c10278240 | - | 201 | 174 | **RNA**→ rplS (PRK05338)Ribosomal_L19 (pfam01245)→ |
| env-810 | JGI20154J14316_10043409 | + | 1626 | 1653 | **RNA**→ rplS (PRK05338)Ribosomal_L19 (pfam01245)→ |
| env-811 | JGI20157J14317_10008721 | - | 382 | 355 | **RNA**→ rplS (PRK05338)Ribosomal_L19 (pfam01245)→ |
| env-812 | 2236876003_126260 | - | 373 | 346 | **RNA**→ rplS (PRK05338)Ribosomal_L19 (pfam01245)→ rplS (PRK05338)Ribosomal_L19 (pfam01245)→ |
| env-813 | JGI20151J14362_10005078 | - | 176 | 149 | **RNA**→ rplS (PRK05338)Ribosomal_L19 (pfam01245)→ |
| env-814 | JGI20156J14371_10014418 | + | 4045 | 4072 | **RNA**→ rplS (PRK05338)Ribosomal_L19 (pfam01245)→ |
| env-815 | JGI20160J14292_10615073 | - | 221 | 194 | **RNA**→ rplS (PRK05338)Ribosomal_L19 (pfam01245)→ |
| env-816 | JGI20158J14315_10346697 | + | 250 | 277 | **RNA**→ rplS (PRK05338)Ribosomal_L19 (pfam01245)→ |
| env-817 | CERM01179909.1 | - | 111 | 84 | **RNA**→ |
| env-818 | JGI24025J20009_10452841 | - | 33 | 6 | **RNA**→ |
| env-819 | LAZR01000052.1 | + | 53070 | 53097 | **RNA**→ rplS (PRK05338)Ribosomal_L19 (pfam01245)→ |
| env-820 | JGI24723J26617_10040665 | + | 256 | 283 | **RNA**→ rplS (PRK05338)Ribosomal_L19 (pfam01245)→ |
| env-821 | KGI_S2_ANT06_2345m_c1000032 | - | 27425 | 27398 | **RNA**→ rplS (PRK05338)Ribosomal_L19 (pfam01245)→ |
| env-822 | KGI_S2_ANT04_2345m_c1056173 | + | 342 | 369 | **RNA**→ rplS (PRK05338)Ribosomal_L19 (pfam01245)→ |
| env-823 | JGI24723J26617_10586359 | + | 61 | 88 | **RNA**→ hypo→ rplS (PRK05338)Ribosomal_L19 (pfam01245)→ |
| Msp-1-1 | NZ_JHZW01000003.1 | + | 1124580 | 1124607 | **RNA**→ rplS (PRK05338)Ribosomal_L19 (pfam01245)→ |
| env-824 | JCVI_SCAF_1101668691466 | - | 995 | 968 | **RNA**→ rplS (PRK05338)Ribosomal_L19 (pfam01245)→ PRK08299 (PRK08299)→ |
| Ksp-1-1 | NC_015496.1 | - | 2413193 | 2413166 | **RNA**→ rplS (PRK05338)Ribosomal_L19 (pfam01245)→ |
| env-825 | JCVI_SCAF_1096627091268 | - | 3048 | 3021 | **RNA**→ rplS (PRK05338)Ribosomal_L19 (pfam01245)→ PRK08299 (PRK08299)→ |
| env-826 | CENF01107965.1 | + | 887 | 914 | **RNA**→ rplS (PRK05338)Ribosomal_L19 (pfam01245)→ |
| env-827 | CENT01102842.1 | - | 285 | 258 | **RNA**→ rplS (PRK05338)Ribosomal_L19 (pfam01245)→ |
| env-828 | CEOF01364937.1 | + | 159 | 186 | **RNA**→ rplS (PRK05338)Ribosomal_L19 (pfam01245)→ PRK08299 (PRK08299)→ |
| env-829 | CESM01052670.1 | + | 1148 | 1175 | **RNA**→ rplS (PRK05338)Ribosomal_L19 (pfam01245)→ PRK08299 (PRK08299)→ |
| env-830 | CETQ01129049.1 | - | 6191 | 6164 | **RNA**→ rplS (PRK05338)Ribosomal_L19 (pfam01245)→ PRK08299 (PRK08299)→ murQ (PRK05441)SIS_Etherase (cd05007)→ |
| env-831 | CEVP01064703.1 | + | 384 | 411 | **RNA**→ rplS (PRK05338)Ribosomal_L19 (pfam01245)→ PRK08299 (PRK08299)→ |
| env-832 | CEWG01096248.1 | - | 610 | 583 | **RNA**→ rplS (PRK05338)Ribosomal_L19 (pfam01245)→ PRK08299 (PRK08299)→ |
| Ddo-1-1 | NZ_JSAQ01000001.1 | - | 3054937 | 3054910 | **RNA**→ rplS (PRK05338)Ribosomal_L19 (pfam01245)→ |
| Dsp-1-1 | NZ_CP009301.1 | - | 3150857 | 3150830 | **RNA**→ rplS (PRK05338)Ribosomal_L19 (pfam01245)→ |
| env-833 | JGI20152J14361_10000826 | - | 19357 | 19330 | **RNA**→ rplS (PRK05338)Ribosomal_L19 (pfam01245)→ Icd (COG0538)PRK08299 (PRK08299)→ |
| env-834 | JGI20151J14362_10000089 | + | 3724 | 3751 | **RNA**→ rplS (PRK05338)Ribosomal_L19 (pfam01245)→ Icd (COG0538)PRK08299 (PRK08299)→ |
| env-835 | JGI20156J14371_10084791 | - | 1014 | 987 | **RNA**→ rplS (PRK05338)Ribosomal_L19 (pfam01245)→ PRK08299 (PRK08299)→ |
| env-836 | JGI20160J14292_10018282 | + | 2194 | 2221 | **RNA**→ rplS (PRK05338)Ribosomal_L19 (pfam01245)→ Icd (COG0538)PRK08299 (PRK08299)→ |
| env-837 | JGI20153J14295_10086476 | + | 211 | 238 | **RNA**→ rplS (PRK05338)Ribosomal_L19 (pfam01245)→ PRK08299 (PRK08299)→ |
| env-838 | JGI20153J14318_10086476 | + | 211 | 238 | **RNA**→ rplS (PRK05338)Ribosomal_L19 (pfam01245)→ PRK08299 (PRK08299)→ |
| env-839 | JGI20159J14440_10043604 | - | 1010 | 983 | **RNA**→ rplS (PRK05338)Ribosomal_L19 (pfam01245)→ PRK08299 (PRK08299)→ PTZ00435 (PTZ00435)→ |
| env-840 | JGI20154J14316_10041093 | + | 2028 | 2055 | **RNA**→ ←hypo |
| env-841 | CERK01035414.1 | + | 22 | 49 | **RNA**→ rplS (PRK05338)Ribosomal_L19 (pfam01245)→ Icd (COG0538)PRK08299 (PRK08299)→ CarbopepD_reg_2 (pfam13715)→ |
| env-842 | JGI20156J14371_10216168 | - | 35 | 8 | **RNA**→ |
| env-843 | JGI20157J14317_10031126 | + | 717 | 744 | **RNA**→ rplS (PRK05338)Ribosomal_L19 (pfam01245)→ Icd (COG0538)PRK08299 (PRK08299)→ |
| env-844 | JGI20155J14468_10062223 | - | 1029 | 1002 | **RNA**→ rplS (PRK05338)Ribosomal_L19 (pfam01245)→ PRK08299 (PRK08299)→ |
| env-845 | JGI20158J14315_10013594 | + | 3165 | 3192 | **RNA**→ rplS (PRK05338)Ribosomal_L19 (pfam01245)→ PRK08299 (PRK08299)→ |
| env-846 | CESO01100995.1 | + | 886 | 913 | **RNA**→ |
| env-847 | JGI20156J14371_10185819 | - | 95 | 68 | **RNA**→ |
| env-848 | JGI20157J14317_10068544 | + | 454 | 481 | **RNA**→ rplS (PRK05338)Ribosomal_L19 (pfam01245)→ PRK08299 (PRK08299)→ |
| env-849 | JGI20155J14468_10063464 | - | 1018 | 991 | **RNA**→ rplS (PRK05338)Ribosomal_L19 (pfam01245)→ PRK08299 (PRK08299)→ |
| env-850 | 2236876010_0174543 | - | 309 | 282 | **RNA**→ rplS (PRK05338)Ribosomal_L19 (pfam01245)→ rplS (PRK05338)Ribosomal_L19 (pfam01245)→ |

| | | | | | |
|---|---|---|---|---|---|
| env-851 | SI47jul10_100m_c1048908 | - | 444 | 417 | **RNA**→ rplS (PRK05338)Ribosomal_L19 (pfam01245)→ |
| env-852 | SI60aug11_100m_c1000269 | - | 14900 | 14873 | **RNA**→ rplS (PRK05338)Ribosomal_L19 (pfam01245)→ Icd (COG0538)PRK08299 (PRK08299)→ CirA (COG1629)PRK10064 (PRK10064)CarbopepD_reg_2 (pfam13715)→ DUF4249 (pfam14054)→ murQ (PRK05441)SIS_Etherase (cd05007)→ hypo→ ThrS (COG0441)thrS (PRK00413)TGS (pfam02824)→ InfC (COG0290)infC (PRK00028)→ rpmI (PRK00172)Ribosomal_L35p (pfam01632)→ rplT (PRK05185)Ribosomal_L20 (pfam00453)→ |
| env-853 | SI60aug11_100m_c1000276 | + | 16471 | 16498 | **RNA**→ rplS (PRK05338)Ribosomal_L19 (pfam01245)→ PRK08299 (PRK08299)→ |
| env-854 | SI34jun09_10m_1000015 | - | 14900 | 14873 | **RNA**→ rplS (PRK05338)Ribosomal_L19 (pfam01245)→ Icd (COG0538)PRK08299 (PRK08299)→ CirA (COG1629)PRK10064 (PRK10064)CarbopepD_reg_2 (pfam13715)→ DUF4249 (pfam14054)→ murQ (PRK05441)SIS_Etherase (cd05007)→ hypo→ ThrS (COG0441)thrS (PRK00413)TGS (pfam02824)→ InfC (COG0290)infC (PRK00028)→ rpmI (PRK00172)Ribosomal_L35p (pfam01632)→ rplT (PRK05185)Ribosomal_L20 (pfam00453)→ |
| env-855 | SI34jun09_120m_1001069 | + | 4978 | 5005 | **RNA**→ rplS (PRK05338)Ribosomal_L19 (pfam01245)→ Icd (COG0538)PRK08299 (PRK08299)→ CirA (COG1629)PRK10064 (PRK10064)CarbopepD_reg_2 (pfam13715)→ DUF4249 (pfam14054)→ murQ (PRK05441)SIS_Etherase (cd05007)→ hypo→ tRNA_SAD (smart00863)ThrS (COG0441)TGS (pfam02824)→ |
| env-856 | SI34jun09_135m_1006065 | - | 2206 | 2179 | **RNA**→ rplS (PRK05338)Ribosomal_L19 (pfam01245)→ Icd (COG0538)PRK08299 (PRK08299)→ CarbopepD_reg_2 (pfam13715)→ |
| env-857 | SI36aug09_100m_1042710 | + | 81 | 108 | **RNA**→ rplS (PRK05338)Ribosomal_L19 (pfam01245)→ |
| env-858 | SI34jun09_100m_1000078 | + | 35369 | 35396 | **RNA**→ rplS (PRK05338)Ribosomal_L19 (pfam01245)→ Icd (COG0538)PRK08299 (PRK08299)→ PRK10064 (PRK10064)Plug (pfam07715)CarbopepD_reg_2 (pfam13715)→ |
| env-859 | SI48aug10_100m_1000414 | - | 12190 | 12163 | **RNA**→ rplS (PRK05338)Ribosomal_L19 (pfam01245)→ Icd (COG0538)PRK08299 (PRK08299)→ CirA (COG1629)PRK10064 (PRK10064)CarbopepD_reg_2 (pfam13715)→ DUF4249 (pfam14054)→ murQ (PRK05441)SIS_Etherase (cd05007)→ hypo→ ThrS (COG0441)thrS (PRK00413)TGS (pfam02824)→ InfC (COG0290)infC (PRK00028)→ rpmI (PRK00172)Ribosomal_L35p (pfam01632)→ rplT (PRK05185)Ribosomal_L20 (pfam00453)→ |
| env-860 | CERW01115635.1 | - | 4803 | 4776 | **RNA**→ hypo→ rplS (PRK05338)Ribosomal_L19 (pfam01245)→ Icd (COG0538)PRK08299 (PRK08299)→ CirA (COG1629)Plug (pfam07715)CarbopepD_reg_2 (pfam13715)OMP_b-brl_3 (pfam14905)→ DUF4249 (pfam14054)→ |
| env-861 | JCVI_READ_1105430340115 | + | 255 | 282 | **RNA**→ rplS (PRK05338)Ribosomal_L19 (pfam01245)→ PRK08299 (PRK08299)→ |
| env-862 | DelMOSum2010_c10759857 | - | 116 | 89 | **RNA**→ hypo→ |
| env-863 | DelMOSum2011_c10081878 | + | 954 | 981 | **RNA**→ rplS (PRK05338)Ribosomal_L19 (pfam01245)→ |
| env-864 | SI48aug10_10m_c1014029 | + | 16 | 43 | **RNA**→ rplS (PRK05338)Ribosomal_L19 (pfam01245)→ PRK08299 (PRK08299)→ |
| env-865 | PML_READ_01299143 | + | 134 | 161 | **RNA**→ |
| env-866 | 2236876010_0240919 | + | 325 | 352 | **RNA**→ |
| env-867 | JGI20157J14317_10001218 | - | 14738 | 14711 | **RNA**→ rplS (PRK05338)Ribosomal_L19 (pfam01245)→ Icd (COG0538)PRK08299 (PRK08299)→ CirA (COG1629)PRK10064 (PRK10064)CarbopepD_reg_2 (pfam13715)→ DUF4249 (pfam14054)→ murQ (PRK05441)SIS_Etherase (cd05007)→ hypo→ ThrS (COG0441)thrS (PRK00413)TGS (pfam02824)→ InfC (COG0290)infC (PRK00028)→ rpmI (PRK00172)Ribosomal_L35p (pfam01632)→ rplT (PRK05185)Ribosomal_L20 (pfam00453)→ |
| env-868 | NCBI_BBAY_READ_1106105285287 | + | 249 | 276 | **RNA**→ rplS (PRK05338)Ribosomal_L19 (pfam01245)→ PRK08299 (PRK08299)→ |
| env-869 | DelMOSum2010_c10181064 | - | 71 | 44 | **RNA**→ |
| env-870 | DelMOSum2010_c10631757 | - | 229 | 202 | **RNA**→ hypo→ rplS (PRK05338)Ribosomal_L19 (pfam01245)→ |
| env-871 | SI36aug09_100m_1082667 | - | 291 | 264 | **RNA**→ rplS (PRK05338)Ribosomal_L19 (pfam01245)→ |
| env-872 | SI34jun09_100m_1043923 | - | 717 | 690 | **RNA**→ rplS (PRK05338)Ribosomal_L19 (pfam01245)→ PRK08299 (PRK08299)→ |
| env-873 | JGI20156J14371_10000105 | - | 20092 | 20065 | **RNA**→ rplS (PRK05338)Ribosomal_L19 (pfam01245)→ Icd (COG0538)PRK08299 (PRK08299)→ |
| env-874 | JGI20160J14292_10261268 | - | 452 | 425 | **RNA**→ rplS (PRK05338)Ribosomal_L19 (pfam01245)→ |
| env-875 | JGI20155J14468_10000942 | - | 20092 | 20065 | **RNA**→ rplS (PRK05338)Ribosomal_L19 (pfam01245)→ Icd (COG0538)PRK08299 (PRK08299)→ |
| env-876 | CERI01041892.1 | - | 67 | 40 | **RNA**→ |
| env-877 | CETY01197871.1 | + | 10782 | 10809 | **RNA**→ rplS (PRK05338)Ribosomal_L19 (pfam01245)→ Icd (COG0538)PRK08299 (PRK08299)→ |
| env-878 | CEWO01125901.1 | - | 186 | 159 | **RNA**→ rplS (PRK05338)Ribosomal_L19 (pfam01245)→ |
| env-879 | F10SI0310a_Sequence0000036835 | - | 83 | 56 | **RNA**→ |
| env-880 | F10SI03100a_Sequence0000055496 | + | 990 | 1017 | **RNA**→ |
| env-881 | CETY01434901.1 | + | 139 | 166 | **RNA**→ rplS (PRK05338)Ribosomal_L19 (pfam01245)→ PRK08299 (PRK08299)→ |
| env-882 | JGI20154J14316_10005236 | + | 6892 | 6919 | **RNA**→ rplS (PRK05338)Ribosomal_L19 (pfam01245)→ Icd (COG0538)PRK08299 (PRK08299)→ |
| env-883 | JGI20155J14468_10106440 | + | 886 | 913 | **RNA**→ |

| | | | | | |
|---|---|---|---|---|---|
| env-884 | SI60aug11_100m_c1103194 | + | 244 | 271 | **RNA**→ rplS (PRK05338)Ribosomal_L19 (pfam01245)→ |
| env-885 | SI48aug10_10m_c1005211 | - | 1208 | 1181 | **RNA**→ rplS (PRK05338)Ribosomal_L19 (pfam01245)→ PRK08299 (PRK08299)→ |
| env-886 | SI34jun09_10m_1014940 | + | 226 | 253 | **RNA**→ rplS (PRK05338)Ribosomal_L19 (pfam01245)→ PRK08299 (PRK08299)→ |
| env-887 | SI34jun09_120m_1058221 | - | 543 | 516 | **RNA**→ rplS (PRK05338)Ribosomal_L19 (pfam01245)→ PRK08299 (PRK08299)→ |
| env-888 | SI34jun09_135m_1193559 | + | 232 | 259 | **RNA**→ |
| env-889 | JGI20151J14362_10577636 | - | 201 | 174 | **RNA**→ rplS (PRK05338)Ribosomal_L19 (pfam01245)→ |
| env-890 | JGI20157J14317_10006145 | - | 8612 | 8585 | **RNA**→ rplS (PRK05338)Ribosomal_L19 (pfam01245)→ Icd (COG0538)PRK08299 (PRK08299)→ |
| env-891 | SI48aug10_100m_1000052 | + | 30997 | 31024 | **RNA**→ rplS (PRK05338)Ribosomal_L19 (pfam01245)→ Icd (COG0538)PRK08299 (PRK08299)→ |
| env-892 | JGI20157J14317_10006052 | - | 4424 | 4397 | **RNA**→ rplS (PRK05338)Ribosomal_L19 (pfam01245)→ Icd (COG0538)PRK08299 (PRK08299)→ |
| env-893 | 2236876001_145660 | - | 390 | 363 | **RNA**→ rplS (PRK05338)Ribosomal_L19 (pfam01245)→ |
| env-894 | JCVI_SCAF_1096627080844 | + | 1234 | 1261 | **RNA**→ rplS (PRK05338)Ribosomal_L19 (pfam01245)→ Icd (COG0538)PRK08299 (PRK08299)→<br>PRK10064 (PRK10064)Plug (pfam07715)CarbopepD_reg_2 (pfam13715)→ |
| env-895 | DelMOWin2010_c10059332 | + | 366 | 393 | **RNA**→ rplS (PRK05338)Ribosomal_L19 (pfam01245)→ PRK08299 (PRK08299)→ |
| env-896 | CENF01410679.1 | + | 457 | 484 | **RNA**→ rplS (PRK05338)Ribosomal_L19 (pfam01245)→ PRK08299 (PRK08299)→ |
| env-897 | CENT01071942.1 | + | 440 | 467 | **RNA**→ rplS (PRK05338)Ribosomal_L19 (pfam01245)→ Icd (COG0538)PRK08299 (PRK08299)→<br>PRK10064 (PRK10064)TonB_dep_Rec (pfam00593)Plug (pfam07715)CarbopepD_reg_2 (pfam13715)OMP_b-brl_3 (pfam1<br>DUF4249 (pfam14054)→ murQ (PRK05441)SIS_Etherase (cd05007)→ hypo→<br>TGS_ThrRS_N (cd01667)ThrS (COG0441)thrS (PRK00413)→ |
| env-898 | CEOP01008385.1 | - | 7910 | 7883 | **RNA**→ rplS (PRK05338)Ribosomal_L19 (pfam01245)→ Icd (COG0538)PRK08299 (PRK08299)→<br>PRK10064 (PRK10064)TonB_dep_Rec (pfam00593)Plug (pfam07715)CarbopepD_reg_2 (pfam13715)OMP_b-brl_3 (pfam1<br>DUF4249 (pfam14054)→ murQ (PRK05441)SIS_Etherase (cd05007)→ hypo→<br>TGS_ThrRS_N (cd01667)ThrS (COG0441)thrS (PRK00413)→ |
| env-899 | CERW01135648.1 | + | 187 | 214 | **RNA**→ rplS (PRK05338)Ribosomal_L19 (pfam01245)→ PRK08299 (PRK08299)→ |
| env-900 | CESF01017371.1 | - | 1161 | 1134 | **RNA**→ rplS (PRK05338)Ribosomal_L19 (pfam01245)→ PRK08299 (PRK08299)→ |
| env-901 | CESM01027070.1 | + | 3699 | 3726 | **RNA**→ rplS (PRK05338)Ribosomal_L19 (pfam01245)→ Icd (COG0538)PRK08299 (PRK08299)→<br>PRK10064 (PRK10064)TonB_dep_Rec (pfam00593)Plug (pfam07715)CarbopepD_reg_2 (pfam13715)OMP_b-brl_3 (pfam1<br>DUF4249 (pfam14054)→ |
| env-902 | CESO01004564.1 | + | 1551 | 1578 | **RNA**→ rplS (PRK05338)Ribosomal_L19 (pfam01245)→ Icd (COG0538)PRK08299 (PRK08299)→<br>PRK10064 (PRK10064)TonB_dep_Rec (pfam00593)Plug (pfam07715)CarbopepD_reg_2 (pfam13715)OMP_b-brl_3 (pfam1<br>DUF4249 (pfam14054)→ murQ (PRK05441)SIS_Etherase (cd05007)→ |
| env-903 | CEVS01180975.1 | - | 1250 | 1223 | **RNA**→ rplS (PRK05338)Ribosomal_L19 (pfam01245)→ PRK08299 (PRK08299)→ |
| env-904 | CEVU01367992.1 | - | 1403 | 1376 | **RNA**→ rplS (PRK05338)Ribosomal_L19 (pfam01245)→ PRK08299 (PRK08299)→ |
| env-905 | CEWG01218791.1 | - | 8217 | 8190 | **RNA**→ rplS (PRK05338)Ribosomal_L19 (pfam01245)→ Icd (COG0538)PRK08299 (PRK08299)→<br>PRK10064 (PRK10064)TonB_dep_Rec (pfam00593)Plug (pfam07715)CarbopepD_reg_2 (pfam13715)OMP_b-brl_3 (pfam1<br>DUF4249 (pfam14054)→ murQ (PRK05441)SIS_Etherase (cd05007)→ hypo→<br>TGS_ThrRS_N (cd01667)ThrS (COG0441)thrS (PRK00413)→ |
| env-906 | NCBI_READ_1112493833240 | + | 25 | 52 | **RNA**→ rplS (PRK05338)Ribosomal_L19 (pfam01245)→ PRK08299 (PRK08299)→ |
| env-907 | CESE01052974.1 | + | 1100 | 1127 | **RNA**→ rplS (PRK05338)Ribosomal_L19 (pfam01245)→ PRK08299 (PRK08299)→ |
| env-908 | CESJ01033277.1 | + | 1112 | 1139 | **RNA**→ rplS (PRK05338)Ribosomal_L19 (pfam01245)→ PRK08299 (PRK08299)→ |
| env-909 | CEST01137492.1 | + | 594 | 621 | **RNA**→ hypo→ |
| env-910 | DelMOSum2011_c10014222 | + | 400 | 427 | **RNA**→ rplS (PRK05338)Ribosomal_L19 (pfam01245)→ Icd (COG0538)PRK08299 (PRK08299)→<br>PRK10064 (PRK10064)TonB_dep_Rec (pfam00593)Plug (pfam07715)CarbopepD_reg_2 (pfam13715)→ |
| env-911 | DelMOSum2011_c10064936 | + | 400 | 427 | **RNA**→ rplS (PRK05338)Ribosomal_L19 (pfam01245)→ PRK08299 (PRK08299)→ |
| env-912 | CESU01016977.1 | - | 4567 | 4540 | **RNA**→ rplS (PRK05338)Ribosomal_L19 (pfam01245)→ Icd (COG0538)PRK08299 (PRK08299)→<br>PRK10064 (PRK10064)TonB_dep_Rec (pfam00593)Plug (pfam07715)CarbopepD_reg_2 (pfam13715)→<br>DUF4249 (pfam14054)→ |
| env-913 | LPjun09P1210m_c1060279 | - | 297 | 270 | **RNA**→ rplS (PRK05338)Ribosomal_L19 (pfam01245)→ |
| env-914 | DelMOSum2010_c10056893 | - | 591 | 564 | **RNA**→ rplS (PRK05338)Ribosomal_L19 (pfam01245)→ PRK08299 (PRK08299)→ |
| env-915 | PML_READ_00864574 | + | 212 | 239 | **RNA**→ |
| env-916 | CEPZ01008967.1 | - | 10616 | 10589 | **RNA**→ rplS (PRK05338)Ribosomal_L19 (pfam01245)→ |
| env-917 | SRS015278_C1831448 | + | 482 | 509 | **RNA**→ rplS (PRK05338)Ribosomal_L19 (pfam01245)→ |
| env-918 | SRS024649_C3604464 | + | 494 | 521 | **RNA**→ |
| Fba-4-1 | NZ_ABHI01000004.1 | + | 89845 | 89872 | **RNA**→ rplS (PRK05338)Ribosomal_L19 (pfam01245)→ rplS (PRK05338)Ribosomal_L19 (pfam01245)→ |
| env-919 | NCBI_BBAY_READ_1105731251167 | - | 848 | 821 | **RNA**→ rplS (PRK05338)Ribosomal_L19 (pfam01245)→ PRK08299 (PRK08299)→ |

| | | | | | |
|---|---|---|---|---|---|
| env-920 | CESI01073316.1 | - | 585 | 558 | **RNA**→ rplS (PRK05338)Ribosomal_L19 (pfam01245)→ |
| env-921 | CESU01070521.1 | - | 784 | 757 | **RNA**→ rplS (PRK05338)Ribosomal_L19 (pfam01245)→ |
| env-922 | Antartic2_1088051 | - | 150 | 123 | **RNA**→ hypo→ |
| env-923 | 2236876011_158566 | + | 32 | 59 | **RNA**→ rplS (PRK05338)Ribosomal_L19 (pfam01245)→ |
| env-924 | 2236876009_152251 | - | 387 | 360 | **RNA**→ rplS (PRK05338)Ribosomal_L19 (pfam01245)→ |
| env-925 | JGI24723J26617_10000165 | - | 51447 | 51420 | **RNA**→ rplS (PRK05338)Ribosomal_L19 (pfam01245)→ |
| env-926 | JGI24024J18818_10020607 | - | 2387 | 2360 | **RNA**→ rplS (PRK05338)Ribosomal_L19 (pfam01245)→ |
| env-927 | LGVF01309355.1 | + | 1747 | 1774 | **RNA**→ rplS (PRK05338)Ribosomal_L19 (pfam01245)→ |
| env-928 | TDF_MC_ARG01_113m_c1056626 | - | 258 | 231 | **RNA**→ hypo→ rplS (PRK05338)Ribosomal_L19 (pfam01245)→ |
| Ffi-1-1 | NZ_AUDM01000017.1 | - | 30024 | 29997 | **RNA**→ rplS (PRK05338)Ribosomal_L19 (pfam01245)→ |
| env-929 | KGI_S1_ANT01_95m_c10018333 | - | 2431 | 2404 | **RNA**→ rplS (PRK05338)Ribosomal_L19 (pfam01245)→ hypo→ |
| env-930 | KGI_S2_ANT04_2345m_c1018541 | + | 769 | 796 | **RNA**→ rplS (PRK05338)Ribosomal_L19 (pfam01245)→ |
| env-931 | KGI_S1_ANT02_95m_c10051843 | - | 1116 | 1089 | **RNA**→ rplS (PRK05338)Ribosomal_L19 (pfam01245)→ |
| Fba-8-1 | NZ_AFPK01000067.1 | + | 24608 | 24635 | **RNA**→ rplS (PRK05338)Ribosomal_L19 (pfam01245)→ rplS (PRK05338)Ribosomal_L19 (pfam01245)→ |
| env-932 | SA_S2_NOR18_50m_1003007 | + | 2084 | 2111 | **RNA**→ rplS (PRK05338)Ribosomal_L19 (pfam01245)→ |
| env-933 | SA_S2_NOR13_50m_c1000705 | + | 3191 | 3218 | **RNA**→ rplS (PRK05338)Ribosomal_L19 (pfam01245)→ |
| env-934 | SA_S1_NOR05_45m_c10004569 | - | 1775 | 1748 | **RNA**→ rplS (PRK05338)Ribosomal_L19 (pfam01245)→ hypo→ |
| env-935 | SA_S1_NOR08_45m_c10000086 | + | 11623 | 11650 | **RNA**→ rplS (PRK05338)Ribosomal_L19 (pfam01245)→ |
| env-936 | SA_S2_NOR15_50m_c10001702 | + | 8902 | 8929 | **RNA**→ rplS (PRK05338)Ribosomal_L19 (pfam01245)→ |
| env-937 | SA_S1_NOR02_45m_c1057368 | + | 341 | 368 | **RNA**→ ←hypo |
| env-938 | JGI20157J14317_10496092 | + | 266 | 293 | **RNA**→ |
| Cca-1-1 | NZ_CDOI01000142.1 | + | 52361 | 52388 | **RNA**→ rplS (PRK05338)Ribosomal_L19 (pfam01245)→ |
| Cca-1-2 | NZ_CDOH01000131.1 | - | 1703 | 1676 | **RNA**→ rplS (PRK05338)Ribosomal_L19 (pfam01245)→ |
| Cca-1-3 | NZ_CDOL01000061.1 | + | 11126 | 11153 | **RNA**→ rplS (PRK05338)Ribosomal_L19 (pfam01245)→ |
| env-939 | JGI25913J50563_1214264 | - | 110 | 83 | **RNA**→ hypo→ |
| env-940 | KVRMV2_100094982 | + | 521 | 548 | **RNA**→ |
| env-941 | KVRMV2_100096620 | + | 400 | 427 | **RNA**→ rplS (PRK05338)Ribosomal_L19 (pfam01245)→ |
| env-942 | KVRMV2_102215955 | + | 253 | 280 | **RNA**→ rplS (PRK05338)Ribosomal_L19 (pfam01245)→ |
| Emi-1-1 | NZ_CP011059.1 | - | 192039 | 192012 | **RNA**→ rplS (PRK05338)Ribosomal_L19 (pfam01245)→ Nudix_Hydrolase_31 (cd04690)NUDIX_4 (pfam14815)→ |
| Wma-1-1 | NZ_CCMH01000053.1 | - | 16198 | 16171 | **RNA**→ rplS (PRK05338)Ribosomal_L19 (pfam01245)→ |
| env-943 | P_2C_Liq_1_UnCty_1237069 | + | 47 | 74 | **RNA**→ rplS (PRK05338)Ribosomal_L19 (pfam01245)→ |
| env-944 | P_1C_Liq_1_UnCty_1029662 | + | 212 | 239 | **RNA**→ rplS (PRK05338)Ribosomal_L19 (pfam01245)→ |
| env-945 | CESF01128812.1 | - | 1808 | 1781 | **RNA**→ rplS (PRK05338)Ribosomal_L19 (pfam01245)→ PRK08299 (PRK08299)→ |
| env-946 | CESO01022412.1 | + | 665 | 692 | **RNA**→ rplS (PRK05338)Ribosomal_L19 (pfam01245)→ PRK08299 (PRK08299)→ |
| env-947 | CEOP01454573.1 | + | 1970 | 1997 | **RNA**→ rplS (PRK05338)Ribosomal_L19 (pfam01245)→ |
| env-948 | CEWO01198114.1 | + | 1577 | 1604 | **RNA**→ rplS (PRK05338)Ribosomal_L19 (pfam01245)→ PRK08299 (PRK08299)→ |
| env-949 | CERI01052802.1 | + | 1424 | 1451 | **RNA**→ rplS (PRK05338)Ribosomal_L19 (pfam01245)→ |
| env-950 | CETY01032138.1 | + | 785 | 812 | **RNA**→ |
| env-951 | JGI20152J14361_10010744 | - | 1575 | 1548 | **RNA**→ rplS (PRK05338)Ribosomal_L19 (pfam01245)→ |
| env-952 | JGI20151J14362_10033063 | + | 947 | 974 | **RNA**→ rplS (PRK05338)Ribosomal_L19 (pfam01245)→ |
| env-953 | JGI20157J14317_10018646 | + | 2505 | 2532 | **RNA**→ rplS (PRK05338)Ribosomal_L19 (pfam01245)→ |
| env-954 | 2236876010_0376568 | + | 129 | 156 | **RNA**→ rplS (PRK05338)Ribosomal_L19 (pfam01245)→ |
| env-955 | 2236876010_0173724 | + | 243 | 270 | **RNA**→ hypo→ |
| env-956 | JGI20156J14371_10012147 | + | 2603 | 2630 | **RNA**→ rplS (PRK05338)Ribosomal_L19 (pfam01245)→ |
| env-957 | 2236876010_0038631 | + | 246 | 273 | **RNA**→ rplS (PRK05338)Ribosomal_L19 (pfam01245)→ hypo→ |
| env-958 | 2236876010_0065632 | - | 228 | 201 | **RNA**→ rplS (PRK05338)Ribosomal_L19 (pfam01245)→ |
| env-959 | DelMOWin2010_c10348025 | - | 98 | 71 | **RNA**→ |
| env-960 | JGI20154J14316_10016038 | + | 2890 | 2917 | **RNA**→ rplS (PRK05338)Ribosomal_L19 (pfam01245)→ |
| env-961 | JGI20160J14292_10020040 | + | 2336 | 2363 | **RNA**→ rplS (PRK05338)Ribosomal_L19 (pfam01245)→ |
| env-962 | JGI20153J14295_10012810 | + | 2889 | 2916 | **RNA**→ rplS (PRK05338)Ribosomal_L19 (pfam01245)→ |
| env-963 | JGI20153J14318_10012810 | + | 2889 | 2916 | **RNA**→ rplS (PRK05338)Ribosomal_L19 (pfam01245)→ |
| env-964 | JGI20159J14440_10003663 | - | 8129 | 8102 | **RNA**→ rplS (PRK05338)Ribosomal_L19 (pfam01245)→ |

| | | | | | |
|---|---|---|---|---|---|
| env-965 | JGI20159J14440_10003889 | + | 6429 | 6456 | **RNA**→ rplS (PRK05338)Ribosomal_L19 (pfam01245)→ |
| env-966 | JGI20155J14468_10099882 | - | 830 | 803 | **RNA**→ rplS (PRK05338)Ribosomal_L19 (pfam01245)→ |
| env-967 | JGI20158J14315_10032857 | - | 673 | 646 | **RNA**→ rplS (PRK05338)Ribosomal_L19 (pfam01245)→ |
| env-968 | 2236876010_0093675 | + | 11 | 38 | **RNA**→ rplS (PRK05338)Ribosomal_L19 (pfam01245)→ |
| env-969 | 2236876010_0167969 | + | 65 | 92 | **RNA**→ rplS (PRK05338)Ribosomal_L19 (pfam01245)→ |
| env-970 | DelMOSum2011_c10093048 | - | 801 | 774 | **RNA**→ rplS (PRK05338)Ribosomal_L19 (pfam01245)→ |
| env-971 | DelMOSum2011_c10001686 | - | 6899 | 6872 | **RNA**→ rplS (PRK05338)Ribosomal_L19 (pfam01245)→ |
| env-972 | SI34jun09_10m_1096433 | - | 156 | 129 | **RNA**→ rplS (PRK05338)Ribosomal_L19 (pfam01245)→ |
| env-973 | CERM01034613.1 | - | 664 | 637 | **RNA**→ rplS (PRK05338)Ribosomal_L19 (pfam01245)→ |
| env-974 | 2236876009_179996 | - | 147 | 120 | **RNA**→ rplS (PRK05338)Ribosomal_L19 (pfam01245)→ |
| env-975 | 2236876010_0260662 | + | 145 | 172 | **RNA**→ |
| env-976 | NCBI_READ_1112749174365 | + | 12 | 39 | **RNA**→ rplS (PRK05338)Ribosomal_L19 (pfam01245)→ |
| env-977 | P_2C_Liq_3_UnCty_100987 | + | 228 | 257 | **RNA**→ rplS (PRK05338)Ribosomal_L19 (pfam01245)→ |
| env-978 | P_2C_Liq_1_UnCty_1005821 | + | 3392 | 3421 | **RNA**→ hypo→ |
| env-979 | P_2C_Liq_2_UnCty_1021870 | + | 790 | 819 | **RNA**→ rplS (PRK05338)Ribosomal_L19 (pfam01245)→ |
| env-980 | JGI20154J14316_10003986 | - | 1590 | 1561 | **RNA**→ rplS (PRK05338)Ribosomal_L19 (pfam01245)→ |
| env-981 | JGI20153J14295_10004614 | + | 8389 | 8418 | **RNA**→ rplS (PRK05338)Ribosomal_L19 (pfam01245)→ |
| env-982 | JGI20153J14318_10004614 | + | 8389 | 8418 | **RNA**→ rplS (PRK05338)Ribosomal_L19 (pfam01245)→ |
| env-983 | JGI20152J14361_10000338 | + | 14595 | 14624 | **RNA**→ rplS (PRK05338)Ribosomal_L19 (pfam01245)→ |
| env-984 | JGI20156J14371_10003171 | + | 2644 | 2673 | **RNA**→ rplS (PRK05338)Ribosomal_L19 (pfam01245)→ |
| env-985 | JGI20154J14316_10025201 | - | 1474 | 1445 | **RNA**→ rplS (PRK05338)Ribosomal_L19 (pfam01245)→ |
| env-986 | JGI20153J14295_10003738 | + | 9965 | 9994 | **RNA**→ |
| env-987 | JGI20153J14318_10003738 | + | 9965 | 9994 | **RNA**→ |
| env-988 | P_2C_Liq_1_UnCty_1000289 | - | 21098 | 21069 | **RNA**→ rplS (PRK05338)Ribosomal_L19 (pfam01245)→ |
| Psp-2-1 | NZ_JHZZ01000001.1 | - | 2913763 | 2913734 | **RNA**→ rplS (PRK05338)Ribosomal_L19 (pfam01245)→ |
| env-989 | CESI01065175.1 | + | 1901 | 1930 | **RNA**→ rplS (PRK05338)Ribosomal_L19 (pfam01245)→ |
| env-990 | CESU01059413.1 | + | 594 | 623 | **RNA**→ rplS (PRK05338)Ribosomal_L19 (pfam01245)→ |
| env-991 | JGI20152J14361_10000044 | - | 80594 | 80565 | **RNA**→ rplS (PRK05338)Ribosomal_L19 (pfam01245)→ |
| env-992 | JGI20153J14295_10523871 | - | 70 | 41 | **RNA**→ ←hypo |
| env-993 | JGI20153J14318_10523871 | - | 70 | 41 | **RNA**→ ←hypo |
| env-994 | DelMOWin2010_c10731323 | + | 65 | 94 | **RNA**→ ←hypo |
| env-995 | JGI20156J14371_10483843 | + | 180 | 209 | **RNA**→ ←hypo |
| env-996 | JGI20159J14440_10053546 | + | 607 | 636 | **RNA**→ rplS (PRK05338)Ribosomal_L19 (pfam01245)→ |
| env-997 | NCBI_READ_1112777322026 | - | 870 | 841 | **RNA**→ rplS (PRK05338)Ribosomal_L19 (pfam01245)→ |
| env-998 | NCBI_READ_1112637402410 | + | 418 | 447 | **RNA**→ rplS (PRK05338)Ribosomal_L19 (pfam01245)→ |
| env-999 | NCBI_READ_1112614318228 | - | 194 | 165 | **RNA**→ rplS (PRK05338)Ribosomal_L19 (pfam01245)→ |
| env-1000 | CESE01039129.1 | + | 14427 | 14456 | **RNA**→ |
| env-1001 | CESJ01285814.1 | - | 2842 | 2813 | **RNA**→ rplS (PRK05338)Ribosomal_L19 (pfam01245)→ |
| env-1002 | CEST01030576.1 | - | 89 | 60 | **RNA**→ |
| Pir-1-1 | NZ_CH724148.1 | + | 36399 | 36428 | **RNA**→ rplS (PRK05338)Ribosomal_L19 (pfam01245)→ |
| env-1003 | BS_KBA_SWE12_21m_c10085874 | + | 203 | 230 | **RNA**→ rplS (PRK05338)Ribosomal_L19 (pfam01245)→ |
| env-1004 | BS_KBB_SWE26_205m_c1058109 | + | 165 | 192 | **RNA**→ rplS (PRK05338)Ribosomal_L19 (pfam01245)→ |
| env-1005 | BS_KBA_SWE21_205m_10102186 | + | 146 | 173 | **RNA**→ rplS (PRK05338)Ribosomal_L19 (pfam01245)→ |
| env-1006 | BS_KBA_SWE02_21m_10115347 | - | 438 | 411 | **RNA**→ rplS (PRK05338)Ribosomal_L19 (pfam01245)→ |
| env-1007 | NCBI_READ_1112494184882 | - | 125 | 96 | **RNA**→ |
| env-1008 | NCBI_READ_1112637830883 | + | 552 | 581 | **RNA**→ rplS (PRK05338)Ribosomal_L19 (pfam01245)→ |
| env-1009 | DelMOSum2010_c10000424 | - | 11560 | 11531 | **RNA**→ rplS (PRK05338)Ribosomal_L19 (pfam01245)→ |
| env-1010 | JGI20157J14317_10004444 | - | 857 | 828 | **RNA**→ rplS (PRK05338)Ribosomal_L19 (pfam01245)→ |
| env-1011 | NCBI_READ_1112493603404 | + | 47 | 76 | **RNA**→ rplS (PRK05338)Ribosomal_L19 (pfam01245)→ |
| env-1012 | NCBI_READ_1112493503141 | + | 144 | 173 | **RNA**→ rplS (PRK05338)Ribosomal_L19 (pfam01245)→ |
| env-1013 | NCBI_READ_1112788476902 | + | 509 | 538 | **RNA**→ rplS (PRK05338)Ribosomal_L19 (pfam01245)→ |
| env-1014 | CEWR01020717.1 | - | 6341 | 6312 | **RNA**→ rplS (PRK05338)Ribosomal_L19 (pfam01245)→ |

| | | | | | |
|---|---|---|---|---|---|
| env-1015 | CESJ01043971.1 | + | 1219 | 1246 | **RNA**→ rplS (PRK05338)Ribosomal_L19 (pfam01245)→ PRK08299 (PRK08299)→ murQ (PRK05441)SIS_Etherase (cd05007)→ hypo→ TGS_ThrRS_N (cd01667)ThrS (COG0441)thrS (PRK00413)→ |
| env-1016 | 2236876002_177911 | - | 239 | 210 | **RNA**→ hypo→ rplS (PRK05338)Ribosomal_L19 (pfam01245)→ |
| env-1017 | DelMOSum2010_c10001649 | - | 15644 | 15615 | **RNA**→ rplS (PRK05338)Ribosomal_L19 (pfam01245)→ |
| env-1018 | JGI20154J14316_10032387 | + | 2473 | 2502 | **RNA**→ ←hypo |
| env-1019 | JGI20153J14295_10012199 | + | 4050 | 4079 | **RNA**→ rplS (PRK05338)Ribosomal_L19 (pfam01245)→ |
| env-1020 | JGI20153J14318_10012199 | + | 4050 | 4079 | **RNA**→ rplS (PRK05338)Ribosomal_L19 (pfam01245)→ |
| env-1021 | SI34jun09_10m_1071064 | + | 169 | 198 | **RNA**→ rplS (PRK05338)Ribosomal_L19 (pfam01245)→ |
| env-1022 | A09P0410a_Sequence0000002007 | - | 383 | 354 | **RNA**→ rplS (PRK05338)Ribosomal_L19 (pfam01245)→ |
| env-1023 | JGI20154J14316_10036828 | + | 484 | 513 | **RNA**→ rplS (PRK05338)Ribosomal_L19 (pfam01245)→ |
| env-1024 | JGI20160J14292_10414066 | - | 250 | 221 | **RNA**→ rplS (PRK05338)Ribosomal_L19 (pfam01245)→ |
| env-1025 | CESI01082352.1 | + | 317 | 346 | **RNA**→ rplS (PRK05338)Ribosomal_L19 (pfam01245)→ |
| env-1026 | JGI20160J14292_10063069 | + | 592 | 621 | **RNA**→ rplS (PRK05338)Ribosomal_L19 (pfam01245)→ |
| env-1027 | CESD01219058.1 | + | 576 | 603 | **RNA**→ rplS (PRK05338)Ribosomal_L19 (pfam01245)→ |
| env-1028 | CETB01155201.1 | + | 708 | 735 | **RNA**→ rplS (PRK05338)Ribosomal_L19 (pfam01245)→ |
| env-1029 | CETH01045262.1 | + | 30526 | 30553 | **RNA**→ rplS (PRK05338)Ribosomal_L19 (pfam01245)→ Icd (COG0538)PRK08299 (PRK08299)→ murQ (PRK05441)SIS_Etherase (cd05007)→ hypo→ TGS_ThrRS_N (cd01667)ThrS (COG0441)thrS (PRK00413)→ |
| env-1030 | CETM01203845.1 | + | 3063 | 3090 | **RNA**→ rplS (PRK05338)Ribosomal_L19 (pfam01245)→ Icd (COG0538)PRK08299 (PRK08299)→ murQ (PRK05441)SIS_Etherase (cd05007)→ hypo→ TGS_ThrRS_N (cd01667)ThrS (COG0441)thrS (PRK00413)→ |
| env-1031 | CETT01281088.1 | + | 569 | 596 | **RNA**→ rplS (PRK05338)Ribosomal_L19 (pfam01245)→ |
| env-1032 | CEUL01311114.1 | - | 456 | 429 | **RNA**→ rplS (PRK05338)Ribosomal_L19 (pfam01245)→ |
| env-1033 | CEUZ01290455.1 | + | 1300 | 1327 | **RNA**→ rplS (PRK05338)Ribosomal_L19 (pfam01245)→ Icd (COG0538)PRK08299 (PRK08299)→ murQ (PRK05441)SIS_Etherase (cd05007)→ hypo→ TGS_ThrRS_N (cd01667)ThrS (COG0441)thrS (PRK00413)→ |
| env-1034 | SI60aug11_100m_c1007508 | - | 2548 | 2519 | **RNA**→ rplS (PRK05338)Ribosomal_L19 (pfam01245)→ |
| env-1035 | JGI20152J14361_10003189 | - | 8383 | 8354 | **RNA**→ rplS (PRK05338)Ribosomal_L19 (pfam01245)→ |
| env-1036 | JGI20153J14295_10000489 | + | 27052 | 27081 | **RNA**→ rplS (PRK05338)Ribosomal_L19 (pfam01245)→ |
| env-1037 | JGI20153J14318_10000489 | + | 27052 | 27081 | **RNA**→ rplS (PRK05338)Ribosomal_L19 (pfam01245)→ |
| Psp-1-1 | NZ_JPDI01000001.1 | - | 2860067 | 2860038 | **RNA**→ rplS (PRK05338)Ribosomal_L19 (pfam01245)→ |
| env-1038 | CEOP01247711.1 | - | 5731 | 5704 | **RNA**→ rplS (PRK05338)Ribosomal_L19 (pfam01245)→ Icd (COG0538)PRK08299 (PRK08299)→ murQ (PRK05441)SIS_Etherase (cd05007)→ hypo→ TGS_ThrRS_N (cd01667)ThrS (COG0441)thrS (PRK00413)→ |
| env-1039 | 2236876001_262712 | - | 39 | 11 | **RNA**→ hypo→ |
| env-1040 | CERB01093551.1 | + | 315 | 342 | **RNA**→ rplS (PRK05338)Ribosomal_L19 (pfam01245)→ |
| env-1041 | CERS01312156.1 | + | 272 | 299 | **RNA**→ rplS (PRK05338)Ribosomal_L19 (pfam01245)→ PRK08299 (PRK08299)→ |
| env-1042 | NCBI_READ_1112290446392 | + | 322 | 351 | **RNA**→ rplS (PRK05338)Ribosomal_L19 (pfam01245)→ |
| env-1043 | NCBI_READ_1112493493141 | + | 227 | 256 | **RNA**→ |
| env-1044 | CESE01277103.1 | - | 107 | 78 | **RNA**→ |
| env-1045 | CEST01047642.1 | + | 9823 | 9852 | **RNA**→ |
| env-1046 | CEWR01079815.1 | + | 9821 | 9850 | **RNA**→ |
| env-1047 | NCBI_READ_1112777250419 | - | 310 | 281 | **RNA**→ rplS (PRK05338)Ribosomal_L19 (pfam01245)→ |
| env-1048 | KGI_S2_ANT05_2345m_c1212107 | - | 177 | 148 | **RNA**→ rplS (PRK05338)Ribosomal_L19 (pfam01245)→ |
| env-1049 | P_2C_Liq_3_UnCty_100705 | - | 3459 | 3430 | **RNA**→ rplS (PRK05338)Ribosomal_L19 (pfam01245)→ |
| env-1050 | P_2C_Liq_1_UnCty_1002384 | - | 5606 | 5577 | **RNA**→ rplS (PRK05338)Ribosomal_L19 (pfam01245)→ |
| env-1051 | P_2C_Liq_2_UnCty_1024292 | - | 447 | 418 | **RNA**→ rplS (PRK05338)Ribosomal_L19 (pfam01245)→ |
| env-1052 | NCBI_READ_1112788537198 | - | 239 | 210 | **RNA**→ rplS (PRK05338)Ribosomal_L19 (pfam01245)→ |
| env-1053 | NCBI_READ_1112493844367 | - | 655 | 626 | **RNA**→ rplS (PRK05338)Ribosomal_L19 (pfam01245)→ |
| env-1054 | NCBI_READ_1112679247542 | + | 278 | 307 | **RNA**→ rplS (PRK05338)Ribosomal_L19 (pfam01245)→ |
| env-1055 | CESI01042215.1 | + | 703 | 732 | **RNA**→ |
| env-1056 | CESJ01095310.1 | - | 463 | 434 | **RNA**→ rplS (PRK05338)Ribosomal_L19 (pfam01245)→ |
| env-1057 | NCBI_READ_1112493752993 | - | 840 | 812 | **RNA**→ rplS (PRK05338)Ribosomal_L19 (pfam01245)→ |
| env-1058 | NCBI_READ_1112777271421 | + | 338 | 367 | **RNA**→ rplS (PRK05338)Ribosomal_L19 (pfam01245)→ |
| Wvi-1-1 | NC_015144.1 | + | 358414 | 358441 | **RNA**→ rplS (PRK05338)Ribosomal_L19 (pfam01245)→ |
| env-1059 | BGMT1_NODE_590531 | - | 606 | 579 | **RNA**→ rplS (PRK05338)Ribosomal_L19 (pfam01245)→ |
| env-1060 | HBC_scaff_10036577 | + | 20118 | 20145 | **RNA**→ rplS (PRK05338)Ribosomal_L19 (pfam01245)→ |

| | | | | | |
|---|---|---|---|---|---|
| env-1061 | HBC_ctgs_1000111 | - | 333 | 306 | RNA→ rplS (PRK05338)Ribosomal_L19 (pfam01245)→ |
| Fbr-1-1 | NC_016001.1 | - | 3443259 | 3443232 | RNA→ rplS (PRK05338)Ribosomal_L19 (pfam01245)→ |
| env-1062 | BS_KBA_SWE02_21m_10304347 | + | 145 | 172 | RNA→ rplS (PRK05338)Ribosomal_L19 (pfam01245)→ |
| env-1063 | P_1C_Liq_1_UnCty_1096570 | - | 282 | 255 | RNA→ rplS (PRK05338)Ribosomal_L19 (pfam01245)→ |
| env-1064 | JGI25910J50241_10412907 | - | 135 | 108 | RNA→ hypo→ |
| env-1065 | JGI25920J50251_10025281 | + | 980 | 1007 | RNA→ rplS (PRK05338)Ribosomal_L19 (pfam01245)→ |
| env-1066 | GXP7IEG01AS1FF | - | 340 | 313 | RNA→ rplS (PRK05338)Ribosomal_L19 (pfam01245)→ |
| env-1067 | GXP7IEG01AL3G3 | + | 149 | 176 | RNA→ |
| env-1068 | ElkS_mat_MD6A_1173344 | + | 220 | 247 | RNA→ |
| env-1069 | JGI11876J14442_10000405 | - | 30539 | 30512 | RNA→ rplS (PRK05338)Ribosomal_L19 (pfam01245)→ |
| Iha-1-1 | NZ_AJJU01000006.1 | - | 7932 | 7905 | RNA→ rplS (PRK05338)Ribosomal_L19 (pfam01245)→ rplS (PRK05338)Ribosomal_L19 (pfam01245)→ |
| env-1070 | CEQG01145189.1 | + | 73521 | 73548 | RNA→ rplS (PRK05338)Ribosomal_L19 (pfam01245)→ |
| env-1071 | CETR01067878.1 | + | 322 | 349 | RNA→ rplS (PRK05338)Ribosomal_L19 (pfam01245)→ |
| env-1072 | CERQ01241836.1 | + | 41 | 68 | RNA→ rplS (PRK05338)Ribosomal_L19 (pfam01245)→ |
| env-1073 | SA_S1_NOR08_45m_c10570168 | + | 216 | 243 | RNA→ |
| env-1074 | SA_S1_NOR08_45m_c10657421 | - | 214 | 187 | RNA→ rplS (PRK05338)Ribosomal_L19 (pfam01245)→ |
| Fjo-1-1 | NC_009441.1 | - | 2640521 | 2640494 | RNA→ rplS (PRK05338)Ribosomal_L19 (pfam01245)→ |
| env-1075 | KGI_S2_ANT06_2345m_c1002444 | + | 3037 | 3064 | RNA→ rplS (PRK05338)Ribosomal_L19 (pfam01245)→ |
| env-1076 | BS_KBA_SWE12_21m_c10001565 | - | 7677 | 7650 | RNA→ rplS (PRK05338)Ribosomal_L19 (pfam01245)→ |
| env-1077 | BS_KBB_SWE26_205m_c1000550 | - | 7615 | 7588 | RNA→ rplS (PRK05338)Ribosomal_L19 (pfam01245)→ |
| env-1078 | BS_KBA_SWE21_205m_10000090 | + | 19045 | 19072 | RNA→ rplS (PRK05338)Ribosomal_L19 (pfam01245)→ |
| env-1079 | BS_KBA_SWE02_21m_10000113 | - | 8549 | 8522 | RNA→ rplS (PRK05338)Ribosomal_L19 (pfam01245)→ |
| env-1080 | SL_9KL_010_SED_10011059 | + | 964 | 991 | RNA→ rplS (PRK05338)Ribosomal_L19 (pfam01245)→ |
| env-1081 | BS_KBA_SWE12_21m_c10000952 | + | 3392 | 3419 | RNA→ ←hypo |
| env-1082 | BS_KBB_SWE26_205m_c1003233 | - | 1828 | 1801 | RNA→ rplS (PRK05338)Ribosomal_L19 (pfam01245)→ |
| env-1083 | BS_KBA_SWE02_21m_10008514 | + | 1890 | 1917 | RNA→ ←hypo |
| env-1084 | LWSO_GLQAYWI01D9JTG | + | 68 | 95 | RNA→ rplS (PRK05338)Ribosomal_L19 (pfam01245)→ |
| Sxi-1-1 | NZ_AUIG01000022.1 | - | 52581 | 52554 | RNA→ rplS (PRK05338)Ribosomal_L19 (pfam01245)→ |
| env-1085 | JGI25910J50241_10309536 | - | 48 | 21 | RNA→ |
| env-1086 | JGI25920J50251_10000309 | - | 8552 | 8525 | RNA→ rplS (PRK05338)Ribosomal_L19 (pfam01245)→ |
| env-1087 | JGI25914J50564_10400042 | + | 203 | 230 | RNA→ ←hypo |
| env-1088 | JGI24770J26754_10001491 | + | 8389 | 8416 | RNA→ rplS (PRK05338)Ribosomal_L19 (pfam01245)→ |
| env-1089 | JGI25917J50250_1002458 | + | 3566 | 3593 | RNA→ rplS (PRK05338)Ribosomal_L19 (pfam01245)→ |
| env-1090 | JGI25914J50564_10000033 | - | 2824 | 2797 | RNA→ rplS (PRK05338)Ribosomal_L19 (pfam01245)→ |
| Gli-1-1 | NZ_JH594606.1 | + | 1012580 | 1012607 | RNA→ rplS (PRK05338)Ribosomal_L19 (pfam01245)→ rplS (PRK05338)Ribosomal_L19 (pfam01245)→ |
| env-1091 | SL_7KL_010_BRINE_10005701 | - | 3752 | 3725 | RNA→ rplS (PRK05338)Ribosomal_L19 (pfam01245)→ |
| env-1092 | JGI25920J50251_10215690 | - | 222 | 195 | RNA→ rplS (PRK05338)Ribosomal_L19 (pfam01245)→ |
| env-1093 | JGI25914J50564_10165301 | - | 359 | 332 | RNA→ rplS (PRK05338)Ribosomal_L19 (pfam01245)→ |
| env-1094 | JGI25914J50564_10198338 | - | 73 | 46 | RNA→ |
| env-1095 | JGI25913J50563_1230681 | - | 286 | 259 | RNA→ rplS (PRK05338)Ribosomal_L19 (pfam01245)→ |
| env-1096 | BS_KBA_SWE21_205m_10014821 | - | 1943 | 1916 | RNA→ rplS (PRK05338)Ribosomal_L19 (pfam01245)→ |
| env-1097 | JGI25912J50252_10330425 | - | 38 | 11 | RNA→ |
| env-1098 | CENH01045038.1 | + | 3 | 30 | RNA→ rplS (PRK05338)Ribosomal_L19 (pfam01245)→ |
| env-1099 | CETA01060825.1 | - | 2104 | 2077 | RNA→ rplS (PRK05338)Ribosomal_L19 (pfam01245)→ |
| env-1100 | CETB01301347.1 | + | 490 | 517 | RNA→ rplS (PRK05338)Ribosomal_L19 (pfam01245)→ |
| env-1101 | CETT01089007.1 | - | 2908 | 2881 | RNA→ rplS (PRK05338)Ribosomal_L19 (pfam01245)→ |
| env-1102 | CEVJ01064547.1 | + | 730 | 757 | RNA→ rplS (PRK05338)Ribosomal_L19 (pfam01245)→ |
| env-1103 | CEWI01163115.1 | + | 358 | 385 | RNA→ rplS (PRK05338)Ribosomal_L19 (pfam01245)→ |
| env-1104 | KGI_S1_ANT01_95m_c10228781 | - | 190 | 163 | RNA→ rplS (PRK05338)Ribosomal_L19 (pfam01245)→ |
| env-1105 | KGI_S1_ANT02_95m_c10005113 | + | 4522 | 4549 | RNA→ rplS (PRK05338)Ribosomal_L19 (pfam01245)→ |
| env-1106 | CERE01034022.1 | + | 347 | 374 | RNA→ rplS (PRK05338)Ribosomal_L19 (pfam01245)→ |
| env-1107 | CENS01098701.1 | - | 541 | 514 | RNA→ rplS (PRK05338)Ribosomal_L19 (pfam01245)→ |

| | | | | | |
|---|---|---|---|---|---|
| env-1108 | CEPZ01082641.1 | - | 112 | 85 | **RNA**→ |
| env-1109 | CEQY01232939.1 | - | 234 | 207 | **RNA**→ rplS (PRK05338)Ribosomal_L19 (pfam01245)→ |
| env-1110 | CERQ01070756.1 | - | 1060 | 1033 | **RNA**→ rplS (PRK05338)Ribosomal_L19 (pfam01245)→ |
| env-1111 | CERU01112183.1 | - | 9021 | 8994 | **RNA**→ rplS (PRK05338)Ribosomal_L19 (pfam01245)→ |
| env-1112 | CESC01027484.1 | + | 19541 | 19568 | **RNA**→ rplS (PRK05338)Ribosomal_L19 (pfam01245)→ |
| env-1113 | CEUN01369745.1 | + | 390 | 417 | **RNA**→ rplS (PRK05338)Ribosomal_L19 (pfam01245)→ |
| Lbl-1-1 | NZ_CH672395.1 | - | 1844571 | 1844544 | **RNA**→ rplS (PRK05338)Ribosomal_L19 (pfam01245)→ |
| Aag-1-1 | NZ_JH621256.1 | - | 89135 | 89108 | **RNA**→ rplS (PRK05338)Ribosomal_L19 (pfam01245)→ |
| Ead-1-1 | NZ_KB907548.1 | - | 10231 | 10204 | **RNA**→ rplS (PRK05338)Ribosomal_L19 (pfam01245)→ |
| env-1114 | TahiMoana_1000007 | - | 24341 | 24314 | **RNA**→ ←hypo |
| env-1115 | topACOD_FV90NF401A1PKH | - | 248 | 221 | **RNA**→ rplS (PRK05338)Ribosomal_L19 (pfam01245)→ |
| Lsp-2-1 | NZ_JYNQ01000001.1 | + | 1019925 | 1019952 | **RNA**→ rplS (PRK05338)Ribosomal_L19 (pfam01245)→ |
| env-1116 | CESE01078901.1 | + | 29392 | 29419 | **RNA**→ rplS (PRK05338)Ribosomal_L19 (pfam01245)→ |
| env-1117 | CEST01016161.1 | + | 29395 | 29422 | **RNA**→ rplS (PRK05338)Ribosomal_L19 (pfam01245)→ |
| Lsp-3-1 | NZ_JHZY01000002.1 | + | 1633784 | 1633811 | **RNA**→ rplS (PRK05338)Ribosomal_L19 (pfam01245)→ |
| env-1118 | CEQG01152154.1 | + | 3057 | 3084 | **RNA**→ rplS (PRK05338)Ribosomal_L19 (pfam01245)→ |
| env-1119 | CEVJ01105358.1 | - | 124 | 97 | **RNA**→ hypo→ |
| env-1120 | CEVO01225740.1 | - | 100 | 73 | **RNA**→ |
| env-1121 | JGI12097J13213_1000092 | + | 32172 | 32199 | **RNA**→ rplS (PRK05338)Ribosomal_L19 (pfam01245)→ |
| env-1122 | JGI11800J13271_1071213 | - | 166 | 139 | **RNA**→ rplS (PRK05338)Ribosomal_L19 (pfam01245)→ |
| env-1123 | CESO01260573.1 | + | 553 | 580 | **RNA**→ rplS (PRK05338)Ribosomal_L19 (pfam01245)→ |
| Lja-1-1 | NZ_JSWF01000016.1 | - | 442 | 415 | **RNA**→ rplS (PRK05338)Ribosomal_L19 (pfam01245)→ |
| Lsp-1-1 | NC_015638.1 | + | 16178 | 16205 | **RNA**→ rplS (PRK05338)Ribosomal_L19 (pfam01245)→ |
| env-1124 | CEPG01053707.1 | + | 643 | 670 | **RNA**→ |
| Pto-1-1 | NC_018721.1 | - | 3532395 | 3532368 | **RNA**→ rplS (PRK05338)Ribosomal_L19 (pfam01245)→ |
| Pgo-1-1 | NZ_APLF01000012.1 | - | 7252 | 7225 | **RNA**→ rplS (PRK05338)Ribosomal_L19 (pfam01245)→ |
| env-1125 | LV_Brine_h2_0102_1076663 | - | 420 | 393 | **RNA**→ rplS (PRK05338)Ribosomal_L19 (pfam01245)→ |
| env-1126 | JGI1221J11331_1011062 | - | 813 | 786 | **RNA**→ rplS (PRK05338)Ribosomal_L19 (pfam01245)→ |
| env-1127 | P_1C_Liq_1_UnCty_1029884 | - | 249 | 222 | **RNA**→ rplS (PRK05338)Ribosomal_L19 (pfam01245)→ |
| env-1128 | P_2C_Liq_2_UnCty_1118043 | + | 295 | 322 | **RNA**→ ←hypo |
| Ptr-1-1 | NZ_KB905345.1 | + | 151414 | 151441 | **RNA**→ rplS (PRK05338)Ribosomal_L19 (pfam01245)→ |
| env-1129 | SL_9KL_010_SED_10069806 | + | 146 | 173 | **RNA**→ rplS (PRK05338)Ribosomal_L19 (pfam01245)→ |
| env-1130 | P_2C_Liq_1_UnCty_1186584 | - | 277 | 250 | **RNA**→ rplS (PRK05338)Ribosomal_L19 (pfam01245)→ |
| env-1131 | P_1C_Liq_2_UnCty_1015821 | + | 305 | 332 | **RNA**→ rplS (PRK05338)Ribosomal_L19 (pfam01245)→ |
| env-1132 | JGI25910J50241_10010778 | - | 306 | 279 | **RNA**→ rplS (PRK05338)Ribosomal_L19 (pfam01245)→ |
| env-1133 | JGI25920J50251_10125176 | + | 96 | 123 | **RNA**→ rplS (PRK05338)Ribosomal_L19 (pfam01245)→ |
| env-1134 | CEPR01006195.1 | + | 270508 | 270535 | **RNA**→ rplS (PRK05338)Ribosomal_L19 (pfam01245)→ |
| env-1135 | CERJ01067812.1 | + | 321 | 348 | **RNA**→ rplS (PRK05338)Ribosomal_L19 (pfam01245)→ |
| env-1136 | JGI25911J50253_10012314 | + | 1397 | 1424 | **RNA**→ rplS (PRK05338)Ribosomal_L19 (pfam01245)→ |
| Gsp-4-1 | NZ_JQLP01000003.1 | - | 9409 | 9382 | **RNA**→ rplS (PRK05338)Ribosomal_L19 (pfam01245)→ |
| env-1137 | JGI24770J26754_10341917 | - | 46 | 19 | **RNA**→ ←hypo |
| env-1138 | JGI24770J26754_10005581 | + | 5539 | 5566 | **RNA**→ rplS (PRK05338)Ribosomal_L19 (pfam01245)→ |
| env-1139 | JGI24723J26617_10022857 | + | 2227 | 2254 | **RNA**→ rplS (PRK05338)Ribosomal_L19 (pfam01245)→ |
| env-1140 | M3P_c10140209 | - | 55 | 28 | **RNA**→ ←hypo |
| env-1141 | JGI1221J11331_1121684 | - | 169 | 142 | **RNA**→ rplS (PRK05338)Ribosomal_L19 (pfam01245)→ |
| env-1142 | JFJP01038398.1 | + | 17693 | 17720 | **RNA**→ rplS (PRK05338)Ribosomal_L19 (pfam01245)→ |
| env-1143 | JGI26054J46599_1167877 | + | 72 | 99 | **RNA**→ rplS (PRK05338)Ribosomal_L19 (pfam01245)→ |
| env-1144 | LWFCAnN_GO09JKT02FQCFO | + | 32 | 59 | **RNA**→ hypo→ rplS (PRK05338)Ribosomal_L19 (pfam01245)→ |
| env-1145 | LM20APR2010E_c18959 | - | 172 | 145 | **RNA**→ rplS (PRK05338)Ribosomal_L19 (pfam01245)→ |
| env-1146 | KGI_S1_ANT02_95m_c10072419 | - | 199 | 172 | **RNA**→ rplS (PRK05338)Ribosomal_L19 (pfam01245)→ |
| env-1147 | JGI1221J11331_1000410 | + | 13452 | 13479 | **RNA**→ rplS (PRK05338)Ribosomal_L19 (pfam01245)→ |
| env-1148 | JGI24770J26754_10209365 | + | 67 | 94 | **RNA**→ rplS (PRK05338)Ribosomal_L19 (pfam01245)→ |

| | | | | | |
|---|---|---|---|---|---|
| env-1149 | LWAeNNiAF_GBUVFP102GIQM4 | - | 107 | 80 | RNA→ ←hypo |
| env-1150 | GXP7IEG01DK1RD | + | 62 | 89 | RNA→ rplS (PRK05338)Ribosomal_L19 (pfam01245)→ |
| env-1151 | TB_FS08_3DRAFT_1076085 | - | 419 | 392 | RNA→ rplS (PRK05338)Ribosomal_L19 (pfam01245)→ |
| env-1152 | KGI_S1_ANT02_95m_c10077220 | + | 767 | 794 | RNA→ rplS (PRK05338)Ribosomal_L19 (pfam01245)→ |
| env-1153 | SR_TTP_S3_1011262 | - | 257 | 230 | RNA→ rplS (PRK05338)Ribosomal_L19 (pfam01245)→ |
| env-1154 | SR_TP_S13sol_1004373 | + | 1564 | 1591 | RNA→ rplS (PRK05338)Ribosomal_L19 (pfam01245)→ |
| env-1155 | CETR01086737.1 | - | 288 | 261 | RNA→ rplS (PRK05338)Ribosomal_L19 (pfam01245)→ |
| env-1156 | CEVJ01064574.1 | - | 35932 | 35905 | RNA→ rplS (PRK05338)Ribosomal_L19 (pfam01245)→ |
| env-1157 | CEVK01071768.1 | - | 2531 | 2504 | RNA→ rplS (PRK05338)Ribosomal_L19 (pfam01245)→ |
| env-1158 | CEVM01085059.1 | - | 146813 | 146786 | RNA→ rplS (PRK05338)Ribosomal_L19 (pfam01245)→ |
| env-1159 | CEVO01080562.1 | - | 4295 | 4268 | RNA→ rplS (PRK05338)Ribosomal_L19 (pfam01245)→ |
| env-1160 | CEVQ01352678.1 | + | 1996 | 2023 | RNA→ |
| env-1161 | JGI12421J11937_10000023 | - | 29464 | 29437 | RNA→ ←hypo |
| env-1162 | JGI25911J50253_10733752 | - | 166 | 139 | RNA→ hypo→ |
| env-1163 | _GVICOHK01DYU3D | - | 65 | 38 | RNA→ hypo→ |
| env-1164 | KGI_S1_ANT02_95m_c10073298 | + | 760 | 787 | RNA→ rplS (PRK05338)Ribosomal_L19 (pfam01245)→ |
| env-1165 | LV_Brine_h2_0102_1044952 | + | 61 | 88 | RNA→ rplS (PRK05338)Ribosomal_L19 (pfam01245)→ |
| env-1166 | PBR_1111292 | + | 3885 | 3912 | RNA→ rplS (PRK05338)Ribosomal_L19 (pfam01245)→ hypo→ hypo→ |
| env-1167 | JGI25910J50241_10023455 | + | 23 | 50 | RNA→ rplS (PRK05338)Ribosomal_L19 (pfam01245)→ |
| env-1168 | LWFCAn_GLO1YSU01DQOFC | - | 166 | 139 | RNA→ hypo→ |
| Fso-1-1 | NZ_AUGO01000009.1 | + | 122444 | 122471 | RNA→ rplS (PRK05338)Ribosomal_L19 (pfam01245)→ |
| env-1169 | TB_PC08_66_10000538 | + | 14971 | 14998 | RNA→ rplS (PRK05338)Ribosomal_L19 (pfam01245)→ |
| env-1170 | TB_GS10_10_10217433 | + | 384 | 411 | RNA→ rplS (PRK05338)Ribosomal_L19 (pfam01245)→ |
| env-1171 | TB_PC08_64_1080822 | - | 480 | 453 | RNA→ rplS (PRK05338)Ribosomal_L19 (pfam01245)→ |
| env-1172 | CESI01032264.1 | + | 1988 | 2015 | RNA→ rplS (PRK05338)Ribosomal_L19 (pfam01245)→ |
| env-1173 | CESU01010912.1 | + | 7776 | 7803 | RNA→ rplS (PRK05338)Ribosomal_L19 (pfam01245)→ |
| env-1174 | Antartic2_1044867 | - | 57 | 30 | RNA→ hypo→ |
| env-1175 | JGI20156J14371_10138076 | - | 258 | 231 | RNA→ rplS (PRK05338)Ribosomal_L19 (pfam01245)→ |
| env-1176 | JGI20154J14316_10002203 | - | 374 | 347 | RNA→ rplS (PRK05338)Ribosomal_L19 (pfam01245)→ |
| env-1177 | JGI20160J14292_10013217 | + | 567 | 594 | RNA→ rplS (PRK05338)Ribosomal_L19 (pfam01245)→ |
| env-1178 | JGI20153J14295_10047621 | - | 358 | 331 | RNA→ rplS (PRK05338)Ribosomal_L19 (pfam01245)→ |
| env-1179 | JGI20153J14318_10047621 | - | 358 | 331 | RNA→ rplS (PRK05338)Ribosomal_L19 (pfam01245)→ |
| env-1180 | JGI20157J14317_10000793 | - | 151 | 124 | RNA→ rplS (PRK05338)Ribosomal_L19 (pfam01245)→ |
| env-1181 | CERM01033640.1 | + | 3672 | 3699 | RNA→ |
| env-1182 | CESO01224126.1 | - | 67 | 40 | RNA→ |
| env-1183 | CERM01182019.1 | + | 636 | 663 | RNA→ rplS (PRK05338)Ribosomal_L19 (pfam01245)→ |
| env-1184 | CESO01277244.1 | - | 422 | 395 | RNA→ rplS (PRK05338)Ribosomal_L19 (pfam01245)→ |
| env-1185 | DelMOSum2010_c10004802 | + | 3737 | 3764 | RNA→ rplS (PRK05338)Ribosomal_L19 (pfam01245)→ |
| env-1186 | LPaug08P2610m_c1071471 | - | 214 | 187 | RNA→ rplS (PRK05338)Ribosomal_L19 (pfam01245)→ |
| env-1187 | JGI20154J14316_10001372 | - | 183 | 156 | RNA→ rplS (PRK05338)Ribosomal_L19 (pfam01245)→ |
| env-1188 | JGI20153J14295_10000027 | - | 965 | 938 | RNA→ rplS (PRK05338)Ribosomal_L19 (pfam01245)→ |
| env-1189 | JGI20153J14318_10000027 | - | 965 | 938 | RNA→ rplS (PRK05338)Ribosomal_L19 (pfam01245)→ |
| env-1190 | JGI20160J14292_10003823 | + | 1395 | 1422 | RNA→ rplS (PRK05338)Ribosomal_L19 (pfam01245)→ |
| env-1191 | DelMOSum2010_c10005021 | - | 5906 | 5879 | RNA→ rplS (PRK05338)Ribosomal_L19 (pfam01245)→ |
| env-1192 | LPaug09P1610m_c1163453 | - | 115 | 88 | RNA→ hypo→ |
| env-1193 | JGI20154J14316_10016259 | - | 2513 | 2486 | RNA→ rplS (PRK05338)Ribosomal_L19 (pfam01245)→ |
| env-1194 | JGI20153J14295_10112779 | - | 627 | 600 | RNA→ rplS (PRK05338)Ribosomal_L19 (pfam01245)→ |
| env-1195 | JGI20153J14318_10112779 | - | 627 | 600 | RNA→ rplS (PRK05338)Ribosomal_L19 (pfam01245)→ |
| env-1196 | JGI20157J14317_10641510 | - | 167 | 140 | RNA→ rplS (PRK05338)Ribosomal_L19 (pfam01245)→ |
| env-1197 | JGI20154J14316_10003287 | + | 9143 | 9170 | RNA→ rplS (PRK05338)Ribosomal_L19 (pfam01245)→ |
| env-1198 | JGI20157J14317_10237281 | - | 352 | 325 | RNA→ rplS (PRK05338)Ribosomal_L19 (pfam01245)→ |
| env-1199 | CESF01069914.1 | + | 3709 | 3736 | RNA→ rplS (PRK05338)Ribosomal_L19 (pfam01245)→ |

| | | | | | |
|---|---|---|---|---|---|
| env-1200 | 3300001197_1005977 | + | 3403 | 3430 | **RNA**→ rplS (PRK05338)Ribosomal_L19 (pfam01245)→ |
| env-1201 | 3300001197_1015663 | + | 4060 | 4087 | **RNA**→ ←monomer_idh (TIGR00178)IDH (pfam03971) |
| env-1202 | JGI24723J26617_10020059 | - | 30 | 3 | **RNA**→ |
| env-1203 | DelMOSum2011_c10017080 | + | 3029 | 3056 | **RNA**→ rplS (PRK05338)Ribosomal_L19 (pfam01245)→ |
| env-1204 | A09P04_1300_Sequence0000012002 | + | 1343 | 1370 | **RNA**→ hypo→ |
| Apa-1-1 | NZ_JACC01000002.1 | + | 197695 | 197722 | **RNA**→ rplS (PRK05338)Ribosomal_L19 (pfam01245)→ |
| env-1205 | SI39nov09_120m_c1067562 | + | 410 | 437 | **RNA**→ rplS (PRK05338)Ribosomal_L19 (pfam01245)→ |
| env-1206 | CESI01135903.1 | - | 37 | 10 | **RNA**→ |
| env-1207 | CETR01240919.1 | + | 43 | 70 | **RNA**→ rplS (PRK05338)Ribosomal_L19 (pfam01245)→ hypo→ |
| env-1208 | JGI24723J26617_10024281 | - | 2550 | 2523 | **RNA**→ rplS (PRK05338)Ribosomal_L19 (pfam01245)→ |
| Fba-6-1 | NZ_AFPB01000181.1 | - | 41364 | 41337 | **RNA**→ rplS (PRK05338)Ribosomal_L19 (pfam01245)→ rplS (PRK05338)Ribosomal_L19 (pfam01245)→ |
| Aat-1-1 | NZ_AQRA01000009.1 | - | 19513 | 19486 | **RNA**→ rplS (PRK05338)Ribosomal_L19 (pfam01245)→ |
| env-1209 | SA_S1_NOR08_45m_c10624926 | - | 92 | 65 | **RNA**→ hypo→ |
| env-1210 | DelMOSum2010_c10001221 | + | 15564 | 15591 | **RNA**→ rplS (PRK05338)Ribosomal_L19 (pfam01245)→ |
| env-1211 | SI34jun09_10m_1058447 | + | 104 | 131 | **RNA**→ rplS (PRK05338)Ribosomal_L19 (pfam01245)→ |
| env-1212 | SI34jun09_100m_1143518 | + | 11 | 38 | **RNA**→ rplS (PRK05338)Ribosomal_L19 (pfam01245)→ |
| env-1213 | CERM01031204.1 | - | 3510 | 3483 | **RNA**→ rplS (PRK05338)Ribosomal_L19 (pfam01245)→ |
| env-1214 | CESO01003542.1 | - | 1110 | 1083 | **RNA**→ rplS (PRK05338)Ribosomal_L19 (pfam01245)→ |
| env-1215 | CEVO01049684.1 | - | 87 | 60 | **RNA**→ |
| env-1216 | 3300001197_1020214 | - | 102 | 75 | **RNA**→ ←hypo |
| env-1217 | JGI24724J26744_10266378 | - | 318 | 291 | **RNA**→ rplS (PRK05338)Ribosomal_L19 (pfam01245)→ |
| env-1218 | JGI24723J26617_10417056 | - | 210 | 183 | **RNA**→ rplS (PRK05338)Ribosomal_L19 (pfam01245)→ |
| Fag-1-1 | NZ_HG315671.1 | + | 2869946 | 2869973 | **RNA**→ rplS (PRK05338)Ribosomal_L19 (pfam01245)→ |
| env-1219 | SA_S1_NOR08_45m_c10036615 | - | 52 | 25 | **RNA**→ |
| env-1220 | CERM01065147.1 | - | 4841 | 4814 | **RNA**→ rplS (PRK05338)Ribosomal_L19 (pfam01245)→ |
| env-1221 | CESU01000186.1 | - | 4638 | 4611 | **RNA**→ rplS (PRK05338)Ribosomal_L19 (pfam01245)→ |
| env-1222 | CEWR01293092.1 | - | 4692 | 4665 | **RNA**→ rplS (PRK05338)Ribosomal_L19 (pfam01245)→ |
| env-1223 | JGI24723J26617_10804348 | - | 30 | 3 | **RNA**→ |
| Ame-1-1 | NZ_JACB01000004.1 | - | 19600 | 19573 | **RNA**→ rplS (PRK05338)Ribosomal_L19 (pfam01245)→ |
| env-1224 | DelMOSum2010_c10203686 | + | 55 | 82 | **RNA**→ rplS (PRK05338)Ribosomal_L19 (pfam01245)→ |
| env-1225 | DelMOWin2010_c10009783 | - | 4961 | 4934 | **RNA**→ rplS (PRK05338)Ribosomal_L19 (pfam01245)→ |
| env-1226 | JGI20158J14315_10383416 | + | 217 | 244 | **RNA**→ rplS (PRK05338)Ribosomal_L19 (pfam01245)→ |
| Fsp-3-1 | NZ_AJXL01000094.1 | - | 14222 | 14195 | **RNA**→ rplS (PRK05338)Ribosomal_L19 (pfam01245)→ |
| env-1227 | KGI_S1_ANT01_95m_c10053628 | + | 1010 | 1037 | **RNA**→ rplS (PRK05338)Ribosomal_L19 (pfam01245)→ |
| env-1228 | KGI_S2_ANT04_2345m_c1008646 | - | 1380 | 1353 | **RNA**→ rplS (PRK05338)Ribosomal_L19 (pfam01245)→ |
| env-1229 | KGI_S1_ANT02_95m_c10015121 | + | 1911 | 1938 | **RNA**→ rplS (PRK05338)Ribosomal_L19 (pfam01245)→ |
| env-1230 | KGI_S1_ANT01_95m_c10161490 | + | 312 | 339 | **RNA**→ rplS (PRK05338)Ribosomal_L19 (pfam01245)→ |
| env-1231 | JGI20157J14317_10035387 | + | 911 | 938 | **RNA**→ rplS (PRK05338)Ribosomal_L19 (pfam01245)→ PRK08299 (PRK08299)→ |
| env-1232 | JGI20157J14317_10082171 | + | 919 | 946 | **RNA**→ rplS (PRK05338)Ribosomal_L19 (pfam01245)→ |
| env-1233 | SRS018157_Baylor_scaffold_84048 | - | 8172 | 8145 | **RNA**→ rplS (PRK05338)Ribosomal_L19 (pfam01245)→ |
| | | | | | DEDDh (cd06127)GIY-YIG_UvrC_Cho (cd10434)EXOIII (smart00479)→ hypo→ |
| env-1234 | SRS051941_LANL_scaffold_40129 | + | 359 | 386 | **RNA**→ rplS (PRK05338)Ribosomal_L19 (pfam01245)→ |
| env-1235 | Buccal_mucosa_LANL_C64921346 | + | 256 | 283 | **RNA**→ |
| env-1236 | SRS052876_LANL_scaffold_53214 | - | 5451 | 5424 | **RNA**→ rplS (PRK05338)Ribosomal_L19 (pfam01245)→ |
| | | | | | DEDDh (cd06127)GIY-YIG_UvrC_Cho (cd10434)EXOIII (smart00479)→ hypo→ |
| env-1237 | SRS015044_WUGC_scaffold_8201 | + | 78 | 105 | **RNA**→ rplS (PRK05338)Ribosomal_L19 (pfam01245)→ |
| env-1238 | SRS063603_LANL_scaffold_47458 | + | 2882 | 2909 | **RNA**→ rplS (PRK05338)Ribosomal_L19 (pfam01245)→ |
| | | | | | DEDDh (cd06127)GIY-YIG_UvrC_Cho (cd10434)EXOIII (smart00479)→ hypo→ |
| env-1239 | SRS024087_C3545532 | + | 415 | 442 | **RNA**→ rplS (PRK05338)Ribosomal_L19 (pfam01245)→ DEDDh (cd06127)EXOIII (smart00479)→ |
| env-1240 | SRS063932_LANL_scaffold_104549 | + | 686 | 713 | **RNA**→ rplS (PRK05338)Ribosomal_L19 (pfam01245)→ |
| | | | | | DEDDh (cd06127)GIY-YIG_UvrC_Cho (cd10434)EXOIII (smart00479)→ hypo→ |
| env-1241 | SRS011126_Baylor_scaffold_17978 | + | 2727 | 2754 | **RNA**→ rplS (PRK05338)Ribosomal_L19 (pfam01245)→ |
| | | | | | DEDDh (cd06127)GIY-YIG_UvrC_Cho (cd10434)EXOIII (smart00479)→ hypo→ |

| | | | | | |
|---|---|---|---|---|---|
| env-1242 | SRS013252_C3667205 | + | 686 | 713 | **RNA**→ rplS (PRK05338)Ribosomal_L19 (pfam01245)→ |
| | | | | | DEDDh (cd06127)GIY-YIG_UvrC_Cho (cd10434)EXOIII (smart00479)→ |
| env-1243 | SRS013533_PGA_scaffold_50908 | + | 1008 | 1035 | **RNA**→ rplS (PRK05338)Ribosomal_L19 (pfam01245)→ |
| env-1244 | SRS015803_C2641419 | - | 493 | 466 | **RNA**→ rplS (PRK05338)Ribosomal_L19 (pfam01245)→ |
| env-1245 | SRS016360_Baylor_scaffold_62601 | + | 2151 | 2178 | **RNA**→ rplS (PRK05338)Ribosomal_L19 (pfam01245)→ |
| | | | | | DEDDh (cd06127)GIY-YIG_UvrC_Cho (cd10434)EXOIII (smart00479)→ hypo→ |
| env-1246 | SRS018665_WUGC_scaffold_58455 | + | 2151 | 2178 | **RNA**→ rplS (PRK05338)Ribosomal_L19 (pfam01245)→ |
| | | | | | DEDDh (cd06127)GIY-YIG_UvrC_Cho (cd10434)EXOIII (smart00479)→ hypo→ |
| env-1247 | SRS019128_WUGC_scaffold_49488 | + | 686 | 713 | **RNA**→ rplS (PRK05338)Ribosomal_L19 (pfam01245)→ |
| env-1248 | SRS024561_LANL_scaffold_36518 | - | 14316 | 14289 | **RNA**→ rplS (PRK05338)Ribosomal_L19 (pfam01245)→ |
| | | | | | DEDDh (cd06127)GIY-YIG_UvrC_Cho (cd10434)EXOIII (smart00479)→ hypo→ |
| env-1249 | SRS045197_C3607834 | - | 1029 | 1002 | **RNA**→ rplS (PRK05338)Ribosomal_L19 (pfam01245)→ DEDDh (cd06127)EXOIII (smart00479)→ |
| env-1250 | SRS047113_C5335430 | - | 293 | 266 | **RNA**→ rplS (PRK05338)Ribosomal_L19 (pfam01245)→ |
| env-1251 | SRS063999_C3543586 | + | 260 | 287 | **RNA**→ rplS (PRK05338)Ribosomal_L19 (pfam01245)→ |
| Csp-6-1 | NZ_KB291246.1 | + | 15364 | 15391 | **RNA**→ rplS (PRK05338)Ribosomal_L19 (pfam01245)→ |
| | | | | | DEDDh (cd06127)GIY-YIG_UvrC_Cho (cd10434)EXOIII (smart00479)→ hypo→ |
| env-1252 | SRS018157_Baylor_scaffold_18962 | + | 6563 | 6590 | **RNA**→ rplS (PRK05338)Ribosomal_L19 (pfam01245)→ |
| env-1253 | JGI24768J34885_10247260 | + | 433 | 460 | **RNA**→ |
| env-1254 | JCVI_SCAF_1101668160045 | + | 894 | 921 | **RNA**→ rplS (PRK05338)Ribosomal_L19 (pfam01245)→ |
| env-1255 | SwRhRL2b_contig_2171902 | + | 1146 | 1173 | **RNA**→ rplS (PRK05338)Ribosomal_L19 (pfam01245)→ |
| env-1256 | PBDCA2_GBB5CE401BK0D1_left | + | 27 | 54 | **RNA**→ |
| Fsu-1-1 | NZ_JATV01000027.1 | + | 3805 | 3832 | **RNA**→ rplS (PRK05338)Ribosomal_L19 (pfam01245)→ |
| env-1257 | JGI25913J50563_1001466 | + | 819 | 846 | **RNA**→ rplS (PRK05338)Ribosomal_L19 (pfam01245)→ |
| env-1258 | 3300000052_140975 | - | 3211 | 3184 | **RNA**→ rplS (PRK05338)Ribosomal_L19 (pfam01245)→ |
| env-1259 | G312J29652_10012531 | - | 1124 | 1097 | **RNA**→ rplS (PRK05338)Ribosomal_L19 (pfam01245)→ |
| env-1260 | KGI_S1_ANT02_95m_c10000047 | - | 11017 | 10990 | **RNA**→ rplS (PRK05338)Ribosomal_L19 (pfam01245)→ |
| env-1261 | JGI24770J26754_10206717 | + | 89 | 116 | **RNA**→ rplS (PRK05338)Ribosomal_L19 (pfam01245)→ |
| env-1262 | TFO_HY__Contig_1126 | + | 12214 | 12241 | **RNA**→ rplS (PRK05338)Ribosomal_L19 (pfam01245)→ |
| Msp-2-1 | NZ_JQNJ01000001.1 | + | 4092805 | 4092832 | **RNA**→ rplS (PRK05338)Ribosomal_L19 (pfam01245)→ |
| env-1263 | P_1C_Sed_3_UnCty_1133928 | - | 236 | 209 | **RNA**→ rplS (PRK05338)Ribosomal_L19 (pfam01245)→ |
| Fco-1-1 | NC_016510.2 | + | 2920503 | 2920530 | **RNA**→ rplS (PRK05338)Ribosomal_L19 (pfam01245)→ |
| Ala-1-1 | NZ_AUMK01000014.1 | - | 41792 | 41765 | **RNA**→ rplS (PRK05338)Ribosomal_L19 (pfam01245)→ |
| env-1264 | JGI24023J19991_10023870 | - | 1433 | 1406 | **RNA**→ rplS (PRK05338)Ribosomal_L19 (pfam01245)→ |
| env-1265 | JGI24724J26744_10048212 | + | 212 | 239 | **RNA**→ rplS (PRK05338)Ribosomal_L19 (pfam01245)→ |
| env-1266 | JGI24723J26617_10001125 | - | 19348 | 19321 | **RNA**→ rplS (PRK05338)Ribosomal_L19 (pfam01245)→ |
| env-1267 | TrichMG_c100001 | - | 14774 | 14747 | **RNA**→ rplS (PRK05338)Ribosomal_L19 (pfam01245)→ |
| Wps-1-1 | NZ_ATMR01000180.1 | + | 83347 | 83374 | **RNA**→ rplS (PRK05338)Ribosomal_L19 (pfam01245)→ |
| env-1268 | AglaG_GBVDVLI01B3NQO | - | 163 | 136 | **RNA**→ rplS (PRK05338)Ribosomal_L19 (pfam01245)→ |
| env-1269 | ElkS_mat_MD6A_1003775 | - | 3249 | 3222 | **RNA**→ ←hypo |
| env-1270 | JGI11876J14442_10004776 | - | 3630 | 3603 | **RNA**→ ←hypo |
| env-1271 | CERE01152256.1 | + | 629 | 656 | **RNA**→ |
| Mze-1-1 | NZ_AULQ01000006.1 | - | 14032 | 14005 | **RNA**→ rplS (PRK05338)Ribosomal_L19 (pfam01245)→ |
| env-1272 | JGI24723J26617_10002074 | - | 4096 | 4069 | **RNA**→ rplS (PRK05338)Ribosomal_L19 (pfam01245)→ |
| Gpo-1-1 | NZ_AUHF01000008.1 | + | 262596 | 262623 | **RNA**→ rplS (PRK05338)Ribosomal_L19 (pfam01245)→ |
| Oma-1-1 | NZ_JADR01000006.1 | - | 9045 | 9018 | **RNA**→ rplS (PRK05338)Ribosomal_L19 (pfam01245)→ |
| env-1273 | wf3_AHAI14839_x1 | + | 659 | 686 | **RNA**→ rplS (PRK05338)Ribosomal_L19 (pfam01245)→ |
| env-1274 | CEGD01004528.1 | - | 427 | 400 | **RNA**→ rplS (PRK05338)Ribosomal_L19 (pfam01245)→ |
| env-1275 | BS_KBA_SWE21_205m_10562486 | - | 229 | 202 | **RNA**→ rplS (PRK05338)Ribosomal_L19 (pfam01245)→ |
| env-1276 | wf3_Contig6796 | + | 1463 | 1490 | **RNA**→ rplS (PRK05338)Ribosomal_L19 (pfam01245)→ |
| env-1277 | CESF01053946.1 | - | 9020 | 8993 | **RNA**→ rplS (PRK05338)Ribosomal_L19 (pfam01245)→ |
| env-1278 | CESO01316884.1 | + | 263 | 290 | **RNA**→ rplS (PRK05338)Ribosomal_L19 (pfam01245)→ |
| env-1279 | JGI24724J26744_10002545 | + | 3020 | 3047 | **RNA**→ rplS (PRK05338)Ribosomal_L19 (pfam01245)→ |
| Fgi-1-1 | NZ_JNCP01000011.1 | + | 2791 | 2818 | **RNA**→ rplS (PRK05338)Ribosomal_L19 (pfam01245)→ |

| | | | | | |
|---|---|---|---|---|---|
| env-1280 | KGI_S1_ANT01_95m_c10010682 | + | 3821 | 3848 | **RNA**→ rplS (PRK05338)Ribosomal_L19 (pfam01245)→ |
| Fde-1-1 | NZ_KE384377.1 | - | 10971 | 10944 | **RNA**→ rplS (PRK05338)Ribosomal_L19 (pfam01245)→ |
| Fsp-8-1 | NZ_JQJY01000004.1 | + | 576152 | 576179 | **RNA**→ rplS (PRK05338)Ribosomal_L19 (pfam01245)→ |
| Ffr-2-1 | NZ_AHKF01000018.1 | - | 29890 | 29863 | **RNA**→ rplS (PRK05338)Ribosomal_L19 (pfam01245)→ rplS (PRK05338)Ribosomal_L19 (pfam01245)→ |
| Fsp-7-1 | NZ_AKZQ01000002.1 | - | 95707 | 95680 | **RNA**→ rplS (PRK05338)Ribosomal_L19 (pfam01245)→ rplS (PRK05338)Ribosomal_L19 (pfam01245)→ |
| Fch-1-1 | NZ_JASY01000001.1 | + | 715110 | 715137 | **RNA**→ rplS (PRK05338)Ribosomal_L19 (pfam01245)→ |
| Fsp-2-1 | NZ_LEKS01000006.1 | - | 39097 | 39070 | **RNA**→ rplS (PRK05338)Ribosomal_L19 (pfam01245)→ |
| Ffr-1-1 | NZ_AUDO01000011.1 | - | 15707 | 15680 | **RNA**→ rplS (PRK05338)Ribosomal_L19 (pfam01245)→ |
| Fre-1-1 | NZ_JPRL01000001.1 | + | 2564906 | 2564933 | **RNA**→ rplS (PRK05338)Ribosomal_L19 (pfam01245)→ |
| env-1281 | SA_S1_NOR08_45m_c10539000 | - | 115 | 88 | **RNA**→ hypo→ |
| Osp-1-1 | NZ_KI912445.1 | - | 8860 | 8833 | **RNA**→ rplS (PRK05338)Ribosomal_L19 (pfam01245)→ |
| Osp-2-1 | NZ_JADT01000012.1 | - | 9076 | 9049 | **RNA**→ rplS (PRK05338)Ribosomal_L19 (pfam01245)→ |
| env-1282 | BS_KBA_SWE12_21m_c10000785 | - | 8390 | 8363 | **RNA**→ ←hypo |
| env-1283 | BS_KBB_SWE26_205m_c1003525 | + | 489 | 516 | **RNA**→ ←hypo |
| env-1284 | BS_KBA_SWE21_205m_10009165 | + | 1230 | 1257 | **RNA**→ ←hypo |
| env-1285 | BS_KBA_SWE02_21m_10000834 | - | 8439 | 8412 | **RNA**→ ←hypo |
| Ssp-2-1 | NZ_JHZX01000001.1 | - | 3745903 | 3745876 | **RNA**→ rplS (PRK05338)Ribosomal_L19 (pfam01245)→ |
| env-1286 | CEUX01158018.1 | - | 198 | 171 | **RNA**→ rplS (PRK05338)Ribosomal_L19 (pfam01245)→ |
| Fsp-4-1 | NZ_JSYM01000007.1 | + | 139958 | 139985 | **RNA**→ rplS (PRK05338)Ribosomal_L19 (pfam01245)→ |
| Fsp-10-1 | NZ_JXQR01000051.1 | + | 159915 | 159942 | **RNA**→ rplS (PRK05338)Ribosomal_L19 (pfam01245)→ |
| env-1287 | KGI_S1_ANT01_95m_c10011075 | + | 3112 | 3139 | **RNA**→ rplS (PRK05338)Ribosomal_L19 (pfam01245)→ |
| env-1288 | KGI_S2_ANT04_2345m_c1004935 | - | 4516 | 4489 | **RNA**→ rplS (PRK05338)Ribosomal_L19 (pfam01245)→ |
| env-1289 | KGI_S1_ANT01_95m_c10002779 | - | 5614 | 5587 | **RNA**→ rplS (PRK05338)Ribosomal_L19 (pfam01245)→ |
| env-1290 | KGI_S2_ANT04_2345m_c1005141 | - | 4576 | 4549 | **RNA**→ rplS (PRK05338)Ribosomal_L19 (pfam01245)→ |
| Gsp-2-1 | NZ_APHJ01000024.1 | + | 94820 | 94847 | **RNA**→ rplS (PRK05338)Ribosomal_L19 (pfam01245)→ |
| Fsp-9-1 | NZ_JARP01000001.1 | - | 168990 | 168963 | **RNA**→ rplS (PRK05338)Ribosomal_L19 (pfam01245)→ |
| env-1291 | KGI_S1_ANT01_95m_c10032372 | + | 345 | 372 | **RNA**→ rplS (PRK05338)Ribosomal_L19 (pfam01245)→ |
| env-1292 | KGI_S2_ANT04_2345m_c1037950 | + | 215 | 242 | **RNA**→ rplS (PRK05338)Ribosomal_L19 (pfam01245)→ |
| env-1293 | ElkS_mat_MD6A_1029448 | - | 765 | 738 | **RNA**→ rplS (PRK05338)Ribosomal_L19 (pfam01245)→ |
| env-1294 | JGI11876J14442_10227469 | - | 312 | 285 | **RNA**→ rplS (PRK05338)Ribosomal_L19 (pfam01245)→ |
| env-1295 | JGI11876J14442_10329633 | - | 316 | 289 | **RNA**→ rplS (PRK05338)Ribosomal_L19 (pfam01245)→ |
| env-1296 | JGI24770J26754_10195704 | - | 676 | 649 | **RNA**→ rplS (PRK05338)Ribosomal_L19 (pfam01245)→ |
| env-1297 | KGI_S1_ANT01_95m_c10062448 | + | 418 | 445 | **RNA**→ rplS (PRK05338)Ribosomal_L19 (pfam01245)→ |
| env-1298 | KGI_S1_ANT01_95m_c10003247 | + | 7447 | 7474 | **RNA**→ rplS (PRK05338)Ribosomal_L19 (pfam01245)→ |
| env-1299 | KGI_S2_ANT04_2345m_c1001347 | + | 8040 | 8067 | **RNA**→ rplS (PRK05338)Ribosomal_L19 (pfam01245)→ |
| env-1300 | KGI_S1_ANT01_95m_c10708988 | + | 201 | 228 | **RNA**→ |
| env-1301 | KGI_S1_ANT01_95m_c10208214 | - | 419 | 392 | **RNA**→ rplS (PRK05338)Ribosomal_L19 (pfam01245)→ |
| env-1302 | JGI24770J26754_10790136 | - | 258 | 231 | **RNA**→ rplS (PRK05338)Ribosomal_L19 (pfam01245)→ |
| env-1303 | LWAnN_GIDYKCY01DFNMC | + | 24 | 51 | **RNA**→ rplS (PRK05338)Ribosomal_L19 (pfam01245)→ |
| Fhi-1-1 | NZ_JPRK01000017.1 | - | 43000 | 42973 | **RNA**→ rplS (PRK05338)Ribosomal_L19 (pfam01245)→ |
| Gsp-3-1 | NZ_AJLT01000006.1 | - | 8930 | 8903 | **RNA**→ rplS (PRK05338)Ribosomal_L19 (pfam01245)→ rplS (PRK05338)Ribosomal_L19 (pfam01245)→ |
| env-1304 | ARMCC_Contig_357991 | + | 153 | 180 | **RNA**→ rplS (PRK05338)Ribosomal_L19 (pfam01245)→ |
| Fsp-6-1 | NZ_AKJZ01000008.1 | + | 64241 | 64268 | **RNA**→ rplS (PRK05338)Ribosomal_L19 (pfam01245)→ rplS (PRK05338)Ribosomal_L19 (pfam01245)→ |
| env-1305 | JGI24770J26754_10441296 | + | 303 | 330 | **RNA**→ |
| env-1306 | JGI25920J50251_10022691 | + | 1368 | 1395 | **RNA**→ rplS (PRK05338)Ribosomal_L19 (pfam01245)→ |
| Gsp-5-1 | NZ_KE384218.1 | - | 1128381 | 1128354 | **RNA**→ rplS (PRK05338)Ribosomal_L19 (pfam01245)→ |
| Fsp-11-1 | NZ_AUEU01000005.1 | - | 10099 | 10072 | **RNA**→ rplS (PRK05338)Ribosomal_L19 (pfam01245)→ |
| Fsp-1-1 | NZ_JQMS01000001.1 | + | 3749524 | 3749551 | **RNA**→ rplS (PRK05338)Ribosomal_L19 (pfam01245)→ |
| env-1307 | KVRMV2_102036113 | - | 475 | 448 | **RNA**→ rplS (PRK05338)Ribosomal_L19 (pfam01245)→ |
| Psp-3-1 | NC_020830.1 | + | 933043 | 933070 | **RNA**→ rplS (PRK05338)Ribosomal_L19 (pfam01245)→ |
| env-1308 | CEOF01395880.1 | - | 44 | 17 | **RNA**→ |
| env-1309 | SI60aug11_200m_c1101747 | - | 140 | 113 | **RNA**→ hypo→ |

| | | | | | |
|---|---|---|---|---|---|
| env-1310 | P_1C_Liq_1_UnCty_1069070 | + | 219 | 246 | **RNA**→ rplS (PRK05338)Ribosomal_L19 (pfam01245)→ |
| Gme-1-1 | NZ_AUHD01000012.1 | - | 8795 | 8768 | **RNA**→ rplS (PRK05338)Ribosomal_L19 (pfam01245)→ |
| env-1311 | KVRMV2_102150733 | + | 448 | 475 | **RNA**→ |
| env-1312 | ElkS_mat_CD2A_1245275 | - | 90 | 63 | **RNA**→ ←hypo |
| env-1313 | JGI11876J14442_10000066 | - | 15974 | 15947 | **RNA**→ rplS (PRK05338)Ribosomal_L19 (pfam01245)→ |
| env-1314 | JGI25317J35169_1088507 | - | 268 | 241 | **RNA**→ rplS (PRK05338)Ribosomal_L19 (pfam01245)→ |
| env-1315 | JGI25321J39296_1089961 | + | 165 | 192 | **RNA**→ rplS (PRK05338)Ribosomal_L19 (pfam01245)→ |
| env-1316 | JGI25321J50212_10001726 | - | 4881 | 4854 | **RNA**→ rplS (PRK05338)Ribosomal_L19 (pfam01245)→ |
| Min-1-1 | NZ_BAEX01000008.1 | - | 31090 | 31063 | **RNA**→ rplS (PRK05338)Ribosomal_L19 (pfam01245)→ rplS (PRK05338)Ribosomal_L19 (pfam01245)→ |
| env-1317 | CEVI01222985.1 | + | 1040 | 1067 | **RNA**→ rplS (PRK05338)Ribosomal_L19 (pfam01245)→ |
| env-1318 | JGI11876J14442_10220788 | + | 165 | 192 | **RNA**→ rplS (PRK05338)Ribosomal_L19 (pfam01245)→ |
| env-1319 | ElkS_mat_MD6A_1191826 | + | 14 | 41 | **RNA**→ rplS (PRK05338)Ribosomal_L19 (pfam01245)→ |
| env-1320 | CENM01024442.1 | - | 460 | 433 | **RNA**→ rplS (PRK05338)Ribosomal_L19 (pfam01245)→ |
| Ssp-1-1 | NZ_JQNQ01000001.1 | - | 3054097 | 3054070 | **RNA**→ rplS (PRK05338)Ribosomal_L19 (pfam01245)→ |
| env-1321 | 3300001605_10299261 | + | 678 | 705 | **RNA**→ rplS (PRK05338)Ribosomal_L19 (pfam01245)→ |
| Fsu-2-1 | NZ_AUCZ01000004.1 | - | 149796 | 149769 | **RNA**→ rplS (PRK05338)Ribosomal_L19 (pfam01245)→ hypo→ monomer_idh (TIGR00178)IDH (pfam03971)→ |
| Fsu-2-2 | NZ_JRLW01000001.1 | - | 149690 | 149663 | **RNA**→ rplS (PRK05338)Ribosomal_L19 (pfam01245)→ hypo→ monomer_idh (TIGR00178)IDH (pfam03971)→ |
| env-1322 | PBR_1002168 | + | 6973 | 7000 | **RNA**→ rplS (PRK05338)Ribosomal_L19 (pfam01245)→ |
| env-1323 | ElkS_mat_CD2A_1001393 | - | 6905 | 6878 | **RNA**→ rplS (PRK05338)Ribosomal_L19 (pfam01245)→ |
| env-1324 | ElkS_mat_CD2A_1002187 | - | 557 | 530 | **RNA**→ rplS (PRK05338)Ribosomal_L19 (pfam01245)→ |
| env-1325 | JGI11876J14442_10004185 | - | 6445 | 6418 | **RNA**→ rplS (PRK05338)Ribosomal_L19 (pfam01245)→ |
| env-1326 | CENR01035721.1 | + | 297 | 324 | **RNA**→ rplS (PRK05338)Ribosomal_L19 (pfam01245)→ |
| env-1327 | 2200738958 | - | 695 | 668 | **RNA**→ rplS (PRK05338)Ribosomal_L19 (pfam01245)→ |
| Mze-2-1 | NZ_AFOE01000017.1 | - | 24032 | 24005 | **RNA**→ rplS (PRK05338)Ribosomal_L19 (pfam01245)→ rplS (PRK05338)Ribosomal_L19 (pfam01245)→ |
| env-1328 | P_2C_Liq_1_UnCty_1043821 | - | 45 | 18 | **RNA**→ |
| Myu-1-1 | NZ_JPFK01000002.1 | + | 125744 | 125771 | **RNA**→ rplS (PRK05338)Ribosomal_L19 (pfam01245)→ |
| Gsa-1-1 | NZ_AUDV01000007.1 | - | 8102 | 8075 | **RNA**→ rplS (PRK05338)Ribosomal_L19 (pfam01245)→ |
| Amu-1-1 | NZ_AUML01000029.1 | + | 59901 | 59928 | **RNA**→ rplS (PRK05338)Ribosomal_L19 (pfam01245)→ |
| Fen-1-1 | NZ_AVCS01000013.1 | + | 115329 | 115356 | **RNA**→ rplS (PRK05338)Ribosomal_L19 (pfam01245)→ |
| Fen-1-2 | NZ_JRLZ01000009.1 | - | 140934 | 140907 | **RNA**→ rplS (PRK05338)Ribosomal_L19 (pfam01245)→ |
| Fli-1-1 | NZ_AVGG01000005.1 | + | 168606 | 168633 | **RNA**→ rplS (PRK05338)Ribosomal_L19 (pfam01245)→ |
| Fca-1-1 | NZ_AVBI01000004.1 | + | 53904 | 53931 | **RNA**→ rplS (PRK05338)Ribosomal_L19 (pfam01245)→ |
| Fca-1-2 | NZ_JRLU01000011.1 | - | 41550 | 41523 | **RNA**→ rplS (PRK05338)Ribosomal_L19 (pfam01245)→ |
| Fsa-1-1 | NZ_AVFO01000024.1 | - | 41542 | 41515 | **RNA**→ rplS (PRK05338)Ribosomal_L19 (pfam01245)→ |
| Fse-1-1 | NZ_JNCA01000006.1 | - | 95236 | 95209 | **RNA**→ rplS (PRK05338)Ribosomal_L19 (pfam01245)→ |
| Fda-1-1 | NZ_AUDK01000004.1 | - | 79117 | 79090 | **RNA**→ rplS (PRK05338)Ribosomal_L19 (pfam01245)→ |
| Dsp-2-1 | NZ_CM001837.1 | - | 2216219 | 2216192 | **RNA**→ rplS (PRK05338)Ribosomal_L19 (pfam01245)→ |
| env-1329 | P_1C_Liq_1_UnCty_1060928 | + | 197 | 224 | **RNA**→ rplS (PRK05338)Ribosomal_L19 (pfam01245)→ |
| env-1330 | P_2C_Liq_1_UnCty_1114980 | - | 96 | 69 | **RNA**→ hypo→ |
| env-1331 | P_2C_Liq_2_UnCty_1069521 | - | 266 | 239 | **RNA**→ rplS (PRK05338)Ribosomal_L19 (pfam01245)→ |
| env-1332 | JGI26524J50256_1000640 | + | 40528 | 40555 | **RNA**→ rplS (PRK05338)Ribosomal_L19 (pfam01245)→ |
| env-1333 | JGI26523J50269_1000212 | - | 2723 | 2696 | **RNA**→ rplS (PRK05338)Ribosomal_L19 (pfam01245)→ |
| env-1334 | JGI26534J51046_1001256 | + | 19707 | 19734 | **RNA**→ rplS (PRK05338)Ribosomal_L19 (pfam01245)→ |
| env-1335 | NCBI_READ_1112777305773 | - | 253 | 226 | **RNA**→ rplS (PRK05338)Ribosomal_L19 (pfam01245)→ |
| env-1336 | PBR_1158914 | + | 213 | 240 | **RNA**→ |
| env-1337 | JGI24768J34885_10254432 | + | 396 | 423 | **RNA**→ rplS (PRK05338)Ribosomal_L19 (pfam01245)→ |
| env-1338 | JGI25913J50563_1000388 | - | 6034 | 6007 | **RNA**→ rplS (PRK05338)Ribosomal_L19 (pfam01245)→ |
| Faq-1-1 | NZ_JRHH01000003.1 | - | 440802 | 440775 | **RNA**→ rplS (PRK05338)Ribosomal_L19 (pfam01245)→ |
| env-1339 | CEUK01343367.1 | - | 1065 | 1038 | **RNA**→ rplS (PRK05338)Ribosomal_L19 (pfam01245)→ |
| env-1340 | JGI24770J26754_10099893 | + | 564 | 591 | **RNA**→ rplS (PRK05338)Ribosomal_L19 (pfam01245)→ |
| env-1341 | JGI25910J50241_10080562 | + | 57 | 84 | **RNA**→ rplS (PRK05338)Ribosomal_L19 (pfam01245)→ |
| env-1342 | CESF01139250.1 | - | 371 | 344 | **RNA**→ rplS (PRK05338)Ribosomal_L19 (pfam01245)→ |

| | | | | | |
|---|---|---|---|---|---|
| Zam-1-1 | NZ_AYXY01000014.1 | - | 3873 | 3846 | **RNA**→ rplS (PRK05338)Ribosomal_L19 (pfam01245)→ |
| Gec-1-1 | NZ_AUHG01000016.1 | - | 9590 | 9563 | **RNA**→ rplS (PRK05338)Ribosomal_L19 (pfam01245)→ |
| env-1343 | CEOS01230303.1 | - | 1375 | 1348 | **RNA**→ rplS (PRK05338)Ribosomal_L19 (pfam01245)→ |
| env-1344 | 2227004913 | + | 161 | 197 | **RNA**→ rplS (PRK05338)Ribosomal_L19 (pfam01245)→ |
| env-1345 | IMNBL1_c0000763 | - | 5212 | 5176 | **RNA**→ rplS (PRK05338)Ribosomal_L19 (pfam01245)→ |
| env-1346 | JGI996J12353_1000635 | - | 5278 | 5242 | **RNA**→ rplS (PRK05338)Ribosomal_L19 (pfam01245)→ |
| env-1347 | TB03JUN2009E__Contig_140630 | - | 66 | 39 | **RNA**→ |
| env-1348 | JGI24770J26754_10070249 | + | 144 | 171 | **RNA**→ rplS (PRK05338)Ribosomal_L19 (pfam01245)→ |
| env-1349 | JGI24025J20009_10007707 | - | 2829 | 2802 | **RNA**→ rplS (PRK05338)Ribosomal_L19 (pfam01245)→ |
| env-1350 | JGI24724J26744_10044739 | - | 1715 | 1688 | **RNA**→ rplS (PRK05338)Ribosomal_L19 (pfam01245)→ |
| env-1351 | TB_GS09_5_10015365 | - | 2511 | 2484 | **RNA**→ rplS (PRK05338)Ribosomal_L19 (pfam01245)→ |
| env-1352 | KGI_S1_ANT01_95m_c10107932 | + | 100 | 127 | **RNA**→ rplS (PRK05338)Ribosomal_L19 (pfam01245)→ |
| env-1353 | SA_S1_NOR08_45m_c10041987 | + | 169 | 196 | **RNA**→ rplS (PRK05338)Ribosomal_L19 (pfam01245)→ |
| env-1354 | KGI_S2_ANT04_2345m_c1369759 | - | 184 | 157 | **RNA**→ rplS (PRK05338)Ribosomal_L19 (pfam01245)→ |
| env-1355 | KGI_S2_ANT05_2345m_c1099771 | + | 146 | 173 | **RNA**→ rplS (PRK05338)Ribosomal_L19 (pfam01245)→ |
| env-1356 | KGI_S1_ANT02_95m_c10047434 | - | 1222 | 1195 | **RNA**→ hypo→ rplS (PRK05338)Ribosomal_L19 (pfam01245)→ |
| env-1357 | SA_S1_NOR05_45m_c10449740 | - | 163 | 136 | **RNA**→ hypo→ rplS (PRK05338)Ribosomal_L19 (pfam01245)→ |
| env-1358 | SA_S2_NOR15_50m_c10770364 | + | 118 | 145 | **RNA**→ rplS (PRK05338)Ribosomal_L19 (pfam01245)→ |
| env-1359 | JGI20156J14371_10179980 | + | 85 | 112 | **RNA**→ rplS (PRK05338)Ribosomal_L19 (pfam01245)→ |
| env-1360 | TDF_MC_ARG01_113m_c1038862 | - | 280 | 253 | **RNA**→ rplS (PRK05338)Ribosomal_L19 (pfam01245)→ |
| env-1361 | CESO01105902.1 | + | 2141 | 2168 | **RNA**→ rplS (PRK05338)Ribosomal_L19 (pfam01245)→ Icd (COG0538)PRK08299 (PRK08299)→<br>murQ (PRK05441)SIS_Etherase (cd05007)→ hypo→ TGS_ThrRS_N (cd01667)ThrS (COG0441)thrS (PRK00413)→ |
| env-1362 | CETY01197370.1 | + | 892 | 919 | **RNA**→ rplS (PRK05338)Ribosomal_L19 (pfam01245)→ Icd (COG0538)PRK08299 (PRK08299)→<br>murQ (PRK05441)SIS_Etherase (cd05007)→ hypo→ TGS_ThrRS_N (cd01667)ThrS (COG0441)thrS (PRK00413)→ |
| env-1363 | CEVX01010802.1 | + | 2158 | 2185 | **RNA**→ |
| env-1364 | DelMOWin2010_c10186470 | + | 427 | 454 | **RNA**→ rplS (PRK05338)Ribosomal_L19 (pfam01245)→ |
| env-1365 | CEWO01359685.1 | + | 828 | 855 | **RNA**→ rplS (PRK05338)Ribosomal_L19 (pfam01245)→ |
| env-1366 | DelMOSum2010_c10245447 | - | 220 | 193 | **RNA**→ rplS (PRK05338)Ribosomal_L19 (pfam01245)→ |
| env-1367 | JCVI_SCAF_1101668030402 | + | 742 | 769 | **RNA**→ rplS (PRK05338)Ribosomal_L19 (pfam01245)→ PRK08299 (PRK08299)→ |
| env-1368 | JCVI_SCAF_1096627111654 | + | 201 | 228 | **RNA**→ rplS (PRK05338)Ribosomal_L19 (pfam01245)→ PRK08299 (PRK08299)→ |
| env-1369 | SA_S1_NOR08_45m_c10116776 | - | 500 | 473 | **RNA**→ rplS (PRK05338)Ribosomal_L19 (pfam01245)→ |
| env-1370 | CENP01239202.1 | - | 14739 | 14712 | **RNA**→ rplS (PRK05338)Ribosomal_L19 (pfam01245)→ PRK08299 (PRK08299)→ |
| env-1371 | CENX01155294.1 | - | 392 | 365 | **RNA**→ rplS (PRK05338)Ribosomal_L19 (pfam01245)→ |
| env-1372 | CEOF01367002.1 | - | 392 | 365 | **RNA**→ rplS (PRK05338)Ribosomal_L19 (pfam01245)→ |
| env-1373 | CEOK01333021.1 | + | 503 | 530 | **RNA**→ rplS (PRK05338)Ribosomal_L19 (pfam01245)→ |
| Jma-1-1 | NZ_JH651379.1 | - | 2994278 | 2994251 | **RNA**→ rplS (PRK05338)Ribosomal_L19 (pfam01245)→ rplS (PRK05338)Ribosomal_L19 (pfam01245)→ |
| env-1374 | SRS043422_LANL_scaffold_17386 | - | 316 | 289 | **RNA**→ rplS (PRK05338)Ribosomal_L19 (pfam01245)→ |
| env-1375 | SRS015947_WUGC_scaffold_22634 | + | 8860 | 8887 | **RNA**→ rplS (PRK05338)Ribosomal_L19 (pfam01245)→ |
| env-1376 | SRS055450_LANL_scaffold_26813 | - | 98 | 71 | **RNA**→ |
| Csp-1-1 | NZ_ABZV01000006.1 | - | 117415 | 117388 | **RNA**→ rplS (PRK05338)Ribosomal_L19 (pfam01245)→ rplS (PRK05338)Ribosomal_L19 (pfam01245)→ |
| env-1377 | KGI_S1_ANT01_95m_c10119168 | - | 553 | 526 | **RNA**→ rplS (PRK05338)Ribosomal_L19 (pfam01245)→ |
| env-1378 | KGI_S2_ANT04_2345m_c1039681 | - | 468 | 441 | **RNA**→ rplS (PRK05338)Ribosomal_L19 (pfam01245)→ |
| env-1379 | KGI_S1_ANT02_95m_c10037526 | + | 1078 | 1105 | **RNA**→ rplS (PRK05338)Ribosomal_L19 (pfam01245)→ |
| env-1380 | KVRMV2_100600626 | + | 510 | 537 | **RNA**→ rplS (PRK05338)Ribosomal_L19 (pfam01245)→ |
| env-1381 | KGI_S2_ANT05_2345m_c1283145 | - | 206 | 179 | **RNA**→ rplS (PRK05338)Ribosomal_L19 (pfam01245)→ |
| env-1382 | KGI_S1_ANT01_95m_c10241621 | - | 446 | 419 | **RNA**→ hypo→ rplS (PRK05338)Ribosomal_L19 (pfam01245)→ |
| env-1383 | SRS015440_WUGC_scaffold_57079 | - | 459 | 432 | **RNA**→ rplS (PRK05338)Ribosomal_L19 (pfam01245)→ rplS (PRK05338)Ribosomal_L19 (pfam01245)→ |
| env-1384 | SRS043018_WUGC_scaffold_66496 | + | 2705 | 2732 | **RNA**→ rplS (PRK05338)Ribosomal_L19 (pfam01245)→ |
| env-1385 | Buccal_mucosa_LANL_C64861354 | + | 291 | 318 | **RNA**→ |
| env-1386 | SRS015470_WUGC_scaffold_44437 | + | 4765 | 4792 | **RNA**→ rplS (PRK05338)Ribosomal_L19 (pfam01245)→ |
| env-1387 | SRS015803_WUGC_scaffold_24800 | - | 3059 | 3032 | **RNA**→ rplS (PRK05338)Ribosomal_L19 (pfam01245)→ |
| env-1388 | SRS018157_Baylor_scaffold_69033 | - | 304 | 277 | **RNA**→ rplS (PRK05338)Ribosomal_L19 (pfam01245)→ |
| env-1389 | SRS019077_WUGC_scaffold_14658 | - | 3232 | 3205 | **RNA**→ rplS (PRK05338)Ribosomal_L19 (pfam01245)→ |

| | | | | | |
|---|---|---|---|---|---|
| env-1390 | SRS045197_C3591711 | + | 1253 | 1280 | **RNA**→ rplS (PRK05338)Ribosomal_L19 (pfam01245)→ |
| env-1391 | SRS047113_LANL_scaffold_83219 | + | 1884 | 1911 | **RNA**→ rplS (PRK05338)Ribosomal_L19 (pfam01245)→ |
| env-1392 | SRS053584_LANL_scaffold_18794 | + | 28521 | 28548 | **RNA**→ rplS (PRK05338)Ribosomal_L19 (pfam01245)→ |
| env-1393 | SRS049318_LANL_scaffold_16971 | + | 18008 | 18035 | **RNA**→ ←MnmE (COG0486)trmE (PRK05291) |
| env-1394 | SRS053917_LANL_scaffold_21642 | - | 195 | 168 | **RNA**→ rplS (PRK05338)Ribosomal_L19 (pfam01245)→ |
| env-1395 | SRS016541_C1605707 | + | 1495 | 1522 | **RNA**→ |
| env-1396 | SRS064329_C2650028 | - | 160 | 133 | **RNA**→ rplS (PRK05338)Ribosomal_L19 (pfam01245)→ |
| env-1397 | SRS013705_Baylor_scaffold_103146 | + | 2218 | 2245 | **RNA**→ |
| env-1398 | SRS014691_C2508076 | + | 212 | 239 | **RNA**→ |
| env-1399 | SRS015899_WUGC_scaffold_20745 | - | 1016 | 989 | **RNA**→ rplS (PRK05338)Ribosomal_L19 (pfam01245)→ DEDDh (cd06127)EXOIII (smart00479)→ |
| env-1400 | SRS016600_C792137 | + | 109 | 136 | **RNA**→ rplS (PRK05338)Ribosomal_L19 (pfam01245)→ |
| env-1401 | SRS018300_Baylor_scaffold_9343 | - | 19627 | 19600 | **RNA**→ rplS (PRK05338)Ribosomal_L19 (pfam01245)→ |
| | | | | | DEDDh (cd06127)GIY-YIG_UvrC_Cho (cd10434)GIYc (smart00465)EXOIII (smart00479)→ hypo→ |
| env-1402 | SRS018394_Baylor_scaffold_38060 | + | 100 | 127 | **RNA**→ DEDDh (cd06127)GIY-YIG_UvrC_Cho (cd10434)GIYc (smart00465)EXOIII (smart00479)→ |
| env-1403 | SRS019077_WUGC_scaffold_36089 | + | 316 | 343 | **RNA**→ rplS (PRK05338)Ribosomal_L19 (pfam01245)→ |
| env-1404 | SRS019591_WUGC_scaffold_29923 | + | 8636 | 8663 | **RNA**→ rplS (PRK05338)Ribosomal_L19 (pfam01245)→ |
| | | | | | DEDDh (cd06127)GIY-YIG_UvrC_Cho (cd10434)GIYc (smart00465)EXOIII (smart00479)→ hypo→ |
| env-1405 | SRS020226_Baylor_scaffold_9915 | - | 98 | 71 | **RNA**→ |
| env-1406 | SRS045197_WUGC_scaffold_16333 | + | 97 | 124 | **RNA**→ rplS (PRK05338)Ribosomal_L19 (pfam01245)→ |
| env-1407 | SRS045715_LANL_scaffold_96337 | - | 197 | 170 | **RNA**→ rplS (PRK05338)Ribosomal_L19 (pfam01245)→ |
| env-1408 | SRS047634_LANL_scaffold_48423 | + | 258 | 285 | **RNA**→ rplS (PRK05338)Ribosomal_L19 (pfam01245)→ |
| | | | | | DEDDh (cd06127)GIY-YIG_UvrC_Cho (cd10434)GIYc (smart00465)EXOIII (smart00479)→ |
| env-1409 | SRS049389_C4597964 | + | 336 | 363 | **RNA**→ rplS (PRK05338)Ribosomal_L19 (pfam01245)→ |
| env-1410 | SRS051378_C1803515 | + | 93 | 120 | **RNA**→ rplS (PRK05338)Ribosomal_L19 (pfam01245)→ |
| env-1411 | SRS052876_LANL_scaffold_22433 | - | 95 | 68 | **RNA**→ |
| env-1412 | SRS054430_C1931853 | - | 145 | 118 | **RNA**→ rplS (PRK05338)Ribosomal_L19 (pfam01245)→ |
| env-1413 | SRS063215_LANL_scaffold_23111 | + | 338 | 365 | **RNA**→ rplS (PRK05338)Ribosomal_L19 (pfam01245)→ |
| | | | | | DEDDh (cd06127)GIY-YIG_UvrC_Cho (cd10434)GIYc (smart00465)EXOIII (smart00479)→ hypo→ |
| Csp-5-1 | NZ_KN390023.1 | + | 11801 | 11828 | **RNA**→ rplS (PRK05338)Ribosomal_L19 (pfam01245)→ |
| | | | | | DEDDh (cd06127)GIY-YIG_UvrC_Cho (cd10434)GIYc (smart00465)EXOIII (smart00479)→ hypo→ |
| env-1414 | SRS014888_C3105885 | + | 94 | 121 | **RNA**→ rplS (PRK05338)Ribosomal_L19 (pfam01245)→ |
| env-1415 | SRS014894_WUGC_scaffold_7097 | - | 2195 | 2168 | **RNA**→ rplS (PRK05338)Ribosomal_L19 (pfam01245)→ |
| env-1416 | SRS015755_WUGC_scaffold_30815 | + | 386 | 413 | **RNA**→ |
| env-1417 | SRS015989_WUGC_scaffold_19659 | - | 95 | 68 | **RNA**→ |
| env-1418 | SRS016575_Baylor_scaffold_9392 | - | 196 | 169 | **RNA**→ rplS (PRK05338)Ribosomal_L19 (pfam01245)→ |
| env-1419 | SRS015762_C3674485 | - | 8573 | 8546 | **RNA**→ rplS (PRK05338)Ribosomal_L19 (pfam01245)→ |
| | | | | | DEDDh (cd06127)GIY-YIG_UvrC_Cho (cd10434)GIYc (smart00465)EXOIII (smart00479)→ hypo→ |
| env-1420 | SRS019122_C5792698 | + | 337 | 364 | **RNA**→ |
| env-1421 | SRS019127_C1676711 | + | 5492 | 5519 | **RNA**→ rplS (PRK05338)Ribosomal_L19 (pfam01245)→ |
| env-1422 | SRS057205_LANL_scaffold_59131 | - | 8746 | 8719 | **RNA**→ rplS (PRK05338)Ribosomal_L19 (pfam01245)→ |
| | | | | | DEDDh (cd06127)GIY-YIG_UvrC_Cho (cd10434)GIYc (smart00465)EXOIII (smart00479)→ hypo→ |
| env-1423 | SRS020862_C943009 | - | 41 | 14 | **RNA**→ |
| env-1424 | SRS023358_LANL_scaffold_6552 | + | 284 | 311 | **RNA**→ |
| Csp-3-1 | NZ_KB291508.1 | - | 173545 | 173518 | **RNA**→ rplS (PRK05338)Ribosomal_L19 (pfam01245)→ |
| | | | | | DEDDh (cd06127)GIY-YIG_UvrC_Cho (cd10434)GIYc (smart00465)EXOIII (smart00479)→ hypo→ |
| env-1425 | SRS050244_LANL_scaffold_96048 | - | 316 | 289 | **RNA**→ rplS (PRK05338)Ribosomal_L19 (pfam01245)→ |
| env-1426 | SRS058336_C2049026 | + | 171 | 198 | **RNA**→ rplS (PRK05338)Ribosomal_L19 (pfam01245)→ |
| Csp-7-1 | NZ_AKFR01000031.1 | - | 313220 | 313193 | **RNA**→ rplS (PRK05338)Ribosomal_L19 (pfam01245)→ |
| | | | | | DEDDh (cd06127)GIY-YIG_UvrC_Cho (cd10434)EXOIII (smart00479)→ hypo→ |
| env-1427 | SRS023930_C2193178 | + | 1147 | 1174 | **RNA**→ rplS (PRK05338)Ribosomal_L19 (pfam01245)→ |
| | | | | | DEDDh (cd06127)GIY-YIG_UvrC_Cho (cd10434)GIYc (smart00465)EXOIII (smart00479)→ |
| env-1428 | SRS013723_Baylor_scaffold_18285 | - | 2329 | 2302 | **RNA**→ rplS (PRK05338)Ribosomal_L19 (pfam01245)→ |
| | | | | | DEDDh (cd06127)GIY-YIG_UvrC_Cho (cd10434)GIYc (smart00465)EXOIII (smart00479)→ hypo→ |
| env-1429 | SRS014888_C3086477 | + | 650 | 677 | **RNA**→ rplS (PRK05338)Ribosomal_L19 (pfam01245)→ |

| | | | | | |
|---|---|---|---|---|---|
| env-1430 | SRS050669_LANL_scaffold_18681 | + | 4293 | 4320 | **RNA**→ rplS (PRK05338)Ribosomal_L19 (pfam01245)→ DEDDh (cd06127)GIY-YIG_UvrC_Cho (cd10434)GIYc (smart00465)EXOIII (smart00479)→ hypo→ |
| env-1431 | SRS016200_C2186320 | + | 683 | 710 | **RNA**→ |
| env-1432 | SRS019128_WUGC_scaffold_9518 | - | 98 | 71 | **RNA**→ |
| env-1433 | SRS022536_LANL_scaffold_19907 | + | 4308 | 4335 | **RNA**→ rplS (PRK05338)Ribosomal_L19 (pfam01245)→ DEDDh (cd06127)GIY-YIG_UvrC_Cho (cd10434)EXOIII (smart00479)→ hypo→ |
| env-1434 | SRS047265_WUGC_scaffold_12298 | + | 1403 | 1430 | **RNA**→ rplS (PRK05338)Ribosomal_L19 (pfam01245)→ |
| Coc-3-1 | NZ_GL573160.1 | + | 1092618 | 1092645 | **RNA**→ rplS (PRK05338)Ribosomal_L19 (pfam01245)→ DEDDh (cd06127)GIY-YIG_UvrC_Cho (cd10434)EXOIII (smart00479)→ hypo→ |
| Coc-4-1 | NZ_AKFV01000020.1 | - | 85462 | 85435 | **RNA**→ rplS (PRK05338)Ribosomal_L19 (pfam01245)→ DEDDh (cd06127)GIY-YIG_UvrC_Cho (cd10434)EXOIII (smart00479)→ hypo→ |
| env-1435 | SRS015064_C1629213 | - | 175 | 148 | **RNA**→ rplS (PRK05338)Ribosomal_L19 (pfam01245)→ |
| env-1436 | SRS015985_C1503945 | - | 334 | 307 | **RNA**→ rplS (PRK05338)Ribosomal_L19 (pfam01245)→ |
| env-1437 | SRS023841_Baylor_scaffold_1736 | + | 704 | 731 | **RNA**→ |
| env-1438 | SRS011098_Baylor_scaffold_6943 | - | 17288 | 17261 | **RNA**→ rplS (PRK05338)Ribosomal_L19 (pfam01245)→ |
| env-1439 | SRS013836_C4761312 | + | 346 | 373 | **RNA**→ rplS (PRK05338)Ribosomal_L19 (pfam01245)→ |
| env-1440 | SRS013949_WUGC_scaffold_10235 | - | 196 | 169 | **RNA**→ rplS (PRK05338)Ribosomal_L19 (pfam01245)→ |
| env-1441 | SRS013950_WUGC_scaffold_18124 | - | 73 | 46 | **RNA**→ |
| env-1442 | SRS015278_WUGC_scaffold_6560 | + | 2113 | 2140 | **RNA**→ rplS (PRK05338)Ribosomal_L19 (pfam01245)→ |
| env-1443 | SRS015899_WUGC_scaffold_28433 | - | 886 | 859 | **RNA**→ rplS (PRK05338)Ribosomal_L19 (pfam01245)→ |
| env-1444 | SRS019073_C2522155 | - | 95 | 68 | **RNA**→ |
| env-1445 | SRS019077_WUGC_scaffold_58916 | - | 190 | 163 | **RNA**→ rplS (PRK05338)Ribosomal_L19 (pfam01245)→ |
| env-1446 | SRS019225_WUGC_scaffold_9244 | - | 22742 | 22715 | **RNA**→ rplS (PRK05338)Ribosomal_L19 (pfam01245)→ |
| env-1447 | SRS024289_LANL_scaffold_9371 | - | 98 | 71 | **RNA**→ |
| Csp-4-1 | NZ_KB291464.1 | - | 78968 | 78941 | **RNA**→ rplS (PRK05338)Ribosomal_L19 (pfam01245)→ |
| env-1448 | SRS011243_Baylor_scaffold_38886 | - | 160 | 133 | **RNA**→ rplS (PRK05338)Ribosomal_L19 (pfam01245)→ |
| env-1449 | SRS013723_Baylor_scaffold_75255 | - | 295 | 268 | **RNA**→ rplS (PRK05338)Ribosomal_L19 (pfam01245)→ |
| env-1450 | SRS015215_WUGC_scaffold_9005 | - | 288 | 261 | **RNA**→ rplS (PRK05338)Ribosomal_L19 (pfam01245)→ |
| env-1451 | SRS015797_WUGC_scaffold_26726 | + | 97 | 124 | **RNA**→ rplS (PRK05338)Ribosomal_L19 (pfam01245)→ |
| env-1452 | SRS016037_WUGC_scaffold_44282 | + | 97 | 124 | **RNA**→ rplS (PRK05338)Ribosomal_L19 (pfam01245)→ |
| env-1453 | SRS019127_WUGC_scaffold_18778 | - | 8687 | 8660 | **RNA**→ rplS (PRK05338)Ribosomal_L19 (pfam01245)→ |
| env-1454 | SRS023926_Baylor_scaffold_61121 | - | 10271 | 10244 | **RNA**→ rplS (PRK05338)Ribosomal_L19 (pfam01245)→ |
| env-1455 | SRS016092_WUGC_scaffold_23364 | + | 5266 | 5293 | **RNA**→ rplS (PRK05338)Ribosomal_L19 (pfam01245)→ |
| env-1456 | SRS013950_C2181950 | - | 72 | 45 | **RNA**→ |
| env-1457 | SRS017304_Baylor_scaffold_31444 | - | 97 | 70 | **RNA**→ |
| env-1458 | SRS055378_LANL_scaffold_18208 | - | 294 | 267 | **RNA**→ rplS (PRK05338)Ribosomal_L19 (pfam01245)→ |
| Coc-2-1 | NZ_JVFB01000050.1 | + | 35837 | 35864 | **RNA**→ rplS (PRK05338)Ribosomal_L19 (pfam01245)→ DEDDh (cd06127)GIY-YIG_UvrC_Cho (cd10434)GIYc (smart00465)EXOIII (smart00479)→ hypo→ |
| Coc-1-1 | NC_013162.1 | + | 98683 | 98710 | **RNA**→ rplS (PRK05338)Ribosomal_L19 (pfam01245)→ DEDDh (cd06127)GIY-YIG_UvrC_Cho (cd10434)EXOIII (smart00479)→ hypo→ |
| env-1459 | SRS022602_Baylor_scaffold_10139 | - | 838 | 811 | **RNA**→ rplS (PRK05338)Ribosomal_L19 (pfam01245)→ DEDDh (cd06127)EXOIII (smart00479)→ |
| env-1460 | SRS019129_C1939915 | + | 361 | 388 | **RNA**→ |
| env-1461 | SRS019225_C1595905 | + | 1474 | 1501 | **RNA**→ rplS (PRK05338)Ribosomal_L19 (pfam01245)→ |
| env-1462 | SRS022530_LANL_scaffold_63608 | - | 33016 | 32989 | **RNA**→ rplS (PRK05338)Ribosomal_L19 (pfam01245)→ |
| env-1463 | SRS063288_LANL_scaffold_25513 | + | 95 | 122 | **RNA**→ rplS (PRK05338)Ribosomal_L19 (pfam01245)→ |
| env-1464 | SRS023595_Baylor_scaffold_90932 | + | 332 | 359 | **RNA**→ |
| env-1465 | SRS063215_C2182525 | + | 130 | 157 | **RNA**→ rplS (PRK05338)Ribosomal_L19 (pfam01245)→ |
| env-1466 | SRS015540_C1108360 | + | 184 | 211 | **RNA**→ rplS (PRK05338)Ribosomal_L19 (pfam01245)→ |
| env-1467 | SRS011098_Baylor_scaffold_10766 | + | 140 | 167 | **RNA**→ rplS (PRK05338)Ribosomal_L19 (pfam01245)→ |
| env-1468 | SRS013836_Baylor_scaffold_6509 | - | 524 | 497 | **RNA**→ rplS (PRK05338)Ribosomal_L19 (pfam01245)→ |
| Csp-9-1 | NZ_KB291222.1 | + | 54393 | 54420 | **RNA**→ rplS (PRK05338)Ribosomal_L19 (pfam01245)→ DEDDh (cd06127)GIY-YIG_UvrC_Cho (cd10434)GIYc (smart00465)EXOIII (smart00479)→ hypo→ |
| env-1469 | SRS015989_WUGC_scaffold_4470 | + | 307 | 334 | **RNA**→ rplS (PRK05338)Ribosomal_L19 (pfam01245)→ DEDDh (cd06127)GIY-YIG_UvrC_Cho (cd10434)GIYc (smart00465)EXOIII (smart00479)→ hypo→ |

| | | | | | |
|---|---|---|---|---|---|
| env-1470 | SRS018145_Baylor_scaffold_33633 | + | 64 | 91 | **RNA**→ rplS (PRK05338)Ribosomal_L19 (pfam01245)→ |
| env-1471 | SRS057692_LANL_scaffold_49475 | + | 78 | 105 | **RNA**→ rplS (PRK05338)Ribosomal_L19 (pfam01245)→ |
| env-1472 | SRS021960_C2154761 | + | 379 | 406 | **RNA**→ |
| env-1473 | SRS024441_LANL_scaffold_52434 | + | 78 | 105 | **RNA**→ rplS (PRK05338)Ribosomal_L19 (pfam01245)→ |
| env-1474 | SRS050628_C1898890 | - | 986 | 959 | **RNA**→ rplS (PRK05338)Ribosomal_L19 (pfam01245)→ DEDDh (cd06127)EXOIII (smart00479)→ |
| Csp-8-1 | NZ_KE150490.1 | - | 376575 | 376548 | **RNA**→ rplS (PRK05338)Ribosomal_L19 (pfam01245)→ |
| | | | | | DEDDh (cd06127)GIY-YIG_UvrC_Cho (cd10434)GIYc (smart00465)EXOIII (smart00479)→ hypo→ |
| Csp-10-1 | NZ_AJZR01000006.1 | + | 50848 | 50875 | **RNA**→ rplS (PRK05338)Ribosomal_L19 (pfam01245)→ rplS (PRK05338)Ribosomal_L19 (pfam01245)→ |
| | | | | | DEDDh (cd06127)GIY-YIG_UvrC_Cho (cd10434)GIYc (smart00465)EXOIII (smart00479)→ |
| | | | | | DEDDh (cd06127)GIY-YIG_UvrC_Cho (cd10434)GIYc (smart00465)EXOIII (smart00479)→ hypo→ hypo→ |
| env-1475 | Buccal_mucosa_LANL_scaffold_29579 | + | 329 | 356 | **RNA**→ rplS (PRK05338)Ribosomal_L19 (pfam01245)→ |
| env-1476 | SRS020222_C1009936 | + | 264 | 291 | **RNA**→ rplS (PRK05338)Ribosomal_L19 (pfam01245)→ |
| env-1477 | SRS014578_WUGC_scaffold_11003 | - | 214 | 187 | **RNA**→ rplS (PRK05338)Ribosomal_L19 (pfam01245)→ |
| env-1478 | SRS016043_WUGC_scaffold_8966 | - | 256 | 229 | **RNA**→ rplS (PRK05338)Ribosomal_L19 (pfam01245)→ |
| env-1479 | SRS018573_WUGC_scaffold_5605 | - | 2301 | 2274 | **RNA**→ rplS (PRK05338)Ribosomal_L19 (pfam01245)→ |
| env-1480 | SRS019591_WUGC_scaffold_18990 | + | 16688 | 16715 | **RNA**→ rplS (PRK05338)Ribosomal_L19 (pfam01245)→ |
| env-1481 | SRS020340_Baylor_scaffold_33924 | - | 288 | 261 | **RNA**→ rplS (PRK05338)Ribosomal_L19 (pfam01245)→ |
| env-1482 | SRS021477_Baylor_scaffold_1231 | + | 5953 | 5980 | **RNA**→ rplS (PRK05338)Ribosomal_L19 (pfam01245)→ |
| env-1483 | SRS023358_LANL_scaffold_44043 | + | 179 | 206 | **RNA**→ rplS (PRK05338)Ribosomal_L19 (pfam01245)→ |
| env-1484 | SRS024021_Baylor_scaffold_8187 | - | 288 | 261 | **RNA**→ rplS (PRK05338)Ribosomal_L19 (pfam01245)→ |
| env-1485 | SRS047100_C2080290 | - | 301 | 274 | **RNA**→ rplS (PRK05338)Ribosomal_L19 (pfam01245)→ |
| Fan-1-1 | NZ_ATTM01000004.1 | + | 10713 | 10740 | **RNA**→ rplS (PRK05338)Ribosomal_L19 (pfam01245)→ |
| Fte-1-1 | NZ_AUDN01000020.1 | - | 101805 | 101778 | **RNA**→ rplS (PRK05338)Ribosomal_L19 (pfam01245)→ |
| env-1486 | SRS017227_Baylor_scaffold_48001 | - | 401 | 374 | **RNA**→ rplS (PRK05338)Ribosomal_L19 (pfam01245)→ |
| env-1487 | SRS055401_LANL_scaffold_34300 | - | 163 | 136 | **RNA**→ rplS (PRK05338)Ribosomal_L19 (pfam01245)→ |
| env-1488 | SRS013949_C3524448 | + | 60 | 87 | **RNA**→ rplS (PRK05338)Ribosomal_L19 (pfam01245)→ |
| env-1489 | SRS063932_LANL_scaffold_105541 | + | 28523 | 28550 | **RNA**→ rplS (PRK05338)Ribosomal_L19 (pfam01245)→ COG4389 (COG4389)SpecificRecomb (pfam10136)→ |
| env-1490 | CEOS01042827.1 | - | 20258 | 20231 | **RNA**→ rplS (PRK05338)Ribosomal_L19 (pfam01245)→ |
| env-1491 | CERQ01126197.1 | + | 4556 | 4583 | **RNA**→ rplS (PRK05338)Ribosomal_L19 (pfam01245)→ |
| env-1492 | CETA01045631.1 | - | 13014 | 12987 | **RNA**→ rplS (PRK05338)Ribosomal_L19 (pfam01245)→ |
| env-1493 | CETB01056197.1 | + | 21881 | 21908 | **RNA**→ rplS (PRK05338)Ribosomal_L19 (pfam01245)→ |
| env-1494 | CETG01362727.1 | - | 665 | 638 | **RNA**→ rplS (PRK05338)Ribosomal_L19 (pfam01245)→ |
| env-1495 | CETR01036094.1 | + | 21881 | 21908 | **RNA**→ rplS (PRK05338)Ribosomal_L19 (pfam01245)→ |
| env-1496 | CETS01044312.1 | + | 9368 | 9395 | **RNA**→ rplS (PRK05338)Ribosomal_L19 (pfam01245)→ |
| env-1497 | CETT01258458.1 | - | 563 | 536 | **RNA**→ rplS (PRK05338)Ribosomal_L19 (pfam01245)→ |
| env-1498 | CEUY01396649.1 | + | 431 | 458 | **RNA**→ rplS (PRK05338)Ribosomal_L19 (pfam01245)→ |
| env-1499 | CEVJ01196187.1 | - | 671 | 644 | **RNA**→ rplS (PRK05338)Ribosomal_L19 (pfam01245)→ |
| env-1500 | CEVK01214642.1 | - | 308 | 281 | **RNA**→ rplS (PRK05338)Ribosomal_L19 (pfam01245)→ |
| env-1501 | CEVM01174021.1 | + | 444 | 471 | **RNA**→ rplS (PRK05338)Ribosomal_L19 (pfam01245)→ |
| env-1502 | CEVO01132965.1 | - | 263 | 236 | **RNA**→ rplS (PRK05338)Ribosomal_L19 (pfam01245)→ |
| env-1503 | CEVQ01096279.1 | - | 594 | 567 | **RNA**→ rplS (PRK05338)Ribosomal_L19 (pfam01245)→ |
| env-1504 | CEWI01238921.1 | - | 396 | 369 | **RNA**→ rplS (PRK05338)Ribosomal_L19 (pfam01245)→ |
| env-1505 | CENM01019596.1 | - | 258 | 231 | **RNA**→ rplS (PRK05338)Ribosomal_L19 (pfam01245)→ |
| env-1506 | JGI24723J26617_10000093 | + | 49411 | 49438 | **RNA**→ rplS (PRK05338)Ribosomal_L19 (pfam01245)→ |
| env-1507 | LGVF01495366.1 | + | 36 | 63 | **RNA**→ rplS (PRK05338)Ribosomal_L19 (pfam01245)→ |
| Mmo-1-1 | NZ_AUHX01000001.1 | - | 133740 | 133713 | **RNA**→ rplS (PRK05338)Ribosomal_L19 (pfam01245)→ |
| env-1508 | CEOS01168860.1 | + | 192 | 219 | **RNA**→ rplS (PRK05338)Ribosomal_L19 (pfam01245)→ |
| env-1509 | CEQE01154469.1 | + | 4 | 31 | **RNA**→ rplS (PRK05338)Ribosomal_L19 (pfam01245)→ |
| env-1510 | CEVM01223312.1 | - | 386 | 359 | **RNA**→ rplS (PRK05338)Ribosomal_L19 (pfam01245)→ |
| env-1511 | CEVF01118313.1 | - | 339 | 312 | **RNA**→ rplS (PRK05338)Ribosomal_L19 (pfam01245)→ |
| env-1512 | TahiMoana_1030440 | - | 764 | 737 | **RNA**→ rplS (PRK05338)Ribosomal_L19 (pfam01245)→ |
| env-1513 | JGI11881J13070_1029301 | + | 172 | 199 | **RNA**→ rplS (PRK05338)Ribosomal_L19 (pfam01245)→ |
| env-1514 | JGI12218J13221_1034272 | - | 80 | 53 | **RNA**→ |

| | | | | | |
|---|---|---|---|---|---|
| env-1515 | JGI11931J13222_1024356 | - | 246 | 219 | **RNA**→ rplS (PRK05338)Ribosomal_L19 (pfam01245)→ |
| env-1516 | CETA01220460.1 | + | 285 | 312 | **RNA**→ rplS (PRK05338)Ribosomal_L19 (pfam01245)→ |
| env-1517 | CETS01220756.1 | + | 255 | 282 | **RNA**→ rplS (PRK05338)Ribosomal_L19 (pfam01245)→ |
| env-1518 | CEVI01118676.1 | - | 855 | 828 | **RNA**→ rplS (PRK05338)Ribosomal_L19 (pfam01245)→ |
| Lsp-4-1 | NZ_JPOL01000003.1 | + | 201753 | 201780 | **RNA**→ rplS (PRK05338)Ribosomal_L19 (pfam01245)→ |
| env-1519 | JGI1221J11331_1178417 | + | 266 | 293 | **RNA**→ |
| env-1520 | SA_S1_NOR08_45m_c10151176 | - | 451 | 424 | **RNA**→ rplS (PRK05338)Ribosomal_L19 (pfam01245)→ |
| env-1521 | SA_S1_NOR08_45m_c10000441 | + | 12622 | 12649 | **RNA**→ rplS (PRK05338)Ribosomal_L19 (pfam01245)→ |
| env-1522 | SA_S2_NOR15_50m_c10000385 | + | 13396 | 13423 | **RNA**→ rplS (PRK05338)Ribosomal_L19 (pfam01245)→ |
| env-1523 | SA_S1_NOR02_45m_c1172796 | - | 185 | 158 | **RNA**→ rplS (PRK05338)Ribosomal_L19 (pfam01245)→ |
| env-1524 | SA_S2_NOR18_50m_1098208 | + | 227 | 254 | **RNA**→ rplS (PRK05338)Ribosomal_L19 (pfam01245)→ |
| env-1525 | SA_S1_NOR05_45m_c10040916 | - | 1024 | 997 | **RNA**→ rplS (PRK05338)Ribosomal_L19 (pfam01245)→ |
| Ama-1-1 | NZ_JACA01000020.1 | + | 179901 | 179928 | **RNA**→ rplS (PRK05338)Ribosomal_L19 (pfam01245)→ |
| env-1526 | JGI25920J50251_10000870 | - | 9504 | 9477 | **RNA**→ rplS (PRK05338)Ribosomal_L19 (pfam01245)→ |
| env-1527 | JGI25909J50240_1100786 | + | 75 | 102 | **RNA**→ rplS (PRK05338)Ribosomal_L19 (pfam01245)→ |
| env-1528 | JGI25911J50253_10030107 | + | 69 | 96 | **RNA**→ rplS (PRK05338)Ribosomal_L19 (pfam01245)→ |
| env-1529 | JGI25912J50252_10144474 | + | 133 | 160 | **RNA**→ rplS (PRK05338)Ribosomal_L19 (pfam01245)→ |
| env-1530 | JGI25908J49247_10036170 | - | 1268 | 1241 | **RNA**→ rplS (PRK05338)Ribosomal_L19 (pfam01245)→ |
| env-1531 | JGI25907J50239_1087629 | + | 148 | 175 | **RNA**→ rplS (PRK05338)Ribosomal_L19 (pfam01245)→ |
| Aca-1-1 | NZ_AUBG01000016.1 | + | 27656 | 27683 | **RNA**→ rplS (PRK05338)Ribosomal_L19 (pfam01245)→ |
| env-1532 | CETR01021789.1 | - | 7980 | 7953 | **RNA**→ rplS (PRK05338)Ribosomal_L19 (pfam01245)→ |
| env-1533 | JGI24723J26617_10120000 | + | 312 | 339 | **RNA**→ rplS (PRK05338)Ribosomal_L19 (pfam01245)→ |
| env-1534 | JGI24724J26744_10003103 | - | 7169 | 7142 | **RNA**→ rplS (PRK05338)Ribosomal_L19 (pfam01245)→ |
| Fsa-2-1 | NZ_JMLU01000004.1 | + | 158055 | 158082 | **RNA**→ rplS (PRK05338)Ribosomal_L19 (pfam01245)→ |
| env-1535 | 3300001592_10082624 | - | 375 | 348 | **RNA**→ rplS (PRK05338)Ribosomal_L19 (pfam01245)→ |
| env-1536 | 3300001605_10000472 | - | 597 | 570 | **RNA**→ rplS (PRK05338)Ribosomal_L19 (pfam01245)→ |
| env-1537 | 3300001592_10006511 | + | 11374 | 11401 | **RNA**→ rplS (PRK05338)Ribosomal_L19 (pfam01245)→ |
| Fba-2-1 | NZ_AAXX01000003.1 | - | 173796 | 173769 | **RNA**→ rplS (PRK05338)Ribosomal_L19 (pfam01245)→ rplS (PRK05338)Ribosomal_L19 (pfam01245)→ |
| Kje-1-1 | NZ_LBMG01000057.1 | + | 26887 | 26914 | **RNA**→ rplS (PRK05338)Ribosomal_L19 (pfam01245)→ |
| Kal-1-1 | NZ_DS544873.1 | + | 1877186 | 1877213 | **RNA**→ rplS (PRK05338)Ribosomal_L19 (pfam01245)→ |
| Kzh-1-1 | NZ_LBMH01000008.1 | - | 5300 | 5273 | **RNA**→ rplS (PRK05338)Ribosomal_L19 (pfam01245)→ |
| env-1538 | TB_FS06_10DRAFT_1000005 | + | 81339 | 81366 | **RNA**→ rplS (PRK05338)Ribosomal_L19 (pfam01245)→ |
| env-1539 | JGI20151J14362_10264229 | - | 319 | 292 | **RNA**→ rplS (PRK05338)Ribosomal_L19 (pfam01245)→ |
| env-1540 | JGI20156J14371_10338308 | - | 271 | 244 | **RNA**→ rplS (PRK05338)Ribosomal_L19 (pfam01245)→ |
| env-1541 | JGI25920J50251_10088380 | + | 139 | 166 | **RNA**→ rplS (PRK05338)Ribosomal_L19 (pfam01245)→ |
| Fin-1-1 | NC_017025.1 | - | 992421 | 992394 | **RNA**→ rplS (PRK05338)Ribosomal_L19 (pfam01245)→ |
| env-1542 | CESJ01019018.1 | - | 182 | 155 | **RNA**→ rplS (PRK05338)Ribosomal_L19 (pfam01245)→ |
| env-1543 | CEWG01277609.1 | - | 481 | 454 | **RNA**→ rplS (PRK05338)Ribosomal_L19 (pfam01245)→ PRK08299 (PRK08299)→ |
| env-1544 | CESM01468209.1 | - | 344 | 317 | **RNA**→ rplS (PRK05338)Ribosomal_L19 (pfam01245)→ |
| env-1545 | 2236876007_0430719 | + | 198 | 225 | **RNA**→ hypo→ |
| env-1546 | CETY01043148.1 | - | 6005 | 5978 | **RNA**→ rplS (PRK05338)Ribosomal_L19 (pfam01245)→ Icd (COG0538)PRK08299 (PRK08299)→ Peptidase_M14NE-CP-C_like (cd11308)CirA (COG1629)TonB-hemlactrns (TIGR01786)CarbopepD_reg_2 (pfam13715)→ DUF4249 (pfam14054)→ murQ (PRK05441)SIS_Etherase (cd05007)→ |
| env-1547 | DelMOSum2010_c10221712 | + | 321 | 348 | **RNA**→ rplS (PRK05338)Ribosomal_L19 (pfam01245)→ |
| env-1548 | CEWO01314910.1 | + | 580 | 607 | **RNA**→ rplS (PRK05338)Ribosomal_L19 (pfam01245)→ |
| env-1549 | JGI20156J14371_10001449 | + | 17641 | 17668 | **RNA**→ rplS (PRK05338)Ribosomal_L19 (pfam01245)→ Icd (COG0538)PRK08299 (PRK08299)→ |
| env-1550 | JGI20155J14468_10020639 | - | 210 | 183 | **RNA**→ rplS (PRK05338)Ribosomal_L19 (pfam01245)→ |
| env-1551 | JGI20160J14292_10134963 | - | 298 | 271 | **RNA**→ rplS (PRK05338)Ribosomal_L19 (pfam01245)→ |
| env-1552 | CERI01264885.1 | + | 831 | 858 | **RNA**→ rplS (PRK05338)Ribosomal_L19 (pfam01245)→ |
| env-1553 | CERM01228874.1 | + | 1239 | 1266 | **RNA**→ rplS (PRK05338)Ribosomal_L19 (pfam01245)→ |
| env-1554 | JGI20156J14371_10182376 | - | 65 | 38 | **RNA**→ |
| env-1555 | 2236876001_539343 | + | 237 | 264 | **RNA**→ rplS (PRK05338)Ribosomal_L19 (pfam01245)→ |
| env-1556 | CESU01013805.1 | + | 6882 | 6909 | **RNA**→ rplS (PRK05338)Ribosomal_L19 (pfam01245)→ |

| | | | | | |
|---|---|---|---|---|---|
| env-1557 | CESI01075687.1 | - | 1194 | 1167 | **RNA**→ rplS (PRK05338)Ribosomal_L19 (pfam01245)→ PRK08299 (PRK08299)→ |
| env-1558 | CEVG01321748.1 | + | 522 | 549 | **RNA**→ rplS (PRK05338)Ribosomal_L19 (pfam01245)→ PRK08299 (PRK08299)→ |
| env-1559 | A09P04_1300_Sequence0000000262 | + | 5688 | 5715 | **RNA**→ rplS (PRK05338)Ribosomal_L19 (pfam01245)→ PRK08299 (PRK08299)→ |
| env-1560 | A09P041000a_Sequence0000000037 | - | 206 | 179 | **RNA**→ rplS (PRK05338)Ribosomal_L19 (pfam01245)→ |
| env-1561 | CENF01216542.1 | + | 866 | 893 | **RNA**→ rplS (PRK05338)Ribosomal_L19 (pfam01245)→ Icd (COG0538)PRK08299 (PRK08299)→ |
| | | | | | PRK10064 (PRK10064)TonB_dep_Rec (pfam00593)Plug (pfam07715)CarbopepD_reg_2 (pfam13715)OMP_b-brl_3 (pfam1 |
| | | | | | DUF4249 (pfam14054)→ murQ (PRK05441)SIS_Etherase (cd05007)→ |
| env-1562 | CETV01076215.1 | + | 63 | 90 | **RNA**→ rplS (PRK05338)Ribosomal_L19 (pfam01245)→ Icd (COG0538)PRK08299 (PRK08299)→ |
| | | | | | murQ (PRK05441)SIS_Etherase (cd05007)→ hypo→ TGS_ThrRS_N (cd01667)thrS (PRK00413)→ |
| env-1563 | JCVI_SCAF_1096627109417 | + | 201 | 228 | **RNA**→ rplS (PRK05338)Ribosomal_L19 (pfam01245)→ Icd (COG0538)PRK08299 (PRK08299)→ |
| env-1564 | SI60aug11_100m_c1000185 | + | 69 | 96 | **RNA**→ rplS (PRK05338)Ribosomal_L19 (pfam01245)→ Icd (COG0538)PRK08299 (PRK08299)→ |
| env-1565 | CESI01023758.1 | - | 17726 | 17699 | **RNA**→ rplS (PRK05338)Ribosomal_L19 (pfam01245)→ Icd (COG0538)PRK08299 (PRK08299)→ |
| | | | | | CirA (COG1629)TonB-hemlactrns (TIGR01786)CarbopepD_reg_2 (pfam13715)→ DUF4249 (pfam14054)→ |
| | | | | | murQ (PRK05441)SIS_Etherase (cd05007)→ hypo→ TGS_ThrRS_N (cd01667)ThrS (COG0441)thrS (PRK00413)→ |
| env-1566 | CESU01036743.1 | + | 15859 | 15886 | **RNA**→ rplS (PRK05338)Ribosomal_L19 (pfam01245)→ PRK08299 (PRK08299)→ |
| env-1567 | CERR01132098.1 | + | 302 | 329 | **RNA**→ rplS (PRK05338)Ribosomal_L19 (pfam01245)→ |
| env-1568 | CEVX01177957.1 | + | 418 | 445 | **RNA**→ ←hypo |
| env-1569 | SI47jul10_100m_c1001824 | - | 4900 | 4873 | **RNA**→ rplS (PRK05338)Ribosomal_L19 (pfam01245)→ Icd (COG0538)PRK08299 (PRK08299)→ |
| env-1570 | SI54feb11_100m_c1021428 | - | 825 | 798 | **RNA**→ rplS (PRK05338)Ribosomal_L19 (pfam01245)→ PRK08299 (PRK08299)→ |
| env-1571 | SI36aug09_135m_c1033193 | + | 185 | 212 | **RNA**→ rplS (PRK05338)Ribosomal_L19 (pfam01245)→ PRK08299 (PRK08299)→ |
| env-1572 | SI47jul10_120m_c1035424 | - | 591 | 564 | **RNA**→ rplS (PRK05338)Ribosomal_L19 (pfam01245)→ PRK08299 (PRK08299)→ |
| env-1573 | SI34jun09_120m_1000013 | + | 68976 | 69003 | **RNA**→ rplS (PRK05338)Ribosomal_L19 (pfam01245)→ Icd (COG0538)PRK08299 (PRK08299)→ |
| env-1574 | SI34jun09_135m_1000008 | - | 90123 | 90096 | **RNA**→ rplS (PRK05338)Ribosomal_L19 (pfam01245)→ Icd (COG0538)PRK08299 (PRK08299)→ |
| env-1575 | SI36aug09_100m_1005555 | - | 1309 | 1282 | **RNA**→ rplS (PRK05338)Ribosomal_L19 (pfam01245)→ PRK08299 (PRK08299)→ |
| env-1576 | SI36aug09_120m_1000042 | + | 41474 | 41501 | **RNA**→ rplS (PRK05338)Ribosomal_L19 (pfam01245)→ Icd (COG0538)PRK08299 (PRK08299)→ |
| env-1577 | SI36aug09_120m_1006408 | - | 625 | 598 | **RNA**→ rplS (PRK05338)Ribosomal_L19 (pfam01245)→ |
| env-1578 | SI34jun09_100m_1013853 | + | 1182 | 1209 | **RNA**→ rplS (PRK05338)Ribosomal_L19 (pfam01245)→ PRK08299 (PRK08299)→ |
| env-1579 | F10SI03120a_Sequence0000008470 | - | 1292 | 1265 | **RNA**→ rplS (PRK05338)Ribosomal_L19 (pfam01245)→ PRK08299 (PRK08299)→ |
| env-1580 | F10SI03135a_Sequence0000028256 | - | 1234 | 1207 | **RNA**→ rplS (PRK05338)Ribosomal_L19 (pfam01245)→ PRK08299 (PRK08299)→ |
| env-1581 | CESE01030659.1 | + | 2760 | 2787 | **RNA**→ rplS (PRK05338)Ribosomal_L19 (pfam01245)→ PRK08299 (PRK08299)→ |
| env-1582 | CEWR01159442.1 | + | 4928 | 4955 | **RNA**→ rplS (PRK05338)Ribosomal_L19 (pfam01245)→ PRK08299 (PRK08299)→ |
| env-1583 | CEOP01456443.1 | - | 1734 | 1707 | **RNA**→ rplS (PRK05338)Ribosomal_L19 (pfam01245)→ Icd (COG0538)PRK08299 (PRK08299)→ |
| env-1584 | CERW01247538.1 | - | 744 | 717 | **RNA**→ rplS (PRK05338)Ribosomal_L19 (pfam01245)→ PRK08299 (PRK08299)→ |
| env-1585 | CEVX01084310.1 | + | 8526 | 8553 | **RNA**→ rplS (PRK05338)Ribosomal_L19 (pfam01245)→ Icd (COG0538)PRK08299 (PRK08299)→ |
| | | | | | PRK10064 (PRK10064)Plug (pfam07715)CarbopepD_reg_2 (pfam13715)→ |
| env-1586 | CEWO01028333.1 | + | 4814 | 4841 | **RNA**→ rplS (PRK05338)Ribosomal_L19 (pfam01245)→ Icd (COG0538)PRK08299 (PRK08299)→ |
| | | | | | PRK10064 (PRK10064)Plug (pfam07715)CarbopepD_reg_2 (pfam13715)→ |
| env-1587 | CETQ01273970.1 | - | 129 | 102 | **RNA**→ hypo→ |
| env-1588 | CETK01030187.1 | + | 60 | 87 | **RNA**→ rplS (PRK05338)Ribosomal_L19 (pfam01245)→ Icd (COG0538)PRK08299 (PRK08299)→ |
| | | | | | CarbopepD_reg_2 (pfam13715)→ |
| env-1589 | CETA01078260.1 | + | 1931 | 1958 | **RNA**→ rplS (PRK05338)Ribosomal_L19 (pfam01245)→ PRK08299 (PRK08299)→ |
| env-1590 | CESM01008740.1 | + | 6469 | 6496 | **RNA**→ rplS (PRK05338)Ribosomal_L19 (pfam01245)→ |
| env-1591 | CEUA01047049.1 | - | 77 | 50 | **RNA**→ |
| env-1592 | CEUB01108690.1 | - | 4211 | 4184 | **RNA**→ rplS (PRK05338)Ribosomal_L19 (pfam01245)→ Icd (COG0538)PRK08299 (PRK08299)→ |
| | | | | | TonB-hemin (TIGR01785)TonB_dep_Rec (pfam00593)Plug (pfam07715)CarbopepD_reg_2 (pfam13715)→ hypo→ |
| env-1593 | CENZ01369401.1 | - | 254 | 227 | **RNA**→ rplS (PRK05338)Ribosomal_L19 (pfam01245)→ |
| env-1594 | CEQF01051053.1 | + | 1217 | 1244 | **RNA**→ rplS (PRK05338)Ribosomal_L19 (pfam01245)→ PRK08299 (PRK08299)→ |
| env-1595 | CERL01234262.1 | + | 134 | 161 | **RNA**→ rplS (PRK05338)Ribosomal_L19 (pfam01245)→ PRK08299 (PRK08299)→ |
| env-1596 | CEQY01193862.1 | - | 1759 | 1732 | **RNA**→ rplS (PRK05338)Ribosomal_L19 (pfam01245)→ Icd (COG0538)PRK08299 (PRK08299)→ |
| env-1597 | CERR01235002.1 | + | 163 | 190 | **RNA**→ rplS (PRK05338)Ribosomal_L19 (pfam01245)→ PRK08299 (PRK08299)→ |
| env-1598 | CESQ01389779.1 | + | 211 | 238 | **RNA**→ rplS (PRK05338)Ribosomal_L19 (pfam01245)→ PRK08299 (PRK08299)→ |
| env-1599 | CETI01152875.1 | + | 187 | 214 | **RNA**→ rplS (PRK05338)Ribosomal_L19 (pfam01245)→ PRK08299 (PRK08299)→ |
| Mth-1-1 | NZ_LDAS01000001.1 | - | 695657 | 695630 | **RNA**→ rplS (PRK05338)Ribosomal_L19 (pfam01245)→ |
| env-1600 | JGI24723J26617_10397431 | + | 150 | 177 | **RNA**→ rplS (PRK05338)Ribosomal_L19 (pfam01245)→ |

| | | | | | |
|---|---|---|---|---|---|
| env-1601 | JGI24724J26744_10000071 | - | 8106 | 8079 | **RNA**→ rplS (PRK05338)Ribosomal_L19 (pfam01245)→ |
| env-1602 | BS_KBA_SWE12_21m_c10564765 | + | 114 | 141 | **RNA**→ rplS (PRK05338)Ribosomal_L19 (pfam01245)→ |
| env-1603 | BS_KBA_SWE02_21m_10243122 | - | 247 | 220 | **RNA**→ rplS (PRK05338)Ribosomal_L19 (pfam01245)→ |
| Bar-1-1 | NZ_AFXZ01000061.1 | - | 9495 | 9468 | **RNA**→ rplS (PRK05338)Ribosomal_L19 (pfam01245)→ rplS (PRK05338)Ribosomal_L19 (pfam01245)→ |
| env-1604 | CESI01155152.1 | - | 1312 | 1285 | **RNA**→ rplS (PRK05338)Ribosomal_L19 (pfam01245)→ PRK08299 (PRK08299)→ |
| env-1605 | JCVI_SCAF_1096627904983 | + | 1367 | 1394 | **RNA**→ rplS (PRK05338)Ribosomal_L19 (pfam01245)→ |
| env-1606 | JGI20152J14361_10239717 | + | 202 | 229 | **RNA**→ |
| env-1607 | CESI01144570.1 | + | 19660 | 19687 | **RNA**→ rplS (PRK05338)Ribosomal_L19 (pfam01245)→ PRK08299 (PRK08299)→ |
| env-1608 | JGI20156J14371_10001677 | + | 8441 | 8468 | **RNA**→ rplS (PRK05338)Ribosomal_L19 (pfam01245)→ Icd (COG0538)PRK08299 (PRK08299)→ |
| | | | | | murQ (PRK05441)SIS (cd04795)→ murQ (PRK05441)SIS_Etherase (cd05007)→ hypo→ |
| | | | | | TGS_ThrRS_N (cd01667)ThrS (COG0441)thrS (PRK00413)→ |
| env-1609 | JGI20160J14292_10002169 | + | 6253 | 6280 | **RNA**→ rplS (PRK05338)Ribosomal_L19 (pfam01245)→ Icd (COG0538)PRK08299 (PRK08299)→ |
| | | | | | murQ (PRK05441)SIS_Etherase (cd05007)→ hypo→ TGS_ThrRS_N (cd01667)ThrS (COG0441)thrS (PRK00413)→ |
| env-1610 | JGI20159J14440_10295666 | + | 144 | 171 | **RNA**→ rplS (PRK05338)Ribosomal_L19 (pfam01245)→ |
| env-1611 | CERI01069492.1 | + | 3368 | 3395 | **RNA**→ rplS (PRK05338)Ribosomal_L19 (pfam01245)→ |
| env-1612 | CESI01307738.1 | - | 4623 | 4596 | **RNA**→ rplS (PRK05338)Ribosomal_L19 (pfam01245)→ Icd (COG0538)PRK08299 (PRK08299)→ |
| | | | | | murQ (PRK05441)SIS_Etherase (cd05007)→ hypo→ TGS_ThrRS_N (cd01667)ThrS (COG0441)thrS (PRK00413)→ |
| env-1613 | CESU01111611.1 | + | 322 | 349 | **RNA**→ rplS (PRK05338)Ribosomal_L19 (pfam01245)→ |
| env-1614 | DelMOSum2010_c10000810 | - | 24769 | 24742 | **RNA**→ rplS (PRK05338)Ribosomal_L19 (pfam01245)→ Icd (COG0538)PRK08299 (PRK08299)→ |
| | | | | | murQ (PRK05441)SIS_Etherase (cd05007)→ hypo→ TGS_ThrRS_N (cd01667)ThrS (COG0441)thrS (PRK00413)→ |
| env-1615 | JGI20160J14292_10000029 | + | 74902 | 74929 | **RNA**→ rplS (PRK05338)Ribosomal_L19 (pfam01245)→ Icd (COG0538)PRK08299 (PRK08299)→ |
| | | | | | murQ (PRK05441)SIS_Etherase (cd05007)→ hypo→ TGS_ThrRS_N (cd01667)ThrS (COG0441)thrS (PRK00413)→ |
| env-1616 | NCBI_BBAY_READ_1105898016080 | - | 698 | 671 | **RNA**→ rplS (PRK05338)Ribosomal_L19 (pfam01245)→ PRK08299 (PRK08299)→ |
| env-1617 | KGI_S1_ANT03_95m_c1092068 | + | 85 | 112 | **RNA**→ rplS (PRK05338)Ribosomal_L19 (pfam01245)→ |
| env-1618 | CENF01157472.1 | + | 115 | 142 | **RNA**→ rplS (PRK05338)Ribosomal_L19 (pfam01245)→ PRK08299 (PRK08299)→ |
| env-1619 | CEOM01030827.1 | - | 7675 | 7648 | **RNA**→ rplS (PRK05338)Ribosomal_L19 (pfam01245)→ Icd (COG0538)PRK08299 (PRK08299)→ |
| env-1620 | CERW01220270.1 | + | 272 | 299 | **RNA**→ rplS (PRK05338)Ribosomal_L19 (pfam01245)→ |
| env-1621 | CESP01276117.1 | - | 1053 | 1026 | **RNA**→ rplS (PRK05338)Ribosomal_L19 (pfam01245)→ PRK08299 (PRK08299)→ |
| env-1622 | CEVP01353816.1 | - | 945 | 918 | **RNA**→ rplS (PRK05338)Ribosomal_L19 (pfam01245)→ PRK08299 (PRK08299)→ |
| env-1623 | CEVS01278067.1 | - | 151 | 124 | **RNA**→ rplS (PRK05338)Ribosomal_L19 (pfam01245)→ |
| env-1624 | TB_GS09_5_10001116 | + | 8952 | 8979 | **RNA**→ ←hypo |
| env-1625 | LGVF01492896.1 | - | 734 | 707 | **RNA**→ rplS (PRK05338)Ribosomal_L19 (pfam01245)→ |
| env-1626 | LGVC01057912.1 | - | 723 | 696 | **RNA**→ rplS (PRK05338)Ribosomal_L19 (pfam01245)→ |
| env-1627 | LGVF01075961.1 | - | 4858 | 4831 | **RNA**→ rplS (PRK05338)Ribosomal_L19 (pfam01245)→ |
| env-1628 | LGVD01061935.1 | + | 3177 | 3204 | **RNA**→ rplS (PRK05338)Ribosomal_L19 (pfam01245)→ |
| env-1629 | JGI24023J19991_10018257 | - | 2685 | 2658 | **RNA**→ rplS (PRK05338)Ribosomal_L19 (pfam01245)→ |
| env-1630 | SA_S1_NOR08_45m_c10644258 | + | 115 | 142 | **RNA**→ rplS (PRK05338)Ribosomal_L19 (pfam01245)→ |
| env-1631 | BS_KBA_SWE12_21m_c10053575 | - | 952 | 925 | **RNA**→ rplS (PRK05338)Ribosomal_L19 (pfam01245)→ |
| env-1632 | BS_KBA_SWE12_21m_c10014505 | - | 2364 | 2337 | **RNA**→ rplS (PRK05338)Ribosomal_L19 (pfam01245)→ |
| env-1633 | BS_KBA_SWE21_205m_10014880 | - | 290 | 263 | **RNA**→ rplS (PRK05338)Ribosomal_L19 (pfam01245)→ |
| env-1634 | BS_KBA_SWE02_21m_10009640 | - | 1361 | 1334 | **RNA**→ rplS (PRK05338)Ribosomal_L19 (pfam01245)→ |
| env-1635 | BS_KBA_SWE12_21m_c10000793 | + | 3415 | 3442 | **RNA**→ rplS (PRK05338)Ribosomal_L19 (pfam01245)→ |
| env-1636 | BS_KBB_SWE26_205m_c1001614 | - | 974 | 947 | **RNA**→ rplS (PRK05338)Ribosomal_L19 (pfam01245)→ |
| env-1637 | BS_KBA_SWE07_21m_c1003990 | - | 996 | 969 | **RNA**→ rplS (PRK05338)Ribosomal_L19 (pfam01245)→ |
| env-1638 | BS_KBA_SWE21_205m_10000010 | + | 4468 | 4495 | **RNA**→ rplS (PRK05338)Ribosomal_L19 (pfam01245)→ |
| env-1639 | BS_KBA_SWE02_21m_10000410 | - | 12530 | 12503 | **RNA**→ rplS (PRK05338)Ribosomal_L19 (pfam01245)→ |
| env-1640 | SA_S1_NOR05_45m_c10006399 | - | 2424 | 2397 | **RNA**→ rplS (PRK05338)Ribosomal_L19 (pfam01245)→ |
| env-1641 | SA_S1_NOR08_45m_c10019755 | - | 2638 | 2611 | **RNA**→ rplS (PRK05338)Ribosomal_L19 (pfam01245)→ |
| env-1642 | SA_S2_NOR15_50m_c10428401 | - | 127 | 100 | **RNA**→ hypo→ |
| env-1643 | SA_S1_NOR02_45m_c1030347 | - | 406 | 379 | **RNA**→ rplS (PRK05338)Ribosomal_L19 (pfam01245)→ |
| env-1644 | SA_S1_NOR05_45m_c10357957 | + | 172 | 199 | **RNA**→ rplS (PRK05338)Ribosomal_L19 (pfam01245)→ |
| env-1645 | SA_S1_NOR08_45m_c10002181 | - | 7235 | 7208 | **RNA**→ rplS (PRK05338)Ribosomal_L19 (pfam01245)→ |
| env-1646 | SA_S2_NOR15_50m_c10098518 | + | 631 | 658 | **RNA**→ rplS (PRK05338)Ribosomal_L19 (pfam01245)→ |

| | | | | | |
|---|---|---|---|---|---|
| env-1647 | SA_S1_NOR02_45m_c1091868 | + | 275 | 302 | **RNA**→ |
| env-1648 | SA_S1_NOR08_45m_c10420270 | + | 193 | 220 | **RNA**→ rplS (PRK05338)Ribosomal_L19 (pfam01245)→ |
| env-1649 | SA_S2_NOR15_50m_c10316835 | - | 257 | 230 | **RNA**→ rplS (PRK05338)Ribosomal_L19 (pfam01245)→ |
| env-1650 | TDF_OR_ARG04_113m_c1012125 | + | 552 | 579 | **RNA**→ rplS (PRK05338)Ribosomal_L19 (pfam01245)→ |
| env-1651 | TDF_MC_ARG01_113m_c1022147 | + | 240 | 267 | **RNA**→ rplS (PRK05338)Ribosomal_L19 (pfam01245)→ |
| env-1652 | TDF_MC_ARG02_113m_c1008450 | - | 506 | 479 | **RNA**→ rplS (PRK05338)Ribosomal_L19 (pfam01245)→ |
| env-1653 | BS_KBA_SWE02_21m_10037748 | - | 1195 | 1168 | **RNA**→ rplS (PRK05338)Ribosomal_L19 (pfam01245)→ |
| env-1654 | 3300000558_10149016 | - | 188 | 161 | **RNA**→ rplS (PRK05338)Ribosomal_L19 (pfam01245)→ |
| env-1655 | 3300001580_10557157 | - | 129 | 102 | **RNA**→ hypo→ |
| env-1656 | JGI24025J20009_10008154 | + | 4607 | 4634 | **RNA**→ rplS (PRK05338)Ribosomal_L19 (pfam01245)→ |
| env-1657 | JGI24023J19991_10004490 | + | 7709 | 7736 | **RNA**→ rplS (PRK05338)Ribosomal_L19 (pfam01245)→ |
| env-1658 | JGI24023J19991_10080503 | - | 958 | 931 | **RNA**→ rplS (PRK05338)Ribosomal_L19 (pfam01245)→ |
| env-1659 | JGI24723J26617_10180956 | + | 346 | 373 | **RNA**→ rplS (PRK05338)Ribosomal_L19 (pfam01245)→ |
| env-1660 | KGI_S1_ANT01_95m_c10580845 | + | 177 | 204 | **RNA**→ |
| env-1661 | SA_S1_NOR05_45m_c10210261 | - | 309 | 282 | **RNA**→ rplS (PRK05338)Ribosomal_L19 (pfam01245)→ |
| env-1662 | SA_S1_NOR08_45m_c10537755 | - | 218 | 191 | **RNA**→ rplS (PRK05338)Ribosomal_L19 (pfam01245)→ |
| env-1663 | BS_KBA_SWE02_21m_10201411 | - | 197 | 170 | **RNA**→ rplS (PRK05338)Ribosomal_L19 (pfam01245)→ |
| env-1664 | JGI24023J19991_10000890 | + | 13567 | 13594 | **RNA**→ rplS (PRK05338)Ribosomal_L19 (pfam01245)→ |
| env-1665 | JGI24023J19991_10066851 | - | 58 | 31 | **RNA**→ |
| env-1666 | JGI24723J26617_10016062 | - | 2589 | 2562 | **RNA**→ rplS (PRK05338)Ribosomal_L19 (pfam01245)→ |
| env-1667 | SA_S2_NOR13_50m_c1002773 | + | 505 | 532 | **RNA**→ rplS (PRK05338)Ribosomal_L19 (pfam01245)→ |
| env-1668 | SA_S1_NOR08_45m_c10050681 | - | 1008 | 981 | **RNA**→ rplS (PRK05338)Ribosomal_L19 (pfam01245)→ |
| env-1669 | SA_S2_NOR18_50m_1016505 | + | 743 | 770 | **RNA**→ rplS (PRK05338)Ribosomal_L19 (pfam01245)→ |
| env-1670 | BS_KBA_SWE12_21m_c10023034 | + | 75 | 102 | **RNA**→ hypo→ rplS (PRK05338)Ribosomal_L19 (pfam01245)→ |
| env-1671 | TDF_MC_ARG01_113m_c1073913 | - | 277 | 250 | **RNA**→ rplS (PRK05338)Ribosomal_L19 (pfam01245)→ |
| env-1672 | SA_S1_NOR05_45m_c10164455 | - | 247 | 220 | **RNA**→ rplS (PRK05338)Ribosomal_L19 (pfam01245)→ |
| env-1673 | SA_S1_NOR08_45m_c10222535 | + | 399 | 426 | **RNA**→ hypo→ |
| env-1674 | KGI_S2_ANT04_2345m_c1080213 | - | 311 | 284 | **RNA**→ rplS (PRK05338)Ribosomal_L19 (pfam01245)→ |
| env-1675 | KGI_S2_ANT05_2345m_c1048372 | + | 258 | 285 | **RNA**→ rplS (PRK05338)Ribosomal_L19 (pfam01245)→ |
| env-1676 | KGI_S1_ANT02_95m_c10148848 | + | 250 | 277 | **RNA**→ rplS (PRK05338)Ribosomal_L19 (pfam01245)→ |
| env-1677 | SA_S1_NOR05_45m_c10033322 | - | 654 | 627 | **RNA**→ rplS (PRK05338)Ribosomal_L19 (pfam01245)→ |
| env-1678 | SA_S2_NOR15_50m_c10038594 | - | 1024 | 997 | **RNA**→ rplS (PRK05338)Ribosomal_L19 (pfam01245)→ |
| env-1679 | SA_S1_NOR02_45m_c1001276 | + | 495 | 522 | **RNA**→ rplS (PRK05338)Ribosomal_L19 (pfam01245)→ |
| env-1680 | DelMOSum2010_c10000022 | + | 68552 | 68579 | **RNA**→ rplS (PRK05338)Ribosomal_L19 (pfam01245)→ |
| env-1681 | JGI20152J14361_10000425 | + | 29939 | 29966 | **RNA**→ rplS (PRK05338)Ribosomal_L19 (pfam01245)→ |
| env-1682 | JGI20156J14371_10000267 | - | 17345 | 17318 | **RNA**→ rplS (PRK05338)Ribosomal_L19 (pfam01245)→ |
| env-1683 | JGI20160J14292_10003446 | - | 6169 | 6142 | **RNA**→ rplS (PRK05338)Ribosomal_L19 (pfam01245)→ |
| env-1684 | JGI20157J14317_10000008 | - | 112600 | 112573 | **RNA**→ rplS (PRK05338)Ribosomal_L19 (pfam01245)→ |
| env-1685 | JGI20159J14440_10247722 | - | 454 | 427 | **RNA**→ rplS (PRK05338)Ribosomal_L19 (pfam01245)→ |
| env-1686 | JGI20155J14468_10011132 | - | 1438 | 1411 | **RNA**→ rplS (PRK05338)Ribosomal_L19 (pfam01245)→ |
| env-1687 | JGI24024J18818_10003851 | - | 4397 | 4370 | **RNA**→ rplS (PRK05338)Ribosomal_L19 (pfam01245)→ |
| env-1688 | JGI24025J20009_10171329 | - | 268 | 241 | **RNA**→ rplS (PRK05338)Ribosomal_L19 (pfam01245)→ |
| env-1689 | JGI24023J19991_10009898 | - | 4045 | 4018 | **RNA**→ rplS (PRK05338)Ribosomal_L19 (pfam01245)→ |
| env-1690 | AMQJ01014829.1 | + | 152 | 179 | **RNA**→ rplS (PRK05338)Ribosomal_L19 (pfam01245)→ |
| env-1691 | BS_KBB_SWE26_205m_c1063562 | - | 562 | 535 | **RNA**→ rplS (PRK05338)Ribosomal_L19 (pfam01245)→ |
| env-1692 | LADL01001103.1 | - | 19831 | 19804 | **RNA**→ rplS (PRK05338)Ribosomal_L19 (pfam01245)→ |
| env-1693 | 3300001605_10098760 | + | 18 | 45 | **RNA**→ rplS (PRK05338)Ribosomal_L19 (pfam01245)→ |
| env-1694 | 3300001605_10728906 | + | 37 | 64 | **RNA**→ rplS (PRK05338)Ribosomal_L19 (pfam01245)→ |
| env-1695 | JGI25319J35699_1005705 | - | 2536 | 2509 | **RNA**→ rplS (PRK05338)Ribosomal_L19 (pfam01245)→ |
| env-1696 | JGI25322J35698_1044001 | + | 313 | 340 | **RNA**→ rplS (PRK05338)Ribosomal_L19 (pfam01245)→ |
| env-1697 | JGI25322J35698_1082485 | - | 304 | 277 | **RNA**→ rplS (PRK05338)Ribosomal_L19 (pfam01245)→ |
| env-1698 | JGI25321J50212_10001262 | + | 10258 | 10285 | **RNA**→ rplS (PRK05338)Ribosomal_L19 (pfam01245)→ |

| | | | | | |
|---|---|---|---|---|---|
| env-1699 | JGI24023J19991_10000724 | - | 16241 | 16214 | **RNA**→ rplS (PRK05338)Ribosomal_L19 (pfam01245)→ |
| env-1700 | JGI24023J19991_10020737 | - | 942 | 915 | **RNA**→ rplS (PRK05338)Ribosomal_L19 (pfam01245)→ |
| env-1701 | JGI24724J26744_10238812 | + | 132 | 159 | **RNA**→ rplS (PRK05338)Ribosomal_L19 (pfam01245)→ |
| env-1702 | GYQO9XB02HASQ4 | + | 72 | 99 | **RNA**→ rplS (PRK05338)Ribosomal_L19 (pfam01245)→ |
| env-1703 | SA_S2_NOR13_50m_c1000423 | - | 7317 | 7290 | **RNA**→ rplS (PRK05338)Ribosomal_L19 (pfam01245)→ |
| env-1704 | SA_S1_NOR05_45m_c10000647 | + | 6814 | 6841 | **RNA**→ rplS (PRK05338)Ribosomal_L19 (pfam01245)→ |
| env-1705 | SA_S1_NOR08_45m_c10010910 | + | 1079 | 1106 | **RNA**→ rplS (PRK05338)Ribosomal_L19 (pfam01245)→ |
| env-1706 | SA_S2_NOR15_50m_c10000830 | - | 7224 | 7197 | **RNA**→ rplS (PRK05338)Ribosomal_L19 (pfam01245)→ |
| env-1707 | SA_S1_NOR02_45m_c1000223 | + | 5930 | 5957 | **RNA**→ rplS (PRK05338)Ribosomal_L19 (pfam01245)→ |
| env-1708 | SA_S2_NOR18_50m_1000296 | + | 7034 | 7061 | **RNA**→ rplS (PRK05338)Ribosomal_L19 (pfam01245)→ |
| env-1709 | SA_S1_NOR08_45m_c10776056 | + | 93 | 120 | **RNA**→ rplS (PRK05338)Ribosomal_L19 (pfam01245)→ |
| env-1710 | SA_S2_NOR15_50m_c10819626 | + | 139 | 166 | **RNA**→ rplS (PRK05338)Ribosomal_L19 (pfam01245)→ |
| env-1711 | KGI_S2_ANT04_2345m_c1093702 | - | 353 | 326 | **RNA**→ rplS (PRK05338)Ribosomal_L19 (pfam01245)→ |
| env-1712 | JGI24024J18818_10091357 | + | 178 | 205 | **RNA**→ rplS (PRK05338)Ribosomal_L19 (pfam01245)→ |
| env-1713 | JGI24023J19991_10004817 | + | 8687 | 8714 | **RNA**→ rplS (PRK05338)Ribosomal_L19 (pfam01245)→ |
| env-1714 | JGI24724J26744_10028319 | + | 646 | 673 | **RNA**→ rplS (PRK05338)Ribosomal_L19 (pfam01245)→ |
| env-1715 | JGI25321J39296_1005427 | - | 539 | 512 | **RNA**→ rplS (PRK05338)Ribosomal_L19 (pfam01245)→ |
| env-1716 | KGI_S1_ANT01_95m_c10099108 | - | 675 | 648 | **RNA**→ rplS (PRK05338)Ribosomal_L19 (pfam01245)→ |
| env-1717 | JGI20152J14361_10116815 | - | 44 | 17 | **RNA**→ |
| env-1718 | CESJ01024341.1 | - | 209 | 182 | **RNA**→ rplS (PRK05338)Ribosomal_L19 (pfam01245)→ |
| env-1719 | CESU01225158.1 | - | 101 | 74 | **RNA**→ ←hypo |
| env-1720 | 2236876010_0042617 | - | 400 | 373 | **RNA**→ ←hypo |
| env-1721 | CESE01142542.1 | - | 615 | 588 | **RNA**→ rplS (PRK05338)Ribosomal_L19 (pfam01245)→ |
| env-1722 | CEST01023914.1 | + | 29031 | 29058 | **RNA**→ rplS (PRK05338)Ribosomal_L19 (pfam01245)→ |
| env-1723 | SI47jul10_100m_c1098063 | + | 194 | 221 | **RNA**→ rplS (PRK05338)Ribosomal_L19 (pfam01245)→ |
| env-1724 | SI60aug11_100m_c1028962 | + | 97 | 124 | **RNA**→ rplS (PRK05338)Ribosomal_L19 (pfam01245)→ |
| env-1725 | SI34jun09_10m_1003941 | - | 2294 | 2267 | **RNA**→ rplS (PRK05338)Ribosomal_L19 (pfam01245)→ |
| env-1726 | SI34jun09_120m_1016406 | + | 1739 | 1766 | **RNA**→ rplS (PRK05338)Ribosomal_L19 (pfam01245)→ |
| env-1727 | SI34jun09_135m_1017351 | + | 105 | 132 | **RNA**→ rplS (PRK05338)Ribosomal_L19 (pfam01245)→ |
| env-1728 | SI34jun09_100m_1011215 | + | 2334 | 2361 | **RNA**→ ←hypo |
| env-1729 | JGI20152J14361_10052984 | - | 284 | 257 | **RNA**→ rplS (PRK05338)Ribosomal_L19 (pfam01245)→ |
| env-1730 | JGI20154J14316_10033520 | - | 1618 | 1591 | **RNA**→ rplS (PRK05338)Ribosomal_L19 (pfam01245)→ |
| env-1731 | JGI20153J14295_10031874 | + | 864 | 891 | **RNA**→ rplS (PRK05338)Ribosomal_L19 (pfam01245)→ |
| env-1732 | JGI20153J14318_10031874 | + | 864 | 891 | **RNA**→ rplS (PRK05338)Ribosomal_L19 (pfam01245)→ |
| env-1733 | SI34jun09_100m_1117558 | - | 60 | 33 | **RNA**→ |
| env-1734 | JGI20154J14316_10134296 | - | 50 | 23 | **RNA**→ ←hypo |
| env-1735 | A09P04500a_Sequence0000006337 | - | 1219 | 1192 | **RNA**→ rplS (PRK05338)Ribosomal_L19 (pfam01245)→ |
| env-1736 | 2236876010_0103866 | - | 296 | 269 | **RNA**→ rplS (PRK05338)Ribosomal_L19 (pfam01245)→ |
| env-1737 | 2236876010_0516017 | - | 121 | 94 | **RNA**→ hypo→ |
| env-1738 | SI48aug10_10m_c1005499 | - | 1214 | 1187 | **RNA**→ rplS (PRK05338)Ribosomal_L19 (pfam01245)→ |
| env-1739 | SI34jun09_100m_1006670 | - | 3438 | 3411 | **RNA**→ rplS (PRK05338)Ribosomal_L19 (pfam01245)→ |
| env-1740 | SI48aug10_100m_1215338 | - | 233 | 206 | **RNA**→ rplS (PRK05338)Ribosomal_L19 (pfam01245)→ |
| env-1741 | JGI20152J14361_10035731 | - | 36 | 9 | **RNA**→ |
| env-1742 | JGI20154J14316_10050212 | + | 864 | 891 | **RNA**→ rplS (PRK05338)Ribosomal_L19 (pfam01245)→ |
| env-1743 | DelMOSum2010_c10404944 | + | 225 | 252 | **RNA**→ rplS (PRK05338)Ribosomal_L19 (pfam01245)→ |
| env-1744 | JGI20152J14361_10057660 | + | 53 | 80 | **RNA**→ rplS (PRK05338)Ribosomal_L19 (pfam01245)→ |
| env-1745 | JGI20160J14292_10015472 | - | 917 | 890 | **RNA**→ rplS (PRK05338)Ribosomal_L19 (pfam01245)→ |
| env-1746 | JGI20153J14295_10017977 | - | 917 | 890 | **RNA**→ rplS (PRK05338)Ribosomal_L19 (pfam01245)→ |
| env-1747 | JGI20153J14318_10017977 | - | 917 | 890 | **RNA**→ rplS (PRK05338)Ribosomal_L19 (pfam01245)→ |
| env-1748 | CESI01016430.1 | - | 9287 | 9260 | **RNA**→ rplS (PRK05338)Ribosomal_L19 (pfam01245)→ |
| env-1749 | CESU01088699.1 | - | 907 | 880 | **RNA**→ rplS (PRK05338)Ribosomal_L19 (pfam01245)→ |
| env-1750 | JGI20160J14292_10015444 | - | 915 | 888 | **RNA**→ rplS (PRK05338)Ribosomal_L19 (pfam01245)→ |

| | | | | | |
|---|---|---|---|---|---|
| env-1751 | JGI20152J14361_10058556 | - | 907 | 880 | **RNA**→ rplS (PRK05338)Ribosomal_L19 (pfam01245)→ |
| env-1752 | JGI20151J14362_10017387 | - | 622 | 595 | **RNA**→ rplS (PRK05338)Ribosomal_L19 (pfam01245)→ |
| env-1753 | JGI20156J14371_10018698 | - | 1553 | 1526 | **RNA**→ rplS (PRK05338)Ribosomal_L19 (pfam01245)→ |
| env-1754 | JGI20153J14295_10043016 | - | 915 | 888 | **RNA**→ rplS (PRK05338)Ribosomal_L19 (pfam01245)→ |
| env-1755 | JGI20153J14318_10043016 | - | 915 | 888 | **RNA**→ rplS (PRK05338)Ribosomal_L19 (pfam01245)→ |
| env-1756 | JGI20157J14317_10051617 | - | 1574 | 1547 | **RNA**→ rplS (PRK05338)Ribosomal_L19 (pfam01245)→ |
| env-1757 | JGI20155J14468_10700974 | - | 81 | 54 | **RNA**→ |
| env-1758 | SA_S1_NOR05_45m_c10078427 | - | 539 | 512 | **RNA**→ rplS (PRK05338)Ribosomal_L19 (pfam01245)→ |
| env-1759 | SA_S1_NOR08_45m_c10045004 | + | 838 | 865 | **RNA**→ rplS (PRK05338)Ribosomal_L19 (pfam01245)→ |
| env-1760 | KGI_S2_ANT05_2345m_c1126697 | + | 69 | 96 | **RNA**→ hypo→ rplS (PRK05338)Ribosomal_L19 (pfam01245)→ |
| env-1761 | DelMOWin2010_c10293423 | - | 119 | 92 | **RNA**→ hypo→ |
| env-1762 | SA_S1_NOR05_45m_c10023438 | + | 31 | 58 | **RNA**→ hypo→ rplS (PRK05338)Ribosomal_L19 (pfam01245)→ |
| env-1763 | SA_S2_NOR15_50m_c10177335 | + | 332 | 359 | **RNA**→ rplS (PRK05338)Ribosomal_L19 (pfam01245)→ |
| env-1764 | wf3_Contig6924 | + | 1855 | 1882 | **RNA**→ rplS (PRK05338)Ribosomal_L19 (pfam01245)→ |
| env-1765 | SA_S2_NOR13_50m_c1013867 | + | 488 | 515 | **RNA**→ rplS (PRK05338)Ribosomal_L19 (pfam01245)→ |
| env-1766 | SA_S1_NOR05_45m_c10010432 | - | 1796 | 1769 | **RNA**→ rplS (PRK05338)Ribosomal_L19 (pfam01245)→ |
| env-1767 | SA_S1_NOR08_45m_c10037082 | - | 634 | 607 | **RNA**→ rplS (PRK05338)Ribosomal_L19 (pfam01245)→ |
| env-1768 | SA_S2_NOR15_50m_c10017076 | - | 2062 | 2035 | **RNA**→ rplS (PRK05338)Ribosomal_L19 (pfam01245)→ |
| env-1769 | SA_S2_NOR15_50m_c10038947 | + | 1097 | 1124 | **RNA**→ rplS (PRK05338)Ribosomal_L19 (pfam01245)→ |
| env-1770 | SA_S2_NOR18_50m_1060527 | - | 230 | 203 | **RNA**→ rplS (PRK05338)Ribosomal_L19 (pfam01245)→ |
| env-1771 | SA_S1_NOR05_45m_c10402627 | + | 248 | 275 | **RNA**→ |
| env-1772 | SA_S1_NOR05_45m_c10000045 | - | 1209 | 1182 | **RNA**→ rplS (PRK05338)Ribosomal_L19 (pfam01245)→ |
| env-1773 | SA_S1_NOR08_45m_c10003536 | - | 7347 | 7320 | **RNA**→ rplS (PRK05338)Ribosomal_L19 (pfam01245)→ |
| env-1774 | SA_S1_NOR08_45m_c10000343 | - | 1310 | 1283 | **RNA**→ rplS (PRK05338)Ribosomal_L19 (pfam01245)→ |
| env-1775 | SA_S2_NOR15_50m_c10001574 | + | 6336 | 6363 | **RNA**→ rplS (PRK05338)Ribosomal_L19 (pfam01245)→ |
| env-1776 | SA_S1_NOR02_45m_c1000618 | - | 359 | 332 | **RNA**→ rplS (PRK05338)Ribosomal_L19 (pfam01245)→ |
| env-1777 | JGI20153J14295_10002993 | + | 9139 | 9166 | **RNA**→ rplS (PRK05338)Ribosomal_L19 (pfam01245)→ hypo→ |
| env-1778 | JGI20153J14318_10002993 | + | 9139 | 9166 | **RNA**→ rplS (PRK05338)Ribosomal_L19 (pfam01245)→ hypo→ |
| env-1779 | TDF_OR_ARG04_113m_c1085059 | + | 254 | 281 | **RNA**→ rplS (PRK05338)Ribosomal_L19 (pfam01245)→ |
| env-1780 | TDF_MC_ARG01_113m_c1019942 | + | 199 | 226 | **RNA**→ rplS (PRK05338)Ribosomal_L19 (pfam01245)→ |
| env-1781 | SA_S1_NOR08_45m_c10002903 | + | 2910 | 2937 | **RNA**→ rplS (PRK05338)Ribosomal_L19 (pfam01245)→ |
| env-1782 | A09P04_1300_Sequence0000007657 | + | 1676 | 1703 | **RNA**→ ←hypo |
| env-1783 | KGI_S1_ANT01_95m_c10287783 | + | 183 | 210 | **RNA**→ rplS (PRK05338)Ribosomal_L19 (pfam01245)→ |
| env-1784 | KGI_S2_ANT06_2345m_c1008730 | + | 1837 | 1864 | **RNA**→ rplS (PRK05338)Ribosomal_L19 (pfam01245)→ |
| env-1785 | KGI_S2_ANT04_2345m_c1006171 | + | 550 | 577 | **RNA**→ rplS (PRK05338)Ribosomal_L19 (pfam01245)→ |
| env-1786 | KGI_S1_ANT03_95m_c1134508 | - | 63 | 36 | **RNA**→ |
| env-1787 | JGI20157J14317_10111054 | - | 633 | 606 | **RNA**→ rplS (PRK05338)Ribosomal_L19 (pfam01245)→ |
| env-1788 | SA_S1_NOR08_45m_c10034502 | - | 1734 | 1707 | **RNA**→ rplS (PRK05338)Ribosomal_L19 (pfam01245)→ |
| env-1789 | SA_S2_NOR15_50m_c10791456 | - | 35 | 8 | **RNA**→ hypo→ |
| env-1790 | SA_S1_NOR05_45m_c10068256 | - | 784 | 757 | **RNA**→ rplS (PRK05338)Ribosomal_L19 (pfam01245)→ |
| env-1791 | SA_S1_NOR02_45m_c1126975 | + | 134 | 161 | **RNA**→ rplS (PRK05338)Ribosomal_L19 (pfam01245)→ |
| env-1792 | JGI24023J19991_10001462 | - | 645 | 618 | **RNA**→ rplS (PRK05338)Ribosomal_L19 (pfam01245)→ |
| env-1793 | TB_FS06_10DRAFT_1000012 | - | 81337 | 81310 | **RNA**→ rplS (PRK05338)Ribosomal_L19 (pfam01245)→ |
| env-1794 | CEQG01202036.1 | - | 389 | 362 | **RNA**→ rplS (PRK05338)Ribosomal_L19 (pfam01245)→ |
| env-1795 | CETB01163705.1 | - | 1198 | 1171 | **RNA**→ rplS (PRK05338)Ribosomal_L19 (pfam01245)→ |
| env-1796 | CETR01130265.1 | - | 4705 | 4678 | **RNA**→ rplS (PRK05338)Ribosomal_L19 (pfam01245)→ |
| env-1797 | CEVI01205692.1 | - | 792 | 765 | **RNA**→ rplS (PRK05338)Ribosomal_L19 (pfam01245)→ |
| env-1798 | CEVK01214383.1 | + | 636 | 663 | **RNA**→ rplS (PRK05338)Ribosomal_L19 (pfam01245)→ |
| env-1799 | CEVM01258184.1 | + | 830 | 857 | **RNA**→ rplS (PRK05338)Ribosomal_L19 (pfam01245)→ |
| env-1800 | CEVO01247352.1 | + | 698 | 725 | **RNA**→ rplS (PRK05338)Ribosomal_L19 (pfam01245)→ |
| env-1801 | CEVQ01122451.1 | - | 320926 | 320899 | **RNA**→ rplS (PRK05338)Ribosomal_L19 (pfam01245)→ |
| env-1802 | KVRMV2_102248759 | - | 166 | 139 | **RNA**→ rplS (PRK05338)Ribosomal_L19 (pfam01245)→ |

| | | | | | |
|---|---|---|---|---|---|
| env-1803 | SA_S1_NOR05_45m_c10338159 | - | 197 | 170 | **RNA**→ rplS (PRK05338)Ribosomal_L19 (pfam01245)→ |
| env-1804 | TB_PC08_66_10120445 | + | 167 | 194 | **RNA**→ rplS (PRK05338)Ribosomal_L19 (pfam01245)→ |
| env-1805 | TB_PC08_3_1025062 | - | 907 | 880 | **RNA**→ rplS (PRK05338)Ribosomal_L19 (pfam01245)→ |
| env-1806 | JGI20156J14371_10038484 | + | 575 | 602 | **RNA**→ rplS (PRK05338)Ribosomal_L19 (pfam01245)→ |
| env-1807 | TB_FS06_10DRAFT_1028770 | + | 1020 | 1047 | **RNA**→ rplS (PRK05338)Ribosomal_L19 (pfam01245)→ |
| env-1808 | TB_FS08_3DRAFT_1011355 | + | 1346 | 1373 | **RNA**→ rplS (PRK05338)Ribosomal_L19 (pfam01245)→ |
| env-1809 | TB_GS10_10_10046912 | - | 1239 | 1212 | **RNA**→ rplS (PRK05338)Ribosomal_L19 (pfam01245)→ |
| env-1810 | TB_PC08_64_1001821 | - | 694 | 667 | **RNA**→ rplS (PRK05338)Ribosomal_L19 (pfam01245)→ |
| env-1811 | TB_GS09_5_10120216 | + | 181 | 208 | **RNA**→ rplS (PRK05338)Ribosomal_L19 (pfam01245)→ |
| env-1812 | SA_S2_NOR13_50m_c1168462 | - | 250 | 223 | **RNA**→ rplS (PRK05338)Ribosomal_L19 (pfam01245)→ |
| env-1813 | F3OLC3J02IOFD0 | + | 181 | 208 | **RNA**→ rplS (PRK05338)Ribosomal_L19 (pfam01245)→ |
| env-1814 | DelMOSum2010_c10132439 | + | 587 | 614 | **RNA**→ rplS (PRK05338)Ribosomal_L19 (pfam01245)→ |
| env-1815 | JGI20154J14316_10553810 | + | 41 | 68 | **RNA**→ rplS (PRK05338)Ribosomal_L19 (pfam01245)→ |
| env-1816 | Y607_1006251 | + | 85823 | 85850 | **RNA**→ rplS (PRK05338)Ribosomal_L19 (pfam01245)→ |
| env-1817 | P_2C_Liq_1_UnCty_1000596 | - | 3980 | 3953 | **RNA**→ rplS (PRK05338)Ribosomal_L19 (pfam01245)→ |
| env-1818 | P_2C_Liq_2_UnCty_1018319 | - | 663 | 636 | **RNA**→ rplS (PRK05338)Ribosomal_L19 (pfam01245)→ |
| env-1819 | CESM01027170.1 | - | 1718 | 1691 | **RNA**→ rplS (PRK05338)Ribosomal_L19 (pfam01245)→ |
| env-1820 | CESO01163408.1 | - | 81 | 54 | **RNA**→ |
| env-1821 | 2236876010_0590763 | + | 140 | 167 | **RNA**→ rplS (PRK05338)Ribosomal_L19 (pfam01245)→ |
| env-1822 | CERM01240744.1 | - | 96 | 69 | **RNA**→ |
| env-1823 | CESO01076655.1 | + | 584 | 611 | **RNA**→ rplS (PRK05338)Ribosomal_L19 (pfam01245)→ |
| env-1824 | SI34jun09_10m_1037670 | - | 665 | 638 | **RNA**→ hypo→ rplS (PRK05338)Ribosomal_L19 (pfam01245)→ |
| env-1825 | JCVI_SCAF_1096627114274 | - | 579 | 552 | **RNA**→ rplS (PRK05338)Ribosomal_L19 (pfam01245)→ |
| env-1826 | KGI_S1_ANT01_95m_c10055806 | - | 1138 | 1111 | **RNA**→ rplS (PRK05338)Ribosomal_L19 (pfam01245)→ |
| env-1827 | KGI_S2_ANT04_2345m_c1015516 | + | 2067 | 2094 | **RNA**→ |
| env-1828 | KGI_S2_ANT05_2345m_c1137445 | - | 257 | 230 | **RNA**→ rplS (PRK05338)Ribosomal_L19 (pfam01245)→ |
| env-1829 | KGI_S1_ANT02_95m_c10013777 | - | 1526 | 1499 | **RNA**→ rplS (PRK05338)Ribosomal_L19 (pfam01245)→ |
| env-1830 | SA_S2_NOR18_50m_1001412 | + | 399 | 426 | **RNA**→ rplS (PRK05338)Ribosomal_L19 (pfam01245)→ |
| env-1831 | JGI24723J26617_10068646 | + | 585 | 612 | **RNA**→ rplS (PRK05338)Ribosomal_L19 (pfam01245)→ |
| env-1832 | SA_S2_NOR13_50m_c1001448 | - | 4743 | 4716 | **RNA**→ rplS (PRK05338)Ribosomal_L19 (pfam01245)→ |
| env-1833 | SA_S1_NOR05_45m_c10001450 | + | 3917 | 3944 | **RNA**→ rplS (PRK05338)Ribosomal_L19 (pfam01245)→ |
| env-1834 | SA_S1_NOR08_45m_c10000307 | + | 15014 | 15041 | **RNA**→ rplS (PRK05338)Ribosomal_L19 (pfam01245)→ |
| env-1835 | SA_S2_NOR15_50m_c10010164 | - | 3285 | 3258 | **RNA**→ rplS (PRK05338)Ribosomal_L19 (pfam01245)→ |
| env-1836 | SA_S1_NOR02_45m_c1019106 | - | 530 | 503 | **RNA**→ rplS (PRK05338)Ribosomal_L19 (pfam01245)→ |
| env-1837 | SA_S1_NOR08_45m_c10000615 | - | 6292 | 6265 | **RNA**→ rplS (PRK05338)Ribosomal_L19 (pfam01245)→ |
| env-1838 | SA_S2_NOR15_50m_c10000097 | + | 10972 | 10999 | **RNA**→ rplS (PRK05338)Ribosomal_L19 (pfam01245)→ |
| env-1839 | SA_S2_NOR15_50m_c10469518 | - | 217 | 190 | **RNA**→ rplS (PRK05338)Ribosomal_L19 (pfam01245)→ |
| env-1840 | JGI24724J26744_11008656 | + | 37 | 64 | **RNA**→ rplS (PRK05338)Ribosomal_L19 (pfam01245)→ |
| env-1841 | TDF_MC_ARG01_113m_c1006717 | + | 93 | 120 | **RNA**→ rplS (PRK05338)Ribosomal_L19 (pfam01245)→ |
| env-1842 | SA_S1_NOR05_45m_c10106893 | + | 70 | 97 | **RNA**→ rplS (PRK05338)Ribosomal_L19 (pfam01245)→ |
| env-1843 | KGI_S1_ANT03_95m_c1004957 | + | 964 | 991 | **RNA**→ rplS (PRK05338)Ribosomal_L19 (pfam01245)→ |
| env-1844 | CESI01143241.1 | - | 582 | 555 | **RNA**→ rplS (PRK05338)Ribosomal_L19 (pfam01245)→ |
| env-1845 | KGI_S1_ANT01_95m_c10121130 | - | 48 | 21 | **RNA**→ |
| env-1846 | TDF_OR_ARG04_113m_c1009977 | - | 584 | 557 | **RNA**→ rplS (PRK05338)Ribosomal_L19 (pfam01245)→ |
| env-1847 | TDF_MC_ARG01_113m_c1004441 | - | 665 | 638 | **RNA**→ rplS (PRK05338)Ribosomal_L19 (pfam01245)→ |
| env-1848 | KGI_S2_ANT04_2345m_c1086413 | + | 531 | 558 | **RNA**→ ←hypo |
| env-1849 | JGI24024J18818_10700231 | + | 18 | 45 | **RNA**→ rplS (PRK05338)Ribosomal_L19 (pfam01245)→ |
| env-1850 | JGI24723J26617_10589038 | + | 80 | 107 | **RNA**→ rplS (PRK05338)Ribosomal_L19 (pfam01245)→ |
| env-1851 | TDF_OR_ARG04_113m_c1145284 | - | 81 | 54 | **RNA**→ ←hypo |
| env-1852 | TDF_MC_ARG01_113m_c1013236 | - | 275 | 248 | **RNA**→ ←hypo |
| env-1853 | CEVE01269246.1 | + | 219 | 246 | **RNA**→ rplS (PRK05338)Ribosomal_L19 (pfam01245)→ |
| env-1854 | CEVF01060294.1 | + | 1089 | 1116 | **RNA**→ rplS (PRK05338)Ribosomal_L19 (pfam01245)→ |

| | | | | | |
|---|---|---|---|---|---|
| Mlu-1-1 | NZ_CP011071.1 | + | 10994 | 11021 | **RNA**→ rplS (PRK05338)Ribosomal_L19 (pfam01245)→ |
| env-1855 | 2236876010_0651002 | - | 265 | 238 | **RNA**→ rplS (PRK05338)Ribosomal_L19 (pfam01245)→ |
| env-1856 | 2236876010_0559523 | + | 257 | 284 | **RNA**→ |
| env-1857 | JGI20154J14316_10006430 | + | 537 | 564 | **RNA**→ rplS (PRK05338)Ribosomal_L19 (pfam01245)→ |
| env-1858 | JGI20153J14295_10000108 | - | 47780 | 47753 | **RNA**→ rplS (PRK05338)Ribosomal_L19 (pfam01245)→ |
| env-1859 | JGI20153J14318_10000108 | - | 47780 | 47753 | **RNA**→ rplS (PRK05338)Ribosomal_L19 (pfam01245)→ |
| env-1860 | JCVI_SCAF_1096627739007 | - | 830 | 803 | **RNA**→ rplS (PRK05338)Ribosomal_L19 (pfam01245)→ PRK08299 (PRK08299)→ |
| env-1861 | CETK01280519.1 | + | 89 | 116 | **RNA**→ rplS (PRK05338)Ribosomal_L19 (pfam01245)→ PRK08299 (PRK08299)→ |
| Ala-2-1 | NZ_AUKX01000041.1 | + | 37352 | 37379 | **RNA**→ rplS (PRK05338)Ribosomal_L19 (pfam01245)→ |
| env-1862 | JGI24723J26617_10101877 | - | 728 | 701 | **RNA**→ rplS (PRK05338)Ribosomal_L19 (pfam01245)→ |
| Ndo-1-1 | NC_020156.1 | + | 117213 | 117240 | **RNA**→ rplS (PRK05338)Ribosomal_L19 (pfam01245)→ |
| env-1863 | DelMOSum2011_c10602551 | - | 154 | 127 | **RNA**→ rplS (PRK05338)Ribosomal_L19 (pfam01245)→ |
| env-1864 | A09P0410a_Sequence0000028955 | + | 1010 | 1037 | **RNA**→ |
| Cal-1-1 | NC_014934.1 | + | 10579 | 10606 | **RNA**→ rplS (PRK05338)Ribosomal_L19 (pfam01245)→ |
| env-1865 | 2236876009_020497 | + | 88 | 115 | **RNA**→ |
| Csp-13-1 | NZ_JUHB01000001.1 | + | 1008351 | 1008378 | **RNA**→ rplS (PRK05338)Ribosomal_L19 (pfam01245)→ |
| Mfo-1-1 | NZ_JQLH01000001.1 | + | 2737717 | 2737744 | **RNA**→ rplS (PRK05338)Ribosomal_L19 (pfam01245)→ |
| Cba-4-1 | NZ_CP009887.1 | + | 4088266 | 4088293 | **RNA**→ rplS (PRK05338)Ribosomal_L19 (pfam01245)→ |
| Cba-2-1 | NZ_CP009976.1 | + | 4128380 | 4128407 | **RNA**→ rplS (PRK05338)Ribosomal_L19 (pfam01245)→ |
| Cba-1-1 | NZ_ATLH01000022.1 | + | 273938 | 273965 | **RNA**→ rplS (PRK05338)Ribosomal_L19 (pfam01245)→ |
| Cba-3-1 | NZ_ATLG01000021.1 | + | 275945 | 275972 | **RNA**→ rplS (PRK05338)Ribosomal_L19 (pfam01245)→ |
| Csp-12-1 | NZ_JQCT01000003.1 | - | 7091 | 7064 | **RNA**→ rplS (PRK05338)Ribosomal_L19 (pfam01245)→ |
| env-1866 | CEQL01331727.1 | - | 1140 | 1113 | **RNA**→ rplS (PRK05338)Ribosomal_L19 (pfam01245)→ PRK08299 (PRK08299)→ |
| env-1867 | BS_KBA_SWE12_21m_c10696970 | + | 162 | 189 | **RNA**→ |
| env-1868 | BS_KBA_SWE02_21m_10122283 | + | 143 | 170 | **RNA**→ rplS (PRK05338)Ribosomal_L19 (pfam01245)→ |
| env-1869 | 3300001605_10642224 | + | 131 | 158 | **RNA**→ rplS (PRK05338)Ribosomal_L19 (pfam01245)→ |
| env-1870 | 3300000558_11705956 | - | 186 | 159 | **RNA**→ rplS (PRK05338)Ribosomal_L19 (pfam01245)→ |
| env-1871 | 3300001605_11328178 | - | 116 | 89 | **RNA**→ hypo→ |
| env-1872 | CENX01111910.1 | + | 8377 | 8404 | **RNA**→ rplS (PRK05338)Ribosomal_L19 (pfam01245)→ Icd (COG0538)PRK08299 (PRK08299)→<br>murQ (PRK05441)SIS_Etherase (cd05007)→ hypo→ |
| env-1873 | CENP01126039.1 | + | 1259 | 1286 | **RNA**→ |
| env-1874 | CEOF01188357.1 | - | 3192 | 3165 | **RNA**→ rplS (PRK05338)Ribosomal_L19 (pfam01245)→ Icd (COG0538)PRK08299 (PRK08299)→<br>murQ (PRK05441)SIS_Etherase (cd05007)→ hypo→ TGS_ThrRS_N (cd01667)ThrS (COG0441)thrS (PRK00413)→ |
| env-1875 | JGI20154J14316_10007303 | - | 5508 | 5481 | **RNA**→ rplS (PRK05338)Ribosomal_L19 (pfam01245)→ |
| env-1876 | JGI20160J14292_10005573 | - | 5508 | 5481 | **RNA**→ rplS (PRK05338)Ribosomal_L19 (pfam01245)→ |
| env-1877 | JGI20153J14295_10005409 | - | 5688 | 5661 | **RNA**→ rplS (PRK05338)Ribosomal_L19 (pfam01245)→ |
| env-1878 | JGI20153J14318_10005409 | - | 5688 | 5661 | **RNA**→ rplS (PRK05338)Ribosomal_L19 (pfam01245)→ |
| env-1879 | JGI20157J14317_10006339 | - | 4204 | 4177 | **RNA**→ rplS (PRK05338)Ribosomal_L19 (pfam01245)→ |
| env-1880 | KGI_S1_ANT01_95m_c10030633 | + | 858 | 885 | **RNA**→ rplS (PRK05338)Ribosomal_L19 (pfam01245)→ |
| env-1881 | SA_S1_NOR08_45m_c10000592 | - | 9103 | 9076 | **RNA**→ rplS (PRK05338)Ribosomal_L19 (pfam01245)→ |
| env-1882 | KGI_S2_ANT04_2345m_c1033268 | + | 162 | 189 | **RNA**→ rplS (PRK05338)Ribosomal_L19 (pfam01245)→ |
| env-1883 | SA_S2_NOR15_50m_c10000395 | - | 9786 | 9759 | **RNA**→ rplS (PRK05338)Ribosomal_L19 (pfam01245)→ |
| env-1884 | KGI_S2_ANT05_2345m_c1037235 | + | 96 | 123 | **RNA**→ rplS (PRK05338)Ribosomal_L19 (pfam01245)→ |
| env-1885 | KGI_S1_ANT02_95m_c10039728 | + | 412 | 439 | **RNA**→ rplS (PRK05338)Ribosomal_L19 (pfam01245)→ |
| env-1886 | JGI24023J19991_10000015 | - | 6676 | 6649 | **RNA**→ rplS (PRK05338)Ribosomal_L19 (pfam01245)→ |
| Orh-1-1 | NC_018016.1 | - | 563396 | 563369 | **RNA**→ rplS (PRK05338)Ribosomal_L19 (pfam01245)→ |
| Orh-3-1 | NZ_CP006828.1 | + | 1411724 | 1411751 | **RNA**→ rplS (PRK05338)Ribosomal_L19 (pfam01245)→ |
| Orh-2-1 | NZ_AXDE01000005.1 | + | 164490 | 164517 | **RNA**→ rplS (PRK05338)Ribosomal_L19 (pfam01245)→ |
| env-1887 | TB_FS06_10DRAFT_1002658 | + | 1277 | 1304 | **RNA**→ rplS (PRK05338)Ribosomal_L19 (pfam01245)→ |
| env-1888 | LGOV01058696.1 | + | 3884 | 3911 | **RNA**→ rplS (PRK05338)Ribosomal_L19 (pfam01245)→ |
| env-1889 | LGVE01220578.1 | - | 1243 | 1216 | **RNA**→ rplS (PRK05338)Ribosomal_L19 (pfam01245)→ |
| env-1890 | LGVF01377133.1 | + | 233 | 260 | **RNA**→ rplS (PRK05338)Ribosomal_L19 (pfam01245)→ |
| env-1891 | topACOD_F95O9CU01CV6MI | + | 310 | 337 | **RNA**→ ←hypo |

| | | | | | |
|---|---|---|---|---|---|
| env-1892 | SRS011243_C2014617 | + | 132 | 159 | **RNA**→ rplS (PRK05338)Ribosomal_L19 (pfam01245)→ |
| env-1893 | SRS013533_PGA_scaffold_14677 | + | 406 | 433 | **RNA**→ rplS (PRK05338)Ribosomal_L19 (pfam01245)→ |
| env-1894 | SRS014888_C3069310 | - | 488 | 461 | **RNA**→ rplS (PRK05338)Ribosomal_L19 (pfam01245)→ |
| env-1895 | SRS015762_C3674271 | - | 5976 | 5949 | **RNA**→ rplS (PRK05338)Ribosomal_L19 (pfam01245)→ |
| env-1896 | SRS019028_C2956184 | + | 301 | 328 | **RNA**→ rplS (PRK05338)Ribosomal_L19 (pfam01245)→ |
| env-1897 | SRS057355_C2324181 | - | 410 | 383 | **RNA**→ rplS (PRK05338)Ribosomal_L19 (pfam01245)→ |
| env-1898 | SRS015893_C2350217 | - | 265 | 238 | **RNA**→ rplS (PRK05338)Ribosomal_L19 (pfam01245)→ |
| env-1899 | SRS016575_Baylor_scaffold_56189 | - | 634 | 607 | **RNA**→ rplS (PRK05338)Ribosomal_L19 (pfam01245)→ |
| env-1900 | SRS015985_WUGC_scaffold_37008 | + | 406 | 433 | **RNA**→ rplS (PRK05338)Ribosomal_L19 (pfam01245)→ |
| env-1901 | SRS013949_WUGC_scaffold_26331 | + | 33 | 60 | **RNA**→ rplS (PRK05338)Ribosomal_L19 (pfam01245)→ |
| env-1902 | SRS015755_WUGC_scaffold_23418 | + | 50 | 77 | **RNA**→ rplS (PRK05338)Ribosomal_L19 (pfam01245)→ |
| env-1903 | SRS015989_WUGC_scaffold_7779 | + | 1237 | 1264 | **RNA**→ rplS (PRK05338)Ribosomal_L19 (pfam01245)→ |
| env-1904 | SRS018665_WUGC_scaffold_42984 | - | 394 | 367 | **RNA**→ rplS (PRK05338)Ribosomal_L19 (pfam01245)→ |
| env-1905 | SRS019077_C2600640 | - | 510 | 483 | **RNA**→ rplS (PRK05338)Ribosomal_L19 (pfam01245)→ |
| env-1906 | SRS019980_Baylor_scaffold_35308 | - | 329 | 302 | **RNA**→ rplS (PRK05338)Ribosomal_L19 (pfam01245)→ |
| env-1907 | SRS021960_Baylor_scaffold_16219 | - | 926 | 899 | **RNA**→ rplS (PRK05338)Ribosomal_L19 (pfam01245)→ |
| env-1908 | SRS022725_C5693545 | + | 21 | 48 | **RNA**→ rplS (PRK05338)Ribosomal_L19 (pfam01245)→ |
| env-1909 | SRS024144_C1769633 | - | 629 | 602 | **RNA**→ rplS (PRK05338)Ribosomal_L19 (pfam01245)→ |
| env-1910 | SRS024447_LANL_scaffold_57217 | + | 716 | 743 | **RNA**→ rplS (PRK05338)Ribosomal_L19 (pfam01245)→ |
| env-1911 | SRS042984_LANL_scaffold_57938 | + | 113 | 140 | **RNA**→ rplS (PRK05338)Ribosomal_L19 (pfam01245)→ |
| env-1912 | SRS043018_WUGC_scaffold_57339 | - | 1047 | 1020 | **RNA**→ rplS (PRK05338)Ribosomal_L19 (pfam01245)→ |
| env-1913 | SRS045197_WUGC_scaffold_67272 | - | 636 | 609 | **RNA**→ rplS (PRK05338)Ribosomal_L19 (pfam01245)→ |
| env-1914 | SRS047100_C2003956 | + | 65 | 92 | **RNA**→ rplS (PRK05338)Ribosomal_L19 (pfam01245)→ |
| env-1915 | SRS047113_LANL_scaffold_55438 | + | 262 | 289 | **RNA**→ rplS (PRK05338)Ribosomal_L19 (pfam01245)→ |
| env-1916 | SRS051378_C1867511 | + | 419 | 446 | **RNA**→ rplS (PRK05338)Ribosomal_L19 (pfam01245)→ |
| env-1917 | SRS053584_LANL_scaffold_44116 | - | 1012 | 985 | **RNA**→ rplS (PRK05338)Ribosomal_L19 (pfam01245)→ |
| env-1918 | SRS019387_C1763302 | + | 725 | 752 | **RNA**→ rplS (PRK05338)Ribosomal_L19 (pfam01245)→ |
| env-1919 | SRS022719_LANL_scaffold_27291 | - | 206354 | 206327 | **RNA**→ rplS (PRK05338)Ribosomal_L19 (pfam01245)→ |
| env-1920 | SRS044662_LANL_scaffold_16481 | + | 271 | 298 | **RNA**→ rplS (PRK05338)Ribosomal_L19 (pfam01245)→ |
| env-1921 | SRS024081_LANL_scaffold_75020 | + | 4020 | 4047 | **RNA**→ rplS (PRK05338)Ribosomal_L19 (pfam01245)→ |
| env-1922 | SRS043663_C4071478 | + | 98 | 125 | **RNA**→ rplS (PRK05338)Ribosomal_L19 (pfam01245)→ |
| env-1923 | SRS058808_LANL_scaffold_27436 | - | 1152 | 1125 | **RNA**→ rplS (PRK05338)Ribosomal_L19 (pfam01245)→ |
| Ran-1-1 | NZ_CP007503.1 | - | 224334 | 224307 | **RNA**→ rplS (PRK05338)Ribosomal_L19 (pfam01245)→ |
| env-1924 | Throat_LANL_C5692256 | - | 55 | 28 | **RNA**→ |
| env-1925 | SRS013945_C1211939 | - | 282 | 255 | **RNA**→ rplS (PRK05338)Ribosomal_L19 (pfam01245)→ |
| env-1926 | SRS014124_WUGC_scaffold_20088 | + | 433 | 460 | **RNA**→ rplS (PRK05338)Ribosomal_L19 (pfam01245)→ |
| env-1927 | SRS050244_C4542361 | + | 282 | 309 | **RNA**→ rplS (PRK05338)Ribosomal_L19 (pfam01245)→ |
| env-1928 | SRS077736_C1862773 | - | 6973 | 6946 | **RNA**→ rplS (PRK05338)Ribosomal_L19 (pfam01245)→ |
| env-1929 | SRS015574_C3403721 | - | 626 | 599 | **RNA**→ rplS (PRK05338)Ribosomal_L19 (pfam01245)→ |
| env-1930 | SRS018394_Baylor_scaffold_2340 | - | 289 | 262 | **RNA**→ rplS (PRK05338)Ribosomal_L19 (pfam01245)→ |
| env-1931 | SRS020226_Baylor_scaffold_48915 | + | 185 | 212 | **RNA**→ rplS (PRK05338)Ribosomal_L19 (pfam01245)→ |
| env-1932 | SRS024580_C3050280 | + | 109 | 136 | **RNA**→ rplS (PRK05338)Ribosomal_L19 (pfam01245)→ |
| Ran-6-1 | NC_017569.1 | + | 2072327 | 2072354 | **RNA**→ rplS (PRK05338)Ribosomal_L19 (pfam01245)→ |
| Ran-8-1 | NZ_AENH01000023.1 | + | 154146 | 154173 | **RNA**→ rplS (PRK05338)Ribosomal_L19 (pfam01245)→ |
| Ran-5-1 | NC_020125.1 | + | 1496908 | 1496935 | **RNA**→ rplS (PRK05338)Ribosomal_L19 (pfam01245)→ |
| Ran-1-2 | NZ_CP007504.1 | - | 234152 | 234125 | **RNA**→ rplS (PRK05338)Ribosomal_L19 (pfam01245)→ |
| Ran-9-1 | NZ_CP007204.1 | - | 224442 | 224415 | **RNA**→ rplS (PRK05338)Ribosomal_L19 (pfam01245)→ |
| env-1933 | SRS064329_C2693709 | + | 819 | 846 | **RNA**→ |
| env-1934 | Contig7224538.1 | + | 301 | 328 | **RNA**→ rplS (PRK05338)Ribosomal_L19 (pfam01245)→ |
| env-1935 | SRS011306_C3614405 | + | 617 | 644 | **RNA**→ rplS (PRK05338)Ribosomal_L19 (pfam01245)→ |
| env-1936 | SRS012279_C2754684 | + | 588 | 615 | **RNA**→ rplS (PRK05338)Ribosomal_L19 (pfam01245)→ |
| env-1937 | SRS014573_C3689455 | - | 453 | 426 | **RNA**→ rplS (PRK05338)Ribosomal_L19 (pfam01245)→ |

| | | | | | |
|---|---|---|---|---|---|
| env-1938 | SRS015434_C5917170 | + | 248 | 275 | **RNA**→ rplS (PRK05338)Ribosomal_L19 (pfam01245)→ |
| env-1939 | SRS015644_C1854550 | + | 125 | 152 | **RNA**→ rplS (PRK05338)Ribosomal_L19 (pfam01245)→ |
| env-1940 | SRS016002_WUGC_scaffold_45788 | + | 233 | 260 | **RNA**→ rplS (PRK05338)Ribosomal_L19 (pfam01245)→ |
| env-1941 | SRS016086_C2526098 | - | 7008 | 6981 | **RNA**→ rplS (PRK05338)Ribosomal_L19 (pfam01245)→ |
| env-1942 | SRS016225_C3019675 | - | 16261 | 16234 | **RNA**→ rplS (PRK05338)Ribosomal_L19 (pfam01245)→ |
| env-1943 | SRS017209_Baylor_scaffold_70466 | - | 6318 | 6291 | **RNA**→ rplS (PRK05338)Ribosomal_L19 (pfam01245)→ |
| env-1944 | SRS018157_Baylor_scaffold_33211 | - | 449 | 422 | **RNA**→ rplS (PRK05338)Ribosomal_L19 (pfam01245)→ |
| env-1945 | SRS018300_C3337187 | + | 271 | 298 | **RNA**→ rplS (PRK05338)Ribosomal_L19 (pfam01245)→ |
| env-1946 | SRS018357_C3935552 | - | 973 | 946 | **RNA**→ rplS (PRK05338)Ribosomal_L19 (pfam01245)→ |
| env-1947 | SRS019071_C5219198 | + | 750 | 777 | **RNA**→ rplS (PRK05338)Ribosomal_L19 (pfam01245)→ |
| env-1948 | SRS019327_C2809319 | - | 4490 | 4463 | **RNA**→ rplS (PRK05338)Ribosomal_L19 (pfam01245)→ |
| env-1949 | SRS019591_WUGC_scaffold_40451 | - | 587 | 560 | **RNA**→ rplS (PRK05338)Ribosomal_L19 (pfam01245)→ |
| env-1950 | SRS022077_Baylor_scaffold_57695 | + | 692 | 719 | **RNA**→ rplS (PRK05338)Ribosomal_L19 (pfam01245)→ |
| env-1951 | SRS022530_LANL_scaffold_35836 | - | 15939 | 15912 | **RNA**→ rplS (PRK05338)Ribosomal_L19 (pfam01245)→ |
| env-1952 | SRS023557_C1767643 | + | 685 | 712 | **RNA**→ rplS (PRK05338)Ribosomal_L19 (pfam01245)→ |
| env-1953 | SRS024015_C2912470 | + | 143 | 170 | **RNA**→ rplS (PRK05338)Ribosomal_L19 (pfam01245)→ |
| env-1954 | SRS024138_C2702299 | + | 249 | 276 | **RNA**→ rplS (PRK05338)Ribosomal_L19 (pfam01245)→ |
| env-1955 | SRS024277_C2782780 | + | 132 | 159 | **RNA**→ rplS (PRK05338)Ribosomal_L19 (pfam01245)→ |
| env-1956 | SRS024441_C3047628 | - | 532 | 505 | **RNA**→ rplS (PRK05338)Ribosomal_L19 (pfam01245)→ |
| env-1957 | SRS042131_C2195835 | - | 173 | 146 | **RNA**→ rplS (PRK05338)Ribosomal_L19 (pfam01245)→ |
| env-1958 | SRS042910_C5619617 | - | 593 | 566 | **RNA**→ rplS (PRK05338)Ribosomal_L19 (pfam01245)→ |
| env-1959 | SRS044373_C2908610 | + | 898 | 925 | **RNA**→ rplS (PRK05338)Ribosomal_L19 (pfam01245)→ |
| env-1960 | SRS049389_WUGC_scaffold_41106 | - | 1464 | 1437 | **RNA**→ rplS (PRK05338)Ribosomal_L19 (pfam01245)→ |
| env-1961 | SRS050669_LANL_scaffold_26170 | - | 6321 | 6294 | **RNA**→ rplS (PRK05338)Ribosomal_L19 (pfam01245)→ |
| env-1962 | SRS054687_C3492060 | - | 1063 | 1036 | **RNA**→ rplS (PRK05338)Ribosomal_L19 (pfam01245)→ |
| env-1963 | SRS056622_C2465234 | - | 126 | 99 | **RNA**→ hypo→ |
| env-1964 | SRS057791_LANL_scaffold_38618 | - | 378 | 351 | **RNA**→ rplS (PRK05338)Ribosomal_L19 (pfam01245)→ |
| env-1965 | SRS058336_LANL_scaffold_49913 | - | 1206 | 1179 | **RNA**→ rplS (PRK05338)Ribosomal_L19 (pfam01245)→ |
| env-1966 | SRS063288_LANL_scaffold_40195 | - | 1268 | 1241 | **RNA**→ rplS (PRK05338)Ribosomal_L19 (pfam01245)→ |
| env-1967 | SRS064423_C2493432 | + | 93 | 120 | **RNA**→ rplS (PRK05338)Ribosomal_L19 (pfam01245)→ |
| env-1968 | SRS011152_Baylor_scaffold_32757 | + | 171 | 198 | **RNA**→ rplS (PRK05338)Ribosomal_L19 (pfam01245)→ |
| env-1969 | SRS011343_C2194933 | + | 209 | 236 | **RNA**→ rplS (PRK05338)Ribosomal_L19 (pfam01245)→ |
| env-1970 | SRS013170_Baylor_scaffold_51649 | + | 154 | 181 | **RNA**→ rplS (PRK05338)Ribosomal_L19 (pfam01245)→ |
| env-1971 | SRS015278_WUGC_scaffold_59883 | + | 397 | 424 | **RNA**→ rplS (PRK05338)Ribosomal_L19 (pfam01245)→ |
| env-1972 | SRS018971_C758067 | + | 179 | 206 | **RNA**→ rplS (PRK05338)Ribosomal_L19 (pfam01245)→ |
| env-1973 | SRS018975_WUGC_scaffold_37778 | - | 742 | 715 | **RNA**→ rplS (PRK05338)Ribosomal_L19 (pfam01245)→ |
| env-1974 | SRS022536_LANL_scaffold_73925 | + | 387 | 414 | **RNA**→ rplS (PRK05338)Ribosomal_L19 (pfam01245)→ |
| env-1975 | SRS023964_C2589681 | - | 1401 | 1374 | **RNA**→ rplS (PRK05338)Ribosomal_L19 (pfam01245)→ |
| env-1976 | SRS051941_C3866993 | - | 218 | 191 | **RNA**→ rplS (PRK05338)Ribosomal_L19 (pfam01245)→ |
| env-1977 | SRS013947_WUGC_scaffold_25354 | - | 1008 | 981 | **RNA**→ rplS (PRK05338)Ribosomal_L19 (pfam01245)→ |
| env-1978 | SRS023930_Baylor_scaffold_9769 | - | 192 | 165 | **RNA**→ rplS (PRK05338)Ribosomal_L19 (pfam01245)→ |
| env-1979 | SRS013723_C3240076 | - | 198 | 171 | **RNA**→ rplS (PRK05338)Ribosomal_L19 (pfam01245)→ |
| env-1980 | SRS015440_WUGC_scaffold_82221 | + | 2031 | 2058 | **RNA**→ rplS (PRK05338)Ribosomal_L19 (pfam01245)→ |
| env-1981 | SRS015470_WUGC_scaffold_52571 | + | 10203 | 10230 | **RNA**→ rplS (PRK05338)Ribosomal_L19 (pfam01245)→ |
| env-1982 | SRS016043_WUGC_scaffold_35871 | - | 521 | 494 | **RNA**→ rplS (PRK05338)Ribosomal_L19 (pfam01245)→ |
| env-1983 | SRS019073_C2517672 | - | 733 | 706 | **RNA**→ rplS (PRK05338)Ribosomal_L19 (pfam01245)→ |
| env-1984 | SRS016200_WUGC_scaffold_14444 | - | 1066 | 1039 | **RNA**→ rplS (PRK05338)Ribosomal_L19 (pfam01245)→ |
| env-1985 | SRS051244_C3506325 | + | 2611 | 2638 | **RNA**→ rplS (PRK05338)Ribosomal_L19 (pfam01245)→ |
| env-1986 | SRS055450_C2078447 | - | 643 | 616 | **RNA**→ rplS (PRK05338)Ribosomal_L19 (pfam01245)→ |
| env-1987 | SRS017439_C3808563 | + | 80 | 107 | **RNA**→ rplS (PRK05338)Ribosomal_L19 (pfam01245)→ |
| env-1988 | SRS019591_C1935122 | + | 1741 | 1768 | **RNA**→ rplS (PRK05338)Ribosomal_L19 (pfam01245)→ |
| env-1989 | SRS020340_Baylor_scaffold_39528 | + | 113 | 140 | **RNA**→ rplS (PRK05338)Ribosomal_L19 (pfam01245)→ |

| | | | | | |
|---|---|---|---|---|---|
| env-1990 | SRS023595_C7179369 | + | 182 | 209 | **RNA**→ rplS (PRK05338)Ribosomal_L19 (pfam01245)→ |
| env-1991 | SRS042643_C2995340 | + | 43 | 70 | **RNA**→ rplS (PRK05338)Ribosomal_L19 (pfam01245)→ |
| env-1992 | SRS057539_C2749897 | + | 2475 | 2502 | **RNA**→ rplS (PRK05338)Ribosomal_L19 (pfam01245)→ |
| env-1993 | SRS014894_WUGC_scaffold_69889 | - | 1011 | 984 | **RNA**→ rplS (PRK05338)Ribosomal_L19 (pfam01245)→ |
| Ran-2-1 | NC_014738.1 | - | 235938 | 235911 | **RNA**→ rplS (PRK05338)Ribosomal_L19 (pfam01245)→ |
| Ran-7-1 | NZ_KB206037.1 | + | 651066 | 651093 | **RNA**→ rplS (PRK05338)Ribosomal_L19 (pfam01245)→ |
| Ran-2-2 | NC_017045.1 | - | 396304 | 396277 | **RNA**→ rplS (PRK05338)Ribosomal_L19 (pfam01245)→ |
| Rco-1-1 | NZ_KB894217.1 | + | 70188 | 70215 | **RNA**→ rplS (PRK05338)Ribosomal_L19 (pfam01245)→ Era (COG1159)era (PRK00089)→ ABC_YhbG (cd03218)AAA (smart00382)BCA_ABC_TP_C (pfam12399)→ PduO_Nterm (TIGR00636)Cob_adeno_trans (pfam01923)→ TPK (cd07995)TPK_B1_binding (smart00983)→ |
| env-1994 | Palatine_Tonsils_LANL_C6201686 | - | 597 | 570 | **RNA**→ rplS (PRK05338)Ribosomal_L19 (pfam01245)→ |
| env-1995 | SRS013252_C3650314 | + | 625 | 652 | **RNA**→ rplS (PRK05338)Ribosomal_L19 (pfam01245)→ |
| env-1996 | SRS015158_WUGC_scaffold_8442 | + | 3252 | 3279 | **RNA**→ rplS (PRK05338)Ribosomal_L19 (pfam01245)→ |
| env-1997 | SRS015797_C2637654 | + | 603 | 630 | **RNA**→ rplS (PRK05338)Ribosomal_L19 (pfam01245)→ |
| env-1998 | SRS019906_C2274015 | + | 1640 | 1667 | **RNA**→ rplS (PRK05338)Ribosomal_L19 (pfam01245)→ |
| env-1999 | SRS022149_LANL_scaffold_18048 | + | 563 | 590 | **RNA**→ |
| env-2000 | SRS047634_LANL_scaffold_86898 | + | 255 | 282 | **RNA**→ rplS (PRK05338)Ribosomal_L19 (pfam01245)→ |
| Ran-3-1 | NZ_CP006649.1 | + | 63395 | 63422 | **RNA**→ rplS (PRK05338)Ribosomal_L19 (pfam01245)→ |
| env-2001 | SRS016541_C1584168 | - | 612 | 585 | **RNA**→ rplS (PRK05338)Ribosomal_L19 (pfam01245)→ |
| env-2002 | SRS023841_C1590187 | + | 410 | 437 | **RNA**→ rplS (PRK05338)Ribosomal_L19 (pfam01245)→ |
| env-2003 | SRS043422_C2629820 | + | 251 | 278 | **RNA**→ rplS (PRK05338)Ribosomal_L19 (pfam01245)→ |
| env-2004 | SRS053917_C3736582 | + | 117 | 144 | **RNA**→ rplS (PRK05338)Ribosomal_L19 (pfam01245)→ |
| env-2005 | SRS012285_Baylor_scaffold_55554 | + | 199 | 226 | **RNA**→ rplS (PRK05338)Ribosomal_L19 (pfam01245)→ |
| env-2006 | SRS013945_C1168989 | - | 161 | 134 | **RNA**→ rplS (PRK05338)Ribosomal_L19 (pfam01245)→ |
| env-2007 | SRS013950_C2108425 | + | 189 | 216 | **RNA**→ rplS (PRK05338)Ribosomal_L19 (pfam01245)→ |
| env-2008 | SRS014473_C1123279 | + | 540 | 567 | **RNA**→ rplS (PRK05338)Ribosomal_L19 (pfam01245)→ |
| env-2009 | SRS014578_WUGC_scaffold_6706 | + | 428 | 455 | **RNA**→ rplS (PRK05338)Ribosomal_L19 (pfam01245)→ |
| env-2010 | SRS015044_WUGC_scaffold_69863 | - | 685 | 658 | **RNA**→ rplS (PRK05338)Ribosomal_L19 (pfam01245)→ |
| env-2011 | SRS015899_C1768151 | - | 210 | 183 | **RNA**→ rplS (PRK05338)Ribosomal_L19 (pfam01245)→ |
| env-2012 | SRS016331_C4139213 | - | 394 | 367 | **RNA**→ rplS (PRK05338)Ribosomal_L19 (pfam01245)→ |
| env-2013 | SRS017227_Baylor_scaffold_32942 | - | 562 | 535 | **RNA**→ rplS (PRK05338)Ribosomal_L19 (pfam01245)→ |
| env-2014 | SRS017691_C5166596 | - | 303 | 276 | **RNA**→ rplS (PRK05338)Ribosomal_L19 (pfam01245)→ |
| env-2015 | SRS019125_C626092 | - | 1070 | 1043 | **RNA**→ rplS (PRK05338)Ribosomal_L19 (pfam01245)→ |
| env-2016 | SRS019129_WUGC_scaffold_11480 | - | 802 | 775 | **RNA**→ rplS (PRK05338)Ribosomal_L19 (pfam01245)→ |
| env-2017 | SRS022083_C1917930 | + | 717 | 744 | **RNA**→ |
| env-2018 | SRS024087_C3403049 | - | 384 | 357 | **RNA**→ rplS (PRK05338)Ribosomal_L19 (pfam01245)→ |
| env-2019 | SRS024649_C3480943 | + | 159 | 186 | **RNA**→ rplS (PRK05338)Ribosomal_L19 (pfam01245)→ |
| env-2020 | SRS043755_WUGC_scaffold_2404 | + | 403 | 430 | **RNA**→ rplS (PRK05338)Ribosomal_L19 (pfam01245)→ |
| env-2021 | SRS047265_C1089128 | + | 277 | 304 | **RNA**→ rplS (PRK05338)Ribosomal_L19 (pfam01245)→ |
| env-2022 | SRS049318_C6415734 | + | 418 | 445 | **RNA**→ rplS (PRK05338)Ribosomal_L19 (pfam01245)→ |
| env-2023 | SRS051930_C3222104 | + | 135 | 162 | **RNA**→ rplS (PRK05338)Ribosomal_L19 (pfam01245)→ |
| env-2024 | SRS052876_LANL_scaffold_28233 | - | 514 | 487 | **RNA**→ rplS (PRK05338)Ribosomal_L19 (pfam01245)→ |
| env-2025 | SRS054430_LANL_scaffold_17686 | - | 3792 | 3765 | **RNA**→ rplS (PRK05338)Ribosomal_L19 (pfam01245)→ |
| env-2026 | SRS055378_C4102388 | - | 904 | 877 | **RNA**→ rplS (PRK05338)Ribosomal_L19 (pfam01245)→ |
| env-2027 | SRS063603_C4216775 | + | 204 | 231 | **RNA**→ rplS (PRK05338)Ribosomal_L19 (pfam01245)→ |
| env-2028 | SRS063932_LANL_scaffold_98887 | - | 303 | 276 | **RNA**→ rplS (PRK05338)Ribosomal_L19 (pfam01245)→ |
| env-2029 | SRS063999_LANL_scaffold_36577 | + | 83 | 110 | **RNA**→ rplS (PRK05338)Ribosomal_L19 (pfam01245)→ |
| env-2030 | SRS064449_C3931012 | + | 199 | 226 | **RNA**→ rplS (PRK05338)Ribosomal_L19 (pfam01245)→ |
| env-2031 | SRS075410_LANL_scaffold_45208 | + | 279 | 306 | **RNA**→ rplS (PRK05338)Ribosomal_L19 (pfam01245)→ |
| env-2032 | SRS021496_C2863013 | + | 18 | 45 | **RNA**→ rplS (PRK05338)Ribosomal_L19 (pfam01245)→ |
| env-2033 | SRS017445_Baylor_scaffold_27071 | - | 1986 | 1959 | **RNA**→ rplS (PRK05338)Ribosomal_L19 (pfam01245)→ |
| env-2034 | CENR01002311.1 | - | 10635 | 10608 | **RNA**→ rplS (PRK05338)Ribosomal_L19 (pfam01245)→ |
| env-2035 | JCVI_SCAF_1096626906398 | - | 1635 | 1608 | **RNA**→ rplS (PRK05338)Ribosomal_L19 (pfam01245)→ |

| | | | | | |
|---|---|---|---|---|---|
| Fge-1-1 | NZ_AUGN01000001.1 | + | 377756 | 377783 | **RNA**→ rplS (PRK05338)Ribosomal_L19 (pfam01245)→ |
| Nma-1-1 | NZ_AP014548.1 | + | 46175 | 46202 | **RNA**→ rplS (PRK05338)Ribosomal_L19 (pfam01245)→ |
| env-2036 | CENJ01235641.1 | - | 101 | 74 | **RNA**→ |
| env-2037 | CEQC01079196.1 | + | 513 | 540 | **RNA**→ rplS (PRK05338)Ribosomal_L19 (pfam01245)→ |
| env-2038 | CEQG01233746.1 | - | 746 | 719 | **RNA**→ rplS (PRK05338)Ribosomal_L19 (pfam01245)→ |
| env-2039 | CEQH01174899.1 | + | 314 | 341 | **RNA**→ rplS (PRK05338)Ribosomal_L19 (pfam01245)→ |
| env-2040 | CEQR01110413.1 | - | 5165 | 5138 | **RNA**→ rplS (PRK05338)Ribosomal_L19 (pfam01245)→ |
| env-2041 | CEQT01267208.1 | - | 212 | 185 | **RNA**→ rplS (PRK05338)Ribosomal_L19 (pfam01245)→ |
| env-2042 | CEQZ01055080.1 | - | 2498 | 2471 | **RNA**→ rplS (PRK05338)Ribosomal_L19 (pfam01245)→ |
| env-2043 | CERB01343087.1 | + | 510 | 537 | **RNA**→ rplS (PRK05338)Ribosomal_L19 (pfam01245)→ |
| env-2044 | CERC01066855.1 | + | 170716 | 170743 | **RNA**→ rplS (PRK05338)Ribosomal_L19 (pfam01245)→ |
| env-2045 | CERD01039641.1 | + | 170722 | 170749 | **RNA**→ rplS (PRK05338)Ribosomal_L19 (pfam01245)→ |
| env-2046 | CERX01145001.1 | + | 1529 | 1556 | **RNA**→ rplS (PRK05338)Ribosomal_L19 (pfam01245)→ |
| env-2047 | JGI20157J14317_10303049 | - | 277 | 250 | **RNA**→ rplS (PRK05338)Ribosomal_L19 (pfam01245)→ |
| env-2048 | DelMOSum2010_c10000073 | - | 56849 | 56822 | **RNA**→ rplS (PRK05338)Ribosomal_L19 (pfam01245)→ |
| env-2049 | A09P0410a_Sequence0000004289 | - | 1199 | 1172 | **RNA**→ rplS (PRK05338)Ribosomal_L19 (pfam01245)→ |
| env-2050 | CESP01013401.1 | + | 29497 | 29523 | **RNA**→ rplS (PRK05338)Ribosomal_L19 (pfam01245)→ Icd (COG0538)PRK08299 (PRK08299)→ |
| | | | | | TonB_dep_Rec (pfam00593)Plug (pfam07715)CarbopepD_reg_2 (pfam13715)OMP_b-brl_3 (pfam14905)→ |
| | | | | | DUF4249 (pfam14054)→ murQ (PRK05441)SIS_Etherase (cd05007)→ |
| env-2051 | CEUM01057183.1 | + | 1335 | 1361 | **RNA**→ rplS (PRK05338)Ribosomal_L19 (pfam01245)→ |
| env-2052 | CESB01235038.1 | + | 467 | 493 | **RNA**→ rplS (PRK05338)Ribosomal_L19 (pfam01245)→ |
| env-2053 | CESZ01095135.1 | - | 719 | 693 | **RNA**→ rplS (PRK05338)Ribosomal_L19 (pfam01245)→ PRK08299 (PRK08299)→ |
| env-2054 | CEOC01268308.1 | - | 738 | 712 | **RNA**→ rplS (PRK05338)Ribosomal_L19 (pfam01245)→ PRK08299 (PRK08299)→ |
| env-2055 | CEQL01119221.1 | - | 829 | 803 | **RNA**→ rplS (PRK05338)Ribosomal_L19 (pfam01245)→ PRK08299 (PRK08299)→ |
| env-2056 | CETL01074928.1 | - | 1402 | 1376 | **RNA**→ rplS (PRK05338)Ribosomal_L19 (pfam01245)→ PRK08299 (PRK08299)→ |
| env-2057 | JCVI_SCAF_1096627050705 | + | 985 | 1011 | **RNA**→ rplS (PRK05338)Ribosomal_L19 (pfam01245)→ Icd (COG0538)PRK08299 (PRK08299)→ |
| | | | | | Peptidase_M14NE-CP-C_like (cd11308)Plug (pfam07715)CarbopepD_reg_2 (pfam13715)→ |
| env-2058 | 85084776 | + | 656 | 682 | **RNA**→ |
| env-2059 | JCVI_READ_1103180508165 | + | 656 | 682 | **RNA**→ |
| env-2060 | CENO01049189.1 | - | 4161 | 4135 | **RNA**→ rplS (PRK05338)Ribosomal_L19 (pfam01245)→ Icd (COG0538)PRK08299 (PRK08299)→ |
| | | | | | Peptidase_M14NE-CP-C_like (cd11308)TonB-hemlactrns (TIGR01786)TonB_dep_Rec (pfam00593)Plug (pfam07715)Carb |
| | | | | | DUF4249 (pfam14054)→ |
| env-2061 | CENQ01101034.1 | + | 138 | 164 | **RNA**→ rplS (PRK05338)Ribosomal_L19 (pfam01245)→ PRK08299 (PRK08299)→ |
| env-2062 | CEOV01119354.1 | + | 225 | 251 | **RNA**→ rplS (PRK05338)Ribosomal_L19 (pfam01245)→ PRK08299 (PRK08299)→ |
| env-2063 | CEPB01008934.1 | + | 225 | 251 | **RNA**→ rplS (PRK05338)Ribosomal_L19 (pfam01245)→ PRK08299 (PRK08299)→ |
| env-2064 | CEQD01038409.1 | + | 1170 | 1196 | **RNA**→ rplS (PRK05338)Ribosomal_L19 (pfam01245)→ Icd (COG0538)PRK08299 (PRK08299)→ |
| | | | | | Peptidase_M14NE-CP-C_like (cd11308)Plug (pfam07715)CarbopepD_reg_2 (pfam13715)→ |
| env-2065 | CETP01090435.1 | - | 265 | 239 | **RNA**→ rplS (PRK05338)Ribosomal_L19 (pfam01245)→ |
| env-2066 | CEUW01136624.1 | + | 327 | 353 | **RNA**→ rplS (PRK05338)Ribosomal_L19 (pfam01245)→ |
| env-2067 | JCVI_SCAF_1096627827830 | - | 1339 | 1315 | **RNA**→ rplS (PRK05338)Ribosomal_L19 (pfam01245)→ |
| env-2068 | CERK01129111.1 | - | 488 | 464 | **RNA**→ rplS (PRK05338)Ribosomal_L19 (pfam01245)→ |
| env-2069 | CEOY01052385.1 | - | 1246 | 1222 | **RNA**→ rplS (PRK05338)Ribosomal_L19 (pfam01245)→ PRK08299 (PRK08299)→ |
| env-2070 | CEWA01020377.1 | - | 206 | 182 | **RNA**→ rplS (PRK05338)Ribosomal_L19 (pfam01245)→ |
| env-2071 | JCVI_SCAF_1101668434874 | - | 68 | 44 | **RNA**→ |
| env-2072 | CEPA01251295.1 | + | 591 | 615 | **RNA**→ |
| env-2073 | CEUM01025643.1 | - | 3227 | 3203 | **RNA**→ rplS (PRK05338)Ribosomal_L19 (pfam01245)→ Icd (COG0538)PRK08299 (PRK08299)→ |
| | | | | | PRK10064 (PRK10064)Plug (pfam07715)CarbopepD_reg_2 (pfam13715)→ |
| env-2074 | CEUR01331033.1 | + | 1016 | 1040 | **RNA**→ rplS (PRK05338)Ribosomal_L19 (pfam01245)→ PRK08299 (PRK08299)→ |
| env-2075 | CEWJ01138850.1 | + | 1316 | 1340 | **RNA**→ rplS (PRK05338)Ribosomal_L19 (pfam01245)→ PRK08299 (PRK08299)→ |
| env-2076 | CEVY01039090.1 | + | 1019 | 1043 | **RNA**→ rplS (PRK05338)Ribosomal_L19 (pfam01245)→ PRK08299 (PRK08299)→ |
| env-2077 | JCVI_SCAF_1101667338675 | - | 229 | 205 | **RNA**→ |
| env-2078 | CEQN01304455.1 | + | 639 | 663 | **RNA**→ |
| env-2079 | CEWH01064632.1 | - | 607 | 583 | **RNA**→ rplS (PRK05338)Ribosomal_L19 (pfam01245)→ PRK08299 (PRK08299)→ |

| | | | | | |
|---|---|---|---|---|---|
| env-2080 | JCVI_SCAF_1096627518681 | - | 1299 | 1275 | **RNA**→ rplS (PRK05338)Ribosomal_L19 (pfam01245)→ |
| env-2081 | JCVI_SCAF_1096628156800 | + | 1484 | 1508 | **RNA**→ rplS (PRK05338)Ribosomal_L19 (pfam01245)→ |
| env-2082 | JCVI_SCAF_1096626505381 | - | 660 | 636 | **RNA**→ rplS (PRK05338)Ribosomal_L19 (pfam01245)→ PRK08299 (PRK08299)→ |
| env-2083 | CEOC01044408.1 | - | 1075 | 1051 | **RNA**→ rplS (PRK05338)Ribosomal_L19 (pfam01245)→ PRK08299 (PRK08299)→ |
| env-2084 | CEOO01047216.1 | - | 229 | 205 | **RNA**→ rplS (PRK05338)Ribosomal_L19 (pfam01245)→ |
| env-2085 | CEOV01066520.1 | + | 633 | 657 | **RNA**→ rplS (PRK05338)Ribosomal_L19 (pfam01245)→ PRK08299 (PRK08299)→ |
| env-2086 | CEOW01181946.1 | - | 83 | 59 | **RNA**→ |
| env-2087 | CEOX01007014.1 | - | 822 | 798 | **RNA**→ rplS (PRK05338)Ribosomal_L19 (pfam01245)→ PRK08299 (PRK08299)→ |
| env-2088 | CEPC01000822.1 | + | 908 | 932 | **RNA**→ rplS (PRK05338)Ribosomal_L19 (pfam01245)→ PRK08299 (PRK08299)→ |
| env-2089 | CEPK01174470.1 | + | 625 | 649 | **RNA**→ rplS (PRK05338)Ribosomal_L19 (pfam01245)→ PRK08299 (PRK08299)→ |
| env-2090 | CEPT01287638.1 | - | 874 | 850 | **RNA**→ rplS (PRK05338)Ribosomal_L19 (pfam01245)→ PRK08299 (PRK08299)→ |
| env-2091 | CEPW01050466.1 | + | 751 | 775 | **RNA**→ |
| env-2092 | CEQM01235848.1 | + | 92 | 116 | **RNA**→ rplS (PRK05338)Ribosomal_L19 (pfam01245)→ PRK08299 (PRK08299)→ |
| env-2093 | CERW01124879.1 | + | 537 | 561 | **RNA**→ rplS (PRK05338)Ribosomal_L19 (pfam01245)→ PRK08299 (PRK08299)→ |
| env-2094 | CETW01029624.1 | + | 2015 | 2039 | **RNA**→ rplS (PRK05338)Ribosomal_L19 (pfam01245)→ PRK08299 (PRK08299)→ |
| env-2095 | CEVL01092845.1 | + | 120 | 144 | **RNA**→ rplS (PRK05338)Ribosomal_L19 (pfam01245)→ PRK08299 (PRK08299)→ |
| env-2096 | CEVZ01428942.1 | - | 1474 | 1450 | **RNA**→ rplS (PRK05338)Ribosomal_L19 (pfam01245)→ PRK08299 (PRK08299)→ |
| env-2097 | JCVI_READ_1108839666479 | + | 178 | 202 | **RNA**→ rplS (PRK05338)Ribosomal_L19 (pfam01245)→ PRK08299 (PRK08299)→ |
| env-2098 | CEQO01202726.1 | + | 82 | 106 | **RNA**→ rplS (PRK05338)Ribosomal_L19 (pfam01245)→ PRK08299 (PRK08299)→ |
| env-2099 | CERB01095802.1 | - | 746 | 722 | **RNA**→ rplS (PRK05338)Ribosomal_L19 (pfam01245)→ PRK08299 (PRK08299)→ |
| env-2100 | CEVN01023152.1 | + | 744 | 768 | **RNA**→ |
| env-2101 | JCVI_READ_1103242406987 | + | 651 | 675 | **RNA**→ rplS (PRK05338)Ribosomal_L19 (pfam01245)→ hypo→ |
| env-2102 | JCVI_SCAF_1096626678284 | - | 913 | 889 | **RNA**→ rplS (PRK05338)Ribosomal_L19 (pfam01245)→ PRK08299 (PRK08299)→ |
| env-2103 | CEVR01021723.1 | - | 210 | 186 | **RNA**→ rplS (PRK05338)Ribosomal_L19 (pfam01245)→ |
| env-2104 | JCVI_SCAF_1096627874827 | + | 442 | 466 | **RNA**→ rplS (PRK05338)Ribosomal_L19 (pfam01245)→ rplS (PRK05338)Ribosomal_L19 (pfam01245)→ |
| env-2105 | JCVI_SCAF_1096627874827 | + | 1299 | 1323 | **RNA**→ rplS (PRK05338)Ribosomal_L19 (pfam01245)→ rplS (PRK05338)Ribosomal_L19 (pfam01245)→ |
| env-2106 | JCVI_SCAF_1101668523738 | - | 116 | 92 | **RNA**→ |
| env-2107 | JCVI_SCAF_1096628400995 | - | 1440 | 1416 | **RNA**→ rplS (PRK05338)Ribosomal_L19 (pfam01245)→ rplS (PRK05338)Ribosomal_L19 (pfam01245)→ |
| | | | | | PRK08299 (PRK08299)→ PRK08299 (PRK08299)→ TrmD (COG0336)trmD (PRK00026)→ |
| | | | | | TrmD (COG0336)trmD (PRK00026)→ |
| env-2108 | JCVI_SCAF_1096628400995 | - | 527 | 503 | **RNA**→ rplS (PRK05338)Ribosomal_L19 (pfam01245)→ rplS (PRK05338)Ribosomal_L19 (pfam01245)→ |
| env-2109 | CEPB01125895.1 | - | 825 | 801 | **RNA**→ rplS (PRK05338)Ribosomal_L19 (pfam01245)→ PRK08299 (PRK08299)→ |
| env-2110 | CETE01078549.1 | - | 790 | 766 | **RNA**→ rplS (PRK05338)Ribosomal_L19 (pfam01245)→ PRK08299 (PRK08299)→ |
| env-2111 | CETH01012318.1 | + | 755 | 779 | **RNA**→ rplS (PRK05338)Ribosomal_L19 (pfam01245)→ Icd (COG0538)PRK08299 (PRK08299)→ |
| env-2112 | CETM01404675.1 | + | 166 | 190 | **RNA**→ rplS (PRK05338)Ribosomal_L19 (pfam01245)→ PRK08299 (PRK08299)→ |
| env-2113 | CETQ01016873.1 | - | 562 | 538 | **RNA**→ rplS (PRK05338)Ribosomal_L19 (pfam01245)→ PRK08299 (PRK08299)→ |
| env-2114 | CEUF01066100.1 | - | 3392 | 3368 | **RNA**→ rplS (PRK05338)Ribosomal_L19 (pfam01245)→ Icd (COG0538)PRK08299 (PRK08299)→ |
| | | | | | PRK10064 (PRK10064)Plug (pfam07715)CarbopepD_reg_2 (pfam13715)OMP_b-brl_3 (pfam14905)→ |
| env-2115 | CEUG01002262.1 | + | 1316 | 1340 | **RNA**→ rplS (PRK05338)Ribosomal_L19 (pfam01245)→ Icd (COG0538)PRK08299 (PRK08299)→ |
| | | | | | PRK10064 (PRK10064)Plug (pfam07715)CarbopepD_reg_2 (pfam13715)→ |
| env-2116 | CEVE01132406.1 | - | 1971 | 1947 | **RNA**→ rplS (PRK05338)Ribosomal_L19 (pfam01245)→ Icd (COG0538)PRK08299 (PRK08299)→ |
| | | | | | Peptidase_M14NE-CP-C_like (cd11308)CarbopepD_reg_2 (pfam13715)→ |
| env-2117 | CEWG01029302.1 | + | 660 | 684 | **RNA**→ rplS (PRK05338)Ribosomal_L19 (pfam01245)→ PRK08299 (PRK08299)→ |
| env-2118 | JGI24820J26691_1003156 | + | 3260 | 3284 | **RNA**→ rplS (PRK05338)Ribosomal_L19 (pfam01245)→ Icd (COG0538)PRK08299 (PRK08299)→ |
| env-2119 | JCVI_READ_1108800196652 | + | 341 | 365 | **RNA**→ rplS (PRK05338)Ribosomal_L19 (pfam01245)→ |
| env-2120 | JCVI_READ_1105333388420 | - | 189 | 165 | **RNA**→ rplS (PRK05338)Ribosomal_L19 (pfam01245)→ |
| env-2121 | CETK01326476.1 | - | 589 | 565 | **RNA**→ rplS (PRK05338)Ribosomal_L19 (pfam01245)→ PRK08299 (PRK08299)→ |
| env-2122 | JCVI_SCAF_1096628212569 | - | 1326 | 1302 | **RNA**→ rplS (PRK05338)Ribosomal_L19 (pfam01245)→ |
| env-2123 | CENO01004142.1 | - | 481 | 457 | **RNA**→ rplS (PRK05338)Ribosomal_L19 (pfam01245)→ |
| Fps-4-1 | NC_009613.3 | - | 1709481 | 1709454 | **RNA**→ rplS (PRK05338)Ribosomal_L19 (pfam01245)→ |
| Fps-1-1 | NZ_CP008878.1 | - | 1965790 | 1965763 | **RNA**→ rplS (PRK05338)Ribosomal_L19 (pfam01245)→ |
| Fps-1-2 | NZ_CP008883.1 | - | 1959068 | 1959041 | **RNA**→ rplS (PRK05338)Ribosomal_L19 (pfam01245)→ |
| Fps-1-3 | NZ_CP008881.1 | - | 1956157 | 1956130 | **RNA**→ rplS (PRK05338)Ribosomal_L19 (pfam01245)→ |

| | | | | | |
|---|---|---|---|---|---|
| Fps-1-4 | NZ_CP007627.1 | + | 1042967 | 1042994 | **RNA**→ rplS (PRK05338)Ribosomal_L19 (pfam01245)→ |
| Fps-2-1 | NZ_CP007206.1 | + | 992155 | 992182 | **RNA**→ rplS (PRK05338)Ribosomal_L19 (pfam01245)→ |
| Fps-3-1 | NZ_CP007207.1 | + | 1881558 | 1881585 | **RNA**→ rplS (PRK05338)Ribosomal_L19 (pfam01245)→ |
| Fps-1-5 | NZ_CP008902.1 | - | 2002522 | 2002495 | **RNA**→ rplS (PRK05338)Ribosomal_L19 (pfam01245)→ |
| Fps-1-6 | NZ_CP010278.1 | - | 2065590 | 2065563 | **RNA**→ rplS (PRK05338)Ribosomal_L19 (pfam01245)→ |
| Fps-1-7 | NZ_CP010275.1 | - | 2106998 | 2106971 | **RNA**→ rplS (PRK05338)Ribosomal_L19 (pfam01245)→ |
| Fps-1-8 | NZ_CP010276.1 | - | 2110820 | 2110793 | **RNA**→ rplS (PRK05338)Ribosomal_L19 (pfam01245)→ |
| Fps-1-9 | NZ_CP010277.1 | - | 2068780 | 2068753 | **RNA**→ rplS (PRK05338)Ribosomal_L19 (pfam01245)→ |
| Fps-1-10 | NZ_CP010274.1 | - | 2108381 | 2108354 | **RNA**→ rplS (PRK05338)Ribosomal_L19 (pfam01245)→ |
| Ran-4-1 | NC_018609.1 | + | 1957039 | 1957066 | **RNA**→ rplS (PRK05338)Ribosomal_L19 (pfam01245)→ |

#### 6.3 Conserved domains

Conserved domains found in protein-coding genes listed in Section 6.2 are shown below, with the first sentence in their description from the Conserved Domain Database (if any). Conserved domains associated with more than one L19-Flavobacteria RNA are assigned a color, while others are shown in gray. The number in parentheses after the colored domain name is the number of occurrences in Section 6.2.

cd00635 (4) Type III Pyridoxal 5-phosphate (PLP)-Dependent Enzymes, YBL036c-like proteins.
cd00834 (35) Beta-ketoacyl-acyl carrier protein (ACP) synthase (KAS), type I and II.
cd01667 (19) TGS _ThrRS_N: ThrRS (threonyl-tRNA Synthetase) is a class II tRNA synthetase that couples threonine to its cognate tRNA.
cd02968 (7) SCO (an acronym for Synthesis of Cytochrome c Oxidase) family; composed of proteins similar to Sco1, a membrane-anchored protein possessing a soluble domain with a TRX fold.
cd03218 (1) ATP-binding cassette component of YhbG transport system.
cd04690 (18) Members of the Nudix hydrolase superfamily catalyze the hydrolysis of NUcleoside DIphosphates linked to other moieties, X.
cd04795 (1) SIS domain.
cd05007 (36) N-acetylmuramic acid 6-phosphate etherase.
cd05233 (1) classical (c) SDRs. [AG]
cd06127 (38) DEDDh 3'-5' exonuclease domain family.
cd07995 (1) Thiamine pyrophosphokinase.
cd10434 (33) Catalytic GIY-YIG domain of nucleotide excision repair endonucleases UvrC, Cho, and similar proteins.
cd11308 (5) Peptidase associated domain: C-terminal domain of M14 N/E carboxypeptidase; putative folding, regulation, or interaction domain.
COG0030 (5) 16S rRNA A1518 and A1519 N6-dimethyltransferase RsmA/KsgA/DIM1 (may also have DNA glycosylase/AP lyase activity) [Translation, ribosomal structure and biogenesis]
COG0290 (4) Translation initiation factor IF-3 [Translation, ribosomal structure and biogenesis]
COG0336 (2) tRNA G37 N-methylase TrmD [Translation, ribosomal structure and biogenesis]
COG0441 (23) Threonyl-tRNA synthetase [Translation, ribosomal structure and biogenesis]
COG0486 (1) tRNA U34 5-carboxymethylaminomethyl modifying GTPase MnmE/TrmE [Translation, ribosomal structure and biogenesis]
COG0525 (3) Valyl-tRNA synthetase [Translation, ribosomal structure and biogenesis]
COG0538 (76) Isocitrate dehydrogenase [Energy production and conversion]
COG0605 (32) Superoxide dismutase [Inorganic ion transport and metabolism]
COG1159 (1) GTPase Era, involved in 16S rRNA processing [Translation, ribosomal structure and biogenesis]
COG1629 (8) Outer membrane receptor proteins, mostly Fe transport [Inorganic ion transport and metabolism]
COG4389 (1) Site-specific recombinase [Replication, recombination and repair]
COG4799 (1) Acetyl-CoA carboxylase, carboxyltransferase component [Lipid transport and metabolism]
pfam00081 (1) Iron/manganese superoxide dismutases, alpha-hairpin domain.
pfam00106 (1) short chain dehydrogenase.
pfam00453 (4) Ribosomal protein L20.
pfam00593 (12) TonB dependent receptor.
pfam01245 (2085) Ribosomal protein L19.
pfam01632 (4) Ribosomal protein L35.
pfam01850 (5) PIN domain.
pfam01923 (1) Cobalamin adenosyltransferase.
pfam02628 (3) Cytochrome oxidase assembly protein.
pfam02824 (5) TGS domain.
pfam03544 (22) Gram-negative bacterial TonB protein C-terminal.
pfam03971 (8) Monomeric isocitrate dehydrogenase.
pfam06245 (7) Protein of unknown function (DUF1015).
pfam07715 (22) TonB-dependent Receptor Plug Domain.
pfam10136 (1) Site-specific recombinase.
pfam12399 (1) Branched-chain amino acid ATP-binding cassette transporter.
pfam13715 (33) CarboxypepD_reg-like domain.
pfam13723 (27) Beta-ketoacyl synthase, N-terminal domain.
pfam14054 (18) Domain of unknown function (DUF4249).
pfam14213 (5) Domain of unknown function (DUF4325).
pfam14815 (1) NUDIX domain.
pfam14905 (10) Outer membrane protein beta-barrel family.
PLN02820 (1) 3-methylcrotonyl-CoA carboxylase, beta chain
PRK00026 (2) tRNA (guanine-N(1)-)-methyltransferase; Reviewed
PRK00028 (4) translation initiation factor IF-3; Reviewed
PRK00089 (1) GTPase Era; Reviewed
PRK00172 (4) 50S ribosomal protein L35; Reviewed

PRK00274 (5) 16S ribosomal RNA methyltransferase KsgA/Dim1 family protein; Reviewed
PRK00413 (23) threonyl-tRNA synthetase; Reviewed
PRK05185 (4) 50S ribosomal protein L20; Provisional
PRK05291 (1) tRNA modification GTPase TrmE; Reviewed
PRK05338 (2085) 50S ribosomal protein L19; Provisional
PRK05441 (37) N-acetylmuramic acid-6-phosphate etherase; Reviewed
PRK05653 (1) 3-ketoacyl-(acyl-carrier-protein) reductase; Validated
PRK05729 (3) valyl-tRNA synthetase; Reviewed
PRK07825 (1) short chain dehydrogenase; Provisional
PRK08299 (224) isocitrate dehydrogenase; Validated
PRK08999 (16) hypothetical protein; Provisional
PRK10064 (20) catecholate siderophore receptor CirA; Provisional
PRK10546 (1) pyrimidine (deoxy)nucleoside triphosphate pyrophosphohydrolase; Provisional
PRK10925 (33) superoxide dismutase; Provisional
PRK13484 (1) putative iron-regulated outer membrane virulence protein; Provisional
PTZ00435 (1) isocitrate dehydrogenase; Provisional
smart00382 (1) ATPases associated with a variety of cellular activities.
smart00465 (18) GIY-YIG type nucleases (URI domain).
smart00479 (38) exonuclease domain in DNA-polymerase alpha and epsilon chain, ribonuclease T and other exonucleases.
smart00863 (1) Threonyl and Alanyl tRNA synthetase second additional domain.
smart00983 (1) Thiamin pyrophosphokinase, vitamin B1 binding domain.
TIGR00044 (4) pyridoxal phosphate enzyme, YggS family. [Unknown function, Enzymes of unknown specificity]
TIGR00178 (8) isocitrate dehydrogenase, NADP-dependent, monomeric type. [Energy metabolism, TCA cycle]
TIGR00636 (1) ATP:cob(I)alamin adenosyltransferase. [Biosynthesis of cofactors, prosthetic groups, and carriers, Heme, porphyrin, and cobalamin]
TIGR01352 (22) TonB family C-terminal domain. [Transport and binding proteins, Cations and iron carrying compounds]
TIGR01785 (1) TonB-dependent heme/hemoglobin receptor family protein. [Transport and binding proteins, Cations and iron carrying compounds, Transport and binding proteins, Porins]
TIGR01786 (3) TonB-dependent hemoglobin/transferrin/lactoferrin receptor family protein. [Transport and binding proteins, Cations and iron carrying compounds, Transport and binding proteins, Porins]
TIGR03150 (8) beta-ketoacyl-acyl-carrier-protein synthase II. [acyl-carrier-protein] synthase 2 (KAS-II, FabF) is involved in the condensation step of fatty acid biosynthesis in which the malonyl donor group is decarboxylated and the resulting carbanion used to attack and extend the acyl group attached to the acyl carrier protein. Most genomes encoding fatty acid biosynthesis contain a number of condensing enzymes, often of all three types: 1, 2 and 3. Synthase 2 is mechanistically related to synthase 1 (KAS-I, FabB) containing a number of absolutely conserved catalytic residues in common. This model is based primarily on genes which are found in apparent operons with other essential genes of fatty acid biosynthesis (GenProp0681). The large gap between the trusted cutoff and the noise cutoff contains many genes which are not found adjacent to genes of the fatty acid pathway in genomes that often also contain a better hit to this model. These genes may be involved in other processes such as polyketide biosyntheses. Some genomes contain more than one above-trusted hit to this model which may result from recent paralogous expansions. Second hits to this model which are not next to other fatty acid biosynthesis genes may be involved in other processes. FabB sequences should fall well below the noise cutoff of this model. [Fatty acid and phospholipid metabolism, Biosynthesis]

#### 6.4 Multiple-sequence alignment

Each L19-Flavobacteria RNA is denoted by an abbreviation (like "Eco-1-1") that refers to a taxonomy given in Section 6.1. The alignment may include sequences containing the RNA motif, as well as flanking sequence. The L19-Flavobacteria RNA itself is denoted by the line underneath marked 5′ and 3′ on either end. Nucleotides in flanking sequences (i.e., not part of the motif) are written in gray letters. Stems of predicted rho-independent transcription terminators, if any, are shaded yellow. (Note: terminator predictions have not been analyzed manually, and many are likely to be false positives. Terminator predictions are those of the RNIE software.) Nucleotides predicted to function (as DNA) as transcription-factor binding sites, if any, are shaded in green. (But note: these predictions are manually annotated, so they might be under-predicted.) Annotated start codons, if any, are shaded green. (Note: start codons are frequently misannotated, especially in environmental samples.) Nucleotides proposed to basepair as part of the consensus structure are shaded in color when they comprise Watson-Crick or G-U pairs. Otherwise they are shaded gray. Conserved stems are also indicated at the bottom of the alignment by angle brackets, where matching < and > denote base-paired columns. Below these angle brackets, the symbol "2" denotes base pairs exhibiting covariation according to the statistically well-founded R-scape method. "1" denotes base pairs exhibiting covariation according to R2R's simplistic method. "0" denotes base pairs that are not observed to mutate and "?" denotes base pairs that have a significant frequency of non-canonical nucleotides for Watson-Crick or G-U pairs (> 5%). Below these base pair annotation is the consensus sequence: "R" = "A" or "G", "Y" = "C" or "U", red nucleotides: nucleotide identity conserved more than 97% of the time, black nucleotides: 90%, gray nucleotides: 75%, red circle (•): nucleotide is present 97% of the time, black circle (•): 90%, gray circle (•): 75%, white circle (◦): 50%. All percentages of sequences just described (e.g. 97% conserved) assume that sequences have been weighted by the GSC algorithm implemented by the Infernal software package.

Duplicate sequences: the following putative homologs are not shown in the alignment because their sequences are identical to a homolog already shown: Cat-1-1, Cba-1-1, Cba-2-1, Cba-3-1, Cca-1-2, Cca-1-3, Cgi-1-1, Cgr-1-1, Coc-1-1, Coc-2-1, Coc-4-1, Csp-1-1, Csp-10-1, Csp-11-1, Csp-12-1, Csp-2-1, Csp-3-1, Csp-4-1, Csp-5-1,

Csp-6-1, Csp-9-1, Ean-1-3, Ean-1-4, Ean-10-1, Ean-12-1, Ean-2-1, Ean-3-1, Ean-4-1, Ean-5-1, Ean-6-1, Ean-7-1, Ean-8-1, Ean-9-1, Eme-2-2, Fca-1-2, Fen-1-2, Fps-1-1, Fps-1-10, Fps-1-2, Fps-1-3, Fps-1-4, Fps-1-5, Fps-1-6, Fps-1-7, Fps-1-8, Fps-1-9, Fps-2-1, Fps-3-1, Fsp-6-1, Fsu-2-2, Gsp-2-1, Jpa-1-2, Jpa-1-3, Lbl-1-1, Mlu-1-1, Nul-1-5, Orh-2-1, Orh-3-1, Psp-1-1, Ran-1-2, Ran-2-2, Ran-5-1, Ran-7-1, Ran-8-1, Ran-9-1, Zpr-1-1, env-100, env-1000, env-1001, env-1002, env-1004, env-1005, env-1006, env-1008, env-101, env-1013, env-1014, env-1018, env-102, env-1020, env-1022, env-1024, env-1028, env-103, env-1030, env-1031, env-1032, env-1033, env-1035, env-1036, env-1037, env-104, env-1041, env-1043, env-1044, env-1045, env-1046, env-105, env-1050, env-1051, env-1054, env-1055, env-1056, env-106, env-1061, env-1065, env-1071, env-1077, env-1078, env-1079, env-108, env-1082, env-1083, env-1087, env-109, env-1090, env-1093, env-1094, env-11, env-110, env-1100, env-1101, env-1102, env-1103, env-1105, env-1108, env-1109, env-111, env-1110, env-1111, env-1112, env-1113, env-1117, env-1119, env-112, env-1120, env-1122, env-1123, env-1126, env-113, env-1133, env-114, env-115, env-1154, env-1156, env-1157, env-1158, env-1159, env-116, env-1160, env-1170, env-1171, env-1173, env-1176, env-1177, env-1178, env-1179, env-1180, env-1184, env-1186, env-1187, env-1188, env-1189, env-119, env-1192, env-1193, env-1194, env-1195, env-1198, env-12, env-1201, env-1204, env-121, env-1211, env-1212, env-1214, env-1218, env-122, env-1226, env-1229, env-1236, env-1238, env-1240, env-1242, env-1243, env-1244, env-1245, env-1246, env-1247, env-1248, env-1249, env-125, env-1250, env-1251, env-1266, env-1270, env-1277, env-1278, env-1279, env-1283, env-1284, env-1285, env-1288, env-129, env-1290, env-1292, env-1294, env-1295, env-1298, env-1299, env-13, env-130, env-131, env-1313, env-1315, env-1316, env-132, env-1324, env-1325, env-133, env-1330, env-1331, env-1333, env-1334, env-1345, env-1346, env-135, env-1350, env-1353, env-1354, env-1355, env-1356, env-1358, env-1362, env-1363, env-1365, env-1371, env-1372, env-1373, env-1376, env-1378, env-1379, env-1386, env-1387, env-1388, env-1389, env-1390, env-1391, env-1392, env-1396, env-1397, env-1398, env-1399, env-14, env-1400, env-1401, env-1402, env-1403, env-1404, env-1405, env-1406, env-1407, env-1408, env-1409, env-1410, env-1411, env-1412, env-1413, env-1415, env-142, env-1420, env-1421, env-1422, env-1426, env-1430, env-1438, env-1439, env-1440, env-1441, env-1442, env-1443, env-1444, env-1445, env-1446, env-1447, env-145, env-1452, env-1453, env-1454, env-1457, env-1458, env-1460, env-1461, env-1463, env-1471, env-1476, env-1477, env-1478, env-1479, env-148, env-1480, env-1481, env-1482, env-1483, env-1484, env-1485, env-1491, env-1492, env-1493, env-1494, env-1495, env-1496, env-1497, env-1498, env-1499, env-15, env-1500, env-1501, env-1502, env-1503, env-1504, env-1506, env-1509, env-151, env-1510, env-1514, env-1515, env-1516, env-1517, env-1518, env-152, env-1522, env-1523, env-1524, env-153, env-1531, env-154, env-1540, env-1548, env-155, env-1550, env-1552, env-1553, env-1560, env-1566, env-1568, env-157, env-1570, env-1571, env-1572, env-1573, env-1574, env-1575, env-1576, env-1577, env-1578, env-1579, env-158, env-1580, env-1584, env-1585, env-1586, env-159, env-1591, env-1592, env-1595, env-1598, env-1599, env-16, env-160, env-1609, env-161, env-1610, env-1613, env-1619, env-162, env-1620, env-1621, env-1622, env-1623, env-1627, env-163, env-1632, env-1633, env-1634, env-1636, env-1637, env-1638, env-1639, env-164, env-1641, env-1642, env-1643, env-1645, env-1646, env-1647, env-1649, env-165, env-1651, env-1652, env-1655, env-166, env-1668, env-1669, env-167, env-1675, env-1676, env-1678, env-1679, env-168, env-1681, env-1682, env-1683, env-1684, env-1685, env-1686, env-1688, env-1689, env-169, env-1691, env-1692, env-1693, env-1694, env-1695, env-1696, env-1697, env-1698, env-17, env-170, env-1704, env-1705, env-1706, env-1707, env-1708, env-171, env-1710, env-1719, env-172, env-1722, env-1724, env-1725, env-1726, env-1727, env-1728, env-1729, env-173, env-1730, env-1731, env-1732, env-1738, env-1739, env-174, env-1740, env-1744, env-1745, env-1746, env-1747, env-1748, env-1749, env-175, env-1753, env-1754, env-1755, env-1756, env-1757, env-1759, env-176, env-1766, env-1767, env-1768, env-1769, env-177, env-1770, env-1773, env-1774, env-1775, env-1776, env-1778, env-1780, env-1784, env-1785, env-1786, env-1795, env-1796, env-1797, env-1798, env-1799, env-18, env-1800, env-1801, env-1808, env-1809, env-181, env-1810, env-1811, env-1827, env-1828, env-1829, env-1833, env-1834, env-1835, env-1836, env-1838, env-1847, env-1852, env-1854, env-1858, env-1859, env-1868, env-187, env-1871, env-1874, env-1876, env-1877, env-1878, env-1879, env-1882, env-1883, env-1884, env-1885, env-1889, env-189, env-1890, env-1895, env-1896, env-1897, env-19, env-190, env-1901, env-1902, env-1903, env-1904, env-1905, env-1906, env-1907, env-1908, env-1909, env-191, env-1910, env-1911, env-1912, env-1913, env-1914, env-1915, env-1916, env-1917, env-192, env-1920, env-1926, env-1927, env-1928, env-193, env-1935, env-1936, env-1937, env-1938, env-1939, env-1940, env-1941, env-1942, env-1943, env-1944, env-1945, env-1946, env-1947, env-1948, env-1949, env-1950, env-1951, env-1952, env-1953, env-1954, env-1955, env-1956, env-1957, env-1958, env-1959, env-196, env-1960, env-1961, env-1962, env-1963, env-1964, env-1965, env-1966, env-1967, env-1969, env-1970, env-1971, env-1972, env-1973, env-1974, env-1975, env-1976, env-1979, env-198, env-1980, env-1981, env-1982, env-1983, env-1985, env-1986, env-199, env-1992, env-20, env-2002, env-2003, env-2004, env-2005, env-2006, env-2007, env-2008, env-2009, env-201, env-2010, env-2011, env-2012, env-2013, env-2014, env-2015, env-2016, env-2017, env-2018, env-2019, env-2020, env-2021, env-2022, env-2023, env-2024, env-2025, env-2026, env-2027, env-2028, env-2029, env-2030, env-2031, env-2037, env-2038, env-2039, env-2040, env-2041, env-2042, env-2043, env-2044, env-2045, env-2046, env-2049, env-205, env-2053, env-2058, env-2059, env-206, env-2060, env-2061, env-2062, env-2063, env-2064, env-2070, env-2072, env-2073, env-2074, env-2075, env-2081, env-2082, env-2083, env-2084, env-2085, env-2086, env-2087, env-2088, env-2089, env-209, env-2090, env-2091, env-2092, env-2093, env-2094, env-2095, env-2096, env-2097, env-2099, env-21, env-2105, env-2106, env-2107, env-2108, env-2109, env-2110, env-2111, env-2112, env-2113, env-2114, env-2115, env-2116, env-2117, env-2118, env-2119, env-2120, env-213, env-22, env-221, env-225, env-228, env-229, env-23, env-230, env-231, env-232, env-233, env-234, env-235, env-236, env-237, env-238, env-239, env-24, env-240, env-241, env-243, env-245, env-25, env-250, env-251, env-252, env-253, env-254, env-255, env-257, env-258, env-259, env-26, env-260, env-261, env-262, env-263, env-264, env-265, env-266, env-267, env-268, env-269, env-270, env-271, env-272, env-276, env-278, env-279, env-28, env-284, env-285, env-286, env-287, env-288, env-289, env-290, env-291, env-292, env-293, env-294, env-295, env-296, env-297, env-298, env-299, env-300, env-301, env-302, env-303, env-304, env-305, env-306, env-307, env-308, env-309, env-311, env-314, env-315, env-316, env-317, env-318, env-319, env-32, env-320, env-321, env-322, env-324, env-328, env-329, env-333, env-334, env-335, env-336, env-337, env-338, env-339, env-340, env-341, env-342, env-343, env-344, env-345, env-346, env-347, env-348, env-349, env-350, env-351, env-352, env-353, env-354, env-355, env-357, env-358, env-359, env-360, env-361, env-362, env-363, env-364, env-365, env-366, env-367, env-368, env-369, env-37, env-370, env-371, env-372, env-373, env-374, env-375, env-376, env-377, env-378, env-379, env-382, env-383, env-384, env-386, env-387, env-388, env-389, env-39, env-397, env-398, env-399, env-408, env-409, env-410, env-415, env-417, env-42, env-423, env-429, env-443, env-444, env-445, env-446, env-447, env-448, env-449, env-450, env-458, env-459, env-465, env-466, env-467, env-468, env-469, env-478, env-48, env-482, env-484, env-488, env-489, env-490, env-491, env-492, env-494, env-495, env-50, env-507, env-511, env-516, env-52, env-528, env-53, env-534, env-539, env-541, env-542, env-543, env-544, env-545, env-546, env-547, env-549, env-550, env-551, env-552, env-553, env-554, env-556, env-560, env-561, env-563, env-564, env-567, env-568, env-569, env-571, env-576, env-582, env-583, env-584, env-586, env-589, env-593, env-594, env-595, env-596, env-597, env-598, env-599, env-600, env-601, env-602, env-603, env-604, env-605, env-606, env-609, env-613, env-618, env-619, env-62, env-620, env-621, env-622, env-624, env-632, env-633, env-634, env-635, env-637, env-638, env-641, env-642, env-643, env-644, env-646, env-647, env-648, env-649, env-650, env-658, env-659, env-66, env-660, env-661, env-662, env-665, env-666, env-674, env-675, env-676, env-681, env-687, env-691, env-696, env-698, env-701, env-705, env-706, env-708, env-709, env-716, env-72, env-721, env-729, env-730, env-731, env-733, env-735, env-736, env-738, env-745, env-749, env-751, env-753, env-754, env-757, env-758, env-759, env-760, env-763, env-768, env-769, env-77, env-770, env-775, env-776, env-777, env-778, env-782, env-786, env-787, env-789, env-792, env-80, env-801, env-802, env-803, env-808, env-814, env-815, env-816, env-817, env-82, env-822, env-827, env-828, env-829, env-83, env-830, env-831, env-832, env-834, env-835, env-836, env-837, env-838, env-84, env-844, env-845, env-846, env-848, env-849, env-85, env-852, env-853, env-854, env-855, env-856, env-857, env-858, env-859, env-861, env-863, env-871, env-872, env-873, env-874, env-875, env-876, env-877, env-878, env-879, env-880, env-885, env-886, env-887, env-888, env-895, env-896, env-897, env-898, env-899, env-900, env-901, env-902, env-903, env-904, env-905, env-908, env-909, env-911, env-912, env-921, env-922, env-930, env-931, env-934, env-935, env-936, env-937, env-94, env-946, env-95, env-952, env-953, env-958, env-96, env-960, env-961, env-962, env-963, env-964, env-965, env-966, env-967, env-97, env-971, env-972, env-973, env-978, env-979, env-98, env-981, env-982, env-984, env-985, env-986, env-987, env-99, env-990, env-993, env-995, env-996

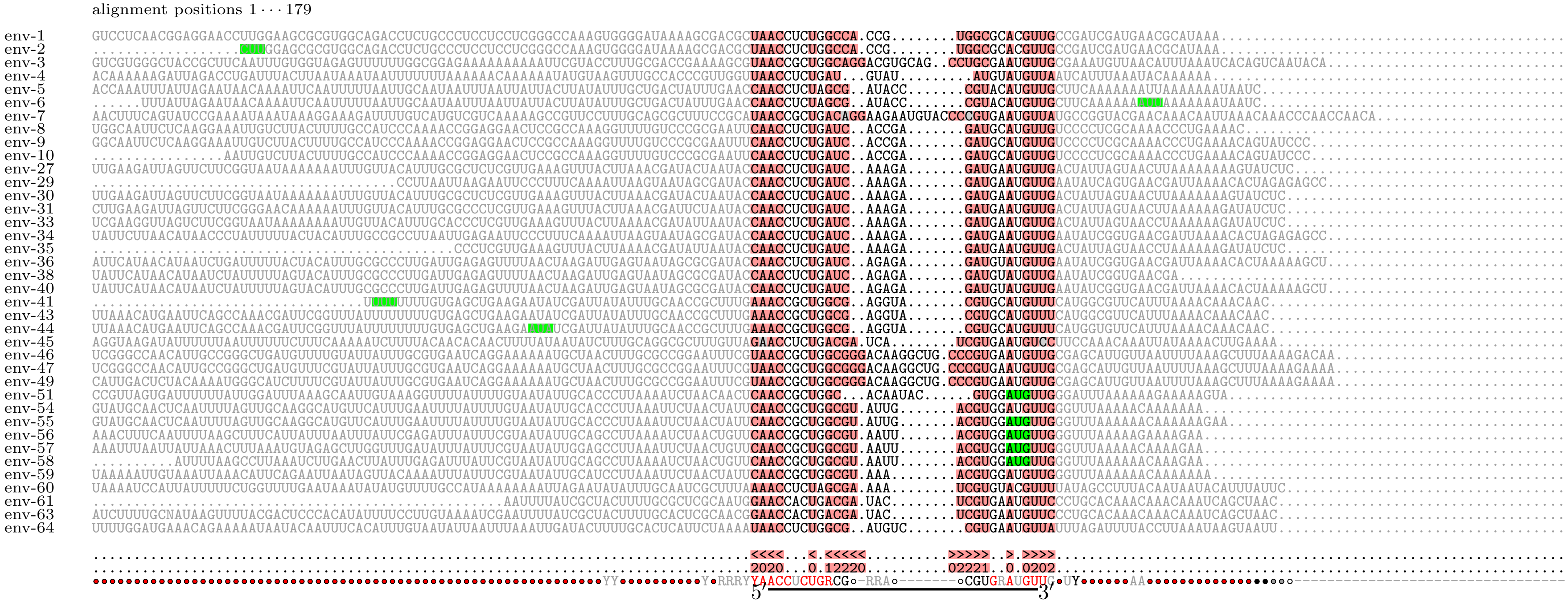

```
env-65    .UGACUUAUUGGGUGAAAAAUAAUUUCAUUAUUCAUUGUACAUUAUCAAUUAUUCAUUAAAUUUGCAGCCAAUUUCAGAUUAACCUCUGGCGA.UGU........UCGUGAACGUUGGUUUGAAUCAACCAUUUUAAAUUUAAUCAA..............................
env-67    AUUGCCUUUUUACUCCUCACUUUCUUCUAAAAAAAUUUUGCCCGAAAGAAAUUUUUAGUACAUUUGCACCCACUUCGGACCAACCUCUGACG..AAACG.......CGUGUAUGUUGCUCUGAUUCAAACUAUUAAUUUUUAGAAAUC............................
Nul-1-1   AGUAGACAUAAACUGCAUUAUUUUCUUUAAAAACAUUUGUUACAGUAAAAGAAGUUUGUAUUUUUGCAGCUCGUAAAAAUUAACCUCUGACGA.UCU........UCGUGAAUGUUACUCUUUACUUAAACAAAUUAAUUCCCAUUCAAC.........................
Fba-1-1   UAAAUCGUUACGACUCAUUUUUUUCCACAAAAACAUUUGUAACAGUAAAAGAAGUUUGUACUUUUGCAGCUCGUAAAAAUUAACCUCUGACGA.UCU........UCGUGAAUGUUACUCUUUACAUAAACAUAUUAAUUCCCAUUCAAC.........................
Nul-1-2   GUAACCAGAAGUAUUCGAUUAUUUUCUUCAAAAUUUUUGUAACAGUAAAAGAAGUUUGUAUUUUCGCAGCUCGUAAAAAUUAACCUCUGACGA.UCU........UCGUGAAUGUUACUCUUUACUUAAACAAAUUAAUUCCCAUUCAAC.........................
Nul-1-3   AAUAGUCAUAAACUGCAUUAUUUUCUUUAAAAACAUUUGUUACAGUAAAAGAAGUUUGUAUUUUUGCAGCUCGUAAAAAUUAACCUCUGACGA.UCU........UCGUGAAUGUUACUCUUUACUUAAACAAAUUAAUUCCCAUUCAAC.........................
Nul-1-4   UAACCAGAAGUAUUCGAUUAUUUUCUUCAAAAAUUUUUGUAACAGUAAAAGAAGUUUGUAUUUUUGCAGCUCGUAAAAAUUAACCUCUGACGA.UCU........UCGUGAAUGUUACUCUUUACUUAAACAAAUUAAUUCCCAUUCAAC.........................
Nul-1-6   UAACCAGAAGUAUUCGAUUAUUUUCUUCAAAAAUUUUUGUAACAGUAAAAGAAGUUUGUAUUUUUGCAGCUCGUAAAAAUUAACCUCUGACGA.UCU........UCGUGAAUGUUACUCUUUACUUAAACAAAUUAAUUCCCAUUUAAC.........................
env-68    AAAUAGUAUUUCGUAUAAAAAUUGCGUAAAAUAUCUUGGAAGUAAUAAUUUAAAGAGUAUUUUUGCAGUCGAUUUAAAAUCAACCUCUGACG..AAGUC......CGUGCAUGUUGGUUUUGACUUAAUCAAAAUAAAAAUCA...............................
Csp-18-1  UGAUUAAAAAAUUAUCAUAUUUAGCAUAUUAAUUAUUUGGCAUAUUCAAAAAAUGUUUUGUACAUUUGCAACCUGUUAUUCAACCUCUGACG..AAGUC......CGUGUAAGUUGCUUAGCCUAAACAUUUUAUUUAAUUAAUA.............................
Csp-15-1  GAACGAAAAAUUUAUUGUAUUUAAUACAUAAAUUGUUUGGCACAUUCAAAAAGUAUUUCUUAUAUUUGCAGCCUGUUAUUCAACCUCUGACG..AAGUC......CGUGUAAGUUGCUUAGCCUAAACAUUUUAUUUAUUAAUA..............................
Csp-16-1  UUUAAUAAAAAUCUUGUAAAAUUUAUUGUGUAAAGUAUUGGCGGUUAGAAGAAUUUGUUUUAUUUUUGCAACCUGUUAACCAACCUCUGACG..AAGUC......CGUGUAAGUUGCUUAGCUUGACAAAAACAAAUUUUUAUUAUAAAA........................
Chi-1-1   UUCUUAAAAAUUCAUUGUAUUUUAAUAAUAAACUGUUUGGAAUAUUUGAAAAAAGAUUUGUAUAUUUGCAACCUGUUAUUCAACCUCUGACG..AAGUC......CGUGUAAGUUGCUUAGCCUUAACAUUUUAUUUUAUUAAUA.............................
env-69    GGGGUGAAAAUUUCAUCAUAUUUAAUAUAAAAUUGUUUGGAAAAUUUCAAAAAUGUUUUGUACAUUUGCAACCUGUUAUUCAACCUCUGACG..AAGUC......CGUGUAAGUUGCUUAGCUUUAACAUUUUAUUUAUUAAUA..............................
Cgr-2-1   AAUGACUGAAAUUCUGCUAUUUUUAAUAAACAAUUGUUUUGAAUAUUUAAAAAAGUAUUUAUAUAUUUGCAACCUGUUAUUCAACCUCUGACG..AAGUC......CGUGUAAGUUGCUUAGCCUAAAUAUUUUAUUUAUUAAUA..............................
Csp-17-1  UAUGAUUGAAAUAAAUUAAAAAUUGUUUUAUAAAGUGUUGGCUAACAGAAAAAUUUAUUUUAUUUUUGCAGUCUGUUAUUCAACCUCUGACG..AAGUC......CGUGUAAGUUGCUUAGCGUAACACAUUUUAUUUUAUUAAUA............................
Fsp-5-1   UUUCUUCAACAUUUCAUAAGAUUUAAUAUUAAUUUGUUUUGAAUAUUUGAAAAUGAUUUGUAUAUUUGCAGCCUGUUAUUCAACCUCUGACG..AAGUC......CGUGUAAGUUGCUUAGCUUUAAACAUUUUAUUUUAAUUAAU............................
env-70    AACCCUUAGAAAGGUAUUUUCUUAUUGUAAAAUUGUUUGGCAAAUUCAAAAAACAUUUCAUAUAUUUGCAGCCUGUUAUUCAACCUCUGACG..AAGUC......CGUGUACGUUGUUUAGCUUGACAAAAAAUUUUUAUUACAAAA...........................
env-71    .....................AUUAUUUCUAAUUUUGCGCCCGCGUUUAAAUUCUUAGGAUUUGAACGAAAAUUGUUUCCCAGCCUCUGGCAA.AAA........UUGUGAAUGCUGUUUAACAAAAACAAUAAUUUAUAGAAGAA.............................
env-73    AGACCAGAUUUGUUAGAAGAUUGAGGUCGGCAAAAGAAUAAUCCUUAAUUUUGCCCCGCUUCAAAAAGAAGUGCGCUCUCCAACCUCUGGCA..AGAGA......UGUUAACGUUGCAGGGCGGUUAAUGCAUUAUUUUUAGGGUU............................
env-74    GUGAUUUUCAUUUUGCAGGCCGAGAGCAAUUUUAAAAAGAAUUUUUAAUUACUUUUGCCCUCCCGUAAGGGACUGCGCGUCAACCUCUGGCA..AAAGA......UGUGAAUGUUGACCCACAGCAAUAAAAACAAAUAAAAAC..............................
env-75    CGAUUUCCACUUUGCGGGUCGCGAGAAGUUCUAAAAAAGAAUUUUUAAUUACUUUUGCCCUCCUGAUUGGAACUGCGUGUCAACCUCUGGCA..AGAGA......UGUGAAUGUUGGCUCACAGCAAUGUAAAUUACUUUAAAC..............................
env-76    CGAUUUCCACUUUGCGGGUCGCGAGCAGUUCUAAAAAAGAAUUUUUAAUUACUUUUGCCCUCCCGAAAGGGACUGCGUGUCAACCUCUGGCA..AGAGA......UGUGAAUGUUGGCUCACAGCAAUGUAAAUUAUUUUAAAC..............................
env-78    ............................................................CUCUGAAAGGGACUGCGUGUCAACCUCUGGCA..AGAGA......UGUGAAUGUUGGCUCACAGCAAUGUAAAUUAUUUUAAAC..............................
env-79    CGAUUUCCACUUUGCAGGUCGCGAGCAGUUCUAAAAAAGAAUUUUUAAUUACUUUUGCCCUCCCGAAAGGGACUGCGUGUCAACCUCUGGCA..AGAGA......UGUGAAUGUUGGCUCACAGCAAUGUAAAUUAUUUUAAAC..............................
env-81    ACUUCCAUUUUGCAGGACGCGAGCAGUUCUAGACAACGAAUUAUUUAUUACUUUUGCCCUCCCGAAAGGGAACUGCGUAUCAACCUCUGGCA..AGAGA......UGUGAAUGUUGGUCCACAGAAAUAGAAAAUAUA.....................................
env-86    ...........................................UUAAUUACUUUUGCCCUCCCGUUAGGGACUGCGCGUCAACCUCUGGCA..AGAGA......UGUGAAUGUUGACCCACAGCAAUAAAAACAAAUAAAAAC..............................
env-87    ACUUCCAUUUUGCAGGACGCGAACAGUUCUAGAGAAAGAAUUAUUUAUUACUUUUGCCCUCCCGAAAGGGAACUGCGUAUCAACCUCUGGCA..AGAGA......UGUGAAUGUUGGUCCACAGAAAUAGAAAAUAUUAUUAUC..............................
env-88    GAUUUCCACUUUGCGGGUCGCGAGCAGUUCUAAAAAAGAAUUUUUAAUUACUUUUGCCCUCCCGAAACGGGACUGCGUGUCAACCUCUGGCA..AGAGA......UGUGAAUGUUGGCUCACAGCAAUGUAAAUUAUUUUAAAC..............................
env-89    GUGAUUUUCAUUUUGCAGGCCGAGAGCAAUUUUAAAAAGAAUUUUUAAUUACUUUUGCCCUCCCGUUAGGUACUGAGCGUCAACCUCUGGCA..AGAGA......UGUGAAUGUUGACCCACAGCAAUAAAAACAAAUAAAAAC..............................
env-90    UGAUUUCCACUUUGCAGGUCGCGAGCAGUUUUAAUCAGAAAUUAUUUACUAAUUUCGCCGUCCCGAAAGGGACUGAGUGUCAACCUCUGGCAA.GAG........UUGUGAAUGUUGGCCCACGGUAAUAGUUAAUUUAUAUACUC.............................
env-91    UGAUUUUCACUUUGCAGGUCGCGAGCAAUUUUAAUCAGAAAUUAUUUACUAAUUUCGCCGUCCCGAAAGGGACUGAGUGUCAACCUCUGGCAA.GAG........UUGUGAAUGUUGGCUCACGGUAAUAGUUAAUUUUAAUACUC.............................
env-92    GAUUUCCACUUUGCAGGUCGCGAGCAAUUUUAAUCAGAAAUUAUUUACUAAUUUUCGCCGUCCCGAAAGGGACUGAGUGUCAACCUCUGGCAA.GAG........UUGUGAAUGUUGGCUCACGGUAAUAGUUAA.........................................
env-93    CGAUUUCCACUUUGCAGGUCGCGAGCAAUUUUAAUCAGAAAUUAUUUACUAAUUUCGCCGUCCCGAAAGGGACUGAGUGUCAACCUCUGGCAA.GAG........UUGUGAAUGUUGGCUCAC......................................................
env-107   UGAUUUCCACUUUGCAGGUCGCGAGCAGUUUUAAUCAGAAAUUAUUUACUAAUUUCGCCGUCCCGAAAGGGACUGAGUGUCAACCUCUGGCAA.GAG........UUGUGAAUGUUGGCCCACGGUAAUAGUUAAUUUUUAUACUC.............................
env-117   UGAUUUUCACUUUGCAGGUCGCGAGCAAUUUUAAUCAGAAAUUAUUUACUAAUUUCGCCGUCCCGAAAGGGACUGAGUGUCAACCUCUGGCAA.GAG........UUGUGAAUGUUG............................................................
env-118   GAUUUCCACUUUGCAGGUCGCGAGCAGUUUUAAUCAGAAAUUAUUUACUAAUUUCGCCGUCCCGAAAGGGGACUGAGUGUCAACCUCUGGCAA.GAG........UUGUGAAUGUUGGCCCACGGUAAUAGUUAAUUUUUAUACUC.............................
env-120   UGAUUUUCACUUUGCAGGUCGCGAGCAGUUUUAAUCAGAAAUUAUUUACUAAUUUCGCCGUCCCGAAAGGGACUGAGUGUCAACCUCUGGCAA.GAG........UUGUGAAUGUUGGCCCACGGUAAUAGUUAAUUUUUAUACUC.............................
env-123   CGAUUUCCACUUUGCAGGUCGCGAGCAAUUUUAAUCAGAAAUUAUUUACUAAUUUCGCCGUCCCGAAAGGGACUGAGUGUCAACCUCUGGCAA.GAG........UUGUGAAUGUUGGCUCACGGUAAUAGUUAAUUUUAAUANUC.............................
env-124   CGAUUUCCACUUUGCAGGUCGAGAGCAAUUUUAAUCAGAAAUUAUUUACUAAUUUCGCCGUCCCGAAAGGGACUGAGUGUCAACCUCUGGCAA.GAG........UUGUGAAUGUUGGCUCACGGUAAUAGUUAAUUUUAAUACUC.............................
env-126   GCUUUAUCAAAAAAGAAUUUUGUUGUACUUUUGCAGCCGAUUUGCUCAUUAAGUAAAAUUAAAGACAAAUCGCUGAUUUUCAACCUCUGGCA..AAAAC......UGUGAAUGUUGUCUUUUAGUUUAUUAAAAAACAAAUAAGAA............................
env-127   GUUUCCUCAAUUAAUGGAUUUGUACUAAUAUUGCACUCCGUUUAACGAAGAAGAAUCAAAUUCGAUAAACAACUGAUAUUCAACCUCUGGCAA.AAA........UUGUGAAUGUUGGUCUUCAGUAAACCUUAAAGUAUCCUUAGC............................
env-128   GGUUUUAUAUUAUUUCGAUUUGUCGUAAUAUUGCGCUCCGUUUACCAAGUCAAAAAAUGAAAAGGUAAGCAACUGAAUUUCAACCUCUGGCAA.AAA........UUGUGAAUGUUGACCUUCAGCAAUUUUAAAGAUUCUGUAAA.............................
Oho-1-1   GUUUCUCCAAUUAAUUGUUUUGUAUUAAUAUUGCAGCCUGUUUACCACCUCUUACAAAAAUGUGAUAAGCAACUGAUAUUCAACCUCUGGCAA.AAA........UUGUGAAUGUUGGCUUUCAGCAAACUAUAAAUUAUCCUUAAG............................
env-134   ACUUUAGUGAUUAGGCUUUUUGUUGUAUUAUUGCAGCCGAUUUGCAACACCCGGAAAACUAAUUGUAAAUCGCUGAUUUUCAACCUCUGGCAA.AAA........UUGUGAAUGUUGUCUUUCGGUUAAUUAAAUUUUUGUUUCAAA............................
env-136   GCGAUUUCCACUUUGCUGUCCGUGAGCAGUUUUAAUUAGAAUUUUUAUCUAUUUUUGCCAUCCCUAAAGGGGCUACGUGUCAACCUCUGGCAA.AAG........UUGUGAAUGUUGGCUCAUAGUGAUUGUAAAUUUUUUAUUCC.............................
env-137   GCGAUUUCCACUUUGCUGUACGCGAGCAGUUUUAAUUAGAAUUUUUAUCUAUUUUUGCCGUCCCCACAGGGACUAAGAGUCAACCUCUGGCAA.AAG........UUGUGAAUGUUGGCUCAUAGUGAUUGUAAAUUUUUUAUACC.............................
env-138   GGCGAUUUCCACUUUGCUGUACGCGAGCAGUUUUAAUUAGAAUUUUUAUCUAUUUUUGCCGUCCCACAGGGACUAAGAGUCAACCUCUGGCAA.AAG........UUGUGAAUGUUGGCUCAUAGUGAUUGUAAAUUUUUUAUACC.............................
env-139   .................GUACGCGAGCAGUUUUAAUUAGAAUUUUUAUCUAUUUUUGCCGUCCCCAAAGGGACUAAGUGUCAACCUCUGGCAA.AAG........UUGUGAAUGUUGGCUCAUAGUGAUUGUAAAUUUUUUAUACC.............................
env-140   GCGAUUUCCACUUUGCUGUCCGUGAGCAGUUUUAAUUAGAAUUUUUAUCUAUUUUUGCCGUCCCUACAGGGGCUAAGUGUCAACCUCUGGCAA.AAG........UUGUGAAUGUUGGCUCAUAGUGAUUGUAAAUUUUUUAUACC.............................
env-141   ....................................................................GCAAGGGUGGCCAACCUCUGACA..AGAGC......UGUGAAUGUUGCCUAUCCUCAAAAUAAAUUUUACUAAGA..............................
env-143   CCCGAUCUUUUGCAGGAUGAUUCUUUGUUAGACAAUUGAUUUAUCGUAAUAUUGCAGCCCCAAAAAUGGGCAAGGUGGCCAACCUCUGACA..AGAGC......UGUGAAUGUUGCCUAUCCUCAAAAUAAAUUUUACUAAGA..............................
env-144   GGACCAGAAACGAUUGAUUAAAAAAGCUAGUCUAUGUUGGUUACAAUAAAAUAAAUAUUAAUUUUGCACGGCAAAGCCAUCAACCUCUGACGA.UAG........UCGUGCAUGUUGCUGUUUUCUAAUCAACUCAUAAAA..................................
env-146   GGACCAGAAACGAUUGAUUAAAAAAGCUAGUCUAUGUUGGUUACAAUAAAAUAAAUAUUAAUUUUGCACGGCAAAGCCAUCAACCUCUGACGA.UAG........UCGUGCAUGUUGCUGUUUUCUAAUCAACUAAUAAAA..................................
Nse-1-1   ACUUAUCAUUCAAUAAAGUUUUUUUUAUUUUAUUUAUUGUACACUUUUAAAUAAGUGUUUAUUUUUGCAGCUCGUUAAAUCAACCUCUGACGA.GGU........UCGUGUAUGUUGUUUUAAUUAAAUAUAAUAAGUCUAUAAAA.............................
env-147   ...........................................................UUUUGCAGGGCGAAAAGAUCAACCGCUGACGA.GGU........UCGUGAAUGUUGGCUUUUUAAAUCAUAAAAGAUCCAAAUU..............................
env-149   UUUUUUUCACUUUUACAGAAUAUUUAGAACCCUCUAUUGUUUAGAUCAAAAUAAAGUUUAAUUUUGCAGGGCGAAAAGAUCAACCGCUGACGA.GGU........UCGUGAAUGUUGACUUUUUAAAUCUUAAAAGAUCCAAAUU..............................
env-150   UUUUUUUCACUUUUACAGAAUAUUUAUAACCCUCUAUUGUUUAGAUCAAAAUAAAGUUUAAUUUUGCAGGGCGAAAAGAUCAACCGCUGACGA.GGU........UCGUGAAUGUUGACUUUUUAAAUCUUAAAAGAUCCAAAUU..............................
env-156   CUUUUGGAAGAAGAUUAAAAUUCUUCAUUAAAACAUGUAAUUAUAAAAAUAAAAUUUAUAAUUUUGCAUCCGUUCUAAAAUAACCUCUGGCG..AGAAC......CGUGAACGUUGUUUUAGCUUGCAUCUUAAAUAUUUAACU..............................
env-178   CUUUUGGAAGAAGAUUAAAAUUCUUCAUUAAAACAUGUAAUUAUAAAAAUAAAAUUUAUAAUUUUGCAUCCGUUCUAAAAUAACCUCUGGCG..AGAAC......CGUGAACGUUGUUUUAGCUUACAUCUUAAAUAUUUAACU..............................
env-179   AGUUAUUCAAGUUUAAUGAAAAAAUGUUGAAAAUAAUUUGUGUAUUUUGAUUAUUUUUCUUUACUUUGCGGAAUGUUAAUCAACCUCUGACG..AGAGC......CGUGAAAGUUGUUUAGCAUUAAUCAUUUUUAUUUAUUAAUA............................
env-180   AGUUAUUCAAGUUUAAUGAAAAAAUGUUGAAAAUAAUUUGUGUAUUUUGAUUAUUUUUCUUUACUUUGCGGACUGUUAAUCAACCUCUGACG..AGAGC......CGUGAAAGUUGUUUAGCGUUAAUCAUUUUUAUUUAUUAAUA............................
env-182   AGUUAUUCAAGUUUAAUGAAAAAAUGUUGAAAAUAAUUUGUGUAUUUUGAUUAUUUUUCUUUACUUUGCGGACUGUUAAUCAACCUCUGACG..AGAGC......CGUGAAAGUUGUUUAGCAUUAAUCAUUUUUAUUUAUUAAUA............................
env-183   AGUUAUUCAAGUUUAAUGAAAAAAUGUUGAAAAUAAUUUGUGUAUUUUGAUUAUUUUUCUUUACUUUGCGGGCUGUUAAUCAACCUCUGACG..AGAGC......CGUGAAAGUUGUUUAGCAUUAAUCAUUUUUAUUUAUUAAUA............................
env-184   AGUUAUUCAAGUUUAAUGAAAAAAUGUUGAAAAUAAUUUGUGUAUUUUGAUUAUUUUUCUUUACUUUGCGGAAUGUUAAUCAACCUCUGACG..AGAGC......CGUGAAAGUUGUUUAGCGUUAAUCAUUUUUAUUUAUUAAUA............................
env-185   AAUGGAUUUCAAUGGGAAUAAAUGCCGCCAACCAUUCUCCGAUUACAGAAAGGUUUAAUUUANNNNNNNNNNNNGUUAAUCAACCUCUGACG..AGAGC......CGUGAAAGUUGUUUAGCGUUAAUCAUUUUUAUUUAUUAAUA............................
env-186   AUAACAAAUUUUAUCUAAAUUUGCACCCGCUUAUAGAGAGUCUUAUUGAUUCUUAGUAAAAAGCUAUAAUUUAAGUUUAUCAACCGCUGACG..AAAAA......CGUGUAUGUUGAUUGAUUAUAAUUUAUUAAAAUCAUCUAAG............................
env-188   ...........GCGAAUAAAAAAUGAUUAUAAAAUUGCACUAAUUUAAUUAUUGAUUAUAUUUGCACUCAAUUUCGACUUAACCUCUGGCG..AUAGA......CGUAAAUGUUAAUUUGAAAAAAUAUAUAAAAAAAAUUACA..............................
env-194   GCGGGGAACGCCCAAGAAAAAAAGUACAUAAAAAGUUGUAACAUUCUAUUAAUUACGUAUUUUUGCACUCGAUUUUACCCCAACCUCUGGCG..AUGUA......CGUGUAUGUUGUUGUAAACAAACCCAUUAAAUUAAAGAGCA............................
env-195   UAAGCAAGAUCAUUGAUUUUAAAAUUUGGAAAAAAACUUGUCAUUUCUAAAUUAAGAACUAUUUUUGCAGCCUGUUAAAACAACCUCUGACGA.GCA........UCGUGAACGUUGUUUUGAUUUAAUUUCAAUAAAGUAGACC..............................
env-197   GGCGUCGCAACGCCCAAAUAAAAAUUCAAAUACUGCUUGGUACUUCUUUAAAAAAACGUAUUUUUGCAUCCGAUUUGAUCCAACCUCUGGCGA.GCA........UCGUGAACGUUGUUUCGAUUAAACCAUAACAAAACGACC...............................
env-200   AAAAAAUUCGUCCAAACUUUUUUAAUUAAGUAAAUUUCUUUGAAAGAAAAAAAAAAUAGUAAAUUCGCAUUCGAUUUGAUAUAACCGCUAACGA.UCA........UCGUGUAUGUUGUAUCAAUUAAAUUAAAAUACUUACAAA...............................
env-202   GAAGAUUAGGCGCUUAAAAAAAAAUAUCAUUUAUCCAUUGUCAAUUAUCAAUUAAAUUCUAUCUUUGCGCCCACAUUAGACCAACCUCUGGCGA.AGU........UCGCGAAUGUUGCUCCGAUUUAAAACCAUAAUUAAAAAAAUUAUC.........................
Rbi-1-1   ACCCCGGGGAGGACCACUAAAUUUGCGCGAUUUUCUUGCCAGUUGGGAUUUAUCGAUUAUUUUUGCACACGAUUUCAGGUCAACCUCUGACG..AGGUA......CGUGCAUGUUGGCCCGAACAGAACAGUAAAGCAAUACC...............................
env-203   ACUUUUGAUUCCGAAGAAAAAUUGCNAAAAAUAUCUUGUAAGUAAUAAUUUAAAGACUAUUUUUGCAUUCGAUUUGAAAUUAACCUCUGACG..AAGUC......CGUGAAUGUUGGUUUUGACUUAAUUAAAAUAAAAAUCA...............................
env-204   ...................................................................CAAACUUCAAAUUAACCACUGGCG..AUAUA......CGUGUAUGUUGAUUUGAAAUUUAAUUGAUUAAACAAAUAGAUU..........................
env-207   ............CCUUGAAUAAUUAGGAAGCACUAUUGUUUAGAUCAAAAUAAAGUUUAAUUUUGCAGGGCGAAAAGAUUAACCACUGACG..AUAUA......CGUGUAUGUUGACUUUUUAAACAAAAAAAGAUCCUAAAU..............................
env-208   .UAAAAUCUAACAUACUUAUAUUUAUUUAAAAUUUUGGUUUGCUAAUAAUUAAAAUUAGAAUAUAUUUGCAACCGCUUAAAAACCUCUGGCGA.AAA........UCGUGCAUGUUCUUUAGCUUUUACAAUAAUACAUUAUAUC..............................
env-210   ..........GUUUGUUAUUUUUUUUAAUUAUCCAUUGUUUAUUAUCAAUUAUUAAUUACAUUUGCGCCCACUUAGACCAACCUCUGGCGA.AAC........UCGUGCAUGUUGCUCUGAUUUAAAACCAUAAUUAAAACAAUUAUC.........................
env-211   UAAAAUCCAUGUUUACAGUUAUUUUAAAUUAAAUAUUAUUUGCUAUAAAAUAAAAUUGAAAUAUAUUUGCACCCGCUUUUAAACCUCUGACGA.UAA........UCGUGAAUGUUUUCUAGCAUUUACAAUAAUACAUUUUAUC..............................
Gsp-1-1   GCCAGAUUUACUUGAUAAAAAUAAUUUGUAAAAAAUUGUAUAUUAAUGAAUAAUGAUUAAUUUUGCAACCGAUUUCAGCUAAACCUCUGACGA.GAA........UCGUGAAUGUUUACUUGAAUUAAUCUAUUAUAAACAUUAAGAA...........................
env-212   AAAAUCCAUUUUUUUUCUGAUUUUGAAUAAAGUAUUGUUUUGUUAUGAAAUAAAAUUAAAAUAUAUUUGCACCCGCUUUAAAACCUCUGACGA.AAA........UCGUGAAUGUUCUAAAGCUUUUUAC.........................................
env-214   ACCUGAUUUAUUGGACAAAAAUUAAUUGUAAAAAAUUGCAGAUUAGCGAAUAAUGAUUAAUUUUGCAUCCGAUUUCAAUUAAACCUCUGACGA.AAA........UCGUGAAUGUUUACUUGGAC..............................................

          ...............................................................................<<<<...<.<<<<<..........>>>>>..>.>>>>..........................................................
          ...............................................................................2020...0.12220..........02221..0.0202..........................................................
          ●●●●●●●●●●●●●●●●●●●●●●●●●●●●●●●●●●●●●●●●●●●●●●●●●●●●●●●●●●●●●●●●●●YY●●●●●●●●●●Y●RRRYYAACCUCUGRCG○-RRA○-------○CGUGRAUGUUG●UY●●●●●●AA●●●●●●●●●●●●●●●●●●○○○---------------------------
                                                                                 5'                                 3'
```

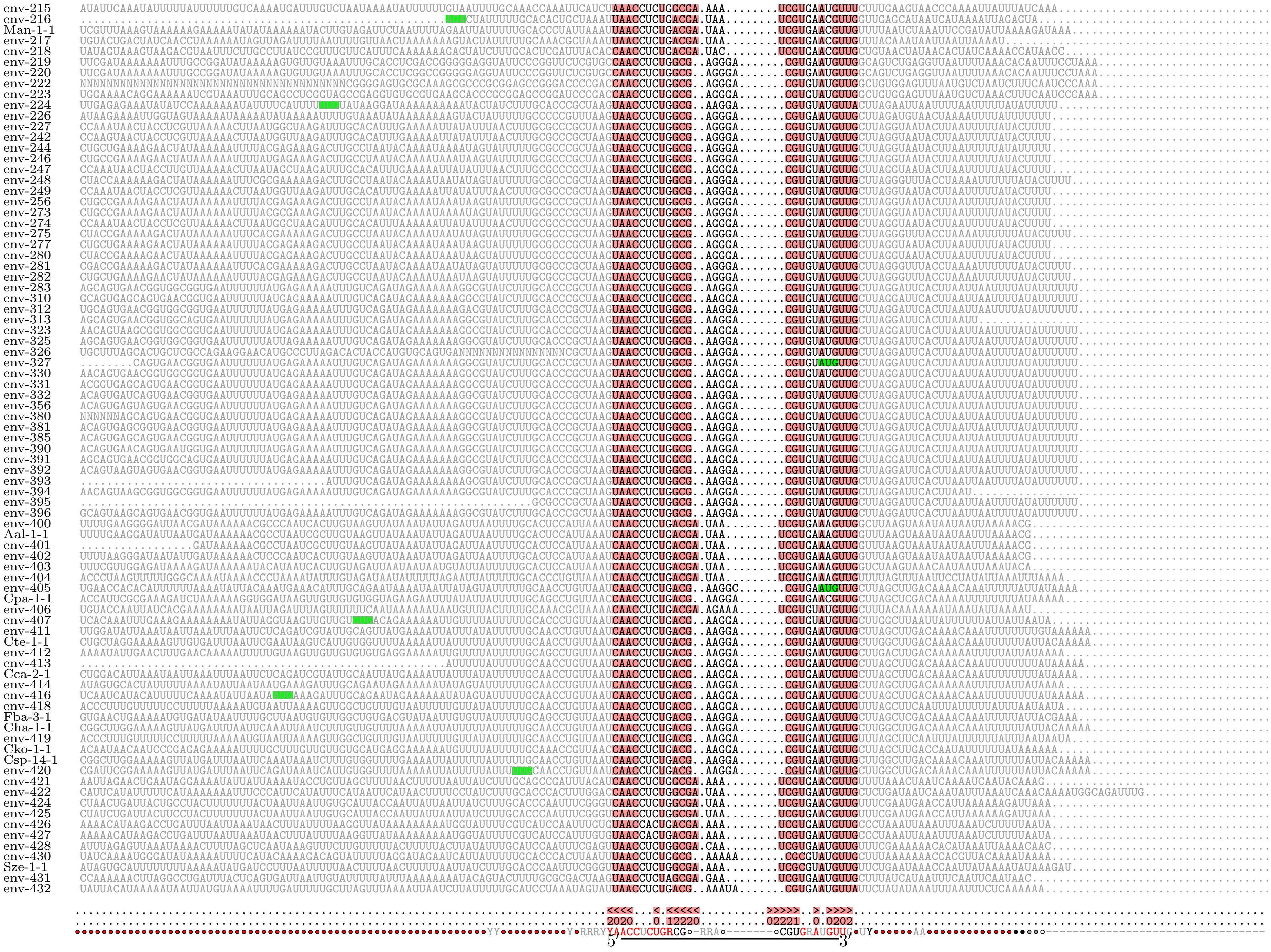
env-215
env-216
Man-1-1
env-217
env-218
env-219
env-220
env-222
env-223
env-224
env-226
env-227
env-242
env-244
env-246
env-247
env-248
env-249
env-256
env-273
env-274
env-275
env-277
env-280
env-281
env-282
env-283
env-310
env-312
env-313
env-323
env-325
env-326
env-327
env-330
env-331
env-332
env-356
env-380
env-381
env-385
env-390
env-391
env-392
env-393
env-394
env-395
env-396
env-400
Aal-1-1
env-401
env-402
env-403
env-404
env-405
Cpa-1-1
env-406
env-407
env-411
Cte-1-1
env-412
env-413
Cca-2-1
env-414
env-416
env-418
Fba-3-1
Cha-1-1
env-419
Cko-1-1
Csp-14-1
env-420
env-421
env-422
env-424
env-425
env-426
env-427
env-428
env-430
Sze-1-1
env-431
env-432
5′
3′

```
env-433   UACCAAAUAUAAAUAAUUAGUUAAUAUUUUGAUUUUUGCGUAGUUUAAAAUUAAUCUUAUUUUUGCAUCCUAAAUAGUAUUAACCUCUGACG..AAAUA.......CGUGAAUGUUAUUCUAUAUAAAUUUAAUUUCUCAAAAAA........................
env-434   UUCCAAAUAUAAAUAAUUAGGUAAAAUUUUGAUUUUUGCUUAGUUUAAAAUUAAUCUUAUUUUUGCAUCCUAAAUAGUAUUAACCUCUGACG..AAAUA.......CGUGAAUGUUAUUCUAUAUAAAUUUAAUUUCUCAAAAAA........................
env-435   UUCCAAAUAUAAAUAAUUAGGUAAUAUUUUGAUUUUUGCGUAGUUUAAAAUUAAUCUUAUUUUUGCAUCCUAAAUAGUAUUAACCUCUGACG..AAAUA.......CGUGAAUGUUAUUCUAUAUAAAUUUAAUUUCUCAAAAAA........................
env-436   UAUUACAUAAAAAUAAUUAUGUAAAAUUUUGAUUUUUGCGUAGUUUAAAAUUAAUCUUAUUUUUGCAUCCUAAAUAGUAUUAACCUCUGACGA.AAU........UCGUGAAUGUUAUUCUAUGUAAACUUAAUUUCUCAAAAA.........................
env-437   UAUUACAUAAAAAUAAUUAUGUAAAAUUUUGAUUUUUGCGUAGUUUAAAAUUAAUCUUAUUUUUGCAUCCUAAAUAGGAUUAACCUCUGACGA.AAU........UCGUGAAUGUUAUUCUUUGUAAACUUAAUUUCUCAAAAA.........................
env-438   .................AUGUGAAAUUUUGAUUUUUGCGUAGUUUAAAAUUAAUCUUAUUUUUGCAUCCUAAAUAGGAUUAACCUCUGACGA.AAU........UCGUGAAUGUUAUUCUAUGUAAACUUAAUUUCUCAAAAA.........................
env-439   UAUUAAGUAAAAAUAAUUAUGUGAAAUUUUGAUUUUUGCGUAGUUUAAAAUUAAUCUUAUUUUUGCAUCCUAAAUAGGAUUAACCUCUGACGA.AAU........UCGUGAAUGUUAUUCUAUGUAAACUUAAUUUCUCAAAAA.........................
env-440   GUUUUAAAUAAAAAAUACUAAUGUAAAAUUUGAUUUUUGCGUAGUUUAAAAUUAAUCUUAUUUUUGCAUCCUAAAUAGAAUUAACCUCUGACGA.AAU........UCGUGAAUGUUACUCUAUGUAAACUUAAUUUCUCAAAAA.........................
env-441   UAUUACAUAAAAAUAAUUAUGUAAAAUUUUGAUUUUUGCUUAGUUUAAAAUUAAUCUUAUUUUUGCAUCCUAAAUAGUAUUAACCUCUGACGA.AAU........UCGUGAAUGUUAUUCUAUGUAAACUUAAUUUCUCAAAAA.........................
env-442   ..............................................UAAAAUUAAUCUUAUUUUUGCAUCCUAAAUAGGAUUAACCUCUGACGA.AAU........UCGUGAAUGUUAUUCUAUGUAAACUUAAUUUCUCAAAAA.........................
env-451   GCAAUGCUUUUAUCACUAUAUAAAAGAAGUUUAACGGUUGCACACUAAAAAAAAAAUAUCUAAUUUUGCAGACCCUAGGAUUAACCGCUGGCGA.UGG........UCGUGAAUGUUGCUCUUAGUUAUUUUUAGCUAAA............................
env-452   GUGUGAACCGAGCAAUUUGAUAAAAUGGUUUAAUGGGUUGCACACUAAAAAAAAAAUAUCUAAUUUUGCAGACCCUAGGUUUAACCGCUGGCGA.UGG........UCGUGAAUGUUGCUCUUAGUCAUUUUUUAGUUAAAAAUC.........................
env-453   GUGUGAACCGAGCAAUUUGAUAAAAUGGUUUUAUGGGUUGCACACUAAAAAAAAAAUAUCUAAUUUUGCAGACCCUAGGUUUAACCGCUGGCGA.UGG........UCGUGAAUGUUGCUCUUAGUCAUUUUUUAGUUAAAAAUC.........................
env-454   GAGAAUACCCUUGCAAUUAGAUAAAAGAGGUUGAAUGGUUGCACACAAAAAAAAAAUAUCUACUUUUGCAACCCUAGGUUCAACCGCUGGCGA.UGG........UCGUGAAUGUUGCUCUUAUUCAUUUUUAGCUAAAAAUC..........................
env-455   .....................AUUAAAAAAAAGAACAGUUGCAAACUAAAAAAAAAAUAUCUAAUUUUGCACCCCUAGGUUCAACCGCUGGCGA.UGG........UCGUGAAUGUUGUUCUUAGUUAUUUUUUAGCUAGAAAUC.........................
env-456   GAUGAUUUUUGAUUGUAUUAAGAUUAAAAAAAAGAACAGUUGCAAACUAAAAAAAAAAUAUCUAAUUUUGCACCCCUAGGCUCAACCGCUGGCGA.UGG........UCGUGAAUGUUGUUCUUAUUUAUUUUUUAGCUAGAAAUC.........................
env-457   ............................................................................GGUUCAACCGCUGGCGA.UGG........UCGUGAAUGUUGCUCUUAGUUAUUUUUAGCUAAAAAUC..........................
env-460   AUGAUUUUUGGUCGUGUUAAGAUUAAAAAAAAAGAACAGUUGCAAACUAAAAAAAAAAUAUCUAAUUUUGCACCCCUAGGUUCAACCGCUGGCGA.UGG........UCGUGAAUGUUGUUCUUAGUUAUUUUUAGCUAAAAUC...........................
env-461   UGAUUAUUGAUUGUAUUAAUAUUAAAAAAAAAAGAACAGUUGCAAACUAAAAAAAAAAUAUCUAAUUUUGCACCCCUAGGUUCAACCGCUGGCGA.UGG........UCGUGAAUGUUGUUCUUAGUUAUUUUUAGCUAAAAUC...........................
env-462   AUGAUUUUUGGUCUUAUUAAGAUUAAAAAAAAAGAACAGUUGCAUACUAAAAAAAAAAUAUCUAAUUUUGCACCCCUAGGUUCAACCGCUGGCGA.UGG........UCGUGAAUGUUGUUCUUAGUUAUUUUUUAGCUAGAAAUC.........................
env-463   UGAUUAUUGAUUGUAUUAAUAUUAAAAAAAAAAGAACAGUUGCAAACUAAAAAAAAAAUAUCUAAUUUUGCACCCCUAGGUUCAACCGCUGGCGA.UGG........UCGUGAAUGUUGUUCUUAGUUAUUUUUUAGCUAGAAAUC.........................
env-464   GAGAACACUCUUGCAAUUAGAUAAAAGGGCUAAACGGUUGCACACUAAAAAAAAAAUAUCUAAUUUUGCAGACCCUAGGUUCAACCGCUGGCGA.UGG........UCGUGAAUGUUGCUCUUAGUUAUUUUUAGCUAAAAAUC..........................
env-470   GAGAACACUCUUGCAAUUAGAUAAAAGGGUUAAACGGUUGCACACUAAAAAAAAAAUAUCUAAUUUUGCAGACCCUAGGUUCAACCGCUGGCGA.UGG........UCGUGAAUGUUGCUCUUAGUUAUUUUUAGCUAAAAAUC..........................
env-471   UGGAUACUGUUGCCAUUUGAUAAAAGGGGUUGAAGGGUUGCACACUAAAAAAAAAAUAUUUAAUUUUGCAAACCCUAGGUUCAACCGCUGGCGA.UGG........UCGUGAAUGUUGCUCUUAGUUAUUUUUAGCUAAUAAUC..........................
env-472   .................................ACGGUUGCACACAAAAAAAAAAAUAUCUAAUUUUGCAAACCCUAGGUUCAACCGCUGGCGA.UGG........UCGUGAAUGUUGCUCUUAGUUAUUUUUAGCUAAAAAUC..........................
env-473   GUAAUACUUUUACUCCUAGAAUAAAGAUUGUUAACUGUUGCACACAAAAAAAAAAAUAUCUAAUUUUGCAGACCCUAGGAUCAACCGCUGGCGA.UGG........UCGUGAAUGUUGCUCUUAGUUAUUUUUAGC................................
env-474   AUGAUUUUUGGUUGUGUUAAGAUUAAAAAAAAAGAACAGUUGCAAACUAAAAAAAAAAUAUCUAAUUUUGCACCCCUAGGUUCAACCGCUGGCGA.UGG........UCGUGAAUGUUGUUCUUAGUUAUUUUUAGCUAAAAUC...........................
env-475   GAUGAUUUUUAGUCGUAUUAAGAUUAAAAAAAAGAACAGUUGCAUACUAAAAAAAAAAUAUCUAAUUUUGCACCCCUAGGUUCAACCGCUGGCGA.UGG........UCGUGAAUGUUGUUCUUAGUUAUUUUUAGCUAGAAAUC..........................
env-476   CUAUCAAAAGCCUAGGAUACAAAACUCUCAUUAGGUAUUGUACUCUAAAAAAAAAAUACAUAGUUUUGCCGACCCUAGGGUCAACCGCUGACGA.UAG........UCGUGAAUGUUGUUCUUAGUUAUUUUUAGCUAAAAAUC..........................
env-477   GAGAAUACUCUUGCAAUUAGAUAAAAGGGCUAAACGGUUGCACACUAAAAAAAAAAUAUCUAAUUUUACAGACCCUAGGUUCAACCGCUGACGA.UGG........UCGUGAAUGUUGCUCCUAGUUAUUUUUAGCUAAAAAUC..........................
env-479   AAAAUACUCUUGCAAUUAGAUAAUAAGGGUUAAACGGUUGCACACAAAAAAAAAAAUAUCUAAUUUUGCAGACCCUAGGUUCAACCGCUGACGA.UGG........UCGUGAAUGUUGCUCUUAGUUAUUUUUAGCUAAAUAUG..........................
env-480   AAACAUUUAGCCCAACAACAAAAAAGAUUUAAUUCUUAGUUUUUUAAUUAAAAAAAAGAAUUAUAUUUGCAACCUUAAAAUCAACCUCUGACGA.UGG........UCGUGAAUGUUGGUUUGUUGAAAACCAUUAAAUAUAAACAA.......................
env-481   CUUCUCAAAUAAAUAAACAAUUGUUGAAAAUCUUUAUUGUGUACUUCAAAAUAAAGUUUAAUUUCGCAAAAAUAAAAGUACAACCUCUGACGA.UGA........UCGUGAAUGUUGGCUUUAAAAACUAUAAAUAAUCUUAUU.........................
env-483   GCAAAAAGGAACACCAUAAAAAACACAACUAUUAAUUGUGCAUUAUUAAUUAUUAAUUAUCUUUGCAGCCAAUUUCGGGUUAACCUCUGGCGA.AAA........UCGUAAAUGUUAUUUCGAGACAACUAUUAAAAAGAUUAAA.......................
env-485   AUAAAAUCAAUUAUUGUUGAUUUGUUAAAUUCUGUAUUGUUUACUGUAAAAUAAAGUUUAUUUUUGCAUUUCGAAAAUGUUAACCUCUGACGA.AAA........UCGUAAAUGUUAAUUUUUAAAAUUAUAUAUCCAAAUU..........................
env-486   ...........................................................................AUAAAUAACCUCUGACGA.AAA........UCGUGAAUGUCAUUUAUCCCAAUACAACAAAAUAAAAU..........................
env-487   AUCCUGAGCUUUAGGAUAAAAAAUCAAAAUAAAUUUAUUGUGGAUUCUAAUUUAUAAACUAUUUUUGCACACUGUUAAAUUAACCUCUGACGA.AAA........UCGUGAAUGUUAGUUCAACAUAAUUGUAAAAUAAAGUA.........................
env-493   AUCCUGAACUUUAGGACAAAAAAUAAAAAUAAAUUUCUUGUGGGUUCUAAUUUAUAAACUAUUUUUGCACACUGUUAAAUUAACCUCUGACGA.AAA........UCGUGAAUGUUAGUUAAACAUAAUUGUAAAAUAAAGUA.........................
env-496   UAAAAAAUGCUUUUUUUUUAUUUUUUUUUAUAAAAAAGGAAAGCAAAAAAUAAAAUAAUUGGUACUUUUGCACUCGCAAUGUAACCUCUGACGA.AAA........UCGUGAAUGUUACUAGCUACAAUUAUUUAAGUAUAUCUA........................
env-497   UAAAAAAUGUUUUUUUUUAUUUUUUUUCAUAAAAAAGGAAAGCAAAAAAUAAAAUAAUUGGUACUUUUGCACUCGCAAUGUAACCUCUGACGA.AAA........UCGUGAAUGUUACUAGCUACAAUUAUUUAAGUAUAUCUA........................
env-498   AUAAAAUCAAUUAUUGUUGAUUUGUUAAAUUCUGUAUUGUUUACUGUAAAAUAAAGUUUAUUUUUGCAAUUCGAAAAUGUUAACCUCUGACGA.AAA........UCGUGAAUGUUAAUUUUUAAAAAUCAUAUAUCCUAAUU.........................
env-499   ACUUAACAAAAUUAAUUAUCAAUUAAUUUCAUUUUUGUGUAGUUUAAAAUUAAUCUUAUUUUUGCAAGCUAAAAUAGUAUUAACCUCUGACGA.AAA........UCGUGAAUGUUAAACUAUACAAUUAAAUUUCUCAAAUA.........................
env-500   AAUAAAUCAAUUAUUGUUGAUUUGUUAAAUUCUGUAUUGUUUACUGUAAAAUAAAGUUUAUUUUUGCAAUUCGAAAAUGUUAACCUCUGACGA.AAA........UCGUGAAUGUUAAUUUUUAAAAAU.......................................
env-501   UAUUAAAUAAAAGUGAAAAAUAAAAUUAAUUUCAUUAUCUAAUCUUCAAAAACUUUUUAUUACAUUUGCACACUCAAAAUUAACCUCUGACGA.AAA........UCGUGAAUGUUAGUUUCUGAAAAAUCAUUAAAAUAAUAAA.......................
env-502   AUAAAAUCAAUUAUUGUUGAUUUGUUAAAUUCUGUAUUGUUUACUGUAAAAUAAAGUUUAUUUUUGCAUUUCAAAAAUGUUAACCUCUGACGA.AAA........UCGUGAAUGUUAAUUUUUAAAAUUAUAUAUCCAAAUU..........................
env-503   AUCCUGAGCUUUAGGAUAAAAAAUCAAAAUAAAUUUCUUGUAGAUUCUAAUUUAUAAACUAUUUUUGCACACUGUUAAAAUAACCUCUGACGA.AAA........UCGUGAAUGUUAGUUCAACAUAAUUGUAAAAUAAAGUA.........................
env-504   ..................................GAUUUUAUUUCCUAAAAUAAAUCAUUAGAUUUGCACUCGCUAAAAAUAACCUCUGACGA.AAA........UCGUGAAUGUUAGUUUAGAAAAUUUUAAAACAAUAAA..........................
env-505   UUCCUGAACUUUUGGAUAAAAAAUCAAAAUAAAUUUCUUGUGGAUUCUAAUUUAUAAACUAUUUUUGCACACUGUUAAAUUAACCUCUGACGA.AAA........UCGUGAAUGUUAGUUCAACAUAAUUGUAAAACAAAGUA.........................
env-506   UAAAAAAUGCUUUUUUUUAUUUUUUUUCAUAAAAAAGGAAAGCAAAAAAUAAAAUAAUUGGUACUUUUGCACUCGCAAUGUAACCUCUGACGA.AAA........UCGUGAAUGUUACUAGCUACAAUUAUUUAAGUAUAUCUA........................
env-508   AAUAAAUCAAUUAUUGUUGAUUUGUUAAAUUCUGUAUUGUUUACUGUAAAAUAAAGUUUAUUUUUGCAAUUCGAAAAUGUUAACCUCUGACGA.AAA........UCGUGAAUGUUAAUUUUUAAAAUUAUAUAUCCAAAUU..........................
env-509   UUAAAAAAUACUUUUUUUUAUUUUUUUCAUAAAAAAGGAAAGCAAAAAAUAAAAUAAUUGGUACUUUUGCACUCGCAAUGUAACCUCUGACGA.AAA........UCGUGAAUGUUACUAGCUACAAUUAUUUAAGUAUAUCUA........................
env-510   AUAAAAUCAAUUAUUGUUGAUUUGUUAAAUUCUGUAUUGUUUACUGUAAAAUAAAGUUUAUUUUUGCAUUUCGAAAAUGUUAACCUCUGACGA.AAA........UCGUGAAUGUUAAUUUUUAAAAUUAUAUAUCCAAACU..........................
env-512   UAGUGCAUAAUGUGAGUUGAAAAAGGAUAAAAUAAUUUUAUUUGGUUUAAAAUAUUUUACUACAUUUGCACACUCAAAAUUAACCUCUGACGA.AAA........UCGUGAAUGUUACUCUUUGAAAAUAAUUAAAUAAUUAAAA.......................
env-513   GAAAAGCCAUAGAAAGAACAAAAAAUAAGACCAGAUUUGUUGGAAGACUAAUCUUUUUAUUAAAUUUGCACUCUCAAAAUUAACCUCUGACGA.AAA........UCGUGAAUGUUAAUUAUUGAAAAACACUAAAAAACAUAAA.......................
env-514   UGGUCCAGAUCAUAUCAAUUAAUUUUUAGGAUCAAGUUGUUUACUUGAAAAUAAAGUUUAAUUUUGCACCACGAAAUGCAUAACCUCUGACGA.AAA........UCGUGAAUGUUAUCUUUUCCAACUCAAUAAAAGAAAAU.........................
env-515   AAGCAUUUGAACAUACCAAAAAAAUUAGACCAGAUUUGUUGGAAGACUAAUAUUUUUGACUAAAUUUGCACACUCAAAAUUAACCUCUGACGA.AAA........UCGUGAAUGUUACUUAUUGAAAAACUAUUAAAAAGUUUAAA......................
env-517   UUCCUGAACUUUAGGGUAAAAAAUCAAAAUAAAUUUCUUGUGGGUUCUAAUUUAUAAACUAUUUUUGCACACUGUUAAAUUAACCUCUGACGA.AAA........UCGUGAAUGUUAGUUAAACAUAAUUGUAAAAUAAAGUA.........................
env-518   AAGCAUUUGAACAUACCAAAAAAAUUAGACCAGAUUUGUUGGAAGACUAAUAUUUUUCAUUAAAUUUGCACACUCAAAAUUAACCUCUGACGA.AAA........UCGUGAAUGUUACUUAUUGAAAAACUAUUAAAAAGUUUAAA......................
env-519   UAAAAAAUGUUUUUUUNUAUUUUUUUUCAUAAAAAAGGAAAGCAAAAAAUAAAAUAAUUGGUACUUUUGCACUCGCAAUGUAACCUCUGACGA.AAA........UCGUGAAUGUUACUAGCUACAAUUAUUUAAGUAUAUCUA........................
env-520   AAUAAAUCAAUUAUUGUUGAUUUGUUAAAUUCUGUAUUGUUUACUGUAAAAUAAAGUUUAUUUUUGCAAUUCGAAAAUGUUAACCUCUGACGA.AAA........UCGUGAAUGUUAAUUUUUAAAAAUCAUAUAUCCUAAUU.........................
env-521   ACUUAAUAAUAUAUAAUUAUCAAUAGUUUCAUUUUUGUGUAGUUUAAAAUUAAUCUUAUUUUUGCAAGCUAAAAUAGUAUUAACCUCUGACGA.AAA........UCGUGAAUGUUAAACUAUUCAAUUAAAUUUCUCAAAUA.........................
env-522   UAAAAAUCAAUUAUUGUUGAUUUGUUAAAUUCUGUAUUGUUUACUGUAAAAUAAAGUUUAUUUUUGCAAUUCGAAAAUGUUAACCUCUGACGA.AAA........UCGUGAAUGUUAAUUUUUAAAAAUCAUAUAUCCUAAUU.........................
env-523   AUCCUGAACAUUAGGUUAAAAAAUCAAAAUAAAUAUCUUGUUGGUUCUAAAUUAUAAACUAUUUUUGCACACUGUUAAAUUAACCUCUGACGA.AAA........UCGUGAAUGUUAGUUCAACAUAAUUGUAAAACAAAGUA.........................
env-524   AUAAAGUCAAUUAUUGAUGAUUUGUUAAAUUCUGUAUUGUUUACUGUAAAAUAAAGUUUACUUUUGCAUUUCGAAAAUGUUAACCUCUGACGA.AAA........UCGUGAAUGUUAAUUUUUUAAAACUAUAUAUCCAAAUU.........................
Tma-1-1   CAAGCUAAUUAUCAACUUAAAAAAAUAUUUCAAUUUUAUUUUAUCAGCAAAAAAAAUAUUUCUAAAUUUGCAAACUCAAAAUUAACCUCUGACGA.AAA........UCGUGAAUGUUAGUUAUUGAAAACCAUAAAAAAUAAAAAA.......................
env-525   UUAAAAAAUGCUUUUUUUUAUUUUUUUUAUAAAAAAGGAAAGCAAAAAAUAAAAUAAUUGGUACUUUUGCACUCGCAAUGUAACCUCUGACGA.AAA........UCGUGAAUGUUACUAGCUACAAUUAUUUAAGUAUAUCUA........................
env-526   AAGCAUUUGAACAUACCAAAAAAAUUAGACCAGAUUUGUUGGAAGACUAAUAUUUUUGAUUAAAUUUGCACACUCAAAAUUAACCUCUGACGA.AAA........UCGUGAAUGUUAAUUUCUGAAAAACAAUUAAAAAGUUUAAA......................
env-527   UGUGCUAAUUAUCAAGCUAAAAAAUAAUCAGUAUUCCUUUUAUCAACUAAAAAAUUAUUAUUAUUUUUGCACUCGCUAAAUUAACCUCUGACGA.AAA........UCGUGAAUGUUAUUUUAGAAAAAAUCUAAAAUUUAAAAG........................
env-529   AAGCCAUAGAAAGAACAAAAAAAAUAAGACCAGAUUUGUUGGAAGACUAAUCUUUUUUAUUAAAUUUGCACUCUCAAAAUUAACCUCUGACGA.AAA........UCGUGAAUGUUAAUUAUUGAAAAACACUAAAAAAACAUAAA......................
env-530   AGACCAGAUCAUAUCAAUUAAUUUUUUAGGAUCAAGUUGUUUACUUGAAAAUAAAGUUUAAUUUUGCACCACGAAAUGCAUAACCUCUGACGA.AAA........UCGUGAAUGUUAUCUUUUCCAACUCAAUAAAAGAAAAU.........................
env-531   CUUAAUAAUAUAUAAUUAUCAGUUAGUUUCAUUUUUGUGUAGUUUAAAAUUAAUCUUAUUUUUGCAAGCUAAAAUAGUAUUAACCUCUGACGA.AAA........UCGUGAAUGUUAAACUAUUCAAUUAAAUUUCUCAAAUA.........................
env-532   AUUAAAAAAACGAAGAGAAAUAAUUUCUUGUAUUGAUAUACAAUUUUAUAAAACUUUUUAUUAAAUUUGCAAACUCAAAAUUAACCUCUGACGA.AAA........UCGUGAAUGUUAGUUUUUAAAAACCAUUAAAAUUAAAAA........................
env-533   AAUAAAUCUAUUAUUGUUGAUUUGUUAAAUUCUGUAUUGUUUACUGUAAAAUAAAGUUUAUUUUUGCAAUUCGAAAAUGUUAACCUCUGACGA.AAA........UCGUGAAUGUUAAUUUUUAAAAAUCAUAUAUCCUAAUU.........................
env-535   GAUUUUUAAAAAUGCUUUUUUAUUUUUUUUAUAAAAAAGGAAAGCAAAAAUAAAAUAAUUGGUACUUUUGCACUCGCAAUGUAACCUCUGACGA.AAA........UCGUGAAUGUUACUAGCUACAAUUAUUUAAGUAUAUCUA........................
env-536   ....................AUUUUUGAGGAUCAAGUGUUUACUUGAAAAUAAAGUUUAAUUUUGCACCACGAAAUGCACAACCUCUGACGA.AAA........UCGUGAAUGUUAUCUUUUCAAACCCAAUAAAAGAAAAU.........................
env-537   UUUUAAAUUUUUAAUAAAAAAAAACACAUACAUUAAUUGGUUGGUUGUUAGAAUAAUAUUAUUUUUGCAGACCAUUAAAUCCAACCUCUGACG..AAAAA.......CGUGAAUGUUGUUUUUAAUUCAUCAAUAAAGUAUAACAUUACU...................
env-538   CUUUUAAAUUUUUAUAAAAAAAAACACAUACAUUAAUUGGUUGAUUGUUAGAAUAAUAUUAUUUUUGCAGACCAUUAAAUCCAACCUCUGACG..AAAAA.......CGUGAAUGUUGUUUUUAAUUCAUCAAUAAAGUAUAACAUUACU...................
env-540   CUUUUAAAUUUUAAUAAAAAAAAACACAUACAUUAAUUGGUUGGUUGUUAGAAUAAUAUUAUUUUUGCAGACCAUUAAAUCCAACCUCUGACG..AAAAA.......CGUGAAUGUUGUUUUUAAUUCAUCAAUAAAGUAUAACAUUACU...................
env-548   CUUUUAAAUUUUUAUAAAAAAAAACACAUACAUUAAUUGGUUGAUUGUUAGAAUAAUAUUAUUUUUGCAGACCAUUAAAUCCAACCUCUGACG..AAAAA.......CGUGAAUGUUGUUUUUAAUUCAUCAAUAAAGUAUAGUAUUACU...................
env-555   GACUUUAAAUUAAUUAAAAAAAACACAAUUUUUUAAUUGGUUGAUUGUUAGAAUAAUAUUAUUUUUGCAGUCCAUUAAAUCCAACCUCUGACG..AAAAA.......CGUGAAUGUUGUUUUUAAUUCAUUAAUAAAGUAUUUCAUUACU...................
env-557   AGAUCAAAUUAAUUAAAAAAAAACACGAUUUUUUAAUUGGUUGAUUGUUAGAAUAAUAUUAUUUUUGCAGUCCAUUAAAUCCAACCUCUGACG..AAAAA.......CGUGAAUGUUGUUUUUAAUUCAUUAAUAAAGUAUUUCAUUACU...................
env-558   UUUAAAUUUUAAUAAAAAAAAACACAUAACAUUUAAUUGGUUGGUUGUUAGAAUAAUAUUAUUUUUGCAGACCAUUAAAUCCAACCUCUGACG..AAAAA.......CGUGAAUGUUGUUUUUAAUUCAUCAAUAAAGUAUAACAUUACU...................
env-559   AAGCCUUAGAAAGAACAAAAAAAAUAAGACCAGAUUUGUUGGAAGAUUAAUCUUUUUUAUUAAAUUUGCACACUCAAAAUCAACCUCUGACGA.AAA........UCGUGAAUGUUGGUCAUUGAAAAACCCAUUAAAAAAAUUAACA.....................

          ....................................................................................<<<<...<.<<<<<..........>>>>>..>.>>>>..................................................
          ....................................................................................2020...0.12220..........02221..0.0202..................................................
          ••••••••••••••••••••••••••••••••••••••••••••••••••••••••••••••••••••••••••YY••••••••••Y•RRRYYAACCUCUGRCG∘-RRA∘-------∘CGUGRAUGUUG•UY••••••AA••••••••••••••••∘∘∘---------------------------
                                                                                          5′──────────────────────────────3′
```

```
env-562    CUUAAAAACGCCUCCUUAAAAAAUAUUUUUAAUACGCUCUUUCUUAUUAAAAAGUUUUUUAUAAAUUUGCAGACUCAAAAUCAACCUCUGACGA.AAA........UCGUGAAUGUUGGUAUUUGUGAAGCAUUUAAAAUAUCAUA..........
env-565    GGCCGAAAAAUAAUAUUAAAAAAUAUUUUCGAUCACCUGUUUUCUAUUAAAAAGUUUUUUAUAAAUUUGCAGACUCAAAAUCAACCUCUGACGA.AAA........UCGUGAAUGUUGGUGUUUGGUCAACCAUUUUAAAAUAUCAUA........
env-566    AAGCCUUAGAAAGAACAAAAAAAAAUAAGACCAGAUUUGUUAAAAGAUUAAUCUUUUUUAUUAAAUUUGCACACUCAAAAUCAACCUCUGACGA.AAA........UCGUGAAUGUUGGUCAUUGAAAAACCCAUUAAAAAAAUUAACA........
env-570    CUUAAAAAACUAAGGUUAAAAAAUAUUUUUAAUACGCUGUUUCUUAUUAAAAAGUUUUUUAUAAAUUUGCAGACUCAAAAUCAACCUCUGACGA.AAA........UCGUGAAUGUUGGUGUUUGGUCAAGUAUUUAAAAUAUCAUA.........
env-572    AAGCCUUAGAAAGAACAAAAAAAAAUAAGACCAGAUUUGUUGAAAGAUUAAUCUUUUUUAUUAAAUUUGCACACUCAAAAUCAACCUCUGACGA.AAA........UCGUGAAUGUUGGUCAUUGAAAAACCCAUUAAAAAAAUUAACA........
env-573    ..................................UUACGCGGUUUCAUAUCAAAAGUUUUUUAUAAAUUUGCAGACUCAAAAUCAACCUCUGACGA.AAA........UCGUGAAUGUUGGUAUUUGAUCAAGUAUUUAAAAUAUCAUAAU.......
Fba-5-1    UUGAAGAGUUACCUAACAAAAAAAUAAAUAUAUUAUUCUUGUAGUUUCUAAAUUAAGAGCUAUUUUUGCACCCGGUUAAUUCAACCUCUGACGA.AAA........UCGUGAAUGUUGAUUGAACAUAAUUAUAAAACAAUAUA...........
env-574    AGCAAUUUUAUUGAAUAAAAAAUUAUUUCAAAUUACUUAUUUCUUAUUUAAAAGUUUUUUAUAAAUUUGCACACUCAAAAUCAACCUCUGACGA.AAA........UCGUGAAUGUUGGUGUUUGAUCAACCAUUUUAAAAUAUCAUA........
env-575    CAUAAAAAAUGAUGCAUUAAAAAAUAUUUUUAAUACGCUGUUUCUUAUUAAAAAGUUUUUUAUAAAUUUGCAGACUCAAAAUCAACCUCUGACGA.AAA........UCGUGAAUGUUGGUAUUUGGUCAAGCAUUUAAAAUAUCAUA.........
env-577    ..................................................UAAGAUAUAACCUAUUUUUGCACACUGUUAAAUCAACCUCUGACGA.AAA........UCGUGAAUGUUGGUCAUUACAUAAUCCUAAAUAAAACAAA.........
env-578    AAGUAAUGUUGAAGUUAUUUAUUAAGACUAAGAAGUUGCCUACUUCAAAAUAAAGUCUAAUUUUGCAAAAAUGAAAUGAACAACCUCUGACGA.AAA........UCGUGAAUGUUGUUACUUUUCAAACAAUAAAAAAAAAUAAU.........
Tme-1-1    AUUUAUUAGAUUAACUAAUAAUCAAUGAUUUAUUUAUUUUUUUUAUCAAAAGAAAAAUUCUAAAUUUGCAGCCACAAAAUCAACCUCUGACGA.AAA........UCGUGAAUGUUGCUUUUAUACAACCAUAAAUAACAAGAAA.........
Zul-1-1    GCAAACGGACAUCUAUUUAAAAAAUAAUUGGAAAUAAUUGUCAGUUCUAAAUUAUAGCGUAUUUUUGCACCCUGUUAAAUCAACCUCUGACGA.AAA........UCGUGAAUGUUGACCAUAACAUAAUCAAUAAAAAAAAUAUC.........
env-579    ..........ACAUUUCAAAAANUUAUCNUAAAUCCCUUGUAGGUUUUAAGAUAUANCNUAUUUUUGCACACUGUUAAAUCAACCUCUGACGA.AAA........UCGUGAAUGUUGGUCAUUACAUAAUCCUAAAUAAAACAAA.........
env-580    CAUAAAAAAUGAUGCAUUAAAAAAUAUUUUUAAUACGCUGUUUCUUAUUAAAAAGUUUUUUAUAAAUUUGCAGACUCAAAAUCAACCUCUGACGA.AAA........UCGUGAAUGUUGGUGUUUGGUCAAGCAUUUAAAAUAUCAUA.........
env-581    CAUAAAAAAUGAUGCAUUAAAAAAUAUUUUUAAUACGCUGUUUCUUAUUAAAAAGUUUUUUAUAAAUUUGCAGACUCAAAAUCAACCUCUGACGA.AAA........UCGUGAAUGUUGGUAUUUGGUCAAGUAUUUAAAAUAUCAUA.........
env-585    CUUAAAACUGAAGCAUUAAAAAAUAUUUUUAAUACGCUGUUUCUUAUUAAAAAGUUUUUUAUAAAUUUGCAGACUCAAAAUCAACCUCUGACGA.AAA........UCGUGAAUGUUGGUGUUUGGUCAAGCAUUUAAAAUAUCAUA.........
env-587    CUUAAAAAGCUAAGGUUAAAAAAUAUUUUUAAUACGCCGUUUCUUAUUAAAAAGUUUUUUAUAAAUUUGCAGACUCAAAAUCAACCUCUGACGA.AAA........UCGUGAAUGUUGGUGUUUGGUCAAGUAUUUAAAAUAUCAUA.........
env-588    CUUGAAAUUGAAGGGUUAAAAAAUAUUUUUAAUAGGGUGUUUCUUAUUAAAAAGUUUUUUAUAAAUUUGCAGACUCAAAAUCAACCUCUGACGA.AAA........UCGUGAAUGUUGGUAUUUGGUCAAGCAUUUAAAAUAUCAUA.........
env-590    GGAUUUAUUGGAAGAGUAAAAAAAAUACAGAUUAUUUUUCUGAUUCAAUAAAAUUUAUUACUUUUGCACCCACAUUUAGACCAACCUCUGACGA.AAA........UCGUGAAUGUUGCUCUGAUAAUCAUUAUUUUAACUUAAUUGUU.......
env-591    AGCAUUUUAGAAUAGAUAAAAAUAAUUUACUCUUAAAGGUUUUAUAUUUAAAAGAAUUUAUUAAAUUUGCAAACUCAAAAUCAACCUCUGACGA.AAA........UCGUGAAUGUUGGUAUUUGAUCAACCAUUAAAAAAAAACAUA.........
env-592    ......................................UUGCCUACUUCAAAAUAAAGUUUAAUUUUGCACCACGAAAUGAACAACCUCUGACGA.AAA........UCGUGAAUGUUGUCUUUUCCAACUCAAUAAAAACAUAAU..........
Zga-1-1    CAUUUUGCCCUAUACCACAAAAAAUCAGUAAAAUAUAUUGCAAAUUCUAAUUUAUAGACUAUUUUUGCACCCUGUUAAAUCAACCUCUGACGA.AAA........UCGUGAAUGUUGACCAUAACAUAAUCAUUAAAACCAUAUC.........
env-607    CUUAAAAAGCUAAGGUUAAAAAAUAUUCUUAAUACGCUGUUUCUUAUUAAAAAGUUUUUUAUAAAUUUGCAAACUCAAAAUCAACCUCUGACGA.AAA........UCGUGAAUGUUGGUGUUUGGUCAAGUAUUUAAAAUAUCAUA.........
env-608    CUUUAAAUUGAAGGGUUAAAAAAUAUUUUUAAUAGGGUGUUUCUUAUUAAAAAGUUUUUUAUAAAUUUGCAGACUCAAAAUCAACCUCUGACGA.AAA........UCGUGAAUGUUGGUAUUUGGUCAAGCAUUUAAAAUAUCAUA.........
env-610    GACAAGUACAUUCCAAAAAAAAAUUUAUCUUAAAUCCCUUGUAGGUUUUAAGAUAUAGCGUAUUUUUGCACCCUGUUAAAUCAACCUCUGACGA.AAA........UCGUGAAUGUUGAUCAUUACAUAAUCCUAAAUAAAACAAC.........
env-611    UCCGUCCGGAUCUUCUUAAAAAAGAAGAAAAAUAAUUUUAUCAUCUUUAGAAUAUUUUAUUAUCUUUGCAAACUCAAAAUCAACCUCUGACGA.AAA........UCGUGAAUGUUGUACAAUUGAAAAAUUUUAAAAAUAAUUUAAUA......
Tov-1-1    UGUACUAAUUAUCAGUAUAAAAAAUAAACAACUUUAAUUUUCAUUCAAAAAAAUAAUCCUUAAUUUGCAGCCACAAAUCAACCUCUGACGA.AAA........UCGUGAAUGUUGCUUUAAACAAUCAUAUAAAUAUCAUUACA.........
env-612    CCCAAUUUAAGUGACUGAAAAUUAUUUCGAAAUACUUAAUUCUAAUUUAAAAGUUUUUUAUAAAUUUGCACACUCAAAAUCAACCUCUGACGA.AAA........UCGUGAAUGUUGAUGAUUGAUCAACCAUUUUAAAAUAUCAUA........
env-614    .........................................................UUUAUUAGCUAUUUUUGCACCCUGUUAAAUCAACCUCUGACGA.AAA........UCGUGAAAGUUGGUCCAAACUAAAUUAAAAAUUAAAGCA..........
Ace-1-1    AUUUUUUCAGAUUCAUAUAUAAAAAAGGAAUAUUCACUUGUAAGUUAGAAAAAAUCCACUAUUUUUGCACUCCAUUAAUUCAACCUCUGACGA.AAA........UCGUGAAAGUUGGCUUAAGUAACUGAUAAUAAAAUACC..........
env-615    ....................................................................................CGCAAUGUAACCUCUGACGA.AAA........UCGUGUAUGUUACUAGCUACAAUUAUUUAAGUAUAACUA.........
env-616    CUAAAUAAUAACUAAUUAUCAAUUAAUUUGAUUUUUGUGUAGUUUAAAAUUAAUCUUAUUUUUGCAAACGAAAAUAGUAUUAACCUCUGACGA.AAA........UCGUGUAUGUUAAA.........................
env-617    UAAAAAGCAGCUUUUUUUGAUUUUUUUCUACAAAAAAGGAAAUCAAAAAUAAAAUAAUUGGUACUUUUGCACUCGCAAUGUAACCUCUGACGA.AAA........UCGUGUAUGUUACUCGCUACAAUUAUUUAAGCAUAUCGA.........
env-623    UUGAAAAGUACUUUUUUUUGAUUUUUUUCUUAAAAAAAGGAAAGUGAAAAUAAAAUAAUUGGUACUUUUGCACUCGCAAUGUAACCUCUGACGA.AAA........UCGUGUAUGUUACUAGCUACAAUUAUUUAAGUAUAUCUA.........
env-625    UAAAAAGCAGCUUUUUUUGAUUUUUUUCUACAAAAAAGGAAAACAAAAAUAAAAUAAUUGGUACUUUUGCACUCGCAAUGUAACCUCUGACGA.AAA........UCGUGUAUGUUACUCGCUACAAUUAUUUAAGCAUAUCGA.........
env-626    UUUUUAAAAAGUUUUUUUUGAUUUUUUUAAACAAAAAAGGAAAAAAAAAAUAAAAUAAUUGGUACUUUUGCACUCGCAAUGUAACCUCUGACGA.AAA........UCGUGUAUGUUACUAGCUACAAUUAUUUAAGUAUAUCUA.........
env-627    UAAAAAGCAGCUUUUUUUUAUUUUUUUCUACAAAAAAGGAAAACAAAAAUAAAAUAAUUGGUACUUUUGCACUCGCAAUGUAACCUCUGACGA.AAA........UCGUGUAUGUUACUCGCUACAAUUAUUUAAGCAUAUCGA.........
env-628    AAUUUUCAAAAUUCAAUCAAAAUAGUGGCCUAAAUAAUUGUCACAUUACAAAAUUUAGGUAUUUUUGCAGCCAAUUAAUUUAACCUCUGGCG..AAAUA.......CGUGUAUGUUGCUUUAGUUUAAUAUUUAAAUUAAUACU.........
env-629    AAUUUUCAAAAUUCAAUCAAAAUAGUGGCCUAAAUAAUUGUCACAUUACAAAAUUUAGGUAUUUUUGCAGCCAAUUAAUCUAACCUCUGGCG..AAAUA.......CGUGUAUGUUGUUUUAAUUUAAUAUUUAAAUUAAUACU.........
env-630    AAUUUUAAUAUUUCAAUCAAAAUAGUGNCCUAAAUAAUUGCCACAUUACAAAAAUUAGGUAUUUUUGCAGCCAAUUAAUUUAACCUCUGGCG..AAAUA.......CGUGUAUGUUACUUUAAUUUAACAUUAUAAAUUAAUACU........
env-631    AAUUUUAAUAUUUCAAUCAAAAUAGUGGCCUAAAUAAUUGCCACAUUACAAAAAUUAGGUAUUUUUGCAGCCAAUUAAUUUAACCUCUGGCG..AAAUA.......CGUGUAUGUUACUUUAAUUUAACAUUAUAAAUUAAUACU........
env-636    AAUUUUAAUAUUUCAUUCAAAAUAGUGACUUAAAUAAUUGCCACAUUACAAAAAUUAGGUAUUUUUGCAUCCAAUUAAUUUAACCUCUGGCG..AAAUA.......CGUGUAUGUUACUUUAAUUUAACAUUAUAAAUUAAUACU........
env-639    AAUUUUAAUAUUUCAUUCAAAAUAGUGGCCUAAAUAAUUGCCACAUUACAAAAAUUAGGUAUUUUUGCAGCCAAUUAAUUUAACCUCUGGCG..AAAUA.......CGUGUAUGUUACUUAAUUUAAUAUUUAUAAAUUAAUACU........
Fsp-12-1   AUUUAUCAGCAACUUAAAAUAAUUUACAAUUUAUAAUUCAUAAUUUACAAUUAUUUUUUACAUUUGCACCCGAAUUAGACUAACCUCUGGCG..AAAUA.......CGUGAAUGUUACUCUAAAUAAACAAUAAUAAAAAUUAUC........
env-640    AAUUUUAAUAUUUCAAUCAAAAUAGUGGCCUAAAUAAUUGCCACAUUACAAAAAUUAGGUAUUUUUGCAGCCAAUUAAUUUAACCUCUGGCG..AAAUA.......CGUGCAUGUUACUUUAAUUUAACAUUAUAAAUUAAUACU........
env-645    AAUUUUCAGUUUUCAAUCAAAAUAGUGGCCUAAAUAAUUGCUACAUAACAAAAAUUAGGUAUUUUUGCAGCCAAUUAAUUUAACCUCUGGCG..AAAUA.......CGUGCAUGUUACUUUGGUUUAACAUUAUAAAUUAAUACU........
env-651    AAAGUCCAAAAAAGUAAAAUAAUUCAAAAUUUAUAAUUUAUAAUUCAAAAUAAUUAUUAAAUUUGCACCCUCAAUUGGAUUAACCUCUGGCG..AAAGA.......CGUGCAUGUUGGUCUGACAUUCAAUAAUUUAAAUCGUAAAUC......
env-652    ACGUGCUUUAGAAAUUUACAAUAAUCGAAAUAAUGAUUUGGAAAAUCAAUAAUUAUCAUUAAUUUUGCAGUCGUAUUGAUUAACCUCUGGCG..AAAGA.......CGUGCAUGUUACUCAAUCUUAAUUAAUUCGAAUUAUUACAA.......
env-653    CCGUGCUUUAGAAAUUUACAAUAAUCGAAAUAAUGAUUUGGAAAAUCAAUAAUUAUCGUUAAUUUUGCAGUCGUAUUGAUUAACCUCUGGCG..AAAGA.......CGUGCAUGUUACUCAAUCUUAAUUAAUUCGAAUUAUUACAA.......
env-654    AAUUUUAAUAUUUCAAUCAAAAUAGUGGCCUAAAUAAUUGCCACAUUACAAAAAUUAGGUAUUUUUGCAGCCAAUUAAUUUAACCUCUGGCG..AAACA.......CGUGCAUGUUACUUUAAUUUAACAUUAUAAAUUAAUACU........
env-655    AUACUUUAAUCAUUGUAGAUUAAAUAAAAAUGAUUAUUUUUGCAGCCGUCUAUACUAUUAAUAUAGGCACAAAUUUAUUCCAACCUCUAGCGA.AAA........UCGUGAAUGUUGUUUUAAAUAUUACAUUAAUAAAAAUUAAA........
env-656    CCAAAAAACUUAGGCCUGAUUUACUCAGUGAUUAAUUGUAUUUUUAUUUAAAAAAAAUACAGUACUUUUGCGCGACUAAGUAACCUCUAGCGA.AAA........UCGUGAAUGUUGCUUUAAUCAUAAUUUCAAUUCAAUAAAC.........
env-657    GCAAAAAGAAACACCAAAAAAAUACAAACAAAAGGUUGUAGAUUAUGAAUUAAUAAUUACUUUUGCACCCAAUUUCGGAUUAACCUCUAGCGA.AAA........UCGUGAAUGUUGUUCUGGAUUAAUUAUAAUUAAAUUUAGAGAU.......
env-663    ACAGUCCCAUUGCAAGCUGAUAAAAUUACUACUGCUAUUGUGUACUCAAAAAAUAAGUUUAAUUUUGCACCACGAAAGAUUAACCACUGACG..AUAAC.......CGUGUAUGUUGGCUUUUUAAACUUAAUAAACUAUCAA..........
env-664    AUAAAUUUAAAAACGUUUCACAUUCCUGAGAAAAAUAAUUUUCCUUGAUUAUAAAAUUAUUAUAUUUGCAAACUUCAAAUUAACCACUGGCG..AUAAC.......CGUGUAUGUUGAUUUGAUUUUAAUCGUUUUAAAAAUUACAUU.......
env-667    ACAGCCCCAUUGCAAGCUGAUAAAAUUGCUACUGCUAUUGUAUACUCAAAAAAUAAGUUUAAUUUUGCACCACAAAAGAUCAACCACUGACG..AUAAC.......CGUGUAUGUUGGCUUUUAAAACUUAAUAAACUAUC...........
env-668    ACAGCCCCAUUGCAAGCUGAUAAAAUUGCUACUGCUAUUGUAUACUCAAAAAAUAAGUUUAAUUUUGCACCACAAAAGAUCAACCACUGACG..AUAAC.......CGUGUAUGUUGGCUUUUAAAACUUAAUAAACUAUCAA.........
env-669    ACAGCCCCAUUGCAAGCUGAUAAAAUUGCUAUUGCUAUUGUAUACUCAAAAAAUAAGUUUAAUUUUGCACCACAAAAGAUCAACCACUGACG..AUAAC.......CGUGUAUGUUGGCUUUUAAAACUUAAUAAACUAUCAU.........
env-670    UAAGACCUGACUUGUUAAGUGAUUAAAUACUAAUCCAAUUUUUCUUUUAAGUAAAAAUAAAGUAUUUUUGCCGAGCAAAACAACCACUGGCGA.AAA........UCGUGAAUGUUGUCUUGCUUAA.......................
env-671    UAAGACCUGACUUGUUAAGUGAUUAAAUACUAAUCCAAUUUUUCUUUUAAGUAAAAAUAAAGUAUUUUUGGCGAGCAAAACAACCACUGGCGA.AAA........UCGUGAAUGUUGUCUUGCUUAAUUUCAAUAUAAAAAUC.........
env-672    UAAGACCUGACUUGUUAAGUGAUUAAAUACUAAUCCAAUUUUUCUUUUAAGUAAAAAUAAAGUAUUUUUGCCGGGCAAAACAACCACUGGCGA.AAA........UCGUGAAUGUUGUUUUGCUUAAUUUCAAUAUAAAAAUC.........
env-673    UAAGACCUGACUUGUUAAGUGAUUAAAUACUAAUCCAAUUUUUCUUUUAAGUAAAAAUAAAGUAUUUUUGCCGGGCAAAACAACCACUGGCGA.AAA........UCGUGAAUGUUGUUUCGCUUAAUUUCAAUAUAAAAAUC.........
env-677    ...................................................................................ACAACCACUGGCGA.AAA........UCGUGAAUGUUGUUUCGCUUAAUUUCAAUAUAAAGAUC.........
env-678    UCAGGCCGUGACUGUUAAGUGAUUAAAUACUAAUCCAAUUUUUCUUAUAAGAAAAAAUAAAGUAGUUGGGCGAGCAAAACAACCACUGGCGA.AAA........UCGUGAAUGUUGUCUUGCUUAAUUUCAAUAUAAAAAUC.........
env-679    .........................................................................UGGCGAGCAAAACAACCACUGGCGA.AAA........UCGUGAAUGUUGUCUUGCUUAAUUUCAAUAUAAAAAUC.........
env-680    GGAAAUUACGCCCUGAUAUUUUUGAUUGAUUUCUUUCUAUAAAUAAAACAAAAAAUGUAUAUUCGCAAUCGAUUUUAAUACAACCACUGACGA.UAA........UCGUGUAUGUUGUAUUAAACUAAACUAAAAUUUUACACA........
env-682    GAACUAAACAAAUUAAACCUGGUAUUAUUAAAUAGUUUUUUUGAUAAAAAAAAAUAGUAAUUUCGCAUUCCAAAUUAUUCAACCACUGACGA.UAA........UCGUGUAUGUUGUGUAACUAAAAUUAAAAUCUUUUCAAUA.......
env-683    ..............................GUGCAGGUAUUGUGUACUCAAAAAAUAAGUUUAAUUUUGCACCACUAAAGAUCAACCACUGACGA.UAA........UCGUGUAUGUUGACUUUUAAAACUAAAUAAAUUAUAAG.........
env-684    CAGCUGCACUAAGGGACUCAGAAAAUUAGUGCGGGUAUUGUAUACUCAAAAAAUAAGUUUAAUUUUGCACCACUAAAGAUCAACCACUGACGA.UAA........UCGUGUAUGUUGACUUUUAAAACUAAAUAAAUUAUCAG.........
env-685    ACAGCUGCACGAAGGACUCAGAAAAUUAGUGUGGGUGUUGUAUACUCAAAAAAUAAGUUUAAUUUUGCAGCACUAAAGAUCAACCACUGACGA.UAA........UCGUGUAUGUUGACUUUUAAAACUAAAUAAAUUAUCAG.........
env-686    GAAGACCAAAUUUACUUGAUGAUUAAAUAUCCACAGUGCUUUUAUGUUAAAUAAAAAAAAAGUACUUUUGUAAAAUAAAGUAACCACUGGCGA.AAA........UCGUGAAUGUUGCUUUUCUUAUAAUUCUAAUGCAACAACA........
Ean-11-1   UUUUUACUAUUACAAUUAUAAAUCAUUCACAAUUUAUUUUGCAGAUUUGAAAAAUUGUUCUAAAUUUGCAAACUGUUAAUCAACCUCUGACG..AUAGA.......CGUGUAUGUUGUUUGAUGUAAUUAUUUAUUUUUAAUUAACA.......
Eme-1-1    UCAAGACCGUACAAUCCAAUUGAAUUAGAAUAAUAAUUUUGCAGAUUUGAAAAAUUGUUCUAAAUUUGCAACCUGUUAAUCAACCUCUGACG..AUAGA.......CGUGUAUGUUGUUUGAUGUAAUUAUUUAUUUUUAAUUAACA.......
Ean-1-1    UCAAGACCGUACAAUCCAAUUGAAUUAGAAUAAUAAUUUUGCAGAUUUGAAAAAUUGUUCUAAAUUUGCAAACUGUUAAUCAACCUCUGACG..AUAGA.......CGUGUAUGUUGUUUGAUGUAAUUAUUUAUUUUUAAUUAACA.......
Ean-1-2    UUUUUACUAUUACAAUUAUAAAUCAUUCACAAUUUAUUUUGCAGAUUUGAAAAAUUGUUCUAAAUUUGCAACCUGUUAAUCAACCUCUGACG..AUAGA.......CGUGUAUGUUGUUUGAUGUAAUUAUUUAUUUUUAAUUAACA.......
Eme-2-1    UGUUUCAGUUUGUUAUGAAUUAUAAUUGAUCAAAUAUUUUGUCAAUAUAGAAAAAUGUUCUAAAUUUGCAUUCCGUUAAUCAACCUCUGACG..AUAGA.......CGUGUAUGUUGUUUGAUGUCAAUUUUUUUAUUAAUUAACA........
env-688    AAAAAAAUUCGUCCAGAUUAUUUAAUUAACUAAAUUUCUCUGAAAGAAAAAAAAAUAGUAAAUUCGCAAUCCAUUUGAUAUAACCGCUGACGA.UAA........UCGUGGAUGUUGUACAAUUUAAAUUAAAAAUUUACAAA.........
env-689    AAGCCUUCUUCUUGAGGAAGAAAAAAAGCAAAAACAUAACUUGUAAACAACAAAAUUAAUUGUACUUUUGCUCCCGCAUUUAACCGCUGACGA.UAA........UCGUGGAUGUUGUAUGCAAGAACCCAUUAAAAUAAAAAC.........
env-690    AAAAAAAUUCGUCCAGAUUUUUUAAUUAACUAAAUUUCUCUGAAAGAAAAAAAAAUAGUAAAUUCGCAGUCCAUUUGAUAUAACCGCUGACGA.UAA........UCGUGGAUGUUGUACAAUUUAAAUUAAAAAUUUACAAA.........
env-692    UCAUCAAAUUCCCCGAACAUUUUUUCACAAUACUAUUGCCGAUUAUUAUUUAUUGCUUACUUUUGCAUCCAAUUUCAUAUUAACCUCUGGCG..AUCUC.......CGUGAAUGUUGGUUUGGAUUAAAUUUUAAAAUUAAAAGAA........
env-693    ..............................AAAUAAUCUCAUUUAUGAAUUAAAUUUUUAUUUUUGCAAACUGAUUUGAAUAACCUCUGGCG..AAAAA.......CGUGAAUGUUGUUUAAGUAUAACCUAAUUAUUAAGUU.........
env-694    ACGCCGUGUGUUACAAAACUUUUUCUAUAAUAAAGUUUCUAGGUUUUACUUUUUACUUAUAUUUGCAGCCAAUUUCGAGUUAACCUCUGGCGA.UCA........UCGCGAAUGUUGUUUCGAAUAAAUACAUAAAAAAACCAUAUAC.......

           ...................................................................................<<<<...<.<<<<<..........>>>>>..>.>>>>.........................................
           ...................................................................................2020...0.12220..........02221..0.0202.........................................
           ●●●●●●●●●●●●●●●●●●●●●●●●●●●●●●●●●●●●●●●●●●●●●●●●●●●●●●●●●●●●●●●●●●●●●●●●YY●●●●●●●●●●Y●RRRYYAACCUCUGRCG○-RRA○-------○CGUGRAUGUUG●UY●●●●●●AA●●●●●●●●●●●●●●●●○○○---------------------------
                                                                                              5'                                   3'
```

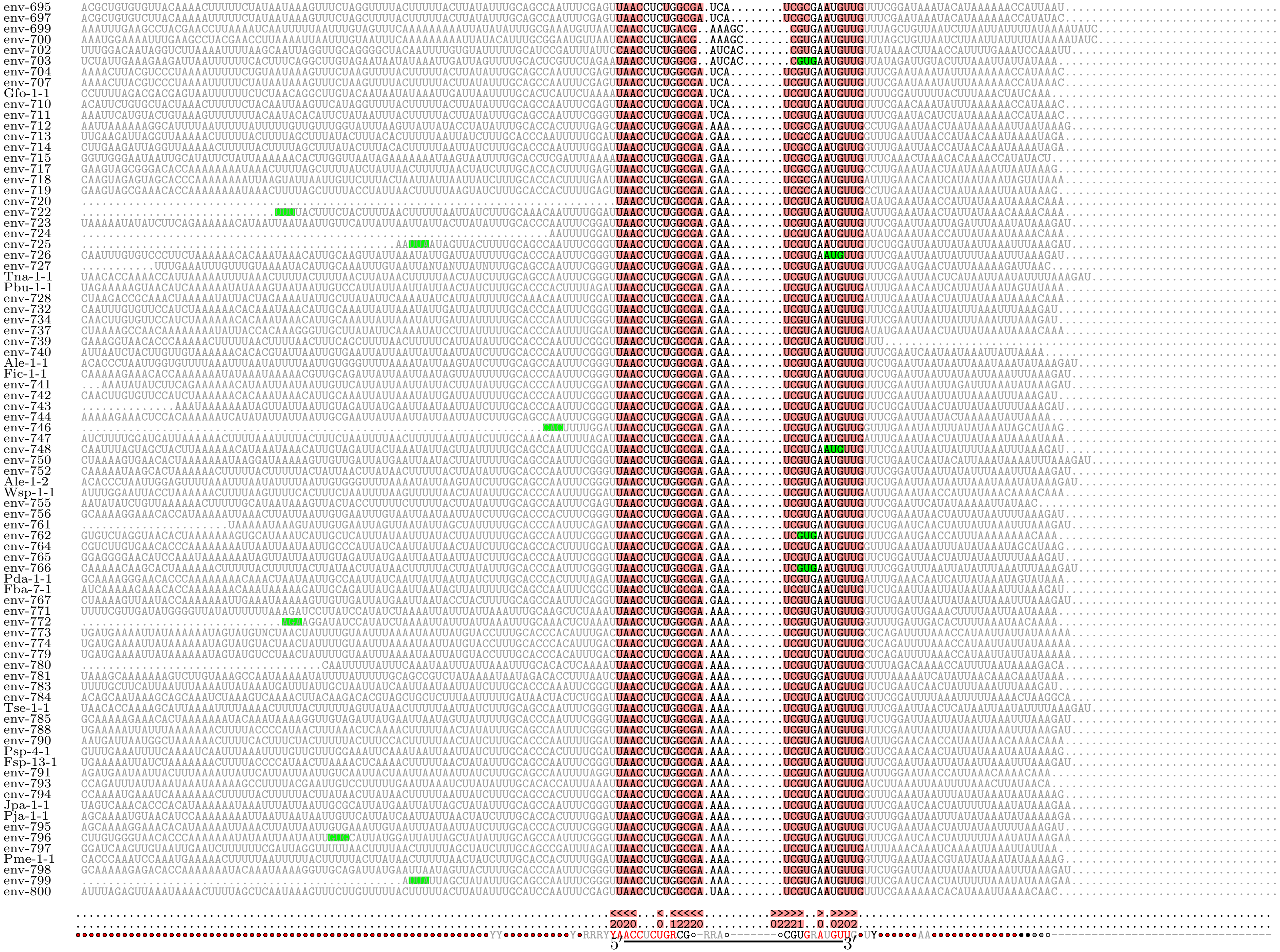

```
env-804   CUAAUUUUUCAUCGUUUAAAAAGUGAAAUUAUUAAUUGCAAGUUAUUUAUAAUUAGUUAUCUUUGCAGCCAAUUUCGAGUUAACCUCUGGCGA.UAA........UCGUGAAUGUUGUUUCGGAUUAAUUAAGAUAAAAAUAAAAGAU..........
env-805   UUGUGCUUUUGAUUUAAAAAAAAAGCUACAAUUCACUUUCUAUAAUUCACUUUUUACUUAUAUUUGCAGCCAAUUUCGAGUUAACCUCUGGCGA.UAA........UCGUGAAUGUUGUUUCGAAUACACUUAUAAAAACCAUAAAC..........
env-806   AAUAUUUUAAGCAUAAAUAAAAGUAUUAUUAAAAAUUGUUCAUUAUCAAUAAUUAAUUAUCUUUGCACCCAAUUUCGGGUUAACCUCUGGCGA.UAA........UCGUGAAUGUUGUUUCGAGUCAACUAUUAAAAAGAUUAAA..........
env-807   UCGGCCUGGGUCUCAAAACUUUUUCUAUAAUAAAGUUUCUGGGAUUUACUUUUUACUUAUAUUUGCAGCCAAUUUCGAGUUAACCUCUGGCGA.UAA........UCGUGAAUGUUGUUUCGAACACACAUAAAAAAACCAUAUAU..........
env-809   GAAAAAGUAUGGUACACAAAAAAUAUAAUUAUUAAUUUGCGCAUUGUUAAUUAUUAAUUAUCUUUGCACCCAAUUUCGGGUUAACCUCUGGCGA.UAA........UCGUGAAUGUUGUUCUGAAUUAAUUAUAAUUAAAUUUAAAGAU..........
env-810   UCGGCCUGGGUCUCAAAACUUUUUCUAUAAUAAAGUUUCUGGGAUUUACUUUUUACUUAUAUUUGCAUCCAAUUUCGAGUUAACCUCUGGCGA.UAA........UCGUGAAUGUUGUUUCGAACACACAUAAAAAAACCAUAUAU..........
env-811   AAUUUGGGCAUGGCUAAAAUAAAUCUACAAUUCACUUUCUAUAAUUCACUUUUUACCUAUAUUUGCAGCCAAUUUCGAGUUAACCUCUGGCGA.UAA........UCGUGAACGUUGUUUCGAAUUCAUAUAAAAAACCCAUAAAC..........
env-812   UAUUAAAGGAAGUCAAAAUAAACUUAUCAAACUAUUGUCUAUUUUGCAAUAAUUCUUUAUUUUUGCACCCAAUUUCGGGUUAACCUCUGGCGA.UAA........UCGUGAACGUUGUUUCGAAAUAAUUAUAAAAUUAAAAC..........
env-813   ........................CUUAUCAAACUAUUGUCUAUUUGCAAUAAUUCUUUAUUUUUGCACCCAAUUUCGGGUUAACCUCUGGCGA.UAA........UCGUGAACGUUGUUUCGAAAUAAUUAUAAAAUUAAAAC..........
env-818   ACGAACAUGGAUUUUUAGUAAAAAGAGCACAAAAUUCUUGUAUGUUUUAAUUUAAGUAUUAAUUUUGCAGCCUGUUAGAAUAACCUCUGACGA.UAA........UCGUGAAUGUUGCUCUA..........
env-819   GCUGAAAUUGAGAAAUAAAUAAAAAGCAUCUAAAUACUUGUAUGUUUUAAUUUAAAUAUUAAUUUUGCACCCUGUUAGAAUAACCUCUGACGA.UAA........UCGUGAAUGUUGCUCUAAUUUAAUUCCUAUUAACUAUACA..........
env-820   ACGAACAUGGAUUUUUAGUAAAAAGAGCACAAAAUUCUUGUAUGUUUUAAUUUAAGUAUUAAUUUUGCAGGCUGUUAGAAUAACCUCUGACGA.UAA........UCGUGAAUGUUGCUCUAAUUUAAUUCCAAUUAACUAUACA..........
env-821   UUUUCUCAUUUUUUAGCUUAAAAAGCAUUAAAAAUACUUGUAGAUUAUAAUUUAAGGAUUAUUUUUGCACCCUGUUAGAAUAACCUCUGACGA.UAA........UCGUGAAUGUUGCUCUAAUUUAAUUCCAAUUAACUAUUCC..........
env-823   .....................AAAAAGCAUAUAAAUACUUGUAUGUUUUAAUUUAAUAUUAAUUUUGCAGCCUGUUAGAAUAACCUCUGACGA.UAA........UCGUGAAUGUUGCUCUAAUUUAAUUCCUAUUUAACUAUACA..........
Msp-1-1   ACAGGAAAUAAGAAAUAAAUAAAAAGCAUAUAAAUACUUGUAUGUUUUAAUUUAAAUAUUAAUUUUGCAGCCUGUUAGAAUAACCUCUGACGA.UAA........UCGUGAAUGUUGCUCUAAUUUAAUUCCUAUUUAACUAUACA..........
env-824   CUACUCAAAUAAAUAAACAAUUGUUUAAAUUCUUUAUUGUAUACUUCAAAAUAAAGUUUAAUUUCGCAAAAAUAAAAGUAUAACCUCUGACGG.UGA........UCGUGAAUGUUGACUUUAAAAACUAUAAAUAAUCUUAUU..........
Ksp-1-1   CGCUUUCGCGAAAGCAAAACCUCUCUACUACUUUUACAUAUUAAAUUAAUACCGCAAGAAAUUCUUGCAACUGAUUAGGCUAACCUCUGACGA.UGU........UCGUGAAUGUUGCCCUAACGUAAAACCACUUUAAUAUCCAUAGAA..........
env-825   CUUCUCAAAUAAAAAAACAAUUGUGAAAUUCUUUAUUGUAUACUUCAAAAUAAAGUUUAAUUUCGCAAAAAUAAAAGUAUAACCUCUGACGA.UGA........UCGUGAAUGUUGGCUUUAAAAACUAUAAAUAAUCUUAUU..........
env-826   CUUCUCAAAUAAAUAAACAAUUGUUGAAAUUCUUUAUUGUAUACUUCAAAAUAAAGUUUAAUUUCGCAAAAAUAAAAGUAUAACCUCUGACGA.UGA........UCGUGAAUGUUGGCUUUAAAAACUAUAAAUAAUCUUAUU..........
Ddo-1-1   UUUCGCGAAAGCGUAAAAAAUAAAAUUAAGACGCACUUUGAACAAUUUAAAACCGCAAGUAAAAUUGCACCCGUUUAGGCUAACCUCUGACGA.UGA........UCGUGAAUGUUGCCCUAACGUAAAACCACUUUAAAAUCCAUAAAA..........
Dsp-1-1   UUUCGCGAAAGCGUAAAGAAUAAAAUUAAGACGCACUUUGAACAACUUAAAACCGCAAGUAAAAUUGCACCCGCUUAGGUUAACCUCUGACGA.UGA........UCGUGAAUGUUGCCCUAACGUAAAACCAAUUUAACAUCCAUAAA..........
env-833   ..AACUUAAGACCAGAUAUUUUUGACUGAUUUCUUUCUAUAAUUUAAAAAAUAAAUGUAAUUUCGCAUUCGAUUUUAUUGUAACCACUGACGA.UAA........UCGUGUACGUUGUAAUAAAUUAAUUUAAAAUAUUACAUA..........
env-839   AAAACUUAAGACCAGAUAUUUUUGACUGAUUUCUUUCUAUAAUUUAAAAAAUAAAUGUAAUUUCGCAUUCGAUUUUAUUGUAACCACUGACGA.UAA........UCGUGUACGUUGUAAUAAAUUAAUUUAAAACUAUUACAUA..........
env-840   AAAACUUAAGACCAGAUAUUUUUGACUGAUUUCUUUCUAUAAUUUAAAAAAUAAAUGUAAUUUCGCAUUCGAUUUUAUUGUAACCACUGACGA.UAA........UCGUGUACGUUGUAAUAAAUUAAUUUAAAACAAACAUA..........
env-841   ............................................................AAUUCGCAAUCGAUUUUGAUAUAACCACUGACGA.UAA........UCGUGUACGUUGUAUUAAAUAAACUUAAAAUCUUACAUA..........
env-842   AAAACUUAAGACCUGAUAUUUUUGACUGAUUUCUUUCUAUAAUUUAAAAAAUAAAUGUAAUUUCGCAUUCGAUUUUAUUGUAACCACUGACGA.UAA........UCGUGUACGUUGUAAUAAA..........
env-843   AAAACUUAAGACCAGAUAUUUUUGACUGAUUUCUUUCUAUAAUUUAAAAAAUAAAUGUAAUUUCGCAUUCGAUUUUAUUGUAACCACUGACGA.UAA........UCGUGUACGUUGUAAUAAAUUAAUUUAAAACAUUACAUA..........
env-847   AAAAAUUAAGACCAGACAUUUUUGACUGAAUUCUUUCUAUAAUUUAAAAAAUAAAUGUAAUUUCGCAGUCGAUUUUAUUGUAACCACUGACGA.UAA........UCGUGUAUGUUGUAAUAAAUUAAUUUAAAACUUUACAUA..........
env-850   AAAAAAACUAAGACCAGAUAUUUUUGACUGAAUUCUUUCUAUAAAUAAAAAAAAAUGUAAUUUCGCAAUCGAUUUUAUUAUAACCACUGACGA.UAA........UCGUGUAUGUUGUAUUAAAUAAUUUAAAAUCUUAUCAAAA..........
env-851   AAAAAAACUAAGACCAGAUAUUUUUGACUGAAUUCUUUCUAUAAAUAAAAAAAAAUGUAAUUUCGCAUUCGAUUUUAUUAUAACCACUGACGA.UAA........UCGUGUAUGUUGUAUUAAAUAAAUUUAAAAUCUUAUCAAA..........
env-860   AAAAAAUUACGACCCGAUAUUUUUGAUUGAUUUCUUUCGAGAAAUAAAAAAAAAAUGUAAUUUCGCAUUCGAUUUUGAUAUAACCACUGACGA.UAA........UCGUGUAUGUUGUAUUAAAUAAACUUAAAAUCUUACAUA..........
env-862   AAAAAAACUAAGACCAGAUAUUUUUGACUGAAUUCUUUCUAAAAAUAAAAAAAAAUGUAAUUUCGCAUUCGAUUUUAUUAUAACCACUGACGA.UAA........UCGUGUAUGUUGUAUUAAAUAAAUUUAAAAUUUUAUCAAA..........
env-864   .....................................................................GCAUUCGAUUUUAUUAUAACCACUGACGA.UAA........UCGUGUAUGUUGUAUUAAAUAAAUUUAAAAUCUUAUCAAA..........
env-865   ACAAAAAACUAAGACCAGAUAUUUUUGACUGAAUUCUUUCUAUAAAUAAAAAAAAAUGUAAUUUCGCAUUCGAUUUUAUUAUAACCACUGACGA.UAA........UCGUGUAUGUUGUAUUAAAUAAUUUAAAAUCUUAUCAAA..........
env-866   AAAAAAACUAAGACCAGAUAUUUUUGACUGAAUUCUUUCUAUAAAUAAAAAAAAAUAGUAAUUUACGCAUUCGAUUUUAUUAUAACCACUGACGA.UAA........UCGUGUAUGUUGUAUUAAAUAAAUUUAAAAUACUUAUCAAA..........
env-867   AAAAAAACUAAGACCAGAUAUUUUUGACUGAAUUCUUUCUAUAAAUAAAAAAAAAUGUAAUUUCGCAUUCGAUUUUAUUAUAACCACUGACGA.UAA........UCGUGUAUGUUGUAUUAAAUAAUUUAAAAUCUUAUCAAA..........
env-868   AAAAAAUUACGACCCGAUAUUUUUGAUUGAUUUCUUUCGAGAAAUAAAAAAAAAAUGUAAUUUCGCAUUCAAUUUUUAUAUAACCACUGACGA.UAA........UCGUGUAUGUUGUAUUAAAUAAACUUAAAAUCUUACAUA..........
env-869   AAAAAACUAAGACCAGAUAUUUUUGACUGAAUUCUUUCUAAAAAUAAAAAAAAAAUGUAAUUUCGCAUUCGAUUUUAUUAUAACCACUGACGA.UAA........UCGUGUAUGUUGUAUUAAAUAAAUUUAAAAUUUUAUCAAA..........
env-870   ..................................................................UAAUUUCGCAUUCGAUUUUAUUAUAACCACUGACGA.UAA........UCGUGGAUGUUGUAAUAAAUUAAUUUAAAAUCUUACACA..........
env-881   GAAAAUUACGCCCUGAUAUUUUUGAUUGAUUUCUUGCGAUAAAUAAAACAAAAAAUGUAUUUCGCAUUCGAUUUUGAUAUAACCACUGACGA.UAA........UCGUGGAUGUUGUAUUAAAUAAUUUAAAAUCUUACACA..........
env-882   AAAUUUAAGACCAGACAUUUUUGACUGAAUUCUUUCUAUAAAUUAAAAAAAAAUGUAAUUUCGCAUUCGAUUUUAUUAUAACCACUGACGA.UAA........UCGUGGAUGUUGUAAUAAAUUAAUUUAAAAUCUUACACA..........
env-883   GAAAAUUACGCCCUAAUAUUUUUGAUUGAUUUCUUUCGAUAAAUAAAACAAAAAAUGUAUUUCGCUUUCGAUUUUGAUAUAACCACUGACGA.UAA........UCGUGGAUGUUGUAUUAAAUAAUUUAAAAUCUUACACA..........
env-884   .....UUAAGACCAGACAUUUUUAGACUGAAAUCUUUCUAUAAAUUAAAAAAAAAUGUAAUUUCGCAUUCGAUUUUAUUAUAACCACUGACGA.UAA........UCGUGGAUGUUGUAAUAAAUUAAUUUAAAAUCUUACACA..........
env-889   .................UUUUUGAUUGAUUUCUUUCGAUAAAUAAAACAAAAAAUGUAUUUCGCAUUCGAUUUUGAUAUAACCACUGACGA.UAA........UCGUGGAUGUUGUAUUAAAUAAUUUAAAAUCUUACACA..........
env-890   AAAUUUAAGACCAGACAUUUUUGACUGAAAUCUUCCUAUAAAUUAAAAAAAAAUGUAAUUUCGCUUUCGAUUUUAUUAUAACCACUGACGA.UAA........UCGUGGAUGUUGUAAUAAUUUAAUUUAAAAUCUUACACA..........
env-891   AAAUUUAAGACCAGACAUUUUUGACUGAAAUCUUUCUAUAAAUUAAAAAAAAAUGUAAUUUCGCAUUCGAUUUUAUUAUAACCACUGACGA.UAA........UCGUGGAUGUUGUAAUAAAUUAAUUUAAAAUCUUACACA..........
env-892   AAAAUUUAAGACCAGACAUUUUUGACUGAAAUCUUUCUAUAAAUUAAAAAAAAAUGUAAUUUCGCAUUCGAUUUUAUUAUAACCACUGACGA.UAA........UCGUGGAUGUUGUAAUAAAUUAAUUUAAAAUCUUACAUA..........
env-893   .........................................CUAUAAAUUAAAAAAAAAUGUAAUUUCGCAUUCGAUUUUAUUAUAACCACUGACGA.UAA........UCGUGGAUGUUGUAAUAAUUAAUUUAAAACCUUACACA..........
env-894   GAAAAUUACGCCCUGAUAUUUUUGAUUGAUUUCUUUCGAUAAAUAAAACAAAAAAUGUAUUUCGCAUUCGAUUUUGAUAUAACCACUGACGA.UAA........UCGUGGAUGUUGUAUUAAAUAAUUUAAAAUCUUACACA..........
env-906   ..................................................................GUAAGGCGUAACGCAUAAUUUCUAUAACCACUGACGA.UAU........UCGUGAAUGUUGUAUCAAUUAAAUUUAAAAUCUUACAUA..........
env-907   AAAAUUAAGACGAUAUAUUUUAAUUAAUUUUUUUAAAAAAUAAAUAAAAAAAAUGUAAUUUCGCAUUAAAAAAUUUAUAUAACCACUGACGA.UAU........UCGUGAAUGUUGUAUCAAUUAAAUUUAAAAUCUUACAUA..........
env-910   AAAAAAACUAAGACCAGAUAUUUUUAACUGAAUUCUUUCUAAAAAUAAAAAAAAAUGUAAUUUCGCAUUCGAUUUUAUUAUAACCACUGACGA.UAA........UCGUGAAUGUUGUACUAAAUAAAUUUAAAAUCUUAUCAAA..........
env-913   ...............................................................................AUUUAUUAUAACCACUGACGA.UAA........UCGUGAAUGUUGUACUAAAUAAAUUUAAAAUCUUAUCAAA..........
env-914   AAAAAAACUAAGACCAGAUAUUUUUGACUGAAUUCUUUCUAUAAAUAAAAAAAANAUGUAAUUUCGCAUUCGAUUUUAUUAUAACCACUGACGA.UAA........UCGUGAAUGUUGUACUAAAUAAAUUUAAAAUCUUAUCAAA..........
env-915   AAAAAACUAAGACCAGAUAUUUUUAACUGAAUUCUUUACUAAAAAAUAAAAAAAAAUGUAAUUUCGCAUUCGAUUUUAUUAUAACCACUGACGA.UAA........UCGUGAAUGUUGUACUAAAUAAAUUUAAAAUCUUAUCAAA..........
env-916   AUUGCAAGAAUUACAAAGAAAAAUAAUUUCAAUUCUAUUUUUAAAGAAAAAAUUAUUUAAAUGUAAAUUUGCAUUGCAACCGCUAAAAUUAACCUCUGACG..AAAGC.......CGUGAAUGUUGUUUAGAUAAAUCCAAAUUUAUUAAAAA..........
env-917   AAUUUCUAUUUCCUAAAAAAAAUAUUUGUAAAAGUUUGGUAGAUAGAAAUAUUCGUCGUACUUUUGCCGCCGAUUUGGUGUAACCUCUGACG..AAAGC.......CGUGAAUGUUGCCCAAUGAUAACAUAAUUUAUUUUUAUUUUUUU..........
env-918   AAUUUCUAUUUCCUAAAAAAAAUAUUUGUAAAAGUUUGGUAGAUAGAAAUAUUCGCCGUACUUUUGCCGCCGAUUUGGUGUAACCUCUGACG..AAAGC.......CGUGAAUGUUGCCCAAUGAUAACAU..........
Fba-4-1   GGAUGAUUAAGUCUUUUAAAAUUAUUAAUUAAUAAUUGCCAAUUACUAAUUAAUAACUAUCUUUGCAGCCAAUUUCGGGUUAACCUCUGGCGA.AAA........UCGUGAACGUUGAUUUGAGAUAAUCAUUAUAAAUAAAACAAA..........
env-919   UAGCAAAAUACAGAGGGAAUAAAAAUUUUUGUUUGGUAUCUUGUUAAAAUAAAAUUAAUUGUACUUUUGCUCCCGCAAUUAACCGCUGACGA.AAA........UCGUGUAUGUUGACUGUAAUGAAUCCAAAAAUUAAAAU..........
env-920   AAAGUAAUGUUGUAGUUAUUUAUUAAGACUAAGAAGUUGCUUACUUCAAAAUAAAGUCUAAUUUUGCAAAAUGAAAUGAAUAACCGCUGACGA.AAA........UCGUGUAUGUUGUCACUUUUCAAACAAUAAAAAAAUAAU..........
env-923   ..................................................UAAAAAAGUCGUAAGUUUGCAACCUCAAAAUUAACCGCUGACGA.AAA........UCGUGUAUGUUGGUUUGUCGAAACCCAUUAAAAUAAUAAGAA..........
env-924   .....................................................................................CAACCUCAAAAUUAACCGCUGACGA.AAA........UCGUGUAUGUUGGUUUGUCGAAACCCAUUAAAAUAAUAAGAA..........
env-925   CCUCGUUUUACUCACCUGUAUUUUUUUUAAUUUACGGCUUGCGCAUUUAAAAAAGUGUUUACAUUUGCAACCUAAAAUUAACCUCUGACGA.AAA........UCGUGUAUGUUGAUUUUUAGCAAUGUUUAAAUCAAUUAUC..........
env-926   AGAAAUUUAUAUGAUCUGAAAAUUAUUUGCUUGUCUUAUUUUUUAUUAAAAAGUUUUUAUUAAAUUUGCAAACUCAAAAUUAACCUCUGACGA.AAA........UCGUGUAUGUUGAUUAUUGAUCAACCAUUUAAAUAAAAUA..........
env-927   AGUAAUUAAAAUGAUCUGAAAAUUAUUUGCUUGUCUUAUUUUUUAUUAAAAACUUUUUAUUAAAUUUGCAAACUCAAAAUUAACCUCUGACGA.AAA........UCGUGUAUGUUGAUUAUUGAUCAACCAUUUAAAUAUAAAUA..........
env-928   ..................UAUUUUUUUUUCAAUCCUGACUUUAAUAAUCAAAACUUUUUUAUAAAUUUGCAAACUCAAAAUCAACCUCUGACGA.AAA........UCGUGUAUGUUGACCAUUGAUCAACCAUUUAAAAUAUCAUA..........
Ffi-1-1   UGAAUAUAAAAAAUCACAAACUUUUUGUAUAAAUUGAUGUAAUUUAAAAAUAAAUUGUAAAUUUGCACCCACAUUUGGACCAACCUCUGACGA.AAA........UCGUGUAUGUUGCUCUGAUAAUCACUAUUUUAACUUAAAUGUU..........
env-929   CCAAUUAUUAAAAGAAUAAUAUUUUUUUAAAUCAUGACUUUGUUAAUCAAAAGUUUUUUAUAAAUUUGCACACUCAAAAUCAACCUCUGACGA.AAA........UCGUGUAUGUUGACGAUUGAUCAACCAUUUUAAAAUAUCAUA..........
Fba-8-1   AGCUUUUAUAGUUUCUAAAAAAAAAUCAAUUGCAAAUACCUAAAAACUAACUACUAAUUAUAUUAUUUGCAGCGCUAAAAUCAACCUCUGACGA.AAA........UCGUGUAUGUUGUUUAGUAGCAAACCAUUCAAAGAACAAA..........
env-932   CCAANUAUUAAAAGAAUAAUAUUUUUUUAAAUCAUGACUUUAUUAAUCAAAAGUUUUUUAUAAAUUUGCACACUCAAAAUCAACCUCUGACGA.AAA........UCGUGUAUGUUGACGAUUGAUCAACCAUUUUAAAAUAUCAUA..........
env-933   CCAAUUAUUAAAAGAAUAAUAUUUUUUUAAAUCAUGACUUUAUUAAUCAAAAGUUUUUUAUAAAUUUGCACACUCAAAAUCAACCUCUGACGA.AAA........UCGUGUAUGUUGACGAUUGAUCAACCAUUUUAAAAUAUCAUA..........
env-938   UUUGCUAAAAUAAAAUAUUUUUAGACGAUAAAAUGUUUUAUUUUCUUUAAAAAAAAGUAUUUUUGCACCCAUUUUGGUCAACCGCUGACG..AAAGA.......CGUGAAUGUUGUACUGGAUA..........
Cca-1-1   AAUACGAAAAUAUGGAAGUAAAAGAAGUAAAAUAAACUUGUAUUUUCAAAAAUAAUAUCUAAAUUUGCACCCGAUUUGGUCAACCUCUGGCG..AAAGA.......CGUGAAUGUUGCCCAAUGAUAACUAAAUUAUAUUUUAAAUUUUU..........
env-939   AAAUUAACGCUUUAUUAUAAAAUUAUUGAAACAAUUAAUUUUGCAUAAUUGAAAAAUAGUAUUACUUUUGCAUCCACUUUAAUCAACCUCUGGCG..AAAGA.......CGUGAAUGUUGCUUGGAUUUAAUGAUUUAUAAUUAACUAUU..........
env-940   UACCACUUAACCAACUAUAAAAAUAAGCCAAAGUAUUAGUUUUAAUUAAAAAAUAUUUUCUAAAUUUGCGCACUCAAAAUCAACCUCUGACG..AAAGA.......CGUGAAUGUUGUUUUAUGAUUAAU..........
env-941   UACCACUUAACCAACUAUAAAAAUAAGCCAAAGUAUUAGUUUUAAUUAAAAAAUAUUUUCUAAAUUUGCGCACUCAAAAUCAACCUCUGACG..AAAGA.......CGUGAAUGUUGUUUUAUGAUUAAUAUUAAAAACCAUUUAAAA..........
env-942   CCUUGCCGAAUUAGAUCAAAAAAAGGCCUCAAAAGGUUUGCGACUAUUGUAAAAAGCACUAUUUUUGCAGUCGAUUUACCCCAACCUCUGGCG..AUAGA.......CGUGAAUGUUGUUGUAAACGUAACAUUAUCAAAUCAAUAGAA..........
Emi-1-1   UUUUUUACUAUACAAUUAAUAAAUCAUUCACAAUUUAUUUUGCAGAUUUGAAAAAUUAUUCUAAAUUUGCAACCUGUUAAUCAACCUCUGACG..AUAGA.......CGUGAAUGUUGUUUGAUGUAAUUAUUUAUUUUUAAUUAACA..........
Wma-1-1   AAUAUUGGAAAGUAAUGAAGAAAACAGCGCAGAAAAUUUGUAAAUAAGAGAAAUAUAAUUAAUUUGCACUCGCAUUGAUUAACCUCUGACG..AAAGA.......CGUGAAAGUUGGUCAAUCUAAAAAAUUGUUUAUUAUUACCAAA..........
env-943   .......................................AUAAUUUUUAUAGAUUAGAAUAAAUUACUAUUUUUGCAAACUAUUAUUAACCUCUGACG..AAGAC.......CGUGAAUGUUAGUAAUAAUUAAAAAAUAAUAAUUAUAAG..........
env-944   GAACAAGAAAUUUAAGACCAGAUUUGUUGGACGAAUAAUUUUAUAGAUUAGAAUAAAUUACUACUUUUGCAAACUAUUAUUAACCUCUGACG..AAGAC.......CGUGAAUGUUAGUAAUAAUUAAAAAAUAAUAAUUAUAAG..........
env-945   CUAUUGAAAUAAGAAAAUAUUUGUUAAUAUUCUUUAUUGUAUACUUAAAAAAUAAAGUUUAAUUUUGCAAAACAAAAAGUUUAACCUCUGACG..AUAAA.......CGUGCAUGUUGACUUUCAAAAUGUUAAAAUAACCCUAUU..........

          ..........................................................................................<<<<...<.<<<<<..........>>>>>..>.>>>>..........................................
          ..........................................................................................2020...0.12220..........02221..0.0202..........................................
          •••••••••••••••••••••••••••••••••••••••••••••••••••••••••••••••YY••••••••••Y•RRRRYYAACCUCUGRCG○-RRA○-------○CGUGRAUGUUG•UY••••••AA•••••••••••••••••••○○○----------------------------
                                                                                      5′                                          3′
```

```
env-947   CUUUUGAAAUAAGAAAAUAUUUGUUAAUAUUCUUUAUUGUAUACUUAAAAAUAAAGUUUAAUUUUGCAAAACAAAAAGUUUAACCUCUGACG..AUAAA.......CGUGCAUGUUGACUUUCAAAAUGUUAAAAUAACCCUAUU...................................
env-948   CUUCUGAAAUAAGAAAAUAUUUGUUAAUAUUCUUUAUUGUAUACUUAAAAAUAAAGUUUAAUUUUGCAAAACAAAAAGUUUAACCUCUGACG..AUAAA.......CGUGCAUGUUGACUUUCAAAAUGUUAAAAUAACCCUAUU...................................
env-949   CUUUUGAAAUAAGAAAAUAUUUGUUAAUAUUCUUUAUUGUAUACUUAAAAAUAAAGUUUAAUUUUGCAAAACAAAAAGUUUAACCUCUGACG..AUAAA.......CGUGCAUGUUGACUUUCAAAAUGUUAAAACAACCCUAUU...................................
env-950   CUUCUGAAAUAAGAAAAUAUUUGUUUAUAUUCUUUAUUGUAUACUUAAAAAUAAAGUUUAAUUUUGCAAAAUAAAAAGUUUAACCUCUGACG..AUAAA.......CGUGUAUGUUGACUUUCAAAAUGUUAAAAUAACCCUAUU...................................
env-951   GCAAAAACUAUCUACCAAAAAAAUAACUUGAAUAGAUUUUAUAUCCUAAAAUAAAUCAUUAGAUUUGCACUCGCUAAAAAUAACCUCUGACGA.AAA........UCGUGAACGUUAGUUUAGAAAAAUUUUAAAAUAAUAAA....................................
env-954   GCAGAAACUAUCUACCAAAAAAAUAACUUGAAUAGAUUUUAUAUCCUAAAAUAAAUCAUUAGAUUUGCAAUCGCUAAAAAUAACCUCUGACGA.AAA........UCGUGAACGUUAGUUUAGAAAAAUUUUAAAACAAUAAA....................................
env-955   CAGAAACUAUCUACCAAAAAAAUAACUUGAAUAGAUUUUAUAUCCUAAAAUAAAUCAUUAGAUUUGCACUCGCUAAAAAUAACCUCUGACGA.AAA........UCGUGAACGUUAGUUUAGAAAAAUUUUAAAACAAUAAA....................................
env-956   GCAAAAACUAUCUACCAAAAAAAUAACUUGAAUAGAUUUUAUAUCCUAAAAUAAAUCAUUAGAUUUGCAUUCGCUAAAAAUAACCUCUGACGA.AAA........UCGUGAACGUUAGUUUAGAAAAAUUUUAAAAUAAUAAA....................................
env-957   GCAGAAACUAUCUACCAAAAAAAUAACUUGAAUAGAUUUUAUAUCCUAAAAUAAAUCAUUAGAUUUGCACUCGCUAAAAAUAACCUCUGACGA.AAA........UCGUGAACGUUAGUUUAGAAAAAUUUUAAAACAAUAAA....................................
env-959   GCAAAACCUAUCUACCAAAAAAAUAACUUGAAUAGAUUUUAUAUCCUAAAAUAAAUCAUUAGAUUUGCACUCGCUAAAAAUAACCUCUGACGA.AAA........UCGUGAACGUUAGUUUAGAAAAAUUUUAAAAUAAUAAA....................................
env-968   ......................................................................CUCGCUAAAAUAACCUCUGACGA.AAA........UCGUGAACGUUAGUUUAGAAAAUUUAAACAAUAAA.......................................
env-969   ...............CCAAAAAAUAACUUGAAUAGAUUUUAUAUCCUAAAAUAAAUCAUUAGAUUUGCACUCGCUAAAAAUAACCUCUGACGA.AAA........UCGUGAACGUUAGUUUAGAAAAUUUUAAAACAAUAAA.....................................
env-970   GCAGAAACUAUCUACCAAAAAAAUAACUUGAAUAGAUUUUAUAUCCUAAAAUAAAUCAUUAGAUUUGCACUCGCUAAAAAUAACCUCUGACGA.AAA........UCGUGAACGUUAGUUUAGAAAAAUUUUAAAACAAUAAA....................................
env-974   GCAGAAACUAUCUACCAAAAAAAUAACUUGAAUAGAUUUUAUAUCCUAAAAUAAAUCAUUAAAUUUGCACUUGCUAAAAAUAACCUCUGACGA.AAA........UCGUGAACGUUAGUUUAGAAAAAUUUUAAAACAAUAAA....................................
env-975   UACUGCAGAAACUAUCUACCAAAAUAACUUGAAUAGAUUUUAUAUCCUAAAAUAAAUCAUUAGAUUUGCACUCGCUAAAAUAACCUCUGACGA.AAA........UCGUGAACGUUAGUUUAGAAAAUUUAAAACAAUAAAAUAGGAAUCUUUAGUAAAAUUUGUACAAGACGAAUUUGU
env-976   ..................................................................UACGCGAGCUAUUACCUCUGACGA.AUA........UCGUGAACGUUGUUUUACAAAAUCAUUAAAUUAAAAAAU..................................
env-977   GCACUGACUAUCAACCCAAAAAAUAAUUAAAAUAAUUGUAUCAACCAAAAAAUAAUUAUACUUUUGCAACCGCUAAAAUUAACCUCUGACGA.AGAAA......UCGUGAAUGUUGCUUUACAAAAUCAAUAAAAUUAAAAGAU................................
env-980   UGCACUGAUUAUCAACCAAAAAAAUAAUCUCAUUUAGUUUUAGUAAUAAAAAAAAGUCCUAUUUUUGCACUCGCUAAAUUAACCUCUGACGA.AGAAA......UCGUGAAUGUUGUUUUACAAAAACAUUAAAUUAAAAU..................................
env-983   UGUACAGAUUAUCAACCAAAAAAAUAAUCUUUAUUAUUUUAGUAAUCAAAAAAAAUCCUAUUUUUGCACUCGCUAAAUUAACCUCUGACGA.AGAAA......UCGUGAAUGUUGUUUUACAAAACUAUUAAAUUAAAAU..................................
env-988   GUACUGACUAUCAACCUAAAAAAUAAUUAAAAUAAUUGUAUCAACAAAAAAAUAAUUAUACUUUUGCAACCGCUAAAUUAACCUCUGACGA.AGAAA......UCGUGAAUGUUGCUUUACAAAAUCAAUUAAAUUAAAAGAU................................
Psp-2-1   GUACUGAUUAUCAACCUAAAAAAUAAAUGCUUUUUAGUUUUAUCAACUGAAAAAAAAUACUAUUUUUGCAACCGCUAAAUUAACCUCUGACGA.AGAAA......UCGUGAAUGUUGCUUUACAACAAAACAUAAACAUUACAU................................
env-989   UAAACUGAUUAUCAAAGAAAAAAUAUUUAUAUAGUUUUAUCAACUAAAAAAAAGUUCUAUUUUUGCACCGCUAAAUUAACCUCUGACGA.AAAAA......UCGUGUAUGUUGUUUUACAAAACAUUAAAAU.....................................
env-991   GCACUAAUUAUCAGUAUAAAAAAUAAUUACCUGAAUCUUUUUUUUUAGUAAAAUAUUUCAUAAAUUUGCAACCUCAAAAUUAACCUCUGACGA.AAAAA......UCGUGAAAGUUGGUUCUUGAAAAACCAUAAAUAAAAUAAAA...............................
env-992   GCACAAAGUAUCACUAUAAAAAAUAAUUACUCGAAUCUUUUUUUAUACUAAAAUAUUUCAUAAAUUUGCAACCUCAAAAUUAACCUCUGACGA.AAAAA......UCGUGAAAGUUGAUCUUGAAAAACCAUAAAUAAAAUAAAA...............................
env-994   ...............UAAAAAAUAAUUACCUGAAUCUUUUUUUUUAGUAAAAUAUUUCAUAAAUUUGCAACCUCAAAAUUAACCUCUGACGA.AAAAA......UCGUGAAAGUUGGUUCUUGAAAAACCAUAAAUAAAAUAAAA...............................
env-997   CUAUUGUACUGACUACAAACUAAAAAAUAGUUGAAUCUUAUUUUUAUCAACUGAAAAAAAGUACUAUUUUUGCAACCGCUAAAUUACCUCUGACGA.AAAUA......UCGUGAACGUUGUUUUACAAAAUCAUUAAAUUAAAAAAU..................................
env-998   GUACUGACUAUCAAACUAAAAAAUAGUUGAAUUUUAUUUUUAUCAACUGAAAAAAAGUACUAUUUUUGCAACCGCUAAAUUAACCUCUGACGA.AAAUA......UCGUGGACGUUGUUUAGGGGGGGGGUUUUUAAAAAAU..................................
env-999   GUACUGACUAUCAAACUAAAAAAUAGUUGAAUUUUAUUUUUAUCAACUGAAAAAAAGUACUAUUUUUGCAAACGCUAAAUUAACCUCUGACGA.AAAUA......UCGUGAACGUUGUUUUACAAAAUCAUUAAAUUAAAAAAU..................................
Pir-1-1   GCACUGACUAUCAAACUAAAAAAUAGUUCAAUUUUAUUUUUAUCAACUAAAAAAAAAAAUACUAUUUUUGCAACCGCUAAAUUAACCUCUGACGA.AAAUA......UCGUGAACGUUGUUUUACACAAUCAAUUAAAUUAAAAAAU...............................
env-1003  AAGCUAUAGAAAGAACUAAAAAAUAAGACCUGAUUUGUUGGAAGACUAAUUUUUUUUAUUAAAUUUGCACUCUUAAAAUUAACCUCUGACGA.AAA........UCGUGAACGUUGUUUUGAAAAACCCAUUAAAAAUUUAACA.................................
env-1007  GCACUGACUAUCAACCUAAAAAAUAGUUGCUUUUAUAUUUUAUCAACUAAAAAAAAGUGCUAUUUUUGCAACCGCUAAAUUAACCUCUGACGA.AGAUA......UCGUGAACGUUGUUUUACAAAAUCAAUUAAAUUAAAAAAU...............................
env-1009  GUACUGACUAUCAACCUAAAAAAUAGUUGCUUUUUAGUUUUAUCAACUAAAAAAAAGUGCUAUUUUUGCAACCGCUAAAUUAACCUCUGACGA.AGAUA......UCGUGAACGUUGUUUUACAAAAUCAAUUAAAUUAGAAAAU...............................
env-1010  GUACUGACUAUCAACCUAAAAAAUAGUUGCUUUUUAGUUUUAUCAACUAAAAAAAAGUGCUAUUUUUGCAACCGCUAAAUUAACCUCUGACGA.AGAUA......UCGUGAACGUUGUUUUACAAAAUCAAUUAAAUUAAAACAU...............................
env-1011  .....................................GAAGGCUGGAGUCACUAAAAAAGUGCUAUUUUUGCAACCGCUAAAUUAACCUCUGACGA.AGAUA......UCGUGAACGUUGUUUUACAAAAUCAUUUUAAAUUAAAAAAU...............................
env-1012  GUACUGACUAUCAACCUAAAAAAUAGUUGCUUUUUAAUUUUAUCAACUAAAAAAAAGUGCUAUUUUUGCAACCGCUAAAUUAACCUCUGACGA.AGAUA......UCGUGAACGUUGUUUUACAAAAUCAUUUUAAAUUAAAAAAU...............................
env-1015  AGUUAGUGAGAUUAAUUUCUAAAAUCAAAAUAGAAGUUGUUUAAUUUAAAAUAAAGUUUAAUUUUGCAGAACUAAAAGAAUAACCUCUGACG..AAAAA.......CGUGAAUGUUGUCUUUCUAAAUUUAUUAUUAUCCAAUU................................
env-1016  AUUGACUCCAAUUAUGAAACAAAAAAUAAUUAAUUUUAGUUUUAUAAUAGAAAAUAAUGACACUUUUGCAACCGCUAAAUUAACCUCUGACGA.AAAAA......UCGUGAAUGUUGUUUUACAAAAUCAAUUAAAUCAAAAGAU...............................
env-1017  UUGCUCCAAUUAUGAAACAAAAAAAUAAUUAAUUUUAGUUUUAUAAUAGAAAAUAAUUACUACUUUUGCAACCGCUAAAUUAACCUCUGACGA.AAAAA......UCGUGAAUGUUGUUUUACAAAAUCAAUUAAAUCAAAAGAU...............................
env-1019  UUGCUCCAAUUAUGAAACAAAAAAAUAAUUAAUUUUAGUUUUAAAAUAGAAAAUAAUUACUACUUUUGCAACCGCUAAAUUAACCUCUGACGA.AAAAA......UCGUGAAUGUUGUUUUACAAAAUCAAUUAAAUCAAAAGAU...............................
env-1021  UUGCUCCAAUUAUGAAACAAAAAAAUAAUUAAUUUUAGUUUUAUAAUAGAAAAUAAUGACUACUUUUGCAACCGCUAAAUUAACCUCUGACGA.AAAAA......UCGUGAAUGUUGUUUUACAAAAUCAAUUAAAUCAAAAGAU...............................
env-1023  UGUACGAACUAUCAAGCAAAAAAAUAAUUAAAUAUAGUUUUAUCAACAAGAAAAUAAUUGUUACUUUUGCAAGCGUUAAAUAACCUCUGACGA.AAAAA......UCGUGAAUGUUGCUUUACAAAAAUCAAUUAAAAUUAAAAGAU.............................
env-1025  CUGCUCCAAUUAUGAAACAAAAAAAUAAUUAAUUUUAGUUUUAUAAUAGAAAAUAAUUACUACUUUUGCAACCGCUAAAUUAACCUCUGACGA.AAAAA......UCGUGAAUGUUGUUUUACAAAAUCAAUUAAAUCAAAAGAU...............................
env-1026  UUGCACCAAUUAUGAAACAAAAAAAUAAUUAAUUUUAGUUUUAUAAUAGAAAAUAAUUACUACUUUUGCAACCGCUAAAUUAACCUCUGACGA.AAAAA......UCGUGAAUGUUGUUUUACAAAAUCAAUUAAAUCAAAAGAU...............................
env-1027  CUUAAUAUAUUAAUAAUUAUCUGCUAUUUCGAUUUUUGCGUAGUUUAAAAUUAAUCUUAUUUUUGCAACCUAAAUAGUAUUAACCUCUGACG..AAAAA.......CGUGAAUGUUACUCUAUAUAAAUUAAAUUUCUCAAAAAA.................................
env-1029  AUUAAAUAAAAAUAAUUACGUAAAAAUUUUGAUUUUUGCGUAGUUUAAAAUUAAUCUUAUUUUUGCAGCCUAAAUAGGUAUUAACCUCUGACG..AAAAA.......CGUGAAUGUUAUUCUAUAUAAAAUAAAUUUGUAAAAAA.................................
env-1034  UGUACUAAUUAUCAAUAUAAAAAAUAAUUAAAUUAAUUAUAUUUUAUCAAUAAAAAAUAAUUAGUACUUUUGCAACCGCUAAAUUAACCUCUGACGA.AAAAA......UCGUGAAUGUUACUUUACAAAAUCAAUAAAUAUUAAAAC..............................
env-1038  CUUAGACCAGAUCUUUUCAAAGAUUAAAUUUAAUUGAAUUUUUCUUUUAAAAAAAAAUAAAGUAUUUUUGCAUUGAAAUUUAACCUCUGACG..AAAAA.......CGUGAAUGUUACUUUUUUAAAAUUUGAAAUAAUUUAUA................................
env-1039  UUAAAAAAUGCUUUUUUUUAUUUUUUUUAUAAAAAAGGAAAGCAAAAAAUAAAAUAAUGGUACUUUUGCACUCGCAAUGUAACCUCUGACGA.AAAA.......UCGUGAAUGUUACUAGCUACAA..............................................
env-1040  CUUAAUAUAUUAAUAAUUAUCAGCUAUUUCGAUUUUUGCAUAGUUUAAAAUUAAUCUUAUUUUUGCAACCUAAAUAUUAUUAACCUCUGACG..AAAAA.......CGUGAAUGUUAUUCUAUAUAAAUUAAAUUUCUCAAAAAA.................................
env-1042  GUACUAAUUAUCAGUAUAAAAAAUAAUUGUUUUUAGUUUUAUCAACAAAAAAUAAUUACUACUUUUGCAAACGCUAAAUUAACCUCUGACGA.AGAAA......UCGUGAACGUUUGUUUUUUUAAAAUAUAAAUAUUAGAU..................................
env-1047  UGUUCUAACUAUCAGGCUAAAAAAUAAUUAGUUUUAGUUUUAUCAAUAAAAAAUAAUUAUACUUUUGCAACCGCUAAAUUAACCUCUGACGA.AGAAA......UCGUGAACGUUACUUUACAAAAAUUAUAAAUAUUAAGAU................................
env-1048  UGUACUAAUUAUCAAGUAAAAAAUAAUUAGUUUUAGUUUUUUCAACAAAAAAUAAUGAUUACUUUUGCAACCGCUAAAUUAACCUCUGACGA.AGAAA......UCGUGAAUGUUACUUUACAAAAAUUAUAAAUAUUACAAU................................
env-1049  GUACUAACUAUCAGUACAAAAAAUAAUUUAAUUUAGUUUUAUCAAUAAAAAAAAGCAUUACUUUUGCAACCGCUAAAUUAACCUCUGACGA.AGAAA......UCGUGAACGUUGUUUUAGAAAAAACUAUUAAAUUAAAAGAU..............................
env-1052  GUACUGACUAUCAACCUAAAAAAUAGUUAGAUUUUAGUUUUAUCAACUAAAAAAAAGUGCUAUUUUUGCAAACGCUAAAUUAACCUCUGACGA.AUAAA......UCGUGAACGUUGUUUUACAAAAUCAAUUAAAAUAAAAAAU..............................
env-1053  UGCUCUAACUAUCAGACUAAAAAAUAAUUAGUUUUGUUUUAGUAACAAAAAAAAAAAUCCUAUUUUUGCACUCGCUAAAUUAACCUCUGACGA.AUAAA......UCGUGAACGUUGUUUUACAAAACCUUUUUAAAUUAAAAGAU.............................
env-1057  UGCUCUAACUAUCAGACUAAAAAAUAAUUAGUUUUGUUUUAGUAACAAAAAAAAAAAUCCUAUUUUUGCACUCGCUAAAUUAAC.UCUGACGA.AUAAA......UCGUGAACGUUGUUUUACAAAACUCAUUAAAUUAAAAAUUUU...........................
env-1058  UGCUCUAACUAUCAGACUAAAAAAUAAUUAGUUUUGUUUUAGUAACAAAAAAAAAAAUCCUAUUUUUGCACUCGCUAAAUUAACCUCUGACGA.AUAAA......UCGUGAACGUUGUUUUUUUUAAACCUAUUAAAUUAAAAGAU.............................
Wvi-1-1   UAUUUAAAAUAAUGUAUGUAUUUUAACUUGUAAACAUUUGUAAAUAACAGAAUUAUAAUUAAUUUUGCACUCGCAUUGAUUAACCUCUGACG..AAAGA.......CGUGAAUGUUGGUCAAUUUAAAUAACUGUUUAUUAUUACUAAA............................
env-1059  .....................................UUUUUGUAAAAUUAAAUAAGAAUGAUUAAUUUUGCAGCCGAUUGAAUAACCUCUGACG..AAAGA.......CGUGAAUGUUAUCAAUUAUAAAUGAUUAUAUUUUUUUAA...............................
env-1060  UUGAGGUUGAUCAUUUCUUAAUUUUUCGUUUGUAUUUUUGUAAAAUUAAAUAAGAAUGAUUAAUUUUGCAGCCGAUUGAAUAACCUCUGACG..AAAGA.......CGUGAAUGUUAUUCAAUUAUAAAUGAUUAUAUUUUUUUAA...............................
Fbr-1-1   AGUAACCUUUAAAUAAUUUUGACACUUAUUUAUUUGUAUAAGUCUGGAUUAAACAGUACUUUUGCACCCAAAUAUUGGACCAACCUCUGGCG..AAACC.......CGUGAAUGUUGAGCUGAUAAAACAUUAAAAUACUAAAAA..............................
env-1062  GCUCAAACAACUUUAAACAAUUUUAAACUUUUUUUAAACAUUUUAAAAAAUUUGUUUUACUUUUGCACCCUCAUUAGACCAACCUCUGGCG..AAACC.......CGUGAAUGUUGCUCUUUUUAAACCAUAAAUAAAAAAUAAA............................
env-1063  ............................................GUAAUUGUAAUUUUUAUGUAAUUUUUGCAACUUCUUAAUCAACCUCUGACG..AUAUA.......CGUGAACGUUGGUUAAGGUAACAGAAAACUAUUUAUA.................................
env-1064  CGAAAAUCCAUUUUCAAAUAUUUUCGAAUUUGAAAUUUGUUUAUUUGAUUUUUUUUAAUACUUUUGCACCCACUUGAAUCAACCUCUGGCG..AUAGA.......CGUGAAUGUUGUUUGAAUCGAAUUUUUUAUUUUUAAAGAAU............................
env-1066  .....................................................................GCUUUUCACUUUGAAUCAACCUCUGGCG..AUAGA.......CGUGAAUGUUGUUUGAAUCGAAUUUAUUAUUUUUAAAACU............................
env-1067  GAAAAUCCUUUCCAAAUAUUUUCUAAUUAGAAAUUUGUUUAUUCGAUUUUUUUAAAAUACUUUUGCACCCACUUUGAAUCAACCUCUGGCG..AUAGA.......CGUGAAUGUUGCUUUGAAUCGAAUUUAUUA.....................................
env-1068  GGAUUUGUUAGAUGAUUGAAAUUUAGAAUUUGUUUUUAUUAAUAAAUAAAAAACACUAGCUUUGCAGCCAAUUUCAGAUUAACCUCUGGCG..AAGUA.......CGUGAAUGUUGAUUUGAAUUAAACACAAUAAACUUAAUAAA...............................
env-1069  GGAUUUGUUAGAUGAUUGAAAUUUAGAAUUUGUUUUUAUUAAUAAAUAAAAAACACUAGCUUUGCAGCCAAUUUCAGAUUAACCUCUGGCG..AAGUA.......CGUGAAUGUUGGUUUGAAUUAAACACAAUAAACUUAAUAAA...............................
Iha-1-1   GAUCUUUUAAAGGAGUAAUCUUUUUAAAUUUUAGGCCAUAUAUAUCUUAAUAUUUAUUAUUUUUGCAUCCGAUUUGGAGUAACCUCUGGCG..AAGUA.......CGUGAAUGUUGCUUUGAAAAAACAAUUAAUAUAUAGCAAA.............................
env-1070  UAGACAAAUAACUACGGAAAAAUAUUGCUUAUAAAGUUGCGGGUAGUGCAAUUUUGUGUAUUUUUGCAACCGAUUUGUUCCAACCUCUGGCG..AAGUA.......CGUGAAUGUUGUUACAAGCUUAACCAUUUUGAAACUGAAACC............................
env-1072  ...........................................GCGUAGUGCAAUUUUGUGUAUUUUUGCAACCGAUUUGUUCCAACCUCUGGCG..AAGUA.......CGUGAAUGUUGUUACAAGCUUAACCAUUUUGAAACUGAAACC............................
env-1073  ACCUGAUUUAUUGGAGUAAAAAAGAUUCGUAAAACAUUGUAGCAAAGCAUAAAUUGUUUAUUUUUGCAACCGAUUUAAUCCAACCUCUGGCG..AAGUA.......CGUGAAUGUUGCAUUGAACUAAUCUUUAGUAAAGAAAUAAC.............................
env-1074  ....GAUUUAUUGAGUAAAAAAGAUUGAUAAAAUAUUGUGACAACAUAUAAAUUGUUUAUUUUUGCACCCGAUUUGAUUCAACCUCUGGCG..AAAUA.......CGUGAAUGUUGUUUUAAACAAACCAAUAUCAAAAUAAACA............................
Fjo-1-1   UUUUAUCAGAUAGUUAAAAUAAUUUUCAAUUUAUAAUUCAUAAUUUAUAAUUAUUUUCUACAUUUGCACCCGAAUUAGACCAACCUCUGGCG..AAAUA.......CGUGAAUGUUGCUUUAUAUAAAACUAUAAUAUAAAAAUUAUC..........................
env-1075  UUUCAAAAAAUAAAUCAUGAAAUUAUAGCUAACCAAUUGCUAAUUAUCAAUAAAUAACUAUCUUUGCACUCAAUUUUGGACCAACCUCUGGCG..AAAUA.......CGUGAAUGUUGCUUUGAAUUACUAAUAAAUAAAAGAAA.............................
env-1076  UUAUUUCUAAAUCCUCCAAAAAAAAUCAAUUAAUUACUUGCAUCAAAGUAGAAUUUACUUAUUUUUGCACCCACUUAAUCCAACCUCUGGCG..AAAUA.......CGUGAAUGUUGUUUUAAUAGAAACUUUAUCAAUUAAUUAAA............................
env-1080  AUGAUCUUAGUUUUUGCAUGAAUAAUUCGAAUUAUUUGUUAGAAUCAAAAAAUAAGAAGUAAUUUUGCACACGCUUAAACCAACCUCUGGCG..AAAUA.......CGUGAAUGUUGCUUUUAGUCUAACACUUAAUAAUUUAUAGAA............................
env-1081  AAAGUAAAAUCAUUUGUAAUCUUUCGACUUUUGACUUGUAACUUUCGACUUUUUAAAAUACUUUUGCACCCCACAUUAGACCAACCUCUGGCG..AAAUC.......CGUGAAUGUUGCUCUUUUUUAAACCAUAAUUAAAAAAAUAAA..........................
env-1084  .............AAUAAAAAUCAUAGACUAUUAAUUUGUUAAUUAUUAAAUAUUAAUUAUUUUUGCACCCUCAUUAGACCAACCUCUGGCG..AAAUC.......CGUGAAUGUUGCUCUGAUUUAAACCAUAAUUAAAAAAUUAUC..........................
Sxi-1-1   UGUCUGAAGAGGAGAAAUAAAUUUACCUCCACCCUUUUAAAAAUAAGAUUUUUUUAUUACUUUUGCACACGUUUCUGAACCAACCUCUGGCG..AAAUC.......CGUGAAUGUUGCUCAGACUUUAUUUUUUAAACCAUUACAGAU............................
env-1085  CGGAUUUACUUGAUGAAAAAAAAUCAUAAUUCAUAAUUCAUAAUUCAUAAUUAUUUUUUAUUUUUGCACGCUCAUUAGACCAACCUCUGGCG..AAAUC.......CGUGAAUGUUGCUCUUUUUAAAACCAUAAUA.....................................
env-1086  UUCUAUUAUAUUUAUUGAUUUUUUUUAGAUUAUCCAUUGUUAAUUAUCAAUUAUUAAUUACAUUUGCACCCACGUUUGACCAACCUCUGGCG..AAAUC.......CGUGAAUGUUGCUCAGAUUUAAAACCAUAAUUAAAAAAAUUAUC.........................
env-1088  UUACUAGUGUAAGGAGAAAAAAUUAUCAAUUAUCCAUUGUUAGUUAUCAAUUAUUUUCUACAUUUGCGCCCACAUUAGACCAACCUCUGGCG..AAAUC.......CGUGAAUGUUGCUCUGACUUAAAAACCAUAAUUAAAAAAAUUAUC.........................

..........................................................................&lt;&lt;&lt;&lt;...&lt;.&lt;&lt;&lt;&lt;&lt;..........&gt;&gt;&gt;&gt;&gt;..&gt;.&gt;&gt;&gt;&gt;...................................
..........................................................................2020...0.12220..........02221..0.0202...................................
••••••••••••••••••••••••••••••••••••••••••••••YY••••••••••Y•RRRYYAACCUCUGRCG○-RRA○-------○CGUGRAUGUUG•UY•••••AA••••••••••••••••○○○-----------------------------
                                                                        5′                         3′
```

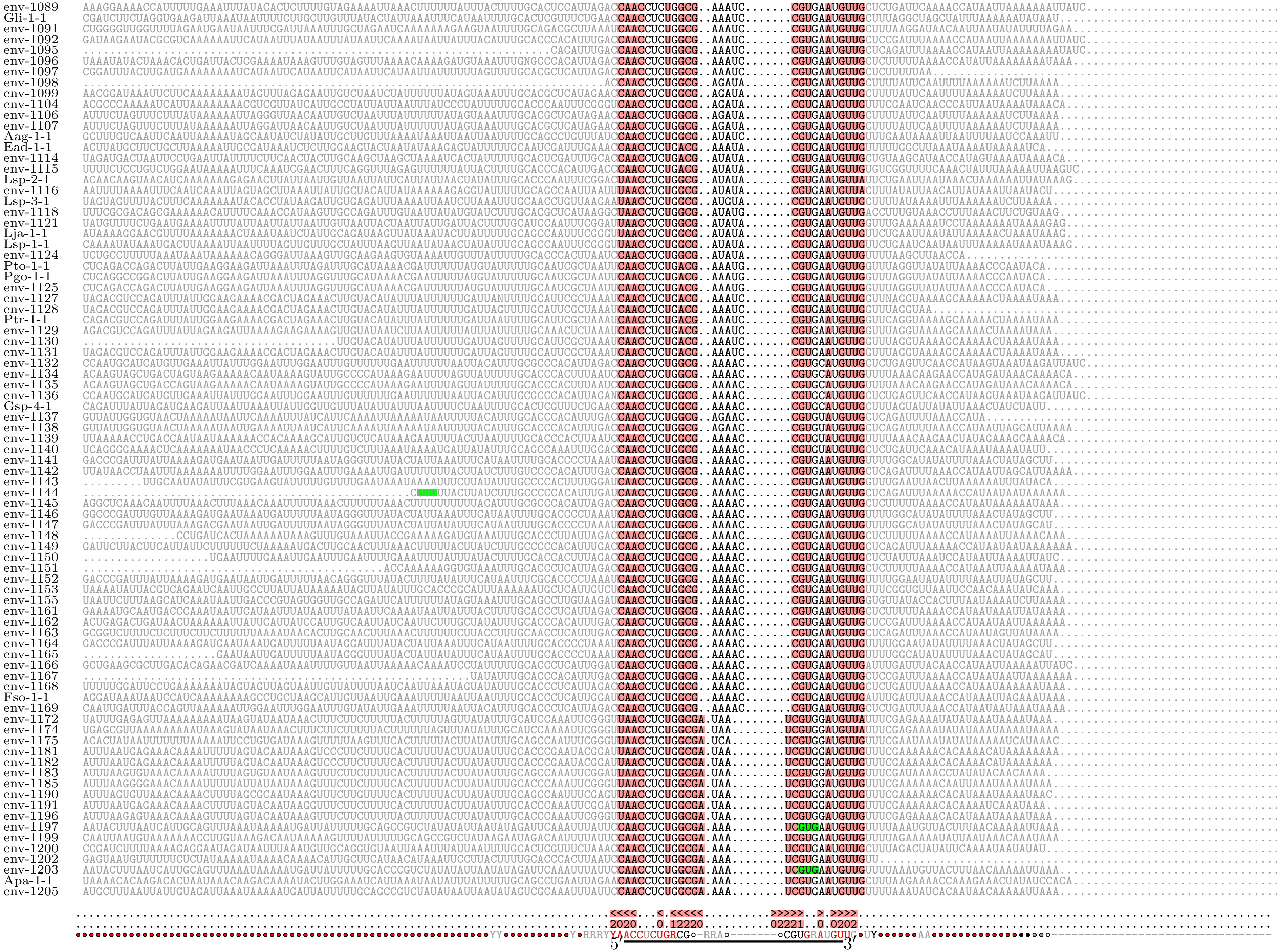
env-1089
Gli-1-1
env-1091
env-1092
env-1095
env-1096
env-1097
env-1098
env-1099
env-1104
env-1106
env-1107
Aag-1-1
Ead-1-1
env-1114
env-1115
Lsp-2-1
env-1116
Lsp-3-1
env-1118
env-1121
Lja-1-1
Lsp-1-1
env-1124
Pto-1-1
Pgo-1-1
env-1125
env-1127
env-1128
Ptr-1-1
env-1129
env-1130
env-1131
env-1132
env-1134
env-1135
env-1136
Gsp-4-1
env-1137
env-1138
env-1139
env-1140
env-1141
env-1142
env-1143
env-1144
env-1145
env-1146
env-1147
env-1148
env-1149
env-1150
env-1151
env-1152
env-1153
env-1155
env-1161
env-1162
env-1163
env-1164
env-1165
env-1166
env-1167
env-1168
Fso-1-1
env-1169
env-1172
env-1174
env-1175
env-1181
env-1182
env-1183
env-1185
env-1190
env-1191
env-1196
env-1197
env-1199
env-1200
env-1202
env-1203
Apa-1-1
env-1205
5'
3'

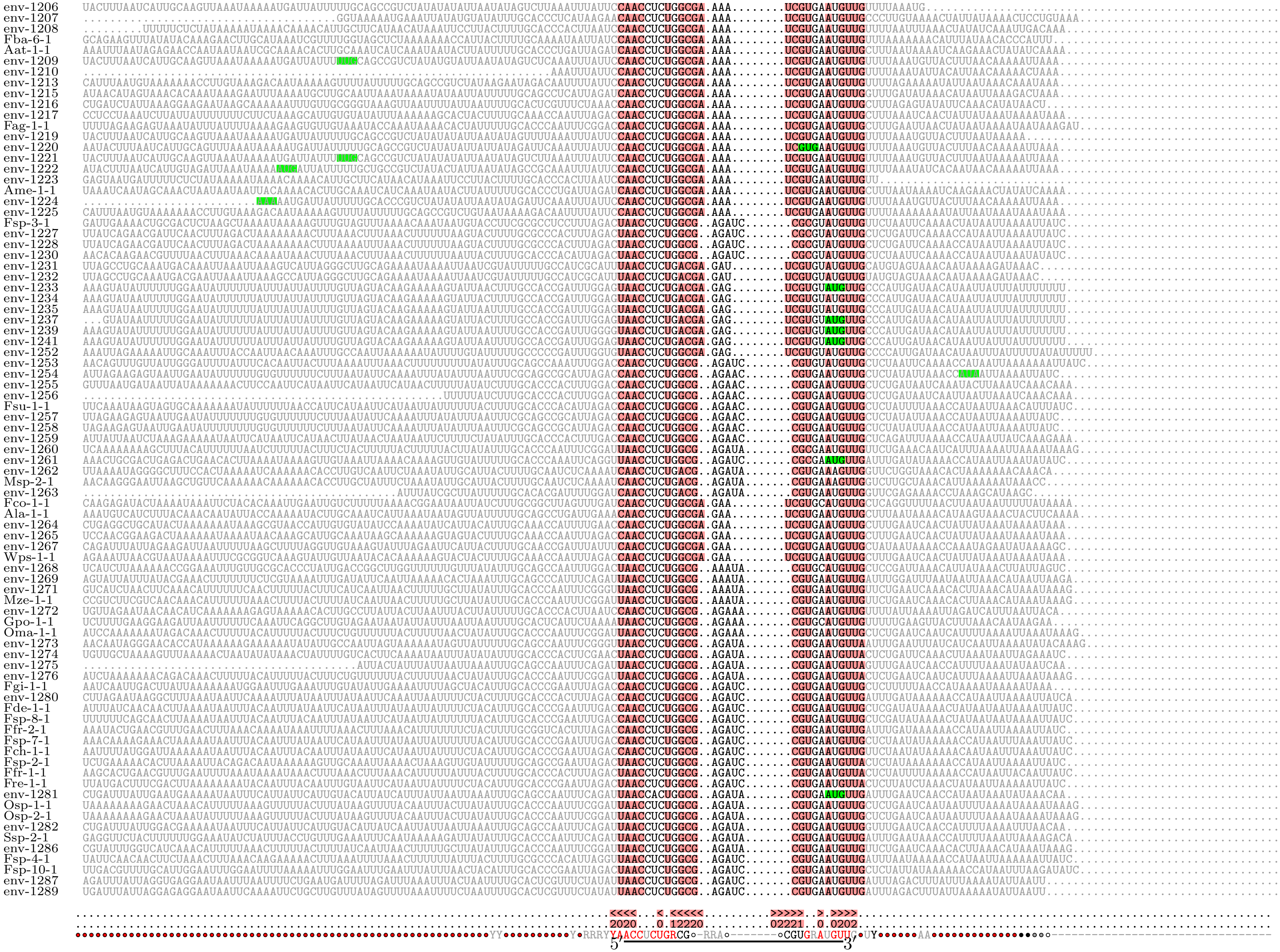
env-1206
env-1207
env-1208
Fba-6-1
Aat-1-1
env-1209
env-1210
env-1213
env-1215
env-1216
env-1217
Fag-1-1
env-1219
env-1220
env-1221
env-1222
env-1223
Ame-1-1
env-1224
env-1225
Fsp-3-1
env-1227
env-1228
env-1230
env-1231
env-1232
env-1233
env-1234
env-1235
env-1237
env-1239
env-1241
env-1252
env-1253
env-1254
env-1255
env-1256
Fsu-1-1
env-1257
env-1258
env-1259
env-1260
env-1261
env-1262
Msp-2-1
env-1263
Fco-1-1
Ala-1-1
env-1264
env-1265
env-1267
Wps-1-1
env-1268
env-1269
env-1271
Mze-1-1
env-1272
Gpo-1-1
Oma-1-1
env-1273
env-1274
env-1275
env-1276
Fgi-1-1
env-1280
Fde-1-1
Fsp-8-1
Ffr-2-1
Fsp-7-1
Fch-1-1
Fsp-2-1
Ffr-1-1
Fre-1-1
env-1281
Osp-1-1
Osp-2-1
env-1282
Ssp-2-1
env-1286
Fsp-4-1
Fsp-10-1
env-1287
env-1289
5′
3′

```
Fsp-9-1   UGGAUUUUCUUUUAUUGGAAUUUGCAAUUUAUUUAUUUUGGAAUUUCAUUUUUUUAAUUACCUUUGCACCCGAAUUAGACUAACCUCUGGCG..AGAUC.......CGUGAAUGUUGCUCUAUAUAAAAACCAUAAUUAUAAGAUAUC....
env-1291  AGAUCUAUUAGGAGAGGAAUAAUUUAAAAAUCUGCUUGUUUAUAGUUUUAAAUUUUAUAAUUUGCACUCGUUUCUAUAUUAACCUCUGGCG..AGAUC.......CGUGAAUGUUGGUUUAGACUUUAUUAAAAAUAUUAAUU....
env-1293  UUAGAAAUAAGUUUUUAAAUUUUCAGUGUUAAAAAUUUAAUAUUCCAUUAAAAACGCUAAAUUUGCACCCGAUUUCAGAUUAACCUCUGGCG..AGAUC.......CGUGAAUGUUGAUUUGAAUAAACAUUAUAAAUAGAAUUACG....
env-1296  ..................................................................CACCCGAAUUAGACUAACCUCUGGCG..AGAUC.......CGUGAAUGUUGCUCUAUAUAAAAAACCAUAAUUAAAGAUAUC....
env-1297  GGAUCUAUUAGGUGAUGAAUAAUCUAAAUUAAGCAAGGUUUAUAAGAUUAAUUUUUAUAAUUUGCACUCGUUUCUAAAUUAACCUCUGGCG..AGAUC.......CGUGAAUGUUGAUUUGGACUUUAUUUAAAUUAUUAAUU....
env-1300  UGAUUUAUUAGGAGAGGAAUAAUUCAAAUUUCUGCUUGUUUAUAGUUUUAAAUUUUAUAAUUUGCACUCGUUUCUAUAUUAACCUCUGGCG..AGAUC.......CGUGAAUGUUGAUUUAGACUUUAUUAAAAAUAUUAAUUAUGGAAU....
env-1301  CCGAUUUAUUAGGAGAAGAUUAAUUUAAAAUUACCUUGUUUAUAGCUUUAAAUUUUAUAAUUUGCACUCGUUUCUAUGUUAACCUCUGGCG..AGAUC.......CGUGAAUGUUGAUUUAGACUAUAUUAAAAAUAUUAAAU....
env-1302  ....................................UUAAACUUUAAACAUUUUUAUCUACUUUUGCAGCCAUAUUAGACUAACCUCUGGCG..AGAUC.......CGUGAAUGUUGAUUUGAUAAAAAACCAUAAUUAUAAUUAUC....
env-1303  .........................................................UUACUUUUGCACCUGAUUUAGACUAACCUCUGGCG..AGAUC.......CGUGAAUGUUGCUCUAAUUAAAACCAUAAUUAAAAAAUUAUC....
Fhi-1-1   AUUUAUCAACAACUUAAAAUAAUUUACAAUUUACAAUUUAUAAUUCACAAUUAUUUUCUACAUUUGCACCCGAAUUAGACUAACCUCUGGCG..AGAUC.......CGUGAAUGUUGCUCUAUAUAAAAAACCAUAAUUAUAAGAUAUC....
Gsp-3-1   CGAUCUUUUAGGAGAAGAUUAAAUUAAGUUUUCUCUUGUUUAUAGUUUUAAAUUAUUUAAUUUGCACUCGUUUCUAUAUUAACCUCUGGCG..AGAUC.......CGUGAAUGUUGAUUUAGACUAUAUUAAAAAUAUUAGAU....
env-1304  UUUUAUCAAAUACUUAAAAUAAUUUAUAAUUUACAAUUUAUAAUUCAUAAUUAUUUUCUACAUUUGCACCCGAAUUAGACUAACCUCUGGCG..AGAUC.......CGUGAAUGUUGCUCUAUAUAAAAACCAUAAUUAUAAGAUAUC....
env-1305  AAAAGUAAAAACAUUAUUAAUCUUUCGACUUUCGACUUGUAACUUUCGACUUUUUUUGUAUUUUGCACCCGAAUUAGACUAACCUCUGGCG..AGAUC.......CGUGAAUGUUGCUCUAUAUAAAACCAUAAUUAUAAGAUAUC....
env-1306  AUGUUUUUACUCUCUAAAAAAAAUAUCAAUAAUACAUUGUUAAUUAUCAAUUAAAUUUUAUAUUUGCGCCCACAUUAGACUAACCUCUGGCG..AGAUC.......CGUGAAUGUUGCUCUUUUUAUAAAACCAUAAUUAAUAAUAUC....
Gsp-5-1   AGAUUUAUUAGGAGAGGAAUAAUUUGAAAUUCUGCUUGUUUAUAGUUUAAAAUUAUCUAGUUUUGCACUCGUUUCUAUAUUAACCUCUGGCG..AGAUC.......CGUGAAUGUUGGUUUAGACUUUAUUAAAAAUAUUAAUU....
Fsp-11-1  UUUACUUUUUUUAUUGGAAUUUGGAAUUUAUUUUUAUUGGAAUUUACUUUUUUUUAAUUACAUUUGCACCCGAAUUAGACUAACCUCUGGCG..AGAUC.......CGUGAAUGUUGCUCUAUUAUAAAACCAUAAUUAUAAGAUAUC....
Fsp-1-1   UAUUCAAGAACUUCUAAACUUUAAACAAGAAAAACUUUAAACUUUAAACUUUUUUAUUUACUUUGCGCCCACAUUAAGUUAACCUCUGGCG..AGAUC.......CGUGAAUGUUGAUUUAAUAAAAACCAUAAUUAAAAAUUAUC....
env-1307  ..................................................AAUUUUUAUAUAAAUUUGCACUCUCAAAAAUAACCUCUGACG..AGAUA.......CGUGUAUGUUGCUUUUGUUCAAUCCUAAUAAAACUAUU....
Psp-3-1   CUCACUAAUUAUCAGCCUAAAAAAUAAACGAAUUAAAUUUCCUCAAUUAAAAAUAAUUACUAAAUUUGCACUCGCUAAAAUAACCUCUGACG..AGAUA.......CGUGAAUGUUGUUUAAGUAAAACUAUAAACUAUUAUAAGAU....
env-1308  GACCUUUUAACAUAAAAAUAUUUGUUGAUAUUUAUCAUUGUUUAGUUUAAAAUAAAGUUUAAUUUUGCACAUUCAAAGAUUAACCUCUGACG..AUAUC.......CGUGAAUGUUAGCUUUUAAAAACUAAU....
env-1309  AACGAAUUUAAUUAUACUUAAAAAAAUGUCUAAUUAAUUUUUAAGAAUAAAAUUAAUUACUAAAUUUGCACUCGCUAAAAUAACCUCUGACG..AUAUC.......CGUGAAUGUUGUUUUUGUAAAACUAUAAAAUAUUAAGAUA....
env-1310  ACGAAGACCCGACUUAUUGGAAGAAUAAAGUUUCAUGCUUGCAUAAUUAUUAUUUUUGUUUACUUUUGCAAUCUCUAAAUUAACCUCUGACG..AGAUC.......CGUGAAUGUUGGUUUAGGUAAUAGAAAAACUAAAAUAAA....
Gme-1-1   CUUGAAAUUUCGAAUUUAAAAUUAUUAAUUUUUAAUUGUUAAUUAUUAAUUAUUCCUUACUUUUGCAUCCAAUUUCAGAUUAACCUCUGACG..AUUUA.......CGUGAAUGUUGGUUUGGAUGAAUCUUAAAAAUUAAAAGAA....
env-1311  AAAAUGAGCUGAUUUGAAAAAAACAAUCGAACAUACUUGCUAGUUUUAAAAAAGGUUUUAAUUUUGCACUCCAUUAAAGUUAACCUCUGACG..AUAUA.......CGUGAAUGUUGGUUU....
env-1312  CAGACGUCCAGAUUUAUUGGAAGAAAUUGAUUAGAAACUUGUACACUUACAUUUUUAAUUAAUUUUGCAUUCGCUAAAUCAACCUCUGACG..AGAUC.......CGUGAAUGUUGGUUUGGAUUAAAGAUAAAUCAAUAUAAA....
env-1314  CUAAAGGGCGUUUCUUCCUAAAAUCCCAUUUUUUACUUGUGUAUCAUCAAAUAUUACUUAAUUUUGCAGUCGAUUAAAAUCAACCGCUGGCG..AGGUC.......CGUGAAUGUUGGUUUUGACAAACCUUUAAAUAUUCAAUUGCGCG....
Min-1-1   ACAGAACUACAAGUAGUUAUUACUAUAAAAUAAUCAUUUGUUAUUCUGUUUUAUUUUUAUAUUUGCACUCGAUUUAGAUCAACCUCUGGCG..AGGUC.......CGUGAAUGUUGUUCUAAGAUUAACAAAUUUAUACAUUAUUACA....
env-1317  UUUGGAUGAAACAGAAAAAUAAUUGCAAUUUCUCAAUUGCAUUAUAAAUUAAAAUUAAUACUUUUGCACUCGUUCUAAAAUAACCUCUGGCG..AGGUC.......CGUGUAUGUUGUUUUGGAUUUUACAUUAAAUAAGCAAUU....
env-1318  UUUGGAUGAAACAGAAAAAUAAUUGCAAUUUCUCAAUUGCAUUAUAAAUUAAAAUUAAUACUUUUGCACUCGUUCUAAAAUAACCUCUGGCG..AGGUC.......CGUGUAUGUUGUUUUGGAUUUUACAUUAAAUAAGAAAUU....
env-1319  ...................................................................ACUCGUUCUAAAAUAACCUCUGGCG..AGGUC.......CGUGUAUGUUGUUUUGGAUUUUACAUUAAAUAAGAAAUU....
env-1320  UUUGGAUGAAACAGAAAAAUAAUUGCAAUUUCUAAGUUGCAUUAUAAAUUAAAAUUAAUACUUUUGCACUCGUUCUAAAAUAACCUCUGGCG..AGGUC.......CGUGAAUGUUGUUUUAGAUUUUACAUUAAAUAAGAAAUU....
Ssp-1-1   UUUGGAUGAAACAGAAAAAUAAUUACAAUUUCUAAGUUGCAUUAUAAAUUAAAAUUAAUACUUUUGCACUCGUUCUAAAAUAACCUCUGGCG..AGGUC.......CGUGAAUGUUGUUUUAGGUUUUACAUUAAAUAAAAUAUU....
env-1321  UUUAGCAUCUGGAUGAAAAUUAUCAAUUAUAAAUUGUUAAUUAUUUAAUUUAUUUUCUUAAUUUGCACCCACUUUGAACCAACCUCUGACG..AUAAC.......CGUGUAUGUUGCUUUGAAUCAAACAUUUAUAAUUACGAUU....
Fsu-2-1   AACCGCGACUGUGACUGAAAACUUCUCAAAUAAAUUUGUGAUUUAAAAUAAAAAUCGUAAUUUCGCAGCCACAUUCGGCCCAACCUCUGACG..AUAAC.......CGUGUAUGUUGCUCUGAUUCAAACAUUUAAUUUAAAACGAUC....
env-1322  AUUAGCAUAAGGAAUAAAAUUUAUCAAUUAUAAAUUGUUAAUUAUAAAUUUAUUUUCUUAAUUUGCACCCACUUUGAACCAACCUCUGACG..AUAAC.......CGUGUAUGUUGCUUUGAAUCAAACAUUUAUAAUUACGAUU....
env-1323  CUAAUUGGAAAUCAAAAUUCAAAAACGCAAAAAACAACUUGUUAUUUAUAAAUAUUGCGUUACUUUUGCAAUCUCAAAAUCAACCUCUGACG..AUAUC.......CGUGUAAGUUGGUCUUGAAAAAUCAUUUAAAAAAAGUACAA....
env-1326  AAUAAUGUCCUUCAAAUUUCAAAAACGCAAAAAUCCGCUUGCUAUUUCUAAAUAUUGCAUUACUUUUGCAAUCUCAAAAUCAACCUCUGACG..AUAUC.......CGUGUAAGUUGGUCUUGAAAAAACUUUAAAAAAUACACAA....
env-1327  CAUCUGCAAUUUUUGAUCAAACAAAAACAAAAAUCUAUUGUUUUAUUAUUAAAUAAUUACUAUUUUGCAACCUGUUAUUCAACCUCUGACG..AUAAC.......CGUGUAAGUUGCUUAGCUUUAACAUUUUAUUUUAUUAAUA....
Mze-2-1   UAAAGCUAAAAACCAAUAAAAAAUCAAGAGAAACGCUUGUCCAAUAAAAAAUUAAAACUAAUUUGCACUCGCAUAUUUACAACCUCUGACG..AUAAC.......CGUGUAAGUUGUUCUUAUUAAACCAAUUAAAAAGAUUAAA....
env-1328  AAGAACAGAUUUUAAUAAAAAUUGAUAGCUAUAUUGAUUAUAGUUUAAUAUUAAUUUUUAUUUUUGCAAACUCAAUUGAAUAACCGCUGGCGA.AAA........UCGUGAAUGUUAUUUAAUGAAAACCAAAU....
Myu-1-1   UUUAUUGGAAGACUAAUUUUUUUUGUUUUUUUAGCCUCUAUGUUUUAACUUUUUAAUUAUCUUUGCAGCCAAUUUCGGAUUAACCUCUGGCGA.AAA........UCGUGAAUGUUACUUCGAUAUAACACUAUAAAAAAUAUUUAACC....
Gsa-1-1   CACAUAAAAACACUUCAAAAAACACAAAAUAAAUAUUGGAUAUUACUAAUAAAUAAGUAUUUUUGCACCCAAUUUCGGGUUAACCUCUGGCGA.AAA........UCGUGAAUGUUAUUCUGAAUAAAACUAUUUAAAUUUUAAAGAU....
Amu-1-1   GAAAAAUAGCCCUUAUAAGAAUUCAAACAUAAAACAUUUGCUAAUCAUUAAAUAAUACUUACUUUGCACCCGAUUAGAUUAACCUCUGGCGA.AAA........UCGUGAAUGUUAUUUUAACAAAACCAAAGAAACUAUAUCAAA....
Fen-1-1   UUCUAUUCUCUAUUUUCCCAAACUCUGUAAAAAAUUUGUAGUUCAAAAUAAAAUUUGUAAUUUUGCACCCACAUUCAGCCCAACCUCUGACG..AUAAC.......CGUGAAUGUUGCUCUGAUUCAAACAUUUAACUUUAACAAUC....
Fli-1-1   AAAAGUUGAUUUCAAACAAAUUCUUUCAAUUUAUUAUUUGCUUUUAAAAUUAUUUCGUAAUUUCGCAGCCUAAUUCAGCCCAACCUCUGACG..AUAAC.......CGUGAAUGUUGCUCUGAUUCAAACAUUAAACAUUAAUAAUC....
Fca-1-1   UGGAGUGGACAGUCAAAAAAACUCACAAUUCAUAAUUCAUAAUUCAUAAUUAUUUAUUAAUUUCGCAGCCAAAUUCAGCCCAACCUCUGACG..AUAAC.......CGUGAAUGUUGCUCUGAUUCAAACAUUUAACAUUAAUAAUC....
Fsa-1-1   AACUGAGACUGAGACUGAAAACUAUCAAAAGAAAUUUGUAGUUUAAAAUAAAAUACGUAAUUUCGCAGCCUAAUUCAGCCCAACCUCUGACG..AUAAC.......CGUGAAUGUUGCUCUGAUUCAAACAUUUAACAUUAAUAAUC....
Fse-1-1   CAUAACUUUUUAACGAAAAUAUUUCAUAACUCAUAACUCAUAACUCAAAACUUUUUUAUACUUUGCACCCACAUUUAAUUAACCUCUGGCG..AUACC.......CGUGAAUGUUGGUUAGAUUUAAAACCAUAAUUAGCAUUUAAA....
Fda-1-1   CAAUUGAUUAUGGAGUUAAAAAAAAUUGGAAUUUGGAAUUUGUAUAUUGAAAAUUUUAAUUACUUUUGCGCCACAUUUAAUUAACCUCUGGCG..AUACC.......CGUGAAUGUUGGUUAGAUUUUAAACCGUAAUUAGCAAUUAAA....
Dsp-2-1   GCUUUCGCGAAAGCAACACUCCACUACAACUUUUACAUACUGAAAUUAAUACCGCAAGAAUUUCUUGCAACUGAUUAGGCUAACCUCUGACG..AUACG.......CGUGAAUGUUGCCCUAAUGUAAAACCACUUUAAUAUCCAUAAA....
env-1329  GAAGACCUGAUUUAUUAGAUGAAUAAAAAACUAAAAAGUUGUAUAAUAAUUAUUUUUAUGUAAUUUUGCAACUUCUUAAUUAACCUCUGACG..AUACC.......CGUGAAUGUUGGUUAAGUUAAUCGAAAAAUUAAAUAAA....
env-1332  UUUCAAGAAAAGUAAGUUUUUUAGCUAUAAAAGACUUGUCUGUUUCGUUAAAAUCAUUAUCUUUGCGCCCAUAUUUGGACUAACCUCUGACG..AUAAC.......CGUGAAUGUUGCUCUGAUUCAAAACAAUUAACAUUAUCUAUC....
env-1335  GAAGUGAUAUUUAACUUAUUUAUUAGUACUAAGAAGUUGCCUACUUCAAAAUAAAGUCUAAUUUUGCAAAACGAAAUAGAUAACCUCUGACG..AAAAC.......CGUGAAUGUUGUCUCUUUUCAAACAAUAAAAAAAAUAAU....
env-1336  CUCCUAACUUAGGAUAAAAAAACUCAUAAUUCAUAACUCAUAAUUCGUAAUUCUUUUUUAUAUUUGCACCCACAUUUGACUAACCUCUGGCG..AAAAC.......CGUGAAUGUUGCUCAGAUUUCAAACCAUAAUAGUCUUAAAA....
env-1337  AACUGAGACUGAAAACUAAAAAUUAUUCAUUAUCCGUUGUCAAUUAUCAAUUCUUUUCUAUAUUUGCACCCACAUUUGACUAACCUCUGGCG..AAAAC.......CGUGAAUGUUGCUCAGAUUUAAAACCAUAAUAAUUAAUUAAA....
env-1338  CUCUAACCUAAUUUUGAAAAGUUUUGGNAUUUGGGAUNUGAAAAUUGAUUUUUUUAAUUAUNUUUGCACCCACAUUUGACUAACCUCUGGCG..AAAAC.......CGUGAAUGUUGCUCAGAUUUCAAACCAUAAUUAGCAUUAAAA....
Faq-1-1   ACUUUUGAUUUGAUAUAAAAAAACUGUCAUUAUUAAUUGUAAAUUAUUAAAUAUUAAUUACAUUUGCACCCACAUUUGACUAACCUCUGGCG..AAAAC.......CGUGAAUGUUGCUCAAGUUUUAAACCAUAAUAGCAUUAAAA....
env-1339  ACCCAAAAUUCAAACAAAAACUUUUCACUUUCUACUUUCUACUUUUAACUUUUUAAUUAUCUUUGCACCCACUUUUGGAUUAACCUCUGGCG..AAAAC.......CGUGAAUGUUGAUUUGAAACAACCAUUAUAAAAGAGAAAAC....
env-1340  AGAUUUAUUGGAUGAAAAAUAAUCANCAAUNCAUUACUAAUAAUUCAUAAUUAUUUUUUACUUUGCACCCACAUUUGACUAACCUCUGGCG..AAAAC.......CGUGAAUGUUGCUCAGAUUUUAAACCAUAAUUAGCAUUAAAA....
env-1341  ........................AUUCAUUAUCCAUUGUCAAUUAUCAAUUCUUUGCUAUAUUUGCACCCACAUUUGACUAACCUCUGGCG..AAAAC.......CGUGAAUGUUGCUCAGAUUCAAAACCAUAAUAAUUAAUUAAA....
env-1342  GGAAGAAUAAUCUUCUUAAAAUUAUUAAUUAUUAAUUGUCAAUUACCAAUUAAUAAUUAUCUUUGCACCCACUUUUGGAUUAACCUCUGGCG..AAAAC.......CGUGAAUGUUGAUUUGAAACAACUAUUAUAAAAGAGAAAAC....
Zam-1-1   ACCCGACCUUCUAAAGAAGAAAUAAUCUCAAACCGAUUGUAUAUUGUAAUUUAUUGAUUAUCUUUGCACUCAUUUCGAAUUAACCUCUGGCG..AUAAC.......CGUGAAUGUUACUUCGAUUAUAACAUUUUAAAAUUUAAUC....
Gec-1-1   CCUUUUAGAGGACGAGUAAAUAUUUCUCUAAUAGGCUUGUAGAAUAAUUUAAAUUGCUUAAUUUGCACUCAUUCUAAAAUAACCUCUGGCG..AUAAC.......CGUGAAUGUUGUUUUGGAUUUUACUUUAAACUAAUACAA....
env-1343  CCUGAUCUUUUAGAUGAAGAUUUUUAAAGCUCAGGUUUGCAGGUAACUAUAAAUUAAUUAAUUUUGCACUCGUUCUAAAAUAACCUCUGGCG..AUAAC.......CGUGAAUGUUGUUUUGGAGAUCAUAUAAACAUAUUAACU....
env-1344  UUUUAAUGCGUUUUCCCUGUGAAAGAAAGAUAAUCAUUUUUUAUUUUGCAAAAAAUGAUUAUCUUUGCGCUCGCUUUACAUAACCUCUGACGAGAAAAGAGUAGCUCGUGAAUGUUAUGUUUCAUUAUUAAACAAUUAAAAGCUAAAAAAAAUC....
env-1347  AAAAAAGAUACUACAACCGAAAACUAAACCUAAAAGUUGUUUAUUUAAAAUAAAGGUGUAAAUUUGCAGCCACAUUUGACUAACCUCUGGCG..AAAAC.......CGUGAAUGUUACUCAGAUUUUAAACCAUAAUUAGCAUUAAAA....
env-1348  UUAUUAGUUUAACAUAAAAUAAUUUGCAAUUCCUAAUUCUUAAUUCAUAAUUAUUUUUUACAUUUGCACCCACAUUUGACUAACCUCUGGCG..AAAAC.......CGUGAAUGUUACUCAGAUUUAAACCAUAAUUAGCAUUAAAA....
env-1349  AAGCUCUUGAGCACACUAAAAAAAAUUAGACCAGAUUUGUUGGAAGACUAAUAUUUUUGAUUAAAUUUGCACACUCAAAAUUAACCUCUGACGA.GAA........UCGUGAAUGUUAAUUUAUGAAAAAUUAAUAAAAAGUUUAAA....
env-1351  UGUAGAGAGCAACCGAAAAAAAACUUGAUAGAAAGUAAAUAAAUUAAAAAUAUUCAUUACAUUUGCACCCGAAUUUAGACUAACCUCUGACGA.GAA........UCGUGAAUGUUACUCUGAUAAAACUAUAAUUAUUAAACCAUU....
env-1352  UCCUAAAAAAAGAAUAAAAAAAUAUUUUAGUUUCUUUUGUGCAAUAAUCAAAAAAAGCAUUAUUUUUGCAAACUCAAAAUUAACCUCUGACGA.GAA........UCGUGAAUGUUAGUUUGUUAAAUCAUUAAAAAUUAAAACA....
env-1357  UCCUAAGAAAAGAAUAAAAAAAUAUUUUAGUUUCUUUUGUGCAAUAAUCAAAAAAAGCAUUAUUUUUGCAAACUCAAAAUUAACCUCUGACGA.GAA........UCGUGAAUGUUAGUUUGUUAAAUCAUUAAAAAUUAAAACA....
env-1359  GCUCUAAUUAUUAGGCAAAAAAAUAAUUCAAGAAGGUUUUAUUUCCCGAAAUAAAUUUCUAAAUUUGCACUCGCUAAAAUUAACCUCUGACGA.GAA........UCGUGAAUGUUACUUUUACUCAAACCCAUUAAAACAUUAAAA....
env-1360  AUUCUAAAAAAAGAAUAAAAAAAUAUUUUAGUUUCUUUUGUGCAAUAAUCAAAAAAAGCAUUAUUUUUGCAAACUCAAAAUUAACCUCUGACGA.GAA........UCGUGAAUGUUAGUUUGUUAAAUCAUUAAAAAUUAAAACA....
env-1361  CUUAGACCAGAUCUUUUCAAAGAUUAACUUUAAUUGAAUUUUUCUUUUAAAAAAAAAAUAAAGUAUUUUUGCAUCAAAAUUUAACCUCUGACG..AGAAC.......CGUGAAUGUUACUUUUUUAAAAUUUGAAAUAAUU....
env-1364  CUUAGACCAGAUCUUUUCAAAGAUUAACUUUAAUUGAAUUUUUCUUUUAAAAAAAAAAUAAAGUAUUUUUGCAUCGAAAUUUAACCUCUGACG..AAAAC.......CGUGAAUGUUACUUUUUUAAAAUUUGAAAUAAUUUAUA....
env-1366  AAAAAAACUAAGACCAGAUAUUUUUNACUGAAUUCUUUCUAAAAAUAAAAAAAANAUGUAAUUUCGCAUUCGAUUUAUUAUAACCACUGACGA.UAA........UCGUGAAUGUUNUACUAAAUAAAUUUAAAAUCUUAUCAAA....
env-1367  AAAAAAACUAAGACCAGAUAUUUUUAACUGAAUUCUUUCUAAAAAUAAAAAAAAAUAUGUAAUUUCGCAUUCGAUUUAUUAUAACCACUGACGA.UAA........UCGUGAAUGUUAUACUAAAUAAAUUUAAAAUCUUAUCAAA....
env-1368  AGAAAAUACGUCCAAAUUUUUUUGAUUAACCAAAUUUCAUUGAAAUAAAAAAAAAUGGUAAAUUCGCAAUCCAAUUGACAUAACCACUGACGA.UAA........UCGUGAAUGUUAUAUCAGUUUAAUUAAAAUCUUACAUA....
env-1369  AACUCAUGUAUUUCGAGCGAAAAAUCAACUAAAAUACUUGUAGGUUUUAAUUUAAGUAUUAAUUUCGCAGCUUGUUAGAAUAACCUCUGACGA.UAA........UCGUGAAUGUUACUCUAAUUUAAUUCCAAUUAACUAUACA....
env-1370  GACCUUUUAAUGUAAAAACAUUUGUUGAUAUUUAUUAUUGUUUAGUUUAAAAUAAAGUUUAAUUUCGCACAAUUAAAGAUUAACCUCUGACGA.UAA........UCGUGAAUGUUAGCUUUUCAAAAUUAAUAUUCCUAAGU....
Jma-1-1   UAAUAUAAUUAUAACAAAAAAUAAACUGUAAAAGGUUGUAUUUUAAUUAAUAAUAAUUAUUUUUGCACCCAAAUUCGGAUUAACCUCUGACGA.UAA........UCGUGAAUGUUACUCUGAAUCAACACAUUAUAAAUAUAGCACA....
env-1374  UAAGAGCUGAAAAUAAAAUAAAAAUGAAUGUACUAUUUGUCAAUUCAAAAAAAAGUAGUACCUUUGUCCCCGAUUUGGAGUAACCUCUGGCGA.AAG........UCGUGAAUGUUGCCCAAUGAUAACAAUAAUUUAUCUAAUUUUUU....
env-1375  UAUCAUAUUCGUUAGGCGACAAAAGUACGAAAAAACUUGCUAAUUCAAAAAAAAAUAGUACCUUUGCCCCCGAUUUGGAGUAACCUCUGGCGA.GAG........UCGUGAAUGUUACCCAAUGAUAACAUAAUUAUUUUAUAUUUUUU....

          ................................................................................<<<<...<.<<<<<..........>>>>>..>.>>>>............................................................
          ................................................................................2020...0.12220..........02221..0.0202............................................................
          ●●●●●●●●●●●●●●●●●●●●●●●●●●●●●●●●●●●●●●●●●●●●●●●●●●●●●●●●●●●●●●●●●●●●●YY●●●●●●●●●●Y●RRRYYAACCUCUGRCG○-RRA○-------○CGUGRAUGUUG●UY●●●●●●AA●●●●●●●●●●●●●●●●○○○-----------------------------
                                                                                  5′                             3′
```

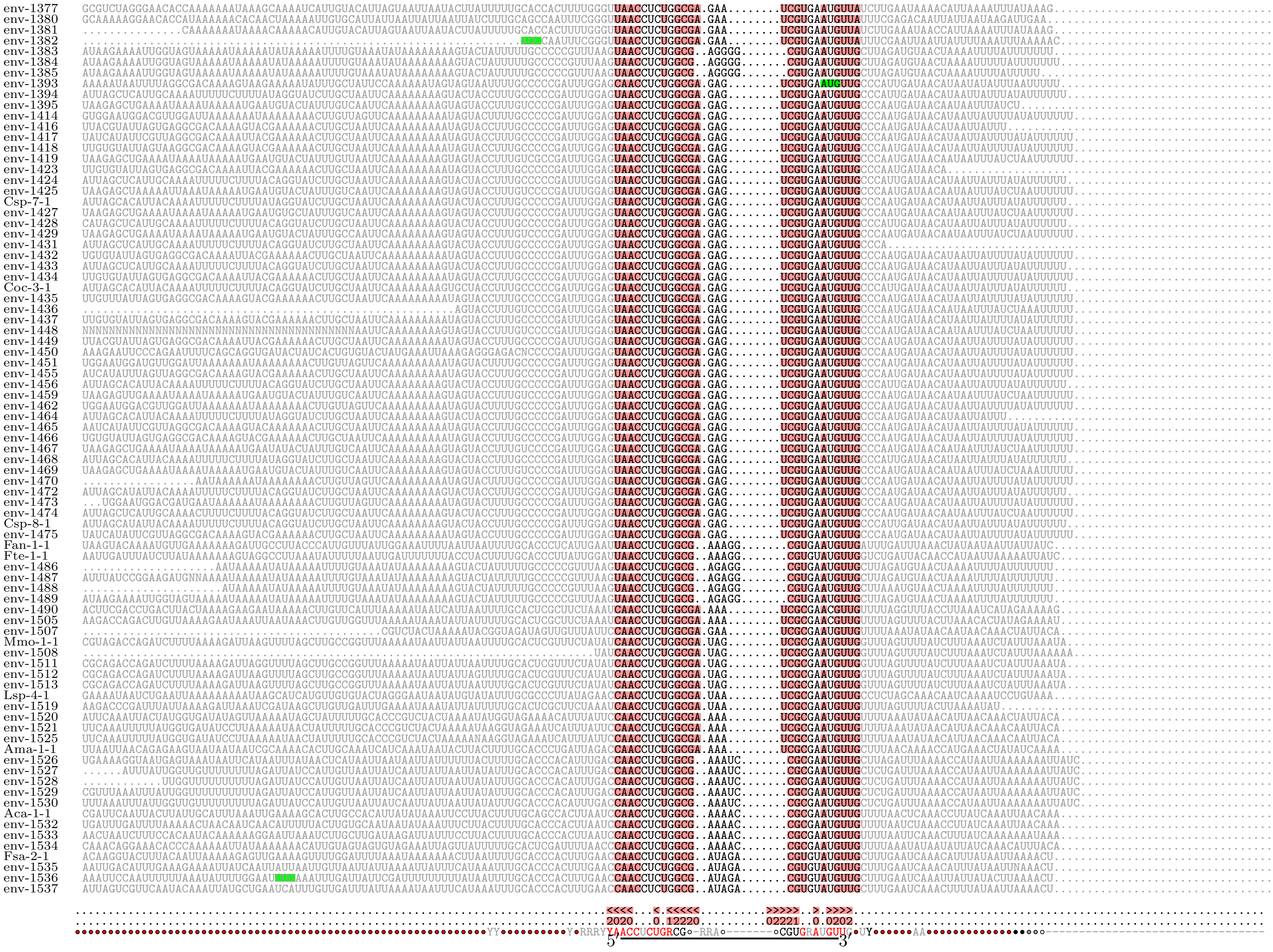
env-1377
env-1380
env-1381
env-1382
env-1383
env-1384
env-1385
env-1393
env-1394
env-1395
env-1414
env-1416
env-1417
env-1418
env-1419
env-1423
env-1424
env-1425
Csp-7-1
env-1427
env-1428
env-1429
env-1431
env-1432
env-1433
env-1434
Coc-3-1
env-1435
env-1436
env-1437
env-1448
env-1449
env-1450
env-1451
env-1455
env-1456
env-1459
env-1462
env-1464
env-1465
env-1466
env-1467
env-1468
env-1469
env-1470
env-1472
env-1473
env-1474
Csp-8-1
env-1475
Fan-1-1
Fte-1-1
env-1486
env-1487
env-1488
env-1489
env-1490
env-1505
env-1507
Mmo-1-1
env-1508
env-1511
env-1512
env-1513
Lsp-4-1
env-1519
env-1520
env-1521
env-1525
Ama-1-1
env-1526
env-1527
env-1528
env-1529
env-1530
Aca-1-1
env-1532
env-1533
env-1534
Fsa-2-1
env-1535
env-1536
env-1537
5′
3′

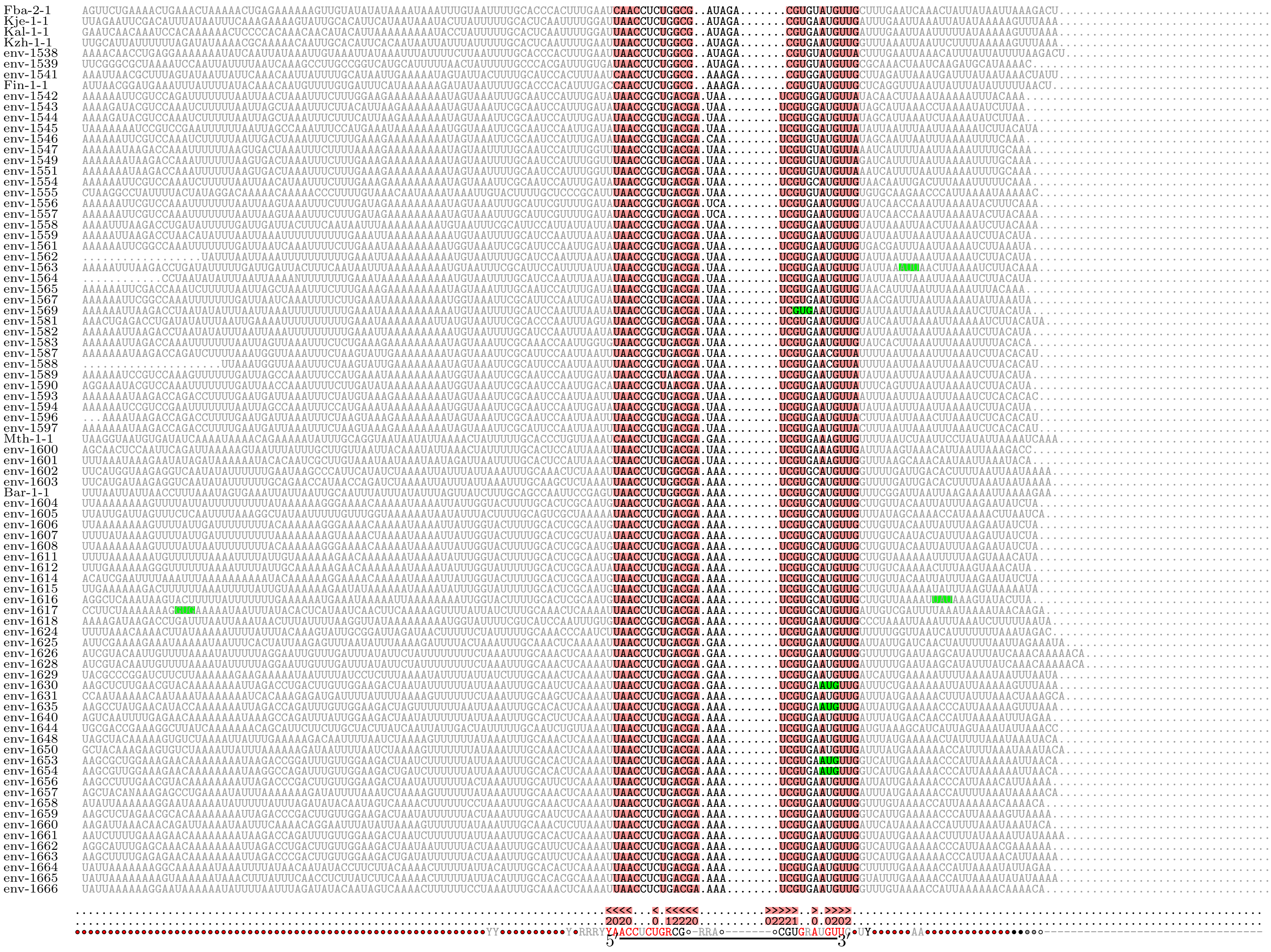

```
env-1667  AAGCUUAUGCGCAUACAAAAAAAAUUAGACCAGAUUUGUUGGAAGACUGAUUUUUUUAGUUAAAUUUGCACUCUUAAAAUUAACCUCUGACGA.AAA........UCGUGAAUGUUGGUUUAUGAAAAACCCAUUAAAAAGUUUAAA........
env-1670  ......UGGAAAGAACAAAAAAAAUAAGGCCAGAUUUGUUGGAAGACUAAUCUUUUUUAUUAAAUUUGCACACUCAAAAUUAACCUCUGACGA.AAA........UCGUGAAUGUUGGUCAUUGAAAAACCCAUUAAAAAAUUAACA........
env-1671  ...........................................................CAUUCUCAAAAUUAACCUCUGACGA.AAA........UCGUGAAUGUUGAUUAUUGAAAAACCCAUUAAAAGUUAAAA........
env-1672  AAGCUUUUGAACGUACAAAAAAAAUUAGACCUGACUUGUUGGAAGACUAAUAUUUUUUACUAAAUUUGCAUUCUCAAAAUUAACCUCUGACGA.AAA........UCGUGAAUGUUGAUCAUUGAAUAACCCAUUAAAUAUUUAAA........
env-1673  AAGCAUUUGAAAGAACAAAAAAAAUUAGACCCGACUUGUUGGAAGACUAAUAUUUUUUACUAAAUUUGCAUUCUCAAAAUUAACCUCUGACGA.AAA........UCGUGAAUGUUGGUAAUUGAAAAAACCCAUUAAACAUUAAAA........
env-1674  UGUGACCGAUAGGCUUAUCAAAAAAACAGCAUUCUUCUUGCUACUUAUCAAUUAUUGCCUAUUUUUGCAAUCUGUUAAAUUAACCUCUGACGA.AAA........UCGUGAAUGUUGAUGUAAAGCAUCAUUAGUAAAUAUUAAACC........
env-1677  AAGCUUAUGCGCAUACAAAAAAAAUUAGGCCAGAUUUGUUGGAAGACUGAUUUUUUUAGUUAAAUUUGCACUCUUAAAAUUAACCUCUGACGA.AAA........UCGUGAAUGUUGGUUUAUGAAAAACCCAUUAAAAAGUUUAAA........
env-1680  ......................................................AAUUUUGCAAAACGGAAUGGAUAACCUCUGACGA.AAA........UCGUGAAUGUUGUCACUUUUCAAACAAUAAAAAAAAUAAC........
env-1687  CAAUCAAAAAAAGUUAAAAAUAAUUUAACUUCUAUCCGUUUUUUAUUAAAAAUAUUUUACUAAAUUUGCAAACUCAAAAUUAACCUCUGACGA.AAA........UCGUGAAUGUUGGUUUUUGAUUAACUAUUUUUAAAUAGAAAAA........
env-1690  ..........................................................AUUUGCACACUCAAAAUUAACCUCUGACGA.AAA........UCGUGAAUGUUGGUCAUUGAAAAACCCAUUAAAAAAUUAACA........
env-1699  UAUUGAAAAAAAGUAAAAAAUAAAUUUUAUUUCAAUAUAUAAUCUUCAAAAACUUUUUAUUACAUUUGCACACUCAAAAUUAACCUCUGACGA.AAA........UCGUGAAUGUUGAUUUUUGAAAACCAUUAAAAUAAUUAAA........
env-1700  UAUUGAAAAAAAGUAAAAAAUAAACUUUAUUUCAAUCUUUAAUCUUCAAAAACUUUUUAUUACAUUUGCACACGCAAAAUUAACCUCUGACGA.AAA........UCGUGAAUGUUGGUAUUUGAAAAACCAUUAAAAUAUAUAAA........
env-1701  AAGCUUUUGAACGUACAAAAAAAAUUAGACCUGACUUGUUGGAAGACUAAUAUUUUUUACUAAAUUUGCAUUCUCAAAAUUAACCUCUGACGA.AAA........UCGUGAAUGUUGGUAAUUGAUAAAACCCAUUAAAAGUUAAAA........
env-1702  .........CAUAACUUCAAAAAUAUCGACAAUCCGCUUGUAGAUUUUAAGAUAUACCGUACUUUUGCGCCCUGUAAUAUUAACCUCUGACGA.AAA........UCGUGAAUGUUGGUGUUCAAAUCACAGUCAAAUAAUAUAUC........
env-1703  AAGCUUUUGAACGUACAAAAAAAAUAAGACCAGACUUGUUGGAAGACUAAUAUUUUUUACUAAAUUUGCAUUCUCAAAAUUAACCUCUGACGA.AAA........UCGUGAAUGUUGAUCAUUGAAAAACCCAUUAAACAUUUAAA........
env-1709  UCCUCAUGAUUACUUUUUAAAAUAAUUCUAAAACAGGAUUUUAUAUUAAAAGUUUUUUAUAAAUUUGCAAACUCUUAAAUUAACCUCUGACGA.AAA........UCGUGAAUGUUGAUUUUUGAAAAACCAUUUUAAAUAAAUACG........
env-1711  AGACCAAGCACAUUCCAAAAAUAUUGUCUUAAAUCCCUUGUAGGUUUUAAGAUAUAACCUAUUUUUGCACAUUGUUAAAUUAACCUCUGACGA.AAA........UCGUGAAUGUUGGUCACUACAUAAUCCUAAAUAAAACAAA........
env-1712  AGCUACAGAAAGAGCCUGAAAAUAUUUAAAAAAAGAUAUUUUAAAUCUAAAAGUUUUUUAUAAAUUUGCAAACUCAAAAUUAACCUCUGACGA.AAA........UCGUGAAUGUUGAUUUAUGAAAAACCAUUUUAAAUAAAAACA........
env-1713  UAUUAAAAAAAGACAGAAAAUAAAAUUAAUNUCCAUAUCUAAUCUUCAAAAACUUUUUAUUACAUUUGCACACUCAAAAUUAACCUCUGACGA.AAA........UCGUGAAUGUUGGUUACUGAAAAAUCAUUAAAACAAUAAAA........
env-1714  AAGCCUUUGAACGUACAAAAAAAAUUAGACCAGACUUGUUGGAAGACUAAUAUUUUUUACUAAAUUUGCAUUCUCAAAAUUAACCUCUGACGA.AAA........UCGUGAAUGUUGAUUAUUGAAAAACCCAUUAAACAUUAAAA........
env-1715  AAGCCUUGGAAAGAACAAAAAAAAUAAGGCCAGAUUUGUUGGAAGACUGAUCUUUUUUAUUAAAUUUGCACACUCAAAAUUAACCUCUGACGA.AAA........UCGUGAAUGUUGGUCAUUGAAAAACCCAUUAAAAAAUUAACA........
env-1716  AAGCUUAUGCGCAUACAAAAAAAAUUAGACCAGAUUUGUUGGAAGACUGAUUUUUUUAGUUAAAUUUGCACUCUUAAAAUUAACCUCUGACGA.AAA........UCGUGAAUGUUGAUUUAUGAAAAACCCAUUAAAAAGUUUAAA........
env-1717  GCAAGAACUAUCAACCAAAAAAAUAACUUCAAUAGGUUUUAUAUCCUAAAAUAAAUUCUUAGAUUUGCACUCGCUAAAAUUAACCUCUGACGA.GAA........UCGUGAAAGUUAGUUGUGGUCAAUUCAA........
env-1718  GCAGAAACUAUCAAUCAAAAAAAUAACUUGAAUAGGUUUUAUUUACUAAAAUAAAUUCUUAAAUUUGCAAUCGCUAAAAUUAACCUCUGACGA.GAA........UCGUGAAAGUUAGUUGUGGCCAACUCAAUAAAACAAAAAA........
env-1720  .................AAAAAAAUAACUUGAAUUAGGUUUAUUUACUAAAUAAAUUCUUAGAUUUGCACUCGCUAAAAUUAACCUCUGACGA.GAA........UCGUGAAAGUUAGUUGUGGUCAAUUCAAUAAAAACAAAAA........
env-1721  GUUUGAACUACCAAACAAAAAAAUAGCUUGAAUAGAUUUUAUUUACUAAAAUAAAUUCAUAGAUUUGCACUCGCUAAAAUUAACCUCUGACGA.GAA........UCGUGAAAGUUAGUUAUGGUAAAUACAUUAAAACAAAAAA........
env-1723  GCAAGAACUAUCAACCAAAAAAAUAACUUGAAUAGGUUUUAUUUACUAAAAUAAAUUCUUAGAUUUGCACUCGCUAAAAUUAACCUCUGACGA.GAA........UCGUGAAAGUUAGUUGUGGUCAAUUCAAUAAAACAAAAAA........
env-1733  GCAAGAACUAUCAACCAAAAAAAUAACUUCAAUAGGUUUUAUAUACUAAAAUAAAUUCUUAGAUUUGCACUCGCUAAAAUUAACCUCUGACGA.GAA........UCGUGAAAGUUAGUUGUGGUCAAUUCAAUAAAACAAAAAA........
env-1734  GCAAGAACUAUCAACCAAAAAAAUAACUUCAAUAGGUUUUAUAUCCUAAAAUAAAUUCUUAGAUUUGCACUCGCUAAAAUUAACCUCUGACGA.GAA........UCGUGAAAGUUAGUUGUGGUCAAUUCAAUAAAAC........
env-1735  GCAAGAACUAUCAACCAAAAAAAUAACUUGAAUAGGUUUUAUUUACUAAAAUAAAUUCUUAGAUUUGCACUCGCUAAAAUUAACCUCUGACGA.GAA........UCGUGAAAGUUAGUUGUGGCCAAUUCAAUAAAACAAAAAA........
env-1736  GUAGAAAAUUAUCAACCAAAAAAAUAACUUAAAUCGGUUUUAUAUCCUAAAUUAAAUUCUUAGAUUUGCACUCGCUAAAAUUAACCUCUGACGA.GAA........UCGUGAAAGUUAGUUCUGGUAAACACCAUAAAUUAUAAUAA........
env-1737  .....................................................AAUUCUUAGAUUUGCACUCGCUAAAAUUAACCUCUGACGA.GAA........UCGUGAAAGUUAGUUGUGGUCAAUUCAAUAAAACAAAAAA........
env-1741  GUAAAGAUUACCAACCAAAAAAAUAACUUGAAUAGGUUUUAUAUCCUAAAUUAAAUUCUUAGAUUUGCACUCGCUAAAAUUAACCUCUGACGA.AAA........UCGUGAAAGUUAGUUAUAGA........
env-1742  GUAAAGAUUACCAACCAAAAAAAUAACUUGAAUAGGUUUUAUAUCCUAAAUUAAAUUCUUAGAUUUGCACUCGCUAAAAUUAACCUCUGACGA.AAA........UCGUGAAAGUUAGUUAUAGAAAAUAUUUUAAAUCAACUAGAA........
env-1743  ...........................UGAAUAGGUUUUAUAUCCUAAAUUAAAUUCUUAGAUUUGCACUCGCUAAAAUUAACCUCUGACGA.AAA........UCGUGAAAGUUAGUUAUAGAAAAUAUUUUAAAUCAACUAGAA........
env-1750  GUAAAGAUUACCAACCGAAAAAAUAACUUGAAUAGGUUUUAUAUCCUAAAUUAAAUUCUUAGAUUUGCACUCGCUAAAAUUAACCUCUGACGA.AAA........UCGUGAAAGUUAGUUAUAGAAAAUAUUUUAAAUUAGAUAUAA........
env-1751  ...............................UAGGUUUUAUAUCCUAAAUUAAAUUCUUAGAUUUGCACUCGCUAAAAUUAACCUCUGACGA.AAA........UCGUGAAAGUUAGUUAUAGAAAAUAUUUUAAAUUAGAUAUAA........
env-1752  GUAAAGAUUACCAACCAAAAAAAUAACUUGAAUAGGUUUUAUAUCCUAAAUUAAAUUCUUAGAUUUGCACUCGCUAAAAUUAACCUCUGACGA.AAA........UCGUGAAAGUUAGUUAUAGAAAAUAUUUUAAAUUAGAUAUAA........
env-1758  UUUUAAAAAAAGGAAAAAAAUAAUUACUCCAGCAAUAUUUAAUCUUCAAAAACUUUUUAUUACAUUUGCACACUCAAAAUUAACCUCUGACGA.AAA........UCGUGAAAGUUGAUUCUUGAAAAACCAUUAAAACAAUUAAA........
env-1760  ............GAGGAAAAUAAUUUCUGNCACUAUUUAUAUUCUUCAAAAACUUUUUAUUACAUUUGCAAACGCAAAAUUAACCUCUGACGA.AAA........UCGUGAAAGUUGAUUCUUGAAAAACCAUUAAAACAUUUAAA........
env-1761  GUAGAAACUAUCAACCAAAAAAAUAACUUGAUUAGGUUUUAUAUCCUAAAUUAAAUUCUUAGAUUUGCACUCGCUAAAAUUAACCUCUGACGA.AAA........UCGUGAAAGUUGAUUAUAGUAAACACAAUAAAAUAUAAUAA........
env-1762  .................................................AACUUUUUAUUACCUUUGCAAACGCAAAAUUAACCUCUGACGA.AAA........UCGUGAAAGUUGAUUAUUGAAAAACUAUUAAAAAAAUCCGAA........
env-1763  UUUUAAAAAAAGGAGAAAAAUAAUUACUCCAGCAAUAUUUAAUCUUCAAAAACUUUUUAUUACAUUUGCACACUCAAAAUUAACCUCUGACGA.AAA........UCGUGAAAGUUGAUUCUUGAAAAACCAUUAAAACAAUUAAA........
env-1764  UAUUGAAUAAAGGAGAAAAAUAAUUUCACUAGCUAUAUAAAUUCUUCAAAAACUUUUCAUUACAUUUGCAAACGCAAAAUUAACCUCUGACGA.AAA........UCGUGAAAGUUGAUUAUUGAAAAAACCAUUAAAACAAUACAA........
env-1765  UAUUAAAUAAAUCAGGAAAAUAAUUUCUGCCACUAUUUAUAUUCUUCAAAAACUUUUUAUUACAUUUGCAAACGCAAAAUUAACCUCUGACGA.AAA........UCGUGAAAGUUGAUUAUUGAAAAACCAUUAAAACAUUUAAA........
env-1771  UAUUAAAUAAAGAAUGAAAAUAAUUUUCAUAGCUAUAUAAAUUCUUCAAAAACUUUUUAUUACCUUUGCAAACGCAAAAUUAACCUCUGACGA.AAA........UCGUGAAAGUUGAUUAUUGAAAAACUAUUAAA........
env-1772  ..............................GCUAUAUAAAUUCUUCAAAAACUUUUUAUUACCUUUGCAAACGCAAAAUUAACCUCUGACGA.AAA........UCGUGAAAGUUGAUUAUUGAAAAACUAUUAAAAAAAUCCAAA........
env-1777  GCUCUAAUUAUCAAGCAAAAAAAUAAUUCAAAUAGAUUUUAUUUACUAAAAUAAAUUCCUAGAUUUGCACUCGCUAAAAUUAACCUCUGACGA.AAA........UCGUGAAAGUUGGUUGUAGUUAACCCAUAAAAUAAAAAA........
env-1779  UUUUAAAAAAAGGAGAAAAAUAAUUUCUACAGCAAUAUUUAAUCUUCAAAAACUUUUUAUUACAUUUGCACACUCAAAAUUAACCUCUGACGA.AAA........UCGUGAAAGUUGAUUCUUGAAAAACCAUUAAAACAAUUAAA........
env-1781  UAUUAAAUAAAGAAUGAAAAUAAUUUUCAUAGCUAUAUAAAUUCUUCAAAAACUUUUUAUUAUCUUUGCAAACGCAAAAUUAACCUCUGACGA.AAA........UCGUGAAAGUUGAUUAUUGAAAAACUAUUAAAAAAAUCCGAA........
env-1782  GCUCUAAUUAUCAAGCAAAAAAAUAAUUCAAAUAGAUUUUAUUUACUAAAAUAAAUUCCUAAAUUUGCACUCGCUAAAAUUAACCUCUGACGA.AAA........UCGUGAAAGUUGGUUGUAGUUAACCCAUAAAAUAAAAAA........
env-1783  UAUUAAAUAAAGGAGGAAAAUAAUUUCUGUCACUAUUUAUAUUCUUCAAAAACUUUUUAUUACAUUUGCAAACGCAAAAUUAACCUCUGACGA.AAA........UCGUGAAAGUUGAUUCUUGAAAAACCAUUAAAACAUUUAAA........
env-1787  AAGUUUAUACUGACUUUGGAUAAAUUCAUUGGUUGUAUUGUGUACUUAAAAAAUAAGUUUAAUUUUGCACGACUAAAGAUCAACCACUGACGA.AAA........UCGUGAAUGUUGGCUUUUAUAAAAAUUAAUUAUUAUAAAG........
env-1788  UUGGAAAAUAGUGAAGAAAAUUAACUCAAGAUAAUAUUUGUUAAAACAAAGUUAUUUUAUAAAUUUGCACACGCUAAAAUCAACCACUGACGA.AAA........UCGUGAAUGUUGCCUUUACAAAACCAUUAAAAAAAAAUACA........
env-1789  UUGGAAAAUAGCGAAGAAAAUUAACUCAAAAUAAUGUUUGUUAAAACAAAGUUAUUUUAUAAAUUUGCACACGCUAAAAUCAACCACUGACGA.AAA........UCGUGAAUGUUGCCUUUAC........
env-1790  ......AAUAGCGAAGAAAAUUAACUUAAAAUAAUGUUUGUUAAAACAAAGUUAUUUUAUAAAUUUGCACACGCUAAAAUCAACCACUGACGA.AAA........UCGUGAAUGUUGCCUUUACAAAACCAUUAAAAUAAAUACA........
env-1791  UUGGAAAAUAGCGAAGAAAAUUAACUCAAAAUAAUGUUUGUUAAAACAAAGUUAUUUUAUAAAUUUGCACACGCUAAAAUCAACCACUGACGA.AAA........UCGUGAAUGUUGCCUUUACAAAACCAUUAAAAUAAAUACA........
env-1792  UAUUCCAAAAGACUCUUUAAUUAUUAAAAAUAGAGGACUGUUAUUUUAAAAUUUUUUUCAUAAAUUUGCAAACUCAAUUUCAACCGCUGACGA.GAA........UCGUGAAUGUUGGUUUUGAAUAUCUAAAAAAUAGUAAA........
env-1793  UUCAAUUAUAAAUUGUAAAAAAUAUUUUUGAUAAAGUAUUGCGGAUUAGAUAACUUUUUCUAUUUUUGCAAACCCAAUCUCAACCGCUGACGA.GAA........UCGUGAAUGUUGCUUUUGGUUAAUCAUUUUUUAAAUAGAC........
env-1794  GUUGGAAUAAAACCGGUCAAAAAAAUCAUAAUUUUACUUGUAGAUUCUAUUAUAUAGACUAAUUUUGCAACCUGUUAAAUCAACCUCUGACGA.GAA........UCGUGAAUGUUGGUCUACGAUAAACCAUAAAUUUUUAUC........
env-1802  AACCAAAUCCGUAACGCAAAAAAACAACCCAACCCUUCUUUUUAAAUUAAAAAAAAGAGCUAUAUUUGCAACCUCAAAAUCAACCUCUGACGA.GAA........UCGUGAAUGUUGGUCAUUGAUAAAACCAUUAAAUAAAAAGAA........
env-1803  CACUCACAAUCAGCGCUUAAAAUAAUAUUAUAUUUUUUUUGCUCAACAAAACAAAUUCCUUACAUUUGCAACCUCAAAAUCAACCUCUGACGA.GAA........UCGUGAAUGUUGGUAUUAGAGAAUCCGUUAAAACAAACAAA........
env-1804  UGGUUGAAUCUACCUCAAAGAAAAAAUAAAUUUAUAUAUUGCAGGAUAAAAUCUUUUUAUUAAUUUUGCAAACUCAAUUUCAACCUCUGACGA.GAA........UCGUGAAUGUUGCUCUUGAUUAAAUAUUUUUUUAUUAAA........
env-1805  AAAUUAAAUGAUUUCAAAGAAAAAAAUAAAUUUAUAUAUUGCAGGAUAAAAUCUUUUUAUUAAUUUUGCAAACUCAAUUUCAACCUCUGACGA.GAA........UCGUGAAUGUUGCUCUUGAUUAAAUAUUUUUUUAUUAAA........
env-1806  AAAUCAAAUGGCAAGAAAUAAUAAUUUAAAAUACUUAUUCAAUUUGUGACAAUAAAAUAUUAUAUUUGCAACCUCAAAAUCAACCUCUGACGA.GAA........UCGUGAAUGUUGGUCAUUGAUAAAACCAUUAAAUAAAAAGAA........
env-1807  UAAAUUAAAUGAUUUCAAAGAAAAAAUAAAUUUAUAUAUUGCAGGAUAAAAUCUUUUUAUUAAUUUUGCAAACUCAAUUUCAACCUCUGACGA.GAA........UCGUGAAUGUUGCUCUUGAUUAAAUAUUUUUUUAUUAAA........
env-1812  ............................................AACAAAACAAAUUCCUUACAUUUGCAACCUCAAAAUCAACCUCUGACGA.GAA........UCGUGAAUGUUGGUAUUAGAGAAUCCGUUAAAACAAACAAA........
env-1813  AGUUGAAAUUACUAUUAUUAAAAAAAAUUAAAUUUAAUAUUGCAGAAUAAAAUUCUUUUUAUUAAUUUUGCAAACUCAAUUUCAACCUCUGACGA.GAA........UCGUGAAUGUUGCUUUUGGUUAAAUAUUUUUUUCAUCAAA........
env-1814  UACCCAGAAAACAGCCCAUAAAAAUAAAGCGAACUGGGGCUUGCAGAAAAUAAAAUUAAUCGUAUUUUUGCCAUCGCAUUUAACCUCUGACGA.AAU........UCGUGUAUGUUGGAUGUAGUAAACCAUAAAAGAUAAAC........
env-1815  ........................................UUGCAGAAAAUAAAAUUAAUCGUAUUUUUGCCAUCGCAUUUAACCUCUGACGA.UAU........UCGUGUAUGUUGGAUGUAGUAAACCAUAAAAGAUAAAC........
env-1816  ACUAUACUCAAUACAAAAUAAAUUCAACCAAUUUAAUUGUAAUUUCAAAAUAAAUUGUAAAUUUGCACCCACAUUUGGACUAACCUCUGACGA.UAA........UCGUGUAUGUUGCUCUGAUAAUCACUAUUUUAACUUAAUUGUU........
env-1817  ACAUUUAAGUUUACUUUUUAUUUUUUUAGAUUUAUUUUGUUGGACUUGAAGAAGUUUGUACUUUUGCCGUCUGUAAAAAAUAACCUCUGACGA.UAA........UCGUGUAUGUUGCUUUUUAGUAAAAACAUUUAAUUCCAAUUUAUU........
env-1818  ACAUUUAAGUUUACUUUUUAUUUUUUUAUAUUUAUUUUGUUGGAAUUGAAGAAGUUUGUACUUUUGCCGUCUGUAAAAAAUAACCUCUGACGA.UAA........UCGUGUAUGUUGCUUUUUAGUAAAAACAUUUAAUUCCAAUUUAUU........
env-1819  AUAAUGUAAAAAAAGUCUUGUAAAGACAAUAAAAAGUUUUAUUUUUGCAGCCGUCUACAAAAUAUUAGACGCUUUUAAUCCAACCUCUGGCGA.AAA........UCGUGGAUGUUGUUUUAAAAAUCAUAUUAACAAAUAAAUAAA........
env-1820  UAAUGUAAAAAAAAGUCUUGUAAAGACAAUAAAAAGUUUUUAUUUUUGCAGCCGUCUAUAAAAUAUUUAGACGCUUUUAAUCCAACCUCUGGCGA.AAA........UCGUGGAUGUUGUUUUAAAGAUCAUAUUAACAAACAAAUAAA........
env-1821  UAAUGCAAGAAAAGUCUUGUAAAGACAAUAAAAAAAAUUUACUUUUGCAGCCGUCUAUAAAAUAAUAGACACCUUUAAUCCAACCUCUGGCGA.AAA........UCGUGGAUGUUGUUUUAAAAAUCAUAUUAACAAACAAUAAA........
env-1822  UAAUGCAAAAAAAGUCUUGUAAAGCCAAUAAAAAUAUUUUACUUUUGCACCCGUCUACAAAAUAAUAGACAACUUUAAUCCAACCUCUGGCGA.AAA........UCGUGGAUGUUGUUUUAAAAAUCAUAUUAACAAACAAAUAAA........
env-1823  UAAUGCAAAAAAAGUCUUGUAAAGCCAAUAAAAAUAUUUUACUUUUGCACCCGUCUAUAAAAUAAUAGACAACUUUAAUCCAACCUCUGGCGA.AAA........UCGUGGAUGUUGUUUUAAAAAUCAUAUUAACAAACAAAUAAA........
env-1824  ..........................................................AAUAAUAGACAACUUUAAUCCAACCUCUGGCGA.AAA........UCGUGGAUGUUGUUUUAAAAAUCAUAUUAACAAACAAAUAAA........
env-1825  ACCUUUUAGAGUAGAAAAUUUUUGAUCGCAUACCUAUUGCCUACUUCGAAAUAAAGUUUAAUUUUGCAGCACCAAAAGAUCAACCUCUGACGA.AAA........UCGUGGAUGUUGACUUUUACAAUCUUAAAAAGAUCCG........
env-1826  GCAUUUAAAAUUGCCAUAAAAUAUUUACAAACCUGCAUAUUAUAAUCAAAUAAUUUUUUAUAAAUUUGCAAACUCAAAAUCAACCUCUGACGA.AAA........UCGUGCAUGUUGGUGAUUGAUCAACCAUUUUAAAAAAUGAUA........
env-1830  GCAUUUAAAAUUGCCAUAAAAUAUUUACAAACCUUCAUAUUAUAAUCAAAUAUUUUUUUAUAAAUUUGCAAACUCAAAAUCAACCUCUGACGA.AAA........UCGUGCAUGUUGGUGAUUGAUCAACCAUUUUAAAAAAUGAUA........
env-1831  UGUUUUCAAACAUCUUAAAAAUAUUUACUAUACAGCUUAUAAUAAUCAAAACUUUUUUUAUAAAUUUGCAAACUCAAAAUCAACCUCUGACGA.AAA........UCGUGCAUGUUGGUGAUUGAUAAACCAUUUUAAAAUAUAAUA........

          ........................................................<<<<...<.<<<<<..........>>>>>..>.>>>>........................................
          ........................................................2020...0.12220..........02221..0.0202........................................
          ●●●●●●●●●●●●●●●●●●●●●●●●●●●●●●●●●●●●●●●●●●●●●●●●●●●●●YY●●●●●●●●●●Y●RRRYYAACCUCUGRCG○-RRA○--------○CGUGRAUGUUG●UY●●●●●●AA●●●●●●●●●●●●●●●●○○○-------------------------------
                                                                                      5′                              3′
```

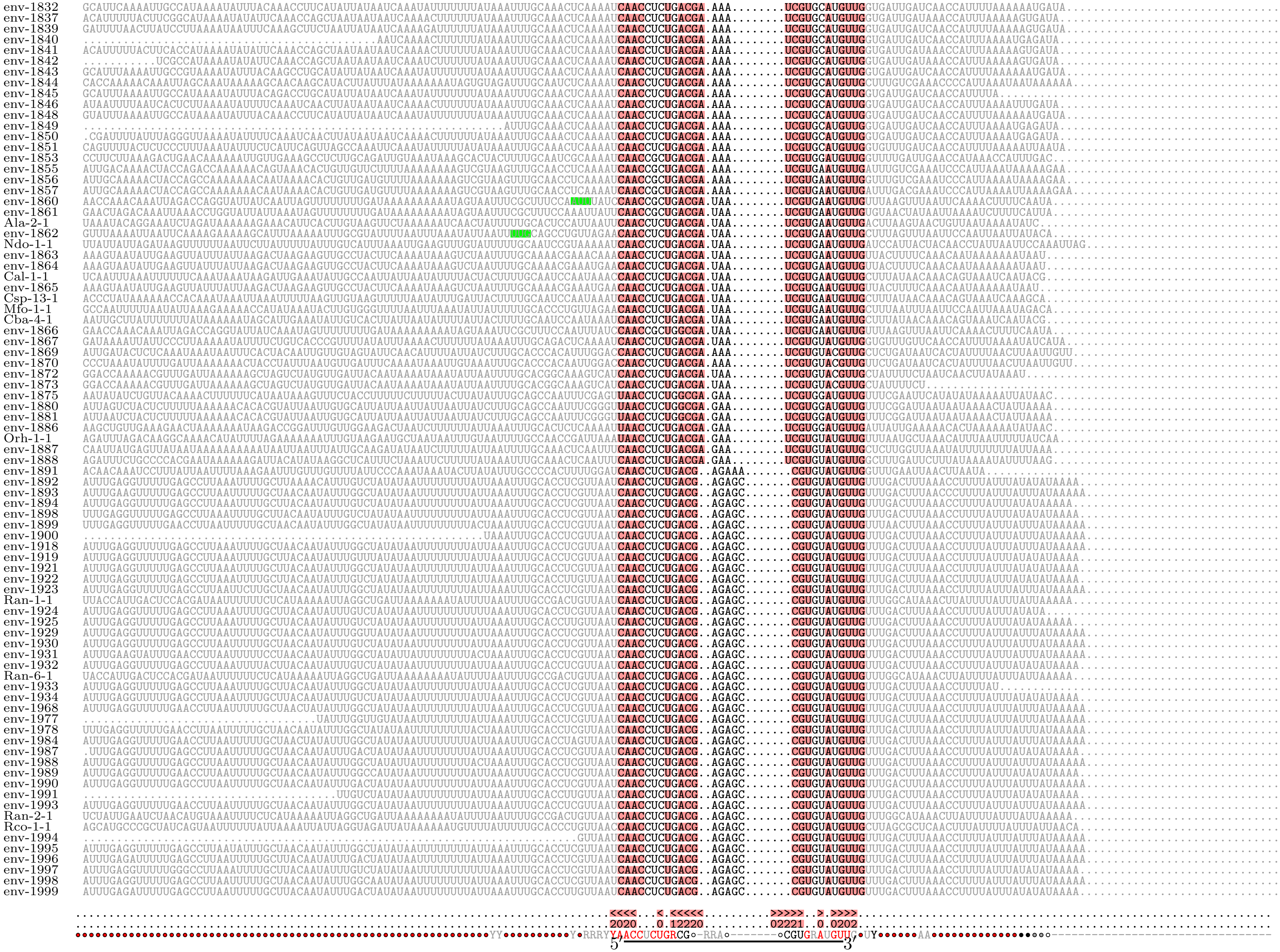
env-1832
env-1837
env-1839
env-1840
env-1841
env-1842
env-1843
env-1844
env-1845
env-1846
env-1848
env-1849
env-1850
env-1851
env-1853
env-1855
env-1856
env-1857
env-1860
env-1861
Ala-2-1
env-1862
Ndo-1-1
env-1863
env-1864
Cal-1-1
env-1865
Csp-13-1
Mfo-1-1
Cba-4-1
env-1866
env-1867
env-1869
env-1870
env-1872
env-1873
env-1875
env-1880
env-1881
env-1886
Orh-1-1
env-1887
env-1888
env-1891
env-1892
env-1893
env-1894
env-1898
env-1899
env-1900
env-1918
env-1919
env-1921
env-1922
env-1923
Ran-1-1
env-1924
env-1925
env-1929
env-1930
env-1931
env-1932
Ran-6-1
env-1933
env-1934
env-1968
env-1977
env-1978
env-1984
env-1987
env-1988
env-1989
env-1990
env-1991
env-1993
Ran-2-1
Rco-1-1
env-1994
env-1995
env-1996
env-1997
env-1998
env-1999
5′
3′

```
env-2000  AUUUGAGGUUUUUGAGCCUUAAUUUUUGCUAACAAUAUUUGGUUAUAUAAUUUUUUUUAUUAAAUUUGCACCUCGUUAAUCAACCUCUGACG..AGAGC.......CGUGUAUGUUGUUUGAUUUUAAACCUUUUAUUUAUUUAUAAAAA...........................
Ran-3-1   UCUAUUGAAUCUGACAUGUAAAUUUUCUCAUAAAAAAUUAGGCUGAUUAAAAAAAAAUAUUUUAAUUUUGCCGACUGUUAAUCAACCUCUGACG..AGAGC.......CGUGUAUGUUGUUUGGCAUAAACUUAUUUUAUUUAUUAAAAAA............................
env-2001  AUUUGAGGUUUUUGAGCCUUAAUUUUUGCUAACAAUAUUUGGCUAUAUAAUUUUUUUUAUUAAAUUUGCACCUCGUUAAUCAACCUCUGACG..AGAGC.......CGUGUAUGUUGUUUGACUUUAAACCUUUUAUUUAUUUAUAAAAA...........................
env-2032  ..............................................................AUUUGCACCUCGUUAAUCAACCUCUGACGA.GAG........UCGUGUAUGUUGUUUGACUUUAAACCUUUUAUUUAUAUAUAAAA............................
env-2033  ..................................AUAUUUGACUAUAUAAUUUUUUUUAUUAAAUUUGCACCUCGUUAAUCAACCUCUGACGA.GAG........UCGUGUAUGUUGUUUGACUUUAAACCUUUUAUUUAUUUAUAAAAA...........................
env-2034  AGUUGUAGAAUAAGAUGUAAUUUCUUCCCACAUCUGUUUGUUCAUUUCAAAAAAAGACUAUCUUUGCACUCGAUUUACACCAACCUCUGACGA.GAC........UCGUGAAUGUUGCUGUAACCAUAACAAUAUCAAAACCAUAACA.............................
env-2035  UAACUUCAUAGAAACUCAGAUAAAUUCUCAGGUUGUAUUGUGUACUAAAAAAAUAAGUUUAUUUUUGCACCACAAAAGAUCAACCACUGACGA.UAC........UCGUGUAUGUUGGCUUUUAAAACUUAAUUAUUAUCAUAG................................
Fge-1-1   ACUAUCGAAUAAUUGUAAAAAACUCAUAAUUUAUAUUUCAUAAUUCAUAAUUAUUUUCUACUUUUGCACCCAUAUUUGACCAACCUCUGACGA.UAC........UCGUGUAUGUUGCUUAGAUUCAACAAGUUAUUUAUAUUAUU..................................
Nma-1-1   UGAUUAUCCAAUAAACAUUCAAAUCCAUAUUAAAUUUUGCAGGAAUAGAUAUUAAUCCUAUUUUUGCAGCUCGUAAAAAUCAACCUCUGACGA.UAC........UCGUGAAUGUUGUUUGUUUACCUAACACAUAAUCUAUUAAG..................................
env-2036  ...........................................................AUAUUAAUUUGCAGCCUGUUAGAACAACCUCUGACGA.UAA........UCGUGAACGUUGCUCUGAUUUAAUUCCAAUUUUAACUAUACA...............................
env-2047  UAAGUGAAGAUUAACAACAUAAAAACAGGUUACGUGGUUGUGUAAAUGAAAAUAAGUUAUAAAUUUGCACCGCGUUAAAAUAACCUCUGGC...UCUCAUU......GUGAAUGUUGUUUUACUAACACUAAUUAACUAAA....................................
env-2048  UAAAUGAAGAUUAACAACAUAAAAACAGGUUACGUGGUUGUGUAAAUGAAAAUAAGUUAUAAAUUUGCACCGCGUUAAAAUAACCUCUGGC...UCUCAUU......GUGAAUGUUGUUUUACUAACACUAAUUAACUAAA....................................
env-2050  AAAACAAAAAAGAUUAGACCUGGAUUGCAUAAAUAAUAAUUUUAUUUUUAAUUCAAAUUGUGUAUGUUUGUGCCCCUAAUUAACCUCUGGAA..CUGU........UUUGAAUGUUGAUUAGACUAAUAAAAAUCAUAUAUU...................................
env-2051  AAAACAAAAAAGAUUAGACCUGGAUUGCUUAAAUAAUAAUUUUAUUUUUGAUUCAAAUUGUGUAUGUUUGUGCCCCUAAUUAACCUCUGGAA..CUGU........UUUGAAUGUUGAUUAGACUAAUAAAAAUCUUAAAUU...................................
env-2052  AAAACAAAAAAGGUUAGACCUGGAUUGCUUAAAAAAUAAUUUUAUUUUUAAUUCAAAUUGUGUAUGUUUGUGCCCCUAAUUAACCUCUGGAA..CUGU........UUUGAAUGUUGAUUAGAUUAAUAAAAAUCUUAAAUU...................................
env-2054  ......................................................................GCCCCUAAUUAACCUCUGGAA..CUGU........UUUGAAUGUUGAUUAGAUUAAUUAAAACCACAAAUU...................................
env-2055  ..................................AAUAAUUUUAUUUUUAAUUCAAAUUGUGUAUGUUUGUGCCCCUAAUUAACCUCUGGAA..CUGU........UUUGAAUGUUGAUUAGACUAAUAAAAAUCUUAAAUU...................................
env-2056  AAAACAAAAAAGAUUAGACCUGGAUUGCUUAAAUAAUAAUUUUAUUUUUAAUUCAAAUUGUGUAUGUUUGUGCCCCUAAUUAACCUCUGGAA..CUGU........UUUGAAUGUUGAUUAGACUAAUAAAAAUCAUAAAUU...................................
env-2057  AAAACAAAAAAGAUUAGACCUGGAUUGCUUAAAUAAUAAUUUUAUUUUUGAUACAAAUUGUGUAUGUUUGUGCCCCUAAUUAACCUCUGGAA..CUGU........UUUGAAUGUUGAUUAGACUAAUAAAAAUCAUAAAUU...................................
env-2065  AAACAAAAAAGAUUAGACCUGGAUUGCUUAAAUAAUAAUUUUAAUUUUUAAUUCAAAUUGUGUAUGUUUGUGCCCCUAAUUAACCUCUGGAA..CUGU........UUUGAAUGUUGAUUAGACUAAUAAAAAUCUUAAAUU...................................
env-2066  AAAACAAAAAAGAUUAGACCUGGAUUGCUUAAGUAAUAAUUUUAUUUUUGAUUCAAAUUGUGUAUGUUUGUGCCCCUAAUUAACCUCUGGAA..CUGU........UUUGAAUGUUGAUUAGACUAAUAAAAAUCUUAAAUU...................................
env-2067  AAACAAAGGUAAUUAAGCCUUAAUACUUAUAAAUAGUUUUUUUUUAACUAUAAAAAUAUGUAAGUUUGCCACCGCUUGAUUAACCUCUGGU...AUUU.........ACGAAUGUUGGUCAUUUUUAAAAUUAAAAUAAA.....................................
env-2068  AAAACAAAGGUAAUUAAGCCUUAAUACAUAUAAAUAGUUUUUUUUUAACUAUAAAAAUAUGUAAGUUUGCCACCGCUUGAUUAACCUCUGGU...AUUU.........ACGAAUGUUGGUCAUUUUUAAAAUUAAAAUAAA.....................................
env-2069  AAAACAAAGGUAAUUAAGCCUUAAUACUUAUAAAUAGUUUUUUUUUAACUAUAAAAAUAUGUAAGUUUGCCACCGCUUGAUUAACCUCUGGU...AUUU.........ACGAAUGUUGGUCAUUUUUAAAAUUAAAAUAAA.....................................
env-2071  AAAACAAAGGUAAUUAAGCCUUAAUAAUUAUAAAUAGUUUUUUUUUAACUAUAAAAAUAUGUAAGUUUGCCACCGCUUGAUUAACCUCUGGU...AUCU.........ACGAAUGUUGGUCAUUUUUAAAAUUAAAAUAAA.....................................
env-2076  AAAACGAAGGUAAUUAAGCCUUAAUACUUAUAAAUAGUUUUUUUUUAACUAUAAAAAUAUGUAAGUUUGCCACCGCUUGAUUAACCUCUGGU...AUCU.........ACGAAUGUUGGUCAUUUUUAAAAUUAAAAUAAA.....................................
env-2077  AAAACAAAGGUAAUUGAGCCUUAAUAGUUAUAAAUAGUUUUUUUUUAACUAUAAAAAUAUGUAAGUUUGCCACCGCUUGAUUAACCUCUGGU...AUCU.........ACGAAUGUUGGUCAUUUUUAAAAUUAAAAUAAA.....................................
env-2078  AAAACAAAGGUAAUUAAGCCUUAAUACAUAUAAAUAGUUUUUUUUUAACUAUAAAAAUAUGUAAGUUUGCCACCGCUUGAUUAACCUCUGGU...AUCU.........ACGAAUGUUGGUCAUUUUUAAAAUA............................................
env-2079  AAAAACAAAGGUAAUUAAGCCUUAAUACUUAUAAAUAGUUUUUUUUUAACUAUAAAAAUAUGUAAGUUUGCCACCGCUUGAUUAACCUCUGGU...AUCU.........ACGAAUGUUGGUCAUUUUUAAAAUUAAAAUAAA.....................................
env-2080  AAAACAAAGGUAAUUAAGCCUUAAUACAUAUAAAUAGUUUUUUUUUAACUAUAAAAAUAUGUAAGUUUGCCACCGCUUGAUUAACCUCUGGU...AUCU.........ACGAAUGUUGGUCAUUUUUAAAAUUAAAAUAAA.....................................
env-2098  AAAACAAAGCUAAUUAAGCCUUAAUACAUGUAAAUAGUUUUUUUUUAACUAUAAAAAUAUGUAAGUUUGCCACCGCUUGAUUAACCUCUGGU...AUCU.........ACGAAUGUUGGUCAUUUUUAAAAUUAAAAUAAA.....................................
env-2100  AAAACAAAGGUAAUUAAGCCUUAAUACAUAUAAAUAGUUUUUUUUUAACUAUAAAAAUAUGUAAGUUUGCCACCGCUUGAUUAACCUCUGGU...AUCU.........ACGAAUGUUGGUCAUUUAUAAAAUUAAAAUAAA.....................................
env-2101  AAAACAAAGGUAAUUAAGCCUUAAUACUUACAAAUAGUUUUUUUUUAACUAUAAAAAUAUGUAAGUUUGCCACCGCUUGAUUAACCUCUGGU...AUCU.........ACGAAUGUUGGUCAUUUUUAAAAUUAAAAUAAA.....................................
env-2102  AAACAAAGGUAAUUAAGCCUUAAUACAUUAUAAAUAGUUUUUUUUUAACUAUAAAAAUAUGUAAGUUUGCCACCGCUUGAUUAACCUCUGGU...AUCU.........ACGAAUGUUGGUCAUUUUUAAAAUUAAAAUAAA.....................................
env-2103  AACAAAGGAAAUUAAGCCUUAAUUACUUUAGAAAUAGUUUUUUUUUAACUUUAAAAAUAUGUAAGUUUGCCUCCGCUUGAUUAACCUCUGGU...AUCU.........ACGAAUGUUGGUCAUUUUAAAAUAAAAUAAA.......................................
env-2104  AAAACAAAGGUAAUUAAGCCUUAAUACUUAUAAAUAGUUUUUUUUUAACUAUAAAAAUAUGUAAGUUUGCCACCGCUUGAUUAACCUCUGGU...AUCU.........ACGAAUGUUGGUCAUUUUUAAAAUUAAAAUAAA.....................................
env-2121  AAAACAAAGGUAAUUAAGCCUUAAUACUUAUAAAUAGUUUUUUUUUAACUAUAAAAAUAUGUAAGUUUGCCACCGCUUGAUUAACCUCUGGU...UUCU.........ACGAAUGUUGGUCAUUUUUAAAAUUAAAAUAAA.....................................
env-2122  AAAACGAAGGUAAUUAAGCCUUAAUACAUAUAAAUAGUUUUUUUUUAACUAUAAAAAUAUGUAAGUUUGCCACCGCUUGAUUAACCUCUGGU...UUCU.........ACGAAUGUUGGUCAUUUAUAAAAUUAAAAUAAA.....................................
env-2123  AAAACAAAGGUAAUUAAGCCUUAAUACAUAUAAAUAGUUUUUUUUUAACUAUAAAAAUAUGUAAGUUUGCCACCGCUUGAUUAACCUCUGGU...UUCU.........ACGAAUGUUGGUCAUUUUUAAAAUUAAAAUAAA.....................................
Fps-4-1   AUGAAACUUUACAGUUAUAAAAAUUAUCAAUUAUUCAUUGUAAAUUAUUAAUAAUUUCUUACUUUUGUGGGCUAUUAGACCAACCUCUGGCG..AGAUC.......CGUGAAUGUUGCUCUGAUUUACAACCAUAAUUAAAUAAUUAUC............................
Ran-4-1   UACCAUUGACUCCACGAUAAUUUUUUCUCAUAAAAAUUAGGCCGAUUAAAAAAAAAUAUUUUAAUUUUGCCGACUGUUAAUCAACCUCUGACG..AGAGC.......CGUGUAUGUUGUUUGGCAUAAACUUAUUUUAUUUAUUAAAA...............................
          ...............................................................................<<<<...<.<<<<<..........>>>>>..>.>>>>...............................................................
          ...............................................................................2020...0.12220..........02221..0.0202...............................................................
          ●●●●●●●●●●●●●●●●●●●●●●●●●●●●●●●●●●●●●●●●●●●●●●●●●●●●●●●●●●●●●●●●●●YY●●●●●●●●●●Y●RRRYYAACCUCUGRCG○-RRA○-------○CGUGRAUGUUG●UY●●●●●●AA●●●●●●●●●●●●●●●●●○○○------------------------------
                                                                                        5′                                  3′
```

alignment positions 180 · · · 190

```
env-1     ........AUG
env-2     ........AUG
env-3     ........AUG
env-4     ........AUG
env-5     ........AUG
env-6     ........AUG
env-7     ........AUG
env-8     ...........
env-9     ........AUG
env-10    ........AUG
env-27    ........AUG
env-29    ........AUG
env-30    ........AUG
env-31    ........AUG
env-33    ........AUG
env-34    ........AUG
env-35    ........AUG
env-36    ........AUG
env-38    ...........
env-40    ........AUG
env-41    ........AUG
env-43    ........AUG
env-44    ........AUG
env-45    ........AUG
env-46    ........AUG
env-47    ........AUG
env-49    ........AUG
env-51    ........AUG
env-54    ........AUG
env-55    ........AUG
env-56    ........AUG
env-57    ........AUG
env-58    ........AUG

          ...........
          ...........
          --------AUG
```

| | |
|---|---|
| env-59 | ........AUG |
| env-60 | ........AUG |
| env-61 | ........AUG |
| env-63 | ........AUG |
| env-64 | ........AUG |
| env-65 | ........AUG |
| env-67 | ........AUG |
| Nul-1-1 | ........AUG |
| Fba-1-1 | ........AUG |
| Nul-1-2 | ........AUG |
| Nul-1-3 | ........AUG |
| Nul-1-4 | ........AUG |
| Nul-1-6 | ........AUG |
| env-68 | ........AUG |
| Csp-18-1 | ........AUG |
| Csp-15-1 | ........AUG |
| Csp-16-1 | ........AUG |
| Chi-1-1 | ........AUG |
| env-69 | ........AUG |
| Cgr-2-1 | ........AUG |
| Csp-17-1 | ........AUG |
| Fsp-5-1 | ........AAU |
| env-70 | ........AUG |
| env-71 | ........AUG |
| env-73 | ........AUG |
| env-74 | ........AUG |
| env-75 | ........AUG |
| env-76 | ........AUG |
| env-78 | ........AUG |
| env-79 | ........AUG |
| env-81 | ........AUG |
| env-86 | ........AUG |
| env-87 | ........AUG |
| env-88 | ........AUG |
| env-89 | ........AUG |
| env-90 | ........AUG |
| env-91 | ........AUG |
| env-92 | ........... |
| env-93 | ........... |
| env-107 | ........AUG |
| env-117 | ........... |
| env-118 | ........AUG |
| env-120 | ........AUG |
| env-123 | ........AUG |
| env-124 | ........AUG |
| env-126 | ........AUG |
| env-127 | ........AUG |
| env-128 | ........AUG |
| Oho-1-1 | ........AUG |
| env-134 | ........AUG |
| env-136 | ........AUG |
| env-137 | ........AUG |
| env-138 | ........AUG |
| env-139 | ........AUG |
| env-140 | ........AUG |
| env-141 | ........AUG |
| env-143 | ........AUG |
| env-144 | ........AUG |
| env-146 | ........AUG |
| Nse-1-1 | ........AUG |
| env-147 | ........AUG |
| env-149 | ........AUG |
| env-150 | ........AUG |
| env-156 | ........AUG |
| env-178 | ........AUG |
| env-179 | ........AUG |
| env-180 | ........AUG |
| env-182 | ........AUG |
| env-183 | ........AUG |
| env-184 | ........AUG |
| env-185 | ........AUG |
| env-186 | ........AUG |
| env-188 | ........AUG |
| env-194 | ........AUG |
| env-195 | ........AUG |
| env-197 | ........AUG |
| env-200 | ........AUG |
| env-202 | ........AUG |
| Rbi-1-1 | ........AUG |
| env-203 | ........AUG |
| env-204 | ........AUG |
| env-207 | ........AUG |
| env-208 | ........AUG |
| env-210 | ........AUG |
| | ........... |
| | ........... |
|  | --------AUG |

```
env-211     ........AUG
Gsp-1-1     ........AUG
env-212     ...........
env-214     ...........
env-215     ........AUG
env-216     ........AUG
Man-1-1     ........AUG
env-217     ........AUG
env-218     ........AUG
env-219     ........AUG
env-220     ........AUG
env-222     ........AUG
env-223     ........AUG
env-224     ........AUG
env-226     ........AUG
env-227     ........AUG
env-242     ........AUG
env-244     ........AUG
env-246     ........AUG
env-247     ........AUG
env-248     ........AUG
env-249     ........AUG
env-256     ........AUG
env-273     ........AUG
env-274     ........AUG
env-275     ........AUG
env-277     ........AUG
env-280     ........AUG
env-281     ........AUG
env-282     ........AUG
env-283     ........AUG
env-310     ........AUG
env-312     ........AUG
env-313     ...........
env-323     ........AUG
env-325     ........AUG
env-326     ........AUG
env-327     ........AUG
env-330     ........AUG
env-331     ........AUG
env-332     ........AUG
env-356     ........AUG
env-380     ........AUG
env-381     ........AUG
env-385     ........AUG
env-390     ........AUG
env-391     ........AUG
env-392     ........AUG
env-393     ........AUG
env-394     ...........
env-395     ........AUG
env-396     ........AUG
env-400     ........AUG
Aal-1-1     ........AUG
env-401     ........AUG
env-402     ........AUG
env-403     ........AUG
env-404     ........AUG
env-405     ........AUG
Cpa-1-1     ........AUG
env-406     ........AUG
env-407     ........AUG
env-411     ........AUG
Cte-1-1     ........AUG
env-412     ........AUG
env-413     ........AUG
Cca-2-1     ........AUG
env-414     ........AUG
env-416     ........AUG
env-418     ........AUG
Fba-3-1     ........AUG
Cha-1-1     ........AUG
env-419     ........AUG
Cko-1-1     ........AUG
Csp-14-1    ........AUG
env-420     ........AUG
env-421     ........AUG
env-422     ........A..
env-424     ........AUG
env-425     ........AUG
env-426     ........AUG
env-427     ........AUG
env-428     ........AUG
env-430     ........AUG

            ...........
            ...........
            --------AUG
```

| | |
|---|---|
| Sze-1-1 | ........AUG |
| env-431 | ........AUG |
| env-432 | ........AUG |
| env-433 | ........AUG |
| env-434 | ........AUG |
| env-435 | ........AUG |
| env-436 | ........AUG |
| env-437 | ........AUG |
| env-438 | ........AUG |
| env-439 | ........AUG |
| env-440 | ........AUG |
| env-441 | ........AUG |
| env-442 | ........AUG |
| env-451 | ........... |
| env-452 | ........AUG |
| env-453 | ........AUG |
| env-454 | ........AUG |
| env-455 | ........AUG |
| env-456 | ........AUG |
| env-457 | ........AUG |
| env-460 | ........AUG |
| env-461 | ........AUG |
| env-462 | ........AUG |
| env-463 | ........AUG |
| env-464 | ........AUG |
| env-470 | ........AUG |
| env-471 | ........AUG |
| env-472 | ........AUG |
| env-473 | ........... |
| env-474 | ........AUG |
| env-475 | ........AUG |
| env-476 | ........AUG |
| env-477 | ........AUG |
| env-479 | ........AUG |
| env-480 | ........AUG |
| env-481 | ........AUG |
| env-483 | ........AUG |
| env-485 | ........AUG |
| env-486 | ........AUG |
| env-487 | ........AUG |
| env-493 | ........AUG |
| env-496 | ........AUG |
| env-497 | ........AUG |
| env-498 | ........AUG |
| env-499 | ........AUG |
| env-500 | ........... |
| env-501 | ........AUG |
| env-502 | ........AUG |
| env-503 | ........AUG |
| env-504 | ........AUG |
| env-505 | ........AUG |
| env-506 | ........AUG |
| env-508 | ........AUG |
| env-509 | ........AUG |
| env-510 | ........AUG |
| env-512 | ........AUG |
| env-513 | ........AUG |
| env-514 | ........AUG |
| env-515 | ........AUG |
| env-517 | ........AUG |
| env-518 | ........AUG |
| env-519 | ........AUG |
| env-520 | ........AUG |
| env-521 | ........AUG |
| env-522 | ........AUG |
| env-523 | ........AUG |
| env-524 | ........AUG |
| Tma-1-1 | ........AUG |
| env-525 | ........AUG |
| env-526 | ........AUG |
| env-527 | ........AUG |
| env-529 | ........AUG |
| env-530 | ........AUG |
| env-531 | ........AUG |
| env-532 | ........AUG |
| env-533 | ........AUG |
| env-535 | ........AUG |
| env-536 | ........AUG |
| env-537 | ........AUG |
| env-538 | ........AUG |
| env-540 | ........AUG |
| env-548 | ........AUG |
| env-555 | ........AUG |
| env-557 | ........AUG |
| | ........... |
| | ........... |
|  | --------AUG |

| | |
|---|---|
| env-558 | ........AUG |
| env-559 | ........AUG |
| env-562 | ........AUG |
| env-565 | ........AUG |
| env-566 | ........AUG |
| env-570 | ........AUG |
| env-572 | ........AUG |
| env-573 | ........... |
| Fba-5-1 | ........AUG |
| env-574 | ........AUG |
| env-575 | ........AUG |
| env-577 | ........AUG |
| env-578 | ........AUG |
| Tme-1-1 | ........AUG |
| Zul-1-1 | ........AUG |
| env-579 | ........AUG |
| env-580 | ........AUG |
| env-581 | ........AUG |
| env-585 | ........AUG |
| env-587 | ........AUG |
| env-588 | ........AUG |
| env-590 | ........AUG |
| env-591 | ........AUG |
| env-592 | ........AUG |
| Zga-1-1 | ........AUG |
| env-607 | ........AUG |
| env-608 | ........AUG |
| env-610 | ........AUG |
| env-611 | ........AUG |
| Tov-1-1 | ........AUG |
| env-612 | ........AUG |
| env-614 | ........AUG |
| Ace-1-1 | ........AUG |
| env-615 | ........AUG |
| env-616 | ........... |
| env-617 | ........AUG |
| env-623 | ........AUG |
| env-625 | ........AUG |
| env-626 | ........AUG |
| env-627 | ........AUG |
| env-628 | ........AUG |
| env-629 | ........AUG |
| env-630 | ........AUG |
| env-631 | ........AUG |
| env-636 | ........AUG |
| env-639 | ........AUG |
| Fsp-12-1 | ........AUG |
| env-640 | ........AUG |
| env-645 | ........AUG |
| env-651 | ........AUG |
| env-652 | ........AUG |
| env-653 | ........AUG |
| env-654 | ........AUG |
| env-655 | ........AUG |
| env-656 | ........AUG |
| env-657 | ........AUG |
| env-663 | ........AUG |
| env-664 | ........AUG |
| env-667 | ........AUA |
| env-668 | ........AUG |
| env-669 | ........AUG |
| env-670 | ........... |
| env-671 | ........AUG |
| env-672 | ........AUG |
| env-673 | ........AUG |
| env-677 | ........AUG |
| env-678 | ........AUG |
| env-679 | ........AUG |
| env-680 | ........AUG |
| env-682 | ........AUG |
| env-683 | ........AUG |
| env-684 | ........AUG |
| env-685 | ........AUG |
| env-686 | ........AUG |
| Ean-11-1 | ........AUG |
| Eme-1-1 | ........AUG |
| Ean-1-1 | ........AUG |
| Ean-1-2 | ........AUG |
| Eme-2-1 | ........AUG |
| env-688 | ........AUG |
| env-689 | ........AUG |
| env-690 | ........AUG |
| env-692 | ........AUG |
| env-693 | ........AUG |
| | ........... |
| | ........... |
|  | --------AUG |

```
env-694    ........AUG
env-695    ........AUG
env-697    ........AUG
env-699    ........AUG
env-700    ........AUG
env-702    ........AUG
env-703    ........AUG
env-704    ........AUG
env-707    ........AUG
Gfo-1-1    ........AAU
env-710    ........AUG
env-711    ........AUG
env-712    ........AUG
env-713    ........AUG
env-714    ........AUG
env-715    ........AUG
env-717    ........AUG
env-718    ........AUG
env-719    ........AUG
env-720    ........AUG
env-722    ........AUG
env-723    ........AUG
env-724    ........AUG
env-725    ........AUG
env-726    ........AUG
env-727    ........AUG
Tna-1-1    ........AUG
Pbu-1-1    ........AUG
env-728    ........AUG
env-732    ........AUG
env-734    ........AUG
env-737    ........AUG
env-739    ...........
env-740    ........AUG
Ale-1-1    ........AUG
Fic-1-1    ........AUG
env-741    ........AUG
env-742    ........AUG
env-743    ........AUG
env-744    ........AUG
env-746    ........AUG
env-747    ........AUG
env-748    ........AUG
env-750    ........AUG
env-752    ........AUG
Ale-1-2    ........AUG
Wsp-1-1    ........AUG
env-755    ........AUG
env-756    ........AUG
env-761    ........AUG
env-762    ........AUG
env-764    ........AUG
env-765    ........AUG
env-766    ........AUG
Pda-1-1    ........AUG
Fba-7-1    ........AUG
env-767    ........AUG
env-771    ........AUG
env-772    ........AUG
env-773    ........AUG
env-774    ........AUG
env-779    ........AUG
env-780    ........AUG
env-781    ........AUG
env-783    ........AUG
env-784    ........AUG
Tse-1-1    ........AUG
env-785    ........AUG
env-788    ........AUG
env-790    ........AUG
Psp-4-1    ........AUG
Fsp-13-1   ........AUG
env-791    ........AUG
env-793    ........AUG
env-794    ........AUG
Jpa-1-1    ........AUG
Pja-1-1    ........UGA
env-795    ........AUG
env-796    ........AUG
env-797    ........AUG
Pme-1-1    ........AUG
env-798    ........AUG
env-799    ........AUG
env-800    ........AUG

           ...........
           ...........
           --------AUG
```

| | |
|---|---|
| env-804 | ........AUG |
| env-805 | ........AUG |
| env-806 | ........AUG |
| env-807 | ........AUG |
| env-809 | ........AUG |
| env-810 | ........AUG |
| env-811 | ........AUG |
| env-812 | ........AUG |
| env-813 | ........AUG |
| env-818 | ........... |
| env-819 | ........AUG |
| env-820 | ........AUG |
| env-821 | ........AUG |
| env-823 | ........AUG |
| Msp-1-1 | ........AUG |
| env-824 | ........AUG |
| Ksp-1-1 | ........AUG |
| env-825 | ........AUG |
| env-826 | ........AUG |
| Ddo-1-1 | ........AUG |
| Dsp-1-1 | ........AUG |
| env-833 | ........AUG |
| env-839 | ........AUG |
| env-840 | ........AUG |
| env-841 | ........AUG |
| env-842 | ........... |
| env-843 | ........AUG |
| env-847 | ........AUG |
| env-850 | ........AUG |
| env-851 | ........AUG |
| env-860 | ........AUG |
| env-862 | ........AUG |
| env-864 | ........AUG |
| env-865 | ........AUG |
| env-866 | ........AUG |
| env-867 | ........AUG |
| env-868 | ........AUG |
| env-869 | ........AUG |
| env-870 | ........AUG |
| env-881 | ........AUG |
| env-882 | ........AUG |
| env-883 | ........AUG |
| env-884 | ........AUG |
| env-889 | ........AUG |
| env-890 | ........AUG |
| env-891 | ........AUG |
| env-892 | ........AUG |
| env-893 | ........AUG |
| env-894 | ........AUG |
| env-906 | ........AUG |
| env-907 | ........AUG |
| env-910 | ........AUG |
| env-913 | ........AUG |
| env-914 | ........AUG |
| env-915 | ........AUG |
| env-916 | ........AUG |
| env-917 | ........AUG |
| env-918 | ........... |
| Fba-4-1 | ........AUG |
| env-919 | ........AUG |
| env-920 | ........AUG |
| env-923 | ........AUG |
| env-924 | ........AUG |
| env-925 | ........AUG |
| env-926 | ........AUG |
| env-927 | ........AUG |
| env-928 | ........AUG |
| Ffi-1-1 | ........AUG |
| env-929 | ........AUG |
| Fba-8-1 | ........AUG |
| env-932 | ........AUG |
| env-933 | ........AUG |
| env-938 | ........... |
| Cca-1-1 | ........AUG |
| env-939 | ........AUG |
| env-940 | ........... |
| env-941 | ........AUG |
| env-942 | ........AUG |
| Emi-1-1 | ........AUG |
| Wma-1-1 | ........AUG |
| env-943 | ........AUG |
| env-944 | ........AUG |
| env-945 | ........AUG |
| env-947 | ........AUG |
| | ........... |
| | ........... |
|  | --------AUG |

| | |
|---|---|
| env-948 | ........AUG |
| env-949 | ........AUG |
| env-950 | ........AUG |
| env-951 | ........AUG |
| env-954 | ........AUG |
| env-955 | ........AUG |
| env-956 | ........AUG |
| env-957 | ........AUG |
| env-959 | ........AUG |
| env-968 | ........AUG |
| env-969 | ........AUG |
| env-970 | ........AUG |
| env-974 | ........AUG |
| env-975 | AACAAGAAAUG |
| env-976 | ........AUG |
| env-977 | ........AUG |
| env-980 | ........AUG |
| env-983 | ........AUG |
| env-988 | ........AUG |
| Psp-2-1 | ........AUG |
| env-989 | ........AUG |
| env-991 | ........AUG |
| env-992 | ........AUG |
| env-994 | ........AUG |
| env-997 | ........AUG |
| env-998 | ........AUG |
| env-999 | ........AUG |
| Pir-1-1 | ........AUG |
| env-1003 | ........AUG |
| env-1007 | ........AUG |
| env-1009 | ........AUG |
| env-1010 | ........AUG |
| env-1011 | ........AUG |
| env-1012 | ........AUG |
| env-1015 | ........AUG |
| env-1016 | ........AUG |
| env-1017 | ........AUG |
| env-1019 | ........AUG |
| env-1021 | ........AUG |
| env-1023 | ........AUG |
| env-1025 | ........AUG |
| env-1026 | ........AUG |
| env-1027 | ........AUG |
| env-1029 | ........AUG |
| env-1034 | ........AUG |
| env-1038 | ........AUG |
| env-1039 | ........... |
| env-1040 | ........AUG |
| env-1042 | ........AUG |
| env-1047 | ........AUG |
| env-1048 | ........AUG |
| env-1049 | ........AUG |
| env-1052 | ........AUG |
| env-1053 | ........AUG |
| env-1057 | ........AUG |
| env-1058 | ........AUG |
| Wvi-1-1 | ........AUG |
| env-1059 | ........AUG |
| env-1060 | ........AUG |
| Fbr-1-1 | ........AUG |
| env-1062 | ........AUG |
| env-1063 | ........AUG |
| env-1064 | ........AUG |
| env-1066 | ........AUG |
| env-1067 | ........... |
| env-1068 | ........AUG |
| env-1069 | ........AUG |
| Iha-1-1 | ........AUG |
| env-1070 | ........AUG |
| env-1072 | ........AUG |
| env-1073 | ........AUG |
| env-1074 | ........AUG |
| Fjo-1-1 | ........AUG |
| env-1075 | ........AUG |
| env-1076 | ........AUG |
| env-1080 | ........AUG |
| env-1081 | ........AUG |
| env-1084 | ........AUG |
| Sxi-1-1 | ........AUG |
| env-1085 | ........... |
| env-1086 | ........AUG |
| env-1088 | ........AUG |
| env-1089 | ........AUG |
| Gli-1-1 | ........AUG |
| | ........... |
| | ........... |
|  | --------AUG |

| | |
|---|---|
| env-1091 | ........AUG |
| env-1092 | ........AUG |
| env-1095 | ........AUG |
| env-1096 | ........AUG |
| env-1097 | ........... |
| env-1098 | ........AUG |
| env-1099 | ........AUG |
| env-1104 | ........AUG |
| env-1106 | ........AUG |
| env-1107 | ........AUG |
| Aag-1-1 | ........AUG |
| Ead-1-1 | ........AUG |
| env-1114 | ........AUG |
| env-1115 | ........AUG |
| Lsp-2-1 | ........AUG |
| env-1116 | ........AUG |
| Lsp-3-1 | ........AUG |
| env-1118 | ........AUG |
| env-1121 | ........AUG |
| Lja-1-1 | ........AUG |
| Lsp-1-1 | ........AUG |
| env-1124 | ........... |
| Pto-1-1 | ........AUG |
| Pgo-1-1 | ........AUG |
| env-1125 | ........AUG |
| env-1127 | ........AUG |
| env-1128 | ........... |
| Ptr-1-1 | ........AUG |
| env-1129 | ........AUG |
| env-1130 | ........AUG |
| env-1131 | ........AUG |
| env-1132 | ........AUG |
| env-1134 | ........AUG |
| env-1135 | ........AUG |
| env-1136 | ........AUG |
| Gsp-4-1 | ........AUG |
| env-1137 | ........... |
| env-1138 | ........AUG |
| env-1139 | ........AUG |
| env-1140 | ........... |
| env-1141 | ........AUG |
| env-1142 | ........AUG |
| env-1143 | ........AUG |
| env-1144 | ........AUG |
| env-1145 | ........AUG |
| env-1146 | ........AUG |
| env-1147 | ........AUG |
| env-1148 | ........AUG |
| env-1149 | ........AUG |
| env-1150 | ........AUG |
| env-1151 | ........AUG |
| env-1152 | ........AUG |
| env-1153 | ........AUG |
| env-1155 | ........AUG |
| env-1161 | ........AUG |
| env-1162 | ........AUG |
| env-1163 | ........AUG |
| env-1164 | ........AUG |
| env-1165 | ........AUG |
| env-1166 | ........AUG |
| env-1167 | ........AUG |
| env-1168 | ........AUG |
| Fso-1-1 | ........AUG |
| env-1169 | ........AUG |
| env-1172 | ........AUG |
| env-1174 | ........... |
| env-1175 | ........AUG |
| env-1181 | ........... |
| env-1182 | ........AUG |
| env-1183 | ........AUG |
| env-1185 | ........AUG |
| env-1190 | ........AUG |
| env-1191 | ........AUG |
| env-1196 | ........AUG |
| env-1197 | ........AUG |
| env-1199 | ........AUG |
| env-1200 | ........AUG |
| env-1202 | ........... |
| env-1203 | ........AUG |
| Apa-1-1 | ........AUG |
| env-1205 | ........AUG |
| env-1206 | ........... |
| env-1207 | ........AUG |
| env-1208 | ........AUG |

...........
...........
--------AUG

| | |
|---|---|
| Fba-6-1 | ........AUG |
| Aat-1-1 | ........AUG |
| env-1209 | ........AUG |
| env-1210 | ........AUG |
| env-1213 | ........AUG |
| env-1215 | ........AUG |
| env-1216 | ........AUG |
| env-1217 | ........AUG |
| Fag-1-1 | ........AUG |
| env-1219 | ........... |
| env-1220 | ........AUG |
| env-1221 | ........AUG |
| env-1222 | ........AUG |
| env-1223 | ........... |
| Ame-1-1 | ........AUG |
| env-1224 | ........AUG |
| env-1225 | ........AUG |
| Fsp-3-1 | ........AUG |
| env-1227 | ........AUG |
| env-1228 | ........AUG |
| env-1230 | ........AUG |
| env-1231 | ........AUG |
| env-1232 | ........AUG |
| env-1233 | ........AUG |
| env-1234 | ........AUG |
| env-1235 | ........AUG |
| env-1237 | ........AUG |
| env-1239 | ........AUG |
| env-1241 | ........AUG |
| env-1252 | ........AUG |
| env-1253 | ........AUG |
| env-1254 | ........AUG |
| env-1255 | ........AUG |
| env-1256 | ........AUG |
| Fsu-1-1 | ........AUG |
| env-1257 | ........AUG |
| env-1258 | ........AUG |
| env-1259 | ........AUG |
| env-1260 | ........AUG |
| env-1261 | ........AUG |
| env-1262 | ........AUG |
| Msp-2-1 | ........AUG |
| env-1263 | ........AUG |
| Fco-1-1 | ........AUG |
| Ala-1-1 | ........AUG |
| env-1264 | ........AUG |
| env-1265 | ........AUG |
| env-1267 | ........AUG |
| Wps-1-1 | ........AUG |
| env-1268 | ........AUG |
| env-1269 | ........UGA |
| env-1271 | ........AUG |
| Mze-1-1 | ........AUG |
| env-1272 | ........AUG |
| Gpo-1-1 | ........AUG |
| Oma-1-1 | ........AUG |
| env-1273 | ........AUG |
| env-1274 | ........AUG |
| env-1275 | ........AUG |
| env-1276 | ........AUG |
| Fgi-1-1 | ........AUG |
| env-1280 | ........UGG |
| Fde-1-1 | ........AUG |
| Fsp-8-1 | ........AUG |
| Ffr-2-1 | ........AUG |
| Fsp-7-1 | ........AUG |
| Fch-1-1 | ........AUG |
| Fsp-2-1 | ........AUG |
| Ffr-1-1 | ........AUG |
| Fre-1-1 | ........AUG |
| env-1281 | ........AUG |
| Osp-1-1 | ........AUG |
| Osp-2-1 | ........AUG |
| env-1282 | ........AUG |
| Ssp-2-1 | ........AUG |
| env-1286 | ........AUG |
| Fsp-4-1 | ........AUG |
| Fsp-10-1 | ........AUG |
| env-1287 | ........AUG |
| env-1289 | ........AUG |
| Fsp-9-1 | ........AUG |
| env-1291 | ........AUG |
| env-1293 | ........AUG |
| env-1296 | ........AUG |
| | ........... |
| | ........... |
|  | --------AUG |

| | |
|---|---|
| env-1297 | ........AUG |
| env-1300 | ........... |
| env-1301 | ........AUG |
| env-1302 | ........AUG |
| env-1303 | ........AUG |
| Fhi-1-1 | ........AUG |
| Gsp-3-1 | ........AUG |
| env-1304 | ........AUG |
| env-1305 | ........AUG |
| env-1306 | ........AUG |
| Gsp-5-1 | ........AUG |
| Fsp-11-1 | ........AUG |
| Fsp-1-1 | ........AUG |
| env-1307 | ........AUG |
| Psp-3-1 | ........AUG |
| env-1308 | ........... |
| env-1309 | ........AUG |
| env-1310 | ........AUG |
| Gme-1-1 | ........AUG |
| env-1311 | ........... |
| env-1312 | ........AUG |
| env-1314 | ........AUG |
| Min-1-1 | ........AUG |
| env-1317 | ........AUG |
| env-1318 | ........AUG |
| env-1319 | ........AUG |
| env-1320 | ........AUG |
| Ssp-1-1 | ........AUG |
| env-1321 | ........AUG |
| Fsu-2-1 | ........AUG |
| env-1322 | ........AUG |
| env-1323 | ........AUG |
| env-1326 | ........AUG |
| env-1327 | ........AUG |
| Mze-2-1 | ........AUG |
| env-1328 | ........... |
| Myu-1-1 | ........AUG |
| Gsa-1-1 | ........AUG |
| Amu-1-1 | ........AUG |
| Fen-1-1 | ........AUG |
| Fli-1-1 | ........AUG |
| Fca-1-1 | ........AUG |
| Fsa-1-1 | ........AUG |
| Fse-1-1 | ........AUG |
| Fda-1-1 | ........AUG |
| Dsp-2-1 | ........AUG |
| env-1329 | ........AUG |
| env-1332 | ........AUG |
| env-1335 | ........AUG |
| env-1336 | ........AUG |
| env-1337 | ........AUG |
| env-1338 | ........AUG |
| Faq-1-1 | ........AUG |
| env-1339 | ........AUG |
| env-1340 | ........AUG |
| env-1341 | ........AUG |
| env-1342 | ........AUG |
| Zam-1-1 | ........AUG |
| Gec-1-1 | ........AUG |
| env-1343 | ........AUG |
| env-1344 | ........AUG |
| env-1347 | ........AUG |
| env-1348 | ........AUG |
| env-1349 | ........AUG |
| env-1351 | ........AUG |
| env-1352 | ........AUG |
| env-1357 | ........AUG |
| env-1359 | ........AUG |
| env-1360 | ........AUG |
| env-1361 | ........... |
| env-1364 | ........AUG |
| env-1366 | ........AUG |
| env-1367 | ........AUG |
| env-1368 | ........AUG |
| env-1369 | ........AUG |
| env-1370 | ........AUG |
| Jma-1-1 | ........AUG |
| env-1374 | ........AUG |
| env-1375 | ........AUG |
| env-1377 | ........AUG |
| env-1380 | ........AUG |
| env-1381 | ........AUG |
| env-1382 | ........AUG |
| env-1383 | ........AUG |
| | ........... |
| | ........... |
|  | --------AUG |

```
env-1384   ........AUG
env-1385   ...........
env-1393   ........AUG
env-1394   ........AUG
env-1395   ...........
env-1414   ........AUG
env-1416   ...........
env-1417   ........AUG
env-1418   ........AUG
env-1419   ........AUG
env-1423   ...........
env-1424   ........AUG
env-1425   ........AUG
Csp-7-1    ........AUG
env-1427   ........AUG
env-1428   ........AUG
env-1429   ........AUG
env-1431   ...........
env-1432   ........AUG
env-1433   ........AUG
env-1434   ........AUG
Coc-3-1    ........AUG
env-1435   ........AUG
env-1436   ........AUG
env-1437   ........AUG
env-1448   ........AUG
env-1449   ........AUG
env-1450   ........AUG
env-1451   ........AUG
env-1455   ........AUG
env-1456   ........AUG
env-1459   ........AUG
env-1462   ........AUG
env-1464   ...........
env-1465   ........AUG
env-1466   ........AUG
env-1467   ........AUG
env-1468   ........AUG
env-1469   ........AUG
env-1470   ........AUG
env-1472   ........AUG
env-1473   ........AUG
env-1474   ........AUG
Csp-8-1    ........AUG
env-1475   ........AUG
Fan-1-1    ........AUG
Fte-1-1    ........AUG
env-1486   ........AUG
env-1487   ........AUG
env-1488   ........AUG
env-1489   ........AUG
env-1490   ........AUG
env-1505   ........AUG
env-1507   ........AUG
Mmo-1-1    ........AUG
env-1508   ........AUG
env-1511   ........AUG
env-1512   ........AUG
env-1513   ........AUG
Lsp-4-1    ........AUG
env-1519   ...........
env-1520   ........AUG
env-1521   ........AUG
env-1525   ........AUG
Ama-1-1    ........AUG
env-1526   ........AUG
env-1527   ........AUG
env-1528   ........AUG
env-1529   ........AUG
env-1530   ........AUG
Aca-1-1    ........AUG
env-1532   ........AUG
env-1533   ........AUG
env-1534   ........AUG
Fsa-2-1    ........AUG
env-1535   ........AUG
env-1536   ........AUG
env-1537   ........AUG
Fba-2-1    ........AUG
Kje-1-1    ........AUG
Kal-1-1    ........AUG
Kzh-1-1    ........AUG
env-1538   ........AUG
env-1539   ........AUG

           ...........
           ...........
           --------AUG
```

```
env-1541   .......AUG
Fin-1-1    .......AUG
env-1542   .......AUG
env-1543   .......AUG
env-1544   .......AUG
env-1545   .......AUG
env-1546   .......AUG
env-1547   .......AUG
env-1549   .......AUG
env-1551   .......AUG
env-1554   .......AUG
env-1555   .......AUG
env-1556   .......AUG
env-1557   .......AUG
env-1558   .......AUG
env-1559   .......AUG
env-1561   .......AUG
env-1562   .......AUG
env-1563   .......AUG
env-1564   .......AUG
env-1565   .......AUG
env-1567   .......AUG
env-1569   .......AUG
env-1581   .......AUG
env-1582   .......AUG
env-1583   .......AUG
env-1587   .......AUG
env-1588   .......AUG
env-1589   .......AUG
env-1590   .......AUG
env-1593   .......AUG
env-1594   .......AUG
env-1596   .......AUG
env-1597   .......AUG
Mth-1-1    .......AUG
env-1600   .......AUG
env-1601   .......AUG
env-1602   .......AUG
env-1603   .......AUG
Bar-1-1    .......AUG
env-1604   .......AUG
env-1605   .......AUG
env-1606   .......AUG
env-1607   .......AUG
env-1608   .......AUG
env-1611   .......AUG
env-1612   .......AUG
env-1614   .......AUG
env-1615   .......AUG
env-1616   .......AUG
env-1617   .......AUG
env-1618   .......AUG
env-1624   .......AUG
env-1625   .......AUG
env-1626   .......AUG
env-1628   .......AUG
env-1629   .......AUG
env-1630   .......AUG
env-1631   .......AUG
env-1635   .......AUG
env-1640   .......AUG
env-1644   .......AUG
env-1648   .......AUG
env-1650   .......AUG
env-1653   .......AUG
env-1654   .......AUG
env-1656   .......AUG
env-1657   .......AUG
env-1658   .......AUG
env-1659   .......AUG
env-1660   .......AUG
env-1661   .......AUG
env-1662   .......AUG
env-1663   .......AUG
env-1664   .......AUG
env-1665   ..........
env-1666   .......AUG
env-1667   .......AUG
env-1670   .......AUG
env-1671   .......AUG
env-1672   .......AUG
env-1673   .......AUG
env-1674   .......AUG
env-1677   .......AUG

           ...........
           ...........
           -------AUG
```

| | |
|---|---|
| env-1680 | ........AUG |
| env-1687 | ........AUG |
| env-1690 | ........AUG |
| env-1699 | ........AUG |
| env-1700 | ........AUG |
| env-1701 | ........AUG |
| env-1702 | ........AUG |
| env-1703 | ........AUG |
| env-1709 | ........AUG |
| env-1711 | ........AUG |
| env-1712 | ........AUG |
| env-1713 | ........AUG |
| env-1714 | ........AUG |
| env-1715 | ........AUG |
| env-1716 | ........AUG |
| env-1717 | ........... |
| env-1718 | ........AUG |
| env-1720 | ........AUG |
| env-1721 | ........AUG |
| env-1723 | ........AUG |
| env-1733 | ........AUG |
| env-1734 | ........... |
| env-1735 | ........AUG |
| env-1736 | ........AUG |
| env-1737 | ........AUG |
| env-1741 | ........... |
| env-1742 | ........AUG |
| env-1743 | ........AUG |
| env-1750 | ........AUG |
| env-1751 | ........AUG |
| env-1752 | ........AUG |
| env-1758 | ........AUG |
| env-1760 | ........AUG |
| env-1761 | ........AUG |
| env-1762 | ........AUG |
| env-1763 | ........AUG |
| env-1764 | ........AUG |
| env-1765 | ........AUG |
| env-1771 | ........... |
| env-1772 | ........AUG |
| env-1777 | ........AUG |
| env-1779 | ........AUG |
| env-1781 | ........AUG |
| env-1782 | ........AUG |
| env-1783 | ........AUG |
| env-1787 | ........AUG |
| env-1788 | ........AUG |
| env-1789 | ........... |
| env-1790 | ........AUG |
| env-1791 | ........AUG |
| env-1792 | ........GAU |
| env-1793 | ........AUG |
| env-1794 | ........AUG |
| env-1802 | ........AUG |
| env-1803 | ........AUG |
| env-1804 | ........AUG |
| env-1805 | ........AUG |
| env-1806 | ........AUG |
| env-1807 | ........AUG |
| env-1812 | ........AUG |
| env-1813 | ........AUG |
| env-1814 | ........AUG |
| env-1815 | ........AUG |
| env-1816 | ........AUG |
| env-1817 | ........AUG |
| env-1818 | ........AUG |
| env-1819 | ........AUG |
| env-1820 | ........AUG |
| env-1821 | ........AUG |
| env-1822 | ........AUG |
| env-1823 | ........AUG |
| env-1824 | ........AUG |
| env-1825 | ........AUG |
| env-1826 | ........AUG |
| env-1830 | ........AUG |
| env-1831 | ........AUG |
| env-1832 | ........AUG |
| env-1837 | ........AUG |
| env-1839 | ........AUG |
| env-1840 | ........AUG |
| env-1841 | ........AUG |
| env-1842 | ........AUG |
| env-1843 | ........AUG |
| env-1844 | ........AUG |
| | ........... |
| | ........... |
|  | --------AUG |

| | |
|---|---|
| env-1845 | ........... |
| env-1846 | ........AUG |
| env-1848 | ........AUG |
| env-1849 | ........AUG |
| env-1850 | ........AUG |
| env-1851 | ........AUG |
| env-1853 | ........AUG |
| env-1855 | ........AUG |
| env-1856 | ........AUG |
| env-1857 | ........AUG |
| env-1860 | ........AUG |
| env-1861 | ........AUG |
| Ala-2-1 | ........AUG |
| env-1862 | ........AUG |
| Ndo-1-1 | ........AUG |
| env-1863 | ........AUG |
| env-1864 | ........AUG |
| Cal-1-1 | ........AUG |
| env-1865 | ........AUG |
| Csp-13-1 | ........AUG |
| Mfo-1-1 | ........AUG |
| Cba-4-1 | ........AUG |
| env-1866 | ........AUG |
| env-1867 | ........AUG |
| env-1869 | ........AUG |
| env-1870 | ........AUG |
| env-1872 | ........AUG |
| env-1873 | ........... |
| env-1875 | ........AUG |
| env-1880 | ........AUG |
| env-1881 | ........AUG |
| env-1886 | ........AUG |
| Orh-1-1 | ........AUG |
| env-1887 | ........AUG |
| env-1888 | ........AUG |
| env-1891 | ........... |
| env-1892 | ........AUG |
| env-1893 | ........AUG |
| env-1894 | ........AUG |
| env-1898 | ........AUG |
| env-1899 | ........AUG |
| env-1900 | ........AUG |
| env-1918 | ........AUG |
| env-1919 | ........AUG |
| env-1921 | ........AUG |
| env-1922 | ........AUG |
| env-1923 | ........AUG |
| Ran-1-1 | ........AUG |
| env-1924 | ........... |
| env-1925 | ........AUG |
| env-1929 | ........AUG |
| env-1930 | ........AUG |
| env-1931 | ........AUG |
| env-1932 | ........AUG |
| Ran-6-1 | ........AUG |
| env-1933 | ........... |
| env-1934 | ........AUG |
| env-1968 | ........AUG |
| env-1977 | ........AUG |
| env-1978 | ........AUG |
| env-1984 | ........AUG |
| env-1987 | ........AUG |
| env-1988 | ........AUG |
| env-1989 | ........AUG |
| env-1990 | ........AUG |
| env-1991 | ........AUG |
| env-1993 | ........AUG |
| Ran-2-1 | ........AUG |
| Rco-1-1 | ........AUG |
| env-1994 | ........AUG |
| env-1995 | ........AUG |
| env-1996 | ........AUG |
| env-1997 | ........AUG |
| env-1998 | ........AUG |
| env-1999 | ........AUG |
| env-2000 | ........AUG |
| Ran-3-1 | ........AUG |
| env-2001 | ........AUG |
| env-2032 | ........AUG |
| env-2033 | ........AUG |
| env-2034 | ........AUG |
| env-2035 | ........AUG |
| Fge-1-1 | ........AUG |
| Nma-1-1 | ........AUG |
| | ........... |
| | ........... |
|  | --------AUG |

```
env-2036   ........AUG
env-2047   ........AUG
env-2048   ........AUG
env-2050   ........AUG
env-2051   ........AUG
env-2052   ........AUG
env-2054   ........AUG
env-2055   ........AUG
env-2056   ........AUG
env-2057   ........AUG
env-2065   ........AUG
env-2066   ........AUG
env-2067   ........AUG
env-2068   ........AUG
env-2069   ........AUG
env-2071   ........AUG
env-2076   ........AUG
env-2077   ........AUG
env-2078   ...........
env-2079   ........AUG
env-2080   ........AUG
env-2098   ........AUG
env-2100   ........AUG
env-2101   ........AUG
env-2102   ........AUG
env-2103   ........AUG
env-2104   ........AUG
env-2121   ........AUG
env-2122   ........AUG
env-2123   ........AUG
Fps-4-1    ........AUG
Ran-4-1    ........AUG
           ...........
           ...........
           --------AUG
```

### 7 L20-Deltaproteobacteria

#### 7.1 Taxa

The taxonomy of each organism containing a putative L20-Deltaproteobacteria RNA is listed, with abbreviations identifying each hit (e.g., "Eco-1-1" and "Eco-1-2" might hypothetically represent two distinct RNAs in *E. coli*). The abbreviations will be used to identify each individual L20-Deltaproteobacteria RNA in Sections 7.2 and 7.4:.

| **abbrev. of hits** | **taxonomy of species** |
|---|---|
| Dac-1-1 | Bacteria Proteobacteria Deltaproteobacteria Desulfuromonadales Desulfuromonadaceae *Desulfuromonas acetoxidans* DSM 684 |
| Dsp-1-1 | Bacteria Proteobacteria Deltaproteobacteria Desulfuromonadales Desulfuromonadaceae *Desulfuromonas sp.* TF |
| Pse-1-1 | Bacteria Proteobacteria Deltaproteobacteria Desulfuromonadales Desulfuromonadaceae *Pelobacter seleniigenes* DSM 18267 |
| Gfe-1-1 | Bacteria Proteobacteria Deltaproteobacteria Desulfuromonadales Geobacteraceae *Geoalkalibacter ferrihydriticus* DSM 17813 |
| Gsu-1-1 | Bacteria Proteobacteria Deltaproteobacteria Desulfuromonadales Geobacteraceae *Geoalkalibacter subterraneus* |
| Gbe-1-1 | Bacteria Proteobacteria Deltaproteobacteria Desulfuromonadales Geobacteraceae *Geobacter bemidjiensis* Bem |
| Gbr-1-1 | Bacteria Proteobacteria Deltaproteobacteria Desulfuromonadales Geobacteraceae *Geobacter bremensis* R1 |
| Glo-1-1 | Bacteria Proteobacteria Deltaproteobacteria Desulfuromonadales Geobacteraceae *Geobacter lovleyi* SZ |
| Gme-1-1 | Bacteria Proteobacteria Deltaproteobacteria Desulfuromonadales Geobacteraceae *Geobacter metallireducens* GS-15 |
| Gme-2-1 | Bacteria Proteobacteria Deltaproteobacteria Desulfuromonadales Geobacteraceae *Geobacter metallireducens* RCH3 |
| Gpi-1-1 | Bacteria Proteobacteria Deltaproteobacteria Desulfuromonadales Geobacteraceae *Geobacter pickeringii* |
| Gso-1-1 | Bacteria Proteobacteria Deltaproteobacteria Desulfuromonadales Geobacteraceae *Geobacter soli* |
| Gsp-1-1 | Bacteria Proteobacteria Deltaproteobacteria Desulfuromonadales Geobacteraceae *Geobacter sp.* FRC-32 |
| Gsp-2-1 | Bacteria Proteobacteria Deltaproteobacteria Desulfuromonadales Geobacteraceae *Geobacter sp.* M18 |
| Gsp-3-1 | Bacteria Proteobacteria Deltaproteobacteria Desulfuromonadales Geobacteraceae *Geobacter sp.* M21 |
| Gsp-4-1 | Bacteria Proteobacteria Deltaproteobacteria Desulfuromonadales Geobacteraceae *Geobacter sp.* OR-1 |
| Gsu-2-1 | Bacteria Proteobacteria Deltaproteobacteria Desulfuromonadales Geobacteraceae *Geobacter sulfurreducens* |
| Gsu-3-1 | Bacteria Proteobacteria Deltaproteobacteria Desulfuromonadales Geobacteraceae *Geobacter sulfurreducens* KN400 |
| Gsu-4-1 | Bacteria Proteobacteria Deltaproteobacteria Desulfuromonadales Geobacteraceae *Geobacter sulfurreducens* PCA |
| Gur-1-1 | Bacteria Proteobacteria Deltaproteobacteria Desulfuromonadales Geobacteraceae *Geobacter uraniireducens* Rf4 |
| Gel-1-1 | Bacteria Proteobacteria Deltaproteobacteria Desulfuromonadales Geobacteraceae *Geopsychrobacter electrodiphilus* DSM 16401 |

| | |
|---|---|
| Pca-1-1 | Bacteria Proteobacteria Deltaproteobacteria Desulfuromonadales Pelobacteraceae *Pelobacter carbinolicus* DSM 2380 |
| Ppr-1-1 | Bacteria Proteobacteria Deltaproteobacteria Desulfuromonadales Pelobacteraceae *Pelobacter propionicus* DSM 2379 |
| env-1 to env-179 | environmental samples |

#### 7.2 Gene contexts

Each L20-Deltaproteobacteria RNA (indicated by "RNA→") is listed. For each hit, the downstream genes predicted to reside in a regulated operon are listed. If the nearest downstream gene is encoding in the opposite strand (and therefore presumed to not be a part of a regulated operon), then that gene is still depicted. Some environmental sequences and some RefSeq entries lack gene annotations, and so no genes are available for such sequences. The direction of each gene is indicated with an arrow (→), and each predicted conserved domain in the gene is named. Conserved domains associated with more than one L20-Deltaproteobacteria RNA are assigned a color; other domains are gray. Information about these conserved domains is given in Section 7.3. The accession of the sequence containing each L20-Deltaproteobacteria RNA is given in the column named "Seq. accession". Accessions beginning with "NC_", "NS_", "NW_" or "NZ_" are contained in RefSeq. Other accession refer to environmental samples. Nucleotide coordinates are given for the 5′ and 3′ boundaries of each L20-Deltaproteobacteria RNA. If the 5′ coordinate is greater than the 3′ coordinate, the RNA is present on the reverse-complement strand of the containing genomic DNA sequence. Each hit is denoted by an abbreviation (like "Eco-1-1") that refers to a taxonomy given in Section 7.1.

| abbrev. | Seq. accession | | 5′ at | 3′ at | genes |
|---|---|---|---|---|---|
| env-1 | 3300000558_10042364 | - | 210 | 132 | RNA→ rplT (PRK05185)Ribosomal_L20 (pfam00453)→ |
| env-2 | KVRMV2_101537624 | - | 193 | 105 | RNA→ rplT (PRK05185)rpl20 (CHL00068)→ |
| env-3 | LGVD01016786.1 | + | 10188 | 10263 | RNA→ rplT (PRK05185)Ribosomal_L20 (pfam00453)→ |
| env-4 | LGVE01096777.1 | + | 810 | 885 | RNA→ rplT (PRK05185)Ribosomal_L20 (pfam00453)→ |
| env-5 | LGVF01151704.1 | - | 2284 | 2209 | RNA→ rplT (PRK05185)Ribosomal_L20 (pfam00453)→ |
| env-6 | LGVF01446591.1 | - | 671 | 594 | RNA→ rplT (PRK05185)Ribosomal_L20 (pfam00453)→ |
| env-7 | JGI24023J19991_10001126 | + | 6175 | 6251 | RNA→ rplT (PRK05185)Ribosomal_L20 (pfam00453)→ |
| Pse-1-1 | NZ_JOMG01000004.1 | + | 547284 | 547360 | RNA→ rplT (PRK05185)Ribosomal_L20 (pfam00453)→ |
| env-8 | M940M_1000089 | - | 2577 | 2501 | RNA→ rplT (PRK05185)Ribosomal_L20 (pfam00453)→ |
| env-9 | M590M1_1000913 | + | 8384 | 8460 | RNA→ rplT (PRK05185)Ribosomal_L20 (pfam00453)→ |
| env-10 | M590M2_1001691 | - | 8449 | 8373 | RNA→ rplT (PRK05185)Ribosomal_L20 (pfam00453)→ |
| env-11 | SA_S1_NOR05_45m_c10229484 | - | 170 | 92 | RNA→ rplT (PRK05185)rpl20 (CHL00068)→ |
| Gel-1-1 | NZ_ARWE01000001.1 | + | 1492573 | 1492658 | RNA→ rplT (PRK05185)Ribosomal_L20 (pfam00453)→ |
| env-12 | KVRMV2_100037767 | - | 14415 | 14351 | RNA→ rplT (PRK05185)Ribosomal_L20 (pfam00453)→ PheS (COG0016)pheS (PRK00488)→ PheT (COG0072)pheT (PRK00629)tRNA_bind_bactPheRS (cd02796)→ |
| env-13 | 3300001605_10126988 | + | 58 | 114 | RNA→ rplT (PRK05185)Ribosomal_L20 (pfam00453)→ PheS (COG0016)pheS (PRK00488)→ syfB (CHL00192)pheT (PRK00629)→ |
| env-14 | 3300001605_10740178 | + | 188 | 244 | RNA→ rplT (PRK05185)Ribosomal_L20 (pfam00453)→ |
| env-15 | 3300001605_10006506 | + | 12971 | 13027 | RNA→ |
| env-16 | 3300001605_10708600 | + | 200 | 256 | RNA→ rplT (PRK05185)Ribosomal_L20 (pfam00453)→ |
| env-17 | LWFCAn_GLO1YSU01EBI9B | + | 3 | 64 | RNA→ hypo→ rplT (PRK05185)Ribosomal_L20 (pfam00453)→ |
| env-18 | lwComb_BHFI11626_g1 | + | 746 | 807 | RNA→ |
| env-19 | 2034067870 | + | 177 | 238 | RNA→ rplT (PRK05185)Ribosomal_L20 (cd07026)→ |
| env-20 | LWFCAn_GLO1YSU01D9AZM | - | 414 | 353 | RNA→ hypo→ rplT (PRK05185)Ribosomal_L20 (pfam00453)→ |
| env-21 | LWFCA_GLZZ6Z001ECB4G | - | 114 | 53 | RNA→ ←hypo |
| env-22 | LWFCAnN_GO09JKT02GDB9I | + | 274 | 335 | RNA→ rplT (PRK05185)Ribosomal_L20 (pfam00453)→ |
| Ppr-1-1 | NC_008609.1 | - | 2488533 | 2488474 | RNA→ rplT (PRK05185)Ribosomal_L20 (pfam00453)→ PheS (COG0016)pheS (PRK00488)→ PheT (COG0072)pheT (PRK00629)tRNA_bind_bactPheRS (cd02796)→ ihfA (PRK00285)IHF_A (cd13835)→ HTH_MlrA-like_sg2 (cd04765)MerR_1 (pfam13411)→ hypo→ hypo→ hypo→ hypo→ hypo→ |
| env-23 | JGI11944J13513_1245443 | - | 94 | 39 | RNA→ |
| env-24 | 3300000032_0015669 | - | 35340 | 35277 | RNA→ rplT (PRK05185)Ribosomal_L20 (pfam00453)→ |
| env-25 | 3300000568_10008200 | + | 165385 | 165448 | RNA→ rplT (PRK05185)Ribosomal_L20 (pfam00453)→ |

| | | | | | |
|---|---|---|---|---|---|
| Gme-1-1 | NC_007517.1 | + | 1582191 | 1582253 | RNA→ rplT (PRK05185)Ribosomal_L20 (pfam00453)→ rplT (PRK05185)Ribosomal_L20 (pfam00453)→ PheS (COG0016)pheS (PRK00488)→ PheS (COG0016)pheS (PRK00488)→ PheT (COG0072)pheT (PRK00629)tRNA_bind_bactPheRS (cd02796)→ PheT (COG0072)pheT (PRK00629)tRNA_bind_bactPheRS (cd02796)→ ihfA (PRK00285)IHF_A (cd13835)→ ihfA (PRK00285)IHF_A (cd13835)→ HTH_MlrA-like_sg2 (cd04765)MerR_1 (pfam13411)→ HTH_MlrA-like_sg2 (cd04765)MerR_1 (pfam13411)→ hypo→ hypo→ hypo→ hypo→ hypo→ SurE (COG0496)surE (PRK00346)→ SurE (COG0496)surE (PRK00346)→ |
| Gme-2-1 | NZ_AGJM01000001.1 | - | 308216 | 308154 | RNA→ rplT (PRK05185)Ribosomal_L20 (pfam00453)→ PheS (COG0016)pheS (PRK00488)→ PheT (COG0072)pheT (PRK00629)tRNA_bind_bactPheRS (cd02796)→ ihfA (PRK00285)IHF_A (cd13835)→ HTH_MlrA-like_sg2 (cd04765)MerR_1 (pfam13411)→ hypo→ hypo→ SurE (COG0496)surE (PRK00346)→ |
| env-26 | JGI12269J14319_10525591 | - | 162 | 102 | RNA→ rplT (PRK05185)rpl20 (CHL00068)→ |
| env-27 | 3300001605_10002885 | + | 13437 | 13493 | RNA→ rplT (PRK05185)Ribosomal_L20 (pfam00453)→ PheS (COG0016)pheS (PRK00488)→ PheT (COG0072)pheT (PRK00629)tRNA_bind_bactPheRS (cd02796)→ hypo→ |
| env-28 | 3300001592_11009265 | + | 13 | 69 | RNA→ rplT (PRK05185)Ribosomal_L20 (pfam00453)→ |
| env-29 | TB_FS08_3DRAFT_1002004 | - | 4554 | 4498 | RNA→ rplT (PRK05185)Ribosomal_L20 (pfam00453)→ PheS (COG0016)pheS (PRK00488)→ PheT (COG0072)pheT (PRK00629)tRNA_bind_bactPheRS (cd02796)→ |
| env-30 | TB_FS06_10DRAFT_1084659 | + | 404 | 460 | RNA→ rplT (PRK05185)Ribosomal_L20 (pfam00453)→ |
| env-31 | TB_PC08_66_10006928 | + | 2535 | 2591 | RNA→ rplT (PRK05185)Ribosomal_L20 (pfam00453)→ PheS (COG0016)pheS (PRK00488)→ PheT (COG0072)pheT (PRK00629)tRNA_bind_bactPheRS (cd02796)→ |
| env-32 | TB_PC08_64_1003822 | - | 2374 | 2318 | RNA→ rplT (PRK05185)Ribosomal_L20 (pfam00453)→ PheS (COG0016)pheS (PRK00488)→ PheT (COG0072)pheT (PRK00629)tRNA_bind_bactPheRS (cd02796)→ |
| env-33 | TB_PC08_3_1055617 | - | 375 | 319 | RNA→ rplT (PRK05185)Ribosomal_L20 (pfam00453)→ |
| env-34 | 3300001580_10000050 | + | 94579 | 94634 | RNA→ rplT (PRK05185)Ribosomal_L20 (pfam00453)→ |
| env-35 | TB_GS10_10_10002215 | + | 3287 | 3343 | RNA→ rplT (PRK05185)Ribosomal_L20 (pfam00453)→ PheS (COG0016)pheS (PRK00488)→ PheT (COG0072)pheT (PRK00629)tRNA_bind_bactPheRS (cd02796)→ |
| env-36 | TB_PC08_64_1182178 | - | 205 | 149 | RNA→ rplT (PRK05185)Ribosomal_L20 (pfam00453)→ |
| env-37 | TB_GS09_5_10173233 | - | 132 | 76 | RNA→ rplT (PRK05185)Ribosomal_L20 (pfam00453)→ |
| env-38 | TB_GS10_10_10001740 | + | 11836 | 11892 | RNA→ rplT (PRK05185)Ribosomal_L20 (pfam00453)→ |
| env-39 | TB_FS08_3DRAFT_1028849 | + | 165 | 221 | RNA→ rplT (PRK05185)Ribosomal_L20 (pfam00453)→ pheS (PRK00488)Phe_tRNA-synt_N (pfam02912)→ |
| env-40 | TB_PC08_64_1167760 | + | 172 | 228 | RNA→ rplT (PRK05185)Ribosomal_L20 (pfam00453)→ |
| env-41 | TB_PC08_66_10223732 | + | 215 | 271 | RNA→ rplT (PRK05185)Ribosomal_L20 (pfam00453)→ |
| env-42 | TB_GS10_10_10049699 | + | 1329 | 1385 | RNA→ rplT (PRK05185)Ribosomal_L20 (pfam00453)→ pheS (PRK00488)→ |
| env-43 | TB_PC08_3_1110535 | + | 168 | 224 | RNA→ rplT (PRK05185)Ribosomal_L20 (pfam00453)→ |
| env-44 | TB_GS10_10_10002500 | - | 11045 | 10989 | RNA→ rplT (PRK05185)Ribosomal_L20 (pfam00453)→ PheS (COG0016)pheS (PRK00488)→ PheT (COG0072)pheT (PRK00629)tRNA_bind_bactPheRS (cd02796)→ S2P-M50_like_1 (cd06158)→ TrpS (COG0180)PRK00927 (PRK00927)→ ScpA (COG1354)scpA (PRK00104)→ scpB (PRK00135)SMC_ScpB (pfam04079)→ |
| env-45 | TB_GS09_5_10420941 | - | 158 | 102 | RNA→ rplT (PRK05185)Ribosomal_L20 (pfam00453)→ |
| env-46 | TB_PC08_66_10333941 | - | 159 | 103 | RNA→ rplT (PRK05185)Ribosomal_L20 (pfam00453)→ |
| env-47 | TB_GS10_10_10234470 | - | 198 | 142 | RNA→ rplT (PRK05185)Ribosomal_L20 (pfam00453)→ |
| env-48 | 3300000568_10942424 | - | 314 | 260 | RNA→ rplT (PRK05185)Ribosomal_L20 (pfam00453)→ |
| env-49 | 3300000032_0003617 | - | 28755 | 28701 | RNA→ rplT (PRK05185)Ribosomal_L20 (pfam00453)→ PheS (COG0016)pheS (PRK00488)→ PheT (COG0072)pheT (PRK00629)tRNA_bind_bactPheRS (cd02796)→ FDX-ACB (smart00896)PheT (COG0072)→ ihfA (PRK00285)IHF_A (cd13835)→ HTH_MlrA-like_sg2 (cd04765)MerR_1 (pfam13411)→ hypo→ |
| env-50 | 3300000568_10174305 | - | 447 | 393 | RNA→ rplT (PRK05185)Ribosomal_L20 (pfam00453)→ |
| env-51 | 3300001567_10000440 | + | 37268 | 37322 | RNA→ rplT (PRK05185)Ribosomal_L20 (pfam00453)→ |
| env-52 | 3300000558_10301824 | + | 217 | 274 | RNA→ rplT (PRK05185)Ribosomal_L20 (pfam00453)→ |
| env-53 | JGI24728J21555_1025623 | - | 728 | 671 | RNA→ rplT (PRK05185)Ribosomal_L20 (pfam00453)→ pheS (PRK00488)Phe_tRNA-synt_N (pfam02912)→ |
| env-54 | 3300001592_10294556 | - | 272 | 218 | RNA→ rplT (PRK05185)Ribosomal_L20 (pfam00453)→ |
| env-55 | 3300001605_10107340 | - | 1573 | 1518 | RNA→ rplT (PRK05185)Ribosomal_L20 (pfam00453)→ PheS (COG0016)pheS (PRK00488)→ |
| env-56 | 3300000501_100057 | + | 10794 | 10848 | RNA→ rplT (PRK05185)Ribosomal_L20 (pfam00453)→ PheS (COG0016)pheS (PRK00488)→ PheT (COG0072)pheT (PRK00629)tRNA_bind_bactPheRS (cd02796)→ |
| env-57 | 3300000558_10291420 | - | 104 | 49 | RNA→ |
| env-58 | 3300000558_11797894 | + | 48 | 103 | RNA→ rplT (PRK05185)Ribosomal_L20 (pfam00453)→ |

| | | | | | |
|---|---|---|---|---|---|
| env-59 | 3300001580_11172117 | - | 105 | 50 | **RNA**→ |
| env-60 | 3300000032_0024462 | - | 15336 | 15280 | **RNA**→ rplT (PRK05185)Ribosomal_L20 (pfam00453)→ PheS (COG0016)pheS (PRK00488)→<br>PheT (COG0072)pheT (PRK00629)tRNA_bind_bactPheRS (cd02796)→ |
| env-61 | JGI24728J21555_1009694 | - | 2660 | 2605 | **RNA**→ rplT (PRK05185)Ribosomal_L20 (pfam00453)→ PheS (COG0016)pheS (PRK00488)→<br>pheT (PRK00629)tRNA_bind_bactPheRS (cd02796)B3_4 (pfam03483)→ |
| env-62 | JGI24728J21555_1023465 | + | 254 | 309 | **RNA**→ rplT (PRK05185)Ribosomal_L20 (pfam00453)→ PheS (COG0016)pheS (PRK00488)→ |
| env-63 | JGI24730J26740_1004602 | - | 3221 | 3166 | **RNA**→ rplT (PRK05185)Ribosomal_L20 (pfam00453)→ PheS (COG0016)pheS (PRK00488)→<br>PheT (COG0072)pheT (PRK00629)tRNA_bind_bactPheRS (cd02796)→ |
| env-64 | JGI24730J26740_1000449 | - | 28604 | 28549 | **RNA**→ rplT (PRK05185)Ribosomal_L20 (pfam00453)→ |
| Glo-1-1 | NC_010814.1 | + | 1993020 | 1993075 | **RNA**→ rplT (PRK05185)Ribosomal_L20 (pfam00453)→ PheS (COG0016)pheS (PRK00488)→<br>PheT (COG0072)pheT (PRK00629)tRNA_bind_bactPheRS (cd02796)→ DUF3373 (pfam11853)→ |
| env-65 | JGI24728J21555_1018455 | + | 1325 | 1380 | **RNA**→ rplT (PRK05185)Ribosomal_L20 (pfam00453)→ pheS (PRK00488)Phe_tRNA-synt_N (pfam02912)→ |
| env-66 | PCEOT_F164E7T01BG9SK | + | 218 | 273 | **RNA**→ rplT (PRK05185)Ribosomal_L20 (pfam00453)→ |
| env-67 | IMNBL1_c0232076 | + | 282 | 341 | **RNA**→ rplT (PRK05185)rpl20 (CHL00068)→ |
| env-68 | JGI994J12354_10020529 | - | 91 | 32 | **RNA**→ ←hypo |
| env-69 | JGI994J12354_10102055 | - | 91 | 32 | **RNA**→ |
| env-70 | JGI24728J21555_1023893 | - | 695 | 633 | **RNA**→ rplT (PRK05185)Ribosomal_L20 (pfam00453)→ pheS (PRK00488)Phe_tRNA-synt_N (pfam02912)→ |
| env-71 | 3300000568_10095751 | - | 3182 | 3123 | **RNA**→ rplT (PRK05185)Ribosomal_L20 (pfam00453)→ |
| env-72 | 3300000568_10166208 | + | 543 | 602 | **RNA**→ rplT (PRK05185)Ribosomal_L20 (pfam00453)→ |
| env-73 | 3300000568_10910092 | - | 126 | 67 | **RNA**→ hypo→ |
| env-74 | 3300001580_10542033 | - | 192 | 133 | **RNA**→ rplT (PRK05185)Ribosomal_L20 (pfam00453)→ |
| env-75 | 3300001567_10003592 | + | 12408 | 12467 | **RNA**→ rplT (PRK05185)Ribosomal_L20 (pfam00453)→ |
| env-76 | 3300000558_10376168 | - | 3122 | 3060 | **RNA**→ rplT (PRK05185)Ribosomal_L20 (pfam00453)→ PheS (COG0016)pheS (PRK00488)→ |
| env-77 | 3300000032_0032030 | - | 654 | 593 | **RNA**→ rplT (PRK05185)Ribosomal_L20 (pfam00453)→ pheS (PRK00488)Phe_tRNA-synt_N (pfam02912)→ |
| env-78 | 3300000568_10567303 | - | 669 | 608 | **RNA**→ rplT (PRK05185)Ribosomal_L20 (pfam00453)→ pheS (PRK00488)Phe_tRNA-synt_N (pfam02912)→ |
| Gso-1-1 | NZ_JXBL01000001.1 | - | 1713733 | 1713668 | **RNA**→ rplT (PRK05185)Ribosomal_L20 (pfam00453)→ PheS (COG0016)pheS (PRK00488)→<br>PheT (COG0072)pheT (PRK00629)tRNA_bind_bactPheRS (cd02796)→ ihfA (PRK00285)IHF_A (cd13835)→<br>HTH_MlrA-like_sg2 (cd04765)MerR_1 (pfam13411)→ hypo→ |
| Gsu-2-1 | NZ_CP010430.1 | - | 4273106 | 4273044 | **RNA**→ rplT (PRK05185)Ribosomal_L20 (pfam00453)→ PheS (COG0016)pheS (PRK00488)→<br>PheT (COG0072)pheT (PRK00629)tRNA_bind_bactPheRS (cd02796)→ ihfA (PRK00285)IHF_A (cd13835)→<br>HTH_MlrA-like_sg2 (cd04765)MerR_1 (pfam13411)→ hypo→ |
| env-79 | ACD53_150 | - | 7941 | 7879 | **RNA**→ rplT (PRK05185)Ribosomal_L20 (pfam00453)→ PheS (COG0016)pheS (PRK00488)→<br>pheT (PRK00629)tRNA_bind_bactPheRS (cd02796)B3_4 (pfam03483)→ PheT (COG0072)pheT (PRK00629)→<br>ihfA (PRK00285)IHF_A (cd13835)→ HTH_MlrA-like_sg2 (cd04765)MerR_1 (pfam13411)→ hypo→ |
| Gsu-4-1 | NC_002939.5 | + | 1664846 | 1664908 | **RNA**→ rplT (PRK05185)Ribosomal_L20 (pfam00453)→ PheS (COG0016)pheS (PRK00488)→<br>PheT (COG0072)pheT (PRK00629)tRNA_bind_bactPheRS (cd02796)→ ihfA (PRK00285)IHF_A (cd13835)→<br>HTH_MlrA-like_sg2 (cd04765)MerR_1 (pfam13411)→ hypo→ hypo→ hypo→ hypo→ |
| Gsu-3-1 | NC_017454.1 | + | 1634500 | 1634562 | **RNA**→ rplT (PRK05185)Ribosomal_L20 (pfam00453)→ PheS (COG0016)pheS (PRK00488)→<br>PheT (COG0072)pheT (PRK00629)tRNA_bind_bactPheRS (cd02796)→ ihfA (PRK00285)IHF_A (cd13835)→<br>HTH_MlrA-like_sg2 (cd04765)MerR_1 (pfam13411)→ hypo→ hypo→ hypo→ hypo→ |
| Gpi-1-1 | NZ_CP009788.1 | - | 1887990 | 1887929 | **RNA**→ rplT (PRK05185)Ribosomal_L20 (pfam00453)→ PheS (COG0016)pheS (PRK00488)→<br>PheT (COG0072)pheT (PRK00629)tRNA_bind_bactPheRS (cd02796)→ ihfA (PRK00285)IHF_A (cd13835)→<br>HTH_MlrA-like_sg2 (cd04765)MerR_1 (pfam13411)→ hypo→ SurE (COG0496)surE (PRK00346)→<br>SurE (COG0496)surE (PRK00346)→ pcm (PRK00312)Pcm (COG2518)→<br>RpoD (COG0568)sigma70-ECF (TIGR02937)→ Apt (COG0503)PRK02304 (PRK02304)→ |
| env-80 | JXWW01164903.1 | - | 235 | 172 | **RNA**→ rplT (PRK05185)Ribosomal_L20 (pfam00453)→ |
| Gsp-4-1 | NZ_BAZF01000009.1 | + | 8601 | 8663 | **RNA**→ rplT (PRK05185)Ribosomal_L20 (pfam00453)→ PheS (COG0016)pheS (PRK00488)→<br>PheT (COG0072)pheT (PRK00629)tRNA_bind_bactPheRS (cd02796)→ ihfA (PRK00285)IHF_A (cd13835)→<br>HTH_MlrA-like_sg2 (cd04765)MerR_1 (pfam13411)→ hypo→ |
| env-81 | 3300001567_10543765 | + | 112 | 171 | **RNA**→ ←hypo |
| env-82 | 3300001567_10545448 | - | 129 | 70 | **RNA**→ ←hypo |
| env-83 | 3300000568_10531113 | + | 84 | 144 | **RNA**→ rplT (PRK05185)Ribosomal_L20 (pfam00453)→ |
| env-84 | 3300000032_0125795 | - | 2173 | 2113 | **RNA**→ rplT (PRK05185)Ribosomal_L20 (pfam00453)→ PheS (COG0016)pheS (PRK00488)→<br>PheT (COG0072)pheT (PRK00629)tRNA_bind_bactPheRS (cd02796)→ |

| | | | | | |
|---|---|---|---|---|---|
| env-85 | 3300000568_10022887 | - | 111929 | 111869 | RNA→ rplT (PRK05185)Ribosomal_L20 (pfam00453)→ PheS (COG0016)pheS (PRK00488)→ PheT (COG0072)pheT (PRK00629)tRNA_bind_bactPheRS (cd02796)→ FDX-ACB (smart00896)PheT (COG0072)→ |
| env-86 | AMWB01069312.1 | - | 684 | 624 | RNA→ rplT (PRK05185)Ribosomal_L20 (pfam00453)→ pheS (PRK00488)Phe_tRNA-synt_N (pfam02912)→ |
| env-87 | TB_GS10_10_10531313 | - | 104 | 42 | RNA→ |
| env-88 | SwBSRL2_contig_2794542 | + | 72 | 133 | RNA→ ←hypo |
| env-89 | 4491734.3.5201783 | - | 124 | 63 | RNA→ ←hypo |
| Gsp-2-1 | NC_014973.1 | - | 2673478 | 2673416 | RNA→ rplT (PRK05185)Ribosomal_L20 (pfam00453)→ PheS (COG0016)pheS (PRK00488)→ |
| env-90 | PRSSGFe2_Sequence0000018884 | + | 1176 | 1238 | RNA→ rplT (PRK05185)Ribosomal_L20 (pfam00453)→ pheS (PRK00488)Phe_tRNA-synt_N (pfam02912)→ |
| Gbr-1-1 | NZ_AUGE01000018.1 | + | 72618 | 72680 | RNA→ rplT (PRK05185)Ribosomal_L20 (pfam00453)→ PheS (COG0016)pheS (PRK00488)→ |
| Gbe-1-1 | NC_011146.1 | + | 2298127 | 2298189 | RNA→ rplT (PRK05185)Ribosomal_L20 (pfam00453)→ PheS (COG0016)pheS (PRK00488)→ |
| Gsp-3-1 | NC_012918.1 | - | 2599031 | 2598969 | RNA→ rplT (PRK05185)Ribosomal_L20 (pfam00453)→ PheS (COG0016)pheS (PRK00488)→ |
| Gsp-1-1 | NC_011979.1 | - | 3716790 | 3716730 | RNA→ rplT (PRK05185)Ribosomal_L20 (pfam00453)→ PheS (COG0016)pheS (PRK00488)→ PheT (COG0072)pheT (PRK00629)tRNA_bind_bactPheRS (cd02796)→ ihfA (PRK00285)IHF_A (cd13835)→ HTH_MlrA-like_sg2 (cd04765)MerR_1 (pfam13411)→ SoxR (COG0789)MerR_1 (pfam13411)→ hypo→ hypo→ hypo→ hypo→ hypo→ |
| env-91 | JGI24731J21663_1002160 | - | 9311 | 9251 | RNA→ rplT (PRK05185)Ribosomal_L20 (pfam00453)→ PheS (COG0016)pheS (PRK00488)→ PheT (COG0072)pheT (PRK00629)tRNA_bind_bactPheRS (cd02796)→ ihfA (PRK00285)IHF_A (cd13835)→ HTH_MlrA-like_sg2 (cd04765)MerR_1 (pfam13411)→ hypo→ |
| Gur-1-1 | NC_009483.1 | - | 3051608 | 3051548 | RNA→ rplT (PRK05185)Ribosomal_L20 (pfam00453)→ PheS (COG0016)pheS (PRK00488)→ PheT (COG0072)pheT (PRK00629)tRNA_bind_bactPheRS (cd02796)→ ihfA (PRK00285)IHF_A (cd13835)→ HTH_MlrA-like_sg2 (cd04765)MerR_1 (pfam13411)→ hypo→ hypo→ hypo→ hypo→ hypo→ hypo→ hypo→ |
| env-92 | LWFCA_GLZZ6Z001BD8DZ | - | 103 | 44 | RNA→ |
| env-93 | PBDCA2_contig38528 | - | 284 | 225 | RNA→ rplT (PRK05185)Ribosomal_L20 (pfam00453)→ |
| env-94 | KVRMV2_101237892 | - | 963 | 877 | RNA→ rplT (PRK05185)Ribosomal_L20 (pfam00453)→ |
| env-95 | LGOV01006858.1 | + | 224 | 303 | RNA→ rplT (PRK05185)Ribosomal_L20 (pfam00453)→ PheS (COG0016)pheS (PRK00488)→ PheT (COG0072)pheT (PRK00629)tRNA_bind_bactPheRS (cd02796)→ ihfA (PRK00285)IHF_A (cd13835)→ HTH_MlrA-like_sg2 (cd04765)MerR_1 (pfam13411)→ SurE (COG0496)surE (PRK00346)→ Pcm (COG2518)Methyltransf_25 (pfam13649)→ YgaA (COG1238)SNARE_assoc (pfam09335)→ LysM (smart00257)NlpD (COG0739)→ RpoD (COG0568)sigma70-ECF (TIGR02937)→ Apt (COG0503)PRK02304 (PRK02304)→ hypo→ hypo→ |
| env-96 | LGVE01160116.1 | - | 2036 | 1957 | RNA→ rplT (PRK05185)Ribosomal_L20 (pfam00453)→ PheS (COG0016)pheS (PRK00488)→ pheT (PRK00629)tRNA_bind_bactPheRS (cd02796)→ |
| env-97 | LGVF01622022.1 | + | 205 | 284 | RNA→ rplT (PRK05185)Ribosomal_L20 (pfam00453)→ PheS (COG0016)pheS (PRK00488)→ |
| env-98 | SL_1KL_011_SED_10290984 | + | 65 | 141 | RNA→ rplT (PRK05185)Ribosomal_L20 (pfam00453)→ |
| env-99 | SL_1KL_011_SED_10360958 | - | 178 | 102 | RNA→ rplT (PRK05185)rpl20 (CHL00068)→ |
| env-100 | LGVD01055386.1 | - | 622 | 544 | RNA→ rplT (PRK05185)Ribosomal_L20 (pfam00453)→ hypo→ |
| env-101 | LGVE01140865.1 | + | 773 | 851 | RNA→ rplT (PRK05185)Ribosomal_L20 (pfam00453)→ PheS (COG0016)pheS (PRK00488)→ |
| env-102 | LGVF01039065.1 | - | 9383 | 9305 | RNA→ rplT (PRK05185)Ribosomal_L20 (pfam00453)→ PheS (COG0016)pheS (PRK00488)→ PheT (COG0072)pheT (PRK00629)tRNA_bind_bactPheRS (cd02796)→ BHL (smart00411)IHF_A (cd13835)→ HTH_MlrA-like_sg2 (cd04765)MerR_1 (pfam13411)→ SurE (COG0496)surE (PRK00346)→ pcm (PRK00312)Pcm (COG2518)→ YgaA (COG1238)SNARE_assoc (pfam09335)→ LysM (smart00257)NlpD (COG0739)→ RpoD (COG0568)sigma70-ECF (TIGR02937)→ Apt (COG0503)PRK02304 (PRK02304)→ |
| env-103 | LGVF01564570.1 | + | 2588 | 2666 | RNA→ rplT (PRK05185)Ribosomal_L20 (pfam00453)→ |
| Gfe-1-1 | NZ_JWJD01000001.1 | + | 251749 | 251826 | RNA→ rplT (PRK05185)Ribosomal_L20 (pfam00453)→ |
| Gsu-1-1 | NZ_CP010311.1 | + | 1689104 | 1689181 | RNA→ rplT (PRK05185)Ribosomal_L20 (pfam00453)→ |
| env-104 | M740_1002203 | - | 1951 | 1857 | RNA→ rplT (PRK05185)Ribosomal_L20 (pfam00453)→ |
| env-105 | M750CN_1001098 | + | 23430 | 23524 | RNA→ rplT (PRK05185)Ribosomal_L20 (pfam00453)→ |
| Dac-1-1 | NZ_AAEW02000011.1 | - | 131799 | 131705 | RNA→ rplT (PRK05185)Ribosomal_L20 (pfam00453)→ rplT (PRK05185)Ribosomal_L20 (pfam00453)→ |
| env-106 | AMWB01069310.1 | - | 7209 | 7132 | RNA→ rplT (PRK05185)Ribosomal_L20 (pfam00453)→ |
| env-107 | 3300000568_10425264 | - | 297 | 221 | RNA→ rplT (PRK05185)Ribosomal_L20 (pfam00453)→ |
| env-108 | 3300000558_10049026 | + | 2911 | 2987 | RNA→ rplT (PRK05185)Ribosomal_L20 (pfam00453)→ |
| env-109 | 3300001580_10052679 | + | 1247 | 1323 | RNA→ rplT (PRK05185)Ribosomal_L20 (pfam00453)→ |

| | | | | | |
|---|---|---|---|---|---|
| env-110 | 2232073740 | - | 430 | 354 | **RNA**→ rplT (PRK05185)Ribosomal_L20 (pfam00453)→ |
| env-111 | 3300000558_10134028 | + | 138 | 216 | **RNA**→ rplT (PRK05185)Ribosomal_L20 (pfam00453)→ |
| env-112 | 3300000558_11394841 | - | 255 | 177 | **RNA**→ rplT (PRK05185)Ribosomal_L20 (pfam00453)→ |
| env-113 | 3300000568_10011739 | - | 718 | 640 | **RNA**→ rplT (PRK05185)Ribosomal_L20 (pfam00453)→ |
| env-114 | 3300001567_10107199 | - | 546 | 468 | **RNA**→ rplT (PRK05185)Ribosomal_L20 (pfam00453)→ |
| Dsp-1-1 | NZ_KI421413.1 | - | 228136 | 228058 | **RNA**→ rplT (PRK05185)Ribosomal_L20 (pfam00453)→ |
| env-115 | 3300001580_10000645 | + | 14984 | 15060 | **RNA**→ rplT (PRK05185)Ribosomal_L20 (pfam00453)→ |
| env-116 | 3300000558_11481507 | - | 595 | 519 | **RNA**→ rplT (PRK05185)Ribosomal_L20 (pfam00453)→ |
| env-117 | 3300000558_11690408 | + | 653 | 728 | **RNA**→ rplT (PRK05185)Ribosomal_L20 (pfam00453)→ |
| Pca-1-1 | NC_007498.2 | + | 1669010 | 1669085 | **RNA**→ rplT (PRK05185)Ribosomal_L20 (pfam00453)→ rplT (PRK05185)Ribosomal_L20 (pfam00453)→ |
| env-118 | 3300001567_10001919 | + | 18381 | 18456 | **RNA**→ rplT (PRK05185)Ribosomal_L20 (pfam00453)→ PheS (COG0016)pheS (PRK00488)→ PheT (COG0072)pheT (PRK00629)tRNA_bind_bactPheRS (cd02796)→ |
| env-119 | 3300000558_10022538 | - | 6809 | 6737 | **RNA**→ rplT (PRK05185)Ribosomal_L20 (pfam00453)→ PheS (COG0016)pheS (PRK00488)→ PheT (COG0072)pheT (PRK00629)tRNA_bind_bactPheRS (cd02796)→ |
| env-120 | 3300000558_10299282 | + | 347 | 425 | **RNA**→ |
| env-121 | SA_S1_NOR05_45m_c10345692 | - | 276 | 199 | **RNA**→ rplT (PRK05185)Ribosomal_L20 (pfam00453)→ |
| env-122 | SA_S1_NOR05_45m_c10511679 | - | 159 | 82 | **RNA**→ rplT (PRK05185)rpl20 (CHL00068)→ |
| env-123 | SA_S2_NOR18_50m_1106132 | - | 249 | 173 | **RNA**→ rplT (PRK05185)Ribosomal_L20 (pfam00453)→ |
| env-124 | SA_S2_NOR13_50m_c1176942 | + | 189 | 265 | **RNA**→ |
| env-125 | SA_S1_NOR05_45m_c10000719 | + | 2825 | 2901 | **RNA**→ rplT (PRK05185)Ribosomal_L20 (pfam00453)→ |
| env-126 | SA_S1_NOR08_45m_c10011655 | + | 1979 | 2055 | **RNA**→ rplT (PRK05185)Ribosomal_L20 (pfam00453)→ |
| env-127 | SA_S1_NOR08_45m_c10015541 | + | 2845 | 2921 | **RNA**→ rplT (PRK05185)Ribosomal_L20 (pfam00453)→ |
| env-128 | SA_S2_NOR15_50m_c10016791 | + | 1291 | 1367 | **RNA**→ rplT (PRK05185)Ribosomal_L20 (pfam00453)→ |
| env-129 | SA_S1_NOR02_45m_c1003376 | - | 909 | 833 | **RNA**→ rplT (PRK05185)Ribosomal_L20 (pfam00453)→ |
| env-130 | SA_S2_NOR15_50m_c10152710 | - | 153 | 77 | **RNA**→ rplT (PRK05185)rpl20 (CHL00068)→ |
| env-131 | SA_S1_NOR05_45m_c10069212 | + | 630 | 706 | **RNA**→ rplT (PRK05185)Ribosomal_L20 (pfam00453)→ |
| env-132 | SA_S1_NOR02_45m_c1134354 | - | 100 | 24 | **RNA**→ ←hypo |
| env-133 | SA_S1_NOR08_45m_c10008401 | - | 164 | 88 | **RNA**→ rplT (PRK05185)rpl20 (CHL00068)→ |
| env-134 | SA_S2_NOR13_50m_c1190193 | + | 137 | 213 | **RNA**→ |
| env-135 | SA_S1_NOR05_45m_c10026777 | - | 566 | 490 | **RNA**→ rplT (PRK05185)Ribosomal_L20 (pfam00453)→ hypo→ |
| env-136 | SA_S1_NOR08_45m_c10014299 | + | 2144 | 2220 | **RNA**→ rplT (PRK05185)Ribosomal_L20 (pfam00453)→ |
| env-137 | SA_S2_NOR15_50m_c10136720 | + | 639 | 715 | **RNA**→ rplT (PRK05185)rpl20 (CHL00068)→ |
| env-138 | SA_S1_NOR02_45m_c1067657 | + | 160 | 236 | **RNA**→ rplT (PRK05185)Ribosomal_L20 (pfam00453)→ |
| env-139 | SA_S2_NOR13_50m_c1078169 | - | 216 | 139 | **RNA**→ rplT (PRK05185)Ribosomal_L20 (pfam00453)→ |
| env-140 | SA_S1_NOR05_45m_c10009856 | + | 2006 | 2083 | **RNA**→ rplT (PRK05185)Ribosomal_L20 (pfam00453)→ |
| env-141 | SA_S1_NOR08_45m_c10000806 | + | 2877 | 2954 | **RNA**→ rplT (PRK05185)Ribosomal_L20 (pfam00453)→ |
| env-142 | SA_S2_NOR15_50m_c10001918 | + | 2889 | 2966 | **RNA**→ rplT (PRK05185)Ribosomal_L20 (pfam00453)→ |
| env-143 | SA_S1_NOR02_45m_c1117957 | + | 138 | 215 | **RNA**→ rplT (PRK05185)rpl20 (CHL00068)→ |
| env-144 | SA_S2_NOR18_50m_1236664 | + | 180 | 257 | **RNA**→ |
| env-145 | SA_S1_NOR05_45m_c10219562 | + | 174 | 251 | **RNA**→ rplT (PRK05185)Ribosomal_L20 (pfam00453)→ |
| env-146 | SA_S2_NOR15_50m_c10184235 | + | 79 | 156 | **RNA**→ rplT (PRK05185)Ribosomal_L20 (pfam00453)→ rplT (PRK05185)Ribosomal_L20 (pfam00453)→ |
| env-147 | SA_S2_NOR15_50m_c10184235 | + | 392 | 469 | **RNA**→ rplT (PRK05185)Ribosomal_L20 (pfam00453)→ rplT (PRK05185)Ribosomal_L20 (pfam00453)→ |
| env-148 | SA_S1_NOR05_45m_c10004739 | + | 2326 | 2403 | **RNA**→ rplT (PRK05185)Ribosomal_L20 (pfam00453)→ |
| env-149 | SA_S1_NOR08_45m_c10004622 | - | 3588 | 3511 | **RNA**→ rplT (PRK05185)Ribosomal_L20 (pfam00453)→ |
| env-150 | SA_S2_NOR15_50m_c10019150 | + | 2898 | 2975 | **RNA**→ rplT (PRK05185)Ribosomal_L20 (pfam00453)→ |
| env-151 | SA_S2_NOR15_50m_c10106824 | - | 231 | 154 | **RNA**→ rplT (PRK05185)Ribosomal_L20 (pfam00453)→ |
| env-152 | SA_S1_NOR02_45m_c1126557 | + | 164 | 241 | **RNA**→ rplT (PRK05185)rpl20 (CHL00068)→ |
| env-153 | ASA120_GJFD58A02GB27U | - | 120 | 43 | **RNA**→ |
| env-154 | BS_KBA_SWE12_21m_c10210524 | + | 169 | 246 | **RNA**→ rplT (PRK05185)Ribosomal_L20 (pfam00453)→ |
| env-155 | BS_KBA_SWE21_205m_10288133 | + | 106 | 183 | **RNA**→ rplT (PRK05185)Ribosomal_L20 (pfam00453)→ |
| env-156 | BS_KBA_SWE02_21m_10654080 | - | 133 | 56 | **RNA**→ |
| env-157 | BS_KBA_SWE12_21m_c10693372 | - | 146 | 69 | **RNA**→ rplT (PRK05185)rpl20 (CHL00068)→ |
| env-158 | BS_KBA_SWE21_205m_10269672 | + | 184 | 261 | **RNA**→ rplT (PRK05185)rpl20 (CHL00068)→ |

| | | | | | |
|---|---|---|---|---|---|
| env-159 | BS_KBA_SWE12_21m_c10454375 | - | 178 | 101 | **RNA**→ rplT (PRK05185)rpl20 (CHL00068)→ |
| env-160 | KGI_S1_ANT01_95m_c10080207 | - | 841 | 763 | **RNA**→ rplT (PRK05185)Ribosomal_L20 (pfam00453)→ pheS (PRK00488)Phe_tRNA-synt_N (pfam02912)→ |
| env-161 | ASLM20b_GM71RDO02F8O89 | + | 134 | 212 | **RNA**→ rplT (PRK05185)Ribosomal_L20 (pfam00453)→ |
| env-162 | ASA120_GJFD58A02J09L8 | - | 396 | 318 | **RNA**→ rplT (PRK05185)Ribosomal_L20 (pfam00453)→ |
| env-163 | SA_S1_NOR08_45m_c10161630 | - | 206 | 128 | **RNA**→ rplT (PRK05185)rpl20 (CHL00068)→ |
| env-164 | SA_S2_NOR15_50m_c10056163 | + | 270 | 348 | **RNA**→ rplT (PRK05185)Ribosomal_L20 (pfam00453)→ hypo→ PheS (COG0016)pheS (PRK00488)→ |
| env-165 | SA_S2_NOR13_50m_c1030706 | - | 679 | 601 | **RNA**→ rplT (PRK05185)Ribosomal_L20 (pfam00453)→ hypo→ |
| | | | | | pheS (PRK00488)Phe_tRNA-synt_N (pfam02912)→ |
| env-166 | SA_S1_NOR08_45m_c10303771 | - | 125 | 47 | **RNA**→ |
| env-167 | SA_S1_NOR05_45m_c10043877 | + | 802 | 880 | **RNA**→ |
| env-168 | SA_S2_NOR15_50m_c10070834 | - | 112 | 34 | **RNA**→ |
| env-169 | SA_S1_NOR05_45m_c10169234 | + | 253 | 331 | **RNA**→ rplT (PRK05185)Ribosomal_L20 (pfam00453)→ |
| env-170 | SA_S1_NOR08_45m_c10013077 | + | 3405 | 3483 | **RNA**→ rplT (PRK05185)Ribosomal_L20 (pfam00453)→ |
| env-171 | SA_S1_NOR08_45m_c10110821 | - | 352 | 274 | **RNA**→ rplT (PRK05185)Ribosomal_L20 (pfam00453)→ |
| env-172 | SA_S2_NOR15_50m_c10446830 | + | 267 | 345 | **RNA**→ |
| env-173 | SA_S2_NOR18_50m_1021448 | + | 190 | 268 | **RNA**→ rplT (PRK05185)Ribosomal_L20 (pfam00453)→ hypo→ |
| | | | | | pheS (PRK00488)Phe_tRNA-synt_N (pfam02912)→ |
| env-174 | LGOV01008591.1 | + | 13588 | 13681 | **RNA**→ rplT (PRK05185)Ribosomal_L20 (pfam00453)→ |
| env-175 | LGVE01183011.1 | + | 6160 | 6253 | **RNA**→ rplT (PRK05185)Ribosomal_L20 (pfam00453)→ |
| env-176 | LGVF01312494.1 | + | 20148 | 20241 | **RNA**→ rplT (PRK05185)Ribosomal_L20 (pfam00453)→ |
| env-177 | 3300001580_10002513 | - | 10447 | 10365 | **RNA**→ rplT (PRK05185)Ribosomal_L20 (pfam00453)→ PheS (COG0016)pheS (PRK00488)→ |
| | | | | | PheT (COG0072)pheT (PRK00629)tRNA_bind_bactPheRS (cd02796)→ ihfA (PRK00285)IHF_A (cd13835)→ |
| | | | | | HTH_MlrA-like_sg2 (cd04765)MerR_1 (pfam13411)→ SurE (COG0496)surE (PRK00346)→ |
| | | | | | Pcm (COG2518)Methyltransf_25 (pfam13649)→ YgaA (COG1238)SNARE_assoc (pfam09335)→ |
| | | | | | LysM (smart00257)LysM (cd00118)Peptidase_M23 (pfam01551)→ RpoD (COG0568)sigma70-ECF (TIGR02937)→ |
| | | | | | Apt (COG0503)PRK02304 (PRK02304)→ hypo→ |
| env-178 | 3300000558_10371857 | - | 2112 | 2038 | **RNA**→ rplT (PRK05185)Ribosomal_L20 (pfam00453)→ PheS (COG0016)pheS (PRK00488)→ |
| env-179 | 3300001580_10030616 | - | 3653 | 3579 | **RNA**→ rplT (PRK05185)Ribosomal_L20 (pfam00453)→ PheS (COG0016)pheS (PRK00488)→ |

#### 7.3 Conserved domains

Conserved domains found in protein-coding genes listed in Section 7.2 are shown below, with the first sentence in their description from the Conserved Domain Database (if any). Conserved domains associated with more than one L20-Deltaproteobacteria RNA are assigned a color, while others are shown in gray. The number in parentheses after the colored domain name is the number of occurrences in Section 7.2.

cd00118 (1) Lysine Motif is a small domain involved in binding peptidoglycan.
cd02796 (35) tRNA-binding-domain-containing prokaryotic phenylalanly tRNA synthetase (PheRS) beta chain.
cd04765 (18) Helix-Turn-Helix DNA binding domain of putative MlrA-like transcription regulators.
cd06158 (1) Uncharacterized homologs of Site-2 protease (S2P), zinc metalloproteases (MEROPS family M50) which cleave transmembrane domains of substrate proteins, regulating intramembrane proteolysis (RIP) of diverse signal transduction mechanisms.
cd07026 (1) Ribosomal protein L20.
cd13835 (18) Alpha subunit of integration host factor (IHFA).
CHL00068 (15) ribosomal protein L20
CHL00192 (1) phenylalanyl-tRNA synthetase beta chain; Provisional
COG0016 (48) Phenylalanyl-tRNA synthetase alpha subunit [Translation, ribosomal structure and biogenesis]
COG0072 (35) Phenylalanyl-tRNA synthetase beta subunit [Translation, ribosomal structure and biogenesis]
COG0180 (1) Tryptophanyl-tRNA synthetase [Translation, ribosomal structure and biogenesis]
COG0496 (8) Broad specificity polyphosphatase and 5'/3'-nucleotidase SurE [Replication, recombination and repair]
COG0503 (4) Adenine/guanine phosphoribosyltransferase or related PRPP-binding protein [Nucleotide transport and metabolism]
COG0568 (4) DNA-directed RNA polymerase, sigma subunit (sigma70/sigma32) [Transcription]
COG0739 (2) Murein DD-endopeptidase MepM and murein hydrolase activator NlpD, contain LysM domain [Cell wall/membrane/envelope biogenesis]
COG0789 (1) DNA-binding transcriptional regulator, MerR family [Transcription]
COG1238 (3) Uncharacterized membrane protein YqaA, SNARE-associated domain [Function unknown]
COG1354 (1) Chromatin segregation and condensation protein Rec8/ScpA/Scc1, kleisin family

[Replication, recombination and repair]
COG2518 (4) Protein-L-isoaspartate O-methyltransferase [Posttranslational modification, protein turnover, chaperones]
pfam00453 (165) Ribosomal protein L20.
pfam01551 (1) Peptidase family M23.
pfam02912 (11) Aminoacyl tRNA synthetase class II, N-terminal domain.
pfam03483 (2) B3/4 domain.
pfam04079 (1) Segregation and condensation complex subunit ScpB.
pfam09335 (3) SNARE associated Golgi protein.
pfam11853 (1) Protein of unknown function (DUF3373).
pfam13411 (19) MerR HTH family regulatory protein.
pfam13649 (2) Methyltransferase domain.
PRK00104 (1) segregation and condensation protein A; Reviewed
PRK00135 (1) segregation and condensation protein B; Reviewed
PRK00285 (17) integration host factor subunit alpha; Reviewed
PRK00312 (2) protein-L-isoaspartate O-methyltransferase; Reviewed
PRK00346 (8) 5'(3')-nucleotidase/polyphosphatase; Provisional
PRK00488 (60) phenylalanyl-tRNA synthetase subunit alpha; Validated
PRK00629 (37) phenylalanyl-tRNA synthetase subunit beta; Reviewed
PRK00927 (1) tryptophanyl-tRNA synthetase; Reviewed
PRK02304 (4) adenine phosphoribosyltransferase; Provisional
PRK05185 (181) 50S ribosomal protein L20; Provisional
smart00257 (3) Lysin motif.
smart00411 (1) bacterial (prokaryotic) histone like domain.
smart00896 (2) Ferredoxin-fold anticodon binding domain.
TIGR02937 (4) RNA polymerase sigma factor, sigma-70 family.

#### 7.4 Multiple-sequence alignment

Each L20-Deltaproteobacteria RNA is denoted by an abbreviation (like "Eco-1-1") that refers to a taxonomy given in Section 7.1. The alignment may include sequences containing the RNA motif, as well as flanking sequence. The L20-Deltaproteobacteria RNA itself is denoted by the line underneath marked 5′ and 3′ on either end. Nucleotides in flanking sequences (i.e., not part of the motif) are written in gray letters. Stems of predicted rho-independent transcription terminators, if any, are shaded yellow. (Note: terminator predictions have not been analyzed manually, and many are likely to be false positives. Terminator predictions are those of the RNIE software.) Nucleotides predicted to function (as DNA) as transcription-factor binding sites, if any, are shaded in green. (But note: these predictions are manually annotated, so they might be under-predicted.) Annotated start codons, if any, are shaded green. (Note: start codons are frequently misannotated, especially in environmental samples.) Nucleotides proposed to basepair as part of the consensus structure are shaded in color when they comprise Watson-Crick or G-U pairs. Otherwise they are shaded gray. Conserved stems are also indicated at the bottom of the alignment by angle brackets, where matching < and > denote base-paired columns. Below these angle brackets, the symbol "2" denotes base pairs exhibiting covariation according to the statistically well-founded R-scape method. "1" denotes base pairs exhibiting covariation according to R2R's simplistic method. "0" denotes base pairs that are not observed to mutate and "?" denotes base pairs that have a significant frequency of non-canonical nucleotides for Watson-Crick or G-U pairs (> 5%). Below these base pair annotation is the consensus sequence: "R" = "A" or "G", "Y" = "C" or "U", red nucleotides: nucleotide identity conserved more than 97% of the time, black nucleotides: 90%, gray nucleotides: 75%, red circle (•): nucleotide is present 97% of the time, black circle (•): 90%, gray circle (•): 75%, white circle (∘): 50%. All percentages of sequences just described (e.g. 97% conserved) assume that sequences have been weighted by the GSC algorithm implemented by the Infernal software package.

Duplicate sequences: the following putative homologs are not shown in the alignment because their sequences are identical to a homolog already shown: Dac-1-1, Gme-2-1, Gsu-3-1, Gsu-4-1, env-10, env-101, env-102, env-105, env-125, env-126, env-127, env-128, env-129, env-132, env-135, env-136, env-137, env-138, env-140, env-141, env-142, env-143, env-144, env-146, env-149, env-150, env-151, env-152, env-155, env-156, env-161, env-164, env-170, env-171, env-172, env-173, env-175, env-176, env-179, env-22, env-31, env-32, env-33, env-36, env-37, env-4, env-40, env-42, env-43, env-45, env-5, env-50, env-51, env-53, env-63, env-66, env-68, env-73, env-74, env-78, env-85, env-96, env-97

alignment positions 1 · · · 77

```
env-1     CGACCGUUGUCUCCAAGGCUGAUGAGCGCAACAUCAGAGAGCUCAUCCCUUACCUGUAAGCUUGACAACUGUUUCAC
env-2     GCGUCAGGGAACCCUGAUCCAUAAAGCAGAUGAGCGUAAUAUUGCGCGGUUGAUUCCCUACUUGUAAAGGGAACUUU
env-3     UCAUAUCGAAGGCUGAUGAGCGCAACAUUGCACAGUUGAUUCCGUAUCUGUAAGGCGGUCUGCCACUGAGGUAGAUU
env-6     .......AAGAGCGAUUCGAAAAAUAUUUCCCAAUUAGUUCCGUAUUUGUAACUUGAUCUGCUGCUUUCUGGCAGUA
env-7     UGUUGCAAAGGCGGAUGCAAAAAAUAUCUCCCAAUUAGUUCCGUAUUUGUAGAUUGACCUACUGUCUUCUGGCAGUG
Pse-1-1   UCUCGUUGCCAAGGCUGAUGCCCGCAACAUGGCUCAAUUGAUUCCUUACCUGUAACACUCGCCUUUCGGGGCGGGUC
env-8     CCUCGUUGCCAAGGCUGAUGCCCGCAACAUGGCUCAAUUGAUUCCCUACAUGUAAUACCCGCCUUUUGAGAUGGGUC
env-9     CCUCGUUGCCAAGGCUGAUGCCCGCAACAUGGCUCAAUUGAUUCCCUACAUGUAAUACCCGCCUUUCGAGAUGGGUC
env-11    CGUGCACAAAGCUGACGAGCGCAACAUUGCGCAACUGAUUCCGUACUUGUAAAAAUUGCCUGUUGGUAGGGCAACAU

          .............................................................................
          .............................................................................
          •••••••••••••••••••••••••••••••••••••••••••••••••••••••••••••••••••••••••••••
```

```
Gel-1-1    GCGUCAAGGAACCCUGGUUCACAAAGCCGACGAGCGUAAUAUUGCAAGGUUGAUCCCCUAUUUGUAGGGAGCUGUAA
env-12     GUCAGAGCACCAUUGUCGACCCGGCAGAUGCUAAGAAUAUAGCCCGUCUGAUCCCUUACAAGUAAGGCUCUAUCGGG
env-13     .....................GUUGAUCAGAAAAACAUCUGUGCUCUGAUUCCGUAUCUGUAAUGACGCUUGUGCGU
env-14     CACGGCGGUACGGUGGCUGCAGUUGAUCAGAAAAAUAUCUGUGCUCUGAUUCCGUACCUGUAAUGACGCUUGUGCGU
env-15     CACGGCGGCACGGUGGCUGCGGUUGAUCAGAAAAACAUCUGUGCUCUGAUUCCGUAUCUGUAAUGACGCUUCUGCGU
env-16     CANGGCGGCACGGUGGCUGCGGUUGAUCAGAAAAACAUCUGUGCUCUGAUUCCGUAUCUGUAAUGACGCUUGUGCGU
env-17     .............................................................................A
env-18     GCCGUUGACCAGAAGAACAUUGCCAGACUGAUUCCGUAUAAAUAAAUUUUCCUCCAGUAAUAGUGGAGGACUGCAUA
env-19     ACCAGAAGAAUAUCGCCCGUUUGAUUCCGUACAAAUAACCAGUUUUCUUUCUCAUUUAUCAUGUGAGAGAACGCAUA
env-20     ..................................................................AGACGCGCAUA
env-21     GAUCAGAAGAACGUUGCCCGGUUGAUUCCGUAUAAGUAACCAGAUUUAUUCGCUCACAGUAUGUGGGUAGAUGCAUA
Ppr-1-1    CUGCGUGGUACUUCCAUGGUGUCCGAUGUGGACCAGAAGAAUAUUUCACGUCUCAUCCCGUACAAGUAAGCGGGACC
env-23     AUACCGUUGCUGCUGUUGAUCAGAAGAACAUCUGCGCACUGAUUCCGUACAUGUAAGUUUGAGCAGCGUGGUUUGCU
env-24     UAACCUGAGGCAGGCCGGCAUGGUUGCCGAUGUUGACCAGAAAAACAUAGCCAGACUUAUCCCCUAUAAAUAGCGGG
env-25     GCAAAGAACAUCGCAAAACUGAUUCCGUAUAUGUAGCUCAGCGGUGACUGGCGCAACGUGCCAGAUCCCUGCAACCA
Gme-1-1    GGGCCAGGAGCCAUGGCUGAUAUCGGGCCGGAGUUAUUCGUUGGUUCGUGUAGCUGUCUCCUGUAUCCGUUUUUAUA
env-26     GGCUUUGCCCGCGAUCGCGCCUGAAGGCGCUCCUGCAAUUUGUGCGGUUAGCCGCCGGAGACAUAAAUACGAUCAGC
env-27     CACCGUUGCUGCCGUUGACCAGAAGAAUAUCUGUGCGUUGAUCCCUUAUAUGUAAUCAGAGCCAGCAACUGCUGUCU
env-28     ..................................................................CCCUGCUGUCU
env-29     GUUGCAUCAGUAGAUCAAAAGAACAUCUGUGCACUGAUUCCAUAUCUUUAAUCCAGGCACAGCCCUGUGCUGUACCG
env-30     GGUAGCAGCCGUUGACCAAAAAAAUAUCUGUGCUUUGAUCCCCUAUUUGUAACCUAGAGGCUGCUUUAGAGUGGUUU
env-34     CACCGUGGCAGCAGUUGAUCACAAAAACAUCUGUGCCUUAAUCCCUUAUAUGUAACCAGAAACAGCAACUGCUGUUU
env-35     GCUGCUGUUGAUCAAAAAAAUAUUUGUGCAUUAAUUCCCUAUAUGUAAAGUGUUGUAUAGGUGCAGCUUUGCUGUGC
env-38     CACUGUGGCAGCAGUUGAUCAUAAAAACAUCUGCGCCUUAAUCCCUUAUAUGUAACCAGAAACAGCAGCAGCUGUUU
env-39     UACGGUGGCUGCAGUGGAUCAAAAAAAUAUUUGUGCUUUGAUUCCCUAUAUGUAGUUAGAAACAACGUCUGUUGUUU
env-41     UACAGUGGCUGCAGUGGAUCAAAAAAAUAUUUCUGCUUUGAUUCCUUAUAUGUAGUCAGAAACAACGCCUGUUGUUU
env-44     CACAGUGGCCGCAGUGGAUCAUAAAAAUAUUUGCGCUUUGAUUCCCUAUAUGUAACCAGAAACAGCAGCUGCUGUUU
env-46     UACCGUGGCUGCAGUGGAUCACAAAAACAUCUGCGCUUUGAUCCCCUAUAUGUAACCAGAAACAGCAGAUGCUGUUU
env-47     CACUGUGGCUGCAGUAGAUCAUAAAAAUAUCUGCGCUUUGAUUCCCUAUCUGUAACCAGAAACAGCCAGUCCUGUUU
env-48     GCAGUUGACCAGAAGAACAUCUGCGCACUGAUUCCUUACCUCGUAACCGUGUAAGGUUAGGCGCACCACUGGUGCGC
env-49     CGCAGUUGACCAGAAGAACAUCUGCGCACUGAUUCCUUACCUGUAACCGUGUAAGGUUAGGCGCACCACUGGUGCGC
env-52     GGUGGAUCAAAAGAACAUCAGCGCACUGAUUCCUUACCUGUAACCGGGUAGGGGCGCUAGGUGCACCGCUGGUGCAC
env-54     GGUGGAUCAAAAGAACAUCAGCGCACUGAUUCCUUACCUGUAAUCAGGUAGGGGCGCUAGGUGCACCUACGGUGCGC
env-55     GUGGAUCAAAAGAACAUCAGCGCACUGAUUCCUUACCUGUAGCCGGGUAGGGGCGCUAGGUGCACCGACUGGUGCAC
env-56     GUGGAUCAAAAGAACAUCAGCGCACUUAUUCCUUACCUGUAACCAGGUAGGGGCGCUAGGUGCACCGUUAGGUGCAC
env-57     GUGGAUCAAAAGAACAUCAGCGCACUGAUUCCUUACCUGUAGCCGGGUAGGGGCGCUAGGUGCACCGACUGGUGCAC
env-58     ...............................CUUACCUGUAGCCGGGUAGGGGCGCUAGGUGCACCGACUGGUGCAC
env-59     GUGGAUCAAAAGAACAUCAGCGCACUGAUUCCUUACCUGUAGCCGGGUAGGGGCGCUAGGUGCACCGACUGGUGCAC
env-60     GUGGAUCAAAAGAACAUCAGCGCACUGAUUCCUUACCUGUAACCGGGUAGGGGCGCUAGGUGCACCGCUAGGUGCAC
env-61     UACCGUUGCCGCUGUGGAUCAAAAAAAUAUCAGUGCUUUGAUUCCGUACAUGUAGUUAGAAACAGCAUCUGCUGUUU
env-62     UACCGUUGCCGCUGUGGAUCAGAAAAACAUCUGUGCUUUGAUUCCGUACAUGUAGUUAGAAACAGCAGCUGCUGUUU
env-64     CACCGUUGCUGCAGUGGACCAGAAAAAUAUCUGCGCCUUGAUUCCGUACAUGUAGUUAGAAACAGCUUUUGCUGUUU
Glo-1-1    CACCGUUGCUGCAGUGGAUCAGAAAAAUAUCUGUGCCUUGAUUCCGUACAUGUAGUUAGAAACAGCUUUUGCUGUUU
env-65     CACCGUUGCUGCAGUGGAUCAGAAAAAUAUCUGUGCCUUGAUUCCGUACAUGUAGUUAGAAACAGCUUUUGCUGUUU
env-67     UGCGGGGUUCCACAAUGGUUUCCGAUGUGGAUCAGAAAAAUAUUUCGCGUCUUAUCCCGUACAAAUAAAAACGGGAA
env-69     UGCGGGGCGUCACUAUGGUGUCUGAUGUGGAUCAAAAAAAUAUUUCACGUCUUAUCCCGUAUAAAUAGAAACGGGAA
env-70     UCAAAAAAAUAUCGCAAGAUUGAUCCCCUAUAAGUAAAAACGGGGCCGGGAUUCAUUCCCGGCACCUGUUUUUCAUA
env-71     GACGCAAAGAACAUCGCAAAAUUGAUUCCGUAUCUAUAGUCUGGGAGUCCGUCCAUUUGUGACAGGGUUCCACCCGA
env-72     GACGCAAAAAAUAUCGCAAAAUUGAUUCCGUAUCUAUAGUCUGGGAGUCCGUCCAUCGGUGACAGGGGUCCACCCGA
env-75     UGACGCCAAGAAUAUCGCAAAGCUGAUUCCCUAUAUGUAAUUUGACCCGCUGGCCUUUGUGCCGGCGUCUCACACGA
env-76     CCUGCGUCAGGGCGGUAUGGUUGCCGACGUUGACCAGAAAAACAUUUCCAAGCUCAUCCCGUACAUGUAACGGGCCA
env-77     UGUUGACGCCAAGAAUAUCGCGAAGCUGAUUCCGUAUAUGUAGUUUGGGUAACGCCCUUGAGGGGGUACCCAGCCAU
Gso-1-1    CGCCGUUGUGGCGGCAGUGGACCACAAGAACAUCGCCAAGCUCAUCCCUUACAUGUAGCGGAGCCUGCUCCGUCAUG
Gsu-2-1    CGCUGUUGUGGCGGCAGUUGAUCACAAGAAUAUUGCCAAGCUUAUCCCUUACAUGUAACGGAGCCUGCUCCGUCAUG
env-79     CGCUGUUGUGGCGGCAGUUGAUCACAAGAAUAUCGCCAAGCUUAUCCCUUACAUGUAACGGAGCCCGCUCCGUCAUG
Gpi-1-1    CGGCGCUUUGGUUGCUGCUGUUGAUCAGAAGAACAUCGCAAAGCUGAUCCCCUACGUGUAGUUAAGGGGCAUGAUUU
env-80     GCCAGGCUGAUUCCGUACAUGUAAUAAGUAAUAAAUAAACGUGAACAGUGAUACUUUCACUGUUCACGUUUUACCAG
Gsp-4-1    GUAAAAGCCGUGAAUGGUGAAAUGUGAUUGUGAAGAAAUAUCUUCACUUUCACUGUUUCCAAUUCACGUUGCAACAG
env-81     AGAUGCGAAGAAUAUCGCAAAGCUGAUUCCGUAUAUGUAGACCUGGUCCCGGCCUCCCAUGGCCGAGGUCCUAACAG
env-82     AGAUGCGAAGAAUAUCGCAAAGCUGAUUCCGUAUAUGUAGACUUGGUCCCGGCCUCCCAUGGCCGAGGUCCUAACAG
env-83     GUAGCACCGGUUGACGCAAAAAACAUCGCCAAGCUGAUUCCUUAUCUUUAAAGUGCAUGAGGCUUGGGCUGUGCGCA
env-84     UGUAUCCCCUGUAGAUGCGAAGAACAUUGCCAGGCUGAUUCCCUACUUGUAGGAAGUGACGGUGACGGCACUACGGA
env-86     UGUAUCGCCGGUAGAUGCGAAGAAUAUUGCCAAGCUGAUUCCUUACUUGUAGGAAGUGACCGCGAUGGCAAUGAGGA
env-87     UGGGCUCUAGGCUUUAGGUAAAAACAGACCCAGAGCCUAAUGCCCAGUGCCUAGAGCCUAGAGCCCGAUUUUUUGAA
env-88     .......AUUCCGUAUAUGUAGAUAAACGUGAAUGUAUGGAAGUCUUAGGCCUUUCAUUAAGUUCACGUCUGACAGU
env-89     ...........GCGGCAUGGUUGCCGAUGUGGAUCAGAAAAACAUUUCCAAGCUUAUCCCGUACAAGUAACGGGGCA
Gsp-2-1    CUGGAGACUGUAAAAACUUGAAAAACCCGGGACUGGGAUUAGGGCUGCGGCGCUACCCCCUCGCCCCUGGUUCACAU
env-90     ACUAGGGACUGGAACACCGGCAUCGAGGUGCGCCUCGGAGCUGUUCGUCUUCAACCGCCGUUUUCGCCCGUUACCGA
Gbr-1-1    GAGACUAGAACCGGAGACUAAAUAGCUAAUAACCUGGACGGGGGCGGAGUUCCCAGCCCCCAGCCCCGGGUUUACCA
Gbe-1-1    GUAACGGGAGGGAGUUGUGCAGGGAUGCUGUGAUUUUUCCAGCCCCUAGCCCCCAGUCCCUAGCCCCGGGUUUACCA
Gsp-3-1    GGAAUGGGACAGGUGGUGCAAGGGAUGCUGAGAUUUUUCCAGCCCCUAGCCCCCAACCCCUAGCCCCGGGUUUACGA
Gsp-1-1    CGGUCGCUGGUCGGCUGUCGGCGGUUGAUAGUUUAAAAACACCGGCGACCACCGUCCAUCGCCUGCCGACCGAUUUA
env-91     GGUCGGUAGUCGGCGGUCGGCGGUCGGAAGUAGAUCAACCAGCGACCACCGACCGCCGACCGCCGACCGGUUUUUAA
Gur-1-1    GUCGGUGGUCGAUAGCAAUGCAUCAGAUUCUUAAUUCACCAUCGACCGGUUACCAUCAACCAUCAACCGAUUUUUAA
env-92     AUCUGGGAUCUGGGACCUGGAAAAACCAGAUUUGUAUUUACCGUUCACAGUUCCCUGUUCACCAUUCACAGCAUUAA
env-93     AACCUUCGUCAGACCGCCAUGGUGGCUGAUGUUGAUCAGAAAAACAUAGCUCGUCUUAUCCCGUACAAAUAGCGGGC
env-94     GCCACUCUCGUGUCCGCUGCAGAUGUGAAAAAUAUCAGUCGACUGAUUCCUUAUCUCUGAUCAGUCGGUUUGACAUU
env-95     UGGUGUCCAAGGCAGACGCUGGCAAUAUUGCCCGUCUCAUCCCUUACAAGUAAUCGGGGCUGAGAGGGCUUGGCUUA
env-98     ..............UCCGGCAAUAUCGCCCAGCUUAUUCCCUACAAGUGACGGCUUGUUUUUGGAGUACCCGUUUCA
env-99     GUUUCCAAAGCGGAUGCCGGCAAUAUCGCCCAGCUUAUUCCUUACAAGUAACUGGUUGUUUUGAAGCACCCGUUUCA
env-100    UGGUGUCCAAGGCAGACGCAGGAAACAUUGCCCGUCUCAUCCCUUACAAGUAAUCGGGGUCGAGUGGGCUUGGCUUA
env-103    CUGCCAAGCGUUUUCGUAAAACCGGUACCGGAAAGAUUCGUCGUAAUAAGGCUUUUACCAGCCAUAUUCNNGGCUUA
Gfe-1-1    GUCGACAGUCGUGACCAUGCCAAUAUCAGCCGCCUAAUUCCUUAUCUCUGAUUGGCAGAUACUGCCUAUCGGCUUAA
Gsu-1-1    CAUCAGUGGCCGCGACGAGAAGAAUGUCAGUCGUCUGAUCCCUUACCUUUAAGGGGGGAUAAAAAUCGGCUUUCUUA

  ...............................................................................
  ...............................................................................
  ...............................................................................
```

```
env-104   CACAAUUGUCGAUGCUGCGGAUCAUAAAAAUAUCAGUUGCCUGAUUCCUUACAAGUAGGGAACGGCAACUCGGCUUA
env-106   GUGCACAAGGCGGAUGCCGGCAACAUCAGUCGCCUGAUUCCUUAUCUGUAACCGGGUAACGAACCAGUUUCGGCUUA
env-107   ......................................................................CGAUUCA
env-108   ACUCUGGUGGCCAAGGUGGACGAAAAAAAUAUCAGUCGGUUGAUUCCCUACAUCUAGGAAUCCGCCAGGGCGAAUCA
env-109   ACUCUGGUGGCCAAGGUGGACGAAAAAAAUAUCAGUCGGUUGAUUCCCUACAUCUAGGAAUCCGCCAGGGCGANUCA
env-110   ....................................CGGUUGAUUCCCUACAUCUAGGAAUCCGCCAGGGCGAAUCA
env-111   UGGUGCACAAGGCUGAUGCCGCCAACAUCAGUCGCCUGAUUCCGUACCUGUAAAAGGGACCAACCAGGAUCGGCUCA
env-112   UGGUGCACAAGGCUGAUGCCGGAAACAUCAGUCGCCUGAUUCCGUACCUGUAAGAGGGACCAACCAGGAUCGGCUCA
env-113   UGGUGCACAAGGCUGAUGCCGCCAACAUCAGUCGCCUGAUUCCGUACCUGUAAAAGGGACCAACCAGGAUCGGCUCA
env-114   UGGUGCACAAGGCUGAUGCCAGCAACAUCAGCCGCCUGAUUCCGUACCUGUAAAAGGGACCAACCAGGAUCGGCUCA
Dsp-1-1   AACCCUUGUGGCCAAGGUGGAUGAAAAGAACAUCAGCCGUUUGAUUCCCUACAUGUAGGUUCAUCCGGCUCGGCUCA
env-115   ACCCUGGUCGCCGCAGUCGAUGCAAAGAACAUCAGCCGUCUGAUUCCCUAUCUCUGAGGAUUCAGUGGCUCGGCCUA
env-116   ..................................GCCGUCUGAUUCCCUAUCUCUGAGGAUUCAGUGGCUCGGCCUA
env-117   GACCCUGGUUGCCGCUGUCGAUGAAAAGAAUAUCAGCCGUCUGAUUCCUUACCUGUAAGGCCUGACGGCUCGGCUUA
Pca-1-1   UUGUCGCAAAAGCUGACCAGAAGAAUAUCAGCCGUCUUAUUCCCUAUAUCUAGGCGUAAGACAGCGGGCCAUGUUUA
env-118   UCGUAGCCAAGGCCGACCACAAAAAUAUCAGCUGUCUUAUCCCUUAUAUUUAAGAGCGAGACGGUGGGUCAUGUUUA
env-119   UCGUUGCCAAGGCUGACCAUAAAAAAUAUCAGCUGUCUUAUCCCUUACAUUUAAGAGCGAGACGGUGGGUUAUAUUUA
env-120   CACCCUGGUCGACAAGGCCGAUGUCAGGAAUAUCCGCUGCCUGAUCCCCUAUCUGUAAGGGCAGGCGGCACCACAAC
env-121   .....................................UGUUUGAUCCCUUACCUCUAAGGGACAGGCAGAAAGCUAC
env-122   GCGACGUCAAAAAUAUCAGCUGUCUGAUCCCUUACCUUUAAGGGAUAGUAGUCAAGGCUGCUAGGUAGCACAUCAAC
env-123   CCUUGGUUGCUAAAUGCGACUCCAAGAAUAUCAGCUGUUUGGUUCCUUACCUUUAAGGACUAGGCGGCAGAUUAACC
env-124   CCUUGGUUGCUAAAUGCGACUCCAAGAAUAUCAGCUGUUUGGUUCCUUACCUUUAAGGACUAGGCGGCAGAUUAACC
env-130   CUUUGGUUGCUAAAUGCGACGCCAAAAACAUCAGCUGUUUGGUCCCUUACCUUUAAGGGCUAGAGAGCAGAUCAACC
env-131   CUUUGGUUGCUAAAUGCGACGCCAAAAACAUCAGCUGUUUGGUCCCUUACCUUUAAGGGCUAGAGAGCAGAUCAACC
env-133   CUUUGGUUGCUAAAUGCGACGCCAAAAACAUCAGCUGUUUGGUCCCUUACCUUUAAGGGCUAGAGAGCAGAUCAACC
env-134   CUGUGGUUGCCAAAUGCGACGUUAAGAACAUCAGCUGCUUGAUUCCUUACCUUUAAGGACAAGUUGGCAGAUUAAUC
env-139   CUUUGGUUGCCAAAUGCGACGCCAAGAACAUCAGCUGUUUGAUCCCUUACCUUUAAGGGCUGGACAGCAGAUCAACC
env-145   CUUUGGUUGCUAAAUGCGACACCAAGAAUAUCAGCUGUUUGGUCCCUUACCUUUAAGGGCUGAAUGGCAGAUCAACA
env-147   CGUGUUCGCAAAAAUGCGACGCCAAGAAUAUCAGCUGUUUAGUCCCUUACCUUUAAGGGCUGGAUGGCAGAUCAACC
env-148   CUUUGGUUGCUAAAUGCGACGCCAAGAAUAUCAGCUGUUUAGUCCCUUACCUUUAAGGGCUGGAUGGCAGAUCAACC
env-153   GACUCUGGUUGACAAGAGCGACGCCAAGAAUAUCAGCUGUCUGAUCCCUUACCUGUAAGGGUUAGCCACCUGUCAAC
env-154   GAUAAGAGCGACGUCAAGAAUAUCCGCUGUUUGAUCCCUUAUCUUUAAGCUUUAAGCUGUAAGGGGUACAGUACAAC
env-157   ACCCUGGUAAACAAGAGCGACGCCAAGAACAUCAGCUGUAUGAUCCCUUACCUUUAAGGACUGUCGGCACAAUAUC
env-158   ACCUUGGUCAAUAAAAGCGACGUCAAGAACAUCAGCUGUUUGAUCCCUUACCUCUAAGGGGUCGGCGGCACAUCAAC
env-159   ACCCUGGUCAAUAAAAGCGACGUCAAGAACAUCAGCUGUUUGAUCCCUUACCUCUAAGGGGUCGGCGGCACAUCAAC
env-160   ACCCUGGUUGACAAGGCCGACGCCAAAAAUAUCCGCUGUCUGAUCCCUUACCUGUAAGGGACGUACAGUAUCCACAC
env-162   ACCCUGGUUGACAAGGCCGACGCCAAGAAUAUCCGCUGUCUGGUUCCUUACCUGUAAGGGACGUACAGUAUCCACAC
env-163   ACCCUGGUUGACAAGUGCGACGCCAAGAAUAUCCGCUGUCUGAUCCCUUACCUGUAAGGGACGUACAGUAUCCACAC
env-165   .................CGACGCCAAGAAUAUCCGCUGUCUGAUCCCUUACCUUUAAGGGACGUACAGUAUCCACAC
env-166   ACCCUGGUUGACAAGUGCGACGCCAAGAAUAUCCGCUGUCUGAUCCCUUAUCUGUAAGGGACGUACAGUAUCCACAC
env-167   ACCCUGGUUGACAAGUGCGACGCCAAGAAUAUCCGCUGUCUGAUCCCUUACCUGUAAGGGACGUNCAGUAUCCACAC
env-168   ACCCUGGUUGACAAGGCCGACGCCAAGAAUAUCCGCUGUCUGAUCCCUUACCUGUAAGGGACGUUCAGUAUCCACAC
env-169   ACCUUGGUUGACAAGGCCGACGCCAAGAAUAUCCGCUGUCUGAUCCCUUACCUGUAAGGGACGUACAGUAUCAACAC
env-174   CGAUUGUUUGUGCCGCAGAUCAUAAAAAUAUCAGUUGCCUGAUCCCUUACAUGUAAGGUUCAUGCAACUCGGCUUUA
env-177   UCGACCGUGGUCCACGCGAGUGACGCCAGGAACAUCAGCCGCCUGAUCCCCUACGUGUAGGCAGGCGGUCGUUUUAC
env-178   GUCGACCCUGGUUCACCCCAGCGAUGUCAGCAGCAUCAGCCGCCUGAUUCCCUACAUGUAACAGGCAUUGCGUACAA
    .............................................................................
    .............................................................................
    •••••••••••••••••••••••••••••••••••••••••••••••••••••••••••••••••••••••••••••
```

alignment positions 78 · · · 216

```
env-1     ACCCCGUGAACUGGGGGG.....AUACGGCUCCCCCUCCAGAUCCGG....................CGACGGCU....UUU.......GGCCGUUG...........ACAAAUUCAACCUAAGGAGAAAGAG...AUG
env-2     UUUCUGUGAACUGUGGGG......ACCAUCUCCCCCUGCAGAUCCAGCAGCGGCAUGUGUCGCAG..AAGAUG......UAUU........CAUUUU...............AAAGAUUAGAAGGAGAAAGUGAG.AUG
env-3     CGGCUGCGAACUGUGGGG......ACGAUCUCCCCCUGCAGAUCCGGCCA..................CGGC.......AAAU.........GCCG..........AAACCAUUUAGAGAGAAGGAGCAAGUGAG.AUG
env-6     CGGUUGUGAACUGUGGGG......ACAAUCUCCCCCUGCAGAUCCAGCCA..................CGGC.......AGUU.........GCCG........AAAAACUAUUCAAUGAGAAGGAGCAAGCGAG.AUG
env-7     AGGUUGUGAACUGUGGGG......ACAAUCUCCCCCUGCAGAUCCAGCCA..................CGGC.......AGAU.........GCCG.........AAAAACUAUUUAUGAGAAGGAGCAAGUGAG.AUG
Pse-1-1   UGGUUGUGAACUGUGGGG......ACA.UCUCCCCCUGCAGAUCCGACCG..................CGGC.......AUAA.........GCCG........AAACAACAUUAUUUGAGAAGGAGCAAGUGAG.AUG
env-8     UGGUUGUGAACUGUGGGG......AUA.UCUCCCCCUGCAGAUCCGACCG..................CGGC.......AUAU.........GCCG........AAACAACAUUAUUUGAGAAGGAGCAAGUGAG.AUG
env-9     UGGUUGUGAACUGUGGGG......AUA.UCUCCCCCUGCAGAUCCGACCG..................CGGC.......AUAU.........GCCG........AAACAACAUUAUUUGAGAAGGAGCAAGUGAG.AUG
env-11    UGGCUGAGAACUGUGGGG......ACAAUCUCCCCCUGUAGAUCCGGCCG..................CGGC.......UUAU.........GCCG.......AAAAACAUACACUUUAUGAAGGAGCAAGAGAG.AUG
Gel-1-1   ACGUUGUGAACUGUGGGG......ACCAUCUCCCCCUGCAGAUCCGGCA..................GCGGU.......CUUA.........ACCGCAAUCUUAAAAUCUUAUUACCCAAGAAGGAGCAAGAGAG.AUG
env-12    UAGUCGUGAACUGGGGGGACAUUUCCAGUCUCCCCCUCCAGAUCCGGCGCAAUACCACAGA.............................................................AGGAGGCAGAU...AUG
env-13    CAUCCCUGAAUCGCGAGG.....AACGG...CCCCUUGCGCAUCGGGCAAA................UUA.........AAAC...........UGA.........................AGGAGUUCAAAG..AUG
env-14    CAUCCCUGAAUCGCGAGG.....AACGG...CCCCUUGCGCAUCGGGCAAA................UUA.........AAAC...........UGA.........................AGGAGUUCAAAG..AUG
env-15    CAUCCCUGAAUCGCGAGG.....AACGG...CCCCUUGCGCAUCGGGCAAA................UUA.........AAAC...........UGA.........................AGGAGUUCAAAG..AUG
env-16    CAUCCCUGAAUCGCGAGG.....AACGG...CCCCUUGCGCAUCGGGCAAA................UUA.........AAAC...........UGA.........................AGGAGUUCAAAG..AUG
env-17    AUGCCAAGAACCAUGAGG....AAAUUGUCUCCCCUUAUGGAUCUGGUAAAUUAACCAAUGA............................................................AGGAGUGCAUUA..AUG
env-18    AUGCCAAGAACCAUGAGG....AAAUUGUCUCCCCUUAUGGAUCUGGUAAAUUAACCAACGA............................................................AGGAG..UGCAUUAAUG
env-19    AUGCCAAGAACCAUGAGG....AAAUUGUCUCCCCUUAUGGAUCUGGCCAAUUCAGCAACGA............................................................AGGAGUAGUUACGUAUG
env-20    AUGCCAUGAACCAUGAGG....AAAUUGUCUCCCCUUGUGGAUCUGGCCAAUUCAGCAACGA............................................................AGGAGUAGUCACGUAUG
env-21    AUGCCAUGAACCAUGAGG....AAAUUGUCUCCCCUUAUGGAUCUGGCCAAUUCAGCAACUA............................................................AGGAGUAGUAACGUAUG
Ppr-1-1   ACGCCAAGAACCACGAGG......ACAGUCUCCCCUUGUGGAUCGGGCAAA................AUU.........CAU............GAU.......................CAAGGAGUAGAAC...AUG
env-23    CAGCCCAGAACCGUGAGG.....AAACGG..CCCCUUGCGGAUCGGGAAU.UCAA..............................................................CCUAAAGGAGUAAGAAG..AUG
env-24    GAGCCAUGAACCGUGAGG....AAAAUGUCUCCCCUUGCGGAUCGGCGAUAAAAAAAAUGACAA..........................................................AGGAGUAGAAGU..AUG
env-25    AGACCGUGAACCGUGAGG.....AAACGUCUCCCCUUGCGGAUCCGGCGAA................AUU.........CAAGCA.........AAU.......................AAAGGAGUAUGU....AUG
Gme-1-1   AGUCCGUGAACCGUGAGG.....AAGCGCCUCCCCUUGCGGAUCCGGCAAA................AUU.........CAUC...........AGU......................AAAAGGAGUAGGUA...AUG
env-26    AAGCUGUGAACCGCGAGG.....AAACGUCUCCCCUUGCGGAUCCGGACCAAUAAAAACC............................................................AAAGGAGUCUGUA...AUG
env-27    CGCCCCAGAACCGUGAGG.....AAACGG..CCCCUUGCGGAUCGGGUACCUAAAACUAA..............................................................AGGAGUAGAAG...AUG
env-28    CGCCCCAGAAUCGUGAGG.....AAACGG..CCCCUUACGGAUCGGGUAACUAUCACAAA..............................................................AGGAGUAGAAG...AUG

          ...<<<.<<.<<<<<<<<...............>>>>>>>>.>>>>>....................<<<<<<<<<<<<......>>>>>>>>>>>>..........................................
          ...112.00..212.201...............102.212..00211....................1221111.1011......1101.1111221..........................................
          •R•CCG•GAACYGYGRGG-----•A••GUCUCCCCYU•CRGAUCYGGY•◦◦◦◦◦◦◦◦◦◦-------------------------------------------------------------◦◦AGGAGYA••••---AUG
           5′                                                                                                                              3′
```

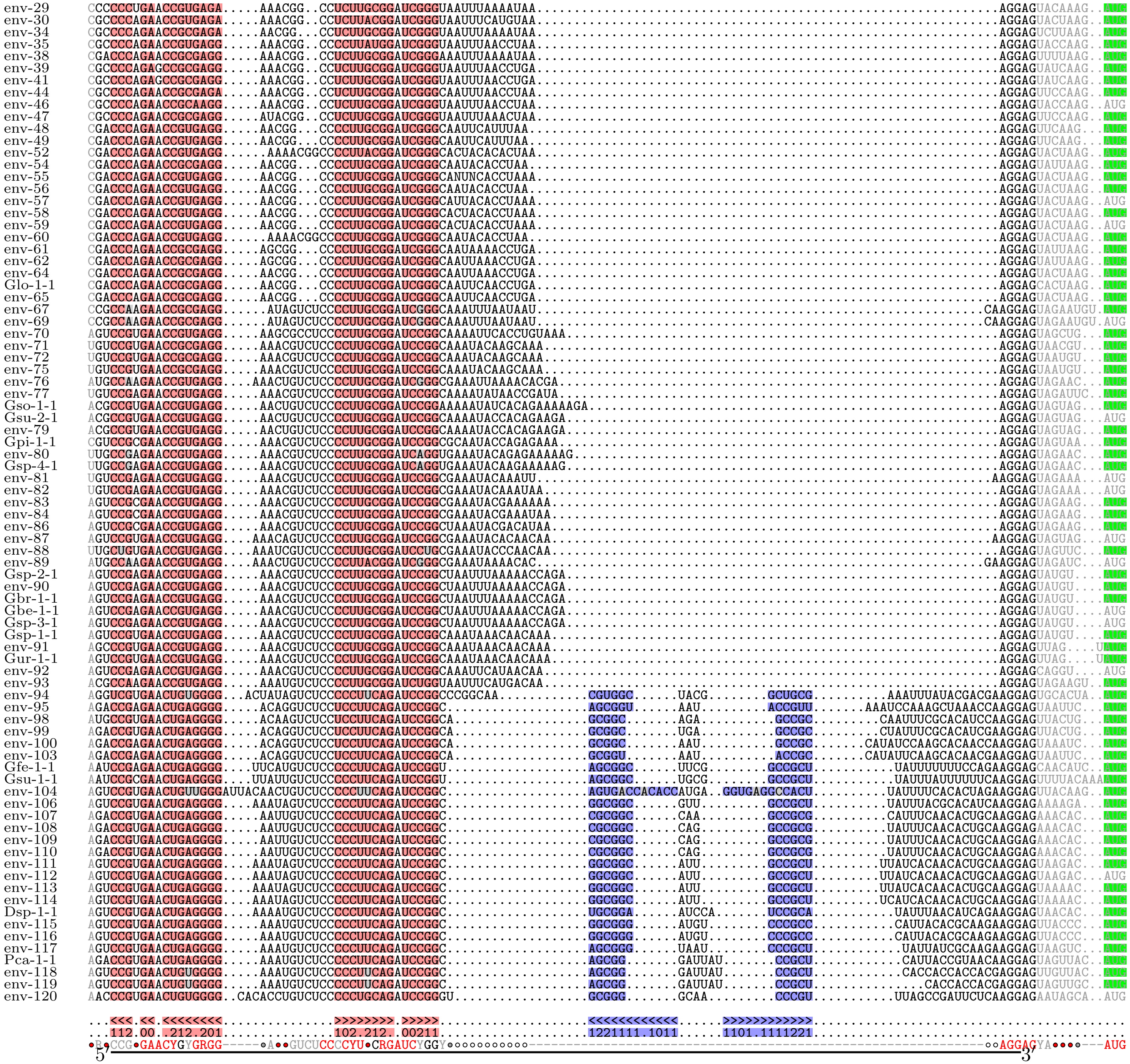
env-29
env-30
env-34
env-35
env-38
env-39
env-41
env-44
env-46
env-47
env-48
env-49
env-52
env-54
env-55
env-56
env-57
env-58
env-59
env-60
env-61
env-62
env-64
Glo-1-1
env-65
env-67
env-69
env-70
env-71
env-72
env-75
env-76
env-77
Gso-1-1
Gsu-2-1
env-79
Gpi-1-1
env-80
Gsp-4-1
env-81
env-82
env-83
env-84
env-86
env-87
env-88
env-89
Gsp-2-1
env-90
Gbr-1-1
Gbe-1-1
Gsp-3-1
Gsp-1-1
env-91
Gur-1-1
env-92
env-93
env-94
env-95
env-98
env-99
env-100
env-103
Gfe-1-1
Gsu-1-1
env-104
env-106
env-107
env-108
env-109
env-110
env-111
env-112
env-113
env-114
Dsp-1-1
env-115
env-116
env-117
Pca-1-1
env-118
env-119
env-120
5'
3'

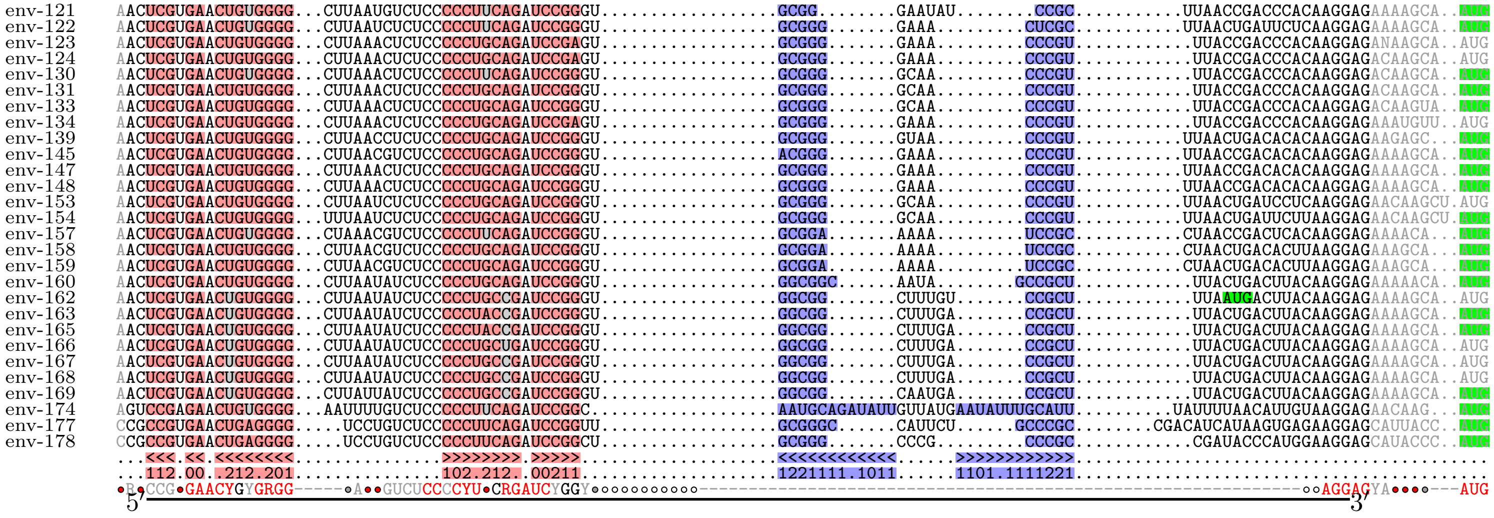

### 8 L25-Gammaproteobacteria

#### 8.1 Taxa

The taxonomy of each organism containing a putative L25-Gammaproteobacteria RNA is listed, with abbreviations identifying each hit (e.g., “Eco-1-1” and “Eco-1-2” might hypothetically represent two distinct RNAs in *E. coli*). The abbreviations will be used to identify each individual L25-Gammaproteobacteria RNA in Sections **??** and 8.2:.

| abbrev. of hits | taxonomy of species |
|---|---|
| Sen-1-1 | AKU_12601Bacteria Proteobacteria Gammaproteobacteria Enterobacteriales Enterobacteriaceae *Salmonella enterica subsp. enterica serovar* Paratyphi A str |
| bsy-1-1 | Bacteria bacteria symbiont BFo1 of Frankliniella occidentalis |
| bsy-2-1 | Bacteria bacteria symbiont BFo2 of Frankliniella occidentalis |
| Baq-1-1 | Bacteria Proteobacteria Gammaproteobacteria Enterobacterales Budviciaceae *Budvicia aquatica* DSM 5075 = ATCC 35567 |
| Lgr-1-1 to Lgr-1-2 | Bacteria Proteobacteria Gammaproteobacteria Enterobacterales Budviciaceae *Leminorella grimontii* ATCC 33999 = DSM 5078 |
| Pfo-1-1 | Bacteria Proteobacteria Gammaproteobacteria Enterobacterales Budviciaceae *Pragia fontium* |
| Bag-1-1 | Bacteria Proteobacteria Gammaproteobacteria Enterobacterales Enterobacteriaceae *Buttiauxella agrestis* |
| Bag-2-1 | Bacteria Proteobacteria Gammaproteobacteria Enterobacterales Enterobacteriaceae *Buttiauxella agrestis* ATCC 33320 |
| Cne-1-1 to Cne-1-3 | Bacteria Proteobacteria Gammaproteobacteria Enterobacterales Enterobacteriaceae *Cedecea neteri* |
| Cam-1-1 to Cam-1-3 | Bacteria Proteobacteria Gammaproteobacteria Enterobacterales Enterobacteriaceae *Citrobacter amalonaticus* |
| Cam-2-1 | Bacteria Proteobacteria Gammaproteobacteria Enterobacterales Enterobacteriaceae *Citrobacter amalonaticus* Y19 |
| Cfa-1-1 | Bacteria Proteobacteria Gammaproteobacteria Enterobacterales Enterobacteriaceae *Citrobacter farmeri* GTC 1319 |
| Cbr-1-1 to Cbr-1-6 | Bacteria Proteobacteria Gammaproteobacteria Enterobacterales Enterobacteriaceae Citrobacter Citrobacter freundii complex *Citrobacter braakii* |
| Cfr-1-1 to Cfr-1-4 | Bacteria Proteobacteria Gammaproteobacteria Enterobacterales Enterobacteriaceae Citrobacter Citrobacter freundii complex *Citrobacter freundii* |
| Cfr-2-1 to Cfr-2-2 | Bacteria Proteobacteria Gammaproteobacteria Enterobacterales Enterobacteriaceae Citrobacter Citrobacter freundii complex *Citrobacter freundii* ATCC 8090 = MTCC 1658 |
| Cfr-3-1 | Bacteria Proteobacteria Gammaproteobacteria Enterobacterales Enterobacteriaceae Citrobacter Citrobacter freundii complex *Citrobacter freundii* CFNIH1 |
| Cfr-4-1 | Bacteria Proteobacteria Gammaproteobacteria Enterobacterales Enterobacteriaceae Citrobacter Citrobacter freundii complex *Citrobacter freundii* MGH 56 |
| Cfr-5-1 | Bacteria Proteobacteria Gammaproteobacteria Enterobacterales Enterobacteriaceae Citrobacter Citrobacter freundii complex *Citrobacter freundii* NBRC 12681 |
| Cfr-6-1 | Bacteria Proteobacteria Gammaproteobacteria Enterobacterales Enterobacteriaceae Citrobacter Citrobacter freundii complex *Citrobacter freundii* RLS1 |

| | |
|---|---|
| Cfr-7-1 | Bacteria Proteobacteria Gammaproteobacteria Enterobacterales Enterobacteriaceae Citrobacter Citrobacter freundii complex *Citrobacter freundii* UCI 31 |
| Cfr-8-1 | Bacteria Proteobacteria Gammaproteobacteria Enterobacterales Enterobacteriaceae Citrobacter Citrobacter freundii complex *Citrobacter freundii* UCI 32 |
| Cse-1-1 | Bacteria Proteobacteria Gammaproteobacteria Enterobacterales Enterobacteriaceae Citrobacter Citrobacter freundii complex *Citrobacter sedlakii* NBRC 105722 |
| Csp-1-1 | Bacteria Proteobacteria Gammaproteobacteria Enterobacterales Enterobacteriaceae Citrobacter Citrobacter freundii complex *Citrobacter sp.* BIDMC107 |
| Csp-2-1 | Bacteria Proteobacteria Gammaproteobacteria Enterobacterales Enterobacteriaceae Citrobacter Citrobacter freundii complex *Citrobacter sp.* BIDMC108 |
| Csp-3-1 | Bacteria Proteobacteria Gammaproteobacteria Enterobacterales Enterobacteriaceae Citrobacter Citrobacter freundii complex *Citrobacter sp.* MGH100 |
| Csp-4-1 | Bacteria Proteobacteria Gammaproteobacteria Enterobacterales Enterobacteriaceae Citrobacter Citrobacter freundii complex *Citrobacter sp.* MGH103 |
| Csp-5-1 | Bacteria Proteobacteria Gammaproteobacteria Enterobacterales Enterobacteriaceae Citrobacter Citrobacter freundii complex *Citrobacter sp.* MGH104 |
| Csp-6-1 | Bacteria Proteobacteria Gammaproteobacteria Enterobacterales Enterobacteriaceae Citrobacter Citrobacter freundii complex *Citrobacter sp.* MGH105 |
| Csp-7-1 | Bacteria Proteobacteria Gammaproteobacteria Enterobacterales Enterobacteriaceae Citrobacter Citrobacter freundii complex *Citrobacter sp.* MGH106 |
| Csp-8-1 | Bacteria Proteobacteria Gammaproteobacteria Enterobacterales Enterobacteriaceae Citrobacter Citrobacter freundii complex *Citrobacter sp.* MGH109 |
| Csp-9-1 | Bacteria Proteobacteria Gammaproteobacteria Enterobacterales Enterobacteriaceae Citrobacter Citrobacter freundii complex *Citrobacter sp.* MGH110 |
| Csp-10-1 | Bacteria Proteobacteria Gammaproteobacteria Enterobacterales Enterobacteriaceae Citrobacter Citrobacter freundii complex *Citrobacter sp.* MGH99 |
| Cwe-1-1 | Bacteria Proteobacteria Gammaproteobacteria Enterobacterales Enterobacteriaceae Citrobacter Citrobacter freundii complex *Citrobacter werkmanii* NBRC 105721 |
| Cko-1-1 to Cko-1-2 | Bacteria Proteobacteria Gammaproteobacteria Enterobacterales Enterobacteriaceae *Citrobacter koseri* |
| Cpa-1-1 | Bacteria Proteobacteria Gammaproteobacteria Enterobacterales Enterobacteriaceae *Citrobacter pasteurii* |
| Cro-1-1 | Bacteria Proteobacteria Gammaproteobacteria Enterobacterales Enterobacteriaceae *Citrobacter rodentium* |
| Cro-2-1 | Bacteria Proteobacteria Gammaproteobacteria Enterobacterales Enterobacteriaceae *Citrobacter rodentium* NBRC 105723 |
| Csp-11-1 | Bacteria Proteobacteria Gammaproteobacteria Enterobacterales Enterobacteriaceae *Citrobacter sp.* JT3 |
| Csp-12-1 | Bacteria Proteobacteria Gammaproteobacteria Enterobacterales Enterobacteriaceae *Citrobacter sp.* MGH 55 |
| Csp-13-1 | Bacteria Proteobacteria Gammaproteobacteria Enterobacterales Enterobacteriaceae *Citrobacter sp.* S-77 |
| Cdu-1-1 | Bacteria Proteobacteria Gammaproteobacteria Enterobacterales Enterobacteriaceae *Cronobacter dublinensis subsp. dublinensis* LMG 23823 |
| Cdu-2-1 | Bacteria Proteobacteria Gammaproteobacteria Enterobacterales Enterobacteriaceae *Cronobacter dublinensis subsp. lactaridi* LMG 23825 |
| Cdu-3-1 | Bacteria Proteobacteria Gammaproteobacteria Enterobacterales Enterobacteriaceae *Cronobacter dublinensis subsp. lausannensis* LMG 23824 |
| Cma-1-1 to Cma-1-5 | Bacteria Proteobacteria Gammaproteobacteria Enterobacterales Enterobacteriaceae *Cronobacter malonaticus* |
| Cma-2-1 | Bacteria Proteobacteria Gammaproteobacteria Enterobacterales Enterobacteriaceae *Cronobacter malonaticus* ENBT0334 |
| Cma-3-1 | Bacteria Proteobacteria Gammaproteobacteria Enterobacterales Enterobacteriaceae *Cronobacter malonaticus* LMG 23826 |
| Cmu-1-1 | Bacteria Proteobacteria Gammaproteobacteria Enterobacterales Enterobacteriaceae *Cronobacter muytjensii* ATCC 51329 |
| Csa-1-1 | Bacteria Proteobacteria Gammaproteobacteria Enterobacterales Enterobacteriaceae *Cronobacter sakazakii* |
| Csa-2-1 | Bacteria Proteobacteria Gammaproteobacteria Enterobacterales Enterobacteriaceae *Cronobacter sakazakii* 2151 |
| Csa-3-1 | Bacteria Proteobacteria Gammaproteobacteria Enterobacterales Enterobacteriaceae *Cronobacter sakazakii* 8399 |
| Csa-4-1 | Bacteria Proteobacteria Gammaproteobacteria Enterobacterales Enterobacteriaceae *Cronobacter sakazakii* E764 |
| Csa-5-1 | Bacteria Proteobacteria Gammaproteobacteria Enterobacterales Enterobacteriaceae *Cronobacter sakazakii* ES35 |
| Csa-6-1 | Bacteria Proteobacteria Gammaproteobacteria Enterobacterales Enterobacteriaceae *Cronobacter sakazakii* ES713 |
| Csa-7-1 | Bacteria Proteobacteria Gammaproteobacteria Enterobacterales Enterobacteriaceae *Cronobacter sakazakii* NBRC 102416 |
| Csa-8-1 | Bacteria Proteobacteria Gammaproteobacteria Enterobacterales Enterobacteriaceae *Cronobacter sakazakii* NCIMB 8272 |
| Ctu-1-1 | Bacteria Proteobacteria Gammaproteobacteria Enterobacterales Enterobacteriaceae *Cronobacter turicensis z*508 |
| Ctu-2-1 | Bacteria Proteobacteria Gammaproteobacteria Enterobacterales Enterobacteriaceae *Cronobacter turicensis z*610 |
| Cun-1-1 to Cun-1-2 | Bacteria Proteobacteria Gammaproteobacteria Enterobacterales Enterobacteriaceae *Cronobacter universalis* NCTC 9529 |
| Eas-1-1 to Eas-1-2 | Bacteria Proteobacteria Gammaproteobacteria Enterobacterales Enterobacteriaceae Enterobacter Enterobacter cloacae complex *Enterobacter asburiae* |
| Eas-2-1 | Bacteria Proteobacteria Gammaproteobacteria Enterobacterales Enterobacteriaceae Enterobacter Enterobacter cloacae complex *Enterobacter asburiae* C1 |
| Eas-3-1 | Bacteria Proteobacteria Gammaproteobacteria Enterobacterales Enterobacteriaceae Enterobacter Enterobacter cloacae complex *Enterobacter asburiae* L1 |
| Eca-1-1 | Bacteria Proteobacteria Gammaproteobacteria Enterobacterales Enterobacteriaceae Enterobacter Enterobacter cloacae complex *Enterobacter cancerogenus* |
| Eca-2-1 | Bacteria Proteobacteria Gammaproteobacteria Enterobacterales Enterobacteriaceae Enterobacter Enterobacter cloacae complex *Enterobacter cancerogenus* YZ1 |
| Ecl-1-1 to Ecl-1-12 | Bacteria Proteobacteria Gammaproteobacteria Enterobacterales Enterobacteriaceae Enterobacter Enterobacter cloacae complex *Enterobacter cloacae* |
| Ecl-2-1 | Bacteria Proteobacteria Gammaproteobacteria Enterobacterales Enterobacteriaceae Enterobacter Enterobacter cloacae complex *Enterobacter cloacae* 'Hoffmann cluster III' MGH 13 |
| Ecl-3-1 | Bacteria Proteobacteria Gammaproteobacteria Enterobacterales Enterobacteriaceae Enterobacter Enterobacter cloacae complex *Enterobacter cloacae* BIDMC 33A |
| Ecl-4-1 | Bacteria Proteobacteria Gammaproteobacteria Enterobacterales Enterobacteriaceae Enterobacter Enterobacter cloacae complex *Enterobacter cloacae* BIDMC 66 |
| Ecl-5-1 | Bacteria Proteobacteria Gammaproteobacteria Enterobacterales Enterobacteriaceae Enterobacter Enterobacter cloacae complex *Enterobacter cloacae* BIDMC 67 |
| Ecl-6-1 | Bacteria Proteobacteria Gammaproteobacteria Enterobacterales Enterobacteriaceae Enterobacter Enterobacter cloacae complex *Enterobacter cloacae* BIDMC 8 |
| Ecl-7-1 | Bacteria Proteobacteria Gammaproteobacteria Enterobacterales Enterobacteriaceae Enterobacter Enterobacter cloacae complex *Enterobacter cloacae* BWH 43 |

| | |
|---|---|
| Ecl-8-1 | Bacteria Proteobacteria Gammaproteobacteria Enterobacterales Enterobacteriaceae Enterobacter Enterobacter cloacae complex *Enterobacter cloacae* CHS 79 |
| Ecl-27-1 | Bacteria Proteobacteria Gammaproteobacteria Enterobacterales Enterobacteriaceae Enterobacter *Enterobacter cloacae complex* 'Hoffmann cluster III' |
| Ecl-28-1 to Ecl-28-6 | Bacteria Proteobacteria Gammaproteobacteria Enterobacterales Enterobacteriaceae Enterobacter *Enterobacter cloacae complex* 'Hoffmann cluster IV' |
| Ecl-29-1 | Bacteria Proteobacteria Gammaproteobacteria Enterobacterales Enterobacteriaceae Enterobacter *Enterobacter cloacae complex sp.* 20432 |
| Ecl-30-1 | Bacteria Proteobacteria Gammaproteobacteria Enterobacterales Enterobacteriaceae Enterobacter *Enterobacter cloacae complex sp.* 35009 |
| Ecl-31-1 | Bacteria Proteobacteria Gammaproteobacteria Enterobacterales Enterobacteriaceae Enterobacter *Enterobacter cloacae complex sp.* 35618 |
| Ecl-32-1 | Bacteria Proteobacteria Gammaproteobacteria Enterobacterales Enterobacteriaceae Enterobacter *Enterobacter cloacae complex sp.* 35651 |
| Ecl-33-1 | Bacteria Proteobacteria Gammaproteobacteria Enterobacterales Enterobacteriaceae Enterobacter *Enterobacter cloacae complex sp.* 35669 |
| Ecl-34-1 | Bacteria Proteobacteria Gammaproteobacteria Enterobacterales Enterobacteriaceae Enterobacter *Enterobacter cloacae complex sp.* 35699 |
| Ecl-35-1 | Bacteria Proteobacteria Gammaproteobacteria Enterobacterales Enterobacteriaceae Enterobacter *Enterobacter cloacae complex sp.* 35731 |
| Ecl-36-1 | Bacteria Proteobacteria Gammaproteobacteria Enterobacterales Enterobacteriaceae Enterobacter *Enterobacter cloacae complex sp.* 35734 |
| Ecl-37-1 | Bacteria Proteobacteria Gammaproteobacteria Enterobacterales Enterobacteriaceae Enterobacter *Enterobacter cloacae complex sp.* 42192 |
| Ecl-38-1 | Bacteria Proteobacteria Gammaproteobacteria Enterobacterales Enterobacteriaceae Enterobacter *Enterobacter cloacae complex sp.* 43444 |
| Ecl-39-1 | Bacteria Proteobacteria Gammaproteobacteria Enterobacterales Enterobacteriaceae Enterobacter *Enterobacter cloacae complex sp.* 44242 |
| Ecl-40-1 | Bacteria Proteobacteria Gammaproteobacteria Enterobacterales Enterobacteriaceae Enterobacter *Enterobacter cloacae complex sp.* 44246 |
| Ecl-41-1 | Bacteria Proteobacteria Gammaproteobacteria Enterobacterales Enterobacteriaceae Enterobacter *Enterobacter cloacae complex sp.* CIDEIMsCOL1 |
| Ecl-42-1 | Bacteria Proteobacteria Gammaproteobacteria Enterobacterales Enterobacteriaceae Enterobacter *Enterobacter cloacae complex sp.* CIDEIMsCOL9 |
| Ecl-43-1 | Bacteria Proteobacteria Gammaproteobacteria Enterobacterales Enterobacteriaceae Enterobacter *Enterobacter cloacae complex sp.* GN02141 |
| Ecl-44-1 | Bacteria Proteobacteria Gammaproteobacteria Enterobacterales Enterobacteriaceae Enterobacter *Enterobacter cloacae complex sp.* GN02208 |
| Ecl-45-1 | Bacteria Proteobacteria Gammaproteobacteria Enterobacterales Enterobacteriaceae Enterobacter *Enterobacter cloacae complex sp.* GN02283 |
| Ecl-46-1 | Bacteria Proteobacteria Gammaproteobacteria Enterobacterales Enterobacteriaceae Enterobacter *Enterobacter cloacae complex sp.* GN02440 |
| Ecl-47-1 | Bacteria Proteobacteria Gammaproteobacteria Enterobacterales Enterobacteriaceae Enterobacter *Enterobacter cloacae complex sp.* GN02548 |
| Ecl-48-1 | Bacteria Proteobacteria Gammaproteobacteria Enterobacterales Enterobacteriaceae Enterobacter *Enterobacter cloacae complex sp.* GN02570 |
| Ecl-49-1 | Bacteria Proteobacteria Gammaproteobacteria Enterobacterales Enterobacteriaceae Enterobacter *Enterobacter cloacae complex sp.* GN02587 |
| Ecl-50-1 | Bacteria Proteobacteria Gammaproteobacteria Enterobacterales Enterobacteriaceae Enterobacter *Enterobacter cloacae complex sp.* GN02692 |
| Ecl-51-1 | Bacteria Proteobacteria Gammaproteobacteria Enterobacterales Enterobacteriaceae Enterobacter *Enterobacter cloacae complex sp.* GN03164 |
| Ecl-9-1 | Bacteria Proteobacteria Gammaproteobacteria Enterobacterales Enterobacteriaceae Enterobacter Enterobacter cloacae complex *Enterobacter cloacae* EC-NIH2 |
| Ecl-10-1 | Bacteria Proteobacteria Gammaproteobacteria Enterobacterales Enterobacteriaceae Enterobacter Enterobacter cloacae complex *Enterobacter cloacae* EC-NIH3 |
| Ecl-11-1 | Bacteria Proteobacteria Gammaproteobacteria Enterobacterales Enterobacteriaceae Enterobacter Enterobacter cloacae complex *Enterobacter cloacae* ECR091 |
| Ecl-12-1 | Bacteria Proteobacteria Gammaproteobacteria Enterobacterales Enterobacteriaceae Enterobacter Enterobacter cloacae complex *Enterobacter cloacae* IIT-BT 08 |
| Ecl-13-1 | Bacteria Proteobacteria Gammaproteobacteria Enterobacterales Enterobacteriaceae Enterobacter Enterobacter cloacae complex *Enterobacter cloacae* JD6301 |
| Ecl-14-1 | Bacteria Proteobacteria Gammaproteobacteria Enterobacterales Enterobacteriaceae Enterobacter Enterobacter cloacae complex *Enterobacter cloacae* JD8715 |
| Ecl-15-1 | Bacteria Proteobacteria Gammaproteobacteria Enterobacterales Enterobacteriaceae Enterobacter Enterobacter cloacae complex *Enterobacter cloacae* MGH 53 |
| Ecl-16-1 | Bacteria Proteobacteria Gammaproteobacteria Enterobacterales Enterobacteriaceae Enterobacter Enterobacter cloacae complex *Enterobacter cloacae* MGH 54 |
| Ecl-17-1 | Bacteria Proteobacteria Gammaproteobacteria Enterobacterales Enterobacteriaceae Enterobacter Enterobacter cloacae complex *Enterobacter cloacae* P101 |
| Ecl-52-1 to Ecl-52-5 | Bacteria Proteobacteria Gammaproteobacteria Enterobacterales Enterobacteriaceae Enterobacter Enterobacter cloacae complex *Enterobacter cloacae subsp. cloacae* |
| Ecl-53-1 | Bacteria Proteobacteria Gammaproteobacteria Enterobacterales Enterobacteriaceae Enterobacter Enterobacter cloacae complex *Enterobacter cloacae subsp. cloacae* SY-70 |
| Ecl-54-1 | Bacteria Proteobacteria Gammaproteobacteria Enterobacterales Enterobacteriaceae Enterobacter Enterobacter cloacae complex *Enterobacter cloacae subsp. dissolvens* |
| Ecl-18-1 | Bacteria Proteobacteria Gammaproteobacteria Enterobacterales Enterobacteriaceae Enterobacter Enterobacter cloacae complex *Enterobacter cloacae* UCI 23 |
| Ecl-19-1 | Bacteria Proteobacteria Gammaproteobacteria Enterobacterales Enterobacteriaceae Enterobacter Enterobacter cloacae complex *Enterobacter cloacae* UCI 24 |
| Ecl-20-1 | Bacteria Proteobacteria Gammaproteobacteria Enterobacterales Enterobacteriaceae Enterobacter Enterobacter cloacae complex *Enterobacter cloacae* UCI 29 |
| Ecl-21-1 | Bacteria Proteobacteria Gammaproteobacteria Enterobacterales Enterobacteriaceae Enterobacter Enterobacter cloacae complex *Enterobacter cloacae* UCI 30 |
| Ecl-22-1 | Bacteria Proteobacteria Gammaproteobacteria Enterobacterales Enterobacteriaceae Enterobacter Enterobacter cloacae complex *Enterobacter cloacae* UCI 35 |

| | |
|---|---|
| Ecl-23-1 | Bacteria Proteobacteria Gammaproteobacteria Enterobacterales Enterobacteriaceae Enterobacter Enterobacter cloacae complex *Enterobacter cloacae* UCI 36 |
| Ecl-24-1 | Bacteria Proteobacteria Gammaproteobacteria Enterobacterales Enterobacteriaceae Enterobacter Enterobacter cloacae complex *Enterobacter cloacae* UCI 39 |
| Ecl-25-1 | Bacteria Proteobacteria Gammaproteobacteria Enterobacterales Enterobacteriaceae Enterobacter Enterobacter cloacae complex *Enterobacter cloacae* UCI 49 |
| Ecl-26-1 | Bacteria Proteobacteria Gammaproteobacteria Enterobacterales Enterobacteriaceae Enterobacter Enterobacter cloacae complex *Enterobacter cloacae* UCI 50 |
| Eho-1-1 to Eho-1-9 | Bacteria Proteobacteria Gammaproteobacteria Enterobacterales Enterobacteriaceae Enterobacter Enterobacter cloacae complex *Enterobacter hormaechei* |
| Eho-2-1 | Bacteria Proteobacteria Gammaproteobacteria Enterobacterales Enterobacteriaceae Enterobacter Enterobacter cloacae complex *Enterobacter hormaechei subsp. hormaechei* |
| Eho-3-1 to Eho-3-2 | Bacteria Proteobacteria Gammaproteobacteria Enterobacterales Enterobacteriaceae Enterobacter Enterobacter cloacae complex *Enterobacter hormaechei subsp. oharae* |
| Eho-4-1 to Eho-4-2 | Bacteria Proteobacteria Gammaproteobacteria Enterobacterales Enterobacteriaceae Enterobacter Enterobacter cloacae complex *Enterobacter hormaechei subsp. steigerwaltii* |
| Eko-1-1 to Eko-1-11 | Bacteria Proteobacteria Gammaproteobacteria Enterobacterales Enterobacteriaceae Enterobacter Enterobacter cloacae complex *Enterobacter kobei* |
| Elu-1-1 to Elu-1-5 | Bacteria Proteobacteria Gammaproteobacteria Enterobacterales Enterobacteriaceae Enterobacter Enterobacter cloacae complex *Enterobacter ludwigii* |
| Esp-1-1 | Bacteria Proteobacteria Gammaproteobacteria Enterobacterales Enterobacteriaceae Enterobacter Enterobacter cloacae complex *Enterobacter sp.* 5-4 |
| Esp-2-1 | Bacteria Proteobacteria Gammaproteobacteria Enterobacterales Enterobacteriaceae Enterobacter Enterobacter cloacae complex *Enterobacter sp.* BIDMC 26 |
| Esp-3-1 | Bacteria Proteobacteria Gammaproteobacteria Enterobacterales Enterobacteriaceae Enterobacter Enterobacter cloacae complex *Enterobacter sp.* BIDMC 27 |
| Esp-4-1 | Bacteria Proteobacteria Gammaproteobacteria Enterobacterales Enterobacteriaceae Enterobacter Enterobacter cloacae complex *Enterobacter sp.* BIDMC 28 |
| Esp-5-1 | Bacteria Proteobacteria Gammaproteobacteria Enterobacterales Enterobacteriaceae Enterobacter Enterobacter cloacae complex *Enterobacter sp.* BIDMC 29 |
| Esp-6-1 | Bacteria Proteobacteria Gammaproteobacteria Enterobacterales Enterobacteriaceae Enterobacter Enterobacter cloacae complex *Enterobacter sp.* BIDMC 30 |
| Esp-7-1 | Bacteria Proteobacteria Gammaproteobacteria Enterobacterales Enterobacteriaceae Enterobacter Enterobacter cloacae complex *Enterobacter sp.* BIDMC100 |
| Esp-8-1 | Bacteria Proteobacteria Gammaproteobacteria Enterobacterales Enterobacteriaceae Enterobacter Enterobacter cloacae complex *Enterobacter sp.* BIDMC109 |
| Esp-9-1 | Bacteria Proteobacteria Gammaproteobacteria Enterobacterales Enterobacteriaceae Enterobacter Enterobacter cloacae complex *Enterobacter sp.* BIDMC87 |
| Esp-10-1 | Bacteria Proteobacteria Gammaproteobacteria Enterobacterales Enterobacteriaceae Enterobacter Enterobacter cloacae complex *Enterobacter sp.* BIDMC93 |
| Esp-11-1 | Bacteria Proteobacteria Gammaproteobacteria Enterobacterales Enterobacteriaceae Enterobacter Enterobacter cloacae complex *Enterobacter sp.* BIDMC94 |
| Esp-12-1 | Bacteria Proteobacteria Gammaproteobacteria Enterobacterales Enterobacteriaceae Enterobacter Enterobacter cloacae complex *Enterobacter sp.* BIDMC99 |
| Esp-13-1 | Bacteria Proteobacteria Gammaproteobacteria Enterobacterales Enterobacteriaceae Enterobacter Enterobacter cloacae complex *Enterobacter sp.* BWH 27 |
| Esp-14-1 | Bacteria Proteobacteria Gammaproteobacteria Enterobacterales Enterobacteriaceae Enterobacter Enterobacter cloacae complex *Enterobacter sp.* BWH 37 |
| Esp-15-1 | Bacteria Proteobacteria Gammaproteobacteria Enterobacterales Enterobacteriaceae Enterobacter Enterobacter cloacae complex *Enterobacter sp.* BWH 39 |
| Esp-16-1 | Bacteria Proteobacteria Gammaproteobacteria Enterobacterales Enterobacteriaceae Enterobacter Enterobacter cloacae complex *Enterobacter sp.* BWH52 |
| Esp-17-1 | Bacteria Proteobacteria Gammaproteobacteria Enterobacterales Enterobacteriaceae Enterobacter Enterobacter cloacae complex *Enterobacter sp.* BWH63 |
| Esp-18-1 | Bacteria Proteobacteria Gammaproteobacteria Enterobacterales Enterobacteriaceae Enterobacter Enterobacter cloacae complex *Enterobacter sp.* BWH64 |
| Esp-19-1 | Bacteria Proteobacteria Gammaproteobacteria Enterobacterales Enterobacteriaceae Enterobacter Enterobacter cloacae complex *Enterobacter sp.* FB |
| Esp-20-1 | Bacteria Proteobacteria Gammaproteobacteria Enterobacterales Enterobacteriaceae Enterobacter Enterobacter cloacae complex *Enterobacter sp.* MGH 1 |
| Esp-21-1 | Bacteria Proteobacteria Gammaproteobacteria Enterobacterales Enterobacteriaceae Enterobacter Enterobacter cloacae complex *Enterobacter sp.* MGH 10 |
| Esp-22-1 | Bacteria Proteobacteria Gammaproteobacteria Enterobacterales Enterobacteriaceae Enterobacter Enterobacter cloacae complex *Enterobacter sp.* MGH 11 |
| Esp-23-1 | Bacteria Proteobacteria Gammaproteobacteria Enterobacterales Enterobacteriaceae Enterobacter Enterobacter cloacae complex *Enterobacter sp.* MGH 12 |
| Esp-24-1 | Bacteria Proteobacteria Gammaproteobacteria Enterobacterales Enterobacteriaceae Enterobacter Enterobacter cloacae complex *Enterobacter sp.* MGH 15 |
| Esp-25-1 | Bacteria Proteobacteria Gammaproteobacteria Enterobacterales Enterobacteriaceae Enterobacter Enterobacter cloacae complex *Enterobacter sp.* MGH 2 |
| Esp-26-1 | Bacteria Proteobacteria Gammaproteobacteria Enterobacterales Enterobacteriaceae Enterobacter Enterobacter cloacae complex *Enterobacter sp.* MGH 3 |
| Esp-27-1 | Bacteria Proteobacteria Gammaproteobacteria Enterobacterales Enterobacteriaceae Enterobacter Enterobacter cloacae complex *Enterobacter sp.* MGH 33 |
| Esp-28-1 | Bacteria Proteobacteria Gammaproteobacteria Enterobacterales Enterobacteriaceae Enterobacter Enterobacter cloacae complex *Enterobacter sp.* MGH 37 |
| Esp-29-1 | Bacteria Proteobacteria Gammaproteobacteria Enterobacterales Enterobacteriaceae Enterobacter Enterobacter cloacae complex *Enterobacter sp.* MGH 4 |
| Esp-30-1 | Bacteria Proteobacteria Gammaproteobacteria Enterobacterales Enterobacteriaceae Enterobacter Enterobacter cloacae complex *Enterobacter sp.* MGH 5 |
| Esp-31-1 | Bacteria Proteobacteria Gammaproteobacteria Enterobacterales Enterobacteriaceae Enterobacter Enterobacter cloacae complex *Enterobacter sp.* MGH 6 |
| Esp-32-1 | Bacteria Proteobacteria Gammaproteobacteria Enterobacterales Enterobacteriaceae Enterobacter Enterobacter cloacae complex *Enterobacter sp.* MGH 7 |
| Esp-33-1 | Bacteria Proteobacteria Gammaproteobacteria Enterobacterales Enterobacteriaceae Enterobacter Enterobacter cloacae complex *Enterobacter sp.* MGH 9 |
| Esp-34-1 | Bacteria Proteobacteria Gammaproteobacteria Enterobacterales Enterobacteriaceae Enterobacter Enterobacter cloacae complex *Enterobacter sp.* MGH119 |
| Esp-35-1 | Bacteria Proteobacteria Gammaproteobacteria Enterobacterales Enterobacteriaceae Enterobacter Enterobacter cloacae complex *Enterobacter sp.* MGH120 |
| Esp-36-1 | Bacteria Proteobacteria Gammaproteobacteria Enterobacterales Enterobacteriaceae Enterobacter Enterobacter cloacae complex *Enterobacter sp.* MGH127 |
| Esp-37-1 | Bacteria Proteobacteria Gammaproteobacteria Enterobacterales Enterobacteriaceae Enterobacter Enterobacter cloacae complex *Enterobacter sp.* MGH128 |

| | |
|---|---|
| Esp-38-1 | Bacteria Proteobacteria Gammaproteobacteria Enterobacterales Enterobacteriaceae Enterobacter Enterobacter cloacae complex *Enterobacter sp.* MGH85 |
| Esp-39-1 | Bacteria Proteobacteria Gammaproteobacteria Enterobacterales Enterobacteriaceae Enterobacter Enterobacter cloacae complex *Enterobacter sp.* MGH86 |
| Esp-40-1 | Bacteria Proteobacteria Gammaproteobacteria Enterobacterales Enterobacteriaceae *Enterobacter sp.* 940_PEND |
| Esp-41-1 | Bacteria Proteobacteria Gammaproteobacteria Enterobacterales Enterobacteriaceae *Enterobacter sp.* BIDMC92 |
| Esp-42-1 | Bacteria Proteobacteria Gammaproteobacteria Enterobacterales Enterobacteriaceae *Enterobacter sp.* Bisph1 |
| Esp-43-1 | Bacteria Proteobacteria Gammaproteobacteria Enterobacterales Enterobacteriaceae *Enterobacter sp.* Bisph2 |
| Esp-44-1 | Bacteria Proteobacteria Gammaproteobacteria Enterobacterales Enterobacteriaceae *Enterobacter sp.* DC1 |
| Esp-45-1 | Bacteria Proteobacteria Gammaproteobacteria Enterobacterales Enterobacteriaceae *Enterobacter sp.* DC3 |
| Esp-46-1 | Bacteria Proteobacteria Gammaproteobacteria Enterobacterales Enterobacteriaceae *Enterobacter sp.* DC4 |
| Esp-47-1 | Bacteria Proteobacteria Gammaproteobacteria Enterobacterales Enterobacteriaceae *Enterobacter sp.* EGD-HP1 |
| Esp-48-1 | Bacteria Proteobacteria Gammaproteobacteria Enterobacterales Enterobacteriaceae *Enterobacter sp.* GN02600 |
| Esp-49-1 | Bacteria Proteobacteria Gammaproteobacteria Enterobacterales Enterobacteriaceae *Enterobacter sp.* T1-1 |
| Esp-50-1 | Bacteria Proteobacteria Gammaproteobacteria Enterobacterales Enterobacteriaceae *Enterobacter sp.* UCD-UG_FMILLET |
| Esp-51-1 | Bacteria Proteobacteria Gammaproteobacteria Enterobacterales Enterobacteriaceae *Enterobacter sp.* ZOR0014 |
| Eba-1-1 | Bacteria Proteobacteria Gammaproteobacteria Enterobacterales *Enterobacteriaceae bacterium* ATCC 29904 |
| Eba-2-1 | Bacteria Proteobacteria Gammaproteobacteria Enterobacterales *Enterobacteriaceae bacterium* B14 |
| Eal-1-1 | Bacteria Proteobacteria Gammaproteobacteria Enterobacterales Enterobacteriaceae *Escherichia albertii* KF1 |
| Eal-2-1 | Bacteria Proteobacteria Gammaproteobacteria Enterobacterales Enterobacteriaceae *Escherichia albertii* NBRC 107761 |
| Eco-1-1 to Eco-1-4 | Bacteria Proteobacteria Gammaproteobacteria Enterobacterales Enterobacteriaceae *Escherichia coli* |
| Eco-2-1 | Bacteria Proteobacteria Gammaproteobacteria Enterobacterales Enterobacteriaceae *Escherichia coli* 1-110-08_S1_C1 |
| Eco-3-1 | Bacteria Proteobacteria Gammaproteobacteria Enterobacterales Enterobacteriaceae *Escherichia coli* 1-110-08_S1_C2 |
| Eco-4-1 | Bacteria Proteobacteria Gammaproteobacteria Enterobacterales Enterobacteriaceae *Escherichia coli* 1-110-08_S1_C3 |
| Eco-5-1 | Bacteria Proteobacteria Gammaproteobacteria Enterobacterales Enterobacteriaceae *Escherichia coli* 1-110-08_S3_C1 |
| Eco-6-1 | Bacteria Proteobacteria Gammaproteobacteria Enterobacterales Enterobacteriaceae *Escherichia coli* 1-110-08_S3_C2 |
| Eco-7-1 | Bacteria Proteobacteria Gammaproteobacteria Enterobacterales Enterobacteriaceae *Escherichia coli* 1-110-08_S3_C3 |
| Eco-8-1 | Bacteria Proteobacteria Gammaproteobacteria Enterobacterales Enterobacteriaceae *Escherichia coli* 1-110-08_S4_C1 |
| Eco-9-1 | Bacteria Proteobacteria Gammaproteobacteria Enterobacterales Enterobacteriaceae *Escherichia coli* 1-110-08_S4_C2 |
| Eco-10-1 | Bacteria Proteobacteria Gammaproteobacteria Enterobacterales Enterobacteriaceae *Escherichia coli* 1-110-08_S4_C3 |
| Eco-11-1 | Bacteria Proteobacteria Gammaproteobacteria Enterobacterales Enterobacteriaceae *Escherichia coli* 1-176-05_S1_C1 |
| Eco-12-1 | Bacteria Proteobacteria Gammaproteobacteria Enterobacterales Enterobacteriaceae *Escherichia coli* 1-176-05_S1_C2 |
| Eco-13-1 | Bacteria Proteobacteria Gammaproteobacteria Enterobacterales Enterobacteriaceae *Escherichia coli* 1-176-05_S1_C3 |
| Eco-14-1 | Bacteria Proteobacteria Gammaproteobacteria Enterobacterales Enterobacteriaceae *Escherichia coli* 1-176-05_S3_C1 |
| Eco-15-1 | Bacteria Proteobacteria Gammaproteobacteria Enterobacterales Enterobacteriaceae *Escherichia coli* 1-176-05_S3_C2 |
| Eco-16-1 | Bacteria Proteobacteria Gammaproteobacteria Enterobacterales Enterobacteriaceae *Escherichia coli* 1-176-05_S4_C1 |
| Eco-17-1 | Bacteria Proteobacteria Gammaproteobacteria Enterobacterales Enterobacteriaceae *Escherichia coli* 1-176-05_S4_C2 |
| Eco-18-1 | Bacteria Proteobacteria Gammaproteobacteria Enterobacterales Enterobacteriaceae *Escherichia coli* 1-176-05_S4_C3 |
| Eco-19-1 | Bacteria Proteobacteria Gammaproteobacteria Enterobacterales Enterobacteriaceae *Escherichia coli* 1-182-04_S1_C1 |
| Eco-20-1 | Bacteria Proteobacteria Gammaproteobacteria Enterobacterales Enterobacteriaceae *Escherichia coli* 1-182-04_S1_C2 |
| Eco-21-1 | Bacteria Proteobacteria Gammaproteobacteria Enterobacterales Enterobacteriaceae *Escherichia coli* 1-182-04_S1_C3 |
| Eco-22-1 | Bacteria Proteobacteria Gammaproteobacteria Enterobacterales Enterobacteriaceae *Escherichia coli* 1-182-04_S3_C1 |
| Eco-23-1 | Bacteria Proteobacteria Gammaproteobacteria Enterobacterales Enterobacteriaceae *Escherichia coli* 1-182-04_S3_C2 |
| Eco-24-1 | Bacteria Proteobacteria Gammaproteobacteria Enterobacterales Enterobacteriaceae *Escherichia coli* 1-182-04_S3_C3 |
| Eco-25-1 | Bacteria Proteobacteria Gammaproteobacteria Enterobacterales Enterobacteriaceae *Escherichia coli* 1-182-04_S4_C1 |
| Eco-26-1 | Bacteria Proteobacteria Gammaproteobacteria Enterobacterales Enterobacteriaceae *Escherichia coli* 1-182-04_S4_C2 |
| Eco-27-1 | Bacteria Proteobacteria Gammaproteobacteria Enterobacterales Enterobacteriaceae *Escherichia coli* 1-182-04_S4_C3 |
| Eco-28-1 | Bacteria Proteobacteria Gammaproteobacteria Enterobacterales Enterobacteriaceae *Escherichia coli* 1-250-04_S1_C1 |
| Eco-29-1 | Bacteria Proteobacteria Gammaproteobacteria Enterobacterales Enterobacteriaceae *Escherichia coli* 1-250-04_S1_C2 |
| Eco-30-1 | Bacteria Proteobacteria Gammaproteobacteria Enterobacterales Enterobacteriaceae *Escherichia coli* 1-250-04_S1_C3 |
| Eco-31-1 | Bacteria Proteobacteria Gammaproteobacteria Enterobacterales Enterobacteriaceae *Escherichia coli* 1-250-04_S3_C1 |
| Eco-32-1 | Bacteria Proteobacteria Gammaproteobacteria Enterobacterales Enterobacteriaceae *Escherichia coli* 1-250-04_S3_C2 |
| Eco-33-1 | Bacteria Proteobacteria Gammaproteobacteria Enterobacterales Enterobacteriaceae *Escherichia coli* 1-250-04_S4_C1 |
| Eco-34-1 | Bacteria Proteobacteria Gammaproteobacteria Enterobacterales Enterobacteriaceae *Escherichia coli* 1-250-04_S4_C2 |
| Eco-35-1 | Bacteria Proteobacteria Gammaproteobacteria Enterobacterales Enterobacteriaceae *Escherichia coli* 1-392-07_S1_C1 |
| Eco-36-1 | Bacteria Proteobacteria Gammaproteobacteria Enterobacterales Enterobacteriaceae *Escherichia coli* 1-392-07_S1_C2 |
| Eco-37-1 | Bacteria Proteobacteria Gammaproteobacteria Enterobacterales Enterobacteriaceae *Escherichia coli* 1-392-07_S3_C1 |
| Eco-38-1 | Bacteria Proteobacteria Gammaproteobacteria Enterobacterales Enterobacteriaceae *Escherichia coli* 1-392-07_S3_C2 |
| Eco-39-1 | Bacteria Proteobacteria Gammaproteobacteria Enterobacterales Enterobacteriaceae *Escherichia coli* 1-392-07_S3_C3 |
| Eco-40-1 | Bacteria Proteobacteria Gammaproteobacteria Enterobacterales Enterobacteriaceae *Escherichia coli* 1-392-07_S4_C1 |
| Eco-41-1 | Bacteria Proteobacteria Gammaproteobacteria Enterobacterales Enterobacteriaceae *Escherichia coli* 1-392-07_S4_C2 |
| Eco-42-1 | Bacteria Proteobacteria Gammaproteobacteria Enterobacterales Enterobacteriaceae *Escherichia coli* 1-392-07_S4_C3 |
| Eco-43-1 | Bacteria Proteobacteria Gammaproteobacteria Enterobacterales Enterobacteriaceae *Escherichia coli* 100329 |
| Eco-44-1 | Bacteria Proteobacteria Gammaproteobacteria Enterobacterales Enterobacteriaceae *Escherichia coli* 103385 |
| Eco-45-1 | Bacteria Proteobacteria Gammaproteobacteria Enterobacterales Enterobacteriaceae *Escherichia coli* 103573 |
| Eco-46-1 | Bacteria Proteobacteria Gammaproteobacteria Enterobacterales Enterobacteriaceae *Escherichia coli* 1047 |
| Eco-47-1 | Bacteria Proteobacteria Gammaproteobacteria Enterobacterales Enterobacteriaceae *Escherichia coli* 10810 |

| | |
|---|---|
| Eco-48-1 | Bacteria Proteobacteria Gammaproteobacteria Enterobacterales Enterobacteriaceae *Escherichia coli* 11117 |
| Eco-49-1 | Bacteria Proteobacteria Gammaproteobacteria Enterobacterales Enterobacteriaceae *Escherichia coli* 112469215-isolate1 |
| Eco-50-1 | Bacteria Proteobacteria Gammaproteobacteria Enterobacterales Enterobacteriaceae *Escherichia coli* 112469218-isolate1 |
| Eco-51-1 | Bacteria Proteobacteria Gammaproteobacteria Enterobacterales Enterobacteriaceae *Escherichia coli* 1240 |
| Eco-52-1 | Bacteria Proteobacteria Gammaproteobacteria Enterobacterales Enterobacteriaceae *Escherichia coli* 1303 |
| Eco-53-1 | Bacteria Proteobacteria Gammaproteobacteria Enterobacterales Enterobacteriaceae *Escherichia coli* 1350 |
| Eco-54-1 | Bacteria Proteobacteria Gammaproteobacteria Enterobacterales Enterobacteriaceae *Escherichia coli* 1365 |
| Eco-55-1 | Bacteria Proteobacteria Gammaproteobacteria Enterobacterales Enterobacteriaceae *Escherichia coli* 148 |
| Eco-56-1 | Bacteria Proteobacteria Gammaproteobacteria Enterobacterales Enterobacteriaceae *Escherichia coli* 1A |
| Eco-57-1 | Bacteria Proteobacteria Gammaproteobacteria Enterobacterales Enterobacteriaceae *Escherichia coli* 2-005-03_S1_C1 |
| Eco-58-1 | Bacteria Proteobacteria Gammaproteobacteria Enterobacterales Enterobacteriaceae *Escherichia coli* 2-005-03_S1_C2 |
| Eco-59-1 | Bacteria Proteobacteria Gammaproteobacteria Enterobacterales Enterobacteriaceae *Escherichia coli* 2-005-03_S1_C3 |
| Eco-60-1 | Bacteria Proteobacteria Gammaproteobacteria Enterobacterales Enterobacteriaceae *Escherichia coli* 2-005-03_S3_C1 |
| Eco-61-1 | Bacteria Proteobacteria Gammaproteobacteria Enterobacterales Enterobacteriaceae *Escherichia coli* 2-005-03_S3_C2 |
| Eco-62-1 | Bacteria Proteobacteria Gammaproteobacteria Enterobacterales Enterobacteriaceae *Escherichia coli* 2-005-03_S3_C3 |
| Eco-63-1 | Bacteria Proteobacteria Gammaproteobacteria Enterobacterales Enterobacteriaceae *Escherichia coli* 2-005-03_S4_C1 |
| Eco-64-1 | Bacteria Proteobacteria Gammaproteobacteria Enterobacterales Enterobacteriaceae *Escherichia coli* 2-005-03_S4_C2 |
| Eco-65-1 | Bacteria Proteobacteria Gammaproteobacteria Enterobacterales Enterobacteriaceae *Escherichia coli* 2-005-03_S4_C3 |
| Eco-66-1 | Bacteria Proteobacteria Gammaproteobacteria Enterobacterales Enterobacteriaceae *Escherichia coli* 2-011-08_S1_C1 |
| Eco-67-1 | Bacteria Proteobacteria Gammaproteobacteria Enterobacterales Enterobacteriaceae *Escherichia coli* 2-011-08_S1_C2 |
| Eco-68-1 | Bacteria Proteobacteria Gammaproteobacteria Enterobacterales Enterobacteriaceae *Escherichia coli* 2-011-08_S1_C3 |
| Eco-69-1 | Bacteria Proteobacteria Gammaproteobacteria Enterobacterales Enterobacteriaceae *Escherichia coli* 2-011-08_S3_C1 |
| Eco-70-1 | Bacteria Proteobacteria Gammaproteobacteria Enterobacterales Enterobacteriaceae *Escherichia coli* 2-011-08_S3_C2 |
| Eco-71-1 | Bacteria Proteobacteria Gammaproteobacteria Enterobacterales Enterobacteriaceae *Escherichia coli* 2-011-08_S3_C3 |
| Eco-72-1 | Bacteria Proteobacteria Gammaproteobacteria Enterobacterales Enterobacteriaceae *Escherichia coli* 2-011-08_S4_C1 |
| Eco-73-1 | Bacteria Proteobacteria Gammaproteobacteria Enterobacterales Enterobacteriaceae *Escherichia coli* 2-011-08_S4_C3 |
| Eco-74-1 | Bacteria Proteobacteria Gammaproteobacteria Enterobacterales Enterobacteriaceae *Escherichia coli* 2-052-05_S1_C1 |
| Eco-75-1 | Bacteria Proteobacteria Gammaproteobacteria Enterobacterales Enterobacteriaceae *Escherichia coli* 2-052-05_S1_C3 |
| Eco-76-1 | Bacteria Proteobacteria Gammaproteobacteria Enterobacterales Enterobacteriaceae *Escherichia coli* 2-052-05_S3_C1 |
| Eco-77-1 | Bacteria Proteobacteria Gammaproteobacteria Enterobacterales Enterobacteriaceae *Escherichia coli* 2-052-05_S3_C2 |
| Eco-78-1 | Bacteria Proteobacteria Gammaproteobacteria Enterobacterales Enterobacteriaceae *Escherichia coli* 2-052-05_S3_C3 |
| Eco-79-1 | Bacteria Proteobacteria Gammaproteobacteria Enterobacterales Enterobacteriaceae *Escherichia coli* 2-052-05_S4_C1 |
| Eco-80-1 | Bacteria Proteobacteria Gammaproteobacteria Enterobacterales Enterobacteriaceae *Escherichia coli* 2-052-05_S4_C2 |
| Eco-81-1 | Bacteria Proteobacteria Gammaproteobacteria Enterobacterales Enterobacteriaceae *Escherichia coli* 2-052-05_S4_C3 |
| Eco-82-1 | Bacteria Proteobacteria Gammaproteobacteria Enterobacterales Enterobacteriaceae *Escherichia coli* 2-156-04_S1_C3 |
| Eco-83-1 | Bacteria Proteobacteria Gammaproteobacteria Enterobacterales Enterobacteriaceae *Escherichia coli* 2-156-04_S3_C1 |
| Eco-84-1 | Bacteria Proteobacteria Gammaproteobacteria Enterobacterales Enterobacteriaceae *Escherichia coli* 2-156-04_S3_C2 |
| Eco-85-1 | Bacteria Proteobacteria Gammaproteobacteria Enterobacterales Enterobacteriaceae *Escherichia coli* 2-156-04_S3_C3 |
| Eco-86-1 | Bacteria Proteobacteria Gammaproteobacteria Enterobacterales Enterobacteriaceae *Escherichia coli* 2-156-04_S4_C1 |
| Eco-87-1 | Bacteria Proteobacteria Gammaproteobacteria Enterobacterales Enterobacteriaceae *Escherichia coli* 2-156-04_S4_C2 |
| Eco-88-1 | Bacteria Proteobacteria Gammaproteobacteria Enterobacterales Enterobacteriaceae *Escherichia coli* 2-156-04_S4_C3 |
| Eco-89-1 | Bacteria Proteobacteria Gammaproteobacteria Enterobacterales Enterobacteriaceae *Escherichia coli* 2-177-06_S1_C1 |
| Eco-90-1 | Bacteria Proteobacteria Gammaproteobacteria Enterobacterales Enterobacteriaceae *Escherichia coli* 2-177-06_S1_C2 |
| Eco-91-1 | Bacteria Proteobacteria Gammaproteobacteria Enterobacterales Enterobacteriaceae *Escherichia coli* 2-177-06_S1_C3 |
| Eco-92-1 | Bacteria Proteobacteria Gammaproteobacteria Enterobacterales Enterobacteriaceae *Escherichia coli* 2-177-06_S3_C1 |
| Eco-93-1 | Bacteria Proteobacteria Gammaproteobacteria Enterobacterales Enterobacteriaceae *Escherichia coli* 2-177-06_S3_C2 |
| Eco-94-1 | Bacteria Proteobacteria Gammaproteobacteria Enterobacterales Enterobacteriaceae *Escherichia coli* 2-177-06_S3_C3 |
| Eco-95-1 | Bacteria Proteobacteria Gammaproteobacteria Enterobacterales Enterobacteriaceae *Escherichia coli* 2-177-06_S4_C1 |
| Eco-96-1 | Bacteria Proteobacteria Gammaproteobacteria Enterobacterales Enterobacteriaceae *Escherichia coli* 2-177-06_S4_C2 |
| Eco-97-1 | Bacteria Proteobacteria Gammaproteobacteria Enterobacterales Enterobacteriaceae *Escherichia coli* 2-177-06_S4_C3 |
| Eco-98-1 | Bacteria Proteobacteria Gammaproteobacteria Enterobacterales Enterobacteriaceae *Escherichia coli* 2-210-07_S1_C2 |
| Eco-99-1 | Bacteria Proteobacteria Gammaproteobacteria Enterobacterales Enterobacteriaceae *Escherichia coli* 2-210-07_S1_C3 |
| Eco-100-1 | Bacteria Proteobacteria Gammaproteobacteria Enterobacterales Enterobacteriaceae *Escherichia coli* 2-210-07_S3_C1 |
| Eco-101-1 | Bacteria Proteobacteria Gammaproteobacteria Enterobacterales Enterobacteriaceae *Escherichia coli* 2-210-07_S3_C2 |
| Eco-102-1 | Bacteria Proteobacteria Gammaproteobacteria Enterobacterales Enterobacteriaceae *Escherichia coli* 2-210-07_S3_C3 |
| Eco-103-1 | Bacteria Proteobacteria Gammaproteobacteria Enterobacterales Enterobacteriaceae *Escherichia coli* 2-210-07_S4_C1 |
| Eco-104-1 | Bacteria Proteobacteria Gammaproteobacteria Enterobacterales Enterobacteriaceae *Escherichia coli* 2-210-07_S4_C2 |
| Eco-105-1 | Bacteria Proteobacteria Gammaproteobacteria Enterobacterales Enterobacteriaceae *Escherichia coli* 2-210-07_S4_C3 |
| Eco-106-1 | Bacteria Proteobacteria Gammaproteobacteria Enterobacterales Enterobacteriaceae *Escherichia coli* 2-222-05_S1_C1 |
| Eco-107-1 | Bacteria Proteobacteria Gammaproteobacteria Enterobacterales Enterobacteriaceae *Escherichia coli* 2-222-05_S1_C2 |
| Eco-108-1 | Bacteria Proteobacteria Gammaproteobacteria Enterobacterales Enterobacteriaceae *Escherichia coli* 2-222-05_S1_C3 |
| Eco-109-1 | Bacteria Proteobacteria Gammaproteobacteria Enterobacterales Enterobacteriaceae *Escherichia coli* 2-222-05_S3_C1 |
| Eco-110-1 | Bacteria Proteobacteria Gammaproteobacteria Enterobacterales Enterobacteriaceae *Escherichia coli* 2-222-05_S3_C2 |
| Eco-111-1 | Bacteria Proteobacteria Gammaproteobacteria Enterobacterales Enterobacteriaceae *Escherichia coli* 2-222-05_S3_C3 |
| Eco-112-1 | Bacteria Proteobacteria Gammaproteobacteria Enterobacterales Enterobacteriaceae *Escherichia coli* 2-222-05_S4_C1 |

| | |
|---|---|
| Eco-113-1 | Bacteria Proteobacteria Gammaproteobacteria Enterobacterales Enterobacteriaceae *Escherichia coli* 2-222-05_S4_C2 |
| Eco-114-1 | Bacteria Proteobacteria Gammaproteobacteria Enterobacterales Enterobacteriaceae *Escherichia coli* 2-222-05_S4_C3 |
| Eco-115-1 | Bacteria Proteobacteria Gammaproteobacteria Enterobacterales Enterobacteriaceae *Escherichia coli* 2-316-03_S1_C1 |
| Eco-116-1 | Bacteria Proteobacteria Gammaproteobacteria Enterobacterales Enterobacteriaceae *Escherichia coli* 2-316-03_S1_C2 |
| Eco-117-1 | Bacteria Proteobacteria Gammaproteobacteria Enterobacterales Enterobacteriaceae *Escherichia coli* 2-316-03_S3_C1 |
| Eco-118-1 | Bacteria Proteobacteria Gammaproteobacteria Enterobacterales Enterobacteriaceae *Escherichia coli* 2-316-03_S3_C2 |
| Eco-119-1 | Bacteria Proteobacteria Gammaproteobacteria Enterobacterales Enterobacteriaceae *Escherichia coli* 2-316-03_S3_C3 |
| Eco-120-1 | Bacteria Proteobacteria Gammaproteobacteria Enterobacterales Enterobacteriaceae *Escherichia coli* 2-316-03_S4_C1 |
| Eco-121-1 | Bacteria Proteobacteria Gammaproteobacteria Enterobacterales Enterobacteriaceae *Escherichia coli* 2-316-03_S4_C2 |
| Eco-122-1 | Bacteria Proteobacteria Gammaproteobacteria Enterobacterales Enterobacteriaceae *Escherichia coli* 2-316-03_S4_C3 |
| Eco-123-1 | Bacteria Proteobacteria Gammaproteobacteria Enterobacterales Enterobacteriaceae *Escherichia coli* 2-427-07_S1_C1 |
| Eco-124-1 | Bacteria Proteobacteria Gammaproteobacteria Enterobacterales Enterobacteriaceae *Escherichia coli* 2-427-07_S1_C2 |
| Eco-125-1 | Bacteria Proteobacteria Gammaproteobacteria Enterobacterales Enterobacteriaceae *Escherichia coli* 2-427-07_S1_C3 |
| Eco-126-1 | Bacteria Proteobacteria Gammaproteobacteria Enterobacterales Enterobacteriaceae *Escherichia coli* 2-427-07_S3_C1 |
| Eco-127-1 | Bacteria Proteobacteria Gammaproteobacteria Enterobacterales Enterobacteriaceae *Escherichia coli* 2-427-07_S3_C3 |
| Eco-128-1 | Bacteria Proteobacteria Gammaproteobacteria Enterobacterales Enterobacteriaceae *Escherichia coli* 2-427-07_S4_C1 |
| Eco-129-1 | Bacteria Proteobacteria Gammaproteobacteria Enterobacterales Enterobacteriaceae *Escherichia coli* 2-427-07_S4_C2 |
| Eco-130-1 | Bacteria Proteobacteria Gammaproteobacteria Enterobacterales Enterobacteriaceae *Escherichia coli* 2-427-07_S4_C3 |
| Eco-131-1 | Bacteria Proteobacteria Gammaproteobacteria Enterobacterales Enterobacteriaceae *Escherichia coli* 2-460-02_S1_C1 |
| Eco-132-1 | Bacteria Proteobacteria Gammaproteobacteria Enterobacterales Enterobacteriaceae *Escherichia coli* 2-460-02_S1_C2 |
| Eco-133-1 | Bacteria Proteobacteria Gammaproteobacteria Enterobacterales Enterobacteriaceae *Escherichia coli* 2-460-02_S1_C3 |
| Eco-134-1 | Bacteria Proteobacteria Gammaproteobacteria Enterobacterales Enterobacteriaceae *Escherichia coli* 2-460-02_S3_C1 |
| Eco-135-1 | Bacteria Proteobacteria Gammaproteobacteria Enterobacterales Enterobacteriaceae *Escherichia coli* 2-460-02_S3_C2 |
| Eco-136-1 | Bacteria Proteobacteria Gammaproteobacteria Enterobacterales Enterobacteriaceae *Escherichia coli* 2-460-02_S3_C3 |
| Eco-137-1 | Bacteria Proteobacteria Gammaproteobacteria Enterobacterales Enterobacteriaceae *Escherichia coli* 2-460-02_S4_C1 |
| Eco-138-1 | Bacteria Proteobacteria Gammaproteobacteria Enterobacterales Enterobacteriaceae *Escherichia coli* 2-460-02_S4_C2 |
| Eco-139-1 | Bacteria Proteobacteria Gammaproteobacteria Enterobacterales Enterobacteriaceae *Escherichia coli* 2-460-02_S4_C3 |
| Eco-140-1 | Bacteria Proteobacteria Gammaproteobacteria Enterobacterales Enterobacteriaceae *Escherichia coli* 2-474-04_S1_C1 |
| Eco-141-1 | Bacteria Proteobacteria Gammaproteobacteria Enterobacterales Enterobacteriaceae *Escherichia coli* 2-474-04_S1_C2 |
| Eco-142-1 | Bacteria Proteobacteria Gammaproteobacteria Enterobacterales Enterobacteriaceae *Escherichia coli* 2-474-04_S3_C1 |
| Eco-143-1 | Bacteria Proteobacteria Gammaproteobacteria Enterobacterales Enterobacteriaceae *Escherichia coli* 2-474-04_S3_C2 |
| Eco-144-1 | Bacteria Proteobacteria Gammaproteobacteria Enterobacterales Enterobacteriaceae *Escherichia coli* 2-474-04_S3_C3 |
| Eco-145-1 | Bacteria Proteobacteria Gammaproteobacteria Enterobacterales Enterobacteriaceae *Escherichia coli* 2-474-04_S4_C1 |
| Eco-146-1 | Bacteria Proteobacteria Gammaproteobacteria Enterobacterales Enterobacteriaceae *Escherichia coli* 2-474-04_S4_C2 |
| Eco-147-1 | Bacteria Proteobacteria Gammaproteobacteria Enterobacterales Enterobacteriaceae *Escherichia coli* 2-474-04_S4_C3 |
| Eco-148-1 | Bacteria Proteobacteria Gammaproteobacteria Enterobacterales Enterobacteriaceae *Escherichia coli* 27A |
| Eco-149-1 | Bacteria Proteobacteria Gammaproteobacteria Enterobacterales Enterobacteriaceae *Escherichia coli* 3-020-07_S1_C1 |
| Eco-150-1 | Bacteria Proteobacteria Gammaproteobacteria Enterobacterales Enterobacteriaceae *Escherichia coli* 3-020-07_S1_C2 |
| Eco-151-1 | Bacteria Proteobacteria Gammaproteobacteria Enterobacterales Enterobacteriaceae *Escherichia coli* 3-020-07_S1_C3 |
| Eco-152-1 | Bacteria Proteobacteria Gammaproteobacteria Enterobacterales Enterobacteriaceae *Escherichia coli* 3-020-07_S3_C1 |
| Eco-153-1 | Bacteria Proteobacteria Gammaproteobacteria Enterobacterales Enterobacteriaceae *Escherichia coli* 3-020-07_S3_C2 |
| Eco-154-1 | Bacteria Proteobacteria Gammaproteobacteria Enterobacterales Enterobacteriaceae *Escherichia coli* 3-020-07_S4_C1 |
| Eco-155-1 | Bacteria Proteobacteria Gammaproteobacteria Enterobacterales Enterobacteriaceae *Escherichia coli* 3-020-07_S4_C2 |
| Eco-156-1 | Bacteria Proteobacteria Gammaproteobacteria Enterobacterales Enterobacteriaceae *Escherichia coli* 3-020-07_S4_C3 |
| Eco-157-1 | Bacteria Proteobacteria Gammaproteobacteria Enterobacterales Enterobacteriaceae *Escherichia coli* 3-073-06_S1_C1 |
| Eco-158-1 | Bacteria Proteobacteria Gammaproteobacteria Enterobacterales Enterobacteriaceae *Escherichia coli* 3-073-06_S1_C2 |
| Eco-159-1 | Bacteria Proteobacteria Gammaproteobacteria Enterobacterales Enterobacteriaceae *Escherichia coli* 3-073-06_S3_C1 |
| Eco-160-1 | Bacteria Proteobacteria Gammaproteobacteria Enterobacterales Enterobacteriaceae *Escherichia coli* 3-073-06_S3_C2 |
| Eco-161-1 | Bacteria Proteobacteria Gammaproteobacteria Enterobacterales Enterobacteriaceae *Escherichia coli* 3-073-06_S4_C1 |
| Eco-162-1 | Bacteria Proteobacteria Gammaproteobacteria Enterobacterales Enterobacteriaceae *Escherichia coli* 3-073-06_S4_C2 |
| Eco-163-1 | Bacteria Proteobacteria Gammaproteobacteria Enterobacterales Enterobacteriaceae *Escherichia coli* 3-073-06_S4_C3 |
| Eco-164-1 | Bacteria Proteobacteria Gammaproteobacteria Enterobacterales Enterobacteriaceae *Escherichia coli* 3-105-05_S1_C1 |
| Eco-165-1 | Bacteria Proteobacteria Gammaproteobacteria Enterobacterales Enterobacteriaceae *Escherichia coli* 3-105-05_S1_C2 |
| Eco-166-1 | Bacteria Proteobacteria Gammaproteobacteria Enterobacterales Enterobacteriaceae *Escherichia coli* 3-105-05_S1_C3 |
| Eco-167-1 | Bacteria Proteobacteria Gammaproteobacteria Enterobacterales Enterobacteriaceae *Escherichia coli* 3-105-05_S3_C1 |
| Eco-168-1 | Bacteria Proteobacteria Gammaproteobacteria Enterobacterales Enterobacteriaceae *Escherichia coli* 3-105-05_S3_C2 |
| Eco-169-1 | Bacteria Proteobacteria Gammaproteobacteria Enterobacterales Enterobacteriaceae *Escherichia coli* 3-105-05_S3_C3 |
| Eco-170-1 | Bacteria Proteobacteria Gammaproteobacteria Enterobacterales Enterobacteriaceae *Escherichia coli* 3-105-05_S4_C1 |
| Eco-171-1 | Bacteria Proteobacteria Gammaproteobacteria Enterobacterales Enterobacteriaceae *Escherichia coli* 3-105-05_S4_C2 |
| Eco-172-1 | Bacteria Proteobacteria Gammaproteobacteria Enterobacterales Enterobacteriaceae *Escherichia coli* 3-105-05_S4_C3 |
| Eco-173-1 | Bacteria Proteobacteria Gammaproteobacteria Enterobacterales Enterobacteriaceae *Escherichia coli* 3-267-03_S1_C1 |
| Eco-174-1 | Bacteria Proteobacteria Gammaproteobacteria Enterobacterales Enterobacteriaceae *Escherichia coli* 3-267-03_S1_C2 |
| Eco-175-1 | Bacteria Proteobacteria Gammaproteobacteria Enterobacterales Enterobacteriaceae *Escherichia coli* 3-267-03_S1_C3 |
| Eco-176-1 | Bacteria Proteobacteria Gammaproteobacteria Enterobacterales Enterobacteriaceae *Escherichia coli* 3-267-03_S3_C1 |
| Eco-177-1 | Bacteria Proteobacteria Gammaproteobacteria Enterobacterales Enterobacteriaceae *Escherichia coli* 3-267-03_S3_C2 |

| | |
|---|---|
| Eco-178-1 | Bacteria Proteobacteria Gammaproteobacteria Enterobacterales Enterobacteriaceae *Escherichia coli* 3-267-03_S4_C1 |
| Eco-179-1 | Bacteria Proteobacteria Gammaproteobacteria Enterobacterales Enterobacteriaceae *Escherichia coli* 3-267-03_S4_C2 |
| Eco-180-1 | Bacteria Proteobacteria Gammaproteobacteria Enterobacterales Enterobacteriaceae *Escherichia coli* 3-373-03_S1_C1 |
| Eco-181-1 | Bacteria Proteobacteria Gammaproteobacteria Enterobacterales Enterobacteriaceae *Escherichia coli* 3-373-03_S1_C2 |
| Eco-182-1 | Bacteria Proteobacteria Gammaproteobacteria Enterobacterales Enterobacteriaceae *Escherichia coli* 3-373-03_S1_C3 |
| Eco-183-1 | Bacteria Proteobacteria Gammaproteobacteria Enterobacterales Enterobacteriaceae *Escherichia coli* 3-373-03_S3_C1 |
| Eco-184-1 | Bacteria Proteobacteria Gammaproteobacteria Enterobacterales Enterobacteriaceae *Escherichia coli* 3-373-03_S3_C2 |
| Eco-185-1 | Bacteria Proteobacteria Gammaproteobacteria Enterobacterales Enterobacteriaceae *Escherichia coli* 3-373-03_S3_C3 |
| Eco-186-1 | Bacteria Proteobacteria Gammaproteobacteria Enterobacterales Enterobacteriaceae *Escherichia coli* 3-373-03_S4_C1 |
| Eco-187-1 | Bacteria Proteobacteria Gammaproteobacteria Enterobacterales Enterobacteriaceae *Escherichia coli* 3-373-03_S4_C2 |
| Eco-188-1 | Bacteria Proteobacteria Gammaproteobacteria Enterobacterales Enterobacteriaceae *Escherichia coli* 3-373-03_S4_C3 |
| Eco-189-1 | Bacteria Proteobacteria Gammaproteobacteria Enterobacterales Enterobacteriaceae *Escherichia coli* 3-475-03_S1_C1 |
| Eco-190-1 | Bacteria Proteobacteria Gammaproteobacteria Enterobacterales Enterobacteriaceae *Escherichia coli* 3-475-03_S1_C2 |
| Eco-191-1 | Bacteria Proteobacteria Gammaproteobacteria Enterobacterales Enterobacteriaceae *Escherichia coli* 3-475-03_S3_C1 |
| Eco-192-1 | Bacteria Proteobacteria Gammaproteobacteria Enterobacterales Enterobacteriaceae *Escherichia coli* 3-475-03_S3_C2 |
| Eco-193-1 | Bacteria Proteobacteria Gammaproteobacteria Enterobacterales Enterobacteriaceae *Escherichia coli* 3-475-03_S4_C1 |
| Eco-194-1 | Bacteria Proteobacteria Gammaproteobacteria Enterobacterales Enterobacteriaceae *Escherichia coli* 3-475-03_S4_C2 |
| Eco-195-1 | Bacteria Proteobacteria Gammaproteobacteria Enterobacterales Enterobacteriaceae *Escherichia coli* 300059 |
| Eco-196-1 | Bacteria Proteobacteria Gammaproteobacteria Enterobacterales Enterobacteriaceae *Escherichia coli* 302053 |
| Eco-197-1 | Bacteria Proteobacteria Gammaproteobacteria Enterobacterales Enterobacteriaceae *Escherichia coli* 320 |
| Eco-198-1 | Bacteria Proteobacteria Gammaproteobacteria Enterobacterales Enterobacteriaceae *Escherichia coli* 328 |
| Eco-199-1 | Bacteria Proteobacteria Gammaproteobacteria Enterobacterales Enterobacteriaceae *Escherichia coli* 38.16 |
| Eco-200-1 | Bacteria Proteobacteria Gammaproteobacteria Enterobacterales Enterobacteriaceae *Escherichia coli* 38.27 |
| Eco-201-1 | Bacteria Proteobacteria Gammaproteobacteria Enterobacterales Enterobacteriaceae *Escherichia coli* 38.34 |
| Eco-202-1 | Bacteria Proteobacteria Gammaproteobacteria Enterobacterales Enterobacteriaceae *Escherichia coli* 38.52 |
| Eco-203-1 | Bacteria Proteobacteria Gammaproteobacteria Enterobacterales Enterobacteriaceae *Escherichia coli* 4-203-08_S1_C1 |
| Eco-204-1 | Bacteria Proteobacteria Gammaproteobacteria Enterobacterales Enterobacteriaceae *Escherichia coli* 4-203-08_S1_C2 |
| Eco-205-1 | Bacteria Proteobacteria Gammaproteobacteria Enterobacterales Enterobacteriaceae *Escherichia coli* 4-203-08_S1_C3 |
| Eco-206-1 | Bacteria Proteobacteria Gammaproteobacteria Enterobacterales Enterobacteriaceae *Escherichia coli* 4-203-08_S3_C1 |
| Eco-207-1 | Bacteria Proteobacteria Gammaproteobacteria Enterobacterales Enterobacteriaceae *Escherichia coli* 4-203-08_S3_C2 |
| Eco-208-1 | Bacteria Proteobacteria Gammaproteobacteria Enterobacterales Enterobacteriaceae *Escherichia coli* 4-203-08_S3_C3 |
| Eco-209-1 | Bacteria Proteobacteria Gammaproteobacteria Enterobacterales Enterobacteriaceae *Escherichia coli* 4-203-08_S4_C2 |
| Eco-210-1 | Bacteria Proteobacteria Gammaproteobacteria Enterobacterales Enterobacteriaceae *Escherichia coli* 4-203-08_S4_C3 |
| Eco-211-1 | Bacteria Proteobacteria Gammaproteobacteria Enterobacterales Enterobacteriaceae *Escherichia coli* 401140 |
| Eco-212-1 | Bacteria Proteobacteria Gammaproteobacteria Enterobacterales Enterobacteriaceae *Escherichia coli* 402290 |
| Eco-213-1 | Bacteria Proteobacteria Gammaproteobacteria Enterobacterales Enterobacteriaceae *Escherichia coli* 435 |
| Eco-214-1 | Bacteria Proteobacteria Gammaproteobacteria Enterobacterales Enterobacteriaceae *Escherichia coli* 4541-1 |
| Eco-215-1 | Bacteria Proteobacteria Gammaproteobacteria Enterobacterales Enterobacteriaceae *Escherichia coli* 4552-1 |
| Eco-216-1 | Bacteria Proteobacteria Gammaproteobacteria Enterobacterales Enterobacteriaceae *Escherichia coli* 5-172-05_S1_C3 |
| Eco-217-1 | Bacteria Proteobacteria Gammaproteobacteria Enterobacterales Enterobacteriaceae *Escherichia coli* 5-172-05_S3_C1 |
| Eco-218-1 | Bacteria Proteobacteria Gammaproteobacteria Enterobacterales Enterobacteriaceae *Escherichia coli* 5-172-05_S3_C3 |
| Eco-219-1 | Bacteria Proteobacteria Gammaproteobacteria Enterobacterales Enterobacteriaceae *Escherichia coli* 5-172-05_S4_C1 |
| Eco-220-1 | Bacteria Proteobacteria Gammaproteobacteria Enterobacterales Enterobacteriaceae *Escherichia coli* 5-172-05_S4_C2 |
| Eco-221-1 | Bacteria Proteobacteria Gammaproteobacteria Enterobacterales Enterobacteriaceae *Escherichia coli* 5-172-05_S4_C3 |
| Eco-222-1 | Bacteria Proteobacteria Gammaproteobacteria Enterobacterales Enterobacteriaceae *Escherichia coli* 5-366-08_S1_C1 |
| Eco-223-1 | Bacteria Proteobacteria Gammaproteobacteria Enterobacterales Enterobacteriaceae *Escherichia coli* 5-366-08_S1_C3 |
| Eco-224-1 | Bacteria Proteobacteria Gammaproteobacteria Enterobacterales Enterobacteriaceae *Escherichia coli* 5-366-08_S3_C1 |
| Eco-225-1 | Bacteria Proteobacteria Gammaproteobacteria Enterobacterales Enterobacteriaceae *Escherichia coli* 5-366-08_S3_C2 |
| Eco-226-1 | Bacteria Proteobacteria Gammaproteobacteria Enterobacterales Enterobacteriaceae *Escherichia coli* 5-366-08_S3_C3 |
| Eco-227-1 | Bacteria Proteobacteria Gammaproteobacteria Enterobacterales Enterobacteriaceae *Escherichia coli* 5-366-08_S4_C1 |
| Eco-228-1 | Bacteria Proteobacteria Gammaproteobacteria Enterobacterales Enterobacteriaceae *Escherichia coli* 5-366-08_S4_C2 |
| Eco-229-1 | Bacteria Proteobacteria Gammaproteobacteria Enterobacterales Enterobacteriaceae *Escherichia coli* 53A |
| Eco-230-1 to Eco-230-2 | Bacteria Proteobacteria Gammaproteobacteria Enterobacterales Enterobacteriaceae *Escherichia coli* 53C |
| Eco-231-1 | Bacteria Proteobacteria Gammaproteobacteria Enterobacterales Enterobacteriaceae *Escherichia coli* 597 |
| Eco-232-1 | Bacteria Proteobacteria Gammaproteobacteria Enterobacterales Enterobacteriaceae *Escherichia coli* 6-175-07_S1_C1 |
| Eco-233-1 | Bacteria Proteobacteria Gammaproteobacteria Enterobacterales Enterobacteriaceae *Escherichia coli* 6-175-07_S1_C2 |
| Eco-234-1 | Bacteria Proteobacteria Gammaproteobacteria Enterobacterales Enterobacteriaceae *Escherichia coli* 6-175-07_S1_C3 |
| Eco-235-1 | Bacteria Proteobacteria Gammaproteobacteria Enterobacterales Enterobacteriaceae *Escherichia coli* 6-175-07_S3_C1 |
| Eco-236-1 | Bacteria Proteobacteria Gammaproteobacteria Enterobacterales Enterobacteriaceae *Escherichia coli* 6-175-07_S3_C2 |
| Eco-237-1 | Bacteria Proteobacteria Gammaproteobacteria Enterobacterales Enterobacteriaceae *Escherichia coli* 6-175-07_S3_C3 |
| Eco-238-1 | Bacteria Proteobacteria Gammaproteobacteria Enterobacterales Enterobacteriaceae *Escherichia coli* 6-175-07_S4_C1 |
| Eco-239-1 | Bacteria Proteobacteria Gammaproteobacteria Enterobacterales Enterobacteriaceae *Escherichia coli* 6-175-07_S4_C2 |
| Eco-240-1 | Bacteria Proteobacteria Gammaproteobacteria Enterobacterales Enterobacteriaceae *Escherichia coli* 6-175-07_S4_C3 |
| Eco-241-1 | Bacteria Proteobacteria Gammaproteobacteria Enterobacterales Enterobacteriaceae *Escherichia coli* 6-319-05_S1_C1 |
| Eco-242-1 | Bacteria Proteobacteria Gammaproteobacteria Enterobacterales Enterobacteriaceae *Escherichia coli* 6-319-05_S1_C2 |

| | |
|---|---|
| Eco-243-1 | Bacteria Proteobacteria Gammaproteobacteria Enterobacterales Enterobacteriaceae *Escherichia coli* 6-319-05_S1_C3 |
| Eco-244-1 | Bacteria Proteobacteria Gammaproteobacteria Enterobacterales Enterobacteriaceae *Escherichia coli* 6-319-05_S3_C1 |
| Eco-245-1 | Bacteria Proteobacteria Gammaproteobacteria Enterobacterales Enterobacteriaceae *Escherichia coli* 6-319-05_S3_C2 |
| Eco-246-1 | Bacteria Proteobacteria Gammaproteobacteria Enterobacterales Enterobacteriaceae *Escherichia coli* 6-319-05_S3_C3 |
| Eco-247-1 | Bacteria Proteobacteria Gammaproteobacteria Enterobacterales Enterobacteriaceae *Escherichia coli* 6-319-05_S4_C2 |
| Eco-248-1 | Bacteria Proteobacteria Gammaproteobacteria Enterobacterales Enterobacteriaceae *Escherichia coli* 6-319-05_S4_C3 |
| Eco-249-1 | Bacteria Proteobacteria Gammaproteobacteria Enterobacterales Enterobacteriaceae *Escherichia coli* 6-537-08_S1_C1 |
| Eco-250-1 | Bacteria Proteobacteria Gammaproteobacteria Enterobacterales Enterobacteriaceae *Escherichia coli* 6-537-08_S1_C2 |
| Eco-251-1 | Bacteria Proteobacteria Gammaproteobacteria Enterobacterales Enterobacteriaceae *Escherichia coli* 6-537-08_S1_C3 |
| Eco-252-1 | Bacteria Proteobacteria Gammaproteobacteria Enterobacterales Enterobacteriaceae *Escherichia coli* 6-537-08_S3_C1 |
| Eco-253-1 | Bacteria Proteobacteria Gammaproteobacteria Enterobacterales Enterobacteriaceae *Escherichia coli* 6-537-08_S3_C2 |
| Eco-254-1 | Bacteria Proteobacteria Gammaproteobacteria Enterobacterales Enterobacteriaceae *Escherichia coli* 6-537-08_S3_C3 |
| Eco-255-1 | Bacteria Proteobacteria Gammaproteobacteria Enterobacterales Enterobacteriaceae *Escherichia coli* 6-537-08_S4_C1 |
| Eco-256-1 | Bacteria Proteobacteria Gammaproteobacteria Enterobacterales Enterobacteriaceae *Escherichia coli* 6-537-08_S4_C2 |
| Eco-257-1 | Bacteria Proteobacteria Gammaproteobacteria Enterobacterales Enterobacteriaceae *Escherichia coli* 606 |
| Eco-258-1 | Bacteria Proteobacteria Gammaproteobacteria Enterobacterales Enterobacteriaceae *Escherichia coli* 668 |
| Eco-259-1 | Bacteria Proteobacteria Gammaproteobacteria Enterobacterales Enterobacteriaceae *Escherichia coli* 681 |
| Eco-260-1 | Bacteria Proteobacteria Gammaproteobacteria Enterobacterales Enterobacteriaceae *Escherichia coli* 7-233-03_S1_C2 |
| Eco-261-1 | Bacteria Proteobacteria Gammaproteobacteria Enterobacterales Enterobacteriaceae *Escherichia coli* 7-233-03_S1_C3 |
| Eco-262-1 | Bacteria Proteobacteria Gammaproteobacteria Enterobacterales Enterobacteriaceae *Escherichia coli* 7-233-03_S3_C1 |
| Eco-263-1 | Bacteria Proteobacteria Gammaproteobacteria Enterobacterales Enterobacteriaceae *Escherichia coli* 7-233-03_S3_C2 |
| Eco-264-1 | Bacteria Proteobacteria Gammaproteobacteria Enterobacterales Enterobacteriaceae *Escherichia coli* 7-233-03_S3_C3 |
| Eco-265-1 | Bacteria Proteobacteria Gammaproteobacteria Enterobacterales Enterobacteriaceae *Escherichia coli* 7-233-03_S4_C1 |
| Eco-266-1 | Bacteria Proteobacteria Gammaproteobacteria Enterobacterales Enterobacteriaceae *Escherichia coli* 7-233-03_S4_C2 |
| Eco-267-1 | Bacteria Proteobacteria Gammaproteobacteria Enterobacterales Enterobacteriaceae *Escherichia coli* 7-233-03_S4_C3 |
| Eco-268-1 | Bacteria Proteobacteria Gammaproteobacteria Enterobacterales Enterobacteriaceae *Escherichia coli* 7996-1 |
| Eco-269-1 | Bacteria Proteobacteria Gammaproteobacteria Enterobacterales Enterobacteriaceae *Escherichia coli* 8-415-05_S1_C1 |
| Eco-270-1 | Bacteria Proteobacteria Gammaproteobacteria Enterobacterales Enterobacteriaceae *Escherichia coli* 8-415-05_S1_C2 |
| Eco-271-1 | Bacteria Proteobacteria Gammaproteobacteria Enterobacterales Enterobacteriaceae *Escherichia coli* 8-415-05_S3_C1 |
| Eco-272-1 | Bacteria Proteobacteria Gammaproteobacteria Enterobacterales Enterobacteriaceae *Escherichia coli* 8-415-05_S3_C2 |
| Eco-273-1 | Bacteria Proteobacteria Gammaproteobacteria Enterobacterales Enterobacteriaceae *Escherichia coli* 8-415-05_S3_C3 |
| Eco-274-1 | Bacteria Proteobacteria Gammaproteobacteria Enterobacterales Enterobacteriaceae *Escherichia coli* 8-415-05_S4_C1 |
| Eco-275-1 | Bacteria Proteobacteria Gammaproteobacteria Enterobacterales Enterobacteriaceae *Escherichia coli* 8-415-05_S4_C2 |
| Eco-276-1 | Bacteria Proteobacteria Gammaproteobacteria Enterobacterales Enterobacteriaceae *Escherichia coli* 8-415-05_S4_C3 |
| Eco-277-1 | Bacteria Proteobacteria Gammaproteobacteria Enterobacterales Enterobacteriaceae *Escherichia coli* 85B |
| Eco-278-1 | Bacteria Proteobacteria Gammaproteobacteria Enterobacterales Enterobacteriaceae *Escherichia coli* 8624 |
| Eco-279-1 | Bacteria Proteobacteria Gammaproteobacteria Enterobacterales Enterobacteriaceae *Escherichia coli* 87A |
| Eco-280-1 | Bacteria Proteobacteria Gammaproteobacteria Enterobacterales Enterobacteriaceae *Escherichia coli* 897 |
| Eco-281-1 | Bacteria Proteobacteria Gammaproteobacteria Enterobacterales Enterobacteriaceae *Escherichia coli* 9.1649 |
| Eco-282-1 | Bacteria Proteobacteria Gammaproteobacteria Enterobacterales Enterobacteriaceae *Escherichia coli* 97.0264 |
| Eco-283-1 | Bacteria Proteobacteria Gammaproteobacteria Enterobacterales Enterobacteriaceae *Escherichia coli* 9B |
| Eco-284-1 | Bacteria Proteobacteria Gammaproteobacteria Enterobacterales Enterobacteriaceae *Escherichia coli* AB42410445-isolate1 |
| Eco-285-1 | Bacteria Proteobacteria Gammaproteobacteria Enterobacterales Enterobacteriaceae *Escherichia coli* AB42554418-isolate1 |
| Eco-286-1 | Bacteria Proteobacteria Gammaproteobacteria Enterobacterales Enterobacteriaceae *Escherichia coli* AB42602061-isolate1 |
| Eco-287-1 | Bacteria Proteobacteria Gammaproteobacteria Enterobacterales Enterobacteriaceae *Escherichia coli* AB43739056-isolate1 |
| Eco-288-1 | Bacteria Proteobacteria Gammaproteobacteria Enterobacterales Enterobacteriaceae *Escherichia coli* ACN001 |
| Eco-289-1 to Eco-289-2 | Bacteria Proteobacteria Gammaproteobacteria Enterobacterales Enterobacteriaceae *Escherichia coli* AD30 |
| Eco-290-1 | Bacteria Proteobacteria Gammaproteobacteria Enterobacterales Enterobacteriaceae *Escherichia coli* APEC IMT5155 |
| Eco-291-1 to Eco-291-2 | Bacteria Proteobacteria Gammaproteobacteria Enterobacterales Enterobacteriaceae *Escherichia coli* ATCC 25922 |
| Eco-292-1 | Bacteria Proteobacteria Gammaproteobacteria Enterobacterales Enterobacteriaceae *Escherichia coli* ATCC 35150 |
| Eco-293-1 | Bacteria Proteobacteria Gammaproteobacteria Enterobacterales Enterobacteriaceae *Escherichia coli* ATCC BAA-2192 |
| Eco-294-1 | Bacteria Proteobacteria Gammaproteobacteria Enterobacterales Enterobacteriaceae *Escherichia coli* ATCC BAA-2193 |
| Eco-295-1 | Bacteria Proteobacteria Gammaproteobacteria Enterobacterales Enterobacteriaceae *Escherichia coli* BIDMC 15 |
| Eco-296-1 | Bacteria Proteobacteria Gammaproteobacteria Enterobacterales Enterobacteriaceae *Escherichia coli* BIDMC 17A |
| Eco-297-1 | Bacteria Proteobacteria Gammaproteobacteria Enterobacterales Enterobacteriaceae *Escherichia coli* BIDMC 17B |
| Eco-298-1 | Bacteria Proteobacteria Gammaproteobacteria Enterobacterales Enterobacteriaceae *Escherichia coli* BIDMC 19A |
| Eco-299-1 | Bacteria Proteobacteria Gammaproteobacteria Enterobacterales Enterobacteriaceae *Escherichia coli* BIDMC 19B |
| Eco-300-1 | Bacteria Proteobacteria Gammaproteobacteria Enterobacterales Enterobacteriaceae *Escherichia coli* BIDMC 20A |
| Eco-301-1 | Bacteria Proteobacteria Gammaproteobacteria Enterobacterales Enterobacteriaceae *Escherichia coli* BIDMC 20B |
| Eco-302-1 | Bacteria Proteobacteria Gammaproteobacteria Enterobacterales Enterobacteriaceae *Escherichia coli* BIDMC 2B |
| Eco-303-1 | Bacteria Proteobacteria Gammaproteobacteria Enterobacterales Enterobacteriaceae *Escherichia coli* BIDMC 3 |
| Eco-304-1 | Bacteria Proteobacteria Gammaproteobacteria Enterobacterales Enterobacteriaceae *Escherichia coli* BIDMC 43a |
| Eco-305-1 | Bacteria Proteobacteria Gammaproteobacteria Enterobacterales Enterobacteriaceae *Escherichia coli* BIDMC 43b |
| Eco-306-1 | Bacteria Proteobacteria Gammaproteobacteria Enterobacterales Enterobacteriaceae *Escherichia coli* BIDMC 49a |
| Eco-307-1 | Bacteria Proteobacteria Gammaproteobacteria Enterobacterales Enterobacteriaceae *Escherichia coli* BIDMC 49b |

| | |
|---|---|
| Eco-308-1 | Bacteria Proteobacteria Gammaproteobacteria Enterobacterales Enterobacteriaceae *Escherichia coli* BIDMC 58 |
| Eco-309-1 | Bacteria Proteobacteria Gammaproteobacteria Enterobacterales Enterobacteriaceae *Escherichia coli* BIDMC 59 |
| Eco-310-1 | Bacteria Proteobacteria Gammaproteobacteria Enterobacterales Enterobacteriaceae *Escherichia coli* BIDMC 6 |
| Eco-311-1 | Bacteria Proteobacteria Gammaproteobacteria Enterobacterales Enterobacteriaceae *Escherichia coli* BIDMC 62 |
| Eco-312-1 | Bacteria Proteobacteria Gammaproteobacteria Enterobacterales Enterobacteriaceae *Escherichia coli* BIDMC 63 |
| Eco-313-1 | Bacteria Proteobacteria Gammaproteobacteria Enterobacterales Enterobacteriaceae *Escherichia coli* BIDMC 64 |
| Eco-314-1 | Bacteria Proteobacteria Gammaproteobacteria Enterobacterales Enterobacteriaceae *Escherichia coli* BIDMC 65 |
| Eco-315-1 | Bacteria Proteobacteria Gammaproteobacteria Enterobacterales Enterobacteriaceae *Escherichia coli* BIDMC 70 |
| Eco-316-1 | Bacteria Proteobacteria Gammaproteobacteria Enterobacterales Enterobacteriaceae *Escherichia coli* BIDMC 71 |
| Eco-317-1 | Bacteria Proteobacteria Gammaproteobacteria Enterobacterales Enterobacteriaceae *Escherichia coli* BIDMC 72 |
| Eco-318-1 | Bacteria Proteobacteria Gammaproteobacteria Enterobacterales Enterobacteriaceae *Escherichia coli* BIDMC 73 |
| Eco-319-1 | Bacteria Proteobacteria Gammaproteobacteria Enterobacterales Enterobacteriaceae *Escherichia coli* BIDMC 74 |
| Eco-320-1 | Bacteria Proteobacteria Gammaproteobacteria Enterobacterales Enterobacteriaceae *Escherichia coli* BIDMC 75 |
| Eco-321-1 | Bacteria Proteobacteria Gammaproteobacteria Enterobacterales Enterobacteriaceae *Escherichia coli* BIDMC 76 |
| Eco-322-1 | Bacteria Proteobacteria Gammaproteobacteria Enterobacterales Enterobacteriaceae *Escherichia coli* BIDMC 77 |
| Eco-323-1 | Bacteria Proteobacteria Gammaproteobacteria Enterobacterales Enterobacteriaceae *Escherichia coli* BIDMC 78 |
| Eco-324-1 | Bacteria Proteobacteria Gammaproteobacteria Enterobacterales Enterobacteriaceae *Escherichia coli* BIDMC 79 |
| Eco-325-1 | Bacteria Proteobacteria Gammaproteobacteria Enterobacterales Enterobacteriaceae *Escherichia coli* BIDMC 82 |
| Eco-326-1 | Bacteria Proteobacteria Gammaproteobacteria Enterobacterales Enterobacteriaceae *Escherichia coli* BIDMC 83 |
| Eco-327-1 | Bacteria Proteobacteria Gammaproteobacteria Enterobacterales Enterobacteriaceae *Escherichia coli* BIDMC 9 |
| Eco-328-1 | Bacteria Proteobacteria Gammaproteobacteria Enterobacterales Enterobacteriaceae *Escherichia coli* BW25113 |
| Eco-329-1 | Bacteria Proteobacteria Gammaproteobacteria Enterobacterales Enterobacteriaceae *Escherichia coli* BWH 34 |
| Eco-330-1 | Bacteria Proteobacteria Gammaproteobacteria Enterobacterales Enterobacteriaceae *Escherichia coli* BWH 40 |
| Eco-331-1 | Bacteria Proteobacteria Gammaproteobacteria Enterobacterales Enterobacteriaceae *Escherichia coli* C1214_90 |
| Eco-332-1 | Bacteria Proteobacteria Gammaproteobacteria Enterobacterales Enterobacteriaceae *Escherichia coli* C1244_91 |
| Eco-333-1 | Bacteria Proteobacteria Gammaproteobacteria Enterobacterales Enterobacteriaceae *Escherichia coli* C12_92 |
| Eco-334-1 | Bacteria Proteobacteria Gammaproteobacteria Enterobacterales Enterobacteriaceae *Escherichia coli* C154_11 |
| Eco-335-1 | Bacteria Proteobacteria Gammaproteobacteria Enterobacterales Enterobacteriaceae *Escherichia coli* C155_11 |
| Eco-336-1 | Bacteria Proteobacteria Gammaproteobacteria Enterobacterales Enterobacteriaceae *Escherichia coli* C157_11 |
| Eco-337-1 | Bacteria Proteobacteria Gammaproteobacteria Enterobacterales Enterobacteriaceae *Escherichia coli* C161_11 |
| Eco-338-1 | Bacteria Proteobacteria Gammaproteobacteria Enterobacterales Enterobacteriaceae *Escherichia coli* C166_11 |
| Eco-339-1 | Bacteria Proteobacteria Gammaproteobacteria Enterobacterales Enterobacteriaceae *Escherichia coli* C170_11 |
| Eco-340-1 | Bacteria Proteobacteria Gammaproteobacteria Enterobacterales Enterobacteriaceae *Escherichia coli* C2139_99 |
| Eco-341-1 | Bacteria Proteobacteria Gammaproteobacteria Enterobacterales Enterobacteriaceae *Escherichia coli* C213_10 |
| Eco-342-1 | Bacteria Proteobacteria Gammaproteobacteria Enterobacterales Enterobacteriaceae *Escherichia coli* C238_91 |
| Eco-343-1 | Bacteria Proteobacteria Gammaproteobacteria Enterobacterales Enterobacteriaceae *Escherichia coli* C260_92 |
| Eco-344-1 | Bacteria Proteobacteria Gammaproteobacteria Enterobacterales Enterobacteriaceae *Escherichia coli* C262_10 |
| Eco-345-1 | Bacteria Proteobacteria Gammaproteobacteria Enterobacterales Enterobacteriaceae *Escherichia coli* C283_09 |
| Eco-346-1 | Bacteria Proteobacteria Gammaproteobacteria Enterobacterales Enterobacteriaceae *Escherichia coli* C295_10 |
| Eco-347-1 | Bacteria Proteobacteria Gammaproteobacteria Enterobacterales Enterobacteriaceae *Escherichia coli* C341_10 |
| Eco-348-1 | Bacteria Proteobacteria Gammaproteobacteria Enterobacterales Enterobacteriaceae *Escherichia coli* C343_08 |
| Eco-349-1 | Bacteria Proteobacteria Gammaproteobacteria Enterobacterales Enterobacteriaceae *Escherichia coli* C347_93 |
| Eco-350-1 | Bacteria Proteobacteria Gammaproteobacteria Enterobacterales Enterobacteriaceae *Escherichia coli* C353_09 |
| Eco-351-1 | Bacteria Proteobacteria Gammaproteobacteria Enterobacterales Enterobacteriaceae *Escherichia coli* C354_03B |
| Eco-352-1 | Bacteria Proteobacteria Gammaproteobacteria Enterobacterales Enterobacteriaceae *Escherichia coli* C40_11 |
| Eco-353-1 | Bacteria Proteobacteria Gammaproteobacteria Enterobacterales Enterobacteriaceae *Escherichia coli* C418_89 |
| Eco-354-1 | Bacteria Proteobacteria Gammaproteobacteria Enterobacterales Enterobacteriaceae *Escherichia coli* C458_10 |
| Eco-355-1 | Bacteria Proteobacteria Gammaproteobacteria Enterobacterales Enterobacteriaceae *Escherichia coli* C488_07 |
| Eco-356-1 | Bacteria Proteobacteria Gammaproteobacteria Enterobacterales Enterobacteriaceae *Escherichia coli* C496_10 |
| Eco-357-1 | Bacteria Proteobacteria Gammaproteobacteria Enterobacterales Enterobacteriaceae *Escherichia coli* C497_10 |
| Eco-358-1 | Bacteria Proteobacteria Gammaproteobacteria Enterobacterales Enterobacteriaceae *Escherichia coli* C527_94 |
| Eco-359-1 | Bacteria Proteobacteria Gammaproteobacteria Enterobacterales Enterobacteriaceae *Escherichia coli* C581_05 |
| Eco-360-1 | Bacteria Proteobacteria Gammaproteobacteria Enterobacterales Enterobacteriaceae *Escherichia coli* C586_05 |
| Eco-361-1 | Bacteria Proteobacteria Gammaproteobacteria Enterobacterales Enterobacteriaceae *Escherichia coli* C58_11 |
| Eco-362-1 | Bacteria Proteobacteria Gammaproteobacteria Enterobacterales Enterobacteriaceae *Escherichia coli* C639_08 |
| Eco-363-1 | Bacteria Proteobacteria Gammaproteobacteria Enterobacterales Enterobacteriaceae *Escherichia coli* C652_10 |
| Eco-364-1 | Bacteria Proteobacteria Gammaproteobacteria Enterobacterales Enterobacteriaceae *Escherichia coli* C654_09 |
| Eco-365-1 | Bacteria Proteobacteria Gammaproteobacteria Enterobacterales Enterobacteriaceae *Escherichia coli* C691-71 (14b) |
| Eco-366-1 | Bacteria Proteobacteria Gammaproteobacteria Enterobacterales Enterobacteriaceae *Escherichia coli* C717_10 |
| Eco-367-1 | Bacteria Proteobacteria Gammaproteobacteria Enterobacterales Enterobacteriaceae *Escherichia coli* C725_88 |
| Eco-368-1 | Bacteria Proteobacteria Gammaproteobacteria Enterobacterales Enterobacteriaceae *Escherichia coli* C732_98 |
| Eco-369-1 | Bacteria Proteobacteria Gammaproteobacteria Enterobacterales Enterobacteriaceae *Escherichia coli* C743_03 |
| Eco-370-1 | Bacteria Proteobacteria Gammaproteobacteria Enterobacterales Enterobacteriaceae *Escherichia coli* C751_03 |
| Eco-371-1 | Bacteria Proteobacteria Gammaproteobacteria Enterobacterales Enterobacteriaceae *Escherichia coli* C78_09C |
| Eco-372-1 | Bacteria Proteobacteria Gammaproteobacteria Enterobacterales Enterobacteriaceae *Escherichia coli* C792_92 |

| | |
|---|---|
| Eco-373-1 | Bacteria Proteobacteria Gammaproteobacteria Enterobacterales Enterobacteriaceae *Escherichia coli* C796_10 |
| Eco-374-1 | Bacteria Proteobacteria Gammaproteobacteria Enterobacterales Enterobacteriaceae *Escherichia coli* C799_92 |
| Eco-375-1 | Bacteria Proteobacteria Gammaproteobacteria Enterobacterales Enterobacteriaceae *Escherichia coli* C79_08 |
| Eco-376-1 | Bacteria Proteobacteria Gammaproteobacteria Enterobacterales Enterobacteriaceae *Escherichia coli* C807_09 |
| Eco-377-1 | Bacteria Proteobacteria Gammaproteobacteria Enterobacterales Enterobacteriaceae *Escherichia coli* C80_08 |
| Eco-378-1 | Bacteria Proteobacteria Gammaproteobacteria Enterobacterales Enterobacteriaceae *Escherichia coli* C824_10 |
| Eco-379-1 | Bacteria Proteobacteria Gammaproteobacteria Enterobacterales Enterobacteriaceae *Escherichia coli* C82_11 |
| Eco-380-1 | Bacteria Proteobacteria Gammaproteobacteria Enterobacterales Enterobacteriaceae *Escherichia coli* C842_97 |
| Eco-381-1 | Bacteria Proteobacteria Gammaproteobacteria Enterobacterales Enterobacteriaceae *Escherichia coli* C844_97 |
| Eco-382-1 | Bacteria Proteobacteria Gammaproteobacteria Enterobacterales Enterobacteriaceae *Escherichia coli* C87_11 |
| Eco-383-1 | Bacteria Proteobacteria Gammaproteobacteria Enterobacterales Enterobacteriaceae *Escherichia coli* C887_10 |
| Eco-384-1 | Bacteria Proteobacteria Gammaproteobacteria Enterobacterales Enterobacteriaceae *Escherichia coli* C900_01 |
| Eco-385-1 | Bacteria Proteobacteria Gammaproteobacteria Enterobacterales Enterobacteriaceae *Escherichia coli* C93_11 |
| Eco-386-1 | Bacteria Proteobacteria Gammaproteobacteria Enterobacterales Enterobacteriaceae *Escherichia coli* C9_92 |
| Eco-387-1 | Bacteria Proteobacteria Gammaproteobacteria Enterobacterales Enterobacteriaceae *Escherichia coli* CHS 68 |
| Eco-388-1 | Bacteria Proteobacteria Gammaproteobacteria Enterobacterales Enterobacteriaceae *Escherichia coli* CHS 69 |
| Eco-389-1 | Bacteria Proteobacteria Gammaproteobacteria Enterobacterales Enterobacteriaceae *Escherichia coli* CHS 77 |
| Eco-390-1 | Bacteria Proteobacteria Gammaproteobacteria Enterobacterales Enterobacteriaceae *Escherichia coli* D6-113.11 |
| Eco-391-1 | Bacteria Proteobacteria Gammaproteobacteria Enterobacterales Enterobacteriaceae *Escherichia coli* DH5[alpha] |
| Eco-392-1 | Bacteria Proteobacteria Gammaproteobacteria Enterobacterales Enterobacteriaceae *Escherichia coli* DSM 30083 = JCM 1649 = ATCC 11775 |
| Eco-393-1 | Bacteria Proteobacteria Gammaproteobacteria Enterobacterales Enterobacteriaceae *Escherichia coli* E1140 |
| Eco-394-1 | Bacteria Proteobacteria Gammaproteobacteria Enterobacterales Enterobacteriaceae *Escherichia coli* E1728 |
| Eco-395-1 | Bacteria Proteobacteria Gammaproteobacteria Enterobacterales Enterobacteriaceae *Escherichia coli* E1777 |
| Eco-396-1 | Bacteria Proteobacteria Gammaproteobacteria Enterobacterales Enterobacteriaceae *Escherichia coli* EC096/10 |
| Eco-397-1 | Bacteria Proteobacteria Gammaproteobacteria Enterobacterales Enterobacteriaceae *Escherichia coli* ECC-1470 |
| Eco-398-1 | Bacteria Proteobacteria Gammaproteobacteria Enterobacterales Enterobacteriaceae *Escherichia coli* ER2796 |
| Eco-399-1 | Bacteria Proteobacteria Gammaproteobacteria Enterobacterales Enterobacteriaceae *Escherichia coli* FAH1 |
| Eco-400-1 | Bacteria Proteobacteria Gammaproteobacteria Enterobacterales Enterobacteriaceae *Escherichia coli* FAH2 |
| Eco-401-1 | Bacteria Proteobacteria Gammaproteobacteria Enterobacterales Enterobacteriaceae *Escherichia coli* FAP1 |
| Eco-402-1 | Bacteria Proteobacteria Gammaproteobacteria Enterobacterales Enterobacteriaceae *Escherichia coli* FAP2 |
| Eco-403-1 | Bacteria Proteobacteria Gammaproteobacteria Enterobacterales Enterobacteriaceae *Escherichia coli* FBH1 |
| Eco-404-1 | Bacteria Proteobacteria Gammaproteobacteria Enterobacterales Enterobacteriaceae *Escherichia coli* FBP1 |
| Eco-405-1 | Bacteria Proteobacteria Gammaproteobacteria Enterobacterales Enterobacteriaceae *Escherichia coli* FCH1 |
| Eco-406-1 | Bacteria Proteobacteria Gammaproteobacteria Enterobacterales Enterobacteriaceae *Escherichia coli* FCP1 |
| Eco-407-1 | Bacteria Proteobacteria Gammaproteobacteria Enterobacterales Enterobacteriaceae *Escherichia coli* G3/10 |
| Eco-408-1 | Bacteria Proteobacteria Gammaproteobacteria Enterobacterales Enterobacteriaceae *Escherichia coli* GSK2022 |
| Eco-409-1 | Bacteria Proteobacteria Gammaproteobacteria Enterobacterales Enterobacteriaceae *Escherichia coli* GSK2023 |
| Eco-410-1 | Bacteria Proteobacteria Gammaproteobacteria Enterobacterales Enterobacteriaceae *Escherichia coli* GSK2024 |
| Eco-411-1 | Bacteria Proteobacteria Gammaproteobacteria Enterobacterales Enterobacteriaceae *Escherichia coli* GSK202B |
| Eco-412-1 | Bacteria Proteobacteria Gammaproteobacteria Enterobacterales Enterobacteriaceae *Escherichia coli* GSK202U |
| Eco-413-1 | Bacteria Proteobacteria Gammaproteobacteria Enterobacterales Enterobacteriaceae *Escherichia coli* IMT8073 |
| Eco-414-1 | Bacteria Proteobacteria Gammaproteobacteria Enterobacterales Enterobacteriaceae *Escherichia coli* IS1 |
| Eco-415-1 | Bacteria Proteobacteria Gammaproteobacteria Enterobacterales Enterobacteriaceae *Escherichia coli* IS25 |
| Eco-416-1 | Bacteria Proteobacteria Gammaproteobacteria Enterobacterales Enterobacteriaceae *Escherichia coli* IS9 |
| Eco-417-1 | Bacteria Proteobacteria Gammaproteobacteria Enterobacterales Enterobacteriaceae *Escherichia coli* JT5 |
| Eco-418-1 | Bacteria Proteobacteria Gammaproteobacteria Enterobacterales Enterobacteriaceae *Escherichia coli* K-12 |
| Eco-419-1 | Bacteria Proteobacteria Gammaproteobacteria Enterobacterales Enterobacteriaceae *Escherichia coli* K02 |
| Eco-420-1 | Bacteria Proteobacteria Gammaproteobacteria Enterobacterales Enterobacteriaceae *Escherichia coli* K1 |
| Eco-421-1 | Bacteria Proteobacteria Gammaproteobacteria Enterobacterales Enterobacteriaceae *Escherichia coli* K2 |
| Eco-422-1 | Bacteria Proteobacteria Gammaproteobacteria Enterobacterales Enterobacteriaceae *Escherichia coli* KLY |
| Eco-423-1 | Bacteria Proteobacteria Gammaproteobacteria Enterobacterales Enterobacteriaceae *Escherichia coli* LAU-EC2 |
| Eco-424-1 | Bacteria Proteobacteria Gammaproteobacteria Enterobacterales Enterobacteriaceae *Escherichia coli* LAU-EC3 |
| Eco-425-1 | Bacteria Proteobacteria Gammaproteobacteria Enterobacterales Enterobacteriaceae *Escherichia coli* LAU-EC4 |
| Eco-426-1 | Bacteria Proteobacteria Gammaproteobacteria Enterobacterales Enterobacteriaceae *Escherichia coli* LAU-EC5 |
| Eco-427-1 | Bacteria Proteobacteria Gammaproteobacteria Enterobacterales Enterobacteriaceae *Escherichia coli* LR09 |
| Eco-428-1 | Bacteria Proteobacteria Gammaproteobacteria Enterobacterales Enterobacteriaceae *Escherichia coli* M1 |
| Eco-429-1 | Bacteria Proteobacteria Gammaproteobacteria Enterobacterales Enterobacteriaceae *Escherichia coli* M10 |
| Eco-430-1 | Bacteria Proteobacteria Gammaproteobacteria Enterobacterales Enterobacteriaceae *Escherichia coli* M11 |
| Eco-431-1 | Bacteria Proteobacteria Gammaproteobacteria Enterobacterales Enterobacteriaceae *Escherichia coli* M12 |
| Eco-432-1 | Bacteria Proteobacteria Gammaproteobacteria Enterobacterales Enterobacteriaceae *Escherichia coli* M13 |
| Eco-433-1 | Bacteria Proteobacteria Gammaproteobacteria Enterobacterales Enterobacteriaceae *Escherichia coli* M14 |
| Eco-434-1 | Bacteria Proteobacteria Gammaproteobacteria Enterobacterales Enterobacteriaceae *Escherichia coli* M15 |
| Eco-435-1 | Bacteria Proteobacteria Gammaproteobacteria Enterobacterales Enterobacteriaceae *Escherichia coli* M16 |
| Eco-436-1 | Bacteria Proteobacteria Gammaproteobacteria Enterobacterales Enterobacteriaceae *Escherichia coli* M17 |
| Eco-437-1 | Bacteria Proteobacteria Gammaproteobacteria Enterobacterales Enterobacteriaceae *Escherichia coli* M18 |

| | |
|---|---|
| Eco-438-1 | Bacteria Proteobacteria Gammaproteobacteria Enterobacterales Enterobacteriaceae *Escherichia coli* M19 |
| Eco-439-1 | Bacteria Proteobacteria Gammaproteobacteria Enterobacterales Enterobacteriaceae *Escherichia coli* M2 |
| Eco-440-1 | Bacteria Proteobacteria Gammaproteobacteria Enterobacterales Enterobacteriaceae *Escherichia coli* M20 |
| Eco-441-1 | Bacteria Proteobacteria Gammaproteobacteria Enterobacterales Enterobacteriaceae *Escherichia coli* M21 |
| Eco-442-1 | Bacteria Proteobacteria Gammaproteobacteria Enterobacterales Enterobacteriaceae *Escherichia coli* M22 |
| Eco-443-1 | Bacteria Proteobacteria Gammaproteobacteria Enterobacterales Enterobacteriaceae *Escherichia coli* M23 |
| Eco-444-1 | Bacteria Proteobacteria Gammaproteobacteria Enterobacterales Enterobacteriaceae *Escherichia coli* M3 |
| Eco-445-1 | Bacteria Proteobacteria Gammaproteobacteria Enterobacterales Enterobacteriaceae *Escherichia coli* M4 |
| Eco-446-1 | Bacteria Proteobacteria Gammaproteobacteria Enterobacterales Enterobacteriaceae *Escherichia coli* M5 |
| Eco-447-1 | Bacteria Proteobacteria Gammaproteobacteria Enterobacterales Enterobacteriaceae *Escherichia coli* M6 |
| Eco-448-1 | Bacteria Proteobacteria Gammaproteobacteria Enterobacterales Enterobacteriaceae *Escherichia coli* M7 |
| Eco-449-1 | Bacteria Proteobacteria Gammaproteobacteria Enterobacterales Enterobacteriaceae *Escherichia coli* M8 |
| Eco-450-1 | Bacteria Proteobacteria Gammaproteobacteria Enterobacterales Enterobacteriaceae *Escherichia coli* M9 |
| Eco-451-1 | Bacteria Proteobacteria Gammaproteobacteria Enterobacterales Enterobacteriaceae *Escherichia coli* MC19 |
| Eco-452-1 | Bacteria Proteobacteria Gammaproteobacteria Enterobacterales Enterobacteriaceae *Escherichia coli* MC21 |
| Eco-453-1 | Bacteria Proteobacteria Gammaproteobacteria Enterobacterales Enterobacteriaceae *Escherichia coli* MC23 |
| Eco-454-1 | Bacteria Proteobacteria Gammaproteobacteria Enterobacterales Enterobacteriaceae *Escherichia coli* MC6002 |
| Eco-455-1 | Bacteria Proteobacteria Gammaproteobacteria Enterobacterales Enterobacteriaceae *Escherichia coli* MC6003 |
| Eco-456-1 | Bacteria Proteobacteria Gammaproteobacteria Enterobacterales Enterobacteriaceae *Escherichia coli* MGH 57 |
| Eco-457-1 | Bacteria Proteobacteria Gammaproteobacteria Enterobacterales Enterobacteriaceae *Escherichia coli* MGH 58 |
| Eco-458-1 | Bacteria Proteobacteria Gammaproteobacteria Enterobacterales Enterobacteriaceae *Escherichia coli* MP1 |
| Eco-459-1 | Bacteria Proteobacteria Gammaproteobacteria Enterobacterales Enterobacteriaceae *Escherichia coli* MRSN 10204 |
| Eco-460-1 | Bacteria Proteobacteria Gammaproteobacteria Enterobacterales Enterobacteriaceae *Escherichia coli* MS499 |
| Eco-461-1 | Bacteria Proteobacteria Gammaproteobacteria Enterobacterales Enterobacteriaceae *Escherichia coli* NB8 |
| Eco-462-1 | Bacteria Proteobacteria Gammaproteobacteria Enterobacterales Enterobacteriaceae *Escherichia coli* NCTC 50110 |
| Eco-463-1 to Eco-463-2 | Bacteria Proteobacteria Gammaproteobacteria Enterobacterales Enterobacteriaceae *Escherichia coli* Nissle 1917 |
| Eco-464-1 | Bacteria Proteobacteria Gammaproteobacteria Enterobacterales Enterobacteriaceae *Escherichia coli* O103:H11 str. 04-3023 |
| Eco-465-1 | Bacteria Proteobacteria Gammaproteobacteria Enterobacterales Enterobacteriaceae *Escherichia coli* O103:H11 str. 2010C-3214 |
| Eco-466-1 | Bacteria Proteobacteria Gammaproteobacteria Enterobacterales Enterobacteriaceae *Escherichia coli* O103:H2 str. 2009C-3279 |
| Eco-467-1 | Bacteria Proteobacteria Gammaproteobacteria Enterobacterales Enterobacteriaceae *Escherichia coli* O103:H2 str. 2010C-4433 |
| Eco-468-1 | Bacteria Proteobacteria Gammaproteobacteria Enterobacterales Enterobacteriaceae *Escherichia coli* O103:H2 str. 2011C-3750 |
| Eco-469-1 | Bacteria Proteobacteria Gammaproteobacteria Enterobacterales Enterobacteriaceae *Escherichia coli* O103:H25 str. 2010C-4529 |
| Eco-470-1 | Bacteria Proteobacteria Gammaproteobacteria Enterobacterales Enterobacteriaceae *Escherichia coli* O104:H21 str. 94-3025 |
| Eco-471-1 | Bacteria Proteobacteria Gammaproteobacteria Enterobacterales Enterobacteriaceae *Escherichia coli* O104:H4 str. 2011EL-1675A |
| Eco-472-1 to Eco-472-2 | Bacteria Proteobacteria Gammaproteobacteria Enterobacterales Enterobacteriaceae *Escherichia coli* O104:H4 str. C227-11 |
| Eco-473-1 | Bacteria Proteobacteria Gammaproteobacteria Enterobacterales Enterobacteriaceae *Escherichia coli* O111:H8 str. 2009C-4126 |
| Eco-474-1 | Bacteria Proteobacteria Gammaproteobacteria Enterobacterales Enterobacteriaceae *Escherichia coli* O111:H8 str. 2009EL-2169 |
| Eco-475-1 | Bacteria Proteobacteria Gammaproteobacteria Enterobacterales Enterobacteriaceae *Escherichia coli* O111:H8 str. 2011C-3453 |
| Eco-476-1 | Bacteria Proteobacteria Gammaproteobacteria Enterobacterales Enterobacteriaceae *Escherichia coli* O111:H8 str. F6627 |
| Eco-477-1 | Bacteria Proteobacteria Gammaproteobacteria Enterobacterales Enterobacteriaceae *Escherichia coli* O111:NM str. 01-3076 |
| Eco-478-1 | Bacteria Proteobacteria Gammaproteobacteria Enterobacterales Enterobacteriaceae *Escherichia coli* O111:NM str. 03-3484 |
| Eco-479-1 | Bacteria Proteobacteria Gammaproteobacteria Enterobacterales Enterobacteriaceae *Escherichia coli* O111:NM str. 04-3211 |
| Eco-480-1 | Bacteria Proteobacteria Gammaproteobacteria Enterobacterales Enterobacteriaceae *Escherichia coli* O111:NM str. 08-4487 |
| Eco-481-1 | Bacteria Proteobacteria Gammaproteobacteria Enterobacterales Enterobacteriaceae *Escherichia coli* O111:NM str. 2009C-4006 |
| Eco-482-1 | Bacteria Proteobacteria Gammaproteobacteria Enterobacterales Enterobacteriaceae *Escherichia coli* O111:NM str. 2009C-4052 |
| Eco-483-1 | Bacteria Proteobacteria Gammaproteobacteria Enterobacterales Enterobacteriaceae *Escherichia coli* O111:NM str. 2010C-3053 |
| Eco-484-1 | Bacteria Proteobacteria Gammaproteobacteria Enterobacterales Enterobacteriaceae *Escherichia coli* O111:NM str. 2010C-3977 |
| Eco-485-1 | Bacteria Proteobacteria Gammaproteobacteria Enterobacterales Enterobacteriaceae *Escherichia coli* O111:NM str. 2010C-4086 |
| Eco-486-1 | Bacteria Proteobacteria Gammaproteobacteria Enterobacterales Enterobacteriaceae *Escherichia coli* O111:NM str. 2010C-4221 |
| Eco-487-1 | Bacteria Proteobacteria Gammaproteobacteria Enterobacterales Enterobacteriaceae *Escherichia coli* O111:NM str. 2010C-4592 |
| Eco-488-1 | Bacteria Proteobacteria Gammaproteobacteria Enterobacterales Enterobacteriaceae *Escherichia coli* O111:NM str. 2010C-4622 |
| Eco-489-1 | Bacteria Proteobacteria Gammaproteobacteria Enterobacterales Enterobacteriaceae *Escherichia coli* O111:NM str. 2010C-4715 |
| Eco-490-1 | Bacteria Proteobacteria Gammaproteobacteria Enterobacterales Enterobacteriaceae *Escherichia coli* O111:NM str. 2010C-4735 |
| Eco-491-1 | Bacteria Proteobacteria Gammaproteobacteria Enterobacterales Enterobacteriaceae *Escherichia coli* O111:NM str. 2010C-4746 |
| Eco-492-1 | Bacteria Proteobacteria Gammaproteobacteria Enterobacterales Enterobacteriaceae *Escherichia coli* O111:NM str. 2010C-4799 |
| Eco-493-1 | Bacteria Proteobacteria Gammaproteobacteria Enterobacterales Enterobacteriaceae *Escherichia coli* O111:NM str. 2010C-4818 |
| Eco-494-1 | Bacteria Proteobacteria Gammaproteobacteria Enterobacterales Enterobacteriaceae *Escherichia coli* O111:NM str. 2011C-3170 |
| Eco-495-1 | Bacteria Proteobacteria Gammaproteobacteria Enterobacterales Enterobacteriaceae *Escherichia coli* O111:NM str. 2011C-3362 |
| Eco-496-1 | Bacteria Proteobacteria Gammaproteobacteria Enterobacterales Enterobacteriaceae *Escherichia coli* O111:NM str. 2011C-3573 |
| Eco-497-1 | Bacteria Proteobacteria Gammaproteobacteria Enterobacterales Enterobacteriaceae *Escherichia coli* O111:NM str. 2011C-3632 |
| Eco-498-1 | Bacteria Proteobacteria Gammaproteobacteria Enterobacterales Enterobacteriaceae *Escherichia coli* O111:NM str. 2011C-3679 |
| Eco-499-1 | Bacteria Proteobacteria Gammaproteobacteria Enterobacterales Enterobacteriaceae *Escherichia coli* O111:NM str. K6722 |
| Eco-500-1 | Bacteria Proteobacteria Gammaproteobacteria Enterobacterales Enterobacteriaceae *Escherichia coli* O111:NM str. K6723 |
| Eco-501-1 | Bacteria Proteobacteria Gammaproteobacteria Enterobacterales Enterobacteriaceae *Escherichia coli* O111:NM str. K6728 |
| Eco-502-1 | Bacteria Proteobacteria Gammaproteobacteria Enterobacterales Enterobacteriaceae *Escherichia coli* O111:NM str. K6890 |

| | |
|---|---|
| Eco-503-1 | Bacteria Proteobacteria Gammaproteobacteria Enterobacterales Enterobacteriaceae *Escherichia coli* O111:NM str. K6895 |
| Eco-504-1 | Bacteria Proteobacteria Gammaproteobacteria Enterobacterales Enterobacteriaceae *Escherichia coli* O111:NM str. K6897 |
| Eco-505-1 | Bacteria Proteobacteria Gammaproteobacteria Enterobacterales Enterobacteriaceae *Escherichia coli* O111:NM str. K6898 |
| Eco-506-1 | Bacteria Proteobacteria Gammaproteobacteria Enterobacterales Enterobacteriaceae *Escherichia coli* O111:NM str. K6904 |
| Eco-507-1 | Bacteria Proteobacteria Gammaproteobacteria Enterobacterales Enterobacteriaceae *Escherichia coli* O111:NM str. K6908 |
| Eco-508-1 | Bacteria Proteobacteria Gammaproteobacteria Enterobacterales Enterobacteriaceae *Escherichia coli* O111:NM str. K6915 |
| Eco-509-1 | Bacteria Proteobacteria Gammaproteobacteria Enterobacterales Enterobacteriaceae *Escherichia coli* O118:H16 str. 06-3256 |
| Eco-510-1 | Bacteria Proteobacteria Gammaproteobacteria Enterobacterales Enterobacteriaceae *Escherichia coli* O118:H16 str. 06-3612 |
| Eco-511-1 | Bacteria Proteobacteria Gammaproteobacteria Enterobacterales Enterobacteriaceae *Escherichia coli* O118:H16 str. 07-4255 |
| Eco-512-1 | Bacteria Proteobacteria Gammaproteobacteria Enterobacterales Enterobacteriaceae *Escherichia coli* O118:H16 str. 08-3651 |
| Eco-513-1 | Bacteria Proteobacteria Gammaproteobacteria Enterobacterales Enterobacteriaceae *Escherichia coli* O118:H16 str. 2009C-4446 |
| Eco-514-1 | Bacteria Proteobacteria Gammaproteobacteria Enterobacterales Enterobacteriaceae *Escherichia coli* O119:H4 str. 03-3458 |
| Eco-515-1 | Bacteria Proteobacteria Gammaproteobacteria Enterobacterales Enterobacteriaceae *Escherichia coli* O121:H19 str. 03-3227 |
| Eco-516-1 | Bacteria Proteobacteria Gammaproteobacteria Enterobacterales Enterobacteriaceae *Escherichia coli* O121:H19 str. 06-3003 |
| Eco-517-1 | Bacteria Proteobacteria Gammaproteobacteria Enterobacterales Enterobacteriaceae *Escherichia coli* O121:H19 str. 06-3822 |
| Eco-518-1 | Bacteria Proteobacteria Gammaproteobacteria Enterobacterales Enterobacteriaceae *Escherichia coli* O121:H19 str. 2009C-4050 |
| Eco-519-1 | Bacteria Proteobacteria Gammaproteobacteria Enterobacterales Enterobacteriaceae *Escherichia coli* O121:H19 str. 2009C-4659 |
| Eco-520-1 | Bacteria Proteobacteria Gammaproteobacteria Enterobacterales Enterobacteriaceae *Escherichia coli* O121:H19 str. 2009C-4750 |
| Eco-521-1 | Bacteria Proteobacteria Gammaproteobacteria Enterobacterales Enterobacteriaceae *Escherichia coli* O121:H19 str. 2009EL1302 |
| Eco-522-1 | Bacteria Proteobacteria Gammaproteobacteria Enterobacterales Enterobacteriaceae *Escherichia coli* O121:H19 str. 2009EL1412 |
| Eco-523-1 | Bacteria Proteobacteria Gammaproteobacteria Enterobacterales Enterobacteriaceae *Escherichia coli* O121:H19 str. 2010C-3609 |
| Eco-524-1 | Bacteria Proteobacteria Gammaproteobacteria Enterobacterales Enterobacteriaceae *Escherichia coli* O121:H19 str. 2010C-3794 |
| Eco-525-1 | Bacteria Proteobacteria Gammaproteobacteria Enterobacterales Enterobacteriaceae *Escherichia coli* O121:H19 str. 2010C-3840 |
| Eco-526-1 | Bacteria Proteobacteria Gammaproteobacteria Enterobacterales Enterobacteriaceae *Escherichia coli* O121:H19 str. 2010C-4254 |
| Eco-527-1 | Bacteria Proteobacteria Gammaproteobacteria Enterobacterales Enterobacteriaceae *Escherichia coli* O121:H19 str. 2010C-4732 |
| Eco-528-1 | Bacteria Proteobacteria Gammaproteobacteria Enterobacterales Enterobacteriaceae *Escherichia coli* O121:H19 str. 2010C-4824 |
| Eco-529-1 | Bacteria Proteobacteria Gammaproteobacteria Enterobacterales Enterobacteriaceae *Escherichia coli* O121:H19 str. 2010C-4966 |
| Eco-530-1 | Bacteria Proteobacteria Gammaproteobacteria Enterobacterales Enterobacteriaceae *Escherichia coli* O121:H19 str. 2010C-4989 |
| Eco-531-1 | Bacteria Proteobacteria Gammaproteobacteria Enterobacterales Enterobacteriaceae *Escherichia coli* O121:H19 str. 2010EL1058 |
| Eco-532-1 | Bacteria Proteobacteria Gammaproteobacteria Enterobacterales Enterobacteriaceae *Escherichia coli* O121:H19 str. 2011C-3072 |
| Eco-533-1 | Bacteria Proteobacteria Gammaproteobacteria Enterobacterales Enterobacteriaceae *Escherichia coli* O121:H19 str. 2011C-3108 |
| Eco-534-1 | Bacteria Proteobacteria Gammaproteobacteria Enterobacterales Enterobacteriaceae *Escherichia coli* O121:H19 str. 2011C-3216 |
| Eco-535-1 | Bacteria Proteobacteria Gammaproteobacteria Enterobacterales Enterobacteriaceae *Escherichia coli* O121:H19 str. 2011C-3500 |
| Eco-536-1 | Bacteria Proteobacteria Gammaproteobacteria Enterobacterales Enterobacteriaceae *Escherichia coli* O121:H19 str. 2011C-3537 |
| Eco-537-1 | Bacteria Proteobacteria Gammaproteobacteria Enterobacterales Enterobacteriaceae *Escherichia coli* O121:H19 str. 2011C-3609 |
| Eco-538-1 | Bacteria Proteobacteria Gammaproteobacteria Enterobacterales Enterobacteriaceae *Escherichia coli* O121:H19 str. F6714 |
| Eco-539-1 | Bacteria Proteobacteria Gammaproteobacteria Enterobacterales Enterobacteriaceae *Escherichia coli* O121:H19 str. K5198 |
| Eco-540-1 | Bacteria Proteobacteria Gammaproteobacteria Enterobacterales Enterobacteriaceae *Escherichia coli* O121:H19 str. K5269 |
| Eco-541-1 | Bacteria Proteobacteria Gammaproteobacteria Enterobacterales Enterobacteriaceae *Escherichia coli* O121:H7 str. 2009C-3299 |
| Eco-542-1 | Bacteria Proteobacteria Gammaproteobacteria Enterobacterales Enterobacteriaceae *Escherichia coli* O123:H11 str. 2009C-3307 |
| Eco-543-1 | Bacteria Proteobacteria Gammaproteobacteria Enterobacterales Enterobacteriaceae *Escherichia coli* O127:H6 str. E2348/69 substr. CVDNalr |
| Eco-544-1 | Bacteria Proteobacteria Gammaproteobacteria Enterobacterales Enterobacteriaceae *Escherichia coli* O127:H6 str. E2348/69 substr. UMD753 |
| Eco-545-1 | Bacteria Proteobacteria Gammaproteobacteria Enterobacterales Enterobacteriaceae *Escherichia coli* O128:H2 str. 2011C-3317 |
| Eco-546-1 | Bacteria Proteobacteria Gammaproteobacteria Enterobacterales Enterobacteriaceae *Escherichia coli* O145:H25 str. 07-3858 |
| Eco-547-1 | Bacteria Proteobacteria Gammaproteobacteria Enterobacterales Enterobacteriaceae *Escherichia coli* O145:H28 str. 2009C-3292 |
| Eco-548-1 to Eco-548-2 | Bacteria Proteobacteria Gammaproteobacteria Enterobacterales Enterobacteriaceae *Escherichia coli* O145:H28 str. 4865/96 |
| Eco-549-1 | Bacteria Proteobacteria Gammaproteobacteria Enterobacterales Enterobacteriaceae *Escherichia coli* O145:H28 str. RM12581 |
| Eco-550-1 | Bacteria Proteobacteria Gammaproteobacteria Enterobacterales Enterobacteriaceae *Escherichia coli* O145:H28 str. RM12761 |
| Eco-551-1 | Bacteria Proteobacteria Gammaproteobacteria Enterobacterales Enterobacteriaceae *Escherichia coli* O145:H28 str. RM13514 |
| Eco-552-1 | Bacteria Proteobacteria Gammaproteobacteria Enterobacterales Enterobacteriaceae *Escherichia coli* O145:H28 str. RM13516 |
| Eco-553-1 | Bacteria Proteobacteria Gammaproteobacteria Enterobacterales Enterobacteriaceae *Escherichia coli* O145:NM str. 06-3484 |
| Eco-554-1 | Bacteria Proteobacteria Gammaproteobacteria Enterobacterales Enterobacteriaceae *Escherichia coli* O145:NM str. 08-4270 |
| Eco-555-1 to Eco-555-2 | Bacteria Proteobacteria Gammaproteobacteria Enterobacterales Enterobacteriaceae *Escherichia coli* O145:NM str. 2010C-3507 |
| Eco-556-1 | Bacteria Proteobacteria Gammaproteobacteria Enterobacterales Enterobacteriaceae *Escherichia coli* O145:NM str. 2010C-3508 |
| Eco-557-1 | Bacteria Proteobacteria Gammaproteobacteria Enterobacterales Enterobacteriaceae *Escherichia coli* O145:NM str. 2010C-3509 |
| Eco-558-1 | Bacteria Proteobacteria Gammaproteobacteria Enterobacterales Enterobacteriaceae *Escherichia coli* O145:NM str. 2010C-3510 |
| Eco-559-1 | Bacteria Proteobacteria Gammaproteobacteria Enterobacterales Enterobacteriaceae *Escherichia coli* O145:NM str. 2010C-3511 |
| Eco-560-1 | Bacteria Proteobacteria Gammaproteobacteria Enterobacterales Enterobacteriaceae *Escherichia coli* O145:NM str. 2010C-3516 |
| Eco-561-1 | Bacteria Proteobacteria Gammaproteobacteria Enterobacterales Enterobacteriaceae *Escherichia coli* O145:NM str. 2010C-3517 |
| Eco-562-1 | Bacteria Proteobacteria Gammaproteobacteria Enterobacterales Enterobacteriaceae *Escherichia coli* O145:NM str. 2010C-3518 |
| Eco-563-1 | Bacteria Proteobacteria Gammaproteobacteria Enterobacterales Enterobacteriaceae *Escherichia coli* O145:NM str. 2010C-3521 |
| Eco-564-1 | Bacteria Proteobacteria Gammaproteobacteria Enterobacterales Enterobacteriaceae *Escherichia coli* O145:NM str. 2010C-3526 |
| Eco-565-1 | Bacteria Proteobacteria Gammaproteobacteria Enterobacterales Enterobacteriaceae *Escherichia coli* O145:NM str. 2010C-4557C2 |
| Eco-566-1 | Bacteria Proteobacteria Gammaproteobacteria Enterobacterales Enterobacteriaceae *Escherichia coli* O146:H21 str. 2010C-3325 |
| Eco-567-1 | Bacteria Proteobacteria Gammaproteobacteria Enterobacterales Enterobacteriaceae *Escherichia coli* O153:H2 str. 2010C-5034 |

| | |
|---|---|
| Eco-568-1 | Bacteria Proteobacteria Gammaproteobacteria Enterobacterales Enterobacteriaceae *Escherichia coli* O156:H25 str. 2011C-3602 |
| Eco-569-1 | Bacteria Proteobacteria Gammaproteobacteria Enterobacterales Enterobacteriaceae *Escherichia coli* O157 str. NCCP15739 |
| Eco-570-1 | Bacteria Proteobacteria Gammaproteobacteria Enterobacterales Enterobacteriaceae *Escherichia coli* O157: str. 2010EL-2044 |
| Eco-571-1 | Bacteria Proteobacteria Gammaproteobacteria Enterobacterales Enterobacteriaceae *Escherichia coli* O157: str. 2010EL-2045 |
| Eco-572-1 | Bacteria Proteobacteria Gammaproteobacteria Enterobacterales Enterobacteriaceae *Escherichia coli* O157:H16 |
| Eco-573-1 | Bacteria Proteobacteria Gammaproteobacteria Enterobacterales Enterobacteriaceae *Escherichia coli* O157:H16 str. 98-3133 |
| Eco-574-1 to Eco-574-3 | Bacteria Proteobacteria Gammaproteobacteria Enterobacterales Enterobacteriaceae *Escherichia coli* O157:H7 |
| Eco-575-1 | Bacteria Proteobacteria Gammaproteobacteria Enterobacterales Enterobacteriaceae *Escherichia coli* O157:H7 str. 06-3745 |
| Eco-576-1 | Bacteria Proteobacteria Gammaproteobacteria Enterobacterales Enterobacteriaceae *Escherichia coli* O157:H7 str. 06-4039 |
| Eco-577-1 | Bacteria Proteobacteria Gammaproteobacteria Enterobacterales Enterobacteriaceae *Escherichia coli* O157:H7 str. 07-3091 |
| Eco-578-1 | Bacteria Proteobacteria Gammaproteobacteria Enterobacterales Enterobacteriaceae *Escherichia coli* O157:H7 str. 07-3391 |
| Eco-579-1 | Bacteria Proteobacteria Gammaproteobacteria Enterobacterales Enterobacteriaceae *Escherichia coli* O157:H7 str. 08-3037 |
| Eco-580-1 | Bacteria Proteobacteria Gammaproteobacteria Enterobacterales Enterobacteriaceae *Escherichia coli* O157:H7 str. 08-3527 |
| Eco-581-1 | Bacteria Proteobacteria Gammaproteobacteria Enterobacterales Enterobacteriaceae *Escherichia coli* O157:H7 str. 08-4169 |
| Eco-582-1 | Bacteria Proteobacteria Gammaproteobacteria Enterobacterales Enterobacteriaceae *Escherichia coli* O157:H7 str. 08-4529 |
| Eco-583-1 | Bacteria Proteobacteria Gammaproteobacteria Enterobacterales Enterobacteriaceae *Escherichia coli* O157:H7 str. 2009C-4258 |
| Eco-584-1 | Bacteria Proteobacteria Gammaproteobacteria Enterobacterales Enterobacteriaceae *Escherichia coli* O157:H7 str. 2009EL1449 |
| Eco-585-1 | Bacteria Proteobacteria Gammaproteobacteria Enterobacterales Enterobacteriaceae *Escherichia coli* O157:H7 str. 2009EL1705 |
| Eco-586-1 | Bacteria Proteobacteria Gammaproteobacteria Enterobacterales Enterobacteriaceae *Escherichia coli* O157:H7 str. 2009EL1913 |
| Eco-587-1 | Bacteria Proteobacteria Gammaproteobacteria Enterobacterales Enterobacteriaceae *Escherichia coli* O157:H7 str. 2009EL2109 |
| Eco-588-1 | Bacteria Proteobacteria Gammaproteobacteria Enterobacterales Enterobacteriaceae *Escherichia coli* O157:H7 str. 2010C-4979C1 |
| Eco-589-1 | Bacteria Proteobacteria Gammaproteobacteria Enterobacterales Enterobacteriaceae *Escherichia coli* O157:H7 str. 2011EL-1107 |
| Eco-590-1 | Bacteria Proteobacteria Gammaproteobacteria Enterobacterales Enterobacteriaceae *Escherichia coli* O157:H7 str. 2011EL-2090 |
| Eco-591-1 | Bacteria Proteobacteria Gammaproteobacteria Enterobacterales Enterobacteriaceae *Escherichia coli* O157:H7 str. 2011EL-2091 |
| Eco-592-1 | Bacteria Proteobacteria Gammaproteobacteria Enterobacterales Enterobacteriaceae *Escherichia coli* O157:H7 str. 2011EL-2092 |
| Eco-593-1 | Bacteria Proteobacteria Gammaproteobacteria Enterobacterales Enterobacteriaceae *Escherichia coli* O157:H7 str. 2011EL-2093 |
| Eco-594-1 | Bacteria Proteobacteria Gammaproteobacteria Enterobacterales Enterobacteriaceae *Escherichia coli* O157:H7 str. 2011EL-2094 |
| Eco-595-1 | Bacteria Proteobacteria Gammaproteobacteria Enterobacterales Enterobacteriaceae *Escherichia coli* O157:H7 str. 2011EL-2096 |
| Eco-596-1 | Bacteria Proteobacteria Gammaproteobacteria Enterobacterales Enterobacteriaceae *Escherichia coli* O157:H7 str. 2011EL-2097 |
| Eco-597-1 | Bacteria Proteobacteria Gammaproteobacteria Enterobacterales Enterobacteriaceae *Escherichia coli* O157:H7 str. 2011EL-2098 |
| Eco-598-1 | Bacteria Proteobacteria Gammaproteobacteria Enterobacterales Enterobacteriaceae *Escherichia coli* O157:H7 str. 2011EL-2099 |
| Eco-599-1 | Bacteria Proteobacteria Gammaproteobacteria Enterobacterales Enterobacteriaceae *Escherichia coli* O157:H7 str. 2011EL-2101 |
| Eco-600-1 | Bacteria Proteobacteria Gammaproteobacteria Enterobacterales Enterobacteriaceae *Escherichia coli* O157:H7 str. 2011EL-2103 |
| Eco-601-1 | Bacteria Proteobacteria Gammaproteobacteria Enterobacterales Enterobacteriaceae *Escherichia coli* O157:H7 str. 2011EL-2104 |
| Eco-602-1 | Bacteria Proteobacteria Gammaproteobacteria Enterobacterales Enterobacteriaceae *Escherichia coli* O157:H7 str. 2011EL-2105 |
| Eco-603-1 | Bacteria Proteobacteria Gammaproteobacteria Enterobacterales Enterobacteriaceae *Escherichia coli* O157:H7 str. 2011EL-2106 |
| Eco-604-1 | Bacteria Proteobacteria Gammaproteobacteria Enterobacterales Enterobacteriaceae *Escherichia coli* O157:H7 str. 2011EL-2107 |
| Eco-605-1 | Bacteria Proteobacteria Gammaproteobacteria Enterobacterales Enterobacteriaceae *Escherichia coli* O157:H7 str. 2011EL-2108 |
| Eco-606-1 | Bacteria Proteobacteria Gammaproteobacteria Enterobacterales Enterobacteriaceae *Escherichia coli* O157:H7 str. 2011EL-2109 |
| Eco-607-1 | Bacteria Proteobacteria Gammaproteobacteria Enterobacterales Enterobacteriaceae *Escherichia coli* O157:H7 str. 2011EL-2111 |
| Eco-608-1 | Bacteria Proteobacteria Gammaproteobacteria Enterobacterales Enterobacteriaceae *Escherichia coli* O157:H7 str. 2011EL-2112 |
| Eco-609-1 | Bacteria Proteobacteria Gammaproteobacteria Enterobacterales Enterobacteriaceae *Escherichia coli* O157:H7 str. 2011EL-2113 |
| Eco-610-1 | Bacteria Proteobacteria Gammaproteobacteria Enterobacterales Enterobacteriaceae *Escherichia coli* O157:H7 str. 2011EL-2114 |
| Eco-611-1 | Bacteria Proteobacteria Gammaproteobacteria Enterobacterales Enterobacteriaceae *Escherichia coli* O157:H7 str. 2011EL-2286 |
| Eco-612-1 | Bacteria Proteobacteria Gammaproteobacteria Enterobacterales Enterobacteriaceae *Escherichia coli* O157:H7 str. 2011EL-2287 |
| Eco-613-1 | Bacteria Proteobacteria Gammaproteobacteria Enterobacterales Enterobacteriaceae *Escherichia coli* O157:H7 str. 2011EL-2288 |
| Eco-614-1 | Bacteria Proteobacteria Gammaproteobacteria Enterobacterales Enterobacteriaceae *Escherichia coli* O157:H7 str. 2011EL-2289 |
| Eco-615-1 | Bacteria Proteobacteria Gammaproteobacteria Enterobacterales Enterobacteriaceae *Escherichia coli* O157:H7 str. 2011EL-2290 |
| Eco-616-1 | Bacteria Proteobacteria Gammaproteobacteria Enterobacterales Enterobacteriaceae *Escherichia coli* O157:H7 str. 2011EL-2312 |
| Eco-617-1 | Bacteria Proteobacteria Gammaproteobacteria Enterobacterales Enterobacteriaceae *Escherichia coli* O157:H7 str. 2011EL-2313 |
| Eco-618-1 | Bacteria Proteobacteria Gammaproteobacteria Enterobacterales Enterobacteriaceae *Escherichia coli* O157:H7 str. EC4127 |
| Eco-619-1 | Bacteria Proteobacteria Gammaproteobacteria Enterobacterales Enterobacteriaceae *Escherichia coli* O157:H7 str. EC4192 |
| Eco-620-1 | Bacteria Proteobacteria Gammaproteobacteria Enterobacterales Enterobacteriaceae *Escherichia coli* O157:H7 str. EC536 |
| Eco-621-1 | Bacteria Proteobacteria Gammaproteobacteria Enterobacterales Enterobacteriaceae *Escherichia coli* O157:H7 str. EDL933 |
| Eco-622-1 | Bacteria Proteobacteria Gammaproteobacteria Enterobacterales Enterobacteriaceae *Escherichia coli* O157:H7 str. F6142 |
| Eco-623-1 | Bacteria Proteobacteria Gammaproteobacteria Enterobacterales Enterobacteriaceae *Escherichia coli* O157:H7 str. F6749 |
| Eco-624-1 | Bacteria Proteobacteria Gammaproteobacteria Enterobacterales Enterobacteriaceae *Escherichia coli* O157:H7 str. F6750 |
| Eco-625-1 | Bacteria Proteobacteria Gammaproteobacteria Enterobacterales Enterobacteriaceae *Escherichia coli* O157:H7 str. F6751 |
| Eco-626-1 | Bacteria Proteobacteria Gammaproteobacteria Enterobacterales Enterobacteriaceae *Escherichia coli* O157:H7 str. F7350 |
| Eco-627-1 | Bacteria Proteobacteria Gammaproteobacteria Enterobacterales Enterobacteriaceae *Escherichia coli* O157:H7 str. F7377 |
| Eco-628-1 | Bacteria Proteobacteria Gammaproteobacteria Enterobacterales Enterobacteriaceae *Escherichia coli* O157:H7 str. F7384 |
| Eco-629-1 | Bacteria Proteobacteria Gammaproteobacteria Enterobacterales Enterobacteriaceae *Escherichia coli* O157:H7 str. F7410 |
| Eco-630-1 | Bacteria Proteobacteria Gammaproteobacteria Enterobacterales Enterobacteriaceae *Escherichia coli* O157:H7 str. F8092B |
| Eco-631-1 | Bacteria Proteobacteria Gammaproteobacteria Enterobacterales Enterobacteriaceae *Escherichia coli* O157:H7 str. G5303 |
| Eco-632-1 | Bacteria Proteobacteria Gammaproteobacteria Enterobacterales Enterobacteriaceae *Escherichia coli* O157:H7 str. H093800014 |

| | |
|---|---|
| Eco-633-1 | Bacteria Proteobacteria Gammaproteobacteria Enterobacterales Enterobacteriaceae *Escherichia coli* O157:H7 str. H2495 |
| Eco-634-1 | Bacteria Proteobacteria Gammaproteobacteria Enterobacterales Enterobacteriaceae *Escherichia coli* O157:H7 str. H2498 |
| Eco-635-1 | Bacteria Proteobacteria Gammaproteobacteria Enterobacterales Enterobacteriaceae *Escherichia coli* O157:H7 str. K1420 |
| Eco-636-1 | Bacteria Proteobacteria Gammaproteobacteria Enterobacterales Enterobacteriaceae *Escherichia coli* O157:H7 str. K1792 |
| Eco-637-1 | Bacteria Proteobacteria Gammaproteobacteria Enterobacterales Enterobacteriaceae *Escherichia coli* O157:H7 str. K1793 |
| Eco-638-1 | Bacteria Proteobacteria Gammaproteobacteria Enterobacterales Enterobacteriaceae *Escherichia coli* O157:H7 str. K1795 |
| Eco-639-1 | Bacteria Proteobacteria Gammaproteobacteria Enterobacterales Enterobacteriaceae *Escherichia coli* O157:H7 str. K1796 |
| Eco-640-1 | Bacteria Proteobacteria Gammaproteobacteria Enterobacterales Enterobacteriaceae *Escherichia coli* O157:H7 str. K1845 |
| Eco-641-1 | Bacteria Proteobacteria Gammaproteobacteria Enterobacterales Enterobacteriaceae *Escherichia coli* O157:H7 str. K1921 |
| Eco-642-1 | Bacteria Proteobacteria Gammaproteobacteria Enterobacterales Enterobacteriaceae *Escherichia coli* O157:H7 str. K1927 |
| Eco-643-1 | Bacteria Proteobacteria Gammaproteobacteria Enterobacterales Enterobacteriaceae *Escherichia coli* O157:H7 str. K2188 |
| Eco-644-1 | Bacteria Proteobacteria Gammaproteobacteria Enterobacterales Enterobacteriaceae *Escherichia coli* O157:H7 str. K2191 |
| Eco-645-1 | Bacteria Proteobacteria Gammaproteobacteria Enterobacterales Enterobacteriaceae *Escherichia coli* O157:H7 str. K2192 |
| Eco-646-1 | Bacteria Proteobacteria Gammaproteobacteria Enterobacterales Enterobacteriaceae *Escherichia coli* O157:H7 str. K2324 |
| Eco-647-1 | Bacteria Proteobacteria Gammaproteobacteria Enterobacterales Enterobacteriaceae *Escherichia coli* O157:H7 str. K2581 |
| Eco-648-1 | Bacteria Proteobacteria Gammaproteobacteria Enterobacterales Enterobacteriaceae *Escherichia coli* O157:H7 str. K2622 |
| Eco-649-1 | Bacteria Proteobacteria Gammaproteobacteria Enterobacterales Enterobacteriaceae *Escherichia coli* O157:H7 str. K2845 |
| Eco-650-1 | Bacteria Proteobacteria Gammaproteobacteria Enterobacterales Enterobacteriaceae *Escherichia coli* O157:H7 str. K2854 |
| Eco-651-1 | Bacteria Proteobacteria Gammaproteobacteria Enterobacterales Enterobacteriaceae *Escherichia coli* O157:H7 str. K4396 |
| Eco-652-1 | Bacteria Proteobacteria Gammaproteobacteria Enterobacterales Enterobacteriaceae *Escherichia coli* O157:H7 str. K4405 |
| Eco-653-1 | Bacteria Proteobacteria Gammaproteobacteria Enterobacterales Enterobacteriaceae *Escherichia coli* O157:H7 str. K4406 |
| Eco-654-1 | Bacteria Proteobacteria Gammaproteobacteria Enterobacterales Enterobacteriaceae *Escherichia coli* O157:H7 str. K4527 |
| Eco-655-1 | Bacteria Proteobacteria Gammaproteobacteria Enterobacterales Enterobacteriaceae *Escherichia coli* O157:H7 str. K5418 |
| Eco-656-1 | Bacteria Proteobacteria Gammaproteobacteria Enterobacterales Enterobacteriaceae *Escherichia coli* O157:H7 str. K5448 |
| Eco-657-1 | Bacteria Proteobacteria Gammaproteobacteria Enterobacterales Enterobacteriaceae *Escherichia coli* O157:H7 str. K5449 |
| Eco-658-1 | Bacteria Proteobacteria Gammaproteobacteria Enterobacterales Enterobacteriaceae *Escherichia coli* O157:H7 str. K5453 |
| Eco-659-1 | Bacteria Proteobacteria Gammaproteobacteria Enterobacterales Enterobacteriaceae *Escherichia coli* O157:H7 str. K5460 |
| Eco-660-1 | Bacteria Proteobacteria Gammaproteobacteria Enterobacterales Enterobacteriaceae *Escherichia coli* O157:H7 str. K5467 |
| Eco-661-1 | Bacteria Proteobacteria Gammaproteobacteria Enterobacterales Enterobacteriaceae *Escherichia coli* O157:H7 str. K5602 |
| Eco-662-1 | Bacteria Proteobacteria Gammaproteobacteria Enterobacterales Enterobacteriaceae *Escherichia coli* O157:H7 str. K5607 |
| Eco-663-1 | Bacteria Proteobacteria Gammaproteobacteria Enterobacterales Enterobacteriaceae *Escherichia coli* O157:H7 str. K5609 |
| Eco-664-1 | Bacteria Proteobacteria Gammaproteobacteria Enterobacterales Enterobacteriaceae *Escherichia coli* O157:H7 str. K5806 |
| Eco-665-1 | Bacteria Proteobacteria Gammaproteobacteria Enterobacterales Enterobacteriaceae *Escherichia coli* O157:H7 str. K5852 |
| Eco-666-1 | Bacteria Proteobacteria Gammaproteobacteria Enterobacterales Enterobacteriaceae *Escherichia coli* O157:H7 str. K6590 |
| Eco-667-1 | Bacteria Proteobacteria Gammaproteobacteria Enterobacterales Enterobacteriaceae *Escherichia coli* O157:H7 str. K6676 |
| Eco-668-1 | Bacteria Proteobacteria Gammaproteobacteria Enterobacterales Enterobacteriaceae *Escherichia coli* O157:H7 str. K6687 |
| Eco-669-1 | Bacteria Proteobacteria Gammaproteobacteria Enterobacterales Enterobacteriaceae *Escherichia coli* O157:H7 str. K7140 |
| Eco-670-1 | Bacteria Proteobacteria Gammaproteobacteria Enterobacterales Enterobacteriaceae *Escherichia coli* O157:H7 str. SS17 |
| Eco-671-1 | Bacteria Proteobacteria Gammaproteobacteria Enterobacterales Enterobacteriaceae *Escherichia coli* O157:H7 str. SS52 |
| Eco-672-1 | Bacteria Proteobacteria Gammaproteobacteria Enterobacterales Enterobacteriaceae *Escherichia coli* O157:H7 str. T1543_06 |
| Eco-673-1 | Bacteria Proteobacteria Gammaproteobacteria Enterobacterales Enterobacteriaceae *Escherichia coli* O157:NM str. 08-4540 |
| Eco-674-1 | Bacteria Proteobacteria Gammaproteobacteria Enterobacterales Enterobacteriaceae *Escherichia coli* O15:H18 str. K1516 |
| Eco-675-1 | Bacteria Proteobacteria Gammaproteobacteria Enterobacterales Enterobacteriaceae *Escherichia coli* O165:H25 str. 2010C-4874 |
| Eco-676-1 | Bacteria Proteobacteria Gammaproteobacteria Enterobacterales Enterobacteriaceae *Escherichia coli* O169:H41 str. F9792 |
| Eco-677-1 | Bacteria Proteobacteria Gammaproteobacteria Enterobacterales Enterobacteriaceae *Escherichia coli* O174:H21 |
| Eco-678-1 | Bacteria Proteobacteria Gammaproteobacteria Enterobacterales Enterobacteriaceae *Escherichia coli* O174:H21 str. 03-3269 |
| Eco-679-1 | Bacteria Proteobacteria Gammaproteobacteria Enterobacterales Enterobacteriaceae *Escherichia coli* O174:H8 str. 04-3038 |
| Eco-680-1 | Bacteria Proteobacteria Gammaproteobacteria Enterobacterales Enterobacteriaceae *Escherichia coli* O177:NM str. 2010C-4558 |
| Eco-681-1 | Bacteria Proteobacteria Gammaproteobacteria Enterobacterales Enterobacteriaceae *Escherichia coli* O25:NM str. E2539C1 |
| Eco-682-1 | Bacteria Proteobacteria Gammaproteobacteria Enterobacterales Enterobacteriaceae *Escherichia coli* O25b:H4-ST131 |
| Eco-683-1 | Bacteria Proteobacteria Gammaproteobacteria Enterobacterales Enterobacteriaceae *Escherichia coli* O26:H1 str. 2009C-4747 |
| Eco-684-1 | Bacteria Proteobacteria Gammaproteobacteria Enterobacterales Enterobacteriaceae *Escherichia coli* O26:H11 str. 03-3500 |
| Eco-685-1 | Bacteria Proteobacteria Gammaproteobacteria Enterobacterales Enterobacteriaceae *Escherichia coli* O26:H11 str. 05-3646 |
| Eco-686-1 | Bacteria Proteobacteria Gammaproteobacteria Enterobacterales Enterobacteriaceae *Escherichia coli* O26:H11 str. 06-3464 |
| Eco-687-1 | Bacteria Proteobacteria Gammaproteobacteria Enterobacterales Enterobacteriaceae *Escherichia coli* O26:H11 str. 2009C-3612 |
| Eco-688-1 | Bacteria Proteobacteria Gammaproteobacteria Enterobacterales Enterobacteriaceae *Escherichia coli* O26:H11 str. 2009C-3689 |
| Eco-689-1 | Bacteria Proteobacteria Gammaproteobacteria Enterobacterales Enterobacteriaceae *Escherichia coli* O26:H11 str. 2009C-3996 |
| Eco-690-1 | Bacteria Proteobacteria Gammaproteobacteria Enterobacterales Enterobacteriaceae *Escherichia coli* O26:H11 str. 2009C-4760 |
| Eco-691-1 | Bacteria Proteobacteria Gammaproteobacteria Enterobacterales Enterobacteriaceae *Escherichia coli* O26:H11 str. 2009C-4826 |
| Eco-692-1 | Bacteria Proteobacteria Gammaproteobacteria Enterobacterales Enterobacteriaceae *Escherichia coli* O26:H11 str. 2010C-3051 |
| Eco-693-1 | Bacteria Proteobacteria Gammaproteobacteria Enterobacterales Enterobacteriaceae *Escherichia coli* O26:H11 str. 2010C-3472 |
| Eco-694-1 | Bacteria Proteobacteria Gammaproteobacteria Enterobacterales Enterobacteriaceae *Escherichia coli* O26:H11 str. 2010C-3871 |
| Eco-695-1 | Bacteria Proteobacteria Gammaproteobacteria Enterobacterales Enterobacteriaceae *Escherichia coli* O26:H11 str. 2010C-3902 |
| Eco-696-1 | Bacteria Proteobacteria Gammaproteobacteria Enterobacterales Enterobacteriaceae *Escherichia coli* O26:H11 str. 2010C-4244 |
| Eco-697-1 | Bacteria Proteobacteria Gammaproteobacteria Enterobacterales Enterobacteriaceae *Escherichia coli* O26:H11 str. 2010C-4430 |

| | |
|---|---|
| Eco-698-1 | Bacteria Proteobacteria Gammaproteobacteria Enterobacterales Enterobacteriaceae *Escherichia coli* O26:H11 str. 2010C-4819 |
| Eco-699-1 | Bacteria Proteobacteria Gammaproteobacteria Enterobacterales Enterobacteriaceae *Escherichia coli* O26:H11 str. 2010C-4834 |
| Eco-700-1 | Bacteria Proteobacteria Gammaproteobacteria Enterobacterales Enterobacteriaceae *Escherichia coli* O26:H11 str. 2010C-5028 |
| Eco-701-1 | Bacteria Proteobacteria Gammaproteobacteria Enterobacterales Enterobacteriaceae *Escherichia coli* O26:H11 str. 2010EL-1699 |
| Eco-702-1 | Bacteria Proteobacteria Gammaproteobacteria Enterobacterales Enterobacteriaceae *Escherichia coli* O26:H11 str. 2011C-3270 |
| Eco-703-1 | Bacteria Proteobacteria Gammaproteobacteria Enterobacterales Enterobacteriaceae *Escherichia coli* O26:H11 str. 2011C-3282 |
| Eco-704-1 | Bacteria Proteobacteria Gammaproteobacteria Enterobacterales Enterobacteriaceae *Escherichia coli* O26:H11 str. 2011C-3387 |
| Eco-705-1 | Bacteria Proteobacteria Gammaproteobacteria Enterobacterales Enterobacteriaceae *Escherichia coli* O26:H11 str. 2011C-3506 |
| Eco-706-1 | Bacteria Proteobacteria Gammaproteobacteria Enterobacterales Enterobacteriaceae *Escherichia coli* O26:H11 str. 2011C-3655 |
| Eco-707-1 | Bacteria Proteobacteria Gammaproteobacteria Enterobacterales Enterobacteriaceae *Escherichia coli* O26:NM str. 2010C-4347 |
| Eco-708-1 | Bacteria Proteobacteria Gammaproteobacteria Enterobacterales Enterobacteriaceae *Escherichia coli* O26:NM str. 2010C-4788 |
| Eco-709-1 | Bacteria Proteobacteria Gammaproteobacteria Enterobacterales Enterobacteriaceae *Escherichia coli* O28ac:NM str. 02-3404 |
| Eco-710-1 | Bacteria Proteobacteria Gammaproteobacteria Enterobacterales Enterobacteriaceae *Escherichia coli* O39:NM str. F8704-2 |
| Eco-711-1 | Bacteria Proteobacteria Gammaproteobacteria Enterobacterales Enterobacteriaceae *Escherichia coli* O45:H2 str. 01-3147 |
| Eco-712-1 | Bacteria Proteobacteria Gammaproteobacteria Enterobacterales Enterobacteriaceae *Escherichia coli* O45:H2 str. 2009C-3686 |
| Eco-713-1 | Bacteria Proteobacteria Gammaproteobacteria Enterobacterales Enterobacteriaceae *Escherichia coli* O45:H2 str. 2009C-4780 |
| Eco-714-1 | Bacteria Proteobacteria Gammaproteobacteria Enterobacterales Enterobacteriaceae *Escherichia coli* O45:H2 str. 2010C-3876 |
| Eco-715-1 | Bacteria Proteobacteria Gammaproteobacteria Enterobacterales Enterobacteriaceae *Escherichia coli* O45:H2 str. 2010C-4211 |
| Eco-716-1 | Bacteria Proteobacteria Gammaproteobacteria Enterobacterales Enterobacteriaceae *Escherichia coli* O55:H7 str. 06-3555 |
| Eco-717-1 | Bacteria Proteobacteria Gammaproteobacteria Enterobacterales Enterobacteriaceae *Escherichia coli* O69:H11 str. 06-3325 |
| Eco-718-1 | Bacteria Proteobacteria Gammaproteobacteria Enterobacterales Enterobacteriaceae *Escherichia coli* O69:H11 str. 07-3763 |
| Eco-719-1 | Bacteria Proteobacteria Gammaproteobacteria Enterobacterales Enterobacteriaceae *Escherichia coli* O69:H11 str. 07-4281 |
| Eco-720-1 | Bacteria Proteobacteria Gammaproteobacteria Enterobacterales Enterobacteriaceae *Escherichia coli* O69:H11 str. 08-4661 |
| Eco-721-1 | Bacteria Proteobacteria Gammaproteobacteria Enterobacterales Enterobacteriaceae *Escherichia coli* O69:H11 str. 2009C-3601 |
| Eco-722-1 | Bacteria Proteobacteria Gammaproteobacteria Enterobacterales Enterobacteriaceae *Escherichia coli* O6:H16 str. 99-3165 |
| Eco-723-1 | Bacteria Proteobacteria Gammaproteobacteria Enterobacterales Enterobacteriaceae *Escherichia coli* O6:H16 str. F5656C1 |
| Eco-724-1 | Bacteria Proteobacteria Gammaproteobacteria Enterobacterales Enterobacteriaceae *Escherichia coli* O6:H16:CFA/II str. B2C |
| Eco-725-1 | Bacteria Proteobacteria Gammaproteobacteria Enterobacterales Enterobacteriaceae *Escherichia coli* O78:H12 str. 00-3279 |
| Eco-726-1 | Bacteria Proteobacteria Gammaproteobacteria Enterobacterales Enterobacteriaceae *Escherichia coli* O79:H7 str. 06-3501 |
| Eco-727-1 | Bacteria Proteobacteria Gammaproteobacteria Enterobacterales Enterobacteriaceae *Escherichia coli* O81:NM str. 02-3012 |
| Eco-728-1 | Bacteria Proteobacteria Gammaproteobacteria Enterobacterales Enterobacteriaceae *Escherichia coli* O86:H34 str. 99-3124 |
| Eco-729-1 to Eco-729-2 | Bacteria Proteobacteria Gammaproteobacteria Enterobacterales Enterobacteriaceae *Escherichia coli* O91 |
| Eco-730-1 | Bacteria Proteobacteria Gammaproteobacteria Enterobacterales Enterobacteriaceae *Escherichia coli* O91:H14 str. 06-3691 |
| Eco-731-1 | Bacteria Proteobacteria Gammaproteobacteria Enterobacterales Enterobacteriaceae *Escherichia coli* O91:H14 str. 2009C-3227 |
| Eco-732-1 | Bacteria Proteobacteria Gammaproteobacteria Enterobacterales Enterobacteriaceae *Escherichia coli* O91:H21 str. 2009C-3740 |
| Eco-733-1 | Bacteria Proteobacteria Gammaproteobacteria Enterobacterales Enterobacteriaceae *Escherichia coli* O91:H21 str. 2009C-4646 |
| Eco-734-1 | Bacteria Proteobacteria Gammaproteobacteria Enterobacterales Enterobacteriaceae *Escherichia coli* O91:NM str. 2009C-3745 |
| Eco-735-1 | Bacteria Proteobacteria Gammaproteobacteria Enterobacterales Enterobacteriaceae *Escherichia coli* PCN033 |
| Eco-736-1 | Bacteria Proteobacteria Gammaproteobacteria Enterobacterales Enterobacteriaceae *Escherichia coli* PCN061 |
| Eco-737-1 | Bacteria Proteobacteria Gammaproteobacteria Enterobacterales Enterobacteriaceae *Escherichia coli* RS218 |
| Eco-738-1 | Bacteria Proteobacteria Gammaproteobacteria Enterobacterales Enterobacteriaceae *Escherichia coli* STEC O174:H2 str. 02-04446 |
| Eco-739-1 | Bacteria Proteobacteria Gammaproteobacteria Enterobacterales Enterobacteriaceae *Escherichia coli* STEC O174:H46 str. I-151 |
| Eco-740-1 | Bacteria Proteobacteria Gammaproteobacteria Enterobacterales Enterobacteriaceae *Escherichia coli* STEC O174:H8 str. 02-07607 |
| Eco-752-1 | Bacteria Proteobacteria Gammaproteobacteria Enterobacterales Enterobacteriaceae *Escherichia coli str.* K-12 substr. MC4100 |
| Eco-753-1 to Eco-753-2 | Bacteria Proteobacteria Gammaproteobacteria Enterobacterales Enterobacteriaceae *Escherichia coli str.* K-12 substr. MG1655 |
| Eco-741-1 | Bacteria Proteobacteria Gammaproteobacteria Enterobacterales Enterobacteriaceae *Escherichia coli* UCI 51 |
| Eco-742-1 | Bacteria Proteobacteria Gammaproteobacteria Enterobacterales Enterobacteriaceae *Escherichia coli* UCI 53 |
| Eco-743-1 | Bacteria Proteobacteria Gammaproteobacteria Enterobacterales Enterobacteriaceae *Escherichia coli* UCI 57 |
| Eco-744-1 | Bacteria Proteobacteria Gammaproteobacteria Enterobacterales Enterobacteriaceae *Escherichia coli* UCI 58 |
| Eco-745-1 | Bacteria Proteobacteria Gammaproteobacteria Enterobacterales Enterobacteriaceae *Escherichia coli* UCI 65 |
| Eco-746-1 | Bacteria Proteobacteria Gammaproteobacteria Enterobacterales Enterobacteriaceae *Escherichia coli* UCI 66 |
| Eco-747-1 | Bacteria Proteobacteria Gammaproteobacteria Enterobacterales Enterobacteriaceae *Escherichia coli* VR50 |
| Eco-750-1 | Bacteria Proteobacteria Gammaproteobacteria Enterobacterales Enterobacteriaceae *Escherichia coli* Wa1 |
| Eco-751-1 | Bacteria Proteobacteria Gammaproteobacteria Enterobacterales Enterobacteriaceae *Escherichia coli* Wa2 |
| Eco-748-1 | Bacteria Proteobacteria Gammaproteobacteria Enterobacterales Enterobacteriaceae *Escherichia coli* WC1 |
| Eco-749-1 | Bacteria Proteobacteria Gammaproteobacteria Enterobacterales Enterobacteriaceae *Escherichia coli* WC2 |
| Ema-1-1 | Bacteria Proteobacteria Gammaproteobacteria Enterobacterales Enterobacteriaceae *Escherichia marmotae* |
| Esp-52-1 | Bacteria Proteobacteria Gammaproteobacteria Enterobacterales Enterobacteriaceae *Escherichia sp.* 3_2_53FAA |
| Esp-53-1 | Bacteria Proteobacteria Gammaproteobacteria Enterobacterales Enterobacteriaceae *Escherichia sp.* TW14182 |
| Esp-54-1 | Bacteria Proteobacteria Gammaproteobacteria Enterobacterales Enterobacteriaceae *Escherichia sp.* TW15838 |
| Fhe-1-1 | Bacteria Proteobacteria Gammaproteobacteria Enterobacterales Enterobacteriaceae *Franconibacter helveticus* |
| Fhe-2-1 | Bacteria Proteobacteria Gammaproteobacteria Enterobacterales Enterobacteriaceae *Franconibacter helveticus* 1159 |
| Fhe-3-1 | Bacteria Proteobacteria Gammaproteobacteria Enterobacterales Enterobacteriaceae *Franconibacter helveticus* 513 |
| Fhe-4-1 | Bacteria Proteobacteria Gammaproteobacteria Enterobacterales Enterobacteriaceae *Franconibacter helveticus* LMG 23732 |
| Fpu-1-1 | Bacteria Proteobacteria Gammaproteobacteria Enterobacterales Enterobacteriaceae *Franconibacter pulveris* |

| | |
|---|---|
| Fpu-2-1 | Bacteria Proteobacteria Gammaproteobacteria Enterobacterales Enterobacteriaceae *Franconibacter pulveris* 1160 |
| Fpu-3-1 | Bacteria Proteobacteria Gammaproteobacteria Enterobacterales Enterobacteriaceae *Franconibacter pulveris* 601 |
| Fpu-4-1 | Bacteria Proteobacteria Gammaproteobacteria Enterobacterales Enterobacteriaceae *Franconibacter pulveris* DSM 19144 |
| Fpu-5-1 | Bacteria Proteobacteria Gammaproteobacteria Enterobacterales Enterobacteriaceae *Franconibacter pulveris* LMG 24059 |
| Kae-1-1 | Bacteria Proteobacteria Gammaproteobacteria Enterobacterales Enterobacteriaceae *Klebsiella aerogenes* |
| Kae-2-1 | Bacteria Proteobacteria Gammaproteobacteria Enterobacterales Enterobacteriaceae *Klebsiella aerogenes* MGH 61 |
| Kae-3-1 | Bacteria Proteobacteria Gammaproteobacteria Enterobacterales Enterobacteriaceae *Klebsiella aerogenes* MGH 62 |
| Kae-4-1 | Bacteria Proteobacteria Gammaproteobacteria Enterobacterales Enterobacteriaceae *Klebsiella aerogenes* MGH 77 |
| Kae-5-1 | Bacteria Proteobacteria Gammaproteobacteria Enterobacterales Enterobacteriaceae *Klebsiella aerogenes* MGH 78 |
| Kae-6-1 | Bacteria Proteobacteria Gammaproteobacteria Enterobacterales Enterobacteriaceae *Klebsiella aerogenes* UCI 15 |
| Kae-7-1 | Bacteria Proteobacteria Gammaproteobacteria Enterobacterales Enterobacteriaceae *Klebsiella aerogenes* UCI 16 |
| Kae-8-1 | Bacteria Proteobacteria Gammaproteobacteria Enterobacterales Enterobacteriaceae *Klebsiella aerogenes* UCI 27 |
| Kae-9-1 | Bacteria Proteobacteria Gammaproteobacteria Enterobacterales Enterobacteriaceae *Klebsiella aerogenes* UCI 28 |
| Kae-10-1 | Bacteria Proteobacteria Gammaproteobacteria Enterobacterales Enterobacteriaceae *Klebsiella aerogenes* UCI 45 |
| Kae-11-1 | Bacteria Proteobacteria Gammaproteobacteria Enterobacterales Enterobacteriaceae *Klebsiella aerogenes* UCI 46 |
| Kae-12-1 | Bacteria Proteobacteria Gammaproteobacteria Enterobacterales Enterobacteriaceae *Klebsiella aerogenes* UCI 47 |
| Kae-13-1 | Bacteria Proteobacteria Gammaproteobacteria Enterobacterales Enterobacteriaceae *Klebsiella aerogenes* UCI 48 |
| Kcf-1-1 | Bacteria Proteobacteria Gammaproteobacteria Enterobacterales Enterobacteriaceae *Klebsiella cf. planticola* B43 |
| Kmi-1-1 to Kmi-1-2 | Bacteria Proteobacteria Gammaproteobacteria Enterobacterales Enterobacteriaceae *Klebsiella michiganensis* |
| Kmi-2-1 | Bacteria Proteobacteria Gammaproteobacteria Enterobacterales Enterobacteriaceae *Klebsiella michiganensis* H1g |
| Kmi-3-1 | Bacteria Proteobacteria Gammaproteobacteria Enterobacterales Enterobacteriaceae *Klebsiella michiganensis* HKOPL1 |
| Kox-1-1 to Kox-1-4 | Bacteria Proteobacteria Gammaproteobacteria Enterobacterales Enterobacteriaceae *Klebsiella oxytoca* |
| Kox-2-1 | Bacteria Proteobacteria Gammaproteobacteria Enterobacterales Enterobacteriaceae *Klebsiella oxytoca* 09-7231 |
| Kox-3-1 | Bacteria Proteobacteria Gammaproteobacteria Enterobacterales Enterobacteriaceae *Klebsiella oxytoca* 10-5244 |
| Kox-4-1 to Kox-4-2 | Bacteria Proteobacteria Gammaproteobacteria Enterobacterales Enterobacteriaceae *Klebsiella oxytoca* 10-5248 |
| Kox-5-1 | Bacteria Proteobacteria Gammaproteobacteria Enterobacterales Enterobacteriaceae *Klebsiella oxytoca* 10-5249 |
| Kox-6-1 | Bacteria Proteobacteria Gammaproteobacteria Enterobacterales Enterobacteriaceae *Klebsiella oxytoca* BRL6-2 |
| Kox-7-1 | Bacteria Proteobacteria Gammaproteobacteria Enterobacterales Enterobacteriaceae *Klebsiella oxytoca* G54 |
| Kox-8-1 | Bacteria Proteobacteria Gammaproteobacteria Enterobacterales Enterobacteriaceae *Klebsiella oxytoca* KA-2 |
| Kox-9-1 | Bacteria Proteobacteria Gammaproteobacteria Enterobacterales Enterobacteriaceae *Klebsiella oxytoca* KONIH1 |
| Kox-10-1 | Bacteria Proteobacteria Gammaproteobacteria Enterobacterales Enterobacteriaceae *Klebsiella oxytoca* OK-1 |
| Kpn-1-1 to Kpn-1-9 | Bacteria Proteobacteria Gammaproteobacteria Enterobacterales Enterobacteriaceae *Klebsiella pneumoniae* |
| Kpn-2-1 | Bacteria Proteobacteria Gammaproteobacteria Enterobacterales Enterobacteriaceae *Klebsiella pneumoniae* 11227-1 |
| Kpn-3-1 to Kpn-3-2 | Bacteria Proteobacteria Gammaproteobacteria Enterobacterales Enterobacteriaceae *Klebsiella pneumoniae* 160_1080 |
| Kpn-4-1 | Bacteria Proteobacteria Gammaproteobacteria Enterobacterales Enterobacteriaceae *Klebsiella pneumoniae* 30660/NJST258_1 |
| Kpn-5-1 | Bacteria Proteobacteria Gammaproteobacteria Enterobacterales Enterobacteriaceae *Klebsiella pneumoniae* 30684/NJST258_2 |
| Kpn-6-1 | Bacteria Proteobacteria Gammaproteobacteria Enterobacterales Enterobacteriaceae *Klebsiella pneumoniae* 4006 |
| Kpn-7-1 | Bacteria Proteobacteria Gammaproteobacteria Enterobacterales Enterobacteriaceae *Klebsiella pneumoniae* 4541-2 |
| Kpn-8-1 | Bacteria Proteobacteria Gammaproteobacteria Enterobacterales Enterobacteriaceae *Klebsiella pneumoniae* 500_1420 |
| Kpn-9-1 | Bacteria Proteobacteria Gammaproteobacteria Enterobacterales Enterobacteriaceae *Klebsiella pneumoniae* 7699 |
| Kpn-10-1 | Bacteria Proteobacteria Gammaproteobacteria Enterobacterales Enterobacteriaceae *Klebsiella pneumoniae* ATCC 25955 |
| Kpn-11-1 | Bacteria Proteobacteria Gammaproteobacteria Enterobacterales Enterobacteriaceae *Klebsiella pneumoniae* BIDMC 1 |
| Kpn-12-1 | Bacteria Proteobacteria Gammaproteobacteria Enterobacterales Enterobacteriaceae *Klebsiella pneumoniae* BIDMC 10 |
| Kpn-13-1 | Bacteria Proteobacteria Gammaproteobacteria Enterobacterales Enterobacteriaceae *Klebsiella pneumoniae* BIDMC 11 |
| Kpn-14-1 | Bacteria Proteobacteria Gammaproteobacteria Enterobacterales Enterobacteriaceae *Klebsiella pneumoniae* BIDMC 12A |
| Kpn-15-1 | Bacteria Proteobacteria Gammaproteobacteria Enterobacterales Enterobacteriaceae *Klebsiella pneumoniae* BIDMC 12B |
| Kpn-16-1 | Bacteria Proteobacteria Gammaproteobacteria Enterobacterales Enterobacteriaceae *Klebsiella pneumoniae* BIDMC 13 |
| Kpn-17-1 | Bacteria Proteobacteria Gammaproteobacteria Enterobacterales Enterobacteriaceae *Klebsiella pneumoniae* BIDMC 14 |
| Kpn-18-1 | Bacteria Proteobacteria Gammaproteobacteria Enterobacterales Enterobacteriaceae *Klebsiella pneumoniae* BIDMC 18A |
| Kpn-19-1 | Bacteria Proteobacteria Gammaproteobacteria Enterobacterales Enterobacteriaceae *Klebsiella pneumoniae* BIDMC 18B |
| Kpn-20-1 | Bacteria Proteobacteria Gammaproteobacteria Enterobacterales Enterobacteriaceae *Klebsiella pneumoniae* BIDMC 18D |
| Kpn-21-1 | Bacteria Proteobacteria Gammaproteobacteria Enterobacterales Enterobacteriaceae *Klebsiella pneumoniae* BIDMC 2A |
| Kpn-22-1 | Bacteria Proteobacteria Gammaproteobacteria Enterobacterales Enterobacteriaceae *Klebsiella pneumoniae* BIDMC 31 |
| Kpn-23-1 | Bacteria Proteobacteria Gammaproteobacteria Enterobacterales Enterobacteriaceae *Klebsiella pneumoniae* BIDMC 32 |
| Kpn-24-1 | Bacteria Proteobacteria Gammaproteobacteria Enterobacterales Enterobacteriaceae *Klebsiella pneumoniae* BIDMC 33B |
| Kpn-25-1 | Bacteria Proteobacteria Gammaproteobacteria Enterobacterales Enterobacteriaceae *Klebsiella pneumoniae* BIDMC 34 |
| Kpn-26-1 | Bacteria Proteobacteria Gammaproteobacteria Enterobacterales Enterobacteriaceae *Klebsiella pneumoniae* BIDMC 4 |
| Kpn-27-1 | Bacteria Proteobacteria Gammaproteobacteria Enterobacterales Enterobacteriaceae *Klebsiella pneumoniae* BIDMC 42a |
| Kpn-28-1 | Bacteria Proteobacteria Gammaproteobacteria Enterobacterales Enterobacteriaceae *Klebsiella pneumoniae* BIDMC 42b |
| Kpn-29-1 | Bacteria Proteobacteria Gammaproteobacteria Enterobacterales Enterobacteriaceae *Klebsiella pneumoniae* BIDMC 45 |
| Kpn-30-1 | Bacteria Proteobacteria Gammaproteobacteria Enterobacterales Enterobacteriaceae *Klebsiella pneumoniae* BIDMC 46a |
| Kpn-31-1 | Bacteria Proteobacteria Gammaproteobacteria Enterobacterales Enterobacteriaceae *Klebsiella pneumoniae* BIDMC 46b |
| Kpn-32-1 | Bacteria Proteobacteria Gammaproteobacteria Enterobacterales Enterobacteriaceae *Klebsiella pneumoniae* BIDMC 47 |
| Kpn-33-1 | Bacteria Proteobacteria Gammaproteobacteria Enterobacterales Enterobacteriaceae *Klebsiella pneumoniae* BIDMC 48 |
| Kpn-34-1 | Bacteria Proteobacteria Gammaproteobacteria Enterobacterales Enterobacteriaceae *Klebsiella pneumoniae* BIDMC 5 |

| | |
|---|---|
| Kpn-35-1 | Bacteria Proteobacteria Gammaproteobacteria Enterobacterales Enterobacteriaceae *Klebsiella pneumoniae* BIDMC 51 |
| Kpn-36-1 | Bacteria Proteobacteria Gammaproteobacteria Enterobacterales Enterobacteriaceae *Klebsiella pneumoniae* BIDMC 52 |
| Kpn-37-1 | Bacteria Proteobacteria Gammaproteobacteria Enterobacterales Enterobacteriaceae *Klebsiella pneumoniae* BIDMC 53 |
| Kpn-38-1 | Bacteria Proteobacteria Gammaproteobacteria Enterobacterales Enterobacteriaceae *Klebsiella pneumoniae* BIDMC 54 |
| Kpn-39-1 | Bacteria Proteobacteria Gammaproteobacteria Enterobacterales Enterobacteriaceae *Klebsiella pneumoniae* BIDMC 55 |
| Kpn-40-1 | Bacteria Proteobacteria Gammaproteobacteria Enterobacterales Enterobacteriaceae *Klebsiella pneumoniae* BIDMC 60 |
| Kpn-41-1 | Bacteria Proteobacteria Gammaproteobacteria Enterobacterales Enterobacteriaceae *Klebsiella pneumoniae* BIDMC 68 |
| Kpn-42-1 | Bacteria Proteobacteria Gammaproteobacteria Enterobacterales Enterobacteriaceae *Klebsiella pneumoniae* BIDMC 69 |
| Kpn-43-1 | Bacteria Proteobacteria Gammaproteobacteria Enterobacterales Enterobacteriaceae *Klebsiella pneumoniae* BIDMC 7A |
| Kpn-44-1 | Bacteria Proteobacteria Gammaproteobacteria Enterobacterales Enterobacteriaceae *Klebsiella pneumoniae* BIDMC 7B |
| Kpn-45-1 | Bacteria Proteobacteria Gammaproteobacteria Enterobacterales Enterobacteriaceae *Klebsiella pneumoniae* BWH 15 |
| Kpn-46-1 | Bacteria Proteobacteria Gammaproteobacteria Enterobacterales Enterobacteriaceae *Klebsiella pneumoniae* BWH 2 |
| Kpn-47-1 | Bacteria Proteobacteria Gammaproteobacteria Enterobacterales Enterobacteriaceae *Klebsiella pneumoniae* BWH 22 |
| Kpn-48-1 | Bacteria Proteobacteria Gammaproteobacteria Enterobacterales Enterobacteriaceae *Klebsiella pneumoniae* BWH 36 |
| Kpn-49-1 | Bacteria Proteobacteria Gammaproteobacteria Enterobacterales Enterobacteriaceae *Klebsiella pneumoniae* BWH 41 |
| Kpn-50-1 | Bacteria Proteobacteria Gammaproteobacteria Enterobacterales Enterobacteriaceae *Klebsiella pneumoniae* BWH 45 |
| Kpn-51-1 | Bacteria Proteobacteria Gammaproteobacteria Enterobacterales Enterobacteriaceae *Klebsiella pneumoniae* BWH 46 |
| Kpn-52-1 | Bacteria Proteobacteria Gammaproteobacteria Enterobacterales Enterobacteriaceae *Klebsiella pneumoniae* BWH 47 |
| Kpn-53-1 | Bacteria Proteobacteria Gammaproteobacteria Enterobacterales Enterobacteriaceae *Klebsiella pneumoniae* BWH 48 |
| Kpn-54-1 | Bacteria Proteobacteria Gammaproteobacteria Enterobacterales Enterobacteriaceae *Klebsiella pneumoniae* CHS 02 |
| Kpn-55-1 | Bacteria Proteobacteria Gammaproteobacteria Enterobacterales Enterobacteriaceae *Klebsiella pneumoniae* CHS 03 |
| Kpn-56-1 | Bacteria Proteobacteria Gammaproteobacteria Enterobacterales Enterobacteriaceae *Klebsiella pneumoniae* CHS 05 |
| Kpn-57-1 | Bacteria Proteobacteria Gammaproteobacteria Enterobacterales Enterobacteriaceae *Klebsiella pneumoniae* CHS 06 |
| Kpn-58-1 | Bacteria Proteobacteria Gammaproteobacteria Enterobacterales Enterobacteriaceae *Klebsiella pneumoniae* CHS 07 |
| Kpn-59-1 | Bacteria Proteobacteria Gammaproteobacteria Enterobacterales Enterobacteriaceae *Klebsiella pneumoniae* CHS 08 |
| Kpn-60-1 | Bacteria Proteobacteria Gammaproteobacteria Enterobacterales Enterobacteriaceae *Klebsiella pneumoniae* CHS 09 |
| Kpn-61-1 | Bacteria Proteobacteria Gammaproteobacteria Enterobacterales Enterobacteriaceae *Klebsiella pneumoniae* CHS 11 |
| Kpn-62-1 | Bacteria Proteobacteria Gammaproteobacteria Enterobacterales Enterobacteriaceae *Klebsiella pneumoniae* CHS 12 |
| Kpn-63-1 | Bacteria Proteobacteria Gammaproteobacteria Enterobacterales Enterobacteriaceae *Klebsiella pneumoniae* CHS 13 |
| Kpn-64-1 | Bacteria Proteobacteria Gammaproteobacteria Enterobacterales Enterobacteriaceae *Klebsiella pneumoniae* CHS 14 |
| Kpn-65-1 | Bacteria Proteobacteria Gammaproteobacteria Enterobacterales Enterobacteriaceae *Klebsiella pneumoniae* CHS 16 |
| Kpn-66-1 | Bacteria Proteobacteria Gammaproteobacteria Enterobacterales Enterobacteriaceae *Klebsiella pneumoniae* CHS 17 |
| Kpn-67-1 | Bacteria Proteobacteria Gammaproteobacteria Enterobacterales Enterobacteriaceae *Klebsiella pneumoniae* CHS 18 |
| Kpn-68-1 | Bacteria Proteobacteria Gammaproteobacteria Enterobacterales Enterobacteriaceae *Klebsiella pneumoniae* CHS 19 |
| Kpn-69-1 | Bacteria Proteobacteria Gammaproteobacteria Enterobacterales Enterobacteriaceae *Klebsiella pneumoniae* CHS 20 |
| Kpn-70-1 | Bacteria Proteobacteria Gammaproteobacteria Enterobacterales Enterobacteriaceae *Klebsiella pneumoniae* CHS 21 |
| Kpn-71-1 | Bacteria Proteobacteria Gammaproteobacteria Enterobacterales Enterobacteriaceae *Klebsiella pneumoniae* CHS 22 |
| Kpn-72-1 | Bacteria Proteobacteria Gammaproteobacteria Enterobacterales Enterobacteriaceae *Klebsiella pneumoniae* CHS 23 |
| Kpn-73-1 | Bacteria Proteobacteria Gammaproteobacteria Enterobacterales Enterobacteriaceae *Klebsiella pneumoniae* CHS 24 |
| Kpn-74-1 | Bacteria Proteobacteria Gammaproteobacteria Enterobacterales Enterobacteriaceae *Klebsiella pneumoniae* CHS 25 |
| Kpn-75-1 | Bacteria Proteobacteria Gammaproteobacteria Enterobacterales Enterobacteriaceae *Klebsiella pneumoniae* CHS 26 |
| Kpn-76-1 | Bacteria Proteobacteria Gammaproteobacteria Enterobacterales Enterobacteriaceae *Klebsiella pneumoniae* CHS 27 |
| Kpn-77-1 | Bacteria Proteobacteria Gammaproteobacteria Enterobacterales Enterobacteriaceae *Klebsiella pneumoniae* CHS 28 |
| Kpn-78-1 | Bacteria Proteobacteria Gammaproteobacteria Enterobacterales Enterobacteriaceae *Klebsiella pneumoniae* CHS 29 |
| Kpn-79-1 | Bacteria Proteobacteria Gammaproteobacteria Enterobacterales Enterobacteriaceae *Klebsiella pneumoniae* CHS 30 |
| Kpn-80-1 | Bacteria Proteobacteria Gammaproteobacteria Enterobacterales Enterobacteriaceae *Klebsiella pneumoniae* CHS 31 |
| Kpn-81-1 | Bacteria Proteobacteria Gammaproteobacteria Enterobacterales Enterobacteriaceae *Klebsiella pneumoniae* CHS 32 |
| Kpn-82-1 | Bacteria Proteobacteria Gammaproteobacteria Enterobacterales Enterobacteriaceae *Klebsiella pneumoniae* CHS 33 |
| Kpn-83-1 | Bacteria Proteobacteria Gammaproteobacteria Enterobacterales Enterobacteriaceae *Klebsiella pneumoniae* CHS 34 |
| Kpn-84-1 | Bacteria Proteobacteria Gammaproteobacteria Enterobacterales Enterobacteriaceae *Klebsiella pneumoniae* CHS 35 |
| Kpn-85-1 | Bacteria Proteobacteria Gammaproteobacteria Enterobacterales Enterobacteriaceae *Klebsiella pneumoniae* CHS 36 |
| Kpn-86-1 | Bacteria Proteobacteria Gammaproteobacteria Enterobacterales Enterobacteriaceae *Klebsiella pneumoniae* CHS 37 |
| Kpn-87-1 | Bacteria Proteobacteria Gammaproteobacteria Enterobacterales Enterobacteriaceae *Klebsiella pneumoniae* CHS 38 |
| Kpn-88-1 | Bacteria Proteobacteria Gammaproteobacteria Enterobacterales Enterobacteriaceae *Klebsiella pneumoniae* CHS 39 |
| Kpn-89-1 | Bacteria Proteobacteria Gammaproteobacteria Enterobacterales Enterobacteriaceae *Klebsiella pneumoniae* CHS 40 |
| Kpn-90-1 | Bacteria Proteobacteria Gammaproteobacteria Enterobacterales Enterobacteriaceae *Klebsiella pneumoniae* CHS 41 |
| Kpn-91-1 | Bacteria Proteobacteria Gammaproteobacteria Enterobacterales Enterobacteriaceae *Klebsiella pneumoniae* CHS 42 |
| Kpn-92-1 | Bacteria Proteobacteria Gammaproteobacteria Enterobacterales Enterobacteriaceae *Klebsiella pneumoniae* CHS 43 |
| Kpn-93-1 | Bacteria Proteobacteria Gammaproteobacteria Enterobacterales Enterobacteriaceae *Klebsiella pneumoniae* CHS 44 |
| Kpn-94-1 | Bacteria Proteobacteria Gammaproteobacteria Enterobacterales Enterobacteriaceae *Klebsiella pneumoniae* CHS 45 |
| Kpn-95-1 | Bacteria Proteobacteria Gammaproteobacteria Enterobacterales Enterobacteriaceae *Klebsiella pneumoniae* CHS 46 |
| Kpn-96-1 | Bacteria Proteobacteria Gammaproteobacteria Enterobacterales Enterobacteriaceae *Klebsiella pneumoniae* CHS 47 |
| Kpn-97-1 | Bacteria Proteobacteria Gammaproteobacteria Enterobacterales Enterobacteriaceae *Klebsiella pneumoniae* CHS 48 |
| Kpn-98-1 | Bacteria Proteobacteria Gammaproteobacteria Enterobacterales Enterobacteriaceae *Klebsiella pneumoniae* CHS 49 |
| Kpn-99-1 | Bacteria Proteobacteria Gammaproteobacteria Enterobacterales Enterobacteriaceae *Klebsiella pneumoniae* CHS 50 |

| | |
|---|---|
| Kpn-100-1 | Bacteria Proteobacteria Gammaproteobacteria Enterobacterales Enterobacteriaceae *Klebsiella pneumoniae* CHS 51 |
| Kpn-101-1 | Bacteria Proteobacteria Gammaproteobacteria Enterobacterales Enterobacteriaceae *Klebsiella pneumoniae* CHS 52 |
| Kpn-102-1 | Bacteria Proteobacteria Gammaproteobacteria Enterobacterales Enterobacteriaceae *Klebsiella pneumoniae* CHS 53 |
| Kpn-103-1 | Bacteria Proteobacteria Gammaproteobacteria Enterobacterales Enterobacteriaceae *Klebsiella pneumoniae* CHS 54 |
| Kpn-104-1 | Bacteria Proteobacteria Gammaproteobacteria Enterobacterales Enterobacteriaceae *Klebsiella pneumoniae* CHS 55 |
| Kpn-105-1 | Bacteria Proteobacteria Gammaproteobacteria Enterobacterales Enterobacteriaceae *Klebsiella pneumoniae* CHS 56 |
| Kpn-106-1 | Bacteria Proteobacteria Gammaproteobacteria Enterobacterales Enterobacteriaceae *Klebsiella pneumoniae* CHS 57 |
| Kpn-107-1 | Bacteria Proteobacteria Gammaproteobacteria Enterobacterales Enterobacteriaceae *Klebsiella pneumoniae* CHS 58 |
| Kpn-108-1 | Bacteria Proteobacteria Gammaproteobacteria Enterobacterales Enterobacteriaceae *Klebsiella pneumoniae* CHS 59 |
| Kpn-109-1 | Bacteria Proteobacteria Gammaproteobacteria Enterobacterales Enterobacteriaceae *Klebsiella pneumoniae* CHS 60 |
| Kpn-110-1 | Bacteria Proteobacteria Gammaproteobacteria Enterobacterales Enterobacteriaceae *Klebsiella pneumoniae* CHS 61 |
| Kpn-111-1 | Bacteria Proteobacteria Gammaproteobacteria Enterobacterales Enterobacteriaceae *Klebsiella pneumoniae* CHS 62 |
| Kpn-112-1 | Bacteria Proteobacteria Gammaproteobacteria Enterobacterales Enterobacteriaceae *Klebsiella pneumoniae* CHS 63 |
| Kpn-113-1 | Bacteria Proteobacteria Gammaproteobacteria Enterobacterales Enterobacteriaceae *Klebsiella pneumoniae* CHS 64 |
| Kpn-114-1 | Bacteria Proteobacteria Gammaproteobacteria Enterobacterales Enterobacteriaceae *Klebsiella pneumoniae* CHS 65 |
| Kpn-115-1 | Bacteria Proteobacteria Gammaproteobacteria Enterobacterales Enterobacteriaceae *Klebsiella pneumoniae* CHS 66 |
| Kpn-116-1 | Bacteria Proteobacteria Gammaproteobacteria Enterobacterales Enterobacteriaceae *Klebsiella pneumoniae* CHS 67 |
| Kpn-117-1 | Bacteria Proteobacteria Gammaproteobacteria Enterobacterales Enterobacteriaceae *Klebsiella pneumoniae* CHS 70 |
| Kpn-118-1 | Bacteria Proteobacteria Gammaproteobacteria Enterobacterales Enterobacteriaceae *Klebsiella pneumoniae* CHS 71 |
| Kpn-119-1 | Bacteria Proteobacteria Gammaproteobacteria Enterobacterales Enterobacteriaceae *Klebsiella pneumoniae* CHS 72 |
| Kpn-120-1 | Bacteria Proteobacteria Gammaproteobacteria Enterobacterales Enterobacteriaceae *Klebsiella pneumoniae* CHS 73 |
| Kpn-121-1 | Bacteria Proteobacteria Gammaproteobacteria Enterobacterales Enterobacteriaceae *Klebsiella pneumoniae* CHS 74 |
| Kpn-122-1 | Bacteria Proteobacteria Gammaproteobacteria Enterobacterales Enterobacteriaceae *Klebsiella pneumoniae* CHS 75 |
| Kpn-123-1 | Bacteria Proteobacteria Gammaproteobacteria Enterobacterales Enterobacteriaceae *Klebsiella pneumoniae* CHS 76 |
| Kpn-124-1 | Bacteria Proteobacteria Gammaproteobacteria Enterobacterales Enterobacteriaceae *Klebsiella pneumoniae* CHS 80 |
| Kpn-125-1 | Bacteria Proteobacteria Gammaproteobacteria Enterobacterales Enterobacteriaceae *Klebsiella pneumoniae* DMC1097 |
| Kpn-126-1 | Bacteria Proteobacteria Gammaproteobacteria Enterobacterales Enterobacteriaceae *Klebsiella pneumoniae* EGD-HP19-C |
| Kpn-127-1 | Bacteria Proteobacteria Gammaproteobacteria Enterobacterales Enterobacteriaceae *Klebsiella pneumoniae* G5-2 |
| Kpn-128-1 | Bacteria Proteobacteria Gammaproteobacteria Enterobacterales Enterobacteriaceae *Klebsiella pneumoniae* HE12 |
| Kpn-129-1 | Bacteria Proteobacteria Gammaproteobacteria Enterobacterales Enterobacteriaceae *Klebsiella pneumoniae* HK787 |
| Kpn-130-1 | Bacteria Proteobacteria Gammaproteobacteria Enterobacterales Enterobacteriaceae *Klebsiella pneumoniae* JT4 |
| Kpn-136-1 | Bacteria Proteobacteria Gammaproteobacteria Enterobacterales Enterobacteriaceae *Klebsiella pneumoniae* Kb140 |
| Kpn-137-1 | Bacteria Proteobacteria Gammaproteobacteria Enterobacterales Enterobacteriaceae *Klebsiella pneumoniae* Kb677 |
| Kpn-131-1 | Bacteria Proteobacteria Gammaproteobacteria Enterobacterales Enterobacteriaceae *Klebsiella pneumoniae* KP1-I |
| Kpn-132-1 | Bacteria Proteobacteria Gammaproteobacteria Enterobacterales Enterobacteriaceae *Klebsiella pneumoniae* KP2-R |
| Kpn-133-1 | Bacteria Proteobacteria Gammaproteobacteria Enterobacterales Enterobacteriaceae *Klebsiella pneumoniae* KP3-S |
| Kpn-134-1 | Bacteria Proteobacteria Gammaproteobacteria Enterobacterales Enterobacteriaceae *Klebsiella pneumoniae* KP4-R |
| Kpn-135-1 | Bacteria Proteobacteria Gammaproteobacteria Enterobacterales Enterobacteriaceae *Klebsiella pneumoniae* KP5-R |
| Kpn-138-1 | Bacteria Proteobacteria Gammaproteobacteria Enterobacterales Enterobacteriaceae *Klebsiella pneumoniae* LAU-KP1 |
| Kpn-139-1 | Bacteria Proteobacteria Gammaproteobacteria Enterobacterales Enterobacteriaceae *Klebsiella pneumoniae* LCT-KP182 |
| Kpn-140-1 | Bacteria Proteobacteria Gammaproteobacteria Enterobacterales Enterobacteriaceae *Klebsiella pneumoniae* LCT-KP289 |
| Kpn-141-1 | Bacteria Proteobacteria Gammaproteobacteria Enterobacterales Enterobacteriaceae *Klebsiella pneumoniae* MGH 29 |
| Kpn-142-1 | Bacteria Proteobacteria Gammaproteobacteria Enterobacterales Enterobacteriaceae *Klebsiella pneumoniae* MGH 35 |
| Kpn-143-1 | Bacteria Proteobacteria Gammaproteobacteria Enterobacterales Enterobacteriaceae *Klebsiella pneumoniae* MGH 39 |
| Kpn-144-1 | Bacteria Proteobacteria Gammaproteobacteria Enterobacterales Enterobacteriaceae *Klebsiella pneumoniae* MGH 43 |
| Kpn-145-1 | Bacteria Proteobacteria Gammaproteobacteria Enterobacterales Enterobacteriaceae *Klebsiella pneumoniae* MGH 45 |
| Kpn-146-1 | Bacteria Proteobacteria Gammaproteobacteria Enterobacterales Enterobacteriaceae *Klebsiella pneumoniae* MGH 47 |
| Kpn-147-1 | Bacteria Proteobacteria Gammaproteobacteria Enterobacterales Enterobacteriaceae *Klebsiella pneumoniae* MGH 51 |
| Kpn-148-1 | Bacteria Proteobacteria Gammaproteobacteria Enterobacterales Enterobacteriaceae *Klebsiella pneumoniae* MGH 52 |
| Kpn-149-1 | Bacteria Proteobacteria Gammaproteobacteria Enterobacterales Enterobacteriaceae *Klebsiella pneumoniae* MGH 60 |
| Kpn-150-1 | Bacteria Proteobacteria Gammaproteobacteria Enterobacterales Enterobacteriaceae *Klebsiella pneumoniae* MGH 63 |
| Kpn-151-1 | Bacteria Proteobacteria Gammaproteobacteria Enterobacterales Enterobacteriaceae *Klebsiella pneumoniae* MGH 64 |
| Kpn-152-1 | Bacteria Proteobacteria Gammaproteobacteria Enterobacterales Enterobacteriaceae *Klebsiella pneumoniae* MGH 65 |
| Kpn-153-1 | Bacteria Proteobacteria Gammaproteobacteria Enterobacterales Enterobacteriaceae *Klebsiella pneumoniae* MGH 66 |
| Kpn-154-1 | Bacteria Proteobacteria Gammaproteobacteria Enterobacterales Enterobacteriaceae *Klebsiella pneumoniae* MGH 67 |
| Kpn-155-1 | Bacteria Proteobacteria Gammaproteobacteria Enterobacterales Enterobacteriaceae *Klebsiella pneumoniae* MGH 69 |
| Kpn-156-1 | Bacteria Proteobacteria Gammaproteobacteria Enterobacterales Enterobacteriaceae *Klebsiella pneumoniae* MGH 70 |
| Kpn-157-1 | Bacteria Proteobacteria Gammaproteobacteria Enterobacterales Enterobacteriaceae *Klebsiella pneumoniae* MGH 71 |
| Kpn-158-1 | Bacteria Proteobacteria Gammaproteobacteria Enterobacterales Enterobacteriaceae *Klebsiella pneumoniae* MGH 72 |
| Kpn-159-1 | Bacteria Proteobacteria Gammaproteobacteria Enterobacterales Enterobacteriaceae *Klebsiella pneumoniae* MGH 73 |
| Kpn-160-1 | Bacteria Proteobacteria Gammaproteobacteria Enterobacterales Enterobacteriaceae *Klebsiella pneumoniae* MGH 74 |
| Kpn-161-1 | Bacteria Proteobacteria Gammaproteobacteria Enterobacterales Enterobacteriaceae *Klebsiella pneumoniae* MGH 75 |
| Kpn-162-1 | Bacteria Proteobacteria Gammaproteobacteria Enterobacterales Enterobacteriaceae *Klebsiella pneumoniae* MGH 79 |
| Kpn-163-1 | Bacteria Proteobacteria Gammaproteobacteria Enterobacterales Enterobacteriaceae *Klebsiella pneumoniae* MRSN 1319 |
| Kpn-164-1 | Bacteria Proteobacteria Gammaproteobacteria Enterobacterales Enterobacteriaceae *Klebsiella pneumoniae* MRSN 2404 |

| | |
|---|---|
| Kpn-165-1 | Bacteria Proteobacteria Gammaproteobacteria Enterobacterales Enterobacteriaceae *Klebsiella pneumoniae* MRSN 3562 |
| Kpn-166-1 | Bacteria Proteobacteria Gammaproteobacteria Enterobacterales Enterobacteriaceae *Klebsiella pneumoniae* MRSN 3852 |
| Kpn-167-1 | Bacteria Proteobacteria Gammaproteobacteria Enterobacterales Enterobacteriaceae *Klebsiella pneumoniae* MRSN 6902 |
| Kpn-168-1 | Bacteria Proteobacteria Gammaproteobacteria Enterobacterales Enterobacteriaceae *Klebsiella pneumoniae* NB60 |
| Kpn-169-1 | Bacteria Proteobacteria Gammaproteobacteria Enterobacterales Enterobacteriaceae *Klebsiella pneumoniae* PR04 |
| Kpn-205-1 to Kpn-205-4 | Bacteria Proteobacteria Gammaproteobacteria Enterobacterales Enterobacteriaceae *Klebsiella pneumoniae subsp. ozaenae* |
| Kpn-206-1 to Kpn-206-7 | Bacteria Proteobacteria Gammaproteobacteria Enterobacterales Enterobacteriaceae *Klebsiella pneumoniae subsp. pneumoniae* |
| Kpn-207-1 | Bacteria Proteobacteria Gammaproteobacteria Enterobacterales Enterobacteriaceae *Klebsiella pneumoniae subsp. pneumoniae* 1158 |
| Kpn-208-1 | Bacteria Proteobacteria Gammaproteobacteria Enterobacterales Enterobacteriaceae *Klebsiella pneumoniae subsp. pneumoniae* BJ1-GA |
| Kpn-209-1 | Bacteria Proteobacteria Gammaproteobacteria Enterobacterales Enterobacteriaceae *Klebsiella pneumoniae subsp. pneumoniae* KKBO-1 |
| Kpn-210-1 | Bacteria Proteobacteria Gammaproteobacteria Enterobacterales Enterobacteriaceae *Klebsiella pneumoniae subsp. pneumoniae* KKBO-4 |
| Kpn-218-1 | Bacteria Proteobacteria Gammaproteobacteria Enterobacterales Enterobacteriaceae *Klebsiella pneumoniae subsp. pneumoniae* Kp13 |
| Kpn-211-1 | Bacteria Proteobacteria Gammaproteobacteria Enterobacterales Enterobacteriaceae *Klebsiella pneumoniae subsp. pneumoniae* KPB-1 |
| Kpn-212-1 | Bacteria Proteobacteria Gammaproteobacteria Enterobacterales Enterobacteriaceae *Klebsiella pneumoniae subsp. pneumoniae* KPB-2 |
| Kpn-213-1 | Bacteria Proteobacteria Gammaproteobacteria Enterobacterales Enterobacteriaceae *Klebsiella pneumoniae subsp. pneumoniae* KPNIH1 |
| Kpn-214-1 | Bacteria Proteobacteria Gammaproteobacteria Enterobacterales Enterobacteriaceae *Klebsiella pneumoniae subsp. pneumoniae* KPNIH10 |
| Kpn-215-1 | Bacteria Proteobacteria Gammaproteobacteria Enterobacterales Enterobacteriaceae *Klebsiella pneumoniae subsp. pneumoniae* KPNIH24 |
| Kpn-216-1 | Bacteria Proteobacteria Gammaproteobacteria Enterobacterales Enterobacteriaceae *Klebsiella pneumoniae subsp. pneumoniae* KPNIH27 |
| Kpn-219-1 | Bacteria Proteobacteria Gammaproteobacteria Enterobacterales Enterobacteriaceae *Klebsiella pneumoniae subsp. pneumoniae* KpQ15 |
| Kpn-220-1 | Bacteria Proteobacteria Gammaproteobacteria Enterobacterales Enterobacteriaceae *Klebsiella pneumoniae subsp. pneumoniae* KpQ24 |
| Kpn-217-1 | Bacteria Proteobacteria Gammaproteobacteria Enterobacterales Enterobacteriaceae *Klebsiella pneumoniae subsp. pneumoniae* KPR0928 |
| Kpn-221-1 | Bacteria Proteobacteria Gammaproteobacteria Enterobacterales Enterobacteriaceae *Klebsiella pneumoniae subsp. pneumoniae* LCT-KP214 |
| Kpn-222-1 | Bacteria Proteobacteria Gammaproteobacteria Enterobacterales Enterobacteriaceae *Klebsiella pneumoniae subsp. pneumoniae* PittNDM01 |
| Kpn-223-1 | Bacteria Proteobacteria Gammaproteobacteria Enterobacterales Enterobacteriaceae *Klebsiella pneumoniae subsp. pneumoniae* SA1 |
| Kpn-224-1 | Bacteria Proteobacteria Gammaproteobacteria Enterobacterales Enterobacteriaceae *Klebsiella pneumoniae subsp. pneumoniae* T69 |
| Kpn-225-1 | Bacteria Proteobacteria Gammaproteobacteria Enterobacterales Enterobacteriaceae *Klebsiella pneumoniae subsp. rhinoscleromatis* SB3432 |
| Kpn-170-1 | Bacteria Proteobacteria Gammaproteobacteria Enterobacterales Enterobacteriaceae *Klebsiella pneumoniae* T2-1-1 |
| Kpn-171-1 | Bacteria Proteobacteria Gammaproteobacteria Enterobacterales Enterobacteriaceae *Klebsiella pneumoniae* T2-1-2 |
| Kpn-172-1 | Bacteria Proteobacteria Gammaproteobacteria Enterobacterales Enterobacteriaceae *Klebsiella pneumoniae* UCI 17 |
| Kpn-173-1 | Bacteria Proteobacteria Gammaproteobacteria Enterobacterales Enterobacteriaceae *Klebsiella pneumoniae* UCI 19 |
| Kpn-174-1 | Bacteria Proteobacteria Gammaproteobacteria Enterobacterales Enterobacteriaceae *Klebsiella pneumoniae* UCI 20 |
| Kpn-175-1 | Bacteria Proteobacteria Gammaproteobacteria Enterobacterales Enterobacteriaceae *Klebsiella pneumoniae* UCI 21 |
| Kpn-176-1 | Bacteria Proteobacteria Gammaproteobacteria Enterobacterales Enterobacteriaceae *Klebsiella pneumoniae* UCI 22 |
| Kpn-177-1 | Bacteria Proteobacteria Gammaproteobacteria Enterobacterales Enterobacteriaceae *Klebsiella pneumoniae* UCI 25 |
| Kpn-178-1 | Bacteria Proteobacteria Gammaproteobacteria Enterobacterales Enterobacteriaceae *Klebsiella pneumoniae* UCI 26 |
| Kpn-179-1 | Bacteria Proteobacteria Gammaproteobacteria Enterobacterales Enterobacteriaceae *Klebsiella pneumoniae* UCI 33 |
| Kpn-180-1 | Bacteria Proteobacteria Gammaproteobacteria Enterobacterales Enterobacteriaceae *Klebsiella pneumoniae* UCI 34 |
| Kpn-181-1 | Bacteria Proteobacteria Gammaproteobacteria Enterobacterales Enterobacteriaceae *Klebsiella pneumoniae* UCI 37 |
| Kpn-182-1 | Bacteria Proteobacteria Gammaproteobacteria Enterobacterales Enterobacteriaceae *Klebsiella pneumoniae* UCI 38 |
| Kpn-183-1 | Bacteria Proteobacteria Gammaproteobacteria Enterobacterales Enterobacteriaceae *Klebsiella pneumoniae* UCI 41 |
| Kpn-184-1 | Bacteria Proteobacteria Gammaproteobacteria Enterobacterales Enterobacteriaceae *Klebsiella pneumoniae* UCI 42 |
| Kpn-185-1 | Bacteria Proteobacteria Gammaproteobacteria Enterobacterales Enterobacteriaceae *Klebsiella pneumoniae* UCI 43 |
| Kpn-186-1 | Bacteria Proteobacteria Gammaproteobacteria Enterobacterales Enterobacteriaceae *Klebsiella pneumoniae* UCI 44 |
| Kpn-187-1 | Bacteria Proteobacteria Gammaproteobacteria Enterobacterales Enterobacteriaceae *Klebsiella pneumoniae* UCI 55 |
| Kpn-188-1 | Bacteria Proteobacteria Gammaproteobacteria Enterobacterales Enterobacteriaceae *Klebsiella pneumoniae* UCI 56 |
| Kpn-189-1 | Bacteria Proteobacteria Gammaproteobacteria Enterobacterales Enterobacteriaceae *Klebsiella pneumoniae* UCI 59 |
| Kpn-190-1 | Bacteria Proteobacteria Gammaproteobacteria Enterobacterales Enterobacteriaceae *Klebsiella pneumoniae* UCI 60 |
| Kpn-191-1 | Bacteria Proteobacteria Gammaproteobacteria Enterobacterales Enterobacteriaceae *Klebsiella pneumoniae* UCI 61 |
| Kpn-192-1 | Bacteria Proteobacteria Gammaproteobacteria Enterobacterales Enterobacteriaceae *Klebsiella pneumoniae* UCI 62 |
| Kpn-193-1 | Bacteria Proteobacteria Gammaproteobacteria Enterobacterales Enterobacteriaceae *Klebsiella pneumoniae* UCI 63 |
| Kpn-194-1 | Bacteria Proteobacteria Gammaproteobacteria Enterobacterales Enterobacteriaceae *Klebsiella pneumoniae* UCI 64 |
| Kpn-195-1 | Bacteria Proteobacteria Gammaproteobacteria Enterobacterales Enterobacteriaceae *Klebsiella pneumoniae* UCI 67 |
| Kpn-196-1 | Bacteria Proteobacteria Gammaproteobacteria Enterobacterales Enterobacteriaceae *Klebsiella pneumoniae* UCI 68 |
| Kpn-197-1 | Bacteria Proteobacteria Gammaproteobacteria Enterobacterales Enterobacteriaceae *Klebsiella pneumoniae* UCICRE 1 |
| Kpn-198-1 | Bacteria Proteobacteria Gammaproteobacteria Enterobacterales Enterobacteriaceae *Klebsiella pneumoniae* UCICRE 13 |
| Kpn-199-1 to Kpn-199-2 | Bacteria Proteobacteria Gammaproteobacteria Enterobacterales Enterobacteriaceae *Klebsiella pneumoniae* UHKPC 52 |
| Kpn-200-1 to Kpn-200-2 | Bacteria Proteobacteria Gammaproteobacteria Enterobacterales Enterobacteriaceae *Klebsiella pneumoniae* UHKPC05 |
| Kpn-201-1 | Bacteria Proteobacteria Gammaproteobacteria Enterobacterales Enterobacteriaceae *Klebsiella pneumoniae* UHKPC07 |
| Kpn-202-1 | Bacteria Proteobacteria Gammaproteobacteria Enterobacterales Enterobacteriaceae *Klebsiella pneumoniae* UHKPC33 |
| Kpn-203-1 to Kpn-203-2 | Bacteria Proteobacteria Gammaproteobacteria Enterobacterales Enterobacteriaceae *Klebsiella pneumoniae* UHKPC45 |
| Kpn-204-1 to Kpn-204-2 | Bacteria Proteobacteria Gammaproteobacteria Enterobacterales Enterobacteriaceae *Klebsiella pneumoniae* VAKPC278 |
| Kqu-1-1 | Bacteria Proteobacteria Gammaproteobacteria Enterobacterales Enterobacteriaceae *Klebsiella quasipneumoniae subsp. quasipneumoniae* |
| Kqu-2-1 to Kqu-2-4 | Bacteria Proteobacteria Gammaproteobacteria Enterobacterales Enterobacteriaceae *Klebsiella quasipneumoniae subsp. similipneumoniae* |
| Ksp-1-1 | Bacteria Proteobacteria Gammaproteobacteria Enterobacterales Enterobacteriaceae *Klebsiella sp.* 10982 |
| Ksp-2-1 | Bacteria Proteobacteria Gammaproteobacteria Enterobacterales Enterobacteriaceae *Klebsiella sp.* AS10 |

| | |
|---|---|
| Ksp-3-1 | Bacteria Proteobacteria Gammaproteobacteria Enterobacterales Enterobacteriaceae *Klebsiella sp.* RIT-PI-d |
| Kva-1-1 to Kva-1-7 | Bacteria Proteobacteria Gammaproteobacteria Enterobacterales Enterobacteriaceae *Klebsiella variicola* |
| Kas-1-1 | Bacteria Proteobacteria Gammaproteobacteria Enterobacterales Enterobacteriaceae *Kluyvera ascorbata* ATCC 33433 |
| Kcr-1-1 | Bacteria Proteobacteria Gammaproteobacteria Enterobacterales Enterobacteriaceae *Kluyvera cryocrescens* |
| Kor-1-1 | Bacteria Proteobacteria Gammaproteobacteria Enterobacterales Enterobacteriaceae *Kosakonia oryzae* |
| Kra-1-1 | Bacteria Proteobacteria Gammaproteobacteria Enterobacterales Enterobacteriaceae *Kosakonia radicincitans* UMEnt01/12 |
| Kra-2-1 | Bacteria Proteobacteria Gammaproteobacteria Enterobacterales Enterobacteriaceae *Kosakonia radicincitans* YD4 |
| Lad-1-1 to Lad-1-2 | Bacteria Proteobacteria Gammaproteobacteria Enterobacterales Enterobacteriaceae *Leclercia adecarboxylata* |
| Lad-2-1 | Bacteria Proteobacteria Gammaproteobacteria Enterobacterales Enterobacteriaceae *Leclercia adecarboxylata* ATCC 23216 = NBRC 102595 |
| Lsp-1-1 | Bacteria Proteobacteria Gammaproteobacteria Enterobacterales Enterobacteriaceae *Leclercia sp.* LK8 |
| Lam-1-1 | Bacteria Proteobacteria Gammaproteobacteria Enterobacterales Enterobacteriaceae *Lelliottia amnigena* CHS 78 |
| Msp-1-1 | Bacteria Proteobacteria Gammaproteobacteria Enterobacterales Enterobacteriaceae *Mangrovibacter sp.* MFB070 |
| Mma-1-1 | Bacteria Proteobacteria Gammaproteobacteria Enterobacterales Enterobacteriaceae *Metakosakonia massiliensis* JC163 |
| Pge-1-1 | Bacteria Proteobacteria Gammaproteobacteria Enterobacterales Enterobacteriaceae *Pluralibacter gergoviae* |
| Pvu-1-1 | Bacteria Proteobacteria Gammaproteobacteria Enterobacterales Enterobacteriaceae *Pseudescherichia vulneris* NBRC 102420 |
| Ror-1-1 | Bacteria Proteobacteria Gammaproteobacteria Enterobacterales Enterobacteriaceae *Raoultella ornithinolytica* |
| Ror-2-1 | Bacteria Proteobacteria Gammaproteobacteria Enterobacterales Enterobacteriaceae *Raoultella ornithinolytica* 2-156-04_S1_C1 |
| Ror-3-1 | Bacteria Proteobacteria Gammaproteobacteria Enterobacterales Enterobacteriaceae *Raoultella ornithinolytica* 2-156-04_S1_C2 |
| Ror-4-1 to Ror-4-2 | Bacteria Proteobacteria Gammaproteobacteria Enterobacterales Enterobacteriaceae *Raoultella ornithinolytica* B6 |
| Rpl-1-1 to Rpl-1-3 | Bacteria Proteobacteria Gammaproteobacteria Enterobacterales Enterobacteriaceae *Raoultella planticola* |
| Rpl-2-1 | Bacteria Proteobacteria Gammaproteobacteria Enterobacterales Enterobacteriaceae *Raoultella planticola* ATCC 33531 |
| Rte-1-1 | Bacteria Proteobacteria Gammaproteobacteria Enterobacterales Enterobacteriaceae *Raoultella terrigena* |
| Sbo-1-1 | Bacteria Proteobacteria Gammaproteobacteria Enterobacterales Enterobacteriaceae *Salmonella bongori serovar* 48:z41:– str. RKS3044 |
| Sen-2-1 to Sen-2-2 | Bacteria Proteobacteria Gammaproteobacteria Enterobacterales Enterobacteriaceae *Salmonella enterica* |
| Sen-3-1 | Bacteria Proteobacteria Gammaproteobacteria Enterobacterales Enterobacteriaceae *Salmonella enterica subsp. arizonae serovar* 62:z36:- str. RKS2983 |
| Sen-4-1 | Bacteria Proteobacteria Gammaproteobacteria Enterobacterales Enterobacteriaceae *Salmonella enterica subsp. enterica* |
| Sen-5-1 to Sen-5-6 | Bacteria Proteobacteria Gammaproteobacteria Enterobacterales Enterobacteriaceae *Salmonella enterica subsp. enterica serovar* 4,12:i:- |
| Sen-6-1 to Sen-6-5 | Bacteria Proteobacteria Gammaproteobacteria Enterobacterales Enterobacteriaceae *Salmonella enterica subsp. enterica serovar* 4,5,12:i- |
| Sen-7-1 to Sen-7-3 | Bacteria Proteobacteria Gammaproteobacteria Enterobacterales Enterobacteriaceae *Salmonella enterica subsp. enterica serovar* 4,[5],12:i:- |
| Sen-8-1 | Bacteria Proteobacteria Gammaproteobacteria Enterobacterales Enterobacteriaceae *Salmonella enterica subsp. enterica serovar* 8,(20):-:z6 |
| Sen-9-1 | Bacteria Proteobacteria Gammaproteobacteria Enterobacterales Enterobacteriaceae *Salmonella enterica subsp. enterica serovar* 9,12:l,v:- str. 94293 |
| Sen-10-1 | Bacteria Proteobacteria Gammaproteobacteria Enterobacterales Enterobacteriaceae *Salmonella enterica subsp. enterica serovar* Abaetetuba str. ATCC 35640 |
| Sen-11-1 | Bacteria Proteobacteria Gammaproteobacteria Enterobacterales Enterobacteriaceae *Salmonella enterica subsp. enterica serovar* Abony str. 0014 |
| Sen-12-1 to Sen-12-3 | Bacteria Proteobacteria Gammaproteobacteria Enterobacterales Enterobacteriaceae *Salmonella enterica subsp. enterica serovar* Agona |
| Sen-13-1 | Bacteria Proteobacteria Gammaproteobacteria Enterobacterales Enterobacteriaceae *Salmonella enterica subsp. enterica serovar* Agona str. 460004 2-1 |
| Sen-14-1 to Sen-14-4 | Bacteria Proteobacteria Gammaproteobacteria Enterobacterales Enterobacteriaceae *Salmonella enterica subsp. enterica serovar* Albany |
| Sen-15-1 to Sen-15-3 | Bacteria Proteobacteria Gammaproteobacteria Enterobacterales Enterobacteriaceae *Salmonella enterica subsp. enterica serovar* Anatum |
| Sen-16-1 | Bacteria Proteobacteria Gammaproteobacteria Enterobacterales Enterobacteriaceae *Salmonella enterica subsp. enterica serovar* Anatum str. 06-0532 |
| Sen-17-1 | Bacteria Proteobacteria Gammaproteobacteria Enterobacterales Enterobacteriaceae *Salmonella enterica subsp. enterica serovar* Anatum str. ATCC BAA-1592 |
| Sen-18-1 | Bacteria Proteobacteria Gammaproteobacteria Enterobacterales Enterobacteriaceae *Salmonella enterica subsp. enterica serovar* Aqua str. NVSL2001 |
| Sen-19-1 | Bacteria Proteobacteria Gammaproteobacteria Enterobacterales Enterobacteriaceae *Salmonella enterica subsp. enterica serovar* Bareilly str. 07-0873 |
| Sen-20-1 | Bacteria Proteobacteria Gammaproteobacteria Enterobacterales Enterobacteriaceae *Salmonella enterica subsp. enterica serovar* Bareilly str. CFSAN000178 |
| Sen-21-1 | Bacteria Proteobacteria Gammaproteobacteria Enterobacterales Enterobacteriaceae *Salmonella enterica subsp. enterica serovar* Bareilly str. CFSAN000179 |
| Sen-22-1 | Bacteria Proteobacteria Gammaproteobacteria Enterobacterales Enterobacteriaceae *Salmonella enterica subsp. enterica serovar* Bareilly str. CFSAN000180 |
| Sen-23-1 | Bacteria Proteobacteria Gammaproteobacteria Enterobacterales Enterobacteriaceae *Salmonella enterica subsp. enterica serovar* Bareilly str. CFSAN000181 |
| Sen-24-1 | Bacteria Proteobacteria Gammaproteobacteria Enterobacterales Enterobacteriaceae *Salmonella enterica subsp. enterica serovar* Bareilly str. CFSAN000182 |
| Sen-25-1 | Bacteria Proteobacteria Gammaproteobacteria Enterobacterales Enterobacteriaceae *Salmonella enterica subsp. enterica serovar* Bareilly str. CFSAN000184 |
| Sen-26-1 | Bacteria Proteobacteria Gammaproteobacteria Enterobacterales Enterobacteriaceae *Salmonella enterica subsp. enterica serovar* Bareilly str. CFSAN000185 |
| Sen-27-1 | Bacteria Proteobacteria Gammaproteobacteria Enterobacterales Enterobacteriaceae *Salmonella enterica subsp. enterica serovar* Bareilly str. CFSAN000186 |
| Sen-28-1 | Bacteria Proteobacteria Gammaproteobacteria Enterobacterales Enterobacteriaceae *Salmonella enterica subsp. enterica serovar* Bareilly str. CFSAN000187 |
| Sen-29-1 | Bacteria Proteobacteria Gammaproteobacteria Enterobacterales Enterobacteriaceae *Salmonella enterica subsp. enterica serovar* Bareilly str. CFSAN000188 |
| Sen-30-1 | Bacteria Proteobacteria Gammaproteobacteria Enterobacterales Enterobacteriaceae *Salmonella enterica subsp. enterica serovar* Bareilly str. CFSAN000190 |
| Sen-31-1 | Bacteria Proteobacteria Gammaproteobacteria Enterobacterales Enterobacteriaceae *Salmonella enterica subsp. enterica serovar* Bareilly str. CFSAN000191 |
| Sen-32-1 | Bacteria Proteobacteria Gammaproteobacteria Enterobacterales Enterobacteriaceae *Salmonella enterica subsp. enterica serovar* Bareilly str. CFSAN000192 |
| Sen-33-1 | Bacteria Proteobacteria Gammaproteobacteria Enterobacterales Enterobacteriaceae *Salmonella enterica subsp. enterica serovar* Bareilly str. CFSAN000193 |
| Sen-34-1 | Bacteria Proteobacteria Gammaproteobacteria Enterobacterales Enterobacteriaceae *Salmonella enterica subsp. enterica serovar* Bareilly str. CFSAN000194 |
| Sen-35-1 | Bacteria Proteobacteria Gammaproteobacteria Enterobacterales Enterobacteriaceae *Salmonella enterica subsp. enterica serovar* Bareilly str. CFSAN000195 |
| Sen-36-1 | Bacteria Proteobacteria Gammaproteobacteria Enterobacterales Enterobacteriaceae *Salmonella enterica subsp. enterica serovar* Bareilly str. CFSAN000196 |
| Sen-37-1 | Bacteria Proteobacteria Gammaproteobacteria Enterobacterales Enterobacteriaceae *Salmonella enterica subsp. enterica serovar* Bareilly str. CFSAN000198 |
| Sen-38-1 | Bacteria Proteobacteria Gammaproteobacteria Enterobacterales Enterobacteriaceae *Salmonella enterica subsp. enterica serovar* Bareilly str. CFSAN000199 |
| Sen-39-1 | Bacteria Proteobacteria Gammaproteobacteria Enterobacterales Enterobacteriaceae *Salmonella enterica subsp. enterica serovar* Bareilly str. CFSAN000201 |
| Sen-40-1 | Bacteria Proteobacteria Gammaproteobacteria Enterobacterales Enterobacteriaceae *Salmonella enterica subsp. enterica serovar* Bareilly str. CFSAN000202 |

| | |
|---|---|
| Sen-41-1 | Bacteria Proteobacteria Gammaproteobacteria Enterobacterales Enterobacteriaceae *Salmonella enterica subsp. enterica serovar* Bareilly str. CFSAN000203 |
| Sen-42-1 | Bacteria Proteobacteria Gammaproteobacteria Enterobacterales Enterobacteriaceae *Salmonella enterica subsp. enterica serovar* Bareilly str. CFSAN000204 |
| Sen-43-1 | Bacteria Proteobacteria Gammaproteobacteria Enterobacterales Enterobacteriaceae *Salmonella enterica subsp. enterica serovar* Bareilly str. CFSAN000205 |
| Sen-44-1 | Bacteria Proteobacteria Gammaproteobacteria Enterobacterales Enterobacteriaceae *Salmonella enterica subsp. enterica serovar* Bareilly str. CFSAN000206 |
| Sen-45-1 | Bacteria Proteobacteria Gammaproteobacteria Enterobacterales Enterobacteriaceae *Salmonella enterica subsp. enterica serovar* Bareilly str. CFSAN000207 |
| Sen-46-1 | Bacteria Proteobacteria Gammaproteobacteria Enterobacterales Enterobacteriaceae *Salmonella enterica subsp. enterica serovar* Bareilly str. CFSAN000208 |
| Sen-47-1 | Bacteria Proteobacteria Gammaproteobacteria Enterobacterales Enterobacteriaceae *Salmonella enterica subsp. enterica serovar* Bareilly str. CFSAN000209 |
| Sen-48-1 | Bacteria Proteobacteria Gammaproteobacteria Enterobacterales Enterobacteriaceae *Salmonella enterica subsp. enterica serovar* Bareilly str. CFSAN000210 |
| Sen-49-1 | Bacteria Proteobacteria Gammaproteobacteria Enterobacterales Enterobacteriaceae *Salmonella enterica subsp. enterica serovar* Bareilly str. CFSAN000211 |
| Sen-50-1 | Bacteria Proteobacteria Gammaproteobacteria Enterobacterales Enterobacteriaceae *Salmonella enterica subsp. enterica serovar* Bareilly str. CFSAN000212 |
| Sen-51-1 | Bacteria Proteobacteria Gammaproteobacteria Enterobacterales Enterobacteriaceae *Salmonella enterica subsp. enterica serovar* Bareilly str. CFSAN000213 |
| Sen-52-1 | Bacteria Proteobacteria Gammaproteobacteria Enterobacterales Enterobacteriaceae *Salmonella enterica subsp. enterica serovar* Bareilly str. CFSAN000214 |
| Sen-53-1 | Bacteria Proteobacteria Gammaproteobacteria Enterobacterales Enterobacteriaceae *Salmonella enterica subsp. enterica serovar* Bareilly str. CFSAN000215 |
| Sen-54-1 | Bacteria Proteobacteria Gammaproteobacteria Enterobacterales Enterobacteriaceae *Salmonella enterica subsp. enterica serovar* Bareilly str. CFSAN000216 |
| Sen-55-1 | Bacteria Proteobacteria Gammaproteobacteria Enterobacterales Enterobacteriaceae *Salmonella enterica subsp. enterica serovar* Bareilly str. CFSAN000217 |
| Sen-56-1 | Bacteria Proteobacteria Gammaproteobacteria Enterobacterales Enterobacteriaceae *Salmonella enterica subsp. enterica serovar* Bareilly str. CFSAN000218 |
| Sen-57-1 | Bacteria Proteobacteria Gammaproteobacteria Enterobacterales Enterobacteriaceae *Salmonella enterica subsp. enterica serovar* Bareilly str. CFSAN000219 |
| Sen-58-1 | Bacteria Proteobacteria Gammaproteobacteria Enterobacterales Enterobacteriaceae *Salmonella enterica subsp. enterica serovar* Bareilly str. CFSAN000220 |
| Sen-59-1 | Bacteria Proteobacteria Gammaproteobacteria Enterobacterales Enterobacteriaceae *Salmonella enterica subsp. enterica serovar* Bareilly str. CFSAN000221 |
| Sen-60-1 | Bacteria Proteobacteria Gammaproteobacteria Enterobacterales Enterobacteriaceae *Salmonella enterica subsp. enterica serovar* Bareilly str. CFSAN000222 |
| Sen-61-1 | Bacteria Proteobacteria Gammaproteobacteria Enterobacterales Enterobacteriaceae *Salmonella enterica subsp. enterica serovar* Bareilly str. CFSAN000223 |
| Sen-62-1 | Bacteria Proteobacteria Gammaproteobacteria Enterobacterales Enterobacteriaceae *Salmonella enterica subsp. enterica serovar* Bareilly str. CFSAN000224 |
| Sen-63-1 | Bacteria Proteobacteria Gammaproteobacteria Enterobacterales Enterobacteriaceae *Salmonella enterica subsp. enterica serovar* Bareilly str. CFSAN000225 |
| Sen-64-1 | Bacteria Proteobacteria Gammaproteobacteria Enterobacterales Enterobacteriaceae *Salmonella enterica subsp. enterica serovar* Bareilly str. CFSAN000227 |
| Sen-65-1 | Bacteria Proteobacteria Gammaproteobacteria Enterobacterales Enterobacteriaceae *Salmonella enterica subsp. enterica serovar* Bareilly str. CFSAN000228 |
| Sen-66-1 | Bacteria Proteobacteria Gammaproteobacteria Enterobacterales Enterobacteriaceae *Salmonella enterica subsp. enterica serovar* Bareilly str. CFSAN000231 |
| Sen-67-1 | Bacteria Proteobacteria Gammaproteobacteria Enterobacterales Enterobacteriaceae *Salmonella enterica subsp. enterica serovar* Bareilly str. CFSAN000235 |
| Sen-68-1 | Bacteria Proteobacteria Gammaproteobacteria Enterobacterales Enterobacteriaceae *Salmonella enterica subsp. enterica serovar* Bareilly str. CFSAN000236 |
| Sen-69-1 | Bacteria Proteobacteria Gammaproteobacteria Enterobacterales Enterobacteriaceae *Salmonella enterica subsp. enterica serovar* Bareilly str. CFSAN000661 |
| Sen-70-1 | Bacteria Proteobacteria Gammaproteobacteria Enterobacterales Enterobacteriaceae *Salmonella enterica subsp. enterica serovar* Bareilly str. CFSAN000662 |
| Sen-71-1 | Bacteria Proteobacteria Gammaproteobacteria Enterobacterales Enterobacteriaceae *Salmonella enterica subsp. enterica serovar* Bareilly str. CFSAN000669 |
| Sen-72-1 | Bacteria Proteobacteria Gammaproteobacteria Enterobacterales Enterobacteriaceae *Salmonella enterica subsp. enterica serovar* Bareilly str. CFSAN000680 |
| Sen-73-1 | Bacteria Proteobacteria Gammaproteobacteria Enterobacterales Enterobacteriaceae *Salmonella enterica subsp. enterica serovar* Bareilly str. CFSAN000700 |
| Sen-74-1 | Bacteria Proteobacteria Gammaproteobacteria Enterobacterales Enterobacteriaceae *Salmonella enterica subsp. enterica serovar* Bareilly str. CFSAN000752 |
| Sen-75-1 | Bacteria Proteobacteria Gammaproteobacteria Enterobacterales Enterobacteriaceae *Salmonella enterica subsp. enterica serovar* Bareilly str. CFSAN000753 |
| Sen-76-1 | Bacteria Proteobacteria Gammaproteobacteria Enterobacterales Enterobacteriaceae *Salmonella enterica subsp. enterica serovar* Bareilly str. CFSAN000754 |
| Sen-77-1 | Bacteria Proteobacteria Gammaproteobacteria Enterobacterales Enterobacteriaceae *Salmonella enterica subsp. enterica serovar* Bareilly str. CFSAN000755 |
| Sen-78-1 | Bacteria Proteobacteria Gammaproteobacteria Enterobacterales Enterobacteriaceae *Salmonella enterica subsp. enterica serovar* Bareilly str. CFSAN000951 |
| Sen-79-1 | Bacteria Proteobacteria Gammaproteobacteria Enterobacterales Enterobacteriaceae *Salmonella enterica subsp. enterica serovar* Bareilly str. CFSAN000952 |
| Sen-80-1 | Bacteria Proteobacteria Gammaproteobacteria Enterobacterales Enterobacteriaceae *Salmonella enterica subsp. enterica serovar* Bareilly str. CFSAN000954 |
| Sen-81-1 | Bacteria Proteobacteria Gammaproteobacteria Enterobacterales Enterobacteriaceae *Salmonella enterica subsp. enterica serovar* Bareilly str. CFSAN000955 |
| Sen-82-1 | Bacteria Proteobacteria Gammaproteobacteria Enterobacterales Enterobacteriaceae *Salmonella enterica subsp. enterica serovar* Bareilly str. CFSAN000956 |
| Sen-83-1 | Bacteria Proteobacteria Gammaproteobacteria Enterobacterales Enterobacteriaceae *Salmonella enterica subsp. enterica serovar* Bareilly str. CFSAN000957 |
| Sen-84-1 | Bacteria Proteobacteria Gammaproteobacteria Enterobacterales Enterobacteriaceae *Salmonella enterica subsp. enterica serovar* Bareilly str. CFSAN000958 |
| Sen-85-1 | Bacteria Proteobacteria Gammaproteobacteria Enterobacterales Enterobacteriaceae *Salmonella enterica subsp. enterica serovar* Bareilly str. CFSAN000959 |
| Sen-86-1 | Bacteria Proteobacteria Gammaproteobacteria Enterobacterales Enterobacteriaceae *Salmonella enterica subsp. enterica serovar* Bareilly str. CFSAN000960 |
| Sen-87-1 | Bacteria Proteobacteria Gammaproteobacteria Enterobacterales Enterobacteriaceae *Salmonella enterica subsp. enterica serovar* Bareilly str. CFSAN000961 |
| Sen-88-1 | Bacteria Proteobacteria Gammaproteobacteria Enterobacterales Enterobacteriaceae *Salmonella enterica subsp. enterica serovar* Bareilly str. CFSAN000962 |
| Sen-89-1 | Bacteria Proteobacteria Gammaproteobacteria Enterobacterales Enterobacteriaceae *Salmonella enterica subsp. enterica serovar* Bareilly str. CFSAN000963 |
| Sen-90-1 | Bacteria Proteobacteria Gammaproteobacteria Enterobacterales Enterobacteriaceae *Salmonella enterica subsp. enterica serovar* Bareilly str. CFSAN000964 |
| Sen-91-1 | Bacteria Proteobacteria Gammaproteobacteria Enterobacterales Enterobacteriaceae *Salmonella enterica subsp. enterica serovar* Bareilly str. CFSAN000965 |
| Sen-92-1 | Bacteria Proteobacteria Gammaproteobacteria Enterobacterales Enterobacteriaceae *Salmonella enterica subsp. enterica serovar* Bareilly str. CFSAN000966 |
| Sen-93-1 | Bacteria Proteobacteria Gammaproteobacteria Enterobacterales Enterobacteriaceae *Salmonella enterica subsp. enterica serovar* Bareilly str. CFSAN000967 |
| Sen-94-1 | Bacteria Proteobacteria Gammaproteobacteria Enterobacterales Enterobacteriaceae *Salmonella enterica subsp. enterica serovar* Bareilly str. CFSAN000968 |
| Sen-95-1 | Bacteria Proteobacteria Gammaproteobacteria Enterobacterales Enterobacteriaceae *Salmonella enterica subsp. enterica serovar* Bareilly str. CFSAN000969 |
| Sen-96-1 | Bacteria Proteobacteria Gammaproteobacteria Enterobacterales Enterobacteriaceae *Salmonella enterica subsp. enterica serovar* Bareilly str. CFSAN000970 |
| Sen-97-1 | Bacteria Proteobacteria Gammaproteobacteria Enterobacterales Enterobacteriaceae *Salmonella enterica subsp. enterica serovar* Bareilly str. CFSAN001087 |
| Sen-98-1 | Bacteria Proteobacteria Gammaproteobacteria Enterobacterales Enterobacteriaceae *Salmonella enterica subsp. enterica serovar* Bareilly str. CFSAN001088 |
| Sen-99-1 | Bacteria Proteobacteria Gammaproteobacteria Enterobacterales Enterobacteriaceae *Salmonella enterica subsp. enterica serovar* Bareilly str. CFSAN001089 |
| Sen-100-1 | Bacteria Proteobacteria Gammaproteobacteria Enterobacterales Enterobacteriaceae *Salmonella enterica subsp. enterica serovar* Bareilly str. CFSAN001090 |
| Sen-101-1 | Bacteria Proteobacteria Gammaproteobacteria Enterobacterales Enterobacteriaceae *Salmonella enterica subsp. enterica serovar* Bareilly str. CFSAN001099 |
| Sen-102-1 | Bacteria Proteobacteria Gammaproteobacteria Enterobacterales Enterobacteriaceae *Salmonella enterica subsp. enterica serovar* Bareilly str. CFSAN001102 |
| Sen-103-1 | Bacteria Proteobacteria Gammaproteobacteria Enterobacterales Enterobacteriaceae *Salmonella enterica subsp. enterica serovar* Bareilly str. CFSAN001105 |
| Sen-104-1 | Bacteria Proteobacteria Gammaproteobacteria Enterobacterales Enterobacteriaceae *Salmonella enterica subsp. enterica serovar* Bareilly str. CFSAN001108 |
| Sen-105-1 | Bacteria Proteobacteria Gammaproteobacteria Enterobacterales Enterobacteriaceae *Salmonella enterica subsp. enterica serovar* Bareilly str. CFSAN001109 |

| | |
|---|---|
| Sen-106-1 | Bacteria Proteobacteria Gammaproteobacteria Enterobacterales Enterobacteriaceae *Salmonella enterica subsp. enterica serovar* Bareilly str. CFSAN001112 |
| Sen-107-1 | Bacteria Proteobacteria Gammaproteobacteria Enterobacterales Enterobacteriaceae *Salmonella enterica subsp. enterica serovar* Bareilly str. CFSAN001115 |
| Sen-108-1 | Bacteria Proteobacteria Gammaproteobacteria Enterobacterales Enterobacteriaceae *Salmonella enterica subsp. enterica serovar* Bareilly str. CFSAN001118 |
| Sen-109-1 | Bacteria Proteobacteria Gammaproteobacteria Enterobacterales Enterobacteriaceae *Salmonella enterica subsp. enterica serovar* Bareilly str. CFSAN001140 |
| Sen-110-1 | Bacteria Proteobacteria Gammaproteobacteria Enterobacterales Enterobacteriaceae *Salmonella enterica subsp. enterica serovar* Bareilly str. CFSAN001591 |
| Sen-111-1 to Sen-111-3 | Bacteria Proteobacteria Gammaproteobacteria Enterobacterales Enterobacteriaceae *Salmonella enterica subsp. enterica serovar* Berta |
| Sen-112-1 to Sen-112-2 | Bacteria Proteobacteria Gammaproteobacteria Enterobacterales Enterobacteriaceae *Salmonella enterica subsp. enterica serovar* Berta str. ATCC 8392 |
| Sen-113-1 | Bacteria Proteobacteria Gammaproteobacteria Enterobacterales Enterobacteriaceae *Salmonella enterica subsp. enterica serovar* Bovismorbificans str. Sal185 |
| Sen-114-1 | Bacteria Proteobacteria Gammaproteobacteria Enterobacterales Enterobacteriaceae *Salmonella enterica subsp. enterica serovar* Bovismorbificans str. Sal609 |
| Sen-115-1 | Bacteria Proteobacteria Gammaproteobacteria Enterobacterales Enterobacteriaceae *Salmonella enterica subsp. enterica serovar* Bovismorbificans str. Sal610 |
| Sen-116-1 | Bacteria Proteobacteria Gammaproteobacteria Enterobacterales Enterobacteriaceae *Salmonella enterica subsp. enterica serovar* Bovismorbificans str. Sal615 |
| Sen-117-1 | Bacteria Proteobacteria Gammaproteobacteria Enterobacterales Enterobacteriaceae *Salmonella enterica subsp. enterica serovar* Bovismorbificans str. Sal616 |
| Sen-118-1 | Bacteria Proteobacteria Gammaproteobacteria Enterobacterales Enterobacteriaceae *Salmonella enterica subsp. enterica serovar* Bovismorbificans str. Sal617 |
| Sen-119-1 | Bacteria Proteobacteria Gammaproteobacteria Enterobacterales Enterobacteriaceae *Salmonella enterica subsp. enterica serovar* Bovismorbificans str. Sal644 |
| Sen-120-1 | Bacteria Proteobacteria Gammaproteobacteria Enterobacterales Enterobacteriaceae *Salmonella enterica subsp. enterica serovar* Bovismorbificans str. Sal676 |
| Sen-121-1 | Bacteria Proteobacteria Gammaproteobacteria Enterobacterales Enterobacteriaceae *Salmonella enterica subsp. enterica serovar* Bovismorbificans str. Sal677 |
| Sen-122-1 | Bacteria Proteobacteria Gammaproteobacteria Enterobacterales Enterobacteriaceae *Salmonella enterica subsp. enterica serovar* Bovismorbificans str. Sal678 |
| Sen-123-1 | Bacteria Proteobacteria Gammaproteobacteria Enterobacterales Enterobacteriaceae *Salmonella enterica subsp. enterica serovar* Bovismorbificans str. Sal679 |
| Sen-124-1 | Bacteria Proteobacteria Gammaproteobacteria Enterobacterales Enterobacteriaceae *Salmonella enterica subsp. enterica serovar* Bovismorbificans str. Sal680 |
| Sen-125-1 | Bacteria Proteobacteria Gammaproteobacteria Enterobacterales Enterobacteriaceae *Salmonella enterica subsp. enterica serovar* Bovismorbificans str. Sal681 |
| Sen-126-1 | Bacteria Proteobacteria Gammaproteobacteria Enterobacterales Enterobacteriaceae *Salmonella enterica subsp. enterica serovar* Bovismorbificans str. Sal682 |
| Sen-127-1 | Bacteria Proteobacteria Gammaproteobacteria Enterobacterales Enterobacteriaceae *Salmonella enterica subsp. enterica serovar* Bovismorbificans str. Sal683 |
| Sen-128-1 to Sen-128-2 | Bacteria Proteobacteria Gammaproteobacteria Enterobacterales Enterobacteriaceae *Salmonella enterica subsp. enterica serovar* Braenderup |
| Sen-129-1 to Sen-129-3 | Bacteria Proteobacteria Gammaproteobacteria Enterobacterales Enterobacteriaceae *Salmonella enterica subsp. enterica serovar* Brandenburg |
| Sen-130-1 | Bacteria Proteobacteria Gammaproteobacteria Enterobacterales Enterobacteriaceae *Salmonella enterica subsp. enterica serovar* Bredeney |
| Sen-131-1 | Bacteria Proteobacteria Gammaproteobacteria Enterobacterales Enterobacteriaceae *Salmonella enterica subsp. enterica serovar* Bredeney str. CF-SAN001080 |
| Sen-132-1 | Bacteria Proteobacteria Gammaproteobacteria Enterobacterales Enterobacteriaceae *Salmonella enterica subsp. enterica serovar* Cerro |
| Sen-133-1 | Bacteria Proteobacteria Gammaproteobacteria Enterobacterales Enterobacteriaceae *Salmonella enterica subsp. enterica serovar* Choleraesuis |
| Sen-134-1 | Bacteria Proteobacteria Gammaproteobacteria Enterobacterales Enterobacteriaceae *Salmonella enterica subsp. enterica serovar* Choleraesuis str. ATCC 10708 |
| Sen-135-1 to Sen-135-2 | Bacteria Proteobacteria Gammaproteobacteria Enterobacterales Enterobacteriaceae *Salmonella enterica subsp. enterica serovar* Cubana str. 76814 |
| Sen-136-1 to Sen-136-6 | Bacteria Proteobacteria Gammaproteobacteria Enterobacterales Enterobacteriaceae *Salmonella enterica subsp. enterica serovar* Derby |
| Sen-137-1 to Sen-137-5 | Bacteria Proteobacteria Gammaproteobacteria Enterobacterales Enterobacteriaceae *Salmonella enterica subsp. enterica serovar* Dublin |
| Sen-138-1 to Sen-138-15 | Bacteria Proteobacteria Gammaproteobacteria Enterobacterales Enterobacteriaceae *Salmonella enterica subsp. enterica serovar* Enteritidis |
| Sen-139-1 | Bacteria Proteobacteria Gammaproteobacteria Enterobacterales Enterobacteriaceae *Salmonella enterica subsp. enterica serovar* Enteritidis str. 18569 |
| Sen-140-1 | Bacteria Proteobacteria Gammaproteobacteria Enterobacterales Enterobacteriaceae *Salmonella enterica subsp. enterica serovar* Enteritidis str. 77-1427 |
| Sen-141-1 | Bacteria Proteobacteria Gammaproteobacteria Enterobacterales Enterobacteriaceae *Salmonella enterica subsp. enterica serovar* Enteritidis str. CDC_2010K_0968 |
| Sen-142-1 | Bacteria Proteobacteria Gammaproteobacteria Enterobacterales Enterobacteriaceae *Salmonella enterica subsp. enterica serovar* Enteritidis str. EC20090641 |
| Sen-143-1 | Bacteria Proteobacteria Gammaproteobacteria Enterobacterales Enterobacteriaceae *Salmonella enterica subsp. enterica serovar* Enteritidis str. EC20090698 |
| Sen-144-1 | Bacteria Proteobacteria Gammaproteobacteria Enterobacterales Enterobacteriaceae *Salmonella enterica subsp. enterica serovar* Enteritidis str. EC20100101 |
| Sen-145-1 | Bacteria Proteobacteria Gammaproteobacteria Enterobacterales Enterobacteriaceae *Salmonella enterica subsp. enterica serovar* Enteritidis str. EC20110221 |
| Sen-146-1 | Bacteria Proteobacteria Gammaproteobacteria Enterobacterales Enterobacteriaceae *Salmonella enterica subsp. enterica serovar* Enteritidis str. EC20110354 |
| Sen-147-1 | Bacteria Proteobacteria Gammaproteobacteria Enterobacterales Enterobacteriaceae *Salmonella enterica subsp. enterica serovar* Enteritidis str. EC20110355 |
| Sen-148-1 | Bacteria Proteobacteria Gammaproteobacteria Enterobacterales Enterobacteriaceae *Salmonella enterica subsp. enterica serovar* Enteritidis str. EC20110356 |
| Sen-149-1 | Bacteria Proteobacteria Gammaproteobacteria Enterobacterales Enterobacteriaceae *Salmonella enterica subsp. enterica serovar* Enteritidis str. EC20110357 |
| Sen-150-1 | Bacteria Proteobacteria Gammaproteobacteria Enterobacterales Enterobacteriaceae *Salmonella enterica subsp. enterica serovar* Enteritidis str. EC20110358 |
| Sen-151-1 | Bacteria Proteobacteria Gammaproteobacteria Enterobacterales Enterobacteriaceae *Salmonella enterica subsp. enterica serovar* Enteritidis str. EC20110359 |
| Sen-152-1 | Bacteria Proteobacteria Gammaproteobacteria Enterobacterales Enterobacteriaceae *Salmonella enterica subsp. enterica serovar* Enteritidis str. EC20110360 |
| Sen-153-1 | Bacteria Proteobacteria Gammaproteobacteria Enterobacterales Enterobacteriaceae *Salmonella enterica subsp. enterica serovar* Enteritidis str. EC20110361 |
| Sen-154-1 | Bacteria Proteobacteria Gammaproteobacteria Enterobacterales Enterobacteriaceae *Salmonella enterica subsp. enterica serovar* Enteritidis str. EC20111095 |
| Sen-155-1 | Bacteria Proteobacteria Gammaproteobacteria Enterobacterales Enterobacteriaceae *Salmonella enterica subsp. enterica serovar* Enteritidis str. EC20111174 |
| Sen-156-1 | Bacteria Proteobacteria Gammaproteobacteria Enterobacterales Enterobacteriaceae *Salmonella enterica subsp. enterica serovar* Enteritidis str. EC20111175 |
| Sen-157-1 | Bacteria Proteobacteria Gammaproteobacteria Enterobacterales Enterobacteriaceae *Salmonella enterica subsp. enterica serovar* Enteritidis str. EC20120002 |
| Sen-158-1 | Bacteria Proteobacteria Gammaproteobacteria Enterobacterales Enterobacteriaceae *Salmonella enterica subsp. enterica serovar* Enteritidis str. EC20120005 |
| Sen-159-1 | Bacteria Proteobacteria Gammaproteobacteria Enterobacterales Enterobacteriaceae *Salmonella enterica subsp. enterica serovar* Enteritidis str. EC20120008 |
| Sen-160-1 | Bacteria Proteobacteria Gammaproteobacteria Enterobacterales Enterobacteriaceae *Salmonella enterica subsp. enterica serovar* Enteritidis str. EC20120916 |
| Sen-161-1 | Bacteria Proteobacteria Gammaproteobacteria Enterobacterales Enterobacteriaceae *Salmonella enterica subsp. enterica serovar* Enteritidis str. EC20121175 |
| Sen-162-1 | Bacteria Proteobacteria Gammaproteobacteria Enterobacterales Enterobacteriaceae *Salmonella enterica subsp. enterica serovar* Enteritidis str. LA5 |
| Sen-163-1 | Bacteria Proteobacteria Gammaproteobacteria Enterobacterales Enterobacteriaceae *Salmonella enterica subsp. enterica serovar* Enteritidis str. SA19940857 |
| Sen-164-1 | Bacteria Proteobacteria Gammaproteobacteria Enterobacterales Enterobacteriaceae *Salmonella enterica subsp. enterica serovar* Gallinarum/pullorum str. FCAV198 |

| | |
|---|---|
| Sen-165-1 | Bacteria Proteobacteria Gammaproteobacteria Enterobacterales Enterobacteriaceae *Salmonella enterica subsp. enterica serovar* Give |
| Sen-166-1 to Sen-166-18 | Bacteria Proteobacteria Gammaproteobacteria Enterobacterales Enterobacteriaceae *Salmonella enterica subsp. enterica serovar* Hadar |
| Sen-167-1 | Bacteria Proteobacteria Gammaproteobacteria Enterobacterales Enterobacteriaceae *Salmonella enterica subsp. enterica serovar* Hadar str. 2012K-0272 |
| Sen-168-1 | Bacteria Proteobacteria Gammaproteobacteria Enterobacterales Enterobacteriaceae *Salmonella enterica subsp. enterica serovar* Hartford str. 2012K-0272 |
| Sen-169-1 to Sen-169-52 | Bacteria Proteobacteria Gammaproteobacteria Enterobacterales Enterobacteriaceae *Salmonella enterica subsp. enterica serovar* Heidelberg |
| Sen-170-1 | Bacteria Proteobacteria Gammaproteobacteria Enterobacterales Enterobacteriaceae *Salmonella enterica subsp. enterica serovar* Heidelberg str. 21381 |
| Sen-171-1 | Bacteria Proteobacteria Gammaproteobacteria Enterobacterales Enterobacteriaceae *Salmonella enterica subsp. enterica serovar* Heidelberg str. 29169 |
| Sen-172-1 | Bacteria Proteobacteria Gammaproteobacteria Enterobacterales Enterobacteriaceae *Salmonella enterica subsp. enterica serovar* Heidelberg str. 32507 |
| Sen-173-1 | Bacteria Proteobacteria Gammaproteobacteria Enterobacterales Enterobacteriaceae *Salmonella enterica subsp. enterica serovar* Heidelberg str. 41567 |
| Sen-174-1 | Bacteria Proteobacteria Gammaproteobacteria Enterobacterales Enterobacteriaceae *Salmonella enterica subsp. enterica serovar* Heidelberg str. 41584 |
| Sen-175-1 | Bacteria Proteobacteria Gammaproteobacteria Enterobacterales Enterobacteriaceae *Salmonella enterica subsp. enterica serovar* Heidelberg str. 579083-10 |
| Sen-176-1 | Bacteria Proteobacteria Gammaproteobacteria Enterobacterales Enterobacteriaceae *Salmonella enterica subsp. enterica serovar* Heidelberg str. 607309-34 |
| Sen-177-1 | Bacteria Proteobacteria Gammaproteobacteria Enterobacterales Enterobacteriaceae *Salmonella enterica subsp. enterica serovar* Heidelberg str. 607309-5 |
| Sen-178-1 | Bacteria Proteobacteria Gammaproteobacteria Enterobacterales Enterobacteriaceae *Salmonella enterica subsp. enterica serovar* Heidelberg str. 622737-21 |
| Sen-179-1 | Bacteria Proteobacteria Gammaproteobacteria Enterobacterales Enterobacteriaceae *Salmonella enterica subsp. enterica serovar* Heidelberg str. 632675-24 |
| Sen-180-1 | Bacteria Proteobacteria Gammaproteobacteria Enterobacterales Enterobacteriaceae *Salmonella enterica subsp. enterica serovar* Heidelberg str. 640151-11 |
| Sen-181-1 | Bacteria Proteobacteria Gammaproteobacteria Enterobacterales Enterobacteriaceae *Salmonella enterica subsp. enterica serovar* Heidelberg str. 670102-3 |
| Sen-182-1 | Bacteria Proteobacteria Gammaproteobacteria Enterobacterales Enterobacteriaceae *Salmonella enterica subsp. enterica serovar* Heidelberg str. 670102-5 |
| Sen-183-1 | Bacteria Proteobacteria Gammaproteobacteria Enterobacterales Enterobacteriaceae *Salmonella enterica subsp. enterica serovar* Heidelberg str. 82-1617 |
| Sen-184-1 | Bacteria Proteobacteria Gammaproteobacteria Enterobacterales Enterobacteriaceae *Salmonella enterica subsp. enterica serovar* Heidelberg str. 84-1004 |
| Sen-185-1 | Bacteria Proteobacteria Gammaproteobacteria Enterobacterales Enterobacteriaceae *Salmonella enterica subsp. enterica serovar* Heidelberg str. 85-0486 |
| Sen-186-1 | Bacteria Proteobacteria Gammaproteobacteria Enterobacterales Enterobacteriaceae *Salmonella enterica subsp. enterica serovar* Heidelberg str. 86-0255 |
| Sen-187-1 | Bacteria Proteobacteria Gammaproteobacteria Enterobacterales Enterobacteriaceae *Salmonella enterica subsp. enterica serovar* Heidelberg str. 87-0208 |
| Sen-188-1 | Bacteria Proteobacteria Gammaproteobacteria Enterobacterales Enterobacteriaceae *Salmonella enterica subsp. enterica serovar* Heidelberg str. 88-0312 |
| Sen-189-1 | Bacteria Proteobacteria Gammaproteobacteria Enterobacterales Enterobacteriaceae *Salmonella enterica subsp. enterica serovar* Heidelberg str. 89-0213 |
| Sen-190-1 | Bacteria Proteobacteria Gammaproteobacteria Enterobacterales Enterobacteriaceae *Salmonella enterica subsp. enterica serovar* Heidelberg str. 92-0138 |
| Sen-191-1 | Bacteria Proteobacteria Gammaproteobacteria Enterobacterales Enterobacteriaceae *Salmonella enterica subsp. enterica serovar* Heidelberg str. 92-0144 |
| Sen-192-1 | Bacteria Proteobacteria Gammaproteobacteria Enterobacterales Enterobacteriaceae *Salmonella enterica subsp. enterica serovar* Heidelberg str. CF-SAN002064 |
| Sen-193-1 | Bacteria Proteobacteria Gammaproteobacteria Enterobacterales Enterobacteriaceae *Salmonella enterica subsp. enterica serovar* Heidelberg str. CF-SAN00323 |
| Sen-194-1 | Bacteria Proteobacteria Gammaproteobacteria Enterobacterales Enterobacteriaceae *Salmonella enterica subsp. enterica serovar* Heidelberg str. CF-SAN00324 |
| Sen-195-1 | Bacteria Proteobacteria Gammaproteobacteria Enterobacterales Enterobacteriaceae *Salmonella enterica subsp. enterica serovar* Heidelberg str. CF-SAN00327 |
| Sen-196-1 | Bacteria Proteobacteria Gammaproteobacteria Enterobacterales Enterobacteriaceae *Salmonella enterica subsp. enterica serovar* Heidelberg str. CVM20752 |
| Sen-197-1 | Bacteria Proteobacteria Gammaproteobacteria Enterobacterales Enterobacteriaceae *Salmonella enterica subsp. enterica serovar* Heidelberg str. CVM24359 |
| Sen-198-1 | Bacteria Proteobacteria Gammaproteobacteria Enterobacterales Enterobacteriaceae *Salmonella enterica subsp. enterica serovar* Heidelberg str. CVM24388 |
| Sen-199-1 | Bacteria Proteobacteria Gammaproteobacteria Enterobacterales Enterobacteriaceae *Salmonella enterica subsp. enterica serovar* Heidelberg str. CVM24391 |
| Sen-200-1 | Bacteria Proteobacteria Gammaproteobacteria Enterobacterales Enterobacteriaceae *Salmonella enterica subsp. enterica serovar* Heidelberg str. N1536 |
| Sen-201-1 | Bacteria Proteobacteria Gammaproteobacteria Enterobacterales Enterobacteriaceae *Salmonella enterica subsp. enterica serovar* Heidelberg str. N15757 |
| Sen-202-1 | Bacteria Proteobacteria Gammaproteobacteria Enterobacterales Enterobacteriaceae *Salmonella enterica subsp. enterica serovar* Heidelberg str. N18393 |
| Sen-203-1 | Bacteria Proteobacteria Gammaproteobacteria Enterobacterales Enterobacteriaceae *Salmonella enterica subsp. enterica serovar* Heidelberg str. N18413 |
| Sen-204-1 | Bacteria Proteobacteria Gammaproteobacteria Enterobacterales Enterobacteriaceae *Salmonella enterica subsp. enterica serovar* Heidelberg str. N18440 |
| Sen-205-1 | Bacteria Proteobacteria Gammaproteobacteria Enterobacterales Enterobacteriaceae *Salmonella enterica subsp. enterica serovar* Heidelberg str. N18453 |
| Sen-206-1 | Bacteria Proteobacteria Gammaproteobacteria Enterobacterales Enterobacteriaceae *Salmonella enterica subsp. enterica serovar* Heidelberg str. N19871 |
| Sen-207-1 | Bacteria Proteobacteria Gammaproteobacteria Enterobacterales Enterobacteriaceae *Salmonella enterica subsp. enterica serovar* Heidelberg str. N19992 |
| Sen-208-1 | Bacteria Proteobacteria Gammaproteobacteria Enterobacterales Enterobacteriaceae *Salmonella enterica subsp. enterica serovar* Heidelberg str. N20134 |
| Sen-209-1 | Bacteria Proteobacteria Gammaproteobacteria Enterobacterales Enterobacteriaceae *Salmonella enterica subsp. enterica serovar* Heidelberg str. N26457 |
| Sen-210-1 | Bacteria Proteobacteria Gammaproteobacteria Enterobacterales Enterobacteriaceae *Salmonella enterica subsp. enterica serovar* Heidelberg str. N29341 |
| Sen-211-1 | Bacteria Proteobacteria Gammaproteobacteria Enterobacterales Enterobacteriaceae *Salmonella enterica subsp. enterica serovar* Heidelberg str. N30678 |
| Sen-212-1 | Bacteria Proteobacteria Gammaproteobacteria Enterobacterales Enterobacteriaceae *Salmonella enterica subsp. enterica serovar* Heidelberg str. N418 |
| Sen-213-1 | Bacteria Proteobacteria Gammaproteobacteria Enterobacterales Enterobacteriaceae *Salmonella enterica subsp. enterica serovar* Heidelberg str. N4630 |
| Sen-214-1 | Bacteria Proteobacteria Gammaproteobacteria Enterobacterales Enterobacteriaceae *Salmonella enterica subsp. enterica serovar* Heidelberg str. N653 |
| Sen-215-1 | Bacteria Proteobacteria Gammaproteobacteria Enterobacterales Enterobacteriaceae *Salmonella enterica subsp. enterica serovar* Heidelberg str. RI-11-013193 |
| Sen-216-1 | Bacteria Proteobacteria Gammaproteobacteria Enterobacterales Enterobacteriaceae *Salmonella enterica subsp. enterica serovar* Heidelberg str. RI-11-014319 |
| Sen-217-1 | Bacteria Proteobacteria Gammaproteobacteria Enterobacterales Enterobacteriaceae *Salmonella enterica subsp. enterica serovar* Heidelberg str. SARA31 |
| Sen-218-1 | Bacteria Proteobacteria Gammaproteobacteria Enterobacterales Enterobacteriaceae *Salmonella enterica subsp. enterica serovar* Heidelberg str. SARA32 |
| Sen-219-1 | Bacteria Proteobacteria Gammaproteobacteria Enterobacterales Enterobacteriaceae *Salmonella enterica subsp. enterica serovar* Heidelberg str. SARA36 |
| Sen-220-1 | Bacteria Proteobacteria Gammaproteobacteria Enterobacterales Enterobacteriaceae *Salmonella enterica subsp. enterica serovar* Heidelberg str. SARA37 |

| | |
|---|---|
| Sen-221-1 | Bacteria Proteobacteria Gammaproteobacteria Enterobacterales Enterobacteriaceae *Salmonella enterica subsp. enterica serovar* Heidelberg str. SARA40 |
| Sen-222-1 | Bacteria Proteobacteria Gammaproteobacteria Enterobacterales Enterobacteriaceae *Salmonella enterica subsp. enterica serovar* Infantis |
| Sen-223-1 to Sen-223-3 | Bacteria Proteobacteria Gammaproteobacteria Enterobacterales Enterobacteriaceae *Salmonella enterica subsp. enterica serovar* Johannesburg |
| Sen-224-1 to Sen-224-52 | Bacteria Proteobacteria Gammaproteobacteria Enterobacterales Enterobacteriaceae *Salmonella enterica subsp. enterica serovar* Kentucky |
| Sen-225-1 to Sen-225-4 | Bacteria Proteobacteria Gammaproteobacteria Enterobacterales Enterobacteriaceae *Salmonella enterica subsp. enterica serovar* Litchfield |
| Sen-226-1 to Sen-226-2 | Bacteria Proteobacteria Gammaproteobacteria Enterobacterales Enterobacteriaceae *Salmonella enterica subsp. enterica serovar* Liverpool |
| Sen-227-1 to Sen-227-2 | Bacteria Proteobacteria Gammaproteobacteria Enterobacterales Enterobacteriaceae *Salmonella enterica subsp. enterica serovar* Livingstone |
| Sen-228-1 | Bacteria Proteobacteria Gammaproteobacteria Enterobacterales Enterobacteriaceae *Salmonella enterica subsp. enterica serovar* Manhattan str. 111113 |
| Sen-229-1 to Sen-229-4 | Bacteria Proteobacteria Gammaproteobacteria Enterobacterales Enterobacteriaceae *Salmonella enterica subsp. enterica serovar* Mbandaka |
| Sen-230-1 | Bacteria Proteobacteria Gammaproteobacteria Enterobacterales Enterobacteriaceae *Salmonella enterica subsp. enterica serovar* Mbandaka str. 2012K-0273 |
| Sen-231-1 | Bacteria Proteobacteria Gammaproteobacteria Enterobacterales Enterobacteriaceae *Salmonella enterica subsp. enterica serovar* Mishmarhaemek |
| Sen-232-1 to Sen-232-5 | Bacteria Proteobacteria Gammaproteobacteria Enterobacterales Enterobacteriaceae *Salmonella enterica subsp. enterica serovar* Montevideo |
| Sen-233-1 | Bacteria Proteobacteria Gammaproteobacteria Enterobacterales Enterobacteriaceae *Salmonella enterica subsp. enterica serovar* Montevideo str. 316111868 |
| Sen-234-1 | Bacteria Proteobacteria Gammaproteobacteria Enterobacterales Enterobacteriaceae *Salmonella enterica subsp. enterica serovar* Montevideo str. 507440-20 |
| Sen-235-1 | Bacteria Proteobacteria Gammaproteobacteria Enterobacterales Enterobacteriaceae *Salmonella enterica subsp. enterica serovar* Montevideo str. USDA-ARS-USMARC-1903 |
| Sen-236-1 to Sen-236-16 | Bacteria Proteobacteria Gammaproteobacteria Enterobacterales Enterobacteriaceae *Salmonella enterica subsp. enterica serovar* Muenchen |
| Sen-237-1 | Bacteria Proteobacteria Gammaproteobacteria Enterobacterales Enterobacteriaceae *Salmonella enterica subsp. enterica serovar* Muenchen str. 08-1125 |
| Sen-238-1 | Bacteria Proteobacteria Gammaproteobacteria Enterobacterales Enterobacteriaceae *Salmonella enterica subsp. enterica serovar* Muenchen str. 2009K-0951 |
| Sen-239-1 | Bacteria Proteobacteria Gammaproteobacteria Enterobacterales Enterobacteriaceae *Salmonella enterica subsp. enterica serovar* Muenster |
| Sen-240-1 | Bacteria Proteobacteria Gammaproteobacteria Enterobacterales Enterobacteriaceae *Salmonella enterica subsp. enterica serovar* Namur str. 05-2929 |
| Sen-241-1 | Bacteria Proteobacteria Gammaproteobacteria Enterobacterales Enterobacteriaceae *Salmonella enterica subsp. enterica serovar* Napoli |
| Sen-242-1 to Sen-242-2 | Bacteria Proteobacteria Gammaproteobacteria Enterobacterales Enterobacteriaceae *Salmonella enterica subsp. enterica serovar* Newport |
| Sen-243-1 | Bacteria Proteobacteria Gammaproteobacteria Enterobacterales Enterobacteriaceae *Salmonella enterica subsp. enterica serovar* Newport str. 36799 |
| Sen-244-1 | Bacteria Proteobacteria Gammaproteobacteria Enterobacterales Enterobacteriaceae *Salmonella enterica subsp. enterica serovar* Newport str. 36801 |
| Sen-245-1 | Bacteria Proteobacteria Gammaproteobacteria Enterobacterales Enterobacteriaceae *Salmonella enterica subsp. enterica serovar* Newport str. 36802 |
| Sen-246-1 | Bacteria Proteobacteria Gammaproteobacteria Enterobacterales Enterobacteriaceae *Salmonella enterica subsp. enterica serovar* Newport str. 36803 |
| Sen-247-1 | Bacteria Proteobacteria Gammaproteobacteria Enterobacterales Enterobacteriaceae *Salmonella enterica subsp. enterica serovar* Newport str. 36804 |
| Sen-248-1 | Bacteria Proteobacteria Gammaproteobacteria Enterobacterales Enterobacteriaceae *Salmonella enterica subsp. enterica serovar* Newport str. 36805 |
| Sen-249-1 | Bacteria Proteobacteria Gammaproteobacteria Enterobacterales Enterobacteriaceae *Salmonella enterica subsp. enterica serovar* Newport str. 36807 |
| Sen-250-1 | Bacteria Proteobacteria Gammaproteobacteria Enterobacterales Enterobacteriaceae *Salmonella enterica subsp. enterica serovar* Newport str. A182RVB |
| Sen-251-1 | Bacteria Proteobacteria Gammaproteobacteria Enterobacterales Enterobacteriaceae *Salmonella enterica subsp. enterica serovar* Newport str. CVM 21538 |
| Sen-252-1 | Bacteria Proteobacteria Gammaproteobacteria Enterobacterales Enterobacteriaceae *Salmonella enterica subsp. enterica serovar* Newport str. CVM 21550 |
| Sen-253-1 | Bacteria Proteobacteria Gammaproteobacteria Enterobacterales Enterobacteriaceae *Salmonella enterica subsp. enterica serovar* Newport str. CVM 22425 |
| Sen-254-1 | Bacteria Proteobacteria Gammaproteobacteria Enterobacterales Enterobacteriaceae *Salmonella enterica subsp. enterica serovar* Newport str. CVM 22462 |
| Sen-255-1 | Bacteria Proteobacteria Gammaproteobacteria Enterobacterales Enterobacteriaceae *Salmonella enterica subsp. enterica serovar* Newport str. CVM 22513 |
| Sen-256-1 | Bacteria Proteobacteria Gammaproteobacteria Enterobacterales Enterobacteriaceae *Salmonella enterica subsp. enterica serovar* Newport str. CVM N1543 |
| Sen-257-1 | Bacteria Proteobacteria Gammaproteobacteria Enterobacterales Enterobacteriaceae *Salmonella enterica subsp. enterica serovar* Newport str. CVM N18486 |
| Sen-258-1 | Bacteria Proteobacteria Gammaproteobacteria Enterobacterales Enterobacteriaceae *Salmonella enterica subsp. enterica serovar* Newport str. DC_10-446 |
| Sen-259-1 | Bacteria Proteobacteria Gammaproteobacteria Enterobacterales Enterobacteriaceae *Salmonella enterica subsp. enterica serovar* Newport str. DC_10-449 |
| Sen-260-1 | Bacteria Proteobacteria Gammaproteobacteria Enterobacterales Enterobacteriaceae *Salmonella enterica subsp. enterica serovar* Newport str. DC_10-450 |
| Sen-261-1 | Bacteria Proteobacteria Gammaproteobacteria Enterobacterales Enterobacteriaceae *Salmonella enterica subsp. enterica serovar* Newport str. MA_10EN1469 |
| Sen-263-1 | Bacteria Proteobacteria Gammaproteobacteria Enterobacterales Enterobacteriaceae *Salmonella enterica subsp. enterica serovar* Newport str. Pond080-2TTA |
| Sen-262-1 | Bacteria Proteobacteria Gammaproteobacteria Enterobacterales Enterobacteriaceae *Salmonella enterica subsp. enterica serovar* Newport str. PRS_2010_0624 |
| Sen-264-1 | Bacteria Proteobacteria Gammaproteobacteria Enterobacterales Enterobacteriaceae *Salmonella enterica subsp. enterica serovar* Newport str. S103RVX |
| Sen-265-1 to Sen-265-2 | Bacteria Proteobacteria Gammaproteobacteria Enterobacterales Enterobacteriaceae *Salmonella enterica subsp. enterica serovar* Newport str. SL254 |
| Sen-266-1 | Bacteria Proteobacteria Gammaproteobacteria Enterobacterales Enterobacteriaceae *Salmonella enterica subsp. enterica serovar* Newport str. USDA-ARS-USMARC-1927 |
| Sen-267-1 | Bacteria Proteobacteria Gammaproteobacteria Enterobacterales Enterobacteriaceae *Salmonella enterica subsp. enterica serovar* Newport str. VA_R100804798 |
| Sen-268-1 | Bacteria Proteobacteria Gammaproteobacteria Enterobacterales Enterobacteriaceae *Salmonella enterica subsp. enterica serovar* Newport str. VA_R100808502 |
| Sen-269-1 | Bacteria Proteobacteria Gammaproteobacteria Enterobacterales Enterobacteriaceae *Salmonella enterica subsp. enterica serovar* Ohio |
| Sen-270-1 to Sen-270-3 | Bacteria Proteobacteria Gammaproteobacteria Enterobacterales Enterobacteriaceae *Salmonella enterica subsp. enterica serovar* Ouakam |
| Sen-271-1 | Bacteria Proteobacteria Gammaproteobacteria Enterobacterales Enterobacteriaceae *Salmonella enterica subsp. enterica serovar* Panama str. ATCC 7378 |
| Sen-272-1 to Sen-272-3 | Bacteria Proteobacteria Gammaproteobacteria Enterobacterales Enterobacteriaceae *Salmonella enterica subsp. enterica serovar* Paratyphi A |
| Sen-273-1 | Bacteria Proteobacteria Gammaproteobacteria Enterobacterales Enterobacteriaceae *Salmonella enterica subsp. enterica serovar* Poona str. 2010K-2244 |
| Sen-274-1 | Bacteria Proteobacteria Gammaproteobacteria Enterobacterales Enterobacteriaceae *Salmonella enterica subsp. enterica serovar* Pullorum |
| Sen-275-1 | Bacteria Proteobacteria Gammaproteobacteria Enterobacterales Enterobacteriaceae *Salmonella enterica subsp. enterica serovar* Pullorum str. ATCC 9120 |
| Sen-276-1 to Sen-276-5 | Bacteria Proteobacteria Gammaproteobacteria Enterobacterales Enterobacteriaceae *Salmonella enterica subsp. enterica serovar* Reading |

| | |
|---|---|
| Sen-277-1 to Sen-277-22 | Bacteria Proteobacteria Gammaproteobacteria Enterobacterales Enterobacteriaceae *Salmonella enterica subsp. enterica serovar* Saintpaul |
| Sen-278-1 to Sen-278-14 | Bacteria Proteobacteria Gammaproteobacteria Enterobacterales Enterobacteriaceae *Salmonella enterica subsp. enterica serovar* Schwarzengrund |
| Sen-279-1 to Sen-279-3 | Bacteria Proteobacteria Gammaproteobacteria Enterobacterales Enterobacteriaceae *Salmonella enterica subsp. enterica serovar* Senftenberg |
| Sen-280-1 | Bacteria Proteobacteria Gammaproteobacteria Enterobacterales Enterobacteriaceae *Salmonella enterica subsp. enterica serovar* Sloterdijk str. ATCC 15791 |
| Sen-281-1 | Bacteria Proteobacteria Gammaproteobacteria Enterobacterales Enterobacteriaceae *Salmonella enterica subsp. enterica serovar* Stanley str. 06-0538 |
| Sen-282-1 | Bacteria Proteobacteria Gammaproteobacteria Enterobacterales Enterobacteriaceae *Salmonella enterica subsp. enterica serovar* Tennessee str. 4535 |
| Sen-283-1 | Bacteria Proteobacteria Gammaproteobacteria Enterobacterales Enterobacteriaceae *Salmonella enterica subsp. enterica serovar* Tennessee str. CDC07-0191 |
| Sen-284-1 | Bacteria Proteobacteria Gammaproteobacteria Enterobacterales Enterobacteriaceae *Salmonella enterica subsp. enterica serovar* Tennessee str. TXSC_TXSC08-19 |
| Sen-285-1 to Sen-285-5 | Bacteria Proteobacteria Gammaproteobacteria Enterobacterales Enterobacteriaceae *Salmonella enterica subsp. enterica serovar* Thompson |
| Sen-286-1 | Bacteria Proteobacteria Gammaproteobacteria Enterobacterales Enterobacteriaceae *Salmonella enterica subsp. enterica serovar* Thompson str. ATCC 8391 |
| Sen-287-1 to Sen-287-1124 | Bacteria Proteobacteria Gammaproteobacteria Enterobacterales Enterobacteriaceae *Salmonella enterica subsp. enterica serovar* Typhi |
| Sen-288-1 | Bacteria Proteobacteria Gammaproteobacteria Enterobacterales Enterobacteriaceae *Salmonella enterica subsp. enterica serovar* Typhi str. CR0063 |
| Sen-289-1 | Bacteria Proteobacteria Gammaproteobacteria Enterobacterales Enterobacteriaceae *Salmonella enterica subsp. enterica serovar* Typhi str. STH2370 |
| Sen-290-1 to Sen-290-7 | Bacteria Proteobacteria Gammaproteobacteria Enterobacterales Enterobacteriaceae *Salmonella enterica subsp. enterica serovar* Typhimurium |
| Sen-291-1 | Bacteria Proteobacteria Gammaproteobacteria Enterobacterales Enterobacteriaceae *Salmonella enterica subsp. enterica serovar* Typhimurium str. 104772 |
| Sen-292-1 | Bacteria Proteobacteria Gammaproteobacteria Enterobacterales Enterobacteriaceae *Salmonella enterica subsp. enterica serovar* Typhimurium str. 108402 |
| Sen-293-1 | Bacteria Proteobacteria Gammaproteobacteria Enterobacterales Enterobacteriaceae *Salmonella enterica subsp. enterica serovar* Typhimurium str. 116045 |
| Sen-294-1 | Bacteria Proteobacteria Gammaproteobacteria Enterobacterales Enterobacteriaceae *Salmonella enterica subsp. enterica serovar* Typhimurium str. 86-0368 |
| Sen-295-1 | Bacteria Proteobacteria Gammaproteobacteria Enterobacterales Enterobacteriaceae *Salmonella enterica subsp. enterica serovar* Typhimurium str. 95799 |
| Sen-296-1 | Bacteria Proteobacteria Gammaproteobacteria Enterobacterales Enterobacteriaceae *Salmonella enterica subsp. enterica serovar* Typhimurium str. 98346 |
| Sen-297-1 | Bacteria Proteobacteria Gammaproteobacteria Enterobacterales Enterobacteriaceae *Salmonella enterica subsp. enterica serovar* Typhimurium str. CDC 2011K-0870 |
| Sen-298-1 | Bacteria Proteobacteria Gammaproteobacteria Enterobacterales Enterobacteriaceae *Salmonella enterica subsp. enterica serovar* Typhimurium str. CDC_2009K1153 |
| Sen-299-1 to Sen-299-330 | Bacteria Proteobacteria Gammaproteobacteria Enterobacterales Enterobacteriaceae *Salmonella enterica subsp. enterica serovar* Typhimurium str. DT104 |
| Sen-300-1 | Bacteria Proteobacteria Gammaproteobacteria Enterobacterales Enterobacteriaceae *Salmonella enterica subsp. enterica serovar* Typhimurium str. L-3553 |
| Sen-301-1 | Bacteria Proteobacteria Gammaproteobacteria Enterobacterales Enterobacteriaceae *Salmonella enterica subsp. enterica serovar* Typhimurium str. LT2-4 |
| Sen-302-1 | Bacteria Proteobacteria Gammaproteobacteria Enterobacterales Enterobacteriaceae *Salmonella enterica subsp. enterica serovar* Typhimurium str. LT2-4_delta.ramA::kan |
| Sen-303-1 | Bacteria Proteobacteria Gammaproteobacteria Enterobacterales Enterobacteriaceae *Salmonella enterica subsp. enterica serovar* Typhimurium str. ST1489 |
| Sen-304-1 | Bacteria Proteobacteria Gammaproteobacteria Enterobacterales Enterobacteriaceae *Salmonella enterica subsp. enterica serovar* Typhimurium str. ST4848 |
| Sen-305-1 to Sen-305-63 | Bacteria Proteobacteria Gammaproteobacteria Enterobacterales Enterobacteriaceae *Salmonella enterica subsp. enterica serovar* Typhimurium var. 5- |
| Sen-306-1 | Bacteria Proteobacteria Gammaproteobacteria Enterobacterales Enterobacteriaceae *Salmonella enterica subsp. enterica serovar* Virchow str. SVQ1 |
| Sen-307-1 to Sen-307-4 | Bacteria Proteobacteria Gammaproteobacteria Enterobacterales Enterobacteriaceae *Salmonella enterica subsp. enterica serovar* Worthington |
| Sen-308-1 | Bacteria Proteobacteria Gammaproteobacteria Enterobacterales Enterobacteriaceae *Salmonella enterica subsp. salamae* |
| Sen-309-1 | Bacteria Proteobacteria Gammaproteobacteria Enterobacterales Enterobacteriaceae *Salmonella enterica subsp. salamae str.* DMA-1 |
| Sbo-2-1 to Sbo-2-10 | Bacteria Proteobacteria Gammaproteobacteria Enterobacterales Enterobacteriaceae *Shigella boydii* |
| Sdy-1-1 | Bacteria Proteobacteria Gammaproteobacteria Enterobacterales Enterobacteriaceae *Shigella dysenteriae* S6205 |
| Sdy-2-1 | Bacteria Proteobacteria Gammaproteobacteria Enterobacterales Enterobacteriaceae *Shigella dysenteriae* SD1D |
| Sfl-1-1 to Sfl-1-2 | Bacteria Proteobacteria Gammaproteobacteria Enterobacterales Enterobacteriaceae *Shigella flexneri* |
| Sfl-2-1 | Bacteria Proteobacteria Gammaproteobacteria Enterobacterales Enterobacteriaceae *Shigella flexneri* 1997005 |
| Sfl-3-1 | Bacteria Proteobacteria Gammaproteobacteria Enterobacterales Enterobacteriaceae *Shigella flexneri* 2000019 |
| Sfl-4-1 | Bacteria Proteobacteria Gammaproteobacteria Enterobacterales Enterobacteriaceae *Shigella flexneri* 2001004 |
| Sfl-5-1 | Bacteria Proteobacteria Gammaproteobacteria Enterobacterales Enterobacteriaceae *Shigella flexneri* 2001025 |
| Sfl-6-1 | Bacteria Proteobacteria Gammaproteobacteria Enterobacterales Enterobacteriaceae *Shigella flexneri* 2001027 |
| Sfl-7-1 | Bacteria Proteobacteria Gammaproteobacteria Enterobacterales Enterobacteriaceae *Shigella flexneri* 2001042 |
| Sfl-8-1 | Bacteria Proteobacteria Gammaproteobacteria Enterobacterales Enterobacteriaceae *Shigella flexneri* 2001044 |
| Sfl-9-1 | Bacteria Proteobacteria Gammaproteobacteria Enterobacterales Enterobacteriaceae *Shigella flexneri* 2001048 |
| Sfl-10-1 | Bacteria Proteobacteria Gammaproteobacteria Enterobacterales Enterobacteriaceae *Shigella flexneri* 2002007 |
| Sfl-11-1 | Bacteria Proteobacteria Gammaproteobacteria Enterobacterales Enterobacteriaceae *Shigella flexneri* 2002021 |
| Sfl-12-1 | Bacteria Proteobacteria Gammaproteobacteria Enterobacterales Enterobacteriaceae *Shigella flexneri* 2002035 |
| Sfl-13-1 | Bacteria Proteobacteria Gammaproteobacteria Enterobacterales Enterobacteriaceae *Shigella flexneri* 2002069 |
| Sfl-14-1 | Bacteria Proteobacteria Gammaproteobacteria Enterobacterales Enterobacteriaceae *Shigella flexneri* 2002091 |
| Sfl-15-1 | Bacteria Proteobacteria Gammaproteobacteria Enterobacterales Enterobacteriaceae *Shigella flexneri* 2002103 |
| Sfl-16-1 | Bacteria Proteobacteria Gammaproteobacteria Enterobacterales Enterobacteriaceae *Shigella flexneri* 2002110 |
| Sfl-17-1 | Bacteria Proteobacteria Gammaproteobacteria Enterobacterales Enterobacteriaceae *Shigella flexneri* 2002127 |
| Sfl-18-1 | Bacteria Proteobacteria Gammaproteobacteria Enterobacterales Enterobacteriaceae *Shigella flexneri* 2002140 |
| Sfl-19-1 | Bacteria Proteobacteria Gammaproteobacteria Enterobacterales Enterobacteriaceae *Shigella flexneri* 2002141 |
| Sfl-20-1 | Bacteria Proteobacteria Gammaproteobacteria Enterobacterales Enterobacteriaceae *Shigella flexneri* 2002142 |
| Sfl-21-1 | Bacteria Proteobacteria Gammaproteobacteria Enterobacterales Enterobacteriaceae *Shigella flexneri* 2003055 |
| Sfl-22-1 | Bacteria Proteobacteria Gammaproteobacteria Enterobacterales Enterobacteriaceae *Shigella flexneri* 2005051 |
| Sfl-23-1 | Bacteria Proteobacteria Gammaproteobacteria Enterobacterales Enterobacteriaceae *Shigella flexneri* 2005AH264 |

| | |
|---|---|
| Sfl-24-1 | Bacteria Proteobacteria Gammaproteobacteria Enterobacterales Enterobacteriaceae *Shigella flexneri* 2005GS061 |
| Sfl-25-1 to Sfl-25-14 | Bacteria Proteobacteria Gammaproteobacteria Enterobacterales Enterobacteriaceae *Shigella flexneri* 2a |
| Sfl-26-1 | Bacteria Proteobacteria Gammaproteobacteria Enterobacterales Enterobacteriaceae *Shigella flexneri* 51575 |
| Sfl-27-1 | Bacteria Proteobacteria Gammaproteobacteria Enterobacterales Enterobacteriaceae *Shigella flexneri* 51576 |
| Sfl-28-1 | Bacteria Proteobacteria Gammaproteobacteria Enterobacterales Enterobacteriaceae *Shigella flexneri* 51577 |
| Sfl-29-1 | Bacteria Proteobacteria Gammaproteobacteria Enterobacterales Enterobacteriaceae *Shigella flexneri* Shi05SX04 |
| Sfl-30-1 | Bacteria Proteobacteria Gammaproteobacteria Enterobacterales Enterobacteriaceae *Shigella flexneri* Shi06AH028 |
| Sfl-31-1 | Bacteria Proteobacteria Gammaproteobacteria Enterobacterales Enterobacteriaceae *Shigella flexneri* Shi06AH091 |
| Sfl-32-1 | Bacteria Proteobacteria Gammaproteobacteria Enterobacterales Enterobacteriaceae *Shigella flexneri* Shi06AH116 |
| Sfl-33-1 | Bacteria Proteobacteria Gammaproteobacteria Enterobacterales Enterobacteriaceae *Shigella flexneri* Shi06AH130 |
| Sfl-34-1 | Bacteria Proteobacteria Gammaproteobacteria Enterobacterales Enterobacteriaceae *Shigella flexneri* Shi06AH135 |
| Sfl-35-1 | Bacteria Proteobacteria Gammaproteobacteria Enterobacterales Enterobacteriaceae *Shigella flexneri* Shi06AH66 |
| Sfl-36-1 | Bacteria Proteobacteria Gammaproteobacteria Enterobacterales Enterobacteriaceae *Shigella flexneri* Shi06GS07 |
| Sfl-37-1 | Bacteria Proteobacteria Gammaproteobacteria Enterobacterales Enterobacteriaceae *Shigella flexneri* Shi06GS37 |
| Sfl-38-1 | Bacteria Proteobacteria Gammaproteobacteria Enterobacterales Enterobacteriaceae *Shigella flexneri* Shi06GS43 |
| Sfl-39-1 | Bacteria Proteobacteria Gammaproteobacteria Enterobacterales Enterobacteriaceae *Shigella flexneri* Shi06GS48 |
| Sfl-40-1 | Bacteria Proteobacteria Gammaproteobacteria Enterobacterales Enterobacteriaceae *Shigella flexneri* Shi06GS55 |
| Sfl-41-1 | Bacteria Proteobacteria Gammaproteobacteria Enterobacterales Enterobacteriaceae *Shigella flexneri* Shi06HN016 |
| Sfl-42-1 | Bacteria Proteobacteria Gammaproteobacteria Enterobacterales Enterobacteriaceae *Shigella flexneri* Shi06HN023 |
| Sfl-43-1 | Bacteria Proteobacteria Gammaproteobacteria Enterobacterales Enterobacteriaceae *Shigella flexneri* Shi06HN091 |
| Sfl-44-1 | Bacteria Proteobacteria Gammaproteobacteria Enterobacterales Enterobacteriaceae *Shigella flexneri* Shi06HN118 |
| Sfl-45-1 | Bacteria Proteobacteria Gammaproteobacteria Enterobacterales Enterobacteriaceae *Shigella flexneri* Shi06HN159 |
| Sfl-46-1 | Bacteria Proteobacteria Gammaproteobacteria Enterobacterales Enterobacteriaceae *Shigella flexneri* Shi06HN244 |
| Sfl-47-1 | Bacteria Proteobacteria Gammaproteobacteria Enterobacterales Enterobacteriaceae *Shigella flexneri* Shi06HN250 |
| Sfl-48-1 | Bacteria Proteobacteria Gammaproteobacteria Enterobacterales Enterobacteriaceae *Shigella flexneri* Shi06HN378 |
| Sfl-49-1 | Bacteria Proteobacteria Gammaproteobacteria Enterobacterales Enterobacteriaceae *Shigella flexneri* Shi06SX36 |
| Sfl-50-1 | Bacteria Proteobacteria Gammaproteobacteria Enterobacterales Enterobacteriaceae *Shigella flexneri* Shi06SX53 |
| Sfl-51-1 to Sfl-51-2 | Bacteria Proteobacteria Gammaproteobacteria Enterobacterales Enterobacteriaceae *Shigella flexneri* Y |
| Sso-1-1 to Sso-1-540 | Bacteria Proteobacteria Gammaproteobacteria Enterobacterales Enterobacteriaceae *Shigella sonnei* |
| Sso-2-1 | Bacteria Proteobacteria Gammaproteobacteria Enterobacterales Enterobacteriaceae *Shigella sonnei* 1DT-1 |
| Sco-1-1 | Bacteria Proteobacteria Gammaproteobacteria Enterobacterales Enterobacteriaceae *Siccibacter colletis* |
| Stu-1-1 | Bacteria Proteobacteria Gammaproteobacteria Enterobacterales Enterobacteriaceae *Siccibacter turicensis* LMG 23730 |
| Tgu-1-1 | Bacteria Proteobacteria Gammaproteobacteria Enterobacterales Enterobacteriaceae *Trabulsiella guamensis* ATCC 49490 |
| Tod-1-1 | Bacteria Proteobacteria Gammaproteobacteria Enterobacterales Enterobacteriaceae *Trabulsiella odontotermitis* |
| Yre-1-1 | Bacteria Proteobacteria Gammaproteobacteria Enterobacterales Enterobacteriaceae *Yokenella regensburgei* ATCC 49455 |
| Eam-1-1 | Bacteria Proteobacteria Gammaproteobacteria Enterobacterales Erwiniaceae *Erwinia amylovora* NBRC 12687 |
| Ebi-1-1 | Bacteria Proteobacteria Gammaproteobacteria Enterobacterales Erwiniaceae *Erwinia billingiae* |
| Ein-1-1 to Ein-1-2 | Bacteria Proteobacteria Gammaproteobacteria Enterobacterales Erwiniaceae *Erwinia iniecta* |
| Ema-2-1 | Bacteria Proteobacteria Gammaproteobacteria Enterobacterales Erwiniaceae *Erwinia mallotivora* |
| Eol-1-1 | Bacteria Proteobacteria Gammaproteobacteria Enterobacterales Erwiniaceae *Erwinia oleae* |
| Esp-55-1 | Bacteria Proteobacteria Gammaproteobacteria Enterobacterales Erwiniaceae *Erwinia sp.* 9145 |
| Etr-1-1 | Bacteria Proteobacteria Gammaproteobacteria Enterobacterales Erwiniaceae *Erwinia tracheiphila* |
| Ety-1-1 | Bacteria Proteobacteria Gammaproteobacteria Enterobacterales Erwiniaceae *Erwinia typographi* |
| Pag-1-1 to Pag-1-6 | Bacteria Proteobacteria Gammaproteobacteria Enterobacterales Erwiniaceae Pantoea Pantoea agglomerans group *Pantoea agglomerans* |
| Pag-2-1 | Bacteria Proteobacteria Gammaproteobacteria Enterobacterales Erwiniaceae Pantoea Pantoea agglomerans group *Pantoea agglomerans* Eh318 |
| Pan-1-1 to Pan-1-5 | Bacteria Proteobacteria Gammaproteobacteria Enterobacterales Erwiniaceae *Pantoea ananatis* |
| Pan-2-1 | Bacteria Proteobacteria Gammaproteobacteria Enterobacterales Erwiniaceae *Pantoea ananatis* BD442 |
| Pan-3-1 to Pan-3-2 | Bacteria Proteobacteria Gammaproteobacteria Enterobacterales Erwiniaceae *Pantoea ananatis* LMG 2665 |
| Pan-4-1 | Bacteria Proteobacteria Gammaproteobacteria Enterobacterales Erwiniaceae *Pantoea ananatis* Sd-1 |
| Pan-5-1 | Bacteria Proteobacteria Gammaproteobacteria Enterobacterales Erwiniaceae *Pantoea anthophila* |
| Pro-1-1 | Bacteria Proteobacteria Gammaproteobacteria Enterobacterales Erwiniaceae *Pantoea rodasii* |
| Prw-1-1 | Bacteria Proteobacteria Gammaproteobacteria Enterobacterales Erwiniaceae *Pantoea rwandensis* |
| Pse-1-1 | Bacteria Proteobacteria Gammaproteobacteria Enterobacterales Erwiniaceae *Pantoea septica* |
| Psp-1-1 | Bacteria Proteobacteria Gammaproteobacteria Enterobacterales Erwiniaceae *Pantoea sp.* 3.5.1 |
| Psp-2-1 | Bacteria Proteobacteria Gammaproteobacteria Enterobacterales Erwiniaceae *Pantoea sp.* 9140 |
| Psp-3-1 | Bacteria Proteobacteria Gammaproteobacteria Enterobacterales Erwiniaceae *Pantoea sp.* BL1 |
| Psp-4-1 | Bacteria Proteobacteria Gammaproteobacteria Enterobacterales Erwiniaceae *Pantoea sp.* CFSAN033090 |
| Psp-5-1 | Bacteria Proteobacteria Gammaproteobacteria Enterobacterales Erwiniaceae *Pantoea sp.* IMH |
| Psp-6-1 | Bacteria Proteobacteria Gammaproteobacteria Enterobacterales Erwiniaceae *Pantoea sp.* MBLJ3 |
| Psp-7-1 | Bacteria Proteobacteria Gammaproteobacteria Enterobacterales Erwiniaceae *Pantoea sp.* NGS-ED-1003 |
| Psp-8-1 | Bacteria Proteobacteria Gammaproteobacteria Enterobacterales Erwiniaceae *Pantoea sp.* PSNIH1 |
| Psp-9-1 | Bacteria Proteobacteria Gammaproteobacteria Enterobacterales Erwiniaceae *Pantoea sp.* PSNIH2 |
| Psp-10-1 | Bacteria Proteobacteria Gammaproteobacteria Enterobacterales Erwiniaceae *Pantoea sp.* RIT-PI-b |
| Psp-11-1 | Bacteria Proteobacteria Gammaproteobacteria Enterobacterales Erwiniaceae *Pantoea sp.* SM3 |
| Pst-1-1 to Pst-1-2 | Bacteria Proteobacteria Gammaproteobacteria Enterobacterales Erwiniaceae *Pantoea stewartii* |

| | |
|---|---|
| Pva-1-1 to Pva-1-3 | Bacteria Proteobacteria Gammaproteobacteria Enterobacterales Erwiniaceae *Pantoea vagans* |
| Pfl-1-1 | Bacteria Proteobacteria Gammaproteobacteria Enterobacterales Erwiniaceae *Phaseolibacter flectens* ATCC 12775 |
| Tmo-1-1 | Bacteria Proteobacteria Gammaproteobacteria Enterobacterales Erwiniaceae *Tatumella morbirosei* |
| Tpt-1-1 to Tpt-1-2 | Bacteria Proteobacteria Gammaproteobacteria Enterobacterales Erwiniaceae *Tatumella ptyseos* ATCC 33301 |
| Tsa-1-1 | Bacteria Proteobacteria Gammaproteobacteria Enterobacterales Erwiniaceae *Tatumella saanichensis* |
| Tsp-1-1 | Bacteria Proteobacteria Gammaproteobacteria Enterobacterales Erwiniaceae *Tatumella sp.* UCD-D_suzukii |
| Ean-1-1 | Bacteria Proteobacteria Gammaproteobacteria Enterobacterales Hafniaceae *Edwardsiella anguillarum* ET070829 |
| Ean-2-1 | Bacteria Proteobacteria Gammaproteobacteria Enterobacterales Hafniaceae *Edwardsiella anguillarum* ET080729 |
| Ean-3-1 | Bacteria Proteobacteria Gammaproteobacteria Enterobacterales Hafniaceae *Edwardsiella anguillarum* ET080813 |
| Ean-4-1 | Bacteria Proteobacteria Gammaproteobacteria Enterobacterales Hafniaceae *Edwardsiella anguillarum* ET081126R |
| Eho-5-1 | Bacteria Proteobacteria Gammaproteobacteria Enterobacterales Hafniaceae *Edwardsiella hoshinae* NBRC 105699 = ATCC 33379 |
| Eic-1-1 | Bacteria Proteobacteria Gammaproteobacteria Enterobacterales Hafniaceae *Edwardsiella ictaluri* ATCC 33202 |
| Epi-1-1 to Epi-1-2 | Bacteria Proteobacteria Gammaproteobacteria Enterobacterales Hafniaceae *Edwardsiella piscicida* |
| Esp-56-1 | Bacteria Proteobacteria Gammaproteobacteria Enterobacterales Hafniaceae *Edwardsiella sp.* EA181011 |
| Eta-1-1 | Bacteria Proteobacteria Gammaproteobacteria Enterobacterales Hafniaceae *Edwardsiella tarda* |
| Eta-2-1 | Bacteria Proteobacteria Gammaproteobacteria Enterobacterales Hafniaceae *Edwardsiella tarda* ATCC 15947 = NBRC 105688 |
| Eta-3-1 | Bacteria Proteobacteria Gammaproteobacteria Enterobacterales Hafniaceae *Edwardsiella tarda* DT |
| Hal-1-1 to Hal-1-4 | Bacteria Proteobacteria Gammaproteobacteria Enterobacterales Hafniaceae *Hafnia alvei* |
| Hal-2-1 | Bacteria Proteobacteria Gammaproteobacteria Enterobacterales Hafniaceae *Hafnia alvei* ATCC 13337 |
| Hal-3-1 | Bacteria Proteobacteria Gammaproteobacteria Enterobacterales Hafniaceae *Hafnia alvei* FB1 |
| Hpa-1-1 to Hpa-1-2 | Bacteria Proteobacteria Gammaproteobacteria Enterobacterales Hafniaceae *Hafnia paralvei* |
| Aen-1-1 | Bacteria Proteobacteria Gammaproteobacteria Enterobacterales Morganellaceae *Arsenophonus endosymbiont str.* Hangzhou of Nilaparvata lugens |
| Ana-1-1 | Bacteria Proteobacteria Gammaproteobacteria Enterobacterales Morganellaceae *Arsenophonus nasoniae* DSM 15247 |
| Mwi-1-1 | Bacteria Proteobacteria Gammaproteobacteria Enterobacterales Morganellaceae *Moellerella wisconsensis* |
| Mmo-1-1 to Mmo-1-8 | Bacteria Proteobacteria Gammaproteobacteria Enterobacterales Morganellaceae *Morganella morganii* |
| Mmo-2-1 | Bacteria Proteobacteria Gammaproteobacteria Enterobacterales Morganellaceae *Morganella morganii* F675 |
| Mmo-3-1 | Bacteria Proteobacteria Gammaproteobacteria Enterobacterales Morganellaceae *Morganella morganii* H1r |
| Mmo-4-1 | Bacteria Proteobacteria Gammaproteobacteria Enterobacterales Morganellaceae *Morganella morganii* IS15 |
| Msp-2-1 | Bacteria Proteobacteria Gammaproteobacteria Enterobacterales Morganellaceae *Morganella sp.* EGD-HP17 |
| Pas-1-1 | Bacteria Proteobacteria Gammaproteobacteria Enterobacterales Morganellaceae *Photorhabdus asymbiotica subsp. australis* DSM 17609 |
| Plu-1-1 to Plu-1-2 | Bacteria Proteobacteria Gammaproteobacteria Enterobacterales Morganellaceae *Photorhabdus luminescens* |
| Plu-2-1 | Bacteria Proteobacteria Gammaproteobacteria Enterobacterales Morganellaceae *Photorhabdus luminescens* BA1 |
| Plu-3-1 | Bacteria Proteobacteria Gammaproteobacteria Enterobacterales Morganellaceae *Photorhabdus luminescens* NBAII H75HRPL105 |
| Plu-4-1 | Bacteria Proteobacteria Gammaproteobacteria Enterobacterales Morganellaceae *Photorhabdus luminescens* NBAII HiPL101 |
| Plu-5-1 | Bacteria Proteobacteria Gammaproteobacteria Enterobacterales Morganellaceae *Photorhabdus luminescens subsp. luminescens* |
| Pte-1-1 | Bacteria Proteobacteria Gammaproteobacteria Enterobacterales Morganellaceae *Photorhabdus temperata subsp. khanii* NC19 |
| Pte-2-1 | Bacteria Proteobacteria Gammaproteobacteria Enterobacterales Morganellaceae *Photorhabdus temperata subsp. temperata* Meg1 |
| Pte-3-1 | Bacteria Proteobacteria Gammaproteobacteria Enterobacterales Morganellaceae *Photorhabdus temperata subsp. thracensis* |
| Pmi-1-1 to Pmi-1-6 | Bacteria Proteobacteria Gammaproteobacteria Enterobacterales Morganellaceae *Proteus mirabilis* |
| Pvu-2-1 | Bacteria Proteobacteria Gammaproteobacteria Enterobacterales Morganellaceae *Proteus vulgaris* |
| Pal-1-1 | Bacteria Proteobacteria Gammaproteobacteria Enterobacterales Morganellaceae *Providencia alcalifaciens* 205/92 |
| Pal-2-1 | Bacteria Proteobacteria Gammaproteobacteria Enterobacterales Morganellaceae *Providencia alcalifaciens* F90-2004 |
| Pal-3-1 | Bacteria Proteobacteria Gammaproteobacteria Enterobacterales Morganellaceae *Providencia alcalifaciens* PAL-1 |
| Pal-4-1 | Bacteria Proteobacteria Gammaproteobacteria Enterobacterales Morganellaceae *Providencia alcalifaciens* PAL-2 |
| Pal-5-1 | Bacteria Proteobacteria Gammaproteobacteria Enterobacterales Morganellaceae *Providencia alcalifaciens* PAL-3 |
| Pal-6-1 | Bacteria Proteobacteria Gammaproteobacteria Enterobacterales Morganellaceae *Providencia alcalifaciens* R90-1475 |
| Pal-7-1 | Bacteria Proteobacteria Gammaproteobacteria Enterobacterales Morganellaceae *Providencia alcalifaciens* RIMD 1656011 |
| Pre-1-1 to Pre-1-2 | Bacteria Proteobacteria Gammaproteobacteria Enterobacterales Morganellaceae *Providencia rettgeri* |
| Pst-2-1 | Bacteria Proteobacteria Gammaproteobacteria Enterobacterales Morganellaceae *Providencia stuartii* |
| Xbo-1-1 | Bacteria Proteobacteria Gammaproteobacteria Enterobacterales Morganellaceae *Xenorhabdus bovienii* |
| Xbo-4-1 | Bacteria Proteobacteria Gammaproteobacteria Enterobacterales Morganellaceae *Xenorhabdus bovienii str. feltiae* Florida |
| Xbo-5-1 | Bacteria Proteobacteria Gammaproteobacteria Enterobacterales Morganellaceae *Xenorhabdus bovienii str. feltiae* France |
| Xbo-6-1 | Bacteria Proteobacteria Gammaproteobacteria Enterobacterales Morganellaceae *Xenorhabdus bovienii str. feltiae* Moldova |
| Xbo-2-1 | Bacteria Proteobacteria Gammaproteobacteria Enterobacterales Morganellaceae *Xenorhabdus bovienii str.* Intermedium |
| Xbo-3-1 | Bacteria Proteobacteria Gammaproteobacteria Enterobacterales Morganellaceae *Xenorhabdus bovienii str.* Jollieti |
| Xbo-7-1 | Bacteria Proteobacteria Gammaproteobacteria Enterobacterales Morganellaceae *Xenorhabdus bovienii str. kraussei* Becker Underwood |
| Xbo-8-1 | Bacteria Proteobacteria Gammaproteobacteria Enterobacterales Morganellaceae *Xenorhabdus bovienii str. kraussei* Quebec |
| Xbo-9-1 | Bacteria Proteobacteria Gammaproteobacteria Enterobacterales Morganellaceae *Xenorhabdus bovienii str. oregonense* |
| Xbo-10-1 | Bacteria Proteobacteria Gammaproteobacteria Enterobacterales Morganellaceae *Xenorhabdus bovienii str. puntauvense* |
| Xca-1-1 | Bacteria Proteobacteria Gammaproteobacteria Enterobacterales Morganellaceae *Xenorhabdus cabanillasii* JM26 |
| Xdo-1-1 | Bacteria Proteobacteria Gammaproteobacteria Enterobacterales Morganellaceae *Xenorhabdus doucetiae* |
| Xgr-1-1 | Bacteria Proteobacteria Gammaproteobacteria Enterobacterales Morganellaceae *Xenorhabdus griffiniae* |
| Xkh-1-1 | Bacteria Proteobacteria Gammaproteobacteria Enterobacterales Morganellaceae *Xenorhabdus khoisanae* |
| Xne-1-1 | Bacteria Proteobacteria Gammaproteobacteria Enterobacterales Morganellaceae *Xenorhabdus nematophila* |
| Xne-2-1 | Bacteria Proteobacteria Gammaproteobacteria Enterobacterales Morganellaceae *Xenorhabdus nematophila* AN6/1 |

| | |
|---|---|
| Xpo-1-1 | Bacteria Proteobacteria Gammaproteobacteria Enterobacterales Morganellaceae *Xenorhabdus poinarii* G6 |
| Xsp-1-1 | Bacteria Proteobacteria Gammaproteobacteria Enterobacterales Morganellaceae *Xenorhabdus sp.* NBAII XenSa04 |
| Xsz-1-1 | Bacteria Proteobacteria Gammaproteobacteria Enterobacterales Morganellaceae *Xenorhabdus szentirmaii* DSM 16338 |
| Bgo-1-1 | Bacteria Proteobacteria Gammaproteobacteria Enterobacterales Pectobacteriaceae *Brenneria goodwinii* |
| Dch-1-1 to Dch-1-2 | Bacteria Proteobacteria Gammaproteobacteria Enterobacterales Pectobacteriaceae *Dickeya chrysanthemi* |
| Dch-2-1 | Bacteria Proteobacteria Gammaproteobacteria Enterobacterales Pectobacteriaceae *Dickeya chrysanthemi* NCPPB 3533 |
| Dch-3-1 | Bacteria Proteobacteria Gammaproteobacteria Enterobacterales Pectobacteriaceae *Dickeya chrysanthemi* NCPPB 402 |
| Dch-4-1 | Bacteria Proteobacteria Gammaproteobacteria Enterobacterales Pectobacteriaceae *Dickeya chrysanthemi* NCPPB 516 |
| Dda-1-1 | Bacteria Proteobacteria Gammaproteobacteria Enterobacterales Pectobacteriaceae *Dickeya dadantii* NCPPB 3537 |
| Dda-2-1 | Bacteria Proteobacteria Gammaproteobacteria Enterobacterales Pectobacteriaceae *Dickeya dadantii* NCPPB 898 |
| Dda-3-1 | Bacteria Proteobacteria Gammaproteobacteria Enterobacterales Pectobacteriaceae *Dickeya dadantii subsp. dieffenbachiae* NCPPB 2976 |
| Ddi-1-1 | Bacteria Proteobacteria Gammaproteobacteria Enterobacterales Pectobacteriaceae *Dickeya dianthicola* IPO 980 |
| Ddi-2-1 | Bacteria Proteobacteria Gammaproteobacteria Enterobacterales Pectobacteriaceae *Dickeya dianthicola* RNS04.9 |
| Dso-1-1 | Bacteria Proteobacteria Gammaproteobacteria Enterobacterales Pectobacteriaceae *Dickeya solani* |
| Dsp-1-1 | Bacteria Proteobacteria Gammaproteobacteria Enterobacterales Pectobacteriaceae *Dickeya sp.* 2B12 |
| Dsp-2-1 | Bacteria Proteobacteria Gammaproteobacteria Enterobacterales Pectobacteriaceae *Dickeya sp.* B16 |
| Dsp-3-1 | Bacteria Proteobacteria Gammaproteobacteria Enterobacterales Pectobacteriaceae *Dickeya sp.* DW 0440 |
| Dsp-4-1 | Bacteria Proteobacteria Gammaproteobacteria Enterobacterales Pectobacteriaceae *Dickeya sp.* MK7 |
| Dsp-5-1 | Bacteria Proteobacteria Gammaproteobacteria Enterobacterales Pectobacteriaceae *Dickeya sp.* NCPPB 3274 |
| Dsp-6-1 | Bacteria Proteobacteria Gammaproteobacteria Enterobacterales Pectobacteriaceae *Dickeya sp.* S1 |
| Dze-1-1 | Bacteria Proteobacteria Gammaproteobacteria Enterobacterales Pectobacteriaceae *Dickeya zeae* CSL RW192 |
| Dze-2-1 | Bacteria Proteobacteria Gammaproteobacteria Enterobacterales Pectobacteriaceae *Dickeya zeae* DZ2Q |
| Dze-3-1 | Bacteria Proteobacteria Gammaproteobacteria Enterobacterales Pectobacteriaceae *Dickeya zeae* EC1 |
| Dze-4-1 | Bacteria Proteobacteria Gammaproteobacteria Enterobacterales Pectobacteriaceae *Dickeya zeae* MK19 |
| Dze-5-1 | Bacteria Proteobacteria Gammaproteobacteria Enterobacterales Pectobacteriaceae *Dickeya zeae* NCPPB 2538 |
| Dze-6-1 | Bacteria Proteobacteria Gammaproteobacteria Enterobacterales Pectobacteriaceae *Dickeya zeae* NCPPB 3531 |
| Lqu-1-1 to Lqu-1-3 | Bacteria Proteobacteria Gammaproteobacteria Enterobacterales Pectobacteriaceae *Lonsdalea quercina subsp. quercina* |
| Pat-1-1 to Pat-1-2 | Bacteria Proteobacteria Gammaproteobacteria Enterobacterales Pectobacteriaceae *Pectobacterium atrosepticum* |
| Pat-2-1 | Bacteria Proteobacteria Gammaproteobacteria Enterobacterales Pectobacteriaceae *Pectobacterium atrosepticum* CFBP 6276 |
| Pat-3-1 | Bacteria Proteobacteria Gammaproteobacteria Enterobacterales Pectobacteriaceae *Pectobacterium atrosepticum* ICMP 1526 |
| Pbe-1-1 to Pbe-1-2 | Bacteria Proteobacteria Gammaproteobacteria Enterobacterales Pectobacteriaceae *Pectobacterium betavasculorum* |
| Pca-1-1 | Bacteria Proteobacteria Gammaproteobacteria Enterobacterales Pectobacteriaceae *Pectobacterium carotovorum* |
| Pca-2-1 | Bacteria Proteobacteria Gammaproteobacteria Enterobacterales Pectobacteriaceae *Pectobacterium carotovorum subsp. actinidiae* |
| Pca-3-1 to Pca-3-15 | Bacteria Proteobacteria Gammaproteobacteria Enterobacterales Pectobacteriaceae *Pectobacterium carotovorum subsp. brasiliense* |
| Pca-4-1 | Bacteria Proteobacteria Gammaproteobacteria Enterobacterales Pectobacteriaceae *Pectobacterium carotovorum subsp. brasiliensis* ICMP 19477 |
| Pca-5-1 to Pca-5-10 | Bacteria Proteobacteria Gammaproteobacteria Enterobacterales Pectobacteriaceae *Pectobacterium carotovorum subsp. carotovorum* |
| Pca-6-1 | Bacteria Proteobacteria Gammaproteobacteria Enterobacterales Pectobacteriaceae *Pectobacterium carotovorum subsp. carotovorum* ICMP 5702 |
| Pca-7-1 | Bacteria Proteobacteria Gammaproteobacteria Enterobacterales Pectobacteriaceae *Pectobacterium carotovorum subsp. carotovorum* UGC32 |
| Pca-8-1 to Pca-8-2 | Bacteria Proteobacteria Gammaproteobacteria Enterobacterales Pectobacteriaceae *Pectobacterium carotovorum subsp. odoriferum* |
| Ppa-1-1 | Bacteria Proteobacteria Gammaproteobacteria Enterobacterales Pectobacteriaceae *Pectobacterium parmentieri* |
| Ppa-2-1 | Bacteria Proteobacteria Gammaproteobacteria Enterobacterales Pectobacteriaceae *Pectobacterium parmentieri* CFIA1002 |
| Ppo-1-1 | Bacteria Proteobacteria Gammaproteobacteria Enterobacterales Pectobacteriaceae *Pectobacterium polaris* |
| Pwa-1-1 to Pwa-1-2 | Bacteria Proteobacteria Gammaproteobacteria Enterobacterales Pectobacteriaceae *Pectobacterium wasabiae* |
| CSo-1-1 | Bacteria Proteobacteria Gammaproteobacteria Enterobacterales Pectobacteriaceae Sodalis *Candidatus* Sodalis pierantonius str. SOPE |
| Spr-1-1 | Bacteria Proteobacteria Gammaproteobacteria Enterobacterales Pectobacteriaceae *Sodalis praecaptivus* |
| Pur-1-1 | Bacteria Proteobacteria Gammaproteobacteria Enterobacterales unclassified Enterobacterales *Phytobacter ursingii* |
| Cmu-2-1 | Bacteria Proteobacteria Gammaproteobacteria Enterobacterales Yersiniaceae *Chania multitudinisentens* RB-25 |
| Rch-1-1 | Bacteria Proteobacteria Gammaproteobacteria Enterobacterales Yersiniaceae *Rouxiella chamberiensis* |
| Sfo-1-1 | Bacteria Proteobacteria Gammaproteobacteria Enterobacterales Yersiniaceae *Serratia fonticola* |
| Sfo-2-1 | Bacteria Proteobacteria Gammaproteobacteria Enterobacterales Yersiniaceae *Serratia fonticola* LMG 7882 |
| Sfo-3-1 | Bacteria Proteobacteria Gammaproteobacteria Enterobacterales Yersiniaceae *Serratia fonticola* UTAD54 |
| Sgr-1-1 | Bacteria Proteobacteria Gammaproteobacteria Enterobacterales Yersiniaceae *Serratia grimesii* |
| Sli-1-1 | Bacteria Proteobacteria Gammaproteobacteria Enterobacterales Yersiniaceae *Serratia liquefaciens* |
| Sli-2-1 | Bacteria Proteobacteria Gammaproteobacteria Enterobacterales Yersiniaceae *Serratia liquefaciens* FK01 |
| Sma-1-1 to Sma-1-4 | Bacteria Proteobacteria Gammaproteobacteria Enterobacterales Yersiniaceae *Serratia marcescens* |
| Sma-2-1 | Bacteria Proteobacteria Gammaproteobacteria Enterobacterales Yersiniaceae *Serratia marcescens* AB42556419-isolate1 |
| Sma-3-1 | Bacteria Proteobacteria Gammaproteobacteria Enterobacterales Yersiniaceae *Serratia marcescens* BIDMC 44 |
| Sma-4-1 | Bacteria Proteobacteria Gammaproteobacteria Enterobacterales Yersiniaceae *Serratia marcescens* BIDMC 50 |
| Sma-5-1 | Bacteria Proteobacteria Gammaproteobacteria Enterobacterales Yersiniaceae *Serratia marcescens* BIDMC 80 |
| Sma-6-1 | Bacteria Proteobacteria Gammaproteobacteria Enterobacterales Yersiniaceae *Serratia marcescens* BIDMC 81 |
| Sma-7-1 | Bacteria Proteobacteria Gammaproteobacteria Enterobacterales Yersiniaceae *Serratia marcescens* H1q |
| Sma-8-1 | Bacteria Proteobacteria Gammaproteobacteria Enterobacterales Yersiniaceae *Serratia marcescens* LCT-SM166 |
| Sma-9-1 | Bacteria Proteobacteria Gammaproteobacteria Enterobacterales Yersiniaceae *Serratia marcescens* LCT-SM262 |
| Sma-10-1 | Bacteria Proteobacteria Gammaproteobacteria Enterobacterales Yersiniaceae *Serratia marcescens* MC458 |
| Sma-11-1 | Bacteria Proteobacteria Gammaproteobacteria Enterobacterales Yersiniaceae *Serratia marcescens* MC459 |

| | |
|---|---|
| Sma-12-1 | Bacteria Proteobacteria Gammaproteobacteria Enterobacterales Yersiniaceae *Serratia marcescens* MC460 |
| Sma-13-1 | Bacteria Proteobacteria Gammaproteobacteria Enterobacterales Yersiniaceae *Serratia marcescens* MC6000 |
| Sma-14-1 | Bacteria Proteobacteria Gammaproteobacteria Enterobacterales Yersiniaceae *Serratia marcescens* MC6001 |
| Sma-15-1 | Bacteria Proteobacteria Gammaproteobacteria Enterobacterales Yersiniaceae *Serratia marcescens* MC620 |
| Sma-16-1 | Bacteria Proteobacteria Gammaproteobacteria Enterobacterales Yersiniaceae *Serratia marcescens* PH1a |
| Sma-17-1 | Bacteria Proteobacteria Gammaproteobacteria Enterobacterales Yersiniaceae *Serratia marcescens* SM39 |
| Sma-18-1 to Sma-18-2 | Bacteria Proteobacteria Gammaproteobacteria Enterobacterales Yersiniaceae *Serratia marcescens subsp. marcescens* |
| Sma-19-1 | Bacteria Proteobacteria Gammaproteobacteria Enterobacterales Yersiniaceae *Serratia marcescens subsp. marcescens* ATCC 13880 |
| Sma-20-1 | Bacteria Proteobacteria Gammaproteobacteria Enterobacterales Yersiniaceae *Serratia marcescens subsp. marcescens* Db11 |
| Sne-1-1 | Bacteria Proteobacteria Gammaproteobacteria Enterobacterales Yersiniaceae *Serratia nematodiphila* DZ0503SBS1 |
| Spl-1-1 | Bacteria Proteobacteria Gammaproteobacteria Enterobacterales Yersiniaceae *Serratia plymuthica* RVH1 |
| Ssp-1-1 | Bacteria Proteobacteria Gammaproteobacteria Enterobacterales Yersiniaceae *Serratia sp.* 506_PEND |
| Ssp-2-1 | Bacteria Proteobacteria Gammaproteobacteria Enterobacterales Yersiniaceae *Serratia sp.* Ag1 |
| Ssp-3-1 | Bacteria Proteobacteria Gammaproteobacteria Enterobacterales Yersiniaceae *Serratia sp.* Ag2 |
| Ssp-4-1 | Bacteria Proteobacteria Gammaproteobacteria Enterobacterales Yersiniaceae *Serratia sp.* DD3 |
| Ssp-5-1 | Bacteria Proteobacteria Gammaproteobacteria Enterobacterales Yersiniaceae *Serratia sp.* FS14 |
| Ssp-6-1 | Bacteria Proteobacteria Gammaproteobacteria Enterobacterales Yersiniaceae *Serratia sp.* H1n |
| Ssp-7-1 | Bacteria Proteobacteria Gammaproteobacteria Enterobacterales Yersiniaceae *Serratia sp.* H1w |
| Ssp-8-1 | Bacteria Proteobacteria Gammaproteobacteria Enterobacterales Yersiniaceae *Serratia sp.* SCBI |
| Ssp-9-1 | Bacteria Proteobacteria Gammaproteobacteria Enterobacterales Yersiniaceae *Serratia sp.* TEL |
| Ssy-1-1 | Bacteria Proteobacteria Gammaproteobacteria Enterobacterales Yersiniaceae *Serratia symbiotica* |
| Sur-1-1 | Bacteria Proteobacteria Gammaproteobacteria Enterobacterales Yersiniaceae *Serratia ureilytica* |
| Yal-1-1 to Yal-1-4 | Bacteria Proteobacteria Gammaproteobacteria Enterobacterales Yersiniaceae *Yersinia aldovae* |
| Yal-2-1 | Bacteria Proteobacteria Gammaproteobacteria Enterobacterales Yersiniaceae *Yersinia aldovae* 670-83 |
| Yal-3-1 | Bacteria Proteobacteria Gammaproteobacteria Enterobacterales Yersiniaceae *Yersinia aleksiciae* |
| Ybe-1-1 | Bacteria Proteobacteria Gammaproteobacteria Enterobacterales Yersiniaceae *Yersinia bercovieri* |
| Yen-1-1 to Yen-1-4 | Bacteria Proteobacteria Gammaproteobacteria Enterobacterales Yersiniaceae *Yersinia enterocolitica* |
| Yen-2-1 | Bacteria Proteobacteria Gammaproteobacteria Enterobacterales Yersiniaceae *Yersinia enterocolitica (type* O:2) str. YE3094/96 |
| Yen-3-1 | Bacteria Proteobacteria Gammaproteobacteria Enterobacterales Yersiniaceae *Yersinia enterocolitica (type* O:5) str. YE53/03 |
| Yen-4-1 | Bacteria Proteobacteria Gammaproteobacteria Enterobacterales Yersiniaceae *Yersinia enterocolitica (type* O:9) str. YE56/03 |
| Yen-5-1 | Bacteria Proteobacteria Gammaproteobacteria Enterobacterales Yersiniaceae *Yersinia enterocolitica subsp. palearctica* 556/8265 |
| Yen-6-1 | Bacteria Proteobacteria Gammaproteobacteria Enterobacterales Yersiniaceae *Yersinia enterocolitica subsp. palearctica* 647/5307 |
| Yen-7-1 | Bacteria Proteobacteria Gammaproteobacteria Enterobacterales Yersiniaceae *Yersinia enterocolitica subsp. palearctica* YO527 |
| Yfr-1-1 to Yfr-1-17 | Bacteria Proteobacteria Gammaproteobacteria Enterobacterales Yersiniaceae *Yersinia frederiksenii* |
| Yfr-2-1 to Yfr-2-2 | Bacteria Proteobacteria Gammaproteobacteria Enterobacterales Yersiniaceae *Yersinia frederiksenii* ATCC 33641 |
| Yfr-3-1 | Bacteria Proteobacteria Gammaproteobacteria Enterobacterales Yersiniaceae *Yersinia frederiksenii* Y225 |
| Yin-1-1 | Bacteria Proteobacteria Gammaproteobacteria Enterobacterales Yersiniaceae *Yersinia intermedia* |
| Ykr-1-1 to Ykr-1-2 | Bacteria Proteobacteria Gammaproteobacteria Enterobacterales Yersiniaceae *Yersinia kristensenii* |
| Yma-1-1 | Bacteria Proteobacteria Gammaproteobacteria Enterobacterales Yersiniaceae *Yersinia massiliensis* |
| Yma-2-1 | Bacteria Proteobacteria Gammaproteobacteria Enterobacterales Yersiniaceae *Yersinia massiliensis* CCUG 53443 |
| Ymo-1-1 to Ymo-1-7 | Bacteria Proteobacteria Gammaproteobacteria Enterobacterales Yersiniaceae *Yersinia mollaretii* |
| Ynu-1-1 | Bacteria Proteobacteria Gammaproteobacteria Enterobacterales Yersiniaceae *Yersinia nurmii* |
| Ype-1-1 to Ype-1-2 | Bacteria Proteobacteria Gammaproteobacteria Enterobacterales Yersiniaceae *Yersinia pekkanenii* |
| Ype-2-1 to Ype-2-9 | Bacteria Proteobacteria Gammaproteobacteria Enterobacterales Yersiniaceae *Yersinia pestis* |
| Ype-3-1 | Bacteria Proteobacteria Gammaproteobacteria Enterobacterales Yersiniaceae *Yersinia pestis* 1045 |
| Ype-4-1 | Bacteria Proteobacteria Gammaproteobacteria Enterobacterales Yersiniaceae *Yersinia pestis* 12 |
| Ype-5-1 | Bacteria Proteobacteria Gammaproteobacteria Enterobacterales Yersiniaceae *Yersinia pestis* 1412 |
| Ype-6-1 | Bacteria Proteobacteria Gammaproteobacteria Enterobacterales Yersiniaceae *Yersinia pestis* 1413 |
| Ype-7-1 | Bacteria Proteobacteria Gammaproteobacteria Enterobacterales Yersiniaceae *Yersinia pestis* 14735 |
| Ype-8-1 | Bacteria Proteobacteria Gammaproteobacteria Enterobacterales Yersiniaceae *Yersinia pestis* 1522 |
| Ype-9-1 | Bacteria Proteobacteria Gammaproteobacteria Enterobacterales Yersiniaceae *Yersinia pestis* 164 |
| Ype-10-1 | Bacteria Proteobacteria Gammaproteobacteria Enterobacterales Yersiniaceae *Yersinia pestis* 1670 |
| Ype-11-1 | Bacteria Proteobacteria Gammaproteobacteria Enterobacterales Yersiniaceae *Yersinia pestis* 2 |
| Ype-12-1 | Bacteria Proteobacteria Gammaproteobacteria Enterobacterales Yersiniaceae *Yersinia pestis* 2944 |
| Ype-13-1 | Bacteria Proteobacteria Gammaproteobacteria Enterobacterales Yersiniaceae *Yersinia pestis* 3067 |
| Ype-14-1 | Bacteria Proteobacteria Gammaproteobacteria Enterobacterales Yersiniaceae *Yersinia pestis* 34008 |
| Ype-15-1 | Bacteria Proteobacteria Gammaproteobacteria Enterobacterales Yersiniaceae *Yersinia pestis* 34202 |
| Ype-16-1 | Bacteria Proteobacteria Gammaproteobacteria Enterobacterales Yersiniaceae *Yersinia pestis* 351001 |
| Ype-17-1 | Bacteria Proteobacteria Gammaproteobacteria Enterobacterales Yersiniaceae *Yersinia pestis* 3770 |
| Ype-18-1 | Bacteria Proteobacteria Gammaproteobacteria Enterobacterales Yersiniaceae *Yersinia pestis* 42095 |
| Ype-19-1 | Bacteria Proteobacteria Gammaproteobacteria Enterobacterales Yersiniaceae *Yersinia pestis* 5 |
| Ype-20-1 | Bacteria Proteobacteria Gammaproteobacteria Enterobacterales Yersiniaceae *Yersinia pestis* 620024 |
| Ype-21-1 | Bacteria Proteobacteria Gammaproteobacteria Enterobacterales Yersiniaceae *Yersinia pestis* 71021 |
| Ype-22-1 | Bacteria Proteobacteria Gammaproteobacteria Enterobacterales Yersiniaceae *Yersinia pestis* 710317 |
| Ype-23-1 | Bacteria Proteobacteria Gammaproteobacteria Enterobacterales Yersiniaceae *Yersinia pestis* 7338 |

| | |
|---|---|
| Ype-24-1 | Bacteria Proteobacteria Gammaproteobacteria Enterobacterales Yersiniaceae *Yersinia pestis* 790 |
| Ype-25-1 | Bacteria Proteobacteria Gammaproteobacteria Enterobacterales Yersiniaceae *Yersinia pestis* 8787 |
| Ype-26-1 | Bacteria Proteobacteria Gammaproteobacteria Enterobacterales Yersiniaceae *Yersinia pestis* 9 |
| Ype-27-1 | Bacteria Proteobacteria Gammaproteobacteria Enterobacterales Yersiniaceae *Yersinia pestis* 945 |
| Ype-28-1 | Bacteria Proteobacteria Gammaproteobacteria Enterobacterales Yersiniaceae *Yersinia pestis* 970754 |
| Ype-29-1 to Ype-29-2 | Bacteria Proteobacteria Gammaproteobacteria Enterobacterales Yersiniaceae *Yersinia pestis* A1122 |
| Ype-30-1 | Bacteria Proteobacteria Gammaproteobacteria Enterobacterales Yersiniaceae *Yersinia pestis* A1973001 |
| Ype-31-1 to Ype-31-2 | Bacteria Proteobacteria Gammaproteobacteria Enterobacterales Yersiniaceae *Yersinia pestis* Angola |
| Ype-32-1 to Ype-32-2 | Bacteria Proteobacteria Gammaproteobacteria Enterobacterales Yersiniaceae *Yersinia pestis* Antiqua |
| Ype-104-1 | Bacteria Proteobacteria Gammaproteobacteria Enterobacterales Yersiniaceae *Yersinia pestis biovar* Orientalis str. AS200901156 |
| Ype-105-1 | Bacteria Proteobacteria Gammaproteobacteria Enterobacterales Yersiniaceae *Yersinia pestis biovar* Orientalis str. AS200901434 |
| Ype-106-1 | Bacteria Proteobacteria Gammaproteobacteria Enterobacterales Yersiniaceae *Yersinia pestis biovar* Orientalis str. AS200901509 |
| Ype-107-1 | Bacteria Proteobacteria Gammaproteobacteria Enterobacterales Yersiniaceae *Yersinia pestis biovar* Orientalis str. AS200901539 |
| Ype-108-1 | Bacteria Proteobacteria Gammaproteobacteria Enterobacterales Yersiniaceae *Yersinia pestis biovar* Orientalis str. AS200902147 |
| Ype-109-1 | Bacteria Proteobacteria Gammaproteobacteria Enterobacterales Yersiniaceae *Yersinia pestis biovar* Orientalis str. BA200901703 |
| Ype-110-1 | Bacteria Proteobacteria Gammaproteobacteria Enterobacterales Yersiniaceae *Yersinia pestis biovar* Orientalis str. BA200901799 |
| Ype-111-1 | Bacteria Proteobacteria Gammaproteobacteria Enterobacterales Yersiniaceae *Yersinia pestis biovar* Orientalis str. BA200901990 |
| Ype-112-1 | Bacteria Proteobacteria Gammaproteobacteria Enterobacterales Yersiniaceae *Yersinia pestis biovar* Orientalis str. BA200902009 |
| Ype-33-1 | Bacteria Proteobacteria Gammaproteobacteria Enterobacterales Yersiniaceae *Yersinia pestis* C1975003 |
| Ype-34-1 | Bacteria Proteobacteria Gammaproteobacteria Enterobacterales Yersiniaceae *Yersinia pestis* CMCC02041 |
| Ype-35-1 | Bacteria Proteobacteria Gammaproteobacteria Enterobacterales Yersiniaceae *Yersinia pestis* CMCC03001 |
| Ype-36-1 | Bacteria Proteobacteria Gammaproteobacteria Enterobacterales Yersiniaceae *Yersinia pestis* CMCC05009 |
| Ype-37-1 | Bacteria Proteobacteria Gammaproteobacteria Enterobacterales Yersiniaceae *Yersinia pestis* CMCC05013 |
| Ype-38-1 | Bacteria Proteobacteria Gammaproteobacteria Enterobacterales Yersiniaceae *Yersinia pestis* CMCC10012 |
| Ype-39-1 | Bacteria Proteobacteria Gammaproteobacteria Enterobacterales Yersiniaceae *Yersinia pestis* CMCC104003 |
| Ype-40-1 | Bacteria Proteobacteria Gammaproteobacteria Enterobacterales Yersiniaceae *Yersinia pestis* CMCC106002 |
| Ype-41-1 | Bacteria Proteobacteria Gammaproteobacteria Enterobacterales Yersiniaceae *Yersinia pestis* CMCC11001 |
| Ype-42-1 | Bacteria Proteobacteria Gammaproteobacteria Enterobacterales Yersiniaceae *Yersinia pestis* CMCC114001 |
| Ype-43-1 | Bacteria Proteobacteria Gammaproteobacteria Enterobacterales Yersiniaceae *Yersinia pestis* CMCC12003 |
| Ype-44-1 | Bacteria Proteobacteria Gammaproteobacteria Enterobacterales Yersiniaceae *Yersinia pestis* CMCC18019 |
| Ype-45-1 | Bacteria Proteobacteria Gammaproteobacteria Enterobacterales Yersiniaceae *Yersinia pestis* CMCC21106 |
| Ype-46-1 | Bacteria Proteobacteria Gammaproteobacteria Enterobacterales Yersiniaceae *Yersinia pestis* CMCC27002 |
| Ype-47-1 | Bacteria Proteobacteria Gammaproteobacteria Enterobacterales Yersiniaceae *Yersinia pestis* CMCC31004 |
| Ype-48-1 | Bacteria Proteobacteria Gammaproteobacteria Enterobacterales Yersiniaceae *Yersinia pestis* CMCC347001 |
| Ype-49-1 | Bacteria Proteobacteria Gammaproteobacteria Enterobacterales Yersiniaceae *Yersinia pestis* CMCC38001 |
| Ype-50-1 | Bacteria Proteobacteria Gammaproteobacteria Enterobacterales Yersiniaceae *Yersinia pestis* CMCC42007 |
| Ype-51-1 | Bacteria Proteobacteria Gammaproteobacteria Enterobacterales Yersiniaceae *Yersinia pestis* CMCC43032 |
| Ype-52-1 | Bacteria Proteobacteria Gammaproteobacteria Enterobacterales Yersiniaceae *Yersinia pestis* CMCC51020 |
| Ype-53-1 | Bacteria Proteobacteria Gammaproteobacteria Enterobacterales Yersiniaceae *Yersinia pestis* CMCC640047 |
| Ype-54-1 | Bacteria Proteobacteria Gammaproteobacteria Enterobacterales Yersiniaceae *Yersinia pestis* CMCC67001 |
| Ype-55-1 | Bacteria Proteobacteria Gammaproteobacteria Enterobacterales Yersiniaceae *Yersinia pestis* CMCC71001 |
| Ype-56-1 | Bacteria Proteobacteria Gammaproteobacteria Enterobacterales Yersiniaceae *Yersinia pestis* CMCC84033 |
| Ype-57-1 | Bacteria Proteobacteria Gammaproteobacteria Enterobacterales Yersiniaceae *Yersinia pestis* CMCC84046 |
| Ype-58-1 | Bacteria Proteobacteria Gammaproteobacteria Enterobacterales Yersiniaceae *Yersinia pestis* CMCC90027 |
| Ype-59-1 | Bacteria Proteobacteria Gammaproteobacteria Enterobacterales Yersiniaceae *Yersinia pestis* CMCC92004 |
| Ype-60-1 | Bacteria Proteobacteria Gammaproteobacteria Enterobacterales Yersiniaceae *Yersinia pestis* CMCC92010 |
| Ype-61-1 | Bacteria Proteobacteria Gammaproteobacteria Enterobacterales Yersiniaceae *Yersinia pestis* CMCC95001 |
| Ype-62-1 | Bacteria Proteobacteria Gammaproteobacteria Enterobacterales Yersiniaceae *Yersinia pestis* CMCC96007 |
| Ype-63-1 | Bacteria Proteobacteria Gammaproteobacteria Enterobacterales Yersiniaceae *Yersinia pestis* CMCC99103 |
| Ype-64-1 | Bacteria Proteobacteria Gammaproteobacteria Enterobacterales Yersiniaceae *Yersinia pestis* CMCCK100001 |
| Ype-65-1 | Bacteria Proteobacteria Gammaproteobacteria Enterobacterales Yersiniaceae *Yersinia pestis* CMCCK110001 |
| Ype-66-1 | Bacteria Proteobacteria Gammaproteobacteria Enterobacterales Yersiniaceae *Yersinia pestis* CMCCN010025 |
| Ype-67-1 | Bacteria Proteobacteria Gammaproteobacteria Enterobacterales Yersiniaceae *Yersinia pestis* D1964001 |
| Ype-68-1 | Bacteria Proteobacteria Gammaproteobacteria Enterobacterales Yersiniaceae *Yersinia pestis* D1964002 |
| Ype-69-1 | Bacteria Proteobacteria Gammaproteobacteria Enterobacterales Yersiniaceae *Yersinia pestis* D1991004 |
| Ype-70-1 | Bacteria Proteobacteria Gammaproteobacteria Enterobacterales Yersiniaceae *Yersinia pestis* E1977001 |
| Ype-71-1 | Bacteria Proteobacteria Gammaproteobacteria Enterobacterales Yersiniaceae *Yersinia pestis* EBD10-058 |
| Ype-72-1 | Bacteria Proteobacteria Gammaproteobacteria Enterobacterales Yersiniaceae *Yersinia pestis* EV76-CN |
| Ype-73-1 | Bacteria Proteobacteria Gammaproteobacteria Enterobacterales Yersiniaceae *Yersinia pestis* F1946001 |
| Ype-74-1 | Bacteria Proteobacteria Gammaproteobacteria Enterobacterales Yersiniaceae *Yersinia pestis* F1954001 |
| Ype-75-1 | Bacteria Proteobacteria Gammaproteobacteria Enterobacterales Yersiniaceae *Yersinia pestis* F1984001 |
| Ype-76-1 | Bacteria Proteobacteria Gammaproteobacteria Enterobacterales Yersiniaceae *Yersinia pestis* G1670A |
| Ype-77-1 | Bacteria Proteobacteria Gammaproteobacteria Enterobacterales Yersiniaceae *Yersinia pestis* G1670E |
| Ype-78-1 | Bacteria Proteobacteria Gammaproteobacteria Enterobacterales Yersiniaceae *Yersinia pestis* H1958004 |
| Ype-79-1 | Bacteria Proteobacteria Gammaproteobacteria Enterobacterales Yersiniaceae *Yersinia pestis* I160001 |

| | |
|---|---|
| Ype-80-1 | Bacteria Proteobacteria Gammaproteobacteria Enterobacterales Yersiniaceae *Yersinia pestis* I1969003 |
| Ype-81-1 | Bacteria Proteobacteria Gammaproteobacteria Enterobacterales Yersiniaceae *Yersinia pestis* I1970005 |
| Ype-82-1 | Bacteria Proteobacteria Gammaproteobacteria Enterobacterales Yersiniaceae *Yersinia pestis* I1991001 |
| Ype-83-1 | Bacteria Proteobacteria Gammaproteobacteria Enterobacterales Yersiniaceae *Yersinia pestis* I1994006 |
| Ype-84-1 | Bacteria Proteobacteria Gammaproteobacteria Enterobacterales Yersiniaceae *Yersinia pestis* I2001001 |
| Ype-85-1 | Bacteria Proteobacteria Gammaproteobacteria Enterobacterales Yersiniaceae *Yersinia pestis* J1963002 |
| Ype-86-1 | Bacteria Proteobacteria Gammaproteobacteria Enterobacterales Yersiniaceae *Yersinia pestis* J1978002 |
| Ype-87-1 | Bacteria Proteobacteria Gammaproteobacteria Enterobacterales Yersiniaceae *Yersinia pestis* K21985002 |
| Ype-88-1 | Bacteria Proteobacteria Gammaproteobacteria Enterobacterales Yersiniaceae *Yersinia pestis* M0000002 |
| Ype-89-1 | Bacteria Proteobacteria Gammaproteobacteria Enterobacterales Yersiniaceae *Yersinia pestis* MGJZ12 |
| Ype-90-1 | Bacteria Proteobacteria Gammaproteobacteria Enterobacterales Yersiniaceae *Yersinia pestis* MGJZ3 |
| Ype-91-1 | Bacteria Proteobacteria Gammaproteobacteria Enterobacterales Yersiniaceae *Yersinia pestis* MGJZ6 |
| Ype-92-1 | Bacteria Proteobacteria Gammaproteobacteria Enterobacterales Yersiniaceae *Yersinia pestis* MGJZ7 |
| Ype-93-1 | Bacteria Proteobacteria Gammaproteobacteria Enterobacterales Yersiniaceae *Yersinia pestis* MGJZ9 |
| Ype-94-1 to Ype-94-2 | Bacteria Proteobacteria Gammaproteobacteria Enterobacterales Yersiniaceae *Yersinia pestis* Pestoides F |
| Ype-95-1 | Bacteria Proteobacteria Gammaproteobacteria Enterobacterales Yersiniaceae *Yersinia pestis* Pestoides G |
| Ype-96-1 | Bacteria Proteobacteria Gammaproteobacteria Enterobacterales Yersiniaceae *Yersinia pestis* SHAN11 |
| Ype-97-1 | Bacteria Proteobacteria Gammaproteobacteria Enterobacterales Yersiniaceae *Yersinia pestis* SHAN12 |
| Ype-113-1 | Bacteria Proteobacteria Gammaproteobacteria Enterobacterales Yersiniaceae *Yersinia pestis str.* Pestoides B |
| Ype-114-1 to Ype-114-2 | Bacteria Proteobacteria Gammaproteobacteria Enterobacterales Yersiniaceae *Yersinia pestis subsp. microtus bv.* Ulegeica |
| Ype-115-1 | Bacteria Proteobacteria Gammaproteobacteria Enterobacterales Yersiniaceae *Yersinia pestis subsp. pestis bv.* Orientalis |
| Ype-98-1 | Bacteria Proteobacteria Gammaproteobacteria Enterobacterales Yersiniaceae *Yersinia pestis* YN1065 |
| Ype-99-1 | Bacteria Proteobacteria Gammaproteobacteria Enterobacterales Yersiniaceae *Yersinia pestis* YN1683 |
| Ype-100-1 | Bacteria Proteobacteria Gammaproteobacteria Enterobacterales Yersiniaceae *Yersinia pestis* YN2179 |
| Ype-101-1 | Bacteria Proteobacteria Gammaproteobacteria Enterobacterales Yersiniaceae *Yersinia pestis* YN2588 |
| Ype-102-1 | Bacteria Proteobacteria Gammaproteobacteria Enterobacterales Yersiniaceae *Yersinia pestis* YN472 |
| Ype-103-1 | Bacteria Proteobacteria Gammaproteobacteria Enterobacterales Yersiniaceae *Yersinia pestis* YN663 |
| Yps-1-1 to Yps-1-4 | Bacteria Proteobacteria Gammaproteobacteria Enterobacterales Yersiniaceae *Yersinia pseudotuberculosis* |
| Yps-2-1 | Bacteria Proteobacteria Gammaproteobacteria Enterobacterales Yersiniaceae *Yersinia pseudotuberculosis* B-6796 |
| Yps-3-1 | Bacteria Proteobacteria Gammaproteobacteria Enterobacterales Yersiniaceae *Yersinia pseudotuberculosis* B-6862 |
| Yps-4-1 | Bacteria Proteobacteria Gammaproteobacteria Enterobacterales Yersiniaceae *Yersinia pseudotuberculosis* B-6863 |
| Yps-5-1 | Bacteria Proteobacteria Gammaproteobacteria Enterobacterales Yersiniaceae *Yersinia pseudotuberculosis* B-6864 |
| Yps-6-1 | Bacteria Proteobacteria Gammaproteobacteria Enterobacterales Yersiniaceae *Yersinia pseudotuberculosis* B-6865 |
| Yps-7-1 | Bacteria Proteobacteria Gammaproteobacteria Enterobacterales Yersiniaceae *Yersinia pseudotuberculosis* B-6866 |
| Yps-8-1 | Bacteria Proteobacteria Gammaproteobacteria Enterobacterales Yersiniaceae *Yersinia pseudotuberculosis* B-7194 |
| Yps-9-1 | Bacteria Proteobacteria Gammaproteobacteria Enterobacterales Yersiniaceae *Yersinia pseudotuberculosis* B-7195 |
| Yps-10-1 to Yps-10-2 | Bacteria Proteobacteria Gammaproteobacteria Enterobacterales Yersiniaceae *Yersinia pseudotuberculosis* IP 32953 |
| Yps-11-1 to Yps-11-2 | Bacteria Proteobacteria Gammaproteobacteria Enterobacterales Yersiniaceae *Yersinia pseudotuberculosis* PB1/+ |
| Yps-13-1 | Bacteria Proteobacteria Gammaproteobacteria Enterobacterales Yersiniaceae *Yersinia pseudotuberculosis str.* PA3606 |
| Yps-12-1 to Yps-12-2 | Bacteria Proteobacteria Gammaproteobacteria Enterobacterales Yersiniaceae *Yersinia pseudotuberculosis* YPIII |
| Yro-1-1 | Bacteria Proteobacteria Gammaproteobacteria Enterobacterales Yersiniaceae *Yersinia rohdei* |
| Yru-1-1 to Yru-1-3 | Bacteria Proteobacteria Gammaproteobacteria Enterobacterales Yersiniaceae *Yersinia ruckeri* |
| Yru-2-1 to Yru-2-2 | Bacteria Proteobacteria Gammaproteobacteria Enterobacterales Yersiniaceae *Yersinia ruckeri* ATCC 29473 |
| Ysi-1-1 | Bacteria Proteobacteria Gammaproteobacteria Enterobacterales Yersiniaceae *Yersinia similis* |
| Ywa-1-1 to Ywa-1-2 | Bacteria Proteobacteria Gammaproteobacteria Enterobacterales Yersiniaceae *Yersinia wautersii* |
| CRe-1-1 | Bacteria Proteobacteria Gammaproteobacteria Enterobacteriales Enterobacteriaceae aphid secondary symbionts *Candidatus* Regiella insecticola LSR1 |
| CRe-2-1 | Bacteria Proteobacteria Gammaproteobacteria Enterobacteriales Enterobacteriaceae aphid secondary symbionts *Candidatus* Regiella insecticola R5.15 |
| Bsp-1-1 | Bacteria Proteobacteria Gammaproteobacteria Enterobacteriales Enterobacteriaceae *Brenneria sp.* EniD312 |
| Cda-1-1 | Bacteria Proteobacteria Gammaproteobacteria Enterobacteriales Enterobacteriaceae *Cedecea davisae* DSM 4568 |
| Cfr-1-1 to Cfr-1-4 | Bacteria Proteobacteria Gammaproteobacteria Enterobacteriales Enterobacteriaceae Citrobacter Citrobacter freundii complex *Citrobacter freundii* |
| Cfr-9-1 | Bacteria Proteobacteria Gammaproteobacteria Enterobacteriales Enterobacteriaceae Citrobacter Citrobacter freundii complex *Citrobacter freundii* 4_7_47CFAA |
| Cfr-2-1 to Cfr-2-2 | Bacteria Proteobacteria Gammaproteobacteria Enterobacteriales Enterobacteriaceae Citrobacter Citrobacter freundii complex *Citrobacter freundii* ATCC 8090 = MTCC 1658 |
| Cfr-10-1 | Bacteria Proteobacteria Gammaproteobacteria Enterobacteriales Enterobacteriaceae Citrobacter Citrobacter freundii complex *Citrobacter freundii* GTC 09479 |
| Cfr-11-1 | Bacteria Proteobacteria Gammaproteobacteria Enterobacteriales Enterobacteriaceae Citrobacter Citrobacter freundii complex *Citrobacter freundii* GTC 09629 |
| Cko-1-1 to Cko-1-2 | Bacteria Proteobacteria Gammaproteobacteria Enterobacteriales Enterobacteriaceae *Citrobacter koseri* |
| Cko-2-1 | Bacteria Proteobacteria Gammaproteobacteria Enterobacteriales Enterobacteriaceae *Citrobacter koseri* ATCC BAA-895 |
| Cro-3-1 | Bacteria Proteobacteria Gammaproteobacteria Enterobacteriales Enterobacteriaceae *Citrobacter rodentium* ICC168 |
| Csp-14-1 | Bacteria Proteobacteria Gammaproteobacteria Enterobacteriales Enterobacteriaceae *Citrobacter sp.* 30_2 |
| Csp-15-1 | Bacteria Proteobacteria Gammaproteobacteria Enterobacteriales Enterobacteriaceae *Citrobacter sp.* A1 |

| | |
|---|---|
| Csp-16-1 | Bacteria Proteobacteria Gammaproteobacteria Enterobacteriales Enterobacteriaceae *Citrobacter sp.* KTE151 |
| Csp-17-1 | Bacteria Proteobacteria Gammaproteobacteria Enterobacteriales Enterobacteriaceae *Citrobacter sp.* KTE30 |
| Csp-18-1 | Bacteria Proteobacteria Gammaproteobacteria Enterobacteriales Enterobacteriaceae *Citrobacter sp.* KTE32 |
| Csp-19-1 | Bacteria Proteobacteria Gammaproteobacteria Enterobacteriales Enterobacteriaceae *Citrobacter sp.* L17 |
| Cyo-1-1 | Bacteria Proteobacteria Gammaproteobacteria Enterobacteriales Enterobacteriaceae *Citrobacter youngae* ATCC 29220 |
| Cco-1-1 | Bacteria Proteobacteria Gammaproteobacteria Enterobacteriales Enterobacteriaceae *Cronobacter condimenti* 1330 |
| Cdu-4-1 | Bacteria Proteobacteria Gammaproteobacteria Enterobacteriales Enterobacteriaceae *Cronobacter dublinensis* 1210 |
| Cdu-5-1 | Bacteria Proteobacteria Gammaproteobacteria Enterobacteriales Enterobacteriaceae *Cronobacter dublinensis* 582 |
| Cma-4-1 | Bacteria Proteobacteria Gammaproteobacteria Enterobacteriales Enterobacteriaceae *Cronobacter malonaticus* 507 |
| Cma-5-1 | Bacteria Proteobacteria Gammaproteobacteria Enterobacteriales Enterobacteriaceae *Cronobacter malonaticus* 681 |
| Csa-9-1 | Bacteria Proteobacteria Gammaproteobacteria Enterobacteriales Enterobacteriaceae *Cronobacter sakazakii* 680 |
| Csa-10-1 | Bacteria Proteobacteria Gammaproteobacteria Enterobacteriales Enterobacteriaceae *Cronobacter sakazakii* 696 |
| Csa-11-1 | Bacteria Proteobacteria Gammaproteobacteria Enterobacteriales Enterobacteriaceae *Cronobacter sakazakii* 701 |
| Csa-12-1 | Bacteria Proteobacteria Gammaproteobacteria Enterobacteriales Enterobacteriaceae *Cronobacter sakazakii* ATCC BAA-894 |
| Csa-13-1 | Bacteria Proteobacteria Gammaproteobacteria Enterobacteriales Enterobacteriaceae *Cronobacter sakazakii* CMCC 45402 |
| Csa-14-1 | Bacteria Proteobacteria Gammaproteobacteria Enterobacteriales Enterobacteriaceae *Cronobacter sakazakii* E899 |
| Csa-15-1 | Bacteria Proteobacteria Gammaproteobacteria Enterobacteriales Enterobacteriaceae *Cronobacter sakazakii* ES15 |
| Csa-16-1 | Bacteria Proteobacteria Gammaproteobacteria Enterobacteriales Enterobacteriaceae *Cronobacter sakazakii* SP291 |
| Ctu-3-1 | Bacteria Proteobacteria Gammaproteobacteria Enterobacteriales Enterobacteriaceae *Cronobacter turicensis* 564 |
| Ctu-4-1 | Bacteria Proteobacteria Gammaproteobacteria Enterobacteriales Enterobacteriaceae *Cronobacter turicensis z*3032 |
| Cun-1-1 to Cun-1-2 | Bacteria Proteobacteria Gammaproteobacteria Enterobacteriales Enterobacteriaceae *Cronobacter universalis* NCTC 9529 |
| Dda-4-1 | Bacteria Proteobacteria Gammaproteobacteria Enterobacteriales Enterobacteriaceae *Dickeya dadantii* 3937 |
| Dda-5-1 | Bacteria Proteobacteria Gammaproteobacteria Enterobacteriales Enterobacteriaceae *Dickeya dadantii* Ech586 |
| Ddi-3-1 | Bacteria Proteobacteria Gammaproteobacteria Enterobacteriales Enterobacteriaceae *Dickeya dianthicola* GBBC 2039 |
| Ddi-4-1 | Bacteria Proteobacteria Gammaproteobacteria Enterobacteriales Enterobacteriaceae *Dickeya dianthicola* NCPPB 3534 |
| Ddi-5-1 | Bacteria Proteobacteria Gammaproteobacteria Enterobacteriales Enterobacteriaceae *Dickeya dianthicola* NCPPB 453 |
| Dpa-1-1 | Bacteria Proteobacteria Gammaproteobacteria Enterobacteriales Enterobacteriaceae *Dickeya paradisiaca* NCPPB 2511 |
| Dso-2-1 | Bacteria Proteobacteria Gammaproteobacteria Enterobacteriales Enterobacteriaceae *Dickeya solani* GBBC 2040 |
| Dso-3-1 | Bacteria Proteobacteria Gammaproteobacteria Enterobacteriales Enterobacteriaceae *Dickeya solani* IPO 2222 |
| Dso-4-1 | Bacteria Proteobacteria Gammaproteobacteria Enterobacteriales Enterobacteriaceae *Dickeya solani* MK10 |
| Dso-5-1 | Bacteria Proteobacteria Gammaproteobacteria Enterobacteriales Enterobacteriaceae *Dickeya solani* MK16 |
| Dso-6-1 | Bacteria Proteobacteria Gammaproteobacteria Enterobacteriales Enterobacteriaceae *Dickeya solani* RNS 08.23.3.1.A |
| Dsp-7-1 | Bacteria Proteobacteria Gammaproteobacteria Enterobacteriales Enterobacteriaceae *Dickeya sp.* CSL RW240 |
| Dsp-8-1 | Bacteria Proteobacteria Gammaproteobacteria Enterobacteriales Enterobacteriaceae *Dickeya sp.* D s0432-1 |
| Dsp-9-1 | Bacteria Proteobacteria Gammaproteobacteria Enterobacteriales Enterobacteriaceae *Dickeya sp.* NCPPB 569 |
| Dze-7-1 | Bacteria Proteobacteria Gammaproteobacteria Enterobacteriales Enterobacteriaceae *Dickeya zeae* MS1 |
| Dze-8-1 | Bacteria Proteobacteria Gammaproteobacteria Enterobacteriales Enterobacteriaceae *Dickeya zeae* NCPPB 3532 |
| Dze-9-1 | Bacteria Proteobacteria Gammaproteobacteria Enterobacteriales Enterobacteriaceae *Dickeya zeae* ZJU1202 |
| Eic-2-1 | Bacteria Proteobacteria Gammaproteobacteria Enterobacteriales Enterobacteriaceae *Edwardsiella ictaluri* 93-146 |
| Eta-4-1 | Bacteria Proteobacteria Gammaproteobacteria Enterobacteriales Enterobacteriaceae *Edwardsiella tarda* ATCC 23685 |
| Eta-5-1 | Bacteria Proteobacteria Gammaproteobacteria Enterobacteriales Enterobacteriaceae *Edwardsiella tarda* C07-087 |
| Eta-6-1 | Bacteria Proteobacteria Gammaproteobacteria Enterobacteriales Enterobacteriaceae *Edwardsiella tarda* EIB202 |
| Eta-7-1 | Bacteria Proteobacteria Gammaproteobacteria Enterobacteriales Enterobacteriaceae *Edwardsiella tarda* FL6-60 |
| Eta-8-1 | Bacteria Proteobacteria Gammaproteobacteria Enterobacteriales Enterobacteriaceae *Edwardsiella tarda* NBRC 105688 |
| Eae-1-1 | Bacteria Proteobacteria Gammaproteobacteria Enterobacteriales Enterobacteriaceae *Enterobacter aerogenes* EA1509E |
| Eae-2-1 | Bacteria Proteobacteria Gammaproteobacteria Enterobacteriales Enterobacteriaceae *Enterobacter aerogenes* FGI35 |
| Eae-3-1 | Bacteria Proteobacteria Gammaproteobacteria Enterobacteriales Enterobacteriaceae *Enterobacter aerogenes* KCTC 2190 |
| Eca-3-1 | Bacteria Proteobacteria Gammaproteobacteria Enterobacteriales Enterobacteriaceae *Enterobacter cancerogenus* ATCC 35316 |
| Eas-1-1 to Eas-1-2 | Bacteria Proteobacteria Gammaproteobacteria Enterobacteriales Enterobacteriaceae Enterobacter Enterobacter cloacae complex *Enterobacter asburiae* |
| Eas-4-1 | Bacteria Proteobacteria Gammaproteobacteria Enterobacteriales Enterobacteriaceae Enterobacter Enterobacter cloacae complex *Enterobacter asburiae* LF7a |
| Ecl-1-1 to Ecl-1-12 | Bacteria Proteobacteria Gammaproteobacteria Enterobacteriales Enterobacteriaceae Enterobacter Enterobacter cloacae complex *Enterobacter cloacae* |
| Ecl-55-1 | Bacteria Proteobacteria Gammaproteobacteria Enterobacteriales Enterobacteriaceae Enterobacter Enterobacter cloacae complex *Enterobacter cloacae* BWH 29 |
| Ecl-56-1 | Bacteria Proteobacteria Gammaproteobacteria Enterobacteriales Enterobacteriaceae Enterobacter Enterobacter cloacae complex *Enterobacter cloacae* BWH 31 |
| Ecl-57-1 | Bacteria Proteobacteria Gammaproteobacteria Enterobacteriales Enterobacteriaceae Enterobacter Enterobacter cloacae complex *Enterobacter cloacae* EC_38VIM1 |
| Ecl-58-1 | Bacteria Proteobacteria Gammaproteobacteria Enterobacteriales Enterobacteriaceae Enterobacter Enterobacter cloacae complex *Enterobacter cloacae* EcWSU1 |
| Ecl-59-1 | Bacteria Proteobacteria Gammaproteobacteria Enterobacteriales Enterobacteriaceae Enterobacter Enterobacter cloacae complex *Enterobacter cloacae* S611 |
| Ecl-60-1 | Bacteria Proteobacteria Gammaproteobacteria Enterobacteriales Enterobacteriaceae Enterobacter Enterobacter cloacae complex *Enterobacter cloacae* SCF1 |
| Ecl-66-1 | Bacteria Proteobacteria Gammaproteobacteria Enterobacteriales Enterobacteriaceae Enterobacter Enterobacter cloacae complex *Enterobacter cloacae str.* Hanford |

| | |
|---|---|
| Ecl-52-1 to Ecl-52-5 | Bacteria Proteobacteria Gammaproteobacteria Enterobacteriales Enterobacteriaceae Enterobacter Enterobacter cloacae complex *Enterobacter cloacae subsp. cloacae* |
| Ecl-67-1 | Bacteria Proteobacteria Gammaproteobacteria Enterobacteriales Enterobacteriaceae Enterobacter Enterobacter cloacae complex *Enterobacter cloacae subsp. cloacae* 08XA1 |
| Ecl-68-1 | Bacteria Proteobacteria Gammaproteobacteria Enterobacteriales Enterobacteriaceae Enterobacter Enterobacter cloacae complex *Enterobacter cloacae subsp. cloacae* ATCC 13047 |
| Ecl-69-1 | Bacteria Proteobacteria Gammaproteobacteria Enterobacteriales Enterobacteriaceae Enterobacter Enterobacter cloacae complex *Enterobacter cloacae subsp. cloacae* ENHKU01 |
| Ecl-70-1 | Bacteria Proteobacteria Gammaproteobacteria Enterobacteriales Enterobacteriaceae Enterobacter Enterobacter cloacae complex *Enterobacter cloacae subsp. dissolvens* SDM |
| Ecl-61-1 | Bacteria Proteobacteria Gammaproteobacteria Enterobacteriales Enterobacteriaceae Enterobacter Enterobacter cloacae complex *Enterobacter cloacae* UCI-CRE 11 |
| Ecl-62-1 | Bacteria Proteobacteria Gammaproteobacteria Enterobacteriales Enterobacteriaceae Enterobacter Enterobacter cloacae complex *Enterobacter cloacae* UCI-CRE 12 |
| Ecl-63-1 | Bacteria Proteobacteria Gammaproteobacteria Enterobacteriales Enterobacteriaceae Enterobacter Enterobacter cloacae complex *Enterobacter cloacae* UCI-CRE 3 |
| Ecl-64-1 | Bacteria Proteobacteria Gammaproteobacteria Enterobacteriales Enterobacteriaceae Enterobacter Enterobacter cloacae complex *Enterobacter cloacae* UCI-CRE 5 |
| Ecl-65-1 | Bacteria Proteobacteria Gammaproteobacteria Enterobacteriales Enterobacteriaceae Enterobacter Enterobacter cloacae complex *Enterobacter cloacae* UCI-CRE 9 |
| Eho-6-1 | Bacteria Proteobacteria Gammaproteobacteria Enterobacteriales Enterobacteriaceae Enterobacter Enterobacter cloacae complex *Enterobacter hormaechei* ATCC 49162 |
| Eho-7-1 | Bacteria Proteobacteria Gammaproteobacteria Enterobacteriales Enterobacteriaceae Enterobacter Enterobacter cloacae complex *Enterobacter hormaechei* YT2 |
| Eho-8-1 | Bacteria Proteobacteria Gammaproteobacteria Enterobacteriales Enterobacteriaceae Enterobacter Enterobacter cloacae complex *Enterobacter hormaechei* YT3 |
| Esp-57-1 | Bacteria Proteobacteria Gammaproteobacteria Enterobacteriales Enterobacteriaceae Enterobacter Enterobacter cloacae complex *Enterobacter sp.* MGH 14 |
| Esp-58-1 | Bacteria Proteobacteria Gammaproteobacteria Enterobacteriales Enterobacteriaceae Enterobacter Enterobacter cloacae complex *Enterobacter sp.* MGH 16 |
| Esp-59-1 | Bacteria Proteobacteria Gammaproteobacteria Enterobacteriales Enterobacteriaceae Enterobacter Enterobacter cloacae complex *Enterobacter sp.* MGH 22 |
| Esp-60-1 | Bacteria Proteobacteria Gammaproteobacteria Enterobacteriales Enterobacteriaceae Enterobacter Enterobacter cloacae complex *Enterobacter sp.* MGH 23 |
| Esp-61-1 | Bacteria Proteobacteria Gammaproteobacteria Enterobacteriales Enterobacteriaceae Enterobacter Enterobacter cloacae complex *Enterobacter sp.* MGH 24 |
| Esp-62-1 | Bacteria Proteobacteria Gammaproteobacteria Enterobacteriales Enterobacteriaceae Enterobacter Enterobacter cloacae complex *Enterobacter sp.* MGH 25 |
| Esp-63-1 | Bacteria Proteobacteria Gammaproteobacteria Enterobacteriales Enterobacteriaceae Enterobacter Enterobacter cloacae complex *Enterobacter sp.* MGH 26 |
| Esp-64-1 | Bacteria Proteobacteria Gammaproteobacteria Enterobacteriales Enterobacteriaceae Enterobacter Enterobacter cloacae complex *Enterobacter sp.* MGH 34 |
| Esp-65-1 | Bacteria Proteobacteria Gammaproteobacteria Enterobacteriales Enterobacteriaceae Enterobacter Enterobacter cloacae complex *Enterobacter sp.* MGH 38 |
| Esp-66-1 | Bacteria Proteobacteria Gammaproteobacteria Enterobacteriales Enterobacteriaceae Enterobacter Enterobacter cloacae complex *Enterobacter sp.* MGH 8 |
| Emo-1-1 | Bacteria Proteobacteria Gammaproteobacteria Enterobacteriales Enterobacteriaceae *Enterobacter mori* LMG 25706 |
| Esp-67-1 | Bacteria Proteobacteria Gammaproteobacteria Enterobacteriales Enterobacteriaceae *Enterobacter sp.* 638 |
| Esp-68-1 | Bacteria Proteobacteria Gammaproteobacteria Enterobacteriales Enterobacteriaceae *Enterobacter sp.* Ag1 |
| Esp-69-1 | Bacteria Proteobacteria Gammaproteobacteria Enterobacteriales Enterobacteriaceae *Enterobacter sp.* E20 |
| Esp-70-1 | Bacteria Proteobacteria Gammaproteobacteria Enterobacteriales Enterobacteriaceae *Enterobacter sp.* GN02454 |
| Esp-71-1 | Bacteria Proteobacteria Gammaproteobacteria Enterobacteriales Enterobacteriaceae *Enterobacter sp.* GN02457 |
| Esp-72-1 | Bacteria Proteobacteria Gammaproteobacteria Enterobacteriales Enterobacteriaceae *Enterobacter sp.* R4-368 |
| Esp-73-1 | Bacteria Proteobacteria Gammaproteobacteria Enterobacteriales Enterobacteriaceae *Enterobacter sp.* SST3 |
| Eba-3-1 | Bacteria Proteobacteria Gammaproteobacteria Enterobacteriales *Enterobacteriaceae bacterium* 9_2_54FAA |
| Eba-4-1 | Bacteria Proteobacteria Gammaproteobacteria Enterobacteriales *Enterobacteriaceae bacterium* LSJC7 |
| Eba-5-1 | Bacteria Proteobacteria Gammaproteobacteria Enterobacteriales *Enterobacteriaceae bacterium strain* FGI 57 |
| Eam-2-1 | Bacteria Proteobacteria Gammaproteobacteria Enterobacteriales Enterobacteriaceae *Erwinia amylovora* 01SFR-BO |
| Eam-3-1 | Bacteria Proteobacteria Gammaproteobacteria Enterobacteriales Enterobacteriaceae *Erwinia amylovora* ACW56400 |
| Eam-4-1 | Bacteria Proteobacteria Gammaproteobacteria Enterobacteriales Enterobacteriaceae *Erwinia amylovora* ATCC 49946 |
| Eam-5-1 | Bacteria Proteobacteria Gammaproteobacteria Enterobacteriales Enterobacteriaceae *Erwinia amylovora* CFBP 1232 |
| Eam-6-1 | Bacteria Proteobacteria Gammaproteobacteria Enterobacteriales Enterobacteriaceae *Erwinia amylovora* CFBP 2585 |
| Eam-7-1 | Bacteria Proteobacteria Gammaproteobacteria Enterobacteriales Enterobacteriaceae *Erwinia amylovora* CFBP1430 |
| Eam-8-1 | Bacteria Proteobacteria Gammaproteobacteria Enterobacteriales Enterobacteriaceae *Erwinia amylovora* Ea266 |
| Eam-9-1 | Bacteria Proteobacteria Gammaproteobacteria Enterobacteriales Enterobacteriaceae *Erwinia amylovora* Ea356 |
| Eam-10-1 | Bacteria Proteobacteria Gammaproteobacteria Enterobacteriales Enterobacteriaceae *Erwinia amylovora* Ea644 |
| Eam-11-1 | Bacteria Proteobacteria Gammaproteobacteria Enterobacteriales Enterobacteriaceae *Erwinia amylovora* LA635 |
| Eam-12-1 | Bacteria Proteobacteria Gammaproteobacteria Enterobacteriales Enterobacteriaceae *Erwinia amylovora* LA636 |
| Eam-13-1 | Bacteria Proteobacteria Gammaproteobacteria Enterobacteriales Enterobacteriaceae *Erwinia amylovora* LA637 |

| | |
|---|---|
| Eam-14-1 | Bacteria Proteobacteria Gammaproteobacteria Enterobacteriales Enterobacteriaceae *Erwinia amylovora* UPN527 |
| Ebi-2-1 | Bacteria Proteobacteria Gammaproteobacteria Enterobacteriales Enterobacteriaceae *Erwinia billingiae* Eb661 |
| Epi-2-1 | Bacteria Proteobacteria Gammaproteobacteria Enterobacteriales Enterobacteriaceae *Erwinia piriflorinigrans* CFBP 5888 |
| Epy-1-1 | Bacteria Proteobacteria Gammaproteobacteria Enterobacteriales Enterobacteriaceae *Erwinia pyrifoliae* DSM 12163 |
| Epy-2-1 | Bacteria Proteobacteria Gammaproteobacteria Enterobacteriales Enterobacteriaceae *Erwinia pyrifoliae* Ep1/96 |
| Esp-74-1 | Bacteria Proteobacteria Gammaproteobacteria Enterobacteriales Enterobacteriaceae *Erwinia sp.* Ejp617 |
| Eta-9-1 | Bacteria Proteobacteria Gammaproteobacteria Enterobacteriales Enterobacteriaceae *Erwinia tasmaniensis* Et1/99 |
| Eto-1-1 | Bacteria Proteobacteria Gammaproteobacteria Enterobacteriales Enterobacteriaceae *Erwinia toletana* DAPP-PG 735 |
| Etr-2-1 | Bacteria Proteobacteria Gammaproteobacteria Enterobacteriales Enterobacteriaceae *Erwinia tracheiphila* PSU-1 |
| Eal-3-1 | Bacteria Proteobacteria Gammaproteobacteria Enterobacteriales Enterobacteriaceae *Escherichia albertii* TW07627 |
| Eal-4-1 | Bacteria Proteobacteria Gammaproteobacteria Enterobacteriales Enterobacteriaceae *Escherichia albertii* TW08933 |
| Eal-5-1 | Bacteria Proteobacteria Gammaproteobacteria Enterobacteriales Enterobacteriaceae *Escherichia albertii* TW15818 |
| Ebl-1-1 | Bacteria Proteobacteria Gammaproteobacteria Enterobacteriales Enterobacteriaceae *Escherichia blattae* DSM 4481 |
| Eco-1-1 to Eco-1-4 | Bacteria Proteobacteria Gammaproteobacteria Enterobacteriales Enterobacteriaceae *Escherichia coli* |
| Eco-754-1 | Bacteria Proteobacteria Gammaproteobacteria Enterobacteriales Enterobacteriaceae *Escherichia coli* 0.1288 |
| Eco-755-1 | Bacteria Proteobacteria Gammaproteobacteria Enterobacteriales Enterobacteriaceae *Escherichia coli* 0.1304 |
| Eco-756-1 | Bacteria Proteobacteria Gammaproteobacteria Enterobacteriales Enterobacteriaceae *Escherichia coli* 042 |
| Eco-757-1 | Bacteria Proteobacteria Gammaproteobacteria Enterobacteriales Enterobacteriaceae *Escherichia coli* 07798 |
| Eco-758-1 | Bacteria Proteobacteria Gammaproteobacteria Enterobacteriales Enterobacteriaceae *Escherichia coli* 08BKT055439 |
| Eco-759-1 | Bacteria Proteobacteria Gammaproteobacteria Enterobacteriales Enterobacteriaceae *Escherichia coli* 08BKT77219 |
| Eco-760-1 | Bacteria Proteobacteria Gammaproteobacteria Enterobacteriales Enterobacteriaceae *Escherichia coli* 09BKT024447 |
| Eco-761-1 | Bacteria Proteobacteria Gammaproteobacteria Enterobacteriales Enterobacteriaceae *Escherichia coli* 09BKT076207 |
| Eco-762-1 | Bacteria Proteobacteria Gammaproteobacteria Enterobacteriales Enterobacteriaceae *Escherichia coli* 09BKT078844 |
| Eco-763-1 | Bacteria Proteobacteria Gammaproteobacteria Enterobacteriales Enterobacteriaceae *Escherichia coli* 1.2264 |
| Eco-764-1 | Bacteria Proteobacteria Gammaproteobacteria Enterobacteriales Enterobacteriaceae *Escherichia coli* 1.2741 |
| Eco-765-1 | Bacteria Proteobacteria Gammaproteobacteria Enterobacteriales Enterobacteriaceae *Escherichia coli* 10.0821 |
| Eco-766-1 | Bacteria Proteobacteria Gammaproteobacteria Enterobacteriales Enterobacteriaceae *Escherichia coli* 10.0833 |
| Eco-767-1 | Bacteria Proteobacteria Gammaproteobacteria Enterobacteriales Enterobacteriaceae *Escherichia coli* 10.0869 |
| Eco-768-1 | Bacteria Proteobacteria Gammaproteobacteria Enterobacteriales Enterobacteriaceae *Escherichia coli* 101-1 |
| Eco-769-1 | Bacteria Proteobacteria Gammaproteobacteria Enterobacteriales Enterobacteriaceae *Escherichia coli* 110957 |
| Eco-770-1 | Bacteria Proteobacteria Gammaproteobacteria Enterobacteriales Enterobacteriaceae *Escherichia coli* 113290 |
| Eco-771-1 | Bacteria Proteobacteria Gammaproteobacteria Enterobacteriales Enterobacteriaceae *Escherichia coli* 113302 |
| Eco-772-1 | Bacteria Proteobacteria Gammaproteobacteria Enterobacteriales Enterobacteriaceae *Escherichia coli* 113303 |
| Eco-773-1 | Bacteria Proteobacteria Gammaproteobacteria Enterobacteriales Enterobacteriaceae *Escherichia coli* 14A |
| Eco-774-1 | Bacteria Proteobacteria Gammaproteobacteria Enterobacteriales Enterobacteriaceae *Escherichia coli* 174750 |
| Eco-775-1 | Bacteria Proteobacteria Gammaproteobacteria Enterobacteriales Enterobacteriaceae *Escherichia coli* 174900 |
| Eco-776-1 | Bacteria Proteobacteria Gammaproteobacteria Enterobacteriales Enterobacteriaceae *Escherichia coli* 178200 |
| Eco-777-1 | Bacteria Proteobacteria Gammaproteobacteria Enterobacteriales Enterobacteriaceae *Escherichia coli* 178850 |
| Eco-778-1 | Bacteria Proteobacteria Gammaproteobacteria Enterobacteriales Enterobacteriaceae *Escherichia coli* 178900 |
| Eco-779-1 | Bacteria Proteobacteria Gammaproteobacteria Enterobacteriales Enterobacteriaceae *Escherichia coli* 179100 |
| Eco-780-1 | Bacteria Proteobacteria Gammaproteobacteria Enterobacteriales Enterobacteriaceae *Escherichia coli* 179550 |
| Eco-781-1 | Bacteria Proteobacteria Gammaproteobacteria Enterobacteriales Enterobacteriaceae *Escherichia coli* 180050 |
| Eco-782-1 | Bacteria Proteobacteria Gammaproteobacteria Enterobacteriales Enterobacteriaceae *Escherichia coli* 180200 |
| Eco-783-1 | Bacteria Proteobacteria Gammaproteobacteria Enterobacteriales Enterobacteriaceae *Escherichia coli* 180600 |
| Eco-784-1 | Bacteria Proteobacteria Gammaproteobacteria Enterobacteriales Enterobacteriaceae *Escherichia coli* 199900.1 |
| Eco-785-1 | Bacteria Proteobacteria Gammaproteobacteria Enterobacteriales Enterobacteriaceae *Escherichia coli* 2.3916 |
| Eco-786-1 | Bacteria Proteobacteria Gammaproteobacteria Enterobacteriales Enterobacteriaceae *Escherichia coli* 2.4168 |
| Eco-787-1 | Bacteria Proteobacteria Gammaproteobacteria Enterobacteriales Enterobacteriaceae *Escherichia coli* 201600.1 |
| Eco-788-1 | Bacteria Proteobacteria Gammaproteobacteria Enterobacteriales Enterobacteriaceae *Escherichia coli* 2362-75 |
| Eco-789-1 | Bacteria Proteobacteria Gammaproteobacteria Enterobacteriales Enterobacteriaceae *Escherichia coli* 2534-86 |
| Eco-790-1 | Bacteria Proteobacteria Gammaproteobacteria Enterobacteriales Enterobacteriaceae *Escherichia coli* 2719100 |
| Eco-791-1 | Bacteria Proteobacteria Gammaproteobacteria Enterobacteriales Enterobacteriaceae *Escherichia coli* 2720900 |
| Eco-792-1 | Bacteria Proteobacteria Gammaproteobacteria Enterobacteriales Enterobacteriaceae *Escherichia coli* 2722950 |
| Eco-793-1 | Bacteria Proteobacteria Gammaproteobacteria Enterobacteriales Enterobacteriaceae *Escherichia coli* 2726800 |
| Eco-794-1 | Bacteria Proteobacteria Gammaproteobacteria Enterobacteriales Enterobacteriaceae *Escherichia coli* 2726950 |
| Eco-795-1 | Bacteria Proteobacteria Gammaproteobacteria Enterobacteriales Enterobacteriaceae *Escherichia coli* 2729250 |
| Eco-796-1 | Bacteria Proteobacteria Gammaproteobacteria Enterobacteriales Enterobacteriaceae *Escherichia coli* 2730350 |
| Eco-797-1 | Bacteria Proteobacteria Gammaproteobacteria Enterobacteriales Enterobacteriaceae *Escherichia coli* 2730450 |
| Eco-798-1 | Bacteria Proteobacteria Gammaproteobacteria Enterobacteriales Enterobacteriaceae *Escherichia coli* 2731150 |
| Eco-799-1 | Bacteria Proteobacteria Gammaproteobacteria Enterobacteriales Enterobacteriaceae *Escherichia coli* 2733950 |
| Eco-800-1 | Bacteria Proteobacteria Gammaproteobacteria Enterobacteriales Enterobacteriaceae *Escherichia coli* 2735000 |
| Eco-801-1 | Bacteria Proteobacteria Gammaproteobacteria Enterobacteriales Enterobacteriaceae *Escherichia coli* 2741950 |
| Eco-802-1 | Bacteria Proteobacteria Gammaproteobacteria Enterobacteriales Enterobacteriaceae *Escherichia coli* 2747800 |
| Eco-803-1 | Bacteria Proteobacteria Gammaproteobacteria Enterobacteriales Enterobacteriaceae *Escherichia coli* 2749250 |
| Eco-804-1 | Bacteria Proteobacteria Gammaproteobacteria Enterobacteriales Enterobacteriaceae *Escherichia coli* 2756500 |

| | |
|---|---|
| Eco-805-1 | Bacteria Proteobacteria Gammaproteobacteria Enterobacteriales Enterobacteriaceae *Escherichia coli* 2762100 |
| Eco-806-1 | Bacteria Proteobacteria Gammaproteobacteria Enterobacteriales Enterobacteriaceae *Escherichia coli* 2770900 |
| Eco-807-1 | Bacteria Proteobacteria Gammaproteobacteria Enterobacteriales Enterobacteriaceae *Escherichia coli* 2780750 |
| Eco-808-1 | Bacteria Proteobacteria Gammaproteobacteria Enterobacteriales Enterobacteriaceae *Escherichia coli* 2785200 |
| Eco-809-1 | Bacteria Proteobacteria Gammaproteobacteria Enterobacteriales Enterobacteriaceae *Escherichia coli* 2788150 |
| Eco-810-1 | Bacteria Proteobacteria Gammaproteobacteria Enterobacteriales Enterobacteriaceae *Escherichia coli* 2845350 |
| Eco-811-1 | Bacteria Proteobacteria Gammaproteobacteria Enterobacteriales Enterobacteriaceae *Escherichia coli* 2845650 |
| Eco-812-1 | Bacteria Proteobacteria Gammaproteobacteria Enterobacteriales Enterobacteriaceae *Escherichia coli* 2846750 |
| Eco-813-1 | Bacteria Proteobacteria Gammaproteobacteria Enterobacteriales Enterobacteriaceae *Escherichia coli* 2848050 |
| Eco-814-1 | Bacteria Proteobacteria Gammaproteobacteria Enterobacteriales Enterobacteriaceae *Escherichia coli* 2850400 |
| Eco-815-1 | Bacteria Proteobacteria Gammaproteobacteria Enterobacteriales Enterobacteriaceae *Escherichia coli* 2850750 |
| Eco-816-1 | Bacteria Proteobacteria Gammaproteobacteria Enterobacteriales Enterobacteriaceae *Escherichia coli* 2851500 |
| Eco-817-1 | Bacteria Proteobacteria Gammaproteobacteria Enterobacteriales Enterobacteriaceae *Escherichia coli* 2853500 |
| Eco-818-1 | Bacteria Proteobacteria Gammaproteobacteria Enterobacteriales Enterobacteriaceae *Escherichia coli* 2854350 |
| Eco-819-1 | Bacteria Proteobacteria Gammaproteobacteria Enterobacteriales Enterobacteriaceae *Escherichia coli* 2860050 |
| Eco-820-1 | Bacteria Proteobacteria Gammaproteobacteria Enterobacteriales Enterobacteriaceae *Escherichia coli* 2860650 |
| Eco-821-1 | Bacteria Proteobacteria Gammaproteobacteria Enterobacteriales Enterobacteriaceae *Escherichia coli* 2861200 |
| Eco-822-1 | Bacteria Proteobacteria Gammaproteobacteria Enterobacteriales Enterobacteriaceae *Escherichia coli* 2862600 |
| Eco-823-1 | Bacteria Proteobacteria Gammaproteobacteria Enterobacteriales Enterobacteriaceae *Escherichia coli* 2864350 |
| Eco-824-1 | Bacteria Proteobacteria Gammaproteobacteria Enterobacteriales Enterobacteriaceae *Escherichia coli* 2865200 |
| Eco-825-1 | Bacteria Proteobacteria Gammaproteobacteria Enterobacteriales Enterobacteriaceae *Escherichia coli* 2866350 |
| Eco-826-1 | Bacteria Proteobacteria Gammaproteobacteria Enterobacteriales Enterobacteriaceae *Escherichia coli* 2866450 |
| Eco-827-1 | Bacteria Proteobacteria Gammaproteobacteria Enterobacteriales Enterobacteriaceae *Escherichia coli* 2866550 |
| Eco-828-1 | Bacteria Proteobacteria Gammaproteobacteria Enterobacteriales Enterobacteriaceae *Escherichia coli* 2866750 |
| Eco-829-1 | Bacteria Proteobacteria Gammaproteobacteria Enterobacteriales Enterobacteriaceae *Escherichia coli* 2867750 |
| Eco-830-1 | Bacteria Proteobacteria Gammaproteobacteria Enterobacteriales Enterobacteriaceae *Escherichia coli* 2871950 |
| Eco-831-1 | Bacteria Proteobacteria Gammaproteobacteria Enterobacteriales Enterobacteriaceae *Escherichia coli* 2872000 |
| Eco-832-1 | Bacteria Proteobacteria Gammaproteobacteria Enterobacteriales Enterobacteriaceae *Escherichia coli* 2872800 |
| Eco-833-1 | Bacteria Proteobacteria Gammaproteobacteria Enterobacteriales Enterobacteriaceae *Escherichia coli* 2875000 |
| Eco-834-1 | Bacteria Proteobacteria Gammaproteobacteria Enterobacteriales Enterobacteriaceae *Escherichia coli* 2875150 |
| Eco-835-1 | Bacteria Proteobacteria Gammaproteobacteria Enterobacteriales Enterobacteriaceae *Escherichia coli* 2886-75 |
| Eco-836-1 | Bacteria Proteobacteria Gammaproteobacteria Enterobacteriales Enterobacteriaceae *Escherichia coli* 3.2303 |
| Eco-837-1 | Bacteria Proteobacteria Gammaproteobacteria Enterobacteriales Enterobacteriaceae *Escherichia coli* 3.2608 |
| Eco-838-1 | Bacteria Proteobacteria Gammaproteobacteria Enterobacteriales Enterobacteriaceae *Escherichia coli* 3.3884 |
| Eco-839-1 | Bacteria Proteobacteria Gammaproteobacteria Enterobacteriales Enterobacteriaceae *Escherichia coli* 3.4870 |
| Eco-840-1 | Bacteria Proteobacteria Gammaproteobacteria Enterobacteriales Enterobacteriaceae *Escherichia coli* 3.4880 |
| Eco-841-1 | Bacteria Proteobacteria Gammaproteobacteria Enterobacteriales Enterobacteriaceae *Escherichia coli* 3003 |
| Eco-842-1 | Bacteria Proteobacteria Gammaproteobacteria Enterobacteriales Enterobacteriaceae *Escherichia coli* 3006 |
| Eco-843-1 | Bacteria Proteobacteria Gammaproteobacteria Enterobacteriales Enterobacteriaceae *Escherichia coli* 3431 |
| Eco-844-1 | Bacteria Proteobacteria Gammaproteobacteria Enterobacteriales Enterobacteriaceae *Escherichia coli* 4.0522 |
| Eco-845-1 | Bacteria Proteobacteria Gammaproteobacteria Enterobacteriales Enterobacteriaceae *Escherichia coli* 4.0967 |
| Eco-846-1 | Bacteria Proteobacteria Gammaproteobacteria Enterobacteriales Enterobacteriaceae *Escherichia coli* 5.0588 |
| Eco-847-1 | Bacteria Proteobacteria Gammaproteobacteria Enterobacteriales Enterobacteriaceae *Escherichia coli* 5.0959 |
| Eco-848-1 | Bacteria Proteobacteria Gammaproteobacteria Enterobacteriales Enterobacteriaceae *Escherichia coli* 5.2239 |
| Eco-849-1 | Bacteria Proteobacteria Gammaproteobacteria Enterobacteriales Enterobacteriaceae *Escherichia coli* 536 |
| Eco-850-1 | Bacteria Proteobacteria Gammaproteobacteria Enterobacteriales Enterobacteriaceae *Escherichia coli* 53638 |
| Eco-851-1 | Bacteria Proteobacteria Gammaproteobacteria Enterobacteriales Enterobacteriaceae *Escherichia coli* 541-1 |
| Eco-852-1 | Bacteria Proteobacteria Gammaproteobacteria Enterobacteriales Enterobacteriaceae *Escherichia coli* 541-15 |
| Eco-853-1 | Bacteria Proteobacteria Gammaproteobacteria Enterobacteriales Enterobacteriaceae *Escherichia coli* 5412 |
| Eco-854-1 | Bacteria Proteobacteria Gammaproteobacteria Enterobacteriales Enterobacteriaceae *Escherichia coli* 55989 |
| Eco-855-1 | Bacteria Proteobacteria Gammaproteobacteria Enterobacteriales Enterobacteriaceae *Escherichia coli* 576-1 |
| Eco-856-1 | Bacteria Proteobacteria Gammaproteobacteria Enterobacteriales Enterobacteriaceae *Escherichia coli* 5905 |
| Eco-857-1 | Bacteria Proteobacteria Gammaproteobacteria Enterobacteriales Enterobacteriaceae *Escherichia coli* 6.0172 |
| Eco-858-1 | Bacteria Proteobacteria Gammaproteobacteria Enterobacteriales Enterobacteriaceae *Escherichia coli* 7.1982 |
| Eco-859-1 | Bacteria Proteobacteria Gammaproteobacteria Enterobacteriales Enterobacteriaceae *Escherichia coli* 75 |
| Eco-860-1 | Bacteria Proteobacteria Gammaproteobacteria Enterobacteriales Enterobacteriaceae *Escherichia coli* 8.0416 |
| Eco-861-1 | Bacteria Proteobacteria Gammaproteobacteria Enterobacteriales Enterobacteriaceae *Escherichia coli* 8.0566 |
| Eco-862-1 | Bacteria Proteobacteria Gammaproteobacteria Enterobacteriales Enterobacteriaceae *Escherichia coli* 8.0569 |
| Eco-863-1 | Bacteria Proteobacteria Gammaproteobacteria Enterobacteriales Enterobacteriaceae *Escherichia coli* 8.0586 |
| Eco-864-1 | Bacteria Proteobacteria Gammaproteobacteria Enterobacteriales Enterobacteriaceae *Escherichia coli* 8.2524 |
| Eco-865-1 | Bacteria Proteobacteria Gammaproteobacteria Enterobacteriales Enterobacteriaceae *Escherichia coli* 83972 |
| Eco-866-1 | Bacteria Proteobacteria Gammaproteobacteria Enterobacteriales Enterobacteriaceae *Escherichia coli* 88.0221 |
| Eco-867-1 | Bacteria Proteobacteria Gammaproteobacteria Enterobacteriales Enterobacteriaceae *Escherichia coli* 88.1042 |
| Eco-868-1 | Bacteria Proteobacteria Gammaproteobacteria Enterobacteriales Enterobacteriaceae *Escherichia coli* 88.1467 |
| Eco-869-1 | Bacteria Proteobacteria Gammaproteobacteria Enterobacteriales Enterobacteriaceae *Escherichia coli* 89.0511 |

| | |
|---|---|
| Eco-870-1 | Bacteria Proteobacteria Gammaproteobacteria Enterobacteriales Enterobacteriaceae *Escherichia coli* 9.0111 |
| Eco-871-1 | Bacteria Proteobacteria Gammaproteobacteria Enterobacteriales Enterobacteriaceae *Escherichia coli* 90.0039 |
| Eco-872-1 | Bacteria Proteobacteria Gammaproteobacteria Enterobacteriales Enterobacteriaceae *Escherichia coli* 90.0091 |
| Eco-873-1 | Bacteria Proteobacteria Gammaproteobacteria Enterobacteriales Enterobacteriaceae *Escherichia coli* 90.2281 |
| Eco-874-1 | Bacteria Proteobacteria Gammaproteobacteria Enterobacteriales Enterobacteriaceae *Escherichia coli* 900105 (10e) |
| Eco-875-1 | Bacteria Proteobacteria Gammaproteobacteria Enterobacteriales Enterobacteriaceae *Escherichia coli* 907357 |
| Eco-876-1 | Bacteria Proteobacteria Gammaproteobacteria Enterobacteriales Enterobacteriaceae *Escherichia coli* 907391 |
| Eco-877-1 | Bacteria Proteobacteria Gammaproteobacteria Enterobacteriales Enterobacteriaceae *Escherichia coli* 907446 |
| Eco-878-1 | Bacteria Proteobacteria Gammaproteobacteria Enterobacteriales Enterobacteriaceae *Escherichia coli* 907672 |
| Eco-879-1 | Bacteria Proteobacteria Gammaproteobacteria Enterobacteriales Enterobacteriaceae *Escherichia coli* 907700 |
| Eco-880-1 | Bacteria Proteobacteria Gammaproteobacteria Enterobacteriales Enterobacteriaceae *Escherichia coli* 907701 |
| Eco-881-1 | Bacteria Proteobacteria Gammaproteobacteria Enterobacteriales Enterobacteriaceae *Escherichia coli* 907710 |
| Eco-882-1 | Bacteria Proteobacteria Gammaproteobacteria Enterobacteriales Enterobacteriaceae *Escherichia coli* 907713 |
| Eco-883-1 | Bacteria Proteobacteria Gammaproteobacteria Enterobacteriales Enterobacteriaceae *Escherichia coli* 907715 |
| Eco-884-1 | Bacteria Proteobacteria Gammaproteobacteria Enterobacteriales Enterobacteriaceae *Escherichia coli* 907779 |
| Eco-885-1 | Bacteria Proteobacteria Gammaproteobacteria Enterobacteriales Enterobacteriaceae *Escherichia coli* 907889 |
| Eco-886-1 | Bacteria Proteobacteria Gammaproteobacteria Enterobacteriales Enterobacteriaceae *Escherichia coli* 907892 |
| Eco-887-1 | Bacteria Proteobacteria Gammaproteobacteria Enterobacteriales Enterobacteriaceae *Escherichia coli* 908519 |
| Eco-888-1 | Bacteria Proteobacteria Gammaproteobacteria Enterobacteriales Enterobacteriaceae *Escherichia coli* 908521 |
| Eco-889-1 | Bacteria Proteobacteria Gammaproteobacteria Enterobacteriales Enterobacteriaceae *Escherichia coli* 908522 |
| Eco-890-1 | Bacteria Proteobacteria Gammaproteobacteria Enterobacteriales Enterobacteriaceae *Escherichia coli* 908524 |
| Eco-891-1 | Bacteria Proteobacteria Gammaproteobacteria Enterobacteriales Enterobacteriaceae *Escherichia coli* 908525 |
| Eco-892-1 | Bacteria Proteobacteria Gammaproteobacteria Enterobacteriales Enterobacteriaceae *Escherichia coli* 908541 |
| Eco-893-1 | Bacteria Proteobacteria Gammaproteobacteria Enterobacteriales Enterobacteriaceae *Escherichia coli* 908555 |
| Eco-894-1 | Bacteria Proteobacteria Gammaproteobacteria Enterobacteriales Enterobacteriaceae *Escherichia coli* 908573 |
| Eco-895-1 | Bacteria Proteobacteria Gammaproteobacteria Enterobacteriales Enterobacteriaceae *Escherichia coli* 908585 |
| Eco-896-1 | Bacteria Proteobacteria Gammaproteobacteria Enterobacteriales Enterobacteriaceae *Escherichia coli* 908616 |
| Eco-897-1 | Bacteria Proteobacteria Gammaproteobacteria Enterobacteriales Enterobacteriaceae *Escherichia coli* 908624 |
| Eco-898-1 | Bacteria Proteobacteria Gammaproteobacteria Enterobacteriales Enterobacteriaceae *Escherichia coli* 908632 |
| Eco-899-1 | Bacteria Proteobacteria Gammaproteobacteria Enterobacteriales Enterobacteriaceae *Escherichia coli* 908658 |
| Eco-900-1 | Bacteria Proteobacteria Gammaproteobacteria Enterobacteriales Enterobacteriaceae *Escherichia coli* 908675 |
| Eco-901-1 | Bacteria Proteobacteria Gammaproteobacteria Enterobacteriales Enterobacteriaceae *Escherichia coli* 908691 |
| Eco-902-1 | Bacteria Proteobacteria Gammaproteobacteria Enterobacteriales Enterobacteriaceae *Escherichia coli* 909945-2 |
| Eco-903-1 | Bacteria Proteobacteria Gammaproteobacteria Enterobacteriales Enterobacteriaceae *Escherichia coli* 909957 |
| Eco-904-1 | Bacteria Proteobacteria Gammaproteobacteria Enterobacteriales Enterobacteriaceae *Escherichia coli* 910096-2 |
| Eco-905-1 | Bacteria Proteobacteria Gammaproteobacteria Enterobacteriales Enterobacteriaceae *Escherichia coli* 93-001 |
| Eco-906-1 | Bacteria Proteobacteria Gammaproteobacteria Enterobacteriales Enterobacteriaceae *Escherichia coli* 93.0055 |
| Eco-907-1 | Bacteria Proteobacteria Gammaproteobacteria Enterobacteriales Enterobacteriaceae *Escherichia coli* 93.0056 |
| Eco-908-1 | Bacteria Proteobacteria Gammaproteobacteria Enterobacteriales Enterobacteriaceae *Escherichia coli* 93.0624 |
| Eco-909-1 | Bacteria Proteobacteria Gammaproteobacteria Enterobacteriales Enterobacteriaceae *Escherichia coli* 94.0618 |
| Eco-910-1 | Bacteria Proteobacteria Gammaproteobacteria Enterobacteriales Enterobacteriaceae *Escherichia coli* 95.0083 |
| Eco-911-1 | Bacteria Proteobacteria Gammaproteobacteria Enterobacteriales Enterobacteriaceae *Escherichia coli* 95.0183 |
| Eco-912-1 | Bacteria Proteobacteria Gammaproteobacteria Enterobacteriales Enterobacteriaceae *Escherichia coli* 95.0941 |
| Eco-913-1 | Bacteria Proteobacteria Gammaproteobacteria Enterobacteriales Enterobacteriaceae *Escherichia coli* 95.0943 |
| Eco-914-1 | Bacteria Proteobacteria Gammaproteobacteria Enterobacteriales Enterobacteriaceae *Escherichia coli* 95.1288 |
| Eco-915-1 | Bacteria Proteobacteria Gammaproteobacteria Enterobacteriales Enterobacteriaceae *Escherichia coli* 95JB1 |
| Eco-916-1 | Bacteria Proteobacteria Gammaproteobacteria Enterobacteriales Enterobacteriaceae *Escherichia coli* 95NR1 |
| Eco-917-1 | Bacteria Proteobacteria Gammaproteobacteria Enterobacteriales Enterobacteriaceae *Escherichia coli* 96.0107 |
| Eco-918-1 | Bacteria Proteobacteria Gammaproteobacteria Enterobacteriales Enterobacteriaceae *Escherichia coli* 96.0109 |
| Eco-919-1 | Bacteria Proteobacteria Gammaproteobacteria Enterobacteriales Enterobacteriaceae *Escherichia coli* 96.0427 |
| Eco-920-1 | Bacteria Proteobacteria Gammaproteobacteria Enterobacteriales Enterobacteriaceae *Escherichia coli* 96.0428 |
| Eco-921-1 | Bacteria Proteobacteria Gammaproteobacteria Enterobacteriales Enterobacteriaceae *Escherichia coli* 96.0497 |
| Eco-922-1 | Bacteria Proteobacteria Gammaproteobacteria Enterobacteriales Enterobacteriaceae *Escherichia coli* 96.0932 |
| Eco-923-1 | Bacteria Proteobacteria Gammaproteobacteria Enterobacteriales Enterobacteriaceae *Escherichia coli* 96.0939 |
| Eco-924-1 | Bacteria Proteobacteria Gammaproteobacteria Enterobacteriales Enterobacteriaceae *Escherichia coli* 96.154 |
| Eco-925-1 | Bacteria Proteobacteria Gammaproteobacteria Enterobacteriales Enterobacteriaceae *Escherichia coli* 97.0003 |
| Eco-926-1 | Bacteria Proteobacteria Gammaproteobacteria Enterobacteriales Enterobacteriaceae *Escherichia coli* 97.0007 |
| Eco-927-1 | Bacteria Proteobacteria Gammaproteobacteria Enterobacteriales Enterobacteriaceae *Escherichia coli* 97.0010 |
| Eco-928-1 | Bacteria Proteobacteria Gammaproteobacteria Enterobacteriales Enterobacteriaceae *Escherichia coli* 97.0246 |
| Eco-929-1 | Bacteria Proteobacteria Gammaproteobacteria Enterobacteriales Enterobacteriaceae *Escherichia coli* 97.0259 |
| Eco-930-1 | Bacteria Proteobacteria Gammaproteobacteria Enterobacteriales Enterobacteriaceae *Escherichia coli* 97.1742 |
| Eco-931-1 | Bacteria Proteobacteria Gammaproteobacteria Enterobacteriales Enterobacteriaceae *Escherichia coli* 99.0670 |
| Eco-932-1 | Bacteria Proteobacteria Gammaproteobacteria Enterobacteriales Enterobacteriaceae *Escherichia coli* 99.0672 |
| Eco-933-1 | Bacteria Proteobacteria Gammaproteobacteria Enterobacteriales Enterobacteriaceae *Escherichia coli* 99.0678 |
| Eco-934-1 | Bacteria Proteobacteria Gammaproteobacteria Enterobacteriales Enterobacteriaceae *Escherichia coli* 99.0713 |

| | |
|---|---|
| Eco-935-1 | Bacteria Proteobacteria Gammaproteobacteria Enterobacteriales Enterobacteriaceae *Escherichia coli* 99.0741 |
| Eco-936-1 | Bacteria Proteobacteria Gammaproteobacteria Enterobacteriales Enterobacteriaceae *Escherichia coli* 99.0814 |
| Eco-937-1 | Bacteria Proteobacteria Gammaproteobacteria Enterobacteriales Enterobacteriaceae *Escherichia coli* 99.0815 |
| Eco-938-1 | Bacteria Proteobacteria Gammaproteobacteria Enterobacteriales Enterobacteriaceae *Escherichia coli* 99.0816 |
| Eco-939-1 | Bacteria Proteobacteria Gammaproteobacteria Enterobacteriales Enterobacteriaceae *Escherichia coli* 99.0839 |
| Eco-940-1 | Bacteria Proteobacteria Gammaproteobacteria Enterobacteriales Enterobacteriaceae *Escherichia coli* 99.0848 |
| Eco-941-1 | Bacteria Proteobacteria Gammaproteobacteria Enterobacteriales Enterobacteriaceae *Escherichia coli* 99.1753 |
| Eco-942-1 | Bacteria Proteobacteria Gammaproteobacteria Enterobacteriales Enterobacteriaceae *Escherichia coli* 99.1762 |
| Eco-943-1 | Bacteria Proteobacteria Gammaproteobacteria Enterobacteriales Enterobacteriaceae *Escherichia coli* 99.1775 |
| Eco-944-1 | Bacteria Proteobacteria Gammaproteobacteria Enterobacteriales Enterobacteriaceae *Escherichia coli* 99.1781 |
| Eco-945-1 | Bacteria Proteobacteria Gammaproteobacteria Enterobacteriales Enterobacteriaceae *Escherichia coli* 99.1793 |
| Eco-946-1 | Bacteria Proteobacteria Gammaproteobacteria Enterobacteriales Enterobacteriaceae *Escherichia coli* 99.1805 |
| Eco-947-1 | Bacteria Proteobacteria Gammaproteobacteria Enterobacteriales Enterobacteriaceae *Escherichia coli* A25922R |
| Eco-948-1 | Bacteria Proteobacteria Gammaproteobacteria Enterobacteriales Enterobacteriaceae *Escherichia coli* A35218R |
| Eco-949-1 | Bacteria Proteobacteria Gammaproteobacteria Enterobacteriales Enterobacteriaceae *Escherichia coli* AA86 |
| Eco-950-1 | Bacteria Proteobacteria Gammaproteobacteria Enterobacteriales Enterobacteriaceae *Escherichia coli* ABU 83972 |
| Eco-289-1 to Eco-289-2 | Bacteria Proteobacteria Gammaproteobacteria Enterobacteriales Enterobacteriaceae *Escherichia coli* AD30 |
| Eco-951-1 | Bacteria Proteobacteria Gammaproteobacteria Enterobacteriales Enterobacteriaceae *Escherichia coli* AI27 |
| Eco-952-1 | Bacteria Proteobacteria Gammaproteobacteria Enterobacteriales Enterobacteriaceae *Escherichia coli* APEC O1 |
| Eco-953-1 | Bacteria Proteobacteria Gammaproteobacteria Enterobacteriales Enterobacteriaceae *Escherichia coli* APEC O78 |
| Eco-954-1 | Bacteria Proteobacteria Gammaproteobacteria Enterobacteriales Enterobacteriaceae *Escherichia coli* ARS4.2123 |
| Eco-291-1 to Eco-291-2 | Bacteria Proteobacteria Gammaproteobacteria Enterobacteriales Enterobacteriaceae *Escherichia coli* ATCC 25922 |
| Eco-955-1 | Bacteria Proteobacteria Gammaproteobacteria Enterobacteriales Enterobacteriaceae *Escherichia coli* ATCC 700728 |
| Eco-956-1 | Bacteria Proteobacteria Gammaproteobacteria Enterobacteriales Enterobacteriaceae *Escherichia coli* ATCC 8739 |
| Eco-957-1 | Bacteria Proteobacteria Gammaproteobacteria Enterobacteriales Enterobacteriaceae *Escherichia coli* ATCC BAA-2196 |
| Eco-958-1 | Bacteria Proteobacteria Gammaproteobacteria Enterobacteriales Enterobacteriaceae *Escherichia coli* ATCC BAA-2209 |
| Eco-959-1 | Bacteria Proteobacteria Gammaproteobacteria Enterobacteriales Enterobacteriaceae *Escherichia coli* ATCC BAA-2215 |
| Eco-960-1 | Bacteria Proteobacteria Gammaproteobacteria Enterobacteriales Enterobacteriaceae *Escherichia coli* ATCC BAA-2219 |
| Eco-961-1 | Bacteria Proteobacteria Gammaproteobacteria Enterobacteriales Enterobacteriaceae *Escherichia coli* B str. REL606 |
| Eco-962-1 | Bacteria Proteobacteria Gammaproteobacteria Enterobacteriales Enterobacteriaceae *Escherichia coli* B088 |
| Eco-963-1 | Bacteria Proteobacteria Gammaproteobacteria Enterobacteriales Enterobacteriaceae *Escherichia coli* B093 |
| Eco-964-1 | Bacteria Proteobacteria Gammaproteobacteria Enterobacteriales Enterobacteriaceae *Escherichia coli* B102 |
| Eco-965-1 | Bacteria Proteobacteria Gammaproteobacteria Enterobacteriales Enterobacteriaceae *Escherichia coli* B103 |
| Eco-966-1 | Bacteria Proteobacteria Gammaproteobacteria Enterobacteriales Enterobacteriaceae *Escherichia coli* B104 |
| Eco-967-1 | Bacteria Proteobacteria Gammaproteobacteria Enterobacteriales Enterobacteriaceae *Escherichia coli* B105 |
| Eco-968-1 | Bacteria Proteobacteria Gammaproteobacteria Enterobacteriales Enterobacteriaceae *Escherichia coli* B106 |
| Eco-969-1 | Bacteria Proteobacteria Gammaproteobacteria Enterobacteriales Enterobacteriaceae *Escherichia coli* B107 |
| Eco-970-1 | Bacteria Proteobacteria Gammaproteobacteria Enterobacteriales Enterobacteriaceae *Escherichia coli* B108 |
| Eco-971-1 | Bacteria Proteobacteria Gammaproteobacteria Enterobacteriales Enterobacteriaceae *Escherichia coli* B109 |
| Eco-972-1 | Bacteria Proteobacteria Gammaproteobacteria Enterobacteriales Enterobacteriaceae *Escherichia coli* B112 |
| Eco-973-1 | Bacteria Proteobacteria Gammaproteobacteria Enterobacteriales Enterobacteriaceae *Escherichia coli* B113 |
| Eco-974-1 | Bacteria Proteobacteria Gammaproteobacteria Enterobacteriales Enterobacteriaceae *Escherichia coli* B114 |
| Eco-975-1 | Bacteria Proteobacteria Gammaproteobacteria Enterobacteriales Enterobacteriaceae *Escherichia coli* B15 |
| Eco-976-1 | Bacteria Proteobacteria Gammaproteobacteria Enterobacteriales Enterobacteriaceae *Escherichia coli* B17 |
| Eco-977-1 | Bacteria Proteobacteria Gammaproteobacteria Enterobacteriales Enterobacteriaceae *Escherichia coli* B171 |
| Eco-978-1 | Bacteria Proteobacteria Gammaproteobacteria Enterobacteriales Enterobacteriaceae *Escherichia coli* B185 |
| Eco-979-1 | Bacteria Proteobacteria Gammaproteobacteria Enterobacteriales Enterobacteriaceae *Escherichia coli* B26-1 |
| Eco-980-1 | Bacteria Proteobacteria Gammaproteobacteria Enterobacteriales Enterobacteriaceae *Escherichia coli* B26-2 |
| Eco-981-1 | Bacteria Proteobacteria Gammaproteobacteria Enterobacteriales Enterobacteriaceae *Escherichia coli* B28-1 |
| Eco-982-1 | Bacteria Proteobacteria Gammaproteobacteria Enterobacteriales Enterobacteriaceae *Escherichia coli* B28-2 |
| Eco-983-1 | Bacteria Proteobacteria Gammaproteobacteria Enterobacteriales Enterobacteriaceae *Escherichia coli* B29-1 |
| Eco-984-1 | Bacteria Proteobacteria Gammaproteobacteria Enterobacteriales Enterobacteriaceae *Escherichia coli* B29-2 |
| Eco-985-1 | Bacteria Proteobacteria Gammaproteobacteria Enterobacteriales Enterobacteriaceae *Escherichia coli* B354 |
| Eco-986-1 | Bacteria Proteobacteria Gammaproteobacteria Enterobacteriales Enterobacteriaceae *Escherichia coli* B36-1 |
| Eco-987-1 | Bacteria Proteobacteria Gammaproteobacteria Enterobacteriales Enterobacteriaceae *Escherichia coli* B36-2 |
| Eco-988-1 | Bacteria Proteobacteria Gammaproteobacteria Enterobacteriales Enterobacteriaceae *Escherichia coli* B40-1 |
| Eco-989-1 | Bacteria Proteobacteria Gammaproteobacteria Enterobacteriales Enterobacteriaceae *Escherichia coli* B40-2 |
| Eco-990-1 | Bacteria Proteobacteria Gammaproteobacteria Enterobacteriales Enterobacteriaceae *Escherichia coli* B41 |
| Eco-991-1 | Bacteria Proteobacteria Gammaproteobacteria Enterobacteriales Enterobacteriaceae *Escherichia coli* B49-2 |
| Eco-992-1 | Bacteria Proteobacteria Gammaproteobacteria Enterobacteriales Enterobacteriaceae *Escherichia coli* B5-2 |
| Eco-993-1 | Bacteria Proteobacteria Gammaproteobacteria Enterobacteriales Enterobacteriaceae *Escherichia coli* B7-1 |
| Eco-994-1 | Bacteria Proteobacteria Gammaproteobacteria Enterobacteriales Enterobacteriaceae *Escherichia coli* B7-2 |
| Eco-995-1 | Bacteria Proteobacteria Gammaproteobacteria Enterobacteriales Enterobacteriaceae *Escherichia coli* B799 |
| Eco-996-1 | Bacteria Proteobacteria Gammaproteobacteria Enterobacteriales Enterobacteriaceae *Escherichia coli* B7A |
| Eco-997-1 | Bacteria Proteobacteria Gammaproteobacteria Enterobacteriales Enterobacteriaceae *Escherichia coli* B83 |

| | |
|---|---|
| Eco-998-1 | Bacteria Proteobacteria Gammaproteobacteria Enterobacteriales Enterobacteriaceae *Escherichia coli* B84 |
| Eco-999-1 | Bacteria Proteobacteria Gammaproteobacteria Enterobacteriales Enterobacteriaceae *Escherichia coli* B85 |
| Eco-1000-1 | Bacteria Proteobacteria Gammaproteobacteria Enterobacteriales Enterobacteriaceae *Escherichia coli* B86 |
| Eco-1001-1 | Bacteria Proteobacteria Gammaproteobacteria Enterobacteriales Enterobacteriaceae *Escherichia coli* B89 |
| Eco-1002-1 | Bacteria Proteobacteria Gammaproteobacteria Enterobacteriales Enterobacteriaceae *Escherichia coli* B90 |
| Eco-1003-1 | Bacteria Proteobacteria Gammaproteobacteria Enterobacteriales Enterobacteriaceae *Escherichia coli* B93 |
| Eco-1004-1 | Bacteria Proteobacteria Gammaproteobacteria Enterobacteriales Enterobacteriaceae *Escherichia coli* B94 |
| Eco-1005-1 | Bacteria Proteobacteria Gammaproteobacteria Enterobacteriales Enterobacteriaceae *Escherichia coli* B95 |
| Eco-1006-1 | Bacteria Proteobacteria Gammaproteobacteria Enterobacteriales Enterobacteriaceae *Escherichia coli* BCE001_MS16 |
| Eco-1007-1 | Bacteria Proteobacteria Gammaproteobacteria Enterobacteriales Enterobacteriaceae *Escherichia coli* BCE002_MS12 |
| Eco-1008-1 | Bacteria Proteobacteria Gammaproteobacteria Enterobacteriales Enterobacteriaceae *Escherichia coli* BCE006_MS-23 |
| Eco-1009-1 | Bacteria Proteobacteria Gammaproteobacteria Enterobacteriales Enterobacteriaceae *Escherichia coli* BCE007_MS-11 |
| Eco-1010-1 | Bacteria Proteobacteria Gammaproteobacteria Enterobacteriales Enterobacteriaceae *Escherichia coli* BCE008_MS-01 |
| Eco-1011-1 | Bacteria Proteobacteria Gammaproteobacteria Enterobacteriales Enterobacteriaceae *Escherichia coli* BCE008_MS-13 |
| Eco-1012-1 | Bacteria Proteobacteria Gammaproteobacteria Enterobacteriales Enterobacteriaceae *Escherichia coli* BCE011_MS-01 |
| Eco-1013-1 | Bacteria Proteobacteria Gammaproteobacteria Enterobacteriales Enterobacteriaceae *Escherichia coli* BCE019_MS-13 |
| Eco-1014-1 | Bacteria Proteobacteria Gammaproteobacteria Enterobacteriales Enterobacteriaceae *Escherichia coli* BCE030_MS-09 |
| Eco-1015-1 | Bacteria Proteobacteria Gammaproteobacteria Enterobacteriales Enterobacteriaceae *Escherichia coli* BCE032_MS-12 |
| Eco-1016-1 | Bacteria Proteobacteria Gammaproteobacteria Enterobacteriales Enterobacteriaceae *Escherichia coli* BCE034_MS-14 |
| Eco-1025-1 | Bacteria Proteobacteria Gammaproteobacteria Enterobacteriales Enterobacteriaceae *Escherichia coli* Bd5610_99 |
| Eco-1017-1 | Bacteria Proteobacteria Gammaproteobacteria Enterobacteriales Enterobacteriaceae *Escherichia coli* BIDMC 19C |
| Eco-1018-1 | Bacteria Proteobacteria Gammaproteobacteria Enterobacteriales Enterobacteriaceae *Escherichia coli* BIDMC 37 |
| Eco-1019-1 | Bacteria Proteobacteria Gammaproteobacteria Enterobacteriales Enterobacteriaceae *Escherichia coli* BIDMC 38 |
| Eco-1020-1 | Bacteria Proteobacteria Gammaproteobacteria Enterobacteriales Enterobacteriaceae *Escherichia coli* BIDMC 39 |
| Eco-1021-1 to Eco-1021-3 | Bacteria Proteobacteria Gammaproteobacteria Enterobacteriales Enterobacteriaceae *Escherichia coli* BL21(DE3) |
| Eco-1022-1 | Bacteria Proteobacteria Gammaproteobacteria Enterobacteriales Enterobacteriaceae *Escherichia coli* BW2952 |
| Eco-1023-1 | Bacteria Proteobacteria Gammaproteobacteria Enterobacteriales Enterobacteriaceae *Escherichia coli* BWH 24 |
| Eco-1024-1 | Bacteria Proteobacteria Gammaproteobacteria Enterobacteriales Enterobacteriaceae *Escherichia coli* BWH 32 |
| Eco-1026-1 | Bacteria Proteobacteria Gammaproteobacteria Enterobacteriales Enterobacteriaceae *Escherichia coli* C-34666 |
| Eco-1027-1 | Bacteria Proteobacteria Gammaproteobacteria Enterobacteriales Enterobacteriaceae *Escherichia coli* CB7326 |
| Eco-1028-1 | Bacteria Proteobacteria Gammaproteobacteria Enterobacteriales Enterobacteriaceae *Escherichia coli* CE418 |
| Eco-1029-1 | Bacteria Proteobacteria Gammaproteobacteria Enterobacteriales Enterobacteriaceae *Escherichia coli* CE516 |
| Eco-1030-1 | Bacteria Proteobacteria Gammaproteobacteria Enterobacteriales Enterobacteriaceae *Escherichia coli* CE549 |
| Eco-1031-1 | Bacteria Proteobacteria Gammaproteobacteria Enterobacteriales Enterobacteriaceae *Escherichia coli* CFT073 |
| Eco-2165-1 | Bacteria Proteobacteria Gammaproteobacteria Enterobacteriales Enterobacteriaceae *Escherichia coli* *chi*7122 |
| Eco-2166-1 | Bacteria Proteobacteria Gammaproteobacteria Enterobacteriales Enterobacteriaceae *Escherichia coli* *clone*A_i1 |
| Eco-1032-1 | Bacteria Proteobacteria Gammaproteobacteria Enterobacteriales Enterobacteriaceae *Escherichia coli* CUMT8 |
| Eco-1033-1 | Bacteria Proteobacteria Gammaproteobacteria Enterobacteriales Enterobacteriaceae *Escherichia coli* DEC10E |
| Eco-1034-1 | Bacteria Proteobacteria Gammaproteobacteria Enterobacteriales Enterobacteriaceae *Escherichia coli* DEC10F |
| Eco-1035-1 | Bacteria Proteobacteria Gammaproteobacteria Enterobacteriales Enterobacteriaceae *Escherichia coli* DEC11A |
| Eco-1036-1 | Bacteria Proteobacteria Gammaproteobacteria Enterobacteriales Enterobacteriaceae *Escherichia coli* DEC11B |
| Eco-1037-1 | Bacteria Proteobacteria Gammaproteobacteria Enterobacteriales Enterobacteriaceae *Escherichia coli* DEC11C |
| Eco-1038-1 | Bacteria Proteobacteria Gammaproteobacteria Enterobacteriales Enterobacteriaceae *Escherichia coli* DEC11D |
| Eco-1039-1 | Bacteria Proteobacteria Gammaproteobacteria Enterobacteriales Enterobacteriaceae *Escherichia coli* DEC11E |
| Eco-1040-1 | Bacteria Proteobacteria Gammaproteobacteria Enterobacteriales Enterobacteriaceae *Escherichia coli* DEC12A |
| Eco-1041-1 | Bacteria Proteobacteria Gammaproteobacteria Enterobacteriales Enterobacteriaceae *Escherichia coli* DEC12B |
| Eco-1042-1 | Bacteria Proteobacteria Gammaproteobacteria Enterobacteriales Enterobacteriaceae *Escherichia coli* DEC12C |
| Eco-1043-1 | Bacteria Proteobacteria Gammaproteobacteria Enterobacteriales Enterobacteriaceae *Escherichia coli* DEC12D |
| Eco-1044-1 | Bacteria Proteobacteria Gammaproteobacteria Enterobacteriales Enterobacteriaceae *Escherichia coli* DEC12E |
| Eco-1045-1 | Bacteria Proteobacteria Gammaproteobacteria Enterobacteriales Enterobacteriaceae *Escherichia coli* DEC13A |
| Eco-1046-1 | Bacteria Proteobacteria Gammaproteobacteria Enterobacteriales Enterobacteriaceae *Escherichia coli* DEC13B |
| Eco-1047-1 | Bacteria Proteobacteria Gammaproteobacteria Enterobacteriales Enterobacteriaceae *Escherichia coli* DEC13C |
| Eco-1048-1 | Bacteria Proteobacteria Gammaproteobacteria Enterobacteriales Enterobacteriaceae *Escherichia coli* DEC13D |
| Eco-1049-1 | Bacteria Proteobacteria Gammaproteobacteria Enterobacteriales Enterobacteriaceae *Escherichia coli* DEC13E |
| Eco-1050-1 | Bacteria Proteobacteria Gammaproteobacteria Enterobacteriales Enterobacteriaceae *Escherichia coli* DEC14A |
| Eco-1051-1 | Bacteria Proteobacteria Gammaproteobacteria Enterobacteriales Enterobacteriaceae *Escherichia coli* DEC14B |
| Eco-1052-1 | Bacteria Proteobacteria Gammaproteobacteria Enterobacteriales Enterobacteriaceae *Escherichia coli* DEC14C |
| Eco-1053-1 | Bacteria Proteobacteria Gammaproteobacteria Enterobacteriales Enterobacteriaceae *Escherichia coli* DEC14D |
| Eco-1054-1 | Bacteria Proteobacteria Gammaproteobacteria Enterobacteriales Enterobacteriaceae *Escherichia coli* DEC15A |
| Eco-1055-1 | Bacteria Proteobacteria Gammaproteobacteria Enterobacteriales Enterobacteriaceae *Escherichia coli* DEC15B |
| Eco-1056-1 | Bacteria Proteobacteria Gammaproteobacteria Enterobacteriales Enterobacteriaceae *Escherichia coli* DEC15C |
| Eco-1057-1 | Bacteria Proteobacteria Gammaproteobacteria Enterobacteriales Enterobacteriaceae *Escherichia coli* DEC15D |
| Eco-1058-1 | Bacteria Proteobacteria Gammaproteobacteria Enterobacteriales Enterobacteriaceae *Escherichia coli* DEC15E |
| Eco-1059-1 | Bacteria Proteobacteria Gammaproteobacteria Enterobacteriales Enterobacteriaceae *Escherichia coli* DEC1A |
| Eco-1060-1 | Bacteria Proteobacteria Gammaproteobacteria Enterobacteriales Enterobacteriaceae *Escherichia coli* DEC1B |

| | |
|---|---|
| Eco-1061-1 | Bacteria Proteobacteria Gammaproteobacteria Enterobacteriales Enterobacteriaceae *Escherichia coli* DEC1C |
| Eco-1062-1 | Bacteria Proteobacteria Gammaproteobacteria Enterobacteriales Enterobacteriaceae *Escherichia coli* DEC1D |
| Eco-1063-1 | Bacteria Proteobacteria Gammaproteobacteria Enterobacteriales Enterobacteriaceae *Escherichia coli* DEC1E |
| Eco-1064-1 | Bacteria Proteobacteria Gammaproteobacteria Enterobacteriales Enterobacteriaceae *Escherichia coli* DEC2B |
| Eco-1065-1 | Bacteria Proteobacteria Gammaproteobacteria Enterobacteriales Enterobacteriaceae *Escherichia coli* DEC2C |
| Eco-1066-1 | Bacteria Proteobacteria Gammaproteobacteria Enterobacteriales Enterobacteriaceae *Escherichia coli* DEC2D |
| Eco-1067-1 | Bacteria Proteobacteria Gammaproteobacteria Enterobacteriales Enterobacteriaceae *Escherichia coli* DEC2E |
| Eco-1068-1 | Bacteria Proteobacteria Gammaproteobacteria Enterobacteriales Enterobacteriaceae *Escherichia coli* DEC3A |
| Eco-1069-1 | Bacteria Proteobacteria Gammaproteobacteria Enterobacteriales Enterobacteriaceae *Escherichia coli* DEC3B |
| Eco-1070-1 | Bacteria Proteobacteria Gammaproteobacteria Enterobacteriales Enterobacteriaceae *Escherichia coli* DEC3C |
| Eco-1071-1 | Bacteria Proteobacteria Gammaproteobacteria Enterobacteriales Enterobacteriaceae *Escherichia coli* DEC3D |
| Eco-1072-1 | Bacteria Proteobacteria Gammaproteobacteria Enterobacteriales Enterobacteriaceae *Escherichia coli* DEC3E |
| Eco-1073-1 | Bacteria Proteobacteria Gammaproteobacteria Enterobacteriales Enterobacteriaceae *Escherichia coli* DEC3F |
| Eco-1074-1 | Bacteria Proteobacteria Gammaproteobacteria Enterobacteriales Enterobacteriaceae *Escherichia coli* DEC4A |
| Eco-1075-1 | Bacteria Proteobacteria Gammaproteobacteria Enterobacteriales Enterobacteriaceae *Escherichia coli* DEC4B |
| Eco-1076-1 | Bacteria Proteobacteria Gammaproteobacteria Enterobacteriales Enterobacteriaceae *Escherichia coli* DEC4C |
| Eco-1077-1 | Bacteria Proteobacteria Gammaproteobacteria Enterobacteriales Enterobacteriaceae *Escherichia coli* DEC4D |
| Eco-1078-1 | Bacteria Proteobacteria Gammaproteobacteria Enterobacteriales Enterobacteriaceae *Escherichia coli* DEC4E |
| Eco-1079-1 | Bacteria Proteobacteria Gammaproteobacteria Enterobacteriales Enterobacteriaceae *Escherichia coli* DEC4F |
| Eco-1080-1 | Bacteria Proteobacteria Gammaproteobacteria Enterobacteriales Enterobacteriaceae *Escherichia coli* DEC5A |
| Eco-1081-1 | Bacteria Proteobacteria Gammaproteobacteria Enterobacteriales Enterobacteriaceae *Escherichia coli* DEC5B |
| Eco-1082-1 | Bacteria Proteobacteria Gammaproteobacteria Enterobacteriales Enterobacteriaceae *Escherichia coli* DEC5C |
| Eco-1083-1 | Bacteria Proteobacteria Gammaproteobacteria Enterobacteriales Enterobacteriaceae *Escherichia coli* DEC5D |
| Eco-1084-1 | Bacteria Proteobacteria Gammaproteobacteria Enterobacteriales Enterobacteriaceae *Escherichia coli* DEC5E |
| Eco-1085-1 | Bacteria Proteobacteria Gammaproteobacteria Enterobacteriales Enterobacteriaceae *Escherichia coli* DEC6A |
| Eco-1086-1 | Bacteria Proteobacteria Gammaproteobacteria Enterobacteriales Enterobacteriaceae *Escherichia coli* DEC6B |
| Eco-1087-1 | Bacteria Proteobacteria Gammaproteobacteria Enterobacteriales Enterobacteriaceae *Escherichia coli* DEC6E |
| Eco-1088-1 | Bacteria Proteobacteria Gammaproteobacteria Enterobacteriales Enterobacteriaceae *Escherichia coli* DEC7A |
| Eco-1089-1 | Bacteria Proteobacteria Gammaproteobacteria Enterobacteriales Enterobacteriaceae *Escherichia coli* DEC7B |
| Eco-1090-1 | Bacteria Proteobacteria Gammaproteobacteria Enterobacteriales Enterobacteriaceae *Escherichia coli* DEC7C |
| Eco-1091-1 | Bacteria Proteobacteria Gammaproteobacteria Enterobacteriales Enterobacteriaceae *Escherichia coli* DEC7D |
| Eco-1092-1 | Bacteria Proteobacteria Gammaproteobacteria Enterobacteriales Enterobacteriaceae *Escherichia coli* DEC7E |
| Eco-1093-1 | Bacteria Proteobacteria Gammaproteobacteria Enterobacteriales Enterobacteriaceae *Escherichia coli* DEC8A |
| Eco-1094-1 | Bacteria Proteobacteria Gammaproteobacteria Enterobacteriales Enterobacteriaceae *Escherichia coli* DEC8B |
| Eco-1095-1 | Bacteria Proteobacteria Gammaproteobacteria Enterobacteriales Enterobacteriaceae *Escherichia coli* DEC8C |
| Eco-1096-1 | Bacteria Proteobacteria Gammaproteobacteria Enterobacteriales Enterobacteriaceae *Escherichia coli* DEC8D |
| Eco-1097-1 | Bacteria Proteobacteria Gammaproteobacteria Enterobacteriales Enterobacteriaceae *Escherichia coli* DEC8E |
| Eco-1098-1 | Bacteria Proteobacteria Gammaproteobacteria Enterobacteriales Enterobacteriaceae *Escherichia coli* DEC9A |
| Eco-1099-1 | Bacteria Proteobacteria Gammaproteobacteria Enterobacteriales Enterobacteriaceae *Escherichia coli* DEC9B |
| Eco-1100-1 | Bacteria Proteobacteria Gammaproteobacteria Enterobacteriales Enterobacteriaceae *Escherichia coli* DEC9C |
| Eco-1101-1 | Bacteria Proteobacteria Gammaproteobacteria Enterobacteriales Enterobacteriaceae *Escherichia coli* DEC9D |
| Eco-1102-1 | Bacteria Proteobacteria Gammaproteobacteria Enterobacteriales Enterobacteriaceae *Escherichia coli* DEC9E |
| Eco-1103-1 to Eco-1103-2 | Bacteria Proteobacteria Gammaproteobacteria Enterobacteriales Enterobacteriaceae *Escherichia coli* DH1 |
| Eco-1104-1 | Bacteria Proteobacteria Gammaproteobacteria Enterobacteriales Enterobacteriaceae *Escherichia coli* E101 |
| Eco-1105-1 | Bacteria Proteobacteria Gammaproteobacteria Enterobacteriales Enterobacteriaceae *Escherichia coli* E110019 |
| Eco-1106-1 | Bacteria Proteobacteria Gammaproteobacteria Enterobacteriales Enterobacteriaceae *Escherichia coli* E1167 |
| Eco-1107-1 | Bacteria Proteobacteria Gammaproteobacteria Enterobacteriales Enterobacteriaceae *Escherichia coli* E128010 |
| Eco-1108-1 | Bacteria Proteobacteria Gammaproteobacteria Enterobacteriales Enterobacteriaceae *Escherichia coli* E1520 |
| Eco-1109-1 | Bacteria Proteobacteria Gammaproteobacteria Enterobacteriales Enterobacteriaceae *Escherichia coli* E22 |
| Eco-1110-1 | Bacteria Proteobacteria Gammaproteobacteria Enterobacteriales Enterobacteriaceae *Escherichia coli* E2265 |
| Eco-1111-1 | Bacteria Proteobacteria Gammaproteobacteria Enterobacteriales Enterobacteriaceae *Escherichia coli* E24377A |
| Eco-1112-1 | Bacteria Proteobacteria Gammaproteobacteria Enterobacteriales Enterobacteriaceae *Escherichia coli* E482 |
| Eco-1113-1 | Bacteria Proteobacteria Gammaproteobacteria Enterobacteriales Enterobacteriaceae *Escherichia coli* EC1734 |
| Eco-1114-1 | Bacteria Proteobacteria Gammaproteobacteria Enterobacteriales Enterobacteriaceae *Escherichia coli* EC1735 |
| Eco-1115-1 | Bacteria Proteobacteria Gammaproteobacteria Enterobacteriales Enterobacteriaceae *Escherichia coli* EC1736 |
| Eco-1116-1 | Bacteria Proteobacteria Gammaproteobacteria Enterobacteriales Enterobacteriaceae *Escherichia coli* EC1737 |
| Eco-1117-1 | Bacteria Proteobacteria Gammaproteobacteria Enterobacteriales Enterobacteriaceae *Escherichia coli* EC1738 |
| Eco-1118-1 | Bacteria Proteobacteria Gammaproteobacteria Enterobacteriales Enterobacteriaceae *Escherichia coli* EC1845 |
| Eco-1119-1 | Bacteria Proteobacteria Gammaproteobacteria Enterobacteriales Enterobacteriaceae *Escherichia coli* EC1846 |
| Eco-1120-1 | Bacteria Proteobacteria Gammaproteobacteria Enterobacteriales Enterobacteriaceae *Escherichia coli* EC1847 |
| Eco-1121-1 | Bacteria Proteobacteria Gammaproteobacteria Enterobacteriales Enterobacteriaceae *Escherichia coli* EC1848 |
| Eco-1122-1 | Bacteria Proteobacteria Gammaproteobacteria Enterobacteriales Enterobacteriaceae *Escherichia coli* EC1849 |
| Eco-1123-1 | Bacteria Proteobacteria Gammaproteobacteria Enterobacteriales Enterobacteriaceae *Escherichia coli* EC1850 |
| Eco-1124-1 | Bacteria Proteobacteria Gammaproteobacteria Enterobacteriales Enterobacteriaceae *Escherichia coli* EC1856 |
| Eco-1125-1 | Bacteria Proteobacteria Gammaproteobacteria Enterobacteriales Enterobacteriaceae *Escherichia coli* EC1862 |

| | |
|---|---|
| Eco-1126-1 | Bacteria Proteobacteria Gammaproteobacteria Enterobacteriales Enterobacteriaceae *Escherichia coli* EC1863 |
| Eco-1127-1 | Bacteria Proteobacteria Gammaproteobacteria Enterobacteriales Enterobacteriaceae *Escherichia coli* EC1864 |
| Eco-1128-1 | Bacteria Proteobacteria Gammaproteobacteria Enterobacteriales Enterobacteriaceae *Escherichia coli* EC1865 |
| Eco-1129-1 | Bacteria Proteobacteria Gammaproteobacteria Enterobacteriales Enterobacteriaceae *Escherichia coli* EC1866 |
| Eco-1130-1 | Bacteria Proteobacteria Gammaproteobacteria Enterobacteriales Enterobacteriaceae *Escherichia coli* EC1868 |
| Eco-1131-1 | Bacteria Proteobacteria Gammaproteobacteria Enterobacteriales Enterobacteriaceae *Escherichia coli* EC1869 |
| Eco-1132-1 | Bacteria Proteobacteria Gammaproteobacteria Enterobacteriales Enterobacteriaceae *Escherichia coli* EC1870 |
| Eco-1133-1 | Bacteria Proteobacteria Gammaproteobacteria Enterobacteriales Enterobacteriaceae *Escherichia coli* EC302/04 |
| Eco-1134-1 | Bacteria Proteobacteria Gammaproteobacteria Enterobacteriales Enterobacteriaceae *Escherichia coli* EC4013 |
| Eco-1135-1 | Bacteria Proteobacteria Gammaproteobacteria Enterobacteriales Enterobacteriaceae *Escherichia coli* EC4100B |
| Eco-1136-1 | Bacteria Proteobacteria Gammaproteobacteria Enterobacteriales Enterobacteriaceae *Escherichia coli* EC4196 |
| Eco-1137-1 | Bacteria Proteobacteria Gammaproteobacteria Enterobacteriales Enterobacteriaceae *Escherichia coli* EC4203 |
| Eco-1138-1 | Bacteria Proteobacteria Gammaproteobacteria Enterobacteriales Enterobacteriaceae *Escherichia coli* EC4402 |
| Eco-1139-1 | Bacteria Proteobacteria Gammaproteobacteria Enterobacteriales Enterobacteriaceae *Escherichia coli* EC4421 |
| Eco-1140-1 | Bacteria Proteobacteria Gammaproteobacteria Enterobacteriales Enterobacteriaceae *Escherichia coli* EC4422 |
| Eco-1141-1 | Bacteria Proteobacteria Gammaproteobacteria Enterobacteriales Enterobacteriaceae *Escherichia coli* EC4436 |
| Eco-1142-1 | Bacteria Proteobacteria Gammaproteobacteria Enterobacteriales Enterobacteriaceae *Escherichia coli* EC4437 |
| Eco-1143-1 | Bacteria Proteobacteria Gammaproteobacteria Enterobacteriales Enterobacteriaceae *Escherichia coli* EC4439 |
| Eco-1144-1 | Bacteria Proteobacteria Gammaproteobacteria Enterobacteriales Enterobacteriaceae *Escherichia coli* EC4448 |
| Eco-1145-1 | Bacteria Proteobacteria Gammaproteobacteria Enterobacteriales Enterobacteriaceae *Escherichia coli* EC96038 |
| Eco-1146-1 | Bacteria Proteobacteria Gammaproteobacteria Enterobacteriales Enterobacteriaceae *Escherichia coli* ECC-Z |
| Eco-1147-1 | Bacteria Proteobacteria Gammaproteobacteria Enterobacteriales Enterobacteriaceae *Escherichia coli* ED1a |
| Eco-1152-1 | Bacteria Proteobacteria Gammaproteobacteria Enterobacteriales Enterobacteriaceae *Escherichia coli* Envira 10/1 |
| Eco-1153-1 | Bacteria Proteobacteria Gammaproteobacteria Enterobacteriales Enterobacteriaceae *Escherichia coli* Envira 8/11 |
| Eco-1148-1 | Bacteria Proteobacteria Gammaproteobacteria Enterobacteriales Enterobacteriaceae *Escherichia coli* EPEC C342-62 |
| Eco-1149-1 | Bacteria Proteobacteria Gammaproteobacteria Enterobacteriales Enterobacteriaceae *Escherichia coli* EPECa12 |
| Eco-1150-1 | Bacteria Proteobacteria Gammaproteobacteria Enterobacteriales Enterobacteriaceae *Escherichia coli* EPECa14 |
| Eco-1151-1 | Bacteria Proteobacteria Gammaproteobacteria Enterobacteriales Enterobacteriaceae *Escherichia coli* ETEC H10407 |
| Eco-1154-1 | Bacteria Proteobacteria Gammaproteobacteria Enterobacteriales Enterobacteriaceae *Escherichia coli* F11 |
| Eco-1155-1 | Bacteria Proteobacteria Gammaproteobacteria Enterobacteriales Enterobacteriaceae *Escherichia coli* FDA504 |
| Eco-1156-1 | Bacteria Proteobacteria Gammaproteobacteria Enterobacteriales Enterobacteriaceae *Escherichia coli* FDA505 |
| Eco-1157-1 | Bacteria Proteobacteria Gammaproteobacteria Enterobacteriales Enterobacteriaceae *Escherichia coli* FDA506 |
| Eco-1158-1 | Bacteria Proteobacteria Gammaproteobacteria Enterobacteriales Enterobacteriaceae *Escherichia coli* FDA507 |
| Eco-1159-1 | Bacteria Proteobacteria Gammaproteobacteria Enterobacteriales Enterobacteriaceae *Escherichia coli* FDA517 |
| Eco-1160-1 | Bacteria Proteobacteria Gammaproteobacteria Enterobacteriales Enterobacteriaceae *Escherichia coli* FRIK1985 |
| Eco-1161-1 | Bacteria Proteobacteria Gammaproteobacteria Enterobacteriales Enterobacteriaceae *Escherichia coli* FRIK1990 |
| Eco-1162-1 | Bacteria Proteobacteria Gammaproteobacteria Enterobacteriales Enterobacteriaceae *Escherichia coli* FRIK1996 |
| Eco-1163-1 | Bacteria Proteobacteria Gammaproteobacteria Enterobacteriales Enterobacteriaceae *Escherichia coli* FRIK1997 |
| Eco-1164-1 | Bacteria Proteobacteria Gammaproteobacteria Enterobacteriales Enterobacteriaceae *Escherichia coli* FRIK1999 |
| Eco-1165-1 | Bacteria Proteobacteria Gammaproteobacteria Enterobacteriales Enterobacteriaceae *Escherichia coli* FRIK2001 |
| Eco-1166-1 | Bacteria Proteobacteria Gammaproteobacteria Enterobacteriales Enterobacteriaceae *Escherichia coli* FRIK523 |
| Eco-1167-1 | Bacteria Proteobacteria Gammaproteobacteria Enterobacteriales Enterobacteriaceae *Escherichia coli* FRIK920 |
| Eco-1168-1 | Bacteria Proteobacteria Gammaproteobacteria Enterobacteriales Enterobacteriaceae *Escherichia coli* FVEC1302 |
| Eco-1169-1 | Bacteria Proteobacteria Gammaproteobacteria Enterobacteriales Enterobacteriaceae *Escherichia coli* FVEC1412 |
| Eco-1170-1 | Bacteria Proteobacteria Gammaproteobacteria Enterobacteriales Enterobacteriaceae *Escherichia coli* G58-1 |
| Eco-1171-1 to Eco-1171-2 | Bacteria Proteobacteria Gammaproteobacteria Enterobacteriales Enterobacteriaceae *Escherichia coli* GM4792 |
| Eco-1172-1 | Bacteria Proteobacteria Gammaproteobacteria Enterobacteriales Enterobacteriaceae *Escherichia coli* H120 |
| Eco-1173-1 | Bacteria Proteobacteria Gammaproteobacteria Enterobacteriales Enterobacteriaceae *Escherichia coli* H252 |
| Eco-1174-1 | Bacteria Proteobacteria Gammaproteobacteria Enterobacteriales Enterobacteriaceae *Escherichia coli* H263 |
| Eco-1175-1 | Bacteria Proteobacteria Gammaproteobacteria Enterobacteriales Enterobacteriaceae *Escherichia coli* H299 |
| Eco-1176-1 | Bacteria Proteobacteria Gammaproteobacteria Enterobacteriales Enterobacteriaceae *Escherichia coli* H30 |
| Eco-1177-1 | Bacteria Proteobacteria Gammaproteobacteria Enterobacteriales Enterobacteriaceae *Escherichia coli* H397 |
| Eco-1178-1 | Bacteria Proteobacteria Gammaproteobacteria Enterobacteriales Enterobacteriaceae *Escherichia coli* H489 |
| Eco-1179-1 | Bacteria Proteobacteria Gammaproteobacteria Enterobacteriales Enterobacteriaceae *Escherichia coli* H494 |
| Eco-1180-1 | Bacteria Proteobacteria Gammaproteobacteria Enterobacteriales Enterobacteriaceae *Escherichia coli* H591 |
| Eco-1181-1 | Bacteria Proteobacteria Gammaproteobacteria Enterobacteriales Enterobacteriaceae *Escherichia coli* H730 |
| Eco-1182-1 | Bacteria Proteobacteria Gammaproteobacteria Enterobacteriales Enterobacteriaceae *Escherichia coli* H736 |
| Eco-1183-1 | Bacteria Proteobacteria Gammaproteobacteria Enterobacteriales Enterobacteriaceae *Escherichia coli* HM26 |
| Eco-1184-1 | Bacteria Proteobacteria Gammaproteobacteria Enterobacteriales Enterobacteriaceae *Escherichia coli* HM27 |
| Eco-1185-1 | Bacteria Proteobacteria Gammaproteobacteria Enterobacteriales Enterobacteriaceae *Escherichia coli* HM46 |
| Eco-1186-1 | Bacteria Proteobacteria Gammaproteobacteria Enterobacteriales Enterobacteriaceae *Escherichia coli* HM605 |
| Eco-1187-1 | Bacteria Proteobacteria Gammaproteobacteria Enterobacteriales Enterobacteriaceae *Escherichia coli* HM65 |
| Eco-1188-1 | Bacteria Proteobacteria Gammaproteobacteria Enterobacteriales Enterobacteriaceae *Escherichia coli* HM69 |
| Eco-1189-1 | Bacteria Proteobacteria Gammaproteobacteria Enterobacteriales Enterobacteriaceae *Escherichia coli* HS |
| Eco-1190-1 | Bacteria Proteobacteria Gammaproteobacteria Enterobacteriales Enterobacteriaceae *Escherichia coli* HVH 1 (4-6876161) |

| | |
|---|---|
| Eco-1191-1 | Bacteria Proteobacteria Gammaproteobacteria Enterobacteriales Enterobacteriaceae *Escherichia coli* HVH 10 (4-6832164) |
| Eco-1192-1 | Bacteria Proteobacteria Gammaproteobacteria Enterobacteriales Enterobacteriaceae *Escherichia coli* HVH 100 (4-2850729) |
| Eco-1193-1 | Bacteria Proteobacteria Gammaproteobacteria Enterobacteriales Enterobacteriaceae *Escherichia coli* HVH 102 (4-6906788) |
| Eco-1194-1 | Bacteria Proteobacteria Gammaproteobacteria Enterobacteriales Enterobacteriaceae *Escherichia coli* HVH 103 (4-5904188) |
| Eco-1195-1 | Bacteria Proteobacteria Gammaproteobacteria Enterobacteriales Enterobacteriaceae *Escherichia coli* HVH 104 (4-6977960) |
| Eco-1196-1 | Bacteria Proteobacteria Gammaproteobacteria Enterobacteriales Enterobacteriaceae *Escherichia coli* HVH 106 (4-6881831) |
| Eco-1197-1 | Bacteria Proteobacteria Gammaproteobacteria Enterobacteriales Enterobacteriaceae *Escherichia coli* HVH 107 (4-5860571) |
| Eco-1198-1 | Bacteria Proteobacteria Gammaproteobacteria Enterobacteriales Enterobacteriaceae *Escherichia coli* HVH 108 (4-6924867) |
| Eco-1199-1 | Bacteria Proteobacteria Gammaproteobacteria Enterobacteriales Enterobacteriaceae *Escherichia coli* HVH 109 (4-6977162) |
| Eco-1200-1 | Bacteria Proteobacteria Gammaproteobacteria Enterobacteriales Enterobacteriaceae *Escherichia coli* HVH 110 (4-6978754) |
| Eco-1201-1 | Bacteria Proteobacteria Gammaproteobacteria Enterobacteriales Enterobacteriaceae *Escherichia coli* HVH 111 (4-7039018) |
| Eco-1202-1 | Bacteria Proteobacteria Gammaproteobacteria Enterobacteriales Enterobacteriaceae *Escherichia coli* HVH 112 (4-5987253) |
| Eco-1203-1 | Bacteria Proteobacteria Gammaproteobacteria Enterobacteriales Enterobacteriaceae *Escherichia coli* HVH 113 (4-7535473) |
| Eco-1204-1 | Bacteria Proteobacteria Gammaproteobacteria Enterobacteriales Enterobacteriaceae *Escherichia coli* HVH 114 (4-7037740) |
| Eco-1205-1 | Bacteria Proteobacteria Gammaproteobacteria Enterobacteriales Enterobacteriaceae *Escherichia coli* HVH 115 (4-4465989) |
| Eco-1206-1 | Bacteria Proteobacteria Gammaproteobacteria Enterobacteriales Enterobacteriaceae *Escherichia coli* HVH 115 (4-4465997) |
| Eco-1207-1 | Bacteria Proteobacteria Gammaproteobacteria Enterobacteriales Enterobacteriaceae *Escherichia coli* HVH 116 (4-6879942) |
| Eco-1208-1 | Bacteria Proteobacteria Gammaproteobacteria Enterobacteriales Enterobacteriaceae *Escherichia coli* HVH 117 (4-6857191) |
| Eco-1209-1 | Bacteria Proteobacteria Gammaproteobacteria Enterobacteriales Enterobacteriaceae *Escherichia coli* HVH 118 (4-7345399) |
| Eco-1210-1 | Bacteria Proteobacteria Gammaproteobacteria Enterobacteriales Enterobacteriaceae *Escherichia coli* HVH 119 (4-6879578) |
| Eco-1211-1 | Bacteria Proteobacteria Gammaproteobacteria Enterobacteriales Enterobacteriaceae *Escherichia coli* HVH 12 (4-7653042) |
| Eco-1212-1 | Bacteria Proteobacteria Gammaproteobacteria Enterobacteriales Enterobacteriaceae *Escherichia coli* HVH 120 (4-6978681) |
| Eco-1213-1 | Bacteria Proteobacteria Gammaproteobacteria Enterobacteriales Enterobacteriaceae *Escherichia coli* HVH 121 (4-6877826) |
| Eco-1214-1 | Bacteria Proteobacteria Gammaproteobacteria Enterobacteriales Enterobacteriaceae *Escherichia coli* HVH 122 (4-6851606) |
| Eco-1215-1 | Bacteria Proteobacteria Gammaproteobacteria Enterobacteriales Enterobacteriaceae *Escherichia coli* HVH 125 (4-2634716) |
| Eco-1216-1 | Bacteria Proteobacteria Gammaproteobacteria Enterobacteriales Enterobacteriaceae *Escherichia coli* HVH 126 (4-6034225) |
| Eco-1217-1 | Bacteria Proteobacteria Gammaproteobacteria Enterobacteriales Enterobacteriaceae *Escherichia coli* HVH 127 (4-7303629) |
| Eco-1218-1 | Bacteria Proteobacteria Gammaproteobacteria Enterobacteriales Enterobacteriaceae *Escherichia coli* HVH 128 (4-7030436) |
| Eco-1219-1 | Bacteria Proteobacteria Gammaproteobacteria Enterobacteriales Enterobacteriaceae *Escherichia coli* HVH 13 (4-7634056) |
| Eco-1220-1 | Bacteria Proteobacteria Gammaproteobacteria Enterobacteriales Enterobacteriaceae *Escherichia coli* HVH 130 (4-7036876) |
| Eco-1221-1 | Bacteria Proteobacteria Gammaproteobacteria Enterobacteriales Enterobacteriaceae *Escherichia coli* HVH 132 (4-6876862) |
| Eco-1222-1 | Bacteria Proteobacteria Gammaproteobacteria Enterobacteriales Enterobacteriaceae *Escherichia coli* HVH 133 (4-4466519) |
| Eco-1223-1 | Bacteria Proteobacteria Gammaproteobacteria Enterobacteriales Enterobacteriaceae *Escherichia coli* HVH 134 (4-6073441) |
| Eco-1224-1 | Bacteria Proteobacteria Gammaproteobacteria Enterobacteriales Enterobacteriaceae *Escherichia coli* HVH 135 (4-4449320) |
| Eco-1225-1 | Bacteria Proteobacteria Gammaproteobacteria Enterobacteriales Enterobacteriaceae *Escherichia coli* HVH 136 (4-5970458) |
| Eco-1226-1 | Bacteria Proteobacteria Gammaproteobacteria Enterobacteriales Enterobacteriaceae *Escherichia coli* HVH 137 (4-2124971) |
| Eco-1227-1 | Bacteria Proteobacteria Gammaproteobacteria Enterobacteriales Enterobacteriaceae *Escherichia coli* HVH 138 (4-6066704) |
| Eco-1228-1 | Bacteria Proteobacteria Gammaproteobacteria Enterobacteriales Enterobacteriaceae *Escherichia coli* HVH 139 (4-3192644) |
| Eco-1229-1 | Bacteria Proteobacteria Gammaproteobacteria Enterobacteriales Enterobacteriaceae *Escherichia coli* HVH 140 (4-5894387) |
| Eco-1230-1 | Bacteria Proteobacteria Gammaproteobacteria Enterobacteriales Enterobacteriaceae *Escherichia coli* HVH 141 (4-5995973) |
| Eco-1231-1 | Bacteria Proteobacteria Gammaproteobacteria Enterobacteriales Enterobacteriaceae *Escherichia coli* HVH 142 (4-5627451) |
| Eco-1232-1 | Bacteria Proteobacteria Gammaproteobacteria Enterobacteriales Enterobacteriaceae *Escherichia coli* HVH 143 (4-5674999) |
| Eco-1233-1 | Bacteria Proteobacteria Gammaproteobacteria Enterobacteriales Enterobacteriaceae *Escherichia coli* HVH 144 (4-4451937) |
| Eco-1234-1 | Bacteria Proteobacteria Gammaproteobacteria Enterobacteriales Enterobacteriaceae *Escherichia coli* HVH 145 (4-5672112) |
| Eco-1235-1 | Bacteria Proteobacteria Gammaproteobacteria Enterobacteriales Enterobacteriaceae *Escherichia coli* HVH 146 (4-3189767) |
| Eco-1236-1 | Bacteria Proteobacteria Gammaproteobacteria Enterobacteriales Enterobacteriaceae *Escherichia coli* HVH 147 (4-5893887) |
| Eco-1237-1 | Bacteria Proteobacteria Gammaproteobacteria Enterobacteriales Enterobacteriaceae *Escherichia coli* HVH 148 (4-3192490) |
| Eco-1238-1 | Bacteria Proteobacteria Gammaproteobacteria Enterobacteriales Enterobacteriaceae *Escherichia coli* HVH 149 (4-4451880) |
| Eco-1239-1 | Bacteria Proteobacteria Gammaproteobacteria Enterobacteriales Enterobacteriaceae *Escherichia coli* HVH 150 (4-3258106) |
| Eco-1240-1 | Bacteria Proteobacteria Gammaproteobacteria Enterobacteriales Enterobacteriaceae *Escherichia coli* HVH 151 (4-5755573) |
| Eco-1241-1 | Bacteria Proteobacteria Gammaproteobacteria Enterobacteriales Enterobacteriaceae *Escherichia coli* HVH 152 (4-3447545) |
| Eco-1242-1 | Bacteria Proteobacteria Gammaproteobacteria Enterobacteriales Enterobacteriaceae *Escherichia coli* HVH 153 (3-9344314) |
| Eco-1243-1 | Bacteria Proteobacteria Gammaproteobacteria Enterobacteriales Enterobacteriaceae *Escherichia coli* HVH 154 (4-5636698) |
| Eco-1244-1 | Bacteria Proteobacteria Gammaproteobacteria Enterobacteriales Enterobacteriaceae *Escherichia coli* HVH 155 (4-4509048) |
| Eco-1245-1 | Bacteria Proteobacteria Gammaproteobacteria Enterobacteriales Enterobacteriaceae *Escherichia coli* HVH 156 (4-3206505) |
| Eco-1246-1 | Bacteria Proteobacteria Gammaproteobacteria Enterobacteriales Enterobacteriaceae *Escherichia coli* HVH 157 (4-3406229) |
| Eco-1247-1 | Bacteria Proteobacteria Gammaproteobacteria Enterobacteriales Enterobacteriaceae *Escherichia coli* HVH 158 (4-3224287) |
| Eco-1248-1 | Bacteria Proteobacteria Gammaproteobacteria Enterobacteriales Enterobacteriaceae *Escherichia coli* HVH 159 (4-5818141) |
| Eco-1249-1 | Bacteria Proteobacteria Gammaproteobacteria Enterobacteriales Enterobacteriaceae *Escherichia coli* HVH 16 (4-7649002) |
| Eco-1250-1 | Bacteria Proteobacteria Gammaproteobacteria Enterobacteriales Enterobacteriaceae *Escherichia coli* HVH 160 (4-5695937) |
| Eco-1251-1 | Bacteria Proteobacteria Gammaproteobacteria Enterobacteriales Enterobacteriaceae *Escherichia coli* HVH 161 (4-3119890) |
| Eco-1252-1 | Bacteria Proteobacteria Gammaproteobacteria Enterobacteriales Enterobacteriaceae *Escherichia coli* HVH 162 (4-5627982) |
| Eco-1253-1 | Bacteria Proteobacteria Gammaproteobacteria Enterobacteriales Enterobacteriaceae *Escherichia coli* HVH 163 (4-4697553) |
| Eco-1254-1 | Bacteria Proteobacteria Gammaproteobacteria Enterobacteriales Enterobacteriaceae *Escherichia coli* HVH 164 (4-5953081) |
| Eco-1255-1 | Bacteria Proteobacteria Gammaproteobacteria Enterobacteriales Enterobacteriaceae *Escherichia coli* HVH 167 (4-6073565) |

| | |
|---|---|
| Eco-1256-1 | Bacteria Proteobacteria Gammaproteobacteria Enterobacteriales Enterobacteriaceae *Escherichia coli* HVH 169 (4-1075578) |
| Eco-1257-1 | Bacteria Proteobacteria Gammaproteobacteria Enterobacteriales Enterobacteriaceae *Escherichia coli* HVH 17 (4-7473087) |
| Eco-1258-1 | Bacteria Proteobacteria Gammaproteobacteria Enterobacteriales Enterobacteriaceae *Escherichia coli* HVH 170 (4-3026949) |
| Eco-1259-1 | Bacteria Proteobacteria Gammaproteobacteria Enterobacteriales Enterobacteriaceae *Escherichia coli* HVH 171 (4-3191958) |
| Eco-1260-1 | Bacteria Proteobacteria Gammaproteobacteria Enterobacteriales Enterobacteriaceae *Escherichia coli* HVH 172 (4-3248542) |
| Eco-1261-1 | Bacteria Proteobacteria Gammaproteobacteria Enterobacteriales Enterobacteriaceae *Escherichia coli* HVH 173 (3-9175482) |
| Eco-1262-1 | Bacteria Proteobacteria Gammaproteobacteria Enterobacteriales Enterobacteriaceae *Escherichia coli* HVH 175 (4-3405184) |
| Eco-1263-1 | Bacteria Proteobacteria Gammaproteobacteria Enterobacteriales Enterobacteriaceae *Escherichia coli* HVH 176 (4-3428664) |
| Eco-1264-1 | Bacteria Proteobacteria Gammaproteobacteria Enterobacteriales Enterobacteriaceae *Escherichia coli* HVH 177 (4-2876612) |
| Eco-1265-1 | Bacteria Proteobacteria Gammaproteobacteria Enterobacteriales Enterobacteriaceae *Escherichia coli* HVH 178 (4-3189163) |
| Eco-1266-1 | Bacteria Proteobacteria Gammaproteobacteria Enterobacteriales Enterobacteriaceae *Escherichia coli* HVH 18 (4-8589585) |
| Eco-1267-1 | Bacteria Proteobacteria Gammaproteobacteria Enterobacteriales Enterobacteriaceae *Escherichia coli* HVH 180 (4-3051617) |
| Eco-1268-1 | Bacteria Proteobacteria Gammaproteobacteria Enterobacteriales Enterobacteriaceae *Escherichia coli* HVH 182 (4-0985554) |
| Eco-1269-1 | Bacteria Proteobacteria Gammaproteobacteria Enterobacteriales Enterobacteriaceae *Escherichia coli* HVH 183 (4-3205932) |
| Eco-1270-1 | Bacteria Proteobacteria Gammaproteobacteria Enterobacteriales Enterobacteriaceae *Escherichia coli* HVH 184 (4-3343286) |
| Eco-1271-1 | Bacteria Proteobacteria Gammaproteobacteria Enterobacteriales Enterobacteriaceae *Escherichia coli* HVH 185 (4-2876639) |
| Eco-1272-1 | Bacteria Proteobacteria Gammaproteobacteria Enterobacteriales Enterobacteriaceae *Escherichia coli* HVH 186 (4-3405044) |
| Eco-1273-1 | Bacteria Proteobacteria Gammaproteobacteria Enterobacteriales Enterobacteriaceae *Escherichia coli* HVH 187 (4-4471660) |
| Eco-1274-1 | Bacteria Proteobacteria Gammaproteobacteria Enterobacteriales Enterobacteriaceae *Escherichia coli* HVH 188 (4-2356988) |
| Eco-1275-1 | Bacteria Proteobacteria Gammaproteobacteria Enterobacteriales Enterobacteriaceae *Escherichia coli* HVH 189 (4-3220125) |
| Eco-1276-1 | Bacteria Proteobacteria Gammaproteobacteria Enterobacteriales Enterobacteriaceae *Escherichia coli* HVH 19 (4-7154984) |
| Eco-1277-1 | Bacteria Proteobacteria Gammaproteobacteria Enterobacteriales Enterobacteriaceae *Escherichia coli* HVH 190 (4-3255514) |
| Eco-1278-1 | Bacteria Proteobacteria Gammaproteobacteria Enterobacteriales Enterobacteriaceae *Escherichia coli* HVH 191 (3-9341900) |
| Eco-1279-1 | Bacteria Proteobacteria Gammaproteobacteria Enterobacteriales Enterobacteriaceae *Escherichia coli* HVH 192 (4-3054470) |
| Eco-1280-1 | Bacteria Proteobacteria Gammaproteobacteria Enterobacteriales Enterobacteriaceae *Escherichia coli* HVH 193 (4-3331423) |
| Eco-1281-1 | Bacteria Proteobacteria Gammaproteobacteria Enterobacteriales Enterobacteriaceae *Escherichia coli* HVH 194 (4-2356805) |
| Eco-1282-1 | Bacteria Proteobacteria Gammaproteobacteria Enterobacteriales Enterobacteriaceae *Escherichia coli* HVH 195 (3-7155360) |
| Eco-1283-1 | Bacteria Proteobacteria Gammaproteobacteria Enterobacteriales Enterobacteriaceae *Escherichia coli* HVH 196 (4-4530470) |
| Eco-1284-1 | Bacteria Proteobacteria Gammaproteobacteria Enterobacteriales Enterobacteriaceae *Escherichia coli* HVH 197 (4-4466217) |
| Eco-1285-1 | Bacteria Proteobacteria Gammaproteobacteria Enterobacteriales Enterobacteriaceae *Escherichia coli* HVH 198 (4-3206106) |
| Eco-1286-1 | Bacteria Proteobacteria Gammaproteobacteria Enterobacteriales Enterobacteriaceae *Escherichia coli* HVH 199 (4-5670322) |
| Eco-1287-1 | Bacteria Proteobacteria Gammaproteobacteria Enterobacteriales Enterobacteriaceae *Escherichia coli* HVH 2 (4-6943160) |
| Eco-1288-1 | Bacteria Proteobacteria Gammaproteobacteria Enterobacteriales Enterobacteriaceae *Escherichia coli* HVH 20 (4-5865042) |
| Eco-1289-1 | Bacteria Proteobacteria Gammaproteobacteria Enterobacteriales Enterobacteriaceae *Escherichia coli* HVH 200 (4-4449924) |
| Eco-1290-1 | Bacteria Proteobacteria Gammaproteobacteria Enterobacteriales Enterobacteriaceae *Escherichia coli* HVH 201 (4-4459431) |
| Eco-1291-1 | Bacteria Proteobacteria Gammaproteobacteria Enterobacteriales Enterobacteriaceae *Escherichia coli* HVH 202 (4-3163997) |
| Eco-1292-1 | Bacteria Proteobacteria Gammaproteobacteria Enterobacteriales Enterobacteriaceae *Escherichia coli* HVH 203 (4-3126218) |
| Eco-1293-1 | Bacteria Proteobacteria Gammaproteobacteria Enterobacteriales Enterobacteriaceae *Escherichia coli* HVH 204 (4-3112802) |
| Eco-1294-1 | Bacteria Proteobacteria Gammaproteobacteria Enterobacteriales Enterobacteriaceae *Escherichia coli* HVH 205 (4-3094677) |
| Eco-1295-1 | Bacteria Proteobacteria Gammaproteobacteria Enterobacteriales Enterobacteriaceae *Escherichia coli* HVH 206 (4-3128229) |
| Eco-1296-1 | Bacteria Proteobacteria Gammaproteobacteria Enterobacteriales Enterobacteriaceae *Escherichia coli* HVH 207 (4-3113221) |
| Eco-1297-1 | Bacteria Proteobacteria Gammaproteobacteria Enterobacteriales Enterobacteriaceae *Escherichia coli* HVH 208 (4-3112292) |
| Eco-1298-1 | Bacteria Proteobacteria Gammaproteobacteria Enterobacteriales Enterobacteriaceae *Escherichia coli* HVH 209 (4-3062651) |
| Eco-1299-1 | Bacteria Proteobacteria Gammaproteobacteria Enterobacteriales Enterobacteriaceae *Escherichia coli* HVH 21 (4-4517873) |
| Eco-1300-1 | Bacteria Proteobacteria Gammaproteobacteria Enterobacteriales Enterobacteriaceae *Escherichia coli* HVH 210 (4-3042480) |
| Eco-1301-1 | Bacteria Proteobacteria Gammaproteobacteria Enterobacteriales Enterobacteriaceae *Escherichia coli* HVH 211 (4-3041891) |
| Eco-1302-1 | Bacteria Proteobacteria Gammaproteobacteria Enterobacteriales Enterobacteriaceae *Escherichia coli* HVH 212 (3-9305343) |
| Eco-1303-1 | Bacteria Proteobacteria Gammaproteobacteria Enterobacteriales Enterobacteriaceae *Escherichia coli* HVH 213 (4-3042928) |
| Eco-1304-1 | Bacteria Proteobacteria Gammaproteobacteria Enterobacteriales Enterobacteriaceae *Escherichia coli* HVH 214 (4-3062198) |
| Eco-1305-1 | Bacteria Proteobacteria Gammaproteobacteria Enterobacteriales Enterobacteriaceae *Escherichia coli* HVH 215 (4-3008371) |
| Eco-1306-1 | Bacteria Proteobacteria Gammaproteobacteria Enterobacteriales Enterobacteriaceae *Escherichia coli* HVH 216 (4-3042952) |
| Eco-1307-1 | Bacteria Proteobacteria Gammaproteobacteria Enterobacteriales Enterobacteriaceae *Escherichia coli* HVH 217 (4-1022806) |
| Eco-1308-1 | Bacteria Proteobacteria Gammaproteobacteria Enterobacteriales Enterobacteriaceae *Escherichia coli* HVH 218 (4-4500903) |
| Eco-1309-1 | Bacteria Proteobacteria Gammaproteobacteria Enterobacteriales Enterobacteriaceae *Escherichia coli* HVH 22 (4-2258986) |
| Eco-1310-1 | Bacteria Proteobacteria Gammaproteobacteria Enterobacteriales Enterobacteriaceae *Escherichia coli* HVH 220 (4-5876842) |
| Eco-1311-1 | Bacteria Proteobacteria Gammaproteobacteria Enterobacteriales Enterobacteriaceae *Escherichia coli* HVH 221 (4-3136817) |
| Eco-1312-1 | Bacteria Proteobacteria Gammaproteobacteria Enterobacteriales Enterobacteriaceae *Escherichia coli* HVH 222 (4-2977443) |
| Eco-1313-1 | Bacteria Proteobacteria Gammaproteobacteria Enterobacteriales Enterobacteriaceae *Escherichia coli* HVH 223 (4-2976528) |
| Eco-1314-1 | Bacteria Proteobacteria Gammaproteobacteria Enterobacteriales Enterobacteriaceae *Escherichia coli* HVH 225 (4-1273116) |
| Eco-1315-1 | Bacteria Proteobacteria Gammaproteobacteria Enterobacteriales Enterobacteriaceae *Escherichia coli* HVH 227 (4-2277670) |
| Eco-1316-1 | Bacteria Proteobacteria Gammaproteobacteria Enterobacteriales Enterobacteriaceae *Escherichia coli* HVH 228 (4-7787030) |
| Eco-1317-1 | Bacteria Proteobacteria Gammaproteobacteria Enterobacteriales Enterobacteriaceae *Escherichia coli* HVH 23 (4-6066488) |
| Eco-1318-1 | Bacteria Proteobacteria Gammaproteobacteria Enterobacteriales Enterobacteriaceae *Escherichia coli* HVH 24 (4-5985145) |
| Eco-1319-1 | Bacteria Proteobacteria Gammaproteobacteria Enterobacteriales Enterobacteriaceae *Escherichia coli* HVH 25 (4-5851939) |
| Eco-1320-1 | Bacteria Proteobacteria Gammaproteobacteria Enterobacteriales Enterobacteriaceae *Escherichia coli* HVH 26 (4-5703913) |

| | |
|---|---|
| Eco-1321-1 | Bacteria Proteobacteria Gammaproteobacteria Enterobacteriales Enterobacteriaceae *Escherichia coli* HVH 27 (4-7449267) |
| Eco-1322-1 | Bacteria Proteobacteria Gammaproteobacteria Enterobacteriales Enterobacteriaceae *Escherichia coli* HVH 28 (4-0907367) |
| Eco-1323-1 | Bacteria Proteobacteria Gammaproteobacteria Enterobacteriales Enterobacteriaceae *Escherichia coli* HVH 29 (4-3418073) |
| Eco-1324-1 | Bacteria Proteobacteria Gammaproteobacteria Enterobacteriales Enterobacteriaceae *Escherichia coli* HVH 3 (4-7276001) |
| Eco-1325-1 | Bacteria Proteobacteria Gammaproteobacteria Enterobacteriales Enterobacteriaceae *Escherichia coli* HVH 30 (4-2661829) |
| Eco-1326-1 | Bacteria Proteobacteria Gammaproteobacteria Enterobacteriales Enterobacteriaceae *Escherichia coli* HVH 31 (4-2602156) |
| Eco-1327-1 | Bacteria Proteobacteria Gammaproteobacteria Enterobacteriales Enterobacteriaceae *Escherichia coli* HVH 32 (4-3773988) |
| Eco-1328-1 | Bacteria Proteobacteria Gammaproteobacteria Enterobacteriales Enterobacteriaceae *Escherichia coli* HVH 33 (4-2174936) |
| Eco-1329-1 | Bacteria Proteobacteria Gammaproteobacteria Enterobacteriales Enterobacteriaceae *Escherichia coli* HVH 35 (4-2962667) |
| Eco-1330-1 | Bacteria Proteobacteria Gammaproteobacteria Enterobacteriales Enterobacteriaceae *Escherichia coli* HVH 36 (4-5675286) |
| Eco-1331-1 | Bacteria Proteobacteria Gammaproteobacteria Enterobacteriales Enterobacteriaceae *Escherichia coli* HVH 37 (4-2773848) |
| Eco-1332-1 | Bacteria Proteobacteria Gammaproteobacteria Enterobacteriales Enterobacteriaceae *Escherichia coli* HVH 38 (4-2774682) |
| Eco-1333-1 | Bacteria Proteobacteria Gammaproteobacteria Enterobacteriales Enterobacteriaceae *Escherichia coli* HVH 39 (4-2679949) |
| Eco-1334-1 | Bacteria Proteobacteria Gammaproteobacteria Enterobacteriales Enterobacteriaceae *Escherichia coli* HVH 4 (4-7276109) |
| Eco-1335-1 | Bacteria Proteobacteria Gammaproteobacteria Enterobacteriales Enterobacteriaceae *Escherichia coli* HVH 40 (4-1219782) |
| Eco-1336-1 | Bacteria Proteobacteria Gammaproteobacteria Enterobacteriales Enterobacteriaceae *Escherichia coli* HVH 41 (4-2677849) |
| Eco-1337-1 | Bacteria Proteobacteria Gammaproteobacteria Enterobacteriales Enterobacteriaceae *Escherichia coli* HVH 42 (4-2100061) |
| Eco-1338-1 | Bacteria Proteobacteria Gammaproteobacteria Enterobacteriales Enterobacteriaceae *Escherichia coli* HVH 43 (4-2173468) |
| Eco-1339-1 | Bacteria Proteobacteria Gammaproteobacteria Enterobacteriales Enterobacteriaceae *Escherichia coli* HVH 44 (4-2298570) |
| Eco-1340-1 | Bacteria Proteobacteria Gammaproteobacteria Enterobacteriales Enterobacteriaceae *Escherichia coli* HVH 45 (4-3129918) |
| Eco-1341-1 | Bacteria Proteobacteria Gammaproteobacteria Enterobacteriales Enterobacteriaceae *Escherichia coli* HVH 46 (4-2758776) |
| Eco-1342-1 | Bacteria Proteobacteria Gammaproteobacteria Enterobacteriales Enterobacteriaceae *Escherichia coli* HVH 48 (4-2658593) |
| Eco-1343-1 | Bacteria Proteobacteria Gammaproteobacteria Enterobacteriales Enterobacteriaceae *Escherichia coli* HVH 5 (4-7148410) |
| Eco-1344-1 | Bacteria Proteobacteria Gammaproteobacteria Enterobacteriales Enterobacteriaceae *Escherichia coli* HVH 50 (4-2593475) |
| Eco-1345-1 | Bacteria Proteobacteria Gammaproteobacteria Enterobacteriales Enterobacteriaceae *Escherichia coli* HVH 51 (4-2172526) |
| Eco-1346-1 | Bacteria Proteobacteria Gammaproteobacteria Enterobacteriales Enterobacteriaceae *Escherichia coli* HVH 53 (4-0631051) |
| Eco-1347-1 | Bacteria Proteobacteria Gammaproteobacteria Enterobacteriales Enterobacteriaceae *Escherichia coli* HVH 55 (4-2646161) |
| Eco-1348-1 | Bacteria Proteobacteria Gammaproteobacteria Enterobacteriales Enterobacteriaceae *Escherichia coli* HVH 56 (4-2153033) |
| Eco-1349-1 | Bacteria Proteobacteria Gammaproteobacteria Enterobacteriales Enterobacteriaceae *Escherichia coli* HVH 58 (4-2839709) |
| Eco-1350-1 | Bacteria Proteobacteria Gammaproteobacteria Enterobacteriales Enterobacteriaceae *Escherichia coli* HVH 59 (4-1119338) |
| Eco-1351-1 | Bacteria Proteobacteria Gammaproteobacteria Enterobacteriales Enterobacteriaceae *Escherichia coli* HVH 6 (3-8296502) |
| Eco-1352-1 | Bacteria Proteobacteria Gammaproteobacteria Enterobacteriales Enterobacteriaceae *Escherichia coli* HVH 61 (4-2736020) |
| Eco-1353-1 | Bacteria Proteobacteria Gammaproteobacteria Enterobacteriales Enterobacteriaceae *Escherichia coli* HVH 63 (4-2542528) |
| Eco-1354-1 | Bacteria Proteobacteria Gammaproteobacteria Enterobacteriales Enterobacteriaceae *Escherichia coli* HVH 65 (4-2262045) |
| Eco-1355-1 | Bacteria Proteobacteria Gammaproteobacteria Enterobacteriales Enterobacteriaceae *Escherichia coli* HVH 68 (4-0888028) |
| Eco-1356-1 | Bacteria Proteobacteria Gammaproteobacteria Enterobacteriales Enterobacteriaceae *Escherichia coli* HVH 69 (4-2837072) |
| Eco-1357-1 | Bacteria Proteobacteria Gammaproteobacteria Enterobacteriales Enterobacteriaceae *Escherichia coli* HVH 7 (4-7315031) |
| Eco-1358-1 | Bacteria Proteobacteria Gammaproteobacteria Enterobacteriales Enterobacteriaceae *Escherichia coli* HVH 70 (4-2963531) |
| Eco-1359-1 | Bacteria Proteobacteria Gammaproteobacteria Enterobacteriales Enterobacteriaceae *Escherichia coli* HVH 73 (4-2393174) |
| Eco-1360-1 | Bacteria Proteobacteria Gammaproteobacteria Enterobacteriales Enterobacteriaceae *Escherichia coli* HVH 74 (4-1034782) |
| Eco-1361-1 | Bacteria Proteobacteria Gammaproteobacteria Enterobacteriales Enterobacteriaceae *Escherichia coli* HVH 76 (4-2538717) |
| Eco-1362-1 | Bacteria Proteobacteria Gammaproteobacteria Enterobacteriales Enterobacteriaceae *Escherichia coli* HVH 77 (4-2605759) |
| Eco-1363-1 | Bacteria Proteobacteria Gammaproteobacteria Enterobacteriales Enterobacteriaceae *Escherichia coli* HVH 78 (4-2735946) |
| Eco-1364-1 | Bacteria Proteobacteria Gammaproteobacteria Enterobacteriales Enterobacteriaceae *Escherichia coli* HVH 79 (4-2512823) |
| Eco-1365-1 | Bacteria Proteobacteria Gammaproteobacteria Enterobacteriales Enterobacteriaceae *Escherichia coli* HVH 80 (4-2428830) |
| Eco-1366-1 | Bacteria Proteobacteria Gammaproteobacteria Enterobacteriales Enterobacteriaceae *Escherichia coli* HVH 82 (4-2209276) |
| Eco-1367-1 | Bacteria Proteobacteria Gammaproteobacteria Enterobacteriales Enterobacteriaceae *Escherichia coli* HVH 83 (4-2051087) |
| Eco-1368-1 | Bacteria Proteobacteria Gammaproteobacteria Enterobacteriales Enterobacteriaceae *Escherichia coli* HVH 84 (4-1021478) |
| Eco-1369-1 | Bacteria Proteobacteria Gammaproteobacteria Enterobacteriales Enterobacteriaceae *Escherichia coli* HVH 85 (4-0792144) |
| Eco-1370-1 | Bacteria Proteobacteria Gammaproteobacteria Enterobacteriales Enterobacteriaceae *Escherichia coli* HVH 86 (4-7026218) |
| Eco-1371-1 | Bacteria Proteobacteria Gammaproteobacteria Enterobacteriales Enterobacteriaceae *Escherichia coli* HVH 87 (4-5977630) |
| Eco-1372-1 | Bacteria Proteobacteria Gammaproteobacteria Enterobacteriales Enterobacteriaceae *Escherichia coli* HVH 88 (4-5854636) |
| Eco-1373-1 | Bacteria Proteobacteria Gammaproteobacteria Enterobacteriales Enterobacteriaceae *Escherichia coli* HVH 89 (4-5885604) |
| Eco-1374-1 | Bacteria Proteobacteria Gammaproteobacteria Enterobacteriales Enterobacteriaceae *Escherichia coli* HVH 9 (4-6942539) |
| Eco-1375-1 | Bacteria Proteobacteria Gammaproteobacteria Enterobacteriales Enterobacteriaceae *Escherichia coli* HVH 90 (4-3191362) |
| Eco-1376-1 | Bacteria Proteobacteria Gammaproteobacteria Enterobacteriales Enterobacteriaceae *Escherichia coli* HVH 91 (4-4638751) |
| Eco-1377-1 | Bacteria Proteobacteria Gammaproteobacteria Enterobacteriales Enterobacteriaceae *Escherichia coli* HVH 92 (4-5930790) |
| Eco-1378-1 | Bacteria Proteobacteria Gammaproteobacteria Enterobacteriales Enterobacteriaceae *Escherichia coli* HVH 95 (4-6074464) |
| Eco-1379-1 | Bacteria Proteobacteria Gammaproteobacteria Enterobacteriales Enterobacteriaceae *Escherichia coli* HVH 96 (4-5934869) |
| Eco-1380-1 | Bacteria Proteobacteria Gammaproteobacteria Enterobacteriales Enterobacteriaceae *Escherichia coli* HVH 98 (4-5799287) |
| Eco-1381-1 | Bacteria Proteobacteria Gammaproteobacteria Enterobacteriales Enterobacteriaceae *Escherichia coli* IAI1 |
| Eco-1382-1 | Bacteria Proteobacteria Gammaproteobacteria Enterobacteriales Enterobacteriaceae *Escherichia coli* IAI39 |
| Eco-1383-1 | Bacteria Proteobacteria Gammaproteobacteria Enterobacteriales Enterobacteriaceae *Escherichia coli* IHE3034 |
| Eco-1384-1 | Bacteria Proteobacteria Gammaproteobacteria Enterobacteriales Enterobacteriaceae *Escherichia coli* IS29 |
| Eco-1385-1 | Bacteria Proteobacteria Gammaproteobacteria Enterobacteriales Enterobacteriaceae *Escherichia coli* IS5 |

| | |
|---|---|
| Eco-1386-1 | Bacteria Proteobacteria Gammaproteobacteria Enterobacteriales Enterobacteriaceae *Escherichia coli* J53 |
| Eco-1387-1 | Bacteria Proteobacteria Gammaproteobacteria Enterobacteriales Enterobacteriaceae *Escherichia coli* J96 |
| Eco-1388-1 | Bacteria Proteobacteria Gammaproteobacteria Enterobacteriales Enterobacteriaceae *Escherichia coli* JB1-95 |
| Eco-1389-1 | Bacteria Proteobacteria Gammaproteobacteria Enterobacteriales Enterobacteriaceae *Escherichia coli* JJ1886 |
| Eco-1390-1 | Bacteria Proteobacteria Gammaproteobacteria Enterobacteriales Enterobacteriaceae *Escherichia coli* Jurua 18/11 |
| Eco-1391-1 | Bacteria Proteobacteria Gammaproteobacteria Enterobacteriales Enterobacteriaceae *Escherichia coli* Jurua 20/10 |
| Eco-1392-1 | Bacteria Proteobacteria Gammaproteobacteria Enterobacteriales Enterobacteriaceae *Escherichia coli* KD1 |
| Eco-1393-1 | Bacteria Proteobacteria Gammaproteobacteria Enterobacteriales Enterobacteriaceae *Escherichia coli* KD2 |
| Eco-1394-1 to Eco-1394-2 | Bacteria Proteobacteria Gammaproteobacteria Enterobacteriales Enterobacteriaceae *Escherichia coli* KO11FL |
| Eco-1395-1 | Bacteria Proteobacteria Gammaproteobacteria Enterobacteriales Enterobacteriaceae *Escherichia coli* KOEGE 10 (25a) |
| Eco-1396-1 | Bacteria Proteobacteria Gammaproteobacteria Enterobacteriales Enterobacteriaceae *Escherichia coli* KOEGE 118 (317a) |
| Eco-1397-1 | Bacteria Proteobacteria Gammaproteobacteria Enterobacteriales Enterobacteriaceae *Escherichia coli* KOEGE 131 (358a) |
| Eco-1398-1 | Bacteria Proteobacteria Gammaproteobacteria Enterobacteriales Enterobacteriaceae *Escherichia coli* KOEGE 3 (4a) |
| Eco-1399-1 | Bacteria Proteobacteria Gammaproteobacteria Enterobacteriales Enterobacteriaceae *Escherichia coli* KOEGE 30 (63a) |
| Eco-1400-1 | Bacteria Proteobacteria Gammaproteobacteria Enterobacteriales Enterobacteriaceae *Escherichia coli* KOEGE 32 (66a) |
| Eco-1401-1 | Bacteria Proteobacteria Gammaproteobacteria Enterobacteriales Enterobacteriaceae *Escherichia coli* KOEGE 33 (68a) |
| Eco-1402-1 | Bacteria Proteobacteria Gammaproteobacteria Enterobacteriales Enterobacteriaceae *Escherichia coli* KOEGE 40 (102a) |
| Eco-1403-1 | Bacteria Proteobacteria Gammaproteobacteria Enterobacteriales Enterobacteriaceae *Escherichia coli* KOEGE 43 (105a) |
| Eco-1404-1 | Bacteria Proteobacteria Gammaproteobacteria Enterobacteriales Enterobacteriaceae *Escherichia coli* KOEGE 44 (106a) |
| Eco-1405-1 | Bacteria Proteobacteria Gammaproteobacteria Enterobacteriales Enterobacteriaceae *Escherichia coli* KOEGE 56 (169a) |
| Eco-1406-1 | Bacteria Proteobacteria Gammaproteobacteria Enterobacteriales Enterobacteriaceae *Escherichia coli* KOEGE 58 (171a) |
| Eco-1407-1 | Bacteria Proteobacteria Gammaproteobacteria Enterobacteriales Enterobacteriaceae *Escherichia coli* KOEGE 61 (174a) |
| Eco-1408-1 | Bacteria Proteobacteria Gammaproteobacteria Enterobacteriales Enterobacteriaceae *Escherichia coli* KOEGE 62 (175a) |
| Eco-1409-1 | Bacteria Proteobacteria Gammaproteobacteria Enterobacteriales Enterobacteriaceae *Escherichia coli* KOEGE 68 (182a) |
| Eco-1410-1 | Bacteria Proteobacteria Gammaproteobacteria Enterobacteriales Enterobacteriaceae *Escherichia coli* KOEGE 7 (16a) |
| Eco-1411-1 | Bacteria Proteobacteria Gammaproteobacteria Enterobacteriales Enterobacteriaceae *Escherichia coli* KOEGE 70 (185a) |
| Eco-1412-1 | Bacteria Proteobacteria Gammaproteobacteria Enterobacteriales Enterobacteriaceae *Escherichia coli* KOEGE 71 (186a) |
| Eco-1413-1 | Bacteria Proteobacteria Gammaproteobacteria Enterobacteriales Enterobacteriaceae *Escherichia coli* KOEGE 73 (195a) |
| Eco-1414-1 | Bacteria Proteobacteria Gammaproteobacteria Enterobacteriales Enterobacteriaceae *Escherichia coli* KOEGE 77 (202a) |
| Eco-1415-1 | Bacteria Proteobacteria Gammaproteobacteria Enterobacteriales Enterobacteriaceae *Escherichia coli* KTE1 |
| Eco-1416-1 | Bacteria Proteobacteria Gammaproteobacteria Enterobacteriales Enterobacteriaceae *Escherichia coli* KTE10 |
| Eco-1417-1 | Bacteria Proteobacteria Gammaproteobacteria Enterobacteriales Enterobacteriaceae *Escherichia coli* KTE100 |
| Eco-1418-1 | Bacteria Proteobacteria Gammaproteobacteria Enterobacteriales Enterobacteriaceae *Escherichia coli* KTE101 |
| Eco-1419-1 | Bacteria Proteobacteria Gammaproteobacteria Enterobacteriales Enterobacteriaceae *Escherichia coli* KTE102 |
| Eco-1420-1 | Bacteria Proteobacteria Gammaproteobacteria Enterobacteriales Enterobacteriaceae *Escherichia coli* KTE103 |
| Eco-1421-1 | Bacteria Proteobacteria Gammaproteobacteria Enterobacteriales Enterobacteriaceae *Escherichia coli* KTE104 |
| Eco-1422-1 | Bacteria Proteobacteria Gammaproteobacteria Enterobacteriales Enterobacteriaceae *Escherichia coli* KTE105 |
| Eco-1423-1 | Bacteria Proteobacteria Gammaproteobacteria Enterobacteriales Enterobacteriaceae *Escherichia coli* KTE106 |
| Eco-1424-1 | Bacteria Proteobacteria Gammaproteobacteria Enterobacteriales Enterobacteriaceae *Escherichia coli* KTE107 |
| Eco-1425-1 | Bacteria Proteobacteria Gammaproteobacteria Enterobacteriales Enterobacteriaceae *Escherichia coli* KTE108 |
| Eco-1426-1 | Bacteria Proteobacteria Gammaproteobacteria Enterobacteriales Enterobacteriaceae *Escherichia coli* KTE109 |
| Eco-1427-1 | Bacteria Proteobacteria Gammaproteobacteria Enterobacteriales Enterobacteriaceae *Escherichia coli* KTE11 |
| Eco-1428-1 | Bacteria Proteobacteria Gammaproteobacteria Enterobacteriales Enterobacteriaceae *Escherichia coli* KTE111 |
| Eco-1429-1 | Bacteria Proteobacteria Gammaproteobacteria Enterobacteriales Enterobacteriaceae *Escherichia coli* KTE112 |
| Eco-1430-1 | Bacteria Proteobacteria Gammaproteobacteria Enterobacteriales Enterobacteriaceae *Escherichia coli* KTE113 |
| Eco-1431-1 | Bacteria Proteobacteria Gammaproteobacteria Enterobacteriales Enterobacteriaceae *Escherichia coli* KTE114 |
| Eco-1432-1 | Bacteria Proteobacteria Gammaproteobacteria Enterobacteriales Enterobacteriaceae *Escherichia coli* KTE115 |
| Eco-1433-1 | Bacteria Proteobacteria Gammaproteobacteria Enterobacteriales Enterobacteriaceae *Escherichia coli* KTE116 |
| Eco-1434-1 | Bacteria Proteobacteria Gammaproteobacteria Enterobacteriales Enterobacteriaceae *Escherichia coli* KTE117 |
| Eco-1435-1 | Bacteria Proteobacteria Gammaproteobacteria Enterobacteriales Enterobacteriaceae *Escherichia coli* KTE118 |
| Eco-1436-1 | Bacteria Proteobacteria Gammaproteobacteria Enterobacteriales Enterobacteriaceae *Escherichia coli* KTE119 |
| Eco-1437-1 | Bacteria Proteobacteria Gammaproteobacteria Enterobacteriales Enterobacteriaceae *Escherichia coli* KTE12 |
| Eco-1438-1 | Bacteria Proteobacteria Gammaproteobacteria Enterobacteriales Enterobacteriaceae *Escherichia coli* KTE120 |
| Eco-1439-1 | Bacteria Proteobacteria Gammaproteobacteria Enterobacteriales Enterobacteriaceae *Escherichia coli* KTE121 |
| Eco-1440-1 | Bacteria Proteobacteria Gammaproteobacteria Enterobacteriales Enterobacteriaceae *Escherichia coli* KTE122 |
| Eco-1441-1 | Bacteria Proteobacteria Gammaproteobacteria Enterobacteriales Enterobacteriaceae *Escherichia coli* KTE123 |
| Eco-1442-1 | Bacteria Proteobacteria Gammaproteobacteria Enterobacteriales Enterobacteriaceae *Escherichia coli* KTE124 |
| Eco-1443-1 | Bacteria Proteobacteria Gammaproteobacteria Enterobacteriales Enterobacteriaceae *Escherichia coli* KTE125 |
| Eco-1444-1 | Bacteria Proteobacteria Gammaproteobacteria Enterobacteriales Enterobacteriaceae *Escherichia coli* KTE126 |
| Eco-1445-1 | Bacteria Proteobacteria Gammaproteobacteria Enterobacteriales Enterobacteriaceae *Escherichia coli* KTE127 |
| Eco-1446-1 | Bacteria Proteobacteria Gammaproteobacteria Enterobacteriales Enterobacteriaceae *Escherichia coli* KTE128 |
| Eco-1447-1 | Bacteria Proteobacteria Gammaproteobacteria Enterobacteriales Enterobacteriaceae *Escherichia coli* KTE129 |
| Eco-1448-1 | Bacteria Proteobacteria Gammaproteobacteria Enterobacteriales Enterobacteriaceae *Escherichia coli* KTE13 |
| Eco-1449-1 | Bacteria Proteobacteria Gammaproteobacteria Enterobacteriales Enterobacteriaceae *Escherichia coli* KTE130 |
| Eco-1450-1 | Bacteria Proteobacteria Gammaproteobacteria Enterobacteriales Enterobacteriaceae *Escherichia coli* KTE131 |

| | |
|---|---|
| Eco-1451-1 | Bacteria Proteobacteria Gammaproteobacteria Enterobacteriales Enterobacteriaceae *Escherichia coli* KTE132 |
| Eco-1452-1 | Bacteria Proteobacteria Gammaproteobacteria Enterobacteriales Enterobacteriaceae *Escherichia coli* KTE133 |
| Eco-1453-1 | Bacteria Proteobacteria Gammaproteobacteria Enterobacteriales Enterobacteriaceae *Escherichia coli* KTE134 |
| Eco-1454-1 | Bacteria Proteobacteria Gammaproteobacteria Enterobacteriales Enterobacteriaceae *Escherichia coli* KTE135 |
| Eco-1455-1 | Bacteria Proteobacteria Gammaproteobacteria Enterobacteriales Enterobacteriaceae *Escherichia coli* KTE136 |
| Eco-1456-1 | Bacteria Proteobacteria Gammaproteobacteria Enterobacteriales Enterobacteriaceae *Escherichia coli* KTE137 |
| Eco-1457-1 | Bacteria Proteobacteria Gammaproteobacteria Enterobacteriales Enterobacteriaceae *Escherichia coli* KTE138 |
| Eco-1458-1 | Bacteria Proteobacteria Gammaproteobacteria Enterobacteriales Enterobacteriaceae *Escherichia coli* KTE139 |
| Eco-1459-1 | Bacteria Proteobacteria Gammaproteobacteria Enterobacteriales Enterobacteriaceae *Escherichia coli* KTE14 |
| Eco-1460-1 | Bacteria Proteobacteria Gammaproteobacteria Enterobacteriales Enterobacteriaceae *Escherichia coli* KTE140 |
| Eco-1461-1 | Bacteria Proteobacteria Gammaproteobacteria Enterobacteriales Enterobacteriaceae *Escherichia coli* KTE141 |
| Eco-1462-1 | Bacteria Proteobacteria Gammaproteobacteria Enterobacteriales Enterobacteriaceae *Escherichia coli* KTE142 |
| Eco-1463-1 | Bacteria Proteobacteria Gammaproteobacteria Enterobacteriales Enterobacteriaceae *Escherichia coli* KTE143 |
| Eco-1464-1 | Bacteria Proteobacteria Gammaproteobacteria Enterobacteriales Enterobacteriaceae *Escherichia coli* KTE144 |
| Eco-1465-1 | Bacteria Proteobacteria Gammaproteobacteria Enterobacteriales Enterobacteriaceae *Escherichia coli* KTE145 |
| Eco-1466-1 | Bacteria Proteobacteria Gammaproteobacteria Enterobacteriales Enterobacteriaceae *Escherichia coli* KTE146 |
| Eco-1467-1 | Bacteria Proteobacteria Gammaproteobacteria Enterobacteriales Enterobacteriaceae *Escherichia coli* KTE147 |
| Eco-1468-1 | Bacteria Proteobacteria Gammaproteobacteria Enterobacteriales Enterobacteriaceae *Escherichia coli* KTE148 |
| Eco-1469-1 | Bacteria Proteobacteria Gammaproteobacteria Enterobacteriales Enterobacteriaceae *Escherichia coli* KTE15 |
| Eco-1470-1 | Bacteria Proteobacteria Gammaproteobacteria Enterobacteriales Enterobacteriaceae *Escherichia coli* KTE150 |
| Eco-1471-1 | Bacteria Proteobacteria Gammaproteobacteria Enterobacteriales Enterobacteriaceae *Escherichia coli* KTE153 |
| Eco-1472-1 | Bacteria Proteobacteria Gammaproteobacteria Enterobacteriales Enterobacteriaceae *Escherichia coli* KTE154 |
| Eco-1473-1 | Bacteria Proteobacteria Gammaproteobacteria Enterobacteriales Enterobacteriaceae *Escherichia coli* KTE155 |
| Eco-1474-1 | Bacteria Proteobacteria Gammaproteobacteria Enterobacteriales Enterobacteriaceae *Escherichia coli* KTE156 |
| Eco-1475-1 | Bacteria Proteobacteria Gammaproteobacteria Enterobacteriales Enterobacteriaceae *Escherichia coli* KTE157 |
| Eco-1476-1 | Bacteria Proteobacteria Gammaproteobacteria Enterobacteriales Enterobacteriaceae *Escherichia coli* KTE158 |
| Eco-1477-1 | Bacteria Proteobacteria Gammaproteobacteria Enterobacteriales Enterobacteriaceae *Escherichia coli* KTE159 |
| Eco-1478-1 | Bacteria Proteobacteria Gammaproteobacteria Enterobacteriales Enterobacteriaceae *Escherichia coli* KTE16 |
| Eco-1479-1 | Bacteria Proteobacteria Gammaproteobacteria Enterobacteriales Enterobacteriaceae *Escherichia coli* KTE160 |
| Eco-1480-1 | Bacteria Proteobacteria Gammaproteobacteria Enterobacteriales Enterobacteriaceae *Escherichia coli* KTE161 |
| Eco-1481-1 | Bacteria Proteobacteria Gammaproteobacteria Enterobacteriales Enterobacteriaceae *Escherichia coli* KTE162 |
| Eco-1482-1 | Bacteria Proteobacteria Gammaproteobacteria Enterobacteriales Enterobacteriaceae *Escherichia coli* KTE163 |
| Eco-1483-1 | Bacteria Proteobacteria Gammaproteobacteria Enterobacteriales Enterobacteriaceae *Escherichia coli* KTE165 |
| Eco-1484-1 | Bacteria Proteobacteria Gammaproteobacteria Enterobacteriales Enterobacteriaceae *Escherichia coli* KTE166 |
| Eco-1485-1 | Bacteria Proteobacteria Gammaproteobacteria Enterobacteriales Enterobacteriaceae *Escherichia coli* KTE167 |
| Eco-1486-1 | Bacteria Proteobacteria Gammaproteobacteria Enterobacteriales Enterobacteriaceae *Escherichia coli* KTE168 |
| Eco-1487-1 | Bacteria Proteobacteria Gammaproteobacteria Enterobacteriales Enterobacteriaceae *Escherichia coli* KTE169 |
| Eco-1488-1 | Bacteria Proteobacteria Gammaproteobacteria Enterobacteriales Enterobacteriaceae *Escherichia coli* KTE17 |
| Eco-1489-1 | Bacteria Proteobacteria Gammaproteobacteria Enterobacteriales Enterobacteriaceae *Escherichia coli* KTE170 |
| Eco-1490-1 | Bacteria Proteobacteria Gammaproteobacteria Enterobacteriales Enterobacteriaceae *Escherichia coli* KTE171 |
| Eco-1491-1 | Bacteria Proteobacteria Gammaproteobacteria Enterobacteriales Enterobacteriaceae *Escherichia coli* KTE172 |
| Eco-1492-1 | Bacteria Proteobacteria Gammaproteobacteria Enterobacteriales Enterobacteriaceae *Escherichia coli* KTE173 |
| Eco-1493-1 | Bacteria Proteobacteria Gammaproteobacteria Enterobacteriales Enterobacteriaceae *Escherichia coli* KTE174 |
| Eco-1494-1 | Bacteria Proteobacteria Gammaproteobacteria Enterobacteriales Enterobacteriaceae *Escherichia coli* KTE175 |
| Eco-1495-1 | Bacteria Proteobacteria Gammaproteobacteria Enterobacteriales Enterobacteriaceae *Escherichia coli* KTE176 |
| Eco-1496-1 | Bacteria Proteobacteria Gammaproteobacteria Enterobacteriales Enterobacteriaceae *Escherichia coli* KTE177 |
| Eco-1497-1 | Bacteria Proteobacteria Gammaproteobacteria Enterobacteriales Enterobacteriaceae *Escherichia coli* KTE178 |
| Eco-1498-1 | Bacteria Proteobacteria Gammaproteobacteria Enterobacteriales Enterobacteriaceae *Escherichia coli* KTE179 |
| Eco-1499-1 | Bacteria Proteobacteria Gammaproteobacteria Enterobacteriales Enterobacteriaceae *Escherichia coli* KTE18 |
| Eco-1500-1 | Bacteria Proteobacteria Gammaproteobacteria Enterobacteriales Enterobacteriaceae *Escherichia coli* KTE180 |
| Eco-1501-1 | Bacteria Proteobacteria Gammaproteobacteria Enterobacteriales Enterobacteriaceae *Escherichia coli* KTE181 |
| Eco-1502-1 | Bacteria Proteobacteria Gammaproteobacteria Enterobacteriales Enterobacteriaceae *Escherichia coli* KTE182 |
| Eco-1503-1 | Bacteria Proteobacteria Gammaproteobacteria Enterobacteriales Enterobacteriaceae *Escherichia coli* KTE183 |
| Eco-1504-1 | Bacteria Proteobacteria Gammaproteobacteria Enterobacteriales Enterobacteriaceae *Escherichia coli* KTE184 |
| Eco-1505-1 | Bacteria Proteobacteria Gammaproteobacteria Enterobacteriales Enterobacteriaceae *Escherichia coli* KTE185 |
| Eco-1506-1 | Bacteria Proteobacteria Gammaproteobacteria Enterobacteriales Enterobacteriaceae *Escherichia coli* KTE186 |
| Eco-1507-1 | Bacteria Proteobacteria Gammaproteobacteria Enterobacteriales Enterobacteriaceae *Escherichia coli* KTE187 |
| Eco-1508-1 | Bacteria Proteobacteria Gammaproteobacteria Enterobacteriales Enterobacteriaceae *Escherichia coli* KTE188 |
| Eco-1509-1 | Bacteria Proteobacteria Gammaproteobacteria Enterobacteriales Enterobacteriaceae *Escherichia coli* KTE189 |
| Eco-1510-1 | Bacteria Proteobacteria Gammaproteobacteria Enterobacteriales Enterobacteriaceae *Escherichia coli* KTE19 |
| Eco-1511-1 | Bacteria Proteobacteria Gammaproteobacteria Enterobacteriales Enterobacteriaceae *Escherichia coli* KTE190 |
| Eco-1512-1 | Bacteria Proteobacteria Gammaproteobacteria Enterobacteriales Enterobacteriaceae *Escherichia coli* KTE191 |
| Eco-1513-1 | Bacteria Proteobacteria Gammaproteobacteria Enterobacteriales Enterobacteriaceae *Escherichia coli* KTE192 |
| Eco-1514-1 | Bacteria Proteobacteria Gammaproteobacteria Enterobacteriales Enterobacteriaceae *Escherichia coli* KTE193 |
| Eco-1515-1 | Bacteria Proteobacteria Gammaproteobacteria Enterobacteriales Enterobacteriaceae *Escherichia coli* KTE194 |

| | |
|---|---|
| Eco-1516-1 | Bacteria Proteobacteria Gammaproteobacteria Enterobacteriales Enterobacteriaceae *Escherichia coli* KTE195 |
| Eco-1517-1 | Bacteria Proteobacteria Gammaproteobacteria Enterobacteriales Enterobacteriaceae *Escherichia coli* KTE196 |
| Eco-1518-1 | Bacteria Proteobacteria Gammaproteobacteria Enterobacteriales Enterobacteriaceae *Escherichia coli* KTE197 |
| Eco-1519-1 | Bacteria Proteobacteria Gammaproteobacteria Enterobacteriales Enterobacteriaceae *Escherichia coli* KTE198 |
| Eco-1520-1 | Bacteria Proteobacteria Gammaproteobacteria Enterobacteriales Enterobacteriaceae *Escherichia coli* KTE199 |
| Eco-1521-1 | Bacteria Proteobacteria Gammaproteobacteria Enterobacteriales Enterobacteriaceae *Escherichia coli* KTE2 |
| Eco-1522-1 | Bacteria Proteobacteria Gammaproteobacteria Enterobacteriales Enterobacteriaceae *Escherichia coli* KTE20 |
| Eco-1523-1 | Bacteria Proteobacteria Gammaproteobacteria Enterobacteriales Enterobacteriaceae *Escherichia coli* KTE200 |
| Eco-1524-1 | Bacteria Proteobacteria Gammaproteobacteria Enterobacteriales Enterobacteriaceae *Escherichia coli* KTE201 |
| Eco-1525-1 | Bacteria Proteobacteria Gammaproteobacteria Enterobacteriales Enterobacteriaceae *Escherichia coli* KTE202 |
| Eco-1526-1 | Bacteria Proteobacteria Gammaproteobacteria Enterobacteriales Enterobacteriaceae *Escherichia coli* KTE203 |
| Eco-1527-1 | Bacteria Proteobacteria Gammaproteobacteria Enterobacteriales Enterobacteriaceae *Escherichia coli* KTE204 |
| Eco-1528-1 | Bacteria Proteobacteria Gammaproteobacteria Enterobacteriales Enterobacteriaceae *Escherichia coli* KTE205 |
| Eco-1529-1 | Bacteria Proteobacteria Gammaproteobacteria Enterobacteriales Enterobacteriaceae *Escherichia coli* KTE206 |
| Eco-1530-1 | Bacteria Proteobacteria Gammaproteobacteria Enterobacteriales Enterobacteriaceae *Escherichia coli* KTE207 |
| Eco-1531-1 | Bacteria Proteobacteria Gammaproteobacteria Enterobacteriales Enterobacteriaceae *Escherichia coli* KTE208 |
| Eco-1532-1 | Bacteria Proteobacteria Gammaproteobacteria Enterobacteriales Enterobacteriaceae *Escherichia coli* KTE209 |
| Eco-1533-1 | Bacteria Proteobacteria Gammaproteobacteria Enterobacteriales Enterobacteriaceae *Escherichia coli* KTE21 |
| Eco-1534-1 | Bacteria Proteobacteria Gammaproteobacteria Enterobacteriales Enterobacteriaceae *Escherichia coli* KTE210 |
| Eco-1535-1 | Bacteria Proteobacteria Gammaproteobacteria Enterobacteriales Enterobacteriaceae *Escherichia coli* KTE211 |
| Eco-1536-1 | Bacteria Proteobacteria Gammaproteobacteria Enterobacteriales Enterobacteriaceae *Escherichia coli* KTE212 |
| Eco-1537-1 | Bacteria Proteobacteria Gammaproteobacteria Enterobacteriales Enterobacteriaceae *Escherichia coli* KTE213 |
| Eco-1538-1 | Bacteria Proteobacteria Gammaproteobacteria Enterobacteriales Enterobacteriaceae *Escherichia coli* KTE214 |
| Eco-1539-1 | Bacteria Proteobacteria Gammaproteobacteria Enterobacteriales Enterobacteriaceae *Escherichia coli* KTE215 |
| Eco-1540-1 | Bacteria Proteobacteria Gammaproteobacteria Enterobacteriales Enterobacteriaceae *Escherichia coli* KTE216 |
| Eco-1541-1 | Bacteria Proteobacteria Gammaproteobacteria Enterobacteriales Enterobacteriaceae *Escherichia coli* KTE217 |
| Eco-1542-1 | Bacteria Proteobacteria Gammaproteobacteria Enterobacteriales Enterobacteriaceae *Escherichia coli* KTE218 |
| Eco-1543-1 | Bacteria Proteobacteria Gammaproteobacteria Enterobacteriales Enterobacteriaceae *Escherichia coli* KTE219 |
| Eco-1544-1 | Bacteria Proteobacteria Gammaproteobacteria Enterobacteriales Enterobacteriaceae *Escherichia coli* KTE22 |
| Eco-1545-1 | Bacteria Proteobacteria Gammaproteobacteria Enterobacteriales Enterobacteriaceae *Escherichia coli* KTE220 |
| Eco-1546-1 | Bacteria Proteobacteria Gammaproteobacteria Enterobacteriales Enterobacteriaceae *Escherichia coli* KTE221 |
| Eco-1547-1 | Bacteria Proteobacteria Gammaproteobacteria Enterobacteriales Enterobacteriaceae *Escherichia coli* KTE222 |
| Eco-1548-1 | Bacteria Proteobacteria Gammaproteobacteria Enterobacteriales Enterobacteriaceae *Escherichia coli* KTE223 |
| Eco-1549-1 | Bacteria Proteobacteria Gammaproteobacteria Enterobacteriales Enterobacteriaceae *Escherichia coli* KTE224 |
| Eco-1550-1 | Bacteria Proteobacteria Gammaproteobacteria Enterobacteriales Enterobacteriaceae *Escherichia coli* KTE225 |
| Eco-1551-1 | Bacteria Proteobacteria Gammaproteobacteria Enterobacteriales Enterobacteriaceae *Escherichia coli* KTE226 |
| Eco-1552-1 | Bacteria Proteobacteria Gammaproteobacteria Enterobacteriales Enterobacteriaceae *Escherichia coli* KTE227 |
| Eco-1553-1 | Bacteria Proteobacteria Gammaproteobacteria Enterobacteriales Enterobacteriaceae *Escherichia coli* KTE228 |
| Eco-1554-1 | Bacteria Proteobacteria Gammaproteobacteria Enterobacteriales Enterobacteriaceae *Escherichia coli* KTE229 |
| Eco-1555-1 | Bacteria Proteobacteria Gammaproteobacteria Enterobacteriales Enterobacteriaceae *Escherichia coli* KTE23 |
| Eco-1556-1 | Bacteria Proteobacteria Gammaproteobacteria Enterobacteriales Enterobacteriaceae *Escherichia coli* KTE230 |
| Eco-1557-1 | Bacteria Proteobacteria Gammaproteobacteria Enterobacteriales Enterobacteriaceae *Escherichia coli* KTE231 |
| Eco-1558-1 | Bacteria Proteobacteria Gammaproteobacteria Enterobacteriales Enterobacteriaceae *Escherichia coli* KTE232 |
| Eco-1559-1 | Bacteria Proteobacteria Gammaproteobacteria Enterobacteriales Enterobacteriaceae *Escherichia coli* KTE233 |
| Eco-1560-1 | Bacteria Proteobacteria Gammaproteobacteria Enterobacteriales Enterobacteriaceae *Escherichia coli* KTE234 |
| Eco-1561-1 | Bacteria Proteobacteria Gammaproteobacteria Enterobacteriales Enterobacteriaceae *Escherichia coli* KTE235 |
| Eco-1562-1 | Bacteria Proteobacteria Gammaproteobacteria Enterobacteriales Enterobacteriaceae *Escherichia coli* KTE236 |
| Eco-1563-1 | Bacteria Proteobacteria Gammaproteobacteria Enterobacteriales Enterobacteriaceae *Escherichia coli* KTE237 |
| Eco-1564-1 | Bacteria Proteobacteria Gammaproteobacteria Enterobacteriales Enterobacteriaceae *Escherichia coli* KTE24 |
| Eco-1565-1 | Bacteria Proteobacteria Gammaproteobacteria Enterobacteriales Enterobacteriaceae *Escherichia coli* KTE240 |
| Eco-1566-1 | Bacteria Proteobacteria Gammaproteobacteria Enterobacteriales Enterobacteriaceae *Escherichia coli* KTE25 |
| Eco-1567-1 | Bacteria Proteobacteria Gammaproteobacteria Enterobacteriales Enterobacteriaceae *Escherichia coli* KTE26 |
| Eco-1568-1 | Bacteria Proteobacteria Gammaproteobacteria Enterobacteriales Enterobacteriaceae *Escherichia coli* KTE27 |
| Eco-1569-1 | Bacteria Proteobacteria Gammaproteobacteria Enterobacteriales Enterobacteriaceae *Escherichia coli* KTE28 |
| Eco-1570-1 | Bacteria Proteobacteria Gammaproteobacteria Enterobacteriales Enterobacteriaceae *Escherichia coli* KTE29 |
| Eco-1571-1 | Bacteria Proteobacteria Gammaproteobacteria Enterobacteriales Enterobacteriaceae *Escherichia coli* KTE3 |
| Eco-1572-1 | Bacteria Proteobacteria Gammaproteobacteria Enterobacteriales Enterobacteriaceae *Escherichia coli* KTE31 |
| Eco-1573-1 | Bacteria Proteobacteria Gammaproteobacteria Enterobacteriales Enterobacteriaceae *Escherichia coli* KTE33 |
| Eco-1574-1 | Bacteria Proteobacteria Gammaproteobacteria Enterobacteriales Enterobacteriaceae *Escherichia coli* KTE34 |
| Eco-1575-1 | Bacteria Proteobacteria Gammaproteobacteria Enterobacteriales Enterobacteriaceae *Escherichia coli* KTE35 |
| Eco-1576-1 | Bacteria Proteobacteria Gammaproteobacteria Enterobacteriales Enterobacteriaceae *Escherichia coli* KTE36 |
| Eco-1577-1 | Bacteria Proteobacteria Gammaproteobacteria Enterobacteriales Enterobacteriaceae *Escherichia coli* KTE37 |
| Eco-1578-1 | Bacteria Proteobacteria Gammaproteobacteria Enterobacteriales Enterobacteriaceae *Escherichia coli* KTE38 |
| Eco-1579-1 | Bacteria Proteobacteria Gammaproteobacteria Enterobacteriales Enterobacteriaceae *Escherichia coli* KTE39 |
| Eco-1580-1 | Bacteria Proteobacteria Gammaproteobacteria Enterobacteriales Enterobacteriaceae *Escherichia coli* KTE4 |

| | |
|---|---|
| Eco-1581-1 | Bacteria Proteobacteria Gammaproteobacteria Enterobacteriales Enterobacteriaceae *Escherichia coli* KTE40 |
| Eco-1582-1 | Bacteria Proteobacteria Gammaproteobacteria Enterobacteriales Enterobacteriaceae *Escherichia coli* KTE41 |
| Eco-1583-1 | Bacteria Proteobacteria Gammaproteobacteria Enterobacteriales Enterobacteriaceae *Escherichia coli* KTE42 |
| Eco-1584-1 | Bacteria Proteobacteria Gammaproteobacteria Enterobacteriales Enterobacteriaceae *Escherichia coli* KTE43 |
| Eco-1585-1 | Bacteria Proteobacteria Gammaproteobacteria Enterobacteriales Enterobacteriaceae *Escherichia coli* KTE44 |
| Eco-1586-1 | Bacteria Proteobacteria Gammaproteobacteria Enterobacteriales Enterobacteriaceae *Escherichia coli* KTE45 |
| Eco-1587-1 | Bacteria Proteobacteria Gammaproteobacteria Enterobacteriales Enterobacteriaceae *Escherichia coli* KTE46 |
| Eco-1588-1 | Bacteria Proteobacteria Gammaproteobacteria Enterobacteriales Enterobacteriaceae *Escherichia coli* KTE47 |
| Eco-1589-1 | Bacteria Proteobacteria Gammaproteobacteria Enterobacteriales Enterobacteriaceae *Escherichia coli* KTE48 |
| Eco-1590-1 | Bacteria Proteobacteria Gammaproteobacteria Enterobacteriales Enterobacteriaceae *Escherichia coli* KTE49 |
| Eco-1591-1 | Bacteria Proteobacteria Gammaproteobacteria Enterobacteriales Enterobacteriaceae *Escherichia coli* KTE5 |
| Eco-1592-1 | Bacteria Proteobacteria Gammaproteobacteria Enterobacteriales Enterobacteriaceae *Escherichia coli* KTE50 |
| Eco-1593-1 | Bacteria Proteobacteria Gammaproteobacteria Enterobacteriales Enterobacteriaceae *Escherichia coli* KTE51 |
| Eco-1594-1 | Bacteria Proteobacteria Gammaproteobacteria Enterobacteriales Enterobacteriaceae *Escherichia coli* KTE52 |
| Eco-1595-1 | Bacteria Proteobacteria Gammaproteobacteria Enterobacteriales Enterobacteriaceae *Escherichia coli* KTE53 |
| Eco-1596-1 | Bacteria Proteobacteria Gammaproteobacteria Enterobacteriales Enterobacteriaceae *Escherichia coli* KTE54 |
| Eco-1597-1 | Bacteria Proteobacteria Gammaproteobacteria Enterobacteriales Enterobacteriaceae *Escherichia coli* KTE55 |
| Eco-1598-1 | Bacteria Proteobacteria Gammaproteobacteria Enterobacteriales Enterobacteriaceae *Escherichia coli* KTE56 |
| Eco-1599-1 | Bacteria Proteobacteria Gammaproteobacteria Enterobacteriales Enterobacteriaceae *Escherichia coli* KTE57 |
| Eco-1600-1 | Bacteria Proteobacteria Gammaproteobacteria Enterobacteriales Enterobacteriaceae *Escherichia coli* KTE58 |
| Eco-1601-1 | Bacteria Proteobacteria Gammaproteobacteria Enterobacteriales Enterobacteriaceae *Escherichia coli* KTE59 |
| Eco-1602-1 | Bacteria Proteobacteria Gammaproteobacteria Enterobacteriales Enterobacteriaceae *Escherichia coli* KTE6 |
| Eco-1603-1 | Bacteria Proteobacteria Gammaproteobacteria Enterobacteriales Enterobacteriaceae *Escherichia coli* KTE60 |
| Eco-1604-1 | Bacteria Proteobacteria Gammaproteobacteria Enterobacteriales Enterobacteriaceae *Escherichia coli* KTE61 |
| Eco-1605-1 | Bacteria Proteobacteria Gammaproteobacteria Enterobacteriales Enterobacteriaceae *Escherichia coli* KTE62 |
| Eco-1606-1 | Bacteria Proteobacteria Gammaproteobacteria Enterobacteriales Enterobacteriaceae *Escherichia coli* KTE63 |
| Eco-1607-1 | Bacteria Proteobacteria Gammaproteobacteria Enterobacteriales Enterobacteriaceae *Escherichia coli* KTE64 |
| Eco-1608-1 | Bacteria Proteobacteria Gammaproteobacteria Enterobacteriales Enterobacteriaceae *Escherichia coli* KTE65 |
| Eco-1609-1 | Bacteria Proteobacteria Gammaproteobacteria Enterobacteriales Enterobacteriaceae *Escherichia coli* KTE66 |
| Eco-1610-1 | Bacteria Proteobacteria Gammaproteobacteria Enterobacteriales Enterobacteriaceae *Escherichia coli* KTE67 |
| Eco-1611-1 | Bacteria Proteobacteria Gammaproteobacteria Enterobacteriales Enterobacteriaceae *Escherichia coli* KTE68 |
| Eco-1612-1 | Bacteria Proteobacteria Gammaproteobacteria Enterobacteriales Enterobacteriaceae *Escherichia coli* KTE69 |
| Eco-1613-1 | Bacteria Proteobacteria Gammaproteobacteria Enterobacteriales Enterobacteriaceae *Escherichia coli* KTE7 |
| Eco-1614-1 | Bacteria Proteobacteria Gammaproteobacteria Enterobacteriales Enterobacteriaceae *Escherichia coli* KTE70 |
| Eco-1615-1 | Bacteria Proteobacteria Gammaproteobacteria Enterobacteriales Enterobacteriaceae *Escherichia coli* KTE71 |
| Eco-1616-1 | Bacteria Proteobacteria Gammaproteobacteria Enterobacteriales Enterobacteriaceae *Escherichia coli* KTE72 |
| Eco-1617-1 | Bacteria Proteobacteria Gammaproteobacteria Enterobacteriales Enterobacteriaceae *Escherichia coli* KTE73 |
| Eco-1618-1 | Bacteria Proteobacteria Gammaproteobacteria Enterobacteriales Enterobacteriaceae *Escherichia coli* KTE74 |
| Eco-1619-1 | Bacteria Proteobacteria Gammaproteobacteria Enterobacteriales Enterobacteriaceae *Escherichia coli* KTE75 |
| Eco-1620-1 | Bacteria Proteobacteria Gammaproteobacteria Enterobacteriales Enterobacteriaceae *Escherichia coli* KTE76 |
| Eco-1621-1 | Bacteria Proteobacteria Gammaproteobacteria Enterobacteriales Enterobacteriaceae *Escherichia coli* KTE77 |
| Eco-1622-1 | Bacteria Proteobacteria Gammaproteobacteria Enterobacteriales Enterobacteriaceae *Escherichia coli* KTE78 |
| Eco-1623-1 | Bacteria Proteobacteria Gammaproteobacteria Enterobacteriales Enterobacteriaceae *Escherichia coli* KTE79 |
| Eco-1624-1 | Bacteria Proteobacteria Gammaproteobacteria Enterobacteriales Enterobacteriaceae *Escherichia coli* KTE8 |
| Eco-1625-1 | Bacteria Proteobacteria Gammaproteobacteria Enterobacteriales Enterobacteriaceae *Escherichia coli* KTE80 |
| Eco-1626-1 | Bacteria Proteobacteria Gammaproteobacteria Enterobacteriales Enterobacteriaceae *Escherichia coli* KTE81 |
| Eco-1627-1 | Bacteria Proteobacteria Gammaproteobacteria Enterobacteriales Enterobacteriaceae *Escherichia coli* KTE82 |
| Eco-1628-1 | Bacteria Proteobacteria Gammaproteobacteria Enterobacteriales Enterobacteriaceae *Escherichia coli* KTE83 |
| Eco-1629-1 | Bacteria Proteobacteria Gammaproteobacteria Enterobacteriales Enterobacteriaceae *Escherichia coli* KTE84 |
| Eco-1630-1 | Bacteria Proteobacteria Gammaproteobacteria Enterobacteriales Enterobacteriaceae *Escherichia coli* KTE85 |
| Eco-1631-1 | Bacteria Proteobacteria Gammaproteobacteria Enterobacteriales Enterobacteriaceae *Escherichia coli* KTE86 |
| Eco-1632-1 | Bacteria Proteobacteria Gammaproteobacteria Enterobacteriales Enterobacteriaceae *Escherichia coli* KTE87 |
| Eco-1633-1 | Bacteria Proteobacteria Gammaproteobacteria Enterobacteriales Enterobacteriaceae *Escherichia coli* KTE88 |
| Eco-1634-1 | Bacteria Proteobacteria Gammaproteobacteria Enterobacteriales Enterobacteriaceae *Escherichia coli* KTE89 |
| Eco-1635-1 | Bacteria Proteobacteria Gammaproteobacteria Enterobacteriales Enterobacteriaceae *Escherichia coli* KTE9 |
| Eco-1636-1 | Bacteria Proteobacteria Gammaproteobacteria Enterobacteriales Enterobacteriaceae *Escherichia coli* KTE90 |
| Eco-1637-1 | Bacteria Proteobacteria Gammaproteobacteria Enterobacteriales Enterobacteriaceae *Escherichia coli* KTE91 |
| Eco-1638-1 | Bacteria Proteobacteria Gammaproteobacteria Enterobacteriales Enterobacteriaceae *Escherichia coli* KTE93 |
| Eco-1639-1 | Bacteria Proteobacteria Gammaproteobacteria Enterobacteriales Enterobacteriaceae *Escherichia coli* KTE94 |
| Eco-1640-1 | Bacteria Proteobacteria Gammaproteobacteria Enterobacteriales Enterobacteriaceae *Escherichia coli* KTE95 |
| Eco-1641-1 | Bacteria Proteobacteria Gammaproteobacteria Enterobacteriales Enterobacteriaceae *Escherichia coli* KTE96 |
| Eco-1642-1 | Bacteria Proteobacteria Gammaproteobacteria Enterobacteriales Enterobacteriaceae *Escherichia coli* KTE97 |
| Eco-1643-1 | Bacteria Proteobacteria Gammaproteobacteria Enterobacteriales Enterobacteriaceae *Escherichia coli* KTE98 |
| Eco-1644-1 | Bacteria Proteobacteria Gammaproteobacteria Enterobacteriales Enterobacteriaceae *Escherichia coli* KTE99 |
| Eco-1645-1 | Bacteria Proteobacteria Gammaproteobacteria Enterobacteriales Enterobacteriaceae *Escherichia coli* LAU-EC10 |

| | |
|---|---|
| Eco-1646-1 | Bacteria Proteobacteria Gammaproteobacteria Enterobacteriales Enterobacteriaceae *Escherichia coli* LAU-EC6 |
| Eco-1647-1 | Bacteria Proteobacteria Gammaproteobacteria Enterobacteriales Enterobacteriaceae *Escherichia coli* LAU-EC7 |
| Eco-1648-1 | Bacteria Proteobacteria Gammaproteobacteria Enterobacteriales Enterobacteriaceae *Escherichia coli* LAU-EC8 |
| Eco-1649-1 | Bacteria Proteobacteria Gammaproteobacteria Enterobacteriales Enterobacteriaceae *Escherichia coli* LAU-EC9 |
| Eco-1650-1 | Bacteria Proteobacteria Gammaproteobacteria Enterobacteriales Enterobacteriaceae *Escherichia coli* LCT-EC106 |
| Eco-1651-1 | Bacteria Proteobacteria Gammaproteobacteria Enterobacteriales Enterobacteriaceae *Escherichia coli* LCT-EC52 |
| Eco-1652-1 | Bacteria Proteobacteria Gammaproteobacteria Enterobacteriales Enterobacteriaceae *Escherichia coli* LCT-EC59 |
| Eco-1653-1 | Bacteria Proteobacteria Gammaproteobacteria Enterobacteriales Enterobacteriaceae *Escherichia coli* LF82 |
| Eco-1654-1 | Bacteria Proteobacteria Gammaproteobacteria Enterobacteriales Enterobacteriaceae *Escherichia coli* LY180 |
| Eco-1655-1 | Bacteria Proteobacteria Gammaproteobacteria Enterobacteriales Enterobacteriaceae *Escherichia coli* M605 |
| Eco-1656-1 | Bacteria Proteobacteria Gammaproteobacteria Enterobacteriales Enterobacteriaceae *Escherichia coli* M718 |
| Eco-1657-1 | Bacteria Proteobacteria Gammaproteobacteria Enterobacteriales Enterobacteriaceae *Escherichia coli* M863 |
| Eco-1658-1 | Bacteria Proteobacteria Gammaproteobacteria Enterobacteriales Enterobacteriaceae *Escherichia coli* M919 |
| Eco-1659-1 | Bacteria Proteobacteria Gammaproteobacteria Enterobacteriales Enterobacteriaceae *Escherichia coli* MA6 |
| Eco-1660-1 | Bacteria Proteobacteria Gammaproteobacteria Enterobacteriales Enterobacteriaceae *Escherichia coli* MP020940.1 |
| Eco-1661-1 | Bacteria Proteobacteria Gammaproteobacteria Enterobacteriales Enterobacteriaceae *Escherichia coli* MP020980.1 |
| Eco-1662-1 | Bacteria Proteobacteria Gammaproteobacteria Enterobacteriales Enterobacteriaceae *Escherichia coli* MP020980.2 |
| Eco-1663-1 | Bacteria Proteobacteria Gammaproteobacteria Enterobacteriales Enterobacteriaceae *Escherichia coli* MP021017.1 |
| Eco-1664-1 | Bacteria Proteobacteria Gammaproteobacteria Enterobacteriales Enterobacteriaceae *Escherichia coli* MP021017.10 |
| Eco-1665-1 | Bacteria Proteobacteria Gammaproteobacteria Enterobacteriales Enterobacteriaceae *Escherichia coli* MP021017.11 |
| Eco-1666-1 | Bacteria Proteobacteria Gammaproteobacteria Enterobacteriales Enterobacteriaceae *Escherichia coli* MP021017.12 |
| Eco-1667-1 | Bacteria Proteobacteria Gammaproteobacteria Enterobacteriales Enterobacteriaceae *Escherichia coli* MP021017.2 |
| Eco-1668-1 | Bacteria Proteobacteria Gammaproteobacteria Enterobacteriales Enterobacteriaceae *Escherichia coli* MP021017.3 |
| Eco-1669-1 | Bacteria Proteobacteria Gammaproteobacteria Enterobacteriales Enterobacteriaceae *Escherichia coli* MP021017.4 |
| Eco-1670-1 | Bacteria Proteobacteria Gammaproteobacteria Enterobacteriales Enterobacteriaceae *Escherichia coli* MP021017.5 |
| Eco-1671-1 | Bacteria Proteobacteria Gammaproteobacteria Enterobacteriales Enterobacteriaceae *Escherichia coli* MP021017.6 |
| Eco-1672-1 | Bacteria Proteobacteria Gammaproteobacteria Enterobacteriales Enterobacteriaceae *Escherichia coli* MP021017.9 |
| Eco-1673-1 | Bacteria Proteobacteria Gammaproteobacteria Enterobacteriales Enterobacteriaceae *Escherichia coli* MP021552.11 |
| Eco-1674-1 | Bacteria Proteobacteria Gammaproteobacteria Enterobacteriales Enterobacteriaceae *Escherichia coli* MP021552.12 |
| Eco-1675-1 | Bacteria Proteobacteria Gammaproteobacteria Enterobacteriales Enterobacteriaceae *Escherichia coli* MP021552.7 |
| Eco-1676-1 | Bacteria Proteobacteria Gammaproteobacteria Enterobacteriales Enterobacteriaceae *Escherichia coli* MP021552.8 |
| Eco-1677-1 | Bacteria Proteobacteria Gammaproteobacteria Enterobacteriales Enterobacteriaceae *Escherichia coli* MP021561.2 |
| Eco-1678-1 | Bacteria Proteobacteria Gammaproteobacteria Enterobacteriales Enterobacteriaceae *Escherichia coli* MP021561.3 |
| Eco-1679-1 | Bacteria Proteobacteria Gammaproteobacteria Enterobacteriales Enterobacteriaceae *Escherichia coli* MP021566.1 |
| Eco-1680-1 | Bacteria Proteobacteria Gammaproteobacteria Enterobacteriales Enterobacteriaceae *Escherichia coli* MS 107-1 |
| Eco-1681-1 | Bacteria Proteobacteria Gammaproteobacteria Enterobacteriales Enterobacteriaceae *Escherichia coli* MS 110-3 |
| Eco-1682-1 | Bacteria Proteobacteria Gammaproteobacteria Enterobacteriales Enterobacteriaceae *Escherichia coli* MS 115-1 |
| Eco-1683-1 | Bacteria Proteobacteria Gammaproteobacteria Enterobacteriales Enterobacteriaceae *Escherichia coli* MS 116-1 |
| Eco-1684-1 | Bacteria Proteobacteria Gammaproteobacteria Enterobacteriales Enterobacteriaceae *Escherichia coli* MS 117-3 |
| Eco-1685-1 | Bacteria Proteobacteria Gammaproteobacteria Enterobacteriales Enterobacteriaceae *Escherichia coli* MS 119-7 |
| Eco-1686-1 | Bacteria Proteobacteria Gammaproteobacteria Enterobacteriales Enterobacteriaceae *Escherichia coli* MS 124-1 |
| Eco-1687-1 | Bacteria Proteobacteria Gammaproteobacteria Enterobacteriales Enterobacteriaceae *Escherichia coli* MS 145-7 |
| Eco-1688-1 | Bacteria Proteobacteria Gammaproteobacteria Enterobacteriales Enterobacteriaceae *Escherichia coli* MS 146-1 |
| Eco-1689-1 | Bacteria Proteobacteria Gammaproteobacteria Enterobacteriales Enterobacteriaceae *Escherichia coli* MS 153-1 |
| Eco-1690-1 | Bacteria Proteobacteria Gammaproteobacteria Enterobacteriales Enterobacteriaceae *Escherichia coli* MS 16-3 |
| Eco-1691-1 | Bacteria Proteobacteria Gammaproteobacteria Enterobacteriales Enterobacteriaceae *Escherichia coli* MS 175-1 |
| Eco-1692-1 | Bacteria Proteobacteria Gammaproteobacteria Enterobacteriales Enterobacteriaceae *Escherichia coli* MS 182-1 |
| Eco-1693-1 | Bacteria Proteobacteria Gammaproteobacteria Enterobacteriales Enterobacteriaceae *Escherichia coli* MS 185-1 |
| Eco-1694-1 | Bacteria Proteobacteria Gammaproteobacteria Enterobacteriales Enterobacteriaceae *Escherichia coli* MS 187-1 |
| Eco-1695-1 | Bacteria Proteobacteria Gammaproteobacteria Enterobacteriales Enterobacteriaceae *Escherichia coli* MS 196-1 |
| Eco-1696-1 | Bacteria Proteobacteria Gammaproteobacteria Enterobacteriales Enterobacteriaceae *Escherichia coli* MS 198-1 |
| Eco-1697-1 | Bacteria Proteobacteria Gammaproteobacteria Enterobacteriales Enterobacteriaceae *Escherichia coli* MS 200-1 |
| Eco-1698-1 | Bacteria Proteobacteria Gammaproteobacteria Enterobacteriales Enterobacteriaceae *Escherichia coli* MS 21-1 |
| Eco-1699-1 | Bacteria Proteobacteria Gammaproteobacteria Enterobacteriales Enterobacteriaceae *Escherichia coli* MS 45-1 |
| Eco-1700-1 | Bacteria Proteobacteria Gammaproteobacteria Enterobacteriales Enterobacteriaceae *Escherichia coli* MS 57-2 |
| Eco-1701-1 | Bacteria Proteobacteria Gammaproteobacteria Enterobacteriales Enterobacteriaceae *Escherichia coli* MS 60-1 |
| Eco-1702-1 | Bacteria Proteobacteria Gammaproteobacteria Enterobacteriales Enterobacteriaceae *Escherichia coli* MS 69-1 |
| Eco-1703-1 | Bacteria Proteobacteria Gammaproteobacteria Enterobacteriales Enterobacteriaceae *Escherichia coli* MS 78-1 |
| Eco-1704-1 | Bacteria Proteobacteria Gammaproteobacteria Enterobacteriales Enterobacteriaceae *Escherichia coli* MS 79-10 |
| Eco-1705-1 | Bacteria Proteobacteria Gammaproteobacteria Enterobacteriales Enterobacteriaceae *Escherichia coli* MS 84-1 |
| Eco-1706-1 | Bacteria Proteobacteria Gammaproteobacteria Enterobacteriales Enterobacteriaceae *Escherichia coli* MS 85-1 |
| Eco-1707-1 | Bacteria Proteobacteria Gammaproteobacteria Enterobacteriales Enterobacteriaceae *Escherichia coli* N1 |
| Eco-1708-1 | Bacteria Proteobacteria Gammaproteobacteria Enterobacteriales Enterobacteriaceae *Escherichia coli* NA114 |
| Eco-1709-1 | Bacteria Proteobacteria Gammaproteobacteria Enterobacteriales Enterobacteriaceae *Escherichia coli* NC101 |
| Eco-1710-1 | Bacteria Proteobacteria Gammaproteobacteria Enterobacteriales Enterobacteriaceae *Escherichia coli* NCCP15647 |

| | |
|---|---|
| Eco-1711-1 to Eco-1711-2 | Bacteria Proteobacteria Gammaproteobacteria Enterobacteriales Enterobacteriaceae *Escherichia coli* NCCP15657 |
| Eco-1712-1 | Bacteria Proteobacteria Gammaproteobacteria Enterobacteriales Enterobacteriaceae *Escherichia coli* NCCP15658 |
| Eco-1713-1 | Bacteria Proteobacteria Gammaproteobacteria Enterobacteriales Enterobacteriaceae *Escherichia coli* NE037 |
| Eco-1714-1 | Bacteria Proteobacteria Gammaproteobacteria Enterobacteriales Enterobacteriaceae *Escherichia coli* NE098 |
| Eco-1715-1 | Bacteria Proteobacteria Gammaproteobacteria Enterobacteriales Enterobacteriaceae *Escherichia coli* NE1487 |
| Eco-463-1 to Eco-463-2 | Bacteria Proteobacteria Gammaproteobacteria Enterobacteriales Enterobacteriaceae *Escherichia coli* Nissle 1917 |
| Eco-1716-1 | Bacteria Proteobacteria Gammaproteobacteria Enterobacteriales Enterobacteriaceae *Escherichia coli* O08 |
| Eco-1717-1 | Bacteria Proteobacteria Gammaproteobacteria Enterobacteriales Enterobacteriaceae *Escherichia coli* O103:H2 str. 12009 |
| Eco-1718-1 | Bacteria Proteobacteria Gammaproteobacteria Enterobacteriales Enterobacteriaceae *Escherichia coli* O103:H2 str. CVM9450 |
| Eco-1719-1 | Bacteria Proteobacteria Gammaproteobacteria Enterobacteriales Enterobacteriaceae *Escherichia coli* O103:H25 str. CVM9340 |
| Eco-1720-1 | Bacteria Proteobacteria Gammaproteobacteria Enterobacteriales Enterobacteriaceae *Escherichia coli* O103:H25 str. NIPH-11060424 |
| Eco-1721-1 | Bacteria Proteobacteria Gammaproteobacteria Enterobacteriales Enterobacteriaceae *Escherichia coli* O104:H4 str. 04-8351 |
| Eco-1722-1 | Bacteria Proteobacteria Gammaproteobacteria Enterobacteriales Enterobacteriaceae *Escherichia coli* O104:H4 str. 09-7901 |
| Eco-1723-1 | Bacteria Proteobacteria Gammaproteobacteria Enterobacteriales Enterobacteriaceae *Escherichia coli* O104:H4 str. 11-02030 |
| Eco-1724-1 | Bacteria Proteobacteria Gammaproteobacteria Enterobacteriales Enterobacteriaceae *Escherichia coli* O104:H4 str. 11-02033-1 |
| Eco-1725-1 | Bacteria Proteobacteria Gammaproteobacteria Enterobacteriales Enterobacteriaceae *Escherichia coli* O104:H4 str. 11-02092 |
| Eco-1726-1 | Bacteria Proteobacteria Gammaproteobacteria Enterobacteriales Enterobacteriaceae *Escherichia coli* O104:H4 str. 11-02093 |
| Eco-1727-1 | Bacteria Proteobacteria Gammaproteobacteria Enterobacteriales Enterobacteriaceae *Escherichia coli* O104:H4 str. 11-02281 |
| Eco-1728-1 | Bacteria Proteobacteria Gammaproteobacteria Enterobacteriales Enterobacteriaceae *Escherichia coli* O104:H4 str. 11-02318 |
| Eco-1729-1 | Bacteria Proteobacteria Gammaproteobacteria Enterobacteriales Enterobacteriaceae *Escherichia coli* O104:H4 str. 11-02913 |
| Eco-1730-1 | Bacteria Proteobacteria Gammaproteobacteria Enterobacteriales Enterobacteriaceae *Escherichia coli* O104:H4 str. 11-03439 |
| Eco-1731-1 | Bacteria Proteobacteria Gammaproteobacteria Enterobacteriales Enterobacteriaceae *Escherichia coli* O104:H4 str. 11-03943 |
| Eco-1732-1 | Bacteria Proteobacteria Gammaproteobacteria Enterobacteriales Enterobacteriaceae *Escherichia coli* O104:H4 str. 11-04080 |
| Eco-1733-1 | Bacteria Proteobacteria Gammaproteobacteria Enterobacteriales Enterobacteriaceae *Escherichia coli* O104:H4 str. 11-3677 |
| Eco-1734-1 | Bacteria Proteobacteria Gammaproteobacteria Enterobacteriales Enterobacteriaceae *Escherichia coli* O104:H4 str. 11-4404 |
| Eco-1735-1 | Bacteria Proteobacteria Gammaproteobacteria Enterobacteriales Enterobacteriaceae *Escherichia coli* O104:H4 str. 11-4522 |
| Eco-1736-1 | Bacteria Proteobacteria Gammaproteobacteria Enterobacteriales Enterobacteriaceae *Escherichia coli* O104:H4 str. 11-4623 |
| Eco-1737-1 | Bacteria Proteobacteria Gammaproteobacteria Enterobacteriales Enterobacteriaceae *Escherichia coli* O104:H4 str. 11-4632 C1 |
| Eco-1738-1 | Bacteria Proteobacteria Gammaproteobacteria Enterobacteriales Enterobacteriaceae *Escherichia coli* O104:H4 str. 11-4632 C2 |
| Eco-1739-1 | Bacteria Proteobacteria Gammaproteobacteria Enterobacteriales Enterobacteriaceae *Escherichia coli* O104:H4 str. 11-4632 C3 |
| Eco-1740-1 | Bacteria Proteobacteria Gammaproteobacteria Enterobacteriales Enterobacteriaceae *Escherichia coli* O104:H4 str. 11-4632 C4 |
| Eco-1741-1 | Bacteria Proteobacteria Gammaproteobacteria Enterobacteriales Enterobacteriaceae *Escherichia coli* O104:H4 str. 11-4632 C5 |
| Eco-1742-1 | Bacteria Proteobacteria Gammaproteobacteria Enterobacteriales Enterobacteriaceae *Escherichia coli* O104:H4 str. 2009EL-2050 |
| Eco-1743-1 | Bacteria Proteobacteria Gammaproteobacteria Enterobacteriales Enterobacteriaceae *Escherichia coli* O104:H4 str. 2009EL-2071 |
| Eco-1744-1 | Bacteria Proteobacteria Gammaproteobacteria Enterobacteriales Enterobacteriaceae *Escherichia coli* O104:H4 str. 2011C-3493 |
| Eco-472-1 to Eco-472-2 | Bacteria Proteobacteria Gammaproteobacteria Enterobacteriales Enterobacteriaceae *Escherichia coli* O104:H4 str. C227-11 |
| Eco-1745-1 | Bacteria Proteobacteria Gammaproteobacteria Enterobacteriales Enterobacteriaceae *Escherichia coli* O104:H4 str. C236-11 |
| Eco-1746-1 | Bacteria Proteobacteria Gammaproteobacteria Enterobacteriales Enterobacteriaceae *Escherichia coli* O104:H4 str. E112/10 |
| Eco-1747-1 | Bacteria Proteobacteria Gammaproteobacteria Enterobacteriales Enterobacteriaceae *Escherichia coli* O104:H4 str. E92/11 |
| Eco-1748-1 | Bacteria Proteobacteria Gammaproteobacteria Enterobacteriales Enterobacteriaceae *Escherichia coli* O104:H4 str. Ec11-4984 |
| Eco-1749-1 | Bacteria Proteobacteria Gammaproteobacteria Enterobacteriales Enterobacteriaceae *Escherichia coli* O104:H4 str. Ec11-4986 |
| Eco-1750-1 | Bacteria Proteobacteria Gammaproteobacteria Enterobacteriales Enterobacteriaceae *Escherichia coli* O104:H4 str. Ec11-4987 |
| Eco-1751-1 | Bacteria Proteobacteria Gammaproteobacteria Enterobacteriales Enterobacteriaceae *Escherichia coli* O104:H4 str. Ec11-4988 |
| Eco-1752-1 | Bacteria Proteobacteria Gammaproteobacteria Enterobacteriales Enterobacteriaceae *Escherichia coli* O104:H4 str. Ec11-5603 |
| Eco-1753-1 | Bacteria Proteobacteria Gammaproteobacteria Enterobacteriales Enterobacteriaceae *Escherichia coli* O104:H4 str. Ec11-5604 |
| Eco-1754-1 | Bacteria Proteobacteria Gammaproteobacteria Enterobacteriales Enterobacteriaceae *Escherichia coli* O104:H4 str. Ec11-6006 |
| Eco-1755-1 | Bacteria Proteobacteria Gammaproteobacteria Enterobacteriales Enterobacteriaceae *Escherichia coli* O104:H4 str. Ec11-9450 |
| Eco-1756-1 | Bacteria Proteobacteria Gammaproteobacteria Enterobacteriales Enterobacteriaceae *Escherichia coli* O104:H4 str. Ec11-9941 |
| Eco-1757-1 | Bacteria Proteobacteria Gammaproteobacteria Enterobacteriales Enterobacteriaceae *Escherichia coli* O104:H4 str. Ec11-9990 |
| Eco-1758-1 | Bacteria Proteobacteria Gammaproteobacteria Enterobacteriales Enterobacteriaceae *Escherichia coli* O104:H4 str. Ec12-0465 |
| Eco-1759-1 | Bacteria Proteobacteria Gammaproteobacteria Enterobacteriales Enterobacteriaceae *Escherichia coli* O104:H4 str. Ec12-0466 |
| Eco-1760-1 | Bacteria Proteobacteria Gammaproteobacteria Enterobacteriales Enterobacteriaceae *Escherichia coli* O104:H4 str. GOS1 |
| Eco-1761-1 | Bacteria Proteobacteria Gammaproteobacteria Enterobacteriales Enterobacteriaceae *Escherichia coli* O104:H4 str. GOS2 |
| Eco-1762-1 | Bacteria Proteobacteria Gammaproteobacteria Enterobacteriales Enterobacteriaceae *Escherichia coli* O104:H4 str. H112180280 |
| Eco-1763-1 | Bacteria Proteobacteria Gammaproteobacteria Enterobacteriales Enterobacteriaceae *Escherichia coli* O104:H4 str. H112180282 |
| Eco-1764-1 | Bacteria Proteobacteria Gammaproteobacteria Enterobacteriales Enterobacteriaceae *Escherichia coli* O104:H4 str. ON2010 |
| Eco-1765-1 | Bacteria Proteobacteria Gammaproteobacteria Enterobacteriales Enterobacteriaceae *Escherichia coli* O104:H4 str. ON2011 |
| Eco-1766-1 | Bacteria Proteobacteria Gammaproteobacteria Enterobacteriales Enterobacteriaceae *Escherichia coli* O104:H4 str. TY-2482 |
| Eco-1767-1 | Bacteria Proteobacteria Gammaproteobacteria Enterobacteriales Enterobacteriaceae *Escherichia coli* O10:K5(L):H4 str. ATCC 23506 |
| Eco-1768-1 | Bacteria Proteobacteria Gammaproteobacteria Enterobacteriales Enterobacteriaceae *Escherichia coli* O111:H- str. 11128 |
| Eco-1769-1 | Bacteria Proteobacteria Gammaproteobacteria Enterobacteriales Enterobacteriaceae *Escherichia coli* O111:H11 str. CFSAN001630 |
| Eco-1770-1 | Bacteria Proteobacteria Gammaproteobacteria Enterobacteriales Enterobacteriaceae *Escherichia coli* O111:H11 str. CVM9455 |
| Eco-1771-1 | Bacteria Proteobacteria Gammaproteobacteria Enterobacteriales Enterobacteriaceae *Escherichia coli* O111:H11 str. CVM9534 |
| Eco-1772-1 | Bacteria Proteobacteria Gammaproteobacteria Enterobacteriales Enterobacteriaceae *Escherichia coli* O111:H11 str. CVM9545 |
| Eco-1773-1 | Bacteria Proteobacteria Gammaproteobacteria Enterobacteriales Enterobacteriaceae *Escherichia coli* O111:H11 str. CVM9553 |

| | |
|---|---|
| Eco-1774-1 | Bacteria Proteobacteria Gammaproteobacteria Enterobacteriales Enterobacteriaceae *Escherichia coli* O111:H8 str. CFSAN001632 |
| Eco-1775-1 | Bacteria Proteobacteria Gammaproteobacteria Enterobacteriales Enterobacteriaceae *Escherichia coli* O111:H8 str. CVM9570 |
| Eco-1776-1 | Bacteria Proteobacteria Gammaproteobacteria Enterobacteriales Enterobacteriaceae *Escherichia coli* O111:H8 str. CVM9574 |
| Eco-1777-1 | Bacteria Proteobacteria Gammaproteobacteria Enterobacteriales Enterobacteriaceae *Escherichia coli* O111:H8 str. CVM9602 |
| Eco-1778-1 | Bacteria Proteobacteria Gammaproteobacteria Enterobacteriales Enterobacteriaceae *Escherichia coli* O111:H8 str. CVM9634 |
| Eco-1779-1 | Bacteria Proteobacteria Gammaproteobacteria Enterobacteriales Enterobacteriaceae *Escherichia coli* O113:H21 str. CL-3 |
| Eco-1780-1 | Bacteria Proteobacteria Gammaproteobacteria Enterobacteriales Enterobacteriaceae *Escherichia coli* O121:H19 str. MT#2 |
| Eco-1781-1 | Bacteria Proteobacteria Gammaproteobacteria Enterobacteriales Enterobacteriaceae *Escherichia coli* O127:H27 str. C43/90 |
| Eco-1782-1 | Bacteria Proteobacteria Gammaproteobacteria Enterobacteriales Enterobacteriaceae *Escherichia coli* O127:H6 str. E2348/69 |
| Eco-548-1 to Eco-548-2 | Bacteria Proteobacteria Gammaproteobacteria Enterobacteriales Enterobacteriaceae *Escherichia coli* O145:H28 str. 4865/96 |
| Eco-1783-1 | Bacteria Proteobacteria Gammaproteobacteria Enterobacteriales Enterobacteriaceae *Escherichia coli* O157 str. NCCP15738 |
| Eco-1784-1 to Eco-1784-2 | Bacteria Proteobacteria Gammaproteobacteria Enterobacteriales Enterobacteriaceae *Escherichia coli* O157:H- str. 493-89 |
| Eco-1785-1 | Bacteria Proteobacteria Gammaproteobacteria Enterobacteriales Enterobacteriaceae *Escherichia coli* O157:H- str. H 2687 |
| Eco-1786-1 | Bacteria Proteobacteria Gammaproteobacteria Enterobacteriales Enterobacteriaceae *Escherichia coli* O157:H43 str. T22 |
| Eco-1787-1 | Bacteria Proteobacteria Gammaproteobacteria Enterobacteriales Enterobacteriaceae *Escherichia coli* O157:H7 EDL933 |
| Eco-1788-1 | Bacteria Proteobacteria Gammaproteobacteria Enterobacteriales Enterobacteriaceae *Escherichia coli* O157:H7 str. 08BKT061141 |
| Eco-1789-1 | Bacteria Proteobacteria Gammaproteobacteria Enterobacteriales Enterobacteriaceae *Escherichia coli* O157:H7 str. 09BKT048303 |
| Eco-1790-1 | Bacteria Proteobacteria Gammaproteobacteria Enterobacteriales Enterobacteriaceae *Escherichia coli* O157:H7 str. 1044 |
| Eco-1791-1 | Bacteria Proteobacteria Gammaproteobacteria Enterobacteriales Enterobacteriaceae *Escherichia coli* O157:H7 str. 1125 |
| Eco-1792-1 | Bacteria Proteobacteria Gammaproteobacteria Enterobacteriales Enterobacteriaceae *Escherichia coli* O157:H7 str. EC1212 |
| Eco-1793-1 | Bacteria Proteobacteria Gammaproteobacteria Enterobacteriales Enterobacteriaceae *Escherichia coli* O157:H7 str. EC4009 |
| Eco-1794-1 | Bacteria Proteobacteria Gammaproteobacteria Enterobacteriales Enterobacteriaceae *Escherichia coli* O157:H7 str. EC4042 |
| Eco-1795-1 | Bacteria Proteobacteria Gammaproteobacteria Enterobacteriales Enterobacteriaceae *Escherichia coli* O157:H7 str. EC4045 |
| Eco-1796-1 | Bacteria Proteobacteria Gammaproteobacteria Enterobacteriales Enterobacteriaceae *Escherichia coli* O157:H7 str. EC4076 |
| Eco-1797-1 | Bacteria Proteobacteria Gammaproteobacteria Enterobacteriales Enterobacteriaceae *Escherichia coli* O157:H7 str. EC4113 |
| Eco-1798-1 | Bacteria Proteobacteria Gammaproteobacteria Enterobacteriales Enterobacteriaceae *Escherichia coli* O157:H7 str. EC4115 |
| Eco-1799-1 | Bacteria Proteobacteria Gammaproteobacteria Enterobacteriales Enterobacteriaceae *Escherichia coli* O157:H7 str. EC4191 |
| Eco-1800-1 | Bacteria Proteobacteria Gammaproteobacteria Enterobacteriales Enterobacteriaceae *Escherichia coli* O157:H7 str. EC4196 |
| Eco-1801-1 | Bacteria Proteobacteria Gammaproteobacteria Enterobacteriales Enterobacteriaceae *Escherichia coli* O157:H7 str. EC4206 |
| Eco-1802-1 | Bacteria Proteobacteria Gammaproteobacteria Enterobacteriales Enterobacteriaceae *Escherichia coli* O157:H7 str. EC4401 |
| Eco-1803-1 | Bacteria Proteobacteria Gammaproteobacteria Enterobacteriales Enterobacteriaceae *Escherichia coli* O157:H7 str. EC4486 |
| Eco-1804-1 | Bacteria Proteobacteria Gammaproteobacteria Enterobacteriales Enterobacteriaceae *Escherichia coli* O157:H7 str. EC4501 |
| Eco-1805-1 | Bacteria Proteobacteria Gammaproteobacteria Enterobacteriales Enterobacteriaceae *Escherichia coli* O157:H7 str. EC508 |
| Eco-1806-1 | Bacteria Proteobacteria Gammaproteobacteria Enterobacteriales Enterobacteriaceae *Escherichia coli* O157:H7 str. EC869 |
| Eco-1807-1 | Bacteria Proteobacteria Gammaproteobacteria Enterobacteriales Enterobacteriaceae *Escherichia coli* O157:H7 str. FRIK2000 |
| Eco-1808-1 | Bacteria Proteobacteria Gammaproteobacteria Enterobacteriales Enterobacteriaceae *Escherichia coli* O157:H7 str. FRIK966 |
| Eco-1809-1 | Bacteria Proteobacteria Gammaproteobacteria Enterobacteriales Enterobacteriaceae *Escherichia coli* O157:H7 str. G5101 |
| Eco-1810-1 | Bacteria Proteobacteria Gammaproteobacteria Enterobacteriales Enterobacteriaceae *Escherichia coli* O157:H7 str. LSU-61 |
| Eco-1811-1 | Bacteria Proteobacteria Gammaproteobacteria Enterobacteriales Enterobacteriaceae *Escherichia coli* O157:H7 str. Sakai |
| Eco-1812-1 | Bacteria Proteobacteria Gammaproteobacteria Enterobacteriales Enterobacteriaceae *Escherichia coli* O157:H7 str. TW14359 |
| Eco-1813-1 | Bacteria Proteobacteria Gammaproteobacteria Enterobacteriales Enterobacteriaceae *Escherichia coli* O157:H7 str. TW14588 |
| Eco-1814-1 | Bacteria Proteobacteria Gammaproteobacteria Enterobacteriales Enterobacteriaceae *Escherichia coli* O26:H11 str. 11368 |
| Eco-1815-1 | Bacteria Proteobacteria Gammaproteobacteria Enterobacteriales Enterobacteriaceae *Escherichia coli* O26:H11 str. 2011C-3274 |
| Eco-1816-1 | Bacteria Proteobacteria Gammaproteobacteria Enterobacteriales Enterobacteriaceae *Escherichia coli* O26:H11 str. CFSAN001629 |
| Eco-1817-1 | Bacteria Proteobacteria Gammaproteobacteria Enterobacteriales Enterobacteriaceae *Escherichia coli* O26:H11 str. CVM10021 |
| Eco-1818-1 | Bacteria Proteobacteria Gammaproteobacteria Enterobacteriales Enterobacteriaceae *Escherichia coli* O26:H11 str. CVM10026 |
| Eco-1819-1 | Bacteria Proteobacteria Gammaproteobacteria Enterobacteriales Enterobacteriaceae *Escherichia coli* O26:H11 str. CVM10030 |
| Eco-1820-1 | Bacteria Proteobacteria Gammaproteobacteria Enterobacteriales Enterobacteriaceae *Escherichia coli* O26:H11 str. CVM10224 |
| Eco-1821-1 | Bacteria Proteobacteria Gammaproteobacteria Enterobacteriales Enterobacteriaceae *Escherichia coli* O26:H11 str. CVM9942 |
| Eco-1822-1 | Bacteria Proteobacteria Gammaproteobacteria Enterobacteriales Enterobacteriaceae *Escherichia coli* O26:H11 str. CVM9952 |
| Eco-1823-1 | Bacteria Proteobacteria Gammaproteobacteria Enterobacteriales Enterobacteriaceae *Escherichia coli* O32:H37 str. P4 |
| Eco-1824-1 | Bacteria Proteobacteria Gammaproteobacteria Enterobacteriales Enterobacteriaceae *Escherichia coli* O45:H2 str. 03-EN-705 |
| Eco-1825-1 | Bacteria Proteobacteria Gammaproteobacteria Enterobacteriales Enterobacteriaceae *Escherichia coli* O55:H7 str. 3256-97 |
| Eco-1826-1 | Bacteria Proteobacteria Gammaproteobacteria Enterobacteriales Enterobacteriaceae *Escherichia coli* O55:H7 str. CB9615 |
| Eco-1827-1 | Bacteria Proteobacteria Gammaproteobacteria Enterobacteriales Enterobacteriaceae *Escherichia coli* O55:H7 str. RM12579 |
| Eco-1828-1 | Bacteria Proteobacteria Gammaproteobacteria Enterobacteriales Enterobacteriaceae *Escherichia coli* O55:H7 str. USDA 5905 |
| Eco-1829-1 | Bacteria Proteobacteria Gammaproteobacteria Enterobacteriales Enterobacteriaceae *Escherichia coli* O5:K4(L):H4 str. ATCC 23502 |
| Eco-1830-1 | Bacteria Proteobacteria Gammaproteobacteria Enterobacteriales Enterobacteriaceae *Escherichia coli* O7:K1 str. CE10 |
| Eco-1831-1 | Bacteria Proteobacteria Gammaproteobacteria Enterobacteriales Enterobacteriaceae *Escherichia coli* O83:H1 str. NRG 857C |
| Eco-1832-1 | Bacteria Proteobacteria Gammaproteobacteria Enterobacteriales Enterobacteriaceae *Escherichia coli* O91:H21 str. B2F1 |
| Eco-1833-1 | Bacteria Proteobacteria Gammaproteobacteria Enterobacteriales Enterobacteriaceae *Escherichia coli* OK1114 |
| Eco-1834-1 | Bacteria Proteobacteria Gammaproteobacteria Enterobacteriales Enterobacteriaceae *Escherichia coli* OK1180 |
| Eco-1835-1 | Bacteria Proteobacteria Gammaproteobacteria Enterobacteriales Enterobacteriaceae *Escherichia coli* OK1357 |
| Eco-1836-1 | Bacteria Proteobacteria Gammaproteobacteria Enterobacteriales Enterobacteriaceae *Escherichia coli* ONT:H33 str. C48/93 |
| Eco-1837-1 | Bacteria Proteobacteria Gammaproteobacteria Enterobacteriales Enterobacteriaceae *Escherichia coli* P0298942.1 |

| | |
|---|---|
| Eco-1838-1 | Bacteria Proteobacteria Gammaproteobacteria Enterobacteriales Enterobacteriaceae *Escherichia coli* P0298942.10 |
| Eco-1839-1 | Bacteria Proteobacteria Gammaproteobacteria Enterobacteriales Enterobacteriaceae *Escherichia coli* P0298942.11 |
| Eco-1840-1 | Bacteria Proteobacteria Gammaproteobacteria Enterobacteriales Enterobacteriaceae *Escherichia coli* P0298942.12 |
| Eco-1841-1 | Bacteria Proteobacteria Gammaproteobacteria Enterobacteriales Enterobacteriaceae *Escherichia coli* P0298942.14 |
| Eco-1842-1 | Bacteria Proteobacteria Gammaproteobacteria Enterobacteriales Enterobacteriaceae *Escherichia coli* P0298942.15 |
| Eco-1843-1 | Bacteria Proteobacteria Gammaproteobacteria Enterobacteriales Enterobacteriaceae *Escherichia coli* P0298942.2 |
| Eco-1844-1 | Bacteria Proteobacteria Gammaproteobacteria Enterobacteriales Enterobacteriaceae *Escherichia coli* P0298942.3 |
| Eco-1845-1 | Bacteria Proteobacteria Gammaproteobacteria Enterobacteriales Enterobacteriaceae *Escherichia coli* P0298942.4 |
| Eco-1846-1 | Bacteria Proteobacteria Gammaproteobacteria Enterobacteriales Enterobacteriaceae *Escherichia coli* P0298942.6 |
| Eco-1847-1 | Bacteria Proteobacteria Gammaproteobacteria Enterobacteriales Enterobacteriaceae *Escherichia coli* P0298942.7 |
| Eco-1848-1 | Bacteria Proteobacteria Gammaproteobacteria Enterobacteriales Enterobacteriaceae *Escherichia coli* P0298942.8 |
| Eco-1849-1 | Bacteria Proteobacteria Gammaproteobacteria Enterobacteriales Enterobacteriaceae *Escherichia coli* P0298942.9 |
| Eco-1850-1 | Bacteria Proteobacteria Gammaproteobacteria Enterobacteriales Enterobacteriaceae *Escherichia coli* P0299438.10 |
| Eco-1851-1 | Bacteria Proteobacteria Gammaproteobacteria Enterobacteriales Enterobacteriaceae *Escherichia coli* P0299438.11 |
| Eco-1852-1 | Bacteria Proteobacteria Gammaproteobacteria Enterobacteriales Enterobacteriaceae *Escherichia coli* P0299438.2 |
| Eco-1853-1 | Bacteria Proteobacteria Gammaproteobacteria Enterobacteriales Enterobacteriaceae *Escherichia coli* P0299438.3 |
| Eco-1854-1 | Bacteria Proteobacteria Gammaproteobacteria Enterobacteriales Enterobacteriaceae *Escherichia coli* P0299438.4 |
| Eco-1855-1 | Bacteria Proteobacteria Gammaproteobacteria Enterobacteriales Enterobacteriaceae *Escherichia coli* P0299438.5 |
| Eco-1856-1 | Bacteria Proteobacteria Gammaproteobacteria Enterobacteriales Enterobacteriaceae *Escherichia coli* P0299438.6 |
| Eco-1857-1 | Bacteria Proteobacteria Gammaproteobacteria Enterobacteriales Enterobacteriaceae *Escherichia coli* P0299438.7 |
| Eco-1858-1 | Bacteria Proteobacteria Gammaproteobacteria Enterobacteriales Enterobacteriaceae *Escherichia coli* P0299438.8 |
| Eco-1859-1 | Bacteria Proteobacteria Gammaproteobacteria Enterobacteriales Enterobacteriaceae *Escherichia coli* P0299438.9 |
| Eco-1860-1 | Bacteria Proteobacteria Gammaproteobacteria Enterobacteriales Enterobacteriaceae *Escherichia coli* P0299483.1 |
| Eco-1861-1 | Bacteria Proteobacteria Gammaproteobacteria Enterobacteriales Enterobacteriaceae *Escherichia coli* P0299483.2 |
| Eco-1862-1 | Bacteria Proteobacteria Gammaproteobacteria Enterobacteriales Enterobacteriaceae *Escherichia coli* P0299483.3 |
| Eco-1863-1 | Bacteria Proteobacteria Gammaproteobacteria Enterobacteriales Enterobacteriaceae *Escherichia coli* P02997067.6 |
| Eco-1864-1 | Bacteria Proteobacteria Gammaproteobacteria Enterobacteriales Enterobacteriaceae *Escherichia coli* P0299917.1 |
| Eco-1865-1 | Bacteria Proteobacteria Gammaproteobacteria Enterobacteriales Enterobacteriaceae *Escherichia coli* P0299917.10 |
| Eco-1866-1 | Bacteria Proteobacteria Gammaproteobacteria Enterobacteriales Enterobacteriaceae *Escherichia coli* P0299917.2 |
| Eco-1867-1 | Bacteria Proteobacteria Gammaproteobacteria Enterobacteriales Enterobacteriaceae *Escherichia coli* P0299917.3 |
| Eco-1868-1 | Bacteria Proteobacteria Gammaproteobacteria Enterobacteriales Enterobacteriaceae *Escherichia coli* P0299917.4 |
| Eco-1869-1 | Bacteria Proteobacteria Gammaproteobacteria Enterobacteriales Enterobacteriaceae *Escherichia coli* P0299917.5 |
| Eco-1870-1 | Bacteria Proteobacteria Gammaproteobacteria Enterobacteriales Enterobacteriaceae *Escherichia coli* P0299917.6 |
| Eco-1871-1 | Bacteria Proteobacteria Gammaproteobacteria Enterobacteriales Enterobacteriaceae *Escherichia coli* P0299917.7 |
| Eco-1872-1 | Bacteria Proteobacteria Gammaproteobacteria Enterobacteriales Enterobacteriaceae *Escherichia coli* P0299917.8 |
| Eco-1873-1 | Bacteria Proteobacteria Gammaproteobacteria Enterobacteriales Enterobacteriaceae *Escherichia coli* P0299917.9 |
| Eco-1874-1 | Bacteria Proteobacteria Gammaproteobacteria Enterobacteriales Enterobacteriaceae *Escherichia coli* P0301867.1 |
| Eco-1875-1 | Bacteria Proteobacteria Gammaproteobacteria Enterobacteriales Enterobacteriaceae *Escherichia coli* P0301867.11 |
| Eco-1876-1 | Bacteria Proteobacteria Gammaproteobacteria Enterobacteriales Enterobacteriaceae *Escherichia coli* P0301867.13 |
| Eco-1877-1 | Bacteria Proteobacteria Gammaproteobacteria Enterobacteriales Enterobacteriaceae *Escherichia coli* P0301867.2 |
| Eco-1878-1 | Bacteria Proteobacteria Gammaproteobacteria Enterobacteriales Enterobacteriaceae *Escherichia coli* P0301867.3 |
| Eco-1879-1 | Bacteria Proteobacteria Gammaproteobacteria Enterobacteriales Enterobacteriaceae *Escherichia coli* P0301867.4 |
| Eco-1880-1 | Bacteria Proteobacteria Gammaproteobacteria Enterobacteriales Enterobacteriaceae *Escherichia coli* P0301867.5 |
| Eco-1881-1 | Bacteria Proteobacteria Gammaproteobacteria Enterobacteriales Enterobacteriaceae *Escherichia coli* P0301867.7 |
| Eco-1882-1 | Bacteria Proteobacteria Gammaproteobacteria Enterobacteriales Enterobacteriaceae *Escherichia coli* P0301867.8 |
| Eco-1883-1 | Bacteria Proteobacteria Gammaproteobacteria Enterobacteriales Enterobacteriaceae *Escherichia coli* P0301904.3 |
| Eco-1884-1 | Bacteria Proteobacteria Gammaproteobacteria Enterobacteriales Enterobacteriaceae *Escherichia coli* P0302293.10 |
| Eco-1885-1 | Bacteria Proteobacteria Gammaproteobacteria Enterobacteriales Enterobacteriaceae *Escherichia coli* P0302293.2 |
| Eco-1886-1 | Bacteria Proteobacteria Gammaproteobacteria Enterobacteriales Enterobacteriaceae *Escherichia coli* P0302293.3 |
| Eco-1887-1 | Bacteria Proteobacteria Gammaproteobacteria Enterobacteriales Enterobacteriaceae *Escherichia coli* P0302293.4 |
| Eco-1888-1 | Bacteria Proteobacteria Gammaproteobacteria Enterobacteriales Enterobacteriaceae *Escherichia coli* P0302293.6 |
| Eco-1889-1 | Bacteria Proteobacteria Gammaproteobacteria Enterobacteriales Enterobacteriaceae *Escherichia coli* P0302293.7 |
| Eco-1890-1 | Bacteria Proteobacteria Gammaproteobacteria Enterobacteriales Enterobacteriaceae *Escherichia coli* P0302293.8 |
| Eco-1891-1 | Bacteria Proteobacteria Gammaproteobacteria Enterobacteriales Enterobacteriaceae *Escherichia coli* P0302293.9 |
| Eco-1892-1 | Bacteria Proteobacteria Gammaproteobacteria Enterobacteriales Enterobacteriaceae *Escherichia coli* P0302308.1 |
| Eco-1893-1 | Bacteria Proteobacteria Gammaproteobacteria Enterobacteriales Enterobacteriaceae *Escherichia coli* P0302308.10 |
| Eco-1894-1 | Bacteria Proteobacteria Gammaproteobacteria Enterobacteriales Enterobacteriaceae *Escherichia coli* P0302308.11 |
| Eco-1895-1 | Bacteria Proteobacteria Gammaproteobacteria Enterobacteriales Enterobacteriaceae *Escherichia coli* P0302308.12 |
| Eco-1896-1 | Bacteria Proteobacteria Gammaproteobacteria Enterobacteriales Enterobacteriaceae *Escherichia coli* P0302308.13 |
| Eco-1897-1 | Bacteria Proteobacteria Gammaproteobacteria Enterobacteriales Enterobacteriaceae *Escherichia coli* P0302308.14 |
| Eco-1898-1 | Bacteria Proteobacteria Gammaproteobacteria Enterobacteriales Enterobacteriaceae *Escherichia coli* P0302308.2 |
| Eco-1899-1 | Bacteria Proteobacteria Gammaproteobacteria Enterobacteriales Enterobacteriaceae *Escherichia coli* P0302308.3 |
| Eco-1900-1 | Bacteria Proteobacteria Gammaproteobacteria Enterobacteriales Enterobacteriaceae *Escherichia coli* P0302308.4 |
| Eco-1901-1 | Bacteria Proteobacteria Gammaproteobacteria Enterobacteriales Enterobacteriaceae *Escherichia coli* P0302308.5 |
| Eco-1902-1 | Bacteria Proteobacteria Gammaproteobacteria Enterobacteriales Enterobacteriaceae *Escherichia coli* P0304777.1 |

| | |
|---|---|
| Eco-1903-1 | Bacteria Proteobacteria Gammaproteobacteria Enterobacteriales Enterobacteriaceae *Escherichia coli* P0304777.10 |
| Eco-1904-1 | Bacteria Proteobacteria Gammaproteobacteria Enterobacteriales Enterobacteriaceae *Escherichia coli* P0304777.11 |
| Eco-1905-1 | Bacteria Proteobacteria Gammaproteobacteria Enterobacteriales Enterobacteriaceae *Escherichia coli* P0304777.12 |
| Eco-1906-1 | Bacteria Proteobacteria Gammaproteobacteria Enterobacteriales Enterobacteriaceae *Escherichia coli* P0304777.13 |
| Eco-1907-1 | Bacteria Proteobacteria Gammaproteobacteria Enterobacteriales Enterobacteriaceae *Escherichia coli* P0304777.14 |
| Eco-1908-1 | Bacteria Proteobacteria Gammaproteobacteria Enterobacteriales Enterobacteriaceae *Escherichia coli* P0304777.15 |
| Eco-1909-1 | Bacteria Proteobacteria Gammaproteobacteria Enterobacteriales Enterobacteriaceae *Escherichia coli* P0304777.2 |
| Eco-1910-1 | Bacteria Proteobacteria Gammaproteobacteria Enterobacteriales Enterobacteriaceae *Escherichia coli* P0304777.3 |
| Eco-1911-1 | Bacteria Proteobacteria Gammaproteobacteria Enterobacteriales Enterobacteriaceae *Escherichia coli* P0304777.4 |
| Eco-1912-1 | Bacteria Proteobacteria Gammaproteobacteria Enterobacteriales Enterobacteriaceae *Escherichia coli* P0304777.5 |
| Eco-1913-1 | Bacteria Proteobacteria Gammaproteobacteria Enterobacteriales Enterobacteriaceae *Escherichia coli* P0304777.7 |
| Eco-1914-1 | Bacteria Proteobacteria Gammaproteobacteria Enterobacteriales Enterobacteriaceae *Escherichia coli* P0304777.8 |
| Eco-1915-1 | Bacteria Proteobacteria Gammaproteobacteria Enterobacteriales Enterobacteriaceae *Escherichia coli* P0304777.9 |
| Eco-1916-1 | Bacteria Proteobacteria Gammaproteobacteria Enterobacteriales Enterobacteriaceae *Escherichia coli* P0304799.3 |
| Eco-1917-1 | Bacteria Proteobacteria Gammaproteobacteria Enterobacteriales Enterobacteriaceae *Escherichia coli* P0304816.1 |
| Eco-1918-1 | Bacteria Proteobacteria Gammaproteobacteria Enterobacteriales Enterobacteriaceae *Escherichia coli* P0304816.10 |
| Eco-1919-1 | Bacteria Proteobacteria Gammaproteobacteria Enterobacteriales Enterobacteriaceae *Escherichia coli* P0304816.11 |
| Eco-1920-1 | Bacteria Proteobacteria Gammaproteobacteria Enterobacteriales Enterobacteriaceae *Escherichia coli* P0304816.12 |
| Eco-1921-1 | Bacteria Proteobacteria Gammaproteobacteria Enterobacteriales Enterobacteriaceae *Escherichia coli* P0304816.13 |
| Eco-1922-1 | Bacteria Proteobacteria Gammaproteobacteria Enterobacteriales Enterobacteriaceae *Escherichia coli* P0304816.14 |
| Eco-1923-1 | Bacteria Proteobacteria Gammaproteobacteria Enterobacteriales Enterobacteriaceae *Escherichia coli* P0304816.15 |
| Eco-1924-1 | Bacteria Proteobacteria Gammaproteobacteria Enterobacteriales Enterobacteriaceae *Escherichia coli* P0304816.2 |
| Eco-1925-1 | Bacteria Proteobacteria Gammaproteobacteria Enterobacteriales Enterobacteriaceae *Escherichia coli* P0304816.3 |
| Eco-1926-1 | Bacteria Proteobacteria Gammaproteobacteria Enterobacteriales Enterobacteriaceae *Escherichia coli* P0304816.4 |
| Eco-1927-1 | Bacteria Proteobacteria Gammaproteobacteria Enterobacteriales Enterobacteriaceae *Escherichia coli* P0304816.5 |
| Eco-1928-1 | Bacteria Proteobacteria Gammaproteobacteria Enterobacteriales Enterobacteriaceae *Escherichia coli* P0304816.6 |
| Eco-1929-1 | Bacteria Proteobacteria Gammaproteobacteria Enterobacteriales Enterobacteriaceae *Escherichia coli* P0304816.7 |
| Eco-1930-1 | Bacteria Proteobacteria Gammaproteobacteria Enterobacteriales Enterobacteriaceae *Escherichia coli* P0304816.8 |
| Eco-1931-1 | Bacteria Proteobacteria Gammaproteobacteria Enterobacteriales Enterobacteriaceae *Escherichia coli* P0304816.9 |
| Eco-1932-1 | Bacteria Proteobacteria Gammaproteobacteria Enterobacteriales Enterobacteriaceae *Escherichia coli* P0305260.1 |
| Eco-1933-1 | Bacteria Proteobacteria Gammaproteobacteria Enterobacteriales Enterobacteriaceae *Escherichia coli* P0305260.10 |
| Eco-1934-1 | Bacteria Proteobacteria Gammaproteobacteria Enterobacteriales Enterobacteriaceae *Escherichia coli* P0305260.11 |
| Eco-1935-1 | Bacteria Proteobacteria Gammaproteobacteria Enterobacteriales Enterobacteriaceae *Escherichia coli* P0305260.12 |
| Eco-1936-1 | Bacteria Proteobacteria Gammaproteobacteria Enterobacteriales Enterobacteriaceae *Escherichia coli* P0305260.13 |
| Eco-1937-1 | Bacteria Proteobacteria Gammaproteobacteria Enterobacteriales Enterobacteriaceae *Escherichia coli* P0305260.15 |
| Eco-1938-1 | Bacteria Proteobacteria Gammaproteobacteria Enterobacteriales Enterobacteriaceae *Escherichia coli* P0305260.2 |
| Eco-1939-1 | Bacteria Proteobacteria Gammaproteobacteria Enterobacteriales Enterobacteriaceae *Escherichia coli* P0305260.3 |
| Eco-1940-1 | Bacteria Proteobacteria Gammaproteobacteria Enterobacteriales Enterobacteriaceae *Escherichia coli* P0305260.4 |
| Eco-1941-1 | Bacteria Proteobacteria Gammaproteobacteria Enterobacteriales Enterobacteriaceae *Escherichia coli* P0305260.5 |
| Eco-1942-1 | Bacteria Proteobacteria Gammaproteobacteria Enterobacteriales Enterobacteriaceae *Escherichia coli* P0305260.6 |
| Eco-1943-1 | Bacteria Proteobacteria Gammaproteobacteria Enterobacteriales Enterobacteriaceae *Escherichia coli* P0305260.7 |
| Eco-1944-1 | Bacteria Proteobacteria Gammaproteobacteria Enterobacteriales Enterobacteriaceae *Escherichia coli* P0305260.8 |
| Eco-1945-1 | Bacteria Proteobacteria Gammaproteobacteria Enterobacteriales Enterobacteriaceae *Escherichia coli* P0305260.9 |
| Eco-2167-1 | Bacteria Proteobacteria Gammaproteobacteria Enterobacteriales Enterobacteriaceae *Escherichia coli p*0305293.1 |
| Eco-2168-1 | Bacteria Proteobacteria Gammaproteobacteria Enterobacteriales Enterobacteriaceae *Escherichia coli p*0305293.10 |
| Eco-2169-1 | Bacteria Proteobacteria Gammaproteobacteria Enterobacteriales Enterobacteriaceae *Escherichia coli p*0305293.11 |
| Eco-2170-1 | Bacteria Proteobacteria Gammaproteobacteria Enterobacteriales Enterobacteriaceae *Escherichia coli p*0305293.12 |
| Eco-2171-1 | Bacteria Proteobacteria Gammaproteobacteria Enterobacteriales Enterobacteriaceae *Escherichia coli p*0305293.13 |
| Eco-2172-1 | Bacteria Proteobacteria Gammaproteobacteria Enterobacteriales Enterobacteriaceae *Escherichia coli p*0305293.14 |
| Eco-2173-1 | Bacteria Proteobacteria Gammaproteobacteria Enterobacteriales Enterobacteriaceae *Escherichia coli p*0305293.15 |
| Eco-2174-1 | Bacteria Proteobacteria Gammaproteobacteria Enterobacteriales Enterobacteriaceae *Escherichia coli p*0305293.2 |
| Eco-2175-1 | Bacteria Proteobacteria Gammaproteobacteria Enterobacteriales Enterobacteriaceae *Escherichia coli p*0305293.3 |
| Eco-2176-1 | Bacteria Proteobacteria Gammaproteobacteria Enterobacteriales Enterobacteriaceae *Escherichia coli p*0305293.4 |
| Eco-2177-1 | Bacteria Proteobacteria Gammaproteobacteria Enterobacteriales Enterobacteriaceae *Escherichia coli p*0305293.5 |
| Eco-2178-1 | Bacteria Proteobacteria Gammaproteobacteria Enterobacteriales Enterobacteriaceae *Escherichia coli p*0305293.6 |
| Eco-2179-1 | Bacteria Proteobacteria Gammaproteobacteria Enterobacteriales Enterobacteriaceae *Escherichia coli p*0305293.7 |
| Eco-2180-1 | Bacteria Proteobacteria Gammaproteobacteria Enterobacteriales Enterobacteriaceae *Escherichia coli p*0305293.8 |
| Eco-2181-1 | Bacteria Proteobacteria Gammaproteobacteria Enterobacteriales Enterobacteriaceae *Escherichia coli p*0305293.9 |
| Eco-1946-1 | Bacteria Proteobacteria Gammaproteobacteria Enterobacteriales Enterobacteriaceae *Escherichia coli* P12b |
| Eco-1947-1 | Bacteria Proteobacteria Gammaproteobacteria Enterobacteriales Enterobacteriaceae *Escherichia coli* P4-96 |
| Eco-1948-1 | Bacteria Proteobacteria Gammaproteobacteria Enterobacteriales Enterobacteriaceae *Escherichia coli* P4-NR |
| Eco-1949-1 | Bacteria Proteobacteria Gammaproteobacteria Enterobacteriales Enterobacteriaceae *Escherichia coli* PA10 |
| Eco-1950-1 | Bacteria Proteobacteria Gammaproteobacteria Enterobacteriales Enterobacteriaceae *Escherichia coli* PA11 |
| Eco-1951-1 | Bacteria Proteobacteria Gammaproteobacteria Enterobacteriales Enterobacteriaceae *Escherichia coli* PA13 |
| Eco-1952-1 | Bacteria Proteobacteria Gammaproteobacteria Enterobacteriales Enterobacteriaceae *Escherichia coli* PA14 |

| | |
|---|---|
| Eco-1953-1 | Bacteria Proteobacteria Gammaproteobacteria Enterobacteriales Enterobacteriaceae *Escherichia coli* PA15 |
| Eco-1954-1 | Bacteria Proteobacteria Gammaproteobacteria Enterobacteriales Enterobacteriaceae *Escherichia coli* PA19 |
| Eco-1955-1 | Bacteria Proteobacteria Gammaproteobacteria Enterobacteriales Enterobacteriaceae *Escherichia coli* PA2 |
| Eco-1956-1 | Bacteria Proteobacteria Gammaproteobacteria Enterobacteriales Enterobacteriaceae *Escherichia coli* PA22 |
| Eco-1957-1 | Bacteria Proteobacteria Gammaproteobacteria Enterobacteriales Enterobacteriaceae *Escherichia coli* PA23 |
| Eco-1958-1 | Bacteria Proteobacteria Gammaproteobacteria Enterobacteriales Enterobacteriaceae *Escherichia coli* PA24 |
| Eco-1959-1 | Bacteria Proteobacteria Gammaproteobacteria Enterobacteriales Enterobacteriaceae *Escherichia coli* PA25 |
| Eco-1960-1 | Bacteria Proteobacteria Gammaproteobacteria Enterobacteriales Enterobacteriaceae *Escherichia coli* PA28 |
| Eco-1961-1 | Bacteria Proteobacteria Gammaproteobacteria Enterobacteriales Enterobacteriaceae *Escherichia coli* PA3 |
| Eco-1962-1 | Bacteria Proteobacteria Gammaproteobacteria Enterobacteriales Enterobacteriaceae *Escherichia coli* PA31 |
| Eco-1963-1 | Bacteria Proteobacteria Gammaproteobacteria Enterobacteriales Enterobacteriaceae *Escherichia coli* PA32 |
| Eco-1964-1 | Bacteria Proteobacteria Gammaproteobacteria Enterobacteriales Enterobacteriaceae *Escherichia coli* PA33 |
| Eco-1965-1 | Bacteria Proteobacteria Gammaproteobacteria Enterobacteriales Enterobacteriaceae *Escherichia coli* PA34 |
| Eco-1966-1 | Bacteria Proteobacteria Gammaproteobacteria Enterobacteriales Enterobacteriaceae *Escherichia coli* PA35 |
| Eco-1967-1 | Bacteria Proteobacteria Gammaproteobacteria Enterobacteriales Enterobacteriaceae *Escherichia coli* PA38 |
| Eco-1968-1 | Bacteria Proteobacteria Gammaproteobacteria Enterobacteriales Enterobacteriaceae *Escherichia coli* PA39 |
| Eco-1969-1 | Bacteria Proteobacteria Gammaproteobacteria Enterobacteriales Enterobacteriaceae *Escherichia coli* PA4 |
| Eco-1970-1 | Bacteria Proteobacteria Gammaproteobacteria Enterobacteriales Enterobacteriaceae *Escherichia coli* PA40 |
| Eco-1971-1 | Bacteria Proteobacteria Gammaproteobacteria Enterobacteriales Enterobacteriaceae *Escherichia coli* PA41 |
| Eco-1972-1 | Bacteria Proteobacteria Gammaproteobacteria Enterobacteriales Enterobacteriaceae *Escherichia coli* PA42 |
| Eco-1973-1 | Bacteria Proteobacteria Gammaproteobacteria Enterobacteriales Enterobacteriaceae *Escherichia coli* PA45 |
| Eco-1974-1 | Bacteria Proteobacteria Gammaproteobacteria Enterobacteriales Enterobacteriaceae *Escherichia coli* PA47 |
| Eco-1975-1 | Bacteria Proteobacteria Gammaproteobacteria Enterobacteriales Enterobacteriaceae *Escherichia coli* PA48 |
| Eco-1976-1 | Bacteria Proteobacteria Gammaproteobacteria Enterobacteriales Enterobacteriaceae *Escherichia coli* PA49 |
| Eco-1977-1 | Bacteria Proteobacteria Gammaproteobacteria Enterobacteriales Enterobacteriaceae *Escherichia coli* PA5 |
| Eco-1978-1 | Bacteria Proteobacteria Gammaproteobacteria Enterobacteriales Enterobacteriaceae *Escherichia coli* PA7 |
| Eco-1979-1 | Bacteria Proteobacteria Gammaproteobacteria Enterobacteriales Enterobacteriaceae *Escherichia coli* PA8 |
| Eco-1980-1 | Bacteria Proteobacteria Gammaproteobacteria Enterobacteriales Enterobacteriaceae *Escherichia coli* PA9 |
| Eco-1981-1 | Bacteria Proteobacteria Gammaproteobacteria Enterobacteriales Enterobacteriaceae *Escherichia coli* PMV-1 |
| Eco-1982-1 | Bacteria Proteobacteria Gammaproteobacteria Enterobacteriales Enterobacteriaceae *Escherichia coli* RN587/1 |
| Eco-1983-1 | Bacteria Proteobacteria Gammaproteobacteria Enterobacteriales Enterobacteriaceae *Escherichia coli* S17 |
| Eco-1984-1 | Bacteria Proteobacteria Gammaproteobacteria Enterobacteriales Enterobacteriaceae *Escherichia coli* SCD1 |
| Eco-1985-1 | Bacteria Proteobacteria Gammaproteobacteria Enterobacteriales Enterobacteriaceae *Escherichia coli* SCD2 |
| Eco-1986-1 | Bacteria Proteobacteria Gammaproteobacteria Enterobacteriales Enterobacteriaceae *Escherichia coli* SCI-07 |
| Eco-1987-1 | Bacteria Proteobacteria Gammaproteobacteria Enterobacteriales Enterobacteriaceae *Escherichia coli* SE11 |
| Eco-1988-1 | Bacteria Proteobacteria Gammaproteobacteria Enterobacteriales Enterobacteriaceae *Escherichia coli* SE15 |
| Eco-1989-1 | Bacteria Proteobacteria Gammaproteobacteria Enterobacteriales Enterobacteriaceae *Escherichia coli* SEPT362 |
| Eco-1990-1 | Bacteria Proteobacteria Gammaproteobacteria Enterobacteriales Enterobacteriaceae *Escherichia coli* SMS-3-5 |
| Eco-1991-1 | Bacteria Proteobacteria Gammaproteobacteria Enterobacteriales Enterobacteriaceae *Escherichia coli* STEC_7v |
| Eco-1992-1 | Bacteria Proteobacteria Gammaproteobacteria Enterobacteriales Enterobacteriaceae *Escherichia coli* STEC_B2F1 |
| Eco-1993-1 | Bacteria Proteobacteria Gammaproteobacteria Enterobacteriales Enterobacteriaceae *Escherichia coli* STEC_C165-02 |
| Eco-1994-1 | Bacteria Proteobacteria Gammaproteobacteria Enterobacteriales Enterobacteriaceae *Escherichia coli* STEC_EH250 |
| Eco-1995-1 | Bacteria Proteobacteria Gammaproteobacteria Enterobacteriales Enterobacteriaceae *Escherichia coli* STEC_MHI813 |
| Eco-1996-1 | Bacteria Proteobacteria Gammaproteobacteria Enterobacteriales Enterobacteriaceae *Escherichia coli* STEC_O31 |
| Eco-1997-1 | Bacteria Proteobacteria Gammaproteobacteria Enterobacteriales Enterobacteriaceae *Escherichia coli* STEC_S1191 |
| Eco-2182-1 | Bacteria Proteobacteria Gammaproteobacteria Enterobacteriales Enterobacteriaceae *Escherichia coli str. 'clone* D i14' |
| Eco-2183-1 | Bacteria Proteobacteria Gammaproteobacteria Enterobacteriales Enterobacteriaceae *Escherichia coli str. 'clone* D i2' |
| Eco-2184-1 | Bacteria Proteobacteria Gammaproteobacteria Enterobacteriales Enterobacteriaceae *Escherichia coli str.* K-12 substr. DH10B |
| Eco-2185-1 | Bacteria Proteobacteria Gammaproteobacteria Enterobacteriales Enterobacteriaceae *Escherichia coli str.* K-12 substr. MDS42 |
| Eco-753-1 to Eco-753-2 | Bacteria Proteobacteria Gammaproteobacteria Enterobacteriales Enterobacteriaceae *Escherichia coli str.* K-12 substr. MG1655 |
| Eco-2186-1 | Bacteria Proteobacteria Gammaproteobacteria Enterobacteriales Enterobacteriaceae *Escherichia coli str.* K-12 substr. MG1655star |
| Eco-2187-1 | Bacteria Proteobacteria Gammaproteobacteria Enterobacteriales Enterobacteriaceae *Escherichia coli str.* K-12 substr. W3110 |
| Eco-1998-1 | Bacteria Proteobacteria Gammaproteobacteria Enterobacteriales Enterobacteriaceae *Escherichia coli* SWW33 |
| Eco-1999-1 | Bacteria Proteobacteria Gammaproteobacteria Enterobacteriales Enterobacteriaceae *Escherichia coli* T1282_01 |
| Eco-2000-1 | Bacteria Proteobacteria Gammaproteobacteria Enterobacteriales Enterobacteriaceae *Escherichia coli* T1840_97 |
| Eco-2001-1 | Bacteria Proteobacteria Gammaproteobacteria Enterobacteriales Enterobacteriaceae *Escherichia coli* T234_00 |
| Eco-2002-1 | Bacteria Proteobacteria Gammaproteobacteria Enterobacteriales Enterobacteriaceae *Escherichia coli* T924_01 |
| Eco-2003-1 | Bacteria Proteobacteria Gammaproteobacteria Enterobacteriales Enterobacteriaceae *Escherichia coli* TA124 |
| Eco-2004-1 | Bacteria Proteobacteria Gammaproteobacteria Enterobacteriales Enterobacteriaceae *Escherichia coli* TA143 |
| Eco-2005-1 | Bacteria Proteobacteria Gammaproteobacteria Enterobacteriales Enterobacteriaceae *Escherichia coli* TA206 |
| Eco-2006-1 | Bacteria Proteobacteria Gammaproteobacteria Enterobacteriales Enterobacteriaceae *Escherichia coli* TA271 |
| Eco-2045-1 | Bacteria Proteobacteria Gammaproteobacteria Enterobacteriales Enterobacteriaceae *Escherichia coli* ThroopD |
| Eco-2007-1 to Eco-2007-2 | Bacteria Proteobacteria Gammaproteobacteria Enterobacteriales Enterobacteriaceae *Escherichia coli* TOP2386 |
| Eco-2008-1 | Bacteria Proteobacteria Gammaproteobacteria Enterobacteriales Enterobacteriaceae *Escherichia coli* TOP2396-1 |
| Eco-2009-1 | Bacteria Proteobacteria Gammaproteobacteria Enterobacteriales Enterobacteriaceae *Escherichia coli* TOP2396-2 |

| | |
|---|---|
| Eco-2010-1 | Bacteria Proteobacteria Gammaproteobacteria Enterobacteriales Enterobacteriaceae *Escherichia coli* TOP2396-3 |
| Eco-2011-1 | Bacteria Proteobacteria Gammaproteobacteria Enterobacteriales Enterobacteriaceae *Escherichia coli* TOP2522-1 |
| Eco-2012-1 to Eco-2012-2 | Bacteria Proteobacteria Gammaproteobacteria Enterobacteriales Enterobacteriaceae *Escherichia coli* TOP2652 |
| Eco-2013-1 | Bacteria Proteobacteria Gammaproteobacteria Enterobacteriales Enterobacteriaceae *Escherichia coli* TOP2662-1 |
| Eco-2014-1 | Bacteria Proteobacteria Gammaproteobacteria Enterobacteriales Enterobacteriaceae *Escherichia coli* TOP2662-2 |
| Eco-2015-1 | Bacteria Proteobacteria Gammaproteobacteria Enterobacteriales Enterobacteriaceae *Escherichia coli* TOP2662-3 |
| Eco-2016-1 | Bacteria Proteobacteria Gammaproteobacteria Enterobacteriales Enterobacteriaceae *Escherichia coli* TOP2662-4 |
| Eco-2017-1 | Bacteria Proteobacteria Gammaproteobacteria Enterobacteriales Enterobacteriaceae *Escherichia coli* TOP382-1 |
| Eco-2018-1 | Bacteria Proteobacteria Gammaproteobacteria Enterobacteriales Enterobacteriaceae *Escherichia coli* TOP382-2 |
| Eco-2019-1 | Bacteria Proteobacteria Gammaproteobacteria Enterobacteriales Enterobacteriaceae *Escherichia coli* TOP382-3 |
| Eco-2020-1 | Bacteria Proteobacteria Gammaproteobacteria Enterobacteriales Enterobacteriaceae *Escherichia coli* TOP550-2 |
| Eco-2021-1 | Bacteria Proteobacteria Gammaproteobacteria Enterobacteriales Enterobacteriaceae *Escherichia coli* TOP550-3 |
| Eco-2022-1 | Bacteria Proteobacteria Gammaproteobacteria Enterobacteriales Enterobacteriaceae *Escherichia coli* TOP550-4 |
| Eco-2023-1 | Bacteria Proteobacteria Gammaproteobacteria Enterobacteriales Enterobacteriaceae *Escherichia coli* TT12B |
| Eco-2024-1 | Bacteria Proteobacteria Gammaproteobacteria Enterobacteriales Enterobacteriaceae *Escherichia coli* TW00353 |
| Eco-2025-1 | Bacteria Proteobacteria Gammaproteobacteria Enterobacteriales Enterobacteriaceae *Escherichia coli* TW06591 |
| Eco-2026-1 | Bacteria Proteobacteria Gammaproteobacteria Enterobacteriales Enterobacteriaceae *Escherichia coli* TW07509 |
| Eco-2027-1 | Bacteria Proteobacteria Gammaproteobacteria Enterobacteriales Enterobacteriaceae *Escherichia coli* TW07793 |
| Eco-2028-1 | Bacteria Proteobacteria Gammaproteobacteria Enterobacteriales Enterobacteriaceae *Escherichia coli* TW07945 |
| Eco-2029-1 | Bacteria Proteobacteria Gammaproteobacteria Enterobacteriales Enterobacteriaceae *Escherichia coli* TW09098 |
| Eco-2030-1 | Bacteria Proteobacteria Gammaproteobacteria Enterobacteriales Enterobacteriaceae *Escherichia coli* TW09109 |
| Eco-2031-1 | Bacteria Proteobacteria Gammaproteobacteria Enterobacteriales Enterobacteriaceae *Escherichia coli* TW09195 |
| Eco-2032-1 | Bacteria Proteobacteria Gammaproteobacteria Enterobacteriales Enterobacteriaceae *Escherichia coli* TW10119 |
| Eco-2033-1 | Bacteria Proteobacteria Gammaproteobacteria Enterobacteriales Enterobacteriaceae *Escherichia coli* TW10246 |
| Eco-2034-1 | Bacteria Proteobacteria Gammaproteobacteria Enterobacteriales Enterobacteriaceae *Escherichia coli* TW10509 |
| Eco-2035-1 | Bacteria Proteobacteria Gammaproteobacteria Enterobacteriales Enterobacteriaceae *Escherichia coli* TW10598 |
| Eco-2036-1 | Bacteria Proteobacteria Gammaproteobacteria Enterobacteriales Enterobacteriaceae *Escherichia coli* TW10722 |
| Eco-2037-1 | Bacteria Proteobacteria Gammaproteobacteria Enterobacteriales Enterobacteriaceae *Escherichia coli* TW10828 |
| Eco-2038-1 | Bacteria Proteobacteria Gammaproteobacteria Enterobacteriales Enterobacteriaceae *Escherichia coli* TW11039 |
| Eco-2039-1 | Bacteria Proteobacteria Gammaproteobacteria Enterobacteriales Enterobacteriaceae *Escherichia coli* TW11681 |
| Eco-2040-1 | Bacteria Proteobacteria Gammaproteobacteria Enterobacteriales Enterobacteriaceae *Escherichia coli* TW14301 |
| Eco-2041-1 | Bacteria Proteobacteria Gammaproteobacteria Enterobacteriales Enterobacteriaceae *Escherichia coli* TW14313 |
| Eco-2042-1 | Bacteria Proteobacteria Gammaproteobacteria Enterobacteriales Enterobacteriaceae *Escherichia coli* TW14425 |
| Eco-2043-1 | Bacteria Proteobacteria Gammaproteobacteria Enterobacteriales Enterobacteriaceae *Escherichia coli* TW15901 |
| Eco-2046-1 | Bacteria Proteobacteria Gammaproteobacteria Enterobacteriales Enterobacteriaceae *Escherichia coli* Tx1686 |
| Eco-2044-1 | Bacteria Proteobacteria Gammaproteobacteria Enterobacteriales Enterobacteriaceae *Escherichia coli* TX1999 |
| Eco-2047-1 | Bacteria Proteobacteria Gammaproteobacteria Enterobacteriales Enterobacteriaceae *Escherichia coli* Tx3800 |
| Eco-2048-1 | Bacteria Proteobacteria Gammaproteobacteria Enterobacteriales Enterobacteriaceae *Escherichia coli* UM146 |
| Eco-2049-1 | Bacteria Proteobacteria Gammaproteobacteria Enterobacteriales Enterobacteriaceae *Escherichia coli* UMEA 3014-1 |
| Eco-2050-1 | Bacteria Proteobacteria Gammaproteobacteria Enterobacteriales Enterobacteriaceae *Escherichia coli* UMEA 3022-1 |
| Eco-2051-1 | Bacteria Proteobacteria Gammaproteobacteria Enterobacteriales Enterobacteriaceae *Escherichia coli* UMEA 3033-1 |
| Eco-2052-1 | Bacteria Proteobacteria Gammaproteobacteria Enterobacteriales Enterobacteriaceae *Escherichia coli* UMEA 3041-1 |
| Eco-2053-1 | Bacteria Proteobacteria Gammaproteobacteria Enterobacteriales Enterobacteriaceae *Escherichia coli* UMEA 3052-1 |
| Eco-2054-1 | Bacteria Proteobacteria Gammaproteobacteria Enterobacteriales Enterobacteriaceae *Escherichia coli* UMEA 3053-1 |
| Eco-2055-1 | Bacteria Proteobacteria Gammaproteobacteria Enterobacteriales Enterobacteriaceae *Escherichia coli* UMEA 3065-1 |
| Eco-2056-1 | Bacteria Proteobacteria Gammaproteobacteria Enterobacteriales Enterobacteriaceae *Escherichia coli* UMEA 3087-1 |
| Eco-2057-1 | Bacteria Proteobacteria Gammaproteobacteria Enterobacteriales Enterobacteriaceae *Escherichia coli* UMEA 3088-1 |
| Eco-2058-1 | Bacteria Proteobacteria Gammaproteobacteria Enterobacteriales Enterobacteriaceae *Escherichia coli* UMEA 3097-1 |
| Eco-2059-1 | Bacteria Proteobacteria Gammaproteobacteria Enterobacteriales Enterobacteriaceae *Escherichia coli* UMEA 3108-1 |
| Eco-2060-1 | Bacteria Proteobacteria Gammaproteobacteria Enterobacteriales Enterobacteriaceae *Escherichia coli* UMEA 3113-1 |
| Eco-2061-1 | Bacteria Proteobacteria Gammaproteobacteria Enterobacteriales Enterobacteriaceae *Escherichia coli* UMEA 3117-1 |
| Eco-2062-1 | Bacteria Proteobacteria Gammaproteobacteria Enterobacteriales Enterobacteriaceae *Escherichia coli* UMEA 3121-1 |
| Eco-2063-1 | Bacteria Proteobacteria Gammaproteobacteria Enterobacteriales Enterobacteriaceae *Escherichia coli* UMEA 3122-1 |
| Eco-2064-1 | Bacteria Proteobacteria Gammaproteobacteria Enterobacteriales Enterobacteriaceae *Escherichia coli* UMEA 3124-1 |
| Eco-2065-1 | Bacteria Proteobacteria Gammaproteobacteria Enterobacteriales Enterobacteriaceae *Escherichia coli* UMEA 3139-1 |
| Eco-2066-1 | Bacteria Proteobacteria Gammaproteobacteria Enterobacteriales Enterobacteriaceae *Escherichia coli* UMEA 3140-1 |
| Eco-2067-1 | Bacteria Proteobacteria Gammaproteobacteria Enterobacteriales Enterobacteriaceae *Escherichia coli* UMEA 3144-1 |
| Eco-2068-1 | Bacteria Proteobacteria Gammaproteobacteria Enterobacteriales Enterobacteriaceae *Escherichia coli* UMEA 3148-1 |
| Eco-2069-1 | Bacteria Proteobacteria Gammaproteobacteria Enterobacteriales Enterobacteriaceae *Escherichia coli* UMEA 3150-1 |
| Eco-2070-1 | Bacteria Proteobacteria Gammaproteobacteria Enterobacteriales Enterobacteriaceae *Escherichia coli* UMEA 3151-1 |
| Eco-2071-1 | Bacteria Proteobacteria Gammaproteobacteria Enterobacteriales Enterobacteriaceae *Escherichia coli* UMEA 3152-1 |
| Eco-2072-1 | Bacteria Proteobacteria Gammaproteobacteria Enterobacteriales Enterobacteriaceae *Escherichia coli* UMEA 3155-1 |
| Eco-2073-1 | Bacteria Proteobacteria Gammaproteobacteria Enterobacteriales Enterobacteriaceae *Escherichia coli* UMEA 3159-1 |
| Eco-2074-1 | Bacteria Proteobacteria Gammaproteobacteria Enterobacteriales Enterobacteriaceae *Escherichia coli* UMEA 3160-1 |
| Eco-2075-1 | Bacteria Proteobacteria Gammaproteobacteria Enterobacteriales Enterobacteriaceae *Escherichia coli* UMEA 3161-1 |

| | |
|---|---|
| Eco-2076-1 | Bacteria Proteobacteria Gammaproteobacteria Enterobacteriales Enterobacteriaceae *Escherichia coli* UMEA 3162-1 |
| Eco-2077-1 | Bacteria Proteobacteria Gammaproteobacteria Enterobacteriales Enterobacteriaceae *Escherichia coli* UMEA 3163-1 |
| Eco-2078-1 | Bacteria Proteobacteria Gammaproteobacteria Enterobacteriales Enterobacteriaceae *Escherichia coli* UMEA 3172-1 |
| Eco-2079-1 | Bacteria Proteobacteria Gammaproteobacteria Enterobacteriales Enterobacteriaceae *Escherichia coli* UMEA 3173-1 |
| Eco-2080-1 | Bacteria Proteobacteria Gammaproteobacteria Enterobacteriales Enterobacteriaceae *Escherichia coli* UMEA 3174-1 |
| Eco-2081-1 | Bacteria Proteobacteria Gammaproteobacteria Enterobacteriales Enterobacteriaceae *Escherichia coli* UMEA 3175-1 |
| Eco-2082-1 | Bacteria Proteobacteria Gammaproteobacteria Enterobacteriales Enterobacteriaceae *Escherichia coli* UMEA 3176-1 |
| Eco-2083-1 | Bacteria Proteobacteria Gammaproteobacteria Enterobacteriales Enterobacteriaceae *Escherichia coli* UMEA 3178-1 |
| Eco-2084-1 | Bacteria Proteobacteria Gammaproteobacteria Enterobacteriales Enterobacteriaceae *Escherichia coli* UMEA 3180-1 |
| Eco-2085-1 | Bacteria Proteobacteria Gammaproteobacteria Enterobacteriales Enterobacteriaceae *Escherichia coli* UMEA 3185-1 |
| Eco-2086-1 | Bacteria Proteobacteria Gammaproteobacteria Enterobacteriales Enterobacteriaceae *Escherichia coli* UMEA 3190-1 |
| Eco-2087-1 | Bacteria Proteobacteria Gammaproteobacteria Enterobacteriales Enterobacteriaceae *Escherichia coli* UMEA 3193-1 |
| Eco-2088-1 | Bacteria Proteobacteria Gammaproteobacteria Enterobacteriales Enterobacteriaceae *Escherichia coli* UMEA 3199-1 |
| Eco-2089-1 | Bacteria Proteobacteria Gammaproteobacteria Enterobacteriales Enterobacteriaceae *Escherichia coli* UMEA 3200-1 |
| Eco-2090-1 | Bacteria Proteobacteria Gammaproteobacteria Enterobacteriales Enterobacteriaceae *Escherichia coli* UMEA 3201-1 |
| Eco-2091-1 | Bacteria Proteobacteria Gammaproteobacteria Enterobacteriales Enterobacteriaceae *Escherichia coli* UMEA 3203-1 |
| Eco-2092-1 | Bacteria Proteobacteria Gammaproteobacteria Enterobacteriales Enterobacteriaceae *Escherichia coli* UMEA 3206-1 |
| Eco-2093-1 | Bacteria Proteobacteria Gammaproteobacteria Enterobacteriales Enterobacteriaceae *Escherichia coli* UMEA 3208-1 |
| Eco-2094-1 | Bacteria Proteobacteria Gammaproteobacteria Enterobacteriales Enterobacteriaceae *Escherichia coli* UMEA 3212-1 |
| Eco-2095-1 | Bacteria Proteobacteria Gammaproteobacteria Enterobacteriales Enterobacteriaceae *Escherichia coli* UMEA 3215-1 |
| Eco-2096-1 | Bacteria Proteobacteria Gammaproteobacteria Enterobacteriales Enterobacteriaceae *Escherichia coli* UMEA 3216-1 |
| Eco-2097-1 | Bacteria Proteobacteria Gammaproteobacteria Enterobacteriales Enterobacteriaceae *Escherichia coli* UMEA 3217-1 |
| Eco-2098-1 | Bacteria Proteobacteria Gammaproteobacteria Enterobacteriales Enterobacteriaceae *Escherichia coli* UMEA 3220-1 |
| Eco-2099-1 | Bacteria Proteobacteria Gammaproteobacteria Enterobacteriales Enterobacteriaceae *Escherichia coli* UMEA 3221-1 |
| Eco-2100-1 | Bacteria Proteobacteria Gammaproteobacteria Enterobacteriales Enterobacteriaceae *Escherichia coli* UMEA 3222-1 |
| Eco-2101-1 | Bacteria Proteobacteria Gammaproteobacteria Enterobacteriales Enterobacteriaceae *Escherichia coli* UMEA 3230-1 |
| Eco-2102-1 | Bacteria Proteobacteria Gammaproteobacteria Enterobacteriales Enterobacteriaceae *Escherichia coli* UMEA 3233-1 |
| Eco-2103-1 | Bacteria Proteobacteria Gammaproteobacteria Enterobacteriales Enterobacteriaceae *Escherichia coli* UMEA 3240-1 |
| Eco-2104-1 | Bacteria Proteobacteria Gammaproteobacteria Enterobacteriales Enterobacteriaceae *Escherichia coli* UMEA 3244-1 |
| Eco-2105-1 | Bacteria Proteobacteria Gammaproteobacteria Enterobacteriales Enterobacteriaceae *Escherichia coli* UMEA 3257-1 |
| Eco-2106-1 | Bacteria Proteobacteria Gammaproteobacteria Enterobacteriales Enterobacteriaceae *Escherichia coli* UMEA 3264-1 |
| Eco-2107-1 | Bacteria Proteobacteria Gammaproteobacteria Enterobacteriales Enterobacteriaceae *Escherichia coli* UMEA 3268-1 |
| Eco-2108-1 | Bacteria Proteobacteria Gammaproteobacteria Enterobacteriales Enterobacteriaceae *Escherichia coli* UMEA 3271-1 |
| Eco-2109-1 | Bacteria Proteobacteria Gammaproteobacteria Enterobacteriales Enterobacteriaceae *Escherichia coli* UMEA 3290-1 |
| Eco-2110-1 | Bacteria Proteobacteria Gammaproteobacteria Enterobacteriales Enterobacteriaceae *Escherichia coli* UMEA 3292-1 |
| Eco-2111-1 | Bacteria Proteobacteria Gammaproteobacteria Enterobacteriales Enterobacteriaceae *Escherichia coli* UMEA 3298-1 |
| Eco-2112-1 | Bacteria Proteobacteria Gammaproteobacteria Enterobacteriales Enterobacteriaceae *Escherichia coli* UMEA 3304-1 |
| Eco-2113-1 | Bacteria Proteobacteria Gammaproteobacteria Enterobacteriales Enterobacteriaceae *Escherichia coli* UMEA 3314-1 |
| Eco-2114-1 | Bacteria Proteobacteria Gammaproteobacteria Enterobacteriales Enterobacteriaceae *Escherichia coli* UMEA 3317-1 |
| Eco-2115-1 | Bacteria Proteobacteria Gammaproteobacteria Enterobacteriales Enterobacteriaceae *Escherichia coli* UMEA 3318-1 |
| Eco-2116-1 | Bacteria Proteobacteria Gammaproteobacteria Enterobacteriales Enterobacteriaceae *Escherichia coli* UMEA 3323-1 |
| Eco-2117-1 | Bacteria Proteobacteria Gammaproteobacteria Enterobacteriales Enterobacteriaceae *Escherichia coli* UMEA 3329-1 |
| Eco-2118-1 | Bacteria Proteobacteria Gammaproteobacteria Enterobacteriales Enterobacteriaceae *Escherichia coli* UMEA 3336-1 |
| Eco-2119-1 | Bacteria Proteobacteria Gammaproteobacteria Enterobacteriales Enterobacteriaceae *Escherichia coli* UMEA 3337-1 |
| Eco-2120-1 | Bacteria Proteobacteria Gammaproteobacteria Enterobacteriales Enterobacteriaceae *Escherichia coli* UMEA 3341-1 |
| Eco-2121-1 | Bacteria Proteobacteria Gammaproteobacteria Enterobacteriales Enterobacteriaceae *Escherichia coli* UMEA 3342-1 |
| Eco-2122-1 | Bacteria Proteobacteria Gammaproteobacteria Enterobacteriales Enterobacteriaceae *Escherichia coli* UMEA 3355-1 |
| Eco-2123-1 | Bacteria Proteobacteria Gammaproteobacteria Enterobacteriales Enterobacteriaceae *Escherichia coli* UMEA 3391-1 |
| Eco-2124-1 | Bacteria Proteobacteria Gammaproteobacteria Enterobacteriales Enterobacteriaceae *Escherichia coli* UMEA 3426-1 |
| Eco-2125-1 | Bacteria Proteobacteria Gammaproteobacteria Enterobacteriales Enterobacteriaceae *Escherichia coli* UMEA 3489-1 |
| Eco-2126-1 | Bacteria Proteobacteria Gammaproteobacteria Enterobacteriales Enterobacteriaceae *Escherichia coli* UMEA 3490-1 |
| Eco-2127-1 | Bacteria Proteobacteria Gammaproteobacteria Enterobacteriales Enterobacteriaceae *Escherichia coli* UMEA 3585-1 |
| Eco-2128-1 | Bacteria Proteobacteria Gammaproteobacteria Enterobacteriales Enterobacteriaceae *Escherichia coli* UMEA 3592-1 |
| Eco-2129-1 | Bacteria Proteobacteria Gammaproteobacteria Enterobacteriales Enterobacteriaceae *Escherichia coli* UMEA 3609-1 |
| Eco-2130-1 | Bacteria Proteobacteria Gammaproteobacteria Enterobacteriales Enterobacteriaceae *Escherichia coli* UMEA 3617-1 |
| Eco-2131-1 | Bacteria Proteobacteria Gammaproteobacteria Enterobacteriales Enterobacteriaceae *Escherichia coli* UMEA 3632-1 |
| Eco-2132-1 | Bacteria Proteobacteria Gammaproteobacteria Enterobacteriales Enterobacteriaceae *Escherichia coli* UMEA 3652-1 |
| Eco-2133-1 | Bacteria Proteobacteria Gammaproteobacteria Enterobacteriales Enterobacteriaceae *Escherichia coli* UMEA 3656-1 |
| Eco-2134-1 | Bacteria Proteobacteria Gammaproteobacteria Enterobacteriales Enterobacteriaceae *Escherichia coli* UMEA 3662-1 |
| Eco-2135-1 | Bacteria Proteobacteria Gammaproteobacteria Enterobacteriales Enterobacteriaceae *Escherichia coli* UMEA 3671-1 |
| Eco-2136-1 | Bacteria Proteobacteria Gammaproteobacteria Enterobacteriales Enterobacteriaceae *Escherichia coli* UMEA 3682-1 |
| Eco-2137-1 | Bacteria Proteobacteria Gammaproteobacteria Enterobacteriales Enterobacteriaceae *Escherichia coli* UMEA 3687-1 |
| Eco-2138-1 | Bacteria Proteobacteria Gammaproteobacteria Enterobacteriales Enterobacteriaceae *Escherichia coli* UMEA 3693-1 |
| Eco-2139-1 | Bacteria Proteobacteria Gammaproteobacteria Enterobacteriales Enterobacteriaceae *Escherichia coli* UMEA 3694-1 |
| Eco-2140-1 | Bacteria Proteobacteria Gammaproteobacteria Enterobacteriales Enterobacteriaceae *Escherichia coli* UMEA 3702-1 |

| | |
|---|---|
| Eco-2141-1 | Bacteria Proteobacteria Gammaproteobacteria Enterobacteriales Enterobacteriaceae *Escherichia coli* UMEA 3703-1 |
| Eco-2142-1 | Bacteria Proteobacteria Gammaproteobacteria Enterobacteriales Enterobacteriaceae *Escherichia coli* UMEA 3705-1 |
| Eco-2143-1 | Bacteria Proteobacteria Gammaproteobacteria Enterobacteriales Enterobacteriaceae *Escherichia coli* UMEA 3707-1 |
| Eco-2144-1 | Bacteria Proteobacteria Gammaproteobacteria Enterobacteriales Enterobacteriaceae *Escherichia coli* UMEA 3718-1 |
| Eco-2145-1 | Bacteria Proteobacteria Gammaproteobacteria Enterobacteriales Enterobacteriaceae *Escherichia coli* UMEA 3805-1 |
| Eco-2146-1 | Bacteria Proteobacteria Gammaproteobacteria Enterobacteriales Enterobacteriaceae *Escherichia coli* UMEA 3821-1 |
| Eco-2147-1 | Bacteria Proteobacteria Gammaproteobacteria Enterobacteriales Enterobacteriaceae *Escherichia coli* UMEA 3834-1 |
| Eco-2148-1 | Bacteria Proteobacteria Gammaproteobacteria Enterobacteriales Enterobacteriaceae *Escherichia coli* UMEA 3889-1 |
| Eco-2149-1 | Bacteria Proteobacteria Gammaproteobacteria Enterobacteriales Enterobacteriaceae *Escherichia coli* UMEA 3893-1 |
| Eco-2150-1 | Bacteria Proteobacteria Gammaproteobacteria Enterobacteriales Enterobacteriaceae *Escherichia coli* UMEA 3899-1 |
| Eco-2151-1 | Bacteria Proteobacteria Gammaproteobacteria Enterobacteriales Enterobacteriaceae *Escherichia coli* UMEA 3955-1 |
| Eco-2152-1 | Bacteria Proteobacteria Gammaproteobacteria Enterobacteriales Enterobacteriaceae *Escherichia coli* UMEA 4075-1 |
| Eco-2153-1 | Bacteria Proteobacteria Gammaproteobacteria Enterobacteriales Enterobacteriaceae *Escherichia coli* UMEA 4076-1 |
| Eco-2154-1 | Bacteria Proteobacteria Gammaproteobacteria Enterobacteriales Enterobacteriaceae *Escherichia coli* UMEA 4207-1 |
| Eco-2155-1 | Bacteria Proteobacteria Gammaproteobacteria Enterobacteriales Enterobacteriaceae *Escherichia coli* UMN026 |
| Eco-2156-1 | Bacteria Proteobacteria Gammaproteobacteria Enterobacteriales Enterobacteriaceae *Escherichia coli* UMNF18 |
| Eco-2157-1 | Bacteria Proteobacteria Gammaproteobacteria Enterobacteriales Enterobacteriaceae *Escherichia coli* UMNK88 |
| Eco-2158-1 | Bacteria Proteobacteria Gammaproteobacteria Enterobacteriales Enterobacteriaceae *Escherichia coli* UTI89 |
| Eco-2159-1 to Eco-2159-2 | Bacteria Proteobacteria Gammaproteobacteria Enterobacteriales Enterobacteriaceae *Escherichia coli* W |
| Eco-2160-1 | Bacteria Proteobacteria Gammaproteobacteria Enterobacteriales Enterobacteriaceae *Escherichia coli* W26 |
| Eco-2161-1 | Bacteria Proteobacteria Gammaproteobacteria Enterobacteriales Enterobacteriaceae *Escherichia coli* WV_060327 |
| Eco-2162-1 | Bacteria Proteobacteria Gammaproteobacteria Enterobacteriales Enterobacteriaceae *Escherichia coli* XH001 |
| Eco-2163-1 | Bacteria Proteobacteria Gammaproteobacteria Enterobacteriales Enterobacteriaceae *Escherichia coli* XH140A |
| Eco-2164-1 | Bacteria Proteobacteria Gammaproteobacteria Enterobacteriales Enterobacteriaceae *Escherichia coli* Xuzhou21 |
| Efe-1-1 to Efe-1-3 | Bacteria Proteobacteria Gammaproteobacteria Enterobacteriales Enterobacteriaceae *Escherichia fergusonii* |
| Efe-2-1 | Bacteria Proteobacteria Gammaproteobacteria Enterobacteriales Enterobacteriaceae *Escherichia fergusonii* ATCC 35469 |
| Efe-3-1 | Bacteria Proteobacteria Gammaproteobacteria Enterobacteriales Enterobacteriaceae *Escherichia fergusonii* B253 |
| Efe-4-1 | Bacteria Proteobacteria Gammaproteobacteria Enterobacteriales Enterobacteriaceae *Escherichia fergusonii* ECD227 |
| Ehe-1-1 | Bacteria Proteobacteria Gammaproteobacteria Enterobacteriales Enterobacteriaceae *Escherichia hermannii* NBRC 105704 |
| Esp-75-1 | Bacteria Proteobacteria Gammaproteobacteria Enterobacteriales Enterobacteriaceae *Escherichia sp.* 1_1_43 |
| Esp-76-1 | Bacteria Proteobacteria Gammaproteobacteria Enterobacteriales Enterobacteriaceae *Escherichia sp.* 4_1_40B |
| Esp-77-1 | Bacteria Proteobacteria Gammaproteobacteria Enterobacteriales Enterobacteriaceae *Escherichia sp.* TW09231 |
| Esp-78-1 | Bacteria Proteobacteria Gammaproteobacteria Enterobacteriales Enterobacteriaceae *Escherichia sp.* TW09276 |
| Esp-79-1 | Bacteria Proteobacteria Gammaproteobacteria Enterobacteriales Enterobacteriaceae *Escherichia sp.* TW09308 |
| Esp-80-1 | Bacteria Proteobacteria Gammaproteobacteria Enterobacteriales Enterobacteriaceae *Escherichia sp.* TW10509 |
| Hal-4-1 | Bacteria Proteobacteria Gammaproteobacteria Enterobacteriales Enterobacteriaceae *Hafnia alvei* ATCC 51873 |
| Kox-1-1 to Kox-1-4 | Bacteria Proteobacteria Gammaproteobacteria Enterobacteriales Enterobacteriaceae *Klebsiella oxytoca* |
| Kox-11-1 | Bacteria Proteobacteria Gammaproteobacteria Enterobacteriales Enterobacteriaceae *Klebsiella oxytoca* 10-5242 |
| Kox-12-1 | Bacteria Proteobacteria Gammaproteobacteria Enterobacteriales Enterobacteriaceae *Klebsiella oxytoca* 10-5243 |
| Kox-13-1 | Bacteria Proteobacteria Gammaproteobacteria Enterobacteriales Enterobacteriaceae *Klebsiella oxytoca* 10-5245 |
| Kox-14-1 | Bacteria Proteobacteria Gammaproteobacteria Enterobacteriales Enterobacteriaceae *Klebsiella oxytoca* 10-5246 |
| Kox-4-1 to Kox-4-2 | Bacteria Proteobacteria Gammaproteobacteria Enterobacteriales Enterobacteriaceae *Klebsiella oxytoca* 10-5248 |
| Kox-15-1 | Bacteria Proteobacteria Gammaproteobacteria Enterobacteriales Enterobacteriaceae *Klebsiella oxytoca* 10-5250 |
| Kox-16-1 | Bacteria Proteobacteria Gammaproteobacteria Enterobacteriales Enterobacteriaceae *Klebsiella oxytoca* 11492-1 |
| Kox-17-1 | Bacteria Proteobacteria Gammaproteobacteria Enterobacteriales Enterobacteriaceae *Klebsiella oxytoca* E718 |
| Kox-18-1 | Bacteria Proteobacteria Gammaproteobacteria Enterobacteriales Enterobacteriaceae *Klebsiella oxytoca* KCTC 1686 |
| Kox-19-1 | Bacteria Proteobacteria Gammaproteobacteria Enterobacteriales Enterobacteriaceae *Klebsiella oxytoca* MGH 28 |
| Kox-20-1 | Bacteria Proteobacteria Gammaproteobacteria Enterobacteriales Enterobacteriaceae *Klebsiella oxytoca* MGH 42 |
| Kpn-1-1 to Kpn-1-9 | Bacteria Proteobacteria Gammaproteobacteria Enterobacteriales Enterobacteriaceae *Klebsiella pneumoniae* |
| Kpn-226-1 | Bacteria Proteobacteria Gammaproteobacteria Enterobacteriales Enterobacteriaceae *Klebsiella pneumoniae* 1162281 |
| Kpn-227-1 | Bacteria Proteobacteria Gammaproteobacteria Enterobacteriales Enterobacteriaceae *Klebsiella pneumoniae* 1191100241 |
| Kpn-228-1 | Bacteria Proteobacteria Gammaproteobacteria Enterobacteriales Enterobacteriaceae *Klebsiella pneumoniae* 120_1020 |
| Kpn-229-1 | Bacteria Proteobacteria Gammaproteobacteria Enterobacteriales Enterobacteriaceae *Klebsiella pneumoniae* 140_1040 |
| Kpn-3-1 to Kpn-3-2 | Bacteria Proteobacteria Gammaproteobacteria Enterobacteriales Enterobacteriaceae *Klebsiella pneumoniae* 160_1080 |
| Kpn-230-1 | Bacteria Proteobacteria Gammaproteobacteria Enterobacteriales Enterobacteriaceae *Klebsiella pneumoniae* 280_1220 |
| Kpn-231-1 | Bacteria Proteobacteria Gammaproteobacteria Enterobacteriales Enterobacteriaceae *Klebsiella pneumoniae* 303K |
| Kpn-232-1 | Bacteria Proteobacteria Gammaproteobacteria Enterobacteriales Enterobacteriaceae *Klebsiella pneumoniae* 342 |
| Kpn-233-1 | Bacteria Proteobacteria Gammaproteobacteria Enterobacteriales Enterobacteriaceae *Klebsiella pneumoniae* 361_1301 |
| Kpn-234-1 | Bacteria Proteobacteria Gammaproteobacteria Enterobacteriales Enterobacteriaceae *Klebsiella pneumoniae* 440_1540 |
| Kpn-235-1 | Bacteria Proteobacteria Gammaproteobacteria Enterobacteriales Enterobacteriaceae *Klebsiella pneumoniae* 540_1460 |
| Kpn-236-1 | Bacteria Proteobacteria Gammaproteobacteria Enterobacteriales Enterobacteriaceae *Klebsiella pneumoniae* 646_1568 |
| Kpn-237-1 | Bacteria Proteobacteria Gammaproteobacteria Enterobacteriales Enterobacteriaceae *Klebsiella pneumoniae* 700603 |
| Kpn-238-1 | Bacteria Proteobacteria Gammaproteobacteria Enterobacteriales Enterobacteriaceae *Klebsiella pneumoniae* ATCC BAA-1705 |
| Kpn-239-1 | Bacteria Proteobacteria Gammaproteobacteria Enterobacteriales Enterobacteriaceae *Klebsiella pneumoniae* ATCC BAA-2146 |
| Kpn-240-1 | Bacteria Proteobacteria Gammaproteobacteria Enterobacteriales Enterobacteriaceae *Klebsiella pneumoniae* BIDMC 12C |

| | |
|---|---|
| Kpn-241-1 | Bacteria Proteobacteria Gammaproteobacteria Enterobacteriales Enterobacteriaceae *Klebsiella pneumoniae* BIDMC 16 |
| Kpn-242-1 | Bacteria Proteobacteria Gammaproteobacteria Enterobacteriales Enterobacteriaceae *Klebsiella pneumoniae* BIDMC 18C |
| Kpn-243-1 | Bacteria Proteobacteria Gammaproteobacteria Enterobacteriales Enterobacteriaceae *Klebsiella pneumoniae* BIDMC 21 |
| Kpn-244-1 | Bacteria Proteobacteria Gammaproteobacteria Enterobacteriales Enterobacteriaceae *Klebsiella pneumoniae* BIDMC 22 |
| Kpn-245-1 | Bacteria Proteobacteria Gammaproteobacteria Enterobacteriales Enterobacteriaceae *Klebsiella pneumoniae* BIDMC 23 |
| Kpn-246-1 | Bacteria Proteobacteria Gammaproteobacteria Enterobacteriales Enterobacteriaceae *Klebsiella pneumoniae* BIDMC 24 |
| Kpn-247-1 | Bacteria Proteobacteria Gammaproteobacteria Enterobacteriales Enterobacteriaceae *Klebsiella pneumoniae* BIDMC 25 |
| Kpn-248-1 | Bacteria Proteobacteria Gammaproteobacteria Enterobacteriales Enterobacteriaceae *Klebsiella pneumoniae* BIDMC 35 |
| Kpn-249-1 | Bacteria Proteobacteria Gammaproteobacteria Enterobacteriales Enterobacteriaceae *Klebsiella pneumoniae* BIDMC 36 |
| Kpn-250-1 | Bacteria Proteobacteria Gammaproteobacteria Enterobacteriales Enterobacteriaceae *Klebsiella pneumoniae* BIDMC 40 |
| Kpn-251-1 | Bacteria Proteobacteria Gammaproteobacteria Enterobacteriales Enterobacteriaceae *Klebsiella pneumoniae* BIDMC 41 |
| Kpn-252-1 | Bacteria Proteobacteria Gammaproteobacteria Enterobacteriales Enterobacteriaceae *Klebsiella pneumoniae* BWH 28 |
| Kpn-253-1 | Bacteria Proteobacteria Gammaproteobacteria Enterobacteriales Enterobacteriaceae *Klebsiella pneumoniae* BWH 30 |
| Kpn-254-1 | Bacteria Proteobacteria Gammaproteobacteria Enterobacteriales Enterobacteriaceae *Klebsiella pneumoniae* CG43 |
| Kpn-255-1 | Bacteria Proteobacteria Gammaproteobacteria Enterobacteriales Enterobacteriaceae *Klebsiella pneumoniae* DMC0526 |
| Kpn-256-1 | Bacteria Proteobacteria Gammaproteobacteria Enterobacteriales Enterobacteriaceae *Klebsiella pneumoniae* DMC0799 |
| Kpn-257-1 | Bacteria Proteobacteria Gammaproteobacteria Enterobacteriales Enterobacteriaceae *Klebsiella pneumoniae* DMC1316 |
| Kpn-324-1 | Bacteria Proteobacteria Gammaproteobacteria Enterobacteriales Enterobacteriaceae *Klebsiella pneumoniae hv*KP1 |
| Kpn-258-1 | Bacteria Proteobacteria Gammaproteobacteria Enterobacteriales Enterobacteriaceae *Klebsiella pneumoniae* JH1 |
| Kpn-259-1 | Bacteria Proteobacteria Gammaproteobacteria Enterobacteriales Enterobacteriaceae *Klebsiella pneumoniae* JHCK1 |
| Kpn-260-1 | Bacteria Proteobacteria Gammaproteobacteria Enterobacteriales Enterobacteriaceae *Klebsiella pneumoniae* JM45 |
| Kpn-261-1 | Bacteria Proteobacteria Gammaproteobacteria Enterobacteriales Enterobacteriaceae *Klebsiella pneumoniae* KCTC 2242 |
| Kpn-262-1 | Bacteria Proteobacteria Gammaproteobacteria Enterobacteriales Enterobacteriaceae *Klebsiella pneumoniae* KP-1 |
| Kpn-263-1 | Bacteria Proteobacteria Gammaproteobacteria Enterobacteriales Enterobacteriaceae *Klebsiella pneumoniae* KP-11 |
| Kpn-264-1 | Bacteria Proteobacteria Gammaproteobacteria Enterobacteriales Enterobacteriaceae *Klebsiella pneumoniae* KP-7 |
| Kpn-265-1 | Bacteria Proteobacteria Gammaproteobacteria Enterobacteriales Enterobacteriaceae *Klebsiella pneumoniae* MGH 17 |
| Kpn-266-1 | Bacteria Proteobacteria Gammaproteobacteria Enterobacteriales Enterobacteriaceae *Klebsiella pneumoniae* MGH 18 |
| Kpn-267-1 | Bacteria Proteobacteria Gammaproteobacteria Enterobacteriales Enterobacteriaceae *Klebsiella pneumoniae* MGH 19 |
| Kpn-268-1 | Bacteria Proteobacteria Gammaproteobacteria Enterobacteriales Enterobacteriaceae *Klebsiella pneumoniae* MGH 20 |
| Kpn-269-1 | Bacteria Proteobacteria Gammaproteobacteria Enterobacteriales Enterobacteriaceae *Klebsiella pneumoniae* MGH 21 |
| Kpn-270-1 | Bacteria Proteobacteria Gammaproteobacteria Enterobacteriales Enterobacteriaceae *Klebsiella pneumoniae* MGH 30 |
| Kpn-271-1 | Bacteria Proteobacteria Gammaproteobacteria Enterobacteriales Enterobacteriaceae *Klebsiella pneumoniae* MGH 31 |
| Kpn-272-1 | Bacteria Proteobacteria Gammaproteobacteria Enterobacteriales Enterobacteriaceae *Klebsiella pneumoniae* MGH 32 |
| Kpn-273-1 | Bacteria Proteobacteria Gammaproteobacteria Enterobacteriales Enterobacteriaceae *Klebsiella pneumoniae* MGH 36 |
| Kpn-274-1 | Bacteria Proteobacteria Gammaproteobacteria Enterobacteriales Enterobacteriaceae *Klebsiella pneumoniae* MGH 40 |
| Kpn-275-1 | Bacteria Proteobacteria Gammaproteobacteria Enterobacteriales Enterobacteriaceae *Klebsiella pneumoniae* MGH 44 |
| Kpn-276-1 | Bacteria Proteobacteria Gammaproteobacteria Enterobacteriales Enterobacteriaceae *Klebsiella pneumoniae* MGH 46 |
| Kpn-277-1 | Bacteria Proteobacteria Gammaproteobacteria Enterobacteriales Enterobacteriaceae *Klebsiella pneumoniae* MGH 48 |
| Kpn-278-1 | Bacteria Proteobacteria Gammaproteobacteria Enterobacteriales Enterobacteriaceae *Klebsiella pneumoniae* MGH 59 |
| Kpn-279-1 | Bacteria Proteobacteria Gammaproteobacteria Enterobacteriales Enterobacteriaceae *Klebsiella pneumoniae* NTUH-K2044 |
| Kpn-205-1 to Kpn-205-4 | Bacteria Proteobacteria Gammaproteobacteria Enterobacteriales Enterobacteriaceae *Klebsiella pneumoniae subsp. ozaenae* |
| Kpn-325-1 | Bacteria Proteobacteria Gammaproteobacteria Enterobacteriales Enterobacteriaceae *Klebsiella pneumoniae subsp. pneumoniae* 1084 |
| Kpn-326-1 | Bacteria Proteobacteria Gammaproteobacteria Enterobacteriales Enterobacteriaceae *Klebsiella pneumoniae subsp. pneumoniae* 12-3578 |
| Kpn-327-1 | Bacteria Proteobacteria Gammaproteobacteria Enterobacteriales Enterobacteriaceae *Klebsiella pneumoniae subsp. pneumoniae* B5055 |
| Kpn-328-1 | Bacteria Proteobacteria Gammaproteobacteria Enterobacteriales Enterobacteriaceae *Klebsiella pneumoniae subsp. pneumoniae* DSM 30104 |
| Kpn-329-1 | Bacteria Proteobacteria Gammaproteobacteria Enterobacteriales Enterobacteriaceae *Klebsiella pneumoniae subsp. pneumoniae* Ecl8 |
| Kpn-330-1 | Bacteria Proteobacteria Gammaproteobacteria Enterobacteriales Enterobacteriaceae *Klebsiella pneumoniae subsp. pneumoniae* HS11286 |
| Kpn-331-1 | Bacteria Proteobacteria Gammaproteobacteria Enterobacteriales Enterobacteriaceae *Klebsiella pneumoniae subsp. pneumoniae* KPNIH11 |
| Kpn-332-1 | Bacteria Proteobacteria Gammaproteobacteria Enterobacteriales Enterobacteriaceae *Klebsiella pneumoniae subsp. pneumoniae* KPNIH12 |
| Kpn-333-1 | Bacteria Proteobacteria Gammaproteobacteria Enterobacteriales Enterobacteriaceae *Klebsiella pneumoniae subsp. pneumoniae* KPNIH14 |
| Kpn-334-1 | Bacteria Proteobacteria Gammaproteobacteria Enterobacteriales Enterobacteriaceae *Klebsiella pneumoniae subsp. pneumoniae* KPNIH16 |
| Kpn-335-1 | Bacteria Proteobacteria Gammaproteobacteria Enterobacteriales Enterobacteriaceae *Klebsiella pneumoniae subsp. pneumoniae* KPNIH17 |
| Kpn-336-1 | Bacteria Proteobacteria Gammaproteobacteria Enterobacteriales Enterobacteriaceae *Klebsiella pneumoniae subsp. pneumoniae* KPNIH18 |
| Kpn-337-1 | Bacteria Proteobacteria Gammaproteobacteria Enterobacteriales Enterobacteriaceae *Klebsiella pneumoniae subsp. pneumoniae* KPNIH19 |
| Kpn-338-1 | Bacteria Proteobacteria Gammaproteobacteria Enterobacteriales Enterobacteriaceae *Klebsiella pneumoniae subsp. pneumoniae* KPNIH2 |
| Kpn-339-1 | Bacteria Proteobacteria Gammaproteobacteria Enterobacteriales Enterobacteriaceae *Klebsiella pneumoniae subsp. pneumoniae* KPNIH20 |
| Kpn-340-1 | Bacteria Proteobacteria Gammaproteobacteria Enterobacteriales Enterobacteriaceae *Klebsiella pneumoniae subsp. pneumoniae* KPNIH21 |
| Kpn-341-1 | Bacteria Proteobacteria Gammaproteobacteria Enterobacteriales Enterobacteriaceae *Klebsiella pneumoniae subsp. pneumoniae* KPNIH22 |
| Kpn-342-1 | Bacteria Proteobacteria Gammaproteobacteria Enterobacteriales Enterobacteriaceae *Klebsiella pneumoniae subsp. pneumoniae* KPNIH23 |
| Kpn-343-1 | Bacteria Proteobacteria Gammaproteobacteria Enterobacteriales Enterobacteriaceae *Klebsiella pneumoniae subsp. pneumoniae* KPNIH4 |
| Kpn-344-1 | Bacteria Proteobacteria Gammaproteobacteria Enterobacteriales Enterobacteriaceae *Klebsiella pneumoniae subsp. pneumoniae* KPNIH5 |
| Kpn-345-1 | Bacteria Proteobacteria Gammaproteobacteria Enterobacteriales Enterobacteriaceae *Klebsiella pneumoniae subsp. pneumoniae* KPNIH6 |
| Kpn-346-1 | Bacteria Proteobacteria Gammaproteobacteria Enterobacteriales Enterobacteriaceae *Klebsiella pneumoniae subsp. pneumoniae* KPNIH7 |
| Kpn-347-1 | Bacteria Proteobacteria Gammaproteobacteria Enterobacteriales Enterobacteriaceae *Klebsiella pneumoniae subsp. pneumoniae* KPNIH8 |
| Kpn-348-1 | Bacteria Proteobacteria Gammaproteobacteria Enterobacteriales Enterobacteriaceae *Klebsiella pneumoniae subsp. pneumoniae* KPNIH9 |

| | |
|---|---|
| Kpn-349-1 | Bacteria Proteobacteria Gammaproteobacteria Enterobacteriales Enterobacteriaceae *Klebsiella pneumoniae subsp. pneumoniae* KpQ3 |
| Kpn-350-1 | Bacteria Proteobacteria Gammaproteobacteria Enterobacteriales Enterobacteriaceae *Klebsiella pneumoniae subsp. pneumoniae* LZ |
| Kpn-351-1 | Bacteria Proteobacteria Gammaproteobacteria Enterobacteriales Enterobacteriaceae *Klebsiella pneumoniae subsp. pneumoniae* MGH 78578 |
| Kpn-352-1 | Bacteria Proteobacteria Gammaproteobacteria Enterobacteriales Enterobacteriaceae *Klebsiella pneumoniae subsp. pneumoniae* MP14 |
| Kpn-353-1 | Bacteria Proteobacteria Gammaproteobacteria Enterobacteriales Enterobacteriaceae *Klebsiella pneumoniae subsp. pneumoniae* ST258-490 |
| Kpn-354-1 | Bacteria Proteobacteria Gammaproteobacteria Enterobacteriales Enterobacteriaceae *Klebsiella pneumoniae subsp. pneumoniae* ST258-K26BO |
| Kpn-355-1 | Bacteria Proteobacteria Gammaproteobacteria Enterobacteriales Enterobacteriaceae *Klebsiella pneumoniae subsp. pneumoniae* ST258-K28BO |
| Kpn-356-1 | Bacteria Proteobacteria Gammaproteobacteria Enterobacteriales Enterobacteriaceae *Klebsiella pneumoniae subsp. pneumoniae* ST512-K30BO |
| Kpn-357-1 | Bacteria Proteobacteria Gammaproteobacteria Enterobacteriales Enterobacteriaceae *Klebsiella pneumoniae subsp. pneumoniae* UKKV901664 |
| Kpn-358-1 | Bacteria Proteobacteria Gammaproteobacteria Enterobacteriales Enterobacteriaceae *Klebsiella pneumoniae subsp. pneumoniae* WGLW1 |
| Kpn-359-1 | Bacteria Proteobacteria Gammaproteobacteria Enterobacteriales Enterobacteriaceae *Klebsiella pneumoniae subsp. pneumoniae* WGLW2 |
| Kpn-360-1 | Bacteria Proteobacteria Gammaproteobacteria Enterobacteriales Enterobacteriaceae *Klebsiella pneumoniae subsp. pneumoniae* WGLW3 |
| Kpn-361-1 | Bacteria Proteobacteria Gammaproteobacteria Enterobacteriales Enterobacteriaceae *Klebsiella pneumoniae subsp. pneumoniae* WGLW5 |
| Kpn-362-1 | Bacteria Proteobacteria Gammaproteobacteria Enterobacteriales Enterobacteriaceae *Klebsiella pneumoniae subsp. rhinoscleromatis* ATCC 13884 |
| Kpn-280-1 | Bacteria Proteobacteria Gammaproteobacteria Enterobacteriales Enterobacteriaceae *Klebsiella pneumoniae* UCICRE 10 |
| Kpn-281-1 | Bacteria Proteobacteria Gammaproteobacteria Enterobacteriales Enterobacteriaceae *Klebsiella pneumoniae* UCICRE 14 |
| Kpn-282-1 | Bacteria Proteobacteria Gammaproteobacteria Enterobacteriales Enterobacteriaceae *Klebsiella pneumoniae* UCICRE 2 |
| Kpn-283-1 | Bacteria Proteobacteria Gammaproteobacteria Enterobacteriales Enterobacteriaceae *Klebsiella pneumoniae* UCICRE 4 |
| Kpn-284-1 | Bacteria Proteobacteria Gammaproteobacteria Enterobacteriales Enterobacteriaceae *Klebsiella pneumoniae* UCICRE 6 |
| Kpn-285-1 | Bacteria Proteobacteria Gammaproteobacteria Enterobacteriales Enterobacteriaceae *Klebsiella pneumoniae* UCICRE 7 |
| Kpn-286-1 | Bacteria Proteobacteria Gammaproteobacteria Enterobacteriales Enterobacteriaceae *Klebsiella pneumoniae* UCICRE 8 |
| Kpn-199-1 to Kpn-199-2 | Bacteria Proteobacteria Gammaproteobacteria Enterobacteriales Enterobacteriaceae *Klebsiella pneumoniae* UHKPC 52 |
| Kpn-287-1 | Bacteria Proteobacteria Gammaproteobacteria Enterobacteriales Enterobacteriaceae *Klebsiella pneumoniae* UHKPC01 |
| Kpn-288-1 | Bacteria Proteobacteria Gammaproteobacteria Enterobacteriales Enterobacteriaceae *Klebsiella pneumoniae* UHKPC02 |
| Kpn-289-1 | Bacteria Proteobacteria Gammaproteobacteria Enterobacteriales Enterobacteriaceae *Klebsiella pneumoniae* UHKPC04 |
| Kpn-200-1 to Kpn-200-2 | Bacteria Proteobacteria Gammaproteobacteria Enterobacteriales Enterobacteriaceae *Klebsiella pneumoniae* UHKPC05 |
| Kpn-290-1 | Bacteria Proteobacteria Gammaproteobacteria Enterobacteriales Enterobacteriaceae *Klebsiella pneumoniae* UHKPC06 |
| Kpn-291-1 | Bacteria Proteobacteria Gammaproteobacteria Enterobacteriales Enterobacteriaceae *Klebsiella pneumoniae* UHKPC09 |
| Kpn-292-1 | Bacteria Proteobacteria Gammaproteobacteria Enterobacteriales Enterobacteriaceae *Klebsiella pneumoniae* UHKPC17 |
| Kpn-293-1 | Bacteria Proteobacteria Gammaproteobacteria Enterobacteriales Enterobacteriaceae *Klebsiella pneumoniae* UHKPC179 |
| Kpn-294-1 | Bacteria Proteobacteria Gammaproteobacteria Enterobacteriales Enterobacteriaceae *Klebsiella pneumoniae* UHKPC18 |
| Kpn-295-1 | Bacteria Proteobacteria Gammaproteobacteria Enterobacteriales Enterobacteriaceae *Klebsiella pneumoniae* UHKPC22 |
| Kpn-296-1 | Bacteria Proteobacteria Gammaproteobacteria Enterobacteriales Enterobacteriaceae *Klebsiella pneumoniae* UHKPC23 |
| Kpn-297-1 | Bacteria Proteobacteria Gammaproteobacteria Enterobacteriales Enterobacteriaceae *Klebsiella pneumoniae* UHKPC24 |
| Kpn-298-1 | Bacteria Proteobacteria Gammaproteobacteria Enterobacteriales Enterobacteriaceae *Klebsiella pneumoniae* UHKPC26 |
| Kpn-299-1 | Bacteria Proteobacteria Gammaproteobacteria Enterobacteriales Enterobacteriaceae *Klebsiella pneumoniae* UHKPC27 |
| Kpn-300-1 | Bacteria Proteobacteria Gammaproteobacteria Enterobacteriales Enterobacteriaceae *Klebsiella pneumoniae* UHKPC28 |
| Kpn-301-1 | Bacteria Proteobacteria Gammaproteobacteria Enterobacteriales Enterobacteriaceae *Klebsiella pneumoniae* UHKPC29 |
| Kpn-302-1 | Bacteria Proteobacteria Gammaproteobacteria Enterobacteriales Enterobacteriaceae *Klebsiella pneumoniae* UHKPC31 |
| Kpn-303-1 | Bacteria Proteobacteria Gammaproteobacteria Enterobacteriales Enterobacteriaceae *Klebsiella pneumoniae* UHKPC32 |
| Kpn-304-1 | Bacteria Proteobacteria Gammaproteobacteria Enterobacteriales Enterobacteriaceae *Klebsiella pneumoniae* UHKPC40 |
| Kpn-203-1 to Kpn-203-2 | Bacteria Proteobacteria Gammaproteobacteria Enterobacteriales Enterobacteriaceae *Klebsiella pneumoniae* UHKPC45 |
| Kpn-305-1 | Bacteria Proteobacteria Gammaproteobacteria Enterobacteriales Enterobacteriaceae *Klebsiella pneumoniae* UHKPC47 |
| Kpn-306-1 | Bacteria Proteobacteria Gammaproteobacteria Enterobacteriales Enterobacteriaceae *Klebsiella pneumoniae* UHKPC48 |
| Kpn-307-1 | Bacteria Proteobacteria Gammaproteobacteria Enterobacteriales Enterobacteriaceae *Klebsiella pneumoniae* UHKPC57 |
| Kpn-308-1 | Bacteria Proteobacteria Gammaproteobacteria Enterobacteriales Enterobacteriaceae *Klebsiella pneumoniae* UHKPC59 |
| Kpn-309-1 | Bacteria Proteobacteria Gammaproteobacteria Enterobacteriales Enterobacteriaceae *Klebsiella pneumoniae* UHKPC61 |
| Kpn-310-1 | Bacteria Proteobacteria Gammaproteobacteria Enterobacteriales Enterobacteriaceae *Klebsiella pneumoniae* UHKPC67 |
| Kpn-311-1 | Bacteria Proteobacteria Gammaproteobacteria Enterobacteriales Enterobacteriaceae *Klebsiella pneumoniae* UHKPC69 |
| Kpn-312-1 | Bacteria Proteobacteria Gammaproteobacteria Enterobacteriales Enterobacteriaceae *Klebsiella pneumoniae* UHKPC77 |
| Kpn-313-1 | Bacteria Proteobacteria Gammaproteobacteria Enterobacteriales Enterobacteriaceae *Klebsiella pneumoniae* UHKPC81 |
| Kpn-314-1 | Bacteria Proteobacteria Gammaproteobacteria Enterobacteriales Enterobacteriaceae *Klebsiella pneumoniae* UHKPC96 |
| Kpn-315-1 | Bacteria Proteobacteria Gammaproteobacteria Enterobacteriales Enterobacteriaceae *Klebsiella pneumoniae* VA360 |
| Kpn-316-1 | Bacteria Proteobacteria Gammaproteobacteria Enterobacteriales Enterobacteriaceae *Klebsiella pneumoniae* VAKPC252 |
| Kpn-317-1 | Bacteria Proteobacteria Gammaproteobacteria Enterobacteriales Enterobacteriaceae *Klebsiella pneumoniae* VAKPC254 |
| Kpn-318-1 | Bacteria Proteobacteria Gammaproteobacteria Enterobacteriales Enterobacteriaceae *Klebsiella pneumoniae* VAKPC269 |
| Kpn-319-1 | Bacteria Proteobacteria Gammaproteobacteria Enterobacteriales Enterobacteriaceae *Klebsiella pneumoniae* VAKPC270 |
| Kpn-320-1 | Bacteria Proteobacteria Gammaproteobacteria Enterobacteriales Enterobacteriaceae *Klebsiella pneumoniae* VAKPC276 |
| Kpn-204-1 to Kpn-204-2 | Bacteria Proteobacteria Gammaproteobacteria Enterobacteriales Enterobacteriaceae *Klebsiella pneumoniae* VAKPC278 |
| Kpn-321-1 | Bacteria Proteobacteria Gammaproteobacteria Enterobacteriales Enterobacteriaceae *Klebsiella pneumoniae* VAKPC280 |
| Kpn-322-1 | Bacteria Proteobacteria Gammaproteobacteria Enterobacteriales Enterobacteriaceae *Klebsiella pneumoniae* VAKPC297 |
| Kpn-323-1 | Bacteria Proteobacteria Gammaproteobacteria Enterobacteriales Enterobacteriaceae *Klebsiella pneumoniae* VAKPC309 |
| Ksp-4-1 | Bacteria Proteobacteria Gammaproteobacteria Enterobacteriales Enterobacteriaceae *Klebsiella sp.* 1_1_55 |
| Ksp-5-1 | Bacteria Proteobacteria Gammaproteobacteria Enterobacteriales Enterobacteriaceae *Klebsiella sp.* 4_1_44FAA |
| Ksp-6-1 | Bacteria Proteobacteria Gammaproteobacteria Enterobacteriales Enterobacteriaceae *Klebsiella sp.* KTE92 |

| | |
|---|---|
| Ksp-7-1 | Bacteria Proteobacteria Gammaproteobacteria Enterobacteriales Enterobacteriaceae *Klebsiella sp.* MS 92-3 |
| Ksp-8-1 | Bacteria Proteobacteria Gammaproteobacteria Enterobacteriales Enterobacteriaceae *Klebsiella sp.* OBRC7 |
| Kva-1-1 to Kva-1-7 | Bacteria Proteobacteria Gammaproteobacteria Enterobacteriales Enterobacteriaceae *Klebsiella variicola* |
| Kva-2-1 | Bacteria Proteobacteria Gammaproteobacteria Enterobacteriales Enterobacteriaceae *Klebsiella variicola* At-22 |
| Era-1-1 | Bacteria Proteobacteria Gammaproteobacteria Enterobacteriales Enterobacteriaceae Kosakonia *Enterobacter radicincitans* DSM 16656 |
| Ksa-1-1 | Bacteria Proteobacteria Gammaproteobacteria Enterobacteriales Enterobacteriaceae *Kosakonia sacchari* SP1 |
| Mmo-1-1 to Mmo-1-8 | Bacteria Proteobacteria Gammaproteobacteria Enterobacteriales Enterobacteriaceae *Morganella morganii* |
| Mmo-5-1 | Bacteria Proteobacteria Gammaproteobacteria Enterobacteriales Enterobacteriaceae *Morganella morganii* SC01 |
| Mmo-6-1 | Bacteria Proteobacteria Gammaproteobacteria Enterobacteriales Enterobacteriaceae *Morganella morganii subsp. morganii* KT |
| Pag-3-1 | Bacteria Proteobacteria Gammaproteobacteria Enterobacteriales Enterobacteriaceae *Pantoea agglomerans* 299R |
| Pag-1-1 to Pag-1-6 | Bacteria Proteobacteria Gammaproteobacteria Enterobacteriales Enterobacteriaceae Pantoea Pantoea agglomerans group *Pantoea agglomerans* |
| Pag-4-1 | Bacteria Proteobacteria Gammaproteobacteria Enterobacteriales Enterobacteriaceae *Pantoea agglomerans* IG1 |
| Pag-5-1 | Bacteria Proteobacteria Gammaproteobacteria Enterobacteriales Enterobacteriaceae *Pantoea agglomerans* Tx10 |
| Pan-1-1 to Pan-1-5 | Bacteria Proteobacteria Gammaproteobacteria Enterobacteriales Enterobacteriaceae *Pantoea ananatis* |
| Pan-6-1 | Bacteria Proteobacteria Gammaproteobacteria Enterobacteriales Enterobacteriaceae *Pantoea ananatis* AJ13355 |
| Pan-7-1 | Bacteria Proteobacteria Gammaproteobacteria Enterobacteriales Enterobacteriaceae *Pantoea ananatis* BRT175 |
| Pan-8-1 | Bacteria Proteobacteria Gammaproteobacteria Enterobacteriales Enterobacteriaceae *Pantoea ananatis* LMG 20103 |
| Pan-9-1 | Bacteria Proteobacteria Gammaproteobacteria Enterobacteriales Enterobacteriaceae *Pantoea ananatis* LMG 5342 |
| Pan-10-1 | Bacteria Proteobacteria Gammaproteobacteria Enterobacteriales Enterobacteriaceae *Pantoea ananatis* PA13 |
| Pdi-1-1 | Bacteria Proteobacteria Gammaproteobacteria Enterobacteriales Enterobacteriaceae *Pantoea dispersa* EGD-AAK13 |
| Psp-12-1 | Bacteria Proteobacteria Gammaproteobacteria Enterobacteriales Enterobacteriaceae *Pantoea sp.* A4 |
| Psp-19-1 | Bacteria Proteobacteria Gammaproteobacteria Enterobacteriales Enterobacteriaceae *Pantoea sp.* aB |
| Psp-13-1 | Bacteria Proteobacteria Gammaproteobacteria Enterobacteriales Enterobacteriaceae *Pantoea sp.* AS-PWVM4 |
| Psp-14-1 | Bacteria Proteobacteria Gammaproteobacteria Enterobacteriales Enterobacteriaceae *Pantoea sp.* At-9b |
| Psp-15-1 | Bacteria Proteobacteria Gammaproteobacteria Enterobacteriales Enterobacteriaceae *Pantoea sp.* GM01 |
| Psp-17-1 | Bacteria Proteobacteria Gammaproteobacteria Enterobacteriales Enterobacteriaceae *Pantoea sp.* Sc1 |
| Psp-16-1 | Bacteria Proteobacteria Gammaproteobacteria Enterobacteriales Enterobacteriaceae *Pantoea sp.* SL1_M5 |
| Psp-18-1 | Bacteria Proteobacteria Gammaproteobacteria Enterobacteriales Enterobacteriaceae *Pantoea sp.* YR343 |
| Pst-3-1 | Bacteria Proteobacteria Gammaproteobacteria Enterobacteriales Enterobacteriaceae *Pantoea stewartii subsp. indologenes* |
| Pst-4-1 | Bacteria Proteobacteria Gammaproteobacteria Enterobacteriales Enterobacteriaceae *Pantoea stewartii subsp. stewartii* DC283 |
| Pva-2-1 | Bacteria Proteobacteria Gammaproteobacteria Enterobacteriales Enterobacteriaceae *Pantoea vagans* C9-1 |
| Pat-4-1 | Bacteria Proteobacteria Gammaproteobacteria Enterobacteriales Enterobacteriaceae *Pectobacterium atrosepticum* SCRI1043 |
| Pca-9-1 | Bacteria Proteobacteria Gammaproteobacteria Enterobacteriales Enterobacteriaceae *Pectobacterium carotovorum subsp. brasiliensis* PBR1692 |
| Pca-10-1 | Bacteria Proteobacteria Gammaproteobacteria Enterobacteriales Enterobacteriaceae *Pectobacterium carotovorum subsp. carotovorum* PC1 |
| Pca-11-1 | Bacteria Proteobacteria Gammaproteobacteria Enterobacteriales Enterobacteriaceae *Pectobacterium carotovorum subsp. carotovorum* PCC21 |
| Pca-12-1 | Bacteria Proteobacteria Gammaproteobacteria Enterobacteriales Enterobacteriaceae *Pectobacterium carotovorum subsp. carotovorum* WPP14 |
| Psp-20-1 | Bacteria Proteobacteria Gammaproteobacteria Enterobacteriales Enterobacteriaceae *Pectobacterium sp.* SCC3193 |
| Pwa-2-1 | Bacteria Proteobacteria Gammaproteobacteria Enterobacteriales Enterobacteriaceae *Pectobacterium wasabiae* CFBP 3304 |
| Pwa-3-1 | Bacteria Proteobacteria Gammaproteobacteria Enterobacteriales Enterobacteriaceae *Pectobacterium wasabiae* WPP163 |
| Pas-2-1 | Bacteria Proteobacteria Gammaproteobacteria Enterobacteriales Enterobacteriaceae *Photorhabdus asymbiotica* |
| Plu-6-1 | Bacteria Proteobacteria Gammaproteobacteria Enterobacteriales Enterobacteriaceae *Photorhabdus luminescens subsp. laumondii* TTO1 |
| Pte-4-1 | Bacteria Proteobacteria Gammaproteobacteria Enterobacteriales Enterobacteriaceae *Photorhabdus temperata* J3 |
| Pte-5-1 | Bacteria Proteobacteria Gammaproteobacteria Enterobacteriales Enterobacteriaceae *Photorhabdus temperata subsp. temperata* M1021 |
| Pst-5-1 | Bacteria Proteobacteria Gammaproteobacteria Enterobacteriales Enterobacteriaceae *Plautia stali symbiont* |
| Pha-1-1 | Bacteria Proteobacteria Gammaproteobacteria Enterobacteriales Enterobacteriaceae *Proteus hauseri* ZMd44 |
| Pmi-1-1 to Pmi-1-6 | Bacteria Proteobacteria Gammaproteobacteria Enterobacteriales Enterobacteriaceae *Proteus mirabilis* |
| Pmi-2-1 | Bacteria Proteobacteria Gammaproteobacteria Enterobacteriales Enterobacteriaceae *Proteus mirabilis* ATCC 29906 |
| Pmi-3-1 | Bacteria Proteobacteria Gammaproteobacteria Enterobacteriales Enterobacteriaceae *Proteus mirabilis* BB2000 |
| Pmi-4-1 | Bacteria Proteobacteria Gammaproteobacteria Enterobacteriales Enterobacteriaceae *Proteus mirabilis* C05028 |
| Pmi-5-1 | Bacteria Proteobacteria Gammaproteobacteria Enterobacteriales Enterobacteriaceae *Proteus mirabilis* HI4320 |
| Pmi-6-1 | Bacteria Proteobacteria Gammaproteobacteria Enterobacteriales Enterobacteriaceae *Proteus mirabilis* PR03 |
| Pmi-7-1 | Bacteria Proteobacteria Gammaproteobacteria Enterobacteriales Enterobacteriaceae *Proteus mirabilis* WGLW4 |
| Pmi-8-1 | Bacteria Proteobacteria Gammaproteobacteria Enterobacteriales Enterobacteriaceae *Proteus mirabilis* WGLW6 |
| Pal-9-1 | Bacteria Proteobacteria Gammaproteobacteria Enterobacteriales Enterobacteriaceae *Providencia alcalifaciens* Dmel2 |
| Pal-8-1 | Bacteria Proteobacteria Gammaproteobacteria Enterobacteriales Enterobacteriaceae *Providencia alcalifaciens* DSM 30120 |
| Pbu-1-1 | Bacteria Proteobacteria Gammaproteobacteria Enterobacteriales Enterobacteriaceae *Providencia burhodogranariea* DSM 19968 |
| Pre-2-1 | Bacteria Proteobacteria Gammaproteobacteria Enterobacteriales Enterobacteriaceae *Providencia rettgeri* Dmel1 |
| Pru-1-1 | Bacteria Proteobacteria Gammaproteobacteria Enterobacteriales Enterobacteriaceae *Providencia rustigianii* DSM 4541 |
| Pst-6-1 | Bacteria Proteobacteria Gammaproteobacteria Enterobacteriales Enterobacteriaceae *Providencia stuartii* ATCC 25827 |
| Pst-7-1 | Bacteria Proteobacteria Gammaproteobacteria Enterobacteriales Enterobacteriaceae *Providencia stuartii* MRSN 2154 |
| Ror-4-1 to Ror-4-2 | Bacteria Proteobacteria Gammaproteobacteria Enterobacteriales Enterobacteriaceae *Raoultella ornithinolytica* B6 |
| Rpl-1-1 to Rpl-1-3 | Bacteria Proteobacteria Gammaproteobacteria Enterobacteriales Enterobacteriaceae *Raoultella planticola* |
| Sbo-3-1 | Bacteria Proteobacteria Gammaproteobacteria Enterobacteriales Enterobacteriaceae *Salmonella bongori* N268-08 |
| Sbo-4-1 | Bacteria Proteobacteria Gammaproteobacteria Enterobacteriales Enterobacteriaceae *Salmonella bongori* NCTC 12419 |
| Sen-2-1 to Sen-2-2 | Bacteria Proteobacteria Gammaproteobacteria Enterobacteriales Enterobacteriaceae *Salmonella enterica* |

| | |
|---|---|
| Sen-310-1 | Bacteria Proteobacteria Gammaproteobacteria Enterobacteriales Enterobacteriaceae *Salmonella enterica subsp. arizonae serovar* 62:z4,z23:– |
| Sen-311-1 | Bacteria Proteobacteria Gammaproteobacteria Enterobacteriales Enterobacteriaceae *Salmonella enterica subsp. diarizonae serovar* 60:r:e,n,x,z15 str. 01-0170 |
| Sen-312-1 | Bacteria Proteobacteria Gammaproteobacteria Enterobacteriales Enterobacteriaceae *Salmonella enterica subsp. enterica serovar* 4,[5],12:i:- str. 08-1700 |
| Sen-313-1 | Bacteria Proteobacteria Gammaproteobacteria Enterobacteriales Enterobacteriaceae *Salmonella enterica subsp. enterica serovar* 4,[5],12:i:- str. 08-1736 |
| Sen-314-1 | Bacteria Proteobacteria Gammaproteobacteria Enterobacteriales Enterobacteriaceae *Salmonella enterica subsp. enterica serovar* 4,[5],12:i:- str. 08-1739 |
| Sen-315-1 | Bacteria Proteobacteria Gammaproteobacteria Enterobacteriales Enterobacteriaceae *Salmonella enterica subsp. enterica serovar* Agona str. 01.O.05 |
| Sen-316-1 | Bacteria Proteobacteria Gammaproteobacteria Enterobacteriales Enterobacteriaceae *Salmonella enterica subsp. enterica serovar* Agona str. 02.O.05 |
| Sen-317-1 | Bacteria Proteobacteria Gammaproteobacteria Enterobacteriales Enterobacteriaceae *Salmonella enterica subsp. enterica serovar* Agona str. 0292 |
| Sen-318-1 | Bacteria Proteobacteria Gammaproteobacteria Enterobacteriales Enterobacteriaceae *Salmonella enterica subsp. enterica serovar* Agona str. 03.O.05 |
| Sen-319-1 | Bacteria Proteobacteria Gammaproteobacteria Enterobacteriales Enterobacteriaceae *Salmonella enterica subsp. enterica serovar* Agona str. 0322 |
| Sen-320-1 | Bacteria Proteobacteria Gammaproteobacteria Enterobacteriales Enterobacteriaceae *Salmonella enterica subsp. enterica serovar* Agona str. 04.O.05 |
| Sen-321-1 | Bacteria Proteobacteria Gammaproteobacteria Enterobacteriales Enterobacteriaceae *Salmonella enterica subsp. enterica serovar* Agona str. 05.O.06 |
| Sen-322-1 | Bacteria Proteobacteria Gammaproteobacteria Enterobacteriales Enterobacteriaceae *Salmonella enterica subsp. enterica serovar* Agona str. 06.O.05 |
| Sen-323-1 | Bacteria Proteobacteria Gammaproteobacteria Enterobacteriales Enterobacteriaceae *Salmonella enterica subsp. enterica serovar* Agona str. 07.O.05 |
| Sen-324-1 | Bacteria Proteobacteria Gammaproteobacteria Enterobacteriales Enterobacteriaceae *Salmonella enterica subsp. enterica serovar* Agona str. 08.A.05 |
| Sen-325-1 | Bacteria Proteobacteria Gammaproteobacteria Enterobacteriales Enterobacteriaceae *Salmonella enterica subsp. enterica serovar* Agona str. 09.F.08 |
| Sen-326-1 | Bacteria Proteobacteria Gammaproteobacteria Enterobacteriales Enterobacteriaceae *Salmonella enterica subsp. enterica serovar* Agona str. 10.A.05 |
| Sen-327-1 | Bacteria Proteobacteria Gammaproteobacteria Enterobacteriales Enterobacteriaceae *Salmonella enterica subsp. enterica serovar* Agona str. 11.A.05 |
| Sen-328-1 | Bacteria Proteobacteria Gammaproteobacteria Enterobacteriales Enterobacteriaceae *Salmonella enterica subsp. enterica serovar* Agona str. 12.A.06 |
| Sen-329-1 | Bacteria Proteobacteria Gammaproteobacteria Enterobacteriales Enterobacteriaceae *Salmonella enterica subsp. enterica serovar* Agona str. 13.E.05 |
| Sen-330-1 | Bacteria Proteobacteria Gammaproteobacteria Enterobacteriales Enterobacteriaceae *Salmonella enterica subsp. enterica serovar* Agona str. 14.E.05 |
| Sen-331-1 | Bacteria Proteobacteria Gammaproteobacteria Enterobacteriales Enterobacteriaceae *Salmonella enterica subsp. enterica serovar* Agona str. 15.H.03 |
| Sen-332-1 | Bacteria Proteobacteria Gammaproteobacteria Enterobacteriales Enterobacteriaceae *Salmonella enterica subsp. enterica serovar* Agona str. 16.H.08 |
| Sen-333-1 | Bacteria Proteobacteria Gammaproteobacteria Enterobacteriales Enterobacteriaceae *Salmonella enterica subsp. enterica serovar* Agona str. 17.H.06 |
| Sen-334-1 | Bacteria Proteobacteria Gammaproteobacteria Enterobacteriales Enterobacteriaceae *Salmonella enterica subsp. enterica serovar* Agona str. 18.H.07 |
| Sen-335-1 | Bacteria Proteobacteria Gammaproteobacteria Enterobacteriales Enterobacteriaceae *Salmonella enterica subsp. enterica serovar* Agona str. 19.F.03 |
| Sen-336-1 | Bacteria Proteobacteria Gammaproteobacteria Enterobacteriales Enterobacteriaceae *Salmonella enterica subsp. enterica serovar* Agona str. 20.H.06 |
| Sen-337-1 | Bacteria Proteobacteria Gammaproteobacteria Enterobacteriales Enterobacteriaceae *Salmonella enterica subsp. enterica serovar* Agona str. 21.H.10 |
| Sen-338-1 | Bacteria Proteobacteria Gammaproteobacteria Enterobacteriales Enterobacteriaceae *Salmonella enterica subsp. enterica serovar* Agona str. 22.H.04 |
| Sen-339-1 | Bacteria Proteobacteria Gammaproteobacteria Enterobacteriales Enterobacteriaceae *Salmonella enterica subsp. enterica serovar* Agona str. 23.F.01 |
| Sen-340-1 | Bacteria Proteobacteria Gammaproteobacteria Enterobacteriales Enterobacteriaceae *Salmonella enterica subsp. enterica serovar* Agona str. 24.H.04 |
| Sen-341-1 | Bacteria Proteobacteria Gammaproteobacteria Enterobacteriales Enterobacteriaceae *Salmonella enterica subsp. enterica serovar* Agona str. 241981 |
| Sen-342-1 | Bacteria Proteobacteria Gammaproteobacteria Enterobacteriales Enterobacteriaceae *Salmonella enterica subsp. enterica serovar* Agona str. 24249 |
| Sen-343-1 | Bacteria Proteobacteria Gammaproteobacteria Enterobacteriales Enterobacteriaceae *Salmonella enterica subsp. enterica serovar* Agona str. 246555-3 |
| Sen-344-1 | Bacteria Proteobacteria Gammaproteobacteria Enterobacteriales Enterobacteriaceae *Salmonella enterica subsp. enterica serovar* Agona str. 26.F.98 |
| Sen-345-1 | Bacteria Proteobacteria Gammaproteobacteria Enterobacteriales Enterobacteriaceae *Salmonella enterica subsp. enterica serovar* Agona str. 266757-1 |
| Sen-346-1 | Bacteria Proteobacteria Gammaproteobacteria Enterobacteriales Enterobacteriaceae *Salmonella enterica subsp. enterica serovar* Agona str. 27.O.98 |
| Sen-347-1 | Bacteria Proteobacteria Gammaproteobacteria Enterobacteriales Enterobacteriaceae *Salmonella enterica subsp. enterica serovar* Agona str. 28.O.08 |
| Sen-348-1 | Bacteria Proteobacteria Gammaproteobacteria Enterobacteriales Enterobacteriaceae *Salmonella enterica subsp. enterica serovar* Agona str. 29.O.08 |
| Sen-349-1 | Bacteria Proteobacteria Gammaproteobacteria Enterobacteriales Enterobacteriaceae *Salmonella enterica subsp. enterica serovar* Agona str. 30.H.04 |
| Sen-350-1 | Bacteria Proteobacteria Gammaproteobacteria Enterobacteriales Enterobacteriaceae *Salmonella enterica subsp. enterica serovar* Agona str. 31.H.09 |
| Sen-351-1 | Bacteria Proteobacteria Gammaproteobacteria Enterobacteriales Enterobacteriaceae *Salmonella enterica subsp. enterica serovar* Agona str. 311387-1 |
| Sen-352-1 | Bacteria Proteobacteria Gammaproteobacteria Enterobacteriales Enterobacteriaceae *Salmonella enterica subsp. enterica serovar* Agona str. 32.A.00 |
| Sen-353-1 | Bacteria Proteobacteria Gammaproteobacteria Enterobacteriales Enterobacteriaceae *Salmonella enterica subsp. enterica serovar* Agona str. 33.A.05 |
| Sen-354-1 | Bacteria Proteobacteria Gammaproteobacteria Enterobacteriales Enterobacteriaceae *Salmonella enterica subsp. enterica serovar* Agona str. 339787 |
| Sen-355-1 | Bacteria Proteobacteria Gammaproteobacteria Enterobacteriales Enterobacteriaceae *Salmonella enterica subsp. enterica serovar* Agona str. 34.H.09 |
| Sen-356-1 | Bacteria Proteobacteria Gammaproteobacteria Enterobacteriales Enterobacteriaceae *Salmonella enterica subsp. enterica serovar* Agona str. 35.H.08 |
| Sen-357-1 | Bacteria Proteobacteria Gammaproteobacteria Enterobacteriales Enterobacteriaceae *Salmonella enterica subsp. enterica serovar* Agona str. 36.H.00 |
| Sen-358-1 | Bacteria Proteobacteria Gammaproteobacteria Enterobacteriales Enterobacteriaceae *Salmonella enterica subsp. enterica serovar* Agona str. 37.F.02 |
| Sen-359-1 | Bacteria Proteobacteria Gammaproteobacteria Enterobacteriales Enterobacteriaceae *Salmonella enterica subsp. enterica serovar* Agona str. 38.O.03 |
| Sen-360-1 | Bacteria Proteobacteria Gammaproteobacteria Enterobacteriales Enterobacteriaceae *Salmonella enterica subsp. enterica serovar* Agona str. 39.O.03 |
| Sen-361-1 | Bacteria Proteobacteria Gammaproteobacteria Enterobacteriales Enterobacteriaceae *Salmonella enterica subsp. enterica serovar* Agona str. 392869-1 |
| Sen-362-1 | Bacteria Proteobacteria Gammaproteobacteria Enterobacteriales Enterobacteriaceae *Salmonella enterica subsp. enterica serovar* Agona str. 392869-2 |
| Sen-363-1 | Bacteria Proteobacteria Gammaproteobacteria Enterobacteriales Enterobacteriaceae *Salmonella enterica subsp. enterica serovar* Agona str. 40.E.08 |
| Sen-364-1 | Bacteria Proteobacteria Gammaproteobacteria Enterobacteriales Enterobacteriaceae *Salmonella enterica subsp. enterica serovar* Agona str. 400095 11 |
| Sen-365-1 | Bacteria Proteobacteria Gammaproteobacteria Enterobacteriales Enterobacteriaceae *Salmonella enterica subsp. enterica serovar* Agona str. 400095 13 |
| Sen-366-1 | Bacteria Proteobacteria Gammaproteobacteria Enterobacteriales Enterobacteriaceae *Salmonella enterica subsp. enterica serovar* Agona str. 400095 14 |
| Sen-367-1 | Bacteria Proteobacteria Gammaproteobacteria Enterobacteriales Enterobacteriaceae *Salmonella enterica subsp. enterica serovar* Agona str. 400095 17 |
| Sen-368-1 | Bacteria Proteobacteria Gammaproteobacteria Enterobacteriales Enterobacteriaceae *Salmonella enterica subsp. enterica serovar* Agona str. 400095 18 |
| Sen-369-1 | Bacteria Proteobacteria Gammaproteobacteria Enterobacteriales Enterobacteriaceae *Salmonella enterica subsp. enterica serovar* Agona str. 409753-6 |
| Sen-370-1 | Bacteria Proteobacteria Gammaproteobacteria Enterobacteriales Enterobacteriaceae *Salmonella enterica subsp. enterica serovar* Agona str. 41.E.09 |
| Sen-371-1 | Bacteria Proteobacteria Gammaproteobacteria Enterobacteriales Enterobacteriaceae *Salmonella enterica subsp. enterica serovar* Agona str. 419639 2-1 |
| Sen-372-1 | Bacteria Proteobacteria Gammaproteobacteria Enterobacteriales Enterobacteriaceae *Salmonella enterica subsp. enterica serovar* Agona str. 42.E.09 |
| Sen-373-1 | Bacteria Proteobacteria Gammaproteobacteria Enterobacteriales Enterobacteriaceae *Salmonella enterica subsp. enterica serovar* Agona str. 43.E.09 |

| | |
|---|---|
| Sen-374-1 | Bacteria Proteobacteria Gammaproteobacteria Enterobacteriales Enterobacteriaceae *Salmonella enterica subsp. enterica serovar* Agona str. 432613 |
| Sen-375-1 | Bacteria Proteobacteria Gammaproteobacteria Enterobacteriales Enterobacteriaceae *Salmonella enterica subsp. enterica serovar* Agona str. 44.E.09 |
| Sen-376-1 | Bacteria Proteobacteria Gammaproteobacteria Enterobacteriales Enterobacteriaceae *Salmonella enterica subsp. enterica serovar* Agona str. 442692 2-4 |
| Sen-377-1 | Bacteria Proteobacteria Gammaproteobacteria Enterobacteriales Enterobacteriaceae *Salmonella enterica subsp. enterica serovar* Agona str. 442692 2-5 |
| Sen-378-1 | Bacteria Proteobacteria Gammaproteobacteria Enterobacteriales Enterobacteriaceae *Salmonella enterica subsp. enterica serovar* Agona str. 447967 1-1 |
| Sen-379-1 | Bacteria Proteobacteria Gammaproteobacteria Enterobacteriales Enterobacteriaceae *Salmonella enterica subsp. enterica serovar* Agona str. 447967 2-1 |
| Sen-380-1 | Bacteria Proteobacteria Gammaproteobacteria Enterobacteriales Enterobacteriaceae *Salmonella enterica subsp. enterica serovar* Agona str. 45.E.09 |
| Sen-381-1 | Bacteria Proteobacteria Gammaproteobacteria Enterobacteriales Enterobacteriaceae *Salmonella enterica subsp. enterica serovar* Agona str. 46.E.09 |
| Sen-382-1 | Bacteria Proteobacteria Gammaproteobacteria Enterobacteriales Enterobacteriaceae *Salmonella enterica subsp. enterica serovar* Agona str. 460004 1-1 |
| Sen-383-1 | Bacteria Proteobacteria Gammaproteobacteria Enterobacteriales Enterobacteriaceae *Salmonella enterica subsp. enterica serovar* Agona str. 467481 |
| Sen-384-1 | Bacteria Proteobacteria Gammaproteobacteria Enterobacteriales Enterobacteriaceae *Salmonella enterica subsp. enterica serovar* Agona str. 47.E.09 |
| Sen-385-1 | Bacteria Proteobacteria Gammaproteobacteria Enterobacteriales Enterobacteriaceae *Salmonella enterica subsp. enterica serovar* Agona str. 48.E.08 |
| Sen-386-1 | Bacteria Proteobacteria Gammaproteobacteria Enterobacteriales Enterobacteriaceae *Salmonella enterica subsp. enterica serovar* Agona str. 49.E.09 |
| Sen-387-1 | Bacteria Proteobacteria Gammaproteobacteria Enterobacteriales Enterobacteriaceae *Salmonella enterica subsp. enterica serovar* Agona str. 50.E.08 |
| Sen-388-1 | Bacteria Proteobacteria Gammaproteobacteria Enterobacteriales Enterobacteriaceae *Salmonella enterica subsp. enterica serovar* Agona str. 51.E.09 |
| Sen-389-1 | Bacteria Proteobacteria Gammaproteobacteria Enterobacteriales Enterobacteriaceae *Salmonella enterica subsp. enterica serovar* Agona str. 52.F.08 |
| Sen-390-1 | Bacteria Proteobacteria Gammaproteobacteria Enterobacteriales Enterobacteriaceae *Salmonella enterica subsp. enterica serovar* Agona str. 53.F.08 |
| Sen-391-1 | Bacteria Proteobacteria Gammaproteobacteria Enterobacteriales Enterobacteriaceae *Salmonella enterica subsp. enterica serovar* Agona str. 54.O.08 |
| Sen-392-1 | Bacteria Proteobacteria Gammaproteobacteria Enterobacteriales Enterobacteriaceae *Salmonella enterica subsp. enterica serovar* Agona str. 55.U.08 |
| Sen-393-1 | Bacteria Proteobacteria Gammaproteobacteria Enterobacteriales Enterobacteriaceae *Salmonella enterica subsp. enterica serovar* Agona str. 557928 |
| Sen-394-1 | Bacteria Proteobacteria Gammaproteobacteria Enterobacteriales Enterobacteriaceae *Salmonella enterica subsp. enterica serovar* Agona str. 56.O.08 |
| Sen-395-1 | Bacteria Proteobacteria Gammaproteobacteria Enterobacteriales Enterobacteriaceae *Salmonella enterica subsp. enterica serovar* Agona str. 57.A.08 |
| Sen-396-1 | Bacteria Proteobacteria Gammaproteobacteria Enterobacteriales Enterobacteriaceae *Salmonella enterica subsp. enterica serovar* Agona str. 58.E.08 |
| Sen-397-1 | Bacteria Proteobacteria Gammaproteobacteria Enterobacteriales Enterobacteriaceae *Salmonella enterica subsp. enterica serovar* Agona str. 59.F.08 |
| Sen-398-1 | Bacteria Proteobacteria Gammaproteobacteria Enterobacteriales Enterobacteriaceae *Salmonella enterica subsp. enterica serovar* Agona str. 60.O.08 |
| Sen-399-1 | Bacteria Proteobacteria Gammaproteobacteria Enterobacteriales Enterobacteriaceae *Salmonella enterica subsp. enterica serovar* Agona str. 61.O.08 |
| Sen-400-1 | Bacteria Proteobacteria Gammaproteobacteria Enterobacteriales Enterobacteriaceae *Salmonella enterica subsp. enterica serovar* Agona str. 62.H.72 |
| Sen-401-1 | Bacteria Proteobacteria Gammaproteobacteria Enterobacteriales Enterobacteriaceae *Salmonella enterica subsp. enterica serovar* Agona str. 620239 |
| Sen-402-1 | Bacteria Proteobacteria Gammaproteobacteria Enterobacteriales Enterobacteriaceae *Salmonella enterica subsp. enterica serovar* Agona str. 63.H.87 |
| Sen-403-1 | Bacteria Proteobacteria Gammaproteobacteria Enterobacteriales Enterobacteriaceae *Salmonella enterica subsp. enterica serovar* Agona str. 632182-2 |
| Sen-404-1 | Bacteria Proteobacteria Gammaproteobacteria Enterobacteriales Enterobacteriaceae *Salmonella enterica subsp. enterica serovar* Agona str. 64.H.00 |
| Sen-405-1 | Bacteria Proteobacteria Gammaproteobacteria Enterobacteriales Enterobacteriaceae *Salmonella enterica subsp. enterica serovar* Agona str. 648586-1 |
| Sen-406-1 | Bacteria Proteobacteria Gammaproteobacteria Enterobacteriales Enterobacteriaceae *Salmonella enterica subsp. enterica serovar* Agona str. 65.H.72 |
| Sen-407-1 | Bacteria Proteobacteria Gammaproteobacteria Enterobacteriales Enterobacteriaceae *Salmonella enterica subsp. enterica serovar* Agona str. 66.F.99 |
| Sen-408-1 | Bacteria Proteobacteria Gammaproteobacteria Enterobacteriales Enterobacteriaceae *Salmonella enterica subsp. enterica serovar* Agona str. 67.H.09 |
| Sen-409-1 | Bacteria Proteobacteria Gammaproteobacteria Enterobacteriales Enterobacteriaceae *Salmonella enterica subsp. enterica serovar* Agona str. 68.U.05 |
| Sen-410-1 | Bacteria Proteobacteria Gammaproteobacteria Enterobacteriales Enterobacteriaceae *Salmonella enterica subsp. enterica serovar* Agona str. 69.H.06 |
| Sen-411-1 | Bacteria Proteobacteria Gammaproteobacteria Enterobacteriales Enterobacteriaceae *Salmonella enterica subsp. enterica serovar* Agona str. 70.E.05 |
| Sen-412-1 | Bacteria Proteobacteria Gammaproteobacteria Enterobacteriales Enterobacteriaceae *Salmonella enterica subsp. enterica serovar* Agona str. 71.E.05 |
| Sen-413-1 | Bacteria Proteobacteria Gammaproteobacteria Enterobacteriales Enterobacteriaceae *Salmonella enterica subsp. enterica serovar* Agona str. 72.A.52 |
| Sen-414-1 | Bacteria Proteobacteria Gammaproteobacteria Enterobacteriales Enterobacteriaceae *Salmonella enterica subsp. enterica serovar* Agona str. 73.H.09 |
| Sen-415-1 | Bacteria Proteobacteria Gammaproteobacteria Enterobacteriales Enterobacteriaceae *Salmonella enterica subsp. enterica serovar* Agona str. ATCC 51957 |
| Sen-416-1 | Bacteria Proteobacteria Gammaproteobacteria Enterobacteriales Enterobacteriaceae *Salmonella enterica subsp. enterica serovar* Agona str. ATCC BAA-707 |
| Sen-417-1 | Bacteria Proteobacteria Gammaproteobacteria Enterobacteriales Enterobacteriaceae *Salmonella enterica subsp. enterica serovar* Agona str. SA-1 |
| Sen-418-1 | Bacteria Proteobacteria Gammaproteobacteria Enterobacteriales Enterobacteriaceae *Salmonella enterica subsp. enterica serovar* Agona str. SA-2 |
| Sen-419-1 | Bacteria Proteobacteria Gammaproteobacteria Enterobacteriales Enterobacteriaceae *Salmonella enterica subsp. enterica serovar* Agona str. SA-3 |
| Sen-420-1 | Bacteria Proteobacteria Gammaproteobacteria Enterobacteriales Enterobacteriaceae *Salmonella enterica subsp. enterica serovar* Agona str. SA-4 |
| Sen-421-1 | Bacteria Proteobacteria Gammaproteobacteria Enterobacteriales Enterobacteriaceae *Salmonella enterica subsp. enterica serovar* Agona str. SA-5 |
| Sen-422-1 | Bacteria Proteobacteria Gammaproteobacteria Enterobacteriales Enterobacteriaceae *Salmonella enterica subsp. enterica serovar* Agona str. SH08SF124 |
| Sen-423-1 | Bacteria Proteobacteria Gammaproteobacteria Enterobacteriales Enterobacteriaceae *Salmonella enterica subsp. enterica serovar* Agona str. SH10GFN094 |
| Sen-424-1 | Bacteria Proteobacteria Gammaproteobacteria Enterobacteriales Enterobacteriaceae *Salmonella enterica subsp. enterica serovar* Agona str. SH11G1113 |
| Sen-425-1 | Bacteria Proteobacteria Gammaproteobacteria Enterobacteriales Enterobacteriaceae *Salmonella enterica subsp. enterica serovar* Albany str. ATCC 51960 |
| Sen-426-1 | Bacteria Proteobacteria Gammaproteobacteria Enterobacteriales Enterobacteriaceae *Salmonella enterica subsp. enterica serovar* Anatum str. CDC 06-0532 |
| Sen-427-1 | Bacteria Proteobacteria Gammaproteobacteria Enterobacteriales Enterobacteriaceae *Salmonella enterica subsp. enterica serovar* Anatum str. USDA 100 |
| Sen-428-1 | Bacteria Proteobacteria Gammaproteobacteria Enterobacteriales Enterobacteriaceae *Salmonella enterica subsp. enterica serovar* Anatum str. USDA-ARS-USMARC-1175 |
| Sen-429-1 | Bacteria Proteobacteria Gammaproteobacteria Enterobacteriales Enterobacteriaceae *Salmonella enterica subsp. enterica serovar* Anatum str. USDA-ARS-USMARC-1735 |
| Sen-430-1 | Bacteria Proteobacteria Gammaproteobacteria Enterobacteriales Enterobacteriaceae *Salmonella enterica subsp. enterica serovar* Baildon str. R6-199 |
| Sen-431-1 | Bacteria Proteobacteria Gammaproteobacteria Enterobacteriales Enterobacteriaceae *Salmonella enterica subsp. enterica serovar* Bareilly str. 06-0784 |
| Sen-432-1 | Bacteria Proteobacteria Gammaproteobacteria Enterobacteriales Enterobacteriaceae *Salmonella enterica subsp. enterica serovar* Bareilly str. 2780 |
| Sen-433-1 | Bacteria Proteobacteria Gammaproteobacteria Enterobacteriales Enterobacteriaceae *Salmonella enterica subsp. enterica serovar* Bareilly str. ATCC 9115 |
| Sen-434-1 | Bacteria Proteobacteria Gammaproteobacteria Enterobacteriales Enterobacteriaceae *Salmonella enterica subsp. enterica serovar* Bareilly str. CFSAN000183 |
| Sen-435-1 | Bacteria Proteobacteria Gammaproteobacteria Enterobacteriales Enterobacteriaceae *Salmonella enterica subsp. enterica serovar* Bareilly str. CFSAN000189 |

| | |
|---|---|
| Sen-436-1 | Bacteria Proteobacteria Gammaproteobacteria Enterobacteriales Enterobacteriaceae *Salmonella enterica subsp. enterica serovar* Bareilly str. CFSAN000197 |
| Sen-437-1 | Bacteria Proteobacteria Gammaproteobacteria Enterobacteriales Enterobacteriaceae *Salmonella enterica subsp. enterica serovar* Bareilly str. CFSAN000200 |
| Sen-112-1 to Sen-112-2 | Bacteria Proteobacteria Gammaproteobacteria Enterobacteriales Enterobacteriaceae *Salmonella enterica subsp. enterica serovar* Berta str. ATCC 8392 |
| Sen-438-1 to Sen-438-7 | Bacteria Proteobacteria Gammaproteobacteria Enterobacteriales Enterobacteriaceae *Salmonella enterica subsp. enterica serovar* Bovismorbificans |
| Sen-439-1 | Bacteria Proteobacteria Gammaproteobacteria Enterobacteriales Enterobacteriaceae *Salmonella enterica subsp. enterica serovar* Bovismorbificans str. 3114 |
| Sen-440-1 | Bacteria Proteobacteria Gammaproteobacteria Enterobacteriales Enterobacteriaceae *Salmonella enterica subsp. enterica serovar* Braenderup str. ATCC BAA-664 |
| Sen-441-1 | Bacteria Proteobacteria Gammaproteobacteria Enterobacteriales Enterobacteriaceae *Salmonella enterica subsp. enterica serovar* Braenderup str. CF-SAN000756 |
| Sen-442-1 | Bacteria Proteobacteria Gammaproteobacteria Enterobacteriales Enterobacteriaceae *Salmonella enterica subsp. enterica serovar* Cerro str. 5569 |
| Sen-443-1 | Bacteria Proteobacteria Gammaproteobacteria Enterobacteriales Enterobacteriaceae *Salmonella enterica subsp. enterica serovar* Cerro str. 818 |
| Sen-444-1 | Bacteria Proteobacteria Gammaproteobacteria Enterobacteriales Enterobacteriaceae *Salmonella enterica subsp. enterica serovar* Cerro str. CFSAN001587 |
| Sen-445-1 | Bacteria Proteobacteria Gammaproteobacteria Enterobacteriales Enterobacteriaceae *Salmonella enterica subsp. enterica serovar* Cerro str. CFSAN001588 |
| Sen-446-1 | Bacteria Proteobacteria Gammaproteobacteria Enterobacteriales Enterobacteriaceae *Salmonella enterica subsp. enterica serovar* Cerro str. CFSAN001589 |
| Sen-447-1 | Bacteria Proteobacteria Gammaproteobacteria Enterobacteriales Enterobacteriaceae *Salmonella enterica subsp. enterica serovar* Cerro str. CFSAN001590 |
| Sen-448-1 | Bacteria Proteobacteria Gammaproteobacteria Enterobacteriales Enterobacteriaceae *Salmonella enterica subsp. enterica serovar* Cerro str. CFSAN001669 |
| Sen-449-1 | Bacteria Proteobacteria Gammaproteobacteria Enterobacteriales Enterobacteriaceae *Salmonella enterica subsp. enterica serovar* Cerro str. CFSAN001670 |
| Sen-450-1 | Bacteria Proteobacteria Gammaproteobacteria Enterobacteriales Enterobacteriaceae *Salmonella enterica subsp. enterica serovar* Cerro str. CFSAN001671 |
| Sen-451-1 | Bacteria Proteobacteria Gammaproteobacteria Enterobacteriales Enterobacteriaceae *Salmonella enterica subsp. enterica serovar* Cerro str. CFSAN001673 |
| Sen-452-1 | Bacteria Proteobacteria Gammaproteobacteria Enterobacteriales Enterobacteriaceae *Salmonella enterica subsp. enterica serovar* Cerro str. CFSAN001674 |
| Sen-453-1 | Bacteria Proteobacteria Gammaproteobacteria Enterobacteriales Enterobacteriaceae *Salmonella enterica subsp. enterica serovar* Cerro str. CFSAN001679 |
| Sen-454-1 | Bacteria Proteobacteria Gammaproteobacteria Enterobacteriales Enterobacteriaceae *Salmonella enterica subsp. enterica serovar* Cerro str. CFSAN001680 |
| Sen-455-1 | Bacteria Proteobacteria Gammaproteobacteria Enterobacteriales Enterobacteriaceae *Salmonella enterica subsp. enterica serovar* Cerro str. CFSAN001681 |
| Sen-456-1 | Bacteria Proteobacteria Gammaproteobacteria Enterobacteriales Enterobacteriaceae *Salmonella enterica subsp. enterica serovar* Cerro str. CFSAN001690 |
| Sen-457-1 | Bacteria Proteobacteria Gammaproteobacteria Enterobacteriales Enterobacteriaceae *Salmonella enterica subsp. enterica serovar* Cerro str. CFSAN001691 |
| Sen-458-1 | Bacteria Proteobacteria Gammaproteobacteria Enterobacteriales Enterobacteriaceae *Salmonella enterica subsp. enterica serovar* Cerro str. CFSAN001692 |
| Sen-459-1 | Bacteria Proteobacteria Gammaproteobacteria Enterobacteriales Enterobacteriaceae *Salmonella enterica subsp. enterica serovar* Cerro str. CFSAN001697 |
| Sen-460-1 | Bacteria Proteobacteria Gammaproteobacteria Enterobacteriales Enterobacteriaceae *Salmonella enterica subsp. enterica serovar* Chester str. ATCC 11997 |
| Sen-135-1 to Sen-135-2 | Bacteria Proteobacteria Gammaproteobacteria Enterobacteriales Enterobacteriaceae *Salmonella enterica subsp. enterica serovar* Cubana str. 76814 |
| Sen-461-1 | Bacteria Proteobacteria Gammaproteobacteria Enterobacteriales Enterobacteriaceae *Salmonella enterica subsp. enterica serovar* Cubana str. CFSAN001083 |
| Sen-462-1 | Bacteria Proteobacteria Gammaproteobacteria Enterobacteriales Enterobacteriaceae *Salmonella enterica subsp. enterica serovar* Cubana str. CFSAN002050 |
| Sen-463-1 | Bacteria Proteobacteria Gammaproteobacteria Enterobacteriales Enterobacteriaceae *Salmonella enterica subsp. enterica serovar* Cubana str. CVM42234 |
| Sen-464-1 | Bacteria Proteobacteria Gammaproteobacteria Enterobacteriales Enterobacteriaceae *Salmonella enterica subsp. enterica serovar* Derby str. 626 |
| Sen-465-1 | Bacteria Proteobacteria Gammaproteobacteria Enterobacteriales Enterobacteriaceae *Salmonella enterica subsp. enterica serovar* Dublin str. DG22 |
| Sen-466-1 | Bacteria Proteobacteria Gammaproteobacteria Enterobacteriales Enterobacteriaceae *Salmonella enterica subsp. enterica serovar* Dublin str. HWS51 |
| Sen-467-1 | Bacteria Proteobacteria Gammaproteobacteria Enterobacteriales Enterobacteriaceae *Salmonella enterica subsp. enterica serovar* Dublin str. SD3246 |
| Sen-468-1 | Bacteria Proteobacteria Gammaproteobacteria Enterobacteriales Enterobacteriaceae *Salmonella enterica subsp. enterica serovar* Dublin str. SL1438 |
| Sen-469-1 | Bacteria Proteobacteria Gammaproteobacteria Enterobacteriales Enterobacteriaceae *Salmonella enterica subsp. enterica serovar* Dublin str. UC16 |
| Sen-470-1 | Bacteria Proteobacteria Gammaproteobacteria Enterobacteriales Enterobacteriaceae *Salmonella enterica subsp. enterica serovar* Eastbourne str. CF-SAN001084 |
| Sen-471-1 | Bacteria Proteobacteria Gammaproteobacteria Enterobacteriales Enterobacteriaceae *Salmonella enterica subsp. enterica serovar* Enteritidis str. 08-0047 |
| Sen-472-1 | Bacteria Proteobacteria Gammaproteobacteria Enterobacteriales Enterobacteriaceae *Salmonella enterica subsp. enterica serovar* Enteritidis str. 08-0128 |
| Sen-473-1 | Bacteria Proteobacteria Gammaproteobacteria Enterobacteriales Enterobacteriaceae *Salmonella enterica subsp. enterica serovar* Enteritidis str. 08-0627 |
| Sen-474-1 | Bacteria Proteobacteria Gammaproteobacteria Enterobacteriales Enterobacteriaceae *Salmonella enterica subsp. enterica serovar* Enteritidis str. 08-1080 |
| Sen-475-1 | Bacteria Proteobacteria Gammaproteobacteria Enterobacteriales Enterobacteriaceae *Salmonella enterica subsp. enterica serovar* Enteritidis str. 10-34587 |
| Sen-476-1 | Bacteria Proteobacteria Gammaproteobacteria Enterobacteriales Enterobacteriaceae *Salmonella enterica subsp. enterica serovar* Enteritidis str. 13-1 |
| Sen-477-1 | Bacteria Proteobacteria Gammaproteobacteria Enterobacteriales Enterobacteriaceae *Salmonella enterica subsp. enterica serovar* Enteritidis str. 13183-1 |
| Sen-478-1 | Bacteria Proteobacteria Gammaproteobacteria Enterobacteriales Enterobacteriaceae *Salmonella enterica subsp. enterica serovar* Enteritidis str. 17927 |
| Sen-479-1 | Bacteria Proteobacteria Gammaproteobacteria Enterobacteriales Enterobacteriaceae *Salmonella enterica subsp. enterica serovar* Enteritidis str. 20037 |
| Sen-480-1 | Bacteria Proteobacteria Gammaproteobacteria Enterobacteriales Enterobacteriaceae *Salmonella enterica subsp. enterica serovar* Enteritidis str. 2009K0958 |
| Sen-481-1 | Bacteria Proteobacteria Gammaproteobacteria Enterobacteriales Enterobacteriaceae *Salmonella enterica subsp. enterica serovar* Enteritidis str. 2009K1651 |
| Sen-482-1 | Bacteria Proteobacteria Gammaproteobacteria Enterobacteriales Enterobacteriaceae *Salmonella enterica subsp. enterica serovar* Enteritidis str. 2009K1726 |
| Sen-483-1 | Bacteria Proteobacteria Gammaproteobacteria Enterobacteriales Enterobacteriaceae *Salmonella enterica subsp. enterica serovar* Enteritidis str. 2010-1237 |
| Sen-484-1 | Bacteria Proteobacteria Gammaproteobacteria Enterobacteriales Enterobacteriaceae *Salmonella enterica subsp. enterica serovar* Enteritidis str. 2010K-0262 |
| Sen-485-1 | Bacteria Proteobacteria Gammaproteobacteria Enterobacteriales Enterobacteriaceae *Salmonella enterica subsp. enterica serovar* Enteritidis str. 2010K-0267 |
| Sen-486-1 | Bacteria Proteobacteria Gammaproteobacteria Enterobacteriales Enterobacteriaceae *Salmonella enterica subsp. enterica serovar* Enteritidis str. 2010K-0271 |
| Sen-487-1 | Bacteria Proteobacteria Gammaproteobacteria Enterobacteriales Enterobacteriaceae *Salmonella enterica subsp. enterica serovar* Enteritidis str. 2010K-0284 |
| Sen-488-1 | Bacteria Proteobacteria Gammaproteobacteria Enterobacteriales Enterobacteriaceae *Salmonella enterica subsp. enterica serovar* Enteritidis str. 2010K-0286 |
| Sen-489-1 | Bacteria Proteobacteria Gammaproteobacteria Enterobacteriales Enterobacteriaceae *Salmonella enterica subsp. enterica serovar* Enteritidis str. 2010K-0669 |
| Sen-490-1 | Bacteria Proteobacteria Gammaproteobacteria Enterobacteriales Enterobacteriaceae *Salmonella enterica subsp. enterica serovar* Enteritidis str. 2010K-2599 |
| Sen-491-1 | Bacteria Proteobacteria Gammaproteobacteria Enterobacteriales Enterobacteriaceae *Salmonella enterica subsp. enterica serovar* Enteritidis str. 22510-1 |
| Sen-492-1 | Bacteria Proteobacteria Gammaproteobacteria Enterobacteriales Enterobacteriaceae *Salmonella enterica subsp. enterica serovar* Enteritidis str. 22704 |
| Sen-493-1 | Bacteria Proteobacteria Gammaproteobacteria Enterobacteriales Enterobacteriaceae *Salmonella enterica subsp. enterica serovar* Enteritidis str. 33944 |
| Sen-494-1 | Bacteria Proteobacteria Gammaproteobacteria Enterobacteriales Enterobacteriaceae *Salmonella enterica subsp. enterica serovar* Enteritidis str. 3402 |

| | |
|---|---|
| Sen-495-1 | Bacteria Proteobacteria Gammaproteobacteria Enterobacteriales Enterobacteriaceae *Salmonella enterica subsp. enterica serovar* Enteritidis str. 436 |
| Sen-496-1 | Bacteria Proteobacteria Gammaproteobacteria Enterobacteriales Enterobacteriaceae *Salmonella enterica subsp. enterica serovar* Enteritidis str. 485549-17 |
| Sen-497-1 | Bacteria Proteobacteria Gammaproteobacteria Enterobacteriales Enterobacteriaceae *Salmonella enterica subsp. enterica serovar* Enteritidis str. 50-3079 |
| Sen-498-1 | Bacteria Proteobacteria Gammaproteobacteria Enterobacteriales Enterobacteriaceae *Salmonella enterica subsp. enterica serovar* Enteritidis str. 50-5646 |
| Sen-499-1 | Bacteria Proteobacteria Gammaproteobacteria Enterobacteriales Enterobacteriaceae *Salmonella enterica subsp. enterica serovar* Enteritidis str. 53-407 |
| Sen-500-1 | Bacteria Proteobacteria Gammaproteobacteria Enterobacteriales Enterobacteriaceae *Salmonella enterica subsp. enterica serovar* Enteritidis str. 543463 22-17 |
| Sen-501-1 | Bacteria Proteobacteria Gammaproteobacteria Enterobacteriales Enterobacteriaceae *Salmonella enterica subsp. enterica serovar* Enteritidis str. 543463 40-18 |
| Sen-502-1 | Bacteria Proteobacteria Gammaproteobacteria Enterobacteriales Enterobacteriaceae *Salmonella enterica subsp. enterica serovar* Enteritidis str. 543463 42-20 |
| Sen-503-1 | Bacteria Proteobacteria Gammaproteobacteria Enterobacteriales Enterobacteriaceae *Salmonella enterica subsp. enterica serovar* Enteritidis str. 561362 1-1 |
| Sen-504-1 | Bacteria Proteobacteria Gammaproteobacteria Enterobacteriales Enterobacteriaceae *Salmonella enterica subsp. enterica serovar* Enteritidis str. 561362 9-7 |
| Sen-505-1 | Bacteria Proteobacteria Gammaproteobacteria Enterobacteriales Enterobacteriaceae *Salmonella enterica subsp. enterica serovar* Enteritidis str. 576709 |
| Sen-506-1 | Bacteria Proteobacteria Gammaproteobacteria Enterobacteriales Enterobacteriaceae *Salmonella enterica subsp. enterica serovar* Enteritidis str. 58-6482 |
| Sen-507-1 | Bacteria Proteobacteria Gammaproteobacteria Enterobacteriales Enterobacteriaceae *Salmonella enterica subsp. enterica serovar* Enteritidis str. 596866-22 |
| Sen-508-1 | Bacteria Proteobacteria Gammaproteobacteria Enterobacteriales Enterobacteriaceae *Salmonella enterica subsp. enterica serovar* Enteritidis str. 596866-70 |
| Sen-509-1 | Bacteria Proteobacteria Gammaproteobacteria Enterobacteriales Enterobacteriaceae *Salmonella enterica subsp. enterica serovar* Enteritidis str. 6.0562-1 |
| Sen-510-1 | Bacteria Proteobacteria Gammaproteobacteria Enterobacteriales Enterobacteriaceae *Salmonella enterica subsp. enterica serovar* Enteritidis str. 607307-2 |
| Sen-511-1 | Bacteria Proteobacteria Gammaproteobacteria Enterobacteriales Enterobacteriaceae *Salmonella enterica subsp. enterica serovar* Enteritidis str. 607307-6 |
| Sen-512-1 | Bacteria Proteobacteria Gammaproteobacteria Enterobacteriales Enterobacteriaceae *Salmonella enterica subsp. enterica serovar* Enteritidis str. 607308-16 |
| Sen-513-1 | Bacteria Proteobacteria Gammaproteobacteria Enterobacteriales Enterobacteriaceae *Salmonella enterica subsp. enterica serovar* Enteritidis str. 607308-19 |
| Sen-514-1 | Bacteria Proteobacteria Gammaproteobacteria Enterobacteriales Enterobacteriaceae *Salmonella enterica subsp. enterica serovar* Enteritidis str. 607308-9 |
| Sen-515-1 | Bacteria Proteobacteria Gammaproteobacteria Enterobacteriales Enterobacteriaceae *Salmonella enterica subsp. enterica serovar* Enteritidis str. 62-1976 |
| Sen-516-1 | Bacteria Proteobacteria Gammaproteobacteria Enterobacteriales Enterobacteriaceae *Salmonella enterica subsp. enterica serovar* Enteritidis str. 622731-39 |
| Sen-517-1 | Bacteria Proteobacteria Gammaproteobacteria Enterobacteriales Enterobacteriaceae *Salmonella enterica subsp. enterica serovar* Enteritidis str. 629163 |
| Sen-518-1 | Bacteria Proteobacteria Gammaproteobacteria Enterobacteriales Enterobacteriaceae *Salmonella enterica subsp. enterica serovar* Enteritidis str. 629164-26 |
| Sen-519-1 | Bacteria Proteobacteria Gammaproteobacteria Enterobacteriales Enterobacteriaceae *Salmonella enterica subsp. enterica serovar* Enteritidis str. 629164-37 |
| Sen-520-1 | Bacteria Proteobacteria Gammaproteobacteria Enterobacteriales Enterobacteriaceae *Salmonella enterica subsp. enterica serovar* Enteritidis str. 635290-58 |
| Sen-521-1 | Bacteria Proteobacteria Gammaproteobacteria Enterobacteriales Enterobacteriaceae *Salmonella enterica subsp. enterica serovar* Enteritidis str. 638970-15 |
| Sen-522-1 | Bacteria Proteobacteria Gammaproteobacteria Enterobacteriales Enterobacteriaceae *Salmonella enterica subsp. enterica serovar* Enteritidis str. 639016-6 |
| Sen-523-1 | Bacteria Proteobacteria Gammaproteobacteria Enterobacteriales Enterobacteriaceae *Salmonella enterica subsp. enterica serovar* Enteritidis str. 639672-46 |
| Sen-524-1 | Bacteria Proteobacteria Gammaproteobacteria Enterobacteriales Enterobacteriaceae *Salmonella enterica subsp. enterica serovar* Enteritidis str. 639672-50 |
| Sen-525-1 | Bacteria Proteobacteria Gammaproteobacteria Enterobacteriales Enterobacteriaceae *Salmonella enterica subsp. enterica serovar* Enteritidis str. 640631 |
| Sen-526-1 | Bacteria Proteobacteria Gammaproteobacteria Enterobacteriales Enterobacteriaceae *Salmonella enterica subsp. enterica serovar* Enteritidis str. 642044 4-1 |
| Sen-527-1 | Bacteria Proteobacteria Gammaproteobacteria Enterobacteriales Enterobacteriaceae *Salmonella enterica subsp. enterica serovar* Enteritidis str. 642044 8-1 |
| Sen-528-1 | Bacteria Proteobacteria Gammaproteobacteria Enterobacteriales Enterobacteriaceae *Salmonella enterica subsp. enterica serovar* Enteritidis str. 642046 4-7 |
| Sen-529-1 | Bacteria Proteobacteria Gammaproteobacteria Enterobacteriales Enterobacteriaceae *Salmonella enterica subsp. enterica serovar* Enteritidis str. 648898 4-5 |
| Sen-530-1 | Bacteria Proteobacteria Gammaproteobacteria Enterobacteriales Enterobacteriaceae *Salmonella enterica subsp. enterica serovar* Enteritidis str. 648899 3-17 |
| Sen-531-1 | Bacteria Proteobacteria Gammaproteobacteria Enterobacteriales Enterobacteriaceae *Salmonella enterica subsp. enterica serovar* Enteritidis str. 648900 1-16 |
| Sen-532-1 | Bacteria Proteobacteria Gammaproteobacteria Enterobacteriales Enterobacteriaceae *Salmonella enterica subsp. enterica serovar* Enteritidis str. 648901 1-17 |
| Sen-533-1 | Bacteria Proteobacteria Gammaproteobacteria Enterobacteriales Enterobacteriaceae *Salmonella enterica subsp. enterica serovar* Enteritidis str. 648901 16-16 |
| Sen-534-1 | Bacteria Proteobacteria Gammaproteobacteria Enterobacteriales Enterobacteriaceae *Salmonella enterica subsp. enterica serovar* Enteritidis str. 648901 39-2 |
| Sen-535-1 | Bacteria Proteobacteria Gammaproteobacteria Enterobacteriales Enterobacteriaceae *Salmonella enterica subsp. enterica serovar* Enteritidis str. 648901 6-18 |
| Sen-536-1 | Bacteria Proteobacteria Gammaproteobacteria Enterobacteriales Enterobacteriaceae *Salmonella enterica subsp. enterica serovar* Enteritidis str. 648902 6-8 |
| Sen-537-1 | Bacteria Proteobacteria Gammaproteobacteria Enterobacteriales Enterobacteriaceae *Salmonella enterica subsp. enterica serovar* Enteritidis str. 648903 1-6 |
| Sen-538-1 | Bacteria Proteobacteria Gammaproteobacteria Enterobacteriales Enterobacteriaceae *Salmonella enterica subsp. enterica serovar* Enteritidis str. 648904 3-6 |
| Sen-539-1 | Bacteria Proteobacteria Gammaproteobacteria Enterobacteriales Enterobacteriaceae *Salmonella enterica subsp. enterica serovar* Enteritidis str. 648905 5-18 |
| Sen-540-1 | Bacteria Proteobacteria Gammaproteobacteria Enterobacteriales Enterobacteriaceae *Salmonella enterica subsp. enterica serovar* Enteritidis str. 653049 13-19 |
| Sen-541-1 | Bacteria Proteobacteria Gammaproteobacteria Enterobacteriales Enterobacteriaceae *Salmonella enterica subsp. enterica serovar* Enteritidis str. 76-2651 |
| Sen-542-1 | Bacteria Proteobacteria Gammaproteobacteria Enterobacteriales Enterobacteriaceae *Salmonella enterica subsp. enterica serovar* Enteritidis str. 77-0424 |
| Sen-543-1 | Bacteria Proteobacteria Gammaproteobacteria Enterobacteriales Enterobacteriaceae *Salmonella enterica subsp. enterica serovar* Enteritidis str. 77-2659 |

| | |
|---|---|
| Sen-544-1 | Bacteria Proteobacteria Gammaproteobacteria Enterobacteriales Enterobacteriaceae *Salmonella enterica subsp. enterica serovar* Enteritidis str. 78-1757 |
| Sen-545-1 | Bacteria Proteobacteria Gammaproteobacteria Enterobacteriales Enterobacteriaceae *Salmonella enterica subsp. enterica serovar* Enteritidis str. 81-2625 |
| Sen-546-1 | Bacteria Proteobacteria Gammaproteobacteria Enterobacteriales Enterobacteriaceae *Salmonella enterica subsp. enterica serovar* Enteritidis str. 8b-1 |
| Sen-547-1 | Bacteria Proteobacteria Gammaproteobacteria Enterobacteriales Enterobacteriaceae *Salmonella enterica subsp. enterica serovar* Enteritidis str. CDC_2010K_0895 |
| Sen-548-1 | Bacteria Proteobacteria Gammaproteobacteria Enterobacteriales Enterobacteriaceae *Salmonella enterica subsp. enterica serovar* Enteritidis str. CDC_2010K_0899 |
| Sen-549-1 | Bacteria Proteobacteria Gammaproteobacteria Enterobacteriales Enterobacteriaceae *Salmonella enterica subsp. enterica serovar* Enteritidis str. CDC_2010K_0956 |
| Sen-550-1 | Bacteria Proteobacteria Gammaproteobacteria Enterobacteriales Enterobacteriaceae *Salmonella enterica subsp. enterica serovar* Enteritidis str. CDC_2010K_1010 |
| Sen-551-1 | Bacteria Proteobacteria Gammaproteobacteria Enterobacteriales Enterobacteriaceae *Salmonella enterica subsp. enterica serovar* Enteritidis str. CDC_2010K_1018 |
| Sen-552-1 | Bacteria Proteobacteria Gammaproteobacteria Enterobacteriales Enterobacteriaceae *Salmonella enterica subsp. enterica serovar* Enteritidis str. CDC_2010K_1441 |
| Sen-553-1 | Bacteria Proteobacteria Gammaproteobacteria Enterobacteriales Enterobacteriaceae *Salmonella enterica subsp. enterica serovar* Enteritidis str. CDC_2010K_1444 |
| Sen-554-1 | Bacteria Proteobacteria Gammaproteobacteria Enterobacteriales Enterobacteriaceae *Salmonella enterica subsp. enterica serovar* Enteritidis str. CDC_2010K_1445 |
| Sen-555-1 | Bacteria Proteobacteria Gammaproteobacteria Enterobacteriales Enterobacteriaceae *Salmonella enterica subsp. enterica serovar* Enteritidis str. CDC_2010K_1455 |
| Sen-556-1 | Bacteria Proteobacteria Gammaproteobacteria Enterobacteriales Enterobacteriaceae *Salmonella enterica subsp. enterica serovar* Enteritidis str. CDC_2010K_1457 |
| Sen-557-1 | Bacteria Proteobacteria Gammaproteobacteria Enterobacteriales Enterobacteriaceae *Salmonella enterica subsp. enterica serovar* Enteritidis str. CDC_2010K_1543 |
| Sen-558-1 | Bacteria Proteobacteria Gammaproteobacteria Enterobacteriales Enterobacteriaceae *Salmonella enterica subsp. enterica serovar* Enteritidis str. CDC_2010K_1558 |
| Sen-559-1 | Bacteria Proteobacteria Gammaproteobacteria Enterobacteriales Enterobacteriaceae *Salmonella enterica subsp. enterica serovar* Enteritidis str. CDC_2010K_1559 |
| Sen-560-1 | Bacteria Proteobacteria Gammaproteobacteria Enterobacteriales Enterobacteriaceae *Salmonella enterica subsp. enterica serovar* Enteritidis str. CDC_2010K_1565 |
| Sen-561-1 | Bacteria Proteobacteria Gammaproteobacteria Enterobacteriales Enterobacteriaceae *Salmonella enterica subsp. enterica serovar* Enteritidis str. CDC_2010K_1566 |
| Sen-562-1 | Bacteria Proteobacteria Gammaproteobacteria Enterobacteriales Enterobacteriaceae *Salmonella enterica subsp. enterica serovar* Enteritidis str. CDC_2010K_1575 |
| Sen-563-1 | Bacteria Proteobacteria Gammaproteobacteria Enterobacteriales Enterobacteriaceae *Salmonella enterica subsp. enterica serovar* Enteritidis str. CDC_2010K_1580 |
| Sen-564-1 | Bacteria Proteobacteria Gammaproteobacteria Enterobacteriales Enterobacteriaceae *Salmonella enterica subsp. enterica serovar* Enteritidis str. CDC_2010K_1594 |
| Sen-565-1 | Bacteria Proteobacteria Gammaproteobacteria Enterobacteriales Enterobacteriaceae *Salmonella enterica subsp. enterica serovar* Enteritidis str. CDC_2010K_1725 |
| Sen-566-1 | Bacteria Proteobacteria Gammaproteobacteria Enterobacteriales Enterobacteriaceae *Salmonella enterica subsp. enterica serovar* Enteritidis str. CDC_2010K_1729 |
| Sen-567-1 | Bacteria Proteobacteria Gammaproteobacteria Enterobacteriales Enterobacteriaceae *Salmonella enterica subsp. enterica serovar* Enteritidis str. CDC_2010K_1745 |
| Sen-568-1 | Bacteria Proteobacteria Gammaproteobacteria Enterobacteriales Enterobacteriaceae *Salmonella enterica subsp. enterica serovar* Enteritidis str. CDC_2010K_1747 |
| Sen-569-1 | Bacteria Proteobacteria Gammaproteobacteria Enterobacteriales Enterobacteriaceae *Salmonella enterica subsp. enterica serovar* Enteritidis str. CDC_2010K_1791 |
| Sen-570-1 | Bacteria Proteobacteria Gammaproteobacteria Enterobacteriales Enterobacteriaceae *Salmonella enterica subsp. enterica serovar* Enteritidis str. CDC_2010K_1795 |
| Sen-571-1 | Bacteria Proteobacteria Gammaproteobacteria Enterobacteriales Enterobacteriaceae *Salmonella enterica subsp. enterica serovar* Enteritidis str. CDC_2010K_1808 |

| | |
|---|---|
| Sen-572-1 | Bacteria Proteobacteria Gammaproteobacteria Enterobacteriales Enterobacteriaceae *Salmonella enterica subsp. enterica serovar* Enteritidis str. CDC_2010K_1810 |
| Sen-573-1 | Bacteria Proteobacteria Gammaproteobacteria Enterobacteriales Enterobacteriaceae *Salmonella enterica subsp. enterica serovar* Enteritidis str. CDC_2010K_1811 |
| Sen-574-1 | Bacteria Proteobacteria Gammaproteobacteria Enterobacteriales Enterobacteriaceae *Salmonella enterica subsp. enterica serovar* Enteritidis str. CDC_2010K_1882 |
| Sen-575-1 | Bacteria Proteobacteria Gammaproteobacteria Enterobacteriales Enterobacteriaceae *Salmonella enterica subsp. enterica serovar* Enteritidis str. CDC_2010K_1884 |
| Sen-576-1 | Bacteria Proteobacteria Gammaproteobacteria Enterobacteriales Enterobacteriaceae *Salmonella enterica subsp. enterica serovar* Enteritidis str. CHS4 |
| Sen-577-1 | Bacteria Proteobacteria Gammaproteobacteria Enterobacteriales Enterobacteriaceae *Salmonella enterica subsp. enterica serovar* Enteritidis str. CHS44 |
| Sen-578-1 | Bacteria Proteobacteria Gammaproteobacteria Enterobacteriales Enterobacteriaceae *Salmonella enterica subsp. enterica serovar* Enteritidis str. CVM_56-3991 |
| Sen-579-1 | Bacteria Proteobacteria Gammaproteobacteria Enterobacteriales Enterobacteriaceae *Salmonella enterica subsp. enterica serovar* Enteritidis str. CVM_69-4941 |
| Sen-580-1 | Bacteria Proteobacteria Gammaproteobacteria Enterobacteriales Enterobacteriaceae *Salmonella enterica subsp. enterica serovar* Enteritidis str. CVM_76-3618 |
| Sen-581-1 | Bacteria Proteobacteria Gammaproteobacteria Enterobacteriales Enterobacteriaceae *Salmonella enterica subsp. enterica serovar* Enteritidis str. CVM_81-2490 |
| Sen-582-1 | Bacteria Proteobacteria Gammaproteobacteria Enterobacteriales Enterobacteriaceae *Salmonella enterica subsp. enterica serovar* Enteritidis str. CVM_N202 |
| Sen-583-1 | Bacteria Proteobacteria Gammaproteobacteria Enterobacteriales Enterobacteriaceae *Salmonella enterica subsp. enterica serovar* Enteritidis str. EC20090135 |
| Sen-584-1 | Bacteria Proteobacteria Gammaproteobacteria Enterobacteriales Enterobacteriaceae *Salmonella enterica subsp. enterica serovar* Enteritidis str. EC20090193 |
| Sen-585-1 | Bacteria Proteobacteria Gammaproteobacteria Enterobacteriales Enterobacteriaceae *Salmonella enterica subsp. enterica serovar* Enteritidis str. EC20090195 |
| Sen-586-1 | Bacteria Proteobacteria Gammaproteobacteria Enterobacteriales Enterobacteriaceae *Salmonella enterica subsp. enterica serovar* Enteritidis str. EC20090332 |
| Sen-587-1 | Bacteria Proteobacteria Gammaproteobacteria Enterobacteriales Enterobacteriaceae *Salmonella enterica subsp. enterica serovar* Enteritidis str. EC20090530 |
| Sen-588-1 | Bacteria Proteobacteria Gammaproteobacteria Enterobacteriales Enterobacteriaceae *Salmonella enterica subsp. enterica serovar* Enteritidis str. EC20090531 |
| Sen-589-1 | Bacteria Proteobacteria Gammaproteobacteria Enterobacteriales Enterobacteriaceae *Salmonella enterica subsp. enterica serovar* Enteritidis str. EC20090884 |
| Sen-590-1 | Bacteria Proteobacteria Gammaproteobacteria Enterobacteriales Enterobacteriaceae *Salmonella enterica subsp. enterica serovar* Enteritidis str. EC20100088 |
| Sen-591-1 | Bacteria Proteobacteria Gammaproteobacteria Enterobacteriales Enterobacteriaceae *Salmonella enterica subsp. enterica serovar* Enteritidis str. EC20100089 |
| Sen-592-1 | Bacteria Proteobacteria Gammaproteobacteria Enterobacteriales Enterobacteriaceae *Salmonella enterica subsp. enterica serovar* Enteritidis str. EC20100100 |
| Sen-593-1 | Bacteria Proteobacteria Gammaproteobacteria Enterobacteriales Enterobacteriaceae *Salmonella enterica subsp. enterica serovar* Enteritidis str. EC20100103 |
| Sen-594-1 | Bacteria Proteobacteria Gammaproteobacteria Enterobacteriales Enterobacteriaceae *Salmonella enterica subsp. enterica serovar* Enteritidis str. EC20100130 |
| Sen-595-1 | Bacteria Proteobacteria Gammaproteobacteria Enterobacteriales Enterobacteriaceae *Salmonella enterica subsp. enterica serovar* Enteritidis str. EC20100131 |
| Sen-596-1 | Bacteria Proteobacteria Gammaproteobacteria Enterobacteriales Enterobacteriaceae *Salmonella enterica subsp. enterica serovar* Enteritidis str. EC20100134 |
| Sen-597-1 | Bacteria Proteobacteria Gammaproteobacteria Enterobacteriales Enterobacteriaceae *Salmonella enterica subsp. enterica serovar* Enteritidis str. EC20100325 |
| Sen-598-1 | Bacteria Proteobacteria Gammaproteobacteria Enterobacteriales Enterobacteriaceae *Salmonella enterica subsp. enterica serovar* Enteritidis str. EC20110222 |
| Sen-599-1 | Bacteria Proteobacteria Gammaproteobacteria Enterobacteriales Enterobacteriaceae *Salmonella enterica subsp. enterica serovar* Enteritidis str. EC20110223 |
| Sen-600-1 | Bacteria Proteobacteria Gammaproteobacteria Enterobacteriales Enterobacteriaceae *Salmonella enterica subsp. enterica serovar* Enteritidis str. EC20110353 |
| Sen-601-1 | Bacteria Proteobacteria Gammaproteobacteria Enterobacteriales Enterobacteriaceae *Salmonella enterica subsp. enterica serovar* Enteritidis str. EC20111510 |
| Sen-602-1 | Bacteria Proteobacteria Gammaproteobacteria Enterobacteriales Enterobacteriaceae *Salmonella enterica subsp. enterica serovar* Enteritidis str. EC20111514 |
| Sen-603-1 | Bacteria Proteobacteria Gammaproteobacteria Enterobacteriales Enterobacteriaceae *Salmonella enterica subsp. enterica serovar* Enteritidis str. EC20111515 |
| Sen-604-1 | Bacteria Proteobacteria Gammaproteobacteria Enterobacteriales Enterobacteriaceae *Salmonella enterica subsp. enterica serovar* Enteritidis str. EC20111554 |
| Sen-605-1 | Bacteria Proteobacteria Gammaproteobacteria Enterobacteriales Enterobacteriaceae *Salmonella enterica subsp. enterica serovar* Enteritidis str. EC20111561 |
| Sen-606-1 | Bacteria Proteobacteria Gammaproteobacteria Enterobacteriales Enterobacteriaceae *Salmonella enterica subsp. enterica serovar* Enteritidis str. EC20111576 |
| Sen-607-1 | Bacteria Proteobacteria Gammaproteobacteria Enterobacteriales Enterobacteriaceae *Salmonella enterica subsp. enterica serovar* Enteritidis str. EC20120003 |
| Sen-608-1 | Bacteria Proteobacteria Gammaproteobacteria Enterobacteriales Enterobacteriaceae *Salmonella enterica subsp. enterica serovar* Enteritidis str. EC20120007 |
| Sen-609-1 | Bacteria Proteobacteria Gammaproteobacteria Enterobacteriales Enterobacteriaceae *Salmonella enterica subsp. enterica serovar* Enteritidis str. EC20120051 |
| Sen-610-1 | Bacteria Proteobacteria Gammaproteobacteria Enterobacteriales Enterobacteriaceae *Salmonella enterica subsp. enterica serovar* Enteritidis str. EC20120200 |
| Sen-611-1 | Bacteria Proteobacteria Gammaproteobacteria Enterobacteriales Enterobacteriaceae *Salmonella enterica subsp. enterica serovar* Enteritidis str. EC20120213 |
| Sen-612-1 | Bacteria Proteobacteria Gammaproteobacteria Enterobacteriales Enterobacteriaceae *Salmonella enterica subsp. enterica serovar* Enteritidis str. EC20120219 |
| Sen-613-1 | Bacteria Proteobacteria Gammaproteobacteria Enterobacteriales Enterobacteriaceae *Salmonella enterica subsp. enterica serovar* Enteritidis str. EC20120229 |
| Sen-614-1 | Bacteria Proteobacteria Gammaproteobacteria Enterobacteriales Enterobacteriaceae *Salmonella enterica subsp. enterica serovar* Enteritidis str. EC20120240 |
| Sen-615-1 | Bacteria Proteobacteria Gammaproteobacteria Enterobacteriales Enterobacteriaceae *Salmonella enterica subsp. enterica serovar* Enteritidis str. EC20120356 |
| Sen-616-1 | Bacteria Proteobacteria Gammaproteobacteria Enterobacteriales Enterobacteriaceae *Salmonella enterica subsp. enterica serovar* Enteritidis str. EC20120469 |
| Sen-617-1 | Bacteria Proteobacteria Gammaproteobacteria Enterobacteriales Enterobacteriaceae *Salmonella enterica subsp. enterica serovar* Enteritidis str. EC20120496 |
| Sen-618-1 | Bacteria Proteobacteria Gammaproteobacteria Enterobacteriales Enterobacteriaceae *Salmonella enterica subsp. enterica serovar* Enteritidis str. EC20120497 |
| Sen-619-1 | Bacteria Proteobacteria Gammaproteobacteria Enterobacteriales Enterobacteriaceae *Salmonella enterica subsp. enterica serovar* Enteritidis str. EC20120498 |
| Sen-620-1 | Bacteria Proteobacteria Gammaproteobacteria Enterobacteriales Enterobacteriaceae *Salmonella enterica subsp. enterica serovar* Enteritidis str. EC20120505 |
| Sen-621-1 | Bacteria Proteobacteria Gammaproteobacteria Enterobacteriales Enterobacteriaceae *Salmonella enterica subsp. enterica serovar* Enteritidis str. EC20120528 |
| Sen-622-1 | Bacteria Proteobacteria Gammaproteobacteria Enterobacteriales Enterobacteriaceae *Salmonella enterica subsp. enterica serovar* Enteritidis str. EC20120544 |
| Sen-623-1 | Bacteria Proteobacteria Gammaproteobacteria Enterobacteriales Enterobacteriaceae *Salmonella enterica subsp. enterica serovar* Enteritidis str. EC20120548 |
| Sen-624-1 | Bacteria Proteobacteria Gammaproteobacteria Enterobacteriales Enterobacteriaceae *Salmonella enterica subsp. enterica serovar* Enteritidis str. EC20120555 |

| | |
|---|---|
| Sen-625-1 | Bacteria Proteobacteria Gammaproteobacteria Enterobacteriales Enterobacteriaceae *Salmonella enterica subsp. enterica serovar* Enteritidis str. EC20120580 |
| Sen-626-1 | Bacteria Proteobacteria Gammaproteobacteria Enterobacteriales Enterobacteriaceae *Salmonella enterica subsp. enterica serovar* Enteritidis str. EC20120581 |
| Sen-627-1 | Bacteria Proteobacteria Gammaproteobacteria Enterobacteriales Enterobacteriaceae *Salmonella enterica subsp. enterica serovar* Enteritidis str. EC20120590 |
| Sen-628-1 | Bacteria Proteobacteria Gammaproteobacteria Enterobacteriales Enterobacteriaceae *Salmonella enterica subsp. enterica serovar* Enteritidis str. EC20120597 |
| Sen-629-1 | Bacteria Proteobacteria Gammaproteobacteria Enterobacteriales Enterobacteriaceae *Salmonella enterica subsp. enterica serovar* Enteritidis str. EC20120686 |
| Sen-630-1 | Bacteria Proteobacteria Gammaproteobacteria Enterobacteriales Enterobacteriaceae *Salmonella enterica subsp. enterica serovar* Enteritidis str. EC20120687 |
| Sen-631-1 | Bacteria Proteobacteria Gammaproteobacteria Enterobacteriales Enterobacteriaceae *Salmonella enterica subsp. enterica serovar* Enteritidis str. EC20120722 |
| Sen-632-1 | Bacteria Proteobacteria Gammaproteobacteria Enterobacteriales Enterobacteriaceae *Salmonella enterica subsp. enterica serovar* Enteritidis str. EC20120734 |
| Sen-633-1 | Bacteria Proteobacteria Gammaproteobacteria Enterobacteriales Enterobacteriaceae *Salmonella enterica subsp. enterica serovar* Enteritidis str. EC20120738 |
| Sen-634-1 | Bacteria Proteobacteria Gammaproteobacteria Enterobacteriales Enterobacteriaceae *Salmonella enterica subsp. enterica serovar* Enteritidis str. EC20120765 |
| Sen-635-1 | Bacteria Proteobacteria Gammaproteobacteria Enterobacteriales Enterobacteriaceae *Salmonella enterica subsp. enterica serovar* Enteritidis str. EC20120773 |
| Sen-636-1 | Bacteria Proteobacteria Gammaproteobacteria Enterobacteriales Enterobacteriaceae *Salmonella enterica subsp. enterica serovar* Enteritidis str. EC20120774 |
| Sen-637-1 | Bacteria Proteobacteria Gammaproteobacteria Enterobacteriales Enterobacteriaceae *Salmonella enterica subsp. enterica serovar* Enteritidis str. EC20120775 |
| Sen-638-1 | Bacteria Proteobacteria Gammaproteobacteria Enterobacteriales Enterobacteriaceae *Salmonella enterica subsp. enterica serovar* Enteritidis str. EC20120776 |
| Sen-639-1 | Bacteria Proteobacteria Gammaproteobacteria Enterobacteriales Enterobacteriaceae *Salmonella enterica subsp. enterica serovar* Enteritidis str. EC20120917 |
| Sen-640-1 | Bacteria Proteobacteria Gammaproteobacteria Enterobacteriales Enterobacteriaceae *Salmonella enterica subsp. enterica serovar* Enteritidis str. EC20120918 |
| Sen-641-1 | Bacteria Proteobacteria Gammaproteobacteria Enterobacteriales Enterobacteriaceae *Salmonella enterica subsp. enterica serovar* Enteritidis str. EC20120925 |
| Sen-642-1 | Bacteria Proteobacteria Gammaproteobacteria Enterobacteriales Enterobacteriaceae *Salmonella enterica subsp. enterica serovar* Enteritidis str. EC20120927 |
| Sen-643-1 | Bacteria Proteobacteria Gammaproteobacteria Enterobacteriales Enterobacteriaceae *Salmonella enterica subsp. enterica serovar* Enteritidis str. EC20120929 |
| Sen-644-1 | Bacteria Proteobacteria Gammaproteobacteria Enterobacteriales Enterobacteriaceae *Salmonella enterica subsp. enterica serovar* Enteritidis str. EC20120963 |
| Sen-645-1 | Bacteria Proteobacteria Gammaproteobacteria Enterobacteriales Enterobacteriaceae *Salmonella enterica subsp. enterica serovar* Enteritidis str. EC20120968 |
| Sen-646-1 | Bacteria Proteobacteria Gammaproteobacteria Enterobacteriales Enterobacteriaceae *Salmonella enterica subsp. enterica serovar* Enteritidis str. EC20120969 |
| Sen-647-1 | Bacteria Proteobacteria Gammaproteobacteria Enterobacteriales Enterobacteriaceae *Salmonella enterica subsp. enterica serovar* Enteritidis str. EC20120970 |
| Sen-648-1 | Bacteria Proteobacteria Gammaproteobacteria Enterobacteriales Enterobacteriaceae *Salmonella enterica subsp. enterica serovar* Enteritidis str. EC20121179 |
| Sen-649-1 | Bacteria Proteobacteria Gammaproteobacteria Enterobacteriales Enterobacteriaceae *Salmonella enterica subsp. enterica serovar* Enteritidis str. EC20121542 |
| Sen-650-1 | Bacteria Proteobacteria Gammaproteobacteria Enterobacteriales Enterobacteriaceae *Salmonella enterica subsp. enterica serovar* Enteritidis str. EC20121671 |
| Sen-651-1 | Bacteria Proteobacteria Gammaproteobacteria Enterobacteriales Enterobacteriaceae *Salmonella enterica subsp. enterica serovar* Enteritidis str. EC20121672 |
| Sen-652-1 | Bacteria Proteobacteria Gammaproteobacteria Enterobacteriales Enterobacteriaceae *Salmonella enterica subsp. enterica serovar* Enteritidis str. EC20121689 |
| Sen-653-1 | Bacteria Proteobacteria Gammaproteobacteria Enterobacteriales Enterobacteriaceae *Salmonella enterica subsp. enterica serovar* Enteritidis str. EC20121744 |
| Sen-654-1 | Bacteria Proteobacteria Gammaproteobacteria Enterobacteriales Enterobacteriaceae *Salmonella enterica subsp. enterica serovar* Enteritidis str. EC20121746 |
| Sen-655-1 | Bacteria Proteobacteria Gammaproteobacteria Enterobacteriales Enterobacteriaceae *Salmonella enterica subsp. enterica serovar* Enteritidis str. EC20121747 |
| Sen-656-1 | Bacteria Proteobacteria Gammaproteobacteria Enterobacteriales Enterobacteriaceae *Salmonella enterica subsp. enterica serovar* Enteritidis str. EC20121748 |
| Sen-657-1 | Bacteria Proteobacteria Gammaproteobacteria Enterobacteriales Enterobacteriaceae *Salmonella enterica subsp. enterica serovar* Enteritidis str. EC20121750 |
| Sen-658-1 | Bacteria Proteobacteria Gammaproteobacteria Enterobacteriales Enterobacteriaceae *Salmonella enterica subsp. enterica serovar* Enteritidis str. EC20121751 |
| Sen-659-1 | Bacteria Proteobacteria Gammaproteobacteria Enterobacteriales Enterobacteriaceae *Salmonella enterica subsp. enterica serovar* Enteritidis str. EC20121753 |
| Sen-660-1 | Bacteria Proteobacteria Gammaproteobacteria Enterobacteriales Enterobacteriaceae *Salmonella enterica subsp. enterica serovar* Enteritidis str. EC20121765 |
| Sen-661-1 | Bacteria Proteobacteria Gammaproteobacteria Enterobacteriales Enterobacteriaceae *Salmonella enterica subsp. enterica serovar* Enteritidis str. EC20121812 |
| Sen-662-1 | Bacteria Proteobacteria Gammaproteobacteria Enterobacteriales Enterobacteriaceae *Salmonella enterica subsp. enterica serovar* Enteritidis str. EC20121825 |
| Sen-663-1 | Bacteria Proteobacteria Gammaproteobacteria Enterobacteriales Enterobacteriaceae *Salmonella enterica subsp. enterica serovar* Enteritidis str. EC20121826 |
| Sen-664-1 | Bacteria Proteobacteria Gammaproteobacteria Enterobacteriales Enterobacteriaceae *Salmonella enterica subsp. enterica serovar* Enteritidis str. EC20121969 |
| Sen-665-1 | Bacteria Proteobacteria Gammaproteobacteria Enterobacteriales Enterobacteriaceae *Salmonella enterica subsp. enterica serovar* Enteritidis str. EC20121970 |
| Sen-666-1 | Bacteria Proteobacteria Gammaproteobacteria Enterobacteriales Enterobacteriaceae *Salmonella enterica subsp. enterica serovar* Enteritidis str. EC20121976 |
| Sen-667-1 | Bacteria Proteobacteria Gammaproteobacteria Enterobacteriales Enterobacteriaceae *Salmonella enterica subsp. enterica serovar* Enteritidis str. EC20121986 |
| Sen-668-1 | Bacteria Proteobacteria Gammaproteobacteria Enterobacteriales Enterobacteriaceae *Salmonella enterica subsp. enterica serovar* Enteritidis str. EC20121989 |
| Sen-669-1 | Bacteria Proteobacteria Gammaproteobacteria Enterobacteriales Enterobacteriaceae *Salmonella enterica subsp. enterica serovar* Enteritidis str. EC20121990 |
| Sen-670-1 | Bacteria Proteobacteria Gammaproteobacteria Enterobacteriales Enterobacteriaceae *Salmonella enterica subsp. enterica serovar* Enteritidis str. EC20122022 |
| Sen-671-1 | Bacteria Proteobacteria Gammaproteobacteria Enterobacteriales Enterobacteriaceae *Salmonella enterica subsp. enterica serovar* Enteritidis str. EC20122026 |
| Sen-672-1 | Bacteria Proteobacteria Gammaproteobacteria Enterobacteriales Enterobacteriaceae *Salmonella enterica subsp. enterica serovar* Enteritidis str. EC20122031 |
| Sen-673-1 | Bacteria Proteobacteria Gammaproteobacteria Enterobacteriales Enterobacteriaceae *Salmonella enterica subsp. enterica serovar* Enteritidis str. EC20122033 |
| Sen-674-1 | Bacteria Proteobacteria Gammaproteobacteria Enterobacteriales Enterobacteriaceae *Salmonella enterica subsp. enterica serovar* Enteritidis str. EC20122045 |
| Sen-675-1 | Bacteria Proteobacteria Gammaproteobacteria Enterobacteriales Enterobacteriaceae *Salmonella enterica subsp. enterica serovar* Enteritidis str. EC20130345 |
| Sen-676-1 | Bacteria Proteobacteria Gammaproteobacteria Enterobacteriales Enterobacteriaceae *Salmonella enterica subsp. enterica serovar* Enteritidis str. EC20130346 |
| Sen-677-1 | Bacteria Proteobacteria Gammaproteobacteria Enterobacteriales Enterobacteriaceae *Salmonella enterica subsp. enterica serovar* Enteritidis str. EC20130348 |
| Sen-678-1 | Bacteria Proteobacteria Gammaproteobacteria Enterobacteriales Enterobacteriaceae *Salmonella enterica subsp. enterica serovar* Enteritidis str. PT23 |
| Sen-679-1 | Bacteria Proteobacteria Gammaproteobacteria Enterobacteriales Enterobacteriaceae *Salmonella enterica subsp. enterica serovar* Enteritidis str. SA19930684 |
| Sen-680-1 | Bacteria Proteobacteria Gammaproteobacteria Enterobacteriales Enterobacteriaceae *Salmonella enterica subsp. enterica serovar* Enteritidis str. SA19942384 |
| Sen-681-1 | Bacteria Proteobacteria Gammaproteobacteria Enterobacteriales Enterobacteriaceae *Salmonella enterica subsp. enterica serovar* Enteritidis str. SA19943269 |
| Sen-682-1 | Bacteria Proteobacteria Gammaproteobacteria Enterobacteriales Enterobacteriaceae *Salmonella enterica subsp. enterica serovar* Enteritidis str. SA19960848 |
| Sen-683-1 | Bacteria Proteobacteria Gammaproteobacteria Enterobacteriales Enterobacteriaceae *Salmonella enterica subsp. enterica serovar* Enteritidis str. SA19961622 |
| Sen-684-1 | Bacteria Proteobacteria Gammaproteobacteria Enterobacteriales Enterobacteriaceae *Salmonella enterica subsp. enterica serovar* Enteritidis str. SA19970510 |
| Sen-685-1 | Bacteria Proteobacteria Gammaproteobacteria Enterobacteriales Enterobacteriaceae *Salmonella enterica subsp. enterica serovar* Enteritidis str. SA19970769 |
| Sen-686-1 | Bacteria Proteobacteria Gammaproteobacteria Enterobacteriales Enterobacteriaceae *Salmonella enterica subsp. enterica serovar* Enteritidis str. SA19971331 |
| Sen-687-1 | Bacteria Proteobacteria Gammaproteobacteria Enterobacteriales Enterobacteriaceae *Salmonella enterica subsp. enterica serovar* Enteritidis str. SA19980677 |
| Sen-688-1 | Bacteria Proteobacteria Gammaproteobacteria Enterobacteriales Enterobacteriaceae *Salmonella enterica subsp. enterica serovar* Enteritidis str. SA19981522 |
| Sen-689-1 | Bacteria Proteobacteria Gammaproteobacteria Enterobacteriales Enterobacteriaceae *Salmonella enterica subsp. enterica serovar* Enteritidis str. SA19981857 |

| | |
|---|---|
| Sen-690-1 | Bacteria Proteobacteria Gammaproteobacteria Enterobacteriales Enterobacteriaceae *Salmonella enterica subsp. enterica serovar* Enteritidis str. SA19982831 |
| Sen-691-1 | Bacteria Proteobacteria Gammaproteobacteria Enterobacteriales Enterobacteriaceae *Salmonella enterica subsp. enterica serovar* Enteritidis str. SA19983126 |
| Sen-692-1 | Bacteria Proteobacteria Gammaproteobacteria Enterobacteriales Enterobacteriaceae *Salmonella enterica subsp. enterica serovar* Enteritidis str. SA19992322 |
| Sen-693-1 | Bacteria Proteobacteria Gammaproteobacteria Enterobacteriales Enterobacteriaceae *Salmonella enterica subsp. enterica serovar* Enteritidis str. SA20083456 |
| Sen-694-1 | Bacteria Proteobacteria Gammaproteobacteria Enterobacteriales Enterobacteriaceae *Salmonella enterica subsp. enterica serovar* Enteritidis str. SA20083636 |
| Sen-695-1 | Bacteria Proteobacteria Gammaproteobacteria Enterobacteriales Enterobacteriaceae *Salmonella enterica subsp. enterica serovar* Enteritidis str. SA20084384 |
| Sen-696-1 | Bacteria Proteobacteria Gammaproteobacteria Enterobacteriales Enterobacteriaceae *Salmonella enterica subsp. enterica serovar* Enteritidis str. SA20090435 |
| Sen-697-1 | Bacteria Proteobacteria Gammaproteobacteria Enterobacteriales Enterobacteriaceae *Salmonella enterica subsp. enterica serovar* Enteritidis str. SA20090877 |
| Sen-698-1 | Bacteria Proteobacteria Gammaproteobacteria Enterobacteriales Enterobacteriaceae *Salmonella enterica subsp. enterica serovar* Enteritidis str. SA20091739 |
| Sen-699-1 | Bacteria Proteobacteria Gammaproteobacteria Enterobacteriales Enterobacteriaceae *Salmonella enterica subsp. enterica serovar* Enteritidis str. SA20092320 |
| Sen-700-1 | Bacteria Proteobacteria Gammaproteobacteria Enterobacteriales Enterobacteriaceae *Salmonella enterica subsp. enterica serovar* Enteritidis str. SA20093266 |
| Sen-701-1 | Bacteria Proteobacteria Gammaproteobacteria Enterobacteriales Enterobacteriaceae *Salmonella enterica subsp. enterica serovar* Enteritidis str. SA20093421 |
| Sen-702-1 | Bacteria Proteobacteria Gammaproteobacteria Enterobacteriales Enterobacteriaceae *Salmonella enterica subsp. enterica serovar* Enteritidis str. SA20093430 |
| Sen-703-1 | Bacteria Proteobacteria Gammaproteobacteria Enterobacteriales Enterobacteriaceae *Salmonella enterica subsp. enterica serovar* Enteritidis str. SA20093538 |
| Sen-704-1 | Bacteria Proteobacteria Gammaproteobacteria Enterobacteriales Enterobacteriaceae *Salmonella enterica subsp. enterica serovar* Enteritidis str. SA20093543 |
| Sen-705-1 | Bacteria Proteobacteria Gammaproteobacteria Enterobacteriales Enterobacteriaceae *Salmonella enterica subsp. enterica serovar* Enteritidis str. SA20093784 |
| Sen-706-1 | Bacteria Proteobacteria Gammaproteobacteria Enterobacteriales Enterobacteriaceae *Salmonella enterica subsp. enterica serovar* Enteritidis str. SA20093788 |
| Sen-707-1 | Bacteria Proteobacteria Gammaproteobacteria Enterobacteriales Enterobacteriaceae *Salmonella enterica subsp. enterica serovar* Enteritidis str. SA20093950 |
| Sen-708-1 | Bacteria Proteobacteria Gammaproteobacteria Enterobacteriales Enterobacteriaceae *Salmonella enterica subsp. enterica serovar* Enteritidis str. SA20093977 |
| Sen-709-1 | Bacteria Proteobacteria Gammaproteobacteria Enterobacteriales Enterobacteriaceae *Salmonella enterica subsp. enterica serovar* Enteritidis str. SA20094079 |
| Sen-710-1 | Bacteria Proteobacteria Gammaproteobacteria Enterobacteriales Enterobacteriaceae *Salmonella enterica subsp. enterica serovar* Enteritidis str. SA20094350 |
| Sen-711-1 | Bacteria Proteobacteria Gammaproteobacteria Enterobacteriales Enterobacteriaceae *Salmonella enterica subsp. enterica serovar* Enteritidis str. SA20094352 |
| Sen-712-1 | Bacteria Proteobacteria Gammaproteobacteria Enterobacteriales Enterobacteriaceae *Salmonella enterica subsp. enterica serovar* Enteritidis str. SA20094383 |
| Sen-713-1 | Bacteria Proteobacteria Gammaproteobacteria Enterobacteriales Enterobacteriaceae *Salmonella enterica subsp. enterica serovar* Enteritidis str. SA20094389 |
| Sen-714-1 | Bacteria Proteobacteria Gammaproteobacteria Enterobacteriales Enterobacteriaceae *Salmonella enterica subsp. enterica serovar* Enteritidis str. SA20094521 |
| Sen-715-1 | Bacteria Proteobacteria Gammaproteobacteria Enterobacteriales Enterobacteriaceae *Salmonella enterica subsp. enterica serovar* Enteritidis str. SA20094642 |
| Sen-716-1 | Bacteria Proteobacteria Gammaproteobacteria Enterobacteriales Enterobacteriaceae *Salmonella enterica subsp. enterica serovar* Enteritidis str. SA20094682 |
| Sen-717-1 | Bacteria Proteobacteria Gammaproteobacteria Enterobacteriales Enterobacteriaceae *Salmonella enterica subsp. enterica serovar* Enteritidis str. SA20094803 |
| Sen-718-1 | Bacteria Proteobacteria Gammaproteobacteria Enterobacteriales Enterobacteriaceae *Salmonella enterica subsp. enterica serovar* Enteritidis str. SA20095309 |
| Sen-719-1 | Bacteria Proteobacteria Gammaproteobacteria Enterobacteriales Enterobacteriaceae *Salmonella enterica subsp. enterica serovar* Enteritidis str. SA20121703 |
| Sen-720-1 | Bacteria Proteobacteria Gammaproteobacteria Enterobacteriales Enterobacteriaceae *Salmonella enterica subsp. enterica serovar* Enteritidis str. SA20123395 |
| Sen-721-1 | Bacteria Proteobacteria Gammaproteobacteria Enterobacteriales Enterobacteriaceae *Salmonella enterica subsp. enterica serovar* Enteritidis str. SARB17 |
| Sen-722-1 | Bacteria Proteobacteria Gammaproteobacteria Enterobacteriales Enterobacteriaceae *Salmonella enterica subsp. enterica serovar* Enteritidis str. SE10 |
| Sen-723-1 | Bacteria Proteobacteria Gammaproteobacteria Enterobacteriales Enterobacteriaceae *Salmonella enterica subsp. enterica serovar* Enteritidis str. SE15-1 |
| Sen-724-1 | Bacteria Proteobacteria Gammaproteobacteria Enterobacteriales Enterobacteriaceae *Salmonella enterica subsp. enterica serovar* Enteritidis str. SE30663 |
| Sen-725-1 | Bacteria Proteobacteria Gammaproteobacteria Enterobacteriales Enterobacteriaceae *Salmonella enterica subsp. enterica serovar* Enteritidis str. SE8a |
| Sen-726-1 | Bacteria Proteobacteria Gammaproteobacteria Enterobacteriales Enterobacteriaceae *Salmonella enterica subsp. enterica serovar* Enteritidis str. SL909 |
| Sen-727-1 | Bacteria Proteobacteria Gammaproteobacteria Enterobacteriales Enterobacteriaceae *Salmonella enterica subsp. enterica serovar* Enteritidis str. SL913 |
| Sen-728-1 | Bacteria Proteobacteria Gammaproteobacteria Enterobacteriales Enterobacteriaceae *Salmonella enterica subsp. enterica serovar* Gallinarum |
| Sen-729-1 | Bacteria Proteobacteria Gammaproteobacteria Enterobacteriales Enterobacteriaceae *Salmonella enterica subsp. enterica serovar* Gallinarum str. 9184 |
| Sen-730-1 | Bacteria Proteobacteria Gammaproteobacteria Enterobacteriales Enterobacteriaceae *Salmonella enterica subsp. enterica serovar* Gallinarum str. SG9 |
| Sen-731-1 | Bacteria Proteobacteria Gammaproteobacteria Enterobacteriales Enterobacteriaceae *Salmonella enterica subsp. enterica serovar* Gallinarum/pullorum str. CDC1983-67 |
| Sen-732-1 | Bacteria Proteobacteria Gammaproteobacteria Enterobacteriales Enterobacteriaceae *Salmonella enterica subsp. enterica serovar* Give str. 564 |
| Sen-733-1 | Bacteria Proteobacteria Gammaproteobacteria Enterobacteriales Enterobacteriaceae *Salmonella enterica subsp. enterica serovar* Give str. S5-487 |
| Sen-734-1 | Bacteria Proteobacteria Gammaproteobacteria Enterobacteriales Enterobacteriaceae *Salmonella enterica subsp. enterica serovar* Give var. 15 str. CF-SAN004343 |
| Sen-735-1 | Bacteria Proteobacteria Gammaproteobacteria Enterobacteriales Enterobacteriaceae *Salmonella enterica subsp. enterica serovar* Hadar str. 2011K-0112 |
| Sen-736-1 | Bacteria Proteobacteria Gammaproteobacteria Enterobacteriales Enterobacteriaceae *Salmonella enterica subsp. enterica serovar* Hadar str. ATCC 51956 |
| Sen-737-1 | Bacteria Proteobacteria Gammaproteobacteria Enterobacteriales Enterobacteriaceae *Salmonella enterica subsp. enterica serovar* Hartford str. 06-0676 |
| Sen-738-1 | Bacteria Proteobacteria Gammaproteobacteria Enterobacteriales Enterobacteriaceae *Salmonella enterica subsp. enterica serovar* Havana str. CFSAN001082 |
| Sen-169-1 to Sen-169-52 | Bacteria Proteobacteria Gammaproteobacteria Enterobacteriales Enterobacteriaceae *Salmonella enterica subsp. enterica serovar* Heidelberg |
| Sen-739-1 | Bacteria Proteobacteria Gammaproteobacteria Enterobacteriales Enterobacteriaceae *Salmonella enterica subsp. enterica serovar* Heidelberg str. 24390 |
| Sen-740-1 | Bacteria Proteobacteria Gammaproteobacteria Enterobacteriales Enterobacteriaceae *Salmonella enterica subsp. enterica serovar* Heidelberg str. 24393 |
| Sen-741-1 | Bacteria Proteobacteria Gammaproteobacteria Enterobacteriales Enterobacteriaceae *Salmonella enterica subsp. enterica serovar* Heidelberg str. 41563 |
| Sen-742-1 | Bacteria Proteobacteria Gammaproteobacteria Enterobacteriales Enterobacteriaceae *Salmonella enterica subsp. enterica serovar* Heidelberg str. 41565 |
| Sen-743-1 | Bacteria Proteobacteria Gammaproteobacteria Enterobacteriales Enterobacteriaceae *Salmonella enterica subsp. enterica serovar* Heidelberg str. 41566 |
| Sen-744-1 | Bacteria Proteobacteria Gammaproteobacteria Enterobacteriales Enterobacteriaceae *Salmonella enterica subsp. enterica serovar* Heidelberg str. 41573 |
| Sen-745-1 | Bacteria Proteobacteria Gammaproteobacteria Enterobacteriales Enterobacteriaceae *Salmonella enterica subsp. enterica serovar* Heidelberg str. 41576 |
| Sen-746-1 | Bacteria Proteobacteria Gammaproteobacteria Enterobacteriales Enterobacteriaceae *Salmonella enterica subsp. enterica serovar* Heidelberg str. 41578 |
| Sen-747-1 | Bacteria Proteobacteria Gammaproteobacteria Enterobacteriales Enterobacteriaceae *Salmonella enterica subsp. enterica serovar* Heidelberg str. 41579 |
| Sen-748-1 | Bacteria Proteobacteria Gammaproteobacteria Enterobacteriales Enterobacteriaceae *Salmonella enterica subsp. enterica serovar* Heidelberg str. 579082-8 |
| Sen-749-1 | Bacteria Proteobacteria Gammaproteobacteria Enterobacteriales Enterobacteriaceae *Salmonella enterica subsp. enterica serovar* Heidelberg str. 579083-19 |
| Sen-750-1 | Bacteria Proteobacteria Gammaproteobacteria Enterobacteriales Enterobacteriaceae *Salmonella enterica subsp. enterica serovar* Heidelberg str. 607310-1 |

| | |
|---|---|
| Sen-751-1 | Bacteria Proteobacteria Gammaproteobacteria Enterobacteriales Enterobacteriaceae *Salmonella enterica subsp. enterica serovar* Heidelberg str. 622737-11 |
| Sen-752-1 | Bacteria Proteobacteria Gammaproteobacteria Enterobacteriales Enterobacteriaceae *Salmonella enterica subsp. enterica serovar* Heidelberg str. 622737-12 |
| Sen-753-1 | Bacteria Proteobacteria Gammaproteobacteria Enterobacteriales Enterobacteriaceae *Salmonella enterica subsp. enterica serovar* Heidelberg str. 622737-14 |
| Sen-754-1 | Bacteria Proteobacteria Gammaproteobacteria Enterobacteriales Enterobacteriaceae *Salmonella enterica subsp. enterica serovar* Heidelberg str. 79-0480 |
| Sen-755-1 | Bacteria Proteobacteria Gammaproteobacteria Enterobacteriales Enterobacteriaceae *Salmonella enterica subsp. enterica serovar* Heidelberg str. 82-2052 |
| Sen-756-1 | Bacteria Proteobacteria Gammaproteobacteria Enterobacteriales Enterobacteriaceae *Salmonella enterica subsp. enterica serovar* Heidelberg str. 83-1068 |
| Sen-757-1 | Bacteria Proteobacteria Gammaproteobacteria Enterobacteriales Enterobacteriaceae *Salmonella enterica subsp. enterica serovar* Heidelberg str. 90-0318 |
| Sen-758-1 | Bacteria Proteobacteria Gammaproteobacteria Enterobacteriales Enterobacteriaceae *Salmonella enterica subsp. enterica serovar* Heidelberg str. CFSAN002069 |
| Sen-759-1 | Bacteria Proteobacteria Gammaproteobacteria Enterobacteriales Enterobacteriaceae *Salmonella enterica subsp. enterica serovar* Heidelberg str. CFSAN00322 |
| Sen-760-1 | Bacteria Proteobacteria Gammaproteobacteria Enterobacteriales Enterobacteriaceae *Salmonella enterica subsp. enterica serovar* Heidelberg str. CFSAN00325 |
| Sen-761-1 | Bacteria Proteobacteria Gammaproteobacteria Enterobacteriales Enterobacteriaceae *Salmonella enterica subsp. enterica serovar* Heidelberg str. CFSAN00326 |
| Sen-762-1 | Bacteria Proteobacteria Gammaproteobacteria Enterobacteriales Enterobacteriaceae *Salmonella enterica subsp. enterica serovar* Heidelberg str. CFSAN00328 |
| Sen-763-1 | Bacteria Proteobacteria Gammaproteobacteria Enterobacteriales Enterobacteriaceae *Salmonella enterica subsp. enterica serovar* Heidelberg str. N1514 |
| Sen-764-1 | Bacteria Proteobacteria Gammaproteobacteria Enterobacteriales Enterobacteriaceae *Salmonella enterica subsp. enterica serovar* Heidelberg str. N189 |
| Sen-765-1 | Bacteria Proteobacteria Gammaproteobacteria Enterobacteriales Enterobacteriaceae *Salmonella enterica subsp. enterica serovar* Heidelberg str. N4403 |
| Sen-766-1 | Bacteria Proteobacteria Gammaproteobacteria Enterobacteriales Enterobacteriaceae *Salmonella enterica subsp. enterica serovar* Heidelberg str. N4496 |
| Sen-767-1 | Bacteria Proteobacteria Gammaproteobacteria Enterobacteriales Enterobacteriaceae *Salmonella enterica subsp. enterica serovar* Heidelberg str. N4541 |
| Sen-768-1 | Bacteria Proteobacteria Gammaproteobacteria Enterobacteriales Enterobacteriaceae *Salmonella enterica subsp. enterica serovar* Heidelberg str. RI-11-013312 |
| Sen-769-1 | Bacteria Proteobacteria Gammaproteobacteria Enterobacteriales Enterobacteriaceae *Salmonella enterica subsp. enterica serovar* Heidelberg str. RI-11-013343 |
| Sen-770-1 | Bacteria Proteobacteria Gammaproteobacteria Enterobacteriales Enterobacteriaceae *Salmonella enterica subsp. enterica serovar* Heidelberg str. RI-11-013374 |
| Sen-771-1 | Bacteria Proteobacteria Gammaproteobacteria Enterobacteriales Enterobacteriaceae *Salmonella enterica subsp. enterica serovar* Heidelberg str. RI-11-013988 |
| Sen-772-1 | Bacteria Proteobacteria Gammaproteobacteria Enterobacteriales Enterobacteriaceae *Salmonella enterica subsp. enterica serovar* Heidelberg str. RI-11-014588 |
| Sen-773-1 | Bacteria Proteobacteria Gammaproteobacteria Enterobacteriales Enterobacteriaceae *Salmonella enterica subsp. enterica serovar* Heidelberg str. SARA 39 |
| Sen-774-1 | Bacteria Proteobacteria Gammaproteobacteria Enterobacteriales Enterobacteriaceae *Salmonella enterica subsp. enterica serovar* Heidelberg str. SARA35 |
| Sen-775-1 | Bacteria Proteobacteria Gammaproteobacteria Enterobacteriales Enterobacteriaceae *Salmonella enterica subsp. enterica serovar* Indiana str. ATCC 51959 |
| Sen-776-1 | Bacteria Proteobacteria Gammaproteobacteria Enterobacteriales Enterobacteriaceae *Salmonella enterica subsp. enterica serovar* Infantis str. 119944 |
| Sen-777-1 | Bacteria Proteobacteria Gammaproteobacteria Enterobacteriales Enterobacteriaceae *Salmonella enterica subsp. enterica serovar* Infantis str. 335-3 |
| Sen-778-1 | Bacteria Proteobacteria Gammaproteobacteria Enterobacteriales Enterobacteriaceae *Salmonella enterica subsp. enterica serovar* Infantis str. SARB27 |
| Sen-779-1 | Bacteria Proteobacteria Gammaproteobacteria Enterobacteriales Enterobacteriaceae *Salmonella enterica subsp. enterica serovar* Inverness str. ATCC 10720 |
| Sen-780-1 | Bacteria Proteobacteria Gammaproteobacteria Enterobacteriales Enterobacteriaceae *Salmonella enterica subsp. enterica serovar* Javiana str. 10721 |
| Sen-781-1 | Bacteria Proteobacteria Gammaproteobacteria Enterobacteriales Enterobacteriaceae *Salmonella enterica subsp. enterica serovar* Javiana str. ATCC BAA-1593 |
| Sen-782-1 | Bacteria Proteobacteria Gammaproteobacteria Enterobacteriales Enterobacteriaceae *Salmonella enterica subsp. enterica serovar* Javiana str. CFSAN001992 |
| Sen-783-1 | Bacteria Proteobacteria Gammaproteobacteria Enterobacteriales Enterobacteriaceae *Salmonella enterica subsp. enterica serovar* Javiana str. PRS_2010_0720 |
| Sen-784-1 | Bacteria Proteobacteria Gammaproteobacteria Enterobacteriales Enterobacteriaceae *Salmonella enterica subsp. enterica serovar* Kentucky str. 0253 |
| Sen-785-1 | Bacteria Proteobacteria Gammaproteobacteria Enterobacteriales Enterobacteriaceae *Salmonella enterica subsp. enterica serovar* Kentucky str. 13562 |
| Sen-786-1 | Bacteria Proteobacteria Gammaproteobacteria Enterobacteriales Enterobacteriaceae *Salmonella enterica subsp. enterica serovar* Kentucky str. 20793 |
| Sen-787-1 | Bacteria Proteobacteria Gammaproteobacteria Enterobacteriales Enterobacteriaceae *Salmonella enterica subsp. enterica serovar* Kentucky str. 22694 |
| Sen-788-1 | Bacteria Proteobacteria Gammaproteobacteria Enterobacteriales Enterobacteriaceae *Salmonella enterica subsp. enterica serovar* Kentucky str. 29166 |
| Sen-789-1 | Bacteria Proteobacteria Gammaproteobacteria Enterobacteriales Enterobacteriaceae *Salmonella enterica subsp. enterica serovar* Kentucky str. 29439 |
| Sen-790-1 | Bacteria Proteobacteria Gammaproteobacteria Enterobacteriales Enterobacteriaceae *Salmonella enterica subsp. enterica serovar* Kentucky str. 5349 |
| Sen-791-1 | Bacteria Proteobacteria Gammaproteobacteria Enterobacteriales Enterobacteriaceae *Salmonella enterica subsp. enterica serovar* Kentucky str. N312 |
| Sen-792-1 | Bacteria Proteobacteria Gammaproteobacteria Enterobacteriales Enterobacteriaceae *Salmonella enterica subsp. enterica serovar* Litchfield str. CFSAN001076 |
| Sen-793-1 | Bacteria Proteobacteria Gammaproteobacteria Enterobacteriales Enterobacteriaceae *Salmonella enterica subsp. enterica serovar* Manhattan str. CFSAN001078 |
| Sen-794-1 | Bacteria Proteobacteria Gammaproteobacteria Enterobacteriales Enterobacteriaceae *Salmonella enterica subsp. enterica serovar* Mbandaka str. 2009K-0807 |

| | |
|---|---|
| Sen-795-1 | Bacteria Proteobacteria Gammaproteobacteria Enterobacteriales Enterobacteriaceae *Salmonella enterica subsp. enterica serovar* Mbandaka str. ATCC 51958 |
| Sen-796-1 | Bacteria Proteobacteria Gammaproteobacteria Enterobacteriales Enterobacteriaceae *Salmonella enterica subsp. enterica serovar* Meleagridis str. 0047 |
| Sen-797-1 | Bacteria Proteobacteria Gammaproteobacteria Enterobacteriales Enterobacteriaceae *Salmonella enterica subsp. enterica serovar* Miami str. 1923 |
| Sen-798-1 | Bacteria Proteobacteria Gammaproteobacteria Enterobacteriales Enterobacteriaceae *Salmonella enterica subsp. enterica serovar* Minnesota str. ATCC 49284 |
| Sen-799-1 | Bacteria Proteobacteria Gammaproteobacteria Enterobacteriales Enterobacteriaceae *Salmonella enterica subsp. enterica serovar* Mississippi str. 2010K-1406 |
| Sen-800-1 | Bacteria Proteobacteria Gammaproteobacteria Enterobacteriales Enterobacteriaceae *Salmonella enterica subsp. enterica serovar* Mississippi str. A4-633 |
| Sen-801-1 | Bacteria Proteobacteria Gammaproteobacteria Enterobacteriales Enterobacteriaceae *Salmonella enterica subsp. enterica serovar* Montevideo str. 19N |
| Sen-802-1 | Bacteria Proteobacteria Gammaproteobacteria Enterobacteriales Enterobacteriaceae *Salmonella enterica subsp. enterica serovar* Montevideo str. 2009083312 |
| Sen-803-1 | Bacteria Proteobacteria Gammaproteobacteria Enterobacteriales Enterobacteriaceae *Salmonella enterica subsp. enterica serovar* Montevideo str. 2009085258 |
| Sen-804-1 | Bacteria Proteobacteria Gammaproteobacteria Enterobacteriales Enterobacteriaceae *Salmonella enterica subsp. enterica serovar* Montevideo str. 29N |
| Sen-805-1 | Bacteria Proteobacteria Gammaproteobacteria Enterobacteriales Enterobacteriaceae *Salmonella enterica subsp. enterica serovar* Montevideo str. 315731156 |
| Sen-806-1 | Bacteria Proteobacteria Gammaproteobacteria Enterobacteriales Enterobacteriaceae *Salmonella enterica subsp. enterica serovar* Montevideo str. 315996572 |
| Sen-807-1 | Bacteria Proteobacteria Gammaproteobacteria Enterobacteriales Enterobacteriaceae *Salmonella enterica subsp. enterica serovar* Montevideo str. 366867 |
| Sen-808-1 | Bacteria Proteobacteria Gammaproteobacteria Enterobacteriales Enterobacteriaceae *Salmonella enterica subsp. enterica serovar* Montevideo str. 413180 |
| Sen-809-1 | Bacteria Proteobacteria Gammaproteobacteria Enterobacteriales Enterobacteriaceae *Salmonella enterica subsp. enterica serovar* Montevideo str. 414877 |
| Sen-810-1 | Bacteria Proteobacteria Gammaproteobacteria Enterobacteriales Enterobacteriaceae *Salmonella enterica subsp. enterica serovar* Montevideo str. 42N |
| Sen-811-1 | Bacteria Proteobacteria Gammaproteobacteria Enterobacteriales Enterobacteriaceae *Salmonella enterica subsp. enterica serovar* Montevideo str. 4441 H |
| Sen-812-1 | Bacteria Proteobacteria Gammaproteobacteria Enterobacteriales Enterobacteriaceae *Salmonella enterica subsp. enterica serovar* Montevideo str. 446600 |
| Sen-813-1 | Bacteria Proteobacteria Gammaproteobacteria Enterobacteriales Enterobacteriaceae *Salmonella enterica subsp. enterica serovar* Montevideo str. 495297-1 |
| Sen-814-1 | Bacteria Proteobacteria Gammaproteobacteria Enterobacteriales Enterobacteriaceae *Salmonella enterica subsp. enterica serovar* Montevideo str. 495297-3 |
| Sen-815-1 | Bacteria Proteobacteria Gammaproteobacteria Enterobacteriales Enterobacteriaceae *Salmonella enterica subsp. enterica serovar* Montevideo str. 495297-4 |
| Sen-816-1 | Bacteria Proteobacteria Gammaproteobacteria Enterobacteriales Enterobacteriaceae *Salmonella enterica subsp. enterica serovar* Montevideo str. 515920-1 |
| Sen-817-1 | Bacteria Proteobacteria Gammaproteobacteria Enterobacteriales Enterobacteriaceae *Salmonella enterica subsp. enterica serovar* Montevideo str. 515920-2 |
| Sen-818-1 | Bacteria Proteobacteria Gammaproteobacteria Enterobacteriales Enterobacteriaceae *Salmonella enterica subsp. enterica serovar* Montevideo str. 531954 |
| Sen-819-1 | Bacteria Proteobacteria Gammaproteobacteria Enterobacteriales Enterobacteriaceae *Salmonella enterica subsp. enterica serovar* Montevideo str. 556150-1 |
| Sen-820-1 | Bacteria Proteobacteria Gammaproteobacteria Enterobacteriales Enterobacteriaceae *Salmonella enterica subsp. enterica serovar* Montevideo str. 556152 |
| Sen-821-1 | Bacteria Proteobacteria Gammaproteobacteria Enterobacteriales Enterobacteriaceae *Salmonella enterica subsp. enterica serovar* Montevideo str. 609458-1 |
| Sen-822-1 | Bacteria Proteobacteria Gammaproteobacteria Enterobacteriales Enterobacteriaceae *Salmonella enterica subsp. enterica serovar* Montevideo str. 609460 |
| Sen-823-1 | Bacteria Proteobacteria Gammaproteobacteria Enterobacteriales Enterobacteriaceae *Salmonella enterica subsp. enterica serovar* Montevideo str. 80959-06 |
| Sen-824-1 | Bacteria Proteobacteria Gammaproteobacteria Enterobacteriales Enterobacteriaceae *Salmonella enterica subsp. enterica serovar* Montevideo str. 81038-01 |
| Sen-825-1 | Bacteria Proteobacteria Gammaproteobacteria Enterobacteriales Enterobacteriaceae *Salmonella enterica subsp. enterica serovar* Montevideo str. 8387 |
| Sen-826-1 | Bacteria Proteobacteria Gammaproteobacteria Enterobacteriales Enterobacteriaceae *Salmonella enterica subsp. enterica serovar* Montevideo str. ATCC BAA710 |
| Sen-827-1 | Bacteria Proteobacteria Gammaproteobacteria Enterobacteriales Enterobacteriaceae *Salmonella enterica subsp. enterica serovar* Montevideo str. CASC_09SCPH15965 |
| Sen-828-1 | Bacteria Proteobacteria Gammaproteobacteria Enterobacteriales Enterobacteriaceae *Salmonella enterica subsp. enterica serovar* Montevideo str. CF-SAN004346 |
| Sen-829-1 | Bacteria Proteobacteria Gammaproteobacteria Enterobacteriales Enterobacteriaceae *Salmonella enterica subsp. enterica serovar* Montevideo str. CT_02035278 |
| Sen-830-1 | Bacteria Proteobacteria Gammaproteobacteria Enterobacteriales Enterobacteriaceae *Salmonella enterica subsp. enterica serovar* Montevideo str. CT_02035318 |
| Sen-831-1 | Bacteria Proteobacteria Gammaproteobacteria Enterobacteriales Enterobacteriaceae *Salmonella enterica subsp. enterica serovar* Montevideo str. CT_02035320 |
| Sen-832-1 | Bacteria Proteobacteria Gammaproteobacteria Enterobacteriales Enterobacteriaceae *Salmonella enterica subsp. enterica serovar* Montevideo str. CT_02035321 |
| Sen-833-1 | Bacteria Proteobacteria Gammaproteobacteria Enterobacteriales Enterobacteriaceae *Salmonella enterica subsp. enterica serovar* Montevideo str. CT_02035327 |
| Sen-834-1 | Bacteria Proteobacteria Gammaproteobacteria Enterobacteriales Enterobacteriaceae *Salmonella enterica subsp. enterica serovar* Montevideo str. FVM_628064 |
| Sen-835-1 | Bacteria Proteobacteria Gammaproteobacteria Enterobacteriales Enterobacteriaceae *Salmonella enterica subsp. enterica serovar* Montevideo str. IA_2009159199 |
| Sen-836-1 | Bacteria Proteobacteria Gammaproteobacteria Enterobacteriales Enterobacteriaceae *Salmonella enterica subsp. enterica serovar* Montevideo str. IA_2010008282 |
| Sen-837-1 | Bacteria Proteobacteria Gammaproteobacteria Enterobacteriales Enterobacteriaceae *Salmonella enterica subsp. enterica serovar* Montevideo str. IA_2010008283 |

| | |
|---|---|
| Sen-838-1 | Bacteria Proteobacteria Gammaproteobacteria Enterobacteriales Enterobacteriaceae *Salmonella enterica subsp. enterica serovar* Montevideo str. IA_2010008284 |
| Sen-839-1 | Bacteria Proteobacteria Gammaproteobacteria Enterobacteriales Enterobacteriaceae *Salmonella enterica subsp. enterica serovar* Montevideo str. IA_2010008285 |
| Sen-840-1 | Bacteria Proteobacteria Gammaproteobacteria Enterobacteriales Enterobacteriaceae *Salmonella enterica subsp. enterica serovar* Montevideo str. IA_2010008286 |
| Sen-841-1 | Bacteria Proteobacteria Gammaproteobacteria Enterobacteriales Enterobacteriaceae *Salmonella enterica subsp. enterica serovar* Montevideo str. IA_2010008287 |
| Sen-842-1 | Bacteria Proteobacteria Gammaproteobacteria Enterobacteriales Enterobacteriaceae *Salmonella enterica subsp. enterica serovar* Montevideo str. LQC 10 |
| Sen-843-1 | Bacteria Proteobacteria Gammaproteobacteria Enterobacteriales Enterobacteriaceae *Salmonella enterica subsp. enterica serovar* Montevideo str. MB101509-0077 |
| Sen-844-1 | Bacteria Proteobacteria Gammaproteobacteria Enterobacteriales Enterobacteriaceae *Salmonella enterica subsp. enterica serovar* Montevideo str. MB102109-0047 |
| Sen-845-1 | Bacteria Proteobacteria Gammaproteobacteria Enterobacteriales Enterobacteriaceae *Salmonella enterica subsp. enterica serovar* Montevideo str. MB110209-0055 |
| Sen-846-1 | Bacteria Proteobacteria Gammaproteobacteria Enterobacteriales Enterobacteriaceae *Salmonella enterica subsp. enterica serovar* Montevideo str. MB111609-0052 |
| Sen-847-1 | Bacteria Proteobacteria Gammaproteobacteria Enterobacteriales Enterobacteriaceae *Salmonella enterica subsp. enterica serovar* Montevideo str. MD_MDA09249507 |
| Sen-848-1 | Bacteria Proteobacteria Gammaproteobacteria Enterobacteriales Enterobacteriaceae *Salmonella enterica subsp. enterica serovar* Montevideo str. NC_MB110209-0054 |
| Sen-849-1 | Bacteria Proteobacteria Gammaproteobacteria Enterobacteriales Enterobacteriaceae *Salmonella enterica subsp. enterica serovar* Montevideo str. OH_2009072675 |
| Sen-850-1 | Bacteria Proteobacteria Gammaproteobacteria Enterobacteriales Enterobacteriaceae *Salmonella enterica subsp. enterica serovar* Montevideo str. SARB30 |
| Sen-851-1 | Bacteria Proteobacteria Gammaproteobacteria Enterobacteriales Enterobacteriaceae *Salmonella enterica subsp. enterica serovar* Montevideo str. SARB31 |
| Sen-852-1 | Bacteria Proteobacteria Gammaproteobacteria Enterobacteriales Enterobacteriaceae *Salmonella enterica subsp. enterica serovar* Montevideo str. USDA-ARS-USMARC-1921 |
| Sen-853-1 | Bacteria Proteobacteria Gammaproteobacteria Enterobacteriales Enterobacteriaceae *Salmonella enterica subsp. enterica serovar* Muenchen str. ATCC 8388 |
| Sen-854-1 | Bacteria Proteobacteria Gammaproteobacteria Enterobacteriales Enterobacteriaceae *Salmonella enterica subsp. enterica serovar* Muenchen str. baa1594 |
| Sen-855-1 | Bacteria Proteobacteria Gammaproteobacteria Enterobacteriales Enterobacteriaceae *Salmonella enterica subsp. enterica serovar* Muenchen str. baa1674 |
| Sen-856-1 | Bacteria Proteobacteria Gammaproteobacteria Enterobacteriales Enterobacteriaceae *Salmonella enterica subsp. enterica serovar* Muenster str. 0315 |
| Sen-857-1 | Bacteria Proteobacteria Gammaproteobacteria Enterobacteriales Enterobacteriaceae *Salmonella enterica subsp. enterica serovar* Muenster str. 420 |
| Sen-858-1 | Bacteria Proteobacteria Gammaproteobacteria Enterobacteriales Enterobacteriaceae *Salmonella enterica subsp. enterica serovar* Muenster str. 660 |
| Sen-859-1 | Bacteria Proteobacteria Gammaproteobacteria Enterobacteriales Enterobacteriaceae *Salmonella enterica subsp. enterica serovar* Muenster str. CFSAN004344 |
| Sen-860-1 | Bacteria Proteobacteria Gammaproteobacteria Enterobacteriales Enterobacteriaceae *Salmonella enterica subsp. enterica serovar* Nchanga str. CFSAN001091 |
| Sen-861-1 | Bacteria Proteobacteria Gammaproteobacteria Enterobacteriales Enterobacteriaceae *Salmonella enterica subsp. enterica serovar* Newport str. #11-2 |
| Sen-862-1 | Bacteria Proteobacteria Gammaproteobacteria Enterobacteriales Enterobacteriaceae *Salmonella enterica subsp. enterica serovar* Newport str. #11-3 |
| Sen-863-1 | Bacteria Proteobacteria Gammaproteobacteria Enterobacteriales Enterobacteriaceae *Salmonella enterica subsp. enterica serovar* Newport str. #11-4 |
| Sen-864-1 | Bacteria Proteobacteria Gammaproteobacteria Enterobacteriales Enterobacteriaceae *Salmonella enterica subsp. enterica serovar* Newport str. 637564_17 |
| Sen-865-1 | Bacteria Proteobacteria Gammaproteobacteria Enterobacteriales Enterobacteriaceae *Salmonella enterica subsp. enterica serovar* Newport str. CDC 2010K-2159 |
| Sen-866-1 | Bacteria Proteobacteria Gammaproteobacteria Enterobacteriales Enterobacteriaceae *Salmonella enterica subsp. enterica serovar* Newport str. CVM 19443 |
| Sen-867-1 | Bacteria Proteobacteria Gammaproteobacteria Enterobacteriales Enterobacteriaceae *Salmonella enterica subsp. enterica serovar* Newport str. CVM 19447 |
| Sen-868-1 | Bacteria Proteobacteria Gammaproteobacteria Enterobacteriales Enterobacteriaceae *Salmonella enterica subsp. enterica serovar* Newport str. CVM 19449 |
| Sen-869-1 | Bacteria Proteobacteria Gammaproteobacteria Enterobacteriales Enterobacteriaceae *Salmonella enterica subsp. enterica serovar* Newport str. CVM 19470 |
| Sen-870-1 | Bacteria Proteobacteria Gammaproteobacteria Enterobacteriales Enterobacteriaceae *Salmonella enterica subsp. enterica serovar* Newport str. CVM 19536 |
| Sen-871-1 | Bacteria Proteobacteria Gammaproteobacteria Enterobacteriales Enterobacteriaceae *Salmonella enterica subsp. enterica serovar* Newport str. CVM 19567 |
| Sen-872-1 | Bacteria Proteobacteria Gammaproteobacteria Enterobacteriales Enterobacteriaceae *Salmonella enterica subsp. enterica serovar* Newport str. CVM 19593 |
| Sen-873-1 | Bacteria Proteobacteria Gammaproteobacteria Enterobacteriales Enterobacteriaceae *Salmonella enterica subsp. enterica serovar* Newport str. CVM 21539 |
| Sen-874-1 | Bacteria Proteobacteria Gammaproteobacteria Enterobacteriales Enterobacteriaceae *Salmonella enterica subsp. enterica serovar* Newport str. CVM 21559 |
| Sen-875-1 | Bacteria Proteobacteria Gammaproteobacteria Enterobacteriales Enterobacteriaceae *Salmonella enterica subsp. enterica serovar* Newport str. CVM 33953 |
| Sen-876-1 | Bacteria Proteobacteria Gammaproteobacteria Enterobacteriales Enterobacteriaceae *Salmonella enterica subsp. enterica serovar* Newport str. CVM 35185 |
| Sen-877-1 | Bacteria Proteobacteria Gammaproteobacteria Enterobacteriales Enterobacteriaceae *Salmonella enterica subsp. enterica serovar* Newport str. CVM 35188 |
| Sen-878-1 | Bacteria Proteobacteria Gammaproteobacteria Enterobacteriales Enterobacteriaceae *Salmonella enterica subsp. enterica serovar* Newport str. CVM 35199 |
| Sen-879-1 | Bacteria Proteobacteria Gammaproteobacteria Enterobacteriales Enterobacteriaceae *Salmonella enterica subsp. enterica serovar* Newport str. CVM 35202 |
| Sen-880-1 | Bacteria Proteobacteria Gammaproteobacteria Enterobacteriales Enterobacteriaceae *Salmonella enterica subsp. enterica serovar* Newport str. CVM 37978 |

| | |
|---|---|
| Sen-881-1 | Bacteria Proteobacteria Gammaproteobacteria Enterobacteriales Enterobacteriaceae *Salmonella enterica subsp. enterica serovar* Newport str. CVM 4176 |
| Sen-882-1 | Bacteria Proteobacteria Gammaproteobacteria Enterobacteriales Enterobacteriaceae *Salmonella enterica subsp. enterica serovar* Newport str. Henan_3 |
| Sen-883-1 | Bacteria Proteobacteria Gammaproteobacteria Enterobacteriales Enterobacteriaceae *Salmonella enterica subsp. enterica serovar* Newport str. JS09102 |
| Sen-884-1 | Bacteria Proteobacteria Gammaproteobacteria Enterobacteriales Enterobacteriaceae *Salmonella enterica subsp. enterica serovar* Newport str. Levine 1 |
| Sen-885-1 | Bacteria Proteobacteria Gammaproteobacteria Enterobacteriales Enterobacteriaceae *Salmonella enterica subsp. enterica serovar* Newport str. Levine 15 |
| Sen-886-1 | Bacteria Proteobacteria Gammaproteobacteria Enterobacteriales Enterobacteriaceae *Salmonella enterica subsp. enterica serovar* Newport str. RI_10P068 |
| Sen-887-1 | Bacteria Proteobacteria Gammaproteobacteria Enterobacteriales Enterobacteriaceae *Salmonella enterica subsp. enterica serovar* Newport str. RI_10P069 |
| Sen-888-1 | Bacteria Proteobacteria Gammaproteobacteria Enterobacteriales Enterobacteriaceae *Salmonella enterica subsp. enterica serovar* Newport str. RI_10P078 |
| Sen-889-1 | Bacteria Proteobacteria Gammaproteobacteria Enterobacteriales Enterobacteriaceae *Salmonella enterica subsp. enterica serovar* Newport str. RI_10P079 |
| Sen-890-1 | Bacteria Proteobacteria Gammaproteobacteria Enterobacteriales Enterobacteriaceae *Salmonella enterica subsp. enterica serovar* Newport str. SH111077 |
| Sen-891-1 | Bacteria Proteobacteria Gammaproteobacteria Enterobacteriales Enterobacteriaceae *Salmonella enterica subsp. enterica serovar* Newport str. Shandong_3 |
| Sen-892-1 | Bacteria Proteobacteria Gammaproteobacteria Enterobacteriales Enterobacteriaceae *Salmonella enterica subsp. enterica serovar* Newport str. USMARC-S3124.1 |
| Sen-893-1 | Bacteria Proteobacteria Gammaproteobacteria Enterobacteriales Enterobacteriaceae *Salmonella enterica subsp. enterica serovar* Newport str. VA_R100506907 |
| Sen-894-1 | Bacteria Proteobacteria Gammaproteobacteria Enterobacteriales Enterobacteriaceae *Salmonella enterica subsp. enterica serovar* Newport str. VA_R100512570 |
| Sen-895-1 | Bacteria Proteobacteria Gammaproteobacteria Enterobacteriales Enterobacteriaceae *Salmonella enterica subsp. enterica serovar* Newport str. VA_R100512572 |
| Sen-896-1 | Bacteria Proteobacteria Gammaproteobacteria Enterobacteriales Enterobacteriaceae *Salmonella enterica subsp. enterica serovar* Newport str. WA_14881 |
| Sen-897-1 | Bacteria Proteobacteria Gammaproteobacteria Enterobacteriales Enterobacteriaceae *Salmonella enterica subsp. enterica serovar* Newport str. WA_14882 |
| Sen-898-1 | Bacteria Proteobacteria Gammaproteobacteria Enterobacteriales Enterobacteriaceae *Salmonella enterica subsp. enterica serovar* Newport str. WA_14885 |
| Sen-899-1 | Bacteria Proteobacteria Gammaproteobacteria Enterobacteriales Enterobacteriaceae *Salmonella enterica subsp. enterica serovar* Newport str. WA_14900 |
| Sen-900-1 | Bacteria Proteobacteria Gammaproteobacteria Enterobacteriales Enterobacteriaceae *Salmonella enterica subsp. enterica serovar* Norwich str. CFSAN001077 |
| Sen-901-1 | Bacteria Proteobacteria Gammaproteobacteria Enterobacteriales Enterobacteriaceae *Salmonella enterica subsp. enterica serovar* Ohio str. CFSAN001079 |
| Sen-902-1 | Bacteria Proteobacteria Gammaproteobacteria Enterobacteriales Enterobacteriaceae *Salmonella enterica subsp. enterica serovar* Oranienburg str. 0250 |
| Sen-903-1 | Bacteria Proteobacteria Gammaproteobacteria Enterobacteriales Enterobacteriaceae *Salmonella enterica subsp. enterica serovar* Oranienburg str. 701 |
| Sen-904-1 | Bacteria Proteobacteria Gammaproteobacteria Enterobacteriales Enterobacteriaceae *Salmonella enterica subsp. enterica serovar* Oranienburg str. S-76 |
| Sen-905-1 | Bacteria Proteobacteria Gammaproteobacteria Enterobacteriales Enterobacteriaceae *Salmonella enterica subsp. enterica serovar* Paratyphi A str. ATCC |
| Sen-906-1 | Bacteria Proteobacteria Gammaproteobacteria Enterobacteriales Enterobacteriaceae *Salmonella enterica subsp. enterica serovar* Paratyphi A str. ATCC 11511 |
| Sen-907-1 | Bacteria Proteobacteria Gammaproteobacteria Enterobacteriales Enterobacteriaceae *Salmonella enterica subsp. enterica serovar* Paratyphi A str. GXS2268 |
| Sen-908-1 | Bacteria Proteobacteria Gammaproteobacteria Enterobacteriales Enterobacteriaceae *Salmonella enterica subsp. enterica serovar* Paratyphi A str. GZ9A00052 |
| Sen-909-1 | Bacteria Proteobacteria Gammaproteobacteria Enterobacteriales Enterobacteriaceae *Salmonella enterica subsp. enterica serovar* Paratyphi A str. JX05-19 |
| Sen-910-1 | Bacteria Proteobacteria Gammaproteobacteria Enterobacteriales Enterobacteriaceae *Salmonella enterica subsp. enterica serovar* Paratyphi A str. YN09620 |
| Sen-911-1 | Bacteria Proteobacteria Gammaproteobacteria Enterobacteriales Enterobacteriaceae *Salmonella enterica subsp. enterica serovar* Paratyphi A str. ZJ98-53 |
| Sen-912-1 | Bacteria Proteobacteria Gammaproteobacteria Enterobacteriales Enterobacteriaceae *Salmonella enterica subsp. enterica serovar* Paratyphi B str. ATCC 10719 |
| Sen-913-1 | Bacteria Proteobacteria Gammaproteobacteria Enterobacteriales Enterobacteriaceae *Salmonella enterica subsp. enterica serovar* Paratyphi B str. ATCC 19940 |
| Sen-914-1 | Bacteria Proteobacteria Gammaproteobacteria Enterobacteriales Enterobacteriaceae *Salmonella enterica subsp. enterica serovar* Paratyphi B str. ATCC 51962 |
| Sen-915-1 | Bacteria Proteobacteria Gammaproteobacteria Enterobacteriales Enterobacteriaceae *Salmonella enterica subsp. enterica serovar* Paratyphi B str. ATCC 8759 |
| Sen-916-1 | Bacteria Proteobacteria Gammaproteobacteria Enterobacteriales Enterobacteriaceae *Salmonella enterica subsp. enterica serovar* Paratyphi B str. ATCC BAA-1585 |
| Sen-917-1 | Bacteria Proteobacteria Gammaproteobacteria Enterobacteriales Enterobacteriaceae *Salmonella enterica subsp. enterica serovar* Paratyphi B str. SARA42 |
| Sen-918-1 | Bacteria Proteobacteria Gammaproteobacteria Enterobacteriales Enterobacteriaceae *Salmonella enterica subsp. enterica serovar* Paratyphi B str. SARA56 |
| Sen-919-1 | Bacteria Proteobacteria Gammaproteobacteria Enterobacteriales Enterobacteriaceae *Salmonella enterica subsp. enterica serovar* Paratyphi B str. SARA61 |
| Sen-920-1 | Bacteria Proteobacteria Gammaproteobacteria Enterobacteriales Enterobacteriaceae *Salmonella enterica subsp. enterica serovar* Paratyphi B str. SARA62 |
| Sen-921-1 | Bacteria Proteobacteria Gammaproteobacteria Enterobacteriales Enterobacteriaceae *Salmonella enterica subsp. enterica serovar* Pomona str. ATCC 10729 |
| Sen-922-1 | Bacteria Proteobacteria Gammaproteobacteria Enterobacteriales Enterobacteriaceae *Salmonella enterica subsp. enterica serovar* Poona str. ATCC BAA-1673 |
| Sen-923-1 | Bacteria Proteobacteria Gammaproteobacteria Enterobacteriales Enterobacteriaceae *Salmonella enterica subsp. enterica serovar* Pullorum str. 13036 |
| Sen-924-1 | Bacteria Proteobacteria Gammaproteobacteria Enterobacteriales Enterobacteriaceae *Salmonella enterica subsp. enterica serovar* Pullorum str. 19945 |
| Sen-925-1 | Bacteria Proteobacteria Gammaproteobacteria Enterobacteriales Enterobacteriaceae *Salmonella enterica subsp. enterica serovar* Pullorum str. S06004 |
| Sen-926-1 | Bacteria Proteobacteria Gammaproteobacteria Enterobacteriales Enterobacteriaceae *Salmonella enterica subsp. enterica serovar* Rissen str. 150 |
| Sen-927-1 | Bacteria Proteobacteria Gammaproteobacteria Enterobacteriales Enterobacteriaceae *Salmonella enterica subsp. enterica serovar* Rubislaw str. ATCC 10717 |

| | |
|---|---|
| Sen-928-1 | Bacteria Proteobacteria Gammaproteobacteria Enterobacteriales Enterobacteriaceae *Salmonella enterica subsp. enterica serovar* Saintpaul str. 9712 |
| Sen-929-1 | Bacteria Proteobacteria Gammaproteobacteria Enterobacteriales Enterobacteriaceae *Salmonella enterica subsp. enterica serovar* Saintpaul str. JO2008 |
| Sen-930-1 | Bacteria Proteobacteria Gammaproteobacteria Enterobacteriales Enterobacteriaceae *Salmonella enterica subsp. enterica serovar* Saintpaul str. S-70 |
| Sen-931-1 | Bacteria Proteobacteria Gammaproteobacteria Enterobacteriales Enterobacteriaceae *Salmonella enterica subsp. enterica serovar* Saintpaul str. SARA26 |
| Sen-932-1 | Bacteria Proteobacteria Gammaproteobacteria Enterobacteriales Enterobacteriaceae *Salmonella enterica subsp. enterica serovar* Senftenberg str. 316235162 |
| Sen-933-1 | Bacteria Proteobacteria Gammaproteobacteria Enterobacteriales Enterobacteriaceae *Salmonella enterica subsp. enterica serovar* Senftenberg str. 361154004 |
| Sen-934-1 | Bacteria Proteobacteria Gammaproteobacteria Enterobacteriales Enterobacteriaceae *Salmonella enterica subsp. enterica serovar* Senftenberg str. 423984-1 |
| Sen-935-1 | Bacteria Proteobacteria Gammaproteobacteria Enterobacteriales Enterobacteriaceae *Salmonella enterica subsp. enterica serovar* Senftenberg str. 423984-2 |
| Sen-936-1 | Bacteria Proteobacteria Gammaproteobacteria Enterobacteriales Enterobacteriaceae *Salmonella enterica subsp. enterica serovar* Senftenberg str. 604314 |
| Sen-937-1 | Bacteria Proteobacteria Gammaproteobacteria Enterobacteriales Enterobacteriaceae *Salmonella enterica subsp. enterica serovar* Senftenberg str. ATCC 43845 |
| Sen-938-1 | Bacteria Proteobacteria Gammaproteobacteria Enterobacteriales Enterobacteriaceae *Salmonella enterica subsp. enterica serovar* Senftenberg str. ATCC 8400 |
| Sen-939-1 | Bacteria Proteobacteria Gammaproteobacteria Enterobacteriales Enterobacteriaceae *Salmonella enterica subsp. enterica serovar* Senftenberg str. NC_MB012510-0038 |
| Sen-940-1 | Bacteria Proteobacteria Gammaproteobacteria Enterobacteriales Enterobacteriaceae *Salmonella enterica subsp. enterica serovar* Senftenberg str. SS209 |
| Sen-941-1 | Bacteria Proteobacteria Gammaproteobacteria Enterobacteriales Enterobacteriaceae *Salmonella enterica subsp. enterica serovar* Soerenga str. 695 |
| Sen-942-1 | Bacteria Proteobacteria Gammaproteobacteria Enterobacteriales Enterobacteriaceae *Salmonella enterica subsp. enterica serovar* Stanley str. ATCC 7308 |
| Sen-943-1 | Bacteria Proteobacteria Gammaproteobacteria Enterobacteriales Enterobacteriaceae *Salmonella enterica subsp. enterica serovar* Stanleyville str. CF-SAN000624 |
| Sen-944-1 | Bacteria Proteobacteria Gammaproteobacteria Enterobacteriales Enterobacteriaceae *Salmonella enterica subsp. enterica serovar* Tallahassee str. 0012 |
| Sen-945-1 | Bacteria Proteobacteria Gammaproteobacteria Enterobacteriales Enterobacteriaceae *Salmonella enterica subsp. enterica serovar* Tennessee str. TXSC_TXSC08-21 |
| Sen-946-1 | Bacteria Proteobacteria Gammaproteobacteria Enterobacteriales Enterobacteriaceae *Salmonella enterica subsp. enterica serovar* Thompson str. 2010K-1863 |
| Sen-947-1 | Bacteria Proteobacteria Gammaproteobacteria Enterobacteriales Enterobacteriaceae *Salmonella enterica subsp. enterica serovar* Thompson str. RM6836 |
| Sen-287-1 to Sen-287-1124 | Bacteria Proteobacteria Gammaproteobacteria Enterobacteriales Enterobacteriaceae *Salmonella enterica subsp. enterica serovar* Typhi |
| Sen-948-1 | Bacteria Proteobacteria Gammaproteobacteria Enterobacteriales Enterobacteriaceae *Salmonella enterica subsp. enterica serovar* Typhi str. BL196 |
| Sen-949-1 | Bacteria Proteobacteria Gammaproteobacteria Enterobacteriales Enterobacteriaceae *Salmonella enterica subsp. enterica serovar* Typhi str. CR0044 |
| Sen-950-1 | Bacteria Proteobacteria Gammaproteobacteria Enterobacteriales Enterobacteriaceae *Salmonella enterica subsp. enterica serovar* Typhi str. P-stx-12 |
| Sen-951-1 | Bacteria Proteobacteria Gammaproteobacteria Enterobacteriales Enterobacteriaceae *Salmonella enterica subsp. enterica serovar* Typhi str. ST0208 |
| Sen-952-1 | Bacteria Proteobacteria Gammaproteobacteria Enterobacteriales Enterobacteriaceae *Salmonella enterica subsp. enterica serovar* Typhi str. Ty21a |
| Sen-953-1 | Bacteria Proteobacteria Gammaproteobacteria Enterobacteriales Enterobacteriaceae *Salmonella enterica subsp. enterica serovar* Typhi str. UJ308A |
| Sen-954-1 | Bacteria Proteobacteria Gammaproteobacteria Enterobacteriales Enterobacteriaceae *Salmonella enterica subsp. enterica serovar* Typhi str. UJ816A |
| Sen-290-1 to Sen-290-7 | Bacteria Proteobacteria Gammaproteobacteria Enterobacteriales Enterobacteriaceae *Salmonella enterica subsp. enterica serovar* Typhimurium |
| Sen-955-1 | Bacteria Proteobacteria Gammaproteobacteria Enterobacteriales Enterobacteriaceae *Salmonella enterica subsp. enterica serovar* Typhimurium str. 14028S |
| Sen-956-1 | Bacteria Proteobacteria Gammaproteobacteria Enterobacteriales Enterobacteriaceae *Salmonella enterica subsp. enterica serovar* Typhimurium str. 34502 |
| Sen-957-1 | Bacteria Proteobacteria Gammaproteobacteria Enterobacteriales Enterobacteriaceae *Salmonella enterica subsp. enterica serovar* Typhimurium str. 35423 |
| Sen-958-1 | Bacteria Proteobacteria Gammaproteobacteria Enterobacteriales Enterobacteriaceae *Salmonella enterica subsp. enterica serovar* Typhimurium str. 36618 |
| Sen-959-1 | Bacteria Proteobacteria Gammaproteobacteria Enterobacteriales Enterobacteriaceae *Salmonella enterica subsp. enterica serovar* Typhimurium str. 798 |
| Sen-960-1 | Bacteria Proteobacteria Gammaproteobacteria Enterobacteriales Enterobacteriaceae *Salmonella enterica subsp. enterica serovar* Typhimurium str. AZ 057 |
| Sen-961-1 | Bacteria Proteobacteria Gammaproteobacteria Enterobacteriales Enterobacteriaceae *Salmonella enterica subsp. enterica serovar* Typhimurium str. CDC_2009K1158 |
| Sen-962-1 | Bacteria Proteobacteria Gammaproteobacteria Enterobacteriales Enterobacteriaceae *Salmonella enterica subsp. enterica serovar* Typhimurium str. CDC_2009K1277 |
| Sen-963-1 | Bacteria Proteobacteria Gammaproteobacteria Enterobacteriales Enterobacteriaceae *Salmonella enterica subsp. enterica serovar* Typhimurium str. CDC_2009K1283 |
| Sen-964-1 | Bacteria Proteobacteria Gammaproteobacteria Enterobacteriales Enterobacteriaceae *Salmonella enterica subsp. enterica serovar* Typhimurium str. CDC_2009K1288 |
| Sen-965-1 | Bacteria Proteobacteria Gammaproteobacteria Enterobacteriales Enterobacteriaceae *Salmonella enterica subsp. enterica serovar* Typhimurium str. D23580 |
| Sen-299-1 to Sen-299-330 | Bacteria Proteobacteria Gammaproteobacteria Enterobacteriales Enterobacteriaceae *Salmonella enterica subsp. enterica serovar* Typhimurium str. DT104 |
| Sen-966-1 | Bacteria Proteobacteria Gammaproteobacteria Enterobacteriales Enterobacteriaceae *Salmonella enterica subsp. enterica serovar* Typhimurium str. DT2 |
| Sen-967-1 | Bacteria Proteobacteria Gammaproteobacteria Enterobacteriales Enterobacteriaceae *Salmonella enterica subsp. enterica serovar* Typhimurium str. SARA13 |
| Sen-968-1 | Bacteria Proteobacteria Gammaproteobacteria Enterobacteriales Enterobacteriaceae *Salmonella enterica subsp. enterica serovar* Typhimurium str. SL1344 |
| Sen-969-1 | Bacteria Proteobacteria Gammaproteobacteria Enterobacteriales Enterobacteriaceae *Salmonella enterica subsp. enterica serovar* Typhimurium str. ST1660/06 |
| Sen-970-1 | Bacteria Proteobacteria Gammaproteobacteria Enterobacteriales Enterobacteriaceae *Salmonella enterica subsp. enterica serovar* Typhimurium str. ST4/74 |
| Sen-971-1 | Bacteria Proteobacteria Gammaproteobacteria Enterobacteriales Enterobacteriaceae *Salmonella enterica subsp. enterica serovar* Typhimurium str. ST4581 |
| Sen-972-1 | Bacteria Proteobacteria Gammaproteobacteria Enterobacteriales Enterobacteriaceae *Salmonella enterica subsp. enterica serovar* Typhimurium str. ST78896 |
| Sen-973-1 | Bacteria Proteobacteria Gammaproteobacteria Enterobacteriales Enterobacteriaceae *Salmonella enterica subsp. enterica serovar* Typhimurium str. STm1 |
| Sen-974-1 | Bacteria Proteobacteria Gammaproteobacteria Enterobacteriales Enterobacteriaceae *Salmonella enterica subsp. enterica serovar* Typhimurium str. STm10 |

| | |
|---|---|
| Sen-975-1 | Bacteria Proteobacteria Gammaproteobacteria Enterobacteriales Enterobacteriaceae *Salmonella enterica subsp. enterica serovar* Typhimurium str. STm11 |
| Sen-976-1 | Bacteria Proteobacteria Gammaproteobacteria Enterobacteriales Enterobacteriaceae *Salmonella enterica subsp. enterica serovar* Typhimurium str. STm12 |
| Sen-977-1 | Bacteria Proteobacteria Gammaproteobacteria Enterobacteriales Enterobacteriaceae *Salmonella enterica subsp. enterica serovar* Typhimurium str. STm2 |
| Sen-978-1 | Bacteria Proteobacteria Gammaproteobacteria Enterobacteriales Enterobacteriaceae *Salmonella enterica subsp. enterica serovar* Typhimurium str. STm3 |
| Sen-979-1 | Bacteria Proteobacteria Gammaproteobacteria Enterobacteriales Enterobacteriaceae *Salmonella enterica subsp. enterica serovar* Typhimurium str. STm4 |
| Sen-980-1 | Bacteria Proteobacteria Gammaproteobacteria Enterobacteriales Enterobacteriaceae *Salmonella enterica subsp. enterica serovar* Typhimurium str. STm5 |
| Sen-981-1 | Bacteria Proteobacteria Gammaproteobacteria Enterobacteriales Enterobacteriaceae *Salmonella enterica subsp. enterica serovar* Typhimurium str. STm6 |
| Sen-982-1 | Bacteria Proteobacteria Gammaproteobacteria Enterobacteriales Enterobacteriaceae *Salmonella enterica subsp. enterica serovar* Typhimurium str. STm7 |
| Sen-983-1 | Bacteria Proteobacteria Gammaproteobacteria Enterobacteriales Enterobacteriaceae *Salmonella enterica subsp. enterica serovar* Typhimurium str. STm8 |
| Sen-984-1 | Bacteria Proteobacteria Gammaproteobacteria Enterobacteriales Enterobacteriaceae *Salmonella enterica subsp. enterica serovar* Typhimurium str. STm9 |
| Sen-985-1 | Bacteria Proteobacteria Gammaproteobacteria Enterobacteriales Enterobacteriaceae *Salmonella enterica subsp. enterica serovar* Typhimurium str. T000240 |
| Sen-986-1 | Bacteria Proteobacteria Gammaproteobacteria Enterobacteriales Enterobacteriaceae *Salmonella enterica subsp. enterica serovar* Typhimurium str. TN061786 |
| Sen-987-1 | Bacteria Proteobacteria Gammaproteobacteria Enterobacteriales Enterobacteriaceae *Salmonella enterica subsp. enterica serovar* Typhimurium str. U288 |
| Sen-988-1 | Bacteria Proteobacteria Gammaproteobacteria Enterobacteriales Enterobacteriaceae *Salmonella enterica subsp. enterica serovar* Typhimurium str. UK-1 |
| Sen-989-1 | Bacteria Proteobacteria Gammaproteobacteria Enterobacteriales Enterobacteriaceae *Salmonella enterica subsp. enterica serovar* Typhimurium str. USDA-ARS-USMARC-1899 |
| Sen-990-1 | Bacteria Proteobacteria Gammaproteobacteria Enterobacteriales Enterobacteriaceae *Salmonella enterica subsp. enterica serovar* Typhimurium var. 5- str. CFSAN001921 |
| Sen-991-1 | Bacteria Proteobacteria Gammaproteobacteria Enterobacteriales Enterobacteriaceae *Salmonella enterica subsp. enterica serovar* Typhimurium var. 5- str. CFSAN004345 |
| Sen-992-1 | Bacteria Proteobacteria Gammaproteobacteria Enterobacteriales Enterobacteriaceae *Salmonella enterica subsp. enterica serovar* Typhimurium var. Copenhagen str. 0084 |
| Sen-993-1 | Bacteria Proteobacteria Gammaproteobacteria Enterobacteriales Enterobacteriaceae *Salmonella enterica subsp. enterica serovar* Uganda str. R8-3404 |
| Sen-994-1 | Bacteria Proteobacteria Gammaproteobacteria Enterobacteriales Enterobacteriaceae *Salmonella enterica subsp. enterica serovar* Urbana str. ATCC 9261 |
| Sen-995-1 | Bacteria Proteobacteria Gammaproteobacteria Enterobacteriales Enterobacteriaceae *Salmonella enterica subsp. enterica serovar* Worthington str. ATCC 9607 |
| Sen-996-1 | Bacteria Proteobacteria Gammaproteobacteria Enterobacteriales Enterobacteriaceae Salmonella Salmonella enterica subsp. entericaserovar Agona *Salmonella enterica subsp. enterica serovar* Agona str. SL483 |
| Sen-997-1 | Bacteria Proteobacteria Gammaproteobacteria Enterobacteriales Enterobacteriaceae Salmonella Salmonella enterica subsp. entericaserovar Choleraesuis *Salmonella enterica subsp. enterica serovar* Choleraesuis str. SCSA50 |
| Sen-998-1 | Bacteria Proteobacteria Gammaproteobacteria Enterobacteriales Enterobacteriaceae Salmonella Salmonella enterica subsp. entericaserovar Dublin *Salmonella enterica subsp. enterica serovar* Dublin str. CT_02021853 |
| Sen-999-1 | Bacteria Proteobacteria Gammaproteobacteria Enterobacteriales Enterobacteriaceae Salmonella Salmonella enterica subsp. entericaserovar Gallinarum/pullorum *Salmonella enterica subsp. enterica serovar* Gallinarum/pullorum str. RKS5078 |
| Sen-1000-1 | Bacteria Proteobacteria Gammaproteobacteria Enterobacteriales Enterobacteriaceae Salmonella Salmonella enterica subsp. entericaserovar Gallinarum *Salmonella enterica subsp. enterica serovar* Gallinarum str. 287/91 |
| Sen-1001-1 | Bacteria Proteobacteria Gammaproteobacteria Enterobacteriales Enterobacteriaceae Salmonella Salmonella enterica subsp. entericaserovar Hadar *Salmonella enterica subsp. enterica serovar* Hadar str. RI_05P066 |
| Sen-1002-1 | Bacteria Proteobacteria Gammaproteobacteria Enterobacteriales Enterobacteriaceae Salmonella Salmonella enterica subsp. entericaserovar Heidelberg *Salmonella enterica subsp. enterica serovar* Heidelberg str. B182 |
| Sen-1003-1 | Bacteria Proteobacteria Gammaproteobacteria Enterobacteriales Enterobacteriaceae Salmonella Salmonella enterica subsp. entericaserovar Heidelberg *Salmonella enterica subsp. enterica serovar* Heidelberg str. SL476 |
| Sen-1004-1 | Bacteria Proteobacteria Gammaproteobacteria Enterobacteriales Enterobacteriaceae Salmonella Salmonella enterica subsp. entericaserovar Heidelberg *Salmonella enterica subsp. enterica serovar* Heidelberg str. SL486 |
| Sen-1005-1 | Bacteria Proteobacteria Gammaproteobacteria Enterobacteriales Enterobacteriaceae Salmonella Salmonella enterica subsp. entericaserovar Kentucky *Salmonella enterica subsp. enterica serovar* Kentucky str. CDC 191 |
| Sen-1006-1 | Bacteria Proteobacteria Gammaproteobacteria Enterobacteriales Enterobacteriaceae Salmonella Salmonella enterica subsp. entericaserovar Kentucky *Salmonella enterica subsp. enterica serovar* Kentucky str. CVM29188 |
| Sen-265-1 to Sen-265-2 | Bacteria Proteobacteria Gammaproteobacteria Enterobacteriales Enterobacteriaceae Salmonella Salmonella enterica subsp. entericaserovar Newport *Salmonella enterica subsp. enterica serovar* Newport str. SL254 |
| Sen-1007-1 | Bacteria Proteobacteria Gammaproteobacteria Enterobacteriales Enterobacteriaceae Salmonella Salmonella enterica subsp. entericaserovar Newport *Salmonella enterica subsp. enterica serovar* Newport str. SL317 |
| Sen-1008-1 | Bacteria Proteobacteria Gammaproteobacteria Enterobacteriales Enterobacteriaceae Salmonella Salmonella enterica subsp. entericaserovar Paratyphi B *Salmonella enterica subsp. enterica serovar* Paratyphi B str. SPB7 |

| | |
|---|---|
| Sen-1009-1 | Bacteria Proteobacteria Gammaproteobacteria Enterobacteriales Enterobacteriaceae Salmonella Salmonella enterica subsp. entericaserovar Saintpaul *Salmonella enterica subsp. enterica serovar* Saintpaul str. SARA23 |
| Sen-1010-1 | Bacteria Proteobacteria Gammaproteobacteria Enterobacteriales Enterobacteriaceae Salmonella Salmonella enterica subsp. entericaserovar Saintpaul *Salmonella enterica subsp. enterica serovar* Saintpaul str. SARA29 |
| Sen-1011-1 | Bacteria Proteobacteria Gammaproteobacteria Enterobacteriales Enterobacteriaceae Salmonella Salmonella enterica subsp. entericaserovar Typhi *Salmonella enterica subsp. enterica serovar* Typhi str. CT18 |
| Sen-1012-1 | Bacteria Proteobacteria Gammaproteobacteria Enterobacteriales Enterobacteriaceae Salmonella Salmonella enterica subsp. entericaserovar Typhi *Salmonella enterica subsp. enterica serovar* Typhi str. E00-7866 |
| Sen-1013-1 | Bacteria Proteobacteria Gammaproteobacteria Enterobacteriales Enterobacteriaceae Salmonella Salmonella enterica subsp. entericaserovar Typhi *Salmonella enterica subsp. enterica serovar* Typhi str. Ty2 |
| Sen-1014-1 | Bacteria Proteobacteria Gammaproteobacteria Enterobacteriales Enterobacteriaceae Salmonella Salmonella enterica subsp. entericaserovar Typhimurium *Salmonella enterica subsp. enterica serovar* Typhimurium str. LT2 |
| Sen-1015-1 | Bacteria Proteobacteria Gammaproteobacteria Enterobacteriales Enterobacteriaceae Salmonella Salmonella enterica subsp. entericaserovar Virchow *Salmonella enterica subsp. enterica serovar* Virchow str. SL491 |
| Sen-1016-1 to Sen-1016-3 | Bacteria Proteobacteria Gammaproteobacteria Enterobacteriales Enterobacteriaceae *Salmonella enterica subsp. houtenae* |
| Sen-1017-1 | Bacteria Proteobacteria Gammaproteobacteria Enterobacteriales Enterobacteriaceae *Salmonella enterica subsp. houtenae serovar* 16:z4,z32:– str. RKS3027 |
| Sen-1018-1 | Bacteria Proteobacteria Gammaproteobacteria Enterobacteriales Enterobacteriaceae *Salmonella enterica subsp. houtenae serovar* 50:g,z51:- str. 01-0133 |
| Sen-1019-1 | Bacteria Proteobacteria Gammaproteobacteria Enterobacteriales Enterobacteriaceae *Salmonella enterica subsp. houtenae str.* ATCC BAA-1581 |
| Sen-1020-1 | Bacteria Proteobacteria Gammaproteobacteria Enterobacteriales Enterobacteriaceae *Salmonella enterica subsp. indica serovar* 6,14,25:z10:1,(2),7 str. 1121 |
| Sen-1021-1 | Bacteria Proteobacteria Gammaproteobacteria Enterobacteriales Enterobacteriaceae *Salmonella enterica subsp. salamae serovar* 58:l,z13,z28:z6 str. 00-0163 |
| Sen-1022-1 | Bacteria Proteobacteria Gammaproteobacteria Enterobacteriales Enterobacteriaceae *Salmonella enterica subsp. salamae str.* 3588/07 |
| Sfo-4-1 | Bacteria Proteobacteria Gammaproteobacteria Enterobacteriales Enterobacteriaceae *Serratia fonticola* AU-AP2C |
| Sli-3-1 | Bacteria Proteobacteria Gammaproteobacteria Enterobacteriales Enterobacteriaceae *Serratia liquefaciens* ATCC 27592 |
| Sma-1-1 to Sma-1-4 | Bacteria Proteobacteria Gammaproteobacteria Enterobacteriales Enterobacteriaceae *Serratia marcescens* |
| Sma-21-1 | Bacteria Proteobacteria Gammaproteobacteria Enterobacteriales Enterobacteriaceae *Serratia marcescens* EGD-HP20 |
| Sma-22-1 | Bacteria Proteobacteria Gammaproteobacteria Enterobacteriales Enterobacteriaceae *Serratia marcescens* FGI94 |
| Sma-23-1 | Bacteria Proteobacteria Gammaproteobacteria Enterobacteriales Enterobacteriaceae *Serratia marcescens* LCT-SM213 |
| Sma-24-1 | Bacteria Proteobacteria Gammaproteobacteria Enterobacteriales Enterobacteriaceae *Serratia marcescens* VGH107 |
| Sma-25-1 | Bacteria Proteobacteria Gammaproteobacteria Enterobacteriales Enterobacteriaceae *Serratia marcescens* W2.3 |
| Sma-26-1 | Bacteria Proteobacteria Gammaproteobacteria Enterobacteriales Enterobacteriaceae *Serratia marcescens* WW4 |
| Sod-1-1 | Bacteria Proteobacteria Gammaproteobacteria Enterobacteriales Enterobacteriaceae *Serratia odorifera* DSM 4582 |
| Spl-2-1 | Bacteria Proteobacteria Gammaproteobacteria Enterobacteriales Enterobacteriaceae *Serratia plymuthica* 4Rx13 |
| Spl-3-1 | Bacteria Proteobacteria Gammaproteobacteria Enterobacteriales Enterobacteriaceae *Serratia plymuthica* A30 |
| Spl-4-1 | Bacteria Proteobacteria Gammaproteobacteria Enterobacteriales Enterobacteriaceae *Serratia plymuthica* AS9 |
| Spl-5-1 | Bacteria Proteobacteria Gammaproteobacteria Enterobacteriales Enterobacteriaceae *Serratia plymuthica* PRI-2C |
| Spl-6-1 | Bacteria Proteobacteria Gammaproteobacteria Enterobacteriales Enterobacteriaceae *Serratia plymuthica* S13 |
| Spr-2-1 | Bacteria Proteobacteria Gammaproteobacteria Enterobacteriales Enterobacteriaceae *Serratia proteamaculans* 568 |
| Ssp-10-1 | Bacteria Proteobacteria Gammaproteobacteria Enterobacteriales Enterobacteriaceae *Serratia sp.* AS12 |
| Ssp-11-1 | Bacteria Proteobacteria Gammaproteobacteria Enterobacteriales Enterobacteriaceae *Serratia sp.* AS13 |
| Ssp-12-1 | Bacteria Proteobacteria Gammaproteobacteria Enterobacteriales Enterobacteriaceae *Serratia sp.* ATCC 39006 |
| Ssp-13-1 | Bacteria Proteobacteria Gammaproteobacteria Enterobacteriales Enterobacteriaceae *Serratia sp.* M24T3 |
| Ssp-14-1 | Bacteria Proteobacteria Gammaproteobacteria Enterobacteriales Enterobacteriaceae *Serratia sp.* S4 |
| Ssy-2-1 | Bacteria Proteobacteria Gammaproteobacteria Enterobacteriales Enterobacteriaceae Serratia *Serratia symbiotica str.* Tucson |
| Sbo-2-1 to Sbo-2-10 | Bacteria Proteobacteria Gammaproteobacteria Enterobacteriales Enterobacteriaceae *Shigella boydii* |
| Sbo-5-1 | Bacteria Proteobacteria Gammaproteobacteria Enterobacteriales Enterobacteriaceae *Shigella boydii* 08-0009 |
| Sbo-6-1 | Bacteria Proteobacteria Gammaproteobacteria Enterobacteriales Enterobacteriaceae *Shigella boydii* 08-0280 |
| Sbo-7-1 | Bacteria Proteobacteria Gammaproteobacteria Enterobacteriales Enterobacteriaceae *Shigella boydii* 08-2671 |
| Sbo-8-1 | Bacteria Proteobacteria Gammaproteobacteria Enterobacteriales Enterobacteriaceae *Shigella boydii* 08-2675 |
| Sbo-9-1 | Bacteria Proteobacteria Gammaproteobacteria Enterobacteriales Enterobacteriaceae *Shigella boydii* 08-6341 |
| Sbo-10-1 | Bacteria Proteobacteria Gammaproteobacteria Enterobacteriales Enterobacteriaceae *Shigella boydii* 09-0344 |
| Sbo-11-1 | Bacteria Proteobacteria Gammaproteobacteria Enterobacteriales Enterobacteriaceae *Shigella boydii* 248-1B |
| Sbo-12-1 | Bacteria Proteobacteria Gammaproteobacteria Enterobacteriales Enterobacteriaceae *Shigella boydii* 4444-74 |
| Sbo-13-1 | Bacteria Proteobacteria Gammaproteobacteria Enterobacteriales Enterobacteriaceae *Shigella boydii* 5216-82 |
| Sbo-14-1 | Bacteria Proteobacteria Gammaproteobacteria Enterobacteriales Enterobacteriaceae *Shigella boydii* ATCC 9905 |
| Sbo-15-1 | Bacteria Proteobacteria Gammaproteobacteria Enterobacteriales Enterobacteriaceae *Shigella boydii* CDC 3083-94 |
| Sbo-16-1 | Bacteria Proteobacteria Gammaproteobacteria Enterobacteriales Enterobacteriaceae *Shigella boydii* S6614 |
| Sbo-17-1 | Bacteria Proteobacteria Gammaproteobacteria Enterobacteriales Enterobacteriaceae *Shigella boydii* S7334 |
| Sbo-18-1 | Bacteria Proteobacteria Gammaproteobacteria Enterobacteriales Enterobacteriaceae *Shigella boydii* Sb227 |
| Sdy-3-1 | Bacteria Proteobacteria Gammaproteobacteria Enterobacteriales Enterobacteriaceae *Shigella dysenteriae* 1012 |
| Sdy-4-1 | Bacteria Proteobacteria Gammaproteobacteria Enterobacteriales Enterobacteriaceae *Shigella dysenteriae* 225-75 |
| Sdy-5-1 | Bacteria Proteobacteria Gammaproteobacteria Enterobacteriales Enterobacteriaceae *Shigella dysenteriae* 4 |
| Sdy-6-1 | Bacteria Proteobacteria Gammaproteobacteria Enterobacteriales Enterobacteriaceae *Shigella dysenteriae* CDC 74-1112 |

| | |
|---|---|
| Sdy-7-1 | Bacteria Proteobacteria Gammaproteobacteria Enterobacteriales Enterobacteriaceae *Shigella dysenteriae* S6554 |
| Sdy-8-1 | Bacteria Proteobacteria Gammaproteobacteria Enterobacteriales Enterobacteriaceae *Shigella dysenteriae* Sd197 |
| Sdy-9-1 | Bacteria Proteobacteria Gammaproteobacteria Enterobacteriales Enterobacteriaceae *Shigella dysenteriae* WRSd3 |
| Sdy-10-1 | Bacteria Proteobacteria Gammaproteobacteria Enterobacteriales Enterobacteriaceae *Shigella dysenteriae* WRSd5 |
| Sfl-1-1 to Sfl-1-2 | Bacteria Proteobacteria Gammaproteobacteria Enterobacteriales Enterobacteriaceae *Shigella flexneri* |
| Sfl-52-1 | Bacteria Proteobacteria Gammaproteobacteria Enterobacteriales Enterobacteriaceae *Shigella flexneri* 2001020 |
| Sfl-53-1 | Bacteria Proteobacteria Gammaproteobacteria Enterobacteriales Enterobacteriaceae *Shigella flexneri* 2002017 |
| Sfl-54-1 | Bacteria Proteobacteria Gammaproteobacteria Enterobacteriales Enterobacteriaceae *Shigella flexneri* 2002028 |
| Sfl-55-1 | Bacteria Proteobacteria Gammaproteobacteria Enterobacteriales Enterobacteriaceae *Shigella flexneri* 2002106 |
| Sfl-56-1 | Bacteria Proteobacteria Gammaproteobacteria Enterobacteriales Enterobacteriaceae *Shigella flexneri* 2003035 |
| Sfl-57-1 | Bacteria Proteobacteria Gammaproteobacteria Enterobacteriales Enterobacteriaceae *Shigella flexneri* 2003036 |
| Sfl-58-1 | Bacteria Proteobacteria Gammaproteobacteria Enterobacteriales Enterobacteriaceae *Shigella flexneri* 2005002 |
| Sfl-59-1 | Bacteria Proteobacteria Gammaproteobacteria Enterobacteriales Enterobacteriaceae *Shigella flexneri* 2005025 |
| Sfl-60-1 | Bacteria Proteobacteria Gammaproteobacteria Enterobacteriales Enterobacteriaceae *Shigella flexneri* 2005184 |
| Sfl-61-1 | Bacteria Proteobacteria Gammaproteobacteria Enterobacteriales Enterobacteriaceae *Shigella flexneri* 2930-71 |
| Sfl-25-1 to Sfl-25-14 | Bacteria Proteobacteria Gammaproteobacteria Enterobacteriales Enterobacteriaceae *Shigella flexneri* 2a |
| Sfl-62-1 | Bacteria Proteobacteria Gammaproteobacteria Enterobacteriales Enterobacteriaceae *Shigella flexneri* 2a str. 2457T |
| Sfl-63-1 | Bacteria Proteobacteria Gammaproteobacteria Enterobacteriales Enterobacteriaceae *Shigella flexneri* 2a str. 301 |
| Sfl-64-1 to Sfl-64-2 | Bacteria Proteobacteria Gammaproteobacteria Enterobacteriales Enterobacteriaceae *Shigella flexneri* 3a |
| Sfl-65-1 | Bacteria Proteobacteria Gammaproteobacteria Enterobacteriales Enterobacteriaceae *Shigella flexneri* 4343-70 |
| Sfl-66-1 | Bacteria Proteobacteria Gammaproteobacteria Enterobacteriales Enterobacteriaceae *Shigella flexneri* 5 str. 8401 |
| Sfl-67-1 | Bacteria Proteobacteria Gammaproteobacteria Enterobacteriales Enterobacteriaceae *Shigella flexneri* 51581 |
| Sfl-68-1 | Bacteria Proteobacteria Gammaproteobacteria Enterobacteriales Enterobacteriaceae *Shigella flexneri* 5a str. M90T |
| Sfl-69-1 | Bacteria Proteobacteria Gammaproteobacteria Enterobacteriales Enterobacteriaceae *Shigella flexneri* 6 |
| Sfl-70-1 | Bacteria Proteobacteria Gammaproteobacteria Enterobacteriales Enterobacteriaceae *Shigella flexneri* CCH060 |
| Sfl-71-1 | Bacteria Proteobacteria Gammaproteobacteria Enterobacteriales Enterobacteriaceae *Shigella flexneri* CDC 796-83 |
| Sfl-72-1 | Bacteria Proteobacteria Gammaproteobacteria Enterobacteriales Enterobacteriaceae *Shigella flexneri* G1663 |
| Sfl-73-1 | Bacteria Proteobacteria Gammaproteobacteria Enterobacteriales Enterobacteriaceae *Shigella flexneri* MT1457 |
| Sfl-74-1 | Bacteria Proteobacteria Gammaproteobacteria Enterobacteriales Enterobacteriaceae *Shigella flexneri* S5644 |
| Sfl-75-1 | Bacteria Proteobacteria Gammaproteobacteria Enterobacteriales Enterobacteriaceae *Shigella flexneri* S5717 |
| Sfl-76-1 | Bacteria Proteobacteria Gammaproteobacteria Enterobacteriales Enterobacteriaceae *Shigella flexneri* S6162 |
| Sfl-77-1 | Bacteria Proteobacteria Gammaproteobacteria Enterobacteriales Enterobacteriaceae *Shigella flexneri* S6585 |
| Sfl-78-1 | Bacteria Proteobacteria Gammaproteobacteria Enterobacteriales Enterobacteriaceae *Shigella flexneri* S6678 |
| Sfl-79-1 | Bacteria Proteobacteria Gammaproteobacteria Enterobacteriales Enterobacteriaceae *Shigella flexneri* S6764 |
| Sfl-80-1 | Bacteria Proteobacteria Gammaproteobacteria Enterobacteriales Enterobacteriaceae *Shigella flexneri* S7737 |
| Sfl-81-1 | Bacteria Proteobacteria Gammaproteobacteria Enterobacteriales Enterobacteriaceae *Shigella flexneri* Shi06GS02 |
| Sfl-82-1 | Bacteria Proteobacteria Gammaproteobacteria Enterobacteriales Enterobacteriaceae *Shigella flexneri* Shi06HN006 |
| Sfl-83-1 | Bacteria Proteobacteria Gammaproteobacteria Enterobacteriales Enterobacteriaceae *Shigella flexneri* Shi06HN344 |
| Sfl-84-1 | Bacteria Proteobacteria Gammaproteobacteria Enterobacteriales Enterobacteriaceae *Shigella flexneri* Shi06HN347 |
| Sfl-85-1 | Bacteria Proteobacteria Gammaproteobacteria Enterobacteriales Enterobacteriaceae *Shigella flexneri* VA-6 |
| Sfl-51-1 to Sfl-51-2 | Bacteria Proteobacteria Gammaproteobacteria Enterobacteriales Enterobacteriaceae *Shigella flexneri* Y |
| Sso-1-1 to Sso-1-540 | Bacteria Proteobacteria Gammaproteobacteria Enterobacteriales Enterobacteriaceae *Shigella sonnei* |
| Sso-3-1 | Bacteria Proteobacteria Gammaproteobacteria Enterobacteriales Enterobacteriaceae *Shigella sonnei* 08-7761 |
| Sso-4-1 | Bacteria Proteobacteria Gammaproteobacteria Enterobacteriales Enterobacteriaceae *Shigella sonnei* 08-7765 |
| Sso-5-1 | Bacteria Proteobacteria Gammaproteobacteria Enterobacteriales Enterobacteriaceae *Shigella sonnei* 09-1032 |
| Sso-6-1 | Bacteria Proteobacteria Gammaproteobacteria Enterobacteriales Enterobacteriaceae *Shigella sonnei* 09-2245 |
| Sso-7-1 | Bacteria Proteobacteria Gammaproteobacteria Enterobacteriales Enterobacteriaceae *Shigella sonnei* 09-4962 |
| Sso-8-1 | Bacteria Proteobacteria Gammaproteobacteria Enterobacteriales Enterobacteriaceae *Shigella sonnei* 3233-85 |
| Sso-9-1 | Bacteria Proteobacteria Gammaproteobacteria Enterobacteriales Enterobacteriaceae *Shigella sonnei* 4822-66 |
| Sso-10-1 | Bacteria Proteobacteria Gammaproteobacteria Enterobacteriales Enterobacteriaceae *Shigella sonnei* 53G |
| Sso-11-1 | Bacteria Proteobacteria Gammaproteobacteria Enterobacteriales Enterobacteriaceae *Shigella sonnei* S6513 |
| Sso-12-1 | Bacteria Proteobacteria Gammaproteobacteria Enterobacteriales Enterobacteriaceae *Shigella sonnei* Ss046 |
| Sso-13-1 | Bacteria Proteobacteria Gammaproteobacteria Enterobacteriales Enterobacteriaceae *Shigella sonnei str.* Moseley |
| Ssp-15-1 | Bacteria Proteobacteria Gammaproteobacteria Enterobacteriales Enterobacteriaceae *Shigella sp.* D9 |
| Sbl-1-1 | Bacteria Proteobacteria Gammaproteobacteria Enterobacteriales Enterobacteriaceae *Shimwellia blattae* DSM 4481 = NBRC 105725 |
| Sgl-1-1 | Bacteria Proteobacteria Gammaproteobacteria Enterobacteriales Enterobacteriaceae *Sodalis glossinidius str. 'morsitans'* |
| Xbo-11-1 | Bacteria Proteobacteria Gammaproteobacteria Enterobacteriales Enterobacteriaceae *Xenorhabdus bovienii* SS-2004 |
| Xne-3-1 | Bacteria Proteobacteria Gammaproteobacteria Enterobacteriales Enterobacteriaceae *Xenorhabdus nematophila* ATCC 19061 |
| Xne-4-1 | Bacteria Proteobacteria Gammaproteobacteria Enterobacteriales Enterobacteriaceae *Xenorhabdus nematophila* F1 |
| Xne-5-1 to Xne-5-2 | Bacteria Proteobacteria Gammaproteobacteria Enterobacteriales Enterobacteriaceae *Xenorhabdus nematophila str.* Anatoliense |
| Xne-6-1 to Xne-6-2 | Bacteria Proteobacteria Gammaproteobacteria Enterobacteriales Enterobacteriaceae *Xenorhabdus nematophila str.* Websteri |
| Yal-4-1 | Bacteria Proteobacteria Gammaproteobacteria Enterobacteriales Enterobacteriaceae *Yersinia aldovae* ATCC 35236 |
| Ybe-2-1 | Bacteria Proteobacteria Gammaproteobacteria Enterobacteriales Enterobacteriaceae *Yersinia bercovieri* ATCC 43970 |
| Yen-8-1 | Bacteria Proteobacteria Gammaproteobacteria Enterobacteriales Enterobacteriaceae *Yersinia enterocolitica (type* O:3) str. YE12/03 |
| Yen-9-1 | Bacteria Proteobacteria Gammaproteobacteria Enterobacteriales Enterobacteriaceae *Yersinia enterocolitica (type* O:5,27) str. YE149/02 |

| | |
|---|---|
| Yen-10-1 | Bacteria Proteobacteria Gammaproteobacteria Enterobacteriales Enterobacteriaceae *Yersinia enterocolitica (type* O:9) str. YE212/02 |
| Yen-11-1 | Bacteria Proteobacteria Gammaproteobacteria Enterobacteriales Enterobacteriaceae *Yersinia enterocolitica* IP 10393 |
| Yen-12-1 | Bacteria Proteobacteria Gammaproteobacteria Enterobacteriales Enterobacteriaceae *Yersinia enterocolitica* IP2222 |
| Yen-13-1 | Bacteria Proteobacteria Gammaproteobacteria Enterobacteriales Enterobacteriaceae *Yersinia enterocolitica* NFO |
| Yen-14-1 | Bacteria Proteobacteria Gammaproteobacteria Enterobacteriales Enterobacteriaceae *Yersinia enterocolitica subsp. enterocolitica* 8081 |
| Yen-15-1 | Bacteria Proteobacteria Gammaproteobacteria Enterobacteriales Enterobacteriaceae *Yersinia enterocolitica subsp. enterocolitica* WA-314 |
| Yen-16-1 | Bacteria Proteobacteria Gammaproteobacteria Enterobacteriales Enterobacteriaceae *Yersinia enterocolitica subsp. palearctica* 105.5R(r) |
| Yen-17-1 | Bacteria Proteobacteria Gammaproteobacteria Enterobacteriales Enterobacteriaceae *Yersinia enterocolitica subsp. palearctica* PhRBD_Ye1 |
| Yen-18-1 | Bacteria Proteobacteria Gammaproteobacteria Enterobacteriales Enterobacteriaceae *Yersinia enterocolitica subsp. palearctica* Y11 |
| Yen-19-1 | Bacteria Proteobacteria Gammaproteobacteria Enterobacteriales Enterobacteriaceae *Yersinia enterocolitica subsp. palearctica* YE-149 |
| Yen-20-1 | Bacteria Proteobacteria Gammaproteobacteria Enterobacteriales Enterobacteriaceae *Yersinia enterocolitica subsp. palearctica* YE-150 |
| Yen-21-1 | Bacteria Proteobacteria Gammaproteobacteria Enterobacteriales Enterobacteriaceae *Yersinia enterocolitica subsp. palearctica* YE-P1 |
| Yen-22-1 | Bacteria Proteobacteria Gammaproteobacteria Enterobacteriales Enterobacteriaceae *Yersinia enterocolitica subsp. palearctica* YE-P4 |
| Yfr-2-1 to Yfr-2-2 | Bacteria Proteobacteria Gammaproteobacteria Enterobacteriales Enterobacteriaceae *Yersinia frederiksenii* ATCC 33641 |
| Yin-2-1 | Bacteria Proteobacteria Gammaproteobacteria Enterobacteriales Enterobacteriaceae *Yersinia intermedia* ATCC 29909 |
| Ykr-2-1 | Bacteria Proteobacteria Gammaproteobacteria Enterobacteriales Enterobacteriaceae *Yersinia kristensenii* ATCC 33638 |
| Ymo-2-1 | Bacteria Proteobacteria Gammaproteobacteria Enterobacteriales Enterobacteriaceae *Yersinia mollaretii* ATCC 43969 |
| Ype-116-1 | Bacteria Proteobacteria Gammaproteobacteria Enterobacteriales Enterobacteriaceae *Yersinia pestis* 113 |
| Ype-117-1 | Bacteria Proteobacteria Gammaproteobacteria Enterobacteriales Enterobacteriaceae *Yersinia pestis* 2330 |
| Ype-118-1 | Bacteria Proteobacteria Gammaproteobacteria Enterobacteriales Enterobacteriaceae *Yersinia pestis* 2501 |
| Ype-119-1 | Bacteria Proteobacteria Gammaproteobacteria Enterobacteriales Enterobacteriaceae *Yersinia pestis* 2504 |
| Ype-120-1 | Bacteria Proteobacteria Gammaproteobacteria Enterobacteriales Enterobacteriaceae *Yersinia pestis* 2506 |
| Ype-121-1 | Bacteria Proteobacteria Gammaproteobacteria Enterobacteriales Enterobacteriaceae *Yersinia pestis* 30017 |
| Ype-122-1 | Bacteria Proteobacteria Gammaproteobacteria Enterobacteriales Enterobacteriaceae *Yersinia pestis* 42013 |
| Ype-123-1 | Bacteria Proteobacteria Gammaproteobacteria Enterobacteriales Enterobacteriaceae *Yersinia pestis* 42091 |
| Ype-124-1 | Bacteria Proteobacteria Gammaproteobacteria Enterobacteriales Enterobacteriaceae *Yersinia pestis* 5761 |
| Ype-125-1 | Bacteria Proteobacteria Gammaproteobacteria Enterobacteriales Enterobacteriaceae *Yersinia pestis* 7 |
| Ype-126-1 | Bacteria Proteobacteria Gammaproteobacteria Enterobacteriales Enterobacteriaceae *Yersinia pestis* 735 |
| Ype-127-1 | Bacteria Proteobacteria Gammaproteobacteria Enterobacteriales Enterobacteriaceae *Yersinia pestis* 780441 |
| Ype-128-1 | Bacteria Proteobacteria Gammaproteobacteria Enterobacteriales Enterobacteriaceae *Yersinia pestis* 91 |
| Ype-29-1 to Ype-29-2 | Bacteria Proteobacteria Gammaproteobacteria Enterobacteriales Enterobacteriaceae *Yersinia pestis* A1122 |
| Ype-129-1 | Bacteria Proteobacteria Gammaproteobacteria Enterobacteriales Enterobacteriaceae *Yersinia pestis* A1956001 |
| Ype-31-1 to Ype-31-2 | Bacteria Proteobacteria Gammaproteobacteria Enterobacteriales Enterobacteriaceae *Yersinia pestis* Angola |
| Ype-32-1 to Ype-32-2 | Bacteria Proteobacteria Gammaproteobacteria Enterobacteriales Enterobacteriaceae *Yersinia pestis* Antiqua |
| Ype-217-1 | Bacteria Proteobacteria Gammaproteobacteria Enterobacteriales Enterobacteriaceae *Yersinia pestis biovar* Antiqua str. B42003004 |
| Ype-218-1 to Ype-218-2 | Bacteria Proteobacteria Gammaproteobacteria Enterobacteriales Enterobacteriaceae *Yersinia pestis biovar* Antiqua str. E1979001 |
| Ype-219-1 | Bacteria Proteobacteria Gammaproteobacteria Enterobacteriales Enterobacteriaceae *Yersinia pestis biovar* Antiqua str. UG05-0454 |
| Ype-220-1 | Bacteria Proteobacteria Gammaproteobacteria Enterobacteriales Enterobacteriaceae *Yersinia pestis biovar* Mediaevalis str. K1973002 |
| Ype-221-1 | Bacteria Proteobacteria Gammaproteobacteria Enterobacteriales Enterobacteriaceae *Yersinia pestis biovar* Medievalis str. Harbin 35 |
| Ype-222-1 | Bacteria Proteobacteria Gammaproteobacteria Enterobacteriales Enterobacteriaceae *Yersinia pestis biovar* Microtus str. 91001 |
| Ype-223-1 | Bacteria Proteobacteria Gammaproteobacteria Enterobacteriales Enterobacteriaceae *Yersinia pestis biovar* Orientalis str. F1991016 |
| Ype-225-1 | Bacteria Proteobacteria Gammaproteobacteria Enterobacteriales Enterobacteriaceae *Yersinia pestis biovar* Orientalis str. India 195 |
| Ype-224-1 | Bacteria Proteobacteria Gammaproteobacteria Enterobacteriales Enterobacteriaceae *Yersinia pestis biovar* Orientalis str. IP275 |
| Ype-226-1 | Bacteria Proteobacteria Gammaproteobacteria Enterobacteriales Enterobacteriaceae *Yersinia pestis biovar* Orientalis str. MG05-1020 |
| Ype-227-1 | Bacteria Proteobacteria Gammaproteobacteria Enterobacteriales Enterobacteriaceae *Yersinia pestis biovar* Orientalis str. PEXU2 |
| Ype-130-1 | Bacteria Proteobacteria Gammaproteobacteria Enterobacteriales Enterobacteriaceae *Yersinia pestis* C1976001 |
| Ype-131-1 | Bacteria Proteobacteria Gammaproteobacteria Enterobacteriales Enterobacteriaceae *Yersinia pestis* C1989001 |
| Ype-132-1 | Bacteria Proteobacteria Gammaproteobacteria Enterobacteriales Enterobacteriaceae *Yersinia pestis* CA88-4125 |
| Ype-133-1 | Bacteria Proteobacteria Gammaproteobacteria Enterobacteriales Enterobacteriaceae *Yersinia pestis* CMCC125002 |
| Ype-134-1 | Bacteria Proteobacteria Gammaproteobacteria Enterobacteriales Enterobacteriaceae *Yersinia pestis* CMCC348002 |
| Ype-135-1 | Bacteria Proteobacteria Gammaproteobacteria Enterobacteriales Enterobacteriaceae *Yersinia pestis* CMCC49003 |
| Ype-136-1 | Bacteria Proteobacteria Gammaproteobacteria Enterobacteriales Enterobacteriaceae *Yersinia pestis* CMCC64001 |
| Ype-137-1 | Bacteria Proteobacteria Gammaproteobacteria Enterobacteriales Enterobacteriaceae *Yersinia pestis* CMCC8211 |
| Ype-138-1 | Bacteria Proteobacteria Gammaproteobacteria Enterobacteriales Enterobacteriaceae *Yersinia pestis* CMCC87001 |
| Ype-139-1 | Bacteria Proteobacteria Gammaproteobacteria Enterobacteriales Enterobacteriaceae *Yersinia pestis* CMCC91090 |
| Ype-140-1 | Bacteria Proteobacteria Gammaproteobacteria Enterobacteriales Enterobacteriaceae *Yersinia pestis* CMCC93014 |
| Ype-141-1 | Bacteria Proteobacteria Gammaproteobacteria Enterobacteriales Enterobacteriaceae *Yersinia pestis* CMCC96001 |
| Ype-142-1 | Bacteria Proteobacteria Gammaproteobacteria Enterobacteriales Enterobacteriaceae *Yersinia pestis* CO92 |
| Ype-143-1 | Bacteria Proteobacteria Gammaproteobacteria Enterobacteriales Enterobacteriaceae *Yersinia pestis* D106004 |
| Ype-144-1 | Bacteria Proteobacteria Gammaproteobacteria Enterobacteriales Enterobacteriaceae *Yersinia pestis* D182038 |
| Ype-145-1 | Bacteria Proteobacteria Gammaproteobacteria Enterobacteriales Enterobacteriaceae *Yersinia pestis* D1982001 |
| Ype-146-1 | Bacteria Proteobacteria Gammaproteobacteria Enterobacteriales Enterobacteriaceae *Yersinia pestis* G1996006 |
| Ype-147-1 | Bacteria Proteobacteria Gammaproteobacteria Enterobacteriales Enterobacteriaceae *Yersinia pestis* G1996010 |
| Ype-148-1 | Bacteria Proteobacteria Gammaproteobacteria Enterobacteriales Enterobacteriaceae *Yersinia pestis* G8786 |
| Ype-149-1 | Bacteria Proteobacteria Gammaproteobacteria Enterobacteriales Enterobacteriaceae *Yersinia pestis* H1959004 |

| | |
|---|---|
| Ype-150-1 | Bacteria Proteobacteria Gammaproteobacteria Enterobacteriales Enterobacteriaceae *Yersinia pestis* INS |
| Ype-151-1 | Bacteria Proteobacteria Gammaproteobacteria Enterobacteriales Enterobacteriaceae *Yersinia pestis* K11973002 |
| Ype-152-1 | Bacteria Proteobacteria Gammaproteobacteria Enterobacteriales Enterobacteriaceae *Yersinia pestis* KIM |
| Ype-153-1 | Bacteria Proteobacteria Gammaproteobacteria Enterobacteriales Enterobacteriaceae *Yersinia pestis* MGJZ11 |
| Ype-154-1 | Bacteria Proteobacteria Gammaproteobacteria Enterobacteriales Enterobacteriaceae *Yersinia pestis* Nepal516 |
| Ype-214-1 | Bacteria Proteobacteria Gammaproteobacteria Enterobacteriales Enterobacteriaceae *Yersinia pestis* Pestoides A |
| Ype-94-1 to Ype-94-2 | Bacteria Proteobacteria Gammaproteobacteria Enterobacteriales Enterobacteriaceae *Yersinia pestis* Pestoides F |
| Ype-155-1 | Bacteria Proteobacteria Gammaproteobacteria Enterobacteriales Enterobacteriaceae *Yersinia pestis* PY-01 |
| Ype-156-1 | Bacteria Proteobacteria Gammaproteobacteria Enterobacteriales Enterobacteriaceae *Yersinia pestis* PY-02 |
| Ype-157-1 | Bacteria Proteobacteria Gammaproteobacteria Enterobacteriales Enterobacteriaceae *Yersinia pestis* PY-03 |
| Ype-158-1 | Bacteria Proteobacteria Gammaproteobacteria Enterobacteriales Enterobacteriaceae *Yersinia pestis* PY-04 |
| Ype-159-1 | Bacteria Proteobacteria Gammaproteobacteria Enterobacteriales Enterobacteriaceae *Yersinia pestis* PY-05 |
| Ype-160-1 | Bacteria Proteobacteria Gammaproteobacteria Enterobacteriales Enterobacteriaceae *Yersinia pestis* PY-06 |
| Ype-161-1 | Bacteria Proteobacteria Gammaproteobacteria Enterobacteriales Enterobacteriaceae *Yersinia pestis* PY-07 |
| Ype-162-1 | Bacteria Proteobacteria Gammaproteobacteria Enterobacteriales Enterobacteriaceae *Yersinia pestis* PY-08 |
| Ype-163-1 | Bacteria Proteobacteria Gammaproteobacteria Enterobacteriales Enterobacteriaceae *Yersinia pestis* PY-09 |
| Ype-164-1 | Bacteria Proteobacteria Gammaproteobacteria Enterobacteriales Enterobacteriaceae *Yersinia pestis* PY-10 |
| Ype-165-1 | Bacteria Proteobacteria Gammaproteobacteria Enterobacteriales Enterobacteriaceae *Yersinia pestis* PY-100 |
| Ype-166-1 | Bacteria Proteobacteria Gammaproteobacteria Enterobacteriales Enterobacteriaceae *Yersinia pestis* PY-101 |
| Ype-167-1 | Bacteria Proteobacteria Gammaproteobacteria Enterobacteriales Enterobacteriaceae *Yersinia pestis* PY-102 |
| Ype-168-1 | Bacteria Proteobacteria Gammaproteobacteria Enterobacteriales Enterobacteriaceae *Yersinia pestis* PY-103 |
| Ype-169-1 | Bacteria Proteobacteria Gammaproteobacteria Enterobacteriales Enterobacteriaceae *Yersinia pestis* PY-11 |
| Ype-170-1 | Bacteria Proteobacteria Gammaproteobacteria Enterobacteriales Enterobacteriaceae *Yersinia pestis* PY-113 |
| Ype-171-1 | Bacteria Proteobacteria Gammaproteobacteria Enterobacteriales Enterobacteriaceae *Yersinia pestis* PY-12 |
| Ype-172-1 | Bacteria Proteobacteria Gammaproteobacteria Enterobacteriales Enterobacteriaceae *Yersinia pestis* PY-13 |
| Ype-173-1 | Bacteria Proteobacteria Gammaproteobacteria Enterobacteriales Enterobacteriaceae *Yersinia pestis* PY-14 |
| Ype-174-1 | Bacteria Proteobacteria Gammaproteobacteria Enterobacteriales Enterobacteriaceae *Yersinia pestis* PY-15 |
| Ype-175-1 | Bacteria Proteobacteria Gammaproteobacteria Enterobacteriales Enterobacteriaceae *Yersinia pestis* PY-16 |
| Ype-176-1 | Bacteria Proteobacteria Gammaproteobacteria Enterobacteriales Enterobacteriaceae *Yersinia pestis* PY-19 |
| Ype-177-1 | Bacteria Proteobacteria Gammaproteobacteria Enterobacteriales Enterobacteriaceae *Yersinia pestis* PY-25 |
| Ype-178-1 | Bacteria Proteobacteria Gammaproteobacteria Enterobacteriales Enterobacteriaceae *Yersinia pestis* PY-29 |
| Ype-179-1 | Bacteria Proteobacteria Gammaproteobacteria Enterobacteriales Enterobacteriaceae *Yersinia pestis* PY-32 |
| Ype-180-1 | Bacteria Proteobacteria Gammaproteobacteria Enterobacteriales Enterobacteriaceae *Yersinia pestis* PY-34 |
| Ype-181-1 | Bacteria Proteobacteria Gammaproteobacteria Enterobacteriales Enterobacteriaceae *Yersinia pestis* PY-36 |
| Ype-182-1 | Bacteria Proteobacteria Gammaproteobacteria Enterobacteriales Enterobacteriaceae *Yersinia pestis* PY-42 |
| Ype-183-1 | Bacteria Proteobacteria Gammaproteobacteria Enterobacteriales Enterobacteriaceae *Yersinia pestis* PY-45 |
| Ype-184-1 | Bacteria Proteobacteria Gammaproteobacteria Enterobacteriales Enterobacteriaceae *Yersinia pestis* PY-46 |
| Ype-185-1 | Bacteria Proteobacteria Gammaproteobacteria Enterobacteriales Enterobacteriaceae *Yersinia pestis* PY-47 |
| Ype-186-1 | Bacteria Proteobacteria Gammaproteobacteria Enterobacteriales Enterobacteriaceae *Yersinia pestis* PY-48 |
| Ype-187-1 | Bacteria Proteobacteria Gammaproteobacteria Enterobacteriales Enterobacteriaceae *Yersinia pestis* PY-52 |
| Ype-188-1 | Bacteria Proteobacteria Gammaproteobacteria Enterobacteriales Enterobacteriaceae *Yersinia pestis* PY-53 |
| Ype-189-1 | Bacteria Proteobacteria Gammaproteobacteria Enterobacteriales Enterobacteriaceae *Yersinia pestis* PY-54 |
| Ype-190-1 | Bacteria Proteobacteria Gammaproteobacteria Enterobacteriales Enterobacteriaceae *Yersinia pestis* PY-55 |
| Ype-191-1 | Bacteria Proteobacteria Gammaproteobacteria Enterobacteriales Enterobacteriaceae *Yersinia pestis* PY-56 |
| Ype-192-1 | Bacteria Proteobacteria Gammaproteobacteria Enterobacteriales Enterobacteriaceae *Yersinia pestis* PY-58 |
| Ype-193-1 | Bacteria Proteobacteria Gammaproteobacteria Enterobacteriales Enterobacteriaceae *Yersinia pestis* PY-59 |
| Ype-194-1 | Bacteria Proteobacteria Gammaproteobacteria Enterobacteriales Enterobacteriaceae *Yersinia pestis* PY-60 |
| Ype-195-1 | Bacteria Proteobacteria Gammaproteobacteria Enterobacteriales Enterobacteriaceae *Yersinia pestis* PY-61 |
| Ype-196-1 | Bacteria Proteobacteria Gammaproteobacteria Enterobacteriales Enterobacteriaceae *Yersinia pestis* PY-63 |
| Ype-197-1 | Bacteria Proteobacteria Gammaproteobacteria Enterobacteriales Enterobacteriaceae *Yersinia pestis* PY-64 |
| Ype-198-1 | Bacteria Proteobacteria Gammaproteobacteria Enterobacteriales Enterobacteriaceae *Yersinia pestis* PY-65 |
| Ype-199-1 | Bacteria Proteobacteria Gammaproteobacteria Enterobacteriales Enterobacteriaceae *Yersinia pestis* PY-66 |
| Ype-200-1 | Bacteria Proteobacteria Gammaproteobacteria Enterobacteriales Enterobacteriaceae *Yersinia pestis* PY-71 |
| Ype-201-1 | Bacteria Proteobacteria Gammaproteobacteria Enterobacteriales Enterobacteriaceae *Yersinia pestis* PY-72 |
| Ype-202-1 | Bacteria Proteobacteria Gammaproteobacteria Enterobacteriales Enterobacteriaceae *Yersinia pestis* PY-76 |
| Ype-203-1 | Bacteria Proteobacteria Gammaproteobacteria Enterobacteriales Enterobacteriaceae *Yersinia pestis* PY-88 |
| Ype-204-1 | Bacteria Proteobacteria Gammaproteobacteria Enterobacteriales Enterobacteriaceae *Yersinia pestis* PY-89 |
| Ype-205-1 | Bacteria Proteobacteria Gammaproteobacteria Enterobacteriales Enterobacteriaceae *Yersinia pestis* PY-90 |
| Ype-206-1 | Bacteria Proteobacteria Gammaproteobacteria Enterobacteriales Enterobacteriaceae *Yersinia pestis* PY-91 |
| Ype-207-1 | Bacteria Proteobacteria Gammaproteobacteria Enterobacteriales Enterobacteriaceae *Yersinia pestis* PY-92 |
| Ype-208-1 | Bacteria Proteobacteria Gammaproteobacteria Enterobacteriales Enterobacteriaceae *Yersinia pestis* PY-93 |
| Ype-209-1 | Bacteria Proteobacteria Gammaproteobacteria Enterobacteriales Enterobacteriaceae *Yersinia pestis* PY-94 |
| Ype-210-1 | Bacteria Proteobacteria Gammaproteobacteria Enterobacteriales Enterobacteriaceae *Yersinia pestis* PY-95 |
| Ype-211-1 | Bacteria Proteobacteria Gammaproteobacteria Enterobacteriales Enterobacteriaceae *Yersinia pestis* PY-96 |
| Ype-212-1 | Bacteria Proteobacteria Gammaproteobacteria Enterobacteriales Enterobacteriaceae *Yersinia pestis* PY-98 |

| | |
|---|---|
| Ype-213-1 | Bacteria Proteobacteria Gammaproteobacteria Enterobacteriales Enterobacteriaceae *Yersinia pestis* PY-99 |
| Ype-215-1 | Bacteria Proteobacteria Gammaproteobacteria Enterobacteriales Enterobacteriaceae *Yersinia pestis* YN2551 |
| Ype-216-1 | Bacteria Proteobacteria Gammaproteobacteria Enterobacteriales Enterobacteriaceae *Yersinia pestis* Z176003 |
| Yps-14-1 | Bacteria Proteobacteria Gammaproteobacteria Enterobacteriales Enterobacteriaceae *Yersinia pseudotuberculosis* IP 31758 |
| Yps-10-1 to Yps-10-2 | Bacteria Proteobacteria Gammaproteobacteria Enterobacteriales Enterobacteriaceae *Yersinia pseudotuberculosis* IP 32953 |
| Yps-15-1 | Bacteria Proteobacteria Gammaproteobacteria Enterobacteriales Enterobacteriaceae *Yersinia pseudotuberculosis* NBRC 105692 |
| Yps-11-1 to Yps-11-2 | Bacteria Proteobacteria Gammaproteobacteria Enterobacteriales Enterobacteriaceae *Yersinia pseudotuberculosis* PB1/+ |
| Yps-12-1 to Yps-12-2 | Bacteria Proteobacteria Gammaproteobacteria Enterobacteriales Enterobacteriaceae *Yersinia pseudotuberculosis* YPIII |
| Yro-2-1 | Bacteria Proteobacteria Gammaproteobacteria Enterobacteriales Enterobacteriaceae *Yersinia rohdei* ATCC 43380 |
| Yru-1-1 to Yru-1-3 | Bacteria Proteobacteria Gammaproteobacteria Enterobacteriales Enterobacteriaceae *Yersinia ruckeri* |
| Yru-2-1 to Yru-2-2 | Bacteria Proteobacteria Gammaproteobacteria Enterobacteriales Enterobacteriaceae *Yersinia ruckeri* ATCC 29473 |
| Yre-2-1 | Bacteria Proteobacteria Gammaproteobacteria Enterobacteriales Enterobacteriaceae *Yokenella regensburgei* ATCC 43003 |
| gpr-1-1 | Bacteria Proteobacteria Gammaproteobacteria gamma proteobacterium WG36 |
| CSc-1-1 | Bacteria Proteobacteria Gammaproteobacteria Orbales Orbaceae *Candidatus* Schmidhempelia bombi str. Bimp |
| Fpe-1-1 | Bacteria Proteobacteria Gammaproteobacteria Orbales Orbaceae *Frischella perrara* |
| Gap-1-1 | Bacteria Proteobacteria Gammaproteobacteria Orbales Orbaceae *Gilliamella apicola* |
| Asu-1-1 | Bacteria Proteobacteria Gammaproteobacteria Pasteurellales Pasteurellaceae *Actinobacillus succinogenes* 130Z |
| Aap-1-1 | Bacteria Proteobacteria Gammaproteobacteria Pasteurellales Pasteurellaceae *Aggregatibacter aphrophilus* |
| Aap-2-1 | Bacteria Proteobacteria Gammaproteobacteria Pasteurellales Pasteurellaceae *Aggregatibacter aphrophilus* ATCC 33389 |
| Aap-3-1 | Bacteria Proteobacteria Gammaproteobacteria Pasteurellales Pasteurellaceae *Aggregatibacter aphrophilus* F0387 |
| Aap-4-1 | Bacteria Proteobacteria Gammaproteobacteria Pasteurellales Pasteurellaceae *Aggregatibacter aphrophilus* NJ8700 |
| Ase-1-1 | Bacteria Proteobacteria Gammaproteobacteria Pasteurellales Pasteurellaceae *Aggregatibacter segnis* ATCC 33393 |
| Gan-1-1 to Gan-1-12 | Bacteria Proteobacteria Gammaproteobacteria Pasteurellales Pasteurellaceae *Gallibacterium anatis* |
| Gan-2-1 | Bacteria Proteobacteria Gammaproteobacteria Pasteurellales Pasteurellaceae *Gallibacterium anatis* 10672-6 |
| Gan-3-1 | Bacteria Proteobacteria Gammaproteobacteria Pasteurellales Pasteurellaceae *Gallibacterium anatis* 12656/12 |
| Gan-4-1 | Bacteria Proteobacteria Gammaproteobacteria Pasteurellales Pasteurellaceae *Gallibacterium anatis* 4895 |
| Gan-5-1 | Bacteria Proteobacteria Gammaproteobacteria Pasteurellales Pasteurellaceae *Gallibacterium anatis* 7990 |
| Gan-6-1 | Bacteria Proteobacteria Gammaproteobacteria Pasteurellales Pasteurellaceae *Gallibacterium anatis* CCM5995 |
| Gan-7-1 | Bacteria Proteobacteria Gammaproteobacteria Pasteurellales Pasteurellaceae *Gallibacterium anatis* DSM 16844 = F 149 |
| Gan-8-1 | Bacteria Proteobacteria Gammaproteobacteria Pasteurellales Pasteurellaceae *Gallibacterium anatis* IPDH697-78 |
| Gan-10-1 | Bacteria Proteobacteria Gammaproteobacteria Pasteurellales Pasteurellaceae *Gallibacterium anatis str.* Avicor |
| Gan-9-1 | Bacteria Proteobacteria Gammaproteobacteria Pasteurellales Pasteurellaceae *Gallibacterium anatis* UMN179 |
| Gge-1-1 | Bacteria Proteobacteria Gammaproteobacteria Pasteurellales Pasteurellaceae *Gallibacterium genomosp.* 1 |
| Gge-2-1 | Bacteria Proteobacteria Gammaproteobacteria Pasteurellales Pasteurellaceae *Gallibacterium genomosp.* 2 |
| Hha-1-1 to Hha-1-2 | Bacteria Proteobacteria Gammaproteobacteria Pasteurellales Pasteurellaceae *Haemophilus haemolyticus* |
| Hha-2-1 | Bacteria Proteobacteria Gammaproteobacteria Pasteurellales Pasteurellaceae *Haemophilus haemolyticus* M21127 |
| Hha-3-1 | Bacteria Proteobacteria Gammaproteobacteria Pasteurellales Pasteurellaceae *Haemophilus haemolyticus* M21621 |
| Hin-1-1 to Hin-1-6 | Bacteria Proteobacteria Gammaproteobacteria Pasteurellales Pasteurellaceae *Haemophilus influenzae* |
| Hin-2-1 | Bacteria Proteobacteria Gammaproteobacteria Pasteurellales Pasteurellaceae *Haemophilus influenzae* 2019 |
| Hin-3-1 | Bacteria Proteobacteria Gammaproteobacteria Pasteurellales Pasteurellaceae *Haemophilus influenzae* 3655 |
| Hin-4-1 | Bacteria Proteobacteria Gammaproteobacteria Pasteurellales Pasteurellaceae *Haemophilus influenzae* 60294N1 |
| Hin-5-1 | Bacteria Proteobacteria Gammaproteobacteria Pasteurellales Pasteurellaceae *Haemophilus influenzae* 7P49H1 |
| Hin-6-1 | Bacteria Proteobacteria Gammaproteobacteria Pasteurellales Pasteurellaceae *Haemophilus influenzae* PittII |
| Hin-7-1 | Bacteria Proteobacteria Gammaproteobacteria Pasteurellales Pasteurellaceae *Haemophilus influenzae* R2866 |
| Hpa-2-1 | Bacteria Proteobacteria Gammaproteobacteria Pasteurellales Pasteurellaceae *Haemophilus parainfluenzae* ATCC 33392 |
| Hsp-1-1 | Bacteria Proteobacteria Gammaproteobacteria Pasteurellales Pasteurellaceae *Haemophilus sp. oral taxon* 851 str. F0397 |
| Mma-2-1 | Bacteria Proteobacteria Gammaproteobacteria Pasteurellales Pasteurellaceae *Mannheimia massilioguelmaensis* |
| Vpa-1-1 to Vpa-1-2 | Bacteria Proteobacteria Gammaproteobacteria Vibrionales Vibrionaceae *Vibrio parahaemolyticus* |
| Sen-1023-1 | CDC07-0191Bacteria Proteobacteria Gammaproteobacteria Enterobacteriales Enterobacteriaceae Salmonella Salmonella enterica subsp. entericaserovar Tennessee *Salmonella enterica subsp. enterica serovar* Tennessee str |
| Sen-1024-1 to Sen-1024-2 | CVM19633Bacteria Proteobacteria Gammaproteobacteria Enterobacteriales Enterobacteriaceae *Salmonella enterica subsp. enterica serovar* Schwarzengrund str |
| Sen-1025-1 | CVM23701Bacteria Proteobacteria Gammaproteobacteria Enterobacteriales Enterobacteriaceae Salmonella Salmonella enterica subsp. entericaserovar 4,[5],12:i:- *Salmonella enterica subsp. enterica serovar* 4,[5],12:i:- str |
| Sen-1026-1 | GA_MM04042433Bacteria Proteobacteria Gammaproteobacteria Enterobacteriales Enterobacteriaceae *Salmonella enterica subsp. enterica serovar* Javiana str |
| Sen-1027-1 | HI_N05-537Bacteria Proteobacteria Gammaproteobacteria Enterobacteriales Enterobacteriaceae Salmonella Salmonella enterica subsp. entericaserovar Weltevreden *Salmonella enterica subsp. enterica serovar* Weltevreden str |
| Sen-1028-1 | P125109Bacteria Proteobacteria Gammaproteobacteria Enterobacteriales Enterobacteriaceae Salmonella Salmonella enterica subsp. entericaserovar Enteritidis *Salmonella enterica subsp. enterica serovar* Enteritidis str |
| Sen-1029-1 | RKS4594Bacteria Proteobacteria Gammaproteobacteria Enterobacteriales Enterobacteriaceae Salmonella Salmonella enterica subsp. entericaserovar Paratyphi C *Salmonella enterica subsp. enterica serovar* Paratyphi C strain |

| | |
|---|---|
| Sen-1030-1 | SC-B67Bacteria Proteobacteria Gammaproteobacteria Enterobacteriales Enterobacteriaceae Salmonella Salmonella enterica subsp. entericaserovar Choleraesuis *Salmonella enterica subsp. enterica serovar* Choleraesuis str |
| Sen-1024-1 to Sen-1024-2 | SL480Bacteria Proteobacteria Gammaproteobacteria Enterobacteriales Enterobacteriaceae *Salmonella enterica subsp. enterica serovar* Schwarzengrund str |
| env-1 to env-370 | environmental samples |

#### 8.2 Multiple-sequence alignment

Each L25-Gammaproteobacteria RNA is denoted by an abbreviation (like "Eco-1-1") that refers to a taxonomy given in Section 8.1. The alignment may include sequences containing the RNA motif, as well as flanking sequence. The L25-Gammaproteobacteria RNA itself is denoted by the line underneath marked 5′ and 3′ on either end. Nucleotides in flanking sequences (i.e., not part of the motif) are written in gray letters. Stems of predicted rho-independent transcription terminators, if any, are shaded yellow. (Note: terminator predictions have not been analyzed manually, and many are likely to be false positives. Terminator predictions are those of the RNIE software.) Nucleotides predicted to function (as DNA) as transcription-factor binding sites, if any, are shaded in green. (But note: these predictions are manually annotated, so they might be under-predicted.) Annotated start codons, if any, are shaded green. (Note: start codons are frequently misannotated, especially in environmental samples.) Nucleotides proposed to basepair as part of the consensus structure are shaded in color when they comprise Watson-Crick or G-U pairs. Otherwise they are shaded gray. Conserved stems are also indicated at the bottom of the alignment by angle brackets, where matching < and > denote base-paired columns. Below these angle brackets, the symbol "2" denotes base pairs exhibiting covariation according to the statistically well-founded R-scape method. "1" denotes base pairs exhibiting covariation according to R2R's simplistic method. "0" denotes base pairs that are not observed to mutate and "?" denotes base pairs that have a significant frequency of non-canonical nucleotides for Watson-Crick or G-U pairs (> 5%). Below these base pair annotation is the consensus sequence: "R" = "A" or "G", "Y" = "C" or "U", red nucleotides: nucleotide identity conserved more than 97% of the time, black nucleotides: 90%, gray nucleotides: 75%, red circle (•): nucleotide is present 97% of the time, black circle (•): 90%, gray circle (•): 75%, white circle (◦): 50%. All percentages of sequences just described (e.g. 97% conserved) assume that sequences have been weighted by the GSC algorithm implemented by the Infernal software package.

Duplicate sequences: the following putative homologs are not shown in the alignment because their sequences are identical to a homolog already shown: Aap-1-1, Aap-2-1, Aap-3-1, Aap-4-1, Ase-1-1, CRe-1-1, Cam-1-1, Cam-1-2, Cam-1-3, Cbr-1-1, Cbr-1-2, Cbr-1-3, Cbr-1-4, Cbr-1-6, Cdu-1-1, Cdu-4-1, Cdu-5-1, Cfr-1-1, Cfr-1-2, Cfr-1-3, Cfr-10-1, Cfr-11-1, Cfr-2-1, Cfr-2-2, Cfr-3-1, Cfr-5-1, Cfr-6-1, Cfr-7-1, Cfr-8-1, Cfr-9-1, Cko-1-1, Cko-2-1, Cma-1-2, Cma-1-3, Cma-1-4, Cma-2-1, Cma-3-1, Cma-5-1, Cro-1-1, Cro-3-1, Csa-1-1, Csa-11-1, Csa-13-1, Csa-14-1, Csa-16-1, Csa-2-1, Csa-3-1, Csa-4-1, Csa-5-1, Csa-6-1, Csa-8-1, Csa-9-1, Csp-1-1, Csp-10-1, Csp-12-1, Csp-14-1, Csp-15-1, Csp-16-1, Csp-19-1, Csp-2-1, Csp-3-1, Csp-4-1, Csp-5-1, Csp-6-1, Csp-7-1, Csp-8-1, Csp-9-1, Ctu-1-1, Ctu-2-1, Ctu-4-1, Cun-1-2, Cyo-1-1, Dch-1-2, Dch-2-1, Dda-4-1, Ddi-1-1, Ddi-2-1, Ddi-4-1, Dso-1-1, Dso-2-1, Dso-3-1, Dso-5-1, Dso-6-1, Dsp-4-1, Dsp-6-1, Dsp-7-1, Dsp-8-1, Dze-3-1, Eae-1-1, Eae-2-1, Eae-3-1, Eal-1-1, Eal-3-1, Eal-4-1, Eal-5-1, Eam-11-1, Eam-12-1, Eam-13-1, Eam-14-1, Eam-2-1, Eam-3-1, Eam-4-1, Eam-5-1, Eam-6-1, Eam-7-1, Eam-8-1, Eam-9-1, Ean-1-1, Ean-2-1, Ean-3-1, Ean-4-1, Eas-1-1, Eas-1-2, Eas-2-1, Eas-3-1, Eba-1-1, Eba-3-1, Eba-5-1, Eca-2-1, Eca-3-1, Ecl-1-1, Ecl-1-10, Ecl-1-11, Ecl-1-12, Ecl-1-2, Ecl-1-3, Ecl-1-4, Ecl-1-5, Ecl-1-6, Ecl-1-7, Ecl-1-8, Ecl-1-9, Ecl-10-1, Ecl-11-1, Ecl-12-1, Ecl-13-1, Ecl-15-1, Ecl-17-1, Ecl-18-1, Ecl-19-1, Ecl-2-1, Ecl-20-1, Ecl-21-1, Ecl-22-1, Ecl-23-1, Ecl-24-1, Ecl-25-1, Ecl-26-1, Ecl-27-1, Ecl-28-1, Ecl-28-2, Ecl-28-3, Ecl-28-4, Ecl-28-5, Ecl-28-6, Ecl-3-1, Ecl-30-1, Ecl-31-1, Ecl-32-1, Ecl-35-1, Ecl-36-1, Ecl-38-1, Ecl-4-1, Ecl-40-1, Ecl-41-1, Ecl-43-1, Ecl-44-1, Ecl-46-1, Ecl-47-1, Ecl-48-1, Ecl-49-1, Ecl-5-1, Ecl-50-1, Ecl-52-2, Ecl-52-4, Ecl-52-5, Ecl-53-1, Ecl-55-1, Ecl-56-1, Ecl-57-1, Ecl-6-1, Ecl-61-1, Ecl-62-1, Ecl-63-1, Ecl-64-1, Ecl-65-1, Ecl-66-1, Ecl-67-1, Ecl-68-1, Ecl-69-1, Ecl-7-1, Ecl-70-1, Ecl-8-1, Ecl-9-1, Eco-1-1, Eco-1-2, Eco-1-3, Eco-1-4, Eco-10-1, Eco-100-1, Eco-1000-1, Eco-1001-1, Eco-1002-1, Eco-1003-1, Eco-1004-1, Eco-1005-1, Eco-1006-1, Eco-1007-1, Eco-1008-1, Eco-1009-1, Eco-101-1, Eco-1010-1, Eco-1011-1, Eco-1012-1, Eco-1013-1, Eco-1014-1, Eco-1015-1, Eco-1016-1, Eco-1017-1, Eco-1018-1, Eco-1019-1, Eco-102-1, Eco-1020-1, Eco-1021-1, Eco-1021-2, Eco-1021-3, Eco-1022-1, Eco-1023-1, Eco-1024-1, Eco-1025-1, Eco-1026-1, Eco-1027-1, Eco-1028-1, Eco-1029-1, Eco-103-1, Eco-1030-1, Eco-1031-1, Eco-1032-1, Eco-1033-1, Eco-1034-1, Eco-1035-1, Eco-1036-1, Eco-1037-1, Eco-1038-1, Eco-1039-1, Eco-104-1, Eco-1040-1, Eco-1041-1, Eco-1042-1, Eco-1043-1, Eco-1044-1, Eco-1045-1, Eco-1046-1, Eco-1047-1, Eco-1048-1, Eco-1049-1, Eco-105-1, Eco-1050-1, Eco-1051-1, Eco-1052-1, Eco-1053-1, Eco-1054-1, Eco-1055-1, Eco-1056-1, Eco-1057-1, Eco-1058-1, Eco-1059-1, Eco-106-1, Eco-1060-1, Eco-1061-1, Eco-1062-1, Eco-1063-1, Eco-1064-1, Eco-1065-1, Eco-1066-1, Eco-1067-1, Eco-1068-1, Eco-1069-1, Eco-107-1, Eco-1070-1, Eco-1071-1, Eco-1072-1, Eco-1073-1, Eco-1074-1, Eco-1075-1, Eco-1076-1, Eco-1077-1, Eco-1078-1, Eco-1079-1, Eco-108-1, Eco-1080-1, Eco-1081-1, Eco-1082-1, Eco-1083-1, Eco-1085-1, Eco-1086-1, Eco-1087-1, Eco-1088-1, Eco-1089-1, Eco-109-1, Eco-1090-1, Eco-1091-1, Eco-1092-1, Eco-1093-1, Eco-1094-1, Eco-1095-1, Eco-1096-1, Eco-1097-1, Eco-1098-1, Eco-1099-1, Eco-11-1, Eco-110-1, Eco-1100-1, Eco-1101-1, Eco-1102-1, Eco-1103-1, Eco-1103-2, Eco-1104-1, Eco-1105-1,

Eco-1106-1, Eco-1107-1, Eco-1108-1, Eco-1109-1, Eco-111-1, Eco-1110-1, Eco-1111-1, Eco-1112-1, Eco-1113-1, Eco-1114-1, Eco-1115-1, Eco-1116-1, Eco-1117-1, Eco-1118-1, Eco-1119-1, Eco-112-1, Eco-1120-1, Eco-1121-1, Eco-1122-1, Eco-1123-1, Eco-1124-1, Eco-1125-1, Eco-1126-1, Eco-1127-1, Eco-1128-1, Eco-1129-1, Eco-113-1, Eco-1130-1, Eco-1131-1, Eco-1132-1, Eco-1133-1, Eco-1134-1, Eco-1135-1, Eco-1136-1, Eco-1137-1, Eco-1138-1, Eco-1139-1, Eco-114-1, Eco-1140-1, Eco-1141-1, Eco-1142-1, Eco-1143-1, Eco-1144-1, Eco-1145-1, Eco-1146-1, Eco-1147-1, Eco-1149-1, Eco-115-1, Eco-1150-1, Eco-1151-1, Eco-1152-1, Eco-1153-1, Eco-1154-1, Eco-1155-1, Eco-1156-1, Eco-1157-1, Eco-1158-1, Eco-1159-1, Eco-116-1, Eco-1160-1, Eco-1161-1, Eco-1162-1, Eco-1163-1, Eco-1164-1, Eco-1165-1, Eco-1166-1, Eco-1167-1, Eco-1168-1, Eco-1169-1, Eco-117-1, Eco-1170-1, Eco-1171-1, Eco-1171-2, Eco-1172-1, Eco-1173-1, Eco-1174-1, Eco-1175-1, Eco-1176-1, Eco-1177-1, Eco-1178-1, Eco-1179-1, Eco-118-1, Eco-1180-1, Eco-1181-1, Eco-1182-1, Eco-1183-1, Eco-1184-1, Eco-1185-1, Eco-1186-1, Eco-1187-1, Eco-1188-1, Eco-1189-1, Eco-119-1, Eco-1190-1, Eco-1191-1, Eco-1192-1, Eco-1193-1, Eco-1194-1, Eco-1195-1, Eco-1196-1, Eco-1197-1, Eco-1198-1, Eco-1199-1, Eco-12-1, Eco-120-1, Eco-1200-1, Eco-1201-1, Eco-1202-1, Eco-1203-1, Eco-1204-1, Eco-1205-1, Eco-1206-1, Eco-1207-1, Eco-1208-1, Eco-1209-1, Eco-121-1, Eco-1210-1, Eco-1211-1, Eco-1212-1, Eco-1213-1, Eco-1214-1, Eco-1215-1, Eco-1216-1, Eco-1217-1, Eco-1218-1, Eco-1219-1, Eco-122-1, Eco-1220-1, Eco-1221-1, Eco-1222-1, Eco-1223-1, Eco-1224-1, Eco-1225-1, Eco-1226-1, Eco-1227-1, Eco-1228-1, Eco-1229-1, Eco-123-1, Eco-1230-1, Eco-1231-1, Eco-1232-1, Eco-1233-1, Eco-1234-1, Eco-1235-1, Eco-1236-1, Eco-1237-1, Eco-1238-1, Eco-1239-1, Eco-124-1, Eco-1240-1, Eco-1241-1, Eco-1242-1, Eco-1243-1, Eco-1244-1, Eco-1245-1, Eco-1246-1, Eco-1247-1, Eco-1248-1, Eco-1249-1, Eco-125-1, Eco-1250-1, Eco-1251-1, Eco-1252-1, Eco-1253-1, Eco-1254-1, Eco-1255-1, Eco-1256-1, Eco-1257-1, Eco-1258-1, Eco-1259-1, Eco-126-1, Eco-1260-1, Eco-1261-1, Eco-1262-1, Eco-1263-1, Eco-1264-1, Eco-1265-1, Eco-1266-1, Eco-1267-1, Eco-1268-1, Eco-1269-1, Eco-127-1, Eco-1270-1, Eco-1271-1, Eco-1272-1, Eco-1273-1, Eco-1274-1, Eco-1275-1, Eco-1276-1, Eco-1277-1, Eco-1278-1, Eco-1279-1, Eco-128-1, Eco-1280-1, Eco-1281-1, Eco-1282-1, Eco-1283-1, Eco-1284-1, Eco-1285-1, Eco-1286-1, Eco-1287-1, Eco-1288-1, Eco-1289-1, Eco-129-1, Eco-1290-1, Eco-1291-1, Eco-1292-1, Eco-1293-1, Eco-1294-1, Eco-1295-1, Eco-1296-1, Eco-1297-1, Eco-1298-1, Eco-1299-1, Eco-13-1, Eco-130-1, Eco-1300-1, Eco-1301-1, Eco-1302-1, Eco-1303-1, Eco-1304-1, Eco-1305-1, Eco-1306-1, Eco-1307-1, Eco-1308-1, Eco-1309-1, Eco-131-1, Eco-1310-1, Eco-1311-1, Eco-1312-1, Eco-1313-1, Eco-1314-1, Eco-1315-1, Eco-1316-1, Eco-1317-1, Eco-1318-1, Eco-1319-1, Eco-132-1, Eco-1320-1, Eco-1321-1, Eco-1322-1, Eco-1323-1, Eco-1324-1, Eco-1325-1, Eco-1326-1, Eco-1327-1, Eco-1328-1, Eco-1329-1, Eco-133-1, Eco-1330-1, Eco-1331-1, Eco-1332-1, Eco-1333-1, Eco-1334-1, Eco-1335-1, Eco-1336-1, Eco-1337-1, Eco-1338-1, Eco-1339-1, Eco-134-1, Eco-1340-1, Eco-1341-1, Eco-1342-1, Eco-1343-1, Eco-1344-1, Eco-1345-1, Eco-1346-1, Eco-1347-1, Eco-1348-1, Eco-1349-1, Eco-135-1, Eco-1350-1, Eco-1351-1, Eco-1352-1, Eco-1353-1, Eco-1354-1, Eco-1355-1, Eco-1356-1, Eco-1357-1, Eco-1358-1, Eco-1359-1, Eco-136-1, Eco-1360-1, Eco-1361-1, Eco-1362-1, Eco-1363-1, Eco-1364-1, Eco-1365-1, Eco-1366-1, Eco-1367-1, Eco-1368-1, Eco-1369-1, Eco-137-1, Eco-1370-1, Eco-1371-1, Eco-1372-1, Eco-1373-1, Eco-1374-1, Eco-1375-1, Eco-1376-1, Eco-1377-1, Eco-1378-1, Eco-1379-1, Eco-138-1, Eco-1380-1, Eco-1381-1, Eco-1382-1, Eco-1383-1, Eco-1384-1, Eco-1385-1, Eco-1386-1, Eco-1387-1, Eco-1388-1, Eco-1389-1, Eco-139-1, Eco-1390-1, Eco-1391-1, Eco-1392-1, Eco-1393-1, Eco-1394-1, Eco-1394-2, Eco-1395-1, Eco-1396-1, Eco-1397-1, Eco-1398-1, Eco-1399-1, Eco-14-1, Eco-140-1, Eco-1400-1, Eco-1401-1, Eco-1402-1, Eco-1403-1, Eco-1404-1, Eco-1405-1, Eco-1406-1, Eco-1407-1, Eco-1408-1, Eco-1409-1, Eco-141-1, Eco-1410-1, Eco-1411-1, Eco-1412-1, Eco-1413-1, Eco-1414-1, Eco-1415-1, Eco-1416-1, Eco-1417-1, Eco-1418-1, Eco-1419-1, Eco-142-1, Eco-1420-1, Eco-1421-1, Eco-1422-1, Eco-1423-1, Eco-1424-1, Eco-1425-1, Eco-1426-1, Eco-1427-1, Eco-1428-1, Eco-1429-1, Eco-143-1, Eco-1430-1, Eco-1431-1, Eco-1432-1, Eco-1433-1, Eco-1434-1, Eco-1435-1, Eco-1436-1, Eco-1437-1, Eco-1438-1, Eco-1439-1, Eco-144-1, Eco-1440-1, Eco-1441-1, Eco-1442-1, Eco-1443-1, Eco-1444-1, Eco-1445-1, Eco-1446-1, Eco-1447-1, Eco-1448-1, Eco-1449-1, Eco-145-1, Eco-1450-1, Eco-1451-1, Eco-1452-1, Eco-1453-1, Eco-1454-1, Eco-1455-1, Eco-1456-1, Eco-1457-1, Eco-1458-1, Eco-1459-1, Eco-146-1, Eco-1460-1, Eco-1461-1, Eco-1462-1, Eco-1463-1, Eco-1464-1, Eco-1465-1, Eco-1466-1, Eco-1467-1, Eco-1468-1, Eco-1469-1, Eco-147-1, Eco-1470-1, Eco-1471-1, Eco-1472-1, Eco-1473-1, Eco-1474-1, Eco-1475-1, Eco-1476-1, Eco-1477-1, Eco-1478-1, Eco-1479-1, Eco-148-1, Eco-1480-1, Eco-1481-1, Eco-1482-1, Eco-1483-1, Eco-1484-1, Eco-1485-1, Eco-1486-1, Eco-1487-1, Eco-1488-1, Eco-1489-1, Eco-149-1, Eco-1490-1, Eco-1491-1, Eco-1492-1, Eco-1493-1, Eco-1494-1, Eco-1495-1, Eco-1496-1, Eco-1497-1, Eco-1498-1, Eco-1499-1, Eco-15-1, Eco-150-1, Eco-1500-1, Eco-1501-1, Eco-1502-1, Eco-1503-1, Eco-1504-1, Eco-1505-1, Eco-1506-1, Eco-1507-1, Eco-1508-1, Eco-1509-1, Eco-151-1, Eco-1510-1, Eco-1511-1, Eco-1512-1, Eco-1513-1, Eco-1514-1, Eco-1515-1, Eco-1516-1, Eco-1517-1, Eco-1518-1, Eco-1519-1, Eco-152-1, Eco-1520-1, Eco-1521-1, Eco-1522-1, Eco-1523-1, Eco-1524-1, Eco-1525-1, Eco-1526-1, Eco-1527-1, Eco-1528-1, Eco-1529-1, Eco-153-1, Eco-1530-1, Eco-1531-1, Eco-1532-1, Eco-1533-1, Eco-1534-1, Eco-1535-1, Eco-1536-1, Eco-1537-1, Eco-1538-1, Eco-1539-1, Eco-154-1, Eco-1540-1, Eco-1541-1, Eco-1542-1, Eco-1543-1, Eco-1544-1, Eco-1545-1, Eco-1546-1, Eco-1547-1, Eco-1548-1, Eco-1549-1, Eco-155-1, Eco-1550-1, Eco-1551-1, Eco-1552-1, Eco-1553-1, Eco-1554-1, Eco-1555-1, Eco-1556-1, Eco-1557-1, Eco-1558-1, Eco-1559-1, Eco-156-1, Eco-1560-1, Eco-1561-1, Eco-1562-1, Eco-1563-1, Eco-1564-1, Eco-1565-1, Eco-1566-1, Eco-1567-1, Eco-1568-1, Eco-1569-1, Eco-157-1, Eco-1570-1, Eco-1571-1, Eco-1572-1, Eco-1573-1, Eco-1574-1, Eco-1575-1, Eco-1576-1, Eco-1577-1, Eco-1578-1, Eco-1579-1, Eco-158-1, Eco-1580-1, Eco-1581-1, Eco-1582-1, Eco-1583-1, Eco-1584-1, Eco-1585-1, Eco-1586-1, Eco-1587-1, Eco-1588-1, Eco-1589-1, Eco-159-1, Eco-1590-1, Eco-1591-1, Eco-1592-1, Eco-1593-1, Eco-1594-1, Eco-1595-1, Eco-1596-1, Eco-1597-1, Eco-1598-1, Eco-1599-1, Eco-16-1, Eco-160-1, Eco-1600-1, Eco-1601-1, Eco-1602-1, Eco-1603-1, Eco-1604-1, Eco-1605-1, Eco-1606-1, Eco-1607-1, Eco-1608-1, Eco-1609-1, Eco-161-1, Eco-1610-1, Eco-1611-1, Eco-1612-1, Eco-1613-1, Eco-1614-1, Eco-1615-1, Eco-1616-1, Eco-1617-1, Eco-1618-1, Eco-1619-1, Eco-162-1, Eco-1620-1, Eco-1621-1, Eco-1622-1, Eco-1623-1, Eco-1624-1, Eco-1625-1, Eco-1626-1, Eco-1627-1, Eco-1628-1, Eco-1629-1, Eco-163-1, Eco-1630-1, Eco-1631-1, Eco-1632-1, Eco-1633-1, Eco-1634-1, Eco-1635-1, Eco-1636-1, Eco-1637-1, Eco-1638-1, Eco-1639-1, Eco-164-1,

Eco-1640-1, Eco-1641-1, Eco-1642-1, Eco-1643-1, Eco-1644-1, Eco-1645-1, Eco-1646-1, Eco-1647-1, Eco-1648-1, Eco-1649-1, Eco-165-1, Eco-1650-1, Eco-1651-1, Eco-1652-1, Eco-1653-1, Eco-1654-1, Eco-1655-1, Eco-1656-1, Eco-1657-1, Eco-1658-1, Eco-1659-1, Eco-166-1, Eco-1660-1, Eco-1661-1, Eco-1662-1, Eco-1663-1, Eco-1664-1, Eco-1665-1, Eco-1666-1, Eco-1667-1, Eco-1668-1, Eco-1669-1, Eco-167-1, Eco-1670-1, Eco-1671-1, Eco-1672-1, Eco-1673-1, Eco-1674-1, Eco-1675-1, Eco-1676-1, Eco-1677-1, Eco-1678-1, Eco-1679-1, Eco-168-1, Eco-1680-1, Eco-1681-1, Eco-1682-1, Eco-1683-1, Eco-1684-1, Eco-1685-1, Eco-1686-1, Eco-1687-1, Eco-1688-1, Eco-1689-1, Eco-169-1, Eco-1690-1, Eco-1691-1, Eco-1692-1, Eco-1693-1, Eco-1694-1, Eco-1695-1, Eco-1696-1, Eco-1697-1, Eco-1698-1, Eco-1699-1, Eco-17-1, Eco-170-1, Eco-1700-1, Eco-1701-1, Eco-1702-1, Eco-1703-1, Eco-1704-1, Eco-1705-1, Eco-1706-1, Eco-1707-1, Eco-1708-1, Eco-1709-1, Eco-171-1, Eco-1710-1, Eco-1711-1, Eco-1711-2, Eco-1712-1, Eco-1713-1, Eco-1714-1, Eco-1715-1, Eco-1716-1, Eco-1717-1, Eco-1718-1, Eco-1719-1, Eco-172-1, Eco-1720-1, Eco-1721-1, Eco-1722-1, Eco-1723-1, Eco-1724-1, Eco-1725-1, Eco-1726-1, Eco-1727-1, Eco-1728-1, Eco-1729-1, Eco-173-1, Eco-1730-1, Eco-1731-1, Eco-1732-1, Eco-1733-1, Eco-1734-1, Eco-1735-1, Eco-1736-1, Eco-1737-1, Eco-1738-1, Eco-1739-1, Eco-1740-1, Eco-1741-1, Eco-1742-1, Eco-1743-1, Eco-1744-1, Eco-1745-1, Eco-1746-1, Eco-1747-1, Eco-1748-1, Eco-1749-1, Eco-175-1, Eco-1750-1, Eco-1751-1, Eco-1752-1, Eco-1753-1, Eco-1754-1, Eco-1755-1, Eco-1756-1, Eco-1757-1, Eco-1758-1, Eco-1759-1, Eco-176-1, Eco-1760-1, Eco-1761-1, Eco-1762-1, Eco-1763-1, Eco-1764-1, Eco-1765-1, Eco-1766-1, Eco-1767-1, Eco-1768-1, Eco-1769-1, Eco-177-1, Eco-1770-1, Eco-1771-1, Eco-1772-1, Eco-1773-1, Eco-1774-1, Eco-1775-1, Eco-1776-1, Eco-1777-1, Eco-1778-1, Eco-1779-1, Eco-178-1, Eco-1780-1, Eco-1781-1, Eco-1782-1, Eco-1783-1, Eco-1784-1, Eco-1784-2, Eco-1785-1, Eco-1786-1, Eco-1787-1, Eco-1788-1, Eco-1789-1, Eco-179-1, Eco-1790-1, Eco-1791-1, Eco-1792-1, Eco-1793-1, Eco-1794-1, Eco-1795-1, Eco-1796-1, Eco-1797-1, Eco-1798-1, Eco-1799-1, Eco-18-1, Eco-180-1, Eco-1800-1, Eco-1801-1, Eco-1802-1, Eco-1803-1, Eco-1804-1, Eco-1805-1, Eco-1806-1, Eco-1807-1, Eco-1808-1, Eco-1809-1, Eco-181-1, Eco-1810-1, Eco-1811-1, Eco-1812-1, Eco-1813-1, Eco-1814-1, Eco-1815-1, Eco-1816-1, Eco-1817-1, Eco-1818-1, Eco-1819-1, Eco-182-1, Eco-1820-1, Eco-1821-1, Eco-1822-1, Eco-1823-1, Eco-1824-1, Eco-1825-1, Eco-1826-1, Eco-1827-1, Eco-1828-1, Eco-1829-1, Eco-183-1, Eco-1830-1, Eco-1831-1, Eco-1832-1, Eco-1833-1, Eco-1834-1, Eco-1835-1, Eco-1836-1, Eco-1837-1, Eco-1838-1, Eco-1839-1, Eco-184-1, Eco-1840-1, Eco-1841-1, Eco-1842-1, Eco-1843-1, Eco-1844-1, Eco-1845-1, Eco-1846-1, Eco-1847-1, Eco-1848-1, Eco-1849-1, Eco-185-1, Eco-1850-1, Eco-1851-1, Eco-1852-1, Eco-1853-1, Eco-1854-1, Eco-1855-1, Eco-1856-1, Eco-1857-1, Eco-1858-1, Eco-1859-1, Eco-186-1, Eco-1860-1, Eco-1861-1, Eco-1862-1, Eco-1863-1, Eco-1864-1, Eco-1865-1, Eco-1866-1, Eco-1867-1, Eco-1868-1, Eco-1869-1, Eco-187-1, Eco-1870-1, Eco-1871-1, Eco-1872-1, Eco-1873-1, Eco-1874-1, Eco-1875-1, Eco-1876-1, Eco-1877-1, Eco-1878-1, Eco-1879-1, Eco-188-1, Eco-1880-1, Eco-1881-1, Eco-1882-1, Eco-1883-1, Eco-1884-1, Eco-1885-1, Eco-1886-1, Eco-1887-1, Eco-1888-1, Eco-1889-1, Eco-189-1, Eco-1890-1, Eco-1891-1, Eco-1892-1, Eco-1893-1, Eco-1894-1, Eco-1895-1, Eco-1896-1, Eco-1897-1, Eco-1898-1, Eco-1899-1, Eco-19-1, Eco-190-1, Eco-1900-1, Eco-1901-1, Eco-1902-1, Eco-1903-1, Eco-1904-1, Eco-1905-1, Eco-1906-1, Eco-1907-1, Eco-1908-1, Eco-1909-1, Eco-191-1, Eco-1910-1, Eco-1911-1, Eco-1912-1, Eco-1913-1, Eco-1914-1, Eco-1915-1, Eco-1916-1, Eco-1917-1, Eco-1918-1, Eco-1919-1, Eco-192-1, Eco-1920-1, Eco-1921-1, Eco-1922-1, Eco-1923-1, Eco-1924-1, Eco-1925-1, Eco-1926-1, Eco-1927-1, Eco-1928-1, Eco-1929-1, Eco-193-1, Eco-1930-1, Eco-1931-1, Eco-1932-1, Eco-1933-1, Eco-1934-1, Eco-1935-1, Eco-1936-1, Eco-1937-1, Eco-1938-1, Eco-1939-1, Eco-194-1, Eco-1940-1, Eco-1941-1, Eco-1942-1, Eco-1943-1, Eco-1944-1, Eco-1945-1, Eco-1946-1, Eco-1947-1, Eco-1948-1, Eco-1949-1, Eco-195-1, Eco-1950-1, Eco-1951-1, Eco-1952-1, Eco-1953-1, Eco-1954-1, Eco-1955-1, Eco-1956-1, Eco-1957-1, Eco-1958-1, Eco-1959-1, Eco-196-1, Eco-1960-1, Eco-1961-1, Eco-1962-1, Eco-1963-1, Eco-1964-1, Eco-1965-1, Eco-1966-1, Eco-1967-1, Eco-1968-1, Eco-1969-1, Eco-197-1, Eco-1970-1, Eco-1971-1, Eco-1972-1, Eco-1973-1, Eco-1974-1, Eco-1975-1, Eco-1976-1, Eco-1977-1, Eco-1978-1, Eco-1979-1, Eco-198-1, Eco-1980-1, Eco-1981-1, Eco-1982-1, Eco-1983-1, Eco-1985-1, Eco-1986-1, Eco-1987-1, Eco-1988-1, Eco-1989-1, Eco-199-1, Eco-1990-1, Eco-1991-1, Eco-1992-1, Eco-1993-1, Eco-1994-1, Eco-1995-1, Eco-1996-1, Eco-1997-1, Eco-1998-1, Eco-1999-1, Eco-2-1, Eco-20-1, Eco-200-1, Eco-2000-1, Eco-2001-1, Eco-2002-1, Eco-2003-1, Eco-2004-1, Eco-2005-1, Eco-2006-1, Eco-2007-1, Eco-2007-2, Eco-2008-1, Eco-2009-1, Eco-201-1, Eco-2010-1, Eco-2011-1, Eco-2012-1, Eco-2012-2, Eco-2013-1, Eco-2014-1, Eco-2015-1, Eco-2016-1, Eco-2017-1, Eco-2018-1, Eco-2019-1, Eco-202-1, Eco-2020-1, Eco-2021-1, Eco-2022-1, Eco-2023-1, Eco-2024-1, Eco-2025-1, Eco-2026-1, Eco-2027-1, Eco-2028-1, Eco-2029-1, Eco-203-1, Eco-2030-1, Eco-2031-1, Eco-2032-1, Eco-2033-1, Eco-2034-1, Eco-2035-1, Eco-2036-1, Eco-2037-1, Eco-2038-1, Eco-2039-1, Eco-204-1, Eco-2040-1, Eco-2041-1, Eco-2042-1, Eco-2043-1, Eco-2044-1, Eco-2045-1, Eco-2046-1, Eco-2047-1, Eco-2048-1, Eco-2049-1, Eco-205-1, Eco-2050-1, Eco-2051-1, Eco-2052-1, Eco-2053-1, Eco-2054-1, Eco-2055-1, Eco-2056-1, Eco-2057-1, Eco-2058-1, Eco-2059-1, Eco-206-1, Eco-2060-1, Eco-2061-1, Eco-2062-1, Eco-2063-1, Eco-2064-1, Eco-2065-1, Eco-2066-1, Eco-2067-1, Eco-2068-1, Eco-2069-1, Eco-207-1, Eco-2070-1, Eco-2071-1, Eco-2072-1, Eco-2073-1, Eco-2074-1, Eco-2075-1, Eco-2076-1, Eco-2077-1, Eco-2078-1, Eco-2079-1, Eco-208-1, Eco-2080-1, Eco-2081-1, Eco-2082-1, Eco-2083-1, Eco-2084-1, Eco-2085-1, Eco-2086-1, Eco-2087-1, Eco-2088-1, Eco-2089-1, Eco-209-1, Eco-2090-1, Eco-2091-1, Eco-2092-1, Eco-2093-1, Eco-2094-1, Eco-2095-1, Eco-2096-1, Eco-2097-1, Eco-2098-1, Eco-2099-1, Eco-21-1, Eco-210-1, Eco-2100-1, Eco-2101-1, Eco-2102-1, Eco-2104-1, Eco-2105-1, Eco-2106-1, Eco-2107-1, Eco-2108-1, Eco-2109-1, Eco-211-1, Eco-2110-1, Eco-2111-1, Eco-2112-1, Eco-2113-1, Eco-2114-1, Eco-2115-1, Eco-2116-1, Eco-2117-1, Eco-2118-1, Eco-2119-1, Eco-212-1, Eco-2120-1, Eco-2121-1, Eco-2122-1, Eco-2123-1, Eco-2124-1, Eco-2125-1, Eco-2126-1, Eco-2127-1, Eco-2128-1, Eco-2129-1, Eco-213-1, Eco-2130-1, Eco-2131-1, Eco-2132-1, Eco-2133-1, Eco-2134-1, Eco-2135-1, Eco-2136-1, Eco-2137-1, Eco-2138-1, Eco-2139-1, Eco-214-1, Eco-2140-1, Eco-2141-1, Eco-2142-1, Eco-2143-1, Eco-2144-1, Eco-2145-1, Eco-2146-1, Eco-2147-1, Eco-2148-1, Eco-2149-1, Eco-215-1, Eco-2150-1, Eco-2151-1, Eco-2152-1, Eco-2153-1, Eco-2154-1, Eco-2155-1, Eco-2156-1, Eco-2157-1, Eco-2158-1, Eco-2159-1, Eco-2159-2, Eco-216-1, Eco-2160-1, Eco-2161-1, Eco-2162-1, Eco-2163-1, Eco-2164-1, Eco-2165-1, Eco-2166-1, Eco-2167-1, Eco-2168-1, Eco-2169-1, Eco-217-1, Eco-2170-1, Eco-2171-1, Eco-2172-1,

Eco-2173-1, Eco-2174-1, Eco-2175-1, Eco-2176-1, Eco-2177-1, Eco-2178-1, Eco-2179-1, Eco-218-1, Eco-2180-1, Eco-2181-1, Eco-2182-1, Eco-2183-1, Eco-2184-1, Eco-2185-1, Eco-2186-1, Eco-2187-1, Eco-219-1, Eco-22-1, Eco-220-1, Eco-221-1, Eco-222-1, Eco-223-1, Eco-224-1, Eco-225-1, Eco-226-1, Eco-227-1, Eco-228-1, Eco-229-1, Eco-23-1, Eco-230-1, Eco-230-2, Eco-231-1, Eco-232-1, Eco-233-1, Eco-234-1, Eco-235-1, Eco-236-1, Eco-237-1, Eco-238-1, Eco-239-1, Eco-24-1, Eco-240-1, Eco-241-1, Eco-242-1, Eco-243-1, Eco-244-1, Eco-245-1, Eco-246-1, Eco-247-1, Eco-248-1, Eco-249-1, Eco-25-1, Eco-250-1, Eco-251-1, Eco-252-1, Eco-253-1, Eco-254-1, Eco-255-1, Eco-256-1, Eco-257-1, Eco-258-1, Eco-259-1, Eco-26-1, Eco-260-1, Eco-261-1, Eco-262-1, Eco-263-1, Eco-264-1, Eco-265-1, Eco-266-1, Eco-267-1, Eco-268-1, Eco-269-1, Eco-27-1, Eco-270-1, Eco-271-1, Eco-272-1, Eco-273-1, Eco-274-1, Eco-275-1, Eco-276-1, Eco-277-1, Eco-278-1, Eco-28-1, Eco-280-1, Eco-281-1, Eco-282-1, Eco-283-1, Eco-284-1, Eco-285-1, Eco-286-1, Eco-287-1, Eco-288-1, Eco-289-1, Eco-289-2, Eco-29-1, Eco-290-1, Eco-291-1, Eco-291-2, Eco-292-1, Eco-293-1, Eco-294-1, Eco-295-1, Eco-296-1, Eco-297-1, Eco-298-1, Eco-299-1, Eco-3-1, Eco-30-1, Eco-300-1, Eco-301-1, Eco-302-1, Eco-303-1, Eco-304-1, Eco-305-1, Eco-306-1, Eco-307-1, Eco-308-1, Eco-309-1, Eco-31-1, Eco-310-1, Eco-311-1, Eco-312-1, Eco-313-1, Eco-314-1, Eco-315-1, Eco-316-1, Eco-317-1, Eco-318-1, Eco-319-1, Eco-32-1, Eco-320-1, Eco-321-1, Eco-322-1, Eco-323-1, Eco-324-1, Eco-325-1, Eco-326-1, Eco-327-1, Eco-328-1, Eco-329-1, Eco-33-1, Eco-330-1, Eco-331-1, Eco-332-1, Eco-333-1, Eco-334-1, Eco-335-1, Eco-336-1, Eco-337-1, Eco-338-1, Eco-339-1, Eco-34-1, Eco-340-1, Eco-341-1, Eco-342-1, Eco-343-1, Eco-344-1, Eco-345-1, Eco-346-1, Eco-347-1, Eco-348-1, Eco-349-1, Eco-35-1, Eco-350-1, Eco-351-1, Eco-352-1, Eco-353-1, Eco-354-1, Eco-355-1, Eco-356-1, Eco-357-1, Eco-358-1, Eco-359-1, Eco-36-1, Eco-360-1, Eco-361-1, Eco-362-1, Eco-363-1, Eco-364-1, Eco-365-1, Eco-366-1, Eco-367-1, Eco-368-1, Eco-369-1, Eco-37-1, Eco-370-1, Eco-371-1, Eco-372-1, Eco-373-1, Eco-374-1, Eco-375-1, Eco-376-1, Eco-377-1, Eco-378-1, Eco-379-1, Eco-38-1, Eco-380-1, Eco-381-1, Eco-382-1, Eco-383-1, Eco-384-1, Eco-385-1, Eco-386-1, Eco-387-1, Eco-388-1, Eco-389-1, Eco-39-1, Eco-390-1, Eco-391-1, Eco-392-1, Eco-393-1, Eco-394-1, Eco-395-1, Eco-396-1, Eco-397-1, Eco-398-1, Eco-399-1, Eco-4-1, Eco-40-1, Eco-400-1, Eco-401-1, Eco-402-1, Eco-403-1, Eco-404-1, Eco-405-1, Eco-406-1, Eco-407-1, Eco-408-1, Eco-409-1, Eco-41-1, Eco-410-1, Eco-411-1, Eco-412-1, Eco-413-1, Eco-414-1, Eco-415-1, Eco-416-1, Eco-417-1, Eco-418-1, Eco-419-1, Eco-42-1, Eco-420-1, Eco-421-1, Eco-422-1, Eco-423-1, Eco-424-1, Eco-425-1, Eco-426-1, Eco-427-1, Eco-428-1, Eco-429-1, Eco-43-1, Eco-430-1, Eco-431-1, Eco-432-1, Eco-433-1, Eco-434-1, Eco-435-1, Eco-436-1, Eco-437-1, Eco-438-1, Eco-439-1, Eco-44-1, Eco-440-1, Eco-441-1, Eco-442-1, Eco-443-1, Eco-444-1, Eco-445-1, Eco-446-1, Eco-447-1, Eco-448-1, Eco-449-1, Eco-45-1, Eco-450-1, Eco-451-1, Eco-452-1, Eco-453-1, Eco-454-1, Eco-455-1, Eco-456-1, Eco-457-1, Eco-458-1, Eco-459-1, Eco-46-1, Eco-460-1, Eco-461-1, Eco-462-1, Eco-463-1, Eco-463-2, Eco-464-1, Eco-465-1, Eco-466-1, Eco-467-1, Eco-468-1, Eco-469-1, Eco-47-1, Eco-470-1, Eco-471-1, Eco-472-1, Eco-472-2, Eco-473-1, Eco-474-1, Eco-475-1, Eco-476-1, Eco-477-1, Eco-478-1, Eco-479-1, Eco-48-1, Eco-480-1, Eco-481-1, Eco-482-1, Eco-483-1, Eco-484-1, Eco-485-1, Eco-486-1, Eco-487-1, Eco-488-1, Eco-489-1, Eco-49-1, Eco-490-1, Eco-491-1, Eco-492-1, Eco-493-1, Eco-494-1, Eco-495-1, Eco-496-1, Eco-497-1, Eco-498-1, Eco-499-1, Eco-5-1, Eco-50-1, Eco-500-1, Eco-501-1, Eco-502-1, Eco-503-1, Eco-504-1, Eco-505-1, Eco-506-1, Eco-507-1, Eco-508-1, Eco-509-1, Eco-51-1, Eco-510-1, Eco-511-1, Eco-512-1, Eco-513-1, Eco-514-1, Eco-515-1, Eco-516-1, Eco-517-1, Eco-518-1, Eco-519-1, Eco-52-1, Eco-520-1, Eco-521-1, Eco-522-1, Eco-523-1, Eco-524-1, Eco-525-1, Eco-526-1, Eco-527-1, Eco-528-1, Eco-529-1, Eco-53-1, Eco-530-1, Eco-531-1, Eco-532-1, Eco-533-1, Eco-534-1, Eco-535-1, Eco-536-1, Eco-537-1, Eco-538-1, Eco-539-1, Eco-54-1, Eco-540-1, Eco-541-1, Eco-542-1, Eco-543-1, Eco-544-1, Eco-545-1, Eco-546-1, Eco-547-1, Eco-548-1, Eco-548-2, Eco-549-1, Eco-55-1, Eco-550-1, Eco-551-1, Eco-552-1, Eco-553-1, Eco-554-1, Eco-555-2, Eco-556-1, Eco-557-1, Eco-558-1, Eco-559-1, Eco-56-1, Eco-560-1, Eco-561-1, Eco-562-1, Eco-563-1, Eco-564-1, Eco-565-1, Eco-566-1, Eco-567-1, Eco-568-1, Eco-569-1, Eco-57-1, Eco-570-1, Eco-571-1, Eco-572-1, Eco-573-1, Eco-574-1, Eco-574-2, Eco-574-3, Eco-575-1, Eco-576-1, Eco-577-1, Eco-578-1, Eco-579-1, Eco-58-1, Eco-580-1, Eco-581-1, Eco-582-1, Eco-583-1, Eco-584-1, Eco-585-1, Eco-586-1, Eco-587-1, Eco-588-1, Eco-589-1, Eco-59-1, Eco-590-1, Eco-591-1, Eco-592-1, Eco-593-1, Eco-594-1, Eco-595-1, Eco-596-1, Eco-597-1, Eco-598-1, Eco-599-1, Eco-6-1, Eco-60-1, Eco-600-1, Eco-601-1, Eco-602-1, Eco-603-1, Eco-604-1, Eco-605-1, Eco-606-1, Eco-607-1, Eco-608-1, Eco-609-1, Eco-61-1, Eco-610-1, Eco-611-1, Eco-612-1, Eco-613-1, Eco-614-1, Eco-615-1, Eco-616-1, Eco-617-1, Eco-618-1, Eco-619-1, Eco-62-1, Eco-620-1, Eco-621-1, Eco-622-1, Eco-623-1, Eco-624-1, Eco-625-1, Eco-626-1, Eco-627-1, Eco-628-1, Eco-629-1, Eco-63-1, Eco-630-1, Eco-631-1, Eco-632-1, Eco-633-1, Eco-634-1, Eco-635-1, Eco-636-1, Eco-637-1, Eco-638-1, Eco-639-1, Eco-64-1, Eco-640-1, Eco-641-1, Eco-642-1, Eco-643-1, Eco-644-1, Eco-645-1, Eco-646-1, Eco-647-1, Eco-648-1, Eco-649-1, Eco-65-1, Eco-650-1, Eco-651-1, Eco-652-1, Eco-653-1, Eco-654-1, Eco-655-1, Eco-656-1, Eco-657-1, Eco-658-1, Eco-659-1, Eco-660-1, Eco-661-1, Eco-662-1, Eco-663-1, Eco-664-1, Eco-665-1, Eco-666-1, Eco-667-1, Eco-668-1, Eco-669-1, Eco-67-1, Eco-670-1, Eco-671-1, Eco-672-1, Eco-673-1, Eco-674-1, Eco-676-1, Eco-677-1, Eco-678-1, Eco-679-1, Eco-68-1, Eco-680-1, Eco-681-1, Eco-682-1, Eco-683-1, Eco-684-1, Eco-685-1, Eco-686-1, Eco-687-1, Eco-688-1, Eco-689-1, Eco-69-1, Eco-690-1, Eco-691-1, Eco-692-1, Eco-693-1, Eco-694-1, Eco-695-1, Eco-696-1, Eco-697-1, Eco-698-1, Eco-699-1, Eco-7-1, Eco-70-1, Eco-700-1, Eco-701-1, Eco-702-1, Eco-703-1, Eco-704-1, Eco-705-1, Eco-706-1, Eco-707-1, Eco-708-1, Eco-709-1, Eco-71-1, Eco-710-1, Eco-711-1, Eco-712-1, Eco-713-1, Eco-714-1, Eco-715-1, Eco-716-1, Eco-717-1, Eco-718-1, Eco-719-1, Eco-72-1, Eco-720-1, Eco-721-1, Eco-722-1, Eco-723-1, Eco-724-1, Eco-725-1, Eco-726-1, Eco-727-1, Eco-728-1, Eco-729-1, Eco-729-2, Eco-73-1, Eco-730-1, Eco-731-1, Eco-732-1, Eco-733-1, Eco-734-1, Eco-735-1, Eco-736-1, Eco-737-1, Eco-738-1, Eco-739-1, Eco-74-1, Eco-740-1, Eco-741-1, Eco-742-1, Eco-743-1, Eco-744-1, Eco-745-1, Eco-746-1, Eco-747-1, Eco-748-1, Eco-749-1, Eco-75-1, Eco-750-1, Eco-751-1, Eco-752-1, Eco-753-1, Eco-753-2, Eco-754-1, Eco-755-1, Eco-756-1, Eco-757-1, Eco-758-1, Eco-759-1, Eco-76-1, Eco-760-1, Eco-761-1, Eco-762-1, Eco-763-1, Eco-765-1, Eco-766-1, Eco-767-1, Eco-768-1, Eco-769-1, Eco-77-1, Eco-770-1, Eco-771-1, Eco-772-1, Eco-773-1, Eco-774-1, Eco-775-1, Eco-776-1, Eco-777-1, Eco-778-1, Eco-779-1, Eco-78-1, Eco-780-1, Eco-781-1, Eco-782-1, Eco-783-1, Eco-784-1, Eco-785-1, Eco-786-1, Eco-787-1, Eco-788-1, Eco-789-1, Eco-79-1, Eco-790-1, Eco-791-1,

Eco-792-1, Eco-793-1, Eco-794-1, Eco-795-1, Eco-796-1, Eco-797-1, Eco-798-1, Eco-799-1, Eco-8-1, Eco-80-1, Eco-800-1, Eco-801-1, Eco-802-1, Eco-803-1, Eco-804-1, Eco-805-1, Eco-806-1, Eco-807-1, Eco-808-1, Eco-809-1, Eco-81-1, Eco-810-1, Eco-811-1, Eco-812-1, Eco-813-1, Eco-814-1, Eco-815-1, Eco-816-1, Eco-817-1, Eco-818-1, Eco-819-1, Eco-82-1, Eco-820-1, Eco-821-1, Eco-822-1, Eco-823-1, Eco-824-1, Eco-825-1, Eco-826-1, Eco-827-1, Eco-828-1, Eco-829-1, Eco-83-1, Eco-830-1, Eco-831-1, Eco-832-1, Eco-833-1, Eco-834-1, Eco-835-1, Eco-836-1, Eco-837-1, Eco-838-1, Eco-839-1, Eco-840-1, Eco-841-1, Eco-842-1, Eco-843-1, Eco-844-1, Eco-845-1, Eco-846-1, Eco-847-1, Eco-848-1, Eco-849-1, Eco-85-1, Eco-850-1, Eco-851-1, Eco-852-1, Eco-853-1, Eco-854-1, Eco-855-1, Eco-856-1, Eco-857-1, Eco-858-1, Eco-859-1, Eco-86-1, Eco-860-1, Eco-861-1, Eco-862-1, Eco-863-1, Eco-864-1, Eco-865-1, Eco-866-1, Eco-867-1, Eco-868-1, Eco-869-1, Eco-87-1, Eco-870-1, Eco-871-1, Eco-872-1, Eco-873-1, Eco-874-1, Eco-875-1, Eco-876-1, Eco-877-1, Eco-878-1, Eco-879-1, Eco-88-1, Eco-880-1, Eco-881-1, Eco-882-1, Eco-883-1, Eco-884-1, Eco-885-1, Eco-886-1, Eco-887-1, Eco-888-1, Eco-889-1, Eco-89-1, Eco-890-1, Eco-891-1, Eco-892-1, Eco-893-1, Eco-894-1, Eco-895-1, Eco-896-1, Eco-897-1, Eco-898-1, Eco-899-1, Eco-9-1, Eco-90-1, Eco-900-1, Eco-901-1, Eco-902-1, Eco-903-1, Eco-904-1, Eco-905-1, Eco-906-1, Eco-907-1, Eco-908-1, Eco-909-1, Eco-91-1, Eco-910-1, Eco-911-1, Eco-912-1, Eco-913-1, Eco-914-1, Eco-915-1, Eco-916-1, Eco-917-1, Eco-918-1, Eco-919-1, Eco-92-1, Eco-920-1, Eco-921-1, Eco-922-1, Eco-923-1, Eco-924-1, Eco-925-1, Eco-926-1, Eco-927-1, Eco-928-1, Eco-929-1, Eco-93-1, Eco-930-1, Eco-931-1, Eco-932-1, Eco-933-1, Eco-934-1, Eco-935-1, Eco-936-1, Eco-937-1, Eco-938-1, Eco-939-1, Eco-94-1, Eco-940-1, Eco-941-1, Eco-942-1, Eco-943-1, Eco-944-1, Eco-945-1, Eco-946-1, Eco-947-1, Eco-948-1, Eco-949-1, Eco-95-1, Eco-950-1, Eco-951-1, Eco-952-1, Eco-953-1, Eco-954-1, Eco-955-1, Eco-956-1, Eco-957-1, Eco-958-1, Eco-959-1, Eco-96-1, Eco-960-1, Eco-961-1, Eco-962-1, Eco-963-1, Eco-964-1, Eco-965-1, Eco-966-1, Eco-967-1, Eco-968-1, Eco-969-1, Eco-97-1, Eco-970-1, Eco-971-1, Eco-972-1, Eco-973-1, Eco-974-1, Eco-975-1, Eco-976-1, Eco-977-1, Eco-978-1, Eco-979-1, Eco-98-1, Eco-980-1, Eco-981-1, Eco-982-1, Eco-983-1, Eco-984-1, Eco-985-1, Eco-986-1, Eco-987-1, Eco-988-1, Eco-989-1, Eco-99-1, Eco-990-1, Eco-991-1, Eco-992-1, Eco-993-1, Eco-994-1, Eco-995-1, Eco-996-1, Eco-997-1, Eco-998-1, Eco-999-1, Efe-1-2, Efe-1-3, Efe-2-1, Efe-3-1, Efe-4-1, Eho-1-1, Eho-1-2, Eho-1-3, Eho-1-4, Eho-1-5, Eho-1-6, Eho-1-7, Eho-1-8, Eho-1-9, Eho-2-1, Eho-3-1, Eho-3-2, Eho-4-1, Eho-4-2, Eho-6-1, Eho-7-1, Eho-8-1, Eic-1-1, Ein-1-2, Eko-1-10, Eko-1-11, Eko-1-2, Eko-1-3, Eko-1-4, Eko-1-5, Eko-1-6, Eko-1-7, Eko-1-9, Elu-1-1, Elu-1-2, Elu-1-3, Elu-1-4, Ema-1-1, Emo-1-1, Epy-1-1, Esp-1-1, Esp-10-1, Esp-11-1, Esp-12-1, Esp-13-1, Esp-15-1, Esp-16-1, Esp-17-1, Esp-2-1, Esp-20-1, Esp-21-1, Esp-22-1, Esp-23-1, Esp-24-1, Esp-25-1, Esp-26-1, Esp-27-1, Esp-28-1, Esp-29-1, Esp-3-1, Esp-30-1, Esp-31-1, Esp-32-1, Esp-33-1, Esp-34-1, Esp-35-1, Esp-36-1, Esp-38-1, Esp-39-1, Esp-4-1, Esp-40-1, Esp-41-1, Esp-44-1, Esp-45-1, Esp-46-1, Esp-47-1, Esp-48-1, Esp-49-1, Esp-5-1, Esp-50-1, Esp-52-1, Esp-54-1, Esp-56-1, Esp-57-1, Esp-58-1, Esp-59-1, Esp-6-1, Esp-60-1, Esp-61-1, Esp-62-1, Esp-63-1, Esp-64-1, Esp-66-1, Esp-67-1, Esp-69-1, Esp-7-1, Esp-70-1, Esp-71-1, Esp-72-1, Esp-73-1, Esp-75-1, Esp-76-1, Esp-8-1, Esp-80-1, Esp-9-1, Eta-2-1, Eta-3-1, Eta-6-1, Eta-7-1, Eta-8-1, Etr-2-1, Fhe-2-1, Fhe-3-1, Fpu-2-1, Fpu-3-1, Fpu-4-1, Gan-1-1, Gan-1-11, Gan-1-12, Gan-1-3, Gan-1-4, Gan-1-7, Gan-1-8, Gan-10-1, Gan-2-1, Gan-4-1, Gan-8-1, Gge-1-1, Gge-2-1, Hal-1-3, Hal-1-4, Hal-2-1, Hal-3-1, Hha-2-1, Hin-1-2, Hin-1-3, Hin-1-4, Hin-1-5, Hin-1-6, Hin-2-1, Hin-3-1, Hin-5-1, Hin-6-1, Hin-7-1, Hpa-1-2, Kae-1-1, Kae-10-1, Kae-11-1, Kae-12-1, Kae-13-1, Kae-4-1, Kae-5-1, Kae-6-1, Kae-7-1, Kae-8-1, Kae-9-1, Kcf-1-1, Kmi-1-1, Kmi-2-1, Kmi-3-1, Kox-1-2, Kox-1-3, Kox-1-4, Kox-10-1, Kox-11-1, Kox-12-1, Kox-13-1, Kox-14-1, Kox-16-1, Kox-17-1, Kox-19-1, Kox-2-1, Kox-20-1, Kox-3-1, Kox-4-1, Kox-5-1, Kox-7-1, Kox-8-1, Kox-9-1, Kpn-1-1, Kpn-1-2, Kpn-1-3, Kpn-1-4, Kpn-1-5, Kpn-1-6, Kpn-1-7, Kpn-1-8, Kpn-1-9, Kpn-10-1, Kpn-100-1, Kpn-101-1, Kpn-102-1, Kpn-103-1, Kpn-104-1, Kpn-105-1, Kpn-106-1, Kpn-107-1, Kpn-108-1, Kpn-109-1, Kpn-11-1, Kpn-110-1, Kpn-111-1, Kpn-112-1, Kpn-113-1, Kpn-114-1, Kpn-115-1, Kpn-116-1, Kpn-117-1, Kpn-118-1, Kpn-119-1, Kpn-12-1, Kpn-120-1, Kpn-121-1, Kpn-122-1, Kpn-123-1, Kpn-124-1, Kpn-125-1, Kpn-126-1, Kpn-127-1, Kpn-128-1, Kpn-129-1, Kpn-13-1, Kpn-130-1, Kpn-131-1, Kpn-132-1, Kpn-133-1, Kpn-134-1, Kpn-135-1, Kpn-136-1, Kpn-137-1, Kpn-138-1, Kpn-139-1, Kpn-14-1, Kpn-140-1, Kpn-141-1, Kpn-142-1, Kpn-143-1, Kpn-144-1, Kpn-145-1, Kpn-146-1, Kpn-147-1, Kpn-148-1, Kpn-149-1, Kpn-15-1, Kpn-150-1, Kpn-151-1, Kpn-152-1, Kpn-153-1, Kpn-154-1, Kpn-155-1, Kpn-156-1, Kpn-157-1, Kpn-158-1, Kpn-159-1, Kpn-16-1, Kpn-160-1, Kpn-161-1, Kpn-162-1, Kpn-163-1, Kpn-164-1, Kpn-165-1, Kpn-166-1, Kpn-167-1, Kpn-168-1, Kpn-17-1, Kpn-170-1, Kpn-171-1, Kpn-172-1, Kpn-173-1, Kpn-174-1, Kpn-175-1, Kpn-176-1, Kpn-177-1, Kpn-178-1, Kpn-179-1, Kpn-18-1, Kpn-180-1, Kpn-181-1, Kpn-182-1, Kpn-183-1, Kpn-184-1, Kpn-185-1, Kpn-186-1, Kpn-187-1, Kpn-188-1, Kpn-189-1, Kpn-19-1, Kpn-190-1, Kpn-191-1, Kpn-192-1, Kpn-193-1, Kpn-195-1, Kpn-196-1, Kpn-197-1, Kpn-198-1, Kpn-199-1, Kpn-199-2, Kpn-2-1, Kpn-20-1, Kpn-200-1, Kpn-200-2, Kpn-201-1, Kpn-202-1, Kpn-203-1, Kpn-203-2, Kpn-204-1, Kpn-204-2, Kpn-205-1, Kpn-205-2, Kpn-205-3, Kpn-205-4, Kpn-206-1, Kpn-206-2, Kpn-206-3, Kpn-206-4, Kpn-206-5, Kpn-206-6, Kpn-206-7, Kpn-207-1, Kpn-208-1, Kpn-209-1, Kpn-21-1, Kpn-210-1, Kpn-211-1, Kpn-212-1, Kpn-213-1, Kpn-214-1, Kpn-215-1, Kpn-216-1, Kpn-217-1, Kpn-218-1, Kpn-22-1, Kpn-220-1, Kpn-221-1, Kpn-222-1, Kpn-223-1, Kpn-224-1, Kpn-225-1, Kpn-227-1, Kpn-228-1, Kpn-229-1, Kpn-23-1, Kpn-230-1, Kpn-231-1, Kpn-232-1, Kpn-233-1, Kpn-234-1, Kpn-235-1, Kpn-236-1, Kpn-237-1, Kpn-238-1, Kpn-239-1, Kpn-24-1, Kpn-240-1, Kpn-241-1, Kpn-242-1, Kpn-243-1, Kpn-244-1, Kpn-245-1, Kpn-246-1, Kpn-247-1, Kpn-248-1, Kpn-249-1, Kpn-25-1, Kpn-250-1, Kpn-251-1, Kpn-252-1, Kpn-253-1, Kpn-254-1, Kpn-255-1, Kpn-256-1, Kpn-257-1, Kpn-258-1, Kpn-259-1, Kpn-26-1, Kpn-260-1, Kpn-261-1, Kpn-262-1, Kpn-263-1, Kpn-264-1, Kpn-265-1, Kpn-266-1, Kpn-267-1, Kpn-268-1, Kpn-269-1, Kpn-27-1, Kpn-270-1, Kpn-271-1, Kpn-272-1, Kpn-273-1, Kpn-274-1, Kpn-275-1, Kpn-276-1, Kpn-277-1, Kpn-278-1, Kpn-279-1, Kpn-28-1, Kpn-280-1, Kpn-282-1, Kpn-283-1, Kpn-284-1, Kpn-285-1, Kpn-286-1, Kpn-287-1, Kpn-288-1, Kpn-289-1, Kpn-29-1, Kpn-290-1, Kpn-291-1, Kpn-292-1, Kpn-293-1, Kpn-294-1, Kpn-295-1, Kpn-296-1, Kpn-297-1, Kpn-298-1, Kpn-299-1, Kpn-3-1, Kpn-3-2, Kpn-30-1, Kpn-300-1, Kpn-301-1, Kpn-302-1, Kpn-303-1, Kpn-304-1, Kpn-305-1, Kpn-306-1, Kpn-307-1, Kpn-308-1, Kpn-309-1, Kpn-31-1, Kpn-310-1, Kpn-311-1, Kpn-312-1, Kpn-313-1, Kpn-314-1,

Kpn-315-1, Kpn-316-1, Kpn-317-1, Kpn-318-1, Kpn-319-1, Kpn-32-1, Kpn-320-1, Kpn-321-1, Kpn-322-1, Kpn-323-1, Kpn-324-1, Kpn-325-1, Kpn-326-1, Kpn-327-1, Kpn-328-1, Kpn-329-1, Kpn-33-1, Kpn-330-1, Kpn-331-1, Kpn-332-1, Kpn-333-1, Kpn-334-1, Kpn-335-1, Kpn-336-1, Kpn-337-1, Kpn-338-1, Kpn-339-1, Kpn-34-1, Kpn-340-1, Kpn-341-1, Kpn-342-1, Kpn-343-1, Kpn-344-1, Kpn-345-1, Kpn-346-1, Kpn-347-1, Kpn-348-1, Kpn-349-1, Kpn-35-1, Kpn-350-1, Kpn-351-1, Kpn-352-1, Kpn-353-1, Kpn-354-1, Kpn-355-1, Kpn-356-1, Kpn-357-1, Kpn-358-1, Kpn-359-1, Kpn-36-1, Kpn-360-1, Kpn-361-1, Kpn-362-1, Kpn-37-1, Kpn-38-1, Kpn-39-1, Kpn-4-1, Kpn-40-1, Kpn-41-1, Kpn-42-1, Kpn-43-1, Kpn-44-1, Kpn-45-1, Kpn-46-1, Kpn-47-1, Kpn-48-1, Kpn-49-1, Kpn-5-1, Kpn-50-1, Kpn-51-1, Kpn-52-1, Kpn-53-1, Kpn-54-1, Kpn-55-1, Kpn-56-1, Kpn-57-1, Kpn-58-1, Kpn-59-1, Kpn-6-1, Kpn-60-1, Kpn-61-1, Kpn-62-1, Kpn-63-1, Kpn-64-1, Kpn-65-1, Kpn-66-1, Kpn-67-1, Kpn-68-1, Kpn-69-1, Kpn-7-1, Kpn-70-1, Kpn-71-1, Kpn-72-1, Kpn-73-1, Kpn-74-1, Kpn-75-1, Kpn-76-1, Kpn-77-1, Kpn-78-1, Kpn-79-1, Kpn-8-1, Kpn-80-1, Kpn-81-1, Kpn-82-1, Kpn-83-1, Kpn-84-1, Kpn-85-1, Kpn-86-1, Kpn-87-1, Kpn-88-1, Kpn-89-1, Kpn-9-1, Kpn-90-1, Kpn-91-1, Kpn-92-1, Kpn-93-1, Kpn-94-1, Kpn-95-1, Kpn-96-1, Kpn-97-1, Kpn-98-1, Kpn-99-1, Kqu-2-1, Kqu-2-2, Kqu-2-3, Kqu-2-4, Kra-1-1, Ksp-4-1, Ksp-5-1, Ksp-6-1, Ksp-7-1, Kva-1-1, Kva-1-2, Kva-1-3, Kva-1-4, Kva-1-6, Kva-1-7, Kva-2-1, Lad-1-2, Lad-2-1, Lam-1-1, Lgr-1-2, Lqu-1-3, Lsp-1-1, Mmo-1-2, Mmo-1-4, Mmo-1-5, Mmo-1-6, Mmo-1-8, Mmo-4-1, Mmo-6-1, Msp-2-1, Pag-1-2, Pag-1-3, Pag-1-4, Pag-2-1, Pal-1-1, Pal-3-1, Pal-4-1, Pal-6-1, Pal-7-1, Pan-1-1, Pan-1-2, Pan-1-4, Pan-1-5, Pan-10-1, Pan-2-1, Pan-3-1, Pan-3-2, Pan-4-1, Pan-6-1, Pan-7-1, Pan-8-1, Pan-9-1, Pat-1-1, Pat-1-2, Pat-2-1, Pat-4-1, Pca-11-1, Pca-12-1, Pca-3-10, Pca-3-11, Pca-3-12, Pca-3-13, Pca-3-14, Pca-3-3, Pca-3-4, Pca-3-5, Pca-3-6, Pca-3-7, Pca-3-8, Pca-3-9, Pca-4-1, Pca-5-4, Pca-5-5, Pca-5-6, Pca-5-8, Pca-8-2, Pdi-1-1, Plu-1-1, Pmi-1-1, Pmi-1-3, Pmi-1-4, Pmi-1-5, Pmi-1-6, Pmi-3-1, Pmi-4-1, Pmi-5-1, Pmi-6-1, Pmi-7-1, Pmi-8-1, Ppa-1-1, Pre-1-2, Prw-1-1, Psp-11-1, Psp-15-1, Psp-16-1, Psp-17-1, Psp-19-1, Psp-2-1, Psp-20-1, Psp-4-1, Psp-6-1, Psp-8-1, Pst-1-2, Pst-2-1, Pst-3-1, Pte-4-1, Pte-5-1, Pva-1-3, Pva-2-1, Pvu-1-1, Pwa-1-1, Pwa-1-2, Ror-2-1, Ror-3-1, Ror-4-1, Ror-4-2, Rpl-1-1, Rpl-1-2, Rpl-1-3, Rpl-2-1, Sbl-1-1, Sbo-1-1, Sbo-10-1, Sbo-11-1, Sbo-12-1, Sbo-14-1, Sbo-15-1, Sbo-16-1, Sbo-17-1, Sbo-18-1, Sbo-2-1, Sbo-2-10, Sbo-2-2, Sbo-2-3, Sbo-2-4, Sbo-2-5, Sbo-2-6, Sbo-2-8, Sbo-2-9, Sbo-4-1, Sbo-5-1, Sbo-6-1, Sbo-7-1, Sbo-8-1, Sbo-9-1, Sdy-1-1, Sdy-10-1, Sdy-2-1, Sdy-3-1, Sdy-4-1, Sdy-5-1, Sdy-6-1, Sdy-7-1, Sdy-9-1, Sen-1-1, Sen-10-1, Sen-100-1, Sen-1000-1, Sen-1001-1, Sen-1002-1, Sen-1003-1, Sen-1004-1, Sen-1005-1, Sen-1006-1, Sen-1008-1, Sen-1009-1, Sen-101-1, Sen-1010-1, Sen-1011-1, Sen-1012-1, Sen-1013-1, Sen-1014-1, Sen-1015-1, Sen-1016-1, Sen-1016-2, Sen-1017-1, Sen-1019-1, Sen-102-1, Sen-1021-1, Sen-1022-1, Sen-1023-1, Sen-1024-1, Sen-1024-2, Sen-1025-1, Sen-1026-1, Sen-1027-1, Sen-1028-1, Sen-1029-1, Sen-103-1, Sen-1030-1, Sen-104-1, Sen-105-1, Sen-106-1, Sen-107-1, Sen-108-1, Sen-109-1, Sen-11-1, Sen-110-1, Sen-111-1, Sen-111-2, Sen-111-3, Sen-112-1, Sen-112-2, Sen-113-1, Sen-114-1, Sen-115-1, Sen-116-1, Sen-117-1, Sen-118-1, Sen-119-1, Sen-12-1, Sen-12-2, Sen-12-3, Sen-120-1, Sen-121-1, Sen-122-1, Sen-123-1, Sen-124-1, Sen-125-1, Sen-126-1, Sen-127-1, Sen-128-1, Sen-128-2, Sen-129-1, Sen-129-2, Sen-129-3, Sen-13-1, Sen-130-1, Sen-131-1, Sen-132-1, Sen-133-1, Sen-134-1, Sen-135-1, Sen-135-2, Sen-136-1, Sen-136-2, Sen-136-3, Sen-136-4, Sen-136-5, Sen-136-6, Sen-137-1, Sen-137-2, Sen-137-3, Sen-137-4, Sen-137-5, Sen-138-1, Sen-138-10, Sen-138-11, Sen-138-12, Sen-138-13, Sen-138-14, Sen-138-15, Sen-138-2, Sen-138-3, Sen-138-4, Sen-138-5, Sen-138-6, Sen-138-7, Sen-138-8, Sen-138-9, Sen-139-1, Sen-14-1, Sen-14-2, Sen-14-3, Sen-14-4, Sen-140-1, Sen-141-1, Sen-142-1, Sen-143-1, Sen-144-1, Sen-145-1, Sen-146-1, Sen-147-1, Sen-148-1, Sen-149-1, Sen-15-1, Sen-15-2, Sen-15-3, Sen-150-1, Sen-151-1, Sen-152-1, Sen-153-1, Sen-154-1, Sen-155-1, Sen-156-1, Sen-157-1, Sen-158-1, Sen-159-1, Sen-16-1, Sen-160-1, Sen-161-1, Sen-162-1, Sen-163-1, Sen-164-1, Sen-165-1, Sen-166-1, Sen-166-10, Sen-166-11, Sen-166-12, Sen-166-13, Sen-166-14, Sen-166-15, Sen-166-16, Sen-166-17, Sen-166-18, Sen-166-2, Sen-166-3, Sen-166-4, Sen-166-5, Sen-166-6, Sen-166-7, Sen-166-8, Sen-166-9, Sen-167-1, Sen-168-1, Sen-169-1, Sen-169-10, Sen-169-11, Sen-169-12, Sen-169-13, Sen-169-14, Sen-169-15, Sen-169-16, Sen-169-17, Sen-169-18, Sen-169-19, Sen-169-2, Sen-169-20, Sen-169-21, Sen-169-22, Sen-169-23, Sen-169-24, Sen-169-25, Sen-169-26, Sen-169-27, Sen-169-28, Sen-169-29, Sen-169-3, Sen-169-30, Sen-169-31, Sen-169-32, Sen-169-33, Sen-169-34, Sen-169-35, Sen-169-36, Sen-169-37, Sen-169-38, Sen-169-39, Sen-169-4, Sen-169-40, Sen-169-41, Sen-169-42, Sen-169-43, Sen-169-44, Sen-169-45, Sen-169-46, Sen-169-47, Sen-169-48, Sen-169-49, Sen-169-5, Sen-169-50, Sen-169-51, Sen-169-52, Sen-169-6, Sen-169-7, Sen-169-8, Sen-169-9, Sen-17-1, Sen-170-1, Sen-171-1, Sen-172-1, Sen-173-1, Sen-174-1, Sen-175-1, Sen-176-1, Sen-177-1, Sen-178-1, Sen-179-1, Sen-18-1, Sen-180-1, Sen-181-1, Sen-182-1, Sen-183-1, Sen-184-1, Sen-185-1, Sen-186-1, Sen-187-1, Sen-188-1, Sen-189-1, Sen-19-1, Sen-190-1, Sen-191-1, Sen-192-1, Sen-193-1, Sen-194-1, Sen-195-1, Sen-196-1, Sen-197-1, Sen-198-1, Sen-199-1, Sen-2-1, Sen-2-2, Sen-20-1, Sen-200-1, Sen-201-1, Sen-202-1, Sen-203-1, Sen-204-1, Sen-205-1, Sen-206-1, Sen-207-1, Sen-208-1, Sen-209-1, Sen-21-1, Sen-210-1, Sen-211-1, Sen-212-1, Sen-213-1, Sen-214-1, Sen-215-1, Sen-216-1, Sen-217-1, Sen-218-1, Sen-219-1, Sen-22-1, Sen-220-1, Sen-221-1, Sen-222-1, Sen-223-2, Sen-223-3, Sen-224-1, Sen-224-10, Sen-224-11, Sen-224-12, Sen-224-13, Sen-224-14, Sen-224-15, Sen-224-16, Sen-224-17, Sen-224-18, Sen-224-19, Sen-224-2, Sen-224-20, Sen-224-21, Sen-224-22, Sen-224-23, Sen-224-24, Sen-224-25, Sen-224-26, Sen-224-27, Sen-224-28, Sen-224-29, Sen-224-3, Sen-224-30, Sen-224-31, Sen-224-32, Sen-224-33, Sen-224-34, Sen-224-35, Sen-224-36, Sen-224-37, Sen-224-38, Sen-224-39, Sen-224-4, Sen-224-40, Sen-224-41, Sen-224-42, Sen-224-43, Sen-224-44, Sen-224-45, Sen-224-46, Sen-224-47, Sen-224-48, Sen-224-49, Sen-224-5, Sen-224-50, Sen-224-51, Sen-224-52, Sen-224-6, Sen-224-7, Sen-224-8, Sen-224-9, Sen-225-2, Sen-225-3, Sen-225-4, Sen-226-2, Sen-227-1, Sen-227-2, Sen-228-1, Sen-229-1, Sen-229-2, Sen-229-3, Sen-229-4, Sen-230-1, Sen-231-1, Sen-232-1, Sen-232-2, Sen-232-3, Sen-232-4, Sen-232-5, Sen-233-1, Sen-234-1, Sen-235-1, Sen-236-1, Sen-236-10, Sen-236-11, Sen-236-12, Sen-236-13, Sen-236-14, Sen-236-15, Sen-236-16, Sen-236-2, Sen-236-3, Sen-236-4, Sen-236-5, Sen-236-6, Sen-236-7, Sen-236-8, Sen-236-9, Sen-237-1, Sen-238-1, Sen-24-1, Sen-240-1, Sen-241-1, Sen-242-1, Sen-242-2, Sen-243-1, Sen-244-1, Sen-245-1, Sen-246-1, Sen-247-1, Sen-248-1, Sen-249-1, Sen-25-1, Sen-250-1, Sen-251-1, Sen-252-1, Sen-253-1, Sen-254-1, Sen-255-1, Sen-256-1, Sen-257-1,

Sen-258-1, Sen-259-1, Sen-26-1, Sen-260-1, Sen-261-1, Sen-262-1, Sen-263-1, Sen-264-1, Sen-265-1, Sen-265-2, Sen-266-1, Sen-267-1, Sen-268-1, Sen-27-1, Sen-270-1, Sen-270-2, Sen-270-3, Sen-271-1, Sen-272-1, Sen-272-2, Sen-272-3, Sen-273-1, Sen-274-1, Sen-275-1, Sen-276-1, Sen-276-2, Sen-276-3, Sen-276-4, Sen-276-5, Sen-277-1, Sen-277-10, Sen-277-11, Sen-277-12, Sen-277-13, Sen-277-14, Sen-277-15, Sen-277-16, Sen-277-17, Sen-277-18, Sen-277-19, Sen-277-2, Sen-277-20, Sen-277-21, Sen-277-22, Sen-277-3, Sen-277-4, Sen-277-5, Sen-277-6, Sen-277-7, Sen-277-8, Sen-277-9, Sen-278-1, Sen-278-10, Sen-278-11, Sen-278-12, Sen-278-13, Sen-278-14, Sen-278-2, Sen-278-3, Sen-278-4, Sen-278-5, Sen-278-6, Sen-278-7, Sen-278-8, Sen-278-9, Sen-279-1, Sen-279-2, Sen-279-3, Sen-28-1, Sen-280-1, Sen-281-1, Sen-282-1, Sen-283-1, Sen-284-1, Sen-285-1, Sen-285-2, Sen-285-3, Sen-285-4, Sen-285-5, Sen-286-1, Sen-287-10, Sen-287-100, Sen-287-1000, Sen-287-1001, Sen-287-1002, Sen-287-1003, Sen-287-1004, Sen-287-1005, Sen-287-1006, Sen-287-1007, Sen-287-1008, Sen-287-1009, Sen-287-101, Sen-287-1010, Sen-287-1011, Sen-287-1012, Sen-287-1013, Sen-287-1014, Sen-287-1015, Sen-287-1016, Sen-287-1017, Sen-287-1018, Sen-287-1019, Sen-287-102, Sen-287-1020, Sen-287-1021, Sen-287-1022, Sen-287-1023, Sen-287-1024, Sen-287-1025, Sen-287-1026, Sen-287-1027, Sen-287-1028, Sen-287-1029, Sen-287-103, Sen-287-1030, Sen-287-1031, Sen-287-1032, Sen-287-1033, Sen-287-1034, Sen-287-1035, Sen-287-1036, Sen-287-1037, Sen-287-1038, Sen-287-1039, Sen-287-104, Sen-287-1040, Sen-287-1041, Sen-287-1042, Sen-287-1043, Sen-287-1044, Sen-287-1045, Sen-287-1046, Sen-287-1047, Sen-287-1048, Sen-287-1049, Sen-287-105, Sen-287-1050, Sen-287-1051, Sen-287-1052, Sen-287-1053, Sen-287-1054, Sen-287-1055, Sen-287-1056, Sen-287-1057, Sen-287-1058, Sen-287-1059, Sen-287-106, Sen-287-1060, Sen-287-1061, Sen-287-1062, Sen-287-1063, Sen-287-1064, Sen-287-1065, Sen-287-1066, Sen-287-1067, Sen-287-1068, Sen-287-1069, Sen-287-107, Sen-287-1070, Sen-287-1071, Sen-287-1072, Sen-287-1073, Sen-287-1074, Sen-287-1075, Sen-287-1076, Sen-287-1077, Sen-287-1078, Sen-287-1079, Sen-287-108, Sen-287-1080, Sen-287-1081, Sen-287-1082, Sen-287-1083, Sen-287-1084, Sen-287-1085, Sen-287-1086, Sen-287-1087, Sen-287-1088, Sen-287-1089, Sen-287-109, Sen-287-1090, Sen-287-1091, Sen-287-1092, Sen-287-1093, Sen-287-1094, Sen-287-1095, Sen-287-1096, Sen-287-1097, Sen-287-1098, Sen-287-1099, Sen-287-11, Sen-287-110, Sen-287-1100, Sen-287-1101, Sen-287-1102, Sen-287-1103, Sen-287-1104, Sen-287-1105, Sen-287-1106, Sen-287-1107, Sen-287-1108, Sen-287-1109, Sen-287-111, Sen-287-1110, Sen-287-1111, Sen-287-1112, Sen-287-1113, Sen-287-1114, Sen-287-1115, Sen-287-1116, Sen-287-1117, Sen-287-1118, Sen-287-1119, Sen-287-112, Sen-287-1120, Sen-287-1121, Sen-287-1122, Sen-287-1123, Sen-287-1124, Sen-287-113, Sen-287-114, Sen-287-115, Sen-287-116, Sen-287-117, Sen-287-118, Sen-287-119, Sen-287-12, Sen-287-120, Sen-287-121, Sen-287-122, Sen-287-123, Sen-287-124, Sen-287-125, Sen-287-126, Sen-287-127, Sen-287-128, Sen-287-129, Sen-287-13, Sen-287-130, Sen-287-131, Sen-287-132, Sen-287-133, Sen-287-134, Sen-287-135, Sen-287-136, Sen-287-137, Sen-287-138, Sen-287-139, Sen-287-14, Sen-287-140, Sen-287-141, Sen-287-142, Sen-287-143, Sen-287-144, Sen-287-145, Sen-287-146, Sen-287-147, Sen-287-148, Sen-287-149, Sen-287-15, Sen-287-150, Sen-287-151, Sen-287-152, Sen-287-153, Sen-287-154, Sen-287-155, Sen-287-156, Sen-287-157, Sen-287-158, Sen-287-159, Sen-287-16, Sen-287-160, Sen-287-161, Sen-287-162, Sen-287-163, Sen-287-164, Sen-287-165, Sen-287-166, Sen-287-167, Sen-287-168, Sen-287-169, Sen-287-17, Sen-287-170, Sen-287-171, Sen-287-172, Sen-287-173, Sen-287-174, Sen-287-175, Sen-287-176, Sen-287-177, Sen-287-178, Sen-287-179, Sen-287-18, Sen-287-180, Sen-287-181, Sen-287-182, Sen-287-183, Sen-287-184, Sen-287-185, Sen-287-186, Sen-287-187, Sen-287-188, Sen-287-189, Sen-287-19, Sen-287-190, Sen-287-191, Sen-287-192, Sen-287-193, Sen-287-194, Sen-287-195, Sen-287-196, Sen-287-197, Sen-287-198, Sen-287-199, Sen-287-2, Sen-287-20, Sen-287-200, Sen-287-201, Sen-287-202, Sen-287-203, Sen-287-204, Sen-287-205, Sen-287-206, Sen-287-207, Sen-287-208, Sen-287-209, Sen-287-21, Sen-287-210, Sen-287-211, Sen-287-212, Sen-287-213, Sen-287-214, Sen-287-215, Sen-287-216, Sen-287-217, Sen-287-218, Sen-287-219, Sen-287-22, Sen-287-220, Sen-287-221, Sen-287-222, Sen-287-223, Sen-287-224, Sen-287-225, Sen-287-226, Sen-287-227, Sen-287-228, Sen-287-229, Sen-287-23, Sen-287-230, Sen-287-231, Sen-287-232, Sen-287-233, Sen-287-234, Sen-287-235, Sen-287-236, Sen-287-237, Sen-287-238, Sen-287-239, Sen-287-24, Sen-287-240, Sen-287-241, Sen-287-242, Sen-287-243, Sen-287-244, Sen-287-245, Sen-287-246, Sen-287-247, Sen-287-248, Sen-287-249, Sen-287-25, Sen-287-250, Sen-287-251, Sen-287-252, Sen-287-253, Sen-287-254, Sen-287-255, Sen-287-256, Sen-287-257, Sen-287-258, Sen-287-259, Sen-287-26, Sen-287-260, Sen-287-261, Sen-287-262, Sen-287-263, Sen-287-264, Sen-287-265, Sen-287-266, Sen-287-267, Sen-287-268, Sen-287-269, Sen-287-27, Sen-287-270, Sen-287-271, Sen-287-272, Sen-287-273, Sen-287-274, Sen-287-275, Sen-287-276, Sen-287-277, Sen-287-278, Sen-287-279, Sen-287-28, Sen-287-280, Sen-287-281, Sen-287-282, Sen-287-283, Sen-287-284, Sen-287-285, Sen-287-286, Sen-287-287, Sen-287-288, Sen-287-289, Sen-287-29, Sen-287-290, Sen-287-291, Sen-287-292, Sen-287-293, Sen-287-294, Sen-287-295, Sen-287-296, Sen-287-297, Sen-287-298, Sen-287-299, Sen-287-3, Sen-287-30, Sen-287-300, Sen-287-301, Sen-287-302, Sen-287-303, Sen-287-304, Sen-287-305, Sen-287-306, Sen-287-307, Sen-287-308, Sen-287-309, Sen-287-31, Sen-287-310, Sen-287-311, Sen-287-312, Sen-287-313, Sen-287-314, Sen-287-315, Sen-287-316, Sen-287-317, Sen-287-318, Sen-287-319, Sen-287-32, Sen-287-320, Sen-287-321, Sen-287-322, Sen-287-323, Sen-287-324, Sen-287-325, Sen-287-326, Sen-287-327, Sen-287-328, Sen-287-329, Sen-287-33, Sen-287-330, Sen-287-331, Sen-287-332, Sen-287-333, Sen-287-334, Sen-287-335, Sen-287-336, Sen-287-337, Sen-287-338, Sen-287-339, Sen-287-34, Sen-287-340, Sen-287-341, Sen-287-342, Sen-287-343, Sen-287-344, Sen-287-345, Sen-287-346, Sen-287-347, Sen-287-348, Sen-287-349, Sen-287-35, Sen-287-350, Sen-287-351, Sen-287-352, Sen-287-353, Sen-287-354, Sen-287-355, Sen-287-356, Sen-287-357, Sen-287-358, Sen-287-359, Sen-287-36, Sen-287-360, Sen-287-361, Sen-287-362, Sen-287-363, Sen-287-364, Sen-287-365, Sen-287-366, Sen-287-367, Sen-287-368, Sen-287-369, Sen-287-37, Sen-287-370, Sen-287-371, Sen-287-372, Sen-287-373, Sen-287-374, Sen-287-375, Sen-287-376, Sen-287-377, Sen-287-378, Sen-287-379, Sen-287-38, Sen-287-380, Sen-287-381, Sen-287-382, Sen-287-383, Sen-287-384, Sen-287-385, Sen-287-386, Sen-287-387, Sen-287-388, Sen-287-389, Sen-287-39, Sen-287-390, Sen-287-391, Sen-287-392, Sen-287-393, Sen-287-394, Sen-287-395, Sen-287-396, Sen-287-397, Sen-287-398, Sen-287-399, Sen-287-4, Sen-287-40, Sen-287-400, Sen-287-401, Sen-287-402, Sen-287-403, Sen-287-404, Sen-287-405, Sen-287-406, Sen-287-407, Sen-287-408, Sen-287-409,

Sen-287-41, Sen-287-410, Sen-287-411, Sen-287-412, Sen-287-413, Sen-287-414, Sen-287-415, Sen-287-416, Sen-287-417, Sen-287-418, Sen-287-419, Sen-287-42, Sen-287-420, Sen-287-421, Sen-287-422, Sen-287-423, Sen-287-424, Sen-287-425, Sen-287-426, Sen-287-427, Sen-287-428, Sen-287-429, Sen-287-43, Sen-287-430, Sen-287-431, Sen-287-432, Sen-287-433, Sen-287-434, Sen-287-435, Sen-287-436, Sen-287-437, Sen-287-438, Sen-287-439, Sen-287-44, Sen-287-440, Sen-287-441, Sen-287-442, Sen-287-443, Sen-287-444, Sen-287-445, Sen-287-446, Sen-287-447, Sen-287-448, Sen-287-449, Sen-287-45, Sen-287-450, Sen-287-451, Sen-287-452, Sen-287-453, Sen-287-454, Sen-287-455, Sen-287-456, Sen-287-457, Sen-287-458, Sen-287-459, Sen-287-46, Sen-287-460, Sen-287-461, Sen-287-462, Sen-287-463, Sen-287-464, Sen-287-465, Sen-287-466, Sen-287-467, Sen-287-468, Sen-287-469, Sen-287-47, Sen-287-470, Sen-287-471, Sen-287-472, Sen-287-473, Sen-287-474, Sen-287-475, Sen-287-476, Sen-287-477, Sen-287-478, Sen-287-479, Sen-287-48, Sen-287-480, Sen-287-481, Sen-287-482, Sen-287-483, Sen-287-484, Sen-287-485, Sen-287-486, Sen-287-487, Sen-287-488, Sen-287-489, Sen-287-49, Sen-287-490, Sen-287-491, Sen-287-492, Sen-287-493, Sen-287-494, Sen-287-495, Sen-287-496, Sen-287-497, Sen-287-498, Sen-287-499, Sen-287-5, Sen-287-50, Sen-287-500, Sen-287-501, Sen-287-502, Sen-287-503, Sen-287-504, Sen-287-505, Sen-287-506, Sen-287-507, Sen-287-508, Sen-287-509, Sen-287-51, Sen-287-510, Sen-287-511, Sen-287-512, Sen-287-513, Sen-287-514, Sen-287-515, Sen-287-516, Sen-287-517, Sen-287-518, Sen-287-519, Sen-287-52, Sen-287-520, Sen-287-521, Sen-287-522, Sen-287-523, Sen-287-524, Sen-287-525, Sen-287-526, Sen-287-527, Sen-287-528, Sen-287-529, Sen-287-53, Sen-287-530, Sen-287-531, Sen-287-532, Sen-287-533, Sen-287-534, Sen-287-535, Sen-287-536, Sen-287-537, Sen-287-538, Sen-287-539, Sen-287-54, Sen-287-540, Sen-287-541, Sen-287-542, Sen-287-543, Sen-287-544, Sen-287-545, Sen-287-546, Sen-287-547, Sen-287-548, Sen-287-549, Sen-287-55, Sen-287-550, Sen-287-551, Sen-287-552, Sen-287-553, Sen-287-554, Sen-287-555, Sen-287-556, Sen-287-557, Sen-287-558, Sen-287-559, Sen-287-56, Sen-287-560, Sen-287-561, Sen-287-562, Sen-287-563, Sen-287-564, Sen-287-565, Sen-287-566, Sen-287-567, Sen-287-568, Sen-287-569, Sen-287-57, Sen-287-570, Sen-287-571, Sen-287-572, Sen-287-573, Sen-287-574, Sen-287-575, Sen-287-576, Sen-287-577, Sen-287-578, Sen-287-579, Sen-287-58, Sen-287-580, Sen-287-581, Sen-287-582, Sen-287-583, Sen-287-584, Sen-287-585, Sen-287-586, Sen-287-587, Sen-287-588, Sen-287-589, Sen-287-59, Sen-287-590, Sen-287-591, Sen-287-592, Sen-287-593, Sen-287-594, Sen-287-595, Sen-287-596, Sen-287-597, Sen-287-598, Sen-287-599, Sen-287-6, Sen-287-60, Sen-287-600, Sen-287-601, Sen-287-602, Sen-287-603, Sen-287-604, Sen-287-605, Sen-287-606, Sen-287-607, Sen-287-608, Sen-287-609, Sen-287-61, Sen-287-610, Sen-287-611, Sen-287-612, Sen-287-613, Sen-287-614, Sen-287-615, Sen-287-616, Sen-287-617, Sen-287-618, Sen-287-619, Sen-287-62, Sen-287-620, Sen-287-621, Sen-287-622, Sen-287-623, Sen-287-624, Sen-287-625, Sen-287-626, Sen-287-627, Sen-287-628, Sen-287-629, Sen-287-63, Sen-287-630, Sen-287-631, Sen-287-632, Sen-287-633, Sen-287-634, Sen-287-635, Sen-287-636, Sen-287-637, Sen-287-638, Sen-287-639, Sen-287-64, Sen-287-640, Sen-287-641, Sen-287-642, Sen-287-643, Sen-287-644, Sen-287-645, Sen-287-646, Sen-287-647, Sen-287-648, Sen-287-649, Sen-287-65, Sen-287-650, Sen-287-651, Sen-287-652, Sen-287-653, Sen-287-654, Sen-287-655, Sen-287-656, Sen-287-657, Sen-287-658, Sen-287-659, Sen-287-66, Sen-287-660, Sen-287-661, Sen-287-662, Sen-287-663, Sen-287-664, Sen-287-665, Sen-287-666, Sen-287-667, Sen-287-668, Sen-287-669, Sen-287-67, Sen-287-670, Sen-287-671, Sen-287-672, Sen-287-673, Sen-287-674, Sen-287-675, Sen-287-676, Sen-287-677, Sen-287-678, Sen-287-679, Sen-287-68, Sen-287-680, Sen-287-681, Sen-287-682, Sen-287-683, Sen-287-684, Sen-287-685, Sen-287-686, Sen-287-687, Sen-287-688, Sen-287-689, Sen-287-69, Sen-287-690, Sen-287-691, Sen-287-692, Sen-287-693, Sen-287-694, Sen-287-695, Sen-287-696, Sen-287-697, Sen-287-698, Sen-287-699, Sen-287-7, Sen-287-70, Sen-287-700, Sen-287-701, Sen-287-702, Sen-287-703, Sen-287-704, Sen-287-705, Sen-287-706, Sen-287-707, Sen-287-708, Sen-287-709, Sen-287-71, Sen-287-710, Sen-287-711, Sen-287-712, Sen-287-713, Sen-287-714, Sen-287-715, Sen-287-716, Sen-287-717, Sen-287-718, Sen-287-719, Sen-287-72, Sen-287-720, Sen-287-721, Sen-287-722, Sen-287-723, Sen-287-724, Sen-287-725, Sen-287-726, Sen-287-727, Sen-287-728, Sen-287-729, Sen-287-73, Sen-287-730, Sen-287-731, Sen-287-732, Sen-287-733, Sen-287-734, Sen-287-735, Sen-287-736, Sen-287-737, Sen-287-738, Sen-287-739, Sen-287-74, Sen-287-740, Sen-287-741, Sen-287-742, Sen-287-743, Sen-287-744, Sen-287-745, Sen-287-746, Sen-287-747, Sen-287-748, Sen-287-749, Sen-287-75, Sen-287-750, Sen-287-751, Sen-287-752, Sen-287-753, Sen-287-754, Sen-287-755, Sen-287-756, Sen-287-757, Sen-287-758, Sen-287-759, Sen-287-76, Sen-287-760, Sen-287-761, Sen-287-762, Sen-287-763, Sen-287-764, Sen-287-765, Sen-287-766, Sen-287-767, Sen-287-768, Sen-287-769, Sen-287-77, Sen-287-770, Sen-287-771, Sen-287-772, Sen-287-773, Sen-287-774, Sen-287-775, Sen-287-776, Sen-287-777, Sen-287-778, Sen-287-779, Sen-287-78, Sen-287-780, Sen-287-781, Sen-287-782, Sen-287-783, Sen-287-784, Sen-287-785, Sen-287-786, Sen-287-787, Sen-287-788, Sen-287-789, Sen-287-79, Sen-287-790, Sen-287-791, Sen-287-792, Sen-287-793, Sen-287-794, Sen-287-795, Sen-287-796, Sen-287-797, Sen-287-798, Sen-287-799, Sen-287-8, Sen-287-80, Sen-287-800, Sen-287-801, Sen-287-802, Sen-287-803, Sen-287-804, Sen-287-805, Sen-287-806, Sen-287-807, Sen-287-808, Sen-287-809, Sen-287-81, Sen-287-810, Sen-287-811, Sen-287-812, Sen-287-813, Sen-287-814, Sen-287-815, Sen-287-816, Sen-287-817, Sen-287-818, Sen-287-819, Sen-287-82, Sen-287-820, Sen-287-821, Sen-287-822, Sen-287-823, Sen-287-824, Sen-287-825, Sen-287-826, Sen-287-827, Sen-287-828, Sen-287-829, Sen-287-83, Sen-287-830, Sen-287-831, Sen-287-832, Sen-287-833, Sen-287-834, Sen-287-835, Sen-287-836, Sen-287-837, Sen-287-838, Sen-287-839, Sen-287-84, Sen-287-840, Sen-287-841, Sen-287-842, Sen-287-843, Sen-287-844, Sen-287-845, Sen-287-846, Sen-287-847, Sen-287-848, Sen-287-849, Sen-287-85, Sen-287-850, Sen-287-851, Sen-287-852, Sen-287-853, Sen-287-854, Sen-287-855, Sen-287-856, Sen-287-857, Sen-287-858, Sen-287-859, Sen-287-86, Sen-287-860, Sen-287-861, Sen-287-862, Sen-287-863, Sen-287-864, Sen-287-865, Sen-287-866, Sen-287-867, Sen-287-868, Sen-287-869, Sen-287-87, Sen-287-870, Sen-287-871, Sen-287-872, Sen-287-873, Sen-287-874, Sen-287-875, Sen-287-876, Sen-287-877, Sen-287-878, Sen-287-879, Sen-287-88, Sen-287-880, Sen-287-881, Sen-287-882, Sen-287-883, Sen-287-884, Sen-287-885, Sen-287-886, Sen-287-887, Sen-287-888, Sen-287-889, Sen-287-89, Sen-287-890, Sen-287-891, Sen-287-892, Sen-287-893, Sen-287-894, Sen-287-895, Sen-287-896, Sen-287-897, Sen-287-898, Sen-287-899, Sen-287-9, Sen-287-90, Sen-287-900, Sen-287-901, Sen-287-902, Sen-287-903, Sen-287-904,

Sen-287-905, Sen-287-906, Sen-287-907, Sen-287-908, Sen-287-909, Sen-287-91, Sen-287-910, Sen-287-911, Sen-287-912, Sen-287-913, Sen-287-914, Sen-287-915, Sen-287-916, Sen-287-917, Sen-287-918, Sen-287-919, Sen-287-92, Sen-287-920, Sen-287-921, Sen-287-922, Sen-287-923, Sen-287-924, Sen-287-925, Sen-287-926, Sen-287-927, Sen-287-928, Sen-287-929, Sen-287-93, Sen-287-930, Sen-287-931, Sen-287-932, Sen-287-933, Sen-287-934, Sen-287-935, Sen-287-936, Sen-287-937, Sen-287-938, Sen-287-939, Sen-287-94, Sen-287-940, Sen-287-941, Sen-287-942, Sen-287-943, Sen-287-944, Sen-287-945, Sen-287-946, Sen-287-947, Sen-287-948, Sen-287-949, Sen-287-95, Sen-287-950, Sen-287-951, Sen-287-952, Sen-287-953, Sen-287-954, Sen-287-955, Sen-287-956, Sen-287-957, Sen-287-958, Sen-287-959, Sen-287-96, Sen-287-960, Sen-287-961, Sen-287-962, Sen-287-963, Sen-287-964, Sen-287-965, Sen-287-966, Sen-287-967, Sen-287-968, Sen-287-969, Sen-287-97, Sen-287-970, Sen-287-971, Sen-287-972, Sen-287-973, Sen-287-974, Sen-287-975, Sen-287-976, Sen-287-977, Sen-287-978, Sen-287-979, Sen-287-98, Sen-287-980, Sen-287-981, Sen-287-982, Sen-287-983, Sen-287-984, Sen-287-985, Sen-287-986, Sen-287-987, Sen-287-988, Sen-287-989, Sen-287-99, Sen-287-990, Sen-287-991, Sen-287-992, Sen-287-993, Sen-287-994, Sen-287-995, Sen-287-996, Sen-287-997, Sen-287-998, Sen-287-999, Sen-288-1, Sen-289-1, Sen-29-1, Sen-290-1, Sen-290-2, Sen-290-3, Sen-290-4, Sen-290-5, Sen-290-6, Sen-290-7, Sen-291-1, Sen-292-1, Sen-293-1, Sen-294-1, Sen-295-1, Sen-296-1, Sen-297-1, Sen-298-1, Sen-299-1, Sen-299-10, Sen-299-100, Sen-299-101, Sen-299-102, Sen-299-103, Sen-299-104, Sen-299-105, Sen-299-106, Sen-299-107, Sen-299-108, Sen-299-109, Sen-299-11, Sen-299-110, Sen-299-111, Sen-299-112, Sen-299-113, Sen-299-114, Sen-299-115, Sen-299-116, Sen-299-117, Sen-299-118, Sen-299-119, Sen-299-12, Sen-299-120, Sen-299-121, Sen-299-122, Sen-299-123, Sen-299-124, Sen-299-125, Sen-299-126, Sen-299-127, Sen-299-128, Sen-299-129, Sen-299-13, Sen-299-130, Sen-299-131, Sen-299-132, Sen-299-133, Sen-299-134, Sen-299-135, Sen-299-136, Sen-299-137, Sen-299-138, Sen-299-139, Sen-299-14, Sen-299-140, Sen-299-141, Sen-299-142, Sen-299-143, Sen-299-144, Sen-299-145, Sen-299-146, Sen-299-147, Sen-299-148, Sen-299-149, Sen-299-15, Sen-299-150, Sen-299-151, Sen-299-152, Sen-299-153, Sen-299-154, Sen-299-155, Sen-299-156, Sen-299-157, Sen-299-158, Sen-299-159, Sen-299-16, Sen-299-160, Sen-299-161, Sen-299-162, Sen-299-163, Sen-299-164, Sen-299-165, Sen-299-166, Sen-299-167, Sen-299-168, Sen-299-169, Sen-299-17, Sen-299-170, Sen-299-171, Sen-299-172, Sen-299-173, Sen-299-174, Sen-299-175, Sen-299-176, Sen-299-177, Sen-299-178, Sen-299-179, Sen-299-18, Sen-299-180, Sen-299-181, Sen-299-182, Sen-299-183, Sen-299-184, Sen-299-185, Sen-299-186, Sen-299-187, Sen-299-188, Sen-299-189, Sen-299-19, Sen-299-190, Sen-299-191, Sen-299-192, Sen-299-193, Sen-299-194, Sen-299-195, Sen-299-196, Sen-299-197, Sen-299-198, Sen-299-199, Sen-299-2, Sen-299-20, Sen-299-200, Sen-299-201, Sen-299-202, Sen-299-203, Sen-299-204, Sen-299-205, Sen-299-206, Sen-299-207, Sen-299-208, Sen-299-209, Sen-299-21, Sen-299-210, Sen-299-211, Sen-299-212, Sen-299-213, Sen-299-214, Sen-299-215, Sen-299-216, Sen-299-217, Sen-299-218, Sen-299-219, Sen-299-22, Sen-299-220, Sen-299-221, Sen-299-222, Sen-299-223, Sen-299-224, Sen-299-225, Sen-299-226, Sen-299-227, Sen-299-228, Sen-299-229, Sen-299-23, Sen-299-230, Sen-299-231, Sen-299-232, Sen-299-233, Sen-299-234, Sen-299-235, Sen-299-236, Sen-299-237, Sen-299-238, Sen-299-239, Sen-299-24, Sen-299-240, Sen-299-241, Sen-299-242, Sen-299-243, Sen-299-244, Sen-299-245, Sen-299-246, Sen-299-247, Sen-299-248, Sen-299-249, Sen-299-25, Sen-299-250, Sen-299-251, Sen-299-252, Sen-299-253, Sen-299-254, Sen-299-255, Sen-299-256, Sen-299-257, Sen-299-258, Sen-299-259, Sen-299-26, Sen-299-260, Sen-299-261, Sen-299-262, Sen-299-263, Sen-299-264, Sen-299-265, Sen-299-266, Sen-299-267, Sen-299-268, Sen-299-269, Sen-299-27, Sen-299-270, Sen-299-271, Sen-299-272, Sen-299-273, Sen-299-274, Sen-299-275, Sen-299-276, Sen-299-277, Sen-299-278, Sen-299-279, Sen-299-28, Sen-299-280, Sen-299-281, Sen-299-282, Sen-299-283, Sen-299-284, Sen-299-285, Sen-299-286, Sen-299-287, Sen-299-288, Sen-299-289, Sen-299-29, Sen-299-290, Sen-299-291, Sen-299-292, Sen-299-293, Sen-299-294, Sen-299-295, Sen-299-296, Sen-299-297, Sen-299-298, Sen-299-299, Sen-299-3, Sen-299-30, Sen-299-300, Sen-299-301, Sen-299-302, Sen-299-303, Sen-299-304, Sen-299-305, Sen-299-306, Sen-299-307, Sen-299-308, Sen-299-309, Sen-299-31, Sen-299-310, Sen-299-311, Sen-299-312, Sen-299-313, Sen-299-314, Sen-299-315, Sen-299-316, Sen-299-317, Sen-299-318, Sen-299-319, Sen-299-32, Sen-299-320, Sen-299-321, Sen-299-322, Sen-299-323, Sen-299-324, Sen-299-325, Sen-299-326, Sen-299-327, Sen-299-328, Sen-299-329, Sen-299-33, Sen-299-330, Sen-299-34, Sen-299-35, Sen-299-36, Sen-299-37, Sen-299-38, Sen-299-39, Sen-299-4, Sen-299-40, Sen-299-41, Sen-299-42, Sen-299-43, Sen-299-44, Sen-299-45, Sen-299-46, Sen-299-47, Sen-299-48, Sen-299-49, Sen-299-5, Sen-299-50, Sen-299-51, Sen-299-52, Sen-299-53, Sen-299-54, Sen-299-55, Sen-299-56, Sen-299-57, Sen-299-58, Sen-299-59, Sen-299-6, Sen-299-60, Sen-299-61, Sen-299-62, Sen-299-63, Sen-299-64, Sen-299-65, Sen-299-66, Sen-299-67, Sen-299-68, Sen-299-69, Sen-299-7, Sen-299-70, Sen-299-71, Sen-299-72, Sen-299-73, Sen-299-74, Sen-299-75, Sen-299-76, Sen-299-77, Sen-299-78, Sen-299-79, Sen-299-8, Sen-299-80, Sen-299-81, Sen-299-82, Sen-299-83, Sen-299-84, Sen-299-85, Sen-299-86, Sen-299-87, Sen-299-88, Sen-299-89, Sen-299-9, Sen-299-90, Sen-299-91, Sen-299-92, Sen-299-93, Sen-299-94, Sen-299-95, Sen-299-96, Sen-299-97, Sen-299-98, Sen-299-99, Sen-30-1, Sen-300-1, Sen-301-1, Sen-302-1, Sen-303-1, Sen-304-1, Sen-305-1, Sen-305-10, Sen-305-11, Sen-305-12, Sen-305-13, Sen-305-14, Sen-305-15, Sen-305-16, Sen-305-17, Sen-305-18, Sen-305-19, Sen-305-2, Sen-305-20, Sen-305-21, Sen-305-22, Sen-305-23, Sen-305-24, Sen-305-25, Sen-305-26, Sen-305-27, Sen-305-28, Sen-305-29, Sen-305-3, Sen-305-30, Sen-305-31, Sen-305-32, Sen-305-33, Sen-305-34, Sen-305-35, Sen-305-36, Sen-305-37, Sen-305-38, Sen-305-39, Sen-305-4, Sen-305-40, Sen-305-41, Sen-305-42, Sen-305-43, Sen-305-44, Sen-305-45, Sen-305-46, Sen-305-47, Sen-305-48, Sen-305-49, Sen-305-5, Sen-305-50, Sen-305-51, Sen-305-52, Sen-305-53, Sen-305-54, Sen-305-55, Sen-305-56, Sen-305-57, Sen-305-58, Sen-305-59, Sen-305-6, Sen-305-60, Sen-305-61, Sen-305-62, Sen-305-63, Sen-305-7, Sen-305-8, Sen-305-9, Sen-306-1, Sen-307-1, Sen-307-2, Sen-307-3, Sen-307-4, Sen-308-1, Sen-309-1, Sen-31-1, Sen-312-1, Sen-313-1, Sen-314-1, Sen-315-1, Sen-316-1, Sen-317-1, Sen-318-1, Sen-319-1, Sen-32-1, Sen-320-1, Sen-321-1, Sen-322-1, Sen-323-1, Sen-324-1, Sen-325-1, Sen-326-1, Sen-327-1, Sen-328-1, Sen-329-1, Sen-33-1, Sen-330-1, Sen-331-1, Sen-332-1, Sen-333-1, Sen-334-1, Sen-335-1, Sen-336-1, Sen-337-1, Sen-338-1, Sen-339-1, Sen-34-1, Sen-340-1, Sen-341-1, Sen-342-1, Sen-343-1, Sen-344-1, Sen-345-1, Sen-346-1, Sen-347-1, Sen-348-1, Sen-349-1, Sen-35-1, Sen-350-1, Sen-351-1, Sen-352-1,

Sen-353-1, Sen-354-1, Sen-355-1, Sen-356-1, Sen-357-1, Sen-358-1, Sen-359-1, Sen-36-1, Sen-360-1, Sen-361-1, Sen-362-1, Sen-363-1, Sen-364-1, Sen-365-1, Sen-366-1, Sen-367-1, Sen-368-1, Sen-369-1, Sen-37-1, Sen-370-1, Sen-371-1, Sen-372-1, Sen-373-1, Sen-374-1, Sen-375-1, Sen-376-1, Sen-377-1, Sen-378-1, Sen-379-1, Sen-38-1, Sen-380-1, Sen-381-1, Sen-382-1, Sen-383-1, Sen-384-1, Sen-385-1, Sen-386-1, Sen-387-1, Sen-388-1, Sen-389-1, Sen-39-1, Sen-390-1, Sen-391-1, Sen-392-1, Sen-393-1, Sen-394-1, Sen-395-1, Sen-396-1, Sen-397-1, Sen-398-1, Sen-399-1, Sen-4-1, Sen-40-1, Sen-400-1, Sen-401-1, Sen-402-1, Sen-403-1, Sen-404-1, Sen-405-1, Sen-406-1, Sen-407-1, Sen-408-1, Sen-409-1, Sen-41-1, Sen-410-1, Sen-411-1, Sen-412-1, Sen-413-1, Sen-414-1, Sen-415-1, Sen-416-1, Sen-417-1, Sen-418-1, Sen-419-1, Sen-42-1, Sen-420-1, Sen-421-1, Sen-422-1, Sen-423-1, Sen-424-1, Sen-425-1, Sen-426-1, Sen-427-1, Sen-428-1, Sen-429-1, Sen-43-1, Sen-430-1, Sen-431-1, Sen-432-1, Sen-433-1, Sen-434-1, Sen-435-1, Sen-436-1, Sen-437-1, Sen-438-4, Sen-438-5, Sen-438-6, Sen-438-7, Sen-439-1, Sen-44-1, Sen-440-1, Sen-441-1, Sen-442-1, Sen-443-1, Sen-444-1, Sen-445-1, Sen-446-1, Sen-447-1, Sen-448-1, Sen-449-1, Sen-45-1, Sen-450-1, Sen-451-1, Sen-452-1, Sen-453-1, Sen-454-1, Sen-455-1, Sen-456-1, Sen-457-1, Sen-458-1, Sen-459-1, Sen-46-1, Sen-460-1, Sen-461-1, Sen-462-1, Sen-463-1, Sen-464-1, Sen-465-1, Sen-466-1, Sen-467-1, Sen-468-1, Sen-469-1, Sen-47-1, Sen-470-1, Sen-471-1, Sen-472-1, Sen-473-1, Sen-474-1, Sen-475-1, Sen-476-1, Sen-477-1, Sen-478-1, Sen-479-1, Sen-48-1, Sen-480-1, Sen-481-1, Sen-482-1, Sen-483-1, Sen-484-1, Sen-485-1, Sen-486-1, Sen-487-1, Sen-488-1, Sen-489-1, Sen-49-1, Sen-490-1, Sen-491-1, Sen-492-1, Sen-493-1, Sen-494-1, Sen-495-1, Sen-496-1, Sen-497-1, Sen-498-1, Sen-499-1, Sen-5-1, Sen-5-2, Sen-5-3, Sen-5-4, Sen-5-5, Sen-5-6, Sen-50-1, Sen-500-1, Sen-501-1, Sen-502-1, Sen-503-1, Sen-504-1, Sen-505-1, Sen-506-1, Sen-507-1, Sen-508-1, Sen-509-1, Sen-51-1, Sen-510-1, Sen-511-1, Sen-512-1, Sen-513-1, Sen-514-1, Sen-515-1, Sen-516-1, Sen-517-1, Sen-518-1, Sen-519-1, Sen-52-1, Sen-520-1, Sen-521-1, Sen-522-1, Sen-523-1, Sen-524-1, Sen-525-1, Sen-526-1, Sen-527-1, Sen-528-1, Sen-529-1, Sen-53-1, Sen-530-1, Sen-531-1, Sen-532-1, Sen-533-1, Sen-534-1, Sen-535-1, Sen-536-1, Sen-537-1, Sen-538-1, Sen-539-1, Sen-54-1, Sen-540-1, Sen-541-1, Sen-542-1, Sen-543-1, Sen-544-1, Sen-545-1, Sen-546-1, Sen-547-1, Sen-548-1, Sen-549-1, Sen-55-1, Sen-550-1, Sen-551-1, Sen-552-1, Sen-553-1, Sen-554-1, Sen-555-1, Sen-556-1, Sen-557-1, Sen-558-1, Sen-559-1, Sen-56-1, Sen-560-1, Sen-561-1, Sen-562-1, Sen-563-1, Sen-564-1, Sen-565-1, Sen-566-1, Sen-567-1, Sen-568-1, Sen-569-1, Sen-57-1, Sen-570-1, Sen-571-1, Sen-572-1, Sen-573-1, Sen-574-1, Sen-575-1, Sen-576-1, Sen-577-1, Sen-578-1, Sen-579-1, Sen-58-1, Sen-580-1, Sen-581-1, Sen-582-1, Sen-583-1, Sen-584-1, Sen-585-1, Sen-586-1, Sen-587-1, Sen-588-1, Sen-589-1, Sen-59-1, Sen-590-1, Sen-591-1, Sen-592-1, Sen-593-1, Sen-594-1, Sen-595-1, Sen-596-1, Sen-597-1, Sen-598-1, Sen-599-1, Sen-6-1, Sen-6-2, Sen-6-3, Sen-6-4, Sen-6-5, Sen-60-1, Sen-600-1, Sen-601-1, Sen-602-1, Sen-603-1, Sen-604-1, Sen-605-1, Sen-606-1, Sen-607-1, Sen-608-1, Sen-609-1, Sen-61-1, Sen-610-1, Sen-611-1, Sen-612-1, Sen-613-1, Sen-614-1, Sen-615-1, Sen-616-1, Sen-617-1, Sen-618-1, Sen-619-1, Sen-62-1, Sen-620-1, Sen-621-1, Sen-622-1, Sen-623-1, Sen-624-1, Sen-625-1, Sen-626-1, Sen-627-1, Sen-628-1, Sen-629-1, Sen-63-1, Sen-630-1, Sen-631-1, Sen-632-1, Sen-633-1, Sen-634-1, Sen-635-1, Sen-636-1, Sen-637-1, Sen-638-1, Sen-639-1, Sen-64-1, Sen-640-1, Sen-641-1, Sen-642-1, Sen-643-1, Sen-644-1, Sen-645-1, Sen-646-1, Sen-647-1, Sen-648-1, Sen-649-1, Sen-65-1, Sen-650-1, Sen-651-1, Sen-652-1, Sen-653-1, Sen-654-1, Sen-655-1, Sen-656-1, Sen-657-1, Sen-658-1, Sen-659-1, Sen-66-1, Sen-660-1, Sen-661-1, Sen-662-1, Sen-663-1, Sen-664-1, Sen-665-1, Sen-666-1, Sen-667-1, Sen-668-1, Sen-669-1, Sen-670-1, Sen-671-1, Sen-672-1, Sen-673-1, Sen-674-1, Sen-675-1, Sen-676-1, Sen-677-1, Sen-678-1, Sen-679-1, Sen-68-1, Sen-680-1, Sen-681-1, Sen-682-1, Sen-683-1, Sen-684-1, Sen-685-1, Sen-686-1, Sen-687-1, Sen-688-1, Sen-689-1, Sen-69-1, Sen-690-1, Sen-691-1, Sen-692-1, Sen-693-1, Sen-694-1, Sen-695-1, Sen-696-1, Sen-697-1, Sen-698-1, Sen-699-1, Sen-7-1, Sen-7-2, Sen-7-3, Sen-70-1, Sen-700-1, Sen-701-1, Sen-702-1, Sen-703-1, Sen-704-1, Sen-705-1, Sen-706-1, Sen-707-1, Sen-708-1, Sen-709-1, Sen-71-1, Sen-710-1, Sen-711-1, Sen-712-1, Sen-713-1, Sen-714-1, Sen-715-1, Sen-716-1, Sen-717-1, Sen-718-1, Sen-719-1, Sen-72-1, Sen-720-1, Sen-721-1, Sen-722-1, Sen-723-1, Sen-724-1, Sen-725-1, Sen-726-1, Sen-727-1, Sen-729-1, Sen-73-1, Sen-730-1, Sen-731-1, Sen-732-1, Sen-733-1, Sen-734-1, Sen-735-1, Sen-736-1, Sen-737-1, Sen-738-1, Sen-739-1, Sen-74-1, Sen-740-1, Sen-741-1, Sen-742-1, Sen-743-1, Sen-744-1, Sen-745-1, Sen-746-1, Sen-747-1, Sen-748-1, Sen-749-1, Sen-75-1, Sen-750-1, Sen-751-1, Sen-752-1, Sen-753-1, Sen-754-1, Sen-755-1, Sen-756-1, Sen-757-1, Sen-758-1, Sen-759-1, Sen-76-1, Sen-760-1, Sen-761-1, Sen-762-1, Sen-763-1, Sen-764-1, Sen-765-1, Sen-766-1, Sen-767-1, Sen-768-1, Sen-769-1, Sen-77-1, Sen-770-1, Sen-771-1, Sen-772-1, Sen-773-1, Sen-774-1, Sen-775-1, Sen-776-1, Sen-777-1, Sen-778-1, Sen-779-1, Sen-78-1, Sen-780-1, Sen-781-1, Sen-782-1, Sen-783-1, Sen-784-1, Sen-785-1, Sen-786-1, Sen-787-1, Sen-788-1, Sen-789-1, Sen-79-1, Sen-790-1, Sen-791-1, Sen-792-1, Sen-793-1, Sen-794-1, Sen-795-1, Sen-796-1, Sen-797-1, Sen-798-1, Sen-799-1, Sen-8-1, Sen-80-1, Sen-800-1, Sen-801-1, Sen-802-1, Sen-803-1, Sen-804-1, Sen-805-1, Sen-806-1, Sen-807-1, Sen-808-1, Sen-809-1, Sen-81-1, Sen-810-1, Sen-811-1, Sen-812-1, Sen-813-1, Sen-814-1, Sen-815-1, Sen-816-1, Sen-817-1, Sen-818-1, Sen-819-1, Sen-82-1, Sen-820-1, Sen-821-1, Sen-822-1, Sen-823-1, Sen-824-1, Sen-825-1, Sen-826-1, Sen-827-1, Sen-829-1, Sen-83-1, Sen-830-1, Sen-831-1, Sen-832-1, Sen-833-1, Sen-835-1, Sen-836-1, Sen-837-1, Sen-838-1, Sen-839-1, Sen-84-1, Sen-840-1, Sen-841-1, Sen-842-1, Sen-843-1, Sen-844-1, Sen-845-1, Sen-846-1, Sen-847-1, Sen-848-1, Sen-849-1, Sen-85-1, Sen-850-1, Sen-851-1, Sen-852-1, Sen-853-1, Sen-854-1, Sen-855-1, Sen-856-1, Sen-857-1, Sen-858-1, Sen-859-1, Sen-86-1, Sen-860-1, Sen-861-1, Sen-862-1, Sen-863-1, Sen-864-1, Sen-865-1, Sen-866-1, Sen-867-1, Sen-868-1, Sen-869-1, Sen-87-1, Sen-870-1, Sen-871-1, Sen-872-1, Sen-873-1, Sen-874-1, Sen-875-1, Sen-876-1, Sen-877-1, Sen-878-1, Sen-879-1, Sen-880-1, Sen-881-1, Sen-882-1, Sen-883-1, Sen-884-1, Sen-885-1, Sen-886-1, Sen-887-1, Sen-888-1, Sen-889-1, Sen-89-1, Sen-890-1, Sen-891-1, Sen-892-1, Sen-893-1, Sen-894-1, Sen-895-1, Sen-896-1, Sen-897-1, Sen-898-1, Sen-899-1, Sen-9-1, Sen-90-1, Sen-900-1, Sen-901-1, Sen-902-1, Sen-903-1, Sen-904-1, Sen-905-1, Sen-906-1, Sen-907-1, Sen-908-1, Sen-909-1, Sen-91-1, Sen-910-1, Sen-911-1, Sen-912-1, Sen-913-1, Sen-914-1, Sen-915-1, Sen-916-1, Sen-917-1, Sen-918-1, Sen-919-1, Sen-92-1, Sen-920-1, Sen-921-1, Sen-922-1, Sen-923-1, Sen-924-1, Sen-925-1, Sen-926-1, Sen-928-1, Sen-929-1, Sen-93-1, Sen-930-1, Sen-931-1, Sen-932-1, Sen-933-1, Sen-934-1, Sen-935-1, Sen-936-1, Sen-937-1, Sen-938-1, Sen-939-1, Sen-94-1, Sen-940-1, Sen-941-1, Sen-942-1,

Sen-943-1, Sen-944-1, Sen-945-1, Sen-946-1, Sen-947-1, Sen-948-1, Sen-949-1, Sen-95-1, Sen-950-1, Sen-951-1, Sen-952-1, Sen-953-1, Sen-954-1, Sen-955-1, Sen-956-1, Sen-957-1, Sen-958-1, Sen-959-1, Sen-96-1, Sen-960-1, Sen-961-1, Sen-962-1, Sen-963-1, Sen-964-1, Sen-965-1, Sen-966-1, Sen-967-1, Sen-968-1, Sen-969-1, Sen-97-1, Sen-970-1, Sen-971-1, Sen-972-1, Sen-973-1, Sen-974-1, Sen-975-1, Sen-976-1, Sen-977-1, Sen-978-1, Sen-979-1, Sen-98-1, Sen-980-1, Sen-981-1, Sen-982-1, Sen-983-1, Sen-984-1, Sen-985-1, Sen-986-1, Sen-987-1, Sen-988-1, Sen-989-1, Sen-99-1, Sen-990-1, Sen-991-1, Sen-992-1, Sen-993-1, Sen-994-1, Sen-995-1, Sen-996-1, Sen-997-1, Sen-998-1, Sen-999-1, Sfl-1-1, Sfl-1-2, Sfl-10-1, Sfl-11-1, Sfl-12-1, Sfl-13-1, Sfl-14-1, Sfl-15-1, Sfl-16-1, Sfl-17-1, Sfl-18-1, Sfl-19-1, Sfl-2-1, Sfl-20-1, Sfl-21-1, Sfl-22-1, Sfl-23-1, Sfl-24-1, Sfl-25-1, Sfl-25-10, Sfl-25-11, Sfl-25-12, Sfl-25-13, Sfl-25-14, Sfl-25-2, Sfl-25-3, Sfl-25-4, Sfl-25-5, Sfl-25-6, Sfl-25-7, Sfl-25-8, Sfl-25-9, Sfl-26-1, Sfl-27-1, Sfl-28-1, Sfl-29-1, Sfl-3-1, Sfl-30-1, Sfl-31-1, Sfl-32-1, Sfl-33-1, Sfl-34-1, Sfl-35-1, Sfl-36-1, Sfl-37-1, Sfl-38-1, Sfl-39-1, Sfl-4-1, Sfl-40-1, Sfl-41-1, Sfl-42-1, Sfl-43-1, Sfl-44-1, Sfl-45-1, Sfl-46-1, Sfl-47-1, Sfl-48-1, Sfl-49-1, Sfl-5-1, Sfl-50-1, Sfl-51-1, Sfl-51-2, Sfl-52-1, Sfl-53-1, Sfl-54-1, Sfl-55-1, Sfl-56-1, Sfl-57-1, Sfl-58-1, Sfl-59-1, Sfl-6-1, Sfl-60-1, Sfl-61-1, Sfl-62-1, Sfl-63-1, Sfl-64-1, Sfl-64-2, Sfl-65-1, Sfl-66-1, Sfl-67-1, Sfl-68-1, Sfl-69-1, Sfl-7-1, Sfl-70-1, Sfl-71-1, Sfl-72-1, Sfl-73-1, Sfl-74-1, Sfl-75-1, Sfl-77-1, Sfl-78-1, Sfl-79-1, Sfl-8-1, Sfl-80-1, Sfl-81-1, Sfl-82-1, Sfl-83-1, Sfl-84-1, Sfl-85-1, Sfl-9-1, Sfo-2-1, Sli-1-1, Sli-3-1, Sma-1-4, Sma-10-1, Sma-11-1, Sma-12-1, Sma-13-1, Sma-15-1, Sma-16-1, Sma-17-1, Sma-2-1, Sma-20-1, Sma-21-1, Sma-23-1, Sma-24-1, Sma-26-1, Sma-3-1, Sma-4-1, Sma-5-1, Sma-6-1, Sma-7-1, Sma-9-1, Spl-3-1, Spl-4-1, Spl-6-1, Sso-1-1, Sso-1-10, Sso-1-100, Sso-1-101, Sso-1-102, Sso-1-103, Sso-1-104, Sso-1-105, Sso-1-106, Sso-1-107, Sso-1-108, Sso-1-109, Sso-1-11, Sso-1-110, Sso-1-111, Sso-1-112, Sso-1-113, Sso-1-114, Sso-1-115, Sso-1-116, Sso-1-117, Sso-1-118, Sso-1-119, Sso-1-12, Sso-1-120, Sso-1-121, Sso-1-122, Sso-1-123, Sso-1-124, Sso-1-125, Sso-1-126, Sso-1-127, Sso-1-128, Sso-1-129, Sso-1-13, Sso-1-130, Sso-1-131, Sso-1-132, Sso-1-133, Sso-1-134, Sso-1-135, Sso-1-136, Sso-1-137, Sso-1-138, Sso-1-139, Sso-1-140, Sso-1-141, Sso-1-142, Sso-1-143, Sso-1-144, Sso-1-145, Sso-1-146, Sso-1-147, Sso-1-148, Sso-1-149, Sso-1-15, Sso-1-150, Sso-1-151, Sso-1-152, Sso-1-153, Sso-1-154, Sso-1-155, Sso-1-156, Sso-1-157, Sso-1-158, Sso-1-159, Sso-1-16, Sso-1-160, Sso-1-161, Sso-1-162, Sso-1-163, Sso-1-164, Sso-1-165, Sso-1-166, Sso-1-167, Sso-1-168, Sso-1-169, Sso-1-17, Sso-1-170, Sso-1-171, Sso-1-172, Sso-1-173, Sso-1-174, Sso-1-175, Sso-1-176, Sso-1-177, Sso-1-178, Sso-1-179, Sso-1-18, Sso-1-180, Sso-1-181, Sso-1-182, Sso-1-183, Sso-1-184, Sso-1-185, Sso-1-186, Sso-1-187, Sso-1-188, Sso-1-189, Sso-1-19, Sso-1-190, Sso-1-191, Sso-1-192, Sso-1-193, Sso-1-194, Sso-1-195, Sso-1-196, Sso-1-197, Sso-1-198, Sso-1-199, Sso-1-20, Sso-1-200, Sso-1-201, Sso-1-202, Sso-1-203, Sso-1-204, Sso-1-205, Sso-1-206, Sso-1-207, Sso-1-208, Sso-1-209, Sso-1-21, Sso-1-210, Sso-1-211, Sso-1-212, Sso-1-213, Sso-1-214, Sso-1-215, Sso-1-216, Sso-1-217, Sso-1-218, Sso-1-219, Sso-1-22, Sso-1-220, Sso-1-221, Sso-1-222, Sso-1-223, Sso-1-224, Sso-1-225, Sso-1-226, Sso-1-227, Sso-1-228, Sso-1-229, Sso-1-23, Sso-1-230, Sso-1-231, Sso-1-232, Sso-1-233, Sso-1-234, Sso-1-235, Sso-1-236, Sso-1-237, Sso-1-238, Sso-1-239, Sso-1-24, Sso-1-240, Sso-1-241, Sso-1-242, Sso-1-243, Sso-1-244, Sso-1-245, Sso-1-246, Sso-1-247, Sso-1-248, Sso-1-249, Sso-1-25, Sso-1-250, Sso-1-251, Sso-1-252, Sso-1-253, Sso-1-254, Sso-1-255, Sso-1-256, Sso-1-257, Sso-1-258, Sso-1-259, Sso-1-26, Sso-1-260, Sso-1-261, Sso-1-262, Sso-1-263, Sso-1-264, Sso-1-265, Sso-1-266, Sso-1-267, Sso-1-268, Sso-1-269, Sso-1-27, Sso-1-270, Sso-1-271, Sso-1-272, Sso-1-273, Sso-1-274, Sso-1-275, Sso-1-276, Sso-1-277, Sso-1-278, Sso-1-279, Sso-1-28, Sso-1-280, Sso-1-281, Sso-1-282, Sso-1-283, Sso-1-284, Sso-1-285, Sso-1-286, Sso-1-287, Sso-1-288, Sso-1-289, Sso-1-29, Sso-1-290, Sso-1-291, Sso-1-292, Sso-1-293, Sso-1-294, Sso-1-295, Sso-1-296, Sso-1-297, Sso-1-298, Sso-1-299, Sso-1-30, Sso-1-300, Sso-1-301, Sso-1-302, Sso-1-303, Sso-1-304, Sso-1-305, Sso-1-306, Sso-1-307, Sso-1-308, Sso-1-309, Sso-1-31, Sso-1-310, Sso-1-311, Sso-1-312, Sso-1-313, Sso-1-314, Sso-1-315, Sso-1-316, Sso-1-317, Sso-1-318, Sso-1-319, Sso-1-32, Sso-1-320, Sso-1-321, Sso-1-322, Sso-1-323, Sso-1-324, Sso-1-325, Sso-1-326, Sso-1-327, Sso-1-328, Sso-1-329, Sso-1-33, Sso-1-330, Sso-1-331, Sso-1-332, Sso-1-333, Sso-1-334, Sso-1-335, Sso-1-336, Sso-1-337, Sso-1-338, Sso-1-339, Sso-1-34, Sso-1-340, Sso-1-341, Sso-1-342, Sso-1-343, Sso-1-344, Sso-1-345, Sso-1-346, Sso-1-347, Sso-1-348, Sso-1-349, Sso-1-35, Sso-1-350, Sso-1-351, Sso-1-352, Sso-1-353, Sso-1-354, Sso-1-355, Sso-1-356, Sso-1-357, Sso-1-358, Sso-1-359, Sso-1-36, Sso-1-360, Sso-1-361, Sso-1-362, Sso-1-363, Sso-1-364, Sso-1-365, Sso-1-366, Sso-1-367, Sso-1-368, Sso-1-369, Sso-1-37, Sso-1-370, Sso-1-371, Sso-1-372, Sso-1-373, Sso-1-374, Sso-1-375, Sso-1-376, Sso-1-377, Sso-1-378, Sso-1-379, Sso-1-38, Sso-1-380, Sso-1-381, Sso-1-382, Sso-1-383, Sso-1-384, Sso-1-385, Sso-1-386, Sso-1-387, Sso-1-388, Sso-1-389, Sso-1-39, Sso-1-390, Sso-1-391, Sso-1-392, Sso-1-393, Sso-1-394, Sso-1-395, Sso-1-396, Sso-1-397, Sso-1-398, Sso-1-399, Sso-1-4, Sso-1-40, Sso-1-400, Sso-1-401, Sso-1-402, Sso-1-403, Sso-1-404, Sso-1-405, Sso-1-406, Sso-1-407, Sso-1-408, Sso-1-409, Sso-1-41, Sso-1-410, Sso-1-411, Sso-1-412, Sso-1-413, Sso-1-414, Sso-1-415, Sso-1-416, Sso-1-417, Sso-1-418, Sso-1-419, Sso-1-42, Sso-1-420, Sso-1-421, Sso-1-422, Sso-1-423, Sso-1-424, Sso-1-425, Sso-1-426, Sso-1-427, Sso-1-428, Sso-1-429, Sso-1-43, Sso-1-430, Sso-1-431, Sso-1-432, Sso-1-433, Sso-1-434, Sso-1-435, Sso-1-436, Sso-1-437, Sso-1-438, Sso-1-439, Sso-1-44, Sso-1-440, Sso-1-441, Sso-1-442, Sso-1-443, Sso-1-444, Sso-1-445, Sso-1-446, Sso-1-447, Sso-1-448, Sso-1-449, Sso-1-45, Sso-1-450, Sso-1-451, Sso-1-452, Sso-1-453, Sso-1-454, Sso-1-455, Sso-1-456, Sso-1-457, Sso-1-458, Sso-1-459, Sso-1-46, Sso-1-460, Sso-1-461, Sso-1-462, Sso-1-463, Sso-1-464, Sso-1-465, Sso-1-466, Sso-1-467, Sso-1-468, Sso-1-469, Sso-1-47, Sso-1-470, Sso-1-471, Sso-1-472, Sso-1-473, Sso-1-474, Sso-1-475, Sso-1-476, Sso-1-477, Sso-1-478, Sso-1-479, Sso-1-48, Sso-1-480, Sso-1-481, Sso-1-482, Sso-1-483, Sso-1-484, Sso-1-485, Sso-1-486, Sso-1-487, Sso-1-488, Sso-1-489, Sso-1-49, Sso-1-490, Sso-1-491, Sso-1-492, Sso-1-493, Sso-1-494, Sso-1-495, Sso-1-496, Sso-1-497, Sso-1-498, Sso-1-499, Sso-1-5, Sso-1-50, Sso-1-500, Sso-1-501, Sso-1-502, Sso-1-503, Sso-1-504, Sso-1-505, Sso-1-506, Sso-1-507, Sso-1-508, Sso-1-509, Sso-1-51, Sso-1-510, Sso-1-511, Sso-1-512, Sso-1-513, Sso-1-514, Sso-1-515, Sso-1-516, Sso-1-517, Sso-1-518, Sso-1-519, Sso-1-52, Sso-1-520, Sso-1-521, Sso-1-522, Sso-1-523, Sso-1-524, Sso-1-525, Sso-1-526, Sso-1-527, Sso-1-528, Sso-1-529, Sso-1-53, Sso-1-530, Sso-1-531, Sso-1-532, Sso-1-533, Sso-1-534, Sso-1-535, Sso-1-536, Sso-1-537, Sso-1-538, Sso-1-54, Sso-1-540, Sso-1-55, Sso-1-56, Sso-1-57, Sso-1-58, Sso-1-59, Sso-1-6, Sso-1-60, Sso-1-61, Sso-1-62, Sso-1-63, Sso-1-64, Sso-1-65, Sso-1-66, Sso-1-67, Sso-1-68, Sso-1-69, Sso-1-7, Sso-1-70, Sso-1-71, Sso-1-72, Sso-1-73, Sso-1-74, Sso-1-75, Sso-1-76, Sso-1-77, Sso-1-78, Sso-1-79,

Sso-1-8, Sso-1-80, Sso-1-81, Sso-1-82, Sso-1-83, Sso-1-84, Sso-1-85, Sso-1-86, Sso-1-87, Sso-1-88, Sso-1-89, Sso-1-9, Sso-1-90, Sso-1-91, Sso-1-92, Sso-1-93, Sso-1-94, Sso-1-95, Sso-1-96, Sso-1-97, Sso-1-98, Sso-1-99, Sso-10-1, Sso-11-1, Sso-12-1, Sso-13-1, Sso-2-1, Sso-3-1, Sso-4-1, Sso-5-1, Sso-6-1, Sso-8-1, Sso-9-1, Ssp-1-1, Ssp-11-1, Ssp-15-1, Ssp-3-1, Ssp-5-1, Ssp-7-1, Stu-1-1, Tmo-1-1, Tpt-1-2, Tsp-1-1, Vpa-1-1, Vpa-1-2, Xbo-1-1, Xbo-10-1, Xbo-3-1, Xbo-4-1, Xbo-5-1, Xbo-6-1, Xbo-7-1, Xne-1-1, Xne-2-1, Xne-4-1, Xne-5-2, Xne-6-1, Xne-6-2, Yal-1-2, Yal-1-4, Yal-2-1, Yal-4-1, Ybe-2-1, Yen-1-1, Yen-1-2, Yen-1-3, Yen-1-4, Yen-10-1, Yen-11-1, Yen-12-1, Yen-13-1, Yen-14-1, Yen-15-1, Yen-16-1, Yen-17-1, Yen-18-1, Yen-19-1, Yen-2-1, Yen-20-1, Yen-22-1, Yen-3-1, Yen-4-1, Yen-5-1, Yen-6-1, Yen-7-1, Yen-8-1, Yen-9-1, Yfr-1-11, Yfr-1-14, Yfr-1-15, Yfr-1-16, Yfr-1-2, Yfr-1-3, Yfr-1-5, Yfr-1-8, Yfr-2-1, Yin-1-1, Yin-2-1, Ykr-1-1, Ykr-1-2, Ymo-1-2, Ymo-1-4, Ymo-1-6, Ymo-1-7, Ymo-2-1, Ype-1-2, Ype-10-1, Ype-100-1, Ype-101-1, Ype-102-1, Ype-103-1, Ype-104-1, Ype-105-1, Ype-106-1, Ype-107-1, Ype-108-1, Ype-109-1, Ype-11-1, Ype-110-1, Ype-111-1, Ype-112-1, Ype-113-1, Ype-114-1, Ype-114-2, Ype-115-1, Ype-116-1, Ype-117-1, Ype-118-1, Ype-119-1, Ype-12-1, Ype-120-1, Ype-121-1, Ype-122-1, Ype-123-1, Ype-124-1, Ype-125-1, Ype-126-1, Ype-127-1, Ype-128-1, Ype-129-1, Ype-13-1, Ype-130-1, Ype-131-1, Ype-133-1, Ype-134-1, Ype-135-1, Ype-136-1, Ype-137-1, Ype-138-1, Ype-139-1, Ype-14-1, Ype-140-1, Ype-141-1, Ype-142-1, Ype-143-1, Ype-144-1, Ype-145-1, Ype-146-1, Ype-147-1, Ype-148-1, Ype-149-1, Ype-15-1, Ype-150-1, Ype-151-1, Ype-152-1, Ype-153-1, Ype-154-1, Ype-155-1, Ype-156-1, Ype-157-1, Ype-158-1, Ype-159-1, Ype-16-1, Ype-160-1, Ype-161-1, Ype-162-1, Ype-163-1, Ype-164-1, Ype-165-1, Ype-166-1, Ype-167-1, Ype-168-1, Ype-169-1, Ype-17-1, Ype-170-1, Ype-171-1, Ype-172-1, Ype-173-1, Ype-174-1, Ype-175-1, Ype-176-1, Ype-177-1, Ype-178-1, Ype-179-1, Ype-18-1, Ype-180-1, Ype-181-1, Ype-182-1, Ype-183-1, Ype-184-1, Ype-185-1, Ype-186-1, Ype-187-1, Ype-188-1, Ype-189-1, Ype-19-1, Ype-190-1, Ype-191-1, Ype-192-1, Ype-193-1, Ype-194-1, Ype-195-1, Ype-196-1, Ype-197-1, Ype-198-1, Ype-199-1, Ype-2-1, Ype-2-2, Ype-2-3, Ype-2-4, Ype-2-5, Ype-2-6, Ype-2-7, Ype-2-8, Ype-2-9, Ype-20-1, Ype-200-1, Ype-201-1, Ype-202-1, Ype-203-1, Ype-204-1, Ype-205-1, Ype-206-1, Ype-207-1, Ype-208-1, Ype-209-1, Ype-21-1, Ype-210-1, Ype-211-1, Ype-212-1, Ype-213-1, Ype-214-1, Ype-215-1, Ype-216-1, Ype-217-1, Ype-218-1, Ype-218-2, Ype-219-1, Ype-22-1, Ype-220-1, Ype-221-1, Ype-222-1, Ype-223-1, Ype-224-1, Ype-225-1, Ype-226-1, Ype-227-1, Ype-23-1, Ype-24-1, Ype-25-1, Ype-26-1, Ype-27-1, Ype-28-1, Ype-29-1, Ype-29-2, Ype-3-1, Ype-30-1, Ype-31-1, Ype-31-2, Ype-32-1, Ype-32-2, Ype-33-1, Ype-34-1, Ype-35-1, Ype-36-1, Ype-37-1, Ype-38-1, Ype-39-1, Ype-4-1, Ype-40-1, Ype-41-1, Ype-42-1, Ype-43-1, Ype-44-1, Ype-45-1, Ype-46-1, Ype-47-1, Ype-48-1, Ype-49-1, Ype-5-1, Ype-50-1, Ype-51-1, Ype-52-1, Ype-53-1, Ype-54-1, Ype-55-1, Ype-56-1, Ype-57-1, Ype-58-1, Ype-59-1, Ype-6-1, Ype-60-1, Ype-61-1, Ype-62-1, Ype-63-1, Ype-64-1, Ype-65-1, Ype-66-1, Ype-67-1, Ype-68-1, Ype-69-1, Ype-7-1, Ype-70-1, Ype-71-1, Ype-72-1, Ype-73-1, Ype-74-1, Ype-75-1, Ype-76-1, Ype-77-1, Ype-78-1, Ype-79-1, Ype-8-1, Ype-80-1, Ype-81-1, Ype-82-1, Ype-83-1, Ype-84-1, Ype-85-1, Ype-86-1, Ype-87-1, Ype-88-1, Ype-89-1, Ype-9-1, Ype-90-1, Ype-91-1, Ype-92-1, Ype-93-1, Ype-94-1, Ype-94-2, Ype-95-1, Ype-96-1, Ype-97-1, Ype-98-1, Ype-99-1, Yps-1-2, Yps-1-3, Yps-1-4, Yps-10-1, Yps-10-2, Yps-11-1, Yps-11-2, Yps-12-1, Yps-12-2, Yps-13-1, Yps-14-1, Yps-2-1, Yps-3-1, Yps-4-1, Yps-5-1, Yps-6-1, Yps-7-1, Yps-8-1, Yps-9-1, Yre-1-1, Yro-1-1, Yru-1-3, Yru-2-1, Ywa-1-2, env-10, env-100, env-102, env-105, env-106, env-107, env-108, env-109, env-11, env-110, env-112, env-113, env-116, env-117, env-118, env-119, env-12, env-120, env-123, env-124, env-128, env-129, env-13, env-130, env-133, env-135, env-137, env-14, env-141, env-145, env-146, env-147, env-15, env-150, env-156, env-16, env-164, env-17, env-176, env-18, env-185, env-186, env-187, env-188, env-19, env-190, env-191, env-193, env-20, env-203, env-206, env-209, env-21, env-215, env-219, env-22, env-220, env-221, env-222, env-223, env-224, env-225, env-226, env-227, env-23, env-231, env-234, env-235, env-238, env-24, env-246, env-247, env-248, env-249, env-25, env-250, env-251, env-253, env-254, env-255, env-256, env-257, env-258, env-259, env-26, env-260, env-261, env-262, env-263, env-267, env-268, env-269, env-27, env-270, env-271, env-272, env-273, env-274, env-275, env-276, env-277, env-278, env-279, env-28, env-280, env-281, env-282, env-283, env-284, env-285, env-286, env-287, env-288, env-289, env-29, env-290, env-291, env-292, env-293, env-294, env-295, env-296, env-297, env-298, env-299, env-3, env-30, env-300, env-301, env-302, env-303, env-304, env-305, env-306, env-307, env-308, env-309, env-31, env-310, env-311, env-312, env-313, env-314, env-315, env-316, env-317, env-318, env-319, env-32, env-320, env-321, env-322, env-323, env-33, env-333, env-338, env-339, env-340, env-341, env-342, env-345, env-352, env-355, env-356, env-36, env-361, env-367, env-369, env-37, env-38, env-39, env-40, env-41, env-42, env-43, env-44, env-45, env-46, env-47, env-48, env-49, env-5, env-51, env-52, env-53, env-56, env-57, env-60, env-61, env-64, env-69, env-81, env-84, env-86, env-87, env-9, env-92, env-93, env-94, env-95, env-96, env-97, env-98, env-99

alignment positions 1 · · · 180

```
env-1     GCGGGUUAUUCUAAUCGUUUUAAGGGAGAUUGCCUAGAAAAAGUUUGCUUUAAUAAAGCCUAUCCUUUAUAAUCCGCGCCCUC.....AAAG..................................................................................UUGGU.
env-2     GCGGAUUAUUCUAAUCGUUUUAAGGGGGAUUGCCUAGAAAAAGUUUGCUUUAAUAAAGUCUAUCCUUUAUAAUCCGCGCCCUC.....AAAG..................................................................................UUGGU.
env-4     GCGAGUUAUUCUAAUCGUUUUAAGGGAGAUUGCCUAGAAAAAGUUUGCUUUAAUAACGUCUAUCCUUUAUAAUCCGCGCCCUC.....AAAG..................................................................................UUGGU.
Mma-2-1   CCAGUCUAAUUUAGUAAUUUAUUUAUACUAAUAUUGUGAUUUUAUUUGAUUUUUUAAAUUGGAUUACUUAUAAUCCACAGGCU.....CAAA..................................................................................AAAGUC
env-6     ........................AAAAAAUUUGCUGUGUAAAUUUGCAUUGUGAGGCUGAAUAUUUUAUAAUCCGUCGCUCU.....C.AAGG................................................................................UUUG..
Hpa-2-1   AUCGCUUUGCCUUGAAAAGCCUAGAAAAAAUUUGCUGUGUAAAUUUGCAUUGUGGGGCUGAAUAUUUUAUAAUCCGUCGCUCU.....C.AAGG................................................................................UUUG..
env-7     AUCGCUUUGCCUUGAAAAGCCUAGAAAAAAUUUGCUGUGUAAAUUUGCAUUGUGGGGCUGAAUAUUUUAUAAUCCGUCGCUCU.....C.AAGG................................................................................UUUG..
env-8     AUCGCUUUGCCUUGAAAAGCCUAGAAAAAAUUUGCUGUGUAAAUUUGCAUUGUGAGGCUGAAUAUUUUAUAAUCCGUCGCUCU.....C.AAGG................................................................................UUUG..
env-34    AUCGCUUUGUCUUGAAAAGCCUAGAAAAAAUUUGCUGUGUAAAUUUGCAUUGUGAGGCUGAAUAUUUUAUAAUCCGUCGCUCU.....C.AAGG................................................................................UUUG..
env-35    AUCGCUUUGCCUUGAAAAGCCUAGAAAAAAUUUGCUGUGUAAAUUUGCAUUGUGGGGCUGAAUAUUUUAUAAUCCGUCGCUCU.....CAAAG.................................................................................UUUGU.
env-50    .............................................................................CGCUCU.....CAAAG.................................................................................UUUGU.
env-54    ............UGAAAAGCCUAGAAAAAAUUUGCUGCGUAAAUUUGCAUUGUGGGGCUGAAUAUUUUAUAAUCCGUCGCUCU.....CAAAG.................................................................................UUUGU.

          ................................................................................<<<.....<<<<<<<<<<<<<<<<<<<<<<<<<<<<<<<<<<<<<<<<<<<<<<<<<<<<<<<<<<<<<<<<<<<<<<<<<<<<<<<<<<<<<<......
          ................................................................................212.....22222.........111....00110.....0.1.111.1111.1.........101...............011....111100......
          ••••••••••••••••••••••••••••••••••••••••YY••UUGYY••••••••••••••••R•UA•AAU•YCGCCCGCU-----U•••○---------------------------------------------------------------------------------•Y••--
```

```
env-55     AUCGCUUUGCCUUGAAAAGCCUAGAAAAAAUUUGCGGUGUAAAUUUGCAUUGUGGGGCUGAAUAUUUUAUAAUCCGUCGCUCU.....CAAAG.................................................................................UUUGU.
env-58     AUCGCUUUGCCUUGAAAAGCCUAGAAAAAAUUUGCGGUGUAAAUUUGCAUUGUGGGGCUGAAUAUUUUAUAAUUCGUCGCUCU.....CAAAG.................................................................................UUUGU.
Hin-1-1    UGGUUCACAUUCUAAUCUUUUCAGGGUAAAAAACCUAGAAAAAGUUUGCUUUGAAAAGAGGGAUAUUCUAUAAUUCGCCGUCU.....C.....AAAGCGU.........................................................................GGCCA.
Hha-1-1    UGCCAAACAUUCUAAUCUUUUCAGGGUGAAAAACCUAGAAAAAGUUUGCUUUGAAAAGCGGGAUAUUCUAUAAUUCGCUGUCU.....C.....AAAGCAU.........................................................................GGCUCA
env-59     UGCUAAGCAUUCUAAUCUUUUCAGGGUAAAAAGCCUAGAAAAAGUUUGCUUUGAAAAGCGGGAUAUUCUAUAAUCCGCCGUCU.....C.....AAAGCAU.........................................................................GGCUCA
Hsp-1-1    CGCUAAACAUUCUAAUCUUUUCAGGGUAAAAAACCUAGAAAAAGUUUGCUUUGAAAAGAGGGUUAUUCUAUAAUUCGCCGUCU.....C......AAAGCAU........................................................................GGCUCA
env-62     UGCUAAGCAUUCUAAUCUUUUCAGGGUAAAAAACCUAGAAAAAGUUUGCUUUGAAAAGAAGGAUAUUCUAUAAUCCGCCGUCU.....C.....AAAGCAU.........................................................................GGCUCA
Hha-1-2    UGCUAAGCAUUCUAAUCUUUUCAGGGUAAAAAACCUAGAAAAAGUUUGCUUUGAAAAAAAGGAUAUUCUAUAAUUCGCCGUCU.....C.....AAAGCAU.........................................................................GGCUCA
env-63     UGCUAAGCAUUCUAAUCUUUUCAGGGUAAAAAACCUAGAAAAAGUUUGCUUUGAAAAGAAGGAUAUUCUAUAAUCCGCCGUCU.....C.....AAAGCAU.........................................................................GGCUCA
Hha-3-1    UGUUAAGUAUUCUAAUCUUUUCAGGGUAAAAAACCUAGAAAAAGUUUGCUUUAAAAAGCGGGAUAUUCUAUAAUUCUCCGUCU.....C.....AAAGCAU.........................................................................GGCUCA
env-65     UGUUAAGUAUUCUAAUCUUUUCAGGGUAAAAAACCUAGAAAAAGUUUGCUUUGAAAAGCGGGAUAUUCUAUAAUUCGCCGUCU.....C.....AAAGCAU.........................................................................GGCUCA
env-66     ....AAGCAUUCUAAUCUUUUCAGGGUAAAAAACCUAGAAAAAGUUUGCUUUGAAAAGAAGGAUAUUCUAUAAUCCGCCGUCU.....C.....AAAGCGU.........................................................................GGCCA.
Hin-4-1    UGGUUCACAUUCUAAUCUUUUCAGGGUAAAAAACCUAGAAAAAGUUUGCUUUGAAAAGAGGGAUAUUCUAUAAUUCGCCGUCU.....C.....AAAGCGU.........................................................................GGCCA.
env-67     UGCUAAGCAUUCUAAUCUUUUCAGGGUAAAAAACCUAGAAAAAGUUUGCUUUGAAAAGAAGGAUAUUCUAUAAUCCGCCGUCU.....C.....AAAGCGU.........................................................................GGCCA.
env-68     UGUUAAGCAUUCUAAUCUUUUCAGGGUAAAAAGCCUAGAAAAAGUUUGCUUUGAAAAGAGGGUUAUUCUAUAAUUCGCCGUCU.....U.....AAAGCAU.........................................................................AGCUCA
env-70     UGCUAAGCAUUCUAAUCUUUUCAGGGUAAAAAGCCUAGAAAAAGUUUGCUUUGAAAAGCGGGAUAUUCUAUAAUCCGCCGUCU.....C.....AAAGCAU.........................................................................GGCUCA
env-71     UGCUAAGCAUUCUAAUCUUUUCAGGGUAAAAAGCCUAGAAAAAGUUUGCUUUGAAAAGAGGGAUAUUCUAUAAUUCUCCCUCU.....C.....AAAGCAU.........................................................................GGCUCA
env-72     UGCUAAACAUUCUAAUCGUUUCAGGGUAAAAAGCCUAGAAAAAGUUUGCUUUGAAAAGAGGGAUAUUCUAUAAUUCUCCGUCU.....C.....AAAGCAU.........................................................................GGCUCA
env-73     CUGUUUUACCUUGGAAUAUCUAGGAUAAAAUUACAGGAUUUAAUUUGCUUUACAAUUGGUAAUUCAUUAUAAUUGCCGACCUC.....AAAGU.................................................................................UUAA..
Asu-1-1    AGCGUUUAUUUUGCCUUUUUUAGCGGUUAAAUACCAGUUUAAAUUUGCUUUCGUAGAAUAAAUUCAUUAUAAUUCACGCCCUC.....AAAGG.................................................................................CUG...
env-74     CCGUUUUUUAGCGUAAUAUCUAGGAAAAUUAACAAGAUUUUCGUUUGCUUUAGCACCACGAAUUCAUUAUAAUGCACCAC.CU.....CAAAGU................................................................................UCAG..
env-75     UGUUUUACCUUGGAAUAUCUAGGAUAAAAUUACAGGAUUUGAUUUGCUUUACAAUCGGUAAUUCAUUAUAAUUGCCGACCUCA.....AAGU..................................................................................UUAG..
env-76     UGUUUUACCUUGGAAUAUCUAGGGUAAAAUUACAGGAUUUGAUUUGCUUUACGAGCAGUAAUUCAUUAUAAUUGCCGACCUCA.....AAG...................................................................................UUCAAU
env-77     CCGUUUUUUAGCGUAAUAUCUAGGAAAAUUAACAAGAUUUUUGUUUGCUUUAGCACCACGAAUUCAUUAUAAUGCACCACCUC.....AAAGU.................................................................................UCAG..
env-78     CCGUUUUUUAGCGUAAUAUCUAGGAAAAUUAACAAGAUUUUCGUUUGCUUUAGCACCACGAAUUCAUUAUAAUGCACCACCUC.....AAAGU.................................................................................UCAG..
env-79     CCGUUUUUGGGCGUAAUAUCUAGGAAAAUUAACAAGAUUUUCGUUUGCUUUAGCGUCAUGGAUUCAUUAUAAUGCACCACCUC.....AAAGU.................................................................................UCAG..
env-80     CCGUUUUGGGGCGUAAUAUCUAGGAAAAUUAACAAGAUUUUUGUUUGCUUUAGCACCACGAAUUCAUUAUAAUGCACCACCUC.....AAAGU.................................................................................UCAG..
env-82     CCGUUUUUUAGCGUAAUAUCUAGGAAAAUUAACAAGAUUUUCGUUUGCUUUAGCACCACGAAUUCAUUAUAAUGCACCACCUC.....AAAGU.................................................................................UCAG..
env-83     CCGUUUUUUGGCGUAAUAUCUAGGAAAAUUAACAAGAUUUUCGUUUGCUUUAGCACCACGAAUUCAUUAUAAUGCACCACCUC.....AAAGU.................................................................................UCAG..
env-85     CCGUUUUUUGGCGUAAUAUCUAGGAAAAUUAACAAGAUUUUCGUUUGCUUUAGCACCACGAAUUCAUUAUAAUGCACCAUCUC.....AAAGU.................................................................................UCAG..
env-88     CCGUUUUUGGGCGUAAUAUCUAGGAAAAUUAACAAGAUUUUUGUUUGCUUUAGCACCACGAAUUCAUUAUAAUGCACCACCUC.....AAAGU.................................................................................UCAG..
env-89     CCGUUUUUGGGCGUAAUAUCUAGGAAAAUUAACAAGAUUUUCGUUUGCUUUAGCGUCAUGAAUUCAUUAUAAUGCACCACCUC.....AAAGU.................................................................................UCAG..
env-90     CCGUUUUUUGGCGUAAUAUCUAGGAAAAUUAACAAGAUUUUCGUUUGCUUUAGCACCAAGAAUUCAUUAUAAUGCACCACCUC.....AAAGU.................................................................................UCAG..
env-91     UGUUUUACCUUGGAAUAUCUAGGGUAAAAUUACAGGAUUUGAUUUGCUUUACGAGCGGUAAUUCAUUAUAAUUGCCGACCUCA.....AAGU..................................................................................UCAA..
env-101    UGUUUUACCUUGGAAUAUCUAGGGUAAAAUUACAGGAUUUGAUUUGCUUUACGAGCAGUAAUUCAUUAUAAUUGCCGACCUCA.....AAGU..................................................................................UCAA..
env-103    UGUUUUACGUUGGAAUAUCUAGGAUAAAAUUACAGGAUUUGAUUUGCUUUACAAUCGGUAAUUCACUAUAAUUACCGACCUCA.....AAGU..................................................................................UUAG..
env-104    UGUUUUACCUUGGAAUAUCUAGGAUAAAAUUACAGGAUUUGAUUUGCUUUACAAUCGGUAAUUCACUAUAAUUACCGACCUCA.....AAGU..................................................................................UUAG..
env-111    UGUUUUACCUUGGAAUAUCUAGGAUAAAAUUACAGGAUUUGAUUUGCUUUACAAUCGGUAAUUCAUUAUAAUUGCCGACCUCA.....AAGU..................................................................................UUAG..
env-114    GCCGGGCCAAUGAGCUGCGCGGGUGAGAAAAGCGGCGAUUUGUAUUGCUGCCGCUACCCUGCUCCGGUAUCAUACCGCCCGCU.....UUCGC.................................................................................CUUGU.
env-115    UGAGCUGCGCGGGUGAGUAAAUAACCUGAAACCGGCUAUUUAUAUUGCUGCAACUGCCCUGCUCCGGUAUCAUACCGCCCGCU.....UUCGC.................................................................................CUUGU.
env-121    UGAGCUGCGCGGGUGAGUAAAUAACCUGAAACCGGCUAUUUAUAUUGCUGCAACUGCCCUGCUCCGGUAUCAUACCGCCCGCU.....UUCGC.................................................................................CUUGU.
Aen-1-1    ACUGAAAUGAAUAUCAACGCCCUAAAAUUAGGACAUCUUUUUCAUUGCUGUAGAGCAUUUUUUUCGCUAAAAUGCCGCCCGCU.....UGAUUAG...............................................................................CAAUCA
Ana-1-1    UUGUUCCUUAAAGGAAUAUAAAUUCCUAAAAUUAGGAACUUUAUUUGCUGUAGCGCGUUUUUUUCGUUAAAAUGCCGCCCGCU.....UGAUUAG...............................................................................CAAUCA
Eba-2-1    GCAACGUAAUAACCAAGUGAACAACGGCACUCACUCGUGUUUUGUUUGCCGCAGCACCAAUACUUCGCUACAAUGCGCCCGCU.....U..CUGC...............................................................................UCUAU.
Dch-3-1    AUGAGCUUACUCAGGUAAGUGAUUCGGGUGAACGAACGCAGCCAACACACCUGCAACUUGAAGUAUGAAGGGUAUGGUUAUUUGAAC..............................................................................................
Sgl-1-1    CGGCAGCGGACAAGUUACGGGCGCCGGAUUUUGUGGUUUCCCGCUUACGCGGCUAACCGGGUUUCGUUUGCUCCGCGCCGGUAAUUCCGGUA..................................................................................AAAUCC
Dsp-3-1    AACGCGACCACGGCCCAAACAGCCGCGCGCGCCAAGGGUUUUAUUUGCCCGCGCCGCCAUAUUCCGCUAGAAUCUCGCCCGCU.....UUUGUUUUCUGG..........................................................................CGG...
Fpe-1-1    GAUAUUCAAUUACCAGAAAUAAUUAUAGGUUAAUCAAUAUUUAAGUUGCGAUACAAGUAAUAAAUCGUUAUAAUUCACACGCU.....UU....................................................................................UAA...
env-122    GAUAUUCAAUUACCAGAGAUAAUUAUAGGUUAAUCAAUAUUUAAGUUGCGAUACGAGUAAUAAAUCGUUAUAAUUCACACGCU.....UU....................................................................................UAA...
Mmo-1-1    CAUAACCCGGAGUGCUGCUUUUUGGGUUUUAUACCGGGUUUCGUUUGCCGUUUGUCCGCAUUUUCGGUAAAAUGCACGCCGCU.....UCACCGGGG.............................................................................GUAA..
Mmo-1-3    CAUAACCCUGAGUGCUGCUUUUUUGGUUUUAUACCGGGUUUCGUUUGCCGUUUGUCCGCAUUUUCGGUAAAAUGCACGCCGCU.....UCACCACGGG............................................................................UAAC..
Mmo-3-1    CCUGACCCGGCGUGCUGCUUUUUGGGUUUUAUGCCGGGUUUCGUUUGCCGUUUGUCCGCAUUUUCGGUAAAAUGCACGCCGCU.....UCAACACGGG............................................................................UAAC..
Mmo-5-1    CAUAACCCUGAGUGCUGCUUUUUUGGUUUUAUACCGGGUUUCGUUUGCCGUUUGUCCGCAUUUUCGGUAAAAUGCACGCCGCU.....UCGCCACGGG............................................................................UAAC..
Mmo-2-1    CAUAACCCUGAGUGCUGCUUUUUUGGUUUUAUACCGGGUUUCGUUUGCCGUUUGUCCGCAUUUUCGGUAAAAUGCACGCCGCU.....UCACCACGGG............................................................................UAAC..
Mmo-1-7    CAUAACCCUGAGUGCUGCUUUUUUUGUUUUAUACCGGGUUUCGUUUGCCGUUUGUCCGCAUUUUCGGUAAAAUGCACGCCGCU.....UCACCACGGG............................................................................UAAC..
Pfl-1-1    GCCAGUGAGGGCCCACAACUUAACACUGCCAACGCCGUUUUUAUUUGCCGCGAACACGAUAAAACGCUAUAAUGCCGCGCGCU.....AUUGC.................................................................................AUGU..
Tpt-1-1    UAAAGCUCACCAGUUGCGCUAAAGCGCCCGCCGGGUGAUUUUAUUUGCCCGGCGUCAGUGAAGCCGUUAUAAUGCCGCGCGCU.....UUUAC.................................................................................AUCCG.
Tsa-1-1    AAAGCCCAUCAAUUGCGCUGAAGAGCGGGCCGGUAGUUUUUUAUUUGCCCGUAGCGGGCUGAGCCGUUAUAAUGCCGCGCGCU.....UUUACA................................................................................UCC...
env-125    GUAAGGCCAAUCAGCUUCGUUAAGCCCUUGUGGCCCGAAUUUAUUUGCCCGCUGCCAGGGAAGCGGCUAUAAUGCCGCGCACU.....UUUAC.................................................................................AUUCG.
Mwi-1-1    GCCGCCACCAACACGUGCUUUGAAUAAUAAACUGUAAUUGUCAUUGCCGCGCUCGCGCUUUUUCGUUAAAAUGCCGCCGCUG.....AGUGAUUUU.U.CG........................................................................CGU...
Spr-1-1    CGCUUCCGCCGCGCCCAUGAAUUACACGGCUGACCGGGUUUUGUUUGCCCCGCGCCGGUAAUUCCGGUAAAAUCCUGCCCGCU.....U.....................................................................................ACUCAC
Eic-2-1    CAGGGGCGUUUUCGCAAGGCGAACCAGCUGCGCGGCUAAUUUAUUUGCCGUCCCGGCGCGGAUCGGCUAGAAUGCCGCCCGCC.....UUGUCG.AGGGU..........................................................................GUUU..
Epi-1-1    CAGGGGCGUUUUCGCAAGGCGAACCAACUGCGCGGCUAAUUUAUUUGCCGUCCCGGCGCGGAUCGGCUAGAAUGCCGCCCGCC.....UUGUCGAGGGU...........................................................................GUCA..
Epi-1-2    CAGGGGCGUUUUCGCAAGGCGAACCAGCUGCGCGGCUAAUUUAUUUGCCGUCCCUGCGCGGAUCGGCUAGAAUGCCGCCCGCC.....UUGUU.AAGGGU..........................................................................GUUA..
Eta-5-1    CAGGGGCGUUUUCGCAAGGCGAACCAGCUGCGCGGCUAAUUUAUUUGCCGUCCCUGCGCGGAUCGGCUAGAAUGCCGCCCGCC.....UUGUU.AAGGGU..........................................................................GUUA..
CSo-1-1    GCGCGCCGCAGCGCCCAUGAAUUUCGCGGCUGACCGGGUUUUGUUUGCCCCGCGCCGGUAAUUCCGGUAAAAUCCUGCCCGCU.....U.....................................................................................ACUCAC
Gan-5-1    GUUACAAAAUUAUCUCGAUUGUUGCAACAAAACGCGCAAUUAACGAUUGCAUUUUCUCGUCAUUUCUCUAUAAUGCGACCGCU.....GA....................................................................................UUUGUU
Gan-7-1    GUUACAAAAUUAUCUCGAUUGUUGCAACAAAACGCGCAAUUAACGAUUGCAUUUUCUCGUCAUUUCUCUAUAAUGCGACCGCU.....GA....................................................................................UUUGUU
Gan-1-2    GUUACAAAAUUAUCUCUAUUGUUGCAACAAAACGCGCAAUUAACGAUUGCAUUUUCUCGUUAUUUCUCUAUAAUGCGACCGCU.....GA....................................................................................UUUGUU
Gan-1-5    GUUACAAAAUUAUCUCGAUUGUUGCAACAAAACGCGCAAUUAACGAUUGCAUUUUCUCGUCAUUUCUCUAUAAUGCGACCGCU.....GA....................................................................................UUUAUU
Gan-1-6    GUUACAAAAUUAUCUCGAUUGUUGCAACAAAACACGCAAUUAACGAUUGCAUUUUCUCGUCAUUUCUCUAUAAUGCGACCGCU.....GA....................................................................................UUUAUU
Gan-6-1    GUUACAAAAUUAUCUCGAUUGUUGCAACAAAAUGCGCAAUUAACGAUUGCAUUUUCUCGUCAUUUCUUUAUAAUGCGACCGCU.....GA....................................................................................UUUAUU
Gan-1-9    GUUACAAAAUUAUCUCGAUUGUUGCAACAAAACGCGCAAUUAACGAUUGCAUUUUCUCGUCAUUUCUCUAUAAUGCGACCGCU.....GA....................................................................................UUUGUU
Gan-1-10   GUUACAAAAUUAUCUCGAUUGUUGCAACAAAACACGCAAUUAACGAUUGCAUUUUCUCGUCAUUUCUCUAUAAUGCGACCGCU.....GA....................................................................................UUUAUU
Gan-3-1    GUUACAAAAUUAUCUCGAUUGUUGCAACAAAACACGCAAUUAACGAUUGCAUUUUCUCGUCAUUUCUCUAUAAUGCGACCGCU.....GA....................................................................................UUUAUU
Gan-9-1    GUUACAAAAUUAUCGCAAUUGUUUAAACAAAACUCUCAAUUAACGAUUGCAUUUUAUCGUCAUUUCUCUAUAAUGCGACAGCU.....GA....................................................................................UUUAUU
CSc-1-1    AUACUUUAUUAAUUAGCUUAAAAAUGACAAGAAAAGAAUUAAGCAUUGCCUCUAUUACUAUAAAUCGUUAUAAUCGCACUGCU.....U.....................................................................................UUU...
env-126    UGCAAUCCAUAUCUGACUUUAUCUUUAUUAAGCUGAUUAUUUGAUUGCGUUUACUAGCAUAAAACUUUAUAAUCCCUAGCGCU.....U.....................................................................................UUUA..
env-127    ............................................................................UAGCGCU.....U.....................................................................................UUUA..
env-131    .....ACGCAAUCAAUAUGAUACAAAGUCGAUAGUAUUAUUUGAUUGCGUUUAAUCGGAUUAAACUUUAUAAUCGCUAGCGCU.....U.....................................................................................UUUA..
env-132    UGCAAUCAAUAUGUGACUUUAUCUUUGUAGAGUUGAUUAUUUGAUUGCGUUUACUAACAUAACACUUUAUAAUCCCUAGCGCU.....U.....................................................................................UUUA..
env-134    UGCAAUCAAUAUGUGACUUUGUCUUUAUUGAGCUGAUUAUUUGAUUGCGUUUACUCACAUAACACAUUAUAAUCCCUAGCGCU.....U.....................................................................................UUUA..
env-136    UGCAAUCAAUAUGUGAUUUUAUCUUUAUAGGAUUGAUUAUUUGAUUGCGUUUACUAGCAUAACACUUUAUAAUCCCUAGCGCU.....U.....................................................................................UUUA..
env-138    UGCAAUCAAUAUGUGAUUUUAUCUUUAUAGGGUUGAUUAUUUGAUUGCGUUUACUAGCAUAACACUUUAUAAUCCCUAGCGCU.....U.....................................................................................UUUA..

           ................................................................................<<<.....<<<<<<<<<<<<<<<<<<<<<<<<<<<<<<<<<<<<<<<<<<<<<<<<<<<<<<<<<<<<<<<<<<<<<<<<<<<<<<<<<<<<<<......
           ................................................................................212.....22222.........111....00110.....0.1.111.1111.1.........101...............011.....111100......
           ••••••••••••••••••••••••••••••••••••••••YY••UUGYY••••••••••••••••R•UA•AAU•YCGCCCGCU-----U∘∘∘∘---------------------------------------------------------------------------------•Y•∘--
```

```
env-139    AUGCAAUUAUGCUUAAUUAGUUCAAUGAUAGAUUGAAUAUUUGAUUGCGUUUAUAUAGCGAAGACUUUAUAAUCGCCUUCGCU.....U...............................UAUA..
Gap-1-1    UGAACACGCAAUCAAUAUGAUACAAAGUCGAUAGUAUUAUUUGAUUGCGUUUAAUUGGAUUAAACUUUAUAAUCGCUAGCGCU.....U...............................UUUA..
env-140    UGUACACGCAAUCAAUAUGAUACAAAGUCGAUAGUAUUAUUUGAUUGCGUUUAAUUGGAUUAAACUUUAUAAUCGCUAGCGCU.....U...............................UUUA..
Dda-1-1    GCUAGCCGCAGCGGUUUUAUUUGCCCGCGCCGCCAUAUUCCGCUAAAAUCUCGCCCGCUUUUAUUGGUAAUAACAACAUCGUU.....A..UG...........................CGAA..
Eho-5-1    CAAGGACGCUAUCGCAAGGCCUACCAGCUGCGCGGCUAAUUUAUUUGCCGUCCCGGCGUAGAUCGGCUAGAAUGCCGCCCGCU.....UUGC............................CGAG..
Eta-1-1    CAAGGACGUUAUCGCAAGGCGUAUCAACUGCGCGGCUAAUUUAUUUGCCGUCCCGGCGCAGAUCGGCUAGAAUGCCGCCCGCU.....UUGC............................CGAG..
Eta-4-1    CAAGGACGUUAUCGCAAGGCGUAUCAACUGCGCGGCUAAUUUAUUUGCCGUCCCGGCGCAGAUCGGCUAGAAUGCCGCCCGCU.....UUGC............................CGAG..
bsy-2-1    UACCAAGGCAGAUUUCGGCGUGCAACGUCCGCGAGUUAGAUUAUUUGCUCUGCCCCCGUUACCCGCUAUAAUGCCGCGCGCU.....UUAU............................CUCA..
env-142    UACCAAGGCCGUUUUCCGCCGCGCGGACUCUGCCAGCUAGAUCAUUUGCCCUCCUACAUUUACCCGCUAUAAUGCCGCGCGCU.....UUG.............................AUUUAA
env-143    UACCAAGGCCGUUUCCGCCGCGCGGACUCUGCCAGCUAGAUCAUUUGCCCUCCCUACAUUUACCCGCUAUAAUGCCGCGCGCU.....UUG.............................AUUUAA
Psp-12-1   CUUUCGUCGCGCCAAUGAGUUGCGCUGAGCGCGGGGCAAACCAUUUGCCCGCGAUACGCAAAGAGGCUAUAAUCCGCCCGCU.....UUUGC...........................UUAU..
Ein-1-1    GGCCGUUUUCGUCGCGCUAAUGAGCUGCGCUGACGCUGUAUUGUUUGCCCGGACCGGCAAACACAGCUAUAAUGCGCCCGCU.....UUUGC...........................UAC...
env-144    GGCCGUUUCCGGCGCGCUAAUGAGUUGCGCUGACGCUGUAUUGUUUGCCCGGGCCGGCAAACACGGCUAUAAUGCCGCCCGCU.....UUUGC...........................UAC...
Psp-9-1    CAGGGACGCUAUCGCCGCGCCAACGAGCUGCGUUGAGCGGGCGUUUGCGUCGACAGGUAAAGAUGGCUAUAAUGCCGCGCGCU.....UUUGU...........................CUG...
Esp-74-1   GGGGCGUUUCCGCCGGGCAAAUGAGCUGCGCUAGCGUUAAUGCAUUGCCCACUGCGGCAAACAUCGUUAUAAUGCCGCCCGCU.....UUUGC...........................UCU...
Eam-10-1   GGGCCGUUUUCGUCGGGCAAAUGAGCUGCGCUAGCGUUAAUACAUUGCCCGCUGCGGCAAACAUCGUUAUAAUGCCGCCCGCU.....UUUGC...........................UAU...
Eam-1-1    GGGCCGUUUCCGUCGGGCAAAUGAGCUGCGCUAGCGUUAAUACAUUGCCCGCUGCGGCAAACAUCGUUAUAAUGCCGCCCGCU.....UUUGC...........................UAU...
Epy-2-1    GGGGCGUUUCCGCCGGGCAAAUGAGCUGCGCUAGCGUUAAUGCAUUGCCCACUGCGGCAAACAUCGUUAUAAUGCCGCCCGCU.....UUUGC...........................UCUGU.
Eto-1-1    GCCGCUUUCGCCGCGCCAACGAGCUGAGUUAACGACGGUAUUGUUUGCCCGCACCGGCAAACGCGGCUAUAAUGCCGCCCGCU.....UUUGC...........................GAAA..
Baq-1-1    GCCGCUAUCGUCGGGCUAAUGAACUGCGCGGAUGACGGUUUUAUUUGCCGUGGGUCGCUAAUUCCGUUAGAAUCCCGCCCGCU.....UUGACUUUUGU.....................UAAUCC
Pfo-1-1    GACGCUACCGCCGAGCCAAUGAACUACGUGGCUAACGGUUUUAUUUGCCGUAGGCGGUGAAUUCCGUUAGAAUCCCGUCCGCU.....U...............................AAA...
Plu-4-1    GGGCAGACGAAUUAUAUUAAUAGGUUCCGAUUGACCUAUUCUAUUUGCUGUAACCGCCAUUUUUCGCUAGAAUGCCGCCCGCU.....U.GAC..ACGAUG...................CUACUA
Pvu-2-1    ACGCUAUCGAAAAGCAGAUACAUUGUAUUAAUCUGGAUUUAUAUUUGCUGUUAGUGGACUUUUUCGUUAAAAUGCGCCCGCU.....UUAUCUA.........................UUUU..
Pha-1-1    UCGCUACCGAAAAGCAGAUACCUUGUAUUAAUCUGGAUUUAUAUUUGCUGUUAGAGGACUUUUUCGUUAAAAUGCGCCCGCU.....UUAUCUA.........................UUUU..
Pmi-1-2    GACGCUUUCGUAAAGCAGAUACCUUAUAUUAAAGUCGAUUUAUAUUUGCUGUUAGAGUGCUUUUUCGUUAAAAUGCGCCCGCU.....UUAUCUG.........................UUUU..
Lgr-1-1    GGCGCUACCGUCGCGCCAACGAGCUGCGCGGAUAACGGUUUUAUUUGCCGUACGGUAAAAUUUCGCUAGAAUCGGCCCCGCU.....U...............................UUAU..
gpr-1-1    AGCUGCGGGGCUAGCGGUUUAAGGCACGAUCAGGGAUUAAUCAUUUGCUGUACGGGCAAUACUUCGGUAACAUACCGCCCGCU.....UUCGC...........................CAUGC.
Rch-1-1    GAGCUGCGCGGCUAGCGAUUAAGCAAAUAUCAGGGAUUAAUCAUUUGCCGUCAGGGCGAUACUUCGGUAACAUACCGCCCGCU.....UUCGC...........................CAUGU.
Ssp-13-1   AGCUACGCGGCUAACGAUUUAAGGCAUUCUCAGGGAUUAAUCAUUUGCCGUACAGGCGAUACUUCGGUAACAUACCGCCCGCU.....UUCGC...........................CAUGU.
env-148    AGCUGCGCGGAUAGCGCUUUAAGGCAUUAUCAGGGAUUAAUCAUUUGCCGUACGGGCGAUACUUCGGUAACAUACCGCCCGCU.....UUCGC...........................CAUGU.
Pru-1-1    UUAAUAUAAAUUAACUUUCUACUUGGCGGAUAGUGAUCUUAAUGAUUGCCCGUUGGGCAUUUUUUCGUUAAACUGCGCCCGCC.....U...UCGUAUAUCUCGC................CUA...
env-149    GCGCUUUCGCCGCGCUAACGAACUGCGCUAAGCCUUAUUAGCGUUUGCCCCUGUAGGGGAUUACCGCUAUAAUGCCGCCCGCU.....UUUGC...........................AUGU..
Pst-1-1    CCGCUUCCGCCGGGCCAAUGAACUCCGCUAGGUCGUAUUAGCGUUUGCCCCGGUGGGGGAUUACGGCUAUAAUGCCGCCCGCU.....UUUGC...........................AUGU..
Pst-4-1    CCGUUUCCGCCGGGCCAAUGAACUCCGCUAGGUCGUAUUAGCGUUUGCCCCGGUGGGGGAUUACGGCUAUAAUGCCGCCCGCU.....UUUGC...........................AUGU..
Bgo-1-1    GCCGGGCAAACGAGUUGCGAGGAUAACUUAUUUCUCGGUUUUAUUUGCCCGACGAACCAUAUUCCGUUAGAAUUCCGUCCGCU.....UUUGC...........................ACC...
Bsp-1-1    CGGCGGGCAAACGAGCUGCGCGGCUAGCUUAUUCACGGUUUUAUUUGCCCGCAGACCCAUAUUCCGCUAGAAUUCAGCCCGCU.....UUUGU...........................AUC...
Dpa-1-1    CGCCGCGCUCAUGAACUGCGCGGCUAACGUCGCCGUGGUUUCGUUUGCCCUACAGGCCAUAUUCCGCUAGAAUUUCGCCCGCU.....UUUGUCCAGUCUG...................UCGUGC
Pal-5-1    GAUGAAUAAAUCAUGCCAAGUCAGAAAGAUAAUGAUCUUAAUGAUUGCCCGCUGGCCGUUUUUUCGUUAAAAUGCGCCCGCC.....U..UUGUAU.GUCUCGG................UUA...
Pal-8-1    GGCGGAUAAACCAUGCCCAAUCAGAAAGAUAAUGAUCUUAAUGAUUGCCCACCCACCGUUUUUUCGUUAAAAUGCUGCCCGCC.....UUUAUAUGUG.UCGCG.................AUAG..
Pal-2-1    GGCGGAUAAACUAUGCCCAAUCAGAAAGAUAAUGAUCUUAAUGAUUGCCCACCCACCGUUUUUUCGUUAAAAUGCUGCCCGCC.....UUUGUAUGUG.UCGCG.................AUAG..
Pal-9-1    GGCGGAUAAACCAUGCCCAAUCAGAAAGAUAAUGAUCUUAAUGAUUGCCCACCCACCGUUUUUUCGUUAAAAUGCUGCCCGCC.....UUUGUAUGUG.UCGCG.................AUAG..
Sma-22-1   CCUGUACUGAUACGCAUCCCGUCCGCCGCUGCGGCAGGUUUCGUUUGCCCUGGCGGCGGCUUUCCGUUAAAAUGCGCCCGCU.....UUACCUGGG.......................UUCG..
env-151    UAGUAUUUUUCUGGCAACUAAUUUGCCAUUUUCCUAACUAGUAUUUGCUGUUCCGCCCGAUUUCCGCUAGAAUGCCAGCCGCU.....UU..............................GUUA..
Dze-7-1    AAUCCCGAUGAGCUUACACAGGUAAGUGAUUCGGGUGAACGAACGCGGCCAACACACCUGCAACUUGAAGUAUGACGGGUAUA.....G..UUAUUU.......................GAAC..
Dze-9-1    UGCAGCGGCCCGAAGGGCGAGGCCCAUUAGAUGGGCCGAGUAACGCAGCCAACACACCUGCAACUUGAAGUAUGACGGGUAUA.....G..UUAUUU.......................GAAC..
Dze-6-1    AAUCCCGAUGAGCUUACUCAGGUAAGUGAUUCGGGUGAACGAACGCGCCAACACACCUGCAACUUGCAGUAUGACGGGUAUA.....G..UUAUUU.......................GAAC..
Dch-1-1    UCGCCGGGCAAACGAACUGCGCGGCUAGCCUUAGGCGGUUUUAUUUGCCCGCACGGCCAUAUUCCGCUAGAAUCUCGCCCGCU.....UUUAUUUCGAAU.AUCA...............ACA...
Dda-5-1    UCGCCGGGCAAACGAACUGCGCGGCUAGCUUUAGGCGGUUUUAUUUGCCCGCACGACCAUAUUCCGCUAGAAUCUCGCCCGCU.....UUUAUUUCGAAUAUCA................ACAU..
Dze-4-1    CCCGAUGAGCUUACACAGGUAAGUGAUUCGGGUGAACGAACGCAGCCAACACACCUGCAACUUGAAGUAUGACGGGUAUAGUU.....A..U............................UUGA..
Dch-4-1    CCCGAUGAGCUUACUCAAGUAAGUGAUUCGGGUGAACGAACGCAGCCAACACACCUGCAACUUGAAGUAUGAAGGGUAUGGUU.....A..U............................UUGA..
Dze-2-1    UCGCCGGGCAAACGAACUGCGCGGCUAGCCUUAGGCGGUUUUAUUUGCCCGCACGGCCAUAUUCCGCUAGAAUCUCGCCCGCU.....UUUAUUUCGAAUAUCA................ACAU..
Dsp-9-1    CGCCGCGCCCAUGAACUGCGCGGCUAAUUCCGCUAUGGUUUUAUUUGCCCGCGCCACCAUAUUCCGCUAGAAUCUCGCCCGCU.....UUUAUUUCGAAUAAC.................CGUAUC
Dze-5-1    UCCCGAUGAGCUUACACAGGUAAGUGAUUCGGGUGAACGAACGCCGCCAACACACCUGCAACUUGAAGUAUGACGGGUAUUGU.....UAUUU...........................GAAC..
Hal-1-1    UCUCUCAGUGGUUAGGCGUUGUAACAAUGAGCGGGCAGAUAUAUUUGCCGUAAGCCCCGUAUUUCGUUAGAAUGCCGCCCGCU.....UUUA............................CUAU..
Hal-1-2    UCUCUCAGCGGUUAGGCGUUGCAACAAUGAGCGGGCAGAUAUAUUUGCCGUAAGCCCCGUAUUUCGUUAGAAUGCCGCCCGCU.....UUUA............................CUAU..
env-152    UCUCUCAGCGGUUAGGCGUUGUAACAAUGAGCGGGCAGAUAUAUUUGCCGUAAGCCCCGUAUUUCGUUAGAAUGCCGCCCGCU.....UUUA............................CUAU..
Hpa-1-1    UUCGCUACGCGGCUAGGUUUUCUUGCCAAAGCGGGCAGAUUUAUUUGCCGUAAGCCCUGUGUUUCGUUAGAAUGCCGCCCGCU.....UUUA............................CUAU..
Hal-4-1    UUCGCUACGCGGCUAGGUUUUCUUGCCAAAGCGGGCAGAUUUAUUUGCCGUAAGCCCUGUGUUUCGUUAGAAUGCCGCCCGCU.....UUUA............................CUAU..
Pst-6-1    CGAGUUGCCGACCAACUCUACUCCUGAUAUUGAUUAGAUUAUGAUUGCUGUAAAGACGGUUUUUCGUUAAAAUGCCGCCCGCU.....U..GGAUGU....UUCGC..............GUCA..
Pst-7-1    CGAGUUGCCGACCAACUCUACUCCUGAUAUUGAUUAGAUUAUGAUUGCUGUAAAGACGGUUUUUCGUUAAAAUGCCGCCCGCU.....U..GGAUGU....UUCGC..............GUCA..
Pre-2-1    CGCGUUGCAGAUUCUCUAUCAUAAAGCGAAAACGAUCAUAAUGAUUGCUGGAUGGCGGAUUUUUCGUUAAAAUGCGCCCGCC.....UUCCGAUAUCUUGUUC.................GCU...
Pre-1-1    CGCGUUGCUGACAGCUCUAUCUUAAAGCUAAGGCGAUCUUAAUGAUUGCUGGAUGGCGGAUUUUUCGUUAAAAUGCGCCCGCC.....UUCAUAUAUCACAUUC.................UCU...
Pbu-1-1    CCAAUAGCUCCUAGCAAUCAUGAGAUUGUGAUUGCGUAGAGGUCAUUUUUCGUUAAAAUACUGCCCGCCUAGGUGUAUCGCU.....UUAGCC..........................AACUUA
Pas-2-1    GAGCAGAUCAAUUAUAUUAAUAUGUCCAAAUGGACGUAUUCAUUUGCUGUAACUACCAUUUUUCGCUAGAAUGCCGCCCGUU.....UAAC............................ACUAC.
Pse-1-1    CCGCUUCCGCCGCGCCAAUGAGCUGCGCUAAGCGGCAGAAUCAUUUGCCCGCUCGCCCAUUACCGCUAUACUGCCGCCCGCU.....UUUGC...........................AUUC..
Psp-7-1    CCGCUUCCGCCGCGCCAAUGAGCUGCGCUAAGCGGCAGAAUCAUUUGCCCGCCCCGCCCAUUACCGCUAUACUGCCGCCCGCU.....UUUGC...........................AUUC..
Pva-1-1    CCGCUUCCGCCGCGCCAAUGAGCUGCGCUAAGCGGCGGAAUCAUUUGCCCGCCCCGCCCAUUACCGCUAUACUGCCGCCCGCU.....UUUGC...........................AUUC..
Msp-1-1    GCAGCCCACCUGGUAAUAGUAAAGAAAGUGGCGCUAACGCUUUAUUUGCCCGAUGGCAGAUUACCCGCUAUAAUGCCGCCCGCU.....UUUGC...........................AUCC..
Eba-4-1    UCACCCCUAUUACCCUGUUAUCCAUGACGUCACCGCAGUUUCGUUGCCCGUUGCUGCGGGUUACGGCUAUAAUGCCGCCCGCU.....CUGC............................AUCC..
Lqu-1-1    CGACGCGCCAAUGAGCUACGCGGCUGAGCGCGCGGCGGUUUUAUUUGCCCGGAGCGCCAGAUUCCGCUAGAAUUCCGCCCGCU.....UUUGC...........................UUCG..
env-153    CGACGCGCCAAUGAGCUACGCGGCUGAGUACGCGGCGGUUUUAUUUGCCCGCAGCGCCAGAUUCCGCUAGAAUUCCGCCCGCU.....UUUGC...........................UUCG..
Lqu-1-2    CGACGCGCCAAUGAGCUACGCGGCUGAGUACGCAGCGGUUUUAUUUGCCCGGAGCGCCAGAUUCCGCUAGAAUUCCGCCCGCU.....UUUGC...........................UUCG..
env-154    NNNNNNNNNNNNNNNNNNNNNNNNNNNNNNNNNNNNNNGGGGGUUUUAUUUGCCCGCAGCGCCAGAUUCCGCUAGAAUUCCGCCCGCU.....UUUGC...........................UUCG..
env-155    ...........CUUGAAGUAUGACGAGUAAUGCUGAGAUUCAUUUGCCCGCAAGGGCAUAUUCCGCUAGAAUUCAGCCCGCU.....UUUG............................UUCG..
Pca-10-1   ACGCACAUGAAACUUGAAGUAUGACGAGUAAUGCUGAGAUUCAUUUGCCCGCAAGGGCAUAUUCCGCUAGAAUUCAGCCCGCU.....UUUG............................UUCG..
Ppo-1-1    ACGCACAUACAACUUGAAGUAUGACGAGUAAUUUUGGGAUUCAUUUGCCCGCAGAGCCAUAUUCCGCUAGAAUUCAGCCCGCU.....UUUG............................UUCG..
Pwa-3-1    ACGUACAUGAAACUUGAAGUAUGACGAGUAAUGUUGGGAUUCAUUUGCCCGCAGAGCCAUAUUCCGCUAGAAUUUAGCCCGCU.....UUUG............................UUCG..
Pwa-2-1    ACGCACAUGAAACUUGAAGUAUGACGAGUAAUGCGGAUUCAUUUGCCCAUGAAGCCAUAUUCCGCUAGAAUUUAGCCCGCU.....UUUG............................UUCG..
Ppa-2-1    ACGCACAUGAAACUUGAAGUAUGACGAGUAAUGUUGGGAUUCAUUUGCCCGCAGAGCCAUAUUCCGCUAGAAUUUAGCCCGCU.....UUUG............................UUCG..
Pbe-1-1    ACACACAUGAAACUUGAAGUAUGACGAGUAAUUUUGGGAUUCAUUUGCCCGCGGAGCCAUAUUCCGCUAGAAUUUAGCCCGCU.....UUUG............................UUCG..
Pca-5-1    CGUCGCGCCAACGAGCUCCGCGGCUAAGUAAUUUUGGGAUUCAUUUGCCCGCGGAGCCAUAUUCCGCUAGAAUUCAGCCCGCU.....UUUG............................UUCG..
Pca-3-1    CGCCGCGCCAACGAGAUCCGCGGCUAAGUAAUUUUGGGAUUCAUUUGCCCGCAGAGCCAUAUUCCGCUAGAAUUCAGCCCGCU.....UUUG............................UUCG..
Pca-3-2    CGCCGCGCCAACGAGAUCCGCGGCUAAGUAAUUUCGGGAUUCAUUUGCCCGCAGAGCCAUAUUCCGCUAGAAUUCAGCCCGCU.....UUUG............................UUCG..
Pca-5-2    CGUCGUGCCAAUGAGCUCCGCGGCUAAGUAAUGCUGAGAUUCAUUUGCCCGCAGAGCCAUAUUCCGCUAGAAUUCAGCCCGCU.....UUUG............................UUCG..

           ................................................................................<<<.....<<<<<<<<<<<<<<<<<<<<<<<<<<<<<<<<<<<<<<<<<<<<<<<<<<<<<<<<<<<<<<<<<<<<<<<<<<<<<<<<<<<<<<<<<<<......
           ................................................................................212.....22222........111....00110.....0.1.111.1111.1.........101...............011.....111100......
           ●●●●●●●●●●●●●●●●●●●●●●●●●●●●●●●●●●●●●●●●●●YY●●UUGYY●●●●●●●●●●●●●●●●●●R●UA●AAU●YCGCCCGCU----U∘∘∘∘-----------------------------------------------------------------------------------●Y●∘--
```

```
Pca-5-3     CGCCGCGCCAACGAGCUCCGCGGCUAAGUAAUUUUGGGAUUCAUUUGCCCGCAGAGCCAUAUUCCGCUAGAAUUCAGCCCGCU.....UUUG...................UUCG..
Pca-5-7     CGCCGCGCCAACGAGAUCCGCGGCUAAGUAAUUUUGGGAUUCAUUUGCCCGCGGAGCCAUAUUCCGCUAGAAUUCAGCCCGCU.....UUUG...................UUCG..
Pca-6-1     CGCCGCGCCAACGAACUCCGCGGCUAAGUAAUUUGGGGAUUCAUUUGCCCAUGUCGCCAUAUUCCGCUAGAAUUCAGCCCGCU.....UUUG...................UUCG..
Pbe-1-2     ACACACAUGAAACUUGAAGUAUGACGAGUAAUUUUGGGAUUCAUUUGCCCGCGGAGCCAUAUUCCGCUAGAAUUUAGCCCGCU.....UUUG...................UUCG..
Pca-5-9     CGCCGCGCCAACGAACUCCGCGGCUAAGUAAUUUUGGGAUUCAUUUGCCCGCAGAGCCAUAUUCCGCUAGAAUUCAGCCCGCU.....UUUG...................UUCG..
Pca-9-1     ACGCACAUGAAACUUGAAGUAUGACGAGUAAUUUUGGGAUUCAUUUGCCCGCGGAGCCAUAUUCCGCUAGAAUUCAGCCCGCU.....UUUG...................UUCG..
Pca-8-1     ACGUACAUGAAACUUGAAGUAUGACGAGUAAUUUUGGGAUUCAUUUGCCCGCAGAGCCAUAUUCCGCUAGAAUUCAGCCCGCU.....UUUG...................UUCG..
Pca-5-10    ACGCACAUGAAACUUGAAGUAUGACGAGUAAUUUUGGGAUUCAUUUGCCCGCAGAGCCAUAUUCCGCUAGAAUUCAGCCCGCU.....UUUG...................UUCG..
Pca-3-15    ACGCACAUGAAACUUGAAGUAUGACGAGUAAUUUUGGGAUUCAUUUGCCCGCGAAGCCAUAUUCCGCUAGAAUUCAGCCCGCU.....UUUG...................UUCG..
Pca-7-1     CGCCGCGCCAAUGAGCUCCGCGGCUAAGUAAUGUUGGGAUUCAUUUGCCCACGAAGCCAUAUUCCGCUAGAAUUUAGCCCGCU.....UUUG...................UUCG..
Pca-1-1     CGGCGCGCCAACGAGCUGCGCGGCUAAGUAAUUUGUGGAUUCAUUUGCCCAUGUCGCCAUAUUCCGCUAGAAUUCAGCCCGCU.....UUUG...................UUUA..
Pca-2-1     CGCCGCGCCAACGAGCUCCGCGGCUAAGUAAUUUGGGGAUUCAUUUGCCCAUGUCGCCAUAUUCCGCUAGAAUUCAGCCCGCU.....UUU....................AUUCAC
Pat-3-1     ACGCACAUGAAACUUGAAGUAUGACGAGUAAUGUUGGGAUUCAUUUGCCCACGAAGCCAUAUUCCGCUAGAAUUUAGCCCGCU.....UUUG...................UCCG..
Pte-1-1     GGGCAGAUGAAUUGUAUUAAUAGGCCCCAAUUGACGUAUUCCAUUUGCUGUCACAACCAUUUUUCGCUAGAAUGCCGCCCGUU.....UAGCG..................CCCG..
Xbo-9-1     GAUCUGUACUGAGCCGAUUAAAUCAGCCCAAAAGUACUUUCUAUUUGCUGUCAUAUAGGUUUUUCGUUAAAAUGCCGCCCGCU.....UAGUCUAG...............UACA..
Plu-6-1     GAGCAGACGAAUUGUACUAAUAGAUUCAGAUUGGCCUAUUCCAUUUGCUGUAACCGCCAUUUUUCGCUAGAAUGCCGCCCGCU.....UGACA.CUAGC............UCUAU.
Plu-2-1     GAGCAGACGAAUUGUACUAAUAGAUUCAGAUUGGCCUAUUCCAUUUGCUGUAACCGCCAUUUUUCGCUAGAAUGCCGCCCGCU.....UAAC...................CUAC..
Xne-3-1     UGUAUUGAGGCCACUUAAGAAACUGGCCGAAAAUAGAUUUAUAUUUGCUGUCAUAUCCGUUUUUCGUUAGAAUGCCGCCCGCU.....UAGCGUAA...............UACA..
Xkh-1-1     UGUACUAAGCUUGCCAAAUAACCUGCCAGAUAGUGCAUUUCUAUUUGCUGUCAUAACGGUUUUUCGUUAGAAUGCCGCCCGCU.....UAACGUAA...............UACA..
Xne-5-1     UGUAUUGAGGCCACUUAAGAAACUGGCCGAAAAUAGAUUUAUAUUUGCUGUCAUAUCCGUUUUUCGUUAGAAUGCCGUCCGCU.....UAGCGUAA...............UACA..
Xgr-1-1     ACCAAACAGGCUAUCCACUAACCAACCGGAAAGCAUAUUUAUAUUUGCUGUCAUAUCGGUUUUUCGCUAGAAUGCCGCCCGCU.....UAACGUGA...............UAAC..
Xdo-1-1     UCUGAAUAGACCAACCACGCACCAACCGGAAAGCAUAUUUAUAUUUGCUGUCAUAUCGGUUUUUCGUUAGAAUGCCGCCCGCU.....UAACGU.................AAUAAC
Xbo-2-1     GAUCUGUACUGAGCCGAUUAAAUCAGCCCAAAAGUACUUUAUAUUUGCUGUCAUAUAGGUUUUUCGUUAAAAUGCCGCCCGCU.....U.....AGUGU............AGU...
Xbo-11-1    GAUCUGUACUGAGCCGAUUAAAUCAGCCCAAAAGUACUUUCUAUUUGCUGUCAUAUAGGUUUUUCGUUAAAAUGCCGCCCGCU.....U.....AGUGU............AGU...
Xbo-8-1     AAUCUGUACUGAGCCGAUUAAAUCAGCCCAAAAGUACUUUCUAUUUGCUGUCAUAUAGGUUUUUCGUUAAAAUGCCGCCCGCU.....U.....AGUGU............AGU...
Pas-1-1     AGCAGAUGAAUUGUAUUAAUAUGUUCAAGAUCGACGUAUUCCAUUUGCUGUAACUGCCAUUUUUCGCUAGAAUGCCGCCCGUU.....UAACG..................CUA...
Xpo-1-1     UGCCAGACAAAUUGCCUGUUAACAAUCUAAAAACAUAUUUAUAUUUGCUGUCAUAUCGGUUUUUCGUUAGAAUGCCGCCCGCU.....CAGUGUGA...............UAUA..
Plu-1-2     GGGCAGACGAAUUAUAUUAAUAGGUUCCGAUUGACCUAUUCUAUUUGCUGUAACCGCCAUUUUUCGCUAGAAUGCCGACCGCU.....UGACA..................CUAUGC
Plu-5-1     AUUAAUAGGAAUCGUACUAAUAGAUUCAGAUUGACCUAUUCCAUUUGCUGUAACCGCCAUUUUUCGCUAGAAUGCCGCCCGCU.....UAAC...................CUAU..
Plu-3-1     GAGCAGACGAAUUAUAUUAAUAGGUUCCGAUUGACCUAUUCUAUUUGCUGUAACCGCCAUUUUUCGCUAGAAUGCCGCCCGCU.....UGACA..................CUAUGC
Pte-2-1     GAGCAGAUGAAUUGUAUUAAUAGGCCCCAAUUGACGUAUUCCAUUUGCUGUCACAACUAUUUUUCGCUAGAAUGCCGCCCGUU.....UAACG..................CCCG..
Pte-3-1     GAGCAGAUGAAUUGUAUUAAUAGGCCCCAAUUGACGUAUUCCAUUUGCUGUCACAACCAUUUUUCGCUAGAAUGCCGCCCGUU.....UAACG..................CCCG..
Xsp-1-1     CCUGUAUUAAGCGGUGUAAUUACUCACAACAAAAUACUUUCUAUUUGCCGUUACAUAGGUUUUUCGUUAAAAUGCCGCCCGCU.....UAGCGU.................AACCCA
Xca-1-1     CCUGUAUUAAGCUGUGUAAUUACUCACGACAAAGUACUUUCUAUUUGCUGUCAUAUAGGUUUUUCGUUAAAAUGCCGCCCGCU.....UAGCGUAA...............UCCA..
Xsz-1-1     UGUACUAAAUCCAGCAACUAAUCGAGCAGAACAUAUAUUUCUAUUUGCUGUUAUAUCGGUUUUUCGUUAGAAUGCCGCCCGCU.....UAGCGUAA...............UACA..
Spl-2-1     GAGCUGUAUUGAUGCCCGCAUCGCGCAGAUUCGGGAGGUUUCCAUUGCCCUGCCGACGGCUUUCCGUUAAAAUGCCGCCCGCU.....UUAGC..UGGG............UUCG..
Sfo-4-1     CGCCGUGCCAAUGAGCUGUAUUAGGCCCGAGCGGCAGGUUUUCAUUGCCCAGCGAGCGGCUUUCCGUUAAAAUGCCGCCCGCU.....U.UAC.CUGGG............UUCG..
Sfo-3-1     CGCCGUGCCAAUGAGCUGUAUUAGCCCCGUGCGGCAGGUUUUCAUUGCCCAGCGAGCGGCUUUCCGUUAAAAUGCCGCCCGCU.....U.UAC.CUGGG............UUCG..
Sfo-1-1     CGCCGUGCCAAUGAGCUGUAUUAAGCCCGUGCGGCAGGUUUUCAUUGCCCAGAGAGCGGCUUUCCGUUAAAAUGCCGCCCGCU.....U.UAC.CUGGG............UUCG..
env-157     UCAAUUGACCUGAGCGAACGGUCCCCAAUCUGGACAGGUUUUCGUUGCCCUGCAGGCGGCUUUCCGUUAAAAUGCCGCGCGCU.....U..UACCUGGG............CCUG..
Sma-1-2     UCAAUUGACCUGAGCGAACGGUCCUCAAUCUGGACAGGUUUUCGUUGCCCUGCGGGCGGCUUUCCGUUAAAAUGCCGCGCGCU.....U..UACCUGGG............CCUG..
Sgr-1-1     GAACUGUACUGAUGUUGGCGGCAGGCACAUUGGGUAGGUUUCCAUUGCCCUGUGGCUGGCUUUCCGUUAAAAUGCCGCCCGCU.....U.UGC.CUGGG............UUCG..
Sma-1-3     UCAGUUGGCCUGAGCGAACGACCCUCAAUCUGGGCAGGUUUUCGUUGCCCUGCAGGCGGCUUUCCGUUAAAAUGCCGCGCGCU.....U.UAC.CUGGG............CCUG..
Sur-1-1     UCAGUUGGCCUGAGCGAACGGCCCUCAAUCUGGGCAGGUUUUCGUUGCCCUGCGUGCGGCUUUCCGUUAAAAUGCCGCGCGCU.....U.UAC.CUGGG............CCUG..
env-158     UCAGUUGGCCUGAGCGAACGGCCCUCAAUCUGGGCAGGUUUUCGUUGCCCUGCAGGCGGCUUUCCGUUAAAAUGCCGCGCGCU.....U.UAC.CUGGG............CCUG..
Sma-14-1    UCAGUUGGCCUGAGCGAACGGCCCUCAAUCUGGGCAGGUUUUCGUUGCCCUGCGGGCGGCUUUCCGUUAAAAUGCCGCGCGCU.....U.UAC.CUGGG............CCUG..
Ssp-8-1     UCAGUUGGCCUGAGCGAACGGCCCUCAAUCUGGGCAGGUUUUCGUUGCCCUGUGGGCGGCUUUCCGUUAAAAUGCCGCGCGCU.....U.UAC.CUGGG............CCUG..
Ssp-9-1     UCAGUUGGCCUGAGCGAACGGCCCCCAAUCUGGGCAGGUUUUCGUUGCCCUGCGGGCGGCUUUCCGUUAAAAUGCCGCGCGCU.....U.UAC.CUGGG............CCUG..
Sne-1-1     UCAAUUGACCUGAGCGAACGGUCCCCAAUCUGGACAGGUUUUCGUUGCCCUACAGGCGGCUUUCCGUUAAAAUGCCGCGCGCU.....U.UAC.UUGGG............CCUG..
Sma-18-1    UCAGUUGGCCUGAGCGAACUGUCCCCAACCUGGGCAGGUUUUCGUUGCCCUGCGGGCGGCUUUCCGUUAAAAUGCCGCGCGCU.....U.UAC.CUGGG............CCUG..
Sma-25-1    UCAAUUGGCCUGAGCGAACUGUCCCCAAUCUGGGCAGGUUUUCAUUGCCCUGCGGAUGGCUUUCCGUUAAAAUGCCGCGCGCU.....U.UAC.CUGGG............CCUG..
Sma-1-1     UCAAUUGGCCUGAGCCAACUGUCCCCAACCUGGGCAGGUUUUCGUUGCCCUGCGGGCGGCUUUCCGUUAAAAUGCCGCGCGCU.....U.UAC.CUGGG............CCUG..
env-159     UCAAUUGGCCUGAGCGAACUGUCCCCAACCUGGGCAGGUUUUCGUUGCCCUGCGGGCGGCUUUCCGUUAAAAUGCCGCGCGCU.....U.UAC.CUGGG............CCUG..
env-160     UCAAUUGGCCUGAGCGAACUGCCGCCAACCCGGGCAGGUUUUCGUUGCCCUGCGGGCGGCUUUCCGUUAAAAUGCCGCGCGCU.....U.UAC.CUGGG............CCUG..
Sli-2-1     UACUCAUCUCCUGGCGCUUACUUAGGUAAGUAACUGGAGUGAGUCAGUGCAGGUAACAACACUGUGACUUGAAAGGCGAC.................................
env-161     AUGAGCUGUACUGAUCUUUGCCCCUUGCAAAUCAGGCAGGUUUCCAUUGCCCUGUGGGCGGCUUUCCGUUAAAAUGCGCCGCU.....UUGC.CUGGG.............UUCG..
Ssp-6-1     CGCCGUGCCAAUGAGUUGUAUUAGGCCUGCGCGGCAGGUUUUGAUUGCCCACGGAGCGGCUUUCCGUUAAAAUGCCGCCCGCU.....U.UAC.CUGGG............UUCG..
Ssp-2-1     CAACUGUAUUAAAGGCAAUAGCACGCACUUGCGGCAGGUUUUCGUUGCCCGCAGAACGGCUUUCCGUUAGAAUGCCGCCCGCU.....U.UGC.CUGGG............UUUA..
Ssy-2-1     GAGCUAUAUUAACCACGGCGGCUAGCGCCUUGGACAGAUUUUCAUUGCCCUGCGGGGGGCAUUCCGUUAGAAUUCCCCCCGCU.....U.UGC.CUGGG............UUCG..
Ssy-1-1     GAGCUAUAUUAACCACGGCGGCUAGCGCCUUGGACAGAUUUUCAUUGCCCUGCGGGGGGCAUUCCGUUAGAAUUCCCCGCGCU.....U.UGC.CUGGG............UUCG..
env-162     UGAGCUGUACUGAUCUUGCCCCUUGCAAAUCAGGCAGGUUUCCAUUGCCCUGUGGGCGGCUUUCCGUUAAAAUGCCGCCCGCU.....U.UGC.CUGGG............UUCG..
env-163     GAGCUGUACUGAUCUUUGCCCCUUGCAAAUCAGGCAGGUUUCCAUUGCCCUGUGGGCGGCUUUCCGUUAAAAUGCCGCCCGCU.....U.UGC.CUGGG............UUCG..
Ssp-14-1    GAGCUGUCUUGAUCCUGACGGCCGGCAAAUCGGGCAGGUUUCCAUUGCCCUGCGGACGGCUUUCCGUUAAACUGCCGCCCGCU.....U.UGC.CUGGG............UUCG..
Cmu-2-1     GCGUUGUAUUAAGACCUGCAACACCCACUUACGGCAGGUUUUCAUUGCCCAGAGAACGGCUUUCCGUUAAAAUGCCGCCCGCU.....U.UGC.CUGGG............UUCG..
Spr-2-1     GAGCUGUCUUGAUCCUGACAGCCGGCAAAUCGGGCAGGUUUCCAUUGCCCUGCGGACGGCUUUCCGUUAAACUGCCGCCCGCU.....U.UGC.CUGGG............UUCG..
Ssp-4-1     AGGUGGCAGCUUGAUUGAGCUUGAGGACAUCAGGCAGGUUUCCAUUGCCCACAGCGCCGCUUUCCGUUAAAAUGCCGCCCGCU.....U.UACU.UGG.............CUAUUU
Spl-1-1     GAGCUGUAUUGAUGCCCGCAUCGCGCAGAUUCGGGAGGUUUCCAUUGCCCUGCCGACGGCUUUCCGUUAAAAUGCCGCCCGCU.....U..UAGCUGGG............UUCG..
Ssp-10-1    GAGCUGUAUUGAUGCCCGCAUCGCGCAGAUUCGGGAGGUUUCCAUUGCCCUGCCGACCGCUUUCCGUUAAAAUGCCGCCCGCU.....U..UAGCUGGG............UUCG..
Sod-1-1     AACUGUAUUGACCGCCAUCCCCCCGCCUGACCGGCAGGUUUUCAUUGCCCUGGCGGUGGCUUUCCGUUAAAAUGCCGCCCGCU.....U.UAC.CCGGG............UCCG..
Spl-5-1     CCAGGCGCCCAGCUUACUCAUCCCCCGGAGCUUACUUAAGUAAGUGACUGAGGUGAGUUAGUGCAGGUAACAACGCUGUGGCU.....U......................GAA...
Ype-132-1   CCCUUGCCUGAGCCGCAUUUGACCAGAGAUAUUCGCAGUUUUAUUUGCUGUCUGAGUGAUAUUUCGCUAGAAUGUCGCCCGCU.....UUACUCUGGG.............UUUA..
Yfr-1-1     AACACCGCUGUAGCGUCAAAUACGCAGGGAAUAUGCGGUAUUGUUUGCUGUCUGGCGGAGACUUCGCUAGAAUAUCGCCCGCU.....UUGCUCUGGG.............UUUA..
Yfr-1-4     AACACCGCUGUAGCGUCAAAUACGCAGGGAAUAUGCGGUAUUGUUUGCUGUCUGGUUGAGACUUCGCUAGAAUAUCGCCCGCU.....UUGCUCUGGG.............UUUA..
Ymo-1-1     CCCUUGCUUAAGCCGCAUUUAGCCUGAGAUUUUCGCAGUUUCAUUUGCUGUUUGGCUGAUAUUUCGCUAGAAUGCCGCCCGCU.....UUGCUCUGGG.............UUUA..
Ymo-1-3     CCCUAGCUUAAGCCGCAUUUAGCCUGAGAUUUUCUCAGUUUCAUUUGCUGUUUGGCCGAUAUUUCGCUAGAAUGCCGCCCGCU.....UUGCUCUGGG.............UUUA..
Yfr-1-6     AACACCGCUGUAGCACCAAGAACGCAGGUUAUAUGCGGUAUUGUUUGCUGUCUGGCUGAGACUUCGCUAGAAUAUCGCCCGCU.....UUGCUCUGGG.............UUUA..
Yfr-1-7     AACACCGCUGUAGCGCCAAGAACGCAGGUUAUAUGCGGUAUUGUUUGCUGUCUGGCUGAGACUUCGCUAGAAUAUCGCCCGCU.....UUGCUCUGGG.............UUUA..
Yen-21-1    CCCUCGCCUAAGCCGCAUUUAGCCUGAUAAUUUCGCAGUUUCGUUUGCUGUCUGCUCAAUACUUCGCUAGAAUGUCGCCCGCU.....UUGCUGUGGG.............UUUA..
Yal-1-1     CCCUUGCCUGAGCCGCAUUUAGCCUGAUAAAUUCGCAGUUUCGUUUGCUGUCUGCCCGAUAUUUCGCUAGAAUGUCGCCCGCU.....UUGCUCUGGG.............UUUA..
Yfr-1-9     AACACUGCCGCGGCGUCAAGUACGUAGGUUAUAUGCGGUAUUGUUUGCUGUCUGGUUGAUACUUCGCUAGAAUAUCGCCCGCU.....UUGCUCUGGG.............UUUA..
Ype-1-1     CCCUCGCCUGAGCCGCAUUCAGCCUGAUAUUUUCGCAGUUUUAUUUGCUGUCUGGUCGAUAUUUCGCUAGAAUGCAGCCCGCU.....UUGCUCUGGG.............UUUA..
Yfr-1-10    AACACCGCUGUAGCGUCAAAUACGCAGGGAAUAUGCGGUAUUGUUUGCUGUCUGGUUGAGACUUCGCUAGAAUAUCGCCCGCU.....UUGCUCUGGG.............UUUA..
Yfr-2-2     AACACUGCCGCGGCGUCAAGUAAGUAGGUUAUAUGCGGUAUUGUUUGCUGUCUGGUUGAUACUUCGCUAGAAUAUCGCCCGCU.....UUGCUCUGGG.............UUUA..
Yfr-1-12    AACACCGCUGUAGCGUCAAAUACGCAGGGAAUAUGCGGUAUUGUUUGCUGUCUGGCGGAGACUUCGCUAGAAUAUCGCCCGCU.....UUGCUCUGGG.............UUUA..

            ................................................................................<<<.....<<<<<<<<<<<<<<<<<<<<<<<<<<<<<<<<<<<<<<<<<<<<<<<<<<<<<<<<<<<<<<<<<<<<<<<<<<<<<<<<<<<<<<<<<<......
            ................................................................................212.....22222.........111....00110.....0.1.111.1111.1.........101...............011.....111100......
            •••••••••••••••••••••••••••••••••••••••YY••UUGYY•••••••••••••••R•UA•AAU•YCGCCCGCU----U∘∘∘∘---------------------------------------------------------------------------•Y••--
```

```
Yal-1-3    CCCUUGCCUGAGCCGCAUUUAGCCUGAUAAAUUCGCAGUUUCGUUUGCUGUCUGCCCGAUAUUUCGCUAGAAUGUCGCCCGCU....UUGCUCUGGG...UUUA..
Ymo-1-5    CCCUCGCUUAAGCCGCAUUUAGCCUGAGAUUUUCGCAGUUUCAUUUGCUGUUUGGCCGAUAUUUCGCUAGAAUGUCGCCCGCU....UUGCACUGGG...UUUA..
Yma-2-1    CCCUCGCCUAAGCCGCAUUUAGCCUGAGAUUUUCGCGGUUUCAUUUGCUGUCGCGCUGAUCUUUCGCUAGAAUGUCGCCCGCU....UUGCUCUGGG...UUUA..
Yma-1-1    CCCUCGCCUAAGCCGCAUUUUGCCUGAGAUUUUCGCGGUUUCAUUUGCUGUCGCGCUGAUCUUUCGCUAGAAUGUCGCCCGCU....UUGCUCUGGG...UUUA..
Yfr-1-13   CCCUCGCCUAAGCCGCAUUUCGCCUGAGAUUUUCGCGGUUUCAUUUGCUGUCGUGCUGAUCUUUCGCUAGAAUGUCGCCCGCU....UUGCUCUGGG...UUUA..
Ybe-1-1    CCCUAGCUUAAGCCGCAUCUAGCCUGAGAUUUUCGCAGUUUUAUUUGCUGUCUGGCUGAUAUUUCGCUAGAAUGUCGCCCGCU....UUGCUCUGGG...UUUA..
Ynu-1-1    CGCUUGGCUAGCGCCUUUCUGCCCUGAGUUAUUUGCGGUUUUGUUUGCUGUCUGGUCGAUAUUUCGCUAGAAUGCCGCCCGCU....UUACUCUGGG...UUCG..
Yru-1-1    CACUGGGUUAGCGCCCUGCGCCCCCAGCUUAUUUGCGGUUUUGUUUGCUGUCUGGGUGAUAUUUCGCUAGAAUGCCGCCCGCU....UUGCUCUGGG...UUCG..
Yru-1-2    CACUGGGUUAGCGUCCUGCGCCCCCAGCUUAUUUGCGGUUUUGUUUGCUGUCUGGGUGAUAUUUCGCUAGAAUGCCGCCCGCU....UUGCUCUGGG...UUCG..
Ssp-12-1   CGCAGAGCGAAUGAACUGCGCGGCUGAUUAAAUGGCGGUUUCAUUUGCCCGCUGGGCCAGAUUCCGAUAGAAUUCGCCCGCU....UUUGU...CUAA..
Pan-5-1    AGCUGCGCUGACGCGGAUGCGCCGCUGGCAGGCGGCUAUAUCGUUUGCUCCCGGCAGGGAUAACCGCUAUACUGCCGCCCGCU....UUUGC...AUGU..
Eas-4-1    GCUUUCGCCGGGCGAAUGAAUUGCGCGGAUAACCGGACUUUUCAUUGAUGUGAAUGGCAUUUGAGUAUAAAAUGCCGCCCGCU....UCAC...AUCC..
env-165    GCUUUCGCCGGGCAAAUGAAUUACGCGGCUGACGGGACUUUUCGUUGAUGUGCGGGGCGUUUGAGUAUAAAAUGCCGCCCGCU....UCAC...AUCC..
Lad-1-1    GCUUCCGCCGCGCCAAUGAAUUGCGGGGAUAACGGGACUUUUCAUUGAUGUAAAAGUCAUUUGAGUAUAAAAUGCCGCCCGCU....UCAC...AUCC..
Ecl-37-1   GCUUCCGUCGCGCCAACGAAUUGCGCGGUUAGCGGGACUUUUCAUUGAUGUGAGGGGCAUUUGAGUAUAAAAUGCCGCCCGCC....UCAC...AUCC..
Esp-37-1   GCUUUCGCCGGGCAAACGAAUUGCGCGGUUAACGGGACUUUUCGUUGAUGCGCGAGCCAUUUGAGUAUAAAAUGCCGCCCGCU....UCAC...AUCC..
Ecl-39-1   GCUUCCGUCGCGCCAACGAAUUGCGCGGGUAGCGGGACUUUUCAUUGAUGUGAGGGGCAUUUGAGUAUAAAAUGCCGCCCGCU....UCAC...AUCC..
env-166    GCUUUCGCCGGGCGAAUGAAUUGCGCGGAUAACCGGACUUUUCAUUGAUGUGAAUGGCAUUUGAGUAUAAAAUGCCGCCCGCU....UCAC...AUCC..
env-167    GUUUUCGUCGCGCCAACGAAUUGCGCGGUUAAGGGGACUUUUCAUUGAUGUGAGGGGCAUUUGAGUAUAAAAUGCCGCCCGCU....UCAU...AUCC..
Elu-1-5    GUUUUCGUCGCGCCAACGAAUUGCGCGGUUAAGGGGACUUUUCAUUGAUGUGAUGGGCAUUUGAGUAUAAAAUGCCGCCCGCU....UCAU...AUCC..
Ecl-58-1   GUUUUCGUCGCGCCAACGAAUUGCGCGGUUAAGGGGACUUUUCAUUGAUGUGAGGGGCGUUUGAGUAUAAAAUGCCGCCCGCU....UCAU...AUCC..
Yre-2-1    CGCUUCCGUCGGGCCAAUGAGUUGCGCUGAAUGUCGCUUUUCCAUUGAUGCGAAAUGCUUUCGCGGCUAUAAUGCCGCCCGCU....UUUGCAU...UCCAU.
env-168    CAGUGGAUGGCUGCGCACCCGCCAGCCUGACGCGGCGGUUUUGAUUGCCGUGGCACGCAUUCGCGGCUAUAAUGCCGCCCGCU....UUUGC...AUUC..
Tod-1-1    AACCGCGACAGCGAAUUGCGCAUUCAGCAGUGGAUGGCUGCGCACCCGCCAGCCUGACGCGGCGGUUUGAUUGCCGNNCGCU....UUUGC...AUUC..
Tgu-1-1    AAGUGGAUGGCGACGCACCCGCCAGCCUGAUGCGGCGGUUUUGAUUGCCGUGGUUUGCAUUCGCGGCUAUAAUGCCGCCCGCU....UUUGC...AUUC..
Psp-5-1    CGCCGCGCCAAUACGCUUCGUUAGCCACCUGCGGUUUAAUACGUUGCCCGCACAGCGAAACCUGGGCUAUAAUGCGCCCCGCU....UUUAC...AGUC..
Ebi-2-1    CCGCCGCGCCAACGAACUUCGUUAAGGCAUCCGCUUUAAUACAUUGCCCGCACAGCCUUAGUUGGGCUAUACUUCGCCCCGCU....UUUGC...UAAU..
Ebi-1-1    CCGCCGCGCCAACGAACUUCGGUAAUGCAUCCGCUUUAAUACAUUGCCCGCACAGCCUUAGUUGGGCUAUACUUCGCCCCGCU....UUUGC...UAAU..
Eol-1-1    CGACGCGCUCACCCGCUUCGUUAGGCACAACGACGUUAAUACAUUGCUCAGCGCUGCCCUCCUGAGCUAUAAUGCGCCCCGCU....UUUGC...CUG...
Esp-55-1   CGACGCGCUCACCCGCUUCGUUAGGCACAACGACGUUAAUACAUUGCUCAGCGCUGCCCUCCUGAGCUAUAAUGCGCCUCGCU....UUUGC...CUG...
env-169    AUCGCCGGGCCAACGAGCUGCGCUAAACGACAUAUUUUAAUCCGUUGCCCGAACGGCCUGAGUUGGGCUAUAAUGCGCCCCGCU....UUUGC...UAAC..
env-170    .......................................CCGUUGCCCGAACGGCCUGAGUUGGGCUAUAAUGCGCCCCGCU....UUUGC...UAAC..
Ety-1-1    CCGCCGGGCCAACGAGCUGCGCUAAACGACAUAUUUUAAUCCGUUGCCCGAACGGCCUGAGUUGGGCUAUAAUGCGCCCCGCU....UUUGC...UAAC..
env-171    ..............AGCUGCGCUAAACGACAUAUUUUAAUCCGUUGCCCGAACGGCCUGAGUUGGGCUAUACUGCGCCCCGCU....UUUGC...UAAC..
env-172    ......................................AAUCCAUUGCCCGUAUGGCCUAAGUUGGCUAUAAUGCGCCCCGCU....UUUGC...UGAC..
Pva-1-2    CCGCGCCAAUGAGCUGCGGUAAUCCCUGCUGGCGGGUAUAUCGUUUGCUACCACCUGGCAUAACGGCUAUACUGCCGCCCGCU....UUUGC...AUGU..
Pro-1-1    CCGCUAUCGCCGUGCCAAUGAAUUGCGUUAAGCGGCAGAAUCAUUUGCCCCGCCCUCUGAAUAUGGCUAUAAUCCCGCCCGCU....UUUGC...AUCC..
env-173    CGCUUCCGUCGGGCAAACGAAUUGCGCGGUUAGCGUACUUUUCGUUGAAGAAAAAGCGAUUCAGGGCUACUAUAUCGCCCGCU....UUAC...AUCC..
env-174    UUCGCCGCGCUAAUGAGCUGCGCGGCUAAGGGUGCGGCUUUUCGUUGCUGCAAACGAUAUUUUCGGCUAUAGUGUCGCCCGCU....UUUGC...AUCC..
env-175    UUUACGUACUACACCCGCUUAGCAGCGUCGGAGAUUUUCAUUUAUUGCCGUUAUCCCCUGAUACGGCUAUAAUGCCGCCCGCU....UUGC...GUCC..
Ehe-1-1    GAUACCGGCGCGCCAACGAAUUACGCGGUUAGCGGCAGUUUUCAUUGAAGCGAACGGCGGUUAUGGCUAUAAUCCCGCCCGCU....UUUGC...AUCC..
Pag-1-1    GGUCCGGCUAACCGGAUGCCGAACCCGGCUGGCGGGUAUAUCGUUUGCUACCACCCGGCAUAACGGCUAUAAUGCCGCCCGCU....UUUGC...AUGU..
env-177    GGUCCGGCUAACCGGAUGCCGAGCCCGGCUGGCGGGUAUAUCGUUUGCUACCACCCGGCAUAACGGCUAUACUGCCGCCCGCU....UUUGC...AUGU..
Pag-1-6    GGUCCGGCUAACCGGAUGCCGAGCCCGGCUGGCGGGUAUAUCGUUUGCUACCACCCGGUAUAACGGCUAUACUGCCGCCCGCU....UUUGC...AUGU..
Pan-1-3    GGUCCGGCUAACCGGAUGCCGAGCCCGGCUGGCGGGUAUAUCGUUUGCUACCAUCCAGCAUAACGGCUAUACUGCCGCCCGCU....UUUGC...AUGU..
Pag-5-1    GGUCCGGCUAACCGGAUGCCGAACCCGGCUGGCGGGUAUAUCGUUUGCUACCAUCCAGCAUAACGGCUAUACUGCCGCCCGCU....UUUGC...AUGU..
Pag-4-1    GGUCCGGCUAACCGGAUGCCGAACCCGGCUGGCGGGUAUAUCGUUUGCUACCACCCGGCAUAACGGCUAUACUGCCGCCCGCU....UUUGC...AUGU..
env-178    ...................UGAACCCGGCGAGCGGGUAUAUCGUUUGCUACCACCNGGCAUAACGGCUAUACUGCCGCCCGCU....UUUGC...AUGU..
env-179    GGCAGGGCUGACCGGAUGCUGAACCCGGCGAGCGGGUAUAUCGUUUGCUACCACCUGGCAUAACGGCUAUACUGCCGCCCGCU....UUUGC...AUGU..
env-180    ...................UGAACCCGGCGAGCGGGUAUAUCGUUUGCUACCACCCGGCAUAACGGCUAUACUGCCGCCCGCU....UUUGC...AUGU..
env-181    ..............................AGCGGGUAUAUCGUUUGCUACCACCCGGCAUAACGGCUAUACUGCCGCCCGCU....UUUGC...AUGU..
Pag-3-1    GUCAGGGCUGACCGGAUGCUGAACCCGGCGAGUGGGUAUAUCGUUUGCUACCACCCGGCAUAACGGCUAUACUGCCGCCCGCU....UUUGC...AUGU..
env-182    GCGAGGGCUGACCGGAUGCUGAACCCGGCGAGCGGGUAUAUCGUUUGCUACCACCCGGCAUAACGGCUAUACUGCCGCCCGCU....UUUGC...AUGU..
Pag-1-5    GGUCCGGCUAACCGGAUGCCGAACCCGGCUGGCGGGUAUAUCGUUUGCUACCACCCGGCAUAACGGCUAUACUGCCGCCCGCU....UUUGC...AUGU..
Psp-1-1    GGUUCGGCUAACCGGAUACCGAACCCGGCUGGCGGGUAUAUCGUUUGCUACCACCUGGCAUAACGGCUAUACUGCCGCCCGCU....UUUGC...AUGU..
env-183    .....................................................UGCCCCGAUCCGGCUAUAAUCCGCCCGCU....UUUGC...AUCC..
env-184    GCGCUUCCGCCGCGCCAAUGAGCUGCGCUAGAUGGCAGAAUCAUUUGCCCCCUGCCCCGAUCCGGCUAUAAUCCGCCCGCU....UUUGC...AUCC..
Eca-1-1    ACGUUAUCGACGCGCGAAUGAGCUGCGUUAAGCGACGGAAUCAUUUGCCCGCGUCUUCUAUUGCGGCUAUAAUGCCGCCCGCU....UUUGC...AUCC..
Psp-3-1    ACGUUAUCGGCGCGCGAAUGAGCUGCGUUAAGCGACGGAAUCAUUUGCCCGCGUCUUCUAUUGCGGCUAUAAUGCCGCCCGCU....UUUGC...AUCC..
env-189    ACGUUAUCGACGCGCGAAUGAGCUGCGUUAAGCGACGGAAUCAUUUGCCCGCGUCUUCUAUUGCGGCUAUAAUGCCGCCCGCU....UUUGC...AUCC..
bsy-1-1    AGGGGCGUUUUCGCCGCGCCAACGAGCUGCGCUGAGUUAUUGCAUUGCCUGGGGGAUGGUAUUCGGCUAUAAUCCCGCCCGCU....UUUG.CCGGGU...UUC...
Epi-2-1    GGGCCGUUACCGCCGGGCAAAUGAGCUGCGCUAACGUUAAUACGUUGCCCGCUGCGGCAAACAUCGUUAUAAUGCCGCCCGCU....UUUGC...UAU...
Eta-9-1    GGGCCGUUUCCGCCGCGCAAAUGAACUGCGCUAGCGUUAAUACAUUGCCCGCUGCAGCAAACAUCGUUAUAAUGCCGCCCGCU....UUUGC...UAU...
env-192    CAUCUUUCCUCCAGGAACGUUUAUGGCUUUAUCUGGCGCGGCAAUUGCCUUCCCCACGGUUCAGGGCUAUAAUGCCGCCCGCU....UUUGC...AUCC..
Kcr-1-1    CACUUGCUGGGCGGUUUCCCCUGCCCGGCAAUUCGAUGUUUUCAUUGCCCGAUAAGGAAUUGGAGGCUAUAAUGCCGCCCGCU....UUCAC...AUCC..
env-194    ...........................GGCUAAAGGGAGUUUUCAUUGAUGUCAGCGCUGAUUACGUAUAAAAUGCCGCCCGCC....UCAC...AUCC..
env-195    CGCUACCGUCGGGCCAAUGAAUUGCGUGGCUAAGGGACUUUUCAUUGAUGUACGGUCUGUUUGAGUAUAAAAUGCCGCCCGCC....UCAC...AUCC..
env-196    GCUACCGCCGGGCGAAUGAAUUGCGUGGCUAAAGGGAGUUUUCAUUGAUGUCAGCGCUGAUUACGUAUAAAAUGCCGCCCGCC....UCAC...AUCC..
Etr-1-1    CCGGGUGAAUGUAGGUGAGCAGUAUCAAGGCGACAUAUAGCCGUUGCCCGCACACCCGAAGUAAGGCUAUAAUCCGCCCCGCU....UUUGC...AACC..
Ema-2-1    ACUUUUCUGACAUCGGCUGGCAGACAGAAACACCGACAUAGCCGUUGCCCGUUUCCCGGAGUAAGGCUAUAAUGCGCCCCGCU....UUUGC...AACC..
env-197    CGCUAUCGCCGGGCCAACGAAUUACGGGGAUAAAGGGACUUUUCAUUGAUGUGCGGACCAUUUGAGUAUAAAAUGCCGCCCUC....UUCAC...AUCC..
env-198    ..........................................................................UGCCGCCCGCU....UCAC...AUCC..
env-199    GCUAUCGCCGGGCUAACGAAUUGCGUGGAUAAAGGGACUUUUCAUUGAUGUGCAGGCCAUUUGAGUAUAAAAUGCCGCCCGCU....UCAC...AUCC..
env-200    GUUUCCGUCGGGCCAAUGAAUUGCGCGGUUAAGGGCACUUUUGAUUGAUGUGACGGGCGUUUGAGUAUAAAAUGCCGCCCGCU....UCAC...AUCC..
env-201    .............................................................................CCGCCCGCU....UUUGC...AUCC..
Ecl-59-1   GCUUUCGCCGCGCGAAUGAAUUGCGCGGCUAAGCGCACUUUUAAUUGAUGUAUCAAUCAUUAGGGGAUAGAAUACCGCCCGCU....UUUGC...AUCC..
env-202    GCUUUCGCCGGGCGCAUGAAUUGCGCGGCUAAGCGCACUUUUGAUUGCUGUAUCAAUCAUUAGGGGAUAGAAUACCGCCCGCU....UUUGC...AUCC..
Eko-1-1    GCUAUCGUCGGGCAAAUGAAUUGCGUGGUUAGCAGGACUUUUCAUUGAUGUGGACAGCGUUUGAGUAUAAAAUGCCGCCCGCU....UCAC...AUCC..
Ecl-33-1   GCUUCCGUCGGGCAAAUGAAUUGCGCGGUUAACGGGACUUUUCAUUGAUGUGCAGGGCAUUUGAGUAUAAAAUGCCGCCCGCU....UCAC...AUCC..
Eko-1-8    GCUAUCGUCGGGCAAACGAAUUGCGUGGUUAGCAGGACUUUUCAUUGAUGUGAGCGGCGUUUGAGUAUAAAAUGCCGCCCGCU....UCAC...AUCC..
Esp-51-1   GCUACCGUCGGGCAAACGAAUUGCGUGGUUAGCAGGACUUUUCAUUGAUGUGAGCGGCGUUUGAGUAUAAAAUGCCGCCCGCU....UCAC...AUCC..
Ecl-45-1   GCUUCCGUCGGGCAAAUGAAUUGCGCGGUUAACGGGACUUUUCAUUGAUGUGCGGGGCAUUUGAGUAUAAAAUGCCGCCCGCU....UCAC...AUCC..
Ecl-52-1   CGUCGCCGACGCCCUAAGCGGUUUUUUUAGUAGGGAUUAAUUCAUUGCCCUGAGGUGGAUUGGGGGCUAUAAUGCCGCCCGCU....UUUGC...AUCC..
Ecl-54-1   GGUCGCCGACGCGCUAAGCGGUUUUUUUAGUGGGGAUUAAUUCAUUGCCCUGAGGUGGAUUGGGGGCUAUAAUGCCGCCCGCU....UUUGC...AUCC..

           ..........................................................................<<<....<<<<<<<<<<<<<<<<<<<<<<<<<<<<<<<<<<<<<<<<<<<<<<<<<<<<<<<<<<<<<<<<<<<<<<<<<<<<<<<<<......
           ..........................................................................212....22222.........111....00110.....0.1.111.1111.1..........101...............011.....111100......
           ●●●●●●●●●●●●●●●●●●●●●●●●●●●●●●●●●●●●●●YY●●UUGYY●●●●●●●●●●●●●●●●R●UA●AAU●YCGCCCGCU----U●●●○------------------------------------------------------------------------●Y●●--
```

```
env-204      CGUCGCCGACGCCCUAAGCGGUUUCUUUAGUGGGGAUUAAUUCAUUGCCCUGAGGUGGAUUGGGGGCUAUAAUGCCGCCCGCU.....UUUGC...AUCC..
Ecl-52-3     CGUCGCCGACGCCCUAAGCGGUUUUUUUAGUGGGGAUUAAUUCAUUGCCCUGAGGUGGAUUGGGGGCUAUAAUGCCGCCCGCU.....UUUGC...AUCC..
env-205      GCUUUCGCCGGGCAAACGAAUUGCGCGGUUAACGGGACUUUUCGUUGAUGCGCGAGCCAUUUGAGUAUAAAAUGCCGCCCGCU.....UCAC....AUCC..
Ecl-51-1     GCUUUCGUCGCGCCAACGAAUUACGCGGUUAGCGGGACUUUUCAUUGAUGUGAGGGGCAUUUGAGUAUAAAAUGCCGCCCGCU.....UCAC....AUCC..
env-207      GCUUUCGUCGCGCCAACGAAUUGCGCGGUUAGCGGGACUUUUCAUUGAUGUGAGGGGCAUUUGAGUAUAAAAUGCCGCCCGCU.....UCAC....AUCC..
env-208      GCUUCCGUCGCGCCAACGAAUUGCGCGGUUAGCGGGACUUUUCAUUGAUGUGAGGGGCAUUUGAGUAUAAAAUGCCGCCCGCU.....UCAC....AUCC..
env-210      ...GGGACUUUUCAUUGAUGUGAGGGGCAUUUGAGUAUAAAAUGCCGCCCGCU.....UCAC....AUCC..
env-211      GCUUCCGUCGCGCCAACGAAUUACGCGGUUAGCGGGACUUUUCAUUGAUGUGAGGGGCAUUUGAGUAUAAAAUGCCGCCCGCU.....UCAC....AUCC..
env-212      GCUUUCGUCGGGCAAAUGAAUUGCGCGGCUAGCGGGACUUUUCAUUGAUGUGAGGGGCAAUUGAGUAUAAAAUGCCGCCCGCU.....UCAC....AUCC..
env-213      GCUUUCGUCGGGCAAAUGAAUUACGCGGCUAGCGGGACUUUUCAUUGAUGUGAGGGGCAAUUGAGUAUAAAAUGCCGCCCGCU.....UCAC....AUCC..
Esp-19-1     GCUUCCGUCGCGCCAACGAAUUGCGCGGUUAGCGGGACUUUUCAUUGAUGUGAGGGGCAUUUGAGUAUAAAAUGCCGCCCGCU.....UCAC....AUCC..
Ecl-34-1     GCUUCCGUCGCGCUAACGAAUUGCGCGGUUAGGGGGACUUUUCAUUGAUGUGCGGGGCAUUUGAGUAUAAAAUGCCGCCCGCU.....UCAC....AUCC..
env-214      GCUUUCGCCGGGCGAAUGAAUUACGGGGUUAGCGGGACUUUUCAUUGAUGUGAAGUGCAUUUGAGUAUAAAAUGCCGCCCGCU.....UCAC....AUCC..
env-216      GCUUUCGCNGGGCGAAUGAAUUACGGGGUUAGCGGGACUUUUCAUUGAUGUGAAGUGCAUUUGAGUAUAAAAUGCCGCCCGCU.....UCAC....AUCC..
Ecl-42-1     GCUUCCGUCGCGCCAACGAAUUGCACGGGUAGCGGGACUUUUCAUUGAUGUGAGGGGCAUUUGAGUAUAAAAUGCCGCCCGCU.....UCAC....AUCC..
env-217      ...UUCGUUGAUGCGCGAGCCGUUUGAGUAUAAAAUGCCGCCCGCU.....UCAC....AUCC..
env-218      GCUUUCGCCGGGCAAACGAAUUGCGCGGUUAACGGGACUUUUCGUUGAUGCGCGAGCCAUUUGAGUAUAAAAUGCCGCCCGCU.....UCAC....AUCC..
env-228      GCUUUCGUCGCGCCAACGAAUUGCGCGGUUAACCGGACUUUUCAUUGAUGUGAGGAGCAUUUGAGUAUAAAAUGCCGCCCGCU.....UCAC....AUCC..
env-229      GCUUCCGUCGCGCCAACGAAUUGCGCGGGUAACGGGACUUUUCAUUGAUGUGAAGGGCAUUUGAGUAUAAAAUGCCGCCCGCU.....UCAC....AUCC..
Ecl-29-1     GCUUCCGUCGCGCCAACGAAUUGCGCGGGUAGCGGGACUUUUCAUUGAUGUGAGGGGCAUUUGAGUAUAAAAUGCCGCCCGCU.....UCAC....AUCC..
Esp-65-1     GCUUUCGCCGGGCAAACGAAUUGCGCGGUUAACGGGACUUUUCGUUGAUGUGCGAGCUAUUUGAGUAUAAAAUGCCGCCCGCU.....UCAC....AUCC..
env-230      ...UUGAUGUGAGGGGCAUUUGAGUAUAAAAUGCCGCCCGCU.....UCAC....AUCC..
env-232      ...GUGAGGGGCAAUUGAGUAUAAAAUGCCGCCCGCU.....UCAC....AUCC..
Ecl-16-1     GCUUCCGUCGCGCCAACGAAUUGCGCGGUUAGCGGGACUUUUCAUUGAUGUGAGGGACAUUUGAGUAUAAAAUGCCGCCCGCU.....UCAC....AUCC..
env-233      GCUUCCGCCGGGCAAAUGAAUUGCGCGGUUAACGGGACUUUUCGUUGAUGUGAGGGGCAUUUGAGUAUAAAAUGCCGCCCGCU.....UCAC....AUCC..
env-236      CGCUAUCGCCGCGCCAAUGAAUUGCGCGGAUAGGCCUGUUUUCAUUGCUGCUGCGGGCAAACGGAUCUAUAAUGCCGCCCGCU.....UUUGC...AUCC..
Sco-1-1      CGUUUCCGGCGCGCCAAUGAAUUGCGCGGCUGAGGCUGUUUUCGUUGCUGCUGCCUGAAAACAAAGCUAUAAUGCCGCCCGCU.....UUUGC...AUCC..
Psp-10-1     CGUUAUCGUCGCGCCAACGAACUGCGCUGAGCGGCGGAAUCAUUUGCCCCUGACACCCCAUUGGCGCUAUAAUGCCGCCCGCU.....UUUGC...AUCC..
Psp-18-1     CGUUAUCGCCGCGCCAAUGAGCUGCGAUAAGCGGCAGAAUCAUUUGCCCCUGCGGCAUCAUUGGGGCUAUAAUGCCGCCCGCU.....UUUGC...AUCC..
env-237      GCUUCCGUCGCGCCAAUGAAUUACGCGGAUAAACGUCCUUUUGAUUGAUACGACGCGCAUUUAAGUAUAAAAUGCCGCCCGCU.....UUCAC...AUCC..
env-239      GCUAUCGCCGUGCCAAUGAAUUACGCGGUUAGGCGCGCUUUUUAUUGAUGUACACGGCAUUAAAGUAUAAAAUGCCGCCCGCU.....UUCAC...AUCC..
Ecl-60-1     GCUUUCGCCGGGCGAAUGAAUUACGCGGAUAGACCCGCUUUUCAUUGCCCUGCCCUGCAAUGAGGGCUAUAAUGCCGCCCGCU.....UUUGC...AUGU..
Csp-13-1     GCUUCCGGCGAGCACAUGAAUUACGUGGUUAAUUGUACUUUUGAUUGAUGUACCAGCCGUUUGAGUAUAGAAUCUCGCCCGCU.....UUUGC...AUAC..
env-240      ...UGUAGUUUCCAUUGCCCCUGUGUGGAUUGAAGGCUAUAAUGCCGCCCGCU.....UUUGC...AUGU..
Cne-1-1      CUUCCGCCGGGCCAAUGAAUUGCGCGGUUAAGCGGUACUUUUCAUUGCCGACAGGGUCAGAUAUGGCUAUAAUGCCGCCCGCU.....UUUGC...AUCC..
env-241      ...GCGGUCAGUAGUACUUUCCAUUGCUGAUAUAGCCAGAUACAGCUAUAAUGCCGCCCGCU.....UUUGC...AUCC..
env-242      ...GCCAGAUACGGCUAUAAUGCCGCCCGCC.....UUUUGC..AUCC..
env-243      GUUUCCGCCGGGCUAAUGAAUUACGCGGUCAGUAGUACUUUCCAUUGCUGAUAUAGCCAGAUACAGCUAUAAUGCCGCCCGCU.....UUUGC...AUCC..
env-244      GUUUCCGCCGGGCUAAUGAAUUACGCGGUCAGUAGUACUUUCCAUUGCUGAUAUAGCCAGAUACAGCUAUAAUGCCGCCCGCU.....UUUGU...CAUCC.
Esp-68-1     CUUCCGCCGGGCCAACGAAUUACGCGGCUAACGGGUACUUUUCAUUGCCGACAGAGCCGGAUACGGCUAUAAUGCCGCCCGCU.....UUUGC...AUCC..
Cne-1-2      CUUCCGCCGGGCCAACGAAUUACGCGGCUAACCGGUACUUUUCAUUGCCGGCAGAGCCGGAUAUGGCUAUAAUGCCGCCCGCU.....UUUGC...AUCC..
Cne-1-3      CUUCCGGCGGGCCAACGAAUUACGCGGCUAACCGGUACUUUUCAUUGCCGGCAGAGCCGGAUACGGCUAUAAUGCCGCCCGCU.....UUUGC...AUCC..
Eco-2103-1   GUUUUCGUCUGGCGCACGAAUUACGCGGUUAAUGGUAGUUUUGAUUGAUGUAUAAGGCGAUUGAGUAUAGAAUCUUGCCCGCU.....UUUGC...AUAC..
Esp-77-1     GUUUUCGUCGGGCACAUGAAUUACGCGGUUAAUAGUAGUUUUGAUUGAUGUAUAAGGCGAUUGAGUAUAGAAUCUCGCCCGCU.....UUUGC...AUAC..
env-245      GUUUUCGUCGGGCGCACGAAUUACGCGGUUAAUGGUAGUUUUGAUUGAUGUAUAAAGCGAUUGAGUAUAGAAUCUCGCCCGCU.....UUUGC...AUAC..
Sso-1-2      NNNNNNNNNNNNNNNNNNNNNNNNNNNNNNNNNNNNNNNNNNNNNNNGAUGUAUAAGGCGAUUGAGUAUAGAAUCUCACCCGCU.....UUUGC...AUAC..
Sso-1-3      NNNNNNNNNNNNNNNNNNNNNNNNNNNNNNNNNNNNNNNNNNNNNNNNNNNNNNNGCGAUUGAGUAUAGAAUCUCACCCGCU.....UUUGC...AUAC..
Eco-555-1    UGGGCGCAGAUUAAGAGGCUACAAUGGACGCCAUGGAGUUUUGAUUGAUGUAUAAGGCGAUUGAGUAUAGAAUCUCGCCCGCU.....UUUGC...AUAC..
env-252      GUUUUCGUCUGGCGCACGAAUUACGCGGUUAAUGGUAGUUUUGAUUGAUGUAUAAGGCGAUUGAGUAUAGAAUCUCGCCCGCU.....UUUGC...AUAC..
Sbo-13-1     GUUUUCGUCGGGCGCACGAAUUACGCGGUUAAUGGUAGUUUUGAUUGAUGUAUAAGGCGAUUGAGUAUAUAAUCUCGCCCGCU.....UUUGC...AUAC..
Eco-174-1    GUUUUCGUCGGGCGCACGAAUUACGUGGUUAAUGGUAGUUUUGAUUGAUGUAUAAGGCGAUUGAGUAUAGAAUCUCGCCCGCU.....UUUGC...AUAC..
env-264      GUUUUCGUCGGGCGCACGAAUUACGCGGUUAAUGGUAGUUUUGAUUGAUGUAUAAGGCGAUUGAGUAUAGAAUAUCGCCCGCU.....UUUGC...AUAC..
env-265      GUCGGGCGCACGAAUUACGCGGUUAAUGGUAGUUUUGANNNNNNNNNNUGUAUAAGGCGAUUGAGUAUAGAAUCUCGCCCGCU.....UUUGC...AUAC..
env-266      GUUUUCGUCGGGCGCACGAAUUACGCGGUUAAUGGUAGUUUUGAUUGAUGUAUAAGGCGAUUGAGUAUAGAAUCUCGCCCGCU.....UUUGC...AUAC..
Eco-1984-1   GUUUUCGUCGGGCGCACGAAUUACGCGGUUAAUGGUAGUUUUGAUUGAUGUAUAAGGCGAUUAAGUAUAGAAUCUCGCCCGCU.....UUUGC...AUAC..
Sso-1-14     GUUUUCGUCGGGCGCACGAAUUACGCGGUUAAUGGUAGUUUUGAUUGAUUUAUAAGGCGAUUGAGUAUAGAAUCUCACCCGCU.....UUUGC...AUAC..
Sso-7-1      GUUUUCGUCGGGCGCACGAAUUACGCGGUUAAUGGUAGUUUUGAUUGAUGUAUAAGGCGAUUGAGUAUAGAAUCUCACCCGCU.....UUUGC...AUAC..
Sso-1-539    NNNNNNNNNNNNNNNNNNNNNNNNNNNNNNNNNNNNNNNNNNNNNNNNNNAUAAGGCGAUUGAGUAUAGAAUCUCACCCGCU.....UUUGC...AUAC..
Esp-78-1     GUUUUCGUCGGGCGCAUGAAUUACGCGGUUAAUAGUAGUUUUGAUUGAUGUAUAAGGCGAUUGAGUAUAGAAUCUCGCCCGCU.....UUUGC...AUAC..
Esp-53-1     GUUUUCGUCGGGCGCAUGAAUUACGCGGUUAAUGGUAGUUUUGAUUGAUGUAUAAGGCGAUUGAGUAUAGAAUCUCGCCCGCU.....UUUGC...AUAC..
Eco-279-1    GUUUUCGUCGGGCGCACGAAUUACGCGGUUAAUGGUAGUUUUGAUUGAUGUAUAAGGCGAUUGAGUAUAGAAUAUUGCCCGCU.....UUUGC...AUAC..
env-324      NNNNNNNNNNNNNNNNNNNNNNNNNNNNNNNNNNNNNNNNNNNNNNNAUGUAUAAAGCGAUUGAGUAUAGAAUCUCGCCCGCU.....UUUGC...AUAC..
env-325      GUUUUCGUCGGGCACACGAAUUACGCGGUUAAUGGUAGUUUUGAUUGAUGUAUAAGGCGAUUGAGUAUAGAAUCUCGCCCGCU.....UUUGC...AUAC..
Eco-1084-1   GUUUUCGUCGGGCGCACGAAUUACGCGGUUAAUGAUAGUUUUGAUUGAUGUAUAAGGCGAUUGAGUAUAGAAUCUCGCCCGCU.....UUUGC...AUAC..
env-326      ...GCCAGGGCCAGAUAUGGCUAUAAUCCCGCCCGCU.....UUUGC...AUCC..
Eal-2-1      GUUAUCGCCGGGCGCAUGAAUUACGCGGUUAAAGGUAGUUUUGAUUGAUGUGAAAGGCGAUUGAGUAUAGAAUCUCGCCCGCU.....UUUGC...AUAC..
Eco-764-1    GUUUUCGUCGGGCGCACGAAUUACGCGGUUAAUGGUAGUUUUAAUUGAUGUGCAAGGCGAUUGAGUAUAGAAUCUCGCCCGCU.....UUUGC...AUAC..
Eco-66-1     GUUUUCGUCGGGCACACGAAUUACGCGGUUAAUGGUAGUUUUAAUUGAUGUGCAAGGCGAUUGAGUAUAGAAUCUCGCCCGCU.....UUUGC...AUAC..
Sfl-76-1     GUUUUCGUCGGGCGCACGAAUUACGCGGUUAAUGGUAGUUUUGAUUGAUGUAUAAGGCGAUUGAGUAUAGAAUCUCGCCCGCU.....UUUGC...AUAC..
env-327      GUUUUCGUCGGGCGCACGAAUUACGCGGUUAAUGGUAGUUUUGAUUGAUGUAUAAGGCGAUUGAGUAUAGAAUCUCGCCCGCU.....UUUGC...AUAC..
Eco-1148-1   GUUUUCGUCGGGCGCACGAAUUACGCGGUUAAUGGUAGUUUUGAUUGAUGUAUAAAGCGAUUGAGUAUAGAAUCUCGCCCGCU.....UUUGC...AUAC..
Sdy-8-1      GUUUUCGUCGGGCGCACGAAUUACGCGGUUAAUGGUAGUUUUGAUUGAUGUAUAAGGCGAUUGAGUAUAGAAUCUCGCCCGCU.....UUUGC...AUAU..
Eco-84-1     GUUUUCGUCGGGCGCACGAAUUACGCGGUUAAUGGUAGUUUUAAUUGAUGUGCAAGGCGAUUGAGUAUAGAAUCUCGCCCGCU.....UUUGC...AUAU..
Cda-1-1      GCUUCCGCCGGGCCAAUGAAUUACGAGGUUAGCGGUACUUUUCAUUGCCGACAUAGCGAGAUAUAGCUAUAAUGCCGCCCGCU.....UUUGC...AUCC..
Bag-1-1      GCUUCCGUCGCGCGAAUGAAUUGCGUGGCUAAAGGUUCUUUUCAUUGAAGCUGGGGCCAGAUAUGGCUAUAAUCCCGCCCGCU.....UUUGC...AUCC..
Bag-2-1      GCUUCCGCCGCGCGAAUGAAUUGCGCGGUUAAAGGUUCUUUUCAUUGAAGCCGAGGCCAGAUAUGGCUAUAAUCCCGCCCGCU.....UUUGC...AUCC..
Esp-42-1     GUUUUCGUCGGGCGCAUGAAUUACGCGGAUAAGUGUGCUUUUAAUUGAUGUAUCGGCUAUUCGAGGAUAGAAUACCGCCCGCU.....UUUGC...AUCC..
env-328      GCAACAAGCCAGGCGCCUGAAAGGGUUAAAGGGGAUUGUUUUCAUUGCCCCACACUCUUGAGAAGGCUAUAAUGCCGCCCGCU.....AUUGC...AUAC..
env-329      ...UUUGGAUGGAUGUACCAGCCGUUUGAGUAUAGAAUCUCGCCCGCU.....UUGGC...AUAC..
Sen-239-1    GCUUCCGCCGGGCGCACGAAUUACGUGGUUAAUUGUACUUUUCAUUGAUGUAACGGCGAUUUGAGUAUAGAAUCCCGCCCGCC.....UUUGC...AUAC..
Sen-728-1    GCUUCCGCCGGGCGCACGAAUUACGUGGUUAAUCGUACUUUUCAUUGAUGUAACGGCGAUUUGAGUAUAGAAUCUCGCCCGCC.....UUUGC...AUGC..
Sen-226-1    GCUUCCGCCGGGCGCACGAAUUACGUGGUUAAUCGUACUUUUCAUUGAUGUAACGGCGAUUUGAGUAUAGAAUCCCGCCCGCC.....UUUGC...AUAC..
Sen-269-1    GCUUCCGCCGGGCGCACGAAUUACGUGGUUAAUUGUACUUUUCAUUGAUGUAACGGCGAUUUGAGUAUAAAAUCCCGCCCGCC.....UUUGC...AUAC..
Sen-225-1    GCUUCCGCCGAGCGCACGAAUUACGUGGUUAAUUGUACUUUUCAUUGAUGUAACGGCGAUUUGAGUAUAGAAUCCCGCCCGCC.....UUUGC...AUAC..

             ...<<<.....<<<<<<<<<<<<<<<<<<<<<<<<<<<<<<<<<<<<<<<<<<<<<<<<<<<<<<<<<<<<<<<<<<<<<<<<<.....
             ...212.....22222........111....00110.....0.1.111.1111.1.........101...............011.....111100......
             •••YY••UUGYY•••R•UA•AAU•YCGCCCGCU-----U••••---•Y•-
```

```
Sen-438-1   GGCGCACGAAUUACGUGGUUAAUUGUACUUUUCAUUGAUGUAACGGCGAUUUNNNNNNNNUUUGAGUAUAGAAUCCCGCCCGCC.....UUUGC................................................................................AUAC..
Sen-287-1   NNNNNNNNNNNNNNNNNNNNNNNNNNNNNNNNNNNNNNNNNNNNNNNNNNNNNNNNNNNNNNNNNNNNNNAAUCCCGCCCGCC.....UUUGC................................................................................AUAC..
Sen-1007-1  GCUUCCGCCGGGCGCACGAAUUACGUGGUUAAUCGUACUUUUCAUUGAUGUAACGGCGAUUUGAGUAUAGAAUCCCGCCCGCC.....UUUGC................................................................................AUAC..
Sen-438-2   UUACGUGGUUAAUUGUACUUUUCAUUGAUGUAACGGCGAUUUNNNNNNNNNNNNNNNNNNNNNNNNNNNNNNNNNNNCCCGCC.....UUUGC................................................................................AUAC..
Sen-23-1    GCUUCCGCCGGGCGCACGAAUUACGUGGUUAAUUGUACUUUUCAUUGAUGUAACGGCGAUUUGAGUAUAGAAUCCCACCCGCC.....UUUGC................................................................................AUAC..
Sen-438-3   GCUUCCGCCGGGCGCACGAAUUACGUGGUUAAUUGUACUUUUCAUUGAUGUAACGGCGAUUUGAGUAUAGAACCCCGCCCGCC.....UUUGC................................................................................AUAC..
Sen-67-1    GCUUCCGCCGGGCGCACGAAUUACGUGGUUAAUUGUACUUUUCAUUGAUGUAACGGCGAUUUGAGUAUAGAAUUCCGCCCGCC.....UUUGC................................................................................AUAC..
Sen-88-1    GCUUCCGCCGGGCGCACGAAUUACGUGGUUAAUUGUACUUUUCAUUGAUGUAACGGCGAUUUGAGUAUAGAAUCCCGCCCGCC.....UUUGC................................................................................AUAC..
Sen-223-1   GCUUCCGCCGGGCGCACGAAUUACGUGGUUAAUUGUACUUUUCAUUGAUGUAACGGCGAUUUGAGUAUAAAAUCCCGCCCGCC.....UUUGC................................................................................AUAC..
Sen-927-1   GCUUCCGCCGGGCGCACGAAUUACGUGGUUAAUUGUACUUUUCAUUGAUGUAACGGCGAUUUGAGUAUAGAAUCCCGCCCGCC.....UUUGC................................................................................AUAC..
Cro-2-1     GCUUCCGCCGGGCGCACGAAUUACGCGGUUAAUCGUAGUUUUGAUUGAUGUGUCCGGCGUUUGAGUAUAGAAUCUCGCCCGCU.....UUUGC................................................................................AUAC..
Cko-1-2     GCUUCCGGCGGGCACAUGAAUUGCGCGGUUAAUCGUACUUUUGAUUGAUGUGAUAGCCAUUUGAGUAUAGAAUCUCGCCCGCU.....UUUGC................................................................................AUAC..
Cam-2-1     GUUUCCGUCGGGCACAUGAAUUACGCGGUUAAUCGUACUUUUGAUUGAUGUAACGGGCGUUUGAGGAUAGAAUCUCGCCCGCU.....UUUGC................................................................................AUAC..
Sen-1020-1  GCUUCCGCCGGGCGCAUGAAUUACGUGGUUAAUCGCACUUUUCAUUGAUGUAACGGCGAUUUGAGUAUAGAAUCUCGCCCGCU.....UUUGC................................................................................AUAC..
env-330     GCUAUCGUCGGGCACAUGAAUUACGCGGUUAAUCGUACUUUUGAUUGAUGUACCAGCCGUUUAAGUAUAGAAUCUCGCCCGCU.....UUUGC................................................................................AUAC..
env-331     GCUUUCGUCGGGCACAUGAAUUACGCGGUUAAUCGUACUUUUGAUUGAUGUACCACCCGUUUGAGUAUAGAAUCUCGCCCGCU.....UUUGC................................................................................AUAC..
Cse-1-1     GCUUUCGCCGGGCGCAUGAAUUGCGCGGUUAAUCGUAGUUUUGAUUGAUGUGCCGGGCGUUUGAGUAUAGAAUCUCGCCCGCU.....UUUGC................................................................................AUAC..
Csp-11-1    GUUUCCGUCGGGCACAUGAAUUACGCGGUUAAUCGUACUUUUGAUUGAUGUAACGGGCGUUUGAGGAUAGAAUCUCGCCCGCU.....UUUGC................................................................................AUAC..
Cwe-1-1     GCUUUCGCCGGGCACAUGAAUUACGCGGUUAAUUGUACUUUUGAUUGAUGUACCAGCCGUUUGAGUAUAGAAUCUCGCCCGCU.....UUUGC................................................................................AUAC..
Sen-310-1   GCUUCCGCCGGGCGCACGAAUUGCGUGGUUAAUCGCACUUUUCAUUGAUGUAACGGCGAUUUGAGUAUAGAAUCUCGCCCGCU.....UUUGC................................................................................AUAC..
Sen-3-1     GCUUCCGCCGGGCGCACGAAUUACGUGGCUAAUCGCACUUUUCAUUGAUGUAACGGCGAUUUGAGUAUAGAAUCUCGCCCGCU.....UUUGC................................................................................AUAC..
Cfr-1-4     GCUUUCGCCGGGCACAUGAAUUACGCGGUUAAUCGUACUUUUGAUUGAUGUACCGACCGUUUGAGUAUAGAAUCUCGCCCGCU.....UUUGC................................................................................AUAC..
Sen-311-1   GCUUCCGCCGGGCGCACGAAUUACGUGGUUAAUCGCACUUUUCAUUGAUGUAACGGCGAUUUGAGUAUAGAAUCUCGCCCGCU.....UUUGC................................................................................AUAC..
Sen-1018-1  GUUUCCGCCGGGCGCACGAAUUACGUGGUUAAUCGCACUUUUCAUUGAUGUAACGGCGAUUUGAGUAUAGAAUCUCGCCCGCU.....UUUGC................................................................................AUAC..
Cbr-1-5     GCUAUCGUCGGGCACAUGAAUUACGCGGUUAGUCGUACUUUUGAUUGAUGUACCAGCCGUUUAAGUAUAGAAUCUCGCCCGCU.....UUUGC................................................................................AUAC..
env-332     GCUUUCGUCGGGCACAUGAAUUACGCGGUUAAUCGUACUUUUGAUUGAUGUACCAGCCGUUUGAGUAUAGAAUCUCGCCCGCU.....UUUGC................................................................................AUAC..
env-334     GCUUCCGCCGGGCACAUGAAUUACGCGGAUAAUCGUACUUUUGAUUGAUGUAACGGGCGUUUGAGUAUAGAAUCUCGCCCGCU.....UUUGC................................................................................AUAC..
Cfa-1-1     GUUUCCGUCGGGCACAUGAAUUACGCGGUUAAUGGUACUUUUGAUUGAUGUAACGGGCGUUUGAGUAUAGAAUCUCGCCCGCU.....UUUGC................................................................................AUAC..
Cpa-1-1     GUUUCCGGCGGGCACAUGAAUUACGCGGUUAAUCGUACUUUUGAUUGAUGUACCAGCCGUUUGAGUAUAGAAUCUCGCCCGCU.....UUUGC................................................................................AUAC..
Csp-17-1    GCUUUCGUCGGGCACAUGAAUUACGCGGUUAAUCGUACUUUUGAUUGAUGUACCAGCUGUUUGAGUAUAGAAUCUCGCCCGCU.....UUUGC................................................................................AUAC..
Cfr-4-1     GCUUUCGCCGGGCACAUGAAUUACGCGGUUAAUCGUACUUUUGAUUGAUAUACCAGCCGUUUGAGUAUAGAAUCUCGCCCGCU.....UUUGC................................................................................AUAC..
Sen-1016-3  CCGUUUCCGCGGGCGCACGAAUUACGUGGUUAAUCGCACUUUUCAUUGAUGUAACGCGAUUUGAGUAUAGAAUCUCGCCCGCU.....UUUGC................................................................................AUAC..
Sbo-3-1     GUUUCCGCCGGGCGUAUGAAUUACGCGGUUAAUCGCACUUUUCAUUGAUGUGGCGAUGAUUUGAGUAUAGAAUCUCGCCCGCU.....UUUGC................................................................................AUAC..
Sen-834-1   GCUUCCGCCGGGCGCACGAAUUACGUGGUUAAUUGUACUUUUCAUUGAUGUAACGGCGAUUUGAGUAUAGAAUCCCGCCCGCC.....UUUGC................................................................................AUAC..
env-335     GUUUCCGUCGGGCACAUGAAUUACGCGGUUAAUGGUACUUUUGAUUGAUGUAACGGGCGUUUGAGUAUAGAAUCUCGCCCGCU.....UUUGC................................................................................AUAC..
Csp-18-1    GCUAUCGUCGGGCACAUGAAUUACGCGGUUAAUCGUACUUUUGAUUGAUGUACCAGCCGUUUAAGUAUAGAAUCUCGCCCGCU.....UUUGC................................................................................AUAC..
Pur-1-1     CGCCGCGCCAAUGAGUUACGCGGCUAAGUUCAUCUCAGUGCGGCUUGUAACAGCGGGGGAUACGGGUAUAAUCCCGCCCGCU.....UUUGC................................................................................AUCC..
Cma-4-1     GGUUCCGCCGCGCCAAUGAAUUACGCGGUUAGCGGCGGUUUUCGUUGCCCCUGCGCGCUCGAAUAGCUAUAAUGCCGCCCGCU.....UUUGC................................................................................AUCC..
Cun-1-1     GGUUCCGCCGCGCCAAUGAAUUACGCGGUUAGCGGCGGUUUUCAUUGCCCCUGCGCGCUCGAAUAGCUAUAAUGCCGCCCGCU.....UUUGC................................................................................AUCC..
Csa-15-1    GGUUCCGCCGCGCCAAUGAAUUACGUGGGUAGCGGCGGUUUUCGUUGCCCCUGCGCGUUCAAAUAGCUAUAAUGCCGCCCGCU.....UUUGC................................................................................AUCC..
Cma-1-1     GGUUCCGCCGCGCCAAUGAAUUACGCGGUUAGCGGCGGUUUUCGUUGCCCCUGCGCGCUCGAAUAGCUAUAAUACCGCCCGCU.....UUUGC................................................................................AUCC..
Csa-7-1     GGUUCCGCCGCGCCAAUGAAUUACGCGGUUAGCGGCGGUUUUCGUUGCCCCUGCGCGCUCGAAUAGCUAUAAUGCCGCCCGCU.....UUUGC................................................................................AUCC..
Csa-10-1    GGUUCCGCCGGGCCAAUGAAUUACGUGGGUAGCGGCGGUUUUCGUUGCCCCUGCGCGCUCAAAUAGCUAUAAUGCCGCCCGCU.....UUUGC................................................................................AUCC..
Ctu-3-1     GCUUCCGGCGCGCCAAUGAAUUACGCGGUUAGCGGCGGUUUUCAUUGCCCGUGCGCGCUCAAAUAGCUAUAAUGCCGCCCGCU.....UUUGC................................................................................AUCC..
Cma-1-5     GAAGGUGACACUUCUGCUUUGCGUUGACAUAGCGGCGGUUUUCGUUGCCCCUGCGCGCUCGAAUAGCUAUAAUGCCGCCCGCU.....UUUGC................................................................................AUCC..
Fpu-1-1     GCGCUGAUAGCCGCCGCGCAGAGGUAACGAAACCGCUGUUUUGAUUGUCACCGGCGUCGUUCGCGGCUAUAAUGCCGCCCGCU.....UUUGC................................................................................AUCC..
Fpu-5-1     GCGCUGAUAGCCGCCGCCCAGAGGUAACGAAACCGCUGUUUUGAUUGUCACCGGCGUCGUUCGCGGCUAUAAUGCCGCCCGCU.....UUUGC................................................................................AUCC..
Fhe-4-1     GCUAUCGCCGGGCUAAUGAAUUGCGCGGCUAAGGGCGGUUUUCGUUGAUGCCGCCGGGCGAUGAAGCUAUAAUGCCGCCCGCU.....UUUGC................................................................................AUCC..
Fhe-1-1     GCUAUCGCCGGGCUAAUGAAUUGCGCGGCUAAGGGCGGUUUUCGUUGCCGCCGCCGGGCGAUGAAGCUAUAAUGCCGCCCGCU.....UUUGC................................................................................AUCC..
Kor-1-1     GUUUCCGGCGGGCACAUGAAUUACGGGGAUAAACGCGCUUUUGAUUGAUGUAUCGAACAUUCAGGGAUAGAAUACCGCCCGCU.....UUUGC................................................................................AUCC..
Ksa-1-1     UCCGGCGGGCACAUGAAUUACGGGGAUAAUAAAUGCGCUUUUGAUUGAUGUAUCGAACAUUCAGGGAUAGAAUACCGCCCGCU.....UUUGC................................................................................AUCC..
Kra-2-1     GCUUCCGGCGGGCAUAUGAAUUGCGUGGGUAAACGUACUUUUGAUUGAUGUAUCGAACGUUCGAGGAUAGAAUACCGCCCGCU.....UUUGC................................................................................AUCC..
Era-1-1     GCUUCCGGCGGGCGCAUGAAUUGCGCGGGUAAACGUACUUUUGAUUGAUGUAUCGAACGUUCGAGGAUAGAAUACCGCCCGCU.....UUUGC................................................................................AUCC..
Cmu-1-1     GCUUCCGGCGCGCUAAUGAAUUACGCGGUUAACGGCGGUUUUCAUUGCCCCUGCGCGCUCAAAUGGCUAUAAUGCCGCCCGCU.....UUUGC................................................................................AUCC..
Cdu-2-1     GCUUCCGGCGCGCUAAUGAAUUACGCGGUUAGCGGCGGUUUUCAUUGCCCGUGCGCGCUCAAAUGGCUAUAAUGCCGCCCGCU.....UUUGC................................................................................AUCC..
Cco-1-1     GCUUCCGUCGCGCCAAUGAAUUACGCGGUUAGCGGCAGUUUUCAUUGCCCGUGCGCGCUCAAAUGGCUAUAAUGCCGCCCGCU.....UUUGC................................................................................AUCC..
Cdu-3-1     GCUUCCGGCGCGCCAAUGAAUUACGCGGUUAGCGGCGGUUUUCAUUGCCCGUGCGCGCUCAAAUGGCUAUAAUGCCGCCCGCU.....UUUGC................................................................................AUCC..
env-336     GCUAUCGCCGGGCCAACGAAUUGCGCGGCUGACGGCCCUUUUGAUUGAUCGAAAAGGCGUUUAAAGGUAAAAUGCCGCCCGCU.....UUCAC................................................................................AUCC..
Kpn-226-1   CGCGCCAACGAGCUGCGUUAAUUACCGGAAAGCUGAGGUUUUGAUUGCCCGGGUGUCGUUUGGCGGCUAUAAUGCCGCCCGCU.....UUUGC................................................................................AUUC..
Kpn-169-1   CGCGCCAACGAGUUGCGUUAAUUACCGGAAAGCUGAGGUUUUGAUUGCCCGGGUGUCGUUUGGCGGCUAUAAUGCCGCCCGCU.....UUUGC................................................................................AUUC..
env-337     CGCGCCAACGAGCUGCGUUAAUUACCGGAAAGCUGAGGUUUUGAUUGCCCGGGUGUCGUUUGGCGGCUAUAAUGCCGCCCGCU.....UUUGC................................................................................AUUC..
Kqu-1-1     CGCGCCAACGAGCUGCGUUAAUUCCCGGAAAGCUGAGGUUUUGAUUGCCCGGGUGUCGUUUGGCGGCUAUAAUGCCGCCCGCU.....UUUGC................................................................................AUUC..
env-343     CGCGCCAACGAGCUGCGUUAAUUAGCGGAAAGCUGAGGUUUUGAUUGCCCGGGUGUCGUUUGGCGGCUAUAAUGCCGCCCGCU.....UUUGC................................................................................AUUC..
env-344     CGCGCCAACGAGCUGCGUUAAUCACCCGAAAGCUGAGGUUUUGAUUGCCCGGGUGUCGUUUGGCGGCUAUAAUGCCGCCCGCU.....UUUGC................................................................................AUUC..
env-346     GCGCCAAUGAGCUGCGUUAAUCCGACCAAAAACUGAGGUUUUGAUUGCCGCUGUGCCGAUUGGCGGCUAUAAUGCCGCCCGCU.....UUUGC................................................................................AUUC..
env-347     CGCGCCAAUGAGCUGCGUUAAUCCGCCAAAAACUGAGGUUUUGAUUGCCGCUGUGCCGAUUGGCGGCUAUAAUGCCGCCCGCU.....UUUGC................................................................................AUUC..
env-348     CGCGCCAACGAGCUGCGUUAAUUACCGGAAAGCUGAGGUUUUGAUUGCCCGGGUGUCGUUUGGCGGCUAUAAUGCCGCCCGCU.....UUUGC................................................................................AUUC..
env-349     CGCGCCAAUGAGCUACGUUAAUCCGCCAAAAACUGAGGUUUUGAUUGCCGCUGUGCCGAUAGGCGGCUAUAAUGCCGCCCGCU.....UUUGC................................................................................AUUC..
env-350     CGCGCUAAUGAGCUACGUUAAUCCGUCAAAAACUGAGGUUUUGAUUGCCGCUGUGCCGAUAGGCGGCUAUAAUGCCGCCCGCU.....UUUGC................................................................................AUUC..
env-351     CGCGCCAAUGAGCUGCGUUAAUCCGCCAAAAACUGAGGUUUUGAUUGCCGCUGUGCCGAUAGGCGGCUAUAAUGCCGCCCGCU.....UUUGC................................................................................AUUC..
env-353     GUCGCGCCAACGAGCUGCGUUAAUCGCCUGCACUGAGGUUUUGAUUGCCGGGGCGCCGAUUGGCGGCUAUAAUGCCGCCCGCU.....UUUGC................................................................................AUUC..
env-354     CGCGCCAAUGAGCUACGUUAAUCCGUCAAAAACUGAGGUUUUGAUUGCCGCUGUGCCGAUAGGCGGCUAUAAUGCCGCCCGCU.....UUUGC................................................................................AUUC..
Ksp-1-1     CGCGCCAACGAGCUGCGUUAAUCAGCCAAAAGCUGAGGUUUUGAUUGCCCGGGUGUCGUUUGGCGGCUAUAAUGCCGCCCGCU.....UUUGC................................................................................AUCC..
Mma-1-1     ACGAGCUGCGCGGCUGACUUUUGCGCCGCCAGCGGCGCUUUUCGUUGCCGCCCGGAAGAUUUUCGGCUAUAGUGUCGCCCGCU.....UUUGC................................................................................AUCC..
Esp-43-1    UUCGCCGCGCCAAUGAGUUACGCGGCUAAGCCAUAGCGGUCCGACUUGUAACAGAGGGGGGAUACGGGUAUAAUCCGCCCGCU.....UUUGC................................................................................AUCC..
Ksp-3-1     GCUACCGUCGCGCCAAUGAAUUACGCGGUUAGUGGUGCUUUUCAUUGCUGUUAAACGCGUACGGGUAUAGAAUACCGCCCGCU.....UUUGC................................................................................ACAC..
Esp-79-1    GUUUUCGUCGGGCGCACGAAUUACGCGGUUAAUGGUAGUUUUGAUUGAUGUGAAAGGCGUUUGAGUAUAGAAUCUCGCCCGCU.....UUUGC................................................................................AUAC..
env-357     ACGUUAUCGGCGCGCCAAUGAGCUGCGCUAGAUGGCAGAAUCAUUUGCCCCUGUGCGCCAAUACGGCUAUAAUGCCGCCCGCU.....UUUGC................................................................................AUCC..
env-358     GACGCUAUCGCCGCGCCAACGAGCUGCGCUAAGCGGCGGAAUCAUUUGCCCCGCUGUGGCAAUCCGGCUAUAAUGCCGCCCGCU.....UUUGC................................................................................AUCC..
Pst-5-1     ACGCUAUCGCCGCGCCAACGAGCGGCGCUAAGCGGCGGAAUCAUUUGCCCCGCUGUGGCAAUCCGGCUAUAAUGCCGCCCGCU.....UUUGC................................................................................AUCC..
env-359     ACGCUAUCGCCGUGCUAACGAGCUGCGCUAAGCGGCGGAAUCAUUUGCCCCGCUGUGGCAAUCCGGCUAUAAUGCCGCCCGCU.....UUUGC................................................................................AUCC..
Psp-14-1    CCGUUUUCGGCGCGCCAACGAGCUGCGAUAAACCACGGAAUCAUUUGCCCGCUGCGGGGAUAGCGGCUAUAAUGCCGCCCGCU.....UUUGC................................................................................AUCC..
Psp-13-1    GCGUUAUCGCCGCGCCAAUGAGCUGCGAUAAGCCACGGAAUCAUUUGCCCGGCGCCGGGAUAGCGGCUAUAAUGCCGCCCGCU.....UUUGC................................................................................AUCC..
Ebl-1-1     UUAACCGGGAGGAUGCAGAAAUCCGCCCCCGGUGGCGCUUUUCGUUGCCCCGCCAGCACAUUGCCGCUAUAAUGCCGCCCGCU.....CUUGC................................................................................AUCC..

            ................................................................................<<<.....<<<<<<<<<<<<<<<<<<<<<<<<<<<<<<<<<<<<<<<<<<<<<<<<<<<<<<<<<<<<<<<<<<<<<<<<<<<<<<<<<......
            ................................................................................212.....22222.........111....00110....0.1.111.1111.1.........101...............011.....111100......
            ••••••••••••••••••••••••••••••••••••••••YY••UUGYY••••••••••••••••R•UA•AAU•YCGCCCGCU-----U••••--------------------------------------------------------------------------------•Y••--
```

```
env-360     CCAGUGGCUGAAUACGCAUCCGGCAGCAUGACUGUACGUUUUGAUUGCCCCACGGAAGAUUGGGGGCUAUAAUGCCGCCCGCU.....UUUGC.......AUCC..
Kas-1-1     AGUCUGCGGCUGCGCUUAACGACCAACUGAAGGUGAGGUUUUCAUUGCCCGUAAGGCAAUAGGCGGCUAUAAUGCCGCCCGCU.....UUCAC.......AUCC..
env-362     AGUCUGCGGCUGCGCUUAACGGCCAACUGAAGGUGAGGUUUUCAUUGCCCGUAAGGCAAUAGGCGGCUAUAAUGCCGCCCGCU.....UUCAC.......AUCC..
env-363     GUUUCCGCCGGGCCAAUGAAUUGCGGGGUUAAAGGGACUUUUGAUUGAUGUAAAGUGCGUUUGAGUAUAAAAAUGCCGCCCGCU.....UGAC........AUCC..
env-364     ...........................................................GUUUGAGUAUAAAAAUGCCGCCCGCU.....UCAU........AUCC..
Rte-1-1     GGGCCAACGAACUGCGCUAAUCGGCAUUGAGACUGAGGUUUUGAUUGCCGUAAUAAUCUCAGGCGGCUAUAAUGCCGCCCGCU.....UUUGC.......AUUC..
Ror-1-1     GGGCCAACGAACUGCGUUAAUCGGCAUUAAGGCUGAGGUUUUGAUUGCCGUAAAAACCUCAGGCGGCUAUAAUGCCGCCCGCU.....UUUGC.......AUUC..
env-365     GGGCAAACGAACUGCGUUAAUCAGCGCCGCGCCUGAGGUUUUCAUUGCCGUCAACGCCAUAGGUGGAUAUAAUGCCGCCCGCU.....UUUGC.......AUUC..
env-366     GGGCAAAUGAACUGCGUUAAUCACCGCCGCGCCUGAGGUUUUCAUUGCCGUCAACGCCAUAGGUGGAUAUAAUGCCGCCCGCU.....UUUGC.......AUUC..
Ksp-8-1     GGGCCAACGAACUGCGCUAAUCGUCCUCACGUCUGAGGUUUUGGUUGCUGCGAACACGUAUGGCGGCUAUAAUGCCGCCCGCU.....UUUGC.......AUUC..
Kmi-1-2     GGGCCAACGAACUGCGCUAAUCGUCCUCACAUCUGAGGUUUUGGUUGCUGCAAACACCUAUGGCGGCUAUAAUGCCGCCCGCU.....UUUGC.......AUUC..
env-368     GGGCCAAUGAAUUGCGCUAAUCCUCCGCGCGCCUGAGGUUUUGGUUGCCGCAAAAACGUAUGGCGGCUAUAAUGCCGCCCGCU.....UUUGC.......AUUC..
Dze-8-1     UCGCCGGGCAAACGAACUGCGCGGCUAGCCUUAGGCGGUUUUAUUUGCCCGCACGGCCAUAUUCCGCUAGAAUCUCGCCCGCU.....UUUAUUUCGAAUAUCAACAUAUAUACCGAAAAUAAUUCGAGUUGCGGG....AAGGCGGCAAGAGAGUGAAUCCCGAUGAGCUUACACAG..
Dda-3-1     UCGCCGGGCAAACGAACUGCGCGGCUAGCCUUAGGCGGUUUUAUUUGCCCGCGCCAUCAUAUUCCGCUAGAAUCUCGCCCGCU.....UUUAUUUCGAAUA.UCA.......ACA...
Dda-2-1     UCGCCGCGCCCAUGAACUGCGCGGCUAAUCUGCGGCGGUUUUAUUUGCCCGCGCCAUCAUAUUCCGCUAGAAUCUCGCCCGCU.....UUUAUUUCGAAUA.UCA.......ACA...
Dze-1-1     UCGCCGGGCAAACGAACUGCGCGGCUAGCCGUAGGCGGUUUUAUUUGCCCGCACGGCCAUAUUCCGCUAGAAUCUCGCCCGCU.....UUUACUUCGAAUAUCAAGAUAUAUACCCAAAAUAAUUCGAGUUGCGGG....AAGGCGGCAAGAGAGUGAAUCCCGAUGAGCUUAC..UCAG..
Ddi-5-1     UCGCCGCGCCCAUGAACUGCGCGGCUAGCCUGCGGCGGUUUUAUUUGCCCGCACCGCCAUAUUCCGUUAAAAUCUGGCCCGCU.....UUUAUUGGUAAUAAC.......AACAUC
Dsp-5-1     UCGCCGCGCCCAUGAGCUACGCGGCUAAUUUGCGGCGGUUUUAUUUGCCCGCGCCGCCAUAUUCCGCUAGAAUCUCGCCCGCU.....UUUAUUUCGAAUAAC.......AGCAUC
Dsp-2-1     UCGCCGCGCCCAUGAGCUACGCGGCUAAUUUGCGGCGGUUUUAUUUGCCCGCGCCGCCAUAUUCCGCUAGAAUCUCGCCCGCU.....UUUAUUUCGAAUAAC.......AGCAUC
Dsp-1-1     CCGCCGUGCCCAUGAGCUACGUGGCUAAUUGGCGGCGGUUUUAUUUGCCCGCGCCGCCAUAUUCCGCUAGAAUCUCGCCCGCU.....UUUAUUU.CGAAUAACG.......GCAU..
Ddi-3-1     UCGCCGCGCCCAUGAACUGCGCGGCUAGUCUGCGGCGGUUUUAUUUGCCCGCGCCGCCAUAUUCCGUUAAAAUCUGGCCCGCU.....U...UUAUUGGUAAU.......AACAAC
env-370     GCGCUAUCGCCGCGCCAACGAGCUGCGCUAAGCGGCGGAAUCAUUUGCCCCGCUGUGGCAAUCCGCUAUAAUGCCCGCCCGCU.....UUUGC.......AUCC..
CRe-2-1     UUCGUGCAUGAUUUUAAACCACUAUUGACACCCUACGGUUUUAUUUGCUGUCUGAUUGAUAUUUCGCUAAAAUCCCGCCUGCU.....U.UGCUUUAGA.......UUAU..
Kpn-281-1   CGCGCCAACGAGCUGCGUUAAUCACCCGAAAGCUGAGGUUUUGAUUGCCCGUGUGUCGUUUGGCGGCUAUAAUGCCGCCCGCU.....UUUGC.......AUUC..
Esp-14-1    GCUUUCGCCGGGCAAACGAAUUGCGUGGUUAACGGGACUUUUCGUUGAUGCGCGAGCCAUUUGAGUAUAAAAAUGCCGCCCGCU.....UCAC........AUCC..
Ywa-1-1     CCCUUGCUUGAGCCGCAUUUGACCAGAGAUAUUCGCAGUUUUAUUUGCUGUCUGAGUGAUAUUUCGCUAGAAUGCCGCCCGCU.....UUACUCUGGG.......UUUA..
Pge-1-1     ACGCUGGGCCACCUUGAGGCUAAAAAGUAGCGCCGCCUUUUCCAUUGCCCGCGCGCCGUUGAAAGCUAUAAUGCCGCCCGCU.....UUUGC.......AUAC..
Pmi-2-1     ACGCUUUCGUAAAGCAGAUACCUUAUAUUAAAGUCGAUUUAUAUUUGCUGUUAGAGUGCUUUUUCGUUAAAAUGCGCCCCGCU.....UUAUCUG.......UUUU..
Kox-1-1     GGGCCAACGAACUACGCUAAUCGCCCUCACGUCUGAGGUUUUGGUUGCUGCAAACACCUAUGGCGGCUAUAAUGCCGCCCGCU.....UUUGC.......AUUC..
Yps-1-1     CCCUUGCCUGAGCCGCAUUUGACCAGAGAUAUUCGCAGUUUUAUUUGCUGUCUGAGUGAUAUUUCGCUAGAAUGUCGCCCGCU.....UUACUCUGGG.......UUUA..
Dso-4-1     AUCGCCGCGCCCAUGAACUGCGCGGCUAGCCGUAGCGGUUUUAUUUGCCCGCGCCGCCAUAUUCCGCUAAAAUCUGGCCCGCU.....U.UUAUU.......GGUAAU
Csa-12-1    AGUUCCGCCGGGCCAAUGAAUUACGCGGAUAGCGGCGGUUUUCGUUGCCCCUGCGCGUUCAAAUAGCUAUAAUGCCGCCCGCU.....UUUGC.......AUCC..
Ecl-14-1    GCUUUCGUCGCGCCAACGAAUUGCGCGGUUAACGGGACUUUUCAUUGAUGUGAGGAGCAUUUGAGUAUAAAAAUGCCGCCCGCU.....UGAC........AUCC..
Yfr-1-17    CCCUUGCCUUAAGCCGCAUUUAGCUGAGAUUUUCGCAGUUUUAUUUGCUGUCUGGCUGAUAUUUCGCUAGAAUGCCGCCCGCU.....UUGCUCUGGG.......UUUA..
Kox-4-2     GGGCCAACGAACUGCGCUAAUCGUCCUCAGGACUGAGGUUUUGGUUGCUGCAAAAACGCAGGGCGGCUAUAAUGCCGCCCGCU.....UUUGC.......AUUC..
Sen-828-1   GCUUCCGCCGGGCGCACGAAUUACGUGGUUAAUCGUACUUUUCAUUGAUGUAACGGCGAUUUGAGUAUAGAAUCUCGCCCGCC.....UUUGC.......AUAC..
Sma-18-2    UCAAUUGACCUGAGCGAACGGUCCCCAAUCUGGACAGGUUUUCGUUGCCCUGCAGGCGGCUUUCCGUUAAAAUGCCGCGCGCU.....U.UAC.CUGGG.......CCUG..
Kae-2-1     CGCGCCAAUGAGCUACGUUAAUCCGUCAAAAACUGAGGUUUUGAUUGCCGCUAUGCCGAUAGGCGGCUAUAAUGCCGCCCGCU.....UUUGC.......AUUC..
Eco-675-1   GUUUUCGUCGGGCGCACGAAUUACGCGGUUAAUGGUAGUUUUGAUUGAUGUAUAAGGCGAUUGAGUAUAGAAUCUCGCCCGCU.....UUUGC.......AUAC..
Yro-2-1     ......................ACGCAGGUUUUGGACAGUAUUGUUUGCUGUCUGGUAGAGACUUCGCUAGAAUAUCGCCCGCU.....UUGCUCUGGG.......UUUA..
Ykr-2-1     CUCUCGCCUAAGCCGCAUUUAGCUGAUAAAUUCGUAGUUUCGUUUGCUGUCUGGUCGAUAUUUCGCUAGAAUGCCGCCCGCU.....UUGCUCUGGG.......UUUA..
Kpn-194-1   CGCGCCAACGAGCUGCGUUAAUUACCGGAAAGCUGAGGUUUUGAUUGCCCGGGUGUCGUUUGGCGGCUAUAAUGCCGUCCGCU.....UUUGC.......AUUC..
Kox-6-1     GGGCCAACGAACUGCGCUAAUCGUCCUCAGGACUGAGGUUUUGGUUGCUGCAAAAACGCAGGGCGGCUAUAAUGCCGCCCGCU.....UUUGC.......AUUC..
Yfr-3-1     CUCUCGCCUAAGCCGCAUUAAGCUGAGAUAAAUUCAUAGUUUCGUUUGCUGUCUGGCCGAUAUUUCGCUAGAAUGUCGCCCGCU.....UUGCUCUGGG.......UUUA..
Kpn-219-1   CGCGCCAACGAGCUGCGUUAAUUACCGGAAAGCUGAGGUUUUGAUUGCCCGGGUGUCGUUUGGCGGCUAUAAUGCCGCCCGCU.....UUUGC.......AUUC..
Yps-15-1    CCCUUGCCUGAGCCGCAUUUGACCGGAGAUAUUCGCAGUUUUAUUUGCUGUCUGAGUGAUAUUUCGCUAGAAUGUCGCCCGCU.....UUACUCUGGG.......UUUA..
Kox-15-1    GGGCCAAUGAACUGCGUUAAUCCUCCGUACGCCUGAGGUUUUGGUUGCCGCAAAAACGUAUGGCGGCUAUAAUGCCGCCCGCU.....UUUGC.......AUUC..
Sma-19-1    UCAAUUGACCUGAGCGAACGGUCCUCAAUCUGGACAGGUUUUCGUUGCCCUGCGGGCGGCUUUCCGUUAAAAUGCCGCGCGCU.....U..UACCUGGG.......CCUG..
Yru-2-2     ACUGGGUUAGCGUCCUGCGCCCCCCAGCUUAUUUGCGGUUUUAUUUGCUGUCUGGUGAUAUUUCGCUAGAAUGCCGCCCGCU.....UUGCUCUGGG.......UUCG..
Sma-8-1     UCAGUUGGCCUGAGCGAAUGGCCCUCAAUCUGGGCAGGUUUUCGUUGCCCUGCGGGCGGCUUUCCGUUAAAAUGCCGCGCGCU.....U.UAC.CUGGG.......CCUG..
Ysi-1-1     CUCUCGCCUGAGCCGCACUUGACCUGCGAGAUCCGCAGUUUUAUUUGCUGUCUGAGCGAUAUUUCGCUAGAAUGCCGCCCGCU.....UUGCUCUGGG.......UUUA..
Kae-3-1     CGAGCCAAUGAGCUACGUUAAUCCGUCAAAAACUGAGGUUUUGAUUGCCGCUGUGCCGAUAGGCGGCUAUAAUGCCGCCCGCU.....UUUGC.......AUUC..
Sbo-2-7     UCAAUUGGCCUGAGCGAACUGCCGCCAACCCGGACAGGUUUUCGUUGCCCUGCGGGCGGCUUUCCGUUAAAAUGCCGCGCGCU.....U.UAC.CUGGG.......CCUG..
Ksp-2-1     GGGCCAACGAACUACGCUAAUCGUCCUCACGUCUGAGGUUUUGGUUGCUGCAAACACCUAUGGCGGCUAUAAUGCCGCCCGCU.....UUUGC.......AUUC..
Yal-3-1     CCCUUGCUUAAGCCGCAUUUAGCCUGAGAUUUUCGCAGUUUUAUUUGCUGUCUGGCUGAUAUUUCGCUAGAAUGUCGCCCGCU.....UUGCUCUGGG.......UUUA..
Kox-18-1    GGGCCAACGAACUGCGCUAAUCGCCCUCACGUCUGAGGUUUUGGUUGCUGCAAACACCUAUGGCGGCUAUAAUGCCGCCCGCU.....UUUGC.......AUUC..
Kva-1-5     CGCGCCAAUGAGCUGCGUUAAUCCGCCAAAAACUGAGGUUUUGAUUGCCGCUGUGCCGAAUGGCGGCUAUAAUGCCGCCCGCU.....UUUGC.......AUUC..
Esp-18-1    GCUUUCGCCGGGCAAACGAAUUGCGCGGUUAACGGGACUUUUCGUUGAUGCGCGAGCCAUUUGAGUAUAAAAAUGCCGCCCGCU.....CCAC........AUCC..
Efe-1-1     GUUUUCGUCGGGCACACGAAUUACGCGGUUAAUGGUAGUUUUAAUUGAUGUGCAAGGCGAUUGAGUAUAGAAUCUCGCCCGCU.....UUUGC.......AUAU..
            ...............................................................................&lt;&lt;&lt;.....&lt;&lt;&lt;&lt;&lt;&lt;&lt;&lt;&lt;&lt;&lt;&lt;&lt;&lt;&lt;&lt;&lt;&lt;&lt;&lt;&lt;&lt;&lt;&lt;&lt;&lt;&lt;&lt;&lt;&lt;&lt;&lt;&lt;&lt;&lt;&lt;&lt;&lt;&lt;&lt;.....
            ...............................................................................212.....22222........111....00110.....0.1.111.1111.1.........101...............011.....111100.....
            ●●●●●●●●●●●●●●●●●●●●●●●●●●●●●●●●●●●●●●●●●YY●●UUGYY●●●●●●●●●●●●●●●R●UA●AAU●YCGCCCGCU----U○○○○------------------------------------●Y○○--
```

alignment positions 181 · · · 360

```
env-1       ...........................................................................................CUUUGAGUCUGUUUAGG..........................................
env-2       ...........................................................................................CUUUGAGUCUGUUUAGG..........................................
env-4       ...........................................................................................CUUUGAGUCUGUUUAGG..........................................
Mma-2-1     ...........................................................................................UUUGAGGCAAAUUUAUG..........................................
env-6       .........................................................................................UCUUUGAGCAAAUUUAGG..........................................
Hpa-2-1     .........................................................................................UCUUUGAGCAAAUUUAGG..........................................
env-7       .........................................................................................UCUUUGAGCAAAUUUAGG..........................................
env-8       .........................................................................................UCUUUGAGCAAAUUUAGG..........................................
env-34      .........................................................................................UCUUUGAGCAAAUUUAGG..........................................
env-35      ..........................................................................................CUUUGAGCAAAUUUAGG..........................................
env-50      ..........................................................................................CUUUGAGCAAAUUUAGG..........................................
env-54      ..........................................................................................CUUUGAGCAAAUUUAGG..........................................
env-55      ..........................................................................................CUUUGAGCAAAUUUAGG..........................................
env-58      ..........................................................................................CUUUGAGCAAAUUUAGG..........................................
Hin-1-1     .................................................................................GUGCUUGUCUUUGAGAACAAUAUUUGG..........................................
Hha-1-1     .................................................................................AUGCUUGUCUUUGAGA..........................GCGA..................UAU..............

            ................&gt;&gt;&gt;&gt;&gt;&gt;&gt;&gt;&gt;&gt;&gt;&gt;&gt;&gt;&gt;&gt;&gt;&gt;&gt;&gt;&gt;&gt;&gt;&gt;&gt;&gt;&gt;&gt;&gt;&gt;&gt;&gt;&gt;&gt;&gt;&gt;&gt;&gt;&gt;&gt;&gt;&gt;&gt;&gt;&gt;&gt;&gt;&gt;&gt;&gt;...........................&lt;&lt;&lt;&lt;&lt;&lt;&lt;&lt;&lt;&lt;&lt;&lt;&lt;&lt;&lt;&lt;&lt;&lt;&lt;&lt;&lt;&lt;&lt;&lt;.........&gt;&gt;&gt;&gt;&gt;&gt;&gt;&gt;&gt;&gt;&gt;&gt;&gt;&gt;
            ................001111.....110...............101.........1.1111.111.1.0.....01100....111.........22222212...........................1111.10.11..100.0..00000.........00000..0.001..
            ----------------------------------------------------------------------------------------------○○○○AAGC●●●●●●●●●●●●------------------------------------------------------
```

```
env-59     ...AUGCUUGUCUUUGAGA...CCAA...UAU...
Hsp-1-1    ...GUGCUUAUUCUUUGAGAACAAUAUUUGG...
env-62     ...AUGCUUGUCUUUGAGAACAAUAUUUCG...
Hha-1-2    ...AUGCUUGUCUUUGAGAACAAUAUUUGG...
env-63     ...AUGCUUGUCUUUGAGAACAAUAUUUGG...
Hha-3-1    ...AUGCUUGUCUUUGAGAACAAUAUUUGG...
env-65     ...AUGCUUGUCUUUGAGAACAAUAUUUGG...
env-66     ...GUGCUUGUCUUUGAGAACAAUAUUUGG...
Hin-4-1    ...GUGCUUGUCUUUGAGAACAAUAUUUGG...
env-67     ...GUGCUUGUCUUUGAGAACAAUAUUUGG...
env-68     ...AUGCUUGUCUUUGAGAACAAUAUUUGG...
env-70     ...AUGCUUGUCUUUGAGA...CCAA...UGU...
env-71     ...AUGCUUGCCUUUGAGAACAAUAUUUGG...
env-72     ...AUGCUUGCCUUUGAGAACAAUAUUUGG...
env-73     ...GCUUUGGGCAAAUUUUUGU...
Asu-1-1    ...UCUUUGGGCAAAUUACGG...
env-74     ...GCUUUGAGCAAAAUUUGUG...
env-75     ...GCUUUGAGCAAUAUUUGU...
env-76     ...CUUUGAGCAAUAUUUGU...
env-77     ...GCUUUGAGCAAAUUUUGUG...
env-78     ...GCUUUGAGCAAAUUUUGUG...
env-79     ...GCUUUGAGCAAAUUUUGUG...
env-80     ...GCUUUGAGCAAAUUUUGUG...
env-82     ...GCUUUGAGCAAAUUUUGUG...
env-83     ...GCUUUGAGCAAAUUUUGUG...
env-85     ...GCUUUGAGCAAAUUUUGUG...
env-88     ...GCUUUGAGCAAAUUUUGUG...
env-89     ...GCUUUGAGCAAAUUUUGUG...
env-90     ...GCUUUGAGCAAAUUUUGUG...
env-91     ...GCUUUGAGCAAUAUUUGU...
env-101    ...GCUUUGAGCAAUAUUUGU...
env-103    ...GCUUUGAGCAAUAUUUGU...
env-104    ...GCUUUGAGCAAUAUUUGU...
env-111    ...GCUUUGAGCAAUAUUUGU...
env-114    ...GUGGAAGCCGCAGUACUCAA...
env-115    ...GUGGAAGCUUGU...AGU...
env-121    ...GUGGAAGCUUGU...AGU...
Aen-1-1    ...CUAAC.AAGUUUUUAUUAUCGGA...
Ana-1-1    ...CUAAC.AAGUUUUUAUUAACAGA...
Eba-2-1    ...GCAGAACGGUUUUAUCGAAU...
Dch-3-1    ...AAAGCGGAUUAUCCUG...
Sgl-1-1    ...UGCCUGCUUAUU...GACA...AGCA...
Dsp-3-1    ...GCAGAAUUCACAGGCACCGUUAAUUCA...
Fpe-1-1    ...AAAGCAACUAACCCUUUG...
env-122    ...AAAGCAACUAACCCUUUG....
Mmo-1-1    ...CCCUGCCGGAGCCAAAUACCGAC...
Mmo-1-3    ...CCCGCUG..AAGCCAAAUACCGAC...
Mmo-3-1    ...CCUGCUG..AAGCCAAAUACCGAC...
Mmo-5-1    ...CCUGCUG..AAGCCAAAUACCGAC...
Mmo-2-1    ...CCUGCUG..AAGCCAAAUACCGAC...
Mmo-1-7    ...CCUGCUG..AAGCCAAAUACCGAC...
Pfl-1-1    ...GUAAUAGUUU...UGG...UUUC...
Tpt-1-1    ...GUAAGAGUGCUUAUGAAUAACCU...
Tsa-1-1    ...UGUGAAGGCGCUUUUGAAUAACCU...
env-125    ...GUAAAAGUACAUU...GG...GAUAA...
Mwi-1-1    ...UGUGCGAACGC..UUAGCCAAAUCUCGAA...
Spr-1-1    ...AAGCACGUUUAUCCUG...
Eic-2-1    ...AGCCUGAGAC.GGGCACACGAUAUCCCG...
Epi-1-1    ...GCCUAAGAC.UGGCGCACGAUAUCCCG...
Epi-1-2    ...ACCU.AAGAC.UGGCGCACGAUAUCCCG...
Eta-5-1    ...ACCU.AAGAC.UGGCGUACGAUAUCCCG...
CSo-1-1    ...AAGCACGUUUAUCCUG...
Gan-5-1    U...UCAGCACCUAUUUUGU...
Gan-7-1    U...UCAGCACCUAUUUUGU...
Gan-1-2    U...UCAGCACCUAUUUUGU...
Gan-1-5    U...UCAGCACCUAUUUUGU...
Gan-1-6    U...UCAGCACCUAUUUUGU...
Gan-6-1    U...UCAGCACCUAUUUUGU...
Gan-1-9    U...UCAGCACCUAUUUUGU...
Gan-1-10   U...UCAGCACCUAUUUUGU...
Gan-3-1    U...UCAGCACCUAUUUUGU...
Gan-9-1    U...UCAGCCCUAUUUUGU...
CSc-1-1    ...AAGCAAAUGUUGGAAUUAUG...
env-126    ...AAGCACUAAUUUUAUAG....
env-127    ...AAGCACUAAUUUUAUAG....
env-131    ...AAGCACCUAUUUUAUAG...
env-132    ...GAGCACUAAUUUUAUAA....
env-134    ...GAGCACUAAUUUUAUAA....
env-136    ...AAGCACUAAUUUUAUAA....
env-138    ...AAGCACUAAUUUUAUAA....
env-139    ...AAGCACCAAUUUGAUAG...
Gap-1-1    ...AAGCACAUAUUUUAUAA....
env-140    ...AAGCACAUAUUUUAUA...
Dda-1-1    ...UAACAAGCAGAGUUAUCAAC...AAGUAUAGUUG...UCAA...C

................>>>>>>>>>>>>>>>>>>>>>>>>>>>>>>>>>>>>>>>>>>>>>>>>>>>>>>>>>>>>>>>>>>>>>>>>>>...<<<<<<<<<<<<<<<<<<<<<<<<.........>>>>>>>>>>>>>>>>
................001111.....110...............101.........1.1111.111.1.0.....01100....111.........22222212...1111.10.11..100.0..00000.........00000..0.001..
----------------------------------------------------------------------------------------------------oooo AAGC ●●ooooooooo----------------------------------------
```

```
Eho-5-1     ...GCAAGGCACACUAUCCCG...
Eta-1-1     ...GCAAGGCACACUAUCCCG...
Eta-4-1     ...GCAAGGCACACUAUCCCG...
bsy-2-1     ...AUAAAGCGUCACUGAAUAAUCCU...
env-142     ...CAAAGCGUCAUUGAAUAAACCU...
env-143     ...CAAAGCGUCAUUGAAUAAACCU...
Psp-12-1    ...GCAUCAGCACAAUCUGAAUCAAUC...
Ein-1-1     ...GCGGAAGC...UGAACG...AUUUAAAC...
env-144     ...GCAGAAGC...UGAACG...AUUUAAAC...
Psp-9-1     ...ACAAGAGCUGAACAAUCCUG...
Esp-74-1    ...GUGAAAGCUGAACAAGACUA...
Eam-10-1    ...GUGAAAGCUGAACAAGACUA...
Eam-1-1     ...GUGAAAGCUGAACAAGACUA...
Epy-2-1     ...GG..AAGCUGAACAAGACUA...
Eto-1-1     ...GCCUGAGCACACUAUAGUACGAACAAUCCAA...
Baq-1-1     ...GCAAGUA...AAGCAGAUUUACUCAA...
Pfo-1-1     ...AAGCAAGAUCGCUCAA...
Plu-4-1     ...UGUUAAAUGUUAAGCCAAAUCUCGAA...
Pvu-2-1     ...UAGAU.AAACCAAAUCUCGAG...
Pha-1-1     ...UAGAU.AAACCAAAUCUCGAG...
Pmi-1-2     ...UAGAU.AAACCAAAUCUCGAG...
Lgr-1-1     ...AAGCAGAACGUAUAAA...
gpr-1-1     ...GCG.AAGCGAAGAUCUAUCA...
Rch-1-1     ...GCG.AAGCGAAGAUCUAUCA...
Ssp-13-1    ...GCG.GAGCGAAGAUCUAUCA...
env-148     ...GCG.AAGCGAGGAUCUAUCA...
Pru-1-1     ...GCGAAAUAAUUUUACAGGGCCAAAUAUCGAA...
env-149     ...GCAUAAGC...GGU...GUUGAAUA...
Pst-1-1     ...GCAUCAGC...GGU...GUUGAAUA...
Pst-4-1     ...GCAUCAGC...GGU...GUUGAAUA...
Bgo-1-1     ...GCAGAAGCUGAAUAACCCAA...
Bsp-1-1     ...GCAAAAGCAGAACAACCCAA...
Dpa-1-1     UAACGCCGC...CAAA.CGGAUACAAGCAAAGUAUCCUG...
Pal-5-1     ...GCGAAAUGAUGUUACAGGGCCAAAUCUCGAA...
Pal-8-1     ...CGAAAUGAUGUUACGAGGCCAAAUCUCGAA...
Pal-2-1     ...CGAAAUGAUGUUACGAGGCCAAAUCUCGAA...
Pal-9-1     ...CGAAAUGAUGUUACGAGGCCAAAUCUCGAA...
Sma-22-1    ...CCCGGGUGGGGU...CGA...AUUA...
env-151     ...AAAGCAGAUUUUCGAA...
Dze-7-1     ...AAAGCGGAUUAUCCUG...
Dze-9-1     ...AAAGCGGAUUAUCCUG...
Dze-6-1     ...AAAGCGGAUUAUCCUG...
Dch-1-1     ...UGGUUAUUUGAACA..AAGC...GGA...UUA...
Dda-5-1     ...UGUUAUUUGAACA..AAGC...GGA...UUA...
Dze-4-1     ...ACAAAGC...GGA...UUA...
Dch-4-1     ...ACAAAGC...GGA...UUA...
Dze-2-1     ...UGUUAUUUGAACA..AAGC...GGA...UUA...
Dsp-9-1     C...GUUAUUUAAACCAAAAGCA...GGG...UUAU...
Dze-5-1     ...AA..AGCGGAUUAUCCUG...
Hal-1-1     ...UAAAAGCAGAAUCUCCUG...
Hal-1-2     ...UAAAAGCAGAAUCUCCUG...
env-152     ...UAAAAGCAGAAUCUCCUG...
Hpa-1-1     ...UAAAAGCAGAAUCUCCUG...
Hal-4-1     ...UAAAAGCAGAAUCUCCUG...
Pst-6-1     ...GCGAAUUACACAUCAACAAGCCAAAUCUCGAA...
Pst-7-1     ...GCGAAUUACACAUCAACAAGCCAAAUCUCGAA...
Pre-2-1     ...GAAUGAAGUAUCCAUAGGCCAAAUCUCGAA...
Pre-1-1     ...GAAUGAAAUAUUCUUAGGCCAAAUCUCGAA...
Pbu-1-1     UUAC...CACU.UAGCCAAAUAUCGAA...
Pas-2-1     ...GUUAAGCCAAAUCUCGAA...
Pse-1-1     ...GCAAUCGCCUUUGAACAACCUG...
Psp-7-1     ...GCAAUCGCCUUUGAACAACCUG...
Pva-1-1     ...GCAAUCGCCUUUGAACAACCUG...
Msp-1-1     ...GUAAAAGCUGAAUAUUUG...
Eba-4-1     ...GCAGAGCAGAAUCUUUG...
Lqu-1-1     ...GCAUAAGCAGAAUCAUCCAA...
env-153     ...GCAUGAGCAGAAUCAUCCAA...
Lqu-1-2     ...GCAUGAGCAGAAUCAUCCAA...
env-154     ...GCAUGAGCAGAAUCAUCCAA...
env-155     ...CAAAAGCUGAAUAACCCCG...
Pca-10-1    ...CAAAAGCUGAAUAACCCCG...
Ppo-1-1     ...CAAAAGCUGAAUAGUCCCA...
Pwa-3-1     ...CAGAAGCUGAAUAACCCAA...
Pwa-2-1     ...CAGAAGCUGAAUAACCCAA...
Ppa-2-1     ...CAGAAGCUGAAUAACCCAA...
Pbe-1-1     ...CAAAAGCUGAAUAACCCAA...
Pca-5-1     ...CAAAAGCUGAAUAACCCAA...
Pca-3-1     ...CAAAAGCUGAAUAACCCAA...
Pca-3-2     ...CAAAAGCUGAAUAACCCAA...
Pca-5-2     ...CAAAAGCUGAAUAACCCAA...
Pca-5-3     ...CAAAAGCUGAAUAACCCAA...
Pca-5-7     ...CAAAAGCUGAAUAACCCAA...
Pca-6-1     ...CAAAAGCUGAAUAACCCAA...
Pbe-1-2     ...CAAAAGCUGAAUAACCCAA...

            ...>>>>>>>>>>>>>>>>>>>>>>>>>>>>>>>>>>>>>>>>>>>>>>>>>>>>>>>>>>>>>>>>>>>>>>>>>>>>>>>>>>>>...<<<<<<<<<<<<<<<<<<<<<<<<........>>>>>>>>>>>>>>>
            ...001111.....110...............101.........1.1111.111.1.0.....01100....111.........22222212...1111.10.11..100.0..00000........00000..0.001..
            ----------oooo AAGC ●●ooooooooo----------
```

```
Pca-5-9      ...CAAAAGCUGAAUAACCCAA...
Pca-9-1      ...CAAAAGCUGAAUAACCCAA...
Pca-8-1      ...CAAAAGCUGAAUAACCCAA...
Pca-5-10     ...CAAAAGCUGAAUAACCCAA...
Pca-3-15     ...CAAAAGCUGAAUAACCCAA...
Pca-7-1      ...CAAAAGCUGAAUAACCCAA...
Pca-1-1      ...CAAAAGCUGAAUAACCCAA...
Pca-2-1      ....AAAAGCUGAAUACCCCAA...
Pat-3-1      ...CAAAAGC...UGG...AUAAC...
Pte-1-1      ...CGUUAAACCAAAUCCCGAA...
Xbo-9-1      ...UUAUAUUAAGCCAAGUCUCGAA...
Plu-6-1      ...GUUAAAUGUUAAGCCAAAUCUCGAA...
Plu-2-1      ...GUUAAGCCAAAUCUCGAA...
Xne-3-1      ...UAAUGUUAAGCCAAGUAUCGAA...
Xkh-1-1      ...UAAUGUUAAGCCAAGUAUCGAA...
Xne-5-1      ...UAAUGUUAAGCCAAGUAUCGAA...
Xgr-1-1      ...UCAUGUUAAGCCAAGUAUCGAA...
Xdo-1-1      CC...AUGUUAAGACAAAUAUCGAA...
Xbo-2-1      ...ACAUUAUAUUAAGCCAAGUCUCGAA...
Xbo-11-1     ...ACAUUAUAUUAAGCCAAGUCUCGAA...
Xbo-8-1      ...ACAUUAUAUUAAGCCAAGUCUCGAA...
Pas-1-1      ...CGUUAAGCCAAAUCUCGAG...
Xpo-1-1      ...UCACGUUAAGCCAAGUAUCGAA...
Plu-1-2      UACUAUGUUAAA...UGUUAAGCCAAAUCUCGAA...
Plu-5-1      ...GUUAAGCCAAAUCUCGAA...
Plu-3-1      UACUAUGUUAAA...UGUUAAGCCAAAUCUCGAA...
Pte-2-1      ...CGUUAAGCCAAAUCUCGAA...
Pte-3-1      ...CGUUAAGCCAAAUCUCGAA...
Xsp-1-1      UA...AUGUUAAGCCAAGUAUCGAA...
Xca-1-1      ...UAAUAUUAAGCCAAGUAUCGAA...
Xsz-1-1      ...UAAUGUUAAGCCAAGUAUCGAA...
Spl-2-1      ...CCCGGUGCUAAAGUGGUUUAUCGAA...
Sfo-4-1      ...CCCAGUGUGAAAGCAGGUUAUCGAG...
Sfo-3-1      ...CCCAGUGUGAAAGCAGGUUAUCGAG...
Sfo-1-1      ...CCCAGUGUGAAAGCAGGUUAUCGAG...
env-157      ...CCCAGCGUUAAAGCGGAUUCCCGAA...
Sma-1-2      ...CCCGGCGUUAAAGCGGAUUUCCGAA...
Sgr-1-1      ...CCCGGCGCUAAAGCA...GGUUAUACCUUAUGCAUUACAGGCUAGG...GUCUGGAAGGCAGC
Sma-1-3      ...CCCAGCGUUAAAGCGGAAUUCCGAA...
Sur-1-1      ...CCCAGCGUUAAAGCGGAAUUCCGAA...
env-158      ...CCCAGCGUUAAAGCGGAAUUCCGAA...
Sma-14-1     ...CCCAGCGUUAAAGCGGAAUUCCGAA...
Ssp-8-1      ...CCCAGCGUUAAAGCGGAAUUCCGAA...
Ssp-9-1      ...CCCAGCGUUAAAGCGGAAUUCCGAA...
Sne-1-1      ...CCCAGCGUUAAAGCGGAUUUCCGAA...
Sma-18-1     ...CCCAGCGUUAAAGCGGGUUUCCGAA...
Sma-25-1     ...CCCAGCGUUAAAGCGGGUUUCCGAA...
Sma-1-1      ...CCCAGCGUUAAAGCGGGUUUCCGAA...
env-159      ...CCCAGCGUUAAAGCGGGUUUCCGAA...
env-160      ...CCCAGCGUUAAAGCGGGUUUCCGAA...
Sli-2-1      ...GGGUAAAUCGAA...
env-161      ...CCCGGUGCUAAGCAGGUUAUCGAA...
Ssp-6-1      ...CCCAGUGUGAAAGCAGGUUAUCGAG...
Ssp-2-1      ...CCCGGUGCAAAAGCAGGUUAUCGAG...
Ssy-2-1      ...CCCGGUGUAGAGGCAGGUUAUCGAA...
Ssy-1-1      ...CCCGGUGUAGAGGCAGGUUAUCGAA...
env-162      ...CCCGGUGCUAAAGCAGGUUAUCGAA...
env-163      ...CCCGGUGCUAAAGCAGGUUAUCGAA...
Ssp-14-1     ...CCCGGUGCUAAAGCAGGUUAUCGAA...
Cmu-2-1      ...CCCGGUGCAAAAGUGGGUCAUCGAA...
Spr-2-1      ...CCCGGUGCUAAAGCA...GGUUAUACCCUACGCAUGACAGGCUGCG...GUCUGGAAGGCCGAC
Ssp-4-1      ...CCAAGGUAAAAGCAGGUUAUCGAG...
Spl-1-1      ...CCCGGUGCUAAAGUGGUUUAUCGAA...
Ssp-10-1     ...CCCGGUGCUAAAGUGGUUUAUCGAA...
Sod-1-1      ...CCCGGCGUAAACGC...GAGUA...AAAC...
Spl-5-1      ...AGGCAA...CGG...GGAAA...
Ype-132-1    ...CCCAGCCA.AAGC...CGA...GUA...
Yfr-1-1      ...CCCAGUCA.AAGC...CGA...GUA...
Yfr-1-4      ...CCCAGUCA.AAGC...CGA...GUA...
Ymo-1-1      ...CCCAAUCA.AAGC...CGA...GUA...
Ymo-1-3      ...CCCAAUCA.AAGC...CGA...GUA...
Yfr-1-6      ...CCCGAUCA.AAGC...CGA...CUA...
Yfr-1-7      ...CCCGAUCA.AAGC...CGA...CUA...
Yen-21-1     ...CCCAAUCA.AAGC...CGA...GUA...
Yal-1-1      ...CCUAAUCA.AAGC...CGA...GUA...
Yfr-1-9      ...CCCAAUCA.AAGC...CGA...GUA...
Ype-1-1      ...CCCAAUCA.AAGC...CGA...GUA...
Yfr-1-10     ...CCCAAUCA.AAGC...CGA...GUA...
Yfr-2-2      ...CCCAAUCA.AAGC...CGA...GUA...
Yfr-1-12     ...CCCAAUCA.AAGC...CGA...GUA...
Yal-1-3      ...CCCAAUCA.AAGC...CGA...GUA...
Ymo-1-5      ...CCCAAUCA.AAGC...CGA...GUA...
Yma-2-1      ...CCCAAUCA.AAGC...CGA...GUA...
Yma-1-1      ...CCCAAUCA.AAGC...CGA...GUA...

             ................>>>>>>>>>>>>>>>>>>>>>>>>>>>>>>>>>>>>>>>>>>>>>>>>>>>>>>>>>>>>>>>>>>>>>>>>>>>>>>>>>>>>>>>>>>>>...<<<<<<<<<<<<<<<<<<<<<<<<........>>>>>>>>>>>>>>
             ................001111.....110...............101.........1.1111.111.1.0.....01100....111.........22222212...1111.10.11..100.0..00000........00000..0.001..
             AAGC
```

| | |
|---|---|
| Yfr-1-13 | CCCAAUCA.AAGC … CGA … GUA |
| Ybe-1-1 | CCCAAUCA.AAGC … CGA … GUA |
| Ynu-1-1 | CCCAAAGU.AAGC … CGA … CUA |
| Yru-1-1 | CCCAAAGU.AAACUGACUAACGAA |
| Yru-1-2 | CCCAAAGU.AAACUGACUAACGAA |
| Ssp-12-1 | ACACAAGC … GGA … AUA |
| Pan-5-1 | GCAUCAGCGGAAUUGAAUAACCUG |
| Eas-4-1 | GUGAGGCAGAACACUCA |
| env-165 | GUGAGGCAGAACACUCA |
| Lad-1-1 | GUGAGGCAGAACACUCA |
| Ecl-37-1 | GUGAGGCAGAACACUCA |
| Esp-37-1 | GUGAGGCAGAACACGCA |
| Ecl-39-1 | GUGAGGCAGAACACUCG |
| env-166 | GUGGGGCAGAACACUCA |
| env-167 | GUGAAGCAGAACACUCA |
| Elu-1-5 | GUGAAGCAGAACACUCA |
| Ecl-58-1 | GUGAAGCAGAACACUCA |
| Yre-2-1 | AUGCA.AAGCAGAUCACUUA |
| env-168 | GCA.AAGCAGAAUAACGG |
| Tod-1-1 | GCA.AAGCAGAAUAACGG |
| Tgu-1-1 | GCA.AAGCAGAAUAACGA |
| Psp-5-1 | GUAAAAGCUGAACACCCCAA |
| Ebi-2-1 | GCAAAAGUUGAACACUCCAA |
| Ebi-1-1 | GCAAAAGUUGAACACUCCAA |
| Eol-1-1 | GCAAAAGCUGAACACUCCCG |
| Esp-55-1 | GCAAAAGCUGAACACUCCCG |
| env-169 | GCAAAAGUUGAACACUCCAA |
| env-170 | GCAAAAGUUGAACACUCCAA |
| Ety-1-1 | GCAAAAGUUGAACACUCCAA |
| env-171 | GCAAAAGUUGAACACUCCAA |
| env-172 | GCAAAAGUUGAACACGCCAA |
| Pva-1-2 | GCAUCAGCGGGAUUGAAUAACCUG |
| Pro-1-1 | GUAAAAGCGUUUGAAUAACCGU |
| env-173 | GUAAAGC … CAG … AAAU |
| env-174 | GCA.AAGCAGAUCACUCG |
| env-175 | GCAGGGCAGAAUCUUCG |
| Ehe-1-1 | GCA.AAGCAGAUAAUCUG |
| Pag-1-1 | GCAUCAGC … GGU … AUUGAAUA |
| env-177 | GCAUCAGC … GGU … AUUGAAUA |
| Pag-1-6 | GCAUCAGC … GGU … AUUGAAUA |
| Pan-1-3 | GCAUCAGC … GGU … AUUGAAUA |
| Pag-5-1 | GCAUCAGC … GGU … AUUGAAUA |
| Pag-4-1 | GCAUCAGC … GGU … AUUGAAUA |
| env-178 | GCAUCAGC … GGU … AUUGAAUA |
| env-179 | GCAUCAGC … GGU … AUUGAAUA |
| env-180 | GCAUCAGC … GGU … AUUGAAUA |
| env-181 | GCAUCAGC … GGU … AUUGAAUA |
| Pag-3-1 | GCAUCAGC … GGU … AUUGAAUA |
| env-182 | GCAUCAGC … GGU … AUUGAAUA |
| Pag-1-5 | GCAUCAGC … GGU … AUUGAAUA |
| Psp-1-1 | GCAUCAGC … GGU … AUUGAAUA |
| env-183 | GCACAAGCGUUUGAAUAACCGU |
| env-184 | GCACAAGCGUUUGAAUAACCGU |
| Eca-1-1 | GUAAAAGCGUAUGAAUAACCG |
| Psp-3-1 | GUAAAAGCAUAUGAAUAACCG |
| env-189 | GUAAAAGCAUAUGAAUAACCG |
| bsy-1-1 | ACCCGGUCA.AAGCUGAACAAUCCAG |
| Epi-2-1 | GCAAAAGCUGAACAAUACUA |
| Eta-9-1 | GCGAAAGCAGAACAAUACCA |
| env-192 | GCA.AAGCAGAACGCUAG |
| Kcr-1-1 | GUG.AAGCAGAUCAAUUG |
| env-194 | GUGAAGCAGAACACCUG |
| env-195 | GUGAAGCAGAACACCUG |
| env-196 | GUGAAGCAGAACACCUG |
| Etr-1-1 | GCACAAGCUGAAAUACCAA |
| Ema-2-1 | GCAGAAGCUGAACCAUUAA |
| env-197 | GUG.AGGCUGAACACUUA |
| env-198 | GUGAGGCAGAACACUUA |
| env-199 | GUGAGGCAGAACACUUA |
| env-200 | GUGAGGCAGAACACUCA |
| env-201 | GCA.AAGCCGAUCAUCUG |
| Ecl-59-1 | GCA.AAGCCGAUCAUCUG |
| env-202 | GCA.AAGCCGAUCAUCUG |
| Eko-1-1 | GUGGGGCAGAACACUCA |
| Ecl-33-1 | GUGGGGCAGAACACUCA |
| Eko-1-8 | GUGGGGCAGAACACUCA |
| Esp-51-1 | GUGGGGCAGAACACUCA |
| Ecl-45-1 | GUGGGGCAGAACACUCA |
| Ecl-52-1 | GCA.AAGCAGAACACUCA |
| Ecl-54-1 | GCA.AAGCAGAACACUCA |
| env-204 | GCA.AAGCAGAACACUCA |
| Ecl-52-3 | GCA.AAGCAGAACACUCA |
| env-205 | GUGAGGCAGAACACUCA |
| Ecl-51-1 | GUGAGGCAGAACACUCA |
| | >>>>>>>>>>>>>>>>>>>>>>>>>>>>>>>>>>>>>>>>>>>>>>>>>>>>>>>>>>>>>>>>>>>>>>>>>>>>>>>>>>>>>>>>>>>> … <<<<<<<<<<<<<<<<<<<<<<<< … >>>>>>>>>>>>>> |
| | 001111.....110...............101.........1.1111.111.1.0.....01100....111.........22222212 … 1111.10.11..100.0..00000 … 00000..0.001.. |
| | ∘∘∘∘AAGC•••••••••••∘ |

```
env-207       GUGAGGCAGAACACUCA
env-208       GUGAGGCAGAACACUCA
env-210       GUGAGGCAGAACACUCA
env-211       GUGAGGCAGAACACUCA
env-212       GUGAGGCAGAACACUCA
env-213       GUGAGGCAGAACACUCA
Esp-19-1      GUGAGGCAGAACACUCA
Ecl-34-1      GUGAGGCAGAACACUCA
env-214       GUGAGGCAGAACACUCA
env-216       GUGAGGCAGAACACUCA
Ecl-42-1      GUGAGGCAGAACACUCA
env-217       GUGAGGCAGAACACUCA
env-218       GUGAGGCAGAACACUCA
env-228       GUGAGGCAGAACACUCA
env-229       GUGAGGCAGAACACUCA
Ecl-29-1      GUGAGGCAGAACACUCA
Esp-65-1      GUGAGGCAGAACACUCA
env-230       GUGAGGCAGAACACUCA
env-232       GUGAGGCAGAACACUCA
Ecl-16-1      GUGAGGCAGAACACUCA
env-233       GUGAGGCAGAACACUCA
env-236      GCA.AAGCAGAACAAUCG
Sco-1-1      GCA.AAGCAGAACAAUCG
Psp-10-1     GUAAAAGCAGUUUGAAUAACCGU
Psp-18-1     GUAAAAGCAGUUUGAAUAACCGU
env-237      GUG.AAGCAGAUCAUUUG
env-239      GUG.AAGCAGAUCAUCUG
Ecl-60-1     GCA.AAGCAGAACACUUA
Csp-13-1     GCA.AAGCAGAUCACUUA
env-240      GCA.AAGCAGAUCACUUA
Cne-1-1      GCA.AAGCAGAAAACUUUA
env-241      GCA.AAGCAGAAAACUUCA
env-242     GCA..AAGCAGAAAACUUCA
env-243      GCA.AAGCAGAAAACUUCA
env-244      GCA.AAGCAGAAAACUUCA
Esp-68-1     GCA.AAGCAGAAAACUUUA
Cne-1-2      GCA.AAGCAGAAAACUUUA
Cne-1-3      GCA.AAGCAGAAAACUUUA
Eco-2103-1   GCA.AAGCAGAUCACUUA
Esp-77-1     GCA.AAGCAGAUCACUUA
env-245      GCA.AAGCAGAUCACUUA
Sso-1-2      GCA.AAGCAGAUCACUUA
Sso-1-3      GCA.AAGCAGAUCACUUA
Eco-555-1    GCA.AAGCAGAUCACUUA
env-252      GCA.AAGCAGAUCACUUA
Sbo-13-1     GCA.AAGCAGAUCACUUA
Eco-174-1    GCA.AAGCAGAUCACUUA
env-264      GCA.AAGCAGAUCACUUA
env-265      GCA.AAGCAGAUCACUUA
env-266      GCA.AAGCAGAUCACUUA
Eco-1984-1   GCA.AAGCAGAUCACUUA
Sso-1-14     GCA.AAGCAGAUCACUUA
Sso-7-1      GCA.AAGCAGAUCACUUA
Sso-1-539    GCA.AAGCAGAUCACUUA
Esp-78-1     GCA.AAGCAGAUCACUUA
Esp-53-1     GCA.AAGCAGAUCACUUA
Eco-279-1    GCA.AAGCAGAUCACUUA
env-324      GCA.AAGCAGAUCACUUA
env-325      GCA.AAGCAGAUCACUUA
Eco-1084-1   GCA.AAGCAGAUCACUUA
env-326      GCA.AAGCAGAUCACUUA
Eal-2-1      GCA.AAGCAGAUCACUUG
Eco-764-1    GCA.AAGCAGAUCAUUUG
Eco-66-1     GCA.AAGCAGAUCAUUUG
Sfl-76-1     GCA.AAGCAGAUCACUUA
env-327      GCA.AAGCAGAUCGCUUA
Eco-1148-1   GCA.AAGCAGAUCACUUA
Sdy-8-1      GCA.AAGCAGAUCACUUA
Eco-84-1     GCA.AAGCAGAUCACUUA
Cda-1-1      GCAAACGCAGAUAACUUU
Bag-1-1      GCA.AAGCAGAUCACUUA
Bag-2-1      GCA.AAGCAGAUCACUUA
Esp-42-1     GCA.AAGCCGAUCAUCUG
env-328      GCA.AAGCAGAUCACUUA
env-329      GCA.AAGCAGAUCACUUA
Sen-239-1    GCA.AAGCAGAUCACUUA
Sen-728-1    GCA.AAGCAGAUCACUUA
Sen-226-1    GCA.AAGUAGAUCACUUA
Sen-269-1    GCA.AAGCAGAUCACUUA
Sen-225-1    GCA.AAGCAGAUCACUUA
Sen-438-1    GCA.AAGCAGAUCACUUA
Sen-287-1    GCA.AAGCAGAUCACUUA
Sen-1007-1   GCA.AAGCAGAUCACUUA
Sen-438-2    GCA.AAGCAGAUCACUUA

................>>>>>>>>>>>>>>>>>>>>>>>>>>>>>>>>>>>>>>>>>>>>>>>>>>>>>>>>>>>>>>>>>>>>>>.........................<<<<<<<<<<<<<<<<<<<<<<.......>>>>>>>>>>>>>
................001111.....110...............101.........1.1111.111.1.0.....01100....111.........22222212.........................1111.10.11..100.0..00000.......00000..0.001..
                                                                                               ○○○○AAGC●●○○○○○○○○○
```

```
Sen-23-1     ...GCA.AAGCAGAUCACUUA...
Sen-438-3    ...GCA.AAGCAGAUCACUUA...
Sen-67-1     ...GCA.AAGCAGAUCACUUA...
Sen-88-1     ...GCA.AAGCAGAUCACUUA...
Sen-223-1    ...GCA.AAGCAGAUCACUUA...
Sen-927-1    ...GCA.AAGCAGAUCACUUA...
Cro-2-1      ...GCA.AAGCAGAUCGCUUA...
Cko-1-2      ...GCA.AAGCAGAUCACUUA...
Cam-2-1      ...GCA.AAGCAGAUCACUUA...
Sen-1020-1   ...GCA.AAGCAGAUCACUUA...
env-330      ...GCA.AAGCAGAUCACUUA...
env-331      ...GCA.AAGCAGAUCACUUA...
Cse-1-1      ...GCA.AAGCAGAUCACUUA...
Csp-11-1     ...GCA.AAGCAGAUCACUUA...
Cwe-1-1      ...GCA.AAGCAGAUCACUUA...
Sen-310-1    ...GCA.AAGCAGAUCACUUA...
Sen-3-1      ...GCA.AAGCAGAUCACUUA...
Cfr-1-4      ...GCA.AAGCAGAUCACUUA...
Sen-311-1    ...GCA.AAGCAGAUCACUUA...
Sen-1018-1   ...GCA.AAGCAGAUCACUUA...
Cbr-1-5      ...GCA.AAGCAGAUCACUUA...
env-332      ...GCA.AAGCAGAUCACUUA...
env-334      ...GCA.AAGCAGAUCACUUA...
Cfa-1-1      ...GCA.AAGCAGAUCACUUA...
Cpa-1-1      ...GCA.AAGCAGAUCACUUA...
Csp-17-1     ...GCA.AAGCAGAUCACUUA...
Cfr-4-1      ...GCA.AAGCAGAUCACUUA...
Sen-1016-3   ...GCA.AAGCAGAUCACUUA...
Sbo-3-1      ...GCA.AAGCAGAUCACUUA...
Sen-834-1    ...GCA.AAGCAGAUCACUUA...
env-335      ...GCA.AAGCAGAUCACUUA...
Csp-18-1     ...GCA.AAGCAGAUCACUUA...
Pur-1-1      ...GCA.AAGCAGAUCACUUG...
Cma-4-1      ...GCA.AAGCCAGUCAUUUG...
Cun-1-1      ...GCA.AAGCCAGUCAUUUG...
Csa-15-1     ...GCA.AAGCCAGUCAUUUG...
Cma-1-1      ...GCA.AAGCCAGUCAUUUG...
Csa-7-1      ...GCA.AAGCCAGUCAUUUG...
Csa-10-1     ...GCA.AAGCCAGUCAUUUG...
Ctu-3-1      ...GCA.AAGCCAGUCAUUUG...
Cma-1-5      ...GCA.AAGCCAGUCAUUUG...
Fpu-1-1      ...GCA.AAGCCGAUCAUCUA...
Fpu-5-1      ...GCA.AAGCCGAUCAUCUA...
Fhe-4-1      ...GCA.AAGCCGAUCAUCUG...
Fhe-1-1      ...GCA.AAGCCGAUCAUCUG...
Kor-1-1      ...GCA.AAGCCGAUCAUCUG...
Ksa-1-1      ...GCA.AAGCCGAUCAUCUG...
Kra-2-1      ...GCA.AAGCCGAUCAUCUG...
Era-1-1      ...GCA.AAGCCGAUCAUCUG...
Cmu-1-1      ...GCA.AAGCCAGUCAUUUA...
Cdu-2-1      ...GCA.AAGCCAGUCAUUCG...
Cco-1-1      ...GCA.AAGCCAGUCAUUCG...
Cdu-3-1      ...GCA.AAGCCAGUCAUUCG...
env-336      ...GUG.AAGCAGAUCAUCUA...
Kpn-226-1    ...GCA.AAGCAGAUAAUCCG...
Kpn-169-1    ...GCA.AAGCAGAUAAUCCG...
env-337      ...GCA.AAGCAGAUAAUCCG...
Kqu-1-1      ...GCA.AAGCAGAUAAUCCG...
env-343      ...GCA.AAGCAGAUAAUCCG...
env-344      ...GCA.AAGCAGAUAAUCCG...
env-346      ...GCA.AAGCAGAUAAUCUG...
env-347      ...GCA.AAGCAGAUAAUCUG...
env-348      ...GCA.AAGCAGAUAAUCUG...
env-349      ...GCA.AAGCAGAUAAUCUG...
env-350      ...GCA.AAGCAGAUAAUCUG...
env-351      ...GCA.AAGCAGAUAAUCUG...
env-353      ...GCA.AAGCAGAUAAUCUG...
env-354      ...GCA.AAGCAGAUAAUCUG...
Ksp-1-1      ...GCA.GAGCAGAUAAUCCG...
Mma-1-1      ...GCA.AAGCAGAUCAUUCG...
Esp-43-1     ...GCA.AAGCAGAUCACUUG...
Ksp-3-1      ...GCA.AAGCAGAUCAAUUA...
Esp-79-1     ...GCA.AAGCAGAUCACUUA...
env-357      ...GUAAAAGCGUUUGAAUAACCGU...
env-358      ...GUAAAAGCGUAUUGAAUAACCGU...
Pst-5-1      ...GUAAAAGCGUAUUGAAUAACCGU...
env-359      ...GUAAAAGCGUAUUGAAUAACCGU...
Psp-14-1     ...GCAAGAGCGUUUGAAUAAUCGU...
Psp-13-1     ...GCAAGAGCGUUUGAAUAAUCGU...
Ebl-1-1      ...GCA.AGGCAGAAAACUUA...
env-360      ...GCA.AAGCAGAACACUCA...
Kas-1-1      ...GUG.AAGCAGAUCAUUUG...
env-362      ...GUG.AAGCAGAUCAUUUG...
env-363      ....GUGAGGCAGAACACUUA...

................>>>>>>>>>>>>>>>>>>>>>>>>>>>>>>>>>>>>>>>>>>>>>>>>>>>>>>>>>>>>>>>>>>>>>>>...........................<<<<<<<<<<<<<<<<<<<<<<<........>>>>>>>>>>>>>
................001111.....110...............101.........1.1111.111.1.0.....01100....111.........22222212...........................1111.10.11..100.0..00000........00000..0.001..
------------------------------------------------------------------------------------------------------------------------------------------------oooooAAGC●●ooooooooooo--------------------------------------------------
```

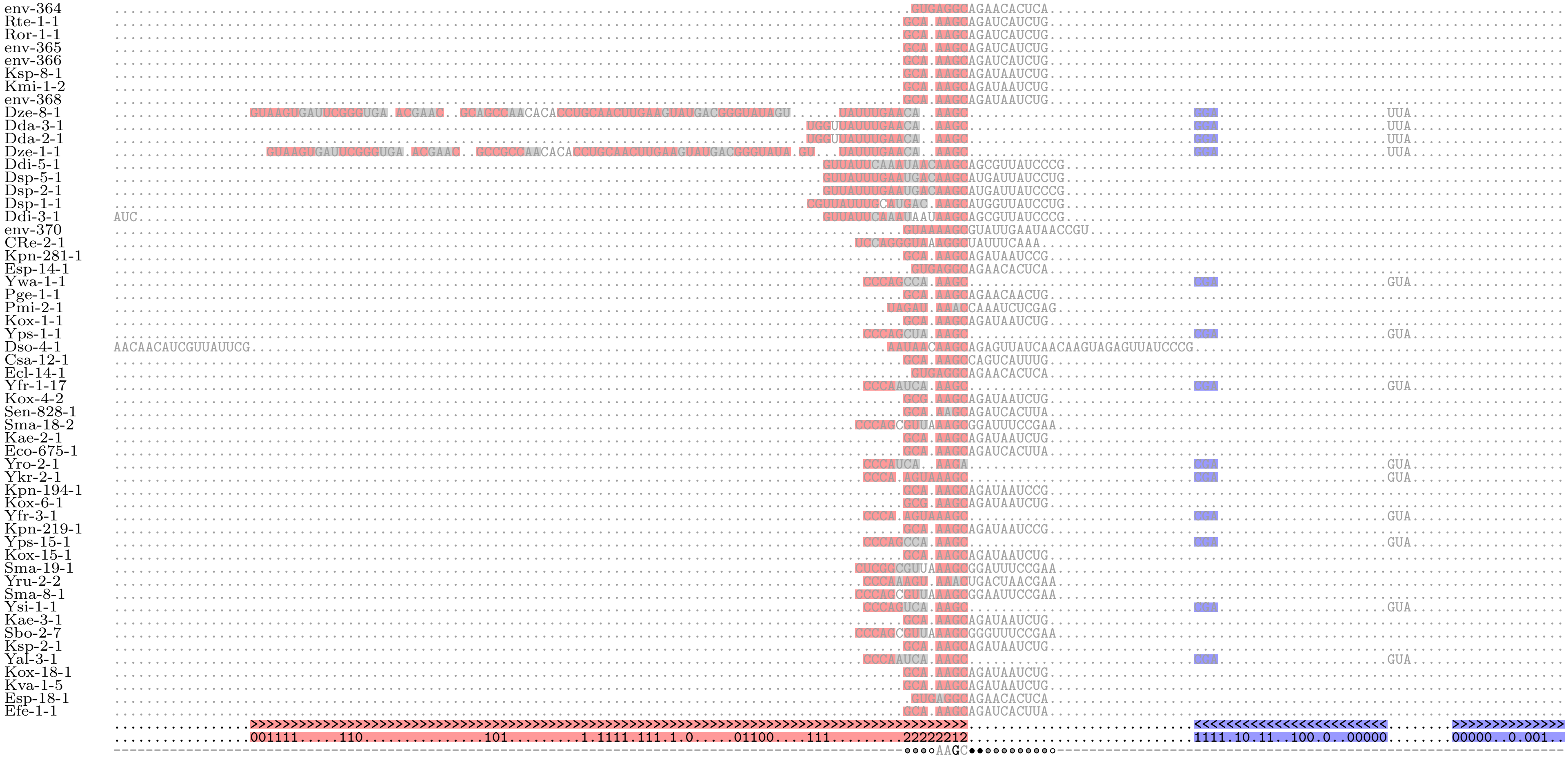

alignment positions 361 · · · 491

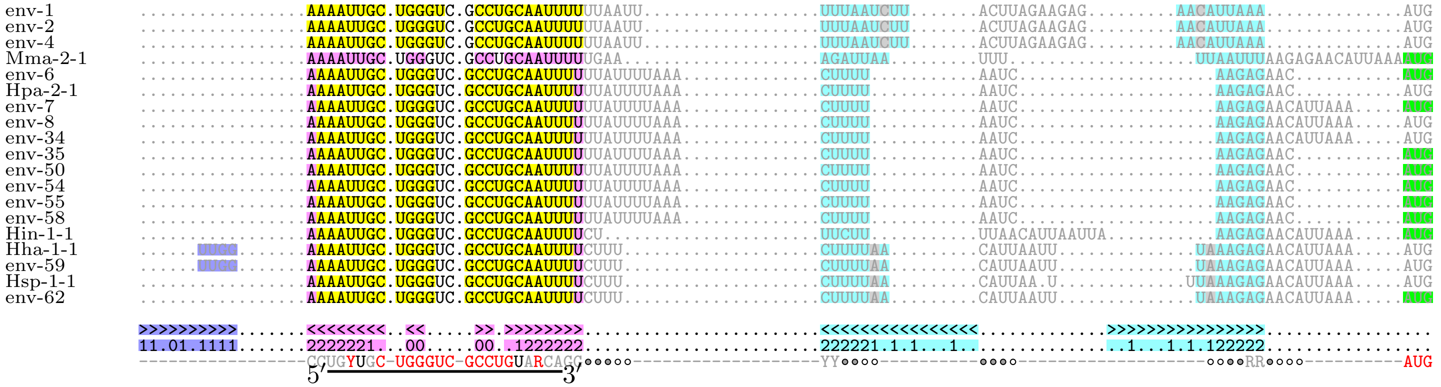

```
Hha-1-2    .................AAAAUUGC.UGGGUC.GCCUGCAAUUUUCUUU...................CUUUUAA.........CAUUAAUU..............UAAAGAGAACAUUAAA.....AUG
env-63     .................AAAAUUGC.UGGGUC.GCCUGCAAUUUUCUUU...................CUUUUAA.........CAUUAAUU..............UAAAGAGAACAUUAAA.....AUG
Hha-3-1    .................AAAAUUGC.UGGGUC.GCCUGCAAUUUUCUUU...................CUUUUAA.........CAUUAAUU..............UAAAGAGAACAUUAAA.....AUG
env-65     .................AAAAUUGC.UGGGUC.GCCUGCAAUUUUCUUU...................CUUUUAA.........CAUUAAUU..............UAAAGAGAACAUUAAA.....AUG
env-66     .................AAAAUUGC.UGGGUC.GCCUGCAAUUUUCUUU...................CUUUUAA.........CAUUAA..............UUUAAAGAGAACAUUAAA.....AUG
Hin-4-1    .................AAAAUUGC.UGGGUC.GCCUGCAAUUUUCUUU...................CUUUUAA.........CAUUAA..............UUUAAAGAGAACAUUAAA.....AUG
env-67     .................AAAAUUGC.UGGGUC.GCCUGCAAUUUUCUUU...................CUUUUAA.........CAUUAA..............UUUAAAGAGAACAUUAAA.....AUG
env-68     .................AAAAUUGC.UGGGUC.GCCUGCAAUUUUCU.....................UUCUUUUAA.......CAUUAAUU............UAAAGAGAACAUUAAA.......AUG
env-70     ......UUGG.......AAAAUUGC.UGGGUC.GCCUGCAAUUUUCU.....................UUCUUUUAA.......CAUUAAUU............UAAAGAGAACAUUAAA.......AUG
env-71     .................AAAAUUGC.UGGGUC.GCCUGCAAUUUUC......................UUUCUUU.........UAACAUUAAUUU...........AAAGAGAACAUUAAA.....AUG
env-72     .................AAAAUUGC.UGGGUC.GCCUGCAAUUUUC......................UUUCUUU.........UAACAUUAAUUU...........AAAGAGAACAUUAAA.....AUG
env-73     .................GGAAUCGC.UGGGUC.GCCUGCGAUUCUUUAA...................UUUUAA..........UCUU....................UAAAAAGAGAACAUUUAAAAUG
Asu-1-1    .................GAAAUUGC.UGGGUC.GCCUGCAAUUUCUGAGAUUAAA.............UCUUA...........CAAUU....................UAAGAGAACAUUAAU...AUG
env-74     .................GAAAUCGC.UGGGUC.GCCUGCGAUUUUUUAAUUUUAA.............UCUUU...........ACAAA....................AGAGAACAUAACA.....AUG
env-75     .................GAAAUCGC.UGGGUC.GCCUGCGAUUUUUUAAUUUUAA.............UCUUU...........ACAAA....................AGAGAACAUAACA.....AUG
env-76     .................GAAAUCGC.UGGGUC.GCCUGCGAUUUUUUAAUUUUAA.............UCUUUU..........A.....................AAGAGAACAUUCAAA.....AUG
env-77     .................GAAAUCGC.UGGGUC.GCCUGCGAUUUUUUAAUUU................UAAUCGUU........ACAAAAGAG.............AAC.AUUACA...........AUG
env-78     .................GAAAUCGC.UGGGUC.GCCUGCGAUUUUUUAAUUUUAA.............UCUUU...........ACA......................AAAGAGAACAUUACA...AUG
env-79     .................GAAAUCGC.UGGGUC.GCCUGCGAUUUUUUAAUUUUAA.............UCUUU...........ACAA.....................AAAGAGAACAUAAAA...AUG
env-80     .................GAAAUCGC.UGGGUC.GCCUGCGAUUUUUUAAUUUUAA.............UCUUU...........ACAAA....................AGAGAACAUAACA.....AUG
env-82     .................GAAAUCGC.UGGGUC.GCCUGCGAUUUUUUAAUUUUAA.............UCUUU...........ACAAA....................AGAGAACAUAACA.....AUG
env-83     .................GAAAUCGC.UGGGUC.GCCUGCGAUUUUUUAAUUUUAA.............UCUUU...........ACAAA....................AGAGAACAUAACA.....AUG
env-85     .................GAAAUCGC.UGGGUC.GCCUGCGAUUUUUUAAUUUUAA.............UCUUU...........ACAAA....................AGAGAACAUAACA.....AUG
env-88     .................GAAAUCGC.UGGGUC.GCCUGCGAUUUUUUAAUUUUAA.............UCUUU...........ACAAA....................AGAGAACAUAACA.....AUG
env-89     .................GAAAUCGC.UGGGUC.GCCUGCGAUUUUUUAAUUUUAA.............UCUUU...........ACAAAA...................AGAGAACAUAACA.....AUG
env-90     .................AAAAUCGC.UGGGUC.GCCUGCGAUUUUUUAAUUUUAA.............UCUUU...........ACAAA....................AGAGAACAUAACA.....AUG
env-91     .................GAAAUCGC.UGGGUC.GCCUGCGAUUUUUUAA...................UUUUAA..........UCUU...................UUAAAGAGAACAUUCAAA..AUG
env-101    .................GAAAUCGC.UGGGUC.GCCUGCGAUUUUUUAA...................UUUUAA..........UCUU...................UUAAAGAGAACAUUCAAA..AUG
env-103    .................GAAAUCGC.UGGGUC.GCCUGCGAUUUUUUAA...................UUUUAA..........UCU....................UUAAAAGAGAACAUUCAAA.AUG
env-104    .................GAAAUCGC.UGGGUC.GCCUGCGAUUUUUUAA...................UUUUAA..........UCU....................UUAAAAGAGAACAUUCAAA.AUG
env-111    .................GAAAUCGC.UGGGUC.GCCUGCGAUUUUUUAA...................UUUUAA..........UCU....................UUAAAAGAGAACAUUCAAA.AUG
env-114    .................CUCGUCAC.UCGGUC.GCAGGUGGCGGGGCACAUUUUAGAAGUAG......UUCU............AAUCG.....................AGAAACAUACA......AUG
env-115    .......AGUCAA....CUCGUCAC.UCGGUC.GCAGGUGGCGGGGCAAAC.................UUUUUUAUA.......................UAUAGAGAAACAUAC.......AUG
env-121    .......AGUCAA....CUCGUCAC.UCGGUC.GCAGGUGGCGGGGCAAA..................CUUU............UUAUAUAU..................AGAGAAACAUAC.....AUG
Aen-1-1    .................UCUGCCAC.UGGGUC.GCCUGUUGCAGAG......................UCUU............UUUAUAUUAU................GAGAACAAAUC......AUG
Ana-1-1    .................UCUGCUAC.UGGGUC.GCCUGUCGCAGAG......................UCUU............UUUACAUUAU................GAGAACAAAUC......AUG
Eba-2-1    ..................CCGCUGC.UGGGUC.GCCUGUAGCGGUAAAU...................UCUUUU..........UGA.....................AAGAGAUAUAAA.......AUG
Dch-3-1    .................CCUGUUGC.UGGGUC.GCCUGUAACAGG.......................GU.CAUUUUAU.....AUAA..............GUAGAGAAAGCA............AUG
Sgl-1-1    ......UGUUUAUCCUGCCUGCUGC.UGGGUC.GCCUGUAGCAGGUUCAA..................UUUCAG..........GUU....................UUGAGAAAAAUU........AUG
Dsp-3-1    .................CCUGUUGC.UGGGUC.GCCUGUAACAGG.......................UUCACUU.........AUCAAA................AAGAGAACGCA..........AUG
Fpe-1-1    .................CCAAUUGC.UGGGUC.GCCUGUGAUUGGU......................UUAUUU..........AACUUAAGAG.............AAAUAAUU............AUG
env-122    .................CCAAUUGC.UGGGUC.GCCUGUGAUUGGU......................UUAUUU..........AACUUAAGAG.............AAAUAAUU............AUG
Mmo-1-1    .................UUUGCUGC.UGGGUC.GCCUGUAGCAGAGAUUU..................UUUUUACUCUU.....GUUA..............AAGAG.AAAAGCU............AUG
Mmo-1-3    .................UUUGCUGC.UGGGUC.GCCUGUAGCAAAGAUUU..................UUUUUACUCUU.....GUUA..............AAGA.GAAAAGUU............AUG
Mmo-3-1    .................UUUGCUGC.UGGGUC.GCCUGUAGCAAAGAUUU..................UUUUUACUCUU.....GUUA..............AAGAG.AAAAGCU............AUG
Mmo-5-1    .................UUUGCUGC.UGGGUC.GCCUGUAGCAAAGAUUU..................UUUUUACUCUU.....GUUA..............AAGAG.AAAAGUU............AUG
Mmo-2-1    .................UUUGCUGC.UGGGUC.GCCUGUAGCAAAGAUUU..................UUUUUACUCUU.....GUUA..............AAGAG.AAAAGUU............AUG
Mmo-1-7    .................UUUGCUGC.UGGGUC.GCCUGUAGCAAAGAUUU..................UUUUUACUCUU.....GUUA..............AAGAG.AAAAGUU............AUG
Pfl-1-1    .......UGAU......UCUGUUGC.UGGGUC.GCCUGUAACAGAAAUUAUUUAAAAGAGA..................................................................AUG
Tpt-1-1    .................UCUGUUGC.UGGGUC.GCCUGUAACAGAAUUACGAAGAGAAAUAAGA...............................................................AUG
Tsa-1-1    .................UCUGUUGC.UGGGUC.GCCUGUAACAGAAUUACCAAGAGAAAUA..................................................................AUG
env-125    ........GCU......UCUGUUGC.UGGGUC.GCCUGUAGCAGAAUUAAAAUGAGAAAUUAAG...............................................................AUG
Mwi-1-1    .................CCUGCUGC.UGGGUC.GCCUGUAGCAGGAUUUUUUA...............UUUUUCUU........UUCUUUAC.............GAGAAAAAAUA...........AUG
Spr-1-1    .................CCUGCUGC.UGGGUC.GCCUGUAGCAGG.......................UUCAA...........UUCCAGGUU...............UUGAGAAAAAUU.......AUG
Eic-2-1    .................UCUGCUGC.UGGGUC.GCCUGUAGCAGGCUUCAUU................UUUUU...........AA.......................AGAGACAGAUCA......AUG
Epi-1-1    .................UCUGCUGC.UGGGUC.GCCUGUAGCAGGCUUCAUU................UUUUU...........AA.......................AGAGACAGAUCA......AUG
Epi-1-2    .................UCUGCUGC.UGGGUC.GCCUGUAGCAGGCU.....................UC.AUUUUU.......UUAA................AGAGAUAGAUCA...........AUG
Eta-5-1    .................UCUGCUGC.UGGGUC.GCCUGUAGCAGGCU.....................UC.AUUUUU.......UUAA................AGAGAUAGAUCA...........AUG
CSo-1-1    .................CCUGCUGC.UGGGUCCGCCUGUAGCAGGUUAAAUUCCA.............GGUUUU..........GAGA...................AAAAUU..............AUG
Gan-5-1    .................CAAGCUAU.UGGGUC.GCCUGUAGCUUG.......................UUUUGUUUAACGACCUUUU...........AGGUCACAACAUAAAGAGAUAUUAAA...AUG
Gan-7-1    .................CAAGCUAC.UGGGUC.GCCUGUAGCUUG.......................UUUUGUUUAACGACCUUUU...........AGGUCACAACAUAAAGAGAUAUUAAA...AUG
Gan-1-2    .................CAAGCUAC.UGGGUC.GCCUGUAGCUUG.......................UUUUGUUUAACGACCUUUU...........AGGUCACAACAUAAAGAGAUAUUAAA...AUG
Gan-1-5    .................CAAGCUAC.UGGGUC.GCCUGUAGCUUG.......................UUUUGUUUAACGACCUUUU...........AGGUCACAACAUAAAGAGAUAUUAAA...AUG
Gan-1-6    .................CAAGCUAC.UGGGUC.GCCUGUAGCUUG.......................UUUUGUUUAACGACCUUUU...........AGGUCACAACAUAAAGAGAUAUUAAA...AUG
Gan-6-1    .................CAAGCUAC.UGGGUC.GCCUGUAGCUUG.......................UUUUGUUUAACGACCUUUU...........AGGUCACAACAUAAAGAGAUAUUAAA...AUG
Gan-1-9    .................CAAGCUAC.UGGGUC.GCCUGUAGCUUG.......................UUUUGUUUAACGACCUUUU...........AGGUCACAACAUAAAGAGAUACUAAA...AUG
Gan-1-10   .................CAAGCUAC.UGGGUC.GCCUGUAGCUUG.......................UUUUGUUUAACGACCUUUU...........AGGUCACAACAUAAAGAGAUACUAAA...AUG
Gan-3-1    .................CAAGCUAC.UGGGUC.GCCUGUAGCUUG.......................UUUUGUUUAACGACCUUUU...........AGGUCACAAUAUAAAGAGAUAUUAAA...AUG
Gan-9-1    .................CAAGCUAC.UGGGUC.GCCUGUAGCUUG.......................UUUUGUCUAACGACC.GUUUA..........GGUCAUAACAUAAAGAGAAUUAAA....AUG
CSc-1-1    .................CCAAUUGC.UGGGUC.GCCUGUAAUUGGUU.....................UUUCU.AUUUUAAAUAGAAACA........UAUUUAAAUAAAGAGACUUAUC......AUG
env-126    .................CCAACUAC.UGGGUC.GCCUGUAAUUGG.......................UUUUUUAUUUG.....AAA...............UAAAUAAGAGAAAUCGAU......AUG
env-127    .................CCAACUAC.UGGGUC.GCCUGUAAUUGG.......................UUUUUUAUUU......GAAAU..............AAAUAAGAGAAAUCAAU......AUG
env-131    .................CCAACUAC.UGGGUC.GCCUGUAAUUGG.......................UUUUUAUUU.......UAAAAU..............AAUUAAGAGAA...........AUG
env-132    .................CCAACUAC.UGGGUC.GCCUGUAAUUGG.......................UUUUUAUUU.......UGAAU...............AAAUAAGAGAAAUCAAU.....AUG
env-134    .................CCAACUAC.UGGGUC.GCCUGUAAUUGG.......................UUUUUAUUU.......UGAAU...............AAAUAAGAGAAAUCAAU.....AUG
env-136    .................CCAACUAC.UGGGUC.GCCUGUAAUUGG.......................UUUUUAUUU.......UGAAU...............AAAUAAGAGAAAUCAAU.....AUG
env-138    .................CCAACUAC.UGGGUC.GCCUGUAAUUGG.......................UUUUUAUUU.......UGAAU...............AAAUAAGAGAAAUCAAU.....AUG
env-139    .................CCAACUAC.UGGGUC.GCCUGUAAUUGGUU.....................UUUAUUUU........AAAAAACUAAGA.........GAAAUAAAC............AUG
Gap-1-1    .................CCAACUAC.UGGGUC.GCCUGUAAUUGGU......................UUUUAUUUU.......AAAAUACUUAAGA.......GAAAUAAAA..............AUG
env-140    .................CCAACUAC.UGGGUC.GCCUGUAAUUGGUU.....................UUUAUUUU........AAAAUAAUUAAGA........GAAAUAAA..............AUG
Dda-1-1    AAGUAUAGUUAUCCCG.CCUGUUGC.UGGGUC.GCCUGUAACAGGUUCA...................UUUUAUC.........AAGUA.................GAGAAAGCA............AUG
Eho-5-1    .................UCUGCUGC.UGGGUC.GCCUGUGGCGGAC......................UUC.AUUUU.......UUUAUC..............GAGAUAGAACC............AUG
Eta-1-1    .................UCUGCUGC.UGGGUC.GCCUGUGGCGGACUUCAUUU...............UUUUAUC...........................GAGAUAGAUCU..........AUG

           >>>>>>>>>>.......<<<<<<<<..<<.....>>.>>>>>>>>.......................<<<<<<<<<<<<<<<<..............>>>>>>>>>>>>>>>>................
           11.01.1111.......2222221...00.....00..1222222.......................222221.1.1...1................1...1.1.122222................
           -----------------CCUGYUGC-UGGGUC-GCCUGUARCAGGooooo------------------YYoooo----------oooo------------------ooooRRoooo----------AUG
                            5'__________________________3'
```

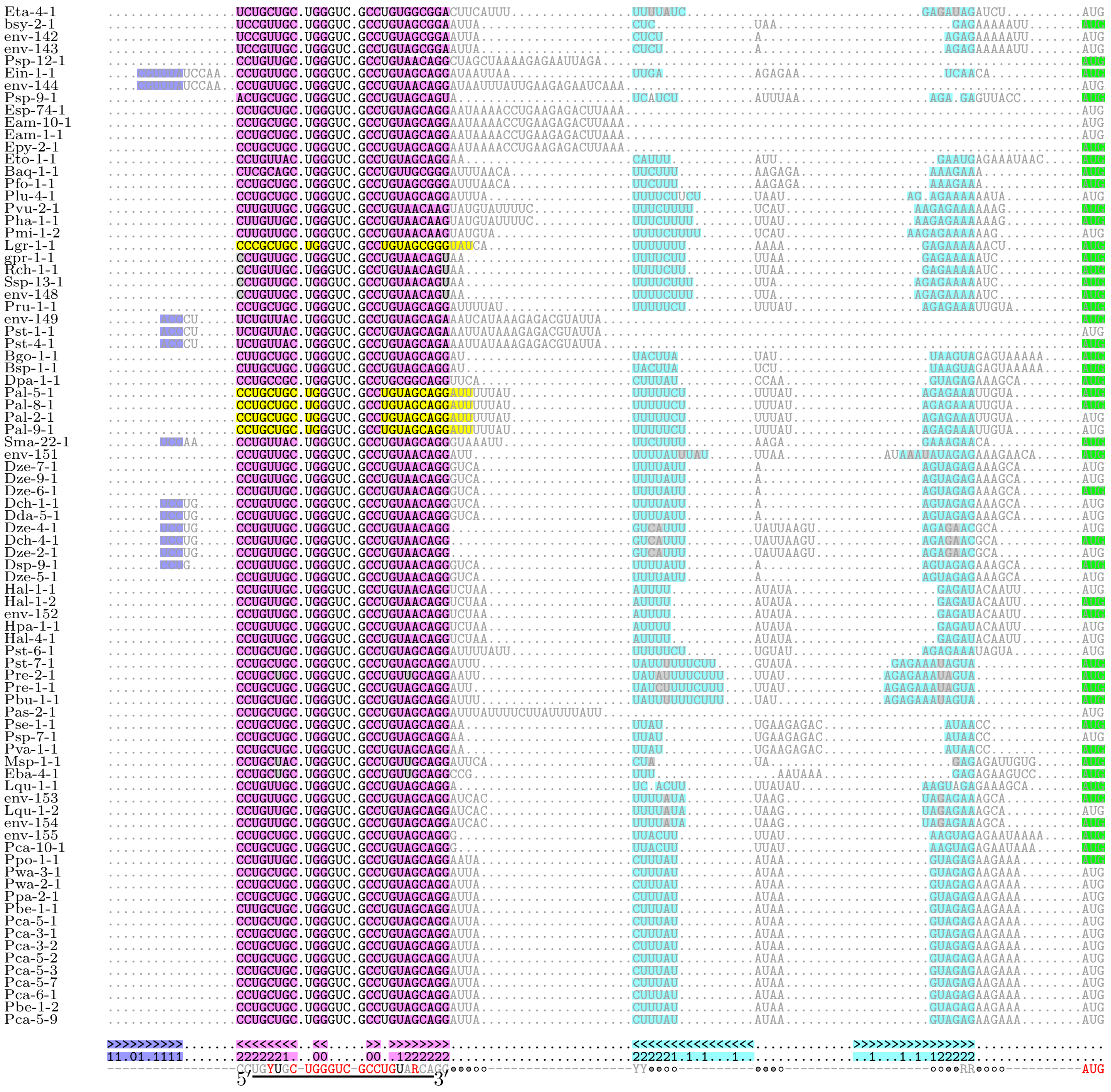
Eta-4-1
bsy-2-1
env-142
env-143
Psp-12-1
Ein-1-1
env-144
Psp-9-1
Esp-74-1
Eam-10-1
Eam-1-1
Epy-2-1
Eto-1-1
Baq-1-1
Pfo-1-1
Plu-4-1
Pvu-2-1
Pha-1-1
Pmi-1-2
Lgr-1-1
gpr-1-1
Rch-1-1
Ssp-13-1
env-148
Pru-1-1
env-149
Pst-1-1
Pst-4-1
Bgo-1-1
Bsp-1-1
Dpa-1-1
Pal-5-1
Pal-8-1
Pal-2-1
Pal-9-1
Sma-22-1
env-151
Dze-7-1
Dze-9-1
Dze-6-1
Dch-1-1
Dda-5-1
Dze-4-1
Dch-4-1
Dze-2-1
Dsp-9-1
Dze-5-1
Hal-1-1
Hal-1-2
env-152
Hpa-1-1
Hal-4-1
Pst-6-1
Pst-7-1
Pre-2-1
Pre-1-1
Pbu-1-1
Pas-2-1
Pse-1-1
Psp-7-1
Pva-1-1
Msp-1-1
Eba-4-1
Lqu-1-1
env-153
Lqu-1-2
env-154
env-155
Pca-10-1
Ppo-1-1
Pwa-3-1
Pwa-2-1
Ppa-2-1
Pbe-1-1
Pca-5-1
Pca-3-1
Pca-3-2
Pca-5-2
Pca-5-3
Pca-5-7
Pca-6-1
Pbe-1-2
Pca-5-9
CCUGYUGC-UGGGUC-GCCUGUARCAGG
YY
RR
AUG
5′
3′

```
Pca-9-1      .................CCUGCUGC.UGGGUC.GCCUGUAGCAGGAUUA....................CUUUAU..........AUAA...................GUAGAGAAGAAA........AUG
Pca-8-1      .................CCUGCUGC.UGGGUC.GCCUGUAGCAGGAUUA....................CUUUAU..........AUAA...................GUAGAGAAGAAA........AUG
Pca-5-10     .................CCUGCUGC.UGGGUC.GCCUGUAGCAGGAUUA....................CUUUAU..........AUAA...................GUAGAGAAGAAA........AUG
Pca-3-15     .................CCUGCUGC.UGGGUC.GCCUGUAGCAGGAUUA....................CUUUAU..........AUAA...................GUAGAGAAGAAA........AUG
Pca-7-1      .................CCUGCUGC.UGGGUC.GCCUGUAGCAGGAUUA....................CUUUAU..........AUAA...................GUAGAGAAGAAA........AUG
Pca-1-1      .................CCUGCUGC.UGGGUC.GCCUGUAGCAGGGUUA....................CUUUAU..........AUAA...................GUAGAGAAAAAA........AUG
Pca-2-1      .................CCUGCUGC.UGGGUC.GCCUGUAGCAGGAUUA....................CUUUAU..........AUAA...................GUAGAGAAGAAA........AUG
Pat-3-1      .......UGAA......CCUGCUGC.UGGGUC.GCCUGUAGCAGGAUUA....................CUUUAU..........AUAA...................GUAGAGAAGAAA........AUG
Pte-1-1      .................CCUGCUGC.UGGGUC.GCCUGUAGCAGGAUUUA...................UUUUCUU.........CUUUA.................AAGAGAAAAAUA.........AUG
Xbo-9-1      .................CCUGCUGC.UGGGUC.GCCUGUAGCAGGAUUUA...................UUUUCUU.........CUUUUACA..............GAGAAAAAAUA..........AUG
Plu-6-1      .................CCUGCUGC.UGGGUC.GCCUGUAGCAGGAUUUA...................UUUUCUU.........CUUA..................AAGAGAAAAAAUA........AUG
Plu-2-1      .................CCUGCUGC.UGGGUC.GCCUGUAGCAGGAUUUA...................UUUUCUU.........CUUAAA................GAGAAAAAACA..........AUG
Xne-3-1      .................CCAGCUGC.UGGGUC.GCCUGUAGCAGGAUUUA...................UUUUCUU.........CUUUUACA..............GAGAAAAAAUA..........AUG
Xkh-1-1      .................CCUGCUGC.UGGGUC.GCCUGUAGCAGGAUUUA...................UUUUCUU.........CUUUUACA..............GAGAAAAAAUA..........AUG
Xne-5-1      .................CCUGCUGC.UGGGUC.GCCUGUAGCAGGAUUUA...................UUUUCUU.........CUUUUACA..............GAGAAAAAAUA..........AUG
Xgr-1-1      .................CCUGCUGC.UGGGUC.GCCUGUAGCAGGAUUUA...................UUUUCUU.........CUUUUACA..............GAGAAAAAAUA..........AUG
Xdo-1-1      .................CCUGCUGC.UGGGUC.GCCUGUAGCAGGAUUUA...................UUUUCUU.........CUUUUACA..............GAGAAAAAAUA..........AUG
Xbo-2-1      .................CCUGCUGC.UGGGUC.GCCUGUAGCAGGAUUUA...................UUUUCUU.........CUUUUACA..............GAGAAAAAAUA..........AUG
Xbo-11-1     .................CCUGCUGC.UGGGUC.GCCUGUAGCAGGAUUUA...................UUUUCUU.........CUUUUACA..............GAGAAAAAAUA..........AUG
Xbo-8-1      .................CCUGCUGC.UGGGUC.GCCUGUAGCAGGAUUUA...................UUUUCUU.........CUUUUACA..............GAGAAAAAAUA..........AUG
Pas-1-1      .................CCUGCUGC.UGGGUC.GCCUGAAGCAGG........................AUUUAUUUUC...CUCUUAAGU.......GAGAAAGAGAAAAAAUA.............AUG
Xpo-1-1      .................CCUGCUGC.UGGGUC.GCCUGUAGCAGG........................AUUUAUUUUCUU....CUUUUC...........AAGAGAAAAAAUA.............AUG
Plu-1-2      .................CCUGCUGC.UGGGUC.GCCUGUAGCAGG........................AUUUAUUUUCUU....CUUA.............AAGAGAAAAAAUA.............AUG
Plu-5-1      .................CCUGCUGC.UGGGUC.GCCUGUAGCAGG........................GUUUAUUUUCUU....CUUA.............AAGAGAAAAAAUA.............AUG
Plu-3-1      .................CCUGCUGC.UGGGUC.GCCUGUAGCAGG........................GUUUACUUUCUU....CUUA.............AAGAGAAAAAACA.............AUG
Pte-2-1      .................CCUGCUGC.UGGGUC.GCCUGUAGCAGG........................AUUUAUUUUCUU....CUUCA............AAGAGAAAAAAUA.............AUG
Pte-3-1      .................CCUGCUGC.UGGGUC.GCCUGUAGCAGG........................AUUUAUUUUCUU....CUUUA............AAGAGAAAAAAUA.............AUG
Xsp-1-1      .................CCUGCUGC.UGGGUC.GCCUGUAGCAGG........................AUUUAUUUUCU.....UCUUUUAU..........AGAGAAAAAAUA.............AUG
Xca-1-1      .................CCUGCUGC.UGGGUC.GCCUGUAGCAGG........................AUUUAUUUUCU.....UCUUUUAU..........AGAGAAAAAAUA.............AUG
Xsz-1-1      .................CCUGCUGC.UGGGUC.GCCUGUAGCAGGAUU.....................UAUUUUCUUCUUU...UCUUAU..........AGAGAA.AAAAUA..............AUG
Spl-2-1      .................CCUGUUAC.UGGGUC.GCCU.UAACAGGGUAAA...................UUUUCUU.........UUAA..................GAGAAAAACA...........AUG
Sfo-4-1      .................CCUGUUAC.UGGGUC.GCCUGUAGCAGGGUAAA...................UUUUCUU.........UUAA..................GAGAAAAAACA..........AUG
Sfo-3-1      .................CCUGUUAC.UGGGUC.GCCUGUAGCAGGGUAAA...................UUUUCUU.........UUAA..................GAGAAAAAACA..........AUG
Sfo-1-1      .................CCUGUUAC.UGGGUC.GCCUGUAGCAGGGUAAA...................UUUUCUU.........UUAA..................GAGAAAAAACA..........AUG
env-157      .................CUUGCUACUUGGGUC.GCCUGUAGCAGGGUAAA...................UUUUCUU.........UUAA..................GAGAAAAAACA..........AUG
Sma-1-2      .................CUUGCUACUUGGGUC.GCCUGUAGCAGGGUAAA...................UUUUCUU.........UUAA..................GAGAAAAAACA..........AUG
Sgr-1-1      GGGUA.AAUGGAA....CUUGCUAC.UGGGUC.GCCUGUAGCAGGGUAAA...................UUUUCUU.........UUAA..................GAGAAAAAACA..........AUG
Sma-1-3      .................CUUGCUACUUGGGUC.GCCUGUAGCAGGGUAAA...................UUUUCUU.........UUAA..................GAGAAAAAACA..........AUG
Sur-1-1      .................CUUGCUACUUGGGUC.GCCUGUAGCAGGGUAAA...................UUUUCUU.........UUAA..................GAGAAAAAACA..........AUG
env-158      .................CUUGCUACUUGGGUC.GCCUGUAGCAGGGUAAA...................UUUUCUU.........UUAA..................GAGAAAAAACA..........AUG
Sma-14-1     .................CUUGCUACUUGGGUC.GCCUGUAGCAGGGUAAA...................UUUUCUU.........UUAA..................GAGAAAAAACA..........AUG
Ssp-8-1      .................CUUGCUACUUGGGUC.GCCUGUAGCAGGGUAAA...................UUUUCUU.........UUAA..................GAGAAAAAACA..........AUG
Ssp-9-1      .................CUUGCUACUUGGGUC.GCCUGUAGCAGGGUAAA...................UUUUCUU.........UUAA..................GAGAAAAAACA..........AUG
Sne-1-1      .................CUUGCUACUUGGGUC.GCCUGUAGCAGGGUAAA...................UUUUCUU.........UUAA..................GAGAAAAAACA..........AUG
Sma-18-1     .................CUUGCUACUUGGGUC.GCCUGUAGCAGGGUAAA...................UUUUCUU.........UUAA..................GAGAAAAAACA..........AUG
Sma-25-1     .................CUUGCUACUUGGGUC.GCCUGUAGCAGGGUAAA...................UUUUCUU.........UUAA..................GAGAAAAAACA..........AUG
Sma-1-1      .................CUUGCUACUUGGGUC.GCCUGUAGCAGGGUAAA...................UUUUCUU.........UUAA..................GAGAAAAAACA..........AUG
env-159      .................CUUGCUACUUGGGUC.GCCUGUAGCAGGGUAAA...................UUUUCUU.........UUAA..................GAGAAAAAACA..........AUG
env-160      .................CUUGCUACUUGGGUC.GCCUGUAGCAGGGUAAA...................UUUUCUU.........UUAA..................GAGAAAAAACA..........AUG
Sli-2-1      .................CUUGCUAC.UGGGUC.GCCUGUAGCAGGGUAAA...................UUUUCUU.........UUAA..................GAGAAAAAACA..........AUG
env-161      .................CUUGCUAC.UGGGUC.GCCUGUAGCAGGUAAU....................UUCUUUU...............................AAGAGAAAAAC..........AUG
Ssp-6-1      .................CCUGUUAC.UGGGUC.GCCUGUAGCAGGGUAAA...................UUUUCUU.........UCAA..................GAGAAAAAACA..........AUG
Ssp-2-1      .................CCUGUUAC.UGGGUC.GCCUGUAGCAGGGUAAA...................UUUUCUU.........UCGA..................GAGAAAAAACA..........AUG
Ssy-2-1      .................CUUGCUAC.UGGGUC.GCCUGUAGCAAAGUAAA...................UUUUCUU.........UUAA..................GAGAAAAAAA...........AUG
Ssy-1-1      .................CUUGCUAC.UGGGUC.GCCUGUAGCAGGGUAAA...................UUUUCUU.........UUAA..................GAGAAAAAAA...........AUG
env-162      .................CUUGCUAC.UGGGUC.GCCUGUAGCAGGGUAAA...................UUUUCUU.........UAA...................GAGAAAAAACA..........AUG
env-163      .................CUUGCUAC.UGGGUC.GCCUGUAGCAGGGUAAA...................UUUUCUU.........UUAA..................GAGAAAAAACA..........AUG
Ssp-14-1     .................CUUGCUAC.UGGGUC.GCCUGUAGCAGGGUAAA...................UUUUCUU.........UUAA..................GAGAAAAAACA..........AUG
Cmu-2-1      .................CUUGUUAC.UGGGUC.GCCUGUAGCAAGGUAAA...................UUUUCUU.........UCUA..................GAGAAAAAACA..........AUG
Spr-2-1      GGGUA.AAUGGAA....CUUGCUAC.UGGGUC.GCCUGUAGCAGGGUAAAU..................UUUCUUU.........UU....................AGAGAAAAAACA.........AUG
Ssp-4-1      .................CCUGUUAC.UGGGUC.GCCUGUAGCAGGAUACA...................UUUUCUU.........AAA...................GAGAAAAAACA..........AUG
Spl-1-1      .................CCUGUUAC.UGGGUC.GCCU.UAACAGGGUAAA...................UUUUCUU.........UUAA..................GAGAAAAAACA..........AUG
Ssp-10-1     .................CCUGUUAC.UGGGUC.GCCU.UAACAGGGUAAA...................UUUUCUU.........UUAA..................GAGAAAAAACA..........AUG
Sod-1-1      .....UGCUUUAUCGAACCUGUUAC.UGGGUC.GCCUGUAGCAGGGUAAA...................UUUUCUU.........UCGA..................GAGAAAAAACA..........AUG
Spl-5-1      .......UGGAA.....CCUGUUAC.UGGGUC.GCCU.UAACAGGGUAAA...................UUUUCUU.........UUAA..................GAGAAAAAACA..........AUG
Ype-132-1    .......UGGAA.....CCUGUUGC.UGGGUC.GCCUGUAGCAGGGUAAA...................UUUUCUU.........UAA...................GAGAAAACAAA..........AUG
Yfr-1-1      .......UGGAA.....CCUGUUGC.UGGGUC.GCCUGUAGCAGGGUAAA...................UUUUCUU.........UAA...................GAGAAAACAAA..........AUG
Yfr-1-4      .......UGGAA.....CCUGUUGC.UGGGUC.GCCUGUAGCAGGGUAAA...................UUUUCUU.........UAA...................GAGAAAACAAA..........AUG
Ymo-1-1      .......UGGAA.....CCUGUUGC.UGGGUC.GCCUGUAGCAGGGUAAA...................UUUUCUU...............................AAGAGAAAACAAA........AUG
Ymo-1-3      .......UGGAA.....CCUGUUGC.UGGGUC.GCCUGUAGCAGGGUAAA...................UUUUCUU...............................AAGAGAAAACAAA........AUG
Yfr-1-6      .......UGGAA.....CCUGUUGC.UGGGUC.GCCUGUAGCAGGGUAAA...................UUUUCUU.........UAA...................GAGAAAACCAA..........AUG
Yfr-1-7      .......UGGAA.....CCUGUUGC.UGGGUC.GCCUGUAGCAGGGUAAA...................UUUUCUU.........UAA...................GAGAAAACCAA..........AUG
Yen-21-1     .......UGGAA.....CCUGUUGC.UGGGUC.GCCUGUAGCAGGGUAAA...................UUUUCUU.........UAA...................GAGAAAACAAA..........AUG
Yal-1-1      .......UGGAA.....CCUGUUGC.UGGGUC.GCCUGUAGCAGGGUAAA...................UUUUCUU.........UAA...................GAGAAAACAAA..........AUG
Yfr-1-9      .......UGGAA.....CCUGUUGC.UGGGUC.GCCUGUAGCAGGGUAAA...................UUUUCUU.........UAA...................GAGAAAACAAA..........AUG
Ype-1-1      .......UGGAA.....CCUGUUGC.UGGGUC.GCCUGUAGCAGGGUAAA...................UUUUCUU.........UAA...................GAGAAAACAAA..........AUG
Yfr-1-10     .......UGGAA.....CCUGUUGC.UGGGUC.GCCUGUAGCAGGGUAAA...................UUUUCUU.........UAA...................GAGAAAACAAA..........AUG
Yfr-2-2      .......UGGAA.....CCUGUUGC.UGGGUC.GCCUGUAGCAGGGUAAA...................UUUUCUU.........UAA...................GAGAAAACAAA..........AUG
Yfr-1-12     .......UGGAA.....CCUGUUGC.UGGGUC.GCCUGUAGCAGGGUAAA...................UUUUCUU.........UAA...................GAGAAAACAAA..........AUG
Yal-1-3      .......UGGAA.....CCUGUUGC.UGGGUC.GCCUGUAGCAGGGUAAA...................UUUUCUU.........UAA...................GAGAAAACAAA..........AUG
Ymo-1-5      .......UGGAA.....CCUGUUGC.UGGGUC.GCCUGUAGCAGGGUAAA...................UUUUCU..........UAAG...................AGAAAACAAA..........AUG
Yma-2-1      .......UGGAA.....CCUGUUGC.UGGGUC.GCCUGUAGCAGGGUAAA...................UUUCUU..........AA.....................GAGAAAAUAAAC........AUG
Yma-1-1      .......UGGAA.....CCUGUUGC.UGGGUC.GCCUGUAGCAGGGUAAA...................UUUCUU..........AA.....................GAGAAAAUAAAC........AUG

             >>>>>>>>>>.......<<<<<<<<..<<.....>>.>>>>>>>>........................<<<<<<<<<<<<<<<<.............>>>>>>>>>>>>>>>>.................
             11.01.1111.......2222221...00.....00..1222222........................222221.1.1...1...............1...1.1.122222.................
             -----------------CCUGYUGC-UGGGUC-GCCUGUARCAGG∘∘∘∘∘-------------------YY∘∘∘∘----------∘∘∘∘-------------------∘∘∘∘RR∘∘∘∘---------AUG
                              5′_________________________3′
```

```
Yfr-1-13  .......UCGAA....CCUGUUGC.UGGGUC.GCCUGUAGCAGGGUAAA...................UUUCUU..........AA.....................GAGAAAAUAAAC........AUG
Ybe-1-1   .......UCGAA....CCUGUUGC.UGGGUC.GCCUGUAGCAGGGUAAA...................UUUCU...........UAAG....................AGAAAACAAA.........AUG
Ynu-1-1   .......UCGAA....CCUGUUGC.UGGGUC.GCCUGUAGCAGGGUAAA...................UUUUCUU.........UUAA..................GAGAAAACAAA..........AUG
Yru-1-1   ................CCUGUUAC.UGGGUC.GCCUGUAGCAGGGUAAA...................UUUUCUU.........UCGA..................GAGAAAACAAA..........AUG
Yru-1-2   ................CCUGUUAC.UGGGUC.GCCUGUAGCAGGGUAAA...................UUUUCUU.........UCGA..................GAGAAAACAAA..........AUG
Ssp-12-1  .......UCCAA....CCUGUUGC.UGGGUC.GCCUGUAGCAGGAUCA....................CUUUUUU.........AAAGUC................GAGAAAGAC............AUG
Pan-5-1   ................CCUGUUAC.UGGGUC.GCCUGUAGCAGGAAUUAAAUAAAGAGACGUAUUA.............................................................AUG
Eas-4-1   ................CCUGCUGC.UGGGUC.GCCUGUAGUAGGGUUUUAAAUAC.............AGA.............GAGAAA....................UCU..............AUG
env-165   ................CCUGCUGC.UGGGUC.GCCUGUAGUAGGGUUUUAAAUAC.............AGA.............GAGAAA....................UCU..............AUG
Lad-1-1   ................CCUGUUGC.UGGGUC.GCCUGUAGCAGGAUUUUAUAUACAGAGAGAAAUCA............................................................AUG
Ecl-37-1  ................CCUGCUGC.UGGGUC.GCCUGUAGUAGGGUUUUAAAUACAGAGAGAAAUCA............................................................AUG
Esp-37-1  ................CCUGCUGC.UGGGUC.GCCUGUAGUAGGGUUUUAAAUACAGAGAGAAAUCA............................................................AUG
Ecl-39-1  ................CCUGCUGC.UGGGUC.GCCUGUAGUAGGGUUUUAAAUACAGAGAGAAAUCA............................................................AUG
env-166   ................CCUGCUGC.UGGGUC.GCCUGUAGCAGGGUUUUAAAUACAGAGAGAAAUCA............................................................AUG
env-167   ................CCUGCUGC.UGGGUC.GCCUGUAGUAGGGUUUAAAAUACAGAGAGAAAUCA............................................................AUG
Elu-1-5   ................CCUGCUGC.UGGGUC.GCCUGUAGUAGGGUUUAAAAUACAGAGAGAAAUCA............................................................AUG
Ecl-58-1  ................CCUGCUGC.UGGGUC.GCCUGUAGUAGGGUUUAAAAUACAGAGAGAAAUCA............................................................AUG
Yre-2-1   ................CCUGCUGC.UGGGUC.GCCUGUAGCAGGAA......................UUAC............AUUAA....................GUAAGAGAGAUAUCA...AUG
env-168   ................UCUGCUGC.UGGGUC.GCCUGUAGCAGAAUUGAAAUAACAGAGAGAUAUCA............................................................AUG
Tod-1-1   ................UCUGCUGC.UGGGUC.GCCUGUAGCAGAAUUGAAAUAACAGAGAGAUAUCA............................................................AUG
Tgu-1-1   ................UCUGCUGC.UGGGUC.GCCUGUAGCAGAAUUGACAUAACAGAGAGAUAUCA............................................................AUG
Psp-5-1   ................CCUGCUGC.UGGGUC.GCCUGUAGCAGGAUAA....................UCA............AAUAAAAGAGAC...............UGAACA...........AUG
Ebi-2-1   ................CCUGUUGC.UGGGUC.GCCUGUAACAGGAUCA....................UCAG............AUAAAGAGA................UUGAACA...........AUG
Ebi-1-1   ................CCUGUUGC.UGGGUC.GCCUGUAACAGGAUCA....................UCAG............AUAAAGAGA................UUGAACA...........AUG
Eol-1-1   ................CCUGCUGC.UGGGUC.GCCUGUAGCAGGAUCA....................UCAG............AUAAAGAGA................UUGAACA...........AUG
Esp-55-1  ................CCUGCUGC.UGGGUC.GCCUGUAGCAGGAUCA....................UCAG............AUAAAGAGA................UUGAACA...........AUG
env-169   ................CCUGCUGC.UGGGUC.GCCUGUAGCAGGA.......................UAAUCAG.........AUAAA..GA...............GAUUGAACA..........AUG
env-170   ................CCUGCUGC.UGGGUC.GCCUGUAGCAGGA.......................UAAUCAG.........AUAAA..GA...............GAUUGAACA..........AUG
Ety-1-1   ................CCUGCUGC.UGGGUC.GCCUGUAGCAGGA.......................UAAUCAG.........AUAAA..GA...............GAUUGAACA..........AUG
env-171   ................CCUGCUGC.UGGGUC.GCCUGUAGCAGGA.......................UAAUC...........AAAUAAAGA...............GAUUGAACA..........AUG
env-172   ................CCUGUUGC.UGGGUC.GCCUGUAGCAGGA.......................UAAUC...........AGAUAAAGA...............GAUUGAACA..........AUG
Pva-1-2   ................CCUGUUAC.UGGGUC.GCCUGUAGCAGGAUUA....................AAUA............AAGAGAUA.................UAUUA.............AUG
Pro-1-1   ................CCUGUUGC.UGGGUC.GCCUGUAGCAGGAACUA...................UUG.............AAGAGAUA..................CAAGA............AUG
env-173   .......CUG......CCUGCUGC.UGGGUC.GCCUGUAGCAGGAAUUAUUGAAGAGAAAUUGAU..............................................................AUG
env-174   ................CCUGUUGC.UGGGUC.GCCUGUAGCAGG........................CAAU............AUUAAAGAGA...............AUUGUC............AUG
env-175   ................CCUGCUGC.UGGGUC.GCCUGUAGCAGGCACUAAUUAAGAGAGAAAAAG..............................................................AUG
Ehe-1-1   ................CCUGUCGC.UGGGUC.GCCUGCGACAGGAUUUAAAGAAGAGAAAAGUUA..............................................................AUG
Pag-1-1   .......ACCAG....CCUGUUAC.UGGGUC.GCCUGUAGCAGGAAUUA...................AAUA............AAGAGAUA.................UAUUA.............AUG
env-177   .......ACCAG....CCUGUUAC.UGGGUC.GCCUGUAGCAGGAAUUA...................AAUA............AAGAGAUA.................UAUUA.............AUG
Pag-1-6   .......ACCAG....CCUGUUAC.UGGGUC.GCCUGUAGCAGGAAUUA...................AAUA............AAGAGAUA.................UAUUA.............AUG
Pan-1-3   .......ACCAG....CCUGUUAC.UGGGUC.GCCUGUAGCAGGAAUUA...................AAUA............AAGAGAUA.................UAUUA.............AUG
Pag-5-1   .......ACCAG....CCUGUUAC.UGGGUC.GCCUGUAGCAGGAAUUA...................AAUA............AAGAGAUA.................UAUUA.............AUG
Pag-4-1   .......ACCAG....CCUGUUAC.UGGGUC.GCCUGUAGCAGGAAUUA...................AAUA............AAGAGAUA.................UAUUA.............AUG
env-178   .......ACCUG....CCUGUUAC.UGGGUC.GCCUGUAGCAGGAUUUA...................AAUA.............AAGAGAUA................UAUUA.............AUG
env-179   .......ACCUG....CCUGUUAC.UGGGUC.GCCUGUAGCAGGAUUUA...................AAUA.............AAGAGAUA................UAUUA.............AUG
env-180   .......ACCUG....CCUGUUAC.UGGGUC.GCCUGUAGCAGGAUUUA...................AAUA.............AAGAGAUA................UAUUA.............AUG
env-181   .......ACCUG....CCUGUUAC.UGGGUC.GCCUGUAGCAGGAUUUA...................AAUA.............AAGAGAUA................UAUUA.............AUG
Pag-3-1   .......ACCUG....CCUGUUAC.UGGGUC.GCCUGUAGCAGGAUUUA...................AAUA.............AAGAGAUA................UAUUA.............AUG
env-182   .......ACCUG....CCUGUUAC.UGGGUC.GCCUGUAGCAGGAUUUA...................AAUA.............AAGAGAUA................UAUUA.............AUG
Pag-1-5   .......ACCUG....CCUGUUAC.UGGGUC.GCCUGUAGCAGGAAUUA...................AAUA.............AAGAGAUA................UAUUA.............AUG
Psp-1-1   .......ACCUG....CCUGUUAC.UGGGUC.GCCUGUAGCAGGAAUUA...................AAUA.............AAGAGAUA................UAUUA.............AUG
env-183   ................CCUGUUGC.UGGGUC.GCCUGUAGCAGGA.......................AUUAUU..........GAAGAGA................AAUAAUG.............AUG
env-184   ................CCUGUUGC.UGGGUC.GCCUGUAGCAGGA.......................AUUAUU..........GAAGAGA................AAUAAUG.............AUG
Eca-1-1   ................UCUGUUGC.UGGGUC.GCCUGUAGCAGAAA......................UGAUU...........GAAGAGA.................AAUCAA.............AUG
Psp-3-1   ................UCUGUUGC.UGGGUC.GCCUGUAGCAGAAA......................UGAUU...........GAAGAGA.................AAUCAA.............AUG
env-189   ................UCUGUUGC.UGGGUC.GCCUGUAGCAGAAA......................UGAUU...........GAAGAGA.................AAUCAA.............AUG
bsy-1-1   ................ACUGCUGC.UGGGUC.GCCUGUAGCAGGAUUAAAAA................UUGA............AGAGAC...................UUAACA............AUG
Epi-2-1   ................CCUGCUGC.UGGGUC.GCCUGUAGCAGGAAUAAAAC................UUGA............AGAGAC...................UUAAA.............AUG
Eta-9-1   ................CCUGCUGC.UGGGUC.GCCUGUAGCAGGAAUAAAAC................UUGA............AGAGAC...................UUAAA.............AUG
env-192   ................CCUGUUGC.UGGGUC.GCCUGUAACAGG........................UGUUAUUU........UAUA.................GAGAGACAUCA...........AUG
Kcr-1-1   ................CCUGCUGC.UGGGUC.GCCUGUAGCAGGAUUUAU..................AUAG............AGAG.....................CUAUCA............AUG
env-194   ................CCUGCUGC.UGGGUC.GCCUGUAGCAGGA.......................UCU.............AUAUAC....................AGAGAGAAAUCA.....AUG
env-195   ................CCUGCUGC.UGGGUC.GCCUGUAGCAGGA.......................UCU.............AUAUAC....................AGAGAGAAAUCA.....AUG
env-196   ................CCUGCUGC.UGGGUC.GCCUGUAGCAGGA.......................UCU.............AUAUAC....................AGAGAGAAAUCA.....AUG
Etr-1-1   ................CCUGUUGC.UGGGUC.GCCUGUAGCAGGAAAAUCAA................UUU.............AA........................GAGACUGAACA......AUG
Ema-2-1   ................CCUGUUGC.UGGGUC.GCCUGUGACAGGAUACUUA.................CUU.............AAA.......................GAGACUGAACA......AUG
env-197   ................CCUGCUGC.UGGGUC.GCCUGUAGCAGGAAUAA...................UAU...........GUACAGAGAGAA................AUAA.............AUG
env-198   ................CCUGCUGC.UGGGUC.GCCUGUAGCAGGAAUAA...................UAU.............GUACAGAG..................AGAAAUCA.........AUG
env-199   ................CCUGCUGC.UGGGUC.GCCUGUAGCAGGAAUAA...................UAU.............GUACAGAG..................AGAAAUCA.........AUG
env-200   ................CCUGCUGC.UGGGUC.GCCUGUAGCAGGACUUA...................UUU.............ACAGAGAG..................AAAUCA...........AUG
env-201   ................CCUGUUGC.UGGGUC.GCCUGUAACAGGAUUUAUUUAA..............GAG.............AGA.......................UUCAA............AUG
Ecl-59-1  ................CCUGUUGC.UGGGUC.GCCUGUAACAGGAUUUAUUUAA..............GAG.............AGA.......................UUCAA............AUG
env-202   ................CCUGUUGC.UGGGUC.GCCUGUAACAGGAUUUAUUUAA..............GAG.............AGA.......................UUCAA............AUG
Eko-1-1   ................CCUGCUGC.UGGGUC.GCCUGUAGUAGGG.......................UUUU............AAAUACAG.................AGAGAAAUCA........AUG
Ecl-33-1  ................CCUGCUGC.UGGGUC.GCCUGUAGUAGGG.......................UUUU............AAAUACAG.................AGAGAAAUCA........AUG
Eko-1-8   ................CCUGCUGC.UGGGUC.GCCUGUAGUAGGG.......................UUUU............AAAUACAG.................AGAGAAAUCA........AUG
Esp-51-1  ................CCUGCUGC.UGGGUC.GCCUGUAGUAGGG.......................UUUU............AAAUACAG.................AGAGAAAUCA........AUG
Ecl-45-1  ................CCUGCUGC.UGGGUC.GCCUGUAGUAGGG.......................UUUU............AAAUACAG.................AGAGAAAUCA........AUG
Ecl-52-1  ................CCUGCUGC.UGGGUC.GCCUGUAGUAGGG.......................UUUUA...........AAUACA..................GAGAGAAAUCA........AUG
Ecl-54-1  ................CCUGCUGC.UGGGUC.GCCUGUAGUAGGG.......................UUUUA...........AAUACA..................GAGAGAAAUCA........AUG
env-204   ................CCUGCUGC.UGGGUC.GCCUGUAGUAGGG.......................UUUUA...........AAUACA..................GAGAGAAAUCA........AUG
Ecl-52-3  ................CCUGCUGC.UGGGUC.GCCUGUAGUAGGG.......................UUUUA...........AAUACA..................GAGAGAAAUCA........AUG
env-205   ................CCUGCUGC.UGGGUC.GCCUGUAGCAGGG.......................UUUUA...........AAUACA..................GAGAGAAAUCA........AUG

          >>>>>>>>>>......<<<<<<<<..<<.....>>.>>>>>>>>........................<<<<<<<<<<<<<<<<.............>>>>>>>>>>>>>>>>................
          11.01.1111......2222221...00.....00..1222222........................222221.1.1...1.............1...1.1.122222................
          ---------------CCUGYUGC-UGGGUC-GCCUGUARCAGG∘∘∘∘∘-----------------YY∘∘∘∘---------∘∘∘∘-----------------∘∘∘∘RR∘∘∘∘---------AUG
                         5′_____________________________3′
```

| | | | | | |
|---|---|---|---|---|---|
| Ecl-51-1 | CCUGCUGC.UGGGUC.GCCUGUAGCAGGG | UUUUA | AAUACA | GAGAGAAAUCA | AUG |
| env-207 | CCUGCUGC.UGGGUC.GCCUGUAGCAGGG | UUUUA | AAUACA | GAGAGAAAUCA | AUG |
| env-208 | CCUGCUGC.UGGGUC.GCCUGUAGCAGGG | UUUUA | AAUACA | GAGAGAAAUCA | AUG |
| env-210 | CCUGCUGC.UGGGUC.GCCUGUAGCAGGG | UUUUA | AAUACA | GAGAGAAAUCA | AUG |
| env-211 | CCUGCUGC.UGGGUC.GCCUGUAGCAGGG | UUUUA | AAUACA | GAGAGAAAUCA | AUG |
| env-212 | CCUGCUGC.UGGGUC.GCCUGUAGUAGGG | UUUUA | AAUACA | GAGAGAAAUCA | AUG |
| env-213 | CCUGCUGC.UGGGUC.GCCUGUAGUAGGG | UUUUA | AAUACA | GAGAGAAAUCA | AUG |
| Esp-19-1 | CCUGCUGC.UGGGUC.GCCUGUAGUAGGG | UUUUA | AAUACA | GAGAGAAAUCA | AUG |
| Ecl-34-1 | CCUGCUGC.UGGGUC.GCCUGUAGUAGGG | UUUUA | AAUACA | GAGAGAAAUCA | AUG |
| env-214 | CCUGCUGC.UGGGUC.GCCUGUAGUAGGG | UUUUA | AAUACA | GAGAGAAAUCA | AUG |
| env-216 | CCUGCUGC.UGGGUC.GCCUGUAGUAGGG | UUUUA | AAUACA | GAGAGAAAUCA | AUG |
| Ecl-42-1 | CCUGCUGC.UGGGUC.GCCUGUAGUAGGG | UUUUA | AAUACA | GAGAGAAAUCA | AUG |
| env-217 | CCUGCUGC.UGGGUC.GCCUGUAGUAGGG | UUUUA | AAUACA | GAGAGAAAUCA | AUG |
| env-218 | CCUGCUGC.UGGGUC.GCCUGUAGUAGGG | UUUUA | AAUACA | GAGAGAAAUCA | AUG |
| env-228 | CCUGCUGC.UGGGUC.GCCUGUAGUAGGG | UUUUA | AAUACA | GAGAGAAAUCA | AUG |
| env-229 | CCUGCUGC.UGGGUC.GCCUGUAGUAGGG | UUUUA | AAUACA | GAGAGAAAUCA | AUG |
| Ecl-29-1 | CCUGCUGC.UGGGUC.GCCUGUAGUAGGG | UUUUA | AAUACA | GAGAGAAAUCA | AUG |
| Esp-65-1 | CCUGCUGC.UGGGUC.GCCUGUAGUAGGG | UUUUA | AAUACA | GAGAGAAAUCA | AUG |
| env-230 | CCUGCUGC.UGGGUC.GCCUGUAGUAGGG | UUUUA | AAUACA | GAGAGAAAUCA | AUG |
| env-232 | CCUGCUGC.UGGGUC.GCCUGUAGUAGGG | UUUUA | AAUACA | GAGAGAAAUCA | AUG |
| Ecl-16-1 | CCUGCUGC.UGGGUC.GCCUGUAGUAGGG | UUUUA | AAUACA | GAGAGAAAUCA | AUG |
| env-233 | CCUGCUGC.UGGGUC.GCCUGUAGUAGGG | UUUUA | AAUACA | GAGAGAAAUCA | AUG |
| env-236 | CCUGUUGC.UGGGUC.GCCUGUAACAGG | UUUUUU | AUUUGAA | GAGAGACAGAA | AUG |
| Sco-1-1 | CCUGUUGC.UGGGUC.GCCUGUAACAGG | UUUUUU | AUUUGAA | GAGAGACAGCA | AUG |
| Psp-10-1 | CCUGUUGC.UGGGUC.GCCUGUAGCAGGAAUUA | UUGA | AGAGAUA | UCAA | AUG |
| Psp-18-1 | CCUGUUGC.UGGGUC.GCCUGUAGCAGGAACUA | UUGA | AGAGACA | UCAA | AUG |
| env-237 | CCUGCUGC.UGGGUC.GCCUGUAGCAGGAAU | AAUU | AAGAGA | GAUUUAA | AUG |
| env-239 | CCUGCUGC.UGGGUC.GCCUGUAGCAGGAAUU | AAUU | AAGAGA | GAUUUAA | AUG |
| Ecl-60-1 | CCUGUUGC.UGGGUC.GCCUGUAACAGGAUUUA | CUUU | AAAG | AGAGACUUCA | AUG |
| Csp-13-1 | CCUGUUGC.UGGGUC.GCCUGUAACAGGA | CUU | AAUUAAGA | GAGAUUUAA | AUG |
| env-240 | CCUGCUGC.UGGGUC.GCCUGUAGCAGGAAUU | AAUA | AAGAGA | UAUUCAA | AUG |
| Cne-1-1 | CCUGCUGC.UGGGUC.GCCUGUAGCAGGCUUA | UUU | AAA | GAGAAACAUUCA | AUG |
| env-241 | CCUGCUGC.UGGGUC.GCCUGUAGCAGG | CUU | AUUUAAA | GAGAAACAGUCA | AUG |
| env-242 | CCUGCUGC.UGGGUC.GCCUGUAGCAGG | CUU | AUUUAAA | GAGAAACAGUCA | AUG |
| env-243 | CCUGCUGC.UGGGUC.G.CUGUAGCAGG | CUU | AUUAAA | GAGAAACAGUCA | AUG |
| env-244 | CCUGCUGC.UGGGUC.GCCUGUAGCAGGCUUA | UUU | AAAGAG | AAACAGUC | AUG |
| Esp-68-1 | CCUGCUGC.UGGGUC.GCCUGUAGCAGGCUUA | UUU | AAAGA | GAAACAGUCA | AUG |
| Cne-1-2 | CCUGCUGC.UGGGUC.GCCUGUAGCAGGCUUA | UUU | AAAGA | GAAACAGUCA | AUG |
| Cne-1-3 | CCUGCUGC.UGGGUC.GCCUGUAGCAGGCUUA | UUU | AAAGA | GAAACAGUCA | AUG |
| Eco-2103-1 | CCUGUUGC.UGGGUC.GCCUGUAGCAGGAUUAA | UUU | AAGA | GAGAAAGAA | AUG |
| Esp-77-1 | CCUGUUGC.UGGGUC.GCCUGUAGCAGGAUUAA | UUU | AAGA | GAGAAAGAA | AUG |
| env-245 | CCUGUUGC.UGGGUC.GCCUGUAGCAGGAUUAA | UUU | AAGA | GAGAAAGAA | AUG |
| Sso-1-2 | CCUGUUGC.UGGGUC.GCCUGUAGCAGGAUUAA | UUU | AAGA | GAGAAAGAA | AUG |
| Sso-1-3 | CCUGUUGC.UGGGUC.GCCUGUAGCAGGAUUAA | UUU | AAGA | GAGAAAGAA | AUG |
| Eco-555-1 | CCUGUUGC.UGGGUC.GCCUGUAGCAGGAUUAA | UUU | AAGA | GAGAAAGAA | AUG |
| env-252 | CCUGUUGC.UGGGUC.GCCUGUAGCAGGAUUAA | UUU | AAGA | GAGAAAGAA | AUG |
| Sbo-13-1 | CCUGUUGC.UGGGUC.GCCUGUAGCAGGAUUAA | UUU | AAGA | GAGAAAGAA | AUG |
| Eco-174-1 | CCUGUUGC.UGGGUC.GCCUGUAGCAGGAUUAA | UUU | AAGA | GAGAAAGAA | AUG |
| env-264 | CCUGUUGC.UGGGUC.GCCUGUAGCAGGAUUAA | UUU | AAGA | GAGAAAGAA | AUG |
| env-265 | CCUGUUGC.UGGGUC.GCCUGUAGCAGGAUUAA | UUU | AAGA | GAGAAAGAA | AUG |
| env-266 | CCUGUUGC.UGGGUC.GCCUGUAGCAGGAUUAA | UUU | AAGA | GAGAAAGAA | AUG |
| Eco-1984-1 | CCUGUUGC.UGGGUC.GCCUGUAGCAGGAUUAA | UUU | AAGA | GAGAAAGAA | AUG |
| Sso-1-14 | CCUGUUGC.UGGGUC.GCCUGUAGCAGGAUUAA | UUU | AAGA | GAGAAAGAA | AUG |
| Sso-7-1 | CCUGUUGC.UGGGUC.GCCUGUAGCAGGAUUAA | UUU | AAGA | GAGAAAGAA | AUG |
| Sso-1-539 | CCUGUUGC.UGGGUC.GCCUGUAGCAGGAUUAA | UUU | AAGA | GAGAAAGAA | AUG |
| Esp-78-1 | CCUGUUGC.UGGGUC.GCCUGUAGCAGGAUUAA | UUU | AAGA | GAGAAAGAA | AUG |
| Esp-53-1 | CCUGUUGC.UGGGUC.GCCUGUAGCAGGAUUAA | UUU | AAGA | GAGAAAGAA | AUG |
| Eco-279-1 | CCUGUUGC.UGGGUC.GCCUGUAGCAGGAUUAA | UUU | AAGA | GAGAAAGAA | AUG |
| env-324 | CCUGUUGC.UGGGUC.GCCUGUAGCAGGAUUAA | UUU | AAGA | GAGAAAGAA | AUG |
| env-325 | CCUGUUGC.UGGGUC.GCCUGUAGCAGGAUUAA | UUU | AAGA | GAGAAAGAA | AUG |
| Eco-1084-1 | CCUGUUGC.UGGGUC.GCCUGUAGCAGGAUUAA | UUU | AAGA | GAGAAAGAA | AUG |
| env-326 | CCUGUUGC.UGGGUC.GCCUGUAACAGGAUUA | UUU | AAGAGA | GAAACAGUUUA | AUG |
| Eal-2-1 | CCUGUUGC.UGGGUC.GCCUGUAGCAGGAUUAA | UUU | AAGAGAG | AAAGAA | AUG |
| Eco-764-1 | CCUGUUGC.UGGGUC.GCCUGUAGCAGGAUUAA | UUU | AAGAGAG | AAAGAA | AUG |
| Eco-66-1 | CCUGUUGC.UGGGUC.GCCUGUAGCAGGAUUAA | UUU | AAGAGAG | AAAGAA | AUG |
| Sfl-76-1 | CUUGUUGC.UGGGUC.GCCUGUAGCAGGAUUAA | UUU | AAGAGAG | AAAGAA | AUG |
| env-327 | CCUGUUGC.UGGGUC.GCCUGUAGCAGGAUUAA | UUU | AAGAGAG | AAAGAA | AUG |
| Eco-1148-1 | CCUGUCGC.UGGGUC.GCCUGUAGCAGGAUUAA | UUU | AAGAGAG | AAAGAA | AUG |
| Sdy-8-1 | CCUGUUGC.UGGGUC.GCCUGUAGCAGGAUUAA | UUU | AAGAGAG | AAAGAA | AUG |
| Eco-84-1 | CCUGUUGC.UGGGUC.GCCUGUAGCAGGAUUAA | UUU | AAGAGAG | AAAGAA | AUG |
| Cda-1-1 | CCUGUUGC.UGGGUC.GCCUGUAGCAGGAGUA | UUU | AAGAGAG | AAACAGUUA | AUG |
| Bag-1-1 | CCUGUUGC.UGGGUC.GCCUGUAACAGGAUUA | UUU | AAGAGAG | AAACAGUUA | AUG |
| Bag-2-1 | CCUGUUGC.UGGGUC.GCCUGUAACAGGAUUA | UUU | AAGAGAG | AAACAGUUA | AUG |
| Esp-42-1 | CCUGUUGC.UGGGUC.GCCUGUAACAGGAAUUA | UUAAA | GAGAGA | UUUAA | AUG |
| env-328 | CCUGCUGC.UGGGUC.GCCUGUAGCAGGAAUUA | UUGAA | GAGAGA | UUCAA | AUG |
| env-329 | CCUGUUGC.UGGUUC.GCCUGUAGCAGGAAUUA | UUAAA | GAGAGA | UUUAA | AUG |
| Sen-239-1 | CCUGUUGC.UGGGUC.GCCUGUAGCAGGAAUUA | UUAAA | GAGAGA | UUUGA | AUG |
| Sen-728-1 | CCUGUUGC.UGGGUC.GCCUGUAGCAGGAAUUA | UUAAA | GAGAGA | UUUAA | AUG |
| Sen-226-1 | CCUGUUGC.UGGGUC.GCCUGUAGCAGGAAUUA | UUAAA | GAGAGA | UUUAA | AUG |
| Sen-269-1 | CCUGUUGC.UGGGUC.GCCUGUAGCAGGAAUUA | UUAAA | GAGAGA | UUUAA | AUG |
| Sen-225-1 | CCUGUUGC.UGGGUC.GCCUGUAGCAGGAAUUA | UUAAA | GAGAGA | UUUAA | AUG |
| Sen-438-1 | CCUGUUGC.UGGGUC.GCCUGUAGCAGGAAUUA | UUAAA | GAGAGA | UUUAA | AUG |
| Sen-287-1 | CCUGUUGC.UGGGUC.GCCUGUAGCAGGAAUUA | UUAAA | GAGAGA | UUUAA | AUG |

```
>>>>>>>>>>.......<<<<<<<<..<<.....>>.>>>>>>>>.........................<<<<<<<<<<<<<<<<.............>>>>>>>>>>>>>>>>.................
11.01.1111.......2222221...00.....00..1222222.........................222221.1.1...1...............1...1.1.122222.................
----------------CCUGYUGC-UGGGUC-GCCUGUARCAGGooooo-----------------YYoooo----------oooo------------------ooooRRoooo----------AUG
                5'                          3'
```

```
Sen-1007-1   ................CCUGUUGC.UGGGUC.GCCUGUAGCAGGAAUUA...................UUAAA...........GAGAGA.................UUUAA.............AUG
Sen-438-2    ................CCUGUUGC.UGGGUC.GCCUGUAGCAGGAAUUA...................UUAAA...........GAGAGA.................UUUAA.............AUG
Sen-23-1     ................CCUGUUGC.UGGGUC.GCCUGUAGCAGGAAUUA...................UUAAA...........GAGAGA.................UUUAA.............AUG
Sen-438-3    ................CCUGUUGC.UGGGUC.GCCUGUAGCAGGAAUUA...................UUAAA...........GAGAGA.................UUUAA.............AUG
Sen-67-1     ................CCUGUUGC.UGGGUC.GCCUGUAGCAGGAAUUA...................UUAAA...........GAGAGA.................UUUAA.............AUG
Sen-88-1     ................CCUGUUGC.UGGGUC.GCCUGUAGCAGGAAUUA...................UUAAA...........GAGAGA.................UUUAA.............AUG
Sen-223-1    ................CCUGUUGC.UGGGUC.GCCUGUAACAGGAAUUA...................UUAAA...........GAGAGA.................UUUAA.............AUG
Sen-927-1    ................CCUGUUGC.UGGGUC.GCCUGUAACAGGAAUUA...................UUAAA...........GAGAGA.................UUUAA.............AUG
Cro-2-1      ................CCUGUUGC.UGGGUC.GCCUGUAGCAGGAAUUA...................UUAAA...........GAGAGA.................UUUAA.............AUG
Cko-1-2      ................CCUGUUGC.UGGGUC.GCCUGUAACAGGAAUUA...................UUUAA...........GAGAGA.................UUUAA.............AUG
Cam-2-1      ................CCUGUUGC.UGGGUC.GCCUGUAACAGGAAUUA...................UUAAA...........GAGAGA.................UUUAA.............AUG
Sen-1020-1   ................CCUGUUGC.UGGGUC.GCCUGUAGCAGGAAUUA...................UUAAA...........GAGAGA.................UUUAA.............AUG
env-330      ................CCUGUUGC.UGGGUC.GCCUGUAGCAGGAAUUA...................UUAAA...........GAGAGA.................UUUAA.............AUG
env-331      ................CCUGUUGC.UGGGUC.GCCUGUAGCAGGAAUUA...................UUAAA...........GAGAGA.................UUUAA.............AUG
Cse-1-1      ................CCUGUUGC.UGGGUC.GCCUGUAGCAGGAAUUA...................UUAAA...........GAGAGA.................UUUAA.............AUG
Csp-11-1     ................CCUGUUGC.UGGGUC.GCCUGUAGCAGGAAUUA...................UUAAA...........GAGAGA.................UUUAA.............AUG
Cwe-1-1      ................CCUGUUGC.UGGGUC.GCCUGUAGCAGGAAUUA...................UUAAA...........GAGAGA.................UUUAA.............AUG
Sen-310-1    ................CCUGUUGC.UGGGUC.GCCUGUAGCAGGAAUUA...................UUAAA...........GAGAGA.................UUUAA.............AUG
Sen-3-1      ................CCUGUUGC.UGGGUC.GCCUGUAGCAGGAAUUA...................UUAAA...........GAGAGA.................UUUAA.............AUG
Cfr-1-4      ................CCUGUUGC.UGGGUC.GCCUGUAGCAGGAAUUA...................UUAAA...........GAGAGA.................UUUAA.............AUG
Sen-311-1    ................CCUGUUGC.UGGGUC.GCCUGUAGCAGGAAUUA...................UUAAA...........GAGAGA.................UUUAA.............AUG
Sen-1018-1   ................CCUGUUGC.UGGGUC.GCCUGUAGCAGGAAUUA...................UUAAA...........GAGAGA.................UUUAA.............AUG
Cbr-1-5      ................CCUGUUGC.UGGGUC.GCCUGUAGCAGGAAUUA...................UUAAA...........GAGAGA.................UUUAA.............AUG
env-332      ................CCUGUUGC.UGGGUC.GCCUGUAGCAGGAAUUA...................UUAAA...........GAGAGA.................UUUAA.............AUG
env-334      ................CCUGUUGC.UGGGUC.GCCUGUAGCAGGAAUUA...................UUAAA...........GAGAGA.................UUUAA.............AUG
Cfa-1-1      ................CCUGUUGC.UGGGUC.GCCUGUAGCAGGAAUUA...................UUAAA...........GAGAGA.................UUUAA.............AUG
Cpa-1-1      ................CCUGUUGC.UGGGUC.GCCUGUAGCAGGAAUUA...................UUAAA...........GAGAGA.................UUUAA.............AUG
Csp-17-1     ................CCUGUUGC.UGGGUC.GCCUGUAGCAGGAAUUA...................UUAAA...........GAGAGA.................UUUAA.............AUG
Cfr-4-1      ................CCUGUUGC.UGGGUC.GCCUGUAGCAGGAAUUA...................UUAAA...........GAGAGA.................UUUAA.............AUG
Sen-1016-3   ................CCUGUUGC.UGGGUC.GCCUGUAGCAGGAAUUA...................UUAAA...........GAGAGA.................UUUAA.............AUG
Sbo-3-1      ................CCUGUUGC.UGGGUC.GCCUGUAGCAGGAAUUA...................UUAAA...........GAGAGA.................UUUAA.............AUG
Sen-834-1    ................CCUGUUGC.UGGGUC.GCCUGUAGCAGGAAUUA...................UUAAA...........AGAGAGA................UUUAA.............AUG
env-335      ................CCUGUUGC.UGGGUC.GCCUGUAGCAGGAAUUA...................UAAA............AGAGAGA..................UUUAA...........AUG
Csp-18-1     ................CCUGUUGC.UGGGUC.GCCUGUAGCAGGAAUUA...................UAAA............AGAGAGA..................UUUAA...........AUG
Pur-1-1      ................CCUGUUGC.UGGGUC.GCCUGUAGCAGGAACUA...................UUGAAG..........AGAG.................UUUUAA.............AUG
Cma-4-1      ................CCUGUUGC.UGGGUC.GCCUGUAACAGGUUUG....................UUU.............AUUA.....................AGAGAG..........AUG
Cun-1-1      ................CCUGUUGC.UGGGUC.GCCUGUAACAGG........................UUUCUUU.........AUUA...............AGAGAGAUGUCA........AUG
Csa-15-1     ................CCUGUUGC.UGGGUC.GCCUGUAACAGG........................UUUCUUU.........AUUA...............AGAGAGAUGUCA........AUG
Cma-1-1      ................CCUGUUGC.UGGGUC.GCCUGUAACAGG........................UUUCUUU.........AUUA...............AGAGAGAUGUCA........AUG
Csa-7-1      ................CCUGUUGC.UGGGUC.GCCUGUAACAGG........................UUUCUUU.........AUUA...............AGAGAGAUGUCA........AUG
Csa-10-1     ................CCUGUUGC.UGGGUC.GCCUGUAACAGG........................UUUCUUU.........AUUA...............AGAGAGAUGUCA........AUG
Ctu-3-1      ................CCUGUUGC.UGGGUC.GCCUGUAACAGG........................UUUCUUU.........AUUA...............AGAGAGAUGUCA........AUG
Cma-1-5      ................CCUGUUGC.UGGGUC.GCCUGUAACAGG........................UUUCUUU.........AUUA...............AGAGAGAUGUCA........AUG
Fpu-1-1      ................CCUGUUGC.UGGGUC.GCCUGUAACAGGAUUA....................UUU.............AUAGAG..................AGACAGUC.........AUG
Fpu-5-1      ................CCUGUUGC.UGGGUC.GCCUGUAACAGGAUUA....................UUU.............AUAGAG..................AGACAGUC.........AUG
Fhe-4-1      ................CCUGUUGC.UGGGUC.GCCUGUAACAGGAUUUA...................UUU.............AUAGA...................GAGACAGUU........AUG
Fhe-1-1      ................CCUGUUGC.UGGGUC.GCCUGUAACAGGAUUUA...................UUU.............AUAGA...................GAGACAGUU........AUG
Kor-1-1      ................CCUGUUGC.UGGGUC.GCCUGUAACAGGAUU.....................AUUU............AAGAG.................AGAUUCAA..........AUG
Ksa-1-1      ................CCUGUUGC.UGGGUC.GCCUGUAACAGGAUU.....................AUUU............AAGAG.................AGAUUCAA..........AUG
Kra-2-1      ................CCUGUUGC.UGGGUC.GCCUGUAACAGGAUUU....................AUUU............AAGAG.................AGAUUUAA..........AUG
Era-1-1      ................CCUGUUGC.UGGGUC.GCCUGUAACAGGAUUU....................AUUU............AAGAG.................AGAUUUAA..........AUG
Cmu-1-1      ................CCUGUUGC.UGGGUC.GCCUGUAACAGGC.......................UUCUUU..........AUUA...............AGAGAGAUGUCA........AUG
Cdu-2-1      ................CCUGUUGC.UGGGUC.GCCUGUAACAGGCUU.....................CUUUAUU...........................AAGAGAG.............AUG
Cco-1-1      ................CCUGUUGC.UGGGUC.GCCUGUAACAGGCUU.....................CUUUAUU...........................AAGAGAG.............AUG
Cdu-3-1      ................CCUGUUGC.UGGGUC.GCCUGUAACAGGCUU.....................CUUUAUU...........................AAGAGAG.............AUG
env-336      ................CCUGUUAC.UGGGUC.GCCUGUAGCAGGAAU.....................AUUU............AAAGAG................AGAUUCAA..........AUG
Kpn-226-1    ................CCUGUUGC.UGGGUC.GCCUGUAGCAGG........................CUUU............AUAUAG................AGAGACAUCA........AUG
Kpn-169-1    ................CCUGUUGC.UGGGUC.GCCUGUAGCAGG........................CUUUAUA...........................UAUAGAGAGACAUCA.....AUG
env-337      ................CCUGUUGC.UGGGUC.GCCUGUAGCAGG........................CUUUAUA...........................UAUAGAGAGACAUCA.....AUG
Kqu-1-1      ................CCUGUUGC.UGGGUC.GCCUGUAGCAGG........................CUUUAUA...........................UAUAGAGAGACAUCA.....AUG
env-343      ................CCUGUUGC.UGGGUC.GCCUGUAGCAGG........................CUUUAUA...........................UAUAGAGAGACAUCA.....AUG
env-344      ................CCUGUUGC.UGGGUC.GCCUGUAGCAGG........................CUUUAUA...........................UAUAGAGAGACAUCA.....AUG
env-346      ................CCUGCAGC.UGGGUC.GCCUGUUGCAGG........................CUUUAUA...........................UAUAGAGAGACAUCA.....AUG
env-347      ................CCUGCAGC.UGGGUC.GCCUGUUGCAGG........................CUUUAUA...........................UAUAGAGAGACAUCA.....AUG
env-348      ................CCUGCAGC.UGGGUC.GCCUGUUGCAGG........................CUUUAUA...........................UAUAGAGAGACAUCA.....AUG
env-349      ................CCUGCUGC.UGGGUC.GCCUGUUGCAGG........................CUUUAUA...........................UAUAGAGAGACAUCA.....AUG
env-350      ................CCUGCUGC.UGGGUC.GCCUGUUGCAGG........................CUUUAUA...........................UAUAGAGAGACAUCA.....AUG
env-351      ................CCUGCUGC.UGGGUC.GCCUGUUGCAGG........................CUUUAUA...........................UAUAGAGAGACAUCA.....AUG
env-353      ................CCUGCUGC.UGGGUC.GCCUGUUGCAGG........................CUUUAUA...........................UAUAGAGAGACAUCA.....AUG
env-354      ................CCUGCUGC.UGGGUC.GCCUGUUGCAGG........................CUUUAUA...........................UAUAGAGAGACAUCA.....AUG
Ksp-1-1      ................CCUGUUGC.UGGGUC.GCCUGUAGCAGG........................CUUUA...........UAUA.................UAGAGAGACAUCA.....AUG
Mma-1-1      ................CCUGUUGC.UGGGUC.GCCUGUAGCAGGCACU....................AUU.............AAAGAG..................AAUUGUC..........AUG
Esp-43-1     ................CCUGUUGC.UGGGUC.GCCUGUAGCAGGAACUAUUUAAGAGAAAUUGAC..................................................................AUG
Ksp-3-1      ................CCUGUUAC.UGGGUC.GCCUGUAGCAGGAUUUAUUUAAGAGAGAAAUUA..................................................................AUG
Esp-79-1     ................CCUGUUGC.UGGGUC.GCCUGUAGCAGGAAUCAAUUAAGAGAGAAAGAA..................................................................AUG
env-357      ................CCUGUUGC.UGGGUC.GCCUGUAGCAGGAACUAUUGAAGAGAU.......................................................................AUG
env-358      ................CCUGUUGC.UGGGUC.GCCUGUAGCAGGAACUAUAGAAGAGAUCUUAAG..................................................................AUG
Pst-5-1      ................CCUGUUGC.UGGGUC.GCCUGUAGCAGGAACUAUAGAAGAGAUCUUAAG..................................................................AUG
env-359      ................CCUGUUGC.UGGGUC.GCCUGUAGCAGGAACUAUAGAAGAGAUCUUAAG..................................................................AUG
Psp-14-1     ................CCUGUUGC.UGGGUC.GCCUGUAGCAGGAAUCAACGAAGAGACAUUAAG..................................................................AUG
Psp-13-1     ................CCUGUUGC.UGGGUC.GCCUGUAGCAGGAAUCAACGAAGAGACAUUAAG..................................................................AUG
Ebl-1-1      ................CCUGUUGC.UGGGUC.GCCUGUGACAGGCACUAUUUAUAGAGAGUUAUCA.................................................................AUG
env-360      ................CCUGCUGC.UGGGUC.GCCUGUAGUAGGGUUUUAAAAUACAGAGAGAAAUCA...............................................................AUG

             >>>>>>>>>>.......<<<<<<<<..<<.....>>.>>>>>>>>........................<<<<<<<<<<<<<<<<.............>>>>>>>>>>>>>>>>................
             11.01.1111.......2222221...00.....00..1222222........................222221.1.1...1...............1...1.1.122222................
             -----------------CCUGYUGC-UGGGUC-GCCUGUARCAGGooooo-----------------YYoooo----------oooo-------------------ooooRRoooo----------AUG
                             5′                            3′
```

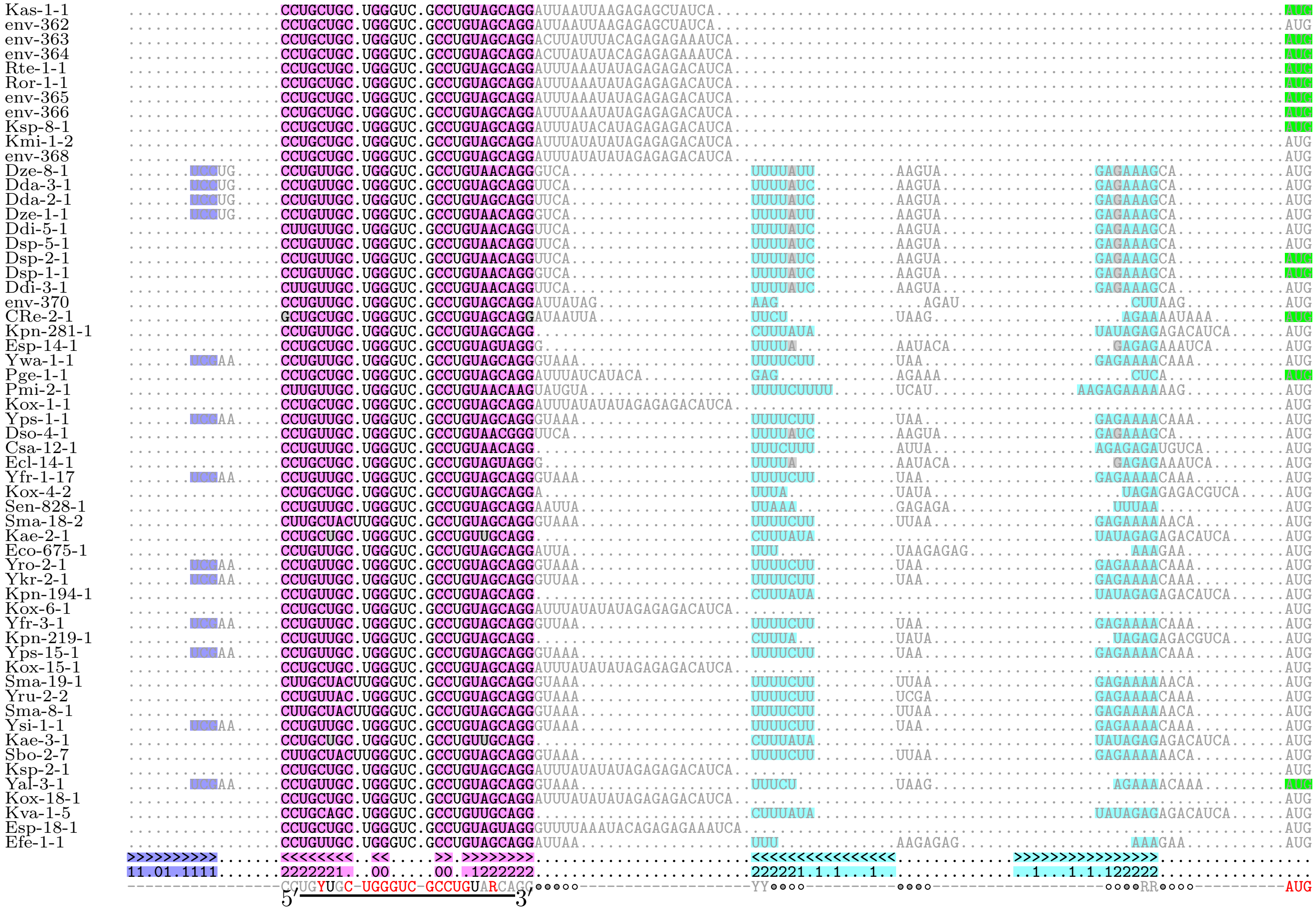

### 9 L31-Actinobacteria

#### 9.1 Taxa

The taxonomy of each organism containing a putative L31-Actinobacteria RNA is listed, with abbreviations identifying each hit (e.g., "Eco-1-1" and "Eco-1-2" might hypothetically represent two distinct RNAs in *E. coli*). The abbreviations will be used to identify each individual L31-Actinobacteria RNA in Sections 9.2 and 9.4:.

| **abbrev. of hits** | **taxonomy of species** |
| --- | --- |
| Ama-1-1 | Bacteria Actinobacteria Actinobacteridae Actinomycetales Actinomycineae Actinomycetaceae *Actinomyces massiliensis* 4401292 |

| | |
|---|---|
| Ama-2-1 | Bacteria Actinobacteria Actinobacteridae Actinomycetales Actinomycineae Actinomycetaceae *Actinomyces massiliensis* F0489 |
| Ane-1-1 | Bacteria Actinobacteria Actinobacteridae Actinomycetales Actinomycineae Actinomycetaceae *Actinomyces neuii* BVS029A5 |
| Vca-1-1 | Bacteria Actinobacteria Actinobacteridae Actinomycetales Actinomycineae Actinomycetaceae *Varibaculum cambriense* DSM 15806 |
| Aha-1-1 | Bacteria Actinobacteria Actinobacteridae Actinomycetales Actinopolysporineae Actinopolysporaceae *Actinopolyspora halophila* DSM 43834 |
| Cac-1-1 | Bacteria Actinobacteria Actinobacteridae Actinomycetales Catenulisporineae Catenulisporaceae *Catenulispora acidiphila* DSM 44928 |
| Gai-1-1 | Bacteria Actinobacteria Actinobacteridae Actinomycetales Corynebacterineae Gordoniaceae *Gordonia aichiensis* NBRC 108223 |
| Gal-1-1 | Bacteria Actinobacteria Actinobacteridae Actinomycetales Corynebacterineae Gordoniaceae *Gordonia alkanivorans* NBRC 16433 |
| Gam-1-1 | Bacteria Actinobacteria Actinobacteridae Actinomycetales Corynebacterineae Gordoniaceae *Gordonia amarae* NBRC 15530 |
| Gam-2-1 | Bacteria Actinobacteria Actinobacteridae Actinomycetales Corynebacterineae Gordoniaceae *Gordonia amicalis* NBRC 100051 = JCM 11271 |
| Gar-1-1 | Bacteria Actinobacteria Actinobacteridae Actinomycetales Corynebacterineae Gordoniaceae *Gordonia araii* NBRC 100433 |
| Gbr-1-1 | Bacteria Actinobacteria Actinobacteridae Actinomycetales Corynebacterineae Gordoniaceae *Gordonia bronchialis* DSM 43247 |
| Gef-1-1 | Bacteria Actinobacteria Actinobacteridae Actinomycetales Corynebacterineae Gordoniaceae *Gordonia effusa* NBRC 100432 |
| Ghi-1-1 to Ghi-1-2 | Bacteria Actinobacteria Actinobacteridae Actinomycetales Corynebacterineae Gordoniaceae *Gordonia hirsuta* DSM 44140 = NBRC 16056 |
| Gkr-1-1 | Bacteria Actinobacteria Actinobacteridae Actinomycetales Corynebacterineae Gordoniaceae *Gordonia kroppenstedtii* DSM 45133 |
| Gma-1-1 | Bacteria Actinobacteria Actinobacteridae Actinomycetales Corynebacterineae Gordoniaceae *Gordonia malaquae* NBRC 108250 |
| Gna-1-1 | Bacteria Actinobacteria Actinobacteridae Actinomycetales Corynebacterineae Gordoniaceae *Gordonia namibiensis* NBRC 108229 |
| Gne-1-1 | Bacteria Actinobacteria Actinobacteridae Actinomycetales Corynebacterineae Gordoniaceae *Gordonia neofelifaecis* NRRL B-59395 |
| Got-1-1 | Bacteria Actinobacteria Actinobacteridae Actinomycetales Corynebacterineae Gordoniaceae *Gordonia otitidis* NBRC 100426 |
| Gpa-1-1 | Bacteria Actinobacteria Actinobacteridae Actinomycetales Corynebacterineae Gordoniaceae *Gordonia paraffinivorans* NBRC 108238 |
| Gpo-1-1 | Bacteria Actinobacteria Actinobacteridae Actinomycetales Corynebacterineae Gordoniaceae *Gordonia polyisoprenivorans* HW436 |
| Gpo-2-1 | Bacteria Actinobacteria Actinobacteridae Actinomycetales Corynebacterineae Gordoniaceae *Gordonia polyisoprenivorans* NBRC 16320 |
| Gpo-3-1 | Bacteria Actinobacteria Actinobacteridae Actinomycetales Corynebacterineae Gordoniaceae *Gordonia polyisoprenivorans* VH2 |
| Grh-1-1 | Bacteria Actinobacteria Actinobacteridae Actinomycetales Corynebacterineae Gordoniaceae *Gordonia rhizosphera* NBRC 16068 |
| Gru-1-1 | Bacteria Actinobacteria Actinobacteridae Actinomycetales Corynebacterineae Gordoniaceae *Gordonia rubripertincta* NBRC 101908 |
| Gsi-1-1 | Bacteria Actinobacteria Actinobacteridae Actinomycetales Corynebacterineae Gordoniaceae *Gordonia sihwensis* NBRC 108236 |
| Gso-1-1 | Bacteria Actinobacteria Actinobacteridae Actinomycetales Corynebacterineae Gordoniaceae *Gordonia soli* NBRC 108243 |
| Gsp-1-1 | Bacteria Actinobacteria Actinobacteridae Actinomycetales Corynebacterineae Gordoniaceae *Gordonia sp.* KTR9 |
| Gsp-2-1 | Bacteria Actinobacteria Actinobacteridae Actinomycetales Corynebacterineae Gordoniaceae *Gordonia sputi* NBRC 100414 |
| Gte-1-1 | Bacteria Actinobacteria Actinobacteridae Actinomycetales Corynebacterineae Gordoniaceae *Gordonia terrae* C-6 |
| Gte-2-1 | Bacteria Actinobacteria Actinobacteridae Actinomycetales Corynebacterineae Gordoniaceae *Gordonia terrae* NBRC 100016 |
| Asu-1-1 | Bacteria Actinobacteria Actinobacteridae Actinomycetales Corynebacterineae Mycobacteriaceae *Amycolicicoccus subflavus* DQS3-9A1 |
| Mab-1-1 to Mab-1-166 | Bacteria Actinobacteria Actinobacteridae Actinomycetales Corynebacterineae Mycobacteriaceae *Mycobacterium abscessus* |
| Mch-1-1 | Bacteria Actinobacteria Actinobacteridae Actinomycetales Corynebacterineae Mycobacteriaceae *Mycobacterium chubuense* NBB4 |
| Mfo-1-1 | Bacteria Actinobacteria Actinobacteridae Actinomycetales Corynebacterineae Mycobacteriaceae *Mycobacterium fortuitum subsp. fortuitum* DSM 46621 |
| Mgi-1-1 | Bacteria Actinobacteria Actinobacteridae Actinomycetales Corynebacterineae Mycobacteriaceae *Mycobacterium gilvum* PYR-GCK |
| Mgi-2-1 | Bacteria Actinobacteria Actinobacteridae Actinomycetales Corynebacterineae Mycobacteriaceae *Mycobacterium gilvum* Spyr1 |
| Mha-1-1 | Bacteria Actinobacteria Actinobacteridae Actinomycetales Corynebacterineae Mycobacteriaceae *Mycobacterium hassiacum* DSM 44199 |
| Mma-1-1 | Bacteria Actinobacteria Actinobacteridae Actinomycetales Corynebacterineae Mycobacteriaceae *Mycobacterium massiliense* CCUG 48898 |
| Mph-1-1 | Bacteria Actinobacteria Actinobacteridae Actinomycetales Corynebacterineae Mycobacteriaceae *Mycobacterium phlei* RIVM601174 |
| Mrh-1-1 | Bacteria Actinobacteria Actinobacteridae Actinomycetales Corynebacterineae Mycobacteriaceae *Mycobacterium rhodesiae* JS60 |
| Mrh-2-1 | Bacteria Actinobacteria Actinobacteridae Actinomycetales Corynebacterineae Mycobacteriaceae *Mycobacterium rhodesiae* NBB3 |
| Msp-1-1 | Bacteria Actinobacteria Actinobacteridae Actinomycetales Corynebacterineae Mycobacteriaceae *Mycobacterium sp.* 141 |
| Msp-2-1 | Bacteria Actinobacteria Actinobacteridae Actinomycetales Corynebacterineae Mycobacteriaceae *Mycobacterium sp.* 155 |
| Msp-3-1 | Bacteria Actinobacteria Actinobacteridae Actinomycetales Corynebacterineae Mycobacteriaceae *Mycobacterium sp.* 360MFTsu5.1 |
| Msp-4-1 | Bacteria Actinobacteria Actinobacteridae Actinomycetales Corynebacterineae Mycobacteriaceae *Mycobacterium sp.* JDM601 |
| Msp-5-1 | Bacteria Actinobacteria Actinobacteridae Actinomycetales Corynebacterineae Mycobacteriaceae *Mycobacterium sp.* JLS |
| Msp-6-1 | Bacteria Actinobacteria Actinobacteridae Actinomycetales Corynebacterineae Mycobacteriaceae *Mycobacterium sp.* KMS |
| Msp-7-1 | Bacteria Actinobacteria Actinobacteridae Actinomycetales Corynebacterineae Mycobacteriaceae *Mycobacterium sp.* MCS |
| Mth-1-1 | Bacteria Actinobacteria Actinobacteridae Actinomycetales Corynebacterineae Mycobacteriaceae *Mycobacterium thermoresistibile* ATCC 19527 |
| Mva-1-1 | Bacteria Actinobacteria Actinobacteridae Actinomycetales Corynebacterineae Mycobacteriaceae *Mycobacterium vaccae* ATCC 25954 |
| Mva-2-1 | Bacteria Actinobacteria Actinobacteridae Actinomycetales Corynebacterineae Mycobacteriaceae *Mycobacterium vanbaalenii* PYR-1 |
| Mab-2-1 | Bacteria Actinobacteria Actinobacteridae Actinomycetales Corynebacterineae Mycobacteriaceae Mycobacterium Mycobacteriumabscessus *Mycobacterium abscessus* 3A-0119-R |
| Mab-3-1 | Bacteria Actinobacteria Actinobacteridae Actinomycetales Corynebacterineae Mycobacteriaceae Mycobacterium Mycobacteriumabscessus *Mycobacterium abscessus* 3A-0122-R |
| Mab-4-1 | Bacteria Actinobacteria Actinobacteridae Actinomycetales Corynebacterineae Mycobacteriaceae Mycobacterium Mycobacteriumabscessus *Mycobacterium abscessus* 3A-0122-S |
| Mab-5-1 | Bacteria Actinobacteria Actinobacteridae Actinomycetales Corynebacterineae Mycobacteriaceae Mycobacterium Mycobacteriumabscessus *Mycobacterium abscessus* 3A-0731 |
| Mab-6-1 | Bacteria Actinobacteria Actinobacteridae Actinomycetales Corynebacterineae Mycobacteriaceae Mycobacterium Mycobacteriumabscessus *Mycobacterium abscessus* 3A-0810-R |

| | |
|---|---|
| Mab-7-1 | Bacteria Actinobacteria Actinobacteridae Actinomycetales Corynebacterineae Mycobacteriaceae Mycobacterium Mycobacteriumabscessus *Mycobacterium abscessus* 3A-0930-R |
| Mab-8-1 | Bacteria Actinobacteria Actinobacteridae Actinomycetales Corynebacterineae Mycobacteriaceae Mycobacterium Mycobacteriumabscessus *Mycobacterium abscessus* 3A-0930-S |
| Mab-9-1 | Bacteria Actinobacteria Actinobacteridae Actinomycetales Corynebacterineae Mycobacteriaceae Mycobacterium Mycobacteriumabscessus *Mycobacterium abscessus* 47J26 |
| Mab-10-1 | Bacteria Actinobacteria Actinobacteridae Actinomycetales Corynebacterineae Mycobacteriaceae Mycobacterium Mycobacteriumabscessus *Mycobacterium abscessus* 4S-0116-R |
| Mab-11-1 | Bacteria Actinobacteria Actinobacteridae Actinomycetales Corynebacterineae Mycobacteriaceae Mycobacterium Mycobacteriumabscessus *Mycobacterium abscessus* 4S-0116-S |
| Mab-12-1 | Bacteria Actinobacteria Actinobacteridae Actinomycetales Corynebacterineae Mycobacteriaceae Mycobacterium Mycobacteriumabscessus *Mycobacterium abscessus* 4S-0206 |
| Mab-13-1 | Bacteria Actinobacteria Actinobacteridae Actinomycetales Corynebacterineae Mycobacteriaceae Mycobacterium Mycobacteriumabscessus *Mycobacterium abscessus* 4S-0303 |
| Mab-14-1 | Bacteria Actinobacteria Actinobacteridae Actinomycetales Corynebacterineae Mycobacteriaceae Mycobacterium Mycobacteriumabscessus *Mycobacterium abscessus* 4S-0726-RA |
| Mab-15-1 | Bacteria Actinobacteria Actinobacteridae Actinomycetales Corynebacterineae Mycobacteriaceae Mycobacterium Mycobacteriumabscessus *Mycobacterium abscessus* 4S-0726-RB |
| Mab-16-1 | Bacteria Actinobacteria Actinobacteridae Actinomycetales Corynebacterineae Mycobacteriaceae Mycobacterium Mycobacteriumabscessus *Mycobacterium abscessus* 5S-0304 |
| Mab-17-1 | Bacteria Actinobacteria Actinobacteridae Actinomycetales Corynebacterineae Mycobacteriaceae Mycobacterium Mycobacteriumabscessus *Mycobacterium abscessus* 5S-0421 |
| Mab-18-1 | Bacteria Actinobacteria Actinobacteridae Actinomycetales Corynebacterineae Mycobacteriaceae Mycobacterium Mycobacteriumabscessus *Mycobacterium abscessus* 5S-0422 |
| Mab-19-1 | Bacteria Actinobacteria Actinobacteridae Actinomycetales Corynebacterineae Mycobacteriaceae Mycobacterium Mycobacteriumabscessus *Mycobacterium abscessus* 5S-0708 |
| Mab-20-1 | Bacteria Actinobacteria Actinobacteridae Actinomycetales Corynebacterineae Mycobacteriaceae Mycobacterium Mycobacteriumabscessus *Mycobacterium abscessus* 5S-0817 |
| Mab-21-1 | Bacteria Actinobacteria Actinobacteridae Actinomycetales Corynebacterineae Mycobacteriaceae Mycobacterium Mycobacteriumabscessus *Mycobacterium abscessus* 5S-0921 |
| Mab-22-1 | Bacteria Actinobacteria Actinobacteridae Actinomycetales Corynebacterineae Mycobacteriaceae Mycobacterium Mycobacteriumabscessus *Mycobacterium abscessus* 5S-1212 |
| Mab-23-1 | Bacteria Actinobacteria Actinobacteridae Actinomycetales Corynebacterineae Mycobacteriaceae Mycobacterium Mycobacteriumabscessus *Mycobacterium abscessus* 5S-1215 |
| Mab-24-1 | Bacteria Actinobacteria Actinobacteridae Actinomycetales Corynebacterineae Mycobacteriaceae Mycobacterium Mycobacteriumabscessus *Mycobacterium abscessus* 6G-0125-R |
| Mab-25-1 | Bacteria Actinobacteria Actinobacteridae Actinomycetales Corynebacterineae Mycobacteriaceae Mycobacterium Mycobacteriumabscessus *Mycobacterium abscessus* 6G-0125-S |
| Mab-26-1 | Bacteria Actinobacteria Actinobacteridae Actinomycetales Corynebacterineae Mycobacteriaceae Mycobacterium Mycobacteriumabscessus *Mycobacterium abscessus* 6G-0212 |
| Mab-27-1 | Bacteria Actinobacteria Actinobacteridae Actinomycetales Corynebacterineae Mycobacteriaceae Mycobacterium Mycobacteriumabscessus *Mycobacterium abscessus* 6G-0728-R |
| Mab-28-1 | Bacteria Actinobacteria Actinobacteridae Actinomycetales Corynebacterineae Mycobacteriaceae Mycobacterium Mycobacteriumabscessus *Mycobacterium abscessus* 6G-0728-S |
| Mab-29-1 | Bacteria Actinobacteria Actinobacteridae Actinomycetales Corynebacterineae Mycobacteriaceae Mycobacterium Mycobacteriumabscessus *Mycobacterium abscessus* 6G-1108 |
| Mab-30-1 | Bacteria Actinobacteria Actinobacteridae Actinomycetales Corynebacterineae Mycobacteriaceae Mycobacterium Mycobacteriumabscessus *Mycobacterium abscessus* 9808 |
| Mab-31-1 | Bacteria Actinobacteria Actinobacteridae Actinomycetales Corynebacterineae Mycobacteriaceae Mycobacterium Mycobacteriumabscessus *Mycobacterium abscessus* M115 |
| Mab-32-1 | Bacteria Actinobacteria Actinobacteridae Actinomycetales Corynebacterineae Mycobacteriaceae Mycobacterium Mycobacteriumabscessus *Mycobacterium abscessus* M139 |

| | |
|---|---|
| Mab-33-1 | Bacteria Actinobacteria Actinobacteridae Actinomycetales Corynebacterineae Mycobacteriaceae Mycobacterium Mycobacteriumabscessus *Mycobacterium abscessus* M148 |
| Mab-34-1 | Bacteria Actinobacteria Actinobacteridae Actinomycetales Corynebacterineae Mycobacteriaceae Mycobacterium Mycobacteriumabscessus *Mycobacterium abscessus* M152 |
| Mab-35-1 | Bacteria Actinobacteria Actinobacteridae Actinomycetales Corynebacterineae Mycobacteriaceae Mycobacterium Mycobacteriumabscessus *Mycobacterium abscessus* M154 |
| Mab-36-1 | Bacteria Actinobacteria Actinobacteridae Actinomycetales Corynebacterineae Mycobacteriaceae Mycobacterium Mycobacteriumabscessus *Mycobacterium abscessus* M156 |
| Mab-37-1 | Bacteria Actinobacteria Actinobacteridae Actinomycetales Corynebacterineae Mycobacteriaceae Mycobacterium Mycobacteriumabscessus *Mycobacterium abscessus* M159 |
| Mab-38-1 | Bacteria Actinobacteria Actinobacteridae Actinomycetales Corynebacterineae Mycobacteriaceae Mycobacterium Mycobacteriumabscessus *Mycobacterium abscessus* M172 |
| Mab-39-1 | Bacteria Actinobacteria Actinobacteridae Actinomycetales Corynebacterineae Mycobacteriaceae Mycobacterium Mycobacteriumabscessus *Mycobacterium abscessus* M93 |
| Mab-40-1 | Bacteria Actinobacteria Actinobacteridae Actinomycetales Corynebacterineae Mycobacteriaceae Mycobacterium Mycobacteriumabscessus *Mycobacterium abscessus* M94 |
| Mab-41-1 | Bacteria Actinobacteria Actinobacteridae Actinomycetales Corynebacterineae Mycobacteriaceae Mycobacterium Mycobacteriumabscessus *Mycobacterium abscessus* MAB_082312_2258 |
| Mab-42-1 | Bacteria Actinobacteria Actinobacteridae Actinomycetales Corynebacterineae Mycobacteriaceae Mycobacterium Mycobacteriumabscessus *Mycobacterium abscessus* MAB_091912_2446 |
| Mab-44-1 | Bacteria Actinobacteria Actinobacteridae Actinomycetales Corynebacterineae Mycobacteriaceae Mycobacterium Mycobacteriumabscessus *Mycobacterium abscessus subsp. bolletii* 1S-151-0930 |
| Mab-45-1 | Bacteria Actinobacteria Actinobacteridae Actinomycetales Corynebacterineae Mycobacteriaceae Mycobacterium Mycobacteriumabscessus *Mycobacterium abscessus subsp. bolletii* 1S-152-0914 |
| Mab-46-1 | Bacteria Actinobacteria Actinobacteridae Actinomycetales Corynebacterineae Mycobacteriaceae Mycobacterium Mycobacteriumabscessus *Mycobacterium abscessus subsp. bolletii* 1S-153-0915 |
| Mab-47-1 | Bacteria Actinobacteria Actinobacteridae Actinomycetales Corynebacterineae Mycobacteriaceae Mycobacterium Mycobacteriumabscessus *Mycobacterium abscessus subsp. bolletii* 1S-154-0310 |
| Mab-48-1 | Bacteria Actinobacteria Actinobacteridae Actinomycetales Corynebacterineae Mycobacteriaceae Mycobacterium Mycobacteriumabscessus *Mycobacterium abscessus subsp. bolletii* 2B-0107 |
| Mab-49-1 | Bacteria Actinobacteria Actinobacteridae Actinomycetales Corynebacterineae Mycobacteriaceae Mycobacterium Mycobacteriumabscessus *Mycobacterium abscessus subsp. bolletii* 2B-0307 |
| Mab-50-1 | Bacteria Actinobacteria Actinobacteridae Actinomycetales Corynebacterineae Mycobacteriaceae Mycobacterium Mycobacteriumabscessus *Mycobacterium abscessus subsp. bolletii* 2B-0626 |
| Mab-51-1 | Bacteria Actinobacteria Actinobacteridae Actinomycetales Corynebacterineae Mycobacteriaceae Mycobacterium Mycobacteriumabscessus *Mycobacterium abscessus subsp. bolletii* 2B-0912-R |
| Mab-52-1 | Bacteria Actinobacteria Actinobacteridae Actinomycetales Corynebacterineae Mycobacteriaceae Mycobacterium Mycobacteriumabscessus *Mycobacterium abscessus subsp. bolletii* 2B-0912-S |
| Mab-53-1 | Bacteria Actinobacteria Actinobacteridae Actinomycetales Corynebacterineae Mycobacteriaceae Mycobacterium Mycobacteriumabscessus *Mycobacterium abscessus subsp. bolletii* 2B-1231 |
| Mab-54-1 | Bacteria Actinobacteria Actinobacteridae Actinomycetales Corynebacterineae Mycobacteriaceae Mycobacterium Mycobacteriumabscessus *Mycobacterium abscessus subsp. bolletii* 50594 |
| Mab-55-1 | Bacteria Actinobacteria Actinobacteridae Actinomycetales Corynebacterineae Mycobacteriaceae Mycobacterium Mycobacteriumabscessus *Mycobacterium abscessus subsp. bolletii* BD |
| Mab-56-1 | Bacteria Actinobacteria Actinobacteridae Actinomycetales Corynebacterineae Mycobacteriaceae Mycobacterium Mycobacteriumabscessus *Mycobacterium abscessus subsp. bolletii* CCUG 48898 = JCM 15300 |
| Mab-57-1 | Bacteria Actinobacteria Actinobacteridae Actinomycetales Corynebacterineae Mycobacteriaceae Mycobacterium Mycobacteriumabscessus *Mycobacterium abscessus subsp. bolletii* CRM-0020 |
| Mab-58-1 | Bacteria Actinobacteria Actinobacteridae Actinomycetales Corynebacterineae Mycobacteriaceae Mycobacterium Mycobacteriumabscessus *Mycobacterium abscessus subsp. bolletii* M18 |
| Mab-43-1 | Bacteria Actinobacteria Actinobacteridae Actinomycetales Corynebacterineae Mycobacteriaceae Mycobacterium Mycobacteriumabscessus *Mycobacterium abscessus* V06705 |

| | |
|---|---|
| Mav-1-1 | Bacteria Actinobacteria Actinobacteridae Actinomycetales Corynebacterineae Mycobacteriaceae Mycobacterium Mycobacteriumavium complex (MAC) *Mycobacterium avium subsp. avium* Env 77 |
| Nas-1-1 | Bacteria Actinobacteria Actinobacteridae Actinomycetales Corynebacterineae Nocardiaceae *Nocardia asteroides* NBRC 15531 |
| Nbr-1-1 | Bacteria Actinobacteria Actinobacteridae Actinomycetales Corynebacterineae Nocardiaceae *Nocardia brasiliensis* ATCC 700358 |
| Ncy-1-1 | Bacteria Actinobacteria Actinobacteridae Actinomycetales Corynebacterineae Nocardiaceae *Nocardia cyriacigeorgica* GUH-2 |
| Nfa-1-1 | Bacteria Actinobacteria Actinobacteridae Actinomycetales Corynebacterineae Nocardiaceae *Nocardia farcinica* IFM 10152 |
| Nsp-1-1 | Bacteria Actinobacteria Actinobacteridae Actinomycetales Corynebacterineae Nocardiaceae *Nocardia sp.* 348MFTsu5.1 |
| Nsp-2-1 | Bacteria Actinobacteria Actinobacteridae Actinomycetales Corynebacterineae Nocardiaceae *Nocardia sp.* BMG111209 |
| Req-1-1 | Bacteria Actinobacteria Actinobacteridae Actinomycetales Corynebacterineae Nocardiaceae *Rhodococcus equi* 103S |
| Req-2-1 | Bacteria Actinobacteria Actinobacteridae Actinomycetales Corynebacterineae Nocardiaceae *Rhodococcus equi* ATCC 33707 |
| Req-3-1 | Bacteria Actinobacteria Actinobacteridae Actinomycetales Corynebacterineae Nocardiaceae *Rhodococcus equi* NBRC 101255 = C 7 |
| Rer-1-1 | Bacteria Actinobacteria Actinobacteridae Actinomycetales Corynebacterineae Nocardiaceae *Rhodococcus erythropolis* CCM2595 |
| Rer-2-1 | Bacteria Actinobacteria Actinobacteridae Actinomycetales Corynebacterineae Nocardiaceae *Rhodococcus erythropolis* DN1 |
| Rer-3-1 | Bacteria Actinobacteria Actinobacteridae Actinomycetales Corynebacterineae Nocardiaceae *Rhodococcus erythropolis* PR4 |
| Rer-4-1 | Bacteria Actinobacteria Actinobacteridae Actinomycetales Corynebacterineae Nocardiaceae *Rhodococcus erythropolis* SK121 |
| Rer-5-1 | Bacteria Actinobacteria Actinobacteridae Actinomycetales Corynebacterineae Nocardiaceae *Rhodococcus erythropolis* XP |
| Rim-1-1 | Bacteria Actinobacteria Actinobacteridae Actinomycetales Corynebacterineae Nocardiaceae *Rhodococcus imtechensis* RKJ300 |
| Rjo-1-1 | Bacteria Actinobacteria Actinobacteridae Actinomycetales Corynebacterineae Nocardiaceae *Rhodococcus jostii* RHA1 |
| Rop-1-1 | Bacteria Actinobacteria Actinobacteridae Actinomycetales Corynebacterineae Nocardiaceae *Rhodococcus opacus* B4 |
| Rop-2-1 | Bacteria Actinobacteria Actinobacteridae Actinomycetales Corynebacterineae Nocardiaceae *Rhodococcus opacus* M213 |
| Rop-3-1 to Rop-3-2 | Bacteria Actinobacteria Actinobacteridae Actinomycetales Corynebacterineae Nocardiaceae *Rhodococcus opacus* PD630 |
| Rpy-1-1 | Bacteria Actinobacteria Actinobacteridae Actinomycetales Corynebacterineae Nocardiaceae *Rhodococcus pyridinivorans* AK37 |
| Rpy-2-1 | Bacteria Actinobacteria Actinobacteridae Actinomycetales Corynebacterineae Nocardiaceae *Rhodococcus pyridinivorans* SB3094 |
| Rqi-1-1 | Bacteria Actinobacteria Actinobacteridae Actinomycetales Corynebacterineae Nocardiaceae *Rhodococcus qingshengii* BKS 20-40 |
| Rrh-1-1 | Bacteria Actinobacteria Actinobacteridae Actinomycetales Corynebacterineae Nocardiaceae *Rhodococcus rhodnii* LMG 5362 |
| Rrh-2-1 | Bacteria Actinobacteria Actinobacteridae Actinomycetales Corynebacterineae Nocardiaceae *Rhodococcus rhodochrous* BKS6-46 |
| Rru-1-1 | Bacteria Actinobacteria Actinobacteridae Actinomycetales Corynebacterineae Nocardiaceae *Rhodococcus ruber* BKS 20-38 |
| Rru-2-1 | Bacteria Actinobacteria Actinobacteridae Actinomycetales Corynebacterineae Nocardiaceae *Rhodococcus ruber* Chol-4 |
| Rsp-1-1 | Bacteria Actinobacteria Actinobacteridae Actinomycetales Corynebacterineae Nocardiaceae *Rhodococcus sp.* 114MFTsu3.1 |
| Rsp-2-1 | Bacteria Actinobacteria Actinobacteridae Actinomycetales Corynebacterineae Nocardiaceae *Rhodococcus sp.* 29MFTsu3.1 |
| Rsp-3-1 | Bacteria Actinobacteria Actinobacteridae Actinomycetales Corynebacterineae Nocardiaceae *Rhodococcus sp.* AW25M09 |
| Rsp-4-1 | Bacteria Actinobacteria Actinobacteridae Actinomycetales Corynebacterineae Nocardiaceae *Rhodococcus sp.* DK17 |
| Rsp-5-1 | Bacteria Actinobacteria Actinobacteridae Actinomycetales Corynebacterineae Nocardiaceae *Rhodococcus sp.* EsD8 |
| Rsp-6-1 | Bacteria Actinobacteria Actinobacteridae Actinomycetales Corynebacterineae Nocardiaceae *Rhodococcus sp.* JVH1 |
| Rsp-7-1 | Bacteria Actinobacteria Actinobacteridae Actinomycetales Corynebacterineae Nocardiaceae *Rhodococcus sp.* P14 |
| Rsp-8-1 | Bacteria Actinobacteria Actinobacteridae Actinomycetales Corynebacterineae Nocardiaceae *Rhodococcus sp.* P27 |
| Rsp-9-1 | Bacteria Actinobacteria Actinobacteridae Actinomycetales Corynebacterineae Nocardiaceae *Rhodococcus sp.* R1101 |
| Rtr-1-1 | Bacteria Actinobacteria Actinobacteridae Actinomycetales Corynebacterineae Nocardiaceae *Rhodococcus triatomae* BKS 15-14 |
| Rwr-1-1 | Bacteria Actinobacteria Actinobacteridae Actinomycetales Corynebacterineae Nocardiaceae *Rhodococcus wratislaviensis* IFP 2016 |
| Sni-1-1 | Bacteria Actinobacteria Actinobacteridae Actinomycetales Corynebacterineae Nocardiaceae *Smaragdicoccus niigatensis* DSM 44881 |
| Tpa-1-1 | Bacteria Actinobacteria Actinobacteridae Actinomycetales Corynebacterineae Tsukamurellaceae *Tsukamurella paurometabola* DSM 20162 |
| Wsp-1-1 | Bacteria Actinobacteria Actinobacteridae Actinomycetales Corynebacterineae Williamsiaceae *Williamsia sp.* D3 |
| Ace-1-1 | Bacteria Actinobacteria Actinobacteridae Actinomycetales Frankineae Acidothermaceae *Acidothermus cellulolyticus* 11B |
| Fal-1-1 | Bacteria Actinobacteria Actinobacteridae Actinomycetales Frankineae Frankiaceae *Frankia alni* ACN14a |
| Fsp-1-1 | Bacteria Actinobacteria Actinobacteridae Actinomycetales Frankineae Frankiaceae *Frankia sp.* BCU110501 |
| Fsp-2-1 | Bacteria Actinobacteria Actinobacteridae Actinomycetales Frankineae Frankiaceae *Frankia sp.* BMG5.12 |
| Fsp-3-1 | Bacteria Actinobacteria Actinobacteridae Actinomycetales Frankineae Frankiaceae *Frankia sp.* CcI3 |
| Fsp-4-1 | Bacteria Actinobacteria Actinobacteridae Actinomycetales Frankineae Frankiaceae *Frankia sp.* CcI6 |
| Fsp-5-1 | Bacteria Actinobacteria Actinobacteridae Actinomycetales Frankineae Frankiaceae *Frankia sp.* EAN1pec |
| Fsp-7-1 | Bacteria Actinobacteria Actinobacteridae Actinomycetales Frankineae Frankiaceae *Frankia sp.* EuI1c |
| Fsp-6-1 | Bacteria Actinobacteria Actinobacteridae Actinomycetales Frankineae Frankiaceae *Frankia sp.* EUN1f |
| Fsp-8-1 | Bacteria Actinobacteria Actinobacteridae Actinomycetales Frankineae Frankiaceae *Frankia sp.* Iso899 |
| Fsp-9-1 | Bacteria Actinobacteria Actinobacteridae Actinomycetales Frankineae Frankiaceae *Frankia sp.* QA3 |
| Fsy-1-1 | Bacteria Actinobacteria Actinobacteridae Actinomycetales Frankineae Frankiaceae *Frankia symbiont of* Datisca glomerata |
| Nmu-1-1 | Bacteria Actinobacteria Actinobacteridae Actinomycetales Frankineae Nakamurellaceae *Nakamurella multipartita* DSM 44233 |
| Spo-1-1 | Bacteria Actinobacteria Actinobacteridae Actinomycetales Frankineae Sporichthyaceae *Sporichthya polymorpha* DSM 43042 |
| Sna-1-1 | Bacteria Actinobacteria Actinobacteridae Actinomycetales Glycomycineae Glycomycetaceae *Stackebrandtia nassauensis* DSM 44728 |
| Bfa-1-1 | Bacteria Actinobacteria Actinobacteridae Actinomycetales Micrococcineae Dermabacteraceae *Brachybacterium faecium* DSM 4810 |
| Bmu-1-1 | Bacteria Actinobacteria Actinobacteridae Actinomycetales Micrococcineae Dermabacteraceae *Brachybacterium muris* UCD-AY4 |
| Bpa-1-1 | Bacteria Actinobacteria Actinobacteridae Actinomycetales Micrococcineae Dermabacteraceae *Brachybacterium paraconglomeratum* LC44 |
| Bsq-1-1 | Bacteria Actinobacteria Actinobacteridae Actinomycetales Micrococcineae Dermabacteraceae *Brachybacterium squillarum* M-6-3 |
| Dsp-1-1 | Bacteria Actinobacteria Actinobacteridae Actinomycetales Micrococcineae Dermabacteraceae *Dermabacter sp.* HFH0086 |
| Afr-1-1 | Bacteria Actinobacteria Actinobacteridae Actinomycetales Micromonosporineae Micromonosporaceae *Actinoplanes friuliensis* DSM 7358 |
| Agl-1-1 | Bacteria Actinobacteria Actinobacteridae Actinomycetales Micromonosporineae Micromonosporaceae *Actinoplanes globisporus* DSM 43857 |
| Ami-1-1 | Bacteria Actinobacteria Actinobacteridae Actinomycetales Micromonosporineae Micromonosporaceae *Actinoplanes missouriensis* 431 |

| | |
|---|---|
| Asp-1-1 | Bacteria Actinobacteria Actinobacteridae Actinomycetales Micromonosporineae Micromonosporaceae *Actinoplanes sp.* N902-109 |
| Asp-2-1 | Bacteria Actinobacteria Actinobacteridae Actinomycetales Micromonosporineae Micromonosporaceae *Actinoplanes sp.* SE50/110 |
| Cko-1-1 | Bacteria Actinobacteria Actinobacteridae Actinomycetales Micromonosporineae Micromonosporaceae *Catelliglobosispora koreensis* DSM 44566 |
| Lal-1-1 | Bacteria Actinobacteria Actinobacteridae Actinomycetales Micromonosporineae Micromonosporaceae *Longispora albida* DSM 44784 |
| Mau-1-1 | Bacteria Actinobacteria Actinobacteridae Actinomycetales Micromonosporineae Micromonosporaceae *Micromonospora aurantiaca* ATCC 27029 |
| Mlu-1-1 | Bacteria Actinobacteria Actinobacteridae Actinomycetales Micromonosporineae Micromonosporaceae *Micromonospora lupini str.* Lupac 08 |
| Msp-8-1 | Bacteria Actinobacteria Actinobacteridae Actinomycetales Micromonosporineae Micromonosporaceae *Micromonospora sp.* ATCC 39149 |
| Msp-9-1 | Bacteria Actinobacteria Actinobacteridae Actinomycetales Micromonosporineae Micromonosporaceae *Micromonospora sp.* CNB394 |
| Msp-10-1 | Bacteria Actinobacteria Actinobacteridae Actinomycetales Micromonosporineae Micromonosporaceae *Micromonospora sp.* L5 |
| Sar-1-1 | Bacteria Actinobacteria Actinobacteridae Actinomycetales Micromonosporineae Micromonosporaceae *Salinispora arenicola* CNH877 |
| Sar-2-1 | Bacteria Actinobacteria Actinobacteridae Actinomycetales Micromonosporineae Micromonosporaceae *Salinispora arenicola* CNH962 |
| Sar-3-1 | Bacteria Actinobacteria Actinobacteridae Actinomycetales Micromonosporineae Micromonosporaceae *Salinispora arenicola* CNP105 |
| Sar-4-1 | Bacteria Actinobacteria Actinobacteridae Actinomycetales Micromonosporineae Micromonosporaceae *Salinispora arenicola* CNP193 |
| Sar-5-1 | Bacteria Actinobacteria Actinobacteridae Actinomycetales Micromonosporineae Micromonosporaceae *Salinispora arenicola* CNR107 |
| Sar-6-1 | Bacteria Actinobacteria Actinobacteridae Actinomycetales Micromonosporineae Micromonosporaceae *Salinispora arenicola* CNS-205 |
| Sar-7-1 | Bacteria Actinobacteria Actinobacteridae Actinomycetales Micromonosporineae Micromonosporaceae *Salinispora arenicola* CNS051 |
| Sar-8-1 | Bacteria Actinobacteria Actinobacteridae Actinomycetales Micromonosporineae Micromonosporaceae *Salinispora arenicola* CNS673 |
| Sar-9-1 | Bacteria Actinobacteria Actinobacteridae Actinomycetales Micromonosporineae Micromonosporaceae *Salinispora arenicola* CNT859 |
| Sar-10-1 | Bacteria Actinobacteria Actinobacteridae Actinomycetales Micromonosporineae Micromonosporaceae *Salinispora arenicola* CNX481 |
| Sar-11-1 | Bacteria Actinobacteria Actinobacteridae Actinomycetales Micromonosporineae Micromonosporaceae *Salinispora arenicola* CNX814 |
| Sar-12-1 | Bacteria Actinobacteria Actinobacteridae Actinomycetales Micromonosporineae Micromonosporaceae *Salinispora arenicola* CNY011 |
| Sar-13-1 | Bacteria Actinobacteria Actinobacteridae Actinomycetales Micromonosporineae Micromonosporaceae *Salinispora arenicola* CNY231 |
| Sar-14-1 | Bacteria Actinobacteria Actinobacteridae Actinomycetales Micromonosporineae Micromonosporaceae *Salinispora arenicola* CNY234 |
| Sar-15-1 | Bacteria Actinobacteria Actinobacteridae Actinomycetales Micromonosporineae Micromonosporaceae *Salinispora arenicola* CNY256 |
| Sar-16-1 | Bacteria Actinobacteria Actinobacteridae Actinomycetales Micromonosporineae Micromonosporaceae *Salinispora arenicola* CNY260 |
| Sar-17-1 | Bacteria Actinobacteria Actinobacteridae Actinomycetales Micromonosporineae Micromonosporaceae *Salinispora arenicola* CNY280 |
| Sar-18-1 | Bacteria Actinobacteria Actinobacteridae Actinomycetales Micromonosporineae Micromonosporaceae *Salinispora arenicola* CNY282 |
| Sar-19-1 | Bacteria Actinobacteria Actinobacteridae Actinomycetales Micromonosporineae Micromonosporaceae *Salinispora arenicola* DSM 45545 |
| Spa-1-1 | Bacteria Actinobacteria Actinobacteridae Actinomycetales Micromonosporineae Micromonosporaceae *Salinispora pacifica* CNQ768 |
| Spa-2-1 | Bacteria Actinobacteria Actinobacteridae Actinomycetales Micromonosporineae Micromonosporaceae *Salinispora pacifica* CNR942 |
| Spa-3-1 | Bacteria Actinobacteria Actinobacteridae Actinomycetales Micromonosporineae Micromonosporaceae *Salinispora pacifica* CNS055 |
| Spa-4-1 | Bacteria Actinobacteria Actinobacteridae Actinomycetales Micromonosporineae Micromonosporaceae *Salinispora pacifica* CNS860 |
| Spa-5-1 | Bacteria Actinobacteria Actinobacteridae Actinomycetales Micromonosporineae Micromonosporaceae *Salinispora pacifica* CNS863 |
| Spa-6-1 | Bacteria Actinobacteria Actinobacteridae Actinomycetales Micromonosporineae Micromonosporaceae *Salinispora pacifica* CNS996 |
| Spa-7-1 | Bacteria Actinobacteria Actinobacteridae Actinomycetales Micromonosporineae Micromonosporaceae *Salinispora pacifica* CNT045 |
| Spa-8-1 | Bacteria Actinobacteria Actinobacteridae Actinomycetales Micromonosporineae Micromonosporaceae *Salinispora pacifica* CNT124 |
| Spa-9-1 | Bacteria Actinobacteria Actinobacteridae Actinomycetales Micromonosporineae Micromonosporaceae *Salinispora pacifica* CNT584 |
| Spa-10-1 | Bacteria Actinobacteria Actinobacteridae Actinomycetales Micromonosporineae Micromonosporaceae *Salinispora pacifica* CNT609 |
| Spa-11-1 | Bacteria Actinobacteria Actinobacteridae Actinomycetales Micromonosporineae Micromonosporaceae *Salinispora pacifica* CNT849 |
| Spa-12-1 | Bacteria Actinobacteria Actinobacteridae Actinomycetales Micromonosporineae Micromonosporaceae *Salinispora pacifica* CNT851 |
| Spa-13-1 | Bacteria Actinobacteria Actinobacteridae Actinomycetales Micromonosporineae Micromonosporaceae *Salinispora pacifica* CNY237 |
| Spa-14-1 | Bacteria Actinobacteria Actinobacteridae Actinomycetales Micromonosporineae Micromonosporaceae *Salinispora pacifica* CNY330 |
| Spa-15-1 | Bacteria Actinobacteria Actinobacteridae Actinomycetales Micromonosporineae Micromonosporaceae *Salinispora pacifica* DSM 45543 |
| Spa-16-1 | Bacteria Actinobacteria Actinobacteridae Actinomycetales Micromonosporineae Micromonosporaceae *Salinispora pacifica* DSM 45544 |
| Spa-17-1 | Bacteria Actinobacteria Actinobacteridae Actinomycetales Micromonosporineae Micromonosporaceae *Salinispora pacifica* DSM 45546 |
| Spa-18-1 | Bacteria Actinobacteria Actinobacteridae Actinomycetales Micromonosporineae Micromonosporaceae *Salinispora pacifica* DSM 45547 |
| Spa-19-1 | Bacteria Actinobacteria Actinobacteridae Actinomycetales Micromonosporineae Micromonosporaceae *Salinispora pacifica* DSM 45548 |
| Spa-20-1 | Bacteria Actinobacteria Actinobacteridae Actinomycetales Micromonosporineae Micromonosporaceae *Salinispora pacifica* DSM 45549 |
| Str-1-1 | Bacteria Actinobacteria Actinobacteridae Actinomycetales Micromonosporineae Micromonosporaceae *Salinispora tropica* CNB-440 |
| Str-2-1 | Bacteria Actinobacteria Actinobacteridae Actinomycetales Micromonosporineae Micromonosporaceae *Salinispora tropica* CNR699 |
| Str-3-1 | Bacteria Actinobacteria Actinobacteridae Actinomycetales Micromonosporineae Micromonosporaceae *Salinispora tropica* CNS416 |
| Vma-1-1 | Bacteria Actinobacteria Actinobacteridae Actinomycetales Micromonosporineae Micromonosporaceae *Verrucosispora maris* AB-18-032 |
| Aal-1-1 | Bacteria Actinobacteria Actinobacteridae Actinomycetales Propionibacterineae Nocardioidaceae *Actinopolymorpha alba* DSM 45243 |
| Ama-3-1 | Bacteria Actinobacteria Actinobacteridae Actinomycetales Propionibacterineae Nocardioidaceae *Aeromicrobium marinum* DSM 15272 |
| Nba-1-1 | Bacteria Actinobacteria Actinobacteridae Actinomycetales Propionibacterineae *Nocardioidaceae bacterium* Broad-1 |
| Nsp-3-1 | Bacteria Actinobacteria Actinobacteridae Actinomycetales Propionibacterineae Nocardioidaceae *Nocardioides sp.* Iso805N |
| Nsp-4-1 | Bacteria Actinobacteria Actinobacteridae Actinomycetales Propionibacterineae Nocardioidaceae *Nocardioides sp.* JS614 |
| Pav-1-1 | Bacteria Actinobacteria Actinobacteridae Actinomycetales Propionibacterineae Propionibacteriaceae *Propionibacterium avidum* 44067 |
| Pav-2-1 | Bacteria Actinobacteria Actinobacteridae Actinomycetales Propionibacterineae Propionibacteriaceae *Propionibacterium avidum* TM16 |
| Psp-1-1 | Bacteria Actinobacteria Actinobacteridae Actinomycetales Propionibacterineae Propionibacteriaceae *Propionibacterium sp.* HGH0353 |
| Psp-2-1 | Bacteria Actinobacteria Actinobacteridae Actinomycetales Propionibacterineae Propionibacteriaceae *Propionibacterium sp.* KPL1838 |
| Psp-3-1 | Bacteria Actinobacteria Actinobacteridae Actinomycetales Propionibacterineae Propionibacteriaceae *Propionibacterium sp.* KPL1852 |
| Psp-4-1 | Bacteria Actinobacteria Actinobacteridae Actinomycetales Propionibacterineae Propionibacteriaceae *Propionibacterium sp.* KPL2000 |
| Psp-5-1 | Bacteria Actinobacteria Actinobacteridae Actinomycetales Propionibacterineae Propionibacteriaceae *Propionibacterium sp.* KPL2005 |
| Ply-1-1 | Bacteria Actinobacteria Actinobacteridae Actinomycetales Propionibacterineae Propionibacteriaceae *Propionimicrobium lymphophilum* ACS-093-V-SCH5 |

| | |
|---|---|
| Ami-2-1 | Bacteria Actinobacteria Actinobacteridae Actinomycetales Pseudonocardineae Actinosynnemataceae *Actinosynnema mirum* DSM 43827 |
| Asp-3-1 | Bacteria Actinobacteria Actinobacteridae Actinomycetales Pseudonocardineae Pseudonocardiaceae *Actinoalloteichus spitiensis* RMV-1378 |
| Aen-1-1 | Bacteria Actinobacteria Actinobacteridae Actinomycetales Pseudonocardineae Pseudonocardiaceae *Actinokineospora enzanensis* DSM 44649 |
| Aaz-1-1 | Bacteria Actinobacteria Actinobacteridae Actinomycetales Pseudonocardineae Pseudonocardiaceae *Amycolatopsis azurea* DSM 43854 |
| Aba-1-1 | Bacteria Actinobacteria Actinobacteridae Actinomycetales Pseudonocardineae Pseudonocardiaceae *Amycolatopsis balhimycina* FH 1894 |
| Abe-1-1 | Bacteria Actinobacteria Actinobacteridae Actinomycetales Pseudonocardineae Pseudonocardiaceae *Amycolatopsis benzoatilytica* AK 16/65 |
| Ade-1-1 | Bacteria Actinobacteria Actinobacteridae Actinomycetales Pseudonocardineae Pseudonocardiaceae *Amycolatopsis decaplanina* DSM 44594 |
| Ame-1-1 | Bacteria Actinobacteria Actinobacteridae Actinomycetales Pseudonocardineae Pseudonocardiaceae *Amycolatopsis mediterranei* RB |
| Ame-2-1 to Ame-2-2 | Bacteria Actinobacteria Actinobacteridae Actinomycetales Pseudonocardineae Pseudonocardiaceae *Amycolatopsis mediterranei* S699 |
| Ame-3-1 | Bacteria Actinobacteria Actinobacteridae Actinomycetales Pseudonocardineae Pseudonocardiaceae *Amycolatopsis mediterranei* U32 |
| Ame-4-1 to Ame-4-2 | Bacteria Actinobacteria Actinobacteridae Actinomycetales Pseudonocardineae Pseudonocardiaceae *Amycolatopsis methanolica* 239 |
| Ani-1-1 | Bacteria Actinobacteria Actinobacteridae Actinomycetales Pseudonocardineae Pseudonocardiaceae *Amycolatopsis nigrescens* CSC17Ta-90 |
| Asp-4-1 | Bacteria Actinobacteria Actinobacteridae Actinomycetales Pseudonocardineae Pseudonocardiaceae *Amycolatopsis sp.* ATCC 39116 |
| Ava-1-1 | Bacteria Actinobacteria Actinobacteridae Actinomycetales Pseudonocardineae Pseudonocardiaceae *Amycolatopsis vancoresmycina* DSM 44592 |
| Pdi-1-1 | Bacteria Actinobacteria Actinobacteridae Actinomycetales Pseudonocardineae Pseudonocardiaceae *Pseudonocardia dioxanivorans* CB1190 |
| Psp-6-1 | Bacteria Actinobacteria Actinobacteridae Actinomycetales Pseudonocardineae Pseudonocardiaceae *Pseudonocardia sp.* P1 |
| Saz-1-1 | Bacteria Actinobacteria Actinobacteridae Actinomycetales Pseudonocardineae Pseudonocardiaceae *Saccharomonospora azurea* NA-128 |
| Saz-2-1 | Bacteria Actinobacteria Actinobacteridae Actinomycetales Pseudonocardineae Pseudonocardiaceae *Saccharomonospora azurea* SZMC 14600 |
| Scy-1-1 | Bacteria Actinobacteria Actinobacteridae Actinomycetales Pseudonocardineae Pseudonocardiaceae *Saccharomonospora cyanea* NA-134 |
| Sgl-1-1 | Bacteria Actinobacteria Actinobacteridae Actinomycetales Pseudonocardineae Pseudonocardiaceae *Saccharomonospora glauca* K62 |
| Sha-1-1 | Bacteria Actinobacteria Actinobacteridae Actinomycetales Pseudonocardineae Pseudonocardiaceae *Saccharomonospora halophila* 8 |
| Sma-1-1 | Bacteria Actinobacteria Actinobacteridae Actinomycetales Pseudonocardineae Pseudonocardiaceae *Saccharomonospora marina* XMU15 |
| Ssa-1-1 | Bacteria Actinobacteria Actinobacteridae Actinomycetales Pseudonocardineae Pseudonocardiaceae *Saccharomonospora saliphila* YIM 90502 |
| Svi-1-1 | Bacteria Actinobacteria Actinobacteridae Actinomycetales Pseudonocardineae Pseudonocardiaceae *Saccharomonospora viridis* DSM 43017 |
| Sxi-1-1 | Bacteria Actinobacteria Actinobacteridae Actinomycetales Pseudonocardineae Pseudonocardiaceae *Saccharomonospora xinjiangensis* XJ-54 |
| Ser-1-1 | Bacteria Actinobacteria Actinobacteridae Actinomycetales Pseudonocardineae Pseudonocardiaceae *Saccharopolyspora erythraea* D |
| Ser-2-1 | Bacteria Actinobacteria Actinobacteridae Actinomycetales Pseudonocardineae Pseudonocardiaceae *Saccharopolyspora erythraea* NRRL 2338 |
| Ssp-1-1 | Bacteria Actinobacteria Actinobacteridae Actinomycetales Pseudonocardineae Pseudonocardiaceae *Saccharopolyspora spinosa* NRRL 18395 |
| Ses-1-1 | Bacteria Actinobacteria Actinobacteridae Actinomycetales Pseudonocardineae Pseudonocardiaceae *Saccharothrix espanaensis* DSM 44229 |
| Sma-2-1 | Bacteria Actinobacteria Actinobacteridae Actinomycetales Pseudonocardineae Pseudonocardiaceae *Sciscionella marina* DSM 45152 |
| Saf-1-1 | Bacteria Actinobacteria Actinobacteridae Actinomycetales Streptomycineae Streptomycetaceae *Streptomyces afghaniensis* 772 |
| Sal-1-1 | Bacteria Actinobacteria Actinobacteridae Actinomycetales Streptomycineae Streptomycetaceae *Streptomyces albus* J1074 |
| Sav-1-1 | Bacteria Actinobacteria Actinobacteridae Actinomycetales Streptomycineae Streptomycetaceae *Streptomyces avermitilis* MA-4680 |
| Sbi-1-1 | Bacteria Actinobacteria Actinobacteridae Actinomycetales Streptomycineae Streptomycetaceae *Streptomyces bingchenggensis* BCW-1 |
| Sbo-1-1 to Sbo-1-2 | Bacteria Actinobacteria Actinobacteridae Actinomycetales Streptomycineae Streptomycetaceae *Streptomyces bottropensis* ATCC 25435 |
| Sca-1-1 | Bacteria Actinobacteria Actinobacteridae Actinomycetales Streptomycineae Streptomycetaceae *Streptomyces canus* 299MFChir4.1 |
| Sca-2-1 to Sca-2-2 | Bacteria Actinobacteria Actinobacteridae Actinomycetales Streptomycineae Streptomycetaceae *Streptomyces cattleya* NRRL 8057 = DSM 46488 |
| Sch-1-1 | Bacteria Actinobacteria Actinobacteridae Actinomycetales Streptomycineae Streptomycetaceae *Streptomyces chartreusis* NRRL 12338 |
| Scl-1-1 to Scl-1-2 | Bacteria Actinobacteria Actinobacteridae Actinomycetales Streptomycineae Streptomycetaceae *Streptomyces clavuligerus* ATCC 27064 |
| Sco-1-1 | Bacteria Actinobacteria Actinobacteridae Actinomycetales Streptomycineae Streptomycetaceae *Streptomyces coelicoflavus* ZG0656 |
| Sco-2-1 | Bacteria Actinobacteria Actinobacteridae Actinomycetales Streptomycineae Streptomycetaceae *Streptomyces coelicolor* A3(2) |
| Sco-3-1 | Bacteria Actinobacteria Actinobacteridae Actinomycetales Streptomycineae Streptomycetaceae *Streptomyces collinus* Tu 365 |
| Sda-1-1 | Bacteria Actinobacteria Actinobacteridae Actinomycetales Streptomycineae Streptomycetaceae *Streptomyces davawensis* JCM 4913 |
| Sfl-1-1 | Bacteria Actinobacteria Actinobacteridae Actinomycetales Streptomycineae Streptomycetaceae *Streptomyces flavogriseus* ATCC 33331 |
| Sfu-1-1 | Bacteria Actinobacteria Actinobacteridae Actinomycetales Streptomycineae Streptomycetaceae *Streptomyces fulvissimus* DSM 40593 |
| Sga-1-1 | Bacteria Actinobacteria Actinobacteridae Actinomycetales Streptomycineae Streptomycetaceae *Streptomyces gancidicus* BKS 13-15 |
| Sgh-1-1 | Bacteria Actinobacteria Actinobacteridae Actinomycetales Streptomycineae Streptomycetaceae *Streptomyces ghanaensis* ATCC 14672 |
| Sgl-2-1 | Bacteria Actinobacteria Actinobacteridae Actinomycetales Streptomycineae Streptomycetaceae *Streptomyces globisporus* C-1027 |
| Sgr-1-1 | Bacteria Actinobacteria Actinobacteridae Actinomycetales Streptomycineae Streptomycetaceae *Streptomyces griseoaurantiacus* M045 |
| Sgr-2-1 | Bacteria Actinobacteria Actinobacteridae Actinomycetales Streptomycineae Streptomycetaceae *Streptomyces griseoflavus* Tu4000 |
| Sgr-4-1 | Bacteria Actinobacteria Actinobacteridae Actinomycetales Streptomycineae Streptomycetaceae *Streptomyces griseus subsp. griseus* NBRC 13350 |
| Sgr-3-1 | Bacteria Actinobacteria Actinobacteridae Actinomycetales Streptomycineae Streptomycetaceae *Streptomyces griseus* XylebKG-1 |
| Shy-1-1 | Bacteria Actinobacteria Actinobacteridae Actinomycetales Streptomycineae Streptomycetaceae *Streptomyces hygroscopicus* ATCC 53653 |
| Shy-2-1 | Bacteria Actinobacteria Actinobacteridae Actinomycetales Streptomycineae Streptomycetaceae *Streptomyces hygroscopicus subsp. jinggangensis* 5008 |
| Shy-3-1 | Bacteria Actinobacteria Actinobacteridae Actinomycetales Streptomycineae Streptomycetaceae *Streptomyces hygroscopicus subsp. jinggangensis* TL01 |
| Sip-1-1 | Bacteria Actinobacteria Actinobacteridae Actinomycetales Streptomycineae Streptomycetaceae *Streptomyces ipomoeae* 91-03 |
| Sli-1-1 | Bacteria Actinobacteria Actinobacteridae Actinomycetales Streptomycineae Streptomycetaceae *Streptomyces lividans* 1326 |
| Sli-2-1 to Sli-2-2 | Bacteria Actinobacteria Actinobacteridae Actinomycetales Streptomycineae Streptomycetaceae *Streptomyces lividans* TK24 |
| Smo-1-1 | Bacteria Actinobacteria Actinobacteridae Actinomycetales Streptomycineae Streptomycetaceae *Streptomyces mobaraensis* NBRC 13819 = DSM 40847 |
| Sni-2-1 | Bacteria Actinobacteria Actinobacteridae Actinomycetales Streptomycineae Streptomycetaceae *Streptomyces niveus* NCIMB 11891 |
| Spr-1-1 | Bacteria Actinobacteria Actinobacteridae Actinomycetales Streptomycineae Streptomycetaceae *Streptomyces pristinaespiralis* ATCC 25486 |
| Spr-2-1 | Bacteria Actinobacteria Actinobacteridae Actinomycetales Streptomycineae Streptomycetaceae *Streptomyces prunicolor* NBRC 13075 |
| Spu-1-1 | Bacteria Actinobacteria Actinobacteridae Actinomycetales Streptomycineae Streptomycetaceae *Streptomyces purpureus* KA281 |
| Sri-1-1 | Bacteria Actinobacteria Actinobacteridae Actinomycetales Streptomycineae Streptomycetaceae *Streptomyces rimosus subsp. rimosus* ATCC 10970 |
| Sro-1-1 | Bacteria Actinobacteria Actinobacteridae Actinomycetales Streptomycineae Streptomycetaceae *Streptomyces roseochromogenes subsp. oscitans* DS 12.976 |

| | |
|---|---|
| Sro-2-1 | Bacteria Actinobacteria Actinobacteridae Actinomycetales Streptomycineae Streptomycetaceae *Streptomyces roseosporus* NRRL 15998 |
| Ssc-1-1 | Bacteria Actinobacteria Actinobacteridae Actinomycetales Streptomycineae Streptomycetaceae *Streptomyces scabiei* 87.22 |
| Ssc-2-1 | Bacteria Actinobacteria Actinobacteridae Actinomycetales Streptomycineae Streptomycetaceae *Streptomyces scabrisporus* DSM 41855 |
| Sso-1-1 | Bacteria Actinobacteria Actinobacteridae Actinomycetales Streptomycineae Streptomycetaceae *Streptomyces somaliensis* DSM 40738 |
| Ssp-2-1 | Bacteria Actinobacteria Actinobacteridae Actinomycetales Streptomycineae Streptomycetaceae *Streptomyces sp.* 303MFCol5.2 |
| Ssp-3-1 | Bacteria Actinobacteria Actinobacteridae Actinomycetales Streptomycineae Streptomycetaceae *Streptomyces sp.* 351MFTsu5.1 |
| Ssp-4-1 | Bacteria Actinobacteria Actinobacteridae Actinomycetales Streptomycineae Streptomycetaceae *Streptomyces sp.* AA0539 |
| Ssp-5-1 | Bacteria Actinobacteria Actinobacteridae Actinomycetales Streptomycineae Streptomycetaceae *Streptomyces sp.* AA4 |
| Ssp-7-1 | Bacteria Actinobacteria Actinobacteridae Actinomycetales Streptomycineae Streptomycetaceae *Streptomyces sp.* Amel2xE9 |
| Ssp-6-1 | Bacteria Actinobacteria Actinobacteridae Actinomycetales Streptomycineae Streptomycetaceae *Streptomyces sp.* ATexAB-D23 |
| Ssp-8-1 | Bacteria Actinobacteria Actinobacteridae Actinomycetales Streptomycineae Streptomycetaceae *Streptomyces sp.* C |
| Ssp-14-1 | Bacteria Actinobacteria Actinobacteridae Actinomycetales Streptomycineae Streptomycetaceae *Streptomyces sp.* CcalMP-8W |
| Ssp-9-1 | Bacteria Actinobacteria Actinobacteridae Actinomycetales Streptomycineae Streptomycetaceae *Streptomyces sp.* CNB091 |
| Ssp-10-1 | Bacteria Actinobacteria Actinobacteridae Actinomycetales Streptomycineae Streptomycetaceae *Streptomyces sp.* CNS615 |
| Ssp-11-1 | Bacteria Actinobacteria Actinobacteridae Actinomycetales Streptomycineae Streptomycetaceae *Streptomyces sp.* CNT302 |
| Ssp-12-1 | Bacteria Actinobacteria Actinobacteridae Actinomycetales Streptomycineae Streptomycetaceae *Streptomyces sp.* CNT372 |
| Ssp-13-1 | Bacteria Actinobacteria Actinobacteridae Actinomycetales Streptomycineae Streptomycetaceae *Streptomyces sp.* CNY228 |
| Ssp-15-1 | Bacteria Actinobacteria Actinobacteridae Actinomycetales Streptomycineae Streptomycetaceae *Streptomyces sp.* DvalAA-83 |
| Ssp-44-1 | Bacteria Actinobacteria Actinobacteridae Actinomycetales Streptomycineae Streptomycetaceae *Streptomyces sp. e*14 |
| Ssp-16-1 | Bacteria Actinobacteria Actinobacteridae Actinomycetales Streptomycineae Streptomycetaceae *Streptomyces sp.* FxanaD5 |
| Ssp-17-1 | Bacteria Actinobacteria Actinobacteridae Actinomycetales Streptomycineae Streptomycetaceae *Streptomyces sp.* GBA 94-10 |
| Ssp-18-1 | Bacteria Actinobacteria Actinobacteridae Actinomycetales Streptomycineae Streptomycetaceae *Streptomyces sp.* HCCB10043 |
| Ssp-19-1 | Bacteria Actinobacteria Actinobacteridae Actinomycetales Streptomycineae Streptomycetaceae *Streptomyces sp.* HGB0020 |
| Ssp-21-1 | Bacteria Actinobacteria Actinobacteridae Actinomycetales Streptomycineae Streptomycetaceae *Streptomyces sp.* HmicA12 |
| Ssp-20-1 | Bacteria Actinobacteria Actinobacteridae Actinomycetales Streptomycineae Streptomycetaceae *Streptomyces sp.* HPH0547 |
| Ssp-22-1 | Bacteria Actinobacteria Actinobacteridae Actinomycetales Streptomycineae Streptomycetaceae *Streptomyces sp.* KhCrAH-244 |
| Ssp-23-1 | Bacteria Actinobacteria Actinobacteridae Actinomycetales Streptomycineae Streptomycetaceae *Streptomyces sp.* KhCrAH-340 |
| Ssp-24-1 | Bacteria Actinobacteria Actinobacteridae Actinomycetales Streptomycineae Streptomycetaceae *Streptomyces sp.* LaPpAH-108 |
| Ssp-25-1 | Bacteria Actinobacteria Actinobacteridae Actinomycetales Streptomycineae Streptomycetaceae *Streptomyces sp.* LaPpAH-165 |
| Ssp-26-1 | Bacteria Actinobacteria Actinobacteridae Actinomycetales Streptomycineae Streptomycetaceae *Streptomyces sp.* LaPpAH-202 |
| Ssp-27-1 | Bacteria Actinobacteria Actinobacteridae Actinomycetales Streptomycineae Streptomycetaceae *Streptomyces sp.* LaPpAH-95 |
| Ssp-28-1 to Ssp-28-2 | Bacteria Actinobacteria Actinobacteridae Actinomycetales Streptomycineae Streptomycetaceae *Streptomyces sp.* Mg1 |
| Ssp-29-1 | Bacteria Actinobacteria Actinobacteridae Actinomycetales Streptomycineae Streptomycetaceae *Streptomyces sp.* PAMC26508 |
| Ssp-31-1 | Bacteria Actinobacteria Actinobacteridae Actinomycetales Streptomycineae Streptomycetaceae *Streptomyces sp.* PsTaAH-124 |
| Ssp-30-1 | Bacteria Actinobacteria Actinobacteridae Actinomycetales Streptomycineae Streptomycetaceae *Streptomyces sp.* PVA 94-07 |
| Ssp-32-1 | Bacteria Actinobacteria Actinobacteridae Actinomycetales Streptomycineae Streptomycetaceae *Streptomyces sp.* R1-NS-10 |
| Ssp-33-1 | Bacteria Actinobacteria Actinobacteridae Actinomycetales Streptomycineae Streptomycetaceae *Streptomyces sp.* S4 |
| Ssp-34-1 | Bacteria Actinobacteria Actinobacteridae Actinomycetales Streptomycineae Streptomycetaceae *Streptomyces sp.* SA3_actF |
| Ssp-35-1 | Bacteria Actinobacteria Actinobacteridae Actinomycetales Streptomycineae Streptomycetaceae *Streptomyces sp.* SA3_actG |
| Ssp-39-1 | Bacteria Actinobacteria Actinobacteridae Actinomycetales Streptomycineae Streptomycetaceae *Streptomyces sp.* ScaeMP-e10 |
| Ssp-40-1 | Bacteria Actinobacteria Actinobacteridae Actinomycetales Streptomycineae Streptomycetaceae *Streptomyces sp.* SirexAA-E |
| Ssp-36-1 | Bacteria Actinobacteria Actinobacteridae Actinomycetales Streptomycineae Streptomycetaceae *Streptomyces sp.* SPB74 |
| Ssp-37-1 | Bacteria Actinobacteria Actinobacteridae Actinomycetales Streptomycineae Streptomycetaceae *Streptomyces sp.* SPB78 |
| Ssp-38-1 | Bacteria Actinobacteria Actinobacteridae Actinomycetales Streptomycineae Streptomycetaceae *Streptomyces sp.* SS |
| Ssp-41-1 | Bacteria Actinobacteria Actinobacteridae Actinomycetales Streptomycineae Streptomycetaceae *Streptomyces sp.* TOR3209 |
| Ssp-42-1 | Bacteria Actinobacteria Actinobacteridae Actinomycetales Streptomycineae Streptomycetaceae *Streptomyces sp.* Tu6071 |
| Ssp-43-1 | Bacteria Actinobacteria Actinobacteridae Actinomycetales Streptomycineae Streptomycetaceae *Streptomyces sp.* W007 |
| Ssu-1-1 | Bacteria Actinobacteria Actinobacteridae Actinomycetales Streptomycineae Streptomycetaceae *Streptomyces sulphureus* DSM 40104 |
| Ssu-2-1 | Bacteria Actinobacteria Actinobacteridae Actinomycetales Streptomycineae Streptomycetaceae *Streptomyces sulphureus* L180 |
| Ssv-1-1 | Bacteria Actinobacteria Actinobacteridae Actinomycetales Streptomycineae Streptomycetaceae *Streptomyces sviceus* ATCC 29083 |
| Sts-1-1 | Bacteria Actinobacteria Actinobacteridae Actinomycetales Streptomycineae Streptomycetaceae *Streptomyces tsukubaensis* NRRL18488 |
| Stu-1-1 | Bacteria Actinobacteria Actinobacteridae Actinomycetales Streptomycineae Streptomycetaceae *Streptomyces turgidiscabies* Car8 |
| Sve-1-1 | Bacteria Actinobacteria Actinobacteridae Actinomycetales Streptomycineae Streptomycetaceae *Streptomyces venezuelae* ATCC 10712 |
| Svi-2-1 | Bacteria Actinobacteria Actinobacteridae Actinomycetales Streptomycineae Streptomycetaceae *Streptomyces violaceusniger* SPC6 |
| Svi-3-1 | Bacteria Actinobacteria Actinobacteridae Actinomycetales Streptomycineae Streptomycetaceae *Streptomyces violaceusniger* Tu 4113 |
| Svi-4-1 | Bacteria Actinobacteria Actinobacteridae Actinomycetales Streptomycineae Streptomycetaceae *Streptomyces viridochromogenes* DSM 40736 |
| Svi-5-1 | Bacteria Actinobacteria Actinobacteridae Actinomycetales Streptomycineae Streptomycetaceae *Streptomyces viridochromogenes* Tue57 |
| Svi-6-1 | Bacteria Actinobacteria Actinobacteridae Actinomycetales Streptomycineae Streptomycetaceae *Streptomyces viridosporus* T7A |
| Svi-7-1 | Bacteria Actinobacteria Actinobacteridae Actinomycetales Streptomycineae Streptomycetaceae *Streptomyces vitaminophilus* DSM 41686 |
| Szi-1-1 | Bacteria Actinobacteria Actinobacteridae Actinomycetales Streptomycineae Streptomycetaceae *Streptomyces zinciresistens* K42 |
| Sba-1-1 | Bacteria Actinobacteria Actinobacteridae Actinomycetales Streptomycineae *Streptomycetaceae bacterium* MP113-05 |
| Nco-1-1 | Bacteria Actinobacteria Actinobacteridae Actinomycetales Streptosporangineae Streptosporangiaceae *Nonomuraea coxensis* DSM 45129 |
| Sro-3-1 | Bacteria Actinobacteria Actinobacteridae Actinomycetales Streptosporangineae Streptosporangiaceae *Streptosporangium roseum* DSM 43021 |
| Tcu-1-1 | Bacteria Actinobacteria Actinobacteridae Actinomycetales Streptosporangineae Thermomonosporaceae *Thermomonospora curvata* DSM 43183 |
| Bas-1-1 | Bacteria Actinobacteria Actinobacteridae Bifidobacteriales Bifidobacteriaceae *Bifidobacterium asteroides* PRL2011 |

| | |
|---|---|
| Blo-1-1 | Bacteria Actinobacteria Actinobacteridae Bifidobacteriales Bifidobacteriaceae *Bifidobacterium longum* D2957 |
| Blo-2-1 | Bacteria Actinobacteria Actinobacteridae Bifidobacteriales Bifidobacteriaceae *Bifidobacterium longum* DJO10A |
| Blo-3-1 | Bacteria Actinobacteria Actinobacteridae Bifidobacteriales Bifidobacteriaceae *Bifidobacterium longum* E18 |
| Blo-4-1 | Bacteria Actinobacteria Actinobacteridae Bifidobacteriales Bifidobacteriaceae *Bifidobacterium longum* NCC2705 |
| Blo-5-1 | Bacteria Actinobacteria Actinobacteridae Bifidobacteriales Bifidobacteriaceae *Bifidobacterium longum subsp. infantis* 157F |
| Blo-6-1 | Bacteria Actinobacteria Actinobacteridae Bifidobacteriales Bifidobacteriaceae *Bifidobacterium longum subsp. infantis* ATCC 15697 |
| Blo-7-1 | Bacteria Actinobacteria Actinobacteridae Bifidobacteriales Bifidobacteriaceae *Bifidobacterium longum subsp. infantis* ATCC 15697 = JCM 1222 |
| Blo-8-1 | Bacteria Actinobacteria Actinobacteridae Bifidobacteriales Bifidobacteriaceae *Bifidobacterium longum subsp. infantis* ATCC 55813 |
| Blo-9-1 | Bacteria Actinobacteria Actinobacteridae Bifidobacteriales Bifidobacteriaceae *Bifidobacterium longum subsp. infantis* CCUG 52486 |
| Blo-10-1 | Bacteria Actinobacteria Actinobacteridae Bifidobacteriales Bifidobacteriaceae *Bifidobacterium longum subsp. longum* 1-6B |
| Blo-11-1 | Bacteria Actinobacteria Actinobacteridae Bifidobacteriales Bifidobacteriaceae *Bifidobacterium longum subsp. longum* 2-2B |
| Blo-12-1 | Bacteria Actinobacteria Actinobacteridae Bifidobacteriales Bifidobacteriaceae *Bifidobacterium longum subsp. longum* 35B |
| Blo-13-1 | Bacteria Actinobacteria Actinobacteridae Bifidobacteriales Bifidobacteriaceae *Bifidobacterium longum subsp. longum* 44B |
| Blo-14-1 | Bacteria Actinobacteria Actinobacteridae Bifidobacteriales Bifidobacteriaceae *Bifidobacterium longum subsp. longum* BBMN68 |
| Blo-15-1 | Bacteria Actinobacteria Actinobacteridae Bifidobacteriales Bifidobacteriaceae *Bifidobacterium longum subsp. longum* CECT 7347 |
| Blo-16-1 | Bacteria Actinobacteria Actinobacteridae Bifidobacteriales Bifidobacteriaceae *Bifidobacterium longum subsp. longum* F8 |
| Blo-17-1 | Bacteria Actinobacteria Actinobacteridae Bifidobacteriales Bifidobacteriaceae *Bifidobacterium longum subsp. longum* JCM 1217 |
| Blo-18-1 | Bacteria Actinobacteria Actinobacteridae Bifidobacteriales Bifidobacteriaceae *Bifidobacterium longum subsp. longum* JDM301 |
| Blo-19-1 | Bacteria Actinobacteria Actinobacteridae Bifidobacteriales Bifidobacteriaceae *Bifidobacterium longum subsp. longum* KACC 91563 |
| Bsp-1-1 | Bacteria Actinobacteria Actinobacteridae Bifidobacteriales Bifidobacteriaceae *Bifidobacterium sp.* 12_1_47BFAA |
| Bth-1-1 | Bacteria Actinobacteria Actinobacteridae Bifidobacteriales Bifidobacteriaceae *Bifidobacterium thermophilum* RBL67 |
| aLL-1-1 | Bacteria Actinobacteria actinobacterium LLX17 |
| Ade-2-1 | Bacteria Actinobacteria Actinomycetales Actinomycetaceae *Actinomyces dentalis* DSM 19115 |
| Age-1-1 | Bacteria Actinobacteria Actinomycetales Actinomycetaceae *Actinomyces gerencseriae* DSM 6844 |
| Apo-1-1 | Bacteria Actinobacteria Actinomycetales Actinomycetaceae *Actinomyces polynesiensis* |
| Asu-2-1 | Bacteria Actinobacteria Actinomycetales Actinomycetaceae *Actinomyces suimastitidis* DSM 15538 |
| Aha-2-1 | Bacteria Actinobacteria Actinomycetales Actinomycetaceae *Arcanobacterium haemolyticum* |
| Bas-2-1 to Bas-2-3 | Bacteria Actinobacteria Bifidobacteriales Bifidobacteriaceae *Bifidobacterium asteroides* |
| Bas-3-1 | Bacteria Actinobacteria Bifidobacteriales Bifidobacteriaceae *Bifidobacterium asteroides* DSM 20089 |
| Bbo-1-1 | Bacteria Actinobacteria Bifidobacteriales Bifidobacteriaceae *Bifidobacterium boum* |
| Bbo-2-1 to Bbo-2-2 | Bacteria Actinobacteria Bifidobacteriales Bifidobacteriaceae *Bifidobacterium boum* DSM 20432 |
| Bco-1-1 | Bacteria Actinobacteria Bifidobacteriales Bifidobacteriaceae *Bifidobacterium coryneforme* |
| Bco-2-1 | Bacteria Actinobacteria Bifidobacteriales Bifidobacteriaceae *Bifidobacterium coryneforme* DSM 20216 |
| Bin-1-1 | Bacteria Actinobacteria Bifidobacteriales Bifidobacteriaceae *Bifidobacterium indicum* LMG 11587 = DSM 20214 |
| Blo-20-1 | Bacteria Actinobacteria Bifidobacteriales Bifidobacteriaceae *Bifidobacterium longum* |
| Blo-21-1 | Bacteria Actinobacteria Bifidobacteriales Bifidobacteriaceae *Bifidobacterium longum* AGR2137 |
| Blo-22-1 to Blo-22-9 | Bacteria Actinobacteria Bifidobacteriales Bifidobacteriaceae *Bifidobacterium longum subsp. infantis* |
| Blo-23-1 | Bacteria Actinobacteria Bifidobacteriales Bifidobacteriaceae *Bifidobacterium longum subsp. infantis* ATCC 15697 = JCM 1222 = DSM 20088 |
| Blo-24-1 | Bacteria Actinobacteria Bifidobacteriales Bifidobacteriaceae *Bifidobacterium longum subsp. infantis* EK3 |
| Blo-25-1 to Blo-25-2 | Bacteria Actinobacteria Bifidobacteriales Bifidobacteriaceae *Bifidobacterium longum subsp. longum* |
| Blo-26-1 | Bacteria Actinobacteria Bifidobacteriales Bifidobacteriaceae *Bifidobacterium longum subsp. longum* 1-5B |
| Blo-27-1 | Bacteria Actinobacteria Bifidobacteriales Bifidobacteriaceae *Bifidobacterium longum subsp. longum* 17-1B |
| Blo-28-1 | Bacteria Actinobacteria Bifidobacteriales Bifidobacteriaceae *Bifidobacterium longum subsp. longum* 7-1B |
| Blo-29-1 | Bacteria Actinobacteria Bifidobacteriales Bifidobacteriaceae *Bifidobacterium longum subsp. longum* 72B |
| Blo-30-1 | Bacteria Actinobacteria Bifidobacteriales Bifidobacteriaceae *Bifidobacterium longum subsp. longum* CMCC P0001 |
| Blo-31-1 | Bacteria Actinobacteria Bifidobacteriales Bifidobacteriaceae *Bifidobacterium longum subsp. longum* EK13 |
| Blo-32-1 | Bacteria Actinobacteria Bifidobacteriales Bifidobacteriaceae *Bifidobacterium longum subsp. longum* EK5 |
| Blo-33-1 | Bacteria Actinobacteria Bifidobacteriales Bifidobacteriaceae *Bifidobacterium longum subsp. longum* GT15 |
| Blo-34-1 | Bacteria Actinobacteria Bifidobacteriales Bifidobacteriaceae *Bifidobacterium longum subsp. suis* |
| Blo-35-1 | Bacteria Actinobacteria Bifidobacteriales Bifidobacteriaceae *Bifidobacterium longum subsp. suis* DSM 20211 |
| Bme-1-1 | Bacteria Actinobacteria Bifidobacteriales Bifidobacteriaceae *Bifidobacterium merycicum* |
| Bme-2-1 | Bacteria Actinobacteria Bifidobacteriales Bifidobacteriaceae *Bifidobacterium merycicum* DSM 6492 |
| Bre-1-1 to Bre-1-2 | Bacteria Actinobacteria Bifidobacteriales Bifidobacteriaceae *Bifidobacterium reuteri* DSM 23975 |
| Bsc-1-1 to Bsc-1-2 | Bacteria Actinobacteria Bifidobacteriales Bifidobacteriaceae *Bifidobacterium scardovii* |
| Bsc-2-1 | Bacteria Actinobacteria Bifidobacteriales Bifidobacteriaceae *Bifidobacterium scardovii* JCM 12489 = DSM 13734 |
| Bsp-2-1 | Bacteria Actinobacteria Bifidobacteriales Bifidobacteriaceae *Bifidobacterium sp.* 7101 |
| Bsp-3-1 | Bacteria Actinobacteria Bifidobacteriales Bifidobacteriaceae *Bifidobacterium sp.* A11 |
| Bsu-1-1 | Bacteria Actinobacteria Bifidobacteriales Bifidobacteriaceae *Bifidobacterium subtile* |
| Bsu-2-1 to Bsu-2-2 | Bacteria Actinobacteria Bifidobacteriales Bifidobacteriaceae *Bifidobacterium subtile* DSM 20096 |
| Bth-2-1 | Bacteria Actinobacteria Bifidobacteriales Bifidobacteriaceae *Bifidobacterium thermacidophilum subsp. porcinum* |
| Bth-3-1 | Bacteria Actinobacteria Bifidobacteriales Bifidobacteriaceae *Bifidobacterium thermacidophilum subsp. porcinum* DSM 17755 |
| Bth-4-1 | Bacteria Actinobacteria Bifidobacteriales Bifidobacteriaceae *Bifidobacterium thermacidophilum subsp. thermacidophilum* |
| Bth-5-1 to Bth-5-2 | Bacteria Actinobacteria Bifidobacteriales Bifidobacteriaceae *Bifidobacterium thermacidophilum subsp. thermacidophilum* DSM 15837 |
| Bth-6-1 | Bacteria Actinobacteria Bifidobacteriales Bifidobacteriaceae *Bifidobacterium thermophilum* |
| Bth-7-1 | Bacteria Actinobacteria Bifidobacteriales Bifidobacteriaceae *Bifidobacterium thermophilum* DSM 20210 |

| | |
|---|---|
| Bth-8-1 | Bacteria Actinobacteria Bifidobacteriales Bifidobacteriaceae *Bifidobacterium thermophilum* DSM 20212 |
| Bts-1-1 | Bacteria Actinobacteria Bifidobacteriales Bifidobacteriaceae *Bifidobacterium tsurumiense* |
| Bts-2-1 to Bts-2-2 | Bacteria Actinobacteria Bifidobacteriales Bifidobacteriaceae *Bifidobacterium tsurumiense* DSM 17777 |
| Aac-1-1 | Bacteria Actinobacteria Catenulisporales Actinospicaceae *Actinospica acidiphila* |
| Aro-1-1 | Bacteria Actinobacteria Catenulisporales Actinospicaceae *Actinospica robiniae* DSM 44927 |
| Gal-2-1 | Bacteria Actinobacteria Corynebacteriales Gordoniaceae *Gordonia alkanivorans* CGMCC 6845 |
| Gal-3-1 | Bacteria Actinobacteria Corynebacteriales Gordoniaceae *Gordonia alkanivorans s*104 |
| Gam-3-1 | Bacteria Actinobacteria Corynebacteriales Gordoniaceae *Gordonia amicalis* CCMA-559 |
| Ghi-1-1 to Ghi-1-2 | Bacteria Actinobacteria Corynebacteriales Gordoniaceae *Gordonia hirsuta* DSM 44140 = NBRC 16056 |
| Gja-1-1 | Bacteria Actinobacteria Corynebacteriales Gordoniaceae *Gordonia jacobaea* |
| Gsh-1-1 | Bacteria Actinobacteria Corynebacteriales Gordoniaceae *Gordonia shandongensis* DSM 45094 |
| Gsi-2-1 | Bacteria Actinobacteria Corynebacteriales Gordoniaceae *Gordonia sihwensis* |
| Gte-3-1 | Bacteria Actinobacteria Corynebacteriales Gordoniaceae *Gordonia terrae* |
| Mab-1-1 to Mab-1-166 | Bacteria Actinobacteria Corynebacteriales Mycobacteriaceae *Mycobacterium abscessus* |
| Mab-59-1 | Bacteria Actinobacteria Corynebacteriales Mycobacteriaceae Mycobacterium *Mycobacterium abscessus* 21 |
| Mab-60-1 | Bacteria Actinobacteria Corynebacteriales Mycobacteriaceae Mycobacterium *Mycobacterium abscessus* CF |
| Mab-61-1 | Bacteria Actinobacteria Corynebacteriales Mycobacteriaceae Mycobacterium *Mycobacterium abscessus* M24 |
| Mab-62-1 | Bacteria Actinobacteria Corynebacteriales Mycobacteriaceae Mycobacterium *Mycobacterium abscessus* MAB_030201_1061 |
| Mab-63-1 | Bacteria Actinobacteria Corynebacteriales Mycobacteriaceae Mycobacterium *Mycobacterium abscessus* MAB_030201_1075 |
| Mab-64-1 | Bacteria Actinobacteria Corynebacteriales Mycobacteriaceae Mycobacterium *Mycobacterium abscessus* MAB_082312_2272 |
| Mab-65-1 | Bacteria Actinobacteria Corynebacteriales Mycobacteriaceae Mycobacterium *Mycobacterium abscessus* MAB_082312_2273 |
| Mab-66-1 | Bacteria Actinobacteria Corynebacteriales Mycobacteriaceae Mycobacterium *Mycobacterium abscessus* MAB_091912_2455 |
| Mab-67-1 | Bacteria Actinobacteria Corynebacteriales Mycobacteriaceae Mycobacterium *Mycobacterium abscessus* MAB_110811_1470 |
| Mab-68-1 | Bacteria Actinobacteria Corynebacteriales Mycobacteriaceae Mycobacterium *Mycobacterium abscessus* MAB_110811_2726 |
| Mab-70-1 to Mab-70-2 | Bacteria Actinobacteria Corynebacteriales Mycobacteriaceae Mycobacterium *Mycobacterium abscessus subsp. abscessus* |
| Mab-71-1 to Mab-71-3 | Bacteria Actinobacteria Corynebacteriales Mycobacteriaceae Mycobacterium *Mycobacterium abscessus subsp. bolletii* |
| Mab-72-1 | Bacteria Actinobacteria Corynebacteriales Mycobacteriaceae Mycobacterium *Mycobacterium abscessus subsp. bolletii* 103 |
| Mab-73-1 | Bacteria Actinobacteria Corynebacteriales Mycobacteriaceae Mycobacterium *Mycobacterium abscessus subsp. bolletii* INCQS 00594 |
| Mab-74-1 | Bacteria Actinobacteria Corynebacteriales Mycobacteriaceae Mycobacterium *Mycobacterium abscessus subsp. massiliense* CCUG 48898 = JCM 15300 |
| Mab-75-1 | Bacteria Actinobacteria Corynebacteriales Mycobacteriaceae Mycobacterium *Mycobacterium abscessus subsp. massiliense str.* GO 06 |
| Mab-69-1 | Bacteria Actinobacteria Corynebacteriales Mycobacteriaceae Mycobacterium *Mycobacterium abscessus* UC22 |
| Mar-1-1 | Bacteria Actinobacteria Corynebacteriales Mycobacteriaceae *Mycobacterium aromaticivorans* JS19b1 = JCM 16368 |
| Mau-2-1 | Bacteria Actinobacteria Corynebacteriales Mycobacteriaceae *Mycobacterium aurum* |
| Mau-3-1 | Bacteria Actinobacteria Corynebacteriales Mycobacteriaceae *Mycobacterium austroafricanum* |
| Mvu-1-1 | Bacteria Actinobacteria Corynebacteriales Mycobacteriaceae Mycobacterium Mycobacterium avium complex (MAC) *Mycobacterium vulneris* |
| Mch-2-1 | Bacteria Actinobacteria Corynebacteriales Mycobacteriaceae *Mycobacterium chelonae* |
| Mch-3-1 | Bacteria Actinobacteria Corynebacteriales Mycobacteriaceae *Mycobacterium chelonae* 1518 |
| Mch-4-1 | Bacteria Actinobacteria Corynebacteriales Mycobacteriaceae *Mycobacterium chlorophenolicum* |
| Mch-5-1 | Bacteria Actinobacteria Corynebacteriales Mycobacteriaceae *Mycobacterium chubuense* |
| Mco-1-1 | Bacteria Actinobacteria Corynebacteriales Mycobacteriaceae *Mycobacterium conceptionense* |
| Mco-2-1 | Bacteria Actinobacteria Corynebacteriales Mycobacteriaceae *Mycobacterium cosmeticum* |
| Mel-1-1 | Bacteria Actinobacteria Corynebacteriales Mycobacteriaceae *Mycobacterium elephantis* |
| Mfa-1-1 | Bacteria Actinobacteria Corynebacteriales Mycobacteriaceae *Mycobacterium farcinogenes* |
| Mfo-2-1 | Bacteria Actinobacteria Corynebacteriales Mycobacteriaceae Mycobacterium Mycobacterium fortuitum complex *Mycobacterium fortuitum* Z58 |
| Mhe-1-1 | Bacteria Actinobacteria Corynebacteriales Mycobacteriaceae *Mycobacterium heraklionense* |
| Mim-1-1 | Bacteria Actinobacteria Corynebacteriales Mycobacteriaceae *Mycobacterium immunogenum* |
| Mka-1-1 | Bacteria Actinobacteria Corynebacteriales Mycobacteriaceae *Mycobacterium kansasii* Z61 |
| Mky-1-1 | Bacteria Actinobacteria Corynebacteriales Mycobacteriaceae *Mycobacterium kyorinense* |
| Mll-1-1 | Bacteria Actinobacteria Corynebacteriales Mycobacteriaceae *Mycobacterium llatzerense* |
| Mma-2-1 | Bacteria Actinobacteria Corynebacteriales Mycobacteriaceae *Mycobacterium mageritense* DSM 44476 = CIP 104973 |
| Mne-1-1 | Bacteria Actinobacteria Corynebacteriales Mycobacteriaceae *Mycobacterium neoaurum* |
| Mne-2-1 | Bacteria Actinobacteria Corynebacteriales Mycobacteriaceae *Mycobacterium neoaurum* ATCC 25795 |
| Mne-3-1 | Bacteria Actinobacteria Corynebacteriales Mycobacteriaceae *Mycobacterium neoaurum* VKM Ac-1815D |
| Mob-1-1 to Mob-1-2 | Bacteria Actinobacteria Corynebacteriales Mycobacteriaceae *Mycobacterium obuense* |
| Mru-1-1 | Bacteria Actinobacteria Corynebacteriales Mycobacteriaceae *Mycobacterium rufum* |
| Mse-1-1 to Mse-1-2 | Bacteria Actinobacteria Corynebacteriales Mycobacteriaceae *Mycobacterium senegalense* |
| Mse-2-1 | Bacteria Actinobacteria Corynebacteriales Mycobacteriaceae *Mycobacterium septicum* DSM 44393 |
| Mse-3-1 to Mse-3-2 | Bacteria Actinobacteria Corynebacteriales Mycobacteriaceae *Mycobacterium setense* |
| Msp-11-1 | Bacteria Actinobacteria Corynebacteriales Mycobacteriaceae *Mycobacterium sp.* EPa45 |
| Msp-12-1 | Bacteria Actinobacteria Corynebacteriales Mycobacteriaceae *Mycobacterium sp.* UM_Kg1 |
| Msp-13-1 | Bacteria Actinobacteria Corynebacteriales Mycobacteriaceae *Mycobacterium sp.* UM_Kg17 |
| Msp-14-1 | Bacteria Actinobacteria Corynebacteriales Mycobacteriaceae *Mycobacterium sp.* UM_Kg27 |
| Msp-15-1 | Bacteria Actinobacteria Corynebacteriales Mycobacteriaceae *Mycobacterium sp.* UM_NYF |
| Msp-16-1 | Bacteria Actinobacteria Corynebacteriales Mycobacteriaceae *Mycobacterium sp.* UM_NZ2 |
| Msp-17-1 | Bacteria Actinobacteria Corynebacteriales Mycobacteriaceae *Mycobacterium sp.* UM_WGJ |

| | |
|---|---|
| Msp-18-1 | Bacteria Actinobacteria Corynebacteriales Mycobacteriaceae *Mycobacterium sp.* UM_WWY |
| Msp-19-1 | Bacteria Actinobacteria Corynebacteriales Mycobacteriaceae *Mycobacterium sp.* UNC280MFTsu5.1 |
| Msp-20-1 | Bacteria Actinobacteria Corynebacteriales Mycobacteriaceae *Mycobacterium sp.* UNC410CL29Cvi84 |
| Msp-21-1 | Bacteria Actinobacteria Corynebacteriales Mycobacteriaceae *Mycobacterium sp.* UNCCL9 |
| Msp-22-1 | Bacteria Actinobacteria Corynebacteriales Mycobacteriaceae *Mycobacterium sp.* URHB0044 |
| Msp-23-1 | Bacteria Actinobacteria Corynebacteriales Mycobacteriaceae *Mycobacterium sp.* URHD0025 |
| Msp-24-1 | Bacteria Actinobacteria Corynebacteriales Mycobacteriaceae *Mycobacterium sp.* VKM Ac-1816D |
| Msp-25-1 | Bacteria Actinobacteria Corynebacteriales Mycobacteriaceae *Mycobacterium sp.* VKM Ac-1817D |
| Mar-2-1 | Bacteria Actinobacteria Corynebacteriales Mycobacteriaceae Mycobacterium Mycobacterium terrae complex *Mycobacterium arupense* |
| Mtu-1-1 | Bacteria Actinobacteria Corynebacteriales Mycobacteriaceae *Mycobacterium tusciae* JS617 |
| Nab-1-1 | Bacteria Actinobacteria Corynebacteriales Nocardiaceae *Nocardia abscessus* NBRC 100374 |
| Nao-1-1 | Bacteria Actinobacteria Corynebacteriales Nocardiaceae *Nocardia aobensis* NBRC 100429 |
| Nar-1-1 | Bacteria Actinobacteria Corynebacteriales Nocardiaceae *Nocardia araoensis* NBRC 100135 |
| Nas-2-1 | Bacteria Actinobacteria Corynebacteriales Nocardiaceae *Nocardia asiatica* NBRC 100129 |
| Nbr-2-1 | Bacteria Actinobacteria Corynebacteriales Nocardiaceae *Nocardia brasiliensis* IFM 10847 |
| Nbr-3-1 | Bacteria Actinobacteria Corynebacteriales Nocardiaceae *Nocardia brasiliensis* NBRC 14402 |
| Nbr-4-1 | Bacteria Actinobacteria Corynebacteriales Nocardiaceae *Nocardia brevicatena* NBRC 12119 |
| Nca-1-1 | Bacteria Actinobacteria Corynebacteriales Nocardiaceae *Nocardia carnea* |
| Nca-2-1 | Bacteria Actinobacteria Corynebacteriales Nocardiaceae *Nocardia carnea* NBRC 14403 |
| Nce-1-1 | Bacteria Actinobacteria Corynebacteriales Nocardiaceae *Nocardia cerradoensis* NBRC 101014 |
| Ncy-2-1 | Bacteria Actinobacteria Corynebacteriales Nocardiaceae *Nocardia cyriacigeorgica* NBRC 100375 |
| Nex-1-1 | Bacteria Actinobacteria Corynebacteriales Nocardiaceae *Nocardia exalbida* NBRC 100660 |
| Nhi-1-1 | Bacteria Actinobacteria Corynebacteriales Nocardiaceae *Nocardia higoensis* NBRC 100133 |
| Nji-1-1 | Bacteria Actinobacteria Corynebacteriales Nocardiaceae *Nocardia jiangxiensis* NBRC 101359 |
| Nni-1-1 | Bacteria Actinobacteria Corynebacteriales Nocardiaceae *Nocardia niigatensis* NBRC 100131 |
| Nno-1-1 | Bacteria Actinobacteria Corynebacteriales Nocardiaceae *Nocardia nova* SH22a |
| Not-1-1 | Bacteria Actinobacteria Corynebacteriales Nocardiaceae *Nocardia otitidiscaviarum* IFM 11049 |
| Not-2-1 | Bacteria Actinobacteria Corynebacteriales Nocardiaceae *Nocardia otitidiscaviarum* NBRC 14405 |
| Npa-1-1 | Bacteria Actinobacteria Corynebacteriales Nocardiaceae *Nocardia paucivorans* NBRC 100373 |
| Npn-1-1 | Bacteria Actinobacteria Corynebacteriales Nocardiaceae *Nocardia pneumoniae* NBRC 100136 |
| Nrh-1-1 to Nrh-1-2 | Bacteria Actinobacteria Corynebacteriales Nocardiaceae *Nocardia rhamnosiphila* NBRC 108938 |
| Nsp-5-1 | Bacteria Actinobacteria Corynebacteriales Nocardiaceae *Nocardia sp.* BMG51109 |
| Nsp-6-1 | Bacteria Actinobacteria Corynebacteriales Nocardiaceae *Nocardia sp.* CNY236 |
| Nsp-7-1 | Bacteria Actinobacteria Corynebacteriales Nocardiaceae *Nocardia sp.* NRRL WC-3656 |
| Nta-1-1 | Bacteria Actinobacteria Corynebacteriales Nocardiaceae *Nocardia takedensis* NBRC 100417 |
| Nte-1-1 | Bacteria Actinobacteria Corynebacteriales Nocardiaceae *Nocardia tenerifensis* NBRC 101015 |
| Nte-2-1 | Bacteria Actinobacteria Corynebacteriales Nocardiaceae *Nocardia testacea* NBRC 100365 |
| Nth-1-1 | Bacteria Actinobacteria Corynebacteriales Nocardiaceae *Nocardia thailandica* NBRC 100428 |
| Ntr-1-1 | Bacteria Actinobacteria Corynebacteriales Nocardiaceae *Nocardia transvalensis* NBRC 15921 |
| Nve-1-1 | Bacteria Actinobacteria Corynebacteriales Nocardiaceae *Nocardia veterana* NBRC 100344 |
| Nvi-1-1 | Bacteria Actinobacteria Corynebacteriales Nocardiaceae *Nocardia vinacea* NBRC 16497 |
| Nvu-1-1 | Bacteria Actinobacteria Corynebacteriales Nocardiaceae *Nocardia vulneris* |
| Rae-1-1 | Bacteria Actinobacteria Corynebacteriales Nocardiaceae *Rhodococcus aetherivorans* |
| Rde-1-1 | Bacteria Actinobacteria Corynebacteriales Nocardiaceae *Rhodococcus defluvii* |
| Rer-6-1 | Bacteria Actinobacteria Corynebacteriales Nocardiaceae *Rhodococcus erythropolis* |
| Rer-7-1 | Bacteria Actinobacteria Corynebacteriales Nocardiaceae *Rhodococcus erythropolis* R138 |
| Rfa-1-1 to Rfa-1-2 | Bacteria Actinobacteria Corynebacteriales Nocardiaceae *Rhodococcus fascians* |
| Rfa-2-1 | Bacteria Actinobacteria Corynebacteriales Nocardiaceae *Rhodococcus fascians* 02-815 |
| Rfa-3-1 | Bacteria Actinobacteria Corynebacteriales Nocardiaceae *Rhodococcus fascians* 02-816c |
| Rfa-4-1 | Bacteria Actinobacteria Corynebacteriales Nocardiaceae *Rhodococcus fascians* 04-516 |
| Rfa-5-1 | Bacteria Actinobacteria Corynebacteriales Nocardiaceae *Rhodococcus fascians* 05-339-1 |
| Rfa-6-1 | Bacteria Actinobacteria Corynebacteriales Nocardiaceae *Rhodococcus fascians* 05-561-1 |
| Rfa-7-1 | Bacteria Actinobacteria Corynebacteriales Nocardiaceae *Rhodococcus fascians* A21d2 |
| Rfa-8-1 | Bacteria Actinobacteria Corynebacteriales Nocardiaceae *Rhodococcus fascians* A25f |
| Rfa-9-1 | Bacteria Actinobacteria Corynebacteriales Nocardiaceae *Rhodococcus fascians* A3b |
| Rfa-10-1 | Bacteria Actinobacteria Corynebacteriales Nocardiaceae *Rhodococcus fascians* A44A |
| Rfa-11-1 | Bacteria Actinobacteria Corynebacteriales Nocardiaceae *Rhodococcus fascians* A73a |
| Rfa-12-1 | Bacteria Actinobacteria Corynebacteriales Nocardiaceae *Rhodococcus fascians* A76 |
| Rfa-13-1 | Bacteria Actinobacteria Corynebacteriales Nocardiaceae *Rhodococcus fascians* A78 |
| Rfa-14-1 | Bacteria Actinobacteria Corynebacteriales Nocardiaceae *Rhodococcus fascians* D188 |
| Rfa-15-1 | Bacteria Actinobacteria Corynebacteriales Nocardiaceae *Rhodococcus fascians* GIC26 |
| Rfa-16-1 | Bacteria Actinobacteria Corynebacteriales Nocardiaceae *Rhodococcus fascians* GIC36 |
| Rfa-17-1 | Bacteria Actinobacteria Corynebacteriales Nocardiaceae *Rhodococcus fascians* LMG 3602 |
| Rfa-18-1 | Bacteria Actinobacteria Corynebacteriales Nocardiaceae *Rhodococcus fascians* LMG 3605 |
| Rfa-19-1 | Bacteria Actinobacteria Corynebacteriales Nocardiaceae *Rhodococcus fascians* LMG 3616 |

| | |
|---|---|
| Rfa-20-1 | Bacteria Actinobacteria Corynebacteriales Nocardiaceae *Rhodococcus fascians* LMG 3625 |
| Rfa-21-1 | Bacteria Actinobacteria Corynebacteriales Nocardiaceae *Rhodococcus fascians* NBRC 12155 = LMG 3623 |
| Rop-4-1 | Bacteria Actinobacteria Corynebacteriales Nocardiaceae *Rhodococcus opacus* |
| Rop-3-1 to Rop-3-2 | Bacteria Actinobacteria Corynebacteriales Nocardiaceae *Rhodococcus opacus* PD630 |
| Rqi-2-1 | Bacteria Actinobacteria Corynebacteriales Nocardiaceae *Rhodococcus qingshengii* |
| Rrh-3-1 | Bacteria Actinobacteria Corynebacteriales Nocardiaceae *Rhodococcus rhodnii* |
| Rrh-4-1 | Bacteria Actinobacteria Corynebacteriales Nocardiaceae *Rhodococcus rhodochrous* |
| Rrh-5-1 | Bacteria Actinobacteria Corynebacteriales Nocardiaceae *Rhodococcus rhodochrous* ATCC 17895 |
| Rrh-6-1 | Bacteria Actinobacteria Corynebacteriales Nocardiaceae *Rhodococcus rhodochrous* ATCC 21198 |
| Rru-3-1 to Rru-3-2 | Bacteria Actinobacteria Corynebacteriales Nocardiaceae *Rhodococcus ruber* |
| Rsp-10-1 | Bacteria Actinobacteria Corynebacteriales Nocardiaceae *Rhodococcus sp.* 311R |
| Rsp-11-1 | Bacteria Actinobacteria Corynebacteriales Nocardiaceae *Rhodococcus sp.* AD45 |
| Rsp-12-1 | Bacteria Actinobacteria Corynebacteriales Nocardiaceae *Rhodococcus sp.* ARP2 |
| Rsp-13-1 | Bacteria Actinobacteria Corynebacteriales Nocardiaceae *Rhodococcus sp.* B7740 |
| Rsp-14-1 | Bacteria Actinobacteria Corynebacteriales Nocardiaceae *Rhodococcus sp.* Chr-9 |
| Rsp-15-1 | Bacteria Actinobacteria Corynebacteriales Nocardiaceae *Rhodococcus sp.* JG-3 |
| Rsp-16-1 | Bacteria Actinobacteria Corynebacteriales Nocardiaceae *Rhodococcus sp.* MEB064 |
| Rsp-22-1 | Bacteria Actinobacteria Corynebacteriales Nocardiaceae *Rhodococcus sp.* p52 |
| Rsp-17-1 | Bacteria Actinobacteria Corynebacteriales Nocardiaceae *Rhodococcus sp.* PML026 |
| Rsp-18-1 | Bacteria Actinobacteria Corynebacteriales Nocardiaceae *Rhodococcus sp.* R04 |
| Rsp-19-1 | Bacteria Actinobacteria Corynebacteriales Nocardiaceae *Rhodococcus sp.* RD6.2 |
| Rsp-20-1 | Bacteria Actinobacteria Corynebacteriales Nocardiaceae *Rhodococcus sp.* UNC23MFCrub1.1 |
| Rsp-21-1 | Bacteria Actinobacteria Corynebacteriales Nocardiaceae *Rhodococcus sp.* UNC363MFTsu5.1 |
| Rwr-2-1 | Bacteria Actinobacteria Corynebacteriales Nocardiaceae *Rhodococcus wratislaviensis* NBRC 100605 |
| Tbi-1-1 | Bacteria Actinobacteria Corynebacteriales *Tomitella biformata* AHU 1821 |
| Tsp-1-1 | Bacteria Actinobacteria Corynebacteriales Tsukamurellaceae *Tsukamurella sp.* 1534 |
| Whe-1-1 | Bacteria Actinobacteria Corynebacteriales Williamsiaceae *Williamsia herbipolensis* |
| Car-1-1 | Bacteria Actinobacteria Frankiales Cryptosporangiaceae *Cryptosporangium arvum* DSM 44712 |
| Fco-1-1 | Bacteria Actinobacteria Frankiales Frankiaceae *Frankia coriariae* |
| Fsa-1-1 | Bacteria Actinobacteria Frankiales Frankiaceae *Frankia saprophytica* |
| Fsp-10-1 | Bacteria Actinobacteria Frankiales Frankiaceae *Frankia sp.* Allo2 |
| Fsp-11-1 | Bacteria Actinobacteria Frankiales Frankiaceae *Frankia sp.* BMG5.23 |
| Fsp-12-1 | Bacteria Actinobacteria Frankiales Frankiaceae *Frankia sp.* CeD |
| Fsp-13-1 | Bacteria Actinobacteria Frankiales Frankiaceae *Frankia sp.* CpI1-S |
| Fsp-14-1 | Bacteria Actinobacteria Frankiales Frankiaceae *Frankia sp.* DC12 |
| Fsp-15-1 | Bacteria Actinobacteria Frankiales Frankiaceae *Frankia sp.* Thr |
| Gar-2-1 | Bacteria Actinobacteria Glycomycetales Glycomycetaceae *Glycomyces arizonensis* DSM 44726 |
| Gsp-3-1 | Bacteria Actinobacteria Glycomycetales Glycomycetaceae *Glycomyces sp.* NRRL B-16210 |
| Gte-4-1 | Bacteria Actinobacteria Glycomycetales Glycomycetaceae *Glycomyces tenuis* |
| Gte-5-1 | Bacteria Actinobacteria Glycomycetales Glycomycetaceae *Glycomyces tenuis* DSM 44171 |
| Hal-1-1 | Bacteria Actinobacteria Glycomycetales Glycomycetaceae *Haloglycomyces albus* DSM 45210 |
| Jal-1-1 | Bacteria Actinobacteria Jiangellales Jiangellaceae *Jiangella alkaliphila* |
| Jga-1-1 | Bacteria Actinobacteria Jiangellales Jiangellaceae *Jiangella gansuensis* DSM 44835 |
| Jmu-1-1 | Bacteria Actinobacteria Jiangellales Jiangellaceae *Jiangella muralis* |
| Cbo-1-1 | Bacteria Actinobacteria Micrococcales Cellulomonadaceae *Cellulomonas bogoriensis* 69B4 = DSM 16987 |
| Bph-1-1 | Bacteria Actinobacteria Micrococcales Dermabacteraceae *Brachybacterium phenoliresistens* |
| Dho-1-1 | Bacteria Actinobacteria Micrococcales Dermabacteraceae *Dermabacter hominis* 1368 |
| Ase-1-1 | Bacteria Actinobacteria Micromonosporales Micromonosporaceae *Actinocatenispora sera* |
| Are-1-1 | Bacteria Actinobacteria Micromonosporales Micromonosporaceae *Actinoplanes rectilineatus* |
| Asu-3-1 | Bacteria Actinobacteria Micromonosporales Micromonosporaceae *Actinoplanes subtropicus* |
| Aut-1-1 | Bacteria Actinobacteria Micromonosporales Micromonosporaceae *Actinoplanes utahensis* |
| Cja-1-1 | Bacteria Actinobacteria Micromonosporales Micromonosporaceae *Catenuloplanes japonicus* |
| Dau-1-1 | Bacteria Actinobacteria Micromonosporales Micromonosporaceae *Dactylosporangium aurantiacum* |
| Mca-1-1 | Bacteria Actinobacteria Micromonosporales Micromonosporaceae *Micromonospora carbonacea* |
| Mch-6-1 | Bacteria Actinobacteria Micromonosporales Micromonosporaceae *Micromonospora chokoriensis* |
| Mgl-1-1 | Bacteria Actinobacteria Micromonosporales Micromonosporaceae *Micromonospora globosa* |
| Mpa-1-1 | Bacteria Actinobacteria Micromonosporales Micromonosporaceae *Micromonospora parva* |
| Mpu-1-1 | Bacteria Actinobacteria Micromonosporales Micromonosporaceae *Micromonospora purpureochromogenes* |
| Msp-26-1 | Bacteria Actinobacteria Micromonosporales Micromonosporaceae *Micromonospora sp.* HK10 |
| Msp-27-1 | Bacteria Actinobacteria Micromonosporales Micromonosporaceae *Micromonospora sp.* M42 |
| Msp-28-1 | Bacteria Actinobacteria Micromonosporales Micromonosporaceae *Micromonospora sp.* RV43 |
| Pan-1-1 | Bacteria Actinobacteria Micromonosporales Micromonosporaceae *Pilimelia anulata* |
| Sar-20-1 | Bacteria Actinobacteria Micromonosporales Micromonosporaceae *Salinispora arenicola* |
| Sar-21-1 | Bacteria Actinobacteria Micromonosporales Micromonosporaceae *Salinispora arenicola* CNB458 |
| Sar-22-1 | Bacteria Actinobacteria Micromonosporales Micromonosporaceae *Salinispora arenicola* CNB527 |

| | |
|---|---|
| Sar-23-1 | Bacteria Actinobacteria Micromonosporales Micromonosporaceae *Salinispora arenicola* CNH646 |
| Sar-24-1 | Bacteria Actinobacteria Micromonosporales Micromonosporaceae *Salinispora arenicola* CNH905 |
| Sar-25-1 | Bacteria Actinobacteria Micromonosporales Micromonosporaceae *Salinispora arenicola* CNH941 |
| Sar-26-1 | Bacteria Actinobacteria Micromonosporales Micromonosporaceae *Salinispora arenicola* CNH963 |
| Sar-27-1 | Bacteria Actinobacteria Micromonosporales Micromonosporaceae *Salinispora arenicola* CNH964 |
| Sar-28-1 | Bacteria Actinobacteria Micromonosporales Micromonosporaceae *Salinispora arenicola* CNQ748 |
| Sar-29-1 | Bacteria Actinobacteria Micromonosporales Micromonosporaceae *Salinispora arenicola* CNR416 |
| Sar-30-1 | Bacteria Actinobacteria Micromonosporales Micromonosporaceae *Salinispora arenicola* CNR425 |
| Sar-31-1 | Bacteria Actinobacteria Micromonosporales Micromonosporaceae *Salinispora arenicola* CNR921 |
| Sar-32-1 | Bacteria Actinobacteria Micromonosporales Micromonosporaceae *Salinispora arenicola* CNS243 |
| Sar-33-1 | Bacteria Actinobacteria Micromonosporales Micromonosporaceae *Salinispora arenicola* CNS299 |
| Sar-34-1 | Bacteria Actinobacteria Micromonosporales Micromonosporaceae *Salinispora arenicola* CNS744 |
| Sar-35-1 | Bacteria Actinobacteria Micromonosporales Micromonosporaceae *Salinispora arenicola* CNT005 |
| Sar-36-1 | Bacteria Actinobacteria Micromonosporales Micromonosporaceae *Salinispora arenicola* CNT798 |
| Sar-37-1 | Bacteria Actinobacteria Micromonosporales Micromonosporaceae *Salinispora arenicola* CNT799 |
| Sar-38-1 | Bacteria Actinobacteria Micromonosporales Micromonosporaceae *Salinispora arenicola* CNT800 |
| Sar-39-1 | Bacteria Actinobacteria Micromonosporales Micromonosporaceae *Salinispora arenicola* CNT850 |
| Sar-40-1 | Bacteria Actinobacteria Micromonosporales Micromonosporaceae *Salinispora arenicola* CNT857 |
| Sar-41-1 | Bacteria Actinobacteria Micromonosporales Micromonosporaceae *Salinispora arenicola* CNX482 |
| Sar-42-1 | Bacteria Actinobacteria Micromonosporales Micromonosporaceae *Salinispora arenicola* CNX508 |
| Sar-43-1 | Bacteria Actinobacteria Micromonosporales Micromonosporaceae *Salinispora arenicola* CNX891 |
| Spa-21-1 | Bacteria Actinobacteria Micromonosporales Micromonosporaceae *Salinispora pacifica* CNR114 |
| Spa-22-1 | Bacteria Actinobacteria Micromonosporales Micromonosporaceae *Salinispora pacifica* CNR894 |
| Spa-23-1 | Bacteria Actinobacteria Micromonosporales Micromonosporaceae *Salinispora pacifica* CNS103 |
| Spa-24-1 | Bacteria Actinobacteria Micromonosporales Micromonosporaceae *Salinispora pacifica* CNS237 |
| Spa-25-1 | Bacteria Actinobacteria Micromonosporales Micromonosporaceae *Salinispora pacifica* CNT001 |
| Spa-26-1 | Bacteria Actinobacteria Micromonosporales Micromonosporaceae *Salinispora pacifica* CNT003 |
| Spa-27-1 | Bacteria Actinobacteria Micromonosporales Micromonosporaceae *Salinispora pacifica* CNT029 |
| Spa-28-1 | Bacteria Actinobacteria Micromonosporales Micromonosporaceae *Salinispora pacifica* CNT084 |
| Spa-29-1 | Bacteria Actinobacteria Micromonosporales Micromonosporaceae *Salinispora pacifica* CNT131 |
| Spa-30-1 | Bacteria Actinobacteria Micromonosporales Micromonosporaceae *Salinispora pacifica* CNT138 |
| Spa-31-1 | Bacteria Actinobacteria Micromonosporales Micromonosporaceae *Salinispora pacifica* CNT148 |
| Spa-32-1 | Bacteria Actinobacteria Micromonosporales Micromonosporaceae *Salinispora pacifica* CNT150 |
| Spa-33-1 | Bacteria Actinobacteria Micromonosporales Micromonosporaceae *Salinispora pacifica* CNT569 |
| Spa-34-1 | Bacteria Actinobacteria Micromonosporales Micromonosporaceae *Salinispora pacifica* CNT603 |
| Spa-35-1 | Bacteria Actinobacteria Micromonosporales Micromonosporaceae *Salinispora pacifica* CNT796 |
| Spa-36-1 | Bacteria Actinobacteria Micromonosporales Micromonosporaceae *Salinispora pacifica* CNT854 |
| Spa-37-1 | Bacteria Actinobacteria Micromonosporales Micromonosporaceae *Salinispora pacifica* CNT855 |
| Spa-38-1 | Bacteria Actinobacteria Micromonosporales Micromonosporaceae *Salinispora pacifica* CNY202 |
| Spa-39-1 | Bacteria Actinobacteria Micromonosporales Micromonosporaceae *Salinispora pacifica* CNY239 |
| Spa-40-1 | Bacteria Actinobacteria Micromonosporales Micromonosporaceae *Salinispora pacifica* CNY331 |
| Str-4-1 | Bacteria Actinobacteria Micromonosporales Micromonosporaceae *Salinispora tropica* CNB476 |
| Str-5-1 | Bacteria Actinobacteria Micromonosporales Micromonosporaceae *Salinispora tropica* CNB536 |
| Str-6-1 | Bacteria Actinobacteria Micromonosporales Micromonosporaceae *Salinispora tropica* CNH898 |
| Str-7-1 | Bacteria Actinobacteria Micromonosporales Micromonosporaceae *Salinispora tropica* CNS197 |
| Str-8-1 | Bacteria Actinobacteria Micromonosporales Micromonosporaceae *Salinispora tropica* CNT250 |
| Str-9-1 | Bacteria Actinobacteria Micromonosporales Micromonosporaceae *Salinispora tropica* CNT261 |
| Str-10-1 | Bacteria Actinobacteria Micromonosporales Micromonosporaceae *Salinispora tropica* CNY012 |
| Nla-1-1 | Bacteria Actinobacteria Nakamurellales Nakamurellaceae *Nakamurella lactea* DSM 19367 |
| Ama-4-1 | Bacteria Actinobacteria Propionibacteriales Nocardioidaceae *Aeromicrobium massiliense* JC14 |
| Msp-29-1 | Bacteria Actinobacteria Propionibacteriales Nocardioidaceae *Marmoricola sp.* URHB0036 |
| Nal-1-1 | Bacteria Actinobacteria Propionibacteriales Nocardioidaceae *Nocardioides alkalitolerans* DSM 16699 |
| Nin-1-1 | Bacteria Actinobacteria Propionibacteriales Nocardioidaceae *Nocardioides insulae* DSM 17944 |
| Nlu-1-1 | Bacteria Actinobacteria Propionibacteriales Nocardioidaceae *Nocardioides luteus* |
| Nsp-8-1 | Bacteria Actinobacteria Propionibacteriales Nocardioidaceae *Nocardioides sp.* UNC345MFTsu5.1 |
| Nsp-9-1 | Bacteria Actinobacteria Propionibacteriales Nocardioidaceae *Nocardioides sp.* URHA0020 |
| Nsp-10-1 | Bacteria Actinobacteria Propionibacteriales Nocardioidaceae *Nocardioides sp.* URHA0032 |
| Psu-1-1 | Bacteria Actinobacteria Propionibacteriales Nocardioidaceae *Propionicicella superfundia* DSM 22317 |
| Ply-2-1 | Bacteria Actinobacteria Propionibacteriales Propionibacteriaceae *Propionimicrobium lymphophilum* DSM 4903 |
| Psp-7-1 | Bacteria Actinobacteria Propionibacteriales Propionibacteriaceae *Propionimicrobium sp.* BV2F7 |
| Acy-1-1 | Bacteria Actinobacteria Pseudonocardiales Pseudonocardiaceae *Actinoalloteichus cyanogriseus* |
| Acy-2-1 | Bacteria Actinobacteria Pseudonocardiales Pseudonocardiaceae *Actinoalloteichus cyanogriseus* DSM 43889 |
| Ain-1-1 | Bacteria Actinobacteria Pseudonocardiales Pseudonocardiaceae *Actinokineospora inagensis* DSM 44258 |
| Asp-5-1 to Asp-5-2 | Bacteria Actinobacteria Pseudonocardiales Pseudonocardiaceae *Actinokineospora spheciospongiae* |
| Aal-2-1 | Bacteria Actinobacteria Pseudonocardiales Pseudonocardiaceae *Allokutzneria albata* |

| | |
|---|---|
| Aha-3-1 | Bacteria Actinobacteria Pseudonocardiales Pseudonocardiaceae *Amycolatopsis halophila* YIM 93223 |
| Aja-1-1 | Bacteria Actinobacteria Pseudonocardiales Pseudonocardiaceae *Amycolatopsis japonica* |
| Aje-1-1 | Bacteria Actinobacteria Pseudonocardiales Pseudonocardiaceae *Amycolatopsis jejuensis* |
| Ame-5-1 to Ame-5-2 | Bacteria Actinobacteria Pseudonocardiales Pseudonocardiaceae *Amycolatopsis mediterranei* |
| Ame-4-1 to Ame-4-2 | Bacteria Actinobacteria Pseudonocardiales Pseudonocardiaceae *Amycolatopsis methanolica* 239 |
| Aor-1-1 | Bacteria Actinobacteria Pseudonocardiales Pseudonocardiaceae *Amycolatopsis orientalis* |
| Aor-2-1 | Bacteria Actinobacteria Pseudonocardiales Pseudonocardiaceae *Amycolatopsis orientalis* DSM 40040 = KCTC 9412 |
| Aor-3-1 | Bacteria Actinobacteria Pseudonocardiales Pseudonocardiaceae *Amycolatopsis orientalis* DSM 43388 |
| Aor-4-1 | Bacteria Actinobacteria Pseudonocardiales Pseudonocardiaceae *Amycolatopsis orientalis* DSM 46075 |
| Ari-1-1 | Bacteria Actinobacteria Pseudonocardiales Pseudonocardiaceae *Amycolatopsis rifamycinica* |
| Asp-6-1 | Bacteria Actinobacteria Pseudonocardiales Pseudonocardiaceae *Amycolatopsis sp.* MJM2582 |
| Ata-1-1 | Bacteria Actinobacteria Pseudonocardiales Pseudonocardiaceae *Amycolatopsis taiwanensis* DSM 45107 |
| Ath-1-1 | Bacteria Actinobacteria Pseudonocardiales Pseudonocardiaceae *Amycolatopsis thermoflava* N1165 |
| Ava-2-1 | Bacteria Actinobacteria Pseudonocardiales Pseudonocardiaceae *Amycolatopsis vancoresmycina* |
| Kar-1-1 | Bacteria Actinobacteria Pseudonocardiales Pseudonocardiaceae *Kibdelosporangium aridum subsp. largum* |
| Ksp-1-1 | Bacteria Actinobacteria Pseudonocardiales Pseudonocardiaceae *Kibdelosporangium sp.* MJ126-NF4 |
| Kal-1-1 | Bacteria Actinobacteria Pseudonocardiales Pseudonocardiaceae *Kutzneria albida* |
| Kal-2-1 | Bacteria Actinobacteria Pseudonocardiales Pseudonocardiaceae *Kutzneria albida* DSM 43870 |
| Ksp-2-1 | Bacteria Actinobacteria Pseudonocardiales Pseudonocardiaceae *Kutzneria sp.* 744 |
| Lae-1-1 to Lae-1-2 | Bacteria Actinobacteria Pseudonocardiales Pseudonocardiaceae *Lechevalieria aerocolonigenes* |
| Lal-2-1 | Bacteria Actinobacteria Pseudonocardiales Pseudonocardiaceae *Lentzea albidocapillata* |
| Pru-1-1 | Bacteria Actinobacteria Pseudonocardiales Pseudonocardiaceae *Prauserella rugosa* |
| Psp-8-1 | Bacteria Actinobacteria Pseudonocardiales Pseudonocardiaceae *Prauserella sp.* Am3 |
| Pac-1-1 | Bacteria Actinobacteria Pseudonocardiales Pseudonocardiaceae *Pseudonocardia acaciae* DSM 45401 |
| Pas-1-1 | Bacteria Actinobacteria Pseudonocardiales Pseudonocardiaceae *Pseudonocardia asaccharolytica* DSM 44247 = NBRC 16224 |
| Pau-1-1 | Bacteria Actinobacteria Pseudonocardiales Pseudonocardiaceae *Pseudonocardia autotrophica* |
| Psp-9-1 | Bacteria Actinobacteria Pseudonocardiales Pseudonocardiaceae *Pseudonocardia sp.* P2 |
| Psp-10-1 | Bacteria Actinobacteria Pseudonocardiales Pseudonocardiaceae *Pseudonocardia spinosispora* DSM 44797 |
| [ir-1-1 | Bacteria Actinobacteria Pseudonocardiales Pseudonocardiaceae Saccharomonospora [Actinopolyspora] iraqiensis IQ-H1 |
| Spa-41-1 | Bacteria Actinobacteria Pseudonocardiales Pseudonocardiaceae *Saccharomonospora paurometabolica* YIM 90007 |
| Ssp-45-1 | Bacteria Actinobacteria Pseudonocardiales Pseudonocardiaceae *Saccharomonospora sp.* CNQ490 |
| Svi-8-1 | Bacteria Actinobacteria Pseudonocardiales Pseudonocardiaceae *Saccharomonospora viridis* |
| Sre-1-1 | Bacteria Actinobacteria Pseudonocardiales Pseudonocardiaceae *Saccharopolyspora rectivirgula* |
| Sre-2-1 | Bacteria Actinobacteria Pseudonocardiales Pseudonocardiaceae *Saccharopolyspora rectivirgula* DSM 43747 |
| Ssp-46-1 | Bacteria Actinobacteria Pseudonocardiales Pseudonocardiaceae *Saccharothrix sp.* NRRL B-16314 |
| Ssy-1-1 | Bacteria Actinobacteria Pseudonocardiales Pseudonocardiaceae *Saccharothrix syringae* |
| Ssp-47-1 | Bacteria Actinobacteria Pseudonocardiales Pseudonocardiaceae *Sciscionella sp.* SE31 |
| Kar-2-1 | Bacteria Actinobacteria Streptomycetales Streptomycetaceae *Kitasatospora arboriphila* |
| Kau-1-1 to Kau-1-2 | Bacteria Actinobacteria Streptomycetales Streptomycetaceae *Kitasatospora aureofaciens* |
| Kaz-1-1 | Bacteria Actinobacteria Streptomycetales Streptomycetaceae *Kitasatospora azatica* KCTC 9699 |
| Sal-2-1 | Bacteria Actinobacteria Streptomycetales Streptomycetaceae *Streptacidiphilus albus* |
| Sal-3-1 | Bacteria Actinobacteria Streptomycetales Streptomycetaceae *Streptacidiphilus albus* JL83 |
| Sje-1-1 | Bacteria Actinobacteria Streptomycetales Streptomycetaceae *Streptacidiphilus jeojiense* |
| Sji-1-1 | Bacteria Actinobacteria Streptomycetales Streptomycetaceae *Streptacidiphilus jiangxiensis* |
| Sru-1-1 | Bacteria Actinobacteria Streptomycetales Streptomycetaceae *Streptacidiphilus rugosus* AM-16 |
| [pa-1-1 | Bacteria Actinobacteria Streptomycetales Streptomycetaceae Streptomyces [Kitasatospora] papulosa |
| Asp-7-1 | Bacteria Actinobacteria Streptomycetales Streptomycetaceae Streptomyces *Actinosporangium sp.* NRRL B-3428 |
| Sac-1-1 | Bacteria Actinobacteria Streptomycetales Streptomycetaceae *Streptomyces achromogenes subsp. achromogenes* |
| Sal-4-1 | Bacteria Actinobacteria Streptomycetales Streptomycetaceae Streptomyces Streptomyces albidoflavus group *Streptomyces albidoflavus* |
| Sal-5-1 | Bacteria Actinobacteria Streptomycetales Streptomycetaceae *Streptomyces alboflavus* |
| Sal-6-1 | Bacteria Actinobacteria Streptomycetales Streptomycetaceae *Streptomyces alboviridis* |
| Sal-7-1 | Bacteria Actinobacteria Streptomycetales Streptomycetaceae *Streptomyces albus* |
| Sal-8-1 to Sal-8-4 | Bacteria Actinobacteria Streptomycetales Streptomycetaceae *Streptomyces albus subsp. albus* |
| Sam-1-1 | Bacteria Actinobacteria Streptomycetales Streptomycetaceae *Streptomyces ambofaciens* ATCC 23877 |
| San-1-1 | Bacteria Actinobacteria Streptomycetales Streptomycetaceae *Streptomyces antibioticus* |
| San-2-1 | Bacteria Actinobacteria Streptomycetales Streptomycetaceae *Streptomyces anulatus* |
| Sat-1-1 | Bacteria Actinobacteria Streptomycetales Streptomycetaceae *Streptomyces atratus* |
| Sat-2-1 | Bacteria Actinobacteria Streptomycetales Streptomycetaceae *Streptomyces atroolivaceus* |
| Sau-1-1 to Sau-1-2 | Bacteria Actinobacteria Streptomycetales Streptomycetaceae *Streptomyces aureocirculatus* |
| Sau-2-1 | Bacteria Actinobacteria Streptomycetales Streptomycetaceae *Streptomyces aureus* |
| Sav-2-1 to Sav-2-2 | Bacteria Actinobacteria Streptomycetales Streptomycetaceae *Streptomyces avermitilis* MA-4680 = NBRC 14893 |
| Sav-3-1 | Bacteria Actinobacteria Streptomycetales Streptomycetaceae *Streptomyces avicenniae* |
| Sba-2-1 | Bacteria Actinobacteria Streptomycetales Streptomycetaceae *Streptomyces baarnensis* |
| Sbi-2-1 to Sbi-2-2 | Bacteria Actinobacteria Streptomycetales Streptomycetaceae *Streptomyces bicolor* |
| Sbi-3-1 to Sbi-3-3 | Bacteria Actinobacteria Streptomycetales Streptomycetaceae *Streptomyces bikiniensis* |

| | |
|---|---|
| Sbr-1-1 | Bacteria Actinobacteria Streptomycetales Streptomycetaceae *Streptomyces brasiliensis* |
| Sca-3-1 | Bacteria Actinobacteria Streptomycetales Streptomycetaceae *Streptomyces caatingaensis* |
| Sca-4-1 to Sca-4-2 | Bacteria Actinobacteria Streptomycetales Streptomycetaceae *Streptomyces californicus* |
| Sca-5-1 to Sca-5-2 | Bacteria Actinobacteria Streptomycetales Streptomycetaceae *Streptomyces capuensis* |
| Sce-1-1 | Bacteria Actinobacteria Streptomycetales Streptomycetaceae *Streptomyces cellulosae* |
| Scy-2-1 | Bacteria Actinobacteria Streptomycetales Streptomycetaceae *Streptomyces cyaneofuscatus* |
| Scy-3-1 | Bacteria Actinobacteria Streptomycetales Streptomycetaceae *Streptomyces cyaneogriseus subsp. noncyanogenus* |
| Sdu-1-1 | Bacteria Actinobacteria Streptomycetales Streptomycetaceae *Streptomyces durhamensis* |
| Ser-3-1 | Bacteria Actinobacteria Streptomycetales Streptomycetaceae *Streptomyces erythrochromogenes* |
| Seu-1-1 | Bacteria Actinobacteria Streptomycetales Streptomycetaceae *Streptomyces europaeiscabiei* |
| Sex-1-1 to Sex-1-2 | Bacteria Actinobacteria Streptomycetales Streptomycetaceae *Streptomyces exfoliatus* |
| Sex-2-1 | Bacteria Actinobacteria Streptomycetales Streptomycetaceae *Streptomyces exfoliatus* DSM 41693 |
| Sfl-2-1 | Bacteria Actinobacteria Streptomycetales Streptomycetaceae *Streptomyces flaveolus* |
| Sfl-3-1 | Bacteria Actinobacteria Streptomycetales Streptomycetaceae *Streptomyces flaveus* |
| Sfl-4-1 | Bacteria Actinobacteria Streptomycetales Streptomycetaceae *Streptomyces flavidovirens* DSM 40150 |
| Sfl-5-1 | Bacteria Actinobacteria Streptomycetales Streptomycetaceae *Streptomyces flavochromogenes* |
| Sfl-6-1 | Bacteria Actinobacteria Streptomycetales Streptomycetaceae *Streptomyces flavotricini* |
| Sfl-7-1 | Bacteria Actinobacteria Streptomycetales Streptomycetaceae *Streptomyces flavovariabilis* |
| Sfl-8-1 | Bacteria Actinobacteria Streptomycetales Streptomycetaceae *Streptomyces flavovirens* |
| Sfl-9-1 | Bacteria Actinobacteria Streptomycetales Streptomycetaceae *Streptomyces floridae* |
| Sfu-2-1 | Bacteria Actinobacteria Streptomycetales Streptomycetaceae *Streptomyces fulvoviolaceus* |
| Sfu-3-1 | Bacteria Actinobacteria Streptomycetales Streptomycetaceae *Streptomyces fulvoviridis* |
| Sga-2-1 | Bacteria Actinobacteria Streptomycetales Streptomycetaceae *Streptomyces galbus* |
| Sgl-3-1 | Bacteria Actinobacteria Streptomycetales Streptomycetaceae *Streptomyces glaucescens* |
| Sgl-4-1 to Sgl-4-2 | Bacteria Actinobacteria Streptomycetales Streptomycetaceae *Streptomyces globisporus subsp. globisporus* |
| Sgr-5-1 | Bacteria Actinobacteria Streptomycetales Streptomycetaceae *Streptomyces graminis* |
| Sgr-6-1 | Bacteria Actinobacteria Streptomycetales Streptomycetaceae *Streptomyces griseoflavus* |
| Sgr-7-1 | Bacteria Actinobacteria Streptomycetales Streptomycetaceae *Streptomyces griseofuscus* |
| Sgr-8-1 | Bacteria Actinobacteria Streptomycetales Streptomycetaceae *Streptomyces griseolus* |
| Sgr-9-1 | Bacteria Actinobacteria Streptomycetales Streptomycetaceae *Streptomyces griseoluteus* |
| Sgr-10-1 | Bacteria Actinobacteria Streptomycetales Streptomycetaceae *Streptomyces griseorubens* |
| Sgr-11-1 | Bacteria Actinobacteria Streptomycetales Streptomycetaceae *Streptomyces griseus* |
| Sgr-12-1 to Sgr-12-9 | Bacteria Actinobacteria Streptomycetales Streptomycetaceae *Streptomyces griseus subsp. griseus* |
| Sgr-13-1 to Sgr-13-5 | Bacteria Actinobacteria Streptomycetales Streptomycetaceae *Streptomyces griseus subsp. rhodochrous* |
| Sha-2-1 | Bacteria Actinobacteria Streptomycetales Streptomycetaceae *Streptomyces halstedii* |
| Shy-4-1 | Bacteria Actinobacteria Streptomycetales Streptomycetaceae *Streptomyces hygroscopicus subsp. hygroscopicus* |
| Sia-1-1 | Bacteria Actinobacteria Streptomycetales Streptomycetaceae *Streptomyces iakyrus* |
| Sir-1-1 | Bacteria Actinobacteria Streptomycetales Streptomycetaceae *Streptomyces iranensis* |
| Ska-1-1 to Ska-1-2 | Bacteria Actinobacteria Streptomycetales Streptomycetaceae *Streptomyces katrae* |
| Sla-1-1 | Bacteria Actinobacteria Streptomycetales Streptomycetaceae *Streptomyces lavendulae* |
| Sla-2-1 to Sla-2-4 | Bacteria Actinobacteria Streptomycetales Streptomycetaceae *Streptomyces lavendulae subsp. lavendulae* |
| Sla-3-1 | Bacteria Actinobacteria Streptomycetales Streptomycetaceae *Streptomyces lavenduligriseus* |
| Sle-1-1 | Bacteria Actinobacteria Streptomycetales Streptomycetaceae *Streptomyces leeuwenhoekii* |
| Sli-2-1 to Sli-2-2 | Bacteria Actinobacteria Streptomycetales Streptomycetaceae *Streptomyces lividans* TK24 |
| Slu-1-1 | Bacteria Actinobacteria Streptomycetales Streptomycetaceae *Streptomyces luteus* |
| Sme-1-1 | Bacteria Actinobacteria Streptomycetales Streptomycetaceae *Streptomyces mediolani* |
| Smi-1-1 | Bacteria Actinobacteria Streptomycetales Streptomycetaceae *Streptomyces mirabilis* |
| Sno-1-1 | Bacteria Actinobacteria Streptomycetales Streptomycetaceae *Streptomyces nodosus* |
| Sod-1-1 | Bacteria Actinobacteria Streptomycetales Streptomycetaceae *Streptomyces odonnellii* |
| Sol-1-1 | Bacteria Actinobacteria Streptomycetales Streptomycetaceae *Streptomyces olindensis* |
| Sol-2-1 to Sol-2-2 | Bacteria Actinobacteria Streptomycetales Streptomycetaceae *Streptomyces olivaceus* |
| Spe-1-1 | Bacteria Actinobacteria Streptomycetales Streptomycetaceae *Streptomyces peruviensis* |
| Spe-2-1 | Bacteria Actinobacteria Streptomycetales Streptomycetaceae *Streptomyces peucetius* |
| Spl-1-1 to Spl-1-2 | Bacteria Actinobacteria Streptomycetales Streptomycetaceae *Streptomyces pluripotens* |
| Spu-2-1 to Spu-2-2 | Bacteria Actinobacteria Streptomycetales Streptomycetaceae *Streptomyces puniceus* |
| Spu-3-1 to Spu-3-2 | Bacteria Actinobacteria Streptomycetales Streptomycetaceae *Streptomyces purpeochromogenes* |
| Sre-3-1 | Bacteria Actinobacteria Streptomycetales Streptomycetaceae *Streptomyces resistomycificus* |
| Sri-2-1 to Sri-2-2 | Bacteria Actinobacteria Streptomycetales Streptomycetaceae *Streptomyces rimosus* |
| Sri-3-1 | Bacteria Actinobacteria Streptomycetales Streptomycetaceae *Streptomyces rimosus* R6-500 |
| Sri-4-1 to Sri-4-20 | Bacteria Actinobacteria Streptomycetales Streptomycetaceae *Streptomyces rimosus subsp. rimosus* |
| Sro-4-1 | Bacteria Actinobacteria Streptomycetales Streptomycetaceae *Streptomyces roseosporus* NRRL 11379 |
| Sro-5-1 | Bacteria Actinobacteria Streptomycetales Streptomycetaceae *Streptomyces roseoverticillatus* |
| Sro-6-1 | Bacteria Actinobacteria Streptomycetales Streptomycetaceae *Streptomyces roseus* |
| Sru-2-1 to Sru-2-2 | Bacteria Actinobacteria Streptomycetales Streptomycetaceae *Streptomyces ruber* |
| Ssc-3-1 to Ssc-3-2 | Bacteria Actinobacteria Streptomycetales Streptomycetaceae *Streptomyces scabiei* |

| | |
|---|---|
| Ssc-4-1 | Bacteria Actinobacteria Streptomycetales Streptomycetaceae *Streptomyces sclerotialus* |
| Ssc-5-1 | Bacteria Actinobacteria Streptomycetales Streptomycetaceae *Streptomyces scopuliridis* RB72 |
| Sse-1-1 | Bacteria Actinobacteria Streptomycetales Streptomycetaceae *Streptomyces seoulensis* |
| Ssp-48-1 | Bacteria Actinobacteria Streptomycetales Streptomycetaceae *Streptomyces sp.* 142MFCol3.1 |
| Ssp-50-1 | Bacteria Actinobacteria Streptomycetales Streptomycetaceae *Streptomyces sp.* AcH 505 |
| Ssp-49-1 | Bacteria Actinobacteria Streptomycetales Streptomycetaceae *Streptomyces sp.* AW19M42 |
| Ssp-51-1 | Bacteria Actinobacteria Streptomycetales Streptomycetaceae *Streptomyces sp.* CNH287 |
| Ssp-52-1 | Bacteria Actinobacteria Streptomycetales Streptomycetaceae *Streptomyces sp.* CNQ431 |
| Ssp-53-1 | Bacteria Actinobacteria Streptomycetales Streptomycetaceae *Streptomyces sp.* CNR698 |
| Ssp-54-1 | Bacteria Actinobacteria Streptomycetales Streptomycetaceae *Streptomyces sp.* CNS606 |
| Ssp-55-1 | Bacteria Actinobacteria Streptomycetales Streptomycetaceae *Streptomyces sp.* CNS654 |
| Ssp-56-1 | Bacteria Actinobacteria Streptomycetales Streptomycetaceae *Streptomyces sp.* CNT360 |
| Ssp-57-1 | Bacteria Actinobacteria Streptomycetales Streptomycetaceae *Streptomyces sp.* DpondAA-B6 |
| Ssp-59-1 | Bacteria Actinobacteria Streptomycetales Streptomycetaceae *Streptomyces sp.* FxanaA7 |
| Ssp-58-1 | Bacteria Actinobacteria Streptomycetales Streptomycetaceae *Streptomyces sp.* FXJ7.023 |
| Ssp-60-1 | Bacteria Actinobacteria Streptomycetales Streptomycetaceae *Streptomyces sp.* GXT6 |
| Ssp-61-1 | Bacteria Actinobacteria Streptomycetales Streptomycetaceae *Streptomyces sp.* HNS054 |
| Ssp-62-1 | Bacteria Actinobacteria Streptomycetales Streptomycetaceae *Streptomyces sp.* JS01 |
| Ssp-63-1 | Bacteria Actinobacteria Streptomycetales Streptomycetaceae *Streptomyces sp.* KE1 |
| Ssp-64-1 | Bacteria Actinobacteria Streptomycetales Streptomycetaceae *Streptomyces sp.* KhCrAH-337 |
| Ssp-65-1 | Bacteria Actinobacteria Streptomycetales Streptomycetaceae *Streptomyces sp.* KhCrAH-40 |
| Ssp-66-1 | Bacteria Actinobacteria Streptomycetales Streptomycetaceae *Streptomyces sp.* M10 |
| Ssp-67-1 | Bacteria Actinobacteria Streptomycetales Streptomycetaceae *Streptomyces sp.* MBT28 |
| Ssp-28-1 to Ssp-28-2 | Bacteria Actinobacteria Streptomycetales Streptomycetaceae *Streptomyces sp.* Mg1 |
| Ssp-68-1 | Bacteria Actinobacteria Streptomycetales Streptomycetaceae *Streptomyces sp.* MNU77 |
| Ssp-69-1 | Bacteria Actinobacteria Streptomycetales Streptomycetaceae *Streptomyces sp.* MUSC 125 |
| Ssp-70-1 | Bacteria Actinobacteria Streptomycetales Streptomycetaceae *Streptomyces sp.* MUSC119T |
| Ssp-71-1 | Bacteria Actinobacteria Streptomycetales Streptomycetaceae *Streptomyces sp.* MUSC136T |
| Ssp-72-1 | Bacteria Actinobacteria Streptomycetales Streptomycetaceae *Streptomyces sp.* MUSC149T |
| Ssp-73-1 | Bacteria Actinobacteria Streptomycetales Streptomycetaceae *Streptomyces sp.* MUSC164 |
| Ssp-74-1 | Bacteria Actinobacteria Streptomycetales Streptomycetaceae *Streptomyces sp.* NBRC 110035 |
| Ssp-75-1 | Bacteria Actinobacteria Streptomycetales Streptomycetaceae *Streptomyces sp.* NRRL B-11253 |
| Ssp-76-1 | Bacteria Actinobacteria Streptomycetales Streptomycetaceae *Streptomyces sp.* NRRL B-12105 |
| Ssp-77-1 | Bacteria Actinobacteria Streptomycetales Streptomycetaceae *Streptomyces sp.* NRRL B-1322 |
| Ssp-78-1 | Bacteria Actinobacteria Streptomycetales Streptomycetaceae *Streptomyces sp.* NRRL B-1347 |
| Ssp-79-1 | Bacteria Actinobacteria Streptomycetales Streptomycetaceae *Streptomyces sp.* NRRL B-1381 |
| Ssp-80-1 | Bacteria Actinobacteria Streptomycetales Streptomycetaceae *Streptomyces sp.* NRRL B-24051 |
| Ssp-81-1 | Bacteria Actinobacteria Streptomycetales Streptomycetaceae *Streptomyces sp.* NRRL B-24720 |
| Ssp-82-1 | Bacteria Actinobacteria Streptomycetales Streptomycetaceae *Streptomyces sp.* NRRL B-3229 |
| Ssp-83-1 | Bacteria Actinobacteria Streptomycetales Streptomycetaceae *Streptomyces sp.* NRRL B-3253 |
| Ssp-84-1 | Bacteria Actinobacteria Streptomycetales Streptomycetaceae *Streptomyces sp.* NRRL F-2202 |
| Ssp-85-1 | Bacteria Actinobacteria Streptomycetales Streptomycetaceae *Streptomyces sp.* NRRL F-2305 |
| Ssp-86-1 | Bacteria Actinobacteria Streptomycetales Streptomycetaceae *Streptomyces sp.* NRRL F-2580 |
| Ssp-87-1 | Bacteria Actinobacteria Streptomycetales Streptomycetaceae *Streptomyces sp.* NRRL F-2664 |
| Ssp-88-1 | Bacteria Actinobacteria Streptomycetales Streptomycetaceae *Streptomyces sp.* NRRL F-2747 |
| Ssp-89-1 | Bacteria Actinobacteria Streptomycetales Streptomycetaceae *Streptomyces sp.* NRRL F-2799 |
| Ssp-90-1 | Bacteria Actinobacteria Streptomycetales Streptomycetaceae *Streptomyces sp.* NRRL F-2890 |
| Ssp-91-1 | Bacteria Actinobacteria Streptomycetales Streptomycetaceae *Streptomyces sp.* NRRL F-3213 |
| Ssp-92-1 | Bacteria Actinobacteria Streptomycetales Streptomycetaceae *Streptomyces sp.* NRRL F-3218 |
| Ssp-93-1 | Bacteria Actinobacteria Streptomycetales Streptomycetaceae *Streptomyces sp.* NRRL F-3273 |
| Ssp-94-1 | Bacteria Actinobacteria Streptomycetales Streptomycetaceae *Streptomyces sp.* NRRL F-3307 |
| Ssp-95-1 | Bacteria Actinobacteria Streptomycetales Streptomycetaceae *Streptomyces sp.* NRRL F-4335 |
| Ssp-96-1 | Bacteria Actinobacteria Streptomycetales Streptomycetaceae *Streptomyces sp.* NRRL F-4428 |
| Ssp-97-1 | Bacteria Actinobacteria Streptomycetales Streptomycetaceae *Streptomyces sp.* NRRL F-4474 |
| Ssp-98-1 | Bacteria Actinobacteria Streptomycetales Streptomycetaceae *Streptomyces sp.* NRRL F-4835 |
| Ssp-99-1 | Bacteria Actinobacteria Streptomycetales Streptomycetaceae *Streptomyces sp.* NRRL F-5008 |
| Ssp-100-1 | Bacteria Actinobacteria Streptomycetales Streptomycetaceae *Streptomyces sp.* NRRL F-5053 |
| Ssp-101-1 | Bacteria Actinobacteria Streptomycetales Streptomycetaceae *Streptomyces sp.* NRRL F-5065 |
| Ssp-102-1 | Bacteria Actinobacteria Streptomycetales Streptomycetaceae *Streptomyces sp.* NRRL F-5126 |
| Ssp-103-1 | Bacteria Actinobacteria Streptomycetales Streptomycetaceae *Streptomyces sp.* NRRL F-5135 |
| Ssp-104-1 | Bacteria Actinobacteria Streptomycetales Streptomycetaceae *Streptomyces sp.* NRRL F-5140 |
| Ssp-105-1 | Bacteria Actinobacteria Streptomycetales Streptomycetaceae *Streptomyces sp.* NRRL F-5193 |
| Ssp-106-1 | Bacteria Actinobacteria Streptomycetales Streptomycetaceae *Streptomyces sp.* NRRL F-525 |
| Ssp-107-1 | Bacteria Actinobacteria Streptomycetales Streptomycetaceae *Streptomyces sp.* NRRL F-5527 |
| Ssp-108-1 | Bacteria Actinobacteria Streptomycetales Streptomycetaceae *Streptomyces sp.* NRRL F-5555 |

| | |
|---|---|
| Ssp-109-1 | Bacteria Actinobacteria Streptomycetales Streptomycetaceae *Streptomyces sp.* NRRL F-5630 |
| Ssp-110-1 | Bacteria Actinobacteria Streptomycetales Streptomycetaceae *Streptomyces sp.* NRRL F-5635 |
| Ssp-111-1 | Bacteria Actinobacteria Streptomycetales Streptomycetaceae *Streptomyces sp.* NRRL F-5639 |
| Ssp-112-1 | Bacteria Actinobacteria Streptomycetales Streptomycetaceae *Streptomyces sp.* NRRL F-5650 |
| Ssp-113-1 | Bacteria Actinobacteria Streptomycetales Streptomycetaceae *Streptomyces sp.* NRRL F-5681 |
| Ssp-114-1 | Bacteria Actinobacteria Streptomycetales Streptomycetaceae *Streptomyces sp.* NRRL F-5702 |
| Ssp-115-1 | Bacteria Actinobacteria Streptomycetales Streptomycetaceae *Streptomyces sp.* NRRL F-5727 |
| Ssp-116-1 | Bacteria Actinobacteria Streptomycetales Streptomycetaceae *Streptomyces sp.* NRRL F-5917 |
| Ssp-117-1 | Bacteria Actinobacteria Streptomycetales Streptomycetaceae *Streptomyces sp.* NRRL F-6628 |
| Ssp-118-1 | Bacteria Actinobacteria Streptomycetales Streptomycetaceae *Streptomyces sp.* NRRL F-6674 |
| Ssp-119-1 | Bacteria Actinobacteria Streptomycetales Streptomycetaceae *Streptomyces sp.* NRRL F-6676 |
| Ssp-120-1 | Bacteria Actinobacteria Streptomycetales Streptomycetaceae *Streptomyces sp.* NRRL F-6677 |
| Ssp-121-1 | Bacteria Actinobacteria Streptomycetales Streptomycetaceae *Streptomyces sp.* NRRL S-1022 |
| Ssp-122-1 | Bacteria Actinobacteria Streptomycetales Streptomycetaceae *Streptomyces sp.* NRRL S-104 |
| Ssp-123-1 | Bacteria Actinobacteria Streptomycetales Streptomycetaceae *Streptomyces sp.* NRRL S-118 |
| Ssp-124-1 | Bacteria Actinobacteria Streptomycetales Streptomycetaceae *Streptomyces sp.* NRRL S-1314 |
| Ssp-125-1 | Bacteria Actinobacteria Streptomycetales Streptomycetaceae *Streptomyces sp.* NRRL S-146 |
| Ssp-126-1 | Bacteria Actinobacteria Streptomycetales Streptomycetaceae *Streptomyces sp.* NRRL S-15 |
| Ssp-127-1 | Bacteria Actinobacteria Streptomycetales Streptomycetaceae *Streptomyces sp.* NRRL S-1824 |
| Ssp-128-1 | Bacteria Actinobacteria Streptomycetales Streptomycetaceae *Streptomyces sp.* NRRL S-1831 |
| Ssp-129-1 | Bacteria Actinobacteria Streptomycetales Streptomycetaceae *Streptomyces sp.* NRRL S-1868 |
| Ssp-130-1 | Bacteria Actinobacteria Streptomycetales Streptomycetaceae *Streptomyces sp.* NRRL S-1896 |
| Ssp-131-1 | Bacteria Actinobacteria Streptomycetales Streptomycetaceae *Streptomyces sp.* NRRL S-237 |
| Ssp-132-1 | Bacteria Actinobacteria Streptomycetales Streptomycetaceae *Streptomyces sp.* NRRL S-241 |
| Ssp-133-1 | Bacteria Actinobacteria Streptomycetales Streptomycetaceae *Streptomyces sp.* NRRL S-244 |
| Ssp-134-1 | Bacteria Actinobacteria Streptomycetales Streptomycetaceae *Streptomyces sp.* NRRL S-31 |
| Ssp-135-1 | Bacteria Actinobacteria Streptomycetales Streptomycetaceae *Streptomyces sp.* NRRL S-325 |
| Ssp-136-1 | Bacteria Actinobacteria Streptomycetales Streptomycetaceae *Streptomyces sp.* NRRL S-340 |
| Ssp-137-1 | Bacteria Actinobacteria Streptomycetales Streptomycetaceae *Streptomyces sp.* NRRL S-37 |
| Ssp-138-1 | Bacteria Actinobacteria Streptomycetales Streptomycetaceae *Streptomyces sp.* NRRL S-378 |
| Ssp-139-1 | Bacteria Actinobacteria Streptomycetales Streptomycetaceae *Streptomyces sp.* NRRL S-455 |
| Ssp-140-1 | Bacteria Actinobacteria Streptomycetales Streptomycetaceae *Streptomyces sp.* NRRL S-474 |
| Ssp-141-1 | Bacteria Actinobacteria Streptomycetales Streptomycetaceae *Streptomyces sp.* NRRL S-475 |
| Ssp-142-1 | Bacteria Actinobacteria Streptomycetales Streptomycetaceae *Streptomyces sp.* NRRL S-481 |
| Ssp-143-1 | Bacteria Actinobacteria Streptomycetales Streptomycetaceae *Streptomyces sp.* NRRL S-515 |
| Ssp-144-1 | Bacteria Actinobacteria Streptomycetales Streptomycetaceae *Streptomyces sp.* NRRL S-575 |
| Ssp-145-1 | Bacteria Actinobacteria Streptomycetales Streptomycetaceae *Streptomyces sp.* NRRL S-623 |
| Ssp-146-1 | Bacteria Actinobacteria Streptomycetales Streptomycetaceae *Streptomyces sp.* NRRL S-646 |
| Ssp-147-1 | Bacteria Actinobacteria Streptomycetales Streptomycetaceae *Streptomyces sp.* NRRL S-813 |
| Ssp-148-1 | Bacteria Actinobacteria Streptomycetales Streptomycetaceae *Streptomyces sp.* NRRL S-87 |
| Ssp-149-1 | Bacteria Actinobacteria Streptomycetales Streptomycetaceae *Streptomyces sp.* NRRL S-920 |
| Ssp-150-1 | Bacteria Actinobacteria Streptomycetales Streptomycetaceae *Streptomyces sp.* NRRL S-98 |
| Ssp-151-1 | Bacteria Actinobacteria Streptomycetales Streptomycetaceae *Streptomyces sp.* NRRL WC-3540 |
| Ssp-152-1 | Bacteria Actinobacteria Streptomycetales Streptomycetaceae *Streptomyces sp.* NRRL WC-3549 |
| Ssp-153-1 | Bacteria Actinobacteria Streptomycetales Streptomycetaceae *Streptomyces sp.* NRRL WC-3626 |
| Ssp-154-1 | Bacteria Actinobacteria Streptomycetales Streptomycetaceae *Streptomyces sp.* NRRL WC-3641 |
| Ssp-155-1 | Bacteria Actinobacteria Streptomycetales Streptomycetaceae *Streptomyces sp.* NRRL WC-3683 |
| Ssp-156-1 | Bacteria Actinobacteria Streptomycetales Streptomycetaceae *Streptomyces sp.* NRRL WC-3702 |
| Ssp-157-1 | Bacteria Actinobacteria Streptomycetales Streptomycetaceae *Streptomyces sp.* NRRL WC-3703 |
| Ssp-158-1 | Bacteria Actinobacteria Streptomycetales Streptomycetaceae *Streptomyces sp.* NRRL WC-3719 |
| Ssp-159-1 | Bacteria Actinobacteria Streptomycetales Streptomycetaceae *Streptomyces sp.* NRRL WC-3725 |
| Ssp-160-1 to Ssp-160-2 | Bacteria Actinobacteria Streptomycetales Streptomycetaceae *Streptomyces sp.* NRRL WC-3744 |
| Ssp-161-1 | Bacteria Actinobacteria Streptomycetales Streptomycetaceae *Streptomyces sp.* NRRL WC-3774 |
| Ssp-162-1 | Bacteria Actinobacteria Streptomycetales Streptomycetaceae *Streptomyces sp.* NRRL WC-3795 |
| Ssp-163-1 | Bacteria Actinobacteria Streptomycetales Streptomycetaceae *Streptomyces sp.* NTK 937 |
| Ssp-164-1 | Bacteria Actinobacteria Streptomycetales Streptomycetaceae *Streptomyces sp.* PCS3-D2 |
| Ssp-165-1 | Bacteria Actinobacteria Streptomycetales Streptomycetaceae *Streptomyces sp.* PRh5 |
| Ssp-166-1 | Bacteria Actinobacteria Streptomycetales Streptomycetaceae *Streptomyces sp.* RSD-27 |
| Ssp-167-1 | Bacteria Actinobacteria Streptomycetales Streptomycetaceae *Streptomyces sp.* SBT349 |
| Ssp-168-1 | Bacteria Actinobacteria Streptomycetales Streptomycetaceae *Streptomyces sp.* SolWspMP-sol2th |
| Ssp-169-1 | Bacteria Actinobacteria Streptomycetales Streptomycetaceae *Streptomyces sp.* TAA040 |
| Ssp-170-1 | Bacteria Actinobacteria Streptomycetales Streptomycetaceae *Streptomyces sp.* TAA204 |
| Ssp-171-1 | Bacteria Actinobacteria Streptomycetales Streptomycetaceae *Streptomyces sp.* TAA486 |
| Ssp-172-1 | Bacteria Actinobacteria Streptomycetales Streptomycetaceae *Streptomyces sp.* Tu 6176 |
| Ssp-173-1 | Bacteria Actinobacteria Streptomycetales Streptomycetaceae *Streptomyces sp.* UNC401CLCol |

| | |
|---|---|
| Ssp-174-1 | Bacteria Actinobacteria Streptomycetales Streptomycetaceae *Streptomyces sp.* URHA0041 |
| Ssp-175-1 | Bacteria Actinobacteria Streptomycetales Streptomycetaceae *Streptomyces sp.* WM6386 |
| Ssp-176-1 | Bacteria Actinobacteria Streptomycetales Streptomycetaceae *Streptomyces sp.* WM6391 |
| Ssp-177-1 | Bacteria Actinobacteria Streptomycetales Streptomycetaceae *Streptomyces sp.* WMMB 714 |
| Sst-1-1 | Bacteria Actinobacteria Streptomycetales Streptomycetaceae *Streptomyces stelliscabiei* |
| Sto-1-1 | Bacteria Actinobacteria Streptomycetales Streptomycetaceae *Streptomyces toyocaensis* |
| Sva-1-1 | Bacteria Actinobacteria Streptomycetales Streptomycetaceae *Streptomyces variegatus* |
| Sva-2-1 to Sva-2-2 | Bacteria Actinobacteria Streptomycetales Streptomycetaceae *Streptomyces varsoviensis* |
| Svi-9-1 | Bacteria Actinobacteria Streptomycetales Streptomycetaceae *Streptomyces vietnamensis* |
| Svi-10-1 | Bacteria Actinobacteria Streptomycetales Streptomycetaceae *Streptomyces vinaceus* |
| Svi-11-1 to Svi-11-2 | Bacteria Actinobacteria Streptomycetales Streptomycetaceae *Streptomyces violaceoruber* |
| Svi-12-1 | Bacteria Actinobacteria Streptomycetales Streptomycetaceae *Streptomyces violaceorubidus* |
| Svi-13-1 to Svi-13-3 | Bacteria Actinobacteria Streptomycetales Streptomycetaceae *Streptomyces virginiae* |
| Svi-14-1 to Svi-14-4 | Bacteria Actinobacteria Streptomycetales Streptomycetaceae *Streptomyces viridochromogenes* |
| Swa-1-1 | Bacteria Actinobacteria Streptomycetales Streptomycetaceae *Streptomyces wadayamensis* |
| Swe-1-1 | Bacteria Actinobacteria Streptomycetales Streptomycetaceae *Streptomyces wedmorensis* |
| Sxa-1-1 | Bacteria Actinobacteria Streptomycetales Streptomycetaceae *Streptomyces xanthophaeus* |
| Sxi-2-1 | Bacteria Actinobacteria Streptomycetales Streptomycetaceae *Streptomyces xiamenensis* |
| Sxy-1-1 | Bacteria Actinobacteria Streptomycetales Streptomycetaceae *Streptomyces xylophagus* |
| Sya-1-1 | Bacteria Actinobacteria Streptomycetales Streptomycetaceae *Streptomyces yangpuensis* |
| Sye-1-1 | Bacteria Actinobacteria Streptomycetales Streptomycetaceae *Streptomyces yeochonensis* CN732 |
| Sye-2-1 | Bacteria Actinobacteria Streptomycetales Streptomycetaceae *Streptomyces yerevanensis* |
| Hcr-1-1 | Bacteria Actinobacteria Streptosporangiales Streptosporangiaceae *Herbidospora cretacea* |
| Mro-1-1 | Bacteria Actinobacteria Streptosporangiales Streptosporangiaceae *Microbispora rosea subsp. nonnitritogenes* |
| Msp-30-1 | Bacteria Actinobacteria Streptosporangiales Streptosporangiaceae *Microbispora sp.* NRRL B-24597 |
| Mgl-2-1 | Bacteria Actinobacteria Streptosporangiales Streptosporangiaceae *Microtetraspora glauca* |
| Nca-3-1 | Bacteria Actinobacteria Streptosporangiales Streptosporangiaceae *Nonomuraea candida* |
| Nsp-11-1 | Bacteria Actinobacteria Streptosporangiales Streptosporangiaceae *Nonomuraea sp.* SBT364 |
| Sam-2-1 | Bacteria Actinobacteria Streptosporangiales Streptosporangiaceae *Streptosporangium amethystogenes* |
| Sro-7-1 | Bacteria Actinobacteria Streptosporangiales Streptosporangiaceae *Streptosporangium roseum* |
| Ama-5-1 | Bacteria Actinobacteria Streptosporangiales Thermomonosporaceae *Actinomadura madurae* LIID-AJ290 |
| Ari-2-1 | Bacteria Actinobacteria Streptosporangiales Thermomonosporaceae *Actinomadura rifamycini* DSM 43936 |
| Dsp-2-1 | Bacteria Firmicutes Clostridia Clostridiales Peptococcaceae *Dehalobacter sp.* FTH1 |
| env-1 to env-666 | environmental samples |

#### 9.2 Gene contexts

Each L31-Actinobacteria RNA (indicated by "RNA→") is listed. For each hit, the downstream genes predicted to reside in a regulated operon are listed. If the nearest downstream gene is encoding in the opposite strand (and therefore presumed to not be a part of a regulated operon), then that gene is still depicted. Some environmental sequences and some RefSeq entries lack gene annotations, and so no genes are available for such sequences. The direction of each gene is indicated with an arrow (→), and each predicted conserved domain in the gene is named. Conserved domains associated with more than one L31-Actinobacteria RNA are assigned a color; other domains are gray. Information about these conserved domains is given in Section 9.3. The accession of the sequence containing each L31-Actinobacteria RNA is given in the column named "Seq. accession". Accessions beginning with "NC_", "NS_", "NW_" or "NZ_" are contained in RefSeq. Other accession refer to environmental samples. Nucleotide coordinates are given for the 5′ and 3′ boundaries of each L31-Actinobacteria RNA. If the 5′ coordinate is greater than the 3′ coordinate, the RNA is present on the reverse-complement strand of the containing genomic DNA sequence. Each hit is denoted by an abbreviation (like "Eco-1-1") that refers to a taxonomy given in Section 9.1.

| abbrev. | Seq. accession | | 5′ at | 3′ at | genes |
|---|---|---|---|---|---|
| env-1 | 3300001239_10346507 | - | 863 | 813 | **RNA→** rpmE (PRK00019)Ribosomal_L31 (pfam01197)→ PCRF (smart00937)PCRF (pfam03462)→ |
| env-2 | topACOD_FV90NF401DDBWP | - | 101 | 65 | **RNA→** |
| Vca-1-1 | NZ_ATUF01000002.1 | - | 402422 | 402378 | **RNA→** rpmE (PRK00019)Ribosomal_L31 (pfam01197)→ hypo→ |
| env-3 | JGI12397J12024_1050070 | + | 67 | 116 | **RNA→** rpmE (PRK00019)Ribosomal_L31 (pfam01197)→ |
| env-4 | JGI24736J21556_1099768 | + | 224 | 283 | **RNA→** rpmE (PRK00019)Ribosomal_L31 (pfam01197)→ |
| env-5 | JGI24739J22299_10173028 | - | 428 | 369 | **RNA→** rpmE (PRK00019)Ribosomal_L31 (pfam01197)→ |
| env-6 | JGI24737J22298_10039126 | - | 417 | 358 | **RNA→** rpmE (PRK00019)Ribosomal_L31 (pfam01197)→ |

| | | | | | |
|---|---|---|---|---|---|
| Cac-1-1 | NC_013131.1 | + | 1360258 | 1360298 | **RNA**→ rpmE (PRK00019)Ribosomal_L31 (pfam01197)→ |
| env-7 | JGI12397J12024_1339528 | + | 56 | 114 | **RNA**→ hypo→ |
| env-8 | LWAnNNiSIP_GDN60OX01COTOZ | + | 264 | 310 | **RNA**→ |
| env-9 | JGI12396J12026_1108615 | + | 2 | 43 | **RNA**→ rpmE (PRK00019)Ribosomal_L31 (pfam01197)→ |
| env-10 | JGI12467J12023_1043457 | + | 377 | 418 | **RNA**→ rpmE (PRK00019)Ribosomal_L31 (pfam01197)→ prfA (PRK00591)PCRF (pfam03462)→ |
| env-11 | JGI12396J12026_1011438 | + | 24 | 84 | **RNA**→ ←hypo |
| env-12 | JGI12273J12029_10084767 | + | 24 | 84 | **RNA**→ rpmE (PRK00019)Ribosomal_L31 (pfam01197)→ |
| env-13 | JGI2065J20421_1025414 | + | 78 | 138 | **RNA**→ rpmE (PRK00019)Ribosomal_L31 (pfam01197)→ PCRF (smart00937)PCRF (pfam03462)→ |
| env-14 | JGI24770J26754_10678085 | + | 122 | 174 | **RNA**→ rpmE (PRK00019)Ribosomal_L31 (pfam01197)→ |
| Apo-1-1 | NZ_CCXH01000177.1 | + | 1390 | 1448 | **RNA**→ rpmE (PRK00019)Ribosomal_L31 (pfam01197)→ |
| env-15 | botACOD_GAKN62C01CC25B | + | 172 | 219 | **RNA**→ rpmE (PRK00019)Ribosomal_L31 (pfam01197)→ |
| env-16 | AglaG_GDN60OX02G4DXO | - | 301 | 242 | **RNA**→ rpmE (PRK00019)Ribosomal_L31 (pfam01197)→ |
| env-17 | JGI12396J12026_1002764 | + | 4 | 65 | **RNA**→ ←hypo |
| env-18 | JGI12396J12026_1094345 | + | 9 | 87 | **RNA**→ rpmE (PRK00019)Ribosomal_L31 (pfam01197)→ |
| Ama-3-1 | NZ_CM001024.1 | + | 677475 | 677520 | **RNA**→ rpmE (PRK00019)Ribosomal_L31 (pfam01197)→ PrfA (COG0216)prfA (PRK00591)→ |
| | | | | | HemK (COG2890)PRK09328 (PRK09328)→ HTH_LUXR (smart00421)CitB (COG2197)→ |
| | | | | | HTH_LUXR (smart00421)CitB (COG2197)→ |
| Nin-1-1 | NZ_KE383929.1 | - | 680299 | 680255 | **RNA**→ rpmE (PRK00019)Ribosomal_L31 (pfam01197)→ |
| env-19 | JGI11800J13271_1023285 | - | 253 | 187 | **RNA**→ rpmE (PRK00019)Ribosomal_L31 (pfam01197)→ |
| env-20 | CEUF01060573.1 | - | 405 | 339 | **RNA**→ rpmE (PRK00019)Ribosomal_L31 (pfam01197)→ |
| Jga-1-1 | NZ_KI911782.1 | + | 1567026 | 1567073 | **RNA**→ rpmE (PRK00019)Ribosomal_L31 (pfam01197)→ |
| Hcr-1-1 | NZ_JODQ01000002.1 | - | 16089 | 16035 | **RNA**→ rpmE (PRK00019)Ribosomal_L31 (pfam01197)→ |
| env-21 | JGI24735J21928_10338363 | - | 327 | 278 | **RNA**→ rpmE (PRK00019)Ribosomal_L31 (pfam01197)→ |
| Asu-2-1 | NZ_KE386848.1 | + | 67014 | 67059 | **RNA**→ rpmE (PRK00019)Ribosomal_L31 (pfam01197)→ |
| | | | | | prfA (PRK00591)RF-1 (pfam00472)PCRF (pfam03462)→ HemK (COG2890)PRK09328 (PRK09328)→ |
| Psp-8-1 | NZ_JTJI01000005.1 | + | 27456 | 27507 | **RNA**→ rpmE (PRK00019)Ribosomal_L31 (pfam01197)→ |
| Pru-1-1 | NZ_JOIJ01000022.1 | + | 44825 | 44876 | **RNA**→ rpmE (PRK00019)Ribosomal_L31 (pfam01197)→ |
| env-22 | JGI25321J39296_1051856 | - | 129 | 78 | **RNA**→ rpmE (PRK00528)rpmE (PRK00019)Ribosomal_L31 (pfam01197)→ |
| env-23 | JGI25321J50212_10048695 | - | 336 | 285 | **RNA**→ rpmE (PRK00019)Ribosomal_L31 (pfam01197)→ |
| env-24 | JGI25317J35169_1083333 | + | 5 | 56 | **RNA**→ rpmE (PRK00019)Ribosomal_L31 (pfam01197)→ |
| env-25 | C688J13580_1096905 | + | 58 | 98 | **RNA**→ hypo→ rpmE (PRK00019)Ribosomal_L31 (pfam01197)→ |
| Sfl-5-1 | NZ_JNZO01000018.1 | + | 362423 | 362470 | **RNA**→ rpmE (PRK00019)Ribosomal_L31 (pfam01197)→ PrfA (COG0216)prfA (PRK00591)→ |
| | | | | | HemK (COG2890)PRK09328 (PRK09328)→ SUA5 (COG0009)TIGR00057 (TIGR00057)→ hypo→ hypo→ hypo→ |
| | | | | | hypo→ GlyA (COG0112)glyA (PRK00011)→ |
| Jmu-1-1 | NZ_KQ434753.1 | - | 67929 | 67882 | **RNA**→ rpmE (PRK00019)Ribosomal_L31 (pfam01197)→ PrfA (COG0216)prfA (PRK00591)→ |
| | | | | | HemK (COG2890)PRK09328 (PRK09328)→ SUA5 (COG0009)TIGR00057 (TIGR00057)→ |
| | | | | | GT_WecA_like (cd06853)mraY (PRK00108)→ hypo→ AtpI2 (COG5336)ATPase_gene1 (pfam09527)→ |
| | | | | | PRK05815 (PRK05815)ATP-synt_A (pfam00119)→ PRK06876 (PRK06876)AtpE (COG0636)→ |
| | | | | | PRK05759 (PRK05759)AtpF (COG0711)→ PRK13430 (PRK13430)OSCP (pfam00213)→ |
| | | | | | AtpA (COG0056)PRK09281 (PRK09281)→ PRK05621 (PRK05621)ATP-synt (pfam00231)→ |
| env-26 | JGI12397J12024_1025645 | + | 23 | 68 | **RNA**→ rpmE (PRK00019)Ribosomal_L31 (pfam01197)→ |
| Ssp-2-1 | NZ_KB911723.1 | - | 80543 | 80496 | **RNA**→ rpmE (PRK00019)Ribosomal_L31 (pfam01197)→ |
| env-27 | FWIROz_GKA24FP02JUPYD | + | 39 | 96 | **RNA**→ rpmE (PRK00019)Ribosomal_L31 (pfam01197)→ |
| env-28 | 3300001592_10007801 | - | 2906 | 2841 | **RNA**→ rpmE (PRK00019)Ribosomal_L31 (pfam01197)→ PrfA (COG0216)prfA (PRK00591)→ |
| | | | | | HemK (COG2890)PRK09328 (PRK09328)→ SUA5 (COG0009)TIGR00057 (TIGR00057)→ |
| Cbo-1-1 | NZ_AXCZ01000025.1 | - | 21537 | 21491 | **RNA**→ rpmE (PRK00019)Ribosomal_L31 (pfam01197)→ |
| | | | | | prfA (PRK00591)RF-1 (pfam00472)PCRF (pfam03462)→ HemK (COG2890)PRK09328 (PRK09328)→ |
| | | | | | SUA5 (COG0009)TIGR00057 (TIGR00057)→ GT_WecA_like (cd06853)mraY (PRK00108)→ hypo→ |
| aLL-1-1 | NZ_KI301992.1 | - | 1665648 | 1665606 | **RNA**→ rpmE (PRK00019)Ribosomal_L31 (pfam01197)→ PrfA (COG0216)prfA (PRK00591)→ |
| | | | | | HemK (COG2890)PRK09328 (PRK09328)→ |
| Msp-29-1 | NZ_KE384490.1 | + | 243863 | 243905 | **RNA**→ rpmE (PRK00019)Ribosomal_L31 (pfam01197)→ |
| Mch-5-1 | NZ_JYNX01000034.1 | - | 42359 | 42308 | **RNA**→ rpmE (PRK00019)Ribosomal_L31 (pfam01197)→ PrfA (COG0216)prfA (PRK00591)→ |
| | | | | | HemK (COG2890)PRK09328 (PRK09328)→ SUA5 (COG0009)TIGR00057 (TIGR00057)→ |
| | | | | | GT_WecA_like (cd06853)mraY (PRK00108)→ |

| | | | | | |
|---|---|---|---|---|---|
| Mch-4-1 | NZ_JYNL01000069.1 | - | 3786 | 3735 | **RNA**→ rpmE (PRK00019)Ribosomal_L31 (pfam01197)→ PrfA (COG0216)prfA (PRK00591)→<br>HemK (COG2890)PRK09328 (PRK09328)→ SUA5 (COG0009)TIGR00057 (TIGR00057)→ hypo→ hypo→ |
| Aha-2-1 | NZ_JUZA01000019.1 | + | 44897 | 44933 | **RNA**→ rpmE (PRK00019)Ribosomal_L31 (pfam01197)→<br>prfA (PRK00591)RF-1 (pfam00472)PCRF (pfam03462)→ hypo→ hypo→<br>SUA5 (COG0009)TIGR00057 (TIGR00057)→ SUA5 (COG0009)TIGR00057 (TIGR00057)→<br>GT_WecA_like (cd06853)mraY (PRK00108)→ hypo→ PRK05815 (PRK05815)ATP-synt_A (pfam00119)→<br>PRK06876 (PRK06876)AtpE (COG0636)→ PRK05759 (PRK05759)AtpF (COG0711)→<br>PRK13430 (PRK13430)OSCP (pfam00213)→ AtpA (COG0056)PRK09281 (PRK09281)→<br>PRK05621 (PRK05621)ATP-synt (pfam00231)→ AtpD (COG0055)PRK09280 (PRK09280)→ |
| Ane-1-1 | NZ_JH815214.1 | + | 304462 | 304498 | **RNA**→ rpmE (PRK00019)Ribosomal_L31 (pfam01197)→<br>prfA (PRK00591)RF-1 (pfam00472)PCRF (pfam03462)→ hypo→ hypo→<br>SUA5 (COG0009)HemK (COG2890)PRK09328 (PRK09328)TIGR00057 (TIGR00057)→<br>SUA5 (COG0009)TIGR00057 (TIGR00057)→ SUA5 (COG0009)TIGR00057 (TIGR00057)→<br>GT_WecA_like (cd06853)mraY (PRK00108)→ hypo→ PRK05815 (PRK05815)ATP-synt_A (pfam00119)→<br>PRK06876 (PRK06876)AtpE (COG0636)→ PRK05759 (PRK05759)AtpF (COG0711)→<br>PRK13430 (PRK13430)OSCP (pfam00213)→ AtpA (COG0056)PRK09281 (PRK09281)→<br>PRK05621 (PRK05621)ATP-synt (pfam00231)→ AtpD (COG0055)PRK09280 (PRK09280)→ |
| env-29 | PRSSGFe2_Sequence0000001542 | - | 7752 | 7691 | **RNA**→ rpmE (PRK00019)Ribosomal_L31 (pfam01197)→ PrfA (COG0216)prfA (PRK00591)→<br>HemK (COG2890)PRK09328 (PRK09328)→ SUA5 (COG0009)TIGR00057 (TIGR00057)→<br>ECA_wecA (TIGR02380)GT_WecA_like (cd06853)→ hypo→ AtpI2 (COG5336)ATPase_gene1 (pfam09527)→<br>PRK05815 (PRK05815)ATP-synt_A (pfam00119)→ PRK07874 (PRK07874)AtpE (COG0636)→<br>PRK05759 (PRK05759)AtpF (COG0711)→ PRK13430 (PRK13430)OSCP (pfam00213)→<br>PRK09281 (PRK09281)ATP-synt_ab_N (pfam02874)→ |
| Sni-1-1 | NZ_AQXZ01000013.1 | - | 17062 | 17017 | **RNA**→ rpmE (PRK00019)Ribosomal_L31 (pfam01197)→ |
| env-30 | JGI24770J26754_10040473 | + | 643 | 688 | **RNA**→ rpmE (PRK00019)Ribosomal_L31 (pfam01197)→ PrfA (COG0216)prfA (PRK00591)→ |
| Asp-2-1 | NC_017803.1 | - | 8255931 | 8255885 | **RNA**→ rpmE (PRK00019)Ribosomal_L31 (pfam01197)→ PrfA (COG0216)prfA (PRK00591)→ |
| env-31 | DCrCPGB_c279856 | - | 114 | 45 | **RNA**→ hypo→ |
| Hal-1-1 | NZ_AZUQ01000001.1 | + | 2107349 | 2107408 | **RNA**→ rpmE (PRK00019)Ribosomal_L31 (pfam01197)→ PrfA (COG0216)prfA (PRK00591)→<br>HemK (COG2890)PRK09328 (PRK09328)→ |
| env-32 | 4491734.3.12108778 | + | 329 | 367 | **RNA**→ rpmE (PRK00019)Ribosomal_L31 (pfam01197)→ |
| Ssp-169-1 | NZ_AUKR01000002.1 | - | 319811 | 319766 | **RNA**→ rpmE (PRK00019)Ribosomal_L31 (pfam01197)→ |
| Dsp-2-1 | NZ_AQYY01000001.1 | + | 1361355 | 1361413 | **RNA**→ rpmE (PRK00019)Ribosomal_L31 (pfam01197)→ PrfA (COG0216)prfA (PRK00591)→<br>HemK (COG2890)PRK09328 (PRK09328)→ SUA5 (COG0009)TIGR00057 (TIGR00057)→<br>GT_WecA_like (cd06853)mraY (PRK00108)→ hypo→ AtpI2 (COG5336)SH3_and_anchor (TIGR04211)→<br>PRK05815 (PRK05815)ATP-synt_A (pfam00119)→ PRK05815 (PRK05815)ATP-synt_A (pfam00119)→<br>PRK07874 (PRK07874)AtpE (COG0636)→ PRK05759 (PRK05759)AtpF (COG0711)→<br>PRK13430 (PRK13430)OSCP (pfam00213)→ AtpA (COG0056)PRK09281 (PRK09281)→<br>PRK05621 (PRK05621)ATP-synt (pfam00231)→ AtpD (COG0055)PRK09280 (PRK09280)→ |
| Mel-1-1 | NZ_LBNO01000049.1 | - | 15168 | 15110 | **RNA**→ rpmE (PRK00019)Ribosomal_L31 (pfam01197)→ PrfA (COG0216)prfA (PRK00591)→<br>HemK (COG2890)PRK09328 (PRK09328)→ SUA5 (COG0009)TIGR00057 (TIGR00057)→<br>GT_WecA_like (cd06853)mraY (PRK00108)→ |
| env-33 | FNTS067_GJ87FRN01EXCYZ | - | 173 | 120 | **RNA**→ hypo→ |
| Car-1-1 | NZ_KK073874.1 | - | 7435021 | 7434972 | **RNA**→ rpmE (PRK00019)Ribosomal_L31 (pfam01197)→ PrfA (COG0216)prfA (PRK00591)→ |
| env-34 | ATED_F4WRAXD01CHLUA | - | 225 | 176 | **RNA**→ rpmE (PRK00019)Ribosomal_L31 (pfam01197)→ |
| Asp-3-1 | NZ_AGVX02000017.1 | - | 538 | 468 | **RNA**→ rpmE (PRK00019)Ribosomal_L31 (pfam01197)→ |
| env-35 | FNTS_GKKEVUV01AVOLL | - | 130 | 92 | **RNA**→ rpmE (PRK00019)Ribosomal_L31 (pfam01197)→ |
| env-36 | FNTS_GKKEVUV01BHC90 | + | 147 | 184 | **RNA**→ |
| Mth-1-1 | NZ_AGVE01000042.1 | - | 531859 | 531799 | **RNA**→ rpmE (PRK00019)Ribosomal_L31 (pfam01197)→ rpmE (PRK00019)Ribosomal_L31 (pfam01197)→ |
| env-37 | JGI24740J21852_10021611 | + | 341 | 395 | **RNA**→ rpmE (PRK00019)Ribosomal_L31 (pfam01197)→ PrfA (COG0216)prfA (PRK00591)→<br>HemK (COG2890)PRK09328 (PRK09328)→ |
| env-38 | JGI24735J21928_10010623 | + | 1057 | 1111 | **RNA**→ rpmE (PRK00019)Ribosomal_L31 (pfam01197)→ PrfA (COG0216)prfA (PRK00591)→<br>HemK (COG2890)PRK09328 (PRK09328)→ |
| env-39 | APMI01059580.1 | - | 1604 | 1551 | **RNA**→ rpmE (PRK00019)Ribosomal_L31 (pfam01197)→ |
| env-40 | JGI12397J12024_1172898 | + | 68 | 120 | **RNA**→ rpmE (PRK00019)Ribosomal_L31 (pfam01197)→ |
| env-41 | JGI12396J12026_1047907 | + | 22 | 72 | **RNA**→ rpmE (PRK00019)Ribosomal_L31 (pfam01197)→ |

| | | | | | |
|---|---|---|---|---|---|
| env-42 | JGI12397J12024_1022031 | + | 22 | 72 | RNA→ rpmE (PRK00019)Ribosomal_L31 (pfam01197)→ |
| Ssp-96-1 | NZ_JYJI01000122.1 | - | 1881 | 1833 | RNA→ rpmE (PRK00019)Ribosomal_L31 (pfam01197)→ |
| Sfl-6-1 | NZ_JNXV01000514.1 | + | 1188 | 1236 | RNA→ rpmE (PRK00019)Ribosomal_L31 (pfam01197)→ |
| Sya-1-1 | NZ_LBMK01000002.1 | + | 777882 | 777930 | RNA→ rpmE (PRK00019)Ribosomal_L31 (pfam01197)→ |
| Ssp-87-1 | NZ_JOFX01000020.1 | + | 50692 | 50740 | RNA→ rpmE (PRK00019)Ribosomal_L31 (pfam01197)→ |
| Ssp-122-1 | NZ_JZWW01000282.1 | - | 1731 | 1683 | RNA→ rpmE (PRK00019)Ribosomal_L31 (pfam01197)→ |
| Ssp-150-1 | NZ_JOJD01000020.1 | - | 213028 | 212980 | RNA→ rpmE (PRK00019)Ribosomal_L31 (pfam01197)→ |
| Svi-13-1 | NZ_JOAK01000005.1 | - | 213389 | 213341 | RNA→ rpmE (PRK00019)Ribosomal_L31 (pfam01197)→ |
| Ssp-97-1 | NZ_JOIB01000042.1 | + | 22450 | 22498 | RNA→ rpmE (PRK00019)Ribosomal_L31 (pfam01197)→ |
| Ssp-148-1 | NZ_JOGB01000028.1 | - | 45090 | 45041 | RNA→ rpmE (PRK00019)Ribosomal_L31 (pfam01197)→ |
| Ssp-88-1 | NZ_JOIS01000002.1 | + | 34345 | 34393 | RNA→ rpmE (PRK00019)Ribosomal_L31 (pfam01197)→ |
| Ssp-164-1 | NZ_JDUZ01000009.1 | + | 57343 | 57391 | RNA→ rpmE (PRK00019)Ribosomal_L31 (pfam01197)→ |
| Svi-13-2 | NZ_LGUV01000348.1 | + | 15469 | 15517 | RNA→ rpmE (PRK00019)Ribosomal_L31 (pfam01197)→ |
| Ssp-95-1 | NZ_JOFY01000017.1 | - | 211039 | 210991 | RNA→ rpmE (PRK00019)Ribosomal_L31 (pfam01197)→ |
| Ssp-132-1 | NZ_JOCY01000015.1 | - | 213281 | 213233 | RNA→ rpmE (PRK00019)Ribosomal_L31 (pfam01197)→ |
| Ssp-28-1 | NZ_DS570390.1 | + | 37435 | 37483 | RNA→ rpmE (PRK00019)Ribosomal_L31 (pfam01197)→ rpmE (PRK00019)Ribosomal_L31 (pfam01197)→ |
| Ssp-28-2 | NZ_CP011664.1 | + | 5363260 | 5363308 | RNA→ rpmE (PRK00019)Ribosomal_L31 (pfam01197)→ |
| Ssp-8-1 | NZ_GG657750.1 | + | 5365177 | 5365225 | RNA→ rpmE (PRK00019)Ribosomal_L31 (pfam01197)→ rpmE (PRK00019)Ribosomal_L31 (pfam01197)→ |
| Ssp-143-1 | NZ_JODE01000026.1 | - | 95012 | 94964 | RNA→ rpmE (PRK00019)Ribosomal_L31 (pfam01197)→ |
| Ssp-144-1 | NZ_JODJ01000019.1 | - | 95012 | 94964 | RNA→ rpmE (PRK00019)Ribosomal_L31 (pfam01197)→ |
| Sla-2-1 | NZ_JOEW01000030.1 | + | 23758 | 23806 | RNA→ rpmE (PRK00019)Ribosomal_L31 (pfam01197)→ |
| Ska-1-1 | NZ_JZWV01000546.1 | + | 4574 | 4622 | RNA→ rpmE (PRK00019)Ribosomal_L31 (pfam01197)→ |
| Sro-6-1 | NZ_LFML01000043.1 | - | 44786 | 44738 | RNA→ rpmE (PRK00019)Ribosomal_L31 (pfam01197)→ |
| Ssp-133-1 | NZ_JOCX01000002.1 | + | 316404 | 316452 | RNA→ rpmE (PRK00019)Ribosomal_L31 (pfam01197)→ |
| Sxa-1-1 | NZ_KL647135.1 | - | 194665 | 194617 | RNA→ rpmE (PRK00019)Ribosomal_L31 (pfam01197)→ |
| Ska-1-2 | NZ_JNZY01000009.1 | - | 192103 | 192055 | RNA→ rpmE (PRK00019)Ribosomal_L31 (pfam01197)→ |
| Ssp-138-1 | NZ_JOBX01000049.1 | - | 61013 | 60965 | RNA→ rpmE (PRK00019)Ribosomal_L31 (pfam01197)→ |
| Ser-3-1 | NZ_JNZG01000011.1 | + | 30135 | 30183 | RNA→ rpmE (PRK00019)Ribosomal_L31 (pfam01197)→ |
| Sla-2-2 | NZ_JOBU01000032.1 | + | 31087 | 31135 | RNA→ rpmE (PRK00019)Ribosomal_L31 (pfam01197)→ |
| Ssp-77-1 | NZ_JOHF01000038.1 | + | 31053 | 31101 | RNA→ rpmE (PRK00019)Ribosomal_L31 (pfam01197)→ |
| Ssp-131-1 | NZ_JODA01000013.1 | - | 129059 | 129011 | RNA→ rpmE (PRK00019)Ribosomal_L31 (pfam01197)→ |
| Ssp-86-1 | NZ_JOIR01000005.1 | + | 53139 | 53187 | RNA→ rpmE (PRK00019)Ribosomal_L31 (pfam01197)→ |
| Sla-1-1 | NZ_JNXL01000021.1 | + | 3212 | 3260 | RNA→ rpmE (PRK00019)Ribosomal_L31 (pfam01197)→ |
| Ssp-166-1 | NZ_JWZS01001028.1 | - | 3788 | 3740 | RNA→ rpmE (PRK00019)Ribosomal_L31 (pfam01197)→ |
| Svi-13-3 | NZ_JNYC01000007.1 | + | 377227 | 377275 | RNA→ rpmE (PRK00019)Ribosomal_L31 (pfam01197)→ |
| Dsp-1-1 | NZ_KE340308.1 | - | 2146886 | 2146842 | RNA→ rpmE (PRK00019)Ribosomal_L31 (pfam01197)→ |
| Dho-1-1 | NZ_KN323183.1 | - | 2148108 | 2148064 | RNA→ rpmE (PRK00019)Ribosomal_L31 (pfam01197)→ |
| Bpa-1-1 | NZ_AGSO01000002.1 | - | 39806 | 39767 | RNA→ rpmE (PRK00019)Ribosomal_L31 (pfam01197)→ rpmE (PRK00019)Ribosomal_L31 (pfam01197)→ |
| Bph-1-1 | NZ_KK069988.1 | + | 25747 | 25790 | RNA→ rpmE (PRK00019)Ribosomal_L31 (pfam01197)→ PrfB (COG1186)prfA (PRK00591)→ |
| | | | | | HemK (COG2890)PRK09328 (PRK09328)→ hypo→ SUA5 (COG0009)TIGR00057 (TIGR00057)→ |
| | | | | | GT_WecA_like (cd06853)mraY (PRK00108)→ hypo→ |
| Bfa-1-1 | NC_013172.1 | - | 2159943 | 2159900 | RNA→ rpmE (PRK00019)Ribosomal_L31 (pfam01197)→ |
| env-43 | 2204806299 | + | 97 | 140 | RNA→ rpmE (PRK00019)Ribosomal_L31 (pfam01197)→ |
| env-44 | 2158018103 | + | 97 | 140 | RNA→ rpmE (PRK00019)Ribosomal_L31 (pfam01197)→ |
| env-45 | 2157611060 | - | 212 | 169 | RNA→ rpmE (PRK00019)Ribosomal_L31 (pfam01197)→ |
| env-46 | 2204808654 | - | 212 | 169 | RNA→ rpmE (PRK00019)Ribosomal_L31 (pfam01197)→ |
| Bmu-1-1 | NZ_AORC01000017.1 | + | 14987 | 15030 | RNA→ rpmE (PRK00019)Ribosomal_L31 (pfam01197)→ |
| | | | | | PCRF (smart00937)RF-1 (pfam00472)PCRF (pfam03462)→ HemK (COG2890)PRK09328 (PRK09328)→ |
| | | | | | PHA03169 (PHA03169)SUA5 (COG0009)TIGR00057 (TIGR00057)→ |
| | | | | | GT_WecA_like (cd06853)mraY (PRK00108)→ hypo→ hypo→ |
| Bsq-1-1 | NZ_AGBX01000007.1 | - | 600878 | 600835 | RNA→ rpmE (PRK00019)Ribosomal_L31 (pfam01197)→ rpmE (PRK00019)Ribosomal_L31 (pfam01197)→ |
| Sba-1-1 | NZ_KI547049.1 | + | 42821 | 42866 | RNA→ rpmE (PRK00019)Ribosomal_L31 (pfam01197)→ |
| env-47 | P3_CLC_Cons55403 | - | 298 | 252 | RNA→ rpmE (PRK00019)Ribosomal_L31 (pfam01197)→ |

| | | | | | |
|---|---|---|---|---|---|
| env-48 | JGI24770J26754_10015274 | + | 2104 | 2148 | **RNA**→ rpmE (PRK00019)Ribosomal_L31 (pfam01197)→ PrfA (COG0216)prfA (PRK00591)→ HemK (COG2890)PRK09328 (PRK09328)→ |
| env-49 | JGI12467J12023_1082663 | + | 17 | 88 | **RNA**→ hypo→ rpmE (PRK00019)Ribosomal_L31 (pfam01197)→ |
| env-50 | 3300001592_10976328 | + | 61 | 104 | **RNA**→ rpmE (PRK00019)Ribosomal_L31 (pfam01197)→ |
| env-51 | 3300001592_10499386 | + | 49 | 92 | **RNA**→ rpmE (PRK00019)Ribosomal_L31 (pfam01197)→ |
| env-52 | JGI12273J12029_10089888 | + | 7 | 85 | **RNA**→ rpmE (PRK00019)Ribosomal_L31 (pfam01197)→ |
| env-53 | JGI12712J15308_10920105 | - | 124 | 57 | **RNA**→ ←hypo |
| env-54 | JGI20163J15578_10158293 | + | 1046 | 1091 | **RNA**→ rpmE (PRK00019)rho (PRK09376)rho_factor (cd01128)Ribosomal_L31 (pfam01197)→ |
| env-55 | JGI12396J12026_1117834 | + | 20 | 67 | **RNA**→ hypo→ |
| env-56 | JGI12630J15595_10565893 | + | 41 | 109 | **RNA**→ rpmE (PRK00019)Ribosomal_L31 (pfam01197)→ |
| env-57 | JGI12396J12026_1055065 | + | 42 | 89 | **RNA**→ rpmE (PRK00019)Ribosomal_L31 (pfam01197)→ prfA (PRK00591)RF-1 (pfam00472)PCRF (pfam03462)→ |
| env-58 | JGI12273J12029_10130660 | + | 34 | 81 | **RNA**→ rpmE (PRK00019)Ribosomal_L31 (pfam01197)→ |
| env-59 | JGI12397J12024_1064796 | + | 44 | 91 | **RNA**→ rpmE (PRK00019)Ribosomal_L31 (pfam01197)→ PCRF (smart00937)PCRF (pfam03462)→ |
| Spo-1-1 | NZ_KB913029.1 | + | 3641099 | 3641144 | **RNA**→ rpmE (PRK00019)Ribosomal_L31 (pfam01197)→ PrfA (COG0216)prfA (PRK00591)→ HemK (COG2890)PRK09328 (PRK09328)→ |
| env-60 | JGI24739J22299_10966797 | + | 95 | 143 | **RNA**→ ←hypo |
| env-61 | JGI26538J50259_1004293 | - | 1226 | 1180 | **RNA**→ rpmE (PRK00019)Ribosomal_L31 (pfam01197)→ PCRF (smart00937)RF-1 (pfam00472)PCRF (pfam03462)→ |
| Ssp-51-1 | NZ_AUKQ01000005.1 | + | 616357 | 616402 | **RNA**→ rpmE (PRK00019)Ribosomal_L31 (pfam01197)→ |
| Sji-1-1 | NZ_BBPN01000044.1 | + | 2914 | 2959 | **RNA**→ rpmE (PRK00019)Ribosomal_L31 (pfam01197)→ |
| Aut-1-1 | NZ_JRTT01000013.1 | + | 144414 | 144459 | **RNA**→ rpmE (PRK00019)Ribosomal_L31 (pfam01197)→ PrfA (COG0216)prfA (PRK00591)→ |
| Mtu-1-1 | NZ_KI912270.1 | - | 156001 | 155939 | **RNA**→ rpmE (PRK00019)Ribosomal_L31 (pfam01197)→ |
| Cko-1-1 | NZ_KB903272.1 | + | 26084 | 26123 | **RNA**→ rpmE (PRK00019)Ribosomal_L31 (pfam01197)→ PrfA (COG0216)prfA (PRK00591)→ |
| Nla-1-1 | NZ_AUFT01000001.1 | - | 90372 | 90294 | **RNA**→ rpmE (PRK00019)Ribosomal_L31 (pfam01197)→ |
| Nsp-9-1 | NZ_JIAR01000025.1 | - | 305751 | 305704 | **RNA**→ rpmE (PRK00019)Ribosomal_L31 (pfam01197)→ PrfA (COG0216)prfA (PRK00591)→ HemK (COG2890)PRK09328 (PRK09328)→ SUA5 (COG0009)TIGR00057 (TIGR00057)→ hypo→ ECA_wecA (TIGR02380)GT_WecA_like (cd06853)→ |
| env-62 | JGI24738J21930_10349179 | - | 206 | 158 | **RNA**→ rpmE (PRK00019)Ribosomal_L31 (pfam01197)→ |
| Afr-1-1 | NC_022657.1 | - | 8170022 | 8169953 | **RNA**→ rpmE (PRK00019)Ribosomal_L31 (pfam01197)→ PrfA (COG0216)prfA (PRK00591)→ |
| env-63 | JGI12397J12024_1210963 | + | 55 | 124 | **RNA**→ ←hypo |
| env-64 | JGI12397J12024_1099088 | + | 55 | 124 | **RNA**→ ←hypo |
| Asu-3-1 | NZ_JOJL01000007.1 | - | 354095 | 354026 | **RNA**→ rpmE (PRK00019)Ribosomal_L31 (pfam01197)→ PrfA (COG0216)prfA (PRK00591)→ |
| env-65 | JGI12397J12024_1237113 | + | 36 | 80 | **RNA**→ ←hypo |
| env-66 | JGI12397J12024_1148967 | + | 11 | 80 | **RNA**→ ←hypo |
| env-67 | JGI12273J12029_10092990 | + | 57 | 97 | **RNA**→ ←hypo |
| env-68 | JGI12272J11983_1168523 | + | 58 | 98 | **RNA**→ rpmE (PRK00019)Ribosomal_L31 (pfam01197)→ |
| env-69 | JGI12271J12027_1090360 | + | 56 | 96 | **RNA**→ ←hypo |
| env-70 | JGI12396J12026_1102605 | + | 43 | 84 | **RNA**→ rpmE (PRK00019)Ribosomal_L31 (pfam01197)→ PCRF (smart00937)RF-1 (pfam00472)PCRF (pfam03462)ABC_tran_CTD (pfam16326)→ |
| env-71 | JGI12396J12026_1237335 | - | 118 | 77 | **RNA**→ ←hypo |
| env-72 | JGI12273J12029_10179347 | + | 43 | 84 | **RNA**→ ←hypo |
| env-73 | JGI12397J12024_1046578 | + | 44 | 85 | **RNA**→ ←hypo |
| env-74 | JGI12273J12029_10030651 | + | 419 | 460 | **RNA**→ rpmE (PRK00019)Ribosomal_L31 (pfam01197)→ PrfA (COG0216)prfA (PRK00591)→ HemK (COG2890)PRK09328 (PRK09328)→ |
| env-75 | JGI12272J11983_1113393 | + | 87 | 128 | **RNA**→ ←hypo |
| env-76 | JGI12467J12023_1025401 | + | 18 | 59 | **RNA**→ ←hypo |
| env-77 | JGI12271J12027_1039259 | + | 18 | 59 | **RNA**→ ←hypo |
| env-78 | JGI12396J12026_1083332 | + | 54 | 95 | **RNA**→ hypo→ rpmE (PRK00019)Ribosomal_L31 (pfam01197)→ |
| env-79 | JGI12397J12024_1046859 | + | 54 | 95 | **RNA**→ hypo→ rpmE (PRK00019)Ribosomal_L31 (pfam01197)→ prfA (PRK00591)PCRF (pfam03462)→ |
| env-80 | JGI12397J12024_1016153 | + | 50 | 91 | **RNA**→ PRK08279 (PRK08279)→ rpmE (PRK00019)Ribosomal_L31 (pfam01197)→ prfA (PRK00591)PCRF (pfam03462)→ |
| Ncy-1-1 | NC_016887.1 | + | 1254812 | 1254859 | **RNA**→ rpmE (PRK00019)Ribosomal_L31 (pfam01197)→ PrfA (COG0216)prfA (PRK00591)→ |
| Ncy-2-1 | NZ_BAFY01000056.1 | + | 16681 | 16728 | **RNA**→ rpmE (PRK00019)Ribosomal_L31 (pfam01197)→ PrfA (COG0216)prfA (PRK00591)→ |

| | | | | | |
|---|---|---|---|---|---|
| Npn-1-1 | NZ_BAGF01000020.1 | + | 89242 | 89291 | **RNA**→ rpmE (PRK00019)Ribosomal_L31 (pfam01197)→ PrfA (COG0216)prfA (PRK00591)→ HemK (COG2890)PRK09328 (PRK09328)→ |
| env-81 | JGI12271J12027_1026620 | + | 102 | 144 | **RNA**→ ←hypo |
| env-82 | JGI12273J12029_10010121 | + | 4 | 46 | **RNA**→ rpmE (PRK00019)Ribosomal_L31 (pfam01197)→ PCRF (smart00937)PCRF (pfam03462)→ |
| env-83 | JGI12397J12024_1000989 | + | 11 | 53 | **RNA**→ ←hypo |
| env-84 | DCrCPGB_c187296 | - | 360 | 318 | **RNA**→ rpmE (PRK00019)Ribosomal_L31 (pfam01197)→ |
| env-85 | JGI12467J12023_1052535 | + | 9 | 51 | **RNA**→ ←hypo |
| env-86 | FNTS_GKKEVUV02H38F2 | - | 160 | 118 | **RNA**→ ←hypo |
| env-87 | JGI12397J12024_1145631 | + | 15 | 60 | **RNA**→ rpmE (PRK00019)Ribosomal_L31 (pfam01197)→ |
| env-88 | JGI24737J22298_10298947 | - | 195 | 153 | **RNA**→ rpmE (PRK00019)Ribosomal_L31 (pfam01197)→ |
| env-89 | JGI24735J21928_10416994 | - | 88 | 46 | **RNA**→ hypo→ |
| Sre-2-1 | NZ_AYJW01000043.1 | - | 1191 | 1142 | **RNA**→ rpmE (PRK00019)Ribosomal_L31 (pfam01197)→ PCRF (smart00937)RF-1 (pfam00472)PCRF (pfam03462)ABC_tran_CTD (pfam16326)→ |
| Sre-1-1 | NZ_JNVU01000048.1 | - | 201384 | 201335 | **RNA**→ rpmE (PRK00019)Ribosomal_L31 (pfam01197)→ PrfA (COG0216)prfA (PRK00591)→ HemK (COG2890)PRK09328 (PRK09328)→ |
| Acy-1-1 | NZ_KL573951.1 | - | 441934 | 441863 | **RNA**→ rpmE (PRK00019)Ribosomal_L31 (pfam01197)→ |
| Acy-2-1 | NZ_AUBJ01000013.1 | - | 277214 | 277143 | **RNA**→ rpmE (PRK00019)Ribosomal_L31 (pfam01197)→ |
| Mrh-2-1 | NC_016604.1 | + | 3820183 | 3820236 | **RNA**→ rpmE (PRK00019)Ribosomal_L31 (pfam01197)→ PrfA (COG0216)prfA (PRK00591)→ HemK (COG2890)PRK09328 (PRK09328)→ SUA5 (COG0009)TIGR00057 (TIGR00057)→ GT_WecA_like (cd06853)mraY (PRK00108)→ |
| env-90 | JGI11758J13082_115022 | + | 103 | 146 | **RNA**→ rpmE (PRK00019)Ribosomal_L31 (pfam01197)→ |
| env-91 | JGI12112J13074_1001942 | - | 1400 | 1357 | **RNA**→ rpmE (PRK00019)Ribosomal_L31 (pfam01197)→ prfA (PRK00591)RF-1 (pfam00472)PCRF (pfam03462)→ |
| env-92 | JGI12060J13090_109431 | - | 402 | 359 | **RNA**→ ←hypo |
| env-93 | CEQG01140696.1 | + | 747 | 790 | **RNA**→ rpmE (PRK00019)Ribosomal_L31 (pfam01197)→ |
| env-94 | JGI12075J13256_1000493 | - | 2999 | 2956 | **RNA**→ rpmE (PRK00019)Ribosomal_L31 (pfam01197)→ PrfA (COG0216)prfA (PRK00591)→ HemK (COG2890)PRK09328 (PRK09328)→ SUA5 (COG0009)TIGR00057 (TIGR00057)→ |
| env-95 | JGI11800J13271_1002272 | - | 358 | 315 | **RNA**→ rpmE (PRK00019)Ribosomal_L31 (pfam01197)→ |
| env-96 | CETS01058554.1 | + | 353 | 396 | **RNA**→ rpmE (PRK00019)Ribosomal_L31 (pfam01197)→ |
| env-97 | SRS016575_C3611077 | + | 309 | 354 | **RNA**→ ←hypo |
| env-98 | SRS020340_Baylor_scaffold_41668 | + | 245 | 290 | **RNA**→ rpmE (PRK00019)Ribosomal_L31 (pfam01197)→ |
| Ade-2-1 | NZ_AUBL01000029.1 | + | 17719 | 17764 | **RNA**→ rpmE (PRK00019)Ribosomal_L31 (pfam01197)→ |
| env-99 | SRS022536_C4437610 | - | 232 | 187 | **RNA**→ rpmE (PRK00019)Ribosomal_L31 (pfam01197)→ |
| env-100 | SRS024144_LANL_scaffold_22347 | + | 53 | 98 | **RNA**→ rpmE (PRK00019)Ribosomal_L31 (pfam01197)→ |
| env-101 | SRS042984_C3945655 | + | 291 | 336 | **RNA**→ rpmE (PRK00019)Ribosomal_L31 (pfam01197)→ |
| env-102 | SRS065099_LANL_scaffold_83872 | - | 432 | 387 | **RNA**→ hypo→ |
| env-103 | SRS017139_Baylor_scaffold_27719 | + | 210 | 255 | **RNA**→ rpmE (PRK00019)Ribosomal_L31 (pfam01197)→ |
| env-104 | SRS024355_LANL_scaffold_54831 | - | 212 | 167 | **RNA**→ rpmE (PRK00019)Ribosomal_L31 (pfam01197)→ |
| env-105 | SRS018443_Baylor_scaffold_61370 | + | 276 | 321 | **RNA**→ rpmE (PRK00019)Ribosomal_L31 (pfam01197)→ |
| env-106 | SRS013252_Baylor_scaffold_60598 | + | 250 | 295 | **RNA**→ rpmE (PRK00019)Ribosomal_L31 (pfam01197)→ |
| env-107 | SRS017691_C5148150 | - | 133 | 88 | **RNA**→ rpmE (PRK00019)rpmE2 (PRK01678)Ribosomal_L31 (pfam01197)→ |
| env-108 | SRS021477_Baylor_scaffold_9272 | + | 137 | 182 | **RNA**→ rpmE (PRK00019)Ribosomal_L31 (pfam01197)→ prfA (PRK00591)PCRF (pfam03462)→ |
| env-109 | SRS024087_C3541330 | - | 796 | 751 | **RNA**→ rpmE (PRK00019)Ribosomal_L31 (pfam01197)→ hypo→ prfA (PRK00591)PCRF (pfam03462)→ |
| env-110 | SRS063932_LANL_scaffold_97365 | + | 249 | 294 | **RNA**→ rpmE (PRK00019)Ribosomal_L31 (pfam01197)→ hypo→ PrfA (COG0216)prfA (PRK00591)→ |
| env-111 | SRS019128_C4186243 | + | 134 | 179 | **RNA**→ rpmE (PRK00019)Ribosomal_L31 (pfam01197)→ |
| Bin-1-1 | NZ_CP006018.1 | - | 1292657 | 1292613 | **RNA**→ rpmE (PRK00019)Ribosomal_L31 (pfam01197)→ prfA (PRK00591)RF-1 (pfam00472)PCRF (pfam03462)→ HemK (COG2890)PRK09328 (PRK09328)→ |
| env-112 | HBC_scaff_10011399 | - | 3944 | 3900 | **RNA**→ rpmE (PRK00019)Ribosomal_L31 (pfam01197)→ prfA (PRK00591)PCRF (pfam03462)→ |
| env-113 | HBC_ctgs_1009601 | + | 1655 | 1699 | **RNA**→ rpmE (PRK00019)Ribosomal_L31 (pfam01197)→ prfA (PRK00591)PCRF (pfam03462)→ PRK05567 (PRK05567)CBS_pair_IMPDH (cd04601)→ |
| Bco-1-1 | NZ_CP007287.1 | - | 1328201 | 1328157 | **RNA**→ rpmE (PRK00019)Ribosomal_L31 (pfam01197)→ prfA (PRK00591)RF-1 (pfam00472)PCRF (pfam03462)→ HemK (COG2890)PRK09328 (PRK09328)→ |
| Bco-2-1 | NZ_JDUF01000003.1 | + | 212969 | 213013 | **RNA**→ rpmE (PRK00019)Ribosomal_L31 (pfam01197)→ prfA (PRK00591)RF-1 (pfam00472)PCRF (pfam03462)→ HemK (COG2890)PRK09328 (PRK09328)→ |

| | | | | | |
|---|---|---|---|---|---|
| env-114 | AF_2010_repII_A01_1248365 | - | 214 | 164 | **RNA**→ ←hypo |
| env-115 | AF_2010_repII_A1_10264337 | - | 138 | 88 | **RNA**→ rpmE (PRK00019)Ribosomal_L31 (pfam01197)→ |
| env-116 | _GX4IIKE02HM9I2 | - | 308 | 260 | **RNA**→ rpmE (PRK00019)Ribosomal_L31 (pfam01197)→ |
| Sro-4-1 | NZ_ABYX02000001.1 | + | 5513786 | 5513833 | **RNA**→ rpmE (PRK00019)Ribosomal_L31 (pfam01197)→ |
| Ssp-18-1 | NZ_AWOQ01000221.1 | + | 3075 | 3122 | **RNA**→ rpmE (PRK00019)Ribosomal_L31 (pfam01197)→ |
| Sro-2-1 | NZ_DS999644.1 | + | 5470394 | 5470441 | **RNA**→ rpmE (PRK00019)Ribosomal_L31 (pfam01197)→ rpmE (PRK00019)Ribosomal_L31 (pfam01197)→ |
| Sgl-4-1 | NZ_JNZK01000005.1 | + | 91932 | 91979 | **RNA**→ rpmE (PRK00019)Ribosomal_L31 (pfam01197)→ |
| Sal-8-1 | NZ_JNWW01000013.1 | + | 257612 | 257659 | **RNA**→ rpmE (PRK00019)Ribosomal_L31 (pfam01197)→ |
| Sgl-2-1 | NZ_JH689271.1 | - | 192859 | 192812 | **RNA**→ rpmE (PRK00019)Ribosomal_L31 (pfam01197)→ |
| Sgr-12-1 | NZ_JOGA01000014.1 | - | 269813 | 269766 | **RNA**→ rpmE (PRK00019)Ribosomal_L31 (pfam01197)→ |
| Ssp-55-1 | NZ_JNLT01000003.1 | - | 25808 | 25761 | **RNA**→ rpmE (PRK00019)Ribosomal_L31 (pfam01197)→ |
| Ssp-62-1 | NZ_JPWW01000002.1 | + | 1728736 | 1728783 | **RNA**→ rpmE (PRK00019)Ribosomal_L31 (pfam01197)→ |
| Ssp-167-1 | NZ_LAVK01000634.1 | - | 420 | 376 | **RNA**→ rpmE (PRK00019)Ribosomal_L31 (pfam01197)→ |
| Sav-3-1 | NZ_JOEK01000002.1 | - | 249434 | 249391 | **RNA**→ rpmE (PRK00019)Ribosomal_L31 (pfam01197)→ |
| Ssp-4-1 | NZ_ALNP01000028.1 | + | 19225 | 19269 | **RNA**→ rpmE (PRK00019)Ribosomal_L31 (pfam01197)→ |
| Sxi-2-1 | NZ_CP009922.2 | + | 4411426 | 4411470 | **RNA**→ rpmE (PRK00019)Ribosomal_L31 (pfam01197)→ |
| Ssp-90-1 | NZ_JOIG01000006.1 | + | 162280 | 162324 | **RNA**→ rpmE (PRK00019)Ribosomal_L31 (pfam01197)→ |
| Jal-1-1 | NZ_LBMC01000041.1 | + | 84087 | 84134 | **RNA**→ rpmE (PRK00019)Ribosomal_L31 (pfam01197)→ PrfA (COG0216)prfA (PRK00591)→ HemK (COG2890)PRK09328 (PRK09328)→ SUA5 (COG0009)TIGR00057 (TIGR00057)→ ECA_wecA (TIGR02380)GT_WecA_like (cd06853)→ hypo→ AtpI2 (COG5336)→ PRK05815 (PRK05815)ATP-synt_A (pfam00119)→ PRK06876 (PRK06876)AtpE (COG0636)→ PRK05759 (PRK05759)AtpF (COG0711)→ PRK13430 (PRK13430)OSCP (pfam00213)→ AtpA (COG0056)PRK09281 (PRK09281)→ PRK05621 (PRK05621)ATP-synt (pfam00231)→ |
| env-117 | JGI20183J14883_110497 | + | 278 | 327 | **RNA**→ ←hypo |
| env-118 | JGI20181J14860_1000476 | + | 1897 | 1946 | **RNA**→ rpmE (PRK00019)Ribosomal_L31 (pfam01197)→ |
| env-119 | JGI20198J14950_1000651 | - | 1554 | 1505 | **RNA**→ rpmE (PRK00019)Ribosomal_L31 (pfam01197)→ |
| env-120 | JGI20200J14955_1000024 | + | 14466 | 14515 | **RNA**→ rpmE (PRK00019)Ribosomal_L31 (pfam01197)→ |
| env-121 | JGI20201J14949_1011227 | + | 406 | 455 | **RNA**→ rpmE (PRK00019)Ribosomal_L31 (pfam01197)→ |
| env-122 | JGI20202J14951_1000077 | + | 2663 | 2712 | **RNA**→ rpmE (PRK00019)Ribosomal_L31 (pfam01197)→ |
| env-123 | JGI20197J15001_1017583 | + | 314 | 363 | **RNA**→ rpmE (PRK00019)Ribosomal_L31 (pfam01197)→ |
| env-124 | JGI20197J15136_1017583 | + | 314 | 363 | **RNA**→ rpmE (PRK00019)Ribosomal_L31 (pfam01197)→ |
| env-125 | R_ear_crease_WUGI_scaffold_53252 | - | 122 | 77 | **RNA**→ |
| Pav-2-1 | NZ_AOUA01000299.1 | + | 6067 | 6112 | **RNA**→ rpmE (PRK00019)Ribosomal_L31 (pfam01197)→ |
| Pav-1-1 | NC_021064.1 | + | 1037423 | 1037468 | **RNA**→ rpmE (PRK00019)Ribosomal_L31 (pfam01197)→ |
| Psp-5-1 | NZ_KI515688.1 | - | 2409959 | 2409914 | **RNA**→ rpmE (PRK00019)Ribosomal_L31 (pfam01197)→ |
| Psp-3-1 | NZ_KI515696.1 | - | 178310 | 178265 | **RNA**→ rpmE (PRK00019)Ribosomal_L31 (pfam01197)→ |
| Psp-2-1 | NZ_KI518468.1 | + | 147059 | 147104 | **RNA**→ rpmE (PRK00019)Ribosomal_L31 (pfam01197)→ |
| Psp-4-1 | NZ_AXMH01000003.1 | - | 663366 | 663321 | **RNA**→ rpmE (PRK00019)Ribosomal_L31 (pfam01197)→ |
| Psp-1-1 | NZ_KE340299.1 | - | 2592352 | 2592307 | **RNA**→ rpmE (PRK00019)Ribosomal_L31 (pfam01197)→ |
| env-126 | JGI12397J12024_1115564 | + | 10 | 52 | **RNA**→ rpmE (PRK00019)Ribosomal_L31 (pfam01197)→ |
| Pac-1-1 | NZ_JIAI01000013.1 | - | 168551 | 168505 | **RNA**→ rpmE (PRK00019)Ribosomal_L31 (pfam01197)→ |
| env-127 | JGI24736J21556_1166172 | + | 166 | 205 | **RNA**→ rpmE (PRK00019)Ribosomal_L31 (pfam01197)→ |
| env-128 | JGI12467J12023_1013674 | + | 485 | 531 | **RNA**→ rpmE (PRK00019)Ribosomal_L31 (pfam01197)→ |
| env-129 | JGI12271J12027_1096703 | + | 1 | 47 | **RNA**→ rpmE (PRK00019)Ribosomal_L31 (pfam01197)→ |
| env-130 | JGI12397J12024_1040638 | + | 77 | 125 | **RNA**→ rpmE (PRK00019)Ribosomal_L31 (pfam01197)→ |
| env-131 | JGI12273J12029_10103667 | + | 517 | 565 | **RNA**→ |
| env-132 | JGI24770J26754_10130366 | - | 170 | 134 | **RNA**→ rpmE (PRK00019)Ribosomal_L31 (pfam01197)→ |
| env-133 | JGI20200J14955_1003644 | + | 1490 | 1528 | **RNA**→ rpmE (PRK00019)Ribosomal_L31 (pfam01197)→ |
| env-134 | JGI20201J14949_1077705 | + | 89 | 127 | **RNA**→ hypo→ rpmE (PRK00019)Ribosomal_L31 (pfam01197)→ |
| env-135 | JGI20202J14951_1033931 | + | 287 | 325 | **RNA**→ ←hypo |
| Bth-2-1 | NZ_JGZS01000003.1 | + | 338005 | 338048 | **RNA**→ rpmE (PRK00019)Ribosomal_L31 (pfam01197)→ |
| Bsc-1-1 | NZ_JULC01000081.1 | + | 2980 | 3024 | **RNA**→ rpmE (PRK00019)Ribosomal_L31 (pfam01197)→ |
| Bth-7-1 | NZ_JDUB01000001.1 | + | 171828 | 171872 | **RNA**→ rpmE (PRK00019)Ribosomal_L31 (pfam01197)→ |

| | | | | | |
|---|---|---|---|---|---|
| Bth-8-1 | NZ_JHWM01000001.1 | - | 74179 | 74133 | **RNA**→ rpmE (PRK00019)Ribosomal_L31 (pfam01197)→ |
| Bth-1-1 | NC_020546.1 | - | 1607274 | 1607228 | **RNA**→ rpmE (PRK00019)Ribosomal_L31 (pfam01197)→ |
| Bth-5-1 | NZ_AUFI01000006.1 | + | 346383 | 346427 | **RNA**→ rpmE (PRK00019)Ribosomal_L31 (pfam01197)→ |
| Bth-4-1 | NZ_JGZT01000005.1 | + | 346830 | 346874 | **RNA**→ rpmE (PRK00019)Ribosomal_L31 (pfam01197)→ |
| Bth-5-2 | NZ_JDTO01000003.1 | + | 250962 | 251006 | **RNA**→ rpmE (PRK00019)Ribosomal_L31 (pfam01197)→ |
| Bth-3-1 | NZ_JDTQ01000009.1 | + | 58022 | 58066 | **RNA**→ rpmE (PRK00019)Ribosomal_L31 (pfam01197)→ |
| Bth-6-1 | NZ_JGZV01000001.1 | - | 350252 | 350208 | **RNA**→ rpmE (PRK00019)Ribosomal_L31 (pfam01197)→ |
| Bbo-2-1 | NZ_JHWO01000001.1 | - | 219911 | 219867 | **RNA**→ rpmE (PRK00019)Ribosomal_L31 (pfam01197)→ |
| Bbo-2-2 | NZ_JDUH01000006.1 | - | 219494 | 219450 | **RNA**→ rpmE (PRK00019)Ribosomal_L31 (pfam01197)→ |
| Bbo-1-1 | NZ_JGYQ01000016.1 | - | 219470 | 219426 | **RNA**→ rpmE (PRK00019)Ribosomal_L31 (pfam01197)→ |
| env-136 | 3300001592_10065837 | - | 1863 | 1818 | **RNA**→ hypo→ rpmE (PRK00019)Ribosomal_L31 (pfam01197)→ |
| Nsp-4-1 | NC_008699.1 | + | 1862060 | 1862105 | **RNA**→ rpmE (PRK00019)Ribosomal_L31 (pfam01197)→ PrfA (COG0216)prfA (PRK00591)→ |
| | | | | | HemK (COG2890)PRK09328 (PRK09328)→ SUA5 (COG0009)TIGR00057 (TIGR00057)→ |
| | | | | | GT_WecA_like (cd06853)mraY (PRK00108)→ |
| env-137 | AP72_2010_repI_A10_1095182 | - | 366 | 312 | **RNA**→ rpmE (PRK00019)Ribosomal_L31 (pfam01197)→ |
| env-138 | JGI12075J13256_1008725 | - | 141 | 76 | **RNA**→ |
| env-139 | CEOS01170558.1 | - | 194 | 129 | **RNA**→ rpmE (PRK00019)Ribosomal_L31 (pfam01197)→ |
| env-140 | JGI12064J13199_110746 | - | 352 | 287 | **RNA**→ rpmE (PRK00019)Ribosomal_L31 (pfam01197)→ |
| env-141 | P3_CLC_Cons13751 | - | 1150 | 1079 | **RNA**→ ←PRK15098 (PRK15098)BglX (COG1472) |
| env-142 | Cons65214 | + | 22 | 93 | **RNA**→ ←PRK15098 (PRK15098)BglX (COG1472) |
| Fsp-8-1 | NZ_ATXA01000002.1 | + | 22334 | 22383 | **RNA**→ rpmE (PRK00019)Ribosomal_L31 (pfam01197)→ |
| env-143 | JGI12635J15846_10174018 | + | 8 | 48 | **RNA**→ hypo→ DUF4387 (pfam14330)→ MhpC (COG0596)menH_SHCHC (TIGR03695)→ |
| Nsp-10-1 | NZ_JIAV01000021.1 | + | 30441 | 30487 | **RNA**→ rpmE (PRK00019)Ribosomal_L31 (pfam01197)→ PrfA (COG0216)prfA (PRK00591)→ |
| | | | | | HemK (COG2890)PRK09328 (PRK09328)→ SUA5 (COG0009)TIGR00057 (TIGR00057)→ |
| | | | | | ECA_wecA (TIGR02380)GT_WecA_like (cd06853)→ |
| env-144 | JGI12273J12029_10078345 | + | 11 | 81 | **RNA**→ rpmE (PRK00019)Ribosomal_L31 (pfam01197)→ |
| Cja-1-1 | NZ_JNXY01000008.1 | + | 8604 | 8652 | **RNA**→ rpmE (PRK00019)Ribosomal_L31 (pfam01197)→ |
| Fsy-1-1 | NC_015656.1 | - | 4523128 | 4523080 | **RNA**→ rpmE (PRK00019)Ribosomal_L31 (pfam01197)→ |
| Fco-1-1 | NZ_JWIO01000003.1 | + | 248178 | 248226 | **RNA**→ rpmE (PRK00019)Ribosomal_L31 (pfam01197)→ |
| env-145 | JGI24770J26754_10245837 | + | 73 | 125 | **RNA**→ rpmE (PRK00019)Ribosomal_L31 (pfam01197)→ prfA (PRK00591)PCRF (pfam03462)→ |
| Ply-1-1 | NZ_KE150269.1 | - | 1488609 | 1488566 | **RNA**→ rpmE (PRK00019)Ribosomal_L31 (pfam01197)→ PrfA (COG0216)prfA (PRK00591)→ |
| | | | | | HemK (COG2890)PRK09328 (PRK09328)→ SUA5 (COG0009)TIGR00057 (TIGR00057)→ |
| | | | | | GT_WecA_like (cd06853)mraY (PRK00108)→ hypo→ AtpI2 (COG5336)ATPase_gene1 (pfam09527)→ |
| | | | | | PRK05815 (PRK05815)ATP-synt_A (pfam00119)→ PRK07874 (PRK07874)AtpE (COG0636)→ |
| | | | | | PRK05759 (PRK05759)AtpF (COG0711)→ AtpH (COG0712)PRK13430 (PRK13430)→ |
| | | | | | AtpA (COG0056)PRK09281 (PRK09281)→ PRK05621 (PRK05621)ATP-synt (pfam00231)→ |
| | | | | | AtpD (COG0055)PRK09280 (PRK09280)→ |
| Psp-7-1 | NZ_AZIS01000003.1 | + | 159103 | 159146 | **RNA**→ rpmE (PRK00019)Ribosomal_L31 (pfam01197)→ PrfA (COG0216)prfA (PRK00591)→ |
| | | | | | HemK (COG2890)PRK09328 (PRK09328)→ SUA5 (COG0009)TIGR00057 (TIGR00057)→ |
| | | | | | GT_WecA_like (cd06853)mraY (PRK00108)→ hypo→ AtpI2 (COG5336)ATPase_gene1 (pfam09527)→ |
| | | | | | PRK05815 (PRK05815)ATP-synt_A (pfam00119)→ PRK07874 (PRK07874)AtpE (COG0636)→ |
| | | | | | PRK05759 (PRK05759)AtpF (COG0711)→ AtpH (COG0712)PRK13430 (PRK13430)→ |
| | | | | | AtpA (COG0056)PRK09281 (PRK09281)→ PRK05621 (PRK05621)ATP-synt (pfam00231)→ |
| | | | | | AtpD (COG0055)PRK09280 (PRK09280)→ |
| Ply-2-1 | NZ_KE384026.1 | - | 390464 | 390421 | **RNA**→ rpmE (PRK00019)Ribosomal_L31 (pfam01197)→ PrfA (COG0216)prfA (PRK00591)→ |
| | | | | | HemK (COG2890)PRK09328 (PRK09328)→ SUA5 (COG0009)TIGR00057 (TIGR00057)→ |
| | | | | | GT_WecA_like (cd06853)mraY (PRK00108)→ hypo→ AtpI2 (COG5336)ATPase_gene1 (pfam09527)→ |
| | | | | | PRK05815 (PRK05815)ATP-synt_A (pfam00119)→ PRK07874 (PRK07874)AtpE (COG0636)→ |
| | | | | | PRK05759 (PRK05759)AtpF (COG0711)→ AtpH (COG0712)PRK13430 (PRK13430)→ |
| | | | | | AtpA (COG0056)PRK09281 (PRK09281)→ PRK05621 (PRK05621)ATP-synt (pfam00231)→ |
| | | | | | AtpD (COG0055)PRK09280 (PRK09280)→ |
| Ssc-2-1 | NZ_KB889571.1 | + | 220 | 266 | **RNA**→ rpmE (PRK00019)Ribosomal_L31 (pfam01197)→ |
| Sri-4-1 | NZ_JNWX01000074.1 | + | 10628 | 10672 | **RNA**→ rpmE (PRK00019)Ribosomal_L31 (pfam01197)→ |
| Sri-2-1 | NZ_JMGX01000008.1 | + | 10567 | 10611 | **RNA**→ rpmE (PRK00019)Ribosomal_L31 (pfam01197)→ |

| | | | | | |
|---|---|---|---|---|---|
| Kau-1-1 | NZ_LGUY01000019.1 | - | 4086 | 4042 | **RNA**→ rpmE (PRK00019)Ribosomal_L31 (pfam01197)→ |
| Ssp-156-1 | NZ_JOCC01000117.1 | - | 4095 | 4051 | **RNA**→ rpmE (PRK00019)Ribosomal_L31 (pfam01197)→ |
| Sri-4-2 | NZ_JOEO01000027.1 | - | 93461 | 93417 | **RNA**→ rpmE (PRK00019)Ribosomal_L31 (pfam01197)→ |
| Sri-4-3 | NZ_JNYK01000063.1 | - | 4106 | 4062 | **RNA**→ rpmE (PRK00019)Ribosomal_L31 (pfam01197)→ |
| Sri-4-4 | NZ_JOBN01000051.1 | - | 4141 | 4097 | **RNA**→ rpmE (PRK00019)Ribosomal_L31 (pfam01197)→ |
| Sri-4-5 | NZ_JOCL01000019.1 | - | 93576 | 93532 | **RNA**→ rpmE (PRK00019)Ribosomal_L31 (pfam01197)→ |
| Sri-4-6 | NZ_JOCR01000115.1 | - | 4074 | 4030 | **RNA**→ rpmE (PRK00019)Ribosomal_L31 (pfam01197)→ |
| Sri-4-7 | NZ_JOCO01000065.1 | + | 10588 | 10632 | **RNA**→ rpmE (PRK00019)Ribosomal_L31 (pfam01197)→ |
| Sri-4-8 | NZ_JOBS01000027.1 | - | 93497 | 93453 | **RNA**→ rpmE (PRK00019)Ribosomal_L31 (pfam01197)→ |
| Ssp-157-1 | NZ_JOCH01000101.1 | - | 4120 | 4076 | **RNA**→ rpmE (PRK00019)Ribosomal_L31 (pfam01197)→ |
| Sca-5-1 | NZ_JNWP01000078.1 | - | 4108 | 4064 | **RNA**→ rpmE (PRK00019)Ribosomal_L31 (pfam01197)→ |
| Sca-5-2 | NZ_JOFM01000099.1 | - | 4116 | 4072 | **RNA**→ rpmE (PRK00019)Ribosomal_L31 (pfam01197)→ |
| Sri-4-9 | NZ_JOBT01000405.1 | + | 10576 | 10620 | **RNA**→ rpmE (PRK00019)Ribosomal_L31 (pfam01197)→ |
| Sri-4-10 | NZ_JOCM01000066.1 | - | 4088 | 4044 | **RNA**→ rpmE (PRK00019)Ribosomal_L31 (pfam01197)→ |
| Sri-4-11 | NZ_JOCP01000071.1 | - | 4099 | 4055 | **RNA**→ rpmE (PRK00019)Ribosomal_L31 (pfam01197)→ |
| Sri-4-12 | NZ_JNYR01000048.1 | - | 4090 | 4046 | **RNA**→ rpmE (PRK00019)Ribosomal_L31 (pfam01197)→ |
| Sla-2-3 | NZ_JOEX01000083.1 | - | 4150 | 4106 | **RNA**→ rpmE (PRK00019)Ribosomal_L31 (pfam01197)→ |
| Sla-2-4 | NZ_JOCN01000068.1 | + | 10603 | 10647 | **RNA**→ rpmE (PRK00019)Ribosomal_L31 (pfam01197)→ |
| Sri-1-1 | NZ_ANSJ01000029.1 | - | 26110 | 26066 | **RNA**→ rpmE (PRK00019)Ribosomal_L31 (pfam01197)→ |
| Sri-3-1 | NZ_JJNO01000011.1 | + | 11204 | 11248 | **RNA**→ rpmE (PRK00019)Ribosomal_L31 (pfam01197)→ |
| Sri-2-2 | NZ_JMGY01000002.1 | + | 188935 | 188979 | **RNA**→ rpmE (PRK00019)Ribosomal_L31 (pfam01197)→ |
| Sri-4-13 | NZ_JOET01000092.1 | - | 4083 | 4039 | **RNA**→ rpmE (PRK00019)Ribosomal_L31 (pfam01197)→ |
| Sgr-6-1 | NZ_LGUW01000086.1 | - | 3993 | 3949 | **RNA**→ rpmE (PRK00019)Ribosomal_L31 (pfam01197)→ |
| Sri-4-14 | NZ_JOBV01000024.1 | + | 10595 | 10639 | **RNA**→ rpmE (PRK00019)Ribosomal_L31 (pfam01197)→ |
| Sri-4-15 | NZ_JOES01000028.1 | - | 85329 | 85285 | **RNA**→ rpmE (PRK00019)Ribosomal_L31 (pfam01197)→ |
| Sri-4-16 | NZ_JOBP01000025.1 | - | 87385 | 87341 | **RNA**→ rpmE (PRK00019)Ribosomal_L31 (pfam01197)→ |
| Sri-4-17 | NZ_JOJJ01000030.1 | + | 10597 | 10641 | **RNA**→ rpmE (PRK00019)Ribosomal_L31 (pfam01197)→ |
| Ssp-75-1 | NZ_JNWN01000030.1 | + | 10583 | 10627 | **RNA**→ rpmE (PRK00019)Ribosomal_L31 (pfam01197)→ |
| Spe-2-1 | NZ_JOCK01000033.1 | + | 10581 | 10625 | **RNA**→ rpmE (PRK00019)Ribosomal_L31 (pfam01197)→ |
| Sri-4-18 | NZ_JOCQ01000154.1 | - | 4106 | 4062 | **RNA**→ rpmE (PRK00019)Ribosomal_L31 (pfam01197)→ |
| Sri-4-19 | NZ_JOBW01000113.1 | + | 10576 | 10620 | **RNA**→ rpmE (PRK00019)Ribosomal_L31 (pfam01197)→ |
| Sri-4-20 | NZ_JOBO01000026.1 | + | 10605 | 10649 | **RNA**→ rpmE (PRK00019)Ribosomal_L31 (pfam01197)→ |
| env-146 | JGI24739J22299_10113857 | + | 487 | 535 | **RNA**→ PRK10959 (PRK10959)→ rpmE (PRK00019)Ribosomal_L31 (pfam01197)→ |
| env-147 | JGI24738J21930_10012013 | - | 456 | 408 | **RNA**→ hypo→ rpmE (PRK00019)Ribosomal_L31 (pfam01197)→ |
| env-148 | JGI24737J22298_10573430 | + | 283 | 331 | **RNA**→ hypo→ |
| Nal-1-1 | NZ_AUFN01000010.1 | - | 437447 | 437399 | **RNA**→ rpmE (PRK00019)Ribosomal_L31 (pfam01197)→ |
| env-149 | JGI25160J50197_1029009 | - | 1165 | 1120 | **RNA**→ rpmE (PRK00019)Ribosomal_L31 (pfam01197)→ |
| Shy-1-1 | NZ_GG657754.1 | - | 3705640 | 3705595 | **RNA**→ rpmE (PRK00019)Ribosomal_L31 (pfam01197)→ rpmE (PRK00019)Ribosomal_L31 (pfam01197)→ |
| Ssp-36-1 | NZ_GG770539.1 | - | 1908349 | 1908302 | **RNA**→ hypo→ rpmE (PRK00019)Ribosomal_L31 (pfam01197)→ |
| Ssp-109-1 | NZ_JOGO01000006.1 | - | 20265 | 20218 | **RNA**→ rpmE (PRK00019)Ribosomal_L31 (pfam01197)→ |
| Ssp-35-1 | NZ_ADXA01000073.1 | - | 20808 | 20761 | **RNA**→ rpmE (PRK00019)Ribosomal_L31 (pfam01197)→ rpmE (PRK00019)Ribosomal_L31 (pfam01197)→ |
| Ssp-37-1 | NZ_GG657742.1 | + | 5431603 | 5431650 | **RNA**→ rpmE (PRK00019)Ribosomal_L31 (pfam01197)→ rpmE (PRK00019)Ribosomal_L31 (pfam01197)→ |
| Ssp-34-1 | NZ_ADXB01000013.1 | - | 18515 | 18468 | **RNA**→ rpmE (PRK00019)Ribosomal_L31 (pfam01197)→ rpmE (PRK00019)Ribosomal_L31 (pfam01197)→ |
| Ssp-42-1 | NZ_CM001165.1 | - | 2277919 | 2277872 | **RNA**→ rpmE (PRK00019)Ribosomal_L31 (pfam01197)→ rpmE (PRK00019)Ribosomal_L31 (pfam01197)→ |
| env-150 | FWIRElOz_GKZ9IRQ02HYXXR | + | 135 | 179 | **RNA**→ rpmE (PRK00019)Ribosomal_L31 (pfam01197)→ |
| Mau-2-1 | NZ_CVQQ01000029.1 | + | 46091 | 46143 | **RNA**→ rpmE (PRK00019)Ribosomal_L31 (pfam01197)→ |
| env-151 | JGI26341J46601_10001885 | - | 1722 | 1672 | **RNA**→ rpmE (PRK00019)Ribosomal_L31 (pfam01197)→ |
| env-152 | JGI26340J50214_10000427 | - | 14824 | 14774 | **RNA**→ rpmE (PRK00019)Ribosomal_L31 (pfam01197)→ |
| env-153 | JGI12397J12024_1078638 | + | 4 | 59 | **RNA**→ ←hypo |
| Psp-10-1 | NZ_KE386835.1 | - | 170013 | 169972 | **RNA**→ rpmE (PRK00019)Ribosomal_L31 (pfam01197)→ |
| env-154 | JGI24739J22299_10349184 | - | 302 | 261 | **RNA**→ rpmE (PRK00019)Ribosomal_L31 (pfam01197)→ |
| env-155 | JGI24737J22298_10666825 | - | 244 | 203 | **RNA**→ rpmE (PRK00019)Ribosomal_L31 (pfam01197)→ |

| | | | | | |
|---|---|---|---|---|---|
| env-156 | FIHLEPW02SY4CH | - | 297 | 256 | **RNA**→ rpmE (PRK00019)Ribosomal_L31 (pfam01197)→ |
| env-157 | JGI20208J14878_1015464 | - | 754 | 701 | **RNA**→ rpmE (PRK00019)Ribosomal_L31 (pfam01197)→ |
| env-158 | JGI20207J14881_1024728 | - | 493 | 439 | **RNA**→ rpmE (PRK00019)Ribosomal_L31 (pfam01197)→ |
| env-159 | JGI20206J14855_1162051 | - | 92 | 38 | **RNA**→ ←hypo |
| env-160 | JGI20208J14878_1000138 | + | 14241 | 14295 | **RNA**→ rpmE (PRK00019)Ribosomal_L31 (pfam01197)→ PrfA (COG0216)prfA (PRK00591)→ |
| | | | | | HemK (COG2890)PRK09328 (PRK09328)→ SUA5 (COG0009)TIGR00057 (TIGR00057)→ |
| | | | | | GT_WecA_like (cd06853)mraY (PRK00108)→ |
| Mha-1-1 | NZ_KB903880.1 | - | 18836 | 18779 | **RNA**→ rpmE (PRK00019)Ribosomal_L31 (pfam01197)→ PrfA (COG0216)prfA (PRK00591)→ |
| | | | | | HemK (COG2890)PRK09328 (PRK09328)→ SUA5 (COG0009)TIGR00057 (TIGR00057)→ |
| | | | | | GT_WecA_like (cd06853)mraY (PRK00108)→ |
| env-161 | JGI12635J15846_10811506 | + | 158 | 208 | **RNA**→ rpmE (PRK00019)Ribosomal_L31 (pfam01197)→ |
| env-162 | JGI12712J15308_10934693 | - | 161 | 94 | **RNA**→ hypo→ |
| env-163 | JGI12271J12027_1083783 | + | 20 | 68 | **RNA**→ hypo→ rpmE (PRK00019)Ribosomal_L31 (pfam01197)→ |
| env-164 | JGI12397J12024_1053925 | + | 25 | 70 | **RNA**→ rpmE (PRK00019)Ribosomal_L31 (pfam01197)→ |
| Asp-5-1 | NZ_AYXG01000169.1 | + | 21786 | 21823 | **RNA**→ efflux_EmrB (TIGR00711)MFS_1 (pfam07690)→ rpmE (PRK00019)Ribosomal_L31 (pfam01197)→ |
| env-165 | JGI24740J21852_10368802 | + | 100 | 168 | **RNA**→ rpmE (PRK00019)Ribosomal_L31 (pfam01197)→ |
| Nlu-1-1 | NZ_JZDQ01000313.1 | + | 1229 | 1274 | **RNA**→ rpmE (PRK00019)Ribosomal_L31 (pfam01197)→ |
| Nba-1-1 | NZ_GL873266.1 | + | 198527 | 198572 | **RNA**→ rpmE (PRK00019)Ribosomal_L31 (pfam01197)→ PrfA (COG0216)prfA (PRK00591)→ |
| Ssp-177-1 | NZ_LAND01000002.1 | - | 44421 | 44376 | **RNA**→ rpmE (PRK00019)Ribosomal_L31 (pfam01197)→ |
| Lal-1-1 | NZ_KB903833.1 | - | 501554 | 501506 | **RNA**→ rpmE (PRK00019)Ribosomal_L31 (pfam01197)→ |
| Gma-1-1 | NZ_BAOP01000039.1 | - | 23950 | 23903 | **RNA**→ rpmE (PRK00019)Ribosomal_L31 (pfam01197)→ |
| Bme-1-1 | NZ_JGZC01000003.1 | + | 96433 | 96480 | **RNA**→ rpmE (PRK00019)Ribosomal_L31 (pfam01197)→ |
| Bme-2-1 | NZ_JDTL01000009.1 | - | 12355 | 12308 | **RNA**→ rpmE (PRK00019)Ribosomal_L31 (pfam01197)→ |
| Bsu-2-1 | NZ_AUFH01000003.1 | - | 180572 | 180527 | **RNA**→ rpmE (PRK00019)Ribosomal_L31 (pfam01197)→ |
| Bsu-1-1 | NZ_JGZR01000005.1 | - | 181538 | 181493 | **RNA**→ rpmE (PRK00019)Ribosomal_L31 (pfam01197)→ |
| Bsu-2-2 | NZ_JDTX01000010.1 | + | 15308 | 15353 | **RNA**→ rpmE (PRK00019)Ribosomal_L31 (pfam01197)→ |
| Bre-1-1 | NZ_JGZK01000017.1 | + | 179460 | 179506 | **RNA**→ rpmE (PRK00019)Ribosomal_L31 (pfam01197)→ |
| Bre-1-2 | NZ_JDUW01000013.1 | + | 107 | 153 | **RNA**→ rpmE (PRK00019)Ribosomal_L31 (pfam01197)→ |
| Blo-34-1 | NZ_JGZA01000001.1 | + | 95476 | 95522 | **RNA**→ rpmE (PRK00019)Ribosomal_L31 (pfam01197)→ |
| Blo-35-1 | NZ_JDUC01000003.1 | + | 20722 | 20768 | **RNA**→ rpmE (PRK00019)Ribosomal_L31 (pfam01197)→ |
| Blo-30-1 | NZ_KE145398.1 | - | 59671 | 59625 | **RNA**→ rpmE (PRK00019)Ribosomal_L31 (pfam01197)→ |
| Blo-18-1 | NC_014169.1 | - | 1896650 | 1896604 | **RNA**→ rpmE (PRK00019)Ribosomal_L31 (pfam01197)→ |
| Blo-20-1 | NZ_CP008885.1 | - | 1899429 | 1899383 | **RNA**→ rpmE (PRK00019)Ribosomal_L31 (pfam01197)→ |
| Blo-21-1 | NZ_ATWX01000002.1 | - | 62575 | 62529 | **RNA**→ rpmE (PRK00019)Ribosomal_L31 (pfam01197)→ |
| env-166 | SRS020869_C3365636 | - | 215 | 169 | **RNA**→ rpmE (PRK00019)Ribosomal_L31 (pfam01197)→ |
| Blo-22-1 | NZ_CCWR01000010.1 | + | 9996 | 10042 | **RNA**→ rpmE (PRK00019)Ribosomal_L31 (pfam01197)→ |
| Blo-22-2 | NZ_CCWS01000010.1 | + | 10024 | 10070 | **RNA**→ rpmE (PRK00019)Ribosomal_L31 (pfam01197)→ |
| Blo-22-3 | NZ_CCWU01000013.1 | + | 9992 | 10038 | **RNA**→ rpmE (PRK00019)Ribosomal_L31 (pfam01197)→ |
| Blo-22-4 | NZ_CCWP01000016.1 | + | 10004 | 10050 | **RNA**→ rpmE (PRK00019)Ribosomal_L31 (pfam01197)→ |
| Blo-22-5 | NZ_CCWO01000014.1 | + | 10001 | 10047 | **RNA**→ rpmE (PRK00019)Ribosomal_L31 (pfam01197)→ |
| Blo-22-6 | NZ_CCWQ01000012.1 | + | 10020 | 10066 | **RNA**→ rpmE (PRK00019)Ribosomal_L31 (pfam01197)→ |
| Blo-22-7 | NZ_CCWN01000013.1 | + | 10016 | 10062 | **RNA**→ rpmE (PRK00019)Ribosomal_L31 (pfam01197)→ |
| Blo-22-8 | NZ_CCWT01000014.1 | + | 10021 | 10067 | **RNA**→ rpmE (PRK00019)Ribosomal_L31 (pfam01197)→ |
| Blo-23-1 | NZ_JDTT01000014.1 | - | 70968 | 70922 | **RNA**→ rpmE (PRK00019)Ribosomal_L31 (pfam01197)→ |
| Blo-6-1 | NC_011593.1 | + | 649451 | 649497 | **RNA**→ rpmE (PRK00019)Ribosomal_L31 (pfam01197)→ |
| Blo-7-1 | NC_017219.1 | + | 649243 | 649289 | **RNA**→ rpmE (PRK00019)Ribosomal_L31 (pfam01197)→ |
| env-167 | BABF01008774.1 | - | 974 | 928 | **RNA**→ rpmE (PRK00019)Ribosomal_L31 (pfam01197)→ |
| env-168 | scaffold29315_2_MH0086 | - | 681 | 635 | **RNA**→ rpmE (PRK00019)Ribosomal_L31 (pfam01197)→ |
| env-169 | SRS018133_C4548354 | - | 305 | 259 | **RNA**→ rpmE (PRK00019)Ribosomal_L31 (pfam01197)→ |
| env-170 | scaffold19704_2_O2.UC-23 | - | 9273 | 9227 | **RNA**→ rpmE (PRK00019)Ribosomal_L31 (pfam01197)→ |
| env-171 | scaffold19269_2_V1.UC-14 | - | 1535 | 1489 | **RNA**→ rpmE (PRK00019)Ribosomal_L31 (pfam01197)→ |
| env-172 | scaffold21097_2_V1.UC-15 | - | 760 | 714 | **RNA**→ rpmE (PRK00019)Ribosomal_L31 (pfam01197)→ |
| env-173 | scaffold17503_1_V1.UC-18 | + | 2099 | 2145 | **RNA**→ rpmE (PRK00019)Ribosomal_L31 (pfam01197)→ |

| | | | | | | |
|---|---|---|---|---|---|---|
| env-174 | scaffold52336_3_V1.UC-6 | + | 189 | 235 | **RNA**→ rpmE (PRK00019)Ribosomal_L31 (pfam01197)→ | |
| env-175 | 4491422.3_NODE_2100 | + | 169 | 215 | **RNA**→ rpmE (PRK00019)Ribosomal_L31 (pfam01197)→ | |
| env-176 | DLF009_scaffold9111_2 | + | 197 | 243 | **RNA**→ rpmE (PRK00019)Ribosomal_L31 (pfam01197)→ | |
| env-177 | DLM001_scaffold39319_11 | - | 1511 | 1465 | **RNA**→ rpmE (PRK00019)Ribosomal_L31 (pfam01197)→ | |
| env-178 | longitudinal_276_85 | - | 13590 | 13544 | **RNA**→ rpmE (PRK00019)Ribosomal_L31 (pfam01197)→ | |
| env-179 | longitudinal_277_74 | - | 54531 | 54485 | **RNA**→ rpmE (PRK00019)Ribosomal_L31 (pfam01197)→ | |
| env-180 | longitudinal_455_238 | + | 17104 | 17150 | **RNA**→ rpmE (PRK00019)Ribosomal_L31 (pfam01197)→ | |
| env-181 | longitudinal_456_166 | + | 17071 | 17117 | **RNA**→ rpmE (PRK00019)Ribosomal_L31 (pfam01197)→ | |
| env-182 | longitudinal_457_211 | + | 244972 | 245018 | **RNA**→ rpmE (PRK00019)Ribosomal_L31 (pfam01197)→ | |
| env-183 | longitudinal_458_181 | + | 17074 | 17120 | **RNA**→ rpmE (PRK00019)Ribosomal_L31 (pfam01197)→ | |
| env-184 | longitudinal_459_243 | + | 17138 | 17184 | **RNA**→ rpmE (PRK00019)Ribosomal_L31 (pfam01197)→ | |
| env-185 | longitudinal_512_194 | - | 163136 | 163090 | **RNA**→ rpmE (PRK00019)Ribosomal_L31 (pfam01197)→ | |
| env-186 | longitudinal_513_220 | + | 245449 | 245495 | **RNA**→ rpmE (PRK00019)Ribosomal_L31 (pfam01197)→ | |
| env-187 | longitudinal_514_219 | + | 245293 | 245339 | **RNA**→ rpmE (PRK00019)Ribosomal_L31 (pfam01197)→ | |
| env-188 | longitudinal_515_8 | + | 17141 | 17187 | **RNA**→ rpmE (PRK00019)Ribosomal_L31 (pfam01197)→ | |
| env-189 | longitudinal_516_182 | - | 163739 | 163693 | **RNA**→ rpmE (PRK00019)Ribosomal_L31 (pfam01197)→ | |
| env-190 | longitudinal_517_226 | - | 112586 | 112540 | **RNA**→ rpmE (PRK00019)Ribosomal_L31 (pfam01197)→ | |
| env-191 | longitudinal_518_210 | - | 116768 | 116722 | **RNA**→ rpmE (PRK00019)Ribosomal_L31 (pfam01197)→ | |
| env-192 | ADJS01003337.1 | - | 8514 | 8468 | **RNA**→ rpmE (PRK00019)Ribosomal_L31 (pfam01197)→ | |
| env-193 | ADJT01002401.1 | + | 120384 | 120430 | **RNA**→ rpmE (PRK00019)Ribosomal_L31 (pfam01197)→ | |
| env-194 | SRS018656_C1658316 | + | 203 | 249 | **RNA**→ rpmE (PRK00019)Ribosomal_L31 (pfam01197)→ | |
| env-195 | SRS020328_C2846861 | - | 160 | 114 | **RNA**→ rpmE (PRK00019)Ribosomal_L31 (pfam01197)→ | |
| Blo-1-1 | NZ_AQGL01000009.1 | - | 343420 | 343374 | **RNA**→ rpmE (PRK00019)Ribosomal_L31 (pfam01197)→ | |
| Blo-22-9 | NZ_CDPT01000001.1 | - | 1421625 | 1421579 | **RNA**→ rpmE (PRK00019)Ribosomal_L31 (pfam01197)→ | |
| Blo-15-1 | NZ_CALH01000107.1 | - | 4653 | 4607 | **RNA**→ rpmE (PRK00019)Ribosomal_L31 (pfam01197)→ | |
| Blo-31-1 | NZ_JNWD01000001.1 | + | 233485 | 233531 | **RNA**→ rpmE (PRK00019)Ribosomal_L31 (pfam01197)→ | |
| Blo-28-1 | NZ_JNVY01000019.1 | + | 152362 | 152408 | **RNA**→ rpmE (PRK00019)Ribosomal_L31 (pfam01197)→ | |
| Blo-24-1 | NZ_JNWB01000013.1 | - | 17648 | 17602 | **RNA**→ rpmE (PRK00019)Ribosomal_L31 (pfam01197)→ | |
| Blo-29-1 | NZ_JNWA01000011.1 | + | 218140 | 218186 | **RNA**→ rpmE (PRK00019)Ribosomal_L31 (pfam01197)→ | |
| Blo-11-1 | NZ_AJTJ01000037.1 | - | 10590 | 10544 | **RNA**→ rpmE (PRK00019)Ribosomal_L31 (pfam01197)→ | |
| Blo-12-1 | NZ_AJTI01000029.1 | + | 4915 | 4961 | **RNA**→ rpmE (PRK00019)Ribosomal_L31 (pfam01197)→ | |
| Blo-26-1 | NZ_JNVX01000017.1 | - | 233903 | 233857 | **RNA**→ rpmE (PRK00019)Ribosomal_L31 (pfam01197)→ | |
| Blo-25-1 | NZ_JGYZ01000008.1 | - | 522649 | 522603 | **RNA**→ rpmE (PRK00019)Ribosomal_L31 (pfam01197)→ | |
| Blo-9-1 | NZ_DS990240.1 | + | 132343 | 132389 | **RNA**→ rpmE (PRK00019)Ribosomal_L31 (pfam01197)→ | rpmE (PRK00019)Ribosomal_L31 (pfam01197)→ |
| Bsp-1-1 | NZ_GL622628.1 | - | 22265 | 22219 | **RNA**→ rpmE (PRK00019)Ribosomal_L31 (pfam01197)→ | |
| Blo-3-1 | NZ_CM002287.1 | - | 1828371 | 1828325 | **RNA**→ rpmE (PRK00019)Ribosomal_L31 (pfam01197)→ | |
| Blo-14-1 | NC_014656.1 | + | 2156860 | 2156906 | **RNA**→ rpmE (PRK00019)Ribosomal_L31 (pfam01197)→ | |
| Blo-17-1 | NC_015067.1 | - | 1854923 | 1854877 | **RNA**→ rpmE (PRK00019)Ribosomal_L31 (pfam01197)→ | |
| Blo-33-1 | NZ_CP006741.1 | - | 1917232 | 1917186 | **RNA**→ rpmE (PRK00019)Ribosomal_L31 (pfam01197)→ | |
| Blo-16-1 | NC_021008.1 | - | 314812 | 314766 | **RNA**→ rpmE (PRK00019)Ribosomal_L31 (pfam01197)→ | |
| Blo-25-2 | NZ_JRWN01000004.1 | + | 171999 | 172045 | **RNA**→ rpmE (PRK00019)Ribosomal_L31 (pfam01197)→ | |
| env-196 | BABC01009804.1 | - | 850 | 804 | **RNA**→ rpmE (PRK00019)Ribosomal_L31 (pfam01197)→ | |
| Blo-19-1 | NC_017221.1 | + | 433187 | 433233 | **RNA**→ rpmE (PRK00019)Ribosomal_L31 (pfam01197)→ | |
| env-197 | hgutS8_s8_172738 | - | 762 | 716 | **RNA**→ rpmE (PRK00019)Ribosomal_L31 (pfam01197)→ | |
| env-198 | BABE01000068.1 | - | 9777 | 9731 | **RNA**→ rpmE (PRK00019)Ribosomal_L31 (pfam01197)→ | |
| env-199 | scaffold8513_5_MH0012 | + | 1213 | 1259 | **RNA**→ rpmE (PRK00019)Ribosomal_L31 (pfam01197)→ | |
| env-200 | DLF013_scaffold22032_2 | - | 754 | 708 | **RNA**→ rpmE (PRK00019)Ribosomal_L31 (pfam01197)→ | |
| env-201 | DLM012_scaffold9385_5 | + | 30093 | 30139 | **RNA**→ rpmE (PRK00019)Ribosomal_L31 (pfam01197)→ | |
| env-202 | DLM018_scaffold48944_2 | - | 1497 | 1451 | **RNA**→ rpmE (PRK00019)Ribosomal_L31 (pfam01197)→ | |
| env-203 | DOF008_scaffold17675_1 | - | 57046 | 57000 | **RNA**→ rpmE (PRK00019)Ribosomal_L31 (pfam01197)→ | |
| env-204 | longitudinal_33_209 | + | 102094 | 102140 | **RNA**→ rpmE (PRK00019)Ribosomal_L31 (pfam01197)→ | |
| env-205 | longitudinal_34_64 | + | 102147 | 102193 | **RNA**→ rpmE (PRK00019)Ribosomal_L31 (pfam01197)→ | |

| | | | | | |
|---|---|---|---|---|---|
| env-206 | longitudinal_35_67 | - | 79278 | 79232 | RNA→ rpmE (PRK00019)Ribosomal_L31 (pfam01197)→ |
| env-207 | longitudinal_36_6 | - | 79301 | 79255 | RNA→ rpmE (PRK00019)Ribosomal_L31 (pfam01197)→ |
| env-208 | longitudinal_37_110 | - | 139321 | 139275 | RNA→ rpmE (PRK00019)Ribosomal_L31 (pfam01197)→ |
| env-209 | longitudinal_59_157 | + | 17124 | 17170 | RNA→ rpmE (PRK00019)Ribosomal_L31 (pfam01197)→ PrfA (COG0216)prfA (PRK00591)→ HemK (COG2890)PRK09328 (PRK09328)→ |
| env-210 | longitudinal_60_167 | + | 21344 | 21390 | RNA→ rpmE (PRK00019)Ribosomal_L31 (pfam01197)→ |
| env-211 | longitudinal_94_39 | - | 19572 | 19526 | RNA→ rpmE (PRK00019)Ribosomal_L31 (pfam01197)→ |
| env-212 | longitudinal_95_34 | + | 21350 | 21396 | RNA→ rpmE (PRK00019)Ribosomal_L31 (pfam01197)→ |
| env-213 | longitudinal_96_7 | + | 102094 | 102140 | RNA→ rpmE (PRK00019)Ribosomal_L31 (pfam01197)→ |
| env-214 | SRS017103_C2706015 | - | 215 | 169 | RNA→ rpmE (PRK00019)Ribosomal_L31 (pfam01197)→ |
| env-215 | SRS052697_LANL_scaffold_51199 | + | 934 | 980 | RNA→ rpmE (PRK00019)Ribosomal_L31 (pfam01197)→ |
| Blo-13-1 | NZ_AJTM01000062.1 | - | 86722 | 86676 | RNA→ rpmE (PRK00019)Ribosomal_L31 (pfam01197)→ |
| Blo-27-1 | NZ_JNVZ01000015.1 | + | 229769 | 229815 | RNA→ rpmE (PRK00019)Ribosomal_L31 (pfam01197)→ |
| Blo-32-1 | NZ_JNWC01000023.1 | - | 297292 | 297246 | RNA→ rpmE (PRK00019)Ribosomal_L31 (pfam01197)→ |
| Blo-10-1 | NZ_AJTF01000083.1 | + | 25032 | 25078 | RNA→ rpmE (PRK00019)Ribosomal_L31 (pfam01197)→ |
| Blo-4-1 | NC_004307.2 | + | 2130767 | 2130813 | RNA→ rpmE (PRK00019)Ribosomal_L31 (pfam01197)→ |
| Blo-8-1 | NZ_GG666850.1 | + | 218460 | 218506 | RNA→ rpmE (PRK00019)Ribosomal_L31 (pfam01197)→ |
| Blo-5-1 | NC_015052.1 | - | 1885479 | 1885433 | RNA→ rpmE (PRK00019)Ribosomal_L31 (pfam01197)→ |
| env-216 | BABF01010041.1 | + | 452 | 498 | RNA→ rpmE (PRK00019)Ribosomal_L31 (pfam01197)→ |
| env-217 | SRS016517_C20400143 | - | 363 | 317 | RNA→ rpmE (PRK00019)Ribosomal_L31 (pfam01197)→ |
| env-218 | SRS016495_LANL_scaffold_979 | + | 781 | 827 | RNA→ |
| Blo-2-1 | NC_010816.1 | + | 2243300 | 2243346 | RNA→ rpmE (PRK00019)Ribosomal_L31 (pfam01197)→ |
| env-219 | JGI24738J21930_10111851 | + | 341 | 398 | RNA→ rpmE (PRK00019)Ribosomal_L31 (pfam01197)→ |
| Ssp-47-1 | NZ_JALM01000005.1 | + | 11373 | 11411 | RNA→ rpmE (PRK00019)Ribosomal_L31 (pfam01197)→ PrfA (COG0216)prfA (PRK00591)→ HemK (COG2890)PRK09328 (PRK09328)→ CpdA (COG1409)→ hypo→ HicB (COG1598)HicB (pfam05534)→ |
| Sma-2-1 | NZ_KB905458.1 | - | 174655 | 174617 | RNA→ rpmE (PRK00019)Ribosomal_L31 (pfam01197)→ PrfA (COG0216)prfA (PRK00591)→ |
| Nrh-1-1 | NZ_JOAJ01000005.1 | + | 675604 | 675651 | RNA→ rpmE (PRK00019)Ribosomal_L31 (pfam01197)→ PrfA (COG0216)prfA (PRK00591)→ |
| Nrh-1-2 | NZ_JOEJ01000005.1 | - | 16032 | 15985 | RNA→ rpmE (PRK00019)Ribosomal_L31 (pfam01197)→ PrfA (COG0216)prfA (PRK00591)→ |
| Nte-2-1 | NZ_BAGJ01000039.1 | + | 21259 | 21306 | RNA→ rpmE (PRK00019)Ribosomal_L31 (pfam01197)→ PrfA (COG0216)prfA (PRK00591)→ HemK (COG2890)PRK09328 (PRK09328)→ HemK (COG2890)PRK09328 (PRK09328)→ |
| Nsp-2-1 | NZ_KB907310.1 | + | 750500 | 750547 | RNA→ rpmE (PRK00019)Ribosomal_L31 (pfam01197)→ PrfA (COG0216)prfA (PRK00591)→ |
| env-220 | JGI12635J15846_10443042 | + | 326 | 380 | RNA→ ←hypo |
| env-221 | JGI12467J12023_1172356 | + | 9 | 51 | RNA→ rpmE (PRK00019)Ribosomal_L31 (pfam01197)→ |
| env-222 | JGI12397J12024_1008566 | + | 21 | 63 | RNA→ rpmE (PRK00019)Ribosomal_L31 (pfam01197)→ PCRF (smart00937)RF-1 (pfam00472)PCRF (pfam03462)→ |
| env-223 | JGI24741J21665_1049030 | + | 81 | 179 | RNA→ rpmE (PRK00019)Ribosomal_L31 (pfam01197)→ prfA (PRK00591)PCRF (pfam03462)→ |
| env-224 | JGI24740J21852_10229407 | + | 87 | 185 | RNA→ hypo→ rpmE (PRK00019)Ribosomal_L31 (pfam01197)→ |
| env-225 | JGI12397J12024_1078760 | + | 13 | 60 | RNA→ rpmE (PRK00019)Ribosomal_L31 (pfam01197)→ |
| env-226 | JGI12396J12026_1036873 | + | 11 | 61 | RNA→ rpmE (PRK00019)Ribosomal_L31 (pfam01197)→ |
| env-227 | FNTS_GKKEVUV01A6HR7 | + | 259 | 297 | RNA→ rpmE (PRK00019)Ribosomal_L31 (pfam01197)→ |
| env-228 | JGI12396J12026_1167238 | + | 15 | 53 | RNA→ rpmE (PRK00019)Ribosomal_L31 (pfam01197)→ |
| env-229 | JGI12396J12026_1310149 | + | 19 | 57 | RNA→ hypo→ rpmE (PRK00019)Ribosomal_L31 (pfam01197)→ |
| env-230 | Incfw_1020660 | + | 204 | 250 | RNA→ rpmE (PRK00019)Ribosomal_L31 (pfam01197)→ |
| env-231 | JGI12273J12029_10368297 | + | 4 | 44 | RNA→ rpmE (PRK00019)Ribosomal_L31 (pfam01197)→ |
| env-232 | JGI24735J21928_10783078 | + | 165 | 212 | RNA→ hypo→ |
| env-233 | JGI12272J11983_1065882 | + | 43 | 88 | RNA→ ←hypo |
| env-234 | JGI12397J12024_1021578 | + | 17 | 62 | RNA→ rpmE (PRK00019)Ribosomal_L31 (pfam01197)→ |
| env-235 | JGI12271J12027_1090917 | + | 17 | 62 | RNA→ rpmE (PRK00019)Ribosomal_L31 (pfam01197)→ |
| env-236 | JGI12273J12029_10052257 | + | 17 | 62 | RNA→ ←hypo |
| Fsp-3-1 | NC_007777.1 | - | 4454117 | 4454072 | RNA→ rpmE (PRK00019)Ribosomal_L31 (pfam01197)→ |
| Fsp-15-1 | NZ_JENI01000016.1 | + | 11880 | 11925 | RNA→ rpmE (PRK00019)Ribosomal_L31 (pfam01197)→ |
| Fsp-11-1 | NZ_JDWE01000027.1 | + | 11908 | 11953 | RNA→ rpmE (PRK00019)Ribosomal_L31 (pfam01197)→ |
| Fsp-12-1 | NZ_JPGU01000036.1 | - | 33938 | 33893 | RNA→ rpmE (PRK00019)Ribosomal_L31 (pfam01197)→ |

| | | | | | |
|---|---|---|---|---|---|
| Fsp-10-1 | NZ_JPHT01000022.1 | - | 73950 | 73905 | **RNA**→ rpmE (PRK00019)Ribosomal_L31 (pfam01197)→ |
| Fsp-4-1 | NZ_AYTZ01000024.1 | + | 10131 | 10176 | **RNA**→ rpmE (PRK00019)Ribosomal_L31 (pfam01197)→ |
| Ssy-1-1 | NZ_JNYO01000029.1 | - | 44929 | 44889 | **RNA**→ rpmE (PRK00019)Ribosomal_L31 (pfam01197)→ |
| Ssp-54-1 | NZ_KE384522.1 | - | 215499 | 215455 | **RNA**→ rpmE (PRK00019)Ribosomal_L31 (pfam01197)→ |
| Ssp-174-1 | NZ_JNIH01000045.1 | + | 12803 | 12848 | **RNA**→ rpmE (PRK00019)Ribosomal_L31 (pfam01197)→ |
| Ssp-32-1 | NZ_BARG01000069.1 | - | 93141 | 93096 | **RNA**→ rpmE (PRK00019)Ribosomal_L31 (pfam01197)→ |
| Sxy-1-1 | NZ_JNWO01000024.1 | - | 100532 | 100487 | **RNA**→ rpmE (PRK00019)Ribosomal_L31 (pfam01197)→ |
| env-237 | JGI12396J12026_1094817 | + | 15 | 73 | **RNA**→ rpmE (PRK00019)Ribosomal_L31 (pfam01197)→ |
| env-238 | JGI12467J12023_1044130 | + | 19 | 77 | **RNA**→ rpmE (PRK00019)Ribosomal_L31 (pfam01197)→ |
| env-239 | JGI12273J12029_10045587 | + | 359 | 415 | **RNA**→ rpmE (PRK00019)Ribosomal_L31 (pfam01197)→ |
| env-240 | JGI24740J21852_10734288 | - | 137 | 91 | **RNA**→ ←hypo |
| Ssp-102-1 | NZ_JOFZ01000002.1 | + | 291142 | 291188 | **RNA**→ rpmE (PRK00019)Ribosomal_L31 (pfam01197)→ |
| Sca-3-1 | NZ_LFXA01000015.1 | + | 32879 | 32925 | **RNA**→ rpmE (PRK00019)Ribosomal_L31 (pfam01197)→ |
| Ghi-1-2 | NZ_ATVK01000007.1 | + | 35312 | 35358 | **RNA**→ rpmE (PRK00019)Ribosomal_L31 (pfam01197)→ |
| Ghi-1-1 | NZ_BANT01000013.1 | + | 35312 | 35358 | **RNA**→ rpmE (PRK00019)Ribosomal_L31 (pfam01197)→ |
| Ssc-5-1 | NZ_JOEI01000009.1 | - | 442082 | 442036 | **RNA**→ rpmE (PRK00019)Ribosomal_L31 (pfam01197)→ |
| Sro-5-1 | NZ_JOFL01000017.1 | + | 10548 | 10594 | **RNA**→ rpmE (PRK00019)Ribosomal_L31 (pfam01197)→ |
| Sva-2-1 | NZ_JOBF01000022.1 | + | 129517 | 129563 | **RNA**→ rpmE (PRK00019)Ribosomal_L31 (pfam01197)→ |
| Sva-2-2 | NZ_JOFN01000033.1 | + | 33993 | 34039 | **RNA**→ rpmE (PRK00019)Ribosomal_L31 (pfam01197)→ |
| Sye-1-1 | NZ_JQNR01000005.1 | + | 2828881 | 2828928 | **RNA**→ rpmE (PRK00019)Ribosomal_L31 (pfam01197)→ |
| env-241 | JGI12397J12024_1054783 | + | 54 | 105 | **RNA**→ rpmE (PRK00019)Ribosomal_L31 (pfam01197)→ PrfA (COG0216)prfA (PRK00591)→ |
| env-242 | JGI12272J11983_1231162 | + | 43 | 94 | **RNA**→ rpmE (PRK00019)Ribosomal_L31 (pfam01197)→ |
| Nex-1-1 | NZ_BAFZ01000027.1 | + | 22758 | 22807 | **RNA**→ rpmE (PRK00019)Ribosomal_L31 (pfam01197)→ PrfA (COG0216)prfA (PRK00591)→ |
| | | | | | HemK (COG2890)PRK09328 (PRK09328)→ |
| Nar-1-1 | NZ_BAFR01000040.1 | + | 1958 | 2005 | **RNA**→ rpmE (PRK00019)Ribosomal_L31 (pfam01197)→ PrfA (COG0216)prfA (PRK00591)→ |
| | | | | | HemK (COG2890)PRK09328 (PRK09328)→ |
| Nab-1-1 | NZ_BAFP01000034.1 | + | 43181 | 43230 | **RNA**→ rpmE (PRK00019)Ribosomal_L31 (pfam01197)→ PrfA (COG0216)prfA (PRK00591)→ |
| | | | | | HemK (COG2890)PRK09328 (PRK09328)→ |
| Mar-1-1 | NZ_JALN02000001.1 | + | 4095907 | 4095961 | **RNA**→ rpmE (PRK00019)Ribosomal_L31 (pfam01197)→ PrfA (COG0216)prfA (PRK00591)→ |
| | | | | | HemK (COG2890)PRK09328 (PRK09328)→ SUA5 (COG0009)TIGR00057 (TIGR00057)→ |
| | | | | | GT_WecA_like (cd06853)mraY (PRK00108)→ |
| env-243 | JGI24770J26754_10202382 | + | 361 | 403 | **RNA**→ rpmE (PRK00019)Ribosomal_L31 (pfam01197)→ |
| env-244 | FWIRElOz_GKZ9IRQ02H6XIE | - | 268 | 216 | **RNA**→ rpmE (PRK00019)Ribosomal_L31 (pfam01197)→ |
| env-245 | JGI26342J46808_1002686 | - | 1270 | 1220 | **RNA**→ rpmE (PRK00019)Ribosomal_L31 (pfam01197)→ |
| env-246 | JGI26344J46810_1010675 | - | 505 | 455 | **RNA**→ rpmE (PRK00019)Ribosomal_L31 (pfam01197)→ |
| env-247 | JGI26347J50199_1039997 | - | 278 | 228 | **RNA**→ rpmE (PRK00019)Ribosomal_L31 (pfam01197)→ |
| Agl-1-1 | NZ_KB903307.1 | - | 53455 | 53386 | **RNA**→ rpmE (PRK00019)Ribosomal_L31 (pfam01197)→ PrfA (COG0216)prfA (PRK00591)→ |
| Msp-15-1 | NZ_LBEW01000064.1 | - | 15115 | 15060 | **RNA**→ rpmE (PRK00019)Ribosomal_L31 (pfam01197)→ PrfA (COG0216)prfA (PRK00591)→ |
| | | | | | HemK (COG2890)PRK09328 (PRK09328)→ SUA5 (COG0009)TIGR00057 (TIGR00057)→ |
| | | | | | GT_WecA_like (cd06853)PRK15119 (PRK15119)→ GT_WecA_like (cd06853)PRK15119 (PRK15119)→ |
| Mco-2-1 | NZ_CCBB010000001.1 | - | 249287 | 249232 | **RNA**→ rpmE (PRK00019)Ribosomal_L31 (pfam01197)→ PrfA (COG0216)prfA (PRK00591)→ |
| | | | | | HemK (COG2890)PRK09328 (PRK09328)→ SUA5 (COG0009)TIGR00057 (TIGR00057)→ |
| | | | | | GT_WecA_like (cd06853)PRK15119 (PRK15119)→ |
| env-248 | JGI12688J13320_109532 | - | 138 | 89 | **RNA**→ rpmE (PRK00019)Ribosomal_L31 (pfam01197)→ |
| env-249 | JGI24735J21928_10093269 | + | 384 | 441 | **RNA**→ rpmE (PRK00019)Ribosomal_L31 (pfam01197)→ prfA (PRK00591)PCRF (pfam03462)→ |
| Nmu-1-1 | NC_013235.1 | + | 2318949 | 2318997 | **RNA**→ rpmE (PRK00019)Ribosomal_L31 (pfam01197)→ |
| env-250 | JGI12635J15846_13193507 | - | 158 | 97 | **RNA**→ hypo→ |
| env-251 | JGI20241J16302_100838 | + | 402 | 463 | **RNA**→ rpmE (PRK00019)Ribosomal_L31 (pfam01197)→ |
| env-252 | JGI20235J16296_1030653 | - | 332 | 271 | **RNA**→ rpmE (PRK00019)Ribosomal_L31 (pfam01197)→ |
| env-253 | C688J13580_1073401 | + | 297 | 353 | **RNA**→ rpmE (PRK00019)rpmE2 (PRK01678)Ribosomal_L31 (pfam01197)→ |
| env-254 | JGI12397J12024_1051954 | + | 41 | 127 | **RNA**→ rpmE (PRK00019)Ribosomal_L31 (pfam01197)→ hypo→ |
| env-255 | C688J13580_1242542 | - | 139 | 84 | **RNA**→ rpmE (PRK00019)Ribosomal_L31 (pfam01197)→ |
| env-256 | JGI12397J12024_1036663 | + | 40 | 82 | **RNA**→ rpmE (PRK00019)Ribosomal_L31 (pfam01197)→ |

| | | | | | |
|---|---|---|---|---|---|
| env-257 | JGI12396J12026_1028942 | + | 39 | 125 | RNA→ rpmE (PRK00019)Ribosomal_L31 (pfam01197)→ prfA (PRK00591)RF-1 (pfam00472)PCRF (pfam03462)→ |
| env-258 | JGI12397J12024_1014281 | + | 48 | 131 | RNA→ rpmE (PRK00019)Ribosomal_L31 (pfam01197)→ prfA (PRK00591)PCRF (pfam03462)→ |
| env-259 | JGI24735J21928_10543660 | - | 202 | 156 | RNA→ ←hypo |
| env-260 | _GVICOHK01AHWYS | - | 222 | 178 | RNA→ rpmE (PRK00019)Ribosomal_L31 (pfam01197)→ |
| env-261 | CTBMPR_c14314 | + | 184 | 228 | RNA→ rpmE (PRK00019)Ribosomal_L31 (pfam01197)→ |
| env-262 | JGI12420J12819_1024840 | + | 306 | 350 | RNA→ rpmE (PRK00019)Ribosomal_L31 (pfam01197)→ |
| env-263 | JGI12397J12024_1056172 | + | 36 | 78 | RNA→ rpmE (PRK00019)Ribosomal_L31 (pfam01197)→ |
| env-264 | JGI24739J22299_10632192 | - | 277 | 228 | RNA→ rpmE (PRK00019)Ribosomal_L31 (pfam01197)→ |
| env-265 | topACOD_FV90NF401CWY9R | + | 385 | 428 | RNA→ |
| env-266 | JGI24735J21928_10186907 | + | 96 | 140 | RNA→ rpmE (PRK00019)Ribosomal_L31 (pfam01197)→ prfA (PRK00591)PCRF (pfam03462)→ |
| Nth-1-1 | NZ_BAGK01000052.1 | + | 1741 | 1791 | RNA→ rpmE (PRK00019)Ribosomal_L31 (pfam01197)→ |
| Pan-1-1 | NZ_JOFP01000001.1 | - | 15114 | 15064 | RNA→ rpmE (PRK00019)Ribosomal_L31 (pfam01197)→ PrfA (COG0216)prfA (PRK00591)→ HemK (COG2890)PRK09328 (PRK09328)→ |
| Nas-1-1 | NZ_BAFO02000037.1 | + | 210935 | 210985 | RNA→ rpmE (PRK00019)Ribosomal_L31 (pfam01197)→ PrfA (COG0216)prfA (PRK00591)→ HemK (COG2890)PRK09328 (PRK09328)→ |
| Sxi-1-1 | NZ_JH636049.1 | + | 3421640 | 3421684 | RNA→ rpmE (PRK00019)Ribosomal_L31 (pfam01197)→ rpmE (PRK00019)Ribosomal_L31 (pfam01197)→ PrfA (COG0216)prfA (PRK00591)→ PrfA (COG0216)prfA (PRK00591)→ hypo→ |
| Ssa-1-1 | NZ_KB912602.1 | - | 65465 | 65418 | RNA→ rpmE (PRK00019)Ribosomal_L31 (pfam01197)→ |
| Mch-1-1 | NC_018027.1 | - | 3984376 | 3984325 | RNA→ rpmE (PRK00019)Ribosomal_L31 (pfam01197)→ PrfA (COG0216)prfA (PRK00591)→ HemK (COG2890)PRK09328 (PRK09328)→ SUA5 (COG0009)TIGR00057 (TIGR00057)→ GT_WecA_like (cd06853)mraY (PRK00108)→ |
| env-267 | SRS015947_C1328584 | - | 203 | 156 | RNA→ rpmE (PRK00019)Ribosomal_L31 (pfam01197)→ |
| env-268 | SRS013170_Baylor_scaffold_89712 | + | 115 | 162 | RNA→ rpmE (PRK00019)Ribosomal_L31 (pfam01197)→ |
| env-269 | SRS015574_WUGC_scaffold_15753 | + | 130 | 177 | RNA→ rpmE (PRK00019)Ribosomal_L31 (pfam01197)→ |
| env-270 | SRS017227_Baylor_scaffold_82921 | + | 846 | 893 | RNA→ |
| env-271 | SRS022149_LANL_scaffold_16970 | + | 119 | 166 | RNA→ rpmE (PRK00019)Ribosomal_L31 (pfam01197)→ |
| env-272 | SRS023964_C2397974 | - | 196 | 149 | RNA→ ←hypo |
| env-273 | SRS055378_LANL_scaffold_81766 | + | 131 | 178 | RNA→ rpmE (PRK00019)Ribosomal_L31 (pfam01197)→ |
| env-274 | SRS044366_WUGC_scaffold_3585 | + | 105 | 152 | RNA→ rpmE (PRK00019)Ribosomal_L31 (pfam01197)→ |
| env-275 | SRS017304_C1861044 | - | 118 | 71 | RNA→ rpmE (PRK00528)rpmE (PRK00019)rpmE2 (PRK01678)Ribosomal_L31 (pfam01197)→ |
| env-276 | SRS024289_C1901398 | + | 126 | 173 | RNA→ rpmE (PRK00019)Ribosomal_L31 (pfam01197)→ |
| env-277 | SRS024649_C3463797 | - | 196 | 149 | RNA→ ←hypo |
| env-278 | SRS058053_C5176642 | - | 196 | 149 | RNA→ ←hypo |
| Age-1-1 | NZ_KE386871.1 | - | 110335 | 110288 | RNA→ rpmE (PRK00019)Ribosomal_L31 (pfam01197)→ |
| Pas-1-1 | NZ_AUII01000018.1 | - | 64638 | 64580 | RNA→ rpmE (PRK00019)Ribosomal_L31 (pfam01197)→ prfA (PRK00591)RF-1 (pfam00472)PCRF (pfam03462)→ HemK (COG2890)PRK09328 (PRK09328)→ SUA5 (COG0009)TIGR00057 (TIGR00057)→ |
| Pdi-1-1 | NC_015312.1 | + | 1759317 | 1759374 | RNA→ rpmE (PRK00019)Ribosomal_L31 (pfam01197)→ PrfA (COG0216)prfA (PRK00591)→ HemK (COG2890)PRK09328 (PRK09328)→ SUA5 (COG0009)TIGR00057 (TIGR00057)→ |
| env-279 | FNTS067_GKA24FP01DWQJ8 | - | 109 | 66 | RNA→ ←hypo |
| env-280 | _GX4IIKE01BKU07 | + | 30 | 75 | RNA→ rpmE (PRK00019)Ribosomal_L31 (pfam01197)→ hypo→ |
| Ame-4-2 | NZ_CP009110.1 | + | 1382283 | 1382334 | RNA→ rpmE (PRK00019)Ribosomal_L31 (pfam01197)→ prfA (PRK00591)RF-1 (pfam00472)PCRF (pfam03462)→ |
| Ame-4-1 | NZ_AQUL01000001.1 | + | 6275366 | 6275417 | RNA→ rpmE (PRK00019)Ribosomal_L31 (pfam01197)→ prfA (PRK00591)RF-1 (pfam00472)PCRF (pfam03462)→ |
| Asp-4-1 | NZ_AFWY03000007.1 | + | 215866 | 215917 | RNA→ rpmE (PRK00019)Ribosomal_L31 (pfam01197)→ prfA (PRK00591)RF-1 (pfam00472)PCRF (pfam03462)→ |
| Ath-1-1 | NZ_KI421511.1 | - | 4345617 | 4345566 | RNA→ rpmE (PRK00019)Ribosomal_L31 (pfam01197)→ prfA (PRK00591)RF-1 (pfam00472)PCRF (pfam03462)→ |
| Sbi-1-1 | NC_016582.1 | - | 4870714 | 4870670 | RNA→ rpmE (PRK00019)Ribosomal_L31 (pfam01197)→ |
| Shy-4-1 | NZ_JOIK01000001.1 | + | 665887 | 665931 | RNA→ rpmE (PRK00019)Ribosomal_L31 (pfam01197)→ |
| Ssp-73-1 | NZ_LAKD01000061.1 | + | 86684 | 86728 | RNA→ rpmE (PRK00019)Ribosomal_L31 (pfam01197)→ |
| Sir-1-1 | NZ_LK022848.1 | - | 4236123 | 4236079 | RNA→ rpmE (PRK00019)Ribosomal_L31 (pfam01197)→ |
| Ssp-165-1 | NZ_JABQ01000013.1 | - | 32688 | 32644 | RNA→ rpmE (PRK00019)Ribosomal_L31 (pfam01197)→ |

| | | | | | |
|---|---|---|---|---|---|
| Svi-3-1 | NC_015957.1 | + | 1549091 | 1549136 | RNA→ rpmE (PRK00019)Ribosomal_L31 (pfam01197)→ |
| env-281 | FNTS_GKKEVUV01CU0U2 | - | 207 | 156 | RNA→ rpmE (PRK00019)Ribosomal_L31 (pfam01197)→ |
| env-282 | JGI24736J21556_1245205 | + | 19 | 73 | RNA→ hypo→ rpmE (PRK00019)Ribosomal_L31 (pfam01197)→ |
| env-283 | JGI20205J14842_1000519 | + | 3917 | 3979 | RNA→ rpmE (PRK00019)Ribosomal_L31 (pfam01197)→ |
| env-284 | JGI20206J14855_1026724 | + | 110 | 172 | RNA→ rpmE (PRK00019)Ribosomal_L31 (pfam01197)→ |
| env-285 | JGI20188J14859_1000358 | + | 3928 | 3990 | RNA→ rpmE (PRK00019)Ribosomal_L31 (pfam01197)→ |
| env-286 | JGI20204J15000_1155101 | + | 129 | 191 | RNA→ |
| env-287 | JGI20204J15135_1155101 | + | 129 | 191 | RNA→ |
| env-288 | JGI12396J12026_1106282 | + | 35 | 77 | RNA→ rpmE (PRK00019)Ribosomal_L31 (pfam01197)→ |
| env-289 | 4491734.3.10488814 | - | 168 | 123 | RNA→ rpmE (PRK00019)Ribosomal_L31 (pfam01197)→ |
| env-290 | JGI20163J15578_11401153 | + | 94 | 155 | RNA→ rpmE (PRK00019)Ribosomal_L31 (pfam01197)→ |
| Gar-1-1 | NZ_BAEE01000076.1 | + | 38199 | 38249 | RNA→ rpmE (PRK00019)Ribosomal_L31 (pfam01197)→ rpmE (PRK00019)Ribosomal_L31 (pfam01197)→ PrfA (COG0216)prfA (PRK00591)→ PrfA (COG0216)prfA (PRK00591)→ HemK (COG2890)PRK09328 (PRK09328)→ HemK (COG2890)PRK09328 (PRK09328)→ SUA5 (COG0009)TIGR00057 (TIGR00057)→ SUA5 (COG0009)TIGR00057 (TIGR00057)→ GlyA (COG0112)glyA (PRK00011)→ GlyA (COG0112)glyA (PRK00011)→ GT_WecA_like (cd06853)mraY (PRK00108)→ GT_WecA_like (cd06853)mraY (PRK00108)→ hypo→ hypo→ |
| Mro-1-1 | NZ_JNZQ01000027.1 | - | 31420 | 31342 | RNA→ rpmE (PRK00019)Ribosomal_L31 (pfam01197)→ |
| env-291 | JGI12271J12027_1030093 | + | 10 | 56 | RNA→ ←hypo |
| env-292 | JGI12273J12029_10015673 | + | 375 | 422 | RNA→ rpmE (PRK00019)Ribosomal_L31 (pfam01197)→ |
| Psu-1-1 | NZ_KE384020.1 | - | 789516 | 789467 | RNA→ rpmE (PRK00019)Ribosomal_L31 (pfam01197)→ PrfA (COG0216)prfA (PRK00591)→ HemK (COG2890)PRK09328 (PRK09328)→ hypo→ SUA5 (COG0009)TIGR00057 (TIGR00057)→ GT_WecA_like (cd06853)PRK15119 (PRK15119)→ hypo→ AtpI2 (COG5336)→ PRK05815 (PRK05815)ATP-synt_A (pfam00119)→ PRK07874 (PRK07874)AtpE (COG0636)→ PRK05759 (PRK05759)AtpF (COG0711)→ AtpH (COG0712)PRK13430 (PRK13430)→ AtpA (COG0056)PRK09281 (PRK09281)→ PRK05621 (PRK05621)ATP-synt (pfam00231)→ AtpD (COG0055)PRK09280 (PRK09280)→ |
| Gne-1-1 | NZ_AEUD01000002.1 | + | 368326 | 368371 | RNA→ rpmE (PRK00019)Ribosomal_L31 (pfam01197)→ rpmE (PRK00019)Ribosomal_L31 (pfam01197)→ |
| Gsi-1-1 | NZ_BANU01000014.1 | - | 24449 | 24403 | RNA→ rpmE (PRK00019)Ribosomal_L31 (pfam01197)→ |
| Gsi-2-1 | NZ_JZDP01000041.1 | - | 24646 | 24600 | RNA→ rpmE (PRK00019)Ribosomal_L31 (pfam01197)→ |
| Nta-1-1 | NZ_BAGG01000023.1 | + | 33157 | 33212 | RNA→ rpmE (PRK00019)Ribosomal_L31 (pfam01197)→ |
| env-293 | JGI24737J22298_11161458 | + | 91 | 139 | RNA→ ←hypo |
| env-294 | JGI12419J13241_1001214 | - | 1703 | 1660 | RNA→ rpmE (PRK00019)Ribosomal_L31 (pfam01197)→ |
| env-295 | JGI24739J22299_10027406 | + | 1671 | 1722 | RNA→ rpmE (PRK00019)Ribosomal_L31 (pfam01197)→ |
| env-296 | JGI24735J21928_10050545 | - | 1004 | 953 | RNA→ rpmE (PRK00019)Ribosomal_L31 (pfam01197)→ PCRF (smart00937)PCRF (pfam03462)→ |
| env-297 | JGI24738J21930_10034417 | - | 910 | 859 | RNA→ rpmE (PRK00019)Ribosomal_L31 (pfam01197)→ PCRF (smart00937)PCRF (pfam03462)→ |
| env-298 | 2044439917 | - | 59 | 9 | RNA→ hypo→ |
| Mob-1-1 | NZ_LAUZ02000011.1 | + | 54331 | 54383 | RNA→ rpmE (PRK00019)Ribosomal_L31 (pfam01197)→ |
| Mob-1-2 | NZ_JYNU01000003.1 | + | 194583 | 194635 | RNA→ rpmE (PRK00019)Ribosomal_L31 (pfam01197)→ |
| env-299 | DCrCPGB_c366807 | + | 142 | 208 | RNA→ rpmE (PRK00019)Ribosomal_L31 (pfam01197)→ |
| Asp-1-1 | NC_021191.1 | - | 8063566 | 8063525 | RNA→ rpmE (PRK00019)Ribosomal_L31 (pfam01197)→ PrfA (COG0216)prfA (PRK00591)→ |
| env-300 | JGI12397J12024_1258028 | + | 36 | 84 | RNA→ rpmE (PRK00019)Ribosomal_L31 (pfam01197)→ |
| env-301 | JGI12396J12026_1152696 | + | 33 | 81 | RNA→ rpmE (PRK00019)Ribosomal_L31 (pfam01197)→ |
| env-302 | JGI12397J12024_1022450 | + | 5 | 53 | RNA→ rpmE (PRK00019)Ribosomal_L31 (pfam01197)→ |
| env-303 | JGI12396J12026_1048802 | + | 4 | 52 | RNA→ rpmE (PRK00019)Ribosomal_L31 (pfam01197)→ hypo→ |
| Nsp-6-1 | NZ_KI519393.1 | - | 15320 | 15272 | RNA→ rpmE (PRK00019)Ribosomal_L31 (pfam01197)→ PrfA (COG0216)prfA (PRK00591)→ HemK (COG2890)PRK09328 (PRK09328)→ HemK (COG2890)PRK09328 (PRK09328)→ SUA5 (COG0009)TIGR00057 (TIGR00057)→ ECA_wecA (TIGR02380)GT_WecA_like (cd06853)→ hypo→ |
| env-304 | JGI12662J13196_1036374 | + | 59 | 111 | RNA→ rpmE (PRK00019)Ribosomal_L31 (pfam01197)→ |
| env-305 | JGI12630J15595_10065573 | - | 185 | 133 | RNA→ rpmE (PRK00019)Ribosomal_L31 (pfam01197)→ |
| env-306 | JGI12701J14581_1053699 | - | 204 | 152 | RNA→ ←hypo |
| env-307 | JGI12635J15846_10023607 | + | 2592 | 2644 | RNA→ rpmE (PRK00019)Ribosomal_L31 (pfam01197)→ |
| env-308 | JGI24739J22299_10989154 | + | 71 | 123 | RNA→ ←hypo |

| | | | | | |
|---|---|---|---|---|---|
| env-309 | JGI24737J22298_10504981 | - | 264 | 212 | RNA→ rpmE (PRK00019)Ribosomal_L31 (pfam01197)→ |
| env-310 | JGI24735J21928_10176247 | + | 442 | 488 | RNA→ rpmE (PRK00019)Ribosomal_L31 (pfam01197)→ |
| env-311 | JGI24738J21930_10548350 | + | 122 | 176 | RNA→ ←hypo |
| env-312 | JGI12654J13325_1015162 | - | 201 | 152 | RNA→ rpmE (PRK00019)Ribosomal_L31 (pfam01197)→ |
| env-313 | JGI12630J15595_10385291 | + | 47 | 96 | RNA→ rpmE (PRK00019)Ribosomal_L31 (pfam01197)→ |
| env-314 | JGI20163J15578_11533527 | + | 79 | 130 | RNA→ rpmE (PRK00019)Ribosomal_L31 (pfam01197)→ |
| Ami-1-1 | NC_017093.1 | - | 7818806 | 7818758 | RNA→ rpmE (PRK00019)Ribosomal_L31 (pfam01197)→ PrfA (COG0216)prfA (PRK00591)→ |
| env-315 | JGI12397J12024_1010539 | + | 13 | 58 | RNA→ rpmE (PRK00019)Ribosomal_L31 (pfam01197)→ |
| env-316 | JGI12271J12027_1149268 | + | 10 | 55 | RNA→ rpmE (PRK00019)Ribosomal_L31 (pfam01197)→ |
| env-317 | JGI12273J12029_10077712 | + | 13 | 58 | RNA→ rpmE (PRK00019)Ribosomal_L31 (pfam01197)→ hypo→ |
| env-318 | DCrCPGB_c531990 | - | 129 | 85 | RNA→ rpmE (PRK00019)Ribosomal_L31 (pfam01197)→ |
| env-319 | JGI12273J12029_10731265 | + | 11 | 55 | RNA→ rpmE (PRK00019)Ribosomal_L31 (pfam01197)→ |
| env-320 | JGI12272J11983_1298237 | + | 5 | 49 | RNA→ rpmE (PRK00019)Ribosomal_L31 (pfam01197)→ |
| env-321 | JGI12397J12024_1066331 | + | 11 | 55 | RNA→ rpmE (PRK00019)Ribosomal_L31 (pfam01197)→ prfA (PRK00591)PCRF (pfam03462)→ |
| Nni-1-1 | NZ_BAGC01000013.1 | + | 24797 | 24839 | RNA→ rpmE (PRK00019)Ribosomal_L31 (pfam01197)→ |
| Nsp-8-1 | NZ_JQJL01000005.1 | + | 229229 | 229271 | RNA→ rpmE (PRK00019)Ribosomal_L31 (pfam01197)→ |
| env-322 | JGI26055J44792_1072012 | + | 221 | 266 | RNA→ rpmE (PRK00019)Ribosomal_L31 (pfam01197)→ |
| Rsp-20-1 | NZ_JMLQ01000015.1 | - | 17505 | 17461 | RNA→ rpmE (PRK00019)Ribosomal_L31 (pfam01197)→ |
| Rsp-16-1 | NZ_JXQS01000036.1 | - | 17102 | 17058 | RNA→ rpmE (PRK00019)Ribosomal_L31 (pfam01197)→ |
| Mpu-1-1 | NZ_JOEU01000058.1 | - | 20946 | 20882 | RNA→ rpmE (PRK00019)Ribosomal_L31 (pfam01197)→ PrfA (COG0216)prfA (PRK00591)→ |
| Msp-27-1 | NZ_KK037233.1 | + | 5461529 | 5461593 | RNA→ rpmE (PRK00019)Ribosomal_L31 (pfam01197)→ PrfA (COG0216)prfA (PRK00591)→ |
| env-323 | JGI24735J21928_10524408 | - | 73 | 31 | RNA→ |
| env-324 | ARMCC_Contig_460253 | + | 48 | 90 | RNA→ rpmE (PRK00019)Ribosomal_L31 (pfam01197)→ |
| Aal-1-1 | NZ_KB892745.1 | + | 20751 | 20792 | RNA→ rpmE (PRK00019)Ribosomal_L31 (pfam01197)→ PrfA (COG0216)prfA (PRK00591)→ |
| | | | | | HemK (COG2890)PRK09328 (PRK09328)→ |
| env-325 | JGI20200J14955_1003645 | - | 246 | 200 | RNA→ rpmE (PRK00019)Ribosomal_L31 (pfam01197)→ |
| env-326 | JGI20200J14955_1019771 | + | 167 | 213 | RNA→ ←hypo |
| env-327 | JGI20200J14955_1023045 | - | 263 | 217 | RNA→ rpmE (PRK00019)Ribosomal_L31 (pfam01197)→ |
| env-328 | JGI20202J14951_1002967 | - | 251 | 205 | RNA→ rpmE (PRK00019)Ribosomal_L31 (pfam01197)→ |
| env-329 | JGI20198J14950_1057835 | + | 78 | 124 | RNA→ ←hypo |
| env-330 | JGI24735J21928_10184830 | - | 549 | 503 | RNA→ rpmE (PRK00019)Ribosomal_L31 (pfam01197)→ |
| env-331 | JGI24739J22299_10397894 | + | 86 | 132 | RNA→ ←hypo |
| env-332 | JGI24737J22298_10271796 | - | 450 | 404 | RNA→ ←hypo |
| env-333 | JGI12467J12023_1177552 | + | 17 | 100 | RNA→ ←hypo |
| env-334 | JGI12272J11983_1096727 | + | 28 | 109 | RNA→ rpmE (PRK00019)Ribosomal_L31 (pfam01197)→ |
| env-335 | JGI24735J21928_10002808 | + | 5639 | 5690 | RNA→ rpmE (PRK00019)Ribosomal_L31 (pfam01197)→ |
| env-336 | JGI24737J22298_10862154 | - | 237 | 199 | RNA→ rpmE (PRK00019)Ribosomal_L31 (pfam01197)→ |
| env-337 | JGI25321J50212_10143090 | + | 305 | 354 | RNA→ rpmE (PRK00019)Ribosomal_L31 (pfam01197)→ |
| env-338 | 4491734.3.365955 | - | 183 | 127 | RNA→ rpmE (PRK00019)Ribosomal_L31 (pfam01197)→ |
| Bts-1-1 | NZ_JGZU01000002.1 | + | 80841 | 80886 | RNA→ ←hypo |
| Bts-2-1 | NZ_JDTR01000003.1 | + | 80645 | 80690 | RNA→ ←hypo |
| Bts-2-2 | NZ_AUCL01000004.1 | + | 80936 | 80981 | RNA→ ←hypo |
| Ama-5-1 | NZ_AWOO02000022.1 | + | 110008 | 110077 | RNA→ rpmE (PRK00019)Ribosomal_L31 (pfam01197)→ |
| env-339 | JGI20163J15578_11194518 | + | 309 | 354 | RNA→ |
| env-340 | JGI12712J15308_10985252 | + | 184 | 237 | RNA→ ←hypo |
| env-341 | botACOD_GAKN62C01DMNDD | + | 189 | 242 | RNA→ rpmE (PRK00019)Ribosomal_L31 (pfam01197)→ |
| env-342 | JGI12420J12819_1025726 | + | 200 | 254 | RNA→ rpmE (PRK00019)Ribosomal_L31 (pfam01197)→ |
| env-343 | JGI12273J12029_10026616 | + | 9 | 57 | RNA→ rpmE (PRK00019)Ribosomal_L31 (pfam01197)→ |
| env-344 | FNTS_GKKEVUV02IK9X6 | - | 344 | 296 | RNA→ rpmE (PRK00019)Ribosomal_L31 (pfam01197)→ |
| env-345 | JGI12397J12024_1004279 | + | 17 | 65 | RNA→ rpmE (PRK00019)Ribosomal_L31 (pfam01197)→ hypo→ |
| Mva-1-1 | NZ_JH814692.1 | + | 273119 | 273170 | RNA→ rpmE (PRK00019)Ribosomal_L31 (pfam01197)→ PrfA (COG0216)prfA (PRK00591)→ |
| | | | | | HemK (COG2890)PRK09328 (PRK09328)→ SUA5 (COG0009)TIGR00057 (TIGR00057)→ |
| | | | | | ECA_wecA (TIGR02380)GT_WecA_like (cd06853)→ |

| | | | | | |
|---|---|---|---|---|---|
| Mru-1-1 | NZ_JROA01000001.1 | - | 4191975 | 4191924 | **RNA**→ rpmE (PRK00019)Ribosomal_L31 (pfam01197)→ PrfA (COG0216)prfA (PRK00591)→ HemK (COG2890)PRK09328 (PRK09328)→ SUA5 (COG0009)TIGR00057 (TIGR00057)→ GT_WecA_like (cd06853)mraY (PRK00108)→ |
| Mva-2-1 | NC_008726.1 | - | 4654670 | 4654618 | **RNA**→ rpmE (PRK00019)Ribosomal_L31 (pfam01197)→ PrfA (COG0216)prfA (PRK00591)→ HemK (COG2890)PRK09328 (PRK09328)→ SUA5 (COG0009)TIGR00057 (TIGR00057)→ GT_WecA_like (cd06853)mraY (PRK00108)→ |
| Mau-3-1 | NZ_HG964453.1 | + | 57934 | 57986 | **RNA**→ rpmE (PRK00019)Ribosomal_L31 (pfam01197)→ PrfA (COG0216)prfA (PRK00591)→ HemK (COG2890)PRK09328 (PRK09328)→ SUA5 (COG0009)TIGR00057 (TIGR00057)→ GT_WecA_like (cd06853)mraY (PRK00108)→ |
| Mgi-1-1 | NC_009338.1 | + | 2399433 | 2399484 | **RNA**→ rpmE (PRK00019)Ribosomal_L31 (pfam01197)→ PrfA (COG0216)prfA (PRK00591)→ HemK (COG2890)PRK09328 (PRK09328)→ SUA5 (COG0009)TIGR00057 (TIGR00057)→ GT_WecA_like (cd06853)mraY (PRK00108)→ |
| Mgi-2-1 | NC_014814.1 | + | 1769516 | 1769567 | **RNA**→ rpmE (PRK00019)Ribosomal_L31 (pfam01197)→ PrfA (COG0216)prfA (PRK00591)→ HemK (COG2890)PRK09328 (PRK09328)→ SUA5 (COG0009)TIGR00057 (TIGR00057)→ GT_WecA_like (cd06853)mraY (PRK00108)→ |
| env-346 | 2200268977 | - | 11805 | 11751 | **RNA**→ rpmE (PRK00019)Ribosomal_L31 (pfam01197)→ |
| env-347 | 2200159822 | - | 2727 | 2673 | **RNA**→ rpmE (PRK00019)Ribosomal_L31 (pfam01197)→ |
| env-348 | JGI1358J11329_10025592 | + | 1331 | 1398 | **RNA**→ rpmE (PRK00019)Ribosomal_L31 (pfam01197)→ |
| Nas-2-1 | NZ_BAFS01000060.1 | + | 2797 | 2847 | **RNA**→ rpmE (PRK00019)Ribosomal_L31 (pfam01197)→ PrfA (COG0216)prfA (PRK00591)→ |
| Mab-65-1 | NZ_JMIA01000002.1 | + | 750325 | 750361 | **RNA**→ rpmE (PRK00019)Ribosomal_L31 (pfam01197)→ PrfA (COG0216)prfA (PRK00591)→ HemK (COG2890)PRK09328 (PRK09328)→ SUA5 (COG0009)TIGR00057 (TIGR00057)→ ECA_wecA (TIGR02380)GT_WecA_like (cd06853)→ |
| Mch-2-1 | NZ_CP010946.1 | + | 1349878 | 1349914 | **RNA**→ rpmE (PRK00019)Ribosomal_L31 (pfam01197)→ PrfA (COG0216)prfA (PRK00591)→ HemK (COG2890)PRK09328 (PRK09328)→ SUA5 (COG0009)TIGR00057 (TIGR00057)→ ECA_wecA (TIGR02380)GT_WecA_like (cd06853)→ |
| env-349 | JGI26345J50200_1206263 | + | 102 | 156 | **RNA**→ rpmE (PRK00019)Ribosomal_L31 (pfam01197)→ |
| env-350 | JGI12396J12026_1038988 | + | 59 | 128 | **RNA**→ ←hypo |
| Fsp-7-1 | NC_014666.1 | + | 1161289 | 1161336 | **RNA**→ rpmE (PRK00019)Ribosomal_L31 (pfam01197)→ PrfA (COG0216)prfA (PRK00591)→ HemK (COG2890)PRK09328 (PRK09328)→ |
| Sca-1-1 | NZ_KB911608.1 | - | 87758 | 87712 | **RNA**→ rpmE (PRK00019)Ribosomal_L31 (pfam01197)→ |
| Ssv-1-1 | NZ_CM000951.1 | + | 6314528 | 6314574 | **RNA**→ rpmE (PRK00019)Ribosomal_L31 (pfam01197)→ hypo→ |
| Ssp-50-1 | NZ_JTIY01000001.1 | - | 828848 | 828802 | **RNA**→ rpmE (PRK00019)Ribosomal_L31 (pfam01197)→ |
| Ssp-19-1 | NZ_KE150433.1 | - | 83516 | 83470 | **RNA**→ rpmE (PRK00019)Ribosomal_L31 (pfam01197)→ |
| Aro-1-1 | NZ_KI632511.1 | - | 5249323 | 5249275 | **RNA**→ rpmE (PRK00019)Ribosomal_L31 (pfam01197)→ |
| Sca-2-1 | NC_016111.1 | + | 4478364 | 4478410 | **RNA**→ rpmE (PRK00019)Ribosomal_L31 (pfam01197)→ |
| Sca-2-2 | NC_017586.1 | + | 4478172 | 4478218 | **RNA**→ rpmE (PRK00019)Ribosomal_L31 (pfam01197)→ |
| Nvi-1-1 | NZ_BAGN01000056.1 | + | 32207 | 32254 | **RNA**→ rpmE (PRK00019)Ribosomal_L31 (pfam01197)→ PrfA (COG0216)prfA (PRK00591)→ HemK (COG2890)PRK09328 (PRK09328)→ |
| env-351 | BICA1_1000552 | + | 2686 | 2734 | **RNA**→ rpmE (PRK00019)Ribosomal_L31 (pfam01197)→ |
| env-352 | JGI12675J13321_109048 | + | 170 | 222 | **RNA**→ rpmE (PRK00019)Ribosomal_L31 (pfam01197)→ |
| env-353 | JGI26342J46808_1016646 | + | 241 | 293 | **RNA**→ rpmE (PRK00019)Ribosomal_L31 (pfam01197)→ |
| env-354 | JGI26345J50200_1290574 | + | 128 | 180 | **RNA**→ |
| env-355 | JGI12713J13577_1075251 | + | 94 | 146 | **RNA**→ rpmE (PRK00019)Ribosomal_L31 (pfam01197)→ |
| env-356 | JGI12669J13542_130180 | - | 221 | 169 | **RNA**→ rpmE (PRK00019)Ribosomal_L31 (pfam01197)→ |
| env-357 | JGI12712J15308_10002766 | - | 597 | 545 | **RNA**→ rpmE (PRK00019)Ribosomal_L31 (pfam01197)→ |
| env-358 | JGI12659J15293_10130778 | - | 420 | 368 | **RNA**→ rpmE (PRK00019)Ribosomal_L31 (pfam01197)→ |
| env-359 | JGI12635J15846_10434338 | + | 340 | 392 | **RNA**→ rpmE (PRK00019)Ribosomal_L31 (pfam01197)→ |
| env-360 | JGI26347J50199_1038113 | - | 440 | 388 | **RNA**→ rpmE (PRK00019)Ribosomal_L31 (pfam01197)→ |
| env-361 | JGI12668J13544_1081006 | - | 183 | 131 | **RNA**→ rpmE (PRK00019)Ribosomal_L31 (pfam01197)→ |
| env-362 | JGI26337J50220_1123253 | + | 72 | 124 | **RNA**→ rpmE (PRK00019)Ribosomal_L31 (pfam01197)→ |
| env-363 | JGI12712J15308_10042764 | + | 278 | 330 | **RNA**→ rpmE (PRK00019)Ribosomal_L31 (pfam01197)→ |
| env-364 | JGI12699J13341_100212 | - | 1510 | 1458 | **RNA**→ rpmE (PRK00019)Ribosomal_L31 (pfam01197)→ |
| env-365 | JGI26338J50219_1041585 | - | 309 | 257 | **RNA**→ rpmE (PRK00019)Ribosomal_L31 (pfam01197)→ |

| | | | | | |
|---|---|---|---|---|---|
| Mim-1-1 | NZ_JXUU01000007.1 | - | 15102 | 15062 | **RNA**→ rpmE (PRK00019)Ribosomal_L31 (pfam01197)→ PrfA (COG0216)prfA (PRK00591)→ HemK (COG2890)PRK09328 (PRK09328)→ SUA5 (COG0009)TIGR00057 (TIGR00057)→ ECA_wecA (TIGR02380)GT_WecA_like (cd06853)→ |
| Mab-73-1 | NZ_AUVF01000002.1 | + | 521591 | 521631 | **RNA**→ rpmE (PRK00019)Ribosomal_L31 (pfam01197)→ PrfA (COG0216)prfA (PRK00591)→ HemK (COG2890)PRK09328 (PRK09328)→ SUA5 (COG0009)TIGR00057 (TIGR00057)→ ECA_wecA (TIGR02380)GT_WecA_like (cd06853)→ |
| Mab-1-2 | NZ_CSZN01000002.1 | - | 459458 | 459418 | **RNA**→ rpmE (PRK00019)Ribosomal_L31 (pfam01197)→ PrfA (COG0216)prfA (PRK00591)→ HemK (COG2890)PRK09328 (PRK09328)→ SUA5 (COG0009)TIGR00057 (TIGR00057)→ ECA_wecA (TIGR02380)GT_WecA_like (cd06853)→ |
| Mab-1-3 | NZ_CSYW01000003.1 | - | 506431 | 506391 | **RNA**→ rpmE (PRK00019)Ribosomal_L31 (pfam01197)→ PrfA (COG0216)prfA (PRK00591)→ HemK (COG2890)PRK09328 (PRK09328)→ SUA5 (COG0009)TIGR00057 (TIGR00057)→ ECA_wecA (TIGR02380)GT_WecA_like (cd06853)→ |
| Mab-1-4 | NZ_CSXQ01000001.1 | - | 309290 | 309250 | **RNA**→ rpmE (PRK00019)Ribosomal_L31 (pfam01197)→ PrfA (COG0216)prfA (PRK00591)→ HemK (COG2890)PRK09328 (PRK09328)→ SUA5 (COG0009)TIGR00057 (TIGR00057)→ ECA_wecA (TIGR02380)GT_WecA_like (cd06853)→ |
| Mab-1-5 | NZ_CSWA01000004.1 | - | 506374 | 506334 | **RNA**→ rpmE (PRK00019)Ribosomal_L31 (pfam01197)→ PrfA (COG0216)prfA (PRK00591)→ HemK (COG2890)PRK09328 (PRK09328)→ SUA5 (COG0009)TIGR00057 (TIGR00057)→ ECA_wecA (TIGR02380)GT_WecA_like (cd06853)→ |
| Mab-1-6 | NZ_CSYH01000003.1 | - | 506327 | 506287 | **RNA**→ rpmE (PRK00019)Ribosomal_L31 (pfam01197)→ PrfA (COG0216)prfA (PRK00591)→ HemK (COG2890)PRK09328 (PRK09328)→ SUA5 (COG0009)TIGR00057 (TIGR00057)→ ECA_wecA (TIGR02380)GT_WecA_like (cd06853)→ |
| Mab-1-7 | NZ_CSZC01000001.1 | - | 742902 | 742862 | **RNA**→ rpmE (PRK00019)Ribosomal_L31 (pfam01197)→ PrfA (COG0216)prfA (PRK00591)→ HemK (COG2890)PRK09328 (PRK09328)→ SUA5 (COG0009)TIGR00057 (TIGR00057)→ ECA_wecA (TIGR02380)GT_WecA_like (cd06853)→ |
| Mab-1-8 | NZ_CSXA01000001.1 | + | 162315 | 162355 | **RNA**→ rpmE (PRK00019)Ribosomal_L31 (pfam01197)→ PrfA (COG0216)prfA (PRK00591)→ HemK (COG2890)PRK09328 (PRK09328)→ SUA5 (COG0009)TIGR00057 (TIGR00057)→ ECA_wecA (TIGR02380)GT_WecA_like (cd06853)→ |
| Mab-1-9 | NZ_CSXG01000001.1 | + | 162314 | 162354 | **RNA**→ rpmE (PRK00019)Ribosomal_L31 (pfam01197)→ PrfA (COG0216)prfA (PRK00591)→ HemK (COG2890)PRK09328 (PRK09328)→ SUA5 (COG0009)TIGR00057 (TIGR00057)→ ECA_wecA (TIGR02380)GT_WecA_like (cd06853)→ |
| Mab-1-10 | NZ_CSWO01000003.1 | - | 506432 | 506392 | **RNA**→ rpmE (PRK00019)Ribosomal_L31 (pfam01197)→ PrfA (COG0216)prfA (PRK00591)→ HemK (COG2890)PRK09328 (PRK09328)→ SUA5 (COG0009)TIGR00057 (TIGR00057)→ ECA_wecA (TIGR02380)GT_WecA_like (cd06853)→ |
| Mab-1-11 | NZ_CSXV01000002.1 | - | 197279 | 197239 | **RNA**→ rpmE (PRK00019)Ribosomal_L31 (pfam01197)→ PrfA (COG0216)prfA (PRK00591)→ HemK (COG2890)PRK09328 (PRK09328)→ SUA5 (COG0009)TIGR00057 (TIGR00057)→ ECA_wecA (TIGR02380)GT_WecA_like (cd06853)→ |
| Mab-1-12 | NZ_CSYV01000002.1 | - | 459141 | 459101 | **RNA**→ rpmE (PRK00019)Ribosomal_L31 (pfam01197)→ PrfA (COG0216)prfA (PRK00591)→ HemK (COG2890)PRK09328 (PRK09328)→ SUA5 (COG0009)TIGR00057 (TIGR00057)→ ECA_wecA (TIGR02380)GT_WecA_like (cd06853)→ |
| Mab-1-13 | NZ_CSXN01000003.1 | - | 506492 | 506452 | **RNA**→ rpmE (PRK00019)Ribosomal_L31 (pfam01197)→ PrfA (COG0216)prfA (PRK00591)→ HemK (COG2890)PRK09328 (PRK09328)→ SUA5 (COG0009)TIGR00057 (TIGR00057)→ ECA_wecA (TIGR02380)GT_WecA_like (cd06853)→ |
| Mab-1-14 | NZ_CSXO01000001.1 | - | 506300 | 506260 | **RNA**→ rpmE (PRK00019)Ribosomal_L31 (pfam01197)→ PrfA (COG0216)prfA (PRK00591)→ HemK (COG2890)PRK09328 (PRK09328)→ SUA5 (COG0009)TIGR00057 (TIGR00057)→ ECA_wecA (TIGR02380)GT_WecA_like (cd06853)→ |
| Mab-1-15 | NZ_CSZL01000002.1 | - | 506309 | 506269 | **RNA**→ rpmE (PRK00019)Ribosomal_L31 (pfam01197)→ PrfA (COG0216)prfA (PRK00591)→ HemK (COG2890)PRK09328 (PRK09328)→ SUA5 (COG0009)TIGR00057 (TIGR00057)→ ECA_wecA (TIGR02380)GT_WecA_like (cd06853)→ |
| Mab-1-16 | NZ_CSYB01000001.1 | - | 309269 | 309229 | **RNA**→ rpmE (PRK00019)Ribosomal_L31 (pfam01197)→ PrfA (COG0216)prfA (PRK00591)→ HemK (COG2890)PRK09328 (PRK09328)→ SUA5 (COG0009)TIGR00057 (TIGR00057)→ ECA_wecA (TIGR02380)GT_WecA_like (cd06853)→ |
| Mab-1-17 | NZ_CSXU01000001.1 | - | 742819 | 742779 | **RNA**→ rpmE (PRK00019)Ribosomal_L31 (pfam01197)→ PrfA (COG0216)prfA (PRK00591)→ HemK (COG2890)PRK09328 (PRK09328)→ SUA5 (COG0009)TIGR00057 (TIGR00057)→ ECA_wecA (TIGR02380)GT_WecA_like (cd06853)→ |
| Mab-1-18 | NZ_CSZA01000001.1 | - | 742884 | 742844 | **RNA**→ rpmE (PRK00019)Ribosomal_L31 (pfam01197)→ PrfA (COG0216)prfA (PRK00591)→ HemK (COG2890)PRK09328 (PRK09328)→ SUA5 (COG0009)TIGR00057 (TIGR00057)→ ECA_wecA (TIGR02380)GT_WecA_like (cd06853)→ |

| | | | | | |
|---|---|---|---|---|---|
| Mab-1-19 | NZ_CSYJ01000002.1 | - | 698905 | 698865 | **RNA**→ rpmE (PRK00019)Ribosomal_L31 (pfam01197)→ PrfA (COG0216)prfA (PRK00591)→ HemK (COG2890)PRK09328 (PRK09328)→ SUA5 (COG0009)TIGR00057 (TIGR00057)→ ECA_wecA (TIGR02380)GT_WecA_like (cd06853)→ |
| Mab-1-20 | NZ_CSUU01000004.1 | + | 10612 | 10652 | **RNA**→ rpmE (PRK00019)Ribosomal_L31 (pfam01197)→ PrfA (COG0216)prfA (PRK00591)→ HemK (COG2890)PRK09328 (PRK09328)→ SUA5 (COG0009)TIGR00057 (TIGR00057)→ ECA_wecA (TIGR02380)GT_WecA_like (cd06853)→ |
| Mab-1-21 | NZ_CSUQ01000003.1 | + | 10596 | 10636 | **RNA**→ rpmE (PRK00019)Ribosomal_L31 (pfam01197)→ PrfA (COG0216)prfA (PRK00591)→ HemK (COG2890)PRK09328 (PRK09328)→ SUA5 (COG0009)TIGR00057 (TIGR00057)→ ECA_wecA (TIGR02380)GT_WecA_like (cd06853)→ |
| Mab-1-22 | NZ_CSUX01000003.1 | + | 475932 | 475972 | **RNA**→ rpmE (PRK00019)Ribosomal_L31 (pfam01197)→ PrfA (COG0216)prfA (PRK00591)→ HemK (COG2890)PRK09328 (PRK09328)→ SUA5 (COG0009)TIGR00057 (TIGR00057)→ ECA_wecA (TIGR02380)GT_WecA_like (cd06853)→ |
| Mab-1-23 | NZ_CSUD01000003.1 | - | 309291 | 309251 | **RNA**→ rpmE (PRK00019)Ribosomal_L31 (pfam01197)→ PrfA (COG0216)prfA (PRK00591)→ HemK (COG2890)PRK09328 (PRK09328)→ SUA5 (COG0009)TIGR00057 (TIGR00057)→ ECA_wecA (TIGR02380)GT_WecA_like (cd06853)→ |
| Mab-1-24 | NZ_CSVO01000002.1 | - | 459138 | 459098 | **RNA**→ rpmE (PRK00019)Ribosomal_L31 (pfam01197)→ PrfA (COG0216)prfA (PRK00591)→ HemK (COG2890)PRK09328 (PRK09328)→ SUA5 (COG0009)TIGR00057 (TIGR00057)→ ECA_wecA (TIGR02380)GT_WecA_like (cd06853)→ |
| Mab-1-25 | NZ_CSVP01000001.1 | - | 459123 | 459083 | **RNA**→ rpmE (PRK00019)Ribosomal_L31 (pfam01197)→ PrfA (COG0216)prfA (PRK00591)→ HemK (COG2890)PRK09328 (PRK09328)→ SUA5 (COG0009)TIGR00057 (TIGR00057)→ ECA_wecA (TIGR02380)GT_WecA_like (cd06853)→ |
| Mab-1-26 | NZ_CSVJ01000002.1 | - | 506338 | 506298 | **RNA**→ rpmE (PRK00019)Ribosomal_L31 (pfam01197)→ PrfA (COG0216)prfA (PRK00591)→ HemK (COG2890)PRK09328 (PRK09328)→ SUA5 (COG0009)TIGR00057 (TIGR00057)→ ECA_wecA (TIGR02380)GT_WecA_like (cd06853)→ |
| Mab-1-27 | NZ_CSYF01000002.1 | - | 506190 | 506150 | **RNA**→ rpmE (PRK00019)Ribosomal_L31 (pfam01197)→ PrfA (COG0216)prfA (PRK00591)→ HemK (COG2890)PRK09328 (PRK09328)→ SUA5 (COG0009)TIGR00057 (TIGR00057)→ ECA_wecA (TIGR02380)GT_WecA_like (cd06853)→ |
| Mab-1-28 | NZ_CSWJ01000002.1 | - | 506432 | 506392 | **RNA**→ rpmE (PRK00019)Ribosomal_L31 (pfam01197)→ PrfA (COG0216)prfA (PRK00591)→ HemK (COG2890)PRK09328 (PRK09328)→ SUA5 (COG0009)TIGR00057 (TIGR00057)→ ECA_wecA (TIGR02380)GT_WecA_like (cd06853)→ |
| Mab-1-29 | NZ_CSWG01000004.1 | - | 309142 | 309102 | **RNA**→ rpmE (PRK00019)Ribosomal_L31 (pfam01197)→ PrfA (COG0216)prfA (PRK00591)→ HemK (COG2890)PRK09328 (PRK09328)→ SUA5 (COG0009)TIGR00057 (TIGR00057)→ ECA_wecA (TIGR02380)GT_WecA_like (cd06853)→ |
| Mab-1-30 | NZ_CSYC01000001.1 | - | 310020 | 309980 | **RNA**→ rpmE (PRK00019)Ribosomal_L31 (pfam01197)→ PrfA (COG0216)prfA (PRK00591)→ HemK (COG2890)PRK09328 (PRK09328)→ SUA5 (COG0009)TIGR00057 (TIGR00057)→ ECA_wecA (TIGR02380)GT_WecA_like (cd06853)→ |
| Mab-1-31 | NZ_CSWT01000001.1 | - | 445981 | 445941 | **RNA**→ rpmE (PRK00019)Ribosomal_L31 (pfam01197)→ PrfA (COG0216)prfA (PRK00591)→ HemK (COG2890)PRK09328 (PRK09328)→ SUA5 (COG0009)TIGR00057 (TIGR00057)→ ECA_wecA (TIGR02380)GT_WecA_like (cd06853)→ |
| Mab-1-32 | NZ_CSXP01000001.1 | - | 1200434 | 1200394 | **RNA**→ rpmE (PRK00019)Ribosomal_L31 (pfam01197)→ PrfA (COG0216)prfA (PRK00591)→ HemK (COG2890)PRK09328 (PRK09328)→ SUA5 (COG0009)TIGR00057 (TIGR00057)→ ECA_wecA (TIGR02380)GT_WecA_like (cd06853)→ |
| Mab-1-33 | NZ_CSXI01000001.1 | + | 244753 | 244793 | **RNA**→ rpmE (PRK00019)Ribosomal_L31 (pfam01197)→ PrfA (COG0216)prfA (PRK00591)→ HemK (COG2890)PRK09328 (PRK09328)→ SUA5 (COG0009)TIGR00057 (TIGR00057)→ ECA_wecA (TIGR02380)GT_WecA_like (cd06853)→ |
| Mab-1-34 | NZ_CSXR01000001.1 | - | 818533 | 818493 | **RNA**→ rpmE (PRK00019)Ribosomal_L31 (pfam01197)→ PrfA (COG0216)prfA (PRK00591)→ HemK (COG2890)PRK09328 (PRK09328)→ SUA5 (COG0009)TIGR00057 (TIGR00057)→ ECA_wecA (TIGR02380)GT_WecA_like (cd06853)→ |
| Mab-1-35 | NZ_CSVD01000002.1 | - | 699015 | 698975 | **RNA**→ rpmE (PRK00019)Ribosomal_L31 (pfam01197)→ PrfA (COG0216)prfA (PRK00591)→ HemK (COG2890)PRK09328 (PRK09328)→ SUA5 (COG0009)TIGR00057 (TIGR00057)→ ECA_wecA (TIGR02380)GT_WecA_like (cd06853)→ |
| Mab-1-36 | NZ_CSVE01000001.1 | + | 148660 | 148700 | **RNA**→ rpmE (PRK00019)Ribosomal_L31 (pfam01197)→ PrfA (COG0216)prfA (PRK00591)→ HemK (COG2890)PRK09328 (PRK09328)→ SUA5 (COG0009)TIGR00057 (TIGR00057)→ ECA_wecA (TIGR02380)GT_WecA_like (cd06853)→ |
| Mab-1-37 | NZ_CSXC01000001.1 | - | 786146 | 786106 | **RNA**→ rpmE (PRK00019)Ribosomal_L31 (pfam01197)→ PrfA (COG0216)prfA (PRK00591)→ HemK (COG2890)PRK09328 (PRK09328)→ SUA5 (COG0009)TIGR00057 (TIGR00057)→ ECA_wecA (TIGR02380)GT_WecA_like (cd06853)→ |

| | | | | | |
|---|---|---|---|---|---|
| Mab-1-38 | NZ_CSUB01000002.1 | - | 622023 | 621983 | **RNA**→ rpmE (PRK00019)Ribosomal_L31 (pfam01197)→ PrfA (COG0216)prfA (PRK00591)→ HemK (COG2890)PRK09328 (PRK09328)→ SUA5 (COG0009)TIGR00057 (TIGR00057)→ ECA_wecA (TIGR02380)GT_WecA_like (cd06853)→ |
| Mab-1-39 | NZ_CSVC01000026.1 | - | 62798 | 62758 | **RNA**→ rpmE (PRK00019)Ribosomal_L31 (pfam01197)→ PrfA (COG0216)prfA (PRK00591)→ HemK (COG2890)PRK09328 (PRK09328)→ SUA5 (COG0009)TIGR00057 (TIGR00057)→ ECA_wecA (TIGR02380)GT_WecA_like (cd06853)→ |
| Mab-1-40 | NZ_CSUE01000001.1 | - | 309387 | 309347 | **RNA**→ rpmE (PRK00019)Ribosomal_L31 (pfam01197)→ PrfA (COG0216)prfA (PRK00591)→ HemK (COG2890)PRK09328 (PRK09328)→ SUA5 (COG0009)TIGR00057 (TIGR00057)→ ECA_wecA (TIGR02380)GT_WecA_like (cd06853)→ |
| Mab-1-41 | NZ_CSUK01000001.1 | + | 382760 | 382800 | **RNA**→ rpmE (PRK00019)Ribosomal_L31 (pfam01197)→ PrfA (COG0216)prfA (PRK00591)→ HemK (COG2890)PRK09328 (PRK09328)→ SUA5 (COG0009)TIGR00057 (TIGR00057)→ ECA_wecA (TIGR02380)GT_WecA_like (cd06853)→ |
| Mab-1-42 | NZ_CSVI01000002.1 | + | 162256 | 162296 | **RNA**→ rpmE (PRK00019)Ribosomal_L31 (pfam01197)→ PrfA (COG0216)prfA (PRK00591)→ HemK (COG2890)PRK09328 (PRK09328)→ SUA5 (COG0009)TIGR00057 (TIGR00057)→ ECA_wecA (TIGR02380)GT_WecA_like (cd06853)→ |
| Mab-1-43 | NZ_CSVH01000001.1 | - | 786306 | 786266 | **RNA**→ rpmE (PRK00019)Ribosomal_L31 (pfam01197)→ PrfA (COG0216)prfA (PRK00591)→ HemK (COG2890)PRK09328 (PRK09328)→ SUA5 (COG0009)TIGR00057 (TIGR00057)→ ECA_wecA (TIGR02380)GT_WecA_like (cd06853)→ |
| Mab-1-44 | NZ_CSTR01000001.1 | + | 245392 | 245432 | **RNA**→ rpmE (PRK00019)Ribosomal_L31 (pfam01197)→ PrfA (COG0216)prfA (PRK00591)→ HemK (COG2890)PRK09328 (PRK09328)→ SUA5 (COG0009)TIGR00057 (TIGR00057)→ ECA_wecA (TIGR02380)GT_WecA_like (cd06853)→ |
| Mab-1-45 | NZ_CSUL01000001.1 | - | 309282 | 309242 | **RNA**→ rpmE (PRK00019)Ribosomal_L31 (pfam01197)→ PrfA (COG0216)prfA (PRK00591)→ HemK (COG2890)PRK09328 (PRK09328)→ SUA5 (COG0009)TIGR00057 (TIGR00057)→ ECA_wecA (TIGR02380)GT_WecA_like (cd06853)→ |
| Mab-1-46 | NZ_CSVW01000021.1 | - | 62685 | 62645 | **RNA**→ rpmE (PRK00019)Ribosomal_L31 (pfam01197)→ PrfA (COG0216)prfA (PRK00591)→ HemK (COG2890)PRK09328 (PRK09328)→ SUA5 (COG0009)TIGR00057 (TIGR00057)→ ECA_wecA (TIGR02380)GT_WecA_like (cd06853)→ |
| Mab-1-47 | NZ_CSUJ01000001.1 | - | 506352 | 506312 | **RNA**→ rpmE (PRK00019)Ribosomal_L31 (pfam01197)→ PrfA (COG0216)prfA (PRK00591)→ HemK (COG2890)PRK09328 (PRK09328)→ SUA5 (COG0009)TIGR00057 (TIGR00057)→ ECA_wecA (TIGR02380)GT_WecA_like (cd06853)→ |
| Mab-1-48 | NZ_CSWY01000003.1 | - | 506245 | 506205 | **RNA**→ rpmE (PRK00019)Ribosomal_L31 (pfam01197)→ PrfA (COG0216)prfA (PRK00591)→ HemK (COG2890)PRK09328 (PRK09328)→ SUA5 (COG0009)TIGR00057 (TIGR00057)→ ECA_wecA (TIGR02380)GT_WecA_like (cd06853)→ |
| Mab-1-49 | NZ_CSTV01000001.1 | - | 309289 | 309249 | **RNA**→ rpmE (PRK00019)Ribosomal_L31 (pfam01197)→ PrfA (COG0216)prfA (PRK00591)→ HemK (COG2890)PRK09328 (PRK09328)→ SUA5 (COG0009)TIGR00057 (TIGR00057)→ ECA_wecA (TIGR02380)GT_WecA_like (cd06853)→ |
| Mab-1-50 | NZ_CSZE01000002.1 | - | 506271 | 506231 | **RNA**→ rpmE (PRK00019)Ribosomal_L31 (pfam01197)→ PrfA (COG0216)prfA (PRK00591)→ HemK (COG2890)PRK09328 (PRK09328)→ SUA5 (COG0009)TIGR00057 (TIGR00057)→ ECA_wecA (TIGR02380)GT_WecA_like (cd06853)→ |
| Mab-1-51 | NZ_CSTG01000003.1 | + | 162313 | 162353 | **RNA**→ rpmE (PRK00019)Ribosomal_L31 (pfam01197)→ PrfA (COG0216)prfA (PRK00591)→ HemK (COG2890)PRK09328 (PRK09328)→ SUA5 (COG0009)TIGR00057 (TIGR00057)→ ECA_wecA (TIGR02380)GT_WecA_like (cd06853)→ |
| Mab-1-52 | NZ_CSVT01000001.1 | - | 685006 | 684966 | **RNA**→ rpmE (PRK00019)Ribosomal_L31 (pfam01197)→ PrfA (COG0216)prfA (PRK00591)→ HemK (COG2890)PRK09328 (PRK09328)→ SUA5 (COG0009)TIGR00057 (TIGR00057)→ ECA_wecA (TIGR02380)GT_WecA_like (cd06853)→ |
| Mab-1-53 | NZ_CSYN01000001.1 | - | 310020 | 309980 | **RNA**→ rpmE (PRK00019)Ribosomal_L31 (pfam01197)→ PrfA (COG0216)prfA (PRK00591)→ HemK (COG2890)PRK09328 (PRK09328)→ SUA5 (COG0009)TIGR00057 (TIGR00057)→ ECA_wecA (TIGR02380)GT_WecA_like (cd06853)→ |
| Mab-1-54 | NZ_CSUV01000003.1 | - | 309253 | 309213 | **RNA**→ rpmE (PRK00019)Ribosomal_L31 (pfam01197)→ PrfA (COG0216)prfA (PRK00591)→ HemK (COG2890)PRK09328 (PRK09328)→ SUA5 (COG0009)TIGR00057 (TIGR00057)→ ECA_wecA (TIGR02380)GT_WecA_like (cd06853)→ |
| Mab-1-55 | NZ_CSYG01000003.1 | - | 415344 | 415304 | **RNA**→ rpmE (PRK00019)Ribosomal_L31 (pfam01197)→ PrfA (COG0216)prfA (PRK00591)→ HemK (COG2890)PRK09328 (PRK09328)→ SUA5 (COG0009)TIGR00057 (TIGR00057)→ ECA_wecA (TIGR02380)GT_WecA_like (cd06853)→ |
| Mab-1-56 | NZ_CSYZ01000001.1 | - | 506610 | 506570 | **RNA**→ rpmE (PRK00019)Ribosomal_L31 (pfam01197)→ PrfA (COG0216)prfA (PRK00591)→ HemK (COG2890)PRK09328 (PRK09328)→ SUA5 (COG0009)TIGR00057 (TIGR00057)→ ECA_wecA (TIGR02380)GT_WecA_like (cd06853)→ |

| | | | | | |
|---|---|---|---|---|---|
| Mab-1-57 | NZ_CSUT01000001.1 | - | 621986 | 621946 | **RNA**→ rpmE (PRK00019)Ribosomal_L31 (pfam01197)→ PrfA (COG0216)prfA (PRK00591)→ HemK (COG2890)PRK09328 (PRK09328)→ SUA5 (COG0009)TIGR00057 (TIGR00057)→ ECA_wecA (TIGR02380)GT_WecA_like (cd06853)→ |
| Mab-1-58 | NZ_CSZI01000001.1 | - | 309973 | 309933 | **RNA**→ rpmE (PRK00019)Ribosomal_L31 (pfam01197)→ PrfA (COG0216)prfA (PRK00591)→ HemK (COG2890)PRK09328 (PRK09328)→ SUA5 (COG0009)TIGR00057 (TIGR00057)→ hypo→ hypo→ |
| Mab-1-59 | NZ_CSUS01000001.1 | - | 353814 | 353774 | **RNA**→ rpmE (PRK00019)Ribosomal_L31 (pfam01197)→ PrfA (COG0216)prfA (PRK00591)→ HemK (COG2890)PRK09328 (PRK09328)→ SUA5 (COG0009)TIGR00057 (TIGR00057)→ ECA_wecA (TIGR02380)GT_WecA_like (cd06853)→ |
| Mab-1-60 | NZ_CSWU01000006.1 | - | 248462 | 248422 | **RNA**→ rpmE (PRK00019)Ribosomal_L31 (pfam01197)→ PrfA (COG0216)prfA (PRK00591)→ HemK (COG2890)PRK09328 (PRK09328)→ SUA5 (COG0009)TIGR00057 (TIGR00057)→ ECA_wecA (TIGR02380)GT_WecA_like (cd06853)→ |
| Mab-1-61 | NZ_CSZU01000001.1 | + | 162316 | 162356 | **RNA**→ rpmE (PRK00019)Ribosomal_L31 (pfam01197)→ PrfA (COG0216)prfA (PRK00591)→ HemK (COG2890)PRK09328 (PRK09328)→ SUA5 (COG0009)TIGR00057 (TIGR00057)→ ECA_wecA (TIGR02380)GT_WecA_like (cd06853)→ |
| Mab-1-62 | NZ_CSWX01000004.1 | - | 506373 | 506333 | **RNA**→ rpmE (PRK00019)Ribosomal_L31 (pfam01197)→ PrfA (COG0216)prfA (PRK00591)→ HemK (COG2890)PRK09328 (PRK09328)→ SUA5 (COG0009)TIGR00057 (TIGR00057)→ ECA_wecA (TIGR02380)GT_WecA_like (cd06853)→ |
| Mab-1-63 | NZ_CSYT01000002.1 | - | 506484 | 506444 | **RNA**→ rpmE (PRK00019)Ribosomal_L31 (pfam01197)→ PrfA (COG0216)prfA (PRK00591)→ HemK (COG2890)PRK09328 (PRK09328)→ SUA5 (COG0009)TIGR00057 (TIGR00057)→ ECA_wecA (TIGR02380)GT_WecA_like (cd06853)→ |
| Mab-1-64 | NZ_CSWQ01000004.1 | - | 506673 | 506633 | **RNA**→ rpmE (PRK00019)Ribosomal_L31 (pfam01197)→ PrfA (COG0216)prfA (PRK00591)→ HemK (COG2890)PRK09328 (PRK09328)→ SUA5 (COG0009)TIGR00057 (TIGR00057)→ ECA_wecA (TIGR02380)GT_WecA_like (cd06853)→ |
| Mab-1-65 | NZ_CSYY01000003.1 | - | 506329 | 506289 | **RNA**→ rpmE (PRK00019)Ribosomal_L31 (pfam01197)→ PrfA (COG0216)prfA (PRK00591)→ HemK (COG2890)PRK09328 (PRK09328)→ SUA5 (COG0009)TIGR00057 (TIGR00057)→ ECA_wecA (TIGR02380)GT_WecA_like (cd06853)→ |
| Mab-1-66 | NZ_CSTZ01000001.1 | + | 476370 | 476410 | **RNA**→ rpmE (PRK00019)Ribosomal_L31 (pfam01197)→ PrfA (COG0216)prfA (PRK00591)→ HemK (COG2890)PRK09328 (PRK09328)→ SUA5 (COG0009)TIGR00057 (TIGR00057)→ ECA_wecA (TIGR02380)GT_WecA_like (cd06853)→ |
| Mab-1-67 | NZ_CSZF01000003.1 | + | 162311 | 162351 | **RNA**→ rpmE (PRK00019)Ribosomal_L31 (pfam01197)→ PrfA (COG0216)prfA (PRK00591)→ HemK (COG2890)PRK09328 (PRK09328)→ SUA5 (COG0009)TIGR00057 (TIGR00057)→ hypo→ |
| Mab-1-68 | NZ_CSXJ01000004.1 | - | 506351 | 506311 | **RNA**→ rpmE (PRK00019)Ribosomal_L31 (pfam01197)→ PrfA (COG0216)prfA (PRK00591)→ HemK (COG2890)PRK09328 (PRK09328)→ SUA5 (COG0009)TIGR00057 (TIGR00057)→ ECA_wecA (TIGR02380)GT_WecA_like (cd06853)→ |
| Mab-1-69 | NZ_CSZV01000002.1 | + | 162295 | 162335 | **RNA**→ rpmE (PRK00019)Ribosomal_L31 (pfam01197)→ PrfA (COG0216)prfA (PRK00591)→ HemK (COG2890)PRK09328 (PRK09328)→ SUA5 (COG0009)TIGR00057 (TIGR00057)→ ECA_wecA (TIGR02380)GT_WecA_like (cd06853)→ |
| Mab-1-70 | NZ_CSTT01000001.1 | + | 930033 | 930073 | **RNA**→ rpmE (PRK00019)Ribosomal_L31 (pfam01197)→ PrfA (COG0216)prfA (PRK00591)→ HemK (COG2890)PRK09328 (PRK09328)→ SUA5 (COG0009)TIGR00057 (TIGR00057)→ ECA_wecA (TIGR02380)GT_WecA_like (cd06853)→ |
| Mab-1-71 | NZ_CSZQ01000006.1 | - | 248596 | 248556 | **RNA**→ rpmE (PRK00019)Ribosomal_L31 (pfam01197)→ PrfA (COG0216)prfA (PRK00591)→ HemK (COG2890)PRK09328 (PRK09328)→ SUA5 (COG0009)TIGR00057 (TIGR00057)→ ECA_wecA (TIGR02380)GT_WecA_like (cd06853)→ |
| Mab-1-72 | NZ_CSXM01000002.1 | - | 501635 | 501595 | **RNA**→ rpmE (PRK00019)Ribosomal_L31 (pfam01197)→ PrfA (COG0216)prfA (PRK00591)→ HemK (COG2890)PRK09328 (PRK09328)→ SUA5 (COG0009)TIGR00057 (TIGR00057)→ ECA_wecA (TIGR02380)GT_WecA_like (cd06853)→ |
| Mab-1-73 | NZ_CSUC01000001.1 | - | 1714344 | 1714304 | **RNA**→ rpmE (PRK00019)Ribosomal_L31 (pfam01197)→ PrfA (COG0216)prfA (PRK00591)→ HemK (COG2890)PRK09328 (PRK09328)→ SUA5 (COG0009)TIGR00057 (TIGR00057)→ ECA_wecA (TIGR02380)GT_WecA_like (cd06853)→ |
| Mab-1-74 | NZ_CSXZ01000001.1 | - | 893217 | 893177 | **RNA**→ rpmE (PRK00019)Ribosomal_L31 (pfam01197)→ PrfA (COG0216)prfA (PRK00591)→ HemK (COG2890)PRK09328 (PRK09328)→ SUA5 (COG0009)TIGR00057 (TIGR00057)→ ECA_wecA (TIGR02380)GT_WecA_like (cd06853)→ |
| Mab-1-75 | NZ_CSVZ01000002.1 | - | 786397 | 786357 | **RNA**→ rpmE (PRK00019)Ribosomal_L31 (pfam01197)→ PrfA (COG0216)prfA (PRK00591)→ HemK (COG2890)PRK09328 (PRK09328)→ SUA5 (COG0009)TIGR00057 (TIGR00057)→ ECA_wecA (TIGR02380)GT_WecA_like (cd06853)→ |

| | | | | | |
|---|---|---|---|---|---|
| Mab-1-76 | NZ_CSUZ01000001.1 | - | 506435 | 506395 | **RNA**→ rpmE (PRK00019)Ribosomal_L31 (pfam01197)→ PrfA (COG0216)prfA (PRK00591)→ HemK (COG2890)PRK09328 (PRK09328)→ SUA5 (COG0009)TIGR00057 (TIGR00057)→ ECA_wecA (TIGR02380)GT_WecA_like (cd06853)→ |
| Mab-1-77 | NZ_CSWM01000002.1 | - | 506367 | 506327 | **RNA**→ rpmE (PRK00019)Ribosomal_L31 (pfam01197)→ PrfA (COG0216)prfA (PRK00591)→ HemK (COG2890)PRK09328 (PRK09328)→ SUA5 (COG0009)TIGR00057 (TIGR00057)→ ECA_wecA (TIGR02380)GT_WecA_like (cd06853)→ |
| Mab-1-78 | NZ_CSYI01000001.1 | - | 248627 | 248587 | **RNA**→ rpmE (PRK00019)Ribosomal_L31 (pfam01197)→ PrfA (COG0216)prfA (PRK00591)→ HemK (COG2890)PRK09328 (PRK09328)→ SUA5 (COG0009)TIGR00057 (TIGR00057)→ ECA_wecA (TIGR02380)GT_WecA_like (cd06853)→ |
| Mab-1-79 | NZ_CSUM01000001.1 | + | 245392 | 245432 | **RNA**→ rpmE (PRK00019)Ribosomal_L31 (pfam01197)→ PrfA (COG0216)prfA (PRK00591)→ HemK (COG2890)PRK09328 (PRK09328)→ SUA5 (COG0009)TIGR00057 (TIGR00057)→ ECA_wecA (TIGR02380)GT_WecA_like (cd06853)→ |
| Mab-1-80 | NZ_CSWP01000004.1 | - | 313093 | 313053 | **RNA**→ rpmE (PRK00019)Ribosomal_L31 (pfam01197)→ PrfA (COG0216)prfA (PRK00591)→ HemK (COG2890)PRK09328 (PRK09328)→ SUA5 (COG0009)TIGR00057 (TIGR00057)→ ECA_wecA (TIGR02380)GT_WecA_like (cd06853)→ |
| Mab-1-81 | NZ_CSYD01000003.1 | - | 506396 | 506356 | **RNA**→ rpmE (PRK00019)Ribosomal_L31 (pfam01197)→ PrfA (COG0216)prfA (PRK00591)→ HemK (COG2890)PRK09328 (PRK09328)→ SUA5 (COG0009)TIGR00057 (TIGR00057)→ ECA_wecA (TIGR02380)GT_WecA_like (cd06853)→ |
| Mab-1-82 | NZ_CSVQ01000001.1 | - | 459269 | 459229 | **RNA**→ rpmE (PRK00019)Ribosomal_L31 (pfam01197)→ PrfA (COG0216)prfA (PRK00591)→ HemK (COG2890)PRK09328 (PRK09328)→ SUA5 (COG0009)TIGR00057 (TIGR00057)→ ECA_wecA (TIGR02380)GT_WecA_like (cd06853)→ |
| Mab-1-83 | NZ_CSZJ01000001.1 | - | 385149 | 385109 | **RNA**→ rpmE (PRK00019)Ribosomal_L31 (pfam01197)→ PrfA (COG0216)prfA (PRK00591)→ HemK (COG2890)PRK09328 (PRK09328)→ SUA5 (COG0009)TIGR00057 (TIGR00057)→ ECA_wecA (TIGR02380)GT_WecA_like (cd06853)→ |
| Mab-1-84 | NZ_CSZB01000001.1 | + | 163125 | 163165 | **RNA**→ rpmE (PRK00019)Ribosomal_L31 (pfam01197)→ PrfA (COG0216)prfA (PRK00591)→ HemK (COG2890)PRK09328 (PRK09328)→ SUA5 (COG0009)TIGR00057 (TIGR00057)→ ECA_wecA (TIGR02380)GT_WecA_like (cd06853)→ |
| Mab-1-85 | NZ_CSWR01000001.1 | + | 244744 | 244784 | **RNA**→ rpmE (PRK00019)Ribosomal_L31 (pfam01197)→ PrfA (COG0216)prfA (PRK00591)→ HemK (COG2890)PRK09328 (PRK09328)→ SUA5 (COG0009)TIGR00057 (TIGR00057)→ ECA_wecA (TIGR02380)GT_WecA_like (cd06853)→ |
| Mab-1-86 | NZ_CSWI01000003.1 | - | 506355 | 506315 | **RNA**→ rpmE (PRK00019)Ribosomal_L31 (pfam01197)→ PrfA (COG0216)prfA (PRK00591)→ HemK (COG2890)PRK09328 (PRK09328)→ SUA5 (COG0009)TIGR00057 (TIGR00057)→ ECA_wecA (TIGR02380)GT_WecA_like (cd06853)→ |
| Mab-1-87 | NZ_CSWV01000002.1 | - | 1045594 | 1045554 | **RNA**→ rpmE (PRK00019)Ribosomal_L31 (pfam01197)→ PrfA (COG0216)prfA (PRK00591)→ HemK (COG2890)PRK09328 (PRK09328)→ SUA5 (COG0009)TIGR00057 (TIGR00057)→ ECA_wecA (TIGR02380)GT_WecA_like (cd06853)→ |
| Mab-1-88 | NZ_CSWW01000002.1 | + | 309303 | 309343 | **RNA**→ rpmE (PRK00019)Ribosomal_L31 (pfam01197)→ PrfA (COG0216)prfA (PRK00591)→ HemK (COG2890)PRK09328 (PRK09328)→ SUA5 (COG0009)TIGR00057 (TIGR00057)→ ECA_wecA (TIGR02380)GT_WecA_like (cd06853)→ |
| Mab-1-89 | NZ_CSZK01000001.1 | - | 445930 | 445890 | **RNA**→ rpmE (PRK00019)Ribosomal_L31 (pfam01197)→ PrfA (COG0216)prfA (PRK00591)→ HemK (COG2890)PRK09328 (PRK09328)→ SUA5 (COG0009)TIGR00057 (TIGR00057)→ ECA_wecA (TIGR02380)GT_WecA_like (cd06853)→ |
| Mab-1-90 | NZ_CSYE01000001.1 | - | 786209 | 786169 | **RNA**→ rpmE (PRK00019)Ribosomal_L31 (pfam01197)→ PrfA (COG0216)prfA (PRK00591)→ HemK (COG2890)PRK09328 (PRK09328)→ SUA5 (COG0009)TIGR00057 (TIGR00057)→ ECA_wecA (TIGR02380)GT_WecA_like (cd06853)→ |
| Mab-1-91 | NZ_CSZW01000003.1 | - | 742881 | 742841 | **RNA**→ rpmE (PRK00019)Ribosomal_L31 (pfam01197)→ PrfA (COG0216)prfA (PRK00591)→ HemK (COG2890)PRK09328 (PRK09328)→ SUA5 (COG0009)TIGR00057 (TIGR00057)→ ECA_wecA (TIGR02380)GT_WecA_like (cd06853)→ |
| Mab-1-92 | NZ_CSXW01000002.1 | - | 506308 | 506268 | **RNA**→ rpmE (PRK00019)Ribosomal_L31 (pfam01197)→ PrfA (COG0216)prfA (PRK00591)→ HemK (COG2890)PRK09328 (PRK09328)→ SUA5 (COG0009)TIGR00057 (TIGR00057)→ ECA_wecA (TIGR02380)GT_WecA_like (cd06853)→ |
| Mab-1-93 | NZ_CSZY01000004.1 | - | 21804 | 21764 | **RNA**→ rpmE (PRK00019)Ribosomal_L31 (pfam01197)→ PrfA (COG0216)prfA (PRK00591)→ HemK (COG2890)PRK09328 (PRK09328)→ SUA5 (COG0009)TIGR00057 (TIGR00057)→ ECA_wecA (TIGR02380)GT_WecA_like (cd06853)→ |
| Mab-1-94 | NZ_CSYP01000001.1 | + | 941699 | 941739 | **RNA**→ rpmE (PRK00019)Ribosomal_L31 (pfam01197)→ PrfA (COG0216)prfA (PRK00591)→ HemK (COG2890)PRK09328 (PRK09328)→ SUA5 (COG0009)TIGR00057 (TIGR00057)→ ECA_wecA (TIGR02380)GT_WecA_like (cd06853)→ |

| | | | | | |
|---|---|---|---|---|---|
| Mab-1-95 | NZ_CSWF01000002.1 | - | 506495 | 506455 | **RNA**→ rpmE (PRK00019)Ribosomal_L31 (pfam01197)→ PrfA (COG0216)prfA (PRK00591)→ HemK (COG2890)PRK09328 (PRK09328)→ SUA5 (COG0009)TIGR00057 (TIGR00057)→ ECA_wecA (TIGR02380)GT_WecA_like (cd06853)→ |
| Mab-1-96 | NZ_CSXY01000002.1 | + | 162311 | 162351 | **RNA**→ rpmE (PRK00019)Ribosomal_L31 (pfam01197)→ PrfA (COG0216)prfA (PRK00591)→ HemK (COG2890)PRK09328 (PRK09328)→ SUA5 (COG0009)TIGR00057 (TIGR00057)→ ECA_wecA (TIGR02380)GT_WecA_like (cd06853)→ |
| Mab-1-97 | NZ_CSZD01000002.1 | + | 162313 | 162353 | **RNA**→ rpmE (PRK00019)Ribosomal_L31 (pfam01197)→ PrfA (COG0216)prfA (PRK00591)→ HemK (COG2890)PRK09328 (PRK09328)→ SUA5 (COG0009)TIGR00057 (TIGR00057)→ ECA_wecA (TIGR02380)GT_WecA_like (cd06853)→ |
| Mab-1-98 | NZ_CSUO01000001.1 | - | 308185 | 308145 | **RNA**→ rpmE (PRK00019)Ribosomal_L31 (pfam01197)→ PrfA (COG0216)prfA (PRK00591)→ HemK (COG2890)PRK09328 (PRK09328)→ SUA5 (COG0009)TIGR00057 (TIGR00057)→ ECA_wecA (TIGR02380)GT_WecA_like (cd06853)→ |
| Mab-1-99 | NZ_CSVK01000004.1 | - | 506337 | 506297 | **RNA**→ rpmE (PRK00019)Ribosomal_L31 (pfam01197)→ PrfA (COG0216)prfA (PRK00591)→ HemK (COG2890)PRK09328 (PRK09328)→ SUA5 (COG0009)TIGR00057 (TIGR00057)→ ECA_wecA (TIGR02380)GT_WecA_like (cd06853)→ |
| Mab-1-100 | NZ_CSYQ01000001.1 | + | 422078 | 422118 | **RNA**→ rpmE (PRK00019)Ribosomal_L31 (pfam01197)→ PrfA (COG0216)prfA (PRK00591)→ HemK (COG2890)PRK09328 (PRK09328)→ SUA5 (COG0009)TIGR00057 (TIGR00057)→ ECA_wecA (TIGR02380)GT_WecA_like (cd06853)→ |
| Mab-1-101 | NZ_CSYX01000001.1 | - | 896926 | 896886 | **RNA**→ rpmE (PRK00019)Ribosomal_L31 (pfam01197)→ PrfA (COG0216)prfA (PRK00591)→ HemK (COG2890)PRK09328 (PRK09328)→ SUA5 (COG0009)TIGR00057 (TIGR00057)→ ECA_wecA (TIGR02380)GT_WecA_like (cd06853)→ |
| Mab-1-102 | NZ_CSXK01000002.1 | - | 506652 | 506612 | **RNA**→ rpmE (PRK00019)Ribosomal_L31 (pfam01197)→ PrfA (COG0216)prfA (PRK00591)→ HemK (COG2890)PRK09328 (PRK09328)→ SUA5 (COG0009)TIGR00057 (TIGR00057)→ ECA_wecA (TIGR02380)GT_WecA_like (cd06853)→ |
| Mab-1-103 | NZ_CSXS01000003.1 | - | 506296 | 506256 | **RNA**→ rpmE (PRK00019)Ribosomal_L31 (pfam01197)→ PrfA (COG0216)prfA (PRK00591)→ HemK (COG2890)PRK09328 (PRK09328)→ SUA5 (COG0009)TIGR00057 (TIGR00057)→ ECA_wecA (TIGR02380)GT_WecA_like (cd06853)→ |
| Mab-1-104 | NZ_CSYA01000001.1 | + | 244753 | 244793 | **RNA**→ rpmE (PRK00019)Ribosomal_L31 (pfam01197)→ PrfA (COG0216)prfA (PRK00591)→ HemK (COG2890)PRK09328 (PRK09328)→ SUA5 (COG0009)TIGR00057 (TIGR00057)→ ECA_wecA (TIGR02380)GT_WecA_like (cd06853)→ |
| Mab-1-105 | NZ_CSWB01000001.1 | - | 506499 | 506459 | **RNA**→ rpmE (PRK00019)Ribosomal_L31 (pfam01197)→ PrfA (COG0216)prfA (PRK00591)→ HemK (COG2890)PRK09328 (PRK09328)→ SUA5 (COG0009)TIGR00057 (TIGR00057)→ ECA_wecA (TIGR02380)GT_WecA_like (cd06853)→ |
| Mab-1-106 | NZ_CSZH01000001.1 | - | 625398 | 625358 | **RNA**→ rpmE (PRK00019)Ribosomal_L31 (pfam01197)→ PrfA (COG0216)prfA (PRK00591)→ HemK (COG2890)PRK09328 (PRK09328)→ SUA5 (COG0009)TIGR00057 (TIGR00057)→ ECA_wecA (TIGR02380)GT_WecA_like (cd06853)→ |
| Mab-1-107 | NZ_CSXE01000001.1 | + | 244753 | 244793 | **RNA**→ rpmE (PRK00019)Ribosomal_L31 (pfam01197)→ PrfA (COG0216)prfA (PRK00591)→ HemK (COG2890)PRK09328 (PRK09328)→ SUA5 (COG0009)TIGR00057 (TIGR00057)→ ECA_wecA (TIGR02380)GT_WecA_like (cd06853)→ |
| Mab-1-108 | NZ_CSYO01000001.1 | - | 309254 | 309214 | **RNA**→ rpmE (PRK00019)Ribosomal_L31 (pfam01197)→ PrfA (COG0216)prfA (PRK00591)→ HemK (COG2890)PRK09328 (PRK09328)→ SUA5 (COG0009)TIGR00057 (TIGR00057)→ ECA_wecA (TIGR02380)GT_WecA_like (cd06853)→ |
| Mab-1-109 | NZ_CSZG01000009.1 | - | 21110 | 21070 | **RNA**→ rpmE (PRK00019)Ribosomal_L31 (pfam01197)→ PrfA (COG0216)prfA (PRK00591)→ HemK (COG2890)PRK09328 (PRK09328)→ SUA5 (COG0009)TIGR00057 (TIGR00057)→ ECA_wecA (TIGR02380)GT_WecA_like (cd06853)→ |
| Mab-1-110 | NZ_CSYR01000003.1 | - | 459464 | 459424 | **RNA**→ rpmE (PRK00019)Ribosomal_L31 (pfam01197)→ PrfA (COG0216)prfA (PRK00591)→ HemK (COG2890)PRK09328 (PRK09328)→ SUA5 (COG0009)TIGR00057 (TIGR00057)→ ECA_wecA (TIGR02380)GT_WecA_like (cd06853)→ |
| Mab-1-111 | NZ_CSZM01000004.1 | - | 132560 | 132520 | **RNA**→ rpmE (PRK00019)Ribosomal_L31 (pfam01197)→ PrfA (COG0216)prfA (PRK00591)→ HemK (COG2890)PRK09328 (PRK09328)→ SUA5 (COG0009)TIGR00057 (TIGR00057)→ ECA_wecA (TIGR02380)GT_WecA_like (cd06853)→ |
| Mab-1-112 | NZ_CSUF01000001.1 | - | 445936 | 445896 | **RNA**→ rpmE (PRK00019)Ribosomal_L31 (pfam01197)→ PrfA (COG0216)prfA (PRK00591)→ HemK (COG2890)PRK09328 (PRK09328)→ SUA5 (COG0009)TIGR00057 (TIGR00057)→ ECA_wecA (TIGR02380)GT_WecA_like (cd06853)→ |
| Mab-1-113 | NZ_CSTF01000001.1 | - | 309258 | 309218 | **RNA**→ rpmE (PRK00019)Ribosomal_L31 (pfam01197)→ PrfA (COG0216)prfA (PRK00591)→ HemK (COG2890)PRK09328 (PRK09328)→ SUA5 (COG0009)TIGR00057 (TIGR00057)→ ECA_wecA (TIGR02380)GT_WecA_like (cd06853)→ |

| | | | | | |
|---|---|---|---|---|---|
| Mab-57-1 | NZ_ATFQ01000015.1 | + | 135878 | 135918 | **RNA**→ rpmE (PRK00019)Ribosomal_L31 (pfam01197)→ PrfA (COG0216)prfA (PRK00591)→ HemK (COG2890)PRK09328 (PRK09328)→ SUA5 (COG0009)TIGR00057 (TIGR00057)→ ECA_wecA (TIGR02380)GT_WecA_like (cd06853)→ |
| Mab-9-1 | NZ_AGQU01000002.1 | + | 364967 | 365007 | **RNA**→ rpmE (PRK00019)Ribosomal_L31 (pfam01197)→ PrfA (COG0216)prfA (PRK00591)→ HemK (COG2890)PRK09328 (PRK09328)→ SUA5 (COG0009)TIGR00057 (TIGR00057)→ ECA_wecA (TIGR02380)GT_WecA_like (cd06853)→ |
| Mab-17-1 | NZ_AKTY01000002.1 | + | 656984 | 657024 | **RNA**→ rpmE (PRK00019)Ribosomal_L31 (pfam01197)→ PrfA (COG0216)prfA (PRK00591)→ HemK (COG2890)PRK09328 (PRK09328)→ SUA5 (COG0009)TIGR00057 (TIGR00057)→ ECA_wecA (TIGR02380)GT_WecA_like (cd06853)→ |
| Mab-38-1 | NZ_AJSE01000006.1 | - | 848090 | 848050 | **RNA**→ rpmE (PRK00019)Ribosomal_L31 (pfam01197)→ PrfA (COG0216)prfA (PRK00591)→ HemK (COG2890)PRK09328 (PRK09328)→ SUA5 (COG0009)TIGR00057 (TIGR00057)→ ECA_wecA (TIGR02380)GT_WecA_like (cd06853)→ |
| Mab-37-1 | NZ_AJSD01000001.1 | + | 321157 | 321197 | **RNA**→ rpmE (PRK00019)Ribosomal_L31 (pfam01197)→ PrfA (COG0216)prfA (PRK00591)→ HemK (COG2890)PRK09328 (PRK09328)→ SUA5 (COG0009)TIGR00057 (TIGR00057)→ ECA_wecA (TIGR02380)GT_WecA_like (cd06853)→ |
| Mab-49-1 | NZ_AKUU01000002.1 | + | 437326 | 437366 | **RNA**→ rpmE (PRK00019)Ribosomal_L31 (pfam01197)→ PrfA (COG0216)prfA (PRK00591)→ HemK (COG2890)PRK09328 (PRK09328)→ SUA5 (COG0009)TIGR00057 (TIGR00057)→ ECA_wecA (TIGR02380)GT_WecA_like (cd06853)→ |
| Mab-45-1 | NZ_AKUJ01000002.1 | + | 812189 | 812229 | **RNA**→ rpmE (PRK00019)Ribosomal_L31 (pfam01197)→ PrfA (COG0216)prfA (PRK00591)→ HemK (COG2890)PRK09328 (PRK09328)→ SUA5 (COG0009)TIGR00057 (TIGR00057)→ ECA_wecA (TIGR02380)GT_WecA_like (cd06853)→ |
| Mab-16-1 | NZ_AKTX01000002.1 | + | 136662 | 136702 | **RNA**→ rpmE (PRK00019)Ribosomal_L31 (pfam01197)→ PrfA (COG0216)prfA (PRK00591)→ HemK (COG2890)PRK09328 (PRK09328)→ SUA5 (COG0009)TIGR00057 (TIGR00057)→ ECA_wecA (TIGR02380)GT_WecA_like (cd06853)→ |
| Mab-52-1 | NZ_AKUW01000004.1 | + | 53562 | 53602 | **RNA**→ rpmE (PRK00019)Ribosomal_L31 (pfam01197)→ PrfA (COG0216)prfA (PRK00591)→ HemK (COG2890)PRK09328 (PRK09328)→ SUA5 (COG0009)TIGR00057 (TIGR00057)→ ECA_wecA (TIGR02380)GT_WecA_like (cd06853)→ |
| Mab-32-1 | NZ_AKVR01000005.1 | - | 756739 | 756699 | **RNA**→ rpmE (PRK00019)Ribosomal_L31 (pfam01197)→ PrfA (COG0216)prfA (PRK00591)→ HemK (COG2890)PRK09328 (PRK09328)→ SUA5 (COG0009)TIGR00057 (TIGR00057)→ ECA_wecA (TIGR02380)GT_WecA_like (cd06853)→ |
| Mab-31-1 | NZ_AJLZ01000010.1 | + | 127082 | 127122 | **RNA**→ rpmE (PRK00019)Ribosomal_L31 (pfam01197)→ PrfA (COG0216)prfA (PRK00591)→ HemK (COG2890)PRK09328 (PRK09328)→ SUA5 (COG0009)TIGR00057 (TIGR00057)→ ECA_wecA (TIGR02380)GT_WecA_like (cd06853)→ |
| Mab-23-1 | NZ_AKUD01000004.1 | - | 490418 | 490378 | **RNA**→ rpmE (PRK00019)Ribosomal_L31 (pfam01197)→ PrfA (COG0216)prfA (PRK00591)→ HemK (COG2890)PRK09328 (PRK09328)→ SUA5 (COG0009)TIGR00057 (TIGR00057)→ ECA_wecA (TIGR02380)GT_WecA_like (cd06853)→ |
| Mab-47-1 | NZ_AKUL01000001.1 | + | 920827 | 920867 | **RNA**→ rpmE (PRK00019)Ribosomal_L31 (pfam01197)→ PrfA (COG0216)prfA (PRK00591)→ HemK (COG2890)PRK09328 (PRK09328)→ SUA5 (COG0009)TIGR00057 (TIGR00057)→ ECA_wecA (TIGR02380)GT_WecA_like (cd06853)→ |
| Mab-22-1 | NZ_AKUC01000004.1 | + | 136640 | 136680 | **RNA**→ rpmE (PRK00019)Ribosomal_L31 (pfam01197)→ PrfA (COG0216)prfA (PRK00591)→ HemK (COG2890)PRK09328 (PRK09328)→ SUA5 (COG0009)TIGR00057 (TIGR00057)→ ECA_wecA (TIGR02380)GT_WecA_like (cd06853)→ |
| Mab-20-1 | NZ_AKUB01000002.1 | + | 136706 | 136746 | **RNA**→ rpmE (PRK00019)Ribosomal_L31 (pfam01197)→ PrfA (COG0216)prfA (PRK00591)→ HemK (COG2890)PRK09328 (PRK09328)→ SUA5 (COG0009)TIGR00057 (TIGR00057)→ ECA_wecA (TIGR02380)GT_WecA_like (cd06853)→ |
| Mab-19-1 | NZ_AKUA01000003.1 | + | 136653 | 136693 | **RNA**→ rpmE (PRK00019)Ribosomal_L31 (pfam01197)→ PrfA (COG0216)prfA (PRK00591)→ HemK (COG2890)PRK09328 (PRK09328)→ SUA5 (COG0009)TIGR00057 (TIGR00057)→ ECA_wecA (TIGR02380)GT_WecA_like (cd06853)→ |
| Mab-48-1 | NZ_AKUN01000002.1 | + | 437379 | 437419 | **RNA**→ rpmE (PRK00019)Ribosomal_L31 (pfam01197)→ PrfA (COG0216)prfA (PRK00591)→ HemK (COG2890)PRK09328 (PRK09328)→ SUA5 (COG0009)TIGR00057 (TIGR00057)→ ECA_wecA (TIGR02380)GT_WecA_like (cd06853)→ |
| Mab-35-1 | NZ_AJMA01000004.1 | - | 313058 | 313018 | **RNA**→ rpmE (PRK00019)Ribosomal_L31 (pfam01197)→ PrfA (COG0216)prfA (PRK00591)→ HemK (COG2890)PRK09328 (PRK09328)→ SUA5 (COG0009)TIGR00057 (TIGR00057)→ ECA_wecA (TIGR02380)GT_WecA_like (cd06853)→ |
| Mab-50-1 | NZ_AKUM01000002.1 | + | 611330 | 611370 | **RNA**→ rpmE (PRK00019)Ribosomal_L31 (pfam01197)→ PrfA (COG0216)prfA (PRK00591)→ HemK (COG2890)PRK09328 (PRK09328)→ SUA5 (COG0009)TIGR00057 (TIGR00057)→ ECA_wecA (TIGR02380)GT_WecA_like (cd06853)→ |

| | | | | | |
|---|---|---|---|---|---|
| Mab-51-1 | NZ_AKUV01000004.1 | + | 437347 | 437387 | **RNA**→ rpmE (PRK00019)Ribosomal_L31 (pfam01197)→ PrfA (COG0216)prfA (PRK00591)→ HemK (COG2890)PRK09328 (PRK09328)→ SUA5 (COG0009)TIGR00057 (TIGR00057)→ ECA_wecA (TIGR02380)GT_WecA_like (cd06853)→ |
| Mab-53-1 | NZ_AKUO01000005.1 | + | 53563 | 53603 | **RNA**→ rpmE (PRK00019)Ribosomal_L31 (pfam01197)→ PrfA (COG0216)prfA (PRK00591)→ HemK (COG2890)PRK09328 (PRK09328)→ SUA5 (COG0009)TIGR00057 (TIGR00057)→ ECA_wecA (TIGR02380)GT_WecA_like (cd06853)→ |
| Mab-44-1 | NZ_AKUI01000002.1 | + | 376605 | 376645 | **RNA**→ rpmE (PRK00019)Ribosomal_L31 (pfam01197)→ PrfA (COG0216)prfA (PRK00591)→ HemK (COG2890)PRK09328 (PRK09328)→ SUA5 (COG0009)TIGR00057 (TIGR00057)→ ECA_wecA (TIGR02380)GT_WecA_like (cd06853)→ |
| Mab-18-1 | NZ_AKTZ01000005.1 | + | 437892 | 437932 | **RNA**→ rpmE (PRK00019)Ribosomal_L31 (pfam01197)→ PrfA (COG0216)prfA (PRK00591)→ HemK (COG2890)PRK09328 (PRK09328)→ SUA5 (COG0009)TIGR00057 (TIGR00057)→ ECA_wecA (TIGR02380)GT_WecA_like (cd06853)→ |
| Mab-36-1 | NZ_AKVU01000008.1 | - | 313101 | 313061 | **RNA**→ rpmE (PRK00019)Ribosomal_L31 (pfam01197)→ PrfA (COG0216)prfA (PRK00591)→ HemK (COG2890)PRK09328 (PRK09328)→ SUA5 (COG0009)TIGR00057 (TIGR00057)→ ECA_wecA (TIGR02380)GT_WecA_like (cd06853)→ |
| Mab-46-1 | NZ_AKUK01000006.1 | + | 53546 | 53586 | **RNA**→ rpmE (PRK00019)Ribosomal_L31 (pfam01197)→ PrfA (COG0216)prfA (PRK00591)→ HemK (COG2890)PRK09328 (PRK09328)→ SUA5 (COG0009)TIGR00057 (TIGR00057)→ ECA_wecA (TIGR02380)GT_WecA_like (cd06853)→ |
| Mab-21-1 | NZ_AKUQ01000005.1 | + | 54178 | 54218 | **RNA**→ rpmE (PRK00019)Ribosomal_L31 (pfam01197)→ PrfA (COG0216)prfA (PRK00591)→ HemK (COG2890)PRK09328 (PRK09328)→ SUA5 (COG0009)TIGR00057 (TIGR00057)→ ECA_wecA (TIGR02380)GT_WecA_like (cd06853)→ |
| Mab-58-1 | NZ_AJSC01000002.1 | - | 671044 | 671004 | **RNA**→ rpmE (PRK00019)Ribosomal_L31 (pfam01197)→ PrfA (COG0216)prfA (PRK00591)→ HemK (COG2890)PRK09328 (PRK09328)→ SUA5 (COG0009)TIGR00057 (TIGR00057)→ ECA_wecA (TIGR02380)GT_WecA_like (cd06853)→ |
| Mab-42-1 | NZ_AYTF01000002.1 | + | 1614438 | 1614478 | **RNA**→ rpmE (PRK00019)Ribosomal_L31 (pfam01197)→ PrfA (COG0216)prfA (PRK00591)→ HemK (COG2890)PRK09328 (PRK09328)→ SUA5 (COG0009)TIGR00057 (TIGR00057)→ hypo→ Rfe (COG0472)→ GT_WecA_like (cd06853)mraY (PRK00108)other_trbK (TIGR04360)→ |
| Mab-64-1 | NZ_JAOW01000004.1 | + | 209627 | 209667 | **RNA**→ rpmE (PRK00019)Ribosomal_L31 (pfam01197)→ PrfA (COG0216)prfA (PRK00591)→ HemK (COG2890)PRK09328 (PRK09328)→ SUA5 (COG0009)TIGR00057 (TIGR00057)→ ECA_wecA (TIGR02380)GT_WecA_like (cd06853)→ |
| Mab-66-1 | NZ_JAOV01000001.1 | + | 1212907 | 1212947 | **RNA**→ rpmE (PRK00019)Ribosomal_L31 (pfam01197)→ PrfA (COG0216)prfA (PRK00591)→ HemK (COG2890)PRK09328 (PRK09328)→ SUA5 (COG0009)TIGR00057 (TIGR00057)→ ECA_wecA (TIGR02380)GT_WecA_like (cd06853)→ |
| Mab-54-1 | NC_021282.1 | + | 1470196 | 1470236 | **RNA**→ rpmE (PRK00019)Ribosomal_L31 (pfam01197)→ PrfA (COG0216)prfA (PRK00591)→ HemK (COG2890)PRK09328 (PRK09328)→ SUA5 (COG0009)TIGR00057 (TIGR00057)→ ECA_wecA (TIGR02380)GT_WecA_like (cd06853)→ |
| Mab-70-1 | NZ_CP009616.1 | - | 3353449 | 3353409 | **RNA**→ rpmE (PRK00019)Ribosomal_L31 (pfam01197)→ PrfA (COG0216)prfA (PRK00591)→ HemK (COG2890)PRK09328 (PRK09328)→ SUA5 (COG0009)TIGR00057 (TIGR00057)→ ECA_wecA (TIGR02380)GT_WecA_like (cd06853)→ |
| Mab-70-2 | NZ_CP009615.1 | - | 3353238 | 3353198 | **RNA**→ rpmE (PRK00019)Ribosomal_L31 (pfam01197)→ PrfA (COG0216)prfA (PRK00591)→ HemK (COG2890)PRK09328 (PRK09328)→ SUA5 (COG0009)TIGR00057 (TIGR00057)→ ECA_wecA (TIGR02380)GT_WecA_like (cd06853)→ |
| Mab-75-1 | NC_018150.2 | - | 3353484 | 3353444 | **RNA**→ rpmE (PRK00019)Ribosomal_L31 (pfam01197)→ PrfA (COG0216)prfA (PRK00591)→ HemK (COG2890)PRK09328 (PRK09328)→ hypo→ hypo→ ECA_wecA (TIGR02380)GT_WecA_like (cd06853)→ |
| Mab-71-1 | NZ_CP009447.1 | - | 3228707 | 3228667 | **RNA**→ rpmE (PRK00019)Ribosomal_L31 (pfam01197)→ PrfA (COG0216)prfA (PRK00591)→ HemK (COG2890)PRK09328 (PRK09328)→ SUA5 (COG0009)TIGR00057 (TIGR00057)→ ECA_wecA (TIGR02380)GT_WecA_like (cd06853)→ |
| Mab-33-1 | NZ_AKVV01000125.1 | + | 9758 | 9798 | **RNA**→ rpmE (PRK00019)Ribosomal_L31 (pfam01197)→ PrfA (COG0216)prfA (PRK00591)→ hypo→ HemK (COG2890)PRK09328 (PRK09328)→ SUA5 (COG0009)TIGR00057 (TIGR00057)→ ECA_wecA (TIGR02380)GT_WecA_like (cd06853)→ |
| Mab-61-1 | NZ_AJLY02000020.1 | + | 60826 | 60866 | **RNA**→ rpmE (PRK00019)Ribosomal_L31 (pfam01197)→ PrfA (COG0216)prfA (PRK00591)→ HemK (COG2890)PRK09328 (PRK09328)→ SUA5 (COG0009)TIGR00057 (TIGR00057)→ ECA_wecA (TIGR02380)GT_WecA_like (cd06853)→ |
| Mab-41-1 | NZ_AYTA01000006.1 | + | 3199797 | 3199837 | **RNA**→ rpmE (PRK00019)Ribosomal_L31 (pfam01197)→ PrfA (COG0216)prfA (PRK00591)→ HemK (COG2890)PRK09328 (PRK09328)→ SUA5 (COG0009)TIGR00057 (TIGR00057)→ ECA_wecA (TIGR02380)GT_WecA_like (cd06853)→ |

| | | | | | |
|---|---|---|---|---|---|
| Mab-55-1 | NZ_AHAS01000006.1 | + | 474490 | 474530 | **RNA**→ rpmE (PRK00019)Ribosomal_L31 (pfam01197)→ PrfA (COG0216)prfA (PRK00591)→ HemK (COG2890)PRK09328 (PRK09328)→ SUA5 (COG0009)TIGR00057 (TIGR00057)→ ECA_wecA (TIGR02380)GT_WecA_like (cd06853)→ |
| Mab-1-114 | NZ_CSZT01000025.1 | - | 39697 | 39657 | **RNA**→ rpmE (PRK00019)Ribosomal_L31 (pfam01197)→ PrfA (COG0216)prfA (PRK00591)→ HemK (COG2890)PRK09328 (PRK09328)→ SUA5 (COG0009)TIGR00057 (TIGR00057)→ ECA_wecA (TIGR02380)GT_WecA_like (cd06853)→ |
| Mab-1-115 | NZ_CSWD01000003.1 | - | 317249 | 317209 | **RNA**→ rpmE (PRK00019)Ribosomal_L31 (pfam01197)→ PrfA (COG0216)prfA (PRK00591)→ HemK (COG2890)PRK09328 (PRK09328)→ SUA5 (COG0009)TIGR00057 (TIGR00057)→ ECA_wecA (TIGR02380)GT_WecA_like (cd06853)→ |
| Mab-1-116 | NZ_CSXL01000001.1 | - | 399190 | 399150 | **RNA**→ rpmE (PRK00019)Ribosomal_L31 (pfam01197)→ PrfA (COG0216)prfA (PRK00591)→ HemK (COG2890)PRK09328 (PRK09328)→ SUA5 (COG0009)TIGR00057 (TIGR00057)→ ECA_wecA (TIGR02380)GT_WecA_like (cd06853)→ |
| Mab-1-117 | NZ_CSXX01000001.1 | - | 143542 | 143502 | **RNA**→ rpmE (PRK00019)Ribosomal_L31 (pfam01197)→ PrfA (COG0216)prfA (PRK00591)→ HemK (COG2890)PRK09328 (PRK09328)→ SUA5 (COG0009)TIGR00057 (TIGR00057)→ ECA_wecA (TIGR02380)GT_WecA_like (cd06853)→ |
| Mab-1-118 | NZ_CSYS01000007.1 | + | 148924 | 148964 | **RNA**→ rpmE (PRK00019)Ribosomal_L31 (pfam01197)→ PrfA (COG0216)prfA (PRK00591)→ HemK (COG2890)PRK09328 (PRK09328)→ SUA5 (COG0009)TIGR00057 (TIGR00057)→ ECA_wecA (TIGR02380)GT_WecA_like (cd06853)→ |
| Mab-1-119 | NZ_CSZR01000007.1 | - | 132076 | 132036 | **RNA**→ rpmE (PRK00019)Ribosomal_L31 (pfam01197)→ PrfA (COG0216)prfA (PRK00591)→ hypo→ hypo→ SUA5 (COG0009)TIGR00057 (TIGR00057)→ ECA_wecA (TIGR02380)GT_WecA_like (cd06853)→ |
| Mab-1-120 | NZ_CSWC01000003.1 | + | 10427 | 10467 | **RNA**→ rpmE (PRK00019)Ribosomal_L31 (pfam01197)→ PrfA (COG0216)prfA (PRK00591)→ HemK (COG2890)PRK09328 (PRK09328)→ SUA5 (COG0009)TIGR00057 (TIGR00057)→ ECA_wecA (TIGR02380)GT_WecA_like (cd06853)→ |
| Mab-1-121 | NZ_CSUY01000001.1 | - | 1112929 | 1112889 | **RNA**→ rpmE (PRK00019)Ribosomal_L31 (pfam01197)→ PrfA (COG0216)prfA (PRK00591)→ HemK (COG2890)PRK09328 (PRK09328)→ SUA5 (COG0009)TIGR00057 (TIGR00057)→ ECA_wecA (TIGR02380)GT_WecA_like (cd06853)→ |
| Mab-1-122 | NZ_CSYU01000006.1 | + | 149331 | 149371 | **RNA**→ rpmE (PRK00019)Ribosomal_L31 (pfam01197)→ PrfA (COG0216)prfA (PRK00591)→ HemK (COG2890)PRK09328 (PRK09328)→ SUA5 (COG0009)TIGR00057 (TIGR00057)→ ECA_wecA (TIGR02380)GT_WecA_like (cd06853)→ |
| Mab-1-123 | NZ_CSYK01000002.1 | - | 294347 | 294307 | **RNA**→ rpmE (PRK00019)Ribosomal_L31 (pfam01197)→ PrfA (COG0216)prfA (PRK00591)→ HemK (COG2890)PRK09328 (PRK09328)→ SUA5 (COG0009)TIGR00057 (TIGR00057)→ ECA_wecA (TIGR02380)GT_WecA_like (cd06853)→ |
| Mab-1-124 | NZ_CSWK01000002.1 | - | 294251 | 294211 | **RNA**→ rpmE (PRK00019)Ribosomal_L31 (pfam01197)→ PrfA (COG0216)prfA (PRK00591)→ HemK (COG2890)PRK09328 (PRK09328)→ SUA5 (COG0009)TIGR00057 (TIGR00057)→ ECA_wecA (TIGR02380)GT_WecA_like (cd06853)→ |
| Mab-1-125 | NZ_CSTX01000003.1 | + | 10449 | 10489 | **RNA**→ rpmE (PRK00019)Ribosomal_L31 (pfam01197)→ PrfA (COG0216)prfA (PRK00591)→ HemK (COG2890)PRK09328 (PRK09328)→ SUA5 (COG0009)TIGR00057 (TIGR00057)→ ECA_wecA (TIGR02380)GT_WecA_like (cd06853)→ |
| Mab-1-126 | NZ_CSUP01000002.1 | - | 294322 | 294282 | **RNA**→ rpmE (PRK00019)Ribosomal_L31 (pfam01197)→ PrfA (COG0216)prfA (PRK00591)→ HemK (COG2890)PRK09328 (PRK09328)→ SUA5 (COG0009)TIGR00057 (TIGR00057)→ ECA_wecA (TIGR02380)GT_WecA_like (cd06853)→ |
| Mab-1-127 | NZ_CSWL01000001.1 | - | 742508 | 742468 | **RNA**→ rpmE (PRK00019)Ribosomal_L31 (pfam01197)→ PrfA (COG0216)prfA (PRK00591)→ HemK (COG2890)PRK09328 (PRK09328)→ SUA5 (COG0009)TIGR00057 (TIGR00057)→ ECA_wecA (TIGR02380)GT_WecA_like (cd06853)→ |
| Mab-1-128 | NZ_CSVB01000002.1 | - | 294465 | 294425 | **RNA**→ rpmE (PRK00019)Ribosomal_L31 (pfam01197)→ PrfA (COG0216)prfA (PRK00591)→ HemK (COG2890)PRK09328 (PRK09328)→ SUA5 (COG0009)TIGR00057 (TIGR00057)→ ECA_wecA (TIGR02380)GT_WecA_like (cd06853)→ |
| Mab-1-129 | NZ_CSTU01000051.1 | - | 20787 | 20747 | **RNA**→ rpmE (PRK00019)Ribosomal_L31 (pfam01197)→ PrfA (COG0216)prfA (PRK00591)→ HemK (COG2890)PRK09328 (PRK09328)→ SUA5 (COG0009)TIGR00057 (TIGR00057)→ ECA_wecA (TIGR02380)GT_WecA_like (cd06853)→ |
| Mab-1-130 | NZ_CSTO01000003.1 | - | 317251 | 317211 | **RNA**→ rpmE (PRK00019)Ribosomal_L31 (pfam01197)→ PrfA (COG0216)prfA (PRK00591)→ HemK (COG2890)PRK09328 (PRK09328)→ SUA5 (COG0009)TIGR00057 (TIGR00057)→ ECA_wecA (TIGR02380)GT_WecA_like (cd06853)→ |
| Mab-1-131 | NZ_CSVA01000001.1 | - | 740453 | 740413 | **RNA**→ rpmE (PRK00019)Ribosomal_L31 (pfam01197)→ PrfA (COG0216)prfA (PRK00591)→ HemK (COG2890)PRK09328 (PRK09328)→ SUA5 (COG0009)TIGR00057 (TIGR00057)→ ECA_wecA (TIGR02380)GT_WecA_like (cd06853)→ |

| | | | | | |
|---|---|---|---|---|---|
| Mab-1-132 | NZ_CSTY01000002.1 | - | 294342 | 294302 | **RNA**→ rpmE (PRK00019)Ribosomal_L31 (pfam01197)→ PrfA (COG0216)prfA (PRK00591)→ HemK (COG2890)PRK09328 (PRK09328)→ SUA5 (COG0009)TIGR00057 (TIGR00057)→ ECA_wecA (TIGR02380)GT_WecA_like (cd06853)→ |
| Mab-1-133 | NZ_CSUG01000005.1 | - | 317113 | 317073 | **RNA**→ rpmE (PRK00019)Ribosomal_L31 (pfam01197)→ PrfA (COG0216)prfA (PRK00591)→ HemK (COG2890)PRK09328 (PRK09328)→ SUA5 (COG0009)TIGR00057 (TIGR00057)→ ECA_wecA (TIGR02380)GT_WecA_like (cd06853)→ |
| Mab-1-134 | NZ_CSVM01000079.1 | - | 10485 | 10445 | **RNA**→ rpmE (PRK00019)Ribosomal_L31 (pfam01197)→ PrfA (COG0216)prfA (PRK00591)→ HemK (COG2890)PRK09328 (PRK09328)→ SUA5 (COG0009)TIGR00057 (TIGR00057)→ ECA_wecA (TIGR02380)GT_WecA_like (cd06853)→ |
| Mab-1-135 | NZ_CSUN01000002.1 | - | 294331 | 294291 | **RNA**→ rpmE (PRK00019)Ribosomal_L31 (pfam01197)→ PrfA (COG0216)prfA (PRK00591)→ HemK (COG2890)PRK09328 (PRK09328)→ SUA5 (COG0009)TIGR00057 (TIGR00057)→ ECA_wecA (TIGR02380)GT_WecA_like (cd06853)→ |
| Mab-1-136 | NZ_CSUI01000009.1 | - | 132037 | 131997 | **RNA**→ rpmE (PRK00019)Ribosomal_L31 (pfam01197)→ PrfA (COG0216)prfA (PRK00591)→ HemK (COG2890)PRK09328 (PRK09328)→ SUA5 (COG0009)TIGR00057 (TIGR00057)→ ECA_wecA (TIGR02380)GT_WecA_like (cd06853)→ |
| Mab-1-137 | NZ_CSVY01000048.1 | - | 20786 | 20746 | **RNA**→ rpmE (PRK00019)Ribosomal_L31 (pfam01197)→ PrfA (COG0216)prfA (PRK00591)→ HemK (COG2890)PRK09328 (PRK09328)→ SUA5 (COG0009)TIGR00057 (TIGR00057)→ ECA_wecA (TIGR02380)GT_WecA_like (cd06853)→ |
| Mab-1-138 | NZ_CSXH01000002.1 | - | 335905 | 335865 | **RNA**→ rpmE (PRK00019)Ribosomal_L31 (pfam01197)→ PrfA (COG0216)prfA (PRK00591)→ HemK (COG2890)PRK09328 (PRK09328)→ SUA5 (COG0009)TIGR00057 (TIGR00057)→ ECA_wecA (TIGR02380)GT_WecA_like (cd06853)→ |
| Mab-1-139 | NZ_CSTS01000003.1 | - | 344480 | 344440 | **RNA**→ rpmE (PRK00019)Ribosomal_L31 (pfam01197)→ PrfA (COG0216)prfA (PRK00591)→ HemK (COG2890)PRK09328 (PRK09328)→ SUA5 (COG0009)TIGR00057 (TIGR00057)→ ECA_wecA (TIGR02380)GT_WecA_like (cd06853)→ |
| Mab-1-140 | NZ_CSUR01000006.1 | - | 299273 | 299233 | **RNA**→ rpmE (PRK00019)Ribosomal_L31 (pfam01197)→ PrfA (COG0216)prfA (PRK00591)→ HemK (COG2890)PRK09328 (PRK09328)→ SUA5 (COG0009)TIGR00057 (TIGR00057)→ ECA_wecA (TIGR02380)GT_WecA_like (cd06853)→ |
| Mab-1-141 | NZ_CSZP01000003.1 | - | 126069 | 126029 | **RNA**→ rpmE (PRK00019)Ribosomal_L31 (pfam01197)→ PrfA (COG0216)prfA (PRK00591)→ HemK (COG2890)PRK09328 (PRK09328)→ SUA5 (COG0009)TIGR00057 (TIGR00057)→ ECA_wecA (TIGR02380)GT_WecA_like (cd06853)→ |
| Mab-1-142 | NZ_CSVV01000003.1 | - | 132014 | 131974 | **RNA**→ rpmE (PRK00019)Ribosomal_L31 (pfam01197)→ PrfA (COG0216)prfA (PRK00591)→ HemK (COG2890)PRK09328 (PRK09328)→ SUA5 (COG0009)TIGR00057 (TIGR00057)→ ECA_wecA (TIGR02380)GT_WecA_like (cd06853)→ |
| Mab-1-143 | NZ_CSVF01000002.1 | - | 294362 | 294322 | **RNA**→ rpmE (PRK00019)Ribosomal_L31 (pfam01197)→ PrfA (COG0216)prfA (PRK00591)→ HemK (COG2890)PRK09328 (PRK09328)→ SUA5 (COG0009)TIGR00057 (TIGR00057)→ ECA_wecA (TIGR02380)GT_WecA_like (cd06853)→ |
| Mab-1-144 | NZ_CSUH01000002.1 | - | 294322 | 294282 | **RNA**→ rpmE (PRK00019)Ribosomal_L31 (pfam01197)→ PrfA (COG0216)prfA (PRK00591)→ HemK (COG2890)PRK09328 (PRK09328)→ SUA5 (COG0009)TIGR00057 (TIGR00057)→ ECA_wecA (TIGR02380)GT_WecA_like (cd06853)→ |
| Mab-1-145 | NZ_CSYM01000002.1 | - | 132500 | 132460 | **RNA**→ rpmE (PRK00019)Ribosomal_L31 (pfam01197)→ PrfA (COG0216)prfA (PRK00591)→ HemK (COG2890)PRK09328 (PRK09328)→ SUA5 (COG0009)TIGR00057 (TIGR00057)→ ECA_wecA (TIGR02380)GT_WecA_like (cd06853)→ |
| Mab-1-146 | NZ_CSVN01000015.1 | - | 132014 | 131974 | **RNA**→ rpmE (PRK00019)Ribosomal_L31 (pfam01197)→ PrfA (COG0216)prfA (PRK00591)→ HemK (COG2890)PRK09328 (PRK09328)→ SUA5 (COG0009)TIGR00057 (TIGR00057)→ ECA_wecA (TIGR02380)GT_WecA_like (cd06853)→ |
| Mab-1-147 | NZ_CSXF01000004.1 | - | 450245 | 450205 | **RNA**→ rpmE (PRK00019)Ribosomal_L31 (pfam01197)→ PrfA (COG0216)prfA (PRK00591)→ HemK (COG2890)PRK09328 (PRK09328)→ SUA5 (COG0009)TIGR00057 (TIGR00057)→ ECA_wecA (TIGR02380)GT_WecA_like (cd06853)→ |
| Mab-1-148 | NZ_CSVL01000002.1 | - | 294322 | 294282 | **RNA**→ rpmE (PRK00019)Ribosomal_L31 (pfam01197)→ PrfA (COG0216)prfA (PRK00591)→ HemK (COG2890)PRK09328 (PRK09328)→ SUA5 (COG0009)TIGR00057 (TIGR00057)→ ECA_wecA (TIGR02380)GT_WecA_like (cd06853)→ |
| Mab-1-149 | NZ_CSTW01000009.1 | - | 132078 | 132038 | **RNA**→ rpmE (PRK00019)Ribosomal_L31 (pfam01197)→ PrfA (COG0216)prfA (PRK00591)→ HemK (COG2890)PRK09328 (PRK09328)→ SUA5 (COG0009)TIGR00057 (TIGR00057)→ ECA_wecA (TIGR02380)GT_WecA_like (cd06853)→ |
| Mab-1-150 | NZ_CSVU01000004.1 | - | 132029 | 131989 | **RNA**→ rpmE (PRK00019)Ribosomal_L31 (pfam01197)→ PrfA (COG0216)prfA (PRK00591)→ HemK (COG2890)PRK09328 (PRK09328)→ SUA5 (COG0009)TIGR00057 (TIGR00057)→ ECA_wecA (TIGR02380)GT_WecA_like (cd06853)→ |

| | | | | | |
|---|---|---|---|---|---|
| Mab-1-151 | NZ_CSUA01000006.1 | - | 299395 | 299355 | **RNA**→ rpmE (PRK00019)Ribosomal_L31 (pfam01197)→ PrfA (COG0216)prfA (PRK00591)→ HemK (COG2890)PRK09328 (PRK09328)→ SUA5 (COG0009)TIGR00057 (TIGR00057)→ ECA_wecA (TIGR02380)GT_WecA_like (cd06853)→ |
| Mab-1-152 | NZ_CSVG01000002.1 | - | 294334 | 294294 | **RNA**→ rpmE (PRK00019)Ribosomal_L31 (pfam01197)→ PrfA (COG0216)prfA (PRK00591)→ HemK (COG2890)PRK09328 (PRK09328)→ SUA5 (COG0009)TIGR00057 (TIGR00057)→ ECA_wecA (TIGR02380)GT_WecA_like (cd06853)→ |
| Mab-1-153 | NZ_CSWS01000002.1 | - | 132041 | 132001 | **RNA**→ rpmE (PRK00019)Ribosomal_L31 (pfam01197)→ PrfA (COG0216)prfA (PRK00591)→ HemK (COG2890)PRK09328 (PRK09328)→ SUA5 (COG0009)TIGR00057 (TIGR00057)→ ECA_wecA (TIGR02380)GT_WecA_like (cd06853)→ |
| Mab-1-154 | NZ_CSUW01000002.1 | - | 143911 | 143871 | **RNA**→ rpmE (PRK00019)Ribosomal_L31 (pfam01197)→ PrfA (COG0216)prfA (PRK00591)→ HemK (COG2890)PRK09328 (PRK09328)→ SUA5 (COG0009)TIGR00057 (TIGR00057)→ ECA_wecA (TIGR02380)GT_WecA_like (cd06853)→ |
| Mab-1-155 | NZ_CSVR01000004.1 | + | 266215 | 266255 | **RNA**→ rpmE (PRK00019)Ribosomal_L31 (pfam01197)→ PrfA (COG0216)prfA (PRK00591)→ HemK (COG2890)PRK09328 (PRK09328)→ SUA5 (COG0009)TIGR00057 (TIGR00057)→ ECA_wecA (TIGR02380)GT_WecA_like (cd06853)→ |
| Mab-1-156 | NZ_CSZO01000004.1 | - | 132026 | 131986 | **RNA**→ rpmE (PRK00019)Ribosomal_L31 (pfam01197)→ PrfA (COG0216)prfA (PRK00591)→ hypo→ hypo→ SUA5 (COG0009)TIGR00057 (TIGR00057)→ ECA_wecA (TIGR02380)GT_WecA_like (cd06853)→ |
| Mab-1-157 | NZ_CSWZ01000003.1 | - | 132152 | 132112 | **RNA**→ rpmE (PRK00019)Ribosomal_L31 (pfam01197)→ PrfA (COG0216)prfA (PRK00591)→ HemK (COG2890)PRK09328 (PRK09328)→ SUA5 (COG0009)TIGR00057 (TIGR00057)→ ECA_wecA (TIGR02380)GT_WecA_like (cd06853)→ |
| Mab-1-158 | NZ_CSWE01000001.1 | + | 657102 | 657142 | **RNA**→ rpmE (PRK00019)Ribosomal_L31 (pfam01197)→ PrfA (COG0216)prfA (PRK00591)→ HemK (COG2890)PRK09328 (PRK09328)→ SUA5 (COG0009)TIGR00057 (TIGR00057)→ ECA_wecA (TIGR02380)GT_WecA_like (cd06853)→ |
| Mab-1-159 | NZ_CSZX01000001.1 | - | 143534 | 143494 | **RNA**→ rpmE (PRK00019)Ribosomal_L31 (pfam01197)→ PrfA (COG0216)prfA (PRK00591)→ HemK (COG2890)PRK09328 (PRK09328)→ SUA5 (COG0009)TIGR00057 (TIGR00057)→ ECA_wecA (TIGR02380)GT_WecA_like (cd06853)→ |
| Mab-1-160 | NZ_CSWN01000005.1 | - | 143616 | 143576 | **RNA**→ rpmE (PRK00019)Ribosomal_L31 (pfam01197)→ PrfA (COG0216)prfA (PRK00591)→ HemK (COG2890)PRK09328 (PRK09328)→ SUA5 (COG0009)TIGR00057 (TIGR00057)→ ECA_wecA (TIGR02380)GT_WecA_like (cd06853)→ |
| Mab-1-161 | NZ_CSXT01000008.1 | - | 144962 | 144922 | **RNA**→ rpmE (PRK00019)Ribosomal_L31 (pfam01197)→ PrfA (COG0216)prfA (PRK00591)→ HemK (COG2890)PRK09328 (PRK09328)→ SUA5 (COG0009)TIGR00057 (TIGR00057)→ ECA_wecA (TIGR02380)GT_WecA_like (cd06853)→ |
| Mab-1-162 | NZ_CSZS01000012.1 | - | 49634 | 49594 | **RNA**→ rpmE (PRK00019)Ribosomal_L31 (pfam01197)→ PrfA (COG0216)prfA (PRK00591)→ HemK (COG2890)PRK09328 (PRK09328)→ SUA5 (COG0009)TIGR00057 (TIGR00057)→ ECA_wecA (TIGR02380)GT_WecA_like (cd06853)→ |
| Mab-1-163 | NZ_CSYL01000003.1 | + | 261973 | 262013 | **RNA**→ rpmE (PRK00019)Ribosomal_L31 (pfam01197)→ PrfA (COG0216)prfA (PRK00591)→ HemK (COG2890)PRK09328 (PRK09328)→ SUA5 (COG0009)TIGR00057 (TIGR00057)→ ECA_wecA (TIGR02380)GT_WecA_like (cd06853)→ |
| Mab-1-164 | NZ_CSWH01000005.1 | - | 299304 | 299264 | **RNA**→ rpmE (PRK00019)Ribosomal_L31 (pfam01197)→ PrfA (COG0216)prfA (PRK00591)→ HemK (COG2890)PRK09328 (PRK09328)→ SUA5 (COG0009)TIGR00057 (TIGR00057)→ ECA_wecA (TIGR02380)GT_WecA_like (cd06853)→ |
| Mab-1-165 | NZ_CSTP01000002.1 | - | 317144 | 317104 | **RNA**→ rpmE (PRK00019)Ribosomal_L31 (pfam01197)→ PrfA (COG0216)prfA (PRK00591)→ HemK (COG2890)PRK09328 (PRK09328)→ SUA5 (COG0009)TIGR00057 (TIGR00057)→ ECA_wecA (TIGR02380)GT_WecA_like (cd06853)→ |
| Mab-43-1 | NZ_AUMY01000010.1 | + | 60813 | 60853 | **RNA**→ rpmE (PRK00019)Ribosomal_L31 (pfam01197)→ PrfA (COG0216)prfA (PRK00591)→ HemK (COG2890)PRK09328 (PRK09328)→ SUA5 (COG0009)TIGR00057 (TIGR00057)→ ECA_wecA (TIGR02380)GT_WecA_like (cd06853)→ |
| Mab-5-1 | NZ_AKVA01000006.1 | + | 237374 | 237414 | **RNA**→ rpmE (PRK00019)Ribosomal_L31 (pfam01197)→ PrfA (COG0216)prfA (PRK00591)→ HemK (COG2890)PRK09328 (PRK09328)→ SUA5 (COG0009)TIGR00057 (TIGR00057)→ ECA_wecA (TIGR02380)GT_WecA_like (cd06853)→ |
| Mab-8-1 | NZ_AKVC01000005.1 | + | 141628 | 141668 | **RNA**→ rpmE (PRK00019)Ribosomal_L31 (pfam01197)→ PrfA (COG0216)prfA (PRK00591)→ HemK (COG2890)PRK09328 (PRK09328)→ SUA5 (COG0009)TIGR00057 (TIGR00057)→ ECA_wecA (TIGR02380)GT_WecA_like (cd06853)→ |
| Mab-14-1 | NZ_AKTV01000003.1 | + | 395176 | 395216 | **RNA**→ rpmE (PRK00019)Ribosomal_L31 (pfam01197)→ PrfA (COG0216)prfA (PRK00591)→ HemK (COG2890)PRK09328 (PRK09328)→ SUA5 (COG0009)TIGR00057 (TIGR00057)→ ECA_wecA (TIGR02380)GT_WecA_like (cd06853)→ |

| | | | | | |
|---|---|---|---|---|---|
| Mab-12-1 | NZ_AKUT01000005.1 | + | 138569 | 138609 | **RNA**→ rpmE (PRK00019)Ribosomal_L31 (pfam01197)→ PrfA (COG0216)prfA (PRK00591)→ HemK (COG2890)PRK09328 (PRK09328)→ SUA5 (COG0009)TIGR00057 (TIGR00057)→ ECA_wecA (TIGR02380)GT_WecA_like (cd06853)→ |
| Mab-11-1 | NZ_AKVE01000002.1 | + | 138525 | 138565 | **RNA**→ rpmE (PRK00019)Ribosomal_L31 (pfam01197)→ PrfA (COG0216)prfA (PRK00591)→ HemK (COG2890)PRK09328 (PRK09328)→ SUA5 (COG0009)TIGR00057 (TIGR00057)→ ECA_wecA (TIGR02380)GT_WecA_like (cd06853)→ |
| Mab-7-1 | NZ_AKVB01000004.1 | + | 141663 | 141703 | **RNA**→ rpmE (PRK00019)Ribosomal_L31 (pfam01197)→ PrfA (COG0216)prfA (PRK00591)→ HemK (COG2890)PRK09328 (PRK09328)→ SUA5 (COG0009)TIGR00057 (TIGR00057)→ ECA_wecA (TIGR02380)GT_WecA_like (cd06853)→ |
| Mab-10-1 | NZ_AKVD01000003.1 | + | 1145427 | 1145467 | **RNA**→ rpmE (PRK00019)Ribosomal_L31 (pfam01197)→ PrfA (COG0216)prfA (PRK00591)→ HemK (COG2890)PRK09328 (PRK09328)→ SUA5 (COG0009)TIGR00057 (TIGR00057)→ ECA_wecA (TIGR02380)GT_WecA_like (cd06853)→ |
| Mab-4-1 | NZ_AKUZ01000006.1 | + | 53482 | 53522 | **RNA**→ rpmE (PRK00019)Ribosomal_L31 (pfam01197)→ PrfA (COG0216)prfA (PRK00591)→ HemK (COG2890)PRK09328 (PRK09328)→ SUA5 (COG0009)TIGR00057 (TIGR00057)→ SUA5 (COG0009)TIGR00057 (TIGR00057)→ ECA_wecA (TIGR02380)GT_WecA_like (cd06853)→ |
| Mab-13-1 | NZ_AKTU01000004.1 | + | 395153 | 395193 | **RNA**→ rpmE (PRK00019)Ribosomal_L31 (pfam01197)→ PrfA (COG0216)prfA (PRK00591)→ HemK (COG2890)PRK09328 (PRK09328)→ SUA5 (COG0009)TIGR00057 (TIGR00057)→ ECA_wecA (TIGR02380)GT_WecA_like (cd06853)→ |
| Mab-6-1 | NZ_AKUP01000009.1 | + | 141650 | 141690 | **RNA**→ rpmE (PRK00019)Ribosomal_L31 (pfam01197)→ PrfA (COG0216)prfA (PRK00591)→ HemK (COG2890)PRK09328 (PRK09328)→ SUA5 (COG0009)TIGR00057 (TIGR00057)→ ECA_wecA (TIGR02380)GT_WecA_like (cd06853)→ |
| Mab-3-1 | NZ_AKUY01000014.1 | + | 53426 | 53466 | **RNA**→ rpmE (PRK00019)Ribosomal_L31 (pfam01197)→ PrfA (COG0216)prfA (PRK00591)→ HemK (COG2890)PRK09328 (PRK09328)→ SUA5 (COG0009)TIGR00057 (TIGR00057)→ ECA_wecA (TIGR02380)GT_WecA_like (cd06853)→ |
| Mab-2-1 | NZ_AKUX01000006.1 | + | 141637 | 141677 | **RNA**→ rpmE (PRK00019)Ribosomal_L31 (pfam01197)→ PrfA (COG0216)prfA (PRK00591)→ HemK (COG2890)PRK09328 (PRK09328)→ SUA5 (COG0009)TIGR00057 (TIGR00057)→ ECA_wecA (TIGR02380)GT_WecA_like (cd06853)→ |
| Mab-27-1 | NZ_AKUS01000005.1 | + | 141661 | 141701 | **RNA**→ rpmE (PRK00019)Ribosomal_L31 (pfam01197)→ PrfA (COG0216)prfA (PRK00591)→ HemK (COG2890)PRK09328 (PRK09328)→ SUA5 (COG0009)TIGR00057 (TIGR00057)→ ECA_wecA (TIGR02380)GT_WecA_like (cd06853)→ |
| Mab-26-1 | NZ_AKUR01000007.1 | + | 141644 | 141684 | **RNA**→ rpmE (PRK00019)Ribosomal_L31 (pfam01197)→ PrfA (COG0216)prfA (PRK00591)→ HemK (COG2890)PRK09328 (PRK09328)→ SUA5 (COG0009)TIGR00057 (TIGR00057)→ ECA_wecA (TIGR02380)GT_WecA_like (cd06853)→ |
| Mab-15-1 | NZ_AKTW01000001.1 | + | 847452 | 847492 | **RNA**→ rpmE (PRK00019)Ribosomal_L31 (pfam01197)→ PrfA (COG0216)prfA (PRK00591)→ HemK (COG2890)PRK09328 (PRK09328)→ SUA5 (COG0009)TIGR00057 (TIGR00057)→ ECA_wecA (TIGR02380)GT_WecA_like (cd06853)→ |
| Mab-40-1 | NZ_AJGG01000034.1 | - | 91946 | 91906 | **RNA**→ rpmE (PRK00019)Ribosomal_L31 (pfam01197)→ PrfA (COG0216)prfA (PRK00591)→ HemK (COG2890)PRK09328 (PRK09328)→ SUA5 (COG0009)TIGR00057 (TIGR00057)→ ECA_wecA (TIGR02380)GT_WecA_like (cd06853)→ |
| Mab-39-1 | NZ_AJGF01000010.1 | + | 316074 | 316114 | **RNA**→ rpmE (PRK00019)Ribosomal_L31 (pfam01197)→ PrfA (COG0216)prfA (PRK00591)→ HemK (COG2890)PRK09328 (PRK09328)→ SUA5 (COG0009)TIGR00057 (TIGR00057)→ ECA_wecA (TIGR02380)GT_WecA_like (cd06853)→ |
| Mab-28-1 | NZ_AKUG01000004.1 | + | 141618 | 141658 | **RNA**→ rpmE (PRK00019)Ribosomal_L31 (pfam01197)→ PrfA (COG0216)prfA (PRK00591)→ HemK (COG2890)PRK09328 (PRK09328)→ SUA5 (COG0009)TIGR00057 (TIGR00057)→ ECA_wecA (TIGR02380)GT_WecA_like (cd06853)→ |
| Mab-34-1 | NZ_AKVT01000011.1 | - | 15076 | 15036 | **RNA**→ rpmE (PRK00019)Ribosomal_L31 (pfam01197)→ PrfA (COG0216)prfA (PRK00591)→ HemK (COG2890)PRK09328 (PRK09328)→ SUA5 (COG0009)TIGR00057 (TIGR00057)→ ECA_wecA (TIGR02380)GT_WecA_like (cd06853)→ |
| Mab-29-1 | NZ_AKUH01000010.1 | + | 53471 | 53511 | **RNA**→ rpmE (PRK00019)Ribosomal_L31 (pfam01197)→ PrfA (COG0216)prfA (PRK00591)→ HemK (COG2890)PRK09328 (PRK09328)→ SUA5 (COG0009)TIGR00057 (TIGR00057)→ ECA_wecA (TIGR02380)GT_WecA_like (cd06853)→ |
| Mab-24-1 | NZ_AKUE01000004.1 | + | 53462 | 53502 | **RNA**→ rpmE (PRK00019)Ribosomal_L31 (pfam01197)→ PrfA (COG0216)prfA (PRK00591)→ HemK (COG2890)PRK09328 (PRK09328)→ SUA5 (COG0009)TIGR00057 (TIGR00057)→ ECA_wecA (TIGR02380)GT_WecA_like (cd06853)→ |
| Mab-25-1 | NZ_AKUF01000006.1 | + | 464209 | 464249 | **RNA**→ rpmE (PRK00019)Ribosomal_L31 (pfam01197)→ PrfA (COG0216)prfA (PRK00591)→ HemK (COG2890)PRK09328 (PRK09328)→ SUA5 (COG0009)TIGR00057 (TIGR00057)→ SUA5 (COG0009)TIGR00057 (TIGR00057)→ ECA_wecA (TIGR02380)GT_WecA_like (cd06853)→ |

| | | | | | |
|---|---|---|---|---|---|
| Mka-1-1 | NZ_JASX01000032.1 | + | 142131 | 142171 | **RNA**→ rpmE (PRK00019)Ribosomal_L31 (pfam01197)→ PrfA (COG0216)prfA (PRK00591)→ HemK (COG2890)PRK09328 (PRK09328)→ SUA5 (COG0009)TIGR00057 (TIGR00057)→ ECA_wecA (TIGR02380)GT_WecA_like (cd06853)other_trbK (TIGR04360)→ |
| Mab-67-1 | NZ_JAOU01000003.1 | + | 1978072 | 1978112 | **RNA**→ rpmE (PRK00019)Ribosomal_L31 (pfam01197)→ PrfA (COG0216)prfA (PRK00591)→ HemK (COG2890)PRK09328 (PRK09328)→ SUA5 (COG0009)TIGR00057 (TIGR00057)→ ECA_wecA (TIGR02380)GT_WecA_like (cd06853)→ |
| Mab-62-1 | NZ_JAOY01000001.1 | + | 3994798 | 3994838 | **RNA**→ rpmE (PRK00019)Ribosomal_L31 (pfam01197)→ PrfA (COG0216)prfA (PRK00591)→ HemK (COG2890)PRK09328 (PRK09328)→ SUA5 (COG0009)TIGR00057 (TIGR00057)→ ECA_wecA (TIGR02380)GT_WecA_like (cd06853)→ |
| Mab-68-1 | NZ_JAOT01000003.1 | + | 174381 | 174421 | **RNA**→ rpmE (PRK00019)Ribosomal_L31 (pfam01197)→ PrfA (COG0216)prfA (PRK00591)→ HemK (COG2890)PRK09328 (PRK09328)→ SUA5 (COG0009)TIGR00057 (TIGR00057)→ ECA_wecA (TIGR02380)GT_WecA_like (cd06853)→ |
| Mch-3-1 | NZ_JAOI01000002.1 | - | 111334 | 111294 | **RNA**→ rpmE (PRK00019)Ribosomal_L31 (pfam01197)→ PrfA (COG0216)prfA (PRK00591)→ HemK (COG2890)PRK09328 (PRK09328)→ SUA5 (COG0009)TIGR00057 (TIGR00057)→ ECA_wecA (TIGR02380)GT_WecA_like (cd06853)→ |
| Mab-1-1 | NC_010397.1 | + | 1446856 | 1446896 | **RNA**→ rpmE (PRK00019)Ribosomal_L31 (pfam01197)→ PrfA (COG0216)prfA (PRK00591)→ HemK (COG2890)PRK09328 (PRK09328)→ SUA5 (COG0009)TIGR00057 (TIGR00057)→ ECA_wecA (TIGR02380)GT_WecA_like (cd06853)→ |
| Mab-60-1 | NZ_HE817967.1 | + | 1449314 | 1449354 | **RNA**→ rpmE (PRK00019)Ribosomal_L31 (pfam01197)→ PrfA (COG0216)prfA (PRK00591)→ HemK (COG2890)PRK09328 (PRK09328)→ SUA5 (COG0009)TIGR00057 (TIGR00057)→ ECA_wecA (TIGR02380)GT_WecA_like (cd06853)→ |
| Mab-30-1 | NZ_KB290571.1 | + | 2284355 | 2284395 | **RNA**→ rpmE (PRK00019)Ribosomal_L31 (pfam01197)→ PrfA (COG0216)prfA (PRK00591)→ HemK (COG2890)PRK09328 (PRK09328)→ SUA5 (COG0009)TIGR00057 (TIGR00057)→ ECA_wecA (TIGR02380)GT_WecA_like (cd06853)→ |
| Mab-1-166 | NZ_LGCJ01000014.1 | + | 53469 | 53509 | **RNA**→ rpmE (PRK00019)Ribosomal_L31 (pfam01197)→ PrfA (COG0216)prfA (PRK00591)→ HemK (COG2890)PRK09328 (PRK09328)→ SUA5 (COG0009)TIGR00057 (TIGR00057)→ ECA_wecA (TIGR02380)GT_WecA_like (cd06853)→ |
| Mab-71-2 | NZ_CP009613.1 | + | 1445609 | 1445649 | **RNA**→ rpmE (PRK00019)Ribosomal_L31 (pfam01197)→ PrfA (COG0216)prfA (PRK00591)→ HemK (COG2890)PRK09328 (PRK09328)→ SUA5 (COG0009)TIGR00057 (TIGR00057)→ ECA_wecA (TIGR02380)GT_WecA_like (cd06853)→ |
| Mab-72-1 | NZ_CP009407.1 | + | 1431380 | 1431420 | **RNA**→ rpmE (PRK00019)Ribosomal_L31 (pfam01197)→ PrfA (COG0216)prfA (PRK00591)→ HemK (COG2890)PRK09328 (PRK09328)→ SUA5 (COG0009)TIGR00057 (TIGR00057)→ ECA_wecA (TIGR02380)GT_WecA_like (cd06853)→ |
| Mab-71-3 | NZ_CP009408.1 | + | 1446345 | 1446385 | **RNA**→ rpmE (PRK00019)Ribosomal_L31 (pfam01197)→ PrfA (COG0216)prfA (PRK00591)→ HemK (COG2890)PRK09328 (PRK09328)→ SUA5 (COG0009)TIGR00057 (TIGR00057)→ ECA_wecA (TIGR02380)GT_WecA_like (cd06853)→ |
| Mab-69-1 | NZ_CP012044.1 | + | 1577915 | 1577955 | **RNA**→ rpmE (PRK00019)Ribosomal_L31 (pfam01197)→ PrfA (COG0216)prfA (PRK00591)→ HemK (COG2890)PRK09328 (PRK09328)→ SUA5 (COG0009)TIGR00057 (TIGR00057)→ ECA_wecA (TIGR02380)GT_WecA_like (cd06853)→ |
| Mab-59-1 | NZ_JAOF01000001.1 | + | 1494986 | 1495026 | **RNA**→ rpmE (PRK00019)Ribosomal_L31 (pfam01197)→ PrfA (COG0216)prfA (PRK00591)→ HemK (COG2890)PRK09328 (PRK09328)→ SUA5 (COG0009)TIGR00057 (TIGR00057)→ ECA_wecA (TIGR02380)GT_WecA_like (cd06853)→ |
| Mab-63-1 | NZ_JAOX01000001.1 | + | 2028893 | 2028933 | **RNA**→ rpmE (PRK00019)Ribosomal_L31 (pfam01197)→ PrfA (COG0216)prfA (PRK00591)→ HemK (COG2890)PRK09328 (PRK09328)→ SUA5 (COG0009)TIGR00057 (TIGR00057)→ ECA_wecA (TIGR02380)GT_WecA_like (cd06853)→ |
| Mfo-2-1 | NZ_JASW01000013.1 | - | 78048 | 78008 | **RNA**→ rpmE (PRK00019)Ribosomal_L31 (pfam01197)→ PrfA (COG0216)prfA (PRK00591)→ HemK (COG2890)PRK09328 (PRK09328)→ SUA5 (COG0009)TIGR00057 (TIGR00057)→ ECA_wecA (TIGR02380)GT_WecA_like (cd06853)other_trbK (TIGR04360)→ |
| Mma-1-1 | NZ_AHAR01000007.1 | + | 396447 | 396487 | **RNA**→ rpmE (PRK00019)Ribosomal_L31 (pfam01197)→ rpmE (PRK00019)Ribosomal_L31 (pfam01197)→ PrfA (COG0216)prfA (PRK00591)→ PrfA (COG0216)prfA (PRK00591)→ HemK (COG2890)PRK09328 (PRK09328)→ HemK (COG2890)RF_mod_PrmC (TIGR03534)→ HemK (COG2890)RF_mod_PrmC (TIGR03534)→ HemK (COG2890)Methyltransf_25 (pfam13649)→ HemK (COG2890)Methyltransf_25 (pfam13649)→ SUA5 (COG0009)TIGR00057 (TIGR00057)→ SUA5 (COG0009)TIGR00057 (TIGR00057)→ ECA_wecA (TIGR02380)GT_WecA_like (cd06853)→ ECA_wecA (TIGR02380)GT_WecA_like (cd06853)→ |

| | | | | | |
|---|---|---|---|---|---|
| Mab-56-1 | NZ_AKVF01000003.1 | + | 670171 | 670211 | **RNA**→ rpmE (PRK00019)Ribosomal_L31 (pfam01197)→ PrfA (COG0216)prfA (PRK00591)→ HemK (COG2890)PRK09328 (PRK09328)→ SUA5 (COG0009)TIGR00057 (TIGR00057)→ ECA_wecA (TIGR02380)GT_WecA_like (cd06853)→ |
| Mab-74-1 | NZ_AP014547.1 | + | 1453421 | 1453461 | **RNA**→ rpmE (PRK00019)Ribosomal_L31 (pfam01197)→ PrfA (COG0216)prfA (PRK00591)→ HemK (COG2890)PRK09328 (PRK09328)→ SUA5 (COG0009)TIGR00057 (TIGR00057)→ ECA_wecA (TIGR02380)GT_WecA_like (cd06853)→ |
| env-366 | JGI12273J12029_10114025 | + | 13 | 58 | **RNA**→ rpmE (PRK00019)Ribosomal_L31 (pfam01197)→ |
| env-367 | JGI12396J12026_1204897 | + | 52 | 134 | **RNA**→ ←hypo |
| env-368 | FNTS_GKKEVUV02IFQEU | - | 183 | 146 | **RNA**→ hypo→ |
| env-369 | JGI20243J16304_127292 | - | 275 | 225 | **RNA**→ ←hypo |
| Tsp-1-1 | NZ_HE997626.1 | - | 24625 | 24578 | **RNA**→ rpmE (PRK00019)Ribosomal_L31 (pfam01197)→ PrfA (COG0216)prfA (PRK00591)→ HemK (COG2890)PRK09328 (PRK09328)→ SUA5 (COG0009)TIGR00057 (TIGR00057)→ |
| Tpa-1-1 | NC_014158.1 | + | 1269385 | 1269433 | **RNA**→ rpmE (PRK00019)Ribosomal_L31 (pfam01197)→ PrfA (COG0216)prfA (PRK00591)→ HemK (COG2890)PRK09328 (PRK09328)→ SUA5 (COG0009)TIGR00057 (TIGR00057)→ |
| Are-1-1 | NZ_JZKF01000016.1 | + | 13468 | 13514 | **RNA**→ rpmE (PRK00019)Ribosomal_L31 (pfam01197)→ PrfA (COG0216)prfA (PRK00591)→ |
| Nca-3-1 | NZ_JOAG01000005.1 | - | 167647 | 167606 | **RNA**→ rpmE (PRK00019)Ribosomal_L31 (pfam01197)→ PrfA (COG0216)prfA (PRK00591)→ HemK (COG2890)PRK09328 (PRK09328)→ SUA5 (COG0009)TIGR00057 (TIGR00057)→ |
| env-370 | botACOD_FV90NF402IKWZZ | + | 231 | 268 | **RNA**→ rpmE (PRK00019)rpmE2 (PRK01678)Ribosomal_L31 (pfam01197)→ |
| env-371 | JGI24735J21928_10271683 | - | 398 | 355 | **RNA**→ hypo→ |
| Mph-1-1 | NZ_AJFJ01000049.1 | - | 15218 | 15165 | **RNA**→ rpmE (PRK00019)Ribosomal_L31 (pfam01197)→ rpmE (PRK00019)Ribosomal_L31 (pfam01197)→ PrfA (COG0216)prfA (PRK00591)→ PrfA (COG0216)prfA (PRK00591)→ HemK (COG2890)PRK09328 (PRK09328)→ HemK (COG2890)PRK09328 (PRK09328)→ SUA5 (COG0009)TIGR00057 (TIGR00057)→ SUA5 (COG0009)TIGR00057 (TIGR00057)→ GT_WecA_like (cd06853)mraY (PRK00108)→ GT_WecA_like (cd06853)mraY (PRK00108)→ |
| env-372 | JGI12635J15846_10020978 | + | 510 | 564 | **RNA**→ rpmE (PRK00019)Ribosomal_L31 (pfam01197)→ |
| env-373 | JGI12682J13319_1038893 | + | 14 | 68 | **RNA**→ ←hypo |
| env-374 | JGI12714J13572_1002355 | - | 413 | 359 | **RNA**→ ←hypo |
| env-375 | JGI12668J13544_1058045 | + | 205 | 259 | **RNA**→ hypo→ |
| env-376 | JGI12712J15308_10190831 | - | 84 | 30 | **RNA**→ ←hypo |
| env-377 | JGI12630J15595_10175159 | - | 290 | 236 | **RNA**→ rpmE (PRK00019)Ribosomal_L31 (pfam01197)→ |
| env-378 | JGI12682J13319_1063338 | + | 232 | 286 | **RNA**→ |
| env-379 | JGI20180J14839_1008125 | + | 143 | 208 | **RNA**→ rpmE (PRK00019)Ribosomal_L31 (pfam01197)→ |
| env-380 | JGI12712J15308_10202108 | - | 470 | 415 | **RNA**→ rpmE (PRK00019)Ribosomal_L31 (pfam01197)→ |
| Sgr-9-1 | NZ_JOBE01000014.1 | - | 122236 | 122189 | **RNA**→ rpmE (PRK00019)Ribosomal_L31 (pfam01197)→ PrfA (COG0216)prfA (PRK00591)→ |
| Sex-1-1 | NZ_JOFB01000019.1 | + | 104773 | 104820 | **RNA**→ rpmE (PRK00019)Ribosomal_L31 (pfam01197)→ PrfA (COG0216)prfA (PRK00591)→ |
| Sex-1-2 | NZ_JNZP01000018.1 | - | 213017 | 212970 | **RNA**→ rpmE (PRK00019)Ribosomal_L31 (pfam01197)→ PrfA (COG0216)prfA (PRK00591)→ |
| Sex-2-1 | NZ_AZSS01000360.1 | - | 24911 | 24864 | **RNA**→ rpmE (PRK00019)Ribosomal_L31 (pfam01197)→ PrfA (COG0216)prfA (PRK00591)→ HemK (COG2890)PRK09328 (PRK09328)→ SUA5 (COG0009)TIGR00057 (TIGR00057)→ etp (PRK11391)Wzb (COG0394)→ |
| Ssp-115-1 | NZ_JOGX01000011.1 | - | 107370 | 107323 | **RNA**→ rpmE (PRK00019)Ribosomal_L31 (pfam01197)→ PrfA (COG0216)prfA (PRK00591)→ HemK (COG2890)PRK09328 (PRK09328)→ SUA5 (COG0009)TIGR00057 (TIGR00057)→ etp (PRK11391)Wzb (COG0394)→ |
| Ssp-105-1 | NZ_JOHZ01000002.1 | + | 13150 | 13197 | **RNA**→ rpmE (PRK00019)Ribosomal_L31 (pfam01197)→ PrfA (COG0216)prfA (PRK00591)→ HemK (COG2890)PRK09328 (PRK09328)→ SUA5 (COG0009)TIGR00057 (TIGR00057)→ etp (PRK11391)Wzb (COG0394)→ |
| Mgl-2-1 | NZ_JOFO01000002.1 | - | 183750 | 183703 | **RNA**→ rpmE (PRK00019)Ribosomal_L31 (pfam01197)→ PrfA (COG0216)prfA (PRK00591)→ HemK (COG2890)PRK09328 (PRK09328)→ SUA5 (COG0009)TIGR00057 (TIGR00057)→ etp (PRK11391)Wzb (COG0394)→ |
| Sgr-12-2 | NZ_JNZT01000101.1 | - | 3778 | 3731 | **RNA**→ rpmE (PRK00019)Ribosomal_L31 (pfam01197)→ PrfA (COG0216)prfA (PRK00591)→ HemK (COG2890)PRK09328 (PRK09328)→ SUA5 (COG0009)TIGR00057 (TIGR00057)→ etp (PRK11391)Wzb (COG0394)→ |
| Sgr-12-3 | NZ_JOEN01000032.1 | - | 52357 | 52310 | **RNA**→ rpmE (PRK00019)Ribosomal_L31 (pfam01197)→ PrfA (COG0216)prfA (PRK00591)→ HemK (COG2890)PRK09328 (PRK09328)→ SUA5 (COG0009)TIGR00057 (TIGR00057)→ etp (PRK11391)Wzb (COG0394)→ |
| Fsp-14-1 | NZ_KQ031391.1 | - | 2026125 | 2026078 | **RNA**→ rpmE (PRK00019)Ribosomal_L31 (pfam01197)→ PrfA (COG0216)prfA (PRK00591)→ HemK (COG2890)PRK09328 (PRK09328)→ |

| | | | | | |
|---|---|---|---|---|---|
| Fsa-1-1 | NZ_KI912266.1 | + | 2655277 | 2655323 | **RNA**→ rpmE (PRK00019)Ribosomal_L31 (pfam01197)→ |
| Gar-2-1 | NZ_AXWO01000001.1 | - | 345857 | 345815 | **RNA**→ rpmE (PRK00019)Ribosomal_L31 (pfam01197)→ PrfA (COG0216)prfA (PRK00591)→ |
| | | | | | HemK (COG2890)PRK09328 (PRK09328)→ SUA5 (COG0009)TIGR00057 (TIGR00057)→ |
| | | | | | LMWPc (smart00226)Wzb (COG0394)→ hypo→ |
| Spa-31-1 | NZ_AZWJ01000013.1 | + | 135671 | 135726 | **RNA**→ rpmE (PRK00019)Ribosomal_L31 (pfam01197)→ |
| Spa-19-1 | NZ_KB900388.1 | - | 2477139 | 2477084 | **RNA**→ rpmE (PRK00019)Ribosomal_L31 (pfam01197)→ rpmE (PRK00019)Ribosomal_L31 (pfam01197)→ |
| Spa-3-1 | NZ_KB894983.1 | - | 51486 | 51431 | **RNA**→ hypo→ rpmE (PRK00019)Ribosomal_L31 (pfam01197)→ |
| Spa-39-1 | NZ_AUKU01000004.1 | - | 50781 | 50726 | **RNA**→ rpmE (PRK00019)Ribosomal_L31 (pfam01197)→ |
| Spa-29-1 | NZ_AZVZ01000001.1 | + | 325608 | 325663 | **RNA**→ rpmE (PRK00019)Ribosomal_L31 (pfam01197)→ |
| Spa-26-1 | NZ_AZWE01000001.1 | - | 50846 | 50791 | **RNA**→ rpmE (PRK00019)Ribosomal_L31 (pfam01197)→ |
| Spa-23-1 | NZ_AZWP01000001.1 | + | 356983 | 357038 | **RNA**→ rpmE (PRK00019)Ribosomal_L31 (pfam01197)→ |
| Spa-22-1 | NZ_AZWN01000007.1 | + | 156487 | 156542 | **RNA**→ rpmE (PRK00019)Ribosomal_L31 (pfam01197)→ |
| Spa-37-1 | NZ_KI911485.1 | - | 160156 | 160101 | **RNA**→ rpmE (PRK00019)Ribosomal_L31 (pfam01197)→ |
| Spa-16-1 | NZ_KB905359.1 | + | 4219288 | 4219343 | **RNA**→ rpmE (PRK00019)Ribosomal_L31 (pfam01197)→ rpmE (PRK00019)Ribosomal_L31 (pfam01197)→ |
| Spa-40-1 | NZ_KB897477.1 | - | 50762 | 50707 | **RNA**→ rpmE (PRK00019)Ribosomal_L31 (pfam01197)→ |
| Spa-14-1 | NZ_KB897006.1 | - | 50602 | 50547 | **RNA**→ rpmE (PRK00019)Ribosomal_L31 (pfam01197)→ rpmE (PRK00019)Ribosomal_L31 (pfam01197)→ |
| Spa-35-1 | NZ_AZWM01000001.1 | - | 50720 | 50665 | **RNA**→ rpmE (PRK00019)Ribosomal_L31 (pfam01197)→ |
| Spa-12-1 | NZ_KB896839.1 | + | 419496 | 419551 | **RNA**→ rpmE (PRK00019)Ribosomal_L31 (pfam01197)→ rpmE (PRK00019)Ribosomal_L31 (pfam01197)→ |
| Spa-25-1 | NZ_AZWC01000002.1 | - | 50728 | 50673 | **RNA**→ rpmE (PRK00019)Ribosomal_L31 (pfam01197)→ |
| Spa-34-1 | NZ_AZVW01000001.1 | - | 50891 | 50836 | **RNA**→ rpmE (PRK00019)Ribosomal_L31 (pfam01197)→ |
| Spa-21-1 | NZ_AZWO01000001.1 | + | 372780 | 372835 | **RNA**→ rpmE (PRK00019)Ribosomal_L31 (pfam01197)→ |
| Spa-1-1 | NZ_KB904824.1 | - | 50851 | 50796 | **RNA**→ rpmE (PRK00019)Ribosomal_L31 (pfam01197)→ rpmE (PRK00019)Ribosomal_L31 (pfam01197)→ |
| Nfa-1-1 | NC_006361.1 | + | 1164117 | 1164159 | **RNA**→ rpmE (PRK00019)Ribosomal_L31 (pfam01197)→ PrfA (COG0216)prfA (PRK00591)→ |
| | | | | | HemK (COG2890)PRK09328 (PRK09328)→ SUA5 (COG0009)TIGR00057 (TIGR00057)→ |
| | | | | | ECA_wecA (TIGR02380)GT_WecA_like (cd06853)→ hypo→ |
| Msp-7-1 | NC_008146.1 | - | 4140648 | 4140593 | **RNA**→ rpmE (PRK00019)Ribosomal_L31 (pfam01197)→ PrfA (COG0216)prfA (PRK00591)→ |
| | | | | | HemK (COG2890)PRK09328 (PRK09328)→ SUA5 (COG0009)TIGR00057 (TIGR00057)→ |
| | | | | | GT_WecA_like (cd06853)mraY (PRK00108)→ |
| Msp-5-1 | NC_009077.1 | - | 4101693 | 4101638 | **RNA**→ rpmE (PRK00019)Ribosomal_L31 (pfam01197)→ PrfA (COG0216)prfA (PRK00591)→ |
| | | | | | HemK (COG2890)PRK09328 (PRK09328)→ SUA5 (COG0009)TIGR00057 (TIGR00057)→ |
| | | | | | GT_WecA_like (cd06853)mraY (PRK00108)→ |
| Msp-6-1 | NC_008705.1 | - | 4175130 | 4175075 | **RNA**→ rpmE (PRK00019)Ribosomal_L31 (pfam01197)→ PrfA (COG0216)prfA (PRK00591)→ |
| | | | | | HemK (COG2890)PRK09328 (PRK09328)→ SUA5 (COG0009)TIGR00057 (TIGR00057)→ |
| | | | | | GT_WecA_like (cd06853)mraY (PRK00108)→ |
| Msp-19-1 | NZ_JQKU01000014.1 | + | 205976 | 206017 | **RNA**→ rpmE (PRK00019)Ribosomal_L31 (pfam01197)→ |
| Mll-1-1 | NZ_JXST01000033.1 | - | 15173 | 15132 | **RNA**→ rpmE (PRK00019)Ribosomal_L31 (pfam01197)→ |
| Msp-3-1 | NZ_KB911477.1 | + | 59673 | 59714 | **RNA**→ rpmE (PRK00019)Ribosomal_L31 (pfam01197)→ PrfA (COG0216)prfA (PRK00591)→ |
| | | | | | HemK (COG2890)PRK09328 (PRK09328)→ SUA5 (COG0009)TIGR00057 (TIGR00057)→ |
| | | | | | GT_WecA_like (cd06853)mraY (PRK00108)→ |
| Msp-20-1 | NZ_JMLM01000001.1 | - | 15820 | 15779 | **RNA**→ rpmE (PRK00019)Ribosomal_L31 (pfam01197)→ PrfA (COG0216)prfA (PRK00591)→ |
| | | | | | HemK (COG2890)PRK09328 (PRK09328)→ SUA5 (COG0009)TIGR00057 (TIGR00057)→ |
| | | | | | GT_WecA_like (cd06853)mraY (PRK00108)→ |
| Nsp-7-1 | NZ_JOJF01000034.1 | - | 15463 | 15396 | **RNA**→ rpmE (PRK00019)Ribosomal_L31 (pfam01197)→ PrfA (COG0216)prfA (PRK00591)→ |
| | | | | | HemK (COG2890)PRK09328 (PRK09328)→ SUA5 (COG0009)TIGR00057 (TIGR00057)→ |
| | | | | | ECA_wecA (TIGR02380)GT_WecA_like (cd06853)→ hypo→ |
| Nve-1-1 | NZ_BAGM01000030.1 | + | 22539 | 22606 | **RNA**→ rpmE (PRK00019)Ribosomal_L31 (pfam01197)→ PrfA (COG0216)prfA (PRK00591)→ |
| | | | | | HemK (COG2890)PRK09328 (PRK09328)→ hypo→ hypo→ SUA5 (COG0009)TIGR00057 (TIGR00057)→ |
| | | | | | ECA_wecA (TIGR02380)GT_WecA_like (cd06853)→ hypo→ |
| Nao-1-1 | NZ_BAFQ01000044.1 | + | 45685 | 45752 | **RNA**→ rpmE (PRK00019)Ribosomal_L31 (pfam01197)→ PrfA (COG0216)prfA (PRK00591)→ |
| | | | | | HemK (COG2890)PRK09328 (PRK09328)→ SUA5 (COG0009)TIGR00057 (TIGR00057)→ |
| | | | | | ECA_wecA (TIGR02380)GT_WecA_like (cd06853)→ hypo→ |
| Nce-1-1 | NZ_BAFW01000042.1 | + | 8980 | 9047 | **RNA**→ rpmE (PRK00019)Ribosomal_L31 (pfam01197)→ PrfA (COG0216)prfA (PRK00591)→ |
| | | | | | HemK (COG2890)PRK09328 (PRK09328)→ SUA5 (COG0009)TIGR00057 (TIGR00057)→ |
| | | | | | ECA_wecA (TIGR02380)GT_WecA_like (cd06853)→ hypo→ |
| env-381 | HBC_scaff_10249466 | + | 482 | 527 | **RNA**→ |

| | | | | | |
|---|---|---|---|---|---|
| env-382 | HBC_scaff_10219859 | - | 180 | 135 | **RNA**→ |
| env-383 | HBC_ctgs_1182059 | - | 180 | 135 | **RNA**→ |
| Bas-2-1 | NZ_KQ033859.1 | - | 419564 | 419519 | **RNA**→ rpmE (PRK00019)Ribosomal_L31 (pfam01197)→ |
| env-384 | HBC_scaff_10141359 | - | 127 | 82 | **RNA**→ hypo→ |
| env-385 | HBC_ctgs_1281286 | - | 127 | 82 | **RNA**→ hypo→ |
| env-386 | HBC_scaff_10035491 | + | 3410 | 3455 | **RNA**→ |
| env-387 | HBC_scaff_10160097 | - | 295 | 250 | **RNA**→ CBS (smart00116)IMPDH (pfam00478)CBS (pfam00571)→ |
| env-388 | HBC_ctgs_1268800 | + | 249 | 294 | **RNA**→ |
| env-389 | HBC_ctgs_1257867 | - | 295 | 250 | **RNA**→ CBS (smart00116)IMPDH (pfam00478)CBS (pfam00571)→ |
| env-390 | HBC_ctgs_1141466 | + | 482 | 527 | **RNA**→ |
| Bas-1-1 | NC_018720.1 | - | 1594861 | 1594816 | **RNA**→ rpmE (PRK00019)Ribosomal_L31 (pfam01197)→ |
| Bas-3-1 | NZ_JDTU01000007.1 | + | 50715 | 50760 | **RNA**→ rpmE (PRK00019)Ribosomal_L31 (pfam01197)→ |
| Bas-2-2 | NZ_KQ033885.1 | - | 1408476 | 1408431 | **RNA**→ rpmE (PRK00019)Ribosomal_L31 (pfam01197)→ |
| Bsp-3-1 | NZ_AWUO01000001.1 | - | 280667 | 280622 | **RNA**→ rpmE (PRK00019)Ribosomal_L31 (pfam01197)→ |
| env-391 | BGMT1_NODE_527181 | - | 9150 | 9105 | **RNA**→ rpmE (PRK00019)Ribosomal_L31 (pfam01197)→ |
| env-392 | HBC_scaff_10281629 | + | 636 | 681 | **RNA**→ |
| env-393 | HBC_ctgs_1093268 | + | 636 | 681 | **RNA**→ |
| env-394 | HBC_scaff_10015902 | - | 112 | 67 | **RNA**→ |
| env-395 | HBC_ctgs_1269501 | + | 204 | 249 | **RNA**→ |
| env-396 | HBC_scaff_10298050 | - | 68 | 23 | **RNA**→ |
| env-397 | HBC_ctgs_1066607 | - | 68 | 23 | **RNA**→ |
| Bsp-2-1 | NZ_AWUN01000004.1 | - | 167539 | 167494 | **RNA**→ rpmE (PRK00019)Ribosomal_L31 (pfam01197)→ |
| Bas-2-3 | NZ_KQ034040.1 | - | 1594375 | 1594330 | **RNA**→ rpmE (PRK00019)Ribosomal_L31 (pfam01197)→ |
| Mma-2-1 | NZ_CCBF010000004.1 | - | 15106 | 15057 | **RNA**→ rpmE (PRK00019)Ribosomal_L31 (pfam01197)→ PrfA (COG0216)prfA (PRK00591)→<br>HemK (COG2890)PRK09328 (PRK09328)→ SUA5 (COG0009)TIGR00057 (TIGR00057)→<br>GT_WecA_like (cd06853)mraY (PRK00108)→ |
| Msp-2-1 | NZ_KB892705.1 | + | 457958 | 457995 | **RNA**→ rpmE (PRK00019)Ribosomal_L31 (pfam01197)→ PrfA (COG0216)prfA (PRK00591)→<br>PrfA (COG0216)prfA (PRK00591)→ HemK (COG2890)PRK09328 (PRK09328)→<br>SUA5 (COG0009)TIGR00057 (TIGR00057)→ GT_WecA_like (cd06853)mraY (PRK00108)→ |
| env-398 | JGI24736J21556_1296931 | - | 194 | 150 | **RNA**→ rpmE (PRK00019)Ribosomal_L31 (pfam01197)→ |
| env-399 | JGI24735J21928_10389060 | + | 23 | 67 | **RNA**→ ←hypo |
| Nji-1-1 | NZ_BAGB01000018.1 | + | 17431 | 17475 | **RNA**→ rpmE (PRK00019)Ribosomal_L31 (pfam01197)→ |
| Nno-1-1 | NZ_CP006850.1 | + | 1393179 | 1393243 | **RNA**→ rpmE (PRK00019)Ribosomal_L31 (pfam01197)→ |
| Nhi-1-1 | NZ_BAGA01000038.1 | + | 57963 | 58015 | **RNA**→ rpmE (PRK00019)Ribosomal_L31 (pfam01197)→ |
| Msp-1-1 | NZ_KB908258.1 | - | 178589 | 178551 | **RNA**→ rpmE (PRK00019)Ribosomal_L31 (pfam01197)→ PrfA (COG0216)prfA (PRK00591)→<br>HemK (COG2890)PRK09328 (PRK09328)→ SUA5 (COG0009)TIGR00057 (TIGR00057)→<br>GT_WecA_like (cd06853)mraY (PRK00108)→ |
| Msp-18-1 | NZ_AUWS01000091.1 | + | 11274 | 11310 | **RNA**→ rpmE (PRK00019)Ribosomal_L31 (pfam01197)→ PrfA (COG0216)prfA (PRK00591)→<br>HemK (COG2890)PRK09328 (PRK09328)→ SUA5 (COG0009)TIGR00057 (TIGR00057)→<br>ECA_wecA (TIGR02380)GT_WecA_like (cd06853)→ |
| Mfo-1-1 | NZ_JH814764.1 | - | 15543 | 15500 | **RNA**→ rpmE (PRK00019)Ribosomal_L31 (pfam01197)→ |
| Msp-25-1 | NZ_CP009914.1 | - | 4482052 | 4482009 | **RNA**→ rpmE (PRK00019)Ribosomal_L31 (pfam01197)→ |
| Mvu-1-1 | NZ_CCBG010000002.1 | + | 2131282 | 2131325 | **RNA**→ rpmE (PRK00019)Ribosomal_L31 (pfam01197)→ PrfA (COG0216)prfA (PRK00591)→<br>HemK (COG2890)PRK09328 (PRK09328)→ SUA5 (COG0009)TIGR00057 (TIGR00057)→<br>GT_WecA_like (cd06853)mraY (PRK00108)→ |
| Mse-1-1 | NZ_LDCO01000024.1 | + | 85147 | 85190 | **RNA**→ rpmE (PRK00019)Ribosomal_L31 (pfam01197)→ |
| Mse-1-2 | NZ_LDPU01000001.1 | + | 5382278 | 5382321 | **RNA**→ rpmE (PRK00019)Ribosomal_L31 (pfam01197)→ |
| Mco-1-1 | NZ_LFOD01000024.1 | + | 84808 | 84851 | **RNA**→ rpmE (PRK00019)Ribosomal_L31 (pfam01197)→ |
| Mfa-1-1 | NZ_HG964484.1 | + | 494652 | 494695 | **RNA**→ rpmE (PRK00019)Ribosomal_L31 (pfam01197)→ |
| Msp-23-1 | NZ_AUEQ01000008.1 | + | 477987 | 478030 | **RNA**→ rpmE (PRK00019)Ribosomal_L31 (pfam01197)→ PrfA (COG0216)prfA (PRK00591)→<br>HemK (COG2890)PRK09328 (PRK09328)→ SUA5 (COG0009)TIGR00057 (TIGR00057)→<br>GT_WecA_like (cd06853)mraY (PRK00108)→ |

| | | | | | |
|---|---|---|---|---|---|
| Mse-2-1 | NZ_HG322952.1 | - | 15319 | 15283 | **RNA**→ rpmE (PRK00019)Ribosomal_L31 (pfam01197)→ PrfA (COG0216)prfA (PRK00591)→ HemK (COG2890)PRK09328 (PRK09328)→ SUA5 (COG0009)TIGR00057 (TIGR00057)→ GT_WecA_like (cd06853)mraY (PRK00108)→ |
| Mse-3-1 | NZ_JTJW01000009.1 | - | 15412 | 15343 | **RNA**→ rpmE (PRK00019)Ribosomal_L31 (pfam01197)→ PrfA (COG0216)prfA (PRK00591)→ HemK (COG2890)PRK09328 (PRK09328)→ SUA5 (COG0009)TIGR00057 (TIGR00057)→ GT_WecA_like (cd06853)mraY (PRK00108)→ |
| Mse-3-2 | NZ_JTLZ01000009.1 | - | 15489 | 15420 | **RNA**→ rpmE (PRK00019)Ribosomal_L31 (pfam01197)→ PrfA (COG0216)prfA (PRK00591)→ HemK (COG2890)PRK09328 (PRK09328)→ SUA5 (COG0009)TIGR00057 (TIGR00057)→ GT_WecA_like (cd06853)mraY (PRK00108)→ |
| Msp-22-1 | NZ_JIAW01000013.1 | + | 210744 | 210800 | **RNA**→ rpmE (PRK00019)Ribosomal_L31 (pfam01197)→ PrfA (COG0216)prfA (PRK00591)→ HemK (COG2890)PRK09328 (PRK09328)→ SUA5 (COG0009)TIGR00057 (TIGR00057)→ |
| Mne-1-1 | NZ_LK021339.1 | + | 855681 | 855737 | **RNA**→ rpmE (PRK00019)Ribosomal_L31 (pfam01197)→ hypo→ hypo→ HemK (COG2890)PRK09328 (PRK09328)→ SUA5 (COG0009)TIGR00057 (TIGR00057)→ GT_WecA_like (cd06853)mraY (PRK00108)→ |
| Mne-2-1 | NZ_JMDW01000012.1 | - | 14937 | 14881 | **RNA**→ rpmE (PRK00019)Ribosomal_L31 (pfam01197)→ PrfA (COG0216)prfA (PRK00591)→ HemK (COG2890)PRK09328 (PRK09328)→ SUA5 (COG0009)TIGR00057 (TIGR00057)→ GT_WecA_like (cd06853)mraY (PRK00108)→ |
| env-400 | botACOD_F64RS5002I4I65 | + | 261 | 317 | **RNA**→ rpmE (PRK00019)Ribosomal_L31 (pfam01197)→ |
| Msp-21-1 | NZ_JQLT01000006.1 | + | 352100 | 352156 | **RNA**→ rpmE (PRK00019)Ribosomal_L31 (pfam01197)→ PrfA (COG0216)prfA (PRK00591)→ HemK (COG2890)PRK09328 (PRK09328)→ SUA5 (COG0009)TIGR00057 (TIGR00057)→ GT_WecA_like (cd06853)mraY (PRK00108)→ |
| Sal-7-1 | NZ_CP010519.1 | - | 2455492 | 2455447 | **RNA**→ rpmE (PRK00019)Ribosomal_L31 (pfam01197)→ |
| Szi-1-1 | NZ_AGBF01000094.1 | - | 21824 | 21778 | **RNA**→ rpmE (PRK00019)Ribosomal_L31 (pfam01197)→ rpmE (PRK00019)Ribosomal_L31 (pfam01197)→ |
| Ssp-17-1 | NZ_CM002271.1 | - | 1971747 | 1971701 | **RNA**→ rpmE (PRK00019)Ribosomal_L31 (pfam01197)→ |
| Ssp-30-1 | NZ_CM002273.1 | - | 2012002 | 2011956 | **RNA**→ rpmE (PRK00019)Ribosomal_L31 (pfam01197)→ |
| Ssp-52-1 | NZ_JTCK01000075.1 | + | 132364 | 132410 | **RNA**→ rpmE (PRK00019)Ribosomal_L31 (pfam01197)→ |
| Ssp-13-1 | NZ_KB898222.1 | - | 216996 | 216950 | **RNA**→ rpmE (PRK00019)Ribosomal_L31 (pfam01197)→ |
| Ssp-66-1 | NZ_KN549149.1 | - | 230060 | 230014 | **RNA**→ rpmE (PRK00019)Ribosomal_L31 (pfam01197)→ |
| Sgr-12-4 | NZ_JOGU01000005.1 | + | 127939 | 127985 | **RNA**→ rpmE (PRK00019)Ribosomal_L31 (pfam01197)→ |
| Ssp-63-1 | NZ_LAYX01000047.1 | - | 37184 | 37138 | **RNA**→ rpmE (PRK00019)Ribosomal_L31 (pfam01197)→ |
| Ssp-33-1 | NZ_FR873700.1 | - | 1935399 | 1935353 | **RNA**→ rpmE (PRK00019)Ribosomal_L31 (pfam01197)→ |
| Ssp-117-1 | NZ_JOHX01000005.1 | - | 52128 | 52082 | **RNA**→ rpmE (PRK00019)Ribosomal_L31 (pfam01197)→ |
| Swa-1-1 | NZ_JHDU01000001.1 | - | 64349 | 64303 | **RNA**→ hypo→ hypo→ |
| Sal-1-1 | NC_020990.1 | - | 1754401 | 1754355 | **RNA**→ rpmE (PRK00019)Ribosomal_L31 (pfam01197)→ |
| Sgr-12-5 | NZ_JNZI01000004.1 | - | 84767 | 84721 | **RNA**→ rpmE (PRK00019)Ribosomal_L31 (pfam01197)→ |
| Sgr-12-6 | NZ_JOGN01000003.1 | - | 52139 | 52093 | **RNA**→ rpmE (PRK00019)Ribosomal_L31 (pfam01197)→ |
| Ssp-83-1 | NZ_JOGQ01000004.1 | + | 154867 | 154913 | **RNA**→ rpmE (PRK00019)Ribosomal_L31 (pfam01197)→ |
| Ssp-26-1 | NZ_KB890732.1 | + | 167437 | 167483 | **RNA**→ rpmE (PRK00019)Ribosomal_L31 (pfam01197)→ |
| Sal-4-1 | NZ_JOII01000005.1 | + | 181272 | 181318 | **RNA**→ rpmE (PRK00019)Ribosomal_L31 (pfam01197)→ |
| Ssp-123-1 | NZ_KL591007.1 | - | 217831 | 217785 | **RNA**→ rpmE (PRK00019)Ribosomal_L31 (pfam01197)→ |
| Sgr-2-1 | NZ_GG657758.1 | - | 2542467 | 2542421 | **RNA**→ rpmE (PRK00019)Ribosomal_L31 (pfam01197)→ rpmE (PRK00019)Ribosomal_L31 (pfam01197)→ |
| Sco-3-1 | NC_021985.1 | + | 5785307 | 5785353 | **RNA**→ rpmE (PRK00019)Ribosomal_L31 (pfam01197)→ |
| Ssp-67-1 | NZ_LARV01000129.1 | + | 3104 | 3150 | **RNA**→ rpmE (PRK00019)Ribosomal_L31 (pfam01197)→ |
| Ssp-98-1 | NZ_KL591912.1 | + | 85133 | 85179 | **RNA**→ rpmE (PRK00019)Ribosomal_L31 (pfam01197)→ |
| Ssp-41-1 | NZ_AGNH01000234.1 | - | 7103 | 7057 | **RNA**→ rpmE (PRK00019)Ribosomal_L31 (pfam01197)→ |
| Ssp-162-1 | NZ_JOCG01000006.1 | + | 231677 | 231723 | **RNA**→ rpmE (PRK00019)Ribosomal_L31 (pfam01197)→ |
| Ssp-137-1 | NZ_JOIZ01000006.1 | + | 380113 | 380159 | **RNA**→ rpmE (PRK00019)Ribosomal_L31 (pfam01197)→ |
| Slu-1-1 | NZ_KN039947.1 | - | 1557714 | 1557668 | **RNA**→ rpmE (PRK00019)Ribosomal_L31 (pfam01197)→ |
| Sga-2-1 | NZ_JRHJ01000022.1 | - | 32900 | 32854 | **RNA**→ rpmE (PRK00019)Ribosomal_L31 (pfam01197)→ PrfA (COG0216)prfA (PRK00591)→ HemK (COG2890)PRK09328 (PRK09328)→ SUA5 (COG0009)TIGR00057 (TIGR00057)→ etp (PRK11391)Wzb (COG0394)→ |
| Sau-2-1 | NZ_JNZD01000033.1 | - | 54400 | 54354 | **RNA**→ rpmE (PRK00019)Ribosomal_L31 (pfam01197)→ |
| Ssp-21-1 | NZ_KB891850.1 | + | 297153 | 297200 | **RNA**→ rpmE (PRK00019)Ribosomal_L31 (pfam01197)→ |

| | | | | | |
|---|---|---|---|---|---|
| Nvu-1-1 | NZ_JNFP01000059.1 | + | 32133 | 32183 | **RNA**→ rpmE (PRK00019)Ribosomal_L31 (pfam01197)→ PrfA (COG0216)prfA (PRK00591)→ HemK (COG2890)PRK09328 (PRK09328)→ SUA5 (COG0009)TIGR00057 (TIGR00057)→ ECA_wecA (TIGR02380)GT_WecA_like (cd06853)→ hypo→ |
| Nbr-3-1 | NZ_BAFT02000053.1 | + | 33276 | 33326 | **RNA**→ rpmE (PRK00019)Ribosomal_L31 (pfam01197)→ PrfA (COG0216)prfA (PRK00591)→ HemK (COG2890)PRK09328 (PRK09328)→ SUA5 (COG0009)TIGR00057 (TIGR00057)→ ECA_wecA (TIGR02380)GT_WecA_like (cd06853)→ hypo→ |
| Nbr-2-1 | NZ_BAUA01000201.1 | + | 29549 | 29599 | **RNA**→ rpmE (PRK00019)Ribosomal_L31 (pfam01197)→ PrfA (COG0216)prfA (PRK00591)→ HemK (COG2890)PRK09328 (PRK09328)→ SUA5 (COG0009)TIGR00057 (TIGR00057)→ ECA_wecA (TIGR02380)GT_WecA_like (cd06853)→ hypo→ |
| Nbr-1-1 | NC_018681.1 | + | 1432741 | 1432791 | **RNA**→ rpmE (PRK00019)Ribosomal_L31 (pfam01197)→ PrfA (COG0216)prfA (PRK00591)→ HemK (COG2890)PRK09328 (PRK09328)→ SUA5 (COG0009)TIGR00057 (TIGR00057)→ ECA_wecA (TIGR02380)GT_WecA_like (cd06853)→ hypo→ |
| Nte-1-1 | NZ_BAGH01000084.1 | + | 1857 | 1906 | **RNA**→ rpmE (PRK00019)Ribosomal_L31 (pfam01197)→ |
| Spr-2-1 | NZ_BARF01000069.1 | + | 97 | 142 | **RNA**→ rpmE (PRK00019)Ribosomal_L31 (pfam01197)→ |
| Ssp-106-1 | NZ_JNXE01000037.1 | + | 11799 | 11844 | **RNA**→ rpmE (PRK00019)Ribosomal_L31 (pfam01197)→ |
| env-401 | JGI24739J22299_10657377 | + | 184 | 230 | **RNA**→ |
| env-402 | JGI24736J21556_1144543 | - | 314 | 268 | **RNA**→ rpmE (PRK00019)Ribosomal_L31 (pfam01197)→ |
| env-403 | JGI24739J22299_10005583 | + | 3113 | 3159 | **RNA**→ rpmE (PRK00019)Ribosomal_L31 (pfam01197)→ |
| env-404 | JGI24737J22298_10014880 | + | 59 | 105 | **RNA**→ rpmE (PRK00019)Ribosomal_L31 (pfam01197)→ |
| env-405 | JGI24735J21928_10026048 | - | 1497 | 1451 | **RNA**→ rpmE (PRK00019)Ribosomal_L31 (pfam01197)→ |
| env-406 | JGI24738J21930_10016695 | + | 1212 | 1258 | **RNA**→ rpmE (PRK00019)Ribosomal_L31 (pfam01197)→ |
| Ssp-146-1 | NZ_JODC01000003.1 | - | 34066 | 34020 | **RNA**→ rpmE (PRK00019)Ribosomal_L31 (pfam01197)→ |
| Smi-1-1 | NZ_KN050746.1 | + | 104683 | 104729 | **RNA**→ rpmE (PRK00019)Ribosomal_L31 (pfam01197)→ |
| Sda-1-1 | NC_020504.1 | - | 3469508 | 3469462 | **RNA**→ rpmE (PRK00019)Ribosomal_L31 (pfam01197)→ |
| Ssp-161-1 | NZ_JOBZ01000107.1 | + | 4806 | 4852 | **RNA**→ rpmE (PRK00019)Ribosomal_L31 (pfam01197)→ |
| Ssp-71-1 | NZ_LBDA01000053.1 | + | 10440 | 10486 | **RNA**→ rpmE (PRK00019)Ribosomal_L31 (pfam01197)→ |
| Sgr-7-1 | NZ_JOFU01000018.1 | - | 82877 | 82831 | **RNA**→ rpmE (PRK00019)Ribosomal_L31 (pfam01197)→ |
| Sgl-3-1 | NZ_CP009438.1 | + | 5215800 | 5215846 | **RNA**→ rpmE (PRK00019)Ribosomal_L31 (pfam01197)→ |
| Ssp-85-1 | NZ_JOFW01000021.1 | + | 12706 | 12752 | **RNA**→ rpmE (PRK00019)Ribosomal_L31 (pfam01197)→ |
| Ssp-121-1 | NZ_JOIU01000003.1 | + | 182946 | 182992 | **RNA**→ rpmE (PRK00019)Ribosomal_L31 (pfam01197)→ |
| Ssp-104-1 | NZ_JOIA01000044.1 | + | 3117 | 3163 | **RNA**→ rpmE (PRK00019)Ribosomal_L31 (pfam01197)→ |
| Ssp-60-1 | NZ_KI601353.1 | - | 78686 | 78640 | **RNA**→ rpmE (PRK00019)Ribosomal_L31 (pfam01197)→ |
| Ssp-72-1 | NZ_LAVA01000109.1 | - | 30407 | 30361 | **RNA**→ rpmE (PRK00019)Ribosomal_L31 (pfam01197)→ |
| Ssp-99-1 | NZ_JOHW01000005.1 | + | 528394 | 528440 | **RNA**→ rpmE (PRK00019)Ribosomal_L31 (pfam01197)→ |
| Ssp-89-1 | NZ_JOIF01000027.1 | + | 9376 | 9423 | **RNA**→ rpmE (PRK00019)Ribosomal_L31 (pfam01197)→ |
| Sbi-2-1 | NZ_JOFS01000024.1 | - | 108312 | 108266 | **RNA**→ rpmE (PRK00019)Ribosomal_L31 (pfam01197)→ |
| Sbi-2-2 | NZ_JOFR01000024.1 | - | 108305 | 108259 | **RNA**→ rpmE (PRK00019)Ribosomal_L31 (pfam01197)→ |
| Ssp-82-1 | NZ_JOGP01000011.1 | - | 82995 | 82949 | **RNA**→ rpmE (PRK00019)Ribosomal_L31 (pfam01197)→ |
| Ssp-3-1 | NZ_KB911781.1 | - | 84488 | 84442 | **RNA**→ rpmE (PRK00019)Ribosomal_L31 (pfam01197)→ |
| Sam-1-1 | NZ_CP012382.1 | + | 5413021 | 5413067 | **RNA**→ rpmE (PRK00019)Ribosomal_L31 (pfam01197)→ |
| Ssp-108-1 | NZ_JOHJ01000018.1 | - | 81408 | 81362 | **RNA**→ rpmE (PRK00019)Ribosomal_L31 (pfam01197)→ |
| Ssp-128-1 | NZ_JOGT01000006.1 | + | 282015 | 282061 | **RNA**→ rpmE (PRK00019)Ribosomal_L31 (pfam01197)→ |
| Ssp-48-1 | NZ_AUKV01000028.1 | + | 12811 | 12857 | **RNA**→ rpmE (PRK00019)Ribosomal_L31 (pfam01197)→ |
| Ssp-124-1 | NZ_JOHU01000015.1 | + | 140224 | 140270 | **RNA**→ rpmE (PRK00019)Ribosomal_L31 (pfam01197)→ |
| Sga-1-1 | NZ_AOHP01000017.1 | + | 10282 | 10328 | **RNA**→ rpmE (PRK00019)Ribosomal_L31 (pfam01197)→ |
| Ssp-110-1 | NZ_JOGJ01000004.1 | - | 81960 | 81914 | **RNA**→ rpmE (PRK00019)Ribosomal_L31 (pfam01197)→ |
| Ssp-140-1 | NZ_KL585965.1 | + | 42291 | 42337 | **RNA**→ rpmE (PRK00019)Ribosomal_L31 (pfam01197)→ |
| Ssp-142-1 | NZ_KL585205.1 | - | 33182 | 33136 | **RNA**→ rpmE (PRK00019)Ribosomal_L31 (pfam01197)→ |
| Ssp-139-1 | NZ_JOCT01000023.1 | + | 96568 | 96614 | **RNA**→ rpmE (PRK00019)Ribosomal_L31 (pfam01197)→ |
| Ssp-107-1 | NZ_JOHL01000041.1 | - | 81671 | 81625 | **RNA**→ rpmE (PRK00019)Ribosomal_L31 (pfam01197)→ |
| Ssp-112-1 | NZ_JOGV01000023.1 | - | 19434 | 19388 | **RNA**→ rpmE (PRK00019)Ribosomal_L31 (pfam01197)→ |
| Svi-5-1 | NZ_AMLP01000083.1 | - | 20043 | 19997 | **RNA**→ hypo→ rpmE (PRK00019)Ribosomal_L31 (pfam01197)→ |
| Ssp-74-1 | NZ_BBNN01000020.1 | + | 195717 | 195763 | **RNA**→ rpmE (PRK00019)Ribosomal_L31 (pfam01197)→ |

| | | | | | |
|---|---|---|---|---|---|
| Sbo-1-1 | NZ_KB405078.1 | - | 426028 | 425982 | RNA→ rpmE (PRK00019)Ribosomal_L31 (pfam01197)→ |
| Sbo-1-2 | NZ_KB911581.1 | + | 1699487 | 1699533 | RNA→ rpmE (PRK00019)Ribosomal_L31 (pfam01197)→ |
| env-407 | JGI12419J13241_1020528 | + | 551 | 597 | RNA→ rpmE (PRK00019)Ribosomal_L31 (pfam01197)→ |
| Ssp-101-1 | NZ_JOHV01000024.1 | - | 31603 | 31557 | RNA→ rpmE (PRK00019)Ribosomal_L31 (pfam01197)→ |
| Ssp-136-1 | NZ_JOIY01000004.1 | - | 81877 | 81831 | RNA→ rpmE (PRK00019)Ribosomal_L31 (pfam01197)→ |
| Sto-1-1 | NZ_JFCB01000017.1 | + | 75109 | 75155 | RNA→ rpmE (PRK00019)Ribosomal_L31 (pfam01197)→ |
| Sch-1-1 | NZ_JH164850.1 | - | 42480 | 42434 | RNA→ rpmE (PRK00019)Ribosomal_L31 (pfam01197)→ |
| Saf-1-1 | NZ_KE354178.1 | + | 20405 | 20451 | RNA→ rpmE (PRK00019)Ribosomal_L31 (pfam01197)→ |
| Sva-1-1 | NZ_JYJH01000019.1 | + | 85068 | 85114 | RNA→ rpmE (PRK00019)Ribosomal_L31 (pfam01197)→ |
| Sfl-7-1 | NZ_JNXD01000019.1 | + | 86558 | 86604 | RNA→ rpmE (PRK00019)Ribosomal_L31 (pfam01197)→ |
| Ssp-154-1 | NZ_KL579021.1 | + | 79524 | 79570 | RNA→ rpmE (PRK00019)Ribosomal_L31 (pfam01197)→ |
| Sol-1-1 | NZ_JJOH01000021.1 | + | 109043 | 109089 | RNA→ rpmE (PRK00019)Ribosomal_L31 (pfam01197)→ |
| Ssp-147-1 | NZ_JODG01000016.1 | - | 75647 | 75601 | RNA→ rpmE (PRK00019)Ribosomal_L31 (pfam01197)→ |
| Sia-1-1 | NZ_JNXI01000026.1 | - | 85292 | 85246 | RNA→ rpmE (PRK00019)Ribosomal_L31 (pfam01197)→ |
| Sro-1-1 | NZ_CM002285.1 | - | 2981947 | 2981901 | RNA→ rpmE (PRK00019)Ribosomal_L31 (pfam01197)→ |
| Scy-3-1 | NZ_CP010849.1 | + | 5238904 | 5238950 | RNA→ rpmE (PRK00019)Ribosomal_L31 (pfam01197)→ |
| Sac-1-1 | NZ_JODT01000015.1 | + | 134233 | 134279 | RNA→ rpmE (PRK00019)Ribosomal_L31 (pfam01197)→ |
| Sla-3-1 | NZ_JOBD01000030.1 | + | 53632 | 53678 | RNA→ rpmE (PRK00019)Ribosomal_L31 (pfam01197)→ |
| Ssp-160-1 | NZ_JNXK01000056.1 | + | 9606 | 9652 | RNA→ rpmE (PRK00019)Ribosomal_L31 (pfam01197)→ |
| San-1-1 | NZ_LGUX01000300.1 | - | 86940 | 86894 | RNA→ rpmE (PRK00019)Ribosomal_L31 (pfam01197)→ |
| Ssp-160-2 | NZ_JOCJ01000056.1 | + | 9610 | 9656 | RNA→ rpmE (PRK00019)Ribosomal_L31 (pfam01197)→ |
| Sfl-3-1 | NZ_JOCU01000056.1 | + | 9567 | 9613 | RNA→ rpmE (PRK00019)Ribosomal_L31 (pfam01197)→ |
| Ssp-94-1 | NZ_JOIC01000059.1 | - | 87663 | 87617 | RNA→ rpmE (PRK00019)Ribosomal_L31 (pfam01197)→ |
| Ssp-159-1 | NZ_JOJH01000041.1 | + | 9637 | 9683 | RNA→ rpmE (PRK00019)Ribosomal_L31 (pfam01197)→ |
| Ssp-130-1 | NZ_JOIV01000034.1 | + | 57899 | 57945 | RNA→ rpmE (PRK00019)Ribosomal_L31 (pfam01197)→ |
| Sfl-2-1 | NZ_JNWV01000050.1 | - | 87704 | 87658 | RNA→ rpmE (PRK00019)Ribosomal_L31 (pfam01197)→ |
| Ssp-155-1 | NZ_JOJG01000055.1 | + | 9473 | 9519 | RNA→ rpmE (PRK00019)Ribosomal_L31 (pfam01197)→ |
| Shy-2-1 | NC_017765.1 | + | 7332293 | 7332339 | RNA→ rpmE (PRK00019)Ribosomal_L31 (pfam01197)→ |
| Shy-3-1 | NC_020895.1 | + | 7026019 | 7026065 | RNA→ rpmE (PRK00019)Ribosomal_L31 (pfam01197)→ |
| Ssp-125-1 | NZ_JOAW01000048.1 | + | 11208 | 11254 | RNA→ rpmE (PRK00019)Ribosomal_L31 (pfam01197)→ |
| Spe-1-1 | NZ_KL575053.1 | + | 113652 | 113698 | RNA→ rpmE (PRK00019)Ribosomal_L31 (pfam01197)→ |
| Ssp-141-1 | NZ_JOJB01000039.1 | - | 22019 | 21973 | RNA→ rpmE (PRK00019)Ribosomal_L31 (pfam01197)→ |
| Svi-14-1 | NZ_LFNT01000024.1 | - | 76388 | 76342 | RNA→ rpmE (PRK00019)Ribosomal_L31 (pfam01197)→ |
| Svi-14-2 | NZ_LGUR01000200.1 | - | 76304 | 76258 | RNA→ rpmE (PRK00019)Ribosomal_L31 (pfam01197)→ |
| Svi-14-3 | NZ_LGUQ01000026.1 | - | 50743 | 50697 | RNA→ rpmE (PRK00019)Ribosomal_L31 (pfam01197)→ |
| Svi-4-1 | NZ_GG657757.1 | + | 6015810 | 6015856 | RNA→ rpmE (PRK00019)Ribosomal_L31 (pfam01197)→ rpmE (PRK00019)Ribosomal_L31 (pfam01197)→ |
| env-408 | JGI25153J46596_10164792 | - | 356 | 310 | RNA→ ←hypo |
| Sol-2-1 | NZ_JOFH01000019.1 | - | 80214 | 80168 | RNA→ rpmE (PRK00019)Ribosomal_L31 (pfam01197)→ |
| Kau-1-2 | NZ_JODU01000015.1 | - | 85909 | 85863 | RNA→ rpmE (PRK00019)Ribosomal_L31 (pfam01197)→ |
| Svi-12-1 | NZ_JODM01000021.1 | + | 27707 | 27753 | RNA→ rpmE (PRK00019)Ribosomal_L31 (pfam01197)→ |
| Sco-1-1 | NZ_AHGS01000015.1 | - | 31365 | 31319 | RNA→ rpmE (PRK00019)Ribosomal_L31 (pfam01197)→ |
| Ssp-58-1 | NZ_APIV01000026.1 | - | 82178 | 82132 | RNA→ rpmE (PRK00019)Ribosomal_L31 (pfam01197)→ |
| Sno-1-1 | NZ_CP009313.1 | + | 5012088 | 5012134 | RNA→ rpmE (PRK00019)Ribosomal_L31 (pfam01197)→ |
| Sgr-1-1 | NZ_AEYX01000041.1 | + | 322182 | 322228 | RNA→ rpmE (PRK00019)Ribosomal_L31 (pfam01197)→ rpmE (PRK00019)Ribosomal_L31 (pfam01197)→ |
| Sce-1-1 | NZ_JOEV01000019.1 | - | 107133 | 107087 | RNA→ rpmE (PRK00019)Ribosomal_L31 (pfam01197)→ |
| Ssp-176-1 | NZ_JXWX01000006.1 | - | 87051 | 87005 | RNA→ rpmE (PRK00019)Ribosomal_L31 (pfam01197)→ |
| Sco-2-1 | NC_003888.3 | + | 5829950 | 5829996 | RNA→ rpmE (PRK00019)Ribosomal_L31 (pfam01197)→ |
| Sli-2-1 | NZ_GG657756.1 | - | 2654278 | 2654232 | RNA→ rpmE (PRK00019)Ribosomal_L31 (pfam01197)→ rpmE (PRK00019)Ribosomal_L31 (pfam01197)→ |
| Sli-1-1 | NZ_CM001889.1 | + | 5849426 | 5849472 | RNA→ rpmE (PRK00019)Ribosomal_L31 (pfam01197)→ |
| Sli-2-2 | NZ_CP009124.1 | - | 2688989 | 2688943 | RNA→ rpmE (PRK00019)Ribosomal_L31 (pfam01197)→ |
| Svi-11-1 | NZ_JOCE01000023.1 | + | 20903 | 20949 | RNA→ rpmE (PRK00019)Ribosomal_L31 (pfam01197)→ |
| Ssp-118-1 | NZ_JNXF01000037.1 | + | 33108 | 33153 | RNA→ rpmE (PRK00019)Ribosomal_L31 (pfam01197)→ |

| | | | | | |
|---|---|---|---|---|---|
| Ssp-120-1 | NZ_JOHP01000066.1 | - | 29002 | 28957 | **RNA**→ rpmE (PRK00019)Ribosomal_L31 (pfam01197)→ |
| Ssp-76-1 | NZ_JNZW01000021.1 | - | 92544 | 92499 | **RNA**→ rpmE (PRK00019)Ribosomal_L31 (pfam01197)→ |
| Ssp-119-1 | NZ_JOHS01000019.1 | + | 33159 | 33204 | **RNA**→ rpmE (PRK00019)Ribosomal_L31 (pfam01197)→ |
| Svi-11-2 | NZ_JODI01000003.1 | + | 207039 | 207085 | **RNA**→ rpmE (PRK00019)Ribosomal_L31 (pfam01197)→ |
| Sgr-12-7 | NZ_JOJE01000033.1 | + | 28750 | 28796 | **RNA**→ rpmE (PRK00019)Ribosomal_L31 (pfam01197)→ |
| Ssc-4-1 | NZ_JODX01000005.1 | - | 199204 | 199158 | **RNA**→ rpmE (PRK00019)Ribosomal_L31 (pfam01197)→ |
| Ssp-69-1 | NZ_JUIG01000017.1 | + | 189710 | 189755 | **RNA**→ rpmE (PRK00019)Ribosomal_L31 (pfam01197)→ |
| Spl-1-1 | NZ_JTDH01000035.1 | + | 15585 | 15630 | **RNA**→ rpmE (PRK00019)Ribosomal_L31 (pfam01197)→ |
| Spl-1-2 | NZ_JUIF01000051.1 | - | 80126 | 80081 | **RNA**→ rpmE (PRK00019)Ribosomal_L31 (pfam01197)→ |
| Ssp-153-1 | NZ_JOCA01000016.1 | + | 83466 | 83511 | **RNA**→ rpmE (PRK00019)Ribosomal_L31 (pfam01197)→ |
| Sye-2-1 | NZ_JNWJ01000028.1 | + | 22848 | 22893 | **RNA**→ rpmE (PRK00019)Ribosomal_L31 (pfam01197)→ |
| Sse-1-1 | NZ_JNXP01000009.1 | + | 359142 | 359189 | **RNA**→ rpmE (PRK00019)Ribosomal_L31 (pfam01197)→ |
| Sfu-2-1 | NZ_JOEY01000004.1 | - | 144771 | 144725 | **RNA**→ rpmE (PRK00019)Ribosomal_L31 (pfam01197)→ |
| Sdu-1-1 | NZ_JNXR01000028.1 | - | 85192 | 85146 | **RNA**→ rpmE (PRK00019)Ribosomal_L31 (pfam01197)→ |
| Sip-1-1 | NZ_AEJC01000389.1 | + | 1739 | 1785 | **RNA**→ rpmE (PRK00019)Ribosomal_L31 (pfam01197)→ |
| Ssc-3-1 | NZ_KL997422.1 | + | 1322338 | 1322384 | **RNA**→ rpmE (PRK00019)Ribosomal_L31 (pfam01197)→ |
| Ssp-59-1 | NZ_LACL01000016.1 | - | 93527 | 93481 | **RNA**→ rpmE (PRK00019)Ribosomal_L31 (pfam01197)→ |
| Ssp-70-1 | NZ_LBMU01000062.1 | - | 24986 | 24941 | **RNA**→ rpmE (PRK00019)Ribosomal_L31 (pfam01197)→ |
| Ssp-24-1 | NZ_KB891987.1 | + | 887555 | 887601 | **RNA**→ rpmE (PRK00019)Ribosomal_L31 (pfam01197)→ |
| Ssp-10-1 | NZ_KB846749.1 | - | 16622 | 16576 | **RNA**→ rpmE (PRK00019)Ribosomal_L31 (pfam01197)→ |
| Ssp-11-1 | NZ_KB898297.1 | - | 18791 | 18745 | **RNA**→ rpmE (PRK00019)Ribosomal_L31 (pfam01197)→ |
| Ssp-53-1 | NZ_KI911521.1 | - | 22367 | 22321 | **RNA**→ rpmE (PRK00019)Ribosomal_L31 (pfam01197)→ |
| Sni-2-1 | NZ_CM002280.1 | - | 2660451 | 2660406 | **RNA**→ rpmE (PRK00019)Ribosomal_L31 (pfam01197)→ |
| Spr-1-1 | NZ_CM000950.1 | - | 2585594 | 2585547 | **RNA**→ hypo→ rpmE (PRK00019)Ribosomal_L31 (pfam01197)→ |
| Ssp-134-1 | NZ_JOCB01000025.1 | - | 28339 | 28293 | **RNA**→ rpmE (PRK00019)Ribosomal_L31 (pfam01197)→ |
| Sau-1-1 | NZ_JOAP01000030.1 | - | 60155 | 60109 | **RNA**→ rpmE (PRK00019)Ribosomal_L31 (pfam01197)→ PrfA (COG0216)prfA (PRK00591)→<br>HemK (COG2890)PRK09328 (PRK09328)→ SUA5 (COG0009)TIGR00057 (TIGR00057)→<br>etp (PRK11391)Wzb (COG0394)→ etp (PRK11391)Wzb (COG0394)→ |
| Sau-1-2 | NZ_JOAR01000022.1 | - | 60248 | 60202 | **RNA**→ rpmE (PRK00019)Ribosomal_L31 (pfam01197)→ PrfA (COG0216)prfA (PRK00591)→<br>HemK (COG2890)PRK09328 (PRK09328)→ SUA5 (COG0009)TIGR00057 (TIGR00057)→<br>etp (PRK11391)Wzb (COG0394)→ etp (PRK11391)Wzb (COG0394)→ |
| Asp-7-1 | NZ_JOGW01000010.1 | + | 24341 | 24387 | **RNA**→ rpmE (PRK00019)Ribosomal_L31 (pfam01197)→ |
| Sal-5-1 | NZ_JNXT01000031.1 | + | 28596 | 28642 | **RNA**→ rpmE (PRK00019)Ribosomal_L31 (pfam01197)→ PrfA (COG0216)prfA (PRK00591)→<br>HemK (COG2890)PRK09328 (PRK09328)→ SUA5 (COG0009)TIGR00057 (TIGR00057)→<br>etp (PRK11391)Wzb (COG0394)→ etp (PRK11391)Wzb (COG0394)→ |
| Sru-2-1 | NZ_JODN01000034.1 | + | 28596 | 28642 | **RNA**→ rpmE (PRK00019)Ribosomal_L31 (pfam01197)→ PrfA (COG0216)prfA (PRK00591)→<br>HemK (COG2890)PRK09328 (PRK09328)→ SUA5 (COG0009)TIGR00057 (TIGR00057)→<br>etp (PRK11391)Wzb (COG0394)→ etp (PRK11391)Wzb (COG0394)→ |
| Sru-2-2 | NZ_JOAQ01000034.1 | + | 28596 | 28642 | **RNA**→ rpmE (PRK00019)Ribosomal_L31 (pfam01197)→ PrfA (COG0216)prfA (PRK00591)→<br>HemK (COG2890)PRK09328 (PRK09328)→ SUA5 (COG0009)TIGR00057 (TIGR00057)→<br>etp (PRK11391)Wzb (COG0394)→ etp (PRK11391)Wzb (COG0394)→ |
| Svi-2-1 | NZ_ASHX01000001.1 | - | 682175 | 682129 | **RNA**→ rpmE (PRK00019)Ribosomal_L31 (pfam01197)→ |
| Stu-1-1 | NZ_AEJB01000521.1 | + | 12742 | 12788 | **RNA**→ rpmE (PRK00019)Ribosomal_L31 (pfam01197)→ |
| Sso-1-1 | NZ_AJJM01000112.1 | + | 4156 | 4202 | **RNA**→ rpmE (PRK00019)Ribosomal_L31 (pfam01197)→ rpmE (PRK00019)Ribosomal_L31 (pfam01197)→ |
| Spu-1-1 | NZ_KB913030.1 | + | 5096632 | 5096678 | **RNA**→ rpmE (PRK00019)Ribosomal_L31 (pfam01197)→ |
| Ssp-12-1 | NZ_KB897776.1 | - | 219651 | 219605 | **RNA**→ rpmE (PRK00019)Ribosomal_L31 (pfam01197)→ |
| Ssp-78-1 | NZ_JOJM01000058.1 | - | 37844 | 37798 | **RNA**→ rpmE (PRK00019)Ribosomal_L31 (pfam01197)→ |
| Ssp-158-1 | NZ_JOCD01000378.1 | - | 4063 | 4017 | **RNA**→ rpmE (PRK00019)Ribosomal_L31 (pfam01197)→ |
| Ssp-79-1 | NZ_JOHG01000018.1 | - | 89216 | 89169 | **RNA**→ rpmE (PRK00019)Ribosomal_L31 (pfam01197)→ |
| Svi-10-1 | NZ_JNYP01000019.1 | - | 145314 | 145267 | **RNA**→ rpmE (PRK00019)Ribosomal_L31 (pfam01197)→ |
| Ssp-92-1 | NZ_JOIP01000013.1 | + | 94481 | 94528 | **RNA**→ rpmE (PRK00019)Ribosomal_L31 (pfam01197)→ |
| Ssp-113-1 | NZ_JOHA01000008.1 | - | 84595 | 84548 | **RNA**→ rpmE (PRK00019)Ribosomal_L31 (pfam01197)→ |
| Ssp-93-1 | NZ_JOIO01000009.1 | + | 238799 | 238846 | **RNA**→ rpmE (PRK00019)Ribosomal_L31 (pfam01197)→ |
| Ssp-114-1 | NZ_JOHD01000020.1 | - | 25512 | 25465 | **RNA**→ rpmE (PRK00019)Ribosomal_L31 (pfam01197)→ |

| | | | | | |
|---|---|---|---|---|---|
| Ssp-84-1 | NZ_JOIH01000014.1 | - | 84509 | 84462 | RNA→ rpmE (PRK00019)Ribosomal_L31 (pfam01197)→ |
| Sgr-13-1 | NZ_JOFF01000006.1 | + | 316771 | 316818 | RNA→ rpmE (PRK00019)Ribosomal_L31 (pfam01197)→ |
| Spu-3-1 | NZ_JOBG01000002.1 | - | 133951 | 133904 | RNA→ rpmE (PRK00019)Ribosomal_L31 (pfam01197)→ |
| Spu-2-1 | NZ_JOFA01000023.1 | + | 12247 | 12294 | RNA→ rpmE (PRK00019)Ribosomal_L31 (pfam01197)→ |
| Sgr-12-8 | NZ_JOIT01000013.1 | - | 26693 | 26646 | RNA→ rpmE (PRK00019)Ribosomal_L31 (pfam01197)→ |
| Sfl-9-1 | NZ_JOAC01000008.1 | - | 83545 | 83498 | RNA→ rpmE (PRK00019)Ribosomal_L31 (pfam01197)→ |
| Spu-2-2 | NZ_JOBQ01000020.1 | + | 12247 | 12294 | RNA→ rpmE (PRK00019)Ribosomal_L31 (pfam01197)→ |
| Spu-3-2 | NZ_JODK01000002.1 | + | 163658 | 163705 | RNA→ rpmE (PRK00019)Ribosomal_L31 (pfam01197)→ |
| Sbr-1-1 | NZ_JNXB01000002.1 | - | 133941 | 133894 | RNA→ rpmE (PRK00019)Ribosomal_L31 (pfam01197)→ |
| Sca-4-1 | NZ_KL573601.1 | - | 30459 | 30412 | RNA→ rpmE (PRK00019)Ribosomal_L31 (pfam01197)→ |
| Sca-4-2 | NZ_KL569933.1 | + | 21968 | 22015 | RNA→ rpmE (PRK00019)Ribosomal_L31 (pfam01197)→ |
| Ssp-151-1 | NZ_JOCW01000015.1 | - | 159046 | 158999 | RNA→ rpmE (PRK00019)Ribosomal_L31 (pfam01197)→ |
| Sgr-13-2 | NZ_LGUZ01000018.1 | - | 139783 | 139736 | RNA→ rpmE (PRK00019)Ribosomal_L31 (pfam01197)→ |
| Sgr-13-3 | NZ_JOFE01000019.1 | - | 139769 | 139722 | RNA→ rpmE (PRK00019)Ribosomal_L31 (pfam01197)→ |
| Sgr-13-4 | NZ_JOFD01000010.1 | + | 270523 | 270570 | RNA→ rpmE (PRK00019)Ribosomal_L31 (pfam01197)→ |
| Sgr-13-5 | NZ_JODO01000012.1 | - | 158845 | 158798 | RNA→ rpmE (PRK00019)Ribosomal_L31 (pfam01197)→ |
| Sat-2-1 | NZ_JNXG01000002.1 | + | 1072335 | 1072381 | RNA→ rpmE (PRK00019)Ribosomal_L31 (pfam01197)→ |
| Sfl-8-1 | NZ_JOAB01000003.1 | - | 225755 | 225709 | RNA→ rpmE (PRK00019)Ribosomal_L31 (pfam01197)→ |
| Ssp-80-1 | NZ_JOAE01000004.1 | + | 218001 | 218047 | RNA→ rpmE (PRK00019)Ribosomal_L31 (pfam01197)→ |
| Ssp-135-1 | NZ_JOIW01000004.1 | - | 216825 | 216779 | RNA→ rpmE (PRK00019)Ribosomal_L31 (pfam01197)→ |
| Sfl-1-1 | NC_016114.1 | - | 2388748 | 2388702 | RNA→ rpmE (PRK00019)Ribosomal_L31 (pfam01197)→ |
| Ssp-29-1 | NC_021055.1 | + | 5042406 | 5042452 | RNA→ rpmE (PRK00019)Ribosomal_L31 (pfam01197)→ |
| [pa-1-1 | NZ_JNYQ01000004.1 | - | 248335 | 248289 | RNA→ rpmE (PRK00019)Ribosomal_L31 (pfam01197)→ |
| Sol-2-2 | NZ_JNWM01000003.1 | + | 1016882 | 1016928 | RNA→ rpmE (PRK00019)Ribosomal_L31 (pfam01197)→ |
| Sfu-3-1 | NZ_JNXH01000004.1 | - | 245003 | 244957 | RNA→ rpmE (PRK00019)Ribosomal_L31 (pfam01197)→ |
| Ssp-126-1 | NZ_KL589835.1 | + | 240377 | 240423 | RNA→ rpmE (PRK00019)Ribosomal_L31 (pfam01197)→ |
| Sgr-11-1 | NZ_JYBE01000037.1 | + | 33072 | 33119 | RNA→ rpmE (PRK00019)Ribosomal_L31 (pfam01197)→ |
| Sme-1-1 | NZ_JOJK01000017.1 | + | 272123 | 272170 | RNA→ rpmE (PRK00019)Ribosomal_L31 (pfam01197)→ |
| Ssp-14-1 | NZ_KB890948.1 | + | 91657 | 91704 | RNA→ rpmE (PRK00019)Ribosomal_L31 (pfam01197)→ |
| Ssp-168-1 | NZ_ARPF01000003.1 | + | 344832 | 344879 | RNA→ rpmE (PRK00019)Ribosomal_L31 (pfam01197)→ |
| Ssp-49-1 | NZ_CBRG010000147.1 | + | 14666 | 14712 | RNA→ rpmE (PRK00019)Ribosomal_L31 (pfam01197)→ |
| Ssp-22-1 | NZ_KB891605.1 | + | 147812 | 147858 | RNA→ rpmE (PRK00019)Ribosomal_L31 (pfam01197)→ |
| Ssp-23-1 | NZ_KB891406.1 | - | 421246 | 421200 | RNA→ rpmE (PRK00019)Ribosomal_L31 (pfam01197)→ |
| Ssp-65-1 | NZ_KI911349.1 | + | 1102284 | 1102330 | RNA→ rpmE (PRK00019)Ribosomal_L31 (pfam01197)→ |
| Ssp-64-1 | NZ_AZVF01000001.1 | - | 420786 | 420740 | RNA→ rpmE (PRK00019)Ribosomal_L31 (pfam01197)→ |
| Ssp-25-1 | NZ_KB891774.1 | + | 108968 | 109014 | RNA→ rpmE (PRK00019)Ribosomal_L31 (pfam01197)→ |
| Ssp-27-1 | NZ_AQWQ01000004.1 | - | 409548 | 409502 | RNA→ rpmE (PRK00019)Ribosomal_L31 (pfam01197)→ |
| Ssp-152-1 | NZ_KL585392.1 | - | 37506 | 37460 | RNA→ rpmE (PRK00019)Ribosomal_L31 (pfam01197)→ rpmE (PRK00019)Ribosomal_L31 (pfam01197)→ |
| Ssp-57-1 | NZ_AZVE01000001.1 | - | 87318 | 87272 | RNA→ rpmE (PRK00019)Ribosomal_L31 (pfam01197)→ |
| Ssp-163-1 | NZ_JJOB01000001.1 | + | 4517158 | 4517204 | RNA→ rpmE (PRK00019)Ribosomal_L31 (pfam01197)→ |
| Ssp-40-1 | NC_015953.1 | + | 5035066 | 5035112 | RNA→ rpmE (PRK00019)Ribosomal_L31 (pfam01197)→ |
| Ssp-15-1 | NZ_ARPI01000001.1 | - | 414568 | 414522 | RNA→ rpmE (PRK00019)Ribosomal_L31 (pfam01197)→ |
| Ssp-6-1 | NZ_KB892001.1 | + | 325904 | 325950 | RNA→ rpmE (PRK00019)Ribosomal_L31 (pfam01197)→ |
| Sha-2-1 | NZ_JOAZ01000017.1 | + | 419289 | 419335 | RNA→ rpmE (PRK00019)Ribosomal_L31 (pfam01197)→ |
| Sgr-8-1 | NZ_JOFC01000033.1 | + | 41970 | 42016 | RNA→ rpmE (PRK00019)Ribosomal_L31 (pfam01197)→ |
| Ssp-43-1 | NZ_AGSW01000051.1 | + | 66077 | 66123 | RNA→ rpmE (PRK00019)Ribosomal_L31 (pfam01197)→ rpmE (PRK00019)Ribosomal_L31 (pfam01197)→ |
| Scy-2-1 | NZ_JOEM01000014.1 | - | 146064 | 146018 | RNA→ rpmE (PRK00019)Ribosomal_L31 (pfam01197)→ |
| Ssp-39-1 | NZ_KB891296.1 | - | 481794 | 481748 | RNA→ rpmE (PRK00019)Ribosomal_L31 (pfam01197)→ |
| Sfu-1-1 | NC_021177.1 | + | 5700829 | 5700875 | RNA→ rpmE (PRK00019)Ribosomal_L31 (pfam01197)→ |
| Sal-6-1 | NZ_JNWU01000016.1 | + | 9488 | 9534 | RNA→ rpmE (PRK00019)Ribosomal_L31 (pfam01197)→ |
| Ssp-145-1 | NZ_JOJC01000009.1 | - | 114395 | 114349 | RNA→ rpmE (PRK00019)Ribosomal_L31 (pfam01197)→ |
| Seu-1-1 | NZ_KQ257779.1 | - | 81970 | 81924 | RNA→ rpmE (PRK00019)Ribosomal_L31 (pfam01197)→ |

| | | | | | |
|---|---|---|---|---|---|
| Sgr-3-1 | NZ_GL877172.1 | - | 2772173 | 2772127 | **RNA**→ rpmE (PRK00019)Ribosomal_L31 (pfam01197)→ rpmE (PRK00019)Ribosomal_L31 (pfam01197)→ |
| Sgr-4-1 | NC_010572.1 | - | 2577363 | 2577317 | **RNA**→ rpmE (PRK00019)Ribosomal_L31 (pfam01197)→ |
| Sgr-12-9 | NZ_JOBR01000007.1 | - | 141549 | 141503 | **RNA**→ rpmE (PRK00019)Ribosomal_L31 (pfam01197)→ |
| Ssp-9-1 | NZ_KB899000.1 | - | 100558 | 100512 | **RNA**→ rpmE (PRK00019)Ribosomal_L31 (pfam01197)→ |
| Ssp-68-1 | NZ_JXUQ01000001.1 | + | 5787746 | 5787792 | **RNA**→ rpmE (PRK00019)Ribosomal_L31 (pfam01197)→ |
| Sal-8-2 | NZ_JOED01000008.1 | + | 96199 | 96245 | **RNA**→ rpmE (PRK00019)Ribosomal_L31 (pfam01197)→ |
| Sba-2-1 | NZ_KL573542.1 | - | 304852 | 304806 | **RNA**→ rpmE (PRK00019)Ribosomal_L31 (pfam01197)→ |
| San-2-1 | NZ_JNXS01000009.1 | - | 156824 | 156778 | **RNA**→ rpmE (PRK00019)Ribosomal_L31 (pfam01197)→ rpmE (PRK00019)Ribosomal_L31 (pfam01197)→ |
| Sod-1-1 | NZ_LATD01000035.1 | + | 9464 | 9510 | **RNA**→ rpmE (PRK00019)Ribosomal_L31 (pfam01197)→ PrfA (COG0216)prfA (PRK00591)→ |
| Ssp-149-1 | NZ_JODF01000062.1 | + | 10265 | 10311 | **RNA**→ rpmE (PRK00019)Ribosomal_L31 (pfam01197)→ |
| Sfl-4-1 | NZ_AUBE01000011.1 | - | 189557 | 189511 | **RNA**→ rpmE (PRK00019)Ribosomal_L31 (pfam01197)→ |
| Ssp-127-1 | NZ_JOGZ01000006.1 | - | 230008 | 229961 | **RNA**→ rpmE (PRK00019)Ribosomal_L31 (pfam01197)→ |
| Ssp-81-1 | NZ_JOJO01000001.1 | + | 783410 | 783457 | **RNA**→ rpmE (PRK00019)Ribosomal_L31 (pfam01197)→ |
| Sat-1-1 | NZ_JQJU01000001.1 | + | 779795 | 779842 | **RNA**→ rpmE (PRK00019)Ribosomal_L31 (pfam01197)→ |
| Ssp-103-1 | NZ_JOHR01000018.1 | - | 2619 | 2572 | **RNA**→ rpmE (PRK00019)Ribosomal_L31 (pfam01197)→ |
| Nsp-3-1 | NZ_KB822580.1 | + | 117055 | 117099 | **RNA**→ rpmE (PRK00019)Ribosomal_L31 (pfam01197)→ PrfA (COG0216)prfA (PRK00591)→ |
| | | | | | HemK (COG2890)PRK09328 (PRK09328)→ SUA5 (COG0009)TIGR00057 (TIGR00057)→ |
| | | | | | GT_WecA_like (cd06853)mraY (PRK00108)→ |
| env-409 | JGI20200J14955_1027293 | - | 434 | 379 | **RNA**→ rpmE (PRK00019)Ribosomal_L31 (pfam01197)→ |
| env-410 | JGI20202J14951_1065050 | + | 200 | 255 | **RNA**→ |
| env-411 | JGI12273J12029_10877295 | + | 59 | 107 | **RNA**→ rpmE (PRK00019)Ribosomal_L31 (pfam01197)→ |
| env-412 | JGI12397J12024_1240506 | + | 83 | 131 | **RNA**→ ←hypo |
| env-413 | JGI12467J12023_1032285 | + | 11 | 56 | **RNA**→ rpmE (PRK00019)Ribosomal_L31 (pfam01197)→ |
| env-414 | JGI12419J13241_1175342 | + | 100 | 140 | **RNA**→ rpmE (PRK00019)Ribosomal_L31 (pfam01197)→ |
| env-415 | JGI12419J13241_1307864 | + | 48 | 89 | **RNA**→ rpmE (PRK00019)Ribosomal_L31 (pfam01197)→ |
| env-416 | 2200077197 | + | 8199 | 8240 | **RNA**→ rpmE (PRK00019)Ribosomal_L31 (pfam01197)→ |
| env-417 | ATED1026_F5B15RW01AGPIR | - | 111 | 72 | **RNA**→ ←hypo |
| env-418 | JGI12635J15846_12820982 | + | 23 | 77 | **RNA**→ hypo→ rpmE (PRK00019)Ribosomal_L31 (pfam01197)→ |
| Svi-14-4 | NZ_LGUP01000177.1 | - | 36425 | 36378 | **RNA**→ rpmE (PRK00019)Ribosomal_L31 (pfam01197)→ PrfA (COG0216)prfA (PRK00591)→ |
| | | | | | HemK (COG2890)PRK09328 (PRK09328)→ SUA5 (COG0009)TIGR00057 (TIGR00057)→ |
| | | | | | etp (PRK11391)Wzb (COG0394)→ GlyA (COG0112)glyA (PRK00011)→ |
| Swe-1-1 | NZ_JNWK01000034.1 | + | 44732 | 44779 | **RNA**→ rpmE (PRK00019)Ribosomal_L31 (pfam01197)→ PrfA (COG0216)prfA (PRK00591)→ |
| | | | | | HemK (COG2890)PRK09328 (PRK09328)→ SUA5 (COG0009)TIGR00057 (TIGR00057)→ |
| | | | | | etp (PRK11391)Wzb (COG0394)→ GlyA (COG0112)glyA (PRK00011)→ |
| Sve-1-1 | NC_018750.1 | + | 5396901 | 5396948 | **RNA**→ rpmE (PRK00019)Ribosomal_L31 (pfam01197)→ PrfA (COG0216)prfA (PRK00591)→ |
| | | | | | HemK (COG2890)PRK09328 (PRK09328)→ SUA5 (COG0009)TIGR00057 (TIGR00057)→ |
| | | | | | etp (PRK11391)Wzb (COG0394)→ GlyA (COG0112)glyA (PRK00011)→ |
| Ssp-38-1 | NZ_AKXV01000009.1 | - | 392719 | 392672 | **RNA**→ rpmE (PRK00019)Ribosomal_L31 (pfam01197)→ PrfA (COG0216)prfA (PRK00591)→ |
| | | | | | HemK (COG2890)PRK09328 (PRK09328)→ SUA5 (COG0009)TIGR00057 (TIGR00057)→ |
| | | | | | etp (PRK11391)Wzb (COG0394)→ GlyA (COG0112)glyA (PRK00011)→ |
| Sbi-3-1 | NZ_JNWL01000038.1 | + | 44138 | 44185 | **RNA**→ rpmE (PRK00019)Ribosomal_L31 (pfam01197)→ PrfA (COG0216)prfA (PRK00591)→ |
| | | | | | HemK (COG2890)PRK09328 (PRK09328)→ SUA5 (COG0009)TIGR00057 (TIGR00057)→ |
| | | | | | etp (PRK11391)Wzb (COG0394)→ |
| Sbi-3-2 | NZ_JOAU01000021.1 | - | 158964 | 158917 | **RNA**→ rpmE (PRK00019)Ribosomal_L31 (pfam01197)→ PrfA (COG0216)prfA (PRK00591)→ |
| | | | | | HemK (COG2890)PRK09328 (PRK09328)→ HemK (COG2890)PRK09328 (PRK09328)→ |
| | | | | | SUA5 (COG0009)TIGR00057 (TIGR00057)→ etp (PRK11391)Wzb (COG0394)→ |
| Sbi-3-3 | NZ_JNXJ01000039.1 | + | 44138 | 44185 | **RNA**→ rpmE (PRK00019)Ribosomal_L31 (pfam01197)→ PrfA (COG0216)prfA (PRK00591)→ |
| | | | | | HemK (COG2890)PRK09328 (PRK09328)→ HemK (COG2890)PRK09328 (PRK09328)→ |
| | | | | | SUA5 (COG0009)TIGR00057 (TIGR00057)→ etp (PRK11391)Wzb (COG0394)→ |
| Sgl-4-2 | NZ_JODW01000008.1 | + | 265005 | 265052 | **RNA**→ rpmE (PRK00019)Ribosomal_L31 (pfam01197)→ PrfA (COG0216)prfA (PRK00591)→ |
| | | | | | HemK (COG2890)PRK09328 (PRK09328)→ SUA5 (COG0009)TIGR00057 (TIGR00057)→ |
| | | | | | etp (PRK11391)Wzb (COG0394)→ |
| Svi-9-1 | NZ_CP010407.1 | + | 5717740 | 5717787 | **RNA**→ rpmE (PRK00019)Ribosomal_L31 (pfam01197)→ PrfA (COG0216)prfA (PRK00591)→ |
| | | | | | HemK (COG2890)PRK09328 (PRK09328)→ SUA5 (COG0009)TIGR00057 (TIGR00057)→ |
| | | | | | etp (PRK11391)Wzb (COG0394)→ |

| | | | | | |
|---|---|---|---|---|---|
| Fsp-2-1 | NZ_KB893751.1 | + | 18012 | 18059 | **RNA**→ rpmE (PRK00019)Ribosomal_L31 (pfam01197)→ |
| Fsp-5-1 | NC_009921.1 | + | 1192038 | 1192085 | **RNA**→ rpmE (PRK00019)Ribosomal_L31 (pfam01197)→ PrfA (COG0216)prfA (PRK00591)→ |
| | | | | | HemK (COG2890)PRK09328 (PRK09328)→ SUA5 (COG0009)TIGR00057 (TIGR00057)→ |
| Fsp-1-1 | NZ_KB891208.1 | + | 394290 | 394337 | **RNA**→ rpmE (PRK00019)Ribosomal_L31 (pfam01197)→ PrfA (COG0216)prfA (PRK00591)→ |
| | | | | | HemK (COG2890)PRK09328 (PRK09328)→ HemK (COG2890)PRK09328 (PRK09328)→ |
| | | | | | SUA5 (COG0009)TIGR00057 (TIGR00057)→ GlyA (COG0112)glyA (PRK00011)→ |
| | | | | | ECA_wecA (TIGR02380)GT_WecA_like (cd06853)→ hypo→ |
| Sgh-1-1 | NZ_DS999641.1 | - | 2595981 | 2595934 | **RNA**→ lytR_cpsA_psr (TIGR00350)LytR_cpsA_psr (pfam03816)→ |
| | | | | | lytR_cpsA_psr (TIGR00350)LytR_cpsA_psr (pfam03816)→ |
| | | | | | lytR_cpsA_psr (TIGR00350)LytR_cpsA_psr (pfam03816)→ rpmE (PRK00019)Ribosomal_L31 (pfam01197)→ |
| | | | | | rpmE (PRK00019)Ribosomal_L31 (pfam01197)→ |
| Svi-6-1 | NZ_JH993790.1 | - | 935516 | 935469 | **RNA**→ lytR_cpsA_psr (TIGR00350)LytR_cpsA_psr (pfam03816)→ |
| | | | | | rpmE (PRK00019)Ribosomal_L31 (pfam01197)→ |
| Ssp-31-1 | NZ_KB892208.1 | + | 277710 | 277757 | **RNA**→ rpmE (PRK00019)Ribosomal_L31 (pfam01197)→ |
| Ssp-172-1 | NZ_KK106988.1 | + | 976736 | 976783 | **RNA**→ rpmE (PRK00019)Ribosomal_L31 (pfam01197)→ |
| Sav-2-1 | NZ_JZJK01000090.1 | + | 1230456 | 1230503 | **RNA**→ rpmE (PRK00019)Ribosomal_L31 (pfam01197)→ |
| Sav-1-1 | NC_003155.4 | - | 3644711 | 3644664 | **RNA**→ rpmE (PRK00019)Ribosomal_L31 (pfam01197)→ |
| Sav-2-2 | NZ_BAVY01000023.1 | - | 106591 | 106544 | **RNA**→ rpmE (PRK00019)Ribosomal_L31 (pfam01197)→ |
| Ssp-7-1 | NZ_KB912967.1 | - | 88385 | 88338 | **RNA**→ rpmE (PRK00019)Ribosomal_L31 (pfam01197)→ |
| Ssp-44-1 | NZ_GG753626.1 | + | 4185762 | 4185809 | **RNA**→ rpmE (PRK00019)Ribosomal_L31 (pfam01197)→ rpmE (PRK00019)Ribosomal_L31 (pfam01197)→ |
| Sgr-10-1 | NZ_KL503830.1 | + | 5116277 | 5116324 | **RNA**→ rpmE (PRK00019)Ribosomal_L31 (pfam01197)→ |
| Aac-1-1 | NZ_JNYX01000011.1 | - | 75643 | 75596 | **RNA**→ rpmE (PRK00019)Ribosomal_L31 (pfam01197)→ |
| Ssp-173-1 | NZ_JMLN01000029.1 | + | 12038 | 12085 | **RNA**→ rpmE (PRK00019)Ribosomal_L31 (pfam01197)→ |
| Ssp-16-1 | NZ_KB891492.1 | - | 84534 | 84487 | **RNA**→ rpmE (PRK00019)Ribosomal_L31 (pfam01197)→ |
| Ssp-61-1 | NZ_LDZX01000024.1 | + | 53213 | 53260 | **RNA**→ rpmE (PRK00019)Ribosomal_L31 (pfam01197)→ |
| Ssp-175-1 | NZ_JXTE01000072.1 | - | 30728 | 30681 | **RNA**→ rpmE (PRK00019)Ribosomal_L31 (pfam01197)→ |
| Fsp-6-1 | NZ_ADGX01000178.1 | + | 1918 | 1965 | **RNA**→ rpmE (PRK00019)Ribosomal_L31 (pfam01197)→ rpmE (PRK00019)Ribosomal_L31 (pfam01197)→ |
| Fsp-13-1 | NZ_JYFN01000018.1 | + | 103666 | 103710 | **RNA**→ rpmE (PRK00019)Ribosomal_L31 (pfam01197)→ |
| Fal-1-1 | NC_008278.1 | - | 6445775 | 6445731 | **RNA**→ rpmE (PRK00019)Ribosomal_L31 (pfam01197)→ |
| Gte-4-1 | NZ_KL571655.1 | + | 204281 | 204341 | **RNA**→ rpmE (PRK00019)Ribosomal_L31 (pfam01197)→ |
| Gte-5-1 | NZ_ATYW01000003.1 | + | 382145 | 382205 | **RNA**→ rpmE (PRK00019)Ribosomal_L31 (pfam01197)→ |
| Mrh-1-1 | NZ_AGIQ01000008.1 | + | 20388 | 20443 | **RNA**→ rpmE (PRK00019)Ribosomal_L31 (pfam01197)→ PrfA (COG0216)prfA (PRK00591)→ |
| | | | | | HemK (COG2890)PRK09328 (PRK09328)→ SUA5 (COG0009)TIGR00057 (TIGR00057)→ |
| | | | | | GT_WecA_like (cd06853)mraY (PRK00108)→ |
| env-419 | JGI24741J21665_1351754 | + | 186 | 229 | **RNA**→ hypo→ |
| env-420 | JGI24740J21852_10109780 | - | 353 | 310 | **RNA**→ rpmE (PRK00019)Ribosomal_L31 (pfam01197)→ |
| env-421 | JGI24741J21665_1395540 | + | 54 | 97 | **RNA**→ rpmE (PRK00019)Ribosomal_L31 (pfam01197)→ |
| env-422 | JGI24735J21928_11010882 | - | 166 | 123 | **RNA**→ rpmE (PRK00019)Ribosomal_L31 (pfam01197)→ |
| env-423 | JGI12269J14319_11006088 | - | 200 | 110 | **RNA**→ rpmE (PRK00019)Ribosomal_L31 (pfam01197)→ |
| env-424 | JGI20186J14852_1074806 | + | 71 | 122 | **RNA**→ rpmE (PRK00019)Ribosomal_L31 (pfam01197)→ |
| env-425 | JGI20188J14859_1014361 | - | 823 | 772 | **RNA**→ rpmE (PRK00019)Ribosomal_L31 (pfam01197)→ |
| env-426 | JGI20190J14840_1016465 | + | 35 | 86 | **RNA**→ rpmE (PRK00019)Ribosomal_L31 (pfam01197)→ |
| env-427 | JGI20208J14878_1005722 | - | 584 | 533 | **RNA**→ rpmE (PRK00019)Ribosomal_L31 (pfam01197)→ |
| Nsp-11-1 | NZ_LAVL01000009.1 | - | 3292 | 3249 | **RNA**→ rpmE (PRK00019)Ribosomal_L31 (pfam01197)→ PrfA (COG0216)prfA (PRK00591)→ |
| | | | | | HemK (COG2890)PRK09328 (PRK09328)→ SUA5 (COG0009)TIGR00057 (TIGR00057)→ hypo→ hypo→ |
| env-428 | JGI26341J46601_10154785 | - | 301 | 238 | **RNA**→ hypo→ |
| env-429 | JGI26340J50214_10419594 | + | 13 | 52 | **RNA**→ rpmE (PRK00019)Ribosomal_L31 (pfam01197)→ |
| env-430 | JGI26340J50214_10140346 | + | 542 | 581 | **RNA**→ |
| env-431 | JGI26341J46601_10019475 | + | 120 | 166 | **RNA**→ rpmE (PRK00019)Ribosomal_L31 (pfam01197)→ PrfA (COG0216)prfA (PRK00591)→ |
| | | | | | HemK (COG2890)PRK09328 (PRK09328)→ |
| env-432 | JGI26339J46600_10188658 | + | 54 | 100 | **RNA**→ rpmE (PRK00019)Ribosomal_L31 (pfam01197)→ |
| env-433 | JGI12712J15308_11072730 | + | 109 | 155 | **RNA**→ rpmE (PRK00019)Ribosomal_L31 (pfam01197)→ |
| env-434 | JGI20133J14441_1007812 | - | 3380 | 3329 | **RNA**→ rpmE (PRK00019)Ribosomal_L31 (pfam01197)→ PrfA (COG0216)prfA (PRK00591)→ |
| | | | | | HemK (COG2890)PRK09328 (PRK09328)→ SUA5 (COG0009)TIGR00057 (TIGR00057)→ Rfe (COG0472)→ |

| | | | | | |
|---|---|---|---|---|---|
| env-435 | 3300001422_101093 | - | 1365 | 1314 | **RNA**→ rpmE (PRK00019)Ribosomal_L31 (pfam01197)→ PrfA (COG0216)prfA (PRK00591)→ |
| Str-2-1 | NZ_KB897247.1 | - | 51465 | 51408 | **RNA**→ rpmE (PRK00019)Ribosomal_L31 (pfam01197)→ |
| Str-1-1 | NC_009380.1 | - | 4173707 | 4173650 | **RNA**→ rpmE (PRK00019)Ribosomal_L31 (pfam01197)→ |
| Str-3-1 | NZ_KB897150.1 | - | 51556 | 51499 | **RNA**→ rpmE (PRK00019)Ribosomal_L31 (pfam01197)→ |
| Str-7-1 | NZ_KI911454.1 | + | 198877 | 198934 | **RNA**→ rpmE (PRK00019)Ribosomal_L31 (pfam01197)→ |
| Str-10-1 | NZ_KK211108.1 | - | 50876 | 50819 | **RNA**→ rpmE (PRK00019)Ribosomal_L31 (pfam01197)→ |
| Str-8-1 | NZ_KK211120.1 | - | 51590 | 51533 | **RNA**→ rpmE (PRK00019)Ribosomal_L31 (pfam01197)→ |
| Str-9-1 | NZ_AUKS01000011.1 | - | 51392 | 51335 | **RNA**→ rpmE (PRK00019)Ribosomal_L31 (pfam01197)→ |
| Sar-29-1 | NZ_AUEJ01000011.1 | - | 51474 | 51417 | **RNA**→ rpmE (PRK00019)Ribosomal_L31 (pfam01197)→ |
| Str-6-1 | NZ_AZXH01000007.1 | - | 51265 | 51208 | **RNA**→ rpmE (PRK00019)Ribosomal_L31 (pfam01197)→ |
| Str-4-1 | NZ_AZXE01000005.1 | - | 51306 | 51249 | **RNA**→ rpmE (PRK00019)Ribosomal_L31 (pfam01197)→ |
| Str-5-1 | NZ_AZXD01000012.1 | - | 51215 | 51158 | **RNA**→ rpmE (PRK00019)Ribosomal_L31 (pfam01197)→ |
| Spa-24-1 | NZ_KE384274.1 | + | 353881 | 353937 | **RNA**→ hypo→ rpmE (PRK00019)Ribosomal_L31 (pfam01197)→ |
| Spa-32-1 | NZ_KI911397.1 | + | 504459 | 504515 | **RNA**→ hypo→ rpmE (PRK00019)Ribosomal_L31 (pfam01197)→ |
| Spa-20-1 | NZ_KB900614.1 | - | 901662 | 901606 | **RNA**→ hypo→ rpmE (PRK00019)Ribosomal_L31 (pfam01197)→ |
| Spa-7-1 | NZ_KB895231.1 | + | 38192 | 38248 | **RNA**→ hypo→ rpmE (PRK00019)Ribosomal_L31 (pfam01197)→ |
| Spa-6-1 | NZ_KB895117.1 | - | 50778 | 50722 | **RNA**→ hypo→ rpmE (PRK00019)Ribosomal_L31 (pfam01197)→ |
| Spa-27-1 | NZ_AZWB01000027.1 | + | 22264 | 22320 | **RNA**→ hypo→ rpmE (PRK00019)Ribosomal_L31 (pfam01197)→ |
| Spa-30-1 | NZ_AZVY01000026.1 | + | 19243 | 19299 | **RNA**→ hypo→ rpmE (PRK00019)Ribosomal_L31 (pfam01197)→ |
| Spa-18-1 | NZ_KB911579.1 | - | 914415 | 914359 | **RNA**→ hypo→ rpmE (PRK00019)Ribosomal_L31 (pfam01197)→ |
| Spa-17-1 | NZ_KB913027.1 | - | 891436 | 891380 | **RNA**→ hypo→ rpmE (PRK00019)Ribosomal_L31 (pfam01197)→ |
| Spa-5-1 | NZ_KB895050.1 | + | 20116 | 20172 | **RNA**→ hypo→ rpmE (PRK00019)Ribosomal_L31 (pfam01197)→ |
| Spa-4-1 | NZ_KB894900.1 | + | 366858 | 366914 | **RNA**→ hypo→ rpmE (PRK00019)Ribosomal_L31 (pfam01197)→ |
| Spa-15-1 | NZ_KB913022.1 | - | 4269611 | 4269555 | **RNA**→ hypo→ rpmE (PRK00019)Ribosomal_L31 (pfam01197)→ |
| Spa-38-1 | NZ_AXVR01000016.1 | + | 40238 | 40293 | **RNA**→ rpmE (PRK00019)Ribosomal_L31 (pfam01197)→ |
| env-436 | JGI24770J26754_10002275 | + | 2336 | 2389 | **RNA**→ rpmE (PRK00019)Ribosomal_L31 (pfam01197)→ |
| Msp-11-1 | NZ_CP011773.1 | - | 4667799 | 4667746 | **RNA**→ rpmE (PRK00019)Ribosomal_L31 (pfam01197)→ PrfA (COG0216)prfA (PRK00591)→<br>HemK (COG2890)PRK09328 (PRK09328)→ SUA5 (COG0009)TIGR00057 (TIGR00057)→<br>GT_WecA_like (cd06853)mraY (PRK00108)→ |
| env-437 | PCEOT_contig21592 | + | 631 | 671 | **RNA**→ rpmE (PRK00019)Ribosomal_L31 (pfam01197)→ prfA (PRK00591)PCRF (pfam03462)→ |
| env-438 | JGI26341J46601_10000390 | + | 11088 | 11139 | **RNA**→ rpmE (PRK00019)Ribosomal_L31 (pfam01197)→ |
| env-439 | JGI26340J50214_10007419 | + | 1129 | 1180 | **RNA**→ rpmE (PRK00019)Ribosomal_L31 (pfam01197)→ |
| env-440 | C688J13580_1085580 | - | 412 | 368 | **RNA**→ hypo→ rpmE (PRK00019)Ribosomal_L31 (pfam01197)→ hypo→ |
| env-441 | JGI26345J50200_1174654 | - | 161 | 109 | **RNA**→ ←hypo |
| env-442 | JGI26341J46601_10705222 | - | 114 | 41 | **RNA**→ hypo→ |
| Gkr-1-1 | NZ_AQYG01000051.1 | - | 16795 | 16754 | **RNA**→ rpmE (PRK00019)Ribosomal_L31 (pfam01197)→ PrfA (COG0216)prfA (PRK00591)→<br>HemK (COG2890)PRK09328 (PRK09328)→ SUA5 (COG0009)TIGR00057 (TIGR00057)→<br>GT_WecA_like (cd06853)mraY (PRK00108)→ GT_WecA_like (cd06853)mraY (PRK00108)→ |
| env-443 | JGI24721J26819_10140833 | - | 944 | 893 | **RNA**→ ←hypo |
| env-444 | JGI24721J44947_10027217 | - | 4805 | 4754 | **RNA**→ rpmE (PRK00019)Ribosomal_L31 (pfam01197)→ PrfA (COG0216)prfA (PRK00591)→<br>HemK (COG2890)PRK09328 (PRK09328)→ SUA5 (COG0009)TIGR00057 (TIGR00057)→<br>GT_WecA_like (cd06853)mraY (PRK00108)→ |
| env-445 | JGI24721J44947_10041083 | - | 3136 | 3085 | **RNA**→ rpmE (PRK00019)Ribosomal_L31 (pfam01197)→ PrfA (COG0216)prfA (PRK00591)→<br>HemK (COG2890)PRK09328 (PRK09328)→ SUA5 (COG0009)TIGR00057 (TIGR00057)→ |
| env-446 | JGI20262J16345_11379 | - | 97 | 56 | **RNA**→ hypo→ |
| env-447 | JGI12272J11983_1061016 | + | 13 | 57 | **RNA**→ rpmE (PRK00019)Ribosomal_L31 (pfam01197)→ |
| env-448 | JGI12467J12023_1038166 | + | 90 | 134 | **RNA**→ rpmE (PRK00019)Ribosomal_L31 (pfam01197)→ |
| env-449 | DCrCPGB_c205975 | - | 57 | 15 | **RNA**→ |
| Mhe-1-1 | NZ_LDPO01000013.1 | - | 14940 | 14896 | **RNA**→ rpmE (PRK00019)Ribosomal_L31 (pfam01197)→ PrfA (COG0216)prfA (PRK00591)→<br>HemK (COG2890)PRK09328 (PRK09328)→ SUA5 (COG0009)TIGR00057 (TIGR00057)→<br>GT_WecA_like (cd06853)mraY (PRK00108)→ |
| Rrh-1-1 | NZ_APMY01000113.1 | + | 67570 | 67615 | **RNA**→ rpmE (PRK00019)Ribosomal_L31 (pfam01197)→ PrfA (COG0216)prfA (PRK00591)→<br>HemK (COG2890)PRK09328 (PRK09328)→ SUA5 (COG0009)TIGR00057 (TIGR00057)→<br>ECA_wecA (TIGR02380)GT_WecA_like (cd06853)→ EPS_sugtrans (TIGR03025)→ hypo→ |

| | | | | | |
|---|---|---|---|---|---|
| Msp-13-1 | NZ_JRML01000027.1 | + | 41320 | 41371 | RNA→ rpmE (PRK00019)Ribosomal_L31 (pfam01197)→ PrfA (COG0216)prfA (PRK00591)→ HemK (COG2890)PRK09328 (PRK09328)→ SUA5 (COG0009)TIGR00057 (TIGR00057)→ GT_WecA_like (cd06853)mraY (PRK00108)→ |
| Mar-2-1 | NZ_LASW02000025.1 | + | 41195 | 41246 | RNA→ rpmE (PRK00019)Ribosomal_L31 (pfam01197)→ PrfA (COG0216)prfA (PRK00591)→ HemK (COG2890)PRK09328 (PRK09328)→ SUA5 (COG0009)TIGR00057 (TIGR00057)→ GT_WecA_like (cd06853)mraY (PRK00108)→ |
| env-450 | 2205805686 | + | 242 | 290 | RNA→ rpmE (PRK00019)Ribosomal_L31 (pfam01197)→ |
| env-451 | 2158117515 | + | 242 | 290 | RNA→ ←hypo |
| Tbi-1-1 | NZ_BAVQ01000020.1 | - | 15062 | 15016 | RNA→ rpmE (PRK00019)Ribosomal_L31 (pfam01197)→ |
| Msp-4-1 | NC_015576.1 | + | 1383199 | 1383245 | RNA→ rpmE (PRK00019)Ribosomal_L31 (pfam01197)→ PrfA (COG0216)prfA (PRK00591)→ HemK (COG2890)PRK09328 (PRK09328)→ SUA5 (COG0009)TIGR00057 (TIGR00057)→ GT_WecA_like (cd06853)mraY (PRK00108)→ |
| Mav-1-1 | NZ_AGAQ01000161.1 | - | 14730 | 14685 | RNA→ rpmE (PRK00019)Ribosomal_L31 (pfam01197)→ PrfA (COG0216)prfA (PRK00591)→ HemK (COG2890)PRK09328 (PRK09328)→ SUA5 (COG0009)TIGR00057 (TIGR00057)→ GT_WecA_like (cd06853)mraY (PRK00108)→ |
| Msp-16-1 | NZ_JRMN01000077.1 | + | 14933 | 14979 | RNA→ rpmE (PRK00019)Ribosomal_L31 (pfam01197)→ PrfA (COG0216)prfA (PRK00591)→ HemK (COG2890)PRK09328 (PRK09328)→ SUA5 (COG0009)TIGR00057 (TIGR00057)→ GT_WecA_like (cd06853)mraY (PRK00108)→ |
| env-452 | JGI24770J26754_10065094 | - | 520 | 475 | RNA→ rpmE (PRK00019)Ribosomal_L31 (pfam01197)→ PCRF (pfam03462)ABC_tran_CTD (pfam16326)→ |
| Msp-12-1 | NZ_JRMK01000124.1 | - | 7553 | 7509 | RNA→ rpmE (PRK00019)Ribosomal_L31 (pfam01197)→ PrfA (COG0216)prfA (PRK00591)→ HemK (COG2890)PRK09328 (PRK09328)→ SUA5 (COG0009)TIGR00057 (TIGR00057)→ GT_WecA_like (cd06853)mraY (PRK00108)→ |
| Msp-14-1 | NZ_JRMM01000086.1 | + | 10717 | 10763 | RNA→ rpmE (PRK00019)Ribosomal_L31 (pfam01197)→ PrfA (COG0216)prfA (PRK00591)→ HemK (COG2890)PRK09328 (PRK09328)→ SUA5 (COG0009)TIGR00057 (TIGR00057)→ GT_WecA_like (cd06853)mraY (PRK00108)→ |
| Msp-17-1 | NZ_AUWR01000034.1 | + | 51013 | 51059 | RNA→ rpmE (PRK00019)Ribosomal_L31 (pfam01197)→ PrfA (COG0216)prfA (PRK00591)→ HemK (COG2890)PRK09328 (PRK09328)→ SUA5 (COG0009)TIGR00057 (TIGR00057)→ GT_WecA_like (cd06853)mraY (PRK00108)→ |
| env-453 | JGI25151J46595_10355844 | + | 12 | 67 | RNA→ rpmE (PRK00019)Ribosomal_L31 (pfam01197)→ |
| Mky-1-1 | NZ_BBKA01000066.1 | - | 14976 | 14925 | RNA→ rpmE (PRK00019)Ribosomal_L31 (pfam01197)→ PrfA (COG0216)prfA (PRK00591)→ hypo→ hypo→ SUA5 (COG0009)TIGR00057 (TIGR00057)→ hypo→ hypo→ |
| Sar-43-1 | NZ_AZWW01000033.1 | + | 1603 | 1659 | RNA→ hypo→ rpmE (PRK00019)Ribosomal_L31 (pfam01197)→ |
| Sar-41-1 | NZ_KI911429.1 | + | 150303 | 150359 | RNA→ hypo→ rpmE (PRK00019)Ribosomal_L31 (pfam01197)→ |
| Sar-42-1 | NZ_KI911487.1 | + | 12550 | 12606 | RNA→ hypo→ rpmE (PRK00019)Ribosomal_L31 (pfam01197)→ |
| Sar-10-1 | NZ_KB896245.1 | - | 51122 | 51066 | RNA→ hypo→ rpmE (PRK00019)Ribosomal_L31 (pfam01197)→ |
| Sar-11-1 | NZ_KB896343.1 | + | 1657 | 1713 | RNA→ hypo→ rpmE (PRK00019)Ribosomal_L31 (pfam01197)→ |
| env-454 | JGI12396J12026_1015431 | + | 45 | 84 | RNA→ hypo→ |
| env-455 | C688J13580_1520551 | - | 158 | 116 | RNA→ rpmE (PRK00019)Ribosomal_L31 (pfam01197)→ |
| env-456 | JGI24737J22298_11100121 | - | 70 | 27 | RNA→ |
| Ase-1-1 | NZ_JOEG01000014.1 | + | 292419 | 292467 | RNA→ rpmE (PRK00019)Ribosomal_L31 (pfam01197)→ |
| Fsp-9-1 | NZ_CM001489.1 | + | 6555193 | 6555238 | RNA→ rpmE (PRK00019)Ribosomal_L31 (pfam01197)→ rpmE (PRK00019)Ribosomal_L31 (pfam01197)→ |
| env-457 | JGI12397J12024_1075610 | + | 11 | 62 | RNA→ rpmE (PRK00019)Ribosomal_L31 (pfam01197)→ |
| env-458 | JGI12273J12029_10446706 | + | 9 | 60 | RNA→ rpmE (PRK00019)Ribosomal_L31 (pfam01197)→ |
| env-459 | JGI12273J12029_10361449 | - | 52 | 1 | RNA→ |
| env-460 | 2044458367 | - | 94 | 11 | RNA→ |
| Spa-33-1 | NZ_AZWQ01000015.1 | + | 224755 | 224811 | RNA→ hypo→ rpmE (PRK00019)Ribosomal_L31 (pfam01197)→ |
| Spa-2-1 | NZ_KB894849.1 | - | 95011 | 94955 | RNA→ hypo→ rpmE (PRK00019)Ribosomal_L31 (pfam01197)→ |
| Vma-1-1 | NC_015434.1 | - | 5587166 | 5587106 | RNA→ rpmE (PRK00019)Ribosomal_L31 (pfam01197)→ PrfA (COG0216)prfA (PRK00591)→ |
| Gbr-1-1 | NC_013441.1 | + | 2009569 | 2009614 | RNA→ rpmE (PRK00019)Ribosomal_L31 (pfam01197)→ |
| Gna-1-1 | NZ_BAHE01000010.1 | - | 25454 | 25409 | RNA→ rpmE (PRK00019)Ribosomal_L31 (pfam01197)→ rpmE (PRK00019)Ribosomal_L31 (pfam01197)→ |
| Gal-1-1 | NZ_BACI01000100.1 | + | 69601 | 69646 | RNA→ rpmE (PRK00019)Ribosomal_L31 (pfam01197)→ rpmE (PRK00019)Ribosomal_L31 (pfam01197)→ |
| Gal-2-1 | NZ_KI629793.1 | + | 441597 | 441642 | RNA→ rpmE (PRK00019)Ribosomal_L31 (pfam01197)→ |
| Gal-3-1 | NZ_HG764859.1 | + | 14350 | 14395 | RNA→ rpmE (PRK00019)Ribosomal_L31 (pfam01197)→ |
| Gam-2-1 | NZ_BANS01000028.1 | + | 75421 | 75466 | RNA→ rpmE (PRK00019)Ribosomal_L31 (pfam01197)→ |
| Gsp-1-1 | NC_018581.1 | + | 2102127 | 2102172 | RNA→ rpmE (PRK00019)Ribosomal_L31 (pfam01197)→ |

| | | | | | |
|---|---|---|---|---|---|
| Gte-1-1 | NZ_AQPW01000013.1 | + | 98992 | 99037 | RNA→ rpmE (PRK00019)Ribosomal_L31 (pfam01197)→ |
| Gpa-1-1 | NZ_BAOQ01000029.1 | + | 64185 | 64230 | RNA→ ←PRK06214 (PRK06214)SiR (cd06199) |
| Gru-1-1 | NZ_BAHB01000043.1 | + | 60751 | 60796 | RNA→ rpmE (PRK00019)Ribosomal_L31 (pfam01197)→ rpmE (PRK00019)Ribosomal_L31 (pfam01197)→ |
| Gam-3-1 | NZ_AWTB01000003.1 | + | 200060 | 200105 | RNA→ rpmE (PRK00019)Ribosomal_L31 (pfam01197)→ |
| Gte-3-1 | NZ_JNXA01000016.1 | - | 25295 | 25250 | RNA→ rpmE (PRK00019)Ribosomal_L31 (pfam01197)→ |
| Gte-2-1 | NZ_BAFD01000064.1 | + | 11988 | 12033 | RNA→ rpmE (PRK00019)Ribosomal_L31 (pfam01197)→ rpmE (PRK00019)Ribosomal_L31 (pfam01197)→ rpmE (PRK00019)Ribosomal_L31 (pfam01197)→ |
| Nsp-1-1 | NZ_KB911522.1 | - | 24205 | 24160 | RNA→ rpmE (PRK00019)Ribosomal_L31 (pfam01197)→ |
| Wsp-1-1 | NZ_AYTE01000018.1 | + | 71375 | 71420 | RNA→ rpmE (PRK00019)Ribosomal_L31 (pfam01197)→ |
| Kar-2-1 | NZ_JNYV01000009.1 | + | 112546 | 112591 | RNA→ rpmE (PRK00019)Ribosomal_L31 (pfam01197)→ |
| Msp-30-1 | NZ_JOAF01000010.1 | + | 112447 | 112492 | RNA→ rpmE (PRK00019)Ribosomal_L31 (pfam01197)→ |
| Gef-1-1 | NZ_BAEH01000017.1 | - | 16990 | 16945 | RNA→ rpmE (PRK00019)Ribosomal_L31 (pfam01197)→ rpmE (PRK00019)Ribosomal_L31 (pfam01197)→ PrfA (COG0216)prfA (PRK00591)→ PrfA (COG0216)prfA (PRK00591)→ HemK (COG2890)PRK09328 (PRK09328)→ HemK (COG2890)PRK09328 (PRK09328)→ HemK (COG2890)PRK09328 (PRK09328)→ SUA5 (COG0009)TIGR00057 (TIGR00057)→ SUA5 (COG0009)TIGR00057 (TIGR00057)→ GlyA (COG0112)glyA (PRK00011)→ GlyA (COG0112)glyA (PRK00011)→ GT_WecA_like (cd06853)mraY (PRK00108)→ GT_WecA_like (cd06853)mraY (PRK00108)→ AnsP (COG1113)AA_permease_2 (pfam13520)→ AnsP (COG1113)AA_permease_2 (pfam13520)→ AnsP (COG1113)AA_permease_2 (pfam13520)→ |
| env-461 | AglaG_F499EG101AZGLU | - | 123 | 78 | RNA→ |
| Gpo-3-1 | NC_016906.1 | + | 2046857 | 2046902 | RNA→ rpmE (PRK00019)Ribosomal_L31 (pfam01197)→ |
| Gpo-2-1 | NZ_BAEI01000018.1 | - | 30702 | 30657 | RNA→ rpmE (PRK00019)Ribosomal_L31 (pfam01197)→ rpmE (PRK00019)Ribosomal_L31 (pfam01197)→ |
| Gpo-1-1 | NZ_ARVZ01000001.1 | - | 3468004 | 3467959 | RNA→ rpmE (PRK00019)Ribosomal_L31 (pfam01197)→ |
| Grh-1-1 | NZ_BAHC01000025.1 | - | 28381 | 28336 | RNA→ rpmE (PRK00019)Ribosomal_L31 (pfam01197)→ rpmE (PRK00019)Ribosomal_L31 (pfam01197)→ |
| env-462 | P3_CLC_Cons143904 | + | 638 | 680 | RNA→ rpmE (PRK00019)Ribosomal_L31 (pfam01197)→ |
| Rde-1-1 | NZ_JPOC01000028.1 | - | 15680 | 15635 | RNA→ rpmE (PRK00019)Ribosomal_L31 (pfam01197)→ |
| Req-3-1 | NZ_APJC01000016.1 | + | 221235 | 221280 | RNA→ rpmE (PRK00019)Ribosomal_L31 (pfam01197)→ |
| Req-1-1 | NC_014659.1 | + | 1850042 | 1850087 | RNA→ rpmE (PRK00019)Ribosomal_L31 (pfam01197)→ |
| Req-2-1 | NZ_CM001149.1 | - | 1501379 | 1501334 | RNA→ rpmE (PRK00019)Ribosomal_L31 (pfam01197)→ |
| env-463 | JGI12060J13090_100106 | + | 585 | 630 | RNA→ rpmE (PRK00019)Ribosomal_L31 (pfam01197)→ |
| env-464 | JGI11829J13087_100888 | + | 1844 | 1889 | RNA→ rpmE (PRK00019)Ribosomal_L31 (pfam01197)→ |
| env-465 | JGI12027J13101_100505 | + | 2020 | 2065 | RNA→ rpmE (PRK00019)Ribosomal_L31 (pfam01197)→ |
| env-466 | JGI12209J13104_110011 | + | 138 | 183 | RNA→ rpmE (PRK00019)Ribosomal_L31 (pfam01197)→ |
| env-467 | JGI12064J13199_100272 | + | 1905 | 1950 | RNA→ rpmE (PRK00019)Ribosomal_L31 (pfam01197)→ |
| env-468 | JGI12075J13256_1000008 | - | 18327 | 18282 | RNA→ rpmE (PRK00019)Ribosomal_L31 (pfam01197)→ |
| env-469 | JGI12097J13213_1000024 | + | 50065 | 50110 | RNA→ rpmE (PRK00019)Ribosomal_L31 (pfam01197)→ |
| env-470 | JGI11758J13082_100031 | - | 5158 | 5113 | RNA→ rpmE (PRK00019)Ribosomal_L31 (pfam01197)→ |
| env-471 | JGI12112J13074_1000205 | - | 1882 | 1837 | RNA→ rpmE (PRK00019)Ribosomal_L31 (pfam01197)→ |
| env-472 | JGI11819J13099_103704 | + | 158 | 203 | RNA→ rpmE (PRK00019)Ribosomal_L31 (pfam01197)→ |
| env-473 | JGI11800J13271_1007055 | - | 1039 | 994 | RNA→ rpmE (PRK00019)Ribosomal_L31 (pfam01197)→ |
| Rfa-3-1 | NZ_JMFE01000037.1 | - | 19407 | 19362 | RNA→ rpmE (PRK00019)Ribosomal_L31 (pfam01197)→ PrfA (COG0216)prfA (PRK00591)→ SHOCT (pfam09851)→ HemK (COG2890)PRK09328 (PRK09328)→ SUA5 (COG0009)TIGR00057 (TIGR00057)→ GlyA (COG0112)glyA (PRK00011)→ ECA_wecA (TIGR02380)GT_WecA_like (cd06853)→ hypo→ |
| Rsp-2-1 | NZ_KB907581.1 | + | 121301 | 121346 | RNA→ rpmE (PRK00019)Ribosomal_L31 (pfam01197)→ PrfA (COG0216)prfA (PRK00591)→ SHOCT (pfam09851)→ HemK (COG2890)PRK09328 (PRK09328)→ HemK (COG2890)PRK09328 (PRK09328)→ SUA5 (COG0009)TIGR00057 (TIGR00057)→ SUA5 (COG0009)TIGR00057 (TIGR00057)→ GlyA (COG0112)glyA (PRK00011)→ ECA_wecA (TIGR02380)GT_WecA_like (cd06853)→ hypo→ |
| Rsp-1-1 | NZ_KB911561.1 | + | 83738 | 83783 | RNA→ rpmE (PRK00019)Ribosomal_L31 (pfam01197)→ PrfA (COG0216)prfA (PRK00591)→ SHOCT (pfam09851)→ HemK (COG2890)PRK09328 (PRK09328)→ SUA5 (COG0009)TIGR00057 (TIGR00057)→ SUA5 (COG0009)TIGR00057 (TIGR00057)→ GlyA (COG0112)glyA (PRK00011)→ ECA_wecA (TIGR02380)GT_WecA_like (cd06853)→ hypo→ |
| Rfa-1-1 | NZ_JOKB01000004.1 | + | 98966 | 99011 | RNA→ rpmE (PRK00019)Ribosomal_L31 (pfam01197)→ PrfA (COG0216)prfA (PRK00591)→ SHOCT (pfam09851)→ HemK (COG2890)PRK09328 (PRK09328)→ SUA5 (COG0009)TIGR00057 (TIGR00057)→ GlyA (COG0112)glyA (PRK00011)→ ECA_wecA (TIGR02380)GT_WecA_like (cd06853)→ hypo→ |

| | | | | | |
|---|---|---|---|---|---|
| Rfa-10-1 | NZ_JMEX01000003.1 | - | 19312 | 19267 | **RNA**→ rpmE (PRK00019)Ribosomal_L31 (pfam01197)→ PrfA (COG0216)prfA (PRK00591)→ SHOCT (pfam09851)→ HemK (COG2890)PRK09328 (PRK09328)→ SUA5 (COG0009)TIGR00057 (TIGR00057)→ GlyA (COG0112)glyA (PRK00011)→ ECA_wecA (TIGR02380)GT_WecA_like (cd06853)→ hypo→ |
| Rfa-11-1 | NZ_JMEW01000005.1 | - | 19530 | 19485 | **RNA**→ rpmE (PRK00019)Ribosomal_L31 (pfam01197)→ PrfA (COG0216)prfA (PRK00591)→ SHOCT (pfam09851)→ HemK (COG2890)PRK09328 (PRK09328)→ SUA5 (COG0009)TIGR00057 (TIGR00057)→ GlyA (COG0112)glyA (PRK00011)→ ECA_wecA (TIGR02380)GT_WecA_like (cd06853)→ hypo→ |
| Rfa-2-1 | NZ_JMFF01000029.1 | + | 83922 | 83967 | **RNA**→ rpmE (PRK00019)Ribosomal_L31 (pfam01197)→ PrfA (COG0216)prfA (PRK00591)→ SHOCT (pfam09851)→ HemK (COG2890)PRK09328 (PRK09328)→ SUA5 (COG0009)TIGR00057 (TIGR00057)→ GlyA (COG0112)glyA (PRK00011)→ ECA_wecA (TIGR02380)GT_WecA_like (cd06853)→ hypo→ |
| Rsp-3-1 | NZ_CAPS01000066.1 | + | 3310 | 3355 | **RNA**→ rpmE (PRK00019)Ribosomal_L31 (pfam01197)→ PrfA (COG0216)prfA (PRK00591)→ |
| Rfa-12-1 | NZ_JMEV01000023.1 | - | 619554 | 619509 | **RNA**→ rpmE (PRK00019)Ribosomal_L31 (pfam01197)→ PrfA (COG0216)prfA (PRK00591)→ |
| Rsp-13-1 | NZ_CP010797.1 | + | 262870 | 262915 | **RNA**→ rpmE (PRK00019)Ribosomal_L31 (pfam01197)→ PrfA (COG0216)prfA (PRK00591)→ |
| Rfa-5-1 | NZ_JMFC01000001.1 | + | 66942 | 66987 | **RNA**→ rpmE (PRK00019)Ribosomal_L31 (pfam01197)→ PrfA (COG0216)prfA (PRK00591)→ |
| Rop-1-1 | NC_012522.1 | - | 1262208 | 1262163 | **RNA**→ rpmE (PRK00019)Ribosomal_L31 (pfam01197)→ PrfA (COG0216)prfA (PRK00591)→ HemK (COG2890)PRK09328 (PRK09328)→ SUA5 (COG0009)TIGR00057 (TIGR00057)→ ECA_wecA (TIGR02380)GT_WecA_like (cd06853)→ hypo→ |
| Rjo-1-1 | NC_008268.1 | - | 1564922 | 1564877 | **RNA**→ rpmE (PRK00019)Ribosomal_L31 (pfam01197)→ PrfA (COG0216)prfA (PRK00591)→ HemK (COG2890)PRK09328 (PRK09328)→ SUA5 (COG0009)TIGR00057 (TIGR00057)→ ECA_wecA (TIGR02380)GT_WecA_like (cd06853)→ hypo→ PRK05815 (PRK05815)ATP-synt_A (pfam00119)→ PRK05815 (PRK05815)ATP-synt_A (pfam00119)→ PRK07874 (PRK07874)AtpE (COG0636)→ PRK05759 (PRK05759)AtpF (COG0711)→ AtpH (COG0712)PRK13430 (PRK13430)→ AtpA (COG0056)PRK09281 (PRK09281)→ AtpG (COG0224)PRK05621 (PRK05621)→ |
| Rop-3-1 | NZ_JH377359.1 | + | 2320813 | 2320858 | **RNA**→ rpmE (PRK00019)Ribosomal_L31 (pfam01197)→ PrfA (COG0216)prfA (PRK00591)→ HemK (COG2890)PRK09328 (PRK09328)→ SUA5 (COG0009)TIGR00057 (TIGR00057)→ ECA_wecA (TIGR02380)GT_WecA_like (cd06853)→ hypo→ PRK05815 (PRK05815)ATP-synt_A (pfam00119)→ PRK05815 (PRK05815)ATP-synt_A (pfam00119)→ PRK07874 (PRK07874)AtpE (COG0636)→ PRK05759 (PRK05759)AtpF (COG0711)→ AtpH (COG0712)PRK13430 (PRK13430)→ AtpA (COG0056)PRK09281 (PRK09281)→ AtpG (COG0224)PRK05621 (PRK05621)→ |
| Rop-3-2 | NZ_CP003949.1 | - | 5793207 | 5793162 | **RNA**→ rpmE (PRK00019)Ribosomal_L31 (pfam01197)→ PrfA (COG0216)prfA (PRK00591)→ HemK (COG2890)PRK09328 (PRK09328)→ SUA5 (COG0009)TIGR00057 (TIGR00057)→ ECA_wecA (TIGR02380)GT_WecA_like (cd06853)→ hypo→ |
| Rim-1-1 | NZ_AJJH01000159.1 | + | 23732 | 23777 | **RNA**→ rpmE (PRK00019)Ribosomal_L31 (pfam01197)→ rpmE (PRK00019)Ribosomal_L31 (pfam01197)→ PrfA (COG0216)prfA (PRK00591)→ PrfA (COG0216)prfA (PRK00591)→ HemK (COG2890)PRK09328 (PRK09328)→ HemK (COG2890)PRK09328 (PRK09328)→ SUA5 (COG0009)TIGR00057 (TIGR00057)→ SUA5 (COG0009)TIGR00057 (TIGR00057)→ ECA_wecA (TIGR02380)GT_WecA_like (cd06853)→ ECA_wecA (TIGR02380)GT_WecA_like (cd06853)→ hypo→ hypo→ |
| Rsp-4-1 | NZ_AJLQ01000087.1 | + | 23811 | 23856 | **RNA**→ rpmE (PRK00019)Ribosomal_L31 (pfam01197)→ PrfA (COG0216)prfA (PRK00591)→ HemK (COG2890)PRK09328 (PRK09328)→ SUA5 (COG0009)TIGR00057 (TIGR00057)→ ECA_wecA (TIGR02380)GT_WecA_like (cd06853)→ hypo→ PRK05815 (PRK05815)ATP-synt_A (pfam00119)→ PRK05815 (PRK05815)ATP-synt_A (pfam00119)→ PRK07874 (PRK07874)AtpE (COG0636)→ PRK05759 (PRK05759)AtpF (COG0711)→ AtpH (COG0712)PRK13430 (PRK13430)→ AtpA (COG0056)PRK09281 (PRK09281)→ AtpG (COG0224)PRK05621 (PRK05621)→ |
| Rsp-6-1 | NZ_AKKP01000036.1 | - | 14697 | 14652 | **RNA**→ ←hypo |
| Rop-2-1 | NZ_AJYC02000166.1 | + | 20908 | 20953 | **RNA**→ rpmE (PRK00019)Ribosomal_L31 (pfam01197)→ PrfA (COG0216)prfA (PRK00591)→ HemK (COG2890)PRK09328 (PRK09328)→ SUA5 (COG0009)TIGR00057 (TIGR00057)→ ECA_wecA (TIGR02380)GT_WecA_like (cd06853)→ hypo→ |
| Rwr-1-1 | NZ_ANIU01000624.1 | - | 15537 | 15492 | **RNA**→ rpmE (PRK00019)Ribosomal_L31 (pfam01197)→ PrfA (COG0216)prfA (PRK00591)→ HemK (COG2890)PRK09328 (PRK09328)→ SUA5 (COG0009)TIGR00057 (TIGR00057)→ ECA_wecA (TIGR02380)GT_WecA_like (cd06853)→ hypo→ |
| Rwr-2-1 | NZ_BAWF01000054.1 | + | 58602 | 58647 | **RNA**→ rpmE (PRK00019)Ribosomal_L31 (pfam01197)→ PrfA (COG0216)prfA (PRK00591)→ HemK (COG2890)PRK09328 (PRK09328)→ SUA5 (COG0009)TIGR00057 (TIGR00057)→ ECA_wecA (TIGR02380)GT_WecA_like (cd06853)→ hypo→ |
| Rfa-21-1 | NZ_JMEN01000002.1 | - | 18231 | 18186 | **RNA**→ rpmE (PRK00019)Ribosomal_L31 (pfam01197)→ PrfA (COG0216)prfA (PRK00591)→ |
| Rfa-1-2 | NZ_LFDS01000014.1 | + | 60916 | 60961 | **RNA**→ rpmE (PRK00019)Ribosomal_L31 (pfam01197)→ PrfA (COG0216)prfA (PRK00591)→ |
| Rsp-15-1 | NZ_KI519398.1 | - | 4412791 | 4412746 | **RNA**→ rpmE (PRK00019)Ribosomal_L31 (pfam01197)→ PrfA (COG0216)prfA (PRK00591)→ |

| | | | | | |
|---|---|---|---|---|---|
| Rsp-17-1 | NZ_KQ031368.1 | - | 244975 | 244930 | RNA→ rpmE (PRK00019)Ribosomal_L31 (pfam01197)→ PrfA (COG0216)prfA (PRK00591)→ |
| Rfa-13-1 | NZ_JMEU01000024.1 | + | 61028 | 61073 | RNA→ rpmE (PRK00019)Ribosomal_L31 (pfam01197)→ PrfA (COG0216)prfA (PRK00591)→ |
| Rfa-14-1 | NZ_JMET01000032.1 | - | 18078 | 18033 | RNA→ rpmE (PRK00019)Ribosomal_L31 (pfam01197)→ PrfA (COG0216)prfA (PRK00591)→ |
| Rfa-6-1 | NZ_JMFB01000009.1 | - | 41568 | 41523 | RNA→ rpmE (PRK00019)Ribosomal_L31 (pfam01197)→ PrfA (COG0216)prfA (PRK00591)→ |
| Rfa-18-1 | NZ_JMEP01000006.1 | - | 18272 | 18227 | RNA→ rpmE (PRK00019)Ribosomal_L31 (pfam01197)→ PrfA (COG0216)prfA (PRK00591)→ |
| Rfa-15-1 | NZ_JMES01000046.1 | + | 65370 | 65415 | RNA→ rpmE (PRK00019)Ribosomal_L31 (pfam01197)→ PrfA (COG0216)prfA (PRK00591)→ |
| Rfa-16-1 | NZ_JMER01000016.1 | - | 18453 | 18408 | RNA→ rpmE (PRK00019)Ribosomal_L31 (pfam01197)→ PrfA (COG0216)prfA (PRK00591)→ |
| Rfa-19-1 | NZ_JMEO01000036.1 | - | 42912 | 42867 | RNA→ rpmE (PRK00019)Ribosomal_L31 (pfam01197)→ PrfA (COG0216)prfA (PRK00591)→ |
| Rfa-17-1 | NZ_JMEQ01000003.1 | - | 18034 | 17989 | RNA→ rpmE (PRK00019)Ribosomal_L31 (pfam01197)→ PrfA (COG0216)prfA (PRK00591)→ |
| Rfa-9-1 | NZ_JMEY01000018.1 | + | 61117 | 61162 | RNA→ rpmE (PRK00019)Ribosomal_L31 (pfam01197)→ PrfA (COG0216)prfA (PRK00591)→ |
| Rfa-4-1 | NZ_JMFD01000009.1 | - | 18100 | 18055 | RNA→ rpmE (PRK00019)Ribosomal_L31 (pfam01197)→ PrfA (COG0216)prfA (PRK00591)→ |
| Rfa-7-1 | NZ_JMFA01000028.1 | - | 18232 | 18187 | RNA→ rpmE (PRK00019)Ribosomal_L31 (pfam01197)→ PrfA (COG0216)prfA (PRK00591)→ |
| Rfa-20-1 | NZ_JMEM01000014.1 | - | 503930 | 503885 | RNA→ rpmE (PRK00019)Ribosomal_L31 (pfam01197)→ PrfA (COG0216)prfA (PRK00591)→ |
| Rfa-8-1 | NZ_JMEZ01000016.1 | + | 635989 | 636034 | RNA→ rpmE (PRK00019)Ribosomal_L31 (pfam01197)→ PrfA (COG0216)prfA (PRK00591)→ |
| Rtr-1-1 | NZ_AODO01000019.1 | - | 15129 | 15084 | RNA→ rpmE (PRK00019)Ribosomal_L31 (pfam01197)→ |
| Rsp-19-1 | NZ_CVQP01000001.1 | + | 953865 | 953910 | RNA→ rpmE (PRK00019)Ribosomal_L31 (pfam01197)→ |
| Rsp-21-1 | NZ_JMLO01000036.1 | + | 39540 | 39585 | RNA→ rpmE (PRK00019)Ribosomal_L31 (pfam01197)→ |
| Sar-21-1 | NZ_AZWI01000036.1 | - | 51029 | 50973 | RNA→ hypo→ rpmE (PRK00019)Ribosomal_L31 (pfam01197)→ |
| Sar-40-1 | NZ_AZWX01000003.1 | - | 51034 | 50978 | RNA→ hypo→ rpmE (PRK00019)Ribosomal_L31 (pfam01197)→ |
| Sar-36-1 | NZ_AZXA01000003.1 | - | 51182 | 51126 | RNA→ hypo→ rpmE (PRK00019)Ribosomal_L31 (pfam01197)→ |
| Sar-38-1 | NZ_AZXG01000012.1 | - | 51140 | 51084 | RNA→ hypo→ rpmE (PRK00019)Ribosomal_L31 (pfam01197)→ |
| Sar-25-1 | NZ_AZWV01000001.1 | + | 346941 | 346997 | RNA→ hypo→ rpmE (PRK00019)Ribosomal_L31 (pfam01197)→ |
| Sar-37-1 | NZ_KI911342.1 | - | 51088 | 51032 | RNA→ hypo→ rpmE (PRK00019)Ribosomal_L31 (pfam01197)→ |
| Sar-39-1 | NZ_KI911438.1 | - | 51088 | 51032 | RNA→ hypo→ rpmE (PRK00019)Ribosomal_L31 (pfam01197)→ |
| Sar-9-1 | NZ_KB898403.1 | - | 50932 | 50876 | RNA→ hypo→ rpmE (PRK00019)Ribosomal_L31 (pfam01197)→ |
| Sar-4-1 | NZ_KB896072.1 | + | 346814 | 346870 | RNA→ hypo→ rpmE (PRK00019)Ribosomal_L31 (pfam01197)→ |
| Spa-11-1 | NZ_KB892476.1 | - | 51107 | 51051 | RNA→ hypo→ rpmE (PRK00019)Ribosomal_L31 (pfam01197)→ |
| Sar-19-1 | NZ_KB913036.1 | - | 2994862 | 2994806 | RNA→ hypo→ rpmE (PRK00019)Ribosomal_L31 (pfam01197)→ |
| Sar-12-1 | NZ_KB896389.1 | + | 143497 | 143553 | RNA→ hypo→ rpmE (PRK00019)Ribosomal_L31 (pfam01197)→ |
| Sar-1-1 | NZ_KB900168.1 | + | 1524 | 1580 | RNA→ hypo→ rpmE (PRK00019)Ribosomal_L31 (pfam01197)→ |
| Sar-27-1 | NZ_JAEY01000041.1 | - | 51135 | 51079 | RNA→ hypo→ rpmE (PRK00019)Ribosomal_L31 (pfam01197)→ |
| Sar-3-1 | NZ_KB896004.1 | + | 346986 | 347042 | RNA→ hypo→ rpmE (PRK00019)Ribosomal_L31 (pfam01197)→ |
| Sar-33-1 | NZ_AUKM01000035.1 | + | 1712 | 1768 | RNA→ hypo→ rpmE (PRK00019)Ribosomal_L31 (pfam01197)→ |
| Sar-6-1 | NC_009953.1 | - | 4579559 | 4579503 | RNA→ hypo→ rpmE (PRK00019)Ribosomal_L31 (pfam01197)→ |
| Sar-32-1 | NZ_KE384262.1 | + | 143944 | 144000 | RNA→ hypo→ rpmE (PRK00019)Ribosomal_L31 (pfam01197)→ |
| Sar-30-1 | NZ_KI519456.1 | - | 51241 | 51185 | RNA→ hypo→ rpmE (PRK00019)Ribosomal_L31 (pfam01197)→ |
| Sar-31-1 | NZ_KI911435.1 | - | 51292 | 51236 | RNA→ hypo→ rpmE (PRK00019)Ribosomal_L31 (pfam01197)→ |
| Sar-16-1 | NZ_KB896701.1 | - | 51106 | 51050 | RNA→ hypo→ rpmE (PRK00019)Ribosomal_L31 (pfam01197)→ |
| Sar-8-1 | NZ_KB892533.1 | + | 146895 | 146951 | RNA→ hypo→ rpmE (PRK00019)Ribosomal_L31 (pfam01197)→ |
| Sar-14-1 | NZ_KB905500.1 | - | 51099 | 51043 | RNA→ hypo→ rpmE (PRK00019)Ribosomal_L31 (pfam01197)→ |
| Sar-34-1 | NZ_KB896161.1 | + | 146321 | 146377 | RNA→ hypo→ rpmE (PRK00019)Ribosomal_L31 (pfam01197)→ |
| Spa-13-1 | NZ_KB896968.1 | + | 1604 | 1660 | RNA→ hypo→ rpmE (PRK00019)Ribosomal_L31 (pfam01197)→ |
| Sar-18-1 | NZ_KB905199.1 | + | 351246 | 351302 | RNA→ hypo→ rpmE (PRK00019)Ribosomal_L31 (pfam01197)→ |
| Sar-5-1 | NZ_KB900324.1 | - | 51168 | 51112 | RNA→ hypo→ rpmE (PRK00019)Ribosomal_L31 (pfam01197)→ |
| Sar-7-1 | NZ_KB895942.1 | - | 51129 | 51073 | RNA→ hypo→ rpmE (PRK00019)Ribosomal_L31 (pfam01197)→ |
| Sar-13-1 | NZ_KB896467.1 | + | 146912 | 146968 | RNA→ hypo→ rpmE (PRK00019)Ribosomal_L31 (pfam01197)→ |
| Sar-15-1 | NZ_KB896531.1 | - | 51113 | 51057 | RNA→ hypo→ rpmE (PRK00019)Ribosomal_L31 (pfam01197)→ |
| Sar-35-1 | NZ_KI911491.1 | - | 51372 | 51316 | RNA→ hypo→ rpmE (PRK00019)Ribosomal_L31 (pfam01197)→ |
| Sar-28-1 | NZ_AZWY01000001.1 | + | 356278 | 356334 | RNA→ hypo→ rpmE (PRK00019)Ribosomal_L31 (pfam01197)→ |
| Sar-26-1 | NZ_KE384230.1 | - | 51394 | 51338 | RNA→ hypo→ rpmE (PRK00019)Ribosomal_L31 (pfam01197)→ |
| Sar-2-1 | NZ_KB900232.1 | - | 51449 | 51393 | RNA→ hypo→ rpmE (PRK00019)Ribosomal_L31 (pfam01197)→ |
| Sar-20-1 | NZ_JNLS01000004.1 | - | 51467 | 51411 | RNA→ hypo→ rpmE (PRK00019)Ribosomal_L31 (pfam01197)→ |

| | | | | | |
|---|---|---|---|---|---|
| Sar-22-1 | NZ_KI911555.1 | - | 50976 | 50920 | **RNA**→ hypo→ rpmE (PRK00019)Ribosomal_L31 (pfam01197)→ |
| Sar-17-1 | NZ_KB896609.1 | - | 51038 | 50982 | **RNA**→ hypo→ rpmE (PRK00019)Ribosomal_L31 (pfam01197)→ |
| Sar-24-1 | NZ_KI911518.1 | + | 208754 | 208810 | **RNA**→ hypo→ rpmE (PRK00019)Ribosomal_L31 (pfam01197)→ |
| Sar-23-1 | NZ_KI911441.1 | + | 352956 | 353012 | **RNA**→ hypo→ rpmE (PRK00019)Ribosomal_L31 (pfam01197)→ |
| env-474 | 2200071588 | + | 82438 | 82519 | **RNA**→ rpmE (PRK00019)Ribosomal_L31 (pfam01197)→ |
| env-475 | JGI12397J12024_1254667 | + | 31 | 93 | **RNA**→ rpmE (PRK00019)Ribosomal_L31 (pfam01197)→ |
| Whe-1-1 | NZ_JXYP01000014.1 | + | 381001 | 381054 | **RNA**→ rpmE (PRK00019)Ribosomal_L31 (pfam01197)→ PrfA (COG0216)prfA (PRK00591)→ hypo→ |
| | | | | | HemK (COG2890)PRK09328 (PRK09328)→ SUA5 (COG0009)TIGR00057 (TIGR00057)→ |
| | | | | | ECA_wecA (TIGR02380)GT_WecA_like (cd06853)→ ECA_wecA (TIGR02380)GT_WecA_like (cd06853)→ |
| Spa-28-1 | NZ_KI911408.1 | - | 162017 | 161962 | **RNA**→ rpmE (PRK00019)Ribosomal_L31 (pfam01197)→ |
| Spa-10-1 | NZ_KB904923.1 | + | 18193 | 18248 | **RNA**→ rpmE (PRK00019)Ribosomal_L31 (pfam01197)→ rpmE (PRK00019)Ribosomal_L31 (pfam01197)→ |
| Spa-36-1 | NZ_AZWR01000034.1 | + | 18288 | 18342 | **RNA**→ rpmE (PRK00019)Ribosomal_L31 (pfam01197)→ |
| Spa-8-1 | NZ_KB895313.1 | - | 50515 | 50461 | **RNA**→ rpmE (PRK00019)Ribosomal_L31 (pfam01197)→ |
| Spa-9-1 | NZ_KB896783.1 | - | 50546 | 50492 | **RNA**→ rpmE (PRK00019)Ribosomal_L31 (pfam01197)→ |
| Dau-1-1 | NZ_JNYJ01000029.1 | - | 93182 | 93146 | **RNA**→ rpmE (PRK00019)Ribosomal_L31 (pfam01197)→ |
| Mlu-1-1 | NZ_HF570108.1 | - | 6128873 | 6128807 | **RNA**→ rpmE (PRK00019)Ribosomal_L31 (pfam01197)→ |
| Mch-6-1 | NZ_JOAN01000006.1 | + | 33922 | 33988 | **RNA**→ rpmE (PRK00019)Ribosomal_L31 (pfam01197)→ |
| Mpa-1-1 | NZ_JNZS01000011.1 | + | 66563 | 66629 | **RNA**→ rpmE (PRK00019)Ribosomal_L31 (pfam01197)→ |
| Rsp-9-1 | NZ_AJVB01000148.1 | + | 7131 | 7176 | **RNA**→ rpmE (PRK00019)Ribosomal_L31 (pfam01197)→ |
| | | | | | PCRF (smart00937)RF-1 (pfam00472)PCRF (pfam03462)→ |
| Rrh-2-1 | NZ_AGVW02000073.1 | + | 58982 | 59027 | **RNA**→ rpmE (PRK00019)Ribosomal_L31 (pfam01197)→ PrfA (COG0216)prfA (PRK00591)→ |
| Rsp-14-1 | NZ_JTIZ01000140.1 | - | 15896 | 15851 | **RNA**→ rpmE (PRK00019)Ribosomal_L31 (pfam01197)→ PrfA (COG0216)prfA (PRK00591)→ |
| Rpy-2-1 | NC_023150.1 | + | 5211019 | 5211064 | **RNA**→ rpmE (PRK00019)Ribosomal_L31 (pfam01197)→ PrfA (COG0216)prfA (PRK00591)→ |
| Rsp-18-1 | NZ_AFAQ01001953.1 | + | 74360 | 74405 | **RNA**→ rpmE (PRK00019)Ribosomal_L31 (pfam01197)→ PrfA (COG0216)prfA (PRK00591)→ |
| Rpy-1-1 | NZ_AHBW01000042.1 | + | 40539 | 40584 | **RNA**→ rpmE (PRK00019)Ribosomal_L31 (pfam01197)→ rpmE (PRK00019)Ribosomal_L31 (pfam01197)→ |
| | | | | | PrfA (COG0216)prfA (PRK00591)→ PrfA (COG0216)prfA (PRK00591)→ |
| Rsp-22-1 | NZ_JPJJ01000011.1 | - | 17491 | 17446 | **RNA**→ rpmE (PRK00019)Ribosomal_L31 (pfam01197)→ PrfA (COG0216)prfA (PRK00591)→ |
| Rsp-5-1 | NZ_CAVJ010000216.1 | - | 15449 | 15404 | **RNA**→ rpmE (PRK00019)Ribosomal_L31 (pfam01197)→ |
| Rae-1-1 | NZ_CP011341.1 | + | 2197037 | 2197082 | **RNA**→ rpmE (PRK00019)Ribosomal_L31 (pfam01197)→ |
| Rrh-6-1 | NZ_AZHI01000044.1 | - | 15673 | 15628 | **RNA**→ rpmE (PRK00019)Ribosomal_L31 (pfam01197)→ |
| Rru-2-1 | NZ_ANGC01000054.1 | + | 15700 | 15745 | **RNA**→ rpmE (PRK00019)Ribosomal_L31 (pfam01197)→ |
| Rru-3-1 | NZ_CCSD01000112.1 | + | 217226 | 217271 | **RNA**→ rpmE (PRK00019)Ribosomal_L31 (pfam01197)→ |
| Rru-3-2 | NZ_LDUF01000007.1 | - | 15446 | 15401 | **RNA**→ rpmE (PRK00019)Ribosomal_L31 (pfam01197)→ |
| Rsp-7-1 | NZ_AJFC01000306.1 | + | 45552 | 45597 | **RNA**→ rpmE (PRK00019)Ribosomal_L31 (pfam01197)→ rpmE (PRK00019)Ribosomal_L31 (pfam01197)→ |
| Rru-1-1 | NZ_AOEX01000028.1 | - | 15828 | 15783 | **RNA**→ rpmE (PRK00019)Ribosomal_L31 (pfam01197)→ |
| Ntr-1-1 | NZ_BAGL01000019.1 | + | 91420 | 91465 | **RNA**→ rpmE (PRK00019)Ribosomal_L31 (pfam01197)→ PrfA (COG0216)prfA (PRK00591)→ |
| | | | | | HemK (COG2890)PRK09328 (PRK09328)→ |
| Not-2-1 | NZ_BAGD01000040.1 | + | 28362 | 28407 | **RNA**→ rpmE (PRK00019)Ribosomal_L31 (pfam01197)→ PrfA (COG0216)prfA (PRK00591)→ |
| | | | | | HemK (COG2890)PRK09328 (PRK09328)→ HemK (COG2890)PRK09328 (PRK09328)→ |
| Not-1-1 | NZ_BATZ01000015.1 | + | 189934 | 189979 | **RNA**→ rpmE (PRK00019)Ribosomal_L31 (pfam01197)→ PrfA (COG0216)prfA (PRK00591)→ |
| | | | | | HemK (COG2890)PRK09328 (PRK09328)→ HemK (COG2890)PRK09328 (PRK09328)→ |
| env-476 | JGI20190J14840_1064086 | + | 179 | 244 | **RNA**→ |
| Msp-26-1 | NZ_KQ058714.1 | + | 43212 | 43277 | **RNA**→ hypo→ rpmE (PRK00019)Ribosomal_L31 (pfam01197)→ |
| env-477 | JGI20205J14842_1097635 | + | 188 | 243 | **RNA**→ hypo→ |
| env-478 | JGI12396J12026_1003816 | + | 12 | 59 | **RNA**→ rpmE (PRK00019)Ribosomal_L31 (pfam01197)→ |
| env-479 | JGI12271J12027_1041588 | + | 10 | 57 | **RNA**→ rpmE (PRK00019)Ribosomal_L31 (pfam01197)→ |
| env-480 | JGI12273J12029_10019447 | + | 10 | 57 | **RNA**→ hypo→ rpmE (PRK00019)Ribosomal_L31 (pfam01197)→ |
| Gsp-3-1 | NZ_JOGR01000006.1 | + | 294303 | 294343 | **RNA**→ rpmE (PRK00019)Ribosomal_L31 (pfam01197)→ |
| | | | | | prfA (PRK00591)RF-1 (pfam00472)PCRF (pfam03462)→ HemK (COG2890)PRK09328 (PRK09328)→ hypo→ |
| | | | | | SUA5 (COG0009)TIGR00057 (TIGR00057)→ LMWPc (smart00226)Wzb (COG0394)→ |
| | | | | | LMWPc (smart00226)LMWPAP (cd16344)→ hypo→ |
| env-481 | JGI24740J21852_10164145 | + | 98 | 134 | **RNA**→ rpmE (PRK00019)Ribosomal_L31 (pfam01197)→ hypo→ |
| env-482 | botACOD_GAKN62C01CSCDR | + | 372 | 436 | **RNA**→ rpmE (PRK00019)Ribosomal_L31 (pfam01197)→ |
| env-483 | topACOD_F95O9CU01EIYTS | + | 236 | 305 | **RNA**→ rpmE (PRK00019)Ribosomal_L31 (pfam01197)→ |

| | | | | | |
|---|---|---|---|---|---|
| env-484 | JGI20263J16346_11281 | - | 78 | 28 | RNA→ hypo→ |
| env-485 | 2210838270 | - | 125 | 37 | RNA→ hypo→ |
| env-486 | 2034117903 | - | 276 | 240 | RNA→ rpmE (PRK00019)Ribosomal_L31 (pfam01197)→ |
| Sam-2-1 | NZ_JOEQ01000004.1 | - | 236796 | 236714 | RNA→ rpmE (PRK00019)Ribosomal_L31 (pfam01197)→ |
| Sro-7-1 | NZ_JOEP01000003.1 | - | 250250 | 250165 | RNA→ rpmE (PRK00019)Ribosomal_L31 (pfam01197)→ PrfA (COG0216)prfA (PRK00591)→ HemK (COG2890)PRK09328 (PRK09328)→ |
| Sro-3-1 | NC_013595.1 | + | 1771338 | 1771423 | RNA→ rpmE (PRK00019)Ribosomal_L31 (pfam01197)→ |
| Mau-1-1 | NC_014391.1 | - | 5788708 | 5788645 | RNA→ rpmE (PRK00019)Ribosomal_L31 (pfam01197)→ PrfA (COG0216)prfA (PRK00591)→ |
| Msp-9-1 | NZ_KB895826.1 | + | 396351 | 396414 | RNA→ hypo→ rpmE (PRK00019)Ribosomal_L31 (pfam01197)→ PrfA (COG0216)prfA (PRK00591)→ hypo→ |
| Msp-10-1 | NC_014815.1 | + | 3318499 | 3318562 | RNA→ rpmE (PRK00019)Ribosomal_L31 (pfam01197)→ PrfA (COG0216)prfA (PRK00591)→ |
| Msp-28-1 | NZ_LEKG01000129.1 | + | 72 | 135 | RNA→ rpmE (PRK00019)Ribosomal_L31 (pfam01197)→ PrfA (COG0216)prfA (PRK00591)→ |
| Mgl-1-1 | NZ_JNZR01000015.1 | - | 22540 | 22477 | RNA→ rpmE (PRK00019)Ribosomal_L31 (pfam01197)→ PrfA (COG0216)prfA (PRK00591)→ |
| env-487 | JGI12698J15209_1012939 | + | 182 | 248 | RNA→ rpmE (PRK00019)Ribosomal_L31 (pfam01197)→ |
| env-488 | JGI12635J15846_10089936 | + | 506 | 573 | RNA→ rpmE (PRK00019)Ribosomal_L31 (pfam01197)→ |
| Msp-8-1 | NZ_GG657738.1 | - | 2230600 | 2230534 | RNA→ rpmE (PRK00019)Ribosomal_L31 (pfam01197)→ rpmE (PRK00019)Ribosomal_L31 (pfam01197)→ |
| env-489 | JGI20206J14855_1044127 | - | 238 | 179 | RNA→ rpmE (PRK00019)Ribosomal_L31 (pfam01197)→ |
| env-490 | JGI20186J14852_1017385 | + | 370 | 429 | RNA→ ←hypo |
| env-491 | JGI20204J15000_1053760 | - | 219 | 160 | RNA→ rpmE (PRK00019)Ribosomal_L31 (pfam01197)→ |
| env-492 | JGI20203J14952_1004251 | - | 245 | 186 | RNA→ rpmE (PRK00019)Ribosomal_L31 (pfam01197)→ |
| env-493 | JGI20204J15135_1053760 | - | 219 | 160 | RNA→ rpmE (PRK00019)Ribosomal_L31 (pfam01197)→ |
| Nco-1-1 | NZ_KB903996.1 | - | 180981 | 180940 | RNA→ rpmE (PRK00019)Ribosomal_L31 (pfam01197)→ |
| env-494 | AP72_2010_repI_A100_1025205 | - | 373 | 326 | RNA→ rpmE (PRK00019)Ribosomal_L31 (pfam01197)→ |
| Mca-1-1 | NZ_JXSX01000002.1 | + | 873352 | 873417 | RNA→ rpmE (PRK00019)Ribosomal_L31 (pfam01197)→ |
| env-495 | JGI20241J16302_105831 | + | 273 | 334 | RNA→ |
| env-496 | JGI12635J15846_10116580 | + | 1117 | 1162 | RNA→ rpmE (PRK00019)Ribosomal_L31 (pfam01197)→ |
| env-497 | JGI12269J14319_10020994 | - | 2767 | 2730 | RNA→ rpmE (PRK00019)Ribosomal_L31 (pfam01197)→ |
| env-498 | JGI12635J15846_13730630 | - | 99 | 50 | RNA→ hypo→ |
| env-499 | JGI12712J15308_10773049 | + | 159 | 209 | RNA→ |
| env-500 | JGI12269J14319_10090585 | - | 241 | 190 | RNA→ rpmE (PRK00019)Ribosomal_L31 (pfam01197)→ |
| env-501 | JGI12269J14319_10493819 | - | 328 | 279 | RNA→ rpmE (PRK00019)Ribosomal_L31 (pfam01197)→ |
| env-502 | JGI20196J14858_1024988 | - | 290 | 239 | RNA→ rpmE (PRK00019)Ribosomal_L31 (pfam01197)→ |
| env-503 | JGI12712J15308_10023311 | + | 1278 | 1329 | RNA→ rpmE (PRK00019)Ribosomal_L31 (pfam01197)→ |
| env-504 | JGI12659J15293_10227025 | + | 221 | 272 | RNA→ rpmE (PRK00019)Ribosomal_L31 (pfam01197)→ |
| env-505 | JGI20188J14859_1052292 | + | 150 | 201 | RNA→ rpmE (PRK00019)Ribosomal_L31 (pfam01197)→ |
| env-506 | JGI20186J14852_1022270 | - | 272 | 221 | RNA→ rpmE (PRK00019)Ribosomal_L31 (pfam01197)→ |
| env-507 | JGI20203J14952_1005414 | + | 1323 | 1374 | RNA→ |
| env-508 | JGI20205J14842_1005807 | - | 759 | 708 | RNA→ rpmE (PRK00019)Ribosomal_L31 (pfam01197)→ |
| env-509 | JGI20186J14852_1008003 | - | 59 | 8 | RNA→ |
| env-510 | JGI20190J14840_1000844 | + | 1262 | 1313 | RNA→ rpmE (PRK00019)Ribosomal_L31 (pfam01197)→ |
| env-511 | JGI20204J15000_1002743 | - | 517 | 466 | RNA→ rpmE (PRK00019)Ribosomal_L31 (pfam01197)→ |
| env-512 | JGI20204J15135_1002743 | - | 517 | 466 | RNA→ rpmE (PRK00019)Ribosomal_L31 (pfam01197)→ |
| env-513 | JGI20183J14883_129429 | - | 281 | 230 | RNA→ rpmE (PRK00019)Ribosomal_L31 (pfam01197)→ |
| env-514 | JGI12635J15846_11642955 | - | 277 | 239 | RNA→ rpmE (PRK00019)Ribosomal_L31 (pfam01197)→ |
| env-515 | C688J13580_1150866 | + | 209 | 247 | RNA→ rpmE (PRK00019)Ribosomal_L31 (pfam01197)→ |
| Nca-2-1 | NZ_BAFV01000016.1 | + | 24171 | 24216 | RNA→ rpmE (PRK00019)Ribosomal_L31 (pfam01197)→ |
| Nca-1-1 | NZ_JNZF01000017.1 | - | 16307 | 16262 | RNA→ rpmE (PRK00019)Ribosomal_L31 (pfam01197)→ |
| Asu-1-1 | NC_015564.1 | - | 3348646 | 3348602 | RNA→ rpmE (PRK00019)Ribosomal_L31 (pfam01197)→ PrfA (COG0216)prfA (PRK00591)→ |
| env-516 | JGI20245J16306_1034373 | + | 145 | 183 | RNA→ rpmE (PRK00019)Ribosomal_L31 (pfam01197)→ |
| env-517 | JGI12635J15846_10593975 | + | 102 | 140 | RNA→ rpmE (PRK00019)Ribosomal_L31 (pfam01197)→ |
| env-518 | JGI26342J46808_1010879 | + | 391 | 429 | RNA→ rpmE (PRK00019)Ribosomal_L31 (pfam01197)→ |
| env-519 | JGI26337J50220_1100613 | - | 177 | 139 | RNA→ hypo→ |
| env-520 | JGI12698J15209_1088486 | + | 118 | 184 | RNA→ |

| | | | | | |
|---|---|---|---|---|---|
| env-521 | JGI12659J15293_10241949 | - | 305 | 265 | **RNA**→ ←hypo |
| env-522 | JGI12630J15595_10661370 | - | 218 | 164 | **RNA**→ hypo→ rpmE (PRK00019)Ribosomal_L31 (pfam01197)→ |
| env-523 | JGI12467J12023_1041632 | + | 41 | 119 | **RNA**→ rpmE (PRK00019)Ribosomal_L31 (pfam01197)→ |
| env-524 | JGI20276J16322_104776 | - | 170 | 120 | **RNA**→ rpmE (PRK00019)Ribosomal_L31 (pfam01197)→ |
| env-525 | JGI20275J16321_100019 | - | 565 | 515 | **RNA**→ rpmE (PRK00019)Ribosomal_L31 (pfam01197)→ PCRF (pfam03462)ABC_tran_CTD (pfam16326)→ |
| env-526 | JGI20176J14865_113912 | - | 189 | 138 | **RNA**→ rpmE (PRK00019)Ribosomal_L31 (pfam01197)→ |
| env-527 | JGI12694J13545_1030456 | + | 50 | 104 | **RNA**→ rpmE (PRK00019)Ribosomal_L31 (pfam01197)→ |
| env-528 | JGI12649J13570_1002003 | - | 2744 | 2690 | **RNA**→ rpmE (PRK00019)Ribosomal_L31 (pfam01197)→ PrfA (COG0216)prfA (PRK00591)→<br>HemK (COG2890)PRK09328 (PRK09328)→ SUA5 (COG0009)TIGR00057 (TIGR00057)→ |
| env-529 | JGI12635J15846_10007700 | - | 7040 | 6986 | **RNA**→ rpmE (PRK00019)Ribosomal_L31 (pfam01197)→ PrfA (COG0216)prfA (PRK00591)→<br>HemK (COG2890)PRK09328 (PRK09328)→ SUA5 (COG0009)TIGR00057 (TIGR00057)→ |
| env-530 | JGI12681J13546_1004659 | - | 463 | 409 | **RNA**→ rpmE (PRK00019)Ribosomal_L31 (pfam01197)→ |
| env-531 | JGI12659J15293_10071495 | + | 208 | 262 | **RNA**→ rpmE (PRK00019)Ribosomal_L31 (pfam01197)→ PrfA (COG0216)prfA (PRK00591)→ |
| env-532 | JGI12659J15293_10081505 | + | 640 | 694 | **RNA**→ |
| env-533 | JGI20235J16296_1010410 | + | 109 | 163 | **RNA**→ rpmE (PRK00019)Ribosomal_L31 (pfam01197)→ |
| env-534 | JGI12635J15846_10479974 | + | 427 | 511 | **RNA**→ rpmE (PRK00019)Ribosomal_L31 (pfam01197)→ |
| env-535 | JGI12648J13191_1102813 | - | 315 | 231 | **RNA**→ rpmE (PRK00019)Ribosomal_L31 (pfam01197)→ |
| env-536 | JGI12270J11330_10000477 | + | 1502 | 1555 | **RNA**→ rpmE (PRK00019)Ribosomal_L31 (pfam01197)→ |
| env-537 | JGI12269J14319_10000786 | + | 1502 | 1555 | **RNA**→ rpmE (PRK00019)Ribosomal_L31 (pfam01197)→ |
| env-538 | 2044437860 | + | 359 | 441 | **RNA**→ |
| Sna-1-1 | NC_013947.1 | + | 1383336 | 1383378 | **RNA**→ rpmE (PRK00019)Ribosomal_L31 (pfam01197)→ |
| env-539 | JGI12269J14319_10084388 | - | 323 | 279 | **RNA**→ rpmE (PRK00019)Ribosomal_L31 (pfam01197)→ |
| env-540 | FIHLEPW02RU19J | - | 259 | 208 | **RNA**→ rpmE (PRK00019)Ribosomal_L31 (pfam01197)→ |
| env-541 | FIHLEPW02PK0H8 | - | 435 | 385 | **RNA**→ hypo→ rpmE (PRK00019)Ribosomal_L31 (pfam01197)→ |
| env-542 | AP72_2010_repI_A10_1090062 | - | 355 | 305 | **RNA**→ hypo→ |
| env-543 | FIHLEPW02TDTYE | - | 192 | 142 | **RNA**→ hypo→ |
| env-544 | AF_2010_repII_A1_10229252 | - | 194 | 144 | **RNA**→ rpmE (PRK00019)Ribosomal_L31 (pfam01197)→ |
| env-545 | AF_2010_repII_A001_10407533 | - | 209 | 159 | **RNA**→ UMPK_like (cd02028)→ |
| env-546 | JGI12712J15308_10689240 | + | 44 | 86 | **RNA**→ hypo→ rpmE (PRK00019)Ribosomal_L31 (pfam01197)→ |
| env-547 | JGI12659J15293_10376808 | + | 47 | 89 | **RNA**→ rpmE (PRK00019)Ribosomal_L31 (pfam01197)→ |
| env-548 | JGI12635J15846_12325035 | + | 141 | 177 | **RNA**→ rpmE (PRK00019)Ribosomal_L31 (pfam01197)→ |
| env-549 | JGI26341J46601_10034216 | + | 1087 | 1123 | **RNA**→ rpmE (PRK00019)Ribosomal_L31 (pfam01197)→ |
| env-550 | JGI26340J50214_10003356 | + | 235 | 271 | **RNA**→ rpmE (PRK00019)Ribosomal_L31 (pfam01197)→ |
| env-551 | JGI12630J15595_10226531 | + | 132 | 168 | **RNA**→ ←hypo |
| env-552 | JGI26346J50198_1083412 | - | 251 | 215 | **RNA**→ rpmE (PRK00019)Ribosomal_L31 (pfam01197)→ |
| env-553 | JGI12419J13241_1069260 | + | 290 | 354 | **RNA**→ rpmE (PRK00019)Ribosomal_L31 (pfam01197)→ |
| env-554 | GBANfinal_FHNL2OP03RHL08 | - | 331 | 267 | **RNA**→ rpmE (PRK00019)Ribosomal_L31 (pfam01197)→ |
| Tcu-1-1 | NC_013510.1 | - | 4476819 | 4476755 | **RNA**→ rpmE (PRK00019)Ribosomal_L31 (pfam01197)→ |
| env-555 | JGI12681J13546_1026784 | - | 235 | 164 | **RNA**→ rpmE (PRK00019)Ribosomal_L31 (pfam01197)→ |
| env-556 | JGI12635J15846_10334590 | - | 676 | 605 | **RNA**→ rpmE (PRK00019)Ribosomal_L31 (pfam01197)→ |
| env-557 | JGI12270J11330_10077884 | + | 1435 | 1471 | **RNA**→ rpmE (PRK00019)Ribosomal_L31 (pfam01197)→ |
| env-558 | JGI12269J14319_10016588 | + | 541 | 577 | **RNA**→ rpmE (PRK00019)Ribosomal_L31 (pfam01197)→ |
| env-559 | JGI12269J14319_10048667 | + | 2404 | 2440 | **RNA**→ rpmE (PRK00019)Ribosomal_L31 (pfam01197)→ |
| env-560 | JGI26340J50214_10285047 | + | 212 | 248 | **RNA**→ rpmE (PRK00019)Ribosomal_L31 (pfam01197)→ |
| env-561 | JGI12631J13338_1101732 | + | 143 | 179 | **RNA**→ ←hypo |
| env-562 | JGI12635J15846_10088123 | - | 1905 | 1869 | **RNA**→ rpmE (PRK00019)Ribosomal_L31 (pfam01197)→ |
| env-563 | JGI12712J15308_10312427 | - | 172 | 136 | **RNA**→ rpmE (PRK00019)Ribosomal_L31 (pfam01197)→ |
| env-564 | JGI12635J15846_10115421 | - | 508 | 472 | **RNA**→ rpmE (PRK00019)Ribosomal_L31 (pfam01197)→ |
| env-565 | JGI26339J46600_10015394 | - | 475 | 439 | **RNA**→ rpmE (PRK00019)Ribosomal_L31 (pfam01197)→ |
| env-566 | JGI26341J46601_10189686 | + | 251 | 287 | **RNA**→ rpmE (PRK00019)Ribosomal_L31 (pfam01197)→ |
| env-567 | JGI26340J50214_10034757 | - | 1226 | 1190 | **RNA**→ rpmE (PRK00019)Ribosomal_L31 (pfam01197)→ |
| env-568 | JGI12270J11330_10005276 | + | 288 | 324 | **RNA**→ rpmE (PRK00019)Ribosomal_L31 (pfam01197)→ |
| env-569 | JGI12269J14319_10440521 | - | 154 | 118 | **RNA**→ rpmE (PRK00019)Ribosomal_L31 (pfam01197)→ |

| | | | | | |
|---|---|---|---|---|---|
| env-570 | JGI26341J46601_10146534 | + | 327 | 363 | RNA→ rpmE (PRK00019)Ribosomal_L31 (pfam01197)→ |
| env-571 | JGI12635J15846_11977336 | - | 238 | 202 | RNA→ rpmE (PRK00019)Ribosomal_L31 (pfam01197)→ |
| env-572 | JGI20272J16309_111800 | - | 126 | 88 | RNA→ ←hypo |
| env-573 | JGI20265J16348_10502 | - | 193 | 155 | RNA→ rpmE (PRK00019)Ribosomal_L31 (pfam01197)→ |
| env-574 | JGI20271J16308_100694 | + | 240 | 278 | RNA→ rpmE (PRK00019)Ribosomal_L31 (pfam01197)→ |
| env-575 | JGI20279J16325_116846 | + | 78 | 116 | RNA→ rpmE (PRK00019)Ribosomal_L31 (pfam01197)→ |
| env-576 | JGI12712J15308_10051662 | - | 98 | 60 | RNA→ hypo→ |
| env-577 | 2200192263 | - | 328 | 245 | RNA→ rpmE (PRK00019)Ribosomal_L31 (pfam01197)→ |
| env-578 | JGI20180J14839_1055467 | - | 123 | 69 | RNA→ rpmE (PRK00528)RpmE (COG0254)rpmE (PRK00019)Ribosomal_L31 (pfam01197)→ |
| env-579 | JGI20249J16330_111215 | + | 99 | 153 | RNA→ |
| env-580 | JGI20235J16296_1001053 | + | 90 | 144 | RNA→ rpmE (PRK00019)Ribosomal_L31 (pfam01197)→ |
| env-581 | JGI20245J16306_1002783 | + | 190 | 244 | RNA→ rpmE (PRK00019)Ribosomal_L31 (pfam01197)→ |
| env-582 | FWIRElOz_GKZ9IRQ02JVM2H | + | 187 | 226 | RNA→ rpmE (PRK00019)Ribosomal_L31 (pfam01197)→ |
| env-583 | FWIRElOz_GKZ9IRQ02I5BP1 | - | 345 | 305 | RNA→ rpmE (PRK00019)Ribosomal_L31 (pfam01197)→ |
| env-584 | JGI24740J21852_10250692 | + | 94 | 133 | RNA→ ←hypo |
| env-585 | JGI24735J21928_10300159 | - | 425 | 386 | RNA→ rpmE (PRK00019)Ribosomal_L31 (pfam01197)→ |
| env-586 | 2044516099 | + | 244 | 282 | RNA→ |
| env-587 | JGI12269J14319_10006866 | + | 697 | 782 | RNA→ rpmE (PRK00019)Ribosomal_L31 (pfam01197)→ PrfA (COG0216)prfA (PRK00591)→<br>HemK (COG2890)PRK09328 (PRK09328)→ SUA5 (COG0009)TIGR00057 (TIGR00057)→<br>GT_WecA_like (cd06853)mraY (PRK00108)PRK07764 (PRK07764)→ |
| env-588 | JGI12269J14319_10074806 | - | 1301 | 1216 | RNA→ rpmE (PRK00019)Ribosomal_L31 (pfam01197)→ |
| env-589 | AP72_2010_repI_A10_1033787 | + | 393 | 430 | RNA→ rpmE (PRK00019)Ribosomal_L31 (pfam01197)→ |
| env-590 | AP72_2010_repI_A100_1084424 | + | 92 | 129 | RNA→ rpmE (PRK00019)Ribosomal_L31 (pfam01197)→ |
| env-591 | FWIRElOz_GKZ9IRQ01DHWYY | + | 73 | 112 | RNA→ rpmE (PRK00019)Ribosomal_L31 (pfam01197)→ |
| env-592 | 2034052008 | - | 318 | 280 | RNA→ hypo→ rpmE (PRK00019)Ribosomal_L31 (pfam01197)→ |
| env-593 | JGI12269J14319_10037047 | + | 1283 | 1334 | RNA→ rpmE (PRK00019)Ribosomal_L31 (pfam01197)→ |
| env-594 | AP72_2010_repI_A10_1129361 | - | 175 | 130 | RNA→ rpmE (PRK00019)Ribosomal_L31 (pfam01197)→ |
| env-595 | AP72_2010_repI_A100_1053081 | + | 201 | 246 | RNA→ rpmE (PRK00019)Ribosomal_L31 (pfam01197)→ |
| env-596 | AP72_2010_repI_A01_1302129 | - | 218 | 173 | RNA→ rpmE (PRK00019)Ribosomal_L31 (pfam01197)→ |
| env-597 | AP72_2010_repI_A10_1217043 | + | 113 | 158 | RNA→ ←hypo |
| env-598 | JGI12631J13338_1003892 | + | 1735 | 1781 | RNA→ rpmE (PRK00019)Ribosomal_L31 (pfam01197)→ |
| env-599 | JGI12635J15846_10008878 | - | 6805 | 6759 | RNA→ rpmE (PRK00019)Ribosomal_L31 (pfam01197)→ |
| env-600 | JGI24738J21930_10757540 | + | 34 | 81 | RNA→ rpmE (PRK00019)Ribosomal_L31 (pfam01197)→ |
| Nbr-4-1 | NZ_BAFU01000023.1 | + | 104700 | 104745 | RNA→ rpmE (PRK00019)Ribosomal_L31 (pfam01197)→ PrfA (COG0216)prfA (PRK00591)→<br>HemK (COG2890)PRK09328 (PRK09328)→ |
| Npa-1-1 | NZ_BAGE01000019.1 | + | 16102 | 16147 | RNA→ rpmE (PRK00019)Ribosomal_L31 (pfam01197)→ |
| env-601 | JGI12272J11983_1372923 | + | 31 | 67 | RNA→ hypo→ rpmE (PRK00019)Ribosomal_L31 (pfam01197)→ |
| env-602 | botACOD_GAKN62C01ECOXR | + | 273 | 311 | RNA→ rpmE (PRK00019)Ribosomal_L31 (pfam01197)→ |
| Gam-1-1 | NZ_BAED01000048.1 | + | 130922 | 130967 | RNA→ rpmE (PRK00019)Ribosomal_L31 (pfam01197)→ rpmE (PRK00019)Ribosomal_L31 (pfam01197)→<br>PrfA (COG0216)prfA (PRK00591)→ PrfA (COG0216)prfA (PRK00591)→<br>HemK (COG2890)PRK09328 (PRK09328)→ HemK (COG2890)PRK09328 (PRK09328)→<br>SUA5 (COG0009)TIGR00057 (TIGR00057)→ SUA5 (COG0009)TIGR00057 (TIGR00057)→<br>GT_WecA_like (cd06853)mraY (PRK00108)→ GT_WecA_like (cd06853)mraY (PRK00108)→ |
| Gso-1-1 | NZ_BANX01000039.1 | - | 25611 | 25566 | RNA→ ←hypo |
| Gsh-1-1 | NZ_AUHE01000013.1 | - | 16550 | 16499 | RNA→ rpmE (PRK00019)Ribosomal_L31 (pfam01197)→ |
| Nsp-5-1 | NZ_JAFQ01000004.1 | + | 3516432 | 3516476 | RNA→ rpmE (PRK00019)Ribosomal_L31 (pfam01197)→ PrfA (COG0216)prfA (PRK00591)→ |
| env-603 | JGI12273J12029_10017424 | + | 27 | 67 | RNA→ hypo→ |
| Ari-2-1 | NZ_AULB01000013.1 | + | 149713 | 149780 | RNA→ rpmE (PRK00019)Ribosomal_L31 (pfam01197)→ |
| env-604 | JGI12272J11983_1083811 | + | 14 | 53 | RNA→ hypo→ rpmE (PRK00019)Ribosomal_L31 (pfam01197)→ |
| env-605 | JGI12467J12023_1087620 | + | 14 | 53 | RNA→ ←hypo |
| env-606 | JGI12397J12024_1060182 | + | 22 | 61 | RNA→ hypo→ rpmE (PRK00019)Ribosomal_L31 (pfam01197)→ |
| env-607 | FNTS_GKKEVUV01BZ0X3 | + | 66 | 105 | RNA→ rpmE (PRK00019)Ribosomal_L31 (pfam01197)→ |
| env-608 | FWIRElOz_GKZ9IRQ02IPUM4 | + | 167 | 206 | RNA→ rpmE (PRK00019)Ribosomal_L31 (pfam01197)→ |

| | | | | | |
|---|---|---|---|---|---|
| Ata-1-1 | NZ_KI912598.1 | + | 15667 | 15704 | **RNA**→ rpmE (PRK00019)Ribosomal_L31 (pfam01197)→ |
| Aha-1-1 | NZ_AQUI01000002.1 | + | 1265681 | 1265725 | **RNA**→ rpmE (PRK00019)Ribosomal_L31 (pfam01197)→ PrfA (COG0216)prfA (PRK00591)→ |
| env-609 | C688J13580_1464814 | - | 190 | 152 | **RNA**→ ←hypo |
| env-610 | SRS065099_LANL_scaffold_20477 | - | 54 | 11 | **RNA**→ |
| env-611 | SRS015064_WUGC_scaffold_2261 | - | 310 | 267 | **RNA**→ rpmE (PRK00019)Ribosomal_L31 (pfam01197)→ |
| env-612 | SRS018443_C3583993 | - | 193 | 150 | **RNA**→ rpmE (PRK00019)Ribosomal_L31 (pfam01197)→ |
| env-613 | SRS011126_C4505688 | + | 336 | 379 | **RNA**→ |
| env-614 | SRS011126_C4505688 | - | 113 | 70 | **RNA**→ hypo→ hypo→ |
| env-615 | SRS015063_C2095482 | + | 120 | 163 | **RNA**→ rpmE (PRK00019)Ribosomal_L31 (pfam01197)→ |
| env-616 | SRS015440_WUGC_scaffold_50169 | + | 447 | 490 | **RNA**→ rpmE (PRK00019)Ribosomal_L31 (pfam01197)→ |
| env-617 | SRS015470_WUGC_scaffold_6374 | - | 193 | 150 | **RNA**→ ←hypo |
| env-618 | SRS015755_WUGC_scaffold_20709 | - | 47 | 4 | **RNA**→ |
| env-619 | SRS015989_C2035914 | - | 193 | 150 | **RNA**→ ←hypo |
| env-620 | SRS016331_C3936733 | + | 121 | 164 | **RNA**→ ←hypo |
| env-621 | SRS016575_Baylor_scaffold_58847 | - | 199 | 156 | **RNA**→ ←hypo |
| env-622 | SRS017139_Baylor_scaffold_68943 | + | 377 | 420 | **RNA**→ ←hypo |
| env-623 | SRS017445_C2964102 | + | 117 | 160 | **RNA**→ rpmE (PRK00019)Ribosomal_L31 (pfam01197)→ |
| env-624 | SRS017691_C4999672 | + | 113 | 156 | **RNA**→ ←hypo |
| env-625 | SRS021477_Baylor_scaffold_64383 | - | 113 | 70 | **RNA**→ |
| env-626 | SRS022536_C4384832 | - | 199 | 156 | **RNA**→ ←hypo |
| env-627 | SRS049268_C4081593 | - | 199 | 156 | **RNA**→ ←hypo |
| env-628 | SRS058053_C5159809 | - | 193 | 150 | **RNA**→ ←hypo |
| Ama-1-1 | NZ_JH711483.1 | + | 455479 | 455522 | **RNA**→ rpmE (PRK00019)Ribosomal_L31 (pfam01197)→ |
| env-629 | SRS011343_Baylor_scaffold_469 | + | 114 | 157 | **RNA**→ ←hypo |
| Pau-1-1 | NZ_JNYD01000008.1 | + | 34980 | 35057 | **RNA**→ rpmE (PRK00019)Ribosomal_L31 (pfam01197)→ PrfA (COG0216)prfA (PRK00591)→ |
| | | | | | HemK (COG2890)PRK09328 (PRK09328)→ hypo→ SUA5 (COG0009)TIGR00057 (TIGR00057)→ |
| | | | | | SUA5 (COG0009)TIGR00057 (TIGR00057)→ |
| Psp-6-1 | NZ_ADUJ01000804.1 | + | 15615 | 15653 | **RNA**→ rpmE (PRK00019)Ribosomal_L31 (pfam01197)→ rpmE (PRK00019)Ribosomal_L31 (pfam01197)→ |
| Ses-1-1 | NC_019673.1 | - | 8141257 | 8141203 | **RNA**→ rpmE (PRK00019)Ribosomal_L31 (pfam01197)→ rpmE (PRK00019)Ribosomal_L31 (pfam01197)→ |
| | | | | | PrfA (COG0216)prfA (PRK00591)→ PrfA (COG0216)prfA (PRK00591)→ |
| | | | | | DAP2 (COG1506)Bile_Hydr_Trans (pfam04775)BAAT_C (pfam08840)→ Lpd (COG1249)PRK07818 (PRK07818)→ |
| | | | | | HemK (COG2890)PRK09328 (PRK09328)→ |
| Ksp-2-1 | NZ_KK037166.1 | + | 7158781 | 7158815 | **RNA**→ rpmE (PRK00019)Ribosomal_L31 (pfam01197)→ |
| Sgl-1-1 | NZ_CM001484.1 | - | 3488024 | 3487984 | **RNA**→ rpmE (PRK00019)Ribosomal_L31 (pfam01197)→ rpmE (PRK00019)Ribosomal_L31 (pfam01197)→ |
| env-630 | JGI24973J35851_1176941 | - | 248 | 206 | **RNA**→ rpmE (PRK00019)Ribosomal_L31 (pfam01197)→ |
| Aba-1-1 | NZ_KB913037.1 | - | 3577594 | 3577551 | **RNA**→ rpmE (PRK00019)Ribosomal_L31 (pfam01197)→ PrfA (COG0216)prfA (PRK00591)→ |
| Ari-1-1 | NZ_JMQI01000011.1 | - | 164510 | 164467 | **RNA**→ rpmE (PRK00019)Ribosomal_L31 (pfam01197)→ PrfA (COG0216)prfA (PRK00591)→ |
| Ava-2-1 | NZ_JNYY01000004.1 | - | 220716 | 220673 | **RNA**→ rpmE (PRK00019)Ribosomal_L31 (pfam01197)→ PrfA (COG0216)prfA (PRK00591)→ |
| Ava-1-1 | NZ_AOUO01000443.1 | - | 35768 | 35725 | **RNA**→ rpmE (PRK00019)Ribosomal_L31 (pfam01197)→ PrfA (COG0216)prfA (PRK00591)→ |
| Aal-2-1 | NZ_JOEF01000005.1 | + | 165358 | 165404 | **RNA**→ rpmE (PRK00019)Ribosomal_L31 (pfam01197)→ |
| env-631 | 2200117070 | + | 1568 | 1604 | **RNA**→ rpmE (PRK00019)Ribosomal_L31 (pfam01197)→ |
| | | | | | prfA (PRK00591)RF-1 (pfam00472)PCRF (pfam03462)→ HemK (COG2890)PRK09328 (PRK09328)→ |
| | | | | | SUA5 (COG0009)TIGR00057 (TIGR00057)→ |
| env-632 | GBANfinal_FHNL2OP04ZB0BC | - | 172 | 136 | **RNA**→ rpmE (PRK00019)Ribosomal_L31 (pfam01197)→ |
| env-633 | JGI24740J21852_10800136 | + | 174 | 210 | **RNA**→ hypo→ |
| env-634 | JGI24737J22298_10394236 | - | 256 | 220 | **RNA**→ rpmE (PRK00019)Ribosomal_L31 (pfam01197)→ |
| env-635 | JGI24735J21928_10110934 | + | 294 | 330 | **RNA**→ rpmE (PRK00019)Ribosomal_L31 (pfam01197)→ PCRF (pfam03462)ABC_tran_CTD (pfam16326)→ |
| env-636 | JGI24738J21930_10339143 | + | 124 | 160 | **RNA**→ ←hypo |
| env-637 | JGI24739J22299_10621000 | + | 75 | 111 | **RNA**→ rpmE (PRK00019)Ribosomal_L31 (pfam01197)→ |
| Lae-1-1 | NZ_JOFI01000046.1 | - | 29517 | 29476 | **RNA**→ rpmE (PRK00019)Ribosomal_L31 (pfam01197)→ PrfA (COG0216)prfA (PRK00591)→ |
| | | | | | DLH (COG0412)Bile_Hydr_Trans (pfam04775)BAAT_C (pfam08840)→ Lpd (COG1249)PRK07818 (PRK07818)→ |
| | | | | | HemK (COG2890)PRK09328 (PRK09328)→ |
| env-638 | SRS053917_LANL_scaffold_22599 | + | 286 | 333 | **RNA**→ ←hypo |

| | | | | | |
|---|---|---|---|---|---|
| env-639 | SRS011255_Baylor_scaffold_25176 | + | 104 | 151 | **RNA**→ rpmE (PRK00019)Ribosomal_L31 (pfam01197)→ |
| env-640 | SRS013252_Baylor_scaffold_19505 | + | 1494 | 1541 | **RNA**→ rpmE (PRK00019)Ribosomal_L31 (pfam01197)→ |
| env-641 | SRS015215_C2366775 | + | 105 | 152 | **RNA**→ rpmE (PRK00019)Ribosomal_L31 (pfam01197)→ |
| env-642 | SRS015278_C1812809 | + | 411 | 458 | **RNA**→ rpmE (PRK00019)Ribosomal_L31 (pfam01197)→ rpmE (PRK00019)Ribosomal_L31 (pfam01197)→ |
| env-643 | SRS015278_C1812809 | - | 206 | 159 | **RNA**→ rpmE (PRK00019)Ribosomal_L31 (pfam01197)→ rpmE (PRK00019)Ribosomal_L31 (pfam01197)→ |
| env-644 | SRS022725_LANL_scaffold_111342 | - | 197 | 150 | **RNA**→ ←hypo |
| env-645 | SRS051930_LANL_scaffold_6579 | - | 323 | 276 | **RNA**→ rpmE (PRK00019)Ribosomal_L31 (pfam01197)→ |
| env-646 | SRS075410_C2925444 | + | 106 | 153 | **RNA**→ rpmE (PRK00019)Ribosomal_L31 (pfam01197)→ |
| Ama-2-1 | NZ_AKFT01000226.1 | - | 10315 | 10268 | **RNA**→ hypo→ rpmE (PRK00019)Ribosomal_L31 (pfam01197)→ |
| Aen-1-1 | NZ_KB894433.1 | + | 15128 | 15170 | **RNA**→ rpmE (PRK00019)Ribosomal_L31 (pfam01197)→ |
| Aha-3-1 | NZ_KI632509.1 | - | 4883378 | 4883317 | **RNA**→ rpmE (PRK00019)Ribosomal_L31 (pfam01197)→ |
| env-647 | JGI24973J35851_1008812 | + | 1167 | 1208 | **RNA**→ rpmE (PRK00019)Ribosomal_L31 (pfam01197)→ prfA (PRK00591)PCRF (pfam03462)→ |
| env-648 | JGI24973J35851_1010391 | - | 1305 | 1264 | **RNA**→ rpmE (PRK00019)Ribosomal_L31 (pfam01197)→<br>prfA (PRK00591)RF-1 (pfam00472)PCRF (pfam03462)→ |
| Ksp-1-1 | NZ_CDME01000015.1 | - | 111616 | 111572 | **RNA**→ rpmE (PRK00019)Ribosomal_L31 (pfam01197)→ PrfA (COG0216)prfA (PRK00591)→<br>HemK (COG2890)PRK09328 (PRK09328)→ |
| Lal-2-1 | NZ_JOEA01000021.1 | + | 93454 | 93495 | **RNA**→ rpmE (PRK00019)Ribosomal_L31 (pfam01197)→ PrfA (COG0216)prfA (PRK00591)→<br>DLH (COG0412)Bile_Hydr_Trans (pfam04775)BAAT_C (pfam08840)→ Lpd (COG1249)PRK07818 (PRK07818)→<br>HemK (COG2890)PRK09328 (PRK09328)→ |
| Lae-1-2 | NZ_JYJG01000134.1 | + | 673 | 714 | **RNA**→ rpmE (PRK00019)Ribosomal_L31 (pfam01197)→ PrfA (COG0216)prfA (PRK00591)→<br>DLH (COG0412)Bile_Hydr_Trans (pfam04775)BAAT_C (pfam08840)→ Lpd (COG1249)PRK07818 (PRK07818)→<br>HemK (COG2890)PRK09328 (PRK09328)→ |
| env-649 | FNTS067_GJ87FRN01CVTQK | - | 106 | 72 | **RNA**→ |
| Psp-9-1 | NZ_AEGE01000437.1 | - | 2109 | 2068 | **RNA**→ rpmE (PRK00019)Ribosomal_L31 (pfam01197)→ PrfA (COG0216)prfA (PRK00591)→ hypo→ hypo→ |
| Aje-1-1 | NZ_JNYZ01000004.1 | - | 127926 | 127889 | **RNA**→ rpmE (PRK00019)Ribosomal_L31 (pfam01197)→ PrfA (COG0216)prfA (PRK00591)→ |
| [ir-1-1 | NZ_KE387135.1 | + | 25080 | 25126 | **RNA**→ rpmE (PRK00019)Ribosomal_L31 (pfam01197)→ PrfA (COG0216)prfA (PRK00591)→<br>HemK (COG2890)PRK09328 (PRK09328)→ SUA5 (COG0009)TIGR00057 (TIGR00057)→<br>ECA_wecA (TIGR02380)GT_WecA_like (cd06853)→ hypo→ |
| Ain-1-1 | NZ_AXWW01000059.1 | - | 14834 | 14796 | **RNA**→ rpmE (PRK00019)Ribosomal_L31 (pfam01197)→ PrfA (COG0216)prfA (PRK00591)→<br>HemK (COG2890)PRK09328 (PRK09328)→ |
| Aja-1-1 | NZ_CP008953.1 | + | 1210103 | 1210138 | **RNA**→ rpmE (PRK00019)Ribosomal_L31 (pfam01197)→ PrfA (COG0216)prfA (PRK00591)→ |
| Asp-6-1 | NZ_JPLW01000003.1 | + | 572205 | 572240 | **RNA**→ rpmE (PRK00019)Ribosomal_L31 (pfam01197)→ PrfA (COG0216)prfA (PRK00591)→ |
| Aaz-1-1 | NZ_ANMG01000003.1 | - | 63867 | 63832 | **RNA**→ rpmE (PRK00019)Ribosomal_L31 (pfam01197)→ PrfA (COG0216)prfA (PRK00591)→ |
| Aor-4-1 | NZ_ASXH01000080.1 | + | 35738 | 35774 | **RNA**→ rpmE (PRK00019)Ribosomal_L31 (pfam01197)→ PrfA (COG0216)prfA (PRK00591)→ |
| Ssp-91-1 | NZ_JOIQ01000021.1 | + | 124197 | 124233 | **RNA**→ rpmE (PRK00019)Ribosomal_L31 (pfam01197)→ PrfA (COG0216)prfA (PRK00591)→ |
| Abe-1-1 | NZ_KB912942.1 | + | 23653 | 23690 | **RNA**→ rpmE (PRK00019)Ribosomal_L31 (pfam01197)→ PrfA (COG0216)prfA (PRK00591)→ |
| Aor-1-1 | NZ_KN912592.1 | + | 508571 | 508607 | **RNA**→ rpmE (PRK00019)Ribosomal_L31 (pfam01197)→ PrfA (COG0216)prfA (PRK00591)→ |
| Aor-3-1 | NZ_ASXG01000124.1 | + | 157002 | 157038 | **RNA**→ rpmE (PRK00019)Ribosomal_L31 (pfam01197)→ PrfA (COG0216)prfA (PRK00591)→ |
| Ade-1-1 | NZ_AOHO01000019.1 | - | 158657 | 158622 | **RNA**→ rpmE (PRK00019)Ribosomal_L31 (pfam01197)→ PrfA (COG0216)prfA (PRK00591)→ |
| Ssp-5-1 | NZ_GG657746.1 | - | 6816029 | 6815993 | **RNA**→ rpmE (PRK00019)Ribosomal_L31 (pfam01197)→ rpmE (PRK00019)Ribosomal_L31 (pfam01197)→<br>PrfA (COG0216)prfA (PRK00591)→ PrfA (COG0216)prfA (PRK00591)→ |
| Aor-2-1 | NZ_ASJB01000091.1 | + | 88996 | 89032 | **RNA**→ rpmE (PRK00019)Ribosomal_L31 (pfam01197)→ PrfA (COG0216)prfA (PRK00591)→ hypo→ |
| env-650 | JGI24737J22298_10153983 | - | 501 | 462 | **RNA**→ rpmE (PRK00019)Ribosomal_L31 (pfam01197)→ prfA (PRK00591)PCRF (pfam03462)→ |
| env-651 | JGI24738J21930_10140397 | - | 125 | 86 | **RNA**→ |
| env-652 | JRHI01000725.1 | + | 25745 | 25791 | **RNA**→ rpmE (PRK00019)Ribosomal_L31 (pfam01197)→ PrfA (COG0216)prfA (PRK00591)→<br>HemK (COG2890)PRK09328 (PRK09328)→ |
| Sha-1-1 | NZ_KB912506.1 | - | 125509 | 125456 | **RNA**→ rpmE (PRK00019)Ribosomal_L31 (pfam01197)→ PrfA (COG0216)prfA (PRK00591)→<br>HemK (COG2890)PRK09328 (PRK09328)→ SUA5 (COG0009)TIGR00057 (TIGR00057)→<br>ECA_wecA (TIGR02380)GT_WecA_like (cd06853)→ hypo→ |
| Spa-41-1 | NZ_KI912213.1 | - | 219937 | 219883 | **RNA**→ rpmE (PRK00019)Ribosomal_L31 (pfam01197)→ PrfA (COG0216)prfA (PRK00591)→<br>HemK (COG2890)PRK09328 (PRK09328)→ SUA5 (COG0009)TIGR00057 (TIGR00057)→<br>ECA_wecA (TIGR02380)GT_WecA_like (cd06853)→ hypo→ |
| Kal-2-1 | NZ_CP007155.1 | - | 8601054 | 8601005 | **RNA**→ rpmE (PRK00019)Ribosomal_L31 (pfam01197)→ |
| Kal-1-1 | NZ_JNYH01000160.1 | - | 17281 | 17232 | **RNA**→ rpmE (PRK00019)Ribosomal_L31 (pfam01197)→ |
| Saz-1-1 | NZ_CM001466.1 | - | 1832387 | 1832343 | **RNA**→ rpmE (PRK00019)Ribosomal_L31 (pfam01197)→ rpmE (PRK00019)Ribosomal_L31 (pfam01197)→ |

| | | | | | |
|---|---|---|---|---|---|
| Saz-2-1 | NZ_AHBX01000091.1 | - | 27128 | 27084 | **RNA**→ rpmE (PRK00019)Ribosomal_L31 (pfam01197)→ |
| Svi-1-1 | NC_013159.1 | - | 3233924 | 3233883 | **RNA**→ rpmE (PRK00019)Ribosomal_L31 (pfam01197)→ |
| Svi-8-1 | NZ_JRZE01000001.1 | - | 241211 | 241170 | **RNA**→ rpmE (PRK00019)Ribosomal_L31 (pfam01197)→ |
| Ssp-45-1 | NZ_AZUM01000002.1 | - | 62099 | 62059 | **RNA**→ rpmE (PRK00019)Ribosomal_L31 (pfam01197)→ PrfA (COG0216)prfA (PRK00591)→ HTH_MerR-like_sg3 (cd01282)HTH_MERR (smart00422)→ HemK (COG2890)PRK09328 (PRK09328)→ SUA5 (COG0009)TIGR00057 (TIGR00057)→ ECA_wecA (TIGR02380)GT_WecA_like (cd06853)→ |
| Kar-1-1 | NZ_JNYM01000248.1 | + | 820 | 862 | **RNA**→ rpmE (PRK00019)Ribosomal_L31 (pfam01197)→ PrfA (COG0216)prfA (PRK00591)→ HemK (COG2890)PRK09328 (PRK09328)→ STE14 (COG2020)→ |
| Ani-1-1 | NZ_ARVW01000001.1 | + | 5123288 | 5123325 | **RNA**→ rpmE (PRK00019)Ribosomal_L31 (pfam01197)→ PrfA (COG0216)prfA (PRK00591)→ |
| Rrh-3-1 | NZ_JOAA01000025.1 | - | 102373 | 102331 | **RNA**→ rpmE (PRK00019)Ribosomal_L31 (pfam01197)→ PrfA (COG0216)prfA (PRK00591)→ HemK (COG2890)PRK09328 (PRK09328)→ |
| Scy-1-1 | NZ_CM001440.1 | - | 4163623 | 4163579 | **RNA**→ rpmE (PRK00019)Ribosomal_L31 (pfam01197)→ rpmE (PRK00019)Ribosomal_L31 (pfam01197)→ PrfA (COG0216)prfA (PRK00591)→ PrfA (COG0216)prfA (PRK00591)→ PrfA (COG0216)prfA (PRK00591)→ |
| env-653 | JGI12271J12027_1317364 | + | 55 | 90 | **RNA**→ ←hypo |
| Ame-3-1 | NC_014318.1 | - | 8567126 | 8567084 | **RNA**→ rpmE (PRK00019)Ribosomal_L31 (pfam01197)→ PrfA (COG0216)prfA (PRK00591)→ |
| Ame-2-1 | NC_017186.1 | - | 8567182 | 8567140 | **RNA**→ rpmE (PRK00019)Ribosomal_L31 (pfam01197)→ PrfA (COG0216)prfA (PRK00591)→ |
| Ame-2-2 | NC_018266.1 | - | 8577099 | 8577057 | **RNA**→ rpmE (PRK00019)Ribosomal_L31 (pfam01197)→ PrfA (COG0216)prfA (PRK00591)→ |
| Ame-1-1 | NC_022116.1 | - | 8577043 | 8577001 | **RNA**→ rpmE (PRK00019)Ribosomal_L31 (pfam01197)→ PrfA (COG0216)prfA (PRK00591)→ |
| Ame-5-1 | NZ_JMQJ01000008.1 | + | 22200 | 22242 | **RNA**→ rpmE (PRK00019)Ribosomal_L31 (pfam01197)→ PrfA (COG0216)prfA (PRK00591)→ |
| Ame-5-2 | NZ_JMQG01000068.1 | - | 49889 | 49847 | **RNA**→ rpmE (PRK00019)Ribosomal_L31 (pfam01197)→ PrfA (COG0216)prfA (PRK00591)→ |
| Ssp-46-1 | NZ_JNXC01000009.1 | + | 134688 | 134733 | **RNA**→ rpmE (PRK00019)Ribosomal_L31 (pfam01197)→ PrfA (COG0216)prfA (PRK00591)→ DLH (COG0412)Bile_Hydr_Trans (pfam04775)BAAT_C (pfam08840)→ Lpd (COG1249)PRK07818 (PRK07818)→ HemK (COG2890)PRK09328 (PRK09328)→ |
| Ami-2-1 | NC_013093.1 | - | 7204512 | 7204468 | **RNA**→ rpmE (PRK00019)Ribosomal_L31 (pfam01197)→ hypo→ hypo→ PrfA (COG0216)prfA (PRK00591)→ |
| Ssp-1-1 | NZ_GL877878.1 | + | 167629 | 167675 | **RNA**→ rpmE (PRK00019)Ribosomal_L31 (pfam01197)→ |
| env-654 | JGI26535J51212_1001019 | + | 71 | 108 | **RNA**→ rpmE (PRK00019)Ribosomal_L31 (pfam01197)→ |
| env-655 | JGI26539J51216_1000587 | - | 26649 | 26612 | **RNA**→ rpmE (PRK00019)Ribosomal_L31 (pfam01197)→ |
| env-656 | JGI26531J51214_10001689 | - | 15226 | 15189 | **RNA**→ rpmE (PRK00019)Ribosomal_L31 (pfam01197)→ |
| Sts-1-1 | NZ_AJSZ01000299.1 | - | 3790 | 3744 | **RNA**→ rpmE (PRK00019)Ribosomal_L31 (pfam01197)→ rpmE (PRK00019)Ribosomal_L31 (pfam01197)→ |
| env-657 | JGI12397J12024_1206537 | + | 4 | 58 | **RNA**→ rpmE (PRK00019)Ribosomal_L31 (pfam01197)→ |
| Rsp-11-1 | NZ_JYOP01000005.1 | - | 14897 | 14852 | **RNA**→ rpmE (PRK00019)Ribosomal_L31 (pfam01197)→ |
| env-658 | JGI11799J14266_104666 | + | 85 | 130 | **RNA**→ ←hypo |
| env-659 | JGI1355J14265_100580 | - | 4687 | 4642 | **RNA**→ rpmE (PRK00019)Ribosomal_L31 (pfam01197)→ |
| env-660 | JGI24972J35695_100056 | - | 16362 | 16317 | **RNA**→ rpmE (PRK00019)Ribosomal_L31 (pfam01197)→ |
| env-661 | JGI24971J35700_100060 | + | 57533 | 57578 | **RNA**→ rpmE (PRK00019)Ribosomal_L31 (pfam01197)→ |
| env-662 | JGI24975J35696_100061 | + | 57533 | 57578 | **RNA**→ rpmE (PRK00019)Ribosomal_L31 (pfam01197)→ |
| Rer-4-1 | NZ_ACNO01000014.1 | + | 120970 | 121015 | **RNA**→ rpmE (PRK00019)Ribosomal_L31 (pfam01197)→ rpmE (PRK00019)Ribosomal_L31 (pfam01197)→ |
| Rqi-1-1 | NZ_AODN01000001.1 | + | 127294 | 127339 | **RNA**→ rpmE (PRK00019)Ribosomal_L31 (pfam01197)→ |
| Rsp-12-1 | NZ_LEKE01000058.1 | - | 16710 | 16665 | **RNA**→ rpmE (PRK00019)Ribosomal_L31 (pfam01197)→ |
| Rrh-5-1 | NZ_ASJJ01000026.1 | + | 157576 | 157621 | **RNA**→ rpmE (PRK00019)Ribosomal_L31 (pfam01197)→ |
| Rsp-10-1 | NZ_CFHW01000012.1 | + | 79898 | 79943 | **RNA**→ rpmE (PRK00019)Ribosomal_L31 (pfam01197)→ |
| Rer-3-1 | NC_012490.1 | - | 4297666 | 4297621 | **RNA**→ rpmE (PRK00019)Ribosomal_L31 (pfam01197)→ hypo→ PrfA (COG0216)prfA (PRK00591)→ HemK (COG2890)PRK09328 (PRK09328)→ SUA5 (COG0009)TIGR00057 (TIGR00057)→ GlyA (COG0112)glyA (PRK00011)→ ECA_wecA (TIGR02380)GT_WecA_like (cd06853)→ ECA_wecA (TIGR02380)GT_WecA_like (cd06853)→ Tim44 (COG4395)→ |
| Rer-1-1 | NC_022115.1 | - | 3994948 | 3994903 | **RNA**→ rpmE (PRK00019)Ribosomal_L31 (pfam01197)→ |
| Rer-7-1 | NZ_CM002793.1 | - | 3935713 | 3935668 | **RNA**→ rpmE (PRK00019)Ribosomal_L31 (pfam01197)→ |
| Rer-6-1 | NZ_CP011295.1 | - | 4024740 | 4024695 | **RNA**→ rpmE (PRK00019)Ribosomal_L31 (pfam01197)→ |
| Rer-5-1 | NZ_AGCF01000002.1 | - | 2284788 | 2284743 | **RNA**→ rpmE (PRK00019)Ribosomal_L31 (pfam01197)→ |
| Rer-2-1 | NZ_AUZK01000027.1 | - | 16252 | 16207 | **RNA**→ rpmE (PRK00019)Ribosomal_L31 (pfam01197)→ |
| Rsp-8-1 | NZ_AVCO01000044.1 | - | 15061 | 15016 | **RNA**→ rpmE (PRK00019)Ribosomal_L31 (pfam01197)→ |
| Rqi-2-1 | NZ_JNCU01000020.1 | + | 109948 | 109993 | **RNA**→ rpmE (PRK00019)Ribosomal_L31 (pfam01197)→ |
| Rop-4-1 | NZ_JOIM01000019.1 | - | 16215 | 16170 | **RNA**→ rpmE (PRK00019)Ribosomal_L31 (pfam01197)→ |
| Rrh-4-1 | NZ_JNWS01000014.1 | + | 192760 | 192805 | **RNA**→ rpmE (PRK00019)Ribosomal_L31 (pfam01197)→ |

| | | | | | |
|---|---|---|---|---|---|
| Ace-1-1 | NC_008578.1 | + | 694535 | 694574 | RNA→ rpmE (PRK00019)Ribosomal_L31 (pfam01197)→ PrfA (COG0216)prfA (PRK00591)→ |
| | | | | | HemK (COG2890)PRK09328 (PRK09328)→ SUA5 (COG0009)TIGR00057 (TIGR00057)→ |
| | | | | | GT_WecA_like (cd06853)mraY (PRK00108)→ |
| Gja-1-1 | NZ_LDTZ01000026.1 | - | 28087 | 28042 | RNA→ rpmE (PRK00019)Ribosomal_L31 (pfam01197)→ PrfA (COG0216)prfA (PRK00591)→ |
| | | | | | HemK (COG2890)PRK09328 (PRK09328)→ SUA5 (COG0009)TIGR00057 (TIGR00057)→ |
| | | | | | GT_WecA_like (cd06853)mraY (PRK00108)→ |
| Gsp-2-1 | NZ_BAFC01000059.1 | - | 28156 | 28111 | RNA→ rpmE (PRK00019)Ribosomal_L31 (pfam01197)→ rpmE (PRK00019)Ribosomal_L31 (pfam01197)→ |
| Got-1-1 | NZ_BAFB01000227.1 | - | 28105 | 28060 | RNA→ rpmE (PRK00019)Ribosomal_L31 (pfam01197)→ rpmE (PRK00019)Ribosomal_L31 (pfam01197)→ |
| Gai-1-1 | NZ_BANR01000032.1 | - | 28002 | 27957 | RNA→ rpmE (PRK00019)Ribosomal_L31 (pfam01197)→ |
| Asp-5-2 | NZ_AYXG01000157.1 | + | 12560 | 12601 | RNA→ rpmE (PRK00019)Ribosomal_L31 (pfam01197)→ PrfA (COG0216)prfA (PRK00591)→ |
| | | | | | HemK (COG2890)PRK09328 (PRK09328)→ |
| env-663 | _GX4IIKE01AVNLA | + | 402 | 450 | RNA→ |
| Sma-1-1 | NZ_CM001439.1 | - | 4262593 | 4262553 | RNA→ rpmE (PRK00019)Ribosomal_L31 (pfam01197)→ rpmE (PRK00019)Ribosomal_L31 (pfam01197)→ |
| | | | | | PrfA (COG0216)prfA (PRK00591)→ PrfA (COG0216)prfA (PRK00591)→ |
| env-664 | JGI26531J51214_10053388 | + | 189 | 235 | RNA→ rpmE (PRK00019)Ribosomal_L31 (pfam01197)→ |
| env-665 | JGI26539J51216_1014970 | + | 457 | 503 | RNA→ rpmE (PRK00019)Ribosomal_L31 (pfam01197)→ |
| Ssu-1-1 | NZ_KB905814.1 | - | 250505 | 250461 | RNA→ rpmE (PRK00019)Ribosomal_L31 (pfam01197)→ |
| Ssu-2-1 | NZ_AJTQ01000139.1 | - | 131331 | 131287 | RNA→ rpmE (PRK00019)Ribosomal_L31 (pfam01197)→ |
| Sje-1-1 | NZ_JOEH01000016.1 | - | 36292 | 36248 | RNA→ rpmE (PRK00019)Ribosomal_L31 (pfam01197)→ |
| Sal-2-1 | NZ_BBPL01000004.1 | - | 265730 | 265686 | RNA→ rpmE (PRK00019)Ribosomal_L31 (pfam01197)→ |
| Sal-3-1 | NZ_JQML01000001.1 | - | 3284063 | 3284019 | RNA→ rpmE (PRK00019)Ribosomal_L31 (pfam01197)→ |
| Sru-1-1 | NZ_JQMJ01000004.1 | - | 6329451 | 6329407 | RNA→ rpmE (PRK00019)Ribosomal_L31 (pfam01197)→ |
| Ssp-100-1 | NZ_JOHT01000012.1 | - | 106860 | 106816 | RNA→ rpmE (PRK00019)Ribosomal_L31 (pfam01197)→ |
| Ssp-129-1 | NZ_JOGD01000005.1 | - | 35544 | 35500 | RNA→ rpmE (PRK00019)Ribosomal_L31 (pfam01197)→ |
| Ssp-171-1 | NZ_AUEV01000006.1 | + | 1894 | 1939 | RNA→ rpmE (PRK00019)Ribosomal_L31 (pfam01197)→ |
| Ssp-170-1 | NZ_KE384281.1 | - | 396059 | 396015 | RNA→ rpmE (PRK00019)Ribosomal_L31 (pfam01197)→ |
| Ssp-56-1 | NZ_AUKO01000001.1 | - | 376807 | 376763 | RNA→ rpmE (PRK00019)Ribosomal_L31 (pfam01197)→ |
| Kaz-1-1 | NZ_JQMO01000003.1 | + | 1403666 | 1403710 | RNA→ rpmE (PRK00019)Ribosomal_L31 (pfam01197)→ |
| Svi-7-1 | NZ_KB904663.1 | + | 33286 | 33330 | RNA→ rpmE (PRK00019)Ribosomal_L31 (pfam01197)→ |
| Smo-1-1 | NZ_AORZ01000143.1 | - | 15463 | 15418 | RNA→ rpmE (PRK00019)Ribosomal_L31 (pfam01197)→ |
| Ama-4-1 | NZ_HE978637.1 | + | 431034 | 431078 | RNA→ rpmE (PRK00019)Ribosomal_L31 (pfam01197)→ PrfA (COG0216)prfA (PRK00591)→ |
| | | | | | HemK (COG2890)PRK09328 (PRK09328)→ YgaA (COG1238)SNARE_assoc (pfam09335)→ |
| | | | | | SUA5 (COG0009)TIGR00057 (TIGR00057)→ ECA_wecA (TIGR02380)GT_WecA_like (cd06853)→ hypo→ hypo→ |
| | | | | | PRK05815 (PRK05815)ATP-synt_A (pfam00119)→ PRK07874 (PRK07874)AtpE (COG0636)→ |
| | | | | | PRK05759 (PRK05759)AtpF (COG0711)→ AtpH (COG0712)PRK13430 (PRK13430)→ |
| | | | | | AtpA (COG0056)PRK09281 (PRK09281)→ PRK05621 (PRK05621)ATP-synt (pfam00231)→ |
| env-666 | JGI24770J26754_10081147 | + | 845 | 890 | RNA→ rpmE (PRK00019)Ribosomal_L31 (pfam01197)→ |
| Bsc-2-1 | NZ_AP012331.1 | + | 769739 | 769783 | RNA→ rpmE (PRK00019)Ribosomal_L31 (pfam01197)→ |
| Bsc-1-2 | NZ_JGZO01000023.1 | - | 265290 | 265246 | RNA→ rpmE (PRK00019)Ribosomal_L31 (pfam01197)→ |
| Ssc-1-1 | NC_013929.1 | - | 3291560 | 3291514 | RNA→ rpmE (PRK00019)Ribosomal_L31 (pfam01197)→ |
| Ssc-3-2 | NZ_KL997517.1 | + | 318375 | 318421 | RNA→ rpmE (PRK00019)Ribosomal_L31 (pfam01197)→ |
| Sle-1-1 | NZ_LN831790.1 | - | 2823297 | 2823251 | RNA→ rpmE (PRK00019)Ribosomal_L31 (pfam01197)→ |
| Scl-1-1 | NZ_CM001015.1 | + | 4930378 | 4930424 | RNA→ rpmE (PRK00019)Ribosomal_L31 (pfam01197)→ |
| Scl-1-2 | NZ_CM000913.1 | + | 4939868 | 4939914 | RNA→ rpmE (PRK00019)Ribosomal_L31 (pfam01197)→ rpmE (PRK00019)Ribosomal_L31 (pfam01197)→ |
| Sre-3-1 | NZ_KL575619.1 | - | 35887 | 35841 | RNA→ rpmE (PRK00019)Ribosomal_L31 (pfam01197)→ |
| Ser-2-1 | NC_009142.1 | - | 7044070 | 7044025 | RNA→ rpmE (PRK00019)Ribosomal_L31 (pfam01197)→ PrfA (COG0216)prfA (PRK00591)→ |
| | | | | | HemK (COG2890)PRK09328 (PRK09328)→ |
| Ser-1-1 | NZ_AVCN01000136.1 | + | 15183 | 15228 | RNA→ rpmE (PRK00019)Ribosomal_L31 (pfam01197)→ PrfA (COG0216)prfA (PRK00591)→ |
| | | | | | HemK (COG2890)PRK09328 (PRK09328)→ |
| Ssp-20-1 | NZ_KE150472.1 | + | 563628 | 563673 | RNA→ rpmE (PRK00019)Ribosomal_L31 (pfam01197)→ |
| Ssp-116-1 | NZ_JOHQ01000026.1 | + | 21477 | 21522 | RNA→ rpmE (PRK00019)Ribosomal_L31 (pfam01197)→ |
| Ssp-111-1 | NZ_JOGK01000004.1 | + | 39032 | 39077 | RNA→ rpmE (PRK00019)Ribosomal_L31 (pfam01197)→ |
| Sgr-5-1 | NZ_JNWT01000006.1 | + | 36815 | 36860 | RNA→ rpmE (PRK00019)Ribosomal_L31 (pfam01197)→ |
| Sal-8-3 | NZ_JODZ01000009.1 | + | 43556 | 43601 | RNA→ rpmE (PRK00019)Ribosomal_L31 (pfam01197)→ |

| | | | | | |
|---|---|---|---|---|---|
| Sal-8-4 | NZ_JODR01000008.1 | + | 36815 | 36860 | **RNA**→ rpmE (PRK00019)Ribosomal_L31 (pfam01197)→ |
| Sst-1-1 | NZ_KQ257875.1 | + | 1320008 | 1320054 | **RNA**→ rpmE (PRK00019)Ribosomal_L31 (pfam01197)→ |
| Mne-3-1 | NC_023036.2 | - | 4254218 | 4254162 | **RNA**→ rpmE (PRK00019)Ribosomal_L31 (pfam01197)→ PrfA (COG0216)prfA (PRK00591)→ HemK (COG2890)PRK09328 (PRK09328)→ SUA5 (COG0009)TIGR00057 (TIGR00057)→ GT_WecA_like (cd06853)mraY (PRK00108)→ |
| Msp-24-1 | NZ_AOHQ01000138.1 | - | 14981 | 14925 | **RNA**→ rpmE (PRK00019)Ribosomal_L31 (pfam01197)→ PrfA (COG0216)prfA (PRK00591)→ HemK (COG2890)PRK09328 (PRK09328)→ SUA5 (COG0009)TIGR00057 (TIGR00057)→ GT_WecA_like (cd06853)mraY (PRK00108)→ |

#### 9.3 Conserved domains

Conserved domains found in protein-coding genes listed in Section 9.2 are shown below, with the first sentence in their description from the Conserved Domain Database (if any). Conserved domains associated with more than one L31-Actinobacteria RNA are assigned a color, while others are shown in gray. The number in parentheses after the colored domain name is the number of occurrences in Section 9.2.

cd01128 (1) Transcription termination factor rho is a bacterial ATP-dependent RNA/DNA helicase.
cd01282 (1) Helix-Turn-Helix DNA binding domain of putative transcription regulators from the MerR superfamily.
cd02028 (1) Uridine monophosphate kinase_like (UMPK_like) is a family of proteins highly similar to the uridine monophosphate kinase (UMPK, EC 2.
cd04601 (1) This cd contains two tandem repeats of the cystathionine beta-synthase (CBS pair) domains in the inosine 5' monophosphate dehydrogenase (IMPDH) protein.
cd06199 (1) Cytochrome p450- like alpha subunits of E.
cd06853 (363) This subfamily contains Escherichia coli WecA, Bacillus subtilis TagO and related proteins.
cd16344 (1) low molecular weight protein arginine phosphatase.
COG0009 (410) tRNA A37 threonylcarbamoyladenosine synthetase subunit TsaC/SUA5/YrdC [Translation, ribosomal structure and biogenesis]
COG0055 (7) FoF1-type ATP synthase, beta subunit [Energy production and conversion]
COG0056 (13) FoF1-type ATP synthase, alpha subunit [Energy production and conversion]
COG0112 (18) Glycine/serine hydroxymethyltransferase [Amino acid transport and metabolism]
COG0216 (536) Protein chain release factor A [Translation, ribosomal structure and biogenesis]
COG0224 (3) FoF1-type ATP synthase, gamma subunit [Energy production and conversion]
COG0254 (1) Ribosomal protein L31 [Translation, ribosomal structure and biogenesis]
COG0394 (28) Protein-tyrosine-phosphatase [Signal transduction mechanisms]
COG0412 (4) Dienelactone hydrolase [Secondary metabolites biosynthesis, transport and catabolism]
COG0472 (2) UDP-N-acetylmuramyl pentapeptide phosphotransferase/UDP-N-acetylglucosamine-1-phosphate transferase [Cell wall/membrane/envelope biogenesis]
COG0596 (1) Pimeloyl-ACP methyl ester carboxylesterase [Coenzyme transport and metabolism, General function prediction only]
COG0636 (14) FoF1-type ATP synthase, membrane subunit c/Archaeal/vacuolar-type H+-ATPase, subunit K [Energy production and conversion]
COG0711 (14) FoF1-type ATP synthase, membrane subunit b or b' [Energy production and conversion]
COG0712 (8) FoF1-type ATP synthase, delta subunit [Energy production and conversion]
COG1113 (3) L-asparagine transporter and related permeases [Amino acid transport and metabolism]
COG1186 (1) Protein chain release factor B [Translation, ribosomal structure and biogenesis]
COG1238 (1) Uncharacterized membrane protein YqaA, SNARE-associated domain [Function unknown]
COG1249 (5) Pyruvate/2-oxoglutarate dehydrogenase complex, dihydrolipoamide dehydrogenase (E3) component or related enzyme [Energy production and conversion]
COG1409 (1) 3',5'-cyclic AMP phosphodiesterase CpdA [Signal transduction mechanisms]
COG1472 (2) Periplasmic beta-glucosidase and related glycosidases [Carbohydrate transport and metabolism]
COG1506 (1) Dipeptidyl aminopeptidase/acylaminoacyl peptidase [Amino acid transport and metabolism]
COG1598 (1) Predicted nuclease of the RNAse H fold, HicB family [Defense mechanisms]
COG2020 (1) Protein-S-isoprenylcysteine O-methyltransferase Ste14 [Posttranslational modification, protein turnover, chaperones]
COG2197 (2) DNA-binding response regulator, NarL/FixJ family, contains REC and HTH domains [Signal transduction mechanisms, Transcription]
COG2890 (456) Methylase of polypeptide chain release factors [Translation, ribosomal structure and biogenesis]
COG4395 (1) Predicted lipid-binding transport protein, Tim44 family [Lipid transport and metabolism]
COG5336 (8) FoF1-type ATP synthase assembly protein I [Energy production and conversion]
pfam00119 (18) ATP synthase A chain.
pfam00213 (6) ATP synthase delta (OSCP) subunit.
pfam00231 (10) ATP synthase.
pfam00472 (24) RF-1 domain.
pfam00478 (2) IMP dehydrogenase / GMP reductase domain.
pfam00571 (2) CBS domain.
pfam01197 (1795) Ribosomal protein L31.
pfam02874 (1) ATP synthase alpha/beta family, beta-barrel domain.
pfam03462 (49) PCRF domain.
pfam03816 (4) Cell envelope-related transcriptional attenuator domain.
pfam04775 (5) Acyl-CoA thioester hydrolase/BAAT N-terminal region.
pfam05534 (1) HicB family.

pfam07690 (1) Major Facilitator Superfamily.
pfam08840 (5) BAAT / Acyl-CoA thioester hydrolase C terminal.
pfam09335 (1) SNARE associated Golgi protein.
pfam09527 (5) Putative F0F1-ATPase subunit Ca2+/Mg2+ transporter.
pfam09851 (7) Short C-terminal domain.
pfam13520 (3) Amino acid permease.
pfam13649 (2) Methyltransferase domain.
pfam14330 (1) Domain of unknown function (DUF4387).
pfam16326 (5) ABC transporter C-terminal domain.
PHA03169 (1) hypothetical protein; Provisional
PRK00011 (18) serine hydroxymethyltransferase; Reviewed
PRK00019 (1795) 50S ribosomal protein L31; Reviewed
PRK00108 (68) phospho-N-acetylmuramoyl-pentapeptide-transferase; Provisional
PRK00528 (3) 50S ribosomal protein L31; Reviewed
PRK00591 (571) peptide chain release factor 1; Validated
PRK01678 (4) 50S ribosomal protein L31 type B; Reviewed
PRK05567 (1) inosine 5'-monophosphate dehydrogenase; Reviewed
PRK05621 (13) F0F1 ATP synthase subunit gamma; Validated
PRK05759 (14) F0F1 ATP synthase subunit B; Validated
PRK05815 (18) F0F1 ATP synthase subunit A; Validated
PRK06214 (1) sulfite reductase; Provisional
PRK06876 (4) F0F1 ATP synthase subunit C; Validated
PRK07764 (1) DNA polymerase III subunits gamma and tau; Validated
PRK07818 (5) dihydrolipoamide dehydrogenase; Reviewed
PRK07874 (10) F0F1 ATP synthase subunit C; Validated
PRK08279 (1) long-chain-acyl-CoA synthetase; Validated
PRK09280 (7) F0F1 ATP synthase subunit beta; Validated
PRK09281 (14) F0F1 ATP synthase subunit alpha; Validated
PRK09328 (452) N5-glutamine S-adenosyl-L-methionine-dependent methyltransferase; Provisional
PRK09376 (1) transcription termination factor Rho; Provisional
PRK10959 (1) outer membrane protein W; Provisional
PRK11391 (26) phosphotyrosine-protein phosphatase; Provisional
PRK13430 (14) F0F1 ATP synthase subunit delta; Provisional
PRK15098 (2) beta-D-glucoside glucohydrolase; Provisional
PRK15119 (4) undecaprenyl-phosphate alpha-N-acetylglucosaminyl 1-phosphate transferase; Provisional
smart00116 (2) Domain in cystathionine beta-synthase and other proteins.
smart00226 (3) Low molecular weight phosphatase family.
smart00421 (2) helix_turn_helix, Lux Regulon.
smart00422 (1) helix_turn_helix, mercury resistance.
smart00937 (12) This domain is found in peptide chain release factors.
TIGR00057 (410) tRNA threonylcarbamoyl adenosine modification protein, Sua5/YciO/YrdC/YwlC family. [Protein synthesis, tRNA and rRNA base modification]
TIGR00350 (4) cell envelope-related function transcriptional attenuator common domain. [Regulatory functions, Other]
TIGR00711 (1) drug resistance transporter, EmrB/QacA subfamily. [Cellular processes, Toxin production and resistance, Transport and binding proteins, Other]
TIGR02380 (291) undecaprenyl-phosphate alpha-N-acetylglucosaminyl 1-phosphatetransferase. [Cell envelope, Biosynthesis and degradation of surface polysaccharides and lipopolysaccharides]
TIGR03025 (1) exopolysaccharide biosynthesis polyprenyl glycosylphosphotransferase.
TIGR03534 (2) protein-(glutamine-N5) methyltransferase, release factor-specific. [LIV]PRx[DE]TE (in Escherichia coli, IPRPDTE) confers specificity for the release factors rather than for ribosomal protein L3. [Protein fate, Protein modification and repair]
TIGR03695 (1) 2-succinyl-6-hydroxy-2,4-cyclohexadiene-1-carboxylate synthase. [Biosynthesis of cofactors, prosthetic groups, and carriers, Menaquinone and ubiquinone]
TIGR04211 (1) SH3 domain protein.
TIGR04360 (3) conjugative transfer region protein TrbK. [Mobile and extrachromosomal element functions, Plasmid functions]

#### 9.4 Multiple-sequence alignment

Each L31-Actinobacteria RNA is denoted by an abbreviation (like "Eco-1-1") that refers to a taxonomy given in Section 9.1. The alignment may include sequences containing the RNA motif, as well as flanking sequence. The L31-Actinobacteria RNA itself is denoted by the line underneath marked 5′ and 3′ on either end. Nucleotides in flanking sequences (i.e., not part of the motif) are written in gray letters. Stems of predicted rho-independent transcription terminators, if any, are shaded yellow. (Note: terminator predictions have not been analyzed manually, and many are likely to be false positives. Terminator predictions are those of the RNIE software.) Nucleotides predicted to function (as DNA) as transcription-factor binding sites, if any, are shaded in green. (But note: these predictions are manually annotated, so they might be under-predicted.) Annotated start codons, if any, are shaded green. (Note: start codons are frequently misannotated, especially in environmental samples.) Nucleotides proposed to basepair as part of the consensus structure are shaded in color when they comprise Watson-Crick or G-U pairs. Otherwise they are shaded gray. Conserved stems are also indicated at the bottom of the alignment by angle brackets, where matching $<$ and $>$ denote base-paired columns. Below these angle brackets, the symbol "2" denotes base pairs exhibiting covariation according to the statistically well-founded R-scape method. "1" denotes base pairs exhibiting covariation according to R2R's simplistic method. "0" denotes base pairs that are not observed to mutate and "?" denotes base pairs that have a significant frequency of non-canonical nucleotides for Watson-Crick or G-U pairs ($> 5\%$). Below these base pair annotation is the consensus sequence: "R" = "A" or "G", "Y" = "C" or "U", red nucleotides: nucleotide identity conserved more than 97% of the time, black nucleotides: 90%, gray nucleotides: 75%, red circle (•): nucleotide is present 97% of the time, black circle (•): 90%, gray circle (•): 75%, white circle (◦): 50%. All percentages of sequences just described (e.g. 97% conserved) assume that sequences have been weighted by the GSC algorithm implemented by the Infernal software package.

Duplicate sequences: the following putative homologs are not shown in the alignment because their sequences are identical to a homolog already shown: Aac-1-1, Acy-2-1, Ade-2-1, Age-1-1, Ama-1-1, Ama-2-1, Ame-1-1, Ame-2-1, Ame-2-2, Ame-4-1, Ame-5-1, Ame-5-2, Ane-1-1, Ava-1-1, Bas-2-1, Bas-3-1, Bbo-1-1, Bbo-2-2, Bco-2-1, Blo-1-1, Blo-10-1, Blo-11-1, Blo-12-1, Blo-13-1, Blo-14-1, Blo-15-1, Blo-16-1, Blo-17-1, Blo-18-1, Blo-2-1, Blo-20-1, Blo-21-1, Blo-22-2, Blo-22-3, Blo-22-4, Blo-22-5, Blo-22-6, Blo-22-7, Blo-22-8, Blo-22-9, Blo-23-1, Blo-24-1, Blo-25-1, Blo-25-2, Blo-26-1, Blo-27-1, Blo-28-1, Blo-29-1, Blo-3-1, Blo-30-1, Blo-31-1, Blo-32-1, Blo-33-1, Blo-35-1, Blo-4-1, Blo-5-1, Blo-6-1, Blo-7-1, Blo-8-1, Blo-9-1, Bme-2-1, Bre-1-2, Bsc-1-2, Bsp-1-1, Bsu-1-1, Bsu-2-2, Bth-1-1, Bth-4-1, Bth-5-2, Bts-2-1, Bts-2-2, Fsp-10-1, Fsp-11-1, Fsp-12-1, Fsp-15-1, Fsp-4-1, Gal-2-1, Gal-3-1, Ghi-1-1, Gpo-1-1, Gpo-2-1, Gte-1-1, Gte-2-1, Gte-5-1, Kal-1-1, Kau-1-1, Mab-1-1, Mab-1-10, Mab-1-100, Mab-1-101, Mab-1-102, Mab-1-103, Mab-1-104, Mab-1-105, Mab-1-106, Mab-1-107, Mab-1-108, Mab-1-109, Mab-1-11, Mab-1-110, Mab-1-111, Mab-1-112, Mab-1-113, Mab-1-115, Mab-1-116, Mab-1-117, Mab-1-118, Mab-1-119, Mab-1-12, Mab-1-120, Mab-1-121, Mab-1-122, Mab-1-123, Mab-1-124, Mab-1-125, Mab-1-126, Mab-1-127, Mab-1-128, Mab-1-129, Mab-1-13, Mab-1-130, Mab-1-131, Mab-1-132, Mab-1-133, Mab-1-134, Mab-1-135, Mab-1-136, Mab-1-137, Mab-1-138, Mab-1-139, Mab-1-14, Mab-1-140, Mab-1-141, Mab-1-142, Mab-1-143, Mab-1-144, Mab-1-145, Mab-1-146, Mab-1-147, Mab-1-148, Mab-1-149, Mab-1-15, Mab-1-150, Mab-1-151, Mab-1-152, Mab-1-153, Mab-1-154, Mab-1-155, Mab-1-156, Mab-1-157, Mab-1-158, Mab-1-159, Mab-1-16, Mab-1-160, Mab-1-161, Mab-1-162, Mab-1-163, Mab-1-164, Mab-1-165, Mab-1-166, Mab-1-17, Mab-1-18, Mab-1-19, Mab-1-2, Mab-1-20, Mab-1-21, Mab-1-22, Mab-1-23, Mab-1-24, Mab-1-25, Mab-1-26, Mab-1-27, Mab-1-28, Mab-1-29, Mab-1-3, Mab-1-30, Mab-1-31, Mab-1-32, Mab-1-33, Mab-1-34, Mab-1-35, Mab-1-36, Mab-1-37, Mab-1-38, Mab-1-39, Mab-1-4, Mab-1-40, Mab-1-41, Mab-1-42, Mab-1-43, Mab-1-44, Mab-1-45, Mab-1-46, Mab-1-47, Mab-1-48, Mab-1-49, Mab-1-5, Mab-1-50, Mab-1-51, Mab-1-52, Mab-1-53, Mab-1-54, Mab-1-55, Mab-1-56, Mab-1-57, Mab-1-58, Mab-1-59, Mab-1-6, Mab-1-60, Mab-1-61, Mab-1-62, Mab-1-63, Mab-1-64, Mab-1-65, Mab-1-66, Mab-1-67, Mab-1-68, Mab-1-69, Mab-1-7, Mab-1-70, Mab-1-71, Mab-1-72, Mab-1-73, Mab-1-74, Mab-1-75, Mab-1-76, Mab-1-77, Mab-1-78, Mab-1-79, Mab-1-8, Mab-1-80, Mab-1-81, Mab-1-82, Mab-1-83, Mab-1-84, Mab-1-85, Mab-1-86, Mab-1-87, Mab-1-88, Mab-1-89, Mab-1-9, Mab-1-90, Mab-1-91, Mab-1-92, Mab-1-93, Mab-1-94, Mab-1-95, Mab-1-96, Mab-1-97, Mab-1-98, Mab-1-99, Mab-10-1, Mab-11-1, Mab-12-1, Mab-13-1, Mab-14-1, Mab-15-1, Mab-16-1, Mab-17-1, Mab-18-1, Mab-19-1, Mab-2-1, Mab-20-1, Mab-21-1, Mab-22-1, Mab-23-1, Mab-24-1, Mab-25-1, Mab-26-1, Mab-27-1, Mab-28-1, Mab-29-1, Mab-3-1, Mab-30-1, Mab-31-1, Mab-32-1, Mab-33-1, Mab-34-1, Mab-35-1, Mab-36-1, Mab-37-1, Mab-38-1, Mab-39-1, Mab-4-1, Mab-40-1, Mab-41-1, Mab-42-1, Mab-43-1, Mab-44-1, Mab-45-1, Mab-46-1, Mab-47-1, Mab-48-1, Mab-49-1, Mab-5-1, Mab-50-1, Mab-51-1, Mab-52-1, Mab-53-1, Mab-54-1, Mab-56-1, Mab-57-1, Mab-58-1, Mab-59-1, Mab-6-1, Mab-60-1, Mab-61-1, Mab-62-1, Mab-63-1, Mab-64-1, Mab-66-1, Mab-67-1, Mab-68-1, Mab-69-1, Mab-7-1, Mab-70-1, Mab-70-2, Mab-71-1, Mab-71-2, Mab-71-3, Mab-72-1, Mab-74-1, Mab-75-1, Mab-8-1, Mab-9-1, Mar-2-1, Mau-3-1, Mch-2-1, Mch-3-1, Mch-4-1, Mco-1-1, Mfa-1-1, Mfo-2-1, Mgi-2-1, Mgl-1-1, Mka-1-1, Mne-2-1, Mse-1-2, Msp-24-1, Msp-25-1, Msp-30-1, Msp-5-1, Msp-6-1, Nas-1-1, Nbr-2-1, Nbr-3-1, Nca-1-1, Nrh-1-2, Pav-1-1, Pav-2-1, Pru-1-1, Psp-2-1, Psp-3-1, Psp-4-1, Psp-5-1, Psp-7-1, Rae-1-1, Req-1-1, Req-2-1, Rer-1-1, Rer-2-1, Rer-3-1, Rer-4-1, Rer-5-1, Rer-6-1, Rer-7-1, Rfa-1-1, Rfa-10-1, Rfa-11-1, Rfa-13-1, Rfa-14-1, Rfa-15-1, Rfa-16-1, Rfa-17-1, Rfa-18-1, Rfa-19-1, Rfa-2-1, Rfa-5-1, Rfa-6-1, Rfa-9-1, Rim-1-1, Rjo-1-1, Rop-2-1, Rop-3-1, Rop-3-2, Rop-4-1, Rpy-1-1, Rpy-2-1, Rqi-1-1, Rqi-2-1, Rrh-5-1, Rrh-6-1, Rru-3-2, Rsp-1-1, Rsp-10-1, Rsp-15-1, Rsp-17-1, Rsp-18-1, Rsp-22-1, Rsp-4-1, Rsp-6-1, Rsp-7-1, Rsp-8-1, Rwr-1-1, Rwr-2-1, Sal-1-1, Sal-3-1, Sal-6-1, Sal-8-3, Sal-8-4, San-1-1, Sar-1-1, Sar-10-1, Sar-12-1, Sar-13-1, Sar-14-1, Sar-15-1, Sar-16-1, Sar-18-1, Sar-19-1, Sar-2-1, Sar-20-1, Sar-23-1, Sar-25-1, Sar-28-1, Sar-29-1, Sar-3-1, Sar-30-1, Sar-31-1, Sar-32-1, Sar-34-1, Sar-35-1, Sar-36-1, Sar-37-1, Sar-38-1, Sar-39-1, Sar-4-1, Sar-41-1, Sar-42-1, Sar-5-1, Sar-6-1, Sar-7-1, Sar-8-1, Sar-9-1, Sau-1-2, Sav-1-1, Sav-2-2, Saz-2-1, Sba-2-1, Sbi-2-2, Sbi-3-2, Sbi-3-3, Sbo-1-2, Sbr-1-1, Sca-2-2, Sca-4-1, Sca-4-2, Sca-5-1, Sca-5-2, Scl-1-2, Sco-2-1, Ser-1-1, Sex-1-1, Sex-1-2, Sfl-2-1, Sfl-3-1, Sfl-6-1, Sfl-7-1, Sfl-9-1, Sfu-3-1, Sga-1-1, Sgl-2-1, Sgr-12-3, Sgr-12-4, Sgr-12-5, Sgr-12-6, Sgr-12-8, Sgr-12-9, Sgr-13-1, Sgr-13-2, Sgr-13-3, Sgr-13-4, Sgr-13-5, Sgr-4-1, Sgr-5-1, Sgr-8-1, Shy-1-1, Shy-3-1, Sla-2-3, Sla-2-4, Sli-1-1, Sli-2-1, Sli-2-2, Sme-1-1, Sol-2-2, Spa-1-1, Spa-10-1, Spa-11-1, Spa-12-1, Spa-13-1, Spa-14-1, Spa-15-1, Spa-16-1, Spa-17-1, Spa-18-1, Spa-19-1, Spa-2-1, Spa-20-1, Spa-21-1, Spa-22-1, Spa-23-1, Spa-26-1, Spa-29-1, Spa-30-1, Spa-34-1, Spa-37-1, Spa-4-1, Spa-40-1, Spa-6-1, Spa-9-1, Spe-2-1, Spl-1-1, Spl-1-2, Spu-2-1, Spu-2-2, Spu-3-1, Spu-3-2, Sre-1-1, Sri-1-1, Sri-2-2, Sri-3-1, Sri-4-10, Sri-4-11, Sri-4-12, Sri-4-13, Sri-4-14, Sri-4-15, Sri-4-17, Sri-4-2, Sri-4-3, Sri-4-4, Sri-4-5, Sri-4-6, Sri-4-7, Sri-4-8, Sri-4-9, Sro-2-1, Sru-2-1, Sru-2-2, Ssc-3-2, Ssp-11-1, Ssp-111-1, Ssp-113-1, Ssp-114-1, Ssp-116-1, Ssp-119-1, Ssp-120-1, Ssp-128-1, Ssp-130-1, Ssp-132-1, Ssp-135-1, Ssp-139-1, Ssp-142-1, Ssp-144-1, Ssp-145-1, Ssp-15-1, Ssp-150-1, Ssp-155-1, Ssp-156-1, Ssp-157-1, Ssp-159-1, Ssp-160-2, Ssp-162-1, Ssp-163-1, Ssp-172-1, Ssp-18-1, Ssp-23-1, Ssp-27-1, Ssp-28-2, Ssp-29-1, Ssp-30-1, Ssp-37-1, Ssp-41-1, Ssp-64-1, Ssp-65-1, Ssp-76-1, Ssp-77-1, Ssp-80-1, Ssp-83-1, Ssp-84-1, Ssp-93-1, Ssp-94-1, Ssp-95-1, Ssp-98-1, Str-10-1, Str-3-1, Str-4-1, Str-5-1, Str-6-1, Str-7-1, Str-8-1, Str-9-1, Sva-2-2, Svi-10-1, Svi-11-1, Svi-13-1, Svi-14-2, Svi-14-3, Svi-6-1, Svi-8-1, Swe-1-1, Tcu-1-1, [pa-1-1, env-100, env-101, env-102, env-106, env-107, env-108, env-109, env-110, env-113, env-118, env-119, env-12, env-120, env-121, env-122, env-123, env-124, env-129, env-134, env-135, env-152, env-155, env-168, env-169, env-171, env-172, env-173, env-174, env-175, env-176, env-177, env-178, env-179, env-180, env-181, env-182, env-183, env-184, env-185, env-186, env-187, env-188, env-189, env-190, env-191, env-192, env-193, env-194, env-195, env-198, env-199, env-200, env-201, env-202, env-203, env-204, env-205, env-206, env-207, env-208, env-209, env-210, env-211, env-212, env-213, env-214, env-215, env-218, env-23, env-234, env-235, env-236, env-246, env-247, env-251, env-269, env-270, env-271, env-272, env-273, env-275, env-276, env-277, env-278, env-284, env-285, env-286, env-287, env-296, env-297, env-305, env-313, env-316, env-326, env-327, env-328, env-332, env-354, env-356, env-357, env-362, env-375, env-376, env-383, env-385, env-387, env-388, env-389, env-392, env-393, env-395, env-397, env-399, env-403, env-404, env-405, env-406, env-420, env-427, env-439, env-44, env-444, env-445, env-451, env-458, env-464, env-465, env-466, env-467, env-468, env-469, env-470, env-471, env-472, env-473, env-480, env-491, env-492, env-493, env-5, env-504, env-507, env-511, env-512, env-519, env-525, env-528, env-529, env-537, env-545, env-550, env-554, env-556, env-559, env-562, env-566, env-567, env-569, env-574, env-575, env-58, env-580, env-581, env-590, env-595, env-599, env-6, env-612, env-613, env-614, env-615, env-616, env-617, env-618, env-619, env-620, env-621, env-622, env-623, env-624, env-625, env-626, env-627, env-628, env-632, env-634, env-635, env-636, env-640, env-641, env-642, env-643, env-644, env-645, env-646, env-655, env-659, env-660, env-661, env-662, env-665, env-71, env-72, env-76, env-77, env-79, env-89, env-91, env-98

```
            alignment positions 1···79

env-1       CAUGCAGAUCCAGAAGACCACCGUUGGACCGUCCAACGGGGCGGGCAACGAGGGCUAGCCGCCCCGCUAGACUUGCCCC
env-2       CGUCCCGGUGAUCCGCGCAUGGGGCAACCCUGAGUGGGAAGAAGCACCCUCACGAUGCCGUUAGACUCGAUGCACACCA
Vca-1-1     GUAGAAGAAAUAAAAGGUCACCCCCACCACAUUUUGCCUGAGAACUGGCGUUAGUAACGUGCAAAGUGGCAUAAUCAAA
env-3       ..............CGUCAGGUCACGUCAGGUCAGGUCAGGUCACGUCUGGUCAGGUCCGGUCUGGCAGACUCGAGCGU
env-4       GAUGGGCGCCGAGGACUGAGCGUCCGACGAGGGAACAUCGCGAUUCGCGGCGGCGUUCGAGCAGGUGGCACAAUCGCCC
Cac-1-1     GGUCCGCGGUGCCGCGCGCCUGCGCCGCACCUCGGACGGGCGUGGAAUAACCACCCCCCGCGGCCCGUUAAGAUGGUUC
env-7       .........................GCCCGAACGGGCACGGAAUUGGCGCUGGGAGCACCGCCCCUGGCAGACUGGACG
env-8       GUCCAGCGACGACGACUGACCCCUGAGGAAUCCGUCCACAGGGGCCUUUUGGCUCAGCCGCCUCGUGGUGGCAUACUUA
env-9       ...............................................................................
env-10      CGUCCGUACGCGCCGGCGGGAAGGUUGGGGGCGUCGCCGACGUUUGGUCCGUACGGUGCGCCCUCUGGCACACUGUCCU
env-11      .........................................................GCUCUGGCACACUGGUCCACGA
env-13      ...UCGCCCUCGGACGGGUAGCGCGGCGCCGGAGGGUCGGACGGUUUGGUUCGGCGGGGCUGAUCGGGCACAAUAGUCC
env-14      AGGCCCGAGCCGCUGUUGAGUCGGUCGGAUCACCCCGCUGCCGCGGUAGCUCCACGACAUGGCAGACUGGGUGACUGGC
Apo-1-1     CGUGGGUCGUGUCGGCCAGUUCACUGCGCCGGGGGCGUACCCGCAUGGACCCGGGGCCGGCGGGUGGCACAAUGCCA
env-15      CAUCGGGAUCUUCGUCUAGCAGCGCCCGGGAGCCGCCCAGGGAAUUGGCCGGCGACCCGGCUUGUUCGUACCAUUGCCA
env-16      CGGAGCGGGUGUCGGACAGGUCACUGCGCCCAGGGCGAAGUCGCAUGGACCGGGGGCCCGUCGAGGUGGCACAAUGCCA
env-17      .............................................................................GC
env-18      ........................................................................GACUGUC
Ama-3-1     CCUGAUCGGACCCGCGCCGGGCCCGGGAACACCCGGUCGGCGAUUUGUGUUGACCCCGUCGACACUGGCACCAUCACUC
Nin-1-1     GGUGCGCGCCCCACGCCGAGGUCGCGCACACCGGCCGGGAAUUGGUGUGCCCCCGAGGGUGUUGACUACUCUUGGUUCG
env-19      GCGUGCGCGUUUAGGCUACUGGCGGCUGACCUGGCAUAAUCGCACCAGGCAUCCGGCCCCAGGGCCGGAACGUCANGAC
env-20      GCGUGCGCGUUUAGGCUACUGGCGGCUGACCUGGCAUAAUCGCACCAGGCAUCCGGCCCCAGGGCCGGAACGUCAUGAC
Jga-1-1     UCCGCUGAUCGUGUUCCCUGGCGGUCGCUGAGCGUCCGCCAGGGAACACGAUCACGGUUCCGGGCGUUUAUACUCACCU
Hcr-1-1     GCAUCGUUCAGGCACGACCCCGGGAAUGCGCGAUGGCCUGGGACGGUUCUCCGAUCUGGCACACUGGUAGCCGAACCGU
env-21      CUCCUCGACCAGCGGCGGGUCUCGGGACCAGGGAAUCUGCGGAUUACGACGCGCGUUGCACCGAGUGGCACAAUCGCCC
Asu-2-1     GGCUAGCAUGUCACCGGUUUCACCCAGCGGUGACGCGCCUGAUAUAGACCGUAAAGGGCGAAGCAUGGCAGAAUAGACG
Psp-8-1     UUCACGGGGUGGGAAUGCCUGGUGGGGGUGGUGCGUUGAAGUGCGUGUCGGGCCGUCUGGCAGAAUUAGCCCGCUGACG
env-22      GGGCGUUUAGGCGACUGGCGGUUGACCUGGCAUAAUGCGUGUCAGUCUGCCGGCCUACGGCCGGGUGGACCGCCGACAC
env-24      ............................................................................CAC
env-25      .......................AUCAUGGAAUGCGGGGCCGCAUCCUCGCGUUGCAUGUCUGGCACACUGGACAACGG
Sfl-5-1     CGCCGUCCGAGGCAGCUCAGGAAUACUUUCGCCGCCACCCCGGUUUGGGGAGAUGCGGCCAGUCCUGGCAGACUGGUAC
Jmu-1-1     GGCGCACAGCAGCACAGCGGCGAACGCGGCGGCCACGUGGCGGUUUGGCUGCGGCGGCCCGCACCUGGAAUACUGAUCU
env-26      ..........................................................GCUCUGGCACACUGGUCCCCU
Ssp-2-1     CGGCCUUCGCGGACCCCGGGGAAUAGAUCCCCCCGCCCCCCGGUUUUGGGGAUGCGGCCAGUCCUGGCAGACUGGUAC
env-27      ..........................................AUUACCUGCAGCGUUGCAGCAGGUGGCACAAUCGCCC
env-28      GCGUCCUCAGUGGCGUCACGGCGCUGAUGGACGCGUCGGGACGGUUUGGUAACUGGCGAGUGAUCAGGCACAAUGGUGG
Cbo-1-1     GCGCGCCCGGGCAGCACGAGGACUGAUCCGGACGGUCGGUCCGGUUAUGGCCCGGACCGACCGUCUGGCAGACUGUCCC
aLL-1-1     CCCGCGGGGCUCCGCCCCGGCCAAGGAAUAGACGGACCCUCGAUUUCCUUGUACCGAGUGCCGAAUGGCACAAUCAAUC
Msp-29-1    AACAGACCGGCACCUGCGAGCGUUGAGCAGGCGUCAAGGUCGAUUCGCCCCCAUGCGGUAGGCGAUGGCACACUUGGAC
Mch-5-1     GUCGGCGCGCAGGAAUAGGGGCCGCCGGGUGCGUGUUUAGGCUGUUGGCGGUUGACCUGGCAUAAUGGGCCGCCGACAC
Aha-2-1     GAUGAGCACGUCGGCCGAUUAGUAGCGCCUGCAGUGAACACUCUCACGUAUGAAUACGUGGAAGAGUAUGGCAUAAUCU
env-29      AGUAGUGAACCGGGCCGGUCCAGGGGGAGGUCCGGCCGUCGAAGUUUGGAUCAGGGGUGCCGAUCGGGCACAAUAGGUC
Sni-1-1     UCGAUGGCUGGCGGUGUUCCACCACAGCCAUCGAUUAACAGCUCAGUGGUCAAAACUGGCAUACUGAGCCGUCGACAAG
env-30      CUGUUAACCCGGUAGGCGGUCGGAGGCCAUGAUUUCGGGACAUCGUGCCCCGCGUGGCAUACUGGAUCGUCGAAUCCAG
Asp-2-1     GUUAUGGCUUGCGAAACCAGUUGCACCUCCUGAUUUCCGGCCGGUACGGGAGGCAUGGCAGACUGGUACAUCGGCCAGU
env-31      CGUUAUCGACCACGGACCGUCGAACGCCUCCUGAUUUCUGAUCAGGUCUUGUGACAUGGCAGACUGGUCCAUCGGCCAG
Hal-1-1     ACGAUCACCGGUUACACUAGUUAGUUGGCGUCAAGAAAGGAUUGACGACCACUGUCCAAGCGCUUCCUCGGAAGCACGA
env-32      GGGCGGCCACUGGCGGGAACACGCAGCGGCCAGCCGGGCGCUGAGCUAGUCGUGCGCGCAUCUGGCACACUUAACCAAC
Ssp-169-1   CCUUCCGGCGGAGCGCGGGGAAUACCACGACCCGCACCCCGGUUUGGGGAGGUGGAGCCAGAAGUGGCAGACUGGUCC
Dsp-2-1     UGGCUGGCGCCGGACAUCGCUCCCGACGGCACGGAUCACCCCGGUUUGGUUACCGCGGUGUGAUCGGGCAAAAUGGUGG
Mel-1-1     UCCGUACCCUGCGGAAUAACGUCGCCGGGGCAGGUGUUUAGGCUGCUGGCGGUCCAACUGGCAUAAUGGACCGUCGACC
env-33      .........................................................GCCGUCCUCUGGCACACUGUUA
Car-1-1     GUCCCUGACCUGUGCGGGAAUGCCGGACCGGUGCGGGCGCGUUUUUCGAUGCGGCAAUACGUCUGACACACUGGACCUC
env-34      UCGGCUUGCGGGCUGGCGCCAGUCCGUUCGCAUCUGAGCCGGGAAUUGGCAACGCGGGGCUUGUUCUGCAUAAUUGAGU
Asp-3-1     CCGGAAACAGAUGACGGCCCUCCCGCGUUGUGGAGGGUGGCGGCGACGCGGUGGGCAUCUGGCACACUGGUUCGCCGAA
env-35      CGGUGCCGGCCCGAGCGGCCGGGACCACCUGGUCGCCACAGCCGGAUGGGACUACUCGGUCUGGCACACUGGUUGACGU
env-36      CGCCGGCGACCGUCAGAGAUCACCGGCAGUCGCCGAGCCAGGCCGCCGGGACUACACCGUCUGGCACACUGGUUGACGU
Mth-1-1     UCCCAGCGGGGAAUAGUCGCCGGUCAUGUGCUGUUUUUCAGGGUCGGCGGUCGAACUGGCAUAAUGGACCGUCGGACAC
env-37      GGUGGACAGGACUCGAUCGACGGUGGGAGGCGGAAUGCCCGGAUUACCUGCGGCGUUGGAGCAGGUGGCACAAUCGCCC
env-38      GGUGGACAGGACUCGAUCGACGGUGGGAGGCGGAAUGCCCGGAUUACCUGCGGCGUUGGAGCAGGUGGCACAAUCGCCC
env-39      GGUGCCGCCGCCUGACCCGAUCGGCCACCGUCGGCAGGCGCUGUUUUCGUGGGGUCGCCGCGGUCUGGCACACUGGUGA
env-40      .............CACCCACCGGCGUCCGCGAGGCACCCCGGAGUGGCGCUGGGAGCACCGCCCCUGGCAGACUGGACG
env-41      ...........................................................UGCUCUGGCAGACUCGAGCG
env-42      ...........................................................UGCUCUGGCAGACUCGAGCG
Ssp-96-1    GAAUAGACGGGGACCCCUUCCCGUUGAAGCAGGCGUCCUCAGAAUUUUGACAGGCGGCCCGGUCCUGGCAGACUGGUCU
Sya-1-1     GAAUAGACCGGGACCCCUUCCCGUUGAAGCAGGCGUCCUCAGAAUUUUGACAGGCGGCCCGGUCCUGGCAGACUGGUCU
Ssp-87-1    GAAUAGAUGCCGACCCCCCUCCCGUUGAGGGAGAUGUCCUCAGAAUUUUGACAGGCAGCCCGGUCCUGGCAGACUGGUCU
Ssp-122-1   GAAUAGAAGCGGACCCCUUCCCGUUGAGGGAGACGUCCUCAGAAUUUUGACAGGCAGCCCGGUCCUGGCAGACUGGUCU

            ...............................................................................
            ...............................................................................
```

```
Ssp-97-1    GAAUAGACCGGGACCCCCUCCCGUUGAGGGUGAUGUCCUCAGAAUUUUGACAGGCAGCCCGGUCCUGGCAGACUGGUCU
Ssp-148-1   AGGAAUAGACCCGGCCCCGUCCCUGUUGAGCAGCACGUCCCGAAUUUUGUAAGGCGGCCCGGUCCUGGCAGACUGGUAC
Ssp-88-1    GAAUAGAUGCCGGACCCCCCUCGUUGAGGGAGGCGUCCUCAGAAUUUUGACAGGCAGCCCGGUCCUGGCAGACUGGUCU
Ssp-164-1   GAAUAGACCCGGCCCCCCUCCCGUUGAGGGUGGUGUCCUCAGAAUUUUGACAGGCGGCCCGGUCCUGGCAGACUGGUCU
Svi-13-2    GAAUAGAUGGCGACCCCUCCCCGUUGAGGAAGGCGUCCUCAGAAUUUUGACAGGCAGCCCGGUCCUGGCAGACUGGUCU
Ssp-28-1    GAAUAGAUGCCGUCCCCGUCCCGUUGAGGGAGGCGUCCUCAGAAUUUUGACAGGCGGCCCGGUCCUGGCAGACUGGUCU
Ssp-8-1     GAAUAGAUGCCGGCCCCCUCCUGUUGAAGGCAGCGUUCCCGGAAUUUUGACAGGCGGCCCGGUCCUGGCAGACUGGUCU
Ssp-143-1   GAAUAGAUGGCGACCCCCUGCCGUUGAGGGAGGCGUCCUCAGAAUUUUGACAGGCAGCCCGGUCCUGGCAGACUGGUCU
Sla-2-1     GAAUAGAUGCCCGCUCCUCCCCGUUGAGGGGAGCGUCCUCAGAAUUUUGACAGGCGGCCCGGUCCUGGCAGACUGGUCU
Ska-1-1     GAAUAGAUGCCGGCCCCACCCCGUUGAGGGGAGCGUUCCGAGAAUUUUGACAGGCGGCCCGGUCCUGGCAGACUGGUCU
Sro-6-1     GAAUAGAUGGCGGACCCCCCUCGUUGAGGGAGGCGUCCUCAGAAUUUUGACAGGCAGCCCGGUCCUGGCAGACUGGUCU
Ssp-133-1   GAAUAGAUGUCGGACCCCCCUCGUUGAGGGAGGCGUCCUCAGAAUUUUGACAGGCAGCCCGGUCCUGGCAGACUGGUCU
Sxa-1-1     GAAUAGAUGCCGGCCCCGUCCCGUUGAGGGAGGCGUCCUCAGAAUUUUGACAGGCAGCCCGGUCCUGGCAGACUGGUCU
Ska-1-2     GAAUAGAUGCCGCCCCCGCCCCGUUGAGGGAGGCGUCCUCAGAAUUUUGACAGGCAGCCCGGUCCUGGCAGACUGGUCU
Ssp-138-1   GAAUAGACCGGGACCCCUUCCCGUUGAAGCAGGCGUCCUCAGAAUUUUGACAGGCGGCCCGGUCCUGGCAGACUGGUCU
Ser-3-1     GAAUAGACCGGCGCCCCCUCCCGUUGAGGGUGAUGUCCUCAGAAUUUUGACAGGCAGCCCGGUCCUGGCAGACUGGUCU
Sla-2-2     GAAUAGACGCGGCCCCCCUCCCGUUGAGGGAGACGUCCUCAGAAUUUUGACAGGCAGCCCGGUCCUGGCAGACUGGUCU
Ssp-131-1   GAAUAGAUGGGGACCCCUUGCCGUUGAGUCAGGCGUCCUCAGAAUUUUGACAGGCAGCCCGGUCCUGGCAGACUGGUCU
Ssp-86-1    GAAUAGAUGGCGGCCCCCCUCCGUUGAGGGAGGCGUCCUCAGAAUUUUGACAGGCAGCCCGGUCCUGGCAGACUGGUCU
Sla-1-1     GAAUAGAUGGCGACCCCCUGCCGUUGAGGGAGGCGUCCUCAGAAUUUUGACAGGCGGCCCGGUCCUGGCAGACUGGUCU
Ssp-166-1   GAAUAGAUGCCGGCCCCACCCCGUUGAGGGGAGCGUUCCGAGAAUUUUGACAGGCGGCCCGGUCCUGGCAGACUGGUCU
Svi-13-3    GAAUAGACGGGGACCCCUUGCCGUUGAGGGAGGCGUCCUCAGAAUUUUGACAGGCAGCCCGGUCCUGGCAGACUGGUCU
Dsp-1-1     CGCCGCUCCGAGCAGGGUCUGUAAGCAGAGUUACAGGCAGCGCACUGGAGCUCGCACGCCCAUUCGUGGGAAACUAAAC
Dho-1-1     CACACUCCGAGCCGCUGCCUGUAAGCAGAGUUACAGGCAGCGCACUGGAGCCAGCAUGCCCAUUCGUGGGAAACUGGAC
Bpa-1-1     CGCCGUCGGCUGCUGCGAUGUUGCGCACGCCGCACCGCGAGCCCUCGCCCCGCCCCGCCCCGGCGUGGCAGACUGUGAC
Bph-1-1     CCGCCUGAGCGGGGAAGGAUCUUCGUCACCCCCGUGCCGUCGGCGUUCCCGCGGCGGCCGCGCUCGUGGCACACUGAAC
Bfa-1-1     UCGGCCUGAGCGGCGCGAUGUUGCGCACGCCGCAUCCCGAUCCCUCGCCCGCAGGGACCGGGGCGUGGCAGACUAUGAC
env-43      ACGGCGCAGGCCCUGCGAUGUUGCGCACGCCGCAUCCGCGGCCCUCGCCCCGCAGGGCUCCGCCGUGGCAGACUGUGAC
env-45      .............................................................GGCGUGGCAGACUGUGAC
env-46      CCCCCGGCUGCUGCGANGUUGCGCACGCCGCACCGCGAGCCCUCGCCCCGCCCCGCCCCGGCGUGGGCAGGACUGUGAC
Bmu-1-1     ACGGAGGUGGGCCACGAGUAUGGCGAACGCCGCACUGCGCUGCCCUGUCCCUUCGGCCCACGACGUGGCACACUGGUGC
Bsq-1-1     CCCUGACGGGUGACCAAGCAGACACCGCGCCGGGCCCGUUCCCCCUUCCCACCUGCGUCGCCGCAUGGCACAAUCGUCC
Sba-1-1     CUCCGGAGGGGCCGUGCGGGAAUACCGGGCGCCGUCACCCCGUUUUGUUCAGGUGCGGCCAGUAGUGGCAGACUGGUAC
env-47      UAGCUCCGCCACCGCCUGACCCGGGGGAACAGACGACCCGGGAUUGGCGUUGAGGAGCUGUCAGAUGGCAGAAUCACCC
env-48      CGGGAACACCACGAUUGGCCCCGGCGUUGGGGGUGGUGGCAGACUUGUCUGUCGGCCCGCGCCCCGUCGGGGUGGCCA
env-49      ................................................................CUCUGGCAGACUGAC
env-50      ....................CCCGGUGGGGUCCGUGGCGCCGGAUUGUGACGAAGGUGGGACCGGUGGCAUGAUAGUCC
env-51      ................................CGUGGCGCCGGAUUGUGACGAAGGUGGGACCGGUGGCAUGAUAGUCC
env-52      ..........................................................................CUGAC
env-53      GCCGACUGCCAUCGGCCAGCGGGAAUGCAGUCUUGUCACAUGUGGUUUACGCGUGUGGCAAACUGAAAGCCGCUUAGUU
env-54      UGCCCGGCCCGCUGCCGGAUUCCGGGGAAUUGACAGUGCGAUGCGCGUGUUCCAUAUCUGGCACACUGGGUAACCCAAU
env-55      .............................................................GCACACUGACAGGUCGAG
env-56      ........................................CGUGGUUUAGGCGUGUGGCAAACUUAAAGCCGCUUAGUU
env-57      ...............................................GCGCGAGGUCCGGGAUGUUGGCACACUAGGCG
env-59      .....................................CUCGCAAUUGGCGCGAGGUCCGGGAUGUUGGCACACUAGGCG
Spo-1-1     CCCGAGACCCCCGAAAUCGCCCGGAAUGUUCGGAGGCGCACCUGCGUUUGGGCAGGUGCGCCAGGACCUGGCAGACUGA
env-60      GGUCCAGCAGACCGGCCUGAAGUAGCCGGGGAACACCGCGAUUGGCUCCGGCGUUACGACGAGUGGCAAACUCUGCCUG
env-61      GGUGCGGCACGCGCGCGGGAACAUCGCGAGCCCGCGGCGUGUUUUGGUGUGCAUGUCAGGGCUCUGGCACACUGAACGG
Ssp-51-1    CCCGCGGCCGCCCCGGGAGGGAAUACCGGCGCCCGCCCCCCGGUUUUGGGGGGUACGGCCAGAAGUGGCAGACUGGUAC
Sji-1-1     GCGCCCACCCGCACAGGGUGGAAUACCUGCGAGGGCAAUGCCGUUUUGGAGCACGGCGACAGUCCUGGCAGACUGGUAC
Aut-1-1     GUUAACCAGCAUGAAUCCAGUUGCGCCUCCUGAUUUCCGGCCGGUACGGGCGGCAUGGCAGACUGGUACAUCGGCCAAG
Mtu-1-1     CCAGGCAGUGCAGGAAUAGCCAGCGCGUUGUGGGCGUUUAGGCGGUUGGCGGUUGACCUGGCAUAAUGGCGCCGACACC
Cko-1-1     GCUGAUCGAGGGGAAACACGGGUAGCAGGGUCGCAUGGAAUAACUUGCGAGUCGAUCCGUUAUAGUUACCGAUCGGCCA
Nla-1-1     GCGACCGGCGGAAUACCCGCCGCCGGUCGCCGGUUGGGAAUGCAGGCGCCGGUCAUGGCACACUUGACCGGCCAUACAC
Nsp-9-1     CCGCGCCCGCUCCGUAGGGGUGCCGGCGCUGGGAAUGUUGUCGAUUACCCAAGCGUUUGGACCCAUGACAGAAUCAUCU
env-62      GGCGACUGAGCGUCGUACGGCGCUCGCACCGGGAAUCCGCAGAUUGGUUCCGGUGUUGCAGCAGAUGGCACAAUCGCCC
Afr-1-1     GUUAUCGACCAAGGACACAGGAACGCCUCCUGAUUUCUGAUCAGGGCAUGUGACAUGGCAGACUGGUCCAUCGGCCAGC
env-63      ..........................UCCUGAUUUCGGAUCAGGGCAGGUGUCGUGGCAGACUGGUCCAUCGGCCAGCA
env-64      ..........................CGCCGAUUUCGCACCGGAGUUACCGGCAUGGCAGACUGGUCCAUCGGCCAGUA
Asu-3-1     UUAUCGACCUUGGAUCAGGAAAGUGGCCUCGAUUUCUUGUCGGGCCGUGUCACAUGGCAGACUGGUCCAUCGGCCAGUA
env-65      .............................................GGCUCCUGCGAGGGAGUCGGUGGCACACUUGGAC
env-66      ......................................................................GACUGGGCU
env-67      ........................GGUCCUGCGGGGCGCGCGGAUUAGCGACGCAGGAACUCCCGCUGGCACACUGAUC
env-68      .......................GGGUCCUGCGGGGCGCGCGGAUUAGCGACGCAGGAACUCCCGCUGGCACACUGAUC
env-69      .........................GUCCUGCGGGGCGCGCGGAUUAGCGACGCAGGAACUCCCGCUGGCACACUGAUC
env-70      ......................................UGAGGAUUCGCGACGCAGCAAGUCCCGCUGGCACACUUCCU
env-73      .....................................GUGAGGAUUCGCGACGCAGCAAGUCCCGCUGGCACACUUCCU
env-74      CACCGCCAGGCGCACCUGUUGUGUCAGCACAGGGUGCGGAGCGAUUCGCAACGCAGCGAGUCCCGCUGGCACACUGCCU
env-75      ...............................................................CGCUGGCACACUGCCU
env-78      ...........................CGUCGAUGCAGCGGCCGUUCUCGGCCGCCGCGCCAGGGCUGGCACACUGCCU
env-80      ...............................GGCGGAAUUGGCGACGCAGCAAGCCCCGCUGGCACACUGCCUUCAGGU
Ncy-1-1     GGGGCGUUCGUACACGUGUCGGUGCAUCUGGCAUACUGGAUCGCCGUGCGUCCGCGCUGUUCGCGGACAAUCCCGCCUA
Ncy-2-1     GGGCGUUCGAUAGACGUGUCGGUGAAUCUGGCAUACUGGAUCGCCGUGCGUCCGCGCUGCUCGCGGACAAUCCCGCCUA
Npn-1-1     GGCGUUUUGGGAGCGUAACGGCAGUUCUGGCAUACUGAACGGUCGUUGCGUCUGCUCAUGUGAGCAGACCUCCCGCCUA
env-81      GGCCUGGCGGCCUGAGCCGCGAGUCGGGAACAAGGCUGCCAGCAGUUGGCGUUGAGCCCAGUGUCGAUGGGACACUGGU
env-82      .............................................................................GU
env-83      ......................................................................GACACUUGU
env-84      AGGCCCUUGCGGCCUGAGGCGAGGCGGGGAACAGGCCGCCAGCAGUUGGCGUUGAGCCUGGUGUCGAUGGGACACUGGU
env-85      ........................................................................CACUUGU
env-86      GAAGGCUCUGGCUGCCUGAGGCGUCGGGGACAAGGCUGCCAGCAGUUGGCGUUGCGCCCAGUGUCGAUGGGACAAUGGU
env-87      ..................................................................GGCAGACUGGACG
env-88      CCCGGGACCGGCGUUCUGAGGAGGUCCGGCGUACGGGGGCGGAUUUCCCCGGUCGGACCACCCAGUGGCACAAUCGUUG
Sre-2-1     CGGAGCUGCCUGGAAUAGCCGGGCGCGUUCCCCGCGUUGAGGUCGAGCAGCCCUGGUCUGGCAAAAUGCCAGAUUGGGU
Acy-1-1     UCGGAAACAGACGGCGACCCCCACGCGUUGUGGAGGGUGGCGGCGACGCGGUGGCCAUCUGGCACACUGGUUCGCCGAA

            ...............................................................................
            ...............................................................................
            ...............................................................................
```

```
Mrh-2-1     UCGCGAUGAGCAGGAAUAGUCGGGCGCUGGCCUGCGUUUAGGCGGUUGGCGGUUGACCUGGCAUAAUGGCGCCGACACC
env-90      CAGGAGUCGGUGGAGGCGAGCGUGGGCGAGCAGCCCCGGAAUUAGCACCGGGGUGCCGGUGUUGGGCACAAUGUCCUGG
env-92      ......................GUGGGCGAGCAGCCCCGGAAUUAGCACCGGGGUGCCGGUGUUGGGCACAAUGUCCUGG
env-93      CAGGAGUCGGUGGAGGCCGGCGUGGGCGAGCAGCCCCGGAAUUAGCACCGGGGUGCCGGUGUUGGGCACAAUGUCCUGG
env-94      CAGGAGUCGGUGGAUGCCGGCGUGGGUGAGCAGCCCCGGAAUUAGCACCGGGGUGCCGGUGUUGGGCACAAUGUCCUGG
env-95      GGCCAGCAAGCGGCGGACGCGACGGGGAGCGGGCCCGGAAUUAGCGGUCGGGUGCCGGUGUUGGGCAUGAUGUCCUGG
env-96      GGCCAGCAGGCGGCGGACGCGACGGGGAGCGGGCCCGGAAUUAGCGGUCGGGUGCCGGUGUUGGGCAUGAUGUCCUGG
env-97      GAUGACGUUGCGGAGCGAGGUGUGCGGUCUCACCGACGGCGUCGUUGUGAGUCGUCGAUUCGCCGUGGCAUAAUCGCCG
env-99      .............................CUCACCGACGGCGUCGUUGUGAGUCGUUGAUCCGCCGUGGCAUAAUCGCCG
env-103     NNNNNNNNNNCUGAGCGAGGUGUGCGGUCUCACCGACGGUGUCGUUGUGAGUCGUCGAUUCGCCGUGGCAUAAUCGCCG
env-104     AAUGACGUUCCUGAGCGAGGUGUGCGGUCUCACCGACGGUGUCGUUGUGAGUCGUUGAUCCGCCGUGGCAUAAUCGCCG
env-105     GAUGGCGUUGCGGAGCGAGGUGUGCGGUCUCACCGACGGCGUCGUUGUGAGUCGUCGAUUCGCCGUGGCAUAAUCGCCG
env-111     GAUGGCGUUGCGGAGCGAGGUGUGCGGUCUCACCGACGGCGUCGUUGUGAGUCGUUGAUUCGCCGUGGCAUAAUCGCCG
Bin-1-1     GGAGUACUGGAAUCUCGGGGAAACCGUUCGGCGUGUGGCGCAAUAGGUUCGGUUCUCCUGAAGGUGCGCUACAAUUGUC
env-112     GGGGUACUGGAAUCUCGGGGAAACCGUUCGGCGUGUGGCGCAAUAGGUUCGGUUCUCCUGAAGGUGCGCUACAAUUGUC
Bco-1-1     GGGGUACUGGAAUCUCGGGGAAGCCGUUCGGCGUGUGGCGCAAUAGGUUCGGUUCUCCUGAAGGUGCGCUACAAUUGUC
env-114     ......CCACGGUCAGCACCUAGGGAAUCUAUGGGGGCCACCGCACGUUGUUGGCUCUGGCACACUACGUAAGGCUGAA
env-115     AGCGGACCACGGUCAGUACCUAGGGAAUCUAUGGGGGCCACCGCACGUUGUUGGCUCUGGCACACUACGUAAGGCUGAA
env-116     GCAGAUCCAGAAGACCACGGUGGCCGAGGCCGAGCACAACAACGGCUCCGCCUGACGUUCUGGCAGAAUGGACCGACAC
Sro-4-1     GGUGCCCGGACGGGCGGCGGGAAUACCAGCGCCCCGCCCCCGGUUUUGGGAGAUACGGCCGGUCCUGGCAGACUGGUAC
Sgl-4-1     GGUGCCCGGCCGGGGGCCGGGAAUACCAGCGCUCCGCCCCCGGUUUUGGGAGAUACGGCCGGUCCUGGCAGACUGGUAC
Sal-8-1     GGUGCACGGCCGGGUGGCGGGAAUACCCGCGCCCCGCCCCCGGUUUUGGGAGAUACGGCCGGUCCUGGCAGACUGGUAC
Sgr-12-1    GGUACACGGCCGGGUGGCGGGAAUACCCGCGCCCCGCCCCCGGUUUUGGGAGAUACGGCCGGUCCUGGCAGACUGGUAC
Ssp-55-1    GGUGCCCGGCCGGGCGGCGGGAAUACCCGCGCCCCGCCCCCGGUUUUGGGAGAUACGGCCGGUCCUGGCAGACUGGUAC
Ssp-62-1    GGUGCACGGCCGGGCGGCGGGAAUAGCUGGGCCCCGCCCCCGGUUUUGGGAGAUACGGCCGGUCCUGGCAGACUGGUAC
Ssp-167-1   GCCGAGCGCGGCACCUCCCGGAAUACCGGGGUGCCCGCCCUCGUUUUGGUGGGCACGCCCAGUGCUGGCAGACUGGACC
Sav-3-1     GGGCCCCGGCGCGGGCCGCGGAAUAACGGGGUCCGCGACCCCGUUUUGGCUGCCGCGACCAGUCCUGGCAGACUGGAGC
Ssp-4-1     CCCGGCCCCGGACAGCGGGGAAUAACGGCCCAGGGCCCCCUCGUUUUGGACGGAGCGGACAGUGCUGGCAGACUGAGCC
Sxi-2-1     CGGGGACGCCUGAGGGCGGGGAAUAACGGUCUGGGCACCCUCGUUUUGGACGGAGCGGACAGUGCUGGCAGACUGAACC
Ssp-90-1    CGGGGACGCCGGAGGGCGGGGAAUAACGGUCUGGGCACCCUCGUUUUGGACGGAGCGGACAGUGCUGGCAGACUGAACC
Jal-1-1     CAACGUUGCAAACCCCCGGCCAACGGUACGAAUCCCGGUCACGGUUUGGCCGGGGCGGCGCGGCCUGGAAUACUGAUCU
env-117     CAGCUACCGCGGGAAUAUCUGCUGCGCUGUUCGGCGUUUCGCCAAUGUCAGGGUGCCUCUGGCACACUGGUCGAUCGAG
env-125     CACGAUCUCGAAGACGACUCCCAGCCAGGACUGACGUCGUCAGGUGGACUGGUGCCGGCCGUUCGGUGUAUCGUCGAAC
Psp-1-1     CACGAUCUCCAAGACGACUCCCAGCCAGGACUGACGUCGCCAGGUGGACCGGUGCCGGCCGUUCGGUGUAUCGUCGAAC
env-126     ........................................................................CACUGAUC
Pac-1-1     CGUCGACGAGUAGGCGCGCCGGGAACACCGACGCGCCGCCCGGCGUUCAAGGCGACGAUGCGACACUGGUCGCGUCAGA
env-127     GCUGGGGGCGGACCGUGGCGUUCCUGGAGCGCGUGCUGGCCGUGGAAUAAGCACCCCCCGACUCCCGUUAAGAUGGUUC
env-128     ................................................................................
env-130     ....CGCGCCGACGACCUGCUCGCAACCGCGCCGGAAUGCCGGCGCGCACCGGCCUGUUGUCGGUCUGGCACACUGGAG
env-131     CUGAGGCGCCGACGACCUGCUCGCAACCGCGCCGGAAUGCCGGCGCGCACCGGCCUGUUGUCGGUCUGGCACACUGGAG
env-132     GCUCAAGCCUGCGGCGUUUGGUCGGCGCAAGCCCCGGAUGGCAGAAUGGGCCCUCGGCCCUGGCCCUCCGGUCGGAGCC
env-133     CCGCGGCCCGUCCUCGGUUUGGAGGGCAUCCCCUCCGGAUGGCAGAAUCGGUCCUUGCCCUGGCCCUCCGGUCGGAGCC
Bth-2-1     UUGAUUGGCUGGUGUCGGCUGCGUUGCUGCUACGCCUUGGGCUCAUUCGCGUGUCGUCCGUAAUGGCGCGUACAAUACU
Bsc-1-1     GGUUCAAGCCCCUGCGGCCCCACUUUCCGCAUGCGCGCCGGGCGCGGCGCAUGUCGUCCUUACCCCGGCGUACAAUCCC
Bth-7-1     UGUAUCAAUCUGCCGGUUCGCGGCGUCGCAUACGCUGUGGGCUCACCAGCGUGUCGUCCGCGACCGCGCGUACAAUACU
Bth-8-1     UGUAUCAAUCUGCCGGUUCGCGGCGUCGCAUACGCUGUGGGCUCACCAGCGUGUCGUCCGCGACCGCGCGUACAAUACU
Bth-5-1     CGCGUUGACGCUGGUGUGUUCCCUUCGGUAUACGCCGUGGGCUCAUCAGCGUGUCGUCCGUAAUGGCGCGUACAAUACU
Bth-3-1     UUGAUUGGCUGGUGUCGGCUGCGUUGCUGCUACGCCUUGGGCUCAUUCGCGUGUCGUCCGUAAUGGCGCGUACAAUACU
Bth-6-1     GGCCCCACUUUUCUUGUGGUUUCGGCGGUUAAGCCGACGGCAUGUUUGGCGUGUCGUCGCCUGUGGCGCGUACAAUACU
Bbo-2-1     GGCCCCACUUUUCUUGUGGUUUCGACGGUUAAGCUGACGGCAUGUUUGGCGUGUCGUCGCCUGUGGCGCGUACAAUACU
env-136     AGGACCGGCCGGACCGGGGAGGGGCCGCUCGGGAACAUCGCGGAUUACCGCGGUGUUCGGGCCCGUGACACAAUCGUGU
Nsp-4-1     GAGCCGCGCCAACGGUUCCGGUCCCGGCAGGGGAACAACGCGGAUUACCGCGGCGUUGGGACCCGUGACACAAUCGUGU
env-137     ...................................CCUUGCGGAUUAUCACCGGGACGCCGUUCCCUGGCACACUAGAUG
env-138     GCGUGCGCGUUUAGGCUACUGGCGGCUGACCUGGCAUAAUCGCACCAGGCAUCCGGCCCCAGGGCCGGAACGUCACGAC
env-139     GCGUGCGCGUUUAGGCUACUGGCGGCUGACCUGGCAUAAUCGCACCAGGCAUCCGGCCCCAGGGCCGGAACGUCAUGAC
env-140     ..............................CCUGGCAUAAUCGCACCAGGCAUCCGGCCCCAGGGCCGGAACGUCACGAC
env-141     GGCCCGUGCUCCCUUGCCAGGUCGUCGUCCAGCCGCCGGACUGACGGCCGAUGUCGUUCGCGGCCGGCCCGGAUAGCAG
env-142     ............................................................GCGGCCGGCCCGGAUAGCAG
Fsp-8-1     CCAUCGGCGAGGGAAUGUGGGAGCUACGCCAAGCGUUUUGAUCAAUGUGCCCUCAGCUGGCACACUGUCUGGUUGACGA
env-143     ..........................................................................ACCGAG
Nsp-10-1    CCCCGGAGCGCACCCCACCGUCCGGGCCCGAGGAACAACGCCGAUUACCCCGGCGUUCUGACCCAUGACACAAUCGUGU
env-144     .......................................................................CAGACUGUC
Cja-1-1     GUGGUUACACCACCCGGAACGGUCACCGUCUCCCGGUGCGCGACCCGCGCCGCGGACAUGGCAGACUGGACCAUCGGCC
Fsy-1-1     GUCCAGGGUUCCCGCGACACCCCCCGCGGGAUGCCCGCCCCGAGGGCGCGCCCGGCUCUGGCAGACUGGAGCGUCGACA
Fco-1-1     CGCCCAGGGUCCCCGCGACACCCCCGCGGGAUGCCCGCCCCGAGGGCGCGCCCGGCUCUGGCAGACUGGAGCGUCGACA
env-145     ........GGGUCGCGGCAGGCGAUACCGGCCGUUUGGGCCAGGGACAUCCGACCUGGCAUAAUCGGUUUCGAUAAACC
Ply-1-1     GACUACCGCUAAUAACGAUUAAGAAAUUCGAUUGCGCCGGUAAUUCGCGUAAGCGUAAAUACUUGCGAUAGCAUUAACC
Ply-2-1     GACUACCGCUAAUAACGAUUAAGAAAUUCGAUUGCGACGGUAAUUCGCGUAAGCGUAAAUACUUGCGAUAGCAUUAACC
Ssc-2-1     GAAUGCCACGACCCCGGUCGUGGUUGGGCGGUACAGACCCGGAUUUGGGUCCGGUCGCCCCCUCCUGGCAGACUGCUCU
Sri-4-1     CGCCCUCCGCAAACCCCCGGGAAUACUUCCCGGACCACCCCGGUUUUGGGAGAUGCGGCCCGUCCUGGCAGACUGGACC
Sri-2-1     CACCCCCCGCAAACCCCCGGGAAUACUUCCCGGACCACCCCGGUUUUGGGAGAUGCGGCCCGUCCUGGCAGACUGGACC
Sgr-6-1     ACCUCCCGGAAACCCCCGGGGAAUACUUCCCGGCCCACCCCGGUUUUGGGAGAUGCGGCCCGUCCUGGCAGACUGGACC
Sri-4-16    CACCCCCAGGAAACCCCCGGGAAUACUUCCCGGACCACCCCGGUUUUGGGAGAUGCGGCCCGUCCUGGCAGACUGGACC
Ssp-75-1    CACCCCCCGGAAACCCCCGGGAAUACUUCCCGGGCCACCCCGGUUUUGGGAGAUGCGGCCCGUCCUGGCAGACUGGACC
Sri-4-18    CGGCCUUCGUUAGGCACCGGGAAUACAUUCCGGACCACCCCGGUUUUGGGAGAUGCGGCCCGUCCUGGCAGACUGGACC
Sri-4-19    CACCCCCCGGAAACCCCCGGGAAUACUUCCCGGACCACCCCGGUUUUGGGAGAUGCGGCCCGUCCUGGCAGACUGGACC
Sri-4-20    CACCCCACGCAAACCCCCGGGAAUACUUCCCGGACCACCCCGGUUUUGGGAGAUGCGGCCCGUCCUGGCAGACUGGACC
env-146     CGCGAAGGCCCGGCGUUGGACCGUCUGCUGGCCGCCACACCGAUUCGCCACCGAGUCGGUCAGGGUGGCAGAAUCACUC
env-147     CGCGAAGGCCCGGCGUUGGACCGUCUGCUGGCCGCCACACCGAUUCGCCUCCGAGUCGGUCAGGGUGGCAGAAUCACUC
env-148     CGCGAAGGCCCGGCGUUGGACCGUCUGCUGGCCGCCACACCGAUUCGCCUCCGAGUCGGUCAGGGUGGCAGAAUCACUC
Nal-1-1     CGAGGGCCGUACGGCCCGCCGCCGGGCUCGGGAAUGCGCGGAUUCGCAUCCGCGUUGAGCCCGGUGGCACAAUUCGUGC
env-149     AGGAUGUCGGACGGGGUCGGGAAUAUCCCGGCCCCACCCCCGGUUUUGGGUGAUGCGGCCAGUGCUGGCAAACUGGAUC
Ssp-36-1    GCGGGCGACCCUCCGUACGGGAAUACGAGUGCCCCCACCCCGGUUUUGGGGGAUGCGGCCAGUGCUGGCAAACUGGUAC
Ssp-109-1   GGCGGGCGCCCCCCGGACGGGAAUACGAGCGCCCCCACCCCGGUUUUGGGGGAUGCGGCCAGUGCUGGCAAACUGGUAC

            ................................................................................
            ................................................................................
            ●●●●●●●●●●●●●●●●●●●●●●●●●●●●●●●●●●●●●●●●●●●●●●●●●●●●●●●●●●●●●●●●●●●●●●●●●●●●●●●●
```

```
Ssp-35-1   GCGGGCGACCCUCCGUACGGGAAUACGAGCGCCCCCACCCCGGUUUUGGGGGAUGCGGCCAGUGCUGGCAAACUGGUCC
Ssp-34-1   GGCGGGCGACCCUCCGUACGGGAAUACGAGCGCCCCCACCCGGUUUUGGGGGAUGCGGCCAGUGCUGGCAAACUGGUCC
Ssp-42-1   GCGGGCGACCCUCCGUACGGGAAUACGAGCGCCCCCACCCCGGUUUUGGGGGAUGCAGCCAGUGCUGGCAAACUGGUCC
env-150    GGCGAGCGGUUAAGCCAGUCGUCCGAAUGAAUGGGCGGCGCGAGGCCCCAGUGGCCGGCAUGGCACACUGGUUGCGCAG
Mau-2-1    GGGCGUUUAGGCUACUGGUGGUUGACCUGGCAUAAUGCAUGUCAGUCUGCCGGCCUCGGCCGGGUGGACCGCCCAACAC
env-151    GUCUCCGUACGGAAUACCUCCGGAGGUGACGGCGUUUGGGCAGGUGGCGGUUGACCUGGCAUAAUUGUCCGCCAGACAC
env-153    ..............................................................................CU
Psp-10-1   CCGGAAUAGCGGGCCGGCACCGGGUGUUCUACGCGGUGUCGGCCCGUUGCACGCUCAAUGGCAGAAUAUGGGCAGACAU
env-154    ....GGCCUGGUCGUCUGACCGCCCGCGCCGGGGCGGGAAUUACGCGGUCAGGACCGGCGUUGUCUAACAUUGAUCCGU
env-156    CGGGCUCGUCGUCUGACCCGAGCUGGCCUCCGGGCGGGAAUUCGCUGGCGCGGACCCGUGUUGACUAGCAUUGAUCCGU
env-157    GGGCGCCGGGAAUCGGGGGUCCCUCACCAGCGUUUAGCCAAGUGUCAAGCCGGUCUGUCACACUCAAGGGCCGGUCAUA
env-158    GGGCGCCGGGAAUCGGGGGUCCGUCAUCGGCGUUUACCACUCCGUCAAGCCGGUCUGUCACACUGAAGGGCCGGUCAUA
env-159    GGCGCCGGGAAUCGGGGGGUCCCUCACCGGCGUUUACCACUCCGUCAAGCCGGUCUGUCACACUGAAGGGCCGGUCAUA
env-160    CCCGAGCGGGAAUCGGGGGUGGGCCAGCGGCGUUUGGCUAUCUGUCAAGCCGGUCUGGCACACUGAAGGGCCGGUCAUA
Mha-1-1    GGGGGAUUCCGGCGCACGCGUACCCGUCGUGCGGCCCGCGGCGCGCCGGAAUCGCGGCCUGGCAUAAUUGAUCCACACC
env-161    CGGUCGAGGCGGGAAUGCGCACCCUCACGCAGGCGUUUAGGGACGUAGCGGCUGACCUGGCAUAAUCGACAGCCGAAGA
env-162    UGACCGCCCGCGUUGCGGGCGGGAAUGCGCACCGGUCACAAGUGGUUAACACGUGUGGCAAACUAAAAGCCGCUUAGUU
env-163    ..............................................................CCAGUGGCACACUUGCGU
env-164    .........................................................CCUGGAAGGAUGGACCCGCACCG
Asp-5-1    GGUCGGCGCCCUGCUGGGCUGCUUCGCGGCGGUCGCCGCCGGGGUCGUCGGGUGGACCUGGCACAAUGGUCGGUCGAAC
env-165    GGCGCGGAGCGCCCGAGGCGGGGAAUGUGGACAUGUCACAAGUGGUUAACGCGUGUGGCAGACUGAAAGCCGCUUAGUU
Nlu-1-1    CGCCAAGAGCUGAGGUCUGUGUGCGCCCUGGGGCCGUCGGGAAUUAUUCGCGGCCCCAGGCGUUCACAUACACUGAAUC
Nba-1-1    UGAGACGACCGGAGUGCGACGAACGUUCUGGGGCCGCCGGGAAUUAUUCGUGGCCCCAGGCGUUCACAUACACUGAAUC
Ssp-177-1  CGGAUCGCGGGGGUAUCGCGGAAUAACGGCCGCCCGCCCCCGGUUUUGGGAGGCACGGCCAGUGAUGGCAGACUGGACC
Lal-1-1    GAACCCGGGUGACCGGCCAGCCGGCCCGGUGGCCCCGAUGCGGCGGGAAUAGCCGCCCCGUGGCAGACUGUUACGUCGG
Gma-1-1    AGGUUAGGCCACCCCAAGACGCUCGCGUCGAGGAGGAUUGGCAGCCGCCCGUGACCUUGUGGAAUAAUCGUUCGGUCAG
Bme-1-1    CGUUCCCGUACCGCUGCGCGUGCCGUCCCGCGGCAUCCCUUCCACGGGCUGCGGUGUCCCAGCCAUCGUCUACCCUUGC
Bsu-2-1    CUCGACGAGGCCAUGCGGAACUACGCCAAGGGCGUCAUGCCGCAGCUAUGCAAGCGUCCGCACAGGUGGCUAUAAUCUG
Bre-1-1    CUGCUCUUUCAAAUGAUGCGAAGGAACGAUUAGCGUGUCUGCAAGACGUAUUCACGUCCAUAUGCACGAGUAUCAUCCC
Blo-34-1   CCCUGCGGCCCCACUCUUUUGCGUCGCCUCCUGCGGCGGCACAGCGCGUCUUCACGUCCAGAUAGGCGCGUACAAUCCC
env-166    CCCUGCGGCCCCACUCUUUUGCGUCGCCUCCUGCGGCGGCACAGCGCGUCUUCACGUCCAGAUAGGCGUGUACAAUCCC
Blo-22-1   CCCUGCGGCCCCACUCUUUUGCGUCGCCUCCUGCGGCGGCACAGCGCGUCUUCACGUCCAGACAGGCGCGUACAAUCCC
env-167    ..........................GUCUCCUGCGGCGGCGUAGCGCGUCUUCACGUCCAGAUAGGCGCGUACAAUCCC
env-170    CCCUGCGGCCCCACUCUUUUGCGUCGUCUCCUGCGGCGGCGCAGCGCGUCUUCACGUCCAGAUAGGCGCGUACAAUCCC
env-196    CCCUGCGGCCCCACUCUUNUGCGUCGUCUCCUGCGGCGGCGCAGCGCGUCUUCACGUCCAGAUAGGCGUGUACAAUCCC
Blo-19-1   CCCUGCGGCCCCACUCUUUUGUGUCGUCUCCUGCGGCGGCGUAGCGCGUCUUCACGUCCAGAUAGGCGCGUACAAUCCC
env-197    CCCUGCGGCCCCACUCUUUUGCGUCGUCUCCUGCGGCGGCGCAGCGCGUCUUCACGUCCAGAUAGGCGUGUACAAUCCC
env-216    CCCUGCGGCCCCACUCUUUUGCGUCGUCUCCUGCGGCGGCGCAGCGCGUCUUCACGUCCAGAUAGGCGCGUAUAAUCCC
env-217    CCCUGCGGCCCCACUCUUUUGCGUCGUCUCCUGCGGCGGCGUAGCGCGUCUUCACGUCCAGAUAGGCGUGUAUAAUCCC
env-219    CGUGCCGCCCUCCGCCGGCCCGCCUCGGUGAGGAAUGCACGGAUUACCCGCGGCGUUGCACCGGGUGGCAGAAUCGCCC
Ssp-47-1   CCGACUGUCGGGAAUGCCUGUUCAUGCAGGUGCGUUGAAAUACCUGCCAACCGGUCUGGCAAACUAUAGGGCUGGACCA
Sma-2-1    CCGACUGUCGGGAAUGCCUGUUCAUGCAGGUGCGUUGAAAUACCUGCCAACCGGUCUGGCAAACUAUAGGGCUGGACCA
Nrh-1-1    GGCGUUGCAGCAGGCAUCGGUGGUUCUGGCAUACUUGACCGCCGUGCGUCUGCGAUUCGCGCAGACAAUCCCGCCUGAA
Nte-2-1    GGCGUUGCAGCGGGCAUCGGUGGUUCUGGCAUACUGGAUCGCCGUGCGUCUGCGAUUCGCGCAGACAAUCCCGCCUGAA
Nsp-2-1    CAGAAGCGUGGGAAGCGCUCUGGCAUACUGGUCCGCUGGUGCGGGCCCAGUCCCCGCGCACCACAACAUCCCCGCCUGA
env-220    AUGGCCGUUGCCUGGCCUCUCAUAUGCUGAGACGGGAUGGCGUGUCCGCCAGGCGUGACAAGGAAUAGCGGCCGCUGCC
env-221    .........................................................................CUGGACG
env-222    .............................................................CGUACUGGCACACUGGACG
env-223    GCCCGGUGGGCUGUGGGAAUGCCCGCAGGCCGACGGGCGUUCGGAAGGUUGUGUGCCGGUCUGGCACACUGGUCACGAC
env-224    GCCCGGUGGGCUAUGGGAAUGCCCGCAGGCCGACGGGCGUUCGGAAGGUUGUGUGCCGGUCUGGCACACUGGUCACGAC
env-225    .....................................................................CACACUCGUAC
env-226    .......................................................................CACUGACAG
env-227    GCGACGAGGACUGACCUGCGGAAUCCCGGUCGCCAGGGCGGCGUUCACCUGCGUGCGCCGCUCUGGCAGACUGGCGC
env-228    ...................................................................GGCAGACUGGCGC
env-229    ...............................................................CUCUGGCAGACUGGCGC
env-230    UGCGCAUGGCCGCCAUCGUAGGGAGCCCCGCGGCACGGGGAAGAUUUCGUCCGGGCCCGCUGUUCUGGCAGACUGUUCC
env-231    ..............................................................................GG
env-232    UGUGCGGGUGUCCGGGUGCUGGAGGGCUCGCGCUGCUCAGCGGUUUUCGAGCAGACGCCCUGGUCUGGCACACUGGUUC
env-233    .................................................................CUGGCACACUGGACG
Fsp-3-1    CGGGGGGCUUCGCCGCAGUGUGGGGACCCUUGCAUCGGCUGGCGGUCGGUGCAUCCGGCCUCUGGCAGACUGGCAGCAG
Ssy-1-1    GACGACCGCCACGGGAACAACCCGGUGCCGGCACGCGUUGCACCGCACGUCAGCCGAUCUGGCACACUGUCGGGUCGAG
Ssp-54-1   GCUGCCCCGGUCCGGGCCGGAAUACCGGACGCCUCCACCCCGUUUUGUUCGUGUGCGGCCGGUAGUGGCAGACUGGUAC
Ssp-174-1  CGCGGAAUCGUACGGGUGGGGAAUUAUCCCGUUCCCGACCUGGUUUUGGGAGAUGCGGCCUGUGCUGGCAGACUGGUCC
Ssp-32-1   AGCCACAACGCACCCGCAGGGAAUAGACGACAGUGACCCCCCGUUUUGGGGGAUGGCCCCAGUCCUGGCAGACUGGUAC
Sxy-1-1    AGCCCCCACGCACCCGCAGGGAAUAGAUCACCUGGGCCCCCCGUUUUGGGAGAUGGCUCCAGUCCUGGCAGACUGGUAC
env-237    ...................................................................GGCACACUGUCCU
env-238    ...............................................................CUCUGGCACACUGUCCU
env-239    UGAGGCCGGCUCCCGCCGGGAAGAUCGAGGGCACCUGCGGCGUUUGGUCCGUACGGUGCGCCCUCUGGCACACUGUCCU
env-240    CUGGCAUACUGGGCCGCUGGUGCGUCCAGCGCAUCCGCGCAGACCAUCCGGCCUCGCCCAGGAGCGGGCCAACGCCUAC
Ssp-102-1  AGGCCCGGUCGGAAUGCCGGGAACAAGGAGUGCAGGACUCCGGUUUUGGGAGAUGCGGACAGUGCUGGCAGACUGGGCC
Sca-3-1    AGCGGCCCCCGGGGCCCCGGGAAUACCGGGCCCCCACCCCCGGUUUUGGGGGAUACGGCCAGGACUGGCAGACUGGUAC
Ghi-1-2    CGAUCUGCAGGUUUCAGAAUAGUCUCGCCGCCGGCGAUUUAGCGGCUGGGGUCGGGCUGUGGAAUAAUUGACCGGUCAG
Ssc-5-1    UAGGUGUGGGGCGGAAUCCGGAAUACUUUCCGCCGCACCCCCGUUUUGGGAGAUACGGCCGGUCCUGGCAGACUGGUAC
Sro-5-1    CCAGGCCCGGAAACCCCCGGGAAUAACCGGCCGCGACCUCCCGUUUUGGGGGAUGCGGCCAGUACUGGCAAACUGGUCC
Sva-2-1    GCCGCCCCGCCGGGCCCCGGGAAUAUCCGGCACCGCCCCCCGGUUUUGGGGUAUGCGGGCAGUACUGGCAAACUGGUCC
Sye-1-1    ACCUGCGACGAUGUCCACGGGAAUGUUCCGGUCCCGGACCUGGUUUUGUCCCAAAGGGCCGGUCCUGGCAGACUGGUCC
env-241    ............................AGCCCGGGUCGCGGCGAUUGGCGCGACGUUCGGGAUGUUGGCACACUAGGCG
env-242    .......................................CGGCGAUUGGCGCGACGUUCGGGAUGUUGGCACACUAGGCG
Nex-1-1    GGCGUUUUGGGAGCGUAACGGCAGUUCUGGCAUACUGAACCGUCGUUGCGUCUGCUCAGGUGAGCAGAGCUCCCGCCUA
Nar-1-1    GGCGUUUUGGGAGCGUAACGGCAGUUCUGGCAUACUGAACGGUCGUUGCGUCUGCUCAGGUGAGCAGAGCUCCCGCCUA
Nab-1-1    GGCGUUUUGGGAGCGUAACGGCAGUUCUGGCAUACUGAACGGUCGUUGCGUCUGCUCAGGUGAGCAGAGCUCCCGCCUA
Mar-1-1    AUCGGCUGAGGAAUAGGCCGAGGCGACCGCGCGUUUGGGCUGGUGACGGUCUACCUGGCAUACUGGAUCGUCGACCACC
env-243    CCCGGUCCGCGCUCAAACCUCCGUUCGGCCGGUUUGGGCAGGUCGACGCCGAACUGGCAUACUGAAAGCCGACCAAACC
env-244    CCGGCCGGGGAAUGUGCGGCGACCAUCGCGCGUUUUGGGAGUACGGCGGCUGACCUGGCAUAAUGGACCGCCGACCACC

           ................................................................................
           ................................................................................
           ●●●●●●●●●●●●●●●●●●●●●●●●●●●●●●●●●●●●●●●●●●●●●●●●●●●●●●●●●●●●●●●●●●●●●●●●●●●●●●●●
```

```
env-245     GGUUAGCGCGGGGAAUGCUCAGCUGCUUGCGCGCGUUUAGGGGGAGAGCGGCUGACCUGGCAUAAUCGACAGCCGAAGC
Agl-1-1     GUUAACCAGACAGGAUCAGGAAAGUCGCCUCGAUUUCUCGCGAGGCCAUGUCACAUGGCAGACUGGUCCAUCGGCCACU
Msp-15-1    CCGCUCGGCGGGGGAAUACCGGCCCGUCCCGGGACGUUUUCGCGGUUGGCGGUUGACCUGGCAUAAUGGACCGCUGACC
Mco-2-1     CCGCACGGCGGGGGAAUACCGGCCCGUCCCGGGACGUUUUCGCGGUUGGCGGCUGACCUGGCAUAAUGGACCGCUGACC
env-248     GGCUAGCUCGGGGAAUGUCCGGCCGCACAUGUGCGUUUAGGGACAUAGCGGCUGAGCUGGCAUAAUGAUCAGCCGAAGU
env-249     UGUGGAAGCGGGACGGAAUAGCCGCCGGUACCCGCGCGUUUAGGAGGUUGGCGGUUGACCUGGCAUAAUGAACCGUCAC
Nmu-1-1     CGCAGCCGCCCCGGAAUAGGCCGCGCGGUCGCGGCGUUUGGCAUAACGACCGUCCGAUCUGGCACACUGGUCGGUCGAG
env-250     GACCGAGACCGGUAAGCGCGCAGGAAAUAUUGACCAGGCGCAAGUGGUUUCAGCGUCUGGCAGACUAGACCGCUAGUGA
env-252     .........................................................GGCAGACUAGACCCGCUAGUGA
env-253     AAACCAACCCGCGCCAUCGUUGGAAUGCACGCAGGGACAUCGAUUGGCGCGCGGUGGAUGCUCCCUGGCAAACUAGAUG
env-254     ........................................CGAGUGGCACUCGGAUGCCGCCCCCUGGCAGACUGGACG
env-255     AAAACACCCCCACGCCACGUUGAGACCGGUGCAGGGACACCGAUUGGCGCCCGGUGGAUGGUCCCUGGCACACUAGUUG
env-256     .........................................CGAGUGGCACUCGAAUGCCGCCCCUGGCACACUGGACG
env-257     ..........................................GAGUGGCACUCGGAUGCCGCCCCUGGCAGACUGGACG
env-258     .................................CAGGGGCCCGAGUGGCACUCGGAUGCCGCCCCUGGCAGACUGGACG
env-259     GGAACCGUGACACGCGCGGGAACAUGAGCGGGCCGGCCGGCGUUUCGACGCACGAGUCCGGUCUCUGGCAGACUGGUCC
env-260     CCCGGGGCACGGCAACGACUGAGUGCUGCCGGACGCCGGGAAGGUCGAGGACCUCCCCGACGUUCUGGCAGACUGUCCC
env-261     CCUCGGCCACUUCGCUCCACCGCACACCGCGAGCCUAGGCUCGAUUUCGGGCUGCGCGCGCGGUCUGGCAGACUGUCCC
env-262     GGUGCGCCGGACGAGCACGCCCGGUGCGGGGCGGGGGCGGAAGAUUUCGAGCCAGCCCGGUGUUCUGGCAGACUGUCCC
env-263     .............................................GGUGGCAGCGGCGCAGUCCCUGGCACACUGGACG
env-264     ...................................GCCCGCGGUUGUGACCGGCGGGUCCGCCUCUGGCACAAUGACCC
env-265     GAGCGGGUCCGAGGCCGCCGAGGAAGGUGGGCGCUGACCCGGAUUUCCGCCCGAUCACGGGGUCGUGGCACACUAGAUG
env-266     UACCGGGGCGGAGGCGGCCGAGGAAGGUGGGCGCUGACCCGGAUUUCCGCCGGAUCGGCGGGUCGUGGCACACUAGAUG
Nth-1-1     GGGGGCGUUUUCGCUCGAUGCGAGGGACCUGGCAUACUGGACCAUCGUGCGUCUGCGCAUCCGCAGACGAUCCCGCCUA
Pan-1-1     GGGGUGUUUUUCGGAGGGUGCGGCCAUAUCUGGCAGAAUGGACCGCUGUGUGUCUGCGGCUUCGCAGACAUCCCGCCUA
Sxi-1-1     AUUAUCCUUGGGAAUAUCGGCCCAUCAGGGUGCGUUUGAACGGGCAGUCUGAGUCUCUGGCAGACUUGUUCCGCUGACG
Ssa-1-1     GCGGCGCGCCCCGCCGGUGCCGGGAAUACGGGCCCCGGCGGGGGCGUUCGCACGGUCUGGCAGACUUGUUCCGCUGACG
Mch-1-1     ACGCGUUUAGGCUGCUGGCGGUUGCCCUGGCAUAAUAAGUGCCAGUCUGCCGGCCUCCGGCCGGGUGGACCGCCGACAC
env-267     CGCGCGAGGUGUCGCGCACGUCUCGGAACCCGAGCAGCGUCGACCAUUGUGCGCCUCACGGGCCGGCGGCAUAAUCACC
env-268     CGCGCGAGGUGUCGCGCACGUCUCGGAACCCGAGCAGCGUCGACCAUUGUGCGCCUCACGGGCCGGCGGCAUAAUCGCC
env-274     CGCGCGAGGUGUCGCGCACGUCUCGGAACCCGAGCAGCGUCGACCAUUGUGCGCCUCACGGGCCGGCGGCAUAAUCACC
Pas-1-1     CCGGGAGCCGAUAACGACUGAGGACGGCUCGGCCGCCGGGAACAGCGCGACGUACCGGAUGGUUCAAUAUGCCGUCAAC
Pdi-1-1     CUGACCUCCCGCACGGGAGUCCGCCACGGGGCCACCGGGAACAGCCGGUGGCCCGUGGUGGUUCAAUUGAGCGUCACGC
env-279     UCGUGGACCCGGGGCCGAAGCUCGCGGGGGUAGACGGCCGGGAAUGUGUCAAGGCCGCGGGGGUUGACUAUCCUGACCC
env-280     ...................................................GCCCGGAUGGUUGAAUGGGCUGUCACGC
Ame-4-2     GUCCAGUACCCGGAAUAUGCGCCGUCCGGGGGGCGUUGACCUGCACGUCAGCGCGUCUGGCAGAAUCAAUCCGCUAACG
Asp-4-1     GUCCCGUACCCGGAAUAUGCGCCGCCCGGGGGGCGUUGACCUGCAUGUCAGCGCGUCUGGCAGAAUCAAUCCGCUAACG
Ath-1-1     GUCCCGUACCCGGAAUAUGCGCCGCCCGGGGGGCGUUGACCUGCACGUCAGCGCGUCUGGCAGAAUCAAUCCGCUAACG
Sbi-1-1     GACCCUCGUCCGAGCCCUGGGAAUAUCUGGGCCCUACCCCCGGUUUUGGGGGAUGCGGCCAGUCCUGGCAGACUGGUUC
Shy-4-1     AGCGUUCGGGGGAGGGCCGGGAAUAUCCCGGUGCGACCCCCGGUUUUGGGGGAUGCGGCCAGUACUGGCAGACUGGAUC
Ssp-73-1    CAGCCUCUUGGGCGGGCCGGGAAUAUCCUGGCCCGCCCCCCGGUUUUGGGAGAUGCGGCCAGUACUGGCAAACUGGAUC
Sir-1-1     CAGCCUCCGGGGCGGGCCGGGAAUAUCCGGGCCCGCCCCCCGGUUUUGGGGGAUGCGGCCAGUACUGGCAAACUGGAUC
Ssp-165-1   CAGCCUCCGGGGCGGGCCGGGAAUAACCGGGCCCGCCCCCCGGUUUUGGGAGAUGCGGCCAGUACUGGCAAACUGGAUC
Svi-3-1     AGCCUCCCGGGAGGGGUCGGGAAUAUCCCGGCCCGACCCCCGGUUUUGGGAGAUGCGGCCAGUACUGGCAAACUGGAUC
env-281     CGAUCCGGCCGCCCCGCGUCGGAUCGCGGUCCGGGCCCCGGGGAACAACCCCGGGACCGGGUGGUUCACUGAUGCGUCA
env-282     ..............................................................CCGCGCCAUCGACGCAA
env-283     GGCCGCCGGAGGUAGCCAGCCGGGAAUCUCCCAGUGCCCGCCGGCGUUACCGCGUCUGGCAGACUGGAAGCCACACCUA
env-288     ..............................................GGCCGGUCGACCGCGAGGUCGCCCCGCUGGAUC
env-289     .........................................................CUCUGGCAGACUGUCAACGGAG
env-290     UGGGCCAGGAGCGCUAGCCGCGGGAAUGCGGGACGGUCAGAUCUGGUUGAGGGGUCUGGCACACUAGAACCUGCGAUUA
Gar-1-1     GGUUGACGCCGGUGGACGGGCAGCGGUCACGAUUUUCCGGCGACGGGACGCGCGUGGCAUACUGGAUCGUCGAAUCAAG
Mro-1-1     GUCACCACGGUGGCCUGCCCAGGAAUAGCCCAUGGCCUGGGAUGGUUGCAGGAUCUGGCACACUGGUAAACCGAACCGC
env-291     .......................................................................CACUGACU
env-292     ACGUGCUCCCGGAGCCCUCAGGCCCGGGAACACGGUCGCAGCGAUUGGCGUUGAGGCGAACGUCGGUGACACACUGUCC
Psu-1-1     GUUCCUCAUGGUGAUCUCGCGGACCACGCCCUGAUCCGGGGAAGUUUUGCGCGGCGGCGCGCGUUUGGCAGAAUGGUCG
Gne-1-1     CGUGGUCGACCUGCCCGCAAGCCCGGCACCUGCACGAUUGGUCGACCGGAGGGGCCCUGUGGAAUAAUUCUCCGGUCAG
Gsi-1-1     UCGCGCUGGAGGACCGCCAGGCCUCGCACCUGCGCGAUUGGCCGGGGUGACCGGGGUUGUGGAAUAAUUCUUCGGUCAG
Gsi-2-1     GUCGCGCUGGAGGACGGCCAGGUCUCGCACCUGCGCGAUUGGCGGGGUGACCGGGGUUGUGGAAUAAUUCUCCGGUCAG
Nta-1-1     CAGGGGCGUUUCGGAAGGUGUCGGCGGAUCUGGCAUACUGGUUCGCCGUGCGUCUGCGCCCCGCAGACAAUCCCGCCUA
env-293     AAAACACCCCCGCCCACCGUUGGGAUCCACGCAGGGGCACCGAUUGGCGUGCGGCGCGCUCUCCCUGGCAGACUAGAUG
env-294     UAGGGGUUGUUGCAAAGCGGGGCUAGGGCGCGGGAUGGCACUCAGCGCCGUGGCAAACUAUGUGGGUACGCGAGACGAC
env-295     CCGCGGCCGCAGGAAUAGCCGCCCCCUGGCACGCGUUUAGGCUGCUGGCGGUUGAGCUGGCAUAAUGGACCGUCGACCC
env-298     GGGCCAGCGAUGGAAUAUCGGCUGCUGCCCGCACGUUUAGGAAGCUGGCGGUUGACCUGGCAUAAUGGACCGCCGACCC
Mob-1-1     CGGGCGCACGUUUAGGCUGUUGGCGGUUGGCCUGGCAUAAUCCAGGGCAGCCUUCCGGCCUCCGGCCGGGUGCCUACAC
Mob-1-2     CGGGCGCGCGUUUAGGCUGUUGGCGGUUGGCCUGGCAUAAUCCAGGGCAGCCUUCCGGCCUCCGGCCGGGUGCCUACAC
env-299     UGCCGUUGUGCCUGCCGGACCGGUCAUGUCCGCCCGCCGUCCAGUGCAGCGGAGCCGCAUGGCAAACUGGUCCAUCGGC
Asp-1-1     UUAUCGACCAAGGACCGGCUGCCGCCUCCUGAUUUCUCCUGGGGGCAAGGCACGUGGCAGACUGGUUCAUCGGCCAGUC
env-300     .............................................GGUGGAAUGCGCACCGCUCCUGGCACACUGGUCG
env-301     ................................................GGAAUGCGCACCGCUCCUGGCACACUGGUCG
env-302     ............................................................................CGU
env-303     .............................................................................GU
Nsp-6-1     GAUGUUUUGAGGGGGUAAUGACGGUUCUGGCAUACUCGAACGUCGUUGCGUCUGCUCGGCCGAGCAGACUUCCCGCCUA
env-304     ......................ACCCCUGGCACGCGUUUAGGCAGCUGGCGGUUGAGCUGGCAUAAUGGACCGCCGACC
env-306     CAGGCCGGCGGAGGAAUAGCUGACCCCUGGCACGCGUUUAGGCAGCUGGCGGUUGAGCUGGCAUAAUGGACCGCCGACC
env-307     CAGGCCGGCGGAGGAAUAGCUGACCCCUGGCACGCGUUUAGGCUGCUGGCGGUUGAGCUGGCAUAAUGGACCGCCGACC
env-308     ..........GGAAUACCCACCCUCUGGCACGCGUUUAGGCUGCUGGCGGUUGAGCUGGCAUAAUGGACCGCCGACCCC
env-309     UGGUACAGGAGGAAUACCCACACUCUGGCACGCGUUUAGGCUGCUGGCGGUUGAGCUGGCAUAAUGGACCGCCGACCCC
env-310     GAAGACGAUGCCGAUGCCGCAGGACUGACCGGCCCCGUCCAGGCCGGAAUGUCCUACCUGCGCAUCGCGUUACCAUCGA
env-311     CGGAUUGCCCGUGGAAUAGCCGUCCCCGUCGGCGCGUUUAGGCUACUAGCGGUUGACCUGGCAUAAUGGGCCGCUGACC
env-312     ..................................GUUUAGGGGCAUAGCGGCUGACCUGGCAUAAUCAACAGCCGAAGU
env-314     ..UUAGCGGGCAACCCCCCCGGGGGAAUUUCCGGUGUGCCAGGCACGUUGUCGCGUCUGGCACACUAGAUACUCGACUA
Ami-1-1     GUUAACCAGGGCGAAUCCGGUUGCGCCUCCUGAUUUCCGGUUGGUACGGCAGGCAUGGCAGACUGGUACAUCGGCCAGU
env-315     .......................................................................ACUGACCC
env-317     ....................................................................CAGACUGACCC

            ...............................................................................
            ...............................................................................
            ●●●●●●●●●●●●●●●●●●●●●●●●●●●●●●●●●●●●●●●●●●●●●●●●●●●●●●●●●●●●●●●●●●●●●●●●●●●●●●●
```

```
env-318     GGUAGUGGCGACGACUGACGACAACGGAAUGUGCCGCCCCGCAUUGGCGUUGUACGCGGCGUCAGUGACAGACUGACCC
env-319     ......................................................................GACUGACCC
env-320     ............................................................................CCC
env-321     ......................................................................GACUGACCC
Nni-1-1     GCCUGUUCAGGACGGUGUCACCCCUCGUGACGGCCACCCGCCCGCGGCGCAUAUACUGUUCCGUCGGCAACCAUGCCUA
Nsp-8-1     UGACCGGCGGCCGACGGCGGGAACAACCUCGCCUCCCCGGCGGUUUGGGGAGGCGAGGGAUCUCGCCGUAAGAUCUCCC
env-322     ACCGGGAACAACUCGGCCGCCGGCCGGGUUGAGACGAUUACCAGCGCGGACAGCUGCUCUGGCAUACUGAUCGGCCGAG
Rsp-20-1    CGACGGAACAAACGUCUCGGCACCGAUGUUGAGACGAUUACCAGCGCGGACAGCUGCUCUGGCAUACUGAUCGGCCGAG
Rsp-16-1    UGAGGAAUACGCAGCCUCCUCACCGACGUUGAGACGAUUACCAGGGCAGAGAGUGGCUCUGGCAUACUGAACGGCCGAG
Mpu-1-1     CGGAGUGUGGGCGGGCGUUGCGGGUGGCGGACCUCCUGUGCCGGGUGGAAGACGCCCGUGGCAGACUGGUCAAUCGGCC
Msp-27-1    GGGAGUGUGGGCGGGCGUUGCGGGUGGCGGACCUCCUGUGCCGGGUGGAAGACGCCCGUGGCAGACUGGUCAAUCGGCC
env-323     GAGCGCCGACCGCUGUUGAGCAGAUGUCACCGCCGGGCUGCGAUUUCCCAGCGCAAUCCUCCGGAUGGCACAAUCGUUG
env-324     .................................CGGGCUGCGAUUUCCCAGCGCAAUCCUCCGGAUGGCACAAUCGUUG
Aal-1-1     UCCGGCGACGAGGACUGACGGAAUUCUCACCGGCCCAGGCCGGUUGAGAGUGCACGUGCCCCAGAUGGCACACUAAUUC
env-325     AGCCGGUUGCCAGGAAUACCCGGCGGCGACCCGCUGUUUGGCUUUGUGACCGGCCCAUCUGGCACACUGGUCGGUCGAG
env-329     ...CGGUUGCCAGGAAUACCCGGCGGCGACCCGCUGUUUGGCUUUGUGACCGGCCCAUCUGGCACACUGGUCGGUCGAG
env-330     ..........................AACCGUCGUUUUGGUCCGGCGGGCACUGCCUGGCAGACUGGUACGGCACACCG
env-331     .............................GGUCGUUUUGGUCCGNCGGGCACUGCCUGGCAGACUGGUACGGCACACCG
env-333     ................................................................CUGUCUGGUUGACGA
env-334     .....................................................CGACUGGCACACUGUCUGGUUGACGA
env-335     GGCCGGAGCGCGGAAUUUUCCUGGUGUGGUCUCGGUUUGGAUGAGCGUGACCAUUCCUGGCACACUGUCUGAUUGACGA
env-336     ...................................GGCAGACUGGACGAUCGGCGCCCAGGCGCUCAUCCAUUCAGGCA
env-337     CGUCAGGGCUUCGGGGACUCCUGAGCCCGCGCAGCGACCGCGAUUGGCAGCGCGGUUGCCGGGAGUGGCAGACUUGGAC
env-338     ...............................................GGCACACUGGAUGUCCCACGGCUGGACCCCAG
Bts-1-1     AUCAGAGUCAACAGAUUGCACAUUCGGUAGUCGGCGUGUUUCUCUACGGGAAAAAGUCGAUAAUCAUGCGUAUAAUCCA
Ama-5-1     CGGCACCGGACGGGCAAGAUCGGGAAUGCCCGGUGACCUCGCCACGUUCGGGUAUCUGGCACACUUGGUGUGCAACCGC
env-339     UGCCCGGCCAGCGGCCGGAUUCCGGGAAUUGACAGUGCCAUGCGCGCGUUCCAUGUCUGGCACACUGGUAAUCCGAAAU
env-340     ACGCCCGGCGCAGGAAUAGCUGACCCCUGGCACACGUUUAGGACGUUGGCGGCUGAGCUGGCAUAAUGGACCGCCGACC
env-341     CACGUUCGCGCGAAAUAAAGCGCCCGUGGCACGCGUUUAGGCUGCUGGCGGUUGAGCUGGCAUAAUGGACCGCCGACCC
env-342     UUGUAGACCGCGGGAAUACCGCCCGCUGCGCACGCGUUUAGGCUGCUGGCGGUUGACCUGGCAUAAUGGCGCUGACACC
env-343     ........................................................................CUGGACC
env-344     ...........................CACCGUCGUGCGCACGGUUAUCUGGCGCAGGUCGGUGUCUGUCACACUGACC
env-345     ................................................................CUGUCACACUUGACC
Mva-1-1     GCGGUGGCUCAGGAAUAGCGGGCCCCGAACGGGCGUUUAGGCUGCUGGCGGUUGACCUGGCAUAAUUGACCGCCGUCAC
Mru-1-1     GUCGGCGCGCAGGAAUAGGGGCCGUCGGGUGCGUGUUUAGGCUGUUGGCGGUUGACCUGGCAUAAUGGACCGCCGACAC
Mva-2-1     GUCUCCGCGCAGGAAUAGUGCGUGCCGAAUGCGCGUUUAGGCUGCUGGCGGUUGACCUGGCAUAAUUGACCGCCGUCAC
Mgi-1-1     UGCGCGUUUAGGCUGCUGGCGGUCGAACUGGCAUAAUCGGUGGCAGUCUGCCGGCCUCAGGCCGGGCGACCGCCGUCAC
env-346     GCGCGGCGGGACACGAGCCAUGGGGUGCGUGUACCUGAGCGCCGGGGGAAUGAGCCCGCGUGGCAGACUGUUGAGACGC
env-347     CUUCCUGGCACGAGCCUGGCCGUGUUGGGUCCACCUGACCAUGGGGGGAAUGAGCCCACGUGGCAGACUGUUGAGACGC
env-348     CGUCGUCGAUCAUCGCGGUGAGCCGGCAGUGAGCCCGGCGGCGGGUCGAGGGCGGGGCGUGGCACACUGGUACGUCGGC
Nas-2-1     GGCGUUUUGGGAGCGUAACGGCAGUUCUGGCAUACUUAACGAUCGUUGCGUCUGCUCAGGUGAGCAGAGCUCCCGCCUA
Mab-65-1    GAAAUCGGGAAUAGUGCGCAGGUCGGCGCCGUUUAGGUUCUACGUAACUGAGCUGGCAUAAUGGCCUCUCAGCCUUUUG
env-349     GUUACGGCCCCGGAAUAGCGGCGCCGCCGGGGGCGUUUCGCAUUGACGGCGGCUGACCUGGCAUAAUCGAUAGCCGACC
env-350     ......................UGCCGCCUGAUUUCGGUUCAGGGUGUGCGACGUGGCAGACUGGUCCAUCGGCCAGCA
Fsp-7-1     AGGCCGGUGUUACAGGGCUGGCGGCGGGCCGUGGGCGCGCCGUUGGCUCGGUCGAAGGUCUGGCAGACUUGGGCUGGUA
Sca-1-1     ACCGCCUGCCCAGUGGCUGGGAAUAGAUCGCGGCGACCCCUCGUUUUGGGGGAUGCGGCCAGUCCUGGCAGACUGGUAC
Ssv-1-1     GCCGUCUGCCGGGGGCUGGGGAAUAGAUCGCGGCGACCCCUCGUUUUGGGGGAUGCGGCCAGUCCUGGCAGACUGGUAC
Ssp-50-1    CUCCGGACAACCGGGCCGCGGAAUAGUCCGGCCCUUCCCCCGGUUUUGGGAGAUACGCCCAGUGCUGGCAGACUGGUCC
Ssp-19-1    GGGCCCCUGCGGGCCGUACGGAAUAGAACACCCGAACCCCCGGUUUUGGGGGAUGGCCCCAGUCCUGGCAGACUGGUAC
Aro-1-1     GUAUACCCACUGAUGUGCGUACGGUCCGCGACGGCGGAAGGGCGGGAAUACCCGGCCUUCCGCCGUUGUAGGAUGGUAC
Sca-2-1     GGGCGGCGGGGGAGCCGGUGGAAUAUCCCGGUAGGUCCUCCGGUUUUGGACAGUACGGCCGGUCCUGGCAAACUGGUCC
Nvi-1-1     GUGUUUUGGGAGCGUAACGGCUGUUCUGGCAUACUGAGCGGUCGUUGCGCCUGCUCCGACGAGCAGGCCUCCCGCCUAA
env-351     ACCCGGCAGGAACCCCGCCGGAAUAGAUCCGGGCCGCCCCCGGUUUUGGGAGAUACGCCCAGUCCUGGCAGACUGGUAC
env-352     GCGCCACCCCGGGAAUGCGCGGGCGGUGGCAGGCGUUUAGGGACACAGCGGCUGAGCUGGCAUAAUCAACAGCCGAAGU
env-353     GCGCCGGCCCGGGAAUGCGCGGACGCUGGCAGGCGUUUAGGGACACAGCGGCUGAGCUGGCAUAAUCAACAGCCGAAGU
env-355     GCGCCGGCCCGGGAAUGCGCGCACGGUGGCGGGCGUUUAGGGACAUAGCGGCUGAGCUGGCAUAAUCAACAGCCGAAGU
env-358     GCGCCGGCCCGGGAAUGCGCGCACGGUGGCNGGCGUUUAGGGACANAGCGGCUGAGCUGGCAUAAUCAACAGCCGAAGU
env-359     GCGCCGGCGCGGGAAUGCGCGGAGCGUGGCAGGCGUUUAGGGGCAUAGCGGCUGAGCUGGCAUAAUCAACAGCCGAAGU
env-360     GCGCCGGCCCGGGAAUGCGCGGACGCCGGCAGGCGUUUAGGGACACAGCGGCUGAGCUGGCAUAAUCAACAGCCGAAGU
env-361     .........CGGGAAUGCGCGGACGCUGGCAGGCGUUUAGGGACACAGCGGCUGAGCUGGCAUAAUCAACAGCCGAAGU
env-363     GCGCCGGCCCGGGAAUGCGCGGACGAGGGCAGGCGUUUAGGGACACAGCGGCUGAGCUGGCAUAAUCAACAGCCGAAGU
env-364     GCGCCGCCCCGGGAAUGCGCGGACGGCGGCAGGCGUUUAGGGACACAGCGGCUGAGCUGGCAUAAUCAACAGCCGAAGU
env-365     ACGCCGGCCCGGGAAUGCGCGGACGCUGGCAGGCGUUUAGGGACACAGCGGCUGAGCUGGCAUAAUCAACAGCCGAAGU
Mim-1-1     AUCGGCCGGGAAUAUCGCGCUGGUCGCCGCCGUUUAGGAUUUGCGUAACUGAGCUGGCAUAAUGGCCUCUCAGCCUUUG
Mab-73-1    CGAGGCCUGGAAUACCGUGCGGGUUGUCGUCGUUUAGGUUCUGCGUGACUGAGCUGGCAUAAUGGCCUCUCAGCCUUUG
Mab-55-1    CGAGGCCUGGAAUACCGUGCGGGUUGUCGUCGUUUAGGUUCUGUGUGACUGAGCUGGCAUAAUGGCCUCUCAGCCUUUG
Mab-1-114   CGAGGCCUGGAAUACCGUGUGGGCUGUCGUCGUUUAGGUUCUGCGUGACUGAGCUGGCAUAAUGGCCUCUCAGCCUUUG
Mma-1-1     CGAGGCCUGGAAUACCGUGCGGGUUGUCGUCGUUUAGGUUCUGCGUGACUGAGCUGGCAUAAUGGCGUCUCAGCCUUUG
env-366     ....................................................................AGAGACUCGUC
env-367     .............................CCUGAUUUCUGAUCCGGAUGUGCGCCAUGGCAGACUGGUCCAUCGGCCAG
env-368     ......................................AAGCCCGUCUGCUGGAUGCCAUGGCAGACUGGUACAUCGGC
env-369     ........................................................................CCGACAU
Tsp-1-1     GAACAGCGGCUUCGCGGACGCCGAUUAACGGCGGGAAUUUCUGCAGACGCGGGACGUUCUGGCAUACUGGAACGUCGGU
Tpa-1-1     CCGACCGGUGACGGGGCCGCGUCGCGGAGCGCGGGAAUUUCUCCGUACGCGGGGCGUUCUGGCAUACUGGAAUGUCGGU
Are-1-1     GUUAUCCAGGGCGUAUCCAGUUGCGCCGCCUGAUUUCCGUCGGGUACGGAGGUCAUGGCAGACUGAUACAUCGGCCAGU
Nca-3-1     GCGGCGUCGUGCCCGGUCGCGGGAAUGCCCGAUGGCCUGGGAUGGUUGCACGAUCUGGCACACUGGUAGCUGACCGUUU
env-370     GAGGCGUCACGGCGAUCUUCGGGAAUGCCGGAUGGCCUGGGGUGGUUGGACGAUCUGGCACACUGGUAGCUGACCAUUU
env-371     GCCCGGUGGGCUGCGGGAAUGCGCGCAGGCCGACGGGCGUUCCGAAGACUGUGUGCCCGUCUGGCACACUGGUCACGAC
Mph-1-1     CCUGCUCUAGCUGGAAUAUCGGGCGCUGGGCCCGCGUUUAGGAACUCGACGGUUGAGCUGGCAUAAUGGACCGUCGACC
env-372     UACUGCCCCCGGAAUAGCGCCACCGCCAGGGGCGUUUCGCAUUGACGGGGGCUGACCUGGCAUAAUCGAUAGCCGACCC
env-373     ...................................................................GAUAGCCGACCC
env-374     UACUGCCCCCGGAAUAGCGCCACCGCCAGGGGCGUUUCGCAUUGACGGGGGCUGACCUGGCAUAAUCGAUAGCCGACCC
env-377     UACUGCCCCCGGAAUAGCGCCACCGCCAGGGGCGUUUCGCAUUGACGGGGGCUGACCUGGCAUAAUCGAUAGCCGACCC
env-378     UACUGCCCCCGGAAUAGCGCCACCGCCAGGGGCGUUUCGCAUUGACGGGGGCUGACCUGGCAUAAUCGAUAGCCGACCC

            ...............................................................................
            ...............................................................................
```

```
env-379     CUCCGGCCGGGCCGUCAGCCGGGAAUCAUCGGAGGCCACGGAAUGUUGGAGGGUCUGGCACACUAGGAAGCCGCGACCA
env-380     .............................CCGGGUGCCCGCGCACGUUACAGCAUCUGGCACACUGGUAGGCCUACGAGA
Sgr-9-1     GGCGUCGUGAGGUAGCCGGGGAAUACUUCCGCCCGGACCCCGGUUUGGGCAGAUGCGGCCAGUCCUGGCAGACUGGUAC
Sex-2-1     UAGCCCCCCGGACAGCCGGGGAAUACUUUCGGCCGGACCCCGGUUUGGGGAGAUGCGGCCGGUCCUGGCAGACUGGUAC
Ssp-115-1   GCCCCUUCGCGGACCCCCGGGAAUACCGGCGCUCCCACCCCGGUUUGGGGAGAUGCGGCCGGUCCUGGCAGACUGGUAC
Ssp-105-1   ACCCCCGGCGGGGCCCGGGGGAAUACCGGCGGUCCCACCCCGGUUUGGGGAGAUGCGGCCAGUCCUGGCAGACUGGUAC
Mgl-2-1     CUGACCCCGCCGGCCCCCGGGAAUACCGGCGCCGCCACCCCGGUUUGGGGAGAUGCGGCCGGUCCUGGCAGACUGGUAC
Sgr-12-2    UAGUCCCCGGGACCACCGCGGAAUAGGACCGGCCCCACCCCGGUUUGGGGAGAUGCGGCCGGUCCUGGCAGACUGGUAC
Fsp-14-1    CAGCCGGUGUUACAGGGCUGGCGGCGGGUCGUGGGCGCGCCGCUGGCCCGGUCGAAGGUCUGGCAAACUUGGGCUGGUA
Fsa-1-1     GGGAUUAGGCGCGGUGCCGGUGGCGUUACGGCACUGGCACCGUGACCCAGUCACCAGGUCUGGCAGACUUGGGCCGGUU
Gar-2-1     AGAGCGGCCUCACGGCCUAUCAUGGUCAAUCGGUGCGCAGGUCGCGCACAUGCCAAGAGCCCUGGACUCCGGGGCCCGA
Spa-31-1    CGACGUUUACGCUGCCAGUGCUGUGGCACGGCCCGCCUGCCAAGCCGACGUGACCCCGUGGCAGACUGGUUCGUCGGCC
Spa-3-1     CGACGUUUACGCUGCCAGUGCCGUGGCGCGGCCCGCCUGCCGAGCCGACGUGACCCCGUGGCAGACUGGUUCGUCGGCC
Spa-39-1    CGACGUUUACGCUGCCAGUGCCGCGGCGCGGCCCGCCUGCCGAGCCAACCUGCCCCCGUGGCAGACUGGUUCGUCGGCC
Spa-35-1    CGACGUUUACGCUGCCGGUGCCGCGGCGCGGCCCGCCUGCGGAGCUAACCUGCCCCCGUGGCAGACUGGUUCGUCGGCC
Spa-25-1    CGACGUUUACGCUGCCGGUGCCGCGGCGCGGCCCGCCUGCCGAGCCAACCUGCCCCCGUGGCAGACUGGUUCGUCGGCC
Nfa-1-1     GGGUGUUUUGGGUGGGUGACGGCGGUUCUGGCAUACUGGAUCGCUGUGCGUCUGCGGCUCGCGCAGACAAUCCCGCCUA
Msp-7-1     UCCGGCCGGAAUAGCCCCGUGCUGCCGCGCGUUUAGGCUGUUGGCGGUUGACCUGGCAUAAUUGACCGCUAGUCACACC
Msp-19-1    CGCUGCCAAGGAAUAGUCACCGGCUCGGCCGCGUUUAGGACCUUGGCGGCUGACCUGGCAUAAUCGACCGCCGAACCCC
Mll-1-1     CGCCGCCAAGGAAUAGUCACCGGCUCGGCCGCGUUUAGGACCUUGGCGGCUGACCUGGCAUAAUCGACCGCCGAACCCC
Msp-3-1     CAGCGCCGAGGAAUAGUCACCGGCCCGCAGGCGUUUAGGACCUUGGCGGCUGACCUGGCAUAAUCGACCGCCGAACCCC
Msp-20-1    AGGUGCCGAGGAAUAGUCACCGGCCCGCAGGCGUUUAGGACCUUGGCGGCUGACCUGGCAUAAUCGACCGCCGAACCCC
Nsp-7-1     GGUGCGUUGUGAGUGCGUAUCGGCGGUUCUGGCAUACUGGACCGUCGGUGCGUCUGCGAUCGGCAGACCAUCCCGCCUA
Nve-1-1     GGGUGCGUUGUGCGUGCAUGUCGGCGGUUUGGCAUACUGGACCGUCGGUGCGUCUGCGAUCGGCAGACCAUCCCGCCUA
Nao-1-1     GGUGCGUUGUGAGUGCGUAUCGGCGGUUCUGGCAUACUGGACCGUCGGUGCGUCUGCGAUCGGCAGACCAUCCCGCCUA
Nce-1-1     GGUGCGUUGUGAGUGCGUAUCGGCGGUUCUGGCAUACUGGACCGUCGGUGCGUCUGCGAUCGGCAGACCAUCCCGCCUA
env-381     GCGGCUCGCCACACCGGUUGGUCGAUUCGGCGUGUGGCGCAAGGGGGUCGUAUUCUCCCCGGGUUGCGCUAGACUAUUA
env-382     GCGGUCCGCCACACCGGUUGGCUGAUUCGGCGUGUGGCGCAAGGGGGUCGUAUUCUCCCCGGGUUGCGCUAGACUAUUA
env-384     UCGACCAGCCACCUCGGUUGAUUGAUUCGGCGUGUGGCGCAAGGGUGCCGUAAUCUCUUCGGGUUGCGCUAGACUAUUA
env-386     .................................................CGUAUUUUCCCCGGGUUGCGCUAGACUAUUA
env-390     GCNGCUCGCCACACCGGUUGNUCGAUUCGGCGUGUGGCGCAAGGGGGUCGUAUUCUCCCCGGGUUGCGCUANACUAUUA
Bas-1-1     CUCGGCUCGCCACACCGGUUGGUGGAUUCGGCGUGUGGCGCAAGGGGUCGUAUUCUCCCCGGGUUGCGCUAGACUAUUA
Bas-2-2     GCUCGACCGGCCCUCGGUUGAUUGAUUCGGCGUGUGGCGCAAGGGUGCCGUAAUCUCUUCGGGUUGCGCUAGACUAUUA
Bsp-3-1     GCGGCUCGCCACACCGGUUGGUCGAUUCGGCGUGUGGCGCAAGGGGGUCGUAUUCUCCCCGGGUUGCGCUAAACUAUUA
env-391     CCGACCAGCCACUCCGGUUGAUGGAUUCGGCGUGUGGCGCAAGGGGAUCGUAUUCUCCCCGGGUUGCGCUAGACUAUUA
env-394     CGACUGGCUGCUCCGGGUUGGUGGAUUCGGCGUGUGGCGCAAGGGGGUCGUAUUUUCCCCGGGUUGCGCUAGACUAUUA
env-396     UCGACCAGCCACUUCGGUUGACGGAUUCGGCGUGUGGCGCAAGGCUGCCGUAAUCUCCCCGGGUUGCGCUAAACUAUUA
Bsp-2-1     UCGGCUCGCCACACCGGUUGGCGGAUUCGGCGUGUGGCGCAAGGGUGCCAUAAUCUCUCCGGGUUGCGCUAGAAUAUUA
Bas-2-3     CGACUGGCUGCUCCGGGUUGGUGGAUUCGGCGUGUGGCGCAAGGGGGUCGUAUUUUCCCCGGGUUGCGCUAGACUAUUA
Mma-2-1     UGGCCGGGCAGGGGAAUACCGCCCCGUCAUCACGCGUUUAGGAUGCUGGCGGUUGACCUGGCAUAAUGGACCGCUGACC
Msp-2-1     ACCGAUGGCCAGGAAUACCGCCGGGUGUGCGCACGUUUAGGCGGCUGGCGGCCGAACUGGCAUAAUGGACCGCCGACCC
env-398     ...........................................................CUUAGUUGGCAACAAUGCCUA
Nji-1-1     AACCGAGCCCGGUGGACCCGGCCCGGAACAAAACCGGGGCGGAUCGGCGUUUACACUGAUUCGUUGGCAACAAUGCCUA
Nno-1-1     GGUGCGUUGUGCGUGCGUAUCGGCGGUUCUGGCAUACUGGACCGUCGGUGCGUCUGCGAUCGGCAGACCAUCCCGCCUA
Nhi-1-1     GUGGUUGUGGGAGGGUGACGACGGUUCUGGCAUACUGGAUCGCCGUGCGUCCGCCGCGUUUCGCGGACAAUCCCGCCUA
Msp-1-1     GACCGAUGGCAAGGAAUACCGUCAGGUAUACACACGUUUAGGCUGCUGGCGGCCGAACUGGCAUAAUGGACCGCCGACC
Msp-18-1    CAUCUGAUGCGUGGAAUAGAGCCUGGCCAGCACGCGUUUAGGCUGCUGGCGGCCGAACUGGCAUAAUGGACCGUCGACC
Mfo-1-1     CAGUAGCGGGCAGGAAUAGUGGCCGGUCAUCCGGCGUUUAGGAGGUUGGCGGUUGACCUGGCAUAAUGGACCGCCGACC
Mvu-1-1     CGGUCCCCGGCAGGAAUAGUGGCCGGUCAUCGCGCGUUUAGGCUGUUGGCGGUUGACCUGGCAUAAUGGACCGCCGACC
Mse-1-1     CAAACGGCGCGGGGAAUAGGCCCCGGUCAUCGGGCGUUUAGGAGGUUGGCGGUUGACCUGGCAUAAUGGACCGCCGACC
Msp-23-1    GGCACCGGCCAGGGAAUAGGCCCCGGUCGUCGGGCGUUUAGGAGGUUGGCGGUUGACCUGGCAUAAUGGACCGCCGACC
Mse-2-1     CGAGUCCGGGCAGGAAUAGCGGCCGGUCAUCGCGCGUUUAGGAGGUUGGCGGUUGACCUGGCAUAAUGGACCGCCGACC
Mse-3-1     GUGACCGGCCGGGGAAUAGGCCCCGGUCAUCCCGCGUUUAGGAAGUUGGCGGUUGACCUGGCAUAAUGGACCGCCGACC
Mse-3-2     GCGACCGGCCGGGGAAUAGCCCCCGGUCAUCCCGCGUUUAGGAAGUUGGCGGUUGACCUGGCAUAAUGGACCGCCGACC
Msp-22-1    GUGACCUCCCAGGAAUAGCCGCCGGUCCGCACGCGUUUAGGAGGUUGGCGGUUGACCUGGCAUAAUGGGCCGCCGAACC
Mne-1-1     GCCGCUGGGCCGGGAAUAGUUGUCGCUGGUCACGCGUUUUCGCAACUGGCGGUUGACCUGGCAUAAUGGACCGUCGACC
env-400     CCCGGCACACCGGGAAUAGUCCGGCUCAAGCACGCGUUUUCGCAACUGGCGGUUGACCUGGCAUAAUGGACCGUCGACC
Msp-21-1    GCCGCUGGGCCGGGAAUAGUUGUCGCUGGUCACGCGUUUUCGCAACUGGCGGUUGACCUGGCAUAAUGGACCGUCGACC
Sal-7-1     GGCCCCUGCGACGGCGCGGGGAAUACCGGGGGCCCCACCCCGGUUUUGGGAGAUGCGGACAGUCCUGGCAGACUGACCC
Szi-1-1     GUGCCCCGGUCCCCGCCCGGGAAUAGAUCACCUCACCCCCCGGUUUGGGAGGAUGCGGUCAGUCCUGGCAGACUGGUAC
Ssp-17-1    CCGGCCGGGCCGGCCUUCGGGAAUACCCAGCGCCGCACCCCGGUUUUGGGAGAUGCGGCGGUCCCUGGCAGACUGGACU
Ssp-52-1    GCCGGCCGGCCGGCCUUCGGGAAUACCCAGCGCCGUACCCCGGUUUUGGGAGAUGCGGCGGUCCCUGGCAGACUGGACU
Ssp-13-1    UCGGUCGGGGGGUCCGUGGGGAAUACCCAGGGGUGUACCCCGGUUUUGGGAGAUGCGGCGGUCCCUGGCAGACUGGACU
Ssp-66-1    GCGAGCUCGGGGUGGCGUGGGAAUACCCGGGGCCGUACCCCGGUUUUGGGAGAUGCGGCGGUCCCUGGCAGACUGGACU
Ssp-63-1    GCCGGCCGGGCUGUCAUGGGGAAUACCCCGGGCCGUACCCCGGUUUUGGGAGAUGCGGCGGUCCCUGGCAGACUGGACU
Ssp-33-1    UCGGUCGGGAGGUCGGUGGGGAAUACCCAGGGGUGUACCCCGGUUUUGGGAGAUGCGGCGGUCCCUGGCAGACUGGACU
Ssp-117-1   GUCGGUCGGGGCGCGGUGGGGAAUACCCAGGGGUGUACCCCGGUUUUGGGAGAUGCGGCGGUCCCUGGCAGACUGGACU
Swa-1-1     UCGGUCGGGGGGUCGGUGGGGAAUACCCAGGGGUGUACCCCGGUUUUGGGAGAUGCGGCGGUCCCUGGCAGACUGGACU
Ssp-26-1    UCGGUCGGGGGGUCGGGGGGGAAUACCCAGGGGUGUACCCCGGUUUUGGGAGAUGCGGCGGUCCCUGGCAGACUGGACU
Sal-4-1     UACCCAGGGGUGUACCCCGGUUAUACCCAGGGGUGUACCCCGGUUUUGGGAGAUGCGGCGGUCCCUGGCAGACUGGACU
Ssp-123-1   GAAGGCCGCCGGCCCGGCCGGAAUAGAUCGGGCCCCACCCCGGUUUUGGGAGAUGCGGCCGGUCCUGGCAGACUGGUAC
Sgr-2-1     AAAUCACCUCCGGACCGCGGGAACAAACGACGGCGGCCCCCGGUUUUGGGAGAAGGCGCCAGUCCUGGCAGACUGGUAC
Sco-3-1     CUUUCGCGCGGAGCCCUCUGGAAUAGAACAAGCGCACCCCCGGUUUGGGUGGAUGGCCCCAGUCCUGGCAGACUGGUAC
Ssp-67-1    AGGGCGUCGUGAACACGGCGGAAUAGAUGACGGCCGCCCCCGGUUUUGGGGGAUGGCGCCAGUCCUGGCAGACUGGACC
Ssp-137-1   AGCCCCCACCGGCCCGCAGGGAAUACACAACGCGAACCCCCGGUUUUGGGUGAUGGCGCCAGUCCUGGCAGACUGGUAA
Slu-1-1     AGGGCGUCGUGAACGCGGGGGAAUAGCCGACGGCAACCCCCGGUUUGGGAAGAGGACACCAGUCCUGGCAAACUGGUAC
Sga-2-1     UCCGCCGGGUGUCGGGGCGGGAAUACCUGCGGGCGCCCCCUGGUUUUGGGAGAUGGCGCCAGUCCUGGCAGACUGGUAC
Sau-2-1     GGCCCCCGUCCCACCUCGGGGAAUAGAUCGCCGCGUCCCCCCGUUUUGGGGGAUACGGCCAGUGCUGGCAAACUGGUAC
Ssp-21-1    GGCGGUGUGCGGGGGCCGGGGAAUAGUUCCGGGCCCACCCCGGUUUUGGGAGAUACGGCCGGUCCUGGCAGACUGGUAC
Nvu-1-1     GUGUUUUGGGAGCGUAGCGACAGUUCUGGCAUACUGAACGGUCGUUGCGUCUGCUCACUCGAGCAGACCUCCCGCCUAA
Nbr-1-1     GUGUUUUGGGAGCGUAGCGACAGUUCUGGCAUACUGAACGGUCGUUGCGUCUGCUCACUCGAGCAGACCUCCCGCCUAA
Nte-1-1     GUGUUUUGGGAGCGUAGGGACUGUUCUGGCAUACUGAACGGUCGUUGCGCCUGCUUCGUCAGAGCAGGGCUCCCGCCUA
Spr-2-1     GCCCCCACCGGCCCGCAGGGGAAUAGACAACGCGAACCCCCGGUUUUGGGAGAUGGCUCCAGUGCUGGCAGACUGGACA
Ssp-106-1   AGCCACAACGCACCCGCGGGGAAUAGACAACAUCAACCCCCCGUUUUGGGAGAUGGCUCCAGUGCUGGCAGACUGGACA

            ................................................................................
            ................................................................................
            ●●●●●●●●●●●●●●●●●●●●●●●●●●●●●●●●●●●●●●●●●●●●●●●●●●●●●●●●●●●●●●●●●●●●●●●●●●●●●●●●
```

```
env-401     GCGAAGCCAAGGGCGUCGUGGAAUAGAUCGCGCUCACCCCCGGUUUUGGGGGAUGGCCCCAGUCCUGGCAGACUGGUAC
env-402     ......................AUAACGGACCCGAACCCCCGGUUUUGGGGGAUGGCCCCAGUCCUGGCAGACUGGUAC
Ssp-146-1   AGCCCCCACCGGCCUGCAGGGAACAAUCUGCAGUCGCCCCCGGUUUUGGGGGAUGGCCCCAGUCCUGGCAGACUGGUAC
Smi-1-1     GGCGUCGUGACCUCGUGAGGGAAUAGACCGGGGCACCCCCUCGUUUUGGGGGAUGCGGCCAGUGCUGGCAGACUGGUAC
Sda-1-1     CGCCGGUUAGGGGUACCCGGGAAUACCUCACCCGCACCCCCCGUUUUGGGGGAUGGCCCCAGUCCUGGCAGACUGGUAC
Ssp-161-1   GGCCGCCCGCCGGGGGCUGGGAAUAGAUCGCGGCGUCCCCUCGUUUUGGGGGAUGCGGCCAGUGCUGGCAGACUGGUAC
Ssp-71-1    GGCCGCCGGACCACCCCGGGGAAUAGAACAGCCGCACCCCCGGUUUUGGGGGAUGGCCCCAGUCCUGGCAGACUGGUAC
Sgr-7-1     ACGGCCGCGGGACCGCCCGGGAAUAGAACAACCGCACCCCCGGUUUGGGGGGAUGGCACCAGUCCUGGCAGACUGGUAC
Sgl-3-1     GGCCCGCGGGGAGCAGUGGGGAACAAACGACCGGCACCCCCGGUUUUGGGAGUAGGCCCCAGUCCUGGCAGACUGGUAC
Ssp-85-1    GCGCCCUCGCCGGCCCGCGGGAAUACCCGGCCACCGCCCCCGGUUUUGGGUGAUGACCCCAGUCCUGGCAGACUGGUGA
Ssp-121-1   GGCCCCCGCCCGCCCGGCCGGAAUAGAACAGCCGCACCCCCGGUUUGGGAGGAUGGCCCCAGUCCUGGCAGACUGGUAC
Ssp-104-1   UAGAGACGGAACCGGCGACGGAAUAGAUCGGAGCGGCCCCCGGUUUUGGGGGAUGGCCCCAGUCCUGGCAGACUGGUAC
Ssp-60-1    GCCUCCCGGGCCCGCCGGCGGAAUAGAACACCCGCACCCCCGGUUUUGGGAGAUGGCACCAGUCCUGGCAGACUGGUAC
Ssp-72-1    GCGGGUCCCGCCCGCGCCGGGAAUAGAUCGCCGAGCCCCCCGGUUUGGGAGGAUGCGGCCAGUCCUGGCAGACUGGUAC
Ssp-99-1    GACCGCGGGACGGACCGCGGGAAUAGAUCACCCCGCACCCCGGUUUUGGGGGAUGGCGCCAGUCCUGGCAGACUGGUAC
Ssp-89-1    AGCCACCCCGCGGGCCCGGGGAAUAGAACAGCCCCACCCCCGGUUUUGGGGGAUGGCGCCAGUCCUGGCAGACUGGUAC
Sbi-2-1     AACCCCUACUCACCUUCGGGGAACAAACAACGGCAACCCCCGGUUUUGGGUGAUGGCCCCAGUCCUGGCAGACUGGUAC
Ssp-82-1    ACCGCUGGCGGGAGCGCUGGGAAUAGAUCGCCGUGACCCCUCGUUUUGGGGGAUGCGGCCAGUGCUGGCAGACUGGUAC
Ssp-3-1     ACCGCUGCCGGGAGCGCUGGGAAUAGAUCACCGCGACCCCUCGUUUUGGGGGAUGCGGCCAGUGCUGGCAGACUGGUAC
Sam-1-1     GCCUCCGAGGGGCCUUCGGGGAAUAGACGGCCCCCGCCCCCGGUUUUGGGGGAUGGCCCCAGUCCUGGCAGACUGGUAC
Ssp-108-1   ACCCGCCGCCGGCCCGCGGGGAAUAGACGGCGCCAACCCCCGGUUUGGGGAGAUGGCACCAGUCCUGGCAGACUGGUUC
Ssp-48-1    CCGGUCCCGGGGCGGCCCGGGAAUAGAUCCUCCACCACCCCGGUUUUGGGGGAUGCGCCCAGUCCUGGCAGACUGGUAC
Ssp-124-1   GAAGCGCGAGGUGGCCCCGGGAAUCUUCCCGGGGCCACCCCCGUUUUGGGAGACGGCGCCAGUCCUGGCAGACUGGUAC
Ssp-110-1   ACCCGCCGCCGGCCCGCGGGGAAUAGACGGCGUCAACCCCCGGUUUGGGGAGAUGGCACCAGUCCUGGCAGACUGGUUC
Ssp-140-1   AACACCCACCGGCCCGCGGGGAACAAACGACGGCAACCCCCGGUUUUGGGAGUAGGCCCCAGUCCUGGCAGACUGGUAC
Ssp-107-1   GAAGCGCGGGGUGGCCCCGGGAAUCUUCCCGGGGCCACCCCCGUUUUGGGAGACGGCGCCAGUCCUGGCAGACUGGUAC
Ssp-112-1   AGGGCGUCGUGAACGCCGGGGAAUAGAUCCCGCCGACCCCCGGUUUGGGGAGAUGGCACCAGUCCUGGCAGACUGGUUC
Svi-5-1     GCCCCCGCUUCCGCCCCUGGGAAUAGAUCACACCGCCCCCCGGUUUUGGGGGAUGCGGUCAGUCCUGGCAGACUGGUAC
Ssp-74-1    AGCCGGCGCCGCCGCGCCGGGAACAAAAGGAGGCGGCCCCCGGUUUGGGGGGAUGGCACCAGUCCUGGCAGACUGGUAC
Sbo-1-1     CGGUGCGAGAAACCCGGUGGGAAUACCUGAGCCCACCCCCUGGUUUGGGUAGAUGGCACCAGUGCUGGCAGACUGGUAC
env-407     GCGGGGCCGCGGACCGCGCGGAAUAGACGGAGGCCGCCCCCGGUUUUGGGGGAUGGGCCCAGUCCUGGCAGACUGGUAC
Ssp-101-1   GGUCCCGGCGGGGCCGUCGGGAAUGCGGGGCCCCACCCCCCGGUUUUGGGAGAGGGCCCCAGUCCUGGCAGACUGGUCC
Ssp-136-1   CCGUGCCGGGGUGCGGCGGGGAAUACUCCGCACCGGCCCCCGGUUUGGGGAGAUGGCCCCAGUCCUGGCAGACUGGUCC
Sto-1-1     ACGCCGGCUAGGGGUGCGGGGAACAAAAGACGGCAACCCCCGGUUUUGGGUGAUGGCCCCAGUCCUGGCAGACUGGUAC
Sch-1-1     AGCCCCCACCGGCCCGCGGGGAACAAACAACGGCAACCCCCGGUUUUGGGGGAUGGCCCCAGUCCUGGCAGACUGGUAC
Saf-1-1     CGGCCGGUAGGAGGCGCGGGGAACAAACGACGGCAACCCCCGGUUUUGGGGGAUGGCCCCAGUCCUGGCAGACUGGUAC
Sva-1-1     ACCCCCACCGGCCCGCGGGGGAAUAGAUCACAGCAACCCCCGGUUUUGGGAGUAGGCCCCAGUCCUGGCAGACUGGUAC
Ssp-154-1   CACUCCCGUCGGCCCGCGGGGAACAAACAACGGCAACCCCCGGUUUUGGGGGAUGGCCCCAGUCCUGGCAGACUGGUAC
Sol-1-1     CACUCCCGUCGGCCCGCGGGGAACAAACAACGGCAACCCCCGGUUUUGGGGGAUGGCCCCAGUCCUGGCAGACUGGUAC
Ssp-147-1   CGACCAGGUGCCCCCGCCGGGAACAAACGACCGCACCCCCCGGUUUGGGAAGAAGGCGCCAGUCCUGGCAGACUGGUAC
Sia-1-1     AGCCACACACGGCCCGCAGGGAACAAACAACGGCAACCCCCGGUUUUGGGAGUAGGCCCCAGUCCUGGCAGACUGGUAC
Sro-1-1     GACCAUUCGCUCACCCGCAGGAAUAGAACAUCCGCACCCCCGGUUUGGGUGGAUGGCCCCAGUCCUGGCAGACUGGUAC
Scy-3-1     CCGCCCCCGCCCCGCCCCGGGAACAAAAGACGGCAACCCCCGGUUUUGGGUGACGGCCCCAGUCCUGGCAGACUGGUAC
Sac-1-1     AGCCGUACUCCCGCCCCCAGGAAUAGAACAACCGCACCCCCGGUUUGGGUGGAUGGCACCAGUCCUGGCAGACUGGUAC
Sla-3-1     ACGCCCCCGCGCACCCCCUGGAAUAGAACGACCGCACCCCCGGUUUGGGUGGAUGGCACCAGUCCUGGCAGACUGGUAC
Ssp-160-1   CCCACCGCUCACCCGAGCCGGAAUAGAACAACCGCACCCCCGGUUUGGGAAGAAGGCACCAGUCCUGGCAGACUGGUAC
Shy-2-1     CCCGUAACUCGCCUGAGCCGGAAUAGAACAACCGCACCCCCGGUUUUGGGGGAUGGCCCCAGUCCUGGCAGACUGGUAC
Ssp-125-1   GGGCGGGGCCGGCCCGCGGGGAAUAGAUCACAACAACCCCCGGUUUUGGGAGUAGGCCCCAGUCCUGGCAGACUGGUAC
Spe-1-1     ACCGCUUCGUGAGCUUCCGGGAAUAGAUCACGGGUCCCCCCGGUUUUGGGGGAUGGCCCCAGUCCUGGCAGACUGGUAC
Ssp-141-1   ACCCCCACCGGCCCGCGGGGGAAUAGAUCACCACAACCCCCGGUUUUGGGAGUAGGCCCCAGUCCUGGCAGACUGGUAC
Svi-14-1    AGCGCUUCGUGAGCUUCAGGGAAUAGAUCACGGGUACCCCCGGUUUUGGGGGAUGGCCCCAGUCCUGGCAGACUGGUAC
Svi-4-1     CCGCUCCACCGGCACGCGGGGAACAAACAACGGCAACCCCCGGUUUUGGGAGUAGGCCCCAGUCCUGGCAGACUGGUAC
env-408     GGGCCCCGGUCACGGCCGGGGAAUAGAUCACUUGUGCCCCCGGUUUUGGGGGAUGCGGCCAGUCCUGGCAGACUGGUAC
Sol-2-1     GCAGCGGGCAGCGGGCGGCGGAAUAGAUCGCGGCCGCCCCCGGUUUUGGAGGGUGGCCCCAGUCCUGGCAGACUGGUAC
Kau-1-2     AGGGCGUCGUCGACACCGGGGAAUAGUUCACGCCGCCCCCCGGUUUUGGGGGAUGGCGCCAGUCCUGGCAGACUGGUAC
Svi-12-1    AGGGCGUCGUCAACACCGGGGAAUAGAUCACCCCGCCCCCCGGUUUUGGGGGAUGGCGCCAGUCCUGGCAGACUGGUAC
Sco-1-1     AGGGCGUCGUGAACGCCGGGGAAUAGUUCACCCCCGCCCCCGGUUUUGGGGGAUGGCCCCAGUCCUGGCAGACUGGUAC
Ssp-58-1    GCGGCGGGCAGCGGGCGGCGGAAUAGAUCGCGGCCGCCCCCGGUUUUGGAGGGUGGCCCCAGUCCUGGCAGACUGGUAC
Sno-1-1     GCGCCGCCGCGGCGGGUCGGGAACAGAUCGGGCCCGCCCCCGGUUUUGGGGGAUACGGCCGGUACUGGCAAACUGGUAC
Sgr-1-1     GGGGGUCGUGGCGGGGACGGAAUAACCGGCCGCUACCCCCCGUUUUGGGGGAUGCGGCCAGUUGCUGGCAGACUGGCAC
Sce-1-1     AGCUCGUCCUGAGCCCGCGGGAAUAGACCACGGCGUCCCCCGGUUUUGGGGGAUGGCGCCAGUCCUGGCAGACUGGUAC
Ssp-176-1   AGGGCGUCGUCAACACCGGGGAAUAGUUCACCCCGCCCCCCGGUUUUGGGGGAUGGCACCAGUCCUGGCAGACUGGUAC
Ssp-118-1   CCCCCACCCCCGGUCGCCGGAAUAACUGGCCGCAACCCCCCGUUUUGGGGGAUGCGGCCAGUUGCUGGCAGACUGGCAC
Svi-11-2    AGCUCGUCCUGAGCCCGCGGGAAUAGACCACGGCGGCCCCCGGUUUUGGGGGAUGGCGCCAGUCCUGGCAGACUGGUAC
Sgr-12-7    GACCCUCGACGACCCACCGGGAACAAACGAGCGCACCCCCUGGUUUUGGGGGAUGGGCCCAGUCCUGGCAGACUGGUAC
Ssc-4-1     ACCCGGGGAGCGCGGGCCCGGAAUAGAUCGCCGGGUCCCCCGGUUUUGGGGGAUGCGGCCAGUACUGGCAGACUGGUAC
Ssp-69-1    AACCACGGACGACCUCGCGGGAAUACCUGACCGUCGCCCCCGGUUUUGGGGGAUGGCGCCAGUCCUGGCAGACUGGUAC
Ssp-153-1   AAACCCCACCGACCUGCGGGGAACAAAGAACGGCAGCCCCCGGUUUUGGGAGAGGACCCCAGUCCUGGCAGACUGGUGA
Sye-2-1     CCACCCCCCUACGGCCAGGGAAUACAAGACCCCGCCCCCCUGUUUUGGGGGAUGACACCAGUCGAUGGCAGACUGGUAC
Sse-1-1     GGGGGCCGCGCGGCCCCCGGGAAUAGAACAGCCCCACCCCCGGUUUUGGGGGAUGGCGCCAGUCCUGGCAGACUGGUAC
Sfu-2-1     AGCCCCCACCGGCCCACGGGGAAUAGAUCCCGGUGACCCCUCGUUUUGGGGGAUGGCCCCAGUCCUGGCAGACUGGUAC
Sdu-1-1     GGUAUGCGGCUCACCCGCGGGAAUAGAACAUCCGCACCCCUGGUUUGGGUGGAUGGCCCCAGUCCUGGCAGACUGGUAC
Sip-1-1     GGGGUCCGGGCGGAGCCGGGGAAUACGUGGGGGUGGCCCCUGGUUUUGGGAGAUGGCGCCAGUCCUGGCAGACUGGUAC
Ssc-3-1     CAGUCAUGGAUCACGGUGGGGAAUACCUGAGCCCACCCCCUGGUUUGGGCAGAUGGCACCAGUGCUGGCAGACUGGUAC
Ssp-59-1    AGCGCGUCGGCAGUCGUGGGAAUACCUGUGCGCACCCCCUGGUUUUGGGAGAUGACGCCAGUGAAUGGCAGACUGGUAC
Ssp-70-1    UGCGACCCCCGACGCCCCCGGAAUAAAGAUCCAAGGUCCCCGGUUUUGGGAGAUGCGGUCGGCCCUGGCAGACUGGACC
Ssp-24-1    AGGCGCCACACACCCCCAGGGAAUAGAAAAGGGACGCCCCCGGUUUUGGGGGAUGGCGCCAGUCCUGGCAGACUGGUAC
Ssp-10-1    CGGCAGGCCCCCGCCGCCGGGAAUACCCGGCGGUGACCCCCGGUUUUGGGAGAUACGGCCGGUCCUGGCAGACUGGUAC
Ssp-53-1    CGGCAGGCCGCCACCGCCGGGAAUACCCGGCGGUGACCCCCGGUUUUGGGAGAUACGGCCGGUCCUGGCAGACUGGUAC
Sni-2-1     CGGACGCGCCCCGUGGCGGCGGAAUAGAUCCGCGGCCACCCCGGUUUUGGAGAUACGGCCGGUCCUGGCAGACUGGAUC
Spr-1-1     AUACCCACUCCGGCCCCCAGGAAUACUUCAGCCCCCACCCCGGUUUUGGGAGAUACGGCCGGUCCUGGCAGACUGGUAC
Ssp-134-1   CGGGCCCCGCGCACCCGGCGGAAUAGAACGGCCGUACCCCCGGUUUGGGAGGAUGGCCCCAGUCCUGGCAGACUGGUAC
Sau-1-1     CCCGCUCCGGGCCGGGCCGGGAAUACUUCCGGGCCCACCCCGGUUUGGGUAGAUGCGGCCAGUCCUGGCAGACUGGUAC
Asp-7-1     ACCUGCGAAAACACCGCCUGGAAUAACUGAGCCCCCAACCCGGUUUUGGGAGAUGCGGCCAGUCCUGGCAAACUGGUAC

            ...............................................................................
            ...............................................................................
            ●●●●●●●●●●●●●●●●●●●●●●●●●●●●●●●●●●●●●●●●●●●●●●●●●●●●●●●●●●●●●●●●●●●●●●●●●●●●●●●
```

```
Sal-5-1     CCCCUCCGGGGUCGCUCCGGGAAUACUUCCGGGCCGACCCCGGUUUGGGCAGAUGCGGCCAGUCCUGGCAGACUGGUAC
Svi-2-1     GAACCGGGGCCGGAACGGCGGAAUACCCGGCGCCCCACCCCGGUUUUGGGAGAUACGGCCGGUCCUGGCAGACUGGUAC
Stu-1-1     CCCCCUCCCGCGCCCUCCGGGAAUAUCCAACCCCCGCCCCCGGUUUUGGGGGAUGCGCCCAGUCCUGGCAGACUGGUAC
Sso-1-1     GUGAUCCGGUGAAAACGGGGGAAUAGCCGCGGCCCCACCCCGGUUUUGGGAGAUACGGCCGGUCCUGGCAGACUGGUAC
Spu-1-1     GAAGUAGAGGCAAGCCGGUGGAAUAGAUCCGGCCACCCCCCGGUUUUGGGAGAUGCGGCCGGUCCUGGCAGACUGGUAC
Ssp-12-1    GCGGAGCAGACGCCGCCGUGGAAUAUCCGGGCCACCACCCCGGUUUUGGGAGAUACGGCCGGUCCUGGCAGACUGGUAC
Ssp-78-1    GGCCCGCGGGAGGGCUCCGGGAAUACUUCCGGGCCCACCCCGGUUUGGGCAGAUGCGGCCGGUCCUGGCAAACUGGUCC
Ssp-158-1   GCCCCGCACCGCCCCCGGUGGAAUACCCGGCGCCCCACCCCGGUUUUGGGAGAUACGGCCGGUCCUGGCAGACUGGUAC
Ssp-79-1    GGUGCCCGCCCCGGGGGCGGGAAUACCUGACGCCCGCCCCCGGUUUUGGGAGAUACGGCCGGUCCUGGCAGACUGGUAC
Ssp-92-1    GGUGCCCGCCCCGGGGGCGGGAAUACCUGACGCCCGCCCCCGGUUUUGGGAGAUACGGCCAGUCCUGGCAGACUGGUAC
Ssp-151-1   GUGCCCGCCCCGGGGGGCGGGAAUACCUGACGCCCGCCCCCGGUUUUGGGAGAUACGGCCAGUCCUGGCAGACUGGUAC
Sat-2-1     AACGUUCCCCGGACGCACGGGAAUACACACGACCGGGUCCUCGUUUUGGGAGAUACGGCCGGUCCUGGCAGACUGGUAC
Sfl-8-1     AACGCUUCCCGGCCCCACGGGAAUACACACGGCCGGGUCCUCGUUUUGGGAGAUACGGCCGGUCCUGGCAGACUGGUAC
Sfl-1-1     AACGCUCCCCGGCCCCACGGGAAUACACACGGCCGGGUCCUCGUUUUGGGAGAUACGGCCGGUCCUGGCAGACUGGUAC
Ssp-126-1   AACGUUCCCCGUACGCACGGGAAUACACACGGCCGGGUCCUCGUUUUGGGAGAUACGGCCGGUCCUGGCAGACUGGUAC
Sgr-11-1    GGUGCACGGCCGGGUGGCGGGAAUACCCGCGCCCCGCCCCCGGUUUUGGGAGAUACGGCCGGUCCUGGCAGACUGGUAC
Ssp-14-1    CACCGGGCGGAGGAGCCCGGGAAUACCUGCGGUCCGCCCCCGGUUUUGGGAGAUACGGCCGGUCCUGGCAGACUGGUAC
Ssp-168-1   CACCGGGCGGAGGGCUCCGGGAAUACCUGCGGUCCGCCCCCGGUUUUGGGAGAUACGGCCGGUCCUGGCAGACUGGUAC
Ssp-49-1    GGGAUGACCGGAGUGCCCGGGAAUAGUUCCGGGCCGACCCCGGUUUUGGGAGAUACGGCCGGUCCUGGCAGACUGGUAC
Ssp-22-1    AAGUGACCAGGCCGGUCCCGGAAUACCCGGCCGUCCUUCCUCGUUUUGGGAGAUACGGCCGGUCCUGGCAGACUGGUAC
Ssp-25-1    GCCCCCGGCCGUGUCCGGGGGAAUACCCGGCCGUCCACCCUGGUUUUGGGAGAUACGGCCGGUCCUGGCAGACUGGUAC
Ssp-152-1   AACGUGCCCGGACCGCCCGGGAAUAGUCACGCCCCCUUCCCCGUUUUGGGAGAUACGGCCGGUCCUGGCAGACUGGUAC
Ssp-57-1    AACGUACCCGGACAGUACGGGAAUAGUCACGUCCCCUUCCUCGUUUUGGGAGAUACGGCCGGUCCUGGCAGACUGGUAC
Ssp-40-1    AACGUACCCGGACAGUACGGGAAUAGUCACGUCCCCCUCCUCGUUUUGGGAGAUACGGCCGGUCCUGGCAGACUGGUCC
Ssp-6-1     CAGUGACCGGGACGGCCGGGGAAUACCCGGCCGUCCACCCCGGUUUUGGGAGAUACGGCCGGUCCUGGCAGACUGGUAC
Sha-2-1     AACGUACCCGGACAGUACGGGAAUAGUCACGUCCCCUUCCUCGUUUUGGGAGAUACGGCCGGUCCUGGCAGACUGGUAC
Ssp-43-1    UCCGUGCGCGGGCACCGGGGGAAUACCUGCGCUCCGCCCCCGGUUUUGGGAGAUACGGCCGGUCCUGGCAGACUGGUAC
Scy-2-1     ACCGGGCGGAAGGGGCCCGGGAAUACCUGCGGCCCGCCCCCGGUUUUGGGAGAUACGGCCGGUCCUGGCAGACUGGUGC
Ssp-39-1    GGCGGGCUGAGGCCCGCGGGGAAUACCUGCGGCCCGCCCCCGGUUUUGGGAGAUACGGCCGGUCCUGGCAGACUGGUAC
Sfu-1-1     CUGGGUGCUCAGGUGUUCGGGAAUACCUGCGGUCCGCCCCCGGUUUUGGGAGAUACGGCCGGUCCUGGCAGACUGGUAC
Seu-1-1     ACCCGUCACGCUCCGUCGGGGAAUACCUGCGCUCCGCCCCCGGUUUUGGGAGAUACGGCCGGUCCUGGCAGACUGGUAC
Sgr-3-1     CCGUGCGCGGCCGGUCGCGGGAAUACCCGCGCCCCGCCCCCGGUUUUGGGAGAUACGGCCGGUCCUGGCAGACUGGUAC
Ssp-9-1     UCCGUGCGCGGGCAACGGGGGAAUACCUGCGCCCCGCCCCCGGUUUUGGGAGAUACGGCCGGUCCUGGCAGACUGGUAC
Ssp-68-1    UCCGUGCGCGGGCACCGGGGGAAUACCUGCGCUCCGUACCCGGUUUUGGGAGAUACGGCCGGUCCUGGCAGACUGGUAC
Sal-8-2     UCCGUGCGCGGAGCGUAGGGGAAUACCUGGCUCCCGUCCCCGGUUUUGGGAGAUACGGCCGGUCCUGGCAGACUGGUAC
San-2-1     CGUGUGCGCGGGGCGUCGGGGAAUACCUGUGCUCCGUCCCCGGUUUUGGGAGAUACGGCCGGUCCUGGCAGACUGGUAC
Sod-1-1     GCGGGCGGCCCGGGAUCAAGGAAUAUCUCCGCCCGGCCCCCGGUUUUGGGAGAUGCGGCCGGUCCUGGCAGACUGGUAC
Ssp-149-1   CCCGGCCCGGAACCCGCCCGGAAUAGAACGGCACCUACCCCCGUUUUGGGAGAUACGGCCGGUCCUGGCAGACUGGUAC
Sfl-4-1     CAGAGCGGGUAGCGGGCGGGGAAUAGUUCCGGCCGGUCCCCGGUUUUGGGAGAUGCGGCCAGUGCUGGCAGACUGGUAC
Ssp-127-1   GAGCGUUCCGCAGUGCGCGGGAAUAGUUGCGGCACGCCCCCGGUUUUGGGAGAUACGGCCGGUCCUGGCAGACUGGUAC
Ssp-81-1    GAACGUUCCGGAACGCUCGGGAAUAGUUGCGGCGCGCCCCCGGUUUUGGGAGAUACGGCCGGUCCUGGCAGACUGGUAC
Sat-1-1     UGUGUUCCCGAACGAUCGGGAAUAGGUGCGGCCCGACCCCGGUUUUGGGAGAUACGGCCGGUCCCUGGCAGACUGGUAC
Ssp-103-1   AGGACGGCGCGCGCCGGGUGGAAUAUCUCCGGGCGUCCCCCCGUUUUGGGAGAUACGGCCGGUCCUGGCAGACUGGUAC
Nsp-3-1     CUGCCUGAACGACGGUCGCCGCGUGACCAGGACCCCCGAGGGAAUUGGUUCGCGGCGGCUGUUGUUCCUAUCAUUGCCC
env-409     .........................................................GCAUACUGGAUCGUCGACCACC
env-410     AGGGGCCCAGGAAUAGGGCCCGGCGCCGACGCGUUUGGGCAGGUGACGGUCAACCUGGCAUACUGGAUCGUCGACCACC
env-411     ......................CACGACUCAGGGACAACCUGAUUUACCACCGGACGGGGAUCAAUGGCAGACUAGACG
env-412     ACAAUGGUGGCCGGCUCGGUGACGAGACUCAGGGACAACCUGAUUUACCACCGGACGGGGAUCAAUGGCAGACUAGACG
env-413     ......................................................................ACACUGUCC
env-414     UGCGCAUCCGCGAUCCGGGGCCGGAAUAGGGACCGGCCUGCGCGGGUUGCAACGUCUGGCACACUGGUAGCCGAACCGC
env-415     .................................CGGCGUGGGUGGGUUGCAACGUCUGGCACACUGGUAGCCGAGCCGC
env-416     GGGCGAUCCGCGGAUCCGGGCCGGAAUACCGACCGGCCUGGGCGGGUUGCAACGUCUGGCACACUGGUAGCCGAGCCGC
env-417     GACUGACGUUCCGGAAUAGAUCACACGCCGCGGGCGUUCACCCGUGGCUAGAGGUCUGGCAUACUAGGUCAGUUCUUAU
env-418     ..........................................................CUGGCACUGGCCACACAUAUA
Svi-14-4    GGGGUCGUGAAGUAACCCGGGAAUACCUCCGGACCGACCCCCGUUUUGGGAGAUACGGCCGGUCCUGGCAGACUGGUAC
Sve-1-1     CGCCCCUGCGGGCAGCCCAGGAAUAGUUCCGGCCCCACCCCGGUUUUGGGAGAUACGGCCGGUCCUGGCAGACUGGUAC
Ssp-38-1    GGGGUCGUGAGGUAACCCGGGAAUACCUCCGGACCGACCCCCGUUUUGGGAGAUACGGCCGGUCCUGGCAGACUGGUAC
Sbi-3-1     GGCGUCGUGAGGUAGGCCGGGAAUACCUCCGGCCCGACCCCGGUUUGGGGAGAUGCGGCCGGUCCUGGCAGACUGGUAC
Sgl-4-2     UAGUCCGCCGCAGGGCCCGGGAAUACCUCCGGACCGGCCCCCGUUUUGGGAGAUGCGGCCGGUCCUGGCAGACUGGUAC
Svi-9-1     CGACCGGCGGACGAGCCCGGGAAUACCUCCGGGCCGACCCCCGUUUUGGGAGAUGCGGCCGGUCCUGGCAGACUGGUAC
Fsp-2-1     GAACACCGGGUCCCGCCGGCGGUGUUGUACCCGGCGGCGGGCCAGGUCCGGUCGGACGGGUCUGGCACACUGGCAAACC
Fsp-5-1     CACCAGGGCCCGGCGGCGGCGUUGCCGUCGGCGGCUGGCUGGUCCGGCCCGGGUCACCGGGUCUGGCACACUGGCGACA
Fsp-1-1     CACCAGGGCCCGGCGGCGGCGUUGUCGUCGGCGGCUGGCUGGUCCGGCCCGGGUCACCGGGUCUGGCACACUGGCGACA
Sgh-1-1     GACCGGCGCCCCACCGACGGGAAUAGAUCGCCCCCACCCCCGGUUUGGGGGAAGGCGCCAGUCCUGGCAGACUGGUAC
Ssp-31-1    CGGGCCGCCUCCGCCGCCGGGAACAAAUGGCCGCACCCCCCGGUUUGGGGAGAAGGCACCAGUCCUGGCAGACUGGUAC
Sav-2-1     AGGCCACGAAAAGCCCUCGGGAAUAGAUCGUCCCCCACCCCGGUUUUGGGGGAUGCGCCCAGUCCUGGCAGACUGGUAC
Ssp-7-1     ACCCUCCCGGCGCGCGCCGGGAAUACCUCGCGCCGCCCCCCGGUUUUGGGGGAUGGCGCCAGUGCUGGCAGACUGGUAC
Ssp-44-1    ACCCUCCCGGCGCCCGCCGGGAAUACCUCGCGCCGCCCCCCGGUUUUGGGGGAUGGCGCCAGUGCUGGCAGACUGGUAC
Sgr-10-1    GGACGCAGGGGUGGCCCCGGGAAUGAUCCCGGGGCCACCCCCGUUUUGGGAGACGGCGCCAGUCCUGGCAGACUGGUAC
Ssp-173-1   GACCGCGAGGUGGCCCCGGGAAUGUUUCCCGGGGCCACCCCCGUUUUGGGAGACGGCCCCAGUCCUGGCAGACUGGUAC
Ssp-16-1    GACCGCGAGGUGGCCCCGGGAAUGUUUCCCGGGGCCACCCCCGUUUUGGGAGACGGCCGCAGUCCUGGCAGACUGGUAC
Ssp-61-1    GGACCGCGGGGUGGCCCCGGGAAUGAUCCCGGGGCCACCCCCGUUUUGGGAGACGGCCCCAGUCCUGGCAGACUGGUAC
Ssp-175-1   GACCCCCGCCGGGGCGCUGGGAAUAGAUCGCCCCGCCUCCCGGUUUUGGGGGAUGCGGCCAGUGCUGGCAGACUGGUAC
Fsp-6-1     GUCGAGAGCUGCUGACCGGGUCCGGUCGGACGGGUCUGGCAGACUGGGAACACCAGCGACACACCAGCGUUACUGGAAC
Fsp-13-1    GGCUGGCUCGUGGGGACGUUGAGCGAGCUGGGAAAGUCACCGGCGGACGUGGCGUCCGCCUCUGGCAGACUGGUGGCAG
Fal-1-1     GGGCGGUCCGCAGCGGCGUUGACGUGGACGGCCAGCGUGCCGGCGGACGUGGCGUCCGCCUCUGGCAGACUGGUGGCAG
Gte-4-1     AGAGCGGCCUCACGGCUUACCAUGGUCAAUCGGUGCGCAGGUCGCGCACACGCCAAGCGCCCUGGAUUCCGGGGCCCGA
Mrh-1-1     GACCUACGCGGAAUAGCCGUUCGCGACGACGCGUUUGGGCUGGUGACGGUCCACCUGGCAUACUGGACCGUCGACCACC
env-419     GAACAAACGGUUCCGGGCCGGCGUUGAGCACCUGUUUGGUCACUGUCGACCCGUCCUGGCACACUGGGGCGGACAACCA
env-421     ...........................GGGUCGUUUUGGUCACUGUCGACCCGUCCUGGCACACUGGGGCGGACAACCA
env-422     GGGAGCGCCGGAAUACCUGGACGGGCGGGGUCGUUUUGGUCACUGUCGACCCGUCCUGGCACACUGGGGCGGACAACCA
env-423     CGGCUCCUAACGCUGUCACGGGGGAAUGCCAGGUCACGGAAAGUGGUUCCAUGAUCUGGCACACUAGAAGCCGCAACCA
env-424     ..........GGGGCGGAACGGGAAUCAGCAGGCCCUCCGGCAUGUUGCAGGGUCUGGCACACUAGUAGCCAUCAGCAG
env-425     CCCCGCGGCCGGGGCGGAACGGGAAUCAGCAGGCCCUCCGGCAUGUUGCAGGGUCUGGCACACUAGUAGCCAUCAGCAG

            ...............................................................................
            ...............................................................................
            ●●●●●●●●●●●●●●●●●●●●●●●●●●●●●●●●●●●●●●●●●●●●●●●●●●●●●●●●●●●●●●●●●●●●●●●●●●●●●●●
```

```
env-426    ..............................................UGCAGGGUCUGGCACACUAGUAGCCAUCAGCAG
Nsp-11-1   GCGCUGCCAUAAGGAUCGGGGAAUGGCCGGUGGCCCGGGCUGGUUGAACCAUCUGGCACACUGGUAGCCGACCGAUUUU
env-428    GCGGCGGCCGCGGCUCAGCCGGGAAUCACCAGGGGCCACGGAACGUUACAACGUCUGGCACACUAGGAAGCCGCGACCA
env-429    ....................................................................AGCCGCUACUA
env-430    GGCCAGGACAGGUAGCACCAGGGAAUGCCUGCCGGACGCACCACGUUACGGCGUCUGGCAGACUAUGAAGCCGCUACUA
env-431    ACUUAGUCCGGGGAAUGCGCGGCUGCCGGCGCGCGUUUAGGGGGAGAGCGGCUGACCUGGCAUAAUCGACAGCCGAAGU
env-432    ...........................GGCGCGCGUUUAGGGGGAGAGCGGCUGACCUGGCAUAAUCGACAGCCGAAGU
env-433    ACUUAGUCCGGGGAAUGCGCGGCUGCCUGCGUGCGUUUAGGGGGAGAGCGGCUGACCUGGCAUAAUCGACAGCCGAAGU
env-434    ACGCCCGCGCGGGAAUGCGUGGGCGACAGCGCGCGUUUAGGGACAUAGCGGCUGAGCUGGCAUAAUCAACAGCCGACGU
env-435    ACGGCCGCGCGGGAAUGCGCAGGCCACGGCGGGCGUUUAUGGACAUAGCGGCUGAGCUGGCAUAAUCGACAGCCGAAGU
Str-2-1    CGGGCCGGCGUUUGCGCUGCCAGCCCCGCAGCCCGCCUGCUGAGCCGACCUGACCCCGUGGCAGACUGGUUAGUCGGCC
Str-1-1    CGGGCCGGCGUUUGCGCUGCCAGCCCCGCAGCCCGCCUGCCGAGCCGACCUGACCCCGUGGCAGACUGGUUAGUCGGCC
Spa-24-1   CGACGUUUGCGCUGCCAGUGGCGCGGCGCGGCCCGAUUUCCGAGCCGAUCUGCCCCCGUGGCAGACUGGUUAGUCGGCC
Spa-32-1   CGACGUUUGCGCUGCCAGUGGCGCGGCGCGGCCCGCUUACCGAGCCGACCUGUCCCCGUGGCAGACUGGUCAGUCGGCC
Spa-7-1    CGACGUUUGCGCUGCCAGUGCCGCAGUAUGGCCCGCCCGCCGAGCCAACCUACCCCCGUGGCAGACUGGUCAGUCGGCC
Spa-27-1   CGACGUUUGCGCUGCCAGUGCCGCAGUAUGGCCCGCCCGCCGAGUCAACCUACCCCCGUGGCAGACUGGUCAGUCGGCC
Spa-5-1    AGACGUUUGCGCUGCCAGUGCCGCAGUAUGGCCCGCCCGCCGAGCCAACCUACCCCCGUGGCAGACUGGUCAGUCGGCC
Spa-38-1   CGACGUUUGCGCUGCCAGUGCCGCGGUUCGGCCCGCCUGCCGAGCCGACCUGCCCCCGUGGCAGACUGGUUAGUCGGCC
env-436    CGCGGCCGGGAAUGUGCAGCGCCCGUCCGGCGUUUUGGGGAUACGGCGGCUGACCUGGCAUAAUUGACCGCCGACCACC
Msp-11-1   AUCGCCUGAGGAAUAGGCCAGGGCGAUCGCGCGUUUGGGCAGGUGAUGGUCCACCUGGCAUACUGGACCGUCGACCACC
env-437    GGGACGAGAUGGGCGACAUCACGAUCGGCUGACGGCCACCCGCCUUGGUCGCGACCACCAGUCGAUGGCACAAUGGACG
env-438    GGUUAGCGCGGGGAAUGCGCGGCGGCCUGCGCGCGUUUAGGGGGAGAGCGGCUGACCUGGCAUAAUCGACAGCCGAAGU
env-440    ...............................GGGGACGGCGGAUUUCGUGCCGGAUGGCGGCGCGUGGCACACUUGUUU
env-441    CCGCCGGCCGCCCGGCCGGCGGAAUGCCAGGUCACGGAAAGUGGUUCCACGGUCUGGCACACUAGAAAGCCGCAACCA
env-442    CCUCACAGCCCGGCCGGGGCGGAAUGCCAGGUCACGGAAAGUGGUUCCAUGGUCUGGCACACUAGAAAGCCGCAACCA
Gkr-1-1    CGAUUGGUCUUCUCGCCGUGGCUUCUGGCAUAAUUGCCCGGUCAACCAGCGGUGGCUUCCGUCGAAGCCGCCACAACCG
env-443    CCGGCAGUCGGGAAUGUGCGCGGGUAUCCCCGCGUUUGGGCUCGAUGUGGUCGAGCUGGCAUAAUCAGCGGUCAACACC
env-446    UAACCCUCCGCACCCCGCGGUUGAGGCAUGCGUCGCAGGGCCGUUUUGGCAACUCAGGGCGGCCCUGGCAGACUGGUAG
env-447    ....................................................................CAGACUGACCC
env-448    GGUAGUGGCGACGAAUGACGGCAACGGAAUGUGCCGCUCAGCAUUGGCGUUGUAUGCGGCGUCAGUGACAGACUGACCC
env-449    GCAGUGGCGACGACUGACCGGUGGCGGAAUGCGCCGCCCGGCAUUGGCGUUGCAGGCAGCGUCAGUGACACACUGACCC
Mhe-1-1    ACUGAUCGAACCCGCGGAAUAGCGGGGGAGCCUCCCGGCGUUUUGGGAGCCGACAGCGUAGCUGGCAUAAUCGAACACC
Rrh-1-1    CGGGGCGCCGCAGCGUUGGGAAACACAGCAGGCGCGAUUGUCGGCCGUCGCCGCUGCUCUGGCAUACUGAACGGCCGAG
Msp-13-1   UCAGGCUCAGCCCCCAGGAAUGCGAACGCCGCUCCCGGCGUUUUGGGGGGCGACAGCGCAGCUGGCAUAAUCAACCACC
env-450    GCUCUGGGUGGAUCCCCAGACUAAGGCGGGGUGCCGGAGUCGGGAAUAAAUCCGGCUCUGGCAUACUUGAGCAGUUGAG
Tbi-1-1    GUAGCGAAUAAGCAGGAUCGAACAAGCGGUAUGCCGGAGUCGGGAACAAAAUCGGCUCUGGCAUACUUGAACAGUUGAG
Msp-4-1    GCGGGCCGGCCCCGCGGAAUGCGCACGACCCUCCCGGCGUUUUGGGGGAGCGCCAGCGCAGCUGGCAUAAUCGAACACC
Mav-1-1    ACUGACCCGGGCCACGGAAUAGCGGGGGAGGCUCCCGACGUUUUGGGAGCCGACAGCGCAGCUGGCAUAAUCGAACACC
Msp-16-1   GACUGACCCGCCCGCGGAAUGCGUGAGCCGCUCCCGCUGUUUUGGGGGAGUGGCAGCGUAGCUGGCAUAAUCGAAAACC
env-452    CUCCAAGACCACGCCGGGAAUGGACAACGACUGAUCGGCGUUUUGAGAGCCGACAGCGCAGCUGGCAUAAUCGCAAACC
Msp-12-1   ACGACUGACCGCUGCGGAAUGCGUGAGCCGCUCCCGCUGUUUUGGGGGAGUGGCAGCGUAGCUGGCAUAAUCGAAAACC
Msp-14-1   GACCCCGCUGACCGCAGGAAUAGCGGGGAGCCUCCCGGCGUUUUGGGAGCCGACAGCGCAGCUGGCAUAAUCGAACACC
Msp-17-1   GACCCCGCUGACCGCAGGAAUAGCGGGGAGCCUCCCGGCGUUUUGGGAGCCGACAGCGCAGCUGGCAUAAUCGAACACC
env-453    .....................................................................GUCGACCACC
Mky-1-1    ACCGCGUCCGGGAAUGCGCGCCGACGUCCCCGCGUUUGGGCUGACGGCGGCUGAGCUGGCAUAAUCAACCGCCAACCCC
Sar-43-1   CGAUGUUCCGGUUGUCGUAUCCGCCGUGCGGCCCGCCUGACGAGCCGACACGCGACCGUGGCACACUGGUCAAUCGGCC
Sar-11-1   CGAUGUUCCGGUUGUCGUAUCCGCCGUGCGGCCCGCCUGACGAGCCGACACGCGACCGUGGCACACUGGUCAAUCGGCC
env-454    ....................................GAUUUCAGAUCGGGACCUGUGACAUGGCAGACUGGUCCAUCGG
env-455    CGACACGCCGCGAAAAUGCCGUGAUCAUGGUCGGGAAGUUGCCGAAGUCCGUGGGUGUUGCACGUCUGGCACACUGGUU
env-456    CAGGGCGAGGAGUAGGGCCGGAGUUCGCCCAAGGAAUCAGCACCGCCCGUCCGGCGUUGCACCUCUGGCACACUGGACA
Ase-1-1    GGCAUAAGUGGCGGGGUGGGGGCGUUACCCAGGCGGUGUGGUUGUGGGAGCACGUGCCACACUGGUACGACGGCCCGUU
Fsp-9-1    GAUGGUCCGGGGGGGCGUUGAAACAGGCUGGGAAGCGUGCCGGCGGACGUGGCGUCCGCCUCUGGCAGACUGGCAGCAG
env-457    ........................................................................CUGUCAG
env-459    AUCGGUCGACCAUAGGGACCCCGGCCGGGAAUGCCGAGCACGGAUUUCGCGCUCUUCCCCGUGCCUGGCACACUGUCAG
env-460    CUGGAACUCAAGCUGGGGCGGGGGAAUGUCUUGUGGCCUGGGAUGGUUGGAGGAUCUGGCACACUGGUAGCCGAACCGU
Spa-33-1   CGGCGUUUGCGCUGCCAGUGCCGCGGCACGGCCCGCCUGCCGAGCCGACCUGUCCCCGUGGCAGACUGGUUAGUCGGCC
Vma-1-1    UGAUGUUGCUACCGGCGACCAGCUGGCGUGGCCCGAUUCCGGGGCCCGCCGAGGGCCAUGGCAGACUGGUCAAUCGGCC
Gbr-1-1    GGCUGGUGACCGAGGAGGAGAUGCACGGUCCGCACGAUUUCUGUUCGUGAGAUCGGGCGUGGCAUACUGAUCGGUCGAG
Gna-1-1    CGUGCCGUUGACCAACCUGCGACCGCUGCGUCCGCGAUUUCCGUCGCGAAGGGCACACAUGGCAUACUGAGCGGUCGAG
Gal-1-1    CCCGCUGUUGACCGACGUGCGACCGCUGCGUCCGCGAUUUCCGUCGCGAACGGCACACAUGGCAUACUGAGCGGUCGAG
Gam-2-1    CAUGCCGUUGACCAACCUGCGGCCGCCGCGUCCACGAUUUCCGUCGUGAACCGCGCACAUGGCAUACUGAGCGGUCGAG
Gsp-1-1    CCCGCCGUUGAGGCACCUGCGACCGCGGUGACCGGGAUUUUUCCCGGUGAUGCCCUACGUGGCAUACUGAUCGGUCGAG
Gpa-1-1    UGCUCUCUGCGACCGCUCGACGGGUGGGUCUCGGCGAUUUCCGUGCCCGAUACCCGACGUGGCAUACUGAGCGGUCGAG
Gru-1-1    CCCGCUGUUGACCGACGUGCGACCGCUGCGUCCGCGAUUUCCGUCGCGAACGGCACACAUGGCAUACUGAGCGGUCGAG
Gam-3-1    CAUGCCGUUGACCAACCUGCGGCCGCUGCGUCCACGAUUUCCGUCGUGAACGGCGCACAUGGCAUACUGAGCGGUCGAG
Gte-3-1    CCCGCCGUUGAGGCAUCUGCGACCGCGGUGAUCGGGAUUUUUCCCGGUGAUGCCCUCCGUGGCAUACUGAUCGGUCGAG
Nsp-1-1    CGCAAUGCAGGCCGUCAGCUCGAUCGCGCGAAGGCGAUUCGCAGGUCGGGAGGCUGUUCUGGCAUACUGAGCGGUCGAG
Wsp-1-1    GAUGUAUGCCCGCCCUCGAUCGCUUGAAUAUUAGCGAUUCGCUGACGUGGUGGAGGCUCUGGCAUACUGAACGGUCGAG
Kar-2-1    GAUGUAUGCCCGCCCUCGAUCGCUUGAAUAUUAGCGAUUCGCUGACGUAGUAGUGGCUCUGGCAUACUGAACGGUCGAG
Gef-1-1    GGUUGAGGCUGCUGAGAUCGCCCAGUGUGUCGACGGAUUGGUAGCAAUACCGCCCGUCGUGGCAUACUGAUCGGUCGAG
env-461    UGAGCCCCUCGAGACCGACGGAUGCGCGUGGUCAGGAUUUCAUUCCCGACCGACCAGCGUGGCAUACUGGUCGGCCGAG
Gpo-3-1    CCCUUCGGUGGGGCACGUCGUCACCGGUCACCGACGAUUUCGGUCCGGUGCGGUACGCGUGGCAUACUGAUCGGUCGAG
Grh-1-1    CCUCGUGGGGUUCGACCGAAGGCGCCGGUAUCGGCGAUUUCGGUGCUGAACGGCCCUCGUGGCAUACUGAGCGGUCGAG
env-462    GGAGUCCUGAGCGGGAACAACUUCGCCGCAGGCACGGUUAGGUCAAACGACAGCCACUCUGGCACACUGAGCGGCUGAG
Rde-1-1    CGGGAAUCAAUCGGGCCGGGGGCGCGUUUAUGAGGAUGCGUCCGCCUCUGGCAGAAUGGGCGAUCGCGUGACAAUCAUG
Req-3-1    CGGGAAUCAUUCGGCGCGGGGGCGCGUUUAUGAGGAAGCGUCCGCCUCUGGCAGAAUGGGCGAUCGCGUGACAACCAUG
env-463    GAAAUACAUAGCCAACUAUGUAUUUCCGCGCACAUGCGCGAAGCGCCGAGGGAACAAUCUGGCAUACUGAACGGCUGAG
Rfa-3-1    CGACAGGGGUUCGUGUUGAAGCCACUGUCGUGAGCGAUUAGCAGGUCGGAUCGCAUAUCUGGCAUACUGAACGGCCGAG
Rsp-2-1    GACAGGGGUUCGUGUUGAAGCCACUGUCGGUGAGCGAUUAGCAGGUCGGAUCGCAUAUCUGGCAUACUGAACGGCCGAG
Rsp-3-1    GCGUUGUAAGCAUCUGUCGGCACAGAUGUACCGGCGAUUAGCAGGUCAGAUGCACGCUCUGGCAUAAUCAACGGCCGAG
Rfa-12-1   GCGUUGUAAGCAUCUGUCGGCACAGAAGUACCGGCGAUUAGCAGGUCAGAUGCACGCUCUGGCAUAAUCAACGGCCGAG
Rsp-13-1   GCGUUGUAAGGAUCUGUCGGCACAGAAGUACCGGCGAUUAGCAGGUCAGAUGCACGCUCUGGCAUAAUCAACGGCCGAG
Rop-1-1    CCACCCGCGUUGACCACACCGCAAGCACAAAUGCCGAUUGGGAUCGCGAACCUGCGAUCUGGCAUACUGAGCGGCUGAG
Rfa-21-1   GCGUUGUAAGCAUCUGUCGGCACAGAAGUACCGACGAUUAGCAGGGCAGAUACACGCUCUGGCAUAAUCAGCGGCCGAG

           ...............................................................................
           ...............................................................................
           ●●●●●●●●●●●●●●●●●●●●●●●●●●●●●●●●●●●●●●●●●●●●●●●●●●●●●●●●●●●●●●●●●●●●●●●●●●●●●●●
```

```
Rfa-1-2    GCGUUGUAAGCAUCUGUCGGCACAGAAGUACCGACGAUUAGCAGGGCAGAUACACGCUCUGGCAUAAUCAACGGCCGAG
Rfa-4-1    GCGUUGUAAGCAUCUGUCGGCACAGAAGUACCGACGAUUAGCAGGGCGGAUGCACGCUCUGGCAUAAUCAACGGCCGAG
Rfa-7-1    GCGUUGUAAGCAUCUGUCGGCACAGAAGUACCGACGAUUAGCAGGGCGGAUGCAUGCUCUGGCAUAAUCAACGGCCGAG
Rfa-20-1   GCGUUGUAAGCAUCUGUCGGCACAGAAGUACCGACGAUUAGUAGGGCGGAUGCACGCUCUGGCAUAAUCAACGGCCGAG
Rfa-8-1    GCGUUGUAAGCAUCUGUCGGCGCAGAAGUACCGACGAUUAGCAGGGCGGAUGCACGCUCUGGCAUAAUCAACGGCCGAG
Rtr-1-1    CGUUUAGGAGAACGUCGGCAGCCUCUGGCAGAAUGAUCGACCGCGCAUCCGGCGCUCGCCGGGUGUUGGAAACACAAUG
Rsp-19-1   CGUUUAGGAGAACGUCGGCAGCCUCUGGCAGAAUGAACGACCGCGCAUCCGGCGUUCGCCGAAUGUUGGAAACUCAAUG
Rsp-21-1   UCGUUAGGACUUGAUGCACGCGUUCUGGCAGAAUGAACGAUCGCGCCCGUGGCACUCGCCGCGUGCGAGCACAACAAUG
Sar-21-1   CGAUGUUCCGGUUGUCGUAUCCGCCGUGUGGCCCGCCUGACGAGCCGACACGCGACCGUGGCACACUGGUCAGUCGGCC
Sar-40-1   CGAUGUUCCGGUUGUCGUAUCCGCCGUGUGGCCCGCCUGACGAGCCGACACGCGACCGUGGCACACUGGUCAAUCGGCC
Sar-27-1   CGAUGUUCCGGUUGUCGUAUCCGCCGUGUGGCCCGCCUGACGAGCCGACACGCGACCGUGGCACACUGGUUAAUCGGCC
Sar-33-1   CGAUGUUCCGGUUGUCGUAUCCGCCGUGCGGCCCGCCUGACGAGCCGACACGCGACCGUGGCACACUGGUCAAUCGGCC
Sar-26-1   CGAUGUUCCGGUUGUCGUAUCCGCCGUGCGGCCCGCCUGACGAGCCGACACGCGACCGUGGCACACUGGUCAAUCGGCC
Sar-22-1   CGGUGUUCCGGUUGUCGUAUCCGCCGUGCGGCCCGCCUGACGAGCCGACACGCGACCGUGGCACACUGGUCAGUCGGCC
Sar-17-1   CGAUGUUCCGGUUGUCGUAUCCGCCGUGUGGCCCGCCUGACGAGCCGACACGCGACCGUGGCACACUGGUCAAUCGGCC
Sar-24-1   CGAUGUUCCGGUUGUCGUAUCCGCCGUGUGGCCCGCCUGACGAGCCGACACGCGACCGUGACACACUGGUCAAUCGGCC
env-474    CAGACCGUGACGUCACGGAUGUGGUGCGCGGCCCGAUUUCGGACCGGCGUGGGCGCCGUGGCACACUGGUCAACCGGCC
env-475    ..................................................CACGCAUGGCACACUGGUCCAUCGGCCAA
Whe-1-1    GCGGCACCGAUUCGGUCCGCAGGCGUCGUUCUGGCAUACUCGUCACGGUUGUGCGCCACCCCGGUGCACAUCAUCACAG
Spa-28-1   CGACGUUUACGCUGCCAGUGCCGCGGCGCGGCCCGCCUGCCGAGCCGACCUGCCCCCGUGGCAGACUGGUUCGUCGGCC
Spa-36-1   CGACGUUUGCGCUGGCAGUGCCGCGGUGUGGCCCGCCUGCCGCGCCAACCCACCCCCGUGGCAGACUGGUUAGUCGGCC
Spa-8-1    CGACGUUUGCGCUGGCAGUGCCGCGGUAUGGCCCGCCUGCCGCGCCAACCCACCCCCGUGGCAGACUGGUUAGUCGGCC
Dau-1-1    AUGCCGGACCCCGGCGCGGCGUUGCAGACACCGGUUGAGGUGCCGUGACCCGCUCAUGGCAGACUUGACCGACGGCCAU
Mlu-1-1    ACGUGUUGACGAUGCCGGACGAGGUGUGCGGCCCGCUUGCCGGGUCGGCUCAGGCCCAUGGCACACUGGUCAAUCGGCC
Mch-6-1    ACGUGUUGACGAUGCCGGACGAGGUGUGCGGCCCGAUUACCGGGUCGGCUCAGGCCCAUGGCACACUGGUCAAUCGGCC
Mpa-1-1    ACGUGUUGACGAUGCCGGACGAGGUGUGCGGCCCGAUUACCGGGUCGGCUCAGGCCCAUGGCACACUGGUCAAUCGGCC
Rsp-9-1    GGGGCACUCGACGACUGACGUCGGCGCCCCGGGGAAUUUCUCCCCGGGGCCCUACGUUUGGCAUAAUCGACAGGCUCAG
Rrh-2-1    GGGGCACUCGACGACUGACGUCGGCGCCCGGGGGAAUUUCUUCCCGGGGCCCUACGUUUGGCAUAAUCGACAGGCUCAG
Rsp-14-1   GGGGCACUCGACGACUGACGUCGGCGCCCCGGGGAAUUUCCUCCCGGGGCCCUACGUUUGGCAUAAUCGGCAGGCUCAG
Rsp-5-1    CGCCGCGGCAGCGGCGCCGCAUGCCCGGGGGCGGAAUUUCUUCCCCCGGGCAUGCGUUUGGCAUAAUCGAACAGCUCAG
Rru-2-1    UCGCCGCGGCCGCGGCGGCGCAUGCCCGGGACGGAAUUUCUCGUCCCGGGCCUGCGUUUGGCAUAAUCGAACAGCUCAG
Rru-3-1    UCGCCGCGGCCGCGGCGGCGCAUGCCCGGGACGGAAUUUCUCGUCCCGGGCCUGCGUUUGGCAUAAUCGAACAGCUCAG
Rru-1-1    UGCCGCAGCCGCGGCGGGGCAUGCCCCGGGACGGAAUUUCUCGUCCCGGGCAUGCGUUUGGCAUAAUCGAACAGCUCAG
Ntr-1-1    UGUUUUGAGUGCGUGUCAGCGGUUCUGGCAUACUGGACCGCUGGUGCGUCUGCUGGUAGAAGGCAGACCAUCCCGCCUA
Not-2-1    CUGGCGCGCACGACCGGAGGGACCGGGAACAAGGCCCCGCGAGGCGGCGUUUAUACUGGUCCGUCGGCAACCAUGCCUA
Not-1-1    CCGGCGCGCACGACCGGAGGGACCGGGAACAAGGCCCCGCGAGGCGGCGUUUAUACUGGUCCGUCGGCAACCAUGCCUA
env-476    GCGGUGUCCGCCGCUCAGCUGGGAAUCUCCAGGGGGCACGGAACGUUACAACGUCUGGCACACUAGGAAGCCGCGACCA
Msp-26-1   GAGCGUUGCGCCUGCCGGACCUGCUGUGUGGGCCGCUCGUCGAGUCGGCGGACGCCCGUGGCACACUGGUCAAUCGGCC
env-477    GUUCCGGAGGCCGGGGGGCGGGGAAUUGCCAGGCCCCGGAGGAUGUUUGAUGGUCUGGCACACUAGUAAGCCGAAACAG
env-478    .....................................................................GAUACUUGUC
env-479    .......................................................................CACUUGUC
Gsp-3-1    UGUCUGAGAGCGGCCUCACGCCUUAAUAUUGAUGCCGGUGCGCAGGUCGCGCACCCACCAUGUGCCCAGACGGGCCCGA
env-481    GGUUGAGAGCGGCCUCACGCCUUAAUAUGGAGAGCUGGUGCGCAGGUCGCGCACCCACUGUGUGCCCGCGAGGGCCUGA
env-482    GCGCCGCCCGUCCAACGAUCGGGAAUGCCCGGUGACCUCGCCGCGUUCCGGUAUCUGGCACACUUGACUGUGCGACCGC
env-483    CCGGCCCGGCCGUCCAAGAUCGGGAAUGACCGGUGACCUCGCCACGUUCAGGUAUCUGGCACACUUGAGGGGCAACCGC
env-484    UCGGCAACUAACGCUGUCACAGGGAAUGCCAGGUCACGGAAAGUGGUUCCAUGGCCUGGCACACUAGAAGCCGCAACCA
env-485    CCGGCCGUUCCAGGCGUACCGGGAAUACCGGAUGGCCUGGACGGUUGACGGAUCUGGCACACUAGUUAGCCGAGCCCGC
env-486    CCGCUGGCCGCACUGGCCAGCGGGAAUUCCAGACCGGGGAAAAUGGUUCCACGGUCUGGCACACUAGAAGCCGCAAUCA
Sam-2-1    GGCCCCGGUCUGUCCAGGGCGGGGAAUGUCUUGUGGCCUGGGAUGGUUGCAGGAUCUGGCACACUGGUAGCCGAACCGC
Sro-7-1    ACAAACGCAGAUCACCUCAGCGGGAAUGUCUUGUGGCCUGGGAUGGUUGGAGCAUCUGGCACACUGGUAGCCGAACCGC
Sro-3-1    CAAACGCAGGUCAUCUCCGCUGGGAAUGUCUUGUGGCCUGGGAUGGUUGGAGCAUCUGGCACACUGGUAGCCGAACCGC
Mau-1-1    UGACGGUGGGCCGGCGUUGGGGGUGGCGGACCUCCUGUGCCGGGUGAAAGAUGCCCGUGGCAGACUGGUCAAUCGGCC
Msp-9-1    UGACGGUGGGCCGGCGUUGGGGGUGGCGGACCUCCUGUGCCGGGUGGAAGACGCCCGUGGCAGACUGGUCAAUCGGCC
Msp-10-1   UGACGGUGGGCCGGCGUUGGGGGUGGCGGACCUCCUGUGCCGGGUGGAAGAUGCCCGUGGCAGACUGGUCAAUCGGCC
Msp-28-1   .........GGCCGGCGUUGGGGGUGGCGGACCUCCUGUGCCGGGUGGAAGAUGCCCGUGGCAGACUGGUCAAUCGGCC
env-487    CAGACGCCACCAGUCCCCGGCGGGAAUCCAGAGCGGUCACGGAGUGUUUAAGCAUCUGGCACACUAGAGGCCGCGACCA
env-488    CACCCUGGGCAUGGACCGAGACCGCUAGCAGCAGGAAAUAUUAACCAGACGUAACUGGUUUGCGCGUCUGGCACACUAG
Msp-8-1    GCACGUUGACGAAGCCGGACCUGCUGUGCGGCCCGCCUGGGGGGUCGGCUCAUGCCCAUGGCACACUGGUCAAUCGGCC
env-489    GACCGAGACCGGUAACGACCGGGAAUUCGCACCCGUCAGACCUGGUUACCAGGUGUGGCAAACUAGGAAGCCGCUAGGA
env-490    GACCGAGACCGGUAACGACCGGGAAUUCGCACCCGUCAGACCUGGUUACCAGGUGUGGCAAACUAGGAAGCCGCUAGGA
Nco-1-1    CGGCGUGCGGGGCUGUGCGCGGGAAUGCGCGAUGGCCUGGGAUGGUUACACGAUCUGGCACACUGGUAGCUGACCAUUU
env-494    ACGGUCGGCANCUGAUCGCGANGGGAAUCGCCAGGCCCCCGCGCGCGUUGUGGGUCUGGCACACUAGUUAGCCGACAAG
Mca-1-1    GCACGUUGACGAAGCCGGACCUGCUGUGCGGCCCGCCCGAAGGGCCGGCUCACGCCCAUGGCACACUGGUCAAUCGGCC
env-495    GCCGGGCUGGGAGGCCCACCGGGAAUGCCGGCCGGGCACUUGUGGUUACAAGUGUCUGGCAGACUGAAAGCCGCACGUA
env-496    ACCACGCUCGGUUCCUGACGGGGAAUGCCAGGUCACGGAAAGUGGUUCCAUGAUCUGGCACACUAGAAAGCCGUAACCA
env-497    AGGCAACUGACGCCGAGGACGGGAAUGCCCGGCCGGGGAAUCCGGUUCCACGGUCUGGCACACUAGAAAGCCGCACCCA
env-498    GCGGCUGACGGUCCCGGACGGGAAUCACCAGGCCCCGCGGUACGUUGCAGGGUCUGGCACACUAGAAAGCCAUCAACAG
env-499    CGGCUGACGGUCCCGGGACGGGAAUCACCAGGCCCCGCAGUACGUUGCAGGGUCUGGCACACUAGAAAGCCAUCAACAG
env-500    CGGCUAACCAAACCACAGGCGGGAAUCACCAGGCCCCCGCGCAUGUUGCAGGGUCUGGCACACUGAAUAGCCAUAACAA
env-501    CAGCUAACCAGACCACAGGCGGGAAUCACCGGGCCCCCGCGCAUGUUGCAGGGCCUGGCACACUGAAUAGCCACCACAA
env-502    CCCGGCCGGCUGGCCAGGCAGCGGGAAUCGACGGGCCUGCCGGCAUGUUGACGUUGUGGCACACUGGUGUGCUGACAAG
env-503    GACCACCGUCGGUUCCUAAGGGGAAUCACCAGGCCCGGUAGCACGUUGCAGGGUCUGGCACACUAUUAGCCAUCAACAG
env-505    GACCACCGUCGGCUCUUAAAGGGAAUCACCAGGCCCGGUAGCACGUUGCAGGGUCUGGCACACUAUUAGCCAUCAACAG
env-506    CCAGCGAGCCGGGGCACAGCGGGAAUCACCAGGCCCGGUAGCACGUUGCAGGGUCUGGCACACUAUCAGCCAUCAACAG
env-508    AGACCACCGUCGGCUCUUAAGGGAAUCACCAGGCCCGGUAGCACGUUGCAGGGUCUGGCACACUAUUAGCCAUCAACAG
env-509    GACCACCGUCGGUUCUUAACGGGAAUCACCAGGCCCGGUAGCACGUUGCAGGGUCUGGCACACUAUUAGCCAUCAACAG
env-510    AGACCACCGUCGGCUCUUAAGGGAAUCACCAGGCCCGGUAGCACGUUGCAGGGUCUGGCACACUAUUAGCCAUCAACAG
env-513    ..................................CCUUCGGCAUGUUGCAGGGUCUGGCACACUGGUAGCCAUCAACAG
env-514    CGAGCGAGGCGAGCAGACUCGGGAAUGCGCGCUGACCCGUUUCGGUUAGCGGGUCUGGCAGUCUAGAUACCUGCGAAUA
env-515    GCCGCGCUCCGUGAAAGCCCGGGAAUGCCAGAUCGGCACAAGUGGUUUCACGGUCUGGCACACUAGAAAGCCGCAACCA
Nca-2-1    AGGCGCGUUUCAGGGGGCGUCGGAGGUUCUGGCAUACUGGACCGCCGUGCGUCUGCGAAUCGCGCAGACUCCCGCCUUA
Asu-1-1    CUACGCUGCCCCGGGAAUAGCGCGGGGCAGCGUGGAGUUGCGGUAAAGCAUGCUGCUUCUGGCAUACUUGAGCGGCUAC
env-516    GCCGACCAAGGCUGGCGCCGAAGGAAUUCCACAUGGUCAAAUCUCGUUCCACGGUUUGACACACUAGAAGCCGCAACCA
env-517    GCCGACCAAGGCNGGCGCCGAAGGAAUUCCACAUGGUCAAAUCUCGUUCCACGGUUUGACACACUAGAAGCCGCAACCA

           ...............................................................................
           ...............................................................................
           •••••••••••••••••••••••••••••••••••••••••••••••••••••••••••••••••••••••••••••••
```

```
env-518    UCGGCAACUAACGCUGUCACAGGGAAUGCCAGGUCACGGAAAGUGGUUCCUUGGUUUGGCACACGAGAAGCCGUAACCA
env-520    CCUCCGGAACCUGUGGCCACGGGGAAUUGUGAGAGGUCACGUCGUGUUACAACGUCUGGCACACUAGAAGCCGCGACCA
env-521    GGUGGGUCAGGACAGGCAAUAGGAAUAUCUGCCGGUCACGAGCGGUUUGCGCGUCUGGCACACUAGAAAGCCGCGACUA
env-522    .....................................................GCUAACGCUGUGCUAGACAACCACGG
env-523    ........................................UCGGGGCGUGUGACAUGGCAGACUGGUCCAUCGGCCAGC
env-524    UCGGCAACUAACGCUGUCACAGGGAAUGCCAGGUCACGGAAAGUGGUUCCAUGGCCUGGCACACUAGAAGCCGCAACCA
env-526    GCCCGCGGCCGGGGCGGGACGGGAAUCAGCAGGCCCUCCGGCAUGUUGCAGGGUCUGGCACACUAGUAGCCAUCAACAG
env-527    ...............................CGGGGCCCAGCGAGCGUUACAGUGGCUGGCACUGGACACCCACUUAUA
env-530    UCGCUUCUGGUGGGCUUUCCGAGGGAAUCCGCGGGGCCCAGCGAGCGUUACAGUGGCUGGCACUGGACACCCACUUAUA
env-531    UCGCUUCUGGUGGGCUUUCCGAGGGAAUCCGCGGGCCCCUGCGAGCGUUACAGUGGCUGGCACUGGACACCCACUUAUA
env-532    CAUGCCCCACCCACCGCGCCCUGGGAAUCCGCGGGCCCCUGCGAGCGUUACAGUGGCUGGCACUGGACACCCAUCUAUA
env-533    CGGCUUUUCGCGGGGAUUCCACGGGAAUCCGCAGGCCCCUGCGAGCGUUACAGUGACUGGCACUGGACACCCACUUAUA
env-534    CUCCUAACCGCCCGCUCACCAGGAAUGCCAGGUCAGCGAAUUUGGUUACACGGUCUGGCACACUAGAAAGCCGCAACCA
env-535    .........................................................................CAACCA
env-536    CUGCCCUGGCCCUCACCUAGCGGGAAUCUUCGGGCGGUGGUUCAUGUUCUGGGUAUGGCACACUGGUGUGCUUACCUAG
env-538    GCUGGAACUCAAGCUGGGGCGGGGAAUGUCUUGUGGCCUGGGAUGGUUGGAGGAUCUGGCACACUGGUAGCCGAACCGU
Sna-1-1    CACGUAGGUCCCAGCGCCGAUGGAAUAGCGCGGCGGGCUAAGAUGUUUCGUCGGCCGCAAGGGCCAAACCCAUACACGA
env-539    GCCGUCCCACCUGGGACGGCGGGAAUGCCAGGUCAGGGAAAUUCGUUUCACGGUCUGGCACACUAGAAGCCCACAACCA
env-540    GGCUAGCACAGCCCCGCCAGAGCUGGGAAUCUUCGCGGCCCACCGCAUGUUUCCCCUCUGGCACACUACGUAAGGCAUA
env-541    ACCACGGUCAGCACGUAGGACCGGGAAUCGUAUGGAGGCCACAGCACGUUGUCGCCUCUGGCACACUACGUAAGGCUGA
env-542    .............UCAGCGCCUAGGGAAUCUAUGGGGGCCACAGCACGUUGUUGGCUCUGGCACACUACGUAAGGCUGA
env-543    ..................................................GUCGCCUCUGGCACACUACGUAAGGCUGA
env-544    GACCACGGUCAGCACGUAGCACCGGGAAUCUAUGGAGGCCACAGCACGUUGUCGCCUCUGGCACACUACGUAAGGCUGA
env-546    .....................................GCCGGCGUUUGCGCGUCUGGCACACUGGAAGUCACAUCCCCA
env-547    ..................................GCCGCCGGCGUUUGCGCGUCUGGCACACUGGAAGUCACAUCCCCA
env-548    UCAUAGCCCCGCUAGAUGCGGGGAAUGCCAGGUCAGCCACAGGUGUUUCAGCAUCUGGCACACUAGAAAGCCGUAACCA
env-549    CCGCUGACCACUCCGGGCAGCGGGAAUGCGGGAUCAGGAAAACUGGUUCCACUGUCUGGCACACUAGAAGCCGCAACCA
env-551    CGCUGCCCCCAUCCAGGAAGCGGGAAUUCGGGACCAGGAAAACUGGUUCCACGGUCUGGCACACUAGAUGCCGCAACCA
env-552    ACCCGAAGACCCCGGACGGCGGGAAUGCCAGGUCACGGAAAGUGGUUCCAUGAUCUGGCACACUAGAAAGCCGCAACUA
env-553    ACGCUGAGCGGCCCCCGAGCGGGGAAUGCCCGGCGACCUCGGUACGUUGUCGCAUCUGGCACACUUGAAGUGCGACAUU
env-555    GCGGGUCGACCAGCACCUUCCCGGGGAAUCGCCACGGCGCCGCCGUGGUUGUAGAUGUGGCACACUGGUGCACCGAUAG
env-557    CCGCUGACCACACCGGACAGCGGGAAUGCGGGAUCAGGAAAACUGGUUCCACUGUCUGGCACACUAGAAGCCGCACCCA
env-558    CCGCUGCCCACGCCGGGCAGCGGGAAUGCGGGAUCAGGAAAACUGGUUCCACUGUCUGGCACACUAGAAGCCGCAACCA
env-560    CCCGCUGACCACUGGGACAGCGGGAAUGCGGGAUCAGGAAAACUGGUUUCACUGUCUGGCACACUAGAAGCCGCAACCA
env-561    CGCGCCCAGCCCCGGGCCGCGGGAAUGCCAGGUCAGUAAAACGAGUUUCACUGUCUGGCACACUAGAAAGCCGCAACCA
env-563    CGCGCCCGGCCCCGGGCCGCGGGAAUGCCAGGUCAGUAAAACGAGUUUCACUGUCUGGCACACUAGAAAGCCGCAACCA
env-564    CGCAGACGCCUGGUAGCGCCGGGAAUGCCAGACCCCGGAAUCUGGUUCCACUGUCUGGCACACUAGAAAGCCGCAAUCA
env-565    CCUGCUGACUCCCGGGACAGCGGGAAUGCGGGAUCAGGAAAACUGGUUCCACUGUCUGGCACACUAGAAGCCGCAACCA
env-568    CCCGCUGACCACUGCGACAGCGGGAAUGCGGGAUCAGGAAAACUGGUUCCACUGUCUGGCACACUAGAAGCCGCAAUCA
env-570    CCGCUGACCAGUCCGGACAGCGGGAAUGCGGGAUCAGGAAAACUGGUUCCACCGUCUGGCACACUAGAAGCCGCAACCA
env-571    GAGCCAGGCUGGCCCGGAGCGGGAAUGCCAGACCACGGAAUCUGGUUCCACUGUCUGGCACACUAGAAAGCCGAAACCA
env-572    UCGGCAACUGACGCUGUAAGAGGGAAUGCCAGGUCACGGAAAGUGGUUCCAUGAUCUGACACACUAAAAGCCGCAACUA
env-573    ...CGGCAACUGACGCUGUACAGGGAAUGGCAGGUCACGGAAAGUGGUUCCAUGAUCUGGCACACUAAAGCCGCAACUA
env-576    GCUCGGCUCCUAACGAUCACCAGGGAAUGCCAGGUCACGGAAAGUGGUUCCAUGGUUUGGCACGCCAAAGCCGCAACCA
env-577    GCCCGGAGACGAUGAGUGUCGUUGGCCGCGACCUGCUGUCGUCGCCGGCACCGGCCCGUGGCAGACUGGGCAAUCGGCC
env-578    UCGGCCGGUCGGCCGGAAGCGGGAAUUACCAGGGCCUGGAACCCGUUUCAGGAUCUGGCACACUGGUAUGCCGAUGAGA
env-579    GGCCCCAGCAGGGCCUGGCCGCGGGAAUCUGCGGGCCGCUGUGAGCGUUACAGCGGCUGGCACUGGACAUCCAUCUAUA
env-582    CCGCUGGCCGGUGCGGCCAGCGGGAAUUCCAGACCGGGAAAAGUGGUUCCACGGUCUGGCACACUAGAAGCCGCAAUCA
env-583    .......................................................................CGCAAUCA
env-584    CCCGCUGGCCGUGAGGCCAGCGGGAAUUCCAGACCGGGAAAAGUGGUUCCACGGUCUGGCACACUAGAAGCCGCAAUCA
env-585    ...................................GGGAAAAGUGGUUCCAUGGUCUGGCACACUAGAAGCCGCAAUCA
env-586    CCGCUGGCCGGUGUGGCCAGCGGGAAUUCCAGACCGGGAAAAGUGGUUCCACGGUCUGGCACACUAGAAGCCGCAAUCA
env-587    UCGGCUCCUGAGCCGAGACAGGGAAUGCCAGGCCGGGGAAUCUGGUUCCACGGUCUGGCACACUAGAAAGCCGCACCCA
env-588    UCGGCUCCUGAGCCGAGACAGGGAAUGCCAGGCCGGGGAAUCUGGUUCCACGGUCUGGCACACUAGAAAGCCGCACCCA
env-589    GAUCGGUAGCUCAGUAGCCGGAGACCGCCGGGGAAUGCGCCAGUUGGCCUCGGUGGUUAGCGAGUCUGGCACACUAUGA
env-591    ........GGUCACGGCCAGCGGGAAUUCCAGACCGGGAAAACUGGUUCCACGGUCUGGCACACUAGAAGCCGCAAUCA
env-592    ..................CCGGAGACCGCCGGGGAAUGCGCAGCCGGCCCCGGCGGUUGGCCGGUCUGGCACACUAUGA
env-593    GACGCGAUCGGGCCGGCUGGCGGGAAUCUCCGGACCGAACUUCACGUUCCCCAUGUGGCACACUUGUGUGCUGACCUAG
env-594    CGCUGGCCGGCCACGGCCAGCGGGAAUUCCAGACCGGGCAAACUGGUUCCACGGUCUGGCACACUAGAAGCCGCAAUCA
env-596    .......CGGCUACGGGCAGCGGGAAUUCCAGACCGGGCAAACUGGUUCCACGGUCUGGCACACUAGAAGCCGCAAUCA
env-597    CGCUGGCCGGUCACGGCCAGCGGGAAUUCCAGACCGGGCAAACUGGUUCCACGGUCUGGCACACUAGAAGCCGCAAUCA
env-598    CUAGGACUGCGUGAUUUCAGGGGGAAUGGCCAGGCACCGGAACACGUUCCAGUGUCUGGCACACUAGAGGCCGUAACUA
env-600    ...............................................CCCUCCGGCUGGCAGACUGAAACGGUACGAGA
Nbr-4-1    GUCCGUCACCGGCCGGCUGGAGCUCGGGCCGCGGAACGAGGCCCGAGACCGGCCCGAGCGAAGGCGGAUGUACGCCUGG
Npa-1-1    GUUCGUCACCGGCCGGUCGAGGCACGGAUCGCGAAGUGCGGUUCGGGCGCGGCCUCAGCGAGGGCGGAGGUACGCCUAG
env-601    ..................................................CUCGGGGUCCCGGAGCGUUCUGUCAGCGU
env-602    CAGGCCGUGCGAUGCCGGUGUUAUGACCUCGUUUUGGUCUGUCCGGCACCGGGCUGGCAGACUGGGACGUCGUAAACCA
Gam-1-1    GUGCCGACUGUGCCCGGCGGACCCGGUUGUCACCGAUUUCAUUCGGGAUGGCCUGUCGUGGCAUAGUGGCAAGGUUGAG
Gso-1-1    GGAGAGAUCGGACCGCUGGUCAGCGGGCUGACCGACGAUUUCGGGGUCGGAGCGGCGCGUGGCAUACUGAGCGGUCGAG
Gsh-1-1    CCGCACCGCGAGGAUCGCUCCGGAGGCGUGGGGGCGAUUGGGCACAGGACCCGAAACUGUGGAAUAAUCGGCCGGUCAG
Nsp-5-1    GCGUUUUGGGAGCGUGUCAGCGGUUCUGGCAUACUGAAUCGCUGGUGCGUCUGCCGAGAGCGGCAGGCCAUCCCGCCUA
env-603    ......................................................GCGACAUGGCAGACUGGUCCAUCGG
Ari-2-1    CGGCGGCUGAAGGGGACGCGGGGGAAUGCCCGGUGACCUCGGCACGUUCUGGUAUCUGGCACACUUGGAGUGCAACCGC
env-604    ...................................................................CACUGGUGGCGC
env-605    ...................................................................CACUGGUGGCGC
env-606    ...........................................................AUCCGUCACACUGGUGGCGC
env-607    ...............GGAACAGGGACGACGCGUAGCGCAUUGACGGACCAGCAGGACCCAUGCGUCACACUGGUGGCGC
env-608    CCGGAGCAGUUCGGCACCGUCACUUCACGGGUCGCCUCACCGCGGCAAGCGGGUCGGCGCCGAACUGCCCCGUCGCACA
Ata-1-1    CCCGCCGCGACGCGGGGGCGCGGGAAUACCCUCCGGUCGUGGCCCGUUGUGUGGUCUGCGAGAAUCAUUCCGCUGAAUG
Aha-1-1    ACCCCUGCGGGAACGGAAUACGUGGUGCGCACCCUGCGUUGCAGGUCAACGGUCUCGGUCUGGCAAAAUCGGGACCCGU
env-609    ......GCGAGGACUGACAGCCACCCGACGCCUGGGGAAUCCGCCGAGGCGUCGGAUGGUUGAAUGGACUGUCACGCAC
env-610    NNNNNNNNNNNNNNNNCAUGUCUCGGAGGCGGGCGUGGCUCGUCCGUUGUGCACUGCAGCGGCGGGAGGCAUAAUCGCC
env-611    UUCGGACGAUGUUGUGCAUGUCUCGGAGGCGGGCGUGGCUCGUCCGUUGUGCACUGCAGCGGCGGGAGGCAUAAUCGCC
env-629    UCUUGGGGAUGUUGUGCAUGUCUCGGAGGCGGGCGUUGCUCCUCCGUUGUGCACUGCAGCGGCGGGAGGCAUAAUCGCC

           ...............................................................................
           ...............................................................................
           ●●●●●●●●●●●●●●●●●●●●●●●●●●●●●●●●●●●●●●●●●●●●●●●●●●●●●●●●●●●●●●●●●●●●●●●●●●●●●●●
```

```
Pau-1-1     GACGACUGACCCGCGCCUCGACGCCCGGCCCCGGUGGGAAUCCCCGCCGGGGCCGGGUGGUUGAAUGGGUCCGUACGCA
Psp-6-1     GGACUAGGUCGCAGGCCACGCACCCGGGCCCCGGUGGGAAUUCCACCGGGGCCCGGGUGGUUGAAUGGUCCGUCAGCUU
Ses-1-1     CCACCAGGCACGGGAACAUCCCAUGACCCCUGGGCGUUGUCACCAGCGUCUACCGCAUCUGGCAGACUGUCAGGUCGAG
Ksp-2-1     GAGUGGUCGACGGGGAACAAACCGACCCACCGGCCUGUUUCAGCAGGUGUCGGGCCAUCUGGCAGAAUGGAUGGUCGAG
Sgl-1-1     CCGGUGUGGGGGAAAUAUCCGCCCGUAAGGGUGCGUUGAAAAGAUCGUCUGCCCCUCUGGCAGACUUGUUCCGCUGACG
env-630     ..........................................CUGAAAUACUGGAGAUGCUCAUGACGCUGCACACUGCA
Aba-1-1     CCUUCCGCGUCGGGAAUAGCUGGUCAGCGGGGUGCGUUCCAGUACCCGUCAGCUCGUCUGGCAGAAUUACCCGCUGAAG
Ari-1-1     AUGGUGGCCUUCGGAAUACCUGGUCAGCGGGGUGCGUUCCAGUACUCGUCUGCCCGUCUGGCAGAAUCACCCGCUGAAG
Ava-2-1     CCCUUCGCGGUGGGAAUAGCUGGUCAGCGGGGUGCGUUCCAGUACACGUCUGCCCGUCUGGCAGAAUUACCCGCUGAAG
Aal-2-1     CCUCCCCGGGCCGUCGAACGGCAGGUGGUCGGGCACAUGCCGACGUCAGGGGAGCGUCUGGCACACUGGUAGGUCGAAG
env-631     UAGCGAGCUCGACUGAGCCGUCCGUCGCCCUCCCCCGUGAGCGCGGGGGAGGGCGCGGUGGUUCAAUGGCAUGACACGG
env-633     AGAGCUCGACUGAGAAACCUCCCCACCCUCCCCCGCGGAAUUCGGGGGAGGGUGCGGUGGUUCAAUGAAGUGAACACGU
env-637     ......CGACUGAGAACCCUCCCCACCCUCCCCCGCGGAAUUCGGGGGAGGGUGCGGUGGUUCAAUGAAGUGAACACGU
Lae-1-1     CCCUCAAGGCCGUGCGGGAGCUCUGGGAACAACCGCGUUUUUGCACGUGUUGAAGGGUCUGGCAAACUAAACGGUCGAG
env-638     NNNNNNNNNNNNNNNNNNNNUCUCGAAGUCUGGCGAUGUUCCUCCGUUGUGCACUGCAGCGGCGGGCGGCAUAAUCGCC
env-639     CCUUGGGGAUGUUGUGCAUAUCUCGAAGUCUGGCGAUGUUCCUCCGUUGUGCACUGCAGCGGCGGGCGGCAUAAUCGCC
Aen-1-1     CGUUCGGGUUCGGGGAAUAGCCCGGUCGCGCCCCGCGUUGCAGCUCACGCCAGCCCGUCUGGCACACUGUCAGGUCGAG
Aha-3-1     GAGGAGUAGCGACGCGGCUCACGCGGGCCCUGCCAUGGUGCGGGGCCCGCGUGUGGCGUCUGGCAGAAUUGAGACGAAG
env-647     GCCCUGAUGAUGCGACGAAGGACCCGGGUUCGAUUCCCGGCACCUCCACCCUGACCGAUCUCAUGAGCUGCACACCGCA
env-648     AUUCCCGGCACCUCCACCACCGCCGCCCUACUCGACCAUGGCUCACGACAGCCAGGGCGAGCAGGGCGGCCUCGCGACA
Ksp-1-1     GGGCCAACGGCGGGAACAAUCCUCCAUGGGGAGGUGUUGAACUGGCUGGUCAGCGCGUCUGGCAAACUAGUAGGUCGAG
Lal-2-1     UGACUUAAGCCACCGAUCAGCCUCGGGAACAACCGCGUUUUUGCACGUGUUGAAGGGUCUGGCAAACUAAAUGGUCGAG
Lae-1-2     CCUGUGAGGUGCUGUGGAGGGCUCGGGAACAACCGCGUUUUUGCUGGUGUUGAAGCGUCUGGCAAACUAAAUGGUCGAG
env-649     ACUCCAGUGGCACGGAAUAUCCCCUGGCAGGGGCAUGUUGAAGGGAUUGCCAGCCCAUCUGGCAGACUGUUAGGUCGAG
Psp-9-1     AGGACUAGUCCGAGCCGCGCCGUCCGGGCCCCGGUGGGAAUUCCACCGGGGCCCGGAUGGUUGAAUGGACCGUCAGCUU
Aje-1-1     GUACAAGGCCCGGGAAUAGCUGGUCACCGGGGUGUGUUGAAGGCCCUGCCGGCUCGUCUGGCAGAAUGACCCGCUGAAG
[ir-1-1     ACGGUUUCCGCCGGGCACCCGGAAUACGGGUCCCGGCGGGGACGUUGUUGUGAGUCUGGCAGACUUGUUCCGCUACACG
Ain-1-1     ACGGCACCGGCCGGGAAUAGCCUGAUGGGGGUGGUCGUUGUACCGGCCGUCAGCGUGUCUGGCAGACUGUCAGGUCGAG
Aja-1-1     ACUCAACGUGUUGGAAUAUGCCGGAGGUGGGGUGCGUUGACGCCACUGCCAGCACAUCUGGCAAAAUCACCCGCUGAAG
Asp-6-1     ACUCAACGUGUUGGAAUAUGCGGGGGCGGGGGUGCGUUGACGCCACUGCCAGCACAUCUGGCAAAAUCACCCGCUGAAG
Aaz-1-1     ACUCAACGUGCUGGAAUAUACCGGGGGAGGGGUGCGUUGACGCCACUGCCAGCACAUCUGGCAAAAUCAUCCGCUGAAG
Aor-4-1     GCGGCCGGCGGCGGAAUACCUGGUCACCGGGGUGUGUUGAAGCCCUUGCCAGCACGUCUGGCAGAAUGACCCGCUGAAG
Ssp-91-1    GCGACCGGCGGCGGAAUACCUGGUCACCGGGGUGUGUUGAAGCCCUUGCCGGCACGUCUGGCAGAAUGACCCGCUGAAG
Abe-1-1     GCGAUCGGCGGCGGAAUACCUGGUCACCGGGGUGUGUUGAAGCCGGUGCCAGCCGGUCUGGCAGAAUGACCCGCUGAAG
Aor-1-1     ACUCAACGUGAUGGAAUAUGCCGGGGGAGGGGUGCGUUAACCCCACUGCCAGCGCAUCUGGCAAAAUCAUCCGCUGAAG
Aor-3-1     GCGAACGGCGGCGGAAUACCUGGUCACCGGGGUGUGUUGAAGCCCUUGCCAGCGCGUCUGGCAGAAUGACCCGCUGAAG
Ade-1-1     ACUCAACGUGCUGGAAUAUGCCGGGGGAGGGGUGCGUUGACGCCACUGCCAGCACAUCUGGCAAAAUCAUCCGCUGAAG
Ssp-5-1     GCCGCGUGCGGCGGAAUACCUGGUCACCGGGGUGUGUUGAAGACCUUGCCGGCACGUCUGGCAGAAUGACCCGCUGAAG
Aor-2-1     ACUCAACGUGAUGGAAUACGCCGGGGGAGGGGUGCGUUAACCCCACUGCCAGCGCAUCUGGCAAAAUCAUCCGCUGAAG
env-650     GCACCCCGCCCGUCGACAGCGGUACGGGGACCUGCCGGGAACAGUAGGCGGGNGNGGAUGGUUGAAUGGACUGUCACGC
env-651     GCACCCCGCCCGUCGACAGCGGUACGGGGACCUGCCGGGAACAGUAGGCGGGCGUGGAUGGUUGAAUGGACUGUCACGC
env-652     UAUCCAGCCCGAGCUGUUGUAAACAUCAGCUCAACGGGCGGGAACAUCGCGGCACCGCUGACGGUUUACACUGCUACGG
Sha-1-1     GAGGGUUUCCGCCGGGUACCCGGAAUACGGGUCCCGGCGGGAACGUUGGAUCCGUCUGGCAGACUUGUUCCGCUACACG
Spa-41-1    ACGGUUUCCGCCGGGUACCCGGAAUACGGGUCCCGGCGGGGACGUUGCUCCGAGUCUGGCAGACUUGUUCCGCUACACG
Kal-2-1     CGAGCGGUCGCCGGGAACAACCGGGUGACCUGGGGCGUUGCAGCCAGAGCCAGUCGAUCUGGCAGAAUUACCGGUCGAG
Saz-1-1     GGCUCGCCGGUGGAAUAGCCACCCGUCAGGGUGCGUUGAAGGACGCGUCUGUUGCUCUGGCAGACUUGUUCCGCUGACG
Svi-1-1     CCGGACGAGAACUGAGCGAGCGGGAACAAGGGCGCACGGAGGAGCGUUGGGCUCUUCUGGCACACUUGAUUCGCUGACG
Ssp-45-1    CGAAGCAGGCGGGAAUACGCGCUCUGCCGGGUGCGUUGCGACUGACGUCGGGGAUUCUGGCACACUUGUUCCGCUGACG
Kar-1-1     CAGGCCACCGCGGGAACAAACCCUCCCCAGGAGGUGUUGUACGAGCUGGUCAGAGCGUCUGGCACACUAGUAGGUCGAG
Ani-1-1     AGGUACGCGCGGGGAAUACCUGCUCACGGGGGUGCGUUGACGCUGGCGUCAGCCCGUCUGGCAGAAUCAUCCGCUGACG
Rrh-3-1     AGCGGGUGAGCGGGAACAAACCUCCCCGAGGAGGUGUUGUACGAGCUGGUCAGCGCGUCUGGCACACUAGUAGGUCGAG
Scy-1-1     UCGUGUCGUGAGGAAUACUCACCCACGAGGGUGCGUUGAAGGCUUCGUCUGUCCCUCUGGCAGACUUGUUCCGCUGACG
env-653     ...........................GGUCCGGGCCCCGGGGAACAACCCCGGGGGCCGGAUGGUUCACUGGAGCGUCA
Ame-3-1     CCUUCCGCGCCGGGAAUAGCUGGUCAGCGGGGUACGUUCCAGUACUCGUCAGCUCGUCUGGCAGAAUUACCCGCUGAAG
Ssp-46-1    ACACCAGCCACGGGGAACAACCGUGCCUGACCUCGCGUUGCAGGGGGUGACGCCCGAUCUGGCACACUGUCGGGUCGAG
Ami-2-1     GACGAGCGCCACGGGAACACCCGAGGAGGUGGACGCGUUGCACCGGGCAGCAGUGGAUCUGGCAGACUGUCAGGUCGAG
Ssp-1-1     CCGAAAGGCCCGCCGGAAUACCGGACGGGCCCCCUGCGUUGCAGGUCACAAGCCCUGGUCUGGCAGAAUGCAGGGUUGA
env-654     ..........GGCACGGCCACCUGCCGAUAGGGCGCGGGAACAUCGGCGGGGGCCGGAUGGUUCAAUGACACGUCACGC
env-656     CGUCGACAGCGGCACGGCCACCUGCCGAUAGGGCGCGGGAACAUCGGCGGGGGCCGGAUGGUUCAAUGACACGUCACGC
Sts-1-1     GCCGUGCCGGAAGCCCCCGGGAAUAGUUCCGCACCGACCCCGGUUUUGGGAGAUACGGCCGGUCCUGGCAGACUGGUUC
env-657     ..............................................................................GG
Rsp-11-1    CGUACAGCCCACUUCCUUCAGAAAGUUCAGACGCGGAUUACCGGCUCUACCACCACGUCUGGCAUACUGAACGGCCGAG
env-658     ACUUCGUACAGCCACCCCGAGGCGUGCGAGACGCGGAUUAUCCGGUGAGGUACCGCAUCUGGCAUACUGAACGGCCGAG
Rsp-12-1    ACUUCGUACAGCCACCCCGAGGCGUGCGAGACGCGGAUUAUCUGGUGAGGUACCGCAUCUGGCAUACUGAACGGCCGAG
Rrh-4-1     CGUACAGCCCACUUCCUUCAGAAAGUUCAGACGCGGAUUACCGGCUCUACCACCACGUCUGGCAUACUGAACGGCCGAG
Ace-1-1     CGGCGCUCCGGUGGCUGGUUCGGCAGCUGGCCGGCGUGCCGGCGGGCAGCCGUCUGGCAGACUGAGAGGCCGGAAGUUA
Gja-1-1     CCGGCGCCCGCGGUCGUCGCCAGCCAGAAGGUUGCGAUUUCCGUCUGGGCCUGGGGCGAUGGCAUACUGAUCGGUCGAG
Gsp-2-1     CCGGCGCCCGCGGUCGUCGCCAACCAGCAGGUUGCGAUUUCCGUCUGGGCCUGGGGCGAUGGCAUACUGAUCGGUCGAG
Got-1-1     ACGACGGGAGCGCCGCGACGUCAUCACGUAGUCGCGAUUUCCGUCAGGGCCUGGCCCGAUGGCAUACUGGUCGGUCGAG
Gai-1-1     GGUUUCCUCGGACAGUAUCGCGGCCGCGAGGCCACGAUUUCCGUGCGGACCCGGCGUGAUGGCAUACUGAUCGGUCGAG
Asp-5-2     CGGUGAGCGGUCGGAAUACUGCAUGGGCCCGGCAGGUUGUACUGACCGUCAGCAUCGUCUGGCACACUGUCAGGUCGAG
env-663     CUCCGGGGGCCGCUAGCUCGCCGUCGCGGUGGGAAUGCCGGGAUUCGCACCGGCGUUGGAUCCGGUGGCACACUUGUGC
Sma-1-1     CUGACCAGACGACCGACGACAGGAACAAGGGCGUGGGCGAGCGCGUUGAAAUGCUCUGGCAGAAUCGCCCGCGUUGACG
env-664     GCCCGGCGCCCGGCCGGCCGAAGACGGCAUCGCGACGAACAGCGUCAUACCCGCUUUUCCGUUCUGGCAUACUGAUGAG
Ssu-1-1     UGACGGCACGGGCCCGGGGGCUGUGCUCGGCGCGAGCACCCGGUUUUGGGGCACGCGGCACCAGGUGGCAAACUUGACC
Ssu-2-1     UGACGGCGCGGCUCCGAUGGCUGCGCCCGGCGCGAGCACCCGGUUUUGGGGCACGCGGCACCAGGUGGCAAACUUGACC
Sje-1-1     GCAGCCCCCGCGCCUAGCAGGAAUACCUGCCGGGCGUCCCCGGUUUUGGGAGAUGCGGCCAGUGCUGGCAGACUGGUAC
Sal-2-1     CGGUGAUCGUGUACACGUCGGAAUAUCCGCGGCCCGUUCCCGGUUUGGGGAGAUGCGGCCAGUGCUGGCAGACUGGUAC
Sru-1-1     AGGGGCGACCUCAGGCCCUGGAAUACCUGCGGGGGGUUCUCGGUUUGGGUGAAUGACGCCGGUCCUGGCAGACUGGUAC
Ssp-100-1   GGCGCCGGGGAGCACCGGGGGAAUACCGUCGUCCGGGCCCCGGUUUUGGGGGGUGCGGUCAGCAGUGGCAAACUUGACC
Ssp-129-1   GGCGCCGGGGAACACCGGGGGAAUACCGUCGUCCGGGCCCCGGUUUUGGGGGGUGCGGUCAGCAGUGGCAAACUUGACC
Ssp-171-1   CCUCCCGGGGGCGCGCACGGGAAUAUCGGGCGCCCGCCCCCGGUUUUGGGAGGUACGGCCAGUGAUGGCAGACUGGACC
Ssp-170-1   CCGGCGGCGCCGGCACGGCGGAAUAAUCACCGUCCGGCCCCGGUUUUGGGAGGUACGGCCGGUGAUGGCAGACUGGACC
Ssp-56-1    CGGGGUGUCGCGGGUCCGGGGAACAAACGGCUUGGCCCCCCGGUUUUGGGGGGUGCGGUCCCCAGUGGCAAACUGGACC

            ................................................................................
            ................................................................................
            ................................................................................
```

```
Kaz-1-1     GCUACCCCCUGAGGCGUGCGGAAUAACCUCGGACCCGCGUCAGUUUGGGAAGAAGCGGUCAGUGCUGGCACACUGGUCC
Svi-7-1     UGAGGUCCCCGGAACGGCGGAAUAACCCCCGGCCGGGACCUGUUUUGAUCCAUGCGUACCUGUACUGGCAAACUGGUGC
Smo-1-1     GCCCCCCGGAAAGAGCCUGGGAAUAACGGGCUCCCGGACCCCGUUUUGGGGAUGCGACCGGUACUGGCAGACUGGUAC
Ama-4-1     CGACAUCGAGAAGCUGCUGCCCGCGGAAUAGAUCCGCCCCCGAUUUCCUUGUAGCGGGUACCACCUGGCAGUAUUGAUC
env-666     UCGAGAAGCUGCUCCAGGCCUGAGGGAAUUUCCCAGCCCUCGAUUUCCUUGAAGAGGGUGCCAACUGGCACAAUCAAUC
Bsc-2-1     GGUUCAAGCCCCUGCGGCCCCACUUUCCGCAUGCGCGCCGGGCGCGGCGCAUGCCGUCCUCACCCCGGCGUACAAUCCC
Ssc-1-1     GGCGGAGGCGGGGCGGCGGGGAAUACCUGAGCCCACCCCCCUGGUUUGGGGAGAUGGCACCAGUGCUGGCAGACUGGUAC
Sle-1-1     CCGGUCCGUUCCCGCCCCGGGAACAAAAGACGGCAACCCCCGGUUUUGGGUGACGGCCCCAGUCCUGGCAGACUGGUAC
Scl-1-1     GCGGCGGCUGAGGGGCCGGGGAAUAGCCGGGCGGUCACCCCGGUUUUGGGAGAUACGGCCGGUCCUGGCAGACUGGUAC
Sre-3-1     AGCCCCCACCGGCCCACGGGGAAUACCUGACCCCCGCCCCCGGUUUUGGGGAUGGCCCCAGUCCUGGCAGACUGGUAC
Ser-2-1     CCGGGUGCGCCGGGAAUAUCCGGGGAGGGCCCCUGCGUUGCAGGUCAGCAGCCUUCGUCUGGCAGAAUGCAAGGUUGGU
Ssp-20-1    CGCCGCAGGCCGGCGCGGCGGAAUAACGGGCGUCGCCCCCCGGUUUUGGGGGGUGCGGCAACCAGUGGCAAACUGGUCC
Sst-1-1     CGGUGCGGGAAACCCGGUGGGAAUACCUGAGCCCACCCCCUGGUUUGGGGAGAUGGCACCAGUGCUGGCAGACUGGUAC
Mne-3-1     GCCGAUCGGCCGGGAAUAGUUGCCGCUGGUCACGCGUUUUCGCAACUGGCGGUUGACCUGGCAUAAUGGACCGUCGACC
            ................................................................................
            ................................................................................
            ●●●●●●●●●●●●●●●●●●●●●●●●●●●●●●●●●●●●●●●●●●●●●●●●●●●●●●●●●●●●●●●●●●●●●●●●●●●●●●●●
```

alignment positions 80 · · · 259

```
env-1       AUCGG.GUCCGGUUCCCGACC.UUCGA......................GCCAU.....................................................UCGAAGGGUCGAUCCGGCG.GCC..................ACGACGAGAAGGAGAACCC
env-2       CCCGG.CUCUGGUUCACAGG.............................CAUG..........................................................CCUGACCCAGCG.GCC...................AUCAACAAAGGGAUCCAU
Vca-1-1     UCCGG.UUCCGGUUCACCUUAUGUA........................ACC.........................................................UGCAUUUGGACCGGC..GCC...............CUGAGAACCUCAAGGAGAAACC
env-3       GCGAC.A.CCGGUUCACGGCGGCAUCC......................AUCCG.....................................................GGGUGCCGACGACCCGGC.UC.................AUCCGAACAGGAGAGCC..
env-4       CUUGG.UUCCGGUUCACCUCAACCCGCUGGUC.................AGCAG................................................GAUCGGCGGC..GAGGACCGGCG.GCC.....................UGAAAGGAAACC..
Cac-1-1     GUCGG.UCCGGUUCACCCCAC............................AGCA.........................................................GUGGAGACCCGGCG.GCC.................ACUCGACCUCAAGGACCGAA
env-7       GCUGGUUCCCGGUUCACGUCG.AGGUGUGC...................GGCUCA..............................................GCGCACCACCGACGACCCGGUG.ACC................GAGACCUCGAGAGGACACC..
env-8       UGUGG.CUCCGGUUCACGGCC..AGC.......................ACCCC...................................................GCUGAGGCCGACCCGGA..GCC...................UCGAAGGAGGAGAUC
env-9       CUUGG.CUCCGGUUCACGAC.............................GCGUGACCA......................................................GUCGACCCGGCG.GCC...............GUCCAUCUCGCGAGGAGACCCG
env-10      CUUGG.CUCCGGUUCACGAC.............................GCGUGACCA......................................................GUCGACCCGGCG.GCC...............GUCCAUCUCGCGAGGAGACCCG
env-11      CCGAG.CUCCGGUUCACGCUGCACCACCGCCCGCC..............GCAC...........................................GGCGGGCA......CGGCGACCCGGCG.GCU...............CCCCCGACCGAGGCACCCGC
env-13      CUUGG.UUCCGGUUCACGCCA.....A...GGAGC..............AGAUCCA........................................GCUCCGCAUCGAAAUGGCGACCCGGU..GCC...................CUCGAAACCAGGAGAAACC
env-14      CACGG.UACCGGUUCCCGUUGGUGCGC......................AAACUCC...................................................GCGC.CCAUCGACCCGGCG.ACC..................ACGAUGAAAGGAACUGCC
Apo-1-1     CUCGG.UCCCGGUUCACCCCCGAG.........................CUCCGGCGUGGGCACCUGC...........................................CCGGGGGACCCGGC..AUC...............CUCGAAUCCUUAAGGAGACACC
env-15      GUCGG.UUCCGGUUCACGUC..GCGC.......................AAGCCC...................................................GCGCUGGACGACCCGGG..ACU...............CUUGAGAGAUGAGGACAUC..
env-16      GACGG.UCCGGUUCACCCCCGAG..........................CACCACGCGUGGGCACCUGC...........................................GCCGGGGGACCCGGC..AUC...............CUCGAAUCCAUAAGGAGACACC
env-17      CCGAG.CUCCGGUUCACGGC.GCCGC.UGUGCA................GAUGG..........................................UGUGCAGGCGGCAGCCGACCCGGCG.GCU...............CUCCCGACCGAGAGGCAGCAC.
env-18      CAGG.CUCCGGUUCACGACGCGG..........................UUCCGAUGCAUUCAGCUUCGUGCUCGCCCUGCGCGAGC..............................CCGC.UCGACCCGGCG.GCC..................ACGACCCUCGAGGAGAUCCC
Ama-3-1     CUUGG.CUCUGGUUCACGG.UGCGC........................AGCC.......................................................GGCAGCCGACCCAGCG.GCC...................CCACGAGAGGACCACC.
Nin-1-1     UUCGG.UUCCGGUUCACGGUCGUGC........................GCUC.......................................................GCAC.ACCGACCCGGC..ACC..................CCUGAUGAGAGGACACC..
env-19      CUCGG.UCCCGGUUCACGCCGCACACUCCUUUCCGGUC...........CAUUC.........................................GGCCGG...GGGGAG...GGCGACCCGG.G.GCC................CCCGAUGGAAGAGGAAGCC..
env-20      CUCGG.UCCCGGUUCACGCCGCACACUCCUUUCCGGUC...........CAUUC.........................................GGCCGG...GGGGAG...GGCGACCCGG.G.GCC................CCCGAUGGAAGAGGAAGCC..
Jga-1-1     GUUGG.UUCCGGUUCACGGUUCGGCA.......................CCC.......................................................UGCCGCACCGACCCGGCG.GCC................CUGAAUGCCCAGGAGUCGC
Hcr-1-1     UGUGG.UUCCGGUUCCCGCC.C......CCUGG................UCU............................................CCAGGCUGACAGAGGCGAUCCGGCG.ACC...............GCGCCCCGGAGAGGAAAAAGG
env-21      CUUGG.UUCCGGUUCACCUCAGCGGC.......................ACCGC....................................................GCGUCUGGGGACCCGGCG.ACU.....................GGAAGGACACC..
Asu-2-1     ACUGG.UUCCGGUUCACCGCGCGGC........................AUGG.....................................................GCCCGGUGGACCCGGC..ACU.............CCUCGAAUCCCUUAAGGAGAACAC
Psp-8-1     UCCGG.CUCCGGUUCACCUUCGCCG........................CAGGAACGA.................................................CGCG.GGAGGACCCGGA..GCC..................ACAGAGAGAGGACACC..
env-22      CUCGG.UUCCGGUUCACGCCGCUGCC.......................UCCCAGGAC.................................................GGGAGG.GCGACCCGGG..GCC..................ACGAUCGAAGAGGAAACC..
env-24      CUCGG.UUCCGGUUCACGCCGCUGCC.......................UCCCAGGAC.................................................GGGAGG.GCGACCCGGG..GCC..................ACGAUCGAAGAGGAAACC..
env-25      CAUGG.UUCCGGUUCACCCGCC...........................GUCC.......................................................GGCGGGACCCGGCG.ACC.................GACAGACGAGGAGCAGCC
Sfl-5-1     GUCGG.CCCCGGUUCACGUUUGCCGCG......................CCC......................................................CGCGG.AAGCGACCCGGA..GCC...............CUCCCGAAACUAGGAGACACC
Jmu-1-1     GCUGG.CUCCGGUUCACGGUGCGACG.......................CCC.......................................................CGUCGUGCUGACCCGGCG.GCC...............CACGACACGCCAGGAGUGCGC
env-26      UCGAG.CACCGGUUCCCGCCCACCGC.......................GCAC.....................................................GC...GGGCGACCCGGCG.GCU...............CCCCGACCGAGGCACAGCC
Ssp-2-1     GUCGG.CUCCGGUUCACGUUGGUCGU.......................AUCCC....................................................GCGA.CAACGACCCGGC..GCC...............CUCCCGAAACUAGGAGACACC
env-27      CUUGG.UUCCGGUUCACCCCAUGACGCCCUCGC................GCC...........................................GCGACGGCA....GGGGACCCGGCG.GCC.....................CGAAAGGAACC..
env-28      GUCGG.UUCCGGUUCACGCCAGGG.........................GGUUCAUCCAGCCCCAUCAGUUUUGC......................................CAUCGGCGACCCGGA..GCC...................CGAGAACAGGAGAAACC
Cbo-1-1     UUCGGUCUGCGGUUCACGCCUGCG.........................AUCAG....................................................CGCAGGCGACCCGGAG.GCC...................CGUCGAUAGGAGAACCA
aLL-1-1     CUUGG.UCCUGGUUCACGACGCGC.........................GCC......................................................GCGCGACGACCCAGC..GCC.....................UCAGAGAGGACACC
Msp-29-1    GCUGG.CUCCGGUUCACGUACCC..........................ACAC......................................................GGUGCGACCCGGCG.GCU..................GUAACCCGAGAGGAAACC..
Mch-5-1     CUCGG.UUCCGGUUCACGCCGCAGCAUCC....................CGACUU..............................................GGGU...UGGCGACCCGGG..GCC..................CACGAUCGAAGAGGAAACC..
Aha-2-1     UUCUG.UUCCGGUUCACCGC.............................UCUG........................................................GCGGACCCGGCG.ACC...............UCGAAUUCCUUAAGGAGAUAAA
env-29      CUCGG.UUCCGGUUCACGCC......GC...GGGGC.............GCAUCCA........................................GCCCCUCAGCAAAGAUGGCGACCCGGU..GCC..................CACGAAACCAGGAGAAAAC
Sni-1-1     UCCGG.UUCCGGUUCACGUGAAGCG........................AAAGAG....................................................CG.UCACGACCCGGCG.ACC.....................AUGAAAGGACACC.
env-30      UCCGG.UUCCGGUUCACACUCGG..........................CAAUCCA...................................................CCGGGUGACCCGGCG.ACC.....................AUGAAAGGACACC.
Asp-2-1     AAGGG.UUCCGGUUCCCGCUCU...........................CACUCAUCAC...............................................AGAGCGACCCGGCG.GCC.................CGCAUCAUCGAGAGGACGCG
env-31      CACGG.UUCCGGUUCCCGC..CCGAG.CUGCCGCUCC............ACCG...........................................GGAGCGGCAGGUCAUGACAGCGACCCGGCG.ACC.................AGACGAUGAGAGGGACCGAC
Hal-1-1     UCCGG.UACCGGUUCACGGUUGGCGUCGCG...................CUCUGUU...............................................CGUGCCGUCGAUCGACCCGGCG.GCC.....................GAAGAAAGGCCCACA.
env-32      UCCGG.CACCGGUUCCCGUG.............................CACCCA........................................................CACGACCCGGCG.GCC...............ACUGAACCGAGAGGCACCACC
Ssp-169-1   GUCGG.UCCCGGUUCACGUCGCACC........................AGCA.......................................................GGUGGGACGACCCGGC..ACC...............CUCCCGAACCUAGGAGACACC
Dsp-2-1     GUCGG.UUCCGGUUCACGCC.......ACGGGGU...............CAAUCCA.........................................GCCCCACCAUCCCGGCGACCCGGC..CCC.....................CGUUAAGGAGAAACC
Mel-1-1     CUCGG.UUCCGGUUCACGCCUCACACCUUCACUG...............AAAU.............................................CGCUGAAGG.....GGCGACCCGGG..UCC..................ACGAUCGAAGAGGAAACC..
env-33      CGAGG.UUCCGGUUCACGG...CGCGCGCC...................CUCC..............................................GGCGCGCGGUCCGACCCGGCG.ACC...............GCCCCUCACGAGGCACAGCC
Car-1-1     AUCGG.UUCCGGUUCACGCUGCGCUCC......................GCCU.....................................................GGAG.GCCGCGACCCGGCG.ACC...............ACGAGAGAGAGGAACC...
env-34      GUUGG.CUCCGGUUCACGUCCGUCCC.......................AGGUCC...................................................GGGA.GGGCGACCCGGCG.GCC.................CCUAUGAGAGGAACCC..
Asp-3-1     CACGG.CUCCGGUUCACC.CCCUCGCCCGGCC...CCGGU.........AGC.......................................ACCGGGCUGGCCAGGC.AGGCCGGACCCGGCG.GCC...............ACGGACGAAGGGACACAC
env-35      ACGAG.CACCGGUUCACGUC.............................AUGUCC....................................................GACGACCCGGCG.GCU...............CCCCAACGAGAGGCACAGCC
env-36      ACGAG.CACCGGUUCACGUC.............................CCUAC.......................................................GACGACCCGGCG.GCU...............CCCCUGACGAGAGGCACAGCC
Mth-1-1     CUCGG.UUCCGGUUCACGUCUCACCGCAGGC.GC...............GCGA.............................................GCGGCCUGAACU..GGCGACCCGGC..ACC..................ACGAUCGAAGAGGACAUC..
env-37      C.UUGGUUCCGGUUCNCCCCGCCGACGUCG...................UGC....................................................CGACG.CGUCGGGGACCCGGCG.ACA.....................GAUAAGGAAAUUCC
env-38      CUUGG.UUCCGGUUCACCCCGCCGACGUCG...................UGC....................................................CGACG.CGUCGGGGACCCGGCG.ACA.....................GAUAAGGAAAUUCC

            ...<<.<.<<<<....<<<<<<<<<<<<<<<<<<<<<<<<<<<<<<<<<........................................>>>>>>>>>>>>>>>>>>>>>>>>>>>>>>>>>..>>>>...>>>..................................
            ...11.2.1220....022222222...........1.....1.111.10.......................................01.111.1.....1..........222222220..0221...211..................................
            ●YCGG-YYCCGGUUCACGYY∘∘∘∘∘---------------------●●●●∘∘-------------------------------------∘∘∘∘●GRCGACCCGGC∘-RCC---------------∘∘∘∘●●●R●●AGGA●●●●∘∘
            5′                                                                                                                      3′
```

```
env-39      GUCGG.CUCCGGUUCACGCU..CGCCCGU....................CCGAU...............................................................GCGGGUGCUGGCGACCGGCG.GCC.....................CUGAACGAGGAGAUGAG
env-40      GCUGG.UUCCGGUUCACGUCACGAGCGCC....................AUC.................................................................GGCGCAAC.GACGACCGGCG.ACC.................GAGACCCCGAGAGGACACC..
env-41      CACGA.CACCGGUUCACGGCGGCACCC......................ACCCG.................................................................GGGUGCCGGCGACCGGC..GUC....................AUCCGAAGAGGAGAGAC.
env-42      CACGA.CACCGGUUCACGGCGGCACCC......................AUCCG.................................................................GGGUGCCGUCGACCGGC..GUC....................AUCCGAAGAGGAGAGAC.
Ssp-96-1    GUCGG.CCCCGGUUCACGCACGUAGC.......................AAUUCG.................................................................GCU.CCUGCGACCGGC..GCC.................CUCCCGAAUCUAGGAGACACC
Sya-1-1     GUCGG.CCCCGGUUCACGCACGUAGC.......................AAUUCG.................................................................GCU.CCUGCGACCGGC..GCC.................CUCCCGAAUCUAGGAGACACC
Ssp-87-1    GUCGG.CCCCGGUUCACGCAAGCAGC.......................AAUUCG.................................................................GCU.CCUGCGACCGGC..GCC.................CUCCCGAAUCUAGGAGACACC
Ssp-122-1   GUCGG.CCCCGGUUCACGCAAGUAGC.......................AAUUCG.................................................................GCU.CCUGCGACCGGC..GCC.................CUCCCGAAUCUAGGAGACACC
Ssp-97-1    GUCGG.CCCCGGUUCACGCAAGUAGC.......................AAUUCG.................................................................GCU.CCUGCGACCGGC..GCC.................CUCCCGAAUCUAGGAGACACC
Ssp-148-1   GUCGG.CUCCGGUUCACGCCGUGGGC.......................AAUUCC.................................................................GCCCCCUGCGACCGGC..GCC.................CUCCCGAUACUAGGAGACACC
Ssp-88-1    GUCGG.CCCCGGUUCACGCAGUGCGC.......................AAUUCG.................................................................GCCC.CUGCGACCGGC..GCC.................CUCCCGAAUCUAGGAGACACC
Ssp-164-1   GUCGG.CCCCGGUUCACGCAGUGCGC.......................AAUUCG.................................................................GCUC.CUGCGACCGGC..GCC.................CUCCCGAAUCUAGGAGACACC
Svi-13-2    GUCGG.CCCCGGUUCACGCAGUGCGC.......................AAUUCG.................................................................GCUC.CUGCGACCGGC..GCC.................CUCCCGAAUCUAGGAGACACC
Ssp-28-1    GUCGG.CCCCGGUUCACGCAGUGCGC.......................AAUUCG.................................................................GCUC.CUGCGACCGGC..GCC.................CUCCCGAAUCUAGGAGACACC
Ssp-8-1     GUCGG.CCCCGGUUCACGCAGUGCGC.......................AAUUCG.................................................................GCUC.CUGCGACCGGC..GCC.................CUCCCGAAUCUAGGAGACACC
Ssp-143-1   GUCGG.CCCCGGUUCACGCAGUGCGC.......................AAUUCG.................................................................GCUC.CUGCGACCGGC..GCC.................CUCCCGAAUCUAGGAGACACC
Sla-2-1     GUCGG.CCCCGGUUCACGCAGUGCGC.......................AAUUCG.................................................................GCUC.CUGCGACCGGC..GCC.................CUCCCGAAUCUAGGAGACACC
Ska-1-1     GUCGG.CCCCGGUUCACGCAGUGCGC.......................AAUUCG.................................................................GCUC.CUGCGACCGGC..GCC.................CUCCCGAAUCUAGGAGACACC
Sro-6-1     GUCGG.CCCCGGUUCACGCAGUGCGC.......................AAUUCG.................................................................GCUC.CUGCGACCGGC..GCC.................CUCCCGAAUCUAGGAGACACC
Ssp-133-1   GUCGG.CCCCGGUUCACGCAGUGCGC.......................AAUUCG.................................................................GCUC.CUGCGACCGGC..GCC.................CUCCCGAAUCUAGGAGACACC
Sxa-1-1     GUCGG.CCCCGGUUCACGCAGUGCGC.......................AAUUCG.................................................................GCUC.CUGCGACCGGC..GCC.................CUCCCGAAUCUAGGAGACACC
Ska-1-2     GUCGG.CCCCGGUUCACGCAGUGCGC.......................AAUUCG.................................................................GCUC.CUGCGACCGGC..GCC.................CUCCCGAAUCUAGGAGACACC
Ssp-138-1   GUCGG.CCCCGGUUCACGCAGUGCGC.......................AAUUCG.................................................................GCUC.CUGCGACCGGC..GCC.................CUCCCGAAUCUAGGAGACACC
Ser-3-1     GUCGG.CCCCGGUUCACGCAGUGCGC.......................AAUUCG.................................................................GCUC.CUGCGACCGGC..GCC.................CUCCCGAAUCUAGGAGACACC
Sla-2-2     GUCGG.CUCCGGUUCACGCAGUGCGC.......................AAUUCG.................................................................GCUC.CUGCGACCGGC..GCC.................CUCCCGAAUCUAGGAGACACC
Ssp-131-1   GUCGG.CCCCGGUUCACGCAGUGCGC.......................AAUUCG.................................................................GCUC.CUGCGACCGGC..GCC.................CUCCCGAAUCUAGGAGACACC
Ssp-86-1    GUCGG.CCCCGGUUCACGCAGUGCGC.......................AAUUCG.................................................................GCUC.CUGCGACCGGC..GCC.................CUCCCGAAUCUAGGAGACACC
Sla-1-1     GUCGG.CCCCGGUUCACGCAGUACGC.......................AAUUCG.................................................................GCUC.CUGCGACCGGC..GCC.................CUCCCGAAUCUAGGAGACACC
Ssp-166-1   GUCGG.CCCCGGUUCACGCAGUACGC.......................AAUUCG.................................................................GCUC.CUGCGACCGGC..GCC.................CUCCCGAAUCUAGGAGACACC
Svi-13-3    GUCGG.CCCCGGUUCACGCAGUACGC.......................AAUUCG.................................................................GCUC.CUGCGACCGGC..GCC.................CUCCCGAAUCUAGGAGACACC
Dsp-1-1     GACGG.UUCUGGUUCACGGC.CCG.........................CACCU....................................................................CGGCGCCGACCAGCG.GCC..................AUGAUCGAAAAGGAGACAUC
Dho-1-1     GACGG.UUCUGGUUCACGGC.CCGC........................ACC.....................................................................GCGGCGCCGACCAGCG.GCC..................AUGAUCGAAAAGGAGACAUC
Bpa-1-1     GUCGG.UUCUGGUUCACGGCGC...........................CCG........................................................................GCACCGACCAGCG.ACC.....................CCGAGCAAGGAGAGAC.
Bph-1-1     GACGG.UUCUGGUUCACGGCGCCC.........................GCC......................................................................GGGCGUCGACCAGCG.ACC.....................CGUGAGAAGGAGACGAG
Bfa-1-1     GUCGG.UUCUGGUUCACGGUGCCC.........................GCC......................................................................GGGCGCCGACCAGCG.ACC.....................CCGAGCAAGGAGAGAC.
env-43      GUCGG.UUCUGGUUCACGGCGCCC.........................GUC......................................................................GGGCGCCGACCAGCG.ACC.....................CCGAGCAAGGAGAGAC.
env-45      GUCGG.UUCUGGUUCACGGCGCCC.........................GCC......................................................................GGGCGCCGACCAGCG.ACC.....................CCGAGCAAGGAGAGAC.
env-46      GUCGG.UUCUGGUUCACGGCGCCC.........................GCC......................................................................GGGCGCCGACCAGCG.ACC.....................CCGAGCAAGGAGAGAC.
Bmu-1-1     GUCGG.UUCUGGUUCACGGCGCCC.........................GCC......................................................................GGGUGCCGACCAGCG.ACC.......................CGAGAAGGAGAGAAG
Bsq-1-1     CUCGG.UUCUGGUUCACGGCACCC.........................GCC......................................................................GGGUGCCGACCAGCG.ACC.....................CGGACGAAGGAGAGAUU
Sba-1-1     GUCGG.CCCCGGUUCACGUGACGC.........................AACUCC...................................................................GUGUCCCGACCGGC..GCC.................CUCCCGAACCUAGGAGCGACC
env-47      GCUGG.UUCCGGUUCACGUC.CGU.........................CUUCAGA..................................................................GCGCGACGACCGGCG.ACC..................UGGAGCCUCGAGGAGAUAGA
env-48      UCUGG.UUCCGGUUCACCCGGCUC.........................GCGA.....................................................................GAGCAGGGACCGGCG.ACC.............CCCAGCGAGAGACUGAGGACACC..
env-49      CCCGG.CUCCGGUUCACGAUGCGGUACCGAUGCAUCCCCGCUCGC....CCUGC..............................................GCGAGC................CCGC.UCGACCGGCG.GCC...................ACGACCUCGAGGAGAUCCC
env-50      GUCGGCCCCCGGUUCACGUGAAC..........................GCCC......................................................................GUUCGCGACCGGCG.GCC......................CCGAAGAGGAGAUUCC
env-51      GUCGGUCCCCGGUUCACGUGAAC..........................GCCC......................................................................GUUCGCGACCGGCG.GCC......................CCGAAGAGGAGAUCGC
env-52      CCCGG.CUCCGGUUCACGAUGCG..........................GUACCGAUGCGUUCCAGCUUCGUGCUCGCCCUGCGCGAGCC..................................CGC.UCGACCGGCG.GCC..................ACGACCCUCGAGGAGAUCCC
env-53      CGCGG.UUCCGGUUCACGUCAUCAAGCGCCAGCGCCU............CAGUGA......................................................GGGCGUCCGCGC.....GACGACCGGCG.GCC.................GCUUGAAGGCAAGGAGCCAGC
env-54      UGCGG.UACCGGUUCCCGGCGGGC.........................GUCAG....................................................................GCUCGCCGACCGGCG.GCC...................GCCAGAACCAGGAGUUAGU
env-55      UCCGG.CUCCGGUUCACGCUCCGCCC.......................ACAU...................................................................GGG.GGAACGACCGGCG.GCC.....................AUCAGAGAGGAUCACC.
env-56      CGCAG.CUCCGGUUCACGUCACAAAGUGCCC..GC...CCC........GUCA....................................................GGGUGAGCGAGGGUAC.....GACGACCGGCG.GCC..................GCUUGAGGCAAGGAGCCAGA
env-57      GCUGG.UUCCGGUUCACGUCGCUGCG.......................CCCG...................................................................CGCAGC.ACGACCGGCG.ACC.................GAGGCCCCGAGAGGACACC..
env-59      GCUGG.UUCCGGUUCACGUCGCUGCG.......................CCCG...................................................................CGCAGC.ACGACCGGCG.ACC.................GAGGCCCCGAGAGGACACC..
Spo-1-1     CCCGG.UUCCGGUUCACGCC.CCGU........................GACCA..................................................................GCGGUGGCGACCGGA..GCC...................AUGAGAAUCAGGAGCGCGU
env-60      UUCGG.UACCGGUUCACGUC.CCGCCC......................ACGG..................................................................GGGCGGCGACGACCGGCG..CC......................UCUGAAAGGACAACC.
env-61      UCGAG.CUCCGGUUCACGUCUGCGC........................AGUC....................................................................GCGCCGACGACCGGCG.GCC.................GCAGCACCGAGAGGAAUCCAA
Ssp-51-1    GUCGG.UUCCGGUUCACGUGGGGC.........................AAAUCC.................................................................GCCCCGCGACCGGC..GCC.................CUCCCGAACCUAGGAGACACC
Sji-1-1     GUCGG.UCCCGGUUCACGGA.CGGC........................AGUCC..................................................................GCCCUUCCGACCGGA..ACC...............CUCCCGCAAACCUAGGAGCAUCC
Aut-1-1     AAGGG.UUCCGGUUCACGCCGCUCC........................AGCA....................................................................GGA.CGGCGACCGGCG.GCC................CGACCAUUCGAGAGGGACCGAG
Mtu-1-1     CUCGG.UUCCGGUUCACGCCGACACCUCC.UGGUCC.............CAAGU.......................................................GGUCCUCGGAGG....GGCGACCGG...GCC.................CACGAUUGAAGAGGAAACC..
Cko-1-1     CCUGG.UACUGGUUCACGCC.............................GCGACAA.......................................................................GGCGACCAGCG.ACC..................GCUUUCUGAGAGGACCCAAG
Nla-1-1     CCAGG.CUCCGGUUCACGGUGCCCGCUUCGAGAGUCGGCCGA.......UCCG...................................................UCGG.CGGCUCA.GCGGCGAGCAGCGACCGGCG.GCC.....................AUCAGAGAGGAACCAGC
Nsp-9-1     GUCGG.CUCCGGUUCACGCCUC.CUC.......................GCAGG..................................................................GAGCGAGGCGACCGGC..GCA.......................UCGAGAGGACACGCA
env-62      CUUGG.CACCGGUUCACGUCACCCGC.......................ACUCA..................................................................GCGGA.GGCGACCGGCG.GCC.......................GUGAAAGGAAACC..
Afr-1-1     AAAGG.UUCCGGUUCACGCU.CGAGU..CG.UCCGCCCU..........AGC.......................................................GGGGCGGUUCGGUGCUGACGGCGACCGGCG.ACC.................ACAUCAUCGAGAGGGACCGAG
env-63      AAAGG.UUCCGGUUCACGCU.CGAGU..CG.UCCGCCCU..........AGC.......................................................GGGGCGGUUCGGUGCUGACGGCGACCGGCG.ACC..................CAUCAUCGAGAGGGACCGAG
env-64      AAAGG.UUCCGGUUCACGC..CCG.AGCC.CACCGCCCU..........GGC.......................................................GGGGCGGCGCGGCGCUGACAGCGACCGGCG.ACC................AGAUCAUCGAGA.GGGACCGAG
Asu-3-1     AAAGG.UUCCGGUUCACGC.CCGAGC.CC..ACCGCCCU..........GGC.......................................................GGGGCGGCGCGGUGCUGACAGCGACCGGCG.ACC.................AGAUCAUCGAGAGGGACCGAG
env-65      GCUGG.CUCCGGUUCACGCACCUC.........................CGCU.....................................................................GAGGUGCGACCGGCG.GCC.................GAGAACCCGAGAGGAACCC..
env-66      GCUGG.UUCCGGUUCACGUCCUAGCCU...GCG.GGCCG..........CCC.......................................................CGGCCACGCGCGAGGCGACGACGACCGGCG.ACC.................CAAGCCCCGAGAGGAAGCC..
env-67      UCAGG.UUCCGGUUCACGUCAUC..........................GCC........................................................................GAUGACGACCGGC..GCC...................GCGUCGGAGAGGACCAUC.
env-68      UCAGG.UUCCGGUUCACGUCAUC..........................GCC........................................................................GAUGACGACCGGC..GCC...................GCUUCGGAGAGGACCAUC.
env-69      UCAGG.UUCCGGUUCACGUCAUC..........................GCC........................................................................GAUGACGACCGGC..GCC...................GCGUCGGAGAGGACCAUC.
env-70      UCUGG.CCCCGGUUCACGUCAUC..........................GCC........................................................................GAUGACGACCGGCG.ACC...................GCUCCGGAGAGGACACC..
env-73      UCUGG.CCCCGGUUCACGUCAUC..........................GCC........................................................................GAUGACGACCGGCG.ACC...................GCUCCGGAGAGGACACC..
env-74      UCAGG.CCCCGGUUCACGUCAUC..........................GCC........................................................................GAUGACGACCGGCG.ACC....................GCUCAGAGAGGACACC..
env-75      UCAGG.CCCCGGUUCACGUCAUC..........................GCC........................................................................GAUGACGACCGGCG.ACC....................GCUCAGAGAGGACACC..
env-78      UCAGG.CCCCGGUUCACGUCAUC..........................GCC........................................................................GAUGACGACCGGCG.ACC...................GCUUCGGAGAGGACAUC..
env-80      UCAGG.UCCCGGUUCACGUCAUC..........................GCC........................................................................GAUGACGACCGGCG.ACC...................AGCUCAGAGAGGACAUC..
Ncy-1-1     GUCGG.CUCCGGUUCACGUUCGACGC.......................UGUCC..................................................................GCG.CGAGCGACCGGC..GCC...................AUUUUCGAGAGGACACCC.
Ncy-2-1     GUCGG.CUCCGGUUCACGUUCGAUGC.......................CAUCC..................................................................GCG.CGAGCGACCGGC..GCC...................AUUUUCGAGAGGACACCC.
Npn-1-1     GUCGG.UUCCGGUUCACGCUCGUCGCG......................AGUU..................................................................CGCG.UGGGCGACCGGCG.GCC..................AUAUCUCGAGAGGACAUCC.
env-81      CGACG.GUCCGGUUCACGCCGUC..........................AGCA......................................................................GACGGCGACCGGCG.CCC..................CCGUAUCGAGAGGACCCGC.

            ...<<.<.<<<<....<<<<<<<<<<<<<<<<<<<<<<<<<<<<<<<<<...............................................>>>>>>>>>>>>>>>>>>>>>>>>>>>>>>>>>..>>>>...>>>......................................
            ...11.2.1220....022222222...........1....1.111.10...............................................01.111.1....1...........222222220..0221...211......................................
            ●YCGG-YYCCGGUUCACGYY●●●○○--------------------------●●●●○---------------------------------------------------------○○○●●GRCGACCCGGC○-RCC-----------------○○●●●●●R●●AGGA●●●●●○
             5′                                                                                                                                      3′
```

```
env-82     CGACG.GUCCGGUUCACGCCGUC..........................AGCA.......................................................................GACGGCGACCGGCG.CCC...................CGUAUCGAGAGGACCCGC.
env-83     CGACG.GUCCGGUUCACGCCGUC..........................AGCA.......................................................................GACGGCGACCGGCG.CCC..................CCGUAUCGAGAGGACCCGC.
env-84     CGACG.GUCCGGUUCACGCCGUC..........................AGCA.......................................................................GACGGCGACCGGCG.CCC..................ACGUAUCGAGAGGACCCGC.
env-85     CGACG.GUCCGGUUCACGCCGUC..........................AGCA.......................................................................GACGGCGACCGGCG.CCC..................CCGUAUGGAGAGGACCCGC.
env-86     CGACG.GUCCGGUUCACGCCGUC..........................AGCA.......................................................................GACGGCGACCGGCG.CCC..................CCGUAUCGAGAGGAUCCGC.
env-87     GCUGG.CUCCGGUUCACGUC.C.CGC.......................ACCC...................................................................GCGAGUGACGACCCGGC..GCC.................GUACGGAGCGAAGGAGCUCGA
env-88     GUCGG.CUCCGGUUCACGCUCGC..........................AGCA.......................................................................GCGGGCGACCGGCG.GCU.................GAGGACCCGAGAGGAAACC..
Sre-2-1    GCUGG.CACCGGUUCACCCGUCGGC........................GGCAAUG..................................................................GCUGGCGGGACCGGCG.GCC....................GAGUAGAGAGGACAAGGC
Acy-1-1    CCCGG.CUCCGGUUCACC.CCCUCACCCGG.CCCCUGGC..........GAGU.......................................................GCCAGGCGACCAGGC.AGGGCGGACCGGCG.GCC...................CAAGAAGGAAGGGACACAC
Mrh-2-1    CUCGG.UCCCGGUUCACGCCGAAACCUCC....................UGGUCAAC.............................................................GGAGG....GGCGACCGGG..GCC.................AACGAUUGAAGAGGAAACC..
env-90     CUCGG.CCCCGGUUCACGCUGCCG.........................CCUC......................................................................CGACGGCGACCGGG..UCC...................AUCGACGAGAGGAACACCC
env-92     CUCGG.CCCCGGUUCACGCUGCCG.........................CCUC......................................................................CGACGGCGACCGGG..UCC...................AUCGACGAGAGGAACACCC
env-93     CUCGG.CCCCGGUUCACGCUGCCG.........................CCUC......................................................................CGACGGCGACCGGG..UCC...................AUCGACGAGAGGAACACCC
env-94     CUCGG.CCCCGGUUCACGCUGCCG.........................CCUC......................................................................CGACGGCGACCGGG..UCC...................AUCGACGAGAGGAACACCC
env-95     CUCGG.CCCCGGUUCACGCUGCCG.........................CCUC......................................................................CGACGGCGACCGGG..UCC...................AUCGACGAGAGGAACACCC
env-96     CUCGG.CCCCGGUUCACGCUGCCG.........................CCUC......................................................................CGACGGCGACCGGG..UCC...................AUCGACGAGAGGAACACCC
env-97     ACCGGUUUCCGGUUCACCUGCGC..........................GGCACCG....................................................................GCGCGGGACCGGG..GCC...............CUCUGAUGGCUUAAGGAGACACC
env-99     ACCGGUUUCCGGUUCACCUGCGC..........................GGCACCG....................................................................GCGCGGGACCGGG..GCC...............CUCUGAUGGCUUAAGGAGACACC
env-103    ACCGGUUUCCGGUUCACCUGCGC..........................GGCACCG....................................................................GCGCGGGACCGGG..GCC...............CUCUGAUGGCUUAAGGAGACACC
env-104    ACCGGUUUCCGGUUCACCUGCGC..........................GGCACCG....................................................................GCGCGGGACCGGG..GCC...............CUCUGAUGGCUUAAGGAGACACC
env-105    ACCGGUUUCCGGUUCACCUGCGC..........................GGCACCG....................................................................GCGCGGGACCGGG..GCC...............CUCUGAUGGCUUAAGGAGACACC
env-111    ACCGGUUUCCGGUUCACCUGCGC..........................GGCACCG....................................................................GCGCGGGACCGGG..GCC...............CUCUGAUGGCUUAAGGAGACACC
Bin-1-1    UCCGG.UUCCGGUUCACGUUGACC.........................AUAGG.....................................................................GGUCGGCGACCGGU..GCC.................CUUUGUCUCCUAGGAGUCCAA
env-112    UCCGG.UUCCGGUUCACGUUGACC.........................AUAGG.....................................................................GGUCGGCGACCGGU..GCC.................CUUUGUCUCCUAGGAGUCCAU
Bco-1-1    UCCGG.UUCCGGUUCACGUUGACC.........................AUAGG.....................................................................GGUCGGCGACCGGU..GCC.................CUUUGUCUCCUAGGAGUCCAA
env-114    AACGG.UUCCGGUUCACGC.CCGUCGG......................AUCCC..................................................................CCGGCGCAGCGACCGGCG.ACC........................AGGCAGGAGUAUUC
env-115    AACGG.UUCCGGUUCACGC.CCGUCGG......................AUCCC..................................................................CCGGCGCAGCGACCGGCG.ACC........................AGGCAGGAGUAUUC
env-116    ACCGG.CUCCGGUUCACGCCCACUGCC......................CUGA...................................................................GGCA..GGGCGACCGGCG.GCU..........CCCGCACCAUGGAUCAAGAGGAUCUGAC
Sro-4-1    GUCGG.CCCCGGUUCACGGCCCC..........................GCAAAUCCGC.................................................................GCGGACGACCGGC..GCC.................CUCCCGAAACUAGGAGACACC
Sgl-4-1    GUCGG.CCCCGGUUCACGGCCCC..........................GCAAAUCCGC.................................................................GCGGACGACCGGC..GCC.................CUCCCGAAACUAGGAGACACC
Sal-8-1    GUCGG.CCCCGGUUCACGGCCCC..........................GCAAAUCCGC.................................................................GCGGACGACCGGC..GCC.................CUCCCGAAACUAGGAGACACC
Sgr-12-1   GUCGG.CCCCGGUUCACGGCCCC..........................GCAAAUCCGC.................................................................GCGGACGACCGGC..GCC.................CUCCCGAAACUAGGAGACACC
Ssp-55-1   GUCGG.CCCCGGUUCACGGCCCC..........................GCAAAUCCGC.................................................................GCGGACGACCGGC..GCC.................CUCCCGAAACUAGGAGACACC
Ssp-62-1   GUCGG.CCCCGGUUCACGGCCCC..........................GCAAAUCCGC.................................................................GCGGACGACCGGC..GCC.................CUCCCGAAACUAGGAGACACC
Ssp-167-1  GUCGG.CCCCGGUUCACGGCGCGC.........................AUCCC.....................................................................GCGUGCCGACCGGC..GCC.................CUCCCGAAACGAGGAGAGACC
Sav-3-1    GUCGG.CCCCGGUUCACGGCACGC.........................GCCC......................................................................GCGUGCCGACCGGC..GCC.................CUCCCGAUCCGAGGAGAGACC
Ssp-4-1    GCCGG.UUCCGGUUCACGGCACGC.........................AUCGG.....................................................................GCGUGCCGACCGGC..GCC................CUCCGAUCAGCAAGGAGAAACC
Sxi-2-1    GCUGG.CUCCGGUUCACGGCACGC.........................AUCGG.....................................................................GUGUGCCGACCGGC..GCC................CUCCAGAACGCAAGGAGAGACC
Ssp-90-1   GCUGG.CUCCGGUUCACGGCACGC.........................AUCGG.....................................................................GCGUGCCGACCGGC..GCC................CUCCAGAACGCAAGGAGAGACC
Jal-1-1    GCUGG.CUCCGGUUCACGUCACGAC........................AUCCC....................................................................GUCGUGCCGACCGGCG.GCC.................CAUGAGACGCCAGGAGUACAG
env-117    CCCGG.CUCCGGUUCACGUC.CCCAGC......................ACCU...................................................................GGUGGGAGACGACCGGCG.GCC..........CGAAAAUCCACGGCUCAG..........
env-125    GCCGG.UUCCGGUUCACGUCCGCAU........................GAUCA....................................................................GU.CGGGCGACCGGG..GCA..................CGUUUGAUUAAGGAGAUCAC
Psp-1-1    GCCGG.UUCCGGUUCACGUCCGCAU........................GAUCA....................................................................GU.CGGGCGACCGGG..GCA..................CGUUUGAUUAAGGAGAUCAC
env-126    GGCGG.UCCCGGUUCACGGCGCC..........................UCCC.......................................................................GGCGGCGACCGGCG.GCC..................ACGUACCGAGAGGAACCC..
Pac-1-1    UUCGG.UUCCGGUUCACCCCCGCC.........................CGCCGA....................................................................GGCGGGGGACCGGCG.ACC....................ACGAUGAGAGGACCCGUA
env-127    GUCGG.UCCCGGUUCACCCCA............................GUGCA........................................................................UGGAGACCGGCG.GCC.................ACUCGACCUCAAGGAGCCGAA
env-128    .GCGG.UUCCGGUUCACGUGCG.CG........................ACCCG....................................................................CGCCGCACGACCGGCG.ACC..................CGACACCCUGAGGAGCACAU
env-130    AGCGG.UUCCGGUUCACGAGC.CCGC.......................AGCCC...................................................................GCGGCGCACGACCGGCG.ACC...................CAGAACCCGAGGAGCACAC
env-131    AGCGG.UUCCGGUUCACGAGC.CCGC.......................AGCCC...................................................................GCGGCGCACGACCGGCG.ACC...................CAGAACCCGAGGAGCACAC
env-132    C.CCG.GUCCGGUUCACGUGC............................CCCG.........................................................................GCACGACCGGCG.CC.....................ACGACAUGAGGAGCACCA
env-133    A.CCG.GUCCGGUUCACGUGCC...........................UCGC........................................................................GGCACGACCGGCG.CC.....................ACGACAUGAGGAGCACC.
Bth-2-1    GACGG.UUCCGGUUCACGCCCGCC.........................AUAGG.....................................................................GGUUGGCGACCGG...GCC..............CCCCCGAAUUCCAUAGGAGACACA
Bsc-1-1    AUAGG.UUCCGGUUCACGCCCGCC.........................AUAGG.....................................................................GGCUGGCGACCGGG..UCC................CCCGAAUUCCAUAGGAGACAUA
Bth-7-1    GACGG.UUCCGGUUCACGCCCGCC.........................AUAGG.....................................................................GGUUGGCGACCGGG..UCC................CCCGAAUUCCAUAGGAGACACA
Bth-8-1    GACGG.UUCCGGUUCACGCCCGCC.........................AUAGG.....................................................................GGCUGGCGACCGGGUCACC..................UGAAUUCCAUAGGAGACACA
Bth-5-1    GACGG.UUCCGGUUCACGCCCGCC.........................AUAGG.....................................................................GGCUGGCGACCGGG..UCC................CCCGAAUUCCAUAGGAGACACA
Bth-3-1    GACGG.UUCCGGUUCACGCCCGCC.........................AUAGG.....................................................................GGCUGGCGACCGGG..UCC................CCCGAAUUCCAUAGGAGACACA
Bth-6-1    CAAGG.UUCCGGUUCACGCCCGCC.........................AUAGG.....................................................................GGCUGGCGACCGGG..UCC................CCCGAAUUCCAUAGGAGACAUA
Bbo-2-1    CGAGG.UUCCGGUUCACGCCCGCC.........................AUAGG.....................................................................GGCUGGCGACCGGG..UCC................CCCGAAUUCCAUAGGAGACAUA
env-136    GCUGG.CUCCGGUUCACGCCUCUC.........................GCAGG.....................................................................GAGGGGCGACCGGCG.GCC......................GAGGAGAGGACACC..
Nsp-4-1    GCUGG.CUCCGGUUCACGGCUCUC.........................GCAGG.....................................................................GAGAGGCGACCGGCG.GCC......................GAUGAGAGGACACC..
env-137    GCUGG.UUCCGGUUCACGUCC..CACGCGC...................ACCA................................................................GCGCGGUACGGGCGACCGGCG.ACU................GAAACCCUCGAGAGGACACC..
env-138    CUCGG.UCCCGGUUCACGCC.GCACACUCCUUUCCGGUC...........CAUUC.......................................................GGCCGG..GGGAG.....GGCGACCGGG..GCC.................CCCGAUGGAAGAGGAAGCC..
env-139    CUCGG.UCCCGGUUCACGCC.GCACACUCCUUUCCGGUC...........CAUUC.......................................................GGCCGG..GGGAG.....GGCGACCGGG..GCC.................CCCGAUGGAAGAGGAAGCC..
env-140    CUCGG.UCCCGGUUCACGCC.GCACACUCCUUUCCGGUC...........CAUUC.......................................................GGCCGG..GGGAG.....GGCGACCGGG..GCC.................CCCGAUGGAAGAGGAAGCC..
env-141    AACGG.UUCCGGUUGACGGCG............................CGAUCGGCAGCACGCCCGCGCCCGUCUUGAGGAGCAC........................................CGCCGACCGGCG.ACCGCCGUUCGCGCCAGAUCGCGGUCCGCAC..........
env-142    AACGG.UUCCGGUUGACGGCG............................CGAUCGGCAGCACGCCCGCGCCCGUCUUGAGGAGCAC........................................CGCCGACCGGCG.ACCGCCGUCGCGCCAGAUCGCGGUCCGCACU..........
Fsp-8-1    CCCGG.UUCCGGUUCACGCUUGUG.........................CAUCAUUAA.................................................................CGCAGGCGACCGGCG.GCC..............AUAAUCGGCACAUA..........
env-143    GUCGGCUUCCUGUUCACGGCG............................ACCGC.......................................................................CGCCGACCAGGCG.ACCGCGACCCAGAUCGCCAAGUACGCCAACC..........
Nsp-10-1   GCUGG.CUCCGGUUCACGCCACCUC........................GUAGG....................................................................GAGG.GGCGACCGGCG.GCC......................GAGGAGAGGACAGC..
env-144    CCCGG.CUCCGGUUCACGAUGCGGUUUCCGUUGCGCUG...........GCUCGAUG....................................................CAGUGC.GCG....CCGC.UCGACCGGCG.GCC..................ACGACCCUCGAGGAGACCCC
Cja-1-1    AGCGG.UUCCGGUUCCCGCCUCGACG.......................AGCC....................................................................CGUCGAGACGACCGGCG.ACC...................ACCCACGAGAGGACCGAGG
Fsy-1-1    CCCGG.CACCGGUUCACGCCCUCGUUC......................AGG....................................................................GGAC.AGGGCGACCGGCG.GCC................AUUUCGGGAAGAAGGAUC....
Fco-1-1    CCCGG.CACCGGUUCACGCCCUCGUUC......................AGG....................................................................GGAC.AGGGCGACCGGCG.GCC................AUUUCGGGAAGAAGGAUC....
env-145    UCAGG.UACCGGUUCACGCCCCCGGGA......................AAUCCA.................................................................UCCAGUCGGCGACCGGCG.ACA....................CAACGCAGAGGACACC..
Ply-1-1    CUCGG.UUCCGGUUCACGCC.............................UUUAUUCGUUU...................................................................GGCGACCGGCG.ACU........................AGACAGGAGAAAUA
Ply-2-1    UUCGG.UUCCGGUUCACGCC.............................UUUAUUCGUUU...................................................................GGCGACCGGCG.ACU........................AGACAGGAGAAAUA
Ssc-2-1    GUCGG.CCCCGGUUCACGUGACGCCC.......................ACUC....................................................................GGGUG.UACGACCGGC..GCC................CUCCCGAACCCAAGGAGACUCC
Sri-4-1    GUCGG.UCCCGGUUCACGUGACGC.........................AAUCA.....................................................................GCGCCACGACCGGA..GCC.................CUCCCGAACCUAGGAGAUUCC
Sri-2-1    GUCGG.UCCCGGUUCACGUGACGC.........................AAUCA.....................................................................GCGCCACGACCGGA..GCC.................CUCCCGAACCUAGGAGAUUCC
Sgr-6-1    GUCGG.UCCCGGUUCACGUGACGC.........................AAUCA.....................................................................GCGCCACGACCGGA..GCC.................CUCCCGAACCUAGGAGAUUCC
Sri-4-16   GUCGG.CCCCGGUUCACGUGACGC.........................AAUCA.....................................................................GCGCCACGACCGGA..GCC.................CUCCCGAACCUAGGAGAUUCC
Ssp-75-1   GUCGG.CCCCGGUUCACGUGACGC.........................AAUCA.....................................................................GCGCCACGACCGGA..GCC.................CUCCCGAACCUAGGAGAUUCC

           ...<<.<.<<<<....<<<<<<<<<<<<<<<<<<<<<<<<<<<<<<<<<................................................>>>>>>>>>>>>>>>>>>>>>>>>>>>>>>>>>..>>>>...>>>......................................
           ...11.2.1220....022222222...........1....1.111.10................................................01.111.1....1...........222222220..0221...211......................................
           ●YCGG-YYCCGGUUCACGYY●●●○○-------------------------------●●●●○-----------------------------------------------------------○○○●●GRCGACCCGGC○-RCG----------------○○●●●●●R●●AGGA●●●●●○○
            5′                                                                                                                                            3′
```

```
Sri-4-18     GUCGG.CCCCGGUUCACGUGACGC..........................AAUCA.....................GCGCCACGACCCGGA..GCC..................CUCCCGAACCUAGGAGAUUCC
Sri-4-19     GUCGG.CCCCGGUUCACGUGACGC..........................AAUCA.....................GCGCCACGACCCGGA..GCC..................CUCCCGAACCUAGGAGAUUCC
Sri-4-20     GUCGG.CCCCGGUUCACGUGACGC..........................AAUCA.....................GCGCCACGACCCGGA..GCC..................CUCCCGAACCUAGGAGAUUCC
env-146      GCUGG.UUCCGGUUCACGUGGGCGUC........................GUAC....................GACGUCGACGACCCGGCG.ACC.....................GUCAUCGAGAGGACACC..
env-147      GCUGG.UUCCGGUUCACGUGGGCGUC........................GUAC....................GACGCCGACGACCCGGCG.ACC.....................GUCAUCGAGAGGACACC..
env-148      GCUGG.UUCCGGUUCACGUGGGCGUC........................GUAC....................GACGCCGACGACCCGGCG.ACC.....................GUCAUCGAGAGGA......
Nal-1-1      GCUGG.UUCCGGUUCACGUGC.GCCC........................AUCCG...................GGGCGGCACGACCCGGCG.ACC.........................GUAGAAGGAGAGCAC
env-149      GUCGG.CCCCGGUUCACGACGUGC..........................AAACCC....................GCUCGUCGACCCGGC..GCC...........CUCCCGAGAAACCAGGAGAAUCC
Ssp-36-1     GUCGG.UCCCGGUUCACGCGGCGCC.........................AUCCCC...................GGCUCCGCGACCCGGC..ACC..................CUCCAGAACCUAGGAGACACC
Ssp-109-1    GUCGG.UCCCGGUUCACGUGGCGCC.........................AUUCCC...................GGCUCCGCGACCCGGC..ACC..................CUCCAGAACCUAGGAGACACC
Ssp-35-1     GUCGG.UCCCGGUUCACGUGGCGCC.........................AUUCCC...................GGCUCCGCGACCCGGC..ACC..................CUCCAGAACCUAGGAGACACC
Ssp-34-1     GUCGG.UCCCGGUUCACGUGGCGCC.........................AUUCCC...................GGCUCCGCGACCCGGC..ACC..................CUCCAGAACCUAGGAGACACC
Ssp-42-1     GUCGG.UCCCGGUUCACGUGGCGCC.........................AUUCCC...................GGCUCCGCGACCCGGC..ACC..................CUCCAGAACCUAGGAGACACC
env-150      CCCGG.UUCCGGUUCACGUCGCGU..........................GAGA......................GCGUGACGACCCGGCG.ACC....................ACCGACGAAAGGACCAGAC
Mau-2-1      CUCGG.UUCCGGUUCACGCCGUAACUCCCCG...................AGA...............CGGGGAG....GGCGACCCGGA..GCC...................ACGAUCGAAGAGGACACC..
env-151      CUUGG.UUCCGGUUCACGCCCCAUCC........................CAGGUCAU..................GGA..GGGCGACCCGGCG.GCC................AGCGAUUGAAGAGGAAACCA.
env-153      CCAGG.CUCCGGUUCACGCCAGCACGACCGCC..................ACGACA...........GGCAC......GGCGACCCGGCG.GCC...................AUGUCUCCCCAGGGAAGGAA
Psp-10-1     UUCGG.CUCCGGUUCACCCGCG............................AAUCC.......................CACGGGACCCGGCG.GCC....................AUGACGAGAGGAACGCAC
env-154      UCCGG.CACCGGUUCACCUCGUGC..........................GUC.......................GCGUGAGGACCCGG...GCC...................CUGAGUGAUGAGGACAUCC.
env-156      UCCGG.CACCGGUUCACCUCGUGC..........................GUC.......................GCGCGAGGACCCGG...GCC...................CUGAGUGAUGAGGACAUCC.
env-157      UGCGG.UUCCGGUUCCCGUCGAU.GCCUC.....................AUAA...............GAGGAAGUUGGCGACCCGGCG.GCC..................GCGACAGCCCAAGGAGAAAAC
env-158      UGCGG.UUCCGGUUCCCGUCGAUGCCCUC.....................AUAA...............GAGGAAGUUGACGACCCGGCG.GCC..................GCGACAGCCCAAGGAGAAAAC
env-159      UGCGG.UUCCGGUUCCCGUCGAUGCCCUC.....................AUAA...............GAGGAAAUUGACGACCCGGCG.GCC..................GCGACAGCCCAAGGAGGAAAC
env-160      UGCGG.UUCCGGUUCCCGUCGAUGCCCUC.....................ACUC...............GAGGAAGUUGGCGACCCGGCG.GCC..................GCGACAGCCCAAGGAGAAGAC
Mha-1-1      CUCGG.UUCCGGUUCACGCCGUCAACUGCCCGA.................ACGAC..........UC.GGCAG....CGGCGACCCGGG..GCC..................CACGAUCGAAGAGGAAACC..
env-161      UACGG.UUCCGGUUCACGUCCGAUCAU.......................CAGCAC.................AUGAA..GGCGACCCGGCG.ACC..................GACGAUUGAAGAGGACACC..
env-162      CGCGG.UUCCGGUUCACGUCAGGCCCUCUAUGA.................GGGCAGCGC......UCCAGGCGGGCGCGACGACCCGGCG.GCC...................GCUUGAGGCAAGGAGCCAGC
env-163      GCUGG.CUCCGGUUCACGCCUUCCGC........................ACCCAC..................GCGU.GGGCGACCCGGC..GCA....................UCGUUUGAGAGGACGACAC
env-164      AUCGG.UUCCGGUUCACCCCG.............................AGCAGCACACC.................CGGGGACCCGGCG.ACC.....................ACCACGAGAGGACCCGAG
Asp-5-1      ACCGG.UUUCGGUUCACCCCG.............................UGA.........................CGGGGACCCGACG.ACC.....................ACCUGACGAGGAGAGAAC
env-165      CGCGG.UUCCGGUUCACGUC..............................AAGCCUCCCUUUUGCAGGCACGCUCCCAUGCGAGCG.......GACGACCCGGCG.GCC...................GCUUGAGGCAAGGAGCCAGA
Nlu-1-1      CUCGG.UGCCGGUUCACGUC.CCGCA........................ACC.................UGCGGCGGCGACCCGGA..ACC................CACUGUCUUGAGAGGAACCUC.
Nba-1-1      CUCGG.UGCCGGUUCACGUC.CCGCA........................ACC.................UGCGGCGGCGACCCGGA..ACC..................CAUGUCUUGAGAGGAACCUC.
Ssp-177-1    GUCGG.CUCCGGUUCACGGAAGG.C.........................AAUCC..................GCCCUUCCGACCCGGC..GCC..................CUCCCGGAACUAGGAGACACC
Lal-1-1      CCAGG.UACCGGUUCACGUC.CCGCG........................AACGG.................CGCGGUGGCGACCCGGCG.ACC...................CCCGUGUGAGAGGACCGCGA
Gma-1-1      UCCGG.UUCCGGUUCACACCGAUGGCA.......................UUC..................UGCC..CGGUGACCCGGCG.ACC....................ACCUUCGAAAGGGACGUA.
Bme-1-1      UGAGG.UUCCGGUUCACGCC.CGCC.........................AACGG..................GGCGAGGCGACCCGGGUCACC...................UGAAUUCCCUAGGAGACACU
Bsu-2-1      UGAGG.UUCCGGUUCACGCC.CGCC.........................AUUGG..................GGCGCGGCGACCCGG.G.UCC...........CCGUUGAAUCCAUAGGAGACAUU
Bre-1-1      UUAGG.UUCCGGUUCACGCUAUGCC.........................AUUGG..................GGCAUGGCGACCCGG.G.UCC..................CCCGAAUCCAUAGGAGACGCA
Blo-34-1     UUAGG.UUCCGGUUCACGUCCUGCC.........................AUAGG..................GGCAUGACGACCCGG.G.UCC..................CCCGAAUCCAUAGGAGACGUA
env-166      UUAGG.UUCCGGUUCACGUCCUGCC.........................AUAGG..................GGCAUGACGACCCGG.G.UCC..................CCCGAAUCCAUAGGAGACGUA
Blo-22-1     UUAGG.UUCCGGUUCACGUCCUGCC.........................AUAGG..................GGCAUGACGACCCGG.G.UCC..................CCCGAAUCCAUAGGAGACGUA
env-167      UUAGG.UUCCGGUUCACGUCUUGCC.........................AUAGG..................GGCAUGACGACCCGG.G.UCC..................CCCGAAUCCAUAGGAGACGUA
env-170      UUAGG.UUCCGGUUCACGUCUUGCC.........................AUAGG..................GGCAUGACGACCCGG.G.UCC..................CCCGAAUCCAUAGGAGACGUA
env-196      UUAGG.UUCCGGUUCACGUCUUGCC.........................AUAGG..................GGCAUGACGACCCGG.G.UCC..................CCCGAAUCCAUAGGAGACGUA
Blo-19-1     UUAGG.UUCCGGUUCACGUCUUGCC.........................AUAGG..................GGCAUGACGACCCGG.G.UCC..................CCCGAAUCCAUAGGAGACGUA
env-197      UUAGG.UUCCGGUUCACGUCUUGCC.........................AUAGG..................GGCAUGACGACCCGG.G.UCC..................CCCGAAUCCAUAGGAGACGUA
env-216      UUAGG.UUCCGGUUCACGUCUUGCC.........................AUAGG..................GGCAUGACGACCCGG.G.UCC..................CCCGAAUCCAUAGGAGACGUA
env-217      UUAGG.UUCCGGUUCACGUCUUGCC.........................AUAGG..................GGCAUGACGACCCGG.G.UCC..................CCCGAAUCCAUAGGAGACGUA
env-219      CUUGG.UUCCGGUUCACCCCGCUGACGUCGUGC.................CGAC............GCGCGAC.....CGGGGACCCGGCG.GCC..........................CGAAAGGAAUCCC.
Ssp-47-1     UCAGG.UUCCGGUUCACCCCG.............................CCAU........................CGGGGACCCGGCG.ACC...................GUCGAUCGAAAGGCGAUGCA
Sma-2-1      UCAGG.UUCCGGUUCACCCCG.............................CUAU........................CGGGGACCCGGCG.ACC...................GUCUAUCGAAAGGCGAUGCA
Nrh-1-1      GUCGG.UUCCGGUUCACGUUCGAGGU........................AGAU....................GCC.CGAGCGACCCGGCG.ACC....................AAUAUCGAGAGGACACCC.
Nte-2-1      GUCGG.UUCCGGUUCACGUUCGGCGC........................AAAG....................GC.CCGAGCGACCCGGCG.ACC....................AAUAUCGAGAGGACACCC.
Nsp-2-1      GUCGG.CUCCGGUUCACGUCCUCGAU........................CGU.....................GUCGAGGGCGACCCGGCG.GCC....................AUUUUCGAAAGGACACCC.
env-220      CUCGG.UUCCGGUUCACGCCCG.AU...ACC...................CACA...............GGUCAUGUACGGGCGACCCGGCG.GCC..................AACGAUUGAAGAGGACACC..
env-221      GCUGG.CUCCGGUUCACGUCCCGC..........................AUCC........................GC.GGACGACCCGGC..GCU..............UCGAGAGCCUUGGAGAAGAC...
env-222      GCUGG.CUCCGGUUCACGUCCCGC..........................AUCC........................GC.GGACGACCCGGC..GCU..............UCGAGAGCCUUGGAGAAGAC...
env-223      CUCGG.CUCCGGUUCACGCACCAACGGGGGGCCAGCCCCC..........CCGUUAACCCCCAAGGCAACACGUACCAAC..........GGGGCAAGCUCCAGG..GGUGCGACCCGGCG.GCC..................GCCAACGAGAGGACCCAUU
env-224      CUCGG.CUCCGGUUCACGCACCAACGGGGGGCCAGCCCCC..........CCGUUAACCCCCAAGGCAACACGUACCAAC..........GGGGCAAGCUCCAGG..GGUGCGACCCGGCG.GCC..................GCCAACGAGAGGACCCAUU
env-225      ACCGG.UUCCGGUUCACGUGCCC.CC........................ACACG.................GGUGGGCACGACCCGGC..GCC.........................AUCACAAGGAGCCCG
env-226      GCCGG.CUCCGGUUCACGUGC.CACCC.......................ACCAG................GGGUGCGCACGACCCGGCG.GCC......................ACGGACAAGGAGCCCCU
env-227      GUCGG.CUCCGGUUCACGUGC.............................GCGA........................GCACGACCCGGCG.GCC..................CCACCCCCA..........
env-228      GUCGG.CUCCGGUUCACGUGC.............................GCGA........................GCACGACCCGGCG.GCC..............CCACCCCCAGUGAGGAGAUCCC
env-229      GCCGG.CUCCGGUUCACGUGC.............................GCGA........................GCACGACCCGGCG.GCC..............CGACCCCCAGUGAGGAGACACC
env-230      GUCGGUCUGCGGUUCACGUGCGCG..........................GCUCG...................CGCGCACGACCCGGAG.GCC.....................CAACCCGAGGAGAACAC
env-231      AGCGG.UUCCGGUUCACGGGC.............................GCGAGA......................GCCCGACCCGGCG.ACC.....................CGACAUCGAGGAGAACC.
env-232      GCUGG.CUCCGGUUCACGUCGUCGCA........................CCA....................UGCACCGACGACCCGGCG.GCC...................GACUACCGCGAGGAGCAUC.
env-233      GCUGG.CUCCGGUUCACGUCG.UGC.........................ACUCA...................GCACCGACGACCCGGC..GCC...........UCGAGAGCCUUGGAGACAUCAC.
Fsp-3-1      ACCGG.CUCCGGUUCACGUCGUC.C.........................ACGC.....................GAGACGACGACCCGGCG.GCC...................ACUUUGAAAGAGGGAUC...
Ssy-1-1      UCCGG.CUCCGGUUCACCUC..............................GUGCAGAA......................GAGGACCCGGCG.GCC....................ACCGAGGAAAGGGACGAGA
Ssp-54-1     GUCGG.CCCCGGUUCACGCGGCGC..........................AAUCC....................GCGACGCGACCCGGC..GCC..................CUCCCGAACCUAGGAGACACC
Ssp-174-1    GCUGG.CCCCGGUUCACGUGACGC..........................AAACCC...................GCGGAGCGACCCGGC..GCC..................CUCCCGAACCUAGGAGCACCC
Ssp-32-1     GUCGG.CCCCGGUUCACGCCCCGC..........................AAACCC...................GCGGAGCGACCCGGC..GCC..................CUCCCGAACCUAGGAGACACC
Sxy-1-1      GUCGG.CCCCGGUUCACGCCCCGC..........................AAACCC...................GCGGAGCGACCCGGC..GCC..................CUCCCGAACCUAGGAGACACC
env-237      CUUGG.CUCCGGUUCACGACGGUGCCUGCGC...................CUGC...............GCGCAGGCCGGGUCGACCCGGCG.GCC..................GCCAAGACCCCGAGGAGACCGC
env-238      CUUGG.CUCCGGUUCACGACGGUGCCUGCGC...................CUGC...............GCGCGGGCCGGGUCGACCCGGCG.GCC..................GCCAAGACCCCGAGGAGACCGC
env-239      CUUGG.CUCCGGUUCACGACGGUGCCUGCG....................CGCG................CGCAGGCCGGGUCGACCCGGCG.GCC..................GCCAAGACCCCGAGGAGACCGC
env-240      AGCGG.UUCCGGUUCACGCACGUGG.........................UCAA.....................CCACGGGCGACCCGGCG.GCC...................GAUUAUCGAGAGGACAUCC.
Ssp-102-1    GCCGG.UCCCGGUUCACGUACGCGC.........................AGUCA....................GCGCGUGCGACCCGGC..ACC..................CUCCCCAGAACUAGGAGAAACC
Sca-3-1      GUCGG.CCCCGGUUCACGGGGGGGC.........................AAUCCC...................GCCCC.UCGACCCGGU..GCC..................CUCCCGAAACUAGGAGAACCC
Ghi-1-2      UCCGG.UUCCGGUUCACAUCGAGGC.........................CCUCC....................GCC.CGAUGACCCGGCG.ACC....................ACAGAGAUUAAGGACCCCCG
Ssc-5-1      GUCGG.UCCCGGUUCACGCACGCGC.........................ACUCC....................GUACGUGCGACCCGGU..GCC.......CUCCCCGCCCCCGCGCGGGG..........

             ...<<.<.<<<<....<<<<<<<<<<<<<<<<<<<<<<<<<<<<<<<<..........................>>>>>>>>>>>>>>>>>>>>>>>>>>>>>>>>..>>>>...>>>...................................
             ...11.2.1220....022222222..........1....1.111.10..........................01.111.1....1..........222222220..0221...211...................................
             •YCGG-YYCCGGUUCACGYY∘∘∘∘∘-------------------•••∘∘----------------------∘∘∘∘∘GRCGACCCGGC∘-RCC-------------------∘∘∘∘•••R••AGGA••••∘∘
             5′                                                                                                    3′
```

```
Sro-5-1    GUCGG.CCCCGGUUCACGUA.AGGC..........AAUCCC..........GCCUCUACGACCCGGA..GCC..........CUCCCGAACCUAGGAGAACCC
Sva-2-1    GCUGG.CCCCGGUUCACGUA.AGGC..........AAUCCC..........GCCUCUACGACCCGGC..GCC..........CUCCCGAACCUUAGGAGUUCCC
Sye-1-1    GCUGG.CCCCGGUUCACGCAUCGGC..........AACCCC..........GCCUGUGCGACCCGGC..GCC..........CUCCGGAACCUAGGAGCACCC
env-241    GCUGG.UUCCGGUUCACGUC...GCUGCG..........CCCG..........CGCAGCCCUGACGACCCGGCG.ACC..........GAGGCCCCGAGAGGAAACC..
env-242    GCUGG.UUCCGGUUCACGUC...GCUGCG..........CCCG..........CGCAGCCCUGACGACCCGGCG.ACC..........GAGGCCCCGAGAGGAAACC..
Nex-1-1    GUCGG.UUCCGGUUCACGCUCAGCGC..........UUGCAC..........GCGC.GGGCGACCCGGCG.ACC..........ACAAUCGAGAGGACAUCC.
Nar-1-1    GUCGG.UUCCGGUUCACGCUUCGCGC..........GUUC..........GCGCG.GGCGACCCGGCG.ACC..........ACAAUCGAGAGGACAUCC.
Nab-1-1    GUCGG.UUCCGGUUCACGCUUCGCGCG..........AAUU..........CGCGUG.GGCGACCCGGCG.ACC..........AAACUCGAGAGGACACCC.
Mar-1-1    UCAGG.UACCGGUUCACGCUCGA..........AUUGCCAGGAAAACAC..........UCGGGCGACCCGGCG.ACC..........AUCGAAUGCAGAGGACACC..
env-243    UCAGG.UACCGGUUCACGUCAGC..........AUUC..........GCUGGCGACCCGGCG.ACC..........AGACCAGAAGAGGACACCA.
env-244    UCAGG.UUCCGGUUCACGUCCAAUACAGC..........AGGUCAG..........GC.....UGGGCGACCCGGCG.GCC..........GACGAUUGAAGAGGACACC..
env-245    UGAGG.UUCCGGUUCACGUCC..........GAUUGCGAGCUUCUCA..........GGGCGACCCGGCG.ACC..........GACGAACGAAGAGGACACC..
Agl-1-1    AAAGG.UUCCGGUUCACGCCCGAGCCCACUGCCCUG..........GCGGGG..........CGGCGCGGUGCAGACA.GCGACCCGGCG.ACC..........AGACCAGCGAGGAGACCGA
Msp-15-1   CUCGG.UUCCGGUUCACGUCCGUACAUCCCC..........AGGUCA..........GGGG.....CGGGCGACCCGGCG.ACC..........AACGAUUGCAGAGGAAAACC.
Mco-2-1    CUCGG.UUCCGGUUCACGUCCGUACAUCCCC..........AGGUCA..........GGGG.....CGGACGACCCGGCG.ACC..........AACGAUUGCAGAGGAAAACC.
env-248    CAGGG.UUCCGGUUCACGUCAGAU..........AUUCAGCAC..........GUUUGGCGACCCGGCG.GCC..........CGCGAAUGAAGAGGACACC..
env-249    CUCGG.UUCCGGUUCACGCUCGA....AGACCU..........UUU..........GGGUCAGGGCUCGGGCGACCCGGCG.ACC..........AACUAUCGAAGAGGACCAAA.
Nmu-1-1    CCCGG.CUCCGGUUCACGCCCGCG.CG..........CAA..........CGACGCGGGCGACCCGGCG.GCC....CCCUGCAAGGCUCUCCCAGGAAGAGGAACGCAC
env-250    UGUGG.UUCCGGUUCACGCCGCGCCCGCCUCGU..........UACC..........ACGAGGCAGGCG.GGCGACCCGGCG.GCC..........GCUUGAGGCAAGGAGCCAAC
env-252    UGUGG.UUCCGGUUCACGCCGCGCCCGCCUCGU..........UACC..........ACGAGGCAGGCG.GGCGACCCGGCG.GCC..........GCUUGAGGCAAGGAGCCAAC
env-253    GCUGG.UUCCGGUUCACGCUCUCGCG..........ACAACACAGGAC..........UGCGGGAACGACCCGGCG.ACC..........GAACCCCUCGAGAGGACACC..
env-254    GCUGG.UUCCGGUUCACGUUCCC..........ACCCGGCAAGGGUGUGACACACACAUCCUGCUGCAGUGCGCCGCACAG..........GGGGACGACCCGGCG.ACC..........GAGACCCCGAGAGGAAACC..
env-255    GCUGG.UUCCGGUUCACGACCCC.....GCUG..........AAGA..........CAGCACCGAGGGGACGACCCGGCG.ACC..........GAACCCGAGAGGACACC..
env-256    GCUGG.UUCCGGUUCACGUCCCC..........ACGA..........GGGGACGACCCGGCG.ACC..........GACACCCCGAGAGGAAACC..
env-257    GCUGG.UUCCGGUUCACGUUCCCG..........AUCAGCAAGGGUGUGACACACACGACCUGCUGCAGUGCGCCGCACA..........CGGGAACGACCCGGCG.ACC..........GAGACCCCGAGAGGAAACC..
env-258    GCUGG.UUCCGGUUCACGUUCCCG..........AUCAGCAGGGUGUGACCACACGACUGCUGCAGUGCGCCGCACA..........CGGGAACGACCCGGCG.ACC..........GAGACCCCGAGAGGAAACC..
env-259    GUUGG.UUCCGGUUCACGUGCCCGC..........ACCC..........GCCCGCACGACCCGGCG.ACC..........AUGAGCCCUGAGGAGUAACA
env-260    CCUGG.UUCCGGUUCACGUGCCCG..........AUCC..........CGUGCACGACCCGGCG.ACC..........GUCUGAUGAGGAGAACCC
env-261    CUUGG.CUCCGGUUCACGUGCGCG..........AACC..........CGUGCACGACCCGGCG.GCC..........CGAGAUCGAGGGAGAGAU
env-262    CUUGG.CUCCGGUUCACGUGCGCA..........AGCC..........UGUCCACGACCCGGCG.GCC..........UCCCAGACGAGGAGAUCCC
env-263    GCUGG.UUCCGGUUCACGUCCCC..........GAUC..........GGUGGCGACCCGGCG.ACC..........UCAUCCCUCGAGAGGACGGAAC
env-264    GCUGG.UUCCGGUUCACGUC.CCGCC..........GUCCGA..........GGCGGCGACGACCCGGCG.ACC..........GUCCCACCGUGAGGAGCACCA
env-265    GCUGG.UUCCGGUUCACGUUCACA..........CCG..........UGUGGGCGACCCGGGG.GCA..........CCUGAUUCCUAGGAGACCUG
env-266    GUCGG.UUCCGGUUCACGUUCACA..........CCG..........UGUGGACGACCCGGGGCACC..........UGAUUCCUAGGAGACCUG
Nth-1-1    GUCGG.UUCCGGUUCACGCAUUCCGC..........GCCCCA..........GCGGCGUGCGACCCGGCG.ACC..........ACGAUCGAGAGGACGCCC.
Pan-1-1    GUCGG.UUCCGGUUCACGUAUCCCGC..........GCUCCA..........GCGGUGUGCGACCCGGCG.ACC..........AUGAUCGAGAGGACGCCC.
Sxi-1-1    UCCGG.CUCCGGUUCACCUCGCGA..........CCGU..........UCGCGGGGAUCCGGCG.GCC..........AGUUGAGAGAGGACACC..
Ssa-1-1    UCCGG.CUCCGGUUCACCUCGUGAC..........ACGGC..........GUCACGGGGAUCCGGCG.GCC..........AUCGAGAGAGGACACC..
Mch-1-1    CUCGG.UUCCGGUUCACGCCGCGACUUCCCG..........GCU..........CGGGAA.....GGCGACCCGGA..GCC..........ACGAUCGAAGAGGAAACC..
env-267    UCCGGCCUCCGGUUCACCCGCAUCGC..........AUUC..........GCG.UGCGGGACCCGGC..GCC..........CUCUGAAUGCUAGGAGAUACC
env-268    UCCGGCCUCCGGUUCACCCGCAUCGC..........AUUC..........GCG.UGCGGGACCCGGC..GCC..........CUCUGAAUGCUAGGAGACACC
env-274    UCCGGCCUCCGGUUCACCCGCAUCGC..........AUUC..........GCG.UGCGGGACCCGGC..GCC..........CUCUGAAUGCUAGGAGACACC
Pas-1-1    CCCGG.UUCCGGUUCACCCUCGGUGCCGCGCUC..........GUC..........GAGC.C.GCACC.AGGGACCCGGCG.ACC..........ACGCGGAGAGGAAGAA..
Pdi-1-1    AGUGG.CUCCGGUUCACCCCUGU.....CGGCG..........AUCC..........CGUCGGCACAGCAGGGGACCCGGCG.GCC..........CUCCCUGGAGAGGAACUUCC
env-279    GUCGG.CUCCGGUUCACGGUCCCC..........AUCG..........GGGGAGCGACCCGGC..GCC..........AUGAGAGGACACC..
env-280    UCCGG.UUCCGGUUCACCCCGCG..........CACAUCAC..........CACGGGGACCCGGC..GCC..........ACUUCGAGAGGAACCCAC
Ame-4-2    UCCGG.CUCCGGUUCACCCCACGUGUCG..........AAGAA..........CGCCGCG..GGGACCCGGCG.GCC..........ACCGAGAGAGGACUCC..
Asp-4-1    UCCGG.CUCCGGUUCACCCCACGUGUCG..........AAGAA..........CGCCGCG..GGGACCCGGCG.GCC..........ACCGAGAGAGGACUCC..
Ath-1-1    UCCGG.CUCCGGUUCACCCCACGUGUCG..........AAGAA..........CGCCGCG..GGGACCCGGCG.GCC..........ACCGAGAGAGGACUCC..
Sbi-1-1    GUCGG.CCCCGGUUCACGUCAGGC..........AACCC..........GCCUGUCGACCCGGU..GCC..........CUCCCGAACCUAGGAGAAUCC
Shy-4-1    GUCGG.UCCCGGUUCACGACAGGC..........AACCC..........GCCCGUCGACCCGGU..GCC..........CUCCCGAACCUAGGAGAUUCC
Ssp-73-1   GUCGG.UCCCGGUUCACGACAGGC..........AACCC..........GCCCGUCGACCCGGU..GCC..........CUCCCGAACCUAGGAGAAUCC
Sir-1-1    GUCGG.UCCCGGUUCACGACAGGC..........AACCC..........GCCCGUCGACCCGGU..GCC..........CUCCCGAACCUAGGAGAUUCC
Ssp-165-1  GUCGG.UCCCGGUUCACGACAGGC..........AAUCC..........GCCCGUCGACCCGGU..GCC..........CUCCCGAACCUAGGAGAUUCC
Svi-3-1    GUCGG.UCCCGGUUCACGACAGGGC..........AUUCC..........GCCC.GUCGACCCGGU..GCC..........CUCCCGAACCUAGGAGAUUCC
env-281    ACGGG.CUCCGGUUCACCCCGG.UCGGU..........GUUCG..........ACCGAUCCGAGGACCCGGC..GCC..........ACAUCGGAAAGGACAUCCC
env-282    CCCGG.CUCCGGUUCACGCCGCUCGACAUG..........GCGA..........CAUGUCGA..GGCGACCCGGCG.GCC..........AUCGGAAGGACCCAG
env-283    UGCGG.UACCGGUUCACGCUCGCUCUGAGGGAC..........CACG..........GUCCUGAGCCGCGAGCGACCCGGCG.ACU..........GCGGAACCAGGAGUCAGG
env-288    CCAGG.CUCCGGUUCACGCCG..........ACAUCCGA..........CGGCGACCCGGCG.GCC..........GACACGAGAGGACACC..
env-289    AACGG.UUCCGGUUCACGGCCUUG..........ACACA..........CGAGGUCGACCCGGCG.GCC..........ACGACGAAAGGUUCACCC
env-290    CGCGG.UACCGGUUCACGCUCGCGCC.CGCGCCC..........CCG..........GGGCGGAAGGCGCU.GCGACCCGGCG.ACC..........GCUGGAGGCAAGGAGCCAUU
Gar-1-1    UCCGG.UUCCGGUUCACACCCCAAGGC..........AAAUUC..........GCCC..GGGUGACCCGGCG.ACC..........AUGAAAGGACACC.
Mro-1-1    AGCGG.UACCGGUUCCCGCAGCCGC..........GCAUCACAGGUGUGCCGAUGCGCGCAGGCCGUGACAG..........GCGAA.GCGACCCGGCG.ACC..........GCGCCUAGCGUUAGGACUAUGA
env-291    CGCGG.UUCCGGUUCACGUCGCGGC..........UUCCC..........GCC.CGGCGACCCGGCG.GCC..........AUCUUCGAGAGGACAUC..
env-292    CGCGG.UUCCGGUUCACGUCGAGGCU..........CCCC..........GGCC.CGACGACCCGGCG.GCC..........AUCUUCGAGAGGACAUC..
Psu-1-1    GUCGG.UGCCGGUUCACGCC......GCCC..........UUCG..........GGGCUUCGAUGGCGACCCGGC..GCC..........UGAGAUGAGGAGAGACG
Gne-1-1    UCCGG.UUCCGGUUCACACCACUGA..........AAA..........UCAGCGGUGACCCGGCG.GCC..........AUGAUGAAGGGACAUCA
Gsi-1-1    UCCGG.UUCCGGUUCACACCACUGA..........AAAA..........UCAGAGGUGACCCGGCG.ACC..........AUGAUGAAGGGACAUCA
Gsi-2-1    UCCGG.UUCCGGUUCACACCACUGA..........AAAA..........UCAGAGGUGACCCGGCG.ACC..........AUGAUGAAGGGACAUCA
Nta-1-1    GUCGG.UUCCGGUUCACGCUCACGGCUCUGC..........UGAC..........GCGGAGC...GGGCGACCCGGCG.ACC..........AUUCUCGAGAGGACACCC.
env-293    GCUGG.UUCCGGUUCACGUCCCGCU..........UCACGC..........GGCGGGACGACCCGGCG.ACC..........GAGACCUCGAGAGGAAACCC.
env-294    GAUGG.UUCCGGUUCACGCCCAGU..........GAAC..........GC.GGGCGACCCGGCG.ACC..........AGAAAGAAGAGGAAC....
env-295    CUCGG.CUCCGGUUCACGCCGACACCUCC..........ACUUCG..........GGAGG...GGCGACCCGG.G.GCC..........CACGAUUGAAGAGGAAACC..
env-298    CUCGG.UUCCGGUUCACGCCGAUACCUCC..........ACUUCG..........GGAG.....GGCGACCCGG.G.GCC..........CACGACUG.........
Mob-1-1    CUCGG.UUCCGGUUCACGCCGAAACCUCCCG..........ACA..........CGGGAGG...GGCGACCCGG.G.GCC..........AACGAAUGAAGAGGAAACC..
Mob-1-2    CUCGG.UUCCGGUUCACGCCGAAACCUCCCG..........ACA..........CGGGAGG...GGCGACCCGGG..GCC..........AACGAAUGAAGAGGAAACC..
env-299    CACGG.UUCCGGUUCACGUCCCGACGGUCGCCCGC..........AGCG..........GCGCCCGGCGCACGGGGCGACCCGGCG.ACC..........ACCAACGAAAGGACCGAGC
Asp-1-1    AAAGG.UUCCGGUUCACGCC..........AGCGGGCAG..........GGCGACCCGGCG.ACC..........UGAUCAUCGAGAGGGACCGAG
env-300    GCUGG.UUCCGGUUCACGUC.CCCGA..........CGAGC..........UCGGGUGGCGACCCGGCG.ACC..........GCAUCCCUCGAGAGGACGGUUC
env-301    GCUGG.UUCCGGUUCACGUC.CCCGA..........CGAGC..........UCGGGUGGCGACCCGGCG.ACC..........GCAUCCCUCGAGAGGACGGUUC
env-302    UCCGG.UUCCGGUUCACGCUGUCCGC..........AUGA..........GAUGACGGCGACCCGGCG.ACC..........ACGAGAGAGGACACC..
env-303    UCCGG.UUCCGGUUCACGCUGUCCGC..........AUGA..........GAUGACGGCGACCCGGCG.ACC..........ACGAGAGAGGACACC..
Nsp-6-1    GUCGG.UUCCGGUUCACGCUCUGCGC..........GCAU..........GUGUUGGGCGACCCGGCG.ACC..........AUAUUCGAGAGGACACCC.

           ...<<.<.<<<<....<<<<<<<<<<<<<<<<<<<<<<<<<<<<<<<<<..........>>>>>>>>>>>>>>>>>>>>>>>>>>>>>>>>>..>>>>...>>>..........
           ...11.2.1220....022222222..........1....1.111.10..........01.111.1....1..........222222220..0221...211..........
           •YCGG-YYCCGGUUCACGYY∘∘∘∘∘-----•••∘∘-----∘∘∘∘∘GRCGACCCGGC∘-RCC-----∘∘∘∘•••R••AGGA•••••∘∘
           5'                                                                                   3'
```

| | | | | |
|---|---|---|---|---|
| env-304 | CUCGG.UUCCGGUUCACGCCGAUACCUCCAC... | ...UUCG... | ...G.GGAGG....GGCGACCGG.G.GCC | ...CACGAGUGAUAGAGGAAACC.. |
| env-306 | CUCGG.UUCCGGUUCACGCCGAUACCUCCAC... | ...UUCG... | ...G.GGAGG....GGCGACCGG.G.GCC | ...CACGAGUGAUAGAGGAAACC.. |
| env-307 | CUCGG.UUCCGGUUCACGCCGAUACCUCCAC... | ...UUCG... | ...G.GGAGG....GGCGACCGG.G.GCC | ....ACGAGUGAUAGAGGAAACC.. |
| env-308 | CUCGG.UUCCGGUUCACGCCGAUACCUCCAC... | ...UUCG... | ...G.GGAGG....GGCGACCGG.G.GCC | ...CACGAUUGAAGAGGAAACC.. |
| env-309 | CUCGG.UUCCGGUUCACGCCGAUACCUCCAC... | ...UUCG... | ...G.GGAGG....GGCGACCGG.G.GCC | ...CACGAUUGAAGAGGAAACC.. |
| env-310 | CACGG.CUCCGGUUCACGCCCGA... | ...UCACCGAU... | ...UCCGGCGACCGGCG.GCC | ...AUCCACCCGAGAGGACACCC. |
| env-311 | UUCGG.UUCCGGUUCACGCUCCAAAGCU... | ...CCAGGUAA... | ...GGCAA..GGGCGACCGGCG.GCC | ...AACGAAUGAAGAGGACACC.. |
| env-312 | UCCGG.UUCCGGUUCACGUCC... | ...GCUUCGUGCUCAUUA... | ...GGGCGACCGGCG.ACC | ...GACGAUAGAAGAGGACAUC.. |
| env-314 | CACGG.UUCCGGUUCACGUCACUUAUGCGC... | ...CGAG... | ...GCGCG....GACGACCGGCG.ACC | ...GCUGAGGCAAGGAGCCAUC |
| Ami-1-1 | AGGGG.UUCCGGUUCACGCCACGUUCC... | ...AGCA... | ...GGAA..UGGCGACCGGCG.ACC | ...CGAUCAUCGAGAGGGACCGAG |
| env-315 | GUCGG.UUCCGGUUCACGUCUCGCG... | ...ACCU... | ...CGC.AGGCGACCGGCG.ACC | ...CUGAGACGAAGGACACUC. |
| env-317 | GUCGG.UUCCGGUUCACGUCUCGCG... | ...ACCU... | ...CGC.AGGCGACCGGCG.ACC | ...CUGAGACGAAGGACACCC. |
| env-318 | GUCGG.UUCCGGUUCACGUCUCGCG... | ...ACCU... | ...CGC.AGGCGACCGGC..GCC | ...CCUGAGACGAAGGACACUC. |
| env-319 | GUCGG.UUCCGGUUCACGUCUCGCG... | ...ACCU... | ...CGC.AGGCGACCGGC..GCC | ...CCUGAGACGAAGGACACUC. |
| env-320 | GUCGG.UUCCGGUUCACGCCUCGCG... | ...ACCU... | ...CGC.AGGCGACCGGC..GCC | ...CCUGAGACGAAGGACACUC. |
| env-321 | GUCGG.CUCCGGUUCACGCCUCGCG... | ...ACCC... | ...UGC.AGGCGACCGGC..GCC | ...CUGAGACGAAGGACACCC. |
| Nni-1-1 | GUCGG.UUCCGGUUCACGUCUCGC... | ...AUC... | ...GCCAGGCGACCGGA..GCC | ...UUUCGAAAGGACAUCC. |
| Nsp-8-1 | GUUGG.CUCCGGUUCACGUC.CGC... | ...ACCCC... | ...GCGAGGCGACCGG...GCC | ...CCUUUGAGAGGACACCC. |
| env-322 | UCCGG.UUCCGGUUCACGCCCACGA... | ...CAG... | ...UCGAGGGCGACCGGCG.ACC | ...ACGAAAGGACACC. |
| Rsp-20-1 | UCCGG.UUCCGGUUCACGCACUCGA... | ...CAG... | ...UCGAG.GCGACCGGCG.ACC | ...AUGAAAGGACACC. |
| Rsp-16-1 | UCCGG.UUCCGGUUCACGCACUCGA... | ...CAG... | ...UCGAG.GCGACCGGCG.ACC | ...AUGAAAGGACACCC |
| Mpu-1-1 | ACCGG.UUCCGGUUCACGCC..CG....AGCCCGUCG... | ...CGGA... | ...CGACGGGGAGCACCGACGGCGACCGGCG.ACC | ...ACCGACGAAAGGACCGAGG |
| Msp-27-1 | ACCGG.UUCCGGUUCACGCC..CGA.GC...CCGUCG... | ...CGGA... | ...CGACGGGGAGCAUCGACGGCGACCGGCG.ACC | ...ACCGACGAAAGGACCGAGG |
| env-323 | GUCGG.CUCCGGUUCACGUCCGC... | ...GCAC... | ...GCGGGCGACCGGCG.GCU | ...GUGAGACCCGAGAGGAAAUC.. |
| env-324 | GUCGG.CUCCGGUUCACGUCCGC... | ...GCAC... | ...GCGGGCGACCGGCG.GCU | ...GUGAGACCCGAGAGGAAAUC.. |
| Aal-1-1 | GCCGG.CUCCGGUUCACGCCCGC... | ...UGCC... | ...GCGGACGACCGGC..GCC | ...AUGAGAGCGGAGGACAGCA. |
| env-325 | CCCGG.CUCCGGUUCACG.CCCAGC... | ...AUCC... | ...GUUGCGGACGACCGGC..GCC | .....ACCGACAAGGGCCUAUCGAAAGAGGAUCCACC |
| env-329 | CCCGG.CUCCGGUUCACG.CCCAGC... | ...AUCC... | ...GUUGCGGACGACCGGC..GCC | .....ACCGACAAGGGCCUAUCGAAAGAGGAUCCACC |
| env-330 | CCCGG.CUCCGGUUCACGUCC... | ...AGCUCGCUGAAA... | ...GGACGACCGGCG.GCC | ...ACCUCGAAAGGGAUCCCA |
| env-331 | CCCGG.CUCCGGUUCACGUCC... | ...AGCUCGCUGANA... | ...GGACGACCGGCG.GCC | ...ACCUCGAAAGGGAUCCCA |
| env-333 | CCCGG.CUCCGGUUCACGUCCUG......UCCCGCGACACCACCG.C.ACCC... | ...ACCC... | ...GUCGGUGGCG.CGCGGGAGUCCCACCAGGGCGACCGGCG.GCC | ...AUUACCGGCGCUG......... |
| env-334 | CCCGG.CUCCGGUUCACGUCCUG......UCCCAUGACACCACC....GCAUCACG... | ...GCAUCACG... | ...GGUGGCGUCC.GGGAGUUCCUCAGGGCGACCGGCG.GCC | ...AUAACCGGCGCCCG......... |
| env-335 | CCCGG.CUCCGGUUCACGUCCGCAAGUGC... | ...GAAAU... | ...GCAC....GGGCGACCGGCG.GCC | ...AUGAUCGGCACCGAACCCAGAAGCG... |
| env-336 | CCCGG.CUCCGGUUCACGUCC... | ...GUAC... | ...GGACGACCGGCG.GCC | ...ACGGAAAGGACCCAC |
| env-337 | ACCGG.CUCCGGUUCACGUCCU.ACGC... | ...AUCC... | ...GCGCGGGACGACCGGCG.GCA | ...CAGGAGAGGACCCC.. |
| env-338 | CCCGG.CUCCGGUUCCCGUCCCCUACCUGCG... | ...UCCG... | ...CGCA..GACGGGGCGACCGGCG.GCC | ...AAUUGACCGAGAGGCACGACC |
| Bts-1-1 | ACAGG.UUCCGGUUCACGCC.CACC... | ...ACAGG... | ...GGUGUGGCGACCGG.G.UCC | ...CCCGAAUUCCAUAGGAGACAAA |
| Ama-5-1 | UGCGG.UACCGGUUCACGUCUCCCACAACUCCCAUGGG... | ...CUUC... | ...CCCAUGCGAGCCU..GGAGGCG.CCCGGCG.ACC | ...GCCUGAGGCGAGGAGAACC. |
| env-339 | UGCGG.UACCGGUUCCCGGCGGGCG... | ...UCA... | ...CGCCCGGCGACCGGCG.ACC | ...GCCAGAACCAGGAGUUAGU |
| env-340 | CUCGG.UUCCGGUUCACGCCGAAUACCUCCAC... | ...UUCG... | ...G.GGAGG.....GGCGACCGG.G.GCC | ...CACGAAUGAAGAGGAAACC.. |
| env-341 | CUCGG.UUCCGGUUCACGCCGAUUACCUCCAC... | ...UUCG... | ...G.GGAGG.....GGCGACCGG.G.GCC | ...CACGAUUGAAGAGGAAACCA. |
| env-342 | CUCGG.UUCCGGUUCACGCCGAAAACCUCCCC... | ...GUCA... | ...GCGGAGG.....GGCGACCGG.G.GCC | ...CUCGAUCGAAUGAGGAAACC.. |
| env-343 | ACCGG.CUCCGGUUCACGUC..CGCGC... | ...ACCC... | ...GCGCCCUGACGACCGGCG.GCC | ...GUCGACCUGAGGAGCACGC |
| env-344 | ACUGG.CUCCGGUUCACGUC..CGCGC... | ...AUCC... | ...GCGCCCUGGCGACCGGCG.GCC | ...GUUGACCCGAGGAGCAUC. |
| env-345 | ACUGG.CUCCGGUUCACGUC..CGCGC... | ...AUCC... | ...GCGCCCUGGCGACCGGCG.GCC | ...GUUGACCCGAGGAGCAUC. |
| Mva-1-1 | CUCGG.UUCCGGUUCACGCCGCAGCCUCCCG... | ...GUA... | ...CGGGAAG....GGCGACCGG...ACC | ...CCACGAUCGAAGAGGAAACC.. |
| Mru-1-1 | CUCGG.UUCCGGUUCACGCCGCAGCAUCCCG... | ...GUA... | ...CGGGAA.....GGCGACCGG.G.GCC | ...CACGAUCGAAGAGGAAACC.. |
| Mva-2-1 | CUCGG.UUCCGGUUCACGCCGCAGCCUCCC... | ...AUCGC... | ...GGGAAG....GGCGACCGG.G.ACC | ...CCACGAUCGAAGAGGAAACC.. |
| Mgi-1-1 | CUCGG.UUCCGGUUCACGCCGCAGCCUCCC... | ...GCUUU... | ...GGGAAG....GGCGACCGG...GCC | ...CACGAUUGAAGAGGAAACC.. |
| env-346 | CCAGG.UACCGGUUCACGCC.CGCCGACG... | ...GCGGGA... | ...UG.CGGCGUGGCGACCGGCG.ACC | ...AACAGAGAGGACGAGGC |
| env-347 | CCAGG.UACCGGUUCACGUC.CGCCGACG... | ...GCGGGA... | ...UG.CGGCGUGGCGACCGGCG.ACC | ...GAGAGGAAGGACAGCGA |
| env-348 | CCAGG.UUCCGGUUCACGCC.CGAGCCGCCCGCUUGA... | ...UGA... | ...UCAAGGCGGC.GCUGACGGCGACCGGCG.ACC | ...AUAUGAUGAAAGGACCAACG |
| Nas-2-1 | GUCGG.UUCCGGUUCACGCUUCGCG... | ...CCAAAUCCA... | ...CGCG.GGCGACCGGCG.ACC | ...AAAAUCGAGAGGACAUCC. |
| Mab-65-1 | AAGGG.UUCCGGUUCACGC... | ...CCACAA... | ...GCGACCGGCG.ACC | ...ACAACACGCAGAGGAAAAGCA |
| env-349 | CUCGG.UCCCGGUUCACGUCUGUUGCUCGCA... | ...GGUCA... | ...UGCG....CAGGCGACCGGCG.GCC | ...ACGAAUGAAGAGGACACC.. |
| env-350 | ACCGG.UUCCGGUUCACGCC...CGAGUCC.ACCGCCCU... | ...GGC... | ...GGGGCGGCGGGGCACUGACGGCGACCGGCG.ACC | ...AGACCAUCGAGAGGACCGAGC |
| Fsp-7-1 | ACAGG.CUCCGGUUCACGCCGUGGCG... | ...ACA... | ...CGCGACGGCGACCGGCG.GCC | ...AACGUGAGAAGAGGGACCACC |
| Sca-1-1 | GUCGG.UCCCGGUUCACGCCUCCGC... | ...AAUCC... | ...GCGAUGGCGACCGGC..ACC | ...CUCCCGGAACUAGGAGACACC |
| Ssv-1-1 | GUCGG.UCCCGGUUCACGCCCCCGC... | ...AAUCC... | ...GCGAUGGCGACCGGC..ACC | ...CUCCCGAAACUAGGAGACACC |
| Ssp-50-1 | GCUGG.CCCCGGUUCACGUACGAGC... | ...AGUCC... | ...GCGCGUGCGACCGGA..GCC | ..CUCCCCGUGAUCCGCGGGGACCUUC.......... |
| Ssp-19-1 | GUCGG.CUCCGGUUCACGUGGCAGC... | ...AAUCC... | ...GCGGCUGCGACCGGC..GCC | ...CUCCCGAAACUAGGAGACACC |
| Aro-1-1 | GUCGG.UCCCGGUUCCCGUCCGCGC... | ...AUCUCC... | ...GCGCCGGCGACCGGCG.GCC | ...CCGAGAACCUAGGAGCACCC |
| Sca-2-1 | GUCGG.CCCCGGUUCACGUCCG.GC... | ...AAUCCC... | ...GCUCGGUCGACCGGU..GCC | ...CUCCCGAACCUAGGAGAACCC |
| Nvi-1-1 | GUCGG.CUCCGGUUCACGUCCUCGC... | ...UAUCU... | ...GCGCGGGCGACCGGCG.GCC | ...AAUAUCGAGAGGACACCC. |
| env-351 | GUCGG.CCCCGGUUCACGUAC.AGGC... | ...AAGCCC... | ...GCCUCGUACGACCGGC..GCC | ...CUCCCGAACCCAGGAGCACCC |
| env-352 | UCAGG.UUCCGGUUCACGUCCUUCGAUG... | ...ACCAGCA... | ...CAUU...GGGCGACCGGCG.ACC | ...GACUACUGAAGAGGACACC.. |
| env-353 | UCAGG.UUCCGGUUCACGUCCUUCGAUG... | ...ACCAGCA... | ...CAUU...GGGCGACCGGCG.ACC | ...GACUACUGAAGAGGACACC.. |
| env-355 | UCAGG.UUCCGGUUCACGUCCUUCGAUG... | ...ACCAGCA... | ...CAUU...GGGCGACCGGCG.ACC | ...GACUACUGAAGAGGACACC.. |
| env-358 | UCAGG.UUCCGGUUCACGUCCUUCGAUG... | ...ACCAGCA... | ...CAUU...GGGCGACCGGCG.ACC | ...GACUACUGAAGAGGACACC.. |
| env-359 | UCAGG.UUCCGGUUCACGUCCUUCGAUG... | ...ACCAGCA... | ...CAUU...GGGCGACCGGCG.ACC | ...GACUACUGAAGAGGACACC.. |
| env-360 | UCAGG.UUCCGGUUCACGUCCUUCGAUG... | ...ACCAGCA... | ...CAUU...GGGCGACCGGCG.ACC | ...GACUACUGAAGAGGACACC.. |
| env-361 | UCAGG.UUCCGGUUCACGUCCUUCGAUG... | ...ACCAGCA... | ...CAUU...GGGCGACCGGCG.ACC | ...GACUACUGAAGAGGACACC.. |
| env-363 | UCAGG.UUCCGGUUCACGUCCUUCGAUG... | ...ACCAGCA... | ...CAUU...GGGCGACCGGCG.ACC | ...GACUACUGAAGAGGACACC.. |
| env-364 | UCAGG.UUCCGGUUCACGUCCUUCGAUG... | ...ACCAGCA... | ...CAUU...GGGCGACCGGCG.ACC | ...GACUACUGAAGAGGACACC.. |
| env-365 | UCAGG.UUCCGGUUCACGUCCUUCGAUG... | ...ACCAGCA... | ...CAUU...GGGCGACCGGCG.ACC | ...GACUACUGAAGAGGACACC.. |
| Mim-1-1 | CAGGG.UUCCGGUUCACGCC... | ...GUAACGCA... | ...GGCGACCGGCG.GCC | ...ACAAACACGAAGAGGAAACACC |
| Mab-73-1 | AAGGG.UUCCGGUUCACGCC... | ...GUGACGCA... | ...GGCGACCGGCG.ACC | ...ACAACACGAAGAGGAAACACC |
| Mab-55-1 | AAGGG.UUCCGGUUCACGCC... | ...GUGACGCA... | ...GGCGACCGGCG.ACC | ...ACAACACGAAGAGGAAACACC |
| Mab-1-114 | AAGGG.UUCCGGUUCACGCC... | ...GUGACGCA... | ...GGCGACCGGCG.ACC | ...ACAACACGAAGAGGAAACACC |
| Mma-1-1 | AAGGG.UUCCGGUUCACGCC... | ...GUGACGCA... | ...GGCGACCGGCG.ACC | ...ACAACACGAAGAGGAAACACC |
| env-366 | GGCGG.UUCCGGUUCACGUCGGCCC... | ...GGCA... | ...GGG.CGGCGACCGGCG.ACC | ...ACGUCACGAGAGGAACCC.. |
| env-367 | CAAGG.UUCCGGUUCACGCC...UGAGC..CACCGCCCUGGGCCA... | ...GAAACG... | ...UGGCG.AGUGCGGUCAAGCGAUGACGGCGACCGGCG.ACC | ...CAUUAUCGAGAGGACCGAGC |
| env-368 | CAAGG.UUCCGGUUCACGUC... | ...GUCAA... | ...GGCGACCGGCG.ACC | ...AUUUUUCCGAGAGGACCGAGC |
| | ...<<.<.<<<<....<<<<<<<<<<<<<<<<<<<<<<<<<<<<<<<<<<<... | ... | ...>>>>>>>>>>>>>>>>>>>>>>>>>>>>>>>>>>..>>>>...>>> | ... |
| | ...11.2.1220....022222222..........1....1.111.10... | ... | ...01.111.1....1..........222222220..0221...211 | ... |
|  | 5′ YCGG-YYCCGGUUCACGYY ---- | ---- | ---- GRCGACCCGGC-RCC 3′ | ---- R AGGA |

```
env-369    UCAGG.UUCCGGUUCACGUCCGAUC........................UUCAGCAC..........................................................GAUUUGGCGACCCGGCG.ACC.................GACUAGUGAAGAGGACACC..
Tsp-1-1    AUGGG.UUCCGGUUCACGGCUCUGGC.......................GAG...............................................................GUCAGAGGCGACCCGGCG.ACC.................ACGAAAGAGAAGGACGGCAC
Tpa-1-1    AUCGG.UUCCGGUUCACGGCCUGGC........................GAUG..............................................................GCCAGGGGCGACCCGGCG.ACC.................ACAAACGAGAAGGAACGGCA
Are-1-1    AGGGG.UUCCGGUUCACGCCAAUC.........................GUGGAA..............................................................GAACGGCGACCCGGCG.ACC.................CGAUCAUCGAGAGGGACCGAG
Nca-3-1    UGCGG.UUCCGGUUCACGCC.............................AUGCGUGAA...............................................................GGCGACCCGGCG.GCC.................GCACCCCGAAAGGACAGCG.
env-370    UGCGG.UUCCGGUUCACGCC.............................ACGGA...................................................................GGCGACCCGGCG.GCC.................GCAGCCCGAAAGGACAGCG.
env-371    CUCGG.CUCCGGUUCACGCC.............................ACGCACGAUCA.............................................................GGCGACCCGGCG.GCC.................GCCAACGAGAGGACCCAUU
Mph-1-1    CUCGG.UUCCGGUUCACGCCUGUCACCUCC...................ACAGAUC...................................................GGAGG.....GGCGACCCGG.G.GCC.................ACGAUCGAAGAGGAAGCC..
env-372    CUCGG.UUCCGGUUCACGUCUGAUGC.......................UCGCAGGUCAC.......................................................GCG.CAGGCGACCCGGCG.GCC.................CACGAAUGAAGAGGAAACU..
env-373    CUCGG.UUCCGGUUCACGUCUGAUGC.......................UCGCAGGUCAC.......................................................GCG.CAGGCGACCCGGCG.GCC.................CACGAAUGAAGAGGAAACU..
env-374    CUCGG.UUCCGGUUCACGUCUGAUGC.......................UCGCAGGUCAC.......................................................GCG.CAGGCGACCCGGCG.GCC.................CAAGAAUGAAGAGGAAACU..
env-377    CUCGG.UUCCGGUUCACGUCUGAUGC.......................UCGCAGGUCAC.......................................................GCG.CAGGCGACCCGGCG.GCC.................CANGAAUGAAGAGGAAACU..
env-378    CUCGG.UUCCGGUUCACGUCUGAUGC.......................UCGCAGGUCAC.......................................................GCG.CAGGCGACCCGGCG.GCC..............................CACGA..........
env-379    CGCGG.UACCGGUUCACGUCACGACGGCCAGCCCC..............GACC............................................GGGGC.CCGGCUAGCGGCGACCCGGCG.ACC.................GCACGAGGCAGGAGUUAAC
env-380    UGCGG.UACCGGUUCACGUC.CCGUGCAGC...................UAGC................................................GCAGCACGGUGGCGACCCGGCG.ACC.................GCGGGAACCAGGAGUUAAA
Sgr-9-1    GUCGG.CCCCGGUUCACGCUCGUCGC.......................AACCC.............................................................GCGGA.AGCGACCCGGA..GCC.................CUCCCGAAACUAGGAGACACC
Sex-2-1    GUCGG.CCCCGGUUCACGUCUUCCGC.......................AAUCC.............................................................GCGG.AGGCGACCCGGU..GCC.................CUCCCGAAACUAGGAGACACC
Ssp-115-1  GUCGG.CCCCGGUUCACGUCCGCCGC.......................AACCC.............................................................GCGGC.GACGACCCGGC..GCC.................CUCCCGGAACUAGGAGACACC
Ssp-105-1  GUCGG.CCCCGGUUCACGUCCGCCGC.......................AACCC.............................................................GCGGC.GACGACCCGGC..GCC.................CUCCCGGAACUAGGAGACACC
Mgl-2-1    GUCGG.CCCCGGUUCACGUCCGCCGC.......................AACCC.............................................................GCGGC.GACGACCCGGU..GCC.................CUCCCGGAACUAGGAGACACC
Sgr-12-2   GUCGG.CCCCGGUUCACGUCCGCCGC.......................AACCC.............................................................GCGGC.GACGACCCGGA..GCC.................CUCCCGGAACUAGGAGACACC
Fsp-14-1   UCAGG.CUCCGGUUCACGUCGUCGC........................GACAU...............................................................GCGGCGGCGACCCGGCG.GCC.................AACGUGAGAAGAGGGACCACC
Fsa-1-1    GGAGG.CUCCGGUUCACGUCGUCGC........................GGCU................................................................GCGACGGCGACCCGGCG.GCC.................AGGUGAGAAGAGGGAAC...
Gar-2-1    UGCGG.UACCGGUUCACGGCCCA..........................GCAA....................................................................UGCGCCGACCCGGCG.ACC.................GAACAGAGAAGAGGGAAACUC
Spa-31-1   ACCGG.UUCCGGUUCACGCCCAUCCGUCG....................UACAC.....................................................CUGCGGCCGGGCGACCCGGCG.ACC.................ACCAACGAGAGGACCGAGG
Spa-3-1    ACCGG.UUCCGGUUCACGCCCAUCCGUCG....................UACAC.....................................................CUGCGGCCGGGCGACCCGGCG.ACC.................ACCAACGAGAGGACCGAGG
Spa-39-1   ACCGG.UUCCGGUUCACGCCCAUCCGUCG....................UACACC....................................................CG.CGGCCGGGCGACCCGGCG.ACC.................ACCAACGAGAGGACCGAGG
Spa-35-1   ACCGG.UUCCGGUUCACGCCCAUCCGUCG....................UACACC....................................................CG.CGGCCGGGCGACCCGGCG.ACC.................ACCAACGAGAGGACCGAGG
Spa-25-1   ACCGG.UUCCGGUUCACGCCCAUCCGUCG....................UACACC....................................................CG.CGGCCGGGCGACCCGGCG.ACC.................ACCAACGAGAGGACCGAGG
Nfa-1-1    GUCGG.UUCCGGUUCACGUUCAC..........................GCCC....................................................................GUGGGCGACCCGGCG.ACC.................AAUAUCGAGAGGACAUCC.
Msp-7-1    CUCGG.UUCCGGUUCACGCCCACCUCA..GC..................GCAG..................................................GUCAUGAGC.GGGCGACCCGGCG.GCC.................CACGAUCGAGAGGACACC..
Msp-19-1   CUCGG.UUCCGGUUCACGUCCGA..........................AUAA....................................................................UCGGGCGACCCGGC..GCC.................AUCGAUCGCAGAGGACC....
Mll-1-1    CUCGG.UUCCGGUUCACGUCCGA..........................AUAA....................................................................UCGGGCGACCCGGC..GCC.................AUCGAUCGCAGAGGACC....
Msp-3-1    CUCGG.UUCCGGUUCACGUCCGA..........................AUUC....................................................................UCGGGCGACCCGGC..GCC.................AUCGAACACAGAGGACC....
Msp-20-1   CUCGG.UUCCGGUUCACGUCCGA..........................AUUC....................................................................UCGGGCGACCCGGC..GCC.................AUCGAACACAGAGGACC....
Nsp-7-1    GUCGG.UUCCGGUUCACGUCCAGCACGGCAGCACGGC............AGAUACA......................................GCCGUGG.GCCG....GGGCGACCCGG.G.GCC.................AGAUCGAGAGGACACCC.
Nve-1-1    GUCGG.UUCCGGUUCACGUCCAUCAAGGCAGCACGGC............AGAUUCC......................................GCCGUGG.GCUC....GGGCGACCCGG.G.GCC.................AUAUCGAGAGGACACCC.
Nao-1-1    GUCGG.UUCCGGUUCACGUCCAUCACGGCAGCACGGC............AGAUACC......................................GCCGUGC.GCGC....GGGCGACCCGG.G.GCC.................AGAUCGAGAGGACACCC.
Nce-1-1    GUCGG.UUCCGGUUCACGUCCAUGACGGCAGCACGGC............AGAUACA......................................GCCGUG.UGCUC....GGGCGACCCGG.G.GCC.................AGAUCGAGAGGACACCC.
env-381    CGCGGUUGCCGGUUCACGUCCGCC.........................ACAGG................................................................GGCGGUCGACCCGG.G.GCC...................................CUU........
env-382    CGCGGUUGCCGGUUCACGUCCGCC.........................ACAGG................................................................GGCGGUCGACCCGG.G.GCC.................CUUGAAUCUAGGAGUAACC
env-384    CGCGGUUGCCGGUUCACGUCCGCC.........................ACAGG................................................................GGCGGUCGACCCGG.G.GCC.................CUUGGAUCUAGGAGUAACC
env-386    CGCGGUUGCCGGUUCACGUCCGCC.........................ACAGG................................................................GGCGGUCGACCCGG.G.GCC.................CUUGAAUCUAGGAGUAACC
env-390    CGCGGUUGCCGGUUCACGUCCGCC.........................ACAGG................................................................GGCGGUCGACCCGG.G.GCC.................................CUU........
Bas-1-1    CGCGGUUGCCGGUUCACGUCCGCC.........................ACAGG................................................................GGCGGUCGACCCGG.G.GCC.................CUUGAAUCUAGGAGUAACC
Bas-2-2    CGCGGUUGCCGGUUCACGUCCGCC.........................ACAGG................................................................GGCGGUCGACCCGG.G.GCC.................CUUGGAUCUAGGAGUAACC
Bsp-3-1    CGCGGUUGCCGGUUCACGUCCGCC.........................ACAGG................................................................GGCGGUCGACCCGG.G.GCC.................CUUGGAUCUAGGAGUAACC
env-391    CGCGGUUGCCGGUUCACGUCCGCC.........................ACAGG................................................................GGCGGUCGACCCGG.G.GCC.................CUUGAAUCUAGGAGUAACC
env-394    UGCGGUUGCCGGUUCACGUCCGCC.........................ACAGG................................................................GGCGGUCGACCCGG.G.GCC.................CUUGAAUCUAGGAGUAACC
env-396    CGCGGUUGCCGGUUCACGUCCGCC.........................ACAGG................................................................GGCGGUCGACCCGG.G.GCC.................CUUGACUCUAGGAGUAACC
Bsp-2-1    CGCGGUUGCCGGUUCACGUCCGCC.........................ACAGG................................................................GGCGGUCGACCCGG.G.GCC.................CUUGGAUCUAGGAGUAACC
Bas-2-3    CGCGGUUACCGGUUCACGUCCGCC.........................ACAGG................................................................GGCGGUCGACCCGG.G.GCC.................CUUGAAUCUAGGAGUAACC
Mma-2-1    CUCGG.UCCCGGUUCACGUCC.CCGCGC.....................AUUA........................................................GAUCGGUGGGCGACCCGG...GCC.................AACGAUGAAGAGGAAACC...
Msp-2-1    CUCGG.UCCCGGUUCACGUCCA...........................UCU.......................................................................UGGGCGACCCGG...GCC.................AACUACUGAAGAGGAACC...
env-398    GUCGG.UUCCGGUUCACGUCCUCA.........................UCUC....................................................................UGUGGGCGACCCGGCG.ACC..........................AUCGAAAGGACAUCC.
Nji-1-1    GUCGG.UUCCGGUUCACGUCCUCA.........................UCUC....................................................................UGUGGGCGACCCGGCG.ACC..........................AUCGAAAGGACAUCC.
Nno-1-1    GUCGG.UUCCGGUUCACGUCCACUACGGCCGCACGGC............ACAUC........................................GCUGUG.GGCC.....GGGCGACCCGG.G.GCC.................AUAUCGAGAGGACACCC.
Nhi-1-1    GUCGG.UCCCGGUUCACGUCCCGAUC.......................ACGAAAAC..........................................................GAUCUGGGCGACCCGGCG.GCC.................AAUAUCGAGAGGACACCC.
Msp-1-1    CUCGG.UCCCGGUUCACGUCC............................ACACGC....................................................................GGGCGACCCGG...GCC.................AACGACUGAAGAGGAACC...
Msp-18-1   CUCGG.UCCCGGUUCACGUCC............................ACAG......................................................................GGGCGACCCGG...GCC.................AACGACUGAAGAGGAACC...
Mfo-1-1    CUCGG.UCCCGGUUCACGUCC............................AUCUGAAUUGC...............................................................GGGCGACCCGG...GCC.................AACGAUUGAAGAGGAACCC..
Mvu-1-1    CUCGG.UCCCGGUUCACGUCC............................AUCUGAAUUGC...............................................................GGGCGACCCGG...GCC.................AACGAUUGAAGAGGAACCC..
Mse-1-1    CUCGG.UCCCGGUUCACGUCC............................AUCUGAAUUGC...............................................................GGGCGACCCGG...GCC.................AACGAUUGAAGAGGAACCC..
Msp-23-1   CUCGG.UCCCGGUUCACGUCC............................AUCCGAAUUGC...............................................................GGGCGACCCGG...GCC.................AACGAUUGAAGAGGAACCC..
Mse-2-1    CUCGG.UCCCGGUUCACGUCC............................GUGC......................................................................GGGCGACCCGG...GCC.................AACGAUUGAAGAGGAACC...
Mse-3-1    CUCGG.UCCCGGUUCACGUCCGCG.UACGUGAGGAGCG...........CAUG............................................CGCUCGAAAUGAUCUGCGGGCGACCCGG...GCC.................AACGAUUGAAGAGGAACCC..
Mse-3-2    CUCGG.UCCCGGUUCACGUCCGCG.UACGUGAGGAGCG...........CAUG............................................CGCUCGAAAUGAUCUGCGGGCGACCCGG...GCC.................AACGAUUGAAGAGGAACCC..
Msp-22-1   CUCGG.UUCCGGUUCACGCUCGAAAGCC.....................AUCAGGUUAG..................................................GGC..UCGGGCGACCAGGCG.ACC.................AACAUUGAAGAGGACACCAGAA
Mne-1-1    CUCGG.UUCCGGUUCACGUUCGA....AGACU.................CCCA..................................................GGUCUGGGCUCGGGCGACCCGGCG.GCC.................AACGAUUGAAGAGGAAAACC.
env-400    CUCGG.UUCCGGUUCACGCUCGA....AGACU.................CCCA..................................................GGUCUGGGCUCGGGCGACCCGGCG.GCC.................AACGAUUGAAGAGGAAAAC.
Msp-21-1   CUCGG.UUCCGGUUCACGCUCGA....AGACU.................CCCA..................................................GGUCUGGGCUCGGGCGACCCGGCG.GCC.................AACGAUUGAAGAGGAAAACC.
Sal-7-1    GUCGG.CCCCGGUUCACGCC.CCGC........................AAUCC...............................................................GCGCAGGCGACCCGGU..GCC.................CUCCCGAGAACACAGGAGAUUCC
Szi-1-1    GUCGG.UCCCGGUUCACGCACCCGU........................AUCCC...............................................................GCGGUGGCGACCCGGU..GCC.................CUCCAUGAACAGUAGGAGAAUCC
Ssp-17-1   GUCGG.CCCCGGUUCACGUGAGCGC........................AAACC...............................................................GCGCGUACGACCCGGC..GCC.................CUCCCGAAACUAGGAGACACC
Ssp-52-1   GUCGG.CCCCGGUUCACGUGAGCGC........................AAACC...............................................................GCGCGUACGACCCGGC..GCC.................CUCCCGAAACUAGGAGACACC
Ssp-13-1   GUCGG.CCCCGGUUCACGUGUGCGC........................AAACC...............................................................GCGUGUGCGACCCGGC..GCC.................CUCCCGAAACUAGGAGACACC
Ssp-66-1   GUCGG.CCCCGGUUCACGUGUGCGC........................AAACC...............................................................GCGUGUGCGACCCGGC..GCC.................CUCCCGAAACUAGGAGACACC
Ssp-63-1   GUCGG.CCCCGGUUCACGUGUGCGC........................AAACC...............................................................GCGUGUGCGACCCGGC..GCC.................CUCCCGAAACUAGGAGACACC
Ssp-33-1   GUCGG.CCCCGGUUCACGUGUGCGC........................AAACC...............................................................GCGUGUGCGACCCGGC..GCC.................CUCCCGAAACUAGGAGACACC
Ssp-117-1  GUCGG.CCCCGGUUCACGUGUGCGC........................AAACC...............................................................GCGUGUGCGACCCGGC..GCC.................CUCCCGAAACUAGGAGACACC
Swa-1-1    GUCGG.CCCCGGUUCACGUGUGCGC........................AAACC...............................................................GCGUGUGCGACCCGGC..GCC.................CUCCCGAAACUAGGAGACACC
Ssp-26-1   GUCGG.CCCCGGUUCACGUGUGCGC........................AAACC...............................................................GCGUGUGCGACCCGGC..GCC.................CUCCCGAAACUAGGAGACACC
Sal-4-1    GUCGG.CCCCGGUUCACGUGUGCGC........................AAACC...............................................................GCGUGUGCGACCCGGC..GCC.................CUCCCGAAACUAGGAGACACC

           ...<<.<.<<<<....<<<<<<<<<<<<<<<<<<<<<<<<<<<<<<<<<<..................................................>>>>>>>>>>>>>>>>>>>>>>>>>>>>>>>>>..>>>>..>>>.......................................
           ...11.2.1220....022222222..........1....1.111.10...................................................01.111.1....1..........222222220..0221...211.......................................
           •YCGG-YYCCGGUUCACGYY•••••--------------------•••••---------------------------------------------------•••••GRCGACCCGGC•-RCC------------------••••••R••AGGA•••••
           5′                                                                                                                                 3′
```

```
Ssp-123-1   GUCGG.CCCCGGUUCACGCACGGGC.......................AUCCC..................................................................GCCCGGGCGACCCGGC..GCC................CUCCCGAAACUAGGAGACACC
Sgr-2-1     GUCGG.CCCCGGUUCACGCUCCCGC.......................ACCGA..................................................................GCGGUGGCGACCCGGA..GCC...............CUCCCGGAACCUAGGAGACACC
Sco-3-1     GUCGG.CUCCGGUUCACGCCUCCGC.......................ACCCC..................................................................GCGAAUGCGACCCGGC..GCC................CUCCCGAAACUAGGAGAACCC
Ssp-67-1    GUCGG.UCCCGGUUCACGCUCCCGC.......................AUCCC..................................................................GCGGAGGCGACCCGGC..ACC................CUCCCGAACCUAGGAGACACC
Ssp-137-1   ACCGG.CCCCGGUUCACGCUCCCGC.......................ACCCC..................................................................GCGGAUGCGACCCGGU..GCC................CUCCCGAACCUAGGAGACACC
Slu-1-1     GUCGG.CUCCGGUUCACGCGCCCGC.......................AUCCC..................................................................GCGGAAGCGACCCGGC..GCC................CUCCCGAACCUAGGAGACACC
Sga-2-1     GUCGG.CUCCGGUUCACGCGCUCGC.......................AUCCC..................................................................GCGGACGCGACCCGGC..GCC................CUCCCGAAACUAGGAGACACC
Sau-2-1     GUCGG.CCCCGGUUCACGCAUCCGC.......................AUCCA..................................................................GCGGCGGCGACCCGGC..GCC................CUCCCGAAACUAGGAGACACC
Ssp-21-1    GUCGG.CUCCGGUUCACGCUCCCGC.......................AAAUCC.................................................................GCGUCUGCGACCCGGC..GCC................CUCCCGAACCUAGGAGACACC
Nvu-1-1     GUCGG.CUCCGGUUCACGUUCCCGCG......................AUUCCG................................................................CGCGUGAGCGACCCGGCG.GCC....................AAUAUCGAGAGGACACCC.
Nbr-1-1     GUCGG.CUCCGGUUCACGUUCCCGCG......................AUUCCA................................................................CGCGCGAGCGACCCGGCG.GCC....................AAUAUCGAGAGGACACCC.
Nte-1-1     GUCGG.CUCCGGUUCACGUUCCCGCG......................AUUCC.................................................................CGCGCGAGCGACCCGGCG.GCC....................AAUAUCGAGAGGACACCC.
Spr-2-1     GUCGG.CCCCGGUUCACGCGACCGC.......................ACCC...................................................................GCGGCAGCGACCCGGC..GCC................CUCCCGAAACUAGGAGACACC
Ssp-106-1   GUCGG.CCCCGGUUCACGCGACCGC.......................ACCC...................................................................GCGGCAGCGACCCGGC..GCC................CUCCCGAAACUAGGAGACACC
env-401     GUCGG.CCCCGGUUCACGCCCCUGC.......................ACCCC..................................................................GCGGGAGCGACCCGGU..GCC................CUCCCGAAACUAGGAGACACC
env-402     GUCGG.CCCCGGUUCACGCCCCUGC.......................ACCCC..................................................................GCGGGAGCGACCCGGC..GCC................CUCCCGAAACUAGGAGACACC
Ssp-146-1   GUCGG.CCCCGGUUCACGCCCCUGC.......................ACCCC..................................................................GCGGGAGCGACCCGGC..GCC................CUCCCGAAACUAGGAGACACC
Smi-1-1     GUCGG.CCCCGGUUCACGCCCCCGC.......................AGCCC..................................................................GCGGGGGCGACCCGGU..GCC................CUCCCGAAACUAGGAGACACC
Sda-1-1     GUCGG.CCCCGGUUCACGCCCCCGC.......................ACCCC..................................................................GCGAUGGCGACCCGGU..GCC................CUCCCGAAACUAGGAGACACC
Ssp-161-1   GUCGG.CCCCGGUUCACGCCCCCGC.......................AGCCC..................................................................GCGAUGGCGACCCGGU..GCC................CUCCCGAAACUAGGAGACACC
Ssp-71-1    GUCGG.CCCCGGUUCACGUCUCCGC.......................AGCCC..................................................................GCGGCGGCGACCCGGC..GCC...............CUCCCGAAAACUAGGAGAAACC
Sgr-7-1     GUCGG.CCCCGGUUCACGUCCCCGC.......................AGCCC..................................................................GCGGCGGCGACCCGGA..GCC...............CUCCCGAAACCUAGGAGAAACC
Sgl-3-1     GUCGG.CCCCGGUUCACGCUGCCGC.......................ACCCC..................................................................GCGGAAGCGACCCGGU..GCC................CUCCCGAACCUAGGAGACACC
Ssp-85-1    GUCGG.CCCCGGUUCACGCUCCCGC.......................ACCCC..................................................................GCGGAGGCGACCCGGC..GCC................CUCCCGAACCUAGGAGACACC
Ssp-121-1   GUCGG.CCCCGGUUCACGCUCCCGC.......................ACCCC..................................................................GCGGCGGCGACCCGGA..GCC...............CUCCCGGAACCUAGGAGACACC
Ssp-104-1   GUCGG.CUCCGGUUCACGUCUCCGC.......................AUCCC..................................................................GCGGUGGCGACCCGGC..GCC................CUCCCGAAACUAGGAGACACC
Ssp-60-1    GUCGG.CCCCGGUUCACGCCCCCGC.......................AGCCC..................................................................GCGGCGGCGACCCGGC..GCC...............CUCCCGAAACCUAGGAGACACC
Ssp-72-1    GUCGG.CCCCGGUUCACGCCUCCGC.......................AUCCC..................................................................GCGGCAGCGACCCGGU..GCC...............CUCCCGAAAUCUAGGAGAACCC
Ssp-99-1    GUCGG.CUCCGGUUCACGCUCCCGU.......................AUCCC..................................................................GCGGCGGCGACCCGGC..GCC................CUCCCGAAACUAGGAGACACC
Ssp-89-1    GUCGG.CCCCGGUUCACGCUCCCGC.......................AUUCCC.................................................................GCGGUGGCGACCCGGC..GCC...............CUCCCGGAACCUAGGAGACACC
Sbi-2-1     GUCGG.CCCCGGUUCACGCUCCCGC.......................AUCCC..................................................................GCGGUUGCGACCCGGC..GCC................CUCCCGAAACUAGGAGACACC
Ssp-82-1    GUCGG.CCCCGGUUCACGCUCCCGC.......................AUCCC..................................................................GCGGUGGCGACCCGGC..GCC................CUCCCGAAACUAGGAGACACC
Ssp-3-1     GUCGG.CCCCGGUUCACGCUCCCGC.......................AUCCC..................................................................GCGGCGGCGACCCGGC..GCC................CUCCCGAAACUAGGAGACACC
Sam-1-1     GUCGG.CUCCGGUUCACGCUCCCGC.......................AUCCC..................................................................GCGGCGGCGACCCGGC..GCC................CUCCCGAACCUAGGAGACACC
Ssp-108-1   GUCGG.CUCCGGUUCACGCUCCCGC.......................AUCCC..................................................................GCGGCGGCGACCCGGC..GCC..............CUCCAGGAACAGUAGGAGACACC
Ssp-48-1    GUCGG.CUCCGGUUCACGCUCCCGC.......................AUCCC..................................................................GCGGCGGCGACCCGGC..GCC................CUCCCGAAACUAGGAGACACC
Ssp-124-1   GUCGG.CUCCGGUUCACGCUCCCGC.......................AUCCC..................................................................GCGGCGGCGACCCGGC..GCC...............CUCCCGAAACCUAGGAGACACC
Ssp-110-1   GUCGG.CUCCGGUUCACGCUCCCGC.......................AUCCC..................................................................GCGGCGGCGACCCGGC..GCC..............CUCCAGGAACAGUAGGAGACACC
Ssp-140-1   GUCGG.CUCCGGUUCACGCUCCCGC.......................AUCCC..................................................................GCGGCGGCGACCCGGC..GCC...............CUCCCGAAACCUAGGAGACACC
Ssp-107-1   GUCGG.CUCCGGUUCACGCUCCCGC.......................AUCCC..................................................................GCGGCGGCGACCCGGC..GCC...............CUCCCGAAACCUAGGAGACACC
Ssp-112-1   GUCGG.CUCCGGUUCACGCUCCCGC.......................AUCCC..................................................................GCGGCGGCGACCCGGC..GCC..............CUCCAGGAACAGUAGGAGACACC
Svi-5-1     GUCGG.CUCCGGUUCACGCUCCCGC.......................AUCCC..................................................................GCGGCGGCGACCCGGC..GCC..............CUCCCGAAACAGUAGGAGACACC
Ssp-74-1    GUCGG.CUCCGGUUCACGCUCCCGC.......................AUCCC..................................................................GCGGCGGCGACCCGGC..GCC................CUCCCGAACCUAGGAGUCACC
Sbo-1-1     GUCGG.CUCCGGUUCACGCCCCCGC.......................AUCCC..................................................................GCGACGGCGACCCGGC..GCC................CUCCCGAACCUAGGAGACACC
env-407     GUCGG.CUCCGGUUCACGCCCCCGC.......................AUCCC..................................................................GCGGCGGCGACCCGGC..GCC...............CUCCCGAAACCUAGGAGACACC
Ssp-101-1   GUCGG.CUCCGGUUCACGUCCCCGC.......................AUCCC..................................................................GCGGCGGCGACCCGGC..GCC................CUCUUGAACCAAGGAGACACC
Ssp-136-1   GUCGG.CCCCGGUUCACGUCCCCGC.......................AUCCC..................................................................GCGGCGGCGACCCGGC..GCC................CUCCCGAACCUAGGAGACACC
Sto-1-1     GUCGG.CCCCGGUUCACGCUCCCGC.......................ACCCC..................................................................GCGGCAGCGACCCGGU..GCC................CUCCCGAACCUAGGAGACACC
Sch-1-1     GUCGG.CCCCGGUUCACGCUCCCGC.......................ACCCC..................................................................GCGGCAGCGACCCGGC..GCC...............CUCCCGAAACCUAGGAGACACC
Saf-1-1     GUCGG.CCCCGGUUCACGCUCCCGC.......................ACCCC..................................................................GCGGCAGCGACCCGGC..GCC...............CUCCCGAAACCUAGGAGACACC
Sva-1-1     GUCGG.CUCCGGUUCACGCUCCCGC.......................ACCCC..................................................................GCGGCAGCGACCCGGC..GCC................CUCCCGAACCUAGGAGACACC
Ssp-154-1   GUCGG.CUCCGGUUCACGCUCCCGC.......................ACCCC..................................................................GCGGCAGCGACCCGGC..GCC...............CUCCCGAAACCUAGGAGACACC
Sol-1-1     GUCGG.CUCCGGUUCACGCUCCCGC.......................ACCCC..................................................................GCGGCAGCGACCCGGC..GCC...............CUCCCGAUACCUAGGAGACACC
Ssp-147-1   GUCGG.CUCCGGUUCACGCUCCCGC.......................ACCCC..................................................................GCGGUGGCGACCCGGC..GCC...............CUCCCGAAACCUAGGAGACACC
Sia-1-1     GUCGG.CUCCGGUUCACGCUCCCGC.......................ACCCC..................................................................GCGGCGGCGACCCGGC..GCC................CUCCCGAACCUAGGAGACACC
Sro-1-1     GUCGG.CUCCGGUUCACGCUCCCGC.......................ACCCC..................................................................GCGGCGGCGACCCGGC..GCC...............CUCCCGAAACCUAGGAGACACC
Scy-3-1     GUCGG.CUCCGGUUCACGCCUCCGC.......................AACCC..................................................................GCGGCUGCGACCCGGC..GCC...............CUCCCGGAACCUAGGAGACACC
Sac-1-1     GUCGG.CUCCGGUUCACGCCUUCGC.......................AUCCC..................................................................GCGGCAGCGACCCGGC..GCC...............CUCCCGAAACCUAGGAGACACC
Sla-3-1     GUCGG.CCCCGGUUCACGCCUCCGC.......................AUCCC..................................................................GCGGCAGCGACCCGGC..GCC...............CUCCCGAAACCUAGGAGACACC
Ssp-160-1   GUCGG.CUCCGGUUCACGCCUCCGC.......................AUCCC..................................................................GCGGCAGCGACCCGGC..GCC...............CUCCCGGAACAUAGGAGACACC
Shy-2-1     GUCGG.CUCCGGUUCACGCCUCCGC.......................AUCCC..................................................................GCGGCAGCGACCCGGC..GCC...............CUCCCGAAACCUAGGAGACACC
Ssp-125-1   GUCGG.CUCCGGUUCACGCUCGCGC.......................ACCCC..................................................................GCGUCAGCGACCCGGC..GCC................CUCCCGAACCUAGGAGACACC
Spe-1-1     GUCGG.CUCCGGUUCACGCUCCCGC.......................ACCCC..................................................................GCGUCAGCGACCCGGC..GCC................CUCCCGAAACUAGGAGACACC
Ssp-141-1   GUCGG.CUCCGGUUCACGCUCCCGC.......................ACCCC..................................................................GCGUCAGCGACCCGGC..GCC................CUCCCGAACCUAGGAGACACC
Svi-14-1    GUCGG.CUCCGGUUCACGCUCCCGC.......................ACCCC..................................................................GCGUCAGCGACCCGGC..GCC................CUCCCGAAACUAGGAGACACC
Svi-4-1     GUCGG.CUCCGGUUCACGCUCCCGC.......................AUCCC..................................................................GCGUCAGCGACCCGGC..GCC...............CUCCCGAAACCUAGGAGACACC
env-408     GUCGG.CUCCGGUUCACGCUCCCGC.......................AUCCC..................................................................GCGUCUGCGACCCGGC..GCC...............CUCCCGAAACCUAGGAGACACC
Sol-2-1     GUCGG.CUCCGGUUCACGCUCCCGC.......................AUCCC..................................................................GCGGCUGCGACCCGGC..GCC..............CUCCCGGAACAGUAGGAGACACC
Kau-1-2     GUCGG.CUCCGGUUCACGCUCCCGC.......................AUCCC..................................................................GCGGCUGCGACCCGGC..GCC..............CUCCCGGAACAGUAGGAGACACC
Svi-12-1    GUCGG.CUCCGGUUCACGCUCCCGC.......................AUCCC..................................................................GCGGCUGCGACCCGGC..GCC..............CUCCCGGAACAGUAGGAGACACC
Sco-1-1     GUCGG.CUCCGGUUCACGCUCCCGC.......................AUCCC..................................................................GCGGCUGCGACCCGGC..GCC..............CUCCCGGAACAGUAGGAGACACC
Ssp-58-1    GUCGG.CUCCGGUUCACGCUCCCGC.......................AUCCC..................................................................GCGGCUGCGACCCGGC..GCC..............CUCCCGGAACAGUAGGAGACACC
Sno-1-1     GUCGG.CUCCGGUUCACGCUCCCGC.......................AUCCC..................................................................GCGGCUGCGACCCGGC..GCC................CUCCCGAACCUAGGAGACACC
Sgr-1-1     GUCGG.CUCCGGUUCACGCCUCCGC.......................AUCCC..................................................................GCGGCUGCGACCCGGC..GCC................CUCCCGAACCUAGGAGACACC
Sce-1-1     GUCGG.CUCCGGUUCACGCUUCCGC.......................AUCCC..................................................................GCGGCUGCGACCCGGC..GCC................CUCCCGAAACUAGGAGACACC
Ssp-176-1   GUCGG.CUCCGGUUCACGCUUCCGC.......................AUCCC..................................................................GCGGCUGCGACCCGGC..GCC..............CUCCCGGAACACUAGGAGACACC
Ssp-118-1   GUCGG.CUCCGGUUCACGC.UCCGC.......................AUCCC..................................................................GCGGCUGCGACCCGGC..GCC................CUCCCGAACCUAGGAGACACC
Svi-11-2    GUCGG.CUCCGGUUCACGCUUCCGC.......................AUCCC..................................................................GCGGCUGCGACCCGGC..GCC................CUCCCGAAACUAGGAGACACC
Sgr-12-7    GUCGG.CUCCGGUUCACGCUUCCGC.......................AUCCC..................................................................GCGGUCGCGACCCGGC..GCC................CUCCCGAAACUAGGAGACACC
Ssc-4-1     GUCGG.CUCCGGUUCACGCUUCCGC.......................AUCCC..................................................................GCGGCCGCGACCCGGC..GCC................CUCCCGAUACUAGGAGACACC
Ssp-69-1    GUCGG.CUCCGGUUCACGCCCCCGC.......................ACCC...................................................................GCGGCGGCGACCCGGC..GCC...............CUCCGAGAACCUAGGAGACACC
Ssp-153-1   GUCGG.CCCCGGUUCACGCCCCCGC.......................ACCC...................................................................GCGGCAGCGACCCGGU..GCC................CUCCCGAACCUAGGAGACACC
Sye-2-1     GUCGG.CCCCGGUUCACGCU.CCGC.......................AGCCC..................................................................GCGGCAGCGACCCGGU..GCC................CUCCCGAAACUAGGAGACACC

            ...<<.<.<<<<....<<<<<<<<<<<<<<<<<<<<<<<<<<<<<<<<<.....................................................>>>>>>>>>>>>>>>>>>>>>>>>>>>>>>>>..>>>>...>>>.......................................
            ...11.2.1220....022222222..........1....1.111.10.....................................................01.111.1....1..........222222220..0221...211.......................................
            •YCGG-YYCCGGUUCACGYY∘∘∘∘∘-----------------------•••∘∘------------------------------------------------∘∘∘∘∘GRCGACCCGGC∘-RCC----------------∘∘∘∘•••R••AGGA•••••∘∘
             5′                                                                                                                                          3′
```

```
Sse-1-1     GUCGG.CUCCGGUUCACGCACCCGC........................AUUCCC......................................................................GCGGUGGCGACCCGGC..GCC................CUCCCGGAACCUAGGAGACACC
Sfu-2-1     GUCGG.CUCCGGUUCACGCCUCCGC........................AAUCC.......................................................................GCGGUUGCGACCCGGC..GCC..................CUCCCGAAACUAGGAGACACC
Sdu-1-1     GUCGG.CUCCGGUUCACGUUCCCGC........................AUCCC.......................................................................GCGGCAGCGACCCGGC..GCC..............CUCCCGAAAUCCUAGGAGACACC
Sip-1-1     GUCGG.CUCCGGUUCACGCUUUCGC........................AGCCC.......................................................................GCGACAGCGACCCGGC..GCC..................CUCCCGAAACUAGGAGACACC
Ssc-3-1     GUCGG.CUCCGGUUCACGCUCUCGC........................AGCCC.......................................................................GCGACAGCGACCCGGC..GCC..................CUCCCGAAACUAGGAGACACC
Ssp-59-1    GUCGG.CUCCGGUUCACGCUCUCGC........................AUCCC.......................................................................GCGGCAGCGACCCGGC..GCC..................CUCCCGAAACUAGGAGACACC
Ssp-70-1    GUCGG.UCCCGGUUCACGCC.CCGC........................AGCCC.......................................................................GCGACGGCGACCCGGU..GCC................CUCCCGAUACCUAGGAGAACCC
Ssp-24-1    GUCGG.CCCCGGUUCACGC.CCCGC........................AAGCCC......................................................................GCGGCAGCGACCCGGC..GCC................CUCCCGGAACCUAGGAGACACC
Ssp-10-1    GUCGG.CCCCGGUUCACGUGCGCGC........................AUCCC.......................................................................GCGCGUGCGACCCGGC..GCC..................CUCCCGGAACUAGGAGAAACC
Ssp-53-1    GUCGG.CCCCGGUUCACGUGCGCGC........................AUCCC.......................................................................GCGCGUGCGACCCGGC..GCC..................CUCCCGGAACUAGGAGAAACC
Sni-2-1     GUCGG.CCCCGGUUCACGUACGAGC........................AUCC........................................................................GCGCGUGCGACCCGGC..GCC..............UUCCUCGCCUGGCGAGG.......
Spr-1-1     GUCGG.CCCCGGUUCACGUACGAGC........................AUCCCC......................................................................GCCCGUGCGACCCGGU..GCC..................CUCCCGAACCUAGGAGACACC
Ssp-134-1   GUCGG.CCCCGGUUCACGCGUCCGC........................AUCCC.......................................................................GCGUCCGCGACCCGGC..GCC................CUCCCGCAACCUAGGAGACACC
Sau-1-1     GUCGG.CCCCGGUUCACGCAUCCGC........................AUCCC.......................................................................GCGCCUGCGACCCGGU..GCC..................CUCCCGAAACUAGGAGACACC
Asp-7-1     GUCGG.CCCCGGUUCACGCAUCCGC........................AUCCC.......................................................................GCGUCUGCGACCCGGU..GCC..................CUCCCGAAACUAGGAGACACC
Sal-5-1     GUCGG.CCCCGGUUCACGCAUCCGC........................AUCCC.......................................................................GCGUCUGCGACCCGGU..GCC..................CUCCCGAAACUAGGAGACACC
Svi-2-1     GUCGG.CCCCGGUUCACGCACCCGC........................AUCCC.......................................................................GCGGGGGCGACCCGGA..GCC..................CUCCCGAAACUAGGAGACACC
Stu-1-1     GUCGG.CCCCGGUUCACGCACCCGC........................AUCCC.......................................................................GCGGAGGCGACCCGGA..GCC..................CUCCCGAAACUAGGAGAACCC
Sso-1-1     GUCGG.CCCCGGUUCACGCACCCGC........................AUCCC.......................................................................GCGGAGGCGACCCGGA..GCC..................CUCCCGAAACUAGGAGACACC
Spu-1-1     GUCGG.CCCCGGUUCACGCACCCGC........................AUCCC.......................................................................GCGCGUGCGACCCGGU..GCC..................CUCCCGAAACUAGGAGACACC
Ssp-12-1    GUCGG.CCCCGGUUCACGCACCCGC........................AUCCC.......................................................................GCGCGUGCGACCCGGC..GCC............CUCCGGAACCAUGGGAGACACC...
Ssp-78-1    GUCGG.CCCCGGUUCACGCACCCGC........................AUCCC.......................................................................GCGCGUGCGACCCGGA..GCC..................CUCCCGAAACUAGGAGACACC
Ssp-158-1   GUCGG.CCCCGGUUCACGCGCCCGC........................AUCCC.......................................................................GCCGACGCGACCCGGA..GCC..................CUCCCGAAACUAGGAGACACC
Ssp-79-1    GUCGG.CCCCGGUUCACGGCCCCGC........................ACUUCC......................................................................GCGUGGACGACCCGGC..GCC..................CUCCCGAAACUAGGAGACACC
Ssp-92-1    GUCGG.CCCCGGUUCACGGCCCCGC........................ACACCC......................................................................GCGUGGACGACCCGGC..GCC..................CUCCCGAAACUAGGAGACACC
Ssp-151-1   GUCGG.CCCCGGUUCACGGCCCCGC........................ACACCC......................................................................GCGUGGACGACCCGGC..GCC..................CUCCCGAAACUAGGAGACACC
Sat-2-1     GUCGG.CCCCGGUUCACGGACGCGC........................AAUCA.......................................................................GCGCGAACGACCCGGA..GCC..................CUCCCGAAUCUAGGAGACACC
Sfl-8-1     GUCGG.UCCCGGUUCACGGACGCGC........................AAUCC.......................................................................GCGCGAACGACCCGGA..ACC..................CUCCCGAAUCUAGGAGACACC
Sfl-1-1     GUCGG.UCCCGGUUCACGGACGCGC........................AAUCC.......................................................................GCGCGAACGACCCGGA..ACC..................CUCCCGAAUCUAGGAGACACC
Ssp-126-1   GUCGG.UCCCGGUUCACGGACGCGC........................AAUCC.......................................................................GCGCGUACGACCCGGA..GCC..................CUCCCGAUACUAGGAGACACC
Sgr-11-1    GUCGG.CCCCGGUUCACGGCCCCGC........................AAAUCC......................................................................GCGCGGACGACCCGGU..GCC..................CUCCCGAAACUAGGAGACACC
Ssp-14-1    GUCGG.CCCCGGUUCACGGCCGCGC........................AAAUCC......................................................................GCGCGGACGACCCGGA..GCC..................CUCCCGAAACUAGGAGACACC
Ssp-168-1   GUCGG.CCCCGGUUCACGGCCGCGC........................AAAUCC......................................................................GCGCGGACGACCCGGA..GCC..................CUCCCGAAACUAGGAGACACC
Ssp-49-1    GUCGG.CCCCGGUUCACGGACACGC........................AAUCC.......................................................................GCGUGUACGACCCGGC..GCC..................CUCCCGAAACUAGGAGACACC
Ssp-22-1    GUCGG.CCCCGGUUCACGGACGCGC........................AAUCC.......................................................................GCGCGUACGACCCGGU..GCC..................CUCCCGAACCUAGGAGACACC
Ssp-25-1    GUCGG.CCCCGGUUCACGGACGCGC........................AAUCC.......................................................................GCGCGUACGACCCGGU..GCC..................CUCCCGAACCUAGGAGACACC
Ssp-152-1   GUCGG.CCCCGGUUCACGGACGCGC........................AAUCC.......................................................................GCGCGUGCGACCCGGC..GCC..................CUCCCGAAUCUAGGAGACACC
Ssp-57-1    GUCGG.CCCCGGUUCACGGACGCGC........................AAUCC.......................................................................GCGCGUGCGACCCGGC..GCC..................CUCCCGAAUCUAGGAGACACC
Ssp-40-1    GUCGG.CCCCGGUUCACGGACGCGC........................AAUCC.......................................................................GCGCGUGCGACCCGGC..GCC..................CUCCCGAAUCUAGGAGACACC
Ssp-6-1     GUCGG.CCCCGGUUCACGGACGCGC........................AAUCC.......................................................................GCGCGUACGACCCGGC..GCC..................CUCCCGAACCUAGGAGAAACC
Sha-2-1     GUCGG.CCCCGGUUCACGGACGCGC........................AAUCC.......................................................................GCGCGUGCGACCCGGA..GCC..................CUCCCGAAUCUAGGAGACACC
Ssp-43-1    GUCGG.CCCCGGUUCACGGCCGCGC........................AAUCC.......................................................................GCGCGGACGACCCGGC..GCC..................CUCCCGAAACUAGGAGACACC
Scy-2-1     GUCGG.CCCCGGUUCACGGCCGCGC........................AAUCC.......................................................................GCGCGGACGACCCGGC..GCC..................CUCCCGAAACUAGGAGACACC
Ssp-39-1    GUCGG.CCCCGGUUCACGGCCGCGC........................AAUCC.......................................................................GCGCGGACGACCCGGC..GCC..................CUCCCGAAACUAGGAGACACC
Sfu-1-1     GUCGG.CCCCGGUUCACGGCCGCGC........................AAUCC.......................................................................GCGCGGACGACCCGGC..GCC..................CUCCCGAAACUAGGAGACACC
Seu-1-1     GUCGG.CCCCGGUUCACGGCCGCGC........................AAUCC.......................................................................GCGCGGACGACCCGGC..GCC..................CUCCCGAAACUAGGAGACACC
Sgr-3-1     GUCGG.CCCCGGUUCACGGCCGCGC........................AAUCC.......................................................................GCGCGGACGACCCGGC..GCC..................CUCCCGAAACUAGGAGACACC
Ssp-9-1     GUCGG.CCCCGGUUCACGGCCGCGC........................AAUCC.......................................................................GCGCGGACGACCCGGC..GCC..................CUCCCGAAACUAGGAGACACC
Ssp-68-1    GUCGG.CCCCGGUUCACGGCCGCGC........................AAUCC.......................................................................GCGCGGACGACCCGGC..GCC..................CUCCCGAAACUAGGAGACACC
Sal-8-2     GUCGG.CCCCGGUUCACGGCCGCGC........................AAUCC.......................................................................GCGCGGACGACCCGGA..GCC..................CUCCCGAAACUAGGAGACACC
San-2-1     GUCGG.CCCCGGUUCACGGCCGCGC........................AAUCC.......................................................................GCGUGGACGACCCGGC..GCC..................CUCCCGAAACUAGGAGACACC
Sod-1-1     GUCGG.CUCCGGUUCACGUACGAGC........................AAUCC.......................................................................GCGCGUGCGACCCGGC..GCC...................CUCCCGAUCUAGGAGAGACC
Ssp-149-1   GUCGG.CCCCGGUUCACGUACGAGC........................AAUCC.......................................................................GCGCGUACGACCCGGC..GCC..................CUCCCGAAACUAGGAGACACC
Sfl-4-1     GUCGG.CCCCGGUUCACGUCCGCGC........................AAUCC.......................................................................GCGCGAGCGACCCGGC..GCC..................CUCCCGAAACUAGGAGUCACC
Ssp-127-1   GUCGG.CCCCGGUUCACGGAUGCGC........................AAAUCC......................................................................GCGCGUACGACCCGGC..GCC..................CUCCCGAACCUAGGAGACACC
Ssp-81-1    GUCGG.CCCCGGUUCACGGAUGCGC........................AAAUCC......................................................................GCGCGUACGACCCGGC..GCC..................CUCCCGAACCUAGGAGACACC
Sat-1-1     GUCGG.CCCCGGUUCACGGAUGCGC........................AAAUCC......................................................................GCGCGUACGACCCGGC..GCC..................CUCCCGAACCUAGGAGACACC
Ssp-103-1   GUCGG.UCCCGGUUCACGUACGCGC........................AACACG......................................................................GCGCGUGCGACCCGGA..GCC..................CUCCCGGAACUAGGAGAAUCC
Nsp-3-1     GUCGG.UUCCGGUUCACGUCU.CGC........................AUCC........................................................................GCGUAGACGACCCGGC..GCC...................CCAAGAGAUGAGGACACC..
env-409     UCCGG.UACCGGUUCACGUCCGCUAUCCAGGGA................UUU.......................................................UCCCGA......GGGCGACCCGGCG.ACC..................GACCAAUGAAGAGGACAUC..
env-410     UCCGG.UACCGGUUCACGUCCGCUAUCCAGGGA................UUU.......................................................UCCCGA......GGGCGACCCGGCG.ACC..................GACCAAUGAAGAGGACAUC..
env-411     GUCGG.UUCCGGUUCACGUCCACGCG.......................CUAA.....................................................................CGCAAGGGCGACCCGGCG.ACU..................GCGACCUCGAGAGGAAGACC.
env-412     GUCGG.UUCCGGUUCACGUCCGCGCG.......................CUAA.....................................................................CGCAAGGGCGACCCGGCG.GCU..................GCGACCUCGAGAGGAAGACC.
env-413     CGCGG.UUCCGGUUCACGUCAGGCG........................CCU.......................................................................CGCCCGACGACCCGGCG.ACC....................ACUUCGGAGAGGACAAGC.
env-414     AGCGG.UACCGGUUCACGUCC............................CGGACC........................................................................GGACGACCCGGCG.ACC..............GCCCGAAUGCGAGAGGAAGAUC.
env-415     AGCGG.UACCGGUUCACGUCCG...........................CUUCC........................................................................CGGACGACCCGGCG.ACC...............GCCCGAUGCGAGAGGAAGAUC.
env-416     AGCGG.UACCGGUUCACGUCCG...........................CCUUC........................................................................CGGACGACCCGGCG.ACC...............GCCCGAUGCGAGAGGAAGAUC.
env-417     AUCGG.UACCGGUUCACGUCAG...........................CCC..........................................................................CUGACGACCCGGCG.ACC...............ACAGCAGAAAGAAGAAAUC...
env-418     UGCGG.UACCGGUUCACGUC.CGUCGC......................GAUAGGCAA.................................................GCGGCGCGGCGACCCGGCG.ACC....................GCCAGAACCAGGAGUCAAC
Svi-14-4    GUCGG.CCCCGGUUCACGCUUGCCGC.......................AACCC......................................................................GCGAC.GGCGACCCGGA..GCC..................CUCCCGAAACUAGGAGACACC
Sve-1-1     GUCGG.CCCCGGUUCACGCUUGCCGC.......................AACCC......................................................................GCGAC.GGCGACCCGGA..GCC..................CUCCCGAAACUAGGAGACACC
Ssp-38-1    GUCGG.CCCCGGUUCACGCUUGCCGC.......................AACCC......................................................................GCGAC.GGCGACCCGGA..GCC..................CUCCCGAAACUAGGAGACACC
Sbi-3-1     GUCGG.CCCCGGUUCACGCAUGCCGC.......................AACCC......................................................................GCGGCG.GCGACCCGGU..GCC................CUCCCGGAACAUAGGAGACACC
Sgl-4-2     GUCGG.CCCCGGUUCACGCUGUCCGC.......................AACCC......................................................................GCGG.CGGCGACCCGGU..GCC..................CUCCCGAAACUAGGAGACACC
Svi-9-1     GUCGG.CCCCGGUUCACGCUGUCCGC.......................AACCC......................................................................GCGG.CGGCGACCCGGA..GCC..................CUCCCGAAACUAGGAGACACC
Fsp-2-1     AUCGG.CUCCGGUUCACGCUGUCCGC.......................GCCC.......................................................................GCGG.CAGCGACCCGGCG.GCC...................ACAUCGGAAGAGGGACC...
Fsp-5-1     ACCGG.CUCCGGUUCACGCCGUCCGC.......................UCCC.......................................................................GCGG.CGGCGACCCGGCG.GCC...................ACAUCGGAAGAGGGAUC...
Fsp-1-1     ACCGG.CUCCGGUUCACGCCGUCCGC.......................UCCC.......................................................................GCGG.UGGCGACCCGGCG.GCC...................ACAUCGAAAGAGGGAUC...
Sgh-1-1     GUCGG.CUCCGGUUCACGCCUGUCGC.......................AUCCC......................................................................GCGGC.GGCGACCCGGC..GCC................CUCCCGGAAUCUAGGAGACACC
Ssp-31-1    GUCGG.CUCCGGUUCACGCCCCCCGC.......................ACCCC......................................................................GCGGC.GGCGACCCGGC..GCC..................CUCCCGAACAUAGGAGACACC
Sav-2-1     GUCGG.CUCCGGUUCACGCCCCCCGC.......................AUCCC......................................................................GCGGC.GGCGACCCGGC..GCC..................CUCCCGAAACUAGGAGACACC
Ssp-7-1     GUCGG.CUCCGGUUCACGCU.UCCGC.......................AUCCC......................................................................GCGGACGGCGACCCGGC..GCC................CUCCCGGAACCUAGGAGACACC
Ssp-44-1    GUCGG.CUCCGGUUCACGCU.UCCGC.......................AUCCC......................................................................GCGGACGGCGACCCGGC..GCC................CUCCCGGAACCUAGGAGACACC

            ...<<.<.<<<<....<<<<<<<<<<<<<<<<<<<<<<<<<<<<<<<<<..................................................>>>>>>>>>>>>>>>>>>>>>>>>>>>>>>>>>..>>>>...>>>.......................................
            ...11.2.1220....022222222...........1....1.111.10..................................................01.111.1....1..........222222220..0221...211.......................................
            •YCGG-YYCCGGUUCACGYY∘∘∘∘∘---------------------•••∘∘-----------------------------------------------------------------∘∘∘∘∘GRCGACCCGGC∘-RCC------------------∘∘∘∘•••R••AGGA•••••∘∘
             5′                                                                                                                                             3′
```

```
Sgr-10-1     GUCGG.CCCGGUUCACGCU.CCCGC........................AUCCC.............................................................GCGGGCGGCGACCCGGC..GCC...............CUCCCGAAACCUAGGAGACACC
Ssp-173-1    GUCGG.CCCGGUUCACGCU.CCCGC........................AUCCC.............................................................GCGGGCGGCGACCCGGC..GCC...............CUCCCGAAACCUAGGAGACACC
Ssp-16-1     GUCGG.CUCCGGUUCACGCU.CCCGC.......................AUCCC.............................................................GCGGGCGGCGACCCGGC..GCC...............CUCCCGAAACCUAGGAGACACC
Ssp-61-1     GUCGG.CUCCGGUUCACGCU.CCCGC.......................AUCCC.............................................................GCGGGCGGCGACCCGGC..GCC...............CUCCCGGAACCUAGGAGACACC
Ssp-175-1    GUCGG.CCCGGUUCACGCC.CCCGC........................AACCC.............................................................GCGGACGGCGACCCGGC..GCC................CUCCCGAAACUAGGAGACACC
Fsp-6-1      AGCGG.CUCCGGUUCACGCCCUCCGC.......................GCCC..............................................................GCGGC.GGCGACCCGGCG.GCC.................ACAUCGGAAGAGGGACC...
Fsp-13-1     AACGG.CUCCGGUUCACGCCCUCC.........................AUCC................................................................GGACGGCGACCCGGCG.GCC.................ACAUCGAAAGAGGGACC...
Fal-1-1      GACGG.CUCCGGUUCACGCCCUCC.........................AUCC................................................................GGACGGCGACCCGGCG.GCC.................ACAUCGAAAGAGGGACC...
Gte-4-1      UGCGG.UACCGGUUCACGGCCC..GCC.CCGCGC...............GAA.......................................................GCGCGUUGGCAGGGGCCGACCCGGCG.ACC..................GAACAGAGAGGGAAACUC
Mrh-1-1      UCAGG.UACCGGUUCACGCUCGAUUACCAGG..................GAAAA........................................................CCUGGU....GGGCGACCCGGCG.ACC..............ACAGAAUGCAGAGGACACC..
env-419      CCCGG.CUCCGGUUCACGCC.............................CGCACGUCCCA..............................................................GGCGACCCGGCG.GCC..................ACCUGAAAGGAUCCAG
env-421      CCCGG.CUCCGGUUCACGCC.............................CGCACGUCCAU..............................................................GGCGACCCGGCG.GCC..................ACCCGAAGGGAUCCAG
env-422      CCCGG.CUCCGGUUCACGCC.............................CGCACGUCCAU..............................................................GGCGACCCGGCG.GCC..................ACCCGAAGGGAUCCAG
env-423      UGCGG.UACCGGUUCACGCUCUGAUGG..CUCUGUUGGCUCCUGGGGGGACGA.........................................CCCCCCAGGCCCC..CGGAAAACCAGAAGGGCGACCCGGCG.ACC...............GCCGAAGGCAGGAGUUUGU
env-424      CGCGG.UACCGGUUCACGUCGCAGCGG......................CAUCA.......................................................GCCGCCCGGCGACCCGGCG.ACC...............GACCAAGGCAGGAGUACAC
env-425      CGCGG.UACCGGUUCACGUCGCAGCGG......................CAUCA.......................................................GCCGCCCGGCGACCCGGCG.ACC...............GACCAAGGCAGGAGUACAC
env-426      CGCGG.UACCGGUUCACGUCGCAGCGG......................CAUCA.......................................................GCCGCCCGGCGACCCGGCG.ACC...............GACCAAGGCAGGAGUACAC
Nsp-11-1     CGCGG.UACCGGUUCCCGCCA.UG.........................AGUU..........................................................CACUGGCGACCCGGCG.ACC...............GCACCCCGAAAGGACAGCC.
env-428      CGCGG.UACCGGUUCACGUCACGAAAGCC.AGC.CC.............CAU.........................................GGUGCUCGGCCA..GUGGCGACCCGGCG.ACC................GCACGAGGCAGGAGUAGAA
env-429      CGCGG.UACCGGUUCACGUCA............................AGUAC..........................................................UGGCGACCCGGCG.ACC...............GCAUGAGGCAAGGAGCCAAC
env-430      CGCGG.UACCGGUUCACGUCA............................AGUAC..........................................................UGGCGACCCGGCG.ACC...............GCAUGAGGCAAGGAGCCAAC
env-431      UGAGG.UUCCGGUUCACGUCC............................AAUAGCUCAUCA....................................................GGGCGACCCGGCG.ACC...............GACGAUUGAAGAGGACAUC..
env-432      UGAGG.UUCCGGUUCACGUCC............................AAUAGCUCAUCA....................................................GGGCGACCCGGCG.ACC...............GACGAUUGAAGAGGACAUC..
env-433      UCAGG.UUCCGGUUCACGUCC............................ACGAACUCAUCA....................................................GGGCGACCCGGCG.ACC...............GACGAUUGAAGAGGACAUC..
env-434      UCAGG.UUCCGGUUCACGUCCGAUU........................UACCAGCAC.....................................................GAUCUGGCGACCCGGCG.ACC...............GACUACUGAAGAGGACACC..
env-435      UCAGG.UUCCGGUUCACGUCCGAUU........................UCUUGUCAC.....................................................GAUCUGGCGACCCGGCG.ACC...............GAUUACUGAAGAGGACACC..
Str-2-1      ACCGG.UUCCGGUUCACGCCCUGUCCGUGG...................UACACC......................................................CCGCGG.CCGGGCGACCCGGCG.ACC...............ACCAACGAGAGGACCGAGG
Str-1-1      ACCGG.UUCCGGUUCACGCCCUGUCCGUGG...................UACACC......................................................CCGCGG.CCGGGCGACCCGGCG.ACC...............ACCAACGAGAGGACCGAGG
Spa-24-1     ACCGG.UUCCGGUUCACGCCCGAUCCGUCG...................UACACC......................................................CG.CGGC.CGGGCGACCCGGCG.ACC...............ACCAACGAGAGGACCGAGG
Spa-32-1     ACCGG.UUCCGGUUCACGCCCGAUCCGUCG...................UACACC......................................................CG.CGGC.CGGGCGACCCGGCG.ACC...............ACCAACGAGAGGACCGAGG
Spa-7-1      ACCGG.UUCCGGUUCACGCCCGAUCCGUUG...................CACAC.......................................................CCACGGC.UGGGCGACCCGGCG.ACC...............ACCAACGAGAGGACCGAGG
Spa-27-1     ACCGG.UUCCGGUUCACGCCCGAUCCGUUG...................CACAC.......................................................CCACGGC.UGGGCGACCCGGCG.ACC...............ACCAACGAAAGGACCGAGG
Spa-5-1      ACCGG.UUCCGGUUCACGCCCGAUCCGUUG...................CACAC.......................................................CCACGGC.UGGGCGACCCGGCG.ACC...............ACCAACGAGAGGACCGAGG
Spa-38-1     ACUGG.UUCCGGUUCACGCCCGAUCCGCG....................UACACC.......................................................CGCGGC.CGGGCGACCCGGCG.ACC...............ACCAACGAGAGGACCGAGG
env-436      UCCGG.UUCCGGUUCACGCCCGAUAUCC.....................CAGGUCA........................................................GGAA..CGGGCGACCCGGCG.GCC..............GACUAUCGAAGAGGACACC..
Msp-11-1     UCAGG.UACCGGUUCACGCUCGA..AUUGC...................CAG.........................................................GUAAAAAUCGGGCGACCCGGCG.ACC..............AUCGAAUGCAGAGGACACC..
env-437      GUCGG.UUCCGGUUCACGCCCC...........................GCG..............................................................GGGGCGACCCGGGGCACC................ACUGAGACAAGGAGAACCC
env-438      UCAGG.UUCCGGUUCACGUCCGAUUAC......................AAGCUUCA.......................................................GUA...GGGCGACCCGGCG.ACC...............ACGACCGAAGAGGACAUC..
env-440      GCUGG.UUCCGGUUCACGUCCGUA.........................CCG...........................................................UGCGGGCGACCCGGGGCACC...............UGAAACCUAGGAGACUGU
env-441      UGCGG.UACCGGUUCACGUUCU.GCUGGC....................UCU.........................................................GCCAGCUAGGGCGACCCGGCG.ACC...............GCCGAAGGCAGGAGUUUGU
env-442      UGCGG.UACCGGUUCACGCUCUGCUGG.CUCUGGGGGG...........ACGA...................................................CCCCCCAGCCCCCCAGCAAGGCGACCCGGCG.ACC...............GCCGAAGGCAGGAGUUUGU
Gkr-1-1      UCCGG.UUCCGGUUCACACCAGC..........................UGC............................................................GCUGGUGACCCGGCG.ACC.....................ACGAAAGGACACC.
env-443      UCCGG.UCCCGGUUCACGUCC.CA.CG......................AGCAGUC......................................................CGAUGCGGGCGACCCGGCG.GCC..............GACGAUAUGAAGAGGGCAUC..
env-446      GCCGG.UCCCGGUUCACGUCCGC..........................ACCC..........................................................GCGG.CGACCCGGCG.GCC................CCGGACCCGAAGGAGCACAU
env-447      GUCGG.UUCCGGUUCACGUGUGCG.........................ACCU..........................................................CGCAGGCGACCCGGCG.ACC...................CUGAGACGAAGGACACCU.
env-448      GUCGG.UUCCGGUUCACGUGUGCG.........................ACCU..........................................................CGCAGGCGACCCGGCG.ACC...................CUGAGACGAAGGACACCU.
env-449      GUCGG.UUCCGGUUCACGUCUGC..........................GACC...........................................................GCAGGCGACCCGGCG.ACC...................CUGAAACGAAGGAC.....
Mhe-1-1      UCAGG.UUCCGGUUCACGUCUGCC.........................ACAU..........................................................GGCAGGCGACCCGGCG.GCC..............GACGAUUGAAGAGGACAUC..
Rrh-1-1      UCCGG.UUCCGGUUCACACCUUCG.........................GCAGG.........................................................CGAAGGUGACCCGGCG.ACC.....................AUCGAAAGGGACACC.
Msp-13-1     UCAGG.UUCCGGUUCACGUUCGCCUCC......................ACUUC......................................................GAAGGCGGGCGACCCGGCG.GCC..............AACGAAUGAAGAGGACAUC..
env-450      UCCGG.UUCCGGUUCACGACUUCGAA.......................AUCC........................................................UUCGAGGCCGACCCGGCG.ACC..................AUUCCCGAGAGGACCCC..
Tbi-1-1      UCCGG.UUCCGGUUCACGACUUCGA........................AACC........................................................UCGAGGCCGACCCGGCG.ACC..................AUUCCCGAGAGGACCCC..
Msp-4-1      UCAGG.UUCCGGUUCACGUCCAGCC........................CACG........................................................GGCGGGGCGACCCGGCG.ACC...............CACGAUCGAAGAGGACACC..
Mav-1-1      UCAGG.UUCCGGUUCACGUCCAGCC........................CAAG........................................................GGC.GGGCGACCCGGCG.ACC...............CACGAUCGAAGAGGACAUC..
Msp-16-1     UCAGG.UUCCGGUUCACGUCCGC..........................CCCACAGC........................................................GCGGGCGACCCGGCG.GCC...............CACGAUUGAAGAGGACACC..
env-452      UCAGG.UUCCGGUUCACGUCCGCC.........................UCGAC.......................................................GGCGGGCGACCCGGCG.GCC...............GACGAUCGAAGAGGACAUC..
Msp-12-1     UCAGG.UUCCGGUUCACGUCCGCC.........................CCAA........................................................GGCGGGCGACCCGGCG.GCC...............GACGAUCGAAGAGGACAUC..
Msp-14-1     UCAGG.UUCCGGUUCACGUCCGCCC........................CGCA.......................................................GGGCGGGCGACCCGGCG.GCC...............GACGAUCGAAGAGGACAUC..
Msp-17-1     UCAGG.UUCCGGUUCACGUCCGCCC........................CGCA.......................................................GGGCGGGCGACCCGGCG.ACC...............GACGAUCGAAGAGGACAUC..
env-453      UCAGG.UUCCGGUUCACGCCC............................GCAAGCCAGGUUCGAACUCCA..............................................GGGCGACCCGGCG.ACC...............ACCUAUCGAAGAGGACACU..
Mky-1-1      UCAGG.UUCCGGUUCACGUCCGAGCAUCG....................GCG.......................................................CGAUGC..GGGCGACCCGGCG.GCC...............GACGAUCGAAGAGGAAAUC..
Sar-43-1     ACUGG.UUCCGGUUCACGUCCGAUC.CG.....................UCGCACAUC..................................................CGCGAUCGGGCGACCCGGCG.ACC...............ACCGACGAGAGGACCGAGG
Sar-11-1     AUUGG.UUCCGGUUCACGUCCGAUC.CG.....................UCGCACAUC..................................................CGCGAUCGGGCGACCCGGCG.ACC...............ACCGACGAGAGGACCGAGG
env-454      CCAGG.UUCCGGUUCACGCCCG...........................GCG................................................................CGGGCGACCCGGCG.ACC..............ACAUCAGUUGAGAGGACCGAGC
env-455      CGCGG.UUCCGGUUCACGCCUGC..........................AGCC............................................................GCAGGCGACCCGGCG.ACC...............CACUUGAGAGC.GGAGCACCC
env-456      GGAGG.UUCCGGUUCACGCCCGC..........................AGUCC..........................................................GCAGGCGACCCGGCG.ACC...................GCAGACAGGAGCAGCU
Ase-1-1      UGAGG.UUCCGGUUCACGCCGUGGCG.......................ACAGG......................................................CGCC.UGGCGACCCGGCG.ACC................UACCUGGAGAGGACCCCC.
Fsp-9-1      ACCGG.CUCCGGUUCACGCCGUUC.........................ACUCC.......................................................GAACGGCGACCCGGCG.GCC................AUAUCGAAAGAGGGACC...
env-457      ACCGG.CUCCGGUUCACGCCUGCACCC......................ACCCG.....................................................GGGUGAGGGCGACCCGGCG.GCC...................ACCGCGAAGGAGCCUCC
env-459      ACCGG.CUCCGGUUCACGCCUGCACCC......................ACCCG.....................................................GGGUGAGGGCGACCCGGCG.GCC.......................................
env-460      AGCGG.UUCCGGUUCCCGCC.....CGCGGC..................ACCCCGGUGUGCUCAGCGCGCAAGCCUCACAG.......................GCGAGAGACAGUGGCGACCCGGCG.ACC..............GCGUUUUGGA..........
Spa-33-1     ACCGG.UUCCGGUUCACGCCCUAUCCGUCGUA.................CACC......................................................UGCGGCC....GGGCGACCCGGCG.ACC...............ACCAACGAGAGGACCGAGG
Vma-1-1      ACCGG.UUCCGGUUCACGCCCCUUGAUCGGCGCGC..............AGCG....................................................GCGCGCCA.....GGGCGACCCGGCG.ACC................ACGACGAGAGGACCGAGG
Gbr-1-1      UCCGG.UUCCGGUUCACACCGUCG.........................AGACA.......................................................CGACGGUGACCCGGCG.ACC....................ACAGAAAGGACAACC
Gna-1-1      UCCGG.UUCCGGUUCACACCGUCG.........................AGAGG.......................................................CGACGGUGACCCGGCG.ACC....................ACAGAAAGGACGCAC
Gal-1-1      UCCGG.UUCCGGUUCACACCGUCG.........................AGAGG.......................................................CGACGGUGACCCGGCG.ACC....................CCAGAAAGGACACCC
Gam-2-1      UCCGG.UUCCGGUUCACACCGUCG.........................AGAGG.......................................................CGACGGUGACCCGGCG.ACC....................ACAGAAAGGACGCAC
Gsp-1-1      UCCGG.UUCCGGUUCACACCGUCG.........................AGAGG.......................................................CGACGGUGACCCGGCG.ACC....................ACAGAAAGGACAGCC
Gpa-1-1      UCCGG.UUCCGGUUCACACCGUCG.........................AGAGG.......................................................CGACGGUGACCCGGCG.ACC....................ACAGAAAGGACACCA
Gru-1-1      UCCGG.UUCCGGUUCACACCGUCG.........................AGAGG.......................................................CGACGGUGACCCGGCG.ACC....................CGAGAAAGGACACCG
Gam-3-1      UCCGG.UUCCGGUUCACACCGUCG.........................ACAGG.......................................................CGACGGUGACCCGGCG.ACC....................AUAGAAAGGACAGCC
Gte-3-1      UCCGG.UUCCGGUUCACACCGUCG.........................ACAAG.......................................................CGACGGUGACCCGGCG.ACC....................ACAGAAAGGACGUAC
Nsp-1-1      UCCGG.UUCCGGUUCACACCUUCG.........................ACUCG.......................................................CGAAGGUGACCCGGCG.ACC.....................AUGAAAGGACACCA

             ...<<.<.<<<<....<<<<<<<<<<<<<<<<<<<<<<<<<<<<<<<<<<...............................>>>>>>>>>>>>>>>>>>>>>>>>>>>>>>>>>..>>>>...>>>......................................
             ...11.2.1220....022222222...........1....1.111.10................................01.111.1....1...........222222220..0221...211......................................
             •YCGG-YYCCGGUUCACGYY○○○○○-----------------------•••••○-------------------------------------------------○○○○○GRCGACCCGGC○-RCC-------------○○○○•••R••AGGA•••••○•
              5′                                                                                                                                  3′
```

```
Wsp-1-1   UCCGG.UUCCGGUUCACACCUUCG.........................AUCCG....................................................................CGAAGGUGACCGGCG.ACC.......................AUUGAAAGGACACCA
Kar-2-1   UCCGG.UUCCGGUUCACACCUUCG.........................AUCCG....................................................................CGAAGGUGACCGGCG.ACC.......................AUUGAAAGGACACCA
Gef-1-1   UCCGG.UUCCGGUUCACACCAUCG.........................AUCCA....................................................................CGAUGGUGACCGGCG.ACC......................AUGAGAAAGGACAUCC
env-461   UCCGG.UUCCGGUUCACACCUUCG.........................AUCCA....................................................................CGAUGGUGACCGGCG.ACC......................ACGAGAAAGGACACCG
Gpo-3-1   UCCGG.UUCCGGUUCACACCGACG.........................AUCCG....................................................................CGUCGGUGACCGGCG.ACC......................ACGAGAAAGGACACCA
Grh-1-1   UCCGG.UUCCGGUUCACACCGUCG.........................AUCCC....................................................................CGACGGUGACCGGCG.ACC.......................AUAGAAAGGACACCA
env-462   UCCGG.UUCCGGUUCACGUCCUC..........................GACU......................................................................GAGGGCGACCGGCG.ACC........................UUGAAGGAAGGGCA
Rde-1-1   UCCGG.UUCCGGUUCACGGCUUCG.........................AAUGG....................................................................CGAGGUCGACCGGCG.ACC.......................AUUGAAAGGACACC.
Req-3-1   UCCGG.UUCCGGUUCACGGCCUCG.........................AACGG....................................................................CGAGGACGACCGGCG.GCC.......................AUCGAAAGGACACC.
env-463   UCCGG.UUCCGGUUCACGUCCUCG.........................AACGG....................................................................CGAGGACGACCGGCG.ACC.......................AUUGAAAGGACACC.
Rfa-3-1   UCCGG.UUCCGGUUCACGUCCUCG.........................AACGG....................................................................CGAGGUCGACCGGCG.ACC......................AUAUGAAAGGACACC.
Rsp-2-1   UCCGG.UUCCGGUUCACGUCCUCG.........................AACGG....................................................................CGAGGUCGACCGGCG.ACC......................AUAUGAAAGGACACC.
Rsp-3-1   UCCGG.UUCCGGUUCACGUCUUCG.........................ACAAG....................................................................CGAAGGCGACCGGCG.ACC...........ACUACGAAAGAACGGUAAGGACACC..
Rfa-12-1  UCCGG.UUCCGGUUCACGUCUUCG.........................AUCGG....................................................................CGAAGACGACCGGCG.ACC............ACUAGAGAGAACGGUAAGGACACC..
Rsp-13-1  UCCGG.UUCCGGUUCACGUCUUCG.........................AUCGG....................................................................CGAAGACGACCGGCG.ACC............ACUAGAGAGAACGGUAAGGACACC..
Rop-1-1   UCCGG.UUCCGGUUCACGUCUUCG.........................AACGG....................................................................CGAAGGCGACCGGCG.ACC.......................AUUGAAAGGACACC.
Rfa-21-1  UCCGG.UUCCGGUUCACGUCUUCG.........................AUCAG....................................................................CGAAGGCGACCGGCG.ACC............AUUACAAAGAACGGUAAGGACACC..
Rfa-1-2   UCCGG.UUCCGGUUCACGUCUUCG.........................AUCAG....................................................................CGAAGGCGACCGGCG.ACC............AUUACAAAGAACGGUAAGGACACC..
Rfa-4-1   UCCGG.UUCCGGUUCACGUCUUCG.........................AUCGG....................................................................CGAAGGCGACCGGCG.ACC............AUUACAAAGAACGGUAAGGACACC..
Rfa-7-1   UCCGG.UUCCGGUUCACGUCUUCG.........................AUCGG....................................................................CGAAGGCGACCGGCG.ACC............AUUACAAAGAACGGUAAGGACACC..
Rfa-20-1  UCCGG.UUCCGGUUCACGUCUUCG.........................AUCGG....................................................................CGAAGGCGACCGGCG.ACC............AUUACAAAGAACGGUAAGGACACC..
Rfa-8-1   UCCGG.UUCCGGUUCACGUCUUCG.........................AUCGG....................................................................CGAAGGCGACCGGCG.ACC............AUUACAAAGAACGGUAAGGACACC..
Rtr-1-1   UCCGG.UUCCGGUUCACGGCUUCG.........................AUCCG....................................................................CGAAGUCGACCGGCG.ACC........................ACGAAAGGACACC.
Rsp-19-1  UCCGG.UUCCGGUUCACGGCUUCG.........................AUUCG....................................................................CGAAGCCGACCGGCG.ACC........................ACGAAAGGACACC.
Rsp-21-1  UCCGG.UUCCGGUUCACGGCUUCG.........................AUUGG....................................................................CGAAGGCGACCGGCG.ACC.......................ACCGAAAGGACACC.
Sar-21-1  ACUGG.UUCCGGUUCACGUCCGAUCCGUCGC..................ACACCC............................................................GCGAC....CGGGCGACCGGCG.ACC...................ACCGACGAGAGGACCGAGG
Sar-40-1  ACUGG.UUCCGGUUCACGUCCGAUCCGUCGC..................ACACCC............................................................GCGAC....CGGGCGACCGGCG.ACC...................ACCGACGAGAGGACCGAGG
Sar-27-1  ACUGG.UUCCGGUUCACGUCCGAUCCGUCGC..................ACACCC............................................................GCGAC....CGGGCGACCGGCG.ACC...................ACCGACGAGAGGACCGAGG
Sar-33-1  ACUGG.UUCCGGUUCACGUCCGAUCCGUCGC..................ACACCC............................................................GCGAC....CGGGCGACCGGCG.ACC...................ACCGACGAGAGGACCGAGG
Sar-26-1  ACUGG.UUCCGGUUCACGUCCGAUCCGUCGC..................ACACCC............................................................GCGAC....CGGGCGACCGGCG.ACC...................AUCGACGAGAGGACCGAGG
Sar-22-1  ACUGG.UUCCGGUUCACGUCCGAUCCGUCGC..................ACACCC............................................................GCGAC....CGGGCGACCGGCG.ACC...................ACCGAGGAGAGGACCGAGG
Sar-17-1  ACUGG.UUCCGGUUCACGUCCGAUCCGUCGC..................ACACCC............................................................GCGAC....UGGGCGACCGGCG.ACC...................ACCGACGAGAGGACCGAGG
Sar-24-1  ACUGG.UUCCGGUUCACGUCCGAUCCGUCGC..................ACACCC............................................................GCGACA....GGGCGACCGGCG.ACC...................ACCGACGAGAGGACCGAGG
env-474   ACCGG.UUCCGGUUCACGCCC....GAGUCCGUGCCCGCCGCG......ACCAUC................................................UGCCGCCGGGU.CGGACGCCGAGGGCGACCGGCG.ACC............AUCGACGAAAGGACCGAGGCGAC...
env-475   CCCGG.UUCCGGUUCACGCC..CG.AGCCC.CCGC..............ACCU..........................................................GCGGUGGGCGCCGACGGCGACCGGCG.ACC...................AUCCUCGAGAGGACCGAGA
Whe-1-1   UCCGG.UUCCGGUUCACACCGCCC.CCGC....................ACAC................................................................GCGGAGAUCGGUGACCGGCG.ACC........................AUGAAAGGACACCA
Spa-28-1  ACCGG.UUCCGGUUCACGCCCAUCUGUCGC...................ACACCC............................................................GCGGCC...GGGCGACCGGCG.ACC...................ACCAACGAGAGGACCGAGG
Spa-36-1  ACCGG.UUCCGGUUCACGCCCAAUCCGUUGC..................ACAC..............................................................GCGGC....UGGGCGACCGGCG.ACC...................ACCAACGAGAGGACCGAGG
Spa-8-1   ACCGG.UUCCGGUUCACGCCCAAUCCGUUGC..................ACAC..............................................................GCGGC....UGGGCGACCGGCG.ACC...................ACCAACGAGAGGACCGAGG
Dau-1-1   CCCGG.UUCCGGUUCACGCC.............................ACAC.........................................................................GGCGACCGGCG.ACC...................ACUGACGAAAGGACCGCGG
Mlu-1-1   ACCGG.UUCCGGUUCACGCC..CG.AGCCCGUCGCGU............ACG.........................................................GCGCGACGAGUGAUGACGGCGACCGGCG.ACC............ACCAACGAGAGGACCGAGGCGAG...
Mch-6-1   ACCGG.UUCCGGUUCACGCC..CG.AGCCCGUCGUGU............ACG.........................................................GCGCGACGAGCGAUGACGGCGACCGGCG.ACC............ACCAACGAGAGGACCGAGGCGAG...
Mpa-1-1   ACCGG.UUCCGGUUCACGCC..CG.AGCCCGUCGCGU............ACG.........................................................GCGCGACGAGCGAUGACGGCGACCGGCG.ACC............ACCAACGAGAGGACCGAGGCGAG...
Rsp-9-1   UCCGG.CUCCGGUUCACGUCUUCG.........................ACCCG....................................................................CGAAGGCGACCGGCG.GCC........................ACGAAAGGACACC.
Rrh-2-1   UCCGG.CUCCGGUUCACGUCUUCG.........................ACCCG....................................................................CGAAGGCGACCGGCG.GCC........................ACGAAAGGACACC.
Rsp-14-1  UCCGG.CUCCGGUUCACGUCUUCG.........................ACCCG....................................................................CGAAGGCGACCGGCG.GCC........................AUGAAAGGACACC.
Rsp-5-1   UCCGG.UCCCGGUUCACGUCUCCG.........................AUCCG....................................................................CGGAGGCGACCGGCG.GCC........................ACGAAAGGACACC.
Rru-2-1   UCCGG.UCCCGGUUCACGUCUCCG.........................AUCCG....................................................................CGGAGGCGACCGGCG.GCC........................ACGAAAGGACACC.
Rru-3-1   UCCGG.UCCCGGUUCACGUCUUCG.........................AUCCG....................................................................CGGAGGCGACCGGCG.GCC........................ACGAAAGGACACC.
Rru-1-1   UCCGG.UCCCGGUUCACGUCUUCG.........................AUCCG....................................................................CGAAGGCGACCGGCG.GCC........................ACGAAAGGACACC.
Ntr-1-1   GUCGG.UUCCGGUUCACGUCCUCG.........................AUCUG....................................................................CGAGGGCGACCGGCG.ACC....................ACAUCGAAAGGACACCC.
Not-2-1   GUCGG.UUCCGGUUCACGUCCUCG.........................GUCGU....................................................................CGAGGGCGACCGGCG.ACC......................AUCGAAAGGACAUCC.
Not-1-1   GUCGG.UUCCGGUUCACGUCCUCG.........................GUCGU....................................................................CGAGGGCGACCGGCG.ACC......................AUCGAAAGGACAUCC.
env-476   CGCGG.UACCGGUUCACGCCACGACGGC.CAGCCCC.............ACAU.........................................................GGAGCCCUGCUA..GCGGCGACCGGCG.ACC...................GCACGAGGCAGGAGUUAAC
Msp-26-1  ACCGG.UUCCGGUUCACGCC..CG..AGUCCCGUCG.............CACCA........................................................CGACGGGACGCCCGACGGCGACCGGCG.ACC...................ACCGACGAAAGGACCGAGG
env-477   UGCGG.UACCGGUUCACGCC.CGGGCGUG....................GCAUUA..............................................................CACGCCCGCGGCGACCGGCG.ACC...................GCCAGAGGCAGGAGUAUUA
env-478   GACGG.UUCCGGUUCACGUCUCCCCG.......................CUUG...................................................................CGGGC.GACGACCGGCG.ACC.................AUGUCACGGAGAGGAUCCCC.
env-479   GACGG.UUCCGGUUCACGUCUCCCCG.......................UUCG...................................................................CGGGC.GGCGACCGGCG.ACC..................ACGUCACGAGAGGAUCACCC
Gsp-3-1   UGCGG.UUCCGGUUCACGGCUA...........................CGCG.......................................................................UAGCCGACCGGCG.ACC.....................CGGAGAAAGGGAAGUC.
env-481   UGCGG.UUCCGGUUCACGGC.............................UGCG.........................................................................GCCGACCGGCG.ACC.....................CGGAGAAAGGGAAGUC.
env-482   UGCGG.UACCGGUUCACGCCCCCCUGU.CUC.CC...............AUGGCCC........................................................GGCCACUGGGUGGAGGCG.CCCGGCG.ACC..................GCCUUAGGCGAGGAGAAUC.
env-483   UGCGG.UACCGGUUCACGCUUCC.CACAUCUUCCACGGG..........UCCG......................................................CCCAUGGGC..GUGCGGAGGCG.CCCGGCG.ACC..................GCCUGAGGCGAGGAGAACCA
env-484   UGCGG.UACCGGUUCACGCUCU.GCUGG.....................UUCU.................................................................CCGGCNAG.GCGACCGGCG.ACC...................GCCGAAGGCAGGAGUUUGU
env-485   AGCGG.UACCGGUUCCCGCC.....CGCGCCUUCCAGGUGU........GCCACGGCGCGCGUGCCAA.....................................GCGGCACGGAGGUGUGAACUCGGCGACCGGCG.ACC................GCGUCUCGCGAGAGGAAAAGAC
env-486   CGCGG.UACCGGUUCCCGCC.............................GAAG.........................................................................GGCGACCGGCG.ACC.................GCUCGAGGCACAGGAGUUGUC
Sam-2-1   AGCGG.UUCCGGUUCCCGCCC.GCGCGCACCUCGGUG............UGCCAGCGCGCAAGCCU...........................................CAUCGGCGAGAGACAGUGGCGACCGGCG.ACC.................GCGUUUUGGAGAGGAAAAGAC
Sro-7-1   AGCGG.UUCCGGUUCCCGCCCGCG.........................CGCACCUCAGUGUGCACACCGCGCGCAAGCCUGACAGGCGAGAGA............................CAGUGGCGACCGGCG.ACC.................GCGCUUUGGAGAGGAACUGAA
Sro-3-1   AGCGG.UUCCGGUUCCCGCCCGCG.........................CGCACCUCAGUGUGCACACCGCGCGCAAGCCUGACAGGCGAGAGA............................CAGUGGCGACCGGCG.ACC.................GCGCUUUGGAGAGGAACUGAA
Mau-1-1   ACCGG.UUCCGGUUCACGCC......CGAGCCCGUCG............UGA........................................................CGACGGGG.CGCACCGACGGCGACCGGCG.ACC...................ACCGACGAAAGGACCGAGG
Msp-9-1   ACCGG.UUCCGGUUCACGCC......CGAGCCCGUCG............UGA........................................................CGACGGGG.CGCACCGACGGCGACCGGCG.ACC...................ACCGACGAAAGGACCGAGG
Msp-10-1  ACCGG.UUCCGGUUCACGCC......CGAGCCCGUCG............UGA........................................................CGACGGGG.CGCACCGACGGCGACCGGCG.ACC...................ACCGACGAAAGGACCGAGG
Msp-28-1  ACCGG.UUCCGGUUCACGCC......CGAGCCCGUCG............UGA........................................................CGACGGGG.CGCACCGACGGCGACCGGCG.ACC...................ACCGACGAAAGGACCGAGG
env-487   CGCGG.UUCCGGUUCACGCCAUACAGGGCCGGCCCGUC...........CCAU.......................................................GACGGGUGA..CC....UGGCGACCGGCG.ACC...................GCACGAGGCAGGAGUAAUC
env-488   AUGGG.UUCCGGUUCACGCCG....CGCCUGCCUCGU............UUCG.......................................................ACGGGGCAGGCGGAAGACGGCGACCGGCG.ACC...................UUUGAGGCAAGGAGCCAAC
Msp-8-1   ACCGG.UUCCGGUUCACGCC...CGAGCCGACGCG..............CGCCA........................................................CGCGUCGGGGCCGCACGGCGACCGGCG.ACC..................ACCAAGCGAGAGGACCGAGG
env-489   CGCGG.UUCCGGUUCACGCCGCGCCUGCCCGU.................UGAA.............................................................ACGGGCAGGCG.GGCGACCGGCG.GCC..................GCUUGAGGCAAGGAGCCAAC
env-490   CGCGG.UUCCGGUUCACGCCGCGCCUGCCCGU.................UGAA.............................................................ACGGGCAGGCG.GGCGACCGGCG.GCC.......................GCUUGAGGC......
Nco-1-1   UGCGG.UUCCGGUUCACGCCAUGC.........................UCA......................................................................GC..GGCGACCGGCG.GCC..................GCAGCCCGAAAGGACAGCG.
env-494   CGCGG.UACCGGUUCACGCC.CGGCG.......................GCAU...................................................................CGCCGCGGCGACCGGCG.ACC....................GACGAGGCAGGAGUAGUG
Mca-1-1   ACCGG.UUCCGGUUCACGCCCG....AUCCCGACACG............CGA.........................................................CGCGUCGGGGCCGCACGGGCGACCGGCG.ACC..................ACCAAGCGAGAGGACCGAGG
env-495   CGCGG.UACCGGUUCACGCCACGAGGGC..CCUGCU.............GAC..........................................................GGCAGGAUGCCC....GGCGACCGGCG.ACC.................GCAUGAUGGCAAGGAGCCACC
env-496   UGCGG.UACCGGUUCACGCUUUGCU........................GAUC....................................................................AGC.AGGCGACCGGCG.ACC...................GCCGAAGGCAGGAGUUUGU
env-497   UGCGG.UACCGGUUCACGCCU............................GAG.........................................................................AGGCGACCGGCG.ACC...................GCUCGAGGCAGGAGUUGUC

          ...<<.<.<<<<....<<<<<<<<<<<<<<<<<<<<<<<<<<<<<<<<<<.................................................>>>>>>>>>>>>>>>>>>>>>>>>>>>>>>>>>..>>>>...>>>......................................
          ...11.2.1220....022222222..........1....1.111.10.................................................01.111.1....1..........222222220..0221...211......................................
          •YCGG-YYCCGGUUCACGYY○○○○○------------------------●●●○○------------------------------------------------○○○○○GRCGACCCGGC○-RCC------------------○○○○●●●R●●AGGA●●●●●○○
          5′                                                                                                                                         3′
```

```
env-498   CGCGG.UACCGGUUCACGCCGGGGCGG......................AGUC...................................................................CCG.CCCGGCGACCGGCG.ACC...................GCCAGAGGCAGGAGUACAA
env-499   CGCGG.UACCGGUUCACGCCGGGGCGG......................AGUCC..................................................................CCG.CCCGGCGACCGGCG.ACC...................GCCAGAGGCAGGAGUACAA
env-500   CGCGG.UACCGGUUCACGCCGGGGCGG......................AUUUA..................................................................CCGGCCCGGCGACCGGCG.ACC...................AGCUGAGGCAGGAGUACAU
env-501   CGCGG.UACCGGUUCACGCCGGGGCGG......................AUUC...................................................................CC.CCCCGGCGACCGGCG.ACC...................AGCUAAGGCAGGAGUACCU
env-502   UGCGG.UACCGGUUCACGUCCGCGGCGC.....................GCA...................................................................GCGCCGCAGGCGACCGGCG.ACC...................GACGGAACCAGGAGAAAC.
env-503   UGCGG.UACCGGUUCACGCCGCGGCGG......................CAUCC..................................................................CCGCCCAGGCGACCGGCG.ACC...................GCCCGAGGCAGGAGUACAA
env-505   UGCGG.UACCGGUUCACGCCGCGGCGG......................CAUCC..................................................................CCGCCCAGGCGACCGGCG.ACC...................GCCAGAGGCAGGAGUACAA
env-506   CGCGG.UACCGGUUCACGCCGCGGCGG......................CAUCC..................................................................CCGCCCAGGCGACCGGCG.ACC...................GCCCGAGGCAGGAGUACAC
env-508   UGCGG.UACCGGUUCACGCCGCGGCGG......................CAUCC..................................................................CCGCCCAGGCGACCGGCG.ACC...................GCCCGAGGCAGGAGUACAA
env-509   UGCGG.UACCGGUUCACGCCGCGGCGG......................CAUCC..................................................................CCGCCCAGGCGACCGGCG.ACC.......................GCCCGAG........
env-510   UGCGG.UACCGGUUCACGCCGCGGCGG......................CAUCC..................................................................CCGCCCAGGCGACCGGCG.ACC...................GCCAGAGGCAGGAGUACAA
env-513   CGCGG.UACCGGUUCACGCCGCGGCGG......................CAUCGU.................................................................CCGCC.CGGCGACCGGCG.ACC...................GACGAAGGCAGGAGUACAC
env-514   CGCGG.UACCGGUUCACGUCC............................UGCG.........................................................................GGACGACCGGCG.ACU..................GCAUGAGGCAAGGAGCCAUC
env-515   CGCGG.UACCGGUUCACGUUC............................GCAA.........................................................................GAGCGACCGGCG.ACC...................GCACGAGGCAGGAGUUGUC
Nca-2-1   GUCGG.UUCCGGUUCACGUUCG...........................GCAAAUAUA...................................................................CGAGCGACCGGCG.ACC...................AAUAUCGAGAGGACAUCC.
Asu-1-1   CGAGG.UACCGGUUCACGUCCAUC.........................GUGG......................................................................GAUGGGCGACCGGCG.ACC.................AGCAAAGGAAGAGGAC.....
env-516   UGCGG.UACCGGUUCACGCC.............................CACAGA........................................................................GGCGACCGGCG.ACC...................GCCCAAGGCAGGAGUUGUC
env-517   UGCGG.UACCGGUUCACGCC.............................CACAGA........................................................................GGCGACCGGCG.ACC...................GCCCAAGGCAGGAGUUGUC
env-518   UGCGG.UACCGGUUCACGCC.............................CGUAAA........................................................................GGCGACCGGCG.ACC...................GCCGAAGGCAGGAGUUUGU
env-520   CGCGG.UUCCGGUUCACGCCACCAAGGGUCGGCCCG.............CCCCAUGA.......................................................CGGG.UGACC....UGGCGACCGGCG.ACC...................GCACGAGGCAGGAGAAAUC
env-521   CGCGG.UACCGGUUCACGUCU............................CGCAGC.......................................................................AGGCGACCGGCG.ACC..................GCUUGAGGCAAGGAGCCAAA
env-522   UGCGG.UACCGGUUCACGUCU.GCCCCG.....................CCAGGCA...............................................................CGGAGCAAGGCGACCGGCG.ACC...................GCCAGAACCAGGAGUCUUG
env-523   AACGG.UUCCGGUUCACGCC..CGA.GCCACCGCCAC...GCUG.....UUCGGC................................................CAGCAUUGUGG.GG.GGCAUUGACGGCGACCGGCG.ACC.................AGACCAUCGAGAGGACCGAGC
env-524   UGCGG.UACCGGUUCACGCUCUGCUGG......................UUCU...................................................................CCGGCAAGGCGACCGGCG.ACC...................GCCGAAGGCAGGAGUUUGU
env-526   UGCGG.UACCGGUUCACGCCGCAGCGGC.....................AUCA..................................................................GCCGCC.CGGCGACCGGCG.ACC...................GACCAAGGCAGGAGUACAC
env-527   UGCGG.UACCGGUUCACGCC.CGCGGCC.....................ACAGGCA...............................................................GGCGGCAAGGCGACCGGCG.ACC..................GCCAAGAACCAGGAGUCACG
env-530   UGCGG.UACCGGUUCACGCC.CGCGGCC.....................ACAGGCA...............................................................GGCGGCAAGGCGACCGGCG.ACC..................GCCAAGAACCAGGAGUCACG
env-531   UGCGG.UACCGGUUCACGCC.CGCGGCC.....................ACAGGCA...............................................................GGCGGCAAGGCGACCGGCG.ACC..................GCCAAGAACCAGGAGUCACG
env-532   UGCGG.UACCGGUUCACGCC.CGCGGCC.....................ACAGGCA...............................................................GGCGGCGAGGCGACCGGCG.ACC..................GGCAAGAACCAGGAG.....
env-533   UGCGG.UACCGGUUCACGCC.CGCGGCC.....................ACAGGCA...............................................................GGUGGCGAGGCGACCGGCG.ACC..................GCCAAGAACCAGGAGUCACG
env-534   UGCGG.UACCGGUUCACGCUCUGUUGGC...UCCUGG.............GGGGACGACCCCCCAGACCCC.........................................CCCGGAAAACCGGGCAGGCGACCGGCG.ACC...................GCUCGAGGCAGGAGUUGUC
env-535   UGCGG.UACCGGUUCACGCUCUGUUGGC...UCCUGG.............GGGGACGACCCCCCAGACCCC.........................................CCCGGAAAACCGGGCAGGCGACCGGCG.ACC...................GCUCGAGGCAGGAGUUGUC
env-536   UGCGG.UUCCGGUUCACGCC.CAGGCGCG....................GUAG.................................................................CGUGCCCAUGGCGACCGGCG.ACC...................GGCGGAACCAGGAGAAUUC
env-538   AGCGG.UUCCGGUUCCCGCCG............................CGCGCACCCCGGUGUGCUCAGCGCGCAAGCCUCACAGGCGAGAGACAG.............................UGGCGACCGGCG.ACC.................GCGUUUUGGAGAGGAAAGGCG
Sna-1-1   UCCGG.UACCGGUUCACGCUGCG..........................CGUG.......................................................................CGCGGCGACCGGCG.ACC...................GAGGACAAGAGGAUGAC..
env-539   UGCGG.UACCGGUUCACGCCCUGC.........................UGGC......................................................................GCGGGGCGACCGGCG.ACC....................GGCGAGGCAGGAGUUUGU
env-540   AGCGG.UACCGGUUCACGCCCCGCCGGA.....................UCC...................................................................UCCGGCGCGGCGACCGGCG.ACC........................AGGCAGGAGUAUUU
env-541   AACGG.UACCGGUUCACGCC.CGCCGGA.....................UCU...................................................................UCCGGCGCGGCGACCGGCG.ACC........................AGGCAGGAGUAAUC
env-542   AACGG.UACCGGUUCACGCC.CGCCGGA.....................UCC...................................................................UCCGGCGCGGCGACCGGCG.ACC........................AGGCAGGAGUAUUC
env-543   AACGG.UACCGGUUCACGCC.CGCCGGA.....................UCC...................................................................UCCGGCGCGGCGACCGGCG.ACC........................AGGCAGGAGUAUUC
env-544   AACGG.UACCGGUUCACGCC.CGCCGGA.....................UCC...................................................................UCCGGCGCGGCGACCGGCG.ACC........................AGGCAGGAGUAUUC
env-546   UGCGG.UACCGGUUCACGCCC.CC.........................UGU.......................................................................GGAGGGCGACCGGCG.ACU....................GCGGAACCAGGAGUCUGA
env-547   UGCGG.UACCGGUUCACGCCC.CC.........................UGC.......................................................................GGAGGGCGACCGGCG.ACU....................GCGGAACCAGGAGUCUGA
env-548   UGCGG.UACCGGUUCACGCU.............................CGAG..........................................................................GGCGACCGGCG.ACC...................GCUCAAGGCAGGAGUUGUC
env-549   UGCGG.UACCGGUUCACGCC.............................CCCG..........................................................................GGCGACCGGCG.ACC...................GCUCAAGGCAGGAGUUGUC
env-551   UGCGG.UACCGGUUCACGCC.............................CGAG..........................................................................GGCGACCGGCG.ACC...................GCUCAAGGCAGGAGUUGUC
env-552   UGCGG.UACCGGUUCACGCC.............................CGCG..........................................................................GGCGACCGGCG.ACC...................GCCGAAGGCAGGAGUUUGU
env-553   CGCGG.UACCGGUUCACGCCGUC..CGGUCGUGCGC..............CGA...........................................................GCGUGCG.CCCCAGAGGGCGACCGGCG.ACC.................GCUCUCAAGCGAGGAGAAGAC
env-555   CGCGG.UACCGGUUCACGCC.CCCGGCGCUCACAGC..............UCAGUAGC......................................................GUUGCGAACGCCGCCCGGCGACCGGCG.ACC...................GCCGGAACCAGGAGAGACA
env-557   UGCGG.UACCGGUUCACGCC.............................CUAG..........................................................................GGCGACCGGCG.ACC...................GCUCGAGGCAGGAGUUGUC
env-558   UGCGG.UACCGGUUCACGCC.............................CUAG..........................................................................GGCGACCGGCG.ACC...................GCUCGAGGCAGGAGUUGUC
env-560   CGCGG.UACCGGUUCACGCC.............................UUCG..........................................................................GGCGACCGGCG.ACC...................GCUCGAGGCAGGAGUUGUC
env-561   UGCGG.UACCGGUUCACGCC.............................UGCG..........................................................................GGCGACCGGCG.ACC...................GCUCAAGGCAGGAGUUGUC
env-563   UGCGG.UACCGGUUCACGCC.............................UGCG..........................................................................GGCGACCGGCG.ACC...................GCUCAAGGCAGGAGUUGUC
env-564   UGCGG.UACCGGUUCACGCC.............................UGCG..........................................................................GGCGACCGGCG.ACC...................GCUCAAGGCAGGAGUUGUC
env-565   CGCGG.UACCGGUUCACGCC.............................CUCG..........................................................................GGCGACCGGCG.ACC...................GCUCGAGGCAGGAGUUGUC
env-568   UGCGG.UACCGGUUCACGCC.............................CUCG..........................................................................GGCGACCGGCG.ACC...................GUUCAAGGCAGGAGUUGUC
env-570   CGCGG.UACCGGUUCACGCC.............................CUCG..........................................................................GGCGACCGGCG.ACC...................GCUCGAGGCAGGAGUUGUC
env-571   UGCGG.UACCGGUUCACGCC.............................UGAG..........................................................................GGCGACCGGCG.ACC...................GCUCAAGGCAGGAGUUGUC
env-572   UGCGG.UACCGGUUCACGUCC............................GUAA.........................................................................GGGCGACCGGCG.ACC...................GCCGAAGGCAGGAGUUUGU
env-573   UGCGG.UACCGGUUCACGCCC............................GUAA.........................................................................GGGCGACCGGCG.ACC...................GCCGAAGGCAGGAGUUUGU
env-576   UGCGG.UACCGGUUCACGCCC............................GUCA.........................................................................GGGCGACCGGCG.ACC....................GCCCAGGCAGGAGUUGUC
env-577   ACCGG.UUCCGGUUCACGCC...UGAGCCCGUCC..GCCGUACGCCC...UGCG................................................GGGCG.GCGGUGGGACGAGC.CGGAUGGCGACCGGCG.ACC...................AGCGAAGAGAGGACCGAGG
env-578   UGCGG.UACCGGUUCACGCC.CGUCGCGG....................GUCUG................................................................CCGCGGCGCGGCGACCGGCG.ACC..................GCCAGAGCCCAGGAGUCACA
env-579   UGCGG.UACCGGUUCACGUC.CGCCGCC.....................CCAGGCA...............................................................GGCGGCGAGGCGACCGGCG.ACU..................GCCAAGAACCAGGAGUCACG
env-582   CGCGG.UACCGGUUCCCGUCCU...........................UCG.........................................................................GGGACGACCGGCG.ACC.................GCUCGAGGCACAGGAGUUGUC
env-583   CGUGG.UACCGGUUCCCGUCCU...........................UCGG........................................................................GGGGCGACCGGCG.ACC.................GCUCGAGGCACAGGAGUUGUC
env-584   CGCGG.UACCGGUUCCCGUCCU...........................UCG.........................................................................GGGGCGACCGGCG.ACC.................GCUCGAGGCACAGGAGUUGUC
env-585   CGCGG.UACCGGUUCCCGUCCU...........................UCG.........................................................................GGGGCGACCGGCG.ACC.................GCUCGAGGCACAGGAGUUGUC
env-586   CGUGG.UACCGGUUCCCGUCC............................UUCG.........................................................................GGGCGACCGGCG.ACC.................GUUCGAGGCACAGGAGUUGUC
env-587   UGCGG.UACCGGUUCACGCCCCGCGUUAGCAGUGCUAGCCCAGGGGG..ACGA................................................CCCCCUGGACC....CCCCGGCGCGGGACGACCGGCG.ACC...................GCUCAAGGCAGGAGUUGUC
env-588   UGCGG.UACCGGUUCACGCCCGGCGUUAACAGUGCUAGCCCAGGGGG..ACGA................................................CCCCCUGGACCC.CC.C..GGCGCGGGACGACCGGCG.ACC...................GCUCAAGGCAGGAGUUCUC
env-589   CGCGG.UACCGGUUCCCGCCU............................GCG..........................................................................AGGCGACCGGCG.ACU..................GCUUGAGACAAGGAGCCAAC
env-591   CGCGG.UACCGGUUCCCGCCCC...........................GCG.........................................................................GGGGCGACCGGCG.ACC.................GCCCGAGGCACAGGAGUUGUC
env-592   CGCGG.UACCGGUUCCCGCCC............................GCGA.........................................................................GGGCGACCGGCG.ACU..................GCUUGAGACAAGGAGCCGAA
env-593   UGCGG.UACCGGUUCACGCCGAUGGCGC.....................GAU...................................................................GCGCCGAUGGCGACCGGCG.ACC....................GAGGAACCAGGAGAAAAC
env-594   CGCGG.UACCGGUUCCCGCC.CGC.........................CAGCGC....................................................................GCGAGGCGACCGGCG.ACC...................GCUCGAGGCAGGAGUUGUC
env-596   CGCGG.UACCGGUUCCCGCC.CGC.........................CAGCGC....................................................................GCGAGGCGACCGGCG.ACC...................GCUCGAGGCAGGAGUUGUC
env-597   CGCGG.UACCGGUUCCCGCC.CGC.........................CAGCGC....................................................................GCGAGGCGACCGGCG.ACC...................GCUCGAGGCAGGAGUUGUC
env-598   UGCGG.UACCGGUUCACGCC.............................ACUUGUUCGAACUA................................................................GGCGACCGGCG.ACC..................GCCCGAAGGCAGGAGUUUGU
env-600   CCCGG.UUCCGGUUCACGUCCAGUC........................CACGU...................................................................GGCUGGGCGACCGGCG.GCC......................ACGAGAAGGGAUCCAU
Nbr-4-1   UCCGG.UUCCGGUUCACGCCCGACG........................CUUG.....................................................................CG.CGGGCGACCGGCG.ACC...................GAUAUCGAGAGGACAUCC.
Npa-1-1   UCCGG.UUCCGGUUCACGUCCGACG........................CUCG.....................................................................CGU.GGGCGACCGGCG.ACC...................GAUAUCGAGAGGACAUCC.

          ...<<.<.<<<<....<<<<<<<<<<<<<<<<<<<<<<<<<<<<<<<<<................................................>>>>>>>>>>>>>>>>>>>>>>>>>>>>>>>>>..>>>>...>>>......................................
          ...11.2.1220....022222222..........1....1.111.10................................................01.111.1....1..........222222220..0221...211......................................
          ●YCGG-YYCCGGUUCACGYY○○○○○------------------------●●●○○------------------------------------------○○○○○GRCGACCCGGC○-RCC-----------------○○○○●●●R●●AGGA●●●●○○
          5′                                                                                                                                 3′
```

| | | | | |
|---|---|---|---|---|
| env-601 | CCCGG.CACCGGUUCACGUC | CUCG | GGCGACCGGCG.GCC | ACUUCUUCAGGAGAUCG |
| env-602 | CCCGG.CUCCGGUUCACGUCG | CGCG | CGGCGACCGGCG.GCC | ACCAGAAGGAUCC.. |
| Gam-1-1 | UCCGG.UACCGGUUCACACCGUCGA | ACA | UCGGCGGUGACCGGCG.ACC | ACCAUCGAAGGGACACCAGGAA..... |
| Gso-1-1 | UCCGG.UACCGGUUCACACCGCCGA | UCG | UCGACGGUGACCGGCG.ACC | ACAGAAAGGACAACC |
| Gsh-1-1 | UCCGG.UUCCGGUUCACACC.CCGAG | UCAGAGUU | CUCGGCGGUGACCGGCG.ACC | AACGAGAAGGACAGAA |
| Nsp-5-1 | GUCGG.UUCCGGUUCACGCCCUCG | AUUU | CGACGGCGACCGGCG.GCC | ACAUCGAAAGGACACCC. |
| env-603 | CCAGG.UUCCGGUUCACGCCC | CGGUGC | GGGCGACCGGCG.ACC | AGACCAUUCGAGAGGACCGAGC |
| Ari-2-1 | UGCGG.UACCGGUUCACGCCUCCCACACCUUCCAUGGU | UCG | GCCAUGC...GUGUCGGGGGCG.CCGGCG.ACC | GCCCCAGGCGAGGAGAAACC |
| env-604 | CCAGG.CGCCGGUUCACGUCC | CGCGC | GGACGACCGGCG.ACC | GCCAGACCCGAGGAGCACCC |
| env-605 | CCAGG.CGCCGGUUCACGUCC | CGCGC | GGACGACCGGCG.ACC | GCCAGACCCGAGGAGCACCG |
| env-606 | CCAGG.CGCCGGUUCACGUCC | CGCGC | GGACGACCGGCG.ACC | GCCAGACCCGAGGAGCACCC |
| env-607 | CCAGG.CGCCGGUUCACGUCC | CGCAC | GGACGACCGGCG.ACC | GCCAGACCCGAGGAGCACCC |
| env-608 | CGCGG.CUCCGGUUCACCUCGG | UCCC | CCGGGGACCGGC..GCC | AUAUUCUCGGAAGGACAUUCC |
| Ata-1-1 | UCCGG.CACCGGUUCACCUCG | CGA | CGAGGACCGGCG.GCC | AGCAAGAGAGGACCCC.. |
| Aha-1-1 | UCCGG.CUCCGGUUCACCUC.UGGU | CGAC | ACCACGAGGACCGGU..GCC | ACGAGGAGAGGAUCGAUC |
| env-609 | ACGGG.CUCCGGUUCACCUCGC | AAC | GCGGGGACCGGC..GCC | ACAUUCUCGGAAGGACAUCCC |
| env-610 | CCGGCCUCCGGUUCACCCGCGG | UGCGC | GCGCGGGACCGGC..GCC | CUCUGG........ |
| env-611 | CCGGCCUCCGGUUCACCCGCGG | UGCGC | GCGCGGGACCGGC..GCC | CUC......... |
| env-629 | CCGGCCUCCGGUUCACCCGCGG | UGCGC | GCGCGGGACCGGC..GCC | CUCUGGAUGCUAGGAGAUACC |
| Pau-1-1 | UCCGG.CUCCGGUUCACCCACAUCC.........GGGGGCU.C | UGCCCCCG | GAAGCCCCCGAACGGCGAGGGGUGUGAGACCGGCG.GCC | GAUCUUCGAGAGGAACAUCC |
| Psp-6-1 | CCCGG.CUCCGGUUCACCCGGC | CCGU | GCGGGACCGGCG.GCC | ACCGAGAGGAGAAAUC |
| Ses-1-1 | UCCGG.CUCCGGUUCACCUCACCCGUGGUU | CCCGC | AACCACC...GAGGACCGGCG.GCC | ACCGAAGAGGGACAAGA |
| Ksp-2-1 | UAAGG.CUCCGGUUCACCC | CACG | GGACCGGCG.GCC | GCCGAUGAGAGGAAACAGC |
| Sgl-1-1 | UCCGG.CUCCGGUUCACCCCG | CGGGUA | CGGGGAUCGGCG.GCC | ACGAGGAGAGGACACC.. |
| env-630 | GCCGG.CUCCGGUUCACCCUGCG | ACGG | CGCGGGGACCGGCG.GCC | ACGAGGAGAGGAACCC.. |
| Aba-1-1 | UCCGG.CGCCGGUUCACCCC.GUGG | GAA | CGACUGGGGACCGGA..GCC | AACGAGAGAGGACACC.. |
| Ari-1-1 | UCCGG.CGCCGGUUCACCCC.GUGG | GAA | CGACUGGGGACCGGA..GCC | AACGAGAGAGGACACC.. |
| Ava-2-1 | UCCGG.CGCCGGUUCACCCC.GUGG | GAA | CGACUGGGGACCGGA..GCC | AACGAGAGAGGACACC.. |
| Aal-2-1 | UCUGG.CUCCGGUUCACCCC..UCGC | UCUC | GCGCACGGGGACCGGCG.GCC | GUCGAGAGAAGAGGACACAG. |
| env-631 | ACCGG.CUCCGGUUCACCCU | GCUC | AGGGACCGGCG.GCC | ACAUCGAGAGGACCCAAC |
| env-633 | ACCGG.CUCCGGUUCACCCU | GCUC | AGGGACCGGCG.GCC | ACUUCGAGAGGACCCAUC |
| env-637 | ACCGG.CUCCGGUUCACCCU | GCUC | AGGGACCGGCG.GCC | ACUUCGAGAGGACCCAUC |
| Lae-1-1 | UCCGG.CUCCGGUUCACCC | GCUCACAAACA | GGGACCGGCG.GCC | AUCCUCUGAAAGGACAAGA |
| env-638 | CCCGGUCUCCGGUUCACCCGCGCUGC | GCUC | GC.GUGCGGGACCGGC..GCC | CUCUGGAUGCUAGGAGAUACC |
| env-639 | CCCGGUCUCCGGUUCACCCGCGCUGC | GCUC | GC.GUGCGGGACCGGC..GCC | CUCUGGAUGCUAGGAGAUACC |
| Aen-1-1 | UCCGG.CUCCGGUUCACCCUGA | UCUGUU | UCGGGGACCGGCG.GCC | ACCAGAGGAGAGGCG |
| Aha-3-1 | UCCGG.CACCGGUUCACCCC..A | CUGGCUCACUUUGGGCCGAGCGUUG | UUCGGGGACCGGCG.GCC | ACAUCAGAAAGGACAACC. |
| env-647 | GCCGG.CUCCGGUUCACCCCG | CUGCUCA | CGGGGACCGGCG.GCC | ACGAGGAGAGGAACCC.. |
| env-648 | GUCUG.CUCCGGUUCACCCCG | CUGCUCA | CGGGGACCGGCG.GCC | ACGAGGAGAGGAACCC.. |
| Ksp-1-1 | UCCGG.CUCCGGUUCACCCCGAAG | AUCA | CGAUGGGGACCGGCG.GCC | ACAACGAGAGGCGAGAUC |
| Lal-2-1 | UCCGG.CUCCGGUUCACCC.GUU | CAUC | AACAGGGACCGGCG.GCC | AUCCUUGAAAGGACAAGA |
| Lae-1-2 | UCCGG.CUCCGGUUCACCC.GUU | CACC | AACUGGGACCGGCG.GCC | AUCCUUGAAAGGACAAGA |
| env-649 | UCCGG.CUCCGGUUCACCC | CUCG | ....GGGACCGGCG.GCC | ACCGAGAAAGGGCAAGAU |
| Psp-9-1 | CCCGG.CACCGGUUCACCCUGGU | CCU | ACCAGGGACCGGCG.GCC | AACGAUGAGGAGAGC |
| Aje-1-1 | UCCGG.CACCGGUUCACCCC | GCAUCA | GGGGACCGGG..GCC | AACGAGAGAGGACACC.. |
| [ir-1-1 | UCCGG.CACCGGUUCACCCCGGUAC | GCCC | GUCCCGGGGAUCCGGCG.GCC | AUCUCCAGAGAGGACACC.. |
| Ain-1-1 | UCCGG.CUCCGGUUCACCCC | GACCCA | GGGGACCGGCG.GCC | ACAUGAGAGAGGCGACCAC |
| Aja-1-1 | UCCGG.CACCGGUUCACCCC | GCUC | GGGGACCGG.G.GCC | AACGAGAGAGGACACC.. |
| Asp-6-1 | UCCGG.CACCGGUUCACCCC | GCUC | GGGGACCGG.G.GCC | AACGAGAGAGGACACC.. |
| Aaz-1-1 | UCCGG.CACCGGUUCACCCC | GAAC | GGGGACCGG.G.GCC | AACGAGAGAGGACACC.. |
| Aor-4-1 | UCCGG.CACCGGUUCACCCC | GUACU | GGGGACCGG.G.GCC | AACGAGAGAGGACACC.. |
| Ssp-91-1 | UCCGG.CACCGGUUCACCCC | GUACU | GGGGACCGG.G.GCC | AACGAGAGAGGACACC.. |
| Abe-1-1 | UCCGG.CACCGGUUCACCCCU | GCAC | AGGGGACCGG.G.GCC | AACGAGAGAGGACACC.. |
| Aor-1-1 | UCCGG.CACCGGUUCACCCC | GCACU | GGGGACCGG.G.GCC | AACGAGAGAGGACACC.. |
| Aor-3-1 | UCCGG.CACCGGUUCACCCC | GCACU | GGGGACCGG.G.GCC | AACGAGAGAGGACACC.. |
| Ade-1-1 | UCCGG.CACCGGUUCACCCC | GCAC | GGGGACCGG.G.GCC | AACGAGAGAGGACACC.. |
| Ssp-5-1 | UCCGG.CACCGGUUCACCCC | GCACU | GGGGACCGG.G.GCC | AACGAGAGAGGACACC.. |
| Aor-2-1 | UCCGG.CACCGGUUCACCCC | GCACU | GGGGACCGG.G.GCC | AACGAGAGAGGACACC.. |
| env-650 | UCCGG.CGCCGGUUCACCCCU | CGACGA | AGGGGACCGGG..GCC | ACAUCGAGAGGAAUCCC |
| env-651 | UCCGG.CGCCGGUUCACCCCU | CGACGA | AGGGGACCGG.G.GCC | ACAUCGAGAGGAAUCCC |
| env-652 | UCCGG.CUCCGGUUCACCCCGCG | UCUGCAUC | CACGGGGACCGGCG.GCC | ACGAAGAGAGGACAUCC. |
| Sha-1-1 | UCCGG.CACCGGUUCACCCCGGCACGCC | CGUUCC | GGCGUCCCGGGGAUCGGC..GCC | AUCUCCAGAGAGGACACC.. |
| Spa-41-1 | UCCGG.CACCGGUUCACCCCGGCACGCC | CGUUCC | GGCGUCCCGGGGAUCCGGC..GCC | AUCUCCAGAGAGGACACC.. |
| Kal-2-1 | UCCGG.CUCCGGUUCACCCCGC...GC | ACGACA | GCACGGCGGGGACCGGCG.GCC | ACCACUGAGAGGACAAGAC |
| Saz-1-1 | UCCGG.CUCCGGUUCACCUCACGC | GGUG | GCGUGGGGAUCGGCG.GCC | AGUUGAGAGAGGACACC.. |
| Svi-1-1 | UCCGG.CUCCGGUUCACCCCGC | UGGUC | GCGGGGACCGGCG.GCC | AUGAGGAGAGGACACC.. |
| Ssp-45-1 | UCCGG.CUCCGGUUCACCCCGC | UGUUC | GCGGGGACCGGC..GCU | ACGAGGAGAGGACACC.. |
| Kar-1-1 | UCCGG.CUCCGGUUCACCCCG | AGUGCGUG | UGGGGACCGGCG.GCC | ACAACGAGAGGCGAGAUC |
| Ani-1-1 | UCCGG.CUCCGGUUCACCCC | AGUUC | GGGGACCGGCG.GCC | ACAGAGAGAGGACACC.. |
| Rrh-3-1 | UCCGG.CUCCGGUUCACCCCG | AGUUCGUG | UGGGGACCGGCG.GCC | ACAACGAGAGGCGAGAUC |
| Scy-1-1 | UCCGG.CUCCGGUUCACCCCACGC | GGUU | GUGUGGGGAUCGGCG.GCC | ACGAGGAGAGGACACC.. |
| env-653 | ACCGG.CUCCGGUUCACCCC | GGUA | ...GGGGACCGGC..GCC | ACACCGGAAAGGACAUCCC |
| Ame-3-1 | UCCGG.CGCCGGUUCACCCC.GU | GGAACG | ACUGGGGACCGGA..GCC | AACGAGAGAGGACACC.. |
| Ssp-46-1 | UCCGG.CUCCGGUUCACCCCG | AAGAACAGCAA | CGGGGACCGGCG.GCC | AACCGAGGAAAGGACGAGA |
| Ami-2-1 | UCCGG.CUCCGGUUCACCCCG.GCU | ACC | AGCGCGGGGACCGGCG.GCC | UCACGAGAAGAACAGGACGAGAC |
| Ssp-1-1 | UCCGG.CACCGGUUCACCGCGCGGC | AGCG | GCCGCGUGGACCGGCG.GCC | AAGCAGAGAGGACGAGCC |
| env-654 | UCCGG.CUCCGGUUCACCCC | UCCCAC | GGGGACCGGC..GCC | ACAUCGAGAGGAACCACC |
| env-656 | UCCGG.CUCCGGUUCACCCC | UCCCAC | GGGGACCGGC..GCC | ACAUCGAGAGGAACCACC |
| Sts-1-1 | GUCGG.CCCCGGUUCACGCGCGUGC | AUUCC | GCCCGUGCGACCGGA..GCC | CUCCCGAAACUAGGAGACACC |
| env-657 | CCAGG.UUCCGGUUCCCGCCCGGGGCGCC | CCAG | GGUGCCGACGGCGACCGGCG.ACC | AGAUCAUCUCGAGAGGACCGACC |
| Rsp-11-1 | UCCGG.UUCCGGUUCACGAUGUCG | AUCGG | CGACGUCGACCGGCG.ACC | AUUGAAAGGACACC. |
| env-658 | UCCGG.UUCCGGUUCACGAUGUCG | AUUGG | CGACGUCGACCGGCG.ACC | AUUGAAAGGACACC. |
| Rsp-12-1 | UCCGG.UUCCGGUUCACGAUGUCG | AUUGG | CGACGUCGACCGGCG.ACC | AUUGAAAGGACACC. |

```
...<<.<.<<<<....<<<<<<<<<<<<<<<<<<<<<<<<<<<<<<<<<<<...>>>>>>>>>>>>>>>>>>>>>>>>>>>>>>>>>..>>>>...>>>......
...11.2.1220....022222222..........1....1.111.10...01.111.1....1..........222222220..0221...211......
YCGG-YYCCGGUUCACGYY ... GRCGACCCGGC-RCC ... R AGGA
5′                                        3′
```

```
Rrh-4-1     UCCGG.UUCCGGUUCACGAUGUCG.........................AUUGG..........................................................................CGACGUCGACCCGGCG.ACC.....................AUUGAAAGGACACC.
Ace-1-1     UCAGG.UACCGGUUCACACCGC...........................CAC.............................................................................GCGGUGACCCGGCG.ACC.......................AGAGGAGACUGC
Gja-1-1     UCCGG.UACCGGUUCACACCGCCG.........................AAUCG..........................................................................CGGCGGUGACCCGGCG.ACC.....................ACAGAAAGGGACACCA
Gsp-2-1     UCCGG.UACCGGUUCACACCGCCG.........................AAUCG..........................................................................CGGCGGUGACCCGGCG.ACC.....................AUAGAAAGGGACACCA
Got-1-1     UCCGG.UACCGGUUCACACCGCCG.........................AAUCG..........................................................................CGGCGGUGACCCGGCG.ACC....................ACGAGAAAGGGACACCA
Gai-1-1     UCCGG.UACCGGUUCACACCGCCG.........................AAUCG..........................................................................CGGCGGUGACCCGGCG.ACC....................ACGAGAAAGGGACACCA
Asp-5-2     CCAGG.CACCGGUUCACCCCAGC..........................UCCA............................................................................GC.GGGGACCCGGCG.GCC..................ACGUGAGAAAGGCGACCAC
env-663     GUCGG.UUCCGGUUCACGUCGUCGC........................AUCCGC........................................................................GCGAGGACGACCCGGGG.GCA......................AGGAGAGGACACCC.
Sma-1-1     UCCGG.CACCGGUUCACCUUCC...........................CGCGU...........................................................................GGA.GGACCCGGCG.GCC...................AUCGAGAGAGGACACC..
env-664     UCGGG.CUCCGGUUCACGUUCGCGC........................GUCC..........................................................................GCGCGGGCGACCCGGCG.GCU..................AAGCGAAAGAGGACUAUCA
Ssu-1-1     GCCGG.UCCCGGUUCACGGGCGGC.........................AACCC..........................................................................GCCUCCCGACCCGGC..GCC...............CUCCCGGACACUAGGAGCAACC
Ssu-2-1     GUCGG.UCCCGGUUCACGGGCGGC.........................AACCC..........................................................................GCCUCCCGACCCGGC..GCC...............CUCCCGGACAUUAGGAGAAACC
Sje-1-1     GUCGG.UCCCGGUUCACGGACGGC.........................AGCCC..........................................................................GCCGCCCGACCCGGA..GCC................CUCCCGAACCUAGGAGCACCC
Sal-2-1     GUCGG.UCCCGGUUCACGAACGGC.........................AACCC..........................................................................GCCGUCCGACCCGGA..GCC................CUCCCGAACCUAGGAGCACCC
Sru-1-1     GUCGG.UCCCGGUUCACGAACGGC.........................AGCCC..........................................................................GCCGUCCGACCCGGU..GCC...............CUCCCAGAACAUAGGAGCAUCC
Ssp-100-1   GUCGG.CCCCGGUUCACGGUGGAC.........................AACCC..........................................................................GUCUCCCGACCCGGC..GCC................CUCCCGAAUCUAGGAGUGACC
Ssp-129-1   GUCGG.CCCCGGUUCACGGUGGAC.........................AACCC..........................................................................GUCUCCCGACCCGGC..GCC................CUCCCGAACCUAGGAGUGACC
Ssp-171-1   GUCGG.CCCCGGUUCACGGCGGAC.........................AAGCCC.........................................................................GUCCUCCGACCCGGU..GCC................CUCCAGGAAAUAGGAGACACC
Ssp-170-1   GUCGG.CCCCGGUUCACGGAGGAC.........................AACCC..........................................................................GUCCCCCGACCCGGU..GCC................CUCCCGGAACUAGGAGACACC
Ssp-56-1    GUCGG.CCCCGGUUCACGGGGGAC.........................AACCC..........................................................................GUCCUUCGACCCGGC..GCC................CUCCCGAACCUAGGAGUGACC
Kaz-1-1     GUCGG.CCCCGGUUCACGUGCGGC.........................AUCCAA.........................................................................GCCGC.CGACCCGGC..GCC...............CUCCCGAUCACUAGGAGAAUCC
Svi-7-1     GUCGG.CCCCGGUUCACGUGCGGC.........................AGCCG..........................................................................GCCGCUCGACCCGGU..GCC..............CUCCCUGACAUUU.GGAGAGCCC
Smo-1-1     GUCGG.CCCCGGUUCACGUACGGC.........................AAUCCC.........................................................................GCCGCACGACCCGGU..GCC................CUCCCGAACCUAGGAGAUCCC
Ama-4-1     CUCGG.UUCUGGUUCACGAGACGC.........................ACUCC..........................................................................GCG.CCCGACCCAGCG.GCC...................GUGACGAGAGGACACCC.
env-666     CUUGG.UUCUGGUUCACGGAUCGC.........................AUCCC..........................................................................GCGAUACGACCCAGCG.ACC.....................GUCGAGAGGACAUC..
Bsc-2-1     AUAGG.UUCCGGUUCACGCCCGCC.........................AUAGG..........................................................................GGCUGGCGACCCGGG..UCC...............CCCGAAUUCCAUAGGAGACAUA
Ssc-1-1     GUCGG.CUCCGGUUCACGCUCCCGC........................AUCCC.........................................................................GCGGCGGCGACCCGGC..GCC................CUCCCGAACCUAGGAGACACC
Sle-1-1     GUCGG.CUCCGGUUCACGCCUCCGC........................AACCC.........................................................................GCGGCUGCGACCCGGC..GCC...............CUCCCGGAACCUAGGAGACACC
Scl-1-1     GUCGG.CCCCGGUUCACGCGCGGAC........................AAUCC.........................................................................GCCCGGGCGACCCGGA..GCC................CUCCCGAAACUAGGAGACACC
Sre-3-1     GUCGG.CUCCGGUUCACGCCUCCGC........................AUCCC.........................................................................GCGGCUGCGACCCGGC..GCC................CUCCCGAAACUAGGAGACACC
Ser-2-1     UCCGG.CACCGGUUCACCGCGCGGC........................GCG...........................................................................GCCGCGUGGACCCGGCG.GCC...................ACGAUGAGAGGACAAGGC
Ssp-20-1    GUCGG.CCCCGGUUCACGGUUGGAC........................AACCC.........................................................................GUCUU.CCGACCCGGC..GCC................CUCCCGAACCUAGGAGUGACC
Sst-1-1     GUCGG.CUCCGGUUCACGCCCUCGC........................AUCCC.........................................................................GCGGCGGCGACCCGGC..GCC................CUCCCGAACCUAGGAGACACC
Mne-3-1     CUCGG.UUCCGGUUCACGCUCGA....AGACU.................CCCA..................................................................GGUCUGGGCUCGGGCGACCCGGCG.GCC................AACGAUUGAAGAGGAAAACC.
            ...<<.<.<<<<....<<<<<<<<<<<<<<<<<<<<<<<<<<<<<<<<<.....................................>>>>>>>>>>>>>>>>>>>>>>>>>>>>>>>>>..>>>>...>>>........................................
            ...11.2.1220....022222222..........1....1.111.10......................................01.111.1....1..........222222220..0221...211........................................
            •YCGG-YYCCGGUUCACGYY•••••------------------------•••••-------------------------------------------------------------------•••••GRCGACCCGGC•-RCC-----------------••••••••R••AGGA••••••
             5'                                                                                                                                              3'
```

alignment positions 260 · · · 272

```
env-1       C.........GUG
env-2       AAAC......AUG
Vca-1-1     ..........AUG
env-3       ..........AUG
env-4       ..........AUG
Cac-1-1     AC........AUG
env-7       ..........AUG
env-8       ..........AUG
env-9       ..........AUG
env-10      ..........AUG
env-11      ..........AUG
env-13      ..........AUG
env-14      GGC.......AUG
Apo-1-1     ..........AUG
env-15      ..........AUG
env-16      ..........AUG
env-17      ..........AUG
env-18      ..........GUG
Ama-3-1     ..........AUG
Nin-1-1     ..........AUG
env-19      ..........AUG
env-20      ..........AUG
Jga-1-1     ..........AUG
Hcr-1-1     U.........AUG
env-21      ..........AUG
Asu-2-1     C.........AUG
Psp-8-1     ..........AUG
env-22      ..........AUG
env-24      ..........AUG
env-25      G.........UGA
Sfl-5-1     ..........UUG
Jmu-1-1     AGC.......AUG
env-26      ..........GUG
Ssp-2-1     ..........UUG
env-27      ..........AUG
env-28      ..........AUG
Cbo-1-1     CU........GUG
aLL-1-1     ..........AUG
Msp-29-1    ..........AUG
Mch-5-1     ..........AUG

            .............
            .............
            ----------RUG
```

```
Aha-2-1      UU........AUG
env-29       ..........AUG
Sni-1-1      ..........AUG
env-30       ..........AUG
Asp-2-1      AGAAGAC...AUG
env-31       CGAC......AUG
Hal-1-1      ..........AUG
env-32       ..........GUG
Ssp-169-1    ..........GUG
Dsp-2-1      ..........AUG
Mel-1-1      ..........AUG
env-33       ..........AUG
Car-1-1      ..........AUG
env-34       ..........AUG
Asp-3-1      CU........GUG
env-35       ..........AUG
env-36       ..........AUG
Mth-1-1      ..........AUG
env-37       ..........AUG
env-38       ..........AUG
env-39       UC........GUG
env-40       ..........AUG
env-41       ..........AUG
env-42       ..........AUG
Ssp-96-1     ..........UUG
Sya-1-1      ..........UUG
Ssp-87-1     ..........UUG
Ssp-122-1    ..........UUG
Ssp-97-1     ..........UUG
Ssp-148-1    ..........UUG
Ssp-88-1     ..........UUG
Ssp-164-1    ..........UUG
Svi-13-2     ..........UUG
Ssp-28-1     ..........UUG
Ssp-8-1      ..........UUG
Ssp-143-1    ..........UUG
Sla-2-1      ..........UUG
Ska-1-1      ..........UUG
Sro-6-1      ..........UUG
Ssp-133-1    ..........UUG
Sxa-1-1      ..........UUG
Ska-1-2      ..........UUG
Ssp-138-1    ..........UUG
Ser-3-1      ..........UUG
Sla-2-2      ..........UUG
Ssp-131-1    ..........UUG
Ssp-86-1     ..........UUG
Sla-1-1      ..........UUG
Ssp-166-1    ..........UUG
Svi-13-3     ..........UUG
Dsp-1-1      AC........AUG
Dho-1-1      AC........AUG
Bpa-1-1      ..........AUG
Bph-1-1      ..........AUG
Bfa-1-1      ..........AUG
env-43       ..........AUG
env-45       ..........AUG
env-46       ..........AUG
Bmu-1-1      CG........AUG
Bsq-1-1      CG........AUG
Sba-1-1      ..........UUG
env-47       CC........GUG
env-48       ..........AUG
env-49       ..........GUG
env-50       C.........GUG
env-51       C.........GUG
env-52       ..........GUG
env-53       AAG.......AUG
env-54       C.........GUG
env-55       ..........AUG
env-56       GAGUA.....AUG
env-57       ..........AUG
env-59       ..........AUG
Spo-1-1      C.........GUG
env-60       ..........AUG
env-61       GA........AUG
Ssp-51-1     UUG.......AAG
Sji-1-1      C.........UUG
Aut-1-1      AAGAC.....AUG
Mtu-1-1      ..........AUG
Cko-1-1      GGCG......AUG
Nla-1-1      U.........AUG
Nsp-9-1      C.........AUG
env-62       ..........AUG

             .............
             .............
             ----------RUG
```

| | |
|---|---|
| Afr-1-1 | AAACC.....AUG |
| env-63 | CAACC.....AUG |
| env-64 | AAGAC.....AUG |
| Asu-3-1 | AGAUA........ |
| env-65 | ..........AUG |
| env-66 | ..........AUG |
| env-67 | ..........AUG |
| env-68 | ..........AUG |
| env-69 | ..........AUG |
| env-70 | ..........AUG |
| env-73 | ..........AUG |
| env-74 | ..........AUG |
| env-75 | ..........AUG |
| env-78 | ..........AUG |
| env-80 | ..........AUG |
| Ncy-1-1 | ..........AUG |
| Ncy-2-1 | ..........AUG |
| Npn-1-1 | ..........AUG |
| env-81 | ..........AAU |
| env-82 | ..........AUG |
| env-83 | ..........AUG |
| env-84 | ..........AUG |
| env-85 | ..........AUG |
| env-86 | ..........AUG |
| env-87 | C.........AUG |
| env-88 | ..........AUG |
| Sre-2-1 | G.........UUG |
| Acy-1-1 | CU........GUG |
| Mrh-2-1 | ..........AUG |
| env-90 | ..........AUG |
| env-92 | ..........AUG |
| env-93 | ..........AUG |
| env-94 | ..........AUG |
| env-95 | ..........AUG |
| env-96 | ..........AUG |
| env-97 | ..........AUG |
| env-99 | ..........AUG |
| env-103 | ..........AUG |
| env-104 | ..........AUG |
| env-105 | ..........AUG |
| env-111 | ..........AUG |
| Bin-1-1 | U.........AUG |
| env-112 | U.........AUG |
| Bco-1-1 | U.........AUG |
| env-114 | ACA.......GUG |
| env-115 | ACA.......GUG |
| env-116 | ..........AUG |
| Sro-4-1 | ..........UUG |
| Sgl-4-1 | ..........UUG |
| Sal-8-1 | ..........UUG |
| Sgr-12-1 | ..........UUG |
| Ssp-55-1 | ..........UUG |
| Ssp-62-1 | ..........UUG |
| Ssp-167-1 | ..........GUG |
| Sav-3-1 | C.........GUG |
| Ssp-4-1 | ..........AUG |
| Sxi-2-1 | ..........AUG |
| Ssp-90-1 | ..........AUG |
| Jal-1-1 | AGC.......AUG |
| env-117 | ..........AUG |
| env-125 | AU........AUG |
| Psp-1-1 | GU........AUG |
| env-126 | ..........AUG |
| Pac-1-1 | A.........GUG |
| env-127 | ..........AUG |
| env-128 | CG........GUG |
| env-130 | ACCC......GUG |
| env-131 | ACCC......GUG |
| env-132 | UC........AUG |
| env-133 | ..........AUG |
| Bth-2-1 | CC........AUG |
| Bsc-1-1 | CC........AUG |
| Bth-7-1 | CC........AUG |
| Bth-8-1 | CC........AUG |
| Bth-5-1 | CC........AUG |
| Bth-3-1 | CC........AUG |
| Bth-6-1 | CC........AUG |
| Bbo-2-1 | CC........AUG |
| env-136 | ..........AUG |
| Nsp-4-1 | ..........AUG |
| env-137 | ..........AUG |
| env-138 | ..........AUG |
| env-139 | ..........AUG |
| env-140 | ..........AUG |
| | ............. |
| | ............. |
|  | ----------RUG |

| | |
|---|---|
| env-141 | ............. |
| env-142 | ............. |
| Fsp-8-1 | ..........AUG |
| env-143 | ............. |
| Nsp-10-1 | ..........AUG |
| env-144 | ..........GUG |
| Cja-1-1 | CAC.......AUG |
| Fsy-1-1 | ..........AUG |
| Fco-1-1 | ..........AUG |
| env-145 | ..........AUG |
| Ply-1-1 | AA........AUG |
| Ply-2-1 | AA........AUG |
| Ssc-2-1 | U.........GUG |
| Sri-4-1 | C.........UUG |
| Sri-2-1 | C.........UUG |
| Sgr-6-1 | C.........UUG |
| Sri-4-16 | C.........UUG |
| Ssp-75-1 | C.........UUG |
| Sri-4-18 | C.........UUG |
| Sri-4-19 | C.........UUG |
| Sri-4-20 | C.........UUG |
| env-146 | ..........AUG |
| env-147 | ..........AUG |
| env-148 | ............. |
| Nal-1-1 | C.........AUG |
| env-149 | C.........UUG |
| Ssp-36-1 | ..........UUG |
| Ssp-109-1 | ..........UUG |
| Ssp-35-1 | ..........UUG |
| Ssp-34-1 | ..........UUG |
| Ssp-42-1 | ..........UUG |
| env-150 | GGC.......AUG |
| Mau-2-1 | ..........AUG |
| env-151 | ..........AUG |
| env-153 | CC........AUG |
| Psp-10-1 | ..........GUG |
| env-154 | ..........AUG |
| env-156 | ..........AUG |
| env-157 | A.........GUG |
| env-158 | A.........GUG |
| env-159 | A.........GUG |
| env-160 | A.........GUG |
| Mha-1-1 | ..........AUG |
| env-161 | ..........AUG |
| env-162 | AAG.......AUG |
| env-163 | CC........AUG |
| env-164 | AGAC......AUG |
| Asp-5-1 | C.........AUG |
| env-165 | GU........AAU |
| Nlu-1-1 | ..........AUG |
| Nba-1-1 | ..........AUG |
| Ssp-177-1 | ..........UUG |
| Lal-1-1 | G.........AUG |
| Gma-1-1 | ..........AUG |
| Bme-1-1 | ..........AUG |
| Bsu-2-1 | ..........AUG |
| Bre-1-1 | A.........AUG |
| Blo-34-1 | AA........AUG |
| env-166 | AA........AUG |
| Blo-22-1 | AA........AUG |
| env-167 | AA........AUG |
| env-170 | AA........AUG |
| env-196 | AA........AUG |
| Blo-19-1 | AA........AUG |
| env-197 | AA........AUG |
| env-216 | AA........AUG |
| env-217 | AA........AUG |
| env-219 | ..........AUG |
| Ssp-47-1 | ..........GUG |
| Sma-2-1 | ..........GUG |
| Nrh-1-1 | ..........AUG |
| Nte-2-1 | ..........AUG |
| Nsp-2-1 | ..........AUG |
| env-220 | ..........AUG |
| env-221 | ..........AUG |
| env-222 | ..........AUG |
| env-223 | C.........GUG |
| env-224 | C.........GUG |
| env-225 | AA........GUG |
| env-226 | C.........GUG |
| env-227 | ..........GUG |
| env-228 | C.........GUG |
| env-229 | U.........GUG |
| env-230 | CU........GUG |
| | ............. |
| | ............. |
|  | ----------RUG |

```
env-231       ..........AUG
env-232       ..........GUG
env-233       ..........AUG
Fsp-3-1       ..........AUG
Ssy-1-1       CG........UUG
Ssp-54-1      ..........UUG
Ssp-174-1     ..........UUG
Ssp-32-1      ..........UUG
Sxy-1-1       ..........UUG
env-237       G.........AUG
env-238       G.........AUG
env-239       G.........AUG
env-240       ..........AUG
Ssp-102-1     C.........UUG
Sca-3-1       ..........UUG
Ghi-1-2       AU........AUG
Ssc-5-1       ..........AUG
Sro-5-1       C.........UUG
Sva-2-1       C.........UUG
Sye-1-1       ..........UUG
env-241       ..........AUG
env-242       ..........AUG
Nex-1-1       ..........AUG
Nar-1-1       ..........AUG
Nab-1-1       ..........AUG
Mar-1-1       ..........AUG
env-243       ..........AUG
env-244       ..........AUG
env-245       ..........AUG
Agl-1-1       GAAAC.....AUG
Msp-15-1      ..........AUG
Mco-2-1       ..........AUG
env-248       ..........AUG
env-249       ..........AUG
Nmu-1-1       G.........AUG
env-250       CG........AUG
env-252       CG........AUG
env-253       ..........AUG
env-254       ..........AUG
env-255       ..........AUG
env-256       ..........AUG
env-257       ..........AUG
env-258       ..........AUG
env-259       GC........GUG
env-260       C.........GUG
env-261       CU........GUG
env-262       U.........GUG
env-263       ACCCCAUC..AUG
env-264       CC........GUG
env-265       ..........UUG
env-266       UU........GUG
Nth-1-1       ..........AUG
Pan-1-1       ..........AUG
Sxi-1-1       ..........AUG
Ssa-1-1       ..........AUG
Mch-1-1       ..........AUG
env-267       ..........AUG
env-268       ..........AUG
env-274       ..........AUG
Pas-1-1       ..........AUG
Pdi-1-1       ..........AUG
env-279       ..........AUG
env-280       ..........GUG
Ame-4-2       ..........AUG
Asp-4-1       ..........AUG
Ath-1-1       ..........AUG
Sbi-1-1       C.........UUG
Shy-4-1       C.........UUG
Ssp-73-1      C.........UUG
Sir-1-1       C.........UUG
Ssp-165-1     C.........UUG
Svi-3-1       C.........UUG
env-281       C.........GUG
env-282       C.........AUG
env-283       AA........AUG
env-288       ..........AUG
env-289       ..........AUG
env-290       C.........AUG
Gar-1-1       ..........AUG
Mro-1-1       C.........AUG
env-291       ..........AUG
env-292       ..........AUG
Psu-1-1       A.........AUG
Gne-1-1       C.........AUG

              .............
              .............
              ----------RUG
```

```
Gsi-1-1     ..........AUG
Gsi-2-1     ..........AUG
Nta-1-1     ..........AUG
env-293     ..........AUG
env-294     ..........AUG
env-295     ..........AUG
env-298     .............
Mob-1-1     ..........AUG
Mob-1-2     ..........AUG
env-299     CUUCC.....AUG
Asp-1-1     CACAC.....AUG
env-300     CACCC.....AUG
env-301     CACCC.....AUG
env-302     ..........AUG
env-303     ..........AUG
Nsp-6-1     ..........AUG
env-304     ..........AUG
env-306     ..........AUG
env-307     ..........AUG
env-308     ..........AUG
env-309     ..........AUG
env-310     ..........AUG
env-311     ..........AUG
env-312     ..........AUG
env-314     AC........AUG
Ami-1-1     UAAAC.....AUG
env-315     ..........AUG
env-317     ..........AUG
env-318     ..........AUG
env-319     ..........AUG
env-320     ..........AUG
env-321     ..........AUG
Nni-1-1     ..........AUG
Nsp-8-1     ..........AUG
env-322     ..........AUG
Rsp-20-1    ..........AUG
Rsp-16-1    ..........AUG
Mpu-1-1     CGAC......AUG
Msp-27-1    CGAC......AUG
env-323     ..........AUG
env-324     ..........AUG
Aal-1-1     ..........GUG
env-325     ..........AUG
env-329     ..........AUG
env-330     CCA.......GUG
env-331     CCA.......GUG
env-333     ..........AUG
env-334     ..........AUG
env-335     ..........GUG
env-336     C.........AUG
env-337     ..........AUG
env-338     ..........AUG
Bts-1-1     CC........AUG
Ama-5-1     ..........AUG
env-339     C.........GUG
env-340     ..........AUG
env-341     ..........AUG
env-342     ..........AUG
env-343     ..........AUG
env-344     ..........AUG
env-345     ..........AUG
Mva-1-1     ..........AUG
Mru-1-1     ..........AUG
Mva-2-1     ..........AUG
Mgi-1-1     ..........AUG
env-346     ..........AUG
env-347     GGC.......AUG
env-348     GCC.......AUG
Nas-2-1     ..........AUG
Mab-65-1    ..........AUG
env-349     ..........AUG
env-350     GACUC.....AUG
Fsp-7-1     ..........AUG
Sca-1-1     ..........UUG
Ssv-1-1     ..........UUG
Ssp-50-1    .............
Ssp-19-1    ..........UUG
Aro-1-1     ..........AUG
Sca-2-1     ..........UUG
Nvi-1-1     ..........AUG
env-351     C.........UUG
env-352     ..........AUG
env-353     ..........AUG
env-355     ..........AUG

            .............
            .............
            ----------RUG
```

```
env-358      ..........AUG
env-359      ..........AUG
env-360      ..........AUG
env-361      ..........AUG
env-363      ..........AUG
env-364      ..........AUG
env-365      ..........AUG
Mim-1-1      ..........AUG
Mab-73-1     ..........AUG
Mab-55-1     ..........AUG
Mab-1-114    ..........AUG
Mma-1-1      ..........AUG
env-366      ..........AUG
env-367      GAC.......AUG
env-368      GAC.......AUG
env-369      ..........AUG
Tsp-1-1      UU........AUG
Tpa-1-1      GAC.......AUG
Are-1-1      AAGAC.....AUG
Nca-3-1      ..........AUG
env-370      ..........AUG
env-371      C.........GUG
Mph-1-1      ..........AUG
env-372      ..........AUG
env-373      ..........AUG
env-374      ..........AUG
env-377      ..........AUG
env-378      ..........AUG
env-379      ..........AUG
env-380      CC........AUG
Sgr-9-1      ..........AUG
Sex-2-1      ..........AUG
Ssp-115-1    ..........AUG
Ssp-105-1    ..........AUG
Mgl-2-1      C.........AUG
Sgr-12-2     ..........UUG
Fsp-14-1     ..........AUG
Fsa-1-1      ..........AUG
Gar-2-1      ..........AUG
Spa-31-1     CGAC......AUG
Spa-3-1      CGAC......AUG
Spa-39-1     CGAC......AUG
Spa-35-1     CGAC......AUG
Spa-25-1     CGAC......AUG
Nfa-1-1      ..........AUG
Msp-7-1      ..........AAU
Msp-19-1     ..........AUG
Mll-1-1      ..........AUG
Msp-3-1      ..........AUG
Msp-20-1     ..........AUG
Nsp-7-1      ..........AUG
Nve-1-1      ..........AUG
Nao-1-1      ..........AUG
Nce-1-1      ..........AUG
env-381      .............
env-382      ..........AUG
env-384      ..........AUG
env-386      ..........AUG
env-390      .............
Bas-1-1      ..........AUG
Bas-2-2      ..........AUG
Bsp-3-1      ..........AUG
env-391      ..........AUG
env-394      ..........AUG
env-396      ..........AUG
Bsp-2-1      ..........AUG
Bas-2-3      ..........AUG
Mma-2-1      ..........AUG
Msp-2-1      ..........AUG
env-398      ..........AUG
Nji-1-1      ..........AUG
Nno-1-1      ..........AUG
Nhi-1-1      ..........AUG
Msp-1-1      ..........AUG
Msp-18-1     ..........AUG
Mfo-1-1      ..........AUG
Mvu-1-1      ..........AUG
Mse-1-1      ..........AUG
Msp-23-1     ..........AUG
Mse-2-1      ..........AUG
Mse-3-1      ..........AUG
Mse-3-2      ..........AUG
Msp-22-1     ..........AUG
Mne-1-1      ..........AUG

             .............
             .............
             ----------RUG
```

```
env-400      .........AUG
Msp-21-1     .........AUG
Sal-7-1      C........UUG
Szi-1-1      C........UUG
Ssp-17-1     .........UUG
Ssp-52-1     .........UUG
Ssp-13-1     .........UUG
Ssp-66-1     .........UUG
Ssp-63-1     .........UUG
Ssp-33-1     .........UUG
Ssp-117-1    .........UUG
Swa-1-1      .........UUG
Ssp-26-1     .........UUG
Sal-4-1      .........UUG
Ssp-123-1    .........UUG
Sgr-2-1      .........UUG
Sco-3-1      C........UUG
Ssp-67-1     .........UUG
Ssp-137-1    .........UUG
Slu-1-1      .........UUG
Sga-2-1      .........UUG
Sau-2-1      .........UUG
Ssp-21-1     .........UUG
Nvu-1-1      .........AUG
Nbr-1-1      .........AUG
Nte-1-1      .........AUG
Spr-2-1      .........UUG
Ssp-106-1    C........UUG
env-401      .........UUG
env-402      .........UUG
Ssp-146-1    .........UUG
Smi-1-1      .........UUG
Sda-1-1      .........UUG
Ssp-161-1    .........UUG
Ssp-71-1     .........UUG
Sgr-7-1      .........UUG
Sgl-3-1      .........UUG
Ssp-85-1     .........UUG
Ssp-121-1    .........UUG
Ssp-104-1    .........UUG
Ssp-60-1     .........UUG
Ssp-72-1     C........UUG
Ssp-99-1     .........UUG
Ssp-89-1     .........UUG
Sbi-2-1      .........UUG
Ssp-82-1     .........UUG
Ssp-3-1      .........UUG
Sam-1-1      .........UUG
Ssp-108-1    .........UUG
Ssp-48-1     .........UUG
Ssp-124-1    .........UUG
Ssp-110-1    .........UUG
Ssp-140-1    .........UUG
Ssp-107-1    .........UUG
Ssp-112-1    .........UUG
Svi-5-1      .........UUG
Ssp-74-1     .........UUG
Sbo-1-1      .........UUG
env-407      .........UUG
Ssp-101-1    .........GUG
Ssp-136-1    .........UUG
Sto-1-1      C........UUG
Sch-1-1      .........UUG
Saf-1-1      .........UUG
Sva-1-1      .........UUG
Ssp-154-1    .........UUG
Sol-1-1      .........UUG
Ssp-147-1    .........UUG
Sia-1-1      .........UUG
Sro-1-1      .........UUG
Scy-3-1      .........UUG
Sac-1-1      .........UUG
Sla-3-1      .........UUG
Ssp-160-1    .........UUG
Shy-2-1      .........UUG
Ssp-125-1    .........UUG
Spe-1-1      .........UUG
Ssp-141-1    .........UUG
Svi-14-1     .........UUG
Svi-4-1      .........UUG
env-408      .........UUG
Sol-2-1      .........UUG
Kau-1-2      .........UUG
Svi-12-1     .........UUG

             ............
             ............
             ---------RUG
```

```
Sco-1-1       ..........UUG
Ssp-58-1      ..........UUG
Sno-1-1       ..........UUG
Sgr-1-1       ..........UUG
Sce-1-1       ..........UUG
Ssp-176-1     ..........UUG
Ssp-118-1     ..........UUG
Svi-11-2      ..........UUG
Sgr-12-7      ..........UUG
Ssc-4-1       ..........UUG
Ssp-69-1      ..........UUG
Ssp-153-1     ..........UUG
Sye-2-1       ..........UUG
Sse-1-1       ..........UUG
Sfu-2-1       ..........UUG
Sdu-1-1       ..........UUG
Sip-1-1       ..........UUG
Ssc-3-1       ..........UUG
Ssp-59-1      ..........UUG
Ssp-70-1      C.........UUG
Ssp-24-1      ..........UUG
Ssp-10-1      C.........UUG
Ssp-53-1      C.........UUG
Sni-2-1       ..........AUG
Spr-1-1       ..........UUG
Ssp-134-1     ..........UUG
Sau-1-1       ..........UUG
Asp-7-1       ..........UUG
Sal-5-1       ..........UUG
Svi-2-1       ..........UUG
Stu-1-1       C.........UUG
Sso-1-1       ..........UUG
Spu-1-1       ..........UUG
Ssp-12-1      ..........UUG
Ssp-78-1      ..........UUG
Ssp-158-1     ..........UUG
Ssp-79-1      ..........UUG
Ssp-92-1      ..........UUG
Ssp-151-1     ..........UUG
Sat-2-1       ..........UUG
Sfl-8-1       ..........UUG
Sfl-1-1       ..........UUG
Ssp-126-1     ..........UUG
Sgr-11-1      ..........UUG
Ssp-14-1      ..........UUG
Ssp-168-1     ..........UUG
Ssp-49-1      C.........UUG
Ssp-22-1      C.........UUG
Ssp-25-1      C.........UUG
Ssp-152-1     ..........UUG
Ssp-57-1      ..........UUG
Ssp-40-1      ..........UUG
Ssp-6-1       C.........UUG
Sha-2-1       ..........UUG
Ssp-43-1      ..........UUG
Scy-2-1       ..........UUG
Ssp-39-1      ..........UUG
Sfu-1-1       ..........UUG
Seu-1-1       ..........UUG
Sgr-3-1       ..........UUG
Ssp-9-1       ..........UUG
Ssp-68-1      ..........UUG
Sal-8-2       ..........UUG
San-2-1       ..........UUG
Sod-1-1       C.........UUG
Ssp-149-1     ..........UUG
Sfl-4-1       ..........UUG
Ssp-127-1     C.........UUG
Ssp-81-1      C.........UUG
Sat-1-1       C.........UUG
Ssp-103-1     C.........UUG
Nsp-3-1       ..........AUG
env-409       ..........AUG
env-410       ..........AUG
env-411       ..........AUG
env-412       ..........AUG
env-413       ..........AUG
env-414       ..........AUG
env-415       ..........AUG
env-416       ..........AUG
env-417       ..........GUG
env-418       GAC.......AUG
Svi-14-4      ..........UUG
Sve-1-1       ..........UUG

              .............
              .............
              ----------RUG
```

```
Ssp-38-1     ..........UUG
Sbi-3-1      ..........UUG
Sgl-4-2      ..........UUG
Svi-9-1      ..........UUG
Fsp-2-1      ..........AUG
Fsp-5-1      ..........AUG
Fsp-1-1      ..........AUG
Sgh-1-1      ..........UUG
Ssp-31-1     ..........UUG
Sav-2-1      ..........UUG
Ssp-7-1      ..........UUG
Ssp-44-1     ..........UUG
Sgr-10-1     ..........UUG
Ssp-173-1    ..........UUG
Ssp-16-1     ..........UUG
Ssp-61-1     ..........UUG
Ssp-175-1    ..........UUG
Fsp-6-1      ..........AUG
Fsp-13-1     ..........AUG
Fal-1-1      ..........AUG
Gte-4-1      ..........AUG
Mrh-1-1      ..........AUG
env-419      ..........AUG
env-421      C.........AUG
env-422      C.........AUG
env-423      CA........AUG
env-424      AAA.......GUG
env-425      AAA.......GUG
env-426      AAA.......GUG
Nsp-11-1     ..........AUG
env-428      C.........GUG
env-429      GACCA.....AUG
env-430      GACCAA.......
env-431      ..........AUG
env-432      ..........AUG
env-433      ..........AUG
env-434      ..........AUG
env-435      ..........AUG
Str-2-1      CGAC......AUG
Str-1-1      CGAC......AUG
Spa-24-1     CGAC......AUG
Spa-32-1     CGAC......AUG
Spa-7-1      CGAC......AUG
Spa-27-1     CGAC......AUG
Spa-5-1      CGAC......AUG
Spa-38-1     CGAC......AUG
env-436      ..........AUG
Msp-11-1     ..........AUG
env-437      C.........AUG
env-438      ..........AUG
env-440      U.........GUG
env-441      CC........AUG
env-442      CAC.......AUG
Gkr-1-1      ..........AUG
env-443      ..........AUG
env-446      C.........GUG
env-447      ..........AUG
env-448      ..........AUG
env-449      .............
Mhe-1-1      ..........AUG
Rrh-1-1      ..........AUG
Msp-13-1     ..........AUG
env-450      ..........AUG
Tbi-1-1      ..........AUG
Msp-4-1      ..........AUG
Mav-1-1      ..........AUG
Msp-16-1     ..........AUG
env-452      ..........AUG
Msp-12-1     ..........AUG
Msp-14-1     ..........AUG
Msp-17-1     ..........AUG
env-453      ..........AUG
Mky-1-1      ..........AUG
Sar-43-1     CGAC......AUG
Sar-11-1     CGAC......AUG
env-454      GACAC.....AUG
env-455      C.........UUG
env-456      ..........GUG
Ase-1-1      ..........AUG
Fsp-9-1      ..........AUG
env-457      C.........GUG
env-459      .............
env-460      .............
Spa-33-1     CGAC......AUG

             .............
             .............
             ----------RUG
```

```
Vma-1-1      CGAC......AUG
Gbr-1-1      ACC.......AUG
Gna-1-1      CAGAC.....AUG
Gal-1-1      AGAU......AUG
Gam-2-1      CAGAC.....AUG
Gsp-1-1      AGAC......AUG
Gpa-1-1      GAC.......AUG
Gru-1-1      AGAU......AUG
Gam-3-1      AGAC......AUG
Gte-3-1      CAGAC.....AUG
Nsp-1-1      GA........AUG
Wsp-1-1      GA........AUG
Kar-2-1      GA........AUG
Gef-1-1      AUC.......AUG
env-461      AGACC.....AUG
Gpo-3-1      GAG.......AUG
Grh-1-1      GAU.......AUG
env-462      CC........AUG
Rde-1-1      ..........AUG
Req-3-1      ..........AUG
env-463      ..........AUG
Rfa-3-1      ..........AUG
Rsp-2-1      ..........AUG
Rsp-3-1      ..........AUG
Rfa-12-1     ..........AUG
Rsp-13-1     ..........AUG
Rop-1-1      ..........AUG
Rfa-21-1     ..........AUG
Rfa-1-2      ..........AUG
Rfa-4-1      ..........AUG
Rfa-7-1      ..........AUG
Rfa-20-1     ..........AUG
Rfa-8-1      ..........AUG
Rtr-1-1      ..........AUG
Rsp-19-1     ..........AUG
Rsp-21-1     ..........AUG
Sar-21-1     CGAC......AUG
Sar-40-1     CGAC......AUG
Sar-27-1     CGAC......AUG
Sar-33-1     CGAC......AUG
Sar-26-1     CGAC......AUG
Sar-22-1     CGAC......AUG
Sar-17-1     CGAC......AUG
Sar-24-1     CGAC......AUG
env-474      ..........AUG
env-475      CAGAC.....AUG
Whe-1-1      G.........AUG
Spa-28-1     CGAC......AUG
Spa-36-1     CGAC......AUG
Spa-8-1      CGAC......AUG
Dau-1-1      CC........AUG
Mlu-1-1      ..........AUG
Mch-6-1      ..........AUG
Mpa-1-1      ..........AUG
Rsp-9-1      ..........AUG
Rrh-2-1      ..........AUG
Rsp-14-1     ..........AUG
Rsp-5-1      ..........AUG
Rru-2-1      ..........AUG
Rru-3-1      ..........AUG
Rru-1-1      ..........AUG
Ntr-1-1      ..........AUG
Not-2-1      ..........AUG
Not-1-1      ..........AUG
env-476      ..........GUG
Msp-26-1     CGAC......AUG
env-477      AC........GUG
env-478      ..........AUG
env-479      ..........AUG
Gsp-3-1      ..........AUG
env-481      ..........AUG
env-482      ..........AUG
env-483      ..........AUG
env-484      CA........AUG
env-485      ..........AUG
env-486      AC........AUG
Sam-2-1      GAC.......AUG
Sro-7-1      GCC.......AUG
Sro-3-1      GCC.......AUG
Mau-1-1      CGAC......AUG
Msp-9-1      CGAC......AUG
Msp-10-1     CGAC......AUG
Msp-28-1     CGAC......AUG
env-487      ..........AUG

             .............
             .............
             ---------RUG
```

| | |
|---|---|
| env-488 | UG........AUG |
| Msp-8-1 | CGAC......AUG |
| env-489 | CCG.......AUG |
| env-490 | ............. |
| Nco-1-1 | ..........AUG |
| env-494 | AA........GUG |
| Mca-1-1 | CGAC......AUG |
| env-495 | UA........AUG |
| env-496 | CC........AUG |
| env-497 | ..........AUG |
| env-498 | A.........GUG |
| env-499 | A.........GUG |
| env-500 | A.........GUG |
| env-501 | A.........GUG |
| env-502 | ..........AUG |
| env-503 | A.........GUG |
| env-505 | A.........GUG |
| env-506 | A.........GUG |
| env-508 | A.........GUG |
| env-509 | ............. |
| env-510 | A.........GUG |
| env-513 | AAA.......GUG |
| env-514 | GAA.......UUG |
| env-515 | CC........AUG |
| Nca-2-1 | ..........AUG |
| Asu-1-1 | ..........AUG |
| env-516 | CC........AUG |
| env-517 | CC........AUG |
| env-518 | CA........AUG |
| env-520 | ..........GUG |
| env-521 | CAA.......GUG |
| env-522 | AC........AUG |
| env-523 | GACUC.....AUG |
| env-524 | CA........AUG |
| env-526 | A.........GUG |
| env-527 | AU........AUG |
| env-530 | AU........AUG |
| env-531 | AU........AUG |
| env-532 | ............. |
| env-533 | AC........AUG |
| env-534 | ..........AUG |
| env-535 | A.........UGA |
| env-536 | A.........UUG |
| env-538 | AC........AUG |
| Sna-1-1 | ..........AUG |
| env-539 | CA........AUG |
| env-540 | AC........GUG |
| env-541 | ACA.......GUG |
| env-542 | ACA.......GUG |
| env-543 | ACA.......GUG |
| env-544 | ACA.......GUG |
| env-546 | A.........AUG |
| env-547 | A.........AUG |
| env-548 | C.........AUG |
| env-549 | AC........AUG |
| env-551 | AC........AUG |
| env-552 | CC........AUG |
| env-553 | U.........UUG |
| env-555 | ..........AUG |
| env-557 | AC........AUG |
| env-558 | AC........AUG |
| env-560 | AC........AUG |
| env-561 | ..........AUG |
| env-563 | ..........AUG |
| env-564 | ..........AUG |
| env-565 | AC........AUG |
| env-568 | AC........AUG |
| env-570 | AU........AUG |
| env-571 | ..........AUG |
| env-572 | CC........AUG |
| env-573 | CC........AUG |
| env-576 | C.........AUG |
| env-577 | CGUU......AUG |
| env-578 | A.........AUG |
| env-579 | AC........AUG |
| env-582 | AC........AUG |
| env-583 | AC........AUG |
| env-584 | AC........AUG |
| env-585 | AC........AUG |
| env-586 | AC........AUG |
| env-587 | ..........AUG |
| env-588 | ..........AUG |
| env-589 | AACAG.....AUG |
| env-591 | AAGAA.....AUG |
| | ............. |
| | ............. |
|  | ----------RUG |

```
env-592     CAGCA.....AUG
env-593     C.........AUG
env-594     ACAC......AUG
env-596     ACAC......AUG
env-597     ACAC......AUG
env-598     C.........AUG
env-600     CCA.......AUG
Nbr-4-1     ..........AUG
Npa-1-1     ..........AUG
env-601     C.........AUG
env-602     ..........AUG
Gam-1-1     ..........AUG
Gso-1-1     ..........AUG
Gsh-1-1     ..........AUG
Nsp-5-1     ..........AUG
env-603     GACUC.....AUG
Ari-2-1     C.........AUG
env-604     ..........AUG
env-605     ..........AUG
env-606     ..........AUG
env-607     ..........AUG
env-608     U.........GUG
Ata-1-1     ..........AUG
Aha-1-1     ..........AUG
env-609     U.........GUG
env-610     ..........AUG
env-611     .............
env-629     U.........AUG
Pau-1-1     C.........AUG
Psp-6-1     CC........AUG
Ses-1-1     CG........UUG
Ksp-2-1     AC........AUG
Sgl-1-1     ..........AUG
env-630     ..........AUG
Aba-1-1     ..........AUG
Ari-1-1     ..........AUG
Ava-2-1     ..........AUG
Aal-2-1     ..........GUG
env-631     ..........GUG
env-633     ..........GUG
env-637     ..........GUG
Lae-1-1     UC........UUG
env-638     U.........AUG
env-639     U.........AUG
Aen-1-1     ACCAGUUCC.GUG
Aha-3-1     ..........AUG
env-647     ..........AUG
env-648     ..........AUG
Ksp-1-1     A.........GUG
Lal-2-1     UC........UUG
Lae-1-2     UC........UUG
env-649     CU........GUG
Psp-9-1     C.........AUG
Aje-1-1     ..........AUG
[ir-1-1     ..........AUG
Ain-1-1     C.........GUG
Aja-1-1     ..........AUG
Asp-6-1     ..........AUG
Aaz-1-1     ..........AUG
Aor-4-1     ..........AUG
Ssp-91-1    ..........AUG
Abe-1-1     ..........AUG
Aor-1-1     ..........AUG
Aor-3-1     ..........AUG
Ade-1-1     ..........AUG
Ssp-5-1     ..........AUG
Aor-2-1     ..........AUG
env-650     ..........GUG
env-651     ..........GUG
env-652     ..........AUG
Sha-1-1     ..........AUG
Spa-41-1    ..........AUG
Kal-2-1     G.........UUG
Saz-1-1     ..........AUG
Svi-1-1     ..........AUG
Ssp-45-1    ..........AUG
Kar-1-1     A.........GUG
Ani-1-1     ..........AUG
Rrh-3-1     A.........GUG
Scy-1-1     ..........AUG
env-653     C.........GUG
Ame-3-1     ..........AUG
Ssp-46-1    CG........UUG
Ami-2-1     G.........UUG

            .............
            .............
            ----------RUG
```

```
Ssp-1-1     G.........UUG
env-654     ..........GUG
env-656     ..........GUG
Sts-1-1     ..........UUG
env-657     AGAUC.....AUG
Rsp-11-1    ..........AUG
env-658     ..........AUG
Rsp-12-1    ..........AUG
Rrh-4-1     ..........AUG
Ace-1-1     G.........UUG
Gja-1-1     GA........AUG
Gsp-2-1     GA........AUG
Got-1-1     GA........AUG
Gai-1-1     GAG.......AUG
Asp-5-2     C.........GUG
env-663     ..........AUG
Sma-1-1     ..........AUG
env-664     C.........AUG
Ssu-1-1     ..........UUG
Ssu-2-1     ..........UUG
Sje-1-1     C.........UUG
Sal-2-1     C.........UUG
Sru-1-1     C.........UUG
Ssp-100-1   ..........UUG
Ssp-129-1   ..........UUG
Ssp-171-1   ..........UUG
Ssp-170-1   C.........UUG
Ssp-56-1    ..........UUG
Kaz-1-1     C.........UUG
Svi-7-1     ..........GUG
Smo-1-1     ..........UUG
Ama-4-1     ..........AUG
env-666     ..........AUG
Bsc-2-1     CC........AUG
Ssc-1-1     ..........UUG
Sle-1-1     ..........UUG
Scl-1-1     ..........UUG
Sre-3-1     ..........UUG
Ser-2-1     G.........UUG
Ssp-20-1    ..........UUG
Sst-1-1     ..........UUG
Mne-3-1     ..........AUG
            .............
            .............
            ----------RUG
```

### 10 L31-Coriobacteria

#### 10.1 Taxa

The taxonomy of each organism containing a putative L31-Coriobacteria RNA is listed, with abbreviations identifying each hit (e.g., "Eco-1-1" and "Eco-1-2" might hypothetically represent two distinct RNAs in *E. coli*). The abbreviations will be used to identify each individual L31-Coriobacteria RNA in Sections 10.2 and 10.4:.

| **abbrev. of hits** | **taxonomy of species** |
|---|---|
| Aeq-1-1 | Bacteria Actinobacteria Coriobacteridae Coriobacteriales Coriobacterineae Coriobacteriaceae *Adlercreutzia equolifaciens* DSM 19450 |
| Ami-1-1 | Bacteria Actinobacteria Coriobacteridae Coriobacteriales Coriobacterineae Coriobacteriaceae *Atopobium minutum* 10063974 |
| Apa-1-1 | Bacteria Actinobacteria Coriobacteridae Coriobacteriales Coriobacterineae Coriobacteriaceae *Atopobium parvulum* DSM 20469 |
| Ari-1-1 | Bacteria Actinobacteria Coriobacteridae Coriobacteriales Coriobacterineae Coriobacteriaceae *Atopobium rimae* ATCC 49626 |
| Asp-1-1 | Bacteria Actinobacteria Coriobacteridae Coriobacteriales Coriobacterineae Coriobacteriaceae *Atopobium sp.* BV3Ac4 |
| Asp-2-1 | Bacteria Actinobacteria Coriobacteridae Coriobacteriales Coriobacterineae Coriobacteriaceae *Atopobium sp. oral taxon* 199 str. F0494 |
| Asp-3-1 | Bacteria Actinobacteria Coriobacteridae Coriobacteriales Coriobacterineae Coriobacteriaceae *Atopobium sp. oral taxon* 810 str. F0209 |
| Ava-1-1 to Ava-1-2 | Bacteria Actinobacteria Coriobacteridae Coriobacteriales Coriobacterineae Coriobacteriaceae *Atopobium vaginae* DSM 15829 |
| Cae-1-1 | Bacteria Actinobacteria Coriobacteridae Coriobacteriales Coriobacterineae Coriobacteriaceae *Collinsella aerofaciens* ATCC 25986 |
| Cin-1-1 | Bacteria Actinobacteria Coriobacteridae Coriobacteriales Coriobacterineae Coriobacteriaceae *Collinsella intestinalis* DSM 13280 |
| Csp-1-1 | Bacteria Actinobacteria Coriobacteridae Coriobacteriales Coriobacterineae Coriobacteriaceae *Collinsella sp.* GD3 |
| Cst-1-1 | Bacteria Actinobacteria Coriobacteridae Coriobacteriales Coriobacterineae Coriobacteriaceae *Collinsella stercoris* DSM 13279 |
| Cta-1-1 | Bacteria Actinobacteria Coriobacteridae Coriobacteriales Coriobacterineae Coriobacteriaceae *Collinsella tanakaei* YIT 12063 |
| Cba-1-1 | Bacteria Actinobacteria Coriobacteridae Coriobacteriales Coriobacterineae *Coriobacteriaceae bacterium* BV3Ac1 |

| | |
|---|---|
| Cba-2-1 | Bacteria Actinobacteria Coriobacteridae Coriobacteriales Coriobacterineae *Coriobacteriaceae bacterium* GD5 |
| Cgl-1-1 | Bacteria Actinobacteria Coriobacteridae Coriobacteriales Coriobacterineae Coriobacteriaceae *Coriobacterium glomerans* PW2 |
| Ele-1-1 | Bacteria Actinobacteria Coriobacteridae Coriobacteriales Coriobacterineae Coriobacteriaceae *Eggerthella lenta* DSM 2243 |
| Esp-1-1 | Bacteria Actinobacteria Coriobacteridae Coriobacteriales Coriobacterineae Coriobacteriaceae *Eggerthella sp.* 1_3_56FAA |
| Esp-2-1 | Bacteria Actinobacteria Coriobacteridae Coriobacteriales Coriobacterineae Coriobacteriaceae *Eggerthella sp.* HGA1 |
| Esp-3-1 | Bacteria Actinobacteria Coriobacteridae Coriobacteriales Coriobacterineae Coriobacteriaceae *Eggerthella sp.* YY7918 |
| Eca-1-1 | Bacteria Actinobacteria Coriobacteridae Coriobacteriales Coriobacterineae Coriobacteriaceae *Enterorhabdus caecimuris* B7 |
| Gpa-1-1 | Bacteria Actinobacteria Coriobacteridae Coriobacteriales Coriobacterineae Coriobacteriaceae *Gordonibacter pamelaeae* 7-10-1-b |
| Opr-1-1 | Bacteria Actinobacteria Coriobacteridae Coriobacteriales Coriobacterineae Coriobacteriaceae *Olsenella profusa* F0195 |
| Osp-1-1 | Bacteria Actinobacteria Coriobacteridae Coriobacteriales Coriobacterineae Coriobacteriaceae *Olsenella sp. oral taxon* 809 str. F0356 |
| Oul-1-1 | Bacteria Actinobacteria Coriobacteridae Coriobacteriales Coriobacterineae Coriobacteriaceae *Olsenella uli* DSM 7084 |
| Sex-1-1 | Bacteria Actinobacteria Coriobacteridae Coriobacteriales Coriobacterineae Coriobacteriaceae *Slackia exigua* ATCC 700122 |
| She-1-1 | Bacteria Actinobacteria Coriobacteridae Coriobacteriales Coriobacterineae Coriobacteriaceae *Slackia heliotrinireducens* DSM 20476 |
| Spi-1-1 | Bacteria Actinobacteria Coriobacteridae Coriobacteriales Coriobacterineae Coriobacteriaceae *Slackia piriformis* YIT 12062 |
| Ssp-1-1 | Bacteria Actinobacteria Coriobacteridae Coriobacteriales Coriobacterineae Coriobacteriaceae *Slackia sp.* CM382 |
| Afo-1-1 | Bacteria Actinobacteria Coriobacteriia Coriobacteriales Atopobiaceae *Atopobium fossor* DSM 15642 |
| Apa-2-1 | Bacteria Actinobacteria Coriobacteriia Coriobacteriales Atopobiaceae *Atopobium parvulum* DNF00906 |
| Asp-4-1 | Bacteria Actinobacteria Coriobacteriia Coriobacteriales Atopobiaceae *Atopobium sp.* BS2 |
| Asp-5-1 | Bacteria Actinobacteria Coriobacteriia Coriobacteriales Atopobiaceae *Atopobium sp.* ICM42b |
| Ava-2-1 | Bacteria Actinobacteria Coriobacteriia Coriobacteriales Atopobiaceae *Atopobium vaginae* |
| Oul-2-1 | Bacteria Actinobacteria Coriobacteriia Coriobacteriales Atopobiaceae *Olsenella uli* MSTE5 |
| Csp-2-1 | Bacteria Actinobacteria Coriobacteriia Coriobacteriales Coriobacteriaceae *Collinsella sp.* 4_8_47FAA |
| Csp-3-1 | Bacteria Actinobacteria Coriobacteriia Coriobacteriales Coriobacteriaceae *Collinsella sp.* MS5 |
| Cba-3-1 | Bacteria Actinobacteria Coriobacteriia Coriobacteriales *Coriobacteriaceae bacterium* 68-1-3 |
| Ema-1-1 | Bacteria Actinobacteria Coriobacteriia Coriobacteriales Coriobacteriaceae *Enorma massiliensis phI* |
| San-1-1 | Bacteria Actinobacteria Coriobacteriia Coriobacteriales Coriobacteriaceae *Senegalimassilia anaerobia* JC110 |
| Ele-2-1 to Ele-2-2 | Bacteria Actinobacteria Coriobacteriia Eggerthellales Eggerthellaceae *Eggerthella lenta* 1_1_60AFAA |
| Emu-1-1 | Bacteria Actinobacteria Coriobacteriia Eggerthellales Eggerthellaceae *Enterorhabdus mucosicola* DSM 19490 |
| env-1 to env-259 | environmental samples |

#### 10.2 Gene contexts

Each L31-Coriobacteria RNA (indicated by "RNA→") is listed. For each hit, the downstream genes predicted to reside in a regulated operon are listed. If the nearest downstream gene is encoding in the opposite strand (and therefore presumed to not be a part of a regulated operon), then that gene is still depicted. Some environmental sequences and some RefSeq entries lack gene annotations, and so no genes are available for such sequences. The direction of each gene is indicated with an arrow (→), and each predicted conserved domain in the gene is named. Conserved domains associated with more than one L31-Coriobacteria RNA are assigned a color; other domains are gray. Information about these conserved domains is given in Section 10.3. The accession of the sequence containing each L31-Coriobacteria RNA is given in the column named "Seq. accession". Accessions beginning with "NC_", "NS_", "NW_" or "NZ_" are contained in RefSeq. Other accession refer to environmental samples. Nucleotide coordinates are given for the 5′ and 3′ boundaries of each L31-Coriobacteria RNA. If the 5′ coordinate is greater than the 3′ coordinate, the RNA is present on the reverse-complement strand of the containing genomic DNA sequence. Each hit is denoted by an abbreviation (like "Eco-1-1") that refers to a taxonomy given in Section 10.1.

| abbrev. | Seq. accession | | 5′ at | 3′ at | genes |
|---|---|---|---|---|---|
| env-1 | JGI24735J21928_10303163 | + | 78 | 111 | **RNA→** rpmE (PRK00528)Ribosomal_L31 (pfam01197)→ RimL (COG1670)Acetyltransf_3 (pfam13302)→ |
| env-2 | AUXO017004512.1 | - | 117 | 84 | **RNA→** rpmE (PRK00019)rpmE2 (PRK01678)Ribosomal_L31 (pfam01197)→ |
| env-3 | AUXO016339607.1 | - | 94 | 61 | **RNA→** rpmE (PRK00019)Ribosomal_L31 (pfam01197)→ |
| env-4 | SRS011111_Baylor_scaffold_9512 | + | 2842 | 2874 | **RNA→** rpmE (PRK00019)Ribosomal_L31 (pfam01197)→ |
| env-5 | SRS023468_LANL_scaffold_7759 | + | 6315 | 6347 | **RNA→** rpmE (PRK00019)Ribosomal_L31 (pfam01197)→ |
| env-6 | SRS023604_Baylor_scaffold_10029 | - | 18845 | 18813 | **RNA→** rpmE (PRK00019)Ribosomal_L31 (pfam01197)→ |
| env-7 | LWSO2_GGWJX9X02HB8PF | + | 220 | 250 | **RNA→** rpmE (PRK00019)Ribosomal_L31 (pfam01197)→ |
| env-8 | 4448818.3_Contig28460 | - | 132 | 94 | **RNA→** ←hypo |
| env-9 | 4448807.3_Contig19481 | - | 164 | 127 | **RNA→** rpmE (PRK00019)rpmE2 (PRK01678)Ribosomal_L31 (pfam01197)→ |
| env-10 | 4448811.3_Contig15515 | - | 234 | 197 | **RNA→** rpmE (PRK00019)Ribosomal_L31 (pfam01197)→ |
| env-11 | 4448816.3_Contig24142 | - | 467 | 430 | **RNA→** rpmE (PRK00019)Ribosomal_L31 (pfam01197)→ |

| | | | | | |
|---|---|---|---|---|---|
| env-12 | 4448820.3_Contig_6849 | - | 894 | 857 | **RNA**→ rpmE (PRK00019)Ribosomal_L31 (pfam01197)→ |
| env-13 | DLM013_scaffold31757_3 | - | 262 | 225 | **RNA**→ rpmE (PRK00019)Ribosomal_L31 (pfam01197)→ |
| Cta-1-1 | NZ_JH126470.1 | + | 134255 | 134292 | **RNA**→ rpmE (PRK00019)Ribosomal_L31 (pfam01197)→ |
| env-14 | HCF12C_23335 | + | 238 | 272 | **RNA**→ rpmE (PRK00019)Ribosomal_L31 (pfam01197)→ |
| env-15 | AUXO018093914.1 | + | 291 | 328 | **RNA**→ rpmE (PRK00019)Ribosomal_L31 (pfam01197)→ |
| env-16 | 4448811.3_Contig11262 | + | 140 | 176 | **RNA**→ ←hypo |
| env-17 | hgutS7_s7_164738 | - | 1067 | 1032 | **RNA**→ rpmE (PRK00019)Ribosomal_L31 (pfam01197)→ |
| env-18 | hgutS7_s7_179880_2 | - | 652 | 617 | **RNA**→ rpmE (PRK00019)Ribosomal_L31 (pfam01197)→ |
| env-19 | scaffold83292_1_MH0041 | - | 442 | 407 | **RNA**→ rpmE (PRK00019)Ribosomal_L31 (pfam01197)→ |
| env-20 | scaffold8232_5_MH0062 | - | 1352 | 1317 | **RNA**→ rpmE (PRK00019)Ribosomal_L31 (pfam01197)→ |
| env-21 | C6781021_1_MH0069 | - | 404 | 369 | **RNA**→ rpmE (PRK00019)Ribosomal_L31 (pfam01197)→ |
| env-22 | scaffold58836_1_MH0085 | + | 477 | 512 | **RNA**→ rpmE (PRK00019)Ribosomal_L31 (pfam01197)→ |
| env-23 | scaffold18205_4_MH0086 | - | 1313 | 1278 | **RNA**→ rpmE (PRK00019)Ribosomal_L31 (pfam01197)→ |
| env-24 | scaffold27957_2_O2.UC-11 | - | 244 | 209 | **RNA**→ rpmE (PRK00019)Ribosomal_L31 (pfam01197)→ |
| env-25 | scaffold61969_3_V1.CD-8 | - | 314 | 279 | **RNA**→ rpmE (PRK00019)Ribosomal_L31 (pfam01197)→ |
| env-26 | scaffold3383_12_V1.UC-15 | - | 531 | 496 | **RNA**→ rpmE (PRK00019)Ribosomal_L31 (pfam01197)→ |
| env-27 | scaffold9272_17_V1.UC-19 | - | 1817 | 1782 | **RNA**→ rpmE (PRK00019)Ribosomal_L31 (pfam01197)→ |
| env-28 | SRS049164_WUGC_scaffold_57754 | - | 167 | 132 | **RNA**→ rpmE (PRK00019)Ribosomal_L31 (pfam01197)→ |
| env-29 | DLF009_scaffold14997_4 | + | 7628 | 7663 | **RNA**→ rpmE (PRK00019)Ribosomal_L31 (pfam01197)→ |
| env-30 | DLM001_scaffold25088_28 | + | 399 | 434 | **RNA**→ rpmE (PRK00019)Ribosomal_L31 (pfam01197)→ |
| env-31 | DLM013_scaffold384_15 | + | 9046 | 9081 | **RNA**→ rpmE (PRK00019)Ribosomal_L31 (pfam01197)→ |
| env-32 | DLM019_scaffold37572_5 | - | 1768 | 1733 | **RNA**→ rpmE (PRK00019)Ribosomal_L31 (pfam01197)→ |
| env-33 | longitudinal_63_388 | + | 48143 | 48178 | **RNA**→ rpmE (PRK00019)Ribosomal_L31 (pfam01197)→ |
| env-34 | longitudinal_64_137 | + | 48145 | 48180 | **RNA**→ rpmE (PRK00019)Ribosomal_L31 (pfam01197)→ |
| env-35 | longitudinal_65_38 | - | 57705 | 57670 | **RNA**→ rpmE (PRK00019)Ribosomal_L31 (pfam01197)→ |
| env-36 | longitudinal_280_54 | + | 6079 | 6114 | **RNA**→ rpmE (PRK00019)Ribosomal_L31 (pfam01197)→ |
| env-37 | longitudinal_281_174 | - | 8873 | 8838 | **RNA**→ rpmE (PRK00019)Ribosomal_L31 (pfam01197)→ |
| env-38 | longitudinal_282_53 | + | 9862 | 9897 | **RNA**→ rpmE (PRK00019)Ribosomal_L31 (pfam01197)→ |
| env-39 | longitudinal_283_45 | + | 6079 | 6114 | **RNA**→ rpmE (PRK00019)Ribosomal_L31 (pfam01197)→ |
| env-40 | longitudinal_284_41 | + | 6076 | 6111 | **RNA**→ rpmE (PRK00019)Ribosomal_L31 (pfam01197)→ |
| env-41 | longitudinal_285_12 | - | 8873 | 8838 | **RNA**→ rpmE (PRK00019)Ribosomal_L31 (pfam01197)→ |
| env-42 | longitudinal_343_62 | + | 6079 | 6114 | **RNA**→ rpmE (PRK00019)Ribosomal_L31 (pfam01197)→ |
| env-43 | longitudinal_381_116 | - | 99840 | 99805 | **RNA**→ rpmE (PRK00019)Ribosomal_L31 (pfam01197)→ |
| env-44 | longitudinal_420_41 | - | 61707 | 61672 | **RNA**→ rpmE (PRK00019)Ribosomal_L31 (pfam01197)→ |
| env-45 | AUXO018332126.1 | + | 54 | 89 | **RNA**→ rpmE (PRK00019)Ribosomal_L31 (pfam01197)→ |
| env-46 | ADJT01003987.1 | + | 3155 | 3190 | **RNA**→ rpmE (PRK00019)Ribosomal_L31 (pfam01197)→ |
| env-47 | SRS012273_C3752464 | + | 1348 | 1383 | **RNA**→ rpmE (PRK00019)Ribosomal_L31 (pfam01197)→ |
| env-48 | SRS013687_Baylor_scaffold_51084 | + | 2102 | 2137 | **RNA**→ rpmE (PRK00019)Ribosomal_L31 (pfam01197)→ |
| env-49 | SRS013951_C2593644 | - | 285 | 250 | **RNA**→ rpmE (PRK00019)Ribosomal_L31 (pfam01197)→ |
| env-50 | SRS014683_C1571330 | + | 245 | 280 | **RNA**→ rpmE (PRK00019)Ribosomal_L31 (pfam01197)→ |
| env-51 | SRS015217_C2166290 | + | 152 | 187 | **RNA**→ rpmE (PRK00019)Ribosomal_L31 (pfam01197)→ |
| env-52 | SRS015782_C3168068 | + | 130 | 165 | **RNA**→ rpmE (PRK00019)Ribosomal_L31 (pfam01197)→ |
| env-53 | SRS017103_C2711990 | - | 528 | 493 | **RNA**→ rpmE (PRK00019)Ribosomal_L31 (pfam01197)→ |
| env-54 | SRS018351_C2783037 | + | 825 | 860 | **RNA**→ rpmE (PRK00019)Ribosomal_L31 (pfam01197)→ |
| env-55 | SRS019030_C1583151 | - | 162 | 127 | **RNA**→ rpmE (PRK00019)Ribosomal_L31 (pfam01197)→ |
| env-56 | SRS020233_C3826123 | - | 273 | 238 | **RNA**→ rpmE (PRK00019)Ribosomal_L31 (pfam01197)→ |
| env-57 | SRS020328_Baylor_scaffold_38533 | + | 5089 | 5124 | **RNA**→ rpmE (PRK00019)Ribosomal_L31 (pfam01197)→ |
| env-58 | SRS020869_C3184531 | + | 109 | 144 | **RNA**→ rpmE (PRK00019)Ribosomal_L31 (pfam01197)→ |
| env-59 | SRS022071_C3051314 | + | 1344 | 1379 | **RNA**→ rpmE (PRK00019)Ribosomal_L31 (pfam01197)→ |
| env-60 | SRS023526_C3010627 | - | 178 | 143 | **RNA**→ rpmE (PRK00019)Ribosomal_L31 (pfam01197)→ |
| env-61 | SRS023914_C2026054 | + | 315 | 350 | **RNA**→ rpmE (PRK00019)Ribosomal_L31 (pfam01197)→ |
| env-62 | SRS047014_C3267765 | - | 281 | 246 | **RNA**→ rpmE (PRK00019)Ribosomal_L31 (pfam01197)→ |

| | | | | | |
|---|---|---|---|---|---|
| env-63 | SRS050422_LANL_scaffold_39298 | - | 288 | 253 | **RNA**→ rpmE (PRK00019)Ribosomal_L31 (pfam01197)→ |
| Cae-1-1 | NZ_AAVN02000005.1 | + | 56643 | 56678 | **RNA**→ rpmE (PRK00019)Ribosomal_L31 (pfam01197)→ rpmE (PRK00019)Ribosomal_L31 (pfam01197)→ hy[illegible] |
| Csp-2-1 | NZ_KN214135.1 | - | 212670 | 212635 | **RNA**→ rpmE (PRK00019)Ribosomal_L31 (pfam01197)→ |
| env-64 | BABA01020564.1 | - | 774 | 739 | **RNA**→ rpmE (PRK00019)Ribosomal_L31 (pfam01197)→ |
| env-65 | 4491422.3_NODE_1607 | - | 337 | 302 | **RNA**→ rpmE (PRK00019)Ribosomal_L31 (pfam01197)→ |
| env-66 | longitudinal_286_70 | - | 8161 | 8126 | **RNA**→ rpmE (PRK00019)Ribosomal_L31 (pfam01197)→ |
| env-67 | longitudinal_287_87 | - | 11143 | 11108 | **RNA**→ rpmE (PRK00019)Ribosomal_L31 (pfam01197)→ |
| env-68 | longitudinal_288_88 | - | 3650 | 3615 | **RNA**→ rpmE (PRK00019)Ribosomal_L31 (pfam01197)→ |
| env-69 | longitudinal_289_88 | - | 8161 | 8126 | **RNA**→ rpmE (PRK00019)Ribosomal_L31 (pfam01197)→ |
| env-70 | longitudinal_290_101 | + | 12604 | 12639 | **RNA**→ rpmE (PRK00019)Ribosomal_L31 (pfam01197)→ |
| env-71 | longitudinal_338_112 | - | 8161 | 8126 | **RNA**→ rpmE (PRK00019)Ribosomal_L31 (pfam01197)→ |
| env-72 | longitudinal_339_157 | + | 12969 | 13004 | **RNA**→ rpmE (PRK00019)Ribosomal_L31 (pfam01197)→ |
| env-73 | longitudinal_340_81 | - | 3439 | 3404 | **RNA**→ rpmE (PRK00019)Ribosomal_L31 (pfam01197)→ |
| env-74 | longitudinal_341_75 | + | 12952 | 12987 | **RNA**→ rpmE (PRK00019)Ribosomal_L31 (pfam01197)→ |
| env-75 | longitudinal_342_144 | + | 12957 | 12992 | **RNA**→ rpmE (PRK00019)Ribosomal_L31 (pfam01197)→ |
| env-76 | SRS014235_WUGC_scaffold_14702 | + | 1430 | 1465 | **RNA**→ rpmE (PRK00019)Ribosomal_L31 (pfam01197)→ |
| env-77 | SRS014979_C3174944 | - | 413 | 378 | **RNA**→ rpmE (PRK00019)Ribosomal_L31 (pfam01197)→ |
| env-78 | SRS017521_Baylor_scaffold_30477 | + | 4003 | 4038 | **RNA**→ rpmE (PRK00019)Ribosomal_L31 (pfam01197)→ |
| env-79 | SRS065504_C3275369 | - | 421 | 386 | **RNA**→ rpmE (PRK00019)Ribosomal_L31 (pfam01197)→ |
| Opr-1-1 | NZ_AWEZ01000030.1 | - | 64182 | 64151 | **RNA**→ rpmE (PRK00019)Ribosomal_L31 (pfam01197)→ |
| Ema-1-1 | NZ_HE978575.1 | + | 590068 | 590102 | **RNA**→ rpmE (PRK00019)Ribosomal_L31 (pfam01197)→ |
| Cba-2-1 | NZ_CAPF01000059.1 | - | 2507 | 2473 | **RNA**→ rpmE (PRK00019)Ribosomal_L31 (pfam01197)→ |
| Csp-1-1 | NZ_CAPI01000012.1 | + | 2962 | 2996 | **RNA**→ rpmE (PRK00019)Ribosomal_L31 (pfam01197)→ |
| Oul-1-1 | NC_014363.1 | + | 119036 | 119067 | **RNA**→ rpmE (PRK00019)Ribosomal_L31 (pfam01197)→ |
| Oul-2-1 | NZ_JANC01000012.1 | - | 45561 | 45530 | **RNA**→ rpmE (PRK00019)Ribosomal_L31 (pfam01197)→ |
| env-80 | AUXO010118135.1 | + | 576 | 609 | **RNA**→ rpmE (PRK00019)Ribosomal_L31 (pfam01197)→ |
| env-81 | 2204694758 | + | 334 | 365 | **RNA**→ |
| Cgl-1-1 | NC_015389.1 | + | 824421 | 824455 | **RNA**→ rpmE (PRK00019)Ribosomal_L31 (pfam01197)→ |
| env-82 | SRS049268_C4052608 | - | 112 | 80 | **RNA**→ hypo→ |
| Ari-1-1 | NZ_ACFE01000002.1 | + | 328239 | 328271 | **RNA**→ rpmE (PRK00019)Ribosomal_L31 (pfam01197)→ rpmE (PRK00019)Ribosomal_L31 (pfam01197)→ |
| Asp-2-1 | NZ_KE150486.1 | - | 1050194 | 1050162 | **RNA**→ rpmE (PRK00019)Ribosomal_L31 (pfam01197)→ |
| env-83 | SRS011086_C2736760 | - | 326 | 294 | **RNA**→ rpmE (PRK00019)Ribosomal_L31 (pfam01197)→ |
| env-84 | SRS017076_Baylor_scaffold_23156 | + | 449 | 481 | **RNA**→ rpmE (PRK00019)Ribosomal_L31 (pfam01197)→ |
| env-85 | SRS075404_C3618531 | + | 64 | 96 | **RNA**→ rpmE (PRK00019)Ribosomal_L31 (pfam01197)→ |
| env-86 | SRS013818_C2768522 | + | 223 | 255 | **RNA**→ rpmE (PRK00019)Ribosomal_L31 (pfam01197)→ |
| env-87 | SRS014271_C2867060 | - | 517 | 485 | **RNA**→ rpmE (PRK00019)Ribosomal_L31 (pfam01197)→ |
| env-88 | SRS015209_WUGC_scaffold_51435 | + | 5240 | 5272 | **RNA**→ rpmE (PRK00019)Ribosomal_L31 (pfam01197)→ |
| env-89 | SRS015272_C3481572 | + | 65 | 97 | **RNA**→ rpmE (PRK00019)Ribosomal_L31 (pfam01197)→ |
| env-90 | SRS015395_C2406090 | + | 1317 | 1349 | **RNA**→ rpmE (PRK00019)Ribosomal_L31 (pfam01197)→ |
| env-91 | SRS015537_C1264717 | + | 921 | 953 | **RNA**→ |
| env-92 | SRS016501_C1505461 | - | 140 | 108 | **RNA**→ hypo→ |
| env-93 | SRS016569_Baylor_scaffold_22226 | - | 109 | 77 | **RNA**→ |
| env-94 | SRS016569_C2401739 | + | 329 | 361 | **RNA**→ |
| env-95 | SRS018739_C3648710 | - | 753 | 721 | **RNA**→ rpmE (PRK00019)Ribosomal_L31 (pfam01197)→ |
| env-96 | SRS020856_C2570764 | + | 315 | 347 | **RNA**→ ←hypo |
| env-97 | SRS022143_WUGC_scaffold_36034 | + | 110 | 142 | **RNA**→ rpmE (PRK00019)Ribosomal_L31 (pfam01197)→ |
| env-98 | SRS023835_Baylor_scaffold_50858 | - | 904 | 872 | **RNA**→ rpmE (PRK00019)Ribosomal_L31 (pfam01197)→ |
| env-99 | SRS024015_Baylor_scaffold_31148 | + | 637 | 669 | **RNA**→ ←hypo |
| env-100 | SRS024277_C2768622 | - | 400 | 368 | **RNA**→ rpmE (PRK00019)Ribosomal_L31 (pfam01197)→ |
| env-101 | SRS042131_WUGC_scaffold_2856 | + | 6 | 38 | **RNA**→ rpmE (PRK00019)Ribosomal_L31 (pfam01197)→ |
| env-102 | SRS042643_WUGC_scaffold_10098 | + | 423 | 455 | **RNA**→ rpmE (PRK00019)Ribosomal_L31 (pfam01197)→ AAA (smart00382)AAA (cd00009)→ |
| env-103 | SRS047210_WUGC_scaffold_60400 | + | 673 | 705 | **RNA**→ rpmE (PRK00019)Ribosomal_L31 (pfam01197)→ |

| | | | | | |
|---|---|---|---|---|---|
| env-104 | SRS047219_WUGC_scaffold_9164 | + | 1127 | 1159 | **RNA**→ rpmE (PRK00019)Ribosomal_L31 (pfam01197)→ |
| env-105 | SRS047824_WUGC_scaffold_52711 | - | 887 | 855 | **RNA**→ rpmE (PRK00019)Ribosomal_L31 (pfam01197)→ |
| env-106 | SRS049147_C4119585 | + | 6 | 38 | **RNA**→ rpmE (PRK00019)Ribosomal_L31 (pfam01197)→ |
| env-107 | SRS049389_C4562091 | - | 221 | 189 | **RNA**→ ←hypo |
| env-108 | SRS056323_LANL_scaffold_14744 | + | 220 | 252 | **RNA**→ rpmE (PRK00019)Ribosomal_L31 (pfam01197)→ |
| env-109 | SRS056622_LANL_scaffold_16043 | - | 1899 | 1867 | **RNA**→ rpmE (PRK00019)Ribosomal_L31 (pfam01197)→ |
| env-110 | SRS057791_C3637891 | + | 69 | 101 | **RNA**→ rpmE (PRK00019)Ribosomal_L31 (pfam01197)→ |
| env-111 | SRS062540_C2336651 | - | 952 | 920 | **RNA**→ rpmE (PRK00019)Ribosomal_L31 (pfam01197)→ |
| env-112 | SRS062544_LANL_scaffold_33267 | + | 527 | 559 | **RNA**→ rpmE (PRK00019)Ribosomal_L31 (pfam01197)→ |
| env-113 | SRS077736_C1828913 | - | 488 | 456 | **RNA**→ rpmE (PRK00019)Ribosomal_L31 (pfam01197)→ |
| Apa-1-1 | NC_013203.1 | + | 128868 | 128900 | **RNA**→ rpmE (PRK00019)Ribosomal_L31 (pfam01197)→ |
| Asp-4-1 | NZ_JDFG01000048.1 | + | 63761 | 63793 | **RNA**→ rpmE (PRK00019)Ribosomal_L31 (pfam01197)→ |
| Asp-5-1 | NZ_JDFH01000035.1 | + | 5736 | 5768 | **RNA**→ rpmE (PRK00019)Ribosomal_L31 (pfam01197)→ |
| env-114 | SRS021496_Baylor_scaffold_7910 | - | 3781 | 3749 | **RNA**→ rpmE (PRK00019)Ribosomal_L31 (pfam01197)→ |
| env-115 | SRS011140_C4988252 | + | 126 | 158 | **RNA**→ rpmE (PRK00019)Ribosomal_L31 (pfam01197)→ |
| env-116 | Buccal_mucosa_LANL_C65146548 | - | 471 | 439 | **RNA**→ hypo→ rpmE (PRK00019)Ribosomal_L31 (pfam01197)→ |
| env-117 | SRS016740_Baylor_scaffold_9629 | + | 219 | 251 | **RNA**→ rpmE (PRK00019)Ribosomal_L31 (pfam01197)→ |
| env-118 | SRS017076_C2662728 | + | 253 | 285 | **RNA**→ rpmE (PRK00019)Ribosomal_L31 (pfam01197)→ |
| env-119 | SRS064329_C2589301 | + | 141 | 173 | **RNA**→ rpmE (PRK00019)Ribosomal_L31 (pfam01197)→ |
| env-120 | Tongue_dor_WUGI_scaffold_55993 | - | 447 | 415 | **RNA**→ rpmE (PRK00019)Ribosomal_L31 (pfam01197)→ |
| env-121 | SRS012279_Baylor_scaffold_57650 | - | 899 | 867 | **RNA**→ rpmE (PRK00019)Ribosomal_L31 (pfam01197)→ |
| env-122 | SRS013234_C2966946 | - | 305 | 273 | **RNA**→ rpmE (PRK00019)Ribosomal_L31 (pfam01197)→ |
| env-123 | SRS013705_C3713865 | - | 413 | 381 | **RNA**→ rpmE (PRK00019)Ribosomal_L31 (pfam01197)→ |
| env-124 | SRS014124_WUGC_scaffold_11711 | - | 938 | 906 | **RNA**→ rpmE (PRK00019)Ribosomal_L31 (pfam01197)→ |
| env-125 | SRS014470_WUGC_scaffold_24370 | - | 840 | 808 | **RNA**→ rpmE (PRK00019)Ribosomal_L31 (pfam01197)→ |
| env-126 | SRS014573_C3542830 | - | 341 | 309 | **RNA**→ rpmE (PRK00019)Ribosomal_L31 (pfam01197)→ |
| env-127 | SRS014689_C1283327 | - | 400 | 368 | **RNA**→ rpmE (PRK00019)Ribosomal_L31 (pfam01197)→ |
| env-128 | SRS015038_C1928591 | + | 27 | 59 | **RNA**→ rpmE (PRK00019)Ribosomal_L31 (pfam01197)→ |
| env-129 | SRS015055_C1676308 | - | 723 | 691 | **RNA**→ rpmE (PRK00019)Ribosomal_L31 (pfam01197)→ |
| env-130 | SRS015057_WUGC_scaffold_9742 | - | 825 | 793 | **RNA**→ rpmE (PRK00019)Ribosomal_L31 (pfam01197)→ |
| env-131 | SRS015434_C5898733 | + | 82 | 114 | **RNA**→ rpmE (PRK00019)Ribosomal_L31 (pfam01197)→ |
| env-132 | SRS015537_WUGC_scaffold_19165 | - | 44 | 12 | **RNA**→ |
| env-133 | SRS015644_WUGC_scaffold_49316 | - | 942 | 910 | **RNA**→ rpmE (PRK00019)Ribosomal_L31 (pfam01197)→ |
| env-134 | SRS015762_C3578704 | - | 639 | 607 | **RNA**→ rpmE (PRK00019)Ribosomal_L31 (pfam01197)→ |
| env-135 | SRS015941_WUGC_scaffold_59228 | - | 663 | 631 | **RNA**→ rpmE (PRK00019)Ribosomal_L31 (pfam01197)→ |
| env-136 | SRS016002_C2556868 | + | 244 | 276 | **RNA**→ rpmE (PRK00019)Ribosomal_L31 (pfam01197)→ |
| env-137 | SRS016086_C2442888 | + | 67 | 99 | **RNA**→ rpmE (PRK00019)Ribosomal_L31 (pfam01197)→ |
| env-138 | SRS016319_WUGC_scaffold_11364 | + | 212 | 244 | **RNA**→ rpmE (PRK00019)Ribosomal_L31 (pfam01197)→ |
| env-139 | SRS016342_Baylor_scaffold_20579 | + | 525 | 557 | **RNA**→ rpmE (PRK00019)Ribosomal_L31 (pfam01197)→ |
| env-140 | SRS017120_C2921280 | - | 413 | 381 | **RNA**→ rpmE (PRK00019)Ribosomal_L31 (pfam01197)→ |
| env-141 | SRS017439_C3698549 | + | 202 | 234 | **RNA**→ |
| env-142 | SRS017713_C3305279 | + | 126 | 158 | **RNA**→ rpmE (PRK00019)Ribosomal_L31 (pfam01197)→ |
| env-143 | SRS017808_Baylor_scaffold_44287 | + | 230 | 262 | **RNA**→ rpmE (PRK00019)Ribosomal_L31 (pfam01197)→ |
| env-144 | SRS018591_WUGC_scaffold_42629 | + | 65 | 97 | **RNA**→ rpmE (PRK00019)Ribosomal_L31 (pfam01197)→ |
| env-145 | SRS018791_WUGC_scaffold_2836 | - | 1590 | 1558 | **RNA**→ rpmE (PRK00019)Ribosomal_L31 (pfam01197)→ |
| env-146 | SRS018969_WUGC_scaffold_53133 | + | 372 | 404 | **RNA**→ rpmE (PRK00019)Ribosomal_L31 (pfam01197)→ |
| env-147 | SRS019022_WUGC_scaffold_39687 | + | 232 | 264 | **RNA**→ rpmE (PRK00019)Ribosomal_L31 (pfam01197)→ |
| env-148 | SRS019045_WUGC_scaffold_40812 | - | 851 | 819 | **RNA**→ rpmE (PRK00019)Ribosomal_L31 (pfam01197)→ |
| env-149 | SRS019071_C5034475 | - | 400 | 368 | **RNA**→ rpmE (PRK00019)Ribosomal_L31 (pfam01197)→ |
| env-150 | SRS019122_C5873801 | - | 732 | 700 | **RNA**→ rpmE (PRK00019)Ribosomal_L31 (pfam01197)→ |
| env-151 | SRS019126_C2212554 | - | 968 | 936 | **RNA**→ rpmE (PRK00019)Ribosomal_L31 (pfam01197)→ |
| env-152 | SRS019389_C2194616 | - | 65 | 33 | **RNA**→ ←hypo |

| | | | | | |
|---|---|---|---|---|---|
| env-153 | SRS019607_WUGC_scaffold_12800 | + | 225 | 257 | **RNA**→ rpmE (PRK00019)Ribosomal_L31 (pfam01197)→ |
| env-154 | SRS019894_WUGC_scaffold_47801 | - | 658 | 626 | **RNA**→ rpmE (PRK00019)Ribosomal_L31 (pfam01197)→ |
| env-155 | SRS020220_Baylor_scaffold_50196 | + | 6 | 38 | **RNA**→ rpmE (PRK00019)Ribosomal_L31 (pfam01197)→ |
| env-156 | SRS020334_Baylor_scaffold_19799 | + | 232 | 264 | **RNA**→ rpmE (PRK00019)Ribosomal_L31 (pfam01197)→ |
| env-157 | SRS021954_Baylor_scaffold_39267 | + | 219 | 251 | **RNA**→ rpmE (PRK00019)Ribosomal_L31 (pfam01197)→ |
| env-158 | SRS022077_Baylor_scaffold_32981 | - | 527 | 495 | **RNA**→ rpmE (PRK00019)Ribosomal_L31 (pfam01197)→ |
| env-159 | SRS022621_C4071744 | + | 67 | 99 | **RNA**→ rpmE (PRK00019)Ribosomal_L31 (pfam01197)→ |
| env-160 | SRS023352_LANL_scaffold_13686 | + | 219 | 251 | **RNA**→ rpmE (PRK00019)Ribosomal_L31 (pfam01197)→ |
| env-161 | SRS023557_C1683448 | - | 223 | 191 | **RNA**→ ←hypo |
| env-162 | SRS023926_Baylor_scaffold_72447 | - | 407 | 375 | **RNA**→ rpmE (PRK00019)Ribosomal_L31 (pfam01197)→ |
| env-163 | SRS024015_C2963407 | - | 44 | 12 | **RNA**→ |
| env-164 | SRS024081_C2590902 | + | 126 | 158 | **RNA**→ rpmE (PRK00019)Ribosomal_L31 (pfam01197)→ |
| env-165 | SRS024138_C2654625 | + | 123 | 155 | **RNA**→ rpmE (PRK00019)Ribosomal_L31 (pfam01197)→ |
| env-166 | SRS024318_C3676033 | + | 65 | 97 | **RNA**→ rpmE (PRK00019)Ribosomal_L31 (pfam01197)→ |
| env-167 | SRS024375_C2434347 | - | 486 | 454 | **RNA**→ rpmE (PRK00019)Ribosomal_L31 (pfam01197)→ |
| env-168 | SRS024580_C3075706 | - | 295 | 263 | **RNA**→ rpmE (PRK00019)Ribosomal_L31 (pfam01197)→ |
| env-169 | SRS042910_C5601812 | - | 747 | 715 | **RNA**→ rpmE (PRK00019)Ribosomal_L31 (pfam01197)→ |
| env-170 | SRS044373_WUGC_scaffold_65551 | + | 486 | 518 | **RNA**→ rpmE (PRK00019)Ribosomal_L31 (pfam01197)→ |
| env-171 | SRS045715_C4791115 | + | 108 | 140 | **RNA**→ rpmE (PRK00019)Ribosomal_L31 (pfam01197)→ |
| env-172 | SRS048791_C3164330 | + | 64 | 96 | **RNA**→ rpmE (PRK00019)Ribosomal_L31 (pfam01197)→ |
| env-173 | SRS050244_LANL_scaffold_45098 | - | 963 | 931 | **RNA**→ rpmE (PRK00019)Ribosomal_L31 (pfam01197)→ |
| env-174 | SRS051791_C1722574 | + | 65 | 97 | **RNA**→ rpmE (PRK00019)Ribosomal_L31 (pfam01197)→ |
| env-175 | SRS053854_C1994549 | + | 108 | 140 | **RNA**→ rpmE (PRK00019)Ribosomal_L31 (pfam01197)→ |
| env-176 | SRS055426_C3209603 | + | 265 | 297 | **RNA**→ |
| env-177 | SRS057205_C3503245 | + | 65 | 97 | **RNA**→ rpmE (PRK00019)Ribosomal_L31 (pfam01197)→ |
| env-178 | SRS057355_C2264004 | + | 154 | 186 | **RNA**→ rpmE (PRK00019)Ribosomal_L31 (pfam01197)→ |
| env-179 | SRS057539_LANL_scaffold_40891 | + | 200 | 232 | **RNA**→ rpmE (PRK00019)Ribosomal_L31 (pfam01197)→ |
| env-180 | SRS058336_C2156394 | + | 459 | 491 | **RNA**→ rpmE (PRK00019)Ribosomal_L31 (pfam01197)→ |
| env-181 | SRS062761_C4283446 | + | 402 | 434 | **RNA**→ rpmE (PRK00019)Ribosomal_L31 (pfam01197)→ |
| env-182 | SRS063193_C2860763 | + | 152 | 184 | **RNA**→ rpmE (PRK00019)Ribosomal_L31 (pfam01197)→ |
| env-183 | SRS064423_LANL_scaffold_34929 | + | 253 | 285 | **RNA**→ rpmE (PRK00019)Ribosomal_L31 (pfam01197)→ |
| env-184 | SRS064774_LANL_scaffold_58893 | - | 1834 | 1802 | **RNA**→ rpmE (PRK00019)Ribosomal_L31 (pfam01197)→ |
| env-185 | SRS065278_LANL_scaffold_41028 | + | 438 | 470 | **RNA**→ rpmE (PRK00019)Ribosomal_L31 (pfam01197)→ |
| Apa-2-1 | NZ_JRND01000030.1 | + | 5302 | 5334 | **RNA**→ rpmE (PRK00019)Ribosomal_L31 (pfam01197)→ |
| Ami-1-1 | NZ_KB822533.1 | + | 353125 | 353157 | **RNA**→ rpmE (PRK00019)Ribosomal_L31 (pfam01197)→ |
| Asp-1-1 | NZ_AWUQ01000004.1 | - | 226500 | 226468 | **RNA**→ rpmE (PRK00019)Ribosomal_L31 (pfam01197)→ hypo→ COG4932 (COG4932)SdrD_B (pfam17210)- |
| Afo-1-1 | NZ_AXXR01000003.1 | - | 295118 | 295086 | **RNA**→ rpmE (PRK00019)Ribosomal_L31 (pfam01197)→ |
| env-186 | SRS023468_C1197102 | + | 689 | 721 | **RNA**→ rpmE (PRK00019)Ribosomal_L31 (pfam01197)→ |
| env-187 | SRS013542_C162841 | - | 662 | 630 | **RNA**→ rpmE (PRK00019)Ribosomal_L31 (pfam01197)→ |
| env-188 | SRS017497_Baylor_scaffold_221 | - | 4581 | 4549 | **RNA**→ rpmE (PRK00019)Ribosomal_L31 (pfam01197)→ |
| Ava-2-1 | NZ_LFWE01000006.1 | - | 17065 | 17033 | **RNA**→ rpmE (PRK00019)Ribosomal_L31 (pfam01197)→ |
| Ava-1-1 | NZ_ADNA01000024.1 | - | 342 | 310 | **RNA**→ hypo→ rpmE (PRK00019)Ribosomal_L31 (pfam01197)→ rpmE (PRK00019)Ribosomal_L31 (pfam011 |
| Ava-1-2 | NZ_ACGK02000004.1 | - | 17226 | 17194 | **RNA**→ ←hypo |
| env-189 | AUXO010832507.1 | - | 84 | 52 | **RNA**→ |
| env-190 | AUXO013415356.1 | + | 198 | 230 | **RNA**→ |
| Csp-3-1 | NZ_CDJG01000012.1 | - | 162765 | 162730 | **RNA**→ rpmE (PRK00019)Ribosomal_L31 (pfam01197)→ |
| env-191 | 4448817.3_Contig32874 | - | 129 | 92 | **RNA**→ rpmE (PRK00019)Ribosomal_L31 (pfam01197)→ |
| env-192 | UnmappedStool_Broad_C253114623 | - | 177 | 145 | **RNA**→ rpmE (PRK00019)Ribosomal_L31 (pfam01197)→ |
| env-193 | SRS012273_C3491547 | - | 170 | 138 | **RNA**→ |
| env-194 | SRS014923_WUGC_scaffold_45412 | + | 37736 | 37768 | **RNA**→ rpmE (PRK00019)Ribosomal_L31 (pfam01197)→ |
| env-195 | SRS015782_C3338422 | - | 62 | 30 | **RNA**→ |
| Cba-3-1 | NZ_CP009302.1 | + | 539611 | 539643 | **RNA**→ rpmE (PRK00019)Ribosomal_L31 (pfam01197)→ |

| | | | | | |
|---|---|---|---|---|---|
| env-196 | longitudinal_131_11 | + | 94369 | 94405 | **RNA**→ rpmE (PRK00019)Ribosomal_L31 (pfam01197)→ |
| env-197 | longitudinal_132_108 | + | 94370 | 94406 | **RNA**→ rpmE (PRK00019)Ribosomal_L31 (pfam01197)→ |
| Cin-1-1 | NZ_GG692710.1 | - | 781470 | 781434 | **RNA**→ rpmE (PRK00019)Ribosomal_L31 (pfam01197)→ |
| env-198 | C5333626_1_V1.UC-13 | - | 179 | 143 | **RNA**→ rpmE (PRK00019)Ribosomal_L31 (pfam01197)→ |
| env-199 | 4448808.3_Contig27735 | + | 205 | 241 | **RNA**→ rpmE (PRK00019)Ribosomal_L31 (pfam01197)→ |
| Cst-1-1 | NZ_DS995474.1 | + | 655651 | 655687 | **RNA**→ rpmE (PRK00019)Ribosomal_L31 (pfam01197)→ |
| env-200 | 4448815.3_Contig8776 | + | 771 | 807 | **RNA**→ rpmE (PRK00019)Ribosomal_L31 (pfam01197)→ |
| env-201 | hgutS7_s7_160723 | + | 1179 | 1211 | **RNA**→ rpmE (PRK00019)Ribosomal_L31 (pfam01197)→ |
| env-202 | scaffold16601_1_MH0086 | + | 702 | 731 | **RNA**→ rpmE (PRK00019)rpmE2 (PRK01678)Ribosomal_L31 (pfam01197)→ |
| env-203 | scaffold88090_1_V1.UC-8 | - | 230 | 201 | **RNA**→ rpmE (PRK00019)Ribosomal_L31 (pfam01197)→ |
| env-204 | scaffold14720_2_V1.UC-9 | + | 1271 | 1300 | **RNA**→ rpmE (PRK00019)Ribosomal_L31 (pfam01197)→ |
| env-205 | UnmappedStool_Broad_C252282338 | + | 322 | 351 | **RNA**→ rpmE (PRK00019)Ribosomal_L31 (pfam01197)→ |
| Aeq-1-1 | NC_022567.1 | - | 1728457 | 1728428 | **RNA**→ rpmE (PRK00019)Ribosomal_L31 (pfam01197)→ |
| Eca-1-1 | NZ_KE159646.1 | - | 1050872 | 1050842 | **RNA**→ rpmE (PRK00019)Ribosomal_L31 (pfam01197)→ |
| Emu-1-1 | NZ_AUGK01000020.1 | + | 30651 | 30681 | **RNA**→ rpmE (PRK00019)Ribosomal_L31 (pfam01197)→ |
| env-206 | scaffold28919_2_O2.UC-11 | - | 258 | 226 | **RNA**→ rpmE (PRK00019)Ribosomal_L31 (pfam01197)→ |
| Esp-3-1 | NC_015738.1 | + | 1175139 | 1175171 | **RNA**→ rpmE (PRK00019)Ribosomal_L31 (pfam01197)→ |
| env-207 | UnmappedStool_Broad_C251071986 | - | 178 | 143 | **RNA**→ rpmE (PRK00019)Ribosomal_L31 (pfam01197)→ |
| env-208 | SRS013687_C2417546 | + | 245 | 280 | **RNA**→ |
| Gpa-1-1 | NC_021021.1 | - | 3191605 | 3191573 | **RNA**→ rpmE (PRK00019)Ribosomal_L31 (pfam01197)→ |
| env-209 | HCF14C_9070 | + | 2040 | 2073 | **RNA**→ rpmE (PRK00019)Ribosomal_L31 (pfam01197)→ |
| env-210 | HCF12C_364780 | + | 265 | 298 | **RNA**→ rpmE (PRK00019)Ribosomal_L31 (pfam01197)→ |
| env-211 | DLF014_scaffold10759_1 | + | 16339 | 16371 | **RNA**→ rpmE (PRK00019)Ribosomal_L31 (pfam01197)→ |
| env-212 | SRS014923_C3176260 | + | 293 | 325 | **RNA**→ rpmE (PRK00019)Ribosomal_L31 (pfam01197)→ |
| env-213 | scaffold261738_2_MH0006 | - | 448 | 417 | **RNA**→ rpmE (PRK00019)Ribosomal_L31 (pfam01197)→ |
| env-214 | scaffold603_48_V1.UC-9 | - | 384 | 353 | **RNA**→ rpmE (PRK00019)Ribosomal_L31 (pfam01197)→ |
| env-215 | 4491403.3_NODE_18899 | + | 21 | 52 | **RNA**→ ←hypo |
| env-216 | scaffold77902_2_O2.UC-20 | - | 409 | 378 | **RNA**→ rpmE (PRK00019)Ribosomal_L31 (pfam01197)→ |
| env-217 | 4491421.3_NODE_60554 | - | 777 | 746 | **RNA**→ rpmE (PRK00019)Ribosomal_L31 (pfam01197)→ |
| env-218 | scaffold30007_3_V1.UC-13 | + | 2845 | 2876 | **RNA**→ ←hypo |
| env-219 | DLM013_scaffold15861_2 | + | 869 | 900 | **RNA**→ rpmE (PRK00019)Ribosomal_L31 (pfam01197)→ |
| env-220 | 4491482.3_NODE_97544 | - | 175288 | 175256 | **RNA**→ rpmE (PRK00019)Ribosomal_L31 (pfam01197)→ |
| env-221 | 4448814.3_Contig25164 | - | 206 | 176 | **RNA**→ rpmE (PRK00019)Ribosomal_L31 (pfam01197)→ |
| Sex-1-1 | NZ_GG700631.1 | - | 393259 | 393228 | **RNA**→ rpmE (PRK00019)Ribosomal_L31 (pfam01197)→ |
| Ssp-1-1 | NZ_ALNO01000001.1 | - | 333235 | 333204 | **RNA**→ rpmE (PRK00019)Ribosomal_L31 (pfam01197)→ rpmE (PRK00019)Ribosomal_L31 (pfam01197)→ |
| Spi-1-1 | NZ_JH815198.1 | + | 276110 | 276143 | **RNA**→ rpmE (PRK00019)Ribosomal_L31 (pfam01197)→ |
| env-222 | AUXO012801284.1 | - | 272 | 242 | **RNA**→ rpmE (PRK00019)Ribosomal_L31 (pfam01197)→ |
| She-1-1 | NC_013165.1 | + | 551648 | 551679 | **RNA**→ rpmE (PRK00019)Ribosomal_L31 (pfam01197)→ |
| env-223 | AUXO013603529.1 | - | 63 | 33 | **RNA**→ hypo→ |
| env-224 | AUXO015651590.1 | - | 285 | 255 | **RNA**→ rpmE (PRK00019)Ribosomal_L31 (pfam01197)→ |
| env-225 | UnmappedStool_Broad_scaffold_663971 | - | 151 | 120 | **RNA**→ rpmE (PRK00019)Ribosomal_L31 (pfam01197)→ |
| env-226 | longitudinal_468_349 | + | 53949 | 53980 | **RNA**→ rpmE (PRK00019)Ribosomal_L31 (pfam01197)→ |
| env-227 | longitudinal_469_265 | + | 204513 | 204544 | **RNA**→ rpmE (PRK00019)Ribosomal_L31 (pfam01197)→ |
| env-228 | longitudinal_470_162 | + | 204460 | 204491 | **RNA**→ rpmE (PRK00019)Ribosomal_L31 (pfam01197)→ |
| env-229 | longitudinal_471_135 | + | 204422 | 204453 | **RNA**→ rpmE (PRK00019)Ribosomal_L31 (pfam01197)→ |
| env-230 | longitudinal_472_50 | + | 103193 | 103224 | **RNA**→ rpmE (PRK00019)Ribosomal_L31 (pfam01197)→ |
| env-231 | longitudinal_473_767 | + | 27039 | 27070 | **RNA**→ rpmE (PRK00019)Ribosomal_L31 (pfam01197)→ |
| env-232 | longitudinal_474_318 | - | 36846 | 36815 | **RNA**→ rpmE (PRK00019)Ribosomal_L31 (pfam01197)→ |
| env-233 | longitudinal_475_209 | - | 8208 | 8177 | **RNA**→ rpmE (PRK00019)Ribosomal_L31 (pfam01197)→ |
| Ele-1-1 | NC_013204.1 | + | 1346970 | 1347001 | **RNA**→ rpmE (PRK00019)Ribosomal_L31 (pfam01197)→ |
| Esp-2-1 | NZ_AEXR01000022.1 | - | 17235 | 17204 | **RNA**→ rpmE (PRK00019)Ribosomal_L31 (pfam01197)→ rpmE (PRK00019)Ribosomal_L31 (pfam01197)→ |
| Esp-1-1 | NZ_GL622579.1 | - | 777790 | 777759 | **RNA**→ rpmE (PRK00019)Ribosomal_L31 (pfam01197)→ |

| | | | | | |
|---|---|---|---|---|---|
| Ele-2-1 | NZ_KN214095.1 | - | 32073 | 32042 | **RNA**→ rpmE (PRK00019)Ribosomal_L31 (pfam01197)→ |
| env-234 | DLF009_scaffold20110_2 | + | 444 | 475 | **RNA**→ rpmE (PRK00019)Ribosomal_L31 (pfam01197)→ |
| env-235 | AUXO017671678.1 | - | 38 | 8 | **RNA**→ |
| env-236 | AUXO015549318.1 | + | 182 | 212 | **RNA**→ rpmE (PRK00019)Ribosomal_L31 (pfam01197)→ |
| env-237 | AUXO011881503.1 | - | 224 | 194 | **RNA**→ rpmE (PRK00019)Ribosomal_L31 (pfam01197)→ |
| San-1-1 | NZ_HE611014.1 | + | 742774 | 742807 | **RNA**→ rpmE (PRK00019)Ribosomal_L31 (pfam01197)→ |
| env-238 | AUXO018607472.1 | - | 131 | 98 | **RNA**→ rpmE (PRK00019)Ribosomal_L31 (pfam01197)→ |
| env-239 | AUXO012093637.1 | + | 87 | 120 | **RNA**→ rpmE (PRK00019)Ribosomal_L31 (pfam01197)→ |
| env-240 | 4448808.3_Contig1066 | + | 3701 | 3732 | **RNA**→ rpmE (PRK00019)Ribosomal_L31 (pfam01197)→ hypo→ |
| env-241 | 4448811.3_Contig16884 | - | 73 | 42 | **RNA**→ ←hypo |
| env-242 | 4448813.3_Contig3373 | + | 962 | 993 | **RNA**→ rpmE (PRK00019)Ribosomal_L31 (pfam01197)→ |
| env-243 | 4448814.3_Contig31153 | - | 302 | 271 | **RNA**→ rpmE (PRK00019)Ribosomal_L31 (pfam01197)→ |
| env-244 | 4448816.3_Contig28865 | - | 364 | 333 | **RNA**→ rpmE (PRK00019)Ribosomal_L31 (pfam01197)→ |
| env-245 | AUXO017402315.1 | - | 978 | 947 | **RNA**→ rpmE (PRK00019)Ribosomal_L31 (pfam01197)→ |
| Cba-1-1 | NZ_AWUP01000003.1 | + | 66691 | 66722 | **RNA**→ rpmE (PRK00019)Ribosomal_L31 (pfam01197)→ |
| Asp-3-1 | NZ_KE952956.1 | + | 5823 | 5854 | **RNA**→ rpmE (PRK00019)Ribosomal_L31 (pfam01197)→ |
| env-246 | 2158020257 | + | 333 | 364 | **RNA**→ ←hypo |
| Osp-1-1 | NZ_JH376563.1 | - | 464047 | 464016 | **RNA**→ rpmE (PRK00019)Ribosomal_L31 (pfam01197)→ |
| env-247 | AUXO013076199.1 | + | 598 | 629 | **RNA**→ rpmE (PRK00019)Ribosomal_L31 (pfam01197)→ |
| env-248 | JGI20225J20221_1001165 | + | 34179 | 34210 | **RNA**→ rpmE (PRK00019)Ribosomal_L31 (pfam01197)→ |
| env-249 | JGI2065J20421_1004925 | + | 1069 | 1100 | **RNA**→ rpmE (PRK00019)Ribosomal_L31 (pfam01197)→ |
| env-250 | SRS024561_C2821916 | + | 707 | 738 | **RNA**→ rpmE (PRK00019)Ribosomal_L31 (pfam01197)→ |
| env-251 | SRS018443_Baylor_scaffold_60373 | - | 2651 | 2620 | **RNA**→ rpmE (PRK00019)rpsP (PRK14521)Ribosomal_L31 (pfam01197)→ |
| env-252 | Supragingival_plaque_LANL_scaffold_407597 | - | 335 | 304 | **RNA**→ rpmE (PRK00019)Ribosomal_L31 (pfam01197)→ |
| env-253 | SRS013836_C4767505 | + | 351 | 382 | **RNA**→ rpmE (PRK00019)Ribosomal_L31 (pfam01197)→ |
| env-254 | SRS047634_C4113595 | + | 242 | 273 | **RNA**→ |
| env-255 | SRS058808_LANL_scaffold_72397 | - | 425 | 394 | **RNA**→ rpmE (PRK00019)Ribosomal_L31 (pfam01197)→ |
| env-256 | HCF12C_292540 | - | 428 | 397 | **RNA**→ rpmE (PRK00019)Ribosomal_L31 (pfam01197)→ |
| env-257 | HCF14C_2_1_1_newblercontig36662 | + | 561 | 592 | **RNA**→ rpmE (PRK00019)Ribosomal_L31 (pfam01197)→ |
| env-258 | HCF12C_352730 | + | 46 | 77 | **RNA**→ rpmE (PRK00019)Ribosomal_L31 (pfam01197)→ |
| env-259 | HCF12C_263188 | - | 719 | 688 | **RNA**→ rpmE (PRK00019)Ribosomal_L31 (pfam01197)→ |
| Ele-2-2 | NZ_KN214097.1 | + | 94244 | 94275 | **RNA**→ ←Transposase_20 (pfam02371) |

#### 10.3 Conserved domains

Conserved domains found in protein-coding genes listed in Section 10.2 are shown below, with the first sentence in their description from the Conserved Domain Database (if any). Conserved domains associated with more than one L31-Coriobacteria RNA are assigned a color, while others are shown in gray. The number in parentheses after the colored domain name is the number of occurrences in Section 10.2.

cd00009 (1) The AAA+ (ATPases Associated with a wide variety of cellular Activities) superfamily represents an ancient group of ATPases belonging to the ASCE (for additional strand, catalytic E) division of the P-loop NTPase fold.
COG1670 (1) Protein N-acetyltransferase, RimJ/RimL family [Translation, ribosomal structure and biogenesis, Posttranslational modification, protein turnover, chaperones]
COG4932 (1) Uncharacterized surface anchored protein [Function unknown]
pfam01197 (277) Ribosomal protein L31.
pfam02371 (1) Transposase IS116/IS110/IS902 family.
pfam13302 (1) Acetyltransferase (GNAT) domain.
pfam17210 (1) SdrD B-like domain.
PRK00019 (276) 50S ribosomal protein L31; Reviewed
PRK00528 (1) 50S ribosomal protein L31; Reviewed
PRK01678 (3) 50S ribosomal protein L31 type B; Reviewed
PRK14521 (1) 30S ribosomal protein S16; Provisional
smart00382 (1) ATPases associated with a variety of cellular activities.

#### 10.4 Multiple-sequence alignment

Each L31-Coriobacteria RNA is denoted by an abbreviation (like "Eco-1-1") that refers to a taxonomy given in Section 10.1. The alignment may include sequences containing the RNA motif, as well as flanking sequence. The L31-Coriobacteria RNA itself is denoted by the line underneath marked 5′ and 3′ on either end. Nucleotides in flanking sequences (i.e., not part of the motif) are written in gray letters. Stems of predicted rho-independent transcription terminators, if any, are shaded yellow. (Note: terminator predictions have not been analyzed manually, and many are likely to be false positives. Terminator predictions are those of the RNIE software.) Nucleotides predicted to function (as DNA) as transcription-factor binding sites, if any, are shaded in green. (But note: these predictions are manually annotated, so they might be under-predicted.) Annotated start codons, if any, are shaded green. (Note: start codons are frequently misannotated, especially in environmental samples.) Nucleotides proposed to basepair as part of the consensus structure are shaded in color when they comprise Watson-Crick or G-U pairs. Otherwise they are shaded gray. Conserved stems are also indicated at the bottom of the alignment by angle brackets, where matching < and > denote base-paired columns. Below these angle brackets, the symbol "2" denotes base pairs exhibiting covariation according to the statistically well-founded R-scape method. "1" denotes base pairs exhibiting covariation according to R2R's simplistic method. "0" denotes base pairs that are not observed to mutate and "?" denotes base pairs that have a significant frequency of non-canonical nucleotides for Watson-Crick or G-U pairs (> 5%). Below these base pair annotation is the consensus sequence: "R" = "A" or "G", "Y" = "C" or "U", red nucleotides: nucleotide identity conserved more than 97% of the time, black nucleotides: 90%, gray nucleotides: 75%, red circle (•): nucleotide is present 97% of the time, black circle (•): 90%, gray circle (•): 75%, white circle (◦): 50%. All percentages of sequences just described (e.g. 97% conserved) assume that sequences have been weighted by the GSC algorithm implemented by the Infernal software package.

Duplicate sequences: the following putative homologs are not shown in the alignment because their sequences are identical to a homolog already shown: Aeq-1-1, Apa-1-1, Apa-2-1, Ari-1-1, Asp-1-1, Asp-2-1, Asp-4-1, Asp-5-1, Ava-1-1, Ava-1-2, Ava-2-1, Cae-1-1, Cin-1-1, Csp-1-1, Csp-2-1, Cst-1-1, Cta-1-1, Ele-1-1, Ele-2-1, Esp-1-1, Esp-2-1, Oul-2-1, Ssp-1-1, env-10, env-100, env-101, env-102, env-103, env-104, env-105, env-106, env-107, env-108, env-109, env-11, env-110, env-111, env-112, env-113, env-117, env-118, env-119, env-12, env-120, env-121, env-122, env-123, env-124, env-125, env-126, env-127, env-128, env-129, env-13, env-130, env-131, env-132, env-133, env-134, env-135, env-136, env-137, env-138, env-139, env-140, env-141, env-142, env-143, env-144, env-145, env-146, env-147, env-148, env-149, env-150, env-151, env-152, env-153, env-154, env-155, env-156, env-157, env-158, env-159, env-160, env-161, env-162, env-163, env-164, env-165, env-166, env-167, env-168, env-169, env-170, env-171, env-172, env-173, env-174, env-175, env-176, env-177, env-178, env-179, env-18, env-180, env-181, env-182, env-183, env-184, env-185, env-187, env-188, env-19, env-193, env-194, env-195, env-197, env-199, env-20, env-203, env-204, env-205, env-208, env-21, env-210, env-215, env-219, env-22, env-226, env-227, env-228, env-229, env-23, env-230, env-231, env-232, env-233, env-239, env-24, env-241, env-242, env-243, env-244, env-245, env-249, env-25, env-252, env-253, env-254, env-255, env-258, env-26, env-27, env-28, env-29, env-30, env-31, env-32, env-33, env-34, env-35, env-36, env-37, env-38, env-39, env-40, env-41, env-42, env-43, env-44, env-45, env-46, env-47, env-48, env-49, env-5, env-50, env-51, env-52, env-53, env-54, env-55, env-56, env-57, env-58, env-59, env-6, env-60, env-61, env-62, env-63, env-65, env-66, env-67, env-68, env-69, env-70, env-71, env-72, env-73, env-74, env-75, env-76, env-77, env-78, env-79, env-84, env-85, env-86, env-87, env-88, env-89, env-90, env-91, env-92, env-93, env-94, env-95, env-96, env-97, env-98, env-99

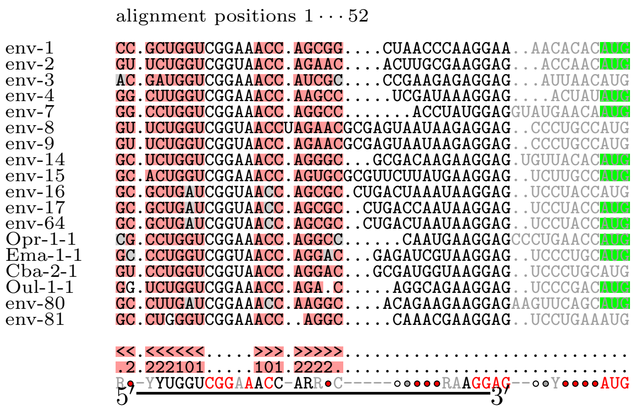

```
Cgl-1-1   GC.UCUGGUCGGAAACC.AGAGC...ACGAUCACAAGGAG..CCCUUGUAUG
env-82    GC.UCUGGUCGGAAACC.AGAGC.....CGAUGUAAGGAG...UCCUGUAUG
env-83    GC.UCUGGUCGGAAACC.AGAGC.....CGAUGUAAGGAG...UCCUGCAUG
env-114   GC.UCUGGUCGGAAACC.AGGGC.....CGAUGUAAGGAG...UUCUGCAUG
env-115   GC.UCUGGUCGGAAACC.AGGGC.....CGAUGUAAGGAG...UCCCGCAUG
env-116   GC.UCUGGUCGGAAACC.AGGGC.....CGAUGUAAGGAG...UCCUGCAUG
Ami-1-1   GC.UCUGGUCGGAAACC.AGAGC.....CGUUAGAAGGAG...UUUUAUAUG
Afo-1-1   GC.UCUGGUCGGAAACC.AGAGU.....CGUUAGAAGGAG....UCCUUAUG
env-186   AG.CCUGAUCGGAAACC.AGGCA.....AACAAGAAGGAG..UCCUGUCAUG
env-189   GC.UCUGGUCGGAAACC.AGGGC.....ACGAAGAAGGAG..UCCUGCUAUG
env-190   GU.CCUGGUCGGAAACC.AGGAC.....ACGAAGAAGGAG..ACCAGAAAUG
Csp-3-1   GU.CCUGGUCGGAAACC.AGGAC..GCGAUUCCAGAGGAG..CCCUGCCAUG
env-191   GC.ACUGGUCGGAAACC.AGUGCGCGUACGAGAAGAGGAG..CCCUGCCAUG
env-192   GCACUUGGUCGGAAACC.AAGGC......ACUAGAAGGAG...ACCAACAUG
Cba-3-1   GA.CUUGGUCGGAAACC.AAGUC.....CUCGAAAAGGAG..AACCCAUAUG
env-196   GU.CCUGGUCGGUAACC.AGGAC.GCGACCAGAAGAGGAG..CCCUGCCAUG
env-198   GU.CCUGGUCGGUAACC.AGGAC.GUGACCAGAAGAGGAG..CCCUGCCAUG
env-200   GU.CCUGGUCGGUAACC.AGGAC.GUGACCAGAAGAGGAG.CCCCUGCCAUG
env-201   GC.CCUGGUCGG.AACC.AGGGC....CGACGAACAGGAG..UCCUGCUAUG
env-202   CC.CUUGGUCGGAAACC.AAGGA........AGUUAGGAG...AACCCAAUG
Eca-1-1   UC.CUUGGUCGGAAACC.AAGGG.......AAGAUAGGAG....AACCCAUG
Emu-1-1   CC.CUUGGUCGGAAACC.AAGGG.......AAGUUAGGAG....AACCCAUG
env-206   AU.CUUGGUCGGAAACC.AAGGU.....AUUAGUGAGGAG....AAUCAAUG
Esp-3-1   GC.CUUGGUCGGAAACC.AAGGC.....AUGUCAAAGGAGAACCAGGAAAUG
env-207   GC.CUUGGUCGGAAACC.AAGGC..ACUCUGUCAAAGGAG..AAUACCAAUG
Gpa-1-1   GC.CUUGGUCGGAAACC.AAGGC.....CUGUCUAAGGAG...AACCAAAUG
env-209   GC.CCUGGUCGGAAACC.AGGGC....CCGAUAAAAGGAG..UCCUGCAAUG
env-211   GU.CCUGGUCGGAAACC.AGGAC.....CGACAAAAGGAG..UCCUGCAAUG
env-212   GC.CUUGGUCGGAAACU.AAGGC.....AUACCAAAGGAG....CAAGAAUG
env-213   GU.CUUGGUCGGAAACC.AAGGC......AUAGUAAGGAG....AACCAAUG
env-214   GC.CUUGGUCGGAAACC.AAGGC......AUAGUAAGGAG....AACCAAUG
env-216   GC.CUUGGUCGGGACCC.AAGGC......AUACGAAGGAG....AACCAAUG
env-217   GC.CUUGGUCGGAAACC.AAGGC......AUACGAAGGAG....AACCAAUG
env-218   GC.CUUGGUCGGAAACC.AAGGC......AUUAGAAGGAG....AACCAAUG
env-220   GU.CUUGGUCGGAAACC.AAGAC.....AAUAAAAAGGAG....AACCAAUG
env-221   GU.CUUGGUCGGAAACC.AAGAC.......CGCAAAGGAG.AAACCAUAAUG
Sex-1-1   GC.CUUGGUCGGAAACC.AAGGC......AUGAAAAGGAG.AACCCAUCAUG
Spi-1-1   GU.CUUGGUCGGAAACC.AAGGC....GCCUGAAAAGGAG.AAACCAACAUG
env-222   GU.CUUGGUCGGAAACC.AAGGC.......AGAAUAGGAG..AACCAACAUG
She-1-1   GU.CUUGGUCGGAAACC.AAGAC......AUGAAAAGGAG.AAACCAUCAUG
env-223   GC.CUUGGUCGGAAACC.AAGGC.......GGAAAAGGAG...AAAACCAUG
env-224   GC.CUUGGUCGGAAACC.AAGGC.......GGAAAAGGAG.AACCAACCAUG
env-225   GC.CUUGGUCGGAAACC.AAGGC......CCGUAAAGGAG..AACCAUAAUG
env-234   GC.CUUGGUCGGAAACC.AAGGC......CCGUGAAGGAG..AACCAUAAUG
env-235   GC.CUUGGUCGGAAACC.AAGGC.......AGAAGAGGAG........AAUG
env-236   GC.CUUGGUCGGAAACC.AAGGC.......AGAAAGGGAG.AACCAGAAAUG
env-237   AC.CUUGGUCGGAAACC.AAGGC.......GGAAUAGGAG......AAAAUG
San-1-1   GC.CUUGGUCGGAAACC.AAGGC....ACGGUAAAAGGAGUACCAGACUAUG
env-238   GC.CCUGGUCGGAAACC.AGGGC....ACGAAAGAAGGAG..UCCCAACAUG
env-240   CG.UCUGGUCGGAAACC.AGACC......UACCAAAGGAG..UCCUGCCAUG
Cba-1-1   UG.CCUGGUCGGAAACC.AGGCC......UACGUUAGGAG....UUGUCAUG
Asp-3-1   CG.CCUGGUCGGAAACC.AGGCC......UACGUUAGGAG....UUGUCAUG
env-246   CG.CCUGGUCGGAAACC.AGGCC......AAACGAAGGAG..UCCUGAAAUG
Osp-1-1   UG.UCUGGUCGGAAACC.AGGCC......GAAAGAAGGAG..UCCUGAAAUG
env-247   CG.CCUGGUCGGAAACC.AGGCC......UACCAAAGGAG..UCCCAGUAUG
env-248   CG.CCUGGUCGGAAACC.AGGCA......AACCAAAGGAG..UCCCGACAUG
env-250   CG.CCUGGUCGGAAACC.AGGCC......AAUCAAGGGAG..UUCCAACAUG
env-251   CG.CCUGGUCGGAAACC.AGGCC......AACCAAGGGAG.UUCCAAACAUG
env-256   CG.CCUGGUCGGAAACC.AGGCC......AACAGAAGGAG....CCAAGAUG
env-257   UG.CCUGGUCGGAAACC.AGGCC......AACAGAAGGAG....UCAAGAUG
env-259   CG.CCUGGUCGGAAACC.AGGCC......AACGUAAGGAG....UCAGCAUG
Ele-2-2   GG.CGGCGCCGGAAGCG.CUGCC......GGAGAAAGGAGACCAGCUAUAUG
          <<.<<<<<<.....>>>.>>>>>.............................
          .2.222101.....101.2222..............................
          R•-YYUGGUCGGAAACC-ARR•C-----∘∘•••RAAGGAG--∘∘Y••••AUG
          5′                                  3′
```

### 11 L31-Firmicutes

#### 11.1 Taxa

The taxonomy of each organism containing a putative L31-Firmicutes RNA is listed, with abbreviations identifying each hit (e.g., "Eco-1-1" and "Eco-1-2" might hypothetically represent two distinct RNAs in *E. coli*). The abbreviations will be used to identify each individual L31-Firmicutes RNA in Sections 11.3 and 11.5:.

| **abbrev. of hits** | **taxonomy of species** |
|---|---|
| Mab-1-1 to Mab-1-2 | Bacteria Actinobacteria Corynebacteriales Mycobacteriaceae *Mycobacterium abscessus* |
| bLF-1-1 | Bacteria bacterium LF-3 |
| Axy-1-1 | Bacteria Firmicutes Bacillales Bacillaceae *Amphibacillus xylanus* NBRC 15112 |
| Afl-1-1 | Bacteria Firmicutes Bacillales Bacillaceae *Anoxybacillus flavithermus* WK1 |
| Bam-1-1 | Bacteria Firmicutes Bacillales Bacillaceae *Bacillus amyloliquefaciens* DSM7 |
| Bam-2-1 | Bacteria Firmicutes Bacillales Bacillaceae *Bacillus amyloliquefaciens* FZB42 |
| Bam-3-1 | Bacteria Firmicutes Bacillales Bacillaceae *Bacillus amyloliquefaciens* LL3 |
| Bam-7-1 | Bacteria Firmicutes Bacillales Bacillaceae *Bacillus amyloliquefaciens subsp. plantarum* CAU B946 |
| Bam-8-1 | Bacteria Firmicutes Bacillales Bacillaceae *Bacillus amyloliquefaciens subsp. plantarum* YAU B9601-Y2 |
| Bam-4-1 | Bacteria Firmicutes Bacillales Bacillaceae *Bacillus amyloliquefaciens* TA208 |
| Bam-5-1 | Bacteria Firmicutes Bacillales Bacillaceae *Bacillus amyloliquefaciens* XH7 |
| Bam-6-1 | Bacteria Firmicutes Bacillales Bacillaceae *Bacillus amyloliquefaciens* Y2 |
| Bat-1-1 | Bacteria Firmicutes Bacillales Bacillaceae *Bacillus atrophaeus* 1942 |
| Baz-1-1 | Bacteria Firmicutes Bacillales Bacillaceae *Bacillus azotoformans* LMG 9581 |
| Bba-1-1 | Bacteria Firmicutes Bacillales Bacillaceae *Bacillus bataviensis* LMG 21833 |
| Bce-1-1 | Bacteria Firmicutes Bacillales Bacillaceae *Bacillus cellulosilyticus* DSM 2522 |
| Bcl-1-1 | Bacteria Firmicutes Bacillales Bacillaceae *Bacillus clausii* KSM-K16 |
| Bco-1-1 | Bacteria Firmicutes Bacillales Bacillaceae *Bacillus coagulans* 2-6 |
| Bco-2-1 | Bacteria Firmicutes Bacillales Bacillaceae *Bacillus coagulans* 36D1 |
| Bco-3-1 | Bacteria Firmicutes Bacillales Bacillaceae *Bacillus coahuilensis m*4-4 |
| Bha-1-1 | Bacteria Firmicutes Bacillales Bacillaceae *Bacillus halodurans* C-125 |
| Bis-1-1 | Bacteria Firmicutes Bacillales Bacillaceae *Bacillus isronensis* B3W22 |
| Bli-1-1 to Bli-1-2 | Bacteria Firmicutes Bacillales Bacillaceae *Bacillus licheniformis* ATCC 14580 (DSM 13) |
| Bma-1-1 | Bacteria Firmicutes Bacillales Bacillaceae *Bacillus macauensis* ZFHKF-1 |
| Bme-1-1 | Bacteria Firmicutes Bacillales Bacillaceae *Bacillus megaterium* DSM 319 |
| Bme-2-1 | Bacteria Firmicutes Bacillales Bacillaceae *Bacillus megaterium* QM B1551 |
| Bme-3-1 | Bacteria Firmicutes Bacillales Bacillaceae *Bacillus megaterium* WSH-002 |
| Bme-4-1 | Bacteria Firmicutes Bacillales Bacillaceae *Bacillus methanolicus* PB1 |
| Bps-1-1 | Bacteria Firmicutes Bacillales Bacillaceae *Bacillus pseudofirmus* OF4 |
| Bps-2-1 | Bacteria Firmicutes Bacillales Bacillaceae *Bacillus pseudomycoides* DSM 12442 |
| Bpu-1-1 | Bacteria Firmicutes Bacillales Bacillaceae *Bacillus pumilus* ATCC 7061 |
| Bpu-2-1 | Bacteria Firmicutes Bacillales Bacillaceae *Bacillus pumilus* SAFR-032 |
| Bse-1-1 | Bacteria Firmicutes Bacillales Bacillaceae *Bacillus selenitireducens* MLS10 |
| Bsp-1-1 | Bacteria Firmicutes Bacillales Bacillaceae *Bacillus sp.* 5B6 |
| Bsp-2-1 | Bacteria Firmicutes Bacillales Bacillaceae *Bacillus sp.* 916 |
| Bsp-3-1 | Bacteria Firmicutes Bacillales Bacillaceae *Bacillus sp.* B14905 |
| Bsp-4-1 | Bacteria Firmicutes Bacillales Bacillaceae *Bacillus sp.* HYC-10 |
| Bsp-5-1 | Bacteria Firmicutes Bacillales Bacillaceae *Bacillus sp.* JS |
| Bsp-6-1 | Bacteria Firmicutes Bacillales Bacillaceae *Bacillus sp.* M 2-6 |
| Bsp-8-1 | Bacteria Firmicutes Bacillales Bacillaceae *Bacillus sp. m*3-13 |
| Bsp-7-1 | Bacteria Firmicutes Bacillales Bacillaceae *Bacillus sp.* NRRL B-14911 |
| Bsu-1-1 | Bacteria Firmicutes Bacillales Bacillaceae *Bacillus subtilis* BSn5 |
| Bsu-2-1 | Bacteria Firmicutes Bacillales Bacillaceae *Bacillus subtilis* QB928 |
| Bsu-3-1 | Bacteria Firmicutes Bacillales Bacillaceae *Bacillus subtilis subsp. spizizenii* ATCC 6633 |
| Bsu-5-1 | Bacteria Firmicutes Bacillales Bacillaceae *Bacillus subtilis subsp. spizizenii str.* W23 |
| Bsu-4-1 | Bacteria Firmicutes Bacillales Bacillaceae *Bacillus subtilis subsp. spizizenii* TU-B-10 |
| Bsu-6-1 to Bsu-6-2 | Bacteria Firmicutes Bacillales Bacillaceae *Bacillus subtilis subsp. subtilis str.* 168 |
| Bsu-7-1 | Bacteria Firmicutes Bacillales Bacillaceae *Bacillus subtilis subsp. subtilis str.* JH642 |
| Bsu-8-1 | Bacteria Firmicutes Bacillales Bacillaceae *Bacillus subtilis subsp. subtilis str.* NCIB 3610 |
| Bsu-9-1 | Bacteria Firmicutes Bacillales Bacillaceae *Bacillus subtilis subsp. subtilis str.* RO-NN-1 |
| Bsu-10-1 | Bacteria Firmicutes Bacillales Bacillaceae *Bacillus subtilis subsp. subtilis str.* SMY |
| Ban-2-1 | Bacteria Firmicutes Bacillales Bacillaceae Bacillus Bacilluscereus group *Bacillus anthracis str.* 'Ames Ancestor' |
| Ban-3-1 | Bacteria Firmicutes Bacillales Bacillaceae Bacillus Bacilluscereus group *Bacillus anthracis str.* A0174 |
| Ban-4-1 | Bacteria Firmicutes Bacillales Bacillaceae Bacillus Bacilluscereus group *Bacillus anthracis str.* A0193 |
| Ban-5-1 | Bacteria Firmicutes Bacillales Bacillaceae Bacillus Bacilluscereus group *Bacillus anthracis str.* A0248 |
| Ban-6-1 | Bacteria Firmicutes Bacillales Bacillaceae Bacillus Bacilluscereus group *Bacillus anthracis str.* A0389 |
| Ban-7-1 | Bacteria Firmicutes Bacillales Bacillaceae Bacillus Bacilluscereus group *Bacillus anthracis str.* A0442 |
| Ban-8-1 | Bacteria Firmicutes Bacillales Bacillaceae Bacillus Bacilluscereus group *Bacillus anthracis str.* A0465 |
| Ban-9-1 | Bacteria Firmicutes Bacillales Bacillaceae Bacillus Bacilluscereus group *Bacillus anthracis str.* A0488 |
| Ban-10-1 | Bacteria Firmicutes Bacillales Bacillaceae Bacillus Bacilluscereus group *Bacillus anthracis str.* A1055 |
| Ban-11-1 | Bacteria Firmicutes Bacillales Bacillaceae Bacillus Bacilluscereus group *Bacillus anthracis str.* Ames |
| Ban-12-1 | Bacteria Firmicutes Bacillales Bacillaceae Bacillus Bacilluscereus group *Bacillus anthracis str.* Australia 94 |
| Ban-13-1 | Bacteria Firmicutes Bacillales Bacillaceae Bacillus Bacilluscereus group *Bacillus anthracis str.* CDC 684 |
| Ban-14-1 | Bacteria Firmicutes Bacillales Bacillaceae Bacillus Bacilluscereus group *Bacillus anthracis str.* CNEVA-9066 |

| | |
|---|---|
| Ban-15-1 | Bacteria Firmicutes Bacillales Bacillaceae Bacillus Bacilluscereus group *Bacillus anthracis str.* H9401 |
| Ban-16-1 | Bacteria Firmicutes Bacillales Bacillaceae Bacillus Bacilluscereus group *Bacillus anthracis str.* Kruger B |
| Ban-17-1 to Ban-17-2 | Bacteria Firmicutes Bacillales Bacillaceae Bacillus Bacilluscereus group *Bacillus anthracis str.* Sterne |
| Ban-18-1 to Ban-18-2 | Bacteria Firmicutes Bacillales Bacillaceae Bacillus Bacilluscereus group *Bacillus anthracis str.* Vollum |
| Ban-19-1 | Bacteria Firmicutes Bacillales Bacillaceae Bacillus Bacilluscereus group *Bacillus anthracis str.* Western North America USA6153 |
| Ban-1-1 | Bacteria Firmicutes Bacillales Bacillaceae Bacillus Bacilluscereus group *Bacillus anthracis* Tsiankovskii-I |
| Bce-2-1 to Bce-2-2 | Bacteria Firmicutes Bacillales Bacillaceae Bacillus Bacilluscereus group *Bacillus cereus* 03BB102 |
| Bce-3-1 | Bacteria Firmicutes Bacillales Bacillaceae Bacillus Bacilluscereus group *Bacillus cereus* 172560W |
| Bce-4-1 | Bacteria Firmicutes Bacillales Bacillaceae Bacillus Bacilluscereus group *Bacillus cereus* 95/8201 |
| Bce-5-1 | Bacteria Firmicutes Bacillales Bacillaceae Bacillus Bacilluscereus group *Bacillus cereus* AH1134 |
| Bce-6-1 | Bacteria Firmicutes Bacillales Bacillaceae Bacillus Bacilluscereus group *Bacillus cereus* AH1271 |
| Bce-7-1 | Bacteria Firmicutes Bacillales Bacillaceae Bacillus Bacilluscereus group *Bacillus cereus* AH1272 |
| Bce-8-1 | Bacteria Firmicutes Bacillales Bacillaceae Bacillus Bacilluscereus group *Bacillus cereus* AH1273 |
| Bce-9-1 | Bacteria Firmicutes Bacillales Bacillaceae Bacillus Bacilluscereus group *Bacillus cereus* AH187 |
| Bce-10-1 | Bacteria Firmicutes Bacillales Bacillaceae Bacillus Bacilluscereus group *Bacillus cereus* AH603 |
| Bce-11-1 | Bacteria Firmicutes Bacillales Bacillaceae Bacillus Bacilluscereus group *Bacillus cereus* AH621 |
| Bce-12-1 | Bacteria Firmicutes Bacillales Bacillaceae Bacillus Bacilluscereus group *Bacillus cereus* AH676 |
| Bce-13-1 | Bacteria Firmicutes Bacillales Bacillaceae Bacillus Bacilluscereus group *Bacillus cereus* AH820 |
| Bce-14-1 to Bce-14-2 | Bacteria Firmicutes Bacillales Bacillaceae Bacillus Bacilluscereus group *Bacillus cereus* ATCC 10876 |
| Bce-15-1 | Bacteria Firmicutes Bacillales Bacillaceae Bacillus Bacilluscereus group *Bacillus cereus* ATCC 10987 |
| Bce-16-1 | Bacteria Firmicutes Bacillales Bacillaceae Bacillus Bacilluscereus group *Bacillus cereus* ATCC 14579 |
| Bce-17-1 to Bce-17-2 | Bacteria Firmicutes Bacillales Bacillaceae Bacillus Bacilluscereus group *Bacillus cereus* ATCC 4342 |
| Bce-18-1 | Bacteria Firmicutes Bacillales Bacillaceae Bacillus Bacilluscereus group *Bacillus cereus* B4264 |
| Bce-19-1 | Bacteria Firmicutes Bacillales Bacillaceae Bacillus Bacilluscereus group *Bacillus cereus* BDRD-Cer4 |
| Bce-20-1 | Bacteria Firmicutes Bacillales Bacillaceae Bacillus Bacilluscereus group *Bacillus cereus* BDRD-ST196 |
| Bce-21-1 | Bacteria Firmicutes Bacillales Bacillaceae Bacillus Bacilluscereus group *Bacillus cereus* BDRD-ST24 |
| Bce-22-1 | Bacteria Firmicutes Bacillales Bacillaceae Bacillus Bacilluscereus group *Bacillus cereus* BDRD-ST26 |
| Bce-23-1 | Bacteria Firmicutes Bacillales Bacillaceae Bacillus Bacilluscereus group *Bacillus cereus* BGSC 6E1 |
| Bce-44-1 | Bacteria Firmicutes Bacillales Bacillaceae Bacillus Bacilluscereus group *Bacillus cereus biovar anthracis str.* CI |
| Bce-24-1 to Bce-24-2 | Bacteria Firmicutes Bacillales Bacillaceae Bacillus Bacilluscereus group *Bacillus cereus* E33L |
| Bce-25-1 | Bacteria Firmicutes Bacillales Bacillaceae Bacillus Bacilluscereus group *Bacillus cereus* F65185 |
| Bce-26-1 | Bacteria Firmicutes Bacillales Bacillaceae Bacillus Bacilluscereus group *Bacillus cereus* F837/76 |
| Bce-27-1 | Bacteria Firmicutes Bacillales Bacillaceae Bacillus Bacilluscereus group *Bacillus cereus* FRI-35 |
| Bce-28-1 to Bce-28-2 | Bacteria Firmicutes Bacillales Bacillaceae Bacillus Bacilluscereus group *Bacillus cereus* G9241 |
| Bce-29-1 | Bacteria Firmicutes Bacillales Bacillaceae Bacillus Bacilluscereus group *Bacillus cereus* G9842 |
| Bce-45-1 | Bacteria Firmicutes Bacillales Bacillaceae Bacillus Bacilluscereus group *Bacillus cereus* $m$1293 |
| Bce-46-1 | Bacteria Firmicutes Bacillales Bacillaceae Bacillus Bacilluscereus group *Bacillus cereus* $m$1550 |
| Bce-30-1 | Bacteria Firmicutes Bacillales Bacillaceae Bacillus Bacilluscereus group *Bacillus cereus* MM3 |
| Bce-31-1 | Bacteria Firmicutes Bacillales Bacillaceae Bacillus Bacilluscereus group *Bacillus cereus* NC7401 |
| Bce-32-1 | Bacteria Firmicutes Bacillales Bacillaceae Bacillus Bacilluscereus group *Bacillus cereus* NVH0597-99 |
| Bce-33-1 | Bacteria Firmicutes Bacillales Bacillaceae Bacillus Bacilluscereus group *Bacillus cereus* Q1 |
| Bce-34-1 | Bacteria Firmicutes Bacillales Bacillaceae Bacillus Bacilluscereus group *Bacillus cereus* R309803 |
| Bce-35-1 | Bacteria Firmicutes Bacillales Bacillaceae Bacillus Bacilluscereus group *Bacillus cereus* Rock1-15 |
| Bce-36-1 | Bacteria Firmicutes Bacillales Bacillaceae Bacillus Bacilluscereus group *Bacillus cereus* Rock1-3 |
| Bce-37-1 | Bacteria Firmicutes Bacillales Bacillaceae Bacillus Bacilluscereus group *Bacillus cereus* Rock3-28 |
| Bce-38-1 | Bacteria Firmicutes Bacillales Bacillaceae Bacillus Bacilluscereus group *Bacillus cereus* Rock3-29 |
| Bce-39-1 | Bacteria Firmicutes Bacillales Bacillaceae Bacillus Bacilluscereus group *Bacillus cereus* Rock3-42 |
| Bce-40-1 | Bacteria Firmicutes Bacillales Bacillaceae Bacillus Bacilluscereus group *Bacillus cereus* Rock3-44 |
| Bce-41-1 | Bacteria Firmicutes Bacillales Bacillaceae Bacillus Bacilluscereus group *Bacillus cereus* Rock4-18 |
| Bce-42-1 | Bacteria Firmicutes Bacillales Bacillaceae Bacillus Bacilluscereus group *Bacillus cereus* Rock4-2 |
| Bce-43-1 | Bacteria Firmicutes Bacillales Bacillaceae Bacillus Bacilluscereus group *Bacillus cereus* W |
| Bcy-1-1 | Bacteria Firmicutes Bacillales Bacillaceae Bacillus Bacilluscereus group *Bacillus cytotoxicus* NVH 391-98 |
| Bmy-1-1 | Bacteria Firmicutes Bacillales Bacillaceae Bacillus Bacilluscereus group *Bacillus mycoides* DSM 2048 |
| Bmy-2-1 | Bacteria Firmicutes Bacillales Bacillaceae Bacillus Bacilluscereus group *Bacillus mycoides* Rock1-4 |
| Bmy-3-1 | Bacteria Firmicutes Bacillales Bacillaceae Bacillus Bacilluscereus group *Bacillus mycoides* Rock3-17 |
| Bth-1-1 | Bacteria Firmicutes Bacillales Bacillaceae Bacillus Bacilluscereus group *Bacillus thuringiensis* BMB171 |
| Bth-2-1 | Bacteria Firmicutes Bacillales Bacillaceae Bacillus Bacilluscereus group *Bacillus thuringiensis* Bt407 |
| Bth-3-1 | Bacteria Firmicutes Bacillales Bacillaceae Bacillus Bacilluscereus group *Bacillus thuringiensis* HD-771 |
| Bth-4-1 | Bacteria Firmicutes Bacillales Bacillaceae Bacillus Bacilluscereus group *Bacillus thuringiensis* IBL 200 |
| Bth-5-1 | Bacteria Firmicutes Bacillales Bacillaceae Bacillus Bacilluscereus group *Bacillus thuringiensis* IBL 4222 |
| Bth-6-1 | Bacteria Firmicutes Bacillales Bacillaceae Bacillus Bacilluscereus group *Bacillus thuringiensis* MC28 |
| Bth-7-1 | Bacteria Firmicutes Bacillales Bacillaceae Bacillus Bacilluscereus group *Bacillus thuringiensis serovar andalousiensis* BGSC 4AW1 |
| Bth-8-1 | Bacteria Firmicutes Bacillales Bacillaceae Bacillus Bacilluscereus group *Bacillus thuringiensis serovar berliner* ATCC 10792 |
| Bth-9-1 | Bacteria Firmicutes Bacillales Bacillaceae Bacillus Bacilluscereus group *Bacillus thuringiensis serovar chinensis* CT-43 |
| Bth-10-1 | Bacteria Firmicutes Bacillales Bacillaceae Bacillus Bacilluscereus group *Bacillus thuringiensis serovar finitimus* YBT-020 |

| | |
|---|---|
| Bth-11-1 | Bacteria Firmicutes Bacillales Bacillaceae Bacillus Bacilluscereus group *Bacillus thuringiensis serovar huazhongensis* BGSC 4BD1 |
| Bth-12-1 | Bacteria Firmicutes Bacillales Bacillaceae Bacillus Bacilluscereus group *Bacillus thuringiensis serovar konkukian str.* 97-27 |
| Bth-13-1 | Bacteria Firmicutes Bacillales Bacillaceae Bacillus Bacilluscereus group *Bacillus thuringiensis serovar kurstaki str.* T03a001 |
| Bth-14-1 | Bacteria Firmicutes Bacillales Bacillaceae Bacillus Bacilluscereus group *Bacillus thuringiensis serovar monterrey* BGSC 4AJ1 |
| Bth-15-1 | Bacteria Firmicutes Bacillales Bacillaceae Bacillus Bacilluscereus group *Bacillus thuringiensis serovar pakistani str.* T13001 |
| Bth-16-1 | Bacteria Firmicutes Bacillales Bacillaceae Bacillus Bacilluscereus group *Bacillus thuringiensis serovar pondicheriensis* BGSC 4BA1 |
| Bth-17-1 | Bacteria Firmicutes Bacillales Bacillaceae Bacillus Bacilluscereus group *Bacillus thuringiensis serovar pulsiensis* BGSC 4CC1 |
| Bth-18-1 | Bacteria Firmicutes Bacillales Bacillaceae Bacillus Bacilluscereus group *Bacillus thuringiensis serovar thuringiensis str.* T01001 |
| Bth-19-1 | Bacteria Firmicutes Bacillales Bacillaceae Bacillus Bacilluscereus group *Bacillus thuringiensis serovar tochigiensis* BGSC 4Y1 |
| Bth-20-1 | Bacteria Firmicutes Bacillales Bacillaceae Bacillus Bacilluscereus group *Bacillus thuringiensis str.* Al Hakam |
| Bwe-1-1 | Bacteria Firmicutes Bacillales Bacillaceae Bacillus Bacilluscereus group *Bacillus weihenstephanensis* KBAB4 |
| Cth-1-1 | Bacteria Firmicutes Bacillales Bacillaceae *Caldalkalibacillus thermarum* TA2.A1 |
| Gka-1-1 | Bacteria Firmicutes Bacillales Bacillaceae *Geobacillus kaustophilus* HTA426 |
| Gsp-1-1 | Bacteria Firmicutes Bacillales Bacillaceae *Geobacillus sp.* C56-T3 |
| Gsp-2-1 | Bacteria Firmicutes Bacillales Bacillaceae *Geobacillus sp.* G11MC16 |
| Gsp-3-1 | Bacteria Firmicutes Bacillales Bacillaceae *Geobacillus sp.* WCH70 |
| Gsp-4-1 | Bacteria Firmicutes Bacillales Bacillaceae *Geobacillus sp.* Y4.1MC1 |
| Gsp-5-1 | Bacteria Firmicutes Bacillales Bacillaceae *Geobacillus sp.* Y412MC10 |
| Gsp-6-1 | Bacteria Firmicutes Bacillales Bacillaceae *Geobacillus sp.* Y412MC52 |
| Gsp-7-1 | Bacteria Firmicutes Bacillales Bacillaceae *Geobacillus sp.* Y412MC61 |
| Gth-1-1 | Bacteria Firmicutes Bacillales Bacillaceae *Geobacillus thermodenitrificans* NG80-2 |
| Gth-2-1 | Bacteria Firmicutes Bacillales Bacillaceae *Geobacillus thermoglucosidasius* C56-YS93 |
| Gth-3-1 | Bacteria Firmicutes Bacillales Bacillaceae *Geobacillus thermoleovorans* CCB_US3_UF5 |
| Hha-1-1 | Bacteria Firmicutes Bacillales Bacillaceae *Halobacillus halophilus* DSM 2266 |
| Lsp-1-1 | Bacteria Firmicutes Bacillales Bacillaceae *Lentibacillus sp.* Grbi |
| Lfu-1-1 | Bacteria Firmicutes Bacillales Bacillaceae *Lysinibacillus fusiformis* ZC1 |
| Oih-1-1 | Bacteria Firmicutes Bacillales Bacillaceae *Oceanobacillus iheyensis* HTE831 |
| Osc-1-1 | Bacteria Firmicutes Bacillales Bacillaceae *Ornithinibacillus scapharcae* TW25 |
| Ssp-1-1 | Bacteria Firmicutes Bacillales Bacillaceae *Salimicrobium sp.* MJ3 |
| Gha-1-1 | Bacteria Firmicutes Bacillales Bacillales Family XI. IncertaeSedis *Gemella haemolysans* ATCC 10379 |
| Ean-1-1 | Bacteria Firmicutes Bacillales Bacillales Family XII. IncertaeSedis *Exiguobacterium antarcticum* B7 |
| Esi-1-1 | Bacteria Firmicutes Bacillales Bacillales Family XII. IncertaeSedis *Exiguobacterium sibiricum* 255-15 |
| Esp-1-1 | Bacteria Firmicutes Bacillales Bacillales Family XII. IncertaeSedis *Exiguobacterium sp.* AT1b |
| Lin-1-1 | Bacteria Firmicutes Bacillales Listeriaceae *Listeria innocua* Clip11262 |
| Liv-1-1 to Liv-1-2 | Bacteria Firmicutes Bacillales Listeriaceae *Listeria ivanovii subsp. ivanovii* PAM 55 |
| Lmo-1-1 | Bacteria Firmicutes Bacillales Listeriaceae *Listeria monocytogenes* 07PF0776 |
| Lmo-2-1 | Bacteria Firmicutes Bacillales Listeriaceae *Listeria monocytogenes* 08-5923 |
| Lmo-3-1 | Bacteria Firmicutes Bacillales Listeriaceae *Listeria monocytogenes* 10403S |
| Lmo-4-1 | Bacteria Firmicutes Bacillales Listeriaceae *Listeria monocytogenes* ATCC 19117 |
| Lmo-5-1 | Bacteria Firmicutes Bacillales Listeriaceae *Listeria monocytogenes* Clip80459 |
| Lmo-6-1 | Bacteria Firmicutes Bacillales Listeriaceae *Listeria monocytogenes* EGD-e |
| Lmo-7-1 | Bacteria Firmicutes Bacillales Listeriaceae *Listeria monocytogenes* F6900 |
| Lmo-13-1 | Bacteria Firmicutes Bacillales Listeriaceae *Listeria monocytogenes* Finland 1998 |
| Lmo-8-1 | Bacteria Firmicutes Bacillales Listeriaceae *Listeria monocytogenes* FSL J1-194 |
| Lmo-9-1 | Bacteria Firmicutes Bacillales Listeriaceae *Listeria monocytogenes* FSL N1-017 |
| Lmo-10-1 | Bacteria Firmicutes Bacillales Listeriaceae *Listeria monocytogenes* FSL N3-165 |
| Lmo-11-1 | Bacteria Firmicutes Bacillales Listeriaceae *Listeria monocytogenes* FSL R2-503 |
| Lmo-12-1 | Bacteria Firmicutes Bacillales Listeriaceae *Listeria monocytogenes* FSL R2-561 |
| Lmo-14-1 | Bacteria Firmicutes Bacillales Listeriaceae *Listeria monocytogenes* HCC23 |
| Lmo-15-1 | Bacteria Firmicutes Bacillales Listeriaceae *Listeria monocytogenes* HPB2262 |
| Lmo-16-1 | Bacteria Firmicutes Bacillales Listeriaceae *Listeria monocytogenes* J0161 |
| Lmo-17-1 | Bacteria Firmicutes Bacillales Listeriaceae *Listeria monocytogenes* J2818 |
| Lmo-18-1 | Bacteria Firmicutes Bacillales Listeriaceae *Listeria monocytogenes* L312 |
| Lmo-19-1 | Bacteria Firmicutes Bacillales Listeriaceae *Listeria monocytogenes* L99 |
| Lmo-20-1 | Bacteria Firmicutes Bacillales Listeriaceae *Listeria monocytogenes* M7 |
| Lmo-29-1 | Bacteria Firmicutes Bacillales Listeriaceae *Listeria monocytogenes serotype* 7 str. SLCC2482 |
| Lmo-21-1 | Bacteria Firmicutes Bacillales Listeriaceae *Listeria monocytogenes* SLCC2372 |
| Lmo-22-1 | Bacteria Firmicutes Bacillales Listeriaceae *Listeria monocytogenes* SLCC2376 |
| Lmo-23-1 | Bacteria Firmicutes Bacillales Listeriaceae *Listeria monocytogenes* SLCC2378 |
| Lmo-24-1 | Bacteria Firmicutes Bacillales Listeriaceae *Listeria monocytogenes* SLCC2479 |
| Lmo-25-1 | Bacteria Firmicutes Bacillales Listeriaceae *Listeria monocytogenes* SLCC2540 |
| Lmo-26-1 | Bacteria Firmicutes Bacillales Listeriaceae *Listeria monocytogenes* SLCC2755 |
| Lmo-27-1 | Bacteria Firmicutes Bacillales Listeriaceae *Listeria monocytogenes* SLCC5850 |
| Lmo-28-1 | Bacteria Firmicutes Bacillales Listeriaceae *Listeria monocytogenes* SLCC7179 |
| Lmo-30-1 | Bacteria Firmicutes Bacillales Listeriaceae *Listeria monocytogenes str.* 1/2a F6854 |

| | |
|---|---|
| Lmo-31-1 | Bacteria Firmicutes Bacillales Listeriaceae *Listeria monocytogenes str.* 4b F2365 |
| Lmo-32-1 | Bacteria Firmicutes Bacillales Listeriaceae *Listeria monocytogenes str.* 4b H7858 |
| Lse-1-1 | Bacteria Firmicutes Bacillales Listeriaceae *Listeria seeligeri serovar* 1/2b str. SLCC3954 |
| Lwe-1-1 | Bacteria Firmicutes Bacillales Listeriaceae *Listeria welshimeri serovar* 6b str. SLCC5334 |
| Lba-1-1 | Bacteria Firmicutes Bacillales *Listeriaceae bacterium* TTU M1-001 |
| Mca-1-1 | Bacteria Firmicutes Bacillales *Macrococcus caseolyticus* JCSC5402 |
| Bbr-1-1 | Bacteria Firmicutes Bacillales Paenibacillaceae *Brevibacillus brevis* NBRC 100599 |
| Bsp-9-1 | Bacteria Firmicutes Bacillales Paenibacillaceae *Brevibacillus sp.* BC25 |
| Bsp-10-1 | Bacteria Firmicutes Bacillales Paenibacillaceae *Brevibacillus sp.* CF112 |
| Pal-1-1 | Bacteria Firmicutes Bacillales Paenibacillaceae *Paenibacillus alvei* DSM 29 |
| Pcu-1-1 | Bacteria Firmicutes Bacillales Paenibacillaceae *Paenibacillus curdlanolyticus* YK9 |
| Pel-1-1 | Bacteria Firmicutes Bacillales Paenibacillaceae *Paenibacillus elgii* B69 |
| Pla-1-1 | Bacteria Firmicutes Bacillales Paenibacillaceae *Paenibacillus lactis* 154 |
| Pla-2-1 | Bacteria Firmicutes Bacillales Paenibacillaceae *Paenibacillus larvae subsp. larvae* BRL-230010 |
| Pmu-1-1 | Bacteria Firmicutes Bacillales Paenibacillaceae *Paenibacillus mucilaginosus* 3016 |
| Pmu-2-1 | Bacteria Firmicutes Bacillales Paenibacillaceae *Paenibacillus mucilaginosus* KNP414 |
| Ppe-1-1 | Bacteria Firmicutes Bacillales Paenibacillaceae *Paenibacillus peoriae* KCTC 3763 |
| Ppo-1-1 | Bacteria Firmicutes Bacillales Paenibacillaceae *Paenibacillus polymyxa* M1 |
| Psp-1-1 | Bacteria Firmicutes Bacillales Paenibacillaceae *Paenibacillus sp.* Aloe-11 |
| Psp-2-1 | Bacteria Firmicutes Bacillales Paenibacillaceae *Paenibacillus sp.* HGF5 |
| Psp-3-1 | Bacteria Firmicutes Bacillales Paenibacillaceae *Paenibacillus sp.* HGF7 |
| Psp-4-1 | Bacteria Firmicutes Bacillales Paenibacillaceae *Paenibacillus sp.* JDR-2 |
| Psp-5-1 | Bacteria Firmicutes Bacillales Paenibacillaceae *Paenibacillus sp. oral taxon* 786 str. D14 |
| Pte-1-1 | Bacteria Firmicutes Bacillales Paenibacillaceae *Paenibacillus terrae* HPL-003 |
| Pvo-1-1 | Bacteria Firmicutes Bacillales Paenibacillaceae *Paenibacillus vortex* V453 |
| Pan-1-1 | Bacteria Firmicutes Bacillales Planococcaceae *Planococcus antarcticus* DSM 14505 |
| Pdo-1-1 | Bacteria Firmicutes Bacillales Planococcaceae *Planococcus donghaensis* MPA1U2 |
| Ssi-1-1 | Bacteria Firmicutes Bacillales Planococcaceae *Solibacillus silvestris* StLB046 |
| Svi-1-1 to Svi-1-2 | Bacteria Firmicutes Bacillales Sporolactobacillaceae *Sporolactobacillus vineae* DSM 21990 = SL153 |
| Sar-1-1 | Bacteria Firmicutes Bacillales *Staphylococcus arlettae* CVD059 |
| Sau-1-1 | Bacteria Firmicutes Bacillales *Staphylococcus aureus* 04-02981 |
| Sau-2-1 | Bacteria Firmicutes Bacillales *Staphylococcus aureus* 08BA02176 |
| Sau-3-1 | Bacteria Firmicutes Bacillales *Staphylococcus aureus* 930918-3 |
| Sau-4-1 | Bacteria Firmicutes Bacillales *Staphylococcus aureus* A10102 |
| Sau-5-1 | Bacteria Firmicutes Bacillales *Staphylococcus aureus* A5937 |
| Sau-6-1 | Bacteria Firmicutes Bacillales *Staphylococcus aureus* A5948 |
| Sau-7-1 | Bacteria Firmicutes Bacillales *Staphylococcus aureus* A6224 |
| Sau-8-1 | Bacteria Firmicutes Bacillales *Staphylococcus aureus* A6300 |
| Sau-9-1 | Bacteria Firmicutes Bacillales *Staphylococcus aureus* A8115 |
| Sau-10-1 | Bacteria Firmicutes Bacillales *Staphylococcus aureus* A8117 |
| Sau-11-1 | Bacteria Firmicutes Bacillales *Staphylococcus aureus* A8796 |
| Sau-12-1 | Bacteria Firmicutes Bacillales *Staphylococcus aureus* A8819 |
| Sau-13-1 | Bacteria Firmicutes Bacillales *Staphylococcus aureus* A9299 |
| Sau-14-1 | Bacteria Firmicutes Bacillales *Staphylococcus aureus* A9635 |
| Sau-15-1 | Bacteria Firmicutes Bacillales *Staphylococcus aureus* A9719 |
| Sau-16-1 | Bacteria Firmicutes Bacillales *Staphylococcus aureus* A9754 |
| Sau-17-1 | Bacteria Firmicutes Bacillales *Staphylococcus aureus* A9763 |
| Sau-18-1 | Bacteria Firmicutes Bacillales *Staphylococcus aureus* A9765 |
| Sau-19-1 | Bacteria Firmicutes Bacillales *Staphylococcus aureus* A9781 |
| Sau-20-1 | Bacteria Firmicutes Bacillales *Staphylococcus aureus* D30 |
| Sau-21-1 | Bacteria Firmicutes Bacillales *Staphylococcus aureus* RF122 |
| Sau-22-1 | Bacteria Firmicutes Bacillales *Staphylococcus aureus subsp. aureus* 11819-97 |
| Sau-23-1 | Bacteria Firmicutes Bacillales *Staphylococcus aureus subsp. aureus* 132 |
| Sau-24-1 | Bacteria Firmicutes Bacillales *Staphylococcus aureus subsp. aureus* 58-424 |
| Sau-25-1 | Bacteria Firmicutes Bacillales *Staphylococcus aureus subsp. aureus* 65-1322 |
| Sau-26-1 | Bacteria Firmicutes Bacillales *Staphylococcus aureus subsp. aureus* 68-397 |
| Sau-27-1 | Bacteria Firmicutes Bacillales *Staphylococcus aureus subsp. aureus* 71193 |
| Sau-28-1 | Bacteria Firmicutes Bacillales *Staphylococcus aureus subsp. aureus* A017934/97 |
| Sau-29-1 | Bacteria Firmicutes Bacillales *Staphylococcus aureus subsp. aureus* ATCC 51811 |
| Sau-30-1 | Bacteria Firmicutes Bacillales *Staphylococcus aureus subsp. aureus* ATCC BAA-39 |
| Sau-31-1 | Bacteria Firmicutes Bacillales *Staphylococcus aureus subsp. aureus* Btn1260 |
| Sau-32-1 | Bacteria Firmicutes Bacillales *Staphylococcus aureus subsp. aureus* C101 |
| Sau-33-1 | Bacteria Firmicutes Bacillales *Staphylococcus aureus subsp. aureus* C160 |
| Sau-34-1 | Bacteria Firmicutes Bacillales *Staphylococcus aureus subsp. aureus* C427 |
| Sau-35-1 | Bacteria Firmicutes Bacillales *Staphylococcus aureus subsp. aureus* COL |

| | |
|---|---|
| Sau-36-1 | Bacteria Firmicutes Bacillales *Staphylococcus aureus subsp. aureus* D139 |
| Sau-37-1 | Bacteria Firmicutes Bacillales *Staphylococcus aureus subsp. aureus* E1410 |
| Sau-38-1 | Bacteria Firmicutes Bacillales *Staphylococcus aureus subsp. aureus* ECT-R 2 |
| Sau-39-1 | Bacteria Firmicutes Bacillales *Staphylococcus aureus subsp. aureus* ED133 |
| Sau-40-1 | Bacteria Firmicutes Bacillales *Staphylococcus aureus subsp. aureus* ED98 |
| Sau-41-1 | Bacteria Firmicutes Bacillales *Staphylococcus aureus subsp. aureus* EMRSA16 |
| Sau-42-1 | Bacteria Firmicutes Bacillales *Staphylococcus aureus subsp. aureus* H19 |
| Sau-43-1 | Bacteria Firmicutes Bacillales *Staphylococcus aureus subsp. aureus* HO 5096 0412 |
| Sau-44-1 | Bacteria Firmicutes Bacillales *Staphylococcus aureus subsp. aureus* JH1 |
| Sau-45-1 | Bacteria Firmicutes Bacillales *Staphylococcus aureus subsp. aureus* JH9 |
| Sau-46-1 | Bacteria Firmicutes Bacillales *Staphylococcus aureus subsp. aureus* JKD6159 |
| Sau-47-1 | Bacteria Firmicutes Bacillales *Staphylococcus aureus subsp. aureus* LGA251 |
| Sau-48-1 | Bacteria Firmicutes Bacillales *Staphylococcus aureus subsp. aureus* M013 |
| Sau-49-1 | Bacteria Firmicutes Bacillales *Staphylococcus aureus subsp. aureus* M1015 |
| Sau-50-1 | Bacteria Firmicutes Bacillales *Staphylococcus aureus subsp. aureus* M809 |
| Sau-51-1 | Bacteria Firmicutes Bacillales *Staphylococcus aureus subsp. aureus* M876 |
| Sau-52-1 | Bacteria Firmicutes Bacillales *Staphylococcus aureus subsp. aureus* M899 |
| Sau-53-1 | Bacteria Firmicutes Bacillales *Staphylococcus aureus subsp. aureus* MN8 |
| Sau-54-1 | Bacteria Firmicutes Bacillales *Staphylococcus aureus subsp. aureus* MR1 |
| Sau-55-1 | Bacteria Firmicutes Bacillales *Staphylococcus aureus subsp. aureus* MRSA252 |
| Sau-56-1 | Bacteria Firmicutes Bacillales *Staphylococcus aureus subsp. aureus* MSHR1132 |
| Sau-57-1 | Bacteria Firmicutes Bacillales *Staphylococcus aureus subsp. aureus* MSSA476 |
| Sau-59-1 to Sau-59-2 | Bacteria Firmicutes Bacillales *Staphylococcus aureus subsp. aureus* Mu3 |
| Sau-60-1 | Bacteria Firmicutes Bacillales *Staphylococcus aureus subsp. aureus* Mu50 |
| Sau-61-1 | Bacteria Firmicutes Bacillales *Staphylococcus aureus subsp. aureus* Mu50-omega |
| Sau-58-1 | Bacteria Firmicutes Bacillales *Staphylococcus aureus subsp. aureus* MW2 |
| Sau-62-1 | Bacteria Firmicutes Bacillales *Staphylococcus aureus subsp. aureus* N315 |
| Sau-63-1 | Bacteria Firmicutes Bacillales *Staphylococcus aureus subsp. aureus* NCTC 8325 |
| Sau-64-1 | Bacteria Firmicutes Bacillales *Staphylococcus aureus subsp. aureus* ST398 |
| Sau-75-1 | Bacteria Firmicutes Bacillales *Staphylococcus aureus subsp. aureus str.* CF-Marseille |
| Sau-76-1 | Bacteria Firmicutes Bacillales *Staphylococcus aureus subsp. aureus str.* JKD6008 |
| Sau-77-1 | Bacteria Firmicutes Bacillales *Staphylococcus aureus subsp. aureus str.* JKD6009 |
| Sau-78-1 | Bacteria Firmicutes Bacillales *Staphylococcus aureus subsp. aureus str.* Newman |
| Sau-65-1 | Bacteria Firmicutes Bacillales *Staphylococcus aureus subsp. aureus* T0131 |
| Sau-66-1 | Bacteria Firmicutes Bacillales *Staphylococcus aureus subsp. aureus* TCH130 |
| Sau-67-1 | Bacteria Firmicutes Bacillales *Staphylococcus aureus subsp. aureus* TCH60 |
| Sau-68-1 | Bacteria Firmicutes Bacillales *Staphylococcus aureus subsp. aureus* TW20 |
| Sau-69-1 | Bacteria Firmicutes Bacillales *Staphylococcus aureus subsp. aureus* USA300_FPR3757 |
| Sau-70-1 | Bacteria Firmicutes Bacillales *Staphylococcus aureus subsp. aureus* USA300_TCH1516 |
| Sau-71-1 | Bacteria Firmicutes Bacillales *Staphylococcus aureus subsp. aureus* USA300_TCH959 |
| Sau-72-1 | Bacteria Firmicutes Bacillales *Staphylococcus aureus subsp. aureus* VC40 |
| Sau-73-1 | Bacteria Firmicutes Bacillales *Staphylococcus aureus subsp. aureus* WBG10049 |
| Sau-74-1 | Bacteria Firmicutes Bacillales *Staphylococcus aureus subsp. aureus* WW2703/97 |
| Sca-1-1 | Bacteria Firmicutes Bacillales *Staphylococcus capitis* SK14 |
| Sca-2-1 | Bacteria Firmicutes Bacillales *Staphylococcus caprae* C87 |
| Sca-3-1 | Bacteria Firmicutes Bacillales *Staphylococcus carnosus subsp. carnosus* TM300 |
| Sep-1-1 | Bacteria Firmicutes Bacillales *Staphylococcus epidermidis* ATCC 12228 |
| Sep-2-1 | Bacteria Firmicutes Bacillales *Staphylococcus epidermidis* BCM-HMP0060 |
| Sep-3-1 | Bacteria Firmicutes Bacillales *Staphylococcus epidermidis* M23864:W1 |
| Sep-4-1 | Bacteria Firmicutes Bacillales *Staphylococcus epidermidis* M23864:W2(grey) |
| Sep-5-1 | Bacteria Firmicutes Bacillales *Staphylococcus epidermidis* RP62A |
| Sep-6-1 | Bacteria Firmicutes Bacillales *Staphylococcus epidermidis* SK135 |
| Sep-7-1 | Bacteria Firmicutes Bacillales *Staphylococcus epidermidis* W23144 |
| Seq-1-1 | Bacteria Firmicutes Bacillales *Staphylococcus equorum subsp. equorum* Mu2 |
| Sha-1-1 | Bacteria Firmicutes Bacillales *Staphylococcus haemolyticus* JCSC1435 |
| Sho-1-1 | Bacteria Firmicutes Bacillales *Staphylococcus hominis* SK119 |
| Sho-2-1 | Bacteria Firmicutes Bacillales *Staphylococcus hominis subsp. hominis* C80 |
| Slu-1-1 | Bacteria Firmicutes Bacillales *Staphylococcus lugdunensis* HKU09-01 |
| Slu-2-1 | Bacteria Firmicutes Bacillales *Staphylococcus lugdunensis* N920143 |
| Sps-1-1 | Bacteria Firmicutes Bacillales *Staphylococcus pseudintermedius* ED99 |
| Sps-2-1 | Bacteria Firmicutes Bacillales *Staphylococcus pseudintermedius* HKU10-03 |
| Ssa-1-1 | Bacteria Firmicutes Bacillales *Staphylococcus saprophyticus subsp. saprophyticus* ATCC 15305 |
| Ssi-2-1 | Bacteria Firmicutes Bacillales *Staphylococcus simiae* CCM 7213 |
| Ssp-2-1 | Bacteria Firmicutes Bacillales *Staphylococcus sp.* OJ82 |
| Swa-1-1 | Bacteria Firmicutes Bacillales *Staphylococcus warneri* L37603 |

| | |
|---|---|
| Apa-1-1 | Bacteria Firmicutes Bacilli Bacillales Bacillaceae *Aeribacillus pallidus* |
| Aha-1-1 | Bacteria Firmicutes Bacilli Bacillales Bacillaceae *Alkalibacillus haloalkaliphilus* C5 |
| Aji-1-1 | Bacteria Firmicutes Bacilli Bacillales Bacillaceae *Amphibacillus jilinensis* Y1 |
| Ama-1-1 | Bacteria Firmicutes Bacilli Bacillales Bacillaceae *Anaerobacillus macyae* |
| Aay-1-1 | Bacteria Firmicutes Bacilli Bacillales Bacillaceae *Anoxybacillus ayderensis* |
| Afl-2-1 | Bacteria Firmicutes Bacilli Bacillales Bacillaceae *Anoxybacillus flavithermus* |
| Afl-3-1 | Bacteria Firmicutes Bacilli Bacillales Bacillaceae *Anoxybacillus flavithermus* AK1 |
| Afl-4-1 | Bacteria Firmicutes Bacilli Bacillales Bacillaceae *Anoxybacillus flavithermus* NBRC 109594 |
| Afl-6-1 | Bacteria Firmicutes Bacilli Bacillales Bacillaceae *Anoxybacillus flavithermus subsp. yunnanensis str.* E13 |
| Afl-5-1 | Bacteria Firmicutes Bacilli Bacillales Bacillaceae *Anoxybacillus flavithermus* TNO-09.006 |
| Age-1-1 to Age-1-2 | Bacteria Firmicutes Bacilli Bacillales Bacillaceae *Anoxybacillus geothermalis* |
| Ago-1-1 | Bacteria Firmicutes Bacilli Bacillales Bacillaceae *Anoxybacillus gonensis* |
| Aka-1-1 | Bacteria Firmicutes Bacilli Bacillales Bacillaceae *Anoxybacillus kamchatkensis* G10 |
| Asp-1-1 | Bacteria Firmicutes Bacilli Bacillales Bacillaceae *Anoxybacillus sp.* BCO1 |
| Asp-2-1 | Bacteria Firmicutes Bacilli Bacillales Bacillaceae *Anoxybacillus sp.* DT3-1 |
| Asp-3-1 | Bacteria Firmicutes Bacilli Bacillales Bacillaceae *Anoxybacillus sp.* KU2-6(11) |
| Asp-4-1 | Bacteria Firmicutes Bacilli Bacillales Bacillaceae *Anoxybacillus sp.* SK3-4 |
| Ate-1-1 | Bacteria Firmicutes Bacilli Bacillales Bacillaceae *Anoxybacillus tepidamans* PS2 |
| Ath-1-1 | Bacteria Firmicutes Bacilli Bacillales Bacillaceae *Anoxybacillus thermarum* |
| Bac-1-1 | Bacteria Firmicutes Bacilli Bacillales Bacillaceae *Bacillus acidiproducens* DSM 23148 |
| Bae-1-1 | Bacteria Firmicutes Bacilli Bacillales Bacillaceae *Bacillus aerophilus* |
| Bak-1-1 | Bacteria Firmicutes Bacilli Bacillales Bacillaceae *Bacillus akibai* JCM 9157 |
| Bal-1-1 | Bacteria Firmicutes Bacilli Bacillales Bacillaceae *Bacillus alcalophilus* ATCC 27647 = CGMCC 1.3604 |
| Bal-2-1 to Bal-2-3 | Bacteria Firmicutes Bacilli Bacillales Bacillaceae *Bacillus altitudinis* |
| Bal-3-1 | Bacteria Firmicutes Bacilli Bacillales Bacillaceae *Bacillus altitudinis* 41KF2b |
| Bal-4-1 to Bal-4-2 | Bacteria Firmicutes Bacilli Bacillales Bacillaceae *Bacillus alveayuensis* |
| Bam-9-1 to Bam-9-3 | Bacteria Firmicutes Bacilli Bacillales Bacillaceae *Bacillus amyloliquefaciens* |
| Bam-10-1 | Bacteria Firmicutes Bacilli Bacillales Bacillaceae *Bacillus amyloliquefaciens* CC178 |
| Bam-11-1 | Bacteria Firmicutes Bacilli Bacillales Bacillaceae *Bacillus amyloliquefaciens* EGD-AQ14 |
| Bam-9-1 to Bam-9-3 | Bacteria Firmicutes Bacilli Bacillales Bacillaceae Bacillus Bacillus amyloliquefaciens group *Bacillus amyloliquefaciens* |
| Bam-15-1 | Bacteria Firmicutes Bacilli Bacillales Bacillaceae Bacillus Bacillus amyloliquefaciens group *Bacillus amyloliquefaciens* EBL11 |
| Bam-16-1 | Bacteria Firmicutes Bacilli Bacillales Bacillaceae Bacillus Bacillus amyloliquefaciens group *Bacillus amyloliquefaciens* HB-26 |
| Bam-17-1 | Bacteria Firmicutes Bacilli Bacillales Bacillaceae Bacillus Bacillus amyloliquefaciens group *Bacillus amyloliquefaciens* KHG19 |
| Bam-18-1 | Bacteria Firmicutes Bacilli Bacillales Bacillaceae Bacillus Bacillus amyloliquefaciens group *Bacillus amyloliquefaciens* Lx-11 |
| Bsi-1-1 | Bacteria Firmicutes Bacilli Bacillales Bacillaceae Bacillus Bacillus amyloliquefaciens group *Bacillus siamensis* |
| Bve-1-1 to Bve-1-4 | Bacteria Firmicutes Bacilli Bacillales Bacillaceae Bacillus Bacillus amyloliquefaciens group *Bacillus velezensis* |
| Bve-2-1 | Bacteria Firmicutes Bacilli Bacillales Bacillaceae Bacillus Bacillus amyloliquefaciens group *Bacillus velezensis* NJN-6 |
| Bve-3-1 | Bacteria Firmicutes Bacilli Bacillales Bacillaceae Bacillus Bacillus amyloliquefaciens group *Bacillus velezensis* SK19.001 |
| Bve-4-1 | Bacteria Firmicutes Bacilli Bacillales Bacillaceae Bacillus Bacillus amyloliquefaciens group *Bacillus velezensis* SQR9 |
| Bve-5-1 | Bacteria Firmicutes Bacilli Bacillales Bacillaceae Bacillus Bacillus amyloliquefaciens group *Bacillus velezensis* TrigoCor1448 |
| Bam-12-1 | Bacteria Firmicutes Bacilli Bacillales Bacillaceae *Bacillus amyloliquefaciens* IT-45 |
| Bam-13-1 | Bacteria Firmicutes Bacilli Bacillales Bacillaceae *Bacillus amyloliquefaciens* LFB112 |
| Bam-19-1 | Bacteria Firmicutes Bacilli Bacillales Bacillaceae *Bacillus amyloliquefaciens subsp. amyloliquefaciens* DC-12 |
| Bam-20-1 | Bacteria Firmicutes Bacilli Bacillales Bacillaceae *Bacillus amyloliquefaciens subsp. plantarum* AS43.3 |
| Bam-21-1 | Bacteria Firmicutes Bacilli Bacillales Bacillaceae *Bacillus amyloliquefaciens subsp. plantarum* M27 |
| Bam-22-1 | Bacteria Firmicutes Bacilli Bacillales Bacillaceae *Bacillus amyloliquefaciens subsp. plantarum* NAU-B3 |
| Bam-23-1 | Bacteria Firmicutes Bacilli Bacillales Bacillaceae *Bacillus amyloliquefaciens subsp. plantarum* UCMB5033 |
| Bam-24-1 | Bacteria Firmicutes Bacilli Bacillales Bacillaceae *Bacillus amyloliquefaciens subsp. plantarum* UCMB5036 |
| Bam-25-1 | Bacteria Firmicutes Bacilli Bacillales Bacillaceae *Bacillus amyloliquefaciens subsp. plantarum* UCMB5113 |
| Bam-14-1 | Bacteria Firmicutes Bacilli Bacillales Bacillaceae *Bacillus amyloliquefaciens* UASWS BA1 |
| Ban-20-1 | Bacteria Firmicutes Bacilli Bacillales Bacillaceae *Bacillus andreraoultii* |
| Baq-1-1 to Baq-1-2 | Bacteria Firmicutes Bacilli Bacillales Bacillaceae *Bacillus aquimaris* |
| Bar-1-1 to Bar-1-5 | Bacteria Firmicutes Bacilli Bacillales Bacillaceae *Bacillus aryabhattai* |
| Bar-2-1 | Bacteria Firmicutes Bacilli Bacillales Bacillaceae *Bacillus aryabhattai* B8W22 |
| Bat-2-1 | Bacteria Firmicutes Bacilli Bacillales Bacillaceae *Bacillus atrophaeus* |
| Bat-3-1 | Bacteria Firmicutes Bacilli Bacillales Bacillaceae *Bacillus atrophaeus* 1013-1 |
| Bat-4-1 | Bacteria Firmicutes Bacilli Bacillales Bacillaceae *Bacillus atrophaeus* 1013-2 |
| Bat-5-1 | Bacteria Firmicutes Bacilli Bacillales Bacillaceae *Bacillus atrophaeus* ATCC 49822-1 |
| Bat-6-1 | Bacteria Firmicutes Bacilli Bacillales Bacillaceae *Bacillus atrophaeus* ATCC 49822-2 |
| Bat-7-1 | Bacteria Firmicutes Bacilli Bacillales Bacillaceae *Bacillus atrophaeus* ATCC 9372-1 |
| Bat-8-1 | Bacteria Firmicutes Bacilli Bacillales Bacillaceae *Bacillus atrophaeus* ATCC 9372-2 |
| Bat-9-1 | Bacteria Firmicutes Bacilli Bacillales Bacillaceae *Bacillus atrophaeus* BACI051-E |
| Bat-10-1 | Bacteria Firmicutes Bacilli Bacillales Bacillaceae *Bacillus atrophaeus* BACI051-N |
| Bat-11-1 | Bacteria Firmicutes Bacilli Bacillales Bacillaceae *Bacillus atrophaeus* C89 |
| Bat-12-1 | Bacteria Firmicutes Bacilli Bacillales Bacillaceae *Bacillus atrophaeus* Detrick-1 |

| | |
|---|---|
| Bat-13-1 | Bacteria Firmicutes Bacilli Bacillales Bacillaceae *Bacillus atrophaeus* Detrick-2 |
| Bat-14-1 | Bacteria Firmicutes Bacilli Bacillales Bacillaceae *Bacillus atrophaeus* Detrick-3 |
| Bat-16-1 | Bacteria Firmicutes Bacilli Bacillales Bacillaceae *Bacillus atrophaeus str.* Dugway |
| Bat-17-1 | Bacteria Firmicutes Bacilli Bacillales Bacillaceae *Bacillus atrophaeus subsp. globigii* |
| Bat-15-1 | Bacteria Firmicutes Bacilli Bacillales Bacillaceae *Bacillus atrophaeus* UCMB-5137 |
| Bau-1-1 | Bacteria Firmicutes Bacilli Bacillales Bacillaceae *Bacillus aurantiacus* DSM 18675 |
| Baz-2-1 | Bacteria Firmicutes Bacilli Bacillales Bacillaceae *Bacillus azotoformans* MEV2011 |
| Bba-2-1 to Bba-2-2 | Bacteria Firmicutes Bacilli Bacillales Bacillaceae *Bacillus badius* |
| Bbo-1-1 | Bacteria Firmicutes Bacilli Bacillales Bacillaceae *Bacillus bogoriensis* ATCC BAA-922 |
| Bbo-2-1 | Bacteria Firmicutes Bacilli Bacillales Bacillaceae *Bacillus bombysepticus str.* Wang |
| Bbo-3-1 | Bacteria Firmicutes Bacilli Bacillales Bacillaceae *Bacillus boroniphilus* JCM 21738 |
| Bbu-1-1 | Bacteria Firmicutes Bacilli Bacillales Bacillaceae *Bacillus butanolivorans* |
| Bca-1-1 | Bacteria Firmicutes Bacilli Bacillales Bacillaceae *Bacillus campisalis* |
| Ban-21-1 to Ban-21-20 | Bacteria Firmicutes Bacilli Bacillales Bacillaceae Bacillus Bacillus cereus group *Bacillus anthracis* |
| Ban-22-1 | Bacteria Firmicutes Bacilli Bacillales Bacillaceae Bacillus Bacillus cereus group *Bacillus anthracis* 52-G |
| Ban-23-1 | Bacteria Firmicutes Bacilli Bacillales Bacillaceae Bacillus Bacillus cereus group *Bacillus anthracis* 8903-G |
| Ban-24-1 | Bacteria Firmicutes Bacilli Bacillales Bacillaceae Bacillus Bacillus cereus group *Bacillus anthracis* 9080-G |
| Ban-25-1 | Bacteria Firmicutes Bacilli Bacillales Bacillaceae Bacillus Bacillus cereus group *Bacillus anthracis* CZC5 |
| Ban-26-1 | Bacteria Firmicutes Bacilli Bacillales Bacillaceae Bacillus Bacillus cereus group *Bacillus anthracis str.* 3154 |
| Ban-27-1 | Bacteria Firmicutes Bacilli Bacillales Bacillaceae Bacillus Bacillus cereus group *Bacillus anthracis str.* 3166 |
| Ban-28-1 | Bacteria Firmicutes Bacilli Bacillales Bacillaceae Bacillus Bacillus cereus group *Bacillus anthracis str.* A16 |
| Ban-29-1 | Bacteria Firmicutes Bacilli Bacillales Bacillaceae Bacillus Bacillus cereus group *Bacillus anthracis str.* A16R |
| Ban-30-1 | Bacteria Firmicutes Bacilli Bacillales Bacillaceae Bacillus Bacillus cereus group *Bacillus anthracis str.* BF1 |
| Ban-31-1 | Bacteria Firmicutes Bacilli Bacillales Bacillaceae Bacillus Bacillus cereus group *Bacillus anthracis str.* Carbosap |
| Ban-32-1 | Bacteria Firmicutes Bacilli Bacillales Bacillaceae Bacillus Bacillus cereus group *Bacillus anthracis str.* Gmb1 |
| Ban-33-1 | Bacteria Firmicutes Bacilli Bacillales Bacillaceae Bacillus Bacillus cereus group *Bacillus anthracis str.* Heroin Ba4599 |
| Ban-35-1 | Bacteria Firmicutes Bacilli Bacillales Bacillaceae Bacillus Bacillus cereus group *Bacillus anthracis str.* Sen2Col2 |
| Ban-36-1 | Bacteria Firmicutes Bacilli Bacillales Bacillaceae Bacillus Bacillus cereus group *Bacillus anthracis str.* Sen3 |
| Ban-17-1 to Ban-17-2 | Bacteria Firmicutes Bacilli Bacillales Bacillaceae Bacillus Bacillus cereus group *Bacillus anthracis str.* Sterne |
| Ban-34-1 | Bacteria Firmicutes Bacilli Bacillales Bacillaceae Bacillus Bacillus cereus group *Bacillus anthracis str.* SVA11 |
| Ban-37-1 | Bacteria Firmicutes Bacilli Bacillales Bacillaceae Bacillus Bacillus cereus group *Bacillus anthracis str.* Turkey32 |
| Ban-38-1 | Bacteria Firmicutes Bacilli Bacillales Bacillaceae Bacillus Bacillus cereus group *Bacillus anthracis str.* V770-NP-1R |
| Ban-39-1 | Bacteria Firmicutes Bacilli Bacillales Bacillaceae Bacillus Bacillus cereus group *Bacillus anthracis str.* V770-NP1-R |
| Ban-18-1 to Ban-18-2 | Bacteria Firmicutes Bacilli Bacillales Bacillaceae Bacillus Bacillus cereus group *Bacillus anthracis str.* Vollum |
| Bce-47-1 to Bce-47-5 | Bacteria Firmicutes Bacilli Bacillales Bacillaceae Bacillus Bacillus cereus group *Bacillus cereus* |
| Bce-2-1 to Bce-2-2 | Bacteria Firmicutes Bacilli Bacillales Bacillaceae Bacillus Bacillus cereus group *Bacillus cereus* 03BB102 |
| Bce-48-1 | Bacteria Firmicutes Bacilli Bacillales Bacillaceae Bacillus Bacillus cereus group *Bacillus cereus* 03BB108 |
| Bce-49-1 | Bacteria Firmicutes Bacilli Bacillales Bacillaceae Bacillus Bacillus cereus group *Bacillus cereus* AND1407 |
| Bce-14-1 to Bce-14-2 | Bacteria Firmicutes Bacilli Bacillales Bacillaceae Bacillus Bacillus cereus group *Bacillus cereus* ATCC 10876 |
| Bce-17-1 to Bce-17-2 | Bacteria Firmicutes Bacilli Bacillales Bacillaceae Bacillus Bacillus cereus group *Bacillus cereus* ATCC 4342 |
| Bce-50-1 | Bacteria Firmicutes Bacilli Bacillales Bacillaceae Bacillus Bacillus cereus group *Bacillus cereus* B5-2 |
| Bce-51-1 | Bacteria Firmicutes Bacilli Bacillales Bacillaceae Bacillus Bacillus cereus group *Bacillus cereus* BAG1O-1 |
| Bce-52-1 | Bacteria Firmicutes Bacilli Bacillales Bacillaceae Bacillus Bacillus cereus group *Bacillus cereus* BAG1O-2 |
| Bce-53-1 | Bacteria Firmicutes Bacilli Bacillales Bacillaceae Bacillus Bacillus cereus group *Bacillus cereus* BAG1O-3 |
| Bce-54-1 | Bacteria Firmicutes Bacilli Bacillales Bacillaceae Bacillus Bacillus cereus group *Bacillus cereus* BAG1X1-1 |
| Bce-55-1 | Bacteria Firmicutes Bacilli Bacillales Bacillaceae Bacillus Bacillus cereus group *Bacillus cereus* BAG1X1-2 |
| Bce-56-1 | Bacteria Firmicutes Bacilli Bacillales Bacillaceae Bacillus Bacillus cereus group *Bacillus cereus* BAG1X1-3 |
| Bce-57-1 | Bacteria Firmicutes Bacilli Bacillales Bacillaceae Bacillus Bacillus cereus group *Bacillus cereus* BAG1X2-1 |
| Bce-58-1 | Bacteria Firmicutes Bacilli Bacillales Bacillaceae Bacillus Bacillus cereus group *Bacillus cereus* BAG1X2-2 |
| Bce-59-1 | Bacteria Firmicutes Bacilli Bacillales Bacillaceae Bacillus Bacillus cereus group *Bacillus cereus* BAG1X2-3 |
| Bce-60-1 | Bacteria Firmicutes Bacilli Bacillales Bacillaceae Bacillus Bacillus cereus group *Bacillus cereus* BAG2O-1 |
| Bce-61-1 | Bacteria Firmicutes Bacilli Bacillales Bacillaceae Bacillus Bacillus cereus group *Bacillus cereus* BAG2O-2 |
| Bce-62-1 | Bacteria Firmicutes Bacilli Bacillales Bacillaceae Bacillus Bacillus cereus group *Bacillus cereus* BAG2O-3 |
| Bce-63-1 | Bacteria Firmicutes Bacilli Bacillales Bacillaceae Bacillus Bacillus cereus group *Bacillus cereus* BAG2X1-1 |
| Bce-64-1 | Bacteria Firmicutes Bacilli Bacillales Bacillaceae Bacillus Bacillus cereus group *Bacillus cereus* BAG2X1-2 |
| Bce-65-1 | Bacteria Firmicutes Bacilli Bacillales Bacillaceae Bacillus Bacillus cereus group *Bacillus cereus* BAG2X1-3 |
| Bce-66-1 | Bacteria Firmicutes Bacilli Bacillales Bacillaceae Bacillus Bacillus cereus group *Bacillus cereus* BAG3O-1 |
| Bce-67-1 | Bacteria Firmicutes Bacilli Bacillales Bacillaceae Bacillus Bacillus cereus group *Bacillus cereus* BAG3O-2 |
| Bce-68-1 | Bacteria Firmicutes Bacilli Bacillales Bacillaceae Bacillus Bacillus cereus group *Bacillus cereus* BAG3X2-1 |
| Bce-69-1 | Bacteria Firmicutes Bacilli Bacillales Bacillaceae Bacillus Bacillus cereus group *Bacillus cereus* BAG3X2-2 |
| Bce-70-1 | Bacteria Firmicutes Bacilli Bacillales Bacillaceae Bacillus Bacillus cereus group *Bacillus cereus* BAG4O-1 |
| Bce-71-1 | Bacteria Firmicutes Bacilli Bacillales Bacillaceae Bacillus Bacillus cereus group *Bacillus cereus* BAG4X12-1 |
| Bce-72-1 | Bacteria Firmicutes Bacilli Bacillales Bacillaceae Bacillus Bacillus cereus group *Bacillus cereus* BAG4X2-1 |
| Bce-73-1 | Bacteria Firmicutes Bacilli Bacillales Bacillaceae Bacillus Bacillus cereus group *Bacillus cereus* BAG5O-1 |
| Bce-74-1 | Bacteria Firmicutes Bacilli Bacillales Bacillaceae Bacillus Bacillus cereus group *Bacillus cereus* BAG5X1-1 |

| | |
|---|---|
| Bce-75-1 | Bacteria Firmicutes Bacilli Bacillales Bacillaceae Bacillus Bacillus cereus group *Bacillus cereus* BAG5X12-1 |
| Bce-76-1 | Bacteria Firmicutes Bacilli Bacillales Bacillaceae Bacillus Bacillus cereus group *Bacillus cereus* BAG5X2-1 |
| Bce-77-1 | Bacteria Firmicutes Bacilli Bacillales Bacillaceae Bacillus Bacillus cereus group *Bacillus cereus* BAG6O-1 |
| Bce-78-1 | Bacteria Firmicutes Bacilli Bacillales Bacillaceae Bacillus Bacillus cereus group *Bacillus cereus* BAG6O-2 |
| Bce-79-1 | Bacteria Firmicutes Bacilli Bacillales Bacillaceae Bacillus Bacillus cereus group *Bacillus cereus* BAG6X1-1 |
| Bce-80-1 | Bacteria Firmicutes Bacilli Bacillales Bacillaceae Bacillus Bacillus cereus group *Bacillus cereus* BAG6X1-2 |
| Bce-81-1 | Bacteria Firmicutes Bacilli Bacillales Bacillaceae Bacillus Bacillus cereus group *Bacillus cereus* BMG1.7 |
| Bce-82-1 | Bacteria Firmicutes Bacilli Bacillales Bacillaceae Bacillus Bacillus cereus group *Bacillus cereus* BtB2-4 |
| Bce-83-1 | Bacteria Firmicutes Bacilli Bacillales Bacillaceae Bacillus Bacillus cereus group *Bacillus cereus* CER057 |
| Bce-84-1 | Bacteria Firmicutes Bacilli Bacillales Bacillaceae Bacillus Bacillus cereus group *Bacillus cereus* CER074 |
| Bce-85-1 | Bacteria Firmicutes Bacilli Bacillales Bacillaceae Bacillus Bacillus cereus group *Bacillus cereus* D17 |
| Bce-24-1 to Bce-24-2 | Bacteria Firmicutes Bacilli Bacillales Bacillaceae Bacillus Bacillus cereus group *Bacillus cereus* E33L |
| Bce-86-1 | Bacteria Firmicutes Bacilli Bacillales Bacillaceae Bacillus Bacillus cereus group *Bacillus cereus* F |
| Bce-28-1 to Bce-28-2 | Bacteria Firmicutes Bacilli Bacillales Bacillaceae Bacillus Bacillus cereus group *Bacillus cereus* G9241 |
| Bce-87-1 | Bacteria Firmicutes Bacilli Bacillales Bacillaceae Bacillus Bacillus cereus group *Bacillus cereus* HD73 |
| Bce-88-1 | Bacteria Firmicutes Bacilli Bacillales Bacillaceae Bacillus Bacillus cereus group *Bacillus cereus* HuA2-1 |
| Bce-89-1 | Bacteria Firmicutes Bacilli Bacillales Bacillaceae Bacillus Bacillus cereus group *Bacillus cereus* HuA2-3 |
| Bce-90-1 | Bacteria Firmicutes Bacilli Bacillales Bacillaceae Bacillus Bacillus cereus group *Bacillus cereus* HuA2-4 |
| Bce-91-1 | Bacteria Firmicutes Bacilli Bacillales Bacillaceae Bacillus Bacillus cereus group *Bacillus cereus* HuA2-9 |
| Bce-92-1 | Bacteria Firmicutes Bacilli Bacillales Bacillaceae Bacillus Bacillus cereus group *Bacillus cereus* HuA3-9 |
| Bce-93-1 | Bacteria Firmicutes Bacilli Bacillales Bacillaceae Bacillus Bacillus cereus group *Bacillus cereus* HuA4-10 |
| Bce-94-1 | Bacteria Firmicutes Bacilli Bacillales Bacillaceae Bacillus Bacillus cereus group *Bacillus cereus* HuB1-1 |
| Bce-95-1 | Bacteria Firmicutes Bacilli Bacillales Bacillaceae Bacillus Bacillus cereus group *Bacillus cereus* HuB13-1 |
| Bce-96-1 | Bacteria Firmicutes Bacilli Bacillales Bacillaceae Bacillus Bacillus cereus group *Bacillus cereus* HuB2-9 |
| Bce-97-1 | Bacteria Firmicutes Bacilli Bacillales Bacillaceae Bacillus Bacillus cereus group *Bacillus cereus* HuB4-10 |
| Bce-98-1 | Bacteria Firmicutes Bacilli Bacillales Bacillaceae Bacillus Bacillus cereus group *Bacillus cereus* HuB4-4 |
| Bce-99-1 | Bacteria Firmicutes Bacilli Bacillales Bacillaceae Bacillus Bacillus cereus group *Bacillus cereus* HuB5-5 |
| Bce-100-1 | Bacteria Firmicutes Bacilli Bacillales Bacillaceae Bacillus Bacillus cereus group *Bacillus cereus* IS075 |
| Bce-101-1 | Bacteria Firmicutes Bacilli Bacillales Bacillaceae Bacillus Bacillus cereus group *Bacillus cereus* IS195 |
| Bce-102-1 | Bacteria Firmicutes Bacilli Bacillales Bacillaceae Bacillus Bacillus cereus group *Bacillus cereus* IS845/00 |
| Bce-103-1 | Bacteria Firmicutes Bacilli Bacillales Bacillaceae Bacillus Bacillus cereus group *Bacillus cereus* ISP2954 |
| Bce-104-1 | Bacteria Firmicutes Bacilli Bacillales Bacillaceae Bacillus Bacillus cereus group *Bacillus cereus* ISP3191 |
| Bce-105-1 | Bacteria Firmicutes Bacilli Bacillales Bacillaceae Bacillus Bacillus cereus group *Bacillus cereus* K-5975c |
| Bce-106-1 | Bacteria Firmicutes Bacilli Bacillales Bacillaceae Bacillus Bacillus cereus group *Bacillus cereus* LCT-BC235 |
| Bce-107-1 | Bacteria Firmicutes Bacilli Bacillales Bacillaceae Bacillus Bacillus cereus group *Bacillus cereus* LCT-BC244 |
| Bce-108-1 | Bacteria Firmicutes Bacilli Bacillales Bacillaceae Bacillus Bacillus cereus group *Bacillus cereus* LCT-BC25 |
| Bce-109-1 | Bacteria Firmicutes Bacilli Bacillales Bacillaceae Bacillus Bacillus cereus group *Bacillus cereus* MC118 |
| Bce-110-1 | Bacteria Firmicutes Bacilli Bacillales Bacillaceae Bacillus Bacillus cereus group *Bacillus cereus* MC67 |
| Bce-111-1 | Bacteria Firmicutes Bacilli Bacillales Bacillaceae Bacillus Bacillus cereus group *Bacillus cereus* MSX-A1 |
| Bce-112-1 | Bacteria Firmicutes Bacilli Bacillales Bacillaceae Bacillus Bacillus cereus group *Bacillus cereus* MSX-A12 |
| Bce-113-1 | Bacteria Firmicutes Bacilli Bacillales Bacillaceae Bacillus Bacillus cereus group *Bacillus cereus* MSX-D12 |
| Bce-147-1 | Bacteria Firmicutes Bacilli Bacillales Bacillaceae Bacillus Bacillus cereus group *Bacillus cereus str.* Schrouff |
| Bce-114-1 | Bacteria Firmicutes Bacilli Bacillales Bacillaceae Bacillus Bacillus cereus group *Bacillus cereus* TIAC219 |
| Bce-115-1 | Bacteria Firmicutes Bacilli Bacillales Bacillaceae Bacillus Bacillus cereus group *Bacillus cereus* VD014 |
| Bce-116-1 | Bacteria Firmicutes Bacilli Bacillales Bacillaceae Bacillus Bacillus cereus group *Bacillus cereus* VD021 |
| Bce-117-1 | Bacteria Firmicutes Bacilli Bacillales Bacillaceae Bacillus Bacillus cereus group *Bacillus cereus* VD022 |
| Bce-118-1 | Bacteria Firmicutes Bacilli Bacillales Bacillaceae Bacillus Bacillus cereus group *Bacillus cereus* VD045 |
| Bce-119-1 | Bacteria Firmicutes Bacilli Bacillales Bacillaceae Bacillus Bacillus cereus group *Bacillus cereus* VD048 |
| Bce-120-1 | Bacteria Firmicutes Bacilli Bacillales Bacillaceae Bacillus Bacillus cereus group *Bacillus cereus* VD078 |
| Bce-121-1 | Bacteria Firmicutes Bacilli Bacillales Bacillaceae Bacillus Bacillus cereus group *Bacillus cereus* VD102 |
| Bce-122-1 | Bacteria Firmicutes Bacilli Bacillales Bacillaceae Bacillus Bacillus cereus group *Bacillus cereus* VD107 |
| Bce-123-1 | Bacteria Firmicutes Bacilli Bacillales Bacillaceae Bacillus Bacillus cereus group *Bacillus cereus* VD115 |
| Bce-124-1 | Bacteria Firmicutes Bacilli Bacillales Bacillaceae Bacillus Bacillus cereus group *Bacillus cereus* VD118 |
| Bce-125-1 | Bacteria Firmicutes Bacilli Bacillales Bacillaceae Bacillus Bacillus cereus group *Bacillus cereus* VD131 |
| Bce-126-1 | Bacteria Firmicutes Bacilli Bacillales Bacillaceae Bacillus Bacillus cereus group *Bacillus cereus* VD133 |
| Bce-127-1 | Bacteria Firmicutes Bacilli Bacillales Bacillaceae Bacillus Bacillus cereus group *Bacillus cereus* VD136 |
| Bce-128-1 | Bacteria Firmicutes Bacilli Bacillales Bacillaceae Bacillus Bacillus cereus group *Bacillus cereus* VD140 |
| Bce-129-1 | Bacteria Firmicutes Bacilli Bacillales Bacillaceae Bacillus Bacillus cereus group *Bacillus cereus* VD142 |
| Bce-130-1 | Bacteria Firmicutes Bacilli Bacillales Bacillaceae Bacillus Bacillus cereus group *Bacillus cereus* VD146 |
| Bce-131-1 | Bacteria Firmicutes Bacilli Bacillales Bacillaceae Bacillus Bacillus cereus group *Bacillus cereus* VD148 |
| Bce-132-1 | Bacteria Firmicutes Bacilli Bacillales Bacillaceae Bacillus Bacillus cereus group *Bacillus cereus* VD154 |
| Bce-133-1 | Bacteria Firmicutes Bacilli Bacillales Bacillaceae Bacillus Bacillus cereus group *Bacillus cereus* VD156 |
| Bce-134-1 | Bacteria Firmicutes Bacilli Bacillales Bacillaceae Bacillus Bacillus cereus group *Bacillus cereus* VD166 |
| Bce-135-1 | Bacteria Firmicutes Bacilli Bacillales Bacillaceae Bacillus Bacillus cereus group *Bacillus cereus* VD169 |
| Bce-136-1 | Bacteria Firmicutes Bacilli Bacillales Bacillaceae Bacillus Bacillus cereus group *Bacillus cereus* VD184 |

| | |
|---|---|
| Bce-137-1 | Bacteria Firmicutes Bacilli Bacillales Bacillaceae Bacillus Bacillus cereus group *Bacillus cereus* VD196 |
| Bce-138-1 | Bacteria Firmicutes Bacilli Bacillales Bacillaceae Bacillus Bacillus cereus group *Bacillus cereus* VD200 |
| Bce-139-1 | Bacteria Firmicutes Bacilli Bacillales Bacillaceae Bacillus Bacillus cereus group *Bacillus cereus* VD214 |
| Bce-140-1 | Bacteria Firmicutes Bacilli Bacillales Bacillaceae Bacillus Bacillus cereus group *Bacillus cereus* VDM006 |
| Bce-141-1 | Bacteria Firmicutes Bacilli Bacillales Bacillaceae Bacillus Bacillus cereus group *Bacillus cereus* VDM019 |
| Bce-142-1 | Bacteria Firmicutes Bacilli Bacillales Bacillaceae Bacillus Bacillus cereus group *Bacillus cereus* VDM021 |
| Bce-143-1 | Bacteria Firmicutes Bacilli Bacillales Bacillaceae Bacillus Bacillus cereus group *Bacillus cereus* VDM022 |
| Bce-144-1 | Bacteria Firmicutes Bacilli Bacillales Bacillaceae Bacillus Bacillus cereus group *Bacillus cereus* VDM034 |
| Bce-145-1 | Bacteria Firmicutes Bacilli Bacillales Bacillaceae Bacillus Bacillus cereus group *Bacillus cereus* VDM053 |
| Bce-146-1 | Bacteria Firmicutes Bacilli Bacillales Bacillaceae Bacillus Bacillus cereus group *Bacillus cereus* VDM062 |
| Bcy-2-1 | Bacteria Firmicutes Bacilli Bacillales Bacillaceae Bacillus Bacillus cereus group *Bacillus cytotoxicus* |
| Bga-1-1 | Bacteria Firmicutes Bacilli Bacillales Bacillaceae Bacillus Bacillus cereus group *Bacillus gaemokensis* |
| Bma-2-1 | Bacteria Firmicutes Bacilli Bacillales Bacillaceae Bacillus Bacillus cereus group *Bacillus manliponensis* |
| Bmy-4-1 to Bmy-4-4 | Bacteria Firmicutes Bacilli Bacillales Bacillaceae Bacillus Bacillus cereus group *Bacillus mycoides* |
| Bmy-5-1 | Bacteria Firmicutes Bacilli Bacillales Bacillaceae Bacillus Bacillus cereus group *Bacillus mycoides* FSL H7-687 |
| Bmy-6-1 | Bacteria Firmicutes Bacilli Bacillales Bacillaceae Bacillus Bacillus cereus group *Bacillus mycoides* FSL R5-860 |
| Bmy-7-1 | Bacteria Firmicutes Bacilli Bacillales Bacillaceae Bacillus Bacillus cereus group *Bacillus mycoides* NBRC 101238 = DSM 11821 |
| Bsp-11-1 | Bacteria Firmicutes Bacilli Bacillales Bacillaceae Bacillus Bacillus cereus group *Bacillus sp.* 7_6_55CFAA_CT2 |
| Bth-21-1 | Bacteria Firmicutes Bacilli Bacillales Bacillaceae Bacillus Bacillus cereus group *Bacillus thuringiensis* |
| Bth-22-1 to Bth-22-2 | Bacteria Firmicutes Bacilli Bacillales Bacillaceae Bacillus Bacillus cereus group *Bacillus thuringiensis* DAR 81934 |
| Bth-23-1 | Bacteria Firmicutes Bacilli Bacillales Bacillaceae Bacillus Bacillus cereus group *Bacillus thuringiensis* DB27 |
| Bth-24-1 | Bacteria Firmicutes Bacilli Bacillales Bacillaceae Bacillus Bacillus cereus group *Bacillus thuringiensis* HD-789 |
| Bth-25-1 | Bacteria Firmicutes Bacilli Bacillales Bacillaceae Bacillus Bacillus cereus group *Bacillus thuringiensis* HD1002 |
| Bth-26-1 | Bacteria Firmicutes Bacilli Bacillales Bacillaceae Bacillus Bacillus cereus group *Bacillus thuringiensis* JM-Mgvxx-63 |
| Bth-27-1 | Bacteria Firmicutes Bacilli Bacillales Bacillaceae Bacillus Bacillus cereus group *Bacillus thuringiensis* LM1212 |
| Bth-28-1 | Bacteria Firmicutes Bacilli Bacillales Bacillaceae Bacillus Bacillus cereus group *Bacillus thuringiensis* Sbt003 |
| Bth-31-1 | Bacteria Firmicutes Bacilli Bacillales Bacillaceae Bacillus Bacillus cereus group *Bacillus thuringiensis serovar aizawai str.* Hu4-2 |
| Bth-32-1 | Bacteria Firmicutes Bacilli Bacillales Bacillaceae Bacillus Bacillus cereus group *Bacillus thuringiensis serovar aizawai str.* Leapi01 |
| Bth-33-1 | Bacteria Firmicutes Bacilli Bacillales Bacillaceae Bacillus Bacillus cereus group *Bacillus thuringiensis serovar galleriae* |
| Bth-34-1 | Bacteria Firmicutes Bacilli Bacillales Bacillaceae Bacillus Bacillus cereus group *Bacillus thuringiensis serovar indiana* |
| Bth-35-1 | Bacteria Firmicutes Bacilli Bacillales Bacillaceae Bacillus Bacillus cereus group *Bacillus thuringiensis serovar israelensis* |
| Bth-36-1 | Bacteria Firmicutes Bacilli Bacillales Bacillaceae Bacillus Bacillus cereus group *Bacillus thuringiensis serovar kurstaki* |
| Bth-37-1 | Bacteria Firmicutes Bacilli Bacillales Bacillaceae Bacillus Bacillus cereus group *Bacillus thuringiensis serovar kurstaki str.* HD-1 |
| Bth-38-1 | Bacteria Firmicutes Bacilli Bacillales Bacillaceae Bacillus Bacillus cereus group *Bacillus thuringiensis serovar kurstaki str.* HD73 |
| Bth-39-1 to Bth-39-2 | Bacteria Firmicutes Bacilli Bacillales Bacillaceae Bacillus Bacillus cereus group *Bacillus thuringiensis serovar kurstaki str.* YBT-1520 |
| Bth-40-1 | Bacteria Firmicutes Bacilli Bacillales Bacillaceae Bacillus Bacillus cereus group *Bacillus thuringiensis serovar mexicanensis* |
| Bth-41-1 | Bacteria Firmicutes Bacilli Bacillales Bacillaceae Bacillus Bacillus cereus group *Bacillus thuringiensis serovar morrisoni* |
| Bth-42-1 | Bacteria Firmicutes Bacilli Bacillales Bacillaceae Bacillus Bacillus cereus group *Bacillus thuringiensis serovar thuringiensis str.* IS5056 |
| Bth-43-1 | Bacteria Firmicutes Bacilli Bacillales Bacillaceae Bacillus Bacillus cereus group *Bacillus thuringiensis serovar tolworthi* NA205-3 |
| Bth-29-1 | Bacteria Firmicutes Bacilli Bacillales Bacillaceae Bacillus Bacillus cereus group *Bacillus thuringiensis* T01-328 |
| Bth-30-1 | Bacteria Firmicutes Bacilli Bacillales Bacillaceae Bacillus Bacillus cereus group *Bacillus thuringiensis* YBT-1518 |
| Bto-1-1 | Bacteria Firmicutes Bacilli Bacillales Bacillaceae Bacillus Bacillus cereus group *Bacillus toyonensis* BCT-7112 |
| Bwe-2-1 | Bacteria Firmicutes Bacilli Bacillales Bacillaceae Bacillus Bacillus cereus group *Bacillus weihenstephanensis* |
| Bch-1-1 | Bacteria Firmicutes Bacilli Bacillales Bacillaceae *Bacillus chagannorensis* DSM 18086 |
| Bci-1-1 | Bacteria Firmicutes Bacilli Bacillales Bacillaceae *Bacillus cihuensis* |
| Bci-2-1 | Bacteria Firmicutes Bacilli Bacillales Bacillaceae *Bacillus circulans* |
| Bcl-2-1 | Bacteria Firmicutes Bacilli Bacillales Bacillaceae *Bacillus clausii* |
| Bco-4-1 to Bco-4-2 | Bacteria Firmicutes Bacilli Bacillales Bacillaceae *Bacillus coagulans* |
| Bco-5-1 | Bacteria Firmicutes Bacilli Bacillales Bacillaceae *Bacillus coagulans* CSIL1 |
| Bco-6-1 | Bacteria Firmicutes Bacilli Bacillales Bacillaceae *Bacillus coagulans* DSM 1 = ATCC 7050 |
| Bco-7-1 | Bacteria Firmicutes Bacilli Bacillales Bacillaceae *Bacillus coagulans* H-1 |
| Bco-8-1 | Bacteria Firmicutes Bacilli Bacillales Bacillaceae *Bacillus coagulans* P38 |
| Bco-9-1 | Bacteria Firmicutes Bacilli Bacillales Bacillaceae *Bacillus coagulans* XZL4 |
| Bco-10-1 | Bacteria Firmicutes Bacilli Bacillales Bacillaceae *Bacillus coagulans* XZL9 |
| Bdi-1-1 | Bacteria Firmicutes Bacilli Bacillales Bacillaceae *Bacillus dielmoensis* |
| Ben-1-1 | Bacteria Firmicutes Bacilli Bacillales Bacillaceae *Bacillus endophyticus* |
| Ben-2-1 | Bacteria Firmicutes Bacilli Bacillales Bacillaceae *Bacillus endophyticus* 2102 |
| Bfi-1-1 | Bacteria Firmicutes Bacilli Bacillales Bacillaceae *Bacillus firmus* |
| Bfi-2-1 | Bacteria Firmicutes Bacilli Bacillales Bacillaceae *Bacillus firmus* DS1 |
| Bfl-1-1 | Bacteria Firmicutes Bacilli Bacillales Bacillaceae *Bacillus flexus* |
| Bfl-2-1 | Bacteria Firmicutes Bacilli Bacillales Bacillaceae *Bacillus flexus* 27Col1.1E |
| Bfl-3-1 | Bacteria Firmicutes Bacilli Bacillales Bacillaceae *Bacillus flexus* T6186-2 |
| Bfo-1-1 | Bacteria Firmicutes Bacilli Bacillales Bacillaceae *Bacillus fordii* DSM 16014 |
| Bgi-1-1 | Bacteria Firmicutes Bacilli Bacillales Bacillaceae *Bacillus ginsengihumi* |
| Bgi-2-1 | Bacteria Firmicutes Bacilli Bacillales Bacillaceae *Bacillus ginsengihumi* J57 |

| | |
|---|---|
| Bgl-1-1 | Bacteria Firmicutes Bacilli Bacillales Bacillaceae *Bacillus glycinifermentans* |
| Bhe-1-1 | Bacteria Firmicutes Bacilli Bacillales Bacillaceae *Bacillus hemicellulosilyticus* JCM 9152 |
| Bin-1-1 | Bacteria Firmicutes Bacilli Bacillales Bacillaceae *Bacillus indicus* |
| Bin-2-1 | Bacteria Firmicutes Bacilli Bacillales Bacillaceae *Bacillus indicus* LMG 22858 |
| Bin-3-1 | Bacteria Firmicutes Bacilli Bacillales Bacillaceae *Bacillus infantis* NRRL B-14911 |
| Bje-1-1 | Bacteria Firmicutes Bacilli Bacillales Bacillaceae *Bacillus jeddahensis* |
| Bko-1-1 | Bacteria Firmicutes Bacilli Bacillales Bacillaceae *Bacillus koreensis* |
| Bkr-1-1 | Bacteria Firmicutes Bacilli Bacillales Bacillaceae *Bacillus kribbensis* DSM 17871 |
| Ble-1-1 | Bacteria Firmicutes Bacilli Bacillales Bacillaceae *Bacillus lehensis* G1 |
| Bli-2-1 to Bli-2-3 | Bacteria Firmicutes Bacilli Bacillales Bacillaceae *Bacillus licheniformis* |
| Bli-3-1 | Bacteria Firmicutes Bacilli Bacillales Bacillaceae *Bacillus licheniformis* 10-1-A |
| Bli-4-1 | Bacteria Firmicutes Bacilli Bacillales Bacillaceae *Bacillus licheniformis* 5-2-D |
| Bli-5-1 | Bacteria Firmicutes Bacilli Bacillales Bacillaceae *Bacillus licheniformis* 9945A |
| Bli-6-1 | Bacteria Firmicutes Bacilli Bacillales Bacillaceae *Bacillus licheniformis* CG-B52 |
| Bli-7-1 | Bacteria Firmicutes Bacilli Bacillales Bacillaceae *Bacillus licheniformis* CGMCC 3963 |
| Bli-8-1 | Bacteria Firmicutes Bacilli Bacillales Bacillaceae *Bacillus licheniformis* F1-1 |
| Bli-9-1 | Bacteria Firmicutes Bacilli Bacillales Bacillaceae *Bacillus licheniformis* F2-1 |
| Bli-10-1 | Bacteria Firmicutes Bacilli Bacillales Bacillaceae *Bacillus licheniformis* S 16 |
| Bli-11-1 | Bacteria Firmicutes Bacilli Bacillales Bacillaceae *Bacillus licheniformis* WX-02 |
| Blo-1-1 | Bacteria Firmicutes Bacilli Bacillales Bacillaceae *Bacillus loiseleuriae* |
| Bma-3-1 | Bacteria Firmicutes Bacilli Bacillales Bacillaceae *Bacillus mannanilyticus* JCM 10596 |
| Bma-4-1 to Bma-4-4 | Bacteria Firmicutes Bacilli Bacillales Bacillaceae *Bacillus marisflavi* |
| Bma-5-1 | Bacteria Firmicutes Bacilli Bacillales Bacillaceae *Bacillus marmarensis* DSM 21297 |
| Bma-6-1 | Bacteria Firmicutes Bacilli Bacillales Bacillaceae *Bacillus massiliogorillae* |
| Bma-7-1 | Bacteria Firmicutes Bacilli Bacillales Bacillaceae *Bacillus massiliosenegalensis* JC6 |
| Bme-5-1 | Bacteria Firmicutes Bacilli Bacillales Bacillaceae *Bacillus megaterium* |
| Bme-6-1 | Bacteria Firmicutes Bacilli Bacillales Bacillaceae *Bacillus megaterium* MSP20.1 |
| Bme-7-1 | Bacteria Firmicutes Bacilli Bacillales Bacillaceae *Bacillus megaterium* NBRC 15308 = ATCC 14581 |
| Bme-8-1 | Bacteria Firmicutes Bacilli Bacillales Bacillaceae *Bacillus megaterium* NCT-2 |
| Bme-9-1 | Bacteria Firmicutes Bacilli Bacillales Bacillaceae *Bacillus megaterium* Q3 |
| Bme-10-1 | Bacteria Firmicutes Bacilli Bacillales Bacillaceae *Bacillus methanolicus* MGA3 |
| Bmo-1-1 | Bacteria Firmicutes Bacilli Bacillales Bacillaceae *Bacillus mojavensis* RO-H-1 = KCTC 3706 |
| Bmo-2-1 | Bacteria Firmicutes Bacilli Bacillales Bacillaceae *Bacillus mojavensis* RRC 101 |
| Bmu-1-1 | Bacteria Firmicutes Bacilli Bacillales Bacillaceae *Bacillus murimartini* |
| Bnd-1-1 | Bacteria Firmicutes Bacilli Bacillales Bacillaceae *Bacillus ndiopicus* |
| Bne-1-1 | Bacteria Firmicutes Bacilli Bacillales Bacillaceae *Bacillus nealsonii* AAU1 |
| Bni-1-1 | Bacteria Firmicutes Bacilli Bacillales Bacillaceae *Bacillus niacini* |
| Boc-1-1 | Bacteria Firmicutes Bacilli Bacillales Bacillaceae *Bacillus oceanisediminis* 2691 |
| Bok-1-1 | Bacteria Firmicutes Bacilli Bacillales Bacillaceae *Bacillus okhensis* |
| Bok-2-1 | Bacteria Firmicutes Bacilli Bacillales Bacillaceae *Bacillus okuhidensis* |
| Bpa-1-1 | Bacteria Firmicutes Bacilli Bacillales Bacillaceae *Bacillus panaciterrae* DSM 19096 |
| Bpa-2-1 | Bacteria Firmicutes Bacilli Bacillales Bacillaceae *Bacillus paralicheniformis* |
| Bpa-3-1 | Bacteria Firmicutes Bacilli Bacillales Bacillaceae *Bacillus paralicheniformis* G-1 |
| Bps-3-1 | Bacteria Firmicutes Bacilli Bacillales Bacillaceae *Bacillus pseudalcaliphilus* |
| Bps-4-1 | Bacteria Firmicutes Bacilli Bacillales Bacillaceae *Bacillus psychrosaccharolyticus* ATCC 23296 |
| Bpu-3-1 to Bpu-3-3 | Bacteria Firmicutes Bacilli Bacillales Bacillaceae *Bacillus pumilus* |
| Bpu-4-1 | Bacteria Firmicutes Bacilli Bacillales Bacillaceae *Bacillus pumilus* BA06 |
| Bpu-5-1 | Bacteria Firmicutes Bacilli Bacillales Bacillaceae *Bacillus pumilus* CCMA-560 |
| Bpu-6-1 | Bacteria Firmicutes Bacilli Bacillales Bacillaceae *Bacillus pumilus* INR7 |
| Bpu-7-1 | Bacteria Firmicutes Bacilli Bacillales Bacillaceae *Bacillus pumilus* S-1 |
| Bru-1-1 | Bacteria Firmicutes Bacilli Bacillales Bacillaceae *Bacillus rubiinfantis* |
| Bsa-1-1 to Bsa-1-3 | Bacteria Firmicutes Bacilli Bacillales Bacillaceae *Bacillus safensis* |
| Bsa-2-1 | Bacteria Firmicutes Bacilli Bacillales Bacillaceae *Bacillus safensis* FO-36b |
| Bsa-3-1 | Bacteria Firmicutes Bacilli Bacillales Bacillaceae *Bacillus safensis* VK |
| Bse-2-1 | Bacteria Firmicutes Bacilli Bacillales Bacillaceae *Bacillus selenatarsenatis* SF-1 |
| Bsi-2-1 | Bacteria Firmicutes Bacilli Bacillales Bacillaceae *Bacillus siamensis* KCTC 13613 |
| Bsi-3-1 | Bacteria Firmicutes Bacilli Bacillales Bacillaceae *Bacillus simplex* BA2H3 |
| Bsm-1-1 | Bacteria Firmicutes Bacilli Bacillales Bacillaceae *Bacillus smithii* |
| Bsm-2-1 | Bacteria Firmicutes Bacilli Bacillales Bacillaceae *Bacillus smithii* 7_3_47FAA |
| Bso-1-1 | Bacteria Firmicutes Bacilli Bacillales Bacillaceae *Bacillus sonorensis* L12 |
| Bso-2-1 | Bacteria Firmicutes Bacilli Bacillales Bacillaceae *Bacillus sonorensis* NBRC 101234 = KCTC 13918 |
| Bsp-12-1 | Bacteria Firmicutes Bacilli Bacillales Bacillaceae *Bacillus sp. (in:* Bacteria) |
| Bsp-13-1 | Bacteria Firmicutes Bacilli Bacillales Bacillaceae *Bacillus sp.* 10403023 |
| Bsp-14-1 | Bacteria Firmicutes Bacilli Bacillales Bacillaceae *Bacillus sp.* 105MF |
| Bsp-15-1 | Bacteria Firmicutes Bacilli Bacillales Bacillaceae *Bacillus sp.* 123MFChir2 |

| | |
|---|---|
| Bsp-16-1 | Bacteria Firmicutes Bacilli Bacillales Bacillaceae *Bacillus sp.* 171095_106 |
| Bsp-17-1 | Bacteria Firmicutes Bacilli Bacillales Bacillaceae *Bacillus sp.* 17376 |
| Bsp-18-1 | Bacteria Firmicutes Bacilli Bacillales Bacillaceae *Bacillus sp.* 1NLA3E |
| Bsp-19-1 | Bacteria Firmicutes Bacilli Bacillales Bacillaceae *Bacillus sp.* 220_BSPC |
| Bsp-20-1 | Bacteria Firmicutes Bacilli Bacillales Bacillaceae *Bacillus sp.* 278922_107 |
| Bsp-21-1 | Bacteria Firmicutes Bacilli Bacillales Bacillaceae *Bacillus sp.* 2_A_57_CT2 |
| Bsp-22-1 | Bacteria Firmicutes Bacilli Bacillales Bacillaceae *Bacillus sp.* 37MA |
| Bsp-23-1 | Bacteria Firmicutes Bacilli Bacillales Bacillaceae *Bacillus sp.* 445_BSPC |
| Bsp-24-1 | Bacteria Firmicutes Bacilli Bacillales Bacillaceae *Bacillus sp.* 522_BSPC |
| Bsp-25-1 | Bacteria Firmicutes Bacilli Bacillales Bacillaceae *Bacillus sp.* 72 |
| Bsp-26-1 | Bacteria Firmicutes Bacilli Bacillales Bacillaceae *Bacillus sp.* 95MFCvi2.1 |
| Bsp-27-1 | Bacteria Firmicutes Bacilli Bacillales Bacillaceae *Bacillus sp.* A053 |
| Bsp-28-1 | Bacteria Firmicutes Bacilli Bacillales Bacillaceae *Bacillus sp.* AP8 |
| Bsp-29-1 | Bacteria Firmicutes Bacilli Bacillales Bacillaceae *Bacillus sp.* Aph1 |
| Bsp-30-1 | Bacteria Firmicutes Bacilli Bacillales Bacillaceae *Bacillus sp.* B-jedd |
| Bsp-31-1 | Bacteria Firmicutes Bacilli Bacillales Bacillaceae *Bacillus sp.* BH072 |
| Bsp-32-1 | Bacteria Firmicutes Bacilli Bacillales Bacillaceae *Bacillus sp.* BSC154 |
| Bsp-33-1 | Bacteria Firmicutes Bacilli Bacillales Bacillaceae *Bacillus sp.* BT1B_CT2 |
| Bsp-34-1 | Bacteria Firmicutes Bacilli Bacillales Bacillaceae *Bacillus sp.* CMAA 1185 |
| Bsp-36-1 | Bacteria Firmicutes Bacilli Bacillales Bacillaceae *Bacillus sp.* Co1-6 |
| Bsp-35-1 | Bacteria Firmicutes Bacilli Bacillales Bacillaceae *Bacillus sp.* CPSM8 |
| Bsp-37-1 | Bacteria Firmicutes Bacilli Bacillales Bacillaceae *Bacillus sp.* EB01 |
| Bsp-38-1 | Bacteria Firmicutes Bacilli Bacillales Bacillaceae *Bacillus sp.* EGD-AK10 |
| Bsp-39-1 | Bacteria Firmicutes Bacilli Bacillales Bacillaceae *Bacillus sp.* FJAT-13831 |
| Bsp-40-1 | Bacteria Firmicutes Bacilli Bacillales Bacillaceae *Bacillus sp.* FJAT-14578 |
| Bsp-41-1 | Bacteria Firmicutes Bacilli Bacillales Bacillaceae *Bacillus sp.* FJAT-26652 |
| Bsp-42-1 | Bacteria Firmicutes Bacilli Bacillales Bacillaceae *Bacillus sp.* FJAT-27231 |
| Bsp-43-1 | Bacteria Firmicutes Bacilli Bacillales Bacillaceae *Bacillus sp.* FJAT-27238 |
| Bsp-44-1 | Bacteria Firmicutes Bacilli Bacillales Bacillaceae *Bacillus sp.* FJAT-27245 |
| Bsp-45-1 | Bacteria Firmicutes Bacilli Bacillales Bacillaceae *Bacillus sp.* FJAT-27251 |
| Bsp-46-1 | Bacteria Firmicutes Bacilli Bacillales Bacillaceae *Bacillus sp.* FJAT-27445 |
| Bsp-47-1 | Bacteria Firmicutes Bacilli Bacillales Bacillaceae *Bacillus sp.* FJAT-27916 |
| Bsp-49-1 | Bacteria Firmicutes Bacilli Bacillales Bacillaceae *Bacillus sp.* GeD10 |
| Bsp-48-1 | Bacteria Firmicutes Bacilli Bacillales Bacillaceae *Bacillus sp.* GO-13 |
| Bsp-50-1 | Bacteria Firmicutes Bacilli Bacillales Bacillaceae *Bacillus sp.* H1a |
| Bsp-51-1 | Bacteria Firmicutes Bacilli Bacillales Bacillaceae *Bacillus sp.* H1m |
| Bsp-52-1 | Bacteria Firmicutes Bacilli Bacillales Bacillaceae *Bacillus sp.* J13 |
| Bsp-53-1 | Bacteria Firmicutes Bacilli Bacillales Bacillaceae *Bacillus sp.* J33 |
| Bsp-54-1 | Bacteria Firmicutes Bacilli Bacillales Bacillaceae *Bacillus sp.* J37 |
| Bsp-55-1 | Bacteria Firmicutes Bacilli Bacillales Bacillaceae *Bacillus sp.* JCA |
| Bsp-56-1 | Bacteria Firmicutes Bacilli Bacillales Bacillaceae *Bacillus sp.* JCM 19045 |
| Bsp-57-1 | Bacteria Firmicutes Bacilli Bacillales Bacillaceae *Bacillus sp.* JCM 19046 |
| Bsp-58-1 | Bacteria Firmicutes Bacilli Bacillales Bacillaceae *Bacillus sp.* JCM 19047 |
| Bsp-59-1 | Bacteria Firmicutes Bacilli Bacillales Bacillaceae *Bacillus sp.* JFL15 |
| Bsp-60-1 | Bacteria Firmicutes Bacilli Bacillales Bacillaceae *Bacillus sp.* KJ-16 |
| Bsp-61-1 | Bacteria Firmicutes Bacilli Bacillales Bacillaceae *Bacillus sp.* KW-12 |
| Bsp-62-1 | Bacteria Firmicutes Bacilli Bacillales Bacillaceae *Bacillus sp.* L1(2012) |
| Bsp-68-1 | Bacteria Firmicutes Bacilli Bacillales Bacillaceae *Bacillus sp.* L_1B0_12 |
| Bsp-69-1 | Bacteria Firmicutes Bacilli Bacillales Bacillaceae *Bacillus sp.* L_1B0_5 |
| Bsp-70-1 | Bacteria Firmicutes Bacilli Bacillales Bacillaceae *Bacillus sp.* L_1B0_8 |
| Bsp-63-1 | Bacteria Firmicutes Bacilli Bacillales Bacillaceae *Bacillus sp.* LK10 |
| Bsp-64-1 | Bacteria Firmicutes Bacilli Bacillales Bacillaceae *Bacillus sp.* LK2 |
| Bsp-65-1 | Bacteria Firmicutes Bacilli Bacillales Bacillaceae *Bacillus sp.* LK7 |
| Bsp-66-1 to Bsp-66-2 | Bacteria Firmicutes Bacilli Bacillales Bacillaceae *Bacillus sp.* LL01 |
| Bsp-67-1 | Bacteria Firmicutes Bacilli Bacillales Bacillaceae *Bacillus sp.* LM 4-2 |
| Bsp-71-1 | Bacteria Firmicutes Bacilli Bacillales Bacillaceae *Bacillus sp.* MB2021 |
| Bsp-72-1 | Bacteria Firmicutes Bacilli Bacillales Bacillaceae *Bacillus sp.* MSP13 |
| Bsp-73-1 | Bacteria Firmicutes Bacilli Bacillales Bacillaceae *Bacillus sp.* MSP5.4 |
| Bsp-74-1 | Bacteria Firmicutes Bacilli Bacillales Bacillaceae *Bacillus sp.* NSP2.1 |
| Bsp-75-1 | Bacteria Firmicutes Bacilli Bacillales Bacillaceae *Bacillus sp.* NSP22.2 |
| Bsp-76-1 | Bacteria Firmicutes Bacilli Bacillales Bacillaceae *Bacillus sp.* NSP9.1 |
| Bsp-77-1 | Bacteria Firmicutes Bacilli Bacillales Bacillaceae *Bacillus sp.* OxB-1 |
| Bsp-78-1 | Bacteria Firmicutes Bacilli Bacillales Bacillaceae *Bacillus sp.* Pc3 |
| Bsp-79-1 | Bacteria Firmicutes Bacilli Bacillales Bacillaceae *Bacillus sp.* REN51N |
| Bsp-80-1 | Bacteria Firmicutes Bacilli Bacillales Bacillaceae *Bacillus sp.* RP1137 |

| | |
|---|---|
| Bsp-81-1 | Bacteria Firmicutes Bacilli Bacillales Bacillaceae *Bacillus sp.* SA1-12 |
| Bsp-82-1 | Bacteria Firmicutes Bacilli Bacillales Bacillaceae *Bacillus sp.* SB47 |
| Bsp-83-1 | Bacteria Firmicutes Bacilli Bacillales Bacillaceae *Bacillus sp.* SB49 |
| Bsp-84-1 | Bacteria Firmicutes Bacilli Bacillales Bacillaceae *Bacillus sp.* SJS |
| Bsp-85-1 | Bacteria Firmicutes Bacilli Bacillales Bacillaceae *Bacillus sp.* TH008 |
| Bsp-86-1 | Bacteria Firmicutes Bacilli Bacillales Bacillaceae *Bacillus sp.* TS-2 |
| Bsp-87-1 | Bacteria Firmicutes Bacilli Bacillales Bacillaceae *Bacillus sp.* UMTAT18 |
| Bsp-88-1 | Bacteria Firmicutes Bacilli Bacillales Bacillaceae *Bacillus sp.* UNC125MFCrub1.1 |
| Bsp-89-1 | Bacteria Firmicutes Bacilli Bacillales Bacillaceae *Bacillus sp.* UNC322MFChir4.1 |
| Bsp-90-1 | Bacteria Firmicutes Bacilli Bacillales Bacillaceae *Bacillus sp.* UNC41MFS5 |
| Bsp-91-1 | Bacteria Firmicutes Bacilli Bacillales Bacillaceae *Bacillus sp.* UNC437CL72CviS29 |
| Bsp-92-1 | Bacteria Firmicutes Bacilli Bacillales Bacillaceae *Bacillus sp.* UNC438CL73TsuS30 |
| Bsp-93-1 | Bacteria Firmicutes Bacilli Bacillales Bacillaceae *Bacillus sp.* UNC69MF |
| Bsp-94-1 | Bacteria Firmicutes Bacilli Bacillales Bacillaceae *Bacillus sp.* URHB0009 |
| Bsp-95-1 | Bacteria Firmicutes Bacilli Bacillales Bacillaceae *Bacillus sp.* WP8 |
| Bsp-96-1 | Bacteria Firmicutes Bacilli Bacillales Bacillaceae *Bacillus sp.* X1(2014) |
| Bsp-97-1 | Bacteria Firmicutes Bacilli Bacillales Bacillaceae *Bacillus sp.* YP1 |
| Bsp-98-1 | Bacteria Firmicutes Bacilli Bacillales Bacillaceae *Bacillus sp.* ZYK |
| Bst-1-1 to Bst-1-6 | Bacteria Firmicutes Bacilli Bacillales Bacillaceae *Bacillus stratosphericus* |
| Bst-2-1 | Bacteria Firmicutes Bacilli Bacillales Bacillaceae *Bacillus stratosphericus* LAMA 585 |
| Bsu-11-1 | Bacteria Firmicutes Bacilli Bacillales Bacillaceae *Bacillus subterraneus* |
| Bsu-12-1 | Bacteria Firmicutes Bacilli Bacillales Bacillaceae *Bacillus subtilis* |
| Bsu-13-1 | Bacteria Firmicutes Bacilli Bacillales Bacillaceae *Bacillus subtilis* B7-s |
| Bsu-14-1 | Bacteria Firmicutes Bacilli Bacillales Bacillaceae *Bacillus subtilis* BEST7003 |
| Bsu-15-1 | Bacteria Firmicutes Bacilli Bacillales Bacillaceae *Bacillus subtilis* E1 |
| Bsu-16-1 | Bacteria Firmicutes Bacilli Bacillales Bacillaceae *Bacillus subtilis* GB03 |
| Bsu-28-1 | Bacteria Firmicutes Bacilli Bacillales Bacillaceae *Bacillus subtilis gt*P20b |
| Bsu-18-1 | Bacteria Firmicutes Bacilli Bacillales Bacillaceae *Bacillus subtilis* Hal1 |
| Bsu-17-1 | Bacteria Firmicutes Bacilli Bacillales Bacillaceae *Bacillus subtilis* HJ5 |
| Bsu-19-1 | Bacteria Firmicutes Bacilli Bacillales Bacillaceae *Bacillus subtilis* KCTC 1028 |
| Bsu-20-1 | Bacteria Firmicutes Bacilli Bacillales Bacillaceae *Bacillus subtilis* MB73/2 |
| Bsu-21-1 | Bacteria Firmicutes Bacilli Bacillales Bacillaceae *Bacillus subtilis* Miyagi-4 |
| Bsu-22-1 | Bacteria Firmicutes Bacilli Bacillales Bacillaceae *Bacillus subtilis* PS216 |
| Bsu-23-1 | Bacteria Firmicutes Bacilli Bacillales Bacillaceae *Bacillus subtilis* PTS-394 |
| Bsu-24-1 | Bacteria Firmicutes Bacilli Bacillales Bacillaceae *Bacillus subtilis* PY79 |
| Bsu-25-1 | Bacteria Firmicutes Bacilli Bacillales Bacillaceae *Bacillus subtilis* QH-1 |
| Bsu-26-1 | Bacteria Firmicutes Bacilli Bacillales Bacillaceae *Bacillus subtilis* S1-4 |
| Bsu-29-1 | Bacteria Firmicutes Bacilli Bacillales Bacillaceae *Bacillus subtilis subsp. inaquosorum* KCTC 13429 |
| Bsu-30-1 | Bacteria Firmicutes Bacilli Bacillales Bacillaceae *Bacillus subtilis subsp. natto* BEST195 |
| Bsu-31-1 | Bacteria Firmicutes Bacilli Bacillales Bacillaceae *Bacillus subtilis subsp. niger* |
| Bsu-32-1 | Bacteria Firmicutes Bacilli Bacillales Bacillaceae *Bacillus subtilis subsp. spizizenii* |
| Bsu-33-1 | Bacteria Firmicutes Bacilli Bacillales Bacillaceae *Bacillus subtilis subsp. spizizenii* DV1-B-1 |
| Bsu-34-1 | Bacteria Firmicutes Bacilli Bacillales Bacillaceae *Bacillus subtilis subsp. spizizenii* RFWG1A3 |
| Bsu-35-1 | Bacteria Firmicutes Bacilli Bacillales Bacillaceae *Bacillus subtilis subsp. spizizenii* RFWG1A4 |
| Bsu-36-1 | Bacteria Firmicutes Bacilli Bacillales Bacillaceae *Bacillus subtilis subsp. spizizenii* RFWG4C10 |
| Bsu-37-1 | Bacteria Firmicutes Bacilli Bacillales Bacillaceae *Bacillus subtilis subsp. spizizenii* RFWG5B15 |
| Bsu-38-1 | Bacteria Firmicutes Bacilli Bacillales Bacillaceae *Bacillus subtilis subsp. subtilis* |
| Bsu-39-1 | Bacteria Firmicutes Bacilli Bacillales Bacillaceae *Bacillus subtilis subsp. subtilis* 6051-HGW |
| Bsu-6-1 to Bsu-6-2 | Bacteria Firmicutes Bacilli Bacillales Bacillaceae *Bacillus subtilis subsp. subtilis str.* 168 |
| Bsu-40-1 | Bacteria Firmicutes Bacilli Bacillales Bacillaceae *Bacillus subtilis subsp. subtilis str.* AG1839 |
| Bsu-41-1 | Bacteria Firmicutes Bacilli Bacillales Bacillaceae *Bacillus subtilis subsp. subtilis str.* AUSI98 |
| Bsu-42-1 | Bacteria Firmicutes Bacilli Bacillales Bacillaceae *Bacillus subtilis subsp. subtilis str.* BAB-1 |
| Bsu-43-1 | Bacteria Firmicutes Bacilli Bacillales Bacillaceae *Bacillus subtilis subsp. subtilis str.* BSP1 |
| Bsu-44-1 | Bacteria Firmicutes Bacilli Bacillales Bacillaceae *Bacillus subtilis subsp. subtilis str.* JH642 substr. AG174 |
| Bsu-45-1 | Bacteria Firmicutes Bacilli Bacillales Bacillaceae *Bacillus subtilis subsp. subtilis str.* MP11 |
| Bsu-46-1 | Bacteria Firmicutes Bacilli Bacillales Bacillaceae *Bacillus subtilis subsp. subtilis str.* MP9 |
| Bsu-47-1 | Bacteria Firmicutes Bacilli Bacillales Bacillaceae *Bacillus subtilis subsp. subtilis str.* OH 131.1 |
| Bsu-48-1 | Bacteria Firmicutes Bacilli Bacillales Bacillaceae *Bacillus subtilis subsp. subtilis str.* SC-8 |
| Bsu-27-1 | Bacteria Firmicutes Bacilli Bacillales Bacillaceae *Bacillus subtilis* XF-1 |
| Bte-1-1 | Bacteria Firmicutes Bacilli Bacillales Bacillaceae *Bacillus tequilensis* KCTC 13622 |
| Bte-2-1 | Bacteria Firmicutes Bacilli Bacillales Bacillaceae *Bacillus testis* |
| Bth-44-1 to Bth-44-5 | Bacteria Firmicutes Bacilli Bacillales Bacillaceae *Bacillus thermoamylovorans* |
| Bva-1-1 | Bacteria Firmicutes Bacilli Bacillales Bacillaceae *Bacillus vallismortis* DV1-F-3 |
| Bvi-1-1 | Bacteria Firmicutes Bacilli Bacillales Bacillaceae *Bacillus vietnamensis* |
| Bvi-2-1 | Bacteria Firmicutes Bacilli Bacillales Bacillaceae *Bacillus vireti* |

| | |
|---|---|
| Bvi-3-1 | Bacteria Firmicutes Bacilli Bacillales Bacillaceae *Bacillus vireti* LMG 21834 |
| Bwa-1-1 | Bacteria Firmicutes Bacilli Bacillales Bacillaceae *Bacillus wakoensis* JCM 9140 |
| Bzh-1-1 | Bacteria Firmicutes Bacilli Bacillales Bacillaceae *Bacillus zhangzhouensis* |
| Cde-1-1 | Bacteria Firmicutes Bacilli Bacillales Bacillaceae *Caldibacillus debilis* DSM 16016 |
| Den-1-1 | Bacteria Firmicutes Bacilli Bacillales Bacillaceae *Domibacillus enclensis* |
| Din-1-1 | Bacteria Firmicutes Bacilli Bacillales Bacillaceae *Domibacillus indicus* |
| Dro-1-1 | Bacteria Firmicutes Bacilli Bacillales Bacillaceae *Domibacillus robiginosus* |
| Dtu-1-1 | Bacteria Firmicutes Bacilli Bacillales Bacillaceae *Domibacillus tundrae* |
| Fge-1-1 | Bacteria Firmicutes Bacilli Bacillales Bacillaceae *Fictibacillus gelatini* DSM 15865 |
| Gca-1-1 | Bacteria Firmicutes Bacilli Bacillales Bacillaceae *Geobacillus caldoxylosilyticus* CIC9 |
| Gic-1-1 | Bacteria Firmicutes Bacilli Bacillales Bacillaceae *Geobacillus icigianus* |
| Gka-2-1 | Bacteria Firmicutes Bacilli Bacillales Bacillaceae *Geobacillus kaustophilus* GBlys |
| Gsp-8-1 | Bacteria Firmicutes Bacilli Bacillales Bacillaceae *Geobacillus sp.* 12AMOR1 |
| Gsp-9-1 | Bacteria Firmicutes Bacilli Bacillales Bacillaceae *Geobacillus sp.* A8 |
| Gsp-10-1 | Bacteria Firmicutes Bacilli Bacillales Bacillaceae *Geobacillus sp.* CAMR5420 |
| Gsp-11-1 | Bacteria Firmicutes Bacilli Bacillales Bacillaceae *Geobacillus sp.* FW23 |
| Gsp-12-1 | Bacteria Firmicutes Bacilli Bacillales Bacillaceae *Geobacillus sp.* GHH01 |
| Gsp-13-1 | Bacteria Firmicutes Bacilli Bacillales Bacillaceae *Geobacillus sp.* JF8 |
| Gsp-14-1 | Bacteria Firmicutes Bacilli Bacillales Bacillaceae *Geobacillus sp.* LC300 |
| Gsp-15-1 | Bacteria Firmicutes Bacilli Bacillales Bacillaceae *Geobacillus sp.* MAS1 |
| Gsp-16-1 | Bacteria Firmicutes Bacilli Bacillales Bacillaceae *Geobacillus sp.* T6 |
| Gsp-17-1 | Bacteria Firmicutes Bacilli Bacillales Bacillaceae *Geobacillus sp.* WSUCF1 |
| Gsp-18-1 | Bacteria Firmicutes Bacilli Bacillales Bacillaceae *Geobacillus sp.* ZGt-1 |
| Gst-1-1 to Gst-1-5 | Bacteria Firmicutes Bacilli Bacillales Bacillaceae *Geobacillus stearothermophilus* |
| Gst-2-1 | Bacteria Firmicutes Bacilli Bacillales Bacillaceae *Geobacillus stearothermophilus* 10 |
| Gst-3-1 | Bacteria Firmicutes Bacilli Bacillales Bacillaceae *Geobacillus stearothermophilus* ATCC 7953 |
| Gsu-1-1 | Bacteria Firmicutes Bacilli Bacillales Bacillaceae *Geobacillus subterraneus* PSS2 |
| Gth-4-1 | Bacteria Firmicutes Bacilli Bacillales Bacillaceae *Geobacillus thermodenitrificans subsp. thermodenitrificans* DSM 465 |
| Gth-5-1 | Bacteria Firmicutes Bacilli Bacillales Bacillaceae *Geobacillus thermoglucosidans* TNO-09.020 |
| Gka-3-1 to Gka-3-2 | Bacteria Firmicutes Bacilli Bacillales Bacillaceae Geobacillus Geobacillus thermoleovorans group *Geobacillus kaustophilus* |
| Gka-4-1 | Bacteria Firmicutes Bacilli Bacillales Bacillaceae Geobacillus Geobacillus thermoleovorans group *Geobacillus kaustophilus* NBRC 102445 |
| Gth-6-1 | Bacteria Firmicutes Bacilli Bacillales Bacillaceae Geobacillus Geobacillus thermoleovorans group *Geobacillus thermocatenulatus* GS-1 |
| Gth-7-1 | Bacteria Firmicutes Bacilli Bacillales Bacillaceae Geobacillus Geobacillus thermoleovorans group *Geobacillus thermoleovorans* B23 |
| Gvu-1-1 | Bacteria Firmicutes Bacilli Bacillales Bacillaceae Geobacillus Geobacillus thermoleovorans group *Geobacillus vulcani* PSS1 |
| Gbo-1-1 | Bacteria Firmicutes Bacilli Bacillales Bacillaceae *Gracilibacillus boraciitolerans* JCM 21714 |
| Gha-2-1 | Bacteria Firmicutes Bacilli Bacillales Bacillaceae *Gracilibacillus halophilus* YIM-C55.5 |
| Gla-1-1 | Bacteria Firmicutes Bacilli Bacillales Bacillaceae *Gracilibacillus lacisalsi* DSM 19029 |
| Hha-2-1 | Bacteria Firmicutes Bacilli Bacillales Bacillaceae *Halalkalibacillus halophilus* DSM 18494 |
| Hda-1-1 | Bacteria Firmicutes Bacilli Bacillales Bacillaceae *Halobacillus dabanensis* |
| Hka-1-1 | Bacteria Firmicutes Bacilli Bacillales Bacillaceae *Halobacillus karajensis* |
| Hku-1-1 | Bacteria Firmicutes Bacilli Bacillales Bacillaceae *Halobacillus kuroshimensis* DSM 18393 |
| Hsp-1-1 | Bacteria Firmicutes Bacilli Bacillales Bacillaceae *Halobacillus sp.* BAB-2008 |
| Hsp-2-1 | Bacteria Firmicutes Bacilli Bacillales Bacillaceae *Halobacillus sp.* BBL2006 |
| Htr-1-1 | Bacteria Firmicutes Bacilli Bacillales Bacillaceae *Halobacillus trueperi* |
| Lbo-1-1 | Bacteria Firmicutes Bacilli Bacillales Bacillaceae *Lysinibacillus boronitolerans* JCM 21713 = 10a = NBRC 103108 |
| Lfu-2-1 | Bacteria Firmicutes Bacilli Bacillales Bacillaceae *Lysinibacillus fusiformis* |
| Lfu-3-1 | Bacteria Firmicutes Bacilli Bacillales Bacillaceae *Lysinibacillus fusiformis* H1k |
| Lfu-4-1 | Bacteria Firmicutes Bacilli Bacillales Bacillaceae *Lysinibacillus fusiformis* ZB2 |
| Lma-1-1 | Bacteria Firmicutes Bacilli Bacillales Bacillaceae *Lysinibacillus manganicus* DSM 26584 |
| Lma-2-1 | Bacteria Firmicutes Bacilli Bacillales Bacillaceae *Lysinibacillus massiliensis* 4400831 = CIP 108448 = CCUG 49529 |
| Lod-1-1 | Bacteria Firmicutes Bacilli Bacillales Bacillaceae *Lysinibacillus odysseyi* 34hs-1 = NBRC 100172 |
| Lsi-1-1 | Bacteria Firmicutes Bacilli Bacillales Bacillaceae *Lysinibacillus sinduriensis* BLB-1 = JCM 15800 |
| Lsp-2-1 | Bacteria Firmicutes Bacilli Bacillales Bacillaceae *Lysinibacillus sp.* A1 |
| Lsp-3-1 | Bacteria Firmicutes Bacilli Bacillales Bacillaceae *Lysinibacillus sp.* BF-4 |
| Lsp-4-1 | Bacteria Firmicutes Bacilli Bacillales Bacillaceae *Lysinibacillus sp.* LK3 |
| Lsp-5-1 to Lsp-5-5 | Bacteria Firmicutes Bacilli Bacillales Bacillaceae *Lysinibacillus sphaericus* |
| Lsp-6-1 | Bacteria Firmicutes Bacilli Bacillales Bacillaceae *Lysinibacillus sphaericus* CBAM5 |
| Lsp-7-1 | Bacteria Firmicutes Bacilli Bacillales Bacillaceae *Lysinibacillus sphaericus* KCTC 3346 |
| Lsp-8-1 | Bacteria Firmicutes Bacilli Bacillales Bacillaceae *Lysinibacillus sphaericus* OT4b.31 |
| Lva-1-1 | Bacteria Firmicutes Bacilli Bacillales Bacillaceae *Lysinibacillus varians* |
| Lxy-1-1 | Bacteria Firmicutes Bacilli Bacillales Bacillaceae *Lysinibacillus xylanilyticus* |
| Oje-1-1 | Bacteria Firmicutes Bacilli Bacillales Bacillaceae *Oceanobacillus jeddahense* |
| Oki-1-1 | Bacteria Firmicutes Bacilli Bacillales Bacillaceae *Oceanobacillus kimchii* X50 |
| Oma-1-1 | Bacteria Firmicutes Bacilli Bacillales Bacillaceae *Oceanobacillus manasiensis* |
| Oon-1-1 | Bacteria Firmicutes Bacilli Bacillales Bacillaceae *Oceanobacillus oncorhynchi* |

| | |
|---|---|
| Opi-1-1 | Bacteria Firmicutes Bacilli Bacillales Bacillaceae *Oceanobacillus picturae* |
| Osp-1-1 | Bacteria Firmicutes Bacilli Bacillales Bacillaceae *Oceanobacillus sp.* Ndiop |
| Oca-1-1 | Bacteria Firmicutes Bacilli Bacillales Bacillaceae *Ornithinibacillus californiensis* |
| Oco-1-1 | Bacteria Firmicutes Bacilli Bacillales Bacillaceae *Ornithinibacillus contaminans* |
| Pca-1-1 | Bacteria Firmicutes Bacilli Bacillales Bacillaceae *Parageobacillus caldoxylosilyticus* NBRC 107762 |
| Pge-1-1 | Bacteria Firmicutes Bacilli Bacillales Bacillaceae *Parageobacillus genomosp.* 1 |
| Pth-1-1 | Bacteria Firmicutes Bacilli Bacillales Bacillaceae *Parageobacillus thermoglucosidasius* |
| Pth-2-1 | Bacteria Firmicutes Bacilli Bacillales Bacillaceae *Parageobacillus thermoglucosidasius* NBRC 107763 |
| Pgl-1-1 | Bacteria Firmicutes Bacilli Bacillales Bacillaceae *Paucisalibacillus globulus* DSM 18846 |
| Psp-6-1 | Bacteria Firmicutes Bacilli Bacillales Bacillaceae *Paucisalibacillus sp.* EB02 |
| Pch-1-1 | Bacteria Firmicutes Bacilli Bacillales Bacillaceae *Pontibacillus chungwhensis* BH030062 |
| Pha-1-1 to Pha-1-2 | Bacteria Firmicutes Bacilli Bacillales Bacillaceae *Pontibacillus halophilus* JSM 076056 = DSM 19796 |
| Pli-1-1 | Bacteria Firmicutes Bacilli Bacillales Bacillaceae *Pontibacillus litoralis* JSM 072002 |
| Pma-1-1 | Bacteria Firmicutes Bacilli Bacillales Bacillaceae *Pontibacillus marinus* BH030004 = DSM 16465 |
| Pya-1-1 | Bacteria Firmicutes Bacilli Bacillales Bacillaceae *Pontibacillus yanchengensis* Y32 |
| Qth-1-1 to Qth-1-2 | Bacteria Firmicutes Bacilli Bacillales Bacillaceae *Quasibacillus thermotolerans* |
| Rma-1-1 | Bacteria Firmicutes Bacilli Bacillales Bacillaceae *Rubeoparvulum massiliense* |
| Sai-1-1 to Sai-1-2 | Bacteria Firmicutes Bacilli Bacillales Bacillaceae *Salinibacillus aidingensis* MSP4 |
| Sko-1-1 | Bacteria Firmicutes Bacilli Bacillales Bacillaceae *Salsuginibacillus kocurii* DSM 18087 |
| Sha-2-1 | Bacteria Firmicutes Bacilli Bacillales Bacillaceae *Sediminibacillus halophilus* NSP9.3 |
| Tai-1-1 | Bacteria Firmicutes Bacilli Bacillales Bacillaceae *Terribacillus aidingensis* |
| Tde-1-1 | Bacteria Firmicutes Bacilli Bacillales Bacillaceae *Thalassobacillus devorans* MSP14 |
| Bba-3-1 | Bacteria Firmicutes Bacilli Bacillales Bacillaceae unclassified Bacillaceae *Bacillaceae bacterium* MTCC 10057 |
| Val-1-1 | Bacteria Firmicutes Bacilli Bacillales Bacillaceae *Virgibacillus alimentarius* |
| Vha-1-1 | Bacteria Firmicutes Bacilli Bacillales Bacillaceae *Virgibacillus halodenitrificans* 1806 |
| Vma-1-1 | Bacteria Firmicutes Bacilli Bacillales Bacillaceae *Virgibacillus massiliensis* |
| Vpa-1-1 | Bacteria Firmicutes Bacilli Bacillales Bacillaceae *Virgibacillus pantothenticus* |
| Vse-1-1 | Bacteria Firmicutes Bacilli Bacillales Bacillaceae *Virgibacillus senegalensis* |
| Vsp-1-1 | Bacteria Firmicutes Bacilli Bacillales Bacillaceae *Virgibacillus sp.* CM-4 |
| Vsp-2-1 | Bacteria Firmicutes Bacilli Bacillales Bacillaceae *Virgibacillus sp.* SK37 |
| Tae-1-1 | Bacteria Firmicutes Bacilli Bacillales Bacillales Family X.Incertae Sedis *Thermicanus aegyptius* DSM 12793 |
| Gbe-1-1 | Bacteria Firmicutes Bacilli Bacillales Bacillales Family XI.Incertae Sedis *Gemella bergeriae* ATCC 700627 |
| Gcu-1-1 | Bacteria Firmicutes Bacilli Bacillales Bacillales Family XI.Incertae Sedis *Gemella cuniculi* DSM 15828 |
| Gha-3-1 | Bacteria Firmicutes Bacilli Bacillales Bacillales Family XI.Incertae Sedis *Gemella haemolysans* M341 |
| Gmo-1-1 | Bacteria Firmicutes Bacilli Bacillales Bacillales Family XI.Incertae Sedis *Gemella morbillorum* M424 |
| Gsa-1-1 | Bacteria Firmicutes Bacilli Bacillales Bacillales Family XI.Incertae Sedis *Gemella sanguinis* ATCC 700632 |
| Gsa-2-1 | Bacteria Firmicutes Bacilli Bacillales Bacillales Family XI.Incertae Sedis *Gemella sanguinis* M325 |
| Eac-1-1 | Bacteria Firmicutes Bacilli Bacillales Bacillales Family XII.Incertae Sedis *Exiguobacterium acetylicum* |
| Eac-2-1 | Bacteria Firmicutes Bacilli Bacillales Bacillales Family XII.Incertae Sedis *Exiguobacterium acetylicum* DSM 20416 |
| Eal-1-1 | Bacteria Firmicutes Bacilli Bacillales Bacillales Family XII.Incertae Sedis *Exiguobacterium alkaliphilum* 12/1 |
| Ean-2-1 | Bacteria Firmicutes Bacilli Bacillales Bacillales Family XII.Incertae Sedis *Exiguobacterium antarcticum* DSM 14480 |
| Eau-1-1 | Bacteria Firmicutes Bacilli Bacillales Bacillales Family XII.Incertae Sedis *Exiguobacterium aurantiacum* DSM 6208 |
| Ech-1-1 | Bacteria Firmicutes Bacilli Bacillales Bacillales Family XII.Incertae Sedis *Exiguobacterium chiriqhucha* |
| Ema-1-1 | Bacteria Firmicutes Bacilli Bacillales Bacillales Family XII.Incertae Sedis *Exiguobacterium marinum* DSM 16307 |
| Eme-1-1 | Bacteria Firmicutes Bacilli Bacillales Bacillales Family XII.Incertae Sedis *Exiguobacterium mexicanum* |
| Eox-1-1 | Bacteria Firmicutes Bacilli Bacillales Bacillales Family XII.Incertae Sedis *Exiguobacterium oxidotolerans* JCM 12280 |
| Epa-1-1 | Bacteria Firmicutes Bacilli Bacillales Bacillales Family XII.Incertae Sedis *Exiguobacterium pavilionensis* RW-2 |
| Esi-2-1 | Bacteria Firmicutes Bacilli Bacillales Bacillales Family XII.Incertae Sedis *Exiguobacterium sibiricum* 7-3 |
| Esp-2-1 | Bacteria Firmicutes Bacilli Bacillales Bacillales Family XII.Incertae Sedis *Exiguobacterium sp.* 8-11-1 |
| Esp-3-1 | Bacteria Firmicutes Bacilli Bacillales Bacillales Family XII.Incertae Sedis *Exiguobacterium sp.* AB2 |
| Esp-4-1 | Bacteria Firmicutes Bacilli Bacillales Bacillales Family XII.Incertae Sedis *Exiguobacterium sp.* JLM-2 |
| Esp-5-1 | Bacteria Firmicutes Bacilli Bacillales Bacillales Family XII.Incertae Sedis *Exiguobacterium sp.* MH3 |
| Esp-6-1 | Bacteria Firmicutes Bacilli Bacillales Bacillales Family XII.Incertae Sedis *Exiguobacterium sp.* NG55 |
| Esp-7-1 | Bacteria Firmicutes Bacilli Bacillales Bacillales Family XII.Incertae Sedis *Exiguobacterium sp.* OS-77 |
| Esp-8-1 | Bacteria Firmicutes Bacilli Bacillales Bacillales Family XII.Incertae Sedis *Exiguobacterium sp.* RIT341 |
| Esp-9-1 | Bacteria Firmicutes Bacilli Bacillales Bacillales Family XII.Incertae Sedis *Exiguobacterium sp.* S17 |
| Esp-10-1 | Bacteria Firmicutes Bacilli Bacillales Bacillales Family XII.Incertae Sedis *Exiguobacterium sp.* ZOR0005 |
| Esp-11-1 | Bacteria Firmicutes Bacilli Bacillales Bacillales Family XII.Incertae Sedis *Exiguobacterium sp.* ZWU0009 |
| Eun-1-1 | Bacteria Firmicutes Bacilli Bacillales Bacillales Family XII.Incertae Sedis *Exiguobacterium undae* 190-11 |
| Eun-2-1 | Bacteria Firmicutes Bacilli Bacillales Bacillales Family XII.Incertae Sedis *Exiguobacterium undae* DSM 14481 |
| Bce-148-1 | Bacteria Firmicutes Bacilli Bacillales *Bhargavaea cecembensis* DSE10 |
| Bca-2-1 | Bacteria Firmicutes Bacilli Bacillales Listeriaceae *Brochothrix campestris* FSL F6-1037 |
| Bth-45-1 to Bth-45-2 | Bacteria Firmicutes Bacilli Bacillales Listeriaceae *Brochothrix thermosphacta* DSM 20171 = FSL F6-1036 |
| Laq-1-1 | Bacteria Firmicutes Bacilli Bacillales Listeriaceae *Listeria aquatica* FSL S10-1188 |
| Lbo-2-1 | Bacteria Firmicutes Bacilli Bacillales Listeriaceae *Listeria booriae* |

| | |
|---|---|
| Lco-1-1 | Bacteria Firmicutes Bacilli Bacillales Listeriaceae *Listeria cornellensis* FSL F6-0969 |
| Lfl-1-1 | Bacteria Firmicutes Bacilli Bacillales Listeriaceae *Listeria fleischmannii* FSL S10-1203 |
| Lfl-2-1 | Bacteria Firmicutes Bacilli Bacillales Listeriaceae *Listeria fleischmannii* LU2006-1 |
| Lfl-3-1 | Bacteria Firmicutes Bacilli Bacillales Listeriaceae *Listeria floridensis* FSL S10-1187 |
| Lgr-1-1 | Bacteria Firmicutes Bacilli Bacillales Listeriaceae *Listeria grandensis* FSL F6-0971 |
| Lgr-2-1 | Bacteria Firmicutes Bacilli Bacillales Listeriaceae *Listeria grayi* DSM 20601 |
| Lgr-3-1 | Bacteria Firmicutes Bacilli Bacillales Listeriaceae *Listeria grayi* FSL F6-1183 |
| Lin-2-1 to Lin-2-3 | Bacteria Firmicutes Bacilli Bacillales Listeriaceae *Listeria innocua* |
| Lin-3-1 | Bacteria Firmicutes Bacilli Bacillales Listeriaceae *Listeria innocua* ATCC 33091 |
| Lin-4-1 | Bacteria Firmicutes Bacilli Bacillales Listeriaceae *Listeria innocua* FSL J1-023 |
| Liv-3-1 | Bacteria Firmicutes Bacilli Bacillales Listeriaceae *Listeria ivanovii subsp. ivanovii* |
| Liv-1-1 to Liv-1-2 | Bacteria Firmicutes Bacilli Bacillales Listeriaceae *Listeria ivanovii subsp. ivanovii* PAM 55 |
| Liv-4-1 to Liv-4-2 | Bacteria Firmicutes Bacilli Bacillales Listeriaceae *Listeria ivanovii subsp. londoniensis* |
| Liv-2-1 | Bacteria Firmicutes Bacilli Bacillales Listeriaceae *Listeria ivanovii* WSLC3009 |
| Lmo-33-1 to Lmo-33-9 | Bacteria Firmicutes Bacilli Bacillales Listeriaceae *Listeria monocytogenes* |
| Lmo-34-1 | Bacteria Firmicutes Bacilli Bacillales Listeriaceae *Listeria monocytogenes* 08-5578 |
| Lmo-35-1 | Bacteria Firmicutes Bacilli Bacillales Listeriaceae *Listeria monocytogenes* 4423 |
| Lmo-36-1 | Bacteria Firmicutes Bacilli Bacillales Listeriaceae *Listeria monocytogenes* 6179 |
| Lmo-37-1 | Bacteria Firmicutes Bacilli Bacillales Listeriaceae *Listeria monocytogenes* CFSAN002349 |
| Lmo-38-1 | Bacteria Firmicutes Bacilli Bacillales Listeriaceae *Listeria monocytogenes* EGD |
| Lmo-39-1 | Bacteria Firmicutes Bacilli Bacillales Listeriaceae *Listeria monocytogenes* FSL F6-684 |
| Lmo-40-1 | Bacteria Firmicutes Bacilli Bacillales Listeriaceae *Listeria monocytogenes* FSL J1-208 |
| Lmo-41-1 | Bacteria Firmicutes Bacilli Bacillales Listeriaceae *Listeria monocytogenes* J1-220 |
| Lmo-42-1 | Bacteria Firmicutes Bacilli Bacillales Listeriaceae *Listeria monocytogenes* J1816 |
| Lmo-44-1 | Bacteria Firmicutes Bacilli Bacillales Listeriaceae *Listeria monocytogenes* Lm19860 |
| Lmo-43-1 | Bacteria Firmicutes Bacilli Bacillales Listeriaceae *Listeria monocytogenes* LM201 |
| Lmo-45-1 | Bacteria Firmicutes Bacilli Bacillales Listeriaceae *Listeria monocytogenes* Lm21045 |
| Lmo-46-1 | Bacteria Firmicutes Bacilli Bacillales Listeriaceae *Listeria monocytogenes* Lm25180 |
| Lmo-47-1 | Bacteria Firmicutes Bacilli Bacillales Listeriaceae *Listeria monocytogenes* Lm26686 |
| Lmo-48-1 | Bacteria Firmicutes Bacilli Bacillales Listeriaceae *Listeria monocytogenes* Lm_1823 |
| Lmo-49-1 | Bacteria Firmicutes Bacilli Bacillales Listeriaceae *Listeria monocytogenes* Lm_1824 |
| Lmo-50-1 | Bacteria Firmicutes Bacilli Bacillales Listeriaceae *Listeria monocytogenes* Lm_1840 |
| Lmo-51-1 | Bacteria Firmicutes Bacilli Bacillales Listeriaceae *Listeria monocytogenes* Lm_1880 |
| Lmo-52-1 | Bacteria Firmicutes Bacilli Bacillales Listeriaceae *Listeria monocytogenes* Lm_1886 |
| Lmo-53-1 | Bacteria Firmicutes Bacilli Bacillales Listeriaceae *Listeria monocytogenes* Lm_1889 |
| Lmo-54-1 | Bacteria Firmicutes Bacilli Bacillales Listeriaceae *Listeria monocytogenes* N53-1 |
| Lmo-55-1 | Bacteria Firmicutes Bacilli Bacillales Listeriaceae *Listeria monocytogenes* QOC1 |
| Lmo-56-1 | Bacteria Firmicutes Bacilli Bacillales Listeriaceae *Listeria monocytogenes* QOC2 |
| Lmo-57-1 | Bacteria Firmicutes Bacilli Bacillales Listeriaceae *Listeria monocytogenes* R479a |
| Lmo-72-1 | Bacteria Firmicutes Bacilli Bacillales Listeriaceae *Listeria monocytogenes serotype* 1/2a str. NCCP No. 15743 |
| Lmo-73-1 | Bacteria Firmicutes Bacilli Bacillales Listeriaceae *Listeria monocytogenes serotype* 4b str. LL195 |
| Lmo-74-1 | Bacteria Firmicutes Bacilli Bacillales Listeriaceae *Listeria monocytogenes serotype* 4bV str. LS542 |
| Lmo-75-1 | Bacteria Firmicutes Bacilli Bacillales Listeriaceae *Listeria monocytogenes serotype* 4bV str. LS642 |
| Lmo-76-1 | Bacteria Firmicutes Bacilli Bacillales Listeriaceae *Listeria monocytogenes serotype* 4bV str. LS643 |
| Lmo-77-1 | Bacteria Firmicutes Bacilli Bacillales Listeriaceae *Listeria monocytogenes serotype* 4bV str. LS644 |
| Lmo-78-1 | Bacteria Firmicutes Bacilli Bacillales Listeriaceae *Listeria monocytogenes serotype* 4bV str. LS645 |
| Lmo-58-1 | Bacteria Firmicutes Bacilli Bacillales Listeriaceae *Listeria monocytogenes* SHL001 |
| Lmo-59-1 | Bacteria Firmicutes Bacilli Bacillales Listeriaceae *Listeria monocytogenes* SHL002 |
| Lmo-60-1 | Bacteria Firmicutes Bacilli Bacillales Listeriaceae *Listeria monocytogenes* SHL004 |
| Lmo-61-1 | Bacteria Firmicutes Bacilli Bacillales Listeriaceae *Listeria monocytogenes* SHL005 |
| Lmo-62-1 | Bacteria Firmicutes Bacilli Bacillales Listeriaceae *Listeria monocytogenes* SHL006 |
| Lmo-63-1 | Bacteria Firmicutes Bacilli Bacillales Listeriaceae *Listeria monocytogenes* SHL007 |
| Lmo-64-1 | Bacteria Firmicutes Bacilli Bacillales Listeriaceae *Listeria monocytogenes* SHL008 |
| Lmo-65-1 | Bacteria Firmicutes Bacilli Bacillales Listeriaceae *Listeria monocytogenes* SHL009 |
| Lmo-66-1 | Bacteria Firmicutes Bacilli Bacillales Listeriaceae *Listeria monocytogenes* SHL010 |
| Lmo-67-1 | Bacteria Firmicutes Bacilli Bacillales Listeriaceae *Listeria monocytogenes* SHL011 |
| Lmo-68-1 | Bacteria Firmicutes Bacilli Bacillales Listeriaceae *Listeria monocytogenes* SHL012 |
| Lmo-69-1 | Bacteria Firmicutes Bacilli Bacillales Listeriaceae *Listeria monocytogenes* SHL013 |
| Lmo-79-1 | Bacteria Firmicutes Bacilli Bacillales Listeriaceae *Listeria monocytogenes str.* Scott A |
| Lmo-70-1 | Bacteria Firmicutes Bacilli Bacillales Listeriaceae *Listeria monocytogenes* WSLC1001 |
| Lmo-71-1 | Bacteria Firmicutes Bacilli Bacillales Listeriaceae *Listeria monocytogenes* WSLC1042 |
| Lne-1-1 | Bacteria Firmicutes Bacilli Bacillales Listeriaceae *Listeria newyorkensis* |
| Lri-1-1 | Bacteria Firmicutes Bacilli Bacillales Listeriaceae *Listeria riparia* FSL S10-1204 |
| Lro-1-1 | Bacteria Firmicutes Bacilli Bacillales Listeriaceae *Listeria rocourtiae* FSL F6-920 |
| Lse-2-1 | Bacteria Firmicutes Bacilli Bacillales Listeriaceae *Listeria seeligeri* |

| | |
|---|---|
| Lse-3-1 | Bacteria Firmicutes Bacilli Bacillales Listeriaceae *Listeria seeligeri* FSL N1-067 |
| Lwe-2-1 | Bacteria Firmicutes Bacilli Bacillales Listeriaceae *Listeria weihenstephanensis* FSL R9-0317 |
| Lba-2-1 | Bacteria Firmicutes Bacilli Bacillales *Listeriaceae bacterium* FSL A5-0209 |
| Aan-1-1 | Bacteria Firmicutes Bacilli Bacillales Paenibacillaceae Aneurinibacillus group *Aneurinibacillus aneurinilyticus* ATCC 12856 |
| Ami-1-1 to Ami-1-3 | Bacteria Firmicutes Bacilli Bacillales Paenibacillaceae Aneurinibacillus group *Aneurinibacillus migulanus* |
| Ate-2-1 | Bacteria Firmicutes Bacilli Bacillales Paenibacillaceae Aneurinibacillus group *Aneurinibacillus terranovensis* DSM 18919 |
| Aty-1-1 | Bacteria Firmicutes Bacilli Bacillales Paenibacillaceae Aneurinibacillus group *Aneurinibacillus tyrosinisolvens* |
| Bag-1-1 | Bacteria Firmicutes Bacilli Bacillales Paenibacillaceae *Brevibacillus agri* 5-2 |
| Bag-2-1 | Bacteria Firmicutes Bacilli Bacillales Paenibacillaceae *Brevibacillus agri* BAB-2500 |
| Bbo-4-1 | Bacteria Firmicutes Bacilli Bacillales Paenibacillaceae *Brevibacillus borstelensis* |
| Bbo-5-1 | Bacteria Firmicutes Bacilli Bacillales Paenibacillaceae *Brevibacillus borstelensis* 3096-7 |
| Bbo-6-1 | Bacteria Firmicutes Bacilli Bacillales Paenibacillaceae *Brevibacillus borstelensis* AK1 |
| Bbo-7-1 | Bacteria Firmicutes Bacilli Bacillales Paenibacillaceae *Brevibacillus borstelensis cifa_chp*40 |
| Bbr-2-1 | Bacteria Firmicutes Bacilli Bacillales Paenibacillaceae *Brevibacillus brevis* |
| Bbr-3-1 | Bacteria Firmicutes Bacilli Bacillales Paenibacillaceae *Brevibacillus brevis* FJAT-0809-GLX |
| Bbr-4-1 | Bacteria Firmicutes Bacilli Bacillales Paenibacillaceae *Brevibacillus brevis* X23 |
| Bfo-2-1 | Bacteria Firmicutes Bacilli Bacillales Paenibacillaceae *Brevibacillus formosus* |
| Bla-1-1 | Bacteria Firmicutes Bacilli Bacillales Paenibacillaceae *Brevibacillus laterosporus* |
| Bla-2-1 | Bacteria Firmicutes Bacilli Bacillales Paenibacillaceae *Brevibacillus laterosporus* DSM 25 |
| Bla-3-1 | Bacteria Firmicutes Bacilli Bacillales Paenibacillaceae *Brevibacillus laterosporus* GI-9 |
| Bla-4-1 | Bacteria Firmicutes Bacilli Bacillales Paenibacillaceae *Brevibacillus laterosporus* LMG 15441 |
| Bla-5-1 | Bacteria Firmicutes Bacilli Bacillales Paenibacillaceae *Brevibacillus laterosporus* PE36 |
| Bpa-4-1 | Bacteria Firmicutes Bacilli Bacillales Paenibacillaceae *Brevibacillus panacihumi* W25 |
| Bre-1-1 | Bacteria Firmicutes Bacilli Bacillales Paenibacillaceae *Brevibacillus reuszeri* |
| Bsp-99-1 | Bacteria Firmicutes Bacilli Bacillales Paenibacillaceae *Brevibacillus sp. ph*R |
| Bth-46-1 | Bacteria Firmicutes Bacilli Bacillales Paenibacillaceae *Brevibacillus thermoruber* 423 |
| Bth-47-1 | Bacteria Firmicutes Bacilli Bacillales Paenibacillaceae *Brevibacillus thermoruber* PM1 |
| Cko-1-1 | Bacteria Firmicutes Bacilli Bacillales Paenibacillaceae *Cohnella kolymensis* |
| Cla-1-1 | Bacteria Firmicutes Bacilli Bacillales Paenibacillaceae *Cohnella laeviribosi* DSM 21336 |
| Cth-2-1 | Bacteria Firmicutes Bacilli Bacillales Paenibacillaceae *Cohnella thermotolerans* DSM 17683 |
| Gma-1-1 | Bacteria Firmicutes Bacilli Bacillales Paenibacillaceae *Gorillibacterium massiliense* |
| Pal-2-1 | Bacteria Firmicutes Bacilli Bacillales Paenibacillaceae *Paenibacillus alginolyticus* DSM 5050 = NBRC 15375 |
| Pal-3-1 | Bacteria Firmicutes Bacilli Bacillales Paenibacillaceae *Paenibacillus algorifonticola* |
| Pal-4-1 | Bacteria Firmicutes Bacilli Bacillales Paenibacillaceae *Paenibacillus alvei* A6-6i-x |
| Pal-5-1 | Bacteria Firmicutes Bacilli Bacillales Paenibacillaceae *Paenibacillus alvei* TS-15 |
| Pan-2-1 | Bacteria Firmicutes Bacilli Bacillales Paenibacillaceae *Paenibacillus antibioticophila* |
| Pas-1-1 | Bacteria Firmicutes Bacilli Bacillales Paenibacillaceae *Paenibacillus assamensis* DSM 18201 |
| Pba-1-1 | Bacteria Firmicutes Bacilli Bacillales Paenibacillaceae *Paenibacillus barengoltzii* G22 |
| Pbe-1-1 | Bacteria Firmicutes Bacilli Bacillales Paenibacillaceae *Paenibacillus beijingensis* |
| Pch-2-1 | Bacteria Firmicutes Bacilli Bacillales Paenibacillaceae *Paenibacillus chitinolyticus* NBRC 15660 |
| Pch-3-1 | Bacteria Firmicutes Bacilli Bacillales Paenibacillaceae *Paenibacillus chondroitinus* |
| Pda-1-1 to Pda-1-3 | Bacteria Firmicutes Bacilli Bacillales Paenibacillaceae *Paenibacillus darwinianus* |
| Peh-1-1 | Bacteria Firmicutes Bacilli Bacillales Paenibacillaceae *Paenibacillus ehimensis* A2 |
| Pgi-1-1 | Bacteria Firmicutes Bacilli Bacillales Paenibacillaceae *Paenibacillus ginsengihumi* DSM 21568 |
| Pgo-1-1 | Bacteria Firmicutes Bacilli Bacillales Paenibacillaceae *Paenibacillus gorillae* |
| Pha-2-1 | Bacteria Firmicutes Bacilli Bacillales Paenibacillaceae *Paenibacillus harenae* DSM 16969 |
| Pla-3-1 | Bacteria Firmicutes Bacilli Bacillales Paenibacillaceae *Paenibacillus larvae* |
| Pla-4-1 | Bacteria Firmicutes Bacilli Bacillales Paenibacillaceae *Paenibacillus larvae subsp. larvae* DSM 25430 |
| Pla-5-1 | Bacteria Firmicutes Bacilli Bacillales Paenibacillaceae *Paenibacillus larvae subsp. larvae* DSM 25719 |
| Pma-2-1 | Bacteria Firmicutes Bacilli Bacillales Paenibacillaceae *Paenibacillus macerans* |
| Pma-3-1 | Bacteria Firmicutes Bacilli Bacillales Paenibacillaceae *Paenibacillus massiliensis* 2301065 = DSM 16942 |
| Pma-4-1 | Bacteria Firmicutes Bacilli Bacillales Paenibacillaceae *Paenibacillus massiliensis* T7 |
| Pmu-3-1 | Bacteria Firmicutes Bacilli Bacillales Paenibacillaceae *Paenibacillus mucilaginosus* K02 |
| Ppa-1-1 | Bacteria Firmicutes Bacilli Bacillales Paenibacillaceae *Paenibacillus panacisoli* DSM 21345 |
| Ppa-2-1 | Bacteria Firmicutes Bacilli Bacillales Paenibacillaceae *Paenibacillus pasadenensis* DSM 19293 |
| Ppe-2-1 | Bacteria Firmicutes Bacilli Bacillales Paenibacillaceae *Paenibacillus peoriae* |
| Ppi-1-1 | Bacteria Firmicutes Bacilli Bacillales Paenibacillaceae *Paenibacillus pini* JCM 16418 |
| Ppi-2-1 | Bacteria Firmicutes Bacilli Bacillales Paenibacillaceae *Paenibacillus pinihumi* DSM 23905 = JCM 16419 |
| Ppo-2-1 to Ppo-2-2 | Bacteria Firmicutes Bacilli Bacillales Paenibacillaceae *Paenibacillus polymyxa* |
| Ppo-3-1 | Bacteria Firmicutes Bacilli Bacillales Paenibacillaceae *Paenibacillus polymyxa* 1-43 |
| Ppo-4-1 | Bacteria Firmicutes Bacilli Bacillales Paenibacillaceae *Paenibacillus polymyxa* A18 |
| Ppo-5-1 | Bacteria Firmicutes Bacilli Bacillales Paenibacillaceae *Paenibacillus polymyxa* ATCC 842 |
| Ppo-6-1 | Bacteria Firmicutes Bacilli Bacillales Paenibacillaceae *Paenibacillus polymyxa* E681 |
| Ppo-7-1 | Bacteria Firmicutes Bacilli Bacillales Paenibacillaceae *Paenibacillus polymyxa* OSY-DF |
| Ppo-8-1 | Bacteria Firmicutes Bacilli Bacillales Paenibacillaceae *Paenibacillus polymyxa* SC2 |

| | |
|---|---|
| Ppo-9-1 | Bacteria Firmicutes Bacilli Bacillales Paenibacillaceae *Paenibacillus polymyxa* SQR-21 |
| Ppo-10-1 | Bacteria Firmicutes Bacilli Bacillales Paenibacillaceae *Paenibacillus polymyxa* TD94 |
| Psa-1-1 | Bacteria Firmicutes Bacilli Bacillales Paenibacillaceae *Paenibacillus sanguinis* 2301083 = DSM 16941 |
| Psp-7-1 | Bacteria Firmicutes Bacilli Bacillales Paenibacillaceae *Paenibacillus sp.* 1-18 |
| Psp-8-1 | Bacteria Firmicutes Bacilli Bacillales Paenibacillaceae *Paenibacillus sp.* 1-49 |
| Psp-9-1 | Bacteria Firmicutes Bacilli Bacillales Paenibacillaceae *Paenibacillus sp.* D9 |
| Psp-10-1 | Bacteria Firmicutes Bacilli Bacillales Paenibacillaceae *Paenibacillus sp.* DMB20 |
| Psp-11-1 | Bacteria Firmicutes Bacilli Bacillales Paenibacillaceae *Paenibacillus sp.* E194 |
| Psp-12-1 | Bacteria Firmicutes Bacilli Bacillales Paenibacillaceae *Paenibacillus sp.* FJAT-27812 |
| Psp-13-1 | Bacteria Firmicutes Bacilli Bacillales Paenibacillaceae *Paenibacillus sp.* FSL H7-689 |
| Psp-14-1 | Bacteria Firmicutes Bacilli Bacillales Paenibacillaceae *Paenibacillus sp.* FSL H8-457 |
| Psp-15-1 | Bacteria Firmicutes Bacilli Bacillales Paenibacillaceae *Paenibacillus sp.* FSL R5-192 |
| Psp-16-1 | Bacteria Firmicutes Bacilli Bacillales Paenibacillaceae *Paenibacillus sp.* FSL R5-808 |
| Psp-17-1 | Bacteria Firmicutes Bacilli Bacillales Paenibacillaceae *Paenibacillus sp.* HGH0039 |
| Psp-18-1 | Bacteria Firmicutes Bacilli Bacillales Paenibacillaceae *Paenibacillus sp.* ICGEB2008 |
| Psp-19-1 | Bacteria Firmicutes Bacilli Bacillales Paenibacillaceae *Paenibacillus sp.* IHBB 10380 |
| Psp-20-1 | Bacteria Firmicutes Bacilli Bacillales Paenibacillaceae *Paenibacillus sp.* J14 |
| Psp-21-1 | Bacteria Firmicutes Bacilli Bacillales Paenibacillaceae *Paenibacillus sp.* JC66 |
| Psp-22-1 | Bacteria Firmicutes Bacilli Bacillales Paenibacillaceae *Paenibacillus sp.* MAEPY1 |
| Psp-23-1 | Bacteria Firmicutes Bacilli Bacillales Paenibacillaceae *Paenibacillus sp.* MAEPY2 |
| Psp-24-1 | Bacteria Firmicutes Bacilli Bacillales Paenibacillaceae *Paenibacillus sp.* P1XP2 |
| Psp-25-1 | Bacteria Firmicutes Bacilli Bacillales Paenibacillaceae *Paenibacillus sp.* P22 |
| Psp-26-1 | Bacteria Firmicutes Bacilli Bacillales Paenibacillaceae *Paenibacillus sp.* PAMC 26794 |
| Psp-27-1 | Bacteria Firmicutes Bacilli Bacillales Paenibacillaceae *Paenibacillus sp.* TCA20 |
| Psp-28-1 | Bacteria Firmicutes Bacilli Bacillales Paenibacillaceae *Paenibacillus sp.* UNC217MF |
| Psp-29-1 | Bacteria Firmicutes Bacilli Bacillales Paenibacillaceae *Paenibacillus sp.* UNC451MF |
| Psp-30-1 | Bacteria Firmicutes Bacilli Bacillales Paenibacillaceae *Paenibacillus sp.* UNCCL52 |
| Psp-31-1 | Bacteria Firmicutes Bacilli Bacillales Paenibacillaceae *Paenibacillus sp.* URHA0014 |
| Psp-32-1 | Bacteria Firmicutes Bacilli Bacillales Paenibacillaceae *Paenibacillus sp.* VKM B-2647 |
| Psp-33-1 | Bacteria Firmicutes Bacilli Bacillales Paenibacillaceae *Paenibacillus sp.* VT-400 |
| Pst-1-1 | Bacteria Firmicutes Bacilli Bacillales Paenibacillaceae *Paenibacillus stellifer* |
| Pta-1-1 | Bacteria Firmicutes Bacilli Bacillales Paenibacillaceae *Paenibacillus taiwanensis* DSM 18679 |
| Pte-2-1 | Bacteria Firmicutes Bacilli Bacillales Paenibacillaceae *Paenibacillus terrae* |
| Pte-3-1 | Bacteria Firmicutes Bacilli Bacillales Paenibacillaceae *Paenibacillus terrigena* DSM 21567 |
| Pty-1-1 | Bacteria Firmicutes Bacilli Bacillales Paenibacillaceae *Paenibacillus tyrfis* |
| Pwy-1-1 | Bacteria Firmicutes Bacilli Bacillales Paenibacillaceae *Paenibacillus wynnii* |
| Tco-1-1 | Bacteria Firmicutes Bacilli Bacillales Paenibacillaceae *Thermobacillus composti* KWC4 |
| Bce-149-1 to Bce-149-2 | Bacteria Firmicutes Bacilli Bacillales Planococcaceae *Bhargavaea cecembensis* |
| Jal-1-1 | Bacteria Firmicutes Bacilli Bacillales Planococcaceae *Jeotgalibacillus alimentarius* |
| Jca-1-1 | Bacteria Firmicutes Bacilli Bacillales Planococcaceae *Jeotgalibacillus campisalis* |
| Jma-1-1 | Bacteria Firmicutes Bacilli Bacillales Planococcaceae *Jeotgalibacillus malaysiensis* |
| Jma-2-1 | Bacteria Firmicutes Bacilli Bacillales Planococcaceae *Jeotgalibacillus marinus* |
| Jso-1-1 | Bacteria Firmicutes Bacilli Bacillales Planococcaceae *Jeotgalibacillus soli* |
| Khu-1-1 | Bacteria Firmicutes Bacilli Bacillales Planococcaceae *Kurthia huakuii* LAM0618 |
| Kma-1-1 | Bacteria Firmicutes Bacilli Bacillales Planococcaceae *Kurthia massiliensis* |
| Ksp-1-1 | Bacteria Firmicutes Bacilli Bacillales Planococcaceae *Kurthia sp.* Dielmo |
| Ksp-2-1 | Bacteria Firmicutes Bacilli Bacillales Planococcaceae *Kurthia sp.* JC8E |
| Psp-34-1 | Bacteria Firmicutes Bacilli Bacillales Planococcaceae *Paenisporosarcina sp.* HGH0030 |
| Psp-35-1 | Bacteria Firmicutes Bacilli Bacillales Planococcaceae *Paenisporosarcina sp.* TG-14 |
| Psp-36-1 | Bacteria Firmicutes Bacilli Bacillales Planococcaceae *Paenisporosarcina sp.* TG20 |
| Pha-3-1 | Bacteria Firmicutes Bacilli Bacillales Planococcaceae *Planococcus halocryophilus* Or1 |
| Pma-5-1 | Bacteria Firmicutes Bacilli Bacillales Planococcaceae *Planococcus massiliensis* |
| Psp-37-1 | Bacteria Firmicutes Bacilli Bacillales Planococcaceae *Planococcus sp.* CAU13 |
| Psp-38-1 | Bacteria Firmicutes Bacilli Bacillales Planococcaceae *Planococcus sp.* L10.15 |
| Psp-39-1 | Bacteria Firmicutes Bacilli Bacillales Planococcaceae *Planococcus sp.* PAMC 21323 |
| Pgl-2-1 | Bacteria Firmicutes Bacilli Bacillales Planococcaceae *Planomicrobium glaciei* |
| Pgl-3-1 | Bacteria Firmicutes Bacilli Bacillales Planococcaceae *Planomicrobium glaciei* CHR43 |
| Sgl-1-1 | Bacteria Firmicutes Bacilli Bacillales Planococcaceae *Sporosarcina globispora* |
| Sko-2-1 | Bacteria Firmicutes Bacilli Bacillales Planococcaceae *Sporosarcina koreensis* |
| Sne-1-1 | Bacteria Firmicutes Bacilli Bacillales Planococcaceae *Sporosarcina newyorkensis* 2681 |
| Ssp-3-1 | Bacteria Firmicutes Bacilli Bacillales Planococcaceae *Sporosarcina sp.* D27 |
| Ssp-4-1 | Bacteria Firmicutes Bacilli Bacillales Planococcaceae *Sporosarcina sp.* EUR3 2.2.2 |
| Ssp-5-1 | Bacteria Firmicutes Bacilli Bacillales Planococcaceae *Sporosarcina sp.* ZBG7A |
| Sur-1-1 | Bacteria Firmicutes Bacilli Bacillales Planococcaceae *Sporosarcina ureae* DSM 2281 |
| Uth-1-1 | Bacteria Firmicutes Bacilli Bacillales Planococcaceae *Ureibacillus thermosphaericus str.* Thermo-BF |

| | |
|---|---|
| Var-1-1 | Bacteria Firmicutes Bacilli Bacillales Planococcaceae *Viridibacillus arenosi* FSL R5-213 |
| Var-2-1 | Bacteria Firmicutes Bacilli Bacillales Planococcaceae *Viridibacillus arvi* |
| Sal-1-1 | Bacteria Firmicutes Bacilli Bacillales *Salinicoccus albus* DSM 19776 |
| Sca-4-1 | Bacteria Firmicutes Bacilli Bacillales *Salinicoccus carnicancri* Crm |
| Sin-1-1 | Bacteria Firmicutes Bacilli Bacillales Sporolactobacillaceae *Sporolactobacillus inulinus* CASD |
| Sla-1-1 | Bacteria Firmicutes Bacilli Bacillales Sporolactobacillaceae *Sporolactobacillus laevolacticus* DSM 442 |
| Ste-1-1 | Bacteria Firmicutes Bacilli Bacillales Sporolactobacillaceae *Sporolactobacillus terrae* DSM 11697 |
| Ste-2-1 | Bacteria Firmicutes Bacilli Bacillales Sporolactobacillaceae *Sporolactobacillus terrae* HKM-1 |
| Svi-1-1 to Svi-1-2 | Bacteria Firmicutes Bacilli Bacillales Sporolactobacillaceae *Sporolactobacillus vineae* DSM 21990 = SL153 |
| Tca-1-1 | Bacteria Firmicutes Bacilli Bacillales Sporolactobacillaceae *Tuberibacillus calidus* DSM 17572 |
| Jma-3-1 | Bacteria Firmicutes Bacilli Bacillales Staphylococcaceae *Jeotgalicoccus marinus* DSM 19772 |
| Jps-1-1 | Bacteria Firmicutes Bacilli Bacillales Staphylococcaceae *Jeotgalicoccus psychrophilus* DSM 19085 |
| Jsa-1-1 | Bacteria Firmicutes Bacilli Bacillales Staphylococcaceae *Jeotgalicoccus saudimassiliensis* |
| Nma-1-1 | Bacteria Firmicutes Bacilli Bacillales Staphylococcaceae *Nosocomiicoccus massiliensis* |
| Sha-3-1 | Bacteria Firmicutes Bacilli Bacillales Staphylococcaceae *Salinicoccus halodurans* |
| Slu-3-1 | Bacteria Firmicutes Bacilli Bacillales Staphylococcaceae *Salinicoccus luteus* DSM 17002 |
| Sro-1-1 | Bacteria Firmicutes Bacilli Bacillales Staphylococcaceae *Salinicoccus roseus* |
| Sse-1-1 | Bacteria Firmicutes Bacilli Bacillales Staphylococcaceae *Salinicoccus sediminis* |
| Ssp-6-1 | Bacteria Firmicutes Bacilli Bacillales Staphylococcaceae *Salinicoccus sp.* YB14-2 |
| Sag-1-1 | Bacteria Firmicutes Bacilli Bacillales Staphylococcaceae *Staphylococcus agnetis* |
| Sar-2-1 | Bacteria Firmicutes Bacilli Bacillales Staphylococcaceae *Staphylococcus argenteus* |
| Sau-79-1 to Sau-79-2 | Bacteria Firmicutes Bacilli Bacillales Staphylococcaceae *Staphylococcus aureus* |
| Sau-80-1 | Bacteria Firmicutes Bacilli Bacillales Staphylococcaceae *Staphylococcus aureus* 07-03450 |
| Sau-81-1 | Bacteria Firmicutes Bacilli Bacillales Staphylococcaceae *Staphylococcus aureus* 07-03451 |
| Sau-82-1 | Bacteria Firmicutes Bacilli Bacillales Staphylococcaceae *Staphylococcus aureus* 08-01059 |
| Sau-83-1 | Bacteria Firmicutes Bacilli Bacillales Staphylococcaceae *Staphylococcus aureus* 08-01062 |
| Sau-84-1 | Bacteria Firmicutes Bacilli Bacillales Staphylococcaceae *Staphylococcus aureus* 08-01084 |
| Sau-85-1 | Bacteria Firmicutes Bacilli Bacillales Staphylococcaceae *Staphylococcus aureus* 08-01085 |
| Sau-86-1 | Bacteria Firmicutes Bacilli Bacillales Staphylococcaceae *Staphylococcus aureus* 08-01229 |
| Sau-87-1 | Bacteria Firmicutes Bacilli Bacillales Staphylococcaceae *Staphylococcus aureus* 08-01728 |
| Sau-88-1 | Bacteria Firmicutes Bacilli Bacillales Staphylococcaceae *Staphylococcus aureus* 08-02906 |
| Sau-89-1 | Bacteria Firmicutes Bacilli Bacillales Staphylococcaceae *Staphylococcus aureus* 08134-6 |
| Sau-90-1 | Bacteria Firmicutes Bacilli Bacillales Staphylococcaceae *Staphylococcus aureus* 08139-6 |
| Sau-91-1 | Bacteria Firmicutes Bacilli Bacillales Staphylococcaceae *Staphylococcus aureus* 08142-8 |
| Sau-92-1 | Bacteria Firmicutes Bacilli Bacillales Staphylococcaceae *Staphylococcus aureus* 08143-5 |
| Sau-93-1 | Bacteria Firmicutes Bacilli Bacillales Staphylococcaceae *Staphylococcus aureus* 09-00736 |
| Sau-94-1 | Bacteria Firmicutes Bacilli Bacillales Staphylococcaceae *Staphylococcus aureus* 09S00475 |
| Sau-95-1 | Bacteria Firmicutes Bacilli Bacillales Staphylococcaceae *Staphylococcus aureus* 09S01694 |
| Sau-96-1 | Bacteria Firmicutes Bacilli Bacillales Staphylococcaceae *Staphylococcus aureus* 1(04GN_04-02-52-07) |
| Sau-97-1 | Bacteria Firmicutes Bacilli Bacillales Staphylococcaceae *Staphylococcus aureus* 10S00488 |
| Sau-98-1 | Bacteria Firmicutes Bacilli Bacillales Staphylococcaceae *Staphylococcus aureus* 10S01493 |
| Sau-99-1 | Bacteria Firmicutes Bacilli Bacillales Staphylococcaceae *Staphylococcus aureus* 1101-1 2010 |
| Sau-100-1 | Bacteria Firmicutes Bacilli Bacillales Staphylococcaceae *Staphylococcus aureus* 1101-1 2011 |
| Sau-101-1 | Bacteria Firmicutes Bacilli Bacillales Staphylococcaceae *Staphylococcus aureus* 1101-2 2010 |
| Sau-102-1 | Bacteria Firmicutes Bacilli Bacillales Staphylococcaceae *Staphylococcus aureus* 1101-2 2011 |
| Sau-103-1 | Bacteria Firmicutes Bacilli Bacillales Staphylococcaceae *Staphylococcus aureus* 110802495 |
| Sau-104-1 | Bacteria Firmicutes Bacilli Bacillales Staphylococcaceae *Staphylococcus aureus* 1110601704 |
| Sau-105-1 | Bacteria Firmicutes Bacilli Bacillales Staphylococcaceae *Staphylococcus aureus* 1110601896 |
| Sau-106-1 | Bacteria Firmicutes Bacilli Bacillales Staphylococcaceae *Staphylococcus aureus* 1110700562 |
| Sau-107-1 | Bacteria Firmicutes Bacilli Bacillales Staphylococcaceae *Staphylococcus aureus* 1110700610 |
| Sau-108-1 | Bacteria Firmicutes Bacilli Bacillales Staphylococcaceae *Staphylococcus aureus* 1110701127 |
| Sau-109-1 | Bacteria Firmicutes Bacilli Bacillales Staphylococcaceae *Staphylococcus aureus* 1110802883 |
| Sau-110-1 | Bacteria Firmicutes Bacilli Bacillales Staphylococcaceae *Staphylococcus aureus* 1110803248 |
| Sau-111-1 | Bacteria Firmicutes Bacilli Bacillales Staphylococcaceae *Staphylococcus aureus* 1110807699 |
| Sau-112-1 | Bacteria Firmicutes Bacilli Bacillales Staphylococcaceae *Staphylococcus aureus* 1110904178 |
| Sau-113-1 | Bacteria Firmicutes Bacilli Bacillales Staphylococcaceae *Staphylococcus aureus* 1111000175 |
| Sau-114-1 | Bacteria Firmicutes Bacilli Bacillales Staphylococcaceae *Staphylococcus aureus* 1111000177 |
| Sau-115-1 | Bacteria Firmicutes Bacilli Bacillales Staphylococcaceae *Staphylococcus aureus* 1111001578 |
| Sau-116-1 | Bacteria Firmicutes Bacilli Bacillales Staphylococcaceae *Staphylococcus aureus* 1111100370 |
| Sau-117-1 | Bacteria Firmicutes Bacilli Bacillales Staphylococcaceae *Staphylococcus aureus* 1111101949 |
| Sau-118-1 | Bacteria Firmicutes Bacilli Bacillales Staphylococcaceae *Staphylococcus aureus* 1111102620 |
| Sau-119-1 | Bacteria Firmicutes Bacilli Bacillales Staphylococcaceae *Staphylococcus aureus* 1111200013 |
| Sau-120-1 | Bacteria Firmicutes Bacilli Bacillales Staphylococcaceae *Staphylococcus aureus* 1111203374 |
| Sau-121-1 | Bacteria Firmicutes Bacilli Bacillales Staphylococcaceae *Staphylococcus aureus* 1111205429 |
| Sau-122-1 | Bacteria Firmicutes Bacilli Bacillales Staphylococcaceae *Staphylococcus aureus* 1111206270 |

| | |
|---|---|
| Sau-123-1 | Bacteria Firmicutes Bacilli Bacillales Staphylococcaceae *Staphylococcus aureus* 11P4 |
| Sau-124-1 | Bacteria Firmicutes Bacilli Bacillales Staphylococcaceae *Staphylococcus aureus* 11P8 |
| Sau-125-1 | Bacteria Firmicutes Bacilli Bacillales Staphylococcaceae *Staphylococcus aureus* 11S00605 |
| Sau-126-1 | Bacteria Firmicutes Bacilli Bacillales Staphylococcaceae *Staphylococcus aureus* 11S00627 |
| Sau-127-1 | Bacteria Firmicutes Bacilli Bacillales Staphylococcaceae *Staphylococcus aureus* 11S01415 |
| Sau-128-1 | Bacteria Firmicutes Bacilli Bacillales Staphylococcaceae *Staphylococcus aureus* 11S01420 |
| Sau-129-1 | Bacteria Firmicutes Bacilli Bacillales Staphylococcaceae *Staphylococcus aureus* 11S01586 |
| Sau-130-1 | Bacteria Firmicutes Bacilli Bacillales Staphylococcaceae *Staphylococcus aureus* 12-03481 |
| Sau-131-1 | Bacteria Firmicutes Bacilli Bacillales Staphylococcaceae *Staphylococcus aureus* 12-ST01564 |
| Sau-132-1 | Bacteria Firmicutes Bacilli Bacillales Staphylococcaceae *Staphylococcus aureus* 12-ST01988 |
| Sau-133-1 | Bacteria Firmicutes Bacilli Bacillales Staphylococcaceae *Staphylococcus aureus* 122 |
| Sau-134-1 | Bacteria Firmicutes Bacilli Bacillales Staphylococcaceae *Staphylococcus aureus* 12464 |
| Sau-135-1 | Bacteria Firmicutes Bacilli Bacillales Staphylococcaceae *Staphylococcus aureus* 12S00040 |
| Sau-136-1 | Bacteria Firmicutes Bacilli Bacillales Staphylococcaceae *Staphylococcus aureus* 12S00579 |
| Sau-137-1 | Bacteria Firmicutes Bacilli Bacillales Staphylococcaceae *Staphylococcus aureus* 12S00881 |
| Sau-138-1 | Bacteria Firmicutes Bacilli Bacillales Staphylococcaceae *Staphylococcus aureus* 12S00972 |
| Sau-139-1 | Bacteria Firmicutes Bacilli Bacillales Staphylococcaceae *Staphylococcus aureus* 12S01153 |
| Sau-140-1 | Bacteria Firmicutes Bacilli Bacillales Staphylococcaceae *Staphylococcus aureus* 12S01399 |
| Sau-141-1 | Bacteria Firmicutes Bacilli Bacillales Staphylococcaceae *Staphylococcus aureus* 13-ST00049 |
| Sau-142-1 | Bacteria Firmicutes Bacilli Bacillales Staphylococcaceae *Staphylococcus aureus* 14(11MN_17-08-66-05) |
| Sau-143-1 | Bacteria Firmicutes Bacilli Bacillales Staphylococcaceae *Staphylococcus aureus* 1484-9 |
| Sau-144-1 | Bacteria Firmicutes Bacilli Bacillales Staphylococcaceae *Staphylococcus aureus* 150211/pool 1 |
| Sau-145-1 | Bacteria Firmicutes Bacilli Bacillales Staphylococcaceae *Staphylococcus aureus* 16(11S1S4-09) |
| Sau-146-1 | Bacteria Firmicutes Bacilli Bacillales Staphylococcaceae *Staphylococcus aureus* 1801-1 2010 |
| Sau-147-1 | Bacteria Firmicutes Bacilli Bacillales Staphylococcaceae *Staphylococcus aureus* 1801-1 2011 |
| Sau-148-1 | Bacteria Firmicutes Bacilli Bacillales Staphylococcaceae *Staphylococcus aureus* 1801-2 2010 |
| Sau-149-1 | Bacteria Firmicutes Bacilli Bacillales Staphylococcaceae *Staphylococcus aureus* 1801-2 2011 |
| Sau-150-1 | Bacteria Firmicutes Bacilli Bacillales Staphylococcaceae *Staphylococcus aureus* 18439-17 |
| Sau-151-1 | Bacteria Firmicutes Bacilli Bacillales Staphylococcaceae *Staphylococcus aureus* 18439-62 |
| Sau-152-1 | Bacteria Firmicutes Bacilli Bacillales Staphylococcaceae *Staphylococcus aureus* 18754-2 |
| Sau-153-1 | Bacteria Firmicutes Bacilli Bacillales Staphylococcaceae *Staphylococcus aureus* 19571 |
| Sau-154-1 | Bacteria Firmicutes Bacilli Bacillales Staphylococcaceae *Staphylococcus aureus* 2(04HN_17-03-52-05) |
| Sau-155-1 | Bacteria Firmicutes Bacilli Bacillales Staphylococcaceae *Staphylococcus aureus* 2009-60-561-1 |
| Sau-156-1 | Bacteria Firmicutes Bacilli Bacillales Staphylococcaceae *Staphylococcus aureus* 2010-60-1240-1 |
| Sau-157-1 | Bacteria Firmicutes Bacilli Bacillales Staphylococcaceae *Staphylococcus aureus* 2010-60-6063-29 |
| Sau-158-1 | Bacteria Firmicutes Bacilli Bacillales Staphylococcaceae *Staphylococcus aureus* 2010-60-6063-39 |
| Sau-159-1 | Bacteria Firmicutes Bacilli Bacillales Staphylococcaceae *Staphylococcus aureus* 2010-60-6511-10 |
| Sau-160-1 | Bacteria Firmicutes Bacilli Bacillales Staphylococcaceae *Staphylococcus aureus* 2010-60-6511-39 |
| Sau-161-1 | Bacteria Firmicutes Bacilli Bacillales Staphylococcaceae *Staphylococcus aureus* 2010-60-6511-5 |
| Sau-162-1 | Bacteria Firmicutes Bacilli Bacillales Staphylococcaceae *Staphylococcus aureus* 2010-60-7626-19 |
| Sau-163-1 | Bacteria Firmicutes Bacilli Bacillales Staphylococcaceae *Staphylococcus aureus* 2011-60-1490-31 |
| Sau-164-1 | Bacteria Firmicutes Bacilli Bacillales Staphylococcaceae *Staphylococcus aureus* 2011-60-2078-5 |
| Sau-165-1 | Bacteria Firmicutes Bacilli Bacillales Staphylococcaceae *Staphylococcus aureus* 2011-60-2256-5 |
| Sau-166-1 | Bacteria Firmicutes Bacilli Bacillales Staphylococcaceae *Staphylococcus aureus* 2011-60-2275-1 |
| Sau-167-1 | Bacteria Firmicutes Bacilli Bacillales Staphylococcaceae *Staphylococcus aureus* 2011-60-2275-7 |
| Sau-168-1 | Bacteria Firmicutes Bacilli Bacillales Staphylococcaceae *Staphylococcus aureus* 22(2K81-5) |
| Sau-169-1 | Bacteria Firmicutes Bacilli Bacillales Staphylococcaceae *Staphylococcus aureus* 22825 |
| Sau-170-1 | Bacteria Firmicutes Bacilli Bacillales Staphylococcaceae *Staphylococcus aureus* 22835 |
| Sau-171-1 | Bacteria Firmicutes Bacilli Bacillales Staphylococcaceae *Staphylococcus aureus* 22837 |
| Sau-172-1 | Bacteria Firmicutes Bacilli Bacillales Staphylococcaceae *Staphylococcus aureus* 22838 |
| Sau-173-1 | Bacteria Firmicutes Bacilli Bacillales Staphylococcaceae *Staphylococcus aureus* 22841 |
| Sau-174-1 | Bacteria Firmicutes Bacilli Bacillales Staphylococcaceae *Staphylococcus aureus* 22843 |
| Sau-175-1 | Bacteria Firmicutes Bacilli Bacillales Staphylococcaceae *Staphylococcus aureus* 22846 |
| Sau-176-1 | Bacteria Firmicutes Bacilli Bacillales Staphylococcaceae *Staphylococcus aureus* 22P79 |
| Sau-177-1 | Bacteria Firmicutes Bacilli Bacillales Staphylococcaceae *Staphylococcus aureus* 23237 |
| Sau-178-1 | Bacteria Firmicutes Bacilli Bacillales Staphylococcaceae *Staphylococcus aureus* 2393-15 |
| Sau-179-1 | Bacteria Firmicutes Bacilli Bacillales Staphylococcaceae *Staphylococcus aureus* 2393-19 |
| Sau-180-1 | Bacteria Firmicutes Bacilli Bacillales Staphylococcaceae *Staphylococcus aureus* 25(2889) |
| Sau-181-1 | Bacteria Firmicutes Bacilli Bacillales Staphylococcaceae *Staphylococcus aureus* 27999-3 |
| Sau-182-1 | Bacteria Firmicutes Bacilli Bacillales Staphylococcaceae *Staphylococcus aureus* 28(18S1K13-24-05) |
| Sau-183-1 | Bacteria Firmicutes Bacilli Bacillales Staphylococcaceae *Staphylococcus aureus* 28(3K2-5) |
| Sau-184-1 | Bacteria Firmicutes Bacilli Bacillales Staphylococcaceae *Staphylococcus aureus* 36P1 |
| Sau-185-1 | Bacteria Firmicutes Bacilli Bacillales Staphylococcaceae *Staphylococcus aureus* 36P5 |
| Sau-186-1 | Bacteria Firmicutes Bacilli Bacillales Staphylococcaceae *Staphylococcus aureus* 37(18S2S5-05) |
| Sau-187-1 | Bacteria Firmicutes Bacilli Bacillales Staphylococcaceae *Staphylococcus aureus* 40P10 |

| | |
|---|---|
| Sau-188-1 | Bacteria Firmicutes Bacilli Bacillales Staphylococcaceae *Staphylococcus aureus* 40P5 1_1 |
| Sau-189-1 | Bacteria Firmicutes Bacilli Bacillales Staphylococcaceae *Staphylococcus aureus* 43P3 |
| Sau-190-1 | Bacteria Firmicutes Bacilli Bacillales Staphylococcaceae *Staphylococcus aureus* 43P8 |
| Sau-191-1 | Bacteria Firmicutes Bacilli Bacillales Staphylococcaceae *Staphylococcus aureus* 44(2608) |
| Sau-192-1 | Bacteria Firmicutes Bacilli Bacillales Staphylococcaceae *Staphylococcus aureus* 45(2607) |
| Sau-193-1 | Bacteria Firmicutes Bacilli Bacillales Staphylococcaceae *Staphylococcus aureus* 47P5 |
| Sau-194-1 | Bacteria Firmicutes Bacilli Bacillales Staphylococcaceae *Staphylococcus aureus* 47P9 |
| Sau-195-1 | Bacteria Firmicutes Bacilli Bacillales Staphylococcaceae *Staphylococcus aureus* 53180-1 |
| Sau-196-1 | Bacteria Firmicutes Bacilli Bacillales Staphylococcaceae *Staphylococcus aureus* 5670-1 |
| Sau-197-1 | Bacteria Firmicutes Bacilli Bacillales Staphylococcaceae *Staphylococcus aureus* 56824-10 |
| Sau-198-1 | Bacteria Firmicutes Bacilli Bacillales Staphylococcaceae *Staphylococcus aureus* 56824-15 |
| Sau-199-1 | Bacteria Firmicutes Bacilli Bacillales Staphylococcaceae *Staphylococcus aureus* 56824-2 |
| Sau-200-1 | Bacteria Firmicutes Bacilli Bacillales Staphylococcaceae *Staphylococcus aureus* 56824-21 |
| Sau-201-1 | Bacteria Firmicutes Bacilli Bacillales Staphylococcaceae *Staphylococcus aureus* 56824-5 |
| Sau-202-1 | Bacteria Firmicutes Bacilli Bacillales Staphylococcaceae *Staphylococcus aureus* 56824-7 |
| Sau-203-1 | Bacteria Firmicutes Bacilli Bacillales Staphylococcaceae *Staphylococcus aureus* 56864-11 |
| Sau-204-1 | Bacteria Firmicutes Bacilli Bacillales Staphylococcaceae *Staphylococcus aureus* 63-D10 |
| Sau-205-1 | Bacteria Firmicutes Bacilli Bacillales Staphylococcaceae *Staphylococcus aureus* 6665-5 |
| Sau-206-1 | Bacteria Firmicutes Bacilli Bacillales Staphylococcaceae *Staphylococcus aureus* 7-B-1 |
| Sau-207-1 | Bacteria Firmicutes Bacilli Bacillales Staphylococcaceae *Staphylococcus aureus* 75495-3 |
| Sau-208-1 | Bacteria Firmicutes Bacilli Bacillales Staphylococcaceae *Staphylococcus aureus* 76669-6 |
| Sau-209-1 | Bacteria Firmicutes Bacilli Bacillales Staphylococcaceae *Staphylococcus aureus* 81070 |
| Sau-210-1 | Bacteria Firmicutes Bacilli Bacillales Staphylococcaceae *Staphylococcus aureus* 81629 |
| Sau-211-1 | Bacteria Firmicutes Bacilli Bacillales Staphylococcaceae *Staphylococcus aureus* 84069-2 |
| Sau-212-1 | Bacteria Firmicutes Bacilli Bacillales Staphylococcaceae *Staphylococcus aureus* 86770-7 |
| Sau-213-1 | Bacteria Firmicutes Bacilli Bacillales Staphylococcaceae *Staphylococcus aureus* 87807-1 |
| Sau-214-1 | Bacteria Firmicutes Bacilli Bacillales Staphylococcaceae *Staphylococcus aureus* 87807-11 |
| Sau-215-1 | Bacteria Firmicutes Bacilli Bacillales Staphylococcaceae *Staphylococcus aureus* 87807-12 |
| Sau-216-1 | Bacteria Firmicutes Bacilli Bacillales Staphylococcaceae *Staphylococcus aureus* 87807-16 |
| Sau-217-1 | Bacteria Firmicutes Bacilli Bacillales Staphylococcaceae *Staphylococcus aureus* 87807-9 |
| Sau-218-1 | Bacteria Firmicutes Bacilli Bacillales Staphylococcaceae *Staphylococcus aureus* 880 |
| Sau-219-1 | Bacteria Firmicutes Bacilli Bacillales Staphylococcaceae *Staphylococcus aureus* 88088-1 |
| Sau-220-1 | Bacteria Firmicutes Bacilli Bacillales Staphylococcaceae *Staphylococcus aureus* 88088-2 |
| Sau-221-1 | Bacteria Firmicutes Bacilli Bacillales Staphylococcaceae *Staphylococcus aureus* 917 |
| Sau-222-1 | Bacteria Firmicutes Bacilli Bacillales Staphylococcaceae *Staphylococcus aureus* 9P9 |
| Sau-223-1 | Bacteria Firmicutes Bacilli Bacillales Staphylococcaceae *Staphylococcus aureus* AGEN6009 |
| Sau-224-1 | Bacteria Firmicutes Bacilli Bacillales Staphylococcaceae *Staphylococcus aureus* AGEN6010 |
| Sau-225-1 | Bacteria Firmicutes Bacilli Bacillales Staphylococcaceae *Staphylococcus aureus* AGEN6033 |
| Sau-226-1 | Bacteria Firmicutes Bacilli Bacillales Staphylococcaceae *Staphylococcus aureus* AGEN6042 |
| Sau-227-1 | Bacteria Firmicutes Bacilli Bacillales Staphylococcaceae *Staphylococcus aureus* AMMC6002 |
| Sau-228-1 | Bacteria Firmicutes Bacilli Bacillales Staphylococcaceae *Staphylococcus aureus* AMMC6003 |
| Sau-229-1 | Bacteria Firmicutes Bacilli Bacillales Staphylococcaceae *Staphylococcus aureus* AMMC6007 |
| Sau-230-1 | Bacteria Firmicutes Bacilli Bacillales Staphylococcaceae *Staphylococcus aureus* AMMC6008 |
| Sau-231-1 | Bacteria Firmicutes Bacilli Bacillales Staphylococcaceae *Staphylococcus aureus* AMMC6011 |
| Sau-232-1 | Bacteria Firmicutes Bacilli Bacillales Staphylococcaceae *Staphylococcus aureus* AMMC6012 |
| Sau-233-1 | Bacteria Firmicutes Bacilli Bacillales Staphylococcaceae *Staphylococcus aureus* AMMC6013 |
| Sau-234-1 | Bacteria Firmicutes Bacilli Bacillales Staphylococcaceae *Staphylococcus aureus* AMMC6014 |
| Sau-235-1 | Bacteria Firmicutes Bacilli Bacillales Staphylococcaceae *Staphylococcus aureus* AMMC6015 |
| Sau-236-1 | Bacteria Firmicutes Bacilli Bacillales Staphylococcaceae *Staphylococcus aureus* AMMC6040 |
| Sau-237-1 | Bacteria Firmicutes Bacilli Bacillales Staphylococcaceae *Staphylococcus aureus* AMMC6050 |
| Sau-238-1 | Bacteria Firmicutes Bacilli Bacillales Staphylococcaceae *Staphylococcus aureus* AMMC6051 |
| Sau-239-1 | Bacteria Firmicutes Bacilli Bacillales Staphylococcaceae *Staphylococcus aureus* AMMC6054 |
| Sau-240-1 | Bacteria Firmicutes Bacilli Bacillales Staphylococcaceae *Staphylococcus aureus* AMMC6068 |
| Sau-241-1 | Bacteria Firmicutes Bacilli Bacillales Staphylococcaceae *Staphylococcus aureus* AMMC6070 |
| Sau-242-1 | Bacteria Firmicutes Bacilli Bacillales Staphylococcaceae *Staphylococcus aureus* AMMC6071 |
| Sau-243-1 | Bacteria Firmicutes Bacilli Bacillales Staphylococcaceae *Staphylococcus aureus* AMMC6072 |
| Sau-244-1 | Bacteria Firmicutes Bacilli Bacillales Staphylococcaceae *Staphylococcus aureus* AMMC6073 |
| Sau-245-1 | Bacteria Firmicutes Bacilli Bacillales Staphylococcaceae *Staphylococcus aureus* AMMC6074 |
| Sau-246-1 | Bacteria Firmicutes Bacilli Bacillales Staphylococcaceae *Staphylococcus aureus* AMMC6075 |
| Sau-247-1 | Bacteria Firmicutes Bacilli Bacillales Staphylococcaceae *Staphylococcus aureus* AMMC6106 |
| Sau-248-1 | Bacteria Firmicutes Bacilli Bacillales Staphylococcaceae *Staphylococcus aureus* AMMC6111 |
| Sau-249-1 | Bacteria Firmicutes Bacilli Bacillales Staphylococcaceae *Staphylococcus aureus* AMMC6114 |
| Sau-250-1 | Bacteria Firmicutes Bacilli Bacillales Staphylococcaceae *Staphylococcus aureus* AMMC6119 |
| Sau-251-1 | Bacteria Firmicutes Bacilli Bacillales Staphylococcaceae *Staphylococcus aureus* BG407 |
| Sau-252-1 | Bacteria Firmicutes Bacilli Bacillales Staphylococcaceae *Staphylococcus aureus* C0626 |

| | |
|---|---|
| Sau-253-1 | Bacteria Firmicutes Bacilli Bacillales Staphylococcaceae *Staphylococcus aureus* C0630 |
| Sau-254-1 | Bacteria Firmicutes Bacilli Bacillales Staphylococcaceae *Staphylococcus aureus* C0637 |
| Sau-255-1 | Bacteria Firmicutes Bacilli Bacillales Staphylococcaceae *Staphylococcus aureus* C0673 |
| Sau-256-1 | Bacteria Firmicutes Bacilli Bacillales Staphylococcaceae *Staphylococcus aureus* C0676 |
| Sau-257-1 | Bacteria Firmicutes Bacilli Bacillales Staphylococcaceae *Staphylococcus aureus* C1655 |
| Sau-258-1 | Bacteria Firmicutes Bacilli Bacillales Staphylococcaceae *Staphylococcus aureus* C1673 |
| Sau-259-1 | Bacteria Firmicutes Bacilli Bacillales Staphylococcaceae *Staphylococcus aureus* C1842 |
| Sau-260-1 | Bacteria Firmicutes Bacilli Bacillales Staphylococcaceae *Staphylococcus aureus* C1891 |
| Sau-261-1 | Bacteria Firmicutes Bacilli Bacillales Staphylococcaceae *Staphylococcus aureus* C1894 |
| Sau-262-1 | Bacteria Firmicutes Bacilli Bacillales Staphylococcaceae *Staphylococcus aureus* C2549 |
| Sau-263-1 | Bacteria Firmicutes Bacilli Bacillales Staphylococcaceae *Staphylococcus aureus* C2679 |
| Sau-264-1 | Bacteria Firmicutes Bacilli Bacillales Staphylococcaceae *Staphylococcus aureus* C2706 |
| Sau-265-1 | Bacteria Firmicutes Bacilli Bacillales Staphylococcaceae *Staphylococcus aureus* C2928 |
| Sau-266-1 | Bacteria Firmicutes Bacilli Bacillales Staphylococcaceae *Staphylococcus aureus* C2930 |
| Sau-267-1 | Bacteria Firmicutes Bacilli Bacillales Staphylococcaceae *Staphylococcus aureus* C2942 |
| Sau-268-1 | Bacteria Firmicutes Bacilli Bacillales Staphylococcaceae *Staphylococcus aureus* C3452 |
| Sau-269-1 | Bacteria Firmicutes Bacilli Bacillales Staphylococcaceae *Staphylococcus aureus* C3489 |
| Sau-270-1 | Bacteria Firmicutes Bacilli Bacillales Staphylococcaceae *Staphylococcus aureus* C3672 |
| Sau-271-1 | Bacteria Firmicutes Bacilli Bacillales Staphylococcaceae *Staphylococcus aureus* C3865 |
| Sau-272-1 | Bacteria Firmicutes Bacilli Bacillales Staphylococcaceae *Staphylococcus aureus* C3965 |
| Sau-273-1 | Bacteria Firmicutes Bacilli Bacillales Staphylococcaceae *Staphylococcus aureus* C4151 |
| Sau-274-1 | Bacteria Firmicutes Bacilli Bacillales Staphylococcaceae *Staphylococcus aureus* C4155 |
| Sau-275-1 | Bacteria Firmicutes Bacilli Bacillales Staphylococcaceae *Staphylococcus aureus* C4668 |
| Sau-276-1 | Bacteria Firmicutes Bacilli Bacillales Staphylococcaceae *Staphylococcus aureus* C5086 |
| Sau-277-1 | Bacteria Firmicutes Bacilli Bacillales Staphylococcaceae *Staphylococcus aureus* C5453 |
| Sau-278-1 | Bacteria Firmicutes Bacilli Bacillales Staphylococcaceae *Staphylococcus aureus* CHAP6022 |
| Sau-279-1 | Bacteria Firmicutes Bacilli Bacillales Staphylococcaceae *Staphylococcus aureus* CHAP6035 |
| Sau-280-1 | Bacteria Firmicutes Bacilli Bacillales Staphylococcaceae *Staphylococcus aureus* CHAP6049 |
| Sau-281-1 | Bacteria Firmicutes Bacilli Bacillales Staphylococcaceae *Staphylococcus aureus* CHAP6102 |
| Sau-307-1 | Bacteria Firmicutes Bacilli Bacillales Staphylococcaceae *Staphylococcus aureus* Chi-10 |
| Sau-308-1 | Bacteria Firmicutes Bacilli Bacillales Staphylococcaceae *Staphylococcus aureus* Chi-4 |
| Sau-309-1 | Bacteria Firmicutes Bacilli Bacillales Staphylococcaceae *Staphylococcus aureus* Chi-8 |
| Sau-282-1 | Bacteria Firmicutes Bacilli Bacillales Staphylococcaceae *Staphylococcus aureus* CHOC6045 |
| Sau-283-1 | Bacteria Firmicutes Bacilli Bacillales Staphylococcaceae *Staphylococcus aureus* CHOC6056 |
| Sau-284-1 | Bacteria Firmicutes Bacilli Bacillales Staphylococcaceae *Staphylococcus aureus* COAS6001 |
| Sau-285-1 | Bacteria Firmicutes Bacilli Bacillales Staphylococcaceae *Staphylococcus aureus* COAS6004 |
| Sau-286-1 | Bacteria Firmicutes Bacilli Bacillales Staphylococcaceae *Staphylococcus aureus* COAS6020 |
| Sau-287-1 | Bacteria Firmicutes Bacilli Bacillales Staphylococcaceae *Staphylococcus aureus* COAS6025 |
| Sau-288-1 | Bacteria Firmicutes Bacilli Bacillales Staphylococcaceae *Staphylococcus aureus* COAS6038 |
| Sau-289-1 | Bacteria Firmicutes Bacilli Bacillales Staphylococcaceae *Staphylococcus aureus* COAS6039 |
| Sau-290-1 | Bacteria Firmicutes Bacilli Bacillales Staphylococcaceae *Staphylococcus aureus* COAS6044 |
| Sau-291-1 | Bacteria Firmicutes Bacilli Bacillales Staphylococcaceae *Staphylococcus aureus* COAS6049 |
| Sau-292-1 | Bacteria Firmicutes Bacilli Bacillales Staphylococcaceae *Staphylococcus aureus* COAS6052 |
| Sau-293-1 | Bacteria Firmicutes Bacilli Bacillales Staphylococcaceae *Staphylococcus aureus* COAS6053 |
| Sau-294-1 | Bacteria Firmicutes Bacilli Bacillales Staphylococcaceae *Staphylococcus aureus* COAS6055 |
| Sau-295-1 | Bacteria Firmicutes Bacilli Bacillales Staphylococcaceae *Staphylococcus aureus* COAS6059 |
| Sau-296-1 | Bacteria Firmicutes Bacilli Bacillales Staphylococcaceae *Staphylococcus aureus* COAS6060 |
| Sau-297-1 | Bacteria Firmicutes Bacilli Bacillales Staphylococcaceae *Staphylococcus aureus* COAS6087 |
| Sau-298-1 | Bacteria Firmicutes Bacilli Bacillales Staphylococcaceae *Staphylococcus aureus* COAS6095 |
| Sau-299-1 | Bacteria Firmicutes Bacilli Bacillales Staphylococcaceae *Staphylococcus aureus* COAS6112 |
| Sau-300-1 | Bacteria Firmicutes Bacilli Bacillales Staphylococcaceae *Staphylococcus aureus* COAS6114 |
| Sau-301-1 | Bacteria Firmicutes Bacilli Bacillales Staphylococcaceae *Staphylococcus aureus* COAS6117 |
| Sau-302-1 | Bacteria Firmicutes Bacilli Bacillales Staphylococcaceae *Staphylococcus aureus* COLL6021 |
| Sau-303-1 | Bacteria Firmicutes Bacilli Bacillales Staphylococcaceae *Staphylococcus aureus* CUHK_BJ2002 |
| Sau-304-1 | Bacteria Firmicutes Bacilli Bacillales Staphylococcaceae *Staphylococcus aureus* CUHK_BJ2007 |
| Sau-305-1 | Bacteria Firmicutes Bacilli Bacillales Staphylococcaceae *Staphylococcus aureus* CUHK_HK1997 |
| Sau-306-1 | Bacteria Firmicutes Bacilli Bacillales Staphylococcaceae *Staphylococcus aureus* CUHK_HK2007 |
| Sau-310-1 | Bacteria Firmicutes Bacilli Bacillales Staphylococcaceae *Staphylococcus aureus* DAR104 |
| Sau-311-1 | Bacteria Firmicutes Bacilli Bacillales Staphylococcaceae *Staphylococcus aureus* DAR1065 |
| Sau-312-1 | Bacteria Firmicutes Bacilli Bacillales Staphylococcaceae *Staphylococcus aureus* DAR1067 |
| Sau-313-1 | Bacteria Firmicutes Bacilli Bacillales Staphylococcaceae *Staphylococcus aureus* DAR1082 |
| Sau-314-1 | Bacteria Firmicutes Bacilli Bacillales Staphylococcaceae *Staphylococcus aureus* DAR1083 |
| Sau-315-1 | Bacteria Firmicutes Bacilli Bacillales Staphylococcaceae *Staphylococcus aureus* DAR1084 |
| Sau-316-1 | Bacteria Firmicutes Bacilli Bacillales Staphylococcaceae *Staphylococcus aureus* DAR1086 |
| Sau-317-1 | Bacteria Firmicutes Bacilli Bacillales Staphylococcaceae *Staphylococcus aureus* DAR1090 |

| | |
|---|---|
| Sau-318-1 | Bacteria Firmicutes Bacilli Bacillales Staphylococcaceae *Staphylococcus aureus* DAR1157 |
| Sau-319-1 | Bacteria Firmicutes Bacilli Bacillales Staphylococcaceae *Staphylococcus aureus* DAR1158 |
| Sau-320-1 | Bacteria Firmicutes Bacilli Bacillales Staphylococcaceae *Staphylococcus aureus* DAR116 |
| Sau-321-1 | Bacteria Firmicutes Bacilli Bacillales Staphylococcaceae *Staphylococcus aureus* DAR1161 |
| Sau-322-1 | Bacteria Firmicutes Bacilli Bacillales Staphylococcaceae *Staphylococcus aureus* DAR1174 |
| Sau-323-1 | Bacteria Firmicutes Bacilli Bacillales Staphylococcaceae *Staphylococcus aureus* DAR1176 |
| Sau-324-1 | Bacteria Firmicutes Bacilli Bacillales Staphylococcaceae *Staphylococcus aureus* DAR1183 |
| Sau-325-1 | Bacteria Firmicutes Bacilli Bacillales Staphylococcaceae *Staphylococcus aureus* DAR1199 |
| Sau-326-1 | Bacteria Firmicutes Bacilli Bacillales Staphylococcaceae *Staphylococcus aureus* DAR1200 |
| Sau-327-1 | Bacteria Firmicutes Bacilli Bacillales Staphylococcaceae *Staphylococcus aureus* DAR126 |
| Sau-328-1 | Bacteria Firmicutes Bacilli Bacillales Staphylococcaceae *Staphylococcus aureus* DAR1275 |
| Sau-329-1 | Bacteria Firmicutes Bacilli Bacillales Staphylococcaceae *Staphylococcus aureus* DAR1276 |
| Sau-330-1 | Bacteria Firmicutes Bacilli Bacillales Staphylococcaceae *Staphylococcus aureus* DAR1278 |
| Sau-331-1 | Bacteria Firmicutes Bacilli Bacillales Staphylococcaceae *Staphylococcus aureus* DAR1280 |
| Sau-332-1 | Bacteria Firmicutes Bacilli Bacillales Staphylococcaceae *Staphylococcus aureus* DAR1291 |
| Sau-333-1 | Bacteria Firmicutes Bacilli Bacillales Staphylococcaceae *Staphylococcus aureus* DAR1293 |
| Sau-334-1 | Bacteria Firmicutes Bacilli Bacillales Staphylococcaceae *Staphylococcus aureus* DAR1295 |
| Sau-335-1 | Bacteria Firmicutes Bacilli Bacillales Staphylococcaceae *Staphylococcus aureus* DAR1310 |
| Sau-336-1 | Bacteria Firmicutes Bacilli Bacillales Staphylococcaceae *Staphylococcus aureus* DAR1311 |
| Sau-337-1 | Bacteria Firmicutes Bacilli Bacillales Staphylococcaceae *Staphylococcus aureus* DAR1313 |
| Sau-338-1 | Bacteria Firmicutes Bacilli Bacillales Staphylococcaceae *Staphylococcus aureus* DAR133 |
| Sau-339-1 | Bacteria Firmicutes Bacilli Bacillales Staphylococcaceae *Staphylococcus aureus* DAR1366 |
| Sau-340-1 | Bacteria Firmicutes Bacilli Bacillales Staphylococcaceae *Staphylococcus aureus* DAR1369 |
| Sau-341-1 | Bacteria Firmicutes Bacilli Bacillales Staphylococcaceae *Staphylococcus aureus* DAR1370 |
| Sau-342-1 | Bacteria Firmicutes Bacilli Bacillales Staphylococcaceae *Staphylococcus aureus* DAR1490 |
| Sau-343-1 | Bacteria Firmicutes Bacilli Bacillales Staphylococcaceae *Staphylococcus aureus* DAR1492 |
| Sau-344-1 | Bacteria Firmicutes Bacilli Bacillales Staphylococcaceae *Staphylococcus aureus* DAR1503 |
| Sau-345-1 | Bacteria Firmicutes Bacilli Bacillales Staphylococcaceae *Staphylococcus aureus* DAR1507 |
| Sau-346-1 | Bacteria Firmicutes Bacilli Bacillales Staphylococcaceae *Staphylococcus aureus* DAR1692 |
| Sau-347-1 | Bacteria Firmicutes Bacilli Bacillales Staphylococcaceae *Staphylococcus aureus* DAR1698 |
| Sau-348-1 | Bacteria Firmicutes Bacilli Bacillales Staphylococcaceae *Staphylococcus aureus* DAR1800 |
| Sau-349-1 | Bacteria Firmicutes Bacilli Bacillales Staphylococcaceae *Staphylococcus aureus* DAR1810 |
| Sau-350-1 | Bacteria Firmicutes Bacilli Bacillales Staphylococcaceae *Staphylococcus aureus* DAR1811 |
| Sau-351-1 | Bacteria Firmicutes Bacilli Bacillales Staphylococcaceae *Staphylococcus aureus* DAR1813 |
| Sau-352-1 | Bacteria Firmicutes Bacilli Bacillales Staphylococcaceae *Staphylococcus aureus* DAR1814 |
| Sau-353-1 | Bacteria Firmicutes Bacilli Bacillales Staphylococcaceae *Staphylococcus aureus* DAR1815 |
| Sau-354-1 | Bacteria Firmicutes Bacilli Bacillales Staphylococcaceae *Staphylococcus aureus* DAR1842 |
| Sau-355-1 | Bacteria Firmicutes Bacilli Bacillales Staphylococcaceae *Staphylococcus aureus* DAR1859 |
| Sau-356-1 | Bacteria Firmicutes Bacilli Bacillales Staphylococcaceae *Staphylococcus aureus* DAR1862 |
| Sau-357-1 | Bacteria Firmicutes Bacilli Bacillales Staphylococcaceae *Staphylococcus aureus* DAR1874 |
| Sau-358-1 | Bacteria Firmicutes Bacilli Bacillales Staphylococcaceae *Staphylococcus aureus* DAR1883 |
| Sau-359-1 | Bacteria Firmicutes Bacilli Bacillales Staphylococcaceae *Staphylococcus aureus* DAR1885 |
| Sau-360-1 | Bacteria Firmicutes Bacilli Bacillales Staphylococcaceae *Staphylococcus aureus* DAR1890 |
| Sau-361-1 | Bacteria Firmicutes Bacilli Bacillales Staphylococcaceae *Staphylococcus aureus* DAR1926 |
| Sau-362-1 | Bacteria Firmicutes Bacilli Bacillales Staphylococcaceae *Staphylococcus aureus* DAR1936 |
| Sau-363-1 | Bacteria Firmicutes Bacilli Bacillales Staphylococcaceae *Staphylococcus aureus* DAR1937 |
| Sau-364-1 | Bacteria Firmicutes Bacilli Bacillales Staphylococcaceae *Staphylococcus aureus* DAR1938 |
| Sau-365-1 | Bacteria Firmicutes Bacilli Bacillales Staphylococcaceae *Staphylococcus aureus* DAR1939 |
| Sau-366-1 | Bacteria Firmicutes Bacilli Bacillales Staphylococcaceae *Staphylococcus aureus* DAR1941 |
| Sau-367-1 | Bacteria Firmicutes Bacilli Bacillales Staphylococcaceae *Staphylococcus aureus* DAR1942 |
| Sau-368-1 | Bacteria Firmicutes Bacilli Bacillales Staphylococcaceae *Staphylococcus aureus* DAR1954 |
| Sau-369-1 | Bacteria Firmicutes Bacilli Bacillales Staphylococcaceae *Staphylococcus aureus* DAR1955 |
| Sau-370-1 | Bacteria Firmicutes Bacilli Bacillales Staphylococcaceae *Staphylococcus aureus* DAR1956 |
| Sau-371-1 | Bacteria Firmicutes Bacilli Bacillales Staphylococcaceae *Staphylococcus aureus* DAR1961 |
| Sau-372-1 | Bacteria Firmicutes Bacilli Bacillales Staphylococcaceae *Staphylococcus aureus* DAR1980 |
| Sau-373-1 | Bacteria Firmicutes Bacilli Bacillales Staphylococcaceae *Staphylococcus aureus* DAR1995 |
| Sau-374-1 | Bacteria Firmicutes Bacilli Bacillales Staphylococcaceae *Staphylococcus aureus* DAR2000 |
| Sau-375-1 | Bacteria Firmicutes Bacilli Bacillales Staphylococcaceae *Staphylococcus aureus* DAR2002 |
| Sau-376-1 | Bacteria Firmicutes Bacilli Bacillales Staphylococcaceae *Staphylococcus aureus* DAR2018 |
| Sau-377-1 | Bacteria Firmicutes Bacilli Bacillales Staphylococcaceae *Staphylococcus aureus* DAR2019 |
| Sau-378-1 | Bacteria Firmicutes Bacilli Bacillales Staphylococcaceae *Staphylococcus aureus* DAR2026 |
| Sau-379-1 | Bacteria Firmicutes Bacilli Bacillales Staphylococcaceae *Staphylococcus aureus* DAR2067 |
| Sau-380-1 | Bacteria Firmicutes Bacilli Bacillales Staphylococcaceae *Staphylococcus aureus* DAR2068 |
| Sau-381-1 | Bacteria Firmicutes Bacilli Bacillales Staphylococcaceae *Staphylococcus aureus* DAR2070 |
| Sau-382-1 | Bacteria Firmicutes Bacilli Bacillales Staphylococcaceae *Staphylococcus aureus* DAR2072 |

| | |
|---|---|
| Sau-383-1 | Bacteria Firmicutes Bacilli Bacillales Staphylococcaceae *Staphylococcus aureus* DAR2075 |
| Sau-384-1 | Bacteria Firmicutes Bacilli Bacillales Staphylococcaceae *Staphylococcus aureus* DAR22 |
| Sau-385-1 | Bacteria Firmicutes Bacilli Bacillales Staphylococcaceae *Staphylococcus aureus* DAR26 |
| Sau-386-1 | Bacteria Firmicutes Bacilli Bacillales Staphylococcaceae *Staphylococcus aureus* DAR3143 |
| Sau-387-1 | Bacteria Firmicutes Bacilli Bacillales Staphylococcaceae *Staphylococcus aureus* DAR3144 |
| Sau-388-1 | Bacteria Firmicutes Bacilli Bacillales Staphylococcaceae *Staphylococcus aureus* DAR3148 |
| Sau-389-1 | Bacteria Firmicutes Bacilli Bacillales Staphylococcaceae *Staphylococcus aureus* DAR3150 |
| Sau-390-1 | Bacteria Firmicutes Bacilli Bacillales Staphylococcaceae *Staphylococcus aureus* DAR3152 |
| Sau-391-1 | Bacteria Firmicutes Bacilli Bacillales Staphylococcaceae *Staphylococcus aureus* DAR3153 |
| Sau-392-1 | Bacteria Firmicutes Bacilli Bacillales Staphylococcaceae *Staphylococcus aureus* DAR3156 |
| Sau-393-1 | Bacteria Firmicutes Bacilli Bacillales Staphylococcaceae *Staphylococcus aureus* DAR3157 |
| Sau-394-1 | Bacteria Firmicutes Bacilli Bacillales Staphylococcaceae *Staphylococcus aureus* DAR3162 |
| Sau-395-1 | Bacteria Firmicutes Bacilli Bacillales Staphylococcaceae *Staphylococcus aureus* DAR3163 |
| Sau-396-1 | Bacteria Firmicutes Bacilli Bacillales Staphylococcaceae *Staphylococcus aureus* DAR3165 |
| Sau-397-1 | Bacteria Firmicutes Bacilli Bacillales Staphylococcaceae *Staphylococcus aureus* DAR3166 |
| Sau-398-1 | Bacteria Firmicutes Bacilli Bacillales Staphylococcaceae *Staphylococcus aureus* DAR3170 |
| Sau-399-1 | Bacteria Firmicutes Bacilli Bacillales Staphylococcaceae *Staphylococcus aureus* DAR3172 |
| Sau-400-1 | Bacteria Firmicutes Bacilli Bacillales Staphylococcaceae *Staphylococcus aureus* DAR3175 |
| Sau-401-1 | Bacteria Firmicutes Bacilli Bacillales Staphylococcaceae *Staphylococcus aureus* DAR3176 |
| Sau-402-1 | Bacteria Firmicutes Bacilli Bacillales Staphylococcaceae *Staphylococcus aureus* DAR3177 |
| Sau-403-1 | Bacteria Firmicutes Bacilli Bacillales Staphylococcaceae *Staphylococcus aureus* DAR3178 |
| Sau-404-1 | Bacteria Firmicutes Bacilli Bacillales Staphylococcaceae *Staphylococcus aureus* DAR3179 |
| Sau-405-1 | Bacteria Firmicutes Bacilli Bacillales Staphylococcaceae *Staphylococcus aureus* DAR3183 |
| Sau-406-1 | Bacteria Firmicutes Bacilli Bacillales Staphylococcaceae *Staphylococcus aureus* DAR3198 |
| Sau-407-1 | Bacteria Firmicutes Bacilli Bacillales Staphylococcaceae *Staphylococcus aureus* DAR3236 |
| Sau-408-1 | Bacteria Firmicutes Bacilli Bacillales Staphylococcaceae *Staphylococcus aureus* DAR3504 |
| Sau-409-1 | Bacteria Firmicutes Bacilli Bacillales Staphylococcaceae *Staphylococcus aureus* DAR3505 |
| Sau-410-1 | Bacteria Firmicutes Bacilli Bacillales Staphylococcaceae *Staphylococcus aureus* DAR3507 |
| Sau-411-1 | Bacteria Firmicutes Bacilli Bacillales Staphylococcaceae *Staphylococcus aureus* DAR3508 |
| Sau-412-1 | Bacteria Firmicutes Bacilli Bacillales Staphylococcaceae *Staphylococcus aureus* DAR3512 |
| Sau-413-1 | Bacteria Firmicutes Bacilli Bacillales Staphylococcaceae *Staphylococcus aureus* DAR3515 |
| Sau-414-1 | Bacteria Firmicutes Bacilli Bacillales Staphylococcaceae *Staphylococcus aureus* DAR3518 |
| Sau-415-1 | Bacteria Firmicutes Bacilli Bacillales Staphylococcaceae *Staphylococcus aureus* DAR3519 |
| Sau-416-1 | Bacteria Firmicutes Bacilli Bacillales Staphylococcaceae *Staphylococcus aureus* DAR3525 |
| Sau-417-1 | Bacteria Firmicutes Bacilli Bacillales Staphylococcaceae *Staphylococcus aureus* DAR3529 |
| Sau-418-1 | Bacteria Firmicutes Bacilli Bacillales Staphylococcaceae *Staphylococcus aureus* DAR3534 |
| Sau-419-1 | Bacteria Firmicutes Bacilli Bacillales Staphylococcaceae *Staphylococcus aureus* DAR3548 |
| Sau-420-1 | Bacteria Firmicutes Bacilli Bacillales Staphylococcaceae *Staphylococcus aureus* DAR3551 |
| Sau-421-1 | Bacteria Firmicutes Bacilli Bacillales Staphylococcaceae *Staphylococcus aureus* DAR3555 |
| Sau-422-1 | Bacteria Firmicutes Bacilli Bacillales Staphylococcaceae *Staphylococcus aureus* DAR3562 |
| Sau-423-1 | Bacteria Firmicutes Bacilli Bacillales Staphylococcaceae *Staphylococcus aureus* DAR3565 |
| Sau-424-1 | Bacteria Firmicutes Bacilli Bacillales Staphylococcaceae *Staphylococcus aureus* DAR3567 |
| Sau-425-1 | Bacteria Firmicutes Bacilli Bacillales Staphylococcaceae *Staphylococcus aureus* DAR3574 |
| Sau-426-1 | Bacteria Firmicutes Bacilli Bacillales Staphylococcaceae *Staphylococcus aureus* DAR3575 |
| Sau-427-1 | Bacteria Firmicutes Bacilli Bacillales Staphylococcaceae *Staphylococcus aureus* DAR3576 |
| Sau-428-1 | Bacteria Firmicutes Bacilli Bacillales Staphylococcaceae *Staphylococcus aureus* DAR3578 |
| Sau-429-1 | Bacteria Firmicutes Bacilli Bacillales Staphylococcaceae *Staphylococcus aureus* DAR3581 |
| Sau-430-1 | Bacteria Firmicutes Bacilli Bacillales Staphylococcaceae *Staphylococcus aureus* DAR3583 |
| Sau-431-1 | Bacteria Firmicutes Bacilli Bacillales Staphylococcaceae *Staphylococcus aureus* DAR3584 |
| Sau-432-1 | Bacteria Firmicutes Bacilli Bacillales Staphylococcaceae *Staphylococcus aureus* DAR3586 |
| Sau-433-1 | Bacteria Firmicutes Bacilli Bacillales Staphylococcaceae *Staphylococcus aureus* DAR3587 |
| Sau-434-1 | Bacteria Firmicutes Bacilli Bacillales Staphylococcaceae *Staphylococcus aureus* DAR3590 |
| Sau-435-1 | Bacteria Firmicutes Bacilli Bacillales Staphylococcaceae *Staphylococcus aureus* DAR3591 |
| Sau-436-1 | Bacteria Firmicutes Bacilli Bacillales Staphylococcaceae *Staphylococcus aureus* DAR3592 |
| Sau-437-1 | Bacteria Firmicutes Bacilli Bacillales Staphylococcaceae *Staphylococcus aureus* DAR3596 |
| Sau-438-1 | Bacteria Firmicutes Bacilli Bacillales Staphylococcaceae *Staphylococcus aureus* DAR3599 |
| Sau-439-1 | Bacteria Firmicutes Bacilli Bacillales Staphylococcaceae *Staphylococcus aureus* DAR3608 |
| Sau-440-1 | Bacteria Firmicutes Bacilli Bacillales Staphylococcaceae *Staphylococcus aureus* DAR3609 |
| Sau-441-1 | Bacteria Firmicutes Bacilli Bacillales Staphylococcaceae *Staphylococcus aureus* DAR3612 |
| Sau-442-1 | Bacteria Firmicutes Bacilli Bacillales Staphylococcaceae *Staphylococcus aureus* DAR3613 |
| Sau-443-1 | Bacteria Firmicutes Bacilli Bacillales Staphylococcaceae *Staphylococcus aureus* DAR3616 |
| Sau-444-1 | Bacteria Firmicutes Bacilli Bacillales Staphylococcaceae *Staphylococcus aureus* DAR3617 |
| Sau-445-1 | Bacteria Firmicutes Bacilli Bacillales Staphylococcaceae *Staphylococcus aureus* DAR3624 |
| Sau-446-1 | Bacteria Firmicutes Bacilli Bacillales Staphylococcaceae *Staphylococcus aureus* DAR3627 |
| Sau-447-1 | Bacteria Firmicutes Bacilli Bacillales Staphylococcaceae *Staphylococcus aureus* DAR3628 |

| | |
|---|---|
| Sau-448-1 | Bacteria Firmicutes Bacilli Bacillales Staphylococcaceae *Staphylococcus aureus* DAR3629 |
| Sau-449-1 | Bacteria Firmicutes Bacilli Bacillales Staphylococcaceae *Staphylococcus aureus* DAR3632 |
| Sau-450-1 | Bacteria Firmicutes Bacilli Bacillales Staphylococcaceae *Staphylococcus aureus* DAR3752 |
| Sau-451-1 | Bacteria Firmicutes Bacilli Bacillales Staphylococcaceae *Staphylococcus aureus* DAR3758 |
| Sau-452-1 | Bacteria Firmicutes Bacilli Bacillales Staphylococcaceae *Staphylococcus aureus* DAR3760 |
| Sau-453-1 | Bacteria Firmicutes Bacilli Bacillales Staphylococcaceae *Staphylococcus aureus* DAR3761 |
| Sau-454-1 | Bacteria Firmicutes Bacilli Bacillales Staphylococcaceae *Staphylococcus aureus* DAR3762 |
| Sau-455-1 | Bacteria Firmicutes Bacilli Bacillales Staphylococcaceae *Staphylococcus aureus* DAR3765 |
| Sau-456-1 | Bacteria Firmicutes Bacilli Bacillales Staphylococcaceae *Staphylococcus aureus* DAR3767 |
| Sau-457-1 | Bacteria Firmicutes Bacilli Bacillales Staphylococcaceae *Staphylococcus aureus* DAR3769 |
| Sau-458-1 | Bacteria Firmicutes Bacilli Bacillales Staphylococcaceae *Staphylococcus aureus* DAR3772 |
| Sau-459-1 | Bacteria Firmicutes Bacilli Bacillales Staphylococcaceae *Staphylococcus aureus* DAR3776 |
| Sau-460-1 | Bacteria Firmicutes Bacilli Bacillales Staphylococcaceae *Staphylococcus aureus* DAR3779 |
| Sau-461-1 | Bacteria Firmicutes Bacilli Bacillales Staphylococcaceae *Staphylococcus aureus* DAR3781 |
| Sau-462-1 | Bacteria Firmicutes Bacilli Bacillales Staphylococcaceae *Staphylococcus aureus* DAR3782 |
| Sau-463-1 | Bacteria Firmicutes Bacilli Bacillales Staphylococcaceae *Staphylococcus aureus* DAR3788 |
| Sau-464-1 | Bacteria Firmicutes Bacilli Bacillales Staphylococcaceae *Staphylococcus aureus* DAR3790 |
| Sau-465-1 | Bacteria Firmicutes Bacilli Bacillales Staphylococcaceae *Staphylococcus aureus* DAR3794 |
| Sau-466-1 | Bacteria Firmicutes Bacilli Bacillales Staphylococcaceae *Staphylococcus aureus* DAR3796 |
| Sau-467-1 | Bacteria Firmicutes Bacilli Bacillales Staphylococcaceae *Staphylococcus aureus* DAR3799 |
| Sau-468-1 | Bacteria Firmicutes Bacilli Bacillales Staphylococcaceae *Staphylococcus aureus* DAR3840 |
| Sau-469-1 | Bacteria Firmicutes Bacilli Bacillales Staphylococcaceae *Staphylococcus aureus* DAR3842 |
| Sau-470-1 | Bacteria Firmicutes Bacilli Bacillales Staphylococcaceae *Staphylococcus aureus* DAR3844 |
| Sau-471-1 | Bacteria Firmicutes Bacilli Bacillales Staphylococcaceae *Staphylococcus aureus* DAR3845 |
| Sau-472-1 | Bacteria Firmicutes Bacilli Bacillales Staphylococcaceae *Staphylococcus aureus* DAR3847 |
| Sau-473-1 | Bacteria Firmicutes Bacilli Bacillales Staphylococcaceae *Staphylococcus aureus* DAR3859 |
| Sau-474-1 | Bacteria Firmicutes Bacilli Bacillales Staphylococcaceae *Staphylococcus aureus* DAR3861 |
| Sau-475-1 | Bacteria Firmicutes Bacilli Bacillales Staphylococcaceae *Staphylococcus aureus* DAR3862 |
| Sau-476-1 | Bacteria Firmicutes Bacilli Bacillales Staphylococcaceae *Staphylococcus aureus* DAR3864 |
| Sau-477-1 | Bacteria Firmicutes Bacilli Bacillales Staphylococcaceae *Staphylococcus aureus* DAR3868 |
| Sau-478-1 | Bacteria Firmicutes Bacilli Bacillales Staphylococcaceae *Staphylococcus aureus* DAR3870 |
| Sau-479-1 | Bacteria Firmicutes Bacilli Bacillales Staphylococcaceae *Staphylococcus aureus* DAR3871 |
| Sau-480-1 | Bacteria Firmicutes Bacilli Bacillales Staphylococcaceae *Staphylococcus aureus* DAR3885 |
| Sau-481-1 | Bacteria Firmicutes Bacilli Bacillales Staphylococcaceae *Staphylococcus aureus* DAR3886 |
| Sau-482-1 | Bacteria Firmicutes Bacilli Bacillales Staphylococcaceae *Staphylococcus aureus* DAR3891 |
| Sau-483-1 | Bacteria Firmicutes Bacilli Bacillales Staphylococcaceae *Staphylococcus aureus* DAR3892 |
| Sau-484-1 | Bacteria Firmicutes Bacilli Bacillales Staphylococcaceae *Staphylococcus aureus* DAR3894 |
| Sau-485-1 | Bacteria Firmicutes Bacilli Bacillales Staphylococcaceae *Staphylococcus aureus* DAR3897 |
| Sau-486-1 | Bacteria Firmicutes Bacilli Bacillales Staphylococcaceae *Staphylococcus aureus* DAR3901 |
| Sau-487-1 | Bacteria Firmicutes Bacilli Bacillales Staphylococcaceae *Staphylococcus aureus* DAR3902 |
| Sau-488-1 | Bacteria Firmicutes Bacilli Bacillales Staphylococcaceae *Staphylococcus aureus* DAR3907 |
| Sau-489-1 | Bacteria Firmicutes Bacilli Bacillales Staphylococcaceae *Staphylococcus aureus* DAR3911 |
| Sau-490-1 | Bacteria Firmicutes Bacilli Bacillales Staphylococcaceae *Staphylococcus aureus* DAR3913 |
| Sau-491-1 | Bacteria Firmicutes Bacilli Bacillales Staphylococcaceae *Staphylococcus aureus* DAR3918 |
| Sau-492-1 | Bacteria Firmicutes Bacilli Bacillales Staphylococcaceae *Staphylococcus aureus* DAR3919 |
| Sau-493-1 | Bacteria Firmicutes Bacilli Bacillales Staphylococcaceae *Staphylococcus aureus* DAR3924 |
| Sau-494-1 | Bacteria Firmicutes Bacilli Bacillales Staphylococcaceae *Staphylococcus aureus* DAR3928 |
| Sau-495-1 | Bacteria Firmicutes Bacilli Bacillales Staphylococcaceae *Staphylococcus aureus* DAR3930 |
| Sau-496-1 | Bacteria Firmicutes Bacilli Bacillales Staphylococcaceae *Staphylococcus aureus* DAR3933 |
| Sau-497-1 | Bacteria Firmicutes Bacilli Bacillales Staphylococcaceae *Staphylococcus aureus* DAR5789 |
| Sau-498-1 | Bacteria Firmicutes Bacilli Bacillales Staphylococcaceae *Staphylococcus aureus* DAR5790 |
| Sau-499-1 | Bacteria Firmicutes Bacilli Bacillales Staphylococcaceae *Staphylococcus aureus* DAR5791 |
| Sau-500-1 | Bacteria Firmicutes Bacilli Bacillales Staphylococcaceae *Staphylococcus aureus* DAR5792 |
| Sau-501-1 | Bacteria Firmicutes Bacilli Bacillales Staphylococcaceae *Staphylococcus aureus* DAR5793 |
| Sau-502-1 | Bacteria Firmicutes Bacilli Bacillales Staphylococcaceae *Staphylococcus aureus* DAR5794 |
| Sau-503-1 | Bacteria Firmicutes Bacilli Bacillales Staphylococcaceae *Staphylococcus aureus* DAR5795 |
| Sau-504-1 | Bacteria Firmicutes Bacilli Bacillales Staphylococcaceae *Staphylococcus aureus* DAR5796 |
| Sau-505-1 | Bacteria Firmicutes Bacilli Bacillales Staphylococcaceae *Staphylococcus aureus* DAR5797 |
| Sau-506-1 | Bacteria Firmicutes Bacilli Bacillales Staphylococcaceae *Staphylococcus aureus* DAR5798 |
| Sau-507-1 | Bacteria Firmicutes Bacilli Bacillales Staphylococcaceae *Staphylococcus aureus* DAR5799 |
| Sau-508-1 | Bacteria Firmicutes Bacilli Bacillales Staphylococcaceae *Staphylococcus aureus* DAR5800 |
| Sau-509-1 | Bacteria Firmicutes Bacilli Bacillales Staphylococcaceae *Staphylococcus aureus* DAR5801 |
| Sau-510-1 | Bacteria Firmicutes Bacilli Bacillales Staphylococcaceae *Staphylococcus aureus* DAR5802 |
| Sau-511-1 | Bacteria Firmicutes Bacilli Bacillales Staphylococcaceae *Staphylococcus aureus* DAR5803 |
| Sau-512-1 | Bacteria Firmicutes Bacilli Bacillales Staphylococcaceae *Staphylococcus aureus* DAR5804 |

| | |
|---|---|
| Sau-513-1 | Bacteria Firmicutes Bacilli Bacillales Staphylococcaceae *Staphylococcus aureus* DAR5805 |
| Sau-514-1 | Bacteria Firmicutes Bacilli Bacillales Staphylococcaceae *Staphylococcus aureus* DAR5806 |
| Sau-515-1 | Bacteria Firmicutes Bacilli Bacillales Staphylococcaceae *Staphylococcus aureus* DAR5807 |
| Sau-516-1 | Bacteria Firmicutes Bacilli Bacillales Staphylococcaceae *Staphylococcus aureus* DAR5808 |
| Sau-517-1 | Bacteria Firmicutes Bacilli Bacillales Staphylococcaceae *Staphylococcus aureus* DAR5809 |
| Sau-518-1 | Bacteria Firmicutes Bacilli Bacillales Staphylococcaceae *Staphylococcus aureus* DAR5810 |
| Sau-519-1 | Bacteria Firmicutes Bacilli Bacillales Staphylococcaceae *Staphylococcus aureus* DAR5811 |
| Sau-520-1 | Bacteria Firmicutes Bacilli Bacillales Staphylococcaceae *Staphylococcus aureus* DAR5812 |
| Sau-521-1 | Bacteria Firmicutes Bacilli Bacillales Staphylococcaceae *Staphylococcus aureus* DAR5813 |
| Sau-522-1 | Bacteria Firmicutes Bacilli Bacillales Staphylococcaceae *Staphylococcus aureus* DAR5814 |
| Sau-523-1 | Bacteria Firmicutes Bacilli Bacillales Staphylococcaceae *Staphylococcus aureus* DAR5815 |
| Sau-524-1 | Bacteria Firmicutes Bacilli Bacillales Staphylococcaceae *Staphylococcus aureus* DAR5816 |
| Sau-525-1 | Bacteria Firmicutes Bacilli Bacillales Staphylococcaceae *Staphylococcus aureus* DAR5817 |
| Sau-526-1 | Bacteria Firmicutes Bacilli Bacillales Staphylococcaceae *Staphylococcus aureus* DAR5819 |
| Sau-527-1 | Bacteria Firmicutes Bacilli Bacillales Staphylococcaceae *Staphylococcus aureus* DAR5820 |
| Sau-528-1 | Bacteria Firmicutes Bacilli Bacillales Staphylococcaceae *Staphylococcus aureus* DAR5821 |
| Sau-529-1 | Bacteria Firmicutes Bacilli Bacillales Staphylococcaceae *Staphylococcus aureus* DAR5822 |
| Sau-530-1 | Bacteria Firmicutes Bacilli Bacillales Staphylococcaceae *Staphylococcus aureus* DAR5823 |
| Sau-531-1 | Bacteria Firmicutes Bacilli Bacillales Staphylococcaceae *Staphylococcus aureus* DAR5824 |
| Sau-532-1 | Bacteria Firmicutes Bacilli Bacillales Staphylococcaceae *Staphylococcus aureus* DAR5825 |
| Sau-533-1 | Bacteria Firmicutes Bacilli Bacillales Staphylococcaceae *Staphylococcus aureus* DAR5826 |
| Sau-534-1 | Bacteria Firmicutes Bacilli Bacillales Staphylococcaceae *Staphylococcus aureus* DAR5827 |
| Sau-535-1 | Bacteria Firmicutes Bacilli Bacillales Staphylococcaceae *Staphylococcus aureus* DAR5828 |
| Sau-536-1 | Bacteria Firmicutes Bacilli Bacillales Staphylococcaceae *Staphylococcus aureus* DAR5829 |
| Sau-537-1 | Bacteria Firmicutes Bacilli Bacillales Staphylococcaceae *Staphylococcus aureus* DAR5830 |
| Sau-538-1 | Bacteria Firmicutes Bacilli Bacillales Staphylococcaceae *Staphylococcus aureus* DAR5831 |
| Sau-539-1 | Bacteria Firmicutes Bacilli Bacillales Staphylococcaceae *Staphylococcus aureus* DAR5832 |
| Sau-540-1 | Bacteria Firmicutes Bacilli Bacillales Staphylococcaceae *Staphylococcus aureus* DAR5833 |
| Sau-541-1 | Bacteria Firmicutes Bacilli Bacillales Staphylococcaceae *Staphylococcus aureus* DAR5834 |
| Sau-542-1 | Bacteria Firmicutes Bacilli Bacillales Staphylococcaceae *Staphylococcus aureus* DAR5835 |
| Sau-543-1 | Bacteria Firmicutes Bacilli Bacillales Staphylococcaceae *Staphylococcus aureus* DAR5836 |
| Sau-544-1 | Bacteria Firmicutes Bacilli Bacillales Staphylococcaceae *Staphylococcus aureus* DAR5837 |
| Sau-545-1 | Bacteria Firmicutes Bacilli Bacillales Staphylococcaceae *Staphylococcus aureus* DAR5838 |
| Sau-546-1 | Bacteria Firmicutes Bacilli Bacillales Staphylococcaceae *Staphylococcus aureus* DAR5839 |
| Sau-547-1 | Bacteria Firmicutes Bacilli Bacillales Staphylococcaceae *Staphylococcus aureus* DAR5840 |
| Sau-548-1 | Bacteria Firmicutes Bacilli Bacillales Staphylococcaceae *Staphylococcus aureus* DAR5841 |
| Sau-549-1 | Bacteria Firmicutes Bacilli Bacillales Staphylococcaceae *Staphylococcus aureus* DAR5842 |
| Sau-550-1 | Bacteria Firmicutes Bacilli Bacillales Staphylococcaceae *Staphylococcus aureus* DAR5843 |
| Sau-551-1 | Bacteria Firmicutes Bacilli Bacillales Staphylococcaceae *Staphylococcus aureus* DAR5844 |
| Sau-552-1 | Bacteria Firmicutes Bacilli Bacillales Staphylococcaceae *Staphylococcus aureus* DAR5845 |
| Sau-553-1 | Bacteria Firmicutes Bacilli Bacillales Staphylococcaceae *Staphylococcus aureus* DAR5846 |
| Sau-554-1 | Bacteria Firmicutes Bacilli Bacillales Staphylococcaceae *Staphylococcus aureus* DAR5847 |
| Sau-555-1 | Bacteria Firmicutes Bacilli Bacillales Staphylococcaceae *Staphylococcus aureus* DAR5848 |
| Sau-556-1 | Bacteria Firmicutes Bacilli Bacillales Staphylococcaceae *Staphylococcus aureus* DAR5849 |
| Sau-557-1 | Bacteria Firmicutes Bacilli Bacillales Staphylococcaceae *Staphylococcus aureus* DAR5850 |
| Sau-558-1 | Bacteria Firmicutes Bacilli Bacillales Staphylococcaceae *Staphylococcus aureus* DAR5851 |
| Sau-559-1 | Bacteria Firmicutes Bacilli Bacillales Staphylococcaceae *Staphylococcus aureus* DAR5852 |
| Sau-560-1 | Bacteria Firmicutes Bacilli Bacillales Staphylococcaceae *Staphylococcus aureus* DAR5853 |
| Sau-561-1 | Bacteria Firmicutes Bacilli Bacillales Staphylococcaceae *Staphylococcus aureus* DAR5854 |
| Sau-562-1 | Bacteria Firmicutes Bacilli Bacillales Staphylococcaceae *Staphylococcus aureus* DAR5855 |
| Sau-563-1 | Bacteria Firmicutes Bacilli Bacillales Staphylococcaceae *Staphylococcus aureus* DAR5856 |
| Sau-564-1 | Bacteria Firmicutes Bacilli Bacillales Staphylococcaceae *Staphylococcus aureus* DAR5857 |
| Sau-565-1 | Bacteria Firmicutes Bacilli Bacillales Staphylococcaceae *Staphylococcus aureus* DAR5858 |
| Sau-566-1 | Bacteria Firmicutes Bacilli Bacillales Staphylococcaceae *Staphylococcus aureus* DAR5859 |
| Sau-567-1 | Bacteria Firmicutes Bacilli Bacillales Staphylococcaceae *Staphylococcus aureus* DAR5860 |
| Sau-568-1 | Bacteria Firmicutes Bacilli Bacillales Staphylococcaceae *Staphylococcus aureus* DAR5861 |
| Sau-569-1 | Bacteria Firmicutes Bacilli Bacillales Staphylococcaceae *Staphylococcus aureus* DAR5862 |
| Sau-570-1 | Bacteria Firmicutes Bacilli Bacillales Staphylococcaceae *Staphylococcus aureus* DAR5863 |
| Sau-571-1 | Bacteria Firmicutes Bacilli Bacillales Staphylococcaceae *Staphylococcus aureus* DAR5864 |
| Sau-572-1 | Bacteria Firmicutes Bacilli Bacillales Staphylococcaceae *Staphylococcus aureus* DAR5865 |
| Sau-573-1 | Bacteria Firmicutes Bacilli Bacillales Staphylococcaceae *Staphylococcus aureus* DAR5866 |
| Sau-574-1 | Bacteria Firmicutes Bacilli Bacillales Staphylococcaceae *Staphylococcus aureus* DAR5867 |
| Sau-575-1 | Bacteria Firmicutes Bacilli Bacillales Staphylococcaceae *Staphylococcus aureus* DAR5868 |
| Sau-576-1 | Bacteria Firmicutes Bacilli Bacillales Staphylococcaceae *Staphylococcus aureus* DAR5869 |
| Sau-577-1 | Bacteria Firmicutes Bacilli Bacillales Staphylococcaceae *Staphylococcus aureus* DAR5870 |

| | |
|---|---|
| Sau-578-1 | Bacteria Firmicutes Bacilli Bacillales Staphylococcaceae *Staphylococcus aureus* DAR5871 |
| Sau-579-1 | Bacteria Firmicutes Bacilli Bacillales Staphylococcaceae *Staphylococcus aureus* DAR5872 |
| Sau-580-1 | Bacteria Firmicutes Bacilli Bacillales Staphylococcaceae *Staphylococcus aureus* DAR5873 |
| Sau-581-1 | Bacteria Firmicutes Bacilli Bacillales Staphylococcaceae *Staphylococcus aureus* DAR5874 |
| Sau-582-1 | Bacteria Firmicutes Bacilli Bacillales Staphylococcaceae *Staphylococcus aureus* DAR5875 |
| Sau-583-1 | Bacteria Firmicutes Bacilli Bacillales Staphylococcaceae *Staphylococcus aureus* DAR5876 |
| Sau-584-1 | Bacteria Firmicutes Bacilli Bacillales Staphylococcaceae *Staphylococcus aureus* DAR5877 |
| Sau-585-1 | Bacteria Firmicutes Bacilli Bacillales Staphylococcaceae *Staphylococcus aureus* DAR5878 |
| Sau-586-1 | Bacteria Firmicutes Bacilli Bacillales Staphylococcaceae *Staphylococcus aureus* DAR5880 |
| Sau-587-1 | Bacteria Firmicutes Bacilli Bacillales Staphylococcaceae *Staphylococcus aureus* DAR5881 |
| Sau-588-1 | Bacteria Firmicutes Bacilli Bacillales Staphylococcaceae *Staphylococcus aureus* DAR5882 |
| Sau-589-1 | Bacteria Firmicutes Bacilli Bacillales Staphylococcaceae *Staphylococcus aureus* DAR5883 |
| Sau-590-1 | Bacteria Firmicutes Bacilli Bacillales Staphylococcaceae *Staphylococcus aureus* DAR5884 |
| Sau-591-1 | Bacteria Firmicutes Bacilli Bacillales Staphylococcaceae *Staphylococcus aureus* DAR5885 |
| Sau-592-1 | Bacteria Firmicutes Bacilli Bacillales Staphylococcaceae *Staphylococcus aureus* DAR5886 |
| Sau-593-1 | Bacteria Firmicutes Bacilli Bacillales Staphylococcaceae *Staphylococcus aureus* DAR5887 |
| Sau-594-1 | Bacteria Firmicutes Bacilli Bacillales Staphylococcaceae *Staphylococcus aureus* DAR5888 |
| Sau-595-1 | Bacteria Firmicutes Bacilli Bacillales Staphylococcaceae *Staphylococcus aureus* DAR5889 |
| Sau-596-1 | Bacteria Firmicutes Bacilli Bacillales Staphylococcaceae *Staphylococcus aureus* DAR5890 |
| Sau-597-1 | Bacteria Firmicutes Bacilli Bacillales Staphylococcaceae *Staphylococcus aureus* DAR5891 |
| Sau-598-1 | Bacteria Firmicutes Bacilli Bacillales Staphylococcaceae *Staphylococcus aureus* DAR5892 |
| Sau-599-1 | Bacteria Firmicutes Bacilli Bacillales Staphylococcaceae *Staphylococcus aureus* DAR5893 |
| Sau-600-1 | Bacteria Firmicutes Bacilli Bacillales Staphylococcaceae *Staphylococcus aureus* DAR5894 |
| Sau-601-1 | Bacteria Firmicutes Bacilli Bacillales Staphylococcaceae *Staphylococcus aureus* DAR5895 |
| Sau-602-1 | Bacteria Firmicutes Bacilli Bacillales Staphylococcaceae *Staphylococcus aureus* DAR5896 |
| Sau-603-1 | Bacteria Firmicutes Bacilli Bacillales Staphylococcaceae *Staphylococcus aureus* DAR5897 |
| Sau-604-1 | Bacteria Firmicutes Bacilli Bacillales Staphylococcaceae *Staphylococcus aureus* DAR5898 |
| Sau-605-1 | Bacteria Firmicutes Bacilli Bacillales Staphylococcaceae *Staphylococcus aureus* DAR948 |
| Sau-606-1 | Bacteria Firmicutes Bacilli Bacillales Staphylococcaceae *Staphylococcus aureus* DAR951 |
| Sau-607-1 | Bacteria Firmicutes Bacilli Bacillales Staphylococcaceae *Staphylococcus aureus* DAR966 |
| Sau-608-1 | Bacteria Firmicutes Bacilli Bacillales Staphylococcaceae *Staphylococcus aureus* DICM09/00997-9HST2 |
| Sau-609-1 | Bacteria Firmicutes Bacilli Bacillales Staphylococcaceae *Staphylococcus aureus* DICM09/01587-13HST |
| Sau-610-1 | Bacteria Firmicutes Bacilli Bacillales Staphylococcaceae *Staphylococcus aureus* F12749 |
| Sau-611-1 | Bacteria Firmicutes Bacilli Bacillales Staphylococcaceae *Staphylococcus aureus* F12753 |
| Sau-612-1 | Bacteria Firmicutes Bacilli Bacillales Staphylococcaceae *Staphylococcus aureus* F12856 |
| Sau-613-1 | Bacteria Firmicutes Bacilli Bacillales Staphylococcaceae *Staphylococcus aureus* F12874 |
| Sau-614-1 | Bacteria Firmicutes Bacilli Bacillales Staphylococcaceae *Staphylococcus aureus* F12915 |
| Sau-615-1 | Bacteria Firmicutes Bacilli Bacillales Staphylococcaceae *Staphylococcus aureus* F12917 |
| Sau-616-1 | Bacteria Firmicutes Bacilli Bacillales Staphylococcaceae *Staphylococcus aureus* F12926_081012 |
| Sau-617-1 | Bacteria Firmicutes Bacilli Bacillales Staphylococcaceae *Staphylococcus aureus* F13022 |
| Sau-618-1 | Bacteria Firmicutes Bacilli Bacillales Staphylococcaceae *Staphylococcus aureus* F13027 |
| Sau-619-1 | Bacteria Firmicutes Bacilli Bacillales Staphylococcaceae *Staphylococcus aureus* F16077 |
| Sau-620-1 | Bacteria Firmicutes Bacilli Bacillales Staphylococcaceae *Staphylococcus aureus* F16104 |
| Sau-621-1 | Bacteria Firmicutes Bacilli Bacillales Staphylococcaceae *Staphylococcus aureus* F16106 |
| Sau-622-1 | Bacteria Firmicutes Bacilli Bacillales Staphylococcaceae *Staphylococcus aureus* F16107 |
| Sau-623-1 | Bacteria Firmicutes Bacilli Bacillales Staphylococcaceae *Staphylococcus aureus* F16108 |
| Sau-624-1 | Bacteria Firmicutes Bacilli Bacillales Staphylococcaceae *Staphylococcus aureus* F16432 |
| Sau-625-1 | Bacteria Firmicutes Bacilli Bacillales Staphylococcaceae *Staphylococcus aureus* F16470 |
| Sau-626-1 | Bacteria Firmicutes Bacilli Bacillales Staphylococcaceae *Staphylococcus aureus* F16533 |
| Sau-627-1 | Bacteria Firmicutes Bacilli Bacillales Staphylococcaceae *Staphylococcus aureus* F19383 |
| Sau-628-1 | Bacteria Firmicutes Bacilli Bacillales Staphylococcaceae *Staphylococcus aureus* F19466 |
| Sau-629-1 | Bacteria Firmicutes Bacilli Bacillales Staphylococcaceae *Staphylococcus aureus* F19470 |
| Sau-630-1 | Bacteria Firmicutes Bacilli Bacillales Staphylococcaceae *Staphylococcus aureus* F19490 |
| Sau-631-1 | Bacteria Firmicutes Bacilli Bacillales Staphylococcaceae *Staphylococcus aureus* F19955 |
| Sau-632-1 | Bacteria Firmicutes Bacilli Bacillales Staphylococcaceae *Staphylococcus aureus* F20105 |
| Sau-633-1 | Bacteria Firmicutes Bacilli Bacillales Staphylococcaceae *Staphylococcus aureus* F20115 |
| Sau-634-1 | Bacteria Firmicutes Bacilli Bacillales Staphylococcaceae *Staphylococcus aureus* F22648 |
| Sau-635-1 | Bacteria Firmicutes Bacilli Bacillales Staphylococcaceae *Staphylococcus aureus* F22893 |
| Sau-636-1 | Bacteria Firmicutes Bacilli Bacillales Staphylococcaceae *Staphylococcus aureus* F22898 |
| Sau-637-1 | Bacteria Firmicutes Bacilli Bacillales Staphylococcaceae *Staphylococcus aureus* F23336 |
| Sau-638-1 | Bacteria Firmicutes Bacilli Bacillales Staphylococcaceae *Staphylococcus aureus* F23347 |
| Sau-639-1 | Bacteria Firmicutes Bacilli Bacillales Staphylococcaceae *Staphylococcus aureus* F23353 |
| Sau-640-1 | Bacteria Firmicutes Bacilli Bacillales Staphylococcaceae *Staphylococcus aureus* F23368 |
| Sau-641-1 | Bacteria Firmicutes Bacilli Bacillales Staphylococcaceae *Staphylococcus aureus* F23585 |
| Sau-642-1 | Bacteria Firmicutes Bacilli Bacillales Staphylococcaceae *Staphylococcus aureus* F23610 |

| | |
|---|---|
| Sau-643-1 | Bacteria Firmicutes Bacilli Bacillales Staphylococcaceae *Staphylococcus aureus* F23613 |
| Sau-644-1 | Bacteria Firmicutes Bacilli Bacillales Staphylococcaceae *Staphylococcus aureus* F25913 |
| Sau-645-1 | Bacteria Firmicutes Bacilli Bacillales Staphylococcaceae *Staphylococcus aureus* F26021 |
| Sau-646-1 | Bacteria Firmicutes Bacilli Bacillales Staphylococcaceae *Staphylococcus aureus* F26032 |
| Sau-647-1 | Bacteria Firmicutes Bacilli Bacillales Staphylococcaceae *Staphylococcus aureus* F26051 |
| Sau-648-1 | Bacteria Firmicutes Bacilli Bacillales Staphylococcaceae *Staphylococcus aureus* F26088 |
| Sau-649-1 | Bacteria Firmicutes Bacilli Bacillales Staphylococcaceae *Staphylococcus aureus* F26858 |
| Sau-650-1 | Bacteria Firmicutes Bacilli Bacillales Staphylococcaceae *Staphylococcus aureus* F26873 |
| Sau-651-1 | Bacteria Firmicutes Bacilli Bacillales Staphylococcaceae *Staphylococcus aureus* F26884 |
| Sau-652-1 | Bacteria Firmicutes Bacilli Bacillales Staphylococcaceae *Staphylococcus aureus* F27233 |
| Sau-653-1 | Bacteria Firmicutes Bacilli Bacillales Staphylococcaceae *Staphylococcus aureus* F27252 |
| Sau-654-1 | Bacteria Firmicutes Bacilli Bacillales Staphylococcaceae *Staphylococcus aureus* F27253 |
| Sau-655-1 | Bacteria Firmicutes Bacilli Bacillales Staphylococcaceae *Staphylococcus aureus* F27254 |
| Sau-656-1 | Bacteria Firmicutes Bacilli Bacillales Staphylococcaceae *Staphylococcus aureus* F29214 |
| Sau-657-1 | Bacteria Firmicutes Bacilli Bacillales Staphylococcaceae *Staphylococcus aureus* F29447 |
| Sau-658-1 | Bacteria Firmicutes Bacilli Bacillales Staphylococcaceae *Staphylococcus aureus* F29450_091412 |
| Sau-659-1 | Bacteria Firmicutes Bacilli Bacillales Staphylococcaceae *Staphylococcus aureus* F29982 |
| Sau-660-1 | Bacteria Firmicutes Bacilli Bacillales Staphylococcaceae *Staphylococcus aureus* F29993 |
| Sau-661-1 | Bacteria Firmicutes Bacilli Bacillales Staphylococcaceae *Staphylococcus aureus* F29997 |
| Sau-662-1 | Bacteria Firmicutes Bacilli Bacillales Staphylococcaceae *Staphylococcus aureus* F31078 |
| Sau-663-1 | Bacteria Firmicutes Bacilli Bacillales Staphylococcaceae *Staphylococcus aureus* F31094 |
| Sau-664-1 | Bacteria Firmicutes Bacilli Bacillales Staphylococcaceae *Staphylococcus aureus* F32119 |
| Sau-665-1 | Bacteria Firmicutes Bacilli Bacillales Staphylococcaceae *Staphylococcus aureus* F32629 |
| Sau-666-1 | Bacteria Firmicutes Bacilli Bacillales Staphylococcaceae *Staphylococcus aureus* F34824 |
| Sau-667-1 | Bacteria Firmicutes Bacilli Bacillales Staphylococcaceae *Staphylococcus aureus* F35307 |
| Sau-668-1 | Bacteria Firmicutes Bacilli Bacillales Staphylococcaceae *Staphylococcus aureus* F36090 |
| Sau-669-1 | Bacteria Firmicutes Bacilli Bacillales Staphylococcaceae *Staphylococcus aureus* F36091 |
| Sau-670-1 | Bacteria Firmicutes Bacilli Bacillales Staphylococcaceae *Staphylococcus aureus* F36097 |
| Sau-671-1 | Bacteria Firmicutes Bacilli Bacillales Staphylococcaceae *Staphylococcus aureus* F36099 |
| Sau-672-1 | Bacteria Firmicutes Bacilli Bacillales Staphylococcaceae *Staphylococcus aureus* F36670 |
| Sau-673-1 | Bacteria Firmicutes Bacilli Bacillales Staphylococcaceae *Staphylococcus aureus* F36683 |
| Sau-674-1 | Bacteria Firmicutes Bacilli Bacillales Staphylococcaceae *Staphylococcus aureus* F36687 |
| Sau-675-1 | Bacteria Firmicutes Bacilli Bacillales Staphylococcaceae *Staphylococcus aureus* F36699 |
| Sau-676-1 | Bacteria Firmicutes Bacilli Bacillales Staphylococcaceae *Staphylococcus aureus* F38211 |
| Sau-677-1 | Bacteria Firmicutes Bacilli Bacillales Staphylococcaceae *Staphylococcus aureus* F40814 |
| Sau-678-1 | Bacteria Firmicutes Bacilli Bacillales Staphylococcaceae *Staphylococcus aureus* F41882 |
| Sau-679-1 | Bacteria Firmicutes Bacilli Bacillales Staphylococcaceae *Staphylococcus aureus* F41886 |
| Sau-680-1 | Bacteria Firmicutes Bacilli Bacillales Staphylococcaceae *Staphylococcus aureus* F41902 |
| Sau-681-1 | Bacteria Firmicutes Bacilli Bacillales Staphylococcaceae *Staphylococcus aureus* F41906 |
| Sau-682-1 | Bacteria Firmicutes Bacilli Bacillales Staphylococcaceae *Staphylococcus aureus* F41913 |
| Sau-683-1 | Bacteria Firmicutes Bacilli Bacillales Staphylococcaceae *Staphylococcus aureus* F42556 |
| Sau-684-1 | Bacteria Firmicutes Bacilli Bacillales Staphylococcaceae *Staphylococcus aureus* F42559 |
| Sau-685-1 | Bacteria Firmicutes Bacilli Bacillales Staphylococcaceae *Staphylococcus aureus* F42568 |
| Sau-686-1 | Bacteria Firmicutes Bacilli Bacillales Staphylococcaceae *Staphylococcus aureus* F42579 |
| Sau-687-1 | Bacteria Firmicutes Bacilli Bacillales Staphylococcaceae *Staphylococcus aureus* F42580 |
| Sau-688-1 | Bacteria Firmicutes Bacilli Bacillales Staphylococcaceae *Staphylococcus aureus* F42582 |
| Sau-689-1 | Bacteria Firmicutes Bacilli Bacillales Staphylococcaceae *Staphylococcus aureus* F43199 |
| Sau-690-1 | Bacteria Firmicutes Bacilli Bacillales Staphylococcaceae *Staphylococcus aureus* F43212 |
| Sau-691-1 | Bacteria Firmicutes Bacilli Bacillales Staphylococcaceae *Staphylococcus aureus* F43221 |
| Sau-692-1 | Bacteria Firmicutes Bacilli Bacillales Staphylococcaceae *Staphylococcus aureus* F43252 |
| Sau-693-1 | Bacteria Firmicutes Bacilli Bacillales Staphylococcaceae *Staphylococcus aureus* F43676 |
| Sau-694-1 | Bacteria Firmicutes Bacilli Bacillales Staphylococcaceae *Staphylococcus aureus* F45300 |
| Sau-695-1 | Bacteria Firmicutes Bacilli Bacillales Staphylococcaceae *Staphylococcus aureus* F45749 |
| Sau-696-1 | Bacteria Firmicutes Bacilli Bacillales Staphylococcaceae *Staphylococcus aureus* F45759 |
| Sau-697-1 | Bacteria Firmicutes Bacilli Bacillales Staphylococcaceae *Staphylococcus aureus* F45779 |
| Sau-698-1 | Bacteria Firmicutes Bacilli Bacillales Staphylococcaceae *Staphylococcus aureus* F46600 |
| Sau-699-1 | Bacteria Firmicutes Bacilli Bacillales Staphylococcaceae *Staphylococcus aureus* F46632 |
| Sau-700-1 | Bacteria Firmicutes Bacilli Bacillales Staphylococcaceae *Staphylococcus aureus* F46780 |
| Sau-701-1 | Bacteria Firmicutes Bacilli Bacillales Staphylococcaceae *Staphylococcus aureus* F46785 |
| Sau-702-1 | Bacteria Firmicutes Bacilli Bacillales Staphylococcaceae *Staphylococcus aureus* F46798 |
| Sau-703-1 | Bacteria Firmicutes Bacilli Bacillales Staphylococcaceae *Staphylococcus aureus* F48420 |
| Sau-704-1 | Bacteria Firmicutes Bacilli Bacillales Staphylococcaceae *Staphylococcus aureus* F48422 |
| Sau-705-1 | Bacteria Firmicutes Bacilli Bacillales Staphylococcaceae *Staphylococcus aureus* F48584 |
| Sau-706-1 | Bacteria Firmicutes Bacilli Bacillales Staphylococcaceae *Staphylococcus aureus* F48603 |
| Sau-707-1 | Bacteria Firmicutes Bacilli Bacillales Staphylococcaceae *Staphylococcus aureus* F48607 |

| | |
|---|---|
| Sau-708-1 | Bacteria Firmicutes Bacilli Bacillales Staphylococcaceae *Staphylococcus aureus* F48611 |
| Sau-709-1 | Bacteria Firmicutes Bacilli Bacillales Staphylococcaceae *Staphylococcus aureus* F48950 |
| Sau-710-1 | Bacteria Firmicutes Bacilli Bacillales Staphylococcaceae *Staphylococcus aureus* F48959 |
| Sau-711-1 | Bacteria Firmicutes Bacilli Bacillales Staphylococcaceae *Staphylococcus aureus* F50076 |
| Sau-712-1 | Bacteria Firmicutes Bacilli Bacillales Staphylococcaceae *Staphylococcus aureus* F50093 |
| Sau-713-1 | Bacteria Firmicutes Bacilli Bacillales Staphylococcaceae *Staphylococcus aureus* F50121 |
| Sau-714-1 | Bacteria Firmicutes Bacilli Bacillales Staphylococcaceae *Staphylococcus aureus* F50126 |
| Sau-715-1 | Bacteria Firmicutes Bacilli Bacillales Staphylococcaceae *Staphylococcus aureus* F52326 |
| Sau-716-1 | Bacteria Firmicutes Bacilli Bacillales Staphylococcaceae *Staphylococcus aureus* F52748 |
| Sau-717-1 | Bacteria Firmicutes Bacilli Bacillales Staphylococcaceae *Staphylococcus aureus* F52754 |
| Sau-718-1 | Bacteria Firmicutes Bacilli Bacillales Staphylococcaceae *Staphylococcus aureus* F53388 |
| Sau-719-1 | Bacteria Firmicutes Bacilli Bacillales Staphylococcaceae *Staphylococcus aureus* F53393 |
| Sau-720-1 | Bacteria Firmicutes Bacilli Bacillales Staphylococcaceae *Staphylococcus aureus* F53399 |
| Sau-721-1 | Bacteria Firmicutes Bacilli Bacillales Staphylococcaceae *Staphylococcus aureus* F53404 |
| Sau-722-1 | Bacteria Firmicutes Bacilli Bacillales Staphylococcaceae *Staphylococcus aureus* F54452 |
| Sau-723-1 | Bacteria Firmicutes Bacilli Bacillales Staphylococcaceae *Staphylococcus aureus* F54461 |
| Sau-724-1 | Bacteria Firmicutes Bacilli Bacillales Staphylococcaceae *Staphylococcus aureus* F54467 |
| Sau-725-1 | Bacteria Firmicutes Bacilli Bacillales Staphylococcaceae *Staphylococcus aureus* F54471 |
| Sau-726-1 | Bacteria Firmicutes Bacilli Bacillales Staphylococcaceae *Staphylococcus aureus* F54494 |
| Sau-727-1 | Bacteria Firmicutes Bacilli Bacillales Staphylococcaceae *Staphylococcus aureus* F55987 |
| Sau-728-1 | Bacteria Firmicutes Bacilli Bacillales Staphylococcaceae *Staphylococcus aureus* F56969 |
| Sau-729-1 | Bacteria Firmicutes Bacilli Bacillales Staphylococcaceae *Staphylococcus aureus* F57021 |
| Sau-730-1 | Bacteria Firmicutes Bacilli Bacillales Staphylococcaceae *Staphylococcus aureus* F57033 |
| Sau-731-1 | Bacteria Firmicutes Bacilli Bacillales Staphylococcaceae *Staphylococcus aureus* F57036 |
| Sau-732-1 | Bacteria Firmicutes Bacilli Bacillales Staphylococcaceae *Staphylococcus aureus* F59156 |
| Sau-733-1 | Bacteria Firmicutes Bacilli Bacillales Staphylococcaceae *Staphylococcus aureus* F59332 |
| Sau-734-1 | Bacteria Firmicutes Bacilli Bacillales Staphylococcaceae *Staphylococcus aureus* F59335 |
| Sau-735-1 | Bacteria Firmicutes Bacilli Bacillales Staphylococcaceae *Staphylococcus aureus* F60346 |
| Sau-736-1 | Bacteria Firmicutes Bacilli Bacillales Staphylococcaceae *Staphylococcus aureus* F60349 |
| Sau-737-1 | Bacteria Firmicutes Bacilli Bacillales Staphylococcaceae *Staphylococcus aureus* F60357 |
| Sau-738-1 | Bacteria Firmicutes Bacilli Bacillales Staphylococcaceae *Staphylococcus aureus* F60360 |
| Sau-739-1 | Bacteria Firmicutes Bacilli Bacillales Staphylococcaceae *Staphylococcus aureus* F62007 |
| Sau-740-1 | Bacteria Firmicutes Bacilli Bacillales Staphylococcaceae *Staphylococcus aureus* F62010 |
| Sau-741-1 | Bacteria Firmicutes Bacilli Bacillales Staphylococcaceae *Staphylococcus aureus* F62013 |
| Sau-742-1 | Bacteria Firmicutes Bacilli Bacillales Staphylococcaceae *Staphylococcus aureus* F63154 |
| Sau-743-1 | Bacteria Firmicutes Bacilli Bacillales Staphylococcaceae *Staphylococcus aureus* F63179 |
| Sau-744-1 | Bacteria Firmicutes Bacilli Bacillales Staphylococcaceae *Staphylococcus aureus* F63687 |
| Sau-745-1 | Bacteria Firmicutes Bacilli Bacillales Staphylococcaceae *Staphylococcus aureus* F63704 |
| Sau-746-1 | Bacteria Firmicutes Bacilli Bacillales Staphylococcaceae *Staphylococcus aureus* F63712 |
| Sau-747-1 | Bacteria Firmicutes Bacilli Bacillales Staphylococcaceae *Staphylococcus aureus* F63730 |
| Sau-748-1 | Bacteria Firmicutes Bacilli Bacillales Staphylococcaceae *Staphylococcus aureus* F63750 |
| Sau-749-1 | Bacteria Firmicutes Bacilli Bacillales Staphylococcaceae *Staphylococcus aureus* F63786 |
| Sau-750-1 | Bacteria Firmicutes Bacilli Bacillales Staphylococcaceae *Staphylococcus aureus* F63787 |
| Sau-751-1 | Bacteria Firmicutes Bacilli Bacillales Staphylococcaceae *Staphylococcus aureus* F64891 |
| Sau-752-1 | Bacteria Firmicutes Bacilli Bacillales Staphylococcaceae *Staphylococcus aureus* F64894 |
| Sau-753-1 | Bacteria Firmicutes Bacilli Bacillales Staphylococcaceae *Staphylococcus aureus* F64896 |
| Sau-754-1 | Bacteria Firmicutes Bacilli Bacillales Staphylococcaceae *Staphylococcus aureus* F64897 |
| Sau-755-1 | Bacteria Firmicutes Bacilli Bacillales Staphylococcaceae *Staphylococcus aureus* F65057 |
| Sau-756-1 | Bacteria Firmicutes Bacilli Bacillales Staphylococcaceae *Staphylococcus aureus* F66300 |
| Sau-757-1 | Bacteria Firmicutes Bacilli Bacillales Staphylococcaceae *Staphylococcus aureus* F66307 |
| Sau-758-1 | Bacteria Firmicutes Bacilli Bacillales Staphylococcaceae *Staphylococcus aureus* F66778 |
| Sau-759-1 | Bacteria Firmicutes Bacilli Bacillales Staphylococcaceae *Staphylococcus aureus* F68118 |
| Sau-760-1 | Bacteria Firmicutes Bacilli Bacillales Staphylococcaceae *Staphylococcus aureus* F70051 |
| Sau-761-1 | Bacteria Firmicutes Bacilli Bacillales Staphylococcaceae *Staphylococcus aureus* F70077 |
| Sau-762-1 | Bacteria Firmicutes Bacilli Bacillales Staphylococcaceae *Staphylococcus aureus* F70893 |
| Sau-763-1 | Bacteria Firmicutes Bacilli Bacillales Staphylococcaceae *Staphylococcus aureus* F71576 |
| Sau-764-1 | Bacteria Firmicutes Bacilli Bacillales Staphylococcaceae *Staphylococcus aureus* F71592 |
| Sau-765-1 | Bacteria Firmicutes Bacilli Bacillales Staphylococcaceae *Staphylococcus aureus* F73187 |
| Sau-766-1 | Bacteria Firmicutes Bacilli Bacillales Staphylococcaceae *Staphylococcus aureus* F73229 |
| Sau-767-1 | Bacteria Firmicutes Bacilli Bacillales Staphylococcaceae *Staphylococcus aureus* F74023 |
| Sau-768-1 | Bacteria Firmicutes Bacilli Bacillales Staphylococcaceae *Staphylococcus aureus* F74035 |
| Sau-769-1 | Bacteria Firmicutes Bacilli Bacillales Staphylococcaceae *Staphylococcus aureus* F74039 |
| Sau-770-1 | Bacteria Firmicutes Bacilli Bacillales Staphylococcaceae *Staphylococcus aureus* F74139 |
| Sau-771-1 | Bacteria Firmicutes Bacilli Bacillales Staphylococcaceae *Staphylococcus aureus* F74220 |
| Sau-772-1 | Bacteria Firmicutes Bacilli Bacillales Staphylococcaceae *Staphylococcus aureus* F74820 |

| | |
|---|---|
| Sau-773-1 | Bacteria Firmicutes Bacilli Bacillales Staphylococcaceae *Staphylococcus aureus* F76218 |
| Sau-774-1 | Bacteria Firmicutes Bacilli Bacillales Staphylococcaceae *Staphylococcus aureus* F76239 |
| Sau-775-1 | Bacteria Firmicutes Bacilli Bacillales Staphylococcaceae *Staphylococcus aureus* F76253 |
| Sau-776-1 | Bacteria Firmicutes Bacilli Bacillales Staphylococcaceae *Staphylococcus aureus* F76996 |
| Sau-777-1 | Bacteria Firmicutes Bacilli Bacillales Staphylococcaceae *Staphylococcus aureus* F77005 |
| Sau-778-1 | Bacteria Firmicutes Bacilli Bacillales Staphylococcaceae *Staphylococcus aureus* F77047 |
| Sau-779-1 | Bacteria Firmicutes Bacilli Bacillales Staphylococcaceae *Staphylococcus aureus* F77709 |
| Sau-780-1 | Bacteria Firmicutes Bacilli Bacillales Staphylococcaceae *Staphylococcus aureus* F77710 |
| Sau-781-1 | Bacteria Firmicutes Bacilli Bacillales Staphylococcaceae *Staphylococcus aureus* F77917 |
| Sau-782-1 | Bacteria Firmicutes Bacilli Bacillales Staphylococcaceae *Staphylococcus aureus* F77919 |
| Sau-783-1 | Bacteria Firmicutes Bacilli Bacillales Staphylococcaceae *Staphylococcus aureus* F77930 |
| Sau-784-1 | Bacteria Firmicutes Bacilli Bacillales Staphylococcaceae *Staphylococcus aureus* F77942 |
| Sau-785-1 | Bacteria Firmicutes Bacilli Bacillales Staphylococcaceae *Staphylococcus aureus* F77960 |
| Sau-786-1 | Bacteria Firmicutes Bacilli Bacillales Staphylococcaceae *Staphylococcus aureus* F77967 |
| Sau-787-1 | Bacteria Firmicutes Bacilli Bacillales Staphylococcaceae *Staphylococcus aureus* F77974 |
| Sau-788-1 | Bacteria Firmicutes Bacilli Bacillales Staphylococcaceae *Staphylococcus aureus* F80349 |
| Sau-789-1 | Bacteria Firmicutes Bacilli Bacillales Staphylococcaceae *Staphylococcus aureus* F80354 |
| Sau-790-1 | Bacteria Firmicutes Bacilli Bacillales Staphylococcaceae *Staphylococcus aureus* F80358 |
| Sau-791-1 | Bacteria Firmicutes Bacilli Bacillales Staphylococcaceae *Staphylococcus aureus* F80375 |
| Sau-792-1 | Bacteria Firmicutes Bacilli Bacillales Staphylococcaceae *Staphylococcus aureus* F80381 |
| Sau-793-1 | Bacteria Firmicutes Bacilli Bacillales Staphylococcaceae *Staphylococcus aureus* F80972 |
| Sau-794-1 | Bacteria Firmicutes Bacilli Bacillales Staphylococcaceae *Staphylococcus aureus* F81035 |
| Sau-795-1 | Bacteria Firmicutes Bacilli Bacillales Staphylococcaceae *Staphylococcus aureus* F81381 |
| Sau-796-1 | Bacteria Firmicutes Bacilli Bacillales Staphylococcaceae *Staphylococcus aureus* F81389 |
| Sau-797-1 | Bacteria Firmicutes Bacilli Bacillales Staphylococcaceae *Staphylococcus aureus* F82568 |
| Sau-798-1 | Bacteria Firmicutes Bacilli Bacillales Staphylococcaceae *Staphylococcus aureus* F82573 |
| Sau-799-1 | Bacteria Firmicutes Bacilli Bacillales Staphylococcaceae *Staphylococcus aureus* F82600 |
| Sau-800-1 | Bacteria Firmicutes Bacilli Bacillales Staphylococcaceae *Staphylococcus aureus* F83251 |
| Sau-801-1 | Bacteria Firmicutes Bacilli Bacillales Staphylococcaceae *Staphylococcus aureus* F83269 |
| Sau-802-1 | Bacteria Firmicutes Bacilli Bacillales Staphylococcaceae *Staphylococcus aureus* F83272 |
| Sau-803-1 | Bacteria Firmicutes Bacilli Bacillales Staphylococcaceae *Staphylococcus aureus* F84093 |
| Sau-804-1 | Bacteria Firmicutes Bacilli Bacillales Staphylococcaceae *Staphylococcus aureus* F84732 |
| Sau-805-1 | Bacteria Firmicutes Bacilli Bacillales Staphylococcaceae *Staphylococcus aureus* F84763 |
| Sau-806-1 | Bacteria Firmicutes Bacilli Bacillales Staphylococcaceae *Staphylococcus aureus* F85184 |
| Sau-807-1 | Bacteria Firmicutes Bacilli Bacillales Staphylococcaceae *Staphylococcus aureus* F85414 |
| Sau-808-1 | Bacteria Firmicutes Bacilli Bacillales Staphylococcaceae *Staphylococcus aureus* F85609 |
| Sau-809-1 | Bacteria Firmicutes Bacilli Bacillales Staphylococcaceae *Staphylococcus aureus* F85660 |
| Sau-810-1 | Bacteria Firmicutes Bacilli Bacillales Staphylococcaceae *Staphylococcus aureus* F85691 |
| Sau-811-1 | Bacteria Firmicutes Bacilli Bacillales Staphylococcaceae *Staphylococcus aureus* F85711 |
| Sau-812-1 | Bacteria Firmicutes Bacilli Bacillales Staphylococcaceae *Staphylococcus aureus* F86379 |
| Sau-813-1 | Bacteria Firmicutes Bacilli Bacillales Staphylococcaceae *Staphylococcus aureus* F87602 |
| Sau-814-1 | Bacteria Firmicutes Bacilli Bacillales Staphylococcaceae *Staphylococcus aureus* F88369 |
| Sau-815-1 | Bacteria Firmicutes Bacilli Bacillales Staphylococcaceae *Staphylococcus aureus* F88375 |
| Sau-816-1 | Bacteria Firmicutes Bacilli Bacillales Staphylococcaceae *Staphylococcus aureus* F88387 |
| Sau-817-1 | Bacteria Firmicutes Bacilli Bacillales Staphylococcaceae *Staphylococcus aureus* F88405 |
| Sau-818-1 | Bacteria Firmicutes Bacilli Bacillales Staphylococcaceae *Staphylococcus aureus* F89046 |
| Sau-819-1 | Bacteria Firmicutes Bacilli Bacillales Staphylococcaceae *Staphylococcus aureus* F89053 |
| Sau-820-1 | Bacteria Firmicutes Bacilli Bacillales Staphylococcaceae *Staphylococcus aureus* F89062 |
| Sau-821-1 | Bacteria Firmicutes Bacilli Bacillales Staphylococcaceae *Staphylococcus aureus* F89073 |
| Sau-822-1 | Bacteria Firmicutes Bacilli Bacillales Staphylococcaceae *Staphylococcus aureus* F89075 |
| Sau-823-1 | Bacteria Firmicutes Bacilli Bacillales Staphylococcaceae *Staphylococcus aureus* F89078 |
| Sau-824-1 | Bacteria Firmicutes Bacilli Bacillales Staphylococcaceae *Staphylococcus aureus* F89441 |
| Sau-825-1 | Bacteria Firmicutes Bacilli Bacillales Staphylococcaceae *Staphylococcus aureus* F89468 |
| Sau-826-1 | Bacteria Firmicutes Bacilli Bacillales Staphylococcaceae *Staphylococcus aureus* F90608 |
| Sau-827-1 | Bacteria Firmicutes Bacilli Bacillales Staphylococcaceae *Staphylococcus aureus* F91072 |
| Sau-828-1 | Bacteria Firmicutes Bacilli Bacillales Staphylococcaceae *Staphylococcus aureus* F91077 |
| Sau-829-1 | Bacteria Firmicutes Bacilli Bacillales Staphylococcaceae *Staphylococcus aureus* F91084 |
| Sau-830-1 | Bacteria Firmicutes Bacilli Bacillales Staphylococcaceae *Staphylococcus aureus* F92047 |
| Sau-831-1 | Bacteria Firmicutes Bacilli Bacillales Staphylococcaceae *Staphylococcus aureus* F92073 |
| Sau-832-1 | Bacteria Firmicutes Bacilli Bacillales Staphylococcaceae *Staphylococcus aureus* F92089 |
| Sau-833-1 | Bacteria Firmicutes Bacilli Bacillales Staphylococcaceae *Staphylococcus aureus* F92105_080312 |
| Sau-834-1 | Bacteria Firmicutes Bacilli Bacillales Staphylococcaceae *Staphylococcus aureus* F92119_080312 |
| Sau-835-1 | Bacteria Firmicutes Bacilli Bacillales Staphylococcaceae *Staphylococcus aureus* F92121 |
| Sau-836-1 | Bacteria Firmicutes Bacilli Bacillales Staphylococcaceae *Staphylococcus aureus* F92125 |
| Sau-837-1 | Bacteria Firmicutes Bacilli Bacillales Staphylococcaceae *Staphylococcus aureus* F92127 |

| | |
|---|---|
| Sau-838-1 | Bacteria Firmicutes Bacilli Bacillales Staphylococcaceae *Staphylococcus aureus* F92128 |
| Sau-839-1 | Bacteria Firmicutes Bacilli Bacillales Staphylococcaceae *Staphylococcus aureus* F92753 |
| Sau-840-1 | Bacteria Firmicutes Bacilli Bacillales Staphylococcaceae *Staphylococcus aureus* F92765 |
| Sau-841-1 | Bacteria Firmicutes Bacilli Bacillales Staphylococcaceae *Staphylococcus aureus* FP_N239 |
| Sau-842-1 | Bacteria Firmicutes Bacilli Bacillales Staphylococcaceae *Staphylococcus aureus* FP_N5203 OX |
| Sau-843-1 | Bacteria Firmicutes Bacilli Bacillales Staphylococcaceae *Staphylococcus aureus* FP_N5208 OX |
| Sau-844-1 | Bacteria Firmicutes Bacilli Bacillales Staphylococcaceae *Staphylococcus aureus* FVRH6001 |
| Sau-845-1 | Bacteria Firmicutes Bacilli Bacillales Staphylococcaceae *Staphylococcus aureus* FVRH6002 |
| Sau-846-1 | Bacteria Firmicutes Bacilli Bacillales Staphylococcaceae *Staphylococcus aureus* FVRH6011 |
| Sau-847-1 | Bacteria Firmicutes Bacilli Bacillales Staphylococcaceae *Staphylococcus aureus* FVRH6021 |
| Sau-848-1 | Bacteria Firmicutes Bacilli Bacillales Staphylococcaceae *Staphylococcus aureus* FVRH6022 |
| Sau-849-1 | Bacteria Firmicutes Bacilli Bacillales Staphylococcaceae *Staphylococcus aureus* FVRH6026 |
| Sau-850-1 | Bacteria Firmicutes Bacilli Bacillales Staphylococcaceae *Staphylococcus aureus* FVRH6035 |
| Sau-851-1 | Bacteria Firmicutes Bacilli Bacillales Staphylococcaceae *Staphylococcus aureus* FVRH6040 |
| Sau-852-1 | Bacteria Firmicutes Bacilli Bacillales Staphylococcaceae *Staphylococcus aureus* FVRH6052 |
| Sau-853-1 | Bacteria Firmicutes Bacilli Bacillales Staphylococcaceae *Staphylococcus aureus* FVRH6065 |
| Sau-854-1 | Bacteria Firmicutes Bacilli Bacillales Staphylococcaceae *Staphylococcus aureus* FVRH6066 |
| Sau-855-1 | Bacteria Firmicutes Bacilli Bacillales Staphylococcaceae *Staphylococcus aureus* FVRH6067 |
| Sau-856-1 | Bacteria Firmicutes Bacilli Bacillales Staphylococcaceae *Staphylococcus aureus* FVRH6074 |
| Sau-857-1 | Bacteria Firmicutes Bacilli Bacillales Staphylococcaceae *Staphylococcus aureus* FVRH6076 |
| Sau-858-1 | Bacteria Firmicutes Bacilli Bacillales Staphylococcaceae *Staphylococcus aureus* FVRH6079 |
| Sau-859-1 | Bacteria Firmicutes Bacilli Bacillales Staphylococcaceae *Staphylococcus aureus* FVRH6118 |
| Sau-860-1 | Bacteria Firmicutes Bacilli Bacillales Staphylococcaceae *Staphylococcus aureus* FVRH6129 |
| Sau-861-1 | Bacteria Firmicutes Bacilli Bacillales Staphylococcaceae *Staphylococcus aureus* FVRH6130 |
| Sau-862-1 | Bacteria Firmicutes Bacilli Bacillales Staphylococcaceae *Staphylococcus aureus* FVRH6131 |
| Sau-863-1 | Bacteria Firmicutes Bacilli Bacillales Staphylococcaceae *Staphylococcus aureus* FVRH6132 |
| Sau-864-1 | Bacteria Firmicutes Bacilli Bacillales Staphylococcaceae *Staphylococcus aureus* FVRH6138 |
| Sau-865-1 | Bacteria Firmicutes Bacilli Bacillales Staphylococcaceae *Staphylococcus aureus* GD2010-052 |
| Sau-866-1 | Bacteria Firmicutes Bacilli Bacillales Staphylococcaceae *Staphylococcus aureus* GD2010-061 |
| Sau-867-1 | Bacteria Firmicutes Bacilli Bacillales Staphylococcaceae *Staphylococcus aureus* GD2010-090 |
| Sau-868-1 | Bacteria Firmicutes Bacilli Bacillales Staphylococcaceae *Staphylococcus aureus* GD2010-102 |
| Sau-869-1 | Bacteria Firmicutes Bacilli Bacillales Staphylococcaceae *Staphylococcus aureus* GD2010-107 |
| Sau-870-1 | Bacteria Firmicutes Bacilli Bacillales Staphylococcaceae *Staphylococcus aureus* GD2010-110 |
| Sau-871-1 | Bacteria Firmicutes Bacilli Bacillales Staphylococcaceae *Staphylococcus aureus* GD2010-112 |
| Sau-872-1 | Bacteria Firmicutes Bacilli Bacillales Staphylococcaceae *Staphylococcus aureus* GD2010-115 |
| Sau-873-1 | Bacteria Firmicutes Bacilli Bacillales Staphylococcaceae *Staphylococcus aureus* GD2010-131 |
| Sau-874-1 | Bacteria Firmicutes Bacilli Bacillales Staphylococcaceae *Staphylococcus aureus* GD2010-147 |
| Sau-875-1 | Bacteria Firmicutes Bacilli Bacillales Staphylococcaceae *Staphylococcus aureus* GD2010-155 |
| Sau-876-1 | Bacteria Firmicutes Bacilli Bacillales Staphylococcaceae *Staphylococcus aureus* GD2010-158 |
| Sau-877-1 | Bacteria Firmicutes Bacilli Bacillales Staphylococcaceae *Staphylococcus aureus* GD2010-159 |
| Sau-878-1 | Bacteria Firmicutes Bacilli Bacillales Staphylococcaceae *Staphylococcus aureus* GD2010-168 |
| Sau-879-1 | Bacteria Firmicutes Bacilli Bacillales Staphylococcaceae *Staphylococcus aureus* GD2010-169 |
| Sau-880-1 | Bacteria Firmicutes Bacilli Bacillales Staphylococcaceae *Staphylococcus aureus* GD2010-177 |
| Sau-881-1 | Bacteria Firmicutes Bacilli Bacillales Staphylococcaceae *Staphylococcus aureus* GGMC6003 |
| Sau-882-1 | Bacteria Firmicutes Bacilli Bacillales Staphylococcaceae *Staphylococcus aureus* GGMC6004 |
| Sau-883-1 | Bacteria Firmicutes Bacilli Bacillales Staphylococcaceae *Staphylococcus aureus* GGMC6008 |
| Sau-884-1 | Bacteria Firmicutes Bacilli Bacillales Staphylococcaceae *Staphylococcus aureus* GGMC6010 |
| Sau-885-1 | Bacteria Firmicutes Bacilli Bacillales Staphylococcaceae *Staphylococcus aureus* GGMC6018 |
| Sau-886-1 | Bacteria Firmicutes Bacilli Bacillales Staphylococcaceae *Staphylococcus aureus* GGMC6024 |
| Sau-887-1 | Bacteria Firmicutes Bacilli Bacillales Staphylococcaceae *Staphylococcus aureus* GGMC6026 |
| Sau-888-1 | Bacteria Firmicutes Bacilli Bacillales Staphylococcaceae *Staphylococcus aureus* GGMC6027 |
| Sau-889-1 | Bacteria Firmicutes Bacilli Bacillales Staphylococcaceae *Staphylococcus aureus* GGMC6036 |
| Sau-890-1 | Bacteria Firmicutes Bacilli Bacillales Staphylococcaceae *Staphylococcus aureus* GGMC6043 |
| Sau-891-1 | Bacteria Firmicutes Bacilli Bacillales Staphylococcaceae *Staphylococcus aureus* GGMC6049 |
| Sau-892-1 | Bacteria Firmicutes Bacilli Bacillales Staphylococcaceae *Staphylococcus aureus* GGMC6053 |
| Sau-893-1 | Bacteria Firmicutes Bacilli Bacillales Staphylococcaceae *Staphylococcus aureus* GGMC6057 |
| Sau-894-1 | Bacteria Firmicutes Bacilli Bacillales Staphylococcaceae *Staphylococcus aureus* H12662 |
| Sau-895-1 | Bacteria Firmicutes Bacilli Bacillales Staphylococcaceae *Staphylococcus aureus* H12818 |
| Sau-896-1 | Bacteria Firmicutes Bacilli Bacillales Staphylococcaceae *Staphylococcus aureus* H12822 |
| Sau-897-1 | Bacteria Firmicutes Bacilli Bacillales Staphylococcaceae *Staphylococcus aureus* H12890 |
| Sau-898-1 | Bacteria Firmicutes Bacilli Bacillales Staphylococcaceae *Staphylococcus aureus* H12893 |
| Sau-899-1 | Bacteria Firmicutes Bacilli Bacillales Staphylococcaceae *Staphylococcus aureus* H12930 |
| Sau-900-1 | Bacteria Firmicutes Bacilli Bacillales Staphylococcaceae *Staphylococcus aureus* H15665 |
| Sau-901-1 | Bacteria Firmicutes Bacilli Bacillales Staphylococcaceae *Staphylococcus aureus* H15899 |
| Sau-902-1 | Bacteria Firmicutes Bacilli Bacillales Staphylococcaceae *Staphylococcus aureus* H15911 |

| | |
|---|---|
| Sau-903-1 | Bacteria Firmicutes Bacilli Bacillales Staphylococcaceae *Staphylococcus aureus* H16097 |
| Sau-904-1 | Bacteria Firmicutes Bacilli Bacillales Staphylococcaceae *Staphylococcus aureus* H18808 |
| Sau-905-1 | Bacteria Firmicutes Bacilli Bacillales Staphylococcaceae *Staphylococcus aureus* H18809 |
| Sau-906-1 | Bacteria Firmicutes Bacilli Bacillales Staphylococcaceae *Staphylococcus aureus* H18811 |
| Sau-907-1 | Bacteria Firmicutes Bacilli Bacillales Staphylococcaceae *Staphylococcus aureus* H18813 |
| Sau-908-1 | Bacteria Firmicutes Bacilli Bacillales Staphylococcaceae *Staphylococcus aureus* H19392 |
| Sau-909-1 | Bacteria Firmicutes Bacilli Bacillales Staphylococcaceae *Staphylococcus aureus* H19398 |
| Sau-910-1 | Bacteria Firmicutes Bacilli Bacillales Staphylococcaceae *Staphylococcus aureus* H19404 |
| Sau-911-1 | Bacteria Firmicutes Bacilli Bacillales Staphylococcaceae *Staphylococcus aureus* H19420 |
| Sau-912-1 | Bacteria Firmicutes Bacilli Bacillales Staphylococcaceae *Staphylococcus aureus* H19476 |
| Sau-913-1 | Bacteria Firmicutes Bacilli Bacillales Staphylococcaceae *Staphylococcus aureus* H19520 |
| Sau-914-1 | Bacteria Firmicutes Bacilli Bacillales Staphylococcaceae *Staphylococcus aureus* H20315 |
| Sau-915-1 | Bacteria Firmicutes Bacilli Bacillales Staphylococcaceae *Staphylococcus aureus* H20321 |
| Sau-916-1 | Bacteria Firmicutes Bacilli Bacillales Staphylococcaceae *Staphylococcus aureus* H20322 |
| Sau-917-1 | Bacteria Firmicutes Bacilli Bacillales Staphylococcaceae *Staphylococcus aureus* H21924 |
| Sau-918-1 | Bacteria Firmicutes Bacilli Bacillales Staphylococcaceae *Staphylococcus aureus* H21935 |
| Sau-919-1 | Bacteria Firmicutes Bacilli Bacillales Staphylococcaceae *Staphylococcus aureus* H22632 |
| Sau-920-1 | Bacteria Firmicutes Bacilli Bacillales Staphylococcaceae *Staphylococcus aureus* H22637 |
| Sau-921-1 | Bacteria Firmicutes Bacilli Bacillales Staphylococcaceae *Staphylococcus aureus* H22656 |
| Sau-922-1 | Bacteria Firmicutes Bacilli Bacillales Staphylococcaceae *Staphylococcus aureus* H23829 |
| Sau-923-1 | Bacteria Firmicutes Bacilli Bacillales Staphylococcaceae *Staphylococcus aureus* H23830 |
| Sau-924-1 | Bacteria Firmicutes Bacilli Bacillales Staphylococcaceae *Staphylococcus aureus* H24637 |
| Sau-925-1 | Bacteria Firmicutes Bacilli Bacillales Staphylococcaceae *Staphylococcus aureus* H24662 |
| Sau-926-1 | Bacteria Firmicutes Bacilli Bacillales Staphylococcaceae *Staphylococcus aureus* H24884 |
| Sau-927-1 | Bacteria Firmicutes Bacilli Bacillales Staphylococcaceae *Staphylococcus aureus* H25869 |
| Sau-928-1 | Bacteria Firmicutes Bacilli Bacillales Staphylococcaceae *Staphylococcus aureus* H25872 |
| Sau-929-1 | Bacteria Firmicutes Bacilli Bacillales Staphylococcaceae *Staphylococcus aureus* H27739 |
| Sau-930-1 | Bacteria Firmicutes Bacilli Bacillales Staphylococcaceae *Staphylococcus aureus* H27761 |
| Sau-931-1 | Bacteria Firmicutes Bacilli Bacillales Staphylococcaceae *Staphylococcus aureus* H27777 |
| Sau-932-1 | Bacteria Firmicutes Bacilli Bacillales Staphylococcaceae *Staphylococcus aureus* H27858 |
| Sau-933-1 | Bacteria Firmicutes Bacilli Bacillales Staphylococcaceae *Staphylococcus aureus* H27862 |
| Sau-934-1 | Bacteria Firmicutes Bacilli Bacillales Staphylococcaceae *Staphylococcus aureus* H27872 |
| Sau-935-1 | Bacteria Firmicutes Bacilli Bacillales Staphylococcaceae *Staphylococcus aureus* H30732 |
| Sau-936-1 | Bacteria Firmicutes Bacilli Bacillales Staphylococcaceae *Staphylococcus aureus* H30737 |
| Sau-937-1 | Bacteria Firmicutes Bacilli Bacillales Staphylococcaceae *Staphylococcus aureus* H31084 |
| Sau-938-1 | Bacteria Firmicutes Bacilli Bacillales Staphylococcaceae *Staphylococcus aureus* H31085 |
| Sau-939-1 | Bacteria Firmicutes Bacilli Bacillales Staphylococcaceae *Staphylococcus aureus* H31086 |
| Sau-940-1 | Bacteria Firmicutes Bacilli Bacillales Staphylococcaceae *Staphylococcus aureus* H31276 |
| Sau-941-1 | Bacteria Firmicutes Bacilli Bacillales Staphylococcaceae *Staphylococcus aureus* H31290 |
| Sau-942-1 | Bacteria Firmicutes Bacilli Bacillales Staphylococcaceae *Staphylococcus aureus* H32126 |
| Sau-943-1 | Bacteria Firmicutes Bacilli Bacillales Staphylococcaceae *Staphylococcus aureus* H32135 |
| Sau-944-1 | Bacteria Firmicutes Bacilli Bacillales Staphylococcaceae *Staphylococcus aureus* H32166 |
| Sau-945-1 | Bacteria Firmicutes Bacilli Bacillales Staphylococcaceae *Staphylococcus aureus* H32175 |
| Sau-946-1 | Bacteria Firmicutes Bacilli Bacillales Staphylococcaceae *Staphylococcus aureus* H32181 |
| Sau-947-1 | Bacteria Firmicutes Bacilli Bacillales Staphylococcaceae *Staphylococcus aureus* H33594 |
| Sau-948-1 | Bacteria Firmicutes Bacilli Bacillales Staphylococcaceae *Staphylococcus aureus* H33602 |
| Sau-949-1 | Bacteria Firmicutes Bacilli Bacillales Staphylococcaceae *Staphylococcus aureus* H33636 |
| Sau-950-1 | Bacteria Firmicutes Bacilli Bacillales Staphylococcaceae *Staphylococcus aureus* H34245 |
| Sau-951-1 | Bacteria Firmicutes Bacilli Bacillales Staphylococcaceae *Staphylococcus aureus* H35283 |
| Sau-952-1 | Bacteria Firmicutes Bacilli Bacillales Staphylococcaceae *Staphylococcus aureus* H37235 |
| Sau-953-1 | Bacteria Firmicutes Bacilli Bacillales Staphylococcaceae *Staphylococcus aureus* H37776 |
| Sau-954-1 | Bacteria Firmicutes Bacilli Bacillales Staphylococcaceae *Staphylococcus aureus* H37781 |
| Sau-955-1 | Bacteria Firmicutes Bacilli Bacillales Staphylococcaceae *Staphylococcus aureus* H38215 |
| Sau-956-1 | Bacteria Firmicutes Bacilli Bacillales Staphylococcaceae *Staphylococcus aureus* H38226 |
| Sau-957-1 | Bacteria Firmicutes Bacilli Bacillales Staphylococcaceae *Staphylococcus aureus* H38239 |
| Sau-958-1 | Bacteria Firmicutes Bacilli Bacillales Staphylococcaceae *Staphylococcus aureus* H40569 |
| Sau-959-1 | Bacteria Firmicutes Bacilli Bacillales Staphylococcaceae *Staphylococcus aureus* H40575 |
| Sau-960-1 | Bacteria Firmicutes Bacilli Bacillales Staphylococcaceae *Staphylococcus aureus* H40583 |
| Sau-961-1 | Bacteria Firmicutes Bacilli Bacillales Staphylococcaceae *Staphylococcus aureus* H41070 |
| Sau-962-1 | Bacteria Firmicutes Bacilli Bacillales Staphylococcaceae *Staphylococcus aureus* H41079 |
| Sau-963-1 | Bacteria Firmicutes Bacilli Bacillales Staphylococcaceae *Staphylococcus aureus* H41109 |
| Sau-964-1 | Bacteria Firmicutes Bacilli Bacillales Staphylococcaceae *Staphylococcus aureus* H41336 |
| Sau-965-1 | Bacteria Firmicutes Bacilli Bacillales Staphylococcaceae *Staphylococcus aureus* H42737 |
| Sau-966-1 | Bacteria Firmicutes Bacilli Bacillales Staphylococcaceae *Staphylococcus aureus* H42759 |
| Sau-967-1 | Bacteria Firmicutes Bacilli Bacillales Staphylococcaceae *Staphylococcus aureus* H43687 |

| | |
|---|---|
| Sau-968-1 | Bacteria Firmicutes Bacilli Bacillales Staphylococcaceae *Staphylococcus aureus* H43690 |
| Sau-969-1 | Bacteria Firmicutes Bacilli Bacillales Staphylococcaceae *Staphylococcus aureus* H43694 |
| Sau-970-1 | Bacteria Firmicutes Bacilli Bacillales Staphylococcaceae *Staphylococcus aureus* H43700 |
| Sau-971-1 | Bacteria Firmicutes Bacilli Bacillales Staphylococcaceae *Staphylococcus aureus* H43986 |
| Sau-972-1 | Bacteria Firmicutes Bacilli Bacillales Staphylococcaceae *Staphylococcus aureus* H43996 |
| Sau-973-1 | Bacteria Firmicutes Bacilli Bacillales Staphylococcaceae *Staphylococcus aureus* H44015 |
| Sau-974-1 | Bacteria Firmicutes Bacilli Bacillales Staphylococcaceae *Staphylococcus aureus* H44713 |
| Sau-975-1 | Bacteria Firmicutes Bacilli Bacillales Staphylococcaceae *Staphylococcus aureus* H45755 |
| Sau-976-1 | Bacteria Firmicutes Bacilli Bacillales Staphylococcaceae *Staphylococcus aureus* H45760 |
| Sau-977-1 | Bacteria Firmicutes Bacilli Bacillales Staphylococcaceae *Staphylococcus aureus* H45765 |
| Sau-978-1 | Bacteria Firmicutes Bacilli Bacillales Staphylococcaceae *Staphylococcus aureus* H47117 |
| Sau-979-1 | Bacteria Firmicutes Bacilli Bacillales Staphylococcaceae *Staphylococcus aureus* H47139 |
| Sau-980-1 | Bacteria Firmicutes Bacilli Bacillales Staphylococcaceae *Staphylococcus aureus* H47169 |
| Sau-981-1 | Bacteria Firmicutes Bacilli Bacillales Staphylococcaceae *Staphylococcus aureus* H47171 |
| Sau-982-1 | Bacteria Firmicutes Bacilli Bacillales Staphylococcaceae *Staphylococcus aureus* H47183 |
| Sau-983-1 | Bacteria Firmicutes Bacilli Bacillales Staphylococcaceae *Staphylococcus aureus* H47224 |
| Sau-984-1 | Bacteria Firmicutes Bacilli Bacillales Staphylococcaceae *Staphylococcus aureus* H47671 |
| Sau-985-1 | Bacteria Firmicutes Bacilli Bacillales Staphylococcaceae *Staphylococcus aureus* H48052 |
| Sau-986-1 | Bacteria Firmicutes Bacilli Bacillales Staphylococcaceae *Staphylococcus aureus* H48054 |
| Sau-987-1 | Bacteria Firmicutes Bacilli Bacillales Staphylococcaceae *Staphylococcus aureus* H48646 |
| Sau-988-1 | Bacteria Firmicutes Bacilli Bacillales Staphylococcaceae *Staphylococcus aureus* H48649 |
| Sau-989-1 | Bacteria Firmicutes Bacilli Bacillales Staphylococcaceae *Staphylococcus aureus* H49831 |
| Sau-990-1 | Bacteria Firmicutes Bacilli Bacillales Staphylococcaceae *Staphylococcus aureus* H49864 |
| Sau-991-1 | Bacteria Firmicutes Bacilli Bacillales Staphylococcaceae *Staphylococcus aureus* H49917 |
| Sau-992-1 | Bacteria Firmicutes Bacilli Bacillales Staphylococcaceae *Staphylococcus aureus* H49922 |
| Sau-993-1 | Bacteria Firmicutes Bacilli Bacillales Staphylococcaceae *Staphylococcus aureus* H50387 |
| Sau-994-1 | Bacteria Firmicutes Bacilli Bacillales Staphylococcaceae *Staphylococcus aureus* H50403 |
| Sau-995-1 | Bacteria Firmicutes Bacilli Bacillales Staphylococcaceae *Staphylococcus aureus* H50411 |
| Sau-996-1 | Bacteria Firmicutes Bacilli Bacillales Staphylococcaceae *Staphylococcus aureus* H51158 |
| Sau-997-1 | Bacteria Firmicutes Bacilli Bacillales Staphylococcaceae *Staphylococcus aureus* H53188 |
| Sau-998-1 | Bacteria Firmicutes Bacilli Bacillales Staphylococcaceae *Staphylococcus aureus* H54403 |
| Sau-999-1 | Bacteria Firmicutes Bacilli Bacillales Staphylococcaceae *Staphylococcus aureus* H56219 |
| Sau-1000-1 | Bacteria Firmicutes Bacilli Bacillales Staphylococcaceae *Staphylococcus aureus* H56227 |
| Sau-1001-1 | Bacteria Firmicutes Bacilli Bacillales Staphylococcaceae *Staphylococcus aureus* H57070 |
| Sau-1002-1 | Bacteria Firmicutes Bacilli Bacillales Staphylococcaceae *Staphylococcus aureus* H57815 |
| Sau-1003-1 | Bacteria Firmicutes Bacilli Bacillales Staphylococcaceae *Staphylococcus aureus* H57821 |
| Sau-1004-1 | Bacteria Firmicutes Bacilli Bacillales Staphylococcaceae *Staphylococcus aureus* H57822 |
| Sau-1005-1 | Bacteria Firmicutes Bacilli Bacillales Staphylococcaceae *Staphylococcus aureus* H57823 |
| Sau-1006-1 | Bacteria Firmicutes Bacilli Bacillales Staphylococcaceae *Staphylococcus aureus* H58165 |
| Sau-1007-1 | Bacteria Firmicutes Bacilli Bacillales Staphylococcaceae *Staphylococcus aureus* H58177 |
| Sau-1008-1 | Bacteria Firmicutes Bacilli Bacillales Staphylococcaceae *Staphylococcus aureus* H58183 |
| Sau-1009-1 | Bacteria Firmicutes Bacilli Bacillales Staphylococcaceae *Staphylococcus aureus* H58191 |
| Sau-1010-1 | Bacteria Firmicutes Bacilli Bacillales Staphylococcaceae *Staphylococcus aureus* H58198 |
| Sau-1011-1 | Bacteria Firmicutes Bacilli Bacillales Staphylococcaceae *Staphylococcus aureus* H59298 |
| Sau-1012-1 | Bacteria Firmicutes Bacilli Bacillales Staphylococcaceae *Staphylococcus aureus* H59303_080912 |
| Sau-1013-1 | Bacteria Firmicutes Bacilli Bacillales Staphylococcaceae *Staphylococcus aureus* H59312_080912 |
| Sau-1014-1 | Bacteria Firmicutes Bacilli Bacillales Staphylococcaceae *Staphylococcus aureus* H59325 |
| Sau-1015-1 | Bacteria Firmicutes Bacilli Bacillales Staphylococcaceae *Staphylococcus aureus* H59329 |
| Sau-1016-1 | Bacteria Firmicutes Bacilli Bacillales Staphylococcaceae *Staphylococcus aureus* H60145 |
| Sau-1017-1 | Bacteria Firmicutes Bacilli Bacillales Staphylococcaceae *Staphylococcus aureus* H61003 |
| Sau-1018-1 | Bacteria Firmicutes Bacilli Bacillales Staphylococcaceae *Staphylococcus aureus* H64299 |
| Sau-1019-1 | Bacteria Firmicutes Bacilli Bacillales Staphylococcaceae *Staphylococcus aureus* H64303 |
| Sau-1020-1 | Bacteria Firmicutes Bacilli Bacillales Staphylococcaceae *Staphylococcus aureus* H64570 |
| Sau-1021-1 | Bacteria Firmicutes Bacilli Bacillales Staphylococcaceae *Staphylococcus aureus* H64941 |
| Sau-1022-1 | Bacteria Firmicutes Bacilli Bacillales Staphylococcaceae *Staphylococcus aureus* H64962 |
| Sau-1023-1 | Bacteria Firmicutes Bacilli Bacillales Staphylococcaceae *Staphylococcus aureus* H64967 |
| Sau-1024-1 | Bacteria Firmicutes Bacilli Bacillales Staphylococcaceae *Staphylococcus aureus* H65794 |
| Sau-1025-1 | Bacteria Firmicutes Bacilli Bacillales Staphylococcaceae *Staphylococcus aureus* H65830 |
| Sau-1026-1 | Bacteria Firmicutes Bacilli Bacillales Staphylococcaceae *Staphylococcus aureus* H65853 |
| Sau-1027-1 | Bacteria Firmicutes Bacilli Bacillales Staphylococcaceae *Staphylococcus aureus* H65856 |
| Sau-1028-1 | Bacteria Firmicutes Bacilli Bacillales Staphylococcaceae *Staphylococcus aureus* H65863 |
| Sau-1029-1 | Bacteria Firmicutes Bacilli Bacillales Staphylococcaceae *Staphylococcus aureus* H65878 |
| Sau-1030-1 | Bacteria Firmicutes Bacilli Bacillales Staphylococcaceae *Staphylococcus aureus* H65881 |
| Sau-1031-1 | Bacteria Firmicutes Bacilli Bacillales Staphylococcaceae *Staphylococcus aureus* H65882 |
| Sau-1032-1 | Bacteria Firmicutes Bacilli Bacillales Staphylococcaceae *Staphylococcus aureus* H65883 |

| | |
|---|---|
| Sau-1033-1 | Bacteria Firmicutes Bacilli Bacillales Staphylococcaceae *Staphylococcus aureus* H67656 |
| Sau-1034-1 | Bacteria Firmicutes Bacilli Bacillales Staphylococcaceae *Staphylococcus aureus* H68244 |
| Sau-1035-1 | Bacteria Firmicutes Bacilli Bacillales Staphylococcaceae *Staphylococcus aureus* H68252 |
| Sau-1036-1 | Bacteria Firmicutes Bacilli Bacillales Staphylococcaceae *Staphylococcus aureus* H68256 |
| Sau-1037-1 | Bacteria Firmicutes Bacilli Bacillales Staphylococcaceae *Staphylococcus aureus* H68996 |
| Sau-1038-1 | Bacteria Firmicutes Bacilli Bacillales Staphylococcaceae *Staphylococcus aureus* H69239 |
| Sau-1039-1 | Bacteria Firmicutes Bacilli Bacillales Staphylococcaceae *Staphylococcus aureus* H70821 |
| Sau-1040-1 | Bacteria Firmicutes Bacilli Bacillales Staphylococcaceae *Staphylococcus aureus* H70826 |
| Sau-1041-1 | Bacteria Firmicutes Bacilli Bacillales Staphylococcaceae *Staphylococcus aureus* H70834 |
| Sau-1042-1 | Bacteria Firmicutes Bacilli Bacillales Staphylococcaceae *Staphylococcus aureus* H71538 |
| Sau-1043-1 | Bacteria Firmicutes Bacilli Bacillales Staphylococcaceae *Staphylococcus aureus* H71553 |
| Sau-1044-1 | Bacteria Firmicutes Bacilli Bacillales Staphylococcaceae *Staphylococcus aureus* H71778 |
| Sau-1045-1 | Bacteria Firmicutes Bacilli Bacillales Staphylococcaceae *Staphylococcus aureus* H71781 |
| Sau-1046-1 | Bacteria Firmicutes Bacilli Bacillales Staphylococcaceae *Staphylococcus aureus* H71785 |
| Sau-1047-1 | Bacteria Firmicutes Bacilli Bacillales Staphylococcaceae *Staphylococcus aureus* H72115 |
| Sau-1048-1 | Bacteria Firmicutes Bacilli Bacillales Staphylococcaceae *Staphylococcus aureus* H72119 |
| Sau-1049-1 | Bacteria Firmicutes Bacilli Bacillales Staphylococcaceae *Staphylococcus aureus* H72129_090612 |
| Sau-1050-1 | Bacteria Firmicutes Bacilli Bacillales Staphylococcaceae *Staphylococcus aureus* H72131 |
| Sau-1051-1 | Bacteria Firmicutes Bacilli Bacillales Staphylococcaceae *Staphylococcus aureus* H72152 |
| Sau-1052-1 | Bacteria Firmicutes Bacilli Bacillales Staphylococcaceae *Staphylococcus aureus* H74056 |
| Sau-1053-1 | Bacteria Firmicutes Bacilli Bacillales Staphylococcaceae *Staphylococcus aureus* H75143 |
| Sau-1054-1 | Bacteria Firmicutes Bacilli Bacillales Staphylococcaceae *Staphylococcus aureus* H75198 |
| Sau-1055-1 | Bacteria Firmicutes Bacilli Bacillales Staphylococcaceae *Staphylococcus aureus* H75421 |
| Sau-1056-1 | Bacteria Firmicutes Bacilli Bacillales Staphylococcaceae *Staphylococcus aureus* H75423 |
| Sau-1057-1 | Bacteria Firmicutes Bacilli Bacillales Staphylococcaceae *Staphylococcus aureus* H75425 |
| Sau-1058-1 | Bacteria Firmicutes Bacilli Bacillales Staphylococcaceae *Staphylococcus aureus* H75430 |
| Sau-1059-1 | Bacteria Firmicutes Bacilli Bacillales Staphylococcaceae *Staphylococcus aureus* H77094 |
| Sau-1060-1 | Bacteria Firmicutes Bacilli Bacillales Staphylococcaceae *Staphylococcus aureus* H77098 |
| Sau-1061-1 | Bacteria Firmicutes Bacilli Bacillales Staphylococcaceae *Staphylococcus aureus* H77102 |
| Sau-1062-1 | Bacteria Firmicutes Bacilli Bacillales Staphylococcaceae *Staphylococcus aureus* H79477 |
| Sau-1063-1 | Bacteria Firmicutes Bacilli Bacillales Staphylococcaceae *Staphylococcus aureus* H79484 |
| Sau-1064-1 | Bacteria Firmicutes Bacilli Bacillales Staphylococcaceae *Staphylococcus aureus* H81433 |
| Sau-1065-1 | Bacteria Firmicutes Bacilli Bacillales Staphylococcaceae *Staphylococcus aureus* H81437 |
| Sau-1066-1 | Bacteria Firmicutes Bacilli Bacillales Staphylococcaceae *Staphylococcus aureus* H81800 |
| Sau-1067-1 | Bacteria Firmicutes Bacilli Bacillales Staphylococcaceae *Staphylococcus aureus* H81809 |
| Sau-1068-1 | Bacteria Firmicutes Bacilli Bacillales Staphylococcaceae *Staphylococcus aureus* H81885 |
| Sau-1069-1 | Bacteria Firmicutes Bacilli Bacillales Staphylococcaceae *Staphylococcus aureus* H81898 |
| Sau-1070-1 | Bacteria Firmicutes Bacilli Bacillales Staphylococcaceae *Staphylococcus aureus* H81901 |
| Sau-1071-1 | Bacteria Firmicutes Bacilli Bacillales Staphylococcaceae *Staphylococcus aureus* H83025 |
| Sau-1072-1 | Bacteria Firmicutes Bacilli Bacillales Staphylococcaceae *Staphylococcus aureus* H83536 |
| Sau-1073-1 | Bacteria Firmicutes Bacilli Bacillales Staphylococcaceae *Staphylococcus aureus* H84489 |
| Sau-1074-1 | Bacteria Firmicutes Bacilli Bacillales Staphylococcaceae *Staphylococcus aureus* H84492 |
| Sau-1075-1 | Bacteria Firmicutes Bacilli Bacillales Staphylococcaceae *Staphylococcus aureus* H84507 |
| Sau-1076-1 | Bacteria Firmicutes Bacilli Bacillales Staphylococcaceae *Staphylococcus aureus* H84956 |
| Sau-1077-1 | Bacteria Firmicutes Bacilli Bacillales Staphylococcaceae *Staphylococcus aureus* H84958 |
| Sau-1078-1 | Bacteria Firmicutes Bacilli Bacillales Staphylococcaceae *Staphylococcus aureus* H84959 |
| Sau-1079-1 | Bacteria Firmicutes Bacilli Bacillales Staphylococcaceae *Staphylococcus aureus* H84973 |
| Sau-1080-1 | Bacteria Firmicutes Bacilli Bacillales Staphylococcaceae *Staphylococcus aureus* H84980 |
| Sau-1081-1 | Bacteria Firmicutes Bacilli Bacillales Staphylococcaceae *Staphylococcus aureus* H85003 |
| Sau-1082-1 | Bacteria Firmicutes Bacilli Bacillales Staphylococcaceae *Staphylococcus aureus* H85008 |
| Sau-1083-1 | Bacteria Firmicutes Bacilli Bacillales Staphylococcaceae *Staphylococcus aureus* H87628 |
| Sau-1084-1 | Bacteria Firmicutes Bacilli Bacillales Staphylococcaceae *Staphylococcus aureus* H87658 |
| Sau-1085-1 | Bacteria Firmicutes Bacilli Bacillales Staphylococcaceae *Staphylococcus aureus* H87668 |
| Sau-1086-1 | Bacteria Firmicutes Bacilli Bacillales Staphylococcaceae *Staphylococcus aureus* H88149 |
| Sau-1087-1 | Bacteria Firmicutes Bacilli Bacillales Staphylococcaceae *Staphylococcus aureus* H88163 |
| Sau-1088-1 | Bacteria Firmicutes Bacilli Bacillales Staphylococcaceae *Staphylococcus aureus* H88170 |
| Sau-1089-1 | Bacteria Firmicutes Bacilli Bacillales Staphylococcaceae *Staphylococcus aureus* H88185 |
| Sau-1090-1 | Bacteria Firmicutes Bacilli Bacillales Staphylococcaceae *Staphylococcus aureus* H88188 |
| Sau-1091-1 | Bacteria Firmicutes Bacilli Bacillales Staphylococcaceae *Staphylococcus aureus* H89939 |
| Sau-1092-1 | Bacteria Firmicutes Bacilli Bacillales Staphylococcaceae *Staphylococcus aureus* H89941 |
| Sau-1093-1 | Bacteria Firmicutes Bacilli Bacillales Staphylococcaceae *Staphylococcus aureus* H90284 |
| Sau-1094-1 | Bacteria Firmicutes Bacilli Bacillales Staphylococcaceae *Staphylococcus aureus* H91384 |
| Sau-1095-1 | Bacteria Firmicutes Bacilli Bacillales Staphylococcaceae *Staphylococcus aureus* HBHO6002 |
| Sau-1096-1 | Bacteria Firmicutes Bacilli Bacillales Staphylococcaceae *Staphylococcus aureus* HBHO6021 |
| Sau-1097-1 | Bacteria Firmicutes Bacilli Bacillales Staphylococcaceae *Staphylococcus aureus* HBHO6022 |

| | |
|---|---|
| Sau-1098-1 | Bacteria Firmicutes Bacilli Bacillales Staphylococcaceae *Staphylococcus aureus* HBHO6024 |
| Sau-1099-1 | Bacteria Firmicutes Bacilli Bacillales Staphylococcaceae *Staphylococcus aureus* HBHO6042 |
| Sau-1100-1 | Bacteria Firmicutes Bacilli Bacillales Staphylococcaceae *Staphylococcus aureus* HBHO6050 |
| Sau-1101-1 | Bacteria Firmicutes Bacilli Bacillales Staphylococcaceae *Staphylococcus aureus* HOAG6005 |
| Sau-1102-1 | Bacteria Firmicutes Bacilli Bacillales Staphylococcaceae *Staphylococcus aureus* HOAG6017 |
| Sau-1103-1 | Bacteria Firmicutes Bacilli Bacillales Staphylococcaceae *Staphylococcus aureus* HOAG6025 |
| Sau-1104-1 | Bacteria Firmicutes Bacilli Bacillales Staphylococcaceae *Staphylococcus aureus* HOAG6034 |
| Sau-1105-1 | Bacteria Firmicutes Bacilli Bacillales Staphylococcaceae *Staphylococcus aureus* HOAG6042 |
| Sau-1106-1 | Bacteria Firmicutes Bacilli Bacillales Staphylococcaceae *Staphylococcus aureus* HOAG6043 |
| Sau-1107-1 | Bacteria Firmicutes Bacilli Bacillales Staphylococcaceae *Staphylococcus aureus* HOAG6047 |
| Sau-1108-1 | Bacteria Firmicutes Bacilli Bacillales Staphylococcaceae *Staphylococcus aureus* HOAG6054 |
| Sau-1109-1 | Bacteria Firmicutes Bacilli Bacillales Staphylococcaceae *Staphylococcus aureus* HOAG6057 |
| Sau-1110-1 | Bacteria Firmicutes Bacilli Bacillales Staphylococcaceae *Staphylococcus aureus* HOAG6059 |
| Sau-1111-1 | Bacteria Firmicutes Bacilli Bacillales Staphylococcaceae *Staphylococcus aureus* HOAG6068 |
| Sau-1112-1 | Bacteria Firmicutes Bacilli Bacillales Staphylococcaceae *Staphylococcus aureus* HOAG6084 |
| Sau-1113-1 | Bacteria Firmicutes Bacilli Bacillales Staphylococcaceae *Staphylococcus aureus* HOAG6092 |
| Sau-1114-1 | Bacteria Firmicutes Bacilli Bacillales Staphylococcaceae *Staphylococcus aureus* HOAG6104 |
| Sau-1115-1 | Bacteria Firmicutes Bacilli Bacillales Staphylococcaceae *Staphylococcus aureus* HSOU6002 |
| Sau-1116-1 | Bacteria Firmicutes Bacilli Bacillales Staphylococcaceae *Staphylococcus aureus* HST-066 |
| Sau-1117-1 | Bacteria Firmicutes Bacilli Bacillales Staphylococcaceae *Staphylococcus aureus* HST-077 |
| Sau-1118-1 | Bacteria Firmicutes Bacilli Bacillales Staphylococcaceae *Staphylococcus aureus* HST-084 |
| Sau-1119-1 | Bacteria Firmicutes Bacilli Bacillales Staphylococcaceae *Staphylococcus aureus* HST-105 |
| Sau-1120-1 | Bacteria Firmicutes Bacilli Bacillales Staphylococcaceae *Staphylococcus aureus* HST-107 |
| Sau-1121-1 | Bacteria Firmicutes Bacilli Bacillales Staphylococcaceae *Staphylococcus aureus* JS046 |
| Sau-1122-1 | Bacteria Firmicutes Bacilli Bacillales Staphylococcaceae *Staphylococcus aureus* KINB6009 |
| Sau-1123-1 | Bacteria Firmicutes Bacilli Bacillales Staphylococcaceae *Staphylococcus aureus* KINB6023 |
| Sau-1124-1 | Bacteria Firmicutes Bacilli Bacillales Staphylococcaceae *Staphylococcus aureus* KINB6024 |
| Sau-1125-1 | Bacteria Firmicutes Bacilli Bacillales Staphylococcaceae *Staphylococcus aureus* KINW6001 |
| Sau-1126-1 | Bacteria Firmicutes Bacilli Bacillales Staphylococcaceae *Staphylococcus aureus* KINW6002 |
| Sau-1127-1 | Bacteria Firmicutes Bacilli Bacillales Staphylococcaceae *Staphylococcus aureus* KINW6003 |
| Sau-1128-1 | Bacteria Firmicutes Bacilli Bacillales Staphylococcaceae *Staphylococcus aureus* KINW6010 |
| Sau-1129-1 | Bacteria Firmicutes Bacilli Bacillales Staphylococcaceae *Staphylococcus aureus* KINW6016 |
| Sau-1130-1 | Bacteria Firmicutes Bacilli Bacillales Staphylococcaceae *Staphylococcus aureus* KINW6019 |
| Sau-1131-1 | Bacteria Firmicutes Bacilli Bacillales Staphylococcaceae *Staphylococcus aureus* KINW6023 |
| Sau-1132-1 | Bacteria Firmicutes Bacilli Bacillales Staphylococcaceae *Staphylococcus aureus* KINW6025 |
| Sau-1133-1 | Bacteria Firmicutes Bacilli Bacillales Staphylococcaceae *Staphylococcus aureus* KINW6027 |
| Sau-1134-1 | Bacteria Firmicutes Bacilli Bacillales Staphylococcaceae *Staphylococcus aureus* KINW6033 |
| Sau-1135-1 | Bacteria Firmicutes Bacilli Bacillales Staphylococcaceae *Staphylococcus aureus* KINW6036 |
| Sau-1136-1 | Bacteria Firmicutes Bacilli Bacillales Staphylococcaceae *Staphylococcus aureus* KINW6040 |
| Sau-1137-1 | Bacteria Firmicutes Bacilli Bacillales Staphylococcaceae *Staphylococcus aureus* KINW6046 |
| Sau-1138-1 | Bacteria Firmicutes Bacilli Bacillales Staphylococcaceae *Staphylococcus aureus* KINW6047 |
| Sau-1139-1 | Bacteria Firmicutes Bacilli Bacillales Staphylococcaceae *Staphylococcus aureus* KINW6048 |
| Sau-1140-1 | Bacteria Firmicutes Bacilli Bacillales Staphylococcaceae *Staphylococcus aureus* KINW6056 |
| Sau-1141-1 | Bacteria Firmicutes Bacilli Bacillales Staphylococcaceae *Staphylococcus aureus* KINW6058 |
| Sau-1142-1 | Bacteria Firmicutes Bacilli Bacillales Staphylococcaceae *Staphylococcus aureus* KINW6061 |
| Sau-1143-1 | Bacteria Firmicutes Bacilli Bacillales Staphylococcaceae *Staphylococcus aureus* KINW6062 |
| Sau-1144-1 | Bacteria Firmicutes Bacilli Bacillales Staphylococcaceae *Staphylococcus aureus* KINW6066 |
| Sau-1145-1 | Bacteria Firmicutes Bacilli Bacillales Staphylococcaceae *Staphylococcus aureus* LAMC0001 |
| Sau-1146-1 | Bacteria Firmicutes Bacilli Bacillales Staphylococcaceae *Staphylococcus aureus* LAMC0010 |
| Sau-1147-1 | Bacteria Firmicutes Bacilli Bacillales Staphylococcaceae *Staphylococcus aureus* LAMC0011 |
| Sau-1148-1 | Bacteria Firmicutes Bacilli Bacillales Staphylococcaceae *Staphylococcus aureus* LAMC0013 |
| Sau-1149-1 | Bacteria Firmicutes Bacilli Bacillales Staphylococcaceae *Staphylococcus aureus* LAMC0016 |
| Sau-1150-1 | Bacteria Firmicutes Bacilli Bacillales Staphylococcaceae *Staphylococcus aureus* LAMC0024 |
| Sau-1151-1 | Bacteria Firmicutes Bacilli Bacillales Staphylococcaceae *Staphylococcus aureus* LAMC0037 |
| Sau-1152-1 | Bacteria Firmicutes Bacilli Bacillales Staphylococcaceae *Staphylococcus aureus* LAMC0047 |
| Sau-1153-1 | Bacteria Firmicutes Bacilli Bacillales Staphylococcaceae *Staphylococcus aureus* LAMC0050 |
| Sau-1154-1 | Bacteria Firmicutes Bacilli Bacillales Staphylococcaceae *Staphylococcus aureus* LAMC6065 |
| Sau-1155-1 | Bacteria Firmicutes Bacilli Bacillales Staphylococcaceae *Staphylococcus aureus* LAMC6066 |
| Sau-1156-1 | Bacteria Firmicutes Bacilli Bacillales Staphylococcaceae *Staphylococcus aureus* LAMC6078 |
| Sau-1157-1 | Bacteria Firmicutes Bacilli Bacillales Staphylococcaceae *Staphylococcus aureus* LAMC6084 |
| Sau-1158-1 | Bacteria Firmicutes Bacilli Bacillales Staphylococcaceae *Staphylococcus aureus* LAMC6094 |
| Sau-1159-1 | Bacteria Firmicutes Bacilli Bacillales Staphylococcaceae *Staphylococcus aureus* LAMC6106 |
| Sau-1160-1 | Bacteria Firmicutes Bacilli Bacillales Staphylococcaceae *Staphylococcus aureus* LAMC6115 |
| Sau-1161-1 | Bacteria Firmicutes Bacilli Bacillales Staphylococcaceae *Staphylococcus aureus* LAMC6117 |
| Sau-1162-1 | Bacteria Firmicutes Bacilli Bacillales Staphylococcaceae *Staphylococcus aureus* LCT-SA67 |

| | |
|---|---|
| Sau-1163-1 | Bacteria Firmicutes Bacilli Bacillales Staphylococcaceae *Staphylococcus aureus* LCT-SAM |
| Sau-1164-1 | Bacteria Firmicutes Bacilli Bacillales Staphylococcaceae *Staphylococcus aureus* LCT-SAO |
| Sau-1165-1 | Bacteria Firmicutes Bacilli Bacillales Staphylococcaceae *Staphylococcus aureus* LPIH6001 |
| Sau-1166-1 | Bacteria Firmicutes Bacilli Bacillales Staphylococcaceae *Staphylococcus aureus* LPIH6004 |
| Sau-1167-1 | Bacteria Firmicutes Bacilli Bacillales Staphylococcaceae *Staphylococcus aureus* LPIH6008 |
| Sau-1168-1 | Bacteria Firmicutes Bacilli Bacillales Staphylococcaceae *Staphylococcus aureus* LPIH6010 |
| Sau-1169-1 | Bacteria Firmicutes Bacilli Bacillales Staphylococcaceae *Staphylococcus aureus* LPIH6011 |
| Sau-1170-1 | Bacteria Firmicutes Bacilli Bacillales Staphylococcaceae *Staphylococcus aureus* LPIH6021 |
| Sau-1171-1 | Bacteria Firmicutes Bacilli Bacillales Staphylococcaceae *Staphylococcus aureus* LysK 1 2010 |
| Sau-1172-1 | Bacteria Firmicutes Bacilli Bacillales Staphylococcaceae *Staphylococcus aureus* LysK 1 2011 |
| Sau-1173-1 | Bacteria Firmicutes Bacilli Bacillales Staphylococcaceae *Staphylococcus aureus* LysK 2 2010 |
| Sau-1174-1 | Bacteria Firmicutes Bacilli Bacillales Staphylococcaceae *Staphylococcus aureus* LysK 2 2011 |
| Sau-1175-1 | Bacteria Firmicutes Bacilli Bacillales Staphylococcaceae *Staphylococcus aureus* Lyso 1 2010 |
| Sau-1176-1 | Bacteria Firmicutes Bacilli Bacillales Staphylococcaceae *Staphylococcus aureus* Lyso 2 2010 |
| Sau-1177-1 | Bacteria Firmicutes Bacilli Bacillales Staphylococcaceae *Staphylococcus aureus* M0002 |
| Sau-1178-1 | Bacteria Firmicutes Bacilli Bacillales Staphylococcaceae *Staphylococcus aureus* M0004 |
| Sau-1179-1 | Bacteria Firmicutes Bacilli Bacillales Staphylococcaceae *Staphylococcus aureus* M0005 |
| Sau-1180-1 | Bacteria Firmicutes Bacilli Bacillales Staphylococcaceae *Staphylococcus aureus* M0007 |
| Sau-1181-1 | Bacteria Firmicutes Bacilli Bacillales Staphylococcaceae *Staphylococcus aureus* M0008 |
| Sau-1182-1 | Bacteria Firmicutes Bacilli Bacillales Staphylococcaceae *Staphylococcus aureus* M0009 |
| Sau-1183-1 | Bacteria Firmicutes Bacilli Bacillales Staphylococcaceae *Staphylococcus aureus* M0010 |
| Sau-1184-1 | Bacteria Firmicutes Bacilli Bacillales Staphylococcaceae *Staphylococcus aureus* M0011 |
| Sau-1185-1 | Bacteria Firmicutes Bacilli Bacillales Staphylococcaceae *Staphylococcus aureus* M0012 |
| Sau-1186-1 | Bacteria Firmicutes Bacilli Bacillales Staphylococcaceae *Staphylococcus aureus* M0013 |
| Sau-1187-1 | Bacteria Firmicutes Bacilli Bacillales Staphylococcaceae *Staphylococcus aureus* M0014 |
| Sau-1188-1 | Bacteria Firmicutes Bacilli Bacillales Staphylococcaceae *Staphylococcus aureus* M0015 |
| Sau-1189-1 | Bacteria Firmicutes Bacilli Bacillales Staphylococcaceae *Staphylococcus aureus* M0016 |
| Sau-1190-1 | Bacteria Firmicutes Bacilli Bacillales Staphylococcaceae *Staphylococcus aureus* M0017 |
| Sau-1191-1 | Bacteria Firmicutes Bacilli Bacillales Staphylococcaceae *Staphylococcus aureus* M0018 |
| Sau-1192-1 | Bacteria Firmicutes Bacilli Bacillales Staphylococcaceae *Staphylococcus aureus* M0021 |
| Sau-1193-1 | Bacteria Firmicutes Bacilli Bacillales Staphylococcaceae *Staphylococcus aureus* M0022 |
| Sau-1194-1 | Bacteria Firmicutes Bacilli Bacillales Staphylococcaceae *Staphylococcus aureus* M0023 |
| Sau-1195-1 | Bacteria Firmicutes Bacilli Bacillales Staphylococcaceae *Staphylococcus aureus* M0024 |
| Sau-1196-1 | Bacteria Firmicutes Bacilli Bacillales Staphylococcaceae *Staphylococcus aureus* M0025 |
| Sau-1197-1 | Bacteria Firmicutes Bacilli Bacillales Staphylococcaceae *Staphylococcus aureus* M0027 |
| Sau-1198-1 | Bacteria Firmicutes Bacilli Bacillales Staphylococcaceae *Staphylococcus aureus* M0028 |
| Sau-1199-1 | Bacteria Firmicutes Bacilli Bacillales Staphylococcaceae *Staphylococcus aureus* M0030 |
| Sau-1200-1 | Bacteria Firmicutes Bacilli Bacillales Staphylococcaceae *Staphylococcus aureus* M0031 |
| Sau-1201-1 | Bacteria Firmicutes Bacilli Bacillales Staphylococcaceae *Staphylococcus aureus* M0032 |
| Sau-1202-1 | Bacteria Firmicutes Bacilli Bacillales Staphylococcaceae *Staphylococcus aureus* M0034 |
| Sau-1203-1 | Bacteria Firmicutes Bacilli Bacillales Staphylococcaceae *Staphylococcus aureus* M0036 |
| Sau-1204-1 | Bacteria Firmicutes Bacilli Bacillales Staphylococcaceae *Staphylococcus aureus* M0037 |
| Sau-1205-1 | Bacteria Firmicutes Bacilli Bacillales Staphylococcaceae *Staphylococcus aureus* M0038 |
| Sau-1206-1 | Bacteria Firmicutes Bacilli Bacillales Staphylococcaceae *Staphylococcus aureus* M0039 |
| Sau-1207-1 | Bacteria Firmicutes Bacilli Bacillales Staphylococcaceae *Staphylococcus aureus* M0040 |
| Sau-1208-1 | Bacteria Firmicutes Bacilli Bacillales Staphylococcaceae *Staphylococcus aureus* M0041 |
| Sau-1209-1 | Bacteria Firmicutes Bacilli Bacillales Staphylococcaceae *Staphylococcus aureus* M0042 |
| Sau-1210-1 | Bacteria Firmicutes Bacilli Bacillales Staphylococcaceae *Staphylococcus aureus* M0043 |
| Sau-1211-1 | Bacteria Firmicutes Bacilli Bacillales Staphylococcaceae *Staphylococcus aureus* M0044 |
| Sau-1212-1 | Bacteria Firmicutes Bacilli Bacillales Staphylococcaceae *Staphylococcus aureus* M0046 |
| Sau-1213-1 | Bacteria Firmicutes Bacilli Bacillales Staphylococcaceae *Staphylococcus aureus* M0047 |
| Sau-1214-1 | Bacteria Firmicutes Bacilli Bacillales Staphylococcaceae *Staphylococcus aureus* M0048 |
| Sau-1215-1 | Bacteria Firmicutes Bacilli Bacillales Staphylococcaceae *Staphylococcus aureus* M0049 |
| Sau-1216-1 | Bacteria Firmicutes Bacilli Bacillales Staphylococcaceae *Staphylococcus aureus* M0050 |
| Sau-1217-1 | Bacteria Firmicutes Bacilli Bacillales Staphylococcaceae *Staphylococcus aureus* M0051 |
| Sau-1218-1 | Bacteria Firmicutes Bacilli Bacillales Staphylococcaceae *Staphylococcus aureus* M0052 |
| Sau-1219-1 | Bacteria Firmicutes Bacilli Bacillales Staphylococcaceae *Staphylococcus aureus* M0054 |
| Sau-1220-1 | Bacteria Firmicutes Bacilli Bacillales Staphylococcaceae *Staphylococcus aureus* M0056 |
| Sau-1221-1 | Bacteria Firmicutes Bacilli Bacillales Staphylococcaceae *Staphylococcus aureus* M0057 |
| Sau-1222-1 | Bacteria Firmicutes Bacilli Bacillales Staphylococcaceae *Staphylococcus aureus* M0058 |
| Sau-1223-1 | Bacteria Firmicutes Bacilli Bacillales Staphylococcaceae *Staphylococcus aureus* M0059 |
| Sau-1224-1 | Bacteria Firmicutes Bacilli Bacillales Staphylococcaceae *Staphylococcus aureus* M0061 |
| Sau-1225-1 | Bacteria Firmicutes Bacilli Bacillales Staphylococcaceae *Staphylococcus aureus* M0062 |
| Sau-1226-1 | Bacteria Firmicutes Bacilli Bacillales Staphylococcaceae *Staphylococcus aureus* M0063 |
| Sau-1227-1 | Bacteria Firmicutes Bacilli Bacillales Staphylococcaceae *Staphylococcus aureus* M0064 |

| | |
|---|---|
| Sau-1228-1 | Bacteria Firmicutes Bacilli Bacillales Staphylococcaceae *Staphylococcus aureus* M0065 |
| Sau-1229-1 | Bacteria Firmicutes Bacilli Bacillales Staphylococcaceae *Staphylococcus aureus* M0067 |
| Sau-1230-1 | Bacteria Firmicutes Bacilli Bacillales Staphylococcaceae *Staphylococcus aureus* M0068 |
| Sau-1231-1 | Bacteria Firmicutes Bacilli Bacillales Staphylococcaceae *Staphylococcus aureus* M0069 |
| Sau-1232-1 | Bacteria Firmicutes Bacilli Bacillales Staphylococcaceae *Staphylococcus aureus* M0070 |
| Sau-1233-1 | Bacteria Firmicutes Bacilli Bacillales Staphylococcaceae *Staphylococcus aureus* M0071 |
| Sau-1234-1 | Bacteria Firmicutes Bacilli Bacillales Staphylococcaceae *Staphylococcus aureus* M0073 |
| Sau-1235-1 | Bacteria Firmicutes Bacilli Bacillales Staphylococcaceae *Staphylococcus aureus* M0074 |
| Sau-1236-1 | Bacteria Firmicutes Bacilli Bacillales Staphylococcaceae *Staphylococcus aureus* M0076 |
| Sau-1237-1 | Bacteria Firmicutes Bacilli Bacillales Staphylococcaceae *Staphylococcus aureus* M0078 |
| Sau-1238-1 | Bacteria Firmicutes Bacilli Bacillales Staphylococcaceae *Staphylococcus aureus* M0079 |
| Sau-1239-1 | Bacteria Firmicutes Bacilli Bacillales Staphylococcaceae *Staphylococcus aureus* M0080 |
| Sau-1240-1 | Bacteria Firmicutes Bacilli Bacillales Staphylococcaceae *Staphylococcus aureus* M0081 |
| Sau-1241-1 | Bacteria Firmicutes Bacilli Bacillales Staphylococcaceae *Staphylococcus aureus* M0082 |
| Sau-1242-1 | Bacteria Firmicutes Bacilli Bacillales Staphylococcaceae *Staphylococcus aureus* M0083 |
| Sau-1243-1 | Bacteria Firmicutes Bacilli Bacillales Staphylococcaceae *Staphylococcus aureus* M0084 |
| Sau-1244-1 | Bacteria Firmicutes Bacilli Bacillales Staphylococcaceae *Staphylococcus aureus* M0086 |
| Sau-1245-1 | Bacteria Firmicutes Bacilli Bacillales Staphylococcaceae *Staphylococcus aureus* M0087 |
| Sau-1246-1 | Bacteria Firmicutes Bacilli Bacillales Staphylococcaceae *Staphylococcus aureus* M0088 |
| Sau-1247-1 | Bacteria Firmicutes Bacilli Bacillales Staphylococcaceae *Staphylococcus aureus* M0089 |
| Sau-1248-1 | Bacteria Firmicutes Bacilli Bacillales Staphylococcaceae *Staphylococcus aureus* M0092 |
| Sau-1249-1 | Bacteria Firmicutes Bacilli Bacillales Staphylococcaceae *Staphylococcus aureus* M0093 |
| Sau-1250-1 | Bacteria Firmicutes Bacilli Bacillales Staphylococcaceae *Staphylococcus aureus* M0094 |
| Sau-1251-1 | Bacteria Firmicutes Bacilli Bacillales Staphylococcaceae *Staphylococcus aureus* M0095 |
| Sau-1252-1 | Bacteria Firmicutes Bacilli Bacillales Staphylococcaceae *Staphylococcus aureus* M0096 |
| Sau-1253-1 | Bacteria Firmicutes Bacilli Bacillales Staphylococcaceae *Staphylococcus aureus* M0097 |
| Sau-1254-1 | Bacteria Firmicutes Bacilli Bacillales Staphylococcaceae *Staphylococcus aureus* M0098 |
| Sau-1255-1 | Bacteria Firmicutes Bacilli Bacillales Staphylococcaceae *Staphylococcus aureus* M0099 |
| Sau-1256-1 | Bacteria Firmicutes Bacilli Bacillales Staphylococcaceae *Staphylococcus aureus* M0100 |
| Sau-1257-1 | Bacteria Firmicutes Bacilli Bacillales Staphylococcaceae *Staphylococcus aureus* M0101 |
| Sau-1258-1 | Bacteria Firmicutes Bacilli Bacillales Staphylococcaceae *Staphylococcus aureus* M0105 |
| Sau-1259-1 | Bacteria Firmicutes Bacilli Bacillales Staphylococcaceae *Staphylococcus aureus* M0107 |
| Sau-1260-1 | Bacteria Firmicutes Bacilli Bacillales Staphylococcaceae *Staphylococcus aureus* M0109 |
| Sau-1261-1 | Bacteria Firmicutes Bacilli Bacillales Staphylococcaceae *Staphylococcus aureus* M0110 |
| Sau-1262-1 | Bacteria Firmicutes Bacilli Bacillales Staphylococcaceae *Staphylococcus aureus* M0111 |
| Sau-1263-1 | Bacteria Firmicutes Bacilli Bacillales Staphylococcaceae *Staphylococcus aureus* M0112 |
| Sau-1264-1 | Bacteria Firmicutes Bacilli Bacillales Staphylococcaceae *Staphylococcus aureus* M0113 |
| Sau-1265-1 | Bacteria Firmicutes Bacilli Bacillales Staphylococcaceae *Staphylococcus aureus* M0114 |
| Sau-1266-1 | Bacteria Firmicutes Bacilli Bacillales Staphylococcaceae *Staphylococcus aureus* M0115 |
| Sau-1267-1 | Bacteria Firmicutes Bacilli Bacillales Staphylococcaceae *Staphylococcus aureus* M0116 |
| Sau-1268-1 | Bacteria Firmicutes Bacilli Bacillales Staphylococcaceae *Staphylococcus aureus* M0117 |
| Sau-1269-1 | Bacteria Firmicutes Bacilli Bacillales Staphylococcaceae *Staphylococcus aureus* M0118 |
| Sau-1270-1 | Bacteria Firmicutes Bacilli Bacillales Staphylococcaceae *Staphylococcus aureus* M0119 |
| Sau-1271-1 | Bacteria Firmicutes Bacilli Bacillales Staphylococcaceae *Staphylococcus aureus* M0120 |
| Sau-1272-1 | Bacteria Firmicutes Bacilli Bacillales Staphylococcaceae *Staphylococcus aureus* M0121 |
| Sau-1273-1 | Bacteria Firmicutes Bacilli Bacillales Staphylococcaceae *Staphylococcus aureus* M0122 |
| Sau-1274-1 | Bacteria Firmicutes Bacilli Bacillales Staphylococcaceae *Staphylococcus aureus* M0123 |
| Sau-1275-1 | Bacteria Firmicutes Bacilli Bacillales Staphylococcaceae *Staphylococcus aureus* M0124 |
| Sau-1276-1 | Bacteria Firmicutes Bacilli Bacillales Staphylococcaceae *Staphylococcus aureus* M0125 |
| Sau-1277-1 | Bacteria Firmicutes Bacilli Bacillales Staphylococcaceae *Staphylococcus aureus* M0126 |
| Sau-1278-1 | Bacteria Firmicutes Bacilli Bacillales Staphylococcaceae *Staphylococcus aureus* M0127 |
| Sau-1279-1 | Bacteria Firmicutes Bacilli Bacillales Staphylococcaceae *Staphylococcus aureus* M0128 |
| Sau-1280-1 | Bacteria Firmicutes Bacilli Bacillales Staphylococcaceae *Staphylococcus aureus* M0130 |
| Sau-1281-1 | Bacteria Firmicutes Bacilli Bacillales Staphylococcaceae *Staphylococcus aureus* M0131 |
| Sau-1282-1 | Bacteria Firmicutes Bacilli Bacillales Staphylococcaceae *Staphylococcus aureus* M0132 |
| Sau-1283-1 | Bacteria Firmicutes Bacilli Bacillales Staphylococcaceae *Staphylococcus aureus* M0133 |
| Sau-1284-1 | Bacteria Firmicutes Bacilli Bacillales Staphylococcaceae *Staphylococcus aureus* M0134 |
| Sau-1285-1 | Bacteria Firmicutes Bacilli Bacillales Staphylococcaceae *Staphylococcus aureus* M0135 |
| Sau-1286-1 | Bacteria Firmicutes Bacilli Bacillales Staphylococcaceae *Staphylococcus aureus* M0136 |
| Sau-1287-1 | Bacteria Firmicutes Bacilli Bacillales Staphylococcaceae *Staphylococcus aureus* M0137 |
| Sau-1288-1 | Bacteria Firmicutes Bacilli Bacillales Staphylococcaceae *Staphylococcus aureus* M0138 |
| Sau-1289-1 | Bacteria Firmicutes Bacilli Bacillales Staphylococcaceae *Staphylococcus aureus* M0139 |
| Sau-1290-1 | Bacteria Firmicutes Bacilli Bacillales Staphylococcaceae *Staphylococcus aureus* M0141 |
| Sau-1291-1 | Bacteria Firmicutes Bacilli Bacillales Staphylococcaceae *Staphylococcus aureus* M0142 |
| Sau-1292-1 | Bacteria Firmicutes Bacilli Bacillales Staphylococcaceae *Staphylococcus aureus* M0143 |

| | |
|---|---|
| Sau-1293-1 | Bacteria Firmicutes Bacilli Bacillales Staphylococcaceae *Staphylococcus aureus* M0145 |
| Sau-1294-1 | Bacteria Firmicutes Bacilli Bacillales Staphylococcaceae *Staphylococcus aureus* M0146 |
| Sau-1295-1 | Bacteria Firmicutes Bacilli Bacillales Staphylococcaceae *Staphylococcus aureus* M0147 |
| Sau-1296-1 | Bacteria Firmicutes Bacilli Bacillales Staphylococcaceae *Staphylococcus aureus* M0148 |
| Sau-1297-1 | Bacteria Firmicutes Bacilli Bacillales Staphylococcaceae *Staphylococcus aureus* M0149 |
| Sau-1298-1 | Bacteria Firmicutes Bacilli Bacillales Staphylococcaceae *Staphylococcus aureus* M0152 |
| Sau-1299-1 | Bacteria Firmicutes Bacilli Bacillales Staphylococcaceae *Staphylococcus aureus* M0153 |
| Sau-1300-1 | Bacteria Firmicutes Bacilli Bacillales Staphylococcaceae *Staphylococcus aureus* M0155 |
| Sau-1301-1 | Bacteria Firmicutes Bacilli Bacillales Staphylococcaceae *Staphylococcus aureus* M0156 |
| Sau-1302-1 | Bacteria Firmicutes Bacilli Bacillales Staphylococcaceae *Staphylococcus aureus* M0157 |
| Sau-1303-1 | Bacteria Firmicutes Bacilli Bacillales Staphylococcaceae *Staphylococcus aureus* M0158 |
| Sau-1304-1 | Bacteria Firmicutes Bacilli Bacillales Staphylococcaceae *Staphylococcus aureus* M0159 |
| Sau-1305-1 | Bacteria Firmicutes Bacilli Bacillales Staphylococcaceae *Staphylococcus aureus* M0160 |
| Sau-1306-1 | Bacteria Firmicutes Bacilli Bacillales Staphylococcaceae *Staphylococcus aureus* M0161 |
| Sau-1307-1 | Bacteria Firmicutes Bacilli Bacillales Staphylococcaceae *Staphylococcus aureus* M0162 |
| Sau-1308-1 | Bacteria Firmicutes Bacilli Bacillales Staphylococcaceae *Staphylococcus aureus* M0163 |
| Sau-1309-1 | Bacteria Firmicutes Bacilli Bacillales Staphylococcaceae *Staphylococcus aureus* M0164 |
| Sau-1310-1 | Bacteria Firmicutes Bacilli Bacillales Staphylococcaceae *Staphylococcus aureus* M0165 |
| Sau-1311-1 | Bacteria Firmicutes Bacilli Bacillales Staphylococcaceae *Staphylococcus aureus* M0166 |
| Sau-1312-1 | Bacteria Firmicutes Bacilli Bacillales Staphylococcaceae *Staphylococcus aureus* M0167 |
| Sau-1313-1 | Bacteria Firmicutes Bacilli Bacillales Staphylococcaceae *Staphylococcus aureus* M0168 |
| Sau-1314-1 | Bacteria Firmicutes Bacilli Bacillales Staphylococcaceae *Staphylococcus aureus* M0169 |
| Sau-1315-1 | Bacteria Firmicutes Bacilli Bacillales Staphylococcaceae *Staphylococcus aureus* M0170 |
| Sau-1316-1 | Bacteria Firmicutes Bacilli Bacillales Staphylococcaceae *Staphylococcus aureus* M0172 |
| Sau-1317-1 | Bacteria Firmicutes Bacilli Bacillales Staphylococcaceae *Staphylococcus aureus* M0174 |
| Sau-1318-1 | Bacteria Firmicutes Bacilli Bacillales Staphylococcaceae *Staphylococcus aureus* M0175 |
| Sau-1319-1 | Bacteria Firmicutes Bacilli Bacillales Staphylococcaceae *Staphylococcus aureus* M0176 |
| Sau-1320-1 | Bacteria Firmicutes Bacilli Bacillales Staphylococcaceae *Staphylococcus aureus* M0178 |
| Sau-1321-1 | Bacteria Firmicutes Bacilli Bacillales Staphylococcaceae *Staphylococcus aureus* M0179 |
| Sau-1322-1 | Bacteria Firmicutes Bacilli Bacillales Staphylococcaceae *Staphylococcus aureus* M0180 |
| Sau-1323-1 | Bacteria Firmicutes Bacilli Bacillales Staphylococcaceae *Staphylococcus aureus* M0181 |
| Sau-1324-1 | Bacteria Firmicutes Bacilli Bacillales Staphylococcaceae *Staphylococcus aureus* M0182 |
| Sau-1325-1 | Bacteria Firmicutes Bacilli Bacillales Staphylococcaceae *Staphylococcus aureus* M0183 |
| Sau-1326-1 | Bacteria Firmicutes Bacilli Bacillales Staphylococcaceae *Staphylococcus aureus* M0184 |
| Sau-1327-1 | Bacteria Firmicutes Bacilli Bacillales Staphylococcaceae *Staphylococcus aureus* M0185 |
| Sau-1328-1 | Bacteria Firmicutes Bacilli Bacillales Staphylococcaceae *Staphylococcus aureus* M0186 |
| Sau-1329-1 | Bacteria Firmicutes Bacilli Bacillales Staphylococcaceae *Staphylococcus aureus* M0187 |
| Sau-1330-1 | Bacteria Firmicutes Bacilli Bacillales Staphylococcaceae *Staphylococcus aureus* M0188 |
| Sau-1331-1 | Bacteria Firmicutes Bacilli Bacillales Staphylococcaceae *Staphylococcus aureus* M0190 |
| Sau-1332-1 | Bacteria Firmicutes Bacilli Bacillales Staphylococcaceae *Staphylococcus aureus* M0191 |
| Sau-1333-1 | Bacteria Firmicutes Bacilli Bacillales Staphylococcaceae *Staphylococcus aureus* M0193 |
| Sau-1334-1 | Bacteria Firmicutes Bacilli Bacillales Staphylococcaceae *Staphylococcus aureus* M0194 |
| Sau-1335-1 | Bacteria Firmicutes Bacilli Bacillales Staphylococcaceae *Staphylococcus aureus* M0195 |
| Sau-1336-1 | Bacteria Firmicutes Bacilli Bacillales Staphylococcaceae *Staphylococcus aureus* M0196 |
| Sau-1337-1 | Bacteria Firmicutes Bacilli Bacillales Staphylococcaceae *Staphylococcus aureus* M0198 |
| Sau-1338-1 | Bacteria Firmicutes Bacilli Bacillales Staphylococcaceae *Staphylococcus aureus* M0199 |
| Sau-1339-1 | Bacteria Firmicutes Bacilli Bacillales Staphylococcaceae *Staphylococcus aureus* M0201 |
| Sau-1340-1 | Bacteria Firmicutes Bacilli Bacillales Staphylococcaceae *Staphylococcus aureus* M0202 |
| Sau-1341-1 | Bacteria Firmicutes Bacilli Bacillales Staphylococcaceae *Staphylococcus aureus* M0203 |
| Sau-1342-1 | Bacteria Firmicutes Bacilli Bacillales Staphylococcaceae *Staphylococcus aureus* M0204 |
| Sau-1343-1 | Bacteria Firmicutes Bacilli Bacillales Staphylococcaceae *Staphylococcus aureus* M0205 |
| Sau-1344-1 | Bacteria Firmicutes Bacilli Bacillales Staphylococcaceae *Staphylococcus aureus* M0207 |
| Sau-1345-1 | Bacteria Firmicutes Bacilli Bacillales Staphylococcaceae *Staphylococcus aureus* M0208 |
| Sau-1346-1 | Bacteria Firmicutes Bacilli Bacillales Staphylococcaceae *Staphylococcus aureus* M0209 |
| Sau-1347-1 | Bacteria Firmicutes Bacilli Bacillales Staphylococcaceae *Staphylococcus aureus* M0211 |
| Sau-1348-1 | Bacteria Firmicutes Bacilli Bacillales Staphylococcaceae *Staphylococcus aureus* M0214 |
| Sau-1349-1 | Bacteria Firmicutes Bacilli Bacillales Staphylococcaceae *Staphylococcus aureus* M0215 |
| Sau-1350-1 | Bacteria Firmicutes Bacilli Bacillales Staphylococcaceae *Staphylococcus aureus* M0218 |
| Sau-1351-1 | Bacteria Firmicutes Bacilli Bacillales Staphylococcaceae *Staphylococcus aureus* M0219 |
| Sau-1352-1 | Bacteria Firmicutes Bacilli Bacillales Staphylococcaceae *Staphylococcus aureus* M0220 |
| Sau-1353-1 | Bacteria Firmicutes Bacilli Bacillales Staphylococcaceae *Staphylococcus aureus* M0222 |
| Sau-1354-1 | Bacteria Firmicutes Bacilli Bacillales Staphylococcaceae *Staphylococcus aureus* M0224 |
| Sau-1355-1 | Bacteria Firmicutes Bacilli Bacillales Staphylococcaceae *Staphylococcus aureus* M0225 |
| Sau-1356-1 | Bacteria Firmicutes Bacilli Bacillales Staphylococcaceae *Staphylococcus aureus* M0226 |
| Sau-1357-1 | Bacteria Firmicutes Bacilli Bacillales Staphylococcaceae *Staphylococcus aureus* M0227 |

| | |
|---|---|
| Sau-1358-1 | Bacteria Firmicutes Bacilli Bacillales Staphylococcaceae *Staphylococcus aureus* M0228 |
| Sau-1359-1 | Bacteria Firmicutes Bacilli Bacillales Staphylococcaceae *Staphylococcus aureus* M0229 |
| Sau-1360-1 | Bacteria Firmicutes Bacilli Bacillales Staphylococcaceae *Staphylococcus aureus* M0230 |
| Sau-1361-1 | Bacteria Firmicutes Bacilli Bacillales Staphylococcaceae *Staphylococcus aureus* M0231 |
| Sau-1362-1 | Bacteria Firmicutes Bacilli Bacillales Staphylococcaceae *Staphylococcus aureus* M0232 |
| Sau-1363-1 | Bacteria Firmicutes Bacilli Bacillales Staphylococcaceae *Staphylococcus aureus* M0234 |
| Sau-1364-1 | Bacteria Firmicutes Bacilli Bacillales Staphylococcaceae *Staphylococcus aureus* M0236 |
| Sau-1365-1 | Bacteria Firmicutes Bacilli Bacillales Staphylococcaceae *Staphylococcus aureus* M0238 |
| Sau-1366-1 | Bacteria Firmicutes Bacilli Bacillales Staphylococcaceae *Staphylococcus aureus* M0241 |
| Sau-1367-1 | Bacteria Firmicutes Bacilli Bacillales Staphylococcaceae *Staphylococcus aureus* M0242 |
| Sau-1368-1 | Bacteria Firmicutes Bacilli Bacillales Staphylococcaceae *Staphylococcus aureus* M0243 |
| Sau-1369-1 | Bacteria Firmicutes Bacilli Bacillales Staphylococcaceae *Staphylococcus aureus* M0244 |
| Sau-1370-1 | Bacteria Firmicutes Bacilli Bacillales Staphylococcaceae *Staphylococcus aureus* M0245 |
| Sau-1371-1 | Bacteria Firmicutes Bacilli Bacillales Staphylococcaceae *Staphylococcus aureus* M0246 |
| Sau-1372-1 | Bacteria Firmicutes Bacilli Bacillales Staphylococcaceae *Staphylococcus aureus* M0247 |
| Sau-1373-1 | Bacteria Firmicutes Bacilli Bacillales Staphylococcaceae *Staphylococcus aureus* M0248 |
| Sau-1374-1 | Bacteria Firmicutes Bacilli Bacillales Staphylococcaceae *Staphylococcus aureus* M0249 |
| Sau-1375-1 | Bacteria Firmicutes Bacilli Bacillales Staphylococcaceae *Staphylococcus aureus* M0251 |
| Sau-1376-1 | Bacteria Firmicutes Bacilli Bacillales Staphylococcaceae *Staphylococcus aureus* M0253 |
| Sau-1377-1 | Bacteria Firmicutes Bacilli Bacillales Staphylococcaceae *Staphylococcus aureus* M0254 |
| Sau-1378-1 | Bacteria Firmicutes Bacilli Bacillales Staphylococcaceae *Staphylococcus aureus* M0255 |
| Sau-1379-1 | Bacteria Firmicutes Bacilli Bacillales Staphylococcaceae *Staphylococcus aureus* M0256 |
| Sau-1380-1 | Bacteria Firmicutes Bacilli Bacillales Staphylococcaceae *Staphylococcus aureus* M0258 |
| Sau-1381-1 | Bacteria Firmicutes Bacilli Bacillales Staphylococcaceae *Staphylococcus aureus* M0259 |
| Sau-1382-1 | Bacteria Firmicutes Bacilli Bacillales Staphylococcaceae *Staphylococcus aureus* M0262 |
| Sau-1383-1 | Bacteria Firmicutes Bacilli Bacillales Staphylococcaceae *Staphylococcus aureus* M0263 |
| Sau-1384-1 | Bacteria Firmicutes Bacilli Bacillales Staphylococcaceae *Staphylococcus aureus* M0264 |
| Sau-1385-1 | Bacteria Firmicutes Bacilli Bacillales Staphylococcaceae *Staphylococcus aureus* M0265 |
| Sau-1386-1 | Bacteria Firmicutes Bacilli Bacillales Staphylococcaceae *Staphylococcus aureus* M0267 |
| Sau-1387-1 | Bacteria Firmicutes Bacilli Bacillales Staphylococcaceae *Staphylococcus aureus* M0268 |
| Sau-1388-1 | Bacteria Firmicutes Bacilli Bacillales Staphylococcaceae *Staphylococcus aureus* M0269 |
| Sau-1389-1 | Bacteria Firmicutes Bacilli Bacillales Staphylococcaceae *Staphylococcus aureus* M0271 |
| Sau-1390-1 | Bacteria Firmicutes Bacilli Bacillales Staphylococcaceae *Staphylococcus aureus* M0272 |
| Sau-1391-1 | Bacteria Firmicutes Bacilli Bacillales Staphylococcaceae *Staphylococcus aureus* M0275 |
| Sau-1392-1 | Bacteria Firmicutes Bacilli Bacillales Staphylococcaceae *Staphylococcus aureus* M0276 |
| Sau-1393-1 | Bacteria Firmicutes Bacilli Bacillales Staphylococcaceae *Staphylococcus aureus* M0277 |
| Sau-1394-1 | Bacteria Firmicutes Bacilli Bacillales Staphylococcaceae *Staphylococcus aureus* M0278 |
| Sau-1395-1 | Bacteria Firmicutes Bacilli Bacillales Staphylococcaceae *Staphylococcus aureus* M0281 |
| Sau-1396-1 | Bacteria Firmicutes Bacilli Bacillales Staphylococcaceae *Staphylococcus aureus* M0283 |
| Sau-1397-1 | Bacteria Firmicutes Bacilli Bacillales Staphylococcaceae *Staphylococcus aureus* M0284 |
| Sau-1398-1 | Bacteria Firmicutes Bacilli Bacillales Staphylococcaceae *Staphylococcus aureus* M0285 |
| Sau-1399-1 | Bacteria Firmicutes Bacilli Bacillales Staphylococcaceae *Staphylococcus aureus* M0286 |
| Sau-1400-1 | Bacteria Firmicutes Bacilli Bacillales Staphylococcaceae *Staphylococcus aureus* M0287 |
| Sau-1401-1 | Bacteria Firmicutes Bacilli Bacillales Staphylococcaceae *Staphylococcus aureus* M0289 |
| Sau-1402-1 | Bacteria Firmicutes Bacilli Bacillales Staphylococcaceae *Staphylococcus aureus* M0290 |
| Sau-1403-1 | Bacteria Firmicutes Bacilli Bacillales Staphylococcaceae *Staphylococcus aureus* M0291 |
| Sau-1404-1 | Bacteria Firmicutes Bacilli Bacillales Staphylococcaceae *Staphylococcus aureus* M0292 |
| Sau-1405-1 | Bacteria Firmicutes Bacilli Bacillales Staphylococcaceae *Staphylococcus aureus* M0293 |
| Sau-1406-1 | Bacteria Firmicutes Bacilli Bacillales Staphylococcaceae *Staphylococcus aureus* M0295 |
| Sau-1407-1 | Bacteria Firmicutes Bacilli Bacillales Staphylococcaceae *Staphylococcus aureus* M0296 |
| Sau-1408-1 | Bacteria Firmicutes Bacilli Bacillales Staphylococcaceae *Staphylococcus aureus* M0297 |
| Sau-1409-1 | Bacteria Firmicutes Bacilli Bacillales Staphylococcaceae *Staphylococcus aureus* M0298 |
| Sau-1410-1 | Bacteria Firmicutes Bacilli Bacillales Staphylococcaceae *Staphylococcus aureus* M0299 |
| Sau-1411-1 | Bacteria Firmicutes Bacilli Bacillales Staphylococcaceae *Staphylococcus aureus* M0300 |
| Sau-1412-1 | Bacteria Firmicutes Bacilli Bacillales Staphylococcaceae *Staphylococcus aureus* M0301 |
| Sau-1413-1 | Bacteria Firmicutes Bacilli Bacillales Staphylococcaceae *Staphylococcus aureus* M0302 |
| Sau-1414-1 | Bacteria Firmicutes Bacilli Bacillales Staphylococcaceae *Staphylococcus aureus* M0303 |
| Sau-1415-1 | Bacteria Firmicutes Bacilli Bacillales Staphylococcaceae *Staphylococcus aureus* M0304 |
| Sau-1416-1 | Bacteria Firmicutes Bacilli Bacillales Staphylococcaceae *Staphylococcus aureus* M0305 |
| Sau-1417-1 | Bacteria Firmicutes Bacilli Bacillales Staphylococcaceae *Staphylococcus aureus* M0307 |
| Sau-1418-1 | Bacteria Firmicutes Bacilli Bacillales Staphylococcaceae *Staphylococcus aureus* M0309 |
| Sau-1419-1 | Bacteria Firmicutes Bacilli Bacillales Staphylococcaceae *Staphylococcus aureus* M0310 |
| Sau-1420-1 | Bacteria Firmicutes Bacilli Bacillales Staphylococcaceae *Staphylococcus aureus* M0311 |
| Sau-1421-1 | Bacteria Firmicutes Bacilli Bacillales Staphylococcaceae *Staphylococcus aureus* M0313 |
| Sau-1422-1 | Bacteria Firmicutes Bacilli Bacillales Staphylococcaceae *Staphylococcus aureus* M0314 |

| | |
|---|---|
| Sau-1423-1 | Bacteria Firmicutes Bacilli Bacillales Staphylococcaceae *Staphylococcus aureus* M0316 |
| Sau-1424-1 | Bacteria Firmicutes Bacilli Bacillales Staphylococcaceae *Staphylococcus aureus* M0317 |
| Sau-1425-1 | Bacteria Firmicutes Bacilli Bacillales Staphylococcaceae *Staphylococcus aureus* M0318 |
| Sau-1426-1 | Bacteria Firmicutes Bacilli Bacillales Staphylococcaceae *Staphylococcus aureus* M0319 |
| Sau-1427-1 | Bacteria Firmicutes Bacilli Bacillales Staphylococcaceae *Staphylococcus aureus* M0320 |
| Sau-1428-1 | Bacteria Firmicutes Bacilli Bacillales Staphylococcaceae *Staphylococcus aureus* M0321 |
| Sau-1429-1 | Bacteria Firmicutes Bacilli Bacillales Staphylococcaceae *Staphylococcus aureus* M0322 |
| Sau-1430-1 | Bacteria Firmicutes Bacilli Bacillales Staphylococcaceae *Staphylococcus aureus* M0323 |
| Sau-1431-1 | Bacteria Firmicutes Bacilli Bacillales Staphylococcaceae *Staphylococcus aureus* M0324 |
| Sau-1432-1 | Bacteria Firmicutes Bacilli Bacillales Staphylococcaceae *Staphylococcus aureus* M0325 |
| Sau-1433-1 | Bacteria Firmicutes Bacilli Bacillales Staphylococcaceae *Staphylococcus aureus* M0327 |
| Sau-1434-1 | Bacteria Firmicutes Bacilli Bacillales Staphylococcaceae *Staphylococcus aureus* M0331 |
| Sau-1435-1 | Bacteria Firmicutes Bacilli Bacillales Staphylococcaceae *Staphylococcus aureus* M0332 |
| Sau-1436-1 | Bacteria Firmicutes Bacilli Bacillales Staphylococcaceae *Staphylococcus aureus* M0333 |
| Sau-1437-1 | Bacteria Firmicutes Bacilli Bacillales Staphylococcaceae *Staphylococcus aureus* M0335 |
| Sau-1438-1 | Bacteria Firmicutes Bacilli Bacillales Staphylococcaceae *Staphylococcus aureus* M0336 |
| Sau-1439-1 | Bacteria Firmicutes Bacilli Bacillales Staphylococcaceae *Staphylococcus aureus* M0337 |
| Sau-1440-1 | Bacteria Firmicutes Bacilli Bacillales Staphylococcaceae *Staphylococcus aureus* M0338 |
| Sau-1441-1 | Bacteria Firmicutes Bacilli Bacillales Staphylococcaceae *Staphylococcus aureus* M0339 |
| Sau-1442-1 | Bacteria Firmicutes Bacilli Bacillales Staphylococcaceae *Staphylococcus aureus* M0341 |
| Sau-1443-1 | Bacteria Firmicutes Bacilli Bacillales Staphylococcaceae *Staphylococcus aureus* M0342 |
| Sau-1444-1 | Bacteria Firmicutes Bacilli Bacillales Staphylococcaceae *Staphylococcus aureus* M0343 |
| Sau-1445-1 | Bacteria Firmicutes Bacilli Bacillales Staphylococcaceae *Staphylococcus aureus* M0344 |
| Sau-1446-1 | Bacteria Firmicutes Bacilli Bacillales Staphylococcaceae *Staphylococcus aureus* M0345 |
| Sau-1447-1 | Bacteria Firmicutes Bacilli Bacillales Staphylococcaceae *Staphylococcus aureus* M0346 |
| Sau-1448-1 | Bacteria Firmicutes Bacilli Bacillales Staphylococcaceae *Staphylococcus aureus* M0348 |
| Sau-1449-1 | Bacteria Firmicutes Bacilli Bacillales Staphylococcaceae *Staphylococcus aureus* M0349 |
| Sau-1450-1 | Bacteria Firmicutes Bacilli Bacillales Staphylococcaceae *Staphylococcus aureus* M0352 |
| Sau-1451-1 | Bacteria Firmicutes Bacilli Bacillales Staphylococcaceae *Staphylococcus aureus* M0353 |
| Sau-1452-1 | Bacteria Firmicutes Bacilli Bacillales Staphylococcaceae *Staphylococcus aureus* M0354 |
| Sau-1453-1 | Bacteria Firmicutes Bacilli Bacillales Staphylococcaceae *Staphylococcus aureus* M0355 |
| Sau-1454-1 | Bacteria Firmicutes Bacilli Bacillales Staphylococcaceae *Staphylococcus aureus* M0356 |
| Sau-1455-1 | Bacteria Firmicutes Bacilli Bacillales Staphylococcaceae *Staphylococcus aureus* M0357 |
| Sau-1456-1 | Bacteria Firmicutes Bacilli Bacillales Staphylococcaceae *Staphylococcus aureus* M0358 |
| Sau-1457-1 | Bacteria Firmicutes Bacilli Bacillales Staphylococcaceae *Staphylococcus aureus* M0359 |
| Sau-1458-1 | Bacteria Firmicutes Bacilli Bacillales Staphylococcaceae *Staphylococcus aureus* M0360 |
| Sau-1459-1 | Bacteria Firmicutes Bacilli Bacillales Staphylococcaceae *Staphylococcus aureus* M0361 |
| Sau-1460-1 | Bacteria Firmicutes Bacilli Bacillales Staphylococcaceae *Staphylococcus aureus* M0362 |
| Sau-1461-1 | Bacteria Firmicutes Bacilli Bacillales Staphylococcaceae *Staphylococcus aureus* M0365 |
| Sau-1462-1 | Bacteria Firmicutes Bacilli Bacillales Staphylococcaceae *Staphylococcus aureus* M0366 |
| Sau-1463-1 | Bacteria Firmicutes Bacilli Bacillales Staphylococcaceae *Staphylococcus aureus* M0368 |
| Sau-1464-1 | Bacteria Firmicutes Bacilli Bacillales Staphylococcaceae *Staphylococcus aureus* M0369 |
| Sau-1465-1 | Bacteria Firmicutes Bacilli Bacillales Staphylococcaceae *Staphylococcus aureus* M0370 |
| Sau-1466-1 | Bacteria Firmicutes Bacilli Bacillales Staphylococcaceae *Staphylococcus aureus* M0371 |
| Sau-1467-1 | Bacteria Firmicutes Bacilli Bacillales Staphylococcaceae *Staphylococcus aureus* M0372 |
| Sau-1468-1 | Bacteria Firmicutes Bacilli Bacillales Staphylococcaceae *Staphylococcus aureus* M0373 |
| Sau-1469-1 | Bacteria Firmicutes Bacilli Bacillales Staphylococcaceae *Staphylococcus aureus* M0376 |
| Sau-1470-1 | Bacteria Firmicutes Bacilli Bacillales Staphylococcaceae *Staphylococcus aureus* M0377 |
| Sau-1471-1 | Bacteria Firmicutes Bacilli Bacillales Staphylococcaceae *Staphylococcus aureus* M0378 |
| Sau-1472-1 | Bacteria Firmicutes Bacilli Bacillales Staphylococcaceae *Staphylococcus aureus* M0379 |
| Sau-1473-1 | Bacteria Firmicutes Bacilli Bacillales Staphylococcaceae *Staphylococcus aureus* M0380 |
| Sau-1474-1 | Bacteria Firmicutes Bacilli Bacillales Staphylococcaceae *Staphylococcus aureus* M0382 |
| Sau-1475-1 | Bacteria Firmicutes Bacilli Bacillales Staphylococcaceae *Staphylococcus aureus* M0383 |
| Sau-1476-1 | Bacteria Firmicutes Bacilli Bacillales Staphylococcaceae *Staphylococcus aureus* M0384 |
| Sau-1477-1 | Bacteria Firmicutes Bacilli Bacillales Staphylococcaceae *Staphylococcus aureus* M0385 |
| Sau-1478-1 | Bacteria Firmicutes Bacilli Bacillales Staphylococcaceae *Staphylococcus aureus* M0386 |
| Sau-1479-1 | Bacteria Firmicutes Bacilli Bacillales Staphylococcaceae *Staphylococcus aureus* M0388 |
| Sau-1480-1 | Bacteria Firmicutes Bacilli Bacillales Staphylococcaceae *Staphylococcus aureus* M0389 |
| Sau-1481-1 | Bacteria Firmicutes Bacilli Bacillales Staphylococcaceae *Staphylococcus aureus* M0390 |
| Sau-1482-1 | Bacteria Firmicutes Bacilli Bacillales Staphylococcaceae *Staphylococcus aureus* M0394 |
| Sau-1483-1 | Bacteria Firmicutes Bacilli Bacillales Staphylococcaceae *Staphylococcus aureus* M0395 |
| Sau-1484-1 | Bacteria Firmicutes Bacilli Bacillales Staphylococcaceae *Staphylococcus aureus* M0397 |
| Sau-1485-1 | Bacteria Firmicutes Bacilli Bacillales Staphylococcaceae *Staphylococcus aureus* M0398 |
| Sau-1486-1 | Bacteria Firmicutes Bacilli Bacillales Staphylococcaceae *Staphylococcus aureus* M0399 |
| Sau-1487-1 | Bacteria Firmicutes Bacilli Bacillales Staphylococcaceae *Staphylococcus aureus* M0400 |

| | |
|---|---|
| Sau-1488-1 | Bacteria Firmicutes Bacilli Bacillales Staphylococcaceae *Staphylococcus aureus* M0402 |
| Sau-1489-1 | Bacteria Firmicutes Bacilli Bacillales Staphylococcaceae *Staphylococcus aureus* M0403 |
| Sau-1490-1 | Bacteria Firmicutes Bacilli Bacillales Staphylococcaceae *Staphylococcus aureus* M0405 |
| Sau-1491-1 | Bacteria Firmicutes Bacilli Bacillales Staphylococcaceae *Staphylococcus aureus* M0407 |
| Sau-1492-1 | Bacteria Firmicutes Bacilli Bacillales Staphylococcaceae *Staphylococcus aureus* M0409 |
| Sau-1493-1 | Bacteria Firmicutes Bacilli Bacillales Staphylococcaceae *Staphylococcus aureus* M0410 |
| Sau-1494-1 | Bacteria Firmicutes Bacilli Bacillales Staphylococcaceae *Staphylococcus aureus* M0411 |
| Sau-1495-1 | Bacteria Firmicutes Bacilli Bacillales Staphylococcaceae *Staphylococcus aureus* M0412 |
| Sau-1496-1 | Bacteria Firmicutes Bacilli Bacillales Staphylococcaceae *Staphylococcus aureus* M0413 |
| Sau-1497-1 | Bacteria Firmicutes Bacilli Bacillales Staphylococcaceae *Staphylococcus aureus* M0414 |
| Sau-1498-1 | Bacteria Firmicutes Bacilli Bacillales Staphylococcaceae *Staphylococcus aureus* M0416 |
| Sau-1499-1 | Bacteria Firmicutes Bacilli Bacillales Staphylococcaceae *Staphylococcus aureus* M0417 |
| Sau-1500-1 | Bacteria Firmicutes Bacilli Bacillales Staphylococcaceae *Staphylococcus aureus* M0418 |
| Sau-1501-1 | Bacteria Firmicutes Bacilli Bacillales Staphylococcaceae *Staphylococcus aureus* M0419 |
| Sau-1502-1 | Bacteria Firmicutes Bacilli Bacillales Staphylococcaceae *Staphylococcus aureus* M0420 |
| Sau-1503-1 | Bacteria Firmicutes Bacilli Bacillales Staphylococcaceae *Staphylococcus aureus* M0421 |
| Sau-1504-1 | Bacteria Firmicutes Bacilli Bacillales Staphylococcaceae *Staphylococcus aureus* M0423 |
| Sau-1505-1 | Bacteria Firmicutes Bacilli Bacillales Staphylococcaceae *Staphylococcus aureus* M0425 |
| Sau-1506-1 | Bacteria Firmicutes Bacilli Bacillales Staphylococcaceae *Staphylococcus aureus* M0426 |
| Sau-1507-1 | Bacteria Firmicutes Bacilli Bacillales Staphylococcaceae *Staphylococcus aureus* M0428 |
| Sau-1508-1 | Bacteria Firmicutes Bacilli Bacillales Staphylococcaceae *Staphylococcus aureus* M0429 |
| Sau-1509-1 | Bacteria Firmicutes Bacilli Bacillales Staphylococcaceae *Staphylococcus aureus* M0430 |
| Sau-1510-1 | Bacteria Firmicutes Bacilli Bacillales Staphylococcaceae *Staphylococcus aureus* M0431 |
| Sau-1511-1 | Bacteria Firmicutes Bacilli Bacillales Staphylococcaceae *Staphylococcus aureus* M0432 |
| Sau-1512-1 | Bacteria Firmicutes Bacilli Bacillales Staphylococcaceae *Staphylococcus aureus* M0433 |
| Sau-1513-1 | Bacteria Firmicutes Bacilli Bacillales Staphylococcaceae *Staphylococcus aureus* M0434 |
| Sau-1514-1 | Bacteria Firmicutes Bacilli Bacillales Staphylococcaceae *Staphylococcus aureus* M0435 |
| Sau-1515-1 | Bacteria Firmicutes Bacilli Bacillales Staphylococcaceae *Staphylococcus aureus* M0437 |
| Sau-1516-1 | Bacteria Firmicutes Bacilli Bacillales Staphylococcaceae *Staphylococcus aureus* M0439 |
| Sau-1517-1 | Bacteria Firmicutes Bacilli Bacillales Staphylococcaceae *Staphylococcus aureus* M0440 |
| Sau-1518-1 | Bacteria Firmicutes Bacilli Bacillales Staphylococcaceae *Staphylococcus aureus* M0441 |
| Sau-1519-1 | Bacteria Firmicutes Bacilli Bacillales Staphylococcaceae *Staphylococcus aureus* M0442 |
| Sau-1520-1 | Bacteria Firmicutes Bacilli Bacillales Staphylococcaceae *Staphylococcus aureus* M0443 |
| Sau-1521-1 | Bacteria Firmicutes Bacilli Bacillales Staphylococcaceae *Staphylococcus aureus* M0444 |
| Sau-1522-1 | Bacteria Firmicutes Bacilli Bacillales Staphylococcaceae *Staphylococcus aureus* M0445 |
| Sau-1523-1 | Bacteria Firmicutes Bacilli Bacillales Staphylococcaceae *Staphylococcus aureus* M0447 |
| Sau-1524-1 | Bacteria Firmicutes Bacilli Bacillales Staphylococcaceae *Staphylococcus aureus* M0448 |
| Sau-1525-1 | Bacteria Firmicutes Bacilli Bacillales Staphylococcaceae *Staphylococcus aureus* M0449 |
| Sau-1526-1 | Bacteria Firmicutes Bacilli Bacillales Staphylococcaceae *Staphylococcus aureus* M0451 |
| Sau-1527-1 | Bacteria Firmicutes Bacilli Bacillales Staphylococcaceae *Staphylococcus aureus* M0452 |
| Sau-1528-1 | Bacteria Firmicutes Bacilli Bacillales Staphylococcaceae *Staphylococcus aureus* M0453 |
| Sau-1529-1 | Bacteria Firmicutes Bacilli Bacillales Staphylococcaceae *Staphylococcus aureus* M0454 |
| Sau-1530-1 | Bacteria Firmicutes Bacilli Bacillales Staphylococcaceae *Staphylococcus aureus* M0456 |
| Sau-1531-1 | Bacteria Firmicutes Bacilli Bacillales Staphylococcaceae *Staphylococcus aureus* M0457 |
| Sau-1532-1 | Bacteria Firmicutes Bacilli Bacillales Staphylococcaceae *Staphylococcus aureus* M0458 |
| Sau-1533-1 | Bacteria Firmicutes Bacilli Bacillales Staphylococcaceae *Staphylococcus aureus* M0459 |
| Sau-1534-1 | Bacteria Firmicutes Bacilli Bacillales Staphylococcaceae *Staphylococcus aureus* M0462 |
| Sau-1535-1 | Bacteria Firmicutes Bacilli Bacillales Staphylococcaceae *Staphylococcus aureus* M0463 |
| Sau-1536-1 | Bacteria Firmicutes Bacilli Bacillales Staphylococcaceae *Staphylococcus aureus* M0464 |
| Sau-1537-1 | Bacteria Firmicutes Bacilli Bacillales Staphylococcaceae *Staphylococcus aureus* M0465 |
| Sau-1538-1 | Bacteria Firmicutes Bacilli Bacillales Staphylococcaceae *Staphylococcus aureus* M0466 |
| Sau-1539-1 | Bacteria Firmicutes Bacilli Bacillales Staphylococcaceae *Staphylococcus aureus* M0469 |
| Sau-1540-1 | Bacteria Firmicutes Bacilli Bacillales Staphylococcaceae *Staphylococcus aureus* M0471 |
| Sau-1541-1 | Bacteria Firmicutes Bacilli Bacillales Staphylococcaceae *Staphylococcus aureus* M0472 |
| Sau-1542-1 | Bacteria Firmicutes Bacilli Bacillales Staphylococcaceae *Staphylococcus aureus* M0473 |
| Sau-1543-1 | Bacteria Firmicutes Bacilli Bacillales Staphylococcaceae *Staphylococcus aureus* M0475 |
| Sau-1544-1 | Bacteria Firmicutes Bacilli Bacillales Staphylococcaceae *Staphylococcus aureus* M0476 |
| Sau-1545-1 | Bacteria Firmicutes Bacilli Bacillales Staphylococcaceae *Staphylococcus aureus* M0481 |
| Sau-1546-1 | Bacteria Firmicutes Bacilli Bacillales Staphylococcaceae *Staphylococcus aureus* M0482 |
| Sau-1547-1 | Bacteria Firmicutes Bacilli Bacillales Staphylococcaceae *Staphylococcus aureus* M0483 |
| Sau-1548-1 | Bacteria Firmicutes Bacilli Bacillales Staphylococcaceae *Staphylococcus aureus* M0484 |
| Sau-1549-1 | Bacteria Firmicutes Bacilli Bacillales Staphylococcaceae *Staphylococcus aureus* M0486 |
| Sau-1550-1 | Bacteria Firmicutes Bacilli Bacillales Staphylococcaceae *Staphylococcus aureus* M0487 |
| Sau-1551-1 | Bacteria Firmicutes Bacilli Bacillales Staphylococcaceae *Staphylococcus aureus* M0488 |
| Sau-1552-1 | Bacteria Firmicutes Bacilli Bacillales Staphylococcaceae *Staphylococcus aureus* M0490 |

| | |
|---|---|
| Sau-1553-1 | Bacteria Firmicutes Bacilli Bacillales Staphylococcaceae *Staphylococcus aureus* M0491 |
| Sau-1554-1 | Bacteria Firmicutes Bacilli Bacillales Staphylococcaceae *Staphylococcus aureus* M0492 |
| Sau-1555-1 | Bacteria Firmicutes Bacilli Bacillales Staphylococcaceae *Staphylococcus aureus* M0495 |
| Sau-1556-1 | Bacteria Firmicutes Bacilli Bacillales Staphylococcaceae *Staphylococcus aureus* M0496 |
| Sau-1557-1 | Bacteria Firmicutes Bacilli Bacillales Staphylococcaceae *Staphylococcus aureus* M0498 |
| Sau-1558-1 | Bacteria Firmicutes Bacilli Bacillales Staphylococcaceae *Staphylococcus aureus* M0499 |
| Sau-1559-1 | Bacteria Firmicutes Bacilli Bacillales Staphylococcaceae *Staphylococcus aureus* M0501 |
| Sau-1560-1 | Bacteria Firmicutes Bacilli Bacillales Staphylococcaceae *Staphylococcus aureus* M0502 |
| Sau-1561-1 | Bacteria Firmicutes Bacilli Bacillales Staphylococcaceae *Staphylococcus aureus* M0503 |
| Sau-1562-1 | Bacteria Firmicutes Bacilli Bacillales Staphylococcaceae *Staphylococcus aureus* M0505 |
| Sau-1563-1 | Bacteria Firmicutes Bacilli Bacillales Staphylococcaceae *Staphylococcus aureus* M0506 |
| Sau-1564-1 | Bacteria Firmicutes Bacilli Bacillales Staphylococcaceae *Staphylococcus aureus* M0508 |
| Sau-1565-1 | Bacteria Firmicutes Bacilli Bacillales Staphylococcaceae *Staphylococcus aureus* M0509 |
| Sau-1566-1 | Bacteria Firmicutes Bacilli Bacillales Staphylococcaceae *Staphylococcus aureus* M0511 |
| Sau-1567-1 | Bacteria Firmicutes Bacilli Bacillales Staphylococcaceae *Staphylococcus aureus* M0512 |
| Sau-1568-1 | Bacteria Firmicutes Bacilli Bacillales Staphylococcaceae *Staphylococcus aureus* M0514 |
| Sau-1569-1 | Bacteria Firmicutes Bacilli Bacillales Staphylococcaceae *Staphylococcus aureus* M0515 |
| Sau-1570-1 | Bacteria Firmicutes Bacilli Bacillales Staphylococcaceae *Staphylococcus aureus* M0516 |
| Sau-1571-1 | Bacteria Firmicutes Bacilli Bacillales Staphylococcaceae *Staphylococcus aureus* M0518 |
| Sau-1572-1 | Bacteria Firmicutes Bacilli Bacillales Staphylococcaceae *Staphylococcus aureus* M0519 |
| Sau-1573-1 | Bacteria Firmicutes Bacilli Bacillales Staphylococcaceae *Staphylococcus aureus* M0520 |
| Sau-1574-1 | Bacteria Firmicutes Bacilli Bacillales Staphylococcaceae *Staphylococcus aureus* M0522 |
| Sau-1575-1 | Bacteria Firmicutes Bacilli Bacillales Staphylococcaceae *Staphylococcus aureus* M0523 |
| Sau-1576-1 | Bacteria Firmicutes Bacilli Bacillales Staphylococcaceae *Staphylococcus aureus* M0524 |
| Sau-1577-1 | Bacteria Firmicutes Bacilli Bacillales Staphylococcaceae *Staphylococcus aureus* M0525 |
| Sau-1578-1 | Bacteria Firmicutes Bacilli Bacillales Staphylococcaceae *Staphylococcus aureus* M0527 |
| Sau-1579-1 | Bacteria Firmicutes Bacilli Bacillales Staphylococcaceae *Staphylococcus aureus* M0530 |
| Sau-1580-1 | Bacteria Firmicutes Bacilli Bacillales Staphylococcaceae *Staphylococcus aureus* M0532 |
| Sau-1581-1 | Bacteria Firmicutes Bacilli Bacillales Staphylococcaceae *Staphylococcus aureus* M0533 |
| Sau-1582-1 | Bacteria Firmicutes Bacilli Bacillales Staphylococcaceae *Staphylococcus aureus* M0534 |
| Sau-1583-1 | Bacteria Firmicutes Bacilli Bacillales Staphylococcaceae *Staphylococcus aureus* M0535 |
| Sau-1584-1 | Bacteria Firmicutes Bacilli Bacillales Staphylococcaceae *Staphylococcus aureus* M0537 |
| Sau-1585-1 | Bacteria Firmicutes Bacilli Bacillales Staphylococcaceae *Staphylococcus aureus* M0538 |
| Sau-1586-1 | Bacteria Firmicutes Bacilli Bacillales Staphylococcaceae *Staphylococcus aureus* M0540 |
| Sau-1587-1 | Bacteria Firmicutes Bacilli Bacillales Staphylococcaceae *Staphylococcus aureus* M0541 |
| Sau-1588-1 | Bacteria Firmicutes Bacilli Bacillales Staphylococcaceae *Staphylococcus aureus* M0542 |
| Sau-1589-1 | Bacteria Firmicutes Bacilli Bacillales Staphylococcaceae *Staphylococcus aureus* M0543 |
| Sau-1590-1 | Bacteria Firmicutes Bacilli Bacillales Staphylococcaceae *Staphylococcus aureus* M0544 |
| Sau-1591-1 | Bacteria Firmicutes Bacilli Bacillales Staphylococcaceae *Staphylococcus aureus* M0545 |
| Sau-1592-1 | Bacteria Firmicutes Bacilli Bacillales Staphylococcaceae *Staphylococcus aureus* M0546 |
| Sau-1593-1 | Bacteria Firmicutes Bacilli Bacillales Staphylococcaceae *Staphylococcus aureus* M0548 |
| Sau-1594-1 | Bacteria Firmicutes Bacilli Bacillales Staphylococcaceae *Staphylococcus aureus* M0549 |
| Sau-1595-1 | Bacteria Firmicutes Bacilli Bacillales Staphylococcaceae *Staphylococcus aureus* M0550 |
| Sau-1596-1 | Bacteria Firmicutes Bacilli Bacillales Staphylococcaceae *Staphylococcus aureus* M0551 |
| Sau-1597-1 | Bacteria Firmicutes Bacilli Bacillales Staphylococcaceae *Staphylococcus aureus* M0552 |
| Sau-1598-1 | Bacteria Firmicutes Bacilli Bacillales Staphylococcaceae *Staphylococcus aureus* M0554 |
| Sau-1599-1 | Bacteria Firmicutes Bacilli Bacillales Staphylococcaceae *Staphylococcus aureus* M0555 |
| Sau-1600-1 | Bacteria Firmicutes Bacilli Bacillales Staphylococcaceae *Staphylococcus aureus* M0556 |
| Sau-1601-1 | Bacteria Firmicutes Bacilli Bacillales Staphylococcaceae *Staphylococcus aureus* M0557 |
| Sau-1602-1 | Bacteria Firmicutes Bacilli Bacillales Staphylococcaceae *Staphylococcus aureus* M0558 |
| Sau-1603-1 | Bacteria Firmicutes Bacilli Bacillales Staphylococcaceae *Staphylococcus aureus* M0559 |
| Sau-1604-1 | Bacteria Firmicutes Bacilli Bacillales Staphylococcaceae *Staphylococcus aureus* M0560 |
| Sau-1605-1 | Bacteria Firmicutes Bacilli Bacillales Staphylococcaceae *Staphylococcus aureus* M0561 |
| Sau-1606-1 | Bacteria Firmicutes Bacilli Bacillales Staphylococcaceae *Staphylococcus aureus* M0563 |
| Sau-1607-1 | Bacteria Firmicutes Bacilli Bacillales Staphylococcaceae *Staphylococcus aureus* M0564 |
| Sau-1608-1 | Bacteria Firmicutes Bacilli Bacillales Staphylococcaceae *Staphylococcus aureus* M0566 |
| Sau-1609-1 | Bacteria Firmicutes Bacilli Bacillales Staphylococcaceae *Staphylococcus aureus* M0567 |
| Sau-1610-1 | Bacteria Firmicutes Bacilli Bacillales Staphylococcaceae *Staphylococcus aureus* M0568 |
| Sau-1611-1 | Bacteria Firmicutes Bacilli Bacillales Staphylococcaceae *Staphylococcus aureus* M0569 |
| Sau-1612-1 | Bacteria Firmicutes Bacilli Bacillales Staphylococcaceae *Staphylococcus aureus* M0570 |
| Sau-1613-1 | Bacteria Firmicutes Bacilli Bacillales Staphylococcaceae *Staphylococcus aureus* M0572 |
| Sau-1614-1 | Bacteria Firmicutes Bacilli Bacillales Staphylococcaceae *Staphylococcus aureus* M0573 |
| Sau-1615-1 | Bacteria Firmicutes Bacilli Bacillales Staphylococcaceae *Staphylococcus aureus* M0577 |
| Sau-1616-1 | Bacteria Firmicutes Bacilli Bacillales Staphylococcaceae *Staphylococcus aureus* M0579 |
| Sau-1617-1 | Bacteria Firmicutes Bacilli Bacillales Staphylococcaceae *Staphylococcus aureus* M0581 |

| | |
|---|---|
| Sau-1618-1 | Bacteria Firmicutes Bacilli Bacillales Staphylococcaceae *Staphylococcus aureus* M0582 |
| Sau-1619-1 | Bacteria Firmicutes Bacilli Bacillales Staphylococcaceae *Staphylococcus aureus* M0583 |
| Sau-1620-1 | Bacteria Firmicutes Bacilli Bacillales Staphylococcaceae *Staphylococcus aureus* M0585 |
| Sau-1621-1 | Bacteria Firmicutes Bacilli Bacillales Staphylococcaceae *Staphylococcus aureus* M0587 |
| Sau-1622-1 | Bacteria Firmicutes Bacilli Bacillales Staphylococcaceae *Staphylococcus aureus* M0588 |
| Sau-1623-1 | Bacteria Firmicutes Bacilli Bacillales Staphylococcaceae *Staphylococcus aureus* M0589 |
| Sau-1624-1 | Bacteria Firmicutes Bacilli Bacillales Staphylococcaceae *Staphylococcus aureus* M0590 |
| Sau-1625-1 | Bacteria Firmicutes Bacilli Bacillales Staphylococcaceae *Staphylococcus aureus* M0591 |
| Sau-1626-1 | Bacteria Firmicutes Bacilli Bacillales Staphylococcaceae *Staphylococcus aureus* M0592 |
| Sau-1627-1 | Bacteria Firmicutes Bacilli Bacillales Staphylococcaceae *Staphylococcus aureus* M0593 |
| Sau-1628-1 | Bacteria Firmicutes Bacilli Bacillales Staphylococcaceae *Staphylococcus aureus* M0594 |
| Sau-1629-1 | Bacteria Firmicutes Bacilli Bacillales Staphylococcaceae *Staphylococcus aureus* M0595 |
| Sau-1630-1 | Bacteria Firmicutes Bacilli Bacillales Staphylococcaceae *Staphylococcus aureus* M0598 |
| Sau-1631-1 | Bacteria Firmicutes Bacilli Bacillales Staphylococcaceae *Staphylococcus aureus* M0599 |
| Sau-1632-1 | Bacteria Firmicutes Bacilli Bacillales Staphylococcaceae *Staphylococcus aureus* M0601 |
| Sau-1633-1 | Bacteria Firmicutes Bacilli Bacillales Staphylococcaceae *Staphylococcus aureus* M0603 |
| Sau-1634-1 | Bacteria Firmicutes Bacilli Bacillales Staphylococcaceae *Staphylococcus aureus* M0604 |
| Sau-1635-1 | Bacteria Firmicutes Bacilli Bacillales Staphylococcaceae *Staphylococcus aureus* M0605 |
| Sau-1636-1 | Bacteria Firmicutes Bacilli Bacillales Staphylococcaceae *Staphylococcus aureus* M0607 |
| Sau-1637-1 | Bacteria Firmicutes Bacilli Bacillales Staphylococcaceae *Staphylococcus aureus* M0608 |
| Sau-1638-1 | Bacteria Firmicutes Bacilli Bacillales Staphylococcaceae *Staphylococcus aureus* M0609 |
| Sau-1639-1 | Bacteria Firmicutes Bacilli Bacillales Staphylococcaceae *Staphylococcus aureus* M0610 |
| Sau-1640-1 | Bacteria Firmicutes Bacilli Bacillales Staphylococcaceae *Staphylococcus aureus* M0611 |
| Sau-1641-1 | Bacteria Firmicutes Bacilli Bacillales Staphylococcaceae *Staphylococcus aureus* M0612 |
| Sau-1642-1 | Bacteria Firmicutes Bacilli Bacillales Staphylococcaceae *Staphylococcus aureus* M0613 |
| Sau-1643-1 | Bacteria Firmicutes Bacilli Bacillales Staphylococcaceae *Staphylococcus aureus* M0615 |
| Sau-1644-1 | Bacteria Firmicutes Bacilli Bacillales Staphylococcaceae *Staphylococcus aureus* M0616 |
| Sau-1645-1 | Bacteria Firmicutes Bacilli Bacillales Staphylococcaceae *Staphylococcus aureus* M0617 |
| Sau-1646-1 | Bacteria Firmicutes Bacilli Bacillales Staphylococcaceae *Staphylococcus aureus* M0618 |
| Sau-1647-1 | Bacteria Firmicutes Bacilli Bacillales Staphylococcaceae *Staphylococcus aureus* M0619 |
| Sau-1648-1 | Bacteria Firmicutes Bacilli Bacillales Staphylococcaceae *Staphylococcus aureus* M0620 |
| Sau-1649-1 | Bacteria Firmicutes Bacilli Bacillales Staphylococcaceae *Staphylococcus aureus* M0621 |
| Sau-1650-1 | Bacteria Firmicutes Bacilli Bacillales Staphylococcaceae *Staphylococcus aureus* M0623 |
| Sau-1651-1 | Bacteria Firmicutes Bacilli Bacillales Staphylococcaceae *Staphylococcus aureus* M0624 |
| Sau-1652-1 | Bacteria Firmicutes Bacilli Bacillales Staphylococcaceae *Staphylococcus aureus* M0625 |
| Sau-1653-1 | Bacteria Firmicutes Bacilli Bacillales Staphylococcaceae *Staphylococcus aureus* M0626 |
| Sau-1654-1 | Bacteria Firmicutes Bacilli Bacillales Staphylococcaceae *Staphylococcus aureus* M0627 |
| Sau-1655-1 | Bacteria Firmicutes Bacilli Bacillales Staphylococcaceae *Staphylococcus aureus* M0630 |
| Sau-1656-1 | Bacteria Firmicutes Bacilli Bacillales Staphylococcaceae *Staphylococcus aureus* M0631 |
| Sau-1657-1 | Bacteria Firmicutes Bacilli Bacillales Staphylococcaceae *Staphylococcus aureus* M0632 |
| Sau-1658-1 | Bacteria Firmicutes Bacilli Bacillales Staphylococcaceae *Staphylococcus aureus* M0634 |
| Sau-1659-1 | Bacteria Firmicutes Bacilli Bacillales Staphylococcaceae *Staphylococcus aureus* M0635 |
| Sau-1660-1 | Bacteria Firmicutes Bacilli Bacillales Staphylococcaceae *Staphylococcus aureus* M0636 |
| Sau-1661-1 | Bacteria Firmicutes Bacilli Bacillales Staphylococcaceae *Staphylococcus aureus* M0637 |
| Sau-1662-1 | Bacteria Firmicutes Bacilli Bacillales Staphylococcaceae *Staphylococcus aureus* M0638 |
| Sau-1663-1 | Bacteria Firmicutes Bacilli Bacillales Staphylococcaceae *Staphylococcus aureus* M0640 |
| Sau-1664-1 | Bacteria Firmicutes Bacilli Bacillales Staphylococcaceae *Staphylococcus aureus* M0642 |
| Sau-1665-1 | Bacteria Firmicutes Bacilli Bacillales Staphylococcaceae *Staphylococcus aureus* M0643 |
| Sau-1666-1 | Bacteria Firmicutes Bacilli Bacillales Staphylococcaceae *Staphylococcus aureus* M0644 |
| Sau-1667-1 | Bacteria Firmicutes Bacilli Bacillales Staphylococcaceae *Staphylococcus aureus* M0645 |
| Sau-1668-1 | Bacteria Firmicutes Bacilli Bacillales Staphylococcaceae *Staphylococcus aureus* M0647 |
| Sau-1669-1 | Bacteria Firmicutes Bacilli Bacillales Staphylococcaceae *Staphylococcus aureus* M0649 |
| Sau-1670-1 | Bacteria Firmicutes Bacilli Bacillales Staphylococcaceae *Staphylococcus aureus* M0650 |
| Sau-1671-1 | Bacteria Firmicutes Bacilli Bacillales Staphylococcaceae *Staphylococcus aureus* M0651 |
| Sau-1672-1 | Bacteria Firmicutes Bacilli Bacillales Staphylococcaceae *Staphylococcus aureus* M0652 |
| Sau-1673-1 | Bacteria Firmicutes Bacilli Bacillales Staphylococcaceae *Staphylococcus aureus* M0653 |
| Sau-1674-1 | Bacteria Firmicutes Bacilli Bacillales Staphylococcaceae *Staphylococcus aureus* M0654 |
| Sau-1675-1 | Bacteria Firmicutes Bacilli Bacillales Staphylococcaceae *Staphylococcus aureus* M0655 |
| Sau-1676-1 | Bacteria Firmicutes Bacilli Bacillales Staphylococcaceae *Staphylococcus aureus* M0656 |
| Sau-1677-1 | Bacteria Firmicutes Bacilli Bacillales Staphylococcaceae *Staphylococcus aureus* M0657 |
| Sau-1678-1 | Bacteria Firmicutes Bacilli Bacillales Staphylococcaceae *Staphylococcus aureus* M0658 |
| Sau-1679-1 | Bacteria Firmicutes Bacilli Bacillales Staphylococcaceae *Staphylococcus aureus* M0659 |
| Sau-1680-1 | Bacteria Firmicutes Bacilli Bacillales Staphylococcaceae *Staphylococcus aureus* M0661 |
| Sau-1681-1 | Bacteria Firmicutes Bacilli Bacillales Staphylococcaceae *Staphylococcus aureus* M0662 |
| Sau-1682-1 | Bacteria Firmicutes Bacilli Bacillales Staphylococcaceae *Staphylococcus aureus* M0664 |

| | |
|---|---|
| Sau-1683-1 | Bacteria Firmicutes Bacilli Bacillales Staphylococcaceae *Staphylococcus aureus* M0666 |
| Sau-1684-1 | Bacteria Firmicutes Bacilli Bacillales Staphylococcaceae *Staphylococcus aureus* M0667 |
| Sau-1685-1 | Bacteria Firmicutes Bacilli Bacillales Staphylococcaceae *Staphylococcus aureus* M0668 |
| Sau-1686-1 | Bacteria Firmicutes Bacilli Bacillales Staphylococcaceae *Staphylococcus aureus* M0669 |
| Sau-1687-1 | Bacteria Firmicutes Bacilli Bacillales Staphylococcaceae *Staphylococcus aureus* M0670 |
| Sau-1688-1 | Bacteria Firmicutes Bacilli Bacillales Staphylococcaceae *Staphylococcus aureus* M0671 |
| Sau-1689-1 | Bacteria Firmicutes Bacilli Bacillales Staphylococcaceae *Staphylococcus aureus* M0672 |
| Sau-1690-1 | Bacteria Firmicutes Bacilli Bacillales Staphylococcaceae *Staphylococcus aureus* M0674 |
| Sau-1691-1 | Bacteria Firmicutes Bacilli Bacillales Staphylococcaceae *Staphylococcus aureus* M0675 |
| Sau-1692-1 | Bacteria Firmicutes Bacilli Bacillales Staphylococcaceae *Staphylococcus aureus* M0677 |
| Sau-1693-1 | Bacteria Firmicutes Bacilli Bacillales Staphylococcaceae *Staphylococcus aureus* M0678 |
| Sau-1694-1 | Bacteria Firmicutes Bacilli Bacillales Staphylococcaceae *Staphylococcus aureus* M0679 |
| Sau-1695-1 | Bacteria Firmicutes Bacilli Bacillales Staphylococcaceae *Staphylococcus aureus* M0680 |
| Sau-1696-1 | Bacteria Firmicutes Bacilli Bacillales Staphylococcaceae *Staphylococcus aureus* M0681 |
| Sau-1697-1 | Bacteria Firmicutes Bacilli Bacillales Staphylococcaceae *Staphylococcus aureus* M0682 |
| Sau-1698-1 | Bacteria Firmicutes Bacilli Bacillales Staphylococcaceae *Staphylococcus aureus* M0683 |
| Sau-1699-1 | Bacteria Firmicutes Bacilli Bacillales Staphylococcaceae *Staphylococcus aureus* M0684 |
| Sau-1700-1 | Bacteria Firmicutes Bacilli Bacillales Staphylococcaceae *Staphylococcus aureus* M0685 |
| Sau-1701-1 | Bacteria Firmicutes Bacilli Bacillales Staphylococcaceae *Staphylococcus aureus* M0686 |
| Sau-1702-1 | Bacteria Firmicutes Bacilli Bacillales Staphylococcaceae *Staphylococcus aureus* M0688 |
| Sau-1703-1 | Bacteria Firmicutes Bacilli Bacillales Staphylococcaceae *Staphylococcus aureus* M0689 |
| Sau-1704-1 | Bacteria Firmicutes Bacilli Bacillales Staphylococcaceae *Staphylococcus aureus* M0690 |
| Sau-1705-1 | Bacteria Firmicutes Bacilli Bacillales Staphylococcaceae *Staphylococcus aureus* M0691 |
| Sau-1706-1 | Bacteria Firmicutes Bacilli Bacillales Staphylococcaceae *Staphylococcus aureus* M0693 |
| Sau-1707-1 | Bacteria Firmicutes Bacilli Bacillales Staphylococcaceae *Staphylococcus aureus* M0696 |
| Sau-1708-1 | Bacteria Firmicutes Bacilli Bacillales Staphylococcaceae *Staphylococcus aureus* M0697 |
| Sau-1709-1 | Bacteria Firmicutes Bacilli Bacillales Staphylococcaceae *Staphylococcus aureus* M0698 |
| Sau-1710-1 | Bacteria Firmicutes Bacilli Bacillales Staphylococcaceae *Staphylococcus aureus* M0699 |
| Sau-1711-1 | Bacteria Firmicutes Bacilli Bacillales Staphylococcaceae *Staphylococcus aureus* M0700 |
| Sau-1712-1 | Bacteria Firmicutes Bacilli Bacillales Staphylococcaceae *Staphylococcus aureus* M0701 |
| Sau-1713-1 | Bacteria Firmicutes Bacilli Bacillales Staphylococcaceae *Staphylococcus aureus* M0702 |
| Sau-1714-1 | Bacteria Firmicutes Bacilli Bacillales Staphylococcaceae *Staphylococcus aureus* M0703 |
| Sau-1715-1 | Bacteria Firmicutes Bacilli Bacillales Staphylococcaceae *Staphylococcus aureus* M0704 |
| Sau-1716-1 | Bacteria Firmicutes Bacilli Bacillales Staphylococcaceae *Staphylococcus aureus* M0706 |
| Sau-1717-1 | Bacteria Firmicutes Bacilli Bacillales Staphylococcaceae *Staphylococcus aureus* M0707 |
| Sau-1718-1 | Bacteria Firmicutes Bacilli Bacillales Staphylococcaceae *Staphylococcus aureus* M0708 |
| Sau-1719-1 | Bacteria Firmicutes Bacilli Bacillales Staphylococcaceae *Staphylococcus aureus* M0709 |
| Sau-1720-1 | Bacteria Firmicutes Bacilli Bacillales Staphylococcaceae *Staphylococcus aureus* M0710 |
| Sau-1721-1 | Bacteria Firmicutes Bacilli Bacillales Staphylococcaceae *Staphylococcus aureus* M0711 |
| Sau-1722-1 | Bacteria Firmicutes Bacilli Bacillales Staphylococcaceae *Staphylococcus aureus* M0713 |
| Sau-1723-1 | Bacteria Firmicutes Bacilli Bacillales Staphylococcaceae *Staphylococcus aureus* M0714 |
| Sau-1724-1 | Bacteria Firmicutes Bacilli Bacillales Staphylococcaceae *Staphylococcus aureus* M0715 |
| Sau-1725-1 | Bacteria Firmicutes Bacilli Bacillales Staphylococcaceae *Staphylococcus aureus* M0716 |
| Sau-1726-1 | Bacteria Firmicutes Bacilli Bacillales Staphylococcaceae *Staphylococcus aureus* M0718 |
| Sau-1727-1 | Bacteria Firmicutes Bacilli Bacillales Staphylococcaceae *Staphylococcus aureus* M0720 |
| Sau-1728-1 | Bacteria Firmicutes Bacilli Bacillales Staphylococcaceae *Staphylococcus aureus* M0721 |
| Sau-1729-1 | Bacteria Firmicutes Bacilli Bacillales Staphylococcaceae *Staphylococcus aureus* M0722 |
| Sau-1730-1 | Bacteria Firmicutes Bacilli Bacillales Staphylococcaceae *Staphylococcus aureus* M0723 |
| Sau-1731-1 | Bacteria Firmicutes Bacilli Bacillales Staphylococcaceae *Staphylococcus aureus* M0724 |
| Sau-1732-1 | Bacteria Firmicutes Bacilli Bacillales Staphylococcaceae *Staphylococcus aureus* M0725 |
| Sau-1733-1 | Bacteria Firmicutes Bacilli Bacillales Staphylococcaceae *Staphylococcus aureus* M0726 |
| Sau-1734-1 | Bacteria Firmicutes Bacilli Bacillales Staphylococcaceae *Staphylococcus aureus* M0727 |
| Sau-1735-1 | Bacteria Firmicutes Bacilli Bacillales Staphylococcaceae *Staphylococcus aureus* M0728 |
| Sau-1736-1 | Bacteria Firmicutes Bacilli Bacillales Staphylococcaceae *Staphylococcus aureus* M0729 |
| Sau-1737-1 | Bacteria Firmicutes Bacilli Bacillales Staphylococcaceae *Staphylococcus aureus* M0730 |
| Sau-1738-1 | Bacteria Firmicutes Bacilli Bacillales Staphylococcaceae *Staphylococcus aureus* M0731 |
| Sau-1739-1 | Bacteria Firmicutes Bacilli Bacillales Staphylococcaceae *Staphylococcus aureus* M0732 |
| Sau-1740-1 | Bacteria Firmicutes Bacilli Bacillales Staphylococcaceae *Staphylococcus aureus* M0733 |
| Sau-1741-1 | Bacteria Firmicutes Bacilli Bacillales Staphylococcaceae *Staphylococcus aureus* M0734 |
| Sau-1742-1 | Bacteria Firmicutes Bacilli Bacillales Staphylococcaceae *Staphylococcus aureus* M0735 |
| Sau-1743-1 | Bacteria Firmicutes Bacilli Bacillales Staphylococcaceae *Staphylococcus aureus* M0736 |
| Sau-1744-1 | Bacteria Firmicutes Bacilli Bacillales Staphylococcaceae *Staphylococcus aureus* M0737 |
| Sau-1745-1 | Bacteria Firmicutes Bacilli Bacillales Staphylococcaceae *Staphylococcus aureus* M0739 |
| Sau-1746-1 | Bacteria Firmicutes Bacilli Bacillales Staphylococcaceae *Staphylococcus aureus* M0741 |
| Sau-1747-1 | Bacteria Firmicutes Bacilli Bacillales Staphylococcaceae *Staphylococcus aureus* M0742 |

| | |
|---|---|
| Sau-1748-1 | Bacteria Firmicutes Bacilli Bacillales Staphylococcaceae *Staphylococcus aureus* M0743 |
| Sau-1749-1 | Bacteria Firmicutes Bacilli Bacillales Staphylococcaceae *Staphylococcus aureus* M0744 |
| Sau-1750-1 | Bacteria Firmicutes Bacilli Bacillales Staphylococcaceae *Staphylococcus aureus* M0745 |
| Sau-1751-1 | Bacteria Firmicutes Bacilli Bacillales Staphylococcaceae *Staphylococcus aureus* M0746 |
| Sau-1752-1 | Bacteria Firmicutes Bacilli Bacillales Staphylococcaceae *Staphylococcus aureus* M0747 |
| Sau-1753-1 | Bacteria Firmicutes Bacilli Bacillales Staphylococcaceae *Staphylococcus aureus* M0749 |
| Sau-1754-1 | Bacteria Firmicutes Bacilli Bacillales Staphylococcaceae *Staphylococcus aureus* M0750 |
| Sau-1755-1 | Bacteria Firmicutes Bacilli Bacillales Staphylococcaceae *Staphylococcus aureus* M0752 |
| Sau-1756-1 | Bacteria Firmicutes Bacilli Bacillales Staphylococcaceae *Staphylococcus aureus* M0753 |
| Sau-1757-1 | Bacteria Firmicutes Bacilli Bacillales Staphylococcaceae *Staphylococcus aureus* M0754 |
| Sau-1758-1 | Bacteria Firmicutes Bacilli Bacillales Staphylococcaceae *Staphylococcus aureus* M0755 |
| Sau-1759-1 | Bacteria Firmicutes Bacilli Bacillales Staphylococcaceae *Staphylococcus aureus* M0757 |
| Sau-1760-1 | Bacteria Firmicutes Bacilli Bacillales Staphylococcaceae *Staphylococcus aureus* M0758 |
| Sau-1761-1 | Bacteria Firmicutes Bacilli Bacillales Staphylococcaceae *Staphylococcus aureus* M0759 |
| Sau-1762-1 | Bacteria Firmicutes Bacilli Bacillales Staphylococcaceae *Staphylococcus aureus* M0760 |
| Sau-1763-1 | Bacteria Firmicutes Bacilli Bacillales Staphylococcaceae *Staphylococcus aureus* M0761 |
| Sau-1764-1 | Bacteria Firmicutes Bacilli Bacillales Staphylococcaceae *Staphylococcus aureus* M0762 |
| Sau-1765-1 | Bacteria Firmicutes Bacilli Bacillales Staphylococcaceae *Staphylococcus aureus* M0763 |
| Sau-1766-1 | Bacteria Firmicutes Bacilli Bacillales Staphylococcaceae *Staphylococcus aureus* M0764 |
| Sau-1767-1 | Bacteria Firmicutes Bacilli Bacillales Staphylococcaceae *Staphylococcus aureus* M0765 |
| Sau-1768-1 | Bacteria Firmicutes Bacilli Bacillales Staphylococcaceae *Staphylococcus aureus* M0766 |
| Sau-1769-1 | Bacteria Firmicutes Bacilli Bacillales Staphylococcaceae *Staphylococcus aureus* M0767 |
| Sau-1770-1 | Bacteria Firmicutes Bacilli Bacillales Staphylococcaceae *Staphylococcus aureus* M0768 |
| Sau-1771-1 | Bacteria Firmicutes Bacilli Bacillales Staphylococcaceae *Staphylococcus aureus* M0771 |
| Sau-1772-1 | Bacteria Firmicutes Bacilli Bacillales Staphylococcaceae *Staphylococcus aureus* M0772 |
| Sau-1773-1 | Bacteria Firmicutes Bacilli Bacillales Staphylococcaceae *Staphylococcus aureus* M0773 |
| Sau-1774-1 | Bacteria Firmicutes Bacilli Bacillales Staphylococcaceae *Staphylococcus aureus* M0774 |
| Sau-1775-1 | Bacteria Firmicutes Bacilli Bacillales Staphylococcaceae *Staphylococcus aureus* M0775 |
| Sau-1776-1 | Bacteria Firmicutes Bacilli Bacillales Staphylococcaceae *Staphylococcus aureus* M0776 |
| Sau-1777-1 | Bacteria Firmicutes Bacilli Bacillales Staphylococcaceae *Staphylococcus aureus* M0777 |
| Sau-1778-1 | Bacteria Firmicutes Bacilli Bacillales Staphylococcaceae *Staphylococcus aureus* M0779 |
| Sau-1779-1 | Bacteria Firmicutes Bacilli Bacillales Staphylococcaceae *Staphylococcus aureus* M0782 |
| Sau-1780-1 | Bacteria Firmicutes Bacilli Bacillales Staphylococcaceae *Staphylococcus aureus* M0783 |
| Sau-1781-1 | Bacteria Firmicutes Bacilli Bacillales Staphylococcaceae *Staphylococcus aureus* M0784 |
| Sau-1782-1 | Bacteria Firmicutes Bacilli Bacillales Staphylococcaceae *Staphylococcus aureus* M0785 |
| Sau-1783-1 | Bacteria Firmicutes Bacilli Bacillales Staphylococcaceae *Staphylococcus aureus* M0786 |
| Sau-1784-1 | Bacteria Firmicutes Bacilli Bacillales Staphylococcaceae *Staphylococcus aureus* M0787 |
| Sau-1785-1 | Bacteria Firmicutes Bacilli Bacillales Staphylococcaceae *Staphylococcus aureus* M0788 |
| Sau-1786-1 | Bacteria Firmicutes Bacilli Bacillales Staphylococcaceae *Staphylococcus aureus* M0789 |
| Sau-1787-1 | Bacteria Firmicutes Bacilli Bacillales Staphylococcaceae *Staphylococcus aureus* M0790 |
| Sau-1788-1 | Bacteria Firmicutes Bacilli Bacillales Staphylococcaceae *Staphylococcus aureus* M0791 |
| Sau-1789-1 | Bacteria Firmicutes Bacilli Bacillales Staphylococcaceae *Staphylococcus aureus* M0793 |
| Sau-1790-1 | Bacteria Firmicutes Bacilli Bacillales Staphylococcaceae *Staphylococcus aureus* M0794 |
| Sau-1791-1 | Bacteria Firmicutes Bacilli Bacillales Staphylococcaceae *Staphylococcus aureus* M0796 |
| Sau-1792-1 | Bacteria Firmicutes Bacilli Bacillales Staphylococcaceae *Staphylococcus aureus* M0798 |
| Sau-1793-1 | Bacteria Firmicutes Bacilli Bacillales Staphylococcaceae *Staphylococcus aureus* M0800 |
| Sau-1794-1 | Bacteria Firmicutes Bacilli Bacillales Staphylococcaceae *Staphylococcus aureus* M0801 |
| Sau-1795-1 | Bacteria Firmicutes Bacilli Bacillales Staphylococcaceae *Staphylococcus aureus* M0802 |
| Sau-1796-1 | Bacteria Firmicutes Bacilli Bacillales Staphylococcaceae *Staphylococcus aureus* M0803 |
| Sau-1797-1 | Bacteria Firmicutes Bacilli Bacillales Staphylococcaceae *Staphylococcus aureus* M0804 |
| Sau-1798-1 | Bacteria Firmicutes Bacilli Bacillales Staphylococcaceae *Staphylococcus aureus* M0805 |
| Sau-1799-1 | Bacteria Firmicutes Bacilli Bacillales Staphylococcaceae *Staphylococcus aureus* M0806 |
| Sau-1800-1 | Bacteria Firmicutes Bacilli Bacillales Staphylococcaceae *Staphylococcus aureus* M0807 |
| Sau-1801-1 | Bacteria Firmicutes Bacilli Bacillales Staphylococcaceae *Staphylococcus aureus* M0808 |
| Sau-1802-1 | Bacteria Firmicutes Bacilli Bacillales Staphylococcaceae *Staphylococcus aureus* M0809 |
| Sau-1803-1 | Bacteria Firmicutes Bacilli Bacillales Staphylococcaceae *Staphylococcus aureus* M0810 |
| Sau-1804-1 | Bacteria Firmicutes Bacilli Bacillales Staphylococcaceae *Staphylococcus aureus* M0811 |
| Sau-1805-1 | Bacteria Firmicutes Bacilli Bacillales Staphylococcaceae *Staphylococcus aureus* M0812 |
| Sau-1806-1 | Bacteria Firmicutes Bacilli Bacillales Staphylococcaceae *Staphylococcus aureus* M0814 |
| Sau-1807-1 | Bacteria Firmicutes Bacilli Bacillales Staphylococcaceae *Staphylococcus aureus* M0816 |
| Sau-1808-1 | Bacteria Firmicutes Bacilli Bacillales Staphylococcaceae *Staphylococcus aureus* M0817 |
| Sau-1809-1 | Bacteria Firmicutes Bacilli Bacillales Staphylococcaceae *Staphylococcus aureus* M0818 |
| Sau-1810-1 | Bacteria Firmicutes Bacilli Bacillales Staphylococcaceae *Staphylococcus aureus* M0819 |
| Sau-1811-1 | Bacteria Firmicutes Bacilli Bacillales Staphylococcaceae *Staphylococcus aureus* M0820 |
| Sau-1812-1 | Bacteria Firmicutes Bacilli Bacillales Staphylococcaceae *Staphylococcus aureus* M0821 |

| | |
|---|---|
| Sau-1813-1 | Bacteria Firmicutes Bacilli Bacillales Staphylococcaceae *Staphylococcus aureus* M0824 |
| Sau-1814-1 | Bacteria Firmicutes Bacilli Bacillales Staphylococcaceae *Staphylococcus aureus* M0826 |
| Sau-1815-1 | Bacteria Firmicutes Bacilli Bacillales Staphylococcaceae *Staphylococcus aureus* M0827 |
| Sau-1816-1 | Bacteria Firmicutes Bacilli Bacillales Staphylococcaceae *Staphylococcus aureus* M0830 |
| Sau-1817-1 | Bacteria Firmicutes Bacilli Bacillales Staphylococcaceae *Staphylococcus aureus* M0832 |
| Sau-1818-1 | Bacteria Firmicutes Bacilli Bacillales Staphylococcaceae *Staphylococcus aureus* M0833 |
| Sau-1819-1 | Bacteria Firmicutes Bacilli Bacillales Staphylococcaceae *Staphylococcus aureus* M0834 |
| Sau-1820-1 | Bacteria Firmicutes Bacilli Bacillales Staphylococcaceae *Staphylococcus aureus* M0836 |
| Sau-1821-1 | Bacteria Firmicutes Bacilli Bacillales Staphylococcaceae *Staphylococcus aureus* M0837 |
| Sau-1822-1 | Bacteria Firmicutes Bacilli Bacillales Staphylococcaceae *Staphylococcus aureus* M0838 |
| Sau-1823-1 | Bacteria Firmicutes Bacilli Bacillales Staphylococcaceae *Staphylococcus aureus* M0839 |
| Sau-1824-1 | Bacteria Firmicutes Bacilli Bacillales Staphylococcaceae *Staphylococcus aureus* M0840 |
| Sau-1825-1 | Bacteria Firmicutes Bacilli Bacillales Staphylococcaceae *Staphylococcus aureus* M0841 |
| Sau-1826-1 | Bacteria Firmicutes Bacilli Bacillales Staphylococcaceae *Staphylococcus aureus* M0842 |
| Sau-1827-1 | Bacteria Firmicutes Bacilli Bacillales Staphylococcaceae *Staphylococcus aureus* M0843 |
| Sau-1828-1 | Bacteria Firmicutes Bacilli Bacillales Staphylococcaceae *Staphylococcus aureus* M0845 |
| Sau-1829-1 | Bacteria Firmicutes Bacilli Bacillales Staphylococcaceae *Staphylococcus aureus* M0846 |
| Sau-1830-1 | Bacteria Firmicutes Bacilli Bacillales Staphylococcaceae *Staphylococcus aureus* M0848 |
| Sau-1831-1 | Bacteria Firmicutes Bacilli Bacillales Staphylococcaceae *Staphylococcus aureus* M0849 |
| Sau-1832-1 | Bacteria Firmicutes Bacilli Bacillales Staphylococcaceae *Staphylococcus aureus* M0850 |
| Sau-1833-1 | Bacteria Firmicutes Bacilli Bacillales Staphylococcaceae *Staphylococcus aureus* M0851 |
| Sau-1834-1 | Bacteria Firmicutes Bacilli Bacillales Staphylococcaceae *Staphylococcus aureus* M0852 |
| Sau-1835-1 | Bacteria Firmicutes Bacilli Bacillales Staphylococcaceae *Staphylococcus aureus* M0853 |
| Sau-1836-1 | Bacteria Firmicutes Bacilli Bacillales Staphylococcaceae *Staphylococcus aureus* M0854 |
| Sau-1837-1 | Bacteria Firmicutes Bacilli Bacillales Staphylococcaceae *Staphylococcus aureus* M0855 |
| Sau-1838-1 | Bacteria Firmicutes Bacilli Bacillales Staphylococcaceae *Staphylococcus aureus* M0857 |
| Sau-1839-1 | Bacteria Firmicutes Bacilli Bacillales Staphylococcaceae *Staphylococcus aureus* M0858 |
| Sau-1840-1 | Bacteria Firmicutes Bacilli Bacillales Staphylococcaceae *Staphylococcus aureus* M0860 |
| Sau-1841-1 | Bacteria Firmicutes Bacilli Bacillales Staphylococcaceae *Staphylococcus aureus* M0861 |
| Sau-1842-1 | Bacteria Firmicutes Bacilli Bacillales Staphylococcaceae *Staphylococcus aureus* M0862 |
| Sau-1843-1 | Bacteria Firmicutes Bacilli Bacillales Staphylococcaceae *Staphylococcus aureus* M0863 |
| Sau-1844-1 | Bacteria Firmicutes Bacilli Bacillales Staphylococcaceae *Staphylococcus aureus* M0864 |
| Sau-1845-1 | Bacteria Firmicutes Bacilli Bacillales Staphylococcaceae *Staphylococcus aureus* M0865 |
| Sau-1846-1 | Bacteria Firmicutes Bacilli Bacillales Staphylococcaceae *Staphylococcus aureus* M0866 |
| Sau-1847-1 | Bacteria Firmicutes Bacilli Bacillales Staphylococcaceae *Staphylococcus aureus* M0867 |
| Sau-1848-1 | Bacteria Firmicutes Bacilli Bacillales Staphylococcaceae *Staphylococcus aureus* M0868 |
| Sau-1849-1 | Bacteria Firmicutes Bacilli Bacillales Staphylococcaceae *Staphylococcus aureus* M0869 |
| Sau-1850-1 | Bacteria Firmicutes Bacilli Bacillales Staphylococcaceae *Staphylococcus aureus* M0870 |
| Sau-1851-1 | Bacteria Firmicutes Bacilli Bacillales Staphylococcaceae *Staphylococcus aureus* M0872 |
| Sau-1852-1 | Bacteria Firmicutes Bacilli Bacillales Staphylococcaceae *Staphylococcus aureus* M0873 |
| Sau-1853-1 | Bacteria Firmicutes Bacilli Bacillales Staphylococcaceae *Staphylococcus aureus* M0874 |
| Sau-1854-1 | Bacteria Firmicutes Bacilli Bacillales Staphylococcaceae *Staphylococcus aureus* M0875 |
| Sau-1855-1 | Bacteria Firmicutes Bacilli Bacillales Staphylococcaceae *Staphylococcus aureus* M0876 |
| Sau-1856-1 | Bacteria Firmicutes Bacilli Bacillales Staphylococcaceae *Staphylococcus aureus* M0878 |
| Sau-1857-1 | Bacteria Firmicutes Bacilli Bacillales Staphylococcaceae *Staphylococcus aureus* M0879 |
| Sau-1858-1 | Bacteria Firmicutes Bacilli Bacillales Staphylococcaceae *Staphylococcus aureus* M0880 |
| Sau-1859-1 | Bacteria Firmicutes Bacilli Bacillales Staphylococcaceae *Staphylococcus aureus* M0882 |
| Sau-1860-1 | Bacteria Firmicutes Bacilli Bacillales Staphylococcaceae *Staphylococcus aureus* M0883 |
| Sau-1861-1 | Bacteria Firmicutes Bacilli Bacillales Staphylococcaceae *Staphylococcus aureus* M0884 |
| Sau-1862-1 | Bacteria Firmicutes Bacilli Bacillales Staphylococcaceae *Staphylococcus aureus* M0885 |
| Sau-1863-1 | Bacteria Firmicutes Bacilli Bacillales Staphylococcaceae *Staphylococcus aureus* M0886 |
| Sau-1864-1 | Bacteria Firmicutes Bacilli Bacillales Staphylococcaceae *Staphylococcus aureus* M0887 |
| Sau-1865-1 | Bacteria Firmicutes Bacilli Bacillales Staphylococcaceae *Staphylococcus aureus* M0888 |
| Sau-1866-1 | Bacteria Firmicutes Bacilli Bacillales Staphylococcaceae *Staphylococcus aureus* M0889 |
| Sau-1867-1 | Bacteria Firmicutes Bacilli Bacillales Staphylococcaceae *Staphylococcus aureus* M0890 |
| Sau-1868-1 | Bacteria Firmicutes Bacilli Bacillales Staphylococcaceae *Staphylococcus aureus* M0891 |
| Sau-1869-1 | Bacteria Firmicutes Bacilli Bacillales Staphylococcaceae *Staphylococcus aureus* M0893 |
| Sau-1870-1 | Bacteria Firmicutes Bacilli Bacillales Staphylococcaceae *Staphylococcus aureus* M0894 |
| Sau-1871-1 | Bacteria Firmicutes Bacilli Bacillales Staphylococcaceae *Staphylococcus aureus* M0895 |
| Sau-1872-1 | Bacteria Firmicutes Bacilli Bacillales Staphylococcaceae *Staphylococcus aureus* M0896 |
| Sau-1873-1 | Bacteria Firmicutes Bacilli Bacillales Staphylococcaceae *Staphylococcus aureus* M0897 |
| Sau-1874-1 | Bacteria Firmicutes Bacilli Bacillales Staphylococcaceae *Staphylococcus aureus* M0898 |
| Sau-1875-1 | Bacteria Firmicutes Bacilli Bacillales Staphylococcaceae *Staphylococcus aureus* M0899 |
| Sau-1876-1 | Bacteria Firmicutes Bacilli Bacillales Staphylococcaceae *Staphylococcus aureus* M0901 |
| Sau-1877-1 | Bacteria Firmicutes Bacilli Bacillales Staphylococcaceae *Staphylococcus aureus* M0902 |

| | |
|---|---|
| Sau-1878-1 | Bacteria Firmicutes Bacilli Bacillales Staphylococcaceae *Staphylococcus aureus* M0903 |
| Sau-1879-1 | Bacteria Firmicutes Bacilli Bacillales Staphylococcaceae *Staphylococcus aureus* M0904 |
| Sau-1880-1 | Bacteria Firmicutes Bacilli Bacillales Staphylococcaceae *Staphylococcus aureus* M0905 |
| Sau-1881-1 | Bacteria Firmicutes Bacilli Bacillales Staphylococcaceae *Staphylococcus aureus* M0906 |
| Sau-1882-1 | Bacteria Firmicutes Bacilli Bacillales Staphylococcaceae *Staphylococcus aureus* M0907 |
| Sau-1883-1 | Bacteria Firmicutes Bacilli Bacillales Staphylococcaceae *Staphylococcus aureus* M0908 |
| Sau-1884-1 | Bacteria Firmicutes Bacilli Bacillales Staphylococcaceae *Staphylococcus aureus* M0909 |
| Sau-1885-1 | Bacteria Firmicutes Bacilli Bacillales Staphylococcaceae *Staphylococcus aureus* M0910 |
| Sau-1886-1 | Bacteria Firmicutes Bacilli Bacillales Staphylococcaceae *Staphylococcus aureus* M0911 |
| Sau-1887-1 | Bacteria Firmicutes Bacilli Bacillales Staphylococcaceae *Staphylococcus aureus* M0913 |
| Sau-1888-1 | Bacteria Firmicutes Bacilli Bacillales Staphylococcaceae *Staphylococcus aureus* M0914 |
| Sau-1889-1 | Bacteria Firmicutes Bacilli Bacillales Staphylococcaceae *Staphylococcus aureus* M0915 |
| Sau-1890-1 | Bacteria Firmicutes Bacilli Bacillales Staphylococcaceae *Staphylococcus aureus* M0916 |
| Sau-1891-1 | Bacteria Firmicutes Bacilli Bacillales Staphylococcaceae *Staphylococcus aureus* M0917 |
| Sau-1892-1 | Bacteria Firmicutes Bacilli Bacillales Staphylococcaceae *Staphylococcus aureus* M0918 |
| Sau-1893-1 | Bacteria Firmicutes Bacilli Bacillales Staphylococcaceae *Staphylococcus aureus* M0919 |
| Sau-1894-1 | Bacteria Firmicutes Bacilli Bacillales Staphylococcaceae *Staphylococcus aureus* M0920 |
| Sau-1895-1 | Bacteria Firmicutes Bacilli Bacillales Staphylococcaceae *Staphylococcus aureus* M0921 |
| Sau-1896-1 | Bacteria Firmicutes Bacilli Bacillales Staphylococcaceae *Staphylococcus aureus* M0922 |
| Sau-1897-1 | Bacteria Firmicutes Bacilli Bacillales Staphylococcaceae *Staphylococcus aureus* M0923 |
| Sau-1898-1 | Bacteria Firmicutes Bacilli Bacillales Staphylococcaceae *Staphylococcus aureus* M0924 |
| Sau-1899-1 | Bacteria Firmicutes Bacilli Bacillales Staphylococcaceae *Staphylococcus aureus* M0928 |
| Sau-1900-1 | Bacteria Firmicutes Bacilli Bacillales Staphylococcaceae *Staphylococcus aureus* M0929 |
| Sau-1901-1 | Bacteria Firmicutes Bacilli Bacillales Staphylococcaceae *Staphylococcus aureus* M0930 |
| Sau-1902-1 | Bacteria Firmicutes Bacilli Bacillales Staphylococcaceae *Staphylococcus aureus* M0931 |
| Sau-1903-1 | Bacteria Firmicutes Bacilli Bacillales Staphylococcaceae *Staphylococcus aureus* M0932 |
| Sau-1904-1 | Bacteria Firmicutes Bacilli Bacillales Staphylococcaceae *Staphylococcus aureus* M0935 |
| Sau-1905-1 | Bacteria Firmicutes Bacilli Bacillales Staphylococcaceae *Staphylococcus aureus* M0936 |
| Sau-1906-1 | Bacteria Firmicutes Bacilli Bacillales Staphylococcaceae *Staphylococcus aureus* M0937 |
| Sau-1907-1 | Bacteria Firmicutes Bacilli Bacillales Staphylococcaceae *Staphylococcus aureus* M0938 |
| Sau-1908-1 | Bacteria Firmicutes Bacilli Bacillales Staphylococcaceae *Staphylococcus aureus* M0939 |
| Sau-1909-1 | Bacteria Firmicutes Bacilli Bacillales Staphylococcaceae *Staphylococcus aureus* M0940 |
| Sau-1910-1 | Bacteria Firmicutes Bacilli Bacillales Staphylococcaceae *Staphylococcus aureus* M0941 |
| Sau-1911-1 | Bacteria Firmicutes Bacilli Bacillales Staphylococcaceae *Staphylococcus aureus* M0942 |
| Sau-1912-1 | Bacteria Firmicutes Bacilli Bacillales Staphylococcaceae *Staphylococcus aureus* M0945 |
| Sau-1913-1 | Bacteria Firmicutes Bacilli Bacillales Staphylococcaceae *Staphylococcus aureus* M0947 |
| Sau-1914-1 | Bacteria Firmicutes Bacilli Bacillales Staphylococcaceae *Staphylococcus aureus* M0948 |
| Sau-1915-1 | Bacteria Firmicutes Bacilli Bacillales Staphylococcaceae *Staphylococcus aureus* M0949 |
| Sau-1916-1 | Bacteria Firmicutes Bacilli Bacillales Staphylococcaceae *Staphylococcus aureus* M0950 |
| Sau-1917-1 | Bacteria Firmicutes Bacilli Bacillales Staphylococcaceae *Staphylococcus aureus* M0951 |
| Sau-1918-1 | Bacteria Firmicutes Bacilli Bacillales Staphylococcaceae *Staphylococcus aureus* M0954 |
| Sau-1919-1 | Bacteria Firmicutes Bacilli Bacillales Staphylococcaceae *Staphylococcus aureus* M0955 |
| Sau-1920-1 | Bacteria Firmicutes Bacilli Bacillales Staphylococcaceae *Staphylococcus aureus* M0956 |
| Sau-1921-1 | Bacteria Firmicutes Bacilli Bacillales Staphylococcaceae *Staphylococcus aureus* M0957 |
| Sau-1922-1 | Bacteria Firmicutes Bacilli Bacillales Staphylococcaceae *Staphylococcus aureus* M0959 |
| Sau-1923-1 | Bacteria Firmicutes Bacilli Bacillales Staphylococcaceae *Staphylococcus aureus* M0960 |
| Sau-1924-1 | Bacteria Firmicutes Bacilli Bacillales Staphylococcaceae *Staphylococcus aureus* M0961 |
| Sau-1925-1 | Bacteria Firmicutes Bacilli Bacillales Staphylococcaceae *Staphylococcus aureus* M0962 |
| Sau-1926-1 | Bacteria Firmicutes Bacilli Bacillales Staphylococcaceae *Staphylococcus aureus* M0965 |
| Sau-1927-1 | Bacteria Firmicutes Bacilli Bacillales Staphylococcaceae *Staphylococcus aureus* M0966 |
| Sau-1928-1 | Bacteria Firmicutes Bacilli Bacillales Staphylococcaceae *Staphylococcus aureus* M0967 |
| Sau-1929-1 | Bacteria Firmicutes Bacilli Bacillales Staphylococcaceae *Staphylococcus aureus* M0968 |
| Sau-1930-1 | Bacteria Firmicutes Bacilli Bacillales Staphylococcaceae *Staphylococcus aureus* M0969 |
| Sau-1931-1 | Bacteria Firmicutes Bacilli Bacillales Staphylococcaceae *Staphylococcus aureus* M0970 |
| Sau-1932-1 | Bacteria Firmicutes Bacilli Bacillales Staphylococcaceae *Staphylococcus aureus* M0971 |
| Sau-1933-1 | Bacteria Firmicutes Bacilli Bacillales Staphylococcaceae *Staphylococcus aureus* M0972 |
| Sau-1934-1 | Bacteria Firmicutes Bacilli Bacillales Staphylococcaceae *Staphylococcus aureus* M0973 |
| Sau-1935-1 | Bacteria Firmicutes Bacilli Bacillales Staphylococcaceae *Staphylococcus aureus* M0974 |
| Sau-1936-1 | Bacteria Firmicutes Bacilli Bacillales Staphylococcaceae *Staphylococcus aureus* M0975 |
| Sau-1937-1 | Bacteria Firmicutes Bacilli Bacillales Staphylococcaceae *Staphylococcus aureus* M0977 |
| Sau-1938-1 | Bacteria Firmicutes Bacilli Bacillales Staphylococcaceae *Staphylococcus aureus* M0979 |
| Sau-1939-1 | Bacteria Firmicutes Bacilli Bacillales Staphylococcaceae *Staphylococcus aureus* M0980 |
| Sau-1940-1 | Bacteria Firmicutes Bacilli Bacillales Staphylococcaceae *Staphylococcus aureus* M0981 |
| Sau-1941-1 | Bacteria Firmicutes Bacilli Bacillales Staphylococcaceae *Staphylococcus aureus* M0982 |
| Sau-1942-1 | Bacteria Firmicutes Bacilli Bacillales Staphylococcaceae *Staphylococcus aureus* M0983 |

| | |
|---|---|
| Sau-1943-1 | Bacteria Firmicutes Bacilli Bacillales Staphylococcaceae *Staphylococcus aureus* M0984 |
| Sau-1944-1 | Bacteria Firmicutes Bacilli Bacillales Staphylococcaceae *Staphylococcus aureus* M0985 |
| Sau-1945-1 | Bacteria Firmicutes Bacilli Bacillales Staphylococcaceae *Staphylococcus aureus* M0986 |
| Sau-1946-1 | Bacteria Firmicutes Bacilli Bacillales Staphylococcaceae *Staphylococcus aureus* M0987 |
| Sau-1947-1 | Bacteria Firmicutes Bacilli Bacillales Staphylococcaceae *Staphylococcus aureus* M0988 |
| Sau-1948-1 | Bacteria Firmicutes Bacilli Bacillales Staphylococcaceae *Staphylococcus aureus* M0989 |
| Sau-1949-1 | Bacteria Firmicutes Bacilli Bacillales Staphylococcaceae *Staphylococcus aureus* M0991 |
| Sau-1950-1 | Bacteria Firmicutes Bacilli Bacillales Staphylococcaceae *Staphylococcus aureus* M0992 |
| Sau-1951-1 | Bacteria Firmicutes Bacilli Bacillales Staphylococcaceae *Staphylococcus aureus* M0993 |
| Sau-1952-1 | Bacteria Firmicutes Bacilli Bacillales Staphylococcaceae *Staphylococcus aureus* M0996 |
| Sau-1953-1 | Bacteria Firmicutes Bacilli Bacillales Staphylococcaceae *Staphylococcus aureus* M0997 |
| Sau-1954-1 | Bacteria Firmicutes Bacilli Bacillales Staphylococcaceae *Staphylococcus aureus* M1000 |
| Sau-1955-1 | Bacteria Firmicutes Bacilli Bacillales Staphylococcaceae *Staphylococcus aureus* M1001 |
| Sau-1956-1 | Bacteria Firmicutes Bacilli Bacillales Staphylococcaceae *Staphylococcus aureus* M1002 |
| Sau-1957-1 | Bacteria Firmicutes Bacilli Bacillales Staphylococcaceae *Staphylococcus aureus* M1003 |
| Sau-1958-1 | Bacteria Firmicutes Bacilli Bacillales Staphylococcaceae *Staphylococcus aureus* M1006 |
| Sau-1959-1 | Bacteria Firmicutes Bacilli Bacillales Staphylococcaceae *Staphylococcus aureus* M1008 |
| Sau-1960-1 | Bacteria Firmicutes Bacilli Bacillales Staphylococcaceae *Staphylococcus aureus* M1009 |
| Sau-1961-1 | Bacteria Firmicutes Bacilli Bacillales Staphylococcaceae *Staphylococcus aureus* M1011 |
| Sau-1962-1 | Bacteria Firmicutes Bacilli Bacillales Staphylococcaceae *Staphylococcus aureus* M1012 |
| Sau-1963-1 | Bacteria Firmicutes Bacilli Bacillales Staphylococcaceae *Staphylococcus aureus* M1013 |
| Sau-1964-1 | Bacteria Firmicutes Bacilli Bacillales Staphylococcaceae *Staphylococcus aureus* M1014 |
| Sau-1965-1 | Bacteria Firmicutes Bacilli Bacillales Staphylococcaceae *Staphylococcus aureus* M1017 |
| Sau-1966-1 | Bacteria Firmicutes Bacilli Bacillales Staphylococcaceae *Staphylococcus aureus* M1018 |
| Sau-1967-1 | Bacteria Firmicutes Bacilli Bacillales Staphylococcaceae *Staphylococcus aureus* M1019 |
| Sau-1968-1 | Bacteria Firmicutes Bacilli Bacillales Staphylococcaceae *Staphylococcus aureus* M1020 |
| Sau-1969-1 | Bacteria Firmicutes Bacilli Bacillales Staphylococcaceae *Staphylococcus aureus* M1021 |
| Sau-1970-1 | Bacteria Firmicutes Bacilli Bacillales Staphylococcaceae *Staphylococcus aureus* M1022 |
| Sau-1971-1 | Bacteria Firmicutes Bacilli Bacillales Staphylococcaceae *Staphylococcus aureus* M1023 |
| Sau-1972-1 | Bacteria Firmicutes Bacilli Bacillales Staphylococcaceae *Staphylococcus aureus* M1024 |
| Sau-1973-1 | Bacteria Firmicutes Bacilli Bacillales Staphylococcaceae *Staphylococcus aureus* M1025 |
| Sau-1974-1 | Bacteria Firmicutes Bacilli Bacillales Staphylococcaceae *Staphylococcus aureus* M1026 |
| Sau-1975-1 | Bacteria Firmicutes Bacilli Bacillales Staphylococcaceae *Staphylococcus aureus* M1027 |
| Sau-1976-1 | Bacteria Firmicutes Bacilli Bacillales Staphylococcaceae *Staphylococcus aureus* M1028 |
| Sau-1977-1 | Bacteria Firmicutes Bacilli Bacillales Staphylococcaceae *Staphylococcus aureus* M1029 |
| Sau-1978-1 | Bacteria Firmicutes Bacilli Bacillales Staphylococcaceae *Staphylococcus aureus* M1030 |
| Sau-1979-1 | Bacteria Firmicutes Bacilli Bacillales Staphylococcaceae *Staphylococcus aureus* M1031 |
| Sau-1980-1 | Bacteria Firmicutes Bacilli Bacillales Staphylococcaceae *Staphylococcus aureus* M1033 |
| Sau-1981-1 | Bacteria Firmicutes Bacilli Bacillales Staphylococcaceae *Staphylococcus aureus* M1035 |
| Sau-1982-1 | Bacteria Firmicutes Bacilli Bacillales Staphylococcaceae *Staphylococcus aureus* M1039 |
| Sau-1983-1 | Bacteria Firmicutes Bacilli Bacillales Staphylococcaceae *Staphylococcus aureus* M1041 |
| Sau-1984-1 | Bacteria Firmicutes Bacilli Bacillales Staphylococcaceae *Staphylococcus aureus* M1042 |
| Sau-1985-1 | Bacteria Firmicutes Bacilli Bacillales Staphylococcaceae *Staphylococcus aureus* M1043 |
| Sau-1986-1 | Bacteria Firmicutes Bacilli Bacillales Staphylococcaceae *Staphylococcus aureus* M1045 |
| Sau-1987-1 | Bacteria Firmicutes Bacilli Bacillales Staphylococcaceae *Staphylococcus aureus* M1046 |
| Sau-1988-1 | Bacteria Firmicutes Bacilli Bacillales Staphylococcaceae *Staphylococcus aureus* M1047 |
| Sau-1989-1 | Bacteria Firmicutes Bacilli Bacillales Staphylococcaceae *Staphylococcus aureus* M1048 |
| Sau-1990-1 | Bacteria Firmicutes Bacilli Bacillales Staphylococcaceae *Staphylococcus aureus* M1049 |
| Sau-1991-1 | Bacteria Firmicutes Bacilli Bacillales Staphylococcaceae *Staphylococcus aureus* M1050 |
| Sau-1992-1 | Bacteria Firmicutes Bacilli Bacillales Staphylococcaceae *Staphylococcus aureus* M1051 |
| Sau-1993-1 | Bacteria Firmicutes Bacilli Bacillales Staphylococcaceae *Staphylococcus aureus* M1052 |
| Sau-1994-1 | Bacteria Firmicutes Bacilli Bacillales Staphylococcaceae *Staphylococcus aureus* M1053 |
| Sau-1995-1 | Bacteria Firmicutes Bacilli Bacillales Staphylococcaceae *Staphylococcus aureus* M1054 |
| Sau-1996-1 | Bacteria Firmicutes Bacilli Bacillales Staphylococcaceae *Staphylococcus aureus* M1055 |
| Sau-1997-1 | Bacteria Firmicutes Bacilli Bacillales Staphylococcaceae *Staphylococcus aureus* M1056 |
| Sau-1998-1 | Bacteria Firmicutes Bacilli Bacillales Staphylococcaceae *Staphylococcus aureus* M1057 |
| Sau-1999-1 | Bacteria Firmicutes Bacilli Bacillales Staphylococcaceae *Staphylococcus aureus* M1058 |
| Sau-2000-1 | Bacteria Firmicutes Bacilli Bacillales Staphylococcaceae *Staphylococcus aureus* M1059 |
| Sau-2001-1 | Bacteria Firmicutes Bacilli Bacillales Staphylococcaceae *Staphylococcus aureus* M1065 |
| Sau-2002-1 | Bacteria Firmicutes Bacilli Bacillales Staphylococcaceae *Staphylococcus aureus* M1066 |
| Sau-2003-1 | Bacteria Firmicutes Bacilli Bacillales Staphylococcaceae *Staphylococcus aureus* M1067 |
| Sau-2004-1 | Bacteria Firmicutes Bacilli Bacillales Staphylococcaceae *Staphylococcus aureus* M1069 |
| Sau-2005-1 | Bacteria Firmicutes Bacilli Bacillales Staphylococcaceae *Staphylococcus aureus* M1070 |
| Sau-2006-1 | Bacteria Firmicutes Bacilli Bacillales Staphylococcaceae *Staphylococcus aureus* M1071 |
| Sau-2007-1 | Bacteria Firmicutes Bacilli Bacillales Staphylococcaceae *Staphylococcus aureus* M1072 |

| | |
|---|---|
| Sau-2008-1 | Bacteria Firmicutes Bacilli Bacillales Staphylococcaceae *Staphylococcus aureus* M1073 |
| Sau-2009-1 | Bacteria Firmicutes Bacilli Bacillales Staphylococcaceae *Staphylococcus aureus* M1074 |
| Sau-2010-1 | Bacteria Firmicutes Bacilli Bacillales Staphylococcaceae *Staphylococcus aureus* M1075 |
| Sau-2011-1 | Bacteria Firmicutes Bacilli Bacillales Staphylococcaceae *Staphylococcus aureus* M1077 |
| Sau-2012-1 | Bacteria Firmicutes Bacilli Bacillales Staphylococcaceae *Staphylococcus aureus* M1079 |
| Sau-2013-1 | Bacteria Firmicutes Bacilli Bacillales Staphylococcaceae *Staphylococcus aureus* M1080 |
| Sau-2014-1 | Bacteria Firmicutes Bacilli Bacillales Staphylococcaceae *Staphylococcus aureus* M1081 |
| Sau-2015-1 | Bacteria Firmicutes Bacilli Bacillales Staphylococcaceae *Staphylococcus aureus* M1082 |
| Sau-2016-1 | Bacteria Firmicutes Bacilli Bacillales Staphylococcaceae *Staphylococcus aureus* M1084 |
| Sau-2017-1 | Bacteria Firmicutes Bacilli Bacillales Staphylococcaceae *Staphylococcus aureus* M1085 |
| Sau-2018-1 | Bacteria Firmicutes Bacilli Bacillales Staphylococcaceae *Staphylococcus aureus* M1086 |
| Sau-2019-1 | Bacteria Firmicutes Bacilli Bacillales Staphylococcaceae *Staphylococcus aureus* M1087 |
| Sau-2020-1 | Bacteria Firmicutes Bacilli Bacillales Staphylococcaceae *Staphylococcus aureus* M1088 |
| Sau-2021-1 | Bacteria Firmicutes Bacilli Bacillales Staphylococcaceae *Staphylococcus aureus* M1089 |
| Sau-2022-1 | Bacteria Firmicutes Bacilli Bacillales Staphylococcaceae *Staphylococcus aureus* M1090 |
| Sau-2023-1 | Bacteria Firmicutes Bacilli Bacillales Staphylococcaceae *Staphylococcus aureus* M1091 |
| Sau-2024-1 | Bacteria Firmicutes Bacilli Bacillales Staphylococcaceae *Staphylococcus aureus* M1094 |
| Sau-2025-1 | Bacteria Firmicutes Bacilli Bacillales Staphylococcaceae *Staphylococcus aureus* M1096 |
| Sau-2026-1 | Bacteria Firmicutes Bacilli Bacillales Staphylococcaceae *Staphylococcus aureus* M1097 |
| Sau-2027-1 | Bacteria Firmicutes Bacilli Bacillales Staphylococcaceae *Staphylococcus aureus* M1098 |
| Sau-2028-1 | Bacteria Firmicutes Bacilli Bacillales Staphylococcaceae *Staphylococcus aureus* M1099 |
| Sau-2029-1 | Bacteria Firmicutes Bacilli Bacillales Staphylococcaceae *Staphylococcus aureus* M1100 |
| Sau-2030-1 | Bacteria Firmicutes Bacilli Bacillales Staphylococcaceae *Staphylococcus aureus* M1101 |
| Sau-2031-1 | Bacteria Firmicutes Bacilli Bacillales Staphylococcaceae *Staphylococcus aureus* M1102 |
| Sau-2032-1 | Bacteria Firmicutes Bacilli Bacillales Staphylococcaceae *Staphylococcus aureus* M1104 |
| Sau-2033-1 | Bacteria Firmicutes Bacilli Bacillales Staphylococcaceae *Staphylococcus aureus* M1105 |
| Sau-2034-1 | Bacteria Firmicutes Bacilli Bacillales Staphylococcaceae *Staphylococcus aureus* M1106 |
| Sau-2035-1 | Bacteria Firmicutes Bacilli Bacillales Staphylococcaceae *Staphylococcus aureus* M1107 |
| Sau-2036-1 | Bacteria Firmicutes Bacilli Bacillales Staphylococcaceae *Staphylococcus aureus* M1108 |
| Sau-2037-1 | Bacteria Firmicutes Bacilli Bacillales Staphylococcaceae *Staphylococcus aureus* M1110 |
| Sau-2038-1 | Bacteria Firmicutes Bacilli Bacillales Staphylococcaceae *Staphylococcus aureus* M1111 |
| Sau-2039-1 | Bacteria Firmicutes Bacilli Bacillales Staphylococcaceae *Staphylococcus aureus* M1112 |
| Sau-2040-1 | Bacteria Firmicutes Bacilli Bacillales Staphylococcaceae *Staphylococcus aureus* M1113 |
| Sau-2041-1 | Bacteria Firmicutes Bacilli Bacillales Staphylococcaceae *Staphylococcus aureus* M1114 |
| Sau-2042-1 | Bacteria Firmicutes Bacilli Bacillales Staphylococcaceae *Staphylococcus aureus* M1115 |
| Sau-2043-1 | Bacteria Firmicutes Bacilli Bacillales Staphylococcaceae *Staphylococcus aureus* M1116 |
| Sau-2044-1 | Bacteria Firmicutes Bacilli Bacillales Staphylococcaceae *Staphylococcus aureus* M1117 |
| Sau-2045-1 | Bacteria Firmicutes Bacilli Bacillales Staphylococcaceae *Staphylococcus aureus* M1120 |
| Sau-2046-1 | Bacteria Firmicutes Bacilli Bacillales Staphylococcaceae *Staphylococcus aureus* M1121 |
| Sau-2047-1 | Bacteria Firmicutes Bacilli Bacillales Staphylococcaceae *Staphylococcus aureus* M1122 |
| Sau-2048-1 | Bacteria Firmicutes Bacilli Bacillales Staphylococcaceae *Staphylococcus aureus* M1123 |
| Sau-2049-1 | Bacteria Firmicutes Bacilli Bacillales Staphylococcaceae *Staphylococcus aureus* M1124 |
| Sau-2050-1 | Bacteria Firmicutes Bacilli Bacillales Staphylococcaceae *Staphylococcus aureus* M1125 |
| Sau-2051-1 | Bacteria Firmicutes Bacilli Bacillales Staphylococcaceae *Staphylococcus aureus* M1127 |
| Sau-2052-1 | Bacteria Firmicutes Bacilli Bacillales Staphylococcaceae *Staphylococcus aureus* M1128 |
| Sau-2053-1 | Bacteria Firmicutes Bacilli Bacillales Staphylococcaceae *Staphylococcus aureus* M1129 |
| Sau-2054-1 | Bacteria Firmicutes Bacilli Bacillales Staphylococcaceae *Staphylococcus aureus* M1130 |
| Sau-2055-1 | Bacteria Firmicutes Bacilli Bacillales Staphylococcaceae *Staphylococcus aureus* M1131 |
| Sau-2056-1 | Bacteria Firmicutes Bacilli Bacillales Staphylococcaceae *Staphylococcus aureus* M1132 |
| Sau-2057-1 | Bacteria Firmicutes Bacilli Bacillales Staphylococcaceae *Staphylococcus aureus* M1134 |
| Sau-2058-1 | Bacteria Firmicutes Bacilli Bacillales Staphylococcaceae *Staphylococcus aureus* M1135 |
| Sau-2059-1 | Bacteria Firmicutes Bacilli Bacillales Staphylococcaceae *Staphylococcus aureus* M1136 |
| Sau-2060-1 | Bacteria Firmicutes Bacilli Bacillales Staphylococcaceae *Staphylococcus aureus* M1137 |
| Sau-2061-1 | Bacteria Firmicutes Bacilli Bacillales Staphylococcaceae *Staphylococcus aureus* M1138 |
| Sau-2062-1 | Bacteria Firmicutes Bacilli Bacillales Staphylococcaceae *Staphylococcus aureus* M1139 |
| Sau-2063-1 | Bacteria Firmicutes Bacilli Bacillales Staphylococcaceae *Staphylococcus aureus* M1140 |
| Sau-2064-1 | Bacteria Firmicutes Bacilli Bacillales Staphylococcaceae *Staphylococcus aureus* M1141 |
| Sau-2065-1 | Bacteria Firmicutes Bacilli Bacillales Staphylococcaceae *Staphylococcus aureus* M1143 |
| Sau-2066-1 | Bacteria Firmicutes Bacilli Bacillales Staphylococcaceae *Staphylococcus aureus* M1144 |
| Sau-2067-1 | Bacteria Firmicutes Bacilli Bacillales Staphylococcaceae *Staphylococcus aureus* M1145 |
| Sau-2068-1 | Bacteria Firmicutes Bacilli Bacillales Staphylococcaceae *Staphylococcus aureus* M1146 |
| Sau-2069-1 | Bacteria Firmicutes Bacilli Bacillales Staphylococcaceae *Staphylococcus aureus* M1147 |
| Sau-2070-1 | Bacteria Firmicutes Bacilli Bacillales Staphylococcaceae *Staphylococcus aureus* M1148 |
| Sau-2071-1 | Bacteria Firmicutes Bacilli Bacillales Staphylococcaceae *Staphylococcus aureus* M1149 |
| Sau-2072-1 | Bacteria Firmicutes Bacilli Bacillales Staphylococcaceae *Staphylococcus aureus* M1150 |

| | |
|---|---|
| Sau-2073-1 | Bacteria Firmicutes Bacilli Bacillales Staphylococcaceae *Staphylococcus aureus* M1151 |
| Sau-2074-1 | Bacteria Firmicutes Bacilli Bacillales Staphylococcaceae *Staphylococcus aureus* M1152 |
| Sau-2075-1 | Bacteria Firmicutes Bacilli Bacillales Staphylococcaceae *Staphylococcus aureus* M1153 |
| Sau-2076-1 | Bacteria Firmicutes Bacilli Bacillales Staphylococcaceae *Staphylococcus aureus* M1154 |
| Sau-2077-1 | Bacteria Firmicutes Bacilli Bacillales Staphylococcaceae *Staphylococcus aureus* M1155 |
| Sau-2078-1 | Bacteria Firmicutes Bacilli Bacillales Staphylococcaceae *Staphylococcus aureus* M1156 |
| Sau-2079-1 | Bacteria Firmicutes Bacilli Bacillales Staphylococcaceae *Staphylococcus aureus* M1157 |
| Sau-2080-1 | Bacteria Firmicutes Bacilli Bacillales Staphylococcaceae *Staphylococcus aureus* M1158 |
| Sau-2081-1 | Bacteria Firmicutes Bacilli Bacillales Staphylococcaceae *Staphylococcus aureus* M1159 |
| Sau-2082-1 | Bacteria Firmicutes Bacilli Bacillales Staphylococcaceae *Staphylococcus aureus* M1160 |
| Sau-2083-1 | Bacteria Firmicutes Bacilli Bacillales Staphylococcaceae *Staphylococcus aureus* M1161 |
| Sau-2084-1 | Bacteria Firmicutes Bacilli Bacillales Staphylococcaceae *Staphylococcus aureus* M1162 |
| Sau-2085-1 | Bacteria Firmicutes Bacilli Bacillales Staphylococcaceae *Staphylococcus aureus* M1163 |
| Sau-2086-1 | Bacteria Firmicutes Bacilli Bacillales Staphylococcaceae *Staphylococcus aureus* M1164 |
| Sau-2087-1 | Bacteria Firmicutes Bacilli Bacillales Staphylococcaceae *Staphylococcus aureus* M1165 |
| Sau-2088-1 | Bacteria Firmicutes Bacilli Bacillales Staphylococcaceae *Staphylococcus aureus* M1166 |
| Sau-2089-1 | Bacteria Firmicutes Bacilli Bacillales Staphylococcaceae *Staphylococcus aureus* M1168 |
| Sau-2090-1 | Bacteria Firmicutes Bacilli Bacillales Staphylococcaceae *Staphylococcus aureus* M1169 |
| Sau-2091-1 | Bacteria Firmicutes Bacilli Bacillales Staphylococcaceae *Staphylococcus aureus* M1171 |
| Sau-2092-1 | Bacteria Firmicutes Bacilli Bacillales Staphylococcaceae *Staphylococcus aureus* M1172 |
| Sau-2093-1 | Bacteria Firmicutes Bacilli Bacillales Staphylococcaceae *Staphylococcus aureus* M1174 |
| Sau-2094-1 | Bacteria Firmicutes Bacilli Bacillales Staphylococcaceae *Staphylococcus aureus* M1175 |
| Sau-2095-1 | Bacteria Firmicutes Bacilli Bacillales Staphylococcaceae *Staphylococcus aureus* M1176 |
| Sau-2096-1 | Bacteria Firmicutes Bacilli Bacillales Staphylococcaceae *Staphylococcus aureus* M1177 |
| Sau-2097-1 | Bacteria Firmicutes Bacilli Bacillales Staphylococcaceae *Staphylococcus aureus* M1178 |
| Sau-2098-1 | Bacteria Firmicutes Bacilli Bacillales Staphylococcaceae *Staphylococcus aureus* M1179 |
| Sau-2099-1 | Bacteria Firmicutes Bacilli Bacillales Staphylococcaceae *Staphylococcus aureus* M118-B |
| Sau-2100-1 | Bacteria Firmicutes Bacilli Bacillales Staphylococcaceae *Staphylococcus aureus* M1180 |
| Sau-2101-1 | Bacteria Firmicutes Bacilli Bacillales Staphylococcaceae *Staphylococcus aureus* M1181 |
| Sau-2102-1 | Bacteria Firmicutes Bacilli Bacillales Staphylococcaceae *Staphylococcus aureus* M1182 |
| Sau-2103-1 | Bacteria Firmicutes Bacilli Bacillales Staphylococcaceae *Staphylococcus aureus* M1183 |
| Sau-2104-1 | Bacteria Firmicutes Bacilli Bacillales Staphylococcaceae *Staphylococcus aureus* M1184 |
| Sau-2105-1 | Bacteria Firmicutes Bacilli Bacillales Staphylococcaceae *Staphylococcus aureus* M1186 |
| Sau-2106-1 | Bacteria Firmicutes Bacilli Bacillales Staphylococcaceae *Staphylococcus aureus* M1187 |
| Sau-2107-1 | Bacteria Firmicutes Bacilli Bacillales Staphylococcaceae *Staphylococcus aureus* M1189 |
| Sau-2108-1 | Bacteria Firmicutes Bacilli Bacillales Staphylococcaceae *Staphylococcus aureus* M1190 |
| Sau-2109-1 | Bacteria Firmicutes Bacilli Bacillales Staphylococcaceae *Staphylococcus aureus* M1191 |
| Sau-2110-1 | Bacteria Firmicutes Bacilli Bacillales Staphylococcaceae *Staphylococcus aureus* M1192 |
| Sau-2111-1 | Bacteria Firmicutes Bacilli Bacillales Staphylococcaceae *Staphylococcus aureus* M1194 |
| Sau-2112-1 | Bacteria Firmicutes Bacilli Bacillales Staphylococcaceae *Staphylococcus aureus* M1195 |
| Sau-2113-1 | Bacteria Firmicutes Bacilli Bacillales Staphylococcaceae *Staphylococcus aureus* M1196 |
| Sau-2114-1 | Bacteria Firmicutes Bacilli Bacillales Staphylococcaceae *Staphylococcus aureus* M1197 |
| Sau-2115-1 | Bacteria Firmicutes Bacilli Bacillales Staphylococcaceae *Staphylococcus aureus* M1200 |
| Sau-2116-1 | Bacteria Firmicutes Bacilli Bacillales Staphylococcaceae *Staphylococcus aureus* M1201 |
| Sau-2117-1 | Bacteria Firmicutes Bacilli Bacillales Staphylococcaceae *Staphylococcus aureus* M1202 |
| Sau-2118-1 | Bacteria Firmicutes Bacilli Bacillales Staphylococcaceae *Staphylococcus aureus* M1203 |
| Sau-2119-1 | Bacteria Firmicutes Bacilli Bacillales Staphylococcaceae *Staphylococcus aureus* M1204 |
| Sau-2120-1 | Bacteria Firmicutes Bacilli Bacillales Staphylococcaceae *Staphylococcus aureus* M1205 |
| Sau-2121-1 | Bacteria Firmicutes Bacilli Bacillales Staphylococcaceae *Staphylococcus aureus* M1206 |
| Sau-2122-1 | Bacteria Firmicutes Bacilli Bacillales Staphylococcaceae *Staphylococcus aureus* M1207 |
| Sau-2123-1 | Bacteria Firmicutes Bacilli Bacillales Staphylococcaceae *Staphylococcus aureus* M1208 |
| Sau-2124-1 | Bacteria Firmicutes Bacilli Bacillales Staphylococcaceae *Staphylococcus aureus* M1209 |
| Sau-2125-1 | Bacteria Firmicutes Bacilli Bacillales Staphylococcaceae *Staphylococcus aureus* M1212 |
| Sau-2126-1 | Bacteria Firmicutes Bacilli Bacillales Staphylococcaceae *Staphylococcus aureus* M1213 |
| Sau-2127-1 | Bacteria Firmicutes Bacilli Bacillales Staphylococcaceae *Staphylococcus aureus* M1214 |
| Sau-2128-1 | Bacteria Firmicutes Bacilli Bacillales Staphylococcaceae *Staphylococcus aureus* M1217 |
| Sau-2129-1 | Bacteria Firmicutes Bacilli Bacillales Staphylococcaceae *Staphylococcus aureus* M1218 |
| Sau-2130-1 | Bacteria Firmicutes Bacilli Bacillales Staphylococcaceae *Staphylococcus aureus* M1219 |
| Sau-2131-1 | Bacteria Firmicutes Bacilli Bacillales Staphylococcaceae *Staphylococcus aureus* M1220 |
| Sau-2132-1 | Bacteria Firmicutes Bacilli Bacillales Staphylococcaceae *Staphylococcus aureus* M1221 |
| Sau-2133-1 | Bacteria Firmicutes Bacilli Bacillales Staphylococcaceae *Staphylococcus aureus* M1222 |
| Sau-2134-1 | Bacteria Firmicutes Bacilli Bacillales Staphylococcaceae *Staphylococcus aureus* M1225 |
| Sau-2135-1 | Bacteria Firmicutes Bacilli Bacillales Staphylococcaceae *Staphylococcus aureus* M1226 |
| Sau-2136-1 | Bacteria Firmicutes Bacilli Bacillales Staphylococcaceae *Staphylococcus aureus* M1227 |
| Sau-2137-1 | Bacteria Firmicutes Bacilli Bacillales Staphylococcaceae *Staphylococcus aureus* M1230 |

| | |
|---|---|
| Sau-2138-1 | Bacteria Firmicutes Bacilli Bacillales Staphylococcaceae *Staphylococcus aureus* M1231 |
| Sau-2139-1 | Bacteria Firmicutes Bacilli Bacillales Staphylococcaceae *Staphylococcus aureus* M1232 |
| Sau-2140-1 | Bacteria Firmicutes Bacilli Bacillales Staphylococcaceae *Staphylococcus aureus* M1233 |
| Sau-2141-1 | Bacteria Firmicutes Bacilli Bacillales Staphylococcaceae *Staphylococcus aureus* M1234 |
| Sau-2142-1 | Bacteria Firmicutes Bacilli Bacillales Staphylococcaceae *Staphylococcus aureus* M1235 |
| Sau-2143-1 | Bacteria Firmicutes Bacilli Bacillales Staphylococcaceae *Staphylococcus aureus* M1236 |
| Sau-2144-1 | Bacteria Firmicutes Bacilli Bacillales Staphylococcaceae *Staphylococcus aureus* M1237 |
| Sau-2145-1 | Bacteria Firmicutes Bacilli Bacillales Staphylococcaceae *Staphylococcus aureus* M1238 |
| Sau-2146-1 | Bacteria Firmicutes Bacilli Bacillales Staphylococcaceae *Staphylococcus aureus* M1239 |
| Sau-2147-1 | Bacteria Firmicutes Bacilli Bacillales Staphylococcaceae *Staphylococcus aureus* M1240 |
| Sau-2148-1 | Bacteria Firmicutes Bacilli Bacillales Staphylococcaceae *Staphylococcus aureus* M1241 |
| Sau-2149-1 | Bacteria Firmicutes Bacilli Bacillales Staphylococcaceae *Staphylococcus aureus* M1242 |
| Sau-2150-1 | Bacteria Firmicutes Bacilli Bacillales Staphylococcaceae *Staphylococcus aureus* M1243 |
| Sau-2151-1 | Bacteria Firmicutes Bacilli Bacillales Staphylococcaceae *Staphylococcus aureus* M1245 |
| Sau-2152-1 | Bacteria Firmicutes Bacilli Bacillales Staphylococcaceae *Staphylococcus aureus* M1246 |
| Sau-2153-1 | Bacteria Firmicutes Bacilli Bacillales Staphylococcaceae *Staphylococcus aureus* M1247 |
| Sau-2154-1 | Bacteria Firmicutes Bacilli Bacillales Staphylococcaceae *Staphylococcus aureus* M1249 |
| Sau-2155-1 | Bacteria Firmicutes Bacilli Bacillales Staphylococcaceae *Staphylococcus aureus* M1250 |
| Sau-2156-1 | Bacteria Firmicutes Bacilli Bacillales Staphylococcaceae *Staphylococcus aureus* M1251 |
| Sau-2157-1 | Bacteria Firmicutes Bacilli Bacillales Staphylococcaceae *Staphylococcus aureus* M1252 |
| Sau-2158-1 | Bacteria Firmicutes Bacilli Bacillales Staphylococcaceae *Staphylococcus aureus* M1254 |
| Sau-2159-1 | Bacteria Firmicutes Bacilli Bacillales Staphylococcaceae *Staphylococcus aureus* M1258 |
| Sau-2160-1 | Bacteria Firmicutes Bacilli Bacillales Staphylococcaceae *Staphylococcus aureus* M1259 |
| Sau-2161-1 | Bacteria Firmicutes Bacilli Bacillales Staphylococcaceae *Staphylococcus aureus* M1260 |
| Sau-2162-1 | Bacteria Firmicutes Bacilli Bacillales Staphylococcaceae *Staphylococcus aureus* M1261 |
| Sau-2163-1 | Bacteria Firmicutes Bacilli Bacillales Staphylococcaceae *Staphylococcus aureus* M1262 |
| Sau-2164-1 | Bacteria Firmicutes Bacilli Bacillales Staphylococcaceae *Staphylococcus aureus* M1263 |
| Sau-2165-1 | Bacteria Firmicutes Bacilli Bacillales Staphylococcaceae *Staphylococcus aureus* M1264 |
| Sau-2166-1 | Bacteria Firmicutes Bacilli Bacillales Staphylococcaceae *Staphylococcus aureus* M1265 |
| Sau-2167-1 | Bacteria Firmicutes Bacilli Bacillales Staphylococcaceae *Staphylococcus aureus* M1267 |
| Sau-2168-1 | Bacteria Firmicutes Bacilli Bacillales Staphylococcaceae *Staphylococcus aureus* M1268 |
| Sau-2169-1 | Bacteria Firmicutes Bacilli Bacillales Staphylococcaceae *Staphylococcus aureus* M1269 |
| Sau-2170-1 | Bacteria Firmicutes Bacilli Bacillales Staphylococcaceae *Staphylococcus aureus* M1270 |
| Sau-2171-1 | Bacteria Firmicutes Bacilli Bacillales Staphylococcaceae *Staphylococcus aureus* M12703 |
| Sau-2172-1 | Bacteria Firmicutes Bacilli Bacillales Staphylococcaceae *Staphylococcus aureus* M12709 |
| Sau-2173-1 | Bacteria Firmicutes Bacilli Bacillales Staphylococcaceae *Staphylococcus aureus* M1271 |
| Sau-2174-1 | Bacteria Firmicutes Bacilli Bacillales Staphylococcaceae *Staphylococcus aureus* M12714 |
| Sau-2175-1 | Bacteria Firmicutes Bacilli Bacillales Staphylococcaceae *Staphylococcus aureus* M12717 |
| Sau-2176-1 | Bacteria Firmicutes Bacilli Bacillales Staphylococcaceae *Staphylococcus aureus* M1272 |
| Sau-2177-1 | Bacteria Firmicutes Bacilli Bacillales Staphylococcaceae *Staphylococcus aureus* M1273 |
| Sau-2178-1 | Bacteria Firmicutes Bacilli Bacillales Staphylococcaceae *Staphylococcus aureus* M1274 |
| Sau-2179-1 | Bacteria Firmicutes Bacilli Bacillales Staphylococcaceae *Staphylococcus aureus* M1276 |
| Sau-2180-1 | Bacteria Firmicutes Bacilli Bacillales Staphylococcaceae *Staphylococcus aureus* M1278 |
| Sau-2181-1 | Bacteria Firmicutes Bacilli Bacillales Staphylococcaceae *Staphylococcus aureus* M1279 |
| Sau-2182-1 | Bacteria Firmicutes Bacilli Bacillales Staphylococcaceae *Staphylococcus aureus* M1280 |
| Sau-2183-1 | Bacteria Firmicutes Bacilli Bacillales Staphylococcaceae *Staphylococcus aureus* M1281 |
| Sau-2184-1 | Bacteria Firmicutes Bacilli Bacillales Staphylococcaceae *Staphylococcus aureus* M1282 |
| Sau-2185-1 | Bacteria Firmicutes Bacilli Bacillales Staphylococcaceae *Staphylococcus aureus* M1283 |
| Sau-2186-1 | Bacteria Firmicutes Bacilli Bacillales Staphylococcaceae *Staphylococcus aureus* M1285 |
| Sau-2187-1 | Bacteria Firmicutes Bacilli Bacillales Staphylococcaceae *Staphylococcus aureus* M1287 |
| Sau-2188-1 | Bacteria Firmicutes Bacilli Bacillales Staphylococcaceae *Staphylococcus aureus* M1288 |
| Sau-2189-1 | Bacteria Firmicutes Bacilli Bacillales Staphylococcaceae *Staphylococcus aureus* M1289 |
| Sau-2190-1 | Bacteria Firmicutes Bacilli Bacillales Staphylococcaceae *Staphylococcus aureus* M1290 |
| Sau-2191-1 | Bacteria Firmicutes Bacilli Bacillales Staphylococcaceae *Staphylococcus aureus* M1292 |
| Sau-2192-1 | Bacteria Firmicutes Bacilli Bacillales Staphylococcaceae *Staphylococcus aureus* M1293 |
| Sau-2193-1 | Bacteria Firmicutes Bacilli Bacillales Staphylococcaceae *Staphylococcus aureus* M1294 |
| Sau-2194-1 | Bacteria Firmicutes Bacilli Bacillales Staphylococcaceae *Staphylococcus aureus* M1295 |
| Sau-2195-1 | Bacteria Firmicutes Bacilli Bacillales Staphylococcaceae *Staphylococcus aureus* M1296 |
| Sau-2196-1 | Bacteria Firmicutes Bacilli Bacillales Staphylococcaceae *Staphylococcus aureus* M1297 |
| Sau-2197-1 | Bacteria Firmicutes Bacilli Bacillales Staphylococcaceae *Staphylococcus aureus* M1298 |
| Sau-2198-1 | Bacteria Firmicutes Bacilli Bacillales Staphylococcaceae *Staphylococcus aureus* M1299 |
| Sau-2199-1 | Bacteria Firmicutes Bacilli Bacillales Staphylococcaceae *Staphylococcus aureus* M1300 |
| Sau-2200-1 | Bacteria Firmicutes Bacilli Bacillales Staphylococcaceae *Staphylococcus aureus* M1301 |
| Sau-2201-1 | Bacteria Firmicutes Bacilli Bacillales Staphylococcaceae *Staphylococcus aureus* M1302 |
| Sau-2202-1 to Sau-2202-2 | Bacteria Firmicutes Bacilli Bacillales Staphylococcaceae *Staphylococcus aureus* M1303 |

| | |
|---|---|
| Sau-2203-1 | Bacteria Firmicutes Bacilli Bacillales Staphylococcaceae *Staphylococcus aureus* M1304 |
| Sau-2204-1 | Bacteria Firmicutes Bacilli Bacillales Staphylococcaceae *Staphylococcus aureus* M13048 |
| Sau-2205-1 | Bacteria Firmicutes Bacilli Bacillales Staphylococcaceae *Staphylococcus aureus* M1305 |
| Sau-2206-1 | Bacteria Firmicutes Bacilli Bacillales Staphylococcaceae *Staphylococcus aureus* M1306 |
| Sau-2207-1 | Bacteria Firmicutes Bacilli Bacillales Staphylococcaceae *Staphylococcus aureus* M13060 |
| Sau-2208-1 | Bacteria Firmicutes Bacilli Bacillales Staphylococcaceae *Staphylococcus aureus* M13065 |
| Sau-2209-1 | Bacteria Firmicutes Bacilli Bacillales Staphylococcaceae *Staphylococcus aureus* M13074 |
| Sau-2210-1 | Bacteria Firmicutes Bacilli Bacillales Staphylococcaceae *Staphylococcus aureus* M1308 |
| Sau-2211-1 | Bacteria Firmicutes Bacilli Bacillales Staphylococcaceae *Staphylococcus aureus* M13099 |
| Sau-2212-1 | Bacteria Firmicutes Bacilli Bacillales Staphylococcaceae *Staphylococcus aureus* M1310 |
| Sau-2213-1 | Bacteria Firmicutes Bacilli Bacillales Staphylococcaceae *Staphylococcus aureus* M1312 |
| Sau-2214-1 | Bacteria Firmicutes Bacilli Bacillales Staphylococcaceae *Staphylococcus aureus* M1313 |
| Sau-2215-1 | Bacteria Firmicutes Bacilli Bacillales Staphylococcaceae *Staphylococcus aureus* M1314 |
| Sau-2216-1 | Bacteria Firmicutes Bacilli Bacillales Staphylococcaceae *Staphylococcus aureus* M1315 |
| Sau-2217-1 | Bacteria Firmicutes Bacilli Bacillales Staphylococcaceae *Staphylococcus aureus* M1316 |
| Sau-2218-1 | Bacteria Firmicutes Bacilli Bacillales Staphylococcaceae *Staphylococcus aureus* M1317 |
| Sau-2219-1 | Bacteria Firmicutes Bacilli Bacillales Staphylococcaceae *Staphylococcus aureus* M1318 |
| Sau-2220-1 | Bacteria Firmicutes Bacilli Bacillales Staphylococcaceae *Staphylococcus aureus* M1319 |
| Sau-2221-1 | Bacteria Firmicutes Bacilli Bacillales Staphylococcaceae *Staphylococcus aureus* M1323 |
| Sau-2222-1 | Bacteria Firmicutes Bacilli Bacillales Staphylococcaceae *Staphylococcus aureus* M1324 |
| Sau-2223-1 | Bacteria Firmicutes Bacilli Bacillales Staphylococcaceae *Staphylococcus aureus* M1325 |
| Sau-2224-1 | Bacteria Firmicutes Bacilli Bacillales Staphylococcaceae *Staphylococcus aureus* M1326 |
| Sau-2225-1 | Bacteria Firmicutes Bacilli Bacillales Staphylococcaceae *Staphylococcus aureus* M1328 |
| Sau-2226-1 | Bacteria Firmicutes Bacilli Bacillales Staphylococcaceae *Staphylococcus aureus* M1329 |
| Sau-2227-1 | Bacteria Firmicutes Bacilli Bacillales Staphylococcaceae *Staphylococcus aureus* M1330 |
| Sau-2228-1 | Bacteria Firmicutes Bacilli Bacillales Staphylococcaceae *Staphylococcus aureus* M1331 |
| Sau-2229-1 | Bacteria Firmicutes Bacilli Bacillales Staphylococcaceae *Staphylococcus aureus* M13316 |
| Sau-2230-1 | Bacteria Firmicutes Bacilli Bacillales Staphylococcaceae *Staphylococcus aureus* M1332 |
| Sau-2231-1 | Bacteria Firmicutes Bacilli Bacillales Staphylococcaceae *Staphylococcus aureus* M1333 |
| Sau-2232-1 | Bacteria Firmicutes Bacilli Bacillales Staphylococcaceae *Staphylococcus aureus* M1334 |
| Sau-2233-1 | Bacteria Firmicutes Bacilli Bacillales Staphylococcaceae *Staphylococcus aureus* M1336 |
| Sau-2234-1 | Bacteria Firmicutes Bacilli Bacillales Staphylococcaceae *Staphylococcus aureus* M1337 |
| Sau-2235-1 | Bacteria Firmicutes Bacilli Bacillales Staphylococcaceae *Staphylococcus aureus* M1338 |
| Sau-2236-1 | Bacteria Firmicutes Bacilli Bacillales Staphylococcaceae *Staphylococcus aureus* M13384 |
| Sau-2237-1 | Bacteria Firmicutes Bacilli Bacillales Staphylococcaceae *Staphylococcus aureus* M1339 |
| Sau-2238-1 | Bacteria Firmicutes Bacilli Bacillales Staphylococcaceae *Staphylococcus aureus* M13403 |
| Sau-2239-1 | Bacteria Firmicutes Bacilli Bacillales Staphylococcaceae *Staphylococcus aureus* M1342 |
| Sau-2240-1 | Bacteria Firmicutes Bacilli Bacillales Staphylococcaceae *Staphylococcus aureus* M1343 |
| Sau-2241-1 | Bacteria Firmicutes Bacilli Bacillales Staphylococcaceae *Staphylococcus aureus* M1344 |
| Sau-2242-1 | Bacteria Firmicutes Bacilli Bacillales Staphylococcaceae *Staphylococcus aureus* M1345 |
| Sau-2243-1 | Bacteria Firmicutes Bacilli Bacillales Staphylococcaceae *Staphylococcus aureus* M1346 |
| Sau-2244-1 | Bacteria Firmicutes Bacilli Bacillales Staphylococcaceae *Staphylococcus aureus* M1347 |
| Sau-2245-1 | Bacteria Firmicutes Bacilli Bacillales Staphylococcaceae *Staphylococcus aureus* M1348 |
| Sau-2246-1 | Bacteria Firmicutes Bacilli Bacillales Staphylococcaceae *Staphylococcus aureus* M1349 |
| Sau-2247-1 | Bacteria Firmicutes Bacilli Bacillales Staphylococcaceae *Staphylococcus aureus* M1350 |
| Sau-2248-1 | Bacteria Firmicutes Bacilli Bacillales Staphylococcaceae *Staphylococcus aureus* M1351 |
| Sau-2249-1 | Bacteria Firmicutes Bacilli Bacillales Staphylococcaceae *Staphylococcus aureus* M1352 |
| Sau-2250-1 | Bacteria Firmicutes Bacilli Bacillales Staphylococcaceae *Staphylococcus aureus* M1353 |
| Sau-2251-1 | Bacteria Firmicutes Bacilli Bacillales Staphylococcaceae *Staphylococcus aureus* M1354 |
| Sau-2252-1 | Bacteria Firmicutes Bacilli Bacillales Staphylococcaceae *Staphylococcus aureus* M1355 |
| Sau-2253-1 | Bacteria Firmicutes Bacilli Bacillales Staphylococcaceae *Staphylococcus aureus* M13559 |
| Sau-2254-1 | Bacteria Firmicutes Bacilli Bacillales Staphylococcaceae *Staphylococcus aureus* M1356 |
| Sau-2255-1 | Bacteria Firmicutes Bacilli Bacillales Staphylococcaceae *Staphylococcus aureus* M1357 |
| Sau-2256-1 | Bacteria Firmicutes Bacilli Bacillales Staphylococcaceae *Staphylococcus aureus* M1358 |
| Sau-2257-1 | Bacteria Firmicutes Bacilli Bacillales Staphylococcaceae *Staphylococcus aureus* M1360 |
| Sau-2258-1 | Bacteria Firmicutes Bacilli Bacillales Staphylococcaceae *Staphylococcus aureus* M1361 |
| Sau-2259-1 | Bacteria Firmicutes Bacilli Bacillales Staphylococcaceae *Staphylococcus aureus* M1362 |
| Sau-2260-1 | Bacteria Firmicutes Bacilli Bacillales Staphylococcaceae *Staphylococcus aureus* M1363 |
| Sau-2261-1 | Bacteria Firmicutes Bacilli Bacillales Staphylococcaceae *Staphylococcus aureus* M1365 |
| Sau-2262-1 | Bacteria Firmicutes Bacilli Bacillales Staphylococcaceae *Staphylococcus aureus* M1366 |
| Sau-2263-1 | Bacteria Firmicutes Bacilli Bacillales Staphylococcaceae *Staphylococcus aureus* M1368 |
| Sau-2264-1 | Bacteria Firmicutes Bacilli Bacillales Staphylococcaceae *Staphylococcus aureus* M1369 |
| Sau-2265-1 | Bacteria Firmicutes Bacilli Bacillales Staphylococcaceae *Staphylococcus aureus* M1370 |
| Sau-2266-1 | Bacteria Firmicutes Bacilli Bacillales Staphylococcaceae *Staphylococcus aureus* M1371 |
| Sau-2267-1 | Bacteria Firmicutes Bacilli Bacillales Staphylococcaceae *Staphylococcus aureus* M1372 |

| | |
|---|---|
| Sau-2268-1 | Bacteria Firmicutes Bacilli Bacillales Staphylococcaceae *Staphylococcus aureus* M1375 |
| Sau-2269-1 | Bacteria Firmicutes Bacilli Bacillales Staphylococcaceae *Staphylococcus aureus* M1376 |
| Sau-2270-1 | Bacteria Firmicutes Bacilli Bacillales Staphylococcaceae *Staphylococcus aureus* M1377 |
| Sau-2271-1 | Bacteria Firmicutes Bacilli Bacillales Staphylococcaceae *Staphylococcus aureus* M1378 |
| Sau-2272-1 | Bacteria Firmicutes Bacilli Bacillales Staphylococcaceae *Staphylococcus aureus* M1379 |
| Sau-2273-1 | Bacteria Firmicutes Bacilli Bacillales Staphylococcaceae *Staphylococcus aureus* M1380 |
| Sau-2274-1 | Bacteria Firmicutes Bacilli Bacillales Staphylococcaceae *Staphylococcus aureus* M1381 |
| Sau-2275-1 | Bacteria Firmicutes Bacilli Bacillales Staphylococcaceae *Staphylococcus aureus* M1382 |
| Sau-2276-1 | Bacteria Firmicutes Bacilli Bacillales Staphylococcaceae *Staphylococcus aureus* M1383 |
| Sau-2277-1 | Bacteria Firmicutes Bacilli Bacillales Staphylococcaceae *Staphylococcus aureus* M1384 |
| Sau-2278-1 | Bacteria Firmicutes Bacilli Bacillales Staphylococcaceae *Staphylococcus aureus* M1385 |
| Sau-2279-1 | Bacteria Firmicutes Bacilli Bacillales Staphylococcaceae *Staphylococcus aureus* M1386 |
| Sau-2280-1 | Bacteria Firmicutes Bacilli Bacillales Staphylococcaceae *Staphylococcus aureus* M1387 |
| Sau-2281-1 | Bacteria Firmicutes Bacilli Bacillales Staphylococcaceae *Staphylococcus aureus* M1388 |
| Sau-2282-1 | Bacteria Firmicutes Bacilli Bacillales Staphylococcaceae *Staphylococcus aureus* M1389 |
| Sau-2283-1 | Bacteria Firmicutes Bacilli Bacillales Staphylococcaceae *Staphylococcus aureus* M1390 |
| Sau-2284-1 | Bacteria Firmicutes Bacilli Bacillales Staphylococcaceae *Staphylococcus aureus* M1391 |
| Sau-2285-1 | Bacteria Firmicutes Bacilli Bacillales Staphylococcaceae *Staphylococcus aureus* M1392 |
| Sau-2286-1 | Bacteria Firmicutes Bacilli Bacillales Staphylococcaceae *Staphylococcus aureus* M1395 |
| Sau-2287-1 | Bacteria Firmicutes Bacilli Bacillales Staphylococcaceae *Staphylococcus aureus* M1396 |
| Sau-2288-1 | Bacteria Firmicutes Bacilli Bacillales Staphylococcaceae *Staphylococcus aureus* M1397 |
| Sau-2289-1 | Bacteria Firmicutes Bacilli Bacillales Staphylococcaceae *Staphylococcus aureus* M1398 |
| Sau-2290-1 | Bacteria Firmicutes Bacilli Bacillales Staphylococcaceae *Staphylococcus aureus* M1399 |
| Sau-2291-1 | Bacteria Firmicutes Bacilli Bacillales Staphylococcaceae *Staphylococcus aureus* M1400 |
| Sau-2292-1 | Bacteria Firmicutes Bacilli Bacillales Staphylococcaceae *Staphylococcus aureus* M1401 |
| Sau-2293-1 | Bacteria Firmicutes Bacilli Bacillales Staphylococcaceae *Staphylococcus aureus* M1402 |
| Sau-2294-1 | Bacteria Firmicutes Bacilli Bacillales Staphylococcaceae *Staphylococcus aureus* M1403 |
| Sau-2295-1 | Bacteria Firmicutes Bacilli Bacillales Staphylococcaceae *Staphylococcus aureus* M1404 |
| Sau-2296-1 | Bacteria Firmicutes Bacilli Bacillales Staphylococcaceae *Staphylococcus aureus* M1406 |
| Sau-2297-1 | Bacteria Firmicutes Bacilli Bacillales Staphylococcaceae *Staphylococcus aureus* M1408 |
| Sau-2298-1 | Bacteria Firmicutes Bacilli Bacillales Staphylococcaceae *Staphylococcus aureus* M1409 |
| Sau-2299-1 | Bacteria Firmicutes Bacilli Bacillales Staphylococcaceae *Staphylococcus aureus* M1410 |
| Sau-2300-1 | Bacteria Firmicutes Bacilli Bacillales Staphylococcaceae *Staphylococcus aureus* M1411 |
| Sau-2301-1 | Bacteria Firmicutes Bacilli Bacillales Staphylococcaceae *Staphylococcus aureus* M1412 |
| Sau-2302-1 | Bacteria Firmicutes Bacilli Bacillales Staphylococcaceae *Staphylococcus aureus* M1413 |
| Sau-2303-1 | Bacteria Firmicutes Bacilli Bacillales Staphylococcaceae *Staphylococcus aureus* M1414 |
| Sau-2304-1 | Bacteria Firmicutes Bacilli Bacillales Staphylococcaceae *Staphylococcus aureus* M1415 |
| Sau-2305-1 | Bacteria Firmicutes Bacilli Bacillales Staphylococcaceae *Staphylococcus aureus* M1416 |
| Sau-2306-1 | Bacteria Firmicutes Bacilli Bacillales Staphylococcaceae *Staphylococcus aureus* M1417 |
| Sau-2307-1 | Bacteria Firmicutes Bacilli Bacillales Staphylococcaceae *Staphylococcus aureus* M1418 |
| Sau-2308-1 | Bacteria Firmicutes Bacilli Bacillales Staphylococcaceae *Staphylococcus aureus* M1420 |
| Sau-2309-1 | Bacteria Firmicutes Bacilli Bacillales Staphylococcaceae *Staphylococcus aureus* M1421 |
| Sau-2310-1 | Bacteria Firmicutes Bacilli Bacillales Staphylococcaceae *Staphylococcus aureus* M1422 |
| Sau-2311-1 | Bacteria Firmicutes Bacilli Bacillales Staphylococcaceae *Staphylococcus aureus* M1423 |
| Sau-2312-1 | Bacteria Firmicutes Bacilli Bacillales Staphylococcaceae *Staphylococcus aureus* M1424 |
| Sau-2313-1 | Bacteria Firmicutes Bacilli Bacillales Staphylococcaceae *Staphylococcus aureus* M1425 |
| Sau-2314-1 | Bacteria Firmicutes Bacilli Bacillales Staphylococcaceae *Staphylococcus aureus* M1426 |
| Sau-2315-1 | Bacteria Firmicutes Bacilli Bacillales Staphylococcaceae *Staphylococcus aureus* M1427 |
| Sau-2316-1 | Bacteria Firmicutes Bacilli Bacillales Staphylococcaceae *Staphylococcus aureus* M1428 |
| Sau-2317-1 | Bacteria Firmicutes Bacilli Bacillales Staphylococcaceae *Staphylococcus aureus* M1429 |
| Sau-2318-1 | Bacteria Firmicutes Bacilli Bacillales Staphylococcaceae *Staphylococcus aureus* M1430 |
| Sau-2319-1 | Bacteria Firmicutes Bacilli Bacillales Staphylococcaceae *Staphylococcus aureus* M1431 |
| Sau-2320-1 | Bacteria Firmicutes Bacilli Bacillales Staphylococcaceae *Staphylococcus aureus* M1432 |
| Sau-2321-1 | Bacteria Firmicutes Bacilli Bacillales Staphylococcaceae *Staphylococcus aureus* M1433 |
| Sau-2322-1 | Bacteria Firmicutes Bacilli Bacillales Staphylococcaceae *Staphylococcus aureus* M1434 |
| Sau-2323-1 | Bacteria Firmicutes Bacilli Bacillales Staphylococcaceae *Staphylococcus aureus* M1435 |
| Sau-2324-1 | Bacteria Firmicutes Bacilli Bacillales Staphylococcaceae *Staphylococcus aureus* M1436 |
| Sau-2325-1 | Bacteria Firmicutes Bacilli Bacillales Staphylococcaceae *Staphylococcus aureus* M1437 |
| Sau-2326-1 | Bacteria Firmicutes Bacilli Bacillales Staphylococcaceae *Staphylococcus aureus* M1438 |
| Sau-2327-1 | Bacteria Firmicutes Bacilli Bacillales Staphylococcaceae *Staphylococcus aureus* M1439 |
| Sau-2328-1 | Bacteria Firmicutes Bacilli Bacillales Staphylococcaceae *Staphylococcus aureus* M1440 |
| Sau-2329-1 | Bacteria Firmicutes Bacilli Bacillales Staphylococcaceae *Staphylococcus aureus* M1441 |
| Sau-2330-1 | Bacteria Firmicutes Bacilli Bacillales Staphylococcaceae *Staphylococcus aureus* M1442 |
| Sau-2331-1 | Bacteria Firmicutes Bacilli Bacillales Staphylococcaceae *Staphylococcus aureus* M1443 |
| Sau-2332-1 | Bacteria Firmicutes Bacilli Bacillales Staphylococcaceae *Staphylococcus aureus* M1444 |

| | |
|---|---|
| Sau-2333-1 | Bacteria Firmicutes Bacilli Bacillales Staphylococcaceae *Staphylococcus aureus* M1445 |
| Sau-2334-1 | Bacteria Firmicutes Bacilli Bacillales Staphylococcaceae *Staphylococcus aureus* M1446 |
| Sau-2335-1 | Bacteria Firmicutes Bacilli Bacillales Staphylococcaceae *Staphylococcus aureus* M1447 |
| Sau-2336-1 | Bacteria Firmicutes Bacilli Bacillales Staphylococcaceae *Staphylococcus aureus* M1448 |
| Sau-2337-1 | Bacteria Firmicutes Bacilli Bacillales Staphylococcaceae *Staphylococcus aureus* M1449 |
| Sau-2338-1 | Bacteria Firmicutes Bacilli Bacillales Staphylococcaceae *Staphylococcus aureus* M1452 |
| Sau-2339-1 | Bacteria Firmicutes Bacilli Bacillales Staphylococcaceae *Staphylococcus aureus* M1453 |
| Sau-2340-1 | Bacteria Firmicutes Bacilli Bacillales Staphylococcaceae *Staphylococcus aureus* M1454 |
| Sau-2341-1 | Bacteria Firmicutes Bacilli Bacillales Staphylococcaceae *Staphylococcus aureus* M1455 |
| Sau-2342-1 | Bacteria Firmicutes Bacilli Bacillales Staphylococcaceae *Staphylococcus aureus* M1456 |
| Sau-2343-1 | Bacteria Firmicutes Bacilli Bacillales Staphylococcaceae *Staphylococcus aureus* M1457 |
| Sau-2344-1 | Bacteria Firmicutes Bacilli Bacillales Staphylococcaceae *Staphylococcus aureus* M1458 |
| Sau-2345-1 | Bacteria Firmicutes Bacilli Bacillales Staphylococcaceae *Staphylococcus aureus* M1459 |
| Sau-2346-1 | Bacteria Firmicutes Bacilli Bacillales Staphylococcaceae *Staphylococcus aureus* M1460 |
| Sau-2347-1 | Bacteria Firmicutes Bacilli Bacillales Staphylococcaceae *Staphylococcus aureus* M1461 |
| Sau-2348-1 | Bacteria Firmicutes Bacilli Bacillales Staphylococcaceae *Staphylococcus aureus* M1464 |
| Sau-2349-1 | Bacteria Firmicutes Bacilli Bacillales Staphylococcaceae *Staphylococcus aureus* M1465 |
| Sau-2350-1 | Bacteria Firmicutes Bacilli Bacillales Staphylococcaceae *Staphylococcus aureus* M1467 |
| Sau-2351-1 | Bacteria Firmicutes Bacilli Bacillales Staphylococcaceae *Staphylococcus aureus* M1468 |
| Sau-2352-1 | Bacteria Firmicutes Bacilli Bacillales Staphylococcaceae *Staphylococcus aureus* M1469 |
| Sau-2353-1 | Bacteria Firmicutes Bacilli Bacillales Staphylococcaceae *Staphylococcus aureus* M1470 |
| Sau-2354-1 | Bacteria Firmicutes Bacilli Bacillales Staphylococcaceae *Staphylococcus aureus* M1471 |
| Sau-2355-1 | Bacteria Firmicutes Bacilli Bacillales Staphylococcaceae *Staphylococcus aureus* M1472 |
| Sau-2356-1 | Bacteria Firmicutes Bacilli Bacillales Staphylococcaceae *Staphylococcus aureus* M1473 |
| Sau-2357-1 | Bacteria Firmicutes Bacilli Bacillales Staphylococcaceae *Staphylococcus aureus* M1474 |
| Sau-2358-1 | Bacteria Firmicutes Bacilli Bacillales Staphylococcaceae *Staphylococcus aureus* M1475 |
| Sau-2359-1 | Bacteria Firmicutes Bacilli Bacillales Staphylococcaceae *Staphylococcus aureus* M1476 |
| Sau-2360-1 | Bacteria Firmicutes Bacilli Bacillales Staphylococcaceae *Staphylococcus aureus* M1477 |
| Sau-2361-1 | Bacteria Firmicutes Bacilli Bacillales Staphylococcaceae *Staphylococcus aureus* M1478 |
| Sau-2362-1 | Bacteria Firmicutes Bacilli Bacillales Staphylococcaceae *Staphylococcus aureus* M1479 |
| Sau-2363-1 | Bacteria Firmicutes Bacilli Bacillales Staphylococcaceae *Staphylococcus aureus* M1480 |
| Sau-2364-1 | Bacteria Firmicutes Bacilli Bacillales Staphylococcaceae *Staphylococcus aureus* M1482 |
| Sau-2365-1 | Bacteria Firmicutes Bacilli Bacillales Staphylococcaceae *Staphylococcus aureus* M1483 |
| Sau-2366-1 | Bacteria Firmicutes Bacilli Bacillales Staphylococcaceae *Staphylococcus aureus* M1484 |
| Sau-2367-1 | Bacteria Firmicutes Bacilli Bacillales Staphylococcaceae *Staphylococcus aureus* M1485 |
| Sau-2368-1 | Bacteria Firmicutes Bacilli Bacillales Staphylococcaceae *Staphylococcus aureus* M1486 |
| Sau-2369-1 | Bacteria Firmicutes Bacilli Bacillales Staphylococcaceae *Staphylococcus aureus* M1487 |
| Sau-2370-1 | Bacteria Firmicutes Bacilli Bacillales Staphylococcaceae *Staphylococcus aureus* M1488 |
| Sau-2371-1 | Bacteria Firmicutes Bacilli Bacillales Staphylococcaceae *Staphylococcus aureus* M1489 |
| Sau-2372-1 | Bacteria Firmicutes Bacilli Bacillales Staphylococcaceae *Staphylococcus aureus* M1490 |
| Sau-2373-1 | Bacteria Firmicutes Bacilli Bacillales Staphylococcaceae *Staphylococcus aureus* M1491 |
| Sau-2374-1 | Bacteria Firmicutes Bacilli Bacillales Staphylococcaceae *Staphylococcus aureus* M1492 |
| Sau-2375-1 | Bacteria Firmicutes Bacilli Bacillales Staphylococcaceae *Staphylococcus aureus* M1493 |
| Sau-2376-1 | Bacteria Firmicutes Bacilli Bacillales Staphylococcaceae *Staphylococcus aureus* M1494 |
| Sau-2377-1 | Bacteria Firmicutes Bacilli Bacillales Staphylococcaceae *Staphylococcus aureus* M1496 |
| Sau-2378-1 | Bacteria Firmicutes Bacilli Bacillales Staphylococcaceae *Staphylococcus aureus* M1497 |
| Sau-2379-1 | Bacteria Firmicutes Bacilli Bacillales Staphylococcaceae *Staphylococcus aureus* M1499 |
| Sau-2380-1 | Bacteria Firmicutes Bacilli Bacillales Staphylococcaceae *Staphylococcus aureus* M1500 |
| Sau-2381-1 | Bacteria Firmicutes Bacilli Bacillales Staphylococcaceae *Staphylococcus aureus* M1501 |
| Sau-2382-1 | Bacteria Firmicutes Bacilli Bacillales Staphylococcaceae *Staphylococcus aureus* M1502 |
| Sau-2383-1 | Bacteria Firmicutes Bacilli Bacillales Staphylococcaceae *Staphylococcus aureus* M1503 |
| Sau-2384-1 | Bacteria Firmicutes Bacilli Bacillales Staphylococcaceae *Staphylococcus aureus* M1504 |
| Sau-2385-1 | Bacteria Firmicutes Bacilli Bacillales Staphylococcaceae *Staphylococcus aureus* M1505 |
| Sau-2386-1 | Bacteria Firmicutes Bacilli Bacillales Staphylococcaceae *Staphylococcus aureus* M1506 |
| Sau-2387-1 | Bacteria Firmicutes Bacilli Bacillales Staphylococcaceae *Staphylococcus aureus* M1507 |
| Sau-2388-1 | Bacteria Firmicutes Bacilli Bacillales Staphylococcaceae *Staphylococcus aureus* M1508 |
| Sau-2389-1 | Bacteria Firmicutes Bacilli Bacillales Staphylococcaceae *Staphylococcus aureus* M1509 |
| Sau-2390-1 | Bacteria Firmicutes Bacilli Bacillales Staphylococcaceae *Staphylococcus aureus* M1511 |
| Sau-2391-1 | Bacteria Firmicutes Bacilli Bacillales Staphylococcaceae *Staphylococcus aureus* M1512 |
| Sau-2392-1 | Bacteria Firmicutes Bacilli Bacillales Staphylococcaceae *Staphylococcus aureus* M1513 |
| Sau-2393-1 | Bacteria Firmicutes Bacilli Bacillales Staphylococcaceae *Staphylococcus aureus* M1514 |
| Sau-2394-1 | Bacteria Firmicutes Bacilli Bacillales Staphylococcaceae *Staphylococcus aureus* M1515 |
| Sau-2395-1 | Bacteria Firmicutes Bacilli Bacillales Staphylococcaceae *Staphylococcus aureus* M1516 |
| Sau-2396-1 | Bacteria Firmicutes Bacilli Bacillales Staphylococcaceae *Staphylococcus aureus* M1518 |
| Sau-2397-1 | Bacteria Firmicutes Bacilli Bacillales Staphylococcaceae *Staphylococcus aureus* M1519 |

| | |
|---|---|
| Sau-2398-1 | Bacteria Firmicutes Bacilli Bacillales Staphylococcaceae *Staphylococcus aureus* M1522 |
| Sau-2399-1 | Bacteria Firmicutes Bacilli Bacillales Staphylococcaceae *Staphylococcus aureus* M1523 |
| Sau-2400-1 | Bacteria Firmicutes Bacilli Bacillales Staphylococcaceae *Staphylococcus aureus* M1524 |
| Sau-2401-1 | Bacteria Firmicutes Bacilli Bacillales Staphylococcaceae *Staphylococcus aureus* M1525 |
| Sau-2402-1 | Bacteria Firmicutes Bacilli Bacillales Staphylococcaceae *Staphylococcus aureus* M1527 |
| Sau-2403-1 | Bacteria Firmicutes Bacilli Bacillales Staphylococcaceae *Staphylococcus aureus* M1528 |
| Sau-2404-1 | Bacteria Firmicutes Bacilli Bacillales Staphylococcaceae *Staphylococcus aureus* M1529 |
| Sau-2405-1 | Bacteria Firmicutes Bacilli Bacillales Staphylococcaceae *Staphylococcus aureus* M1530 |
| Sau-2406-1 | Bacteria Firmicutes Bacilli Bacillales Staphylococcaceae *Staphylococcus aureus* M1532 |
| Sau-2407-1 | Bacteria Firmicutes Bacilli Bacillales Staphylococcaceae *Staphylococcus aureus* M1534 |
| Sau-2408-1 | Bacteria Firmicutes Bacilli Bacillales Staphylococcaceae *Staphylococcus aureus* M1535 |
| Sau-2409-1 | Bacteria Firmicutes Bacilli Bacillales Staphylococcaceae *Staphylococcus aureus* M1536 |
| Sau-2410-1 | Bacteria Firmicutes Bacilli Bacillales Staphylococcaceae *Staphylococcus aureus* M1538 |
| Sau-2411-1 | Bacteria Firmicutes Bacilli Bacillales Staphylococcaceae *Staphylococcus aureus* M1539 |
| Sau-2412-1 | Bacteria Firmicutes Bacilli Bacillales Staphylococcaceae *Staphylococcus aureus* M1541 |
| Sau-2413-1 | Bacteria Firmicutes Bacilli Bacillales Staphylococcaceae *Staphylococcus aureus* M1542 |
| Sau-2414-1 | Bacteria Firmicutes Bacilli Bacillales Staphylococcaceae *Staphylococcus aureus* M1543 |
| Sau-2415-1 | Bacteria Firmicutes Bacilli Bacillales Staphylococcaceae *Staphylococcus aureus* M1545 |
| Sau-2416-1 | Bacteria Firmicutes Bacilli Bacillales Staphylococcaceae *Staphylococcus aureus* M1547 |
| Sau-2417-1 | Bacteria Firmicutes Bacilli Bacillales Staphylococcaceae *Staphylococcus aureus* M1548 |
| Sau-2418-1 | Bacteria Firmicutes Bacilli Bacillales Staphylococcaceae *Staphylococcus aureus* M1549 |
| Sau-2419-1 | Bacteria Firmicutes Bacilli Bacillales Staphylococcaceae *Staphylococcus aureus* M1550 |
| Sau-2420-1 | Bacteria Firmicutes Bacilli Bacillales Staphylococcaceae *Staphylococcus aureus* M1551 |
| Sau-2421-1 | Bacteria Firmicutes Bacilli Bacillales Staphylococcaceae *Staphylococcus aureus* M1552 |
| Sau-2422-1 | Bacteria Firmicutes Bacilli Bacillales Staphylococcaceae *Staphylococcus aureus* M1553 |
| Sau-2423-1 | Bacteria Firmicutes Bacilli Bacillales Staphylococcaceae *Staphylococcus aureus* M1554 |
| Sau-2424-1 | Bacteria Firmicutes Bacilli Bacillales Staphylococcaceae *Staphylococcus aureus* M1555 |
| Sau-2425-1 | Bacteria Firmicutes Bacilli Bacillales Staphylococcaceae *Staphylococcus aureus* M1557 |
| Sau-2426-1 | Bacteria Firmicutes Bacilli Bacillales Staphylococcaceae *Staphylococcus aureus* M1558 |
| Sau-2427-1 | Bacteria Firmicutes Bacilli Bacillales Staphylococcaceae *Staphylococcus aureus* M1559 |
| Sau-2428-1 | Bacteria Firmicutes Bacilli Bacillales Staphylococcaceae *Staphylococcus aureus* M1560 |
| Sau-2429-1 | Bacteria Firmicutes Bacilli Bacillales Staphylococcaceae *Staphylococcus aureus* M1561 |
| Sau-2430-1 | Bacteria Firmicutes Bacilli Bacillales Staphylococcaceae *Staphylococcus aureus* M1562 |
| Sau-2431-1 | Bacteria Firmicutes Bacilli Bacillales Staphylococcaceae *Staphylococcus aureus* M1566 |
| Sau-2432-1 | Bacteria Firmicutes Bacilli Bacillales Staphylococcaceae *Staphylococcus aureus* M1567 |
| Sau-2433-1 | Bacteria Firmicutes Bacilli Bacillales Staphylococcaceae *Staphylococcus aureus* M1569 |
| Sau-2434-1 | Bacteria Firmicutes Bacilli Bacillales Staphylococcaceae *Staphylococcus aureus* M1570 |
| Sau-2435-1 | Bacteria Firmicutes Bacilli Bacillales Staphylococcaceae *Staphylococcus aureus* M1571 |
| Sau-2436-1 | Bacteria Firmicutes Bacilli Bacillales Staphylococcaceae *Staphylococcus aureus* M1572 |
| Sau-2437-1 | Bacteria Firmicutes Bacilli Bacillales Staphylococcaceae *Staphylococcus aureus* M1573 |
| Sau-2438-1 | Bacteria Firmicutes Bacilli Bacillales Staphylococcaceae *Staphylococcus aureus* M1574 |
| Sau-2439-1 | Bacteria Firmicutes Bacilli Bacillales Staphylococcaceae *Staphylococcus aureus* M1575 |
| Sau-2440-1 | Bacteria Firmicutes Bacilli Bacillales Staphylococcaceae *Staphylococcus aureus* M1576 |
| Sau-2441-1 | Bacteria Firmicutes Bacilli Bacillales Staphylococcaceae *Staphylococcus aureus* M1577 |
| Sau-2442-1 | Bacteria Firmicutes Bacilli Bacillales Staphylococcaceae *Staphylococcus aureus* M1579 |
| Sau-2443-1 | Bacteria Firmicutes Bacilli Bacillales Staphylococcaceae *Staphylococcus aureus* M1580 |
| Sau-2444-1 | Bacteria Firmicutes Bacilli Bacillales Staphylococcaceae *Staphylococcus aureus* M16220 |
| Sau-2445-1 | Bacteria Firmicutes Bacilli Bacillales Staphylococcaceae *Staphylococcus aureus* M16438 |
| Sau-2446-1 | Bacteria Firmicutes Bacilli Bacillales Staphylococcaceae *Staphylococcus aureus* M16486 |
| Sau-2447-1 | Bacteria Firmicutes Bacilli Bacillales Staphylococcaceae *Staphylococcus aureus* M16552 |
| Sau-2448-1 | Bacteria Firmicutes Bacilli Bacillales Staphylococcaceae *Staphylococcus aureus* M16586 |
| Sau-2449-1 | Bacteria Firmicutes Bacilli Bacillales Staphylococcaceae *Staphylococcus aureus* M16597 |
| Sau-2450-1 | Bacteria Firmicutes Bacilli Bacillales Staphylococcaceae *Staphylococcus aureus* M17027 |
| Sau-2451-1 | Bacteria Firmicutes Bacilli Bacillales Staphylococcaceae *Staphylococcus aureus* M17033 |
| Sau-2452-1 | Bacteria Firmicutes Bacilli Bacillales Staphylococcaceae *Staphylococcus aureus* M17074 |
| Sau-2453-1 | Bacteria Firmicutes Bacilli Bacillales Staphylococcaceae *Staphylococcus aureus* M17299 |
| Sau-2454-1 | Bacteria Firmicutes Bacilli Bacillales Staphylococcaceae *Staphylococcus aureus* M17301 |
| Sau-2455-1 | Bacteria Firmicutes Bacilli Bacillales Staphylococcaceae *Staphylococcus aureus* M20180 |
| Sau-2456-1 | Bacteria Firmicutes Bacilli Bacillales Staphylococcaceae *Staphylococcus aureus* M20198 |
| Sau-2457-1 | Bacteria Firmicutes Bacilli Bacillales Staphylococcaceae *Staphylococcus aureus* M20676 |
| Sau-2458-1 | Bacteria Firmicutes Bacilli Bacillales Staphylococcaceae *Staphylococcus aureus* M20726 |
| Sau-2459-1 | Bacteria Firmicutes Bacilli Bacillales Staphylococcaceae *Staphylococcus aureus* M20894 |
| Sau-2460-1 | Bacteria Firmicutes Bacilli Bacillales Staphylococcaceae *Staphylococcus aureus* M20913 |
| Sau-2461-1 | Bacteria Firmicutes Bacilli Bacillales Staphylococcaceae *Staphylococcus aureus* M20916 |
| Sau-2462-1 | Bacteria Firmicutes Bacilli Bacillales Staphylococcaceae *Staphylococcus aureus* M24390 |

| | |
|---|---|
| Sau-2463-1 | Bacteria Firmicutes Bacilli Bacillales Staphylococcaceae *Staphylococcus aureus* M24400 |
| Sau-2464-1 | Bacteria Firmicutes Bacilli Bacillales Staphylococcaceae *Staphylococcus aureus* M24416 |
| Sau-2465-1 | Bacteria Firmicutes Bacilli Bacillales Staphylococcaceae *Staphylococcus aureus* M24418 |
| Sau-2466-1 | Bacteria Firmicutes Bacilli Bacillales Staphylococcaceae *Staphylococcus aureus* M24419 |
| Sau-2467-1 | Bacteria Firmicutes Bacilli Bacillales Staphylococcaceae *Staphylococcus aureus* M24487 |
| Sau-2468-1 | Bacteria Firmicutes Bacilli Bacillales Staphylococcaceae *Staphylococcus aureus* M25139 |
| Sau-2469-1 | Bacteria Firmicutes Bacilli Bacillales Staphylococcaceae *Staphylococcus aureus* M267-1 |
| Sau-2470-1 | Bacteria Firmicutes Bacilli Bacillales Staphylococcaceae *Staphylococcus aureus* M27449 |
| Sau-2471-1 | Bacteria Firmicutes Bacilli Bacillales Staphylococcaceae *Staphylococcus aureus* M28112 |
| Sau-2472-1 | Bacteria Firmicutes Bacilli Bacillales Staphylococcaceae *Staphylococcus aureus* M28485 |
| Sau-2473-1 | Bacteria Firmicutes Bacilli Bacillales Staphylococcaceae *Staphylococcus aureus* M28487 |
| Sau-2474-1 | Bacteria Firmicutes Bacilli Bacillales Staphylococcaceae *Staphylococcus aureus* M29068 |
| Sau-2475-1 | Bacteria Firmicutes Bacilli Bacillales Staphylococcaceae *Staphylococcus aureus* M31588 |
| Sau-2476-1 | Bacteria Firmicutes Bacilli Bacillales Staphylococcaceae *Staphylococcus aureus* M32567 |
| Sau-2477-1 | Bacteria Firmicutes Bacilli Bacillales Staphylococcaceae *Staphylococcus aureus* M32669 |
| Sau-2478-1 | Bacteria Firmicutes Bacilli Bacillales Staphylococcaceae *Staphylococcus aureus* M32679 |
| Sau-2479-1 | Bacteria Firmicutes Bacilli Bacillales Staphylococcaceae *Staphylococcus aureus* M32688 |
| Sau-2480-1 | Bacteria Firmicutes Bacilli Bacillales Staphylococcaceae *Staphylococcus aureus* M35158 |
| Sau-2481-1 | Bacteria Firmicutes Bacilli Bacillales Staphylococcaceae *Staphylococcus aureus* M357 |
| Sau-2482-1 | Bacteria Firmicutes Bacilli Bacillales Staphylococcaceae *Staphylococcus aureus* M35933 |
| Sau-2483-1 | Bacteria Firmicutes Bacilli Bacillales Staphylococcaceae *Staphylococcus aureus* M35938 |
| Sau-2484-1 | Bacteria Firmicutes Bacilli Bacillales Staphylococcaceae *Staphylococcus aureus* M35954 |
| Sau-2485-1 | Bacteria Firmicutes Bacilli Bacillales Staphylococcaceae *Staphylococcus aureus* M35955 |
| Sau-2486-1 | Bacteria Firmicutes Bacilli Bacillales Staphylococcaceae *Staphylococcus aureus* M35965 |
| Sau-2487-1 | Bacteria Firmicutes Bacilli Bacillales Staphylococcaceae *Staphylococcus aureus* M36077 |
| Sau-2488-1 | Bacteria Firmicutes Bacilli Bacillales Staphylococcaceae *Staphylococcus aureus* M36089 |
| Sau-2489-1 | Bacteria Firmicutes Bacilli Bacillales Staphylococcaceae *Staphylococcus aureus* M39260 |
| Sau-2490-1 | Bacteria Firmicutes Bacilli Bacillales Staphylococcaceae *Staphylococcus aureus* M39265 |
| Sau-2491-1 | Bacteria Firmicutes Bacilli Bacillales Staphylococcaceae *Staphylococcus aureus* M39274 |
| Sau-2492-1 | Bacteria Firmicutes Bacilli Bacillales Staphylococcaceae *Staphylococcus aureus* M39841 |
| Sau-2493-1 | Bacteria Firmicutes Bacilli Bacillales Staphylococcaceae *Staphylococcus aureus* M41648 |
| Sau-2494-1 | Bacteria Firmicutes Bacilli Bacillales Staphylococcaceae *Staphylococcus aureus* M42184 |
| Sau-2495-1 | Bacteria Firmicutes Bacilli Bacillales Staphylococcaceae *Staphylococcus aureus* M42212 |
| Sau-2496-1 | Bacteria Firmicutes Bacilli Bacillales Staphylococcaceae *Staphylococcus aureus* M42485 |
| Sau-2497-1 | Bacteria Firmicutes Bacilli Bacillales Staphylococcaceae *Staphylococcus aureus* M42496 |
| Sau-2498-1 | Bacteria Firmicutes Bacilli Bacillales Staphylococcaceae *Staphylococcus aureus* M42507 |
| Sau-2499-1 | Bacteria Firmicutes Bacilli Bacillales Staphylococcaceae *Staphylococcus aureus* M42514 |
| Sau-2500-1 | Bacteria Firmicutes Bacilli Bacillales Staphylococcaceae *Staphylococcus aureus* M42518 |
| Sau-2501-1 | Bacteria Firmicutes Bacilli Bacillales Staphylococcaceae *Staphylococcus aureus* M44986 |
| Sau-2502-1 | Bacteria Firmicutes Bacilli Bacillales Staphylococcaceae *Staphylococcus aureus* M44987 |
| Sau-2503-1 | Bacteria Firmicutes Bacilli Bacillales Staphylococcaceae *Staphylococcus aureus* M44988 |
| Sau-2504-1 | Bacteria Firmicutes Bacilli Bacillales Staphylococcaceae *Staphylococcus aureus* M45245 |
| Sau-2505-1 | Bacteria Firmicutes Bacilli Bacillales Staphylococcaceae *Staphylococcus aureus* M457-1 |
| Sau-2506-1 | Bacteria Firmicutes Bacilli Bacillales Staphylococcaceae *Staphylococcus aureus* M45704 |
| Sau-2507-1 | Bacteria Firmicutes Bacilli Bacillales Staphylococcaceae *Staphylococcus aureus* M45726 |
| Sau-2508-1 | Bacteria Firmicutes Bacilli Bacillales Staphylococcaceae *Staphylococcus aureus* M46058 |
| Sau-2509-1 | Bacteria Firmicutes Bacilli Bacillales Staphylococcaceae *Staphylococcus aureus* M46066 |
| Sau-2510-1 | Bacteria Firmicutes Bacilli Bacillales Staphylococcaceae *Staphylococcus aureus* M46070 |
| Sau-2511-1 | Bacteria Firmicutes Bacilli Bacillales Staphylococcaceae *Staphylococcus aureus* M46077 |
| Sau-2512-1 | Bacteria Firmicutes Bacilli Bacillales Staphylococcaceae *Staphylococcus aureus* M49455 |
| Sau-2513-1 | Bacteria Firmicutes Bacilli Bacillales Staphylococcaceae *Staphylococcus aureus* M49474 |
| Sau-2514-1 | Bacteria Firmicutes Bacilli Bacillales Staphylococcaceae *Staphylococcus aureus* M49501 |
| Sau-2515-1 | Bacteria Firmicutes Bacilli Bacillales Staphylococcaceae *Staphylococcus aureus* M49691 |
| Sau-2516-1 | Bacteria Firmicutes Bacilli Bacillales Staphylococcaceae *Staphylococcus aureus* M520-1 |
| Sau-2517-1 | Bacteria Firmicutes Bacilli Bacillales Staphylococcaceae *Staphylococcus aureus* M52055 |
| Sau-2518-1 | Bacteria Firmicutes Bacilli Bacillales Staphylococcaceae *Staphylococcus aureus* M52622 |
| Sau-2519-1 | Bacteria Firmicutes Bacilli Bacillales Staphylococcaceae *Staphylococcus aureus* M53327 |
| Sau-2520-1 | Bacteria Firmicutes Bacilli Bacillales Staphylococcaceae *Staphylococcus aureus* M53356 |
| Sau-2521-1 | Bacteria Firmicutes Bacilli Bacillales Staphylococcaceae *Staphylococcus aureus* M54162 |
| Sau-2522-1 | Bacteria Firmicutes Bacilli Bacillales Staphylococcaceae *Staphylococcus aureus* M55878 |
| Sau-2523-1 | Bacteria Firmicutes Bacilli Bacillales Staphylococcaceae *Staphylococcus aureus* M55890 |
| Sau-2524-1 | Bacteria Firmicutes Bacilli Bacillales Staphylococcaceae *Staphylococcus aureus* M55902 |
| Sau-2525-1 | Bacteria Firmicutes Bacilli Bacillales Staphylococcaceae *Staphylococcus aureus* M56985 |
| Sau-2526-1 | Bacteria Firmicutes Bacilli Bacillales Staphylococcaceae *Staphylococcus aureus* M57729 |
| Sau-2527-1 | Bacteria Firmicutes Bacilli Bacillales Staphylococcaceae *Staphylococcus aureus* M57746 |

| | |
|---|---|
| Sau-2528-1 | Bacteria Firmicutes Bacilli Bacillales Staphylococcaceae *Staphylococcus aureus* M59248 |
| Sau-2529-1 | Bacteria Firmicutes Bacilli Bacillales Staphylococcaceae *Staphylococcus aureus* M59255 |
| Sau-2530-1 | Bacteria Firmicutes Bacilli Bacillales Staphylococcaceae *Staphylococcus aureus* M59257 |
| Sau-2531-1 | Bacteria Firmicutes Bacilli Bacillales Staphylococcaceae *Staphylococcus aureus* M59498 |
| Sau-2532-1 | Bacteria Firmicutes Bacilli Bacillales Staphylococcaceae *Staphylococcus aureus* M62699 |
| Sau-2533-1 | Bacteria Firmicutes Bacilli Bacillales Staphylococcaceae *Staphylococcus aureus* M62794 |
| Sau-2534-1 | Bacteria Firmicutes Bacilli Bacillales Staphylococcaceae *Staphylococcus aureus* M64056 |
| Sau-2535-1 | Bacteria Firmicutes Bacilli Bacillales Staphylococcaceae *Staphylococcus aureus* M64164 |
| Sau-2536-1 | Bacteria Firmicutes Bacilli Bacillales Staphylococcaceae *Staphylococcus aureus* M64191 |
| Sau-2537-1 | Bacteria Firmicutes Bacilli Bacillales Staphylococcaceae *Staphylococcus aureus* M65003 |
| Sau-2538-1 | Bacteria Firmicutes Bacilli Bacillales Staphylococcaceae *Staphylococcus aureus* M66407 |
| Sau-2539-1 | Bacteria Firmicutes Bacilli Bacillales Staphylococcaceae *Staphylococcus aureus* M66410 |
| Sau-2540-1 | Bacteria Firmicutes Bacilli Bacillales Staphylococcaceae *Staphylococcus aureus* M66468 |
| Sau-2541-1 | Bacteria Firmicutes Bacilli Bacillales Staphylococcaceae *Staphylococcus aureus* M66661 |
| Sau-2542-1 | Bacteria Firmicutes Bacilli Bacillales Staphylococcaceae *Staphylococcus aureus* M66673 |
| Sau-2543-1 | Bacteria Firmicutes Bacilli Bacillales Staphylococcaceae *Staphylococcus aureus* M67290 |
| Sau-2544-1 | Bacteria Firmicutes Bacilli Bacillales Staphylococcaceae *Staphylococcus aureus* M67929_020413 |
| Sau-2545-1 | Bacteria Firmicutes Bacilli Bacillales Staphylococcaceae *Staphylococcus aureus* M67938 |
| Sau-2546-1 | Bacteria Firmicutes Bacilli Bacillales Staphylococcaceae *Staphylococcus aureus* M67953 |
| Sau-2547-1 | Bacteria Firmicutes Bacilli Bacillales Staphylococcaceae *Staphylococcus aureus* M67957 |
| Sau-2548-1 | Bacteria Firmicutes Bacilli Bacillales Staphylococcaceae *Staphylococcus aureus* M68467 |
| Sau-2549-1 | Bacteria Firmicutes Bacilli Bacillales Staphylococcaceae *Staphylococcus aureus* M68475 |
| Sau-2550-1 | Bacteria Firmicutes Bacilli Bacillales Staphylococcaceae *Staphylococcus aureus* M70006 |
| Sau-2551-1 | Bacteria Firmicutes Bacilli Bacillales Staphylococcaceae *Staphylococcus aureus* M70013 |
| Sau-2552-1 | Bacteria Firmicutes Bacilli Bacillales Staphylococcaceae *Staphylococcus aureus* M70017 |
| Sau-2553-1 | Bacteria Firmicutes Bacilli Bacillales Staphylococcaceae *Staphylococcus aureus* M70018 |
| Sau-2554-1 | Bacteria Firmicutes Bacilli Bacillales Staphylococcaceae *Staphylococcus aureus* M70073 |
| Sau-2555-1 | Bacteria Firmicutes Bacilli Bacillales Staphylococcaceae *Staphylococcus aureus* M70099 |
| Sau-2556-1 | Bacteria Firmicutes Bacilli Bacillales Staphylococcaceae *Staphylococcus aureus* M70590 |
| Sau-2557-1 | Bacteria Firmicutes Bacilli Bacillales Staphylococcaceae *Staphylococcus aureus* M71813 |
| Sau-2558-1 | Bacteria Firmicutes Bacilli Bacillales Staphylococcaceae *Staphylococcus aureus* M72054 |
| Sau-2559-1 | Bacteria Firmicutes Bacilli Bacillales Staphylococcaceae *Staphylococcus aureus* M72087 |
| Sau-2560-1 | Bacteria Firmicutes Bacilli Bacillales Staphylococcaceae *Staphylococcus aureus* M72095_021113 |
| Sau-2561-1 | Bacteria Firmicutes Bacilli Bacillales Staphylococcaceae *Staphylococcus aureus* M72097 |
| Sau-2562-1 | Bacteria Firmicutes Bacilli Bacillales Staphylococcaceae *Staphylococcus aureus* M73702 |
| Sau-2563-1 | Bacteria Firmicutes Bacilli Bacillales Staphylococcaceae *Staphylococcus aureus* M73703 |
| Sau-2564-1 | Bacteria Firmicutes Bacilli Bacillales Staphylococcaceae *Staphylococcus aureus* M73849 |
| Sau-2565-1 | Bacteria Firmicutes Bacilli Bacillales Staphylococcaceae *Staphylococcus aureus* M74952 |
| Sau-2566-1 | Bacteria Firmicutes Bacilli Bacillales Staphylococcaceae *Staphylococcus aureus* M77202 |
| Sau-2567-1 | Bacteria Firmicutes Bacilli Bacillales Staphylococcaceae *Staphylococcus aureus* M77210 |
| Sau-2568-1 | Bacteria Firmicutes Bacilli Bacillales Staphylococcaceae *Staphylococcus aureus* M77225 |
| Sau-2569-1 | Bacteria Firmicutes Bacilli Bacillales Staphylococcaceae *Staphylococcus aureus* M77229 |
| Sau-2570-1 | Bacteria Firmicutes Bacilli Bacillales Staphylococcaceae *Staphylococcus aureus* M77238 |
| Sau-2571-1 | Bacteria Firmicutes Bacilli Bacillales Staphylococcaceae *Staphylococcus aureus* M79254 |
| Sau-2572-1 | Bacteria Firmicutes Bacilli Bacillales Staphylococcaceae *Staphylococcus aureus* M79256 |
| Sau-2573-1 | Bacteria Firmicutes Bacilli Bacillales Staphylococcaceae *Staphylococcus aureus* M79654 |
| Sau-2574-1 | Bacteria Firmicutes Bacilli Bacillales Staphylococcaceae *Staphylococcus aureus* M79664 |
| Sau-2575-1 | Bacteria Firmicutes Bacilli Bacillales Staphylococcaceae *Staphylococcus aureus* M79667 |
| Sau-2576-1 | Bacteria Firmicutes Bacilli Bacillales Staphylococcaceae *Staphylococcus aureus* M79671 |
| Sau-2577-1 | Bacteria Firmicutes Bacilli Bacillales Staphylococcaceae *Staphylococcus aureus* M79673 |
| Sau-2578-1 | Bacteria Firmicutes Bacilli Bacillales Staphylococcaceae *Staphylococcus aureus* M79684 |
| Sau-2579-1 | Bacteria Firmicutes Bacilli Bacillales Staphylococcaceae *Staphylococcus aureus* M79686 |
| Sau-2580-1 | Bacteria Firmicutes Bacilli Bacillales Staphylococcaceae *Staphylococcus aureus* M80329 |
| Sau-2581-1 | Bacteria Firmicutes Bacilli Bacillales Staphylococcaceae *Staphylococcus aureus* M80333 |
| Sau-2582-1 | Bacteria Firmicutes Bacilli Bacillales Staphylococcaceae *Staphylococcus aureus* M80979 |
| Sau-2583-1 | Bacteria Firmicutes Bacilli Bacillales Staphylococcaceae *Staphylococcus aureus* M80992 |
| Sau-2584-1 | Bacteria Firmicutes Bacilli Bacillales Staphylococcaceae *Staphylococcus aureus* M81424 |
| Sau-2585-1 | Bacteria Firmicutes Bacilli Bacillales Staphylococcaceae *Staphylococcus aureus* M81490 |
| Sau-2586-1 | Bacteria Firmicutes Bacilli Bacillales Staphylococcaceae *Staphylococcus aureus* M81493 |
| Sau-2587-1 | Bacteria Firmicutes Bacilli Bacillales Staphylococcaceae *Staphylococcus aureus* M83332 |
| Sau-2588-1 | Bacteria Firmicutes Bacilli Bacillales Staphylococcaceae *Staphylococcus aureus* M85201 |
| Sau-2589-1 | Bacteria Firmicutes Bacilli Bacillales Staphylococcaceae *Staphylococcus aureus* M86940 |
| Sau-2590-1 | Bacteria Firmicutes Bacilli Bacillales Staphylococcaceae *Staphylococcus aureus* M86943 |
| Sau-2591-1 | Bacteria Firmicutes Bacilli Bacillales Staphylococcaceae *Staphylococcus aureus* M87641 |
| Sau-2592-1 | Bacteria Firmicutes Bacilli Bacillales Staphylococcaceae *Staphylococcus aureus* M87665 |

| | |
|---|---|
| Sau-2593-1 | Bacteria Firmicutes Bacilli Bacillales Staphylococcaceae *Staphylococcus aureus* M90383 |
| Sau-2594-1 | Bacteria Firmicutes Bacilli Bacillales Staphylococcaceae *Staphylococcus aureus* M90388 |
| Sau-2595-1 | Bacteria Firmicutes Bacilli Bacillales Staphylococcaceae *Staphylococcus aureus* M90396 |
| Sau-2596-1 | Bacteria Firmicutes Bacilli Bacillales Staphylococcaceae *Staphylococcus aureus* M91141 |
| Sau-2597-1 | Bacteria Firmicutes Bacilli Bacillales Staphylococcaceae *Staphylococcus aureus* M91155 |
| Sau-2598-1 | Bacteria Firmicutes Bacilli Bacillales Staphylococcaceae *Staphylococcus aureus* M91160 |
| Sau-2599-1 | Bacteria Firmicutes Bacilli Bacillales Staphylococcaceae *Staphylococcus aureus* M91652 |
| Sau-2600-1 | Bacteria Firmicutes Bacilli Bacillales Staphylococcaceae *Staphylococcus aureus* M92071 |
| Sau-2601-1 | Bacteria Firmicutes Bacilli Bacillales Staphylococcaceae *Staphylococcus aureus* M92086 |
| Sau-2602-1 | Bacteria Firmicutes Bacilli Bacillales Staphylococcaceae *Staphylococcus aureus* MISS6007 |
| Sau-2603-1 | Bacteria Firmicutes Bacilli Bacillales Staphylococcaceae *Staphylococcus aureus* MISS6012 |
| Sau-2604-1 | Bacteria Firmicutes Bacilli Bacillales Staphylococcaceae *Staphylococcus aureus* MISS6017 |
| Sau-2605-1 | Bacteria Firmicutes Bacilli Bacillales Staphylococcaceae *Staphylococcus aureus* MISS6019 |
| Sau-2606-1 | Bacteria Firmicutes Bacilli Bacillales Staphylococcaceae *Staphylococcus aureus* MISS6020 |
| Sau-2607-1 | Bacteria Firmicutes Bacilli Bacillales Staphylococcaceae *Staphylococcus aureus* MISS6027 |
| Sau-2608-1 | Bacteria Firmicutes Bacilli Bacillales Staphylococcaceae *Staphylococcus aureus* MISS6033 |
| Sau-2609-1 | Bacteria Firmicutes Bacilli Bacillales Staphylococcaceae *Staphylococcus aureus* MISS6037 |
| Sau-2610-1 | Bacteria Firmicutes Bacilli Bacillales Staphylococcaceae *Staphylococcus aureus* MISS6042 |
| Sau-2611-1 | Bacteria Firmicutes Bacilli Bacillales Staphylococcaceae *Staphylococcus aureus* MISS6048 |
| Sau-2612-1 | Bacteria Firmicutes Bacilli Bacillales Staphylococcaceae *Staphylococcus aureus* MISS6052 |
| Sau-2613-1 | Bacteria Firmicutes Bacilli Bacillales Staphylococcaceae *Staphylococcus aureus* MISS6057 |
| Sau-2614-1 | Bacteria Firmicutes Bacilli Bacillales Staphylococcaceae *Staphylococcus aureus* MISS6078 |
| Sau-2615-1 | Bacteria Firmicutes Bacilli Bacillales Staphylococcaceae *Staphylococcus aureus* MISS6080 |
| Sau-2616-1 | Bacteria Firmicutes Bacilli Bacillales Staphylococcaceae *Staphylococcus aureus* MISS6092 |
| Sau-2617-1 | Bacteria Firmicutes Bacilli Bacillales Staphylococcaceae *Staphylococcus aureus* MISS6103 |
| Sau-2618-1 | Bacteria Firmicutes Bacilli Bacillales Staphylococcaceae *Staphylococcus aureus* MISS6110 |
| Sau-2619-1 | Bacteria Firmicutes Bacilli Bacillales Staphylococcaceae *Staphylococcus aureus* MISS6119 |
| Sau-2620-1 | Bacteria Firmicutes Bacilli Bacillales Staphylococcaceae *Staphylococcus aureus* MISS6125 |
| Sau-2621-1 | Bacteria Firmicutes Bacilli Bacillales Staphylococcaceae *Staphylococcus aureus* MISS6131 |
| Sau-2622-1 | Bacteria Firmicutes Bacilli Bacillales Staphylococcaceae *Staphylococcus aureus* MISS6132 |
| Sau-2623-1 | Bacteria Firmicutes Bacilli Bacillales Staphylococcaceae *Staphylococcus aureus* MISS6134 |
| Sau-2624-1 | Bacteria Firmicutes Bacilli Bacillales Staphylococcaceae *Staphylococcus aureus* MISS6176 |
| Sau-2625-1 | Bacteria Firmicutes Bacilli Bacillales Staphylococcaceae *Staphylococcus aureus* MM66 |
| Sau-2626-1 | Bacteria Firmicutes Bacilli Bacillales Staphylococcaceae *Staphylococcus aureus* MRSA-118 |
| Sau-2627-1 | Bacteria Firmicutes Bacilli Bacillales Staphylococcaceae *Staphylococcus aureus* MRSA-136 |
| Sau-2628-1 | Bacteria Firmicutes Bacilli Bacillales Staphylococcaceae *Staphylococcus aureus* MRSA_CVM43477 |
| Sau-2629-1 | Bacteria Firmicutes Bacilli Bacillales Staphylococcaceae *Staphylococcus aureus* MRSN 2761 |
| Sau-2630-1 | Bacteria Firmicutes Bacilli Bacillales Staphylococcaceae *Staphylococcus aureus* MRSN 8611 |
| Sau-2631-1 | Bacteria Firmicutes Bacilli Bacillales Staphylococcaceae *Staphylococcus aureus* MRSN 8613 |
| Sau-2632-1 | Bacteria Firmicutes Bacilli Bacillales Staphylococcaceae *Staphylococcus aureus* MSSA-123 |
| Sau-2633-1 | Bacteria Firmicutes Bacilli Bacillales Staphylococcaceae *Staphylococcus aureus* MSSA-37 |
| Sau-2634-1 | Bacteria Firmicutes Bacilli Bacillales Staphylococcaceae *Staphylococcus aureus* MSSA-47 |
| Sau-2635-1 | Bacteria Firmicutes Bacilli Bacillales Staphylococcaceae *Staphylococcus aureus* MSSA-93 |
| Sau-2636-1 | Bacteria Firmicutes Bacilli Bacillales Staphylococcaceae *Staphylococcus aureus* N0426 |
| Sau-2637-1 | Bacteria Firmicutes Bacilli Bacillales Staphylococcaceae *Staphylococcus aureus* N1343 |
| Sau-2638-1 | Bacteria Firmicutes Bacilli Bacillales Staphylococcaceae *Staphylococcus aureus* OCMM6002 |
| Sau-2639-1 | Bacteria Firmicutes Bacilli Bacillales Staphylococcaceae *Staphylococcus aureus* OCMM6003 |
| Sau-2640-1 | Bacteria Firmicutes Bacilli Bacillales Staphylococcaceae *Staphylococcus aureus* OCMM6004 |
| Sau-2641-1 | Bacteria Firmicutes Bacilli Bacillales Staphylococcaceae *Staphylococcus aureus* OCMM6015 |
| Sau-2642-1 | Bacteria Firmicutes Bacilli Bacillales Staphylococcaceae *Staphylococcus aureus* OCMM6017 |
| Sau-2643-1 | Bacteria Firmicutes Bacilli Bacillales Staphylococcaceae *Staphylococcus aureus* OCMM6035 |
| Sau-2644-1 | Bacteria Firmicutes Bacilli Bacillales Staphylococcaceae *Staphylococcus aureus* OCMM6037 |
| Sau-2645-1 | Bacteria Firmicutes Bacilli Bacillales Staphylococcaceae *Staphylococcus aureus* OCMM6045 |
| Sau-2646-1 | Bacteria Firmicutes Bacilli Bacillales Staphylococcaceae *Staphylococcus aureus* OCMM6046 |
| Sau-2647-1 | Bacteria Firmicutes Bacilli Bacillales Staphylococcaceae *Staphylococcus aureus* OCMM6047 |
| Sau-2648-1 | Bacteria Firmicutes Bacilli Bacillales Staphylococcaceae *Staphylococcus aureus* OCMM6048 |
| Sau-2649-1 | Bacteria Firmicutes Bacilli Bacillales Staphylococcaceae *Staphylococcus aureus* OCMM6049 |
| Sau-2650-1 | Bacteria Firmicutes Bacilli Bacillales Staphylococcaceae *Staphylococcus aureus* OCMM6052 |
| Sau-2651-1 | Bacteria Firmicutes Bacilli Bacillales Staphylococcaceae *Staphylococcus aureus* OCMM6057 |
| Sau-2652-1 | Bacteria Firmicutes Bacilli Bacillales Staphylococcaceae *Staphylococcus aureus* OCMM6065 |
| Sau-2653-1 | Bacteria Firmicutes Bacilli Bacillales Staphylococcaceae *Staphylococcus aureus* OCMM6066 |
| Sau-2654-1 | Bacteria Firmicutes Bacilli Bacillales Staphylococcaceae *Staphylococcus aureus* OCMM6067 |
| Sau-2655-1 | Bacteria Firmicutes Bacilli Bacillales Staphylococcaceae *Staphylococcus aureus* OCMM6071 |
| Sau-2656-1 | Bacteria Firmicutes Bacilli Bacillales Staphylococcaceae *Staphylococcus aureus* OCMM6075 |
| Sau-2657-1 | Bacteria Firmicutes Bacilli Bacillales Staphylococcaceae *Staphylococcus aureus* OCMM6085 |

| | |
|---|---|
| Sau-2658-1 | Bacteria Firmicutes Bacilli Bacillales Staphylococcaceae *Staphylococcus aureus* OCMM6090 |
| Sau-2659-1 | Bacteria Firmicutes Bacilli Bacillales Staphylococcaceae *Staphylococcus aureus* OCMM6095 |
| Sau-2660-1 | Bacteria Firmicutes Bacilli Bacillales Staphylococcaceae *Staphylococcus aureus* OCMM6096 |
| Sau-2661-1 | Bacteria Firmicutes Bacilli Bacillales Staphylococcaceae *Staphylococcus aureus* OCMM6102 |
| Sau-2662-1 | Bacteria Firmicutes Bacilli Bacillales Staphylococcaceae *Staphylococcus aureus* OCMM6103 |
| Sau-2663-1 | Bacteria Firmicutes Bacilli Bacillales Staphylococcaceae *Staphylococcus aureus* OCMM6109 |
| Sau-2664-1 | Bacteria Firmicutes Bacilli Bacillales Staphylococcaceae *Staphylococcus aureus* OCMM6110 |
| Sau-2665-1 | Bacteria Firmicutes Bacilli Bacillales Staphylococcaceae *Staphylococcus aureus* OCMM6114 |
| Sau-2666-1 | Bacteria Firmicutes Bacilli Bacillales Staphylococcaceae *Staphylococcus aureus* OCMM6115 |
| Sau-2667-1 | Bacteria Firmicutes Bacilli Bacillales Staphylococcaceae *Staphylococcus aureus* OCMM6117 |
| Sau-2668-1 | Bacteria Firmicutes Bacilli Bacillales Staphylococcaceae *Staphylococcus aureus* OCMM6121 |
| Sau-2669-1 | Bacteria Firmicutes Bacilli Bacillales Staphylococcaceae *Staphylococcus aureus* OCMM6123 |
| Sau-2670-1 | Bacteria Firmicutes Bacilli Bacillales Staphylococcaceae *Staphylococcus aureus* OCMM6124 |
| Sau-2671-1 | Bacteria Firmicutes Bacilli Bacillales Staphylococcaceae *Staphylococcus aureus* OCMM6125 |
| Sau-2672-1 | Bacteria Firmicutes Bacilli Bacillales Staphylococcaceae *Staphylococcus aureus* OCMM6129 |
| Sau-2673-1 | Bacteria Firmicutes Bacilli Bacillales Staphylococcaceae *Staphylococcus aureus* OCMM6137 |
| Sau-2674-1 | Bacteria Firmicutes Bacilli Bacillales Staphylococcaceae *Staphylococcus aureus* OCMM6140 |
| Sau-2675-1 | Bacteria Firmicutes Bacilli Bacillales Staphylococcaceae *Staphylococcus aureus* P0218 |
| Sau-2676-1 | Bacteria Firmicutes Bacilli Bacillales Staphylococcaceae *Staphylococcus aureus* PA11 |
| Sau-2677-1 | Bacteria Firmicutes Bacilli Bacillales Staphylococcaceae *Staphylococcus aureus* PA57 |
| Sau-2678-1 | Bacteria Firmicutes Bacilli Bacillales Staphylococcaceae *Staphylococcus aureus* PLAC6004 |
| Sau-2679-1 | Bacteria Firmicutes Bacilli Bacillales Staphylococcaceae *Staphylococcus aureus* PLAC6012 |
| Sau-2680-1 | Bacteria Firmicutes Bacilli Bacillales Staphylococcaceae *Staphylococcus aureus* PLAC6017 |
| Sau-2681-1 | Bacteria Firmicutes Bacilli Bacillales Staphylococcaceae *Staphylococcus aureus* PLAC6019 |
| Sau-2682-1 | Bacteria Firmicutes Bacilli Bacillales Staphylococcaceae *Staphylococcus aureus* PLAC6024 |
| Sau-2683-1 | Bacteria Firmicutes Bacilli Bacillales Staphylococcaceae *Staphylococcus aureus* PLAC6043 |
| Sau-2684-1 | Bacteria Firmicutes Bacilli Bacillales Staphylococcaceae *Staphylococcus aureus* PLAC6044 |
| Sau-2685-1 | Bacteria Firmicutes Bacilli Bacillales Staphylococcaceae *Staphylococcus aureus* PLAC6055 |
| Sau-2686-1 | Bacteria Firmicutes Bacilli Bacillales Staphylococcaceae *Staphylococcus aureus* PLAC6058 |
| Sau-2687-1 | Bacteria Firmicutes Bacilli Bacillales Staphylococcaceae *Staphylococcus aureus* R0294 |
| Sau-2688-1 | Bacteria Firmicutes Bacilli Bacillales Staphylococcaceae *Staphylococcus aureus* R0353 |
| Sau-2689-1 | Bacteria Firmicutes Bacilli Bacillales Staphylococcaceae *Staphylococcus aureus* R0357 |
| Sau-2690-1 | Bacteria Firmicutes Bacilli Bacillales Staphylococcaceae *Staphylococcus aureus* R0487 |
| Sau-2691-1 | Bacteria Firmicutes Bacilli Bacillales Staphylococcaceae *Staphylococcus aureus* R0545 |
| Sau-2692-1 | Bacteria Firmicutes Bacilli Bacillales Staphylococcaceae *Staphylococcus aureus* R0611 |
| Sau-2693-1 | Bacteria Firmicutes Bacilli Bacillales Staphylococcaceae *Staphylococcus aureus* R0615 |
| Sau-2694-1 | Bacteria Firmicutes Bacilli Bacillales Staphylococcaceae *Staphylococcus aureus* Rd.290 |
| Sau-2695-1 | Bacteria Firmicutes Bacilli Bacillales Staphylococcaceae *Staphylococcus aureus* Rd.3 |
| Sau-2696-1 | Bacteria Firmicutes Bacilli Bacillales Staphylococcaceae *Staphylococcus aureus* Rd.40 |
| Sau-2697-1 | Bacteria Firmicutes Bacilli Bacillales Staphylococcaceae *Staphylococcus aureus* Rd.51 |
| Sau-2698-1 | Bacteria Firmicutes Bacilli Bacillales Staphylococcaceae *Staphylococcus aureus* Rd.545 |
| Sau-2699-1 | Bacteria Firmicutes Bacilli Bacillales Staphylococcaceae *Staphylococcus aureus* Rd.60 |
| Sau-2700-1 | Bacteria Firmicutes Bacilli Bacillales Staphylococcaceae *Staphylococcus aureus* Rd.614 |
| Sau-2701-1 | Bacteria Firmicutes Bacilli Bacillales Staphylococcaceae *Staphylococcus aureus* Rd.9 |
| Sau-2702-1 | Bacteria Firmicutes Bacilli Bacillales Staphylococcaceae *Staphylococcus aureus* S0924 |
| Sau-2703-1 | Bacteria Firmicutes Bacilli Bacillales Staphylococcaceae *Staphylococcus aureus* S11444 |
| Sau-2704-1 | Bacteria Firmicutes Bacilli Bacillales Staphylococcaceae *Staphylococcus aureus* S1475 |
| Sau-2705-1 | Bacteria Firmicutes Bacilli Bacillales Staphylococcaceae *Staphylococcus aureus* S29036 |
| Sau-2706-1 | Bacteria Firmicutes Bacilli Bacillales Staphylococcaceae *Staphylococcus aureus* S30087 |
| Sau-2707-1 | Bacteria Firmicutes Bacilli Bacillales Staphylococcaceae *Staphylococcus aureus* S39407 |
| Sau-2708-1 | Bacteria Firmicutes Bacilli Bacillales Staphylococcaceae *Staphylococcus aureus* S39411 |
| Sau-2709-1 | Bacteria Firmicutes Bacilli Bacillales Staphylococcaceae *Staphylococcus aureus* S56450 |
| Sau-2710-1 | Bacteria Firmicutes Bacilli Bacillales Staphylococcaceae *Staphylococcus aureus* S56_POEL |
| Sau-2711-1 | Bacteria Firmicutes Bacilli Bacillales Staphylococcaceae *Staphylococcus aureus* S62_POEL |
| Sau-2712-1 | Bacteria Firmicutes Bacilli Bacillales Staphylococcaceae *Staphylococcus aureus* S63_POEL |
| Sau-2713-1 | Bacteria Firmicutes Bacilli Bacillales Staphylococcaceae *Staphylococcus aureus* S64_POEL |
| Sau-2714-1 | Bacteria Firmicutes Bacilli Bacillales Staphylococcaceae *Staphylococcus aureus* S69_POEL |
| Sau-2715-1 | Bacteria Firmicutes Bacilli Bacillales Staphylococcaceae *Staphylococcus aureus* S72396 |
| Sau-2716-1 | Bacteria Firmicutes Bacilli Bacillales Staphylococcaceae *Staphylococcus aureus* SA0077 |
| Sau-2717-1 | Bacteria Firmicutes Bacilli Bacillales Staphylococcaceae *Staphylococcus aureus* SA10 |
| Sau-2718-1 | Bacteria Firmicutes Bacilli Bacillales Staphylococcaceae *Staphylococcus aureus* SA10-LAU |
| Sau-2719-1 | Bacteria Firmicutes Bacilli Bacillales Staphylococcaceae *Staphylococcus aureus* SA11-LAU |
| Sau-2720-1 | Bacteria Firmicutes Bacilli Bacillales Staphylococcaceae *Staphylococcus aureus* SA12-LAU |
| Sau-2721-1 | Bacteria Firmicutes Bacilli Bacillales Staphylococcaceae *Staphylococcus aureus* SA3-LAU |
| Sau-2722-1 | Bacteria Firmicutes Bacilli Bacillales Staphylococcaceae *Staphylococcus aureus* SA5-LAU |

| | |
|---|---|
| Sau-2723-1 | Bacteria Firmicutes Bacilli Bacillales Staphylococcaceae *Staphylococcus aureus* SA6-LAU |
| Sau-2724-1 | Bacteria Firmicutes Bacilli Bacillales Staphylococcaceae *Staphylococcus aureus* SA7-LAU |
| Sau-2725-1 | Bacteria Firmicutes Bacilli Bacillales Staphylococcaceae *Staphylococcus aureus* SA8-LAU |
| Sau-2726-1 | Bacteria Firmicutes Bacilli Bacillales Staphylococcaceae *Staphylococcus aureus* SA9-LAU |
| Sau-2727-1 | Bacteria Firmicutes Bacilli Bacillales Staphylococcaceae *Staphylococcus aureus* SARM C4155 |
| Sau-2728-1 | Bacteria Firmicutes Bacilli Bacillales Staphylococcaceae *Staphylococcus aureus* SARM C5621 |
| Sau-2820-1 | Bacteria Firmicutes Bacilli Bacillales Staphylococcaceae *Staphylococcus aureus* Sau 194 |
| Sau-2821-1 | Bacteria Firmicutes Bacilli Bacillales Staphylococcaceae *Staphylococcus aureus* Sau 44 |
| Sau-2822-1 | Bacteria Firmicutes Bacilli Bacillales Staphylococcaceae *Staphylococcus aureus* Sau 46 |
| Sau-2823-1 | Bacteria Firmicutes Bacilli Bacillales Staphylococcaceae *Staphylococcus aureus* Sau 47 |
| Sau-2824-1 | Bacteria Firmicutes Bacilli Bacillales Staphylococcaceae *Staphylococcus aureus* Sau 74 |
| Sau-2825-1 | Bacteria Firmicutes Bacilli Bacillales Staphylococcaceae *Staphylococcus aureus* Sau 93 |
| Sau-2826-1 | Bacteria Firmicutes Bacilli Bacillales Staphylococcaceae *Staphylococcus aureus* Sau 95 |
| Sau-2827-1 | Bacteria Firmicutes Bacilli Bacillales Staphylococcaceae *Staphylococcus aureus* Sau 98 |
| Sau-2828-1 | Bacteria Firmicutes Bacilli Bacillales Staphylococcaceae *Staphylococcus aureus* Sau 99 |
| Sau-2729-1 | Bacteria Firmicutes Bacilli Bacillales Staphylococcaceae *Staphylococcus aureus* SCLE6003 |
| Sau-2730-1 | Bacteria Firmicutes Bacilli Bacillales Staphylococcaceae *Staphylococcus aureus* SCLE6005 |
| Sau-2731-1 | Bacteria Firmicutes Bacilli Bacillales Staphylococcaceae *Staphylococcus aureus* SCLE6010 |
| Sau-2732-1 | Bacteria Firmicutes Bacilli Bacillales Staphylococcaceae *Staphylococcus aureus* SCLE6011 |
| Sau-2733-1 | Bacteria Firmicutes Bacilli Bacillales Staphylococcaceae *Staphylococcus aureus* SCLE6012 |
| Sau-2734-1 | Bacteria Firmicutes Bacilli Bacillales Staphylococcaceae *Staphylococcus aureus* SCLE6029 |
| Sau-2735-1 | Bacteria Firmicutes Bacilli Bacillales Staphylococcaceae *Staphylococcus aureus* SCLE6032 |
| Sau-2736-1 | Bacteria Firmicutes Bacilli Bacillales Staphylococcaceae *Staphylococcus aureus* SCLE6035 |
| Sau-2737-1 | Bacteria Firmicutes Bacilli Bacillales Staphylococcaceae *Staphylococcus aureus* SCLE6040 |
| Sau-2738-1 | Bacteria Firmicutes Bacilli Bacillales Staphylococcaceae *Staphylococcus aureus* SCLE6100 |
| Sau-2739-1 | Bacteria Firmicutes Bacilli Bacillales Staphylococcaceae *Staphylococcus aureus* SCOA6006 |
| Sau-2740-1 | Bacteria Firmicutes Bacilli Bacillales Staphylococcaceae *Staphylococcus aureus* SCOA6008 |
| Sau-2741-1 | Bacteria Firmicutes Bacilli Bacillales Staphylococcaceae *Staphylococcus aureus* SCOA6009 |
| Sau-2742-1 | Bacteria Firmicutes Bacilli Bacillales Staphylococcaceae *Staphylococcus aureus* SCOA6011 |
| Sau-2743-1 | Bacteria Firmicutes Bacilli Bacillales Staphylococcaceae *Staphylococcus aureus* SCOA6012 |
| Sau-2744-1 | Bacteria Firmicutes Bacilli Bacillales Staphylococcaceae *Staphylococcus aureus* SCOA6048 |
| Sau-2745-1 | Bacteria Firmicutes Bacilli Bacillales Staphylococcaceae *Staphylococcus aureus* SJOS6001 |
| Sau-2746-1 | Bacteria Firmicutes Bacilli Bacillales Staphylococcaceae *Staphylococcus aureus* SJOS6002 |
| Sau-2747-1 | Bacteria Firmicutes Bacilli Bacillales Staphylococcaceae *Staphylococcus aureus* SJOS6003 |
| Sau-2748-1 | Bacteria Firmicutes Bacilli Bacillales Staphylococcaceae *Staphylococcus aureus* SJOS6004 |
| Sau-2749-1 | Bacteria Firmicutes Bacilli Bacillales Staphylococcaceae *Staphylococcus aureus* SJOS6016 |
| Sau-2750-1 | Bacteria Firmicutes Bacilli Bacillales Staphylococcaceae *Staphylococcus aureus* SJOS6022 |
| Sau-2751-1 | Bacteria Firmicutes Bacilli Bacillales Staphylococcaceae *Staphylococcus aureus* SJOS6027 |
| Sau-2752-1 | Bacteria Firmicutes Bacilli Bacillales Staphylococcaceae *Staphylococcus aureus* SJOS6050 |
| Sau-2753-1 | Bacteria Firmicutes Bacilli Bacillales Staphylococcaceae *Staphylococcus aureus* SJOS6051 |
| Sau-2754-1 | Bacteria Firmicutes Bacilli Bacillales Staphylococcaceae *Staphylococcus aureus* SJOS6052 |
| Sau-2755-1 | Bacteria Firmicutes Bacilli Bacillales Staphylococcaceae *Staphylococcus aureus* SJOS6053 |
| Sau-2756-1 | Bacteria Firmicutes Bacilli Bacillales Staphylococcaceae *Staphylococcus aureus* SJOS6061 |
| Sau-2757-1 | Bacteria Firmicutes Bacilli Bacillales Staphylococcaceae *Staphylococcus aureus* SJOS6071 |
| Sau-2758-1 | Bacteria Firmicutes Bacilli Bacillales Staphylococcaceae *Staphylococcus aureus* SJOS6072 |
| Sau-2759-1 | Bacteria Firmicutes Bacilli Bacillales Staphylococcaceae *Staphylococcus aureus* SJOS6076 |
| Sau-2760-1 | Bacteria Firmicutes Bacilli Bacillales Staphylococcaceae *Staphylococcus aureus* SJOS6095 |
| Sau-2761-1 | Bacteria Firmicutes Bacilli Bacillales Staphylococcaceae *Staphylococcus aureus* SJOS6098 |
| Sau-2762-1 | Bacteria Firmicutes Bacilli Bacillales Staphylococcaceae *Staphylococcus aureus* SJOS6100 |
| Sau-2763-1 | Bacteria Firmicutes Bacilli Bacillales Staphylococcaceae *Staphylococcus aureus* SJOS6105 |
| Sau-2764-1 | Bacteria Firmicutes Bacilli Bacillales Staphylococcaceae *Staphylococcus aureus* SJOS6106 |
| Sau-2765-1 | Bacteria Firmicutes Bacilli Bacillales Staphylococcaceae *Staphylococcus aureus* SJOS6118 |
| Sau-2766-1 | Bacteria Firmicutes Bacilli Bacillales Staphylococcaceae *Staphylococcus aureus* SJOS6123 |
| Sau-2767-1 | Bacteria Firmicutes Bacilli Bacillales Staphylococcaceae *Staphylococcus aureus* SJOS6124 |
| Sau-2768-1 | Bacteria Firmicutes Bacilli Bacillales Staphylococcaceae *Staphylococcus aureus* SJOS6125 |
| Sau-2769-1 | Bacteria Firmicutes Bacilli Bacillales Staphylococcaceae *Staphylococcus aureus* SJOS6126 |
| Sau-2770-1 | Bacteria Firmicutes Bacilli Bacillales Staphylococcaceae *Staphylococcus aureus* SJOS6139 |
| Sau-2771-1 | Bacteria Firmicutes Bacilli Bacillales Staphylococcaceae *Staphylococcus aureus* SJOS6144 |
| Sau-2772-1 | Bacteria Firmicutes Bacilli Bacillales Staphylococcaceae *Staphylococcus aureus* SJOS6145 |
| Sau-2773-1 | Bacteria Firmicutes Bacilli Bacillales Staphylococcaceae *Staphylococcus aureus* SJUD6010 |
| Sau-2774-1 | Bacteria Firmicutes Bacilli Bacillales Staphylococcaceae *Staphylococcus aureus* SJUD6014 |
| Sau-2775-1 | Bacteria Firmicutes Bacilli Bacillales Staphylococcaceae *Staphylococcus aureus* SJUD6015 |
| Sau-2776-1 | Bacteria Firmicutes Bacilli Bacillales Staphylococcaceae *Staphylococcus aureus* SJUD6018 |
| Sau-2777-1 | Bacteria Firmicutes Bacilli Bacillales Staphylococcaceae *Staphylococcus aureus* SJUD6021 |
| Sau-2778-1 | Bacteria Firmicutes Bacilli Bacillales Staphylococcaceae *Staphylococcus aureus* SJUD6022 |

| | |
|---|---|
| Sau-2779-1 | Bacteria Firmicutes Bacilli Bacillales Staphylococcaceae *Staphylococcus aureus* SJUD6028 |
| Sau-2780-1 | Bacteria Firmicutes Bacilli Bacillales Staphylococcaceae *Staphylococcus aureus* SJUD6040 |
| Sau-2781-1 | Bacteria Firmicutes Bacilli Bacillales Staphylococcaceae *Staphylococcus aureus* SJUD6041 |
| Sau-2782-1 | Bacteria Firmicutes Bacilli Bacillales Staphylococcaceae *Staphylococcus aureus* SJUD6053 |
| Sau-2783-1 | Bacteria Firmicutes Bacilli Bacillales Staphylococcaceae *Staphylococcus aureus* SJUD6056 |
| Sau-2784-1 | Bacteria Firmicutes Bacilli Bacillales Staphylococcaceae *Staphylococcus aureus* SJUD6063 |
| Sau-2785-1 | Bacteria Firmicutes Bacilli Bacillales Staphylococcaceae *Staphylococcus aureus* SJUD6064 |
| Sau-2786-1 | Bacteria Firmicutes Bacilli Bacillales Staphylococcaceae *Staphylococcus aureus* SJUD6072 |
| Sau-2787-1 | Bacteria Firmicutes Bacilli Bacillales Staphylococcaceae *Staphylococcus aureus* SJUD6074 |
| Sau-2788-1 | Bacteria Firmicutes Bacilli Bacillales Staphylococcaceae *Staphylococcus aureus* SJUD6085 |
| Sau-2789-1 | Bacteria Firmicutes Bacilli Bacillales Staphylococcaceae *Staphylococcus aureus* SJUD6086 |
| Sau-2790-1 | Bacteria Firmicutes Bacilli Bacillales Staphylococcaceae *Staphylococcus aureus* SJUD6087 |
| Sau-2791-1 | Bacteria Firmicutes Bacilli Bacillales Staphylococcaceae *Staphylococcus aureus* SJUD6098 |
| Sau-2792-1 | Bacteria Firmicutes Bacilli Bacillales Staphylococcaceae *Staphylococcus aureus* SJUD6101 |
| Sau-2793-1 | Bacteria Firmicutes Bacilli Bacillales Staphylococcaceae *Staphylococcus aureus* SJUD6104 |
| Sau-2794-1 | Bacteria Firmicutes Bacilli Bacillales Staphylococcaceae *Staphylococcus aureus* SJUD6107 |
| Sau-2795-1 | Bacteria Firmicutes Bacilli Bacillales Staphylococcaceae *Staphylococcus aureus* SJUD6109 |
| Sau-2796-1 | Bacteria Firmicutes Bacilli Bacillales Staphylococcaceae *Staphylococcus aureus* SJUD6110 |
| Sau-2797-1 | Bacteria Firmicutes Bacilli Bacillales Staphylococcaceae *Staphylococcus aureus* SJUD6111 |
| Sau-2798-1 | Bacteria Firmicutes Bacilli Bacillales Staphylococcaceae *Staphylococcus aureus* SJUD6114 |
| Sau-2799-1 | Bacteria Firmicutes Bacilli Bacillales Staphylococcaceae *Staphylococcus aureus* SJUD6118 |
| Sau-2800-1 | Bacteria Firmicutes Bacilli Bacillales Staphylococcaceae *Staphylococcus aureus* SJUD6120 |
| Sau-2801-1 | Bacteria Firmicutes Bacilli Bacillales Staphylococcaceae *Staphylococcus aureus* SJUD6124 |
| Sau-2802-1 | Bacteria Firmicutes Bacilli Bacillales Staphylococcaceae *Staphylococcus aureus* SMMC0005 |
| Sau-2803-1 | Bacteria Firmicutes Bacilli Bacillales Staphylococcaceae *Staphylococcus aureus* SMMC6007 |
| Sau-2804-1 | Bacteria Firmicutes Bacilli Bacillales Staphylococcaceae *Staphylococcus aureus* SMMC6011 |
| Sau-2805-1 | Bacteria Firmicutes Bacilli Bacillales Staphylococcaceae *Staphylococcus aureus* SMMC6014 |
| Sau-2806-1 | Bacteria Firmicutes Bacilli Bacillales Staphylococcaceae *Staphylococcus aureus* SMMC6016 |
| Sau-2807-1 | Bacteria Firmicutes Bacilli Bacillales Staphylococcaceae *Staphylococcus aureus* SMMC6022 |
| Sau-2808-1 | Bacteria Firmicutes Bacilli Bacillales Staphylococcaceae *Staphylococcus aureus* SMMC6030 |
| Sau-2809-1 | Bacteria Firmicutes Bacilli Bacillales Staphylococcaceae *Staphylococcus aureus* SMMC6034 |
| Sau-2810-1 | Bacteria Firmicutes Bacilli Bacillales Staphylococcaceae *Staphylococcus aureus* SMMC6037 |
| Sau-2811-1 | Bacteria Firmicutes Bacilli Bacillales Staphylococcaceae *Staphylococcus aureus* SMMC6038 |
| Sau-2812-1 | Bacteria Firmicutes Bacilli Bacillales Staphylococcaceae *Staphylococcus aureus* SMMC6051 |
| Sau-2813-1 | Bacteria Firmicutes Bacilli Bacillales Staphylococcaceae *Staphylococcus aureus* SMMC6067 |
| Sau-2814-1 | Bacteria Firmicutes Bacilli Bacillales Staphylococcaceae *Staphylococcus aureus* SMMC6076 |
| Sau-2815-1 | Bacteria Firmicutes Bacilli Bacillales Staphylococcaceae *Staphylococcus aureus* SMMC6085 |
| Sau-2816-1 | Bacteria Firmicutes Bacilli Bacillales Staphylococcaceae *Staphylococcus aureus* SMMC6087 |
| Sau-2817-1 | Bacteria Firmicutes Bacilli Bacillales Staphylococcaceae *Staphylococcus aureus* SMMC6097 |
| Sau-2818-1 | Bacteria Firmicutes Bacilli Bacillales Staphylococcaceae *Staphylococcus aureus* SMMC6108 |
| Sau-2819-1 | Bacteria Firmicutes Bacilli Bacillales Staphylococcaceae *Staphylococcus aureus* SMMC6121 |
| Sau-3721-1 | Bacteria Firmicutes Bacilli Bacillales Staphylococcaceae *Staphylococcus aureus subsp. anaerobius* |
| Sau-3722-1 to Sau-3722-6 | Bacteria Firmicutes Bacilli Bacillales Staphylococcaceae *Staphylococcus aureus subsp. aureus* |
| Sau-3723-1 | Bacteria Firmicutes Bacilli Bacillales Staphylococcaceae *Staphylococcus aureus subsp. aureus* 21205 |
| Sau-3724-1 | Bacteria Firmicutes Bacilli Bacillales Staphylococcaceae *Staphylococcus aureus subsp. aureus* 21230 |
| Sau-3725-1 | Bacteria Firmicutes Bacilli Bacillales Staphylococcaceae *Staphylococcus aureus subsp. aureus* 21239 |
| Sau-3726-1 | Bacteria Firmicutes Bacilli Bacillales Staphylococcaceae *Staphylococcus aureus subsp. aureus* 21251 |
| Sau-3727-1 | Bacteria Firmicutes Bacilli Bacillales Staphylococcaceae *Staphylococcus aureus subsp. aureus* 21267 |
| Sau-3728-1 | Bacteria Firmicutes Bacilli Bacillales Staphylococcaceae *Staphylococcus aureus subsp. aureus* 21275 |
| Sau-3729-1 | Bacteria Firmicutes Bacilli Bacillales Staphylococcaceae *Staphylococcus aureus subsp. aureus* 21304 |
| Sau-3730-1 | Bacteria Firmicutes Bacilli Bacillales Staphylococcaceae *Staphylococcus aureus subsp. aureus* 21311 |
| Sau-3731-1 | Bacteria Firmicutes Bacilli Bacillales Staphylococcaceae *Staphylococcus aureus subsp. aureus* 21314 |
| Sau-3732-1 | Bacteria Firmicutes Bacilli Bacillales Staphylococcaceae *Staphylococcus aureus subsp. aureus* 21320 |
| Sau-3733-1 | Bacteria Firmicutes Bacilli Bacillales Staphylococcaceae *Staphylococcus aureus subsp. aureus* 21321 |
| Sau-3734-1 | Bacteria Firmicutes Bacilli Bacillales Staphylococcaceae *Staphylococcus aureus subsp. aureus* 21337 |
| Sau-3735-1 | Bacteria Firmicutes Bacilli Bacillales Staphylococcaceae *Staphylococcus aureus subsp. aureus* 21338 |
| Sau-3736-1 | Bacteria Firmicutes Bacilli Bacillales Staphylococcaceae *Staphylococcus aureus subsp. aureus* 300-169 |
| Sau-3737-1 | Bacteria Firmicutes Bacilli Bacillales Staphylococcaceae *Staphylococcus aureus subsp. aureus* 301-188 |
| Sau-3738-1 | Bacteria Firmicutes Bacilli Bacillales Staphylococcaceae *Staphylococcus aureus subsp. aureus* CO-06 |
| Sau-3739-1 | Bacteria Firmicutes Bacilli Bacillales Staphylococcaceae *Staphylococcus aureus subsp. aureus* CO-30 |
| Sau-3740-1 | Bacteria Firmicutes Bacilli Bacillales Staphylococcaceae *Staphylococcus aureus subsp. aureus* CO-41 |
| Sau-3741-1 | Bacteria Firmicutes Bacilli Bacillales Staphylococcaceae *Staphylococcus aureus subsp. aureus* CO-49 |
| Sau-3742-1 | Bacteria Firmicutes Bacilli Bacillales Staphylococcaceae *Staphylococcus aureus subsp. aureus* CO-99 |
| Sau-3743-1 to Sau-3743-2 | Bacteria Firmicutes Bacilli Bacillales Staphylococcaceae *Staphylococcus aureus subsp. aureus* DSM 20231 |
| Sau-3744-1 | Bacteria Firmicutes Bacilli Bacillales Staphylococcaceae *Staphylococcus aureus subsp. aureus* H29 |

| | |
|---|---|
| Sau-3745-1 | Bacteria Firmicutes Bacilli Bacillales Staphylococcaceae *Staphylococcus aureus subsp. aureus* MRSA_PR1 |
| Sau-59-1 to Sau-59-2 | Bacteria Firmicutes Bacilli Bacillales Staphylococcaceae *Staphylococcus aureus subsp. aureus* Mu3 |
| Sau-3746-1 | Bacteria Firmicutes Bacilli Bacillales Staphylococcaceae *Staphylococcus aureus subsp. aureus* PB32 |
| Sau-3747-1 | Bacteria Firmicutes Bacilli Bacillales Staphylococcaceae *Staphylococcus aureus subsp. aureus* S0460 |
| Sau-3748-1 | Bacteria Firmicutes Bacilli Bacillales Staphylococcaceae *Staphylococcus aureus subsp. aureus* S0462 |
| Sau-3749-1 | Bacteria Firmicutes Bacilli Bacillales Staphylococcaceae *Staphylococcus aureus subsp. aureus* S1800 |
| Sau-3750-1 | Bacteria Firmicutes Bacilli Bacillales Staphylococcaceae *Staphylococcus aureus subsp. aureus* S1805 |
| Sau-3751-1 | Bacteria Firmicutes Bacilli Bacillales Staphylococcaceae *Staphylococcus aureus subsp. aureus* S2395 |
| Sau-3752-1 | Bacteria Firmicutes Bacilli Bacillales Staphylococcaceae *Staphylococcus aureus subsp. aureus* S2396 |
| Sau-3753-1 | Bacteria Firmicutes Bacilli Bacillales Staphylococcaceae *Staphylococcus aureus subsp. aureus* S2397 |
| Sau-3754-1 | Bacteria Firmicutes Bacilli Bacillales Staphylococcaceae *Staphylococcus aureus subsp. aureus* S2398 |
| Sau-3755-1 | Bacteria Firmicutes Bacilli Bacillales Staphylococcaceae *Staphylococcus aureus subsp. aureus* SA268 |
| Sau-3756-1 | Bacteria Firmicutes Bacilli Bacillales Staphylococcaceae *Staphylococcus aureus subsp. aureus* SK1585 |
| Sau-3757-1 | Bacteria Firmicutes Bacilli Bacillales Staphylococcaceae *Staphylococcus aureus subsp. aureus* ST772-MRSA-V |
| Sau-2829-1 | Bacteria Firmicutes Bacilli Bacillales Staphylococcaceae *Staphylococcus aureus* T12805 |
| Sau-2830-1 | Bacteria Firmicutes Bacilli Bacillales Staphylococcaceae *Staphylococcus aureus* T12818 |
| Sau-2831-1 | Bacteria Firmicutes Bacilli Bacillales Staphylococcaceae *Staphylococcus aureus* T12849 |
| Sau-2832-1 | Bacteria Firmicutes Bacilli Bacillales Staphylococcaceae *Staphylococcus aureus* T12923 |
| Sau-2833-1 | Bacteria Firmicutes Bacilli Bacillales Staphylococcaceae *Staphylococcus aureus* T13096 |
| Sau-2834-1 | Bacteria Firmicutes Bacilli Bacillales Staphylococcaceae *Staphylococcus aureus* T13125 |
| Sau-2835-1 | Bacteria Firmicutes Bacilli Bacillales Staphylococcaceae *Staphylococcus aureus* T13126 |
| Sau-2836-1 | Bacteria Firmicutes Bacilli Bacillales Staphylococcaceae *Staphylococcus aureus* T13127 |
| Sau-2837-1 | Bacteria Firmicutes Bacilli Bacillales Staphylococcaceae *Staphylococcus aureus* T13128 |
| Sau-2838-1 | Bacteria Firmicutes Bacilli Bacillales Staphylococcaceae *Staphylococcus aureus* T15889 |
| Sau-2839-1 | Bacteria Firmicutes Bacilli Bacillales Staphylococcaceae *Staphylococcus aureus* T15898 |
| Sau-2840-1 | Bacteria Firmicutes Bacilli Bacillales Staphylococcaceae *Staphylococcus aureus* T15928 |
| Sau-2841-1 | Bacteria Firmicutes Bacilli Bacillales Staphylococcaceae *Staphylococcus aureus* T16117 |
| Sau-2842-1 | Bacteria Firmicutes Bacilli Bacillales Staphylococcaceae *Staphylococcus aureus* T16281 |
| Sau-2843-1 | Bacteria Firmicutes Bacilli Bacillales Staphylococcaceae *Staphylococcus aureus* T16600 |
| Sau-2844-1 | Bacteria Firmicutes Bacilli Bacillales Staphylococcaceae *Staphylococcus aureus* T16615 |
| Sau-2845-1 | Bacteria Firmicutes Bacilli Bacillales Staphylococcaceae *Staphylococcus aureus* T16619 |
| Sau-2846-1 | Bacteria Firmicutes Bacilli Bacillales Staphylococcaceae *Staphylococcus aureus* T16664 |
| Sau-2847-1 | Bacteria Firmicutes Bacilli Bacillales Staphylococcaceae *Staphylococcus aureus* T16669 |
| Sau-2848-1 | Bacteria Firmicutes Bacilli Bacillales Staphylococcaceae *Staphylococcus aureus* T16676 |
| Sau-2849-1 | Bacteria Firmicutes Bacilli Bacillales Staphylococcaceae *Staphylococcus aureus* T17131 |
| Sau-2850-1 | Bacteria Firmicutes Bacilli Bacillales Staphylococcaceae *Staphylococcus aureus* T17141 |
| Sau-2851-1 | Bacteria Firmicutes Bacilli Bacillales Staphylococcaceae *Staphylococcus aureus* T18851 |
| Sau-2852-1 | Bacteria Firmicutes Bacilli Bacillales Staphylococcaceae *Staphylococcus aureus* T18865 |
| Sau-2853-1 | Bacteria Firmicutes Bacilli Bacillales Staphylococcaceae *Staphylococcus aureus* T20118 |
| Sau-2854-1 | Bacteria Firmicutes Bacilli Bacillales Staphylococcaceae *Staphylococcus aureus* T20142 |
| Sau-2855-1 | Bacteria Firmicutes Bacilli Bacillales Staphylococcaceae *Staphylococcus aureus* T22051 |
| Sau-2856-1 | Bacteria Firmicutes Bacilli Bacillales Staphylococcaceae *Staphylococcus aureus* T22071_080712 |
| Sau-2857-1 | Bacteria Firmicutes Bacilli Bacillales Staphylococcaceae *Staphylococcus aureus* T22073 |
| Sau-2858-1 | Bacteria Firmicutes Bacilli Bacillales Staphylococcaceae *Staphylococcus aureus* T22074 |
| Sau-2859-1 | Bacteria Firmicutes Bacilli Bacillales Staphylococcaceae *Staphylococcus aureus* T23068 |
| Sau-2860-1 | Bacteria Firmicutes Bacilli Bacillales Staphylococcaceae *Staphylococcus aureus* T23868 |
| Sau-2861-1 | Bacteria Firmicutes Bacilli Bacillales Staphylococcaceae *Staphylococcus aureus* T24515 |
| Sau-2862-1 | Bacteria Firmicutes Bacilli Bacillales Staphylococcaceae *Staphylococcus aureus* T25283 |
| Sau-2863-1 | Bacteria Firmicutes Bacilli Bacillales Staphylococcaceae *Staphylococcus aureus* T25611 |
| Sau-2864-1 | Bacteria Firmicutes Bacilli Bacillales Staphylococcaceae *Staphylococcus aureus* T26751 |
| Sau-2865-1 | Bacteria Firmicutes Bacilli Bacillales Staphylococcaceae *Staphylococcus aureus* T28640 |
| Sau-2866-1 | Bacteria Firmicutes Bacilli Bacillales Staphylococcaceae *Staphylococcus aureus* T28649 |
| Sau-2867-1 | Bacteria Firmicutes Bacilli Bacillales Staphylococcaceae *Staphylococcus aureus* T28653 |
| Sau-2868-1 | Bacteria Firmicutes Bacilli Bacillales Staphylococcaceae *Staphylococcus aureus* T28847 |
| Sau-2869-1 | Bacteria Firmicutes Bacilli Bacillales Staphylococcaceae *Staphylococcus aureus* T28854 |
| Sau-2870-1 | Bacteria Firmicutes Bacilli Bacillales Staphylococcaceae *Staphylococcus aureus* T30033 |
| Sau-2871-1 | Bacteria Firmicutes Bacilli Bacillales Staphylococcaceae *Staphylococcus aureus* T30565 |
| Sau-2872-1 | Bacteria Firmicutes Bacilli Bacillales Staphylococcaceae *Staphylococcus aureus* T30569 |
| Sau-2873-1 | Bacteria Firmicutes Bacilli Bacillales Staphylococcaceae *Staphylococcus aureus* T31892 |
| Sau-2874-1 | Bacteria Firmicutes Bacilli Bacillales Staphylococcaceae *Staphylococcus aureus* T32070 |
| Sau-2875-1 | Bacteria Firmicutes Bacilli Bacillales Staphylococcaceae *Staphylococcus aureus* T32086 |
| Sau-2876-1 | Bacteria Firmicutes Bacilli Bacillales Staphylococcaceae *Staphylococcus aureus* T32092 |
| Sau-2877-1 | Bacteria Firmicutes Bacilli Bacillales Staphylococcaceae *Staphylococcus aureus* T33139 |
| Sau-2878-1 | Bacteria Firmicutes Bacilli Bacillales Staphylococcaceae *Staphylococcus aureus* T33147 |
| Sau-2879-1 | Bacteria Firmicutes Bacilli Bacillales Staphylococcaceae *Staphylococcus aureus* T34011 |

| | |
|---|---|
| Sau-2880-1 | Bacteria Firmicutes Bacilli Bacillales Staphylococcaceae *Staphylococcus aureus* T34036 |
| Sau-2881-1 | Bacteria Firmicutes Bacilli Bacillales Staphylococcaceae *Staphylococcus aureus* T34696 |
| Sau-2882-1 | Bacteria Firmicutes Bacilli Bacillales Staphylococcaceae *Staphylococcus aureus* T34703 |
| Sau-2883-1 | Bacteria Firmicutes Bacilli Bacillales Staphylococcaceae *Staphylococcus aureus* T35594 |
| Sau-2884-1 | Bacteria Firmicutes Bacilli Bacillales Staphylococcaceae *Staphylococcus aureus* T35596 |
| Sau-2885-1 | Bacteria Firmicutes Bacilli Bacillales Staphylococcaceae *Staphylococcus aureus* T35604 |
| Sau-2886-1 | Bacteria Firmicutes Bacilli Bacillales Staphylococcaceae *Staphylococcus aureus* T35615 |
| Sau-2887-1 | Bacteria Firmicutes Bacilli Bacillales Staphylococcaceae *Staphylococcus aureus* T35622 |
| Sau-2888-1 | Bacteria Firmicutes Bacilli Bacillales Staphylococcaceae *Staphylococcus aureus* T36111 |
| Sau-2889-1 | Bacteria Firmicutes Bacilli Bacillales Staphylococcaceae *Staphylococcus aureus* T36127 |
| Sau-2890-1 | Bacteria Firmicutes Bacilli Bacillales Staphylococcaceae *Staphylococcus aureus* T37840 |
| Sau-2891-1 | Bacteria Firmicutes Bacilli Bacillales Staphylococcaceae *Staphylococcus aureus* T37849 |
| Sau-2892-1 | Bacteria Firmicutes Bacilli Bacillales Staphylococcaceae *Staphylococcus aureus* T39082 |
| Sau-2893-1 | Bacteria Firmicutes Bacilli Bacillales Staphylococcaceae *Staphylococcus aureus* T39093 |
| Sau-2894-1 | Bacteria Firmicutes Bacilli Bacillales Staphylococcaceae *Staphylococcus aureus* T39102 |
| Sau-2895-1 | Bacteria Firmicutes Bacilli Bacillales Staphylococcaceae *Staphylococcus aureus* T39105 |
| Sau-2896-1 | Bacteria Firmicutes Bacilli Bacillales Staphylococcaceae *Staphylococcus aureus* T39328 |
| Sau-2897-1 | Bacteria Firmicutes Bacilli Bacillales Staphylococcaceae *Staphylococcus aureus* T39363 |
| Sau-2898-1 | Bacteria Firmicutes Bacilli Bacillales Staphylococcaceae *Staphylococcus aureus* T40867 |
| Sau-2899-1 | Bacteria Firmicutes Bacilli Bacillales Staphylococcaceae *Staphylococcus aureus* T41081 |
| Sau-2900-1 | Bacteria Firmicutes Bacilli Bacillales Staphylococcaceae *Staphylococcus aureus* T42547 |
| Sau-2901-1 | Bacteria Firmicutes Bacilli Bacillales Staphylococcaceae *Staphylococcus aureus* T42550 |
| Sau-2902-1 | Bacteria Firmicutes Bacilli Bacillales Staphylococcaceae *Staphylococcus aureus* T42722 |
| Sau-2903-1 | Bacteria Firmicutes Bacilli Bacillales Staphylococcaceae *Staphylococcus aureus* T44444 |
| Sau-2904-1 | Bacteria Firmicutes Bacilli Bacillales Staphylococcaceae *Staphylococcus aureus* T44461 |
| Sau-2905-1 | Bacteria Firmicutes Bacilli Bacillales Staphylococcaceae *Staphylococcus aureus* T44811 |
| Sau-2906-1 | Bacteria Firmicutes Bacilli Bacillales Staphylococcaceae *Staphylococcus aureus* T44830 |
| Sau-2907-1 | Bacteria Firmicutes Bacilli Bacillales Staphylococcaceae *Staphylococcus aureus* T44834 |
| Sau-2908-1 | Bacteria Firmicutes Bacilli Bacillales Staphylococcaceae *Staphylococcus aureus* T45836 |
| Sau-2909-1 | Bacteria Firmicutes Bacilli Bacillales Staphylococcaceae *Staphylococcus aureus* T45839 |
| Sau-2910-1 | Bacteria Firmicutes Bacilli Bacillales Staphylococcaceae *Staphylococcus aureus* T45940 |
| Sau-2911-1 | Bacteria Firmicutes Bacilli Bacillales Staphylococcaceae *Staphylococcus aureus* T45945 |
| Sau-2912-1 | Bacteria Firmicutes Bacilli Bacillales Staphylococcaceae *Staphylococcus aureus* T45974 |
| Sau-2913-1 | Bacteria Firmicutes Bacilli Bacillales Staphylococcaceae *Staphylococcus aureus* T47161 |
| Sau-2914-1 | Bacteria Firmicutes Bacilli Bacillales Staphylococcaceae *Staphylococcus aureus* T47181 |
| Sau-2915-1 | Bacteria Firmicutes Bacilli Bacillales Staphylococcaceae *Staphylococcus aureus* T49198 |
| Sau-2916-1 | Bacteria Firmicutes Bacilli Bacillales Staphylococcaceae *Staphylococcus aureus* T49201 |
| Sau-2917-1 | Bacteria Firmicutes Bacilli Bacillales Staphylococcaceae *Staphylococcus aureus* T49441 |
| Sau-2918-1 | Bacteria Firmicutes Bacilli Bacillales Staphylococcaceae *Staphylococcus aureus* T49833 |
| Sau-2919-1 | Bacteria Firmicutes Bacilli Bacillales Staphylococcaceae *Staphylococcus aureus* T51001 |
| Sau-2920-1 | Bacteria Firmicutes Bacilli Bacillales Staphylococcaceae *Staphylococcus aureus* T51022 |
| Sau-2921-1 | Bacteria Firmicutes Bacilli Bacillales Staphylococcaceae *Staphylococcus aureus* T52642 |
| Sau-2922-1 | Bacteria Firmicutes Bacilli Bacillales Staphylococcaceae *Staphylococcus aureus* T52649 |
| Sau-2923-1 | Bacteria Firmicutes Bacilli Bacillales Staphylococcaceae *Staphylococcus aureus* T54280 |
| Sau-2924-1 | Bacteria Firmicutes Bacilli Bacillales Staphylococcaceae *Staphylococcus aureus* T54283 |
| Sau-2925-1 | Bacteria Firmicutes Bacilli Bacillales Staphylococcaceae *Staphylococcus aureus* T54294 |
| Sau-2926-1 | Bacteria Firmicutes Bacilli Bacillales Staphylococcaceae *Staphylococcus aureus* T55950 |
| Sau-2927-1 | Bacteria Firmicutes Bacilli Bacillales Staphylococcaceae *Staphylococcus aureus* T55954 |
| Sau-2928-1 | Bacteria Firmicutes Bacilli Bacillales Staphylococcaceae *Staphylococcus aureus* T55959 |
| Sau-2929-1 | Bacteria Firmicutes Bacilli Bacillales Staphylococcaceae *Staphylococcus aureus* T55960 |
| Sau-2930-1 | Bacteria Firmicutes Bacilli Bacillales Staphylococcaceae *Staphylococcus aureus* T56203 |
| Sau-2931-1 | Bacteria Firmicutes Bacilli Bacillales Staphylococcaceae *Staphylococcus aureus* T57397 |
| Sau-2932-1 | Bacteria Firmicutes Bacilli Bacillales Staphylococcaceae *Staphylococcus aureus* T57402 |
| Sau-2933-1 | Bacteria Firmicutes Bacilli Bacillales Staphylococcaceae *Staphylococcus aureus* T57415 |
| Sau-2934-1 | Bacteria Firmicutes Bacilli Bacillales Staphylococcaceae *Staphylococcus aureus* T59607 |
| Sau-2935-1 | Bacteria Firmicutes Bacilli Bacillales Staphylococcaceae *Staphylococcus aureus* T59609 |
| Sau-2936-1 | Bacteria Firmicutes Bacilli Bacillales Staphylococcaceae *Staphylococcus aureus* T59618 |
| Sau-2937-1 | Bacteria Firmicutes Bacilli Bacillales Staphylococcaceae *Staphylococcus aureus* T59813 |
| Sau-2938-1 | Bacteria Firmicutes Bacilli Bacillales Staphylococcaceae *Staphylococcus aureus* T60795 |
| Sau-2939-1 | Bacteria Firmicutes Bacilli Bacillales Staphylococcaceae *Staphylococcus aureus* T60805 |
| Sau-2940-1 | Bacteria Firmicutes Bacilli Bacillales Staphylococcaceae *Staphylococcus aureus* T62328 |
| Sau-2941-1 | Bacteria Firmicutes Bacilli Bacillales Staphylococcaceae *Staphylococcus aureus* T62346 |
| Sau-2942-1 | Bacteria Firmicutes Bacilli Bacillales Staphylococcaceae *Staphylococcus aureus* T62848 |
| Sau-2943-1 | Bacteria Firmicutes Bacilli Bacillales Staphylococcaceae *Staphylococcus aureus* T62854 |
| Sau-2944-1 | Bacteria Firmicutes Bacilli Bacillales Staphylococcaceae *Staphylococcus aureus* T62976 |

| | |
|---|---|
| Sau-2945-1 | Bacteria Firmicutes Bacilli Bacillales Staphylococcaceae *Staphylococcus aureus* T63897 |
| Sau-2946-1 | Bacteria Firmicutes Bacilli Bacillales Staphylococcaceae *Staphylococcus aureus* T63898 |
| Sau-2947-1 | Bacteria Firmicutes Bacilli Bacillales Staphylococcaceae *Staphylococcus aureus* T64318 |
| Sau-2948-1 | Bacteria Firmicutes Bacilli Bacillales Staphylococcaceae *Staphylococcus aureus* T66243 |
| Sau-2949-1 | Bacteria Firmicutes Bacilli Bacillales Staphylococcaceae *Staphylococcus aureus* T66282 |
| Sau-2950-1 | Bacteria Firmicutes Bacilli Bacillales Staphylococcaceae *Staphylococcus aureus* T66286 |
| Sau-2951-1 | Bacteria Firmicutes Bacilli Bacillales Staphylococcaceae *Staphylococcus aureus* T66292 |
| Sau-2952-1 | Bacteria Firmicutes Bacilli Bacillales Staphylococcaceae *Staphylococcus aureus* T66622 |
| Sau-2953-1 | Bacteria Firmicutes Bacilli Bacillales Staphylococcaceae *Staphylococcus aureus* T66632 |
| Sau-2954-1 | Bacteria Firmicutes Bacilli Bacillales Staphylococcaceae *Staphylococcus aureus* T67319 |
| Sau-2955-1 | Bacteria Firmicutes Bacilli Bacillales Staphylococcaceae *Staphylococcus aureus* T67327 |
| Sau-2956-1 | Bacteria Firmicutes Bacilli Bacillales Staphylococcaceae *Staphylococcus aureus* T69686 |
| Sau-2957-1 | Bacteria Firmicutes Bacilli Bacillales Staphylococcaceae *Staphylococcus aureus* T69703 |
| Sau-2958-1 | Bacteria Firmicutes Bacilli Bacillales Staphylococcaceae *Staphylococcus aureus* T69716 |
| Sau-2959-1 | Bacteria Firmicutes Bacilli Bacillales Staphylococcaceae *Staphylococcus aureus* T69721 |
| Sau-2960-1 | Bacteria Firmicutes Bacilli Bacillales Staphylococcaceae *Staphylococcus aureus* T70161 |
| Sau-2961-1 | Bacteria Firmicutes Bacilli Bacillales Staphylococcaceae *Staphylococcus aureus* T70944 |
| Sau-2962-1 | Bacteria Firmicutes Bacilli Bacillales Staphylococcaceae *Staphylococcus aureus* T71868 |
| Sau-2963-1 | Bacteria Firmicutes Bacilli Bacillales Staphylococcaceae *Staphylococcus aureus* T71872 |
| Sau-2964-1 | Bacteria Firmicutes Bacilli Bacillales Staphylococcaceae *Staphylococcus aureus* T71888 |
| Sau-2965-1 | Bacteria Firmicutes Bacilli Bacillales Staphylococcaceae *Staphylococcus aureus* T72814 |
| Sau-2966-1 | Bacteria Firmicutes Bacilli Bacillales Staphylococcaceae *Staphylococcus aureus* T72838 |
| Sau-2967-1 | Bacteria Firmicutes Bacilli Bacillales Staphylococcaceae *Staphylococcus aureus* T72876 |
| Sau-2968-1 | Bacteria Firmicutes Bacilli Bacillales Staphylococcaceae *Staphylococcus aureus* T72889 |
| Sau-2969-1 | Bacteria Firmicutes Bacilli Bacillales Staphylococcaceae *Staphylococcus aureus* T74005 |
| Sau-2970-1 | Bacteria Firmicutes Bacilli Bacillales Staphylococcaceae *Staphylococcus aureus* T74060 |
| Sau-2971-1 | Bacteria Firmicutes Bacilli Bacillales Staphylococcaceae *Staphylococcus aureus* T74091 |
| Sau-2972-1 | Bacteria Firmicutes Bacilli Bacillales Staphylococcaceae *Staphylococcus aureus* T74141 |
| Sau-2973-1 | Bacteria Firmicutes Bacilli Bacillales Staphylococcaceae *Staphylococcus aureus* T74154 |
| Sau-2974-1 | Bacteria Firmicutes Bacilli Bacillales Staphylococcaceae *Staphylococcus aureus* T74602 |
| Sau-2975-1 | Bacteria Firmicutes Bacilli Bacillales Staphylococcaceae *Staphylococcus aureus* T76010 |
| Sau-2976-1 | Bacteria Firmicutes Bacilli Bacillales Staphylococcaceae *Staphylococcus aureus* T76018 |
| Sau-2977-1 | Bacteria Firmicutes Bacilli Bacillales Staphylococcaceae *Staphylococcus aureus* T76030 |
| Sau-2978-1 | Bacteria Firmicutes Bacilli Bacillales Staphylococcaceae *Staphylococcus aureus* T77565 |
| Sau-2979-1 | Bacteria Firmicutes Bacilli Bacillales Staphylococcaceae *Staphylococcus aureus* T77618 |
| Sau-2980-1 | Bacteria Firmicutes Bacilli Bacillales Staphylococcaceae *Staphylococcus aureus* T78544 |
| Sau-2981-1 | Bacteria Firmicutes Bacilli Bacillales Staphylococcaceae *Staphylococcus aureus* T79159 |
| Sau-2982-1 | Bacteria Firmicutes Bacilli Bacillales Staphylococcaceae *Staphylococcus aureus* T81559 |
| Sau-2983-1 | Bacteria Firmicutes Bacilli Bacillales Staphylococcaceae *Staphylococcus aureus* T82627 |
| Sau-2984-1 | Bacteria Firmicutes Bacilli Bacillales Staphylococcaceae *Staphylococcus aureus* T83543 |
| Sau-2985-1 | Bacteria Firmicutes Bacilli Bacillales Staphylococcaceae *Staphylococcus aureus* T83563 |
| Sau-2986-1 | Bacteria Firmicutes Bacilli Bacillales Staphylococcaceae *Staphylococcus aureus* T83705 |
| Sau-2987-1 | Bacteria Firmicutes Bacilli Bacillales Staphylococcaceae *Staphylococcus aureus* T83715 |
| Sau-2988-1 | Bacteria Firmicutes Bacilli Bacillales Staphylococcaceae *Staphylococcus aureus* T84592 |
| Sau-2989-1 | Bacteria Firmicutes Bacilli Bacillales Staphylococcaceae *Staphylococcus aureus* T84594 |
| Sau-2990-1 | Bacteria Firmicutes Bacilli Bacillales Staphylococcaceae *Staphylococcus aureus* T84595 |
| Sau-2991-1 | Bacteria Firmicutes Bacilli Bacillales Staphylococcaceae *Staphylococcus aureus* T86006 |
| Sau-2992-1 | Bacteria Firmicutes Bacilli Bacillales Staphylococcaceae *Staphylococcus aureus* T86045 |
| Sau-2993-1 | Bacteria Firmicutes Bacilli Bacillales Staphylococcaceae *Staphylococcus aureus* T86048 |
| Sau-2994-1 | Bacteria Firmicutes Bacilli Bacillales Staphylococcaceae *Staphylococcus aureus* T86055 |
| Sau-2995-1 | Bacteria Firmicutes Bacilli Bacillales Staphylococcaceae *Staphylococcus aureus* T86574 |
| Sau-2996-1 | Bacteria Firmicutes Bacilli Bacillales Staphylococcaceae *Staphylococcus aureus* T86792 |
| Sau-2997-1 | Bacteria Firmicutes Bacilli Bacillales Staphylococcaceae *Staphylococcus aureus* T86825 |
| Sau-2998-1 | Bacteria Firmicutes Bacilli Bacillales Staphylococcaceae *Staphylococcus aureus* T87526 |
| Sau-2999-1 | Bacteria Firmicutes Bacilli Bacillales Staphylococcaceae *Staphylococcus aureus* T87965 |
| Sau-3000-1 | Bacteria Firmicutes Bacilli Bacillales Staphylococcaceae *Staphylococcus aureus* T87972 |
| Sau-3001-1 | Bacteria Firmicutes Bacilli Bacillales Staphylococcaceae *Staphylococcus aureus* T89463 |
| Sau-3002-1 | Bacteria Firmicutes Bacilli Bacillales Staphylococcaceae *Staphylococcus aureus* T89467 |
| Sau-3003-1 | Bacteria Firmicutes Bacilli Bacillales Staphylococcaceae *Staphylococcus aureus* T89906 |
| Sau-3004-1 | Bacteria Firmicutes Bacilli Bacillales Staphylococcaceae *Staphylococcus aureus* T92876 |
| Sau-3005-1 | Bacteria Firmicutes Bacilli Bacillales Staphylococcaceae *Staphylococcus aureus* T92889 |
| Sau-3008-1 | Bacteria Firmicutes Bacilli Bacillales Staphylococcaceae *Staphylococcus aureus* Tur-12 |
| Sau-3009-1 | Bacteria Firmicutes Bacilli Bacillales Staphylococcaceae *Staphylococcus aureus* Tur-15 |
| Sau-3010-1 | Bacteria Firmicutes Bacilli Bacillales Staphylococcaceae *Staphylococcus aureus* Tur-16 |
| Sau-3011-1 | Bacteria Firmicutes Bacilli Bacillales Staphylococcaceae *Staphylococcus aureus* Tur-20 |

| | |
|---|---|
| Sau-3012-1 | Bacteria Firmicutes Bacilli Bacillales Staphylococcaceae *Staphylococcus aureus* Tur-22 |
| Sau-3013-1 | Bacteria Firmicutes Bacilli Bacillales Staphylococcaceae *Staphylococcus aureus* Tur-4 |
| Sau-3014-1 | Bacteria Firmicutes Bacilli Bacillales Staphylococcaceae *Staphylococcus aureus* Tur-5 |
| Sau-3015-1 | Bacteria Firmicutes Bacilli Bacillales Staphylococcaceae *Staphylococcus aureus* Tur-6 |
| Sau-3006-1 | Bacteria Firmicutes Bacilli Bacillales Staphylococcaceae *Staphylococcus aureus* TUSH6001 |
| Sau-3007-1 | Bacteria Firmicutes Bacilli Bacillales Staphylococcaceae *Staphylococcus aureus* TUSH6007 |
| Sau-3016-1 | Bacteria Firmicutes Bacilli Bacillales Staphylococcaceae *Staphylococcus aureus* UB 08-116 |
| Sau-3017-1 | Bacteria Firmicutes Bacilli Bacillales Staphylococcaceae *Staphylococcus aureus* UB 08-172 |
| Sau-3018-1 | Bacteria Firmicutes Bacilli Bacillales Staphylococcaceae *Staphylococcus aureus* UCIM6009 |
| Sau-3019-1 | Bacteria Firmicutes Bacilli Bacillales Staphylococcaceae *Staphylococcus aureus* UCIM6014 |
| Sau-3020-1 | Bacteria Firmicutes Bacilli Bacillales Staphylococcaceae *Staphylococcus aureus* UCIM6015 |
| Sau-3021-1 | Bacteria Firmicutes Bacilli Bacillales Staphylococcaceae *Staphylococcus aureus* UCIM6018 |
| Sau-3022-1 | Bacteria Firmicutes Bacilli Bacillales Staphylococcaceae *Staphylococcus aureus* UCIM6019 |
| Sau-3023-1 | Bacteria Firmicutes Bacilli Bacillales Staphylococcaceae *Staphylococcus aureus* UCIM6034 |
| Sau-3024-1 | Bacteria Firmicutes Bacilli Bacillales Staphylococcaceae *Staphylococcus aureus* UCIM6042 |
| Sau-3025-1 | Bacteria Firmicutes Bacilli Bacillales Staphylococcaceae *Staphylococcus aureus* UCIM6047 |
| Sau-3026-1 | Bacteria Firmicutes Bacilli Bacillales Staphylococcaceae *Staphylococcus aureus* UCIM6049 |
| Sau-3027-1 | Bacteria Firmicutes Bacilli Bacillales Staphylococcaceae *Staphylococcus aureus* UCIM6050 |
| Sau-3028-1 | Bacteria Firmicutes Bacilli Bacillales Staphylococcaceae *Staphylococcus aureus* UCIM6053 |
| Sau-3029-1 | Bacteria Firmicutes Bacilli Bacillales Staphylococcaceae *Staphylococcus aureus* UCIM6055 |
| Sau-3030-1 | Bacteria Firmicutes Bacilli Bacillales Staphylococcaceae *Staphylococcus aureus* UCIM6056 |
| Sau-3031-1 | Bacteria Firmicutes Bacilli Bacillales Staphylococcaceae *Staphylococcus aureus* UCIM6057 |
| Sau-3032-1 | Bacteria Firmicutes Bacilli Bacillales Staphylococcaceae *Staphylococcus aureus* UCIM6065 |
| Sau-3033-1 | Bacteria Firmicutes Bacilli Bacillales Staphylococcaceae *Staphylococcus aureus* UCIM6066 |
| Sau-3034-1 | Bacteria Firmicutes Bacilli Bacillales Staphylococcaceae *Staphylococcus aureus* UCIM6069 |
| Sau-3035-1 | Bacteria Firmicutes Bacilli Bacillales Staphylococcaceae *Staphylococcus aureus* UCIM6070 |
| Sau-3036-1 | Bacteria Firmicutes Bacilli Bacillales Staphylococcaceae *Staphylococcus aureus* UCIM6075 |
| Sau-3037-1 | Bacteria Firmicutes Bacilli Bacillales Staphylococcaceae *Staphylococcus aureus* UCIM6076 |
| Sau-3038-1 | Bacteria Firmicutes Bacilli Bacillales Staphylococcaceae *Staphylococcus aureus* UCIM6077 |
| Sau-3039-1 | Bacteria Firmicutes Bacilli Bacillales Staphylococcaceae *Staphylococcus aureus* UCIM6080 |
| Sau-3040-1 | Bacteria Firmicutes Bacilli Bacillales Staphylococcaceae *Staphylococcus aureus* UCIM6081 |
| Sau-3041-1 | Bacteria Firmicutes Bacilli Bacillales Staphylococcaceae *Staphylococcus aureus* UCIM6084 |
| Sau-3042-1 | Bacteria Firmicutes Bacilli Bacillales Staphylococcaceae *Staphylococcus aureus* UCIM6085 |
| Sau-3043-1 | Bacteria Firmicutes Bacilli Bacillales Staphylococcaceae *Staphylococcus aureus* UCIM6101 |
| Sau-3044-1 | Bacteria Firmicutes Bacilli Bacillales Staphylococcaceae *Staphylococcus aureus* UCIM6104 |
| Sau-3045-1 | Bacteria Firmicutes Bacilli Bacillales Staphylococcaceae *Staphylococcus aureus* UCIM6105 |
| Sau-3046-1 | Bacteria Firmicutes Bacilli Bacillales Staphylococcaceae *Staphylococcus aureus* UCIM6106 |
| Sau-3047-1 | Bacteria Firmicutes Bacilli Bacillales Staphylococcaceae *Staphylococcus aureus* UCIM6115 |
| Sau-3048-1 | Bacteria Firmicutes Bacilli Bacillales Staphylococcaceae *Staphylococcus aureus* UCIM6119 |
| Sau-3049-1 | Bacteria Firmicutes Bacilli Bacillales Staphylococcaceae *Staphylococcus aureus* UCIM6123 |
| Sau-3050-1 | Bacteria Firmicutes Bacilli Bacillales Staphylococcaceae *Staphylococcus aureus* UCIM6126 |
| Sau-3051-1 | Bacteria Firmicutes Bacilli Bacillales Staphylococcaceae *Staphylococcus aureus* UCIM6134 |
| Sau-3052-1 | Bacteria Firmicutes Bacilli Bacillales Staphylococcaceae *Staphylococcus aureus* UCIM6144 |
| Sau-3053-1 | Bacteria Firmicutes Bacilli Bacillales Staphylococcaceae *Staphylococcus aureus* UCIM6145 |
| Sau-3054-1 | Bacteria Firmicutes Bacilli Bacillales Staphylococcaceae *Staphylococcus aureus* UCIM6146 |
| Sau-3055-1 | Bacteria Firmicutes Bacilli Bacillales Staphylococcaceae *Staphylococcus aureus* UCIM6147 |
| Sau-3056-1 | Bacteria Firmicutes Bacilli Bacillales Staphylococcaceae *Staphylococcus aureus* USA300-ISMMS1 |
| Sau-3057-1 | Bacteria Firmicutes Bacilli Bacillales Staphylococcaceae *Staphylococcus aureus* USA7 |
| Sau-3058-1 | Bacteria Firmicutes Bacilli Bacillales Staphylococcaceae *Staphylococcus aureus* USA_1 |
| Sau-3059-1 | Bacteria Firmicutes Bacilli Bacillales Staphylococcaceae *Staphylococcus aureus* USA_11 |
| Sau-3060-1 | Bacteria Firmicutes Bacilli Bacillales Staphylococcaceae *Staphylococcus aureus* USA_12 |
| Sau-3061-1 | Bacteria Firmicutes Bacilli Bacillales Staphylococcaceae *Staphylococcus aureus* USA_15 |
| Sau-3062-1 | Bacteria Firmicutes Bacilli Bacillales Staphylococcaceae *Staphylococcus aureus* USA_6 |
| Sau-3063-1 | Bacteria Firmicutes Bacilli Bacillales Staphylococcaceae *Staphylococcus aureus* USA_8 |
| Sau-3064-1 | Bacteria Firmicutes Bacilli Bacillales Staphylococcaceae *Staphylococcus aureus* USA_9 |
| Sau-3065-1 | Bacteria Firmicutes Bacilli Bacillales Staphylococcaceae *Staphylococcus aureus* VE07/03048STA |
| Sau-3066-1 | Bacteria Firmicutes Bacilli Bacillales Staphylococcaceae *Staphylococcus aureus* VE08/02126ST1 |
| Sau-3067-1 | Bacteria Firmicutes Bacilli Bacillales Staphylococcaceae *Staphylococcus aureus* VE08/02242ST1 |
| Sau-3068-1 | Bacteria Firmicutes Bacilli Bacillales Staphylococcaceae *Staphylococcus aureus* VE08/02244ST1 |
| Sau-3069-1 | Bacteria Firmicutes Bacilli Bacillales Staphylococcaceae *Staphylococcus aureus* VET0050R |
| Sau-3070-1 | Bacteria Firmicutes Bacilli Bacillales Staphylococcaceae *Staphylococcus aureus* VET0051R |
| Sau-3071-1 | Bacteria Firmicutes Bacilli Bacillales Staphylococcaceae *Staphylococcus aureus* VET0052R |
| Sau-3072-1 | Bacteria Firmicutes Bacilli Bacillales Staphylococcaceae *Staphylococcus aureus* VET0053R |
| Sau-3073-1 | Bacteria Firmicutes Bacilli Bacillales Staphylococcaceae *Staphylococcus aureus* VET0054R |
| Sau-3074-1 | Bacteria Firmicutes Bacilli Bacillales Staphylococcaceae *Staphylococcus aureus* VET0055R |

| | |
|---|---|
| Sau-3075-1 | Bacteria Firmicutes Bacilli Bacillales Staphylococcaceae *Staphylococcus aureus* VET0057R |
| Sau-3076-1 | Bacteria Firmicutes Bacilli Bacillales Staphylococcaceae *Staphylococcus aureus* VET0058R |
| Sau-3077-1 | Bacteria Firmicutes Bacilli Bacillales Staphylococcaceae *Staphylococcus aureus* VET0059R |
| Sau-3078-1 | Bacteria Firmicutes Bacilli Bacillales Staphylococcaceae *Staphylococcus aureus* VET0060R |
| Sau-3079-1 | Bacteria Firmicutes Bacilli Bacillales Staphylococcaceae *Staphylococcus aureus* VET0061R |
| Sau-3080-1 | Bacteria Firmicutes Bacilli Bacillales Staphylococcaceae *Staphylococcus aureus* VET0062R |
| Sau-3081-1 | Bacteria Firmicutes Bacilli Bacillales Staphylococcaceae *Staphylococcus aureus* VET0063R |
| Sau-3082-1 | Bacteria Firmicutes Bacilli Bacillales Staphylococcaceae *Staphylococcus aureus* VET0065R |
| Sau-3083-1 | Bacteria Firmicutes Bacilli Bacillales Staphylococcaceae *Staphylococcus aureus* VET0066R |
| Sau-3084-1 | Bacteria Firmicutes Bacilli Bacillales Staphylococcaceae *Staphylococcus aureus* VET0067R |
| Sau-3085-1 | Bacteria Firmicutes Bacilli Bacillales Staphylococcaceae *Staphylococcus aureus* VET0069R |
| Sau-3086-1 | Bacteria Firmicutes Bacilli Bacillales Staphylococcaceae *Staphylococcus aureus* VET0070R |
| Sau-3087-1 | Bacteria Firmicutes Bacilli Bacillales Staphylococcaceae *Staphylococcus aureus* VET0072R |
| Sau-3088-1 | Bacteria Firmicutes Bacilli Bacillales Staphylococcaceae *Staphylococcus aureus* VET0073R |
| Sau-3089-1 | Bacteria Firmicutes Bacilli Bacillales Staphylococcaceae *Staphylococcus aureus* VET0075R |
| Sau-3090-1 | Bacteria Firmicutes Bacilli Bacillales Staphylococcaceae *Staphylococcus aureus* VET0076R |
| Sau-3091-1 | Bacteria Firmicutes Bacilli Bacillales Staphylococcaceae *Staphylococcus aureus* VET0077R |
| Sau-3092-1 | Bacteria Firmicutes Bacilli Bacillales Staphylococcaceae *Staphylococcus aureus* VET0078R |
| Sau-3093-1 | Bacteria Firmicutes Bacilli Bacillales Staphylococcaceae *Staphylococcus aureus* VET0080R |
| Sau-3094-1 | Bacteria Firmicutes Bacilli Bacillales Staphylococcaceae *Staphylococcus aureus* VET0081R |
| Sau-3095-1 | Bacteria Firmicutes Bacilli Bacillales Staphylococcaceae *Staphylococcus aureus* VET0083R |
| Sau-3096-1 | Bacteria Firmicutes Bacilli Bacillales Staphylococcaceae *Staphylococcus aureus* VET0084R |
| Sau-3097-1 | Bacteria Firmicutes Bacilli Bacillales Staphylococcaceae *Staphylococcus aureus* VET0085R |
| Sau-3098-1 | Bacteria Firmicutes Bacilli Bacillales Staphylococcaceae *Staphylococcus aureus* VET0086R |
| Sau-3099-1 | Bacteria Firmicutes Bacilli Bacillales Staphylococcaceae *Staphylococcus aureus* VET0087R |
| Sau-3100-1 | Bacteria Firmicutes Bacilli Bacillales Staphylococcaceae *Staphylococcus aureus* VET0088R |
| Sau-3101-1 | Bacteria Firmicutes Bacilli Bacillales Staphylococcaceae *Staphylococcus aureus* VET0089R |
| Sau-3102-1 | Bacteria Firmicutes Bacilli Bacillales Staphylococcaceae *Staphylococcus aureus* VET0090R |
| Sau-3103-1 | Bacteria Firmicutes Bacilli Bacillales Staphylococcaceae *Staphylococcus aureus* VET0091R |
| Sau-3104-1 | Bacteria Firmicutes Bacilli Bacillales Staphylococcaceae *Staphylococcus aureus* VET0092R |
| Sau-3105-1 | Bacteria Firmicutes Bacilli Bacillales Staphylococcaceae *Staphylococcus aureus* VET0093R |
| Sau-3106-1 | Bacteria Firmicutes Bacilli Bacillales Staphylococcaceae *Staphylococcus aureus* VET0094R |
| Sau-3107-1 | Bacteria Firmicutes Bacilli Bacillales Staphylococcaceae *Staphylococcus aureus* VET0095R |
| Sau-3108-1 | Bacteria Firmicutes Bacilli Bacillales Staphylococcaceae *Staphylococcus aureus* VET0097R |
| Sau-3109-1 | Bacteria Firmicutes Bacilli Bacillales Staphylococcaceae *Staphylococcus aureus* VET0098R |
| Sau-3110-1 | Bacteria Firmicutes Bacilli Bacillales Staphylococcaceae *Staphylococcus aureus* VET0099R |
| Sau-3111-1 | Bacteria Firmicutes Bacilli Bacillales Staphylococcaceae *Staphylococcus aureus* VET0101R |
| Sau-3112-1 | Bacteria Firmicutes Bacilli Bacillales Staphylococcaceae *Staphylococcus aureus* VET0102R |
| Sau-3113-1 | Bacteria Firmicutes Bacilli Bacillales Staphylococcaceae *Staphylococcus aureus* VET0103R |
| Sau-3114-1 | Bacteria Firmicutes Bacilli Bacillales Staphylococcaceae *Staphylococcus aureus* VET0104R |
| Sau-3115-1 | Bacteria Firmicutes Bacilli Bacillales Staphylococcaceae *Staphylococcus aureus* VET0105R |
| Sau-3116-1 | Bacteria Firmicutes Bacilli Bacillales Staphylococcaceae *Staphylococcus aureus* VET0107R |
| Sau-3117-1 | Bacteria Firmicutes Bacilli Bacillales Staphylococcaceae *Staphylococcus aureus* VET0110R |
| Sau-3118-1 | Bacteria Firmicutes Bacilli Bacillales Staphylococcaceae *Staphylococcus aureus* VET0111R |
| Sau-3119-1 | Bacteria Firmicutes Bacilli Bacillales Staphylococcaceae *Staphylococcus aureus* VET0112R |
| Sau-3120-1 | Bacteria Firmicutes Bacilli Bacillales Staphylococcaceae *Staphylococcus aureus* VET0113R |
| Sau-3121-1 | Bacteria Firmicutes Bacilli Bacillales Staphylococcaceae *Staphylococcus aureus* VET0114R |
| Sau-3122-1 | Bacteria Firmicutes Bacilli Bacillales Staphylococcaceae *Staphylococcus aureus* VET0115R |
| Sau-3123-1 | Bacteria Firmicutes Bacilli Bacillales Staphylococcaceae *Staphylococcus aureus* VET0116R |
| Sau-3124-1 | Bacteria Firmicutes Bacilli Bacillales Staphylococcaceae *Staphylococcus aureus* VET0117R |
| Sau-3125-1 | Bacteria Firmicutes Bacilli Bacillales Staphylococcaceae *Staphylococcus aureus* VET0119R |
| Sau-3126-1 | Bacteria Firmicutes Bacilli Bacillales Staphylococcaceae *Staphylococcus aureus* VET0120R |
| Sau-3127-1 | Bacteria Firmicutes Bacilli Bacillales Staphylococcaceae *Staphylococcus aureus* VET0123R |
| Sau-3128-1 | Bacteria Firmicutes Bacilli Bacillales Staphylococcaceae *Staphylococcus aureus* VET0124R |
| Sau-3129-1 | Bacteria Firmicutes Bacilli Bacillales Staphylococcaceae *Staphylococcus aureus* VET0125R |
| Sau-3130-1 | Bacteria Firmicutes Bacilli Bacillales Staphylococcaceae *Staphylococcus aureus* VET0126R |
| Sau-3131-1 | Bacteria Firmicutes Bacilli Bacillales Staphylococcaceae *Staphylococcus aureus* VET0127R |
| Sau-3132-1 | Bacteria Firmicutes Bacilli Bacillales Staphylococcaceae *Staphylococcus aureus* VET0128R |
| Sau-3133-1 | Bacteria Firmicutes Bacilli Bacillales Staphylococcaceae *Staphylococcus aureus* VET0129R |
| Sau-3134-1 | Bacteria Firmicutes Bacilli Bacillales Staphylococcaceae *Staphylococcus aureus* VET0130R |
| Sau-3135-1 | Bacteria Firmicutes Bacilli Bacillales Staphylococcaceae *Staphylococcus aureus* VET0132R |
| Sau-3136-1 | Bacteria Firmicutes Bacilli Bacillales Staphylococcaceae *Staphylococcus aureus* VET0133R |
| Sau-3137-1 | Bacteria Firmicutes Bacilli Bacillales Staphylococcaceae *Staphylococcus aureus* VET0134R |
| Sau-3138-1 | Bacteria Firmicutes Bacilli Bacillales Staphylococcaceae *Staphylococcus aureus* VET0136R |
| Sau-3139-1 | Bacteria Firmicutes Bacilli Bacillales Staphylococcaceae *Staphylococcus aureus* VET0140R |

| | |
|---|---|
| Sau-3140-1 | Bacteria Firmicutes Bacilli Bacillales Staphylococcaceae *Staphylococcus aureus* VET0141R |
| Sau-3141-1 | Bacteria Firmicutes Bacilli Bacillales Staphylococcaceae *Staphylococcus aureus* VET0148R |
| Sau-3142-1 | Bacteria Firmicutes Bacilli Bacillales Staphylococcaceae *Staphylococcus aureus* VET0150R |
| Sau-3143-1 | Bacteria Firmicutes Bacilli Bacillales Staphylococcaceae *Staphylococcus aureus* VET0151R |
| Sau-3144-1 | Bacteria Firmicutes Bacilli Bacillales Staphylococcaceae *Staphylococcus aureus* VET0152R |
| Sau-3145-1 | Bacteria Firmicutes Bacilli Bacillales Staphylococcaceae *Staphylococcus aureus* VET0154R |
| Sau-3146-1 | Bacteria Firmicutes Bacilli Bacillales Staphylococcaceae *Staphylococcus aureus* VET0155R |
| Sau-3147-1 | Bacteria Firmicutes Bacilli Bacillales Staphylococcaceae *Staphylococcus aureus* VET0156R |
| Sau-3148-1 | Bacteria Firmicutes Bacilli Bacillales Staphylococcaceae *Staphylococcus aureus* VET0157R |
| Sau-3149-1 | Bacteria Firmicutes Bacilli Bacillales Staphylococcaceae *Staphylococcus aureus* VET0158R |
| Sau-3150-1 | Bacteria Firmicutes Bacilli Bacillales Staphylococcaceae *Staphylococcus aureus* VET0159R |
| Sau-3151-1 | Bacteria Firmicutes Bacilli Bacillales Staphylococcaceae *Staphylococcus aureus* VET0161R |
| Sau-3152-1 | Bacteria Firmicutes Bacilli Bacillales Staphylococcaceae *Staphylococcus aureus* VET0162R |
| Sau-3153-1 | Bacteria Firmicutes Bacilli Bacillales Staphylococcaceae *Staphylococcus aureus* VET0164R |
| Sau-3154-1 | Bacteria Firmicutes Bacilli Bacillales Staphylococcaceae *Staphylococcus aureus* VET0165R |
| Sau-3155-1 | Bacteria Firmicutes Bacilli Bacillales Staphylococcaceae *Staphylococcus aureus* VET0166R |
| Sau-3156-1 | Bacteria Firmicutes Bacilli Bacillales Staphylococcaceae *Staphylococcus aureus* VET0167R |
| Sau-3157-1 | Bacteria Firmicutes Bacilli Bacillales Staphylococcaceae *Staphylococcus aureus* VET0171R |
| Sau-3158-1 | Bacteria Firmicutes Bacilli Bacillales Staphylococcaceae *Staphylococcus aureus* VET0176R |
| Sau-3159-1 | Bacteria Firmicutes Bacilli Bacillales Staphylococcaceae *Staphylococcus aureus* VET0177R |
| Sau-3160-1 | Bacteria Firmicutes Bacilli Bacillales Staphylococcaceae *Staphylococcus aureus* VET0178R |
| Sau-3161-1 | Bacteria Firmicutes Bacilli Bacillales Staphylococcaceae *Staphylococcus aureus* VET0179R |
| Sau-3162-1 | Bacteria Firmicutes Bacilli Bacillales Staphylococcaceae *Staphylococcus aureus* VET0180R |
| Sau-3163-1 | Bacteria Firmicutes Bacilli Bacillales Staphylococcaceae *Staphylococcus aureus* VET0183R |
| Sau-3164-1 | Bacteria Firmicutes Bacilli Bacillales Staphylococcaceae *Staphylococcus aureus* VET0184R |
| Sau-3165-1 | Bacteria Firmicutes Bacilli Bacillales Staphylococcaceae *Staphylococcus aureus* VET0188R |
| Sau-3166-1 | Bacteria Firmicutes Bacilli Bacillales Staphylococcaceae *Staphylococcus aureus* VET0189R |
| Sau-3167-1 | Bacteria Firmicutes Bacilli Bacillales Staphylococcaceae *Staphylococcus aureus* VET0190R |
| Sau-3168-1 | Bacteria Firmicutes Bacilli Bacillales Staphylococcaceae *Staphylococcus aureus* VET0191R |
| Sau-3169-1 | Bacteria Firmicutes Bacilli Bacillales Staphylococcaceae *Staphylococcus aureus* VET0192R |
| Sau-3170-1 | Bacteria Firmicutes Bacilli Bacillales Staphylococcaceae *Staphylococcus aureus* VET0193R |
| Sau-3171-1 | Bacteria Firmicutes Bacilli Bacillales Staphylococcaceae *Staphylococcus aureus* VET0194R |
| Sau-3172-1 | Bacteria Firmicutes Bacilli Bacillales Staphylococcaceae *Staphylococcus aureus* VET0195R |
| Sau-3173-1 | Bacteria Firmicutes Bacilli Bacillales Staphylococcaceae *Staphylococcus aureus* VET0197R |
| Sau-3174-1 | Bacteria Firmicutes Bacilli Bacillales Staphylococcaceae *Staphylococcus aureus* VET0198R |
| Sau-3175-1 | Bacteria Firmicutes Bacilli Bacillales Staphylococcaceae *Staphylococcus aureus* VET0200R |
| Sau-3176-1 | Bacteria Firmicutes Bacilli Bacillales Staphylococcaceae *Staphylococcus aureus* VET0203R |
| Sau-3177-1 | Bacteria Firmicutes Bacilli Bacillales Staphylococcaceae *Staphylococcus aureus* VET0212R |
| Sau-3178-1 | Bacteria Firmicutes Bacilli Bacillales Staphylococcaceae *Staphylococcus aureus* VET0213R |
| Sau-3179-1 | Bacteria Firmicutes Bacilli Bacillales Staphylococcaceae *Staphylococcus aureus* VET0214R |
| Sau-3180-1 | Bacteria Firmicutes Bacilli Bacillales Staphylococcaceae *Staphylococcus aureus* VET0215R |
| Sau-3181-1 | Bacteria Firmicutes Bacilli Bacillales Staphylococcaceae *Staphylococcus aureus* VET0216R |
| Sau-3182-1 | Bacteria Firmicutes Bacilli Bacillales Staphylococcaceae *Staphylococcus aureus* VET0217R |
| Sau-3183-1 | Bacteria Firmicutes Bacilli Bacillales Staphylococcaceae *Staphylococcus aureus* VET0218R |
| Sau-3184-1 | Bacteria Firmicutes Bacilli Bacillales Staphylococcaceae *Staphylococcus aureus* VET0219R |
| Sau-3185-1 | Bacteria Firmicutes Bacilli Bacillales Staphylococcaceae *Staphylococcus aureus* VET0221R |
| Sau-3186-1 | Bacteria Firmicutes Bacilli Bacillales Staphylococcaceae *Staphylococcus aureus* VET0222R |
| Sau-3187-1 | Bacteria Firmicutes Bacilli Bacillales Staphylococcaceae *Staphylococcus aureus* VET0224R |
| Sau-3188-1 | Bacteria Firmicutes Bacilli Bacillales Staphylococcaceae *Staphylococcus aureus* VET0227R |
| Sau-3189-1 | Bacteria Firmicutes Bacilli Bacillales Staphylococcaceae *Staphylococcus aureus* VET0228R |
| Sau-3190-1 | Bacteria Firmicutes Bacilli Bacillales Staphylococcaceae *Staphylococcus aureus* VET0229R |
| Sau-3191-1 | Bacteria Firmicutes Bacilli Bacillales Staphylococcaceae *Staphylococcus aureus* VET0230R |
| Sau-3192-1 | Bacteria Firmicutes Bacilli Bacillales Staphylococcaceae *Staphylococcus aureus* VET0232R |
| Sau-3193-1 | Bacteria Firmicutes Bacilli Bacillales Staphylococcaceae *Staphylococcus aureus* VET0235R |
| Sau-3194-1 | Bacteria Firmicutes Bacilli Bacillales Staphylococcaceae *Staphylococcus aureus* VET0236R |
| Sau-3195-1 | Bacteria Firmicutes Bacilli Bacillales Staphylococcaceae *Staphylococcus aureus* VET0237R |
| Sau-3196-1 | Bacteria Firmicutes Bacilli Bacillales Staphylococcaceae *Staphylococcus aureus* VET0241R |
| Sau-3197-1 | Bacteria Firmicutes Bacilli Bacillales Staphylococcaceae *Staphylococcus aureus* VET0243R |
| Sau-3198-1 | Bacteria Firmicutes Bacilli Bacillales Staphylococcaceae *Staphylococcus aureus* VET0244R |
| Sau-3199-1 | Bacteria Firmicutes Bacilli Bacillales Staphylococcaceae *Staphylococcus aureus* VET0247R |
| Sau-3200-1 | Bacteria Firmicutes Bacilli Bacillales Staphylococcaceae *Staphylococcus aureus* VET0249R |
| Sau-3201-1 | Bacteria Firmicutes Bacilli Bacillales Staphylococcaceae *Staphylococcus aureus* VET0251R |
| Sau-3202-1 | Bacteria Firmicutes Bacilli Bacillales Staphylococcaceae *Staphylococcus aureus* VET0252R |
| Sau-3203-1 | Bacteria Firmicutes Bacilli Bacillales Staphylococcaceae *Staphylococcus aureus* VET0253R |
| Sau-3204-1 | Bacteria Firmicutes Bacilli Bacillales Staphylococcaceae *Staphylococcus aureus* VET0254R |

| | |
|---|---|
| Sau-3205-1 | Bacteria Firmicutes Bacilli Bacillales Staphylococcaceae *Staphylococcus aureus* VET0256R |
| Sau-3206-1 | Bacteria Firmicutes Bacilli Bacillales Staphylococcaceae *Staphylococcus aureus* VET0259R |
| Sau-3207-1 | Bacteria Firmicutes Bacilli Bacillales Staphylococcaceae *Staphylococcus aureus* VET0260R |
| Sau-3208-1 | Bacteria Firmicutes Bacilli Bacillales Staphylococcaceae *Staphylococcus aureus* VET0261R |
| Sau-3209-1 | Bacteria Firmicutes Bacilli Bacillales Staphylococcaceae *Staphylococcus aureus* VET0263R |
| Sau-3210-1 | Bacteria Firmicutes Bacilli Bacillales Staphylococcaceae *Staphylococcus aureus* VET0267R |
| Sau-3211-1 | Bacteria Firmicutes Bacilli Bacillales Staphylococcaceae *Staphylococcus aureus* VET0268R |
| Sau-3212-1 | Bacteria Firmicutes Bacilli Bacillales Staphylococcaceae *Staphylococcus aureus* VET0269R |
| Sau-3213-1 | Bacteria Firmicutes Bacilli Bacillales Staphylococcaceae *Staphylococcus aureus* VET0270R |
| Sau-3214-1 | Bacteria Firmicutes Bacilli Bacillales Staphylococcaceae *Staphylococcus aureus* VET0271R |
| Sau-3215-1 | Bacteria Firmicutes Bacilli Bacillales Staphylococcaceae *Staphylococcus aureus* VET0275R |
| Sau-3216-1 | Bacteria Firmicutes Bacilli Bacillales Staphylococcaceae *Staphylococcus aureus* VET0278R |
| Sau-3217-1 | Bacteria Firmicutes Bacilli Bacillales Staphylococcaceae *Staphylococcus aureus* VET0279R |
| Sau-3218-1 | Bacteria Firmicutes Bacilli Bacillales Staphylococcaceae *Staphylococcus aureus* VET0280R |
| Sau-3219-1 | Bacteria Firmicutes Bacilli Bacillales Staphylococcaceae *Staphylococcus aureus* VET0282R |
| Sau-3220-1 | Bacteria Firmicutes Bacilli Bacillales Staphylococcaceae *Staphylococcus aureus* VET0283R |
| Sau-3221-1 | Bacteria Firmicutes Bacilli Bacillales Staphylococcaceae *Staphylococcus aureus* VET0284R |
| Sau-3222-1 | Bacteria Firmicutes Bacilli Bacillales Staphylococcaceae *Staphylococcus aureus* VET0286R |
| Sau-3223-1 | Bacteria Firmicutes Bacilli Bacillales Staphylococcaceae *Staphylococcus aureus* VET0287R |
| Sau-3224-1 | Bacteria Firmicutes Bacilli Bacillales Staphylococcaceae *Staphylococcus aureus* VET0288R |
| Sau-3225-1 | Bacteria Firmicutes Bacilli Bacillales Staphylococcaceae *Staphylococcus aureus* VET0289R |
| Sau-3226-1 | Bacteria Firmicutes Bacilli Bacillales Staphylococcaceae *Staphylococcus aureus* VET0290R |
| Sau-3227-1 | Bacteria Firmicutes Bacilli Bacillales Staphylococcaceae *Staphylococcus aureus* VET0291R |
| Sau-3228-1 | Bacteria Firmicutes Bacilli Bacillales Staphylococcaceae *Staphylococcus aureus* VET0292R |
| Sau-3229-1 | Bacteria Firmicutes Bacilli Bacillales Staphylococcaceae *Staphylococcus aureus* VET0293R |
| Sau-3230-1 | Bacteria Firmicutes Bacilli Bacillales Staphylococcaceae *Staphylococcus aureus* VET0294R |
| Sau-3231-1 | Bacteria Firmicutes Bacilli Bacillales Staphylococcaceae *Staphylococcus aureus* VET0295R |
| Sau-3232-1 | Bacteria Firmicutes Bacilli Bacillales Staphylococcaceae *Staphylococcus aureus* VET0296R |
| Sau-3233-1 | Bacteria Firmicutes Bacilli Bacillales Staphylococcaceae *Staphylococcus aureus* VET0297R |
| Sau-3234-1 | Bacteria Firmicutes Bacilli Bacillales Staphylococcaceae *Staphylococcus aureus* VET0298R |
| Sau-3235-1 | Bacteria Firmicutes Bacilli Bacillales Staphylococcaceae *Staphylococcus aureus* VET0299R |
| Sau-3236-1 | Bacteria Firmicutes Bacilli Bacillales Staphylococcaceae *Staphylococcus aureus* VET0300R |
| Sau-3237-1 | Bacteria Firmicutes Bacilli Bacillales Staphylococcaceae *Staphylococcus aureus* VET0301R |
| Sau-3238-1 | Bacteria Firmicutes Bacilli Bacillales Staphylococcaceae *Staphylococcus aureus* VET0302R |
| Sau-3239-1 | Bacteria Firmicutes Bacilli Bacillales Staphylococcaceae *Staphylococcus aureus* VET0303R |
| Sau-3240-1 | Bacteria Firmicutes Bacilli Bacillales Staphylococcaceae *Staphylococcus aureus* VET0304R |
| Sau-3241-1 | Bacteria Firmicutes Bacilli Bacillales Staphylococcaceae *Staphylococcus aureus* VET0305R |
| Sau-3242-1 | Bacteria Firmicutes Bacilli Bacillales Staphylococcaceae *Staphylococcus aureus* VET0306R |
| Sau-3243-1 | Bacteria Firmicutes Bacilli Bacillales Staphylococcaceae *Staphylococcus aureus* VET0307R |
| Sau-3244-1 | Bacteria Firmicutes Bacilli Bacillales Staphylococcaceae *Staphylococcus aureus* VET0308R |
| Sau-3245-1 | Bacteria Firmicutes Bacilli Bacillales Staphylococcaceae *Staphylococcus aureus* VET0309R |
| Sau-3246-1 | Bacteria Firmicutes Bacilli Bacillales Staphylococcaceae *Staphylococcus aureus* VET0312R |
| Sau-3247-1 | Bacteria Firmicutes Bacilli Bacillales Staphylococcaceae *Staphylococcus aureus* VET0313R |
| Sau-3248-1 | Bacteria Firmicutes Bacilli Bacillales Staphylococcaceae *Staphylococcus aureus* VET0314R |
| Sau-3249-1 | Bacteria Firmicutes Bacilli Bacillales Staphylococcaceae *Staphylococcus aureus* VET0315R |
| Sau-3250-1 | Bacteria Firmicutes Bacilli Bacillales Staphylococcaceae *Staphylococcus aureus* VET0316R |
| Sau-3251-1 | Bacteria Firmicutes Bacilli Bacillales Staphylococcaceae *Staphylococcus aureus* VET0317R |
| Sau-3252-1 | Bacteria Firmicutes Bacilli Bacillales Staphylococcaceae *Staphylococcus aureus* VET0318R |
| Sau-3253-1 | Bacteria Firmicutes Bacilli Bacillales Staphylococcaceae *Staphylococcus aureus* VET0319R |
| Sau-3254-1 | Bacteria Firmicutes Bacilli Bacillales Staphylococcaceae *Staphylococcus aureus* VET0320R |
| Sau-3255-1 | Bacteria Firmicutes Bacilli Bacillales Staphylococcaceae *Staphylococcus aureus* VET0322R |
| Sau-3256-1 | Bacteria Firmicutes Bacilli Bacillales Staphylococcaceae *Staphylococcus aureus* VET0323R |
| Sau-3257-1 | Bacteria Firmicutes Bacilli Bacillales Staphylococcaceae *Staphylococcus aureus* VET0325R |
| Sau-3258-1 | Bacteria Firmicutes Bacilli Bacillales Staphylococcaceae *Staphylococcus aureus* VET0326R |
| Sau-3259-1 | Bacteria Firmicutes Bacilli Bacillales Staphylococcaceae *Staphylococcus aureus* VET0329R |
| Sau-3260-1 | Bacteria Firmicutes Bacilli Bacillales Staphylococcaceae *Staphylococcus aureus* VET0331R |
| Sau-3261-1 | Bacteria Firmicutes Bacilli Bacillales Staphylococcaceae *Staphylococcus aureus* VET0332R |
| Sau-3262-1 | Bacteria Firmicutes Bacilli Bacillales Staphylococcaceae *Staphylococcus aureus* VET0333R |
| Sau-3263-1 | Bacteria Firmicutes Bacilli Bacillales Staphylococcaceae *Staphylococcus aureus* VET0335R |
| Sau-3264-1 | Bacteria Firmicutes Bacilli Bacillales Staphylococcaceae *Staphylococcus aureus* VET0336R |
| Sau-3265-1 | Bacteria Firmicutes Bacilli Bacillales Staphylococcaceae *Staphylococcus aureus* VET0337R |
| Sau-3266-1 | Bacteria Firmicutes Bacilli Bacillales Staphylococcaceae *Staphylococcus aureus* VET0338R |
| Sau-3267-1 | Bacteria Firmicutes Bacilli Bacillales Staphylococcaceae *Staphylococcus aureus* VET0339R |
| Sau-3268-1 | Bacteria Firmicutes Bacilli Bacillales Staphylococcaceae *Staphylococcus aureus* VET0340R |
| Sau-3269-1 | Bacteria Firmicutes Bacilli Bacillales Staphylococcaceae *Staphylococcus aureus* VET0341R |

| | |
|---|---|
| Sau-3270-1 | Bacteria Firmicutes Bacilli Bacillales Staphylococcaceae *Staphylococcus aureus* VET0342R |
| Sau-3271-1 | Bacteria Firmicutes Bacilli Bacillales Staphylococcaceae *Staphylococcus aureus* VET0343R |
| Sau-3272-1 | Bacteria Firmicutes Bacilli Bacillales Staphylococcaceae *Staphylococcus aureus* VET0352R |
| Sau-3273-1 | Bacteria Firmicutes Bacilli Bacillales Staphylococcaceae *Staphylococcus aureus* VET0353R |
| Sau-3274-1 | Bacteria Firmicutes Bacilli Bacillales Staphylococcaceae *Staphylococcus aureus* VET0354R |
| Sau-3275-1 | Bacteria Firmicutes Bacilli Bacillales Staphylococcaceae *Staphylococcus aureus* VET0357R |
| Sau-3276-1 | Bacteria Firmicutes Bacilli Bacillales Staphylococcaceae *Staphylococcus aureus* VET0358R |
| Sau-3277-1 | Bacteria Firmicutes Bacilli Bacillales Staphylococcaceae *Staphylococcus aureus* VET0359R |
| Sau-3278-1 | Bacteria Firmicutes Bacilli Bacillales Staphylococcaceae *Staphylococcus aureus* VET0360R |
| Sau-3279-1 | Bacteria Firmicutes Bacilli Bacillales Staphylococcaceae *Staphylococcus aureus* VET0361R |
| Sau-3280-1 | Bacteria Firmicutes Bacilli Bacillales Staphylococcaceae *Staphylococcus aureus* VET0363R |
| Sau-3281-1 | Bacteria Firmicutes Bacilli Bacillales Staphylococcaceae *Staphylococcus aureus* VET0366R |
| Sau-3282-1 | Bacteria Firmicutes Bacilli Bacillales Staphylococcaceae *Staphylococcus aureus* VET0367R |
| Sau-3283-1 | Bacteria Firmicutes Bacilli Bacillales Staphylococcaceae *Staphylococcus aureus* VET0368R |
| Sau-3284-1 | Bacteria Firmicutes Bacilli Bacillales Staphylococcaceae *Staphylococcus aureus* VET0369R |
| Sau-3285-1 | Bacteria Firmicutes Bacilli Bacillales Staphylococcaceae *Staphylococcus aureus* VET0373R |
| Sau-3286-1 | Bacteria Firmicutes Bacilli Bacillales Staphylococcaceae *Staphylococcus aureus* VET0376R |
| Sau-3287-1 | Bacteria Firmicutes Bacilli Bacillales Staphylococcaceae *Staphylococcus aureus* VET0382R |
| Sau-3288-1 | Bacteria Firmicutes Bacilli Bacillales Staphylococcaceae *Staphylococcus aureus* VET0383R |
| Sau-3289-1 | Bacteria Firmicutes Bacilli Bacillales Staphylococcaceae *Staphylococcus aureus* VET0384R |
| Sau-3290-1 | Bacteria Firmicutes Bacilli Bacillales Staphylococcaceae *Staphylococcus aureus* VET0387R |
| Sau-3291-1 | Bacteria Firmicutes Bacilli Bacillales Staphylococcaceae *Staphylococcus aureus* VET0388R |
| Sau-3292-1 | Bacteria Firmicutes Bacilli Bacillales Staphylococcaceae *Staphylococcus aureus* VET0389R |
| Sau-3293-1 | Bacteria Firmicutes Bacilli Bacillales Staphylococcaceae *Staphylococcus aureus* VET0390R |
| Sau-3294-1 | Bacteria Firmicutes Bacilli Bacillales Staphylococcaceae *Staphylococcus aureus* VET0393R |
| Sau-3295-1 | Bacteria Firmicutes Bacilli Bacillales Staphylococcaceae *Staphylococcus aureus* VET0394R |
| Sau-3296-1 | Bacteria Firmicutes Bacilli Bacillales Staphylococcaceae *Staphylococcus aureus* VET0395R |
| Sau-3297-1 | Bacteria Firmicutes Bacilli Bacillales Staphylococcaceae *Staphylococcus aureus* VET0396R |
| Sau-3298-1 | Bacteria Firmicutes Bacilli Bacillales Staphylococcaceae *Staphylococcus aureus* VET0399R |
| Sau-3299-1 | Bacteria Firmicutes Bacilli Bacillales Staphylococcaceae *Staphylococcus aureus* VET0400R |
| Sau-3300-1 | Bacteria Firmicutes Bacilli Bacillales Staphylococcaceae *Staphylococcus aureus* VET0401R |
| Sau-3301-1 | Bacteria Firmicutes Bacilli Bacillales Staphylococcaceae *Staphylococcus aureus* VET0402R |
| Sau-3302-1 | Bacteria Firmicutes Bacilli Bacillales Staphylococcaceae *Staphylococcus aureus* VET0403R |
| Sau-3303-1 | Bacteria Firmicutes Bacilli Bacillales Staphylococcaceae *Staphylococcus aureus* VET0406R |
| Sau-3304-1 | Bacteria Firmicutes Bacilli Bacillales Staphylococcaceae *Staphylococcus aureus* VET0407R |
| Sau-3305-1 | Bacteria Firmicutes Bacilli Bacillales Staphylococcaceae *Staphylococcus aureus* VET0410R |
| Sau-3306-1 | Bacteria Firmicutes Bacilli Bacillales Staphylococcaceae *Staphylococcus aureus* VET0411R |
| Sau-3307-1 | Bacteria Firmicutes Bacilli Bacillales Staphylococcaceae *Staphylococcus aureus* VET0412R |
| Sau-3308-1 | Bacteria Firmicutes Bacilli Bacillales Staphylococcaceae *Staphylococcus aureus* VET0413R |
| Sau-3309-1 | Bacteria Firmicutes Bacilli Bacillales Staphylococcaceae *Staphylococcus aureus* VET0414R |
| Sau-3310-1 | Bacteria Firmicutes Bacilli Bacillales Staphylococcaceae *Staphylococcus aureus* VET0415R |
| Sau-3311-1 | Bacteria Firmicutes Bacilli Bacillales Staphylococcaceae *Staphylococcus aureus* VET0417R |
| Sau-3312-1 | Bacteria Firmicutes Bacilli Bacillales Staphylococcaceae *Staphylococcus aureus* VET0418R |
| Sau-3313-1 | Bacteria Firmicutes Bacilli Bacillales Staphylococcaceae *Staphylococcus aureus* VET0421R |
| Sau-3314-1 | Bacteria Firmicutes Bacilli Bacillales Staphylococcaceae *Staphylococcus aureus* VET0422R |
| Sau-3315-1 | Bacteria Firmicutes Bacilli Bacillales Staphylococcaceae *Staphylococcus aureus* VET0423R |
| Sau-3316-1 | Bacteria Firmicutes Bacilli Bacillales Staphylococcaceae *Staphylococcus aureus* VET0424R |
| Sau-3317-1 | Bacteria Firmicutes Bacilli Bacillales Staphylococcaceae *Staphylococcus aureus* VET0425R |
| Sau-3318-1 | Bacteria Firmicutes Bacilli Bacillales Staphylococcaceae *Staphylococcus aureus* VET0426R |
| Sau-3319-1 | Bacteria Firmicutes Bacilli Bacillales Staphylococcaceae *Staphylococcus aureus* VET0427R |
| Sau-3320-1 | Bacteria Firmicutes Bacilli Bacillales Staphylococcaceae *Staphylococcus aureus* VET0428R |
| Sau-3321-1 | Bacteria Firmicutes Bacilli Bacillales Staphylococcaceae *Staphylococcus aureus* VET0431R |
| Sau-3322-1 | Bacteria Firmicutes Bacilli Bacillales Staphylococcaceae *Staphylococcus aureus* VET0432R |
| Sau-3323-1 | Bacteria Firmicutes Bacilli Bacillales Staphylococcaceae *Staphylococcus aureus* VET0433R |
| Sau-3324-1 | Bacteria Firmicutes Bacilli Bacillales Staphylococcaceae *Staphylococcus aureus* VET0434R |
| Sau-3325-1 | Bacteria Firmicutes Bacilli Bacillales Staphylococcaceae *Staphylococcus aureus* VET0435R |
| Sau-3326-1 | Bacteria Firmicutes Bacilli Bacillales Staphylococcaceae *Staphylococcus aureus* VET0436R |
| Sau-3327-1 | Bacteria Firmicutes Bacilli Bacillales Staphylococcaceae *Staphylococcus aureus* VET0437R |
| Sau-3328-1 | Bacteria Firmicutes Bacilli Bacillales Staphylococcaceae *Staphylococcus aureus* VET0438R |
| Sau-3329-1 | Bacteria Firmicutes Bacilli Bacillales Staphylococcaceae *Staphylococcus aureus* VET0439R |
| Sau-3330-1 | Bacteria Firmicutes Bacilli Bacillales Staphylococcaceae *Staphylococcus aureus* VET0440R |
| Sau-3331-1 | Bacteria Firmicutes Bacilli Bacillales Staphylococcaceae *Staphylococcus aureus* VET0442R |
| Sau-3332-1 | Bacteria Firmicutes Bacilli Bacillales Staphylococcaceae *Staphylococcus aureus* VET0443R |
| Sau-3333-1 | Bacteria Firmicutes Bacilli Bacillales Staphylococcaceae *Staphylococcus aureus* VET0444R |
| Sau-3334-1 | Bacteria Firmicutes Bacilli Bacillales Staphylococcaceae *Staphylococcus aureus* VET0446R |

| | |
|---|---|
| Sau-3335-1 | Bacteria Firmicutes Bacilli Bacillales Staphylococcaceae *Staphylococcus aureus* VET0451R |
| Sau-3336-1 | Bacteria Firmicutes Bacilli Bacillales Staphylococcaceae *Staphylococcus aureus* VET0452R |
| Sau-3337-1 | Bacteria Firmicutes Bacilli Bacillales Staphylococcaceae *Staphylococcus aureus* VET0454R |
| Sau-3338-1 | Bacteria Firmicutes Bacilli Bacillales Staphylococcaceae *Staphylococcus aureus* VET0455R |
| Sau-3339-1 | Bacteria Firmicutes Bacilli Bacillales Staphylococcaceae *Staphylococcus aureus* VET0456R |
| Sau-3340-1 | Bacteria Firmicutes Bacilli Bacillales Staphylococcaceae *Staphylococcus aureus* VET0459R |
| Sau-3341-1 | Bacteria Firmicutes Bacilli Bacillales Staphylococcaceae *Staphylococcus aureus* VET0460R |
| Sau-3342-1 | Bacteria Firmicutes Bacilli Bacillales Staphylococcaceae *Staphylococcus aureus* VET0462R |
| Sau-3343-1 | Bacteria Firmicutes Bacilli Bacillales Staphylococcaceae *Staphylococcus aureus* VET0463R |
| Sau-3344-1 | Bacteria Firmicutes Bacilli Bacillales Staphylococcaceae *Staphylococcus aureus* VET0464R |
| Sau-3345-1 | Bacteria Firmicutes Bacilli Bacillales Staphylococcaceae *Staphylococcus aureus* VET0465R |
| Sau-3346-1 | Bacteria Firmicutes Bacilli Bacillales Staphylococcaceae *Staphylococcus aureus* VET0467R |
| Sau-3347-1 | Bacteria Firmicutes Bacilli Bacillales Staphylococcaceae *Staphylococcus aureus* VET0468R |
| Sau-3348-1 | Bacteria Firmicutes Bacilli Bacillales Staphylococcaceae *Staphylococcus aureus* VET0469R |
| Sau-3349-1 | Bacteria Firmicutes Bacilli Bacillales Staphylococcaceae *Staphylococcus aureus* VET0470R |
| Sau-3350-1 | Bacteria Firmicutes Bacilli Bacillales Staphylococcaceae *Staphylococcus aureus* VET0471R |
| Sau-3351-1 | Bacteria Firmicutes Bacilli Bacillales Staphylococcaceae *Staphylococcus aureus* VET0472R |
| Sau-3352-1 | Bacteria Firmicutes Bacilli Bacillales Staphylococcaceae *Staphylococcus aureus* VET0473R |
| Sau-3353-1 | Bacteria Firmicutes Bacilli Bacillales Staphylococcaceae *Staphylococcus aureus* VET0478R |
| Sau-3354-1 | Bacteria Firmicutes Bacilli Bacillales Staphylococcaceae *Staphylococcus aureus* VET0481R |
| Sau-3355-1 | Bacteria Firmicutes Bacilli Bacillales Staphylococcaceae *Staphylococcus aureus* VET0485R |
| Sau-3356-1 | Bacteria Firmicutes Bacilli Bacillales Staphylococcaceae *Staphylococcus aureus* VET0486R |
| Sau-3357-1 | Bacteria Firmicutes Bacilli Bacillales Staphylococcaceae *Staphylococcus aureus* VET0488R |
| Sau-3358-1 | Bacteria Firmicutes Bacilli Bacillales Staphylococcaceae *Staphylococcus aureus* VET0489R |
| Sau-3359-1 | Bacteria Firmicutes Bacilli Bacillales Staphylococcaceae *Staphylococcus aureus* VET0519S |
| Sau-3360-1 | Bacteria Firmicutes Bacilli Bacillales Staphylococcaceae *Staphylococcus aureus* VET0596R |
| Sau-3361-1 | Bacteria Firmicutes Bacilli Bacillales Staphylococcaceae *Staphylococcus aureus* VET0612S |
| Sau-3362-1 | Bacteria Firmicutes Bacilli Bacillales Staphylococcaceae *Staphylococcus aureus* VET0632S |
| Sau-3363-1 | Bacteria Firmicutes Bacilli Bacillales Staphylococcaceae *Staphylococcus aureus* VET0645S |
| Sau-3364-1 | Bacteria Firmicutes Bacilli Bacillales Staphylococcaceae *Staphylococcus aureus* VET0657S |
| Sau-3365-1 | Bacteria Firmicutes Bacilli Bacillales Staphylococcaceae *Staphylococcus aureus* VET0678S |
| Sau-3366-1 | Bacteria Firmicutes Bacilli Bacillales Staphylococcaceae *Staphylococcus aureus* VET0679S |
| Sau-3367-1 | Bacteria Firmicutes Bacilli Bacillales Staphylococcaceae *Staphylococcus aureus* VET0765S |
| Sau-3368-1 | Bacteria Firmicutes Bacilli Bacillales Staphylococcaceae *Staphylococcus aureus* VET0779S |
| Sau-3369-1 | Bacteria Firmicutes Bacilli Bacillales Staphylococcaceae *Staphylococcus aureus* VET0787S |
| Sau-3370-1 | Bacteria Firmicutes Bacilli Bacillales Staphylococcaceae *Staphylococcus aureus* VET0798S |
| Sau-3371-1 | Bacteria Firmicutes Bacilli Bacillales Staphylococcaceae *Staphylococcus aureus* VET0809S |
| Sau-3372-1 | Bacteria Firmicutes Bacilli Bacillales Staphylococcaceae *Staphylococcus aureus* VET0810S |
| Sau-3373-1 | Bacteria Firmicutes Bacilli Bacillales Staphylococcaceae *Staphylococcus aureus* VET0889S |
| Sau-3374-1 | Bacteria Firmicutes Bacilli Bacillales Staphylococcaceae *Staphylococcus aureus* VET0919S |
| Sau-3375-1 | Bacteria Firmicutes Bacilli Bacillales Staphylococcaceae *Staphylococcus aureus* VET0920S |
| Sau-3376-1 | Bacteria Firmicutes Bacilli Bacillales Staphylococcaceae *Staphylococcus aureus* VET1020S |
| Sau-3377-1 | Bacteria Firmicutes Bacilli Bacillales Staphylococcaceae *Staphylococcus aureus* VET1024S |
| Sau-3378-1 | Bacteria Firmicutes Bacilli Bacillales Staphylococcaceae *Staphylococcus aureus* VET1025S |
| Sau-3379-1 | Bacteria Firmicutes Bacilli Bacillales Staphylococcaceae *Staphylococcus aureus* VET1026S |
| Sau-3380-1 | Bacteria Firmicutes Bacilli Bacillales Staphylococcaceae *Staphylococcus aureus* VET1035S |
| Sau-3381-1 | Bacteria Firmicutes Bacilli Bacillales Staphylococcaceae *Staphylococcus aureus* VET1048S |
| Sau-3382-1 | Bacteria Firmicutes Bacilli Bacillales Staphylococcaceae *Staphylococcus aureus* VET1052S |
| Sau-3383-1 | Bacteria Firmicutes Bacilli Bacillales Staphylococcaceae *Staphylococcus aureus* VET1054S |
| Sau-3384-1 | Bacteria Firmicutes Bacilli Bacillales Staphylococcaceae *Staphylococcus aureus* VET1068S |
| Sau-3385-1 | Bacteria Firmicutes Bacilli Bacillales Staphylococcaceae *Staphylococcus aureus* VET1088S |
| Sau-3386-1 | Bacteria Firmicutes Bacilli Bacillales Staphylococcaceae *Staphylococcus aureus* VET1103S |
| Sau-3387-1 | Bacteria Firmicutes Bacilli Bacillales Staphylococcaceae *Staphylococcus aureus* VET1104S |
| Sau-3388-1 | Bacteria Firmicutes Bacilli Bacillales Staphylococcaceae *Staphylococcus aureus* VET1168S |
| Sau-3389-1 | Bacteria Firmicutes Bacilli Bacillales Staphylococcaceae *Staphylococcus aureus* VET1181S |
| Sau-3390-1 | Bacteria Firmicutes Bacilli Bacillales Staphylococcaceae *Staphylococcus aureus* VET1245S |
| Sau-3391-1 | Bacteria Firmicutes Bacilli Bacillales Staphylococcaceae *Staphylococcus aureus* VET1368S |
| Sau-3392-1 | Bacteria Firmicutes Bacilli Bacillales Staphylococcaceae *Staphylococcus aureus* VET1373S |
| Sau-3393-1 | Bacteria Firmicutes Bacilli Bacillales Staphylococcaceae *Staphylococcus aureus* VET1411S |
| Sau-3394-1 | Bacteria Firmicutes Bacilli Bacillales Staphylococcaceae *Staphylococcus aureus* VET1419S |
| Sau-3395-1 | Bacteria Firmicutes Bacilli Bacillales Staphylococcaceae *Staphylococcus aureus* VET1422S |
| Sau-3396-1 | Bacteria Firmicutes Bacilli Bacillales Staphylococcaceae *Staphylococcus aureus* VET1434S |
| Sau-3397-1 | Bacteria Firmicutes Bacilli Bacillales Staphylococcaceae *Staphylococcus aureus* VET1468S |
| Sau-3398-1 | Bacteria Firmicutes Bacilli Bacillales Staphylococcaceae *Staphylococcus aureus* VET1495S |
| Sau-3399-1 | Bacteria Firmicutes Bacilli Bacillales Staphylococcaceae *Staphylococcus aureus* VET1499S |

| | |
|---|---|
| Sau-3400-1 | Bacteria Firmicutes Bacilli Bacillales Staphylococcaceae *Staphylococcus aureus* VET1515S |
| Sau-3401-1 | Bacteria Firmicutes Bacilli Bacillales Staphylococcaceae *Staphylococcus aureus* VET1518S |
| Sau-3402-1 | Bacteria Firmicutes Bacilli Bacillales Staphylococcaceae *Staphylococcus aureus* VET1519S |
| Sau-3403-1 | Bacteria Firmicutes Bacilli Bacillales Staphylococcaceae *Staphylococcus aureus* VET1526S |
| Sau-3404-1 | Bacteria Firmicutes Bacilli Bacillales Staphylococcaceae *Staphylococcus aureus* VET1646S |
| Sau-3405-1 | Bacteria Firmicutes Bacilli Bacillales Staphylococcaceae *Staphylococcus aureus* VET1653S |
| Sau-3406-1 | Bacteria Firmicutes Bacilli Bacillales Staphylococcaceae *Staphylococcus aureus* VET1727S |
| Sau-3407-1 | Bacteria Firmicutes Bacilli Bacillales Staphylococcaceae *Staphylococcus aureus* VET1738S |
| Sau-3408-1 | Bacteria Firmicutes Bacilli Bacillales Staphylococcaceae *Staphylococcus aureus* VET1779S |
| Sau-3409-1 | Bacteria Firmicutes Bacilli Bacillales Staphylococcaceae *Staphylococcus aureus* VET1816S |
| Sau-3410-1 | Bacteria Firmicutes Bacilli Bacillales Staphylococcaceae *Staphylococcus aureus* VET1817S |
| Sau-3411-1 | Bacteria Firmicutes Bacilli Bacillales Staphylococcaceae *Staphylococcus aureus* VET1822S |
| Sau-3412-1 | Bacteria Firmicutes Bacilli Bacillales Staphylococcaceae *Staphylococcus aureus* VET1831R |
| Sau-3413-1 | Bacteria Firmicutes Bacilli Bacillales Staphylococcaceae *Staphylococcus aureus* VET1832R |
| Sau-3414-1 | Bacteria Firmicutes Bacilli Bacillales Staphylococcaceae *Staphylococcus aureus* VET1833R |
| Sau-3415-1 | Bacteria Firmicutes Bacilli Bacillales Staphylococcaceae *Staphylococcus aureus* VET1834R |
| Sau-3416-1 | Bacteria Firmicutes Bacilli Bacillales Staphylococcaceae *Staphylococcus aureus* VET1835R |
| Sau-3417-1 | Bacteria Firmicutes Bacilli Bacillales Staphylococcaceae *Staphylococcus aureus* VET1838R |
| Sau-3418-1 | Bacteria Firmicutes Bacilli Bacillales Staphylococcaceae *Staphylococcus aureus* VET1839R |
| Sau-3419-1 | Bacteria Firmicutes Bacilli Bacillales Staphylococcaceae *Staphylococcus aureus* VET1842R |
| Sau-3420-1 | Bacteria Firmicutes Bacilli Bacillales Staphylococcaceae *Staphylococcus aureus* VET1843R |
| Sau-3421-1 | Bacteria Firmicutes Bacilli Bacillales Staphylococcaceae *Staphylococcus aureus* VET1844R |
| Sau-3422-1 | Bacteria Firmicutes Bacilli Bacillales Staphylococcaceae *Staphylococcus aureus* VET1845R |
| Sau-3423-1 | Bacteria Firmicutes Bacilli Bacillales Staphylococcaceae *Staphylococcus aureus* VET1846R |
| Sau-3424-1 | Bacteria Firmicutes Bacilli Bacillales Staphylococcaceae *Staphylococcus aureus* VET1848R |
| Sau-3425-1 | Bacteria Firmicutes Bacilli Bacillales Staphylococcaceae *Staphylococcus aureus* VET1849R |
| Sau-3426-1 | Bacteria Firmicutes Bacilli Bacillales Staphylococcaceae *Staphylococcus aureus* VET1850R |
| Sau-3427-1 | Bacteria Firmicutes Bacilli Bacillales Staphylococcaceae *Staphylococcus aureus* VET1851R |
| Sau-3428-1 | Bacteria Firmicutes Bacilli Bacillales Staphylococcaceae *Staphylococcus aureus* VET1852R |
| Sau-3429-1 | Bacteria Firmicutes Bacilli Bacillales Staphylococcaceae *Staphylococcus aureus* VET1853R |
| Sau-3430-1 | Bacteria Firmicutes Bacilli Bacillales Staphylococcaceae *Staphylococcus aureus* VET1854R |
| Sau-3431-1 | Bacteria Firmicutes Bacilli Bacillales Staphylococcaceae *Staphylococcus aureus* VET1855R |
| Sau-3432-1 | Bacteria Firmicutes Bacilli Bacillales Staphylococcaceae *Staphylococcus aureus* VET1856R |
| Sau-3433-1 | Bacteria Firmicutes Bacilli Bacillales Staphylococcaceae *Staphylococcus aureus* VET1858R |
| Sau-3434-1 | Bacteria Firmicutes Bacilli Bacillales Staphylococcaceae *Staphylococcus aureus* VET1859R |
| Sau-3435-1 | Bacteria Firmicutes Bacilli Bacillales Staphylococcaceae *Staphylococcus aureus* VET1860R |
| Sau-3436-1 | Bacteria Firmicutes Bacilli Bacillales Staphylococcaceae *Staphylococcus aureus* VET1861R |
| Sau-3437-1 | Bacteria Firmicutes Bacilli Bacillales Staphylococcaceae *Staphylococcus aureus* VET1865R |
| Sau-3438-1 | Bacteria Firmicutes Bacilli Bacillales Staphylococcaceae *Staphylococcus aureus* VET1866R |
| Sau-3439-1 | Bacteria Firmicutes Bacilli Bacillales Staphylococcaceae *Staphylococcus aureus* VET1867R |
| Sau-3440-1 | Bacteria Firmicutes Bacilli Bacillales Staphylococcaceae *Staphylococcus aureus* VET1868R |
| Sau-3441-1 | Bacteria Firmicutes Bacilli Bacillales Staphylococcaceae *Staphylococcus aureus* VET1869R |
| Sau-3442-1 | Bacteria Firmicutes Bacilli Bacillales Staphylococcaceae *Staphylococcus aureus* VET1871R |
| Sau-3443-1 | Bacteria Firmicutes Bacilli Bacillales Staphylococcaceae *Staphylococcus aureus* VET1872R |
| Sau-3444-1 | Bacteria Firmicutes Bacilli Bacillales Staphylococcaceae *Staphylococcus aureus* VET1873R |
| Sau-3445-1 | Bacteria Firmicutes Bacilli Bacillales Staphylococcaceae *Staphylococcus aureus* VET1875R |
| Sau-3446-1 | Bacteria Firmicutes Bacilli Bacillales Staphylococcaceae *Staphylococcus aureus* VET1876R |
| Sau-3447-1 | Bacteria Firmicutes Bacilli Bacillales Staphylococcaceae *Staphylococcus aureus* VET1877R |
| Sau-3448-1 | Bacteria Firmicutes Bacilli Bacillales Staphylococcaceae *Staphylococcus aureus* VET1880R |
| Sau-3449-1 | Bacteria Firmicutes Bacilli Bacillales Staphylococcaceae *Staphylococcus aureus* VET1884R |
| Sau-3450-1 | Bacteria Firmicutes Bacilli Bacillales Staphylococcaceae *Staphylococcus aureus* VET1886R |
| Sau-3451-1 | Bacteria Firmicutes Bacilli Bacillales Staphylococcaceae *Staphylococcus aureus* VET1890R |
| Sau-3452-1 | Bacteria Firmicutes Bacilli Bacillales Staphylococcaceae *Staphylococcus aureus* VET1892R |
| Sau-3453-1 | Bacteria Firmicutes Bacilli Bacillales Staphylococcaceae *Staphylococcus aureus* VET1893R |
| Sau-3454-1 | Bacteria Firmicutes Bacilli Bacillales Staphylococcaceae *Staphylococcus aureus* VET1895R |
| Sau-3455-1 | Bacteria Firmicutes Bacilli Bacillales Staphylococcaceae *Staphylococcus aureus* VET1896R |
| Sau-3456-1 | Bacteria Firmicutes Bacilli Bacillales Staphylococcaceae *Staphylococcus aureus* VET1897R |
| Sau-3457-1 | Bacteria Firmicutes Bacilli Bacillales Staphylococcaceae *Staphylococcus aureus* VET1898R |
| Sau-3458-1 | Bacteria Firmicutes Bacilli Bacillales Staphylococcaceae *Staphylococcus aureus* VET1899R |
| Sau-3459-1 | Bacteria Firmicutes Bacilli Bacillales Staphylococcaceae *Staphylococcus aureus* VET1901R |
| Sau-3460-1 | Bacteria Firmicutes Bacilli Bacillales Staphylococcaceae *Staphylococcus aureus* VET1904R |
| Sau-3461-1 | Bacteria Firmicutes Bacilli Bacillales Staphylococcaceae *Staphylococcus aureus* VET1905R |
| Sau-3462-1 | Bacteria Firmicutes Bacilli Bacillales Staphylococcaceae *Staphylococcus aureus* VET1906R |
| Sau-3463-1 | Bacteria Firmicutes Bacilli Bacillales Staphylococcaceae *Staphylococcus aureus* VET1907R |
| Sau-3464-1 | Bacteria Firmicutes Bacilli Bacillales Staphylococcaceae *Staphylococcus aureus* VET1908R |

| | |
|---|---|
| Sau-3465-1 | Bacteria Firmicutes Bacilli Bacillales Staphylococcaceae *Staphylococcus aureus* VET1909R |
| Sau-3466-1 | Bacteria Firmicutes Bacilli Bacillales Staphylococcaceae *Staphylococcus aureus* VET1910R |
| Sau-3467-1 | Bacteria Firmicutes Bacilli Bacillales Staphylococcaceae *Staphylococcus aureus* VET1911R |
| Sau-3468-1 | Bacteria Firmicutes Bacilli Bacillales Staphylococcaceae *Staphylococcus aureus* VET1912R |
| Sau-3469-1 | Bacteria Firmicutes Bacilli Bacillales Staphylococcaceae *Staphylococcus aureus* VET1913R |
| Sau-3470-1 | Bacteria Firmicutes Bacilli Bacillales Staphylococcaceae *Staphylococcus aureus* VET1914R |
| Sau-3471-1 | Bacteria Firmicutes Bacilli Bacillales Staphylococcaceae *Staphylococcus aureus* VET1915R |
| Sau-3472-1 | Bacteria Firmicutes Bacilli Bacillales Staphylococcaceae *Staphylococcus aureus* VET1917S |
| Sau-3473-1 | Bacteria Firmicutes Bacilli Bacillales Staphylococcaceae *Staphylococcus aureus* VET1918S |
| Sau-3474-1 | Bacteria Firmicutes Bacilli Bacillales Staphylococcaceae *Staphylococcus aureus* VET1923S |
| Sau-3475-1 | Bacteria Firmicutes Bacilli Bacillales Staphylococcaceae *Staphylococcus aureus* VET1949S |
| Sau-3476-1 | Bacteria Firmicutes Bacilli Bacillales Staphylococcaceae *Staphylococcus aureus* VET1950S |
| Sau-3477-1 | Bacteria Firmicutes Bacilli Bacillales Staphylococcaceae *Staphylococcus aureus* VET1956S |
| Sau-3478-1 | Bacteria Firmicutes Bacilli Bacillales Staphylococcaceae *Staphylococcus aureus* VET1959S |
| Sau-3479-1 | Bacteria Firmicutes Bacilli Bacillales Staphylococcaceae *Staphylococcus aureus* W12461 |
| Sau-3480-1 | Bacteria Firmicutes Bacilli Bacillales Staphylococcaceae *Staphylococcus aureus* W12538 |
| Sau-3481-1 | Bacteria Firmicutes Bacilli Bacillales Staphylococcaceae *Staphylococcus aureus* W12582 |
| Sau-3482-1 | Bacteria Firmicutes Bacilli Bacillales Staphylococcaceae *Staphylococcus aureus* W12583 |
| Sau-3483-1 | Bacteria Firmicutes Bacilli Bacillales Staphylococcaceae *Staphylococcus aureus* W12586 |
| Sau-3484-1 | Bacteria Firmicutes Bacilli Bacillales Staphylococcaceae *Staphylococcus aureus* W12628 |
| Sau-3485-1 | Bacteria Firmicutes Bacilli Bacillales Staphylococcaceae *Staphylococcus aureus* W13012_022013 |
| Sau-3486-1 | Bacteria Firmicutes Bacilli Bacillales Staphylococcaceae *Staphylococcus aureus* W15951 |
| Sau-3487-1 | Bacteria Firmicutes Bacilli Bacillales Staphylococcaceae *Staphylococcus aureus* W15974 |
| Sau-3488-1 | Bacteria Firmicutes Bacilli Bacillales Staphylococcaceae *Staphylococcus aureus* W15976 |
| Sau-3489-1 | Bacteria Firmicutes Bacilli Bacillales Staphylococcaceae *Staphylococcus aureus* W15990 |
| Sau-3490-1 | Bacteria Firmicutes Bacilli Bacillales Staphylococcaceae *Staphylococcus aureus* W15995 |
| Sau-3491-1 | Bacteria Firmicutes Bacilli Bacillales Staphylococcaceae *Staphylococcus aureus* W15997 |
| Sau-3492-1 | Bacteria Firmicutes Bacilli Bacillales Staphylococcaceae *Staphylococcus aureus* W16001 |
| Sau-3493-1 | Bacteria Firmicutes Bacilli Bacillales Staphylococcaceae *Staphylococcus aureus* W16013 |
| Sau-3494-1 | Bacteria Firmicutes Bacilli Bacillales Staphylococcaceae *Staphylococcus aureus* W16028 |
| Sau-3495-1 | Bacteria Firmicutes Bacilli Bacillales Staphylococcaceae *Staphylococcus aureus* W18442 |
| Sau-3496-1 | Bacteria Firmicutes Bacilli Bacillales Staphylococcaceae *Staphylococcus aureus* W18443 |
| Sau-3497-1 | Bacteria Firmicutes Bacilli Bacillales Staphylococcaceae *Staphylococcus aureus* W19220 |
| Sau-3498-1 | Bacteria Firmicutes Bacilli Bacillales Staphylococcaceae *Staphylococcus aureus* W19225 |
| Sau-3499-1 | Bacteria Firmicutes Bacilli Bacillales Staphylococcaceae *Staphylococcus aureus* W19256 |
| Sau-3500-1 | Bacteria Firmicutes Bacilli Bacillales Staphylococcaceae *Staphylococcus aureus* W19270 |
| Sau-3501-1 | Bacteria Firmicutes Bacilli Bacillales Staphylococcaceae *Staphylococcus aureus* W19276 |
| Sau-3502-1 | Bacteria Firmicutes Bacilli Bacillales Staphylococcaceae *Staphylococcus aureus* W20096 |
| Sau-3503-1 | Bacteria Firmicutes Bacilli Bacillales Staphylococcaceae *Staphylococcus aureus* W20433 |
| Sau-3504-1 | Bacteria Firmicutes Bacilli Bacillales Staphylococcaceae *Staphylococcus aureus* W21472 |
| Sau-3505-1 | Bacteria Firmicutes Bacilli Bacillales Staphylococcaceae *Staphylococcus aureus* W21479 |
| Sau-3506-1 | Bacteria Firmicutes Bacilli Bacillales Staphylococcaceae *Staphylococcus aureus* W21932 |
| Sau-3507-1 | Bacteria Firmicutes Bacilli Bacillales Staphylococcaceae *Staphylococcus aureus* W21940 |
| Sau-3508-1 | Bacteria Firmicutes Bacilli Bacillales Staphylococcaceae *Staphylococcus aureus* W21945 |
| Sau-3509-1 | Bacteria Firmicutes Bacilli Bacillales Staphylococcaceae *Staphylococcus aureus* W23207 |
| Sau-3510-1 | Bacteria Firmicutes Bacilli Bacillales Staphylococcaceae *Staphylococcus aureus* W23214 |
| Sau-3511-1 | Bacteria Firmicutes Bacilli Bacillales Staphylococcaceae *Staphylococcus aureus* W23225 |
| Sau-3512-1 | Bacteria Firmicutes Bacilli Bacillales Staphylococcaceae *Staphylococcus aureus* W24177 |
| Sau-3513-1 | Bacteria Firmicutes Bacilli Bacillales Staphylococcaceae *Staphylococcus aureus* W24216 |
| Sau-3514-1 | Bacteria Firmicutes Bacilli Bacillales Staphylococcaceae *Staphylococcus aureus* W25097 |
| Sau-3515-1 | Bacteria Firmicutes Bacilli Bacillales Staphylococcaceae *Staphylococcus aureus* W25787 |
| Sau-3516-1 | Bacteria Firmicutes Bacilli Bacillales Staphylococcaceae *Staphylococcus aureus* W25797 |
| Sau-3517-1 | Bacteria Firmicutes Bacilli Bacillales Staphylococcaceae *Staphylococcus aureus* W25799 |
| Sau-3518-1 | Bacteria Firmicutes Bacilli Bacillales Staphylococcaceae *Staphylococcus aureus* W25814 |
| Sau-3519-1 | Bacteria Firmicutes Bacilli Bacillales Staphylococcaceae *Staphylococcus aureus* W27852 |
| Sau-3520-1 | Bacteria Firmicutes Bacilli Bacillales Staphylococcaceae *Staphylococcus aureus* W28469 |
| Sau-3521-1 | Bacteria Firmicutes Bacilli Bacillales Staphylococcaceae *Staphylococcus aureus* W28921 |
| Sau-3522-1 | Bacteria Firmicutes Bacilli Bacillales Staphylococcaceae *Staphylococcus aureus* W29625 |
| Sau-3523-1 | Bacteria Firmicutes Bacilli Bacillales Staphylococcaceae *Staphylococcus aureus* W29641 |
| Sau-3524-1 | Bacteria Firmicutes Bacilli Bacillales Staphylococcaceae *Staphylococcus aureus* W29651 |
| Sau-3525-1 | Bacteria Firmicutes Bacilli Bacillales Staphylococcaceae *Staphylococcus aureus* W31852 |
| Sau-3526-1 | Bacteria Firmicutes Bacilli Bacillales Staphylococcaceae *Staphylococcus aureus* W31862 |
| Sau-3527-1 | Bacteria Firmicutes Bacilli Bacillales Staphylococcaceae *Staphylococcus aureus* W31936 |
| Sau-3528-1 | Bacteria Firmicutes Bacilli Bacillales Staphylococcaceae *Staphylococcus aureus* W31947 |
| Sau-3529-1 | Bacteria Firmicutes Bacilli Bacillales Staphylococcaceae *Staphylococcus aureus* W31983 |

| | |
|---|---|
| Sau-3530-1 | Bacteria Firmicutes Bacilli Bacillales Staphylococcaceae *Staphylococcus aureus* W32014 |
| Sau-3531-1 | Bacteria Firmicutes Bacilli Bacillales Staphylococcaceae *Staphylococcus aureus* W33563 |
| Sau-3532-1 | Bacteria Firmicutes Bacilli Bacillales Staphylococcaceae *Staphylococcus aureus* W33578 |
| Sau-3533-1 | Bacteria Firmicutes Bacilli Bacillales Staphylococcaceae *Staphylococcus aureus* W33592 |
| Sau-3534-1 | Bacteria Firmicutes Bacilli Bacillales Staphylococcaceae *Staphylococcus aureus* W33796 |
| Sau-3535-1 | Bacteria Firmicutes Bacilli Bacillales Staphylococcaceae *Staphylococcus aureus* W34837 |
| Sau-3536-1 | Bacteria Firmicutes Bacilli Bacillales Staphylococcaceae *Staphylococcus aureus* W34839 |
| Sau-3537-1 | Bacteria Firmicutes Bacilli Bacillales Staphylococcaceae *Staphylococcus aureus* W34841 |
| Sau-3538-1 | Bacteria Firmicutes Bacilli Bacillales Staphylococcaceae *Staphylococcus aureus* W34848 |
| Sau-3539-1 | Bacteria Firmicutes Bacilli Bacillales Staphylococcaceae *Staphylococcus aureus* W35325 |
| Sau-3540-1 | Bacteria Firmicutes Bacilli Bacillales Staphylococcaceae *Staphylococcus aureus* W35356 |
| Sau-3541-1 | Bacteria Firmicutes Bacilli Bacillales Staphylococcaceae *Staphylococcus aureus* W35455 |
| Sau-3542-1 | Bacteria Firmicutes Bacilli Bacillales Staphylococcaceae *Staphylococcus aureus* W36553 |
| Sau-3543-1 | Bacteria Firmicutes Bacilli Bacillales Staphylococcaceae *Staphylococcus aureus* W36570 |
| Sau-3544-1 | Bacteria Firmicutes Bacilli Bacillales Staphylococcaceae *Staphylococcus aureus* W38532 |
| Sau-3545-1 | Bacteria Firmicutes Bacilli Bacillales Staphylococcaceae *Staphylococcus aureus* W38597 |
| Sau-3546-1 | Bacteria Firmicutes Bacilli Bacillales Staphylococcaceae *Staphylococcus aureus* W39821 |
| Sau-3547-1 | Bacteria Firmicutes Bacilli Bacillales Staphylococcaceae *Staphylococcus aureus* W39826 |
| Sau-3548-1 | Bacteria Firmicutes Bacilli Bacillales Staphylococcaceae *Staphylococcus aureus* W39902 |
| Sau-3549-1 | Bacteria Firmicutes Bacilli Bacillales Staphylococcaceae *Staphylococcus aureus* W41757 |
| Sau-3550-1 | Bacteria Firmicutes Bacilli Bacillales Staphylococcaceae *Staphylococcus aureus* W41765 |
| Sau-3551-1 | Bacteria Firmicutes Bacilli Bacillales Staphylococcaceae *Staphylococcus aureus* W41914 |
| Sau-3552-1 | Bacteria Firmicutes Bacilli Bacillales Staphylococcaceae *Staphylococcus aureus* W41918 |
| Sau-3553-1 | Bacteria Firmicutes Bacilli Bacillales Staphylococcaceae *Staphylococcus aureus* W42196 |
| Sau-3554-1 | Bacteria Firmicutes Bacilli Bacillales Staphylococcaceae *Staphylococcus aureus* W42255 |
| Sau-3555-1 | Bacteria Firmicutes Bacilli Bacillales Staphylococcaceae *Staphylococcus aureus* W42270 |
| Sau-3556-1 | Bacteria Firmicutes Bacilli Bacillales Staphylococcaceae *Staphylococcus aureus* W42298_110712 |
| Sau-3557-1 | Bacteria Firmicutes Bacilli Bacillales Staphylococcaceae *Staphylococcus aureus* W42304 |
| Sau-3558-1 | Bacteria Firmicutes Bacilli Bacillales Staphylococcaceae *Staphylococcus aureus* W42929 |
| Sau-3559-1 | Bacteria Firmicutes Bacilli Bacillales Staphylococcaceae *Staphylococcus aureus* W42932 |
| Sau-3560-1 | Bacteria Firmicutes Bacilli Bacillales Staphylococcaceae *Staphylococcus aureus* W45421 |
| Sau-3561-1 | Bacteria Firmicutes Bacilli Bacillales Staphylococcaceae *Staphylococcus aureus* W45747 |
| Sau-3562-1 | Bacteria Firmicutes Bacilli Bacillales Staphylococcaceae *Staphylococcus aureus* W45750 |
| Sau-3563-1 | Bacteria Firmicutes Bacilli Bacillales Staphylococcaceae *Staphylococcus aureus* W45755_111412 |
| Sau-3564-1 | Bacteria Firmicutes Bacilli Bacillales Staphylococcaceae *Staphylococcus aureus* W45757 |
| Sau-3565-1 | Bacteria Firmicutes Bacilli Bacillales Staphylococcaceae *Staphylococcus aureus* W45758 |
| Sau-3566-1 | Bacteria Firmicutes Bacilli Bacillales Staphylococcaceae *Staphylococcus aureus* W46839 |
| Sau-3567-1 | Bacteria Firmicutes Bacilli Bacillales Staphylococcaceae *Staphylococcus aureus* W46883 |
| Sau-3568-1 | Bacteria Firmicutes Bacilli Bacillales Staphylococcaceae *Staphylococcus aureus* W48872 |
| Sau-3569-1 | Bacteria Firmicutes Bacilli Bacillales Staphylococcaceae *Staphylococcus aureus* W48906 |
| Sau-3570-1 | Bacteria Firmicutes Bacilli Bacillales Staphylococcaceae *Staphylococcus aureus* W48969 |
| Sau-3571-1 | Bacteria Firmicutes Bacilli Bacillales Staphylococcaceae *Staphylococcus aureus* W48973 |
| Sau-3572-1 | Bacteria Firmicutes Bacilli Bacillales Staphylococcaceae *Staphylococcus aureus* W50092 |
| Sau-3573-1 | Bacteria Firmicutes Bacilli Bacillales Staphylococcaceae *Staphylococcus aureus* W50101 |
| Sau-3574-1 | Bacteria Firmicutes Bacilli Bacillales Staphylococcaceae *Staphylococcus aureus* W50110 |
| Sau-3575-1 | Bacteria Firmicutes Bacilli Bacillales Staphylococcaceae *Staphylococcus aureus* W51776 |
| Sau-3576-1 | Bacteria Firmicutes Bacilli Bacillales Staphylococcaceae *Staphylococcus aureus* W52104 |
| Sau-3577-1 | Bacteria Firmicutes Bacilli Bacillales Staphylococcaceae *Staphylococcus aureus* W52117 |
| Sau-3578-1 | Bacteria Firmicutes Bacilli Bacillales Staphylococcaceae *Staphylococcus aureus* W52120 |
| Sau-3579-1 | Bacteria Firmicutes Bacilli Bacillales Staphylococcaceae *Staphylococcus aureus* W52294 |
| Sau-3580-1 | Bacteria Firmicutes Bacilli Bacillales Staphylococcaceae *Staphylococcus aureus* W52298 |
| Sau-3581-1 | Bacteria Firmicutes Bacilli Bacillales Staphylococcaceae *Staphylococcus aureus* W53395 |
| Sau-3582-1 | Bacteria Firmicutes Bacilli Bacillales Staphylococcaceae *Staphylococcus aureus* W55442 |
| Sau-3583-1 | Bacteria Firmicutes Bacilli Bacillales Staphylococcaceae *Staphylococcus aureus* W55446 |
| Sau-3584-1 | Bacteria Firmicutes Bacilli Bacillales Staphylococcaceae *Staphylococcus aureus* W55451 |
| Sau-3585-1 | Bacteria Firmicutes Bacilli Bacillales Staphylococcaceae *Staphylococcus aureus* W55536 |
| Sau-3586-1 | Bacteria Firmicutes Bacilli Bacillales Staphylococcaceae *Staphylococcus aureus* W56227 |
| Sau-3587-1 | Bacteria Firmicutes Bacilli Bacillales Staphylococcaceae *Staphylococcus aureus* W56243 |
| Sau-3588-1 | Bacteria Firmicutes Bacilli Bacillales Staphylococcaceae *Staphylococcus aureus* W56246 |
| Sau-3589-1 | Bacteria Firmicutes Bacilli Bacillales Staphylococcaceae *Staphylococcus aureus* W56912 |
| Sau-3590-1 | Bacteria Firmicutes Bacilli Bacillales Staphylococcaceae *Staphylococcus aureus* W57390 |
| Sau-3591-1 | Bacteria Firmicutes Bacilli Bacillales Staphylococcaceae *Staphylococcus aureus* W58473 |
| Sau-3592-1 | Bacteria Firmicutes Bacilli Bacillales Staphylococcaceae *Staphylococcus aureus* W60060 |
| Sau-3593-1 | Bacteria Firmicutes Bacilli Bacillales Staphylococcaceae *Staphylococcus aureus* W60774 |
| Sau-3594-1 | Bacteria Firmicutes Bacilli Bacillales Staphylococcaceae *Staphylococcus aureus* W60784 |

| | |
|---|---|
| Sau-3595-1 | Bacteria Firmicutes Bacilli Bacillales Staphylococcaceae *Staphylococcus aureus* W60801 |
| Sau-3596-1 | Bacteria Firmicutes Bacilli Bacillales Staphylococcaceae *Staphylococcus aureus* W62344 |
| Sau-3597-1 | Bacteria Firmicutes Bacilli Bacillales Staphylococcaceae *Staphylococcus aureus* W62354 |
| Sau-3598-1 | Bacteria Firmicutes Bacilli Bacillales Staphylococcaceae *Staphylococcus aureus* W62934 |
| Sau-3599-1 | Bacteria Firmicutes Bacilli Bacillales Staphylococcaceae *Staphylococcus aureus* W62972 |
| Sau-3600-1 | Bacteria Firmicutes Bacilli Bacillales Staphylococcaceae *Staphylococcus aureus* W64921 |
| Sau-3601-1 | Bacteria Firmicutes Bacilli Bacillales Staphylococcaceae *Staphylococcus aureus* W64953 |
| Sau-3602-1 | Bacteria Firmicutes Bacilli Bacillales Staphylococcaceae *Staphylococcus aureus* W64958 |
| Sau-3603-1 | Bacteria Firmicutes Bacilli Bacillales Staphylococcaceae *Staphylococcus aureus* W65865 |
| Sau-3604-1 | Bacteria Firmicutes Bacilli Bacillales Staphylococcaceae *Staphylococcus aureus* W65885 |
| Sau-3605-1 | Bacteria Firmicutes Bacilli Bacillales Staphylococcaceae *Staphylococcus aureus* W65904 |
| Sau-3606-1 | Bacteria Firmicutes Bacilli Bacillales Staphylococcaceae *Staphylococcus aureus* W65910 |
| Sau-3607-1 | Bacteria Firmicutes Bacilli Bacillales Staphylococcaceae *Staphylococcus aureus* W68479 |
| Sau-3608-1 | Bacteria Firmicutes Bacilli Bacillales Staphylococcaceae *Staphylococcus aureus* W70568 |
| Sau-3609-1 | Bacteria Firmicutes Bacilli Bacillales Staphylococcaceae *Staphylococcus aureus* W70590 |
| Sau-3610-1 | Bacteria Firmicutes Bacilli Bacillales Staphylococcaceae *Staphylococcus aureus* W70599 |
| Sau-3611-1 | Bacteria Firmicutes Bacilli Bacillales Staphylococcaceae *Staphylococcus aureus* W70659 |
| Sau-3612-1 | Bacteria Firmicutes Bacilli Bacillales Staphylococcaceae *Staphylococcus aureus* W73060 |
| Sau-3613-1 | Bacteria Firmicutes Bacilli Bacillales Staphylococcaceae *Staphylococcus aureus* W73592 |
| Sau-3614-1 | Bacteria Firmicutes Bacilli Bacillales Staphylococcaceae *Staphylococcus aureus* W73601 |
| Sau-3615-1 | Bacteria Firmicutes Bacilli Bacillales Staphylococcaceae *Staphylococcus aureus* W73649 |
| Sau-3616-1 | Bacteria Firmicutes Bacilli Bacillales Staphylococcaceae *Staphylococcus aureus* W73676 |
| Sau-3617-1 | Bacteria Firmicutes Bacilli Bacillales Staphylococcaceae *Staphylococcus aureus* W73738 |
| Sau-3618-1 | Bacteria Firmicutes Bacilli Bacillales Staphylococcaceae *Staphylococcus aureus* W75098 |
| Sau-3619-1 | Bacteria Firmicutes Bacilli Bacillales Staphylococcaceae *Staphylococcus aureus* W75113 |
| Sau-3620-1 | Bacteria Firmicutes Bacilli Bacillales Staphylococcaceae *Staphylococcus aureus* W75119 |
| Sau-3621-1 | Bacteria Firmicutes Bacilli Bacillales Staphylococcaceae *Staphylococcus aureus* W75132 |
| Sau-3622-1 | Bacteria Firmicutes Bacilli Bacillales Staphylococcaceae *Staphylococcus aureus* W75229 |
| Sau-3623-1 | Bacteria Firmicutes Bacilli Bacillales Staphylococcaceae *Staphylococcus aureus* W75234 |
| Sau-3624-1 | Bacteria Firmicutes Bacilli Bacillales Staphylococcaceae *Staphylococcus aureus* W75242 |
| Sau-3625-1 | Bacteria Firmicutes Bacilli Bacillales Staphylococcaceae *Staphylococcus aureus* W75262 |
| Sau-3626-1 | Bacteria Firmicutes Bacilli Bacillales Staphylococcaceae *Staphylococcus aureus* W75539 |
| Sau-3627-1 | Bacteria Firmicutes Bacilli Bacillales Staphylococcaceae *Staphylococcus aureus* W76127 |
| Sau-3628-1 | Bacteria Firmicutes Bacilli Bacillales Staphylococcaceae *Staphylococcus aureus* W76930 |
| Sau-3629-1 | Bacteria Firmicutes Bacilli Bacillales Staphylococcaceae *Staphylococcus aureus* W76994_072512 |
| Sau-3630-1 | Bacteria Firmicutes Bacilli Bacillales Staphylococcaceae *Staphylococcus aureus* W76995 |
| Sau-3631-1 | Bacteria Firmicutes Bacilli Bacillales Staphylococcaceae *Staphylococcus aureus* W79328 |
| Sau-3632-1 | Bacteria Firmicutes Bacilli Bacillales Staphylococcaceae *Staphylococcus aureus* W79968 |
| Sau-3633-1 | Bacteria Firmicutes Bacilli Bacillales Staphylococcaceae *Staphylococcus aureus* W79990_080112 |
| Sau-3634-1 | Bacteria Firmicutes Bacilli Bacillales Staphylococcaceae *Staphylococcus aureus* W80011 |
| Sau-3635-1 | Bacteria Firmicutes Bacilli Bacillales Staphylococcaceae *Staphylococcus aureus* W80013 |
| Sau-3636-1 | Bacteria Firmicutes Bacilli Bacillales Staphylococcaceae *Staphylococcus aureus* W80041 |
| Sau-3637-1 | Bacteria Firmicutes Bacilli Bacillales Staphylococcaceae *Staphylococcus aureus* W80042 |
| Sau-3638-1 | Bacteria Firmicutes Bacilli Bacillales Staphylococcaceae *Staphylococcus aureus* W81816 |
| Sau-3639-1 | Bacteria Firmicutes Bacilli Bacillales Staphylococcaceae *Staphylococcus aureus* W82215 |
| Sau-3640-1 | Bacteria Firmicutes Bacilli Bacillales Staphylococcaceae *Staphylococcus aureus* W82228 |
| Sau-3641-1 | Bacteria Firmicutes Bacilli Bacillales Staphylococcaceae *Staphylococcus aureus* W82238 |
| Sau-3642-1 | Bacteria Firmicutes Bacilli Bacillales Staphylococcaceae *Staphylococcus aureus* W82239 |
| Sau-3643-1 | Bacteria Firmicutes Bacilli Bacillales Staphylococcaceae *Staphylococcus aureus* W82303 |
| Sau-3644-1 | Bacteria Firmicutes Bacilli Bacillales Staphylococcaceae *Staphylococcus aureus* W82310 |
| Sau-3645-1 | Bacteria Firmicutes Bacilli Bacillales Staphylococcaceae *Staphylococcus aureus* W83133 |
| Sau-3646-1 | Bacteria Firmicutes Bacilli Bacillales Staphylococcaceae *Staphylococcus aureus* W83137 |
| Sau-3647-1 | Bacteria Firmicutes Bacilli Bacillales Staphylococcaceae *Staphylococcus aureus* W83140 |
| Sau-3648-1 | Bacteria Firmicutes Bacilli Bacillales Staphylococcaceae *Staphylococcus aureus* W85047 |
| Sau-3649-1 | Bacteria Firmicutes Bacilli Bacillales Staphylococcaceae *Staphylococcus aureus* W85326 |
| Sau-3650-1 | Bacteria Firmicutes Bacilli Bacillales Staphylococcaceae *Staphylococcus aureus* W85432 |
| Sau-3651-1 | Bacteria Firmicutes Bacilli Bacillales Staphylococcaceae *Staphylococcus aureus* W85440 |
| Sau-3652-1 | Bacteria Firmicutes Bacilli Bacillales Staphylococcaceae *Staphylococcus aureus* W85806_020613 |
| Sau-3653-1 | Bacteria Firmicutes Bacilli Bacillales Staphylococcaceae *Staphylococcus aureus* W85807 |
| Sau-3654-1 | Bacteria Firmicutes Bacilli Bacillales Staphylococcaceae *Staphylococcus aureus* W85810 |
| Sau-3655-1 | Bacteria Firmicutes Bacilli Bacillales Staphylococcaceae *Staphylococcus aureus* W85837 |
| Sau-3656-1 | Bacteria Firmicutes Bacilli Bacillales Staphylococcaceae *Staphylococcus aureus* W86059 |
| Sau-3657-1 | Bacteria Firmicutes Bacilli Bacillales Staphylococcaceae *Staphylococcus aureus* W86063 |
| Sau-3658-1 | Bacteria Firmicutes Bacilli Bacillales Staphylococcaceae *Staphylococcus aureus* W88551 |
| Sau-3659-1 | Bacteria Firmicutes Bacilli Bacillales Staphylococcaceae *Staphylococcus aureus* W88555 |

| | |
|---|---|
| Sau-3660-1 | Bacteria Firmicutes Bacilli Bacillales Staphylococcaceae *Staphylococcus aureus* W89268 |
| Sau-3661-1 | Bacteria Firmicutes Bacilli Bacillales Staphylococcaceae *Staphylococcus aureus* W89577 |
| Sau-3662-1 | Bacteria Firmicutes Bacilli Bacillales Staphylococcaceae *Staphylococcus aureus* W89579 |
| Sau-3663-1 | Bacteria Firmicutes Bacilli Bacillales Staphylococcaceae *Staphylococcus aureus* W89581 |
| Sau-3664-1 | Bacteria Firmicutes Bacilli Bacillales Staphylococcaceae *Staphylococcus aureus* W91963 |
| Sau-3665-1 | Bacteria Firmicutes Bacilli Bacillales Staphylococcaceae *Staphylococcus aureus* W92052 |
| Sau-3666-1 | Bacteria Firmicutes Bacilli Bacillales Staphylococcaceae *Staphylococcus aureus* W92100 |
| Sau-3667-1 | Bacteria Firmicutes Bacilli Bacillales Staphylococcaceae *Staphylococcus aureus* W92106 |
| Sau-3668-1 | Bacteria Firmicutes Bacilli Bacillales Staphylococcaceae *Staphylococcus aureus* WAMC6003 |
| Sau-3669-1 | Bacteria Firmicutes Bacilli Bacillales Staphylococcaceae *Staphylococcus aureus* WAMC6010 |
| Sau-3670-1 | Bacteria Firmicutes Bacilli Bacillales Staphylococcaceae *Staphylococcus aureus* WAMC6013 |
| Sau-3671-1 | Bacteria Firmicutes Bacilli Bacillales Staphylococcaceae *Staphylococcus aureus* WAMC6030 |
| Sau-3672-1 | Bacteria Firmicutes Bacilli Bacillales Staphylococcaceae *Staphylococcus aureus* WAMC6043 |
| Sau-3673-1 | Bacteria Firmicutes Bacilli Bacillales Staphylococcaceae *Staphylococcus aureus* WAMC6046 |
| Sau-3674-1 | Bacteria Firmicutes Bacilli Bacillales Staphylococcaceae *Staphylococcus aureus* WAMC6049 |
| Sau-3675-1 | Bacteria Firmicutes Bacilli Bacillales Staphylococcaceae *Staphylococcus aureus* WAMC6055 |
| Sau-3676-1 | Bacteria Firmicutes Bacilli Bacillales Staphylococcaceae *Staphylococcus aureus* WAMC6060 |
| Sau-3677-1 | Bacteria Firmicutes Bacilli Bacillales Staphylococcaceae *Staphylococcus aureus* WAMC6062 |
| Sau-3678-1 | Bacteria Firmicutes Bacilli Bacillales Staphylococcaceae *Staphylococcus aureus* WAMC6071 |
| Sau-3679-1 | Bacteria Firmicutes Bacilli Bacillales Staphylococcaceae *Staphylococcus aureus* WAMC6080 |
| Sau-3680-1 | Bacteria Firmicutes Bacilli Bacillales Staphylococcaceae *Staphylococcus aureus* WAMC6081 |
| Sau-3681-1 | Bacteria Firmicutes Bacilli Bacillales Staphylococcaceae *Staphylococcus aureus* WAMC6086 |
| Sau-3682-1 | Bacteria Firmicutes Bacilli Bacillales Staphylococcaceae *Staphylococcus aureus* WAMC6102 |
| Sau-3683-1 | Bacteria Firmicutes Bacilli Bacillales Staphylococcaceae *Staphylococcus aureus* WAMC6108 |
| Sau-3684-1 | Bacteria Firmicutes Bacilli Bacillales Staphylococcaceae *Staphylococcus aureus* WMCA6006 |
| Sau-3685-1 | Bacteria Firmicutes Bacilli Bacillales Staphylococcaceae *Staphylococcus aureus* WMCA6015 |
| Sau-3686-1 | Bacteria Firmicutes Bacilli Bacillales Staphylococcaceae *Staphylococcus aureus* WMCA6024 |
| Sau-3687-1 | Bacteria Firmicutes Bacilli Bacillales Staphylococcaceae *Staphylococcus aureus* WMCA6025 |
| Sau-3688-1 | Bacteria Firmicutes Bacilli Bacillales Staphylococcaceae *Staphylococcus aureus* WMCA6040 |
| Sau-3689-1 | Bacteria Firmicutes Bacilli Bacillales Staphylococcaceae *Staphylococcus aureus* WMCA6041 |
| Sau-3690-1 | Bacteria Firmicutes Bacilli Bacillales Staphylococcaceae *Staphylococcus aureus* WMCA6084 |
| Sau-3691-1 | Bacteria Firmicutes Bacilli Bacillales Staphylococcaceae *Staphylococcus aureus* WMCA6086 |
| Sau-3692-1 | Bacteria Firmicutes Bacilli Bacillales Staphylococcaceae *Staphylococcus aureus* WMCA6091 |
| Sau-3693-1 | Bacteria Firmicutes Bacilli Bacillales Staphylococcaceae *Staphylococcus aureus* WMCA6092 |
| Sau-3694-1 | Bacteria Firmicutes Bacilli Bacillales Staphylococcaceae *Staphylococcus aureus* WMCA6100 |
| Sau-3695-1 | Bacteria Firmicutes Bacilli Bacillales Staphylococcaceae *Staphylococcus aureus* WMCA6123 |
| Sau-3696-1 | Bacteria Firmicutes Bacilli Bacillales Staphylococcaceae *Staphylococcus aureus* WMCA6126 |
| Sau-3697-1 | Bacteria Firmicutes Bacilli Bacillales Staphylococcaceae *Staphylococcus aureus* WMCA6180 |
| Sau-3698-1 | Bacteria Firmicutes Bacilli Bacillales Staphylococcaceae *Staphylococcus aureus* WMCS6001 |
| Sau-3699-1 | Bacteria Firmicutes Bacilli Bacillales Staphylococcaceae *Staphylococcus aureus* WMCS6011 |
| Sau-3700-1 | Bacteria Firmicutes Bacilli Bacillales Staphylococcaceae *Staphylococcus aureus* WMCS6016 |
| Sau-3701-1 | Bacteria Firmicutes Bacilli Bacillales Staphylococcaceae *Staphylococcus aureus* WMCS6019 |
| Sau-3702-1 | Bacteria Firmicutes Bacilli Bacillales Staphylococcaceae *Staphylococcus aureus* WMCS6038 |
| Sau-3703-1 | Bacteria Firmicutes Bacilli Bacillales Staphylococcaceae *Staphylococcus aureus* WMCS6039 |
| Sau-3704-1 | Bacteria Firmicutes Bacilli Bacillales Staphylococcaceae *Staphylococcus aureus* WMCS6047 |
| Sau-3705-1 | Bacteria Firmicutes Bacilli Bacillales Staphylococcaceae *Staphylococcus aureus* WMCS6055 |
| Sau-3706-1 | Bacteria Firmicutes Bacilli Bacillales Staphylococcaceae *Staphylococcus aureus* WMCS6079 |
| Sau-3707-1 | Bacteria Firmicutes Bacilli Bacillales Staphylococcaceae *Staphylococcus aureus* WMCS6087 |
| Sau-3708-1 | Bacteria Firmicutes Bacilli Bacillales Staphylococcaceae *Staphylococcus aureus* WMCS6114 |
| Sau-3709-1 | Bacteria Firmicutes Bacilli Bacillales Staphylococcaceae *Staphylococcus aureus* WMCS6115 |
| Sau-3710-1 | Bacteria Firmicutes Bacilli Bacillales Staphylococcaceae *Staphylococcus aureus* ZTA09/03576-8HST |
| Sau-3711-1 | Bacteria Firmicutes Bacilli Bacillales Staphylococcaceae *Staphylococcus aureus* ZTA09/03576-9HST |
| Sau-3712-1 | Bacteria Firmicutes Bacilli Bacillales Staphylococcaceae *Staphylococcus aureus* ZTA09/03734-9HSA |
| Sau-3713-1 | Bacteria Firmicutes Bacilli Bacillales Staphylococcaceae *Staphylococcus aureus* ZTA09/03739-9HSA |
| Sau-3714-1 | Bacteria Firmicutes Bacilli Bacillales Staphylococcaceae *Staphylococcus aureus* ZTA09/03745-9HSA |
| Sau-3715-1 | Bacteria Firmicutes Bacilli Bacillales Staphylococcaceae *Staphylococcus aureus* ZTA10/00045-9HST |
| Sau-3716-1 | Bacteria Firmicutes Bacilli Bacillales Staphylococcaceae *Staphylococcus aureus* ZTA10/00047-8HST |
| Sau-3717-1 | Bacteria Firmicutes Bacilli Bacillales Staphylococcaceae *Staphylococcus aureus* ZTA10/00058-8HST |
| Sau-3718-1 | Bacteria Firmicutes Bacilli Bacillales Staphylococcaceae *Staphylococcus aureus* ZTA10/02412-8HSA |
| Sau-3719-1 | Bacteria Firmicutes Bacilli Bacillales Staphylococcaceae *Staphylococcus aureus* ZTA11/00189-8HSA |
| Sau-3720-1 | Bacteria Firmicutes Bacilli Bacillales Staphylococcaceae *Staphylococcus aureus* ZTA11/03130-3ST |
| Sca-5-1 to Sca-5-21 | Bacteria Firmicutes Bacilli Bacillales Staphylococcaceae *Staphylococcus capitis* |
| Sca-6-1 | Bacteria Firmicutes Bacilli Bacillales Staphylococcaceae *Staphylococcus capitis subsp. capitis* |
| Sca-7-1 | Bacteria Firmicutes Bacilli Bacillales Staphylococcaceae *Staphylococcus caprae* |
| Sca-8-1 | Bacteria Firmicutes Bacilli Bacillales Staphylococcaceae *Staphylococcus carnosus subsp. utilis* |

| | |
|---|---|
| Sch-1-1 | Bacteria Firmicutes Bacilli Bacillales Staphylococcaceae *Staphylococcus chromogenes* MU 970 |
| Sco-1-1 | Bacteria Firmicutes Bacilli Bacillales Staphylococcaceae *Staphylococcus cohnii hu*-01 |
| Sco-2-1 to Sco-2-3 | Bacteria Firmicutes Bacilli Bacillales Staphylococcaceae *Staphylococcus cohnii subsp. cohnii* |
| Sco-3-1 | Bacteria Firmicutes Bacilli Bacillales Staphylococcaceae *Staphylococcus condimenti* |
| Sep-8-1 to Sep-8-3 | Bacteria Firmicutes Bacilli Bacillales Staphylococcaceae *Staphylococcus epidermidis* |
| Sep-9-1 | Bacteria Firmicutes Bacilli Bacillales Staphylococcaceae *Staphylococcus epidermidis* AG42 |
| Sep-10-1 | Bacteria Firmicutes Bacilli Bacillales Staphylococcaceae *Staphylococcus epidermidis* M0026 |
| Sep-11-1 | Bacteria Firmicutes Bacilli Bacillales Staphylococcaceae *Staphylococcus epidermidis* PM221 |
| Sep-12-1 | Bacteria Firmicutes Bacilli Bacillales Staphylococcaceae *Staphylococcus epidermidis* UC7032 |
| Sep-13-1 | Bacteria Firmicutes Bacilli Bacillales Staphylococcaceae *Staphylococcus epidermidis* VCU013 |
| Sep-14-1 | Bacteria Firmicutes Bacilli Bacillales Staphylococcaceae *Staphylococcus epidermidis* VCU014 |
| Sep-15-1 | Bacteria Firmicutes Bacilli Bacillales Staphylococcaceae *Staphylococcus epidermidis* VCU036 |
| Sep-16-1 | Bacteria Firmicutes Bacilli Bacillales Staphylococcaceae *Staphylococcus epidermidis* VCU050 |
| Sep-17-1 | Bacteria Firmicutes Bacilli Bacillales Staphylococcaceae *Staphylococcus epidermidis* VCU111 |
| Seq-2-1 | Bacteria Firmicutes Bacilli Bacillales Staphylococcaceae *Staphylococcus equorum subsp. equorum* |
| Sga-1-1 | Bacteria Firmicutes Bacilli Bacillales Staphylococcaceae *Staphylococcus gallinarum* |
| Sha-4-1 to Sha-4-119 | Bacteria Firmicutes Bacilli Bacillales Staphylococcaceae *Staphylococcus haemolyticus* |
| Sha-5-1 | Bacteria Firmicutes Bacilli Bacillales Staphylococcaceae *Staphylococcus haemolyticus* DNF00585 |
| Sho-3-1 to Sho-3-17 | Bacteria Firmicutes Bacilli Bacillales Staphylococcaceae *Staphylococcus hominis* |
| Shy-1-1 | Bacteria Firmicutes Bacilli Bacillales Staphylococcaceae *Staphylococcus hyicus* |
| Slu-4-1 | Bacteria Firmicutes Bacilli Bacillales Staphylococcaceae *Staphylococcus lugdunensis* UCIM6116 |
| Slu-5-1 | Bacteria Firmicutes Bacilli Bacillales Staphylococcaceae *Staphylococcus lugdunensis* VCU148 |
| Slu-6-1 | Bacteria Firmicutes Bacilli Bacillales Staphylococcaceae *Staphylococcus lugdunensis* VCU150 |
| Smi-1-1 | Bacteria Firmicutes Bacilli Bacillales Staphylococcaceae *Staphylococcus microti* |
| Spa-1-1 to Spa-1-2 | Bacteria Firmicutes Bacilli Bacillales Staphylococcaceae *Staphylococcus pasteuri* |
| Spe-1-1 to Spe-1-2 | Bacteria Firmicutes Bacilli Bacillales Staphylococcaceae *Staphylococcus pettenkoferi* |
| Sps-3-1 | Bacteria Firmicutes Bacilli Bacillales Staphylococcaceae *Staphylococcus pseudintermedius* E140 |
| Ssa-2-1 to Ssa-2-3 | Bacteria Firmicutes Bacilli Bacillales Staphylococcaceae *Staphylococcus saprophyticus* |
| Ssc-1-1 to Ssc-1-4 | Bacteria Firmicutes Bacilli Bacillales Staphylococcaceae *Staphylococcus schleiferi* |
| Ssc-2-1 to Ssc-2-3 | Bacteria Firmicutes Bacilli Bacillales Staphylococcaceae *Staphylococcus schweitzeri* |
| Ssc-3-1 | Bacteria Firmicutes Bacilli Bacillales Staphylococcaceae *Staphylococcus sciuri subsp. sciuri* |
| Ssc-4-1 | Bacteria Firmicutes Bacilli Bacillales Staphylococcaceae *Staphylococcus sciuri subsp. sciuri* Z8 |
| Ssp-7-1 | Bacteria Firmicutes Bacilli Bacillales Staphylococcaceae *Staphylococcus sp.* M0480 |
| Ssp-8-1 | Bacteria Firmicutes Bacilli Bacillales Staphylococcaceae *Staphylococcus sp.* TE8 |
| Ssp-9-1 | Bacteria Firmicutes Bacilli Bacillales Staphylococcaceae *Staphylococcus sp.* URHA0057 |
| Ssp-10-1 | Bacteria Firmicutes Bacilli Bacillales Staphylococcaceae *Staphylococcus sp.* ZWU0021 |
| Ssu-1-1 | Bacteria Firmicutes Bacilli Bacillales Staphylococcaceae *Staphylococcus succinus subsp. succinus* |
| Swa-2-1 to Swa-2-6 | Bacteria Firmicutes Bacilli Bacillales Staphylococcaceae *Staphylococcus warneri* |
| Swa-3-1 | Bacteria Firmicutes Bacilli Bacillales Staphylococcaceae *Staphylococcus warneri* Lyso 1 2011 |
| Swa-4-1 | Bacteria Firmicutes Bacilli Bacillales Staphylococcaceae *Staphylococcus warneri* Lyso 2 2011 |
| Sxy-1-1 | Bacteria Firmicutes Bacilli Bacillales Staphylococcaceae *Staphylococcus xylosus* |
| Sxy-2-1 | Bacteria Firmicutes Bacilli Bacillales Staphylococcaceae *Staphylococcus xylosus* DMB3-Bh1 |
| Sau-79-1 to Sau-79-2 | Bacteria Firmicutes Bacilli Bacillales *Staphylococcus aureus* |
| Sau-3758-1 | Bacteria Firmicutes Bacilli Bacillales *Staphylococcus aureus* 16K |
| Sau-3759-1 | Bacteria Firmicutes Bacilli Bacillales *Staphylococcus aureus* 3957 |
| Sau-3760-1 | Bacteria Firmicutes Bacilli Bacillales *Staphylococcus aureus* B147830 |
| Sau-3761-1 | Bacteria Firmicutes Bacilli Bacillales *Staphylococcus aureus* B40723 |
| Sau-3762-1 | Bacteria Firmicutes Bacilli Bacillales *Staphylococcus aureus* B40950 |
| Sau-3763-1 | Bacteria Firmicutes Bacilli Bacillales *Staphylococcus aureus* B53639 |
| Sau-3764-1 | Bacteria Firmicutes Bacilli Bacillales *Staphylococcus aureus* Bmb9393 |
| Sau-3765-1 | Bacteria Firmicutes Bacilli Bacillales *Staphylococcus aureus* CA-347 |
| Sau-3766-1 | Bacteria Firmicutes Bacilli Bacillales *Staphylococcus aureus* CN79 |
| Sau-3767-1 | Bacteria Firmicutes Bacilli Bacillales *Staphylococcus aureus* HI010 |
| Sau-3768-1 | Bacteria Firmicutes Bacilli Bacillales *Staphylococcus aureus* HI010B |
| Sau-3769-1 | Bacteria Firmicutes Bacilli Bacillales *Staphylococcus aureus* HI013 |
| Sau-3770-1 | Bacteria Firmicutes Bacilli Bacillales *Staphylococcus aureus* HI022 |
| Sau-3771-1 | Bacteria Firmicutes Bacilli Bacillales *Staphylococcus aureus* HI049 |
| Sau-3772-1 | Bacteria Firmicutes Bacilli Bacillales *Staphylococcus aureus* HI049B |
| Sau-3773-1 | Bacteria Firmicutes Bacilli Bacillales *Staphylococcus aureus* HI049C |
| Sau-3774-1 | Bacteria Firmicutes Bacilli Bacillales *Staphylococcus aureus* HI111 |
| Sau-3775-1 | Bacteria Firmicutes Bacilli Bacillales *Staphylococcus aureus* HI168 |
| Sau-3776-1 | Bacteria Firmicutes Bacilli Bacillales *Staphylococcus aureus* HIF003_B2N-C |
| Sau-3777-1 | Bacteria Firmicutes Bacilli Bacillales *Staphylococcus aureus* KLT6 |
| Sau-3778-1 | Bacteria Firmicutes Bacilli Bacillales *Staphylococcus aureus* KT/314250 |
| Sau-3779-1 | Bacteria Firmicutes Bacilli Bacillales *Staphylococcus aureus* KT/Y21 |

| | |
|---|---|
| Sau-3780-1 | Bacteria Firmicutes Bacilli Bacillales *Staphylococcus aureus* LVP2 |
| Sau-3781-1 | Bacteria Firmicutes Bacilli Bacillales *Staphylococcus aureus* LVP5 |
| Sau-3782-1 | Bacteria Firmicutes Bacilli Bacillales *Staphylococcus aureus* M0001 |
| Sau-3783-1 | Bacteria Firmicutes Bacilli Bacillales *Staphylococcus aureus* M0006 |
| Sau-3784-1 | Bacteria Firmicutes Bacilli Bacillales *Staphylococcus aureus* M0029 |
| Sau-3785-1 | Bacteria Firmicutes Bacilli Bacillales *Staphylococcus aureus* M0035 |
| Sau-3786-1 | Bacteria Firmicutes Bacilli Bacillales *Staphylococcus aureus* M0045 |
| Sau-3787-1 | Bacteria Firmicutes Bacilli Bacillales *Staphylococcus aureus* M0055 |
| Sau-3788-1 | Bacteria Firmicutes Bacilli Bacillales *Staphylococcus aureus* M0060 |
| Sau-3789-1 | Bacteria Firmicutes Bacilli Bacillales *Staphylococcus aureus* M0066 |
| Sau-3790-1 | Bacteria Firmicutes Bacilli Bacillales *Staphylococcus aureus* M0075 |
| Sau-3791-1 | Bacteria Firmicutes Bacilli Bacillales *Staphylococcus aureus* M0077 |
| Sau-3792-1 | Bacteria Firmicutes Bacilli Bacillales *Staphylococcus aureus* M0102 |
| Sau-3793-1 | Bacteria Firmicutes Bacilli Bacillales *Staphylococcus aureus* M0103 |
| Sau-3794-1 | Bacteria Firmicutes Bacilli Bacillales *Staphylococcus aureus* M0104 |
| Sau-3795-1 | Bacteria Firmicutes Bacilli Bacillales *Staphylococcus aureus* M0108 |
| Sau-3796-1 | Bacteria Firmicutes Bacilli Bacillales *Staphylococcus aureus* M0144 |
| Sau-3797-1 | Bacteria Firmicutes Bacilli Bacillales *Staphylococcus aureus* M0150 |
| Sau-3798-1 | Bacteria Firmicutes Bacilli Bacillales *Staphylococcus aureus* M0154 |
| Sau-3799-1 | Bacteria Firmicutes Bacilli Bacillales *Staphylococcus aureus* M0171 |
| Sau-3800-1 | Bacteria Firmicutes Bacilli Bacillales *Staphylococcus aureus* M0173 |
| Sau-3801-1 | Bacteria Firmicutes Bacilli Bacillales *Staphylococcus aureus* M0177 |
| Sau-3802-1 | Bacteria Firmicutes Bacilli Bacillales *Staphylococcus aureus* M0192 |
| Sau-3803-1 | Bacteria Firmicutes Bacilli Bacillales *Staphylococcus aureus* M0197 |
| Sau-3804-1 | Bacteria Firmicutes Bacilli Bacillales *Staphylococcus aureus* M0200 |
| Sau-3805-1 | Bacteria Firmicutes Bacilli Bacillales *Staphylococcus aureus* M0210 |
| Sau-3806-1 | Bacteria Firmicutes Bacilli Bacillales *Staphylococcus aureus* M0212 |
| Sau-3807-1 | Bacteria Firmicutes Bacilli Bacillales *Staphylococcus aureus* M0213 |
| Sau-3808-1 | Bacteria Firmicutes Bacilli Bacillales *Staphylococcus aureus* M0216 |
| Sau-3809-1 | Bacteria Firmicutes Bacilli Bacillales *Staphylococcus aureus* M0221 |
| Sau-3810-1 | Bacteria Firmicutes Bacilli Bacillales *Staphylococcus aureus* M0235 |
| Sau-3811-1 | Bacteria Firmicutes Bacilli Bacillales *Staphylococcus aureus* M0237 |
| Sau-3812-1 | Bacteria Firmicutes Bacilli Bacillales *Staphylococcus aureus* M0239 |
| Sau-3813-1 | Bacteria Firmicutes Bacilli Bacillales *Staphylococcus aureus* M0240 |
| Sau-3814-1 | Bacteria Firmicutes Bacilli Bacillales *Staphylococcus aureus* M0250 |
| Sau-3815-1 | Bacteria Firmicutes Bacilli Bacillales *Staphylococcus aureus* M0252 |
| Sau-3816-1 | Bacteria Firmicutes Bacilli Bacillales *Staphylococcus aureus* M0270 |
| Sau-3817-1 | Bacteria Firmicutes Bacilli Bacillales *Staphylococcus aureus* M0273 |
| Sau-3818-1 | Bacteria Firmicutes Bacilli Bacillales *Staphylococcus aureus* M0279 |
| Sau-3819-1 | Bacteria Firmicutes Bacilli Bacillales *Staphylococcus aureus* M0280 |
| Sau-3820-1 | Bacteria Firmicutes Bacilli Bacillales *Staphylococcus aureus* M0288 |
| Sau-3821-1 | Bacteria Firmicutes Bacilli Bacillales *Staphylococcus aureus* M0294 |
| Sau-3822-1 | Bacteria Firmicutes Bacilli Bacillales *Staphylococcus aureus* M0306 |
| Sau-3823-1 | Bacteria Firmicutes Bacilli Bacillales *Staphylococcus aureus* M0312 |
| Sau-3824-1 | Bacteria Firmicutes Bacilli Bacillales *Staphylococcus aureus* M0326 |
| Sau-3825-1 | Bacteria Firmicutes Bacilli Bacillales *Staphylococcus aureus* M0328 |
| Sau-3826-1 | Bacteria Firmicutes Bacilli Bacillales *Staphylococcus aureus* M0329 |
| Sau-3827-1 | Bacteria Firmicutes Bacilli Bacillales *Staphylococcus aureus* M0330 |
| Sau-3828-1 | Bacteria Firmicutes Bacilli Bacillales *Staphylococcus aureus* M0334 |
| Sau-3829-1 | Bacteria Firmicutes Bacilli Bacillales *Staphylococcus aureus* M0340 |
| Sau-3830-1 | Bacteria Firmicutes Bacilli Bacillales *Staphylococcus aureus* M0347 |
| Sau-3831-1 | Bacteria Firmicutes Bacilli Bacillales *Staphylococcus aureus* M0350 |
| Sau-3832-1 | Bacteria Firmicutes Bacilli Bacillales *Staphylococcus aureus* M0351 |
| Sau-3833-1 | Bacteria Firmicutes Bacilli Bacillales *Staphylococcus aureus* M0363 |
| Sau-3834-1 | Bacteria Firmicutes Bacilli Bacillales *Staphylococcus aureus* M0364 |
| Sau-3835-1 | Bacteria Firmicutes Bacilli Bacillales *Staphylococcus aureus* M0367 |
| Sau-3836-1 | Bacteria Firmicutes Bacilli Bacillales *Staphylococcus aureus* M0374 |
| Sau-3837-1 | Bacteria Firmicutes Bacilli Bacillales *Staphylococcus aureus* M0375 |
| Sau-3838-1 | Bacteria Firmicutes Bacilli Bacillales *Staphylococcus aureus* M0391 |
| Sau-3839-1 | Bacteria Firmicutes Bacilli Bacillales *Staphylococcus aureus* M0396 |
| Sau-3840-1 | Bacteria Firmicutes Bacilli Bacillales *Staphylococcus aureus* M0404 |
| Sau-3841-1 | Bacteria Firmicutes Bacilli Bacillales *Staphylococcus aureus* M0406 |
| Sau-3842-1 | Bacteria Firmicutes Bacilli Bacillales *Staphylococcus aureus* M0408 |
| Sau-3843-1 | Bacteria Firmicutes Bacilli Bacillales *Staphylococcus aureus* M0415 |
| Sau-3844-1 | Bacteria Firmicutes Bacilli Bacillales *Staphylococcus aureus* M0424 |

| | |
|---|---|
| Sau-3845-1 | Bacteria Firmicutes Bacilli Bacillales *Staphylococcus aureus* M0427 |
| Sau-3846-1 | Bacteria Firmicutes Bacilli Bacillales *Staphylococcus aureus* M0438 |
| Sau-3847-1 | Bacteria Firmicutes Bacilli Bacillales *Staphylococcus aureus* M0450 |
| Sau-3848-1 | Bacteria Firmicutes Bacilli Bacillales *Staphylococcus aureus* M0455 |
| Sau-3849-1 | Bacteria Firmicutes Bacilli Bacillales *Staphylococcus aureus* M0460 |
| Sau-3850-1 | Bacteria Firmicutes Bacilli Bacillales *Staphylococcus aureus* M0467 |
| Sau-3851-1 | Bacteria Firmicutes Bacilli Bacillales *Staphylococcus aureus* M0468 |
| Sau-3852-1 | Bacteria Firmicutes Bacilli Bacillales *Staphylococcus aureus* M0478 |
| Sau-3853-1 | Bacteria Firmicutes Bacilli Bacillales *Staphylococcus aureus* M0489 |
| Sau-3854-1 | Bacteria Firmicutes Bacilli Bacillales *Staphylococcus aureus* M0493 |
| Sau-3855-1 | Bacteria Firmicutes Bacilli Bacillales *Staphylococcus aureus* M0494 |
| Sau-3856-1 | Bacteria Firmicutes Bacilli Bacillales *Staphylococcus aureus* M0510 |
| Sau-3857-1 | Bacteria Firmicutes Bacilli Bacillales *Staphylococcus aureus* M0513 |
| Sau-3858-1 | Bacteria Firmicutes Bacilli Bacillales *Staphylococcus aureus* M0528 |
| Sau-3859-1 | Bacteria Firmicutes Bacilli Bacillales *Staphylococcus aureus* M0529 |
| Sau-3860-1 | Bacteria Firmicutes Bacilli Bacillales *Staphylococcus aureus* M0531 |
| Sau-3861-1 | Bacteria Firmicutes Bacilli Bacillales *Staphylococcus aureus* M0536 |
| Sau-3862-1 | Bacteria Firmicutes Bacilli Bacillales *Staphylococcus aureus* M0539 |
| Sau-3863-1 | Bacteria Firmicutes Bacilli Bacillales *Staphylococcus aureus* M0547 |
| Sau-3864-1 | Bacteria Firmicutes Bacilli Bacillales *Staphylococcus aureus* M0562 |
| Sau-3865-1 | Bacteria Firmicutes Bacilli Bacillales *Staphylococcus aureus* M0565 |
| Sau-3866-1 | Bacteria Firmicutes Bacilli Bacillales *Staphylococcus aureus* M0571 |
| Sau-3867-1 | Bacteria Firmicutes Bacilli Bacillales *Staphylococcus aureus* M0580 |
| Sau-3868-1 | Bacteria Firmicutes Bacilli Bacillales *Staphylococcus aureus* M0584 |
| Sau-3869-1 | Bacteria Firmicutes Bacilli Bacillales *Staphylococcus aureus* M0586 |
| Sau-3870-1 | Bacteria Firmicutes Bacilli Bacillales *Staphylococcus aureus* M0602 |
| Sau-3871-1 | Bacteria Firmicutes Bacilli Bacillales *Staphylococcus aureus* M0622 |
| Sau-3872-1 | Bacteria Firmicutes Bacilli Bacillales *Staphylococcus aureus* M0628 |
| Sau-3873-1 | Bacteria Firmicutes Bacilli Bacillales *Staphylococcus aureus* M0633 |
| Sau-3874-1 | Bacteria Firmicutes Bacilli Bacillales *Staphylococcus aureus* M0646 |
| Sau-3875-1 | Bacteria Firmicutes Bacilli Bacillales *Staphylococcus aureus* M0648 |
| Sau-3876-1 | Bacteria Firmicutes Bacilli Bacillales *Staphylococcus aureus* M0660 |
| Sau-3877-1 | Bacteria Firmicutes Bacilli Bacillales *Staphylococcus aureus* M0673 |
| Sau-3878-1 | Bacteria Firmicutes Bacilli Bacillales *Staphylococcus aureus* M0676 |
| Sau-3879-1 | Bacteria Firmicutes Bacilli Bacillales *Staphylococcus aureus* M0687 |
| Sau-3880-1 | Bacteria Firmicutes Bacilli Bacillales *Staphylococcus aureus* M0692 |
| Sau-3881-1 | Bacteria Firmicutes Bacilli Bacillales *Staphylococcus aureus* M0695 |
| Sau-3882-1 | Bacteria Firmicutes Bacilli Bacillales *Staphylococcus aureus* M0719 |
| Sau-3883-1 | Bacteria Firmicutes Bacilli Bacillales *Staphylococcus aureus* M0769 |
| Sau-3884-1 | Bacteria Firmicutes Bacilli Bacillales *Staphylococcus aureus* M0770 |
| Sau-3885-1 | Bacteria Firmicutes Bacilli Bacillales *Staphylococcus aureus* M0780 |
| Sau-3886-1 | Bacteria Firmicutes Bacilli Bacillales *Staphylococcus aureus* M0792 |
| Sau-3887-1 | Bacteria Firmicutes Bacilli Bacillales *Staphylococcus aureus* M0799 |
| Sau-3888-1 | Bacteria Firmicutes Bacilli Bacillales *Staphylococcus aureus* M0822 |
| Sau-3889-1 | Bacteria Firmicutes Bacilli Bacillales *Staphylococcus aureus* M0823 |
| Sau-3890-1 | Bacteria Firmicutes Bacilli Bacillales *Staphylococcus aureus* M0831 |
| Sau-3891-1 | Bacteria Firmicutes Bacilli Bacillales *Staphylococcus aureus* M0844 |
| Sau-3892-1 | Bacteria Firmicutes Bacilli Bacillales *Staphylococcus aureus* M0871 |
| Sau-3893-1 | Bacteria Firmicutes Bacilli Bacillales *Staphylococcus aureus* M0877 |
| Sau-3894-1 | Bacteria Firmicutes Bacilli Bacillales *Staphylococcus aureus* M0892 |
| Sau-3895-1 | Bacteria Firmicutes Bacilli Bacillales *Staphylococcus aureus* M0900 |
| Sau-3896-1 | Bacteria Firmicutes Bacilli Bacillales *Staphylococcus aureus* M0927 |
| Sau-3897-1 | Bacteria Firmicutes Bacilli Bacillales *Staphylococcus aureus* M0934 |
| Sau-3898-1 | Bacteria Firmicutes Bacilli Bacillales *Staphylococcus aureus* M0943 |
| Sau-3899-1 | Bacteria Firmicutes Bacilli Bacillales *Staphylococcus aureus* M0944 |
| Sau-3900-1 | Bacteria Firmicutes Bacilli Bacillales *Staphylococcus aureus* M0946 |
| Sau-3901-1 | Bacteria Firmicutes Bacilli Bacillales *Staphylococcus aureus* M0953 |
| Sau-3902-1 | Bacteria Firmicutes Bacilli Bacillales *Staphylococcus aureus* M0964 |
| Sau-3903-1 | Bacteria Firmicutes Bacilli Bacillales *Staphylococcus aureus* M0978 |
| Sau-3904-1 | Bacteria Firmicutes Bacilli Bacillales *Staphylococcus aureus* M0994 |
| Sau-3905-1 | Bacteria Firmicutes Bacilli Bacillales *Staphylococcus aureus* M0998 |
| Sau-3906-1 | Bacteria Firmicutes Bacilli Bacillales *Staphylococcus aureus* M0999 |
| Sau-3907-1 | Bacteria Firmicutes Bacilli Bacillales *Staphylococcus aureus* M1 |
| Sau-3908-1 | Bacteria Firmicutes Bacilli Bacillales *Staphylococcus aureus* M1007 |
| Sau-3909-1 | Bacteria Firmicutes Bacilli Bacillales *Staphylococcus aureus* M1010 |

| | |
|---|---|
| Sau-3910-1 | Bacteria Firmicutes Bacilli Bacillales *Staphylococcus aureus* M1015 |
| Sau-3911-1 | Bacteria Firmicutes Bacilli Bacillales *Staphylococcus aureus* M1016 |
| Sau-3912-1 | Bacteria Firmicutes Bacilli Bacillales *Staphylococcus aureus* M1034 |
| Sau-3913-1 | Bacteria Firmicutes Bacilli Bacillales *Staphylococcus aureus* M1036 |
| Sau-3914-1 | Bacteria Firmicutes Bacilli Bacillales *Staphylococcus aureus* M1037 |
| Sau-3915-1 | Bacteria Firmicutes Bacilli Bacillales *Staphylococcus aureus* M1044 |
| Sau-3916-1 | Bacteria Firmicutes Bacilli Bacillales *Staphylococcus aureus* M1060 |
| Sau-3917-1 | Bacteria Firmicutes Bacilli Bacillales *Staphylococcus aureus* M1061 |
| Sau-3918-1 | Bacteria Firmicutes Bacilli Bacillales *Staphylococcus aureus* M1062 |
| Sau-3919-1 | Bacteria Firmicutes Bacilli Bacillales *Staphylococcus aureus* M1063 |
| Sau-3920-1 | Bacteria Firmicutes Bacilli Bacillales *Staphylococcus aureus* M1064 |
| Sau-3921-1 | Bacteria Firmicutes Bacilli Bacillales *Staphylococcus aureus* M1068 |
| Sau-3922-1 | Bacteria Firmicutes Bacilli Bacillales *Staphylococcus aureus* M1076 |
| Sau-3923-1 | Bacteria Firmicutes Bacilli Bacillales *Staphylococcus aureus* M1078 |
| Sau-3924-1 | Bacteria Firmicutes Bacilli Bacillales *Staphylococcus aureus* M1083 |
| Sau-3925-1 | Bacteria Firmicutes Bacilli Bacillales *Staphylococcus aureus* M1092 |
| Sau-3926-1 | Bacteria Firmicutes Bacilli Bacillales *Staphylococcus aureus* M1093 |
| Sau-3927-1 | Bacteria Firmicutes Bacilli Bacillales *Staphylococcus aureus* M1095 |
| Sau-3928-1 | Bacteria Firmicutes Bacilli Bacillales *Staphylococcus aureus* M1103 |
| Sau-3929-1 | Bacteria Firmicutes Bacilli Bacillales *Staphylococcus aureus* M1109 |
| Sau-3930-1 | Bacteria Firmicutes Bacilli Bacillales *Staphylococcus aureus* M1119 |
| Sau-3931-1 | Bacteria Firmicutes Bacilli Bacillales *Staphylococcus aureus* M1126 |
| Sau-3932-1 | Bacteria Firmicutes Bacilli Bacillales *Staphylococcus aureus* M1142 |
| Sau-3933-1 | Bacteria Firmicutes Bacilli Bacillales *Staphylococcus aureus* M1167 |
| Sau-3934-1 | Bacteria Firmicutes Bacilli Bacillales *Staphylococcus aureus* M1170 |
| Sau-3935-1 | Bacteria Firmicutes Bacilli Bacillales *Staphylococcus aureus* M1188 |
| Sau-3936-1 | Bacteria Firmicutes Bacilli Bacillales *Staphylococcus aureus* M1193 |
| Sau-3937-1 | Bacteria Firmicutes Bacilli Bacillales *Staphylococcus aureus* M1198 |
| Sau-3938-1 | Bacteria Firmicutes Bacilli Bacillales *Staphylococcus aureus* M1199 |
| Sau-3939-1 | Bacteria Firmicutes Bacilli Bacillales *Staphylococcus aureus* M1215 |
| Sau-3940-1 | Bacteria Firmicutes Bacilli Bacillales *Staphylococcus aureus* M1216 |
| Sau-3941-1 | Bacteria Firmicutes Bacilli Bacillales *Staphylococcus aureus* M1223 |
| Sau-3942-1 | Bacteria Firmicutes Bacilli Bacillales *Staphylococcus aureus* M1224 |
| Sau-3943-1 | Bacteria Firmicutes Bacilli Bacillales *Staphylococcus aureus* M1228 |
| Sau-3944-1 | Bacteria Firmicutes Bacilli Bacillales *Staphylococcus aureus* M1229 |
| Sau-3945-1 | Bacteria Firmicutes Bacilli Bacillales *Staphylococcus aureus* M1244 |
| Sau-3946-1 | Bacteria Firmicutes Bacilli Bacillales *Staphylococcus aureus* M1248 |
| Sau-3947-1 | Bacteria Firmicutes Bacilli Bacillales *Staphylococcus aureus* M1253 |
| Sau-3948-1 | Bacteria Firmicutes Bacilli Bacillales *Staphylococcus aureus* M1255 |
| Sau-3949-1 | Bacteria Firmicutes Bacilli Bacillales *Staphylococcus aureus* M1256 |
| Sau-3950-1 | Bacteria Firmicutes Bacilli Bacillales *Staphylococcus aureus* M1257 |
| Sau-3951-1 | Bacteria Firmicutes Bacilli Bacillales *Staphylococcus aureus* M1275 |
| Sau-3952-1 | Bacteria Firmicutes Bacilli Bacillales *Staphylococcus aureus* M1277 |
| Sau-3953-1 | Bacteria Firmicutes Bacilli Bacillales *Staphylococcus aureus* M1286 |
| Sau-3954-1 | Bacteria Firmicutes Bacilli Bacillales *Staphylococcus aureus* M1291 |
| Sau-3955-1 | Bacteria Firmicutes Bacilli Bacillales *Staphylococcus aureus* M1309 |
| Sau-3956-1 | Bacteria Firmicutes Bacilli Bacillales *Staphylococcus aureus* M1311 |
| Sau-3957-1 | Bacteria Firmicutes Bacilli Bacillales *Staphylococcus aureus* M1320 |
| Sau-3958-1 | Bacteria Firmicutes Bacilli Bacillales *Staphylococcus aureus* M1321 |
| Sau-3959-1 | Bacteria Firmicutes Bacilli Bacillales *Staphylococcus aureus* M1322 |
| Sau-3960-1 | Bacteria Firmicutes Bacilli Bacillales *Staphylococcus aureus* M1359 |
| Sau-3961-1 | Bacteria Firmicutes Bacilli Bacillales *Staphylococcus aureus* M1367 |
| Sau-3962-1 | Bacteria Firmicutes Bacilli Bacillales *Staphylococcus aureus* M1373 |
| Sau-3963-1 | Bacteria Firmicutes Bacilli Bacillales *Staphylococcus aureus* M1374 |
| Sau-3964-1 | Bacteria Firmicutes Bacilli Bacillales *Staphylococcus aureus* M1394 |
| Sau-3965-1 | Bacteria Firmicutes Bacilli Bacillales *Staphylococcus aureus* M1405 |
| Sau-3966-1 | Bacteria Firmicutes Bacilli Bacillales *Staphylococcus aureus* M1407 |
| Sau-3967-1 | Bacteria Firmicutes Bacilli Bacillales *Staphylococcus aureus* M1450 |
| Sau-3968-1 | Bacteria Firmicutes Bacilli Bacillales *Staphylococcus aureus* M1451 |
| Sau-3969-1 | Bacteria Firmicutes Bacilli Bacillales *Staphylococcus aureus* M1462 |
| Sau-3970-1 | Bacteria Firmicutes Bacilli Bacillales *Staphylococcus aureus* M1463 |
| Sau-3971-1 | Bacteria Firmicutes Bacilli Bacillales *Staphylococcus aureus* M1466 |
| Sau-3972-1 | Bacteria Firmicutes Bacilli Bacillales *Staphylococcus aureus* M1481 |
| Sau-3973-1 | Bacteria Firmicutes Bacilli Bacillales *Staphylococcus aureus* M1510 |
| Sau-3974-1 | Bacteria Firmicutes Bacilli Bacillales *Staphylococcus aureus* M1520 |

| | |
|---|---|
| Sau-3975-1 | Bacteria Firmicutes Bacilli Bacillales *Staphylococcus aureus* M1521 |
| Sau-3976-1 | Bacteria Firmicutes Bacilli Bacillales *Staphylococcus aureus* M1531 |
| Sau-3977-1 | Bacteria Firmicutes Bacilli Bacillales *Staphylococcus aureus* M1533 |
| Sau-3978-1 | Bacteria Firmicutes Bacilli Bacillales *Staphylococcus aureus* M1544 |
| Sau-3979-1 | Bacteria Firmicutes Bacilli Bacillales *Staphylococcus aureus* M1556 |
| Sau-3980-1 | Bacteria Firmicutes Bacilli Bacillales *Staphylococcus aureus* M1563 |
| Sau-3981-1 | Bacteria Firmicutes Bacilli Bacillales *Staphylococcus aureus* M1565 |
| Sau-3982-1 | Bacteria Firmicutes Bacilli Bacillales *Staphylococcus aureus* M1578 |
| Sau-3983-1 | Bacteria Firmicutes Bacilli Bacillales *Staphylococcus aureus* M2 |
| Sau-3984-1 | Bacteria Firmicutes Bacilli Bacillales *Staphylococcus aureus* NN54 |
| Sau-3985-1 | Bacteria Firmicutes Bacilli Bacillales *Staphylococcus aureus* O11 |
| Sau-3986-1 | Bacteria Firmicutes Bacilli Bacillales *Staphylococcus aureus* O46 |
| Sau-3987-1 | Bacteria Firmicutes Bacilli Bacillales *Staphylococcus aureus* PM1 |
| Sau-3988-1 | Bacteria Firmicutes Bacilli Bacillales *Staphylococcus aureus* S1 |
| Sau-3989-1 | Bacteria Firmicutes Bacilli Bacillales *Staphylococcus aureus* S100 |
| Sau-3990-1 | Bacteria Firmicutes Bacilli Bacillales *Staphylococcus aureus* S123 |
| Sau-3991-1 | Bacteria Firmicutes Bacilli Bacillales *Staphylococcus aureus* S130 |
| Sau-3992-1 | Bacteria Firmicutes Bacilli Bacillales *Staphylococcus aureus* S94 |
| Sau-3993-1 | Bacteria Firmicutes Bacilli Bacillales *Staphylococcus aureus* SA16 |
| Sau-3994-1 | Bacteria Firmicutes Bacilli Bacillales *Staphylococcus aureus* SA_ST125_MupR |
| Sau-3722-1 to Sau-3722-6 | Bacteria Firmicutes Bacilli Bacillales *Staphylococcus aureus subsp. aureus* |
| Sau-3996-1 | Bacteria Firmicutes Bacilli Bacillales *Staphylococcus aureus subsp. aureus* 06BA18369 |
| Sau-3997-1 | Bacteria Firmicutes Bacilli Bacillales *Staphylococcus aureus subsp. aureus* 091751 |
| Sau-3998-1 | Bacteria Firmicutes Bacilli Bacillales *Staphylococcus aureus subsp. aureus* 103564 |
| Sau-3999-1 | Bacteria Firmicutes Bacilli Bacillales *Staphylococcus aureus subsp. aureus* 112808A |
| Sau-4000-1 | Bacteria Firmicutes Bacilli Bacillales *Staphylococcus aureus subsp. aureus* 118 |
| Sau-4001-1 | Bacteria Firmicutes Bacilli Bacillales *Staphylococcus aureus subsp. aureus* 120 |
| Sau-4002-1 | Bacteria Firmicutes Bacilli Bacillales *Staphylococcus aureus subsp. aureus* 122051 |
| Sau-4003-1 | Bacteria Firmicutes Bacilli Bacillales *Staphylococcus aureus subsp. aureus* 21172 |
| Sau-4004-1 | Bacteria Firmicutes Bacilli Bacillales *Staphylococcus aureus subsp. aureus* 21178 |
| Sau-4005-1 | Bacteria Firmicutes Bacilli Bacillales *Staphylococcus aureus subsp. aureus* 21189 |
| Sau-4006-1 | Bacteria Firmicutes Bacilli Bacillales *Staphylococcus aureus subsp. aureus* 21193 |
| Sau-4007-1 | Bacteria Firmicutes Bacilli Bacillales *Staphylococcus aureus subsp. aureus* 21194 |
| Sau-4008-1 | Bacteria Firmicutes Bacilli Bacillales *Staphylococcus aureus subsp. aureus* 21195 |
| Sau-4009-1 | Bacteria Firmicutes Bacilli Bacillales *Staphylococcus aureus subsp. aureus* 21196 |
| Sau-4010-1 | Bacteria Firmicutes Bacilli Bacillales *Staphylococcus aureus subsp. aureus* 21200 |
| Sau-4011-1 | Bacteria Firmicutes Bacilli Bacillales *Staphylococcus aureus subsp. aureus* 21201 |
| Sau-4012-1 | Bacteria Firmicutes Bacilli Bacillales *Staphylococcus aureus subsp. aureus* 21202 |
| Sau-4013-1 | Bacteria Firmicutes Bacilli Bacillales *Staphylococcus aureus subsp. aureus* 21209 |
| Sau-4014-1 | Bacteria Firmicutes Bacilli Bacillales *Staphylococcus aureus subsp. aureus* 21232 |
| Sau-4015-1 | Bacteria Firmicutes Bacilli Bacillales *Staphylococcus aureus subsp. aureus* 21235 |
| Sau-4016-1 | Bacteria Firmicutes Bacilli Bacillales *Staphylococcus aureus subsp. aureus* 21236 |
| Sau-4017-1 | Bacteria Firmicutes Bacilli Bacillales *Staphylococcus aureus subsp. aureus* 21252 |
| Sau-4018-1 | Bacteria Firmicutes Bacilli Bacillales *Staphylococcus aureus subsp. aureus* 21259 |
| Sau-4019-1 | Bacteria Firmicutes Bacilli Bacillales *Staphylococcus aureus subsp. aureus* 21262 |
| Sau-4020-1 | Bacteria Firmicutes Bacilli Bacillales *Staphylococcus aureus subsp. aureus* 21264 |
| Sau-4021-1 | Bacteria Firmicutes Bacilli Bacillales *Staphylococcus aureus subsp. aureus* 21266 |
| Sau-4022-1 | Bacteria Firmicutes Bacilli Bacillales *Staphylococcus aureus subsp. aureus* 21269 |
| Sau-4023-1 | Bacteria Firmicutes Bacilli Bacillales *Staphylococcus aureus subsp. aureus* 21272 |
| Sau-4024-1 | Bacteria Firmicutes Bacilli Bacillales *Staphylococcus aureus subsp. aureus* 21282 |
| Sau-4025-1 | Bacteria Firmicutes Bacilli Bacillales *Staphylococcus aureus subsp. aureus* 21283 |
| Sau-4026-1 | Bacteria Firmicutes Bacilli Bacillales *Staphylococcus aureus subsp. aureus* 21305 |
| Sau-4027-1 | Bacteria Firmicutes Bacilli Bacillales *Staphylococcus aureus subsp. aureus* 21310 |
| Sau-4028-1 | Bacteria Firmicutes Bacilli Bacillales *Staphylococcus aureus subsp. aureus* 21331 |
| Sau-4029-1 | Bacteria Firmicutes Bacilli Bacillales *Staphylococcus aureus subsp. aureus* 21333 |
| Sau-4030-1 | Bacteria Firmicutes Bacilli Bacillales *Staphylococcus aureus subsp. aureus* 21334 |
| Sau-4031-1 | Bacteria Firmicutes Bacilli Bacillales *Staphylococcus aureus subsp. aureus* 21340 |
| Sau-4032-1 | Bacteria Firmicutes Bacilli Bacillales *Staphylococcus aureus subsp. aureus* 21342 |
| Sau-4033-1 | Bacteria Firmicutes Bacilli Bacillales *Staphylococcus aureus subsp. aureus* 21343 |
| Sau-4034-1 | Bacteria Firmicutes Bacilli Bacillales *Staphylococcus aureus subsp. aureus* 21345 |
| Sau-4035-1 | Bacteria Firmicutes Bacilli Bacillales *Staphylococcus aureus subsp. aureus* 333 |
| Sau-4036-1 | Bacteria Firmicutes Bacilli Bacillales *Staphylococcus aureus subsp. aureus* 3989 |
| Sau-4037-1 | Bacteria Firmicutes Bacilli Bacillales *Staphylococcus aureus subsp. aureus* 55/2053 |
| Sau-4038-1 | Bacteria Firmicutes Bacilli Bacillales *Staphylococcus aureus subsp. aureus* 6850 |
| Sau-4039-1 | Bacteria Firmicutes Bacilli Bacillales *Staphylococcus aureus subsp. aureus* CBD-635 |

| | |
|---|---|
| Sau-4040-1 | Bacteria Firmicutes Bacilli Bacillales *Staphylococcus aureus subsp. aureus* CGS00 |
| Sau-4041-1 | Bacteria Firmicutes Bacilli Bacillales *Staphylococcus aureus subsp. aureus* CGS01 |
| Sau-4042-1 | Bacteria Firmicutes Bacilli Bacillales *Staphylococcus aureus subsp. aureus* CGS03 |
| Sau-4043-1 | Bacteria Firmicutes Bacilli Bacillales *Staphylococcus aureus subsp. aureus* CIG1057 |
| Sau-4044-1 | Bacteria Firmicutes Bacilli Bacillales *Staphylococcus aureus subsp. aureus* CIG1096 |
| Sau-4045-1 | Bacteria Firmicutes Bacilli Bacillales *Staphylococcus aureus subsp. aureus* CIG1114 |
| Sau-4046-1 | Bacteria Firmicutes Bacilli Bacillales *Staphylococcus aureus subsp. aureus* CIG1150 |
| Sau-4047-1 | Bacteria Firmicutes Bacilli Bacillales *Staphylococcus aureus subsp. aureus* CIG1165 |
| Sau-4048-1 | Bacteria Firmicutes Bacilli Bacillales *Staphylococcus aureus subsp. aureus* CIG1176 |
| Sau-4049-1 | Bacteria Firmicutes Bacilli Bacillales *Staphylococcus aureus subsp. aureus* CIG1213 |
| Sau-4050-1 | Bacteria Firmicutes Bacilli Bacillales *Staphylococcus aureus subsp. aureus* CIG1214 |
| Sau-4051-1 | Bacteria Firmicutes Bacilli Bacillales *Staphylococcus aureus subsp. aureus* CIG1233 |
| Sau-4052-1 | Bacteria Firmicutes Bacilli Bacillales *Staphylococcus aureus subsp. aureus* CIG1242 |
| Sau-4053-1 | Bacteria Firmicutes Bacilli Bacillales *Staphylococcus aureus subsp. aureus* CIG1267 |
| Sau-4054-1 | Bacteria Firmicutes Bacilli Bacillales *Staphylococcus aureus subsp. aureus* CIG149 |
| Sau-4055-1 | Bacteria Firmicutes Bacilli Bacillales *Staphylococcus aureus subsp. aureus* CIG1500 |
| Sau-4056-1 | Bacteria Firmicutes Bacilli Bacillales *Staphylococcus aureus subsp. aureus* CIG1524 |
| Sau-4057-1 | Bacteria Firmicutes Bacilli Bacillales *Staphylococcus aureus subsp. aureus* CIG1605 |
| Sau-4058-1 | Bacteria Firmicutes Bacilli Bacillales *Staphylococcus aureus subsp. aureus* CIG1612 |
| Sau-4059-1 | Bacteria Firmicutes Bacilli Bacillales *Staphylococcus aureus subsp. aureus* CIG1750 |
| Sau-4060-1 | Bacteria Firmicutes Bacilli Bacillales *Staphylococcus aureus subsp. aureus* CIG1769 |
| Sau-4061-1 | Bacteria Firmicutes Bacilli Bacillales *Staphylococcus aureus subsp. aureus* CIG1770 |
| Sau-4062-1 | Bacteria Firmicutes Bacilli Bacillales *Staphylococcus aureus subsp. aureus* CIG1835 |
| Sau-4063-1 | Bacteria Firmicutes Bacilli Bacillales *Staphylococcus aureus subsp. aureus* CIG2018 |
| Sau-4064-1 | Bacteria Firmicutes Bacilli Bacillales *Staphylococcus aureus subsp. aureus* CIG290 |
| Sau-4065-1 | Bacteria Firmicutes Bacilli Bacillales *Staphylococcus aureus subsp. aureus* CIG547 |
| Sau-4066-1 | Bacteria Firmicutes Bacilli Bacillales *Staphylococcus aureus subsp. aureus* CIGC128 |
| Sau-4067-1 | Bacteria Firmicutes Bacilli Bacillales *Staphylococcus aureus subsp. aureus* CIGC340D |
| Sau-4068-1 | Bacteria Firmicutes Bacilli Bacillales *Staphylococcus aureus subsp. aureus* CIGC341D |
| Sau-4069-1 | Bacteria Firmicutes Bacilli Bacillales *Staphylococcus aureus subsp. aureus* CIGC345D |
| Sau-4070-1 | Bacteria Firmicutes Bacilli Bacillales *Staphylococcus aureus subsp. aureus* CIGC348 |
| Sau-4071-1 | Bacteria Firmicutes Bacilli Bacillales *Staphylococcus aureus subsp. aureus* CIGC93 |
| Sau-4072-1 | Bacteria Firmicutes Bacilli Bacillales *Staphylococcus aureus subsp. aureus* CM05 |
| Sau-4073-1 | Bacteria Firmicutes Bacilli Bacillales *Staphylococcus aureus subsp. aureus* CN1 |
| Sau-4074-1 | Bacteria Firmicutes Bacilli Bacillales *Staphylococcus aureus subsp. aureus* CO-08 |
| Sau-4075-1 | Bacteria Firmicutes Bacilli Bacillales *Staphylococcus aureus subsp. aureus* CO-23 |
| Sau-4076-1 | Bacteria Firmicutes Bacilli Bacillales *Staphylococcus aureus subsp. aureus* CO-85 |
| Sau-4077-1 | Bacteria Firmicutes Bacilli Bacillales *Staphylococcus aureus subsp. aureus* CO-86 |
| Sau-4078-1 | Bacteria Firmicutes Bacilli Bacillales *Staphylococcus aureus subsp. aureus* CO-98 |
| Sau-4079-1 | Bacteria Firmicutes Bacilli Bacillales *Staphylococcus aureus subsp. aureus* DR10 |
| Sau-3743-1 to Sau-3743-2 | Bacteria Firmicutes Bacilli Bacillales *Staphylococcus aureus subsp. aureus* DSM 20231 |
| Sau-4080-1 | Bacteria Firmicutes Bacilli Bacillales *Staphylococcus aureus subsp. aureus* GR1 |
| Sau-4081-1 | Bacteria Firmicutes Bacilli Bacillales *Staphylococcus aureus subsp. aureus* IS-105 |
| Sau-4082-1 | Bacteria Firmicutes Bacilli Bacillales *Staphylococcus aureus subsp. aureus* IS-111 |
| Sau-4083-1 | Bacteria Firmicutes Bacilli Bacillales *Staphylococcus aureus subsp. aureus* IS-122 |
| Sau-4084-1 | Bacteria Firmicutes Bacilli Bacillales *Staphylococcus aureus subsp. aureus* IS-157 |
| Sau-4085-1 | Bacteria Firmicutes Bacilli Bacillales *Staphylococcus aureus subsp. aureus* IS-160 |
| Sau-4086-1 | Bacteria Firmicutes Bacilli Bacillales *Staphylococcus aureus subsp. aureus* IS-24 |
| Sau-4087-1 | Bacteria Firmicutes Bacilli Bacillales *Staphylococcus aureus subsp. aureus* IS-3 |
| Sau-4088-1 | Bacteria Firmicutes Bacilli Bacillales *Staphylococcus aureus subsp. aureus* IS-88 |
| Sau-4089-1 | Bacteria Firmicutes Bacilli Bacillales *Staphylococcus aureus subsp. aureus* IS-91 |
| Sau-4090-1 | Bacteria Firmicutes Bacilli Bacillales *Staphylococcus aureus subsp. aureus* IS-99 |
| Sau-4091-1 | Bacteria Firmicutes Bacilli Bacillales *Staphylococcus aureus subsp. aureus* IS-M |
| Sau-4092-1 | Bacteria Firmicutes Bacilli Bacillales *Staphylococcus aureus subsp. aureus* KPL1828 |
| Sau-4093-1 | Bacteria Firmicutes Bacilli Bacillales *Staphylococcus aureus subsp. aureus* KPL1845 |
| Sau-4094-1 | Bacteria Firmicutes Bacilli Bacillales *Staphylococcus aureus subsp. aureus* LCT-SA112 |
| Sau-4095-1 | Bacteria Firmicutes Bacilli Bacillales *Staphylococcus aureus subsp. aureus* MRGR3 |
| Sau-4096-1 | Bacteria Firmicutes Bacilli Bacillales *Staphylococcus aureus subsp. aureus* MRSA131 |
| Sau-4097-1 | Bacteria Firmicutes Bacilli Bacillales *Staphylococcus aureus subsp. aureus* MRSA177 |
| Sau-4098-1 | Bacteria Firmicutes Bacilli Bacillales *Staphylococcus aureus subsp. aureus* NN50 |
| Sau-4099-1 | Bacteria Firmicutes Bacilli Bacillales *Staphylococcus aureus subsp. aureus* PSP1996 |
| Sau-4100-1 | Bacteria Firmicutes Bacilli Bacillales *Staphylococcus aureus subsp. aureus* RN4220 |
| Sau-4101-1 | Bacteria Firmicutes Bacilli Bacillales *Staphylococcus aureus subsp. aureus* SA40 |
| Sau-4102-1 | Bacteria Firmicutes Bacilli Bacillales *Staphylococcus aureus subsp. aureus* SA957 |
| Sau-4103-1 to Sau-4103-8 | Bacteria Firmicutes Bacilli Bacillales *Staphylococcus aureus subsp. aureus* ST228 |

| | |
|---|---|
| Sau-4121-1 | Bacteria Firmicutes Bacilli Bacillales *Staphylococcus aureus subsp. aureus str.* Newbould 305 |
| Sau-4105-1 | Bacteria Firmicutes Bacilli Bacillales *Staphylococcus aureus subsp. aureus* Tager 104 |
| Sau-4104-1 | Bacteria Firmicutes Bacilli Bacillales *Staphylococcus aureus subsp. aureus* TCH70 |
| Sau-4106-1 | Bacteria Firmicutes Bacilli Bacillales *Staphylococcus aureus subsp. aureus* VCU006 |
| Sau-4107-1 | Bacteria Firmicutes Bacilli Bacillales *Staphylococcus aureus subsp. aureus* VH60 |
| Sau-4108-1 | Bacteria Firmicutes Bacilli Bacillales *Staphylococcus aureus subsp. aureus* VRS1 |
| Sau-4109-1 | Bacteria Firmicutes Bacilli Bacillales *Staphylococcus aureus subsp. aureus* VRS10 |
| Sau-4110-1 | Bacteria Firmicutes Bacilli Bacillales *Staphylococcus aureus subsp. aureus* VRS11a |
| Sau-4111-1 | Bacteria Firmicutes Bacilli Bacillales *Staphylococcus aureus subsp. aureus* VRS11b |
| Sau-4112-1 | Bacteria Firmicutes Bacilli Bacillales *Staphylococcus aureus subsp. aureus* VRS2 |
| Sau-4113-1 | Bacteria Firmicutes Bacilli Bacillales *Staphylococcus aureus subsp. aureus* VRS3a |
| Sau-4114-1 | Bacteria Firmicutes Bacilli Bacillales *Staphylococcus aureus subsp. aureus* VRS4 |
| Sau-4115-1 | Bacteria Firmicutes Bacilli Bacillales *Staphylococcus aureus subsp. aureus* VRS5 |
| Sau-4116-1 | Bacteria Firmicutes Bacilli Bacillales *Staphylococcus aureus subsp. aureus* VRS6 |
| Sau-4117-1 | Bacteria Firmicutes Bacilli Bacillales *Staphylococcus aureus subsp. aureus* VRS7 |
| Sau-4118-1 | Bacteria Firmicutes Bacilli Bacillales *Staphylococcus aureus subsp. aureus* VRS8 |
| Sau-4119-1 | Bacteria Firmicutes Bacilli Bacillales *Staphylococcus aureus subsp. aureus* VRS9 |
| Sau-4120-1 | Bacteria Firmicutes Bacilli Bacillales *Staphylococcus aureus subsp. aureus* Z172 |
| Sau-3995-1 | Bacteria Firmicutes Bacilli Bacillales *Staphylococcus aureus* VH221 |
| Sca-9-1 | Bacteria Firmicutes Bacilli Bacillales *Staphylococcus capitis* CR01 |
| Sca-10-1 | Bacteria Firmicutes Bacilli Bacillales *Staphylococcus capitis* QN1 |
| Sca-11-1 | Bacteria Firmicutes Bacilli Bacillales *Staphylococcus capitis* VCU116 |
| Sde-1-1 | Bacteria Firmicutes Bacilli Bacillales *Staphylococcus delphini* 8086 |
| Sep-8-1 to Sep-8-3 | Bacteria Firmicutes Bacilli Bacillales *Staphylococcus epidermidis* |
| Sep-18-1 | Bacteria Firmicutes Bacilli Bacillales *Staphylococcus epidermidis* 12142587 |
| Sep-19-1 | Bacteria Firmicutes Bacilli Bacillales *Staphylococcus epidermidis* 14.1.R1.SE |
| Sep-20-1 | Bacteria Firmicutes Bacilli Bacillales *Staphylococcus epidermidis* APO27 |
| Sep-21-1 | Bacteria Firmicutes Bacilli Bacillales *Staphylococcus epidermidis* APO35 |
| Sep-22-1 | Bacteria Firmicutes Bacilli Bacillales *Staphylococcus epidermidis* AU12-03 |
| Sep-23-1 | Bacteria Firmicutes Bacilli Bacillales *Staphylococcus epidermidis* BVS058A4 |
| Sep-24-1 | Bacteria Firmicutes Bacilli Bacillales *Staphylococcus epidermidis* CIM28 |
| Sep-25-1 | Bacteria Firmicutes Bacilli Bacillales *Staphylococcus epidermidis* CIM37 |
| Sep-26-1 | Bacteria Firmicutes Bacilli Bacillales *Staphylococcus epidermidis* CIM40 |
| Sep-27-1 | Bacteria Firmicutes Bacilli Bacillales *Staphylococcus epidermidis* E13A |
| Sep-28-1 | Bacteria Firmicutes Bacilli Bacillales *Staphylococcus epidermidis* FRI909 |
| Sep-29-1 | Bacteria Firmicutes Bacilli Bacillales *Staphylococcus epidermidis* IS-250 |
| Sep-30-1 | Bacteria Firmicutes Bacilli Bacillales *Staphylococcus epidermidis* IS-K |
| Sep-31-1 | Bacteria Firmicutes Bacilli Bacillales *Staphylococcus epidermidis* M0881 |
| Sep-32-1 | Bacteria Firmicutes Bacilli Bacillales *Staphylococcus epidermidis* MC16 |
| Sep-33-1 | Bacteria Firmicutes Bacilli Bacillales *Staphylococcus epidermidis* MC19 |
| Sep-34-1 | Bacteria Firmicutes Bacilli Bacillales *Staphylococcus epidermidis* MC28 |
| Sep-35-1 | Bacteria Firmicutes Bacilli Bacillales *Staphylococcus epidermidis* NIH04003 |
| Sep-36-1 | Bacteria Firmicutes Bacilli Bacillales *Staphylococcus epidermidis* NIH04008 |
| Sep-37-1 | Bacteria Firmicutes Bacilli Bacillales *Staphylococcus epidermidis* NIH05001 |
| Sep-38-1 | Bacteria Firmicutes Bacilli Bacillales *Staphylococcus epidermidis* NIH05003 |
| Sep-39-1 | Bacteria Firmicutes Bacilli Bacillales *Staphylococcus epidermidis* NIH05005 |
| Sep-40-1 | Bacteria Firmicutes Bacilli Bacillales *Staphylococcus epidermidis* NIH051475 |
| Sep-41-1 | Bacteria Firmicutes Bacilli Bacillales *Staphylococcus epidermidis* NIH051668 |
| Sep-42-1 | Bacteria Firmicutes Bacilli Bacillales *Staphylococcus epidermidis* NIH06004 |
| Sep-43-1 | Bacteria Firmicutes Bacilli Bacillales *Staphylococcus epidermidis* NIH08001 |
| Sep-44-1 | Bacteria Firmicutes Bacilli Bacillales *Staphylococcus epidermidis* NIHLM001 |
| Sep-45-1 | Bacteria Firmicutes Bacilli Bacillales *Staphylococcus epidermidis* NIHLM003 |
| Sep-46-1 | Bacteria Firmicutes Bacilli Bacillales *Staphylococcus epidermidis* NIHLM008 |
| Sep-47-1 | Bacteria Firmicutes Bacilli Bacillales *Staphylococcus epidermidis* NIHLM015 |
| Sep-48-1 | Bacteria Firmicutes Bacilli Bacillales *Staphylococcus epidermidis* NIHLM018 |
| Sep-49-1 | Bacteria Firmicutes Bacilli Bacillales *Staphylococcus epidermidis* NIHLM020 |
| Sep-50-1 | Bacteria Firmicutes Bacilli Bacillales *Staphylococcus epidermidis* NIHLM021 |
| Sep-51-1 | Bacteria Firmicutes Bacilli Bacillales *Staphylococcus epidermidis* NIHLM023 |
| Sep-52-1 | Bacteria Firmicutes Bacilli Bacillales *Staphylococcus epidermidis* NIHLM031 |
| Sep-53-1 | Bacteria Firmicutes Bacilli Bacillales *Staphylococcus epidermidis* NIHLM037 |
| Sep-54-1 | Bacteria Firmicutes Bacilli Bacillales *Staphylococcus epidermidis* NIHLM039 |
| Sep-55-1 | Bacteria Firmicutes Bacilli Bacillales *Staphylococcus epidermidis* NIHLM040 |
| Sep-56-1 | Bacteria Firmicutes Bacilli Bacillales *Staphylococcus epidermidis* NIHLM049 |
| Sep-57-1 | Bacteria Firmicutes Bacilli Bacillales *Staphylococcus epidermidis* NIHLM053 |
| Sep-58-1 | Bacteria Firmicutes Bacilli Bacillales *Staphylococcus epidermidis* NIHLM057 |

| | |
|---|---|
| Sep-59-1 | Bacteria Firmicutes Bacilli Bacillales *Staphylococcus epidermidis* NIHLM061 |
| Sep-60-1 | Bacteria Firmicutes Bacilli Bacillales *Staphylococcus epidermidis* NIHLM067 |
| Sep-61-1 | Bacteria Firmicutes Bacilli Bacillales *Staphylococcus epidermidis* NIHLM087 |
| Sep-62-1 | Bacteria Firmicutes Bacilli Bacillales *Staphylococcus epidermidis* NIHLM088 |
| Sep-63-1 | Bacteria Firmicutes Bacilli Bacillales *Staphylococcus epidermidis* NIHLM095 |
| Sep-64-1 | Bacteria Firmicutes Bacilli Bacillales *Staphylococcus epidermidis* Scl19 |
| Sep-65-1 | Bacteria Firmicutes Bacilli Bacillales *Staphylococcus epidermidis* Scl22 |
| Sep-66-1 | Bacteria Firmicutes Bacilli Bacillales *Staphylococcus epidermidis* Scl25 |
| Sep-67-1 | Bacteria Firmicutes Bacilli Bacillales *Staphylococcus epidermidis* Scl31 |
| Sep-68-1 | Bacteria Firmicutes Bacilli Bacillales *Staphylococcus epidermidis* VCU028 |
| Sep-69-1 | Bacteria Firmicutes Bacilli Bacillales *Staphylococcus epidermidis* VCU037 |
| Sep-70-1 | Bacteria Firmicutes Bacilli Bacillales *Staphylococcus epidermidis* VCU041 |
| Sep-71-1 | Bacteria Firmicutes Bacilli Bacillales *Staphylococcus epidermidis* VCU045 |
| Sep-72-1 | Bacteria Firmicutes Bacilli Bacillales *Staphylococcus epidermidis* VCU065 |
| Sep-73-1 | Bacteria Firmicutes Bacilli Bacillales *Staphylococcus epidermidis* VCU071 |
| Sep-74-1 | Bacteria Firmicutes Bacilli Bacillales *Staphylococcus epidermidis* VCU081 |
| Sep-75-1 | Bacteria Firmicutes Bacilli Bacillales *Staphylococcus epidermidis* VCU105 |
| Sep-76-1 | Bacteria Firmicutes Bacilli Bacillales *Staphylococcus epidermidis* VCU109 |
| Sep-77-1 | Bacteria Firmicutes Bacilli Bacillales *Staphylococcus epidermidis* VCU117 |
| Sep-78-1 | Bacteria Firmicutes Bacilli Bacillales *Staphylococcus epidermidis* VCU118 |
| Sep-79-1 | Bacteria Firmicutes Bacilli Bacillales *Staphylococcus epidermidis* VCU120 |
| Sep-80-1 | Bacteria Firmicutes Bacilli Bacillales *Staphylococcus epidermidis* VCU123 |
| Sep-81-1 | Bacteria Firmicutes Bacilli Bacillales *Staphylococcus epidermidis* VCU125 |
| Sep-82-1 | Bacteria Firmicutes Bacilli Bacillales *Staphylococcus epidermidis* VCU126 |
| Sep-83-1 | Bacteria Firmicutes Bacilli Bacillales *Staphylococcus epidermidis* VCU127 |
| Sep-84-1 | Bacteria Firmicutes Bacilli Bacillales *Staphylococcus epidermidis* VCU128 |
| Sep-85-1 | Bacteria Firmicutes Bacilli Bacillales *Staphylococcus epidermidis* VCU129 |
| Sep-86-1 | Bacteria Firmicutes Bacilli Bacillales *Staphylococcus epidermidis* VCU144 |
| Sep-87-1 | Bacteria Firmicutes Bacilli Bacillales *Staphylococcus epidermidis* WI05 |
| Sep-88-1 | Bacteria Firmicutes Bacilli Bacillales *Staphylococcus epidermidis* WI09 |
| Seq-3-1 | Bacteria Firmicutes Bacilli Bacillales *Staphylococcus equorum* UMC-CNS-924 |
| Sha-4-1 to Sha-4-119 | Bacteria Firmicutes Bacilli Bacillales *Staphylococcus haemolyticus* |
| Sha-6-1 | Bacteria Firmicutes Bacilli Bacillales *Staphylococcus haemolyticus* R1P1 |
| Sho-5-1 | Bacteria Firmicutes Bacilli Bacillales *Staphylococcus hominis subsp. hominis* ZBW5 |
| Sho-4-1 | Bacteria Firmicutes Bacilli Bacillales *Staphylococcus hominis* VCU122 |
| Sin-2-1 | Bacteria Firmicutes Bacilli Bacillales *Staphylococcus intermedius* NCTC 11048 |
| Slu-7-1 | Bacteria Firmicutes Bacilli Bacillales *Staphylococcus lugdunensis* ACS-027-V-Sch2 |
| Slu-8-1 | Bacteria Firmicutes Bacilli Bacillales *Staphylococcus lugdunensis* M23590 |
| Slu-9-1 | Bacteria Firmicutes Bacilli Bacillales *Staphylococcus lugdunensis* VCU139 |
| Sma-1-1 | Bacteria Firmicutes Bacilli Bacillales *Staphylococcus massiliensis* CCUG 55927 |
| Sma-2-1 | Bacteria Firmicutes Bacilli Bacillales *Staphylococcus massiliensis* S46 |
| Spa-2-1 | Bacteria Firmicutes Bacilli Bacillales *Staphylococcus pasteuri* SP1 |
| Spe-2-1 | Bacteria Firmicutes Bacilli Bacillales *Staphylococcus pettenkoferi* VCU012 |
| Ssa-3-1 | Bacteria Firmicutes Bacilli Bacillales *Staphylococcus saprophyticus subsp. saprophyticus* KACC 16562 |
| Ssi-3-1 | Bacteria Firmicutes Bacilli Bacillales *Staphylococcus simulans* ACS-120-V-Sch1 |
| Ssi-4-1 | Bacteria Firmicutes Bacilli Bacillales *Staphylococcus simulans* UMC-CNS-990 |
| Ssp-11-1 | Bacteria Firmicutes Bacilli Bacillales *Staphylococcus sp.* AL1 |
| Ssp-12-1 | Bacteria Firmicutes Bacilli Bacillales *Staphylococcus sp.* E463 |
| Ssp-13-1 | Bacteria Firmicutes Bacilli Bacillales *Staphylococcus sp.* EGD-HP3 |
| Ssp-14-1 | Bacteria Firmicutes Bacilli Bacillales *Staphylococcus sp.* HGB0015 |
| Svi-2-1 | Bacteria Firmicutes Bacilli Bacillales *Staphylococcus vitulinus* F1028 |
| Swa-5-1 | Bacteria Firmicutes Bacilli Bacillales *Staphylococcus warneri* A487 |
| Swa-6-1 | Bacteria Firmicutes Bacilli Bacillales *Staphylococcus warneri* SG1 |
| Swa-7-1 | Bacteria Firmicutes Bacilli Bacillales *Staphylococcus warneri* VCU121 |
| Sxy-3-1 | Bacteria Firmicutes Bacilli Bacillales *Staphylococcus xylosus* NJ |
| Bba-4-1 | Bacteria Firmicutes *Bacilli bacterium* VT-13-104 |
| Fso-1-1 | Bacteria Firmicutes Bacilli Lactobacillales Aerococcaceae *Facklamia sourekii* ATCC 700629 |
| Aot-1-1 | Bacteria Firmicutes Bacilli Lactobacillales Carnobacteriaceae *Alloiococcus otitis* ATCC 51267 |
| Aph-1-1 | Bacteria Firmicutes Bacilli Lactobacillales Carnobacteriaceae *Atopobacter phocae* ATCC BAA-285 |
| Ata-1-1 | Bacteria Firmicutes Bacilli Lactobacillales Carnobacteriaceae *Atopococcus tabaci* DSM 17538 |
| Cal-1-1 | Bacteria Firmicutes Bacilli Lactobacillales Carnobacteriaceae *Carnobacterium alterfunditum* DSM 5972 |
| Cdi-1-1 | Bacteria Firmicutes Bacilli Lactobacillales Carnobacteriaceae *Carnobacterium divergens* DSM 20623 |
| Cfu-1-1 | Bacteria Firmicutes Bacilli Lactobacillales Carnobacteriaceae *Carnobacterium funditum* DSM 5970 |
| Cga-1-1 | Bacteria Firmicutes Bacilli Lactobacillales Carnobacteriaceae *Carnobacterium gallinarum* DSM 4847 |
| Cin-1-1 | Bacteria Firmicutes Bacilli Lactobacillales Carnobacteriaceae *Carnobacterium inhibens subsp. inhibens* DSM 13024 |

| | |
|---|---|
| Cje-1-1 | Bacteria Firmicutes Bacilli Lactobacillales Carnobacteriaceae *Carnobacterium jeotgali* |
| Cje-2-1 | Bacteria Firmicutes Bacilli Lactobacillales Carnobacteriaceae *Carnobacterium jeotgali* MS3 |
| Cma-1-1 | Bacteria Firmicutes Bacilli Lactobacillales Carnobacteriaceae *Carnobacterium maltaromaticum* |
| Cma-2-1 | Bacteria Firmicutes Bacilli Lactobacillales Carnobacteriaceae *Carnobacterium maltaromaticum* DSM 20342 |
| Cma-3-1 | Bacteria Firmicutes Bacilli Lactobacillales Carnobacteriaceae *Carnobacterium maltaromaticum* LMA28 |
| Cmo-1-1 | Bacteria Firmicutes Bacilli Lactobacillales Carnobacteriaceae *Carnobacterium mobile* DSM 4848 |
| Cpl-1-1 | Bacteria Firmicutes Bacilli Lactobacillales Carnobacteriaceae *Carnobacterium pleistocenium* FTR1 |
| Csp-1-1 | Bacteria Firmicutes Bacilli Lactobacillales Carnobacteriaceae *Carnobacterium sp.* WN1359 |
| Csp-2-1 | Bacteria Firmicutes Bacilli Lactobacillales Carnobacteriaceae *Carnobacterium sp.* WN1374 |
| Csp-3-1 | Bacteria Firmicutes Bacilli Lactobacillales Carnobacteriaceae *Carnobacterium sp.* ZWU0011 |
| Gel-1-1 | Bacteria Firmicutes Bacilli Lactobacillales Carnobacteriaceae *Granulicatella elegans* ATCC 700633 |
| Gsp-19-1 | Bacteria Firmicutes Bacilli Lactobacillales Carnobacteriaceae *Granulicatella sp.* 572.rep1_STHE |
| Gsp-20-1 | Bacteria Firmicutes Bacilli Lactobacillales Carnobacteriaceae *Granulicatella sp.* 572.rep2_STHE |
| Lna-1-1 | Bacteria Firmicutes Bacilli Lactobacillales Carnobacteriaceae *Lacticigenium naphtae* DSM 19658 |
| Bse-3-1 | Bacteria Firmicutes Bacilli Lactobacillales Enterococcaceae *Bavariicoccus seileri* DSM 19936 |
| Cma-4-1 | Bacteria Firmicutes Bacilli Lactobacillales Enterococcaceae *Catellicoccus marimammalium* M35/04/3 |
| Eas-1-1 | Bacteria Firmicutes Bacilli Lactobacillales Enterococcaceae *Enterococcus asini* ATCC 700915 |
| Eav-1-1 | Bacteria Firmicutes Bacilli Lactobacillales Enterococcaceae *Enterococcus avium* |
| Eav-2-1 to Eav-2-2 | Bacteria Firmicutes Bacilli Lactobacillales Enterococcaceae *Enterococcus avium* ATCC 14025 |
| Eca-1-1 | Bacteria Firmicutes Bacilli Lactobacillales Enterococcaceae *Enterococcus caccae* ATCC BAA-1240 |
| Eca-2-1 | Bacteria Firmicutes Bacilli Lactobacillales Enterococcaceae *Enterococcus casseliflavus* 14-MB-W-14 |
| Eca-3-1 | Bacteria Firmicutes Bacilli Lactobacillales Enterococcaceae *Enterococcus casseliflavus* ATCC 12755 |
| Eca-4-1 | Bacteria Firmicutes Bacilli Lactobacillales Enterococcaceae *Enterococcus casseliflavus* EC20 |
| Ece-1-1 to Ece-1-7 | Bacteria Firmicutes Bacilli Lactobacillales Enterococcaceae *Enterococcus cecorum* |
| Ece-2-1 | Bacteria Firmicutes Bacilli Lactobacillales Enterococcaceae *Enterococcus cecorum* DSM 20682 = ATCC 43198 |
| Eco-1-1 | Bacteria Firmicutes Bacilli Lactobacillales Enterococcaceae *Enterococcus columbae* DSM 7374 = ATCC 51263 |
| Edi-1-1 | Bacteria Firmicutes Bacilli Lactobacillales Enterococcaceae *Enterococcus dispar* ATCC 51266 |
| Edu-1-1 to Edu-1-2 | Bacteria Firmicutes Bacilli Lactobacillales Enterococcaceae *Enterococcus durans* |
| Edu-2-1 to Edu-2-2 | Bacteria Firmicutes Bacilli Lactobacillales Enterococcaceae *Enterococcus durans* ATCC 6056 |
| Edu-3-1 | Bacteria Firmicutes Bacilli Lactobacillales Enterococcaceae *Enterococcus durans* FB129-CNAB-4 |
| Edu-4-1 | Bacteria Firmicutes Bacilli Lactobacillales Enterococcaceae *Enterococcus durans* IPLA 655 |
| Efa-1-1 to Efa-1-87 | Bacteria Firmicutes Bacilli Lactobacillales Enterococcaceae *Enterococcus faecalis* |
| Efa-2-1 | Bacteria Firmicutes Bacilli Lactobacillales Enterococcaceae *Enterococcus faecalis* 02-MB-BW-10 |
| Efa-3-1 | Bacteria Firmicutes Bacilli Lactobacillales Enterococcaceae *Enterococcus faecalis* 02-MB-P-10 |
| Efa-4-1 | Bacteria Firmicutes Bacilli Lactobacillales Enterococcaceae *Enterococcus faecalis* 06-MB-DW-09 |
| Efa-5-1 | Bacteria Firmicutes Bacilli Lactobacillales Enterococcaceae *Enterococcus faecalis* 06-MB-S-04 |
| Efa-6-1 | Bacteria Firmicutes Bacilli Lactobacillales Enterococcaceae *Enterococcus faecalis* 06-MB-S-10 |
| Efa-7-1 | Bacteria Firmicutes Bacilli Lactobacillales Enterococcaceae *Enterococcus faecalis* 10244 |
| Efa-8-1 | Bacteria Firmicutes Bacilli Lactobacillales Enterococcaceae *Enterococcus faecalis* 13-SD-W-01 |
| Efa-9-1 | Bacteria Firmicutes Bacilli Lactobacillales Enterococcaceae *Enterococcus faecalis* 20-SD-BW-06 |
| Efa-10-1 | Bacteria Firmicutes Bacilli Lactobacillales Enterococcaceae *Enterococcus faecalis* 20-SD-BW-08 |
| Efa-11-1 | Bacteria Firmicutes Bacilli Lactobacillales Enterococcaceae *Enterococcus faecalis* 20.SD.W.06 |
| Efa-12-1 | Bacteria Firmicutes Bacilli Lactobacillales Enterococcaceae *Enterococcus faecalis* 599 |
| Efa-13-1 | Bacteria Firmicutes Bacilli Lactobacillales Enterococcaceae *Enterococcus faecalis* 918 |
| Efa-14-1 | Bacteria Firmicutes Bacilli Lactobacillales Enterococcaceae *Enterococcus faecalis* ATCC 10100 |
| Efa-15-1 | Bacteria Firmicutes Bacilli Lactobacillales Enterococcaceae *Enterococcus faecalis* ATCC 19433 |
| Efa-16-1 | Bacteria Firmicutes Bacilli Lactobacillales Enterococcaceae *Enterococcus faecalis* ATCC 27275 |
| Efa-17-1 | Bacteria Firmicutes Bacilli Lactobacillales Enterococcaceae *Enterococcus faecalis* ATCC 27959 |
| Efa-18-1 to Efa-18-2 | Bacteria Firmicutes Bacilli Lactobacillales Enterococcaceae *Enterococcus faecalis* ATCC 29200 |
| Efa-19-1 | Bacteria Firmicutes Bacilli Lactobacillales Enterococcaceae *Enterococcus faecalis* ATCC 29212 |
| Efa-20-1 | Bacteria Firmicutes Bacilli Lactobacillales Enterococcaceae *Enterococcus faecalis* ATCC 35038 |
| Efa-21-1 | Bacteria Firmicutes Bacilli Lactobacillales Enterococcaceae *Enterococcus faecalis* ATCC 6055 |
| Efa-22-1 | Bacteria Firmicutes Bacilli Lactobacillales Enterococcaceae *Enterococcus faecalis* AZ19 |
| Efa-23-1 | Bacteria Firmicutes Bacilli Lactobacillales Enterococcaceae *Enterococcus faecalis* B83616-1 |
| Efa-24-1 | Bacteria Firmicutes Bacilli Lactobacillales Enterococcaceae *Enterococcus faecalis* BM4539 |
| Efa-25-1 | Bacteria Firmicutes Bacilli Lactobacillales Enterococcaceae *Enterococcus faecalis* BM4654 |
| Efa-26-1 | Bacteria Firmicutes Bacilli Lactobacillales Enterococcaceae *Enterococcus faecalis* D811610-10 |
| Efa-27-1 | Bacteria Firmicutes Bacilli Lactobacillales Enterococcaceae *Enterococcus faecalis* DAPTO 512 |
| Efa-28-1 | Bacteria Firmicutes Bacilli Lactobacillales Enterococcaceae *Enterococcus faecalis* DAPTO 516 |
| Efa-29-1 | Bacteria Firmicutes Bacilli Lactobacillales Enterococcaceae *Enterococcus faecalis* DENG1 |
| Efa-30-1 | Bacteria Firmicutes Bacilli Lactobacillales Enterococcaceae *Enterococcus faecalis* E12 |
| Efa-47-1 | Bacteria Firmicutes Bacilli Lactobacillales Enterococcaceae *Enterococcus faecalis* Efa HS0914 |
| Efa-48-1 | Bacteria Firmicutes Bacilli Lactobacillales Enterococcaceae *Enterococcus faecalis* EnGen0058 |
| Efa-49-1 | Bacteria Firmicutes Bacilli Lactobacillales Enterococcaceae *Enterococcus faecalis* EnGen0059 |
| Efa-50-1 | Bacteria Firmicutes Bacilli Lactobacillales Enterococcaceae *Enterococcus faecalis* EnGen0060 |

| | |
|---|---|
| Efa-51-1 | Bacteria Firmicutes Bacilli Lactobacillales Enterococcaceae *Enterococcus faecalis* EnGen0061 |
| Efa-52-1 | Bacteria Firmicutes Bacilli Lactobacillales Enterococcaceae *Enterococcus faecalis* EnGen0062 |
| Efa-53-1 | Bacteria Firmicutes Bacilli Lactobacillales Enterococcaceae *Enterococcus faecalis* EnGen0063 |
| Efa-54-1 | Bacteria Firmicutes Bacilli Lactobacillales Enterococcaceae *Enterococcus faecalis* EnGen0064 |
| Efa-55-1 | Bacteria Firmicutes Bacilli Lactobacillales Enterococcaceae *Enterococcus faecalis* EnGen0065 |
| Efa-56-1 | Bacteria Firmicutes Bacilli Lactobacillales Enterococcaceae *Enterococcus faecalis* EnGen0066 |
| Efa-57-1 | Bacteria Firmicutes Bacilli Lactobacillales Enterococcaceae *Enterococcus faecalis* EnGen0067 |
| Efa-58-1 | Bacteria Firmicutes Bacilli Lactobacillales Enterococcaceae *Enterococcus faecalis* EnGen0068 |
| Efa-59-1 | Bacteria Firmicutes Bacilli Lactobacillales Enterococcaceae *Enterococcus faecalis* EnGen0069 |
| Efa-60-1 | Bacteria Firmicutes Bacilli Lactobacillales Enterococcaceae *Enterococcus faecalis* EnGen0070 |
| Efa-61-1 | Bacteria Firmicutes Bacilli Lactobacillales Enterococcaceae *Enterococcus faecalis* EnGen0071 |
| Efa-62-1 | Bacteria Firmicutes Bacilli Lactobacillales Enterococcaceae *Enterococcus faecalis* EnGen0072 |
| Efa-63-1 | Bacteria Firmicutes Bacilli Lactobacillales Enterococcaceae *Enterococcus faecalis* EnGen0073 |
| Efa-64-1 | Bacteria Firmicutes Bacilli Lactobacillales Enterococcaceae *Enterococcus faecalis* EnGen0074 |
| Efa-65-1 | Bacteria Firmicutes Bacilli Lactobacillales Enterococcaceae *Enterococcus faecalis* EnGen0075 |
| Efa-66-1 | Bacteria Firmicutes Bacilli Lactobacillales Enterococcaceae *Enterococcus faecalis* EnGen0076 |
| Efa-67-1 | Bacteria Firmicutes Bacilli Lactobacillales Enterococcaceae *Enterococcus faecalis* EnGen0078 |
| Efa-68-1 | Bacteria Firmicutes Bacilli Lactobacillales Enterococcaceae *Enterococcus faecalis* EnGen0079 |
| Efa-69-1 | Bacteria Firmicutes Bacilli Lactobacillales Enterococcaceae *Enterococcus faecalis* EnGen0080 |
| Efa-70-1 | Bacteria Firmicutes Bacilli Lactobacillales Enterococcaceae *Enterococcus faecalis* EnGen0081 |
| Efa-71-1 | Bacteria Firmicutes Bacilli Lactobacillales Enterococcaceae *Enterococcus faecalis* EnGen0082 |
| Efa-72-1 | Bacteria Firmicutes Bacilli Lactobacillales Enterococcaceae *Enterococcus faecalis* EnGen0083 |
| Efa-73-1 | Bacteria Firmicutes Bacilli Lactobacillales Enterococcaceae *Enterococcus faecalis* EnGen0084 |
| Efa-74-1 | Bacteria Firmicutes Bacilli Lactobacillales Enterococcaceae *Enterococcus faecalis* EnGen0085 |
| Efa-75-1 | Bacteria Firmicutes Bacilli Lactobacillales Enterococcaceae *Enterococcus faecalis* EnGen0086 |
| Efa-76-1 | Bacteria Firmicutes Bacilli Lactobacillales Enterococcaceae *Enterococcus faecalis* EnGen0087 |
| Efa-77-1 | Bacteria Firmicutes Bacilli Lactobacillales Enterococcaceae *Enterococcus faecalis* EnGen0088 |
| Efa-78-1 | Bacteria Firmicutes Bacilli Lactobacillales Enterococcaceae *Enterococcus faecalis* EnGen0089 |
| Efa-79-1 | Bacteria Firmicutes Bacilli Lactobacillales Enterococcaceae *Enterococcus faecalis* EnGen0090 |
| Efa-80-1 | Bacteria Firmicutes Bacilli Lactobacillales Enterococcaceae *Enterococcus faecalis* EnGen0091 |
| Efa-81-1 | Bacteria Firmicutes Bacilli Lactobacillales Enterococcaceae *Enterococcus faecalis* EnGen0092 |
| Efa-82-1 | Bacteria Firmicutes Bacilli Lactobacillales Enterococcaceae *Enterococcus faecalis* EnGen0093 |
| Efa-83-1 | Bacteria Firmicutes Bacilli Lactobacillales Enterococcaceae *Enterococcus faecalis* EnGen0094 |
| Efa-84-1 | Bacteria Firmicutes Bacilli Lactobacillales Enterococcaceae *Enterococcus faecalis* EnGen0095 |
| Efa-85-1 | Bacteria Firmicutes Bacilli Lactobacillales Enterococcaceae *Enterococcus faecalis* EnGen0096 |
| Efa-86-1 | Bacteria Firmicutes Bacilli Lactobacillales Enterococcaceae *Enterococcus faecalis* EnGen0097 |
| Efa-87-1 | Bacteria Firmicutes Bacilli Lactobacillales Enterococcaceae *Enterococcus faecalis* EnGen0098 |
| Efa-88-1 | Bacteria Firmicutes Bacilli Lactobacillales Enterococcaceae *Enterococcus faecalis* EnGen0099 |
| Efa-89-1 | Bacteria Firmicutes Bacilli Lactobacillales Enterococcaceae *Enterococcus faecalis* EnGen0100 |
| Efa-90-1 | Bacteria Firmicutes Bacilli Lactobacillales Enterococcaceae *Enterococcus faecalis* EnGen0101 |
| Efa-91-1 | Bacteria Firmicutes Bacilli Lactobacillales Enterococcaceae *Enterococcus faecalis* EnGen0102 |
| Efa-92-1 | Bacteria Firmicutes Bacilli Lactobacillales Enterococcaceae *Enterococcus faecalis* EnGen0103 |
| Efa-93-1 | Bacteria Firmicutes Bacilli Lactobacillales Enterococcaceae *Enterococcus faecalis* EnGen0104 |
| Efa-94-1 | Bacteria Firmicutes Bacilli Lactobacillales Enterococcaceae *Enterococcus faecalis* EnGen0105 |
| Efa-95-1 | Bacteria Firmicutes Bacilli Lactobacillales Enterococcaceae *Enterococcus faecalis* EnGen0106 |
| Efa-96-1 | Bacteria Firmicutes Bacilli Lactobacillales Enterococcaceae *Enterococcus faecalis* EnGen0107 |
| Efa-97-1 | Bacteria Firmicutes Bacilli Lactobacillales Enterococcaceae *Enterococcus faecalis* EnGen0108 |
| Efa-98-1 | Bacteria Firmicutes Bacilli Lactobacillales Enterococcaceae *Enterococcus faecalis* EnGen0109 |
| Efa-99-1 | Bacteria Firmicutes Bacilli Lactobacillales Enterococcaceae *Enterococcus faecalis* EnGen0110 |
| Efa-100-1 | Bacteria Firmicutes Bacilli Lactobacillales Enterococcaceae *Enterococcus faecalis* EnGen0111 |
| Efa-101-1 | Bacteria Firmicutes Bacilli Lactobacillales Enterococcaceae *Enterococcus faecalis* EnGen0112 |
| Efa-102-1 | Bacteria Firmicutes Bacilli Lactobacillales Enterococcaceae *Enterococcus faecalis* EnGen0113 |
| Efa-103-1 | Bacteria Firmicutes Bacilli Lactobacillales Enterococcaceae *Enterococcus faecalis* EnGen0114 |
| Efa-104-1 | Bacteria Firmicutes Bacilli Lactobacillales Enterococcaceae *Enterococcus faecalis* EnGen0115 |
| Efa-105-1 | Bacteria Firmicutes Bacilli Lactobacillales Enterococcaceae *Enterococcus faecalis* EnGen0116 |
| Efa-106-1 | Bacteria Firmicutes Bacilli Lactobacillales Enterococcaceae *Enterococcus faecalis* EnGen0117 |
| Efa-107-1 | Bacteria Firmicutes Bacilli Lactobacillales Enterococcaceae *Enterococcus faecalis* EnGen0118 |
| Efa-108-1 | Bacteria Firmicutes Bacilli Lactobacillales Enterococcaceae *Enterococcus faecalis* EnGen0119 |
| Efa-109-1 | Bacteria Firmicutes Bacilli Lactobacillales Enterococcaceae *Enterococcus faecalis* EnGen0120 |
| Efa-110-1 | Bacteria Firmicutes Bacilli Lactobacillales Enterococcaceae *Enterococcus faecalis* EnGen0194 |
| Efa-111-1 | Bacteria Firmicutes Bacilli Lactobacillales Enterococcaceae *Enterococcus faecalis* EnGen0195 |
| Efa-112-1 | Bacteria Firmicutes Bacilli Lactobacillales Enterococcaceae *Enterococcus faecalis* EnGen0196 |
| Efa-113-1 | Bacteria Firmicutes Bacilli Lactobacillales Enterococcaceae *Enterococcus faecalis* EnGen0197 |
| Efa-114-1 | Bacteria Firmicutes Bacilli Lactobacillales Enterococcaceae *Enterococcus faecalis* EnGen0198 |
| Efa-115-1 | Bacteria Firmicutes Bacilli Lactobacillales Enterococcaceae *Enterococcus faecalis* EnGen0199 |

| | |
|---|---|
| Efa-116-1 | Bacteria Firmicutes Bacilli Lactobacillales Enterococcaceae *Enterococcus faecalis* EnGen0200 |
| Efa-117-1 | Bacteria Firmicutes Bacilli Lactobacillales Enterococcaceae *Enterococcus faecalis* EnGen0201 |
| Efa-118-1 | Bacteria Firmicutes Bacilli Lactobacillales Enterococcaceae *Enterococcus faecalis* EnGen0202 |
| Efa-119-1 | Bacteria Firmicutes Bacilli Lactobacillales Enterococcaceae *Enterococcus faecalis* EnGen0203 |
| Efa-120-1 | Bacteria Firmicutes Bacilli Lactobacillales Enterococcaceae *Enterococcus faecalis* EnGen0204 |
| Efa-121-1 | Bacteria Firmicutes Bacilli Lactobacillales Enterococcaceae *Enterococcus faecalis* EnGen0205 |
| Efa-122-1 | Bacteria Firmicutes Bacilli Lactobacillales Enterococcaceae *Enterococcus faecalis* EnGen0206 |
| Efa-123-1 | Bacteria Firmicutes Bacilli Lactobacillales Enterococcaceae *Enterococcus faecalis* EnGen0207 |
| Efa-124-1 | Bacteria Firmicutes Bacilli Lactobacillales Enterococcaceae *Enterococcus faecalis* EnGen0208 |
| Efa-125-1 | Bacteria Firmicutes Bacilli Lactobacillales Enterococcaceae *Enterococcus faecalis* EnGen0209 |
| Efa-126-1 | Bacteria Firmicutes Bacilli Lactobacillales Enterococcaceae *Enterococcus faecalis* EnGen0210 |
| Efa-127-1 | Bacteria Firmicutes Bacilli Lactobacillales Enterococcaceae *Enterococcus faecalis* EnGen0211 |
| Efa-128-1 | Bacteria Firmicutes Bacilli Lactobacillales Enterococcaceae *Enterococcus faecalis* EnGen0212 |
| Efa-129-1 | Bacteria Firmicutes Bacilli Lactobacillales Enterococcaceae *Enterococcus faecalis* EnGen0213 |
| Efa-130-1 | Bacteria Firmicutes Bacilli Lactobacillales Enterococcaceae *Enterococcus faecalis* EnGen0214 |
| Efa-131-1 | Bacteria Firmicutes Bacilli Lactobacillales Enterococcaceae *Enterococcus faecalis* EnGen0215 |
| Efa-132-1 | Bacteria Firmicutes Bacilli Lactobacillales Enterococcaceae *Enterococcus faecalis* EnGen0216 |
| Efa-133-1 | Bacteria Firmicutes Bacilli Lactobacillales Enterococcaceae *Enterococcus faecalis* EnGen0217 |
| Efa-134-1 | Bacteria Firmicutes Bacilli Lactobacillales Enterococcaceae *Enterococcus faecalis* EnGen0218 |
| Efa-135-1 | Bacteria Firmicutes Bacilli Lactobacillales Enterococcaceae *Enterococcus faecalis* EnGen0219 |
| Efa-136-1 | Bacteria Firmicutes Bacilli Lactobacillales Enterococcaceae *Enterococcus faecalis* EnGen0220 |
| Efa-137-1 | Bacteria Firmicutes Bacilli Lactobacillales Enterococcaceae *Enterococcus faecalis* EnGen0221 |
| Efa-138-1 | Bacteria Firmicutes Bacilli Lactobacillales Enterococcaceae *Enterococcus faecalis* EnGen0222 |
| Efa-139-1 | Bacteria Firmicutes Bacilli Lactobacillales Enterococcaceae *Enterococcus faecalis* EnGen0223 |
| Efa-140-1 | Bacteria Firmicutes Bacilli Lactobacillales Enterococcaceae *Enterococcus faecalis* EnGen0224 |
| Efa-141-1 | Bacteria Firmicutes Bacilli Lactobacillales Enterococcaceae *Enterococcus faecalis* EnGen0225 |
| Efa-142-1 | Bacteria Firmicutes Bacilli Lactobacillales Enterococcaceae *Enterococcus faecalis* EnGen0226 |
| Efa-143-1 | Bacteria Firmicutes Bacilli Lactobacillales Enterococcaceae *Enterococcus faecalis* EnGen0228 |
| Efa-144-1 | Bacteria Firmicutes Bacilli Lactobacillales Enterococcaceae *Enterococcus faecalis* EnGen0231 |
| Efa-145-1 | Bacteria Firmicutes Bacilli Lactobacillales Enterococcaceae *Enterococcus faecalis* EnGen0232 |
| Efa-146-1 | Bacteria Firmicutes Bacilli Lactobacillales Enterococcaceae *Enterococcus faecalis* EnGen0233 |
| Efa-147-1 | Bacteria Firmicutes Bacilli Lactobacillales Enterococcaceae *Enterococcus faecalis* EnGen0234 |
| Efa-148-1 | Bacteria Firmicutes Bacilli Lactobacillales Enterococcaceae *Enterococcus faecalis* EnGen0235 |
| Efa-149-1 | Bacteria Firmicutes Bacilli Lactobacillales Enterococcaceae *Enterococcus faecalis* EnGen0237 |
| Efa-150-1 | Bacteria Firmicutes Bacilli Lactobacillales Enterococcaceae *Enterococcus faecalis* EnGen0238 |
| Efa-151-1 | Bacteria Firmicutes Bacilli Lactobacillales Enterococcaceae *Enterococcus faecalis* EnGen0239 |
| Efa-152-1 | Bacteria Firmicutes Bacilli Lactobacillales Enterococcaceae *Enterococcus faecalis* EnGen0240 |
| Efa-153-1 | Bacteria Firmicutes Bacilli Lactobacillales Enterococcaceae *Enterococcus faecalis* EnGen0241 |
| Efa-154-1 | Bacteria Firmicutes Bacilli Lactobacillales Enterococcaceae *Enterococcus faecalis* EnGen0242 |
| Efa-155-1 | Bacteria Firmicutes Bacilli Lactobacillales Enterococcaceae *Enterococcus faecalis* EnGen0243 |
| Efa-156-1 | Bacteria Firmicutes Bacilli Lactobacillales Enterococcaceae *Enterococcus faecalis* EnGen0244 |
| Efa-157-1 | Bacteria Firmicutes Bacilli Lactobacillales Enterococcaceae *Enterococcus faecalis* EnGen0245 |
| Efa-158-1 | Bacteria Firmicutes Bacilli Lactobacillales Enterococcaceae *Enterococcus faecalis* EnGen0246 |
| Efa-159-1 | Bacteria Firmicutes Bacilli Lactobacillales Enterococcaceae *Enterococcus faecalis* EnGen0247 |
| Efa-160-1 | Bacteria Firmicutes Bacilli Lactobacillales Enterococcaceae *Enterococcus faecalis* EnGen0248 |
| Efa-161-1 | Bacteria Firmicutes Bacilli Lactobacillales Enterococcaceae *Enterococcus faecalis* EnGen0249 |
| Efa-162-1 | Bacteria Firmicutes Bacilli Lactobacillales Enterococcaceae *Enterococcus faecalis* EnGen0250 |
| Efa-163-1 | Bacteria Firmicutes Bacilli Lactobacillales Enterococcaceae *Enterococcus faecalis* EnGen0251 |
| Efa-164-1 | Bacteria Firmicutes Bacilli Lactobacillales Enterococcaceae *Enterococcus faecalis* EnGen0252 |
| Efa-165-1 | Bacteria Firmicutes Bacilli Lactobacillales Enterococcaceae *Enterococcus faecalis* EnGen0279 |
| Efa-166-1 | Bacteria Firmicutes Bacilli Lactobacillales Enterococcaceae *Enterococcus faecalis* EnGen0280 |
| Efa-167-1 | Bacteria Firmicutes Bacilli Lactobacillales Enterococcaceae *Enterococcus faecalis* EnGen0281 |
| Efa-168-1 | Bacteria Firmicutes Bacilli Lactobacillales Enterococcaceae *Enterococcus faecalis* EnGen0282 |
| Efa-169-1 | Bacteria Firmicutes Bacilli Lactobacillales Enterococcaceae *Enterococcus faecalis* EnGen0283 |
| Efa-170-1 | Bacteria Firmicutes Bacilli Lactobacillales Enterococcaceae *Enterococcus faecalis* EnGen0284 |
| Efa-171-1 | Bacteria Firmicutes Bacilli Lactobacillales Enterococcaceae *Enterococcus faecalis* EnGen0285 |
| Efa-172-1 | Bacteria Firmicutes Bacilli Lactobacillales Enterococcaceae *Enterococcus faecalis* EnGen0286 |
| Efa-173-1 | Bacteria Firmicutes Bacilli Lactobacillales Enterococcaceae *Enterococcus faecalis* EnGen0287 |
| Efa-174-1 | Bacteria Firmicutes Bacilli Lactobacillales Enterococcaceae *Enterococcus faecalis* EnGen0289 |
| Efa-175-1 | Bacteria Firmicutes Bacilli Lactobacillales Enterococcaceae *Enterococcus faecalis* EnGen0290 |
| Efa-176-1 | Bacteria Firmicutes Bacilli Lactobacillales Enterococcaceae *Enterococcus faecalis* EnGen0291 |
| Efa-177-1 | Bacteria Firmicutes Bacilli Lactobacillales Enterococcaceae *Enterococcus faecalis* EnGen0293 |
| Efa-178-1 | Bacteria Firmicutes Bacilli Lactobacillales Enterococcaceae *Enterococcus faecalis* EnGen0294 |
| Efa-179-1 | Bacteria Firmicutes Bacilli Lactobacillales Enterococcaceae *Enterococcus faecalis* EnGen0295 |
| Efa-180-1 | Bacteria Firmicutes Bacilli Lactobacillales Enterococcaceae *Enterococcus faecalis* EnGen0297 |

| | |
|---|---|
| Efa-181-1 | Bacteria Firmicutes Bacilli Lactobacillales Enterococcaceae *Enterococcus faecalis* EnGen0298 |
| Efa-182-1 | Bacteria Firmicutes Bacilli Lactobacillales Enterococcaceae *Enterococcus faecalis* EnGen0299 |
| Efa-183-1 | Bacteria Firmicutes Bacilli Lactobacillales Enterococcaceae *Enterococcus faecalis* EnGen0300 |
| Efa-184-1 | Bacteria Firmicutes Bacilli Lactobacillales Enterococcaceae *Enterococcus faecalis* EnGen0301 |
| Efa-185-1 | Bacteria Firmicutes Bacilli Lactobacillales Enterococcaceae *Enterococcus faecalis* EnGen0302 |
| Efa-186-1 | Bacteria Firmicutes Bacilli Lactobacillales Enterococcaceae *Enterococcus faecalis* EnGen0303 |
| Efa-187-1 | Bacteria Firmicutes Bacilli Lactobacillales Enterococcaceae *Enterococcus faecalis* EnGen0304 |
| Efa-188-1 | Bacteria Firmicutes Bacilli Lactobacillales Enterococcaceae *Enterococcus faecalis* EnGen0305 |
| Efa-189-1 | Bacteria Firmicutes Bacilli Lactobacillales Enterococcaceae *Enterococcus faecalis* EnGen0306 |
| Efa-190-1 | Bacteria Firmicutes Bacilli Lactobacillales Enterococcaceae *Enterococcus faecalis* EnGen0307 |
| Efa-191-1 | Bacteria Firmicutes Bacilli Lactobacillales Enterococcaceae *Enterococcus faecalis* EnGen0308 |
| Efa-192-1 | Bacteria Firmicutes Bacilli Lactobacillales Enterococcaceae *Enterococcus faecalis* EnGen0310 |
| Efa-193-1 | Bacteria Firmicutes Bacilli Lactobacillales Enterococcaceae *Enterococcus faecalis* EnGen0310 = MMH594 |
| Efa-194-1 | Bacteria Firmicutes Bacilli Lactobacillales Enterococcaceae *Enterococcus faecalis* EnGen0311 |
| Efa-195-1 | Bacteria Firmicutes Bacilli Lactobacillales Enterococcaceae *Enterococcus faecalis* EnGen0326 |
| Efa-196-1 | Bacteria Firmicutes Bacilli Lactobacillales Enterococcaceae *Enterococcus faecalis* EnGen0327 |
| Efa-197-1 | Bacteria Firmicutes Bacilli Lactobacillales Enterococcaceae *Enterococcus faecalis* EnGen0328 |
| Efa-198-1 | Bacteria Firmicutes Bacilli Lactobacillales Enterococcaceae *Enterococcus faecalis* EnGen0329 |
| Efa-199-1 | Bacteria Firmicutes Bacilli Lactobacillales Enterococcaceae *Enterococcus faecalis* EnGen0330 |
| Efa-200-1 | Bacteria Firmicutes Bacilli Lactobacillales Enterococcaceae *Enterococcus faecalis* EnGen0331 |
| Efa-201-1 | Bacteria Firmicutes Bacilli Lactobacillales Enterococcaceae *Enterococcus faecalis* EnGen0332 |
| Efa-202-1 | Bacteria Firmicutes Bacilli Lactobacillales Enterococcaceae *Enterococcus faecalis* EnGen0333 |
| Efa-203-1 | Bacteria Firmicutes Bacilli Lactobacillales Enterococcaceae *Enterococcus faecalis* EnGen0334 |
| Efa-204-1 | Bacteria Firmicutes Bacilli Lactobacillales Enterococcaceae *Enterococcus faecalis* EnGen0335 |
| Efa-205-1 | Bacteria Firmicutes Bacilli Lactobacillales Enterococcaceae *Enterococcus faecalis* EnGen0336 |
| Efa-206-1 | Bacteria Firmicutes Bacilli Lactobacillales Enterococcaceae *Enterococcus faecalis* EnGen0337 |
| Efa-207-1 | Bacteria Firmicutes Bacilli Lactobacillales Enterococcaceae *Enterococcus faecalis* EnGen0338 |
| Efa-208-1 | Bacteria Firmicutes Bacilli Lactobacillales Enterococcaceae *Enterococcus faecalis* EnGen0339 |
| Efa-209-1 | Bacteria Firmicutes Bacilli Lactobacillales Enterococcaceae *Enterococcus faecalis* EnGen0340 |
| Efa-210-1 | Bacteria Firmicutes Bacilli Lactobacillales Enterococcaceae *Enterococcus faecalis* EnGen0341 |
| Efa-211-1 | Bacteria Firmicutes Bacilli Lactobacillales Enterococcaceae *Enterococcus faecalis* EnGen0342 |
| Efa-212-1 | Bacteria Firmicutes Bacilli Lactobacillales Enterococcaceae *Enterococcus faecalis* EnGen0344 |
| Efa-213-1 | Bacteria Firmicutes Bacilli Lactobacillales Enterococcaceae *Enterococcus faecalis* EnGen0345 |
| Efa-214-1 | Bacteria Firmicutes Bacilli Lactobacillales Enterococcaceae *Enterococcus faecalis* EnGen0346 |
| Efa-215-1 | Bacteria Firmicutes Bacilli Lactobacillales Enterococcaceae *Enterococcus faecalis* EnGen0348 |
| Efa-216-1 | Bacteria Firmicutes Bacilli Lactobacillales Enterococcaceae *Enterococcus faecalis* EnGen0350 |
| Efa-217-1 | Bacteria Firmicutes Bacilli Lactobacillales Enterococcaceae *Enterococcus faecalis* EnGen0351 |
| Efa-218-1 | Bacteria Firmicutes Bacilli Lactobacillales Enterococcaceae *Enterococcus faecalis* EnGen0352 |
| Efa-219-1 | Bacteria Firmicutes Bacilli Lactobacillales Enterococcaceae *Enterococcus faecalis* EnGen0354 |
| Efa-220-1 | Bacteria Firmicutes Bacilli Lactobacillales Enterococcaceae *Enterococcus faecalis* EnGen0355 |
| Efa-221-1 | Bacteria Firmicutes Bacilli Lactobacillales Enterococcaceae *Enterococcus faecalis* EnGen0356 |
| Efa-222-1 | Bacteria Firmicutes Bacilli Lactobacillales Enterococcaceae *Enterococcus faecalis* EnGen0357 |
| Efa-223-1 | Bacteria Firmicutes Bacilli Lactobacillales Enterococcaceae *Enterococcus faecalis* EnGen0358 |
| Efa-224-1 | Bacteria Firmicutes Bacilli Lactobacillales Enterococcaceae *Enterococcus faecalis* EnGen0359 |
| Efa-225-1 | Bacteria Firmicutes Bacilli Lactobacillales Enterococcaceae *Enterococcus faecalis* EnGen0360 |
| Efa-226-1 | Bacteria Firmicutes Bacilli Lactobacillales Enterococcaceae *Enterococcus faecalis* EnGen0361 |
| Efa-227-1 | Bacteria Firmicutes Bacilli Lactobacillales Enterococcaceae *Enterococcus faecalis* EnGen0362 |
| Efa-228-1 | Bacteria Firmicutes Bacilli Lactobacillales Enterococcaceae *Enterococcus faecalis* EnGen0363 |
| Efa-229-1 | Bacteria Firmicutes Bacilli Lactobacillales Enterococcaceae *Enterococcus faecalis* EnGen0364 |
| Efa-230-1 | Bacteria Firmicutes Bacilli Lactobacillales Enterococcaceae *Enterococcus faecalis* EnGen0365 |
| Efa-231-1 | Bacteria Firmicutes Bacilli Lactobacillales Enterococcaceae *Enterococcus faecalis* EnGen0366 |
| Efa-232-1 | Bacteria Firmicutes Bacilli Lactobacillales Enterococcaceae *Enterococcus faecalis* EnGen0367 |
| Efa-233-1 | Bacteria Firmicutes Bacilli Lactobacillales Enterococcaceae *Enterococcus faecalis* EnGen0368 |
| Efa-234-1 | Bacteria Firmicutes Bacilli Lactobacillales Enterococcaceae *Enterococcus faecalis* EnGen0369 |
| Efa-235-1 | Bacteria Firmicutes Bacilli Lactobacillales Enterococcaceae *Enterococcus faecalis* EnGen0370 |
| Efa-236-1 | Bacteria Firmicutes Bacilli Lactobacillales Enterococcaceae *Enterococcus faecalis* EnGen0374 |
| Efa-237-1 | Bacteria Firmicutes Bacilli Lactobacillales Enterococcaceae *Enterococcus faecalis* EnGen0400 |
| Efa-238-1 | Bacteria Firmicutes Bacilli Lactobacillales Enterococcaceae *Enterococcus faecalis* EnGen0401 |
| Efa-239-1 | Bacteria Firmicutes Bacilli Lactobacillales Enterococcaceae *Enterococcus faecalis* EnGen0402 |
| Efa-240-1 | Bacteria Firmicutes Bacilli Lactobacillales Enterococcaceae *Enterococcus faecalis* EnGen0403 |
| Efa-241-1 | Bacteria Firmicutes Bacilli Lactobacillales Enterococcaceae *Enterococcus faecalis* EnGen0404 |
| Efa-242-1 | Bacteria Firmicutes Bacilli Lactobacillales Enterococcaceae *Enterococcus faecalis* EnGen0405 |
| Efa-243-1 | Bacteria Firmicutes Bacilli Lactobacillales Enterococcaceae *Enterococcus faecalis* EnGen0406 |
| Efa-244-1 | Bacteria Firmicutes Bacilli Lactobacillales Enterococcaceae *Enterococcus faecalis* EnGen0407 |
| Efa-245-1 | Bacteria Firmicutes Bacilli Lactobacillales Enterococcaceae *Enterococcus faecalis* EnGen0408 |

| | |
|---|---|
| Efa-246-1 | Bacteria Firmicutes Bacilli Lactobacillales Enterococcaceae *Enterococcus faecalis* EnGen0409 |
| Efa-247-1 | Bacteria Firmicutes Bacilli Lactobacillales Enterococcaceae *Enterococcus faecalis* EnGen0410 |
| Efa-248-1 | Bacteria Firmicutes Bacilli Lactobacillales Enterococcaceae *Enterococcus faecalis* EnGen0411 |
| Efa-249-1 | Bacteria Firmicutes Bacilli Lactobacillales Enterococcaceae *Enterococcus faecalis* EnGen0412 |
| Efa-250-1 | Bacteria Firmicutes Bacilli Lactobacillales Enterococcaceae *Enterococcus faecalis* EnGen0413 |
| Efa-251-1 | Bacteria Firmicutes Bacilli Lactobacillales Enterococcaceae *Enterococcus faecalis* EnGen0414 |
| Efa-252-1 | Bacteria Firmicutes Bacilli Lactobacillales Enterococcaceae *Enterococcus faecalis* EnGen0415 |
| Efa-253-1 | Bacteria Firmicutes Bacilli Lactobacillales Enterococcaceae *Enterococcus faecalis* EnGen0416 |
| Efa-254-1 | Bacteria Firmicutes Bacilli Lactobacillales Enterococcaceae *Enterococcus faecalis* EnGen0417 |
| Efa-255-1 | Bacteria Firmicutes Bacilli Lactobacillales Enterococcaceae *Enterococcus faecalis* EnGen0418 |
| Efa-256-1 | Bacteria Firmicutes Bacilli Lactobacillales Enterococcaceae *Enterococcus faecalis* EnGen0419 |
| Efa-257-1 | Bacteria Firmicutes Bacilli Lactobacillales Enterococcaceae *Enterococcus faecalis* EnGen0420 |
| Efa-258-1 | Bacteria Firmicutes Bacilli Lactobacillales Enterococcaceae *Enterococcus faecalis* EnGen0421 |
| Efa-259-1 | Bacteria Firmicutes Bacilli Lactobacillales Enterococcaceae *Enterococcus faecalis* EnGen0422 |
| Efa-260-1 | Bacteria Firmicutes Bacilli Lactobacillales Enterococcaceae *Enterococcus faecalis* EnGen0423 |
| Efa-261-1 | Bacteria Firmicutes Bacilli Lactobacillales Enterococcaceae *Enterococcus faecalis* EnGen0424 |
| Efa-262-1 | Bacteria Firmicutes Bacilli Lactobacillales Enterococcaceae *Enterococcus faecalis* EnGen0425 |
| Efa-263-1 | Bacteria Firmicutes Bacilli Lactobacillales Enterococcaceae *Enterococcus faecalis* EnGen0426 |
| Efa-264-1 | Bacteria Firmicutes Bacilli Lactobacillales Enterococcaceae *Enterococcus faecalis* EnGen0427 |
| Efa-31-1 | Bacteria Firmicutes Bacilli Lactobacillales Enterococcaceae *Enterococcus faecalis* ERV103 |
| Efa-32-1 | Bacteria Firmicutes Bacilli Lactobacillales Enterococcaceae *Enterococcus faecalis* ERV116 |
| Efa-33-1 | Bacteria Firmicutes Bacilli Lactobacillales Enterococcaceae *Enterococcus faecalis* ERV129 |
| Efa-34-1 | Bacteria Firmicutes Bacilli Lactobacillales Enterococcaceae *Enterococcus faecalis* ERV25 |
| Efa-35-1 | Bacteria Firmicutes Bacilli Lactobacillales Enterococcaceae *Enterococcus faecalis* ERV31 |
| Efa-36-1 | Bacteria Firmicutes Bacilli Lactobacillales Enterococcaceae *Enterococcus faecalis* ERV37 |
| Efa-37-1 | Bacteria Firmicutes Bacilli Lactobacillales Enterococcaceae *Enterococcus faecalis* ERV41 |
| Efa-38-1 | Bacteria Firmicutes Bacilli Lactobacillales Enterococcaceae *Enterococcus faecalis* ERV62 |
| Efa-39-1 | Bacteria Firmicutes Bacilli Lactobacillales Enterococcaceae *Enterococcus faecalis* ERV63 |
| Efa-40-1 | Bacteria Firmicutes Bacilli Lactobacillales Enterococcaceae *Enterococcus faecalis* ERV65 |
| Efa-41-1 | Bacteria Firmicutes Bacilli Lactobacillales Enterococcaceae *Enterococcus faecalis* ERV68 |
| Efa-42-1 | Bacteria Firmicutes Bacilli Lactobacillales Enterococcaceae *Enterococcus faecalis* ERV72 |
| Efa-43-1 | Bacteria Firmicutes Bacilli Lactobacillales Enterococcaceae *Enterococcus faecalis* ERV73 |
| Efa-44-1 | Bacteria Firmicutes Bacilli Lactobacillales Enterococcaceae *Enterococcus faecalis* ERV81 |
| Efa-45-1 | Bacteria Firmicutes Bacilli Lactobacillales Enterococcaceae *Enterococcus faecalis* ERV85 |
| Efa-46-1 | Bacteria Firmicutes Bacilli Lactobacillales Enterococcaceae *Enterococcus faecalis* ERV93 |
| Efa-265-1 | Bacteria Firmicutes Bacilli Lactobacillales Enterococcaceae *Enterococcus faecalis* F01966 |
| Efa-266-1 | Bacteria Firmicutes Bacilli Lactobacillales Enterococcaceae *Enterococcus faecalis* FL2 |
| Efa-267-1 | Bacteria Firmicutes Bacilli Lactobacillales Enterococcaceae *Enterococcus faecalis* GA2 |
| Efa-268-1 | Bacteria Firmicutes Bacilli Lactobacillales Enterococcaceae *Enterococcus faecalis* GAN13 |
| Efa-269-1 | Bacteria Firmicutes Bacilli Lactobacillales Enterococcaceae *Enterococcus faecalis* JH2-2 |
| Efa-270-1 | Bacteria Firmicutes Bacilli Lactobacillales Enterococcaceae *Enterococcus faecalis* KI-6-1-110608-1 |
| Efa-271-1 | Bacteria Firmicutes Bacilli Lactobacillales Enterococcaceae *Enterococcus faecalis* KS19 |
| Efa-272-1 | Bacteria Firmicutes Bacilli Lactobacillales Enterococcaceae *Enterococcus faecalis* LA3B-2 |
| Efa-273-1 | Bacteria Firmicutes Bacilli Lactobacillales Enterococcaceae *Enterococcus faecalis* M7 |
| Efa-274-1 | Bacteria Firmicutes Bacilli Lactobacillales Enterococcaceae *Enterococcus faecalis* MA1 |
| Efa-275-1 | Bacteria Firmicutes Bacilli Lactobacillales Enterococcaceae *Enterococcus faecalis* MD6 |
| Efa-276-1 | Bacteria Firmicutes Bacilli Lactobacillales Enterococcaceae *Enterococcus faecalis* MN16 |
| Efa-278-1 | Bacteria Firmicutes Bacilli Lactobacillales Enterococcaceae *Enterococcus faecalis* MTmid8 |
| Efa-277-1 | Bacteria Firmicutes Bacilli Lactobacillales Enterococcaceae *Enterococcus faecalis* MTUP9 |
| Efa-279-1 | Bacteria Firmicutes Bacilli Lactobacillales Enterococcaceae *Enterococcus faecalis* NJ44 |
| Efa-280-1 | Bacteria Firmicutes Bacilli Lactobacillales Enterococcaceae *Enterococcus faecalis* NY9 |
| Efa-281-1 | Bacteria Firmicutes Bacilli Lactobacillales Enterococcaceae *Enterococcus faecalis* OG1X |
| Efa-282-1 | Bacteria Firmicutes Bacilli Lactobacillales Enterococcaceae *Enterococcus faecalis* R508 |
| Efa-283-1 | Bacteria Firmicutes Bacilli Lactobacillales Enterococcaceae *Enterococcus faecalis* RP2S-4 |
| Efa-284-1 | Bacteria Firmicutes Bacilli Lactobacillales Enterococcaceae *Enterococcus faecalis* SLO2C-1 |
| Efa-312-1 | Bacteria Firmicutes Bacilli Lactobacillales Enterococcaceae *Enterococcus faecalis str.* Symbioflor 1 |
| Efa-285-1 | Bacteria Firmicutes Bacilli Lactobacillales Enterococcaceae *Enterococcus faecalis* TX0012 |
| Efa-286-1 | Bacteria Firmicutes Bacilli Lactobacillales Enterococcaceae *Enterococcus faecalis* TX0017 |
| Efa-287-1 | Bacteria Firmicutes Bacilli Lactobacillales Enterococcaceae *Enterococcus faecalis* TX0027 |
| Efa-288-1 | Bacteria Firmicutes Bacilli Lactobacillales Enterococcaceae *Enterococcus faecalis* TX0031 |
| Efa-289-1 | Bacteria Firmicutes Bacilli Lactobacillales Enterococcaceae *Enterococcus faecalis* TX0043 |
| Efa-290-1 | Bacteria Firmicutes Bacilli Lactobacillales Enterococcaceae *Enterococcus faecalis* TX0102 |
| Efa-291-1 | Bacteria Firmicutes Bacilli Lactobacillales Enterococcaceae *Enterococcus faecalis* TX0109 |
| Efa-292-1 | Bacteria Firmicutes Bacilli Lactobacillales Enterococcaceae *Enterococcus faecalis* TX0309A |
| Efa-293-1 | Bacteria Firmicutes Bacilli Lactobacillales Enterococcaceae *Enterococcus faecalis* TX0309B |

| | |
|---|---|
| Efa-294-1 | Bacteria Firmicutes Bacilli Lactobacillales Enterococcaceae *Enterococcus faecalis* TX0312 |
| Efa-295-1 | Bacteria Firmicutes Bacilli Lactobacillales Enterococcaceae *Enterococcus faecalis* TX0411 |
| Efa-296-1 | Bacteria Firmicutes Bacilli Lactobacillales Enterococcaceae *Enterococcus faecalis* TX0470 |
| Efa-297-1 | Bacteria Firmicutes Bacilli Lactobacillales Enterococcaceae *Enterococcus faecalis* TX0630 |
| Efa-298-1 | Bacteria Firmicutes Bacilli Lactobacillales Enterococcaceae *Enterococcus faecalis* TX0645 |
| Efa-299-1 | Bacteria Firmicutes Bacilli Lactobacillales Enterococcaceae *Enterococcus faecalis* TX0860 |
| Efa-300-1 | Bacteria Firmicutes Bacilli Lactobacillales Enterococcaceae *Enterococcus faecalis* TX1302 |
| Efa-301-1 | Bacteria Firmicutes Bacilli Lactobacillales Enterococcaceae *Enterococcus faecalis* TX1341 |
| Efa-302-1 | Bacteria Firmicutes Bacilli Lactobacillales Enterococcaceae *Enterococcus faecalis* TX1342 |
| Efa-303-1 | Bacteria Firmicutes Bacilli Lactobacillales Enterococcaceae *Enterococcus faecalis* TX1346 |
| Efa-304-1 | Bacteria Firmicutes Bacilli Lactobacillales Enterococcaceae *Enterococcus faecalis* TX2137 |
| Efa-305-1 | Bacteria Firmicutes Bacilli Lactobacillales Enterococcaceae *Enterococcus faecalis* TX2141 |
| Efa-306-1 | Bacteria Firmicutes Bacilli Lactobacillales Enterococcaceae *Enterococcus faecalis* TX4000 |
| Efa-307-1 | Bacteria Firmicutes Bacilli Lactobacillales Enterococcaceae *Enterococcus faecalis* TX4244 |
| Efa-308-1 | Bacteria Firmicutes Bacilli Lactobacillales Enterococcaceae *Enterococcus faecalis* UP2S-6 |
| Efa-309-1 to Efa-309-3 | Bacteria Firmicutes Bacilli Lactobacillales Enterococcaceae *Enterococcus faecalis* V583 |
| Efa-310-1 | Bacteria Firmicutes Bacilli Lactobacillales Enterococcaceae *Enterococcus faecalis* VC1B-1 |
| Efa-311-1 | Bacteria Firmicutes Bacilli Lactobacillales Enterococcaceae *Enterococcus faecalis* WKS-26-18-2 |
| Efa-313-1 | Bacteria Firmicutes Bacilli Lactobacillales Enterococcaceae *Enterococcus faecium* |
| Efa-314-1 | Bacteria Firmicutes Bacilli Lactobacillales Enterococcaceae *Enterococcus faecium* 10/96A |
| Efa-315-1 | Bacteria Firmicutes Bacilli Lactobacillales Enterococcaceae *Enterococcus faecium* 13.SD.W.09 |
| Efa-316-1 | Bacteria Firmicutes Bacilli Lactobacillales Enterococcaceae *Enterococcus faecium* 503 |
| Efa-317-1 | Bacteria Firmicutes Bacilli Lactobacillales Enterococcaceae *Enterococcus faecium* 504 |
| Efa-318-1 | Bacteria Firmicutes Bacilli Lactobacillales Enterococcaceae *Enterococcus faecium* 505 |
| Efa-319-1 | Bacteria Firmicutes Bacilli Lactobacillales Enterococcaceae *Enterococcus faecium* 506 |
| Efa-320-1 | Bacteria Firmicutes Bacilli Lactobacillales Enterococcaceae *Enterococcus faecium* 509 |
| Efa-321-1 | Bacteria Firmicutes Bacilli Lactobacillales Enterococcaceae *Enterococcus faecium* 510 |
| Efa-322-1 | Bacteria Firmicutes Bacilli Lactobacillales Enterococcaceae *Enterococcus faecium* 511 |
| Efa-323-1 | Bacteria Firmicutes Bacilli Lactobacillales Enterococcaceae *Enterococcus faecium* 513 |
| Efa-324-1 | Bacteria Firmicutes Bacilli Lactobacillales Enterococcaceae *Enterococcus faecium* 514 |
| Efa-325-1 | Bacteria Firmicutes Bacilli Lactobacillales Enterococcaceae *Enterococcus faecium* 515 |
| Efa-326-1 | Bacteria Firmicutes Bacilli Lactobacillales Enterococcaceae *Enterococcus faecium* ATCC 8459 |
| Efa-327-1 | Bacteria Firmicutes Bacilli Lactobacillales Enterococcaceae *Enterococcus faecium* Aus0085 |
| Efa-328-1 | Bacteria Firmicutes Bacilli Lactobacillales Enterococcaceae *Enterococcus faecium* BM4538 |
| Efa-329-1 | Bacteria Firmicutes Bacilli Lactobacillales Enterococcaceae *Enterococcus faecium* C1904 |
| Efa-330-1 | Bacteria Firmicutes Bacilli Lactobacillales Enterococcaceae *Enterococcus faecium* C309 |
| Efa-331-1 | Bacteria Firmicutes Bacilli Lactobacillales Enterococcaceae *Enterococcus faecium* C497 |
| Efa-332-1 | Bacteria Firmicutes Bacilli Lactobacillales Enterococcaceae *Enterococcus faecium* C621 |
| Efa-333-1 | Bacteria Firmicutes Bacilli Lactobacillales Enterococcaceae *Enterococcus faecium* C68 |
| Efa-334-1 | Bacteria Firmicutes Bacilli Lactobacillales Enterococcaceae *Enterococcus faecium* E1165 |
| Efa-335-1 | Bacteria Firmicutes Bacilli Lactobacillales Enterococcaceae *Enterococcus faecium* E155 |
| Efa-336-1 | Bacteria Firmicutes Bacilli Lactobacillales Enterococcaceae *Enterococcus faecium* E417 |
| Efa-337-1 | Bacteria Firmicutes Bacilli Lactobacillales Enterococcaceae *Enterococcus faecium* E422 |
| Efa-338-1 | Bacteria Firmicutes Bacilli Lactobacillales Enterococcaceae *Enterococcus faecium* E4452 |
| Efa-339-1 | Bacteria Firmicutes Bacilli Lactobacillales Enterococcaceae *Enterococcus faecium* E4453 |
| Efa-340-1 | Bacteria Firmicutes Bacilli Lactobacillales Enterococcaceae *Enterococcus faecium* E525 |
| Efa-350-1 | Bacteria Firmicutes Bacilli Lactobacillales Enterococcaceae *Enterococcus faecium* EnGen0001 |
| Efa-351-1 | Bacteria Firmicutes Bacilli Lactobacillales Enterococcaceae *Enterococcus faecium* EnGen0002 |
| Efa-352-1 | Bacteria Firmicutes Bacilli Lactobacillales Enterococcaceae *Enterococcus faecium* EnGen0003 |
| Efa-353-1 | Bacteria Firmicutes Bacilli Lactobacillales Enterococcaceae *Enterococcus faecium* EnGen0004 |
| Efa-354-1 | Bacteria Firmicutes Bacilli Lactobacillales Enterococcaceae *Enterococcus faecium* EnGen0005 |
| Efa-355-1 | Bacteria Firmicutes Bacilli Lactobacillales Enterococcaceae *Enterococcus faecium* EnGen0006 |
| Efa-356-1 | Bacteria Firmicutes Bacilli Lactobacillales Enterococcaceae *Enterococcus faecium* EnGen0007 |
| Efa-357-1 | Bacteria Firmicutes Bacilli Lactobacillales Enterococcaceae *Enterococcus faecium* EnGen0008 |
| Efa-358-1 | Bacteria Firmicutes Bacilli Lactobacillales Enterococcaceae *Enterococcus faecium* EnGen0009 |
| Efa-359-1 | Bacteria Firmicutes Bacilli Lactobacillales Enterococcaceae *Enterococcus faecium* EnGen0010 |
| Efa-360-1 | Bacteria Firmicutes Bacilli Lactobacillales Enterococcaceae *Enterococcus faecium* EnGen0011 |
| Efa-361-1 | Bacteria Firmicutes Bacilli Lactobacillales Enterococcaceae *Enterococcus faecium* EnGen0012 |
| Efa-362-1 | Bacteria Firmicutes Bacilli Lactobacillales Enterococcaceae *Enterococcus faecium* EnGen0013 |
| Efa-363-1 | Bacteria Firmicutes Bacilli Lactobacillales Enterococcaceae *Enterococcus faecium* EnGen0014 |
| Efa-364-1 | Bacteria Firmicutes Bacilli Lactobacillales Enterococcaceae *Enterococcus faecium* EnGen0015 |
| Efa-365-1 | Bacteria Firmicutes Bacilli Lactobacillales Enterococcaceae *Enterococcus faecium* EnGen0016 |
| Efa-366-1 | Bacteria Firmicutes Bacilli Lactobacillales Enterococcaceae *Enterococcus faecium* EnGen0017 |
| Efa-367-1 | Bacteria Firmicutes Bacilli Lactobacillales Enterococcaceae *Enterococcus faecium* EnGen0018 |
| Efa-368-1 | Bacteria Firmicutes Bacilli Lactobacillales Enterococcaceae *Enterococcus faecium* EnGen0019 |

| | |
|---|---|
| Efa-369-1 | Bacteria Firmicutes Bacilli Lactobacillales Enterococcaceae *Enterococcus faecium* EnGen0020 |
| Efa-370-1 | Bacteria Firmicutes Bacilli Lactobacillales Enterococcaceae *Enterococcus faecium* EnGen0021 |
| Efa-371-1 | Bacteria Firmicutes Bacilli Lactobacillales Enterococcaceae *Enterococcus faecium* EnGen0022 |
| Efa-372-1 | Bacteria Firmicutes Bacilli Lactobacillales Enterococcaceae *Enterococcus faecium* EnGen0024 |
| Efa-373-1 | Bacteria Firmicutes Bacilli Lactobacillales Enterococcaceae *Enterococcus faecium* EnGen0025 |
| Efa-374-1 | Bacteria Firmicutes Bacilli Lactobacillales Enterococcaceae *Enterococcus faecium* EnGen0026 |
| Efa-375-1 | Bacteria Firmicutes Bacilli Lactobacillales Enterococcaceae *Enterococcus faecium* EnGen0027 |
| Efa-376-1 | Bacteria Firmicutes Bacilli Lactobacillales Enterococcaceae *Enterococcus faecium* EnGen0028 |
| Efa-377-1 | Bacteria Firmicutes Bacilli Lactobacillales Enterococcaceae *Enterococcus faecium* EnGen0029 |
| Efa-378-1 | Bacteria Firmicutes Bacilli Lactobacillales Enterococcaceae *Enterococcus faecium* EnGen0030 |
| Efa-379-1 | Bacteria Firmicutes Bacilli Lactobacillales Enterococcaceae *Enterococcus faecium* EnGen0031 |
| Efa-380-1 | Bacteria Firmicutes Bacilli Lactobacillales Enterococcaceae *Enterococcus faecium* EnGen0032 |
| Efa-381-1 | Bacteria Firmicutes Bacilli Lactobacillales Enterococcaceae *Enterococcus faecium* EnGen0033 |
| Efa-382-1 | Bacteria Firmicutes Bacilli Lactobacillales Enterococcaceae *Enterococcus faecium* EnGen0034 |
| Efa-383-1 | Bacteria Firmicutes Bacilli Lactobacillales Enterococcaceae *Enterococcus faecium* EnGen0035 |
| Efa-384-1 | Bacteria Firmicutes Bacilli Lactobacillales Enterococcaceae *Enterococcus faecium* EnGen0036 |
| Efa-385-1 | Bacteria Firmicutes Bacilli Lactobacillales Enterococcaceae *Enterococcus faecium* EnGen0038 |
| Efa-386-1 | Bacteria Firmicutes Bacilli Lactobacillales Enterococcaceae *Enterococcus faecium* EnGen0039 |
| Efa-387-1 | Bacteria Firmicutes Bacilli Lactobacillales Enterococcaceae *Enterococcus faecium* EnGen0040 |
| Efa-388-1 | Bacteria Firmicutes Bacilli Lactobacillales Enterococcaceae *Enterococcus faecium* EnGen0041 |
| Efa-389-1 | Bacteria Firmicutes Bacilli Lactobacillales Enterococcaceae *Enterococcus faecium* EnGen0042 |
| Efa-390-1 | Bacteria Firmicutes Bacilli Lactobacillales Enterococcaceae *Enterococcus faecium* EnGen0043 |
| Efa-391-1 | Bacteria Firmicutes Bacilli Lactobacillales Enterococcaceae *Enterococcus faecium* EnGen0044 |
| Efa-392-1 | Bacteria Firmicutes Bacilli Lactobacillales Enterococcaceae *Enterococcus faecium* EnGen0045 |
| Efa-393-1 | Bacteria Firmicutes Bacilli Lactobacillales Enterococcaceae *Enterococcus faecium* EnGen0046 |
| Efa-394-1 | Bacteria Firmicutes Bacilli Lactobacillales Enterococcaceae *Enterococcus faecium* EnGen0047 |
| Efa-395-1 | Bacteria Firmicutes Bacilli Lactobacillales Enterococcaceae *Enterococcus faecium* EnGen0048 |
| Efa-396-1 | Bacteria Firmicutes Bacilli Lactobacillales Enterococcaceae *Enterococcus faecium* EnGen0049 |
| Efa-397-1 | Bacteria Firmicutes Bacilli Lactobacillales Enterococcaceae *Enterococcus faecium* EnGen0050 |
| Efa-398-1 | Bacteria Firmicutes Bacilli Lactobacillales Enterococcaceae *Enterococcus faecium* EnGen0051 |
| Efa-399-1 | Bacteria Firmicutes Bacilli Lactobacillales Enterococcaceae *Enterococcus faecium* EnGen0052 |
| Efa-400-1 | Bacteria Firmicutes Bacilli Lactobacillales Enterococcaceae *Enterococcus faecium* EnGen0054 |
| Efa-401-1 | Bacteria Firmicutes Bacilli Lactobacillales Enterococcaceae *Enterococcus faecium* EnGen0056 |
| Efa-402-1 | Bacteria Firmicutes Bacilli Lactobacillales Enterococcaceae *Enterococcus faecium* EnGen0057 |
| Efa-403-1 | Bacteria Firmicutes Bacilli Lactobacillales Enterococcaceae *Enterococcus faecium* EnGen0124 |
| Efa-404-1 | Bacteria Firmicutes Bacilli Lactobacillales Enterococcaceae *Enterococcus faecium* EnGen0125 |
| Efa-405-1 | Bacteria Firmicutes Bacilli Lactobacillales Enterococcaceae *Enterococcus faecium* EnGen0126 |
| Efa-406-1 | Bacteria Firmicutes Bacilli Lactobacillales Enterococcaceae *Enterococcus faecium* EnGen0127 |
| Efa-407-1 | Bacteria Firmicutes Bacilli Lactobacillales Enterococcaceae *Enterococcus faecium* EnGen0128 |
| Efa-408-1 | Bacteria Firmicutes Bacilli Lactobacillales Enterococcaceae *Enterococcus faecium* EnGen0129 |
| Efa-409-1 | Bacteria Firmicutes Bacilli Lactobacillales Enterococcaceae *Enterococcus faecium* EnGen0130 |
| Efa-410-1 | Bacteria Firmicutes Bacilli Lactobacillales Enterococcaceae *Enterococcus faecium* EnGen0131 |
| Efa-411-1 | Bacteria Firmicutes Bacilli Lactobacillales Enterococcaceae *Enterococcus faecium* EnGen0132 |
| Efa-412-1 | Bacteria Firmicutes Bacilli Lactobacillales Enterococcaceae *Enterococcus faecium* EnGen0133 |
| Efa-413-1 | Bacteria Firmicutes Bacilli Lactobacillales Enterococcaceae *Enterococcus faecium* EnGen0134 |
| Efa-414-1 | Bacteria Firmicutes Bacilli Lactobacillales Enterococcaceae *Enterococcus faecium* EnGen0135 |
| Efa-415-1 | Bacteria Firmicutes Bacilli Lactobacillales Enterococcaceae *Enterococcus faecium* EnGen0136 |
| Efa-416-1 | Bacteria Firmicutes Bacilli Lactobacillales Enterococcaceae *Enterococcus faecium* EnGen0137 |
| Efa-417-1 | Bacteria Firmicutes Bacilli Lactobacillales Enterococcaceae *Enterococcus faecium* EnGen0138 |
| Efa-418-1 | Bacteria Firmicutes Bacilli Lactobacillales Enterococcaceae *Enterococcus faecium* EnGen0140 |
| Efa-419-1 | Bacteria Firmicutes Bacilli Lactobacillales Enterococcaceae *Enterococcus faecium* EnGen0142 |
| Efa-420-1 | Bacteria Firmicutes Bacilli Lactobacillales Enterococcaceae *Enterococcus faecium* EnGen0145 |
| Efa-421-1 | Bacteria Firmicutes Bacilli Lactobacillales Enterococcaceae *Enterococcus faecium* EnGen0146 |
| Efa-422-1 | Bacteria Firmicutes Bacilli Lactobacillales Enterococcaceae *Enterococcus faecium* EnGen0147 |
| Efa-423-1 | Bacteria Firmicutes Bacilli Lactobacillales Enterococcaceae *Enterococcus faecium* EnGen0148 |
| Efa-424-1 | Bacteria Firmicutes Bacilli Lactobacillales Enterococcaceae *Enterococcus faecium* EnGen0149 |
| Efa-425-1 | Bacteria Firmicutes Bacilli Lactobacillales Enterococcaceae *Enterococcus faecium* EnGen0150 |
| Efa-426-1 | Bacteria Firmicutes Bacilli Lactobacillales Enterococcaceae *Enterococcus faecium* EnGen0151 |
| Efa-427-1 | Bacteria Firmicutes Bacilli Lactobacillales Enterococcaceae *Enterococcus faecium* EnGen0152 |
| Efa-428-1 | Bacteria Firmicutes Bacilli Lactobacillales Enterococcaceae *Enterococcus faecium* EnGen0153 |
| Efa-429-1 | Bacteria Firmicutes Bacilli Lactobacillales Enterococcaceae *Enterococcus faecium* EnGen0155 |
| Efa-430-1 | Bacteria Firmicutes Bacilli Lactobacillales Enterococcaceae *Enterococcus faecium* EnGen0156 |
| Efa-431-1 | Bacteria Firmicutes Bacilli Lactobacillales Enterococcaceae *Enterococcus faecium* EnGen0158 |
| Efa-432-1 | Bacteria Firmicutes Bacilli Lactobacillales Enterococcaceae *Enterococcus faecium* EnGen0159 |
| Efa-433-1 | Bacteria Firmicutes Bacilli Lactobacillales Enterococcaceae *Enterococcus faecium* EnGen0160 |

| | |
|---|---|
| Efa-434-1 | Bacteria Firmicutes Bacilli Lactobacillales Enterococcaceae *Enterococcus faecium* EnGen0161 |
| Efa-435-1 | Bacteria Firmicutes Bacilli Lactobacillales Enterococcaceae *Enterococcus faecium* EnGen0162 |
| Efa-436-1 | Bacteria Firmicutes Bacilli Lactobacillales Enterococcaceae *Enterococcus faecium* EnGen0163 |
| Efa-437-1 | Bacteria Firmicutes Bacilli Lactobacillales Enterococcaceae *Enterococcus faecium* EnGen0164 |
| Efa-438-1 | Bacteria Firmicutes Bacilli Lactobacillales Enterococcaceae *Enterococcus faecium* EnGen0165 |
| Efa-439-1 | Bacteria Firmicutes Bacilli Lactobacillales Enterococcaceae *Enterococcus faecium* EnGen0166 |
| Efa-440-1 | Bacteria Firmicutes Bacilli Lactobacillales Enterococcaceae *Enterococcus faecium* EnGen0167 |
| Efa-441-1 | Bacteria Firmicutes Bacilli Lactobacillales Enterococcaceae *Enterococcus faecium* EnGen0168 |
| Efa-442-1 | Bacteria Firmicutes Bacilli Lactobacillales Enterococcaceae *Enterococcus faecium* EnGen0169 |
| Efa-443-1 | Bacteria Firmicutes Bacilli Lactobacillales Enterococcaceae *Enterococcus faecium* EnGen0170 |
| Efa-444-1 | Bacteria Firmicutes Bacilli Lactobacillales Enterococcaceae *Enterococcus faecium* EnGen0171 |
| Efa-445-1 | Bacteria Firmicutes Bacilli Lactobacillales Enterococcaceae *Enterococcus faecium* EnGen0172 |
| Efa-446-1 | Bacteria Firmicutes Bacilli Lactobacillales Enterococcaceae *Enterococcus faecium* EnGen0173 |
| Efa-447-1 | Bacteria Firmicutes Bacilli Lactobacillales Enterococcaceae *Enterococcus faecium* EnGen0174 |
| Efa-448-1 | Bacteria Firmicutes Bacilli Lactobacillales Enterococcaceae *Enterococcus faecium* EnGen0175 |
| Efa-449-1 | Bacteria Firmicutes Bacilli Lactobacillales Enterococcaceae *Enterococcus faecium* EnGen0176 |
| Efa-450-1 | Bacteria Firmicutes Bacilli Lactobacillales Enterococcaceae *Enterococcus faecium* EnGen0177 |
| Efa-451-1 | Bacteria Firmicutes Bacilli Lactobacillales Enterococcaceae *Enterococcus faecium* EnGen0178 |
| Efa-452-1 | Bacteria Firmicutes Bacilli Lactobacillales Enterococcaceae *Enterococcus faecium* EnGen0179 |
| Efa-453-1 | Bacteria Firmicutes Bacilli Lactobacillales Enterococcaceae *Enterococcus faecium* EnGen0180 |
| Efa-454-1 | Bacteria Firmicutes Bacilli Lactobacillales Enterococcaceae *Enterococcus faecium* EnGen0181 |
| Efa-455-1 | Bacteria Firmicutes Bacilli Lactobacillales Enterococcaceae *Enterococcus faecium* EnGen0182 |
| Efa-456-1 | Bacteria Firmicutes Bacilli Lactobacillales Enterococcaceae *Enterococcus faecium* EnGen0183 |
| Efa-457-1 | Bacteria Firmicutes Bacilli Lactobacillales Enterococcaceae *Enterococcus faecium* EnGen0184 |
| Efa-458-1 | Bacteria Firmicutes Bacilli Lactobacillales Enterococcaceae *Enterococcus faecium* EnGen0185 |
| Efa-459-1 | Bacteria Firmicutes Bacilli Lactobacillales Enterococcaceae *Enterococcus faecium* EnGen0186 |
| Efa-460-1 | Bacteria Firmicutes Bacilli Lactobacillales Enterococcaceae *Enterococcus faecium* EnGen0187 |
| Efa-461-1 | Bacteria Firmicutes Bacilli Lactobacillales Enterococcaceae *Enterococcus faecium* EnGen0188 |
| Efa-462-1 | Bacteria Firmicutes Bacilli Lactobacillales Enterococcaceae *Enterococcus faecium* EnGen0189 |
| Efa-463-1 | Bacteria Firmicutes Bacilli Lactobacillales Enterococcaceae *Enterococcus faecium* EnGen0190 |
| Efa-464-1 | Bacteria Firmicutes Bacilli Lactobacillales Enterococcaceae *Enterococcus faecium* EnGen0191 |
| Efa-465-1 | Bacteria Firmicutes Bacilli Lactobacillales Enterococcaceae *Enterococcus faecium* EnGen0192 |
| Efa-466-1 | Bacteria Firmicutes Bacilli Lactobacillales Enterococcaceae *Enterococcus faecium* EnGen0193 |
| Efa-467-1 | Bacteria Firmicutes Bacilli Lactobacillales Enterococcaceae *Enterococcus faecium* EnGen0253 |
| Efa-468-1 | Bacteria Firmicutes Bacilli Lactobacillales Enterococcaceae *Enterococcus faecium* EnGen0254 |
| Efa-469-1 | Bacteria Firmicutes Bacilli Lactobacillales Enterococcaceae *Enterococcus faecium* EnGen0255 |
| Efa-470-1 | Bacteria Firmicutes Bacilli Lactobacillales Enterococcaceae *Enterococcus faecium* EnGen0256 |
| Efa-471-1 | Bacteria Firmicutes Bacilli Lactobacillales Enterococcaceae *Enterococcus faecium* EnGen0257 |
| Efa-472-1 | Bacteria Firmicutes Bacilli Lactobacillales Enterococcaceae *Enterococcus faecium* EnGen0258 |
| Efa-473-1 | Bacteria Firmicutes Bacilli Lactobacillales Enterococcaceae *Enterococcus faecium* EnGen0259 |
| Efa-474-1 | Bacteria Firmicutes Bacilli Lactobacillales Enterococcaceae *Enterococcus faecium* EnGen0260 |
| Efa-475-1 | Bacteria Firmicutes Bacilli Lactobacillales Enterococcaceae *Enterococcus faecium* EnGen0261 |
| Efa-476-1 | Bacteria Firmicutes Bacilli Lactobacillales Enterococcaceae *Enterococcus faecium* EnGen0262 |
| Efa-477-1 | Bacteria Firmicutes Bacilli Lactobacillales Enterococcaceae *Enterococcus faecium* EnGen0263 |
| Efa-478-1 | Bacteria Firmicutes Bacilli Lactobacillales Enterococcaceae *Enterococcus faecium* EnGen0264 |
| Efa-479-1 | Bacteria Firmicutes Bacilli Lactobacillales Enterococcaceae *Enterococcus faecium* EnGen0265 |
| Efa-480-1 | Bacteria Firmicutes Bacilli Lactobacillales Enterococcaceae *Enterococcus faecium* EnGen0267 |
| Efa-481-1 | Bacteria Firmicutes Bacilli Lactobacillales Enterococcaceae *Enterococcus faecium* EnGen0312 |
| Efa-482-1 | Bacteria Firmicutes Bacilli Lactobacillales Enterococcaceae *Enterococcus faecium* EnGen0313 |
| Efa-483-1 | Bacteria Firmicutes Bacilli Lactobacillales Enterococcaceae *Enterococcus faecium* EnGen0314 |
| Efa-484-1 | Bacteria Firmicutes Bacilli Lactobacillales Enterococcaceae *Enterococcus faecium* EnGen0315 |
| Efa-485-1 | Bacteria Firmicutes Bacilli Lactobacillales Enterococcaceae *Enterococcus faecium* EnGen0316 |
| Efa-486-1 | Bacteria Firmicutes Bacilli Lactobacillales Enterococcaceae *Enterococcus faecium* EnGen0317 |
| Efa-487-1 | Bacteria Firmicutes Bacilli Lactobacillales Enterococcaceae *Enterococcus faecium* EnGen0318 |
| Efa-488-1 | Bacteria Firmicutes Bacilli Lactobacillales Enterococcaceae *Enterococcus faecium* EnGen0319 |
| Efa-489-1 | Bacteria Firmicutes Bacilli Lactobacillales Enterococcaceae *Enterococcus faecium* EnGen0320 |
| Efa-490-1 | Bacteria Firmicutes Bacilli Lactobacillales Enterococcaceae *Enterococcus faecium* EnGen0321 |
| Efa-491-1 | Bacteria Firmicutes Bacilli Lactobacillales Enterococcaceae *Enterococcus faecium* EnGen0322 |
| Efa-492-1 | Bacteria Firmicutes Bacilli Lactobacillales Enterococcaceae *Enterococcus faecium* EnGen0323 |
| Efa-493-1 | Bacteria Firmicutes Bacilli Lactobacillales Enterococcaceae *Enterococcus faecium* EnGen0371 |
| Efa-494-1 | Bacteria Firmicutes Bacilli Lactobacillales Enterococcaceae *Enterococcus faecium* EnGen0372 |
| Efa-495-1 | Bacteria Firmicutes Bacilli Lactobacillales Enterococcaceae *Enterococcus faecium* EnGen0375 |
| Efa-496-1 to Efa-496-2 | Bacteria Firmicutes Bacilli Lactobacillales Enterococcaceae *Enterococcus faecium* EnGen0376 |
| Efa-497-1 to Efa-497-2 | Bacteria Firmicutes Bacilli Lactobacillales Enterococcaceae *Enterococcus faecium* EnGen0377 |
| Efa-341-1 | Bacteria Firmicutes Bacilli Lactobacillales Enterococcaceae *Enterococcus faecium* ERV1 |

| | |
|---|---|
| Efa-342-1 | Bacteria Firmicutes Bacilli Lactobacillales Enterococcaceae *Enterococcus faecium* ERV102 |
| Efa-343-1 | Bacteria Firmicutes Bacilli Lactobacillales Enterococcaceae *Enterococcus faecium* ERV161 |
| Efa-344-1 | Bacteria Firmicutes Bacilli Lactobacillales Enterococcaceae *Enterococcus faecium* ERV165 |
| Efa-345-1 | Bacteria Firmicutes Bacilli Lactobacillales Enterococcaceae *Enterococcus faecium* ERV168 |
| Efa-346-1 | Bacteria Firmicutes Bacilli Lactobacillales Enterococcaceae *Enterococcus faecium* ERV26 |
| Efa-347-1 | Bacteria Firmicutes Bacilli Lactobacillales Enterococcaceae *Enterococcus faecium* ERV38 |
| Efa-348-1 | Bacteria Firmicutes Bacilli Lactobacillales Enterococcaceae *Enterococcus faecium* ERV69 |
| Efa-349-1 | Bacteria Firmicutes Bacilli Lactobacillales Enterococcaceae *Enterococcus faecium* ERV99 |
| Efa-498-1 | Bacteria Firmicutes Bacilli Lactobacillales Enterococcaceae *Enterococcus faecium* LA4B-2 |
| Efa-499-1 | Bacteria Firmicutes Bacilli Lactobacillales Enterococcaceae *Enterococcus faecium* LCT-EF128 |
| Efa-500-1 | Bacteria Firmicutes Bacilli Lactobacillales Enterococcaceae *Enterococcus faecium* LCT-EF20 |
| Efa-501-1 | Bacteria Firmicutes Bacilli Lactobacillales Enterococcaceae *Enterococcus faecium* LCT-EF258 |
| Efa-502-1 | Bacteria Firmicutes Bacilli Lactobacillales Enterococcaceae *Enterococcus faecium* LCT-EF297 |
| Efa-503-1 | Bacteria Firmicutes Bacilli Lactobacillales Enterococcaceae *Enterococcus faecium* LCT-EF301 |
| Efa-504-1 | Bacteria Firmicutes Bacilli Lactobacillales Enterococcaceae *Enterococcus faecium* LCT-EF90 |
| Efa-505-1 | Bacteria Firmicutes Bacilli Lactobacillales Enterococcaceae *Enterococcus faecium* MRSN 11639 |
| Efa-506-1 | Bacteria Firmicutes Bacilli Lactobacillales Enterococcaceae *Enterococcus faecium* MRSN 3418 |
| Efa-507-1 | Bacteria Firmicutes Bacilli Lactobacillales Enterococcaceae *Enterococcus faecium* NEF1 |
| Efa-508-1 | Bacteria Firmicutes Bacilli Lactobacillales Enterococcaceae *Enterococcus faecium* NRRL B-2354 |
| Efa-509-1 | Bacteria Firmicutes Bacilli Lactobacillales Enterococcaceae *Enterococcus faecium* OC2A-1 |
| Efa-510-1 | Bacteria Firmicutes Bacilli Lactobacillales Enterococcaceae *Enterococcus faecium* P1123 |
| Efa-511-1 | Bacteria Firmicutes Bacilli Lactobacillales Enterococcaceae *Enterococcus faecium* P1137 |
| Efa-512-1 | Bacteria Firmicutes Bacilli Lactobacillales Enterococcaceae *Enterococcus faecium* P1139 |
| Efa-513-1 | Bacteria Firmicutes Bacilli Lactobacillales Enterococcaceae *Enterococcus faecium* P1140 |
| Efa-514-1 | Bacteria Firmicutes Bacilli Lactobacillales Enterococcaceae *Enterococcus faecium* P1190 |
| Efa-515-1 | Bacteria Firmicutes Bacilli Lactobacillales Enterococcaceae *Enterococcus faecium* P1986 |
| Efa-516-1 | Bacteria Firmicutes Bacilli Lactobacillales Enterococcaceae *Enterococcus faecium* R446 |
| Efa-517-1 | Bacteria Firmicutes Bacilli Lactobacillales Enterococcaceae *Enterococcus faecium* R494 |
| Efa-518-1 | Bacteria Firmicutes Bacilli Lactobacillales Enterococcaceae *Enterococcus faecium* R496 |
| Efa-519-1 | Bacteria Firmicutes Bacilli Lactobacillales Enterococcaceae *Enterococcus faecium* R497 |
| Efa-520-1 | Bacteria Firmicutes Bacilli Lactobacillales Enterococcaceae *Enterococcus faecium* R499 |
| Efa-521-1 | Bacteria Firmicutes Bacilli Lactobacillales Enterococcaceae *Enterococcus faecium* R501 |
| Efa-522-1 | Bacteria Firmicutes Bacilli Lactobacillales Enterococcaceae *Enterococcus faecium* S447 |
| Efa-523-1 | Bacteria Firmicutes Bacilli Lactobacillales Enterococcaceae *Enterococcus faecium* SB2C-2 |
| Efa-524-1 | Bacteria Firmicutes Bacilli Lactobacillales Enterococcaceae *Enterococcus faecium* SD1C-2 |
| Efa-525-1 | Bacteria Firmicutes Bacilli Lactobacillales Enterococcaceae *Enterococcus faecium* SD2A-2 |
| Efa-526-1 | Bacteria Firmicutes Bacilli Lactobacillales Enterococcaceae *Enterococcus faecium* SD3B-2 |
| Efa-527-1 | Bacteria Firmicutes Bacilli Lactobacillales Enterococcaceae *Enterococcus faecium* T110 |
| Efa-528-1 | Bacteria Firmicutes Bacilli Lactobacillales Enterococcaceae *Enterococcus faecium* TX0082 |
| Efa-529-1 | Bacteria Firmicutes Bacilli Lactobacillales Enterococcaceae *Enterococcus faecium* TX0133A |
| Efa-532-1 | Bacteria Firmicutes Bacilli Lactobacillales Enterococcaceae *Enterococcus faecium* TX0133a01 |
| Efa-533-1 | Bacteria Firmicutes Bacilli Lactobacillales Enterococcaceae *Enterococcus faecium* TX0133a04 |
| Efa-530-1 | Bacteria Firmicutes Bacilli Lactobacillales Enterococcaceae *Enterococcus faecium* TX0133B |
| Efa-531-1 | Bacteria Firmicutes Bacilli Lactobacillales Enterococcaceae *Enterococcus faecium* TX0133C |
| Efa-534-1 | Bacteria Firmicutes Bacilli Lactobacillales Enterococcaceae *Enterococcus faecium* TX1337RF |
| Efa-535-1 | Bacteria Firmicutes Bacilli Lactobacillales Enterococcaceae *Enterococcus faecium* UC10237 |
| Efa-536-1 | Bacteria Firmicutes Bacilli Lactobacillales Enterococcaceae *Enterococcus faecium* UC7251 |
| Efa-537-1 | Bacteria Firmicutes Bacilli Lactobacillales Enterococcaceae *Enterococcus faecium* UC7256 |
| Efa-538-1 | Bacteria Firmicutes Bacilli Lactobacillales Enterococcaceae *Enterococcus faecium* UC7267 |
| Efa-539-1 | Bacteria Firmicutes Bacilli Lactobacillales Enterococcaceae *Enterococcus faecium* UC8668 |
| Efa-540-1 | Bacteria Firmicutes Bacilli Lactobacillales Enterococcaceae *Enterococcus faecium* UC8733 |
| Efa-541-1 | Bacteria Firmicutes Bacilli Lactobacillales Enterococcaceae *Enterococcus faecium* V689 |
| Efa-542-1 | Bacteria Firmicutes Bacilli Lactobacillales Enterococcaceae *Enterococcus faecium* VRE0576 |
| Efa-543-1 | Bacteria Firmicutes Bacilli Lactobacillales Enterococcaceae *Enterococcus faecium* VRE1044 |
| Efa-544-1 | Bacteria Firmicutes Bacilli Lactobacillales Enterococcaceae *Enterococcus faecium* VRE1261 |
| Efa-545-1 | Bacteria Firmicutes Bacilli Lactobacillales Enterococcaceae *Enterococcus faecium* VSE1036 |
| Efl-1-1 | Bacteria Firmicutes Bacilli Lactobacillales Enterococcaceae *Enterococcus flavescens* ATCC 49996 |
| Ega-1-1 to Ega-1-2 | Bacteria Firmicutes Bacilli Lactobacillales Enterococcaceae *Enterococcus gallinarum* |
| Egi-1-1 | Bacteria Firmicutes Bacilli Lactobacillales Enterococcaceae *Enterococcus gilvus* ATCC BAA-350 |
| Eha-1-1 | Bacteria Firmicutes Bacilli Lactobacillales Enterococcaceae *Enterococcus haemoperoxidus* ATCC BAA-382 |
| Ehi-1-1 to Ehi-1-2 | Bacteria Firmicutes Bacilli Lactobacillales Enterococcaceae *Enterococcus hirae* |
| Ehi-2-1 to Ehi-2-2 | Bacteria Firmicutes Bacilli Lactobacillales Enterococcaceae *Enterococcus hirae* ATCC 9790 |
| Eit-1-1 | Bacteria Firmicutes Bacilli Lactobacillales Enterococcaceae *Enterococcus italicus* DSM 15952 |
| Ema-2-1 | Bacteria Firmicutes Bacilli Lactobacillales Enterococcaceae *Enterococcus malodoratus* ATCC 43197 |
| Ema-3-1 | Bacteria Firmicutes Bacilli Lactobacillales Enterococcaceae *Enterococcus massiliensis* |

| | |
|---|---|
| Emo-1-1 | Bacteria Firmicutes Bacilli Lactobacillales Enterococcaceae *Enterococcus moraviensis* ATCC BAA-383 |
| Emu-1-1 | Bacteria Firmicutes Bacilli Lactobacillales Enterococcaceae *Enterococcus mundtii* ATCC 882 |
| Emu-4-1 | Bacteria Firmicutes Bacilli Lactobacillales Enterococcaceae *Enterococcus mundtii crl*1656 |
| Emu-2-1 | Bacteria Firmicutes Bacilli Lactobacillales Enterococcaceae *Enterococcus mundtii* CRL35 |
| Emu-3-1 | Bacteria Firmicutes Bacilli Lactobacillales Enterococcaceae *Enterococcus mundtii* QU 25 |
| Epa-2-1 | Bacteria Firmicutes Bacilli Lactobacillales Enterococcaceae *Enterococcus pallens* ATCC BAA-351 |
| Eph-1-1 | Bacteria Firmicutes Bacilli Lactobacillales Enterococcaceae *Enterococcus phoeniculicola* ATCC BAA-412 |
| Era-1-1 | Bacteria Firmicutes Bacilli Lactobacillales Enterococcaceae *Enterococcus raffinosus* ATCC 49464 |
| Esa-1-1 | Bacteria Firmicutes Bacilli Lactobacillales Enterococcaceae *Enterococcus saccharolyticus* 30_1 |
| Esa-2-1 to Esa-2-2 | Bacteria Firmicutes Bacilli Lactobacillales Enterococcaceae *Enterococcus saccharolyticus* ATCC 43076 |
| Esp-12-1 | Bacteria Firmicutes Bacilli Lactobacillales Enterococcaceae *Enterococcus sp.* 1140_ESPC |
| Esp-13-1 | Bacteria Firmicutes Bacilli Lactobacillales Enterococcaceae *Enterococcus sp.* 255_ESPC |
| Esp-14-1 | Bacteria Firmicutes Bacilli Lactobacillales Enterococcaceae *Enterococcus sp.* C1 |
| Esp-15-1 | Bacteria Firmicutes Bacilli Lactobacillales Enterococcaceae *Enterococcus sp.* GMD3E |
| Esp-16-1 | Bacteria Firmicutes Bacilli Lactobacillales Enterococcaceae *Enterococcus sp.* GMD5E |
| Esp-17-1 | Bacteria Firmicutes Bacilli Lactobacillales Enterococcaceae *Enterococcus sp.* TR |
| Esu-1-1 | Bacteria Firmicutes Bacilli Lactobacillales Enterococcaceae *Enterococcus sulfureus* ATCC 49903 |
| Evi-1-1 | Bacteria Firmicutes Bacilli Lactobacillales Enterococcaceae *Enterococcus villorum* ATCC 700913 |
| Mpl-1-1 to Mpl-1-11 | Bacteria Firmicutes Bacilli Lactobacillales Enterococcaceae *Melissococcus plutonius* |
| Tmu-1-1 | Bacteria Firmicutes Bacilli Lactobacillales Enterococcaceae *Tetragenococcus muriaticus* 3MR10-3 |
| Tmu-2-1 | Bacteria Firmicutes Bacilli Lactobacillales Enterococcaceae *Tetragenococcus muriaticus* DSM 15685 |
| Tmu-3-1 | Bacteria Firmicutes Bacilli Lactobacillales Enterococcaceae *Tetragenococcus muriaticus* PMC-11-5 |
| Vlu-1-1 | Bacteria Firmicutes Bacilli Lactobacillales Enterococcaceae *Vagococcus lutrae* LBD1 |
| Lac-1-1 | Bacteria Firmicutes Bacilli Lactobacillales Lactobacillaceae *Lactobacillus acidophilus* |
| Lac-2-1 | Bacteria Firmicutes Bacilli Lactobacillales Lactobacillaceae *Lactobacillus acidophilus* CFH |
| Lac-3-1 | Bacteria Firmicutes Bacilli Lactobacillales Lactobacillaceae *Lactobacillus acidophilus* CIP 76.13 |
| Lac-4-1 | Bacteria Firmicutes Bacilli Lactobacillales Lactobacillaceae *Lactobacillus acidophilus* CIRM-BIA 442 |
| Lac-5-1 | Bacteria Firmicutes Bacilli Lactobacillales Lactobacillaceae *Lactobacillus acidophilus* CIRM-BIA 445 |
| Lac-6-1 | Bacteria Firmicutes Bacilli Lactobacillales Lactobacillaceae *Lactobacillus acidophilus* DSM 20079 = JCM 1132 = NBRC 13951 |
| Lac-7-1 | Bacteria Firmicutes Bacilli Lactobacillales Lactobacillaceae *Lactobacillus acidophilus* DSM 20242 |
| Lac-8-1 | Bacteria Firmicutes Bacilli Lactobacillales Lactobacillaceae *Lactobacillus acidophilus* DSM 9126 |
| Lac-9-1 | Bacteria Firmicutes Bacilli Lactobacillales Lactobacillaceae *Lactobacillus acidophilus* La-14 |
| Lag-1-1 | Bacteria Firmicutes Bacilli Lactobacillales Lactobacillaceae *Lactobacillus agilis* |
| Lan-1-1 | Bacteria Firmicutes Bacilli Lactobacillales Lactobacillaceae *Lactobacillus animalis* |
| Lan-2-1 | Bacteria Firmicutes Bacilli Lactobacillales Lactobacillaceae *Lactobacillus animalis* KCTC 3501 = DSM 20602 |
| Lap-1-1 | Bacteria Firmicutes Bacilli Lactobacillales Lactobacillaceae *Lactobacillus apis* |
| Lap-2-1 | Bacteria Firmicutes Bacilli Lactobacillales Lactobacillaceae *Lactobacillus apodemi* DSM 16634 = JCM 16172 |
| Lbr-1-1 to Lbr-1-6 | Bacteria Firmicutes Bacilli Lactobacillales Lactobacillaceae *Lactobacillus brevis* |
| Lbr-2-1 | Bacteria Firmicutes Bacilli Lactobacillales Lactobacillaceae *Lactobacillus brevis* AG48 |
| Lbr-3-1 | Bacteria Firmicutes Bacilli Lactobacillales Lactobacillaceae *Lactobacillus brevis* ATCC 14869 = DSM 20054 |
| Lbr-4-1 | Bacteria Firmicutes Bacilli Lactobacillales Lactobacillaceae *Lactobacillus brevis* EW |
| Lbr-5-1 | Bacteria Firmicutes Bacilli Lactobacillales Lactobacillaceae *Lactobacillus brevis* KB290 |
| Lca-1-1 to Lca-1-6 | Bacteria Firmicutes Bacilli Lactobacillales Lactobacillaceae *Lactobacillus casei* |
| Lca-2-1 | Bacteria Firmicutes Bacilli Lactobacillales Lactobacillaceae *Lactobacillus casei* 12A |
| Lca-3-1 | Bacteria Firmicutes Bacilli Lactobacillales Lactobacillaceae *Lactobacillus casei* 21/1 |
| Lca-4-1 | Bacteria Firmicutes Bacilli Lactobacillales Lactobacillaceae *Lactobacillus casei* 32G |
| Lca-5-1 | Bacteria Firmicutes Bacilli Lactobacillales Lactobacillaceae *Lactobacillus casei* 5b |
| Lca-6-1 to Lca-6-2 | Bacteria Firmicutes Bacilli Lactobacillales Lactobacillaceae *Lactobacillus casei* A2-362 |
| Lca-7-1 | Bacteria Firmicutes Bacilli Lactobacillales Lactobacillaceae *Lactobacillus casei* CRF28 |
| Lca-9-1 | Bacteria Firmicutes Bacilli Lactobacillales Lactobacillaceae *Lactobacillus casei* Lc-10 |
| Lca-10-1 | Bacteria Firmicutes Bacilli Lactobacillales Lactobacillaceae *Lactobacillus casei* LcA |
| Lca-11-1 | Bacteria Firmicutes Bacilli Lactobacillales Lactobacillaceae *Lactobacillus casei* LcY |
| Lca-8-1 | Bacteria Firmicutes Bacilli Lactobacillales Lactobacillaceae *Lactobacillus casei* LOCK919 |
| Lca-12-1 | Bacteria Firmicutes Bacilli Lactobacillales Lactobacillaceae *Lactobacillus casei* Lpc-37 |
| Lca-13-1 | Bacteria Firmicutes Bacilli Lactobacillales Lactobacillaceae *Lactobacillus casei* M36 |
| Lca-18-1 | Bacteria Firmicutes Bacilli Lactobacillales Lactobacillaceae *Lactobacillus casei str.* Zhang |
| Lca-19-1 | Bacteria Firmicutes Bacilli Lactobacillales Lactobacillaceae *Lactobacillus casei subsp. casei* ATCC 393 |
| Lca-14-1 | Bacteria Firmicutes Bacilli Lactobacillales Lactobacillaceae *Lactobacillus casei* T71499 |
| Lca-15-1 | Bacteria Firmicutes Bacilli Lactobacillales Lactobacillaceae *Lactobacillus casei* UCD174 |
| Lca-16-1 | Bacteria Firmicutes Bacilli Lactobacillales Lactobacillaceae *Lactobacillus casei* UW1 |
| Lca-17-1 to Lca-17-2 | Bacteria Firmicutes Bacilli Lactobacillales Lactobacillaceae *Lactobacillus casei* UW4 |
| Lce-1-1 | Bacteria Firmicutes Bacilli Lactobacillales Lactobacillaceae *Lactobacillus ceti* DSM 22408 |
| Lco-2-1 | Bacteria Firmicutes Bacilli Lactobacillales Lactobacillaceae *Lactobacillus composti* DSM 18527 = JCM 14202 |
| Lcr-1-1 | Bacteria Firmicutes Bacilli Lactobacillales Lactobacillaceae *Lactobacillus crispatus* EM-LC1 |
| Lcr-2-1 | Bacteria Firmicutes Bacilli Lactobacillales Lactobacillaceae *Lactobacillus crispatus* FB049-03 |

| | |
|---|---|
| Lcr-3-1 | Bacteria Firmicutes Bacilli Lactobacillales Lactobacillaceae *Lactobacillus crispatus* FB077-07 |
| Lcr-4-1 | Bacteria Firmicutes Bacilli Lactobacillales Lactobacillaceae *Lactobacillus crispatus* MV-1A-US |
| Lcr-5-1 | Bacteria Firmicutes Bacilli Lactobacillales Lactobacillaceae *Lactobacillus crispatus* SJ-3C-US |
| Lcu-1-1 | Bacteria Firmicutes Bacilli Lactobacillales Lactobacillaceae *Lactobacillus curieae* |
| Leq-1-1 | Bacteria Firmicutes Bacilli Lactobacillales Lactobacillaceae *Lactobacillus equi* DPC 6820 |
| Leq-2-1 | Bacteria Firmicutes Bacilli Lactobacillales Lactobacillaceae *Lactobacillus equi* DSM 15833 = JCM 10991 |
| Lfa-1-1 | Bacteria Firmicutes Bacilli Lactobacillales Lactobacillaceae *Lactobacillus fabifermentans* DSM 21115 |
| Lfa-2-1 | Bacteria Firmicutes Bacilli Lactobacillales Lactobacillaceae *Lactobacillus fabifermentans* T30PCM01 |
| Lfa-3-1 | Bacteria Firmicutes Bacilli Lactobacillales Lactobacillaceae *Lactobacillus farciminis* KCTC 3681 = DSM 20184 |
| Lfa-4-1 | Bacteria Firmicutes Bacilli Lactobacillales Lactobacillaceae *Lactobacillus farraginis* DSM 18382 = JCM 14108 |
| Lfe-1-1 to Lfe-1-4 | Bacteria Firmicutes Bacilli Lactobacillales Lactobacillaceae *Lactobacillus fermentum* |
| Lfe-2-1 | Bacteria Firmicutes Bacilli Lactobacillales Lactobacillaceae *Lactobacillus fermentum* 3872 |
| Lfe-3-1 | Bacteria Firmicutes Bacilli Lactobacillales Lactobacillaceae *Lactobacillus fermentum* F-6 |
| Lfe-4-1 | Bacteria Firmicutes Bacilli Lactobacillales Lactobacillaceae *Lactobacillus fermentum* FTDC8312 |
| Lfe-5-1 | Bacteria Firmicutes Bacilli Lactobacillales Lactobacillaceae *Lactobacillus fermentum* L930BB |
| Lfe-6-1 | Bacteria Firmicutes Bacilli Lactobacillales Lactobacillaceae *Lactobacillus fermentum* Lf1 |
| Lfe-7-1 | Bacteria Firmicutes Bacilli Lactobacillales Lactobacillaceae *Lactobacillus fermentum* MTCC 8711 |
| Lfe-8-1 | Bacteria Firmicutes Bacilli Lactobacillales Lactobacillaceae *Lactobacillus fermentum* NB-22 |
| Lga-1-1 | Bacteria Firmicutes Bacilli Lactobacillales Lactobacillaceae *Lactobacillus gallinarum* DSM 10532 = JCM 2011 |
| Lga-2-1 to Lga-2-6 | Bacteria Firmicutes Bacilli Lactobacillales Lactobacillaceae *Lactobacillus gasseri* |
| Lga-3-1 | Bacteria Firmicutes Bacilli Lactobacillales Lactobacillaceae *Lactobacillus gasseri* 130918 |
| Lga-4-1 | Bacteria Firmicutes Bacilli Lactobacillales Lactobacillaceae *Lactobacillus gasseri* 2016 |
| Lga-5-1 | Bacteria Firmicutes Bacilli Lactobacillales Lactobacillaceae *Lactobacillus gasseri* ATCC 33323 = JCM 1131 |
| Lga-6-1 | Bacteria Firmicutes Bacilli Lactobacillales Lactobacillaceae *Lactobacillus gasseri* CECT 5714 |
| Lga-7-1 | Bacteria Firmicutes Bacilli Lactobacillales Lactobacillaceae *Lactobacillus gasseri* JV-V03 |
| Lga-8-1 | Bacteria Firmicutes Bacilli Lactobacillales Lactobacillaceae *Lactobacillus gasseri* K7 |
| Lga-9-1 | Bacteria Firmicutes Bacilli Lactobacillales Lactobacillaceae *Lactobacillus gasseri* SJ-9E-US |
| Lga-10-1 | Bacteria Firmicutes Bacilli Lactobacillales Lactobacillaceae *Lactobacillus gasseri* SV-16A-US |
| Lgi-1-1 | Bacteria Firmicutes Bacilli Lactobacillales Lactobacillaceae *Lactobacillus ginsenosidimutans* |
| Lha-1-1 | Bacteria Firmicutes Bacilli Lactobacillales Lactobacillaceae *Lactobacillus hamsteri* DSM 5661 = JCM 6256 |
| Lha-2-1 | Bacteria Firmicutes Bacilli Lactobacillales Lactobacillaceae *Lactobacillus harbinensis* DSM 16991 |
| Lha-3-1 | Bacteria Firmicutes Bacilli Lactobacillales Lactobacillaceae *Lactobacillus hayakitensis* DSM 18933 = JCM 14209 |
| Lhe-1-1 | Bacteria Firmicutes Bacilli Lactobacillales Lactobacillaceae *Lactobacillus heilongjiangensis* |
| Lhe-2-1 | Bacteria Firmicutes Bacilli Lactobacillales Lactobacillaceae *Lactobacillus helsingborgensis* |
| Lhe-3-1 to Lhe-3-2 | Bacteria Firmicutes Bacilli Lactobacillales Lactobacillaceae *Lactobacillus helveticus* |
| Lhe-4-1 | Bacteria Firmicutes Bacilli Lactobacillales Lactobacillaceae *Lactobacillus helveticus* CIRM-BIA 101 |
| Lhe-5-1 | Bacteria Firmicutes Bacilli Lactobacillales Lactobacillaceae *Lactobacillus helveticus* CIRM-BIA 103 |
| Lhe-6-1 | Bacteria Firmicutes Bacilli Lactobacillales Lactobacillaceae *Lactobacillus helveticus* CIRM-BIA 104 |
| Lhe-7-1 | Bacteria Firmicutes Bacilli Lactobacillales Lactobacillaceae *Lactobacillus helveticus* CIRM-BIA 951 |
| Lhe-8-1 | Bacteria Firmicutes Bacilli Lactobacillales Lactobacillaceae *Lactobacillus helveticus* CIRM-BIA 953 |
| Lhe-9-1 | Bacteria Firmicutes Bacilli Lactobacillales Lactobacillaceae *Lactobacillus helveticus* CNRZ32 |
| Lhe-10-1 | Bacteria Firmicutes Bacilli Lactobacillales Lactobacillaceae *Lactobacillus helveticus* H9 |
| Lhe-11-1 | Bacteria Firmicutes Bacilli Lactobacillales Lactobacillaceae *Lactobacillus herbarum* |
| Lho-1-1 | Bacteria Firmicutes Bacilli Lactobacillales Lactobacillaceae *Lactobacillus hokkaidonensis* JCM 18461 |
| Lin-5-1 | Bacteria Firmicutes Bacilli Lactobacillales Lactobacillaceae *Lactobacillus iners* ATCC 55195 |
| Lin-6-1 | Bacteria Firmicutes Bacilli Lactobacillales Lactobacillaceae *Lactobacillus ingluviei str.* Autruche 4 |
| Ljo-1-1 | Bacteria Firmicutes Bacilli Lactobacillales Lactobacillaceae *Lactobacillus johnsonii* 16 |
| Ljo-2-1 | Bacteria Firmicutes Bacilli Lactobacillales Lactobacillaceae *Lactobacillus johnsonii* N6.2 |
| Ljo-3-1 | Bacteria Firmicutes Bacilli Lactobacillales Lactobacillaceae *Lactobacillus johnsonii pf*01 |
| Lke-1-1 | Bacteria Firmicutes Bacilli Lactobacillales Lactobacillaceae *Lactobacillus kefiranofaciens subsp. kefiranofaciens* DSM 5016 = JCM 6985 |
| Lki-1-1 | Bacteria Firmicutes Bacilli Lactobacillales Lactobacillaceae *Lactobacillus kimbladii* |
| Lki-2-1 | Bacteria Firmicutes Bacilli Lactobacillales Lactobacillaceae *Lactobacillus kisonensis* F0435 |
| Lki-3-1 | Bacteria Firmicutes Bacilli Lactobacillales Lactobacillaceae *Lactobacillus kitasatonis* DSM 16761 = JCM 1039 |
| Lko-1-1 | Bacteria Firmicutes Bacilli Lactobacillales Lactobacillaceae *Lactobacillus koreensis* |
| Lku-1-1 | Bacteria Firmicutes Bacilli Lactobacillales Lactobacillaceae *Lactobacillus kullabergensis* |
| Lku-2-1 to Lku-2-2 | Bacteria Firmicutes Bacilli Lactobacillales Lactobacillaceae *Lactobacillus kunkeei* |
| Lku-3-1 | Bacteria Firmicutes Bacilli Lactobacillales Lactobacillaceae *Lactobacillus kunkeei* EFB6 |
| Lma-3-1 to Lma-3-2 | Bacteria Firmicutes Bacilli Lactobacillales Lactobacillaceae *Lactobacillus mali* KCTC 3596 = DSM 20444 |
| Lme-1-1 | Bacteria Firmicutes Bacilli Lactobacillales Lactobacillaceae *Lactobacillus melliventris* |
| Lmu-1-1 to Lmu-1-2 | Bacteria Firmicutes Bacilli Lactobacillales Lactobacillaceae *Lactobacillus mucosae* |
| Lmu-2-1 | Bacteria Firmicutes Bacilli Lactobacillales Lactobacillaceae *Lactobacillus mucosae* LM1 |
| Lmu-3-1 | Bacteria Firmicutes Bacilli Lactobacillales Lactobacillaceae *Lactobacillus murinus* ASF361 |
| Lna-2-1 | Bacteria Firmicutes Bacilli Lactobacillales Lactobacillaceae *Lactobacillus namurensis str.* Chizuka 01 |
| Lno-1-1 | Bacteria Firmicutes Bacilli Lactobacillales Lactobacillaceae *Lactobacillus nodensis* DSM 19682 = JCM 14932 = NBRC 107160 |
| Lor-1-1 | Bacteria Firmicutes Bacilli Lactobacillales Lactobacillaceae *Lactobacillus oris* F0423 |

| | |
|---|---|
| Lor-2-1 | Bacteria Firmicutes Bacilli Lactobacillales Lactobacillaceae *Lactobacillus oryzae* JCM 18671 |
| Lot-1-1 | Bacteria Firmicutes Bacilli Lactobacillales Lactobacillaceae *Lactobacillus otakiensis* JCM 15040 |
| Lpa-1-1 | Bacteria Firmicutes Bacilli Lactobacillales Lactobacillaceae *Lactobacillus panis* DSM 6035 |
| Lpa-2-1 | Bacteria Firmicutes Bacilli Lactobacillales Lactobacillaceae *Lactobacillus parabrevis* ATCC 53295 |
| Lpa-3-1 to Lpa-3-2 | Bacteria Firmicutes Bacilli Lactobacillales Lactobacillaceae *Lactobacillus paracasei* |
| Lpa-4-1 | Bacteria Firmicutes Bacilli Lactobacillales Lactobacillaceae *Lactobacillus paracasei* COM0101 |
| Lpa-5-1 | Bacteria Firmicutes Bacilli Lactobacillales Lactobacillaceae *Lactobacillus paracasei* N1115 |
| Lpa-6-1 | Bacteria Firmicutes Bacilli Lactobacillales Lactobacillaceae *Lactobacillus paracasei* NRIC 0644 |
| Lpa-7-1 | Bacteria Firmicutes Bacilli Lactobacillales Lactobacillaceae *Lactobacillus paracasei* NRIC 1917 |
| Lpa-8-1 | Bacteria Firmicutes Bacilli Lactobacillales Lactobacillaceae *Lactobacillus paracasei* NRIC 1981 |
| Lpa-9-1 | Bacteria Firmicutes Bacilli Lactobacillales Lactobacillaceae *Lactobacillus paracasei subsp. paracasei* 8700:2 |
| Lpa-10-1 | Bacteria Firmicutes Bacilli Lactobacillales Lactobacillaceae *Lactobacillus paracasei subsp. paracasei* CNCM I-2877 |
| Lpa-11-1 | Bacteria Firmicutes Bacilli Lactobacillales Lactobacillaceae *Lactobacillus paracasei subsp. paracasei* CNCM I-4270 |
| Lpa-12-1 | Bacteria Firmicutes Bacilli Lactobacillales Lactobacillaceae *Lactobacillus paracasei subsp. paracasei* CNCM I-4649 |
| Lpa-13-1 | Bacteria Firmicutes Bacilli Lactobacillales Lactobacillaceae *Lactobacillus paracasei subsp. paracasei* JCM 8130 |
| Lpa-14-1 | Bacteria Firmicutes Bacilli Lactobacillales Lactobacillaceae *Lactobacillus paracasei subsp. paracasei* Lpp120 |
| Lpa-15-1 | Bacteria Firmicutes Bacilli Lactobacillales Lactobacillaceae *Lactobacillus paracasei subsp. paracasei* Lpp122 |
| Lpa-16-1 | Bacteria Firmicutes Bacilli Lactobacillales Lactobacillaceae *Lactobacillus paracasei subsp. paracasei* Lpp125 |
| Lpa-17-1 | Bacteria Firmicutes Bacilli Lactobacillales Lactobacillaceae *Lactobacillus paracasei subsp. paracasei* Lpp14 |
| Lpa-18-1 | Bacteria Firmicutes Bacilli Lactobacillales Lactobacillaceae *Lactobacillus paracasei subsp. paracasei* Lpp17 |
| Lpa-19-1 | Bacteria Firmicutes Bacilli Lactobacillales Lactobacillaceae *Lactobacillus paracasei subsp. paracasei* Lpp189 |
| Lpa-20-1 | Bacteria Firmicutes Bacilli Lactobacillales Lactobacillaceae *Lactobacillus paracasei subsp. paracasei* Lpp219 |
| Lpa-21-1 | Bacteria Firmicutes Bacilli Lactobacillales Lactobacillaceae *Lactobacillus paracasei subsp. paracasei* Lpp22 |
| Lpa-22-1 | Bacteria Firmicutes Bacilli Lactobacillales Lactobacillaceae *Lactobacillus paracasei subsp. paracasei* Lpp221 |
| Lpa-23-1 | Bacteria Firmicutes Bacilli Lactobacillales Lactobacillaceae *Lactobacillus paracasei subsp. paracasei* Lpp223 |
| Lpa-24-1 | Bacteria Firmicutes Bacilli Lactobacillales Lactobacillaceae *Lactobacillus paracasei subsp. paracasei* Lpp225 |
| Lpa-25-1 | Bacteria Firmicutes Bacilli Lactobacillales Lactobacillaceae *Lactobacillus paracasei subsp. paracasei* Lpp226 |
| Lpa-26-1 | Bacteria Firmicutes Bacilli Lactobacillales Lactobacillaceae *Lactobacillus paracasei subsp. paracasei* Lpp227 |
| Lpa-27-1 | Bacteria Firmicutes Bacilli Lactobacillales Lactobacillaceae *Lactobacillus paracasei subsp. paracasei* Lpp228 |
| Lpa-28-1 | Bacteria Firmicutes Bacilli Lactobacillales Lactobacillaceae *Lactobacillus paracasei subsp. paracasei* Lpp229 |
| Lpa-29-1 | Bacteria Firmicutes Bacilli Lactobacillales Lactobacillaceae *Lactobacillus paracasei subsp. paracasei* Lpp230 |
| Lpa-30-1 | Bacteria Firmicutes Bacilli Lactobacillales Lactobacillaceae *Lactobacillus paracasei subsp. paracasei* Lpp37 |
| Lpa-31-1 | Bacteria Firmicutes Bacilli Lactobacillales Lactobacillaceae *Lactobacillus paracasei subsp. paracasei* Lpp43 |
| Lpa-32-1 | Bacteria Firmicutes Bacilli Lactobacillales Lactobacillaceae *Lactobacillus paracasei subsp. paracasei* Lpp46 |
| Lpa-33-1 | Bacteria Firmicutes Bacilli Lactobacillales Lactobacillaceae *Lactobacillus paracasei subsp. paracasei* Lpp49 |
| Lpa-34-1 | Bacteria Firmicutes Bacilli Lactobacillales Lactobacillaceae *Lactobacillus paracasei subsp. paracasei* Lpp7 |
| Lpa-35-1 | Bacteria Firmicutes Bacilli Lactobacillales Lactobacillaceae *Lactobacillus paracasei subsp. paracasei* Lpp70 |
| Lpa-36-1 | Bacteria Firmicutes Bacilli Lactobacillales Lactobacillaceae *Lactobacillus paracasei subsp. paracasei* Lpp74 |
| Lpa-37-1 | Bacteria Firmicutes Bacilli Lactobacillales Lactobacillaceae *Lactobacillus paracasei subsp. tolerans* Lpl14 |
| Lpa-38-1 | Bacteria Firmicutes Bacilli Lactobacillales Lactobacillaceae *Lactobacillus paracasei subsp. tolerans* Lpl7 |
| Lpa-39-1 | Bacteria Firmicutes Bacilli Lactobacillales Lactobacillaceae *Lactobacillus parafarraginis* F0439 |
| Lpa-40-1 | Bacteria Firmicutes Bacilli Lactobacillales Lactobacillaceae *Lactobacillus paralimentarius* DSM 13238 = JCM 10415 |
| Lpa-41-1 | Bacteria Firmicutes Bacilli Lactobacillales Lactobacillaceae *Lactobacillus paraplantarum* |
| Lpe-1-1 | Bacteria Firmicutes Bacilli Lactobacillales Lactobacillaceae *Lactobacillus pentosus* |
| Lpe-2-1 | Bacteria Firmicutes Bacilli Lactobacillales Lactobacillaceae *Lactobacillus pentosus* KCA1 |
| Lpl-1-1 | Bacteria Firmicutes Bacilli Lactobacillales Lactobacillaceae *Lactobacillus plantarum* |
| Lpl-2-1 | Bacteria Firmicutes Bacilli Lactobacillales Lactobacillaceae *Lactobacillus plantarum* 16 |
| Lpl-3-1 | Bacteria Firmicutes Bacilli Lactobacillales Lactobacillaceae *Lactobacillus plantarum* 19L3 |
| Lpl-4-1 | Bacteria Firmicutes Bacilli Lactobacillales Lactobacillaceae *Lactobacillus plantarum* 2165 |
| Lpl-5-1 | Bacteria Firmicutes Bacilli Lactobacillales Lactobacillaceae *Lactobacillus plantarum* 4_3 |
| Lpl-6-1 | Bacteria Firmicutes Bacilli Lactobacillales Lactobacillaceae *Lactobacillus plantarum* AY01 |
| Lpl-7-1 | Bacteria Firmicutes Bacilli Lactobacillales Lactobacillaceae *Lactobacillus plantarum* CMPG5300 |
| Lpl-8-1 | Bacteria Firmicutes Bacilli Lactobacillales Lactobacillaceae *Lactobacillus plantarum* EGD-AQ4 |
| Lpl-9-1 | Bacteria Firmicutes Bacilli Lactobacillales Lactobacillaceae *Lactobacillus plantarum* IPLA88 |
| Lpl-10-1 | Bacteria Firmicutes Bacilli Lactobacillales Lactobacillaceae *Lactobacillus plantarum* LP91 |
| Lpl-14-1 | Bacteria Firmicutes Bacilli Lactobacillales Lactobacillaceae *Lactobacillus plantarum subsp. plantarum* |
| Lpl-15-1 | Bacteria Firmicutes Bacilli Lactobacillales Lactobacillaceae *Lactobacillus plantarum subsp. plantarum* ATCC 14917 |
| Lpl-16-1 | Bacteria Firmicutes Bacilli Lactobacillales Lactobacillaceae *Lactobacillus plantarum subsp. plantarum* ATCC 14917 = JCM 1149 = CGMCC 1.2437 |
| Lpl-17-1 | Bacteria Firmicutes Bacilli Lactobacillales Lactobacillaceae *Lactobacillus plantarum subsp. plantarum* NC8 |
| Lpl-18-1 | Bacteria Firmicutes Bacilli Lactobacillales Lactobacillaceae *Lactobacillus plantarum subsp. plantarum* P-8 |
| Lpl-11-1 | Bacteria Firmicutes Bacilli Lactobacillales Lactobacillaceae *Lactobacillus plantarum* UCMA 3037 |
| Lpl-12-1 | Bacteria Firmicutes Bacilli Lactobacillales Lactobacillaceae *Lactobacillus plantarum* WJL |
| Lpl-13-1 | Bacteria Firmicutes Bacilli Lactobacillales Lactobacillaceae *Lactobacillus plantarum* ZJ316 |
| Lpo-1-1 | Bacteria Firmicutes Bacilli Lactobacillales Lactobacillaceae *Lactobacillus pobuzihii* E100301 |
| Lre-1-1 | Bacteria Firmicutes Bacilli Lactobacillales Lactobacillaceae *Lactobacillus reuteri* |

| | |
|---|---|
| Lre-2-1 | Bacteria Firmicutes Bacilli Lactobacillales Lactobacillaceae *Lactobacillus reuteri* ATCC 53608 |
| Lre-3-1 | Bacteria Firmicutes Bacilli Lactobacillales Lactobacillaceae *Lactobacillus reuteri* I5007 |
| Lre-5-1 | Bacteria Firmicutes Bacilli Lactobacillales Lactobacillaceae *Lactobacillus reuteri lpuph* |
| Lre-6-1 | Bacteria Firmicutes Bacilli Lactobacillales Lactobacillaceae *Lactobacillus reuteri mlc*3 |
| Lre-4-1 | Bacteria Firmicutes Bacilli Lactobacillales Lactobacillaceae *Lactobacillus reuteri* TD1 |
| Lrh-1-1 to Lrh-1-28 | Bacteria Firmicutes Bacilli Lactobacillales Lactobacillaceae *Lactobacillus rhamnosus* |
| Lrh-2-1 | Bacteria Firmicutes Bacilli Lactobacillales Lactobacillaceae *Lactobacillus rhamnosus* 51B |
| Lrh-3-1 | Bacteria Firmicutes Bacilli Lactobacillales Lactobacillaceae *Lactobacillus rhamnosus* ATCC 21052 |
| Lrh-4-1 | Bacteria Firmicutes Bacilli Lactobacillales Lactobacillaceae *Lactobacillus rhamnosus* CASL |
| Lrh-5-1 | Bacteria Firmicutes Bacilli Lactobacillales Lactobacillaceae *Lactobacillus rhamnosus* CRL1505 |
| Lrh-6-1 | Bacteria Firmicutes Bacilli Lactobacillales Lactobacillaceae *Lactobacillus rhamnosus* E800 |
| Lrh-7-1 | Bacteria Firmicutes Bacilli Lactobacillales Lactobacillaceae *Lactobacillus rhamnosus* K32 |
| Lrh-8-1 | Bacteria Firmicutes Bacilli Lactobacillales Lactobacillaceae *Lactobacillus rhamnosus* L31 |
| Lrh-9-1 | Bacteria Firmicutes Bacilli Lactobacillales Lactobacillaceae *Lactobacillus rhamnosus* L34 |
| Lrh-10-1 | Bacteria Firmicutes Bacilli Lactobacillales Lactobacillaceae *Lactobacillus rhamnosus* L35 |
| Lrh-11-1 | Bacteria Firmicutes Bacilli Lactobacillales Lactobacillaceae *Lactobacillus rhamnosus* LOCK900 |
| Lrh-12-1 | Bacteria Firmicutes Bacilli Lactobacillales Lactobacillaceae *Lactobacillus rhamnosus* LOCK908 |
| Lrh-13-1 | Bacteria Firmicutes Bacilli Lactobacillales Lactobacillaceae *Lactobacillus rhamnosus* LR231 |
| Lrh-14-1 | Bacteria Firmicutes Bacilli Lactobacillales Lactobacillaceae *Lactobacillus rhamnosus* LRHMDP2 |
| Lrh-15-1 | Bacteria Firmicutes Bacilli Lactobacillales Lactobacillaceae *Lactobacillus rhamnosus* LRHMDP3 |
| Lrh-16-1 | Bacteria Firmicutes Bacilli Lactobacillales Lactobacillaceae *Lactobacillus rhamnosus* PEL5 |
| Lrh-17-1 | Bacteria Firmicutes Bacilli Lactobacillales Lactobacillaceae *Lactobacillus rhamnosus* PEL6 |
| Lrh-18-1 | Bacteria Firmicutes Bacilli Lactobacillales Lactobacillaceae *Lactobacillus rhamnosus* R0011 |
| Lro-2-1 to Lro-2-2 | Bacteria Firmicutes Bacilli Lactobacillales Lactobacillaceae *Lactobacillus rossiae* DSM 15814 |
| Lru-1-1 to Lru-1-2 | Bacteria Firmicutes Bacilli Lactobacillales Lactobacillaceae *Lactobacillus ruminis* |
| Lru-2-1 | Bacteria Firmicutes Bacilli Lactobacillales Lactobacillaceae *Lactobacillus ruminis* ATCC 25644 |
| Lru-3-1 | Bacteria Firmicutes Bacilli Lactobacillales Lactobacillaceae *Lactobacillus ruminis* DPC 6832 |
| Lru-4-1 | Bacteria Firmicutes Bacilli Lactobacillales Lactobacillaceae *Lactobacillus ruminis* S23 |
| Lsa-1-1 | Bacteria Firmicutes Bacilli Lactobacillales Lactobacillaceae *Lactobacillus saerimneri* 30a |
| Lsa-2-1 | Bacteria Firmicutes Bacilli Lactobacillales Lactobacillaceae *Lactobacillus saerimneri* DSM 16049 |
| Lsa-3-1 | Bacteria Firmicutes Bacilli Lactobacillales Lactobacillaceae *Lactobacillus salivarius* |
| Lsa-7-1 | Bacteria Firmicutes Bacilli Lactobacillales Lactobacillaceae *Lactobacillus salivarius cp*400 |
| Lsa-4-1 | Bacteria Firmicutes Bacilli Lactobacillales Lactobacillaceae *Lactobacillus salivarius* GJ-24 |
| Lsa-5-1 | Bacteria Firmicutes Bacilli Lactobacillales Lactobacillaceae *Lactobacillus salivarius* NIAS840 |
| Lsa-6-1 | Bacteria Firmicutes Bacilli Lactobacillales Lactobacillaceae *Lactobacillus salivarius* SMXD51 |
| Lsa-8-1 | Bacteria Firmicutes Bacilli Lactobacillales Lactobacillaceae *Lactobacillus salivarius str.* Ren |
| Lsh-1-1 | Bacteria Firmicutes Bacilli Lactobacillales Lactobacillaceae *Lactobacillus shenzhenensis* LY-73 |
| Lsp-9-1 | Bacteria Firmicutes Bacilli Lactobacillales Lactobacillaceae *Lactobacillus sp.* 7_1_47FAA |
| Lsp-10-1 | Bacteria Firmicutes Bacilli Lactobacillales Lactobacillaceae *Lactobacillus sp.* ASF360 |
| Lsp-11-1 | Bacteria Firmicutes Bacilli Lactobacillales Lactobacillaceae *Lactobacillus sp.* FMNP02 |
| Lsp-12-1 | Bacteria Firmicutes Bacilli Lactobacillales Lactobacillaceae *Lactobacillus sp.* OTU4228 |
| Lsp-13-1 | Bacteria Firmicutes Bacilli Lactobacillales Lactobacillaceae *Lactobacillus sp. wk*B10 |
| Lsp-14-1 | Bacteria Firmicutes Bacilli Lactobacillales Lactobacillaceae *Lactobacillus sp. wk*B8 |
| Lsp-15-1 | Bacteria Firmicutes Bacilli Lactobacillales Lactobacillaceae *Lactobacillus spicheri* |
| Lsu-1-1 | Bacteria Firmicutes Bacilli Lactobacillales Lactobacillaceae *Lactobacillus sucicola* DSM 21376 = JCM 15457 |
| Lvi-1-1 | Bacteria Firmicutes Bacilli Lactobacillales Lactobacillaceae *Lactobacillus vini* JP7.8.9 |
| Lwa-1-1 | Bacteria Firmicutes Bacilli Lactobacillales Lactobacillaceae *Lactobacillus wasatchensis* |
| Pac-1-1 | Bacteria Firmicutes Bacilli Lactobacillales Lactobacillaceae *Pediococcus acidilactici* D3 |
| Pac-3-1 | Bacteria Firmicutes Bacilli Lactobacillales Lactobacillaceae Pediococcus Pediococcus acidilactici group *Pediococcus acidilactici* AGR20 |
| Pac-2-1 | Bacteria Firmicutes Bacilli Lactobacillales Lactobacillaceae *Pediococcus acidilactici* MA18/5M |
| Pda-2-1 | Bacteria Firmicutes Bacilli Lactobacillales Lactobacillaceae *Pediococcus damnosus* LMG 28219 |
| Plo-1-1 | Bacteria Firmicutes Bacilli Lactobacillales Lactobacillaceae *Pediococcus lolii* NGRI 0510Q |
| Ppe-3-1 | Bacteria Firmicutes Bacilli Lactobacillales Lactobacillaceae *Pediococcus pentosaceus* CGMCC 7049 |
| Ppe-4-1 | Bacteria Firmicutes Bacilli Lactobacillales Lactobacillaceae *Pediococcus pentosaceus* IE-3 |
| Ppe-5-1 | Bacteria Firmicutes Bacilli Lactobacillales Lactobacillaceae *Pediococcus pentosaceus* SL4 |
| Lci-1-1 | Bacteria Firmicutes Bacilli Lactobacillales *Leuconostoc citreum* LBAE C10 |
| Lci-2-1 | Bacteria Firmicutes Bacilli Lactobacillales *Leuconostoc citreum* LBAE C11 |
| Lci-3-1 | Bacteria Firmicutes Bacilli Lactobacillales *Leuconostoc citreum* LBAE E16 |
| Lci-4-1 to Lci-4-3 | Bacteria Firmicutes Bacilli Lactobacillales Leuconostocaceae *Leuconostoc citreum* |
| Wci-1-1 to Wci-1-3 | Bacteria Firmicutes Bacilli Lactobacillales Leuconostocaceae *Weissella cibaria* |
| Whe-1-1 | Bacteria Firmicutes Bacilli Lactobacillales Leuconostocaceae *Weissella hellenica* |
| Spn-1-1 to Spn-1-8 | Bacteria Firmicutes Bacilli Lactobacillales Streptococcaceae *Streptococcus pneumoniae* |
| Ssp-15-1 | Bacteria Firmicutes Bacilli Lactobacillales Streptococcaceae *Streptococcus sp.* 263_SSPC |
| Wko-1-1 | Bacteria Firmicutes Bacilli Lactobacillales *Weissella koreensis* KCTC 3621 |
| Bma-8-1 | Bacteria Firmicutes Clostridia Clostridiales Clostridiaceae *Beduini massiliensis* |

| | |
|---|---|
| Cul-1-1 | Bacteria Firmicutes Clostridia Clostridiales Clostridiaceae *Clostridium ultunense* Esp |
| Cba-1-1 | Bacteria Firmicutes Clostridia *Clostridiales bacterium* VE202-01 |
| Esp-18-1 | Bacteria Firmicutes Clostridia Clostridiales Eubacteriaceae *Eubacterium sp.* 3_1_31 |
| Bsp-100-1 | Bacteria Firmicutes Clostridia Clostridiales Lachnospiraceae *Butyrivibrio sp.* MC2021 |
| Lba-3-1 | Bacteria Firmicutes Clostridia Clostridiales *Lachnospiraceae bacterium* A2 |
| Cmi-1-1 | Bacteria Firmicutes Erysipelotrichi Erysipelotrichales Erysipelotrichaceae *Catenibacterium mitsuokai* DSM 15897 |
| Csp-4-1 | Bacteria Firmicutes Erysipelotrichi Erysipelotrichales Erysipelotrichaceae *Clostridium spiroforme* DSM 1552 |
| Edo-1-1 | Bacteria Firmicutes Erysipelotrichi Erysipelotrichales Erysipelotrichaceae *Eubacterium dolichum* DSM 3991 |
| Hfi-1-1 | Bacteria Firmicutes Erysipelotrichi Erysipelotrichales Erysipelotrichaceae *Holdemania filiformis* DSM 12042 |
| Tsp-1-1 | Bacteria Firmicutes Erysipelotrichi Erysipelotrichales Erysipelotrichaceae *Turicibacter sp.* HGF1 |
| Tsp-2-1 | Bacteria Firmicutes Erysipelotrichi Erysipelotrichales Erysipelotrichaceae *Turicibacter sp.* PC909 |
| Ast-1-1 | Bacteria Firmicutes Erysipelotrichia Erysipelotrichales Erysipelotrichaceae *Allobaculum stercoricanis* DSM 13633 |
| CSt-1-1 | Bacteria Firmicutes Erysipelotrichia Erysipelotrichales Erysipelotrichaceae *Candidatus* Stoquefichus massiliensis AP9 |
| CSt-2-1 | Bacteria Firmicutes Erysipelotrichia Erysipelotrichales Erysipelotrichaceae *Candidatus* Stoquefichus sp. SB1 |
| Csp-5-1 | Bacteria Firmicutes Erysipelotrichia Erysipelotrichales Erysipelotrichaceae *Coprobacillus sp.* 29_1 |
| Csp-6-1 | Bacteria Firmicutes Erysipelotrichia Erysipelotrichales Erysipelotrichaceae *Coprobacillus sp.* 8_1_38FAA |
| Csp-7-1 | Bacteria Firmicutes Erysipelotrichia Erysipelotrichales Erysipelotrichaceae *Coprobacillus sp.* D6 |
| Eca-5-1 | Bacteria Firmicutes Erysipelotrichia Erysipelotrichales Erysipelotrichaceae *Eggerthia catenaformis* OT 569 = DSM 20559 |
| [sa-1-1 | Bacteria Firmicutes Erysipelotrichia Erysipelotrichales Erysipelotrichaceae Erysipelatoclostridium [Clostridium] saccharogumia DSM 17460 |
| Eba-1-1 | Bacteria Firmicutes Erysipelotrichia Erysipelotrichales *Erysipelotrichaceae bacterium* 5_2_54FAA |
| Hma-1-1 | Bacteria Firmicutes Erysipelotrichia Erysipelotrichales Erysipelotrichaceae *Holdemania massiliensis* AP2 |
| Kvi-1-1 | Bacteria Firmicutes Erysipelotrichia Erysipelotrichales Erysipelotrichaceae *Kandleria vitulina* DSM 20405 |
| Kvi-2-1 | Bacteria Firmicutes Erysipelotrichia Erysipelotrichales Erysipelotrichaceae *Kandleria vitulina* MC3001 |
| Kvi-3-1 | Bacteria Firmicutes Erysipelotrichia Erysipelotrichales Erysipelotrichaceae *Kandleria vitulina* WCE2011 |
| Fba-1-1 | Bacteria *Firmicutes bacterium* M10-2 |
| Cma-5-1 | Bacteria Firmicutes Lactobacillales Carnobacteriaceae *Carnobacterium maltaromaticum* ATCC 35586 |
| Csp-8-1 | Bacteria Firmicutes Lactobacillales Carnobacteriaceae *Carnobacterium sp.* 17-4 |
| Csp-9-1 | Bacteria Firmicutes Lactobacillales Carnobacteriaceae *Carnobacterium sp.* AT7 |
| Gad-1-1 | Bacteria Firmicutes Lactobacillales Carnobacteriaceae *Granulicatella adiacens* ATCC 49175 |
| Eca-6-1 | Bacteria Firmicutes Lactobacillales Enterococcaceae *Enterococcus casseliflavus* EC10 |
| Eca-7-1 | Bacteria Firmicutes Lactobacillales Enterococcaceae *Enterococcus casseliflavus* EC30 |
| Efa-546-1 | Bacteria Firmicutes Lactobacillales Enterococcaceae *Enterococcus faecalis* AR01/DG |
| Efa-18-1 to Efa-18-2 | Bacteria Firmicutes Lactobacillales Enterococcaceae *Enterococcus faecalis* ATCC 29200 |
| Efa-547-1 | Bacteria Firmicutes Lactobacillales Enterococcaceae *Enterococcus faecalis* ATCC 4200 |
| Efa-548-1 | Bacteria Firmicutes Lactobacillales Enterococcaceae *Enterococcus faecalis* CH188 |
| Efa-549-1 | Bacteria Firmicutes Lactobacillales Enterococcaceae *Enterococcus faecalis* D32 |
| Efa-550-1 | Bacteria Firmicutes Lactobacillales Enterococcaceae *Enterococcus faecalis* D6 |
| Efa-551-1 | Bacteria Firmicutes Lactobacillales Enterococcaceae *Enterococcus faecalis* DS5 |
| Efa-552-1 | Bacteria Firmicutes Lactobacillales Enterococcaceae *Enterococcus faecalis* E1Sol |
| Efa-553-1 | Bacteria Firmicutes Lactobacillales Enterococcaceae *Enterococcus faecalis* Fly1 |
| Efa-554-1 | Bacteria Firmicutes Lactobacillales Enterococcaceae *Enterococcus faecalis* HH22 |
| Efa-555-1 | Bacteria Firmicutes Lactobacillales Enterococcaceae *Enterococcus faecalis* HIP11704 |
| Efa-556-1 | Bacteria Firmicutes Lactobacillales Enterococcaceae *Enterococcus faecalis* JH1 |
| Efa-557-1 | Bacteria Firmicutes Lactobacillales Enterococcaceae *Enterococcus faecalis* Merz96 |
| Efa-558-1 | Bacteria Firmicutes Lactobacillales Enterococcaceae *Enterococcus faecalis* OG1RF |
| Efa-559-1 | Bacteria Firmicutes Lactobacillales Enterococcaceae *Enterococcus faecalis* PC1.1 |
| Efa-560-1 | Bacteria Firmicutes Lactobacillales Enterococcaceae *Enterococcus faecalis* R712 |
| Efa-561-1 | Bacteria Firmicutes Lactobacillales Enterococcaceae *Enterococcus faecalis* S613 |
| Efa-562-1 | Bacteria Firmicutes Lactobacillales Enterococcaceae *Enterococcus faecalis* T1 |
| Efa-563-1 | Bacteria Firmicutes Lactobacillales Enterococcaceae *Enterococcus faecalis* T11 |
| Efa-564-1 | Bacteria Firmicutes Lactobacillales Enterococcaceae *Enterococcus faecalis* T2 |
| Efa-565-1 | Bacteria Firmicutes Lactobacillales Enterococcaceae *Enterococcus faecalis* T3 |
| Efa-566-1 | Bacteria Firmicutes Lactobacillales Enterococcaceae *Enterococcus faecalis* T8 |
| Efa-567-1 | Bacteria Firmicutes Lactobacillales Enterococcaceae *Enterococcus faecalis* TUSoD Ef11 |
| Efa-568-1 | Bacteria Firmicutes Lactobacillales Enterococcaceae *Enterococcus faecalis* TX0104 |
| Efa-569-1 | Bacteria Firmicutes Lactobacillales Enterococcaceae *Enterococcus faecalis* TX0855 |
| Efa-570-1 | Bacteria Firmicutes Lactobacillales Enterococcaceae *Enterococcus faecalis* TX1322 |
| Efa-571-1 | Bacteria Firmicutes Lactobacillales Enterococcaceae *Enterococcus faecalis* TX2134 |
| Efa-572-1 | Bacteria Firmicutes Lactobacillales Enterococcaceae *Enterococcus faecalis* TX4248 |
| Efa-309-1 to Efa-309-3 | Bacteria Firmicutes Lactobacillales Enterococcaceae *Enterococcus faecalis* V583 |
| Efa-573-1 | Bacteria Firmicutes Lactobacillales Enterococcaceae *Enterococcus faecalis* X98 |
| Efa-574-1 | Bacteria Firmicutes Lactobacillales Enterococcaceae *Enterococcus faecium* 1,141,733 |
| Efa-575-1 | Bacteria Firmicutes Lactobacillales Enterococcaceae *Enterococcus faecium* 1,230,933 |
| Efa-576-1 | Bacteria Firmicutes Lactobacillales Enterococcaceae *Enterococcus faecium* 1,231,408 |
| Efa-577-1 | Bacteria Firmicutes Lactobacillales Enterococcaceae *Enterococcus faecium* 1,231,410 |

| | |
|---|---|
| Efa-578-1 | Bacteria Firmicutes Lactobacillales Enterococcaceae *Enterococcus faecium* 1,231,501 |
| Efa-579-1 | Bacteria Firmicutes Lactobacillales Enterococcaceae *Enterococcus faecium* 1,231,502 |
| Efa-580-1 | Bacteria Firmicutes Lactobacillales Enterococcaceae *Enterococcus faecium* Aus0004 |
| Efa-581-1 | Bacteria Firmicutes Lactobacillales Enterococcaceae *Enterococcus faecium* Com12 |
| Efa-582-1 | Bacteria Firmicutes Lactobacillales Enterococcaceae *Enterococcus faecium* Com15 |
| Efa-583-1 | Bacteria Firmicutes Lactobacillales Enterococcaceae *Enterococcus faecium* D344SRF |
| Efa-584-1 | Bacteria Firmicutes Lactobacillales Enterococcaceae *Enterococcus faecium* DO |
| Efa-585-1 | Bacteria Firmicutes Lactobacillales Enterococcaceae *Enterococcus faecium* E1039 |
| Efa-586-1 | Bacteria Firmicutes Lactobacillales Enterococcaceae *Enterococcus faecium* E1071 |
| Efa-587-1 | Bacteria Firmicutes Lactobacillales Enterococcaceae *Enterococcus faecium* E1162 |
| Efa-588-1 | Bacteria Firmicutes Lactobacillales Enterococcaceae *Enterococcus faecium* E1636 |
| Efa-589-1 | Bacteria Firmicutes Lactobacillales Enterococcaceae *Enterococcus faecium* E1679 |
| Efa-590-1 | Bacteria Firmicutes Lactobacillales Enterococcaceae *Enterococcus faecium* E980 |
| Efa-591-1 | Bacteria Firmicutes Lactobacillales Enterococcaceae *Enterococcus faecium* PC4.1 |
| Efa-592-1 | Bacteria Firmicutes Lactobacillales Enterococcaceae *Enterococcus faecium* TC 6 |
| Efa-593-1 | Bacteria Firmicutes Lactobacillales Enterococcaceae *Enterococcus faecium* TX1330 |
| Efa-594-1 | Bacteria Firmicutes Lactobacillales Enterococcaceae *Enterococcus faecium* U0317 |
| Ega-2-1 | Bacteria Firmicutes Lactobacillales Enterococcaceae *Enterococcus gallinarum* EG2 |
| Ehi-2-1 to Ehi-2-2 | Bacteria Firmicutes Lactobacillales Enterococcaceae *Enterococcus hirae* ATCC 9790 |
| Esp-19-1 | Bacteria Firmicutes Lactobacillales Enterococcaceae *Enterococcus sp.* GMD1E |
| Esp-20-1 | Bacteria Firmicutes Lactobacillales Enterococcaceae *Enterococcus sp.* GMD2E |
| Esp-21-1 | Bacteria Firmicutes Lactobacillales Enterococcaceae *Enterococcus sp.* GMD4E |
| Mpl-2-1 | Bacteria Firmicutes Lactobacillales Enterococcaceae *Melissococcus plutonius* ATCC 35311 |
| Mpl-3-1 | Bacteria Firmicutes Lactobacillales Enterococcaceae *Melissococcus plutonius* DAT561 |
| Tha-1-1 | Bacteria Firmicutes Lactobacillales Enterococcaceae *Tetragenococcus halophilus* NBRC 12172 |
| Lac-10-1 | Bacteria Firmicutes Lactobacillales Lactobacillaceae *Lactobacillus acidipiscis* KCTC 13900 |
| Lac-11-1 | Bacteria Firmicutes Lactobacillales Lactobacillaceae *Lactobacillus acidophilus* 30SC |
| Lac-12-1 | Bacteria Firmicutes Lactobacillales Lactobacillaceae *Lactobacillus acidophilus* ATCC 4796 |
| Lac-13-1 | Bacteria Firmicutes Lactobacillales Lactobacillaceae *Lactobacillus acidophilus* NCFM |
| Lam-1-1 | Bacteria Firmicutes Lactobacillales Lactobacillaceae *Lactobacillus amylolyticus* DSM 11664 |
| Lam-2-1 | Bacteria Firmicutes Lactobacillales Lactobacillaceae *Lactobacillus amylovorus* GRL 1112 |
| Lam-3-1 | Bacteria Firmicutes Lactobacillales Lactobacillaceae *Lactobacillus amylovorus* GRL1118 |
| Lan-3-1 | Bacteria Firmicutes Lactobacillales Lactobacillaceae *Lactobacillus antri* DSM 16041 |
| Lbr-6-1 | Bacteria Firmicutes Lactobacillales Lactobacillaceae *Lactobacillus brevis* ATCC 367 |
| Lbr-7-1 | Bacteria Firmicutes Lactobacillales Lactobacillaceae *Lactobacillus brevis subsp. gravesensis* ATCC 27305 |
| Lbu-1-1 | Bacteria Firmicutes Lactobacillales Lactobacillaceae *Lactobacillus buchneri* ATCC 11577 |
| Lbu-2-1 | Bacteria Firmicutes Lactobacillales Lactobacillaceae *Lactobacillus buchneri* CD034 |
| Lbu-3-1 | Bacteria Firmicutes Lactobacillales Lactobacillaceae *Lactobacillus buchneri* NRRL B-30929 |
| Lca-20-1 | Bacteria Firmicutes Lactobacillales Lactobacillaceae *Lactobacillus casei* ATCC 334 |
| Lca-21-1 | Bacteria Firmicutes Lactobacillales Lactobacillaceae *Lactobacillus casei* BD-II |
| Lca-22-1 | Bacteria Firmicutes Lactobacillales Lactobacillaceae *Lactobacillus casei* BL23 |
| Lca-23-1 | Bacteria Firmicutes Lactobacillales Lactobacillaceae *Lactobacillus casei* LC2W |
| Lca-24-1 | Bacteria Firmicutes Lactobacillales Lactobacillaceae *Lactobacillus casei* W56 |
| Lco-3-1 | Bacteria Firmicutes Lactobacillales Lactobacillaceae *Lactobacillus coleohominis* 101-4-CHN |
| Lcr-6-1 | Bacteria Firmicutes Lactobacillales Lactobacillaceae *Lactobacillus crispatus* 125-2-CHN |
| Lcr-7-1 | Bacteria Firmicutes Lactobacillales Lactobacillaceae *Lactobacillus crispatus* 214-1 |
| Lcr-8-1 | Bacteria Firmicutes Lactobacillales Lactobacillaceae *Lactobacillus crispatus* CTV-05 |
| Lcr-9-1 | Bacteria Firmicutes Lactobacillales Lactobacillaceae *Lactobacillus crispatus* JV-V01 |
| Lcr-10-1 | Bacteria Firmicutes Lactobacillales Lactobacillaceae *Lactobacillus crispatus* MV-3A-US |
| Lcr-11-1 | Bacteria Firmicutes Lactobacillales Lactobacillaceae *Lactobacillus crispatus* ST1 |
| Lfe-9-1 | Bacteria Firmicutes Lactobacillales Lactobacillaceae *Lactobacillus fermentum* 28-3-CHN |
| Lfe-10-1 | Bacteria Firmicutes Lactobacillales Lactobacillaceae *Lactobacillus fermentum* ATCC 14931 |
| Lfe-11-1 | Bacteria Firmicutes Lactobacillales Lactobacillaceae *Lactobacillus fermentum* CECT 5716 |
| Lfe-12-1 | Bacteria Firmicutes Lactobacillales Lactobacillaceae *Lactobacillus fermentum* IFO 3956 |
| Lga-11-1 | Bacteria Firmicutes Lactobacillales Lactobacillaceae *Lactobacillus gasseri* 202-4 |
| Lga-12-1 | Bacteria Firmicutes Lactobacillales Lactobacillaceae *Lactobacillus gasseri* ATCC 33323 |
| Lga-13-1 | Bacteria Firmicutes Lactobacillales Lactobacillaceae *Lactobacillus gastricus* PS3 |
| Lgi-2-1 | Bacteria Firmicutes Lactobacillales Lactobacillaceae *Lactobacillus gigeriorum* CRBIP 24.85 |
| Lhe-12-1 | Bacteria Firmicutes Lactobacillales Lactobacillaceae *Lactobacillus helveticus* DPC 4571 |
| Lhe-13-1 | Bacteria Firmicutes Lactobacillales Lactobacillaceae *Lactobacillus helveticus* DSM 20075 |
| Lhe-14-1 | Bacteria Firmicutes Lactobacillales Lactobacillaceae *Lactobacillus helveticus* H10 |
| Lhe-15-1 | Bacteria Firmicutes Lactobacillales Lactobacillaceae *Lactobacillus helveticus* R0052 |
| Lhi-1-1 | Bacteria Firmicutes Lactobacillales Lactobacillaceae *Lactobacillus hilgardii* ATCC 8290 |
| Lho-2-1 | Bacteria Firmicutes Lactobacillales Lactobacillaceae *Lactobacillus hominis* CRBIP 24.179 |
| Lin-7-1 | Bacteria Firmicutes Lactobacillales Lactobacillaceae *Lactobacillus iners* AB-1 |

| | |
|---|---|
| Lin-8-1 | Bacteria Firmicutes Lactobacillales Lactobacillaceae *Lactobacillus iners* DSM 13335 |
| Lin-13-1 | Bacteria Firmicutes Lactobacillales Lactobacillaceae *Lactobacillus iners* LactinV 01V1-a |
| Lin-14-1 | Bacteria Firmicutes Lactobacillales Lactobacillaceae *Lactobacillus iners* LactinV 03V1-b |
| Lin-15-1 | Bacteria Firmicutes Lactobacillales Lactobacillaceae *Lactobacillus iners* LactinV 09V1-c |
| Lin-16-1 | Bacteria Firmicutes Lactobacillales Lactobacillaceae *Lactobacillus iners* LactinV 11V1-d |
| Lin-9-1 | Bacteria Firmicutes Lactobacillales Lactobacillaceae *Lactobacillus iners* LEAF 2052A-d |
| Lin-10-1 | Bacteria Firmicutes Lactobacillales Lactobacillaceae *Lactobacillus iners* LEAF 2053A-b |
| Lin-11-1 | Bacteria Firmicutes Lactobacillales Lactobacillaceae *Lactobacillus iners* LEAF 2062A-h1 |
| Lin-12-1 | Bacteria Firmicutes Lactobacillales Lactobacillaceae *Lactobacillus iners* LEAF 3008A-a |
| Lin-17-1 | Bacteria Firmicutes Lactobacillales Lactobacillaceae *Lactobacillus iners* SPIN 1401G |
| Lin-18-1 | Bacteria Firmicutes Lactobacillales Lactobacillaceae *Lactobacillus iners* SPIN 2503V10-D |
| Lin-19-1 | Bacteria Firmicutes Lactobacillales Lactobacillaceae *Lactobacillus iners* UPII 143-D |
| Lin-20-1 | Bacteria Firmicutes Lactobacillales Lactobacillaceae *Lactobacillus iners* UPII 60-B |
| Ljo-4-1 | Bacteria Firmicutes Lactobacillales Lactobacillaceae *Lactobacillus johnsonii* ATCC 33200 |
| Ljo-5-1 | Bacteria Firmicutes Lactobacillales Lactobacillaceae *Lactobacillus johnsonii* DPC 6026 |
| Ljo-6-1 | Bacteria Firmicutes Lactobacillales Lactobacillaceae *Lactobacillus johnsonii* FI9785 |
| Ljo-7-1 | Bacteria Firmicutes Lactobacillales Lactobacillaceae *Lactobacillus johnsonii* NCC 533 |
| Lke-2-1 | Bacteria Firmicutes Lactobacillales Lactobacillaceae *Lactobacillus kefiranofaciens* ZW3 |
| Lma-4-1 | Bacteria Firmicutes Lactobacillales Lactobacillaceae *Lactobacillus malefermentans* KCTC 3548 |
| Lma-3-1 to Lma-3-2 | Bacteria Firmicutes Lactobacillales Lactobacillaceae *Lactobacillus mali* KCTC 3596 = DSM 20444 |
| Lor-3-1 | Bacteria Firmicutes Lactobacillales Lactobacillaceae *Lactobacillus oris* PB013-T2-3 |
| Lpa-42-1 | Bacteria Firmicutes Lactobacillales Lactobacillaceae *Lactobacillus paracasei subsp. paracasei* ATCC 25302 |
| Lpl-19-1 | Bacteria Firmicutes Lactobacillales Lactobacillaceae *Lactobacillus plantarum* JDM1 |
| Lpl-21-1 | Bacteria Firmicutes Lactobacillales Lactobacillaceae *Lactobacillus plantarum subsp. plantarum* ST-III |
| Lpl-20-1 | Bacteria Firmicutes Lactobacillales Lactobacillaceae *Lactobacillus plantarum* WCFS1 |
| Lre-7-1 | Bacteria Firmicutes Lactobacillales Lactobacillaceae *Lactobacillus reuteri* 100-23 |
| Lre-8-1 | Bacteria Firmicutes Lactobacillales Lactobacillaceae *Lactobacillus reuteri* CF48-3A |
| Lre-9-1 | Bacteria Firmicutes Lactobacillales Lactobacillaceae *Lactobacillus reuteri* DSM 20016 |
| Lre-10-1 | Bacteria Firmicutes Lactobacillales Lactobacillaceae *Lactobacillus reuteri* JCM 1112 |
| Lre-11-1 | Bacteria Firmicutes Lactobacillales Lactobacillaceae *Lactobacillus reuteri* MM2-3 |
| Lre-12-1 | Bacteria Firmicutes Lactobacillales Lactobacillaceae *Lactobacillus reuteri* MM4-1A |
| Lre-13-1 | Bacteria Firmicutes Lactobacillales Lactobacillaceae *Lactobacillus reuteri* SD2112 |
| Lrh-19-1 | Bacteria Firmicutes Lactobacillales Lactobacillaceae *Lactobacillus rhamnosus* ATCC 8530 |
| Lrh-20-1 to Lrh-20-2 | Bacteria Firmicutes Lactobacillales Lactobacillaceae *Lactobacillus rhamnosus* GG |
| Lrh-21-1 | Bacteria Firmicutes Lactobacillales Lactobacillaceae *Lactobacillus rhamnosus* HN001 |
| Lrh-23-1 | Bacteria Firmicutes Lactobacillales Lactobacillaceae *Lactobacillus rhamnosus* Lc 705 |
| Lrh-22-1 | Bacteria Firmicutes Lactobacillales Lactobacillaceae *Lactobacillus rhamnosus* LMS2-1 |
| Lru-5-1 | Bacteria Firmicutes Lactobacillales Lactobacillaceae *Lactobacillus ruminis* ATCC 27782 |
| Lru-6-1 | Bacteria Firmicutes Lactobacillales Lactobacillaceae *Lactobacillus ruminis* SPM0211 |
| Lsa-9-1 | Bacteria Firmicutes Lactobacillales Lactobacillaceae *Lactobacillus salivarius* ACS-116-V-Col5a |
| Lsa-10-1 | Bacteria Firmicutes Lactobacillales Lactobacillaceae *Lactobacillus salivarius* ATCC 11741 |
| Lsa-11-1 | Bacteria Firmicutes Lactobacillales Lactobacillaceae *Lactobacillus salivarius* CECT 5713 |
| Lsa-12-1 | Bacteria Firmicutes Lactobacillales Lactobacillaceae *Lactobacillus salivarius* UCC118 |
| Lsa-13-1 | Bacteria Firmicutes Lactobacillales Lactobacillaceae *Lactobacillus sanfranciscensis* TMW 1.1304 |
| Lsu-2-1 | Bacteria Firmicutes Lactobacillales Lactobacillaceae *Lactobacillus suebicus* KCTC 3549 |
| Lul-1-1 | Bacteria Firmicutes Lactobacillales Lactobacillaceae *Lactobacillus ultunensis* DSM 16047 |
| Lva-2-1 | Bacteria Firmicutes Lactobacillales Lactobacillaceae *Lactobacillus vaginalis* ATCC 49540 |
| Lve-1-1 | Bacteria Firmicutes Lactobacillales Lactobacillaceae *Lactobacillus versmoldensis* KCTC 3814 |
| Lvi-2-1 | Bacteria Firmicutes Lactobacillales Lactobacillaceae *Lactobacillus vini* DSM 20605 |
| Lze-1-1 | Bacteria Firmicutes Lactobacillales Lactobacillaceae *Lactobacillus zeae* KCTC 3804 |
| Pac-4-1 | Bacteria Firmicutes Lactobacillales Lactobacillaceae *Pediococcus acidilactici* 7_4 |
| Pac-5-1 | Bacteria Firmicutes Lactobacillales Lactobacillaceae *Pediococcus acidilactici* DSM 20284 |
| Pcl-1-1 | Bacteria Firmicutes Lactobacillales Lactobacillaceae *Pediococcus claussenii* ATCC BAA-344 |
| Ppe-6-1 | Bacteria Firmicutes Lactobacillales Lactobacillaceae *Pediococcus pentosaceus* ATCC 25745 |
| Lca-25-1 | Bacteria Firmicutes Lactobacillales *Leuconostoc carnosum* JB16 |
| Lci-5-1 | Bacteria Firmicutes Lactobacillales *Leuconostoc citreum* KM20 |
| Lki-4-1 | Bacteria Firmicutes Lactobacillales *Leuconostoc kimchii* IMSNU 11154 |
| Lsp-16-1 | Bacteria Firmicutes Lactobacillales *Leuconostoc sp.* C2 |
| Wci-2-1 | Bacteria Firmicutes Lactobacillales *Weissella cibaria* KACC 11862 |
| Wko-2-1 | Bacteria Firmicutes Lactobacillales *Weissella koreensis* KACC 15510 |
| Hco-1-1 | Bacteria Haloplasmatales Haloplasmataceae *Haloplasma contractile* SSD-17B |
| Ecl-1-1 | Bacteria Proteobacteria Gammaproteobacteria Enterobacteriales Enterobacteriaceae Enterobacter Enterobacter cloacae complex *Enterobacter cloacae* |
| Psp-40-1 | Bacteria Proteobacteria Gammaproteobacteria Pseudomonadales Pseudomonadaceae *Pseudomonas sp.* EGD-AK9 |
| Xca-1-1 | Bacteria Proteobacteria Gammaproteobacteria Xanthomonadales Xanthomonadaceae *Xanthomonas campestris pv. musacearum* NCPPB 2005 |
| Xca-2-1 | Bacteria Proteobacteria Gammaproteobacteria Xanthomonadales Xanthomonadaceae *Xanthomonas campestris pv. musacearum* NCPPB 4379 |

| | |
|---|---|
| Xca-3-1 | Bacteria Proteobacteria Gammaproteobacteria Xanthomonadales Xanthomonadaceae *Xanthomonas campestris pv. musacearum* NCPPB 4380 |
| Xca-4-1 | Bacteria Proteobacteria Gammaproteobacteria Xanthomonadales Xanthomonadaceae *Xanthomonas campestris pv. musacearum* NCPPB 4392 |
| Xca-5-1 | Bacteria Proteobacteria Gammaproteobacteria Xanthomonadales Xanthomonadaceae *Xanthomonas campestris pv. musacearum* NCPPB 4394 |
| Xor-1-1 | Bacteria Proteobacteria Gammaproteobacteria Xanthomonadales Xanthomonadaceae *Xanthomonas oryzae pv. oryzae* PXO99A |
| Xva-1-1 | Bacteria Proteobacteria Gammaproteobacteria Xanthomonadales Xanthomonadaceae *Xanthomonas vasicola pv. vasculorum* NCPPB 1326 |
| Xva-2-1 | Bacteria Proteobacteria Gammaproteobacteria Xanthomonadales Xanthomonadaceae *Xanthomonas vasicola pv. vasculorum* NCPPB 1381 |
| Xva-3-1 | Bacteria Proteobacteria Gammaproteobacteria Xanthomonadales Xanthomonadaceae *Xanthomonas vasicola pv. vasculorum* NCPPB 206 |
| Agr-1-1 | Bacteria Tenericutes Mollicutes Acholeplasmatales Acholeplasmataceae *Acholeplasma granularum* ATCC 19168 |
| Aoc-1-1 | Bacteria Tenericutes Mollicutes Acholeplasmatales Acholeplasmataceae *Acholeplasma oculi* |
| env-1 to env-1514 | environmental samples |

#### 11.2 Notes

Superscript numbers are used to annotate any L31-Firmicutes RNA that has special characteristics (described below). These numeric annotations will be used in Sections 11.3 and 11.5:. The meaning of the numbers are as follows.

[1] A different alignment would have a somewhat stronger hairpin, but would somewhat lose conserved nucleotides. It's unclear which alignment of this sequence is better.

#### 11.3 Gene contexts

Each L31-Firmicutes RNA (indicated by "RNA→") is listed. For each hit, the downstream genes predicted to reside in a regulated operon are listed. If the nearest downstream gene is encoding in the opposite strand (and therefore presumed to not be a part of a regulated operon), then that gene is still depicted. Some environmental sequences and some RefSeq entries lack gene annotations, and so no genes are available for such sequences. The direction of each gene is indicated with an arrow (→), and each predicted conserved domain in the gene is named. Conserved domains associated with more than one L31-Firmicutes RNA are assigned a color; other domains are gray. Information about these conserved domains is given in Section 11.4. The accession of the sequence containing each L31-Firmicutes RNA is given in the column named "Seq. accession". Accessions beginning with "NC_", "NS_", "NW_" or "NZ_" are contained in RefSeq. Other accession refer to environmental samples. Nucleotide coordinates are given for the 5′ and 3′ boundaries of each L31-Firmicutes RNA. If the 5′ coordinate is greater than the 3′ coordinate, the RNA is present on the reverse-complement strand of the containing genomic DNA sequence. Each hit is denoted by an abbreviation (like "Eco-1-1") that refers to a taxonomy given in Section 11.1. (Superscript numbers refer to annotations listed in Section 11.2)

| abbrev. | Seq. accession | | 5′ at | 3′ at | genes |
|---|---|---|---|---|---|
| env-1 | SL_5KL_010_SED_10059915 | - | 521 | 483 | **RNA**→ hypo→ |
| env-2 | 2200881906 | - | 21321 | 21291 | **RNA**→ HXXSHH (pfam07586)→ |
| env-3 | BISONR_C7447 | + | 805 | 839 | **RNA**→ hypo→ |
| env-4 | JGI24025J20009_10001343 | + | 4419 | 4451 | **RNA**→ ←REC (smart00448)Trans_reg_C (smart00862)OmpR (COG0745) |
| env-5 | JGI24724J26744_10982735 | + | 117 | 149 | **RNA**→ ←Trans_reg_C (smart00862)trans_reg_C (cd00383)Trans_reg_C (pfam00486) |
| env-6 | JGI994J12354_10174172 | + | 602 | 631 | **RNA**→ int (PHA02601)→ |
| Cmo-1-1 | NZ_JQMR01000001.1 | - | 1464079 | 1464047 | **RNA**→ rpmE2 (PRK01678)Ribosomal_L31 (pfam01197)→ |
| env-7 | ASHM260b_GM9OZ8V02GSY54 | + | 254 | 292 | **RNA**→ hypo→ |
| env-8 | 08_19_CONTROL_DNA_scaffold7482_7 | - | 8235 | 8202 | **RNA**→ rpmE (PRK00019)Ribosomal_L31 (pfam01197)→ |
| Ast-1-1 | NZ_KB913034.1 | - | 1959703 | 1959670 | **RNA**→ rpmE (PRK00019)Ribosomal_L31 (pfam01197)→ |
| env-9 | JGI24713J26584_10168935 | + | 112 | 146 | **RNA**→ PurM (COG0150)PRK05385 (PRK05385)→ |
| env-10 | SL_1KL_011_SED_10415685 | - | 130 | 99 | **RNA**→ hypo→ |
| env-11 | SL_9KL_010_SED_10025487 | + | 283 | 314 | **RNA**→ dnaG (PRK05667)USP_Like (cd00293)→ |
| env-12 | SL_7KL_010_BRINE_10007095 | + | 279 | 310 | **RNA**→ dnaG (PRK05667)USP_Like (cd00293)→ |
| env-13 | HCF14C_2_1_1_newblercontig17407 | + | 809 | 842 | **RNA**→ rpmE (PRK00019)Ribosomal_L31 (pfam01197)→ |

| | | | | | |
|---|---|---|---|---|---|
| env-14 | HCF12C_35575 | - | 214 | 181 | **RNA**→ rpmE (PRK00019)Ribosomal_L31 (pfam01197)→ |
| env-15 | scaffold5088_2_V1.CD-11 | - | 8404 | 8367 | **RNA**→ hypo→ |
| env-16 | RUMENNODE_3996658_1 | - | 12142 | 12105 | **RNA**→ rpmE (PRK00019)Ribosomal_L31 (pfam01197)→ RhaT (COG0697)→ |
| | | | | | ArcA (COG2235)PRK01388 (PRK01388)→ PRK02102 (PRK02102)ArgF (COG0078)→ |
| | | | | | MetG (COG0143)PRK11893 (PRK11893)→ |
| env-17 | topACOD_FV90NF401C3W26 | - | 154 | 126 | **RNA**→ hypo→ |
| env-18 | SRS016335_C5226868 | - | 1031 | 997 | **RNA**→ ←hypo |
| env-19 | RUMENNODE_2334914_25000 | - | 39974 | 39943 | **RNA**→ hypo→ |
| env-20 | JGI12341J12878_1000327 | - | 8875 | 8843 | **RNA**→ rpmE (PRK00019)Ribosomal_L31 (pfam01197)→ |
| env-21 | JGI12298J13902_1000429 | + | 6154 | 6186 | **RNA**→ rpmE (PRK00019)Ribosomal_L31 (pfam01197)→ |
| env-22 | JGI12298J13902_1007687 | + | 173 | 205 | **RNA**→ ←hypo |
| Cde-1-1 | NZ_KB912881.1 | - | 64384 | 64352 | **RNA**→ rpmE (PRK00019)Ribosomal_L31 (pfam01197)→ |
| Psp-27-1 | NZ_BBIW01000023.1 | - | 4864 | 4830 | **RNA**→ rpmE (PRK00019)Ribosomal_L31 (pfam01197)→ |
| Lsp-1-1 | NZ_AGAV01000016.1 | + | 19061 | 19094 | **RNA**→ rpmE2 (PRK01678)Ribosomal_L31 (pfam01197)→ rpmE2 (PRK01678)Ribosomal_L31 (pfam01197 |
| Xor-1-1 | NC_010717.2 | + | 5213900 | 5213934 | **RNA**→ HSDR_N (pfam04313)HSDR_N_2 (pfam13588)→ |
| Xva-3-1 | NZ_AKBM01000002.1 | + | 13187 | 13221 | **RNA**→ HSDR_N (pfam04313)HSDR_N_2 (pfam13588)→ |
| Xva-2-1 | NZ_KB373231.1 | - | 59457 | 59423 | **RNA**→ HSDR_N (pfam04313)HSDR_N_2 (pfam13588)→ |
| Xca-5-1 | NZ_KB373047.1 | + | 93162 | 93196 | **RNA**→ HSDR_N (pfam04313)HSDR_N_2 (pfam13588)→ |
| Xca-2-1 | NZ_KB372855.1 | - | 57748 | 57714 | **RNA**→ HSDR_N (pfam04313)HSDR_N_2 (pfam13588)→ |
| Xca-4-1 | NZ_KB373012.1 | - | 34001 | 33967 | **RNA**→ HSDR_N (pfam04313)HSDR_N_2 (pfam13588)→ |
| Xca-3-1 | NZ_KB372889.1 | - | 57694 | 57660 | **RNA**→ HSDR_N (pfam04313)HSDR_N_2 (pfam13588)→ |
| Xva-1-1 | NZ_KB373142.1 | - | 59417 | 59383 | **RNA**→ HSDR_N (pfam04313)HSDR_N_2 (pfam13588)→ |
| Xca-1-1 | NZ_AKBE01000040.1 | + | 196 | 230 | **RNA**→ HSDR_N (pfam04313)HSDR_N_2 (pfam13588)→ |
| env-23 | JGI24721J26819_10249650 | + | 591 | 632 | **RNA**→ hypo→ |
| env-24 | JGI24721J44947_10005316 | + | 2145 | 2186 | **RNA**→ vWFA (cd00198)→ hypo→ hypo→ AAA (smart00382)SpoVK (COG0464)CDC48 (TIGR01243)→ |
| | | | | | NHT_00031 (TIGR04181)AHBA_syn (cd00616)→ CesT (pfam05932)YbjN (pfam10722)FtsA (pfam14450)→ |
| env-25 | 2016923541 | + | 557 | 598 | **RNA**→ hypo→ |
| env-26 | JGI20130J15003_1005706 | + | 4068 | 4109 | **RNA**→ vWFA (cd00198)→ hypo→ hypo→ AAA (smart00382)SpoVK (COG0464)CDC48 (TIGR01243)→ |
| env-27 | JGI20130J15133_1005706 | + | 4068 | 4109 | **RNA**→ vWFA (cd00198)→ hypo→ hypo→ AAA (smart00382)SpoVK (COG0464)CDC48 (TIGR01243)→ |
| env-28 | SRS015985_C1514630 | + | 276 | 309 | **RNA**→ rpmE (PRK00019)rpmE2 (PRK01678)L31 (TIGR00105)Ribosomal_L31 (pfam01197)→ |
| env-29 | SRS018443_C3751049 | + | 219 | 252 | **RNA**→ rpmE2 (PRK01678)Ribosomal_L31 (pfam01197)→ |
| env-30 | Subgingival_plaque_LANL_C11194098 | - | 247 | 214 | **RNA**→ rpmE2 (PRK01678)Ribosomal_L31 (pfam01197)→ |
| env-31 | SRS011152_C4313074 | - | 268 | 235 | **RNA**→ rpmE2 (PRK01678)Ribosomal_L31 (pfam01197)→ |
| env-32 | SRS013533_C5440500 | - | 268 | 235 | **RNA**→ rpmE2 (PRK01678)Ribosomal_L31 (pfam01197)→ |
| env-33 | SRS013723_C3407064 | - | 247 | 214 | **RNA**→ rpmE2 (PRK01678)Ribosomal_L31 (pfam01197)→ |
| env-34 | SRS013945_WUGC_scaffold_2673 | - | 368 | 335 | **RNA**→ rpmE2 (PRK01678)Ribosomal_L31 (pfam01197)→ |
| env-35 | SRS013949_C3416715 | + | 114 | 147 | **RNA**→ rpmE2 (PRK01678)Ribosomal_L31 (pfam01197)→ |
| env-36 | SRS014690_C1785810 | + | 89 | 122 | **RNA**→ rpmE2 (PRK01678)Ribosomal_L31 (pfam01197)→ |
| env-37 | SRS015044_WUGC_scaffold_37694 | - | 255 | 222 | **RNA**→ rpmE2 (PRK01678)Ribosomal_L31 (pfam01197)→ |
| env-38 | SRS015470_C3354823 | + | 848 | 881 | **RNA**→ rpmE2 (PRK01678)Ribosomal_L31 (pfam01197)→ |
| env-39 | SRS015755_C1607169 | + | 161 | 194 | **RNA**→ rpmE2 (PRK01678)Ribosomal_L31 (pfam01197)→ |
| env-40 | SRS015803_C2579404 | - | 268 | 235 | **RNA**→ rpmE2 (PRK01678)Ribosomal_L31 (pfam01197)→ |
| env-41 | SRS015947_WUGC_scaffold_25971 | - | 575 | 542 | **RNA**→ rpmE2 (PRK01678)Ribosomal_L31 (pfam01197)→ |
| env-42 | SRS015989_C2127565 | + | 350 | 383 | **RNA**→ rpmE2 (PRK01678)Ribosomal_L31 (pfam01197)→ |
| env-43 | SRS016200_C2226629 | - | 247 | 214 | **RNA**→ rpmE2 (PRK01678)Ribosomal_L31 (pfam01197)→ |
| env-44 | SRS016360_Baylor_scaffold_38732 | - | 406 | 373 | **RNA**→ rpmE2 (PRK01678)Ribosomal_L31 (pfam01197)→ |
| env-45 | SRS017139_Baylor_scaffold_102199 | + | 117 | 150 | **RNA**→ rpmE2 (PRK01678)Ribosomal_L31 (pfam01197)→ |
| env-46 | SRS017227_Baylor_scaffold_86694 | + | 354 | 387 | **RNA**→ rpmE2 (PRK01678)Ribosomal_L31 (pfam01197)→ |
| env-47 | SRS018573_C2469306 | - | 223 | 190 | **RNA**→ rpmE2 (PRK01678)Ribosomal_L31 (pfam01197)→ |
| env-48 | SRS018665_C3678853 | + | 290 | 323 | **RNA**→ rpmE2 (PRK01678)Ribosomal_L31 (pfam01197)→ |
| env-49 | SRS019029_C3197161 | + | 182 | 215 | **RNA**→ rpmE2 (PRK01678)Ribosomal_L31 (pfam01197)→ |
| env-50 | SRS019225_C1506735 | + | 84 | 117 | **RNA**→ rpmE2 (PRK01678)Ribosomal_L31 (pfam01197)→ |
| env-51 | SRS019387_C1744036 | - | 62 | 29 | **RNA**→ |

| | | | | | |
|---|---|---|---|---|---|
| env-52 | SRS019591_WUGC_scaffold_36465 | - | 268 | 235 | **RNA**→ rpmE2 (PRK01678)Ribosomal_L31 (pfam01197)→ |
| env-53 | SRS019980_C3067094 | + | 4283 | 4316 | **RNA**→ rpmE2 (PRK01678)Ribosomal_L31 (pfam01197)→ |
| env-54 | SRS020226_C3471976 | - | 268 | 235 | **RNA**→ rpmE2 (PRK01678)Ribosomal_L31 (pfam01197)→ |
| env-55 | SRS020340_Baylor_scaffold_9891 | + | 1969 | 2002 | **RNA**→ rpmE2 (PRK01678)Ribosomal_L31 (pfam01197)→ |
| env-56 | SRS022725_LANL_scaffold_22307 | + | 709 | 742 | **RNA**→ rpmE2 (PRK01678)Ribosomal_L31 (pfam01197)→ |
| env-57 | SRS023358_C3441410 | + | 264 | 297 | **RNA**→ rpmE2 (PRK01678)Ribosomal_L31 (pfam01197)→ |
| env-58 | SRS023595_C6937803 | - | 247 | 214 | **RNA**→ rpmE2 (PRK01678)Ribosomal_L31 (pfam01197)→ |
| env-59 | SRS023938_Baylor_scaffold_23028 | - | 14585 | 14552 | **RNA**→ rpmE2 (PRK01678)Ribosomal_L31 (pfam01197)→ |
| env-60 | SRS024021_C2019462 | + | 712 | 745 | **RNA**→ rpmE2 (PRK01678)Ribosomal_L31 (pfam01197)→ |
| env-61 | SRS024355_C4162431 | - | 268 | 235 | **RNA**→ rpmE2 (PRK01678)Ribosomal_L31 (pfam01197)→ |
| env-62 | SRS024447_LANL_scaffold_46821 | + | 2480 | 2513 | **RNA**→ rpmE2 (PRK01678)Ribosomal_L31 (pfam01197)→ |
| env-63 | SRS024649_C3520887 | - | 189 | 156 | **RNA**→ rpmE2 (PRK01678)Ribosomal_L31 (pfam01197)→ |
| env-64 | SRS043772_C1744867 | + | 214 | 247 | **RNA**→ rpmE2 (PRK01678)Ribosomal_L31 (pfam01197)→ |
| env-65 | SRS045197_C3371889 | - | 247 | 214 | **RNA**→ rpmE2 (PRK01678)Ribosomal_L31 (pfam01197)→ |
| env-66 | SRS045313_C1320319 | - | 229 | 196 | **RNA**→ rpmE2 (PRK01678)Ribosomal_L31 (pfam01197)→ |
| env-67 | SRS047100_C2004330 | + | 101 | 134 | **RNA**→ rpmE2 (PRK01678)Ribosomal_L31 (pfam01197)→ |
| env-68 | SRS047113_C5226893 | + | 186 | 219 | **RNA**→ rpmE2 (PRK01678)Ribosomal_L31 (pfam01197)→ |
| env-69 | SRS051930_C3120842 | - | 245 | 212 | **RNA**→ rpmE2 (PRK01678)Ribosomal_L31 (pfam01197)→ |
| env-70 | SRS051941_C3883544 | + | 96 | 129 | **RNA**→ rpmE2 (PRK01678)Ribosomal_L31 (pfam01197)→ |
| env-71 | SRS052876_C2340555 | + | 941 | 974 | **RNA**→ rpmE2 (PRK01678)Ribosomal_L31 (pfam01197)→ |
| env-72 | SRS063215_C2148515 | - | 183 | 150 | **RNA**→ rpmE2 (PRK01678)Ribosomal_L31 (pfam01197)→ |
| Gsp-20-1 | NZ_JVBN01000045.1 | - | 69014 | 68981 | **RNA**→ rpmE2 (PRK01678)Ribosomal_L31 (pfam01197)→ |
| Gsp-19-1 | NZ_JVBO01000013.1 | - | 39564 | 39531 | **RNA**→ rpmE2 (PRK01678)Ribosomal_L31 (pfam01197)→ |
| Gad-1-1 | NZ_GG694015.1 | + | 59433 | 59466 | **RNA**→ rpmE2 (PRK01678)Ribosomal_L31 (pfam01197)→ |
| env-73 | Palatine_Tonsils_LANL_C6167790 | - | 268 | 235 | **RNA**→ rpmE2 (PRK01678)Ribosomal_L31 (pfam01197)→ |
| env-74 | SRS011086_C2691210 | - | 244 | 211 | **RNA**→ rpmE2 (PRK01678)Ribosomal_L31 (pfam01197)→ |
| env-75 | SRS014692_C1451880 | - | 214 | 181 | **RNA**→ rpmE2 (PRK01678)Ribosomal_L31 (pfam01197)→ |
| env-76 | SRS016740_Baylor_scaffold_14962 | - | 415 | 382 | **RNA**→ rpmE2 (PRK01678)Ribosomal_L31 (pfam01197)→ |
| env-77 | SRS017076_Baylor_scaffold_28499 | + | 465 | 498 | **RNA**→ rpmE2 (PRK01678)Ribosomal_L31 (pfam01197)→ |
| env-78 | SRS043239_C1988916 | + | 172 | 205 | **RNA**→ rpmE2 (PRK01678)Ribosomal_L31 (pfam01197)→ |
| env-79 | SRS062878_LANL_scaffold_15918 | + | 3373 | 3406 | **RNA**→ rpmE2 (PRK01678)Ribosomal_L31 (pfam01197)→ |
| env-80 | SRS064329_LANL_scaffold_17894 | + | 446 | 479 | **RNA**→ rpmE2 (PRK01678)Ribosomal_L31 (pfam01197)→ |
| env-81 | SRS075404_C3523540 | - | 268 | 235 | **RNA**→ rpmE2 (PRK01678)Ribosomal_L31 (pfam01197)→ |
| env-82 | Throat_LANL_C5742344 | + | 171 | 204 | **RNA**→ rpmE2 (PRK01678)Ribosomal_L31 (pfam01197)→ |
| env-83 | Contig123769.1 | - | 237 | 204 | **RNA**→ rpmE2 (PRK01678)Ribosomal_L31 (pfam01197)→ |
| env-84 | SRS011243_C2055318 | + | 1110 | 1143 | **RNA**→ rpmE2 (PRK01678)Ribosomal_L31 (pfam01197)→ |
| env-85 | SRS011306_Baylor_scaffold_38054 | - | 414 | 381 | **RNA**→ rpmE2 (PRK01678)Ribosomal_L31 (pfam01197)→ |
| env-86 | SRS013164_C2328665 | - | 268 | 235 | **RNA**→ rpmE2 (PRK01678)Ribosomal_L31 (pfam01197)→ |
| env-87 | SRS013234_Baylor_scaffold_22883 | + | 896 | 929 | **RNA**→ rpmE2 (PRK01678)Ribosomal_L31 (pfam01197)→ |
| env-88 | SRS013502_C2111881 | - | 244 | 211 | **RNA**→ rpmE2 (PRK01678)Ribosomal_L31 (pfam01197)→ |
| env-89 | SRS013818_C2698254 | - | 268 | 235 | **RNA**→ rpmE2 (PRK01678)Ribosomal_L31 (pfam01197)→ |
| env-90 | SRS013879_C2728982 | + | 120 | 153 | **RNA**→ rpmE2 (PRK01678)Ribosomal_L31 (pfam01197)→ |
| env-91 | SRS013947_C1595611 | + | 309 | 342 | **RNA**→ rpmE2 (PRK01678)Ribosomal_L31 (pfam01197)→ |
| env-92 | SRS013948_C1493527 | + | 350 | 383 | **RNA**→ rpmE2 (PRK01678)Ribosomal_L31 (pfam01197)→ |
| env-93 | SRS014124_WUGC_scaffold_3490 | - | 2369 | 2336 | **RNA**→ rpmE2 (PRK01678)Ribosomal_L31 (pfam01197)→ |
| env-94 | SRS014271_C2843132 | + | 377 | 410 | **RNA**→ rpmE2 (PRK01678)Ribosomal_L31 (pfam01197)→ |
| env-95 | SRS014573_WUGC_scaffold_16789 | - | 448 | 415 | **RNA**→ rpmE2 (PRK01678)Ribosomal_L31 (pfam01197)→ |
| env-96 | SRS014684_WUGC_scaffold_42590 | + | 348 | 381 | **RNA**→ rpmE2 (PRK01678)Ribosomal_L31 (pfam01197)→ |
| env-97 | SRS014888_C3007517 | - | 268 | 235 | **RNA**→ rpmE2 (PRK01678)Ribosomal_L31 (pfam01197)→ |
| env-98 | SRS015038_C1904815 | + | 588 | 621 | **RNA**→ rpmE2 (PRK01678)Ribosomal_L31 (pfam01197)→ |
| env-99 | SRS015057_C3337539 | + | 168 | 201 | **RNA**→ rpmE2 (PRK01678)Ribosomal_L31 (pfam01197)→ |
| env-100 | SRS015272_C3470509 | - | 244 | 211 | **RNA**→ rpmE2 (PRK01678)Ribosomal_L31 (pfam01197)→ |

| | | | | | |
|---|---|---|---|---|---|
| env-101 | SRS015762_WUGC_scaffold_27214 | - | 414 | 381 | **RNA**→ rpmE2 (PRK01678)Ribosomal_L31 (pfam01197)→ |
| env-102 | SRS015893_C2300612 | + | 271 | 304 | **RNA**→ rpmE2 (PRK01678)Ribosomal_L31 (pfam01197)→ |
| env-103 | SRS015941_C3418565 | - | 268 | 235 | **RNA**→ rpmE2 (PRK01678)Ribosomal_L31 (pfam01197)→ |
| env-104 | SRS016002_WUGC_scaffold_52168 | + | 5156 | 5189 | **RNA**→ rpmE2 (PRK01678)Ribosomal_L31 (pfam01197)→ |
| env-105 | SRS016037_C1931603 | - | 241 | 208 | **RNA**→ rpmE2 (PRK01678)Ribosomal_L31 (pfam01197)→ |
| env-106 | SRS016086_WUGC_scaffold_11609 | + | 396 | 429 | **RNA**→ rpmE2 (PRK01678)Ribosomal_L31 (pfam01197)→ |
| env-107 | SRS016225_C2969359 | - | 268 | 235 | **RNA**→ rpmE2 (PRK01678)Ribosomal_L31 (pfam01197)→ |
| env-108 | SRS016319_WUGC_scaffold_44666 | + | 293 | 326 | **RNA**→ rpmE2 (PRK01678)Ribosomal_L31 (pfam01197)→ |
| env-109 | SRS016342_Baylor_scaffold_17442 | + | 168 | 201 | **RNA**→ rpmE2 (PRK01678)Ribosomal_L31 (pfam01197)→ |
| env-110 | SRS016501_LANL_scaffold_6156 | - | 521 | 488 | **RNA**→ rpmE2 (PRK01678)Ribosomal_L31 (pfam01197)→ |
| env-111 | SRS016529_Baylor_scaffold_15023 | + | 368 | 401 | **RNA**→ rpmE2 (PRK01678)Ribosomal_L31 (pfam01197)→ |
| env-112 | SRS016569_C2383218 | - | 244 | 211 | **RNA**→ rpmE2 (PRK01678)Ribosomal_L31 (pfam01197)→ |
| env-113 | SRS017209_Baylor_scaffold_58383 | - | 268 | 235 | **RNA**→ rpmE2 (PRK01678)Ribosomal_L31 (pfam01197)→ |
| env-114 | SRS017439_Baylor_scaffold_37092 | + | 293 | 326 | **RNA**→ rpmE2 (PRK01678)Ribosomal_L31 (pfam01197)→ |
| env-115 | SRS017713_C3280511 | - | 268 | 235 | **RNA**→ rpmE2 (PRK01678)Ribosomal_L31 (pfam01197)→ |
| env-116 | SRS018145_Baylor_scaffold_27253 | + | 848 | 881 | **RNA**→ rpmE2 (PRK01678)Ribosomal_L31 (pfam01197)→ |
| env-117 | SRS018300_C3233287 | - | 226 | 193 | **RNA**→ rpmE2 (PRK01678)Ribosomal_L31 (pfam01197)→ |
| env-118 | SRS018357_Baylor_scaffold_42581 | + | 135 | 168 | **RNA**→ rpmE2 (PRK01678)Ribosomal_L31 (pfam01197)→ |
| env-119 | SRS018439_Baylor_scaffold_58366 | + | 168 | 201 | **RNA**→ rpmE2 (PRK01678)Ribosomal_L31 (pfam01197)→ |
| env-120 | SRS018739_C3514291 | + | 146 | 179 | **RNA**→ rpmE2 (PRK01678)Ribosomal_L31 (pfam01197)→ |
| env-121 | SRS018969_C2923925 | + | 368 | 401 | **RNA**→ rpmE2 (PRK01678)Ribosomal_L31 (pfam01197)→ |
| env-122 | SRS019026_C1883714 | - | 268 | 235 | **RNA**→ rpmE2 (PRK01678)Ribosomal_L31 (pfam01197)→ |
| env-123 | SRS019027_C1899173 | + | 198 | 231 | **RNA**→ rpmE2 (PRK01678)Ribosomal_L31 (pfam01197)→ |
| env-124 | SRS019045_C2664370 | - | 268 | 235 | **RNA**→ rpmE2 (PRK01678)Ribosomal_L31 (pfam01197)→ |
| env-125 | SRS019126_WUGC_scaffold_19544 | - | 439 | 406 | **RNA**→ rpmE2 (PRK01678)Ribosomal_L31 (pfam01197)→ |
| env-126 | SRS019127_C1552693 | + | 350 | 383 | **RNA**→ |
| env-127 | SRS019219_WUGC_scaffold_61972 | - | 268 | 235 | **RNA**→ rpmE2 (PRK01678)Ribosomal_L31 (pfam01197)→ |
| env-128 | SRS019327_C2685469 | + | 198 | 231 | **RNA**→ rpmE2 (PRK01678)Ribosomal_L31 (pfam01197)→ |
| env-129 | SRS019389_C2182350 | - | 268 | 235 | **RNA**→ rpmE2 (PRK01678)Ribosomal_L31 (pfam01197)→ |
| env-130 | SRS019607_WUGC_scaffold_1340 | - | 345 | 312 | **RNA**→ rpmE2 (PRK01678)Ribosomal_L31 (pfam01197)→ |
| env-131 | SRS019894_WUGC_scaffold_51316 | + | 473 | 506 | **RNA**→ rpmE2 (PRK01678)Ribosomal_L31 (pfam01197)→ |
| env-132 | SRS019974_Baylor_scaffold_9326 | + | 378 | 411 | **RNA**→ rpmE2 (PRK01678)Ribosomal_L31 (pfam01197)→ |
| env-133 | SRS020334_Baylor_scaffold_42522 | + | 168 | 201 | **RNA**→ rpmE2 (PRK01678)Ribosomal_L31 (pfam01197)→ |
| env-134 | SRS020856_Baylor_scaffold_27114 | - | 452 | 419 | **RNA**→ rpmE2 (PRK01678)Ribosomal_L31 (pfam01197)→ |
| env-135 | SRS021496_Baylor_scaffold_12527 | + | 531 | 564 | **RNA**→ rpmE2 (PRK01678)Ribosomal_L31 (pfam01197)→ |
| env-136 | SRS021954_Baylor_scaffold_16680 | - | 268 | 235 | **RNA**→ rpmE2 (PRK01678)Ribosomal_L31 (pfam01197)→ |
| env-137 | SRS022077_Baylor_scaffold_52875 | + | 492 | 525 | **RNA**→ rpmE2 (PRK01678)Ribosomal_L31 (pfam01197)→ |
| env-138 | SRS022143_WUGC_scaffold_12069 | - | 406 | 373 | **RNA**→ rpmE2 (PRK01678)Ribosomal_L31 (pfam01197)→ |
| env-139 | SRS022621_Baylor_scaffold_80034 | - | 244 | 211 | **RNA**→ rpmE2 (PRK01678)Ribosomal_L31 (pfam01197)→ |
| env-140 | SRS023352_C4538890 | - | 268 | 235 | **RNA**→ rpmE2 (PRK01678)Ribosomal_L31 (pfam01197)→ |
| env-141 | SRS023557_Baylor_scaffold_4832 | - | 347 | 314 | **RNA**→ rpmE2 (PRK01678)Ribosomal_L31 (pfam01197)→ |
| env-142 | SRS023617_C3098017 | - | 244 | 211 | **RNA**→ rpmE2 (PRK01678)Ribosomal_L31 (pfam01197)→ |
| env-143 | SRS023835_Baylor_scaffold_35031 | + | 515 | 548 | **RNA**→ rpmE2 (PRK01678)Ribosomal_L31 (pfam01197)→ |
| env-144 | SRS023926_C2767560 | - | 244 | 211 | **RNA**→ rpmE2 (PRK01678)Ribosomal_L31 (pfam01197)→ |
| env-145 | SRS023958_Baylor_scaffold_30996 | + | 299 | 332 | **RNA**→ rpmE2 (PRK01678)Ribosomal_L31 (pfam01197)→ |
| env-146 | SRS024015_Baylor_scaffold_18521 | + | 1374 | 1407 | **RNA**→ rpmE2 (PRK01678)Ribosomal_L31 (pfam01197)→ |
| env-147 | SRS024081_C2462482 | - | 244 | 211 | **RNA**→ rpmE2 (PRK01678)Ribosomal_L31 (pfam01197)→ |
| env-148 | SRS024138_C2623584 | + | 135 | 168 | **RNA**→ rpmE2 (PRK01678)Ribosomal_L31 (pfam01197)→ |
| env-149 | SRS024277_LANL_scaffold_21966 | + | 716 | 749 | **RNA**→ rpmE2 (PRK01678)Ribosomal_L31 (pfam01197)→ |
| env-150 | SRS024375_C2363158 | + | 113 | 146 | **RNA**→ rpmE2 (PRK01678)Ribosomal_L31 (pfam01197)→ |
| env-151 | SRS024580_LANL_scaffold_27268 | + | 614 | 647 | **RNA**→ rpmE2 (PRK01678)Ribosomal_L31 (pfam01197)→ |
| env-152 | SRS024637_LANL_scaffold_24099 | - | 453 | 420 | **RNA**→ rpmE2 (PRK01678)Ribosomal_L31 (pfam01197)→ |

| | | | | | |
|---|---|---|---|---|---|
| env-153 | SRS042643_C3064743 | - | 268 | 235 | **RNA**→ rpmE2 (PRK01678)Ribosomal_L31 (pfam01197)→ |
| env-154 | SRS042910_C5549436 | + | 368 | 401 | **RNA**→ |
| env-155 | SRS044486_C2150381 | + | 120 | 153 | **RNA**→ rpmE2 (PRK01678)Ribosomal_L31 (pfam01197)→ |
| env-156 | SRS045127_LANL_scaffold_25618 | + | 1248 | 1281 | **RNA**→ rpmE2 (PRK01678)Ribosomal_L31 (pfam01197)→ |
| env-157 | SRS045715_LANL_scaffold_6690 | + | 225 | 258 | **RNA**→ rpmE2 (PRK01678)Ribosomal_L31 (pfam01197)→ |
| env-158 | SRS048411_C6826029 | - | 214 | 181 | **RNA**→ rpmE2 (PRK01678)Ribosomal_L31 (pfam01197)→ |
| env-159 | SRS048791_LANL_scaffold_26361 | + | 823 | 856 | **RNA**→ rpmE2 (PRK01678)Ribosomal_L31 (pfam01197)→ |
| env-160 | SRS049147_LANL_scaffold_33494 | + | 596 | 629 | **RNA**→ rpmE2 (PRK01678)Ribosomal_L31 (pfam01197)→ |
| env-161 | SRS049389_C4709343 | - | 268 | 235 | **RNA**→ rpmE2 (PRK01678)Ribosomal_L31 (pfam01197)→ |
| env-162 | SRS050029_C709046 | - | 268 | 235 | **RNA**→ rpmE2 (PRK01678)Ribosomal_L31 (pfam01197)→ |
| env-163 | SRS050244_LANL_scaffold_4525 | - | 353 | 320 | **RNA**→ rpmE2 (PRK01678)Ribosomal_L31 (pfam01197)→ |
| env-164 | SRS050669_C2638407 | + | 270 | 303 | **RNA**→ rpmE2 (PRK01678)Ribosomal_L31 (pfam01197)→ |
| env-165 | SRS052227_LANL_scaffold_51410 | + | 309 | 342 | **RNA**→ rpmE2 (PRK01678)Ribosomal_L31 (pfam01197)→ |
| env-166 | SRS053854_LANL_scaffold_49593 | + | 459 | 492 | **RNA**→ rpmE2 (PRK01678)Ribosomal_L31 (pfam01197)→ |
| env-167 | SRS054687_LANL_scaffold_44325 | - | 268 | 235 | **RNA**→ rpmE2 (PRK01678)Ribosomal_L31 (pfam01197)→ |
| env-168 | SRS055426_C3289545 | + | 1043 | 1076 | **RNA**→ rpmE2 (PRK01678)Ribosomal_L31 (pfam01197)→ |
| env-169 | SRS056323_LANL_scaffold_27832 | + | 486 | 519 | **RNA**→ rpmE2 (PRK01678)Ribosomal_L31 (pfam01197)→ |
| env-170 | SRS056622_LANL_scaffold_18765 | - | 268 | 235 | **RNA**→ rpmE2 (PRK01678)Ribosomal_L31 (pfam01197)→ |
| env-171 | SRS057355_C2278348 | - | 268 | 235 | **RNA**→ rpmE2 (PRK01678)Ribosomal_L31 (pfam01197)→ |
| env-172 | SRS057539_LANL_scaffold_15960 | + | 387 | 420 | **RNA**→ rpmE2 (PRK01678)Ribosomal_L31 (pfam01197)→ |
| env-173 | SRS057692_LANL_scaffold_21980 | + | 239 | 272 | **RNA**→ rpmE2 (PRK01678)Ribosomal_L31 (pfam01197)→ |
| env-174 | SRS057791_C3542302 | + | 165 | 198 | **RNA**→ rpmE2 (PRK01678)Ribosomal_L31 (pfam01197)→ |
| env-175 | SRS058336_LANL_scaffold_61029 | - | 4000 | 3967 | **RNA**→ rpmE2 (PRK01678)Ribosomal_L31 (pfam01197)→ |
| env-176 | SRS062540_LANL_scaffold_56123 | + | 317 | 350 | **RNA**→ rpmE2 (PRK01678)Ribosomal_L31 (pfam01197)→ |
| env-177 | SRS062544_C2960130 | + | 486 | 519 | **RNA**→ rpmE2 (PRK01678)Ribosomal_L31 (pfam01197)→ |
| env-178 | SRS062761_C4244060 | + | 411 | 444 | **RNA**→ rpmE2 (PRK01678)Ribosomal_L31 (pfam01197)→ |
| env-179 | SRS063193_LANL_scaffold_32922 | + | 1090 | 1123 | **RNA**→ rpmE2 (PRK01678)Ribosomal_L31 (pfam01197)→ |
| env-180 | SRS063287_LANL_scaffold_18471 | + | 811 | 844 | **RNA**→ rpmE2 (PRK01678)Ribosomal_L31 (pfam01197)→ |
| env-181 | SRS064423_LANL_scaffold_27210 | + | 531 | 564 | **RNA**→ rpmE2 (PRK01678)Ribosomal_L31 (pfam01197)→ |
| env-182 | SRS064774_C2766594 | - | 244 | 211 | **RNA**→ rpmE2 (PRK01678)Ribosomal_L31 (pfam01197)→ |
| env-183 | SRS065278_LANL_scaffold_64678 | + | 186 | 219 | **RNA**→ rpmE2 (PRK01678)Ribosomal_L31 (pfam01197)→ |
| env-184 | SRS077736_C1812325 | + | 270 | 303 | **RNA**→ rpmE2 (PRK01678)Ribosomal_L31 (pfam01197)→ |
| Cma-1-1 | NZ_JUUF01000442.1 | - | 723 | 690 | **RNA**→ rpmE2 (PRK01678)Ribosomal_L31 (pfam01197)→ |
| Ssp-15-1 | NZ_JVNU01000059.1 | - | 61303 | 61270 | **RNA**→ rpmE2 (PRK01678)Ribosomal_L31 (pfam01197)→ |
| env-185 | SRS044373_WUGC_scaffold_4958 | - | 268 | 235 | **RNA**→ rpmE2 (PRK01678)Ribosomal_L31 (pfam01197)→ |
| env-186 | SRS011140_Baylor_scaffold_5150 | - | 244 | 211 | **RNA**→ rpmE2 (PRK01678)Ribosomal_L31 (pfam01197)→ |
| env-187 | SRS012279_Baylor_scaffold_62508 | + | 168 | 201 | **RNA**→ rpmE2 (PRK01678)Ribosomal_L31 (pfam01197)→ |
| env-188 | SRS014470_WUGC_scaffold_48055 | + | 444 | 477 | **RNA**→ rpmE2 (PRK01678)Ribosomal_L31 (pfam01197)→ |
| env-189 | SRS014689_C1291827 | + | 299 | 332 | **RNA**→ rpmE2 (PRK01678)Ribosomal_L31 (pfam01197)→ |
| env-190 | SRS015209_WUGC_scaffold_29650 | + | 114 | 147 | **RNA**→ rpmE2 (PRK01678)Ribosomal_L31 (pfam01197)→ |
| env-191 | SRS015395_WUGC_scaffold_12946 | - | 404 | 371 | **RNA**→ rpmE2 (PRK01678)Ribosomal_L31 (pfam01197)→ |
| env-192 | SRS015434_C5741645 | + | 153 | 186 | **RNA**→ rpmE2 (PRK01678)Ribosomal_L31 (pfam01197)→ |
| env-193 | SRS015537_WUGC_scaffold_11805 | + | 344 | 377 | **RNA**→ rpmE2 (PRK01678)Ribosomal_L31 (pfam01197)→ |
| env-194 | SRS015644_C1875711 | + | 241 | 274 | **RNA**→ rpmE2 (PRK01678)Ribosomal_L31 (pfam01197)→ |
| env-195 | SRS015797_C2623703 | - | 268 | 235 | **RNA**→ rpmE2 (PRK01678)Ribosomal_L31 (pfam01197)→ |
| env-196 | SRS017533_Baylor_scaffold_28274 | + | 359 | 392 | **RNA**→ rpmE2 (PRK01678)Ribosomal_L31 (pfam01197)→ |
| env-197 | SRS017808_Baylor_scaffold_52314 | + | 368 | 401 | **RNA**→ rpmE2 (PRK01678)Ribosomal_L31 (pfam01197)→ |
| env-198 | SRS018591_WUGC_scaffold_33617 | + | 344 | 377 | **RNA**→ rpmE2 (PRK01678)Ribosomal_L31 (pfam01197)→ |
| env-199 | SRS018791_C2231902 | + | 258 | 291 | **RNA**→ rpmE2 (PRK01678)Ribosomal_L31 (pfam01197)→ |
| env-200 | SRS020220_Baylor_scaffold_50081 | - | 244 | 211 | **RNA**→ rpmE2 (PRK01678)Ribosomal_L31 (pfam01197)→ |
| env-201 | SRS022530_C2101381 | + | 194 | 227 | **RNA**→ rpmE2 (PRK01678)Ribosomal_L31 (pfam01197)→ |
| env-202 | SRS022719_C2849471 | - | 268 | 235 | **RNA**→ rpmE2 (PRK01678)Ribosomal_L31 (pfam01197)→ |

| | | | | | |
|---|---|---|---|---|---|
| env-203 | SRS024318_LANL_scaffold_10943 | - | 268 | 235 | **RNA**→ rpmE2 (PRK01678)Ribosomal_L31 (pfam01197)→ |
| env-204 | SRS024441_LANL_scaffold_48378 | + | 368 | 401 | **RNA**→ rpmE2 (PRK01678)Ribosomal_L31 (pfam01197)→ |
| env-205 | SRS042131_C2145624 | + | 93 | 126 | **RNA**→ rpmE2 (PRK01678)Ribosomal_L31 (pfam01197)→ |
| env-206 | SRS047210_WUGC_scaffold_39313 | + | 344 | 377 | **RNA**→ rpmE2 (PRK01678)Ribosomal_L31 (pfam01197)→ |
| env-207 | SRS053603_LANL_scaffold_60119 | + | 368 | 401 | **RNA**→ rpmE2 (PRK01678)Ribosomal_L31 (pfam01197)→ |
| env-208 | SRS063288_LANL_scaffold_41825 | - | 431 | 398 | **RNA**→ rpmE2 (PRK01678)Ribosomal_L31 (pfam01197)→ hypo→ |
| env-209 | SRS011126_C4392817 | + | 93 | 126 | **RNA**→ rpmE2 (PRK01678)Ribosomal_L31 (pfam01197)→ |
| env-210 | SRS016092_WUGC_scaffold_23228 | + | 983 | 1016 | **RNA**→ rpmE2 (PRK01678)Ribosomal_L31 (pfam01197)→ |
| env-211 | SRS022536_LANL_scaffold_79011 | - | 827 | 794 | **RNA**→ rpmE2 (PRK01678)Ribosomal_L31 (pfam01197)→ |
| env-212 | SRS065099_LANL_scaffold_8242 | + | 652 | 685 | **RNA**→ rpmE2 (PRK01678)Ribosomal_L31 (pfam01197)→ |
| env-213 | SRS051791_LANL_scaffold_16929 | + | 1106 | 1139 | **RNA**→ rpmE2 (PRK01678)Ribosomal_L31 (pfam01197)→ |
| Eas-1-1 | NZ_KB946293.1 | + | 500548 | 500580 | **RNA**→ rpmE2 (PRK01678)Ribosomal_L31 (pfam01197)→ |
| env-214 | JGI12273J12029_10202314 | + | 41 | 80 | **RNA**→ ←hypo |
| Pch-2-1 | NZ_BBJT01000019.1 | + | 57434 | 57468 | **RNA**→ ←hypo |
| Psp-3-1 | NZ_AFDH01000010.1 | + | 58110 | 58144 | **RNA**→ ←hypo |
| Psp-17-1 | NZ_KE150412.1 | - | 2575869 | 2575835 | **RNA**→ rpmE (PRK00019)Ribosomal_L31 (pfam01197)→ |
| Wci-1-1 | NZ_JWHV01000015.1 | + | 4485 | 4517 | **RNA**→ rpmE2 (PRK01678)Ribosomal_L31 (pfam01197)→ |
| Wci-1-2 | NZ_JWHU01000017.1 | - | 4782 | 4750 | **RNA**→ rpmE2 (PRK01678)Ribosomal_L31 (pfam01197)→ |
| env-215 | BS_KBA_SWE02_21m_10495365 | - | 109 | 79 | **RNA**→ rpmE (PRK00528)rpmE (PRK00019)rpmE2 (PRK01678)Ribosomal_L31 (pfam01197)→ |
| env-216 | 08_07_GOS_DNA_scaffold115360_2 | - | 274 | 241 | **RNA**→ rpmE (PRK00019)Ribosomal_L31 (pfam01197)→ |
| env-217 | 08_19_CONTROL_DNA_scaffold125820_3 | - | 76 | 43 | **RNA**→ |
| env-218 | 4491412.3_NODE_32910 | - | 173 | 140 | **RNA**→ rpmE (PRK00019)Ribosomal_L31 (pfam01197)→ |
| env-219 | JGI24709J26583_10035806 | + | 1098 | 1132 | **RNA**→ rpmE (PRK00019)Ribosomal_L31 (pfam01197)→ Tdk (COG1435)PRK04296 (PRK04296)→ PrfA (COG0216)prfA (PRK00591)→ |
| env-220 | JGI24707J26582_10000007 | + | 1235 | 1269 | **RNA**→ rpmE (PRK00019)Ribosomal_L31 (pfam01197)→ Tdk (COG1435)PRK04296 (PRK04296)→ PrfA (COG0216)prfA (PRK00591)→ HemK (COG2890)PRK09328 (PRK09328)→ |
| env-221 | JGI24708J26588_10003174 | - | 10819 | 10785 | **RNA**→ rpmE (PRK00019)Ribosomal_L31 (pfam01197)→ Tdk (COG1435)PRK04296 (PRK04296)→ PrfA (COG0216)prfA (PRK00591)→ HemK (COG2890)PRK09328 (PRK09328)→ |
| env-222 | JGI24710J26742_10050102 | - | 660 | 626 | **RNA**→ rpmE (PRK00019)Ribosomal_L31 (pfam01197)→ Tdk (COG1435)PRK04296 (PRK04296)→ |
| env-223 | JGI24500J29687_10037880 | + | 135 | 169 | **RNA**→ rpmE (PRK00019)Ribosomal_L31 (pfam01197)→ Tdk (COG1435)PRK04296 (PRK04296)→ PrfA (COG0216)prfA (PRK00591)→ HemK (COG2890)PRK09328 (PRK09328)→ |
| env-224 | JGI24711J26586_10152175 | + | 278 | 312 | **RNA**→ rpmE (PRK00019)Ribosomal_L31 (pfam01197)→ |
| env-225 | AUXO011483282.1 | - | 168 | 141 | **RNA**→ rpmE (PRK00019)Ribosomal_L31 (pfam01197)→ |
| env-226 | AUXO010436543.1 | + | 860 | 887 | **RNA**→ rpmE (PRK00019)Ribosomal_L31 (pfam01197)→ |
| Epa-2-1 | NZ_KB946300.1 | + | 1021113 | 1021145 | **RNA**→ rpmE2 (PRK01678)Ribosomal_L31 (pfam01197)→ |
| env-227 | JGI24025J20009_10344154 | - | 59 | 26 | **RNA**→ |
| env-228 | BARV01035863.1 | + | 366 | 399 | **RNA**→ hypo→ |
| env-229 | UnmappedStool_Broad_scaffold_766897 | + | 949 | 975 | **RNA**→ AprE (COG1404)T7SS_mycosin (TIGR03921)→ |
| env-230 | longitudinal_143_760 | - | 43663 | 43630 | **RNA**→ rpmE2 (PRK01678)Ribosomal_L31 (pfam01197)→ |
| env-231 | longitudinal_144_2080 | - | 43746 | 43713 | **RNA**→ rpmE2 (PRK01678)Ribosomal_L31 (pfam01197)→ |
| env-232 | longitudinal_145_864 | - | 2639 | 2606 | **RNA**→ rpmE2 (PRK01678)Ribosomal_L31 (pfam01197)→ |
| Lca-1-1 | NZ_JXNV01000006.1 | + | 54157 | 54190 | **RNA**→ rpmE2 (PRK01678)Ribosomal_L31 (pfam01197)→ |
| Lca-17-1 | NZ_AQVS01000053.1 | - | 10357 | 10324 | **RNA**→ rpmE2 (PRK01678)Ribosomal_L31 (pfam01197)→ |
| Lpa-33-1 | NZ_ANKJ01000022.1 | + | 2648 | 2681 | **RNA**→ rpmE2 (PRK01678)Ribosomal_L31 (pfam01197)→ |
| Lpa-29-1 | NZ_ANKY01000024.1 | + | 53357 | 53390 | **RNA**→ rpmE2 (PRK01678)Ribosomal_L31 (pfam01197)→ |
| Lpa-14-1 | NZ_ANMK01000192.1 | - | 3633 | 3600 | **RNA**→ rpmE2 (PRK01678)Ribosomal_L31 (pfam01197)→ |
| Lpa-24-1 | NZ_ANMM01000002.1 | - | 44657 | 44624 | **RNA**→ rpmE2 (PRK01678)Ribosomal_L31 (pfam01197)→ |
| Lpa-32-1 | NZ_ANMJ01000006.1 | + | 1533 | 1566 | **RNA**→ rpmE2 (PRK01678)Ribosomal_L31 (pfam01197)→ |
| Lpa-25-1 | NZ_ANMN01000105.1 | - | 46066 | 46033 | **RNA**→ rpmE2 (PRK01678)Ribosomal_L31 (pfam01197)→ |
| Lpa-37-1 | NZ_ANKB01000003.1 | - | 44594 | 44561 | **RNA**→ rpmE2 (PRK01678)Ribosomal_L31 (pfam01197)→ |
| Lpa-36-1 | NZ_ANJU01000008.1 | + | 58114 | 58147 | **RNA**→ rpmE2 (PRK01678)Ribosomal_L31 (pfam01197)→ |
| Lpa-27-1 | NZ_ANKA01000008.1 | - | 29182 | 29149 | **RNA**→ rpmE2 (PRK01678)Ribosomal_L31 (pfam01197)→ |
| Lpa-38-1 | NZ_ANKV01000009.1 | - | 44491 | 44458 | **RNA**→ rpmE2 (PRK01678)Ribosomal_L31 (pfam01197)→ |

| | | | | | |
|---|---|---|---|---|---|
| Lpa-15-1 | NZ_ANKW01000003.1 | + | 99166 | 99199 | **RNA**→ rpmE2 (PRK01678)Ribosomal_L31 (pfam01197)→ |
| Lpa-23-1 | NZ_ANML01000015.1 | + | 50153 | 50186 | **RNA**→ rpmE2 (PRK01678)Ribosomal_L31 (pfam01197)→ |
| Lpa-16-1 | NZ_ANKM01000001.1 | - | 46054 | 46021 | **RNA**→ rpmE2 (PRK01678)Ribosomal_L31 (pfam01197)→ |
| Lpa-19-1 | NZ_ANJY01000018.1 | - | 13430 | 13397 | **RNA**→ rpmE2 (PRK01678)Ribosomal_L31 (pfam01197)→ |
| Lpa-21-1 | NZ_ANMI01000231.1 | + | 4976 | 5009 | **RNA**→ rpmE2 (PRK01678)Ribosomal_L31 (pfam01197)→ |
| Lpa-18-1 | NZ_ANMH01000022.1 | + | 1835 | 1868 | **RNA**→ rpmE2 (PRK01678)Ribosomal_L31 (pfam01197)→ |
| Lpa-31-1 | NZ_ANKI01000072.1 | + | 10689 | 10722 | **RNA**→ rpmE2 (PRK01678)Ribosomal_L31 (pfam01197)→ |
| Lpa-35-1 | NZ_ANLF01000101.1 | + | 3217 | 3250 | **RNA**→ rpmE2 (PRK01678)Ribosomal_L31 (pfam01197)→ |
| Lpa-20-1 | NZ_ANJS01000007.1 | + | 49686 | 49719 | **RNA**→ rpmE2 (PRK01678)Ribosomal_L31 (pfam01197)→ |
| Lpa-10-1 | NZ_ANLE01000428.1 | - | 11217 | 11184 | **RNA**→ rpmE2 (PRK01678)Ribosomal_L31 (pfam01197)→ |
| Lpa-12-1 | NZ_ANKH01000172.1 | - | 1356 | 1323 | **RNA**→ rpmE2 (PRK01678)Ribosomal_L31 (pfam01197)→ |
| Lpa-30-1 | NZ_ANKG01000146.1 | + | 1496 | 1529 | **RNA**→ rpmE2 (PRK01678)Ribosomal_L31 (pfam01197)→ |
| Lpa-28-1 | NZ_ANJT01000272.1 | + | 1638 | 1671 | **RNA**→ rpmE2 (PRK01678)Ribosomal_L31 (pfam01197)→ |
| Lpa-17-1 | NZ_ANJZ01000255.1 | + | 27968 | 28001 | **RNA**→ rpmE2 (PRK01678)Ribosomal_L31 (pfam01197)→ |
| Lpa-34-1 | NZ_ANJV01000073.1 | - | 302 | 269 | **RNA**→ rpmE2 (PRK01678)Ribosomal_L31 (pfam01197)→ |
| Lpa-22-1 | NZ_ANKF01000041.1 | + | 18918 | 18951 | **RNA**→ rpmE2 (PRK01678)Ribosomal_L31 (pfam01197)→ |
| Lpa-11-1 | NZ_ANJX01000256.1 | - | 27031 | 26998 | **RNA**→ rpmE2 (PRK01678)Ribosomal_L31 (pfam01197)→ |
| Lpa-26-1 | NZ_ANKK01000559.1 | + | 21036 | 21069 | **RNA**→ rpmE2 (PRK01678)Ribosomal_L31 (pfam01197)→ |
| Lpa-6-1 | NZ_BAYM01000151.1 | - | 43683 | 43650 | **RNA**→ rpmE2 (PRK01678)Ribosomal_L31 (pfam01197)→ |
| Lpa-7-1 | NZ_BAYN01000089.1 | + | 23335 | 23368 | **RNA**→ rpmE2 (PRK01678)Ribosomal_L31 (pfam01197)→ |
| Lpa-8-1 | NZ_BAYO01000435.1 | - | 618 | 585 | **RNA**→ rpmE2 (PRK01678)Ribosomal_L31 (pfam01197)→ |
| Lca-5-1 | NZ_AUYM01000001.1 | + | 51678 | 51711 | **RNA**→ rpmE2 (PRK01678)Ribosomal_L31 (pfam01197)→ |
| Lca-1-2 | NZ_JVUB01000117.1 | + | 92912 | 92945 | **RNA**→ rpmE2 (PRK01678)Ribosomal_L31 (pfam01197)→ |
| Lca-1-3 | NZ_JVUA01000053.1 | - | 246 | 213 | **RNA**→ rpmE2 (PRK01678)Ribosomal_L31 (pfam01197)→ |
| Lca-1-4 | NZ_JUQW01000038.1 | + | 57117 | 57150 | **RNA**→ rpmE2 (PRK01678)Ribosomal_L31 (pfam01197)→ |
| Lca-12-1 | NZ_AFYU01000056.1 | - | 16249 | 16216 | **RNA**→ rpmE2 (PRK01678)Ribosomal_L31 (pfam01197)→ |
| Lca-14-1 | NZ_AFYP01000041.1 | - | 44459 | 44426 | **RNA**→ rpmE2 (PRK01678)Ribosomal_L31 (pfam01197)→ |
| Lca-15-1 | NZ_AFYQ01000094.1 | - | 7962 | 7929 | **RNA**→ rpmE2 (PRK01678)Ribosomal_L31 (pfam01197)→ |
| Lca-16-1 | NZ_AFYR01000102.1 | - | 44152 | 44119 | **RNA**→ rpmE2 (PRK01678)Ribosomal_L31 (pfam01197)→ |
| Lca-17-2 | NZ_AFYS01000052.1 | - | 106073 | 106040 | **RNA**→ rpmE2 (PRK01678)Ribosomal_L31 (pfam01197)→ |
| Lca-9-1 | NZ_AFYT01000048.1 | - | 11682 | 11649 | **RNA**→ rpmE2 (PRK01678)Ribosomal_L31 (pfam01197)→ |
| Lca-13-1 | NZ_AFYO01000057.1 | - | 34209 | 34176 | **RNA**→ rpmE2 (PRK01678)Ribosomal_L31 (pfam01197)→ |
| Lca-3-1 | NZ_AFYK01000043.1 | - | 38236 | 38203 | **RNA**→ rpmE2 (PRK01678)Ribosomal_L31 (pfam01197)→ |
| Lpa-4-1 | NZ_BAGT01000089.1 | - | 46077 | 46044 | **RNA**→ rpmE2 (PRK01678)Ribosomal_L31 (pfam01197)→ |
| Lca-20-1 | NC_008526.1 | - | 2550073 | 2550040 | **RNA**→ rpmE2 (PRK01678)Ribosomal_L31 (pfam01197)→ |
| Lpa-42-1 | NZ_GG670156.1 | + | 291360 | 291393 | **RNA**→ rpmE2 (PRK01678)Ribosomal_L31 (pfam01197)→ |
| Lca-11-1 | NZ_CM001848.2 | - | 2654316 | 2654283 | **RNA**→ rpmE2 (PRK01678)Ribosomal_L31 (pfam01197)→ |
| Lca-22-1 | NC_010999.1 | - | 2690859 | 2690826 | **RNA**→ rpmE2 (PRK01678)Ribosomal_L31 (pfam01197)→ |
| Lca-21-1 | NC_017474.1 | - | 2681824 | 2681791 | **RNA**→ rpmE2 (PRK01678)Ribosomal_L31 (pfam01197)→ |
| Lca-23-1 | NC_017473.1 | - | 2650937 | 2650904 | **RNA**→ rpmE2 (PRK01678)Ribosomal_L31 (pfam01197)→ |
| Lpa-3-1 | NZ_CP012187.1 | - | 2514242 | 2514209 | **RNA**→ rpmE2 (PRK01678)Ribosomal_L31 (pfam01197)→ |
| Lca-24-1 | NC_018641.1 | - | 2687463 | 2687430 | **RNA**→ rpmE2 (PRK01678)Ribosomal_L31 (pfam01197)→ |
| Lca-8-1 | NC_021721.1 | - | 2731500 | 2731467 | **RNA**→ rpmE2 (PRK01678)Ribosomal_L31 (pfam01197)→ |
| Lpa-9-1 | NC_022112.1 | - | 2544386 | 2544353 | **RNA**→ rpmE2 (PRK01678)Ribosomal_L31 (pfam01197)→ |
| Lpa-5-1 | NZ_CP007122.1 | - | 2486141 | 2486108 | **RNA**→ rpmE2 (PRK01678)Ribosomal_L31 (pfam01197)→ |
| Lpa-13-1 | NZ_AP012541.1 | - | 2658289 | 2658256 | **RNA**→ rpmE2 (PRK01678)Ribosomal_L31 (pfam01197)→ |
| Lca-18-1 | NC_014334.2 | - | 2478752 | 2478719 | **RNA**→ rpmE2 (PRK01678)Ribosomal_L31 (pfam01197)→ |
| Lca-2-1 | NZ_CP006690.1 | - | 2512933 | 2512900 | **RNA**→ rpmE2 (PRK01678)Ribosomal_L31 (pfam01197)→ |
| Lca-10-1 | NZ_CM001861.1 | - | 2680824 | 2680791 | **RNA**→ rpmE2 (PRK01678)Ribosomal_L31 (pfam01197)→ |
| Lpa-3-2 | NZ_CP012148.1 | - | 2678238 | 2678205 | **RNA**→ rpmE2 (PRK01678)Ribosomal_L31 (pfam01197)→ |
| Lca-4-1 | NZ_AFYL01000034.1 | - | 39968 | 39935 | **RNA**→ rpmE2 (PRK01678)Ribosomal_L31 (pfam01197)→ |
| Lca-6-1 | NZ_AFYM01000082.1 | - | 5581 | 5548 | **RNA**→ rpmE2 (PRK01678)Ribosomal_L31 (pfam01197)→ |

| | | | | | | |
|---|---|---|---|---|---|---|
| Lca-7-1 | NZ_AFYN01000040.1 | - | 56503 | 56470 | **RNA**→ | rpmE2 (PRK01678)Ribosomal_L31 (pfam01197)→ |
| Lca-6-2 | NZ_AZOE01000015.1 | - | 5466 | 5433 | **RNA**→ | rpmE2 (PRK01678)Ribosomal_L31 (pfam01197)→ |
| Lze-1-1 | NZ_BACQ01000037.1 | + | 45109 | 45142 | **RNA**→ | rpmE2 (PRK01678)Ribosomal_L31 (pfam01197)→ rpmE2 (PRK01678)Ribosomal_L31 (pfam01197 |
| Lca-1-5 | NZ_LCUN01000013.1 | + | 753391 | 753424 | **RNA**→ | rpmE2 (PRK01678)Ribosomal_L31 (pfam01197)→ |
| Lsp-11-1 | NZ_KN714896.1 | + | 1114375 | 1114408 | **RNA**→ | rpmE2 (PRK01678)Ribosomal_L31 (pfam01197)→ |
| Lca-19-1 | NZ_AP012544.1 | - | 2516623 | 2516590 | **RNA**→ | rpmE2 (PRK01678)Ribosomal_L31 (pfam01197)→ |
| Lrh-14-1 | NZ_AMQW01000012.1 | - | 9617 | 9584 | **RNA**→ | rpmE2 (PRK01678)Ribosomal_L31 (pfam01197)→ |
| Lrh-15-1 | NZ_AMQX01000009.1 | + | 73746 | 73779 | **RNA**→ | rpmE2 (PRK01678)Ribosomal_L31 (pfam01197)→ |
| Lrh-1-1 | NZ_JUWG01000100.1 | - | 9704 | 9671 | **RNA**→ | rpmE2 (PRK01678)Ribosomal_L31 (pfam01197)→ |
| Lrh-1-2 | NZ_JUWQ01000064.1 | + | 8431 | 8464 | **RNA**→ | rpmE2 (PRK01678)Ribosomal_L31 (pfam01197)→ |
| env-233 | longitudinal_146_787 | + | 90575 | 90608 | **RNA**→ | rpmE2 (PRK01678)Ribosomal_L31 (pfam01197)→ |
| Lrh-21-1 | NZ_ABWJ01000022.1 | - | 40553 | 40520 | **RNA**→ | rpmE2 (PRK01678)Ribosomal_L31 (pfam01197)→ rpmE2 (PRK01678)Ribosomal_L31 (pfam01197 |
| Lrh-1-3 | NZ_LFNB01000004.1 | - | 40563 | 40530 | **RNA**→ | rpmE2 (PRK01678)Ribosomal_L31 (pfam01197)→ |
| Lrh-1-4 | NZ_JUIL01000391.1 | + | 6444 | 6477 | **RNA**→ | rpmE2 (PRK01678)Ribosomal_L31 (pfam01197)→ |
| Lrh-1-5 | NZ_JVPR01000084.1 | + | 1525 | 1558 | **RNA**→ | rpmE2 (PRK01678)Ribosomal_L31 (pfam01197)→ |
| Lrh-1-6 | NZ_JVQV01000076.1 | + | 3853 | 3886 | **RNA**→ | rpmE2 (PRK01678)Ribosomal_L31 (pfam01197)→ |
| Lrh-1-7 | NZ_JVIZ01000364.1 | - | 40597 | 40564 | **RNA**→ | rpmE2 (PRK01678)Ribosomal_L31 (pfam01197)→ |
| Lrh-1-8 | NZ_JULF01000226.1 | - | 3136 | 3103 | **RNA**→ | rpmE2 (PRK01678)Ribosomal_L31 (pfam01197)→ |
| Lrh-1-9 | NZ_JPZB01000036.1 | - | 4961 | 4928 | **RNA**→ | rpmE2 (PRK01678)Ribosomal_L31 (pfam01197)→ |
| Lrh-16-1 | NZ_JDFQ01000007.1 | + | 73438 | 73471 | **RNA**→ | rpmE2 (PRK01678)Ribosomal_L31 (pfam01197)→ |
| Lrh-17-1 | NZ_JDFR01000003.1 | - | 40564 | 40531 | **RNA**→ | rpmE2 (PRK01678)Ribosomal_L31 (pfam01197)→ |
| Lrh-6-1 | NZ_JDRW01000008.1 | + | 93238 | 93271 | **RNA**→ | rpmE2 (PRK01678)Ribosomal_L31 (pfam01197)→ |
| Lrh-20-1 | NC_013198.1 | - | 2611055 | 2611022 | **RNA**→ | rpmE2 (PRK01678)Ribosomal_L31 (pfam01197)→ |
| Lrh-20-2 | NC_017482.1 | - | 2606025 | 2605992 | **RNA**→ | rpmE2 (PRK01678)Ribosomal_L31 (pfam01197)→ |
| Lrh-1-10 | NZ_JTDC01000034.1 | + | 23421 | 23454 | **RNA**→ | rpmE2 (PRK01678)Ribosomal_L31 (pfam01197)→ |
| Lrh-1-11 | NZ_JWHC01000058.1 | - | 565 | 532 | **RNA**→ | rpmE2 (PRK01678)Ribosomal_L31 (pfam01197)→ |
| Lrh-1-12 | NZ_JYCS01000044.1 | + | 1501 | 1534 | **RNA**→ | rpmE2 (PRK01678)Ribosomal_L31 (pfam01197)→ |
| Lrh-5-1 | NZ_ATBI01000016.1 | - | 42313 | 42280 | **RNA**→ | rpmE2 (PRK01678)Ribosomal_L31 (pfam01197)→ |
| Lrh-1-13 | NZ_LAZE01000069.1 | - | 88154 | 88121 | **RNA**→ | rpmE2 (PRK01678)Ribosomal_L31 (pfam01197)→ |
| Lrh-1-14 | NZ_JUIH01000135.1 | - | 7237 | 7204 | **RNA**→ | rpmE2 (PRK01678)Ribosomal_L31 (pfam01197)→ |
| Lrh-1-15 | NZ_LFNA01000007.1 | - | 565 | 532 | **RNA**→ | rpmE2 (PRK01678)Ribosomal_L31 (pfam01197)→ |
| Lrh-1-16 | NZ_JUII01000291.1 | - | 7276 | 7243 | **RNA**→ | rpmE2 (PRK01678)Ribosomal_L31 (pfam01197)→ |
| Lrh-1-17 | NZ_JUIK01000056.1 | - | 6917 | 6884 | **RNA**→ | rpmE2 (PRK01678)Ribosomal_L31 (pfam01197)→ |
| Lrh-1-18 | NZ_JVLT01000012.1 | + | 180026 | 180059 | **RNA**→ | rpmE2 (PRK01678)Ribosomal_L31 (pfam01197)→ |
| Lrh-1-19 | NZ_JVDP01000093.1 | - | 4098 | 4065 | **RNA**→ | rpmE2 (PRK01678)Ribosomal_L31 (pfam01197)→ |
| Lrh-1-20 | NZ_JVCX01000037.1 | + | 27316 | 27349 | **RNA**→ | rpmE2 (PRK01678)Ribosomal_L31 (pfam01197)→ |
| Lrh-1-21 | NZ_JUTS01000037.1 | + | 27414 | 27447 | **RNA**→ | rpmE2 (PRK01678)Ribosomal_L31 (pfam01197)→ |
| Lrh-1-22 | NZ_JUPX01000230.1 | + | 1814 | 1847 | **RNA**→ | rpmE2 (PRK01678)Ribosomal_L31 (pfam01197)→ |
| Lrh-1-23 | NZ_JUTB01000107.1 | + | 26945 | 26978 | **RNA**→ | rpmE2 (PRK01678)Ribosomal_L31 (pfam01197)→ |
| Lrh-1-24 | NZ_JUPA01000162.1 | - | 40691 | 40658 | **RNA**→ | rpmE2 (PRK01678)Ribosomal_L31 (pfam01197)→ |
| Lrh-1-25 | NZ_JUMP01000096.1 | - | 3386 | 3353 | **RNA**→ | rpmE2 (PRK01678)Ribosomal_L31 (pfam01197)→ |
| Lrh-1-26 | NZ_JUON01000171.1 | + | 1814 | 1847 | **RNA**→ | rpmE2 (PRK01678)Ribosomal_L31 (pfam01197)→ |
| Lrh-1-27 | NZ_JUMO01000067.1 | - | 9622 | 9589 | **RNA**→ | rpmE2 (PRK01678)Ribosomal_L31 (pfam01197)→ |
| Lrh-1-28 | NZ_JUKV01000061.1 | - | 3386 | 3353 | **RNA**→ | rpmE2 (PRK01678)Ribosomal_L31 (pfam01197)→ |
| Lrh-18-1 | NZ_AGKC01000002.1 | - | 40627 | 40594 | **RNA**→ | rpmE2 (PRK01678)Ribosomal_L31 (pfam01197)→ |
| Lrh-8-1 | NZ_AYTQ01000044.1 | - | 40961 | 40928 | **RNA**→ | rpmE2 (PRK01678)Ribosomal_L31 (pfam01197)→ |
| Lrh-2-1 | NZ_JMSI01000003.1 | + | 364620 | 364653 | **RNA**→ | rpmE2 (PRK01678)Ribosomal_L31 (pfam01197)→ |
| Lrh-7-1 | NZ_JNNV01000002.1 | + | 523226 | 523259 | **RNA**→ | rpmE2 (PRK01678)Ribosomal_L31 (pfam01197)→ |
| Lrh-13-1 | NZ_AZHJ01000147.1 | - | 296 | 263 | **RNA**→ | rpmE2 (PRK01678)Ribosomal_L31 (pfam01197)→ |
| Lrh-4-1 | NZ_JH159058.1 | + | 1505 | 1538 | **RNA**→ | rpmE2 (PRK01678)Ribosomal_L31 (pfam01197)→ |
| Lrh-3-1 | NZ_JH379496.1 | + | 1535 | 1568 | **RNA**→ | rpmE2 (PRK01678)Ribosomal_L31 (pfam01197)→ |
| Lrh-23-1 | NC_013199.1 | - | 2590593 | 2590560 | **RNA**→ | rpmE2 (PRK01678)Ribosomal_L31 (pfam01197)→ |

| | | | | | |
|---|---|---|---|---|---|
| Lrh-19-1 | NC_017491.1 | - | 2582498 | 2582465 | **RNA**→ rpmE2 (PRK01678)Ribosomal_L31 (pfam01197)→ |
| Lrh-12-1 | NC_021725.1 | - | 2608939 | 2608906 | **RNA**→ rpmE2 (PRK01678)Ribosomal_L31 (pfam01197)→ |
| Lrh-11-1 | NC_021723.1 | - | 2514116 | 2514083 | **RNA**→ rpmE2 (PRK01678)Ribosomal_L31 (pfam01197)→ |
| Lrh-22-1 | NZ_GG692961.1 | + | 684435 | 684468 | **RNA**→ rpmE2 (PRK01678)Ribosomal_L31 (pfam01197)→ |
| Lrh-10-1 | NZ_AYTP01000042.1 | + | 63172 | 63205 | **RNA**→ rpmE2 (PRK01678)Ribosomal_L31 (pfam01197)→ |
| Lrh-9-1 | NZ_AYTR01000028.1 | - | 40856 | 40823 | **RNA**→ rpmE2 (PRK01678)Ribosomal_L31 (pfam01197)→ |
| Lca-1-6 | NZ_JUPZ01000004.1 | + | 21466 | 21499 | **RNA**→ rpmE2 (PRK01678)Ribosomal_L31 (pfam01197)→ |
| Fba-1-1 | NZ_KE159698.1 | - | 41156 | 41122 | **RNA**→ rpmE (PRK00019)Ribosomal_L31 (pfam01197)→ PRK07681 (PRK07681)AspB (COG0436)→ hyp |
| env-234 | UnmappedStool_Broad_scaffold_36047 | + | 3764 | 3797 | **RNA**→ rpmE (PRK00019)Ribosomal_L31 (pfam01197)→ |
| env-235 | SRS017521_C2731538 | - | 110 | 77 | **RNA**→ rpmE (PRK00019)Ribosomal_L31 (pfam01197)→ |
| env-236 | SRS019601_WUGC_scaffold_47824 | - | 1213 | 1180 | **RNA**→ rpmE (PRK00019)Ribosomal_L31 (pfam01197)→ |
| Esp-18-1 | NZ_JH594450.1 | + | 256315 | 256348 | **RNA**→ rpmE (PRK00019)Ribosomal_L31 (pfam01197)→ |
| Eba-1-1 | NZ_KI391958.1 | + | 308940 | 308973 | **RNA**→ rpmE (PRK00019)Ribosomal_L31 (pfam01197)→ |
| env-237 | AUXO015079216.1 | - | 64 | 31 | **RNA**→ hypo→ |
| env-238 | AUXO018658602.1 | + | 131 | 164 | **RNA**→ rpmE (PRK00019)Ribosomal_L31 (pfam01197)→ |
| env-239 | SRS049318_C6167461 | - | 259 | 226 | **RNA**→ PRK05452 (PRK05452)Rubredoxin (pfam00301)→ |
| env-240 | JGI24724J26744_10023391 | + | 2361 | 2388 | **RNA**→ hypo→ |
| env-241 | 3300000052_087558 | + | 1786 | 1819 | **RNA**→ rpmE (PRK00019)Ribosomal_L31 (pfam01197)→ hypo→ |
| Pwy-1-1 | NZ_JQCR01000002.1 | - | 1896394 | 1896362 | **RNA**→ rpmE (PRK00019)Ribosomal_L31 (pfam01197)→ |
| Bsp-41-1 | NZ_LIGF01000001.1 | + | 2484836 | 2484867 | **RNA**→ rpmE2 (PRK01678)Ribosomal_L31 (pfam01197)→ |
| env-242 | 3300001580_10921771 | + | 174 | 205 | **RNA**→ rpmE (PRK00019)Ribosomal_L31 (pfam01197)→ |
| env-243 | 3300001580_11398849 | + | 190 | 221 | **RNA**→ |
| env-244 | 2227341862 | - | 257 | 226 | **RNA**→ rpmE (PRK00019)Ribosomal_L31 (pfam01197)→ |
| env-245 | JGI994J12354_10052525 | + | 1979 | 2010 | **RNA**→ ←hypo |
| env-246 | KGI_S1_ANT02_95m_c10164418 | - | 454 | 421 | **RNA**→ rpmE2 (PRK01678)Ribosomal_L31 (pfam01197)→ |
| Cal-1-1 | NZ_JQLG01000004.1 | + | 1333377 | 1333410 | **RNA**→ rpmE2 (PRK01678)Ribosomal_L31 (pfam01197)→ |
| env-247 | KGI_S1_ANT01_95m_c10000058 | + | 18408 | 18441 | **RNA**→ rpmE2 (PRK01678)Ribosomal_L31 (pfam01197)→ |
| env-248 | KGI_S2_ANT05_2345m_c1001626 | - | 5035 | 5002 | **RNA**→ rpmE2 (PRK01678)Ribosomal_L31 (pfam01197)→ |
| Cpl-1-1 | NZ_JQLQ01000002.1 | - | 2377756 | 2377723 | **RNA**→ rpmE2 (PRK01678)Ribosomal_L31 (pfam01197)→ hypo→ |
| Csp-1-1 | NC_022606.1 | + | 216680 | 216713 | **RNA**→ rpmE2 (PRK01678)Ribosomal_L31 (pfam01197)→ |
| Cin-1-1 | NZ_JQIV01000006.1 | + | 1981645 | 1981678 | **RNA**→ rpmE2 (PRK01678)Ribosomal_L31 (pfam01197)→ |
| Csp-9-1 | NZ_ABHH01000004.1 | - | 75193 | 75160 | **RNA**→ rpmE2 (PRK01678)Ribosomal_L31 (pfam01197)→ rpmE2 (PRK01678)Ribosomal_L31 (pfam01197 |
| Cje-1-1 | NZ_JEMH01000028.1 | + | 73321 | 73354 | **RNA**→ rpmE2 (PRK01678)Ribosomal_L31 (pfam01197)→ |
| Cje-2-1 | NZ_JQNF01000010.1 | + | 922204 | 922237 | **RNA**→ rpmE2 (PRK01678)Ribosomal_L31 (pfam01197)→ |
| Pas-1-1 | NZ_AULU01000019.1 | + | 60564 | 60597 | **RNA**→ rpmE (PRK00019)Ribosomal_L31 (pfam01197)→ |
| env-249 | AUXO012865815.1 | - | 7672 | 7641 | **RNA**→ rpmE (PRK00019)Ribosomal_L31 (pfam01197)→ U3_snoRNA_assoc (pfam08297)→ |
| | | | | | PRK05571 (PRK05571)LacAB_rpiB (pfam02502)→ DUF1146 (pfam06612)→ |
| | | | | | PRK11649 (PRK11649)Peptidase_M23 (pfam01551)→ MreB_like (cd10225)PRK13930 (PRK13930)→ |
| | | | | | ECA_wecA (TIGR02380)GT_WecA_like (cd06853)→ LytTR (smart00850)LytTR (pfam04397)→ |
| env-250 | RUMENNODE_2681326_1 | + | 10151 | 10184 | **RNA**→ rpmE (PRK00019)Ribosomal_L31 (pfam01197)→ |
| env-251 | 2204978118 | - | 261 | 228 | **RNA**→ rpmE2 (PRK01678)Ribosomal_L31 (pfam01197)→ |
| env-252 | 2157636633 | - | 261 | 228 | **RNA**→ rpmE2 (PRK01678)Ribosomal_L31 (pfam01197)→ |
| Lbu-2-1 | NC_018610.1 | - | 1624612 | 1624579 | **RNA**→ rpmE2 (PRK01678)Ribosomal_L31 (pfam01197)→ |
| Tca-1-1 | NZ_AUMM01000004.1 | + | 100153 | 100187 | **RNA**→ rpmE (PRK00019)Ribosomal_L31 (pfam01197)→ |
| Sin-1-1 | NZ_AFVQ02000220.1 | - | 17111 | 17077 | **RNA**→ rpmE2 (PRK01678)Ribosomal_L31 (pfam01197)→ |
| Sla-1-1 | NZ_AWTC01000005.1 | - | 157176 | 157144 | **RNA**→ rpmE2 (PRK01678)Ribosomal_L31 (pfam01197)→ |
| Svi-1-2 | NZ_KB899046.1 | - | 55946 | 55912 | **RNA**→ rpmE2 (PRK01678)Ribosomal_L31 (pfam01197)→ |
| Svi-1-1 | NZ_BAEY01000002.1 | - | 55800 | 55766 | **RNA**→ rpmE2 (PRK01678)Ribosomal_L31 (pfam01197)→ rpmE2 (PRK01678)Ribosomal_L31 (pfam01197 |
| Ste-2-1 | NZ_JAQS01000032.1 | + | 2004 | 2038 | **RNA**→ rpmE2 (PRK01678)Ribosomal_L31 (pfam01197)→ hypo→ Tdk (COG1435)PRK04296 (PRK04296 |
| Ste-1-1 | NZ_JFZC01000099.1 | + | 3894 | 3928 | **RNA**→ rpmE2 (PRK01678)Ribosomal_L31 (pfam01197)→ hypo→ Tdk (COG1435)PRK04296 (PRK04296 |
| Pma-3-1 | NZ_KB898207.1 | - | 5632 | 5598 | **RNA**→ rpmE (PRK00019)Ribosomal_L31 (pfam01197)→ |
| Pma-4-1 | NZ_ASSE01000237.1 | + | 58736 | 58770 | **RNA**→ rpmE (PRK00019)Ribosomal_L31 (pfam01197)→ |
| Ppa-1-1 | NZ_AUFO01000034.1 | - | 5491 | 5457 | **RNA**→ rpmE (PRK00019)Ribosomal_L31 (pfam01197)→ |

| | | | | | |
|---|---|---|---|---|---|
| Hco-1-1 | NZ_AFNU02000004.1 | + | 304522 | 304555 | **RNA**→ rpmE (PRK00019)Ribosomal_L31 (pfam01197)→ |
| Esp-4-1 | NZ_LDNY01000001.1 | + | 240696 | 240729 | **RNA**→ rpmE2 (PRK01678)Ribosomal_L31 (pfam01197)→ NhaP (COG0025)PRK05326 (PRK05326)→ |
| Esp-1-1 | NC_012673.1 | - | 1311255 | 1311222 | **RNA**→ rpmE2 (PRK01678)Ribosomal_L31 (pfam01197)→ NhaP (COG0025)PRK05326 (PRK05326)→ |
| Esp-6-1 | NZ_JPOD01000003.1 | + | 360593 | 360626 | **RNA**→ rpmE2 (PRK01678)Ribosomal_L31 (pfam01197)→ NhaP (COG0025)PRK05326 (PRK05326)→ |
| Ema-1-1 | NZ_KK211189.1 | + | 798722 | 798755 | **RNA**→ rpmE2 (PRK01678)Ribosomal_L31 (pfam01197)→ NhaP (COG0025)PRK05326 (PRK05326)→ |
| Eme-1-1 | NZ_JQGI01000001.1 | - | 1863165 | 1863132 | **RNA**→ rpmE2 (PRK01678)Ribosomal_L31 (pfam01197)→ NhaP (COG0025)PRK05326 (PRK05326)→ |
| Esp-2-1 | NZ_ATKK01000002.1 | - | 493876 | 493843 | **RNA**→ rpmE2 (PRK01678)Ribosomal_L31 (pfam01197)→ |
| Esp-10-1 | NZ_JTLB01000028.1 | - | 13345 | 13312 | **RNA**→ rpmE2 (PRK01678)Ribosomal_L31 (pfam01197)→ |
| Esp-3-1 | NZ_JNAA01000057.1 | - | 81113 | 81080 | **RNA**→ rpmE2 (PRK01678)Ribosomal_L31 (pfam01197)→ |
| Eal-1-1 | NZ_JJMW01000013.1 | + | 23661 | 23694 | **RNA**→ rpmE2 (PRK01678)Ribosomal_L31 (pfam01197)→ |
| Eau-1-1 | NZ_JNIQ01000001.1 | + | 873212 | 873245 | **RNA**→ rpmE2 (PRK01678)Ribosomal_L31 (pfam01197)→ |
| Ech-1-1 | NZ_JNIP01000001.1 | + | 361382 | 361419 | **RNA**→ rpmE2 (PRK01678)Ribosomal_L31 (pfam01197)→ |
| Epa-1-1 | NZ_ATCL01000014.1 | + | 231351 | 231388 | **RNA**→ rpmE2 (PRK01678)Ribosomal_L31 (pfam01197)→ |
| Esp-9-1 | NZ_ASXD01000010.1 | - | 15774 | 15737 | **RNA**→ rpmE2 (PRK01678)Ribosomal_L31 (pfam01197)→ NhaP (COG0025)PRK05326 (PRK05326)→ |
| | | | | | Tdk (COG1435)PRK04296 (PRK04296)→ |
| env-253 | 3300000558_10315482 | - | 274 | 246 | **RNA**→ rpmE (PRK00019)Ribosomal_L31 (pfam01197)→ |
| Lba-3-1 | NZ_KE159636.1 | - | 2955335 | 2955279 | **RNA**→ ArgC (COG0002)argC (PRK00436)→ ArgC (COG0002)argC (PRK00436)→ |
| | | | | | RimI (COG0456)Acetyltransf_7 (pfam13508)→ argJ (PRK05388)ArgJ (pfam01960)→ hypo→ |
| | | | | | PRK00942 (PRK00942)AAK_NAGK-C (cd04250)→ |
| env-254 | JGI24712J26585_10086008 | - | 716 | 686 | **RNA**→ rpmE (PRK00019)Ribosomal_L31 (pfam01197)→ Tdk (COG1435)PRK04296 (PRK04296)→ |
| env-255 | JGI24711J26586_10149403 | - | 518 | 488 | **RNA**→ rpmE (PRK00019)Ribosomal_L31 (pfam01197)→ Tdk (COG1435)PRK04296 (PRK04296)→ |
| env-256 | JGI24710J26742_10010214 | - | 6368 | 6338 | **RNA**→ rpmE (PRK00019)Ribosomal_L31 (pfam01197)→ Tdk (COG1435)PRK04296 (PRK04296)→ |
| | | | | | TadA (COG0590)MafB19-deam (pfam14437)→ hypo→ PRK10351 (PRK10351)ACPS (pfam01648)→ hypo- |
| | | | | | Peptidase_S9_N (pfam02897)→ TktA (COG0021)PRK05899 (PRK05899)→ |
| | | | | | MnmG (COG0445)PRK05192 (PRK05192)→ |
| env-257 | JGI24501J29690_1009900 | + | 30 | 60 | **RNA**→ rpmE (PRK00019)Ribosomal_L31 (pfam01197)→ Tdk (COG1435)PRK04296 (PRK04296)→ |
| | | | | | TadA (COG0590)MafB19-deam (pfam14437)→ hypo→ pantethn_trn (TIGR00556)ACPS (pfam01648)→ |
| env-258 | JGI24709J26583_10005562 | - | 7770 | 7740 | **RNA**→ rpmE (PRK00019)Ribosomal_L31 (pfam01197)→ Tdk (COG1435)PRK04296 (PRK04296)→ |
| | | | | | TadA (COG0590)MafB19-deam (pfam14437)→ hypo→ PRK10351 (PRK10351)ACPS (pfam01648)→ hypo- |
| | | | | | Peptidase_S9_N (pfam02897)→ TktA (COG0021)PRK05899 (PRK05899)→ |
| | | | | | MnmG (COG0445)PRK05192 (PRK05192)→ RsmG (COG0357)gidB (PRK00107)→ |
| env-259 | JGI24710J26742_10005602 | - | 43 | 13 | **RNA**→ |
| env-260 | JGI24500J29687_10005571 | + | 1219 | 1249 | **RNA**→ rpmE (PRK00019)Ribosomal_L31 (pfam01197)→ Tdk (COG1435)PRK04296 (PRK04296)→ |
| | | | | | TadA (COG0590)MafB19-deam (pfam14437)→ |
| env-261 | JGI24501J29690_1029221 | + | 1220 | 1250 | **RNA**→ |
| env-262 | JGI24710J26742_10016493 | + | 1755 | 1785 | **RNA**→ rpmE (PRK00019)Ribosomal_L31 (pfam01197)→ Tdk (COG1435)PRK04296 (PRK04296)→ |
| | | | | | TadA (COG0590)MafB19-deam (pfam14437)→ hypo→ PRK10351 (PRK10351)ACPS (pfam01648)→ hypo- |
| | | | | | Peptidase_S9_N (pfam02897)→ |
| env-263 | JGI24500J29687_10031739 | + | 4985 | 5015 | **RNA**→ |
| env-264 | JGI24502J29692_10005043 | + | 3736 | 3766 | **RNA**→ rpmE (PRK00019)Ribosomal_L31 (pfam01197)→ |
| env-265 | JGI24501J29690_1033374 | + | 4316 | 4346 | **RNA**→ |
| env-266 | JGI24503J29689_10039384 | + | 6502 | 6532 | **RNA**→ rpmE (PRK00019)Ribosomal_L31 (pfam01197)→ |
| env-267 | AUXO014528965.1 | + | 1985 | 2017 | **RNA**→ rpmE (PRK00019)Ribosomal_L31 (pfam01197)→ |
| env-268 | IMNBL1_c0363154 | + | 140 | 173 | **RNA**→ rpmE (PRK00019)Ribosomal_L31 (pfam01197)→ |
| Lki-4-1 | NC_014136.1 | + | 1067907 | 1067938 | **RNA**→ rpmE2 (PRK01678)Ribosomal_L31 (pfam01197)→ |
| Lsp-16-1 | NC_015734.1 | - | 1357217 | 1357186 | **RNA**→ rpmE2 (PRK01678)Ribosomal_L31 (pfam01197)→ |
| Gel-1-1 | NZ_KI391971.1 | - | 1673422 | 1673389 | **RNA**→ rpmE2 (PRK01678)Ribosomal_L31 (pfam01197)→ |
| env-269 | SRS043422_C2569170 | - | 275 | 242 | **RNA**→ rpmE2 (PRK01678)Ribosomal_L31 (pfam01197)→ |
| env-270 | Keratinized_gingiva_LANL_C5703483 | - | 63 | 30 | **RNA**→ |
| env-271 | SRS011310_C881309 | + | 999 | 1032 | **RNA**→ |
| env-272 | SRS014687_WUGC_scaffold_8481 | + | 7722 | 7755 | **RNA**→ rpmE2 (PRK01678)Ribosomal_L31 (pfam01197)→ |
| env-273 | SRS017025_Baylor_scaffold_6778 | - | 376 | 343 | **RNA**→ rpmE2 (PRK01678)Ribosomal_L31 (pfam01197)→ |
| env-274 | SRS019329_WUGC_scaffold_4099 | + | 368 | 401 | **RNA**→ |
| env-275 | SRS022532_LANL_scaffold_5089 | - | 4704 | 4671 | **RNA**→ rpmE2 (PRK01678)Ribosomal_L31 (pfam01197)→ |

| | | | | | |
|---|---|---|---|---|---|
| env-276 | SRS045049_C1523482 | + | 219 | 252 | RNA→ rpmE2 (PRK01678)Ribosomal_L31 (pfam01197)→ |
| env-277 | SRS045262_C1292262 | - | 269 | 236 | RNA→ rpmE2 (PRK01678)Ribosomal_L31 (pfam01197)→ |
| env-278 | SRS046686_C925134 | + | 3676 | 3709 | RNA→ rpmE2 (PRK01678)Ribosomal_L31 (pfam01197)→ |
| env-279 | SRS063272_LANL_scaffold_6091 | + | 2727 | 2760 | RNA→ rpmE2 (PRK01678)Ribosomal_L31 (pfam01197)→ |
| env-280 | SRS024281_C971068 | + | 203 | 236 | RNA→ rpmE2 (PRK01678)Ribosomal_L31 (pfam01197)→ |
| env-281 | SRS015040_WUGC_scaffold_8734 | + | 814 | 847 | RNA→ rpmE (PRK00019)rpmE2 (PRK01678)Ribosomal_L31 (pfam01197)→ |
| env-282 | SRS057022_LANL_scaffold_6226 | - | 269 | 236 | RNA→ rpmE2 (PRK01678)Ribosomal_L31 (pfam01197)→ |
| env-283 | SRS064809_C825416 | - | 269 | 236 | RNA→ rpmE2 (PRK01678)Ribosomal_L31 (pfam01197)→ |
| env-284 | SRS062878_LANL_scaffold_75258 | + | 1643 | 1676 | RNA→ rpmE2 (PRK01678)Ribosomal_L31 (pfam01197)→ |
| env-285 | SRS013946_WUGC_scaffold_12274 | + | 714 | 747 | RNA→ rpmE2 (PRK01678)Ribosomal_L31 (pfam01197)→ |
| env-286 | SRS019125_C601743 | + | 481 | 514 | RNA→ rpmE2 (PRK01678)Ribosomal_L31 (pfam01197)→ |
| env-287 | SRS013506_C1062755 | + | 32 | 65 | RNA→ rpmE2 (PRK01678)Ribosomal_L31 (pfam01197)→ |
| env-288 | SRS017215_C2571075 | - | 275 | 242 | RNA→ rpmE2 (PRK01678)Ribosomal_L31 (pfam01197)→ |
| env-289 | SRS019976_C721550 | - | 269 | 236 | RNA→ rpmE2 (PRK01678)Ribosomal_L31 (pfam01197)→ |
| env-290 | JGI994J12354_10282062 | - | 254 | 221 | RNA→ rpmE (PRK00019)Ribosomal_L31 (pfam01197)→ |
| env-291 | AUXO015979915.1 | + | 3013 | 3043 | RNA→ rpmE (PRK00019)Ribosomal_L31 (pfam01197)→ |
| Lsa-13-1 | NC_015978.1 | + | 397684 | 397715 | RNA→ rpmE2 (PRK01678)Ribosomal_L31 (pfam01197)→ |
| Cla-1-1 | NZ_KB899855.1 | - | 6693 | 6660 | RNA→ rpmE (PRK00019)Ribosomal_L31 (pfam01197)→ hypo→ |
| env-292 | JGI26057J44791_1087402 | - | 384 | 351 | RNA→ rpmE (PRK00019)Ribosomal_L31 (pfam01197)→ |
| Cth-2-1 | NZ_AUCP01000036.1 | - | 6324 | 6291 | RNA→ rpmE (PRK00019)Ribosomal_L31 (pfam01197)→ rpmE (PRK00019)Ribosomal_L31 (pfam01197)→ |
| env-293 | 2200552946 | - | 269 | 235 | RNA→ rpmE (PRK00019)Ribosomal_L31 (pfam01197)→ |
| env-294 | SRS043422_LANL_scaffold_14382 | - | 926 | 891 | RNA→ rpmE2 (PRK01678)Ribosomal_L31 (pfam01197)→ Tdk (COG1435)PRK04296 (PRK04296)→ |
| env-295 | SRS013946_WUGC_scaffold_5190 | + | 2187 | 2222 | RNA→ rpmE2 (PRK01678)Ribosomal_L31 (pfam01197)→ Tdk (COG1435)PRK04296 (PRK04296)→ |
| | | | | | PrfA (COG0216)prfA (PRK00591)→ HemK (COG2890)PRK09328 (PRK09328)→ |
| env-296 | SRS019125_C619186 | - | 154 | 119 | RNA→ rpmE2 (PRK01678)Ribosomal_L31 (pfam01197)→ |
| env-297 | SRS019591_WUGC_scaffold_32512 | + | 94 | 129 | RNA→ rpmE2 (PRK01678)Ribosomal_L31 (pfam01197)→ |
| env-298 | SRS042984_C4030266 | - | 687 | 652 | RNA→ rpmE2 (PRK01678)Ribosomal_L31 (pfam01197)→ |
| Gmo-1-1 | NZ_GL622607.1 | - | 253537 | 253502 | RNA→ rpmE2 (PRK01678)Ribosomal_L31 (pfam01197)→ |
| env-299 | SRS013723_C3219578 | - | 291 | 256 | RNA→ rpmE2 (PRK01678)Ribosomal_L31 (pfam01197)→ |
| env-300 | SRS047265_WUGC_scaffold_11943 | + | 10 | 45 | RNA→ hypo→ |
| Spa-1-1 | NZ_JUOB01000065.1 | + | 14924 | 14959 | RNA→ rpmE2 (PRK01678)Ribosomal_L31 (pfam01197)→ Tdk (COG1435)PRK04296 (PRK04296)→ |
| | | | | | PrfA (COG0216)prfA (PRK00591)→ HemK (COG2890)PRK09328 (PRK09328)→ |
| env-301 | SRS024017_C910720 | + | 74 | 109 | RNA→ rpmE2 (PRK01678)Ribosomal_L31 (pfam01197)→ |
| env-302 | SRS017088_Baylor_scaffold_355 | - | 773 | 738 | RNA→ rpmE2 (PRK01678)Ribosomal_L31 (pfam01197)→ Tdk (COG1435)PRK04296 (PRK04296)→ |
| env-303 | SRS011086_WUGC_scaffold_19008 | - | 3516 | 3481 | RNA→ rpmE2 (PRK01678)Ribosomal_L31 (pfam01197)→ Tdk (COG1435)PRK04296 (PRK04296)→ |
| env-304 | SRS062878_LANL_scaffold_75449 | + | 78 | 113 | RNA→ rpmE2 (PRK01678)Ribosomal_L31 (pfam01197)→ |
| env-305 | SRS064329_LANL_scaffold_50995 | - | 917 | 882 | RNA→ rpmE2 (PRK01678)Ribosomal_L31 (pfam01197)→ Tdk (COG1435)PRK04296 (PRK04296)→ |
| env-306 | SRS075404_C3430783 | - | 262 | 227 | RNA→ rpmE2 (PRK01678)Ribosomal_L31 (pfam01197)→ |
| env-307 | Contig7283290.1 | + | 2435 | 2470 | RNA→ rpmE2 (PRK01678)Ribosomal_L31 (pfam01197)→ Tdk (COG1435)PRK04296 (PRK04296)→ |
| | | | | | PrfA (COG0216)prfA (PRK00591)→ |
| env-308 | Tongue_dor_WUGI_scaffold_7283291 | + | 2435 | 2470 | RNA→ rpmE2 (PRK01678)Ribosomal_L31 (pfam01197)→ Tdk (COG1435)PRK04296 (PRK04296)→ |
| | | | | | PrfA (COG0216)prfA (PRK00591)→ |
| env-309 | SRS011140_Baylor_scaffold_32790 | + | 1053 | 1088 | RNA→ rpmE2 (PRK01678)Ribosomal_L31 (pfam01197)→ |
| env-310 | SRS011306_Baylor_scaffold_64442 | - | 2134 | 2099 | RNA→ rpmE2 (PRK01678)Ribosomal_L31 (pfam01197)→ Tdk (COG1435)PRK04296 (PRK04296)→ |
| | | | | | PrfA (COG0216)prfA (PRK00591)→ hypo→ |
| env-311 | SRS013164_Baylor_scaffold_712 | - | 2890 | 2855 | RNA→ rpmE2 (PRK01678)Ribosomal_L31 (pfam01197)→ Tdk (COG1435)PRK04296 (PRK04296)→ |
| | | | | | PrfA (COG0216)prfA (PRK00591)→ HemK (COG2890)PRK09328 (PRK09328)→ |
| env-312 | SRS013502_C2198940 | + | 2237 | 2272 | RNA→ rpmE2 (PRK01678)Ribosomal_L31 (pfam01197)→ |
| env-313 | SRS013879_WUGC_scaffold_47221 | + | 203 | 238 | RNA→ rpmE2 (PRK01678)Ribosomal_L31 (pfam01197)→ Tdk (COG1435)PRK04296 (PRK04296)→ |
| | | | | | PrfA (COG0216)prfA (PRK00591)→ |
| env-314 | SRS013947_C1620328 | - | 850 | 815 | RNA→ rpmE2 (PRK01678)Ribosomal_L31 (pfam01197)→ Tdk (COG1435)PRK04296 (PRK04296)→ |
| env-315 | SRS013948_C1539802 | + | 2096 | 2131 | RNA→ rpmE2 (PRK01678)Ribosomal_L31 (pfam01197)→ |
| env-316 | SRS014124_WUGC_scaffold_51107 | - | 1607 | 1572 | RNA→ rpmE2 (PRK01678)Ribosomal_L31 (pfam01197)→ Tdk (COG1435)PRK04296 (PRK04296)→ |
| | | | | | PrfA (COG0216)prfA (PRK00591)→ PCRF (smart00937)PCRF (pfam03462)→ |

| | | | | | |
|---|---|---|---|---|---|
| env-317 | SRS014271_C2958958 | + | 1617 | 1652 | **RNA**→ rpmE2 (PRK01678)Ribosomal_L31 (pfam01197)→ Tdk (COG1435)PRK04296 (PRK04296)→ PrfA (COG0216)prfA (PRK00591)→ |
| env-318 | SRS014470_C3747072 | + | 118 | 153 | **RNA**→ rpmE2 (PRK01678)Ribosomal_L31 (pfam01197)→ Tdk (COG1435)PRK04296 (PRK04296)→ PCRF (smart00937)PrfA (COG0216)→ |
| env-319 | SRS014573_WUGC_scaffold_12834 | + | 2379 | 2414 | **RNA**→ rpmE2 (PRK01678)Ribosomal_L31 (pfam01197)→ Tdk (COG1435)PRK04296 (PRK04296)→ PrfA (COG0216)prfA (PRK00591)→ HemK (COG2890)PRK09328 (PRK09328)→ |
| env-320 | SRS014684_WUGC_scaffold_38221 | + | 2822 | 2857 | **RNA**→ |
| env-321 | SRS014888_WUGC_scaffold_15364 | - | 262 | 227 | **RNA**→ rpmE2 (PRK01678)Ribosomal_L31 (pfam01197)→ |
| env-322 | SRS015038_C1778554 | + | 200 | 235 | **RNA**→ |
| env-323 | SRS015057_C3433924 | - | 538 | 503 | **RNA**→ rpmE2 (PRK01678)Ribosomal_L31 (pfam01197)→ Tdk (COG1435)PRK04296 (PRK04296)→ |
| env-324 | SRS015209_C1830245 | + | 66 | 101 | **RNA**→ rpmE2 (PRK01678)Ribosomal_L31 (pfam01197)→ Tdk (COG1435)PRK04296 (PRK04296)→ |
| env-325 | SRS015395_C2370059 | + | 417 | 452 | **RNA**→ rpmE2 (PRK01678)Ribosomal_L31 (pfam01197)→ |
| env-326 | SRS015644_WUGC_scaffold_3189 | - | 3443 | 3408 | **RNA**→ rpmE2 (PRK01678)Ribosomal_L31 (pfam01197)→ Tdk (COG1435)PRK04296 (PRK04296)→ PrfA (COG0216)prfA (PRK00591)→ HemK (COG2890)PRK09328 (PRK09328)→ |
| env-327 | SRS015762_WUGC_scaffold_41049 | + | 1448 | 1483 | **RNA**→ rpmE2 (PRK01678)Ribosomal_L31 (pfam01197)→ Tdk (COG1435)PRK04296 (PRK04296)→ |
| env-328 | SRS015893_C2246048 | + | 233 | 268 | **RNA**→ rpmE2 (PRK01678)Ribosomal_L31 (pfam01197)→ |
| env-329 | SRS015941_WUGC_scaffold_75574 | - | 3643 | 3608 | **RNA**→ rpmE2 (PRK01678)Ribosomal_L31 (pfam01197)→ Tdk (COG1435)PRK04296 (PRK04296)→ PrfA (COG0216)prfA (PRK00591)→ HemK (COG2890)PRK09328 (PRK09328)→ |
| env-330 | SRS016002_C2451425 | - | 191 | 156 | **RNA**→ rpmE2 (PRK01678)Ribosomal_L31 (pfam01197)→ |
| env-331 | SRS016037_WUGC_scaffold_43287 | + | 2263 | 2298 | **RNA**→ rpmE2 (PRK01678)Ribosomal_L31 (pfam01197)→ Tdk (COG1435)PRK04296 (PRK04296)→ PCRF (smart00937)PrfA (COG0216)→ |
| env-332 | SRS016319_C4011283 | + | 1508 | 1543 | **RNA**→ rpmE2 (PRK01678)Ribosomal_L31 (pfam01197)→ |
| env-333 | SRS016501_LANL_scaffold_17609 | + | 16039 | 16074 | **RNA**→ rpmE2 (PRK01678)Ribosomal_L31 (pfam01197)→ Tdk (COG1435)PRK04296 (PRK04296)→ PrfA (COG0216)prfA (PRK00591)→ HemK (COG2890)Methyltransf_25 (pfam13649)→ HemK (COG2890)PRK09328 (PRK09328)→ |
| env-334 | SRS016529_Baylor_scaffold_10943 | - | 3202 | 3167 | **RNA**→ rpmE2 (PRK01678)Ribosomal_L31 (pfam01197)→ Tdk (COG1435)PRK04296 (PRK04296)→ PrfA (COG0216)prfA (PRK00591)→ HemK (COG2890)PRK09328 (PRK09328)→ |
| env-335 | SRS016569_Baylor_scaffold_31963 | + | 118 | 153 | **RNA**→ rpmE2 (PRK01678)Ribosomal_L31 (pfam01197)→ Tdk (COG1435)PRK04296 (PRK04296)→ PrfA (COG0216)prfA (PRK00591)→ HemK (COG2890)PRK09328 (PRK09328)→ |
| env-336 | SRS017533_Baylor_scaffold_37499 | - | 805 | 770 | **RNA**→ rpmE2 (PRK01678)Ribosomal_L31 (pfam01197)→ Tdk (COG1435)PRK04296 (PRK04296)→ |
| env-337 | SRS017713_C3328455 | + | 417 | 452 | **RNA**→ rpmE2 (PRK01678)Ribosomal_L31 (pfam01197)→ |
| env-338 | SRS018300_Baylor_scaffold_22706 | + | 2963 | 2998 | **RNA**→ rpmE2 (PRK01678)Ribosomal_L31 (pfam01197)→ Tdk (COG1435)PRK04296 (PRK04296)→ |
| env-339 | SRS018357_C3945649 | - | 294 | 259 | **RNA**→ rpmE2 (PRK01678)Ribosomal_L31 (pfam01197)→ |
| env-340 | SRS018591_WUGC_scaffold_4474 | + | 2628 | 2663 | **RNA**→ rpmE2 (PRK01678)Ribosomal_L31 (pfam01197)→ Tdk (COG1435)PRK04296 (PRK04296)→ PrfA (COG0216)prfA (PRK00591)→ hypo→ |
| env-341 | SRS018739_C3708741 | - | 3712 | 3677 | **RNA**→ rpmE2 (PRK01678)Ribosomal_L31 (pfam01197)→ Tdk (COG1435)PRK04296 (PRK04296)→ PrfA (COG0216)prfA (PRK00591)→ HemK (COG2890)PRK09328 (PRK09328)→ |
| env-342 | SRS018969_C2941119 | - | 810 | 775 | **RNA**→ rpmE2 (PRK01678)Ribosomal_L31 (pfam01197)→ Tdk (COG1435)PRK04296 (PRK04296)→ |
| env-343 | SRS019022_WUGC_scaffold_46130 | + | 7621 | 7656 | **RNA**→ rpmE2 (PRK01678)Ribosomal_L31 (pfam01197)→ Tdk (COG1435)PRK04296 (PRK04296)→ PrfA (COG0216)prfA (PRK00591)→ hypo→ |
| env-344 | SRS019045_WUGC_scaffold_60915 | + | 699 | 734 | **RNA**→ rpmE2 (PRK01678)Ribosomal_L31 (pfam01197)→ Tdk (COG1435)PRK04296 (PRK04296)→ PrfA (COG0216)prfA (PRK00591)→ |
| env-345 | SRS019120_C1292577 | - | 263 | 228 | **RNA**→ rpmE2 (PRK01678)Ribosomal_L31 (pfam01197)→ |
| env-346 | SRS019219_WUGC_scaffold_73816 | - | 1354 | 1319 | **RNA**→ rpmE2 (PRK01678)Ribosomal_L31 (pfam01197)→ Tdk (COG1435)PRK04296 (PRK04296)→ PCRF (smart00937)PrfA (COG0216)→ |
| env-347 | SRS019327_WUGC_scaffold_24797 | - | 2643 | 2608 | **RNA**→ rpmE2 (PRK01678)Ribosomal_L31 (pfam01197)→ Tdk (COG1435)PRK04296 (PRK04296)→ PCRF (smart00937)RF-1 (pfam00472)PCRF (pfam03462)→ PrfA (COG0216)prfA (PRK00591)→ HemK (COG2890)PRK09328 (PRK09328)→ |
| env-348 | SRS019389_C2216180 | - | 368 | 333 | **RNA**→ rpmE2 (PRK01678)Ribosomal_L31 (pfam01197)→ |
| env-349 | SRS019607_C2754939 | + | 1076 | 1111 | **RNA**→ rpmE2 (PRK01678)Ribosomal_L31 (pfam01197)→ Tdk (COG1435)PRK04296 (PRK04296)→ PCRF (smart00937)PrfA (COG0216)→ |
| env-350 | SRS019894_C3902787 | + | 118 | 153 | **RNA**→ rpmE2 (PRK01678)Ribosomal_L31 (pfam01197)→ Tdk (COG1435)PRK04296 (PRK04296)→ |
| env-351 | SRS019974_Baylor_scaffold_34514 | - | 187 | 152 | **RNA**→ rpmE2 (PRK01678)Ribosomal_L31 (pfam01197)→ |
| env-352 | SRS020334_C2666501 | - | 1354 | 1319 | **RNA**→ rpmE2 (PRK01678)Ribosomal_L31 (pfam01197)→ Tdk (COG1435)PRK04296 (PRK04296)→ PCRF (smart00937)PrfA (COG0216)→ |
| env-353 | SRS020856_C2525502 | + | 118 | 153 | **RNA**→ rpmE2 (PRK01678)Ribosomal_L31 (pfam01197)→ |

| | | | | | |
|---|---|---|---|---|---|
| env-354 | SRS022077_Baylor_scaffold_56764 | - | 2255 | 2220 | RNA→ rpmE2 (PRK01678)Ribosomal_L31 (pfam01197)→ Tdk (COG1435)PRK04296 (PRK04296)→ PrfA (COG0216)prfA (PRK00591)→ hypo→ |
| env-355 | SRS022530_LANL_scaffold_14562 | - | 5660 | 5625 | RNA→ rpmE2 (PRK01678)Ribosomal_L31 (pfam01197)→ Tdk (COG1435)PRK04296 (PRK04296)→ PrfA (COG0216)prfA (PRK00591)→ HemK (COG2890)PRK09328 (PRK09328)→ |
| env-356 | SRS022719_LANL_scaffold_70646 | + | 27983 | 28018 | RNA→ rpmE2 (PRK01678)Ribosomal_L31 (pfam01197)→ Tdk (COG1435)PRK04296 (PRK04296)→ PrfA (COG0216)prfA (PRK00591)→ HemK (COG2890)PRK09328 (PRK09328)→ |
| env-357 | SRS023557_Baylor_scaffold_14708 | + | 229 | 264 | RNA→ rpmE2 (PRK01678)Ribosomal_L31 (pfam01197)→ Tdk (COG1435)PRK04296 (PRK04296)→ PrfA (COG0216)prfA (PRK00591)→ HemK (COG2890)PRK09328 (PRK09328)→ |
| env-358 | SRS023617_Baylor_scaffold_71319 | - | 3925 | 3890 | RNA→ rpmE2 (PRK01678)Ribosomal_L31 (pfam01197)→ Tdk (COG1435)PRK04296 (PRK04296)→ PrfA (COG0216)prfA (PRK00591)→ HemK (COG2890)PRK09328 (PRK09328)→ |
| env-359 | SRS024015_Baylor_scaffold_40979 | - | 1229 | 1194 | RNA→ rpmE2 (PRK01678)Ribosomal_L31 (pfam01197)→ Tdk (COG1435)PRK04296 (PRK04296)→ PrfA (COG0216)prfA (PRK00591)→ |
| env-360 | SRS024081_LANL_scaffold_2024 | + | 46179 | 46214 | RNA→ rpmE2 (PRK01678)Ribosomal_L31 (pfam01197)→ Tdk (COG1435)PRK04296 (PRK04296)→ PrfA (COG0216)prfA (PRK00591)→ HemK (COG2890)PRK09328 (PRK09328)→ |
| env-361 | SRS024277_LANL_scaffold_37737 | - | 2929 | 2894 | RNA→ rpmE2 (PRK01678)Ribosomal_L31 (pfam01197)→ Tdk (COG1435)PRK04296 (PRK04296)→ PrfA (COG0216)prfA (PRK00591)→ HemK (COG2890)PRK09328 (PRK09328)→ |
| env-362 | SRS024375_LANL_scaffold_1816 | - | 937 | 902 | RNA→ rpmE2 (PRK01678)Ribosomal_L31 (pfam01197)→ Tdk (COG1435)PRK04296 (PRK04296)→ |
| env-363 | SRS024580_C3090129 | - | 224 | 189 | RNA→ rpmE2 (PRK01678)Ribosomal_L31 (pfam01197)→ |
| env-364 | SRS042131_WUGC_scaffold_29424 | + | 4098 | 4133 | RNA→ rpmE2 (PRK01678)Ribosomal_L31 (pfam01197)→ Tdk (COG1435)PRK04296 (PRK04296)→ |
| env-365 | SRS042643_C3133050 | + | 468 | 503 | RNA→ rpmE2 (PRK01678)Ribosomal_L31 (pfam01197)→ Tdk (COG1435)PRK04296 (PRK04296)→ PrfA (COG0216)prfA (PRK00591)→ HemK (COG2890)PRK09328 (PRK09328)→ |
| env-366 | SRS042910_C5598748 | + | 491 | 526 | RNA→ rpmE2 (PRK01678)Ribosomal_L31 (pfam01197)→ |
| env-367 | SRS043663_LANL_scaffold_3211 | + | 5442 | 5477 | RNA→ rpmE2 (PRK01678)Ribosomal_L31 (pfam01197)→ Tdk (COG1435)PRK04296 (PRK04296)→ PrfA (COG0216)prfA (PRK00591)→ |
| env-368 | SRS044662_LANL_scaffold_39333 | + | 2021 | 2056 | RNA→ rpmE2 (PRK01678)Ribosomal_L31 (pfam01197)→ Tdk (COG1435)PRK04296 (PRK04296)→ PrfA (COG0216)prfA (PRK00591)→ HemK (COG2890)PRK09328 (PRK09328)→ |
| env-369 | SRS045127_LANL_scaffold_35833 | + | 349 | 384 | RNA→ rpmE2 (PRK01678)Ribosomal_L31 (pfam01197)→ Tdk (COG1435)PRK04296 (PRK04296)→ |
| env-370 | SRS045715_LANL_scaffold_77986 | - | 3368 | 3333 | RNA→ rpmE2 (PRK01678)Ribosomal_L31 (pfam01197)→ Tdk (COG1435)PRK04296 (PRK04296)→ PrfA (COG0216)prfA (PRK00591)→ HemK (COG2890)PRK09328 (PRK09328)→ |
| env-371 | SRS047210_C3329896 | + | 1570 | 1605 | RNA→ rpmE2 (PRK01678)Ribosomal_L31 (pfam01197)→ Tdk (COG1435)PRK04296 (PRK04296)→ |
| env-372 | SRS048411_C6946699 | - | 702 | 667 | RNA→ rpmE2 (PRK01678)Ribosomal_L31 (pfam01197)→ Tdk (COG1435)PRK04296 (PRK04296)→ |
| env-373 | SRS048791_C3209121 | + | 266 | 301 | RNA→ rpmE2 (PRK01678)Ribosomal_L31 (pfam01197)→ |
| env-374 | SRS049147_C4027824 | - | 262 | 227 | RNA→ rpmE2 (PRK01678)Ribosomal_L31 (pfam01197)→ |
| env-375 | SRS049389_C4719865 | - | 154 | 119 | RNA→ rpmE2 (PRK01678)Ribosomal_L31 (pfam01197)→ |
| env-376 | SRS050244_LANL_scaffold_70126 | + | 6532 | 6567 | RNA→ rpmE2 (PRK01678)Ribosomal_L31 (pfam01197)→ Tdk (COG1435)PRK04296 (PRK04296)→ PCRF (smart00937)PrfA (COG0216)→ |
| env-377 | SRS050669_LANL_scaffold_8194 | - | 943 | 908 | RNA→ rpmE2 (PRK01678)Ribosomal_L31 (pfam01197)→ Tdk (COG1435)PRK04296 (PRK04296)→ |
| env-378 | SRS051791_LANL_scaffold_43760 | + | 9622 | 9657 | RNA→ rpmE2 (PRK01678)Ribosomal_L31 (pfam01197)→ Tdk (COG1435)PRK04296 (PRK04296)→ PrfA (COG0216)prfA (PRK00591)→ HemK (COG2890)PRK09328 (PRK09328)→ |
| env-379 | SRS052227_LANL_scaffold_52022 | - | 2929 | 2894 | RNA→ rpmE2 (PRK01678)Ribosomal_L31 (pfam01197)→ Tdk (COG1435)PRK04296 (PRK04296)→ PrfA (COG0216)prfA (PRK00591)→ HemK (COG2890)PRK09328 (PRK09328)→ |
| env-380 | SRS054687_C3486600 | + | 417 | 452 | RNA→ rpmE2 (PRK01678)Ribosomal_L31 (pfam01197)→ Tdk (COG1435)PRK04296 (PRK04296)→ |
| env-381 | SRS055426_LANL_scaffold_55091 | - | 2268 | 2233 | RNA→ rpmE2 (PRK01678)Ribosomal_L31 (pfam01197)→ Tdk (COG1435)PRK04296 (PRK04296)→ PrfA (COG0216)prfA (PRK00591)→ hypo→ |
| env-382 | SRS056622_LANL_scaffold_45323 | + | 692 | 727 | RNA→ rpmE2 (PRK01678)Ribosomal_L31 (pfam01197)→ Tdk (COG1435)PRK04296 (PRK04296)→ PrfA (COG0216)prfA (PRK00591)→ HemK (COG2890)PRK09328 (PRK09328)→ |
| env-383 | SRS057205_LANL_scaffold_27098 | + | 2436 | 2471 | RNA→ rpmE2 (PRK01678)Ribosomal_L31 (pfam01197)→ |
| env-384 | SRS057539_LANL_scaffold_62110 | + | 203 | 238 | RNA→ rpmE2 (PRK01678)Ribosomal_L31 (pfam01197)→ Tdk (COG1435)PRK04296 (PRK04296)→ |
| env-385 | SRS057791_LANL_scaffold_34993 | + | 1095 | 1130 | RNA→ rpmE2 (PRK01678)Ribosomal_L31 (pfam01197)→ Tdk (COG1435)PRK04296 (PRK04296)→ PrfA (COG0216)prfA (PRK00591)→ HemK (COG2890)PRK09328 (PRK09328)→ |
| env-386 | SRS058336_C1967939 | - | 245 | 210 | RNA→ rpmE2 (PRK01678)Ribosomal_L31 (pfam01197)→ |
| env-387 | SRS062544_C2916403 | + | 186 | 221 | RNA→ rpmE2 (PRK01678)Ribosomal_L31 (pfam01197)→ |
| env-388 | SRS062761_LANL_scaffold_51063 | - | 2240 | 2205 | RNA→ rpmE2 (PRK01678)Ribosomal_L31 (pfam01197)→ Tdk (COG1435)PRK04296 (PRK04296)→ PrfA (COG0216)prfA (PRK00591)→ hypo→ |
| env-389 | SRS063193_LANL_scaffold_65781 | + | 1299 | 1334 | RNA→ rpmE2 (PRK01678)Ribosomal_L31 (pfam01197)→ Tdk (COG1435)PRK04296 (PRK04296)→ PrfA (COG0216)prfA (PRK00591)→ hypo→ |

| | | | | | |
|---|---|---|---|---|---|
| env-390 | SRS063288_C2613515 | - | 1918 | 1883 | **RNA**→ rpmE2 (PRK01678)Ribosomal_L31 (pfam01197)→ Tdk (COG1435)PRK04296 (PRK04296)→ PrfA (COG0216)prfA (PRK00591)→ |
| env-391 | SRS064423_LANL_scaffold_69154 | - | 1354 | 1319 | **RNA**→ rpmE2 (PRK01678)Ribosomal_L31 (pfam01197)→ Tdk (COG1435)PRK04296 (PRK04296)→ PCRF (smart00937)PrfA (COG0216)→ |
| env-392 | SRS064774_C2890698 | - | 627 | 592 | **RNA**→ rpmE2 (PRK01678)Ribosomal_L31 (pfam01197)→ Tdk (COG1435)PRK04296 (PRK04296)→ |
| env-393 | SRS065278_C2548946 | + | 1202 | 1237 | **RNA**→ rpmE2 (PRK01678)Ribosomal_L31 (pfam01197)→ |
| env-394 | SRS077736_LANL_scaffold_17623 | + | 229 | 264 | **RNA**→ rpmE2 (PRK01678)Ribosomal_L31 (pfam01197)→ Tdk (COG1435)PRK04296 (PRK04296)→ PrfA (COG0216)prfA (PRK00591)→ HemK (COG2890)Methyltransf_25 (pfam13649)→ |
| Gsa-2-1 | NZ_ACRY02000001.1 | + | 10104 | 10139 | **RNA**→ rpmE2 (PRK01678)Ribosomal_L31 (pfam01197)→ Tdk (COG1435)PRK04296 (PRK04296)→ PrfA (COG0216)prfA (PRK00591)→ hypo→ HemK (COG2890)PRK09328 (PRK09328)→ |
| env-395 | SRS015540_C1145684 | - | 441 | 406 | **RNA**→ rpmE2 (PRK01678)Ribosomal_L31 (pfam01197)→ Tdk (COG1435)PRK04296 (PRK04296)→ |
| env-396 | SRS015985_WUGC_scaffold_1374 | + | 106 | 141 | **RNA**→ rpmE2 (PRK01678)Ribosomal_L31 (pfam01197)→ |
| env-397 | SRS062878_LANL_scaffold_77532 | + | 103 | 138 | **RNA**→ rpmE2 (PRK01678)Ribosomal_L31 (pfam01197)→ |
| env-398 | SRS014686_C893659 | - | 161 | 126 | **RNA**→ rpmE2 (PRK01678)Ribosomal_L31 (pfam01197)→ |
| env-399 | SRS014690_WUGC_scaffold_25959 | - | 648 | 613 | **RNA**→ rpmE2 (PRK01678)Ribosomal_L31 (pfam01197)→ Tdk (COG1435)PRK04296 (PRK04296)→ |
| env-400 | SRS015799_C834453 | + | 101 | 136 | **RNA**→ rpmE2 (PRK01678)Ribosomal_L31 (pfam01197)→ |
| env-401 | SRS015895_C570099 | - | 828 | 793 | **RNA**→ rpmE2 (PRK01678)Ribosomal_L31 (pfam01197)→ Tdk (COG1435)PRK04296 (PRK04296)→ |
| env-402 | SRS015989_C2178125 | + | 3039 | 3074 | **RNA**→ rpmE2 (PRK01678)Ribosomal_L31 (pfam01197)→ Tdk (COG1435)PRK04296 (PRK04296)→ |
| env-403 | SRS017025_C1697043 | - | 417 | 382 | **RNA**→ rpmE2 (PRK01678)Ribosomal_L31 (pfam01197)→ |
| env-404 | SRS017215_Baylor_scaffold_6391 | + | 667 | 702 | **RNA**→ rpmE2 (PRK01678)Ribosomal_L31 (pfam01197)→ Tdk (COG1435)PRK04296 (PRK04296)→ |
| env-405 | SRS018971_WUGC_scaffold_9088 | + | 2824 | 2859 | **RNA**→ rpmE2 (PRK01678)Ribosomal_L31 (pfam01197)→ Tdk (COG1435)PRK04296 (PRK04296)→ PrfA (COG0216)prfA (PRK00591)→ HemK (COG2890)PRK09328 (PRK09328)→ |
| env-406 | SRS019587_C1062782 | + | 2489 | 2524 | **RNA**→ rpmE2 (PRK01678)Ribosomal_L31 (pfam01197)→ Tdk (COG1435)PRK04296 (PRK04296)→ |
| env-407 | SRS022625_Baylor_scaffold_8635 | + | 256 | 291 | **RNA**→ rpmE2 (PRK01678)Ribosomal_L31 (pfam01197)→ |
| env-408 | SRS023987_Baylor_scaffold_8935 | + | 101 | 136 | **RNA**→ rpmE2 (PRK01678)Ribosomal_L31 (pfam01197)→ |
| env-409 | SRS024447_LANL_scaffold_55871 | + | 3040 | 3075 | **RNA**→ rpmE2 (PRK01678)Ribosomal_L31 (pfam01197)→ Tdk (COG1435)PRK04296 (PRK04296)→ PrfA (COG0216)prfA (PRK00591)→ |
| env-410 | SRS024470_C1034718 | - | 291 | 256 | **RNA**→ rpmE2 (PRK01678)Ribosomal_L31 (pfam01197)→ |
| env-411 | SRS043755_C2262638 | + | 181 | 216 | **RNA**→ rpmE2 (PRK01678)Ribosomal_L31 (pfam01197)→ |
| env-412 | SRS050029_WUGC_scaffold_8539 | - | 2986 | 2951 | **RNA**→ rpmE2 (PRK01678)Ribosomal_L31 (pfam01197)→ Tdk (COG1435)PRK04296 (PRK04296)→ PrfA (COG0216)prfA (PRK00591)→ HemK (COG2890)PRK09328 (PRK09328)→ |
| env-413 | SRS054776_C1596217 | + | 158 | 193 | **RNA**→ rpmE2 (PRK01678)Ribosomal_L31 (pfam01197)→ |
| env-414 | SRS056892_LANL_scaffold_10358 | - | 1175 | 1140 | **RNA**→ rpmE2 (PRK01678)Ribosomal_L31 (pfam01197)→ Tdk (COG1435)PRK04296 (PRK04296)→ PrfA (COG0216)prfA (PRK00591)→ |
| env-415 | SRS063272_C727711 | + | 103 | 138 | **RNA**→ rpmE2 (PRK01678)Ribosomal_L31 (pfam01197)→ |
| Gha-3-1 | NZ_GL883582.1 | + | 200120 | 200155 | **RNA**→ rpmE2 (PRK01678)Ribosomal_L31 (pfam01197)→ Tdk (COG1435)PRK04296 (PRK04296)→ PrfA (COG0216)prfA (PRK00591)→ HemK (COG2890)PRK09328 (PRK09328)→ hypo→ |
| env-416 | Keratinized_gingiva_LANL_scaffold_12580 | - | 518 | 483 | **RNA**→ rpmE2 (PRK01678)Ribosomal_L31 (pfam01197)→ Tdk (COG1435)PRK04296 (PRK04296)→ |
| env-417 | SRS020222_C1000314 | + | 31 | 66 | **RNA**→ rpmE2 (PRK01678)Ribosomal_L31 (pfam01197)→ |
| env-418 | SRS021986_Baylor_scaffold_5911 | + | 3030 | 3065 | **RNA**→ rpmE2 (PRK01678)Ribosomal_L31 (pfam01197)→ |
| env-419 | SRS024281_C990110 | - | 3626 | 3591 | **RNA**→ rpmE2 (PRK01678)Ribosomal_L31 (pfam01197)→ Tdk (COG1435)PRK04296 (PRK04296)→ PrfA (COG0216)prfA (PRK00591)→ HemK (COG2890)PRK09328 (PRK09328)→ |
| env-420 | SRS055495_C1103822 | + | 223 | 258 | **RNA**→ rpmE2 (PRK01678)Ribosomal_L31 (pfam01197)→ |
| env-421 | SRS056157_LANL_scaffold_3130 | - | 22443 | 22408 | **RNA**→ rpmE2 (PRK01678)Ribosomal_L31 (pfam01197)→ Tdk (COG1435)PRK04296 (PRK04296)→ PrfA (COG0216)prfA (PRK00591)→ HemK (COG2890)PRK09328 (PRK09328)→ |
| env-422 | SRS011247_Baylor_scaffold_24154 | + | 56 | 91 | **RNA**→ rpmE2 (PRK01678)Ribosomal_L31 (pfam01197)→ Tdk (COG1435)PRK04296 (PRK04296)→ PCRF (smart00937)PCRF (pfam03462)→ |
| env-423 | SRS013506_C1071539 | - | 305 | 270 | **RNA**→ rpmE2 (PRK01678)Ribosomal_L31 (pfam01197)→ |
| env-424 | SRS013711_C1759566 | + | 247 | 282 | **RNA**→ rpmE2 (PRK01678)Ribosomal_L31 (pfam01197)→ |
| env-425 | SRS013947_WUGC_scaffold_13565 | + | 157 | 192 | **RNA**→ rpmE2 (PRK01678)Ribosomal_L31 (pfam01197)→ Tdk (COG1435)PRK04296 (PRK04296)→ |
| env-426 | SRS014687_WUGC_scaffold_3449 | - | 286 | 251 | **RNA**→ rpmE2 (PRK01678)Ribosomal_L31 (pfam01197)→ |
| env-427 | SRS014691_C2541310 | + | 149 | 184 | **RNA**→ rpmE2 (PRK01678)Ribosomal_L31 (pfam01197)→ |
| env-428 | SRS015040_WUGC_scaffold_9073 | + | 246 | 281 | **RNA**→ rpmE2 (PRK01678)Ribosomal_L31 (pfam01197)→ |
| env-429 | SRS015154_C749644 | + | 103 | 138 | **RNA**→ rpmE2 (PRK01678)Ribosomal_L31 (pfam01197)→ Tdk (COG1435)PRK04296 (PRK04296)→ |
| env-430 | SRS015374_C303312 | + | 108 | 143 | **RNA**→ rpmE2 (PRK01678)Ribosomal_L31 (pfam01197)→ |

| | | | | | |
|---|---|---|---|---|---|
| env-431 | SRS015378_C1036454 | + | 558 | 593 | **RNA**→ rpmE2 (PRK01678)Ribosomal_L31 (pfam01197)→ |
| env-432 | SRS015436_C1704957 | - | 351 | 316 | **RNA**→ rpmE2 (PRK01678)Ribosomal_L31 (pfam01197)→ |
| env-433 | SRS015440_WUGC_scaffold_48560 | - | 6395 | 6360 | **RNA**→ rpmE2 (PRK01678)Ribosomal_L31 (pfam01197)→ Tdk (COG1435)PRK04296 (PRK04296)→ |
| | | | | | PrfA (COG0216)prfA (PRK00591)→ HemK (COG2890)PRK09328 (PRK09328)→ |
| env-434 | SRS015470_WUGC_scaffold_63490 | - | 348 | 313 | **RNA**→ rpmE2 (PRK01678)Ribosomal_L31 (pfam01197)→ |
| env-435 | SRS015745_WUGC_scaffold_6005 | - | 482 | 447 | **RNA**→ rpmE2 (PRK01678)Ribosomal_L31 (pfam01197)→ Tdk (COG1435)PRK04296 (PRK04296)→ |
| env-436 | SRS016196_C484943 | + | 277 | 312 | **RNA**→ rpmE2 (PRK01678)Ribosomal_L31 (pfam01197)→ Tdk (COG1435)PRK04296 (PRK04296)→ |
| env-437 | SRS016600_Baylor_scaffold_2683 | + | 3030 | 3065 | **RNA**→ rpmE2 (PRK01678)Ribosomal_L31 (pfam01197)→ Tdk (COG1435)PRK04296 (PRK04296)→ |
| | | | | | PrfA (COG0216)prfA (PRK00591)→ HemK (COG2890)PRK09328 (PRK09328)→ |
| env-438 | SRS017013_Baylor_scaffold_6676 | - | 592 | 557 | **RNA**→ rpmE2 (PRK01678)Ribosomal_L31 (pfam01197)→ |
| env-439 | SRS018329_C1551846 | + | 51 | 86 | **RNA**→ rpmE2 (PRK01678)Ribosomal_L31 (pfam01197)→ |
| env-440 | SRS018359_C1220600 | - | 348 | 313 | **RNA**→ rpmE2 (PRK01678)Ribosomal_L31 (pfam01197)→ |
| env-441 | SRS019073_C2443412 | - | 309 | 274 | **RNA**→ rpmE2 (PRK01678)Ribosomal_L31 (pfam01197)→ |
| env-442 | SRS019125_C619240 | - | 154 | 119 | **RNA**→ rpmE2 (PRK01678)Ribosomal_L31 (pfam01197)→ |
| env-443 | SRS019126_C2186810 | + | 232 | 267 | **RNA**→ rpmE2 (PRK01678)Ribosomal_L31 (pfam01197)→ Tdk (COG1435)PRK04296 (PRK04296)→ |
| env-444 | SRS019221_C295755 | + | 3024 | 3059 | **RNA**→ rpmE2 (PRK01678)Ribosomal_L31 (pfam01197)→ Tdk (COG1435)PRK04296 (PRK04296)→ |
| env-445 | SRS019976_C716661 | + | 59 | 94 | **RNA**→ rpmE2 (PRK01678)Ribosomal_L31 (pfam01197)→ Tdk (COG1435)PRK04296 (PRK04296)→ |
| env-446 | SRS020858_Baylor_scaffold_5085 | + | 3032 | 3067 | **RNA**→ rpmE2 (PRK01678)Ribosomal_L31 (pfam01197)→ Tdk (COG1435)PRK04296 (PRK04296)→ |
| | | | | | PrfA (COG0216)prfA (PRK00591)→ HemK (COG2890)PRK09328 (PRK09328)→ |
| env-447 | SRS021473_C876560 | - | 603 | 568 | **RNA**→ rpmE2 (PRK01678)Ribosomal_L31 (pfam01197)→ Tdk (COG1435)PRK04296 (PRK04296)→ |
| env-448 | SRS022079_C372086 | - | 276 | 241 | **RNA**→ rpmE2 (PRK01678)Ribosomal_L31 (pfam01197)→ |
| env-449 | SRS022145_C598868 | - | 166 | 131 | **RNA**→ rpmE2 (PRK01678)Ribosomal_L31 (pfam01197)→ |
| env-450 | SRS022532_LANL_scaffold_6630 | - | 1673 | 1638 | **RNA**→ rpmE2 (PRK01678)Ribosomal_L31 (pfam01197)→ Tdk (COG1435)PRK04296 (PRK04296)→ |
| | | | | | Tdk (COG1435)PRK04296 (PRK04296)→ hypo→ PrfA (COG0216)prfA (PRK00591)→ |
| env-451 | SRS022721_LANL_scaffold_429 | + | 210 | 245 | **RNA**→ rpmE2 (PRK01678)Ribosomal_L31 (pfam01197)→ |
| env-452 | SRS023837_C579087 | - | 296 | 261 | **RNA**→ rpmE2 (PRK01678)Ribosomal_L31 (pfam01197)→ |
| env-453 | SRS024140_C1189075 | + | 4828 | 4863 | **RNA**→ rpmE2 (PRK01678)Ribosomal_L31 (pfam01197)→ |
| env-454 | SRS024377_LANL_scaffold_4712 | - | 1160 | 1125 | **RNA**→ rpmE2 (PRK01678)Ribosomal_L31 (pfam01197)→ Tdk (COG1435)PRK04296 (PRK04296)→ |
| | | | | | PrfA (COG0216)prfA (PRK00591)→ |
| env-455 | SRS042457_C486662 | - | 351 | 316 | **RNA**→ rpmE2 (PRK01678)Ribosomal_L31 (pfam01197)→ |
| env-456 | SRS045254_C914869 | - | 3935 | 3900 | **RNA**→ rpmE2 (PRK01678)Ribosomal_L31 (pfam01197)→ Tdk (COG1435)PRK04296 (PRK04296)→ |
| | | | | | PrfA (COG0216)prfA (PRK00591)→ HemK (COG2890)PRK09328 (PRK09328)→ |
| env-457 | SRS045978_C985323 | + | 72 | 107 | **RNA**→ rpmE2 (PRK01678)Ribosomal_L31 (pfam01197)→ |
| env-458 | SRS050628_LANL_scaffold_17394 | + | 458 | 493 | **RNA**→ rpmE2 (PRK01678)Ribosomal_L31 (pfam01197)→ |
| env-459 | SRS051116_C251615 | + | 534 | 569 | **RNA**→ rpmE2 (PRK01678)Ribosomal_L31 (pfam01197)→ |
| env-460 | SRS052668_LANL_scaffold_27772 | + | 2906 | 2941 | **RNA**→ rpmE2 (PRK01678)Ribosomal_L31 (pfam01197)→ Tdk (COG1435)PRK04296 (PRK04296)→ |
| | | | | | PrfA (COG0216)prfA (PRK00591)→ |
| env-461 | SRS052874_LANL_scaffold_5089 | - | 290 | 255 | **RNA**→ hypo→ |
| env-462 | SRS054569_LANL_scaffold_12877 | - | 354 | 319 | **RNA**→ rpmE2 (PRK01678)Ribosomal_L31 (pfam01197)→ |
| env-463 | SRS058105_LANL_scaffold_13652 | + | 469 | 504 | **RNA**→ rpmE2 (PRK01678)Ribosomal_L31 (pfam01197)→ |
| env-464 | SRS064809_C824160 | - | 603 | 568 | **RNA**→ rpmE2 (PRK01678)Ribosomal_L31 (pfam01197)→ Tdk (COG1435)PRK04296 (PRK04296)→ |
| env-465 | SRS075406_C835742 | - | 348 | 313 | **RNA**→ rpmE2 (PRK01678)Ribosomal_L31 (pfam01197)→ |
| env-466 | SRS077738_LANL_scaffold_4679 | - | 647 | 612 | **RNA**→ rpmE2 (PRK01678)Ribosomal_L31 (pfam01197)→ Tdk (COG1435)PRK04296 (PRK04296)→ |
| Gha-1-1 | NZ_ACDZ02000014.1 | - | 557531 | 557496 | **RNA**→ rpmE2 (PRK01678)Ribosomal_L31 (pfam01197)→ rpmE2 (PRK01678)Ribosomal_L31 (pfam01197 |
| | | | | | Tdk (COG1435)PRK04296 (PRK04296)→ Tdk (COG1435)PRK04296 (PRK04296)→ |
| | | | | | PrfA (COG0216)prfA (PRK00591)→ PrfA (COG0216)prfA (PRK00591)→ |
| | | | | | HemK (COG2890)PRK09328 (PRK09328)→ HemK (COG2890)PRK09328 (PRK09328)→ |
| env-467 | SRS053917_LANL_scaffold_15970 | - | 444 | 409 | **RNA**→ rpmE2 (PRK01678)Ribosomal_L31 (pfam01197)→ |
| env-468 | Subgingival_plaque_LANL_C11235616 | - | 423 | 388 | **RNA**→ rpmE2 (PRK01678)Ribosomal_L31 (pfam01197)→ |
| env-469 | SRS012285_Baylor_scaffold_61830 | - | 316 | 281 | **RNA**→ rpmE2 (PRK01678)Ribosomal_L31 (pfam01197)→ |
| env-470 | SRS013252_C3418322 | - | 215 | 180 | **RNA**→ rpmE2 (PRK01678)Ribosomal_L31 (pfam01197)→ |
| env-471 | SRS015574_C3437002 | + | 238 | 273 | **RNA**→ rpmE2 (PRK01678)Ribosomal_L31 (pfam01197)→ |
| env-472 | SRS022725_C5893417 | - | 3760 | 3725 | **RNA**→ rpmE2 (PRK01678)Ribosomal_L31 (pfam01197)→ |
| env-473 | SRS047634_C4297156 | - | 346 | 311 | **RNA**→ rpmE2 (PRK01678)Ribosomal_L31 (pfam01197)→ |

| | | | | | |
|---|---|---|---|---|---|
| env-474 | SRS049268_C4258038 | - | 364 | 329 | RNA→ rpmE2 (PRK01678)Ribosomal_L31 (pfam01197)→ |
| env-475 | SRS049318_LANL_scaffold_177316 | - | 852 | 817 | RNA→ rpmE2 (PRK01678)Ribosomal_L31 (pfam01197)→ |
| env-476 | SRS049389_C4591629 | - | 154 | 119 | RNA→ rpmE2 (PRK01678)Ribosomal_L31 (pfam01197)→ |
| env-477 | SRS051244_C3395339 | - | 351 | 316 | RNA→ rpmE2 (PRK01678)Ribosomal_L31 (pfam01197)→ |
| env-478 | SRS063932_C3959092 | - | 4053 | 4018 | RNA→ rpmE2 (PRK01678)Ribosomal_L31 (pfam01197)→ |
| env-479 | SRS014126_C559610 | - | 349 | 314 | RNA→ rpmE2 (PRK01678)Ribosomal_L31 (pfam01197)→ |
| env-480 | SRS015060_C1555206 | - | 905 | 870 | RNA→ rpmE2 (PRK01678)Ribosomal_L31 (pfam01197)→ Tdk (COG1435)PRK04296 (PRK04296)→ |
| env-481 | SRS022536_LANL_scaffold_96129 | - | 893 | 858 | RNA→ rpmE2 (PRK01678)Ribosomal_L31 (pfam01197)→ |
| env-482 | SRS017511_C3422862 | + | 34 | 69 | RNA→ rpmE2 (PRK01678)Ribosomal_L31 (pfam01197)→ |
| env-483 | SRS015537_C1260669 | + | 407 | 442 | RNA→ rpmE2 (PRK01678)Ribosomal_L31 (pfam01197)→ |
| env-484 | SRS019126_WUGC_scaffold_4752 | - | 943 | 908 | RNA→ rpmE2 (PRK01678)Ribosomal_L31 (pfam01197)→ Tdk (COG1435)PRK04296 (PRK04296)→ |
| env-485 | SRS020220_C3566657 | - | 510 | 475 | RNA→ rpmE2 (PRK01678)Ribosomal_L31 (pfam01197)→ Tdk (COG1435)PRK04296 (PRK04296)→ |
| env-486 | SRS024441_LANL_scaffold_37855 | + | 2333 | 2368 | RNA→ rpmE2 (PRK01678)Ribosomal_L31 (pfam01197)→ Tdk (COG1435)PRK04296 (PRK04296)→ |
| | | | | | PrfA (COG0216)prfA (PRK00591)→ HemK (COG2890)PRK09328 (PRK09328)→ |
| Gsa-1-1 | NZ_JNJO01000009.1 | - | 39706 | 39671 | RNA→ rpmE2 (PRK01678)Ribosomal_L31 (pfam01197)→ Tdk (COG1435)PRK04296 (PRK04296)→ |
| | | | | | PrfA (COG0216)prfA (PRK00591)→ HemK (COG2890)PRK09328 (PRK09328)→ |
| env-487 | SRS019387_C1674147 | - | 222 | 187 | RNA→ rpmE2 (PRK01678)Ribosomal_L31 (pfam01197)→ |
| env-488 | SRS043676_WUGC_scaffold_7376 | - | 288 | 253 | RNA→ |
| env-489 | SRS045262_WUGC_scaffold_5583 | - | 417 | 382 | RNA→ rpmE2 (PRK01678)Ribosomal_L31 (pfam01197)→ |
| env-490 | SRS057022_LANL_scaffold_32496 | - | 164 | 129 | RNA→ rpmE2 (PRK01678)Ribosomal_L31 (pfam01197)→ |
| Gcu-1-1 | NZ_KE384431.1 | + | 63435 | 63471 | RNA→ rpmE2 (PRK01678)Ribosomal_L31 (pfam01197)→ |
| Gbe-1-1 | NZ_KI271855.1 | + | 37507 | 37542 | RNA→ rpmE2 (PRK01678)Ribosomal_L31 (pfam01197)→ |
| env-491 | AUXO018174862.1 | - | 1094 | 1063 | RNA→ rpmE (PRK00019)Ribosomal_L31 (pfam01197)→ PRK05571 (PRK05571)LacAB_rpiB (pfam02502 |
| | | | | | int_mem_ywzB (TIGR02327)DUF1146 (pfam06612)→ hypo→ |
| env-492 | NLM004_scaffold46339_1 | - | 437 | 405 | RNA→ rpmE (PRK00019)Ribosomal_L31 (pfam01197)→ YycJ-like_MBL-fold (cd07733)→ |
| env-493 | SRS012273_C3440542 | + | 90 | 122 | RNA→ rpmE (PRK00019)Ribosomal_L31 (pfam01197)→ |
| env-494 | scaffold33486_3_MH0073 | - | 380 | 348 | RNA→ rpmE (PRK00019)Ribosomal_L31 (pfam01197)→ YycJ-like_MBL-fold (cd07733)→ |
| env-495 | NOF010_scaffold2338_8 | - | 578 | 546 | RNA→ rpmE (PRK00019)Ribosomal_L31 (pfam01197)→ |
| | | | | | Lactamase_B (smart00849)metallo-hydrolase-like_MBL-fold (cd06262)→ |
| env-496 | NOF012_C785263_1 | - | 175 | 143 | RNA→ rpmE (PRK00019)Ribosomal_L31 (pfam01197)→ |
| env-497 | NOM002_scaffold12570_2 | - | 538 | 506 | RNA→ rpmE (PRK00019)Ribosomal_L31 (pfam01197)→ |
| | | | | | Lactamase_B (smart00849)metallo-hydrolase-like_MBL-fold (cd06262)→ |
| env-498 | NOM028_scaffold34027_1 | - | 2866 | 2834 | RNA→ rpmE (PRK00019)Ribosomal_L31 (pfam01197)→ Lactamase_B (smart00849)PhnP (COG1235)→ |
| | | | | | PRK00103 (PRK00103)SPOUT_MTase (pfam02590)→ hypo→ PRK15482 (PRK15482)RpiR (COG1737)→ |
| env-499 | 4491401.3_NODE_43820 | - | 1176 | 1144 | RNA→ rpmE (PRK00019)Ribosomal_L31 (pfam01197)→ |
| env-500 | SRS015854_C5461434 | + | 375 | 407 | RNA→ rpmE (PRK00019)Ribosomal_L31 (pfam01197)→ |
| Csp-6-1 | NZ_KQ235810.1 | - | 156185 | 156153 | RNA→ rpmE (PRK00019)Ribosomal_L31 (pfam01197)→ Lactamase_B (smart00849)PhnP (COG1235)→ |
| | | | | | PRK00103 (PRK00103)SPOUT_MTase (pfam02590)→ |
| env-501 | NLM032_scaffold64463_2 | + | 2904 | 2936 | RNA→ rpmE (PRK00019)Ribosomal_L31 (pfam01197)→ Lactamase_B (smart00849)PhnP (COG1235)→ |
| | | | | | PRK00103 (PRK00103)SPOUT_MTase (pfam02590)→ hypo→ PRK15482 (PRK15482)RpiR (COG1737)→ |
| env-502 | BABG01003555.1 | - | 601 | 569 | RNA→ rpmE (PRK00019)Ribosomal_L31 (pfam01197)→ |
| | | | | | Lactamase_B (smart00849)metallo-hydrolase-like_MBL-fold (cd06262)→ |
| env-503 | NLM027_scaffold7634_1 | + | 182 | 214 | RNA→ rpmE (PRK00019)Ribosomal_L31 (pfam01197)→ Lactamase_B (smart00849)PhnP (COG1235)→ |
| | | | | | PRK00103 (PRK00103)SPOUT_MTase (pfam02590)→ |
| env-504 | NOF005_scaffold9442_3 | + | 1100 | 1132 | RNA→ rpmE (PRK00019)Ribosomal_L31 (pfam01197)→ |
| env-505 | NOM009_scaffold48146_7 | + | 6961 | 6993 | RNA→ rpmE (PRK00019)Ribosomal_L31 (pfam01197)→ Lactamase_B (smart00849)PhnP (COG1235)→ |
| | | | | | PRK00103 (PRK00103)SPOUT_MTase (pfam02590)→ hypo→ PRK15482 (PRK15482)RpiR (COG1737)→ |
| env-506 | SRS020233_C4059603 | - | 223 | 191 | RNA→ rpmE (PRK00019)Ribosomal_L31 (pfam01197)→ |
| [sa-1-1 | NZ_JMLH01000001.1 | - | 51630 | 51598 | RNA→ rpmE (PRK00019)Ribosomal_L31 (pfam01197)→ Lactamase_B (smart00849)PhnP (COG1235)→ |
| | | | | | PRK00103 (PRK00103)SPOUT_MTase (pfam02590)→ |
| Cba-1-1 | NZ_BAHP02000099.1 | - | 13426 | 13394 | RNA→ rpmE (PRK00019)Ribosomal_L31 (pfam01197)→ Lactamase_B (smart00849)PhnP (COG1235)→ |
| | | | | | PRK00103 (PRK00103)SPOUT_MTase (pfam02590)→ |
| env-507 | SRS014313_WUGC_scaffold_17457 | + | 2437 | 2470 | RNA→ rpmE (PRK00019)Ribosomal_L31 (pfam01197)→ |
| env-508 | SRS016517_C20315060 | + | 178 | 211 | RNA→ rpmE (PRK00019)Ribosomal_L31 (pfam01197)→ |
| env-509 | SRS011586_C3323726 | + | 256 | 289 | RNA→ rpmE (PRK00019)rpmE2 (PRK01678)Ribosomal_L31 (pfam01197)→ |

| | | | | | |
|---|---|---|---|---|---|
| env-510 | SRS016267_C4307118 | + | 16 | 49 | **RNA**→ rpmE (PRK00019)Ribosomal_L31 (pfam01197)→ |
| env-511 | SRS016495_C1903365 | + | 841 | 874 | **RNA**→ hypo→ |
| env-512 | SRS019601_C1887446 | - | 239 | 206 | **RNA**→ rpmE (PRK00019)Ribosomal_L31 (pfam01197)→ |
| Csp-4-1 | NZ_DS562854.1 | - | 119936 | 119903 | **RNA**→ rpmE (PRK00019)Ribosomal_L31 (pfam01197)→ rpmE (PRK00019)Ribosomal_L31 (pfam01197)– |
| env-513 | AKZA01000288.1 | + | 213 | 245 | **RNA**→ rpmE2 (PRK01678)Ribosomal_L31 (pfam01197)→ |
| Eca-6-1 | NZ_GG692817.1 | + | 313634 | 313666 | **RNA**→ rpmE2 (PRK01678)Ribosomal_L31 (pfam01197)→ rpmE2 (PRK01678)Ribosomal_L31 (pfam01197 |
| Eca-7-1 | NZ_GG670387.1 | + | 312337 | 312369 | **RNA**→ rpmE2 (PRK01678)Ribosomal_L31 (pfam01197)→ rpmE2 (PRK01678)Ribosomal_L31 (pfam01197 |
| Eca-4-1 | NC_020995.1 | - | 2799854 | 2799822 | **RNA**→ rpmE2 (PRK01678)Ribosomal_L31 (pfam01197)→ |
| Efl-1-1 | NZ_KB946265.1 | - | 427638 | 427606 | **RNA**→ rpmE2 (PRK01678)Ribosomal_L31 (pfam01197)→ |
| Eca-2-1 | NZ_KE350248.1 | - | 72485 | 72453 | **RNA**→ rpmE2 (PRK01678)Ribosomal_L31 (pfam01197)→ |
| Eca-3-1 | NZ_GL872323.1 | + | 1257066 | 1257098 | **RNA**→ rpmE2 (PRK01678)Ribosomal_L31 (pfam01197)→ |
| Efa-315-1 | NZ_KE350268.1 | - | 261980 | 261948 | **RNA**→ rpmE2 (PRK01678)Ribosomal_L31 (pfam01197)→ |
| Efa-4-1 | NZ_KE351443.1 | - | 174912 | 174880 | **RNA**→ rpmE2 (PRK01678)Ribosomal_L31 (pfam01197)→ |
| Esp-14-1 | NZ_AKKS01000010.1 | + | 102506 | 102538 | **RNA**→ rpmE2 (PRK01678)Ribosomal_L31 (pfam01197)→ |
| env-514 | AUXO014118027.1 | + | 50137 | 50167 | **RNA**→ rpmE (PRK00019)Ribosomal_L31 (pfam01197)→ TadA (COG0590)MafB19-deam (pfam14437)→ I<br>hypo→ LemA (pfam04011)→ |
| Hma-1-1 | NZ_HE998568.1 | - | 708686 | 708652 | **RNA**→ rpmE (PRK00019)Ribosomal_L31 (pfam01197)→ |
| env-515 | SRS054590_LANL_scaffold_6972 | - | 112 | 78 | **RNA**→ rpmE (PRK00019)rpmE2 (PRK01678)Ribosomal_L31 (pfam01197)→ |
| Hfi-1-1 | NZ_GG657553.1 | - | 397887 | 397853 | **RNA**→ rpmE (PRK00019)Ribosomal_L31 (pfam01197)→ |
| env-516 | SRS015217_WUGC_scaffold_37705 | + | 9104 | 9137 | **RNA**→ rpmE (PRK00019)Ribosomal_L31 (pfam01197)→ Tdk (COG1435)PRK04296 (PRK04296)→<br>NDK (smart00562)NDK (pfam00334)→ |
| Edo-1-1 | NZ_DS483478.1 | - | 12891 | 12858 | **RNA**→ rpmE (PRK00019)Ribosomal_L31 (pfam01197)→ Tdk (COG1435)PRK04296 (PRK04296)→<br>NDK (smart00562)NDK (pfam00334)→ |
| env-517 | AUXO014926912.1 | + | 1177 | 1210 | **RNA**→ |
| Rma-1-1 | NZ_CVPE01000005.1 | - | 403752 | 403721 | **RNA**→ rpmE (PRK00019)Ribosomal_L31 (pfam01197)→ |
| env-518 | JGI12273J12029_10010743 | + | 2725 | 2769 | **RNA**→ rpmE (PRK00019)Ribosomal_L31 (pfam01197)→ |
| env-519 | 2200826318 | - | 563 | 529 | **RNA**→ ←hypo |
| Lpa-39-1 | NZ_JH414991.1 | + | 1619 | 1652 | **RNA**→ rpmE2 (PRK01678)Ribosomal_L31 (pfam01197)→ |
| env-520 | HBC_scaff_10134855 | - | 119 | 87 | **RNA**→ rpmE (PRK00019)rpmE2 (PRK01678)Ribosomal_L31 (pfam01197)→ |
| env-521 | HBC_ctgs_1288573 | - | 119 | 87 | **RNA**→ rpmE (PRK00019)rpmE2 (PRK01678)Ribosomal_L31 (pfam01197)→ |
| Lku-2-1 | NZ_JPYX01000004.1 | - | 23057 | 23025 | **RNA**→ rpmE2 (PRK01678)Ribosomal_L31 (pfam01197)→ |
| Lku-2-2 | NZ_JPUI01000040.1 | + | 29050 | 29082 | **RNA**→ rpmE2 (PRK01678)Ribosomal_L31 (pfam01197)→ |
| Lku-3-1 | NZ_AZBY01000009.1 | - | 22976 | 22944 | **RNA**→ rpmE2 (PRK01678)Ribosomal_L31 (pfam01197)→ |
| env-522 | AUXO010035532.1 | - | 150 | 121 | **RNA**→ rpmE (PRK00019)Ribosomal_L31 (pfam01197)→ |
| env-523 | AUXO010163029.1 | - | 150 | 121 | **RNA**→ rpmE (PRK00019)Ribosomal_L31 (pfam01197)→ |
| env-524 | JGI24502J29692_10015897 | + | 1091 | 1123 | **RNA**→ rpmE (PRK00019)Ribosomal_L31 (pfam01197)→ |
| env-525 | JGI24503J29689_10022200 | + | 1121 | 1153 | **RNA**→ rpmE (PRK00019)Ribosomal_L31 (pfam01197)→ |
| env-526 | JGI24500J29687_10017177 | + | 1046 | 1078 | **RNA**→ rpmE (PRK00019)Ribosomal_L31 (pfam01197)→ |
| env-527 | SRS014979_C3222903 | + | 585 | 613 | **RNA**→ NrdG (COG0602)Fer4_12 (pfam13353)→ Dut (COG0756)dut (PRK00601)→ |
| env-528 | SRS065504_LANL_scaffold_46983 | - | 2440 | 2412 | **RNA**→ NrdG (COG0602)Fer4_12 (pfam13353)→ Dut (COG0756)dut (PRK00601)→ |
| env-529 | scaffold1558_1_MH0054 | + | 596 | 624 | **RNA**→ NrdG (COG0602)Fer4_12 (pfam13353)→ |
| env-530 | scaffold31141_5_O2.UC-19 | - | 124 | 96 | **RNA**→ NrdG (COG0602)Fer4_12 (pfam13353)→ Dut (COG0756)dut (PRK00601)→ |
| env-531 | SRS017821_C2327271 | + | 300 | 328 | **RNA**→ NrdG (COG0602)Fer4_12 (pfam13353)→ Dut (COG0756)dut (PRK00601)→ |
| env-532 | SRS024435_C3071163 | - | 138 | 110 | **RNA**→ NrdG (COG0602)Fer4_12 (pfam13353)→ Dut (COG0756)dut (PRK00601)→ |
| env-533 | scaffold53311_1_MH0066 | - | 2650 | 2622 | **RNA**→ NrdG (COG0602)Fer4_12 (pfam13353)→ Dut (COG0756)dut (PRK00601)→ |
| env-534 | scaffold65125_1_V1.UC-10 | + | 484 | 512 | **RNA**→ NrdG (COG0602)Fer4_12 (pfam13353)→ trimeric_dUTPase (cd07557)dut (PRK00601)→ |
| env-535 | SRS014235_C3411809 | - | 244 | 216 | **RNA**→ NrdG (COG0602)Fer4_12 (pfam13353)→ trimeric_dUTPase (cd07557)dut (PRK00601)→ |
| env-536 | scaffold3439_3_MH0002 | - | 672 | 644 | **RNA**→ NrdG (COG0602)Fer4_12 (pfam13353)→ Dut (COG0756)dut (PRK00601)→ |
| env-537 | C6319386_1_MH0064 | + | 429 | 457 | **RNA**→ NrdG (COG0602)Fer4_12 (pfam13353)→ trimeric_dUTPase (cd07557)dut (PRK00601)→ |
| env-538 | scaffold208933_1_MH0006 | - | 2939 | 2911 | **RNA**→ NrdG (COG0602)Fer4_12 (pfam13353)→ Dut (COG0756)dut (PRK00601)→ |
| env-539 | UnmappedStool_Broad_scaffold_489424 | + | 400 | 428 | **RNA**→ NrdG (COG0602)Fer4_12 (pfam13353)→ |
| env-540 | SRS011586_WUGC_scaffold_28380 | + | 890 | 918 | **RNA**→ NrdG (COG0602)Fer4_12 (pfam13353)→ dut (PRK00601)→ |
| env-541 | SRS012273_C3719647 | - | 681 | 653 | **RNA**→ NrdG (COG0602)Fer4_12 (pfam13353)→ Dut (COG0756)dut (PRK00601)→ |

| | | | | | |
|---|---|---|---|---|---|
| env-542 | SRS014613_WUGC_scaffold_29094 | - | 2441 | 2413 | **RNA**→ NrdG (COG0602)Fer4_12 (pfam13353)→ Dut (COG0756)dut (PRK00601)→ |
| env-543 | SRS015782_C3240512 | - | 148 | 120 | **RNA**→ NrdG (COG0602)Fer4_12 (pfam13353)→ Dut (COG0756)dut (PRK00601)→ |
| env-544 | SRS020869_Baylor_scaffold_15611 | + | 608 | 636 | **RNA**→ NrdG (COG0602)Fer4_12 (pfam13353)→ trimeric_dUTPase (cd07557)dut (PRK00601)→ |
| env-545 | SRS022071_C3053836 | - | 2440 | 2412 | **RNA**→ NrdG (COG0602)Fer4_12 (pfam13353)→ Dut (COG0756)dut (PRK00601)→ |
| env-546 | SRS047014_C3437135 | - | 64 | 36 | **RNA**→ NrdG (COG0602)Fer4_12 (pfam13353)→ |
| env-547 | SRS064276_LANL_scaffold_22062 | - | 2440 | 2412 | **RNA**→ NrdG (COG0602)Fer4_12 (pfam13353)→ Dut (COG0756)dut (PRK00601)→ |
| Fso-1-1 | NZ_KI912400.1 | + | 87614 | 87648 | **RNA**→ rpmE2 (PRK01678)Ribosomal_L31 (pfam01197)→ |
| env-548 | HCF12C_316963 | + | 338 | 369 | **RNA**→ rpmE (PRK00019)Ribosomal_L31 (pfam01197)→ secG (TIGR00810)SecG (pfam03840)→ |
| env-549 | HCF12C_1_1_newblercontig523487 | - | 64 | 33 | **RNA**→ hypo→ |
| env-550 | JGI24724J26744_10133593 | + | 70 | 102 | **RNA**→ rpmE (PRK00019)Ribosomal_L31 (pfam01197)→ Tdk (COG1435)PRK04296 (PRK04296)→ |
| Psp-21-1 | NZ_HE610963.1 | + | 234443 | 234480 | **RNA**→ rpmE (PRK00019)Ribosomal_L31 (pfam01197)→ rpmE (PRK00019)Ribosomal_L31 (pfam01197)→ |
| env-551 | AUXO014364427.1 | - | 119 | 87 | **RNA**→ rpmE (PRK00019)rpmE2 (PRK01678)Ribosomal_L31 (pfam01197)→ |
| env-552 | AUXO016753630.1 | + | 178 | 210 | **RNA**→ rpmE (PRK00019)Ribosomal_L31 (pfam01197)→ Aes (COG0657)Abhydrolase_3 (pfam07859)→ |
| env-553 | RUMENNODE_4188936_197001 | - | 14624 | 14592 | **RNA**→ ←F420_ligase (pfam01996) |
| env-554 | RUMENNODE_4260751_1 | + | 37231 | 37263 | **RNA**→ rpmE (PRK00019)Ribosomal_L31 (pfam01197)→ |
| env-555 | RUMENNODE_4260751_1 | + | 24591 | 24623 | **RNA**→ rpmE (PRK00019)Ribosomal_L31 (pfam01197)→ |
| Ssp-3-1 | NZ_AZUC01000056.1 | + | 43289 | 43319 | **RNA**→ rpmE (PRK00019)Ribosomal_L31 (pfam01197)→ |
| Ssp-5-1 | NZ_JWIB01000052.1 | - | 9860 | 9830 | **RNA**→ rpmE (PRK00019)Ribosomal_L31 (pfam01197)→ |
| env-556 | JGI24024J18818_10001192 | - | 358 | 328 | **RNA**→ rpmE (PRK00019)Ribosomal_L31 (pfam01197)→ |
| env-557 | JGI24024J18818_10116008 | + | 495 | 525 | **RNA**→ rpmE (PRK00019)Ribosomal_L31 (pfam01197)→ |
| env-558 | JGI2065J20421_1018486 | - | 780 | 747 | **RNA**→ rpmE (PRK00019)Ribosomal_L31 (pfam01197)→ hypo→ |
| env-559 | JGI26050J44790_1028750 | + | 320 | 352 | **RNA**→ |
| env-560 | JGI26051J44788_1026204 | - | 292 | 260 | **RNA**→ rpmE2 (PRK01678)Ribosomal_L31 (pfam01197)→ |
| env-561 | JGI26053J44789_1023599 | - | 308 | 276 | **RNA**→ rpmE2 (PRK01678)Ribosomal_L31 (pfam01197)→ |
| env-562 | JGI26052J44787_1030737 | + | 185 | 217 | **RNA**→ rpmE2 (PRK01678)Ribosomal_L31 (pfam01197)→ |
| env-563 | JGI26055J44792_1008714 | + | 1514 | 1546 | **RNA**→ rpmE2 (PRK01678)Ribosomal_L31 (pfam01197)→ |
| env-564 | JGI26057J44791_1000928 | - | 1554 | 1522 | **RNA**→ rpmE2 (PRK01678)Ribosomal_L31 (pfam01197)→ |
| env-565 | JGI26054J46599_1005137 | + | 1615 | 1647 | **RNA**→ rpmE2 (PRK01678)Ribosomal_L31 (pfam01197)→ |
| Wci-1-3 | NZ_JWHT01000082.1 | + | 3980 | 4012 | **RNA**→ rpmE2 (PRK01678)Ribosomal_L31 (pfam01197)→ |
| Wci-2-1 | NZ_AEKT01000044.1 | - | 80525 | 80493 | **RNA**→ rpmE2 (PRK01678)Ribosomal_L31 (pfam01197)→ rpmE2 (PRK01678)Ribosomal_L31 (pfam01197 |
| Whe-1-1 | NZ_BBIK01000004.1 | + | 62770 | 62802 | **RNA**→ rpmE2 (PRK01678)Ribosomal_L31 (pfam01197)→ |
| env-566 | BMHBC_2_3182488 | - | 96 | 63 | **RNA**→ hypo→ |
| env-567 | BMHBC_2_4390494 | - | 96 | 63 | **RNA**→ hypo→ |
| env-568 | BMHBC_2_6409884 | + | 19 | 52 | **RNA**→ |
| env-569 | BMHBC_2_1969890 | + | 19 | 52 | **RNA**→ |
| env-570 | BMHBC_2_1356989 | + | 19 | 52 | **RNA**→ |
| env-571 | BMHBC_2_7096089 | + | 19 | 52 | **RNA**→ |
| env-572 | BMHBC_2_3756228 | - | 96 | 63 | **RNA**→ hypo→ |
| env-573 | BMHBC_2_5769464 | - | 96 | 63 | **RNA**→ hypo→ |
| env-574 | BMHBC_2_5120317 | - | 96 | 63 | **RNA**→ hypo→ |
| env-575 | BMHBC_2_2565350 | - | 96 | 63 | **RNA**→ hypo→ |
| [1]env-576 | 3300001422_104017 | - | 1678 | 1638 | **RNA**→ ←COG5421 (COG5421) |
| env-577 | C927572_1 | - | 10779 | 10748 | **RNA**→ ←hypo |
| Lci-4-1 | NZ_CCNH01000002.1 | + | 172817 | 172848 | **RNA**→ rpmE2 (PRK01678)Ribosomal_L31 (pfam01197)→ |
| Lci-4-2 | NZ_CCNG01000004.1 | + | 146295 | 146326 | **RNA**→ rpmE2 (PRK01678)Ribosomal_L31 (pfam01197)→ |
| Lci-4-3 | NZ_JVUU01000031.1 | - | 65028 | 64997 | **RNA**→ rpmE2 (PRK01678)Ribosomal_L31 (pfam01197)→ |
| Lci-5-1 | NC_010471.1 | + | 373008 | 373039 | **RNA**→ rpmE2 (PRK01678)Ribosomal_L31 (pfam01197)→ |
| Lci-2-1 | NZ_CAGF01000011.1 | - | 59597 | 59566 | **RNA**→ rpmE2 (PRK01678)Ribosomal_L31 (pfam01197)→ |
| Lci-1-1 | NZ_CAGE01000004.1 | - | 59603 | 59572 | **RNA**→ rpmE2 (PRK01678)Ribosomal_L31 (pfam01197)→ |
| Lci-3-1 | NZ_CAGG01000003.1 | - | 59603 | 59572 | **RNA**→ rpmE2 (PRK01678)Ribosomal_L31 (pfam01197)→ |
| Lca-25-1 | NC_018673.1 | - | 1311252 | 1311221 | **RNA**→ rpmE2 (PRK01678)Ribosomal_L31 (pfam01197)→ |
| env-578 | F76GVBA01AN3WN | - | 315 | 284 | **RNA**→ rpmE2 (PRK01678)Ribosomal_L31 (pfam01197)→ |

| | | | | | |
|---|---|---|---|---|---|
| env-579 | AUXO011646564.1 | + | 63 | 96 | **RNA**→ |
| Psa-1-1 | NZ_KB895396.1 | - | 5948 | 5915 | **RNA**→ rpmE (PRK00019)Ribosomal_L31 (pfam01197)→ |
| env-580 | AUXO010493877.1 | - | 441 | 409 | **RNA**→ rpmE (PRK00019)Ribosomal_L31 (pfam01197)→ MetG (COG0143)PRK12267 (PRK12267)→ |
| env-581 | JGI12273J12029_10023051 | + | 247 | 288 | **RNA**→ rpmE (PRK00019)Ribosomal_L31 (pfam01197)→ |
| env-582 | scaffold70953_1_MH0035 | - | 9668 | 9638 | **RNA**→ rpmE (PRK00019)Ribosomal_L31 (pfam01197)→ Tdk (COG1435)PRK04296 (PRK04296)→ TadA (COG0590)MafB19-deam (pfam14437)→ hypo→ hypo→ hypo→ hypo→ |
| env-583 | SRS022609_Baylor_scaffold_81875 | - | 4601 | 4571 | **RNA**→ rpmE (PRK00019)Ribosomal_L31 (pfam01197)→ Tdk (COG1435)PRK04296 (PRK04296)→ TadA (COG0590)MafB19-deam (pfam14437)→ hypo→ hypo→ hypo→ hypo→ hypo→ |
| env-584 | scaffold175278_1_MH0012 | - | 648 | 618 | **RNA**→ rpmE (PRK00019)Ribosomal_L31 (pfam01197)→ Tdk (COG1435)PRK04296 (PRK04296)→ |
| env-585 | C3957884_1_MH0040 | + | 1247 | 1277 | **RNA**→ rpmE (PRK00019)Ribosomal_L31 (pfam01197)→ Tdk (COG1435)PRK04296 (PRK04296)→ |
| env-586 | scaffold111154_1_MH0065 | + | 2910 | 2940 | **RNA**→ rpmE (PRK00019)Ribosomal_L31 (pfam01197)→ Tdk (COG1435)PRK04296 (PRK04296)→ TadA (COG0590)MafB19-deam (pfam14437)→ hypo→ hypo→ hypo→ hypo→ |
| env-587 | 4491423.3_NODE_1010 | - | 11026 | 10996 | **RNA**→ rpmE (PRK00019)Ribosomal_L31 (pfam01197)→ Tdk (COG1435)PRK04296 (PRK04296)→ TadA (COG0590)MafB19-deam (pfam14437)→ hypo→ hypo→ hypo→ hypo→ hypo→ |
| env-588 | scaffold73462_1_MH0044 | - | 3763 | 3733 | **RNA**→ rpmE (PRK00019)Ribosomal_L31 (pfam01197)→ Tdk (COG1435)PRK04296 (PRK04296)→ TadA (COG0590)MafB19-deam (pfam14437)→ hypo→ hypo→ hypo→ hypo→ |
| env-589 | scaffold17516_6_MH0066 | + | 1479 | 1509 | **RNA**→ rpmE (PRK00019)Ribosomal_L31 (pfam01197)→ Tdk (COG1435)PRK04296 (PRK04296)→ TadA (COG0590)MafB19-deam (pfam14437)→ hypo→ hypo→ hypo→ hypo→ |
| env-590 | scaffold114987_1_MH0070 | - | 572 | 542 | **RNA**→ rpmE (PRK00019)Ribosomal_L31 (pfam01197)→ Tdk (COG1435)PRK04296 (PRK04296)→ |
| env-591 | scaffold24028_2_MH0076 | + | 2809 | 2839 | **RNA**→ rpmE (PRK00019)Ribosomal_L31 (pfam01197)→ Tdk (COG1435)PRK04296 (PRK04296)→ TadA (COG0590)MafB19-deam (pfam14437)→ hypo→ hypo→ hypo→ hypo→ |
| env-592 | C3731285_1_MH0081 | - | 958 | 928 | **RNA**→ rpmE (PRK00019)Ribosomal_L31 (pfam01197)→ Tdk (COG1435)PRK04296 (PRK04296)→ hypo[illegible] |
| env-593 | SRS015663_C3126280 | + | 4752 | 4782 | **RNA**→ rpmE (PRK00019)Ribosomal_L31 (pfam01197)→ Tdk (COG1435)PRK04296 (PRK04296)→ TadA (COG0590)MafB19-deam (pfam14437)→ |
| env-594 | SRS019582_WUGC_scaffold_86237 | - | 1630 | 1600 | **RNA**→ rpmE (PRK00019)Ribosomal_L31 (pfam01197)→ Tdk (COG1435)PRK04296 (PRK04296)→ PRK10860 (PRK10860)TadA (COG0590)→ hypo→ hypo→ hypo→ |
| env-595 | SRS051031_LANL_scaffold_20891 | - | 3442 | 3412 | **RNA**→ rpmE (PRK00019)Ribosomal_L31 (pfam01197)→ Tdk (COG1435)PRK04296 (PRK04296)→ PRK10860 (PRK10860)TadA (COG0590)→ hypo→ hypo→ hypo→ hypo→ hypo→ |
| Cko-1-1 | NZ_JXAL01000029.1 | + | 95618 | 95652 | **RNA**→ rpmE (PRK00019)Ribosomal_L31 (pfam01197)→ spore_YhcN_YlaJ (TIGR02898)→ hypo→ hypo[illegible] hypo→ hypo→ |
| env-596 | CTBMPR_c3813 | - | 1882 | 1848 | **RNA**→ rpmE (PRK00019)Ribosomal_L31 (pfam01197)→ hypo→ |
| env-597 | 2200758960 | - | 2614 | 2581 | **RNA**→ rpmE (PRK00019)Ribosomal_L31 (pfam01197)→ hypo→ hypo→ hypo→ hypo→ hypo→ DnaX (COG2812)PRK05563 (PRK05563)→ |
| env-598 | CTBMPR_c24234 | + | 49 | 82 | **RNA**→ rpmE (PRK00019)Ribosomal_L31 (pfam01197)→ |
| Pta-1-1 | NZ_AULE01000002.1 | - | 8222 | 8189 | **RNA**→ rpmE (PRK00019)Ribosomal_L31 (pfam01197)→ |
| Psp-11-1 | NZ_JMDX01000001.1 | - | 7605 | 7572 | **RNA**→ rpmE (PRK00019)Ribosomal_L31 (pfam01197)→ |
| Pal-4-1 | NZ_ATMS01000017.1 | - | 7605 | 7572 | **RNA**→ rpmE (PRK00019)Ribosomal_L31 (pfam01197)→ |
| Psp-28-1 | NZ_JMLT01000022.1 | + | 85290 | 85323 | **RNA**→ rpmE (PRK00019)Ribosomal_L31 (pfam01197)→ |
| env-599 | JGI25151J46595_10251769 | - | 302 | 269 | **RNA**→ rpmE (PRK00019)Ribosomal_L31 (pfam01197)→ |
| Pal-1-1 | NZ_AMBZ01000002.1 | + | 805774 | 805807 | **RNA**→ rpmE (PRK00019)Ribosomal_L31 (pfam01197)→ rpmE (PRK00019)Ribosomal_L31 (pfam01197)[illegible] |
| Pal-5-1 | NZ_ATMT01000041.1 | + | 2855 | 2888 | **RNA**→ rpmE (PRK00019)Ribosomal_L31 (pfam01197)→ |
| Pte-3-1 | NZ_KB895427.1 | + | 72449 | 72482 | **RNA**→ rpmE (PRK00019)Ribosomal_L31 (pfam01197)→ |
| env-600 | JGI25151J46595_10020550 | - | 1564 | 1531 | **RNA**→ rpmE (PRK00019)Ribosomal_L31 (pfam01197)→ |
| Psp-32-1 | NZ_JXAK01000056.1 | - | 6498 | 6465 | **RNA**→ rpmE (PRK00019)Ribosomal_L31 (pfam01197)→ |
| env-601 | JGI12341J12878_1004125 | + | 582 | 615 | **RNA**→ rpmE (PRK00019)Ribosomal_L31 (pfam01197)→ hypo→ hypo→ hypo→ |
| env-602 | sg4i_contig13910 | + | 382 | 415 | **RNA**→ rpmE (PRK00019)Ribosomal_L31 (pfam01197)→ |
| env-603 | sg4i_contig20097 | + | 2292 | 2325 | **RNA**→ ←hypo |
| env-604 | JGI12341J12878_1005421 | - | 1095 | 1062 | **RNA**→ rpmE (PRK00019)Ribosomal_L31 (pfam01197)→ hypo→ hypo→ hypo→ |
| env-605 | sgz454il_contig08209 | + | 30513 | 30546 | **RNA**→ rpmE (PRK00019)Ribosomal_L31 (pfam01197)→ |
| env-606 | 2200252324 | - | 7897 | 7866 | **RNA**→ rpmE (PRK00019)Ribosomal_L31 (pfam01197)→ DUF4969 (pfam16339)→ |
| env-607 | GBANfinal_FHNL2OP04XNKIW | - | 208 | 170 | **RNA**→ rpmE (PRK00019)Ribosomal_L31 (pfam01197)→ |
| env-608 | JGI25151J46595_10336534 | + | 21 | 56 | **RNA**→ ←hypo |
| Pan-2-1 | NZ_HG005146.1 | + | 43659 | 43692 | **RNA**→ rpmE (PRK00019)Ribosomal_L31 (pfam01197)→ |
| env-609 | 2200180934 | + | 143 | 179 | **RNA**→ ←hypo |
| Pte-2-1 | NZ_JTHP01000028.1 | + | 65669 | 65702 | **RNA**→ rpmE (PRK00019)Ribosomal_L31 (pfam01197)→ |

| | | | | | |
|---|---|---|---|---|---|
| Psp-7-1 | NZ_ASSB01000327.1 | + | 24049 | 24082 | **RNA**→ rpmE (PRK00019)Ribosomal_L31 (pfam01197)→ |
| Psp-1-1 | NZ_JH601038.1 | + | 56254 | 56287 | **RNA**→ rpmE (PRK00019)Ribosomal_L31 (pfam01197)→ rpmE (PRK00019)Ribosomal_L31 (pfam01197)– |
| Ppe-1-1 | NZ_AGFX01000014.1 | + | 59100 | 59133 | **RNA**→ rpmE (PRK00019)Ribosomal_L31 (pfam01197)→ rpmE (PRK00019)Ribosomal_L31 (pfam01197)– |
| Ppo-3-1 | NZ_ASRZ01000123.1 | + | 30332 | 30365 | **RNA**→ rpmE (PRK00019)Ribosomal_L31 (pfam01197)→ |
| Ppo-4-1 | NZ_JWJJ01000013.1 | - | 4811 | 4778 | **RNA**→ rpmE (PRK00019)Ribosomal_L31 (pfam01197)→ |
| Ppo-7-1 | NZ_AIPP01000115.1 | + | 55583 | 55616 | **RNA**→ rpmE (PRK00019)Ribosomal_L31 (pfam01197)→ |
| Psp-18-1 | NZ_AMQU01000013.1 | + | 55780 | 55813 | **RNA**→ rpmE (PRK00019)Ribosomal_L31 (pfam01197)→ |
| Ppo-2-1 | NZ_CP009909.1 | + | 3174641 | 3174674 | **RNA**→ rpmE (PRK00019)Ribosomal_L31 (pfam01197)→ |
| Ppo-9-1 | NZ_CP006872.1 | + | 173664 | 173697 | **RNA**→ rpmE (PRK00019)Ribosomal_L31 (pfam01197)→ |
| Ppo-5-1 | NZ_GL905390.1 | + | 151743 | 151776 | **RNA**→ rpmE (PRK00019)Ribosomal_L31 (pfam01197)→ |
| Ppo-1-1 | NC_017542.1 | + | 175402 | 175435 | **RNA**→ rpmE (PRK00019)Ribosomal_L31 (pfam01197)→ |
| Ppo-2-2 | NZ_CP010268.1 | - | 2491086 | 2491053 | **RNA**→ rpmE (PRK00019)Ribosomal_L31 (pfam01197)→ |
| Ppo-8-1 | NC_014622.2 | + | 176703 | 176736 | **RNA**→ rpmE (PRK00019)Ribosomal_L31 (pfam01197)→ |
| Psp-8-1 | NZ_ASRY01000124.1 | + | 19295 | 19328 | **RNA**→ rpmE (PRK00019)Ribosomal_L31 (pfam01197)→ |
| Pte-1-1 | NC_016641.1 | + | 1753265 | 1753298 | **RNA**→ rpmE (PRK00019)Ribosomal_L31 (pfam01197)→ |
| Psp-30-1 | NZ_JMLR01000019.1 | - | 4809 | 4776 | **RNA**→ rpmE (PRK00019)Ribosomal_L31 (pfam01197)→ |
| Ppe-2-1 | NZ_CP011512.1 | + | 178280 | 178313 | **RNA**→ rpmE (PRK00019)Ribosomal_L31 (pfam01197)→ |
| Ppo-6-1 | NC_014483.2 | + | 176239 | 176272 | **RNA**→ rpmE (PRK00019)Ribosomal_L31 (pfam01197)→ |
| Ppo-10-1 | NZ_ASSA01000052.1 | - | 4575 | 4542 | **RNA**→ rpmE (PRK00019)Ribosomal_L31 (pfam01197)→ |
| env-610 | GI4A6AL01D1VT0 | - | 327 | 303 | **RNA**→ rplB (PRK09374)Ribosomal_L2_C (pfam03947)→ |
| Ppi-1-1 | NZ_BAVZ01000024.1 | + | 39507 | 39540 | **RNA**→ rpmE (PRK00019)Ribosomal_L31 (pfam01197)→ |
| Psp-24-1 | NZ_JRNV01000066.1 | + | 20032 | 20065 | **RNA**→ rpmE (PRK00019)Ribosomal_L31 (pfam01197)→ |
| Psp-19-1 | NZ_CP010976.1 | + | 5003103 | 5003136 | **RNA**→ rpmE (PRK00019)Ribosomal_L31 (pfam01197)→ |
| env-611 | MA40A_GDRVJ9S02I8S30_right | + | 185 | 218 | **RNA**→ rpmE (PRK00019)Ribosomal_L31 (pfam01197)→ |
| env-612 | JGI25162J39368_1010853 | - | 531 | 498 | **RNA**→ rpmE (PRK00019)Ribosomal_L31 (pfam01197)→ |
| env-613 | CLOF_F4EQGSL02GBV14 | - | 158 | 125 | **RNA**→ rpmE (PRK00019)Ribosomal_L31 (pfam01197)→ |
| Psp-33-1 | NZ_LELF01000050.1 | - | 4969 | 4936 | **RNA**→ rpmE (PRK00019)Ribosomal_L31 (pfam01197)→ |
| Psp-13-1 | NZ_ASPU01000009.1 | - | 4594 | 4561 | **RNA**→ rpmE (PRK00019)Ribosomal_L31 (pfam01197)→ |
| Psp-15-1 | NZ_ASPR01000001.1 | + | 36319 | 36352 | **RNA**→ rpmE (PRK00019)Ribosomal_L31 (pfam01197)→ |
| Psp-26-1 | NZ_ANHX01000073.1 | - | 4568 | 4535 | **RNA**→ rpmE (PRK00019)Ribosomal_L31 (pfam01197)→ |
| Psp-23-1 | NZ_AWUK01000005.1 | - | 4782 | 4749 | **RNA**→ rpmE (PRK00019)Ribosomal_L31 (pfam01197)→ |
| Psp-22-1 | NZ_AWUJ01000041.1 | + | 47513 | 47546 | **RNA**→ rpmE (PRK00019)Ribosomal_L31 (pfam01197)→ |
| Psp-2-1 | NZ_AEXS01000174.1 | - | 4730 | 4697 | **RNA**→ rpmE (PRK00019)Ribosomal_L31 (pfam01197)→ rpmE (PRK00019)Ribosomal_L31 (pfam01197)– rpmE (PRK00019)Ribosomal_L31 (pfam01197)→ |
| Pvo-1-1 | NZ_ADHJ01000005.1 | + | 51394 | 51427 | **RNA**→ rpmE (PRK00019)Ribosomal_L31 (pfam01197)→ rpmE (PRK00019)Ribosomal_L31 (pfam01197)– |
| Psp-16-1 | NZ_ASPT01000039.1 | + | 51294 | 51327 | **RNA**→ rpmE (PRK00019)Ribosomal_L31 (pfam01197)→ |
| Gsp-5-1 | NC_013406.1 | + | 153054 | 153087 | **RNA**→ rpmE (PRK00019)Ribosomal_L31 (pfam01197)→ |
| Psp-14-1 | NZ_ASPW01000021.1 | - | 4809 | 4776 | **RNA**→ rpmE (PRK00019)Ribosomal_L31 (pfam01197)→ |
| Psp-10-1 | NZ_LAZU01000035.1 | + | 58827 | 58860 | **RNA**→ rpmE (PRK00019)Ribosomal_L31 (pfam01197)→ |
| Bsp-52-1 | NZ_JAEO01000062.1 | - | 4215 | 4182 | **RNA**→ rpmE (PRK00019)Ribosomal_L31 (pfam01197)→ |
| Pla-1-1 | NZ_AGIP01000028.1 | - | 5136 | 5103 | **RNA**→ rpmE (PRK00019)Ribosomal_L31 (pfam01197)→ rpmE (PRK00019)Ribosomal_L31 (pfam01197)– rpmE (PRK00019)Ribosomal_L31 (pfam01197)→ |
| env-614 | CLOF_F4EQGSL02H5UCR | - | 445 | 412 | **RNA**→ rpmE (PRK00019)Ribosomal_L31 (pfam01197)→ |
| Pst-1-1 | NZ_CP009286.1 | + | 179434 | 179466 | **RNA**→ rpmE (PRK00019)Ribosomal_L31 (pfam01197)→ |
| Psp-20-1 | NZ_JADQ01000029.1 | + | 50634 | 50667 | **RNA**→ rpmE (PRK00019)Ribosomal_L31 (pfam01197)→ |
| env-615 | 2200248758 | - | 316 | 283 | **RNA**→ rpmE (PRK00019)Ribosomal_L31 (pfam01197)→ |
| Pba-1-1 | NZ_KE159656.1 | - | 11310 | 11277 | **RNA**→ rpmE (PRK00019)Ribosomal_L31 (pfam01197)→ |
| Psp-5-1 | NZ_GG695996.1 | + | 45052 | 45085 | **RNA**→ rpmE (PRK00019)Ribosomal_L31 (pfam01197)→ rpmE (PRK00019)Ribosomal_L31 (pfam01197)– |
| Pma-2-1 | NZ_KN125580.1 | - | 10211 | 10177 | **RNA**→ rpmE (PRK00019)Ribosomal_L31 (pfam01197)→ |
| Pbe-1-1 | NZ_CP011058.1 | + | 2435438 | 2435472 | **RNA**→ rpmE (PRK00019)Ribosomal_L31 (pfam01197)→ |
| Pha-2-1 | NZ_AULV01000020.1 | - | 5328 | 5295 | **RNA**→ rpmE (PRK00019)Ribosomal_L31 (pfam01197)→ YscJ_FliF_C (pfam08345)→ |
| Ppi-2-1 | NZ_AULX01000026.1 | + | 122523 | 122557 | **RNA**→ rpmE (PRK00019)Ribosomal_L31 (pfam01197)→ hypo→ hypo→ hypo→ hypo→ hypo→ |
| env-616 | sgz454il_contig00281 | + | 26477 | 26515 | **RNA**→ rpmE (PRK00019)Ribosomal_L31 (pfam01197)→ hypo→ |

| | | | | | |
|---|---|---|---|---|---|
| env-617 | JGI12341J12878_1005803 | + | 718 | 755 | RNA→ rpmE (PRK00019)Ribosomal_L31 (pfam01197)→ hypo→ |
| env-618 | JGI12298J13902_1003093 | + | 1096 | 1133 | RNA→ rpmE (PRK00019)Ribosomal_L31 (pfam01197)→ FieF (COG0053)ZT_dimer (pfam16916)→ hypo– hypo→ |
| Psp-12-1 | NZ_LIGH01000001.1 | - | 1890069 | 1890036 | RNA→ rpmE (PRK00019)Ribosomal_L31 (pfam01197)→ hypo→ |
| env-619 | 2200327300 | + | 8808 | 8844 | RNA→ rpmE (PRK00019)Ribosomal_L31 (pfam01197)→ hypo→ |
| env-620 | 2200095100 | + | 989 | 1025 | RNA→ rpmE (PRK00019)Ribosomal_L31 (pfam01197)→ |
| env-621 | JGI12341J12878_1000292 | - | 5025 | 4989 | RNA→ rpmE (PRK00019)Ribosomal_L31 (pfam01197)→ hypo→ |
| env-622 | JGI12298J13902_1000152 | - | 5280 | 5244 | RNA→ rpmE (PRK00019)Ribosomal_L31 (pfam01197)→ hypo→ |
| Tco-1-1 | NC_019897.1 | - | 3756453 | 3756415 | RNA→ rpmE (PRK00019)Ribosomal_L31 (pfam01197)→ hypo→ |
| env-623 | 2200093379 | + | 51 | 89 | RNA→ rpmE (PRK00019)Ribosomal_L31 (pfam01197)→ hypo→ hypo→ hypo→ hypo→ |
| Pal-3-1 | NZ_KQ040784.1 | - | 5386 | 5353 | RNA→ rpmE (PRK00019)Ribosomal_L31 (pfam01197)→ hypo→ |
| Pgo-1-1 | NZ_CBVJ010000144.1 | + | 40238 | 40271 | RNA→ rpmE (PRK00019)Ribosomal_L31 (pfam01197)→ hypo→ hypo→ hypo→ hypo→ hypo→ |
| Psp-4-1 | NC_012914.1 | + | 153732 | 153765 | RNA→ rpmE (PRK00019)Ribosomal_L31 (pfam01197)→ hypo→ hypo→ hypo→ hypo→ hypo→ hypo→ h hypo→ |
| env-624 | 2200533952 | - | 5624 | 5591 | RNA→ ←hypo |
| Pda-1-1 | NZ_KK082278.1 | + | 22447 | 22481 | RNA→ rpmE (PRK00019)Ribosomal_L31 (pfam01197)→ hypo→ |
| Pda-1-2 | NZ_KK082163.1 | - | 5621 | 5587 | RNA→ rpmE (PRK00019)Ribosomal_L31 (pfam01197)→ hypo→ |
| Pda-1-3 | NZ_KK082388.1 | - | 5525 | 5491 | RNA→ rpmE (PRK00019)Ribosomal_L31 (pfam01197)→ hypo→ |
| Ppa-2-1 | NZ_KE383861.1 | - | 5568 | 5536 | RNA→ rpmE (PRK00019)Ribosomal_L31 (pfam01197)→ hypo→ |
| Psp-25-1 | NZ_CBRA020000271.1 | + | 1490 | 1523 | RNA→ rpmE (PRK00019)Ribosomal_L31 (pfam01197)→ hypo→ hypo→ hypo→ hypo→ hypo→ hypo→ h DnaX (COG2812)PRK05563 (PRK05563)→ PRK00153 (PRK00153)YbaB_DNA_bd (pfam02575)→ |
| Psp-9-1 | NZ_JZEJ01000001.1 | + | 3977749 | 3977782 | RNA→ rpmE (PRK00019)Ribosomal_L31 (pfam01197)→ hypo→ hypo→ hypo→ hypo→ hypo→ hypo→ h |
| env-625 | 2200249840 | - | 249 | 211 | RNA→ rpmE (PRK00019)Ribosomal_L31 (pfam01197)→ |
| Gma-1-1 | NZ_CBQR020000100.1 | - | 8782 | 8748 | RNA→ rpmE (PRK00019)Ribosomal_L31 (pfam01197)→ |
| env-626 | CTBMPR_c370 | - | 7039 | 7005 | RNA→ rpmE (PRK00019)Ribosomal_L31 (pfam01197)→ A_NRPS_TlmIV_like (cd12114)Spore_YhcN_YlaJ (pfam09580)→ |
| env-627 | 2200234285 | + | 2038 | 2072 | RNA→ rpmE (PRK00019)Ribosomal_L31 (pfam01197)→ |
| env-628 | 2204944873 | + | 141 | 172 | RNA→ rpmE (PRK00019)Ribosomal_L31 (pfam01197)→ |
| env-629 | 2158374351 | + | 138 | 169 | RNA→ rpmE (PRK00019)Ribosomal_L31 (pfam01197)→ |
| Bma-3-1 | NZ_BAMO01000003.1 | - | 61835 | 61804 | RNA→ rpmE (PRK00019)Ribosomal_L31 (pfam01197)→ |
| env-630 | MA55A_contig00036 | + | 7249 | 7280 | RNA→ rpmE (PRK00019)Ribosomal_L31 (pfam01197)→ |
| Bau-1-1 | NZ_KE386943.1 | - | 264511 | 264481 | RNA→ rpmE2 (PRK01678)Ribosomal_L31 (pfam01197)→ |
| Bne-1-1 | NZ_ASRU01000098.1 | - | 9343 | 9313 | RNA→ rpmE (PRK00019)Ribosomal_L31 (pfam01197)→ |
| Bci-2-1 | NZ_LDPH01000003.1 | - | 34497 | 34467 | RNA→ rpmE (PRK00019)Ribosomal_L31 (pfam01197)→ |
| env-631 | Amiter_FTLR0DA01DQ0VC | + | 426 | 458 | RNA→ |
| env-632 | Amiter_contig95164 | + | 10 | 42 | RNA→ rpmE (PRK00019)Ribosomal_L31 (pfam01197)→ |
| Eca-5-1 | NZ_KB446648.1 | + | 222167 | 222200 | RNA→ rpmE (PRK00019)Ribosomal_L31 (pfam01197)→ |
| CSt-2-1 | NZ_LN852696.1 | + | 451428 | 451461 | RNA→ rpmE (PRK00019)Ribosomal_L31 (pfam01197)→ |
| env-633 | AUXO014461646.1 | + | 918 | 951 | RNA→ rpmE (PRK00019)Ribosomal_L31 (pfam01197)→ |
| Kvi-2-1 | NZ_KK211400.1 | + | 200045 | 200078 | RNA→ rpmE (PRK00019)Ribosomal_L31 (pfam01197)→ |
| Kvi-1-1 | NZ_KL370857.1 | + | 165470 | 165503 | RNA→ rpmE (PRK00019)Ribosomal_L31 (pfam01197)→ |
| Kvi-3-1 | NZ_JHXA01000019.1 | - | 17181 | 17148 | RNA→ rpmE (PRK00019)Ribosomal_L31 (pfam01197)→ |
| env-634 | 2227236341 | + | 1931 | 1964 | RNA→ rpmE (PRK00019)Ribosomal_L31 (pfam01197)→ |
| env-635 | SRS050422_C2834638 | + | 269 | 301 | RNA→ rpmE (PRK00019)Ribosomal_L31 (pfam01197)→ Lactamase_B (smart00849)PhnP (COG1235)→ PRK00103 (PRK00103)SPOUT_MTase (pfam02590)→ |
| env-636 | C2491656_1_MH0048 | - | 367 | 335 | RNA→ rpmE (PRK00019)Ribosomal_L31 (pfam01197)→ hypo→ |
| env-637 | scaffold20746_3_MH0080 | - | 184 | 152 | RNA→ rpmE (PRK00019)Ribosomal_L31 (pfam01197)→ |
| env-638 | scaffold62983_2_V1.UC-8 | + | 8996 | 9028 | RNA→ rpmE (PRK00019)Ribosomal_L31 (pfam01197)→ Lactamase_B (smart00849)PhnP (COG1235)→ PRK00103 (PRK00103)SPOUT_MTase (pfam02590)→ |
| env-639 | NOM005_C709845_1 | - | 2708 | 2676 | RNA→ rpmE (PRK00019)Ribosomal_L31 (pfam01197)→ Lactamase_B (smart00849)PhnP (COG1235)→ PRK00103 (PRK00103)SPOUT_MTase (pfam02590)→ hypo→ PRK15482 (PRK15482)RpiR (COG1737)→ |
| env-640 | SRS011271_C3070664 | + | 129 | 161 | RNA→ rpmE (PRK00019)Ribosomal_L31 (pfam01197)→ Lactamase_B (smart00849)metallo-hydrolase-like_MBL-fold (cd06262)→ |
| env-641 | SRS013951_C2550116 | - | 63 | 31 | RNA→ |

| | | | | | |
|---|---|---|---|---|---|
| env-642 | SRS015663_C3017608 | - | 163 | 131 | **RNA**→ rpmE (PRK00019)Ribosomal_L31 (pfam01197)→ |
| env-643 | SRS016095_WUGC_scaffold_38231 | + | 6582 | 6614 | **RNA**→ |
| env-644 | SRS048164_C2096952 | + | 94 | 126 | **RNA**→ rpmE (PRK00019)Ribosomal_L31 (pfam01197)→ |
| env-645 | SRS051031_LANL_scaffold_12183 | - | 370 | 338 | **RNA**→ rpmE (PRK00019)Ribosomal_L31 (pfam01197)→ YycJ-like_MBL-fold (cd07733)→ |
| env-646 | SRS058723_C6768793 | - | 194 | 162 | **RNA**→ rpmE (PRK00019)Ribosomal_L31 (pfam01197)→ |
| env-647 | SRS063985_C2384154 | - | 226 | 194 | **RNA**→ rpmE (PRK00019)Ribosomal_L31 (pfam01197)→ |
| bLF-1-1 | NZ_CCMM01000012.1 | - | 232933 | 232901 | **RNA**→ rpmE (PRK00019)Ribosomal_L31 (pfam01197)→ Lactamase_B (smart00849)PhnP (COG1235)→ |
| | | | | | PRK00103 (PRK00103)SPOUT_MTase (pfam02590)→ |
| env-648 | SL_9KL_010_SED_10062519 | + | 317 | 344 | **RNA**→ rpmE (PRK00019)Ribosomal_L31 (pfam01197)→ |
| env-649 | SL_3KL_010_SED_10161252 | + | 369 | 396 | **RNA**→ rpmE (PRK00019)Ribosomal_L31 (pfam01197)→ Tdk (COG1435)PRK04296 (PRK04296)→ |
| env-650 | JGI24500J29687_10038433 | + | 5406 | 5434 | **RNA**→ rpmE (PRK00019)Ribosomal_L31 (pfam01197)→ Tdk (COG1435)PRK04296 (PRK04296)→ |
| | | | | | TadA (COG0590)MafB19-deam (pfam14437)→ |
| env-651 | JGI24502J29692_10023353 | + | 2154 | 2182 | **RNA**→ rpmE (PRK00019)Ribosomal_L31 (pfam01197)→ Tdk (COG1435)PRK04296 (PRK04296)→ |
| | | | | | TadA (COG0590)MafB19-deam (pfam14437)→ hypo→ hypo→ TktA (COG0021)PRK05899 (PRK05899)→ |
| | | | | | MnmG (COG0445)PRK05192 (PRK05192)→ RsmG (COG0357)gidB (PRK00107)→ |
| | | | | | ParB (smart00470)Spo0J (COG1475)→ DUF951 (pfam06107)→ |
| env-652 | JGI24501J29690_1029513 | + | 2144 | 2172 | **RNA**→ rpmE (PRK00019)Ribosomal_L31 (pfam01197)→ Tdk (COG1435)PRK04296 (PRK04296)→ |
| | | | | | TadA (COG0590)MafB19-deam (pfam14437)→ |
| env-653 | JGI24503J29689_10024265 | + | 9945 | 9973 | **RNA**→ rpmE (PRK00019)Ribosomal_L31 (pfam01197)→ Tdk (COG1435)PRK04296 (PRK04296)→ |
| | | | | | TadA (COG0590)MafB19-deam (pfam14437)→ |
| env-654 | JGI24499J29688_1021746 | + | 118 | 146 | **RNA**→ rpmE (PRK00019)Ribosomal_L31 (pfam01197)→ |
| env-655 | RUMENNODE_1702127_1 | - | 23333 | 23300 | **RNA**→ rpmE (PRK00019)Ribosomal_L31 (pfam01197)→ |
| env-656 | RUMENNODE_3744369_1 | - | 1562 | 1528 | **RNA**→ rpmE (PRK00019)Ribosomal_L31 (pfam01197)→ hypo→ hypo→ 2A78 (TIGR00950)EamA (pfam0 |
| | | | | | MetG (COG0143)PRK12267 (PRK12267)→ |
| env-657 | AUXO015798099.1 | + | 1486 | 1520 | **RNA**→ rpmE (PRK00019)Ribosomal_L31 (pfam01197)→ |
| env-658 | RUMENNODE_3527277_1 | + | 1709 | 1743 | **RNA**→ (big gap) RhaT (COG0697)→ ArcA (COG2235)PRK01388 (PRK01388)→ |
| | | | | | ArcA (COG2235)PRK01388 (PRK01388)→ ArcA (COG2235)PRK01388 (PRK01388)→ |
| env-659 | RUMENNODE_3876952_1 | - | 3758 | 3725 | **RNA**→ rpmE (PRK00019)Ribosomal_L31 (pfam01197)→ |
| env-660 | 2211729444 | + | 222 | 255 | **RNA**→ rpmE (PRK00019)Ribosomal_L31 (pfam01197)→ |
| env-661 | 2156473860 | + | 222 | 255 | **RNA**→ rpmE (PRK00019)Ribosomal_L31 (pfam01197)→ |
| env-662 | RUMENNODE_3728468_1 | - | 2906 | 2873 | **RNA**→ rpmE (PRK00019)Ribosomal_L31 (pfam01197)→ EamA (pfam00892)→ |
| | | | | | DegV (TIGR00762)DegV (pfam02645)→ Hydrolase_4 (pfam12146)→ |
| env-663 | RUMENNODE_3917021_13000 | - | 6817 | 6784 | **RNA**→ rpmE (PRK00019)Ribosomal_L31 (pfam01197)→ |
| env-664 | AUXO018183523.1 | - | 52 | 19 | **RNA**→ |
| env-665 | JGI24707J26582_10000610 | + | 18900 | 18931 | **RNA**→ rpmE (PRK00019)Ribosomal_L31 (pfam01197)→ |
| env-666 | JGI24708J26588_10001567 | - | 9771 | 9740 | **RNA**→ rpmE (PRK00019)Ribosomal_L31 (pfam01197)→ |
| env-667 | JGI24712J26585_10002850 | - | 486 | 455 | **RNA**→ rpmE (PRK00019)Ribosomal_L31 (pfam01197)→ |
| env-668 | JGI24712J26585_10002883 | - | 486 | 455 | **RNA**→ rpmE (PRK00019)Ribosomal_L31 (pfam01197)→ |
| env-669 | JGI24711J26586_10044882 | + | 779 | 810 | **RNA**→ rpmE (PRK00019)Ribosomal_L31 (pfam01197)→ |
| env-670 | JGI24709J26583_10001041 | - | 9926 | 9895 | **RNA**→ rpmE (PRK00019)Ribosomal_L31 (pfam01197)→ |
| env-671 | JGI24710J26742_10005051 | + | 8426 | 8457 | **RNA**→ rpmE (PRK00019)Ribosomal_L31 (pfam01197)→ |
| env-672 | JGI24500J29687_10039037 | + | 2191 | 2222 | **RNA**→ rpmE (PRK00019)Ribosomal_L31 (pfam01197)→ |
| env-673 | JGI24502J29692_10024412 | + | 325 | 356 | **RNA**→ rpmE (PRK00019)Ribosomal_L31 (pfam01197)→ |
| env-674 | JGI24501J29690_1058253 | + | 141 | 172 | **RNA**→ rpmE (PRK00019)Ribosomal_L31 (pfam01197)→ |
| env-675 | JGI24503J29689_10041332 | + | 125 | 156 | **RNA**→ rpmE (PRK00019)Ribosomal_L31 (pfam01197)→ |
| env-676 | JGI24712J26585_10116731 | - | 699 | 670 | **RNA**→ rpmE (PRK00019)Ribosomal_L31 (pfam01197)→ |
| env-677 | JGI24502J29692_10067855 | + | 403 | 432 | **RNA**→ rpmE (PRK00019)Ribosomal_L31 (pfam01197)→ |
| env-678 | AUXO010821195.1 | + | 270 | 300 | **RNA**→ rpmE (PRK00019)Ribosomal_L31 (pfam01197)→ |
| env-679 | RUMENNODE_4230474_1 | - | 923 | 893 | **RNA**→ rpmE (PRK00019)Ribosomal_L31 (pfam01197)→ PRK05571 (PRK05571)LacAB_rpiB (pfam02502 |
| | | | | | hypo→ int_mem_ywzB (TIGR02327)DUF1146 (pfam06612)→ |
| env-680 | AUXO015533467.1 | - | 756 | 724 | **RNA**→ rpmE (PRK00019)Ribosomal_L31 (pfam01197)→ PRK05571 (PRK05571)LacAB_rpiB (pfam02502 |
| env-681 | AUXO014634809.1 | + | 2187 | 2219 | **RNA**→ rpmE (PRK00019)Ribosomal_L31 (pfam01197)→ |
| Lga-2-1 | NZ_JVEV01000034.1 | + | 23868 | 23900 | **RNA**→ rpmE2 (PRK01678)Ribosomal_L31 (pfam01197)→ |
| Lho-2-1 | NZ_CAKE01000002.1 | - | 165569 | 165537 | **RNA**→ rpmE2 (PRK01678)Ribosomal_L31 (pfam01197)→ rpmE2 (PRK01678)Ribosomal_L31 (pfam01197 |

| | | | | | |
|---|---|---|---|---|---|
| env-682 | UnmappedStool_Broad_C251615744 | - | 374 | 333 | **RNA**→ ←COG4603 (COG4603) |
| env-683 | SRS011239_C2568925 | - | 248 | 207 | **RNA**→ ←COG4603 (COG4603) |
| env-684 | JGI24707J26582_10604531 | - | 183 | 149 | **RNA**→ rpmE2 (PRK01678)Ribosomal_L31 (pfam01197)→ |
| Bse-3-1 | NZ_ATXL01000033.1 | - | 11293 | 11261 | **RNA**→ RpmE (COG0254)rpmE2 (PRK01678)→ |
| Aph-1-1 | NZ_JAGN01000024.1 | - | 682 | 650 | **RNA**→ rpmE2 (PRK01678)Ribosomal_L31 (pfam01197)→ |
| env-685 | scaffold9639_3_MH0012 | + | 1656 | 1677 | **RNA**→ biotinyl_domain (cd06850)DRE_TIM_PC_TC_5S (cd07937)AccC (COG0439)PRK12999 (PRK12999)PYC_O hypo→ |
| env-686 | 3300000558_10113212 | + | 2767 | 2797 | **RNA**→ rpmE (PRK00019)Ribosomal_L31 (pfam01197)→ |
| env-687 | 3300001580_11274066 | - | 128 | 98 | **RNA**→ rpmE (PRK00019)Ribosomal_L31 (pfam01197)→ |
| Lbu-1-1 | NZ_GG669719.1 | + | 45475 | 45508 | **RNA**→ rpmE2 (PRK01678)Ribosomal_L31 (pfam01197)→ |
| Lhi-1-1 | NZ_GG669998.1 | + | 89473 | 89506 | **RNA**→ rpmE2 (PRK01678)Ribosomal_L31 (pfam01197)→ |
| Lbr-7-1 | NZ_GG669611.1 | + | 46598 | 46631 | **RNA**→ rpmE2 (PRK01678)Ribosomal_L31 (pfam01197)→ |
| Lfa-4-1 | NZ_BAKI01000010.1 | + | 39794 | 39827 | **RNA**→ rpmE2 (PRK01678)Ribosomal_L31 (pfam01197)→ |
| Lcu-1-1 | NZ_JTAL01000005.1 | - | 26638 | 26605 | **RNA**→ rpmE2 (PRK01678)Ribosomal_L31 (pfam01197)→ |
| Lki-2-1 | NZ_JH591051.1 | + | 2889 | 2922 | **RNA**→ rpmE2 (PRK01678)Ribosomal_L31 (pfam01197)→ |
| Lot-1-1 | NZ_BASH01000008.1 | + | 45034 | 45067 | **RNA**→ rpmE2 (PRK01678)Ribosomal_L31 (pfam01197)→ |
| env-688 | 2205249974 | - | 100 | 67 | **RNA**→ hypo→ |
| env-689 | 2204581147 | - | 204 | 171 | **RNA**→ rpmE2 (PRK01678)Ribosomal_L31 (pfam01197)→ |
| env-690 | 2158547212 | - | 204 | 171 | **RNA**→ rpmE2 (PRK01678)Ribosomal_L31 (pfam01197)→ |
| Lbu-3-1 | NC_015428.1 | - | 1626594 | 1626561 | **RNA**→ rpmE2 (PRK01678)Ribosomal_L31 (pfam01197)→ |
| Ata-1-1 | NZ_AUCD01000015.1 | + | 29931 | 29963 | **RNA**→ rpmE2 (PRK01678)Ribosomal_L31 (pfam01197)→ |
| env-691 | 2200849697 | + | 2519 | 2552 | **RNA**→ rpmE (PRK00019)Ribosomal_L31 (pfam01197)→ spore_YhcN_YlaJ (TIGR02898)→ hypo→ hypo→ |
| Pcu-1-1 | NZ_AEDD01000010.1 | - | 172375 | 172342 | **RNA**→ rpmE (PRK00019)Ribosomal_L31 (pfam01197)→ rpmE (PRK00019)Ribosomal_L31 (pfam01197)→ hypo→ hypo→ |
| env-692 | SRS011111_C511282 | + | 1560 | 1591 | **RNA**→ rpmE2 (PRK01678)Ribosomal_L31 (pfam01197)→ |
| env-693 | SRS023604_C786152 | - | 5199 | 5168 | **RNA**→ rpmE2 (PRK01678)Ribosomal_L31 (pfam01197)→ |
| env-694 | SRS015072_WUGC_scaffold_306 | + | 2849 | 2880 | **RNA**→ rpmE2 (PRK01678)Ribosomal_L31 (pfam01197)→ |
| env-695 | SRS015073_C87470 | - | 246 | 215 | **RNA**→ rpmE2 (PRK01678)Ribosomal_L31 (pfam01197)→ |
| env-696 | SRS015225_WUGC_scaffold_40 | + | 4907 | 4938 | **RNA**→ rpmE2 (PRK01678)Ribosomal_L31 (pfam01197)→ |
| env-697 | SRS042428_WUGC_scaffold_125 | + | 5251 | 5282 | **RNA**→ rpmE2 (PRK01678)Ribosomal_L31 (pfam01197)→ |
| env-698 | SRS044742_C45202 | + | 404 | 435 | **RNA**→ |
| env-699 | SRS049237_WUGC_scaffold_364 | - | 44771 | 44740 | **RNA**→ rpmE2 (PRK01678)Ribosomal_L31 (pfam01197)→ |
| env-700 | SRS051505_C116585 | - | 5949 | 5918 | **RNA**→ rpmE2 (PRK01678)Ribosomal_L31 (pfam01197)→ |
| env-701 | SRS052756_LANL_scaffold_202 | + | 30811 | 30842 | **RNA**→ rpmE2 (PRK01678)Ribosomal_L31 (pfam01197)→ |
| env-702 | SRS054962_LANL_scaffold_154 | + | 7561 | 7592 | **RNA**→ rpmE2 (PRK01678)Ribosomal_L31 (pfam01197)→ |
| env-703 | SRS055298_LANL_scaffold_1060 | + | 56113 | 56144 | **RNA**→ rpmE2 (PRK01678)Ribosomal_L31 (pfam01197)→ |
| Lin-7-1 | NZ_ADHG01000001.1 | - | 204612 | 204581 | **RNA**→ rpmE2 (PRK01678)Ribosomal_L31 (pfam01197)→ rpmE2 (PRK01678)Ribosomal_L31 (pfam01197 |
| Lin-11-1 | NZ_AEKJ01000015.1 | - | 74795 | 74764 | **RNA**→ rpmE2 (PRK01678)Ribosomal_L31 (pfam01197)→ rpmE2 (PRK01678)Ribosomal_L31 (pfam01197 |
| Lin-12-1 | NZ_AEKK01000016.1 | - | 44828 | 44797 | **RNA**→ rpmE2 (PRK01678)Ribosomal_L31 (pfam01197)→ rpmE2 (PRK01678)Ribosomal_L31 (pfam01197 |
| Lin-9-1 | NZ_AEKI01000026.1 | - | 46443 | 46412 | **RNA**→ rpmE2 (PRK01678)Ribosomal_L31 (pfam01197)→ rpmE2 (PRK01678)Ribosomal_L31 (pfam01197 |
| Lin-15-1 | NZ_AEHO01000007.1 | - | 160441 | 160410 | **RNA**→ rpmE2 (PRK01678)Ribosomal_L31 (pfam01197)→ rpmE2 (PRK01678)Ribosomal_L31 (pfam01197 |
| Lin-14-1 | NZ_AEHP01000054.1 | - | 71416 | 71385 | **RNA**→ rpmE2 (PRK01678)Ribosomal_L31 (pfam01197)→ rpmE2 (PRK01678)Ribosomal_L31 (pfam01197 |
| Lin-18-1 | NZ_AEHR01000018.1 | - | 44815 | 44784 | **RNA**→ rpmE2 (PRK01678)Ribosomal_L31 (pfam01197)→ rpmE2 (PRK01678)Ribosomal_L31 (pfam01197 |
| Lin-20-1 | NZ_AEXK01000022.1 | - | 74609 | 74578 | **RNA**→ rpmE2 (PRK01678)Ribosomal_L31 (pfam01197)→ rpmE2 (PRK01678)Ribosomal_L31 (pfam01197 |
| Lin-17-1 | NZ_AEXP01000019.1 | - | 175111 | 175080 | **RNA**→ rpmE2 (PRK01678)Ribosomal_L31 (pfam01197)→ rpmE2 (PRK01678)Ribosomal_L31 (pfam01197 |
| Lin-8-1 | NZ_GG700805.1 | + | 256231 | 256262 | **RNA**→ rpmE2 (PRK01678)Ribosomal_L31 (pfam01197)→ |
| Lin-5-1 | NZ_GL622333.1 | - | 1005713 | 1005682 | **RNA**→ rpmE2 (PRK01678)Ribosomal_L31 (pfam01197)→ |
| Lin-10-1 | NZ_AEKH01000010.1 | - | 74730 | 74699 | **RNA**→ rpmE2 (PRK01678)Ribosomal_L31 (pfam01197)→ rpmE2 (PRK01678)Ribosomal_L31 (pfam01197 |
| env-704 | SRS016105_WUGC_scaffold_100 | - | 158089 | 158058 | **RNA**→ rpmE2 (PRK01678)Ribosomal_L31 (pfam01197)→ |
| env-705 | SRS023468_LANL_scaffold_19280 | + | 4225 | 4256 | **RNA**→ rpmE2 (PRK01678)Ribosomal_L31 (pfam01197)→ |
| env-706 | SRS016111_WUGC_scaffold_61 | - | 45180 | 45149 | **RNA**→ rpmE2 (PRK01678)Ribosomal_L31 (pfam01197)→ |
| env-707 | SRS022545_LANL_scaffold_139 | - | 1695 | 1664 | **RNA**→ rpmE2 (PRK01678)Ribosomal_L31 (pfam01197)→ |

| | | | | | |
|---|---|---|---|---|---|
| env-708 | SRS050184_LANL_scaffold_130 | - | 17992 | 17961 | **RNA**→ rpmE2 (PRK01678)Ribosomal_L31 (pfam01197)→ |
| env-709 | SRS064704_LANL_scaffold_211 | - | 17993 | 17962 | **RNA**→ rpmE2 (PRK01678)Ribosomal_L31 (pfam01197)→ |
| Lin-16-1 | NZ_AEHN01000014.1 | - | 44898 | 44867 | **RNA**→ rpmE2 (PRK01678)Ribosomal_L31 (pfam01197)→ rpmE2 (PRK01678)Ribosomal_L31 (pfam01197 |
| Lin-19-1 | NZ_AEXJ01000019.1 | - | 44735 | 44704 | **RNA**→ rpmE2 (PRK01678)Ribosomal_L31 (pfam01197)→ rpmE2 (PRK01678)Ribosomal_L31 (pfam01197 |
| Lsp-9-1 | NZ_JH164952.1 | - | 574321 | 574290 | **RNA**→ rpmE2 (PRK01678)Ribosomal_L31 (pfam01197)→ |
| Lin-13-1 | NZ_AEHQ01000072.1 | - | 7349 | 7318 | **RNA**→ rpmE2 (PRK01678)Ribosomal_L31 (pfam01197)→ rpmE2 (PRK01678)Ribosomal_L31 (pfam01197 |
| env-710 | 2205772966 | + | 552 | 584 | **RNA**→ rpmE2 (PRK01678)Ribosomal_L31 (pfam01197)→ |
| env-711 | 2157969722 | + | 325 | 357 | **RNA**→ rpmE2 (PRK01678)Ribosomal_L31 (pfam01197)→ |
| env-712 | SRS075419_C110109 | - | 966 | 934 | **RNA**→ rpmE2 (PRK01678)Ribosomal_L31 (pfam01197)→ |
| env-713 | SRS019379_C28234 | + | 18298 | 18330 | **RNA**→ rpmE2 (PRK01678)Ribosomal_L31 (pfam01197)→ |
| env-714 | SRS022734_LANL_scaffold_274 | + | 10950 | 10982 | **RNA**→ rpmE2 (PRK01678)Ribosomal_L31 (pfam01197)→ |
| env-715 | SRS023850_C136155 | + | 237 | 269 | **RNA**→ rpmE2 (PRK01678)Ribosomal_L31 (pfam01197)→ |
| env-716 | SRS024428_C55026 | + | 18356 | 18388 | **RNA**→ rpmE2 (PRK01678)Ribosomal_L31 (pfam01197)→ |
| env-717 | SRS047335_WUGC_scaffold_762 | + | 4747 | 4779 | **RNA**→ rpmE2 (PRK01678)Ribosomal_L31 (pfam01197)→ |
| env-718 | SRS052620_C35955 | + | 10950 | 10982 | **RNA**→ rpmE2 (PRK01678)Ribosomal_L31 (pfam01197)→ |
| env-719 | SRS062520_LANL_scaffold_96 | - | 17372 | 17340 | **RNA**→ rpmE2 (PRK01678)Ribosomal_L31 (pfam01197)→ |
| Lga-2-2 | NZ_JVOU01000020.1 | - | 137321 | 137289 | **RNA**→ rpmE2 (PRK01678)Ribosomal_L31 (pfam01197)→ |
| Lga-2-3 | NZ_JVOL01000001.1 | - | 22205 | 22173 | **RNA**→ rpmE2 (PRK01678)Ribosomal_L31 (pfam01197)→ |
| Lga-2-4 | NZ_JUTQ01000039.1 | - | 22209 | 22177 | **RNA**→ rpmE2 (PRK01678)Ribosomal_L31 (pfam01197)→ |
| Lga-12-1 | NC_008530.1 | + | 296324 | 296356 | **RNA**→ rpmE2 (PRK01678)Ribosomal_L31 (pfam01197)→ |
| Lga-11-1 | NZ_ACOZ01000042.1 | + | 4725 | 4757 | **RNA**→ rpmE2 (PRK01678)Ribosomal_L31 (pfam01197)→ rpmE2 (PRK01678)Ribosomal_L31 (pfam01197 |
| Lga-9-1 | NZ_KN050667.1 | - | 187618 | 187586 | **RNA**→ rpmE2 (PRK01678)Ribosomal_L31 (pfam01197)→ |
| Lga-10-1 | NZ_KN050675.1 | - | 196087 | 196055 | **RNA**→ rpmE2 (PRK01678)Ribosomal_L31 (pfam01197)→ |
| Lga-4-1 | NZ_AUUE01000002.1 | + | 86255 | 86287 | **RNA**→ rpmE2 (PRK01678)Ribosomal_L31 (pfam01197)→ |
| Lga-5-1 | NZ_BALQ01000001.1 | + | 396473 | 396505 | **RNA**→ rpmE2 (PRK01678)Ribosomal_L31 (pfam01197)→ |
| Lga-6-1 | NZ_AKFQ01000002.1 | - | 180385 | 180353 | **RNA**→ rpmE2 (PRK01678)Ribosomal_L31 (pfam01197)→ rpmE2 (PRK01678)Ribosomal_L31 (pfam01197 |
| Ljo-3-1 | NZ_AFQJ01000004.1 | + | 1347121 | 1347153 | **RNA**→ rpmE2 (PRK01678)Ribosomal_L31 (pfam01197)→ |
| Ljo-5-1 | NC_017477.1 | + | 311419 | 311451 | **RNA**→ rpmE2 (PRK01678)Ribosomal_L31 (pfam01197)→ |
| Ljo-6-1 | NC_013504.1 | + | 280910 | 280942 | **RNA**→ rpmE2 (PRK01678)Ribosomal_L31 (pfam01197)→ |
| Ljo-2-1 | NC_022909.1 | + | 308188 | 308220 | **RNA**→ rpmE2 (PRK01678)Ribosomal_L31 (pfam01197)→ |
| env-720 | scaffold23403_2_V1.CD-11 | + | 274 | 306 | **RNA**→ rpmE2 (PRK01678)Ribosomal_L31 (pfam01197)→ |
| Ljo-7-1 | NC_005362.1 | + | 301703 | 301735 | **RNA**→ rpmE2 (PRK01678)Ribosomal_L31 (pfam01197)→ |
| env-721 | SRS011269_Baylor_scaffold_351 | - | 274 | 242 | **RNA**→ rpmE2 (PRK01678)Ribosomal_L31 (pfam01197)→ |
| Lga-2-5 | NZ_JUKW01000060.1 | + | 23407 | 23439 | **RNA**→ rpmE2 (PRK01678)Ribosomal_L31 (pfam01197)→ |
| Lga-7-1 | NZ_GL379581.1 | - | 187468 | 187436 | **RNA**→ rpmE2 (PRK01678)Ribosomal_L31 (pfam01197)→ |
| Lga-3-1 | NZ_CP006809.1 | + | 312647 | 312679 | **RNA**→ rpmE2 (PRK01678)Ribosomal_L31 (pfam01197)→ |
| Lga-8-1 | NZ_KL402718.1 | + | 399124 | 399156 | **RNA**→ rpmE2 (PRK01678)Ribosomal_L31 (pfam01197)→ |
| Ljo-1-1 | NZ_LIGY01000084.1 | - | 24710 | 24678 | **RNA**→ rpmE2 (PRK01678)Ribosomal_L31 (pfam01197)→ |
| env-722 | DOF012_scaffold379_3 | + | 2956 | 2988 | **RNA**→ rpmE2 (PRK01678)Ribosomal_L31 (pfam01197)→ |
| Lga-2-6 | NZ_JVGG01000050.1 | + | 18423 | 18455 | **RNA**→ rpmE2 (PRK01678)Ribosomal_L31 (pfam01197)→ |
| Ljo-4-1 | NZ_GG670121.1 | - | 182612 | 182580 | **RNA**→ rpmE2 (PRK01678)Ribosomal_L31 (pfam01197)→ |
| Lgi-2-1 | NZ_CAKC01000030.1 | + | 8283 | 8315 | **RNA**→ rpmE2 (PRK01678)Ribosomal_L31 (pfam01197)→ rpmE2 (PRK01678)Ribosomal_L31 (pfam01197 |
| Lsp-10-1 | NZ_KB822435.1 | - | 181390 | 181358 | **RNA**→ rpmE2 (PRK01678)Ribosomal_L31 (pfam01197)→ |
| Lcr-11-1 | NC_014106.1 | + | 276225 | 276256 | **RNA**→ rpmE2 (PRK01678)Ribosomal_L31 (pfam01197)→ |
| env-723 | SRS056796_LANL_scaffold_1239 | - | 1183 | 1152 | **RNA**→ rpmE2 (PRK01678)Ribosomal_L31 (pfam01197)→ |
| env-724 | DLM012_C404467_1 | - | 274 | 243 | **RNA**→ rpmE2 (PRK01678)Ribosomal_L31 (pfam01197)→ |
| env-725 | SRS015425_WUGC_scaffold_461 | + | 3773 | 3804 | **RNA**→ rpmE2 (PRK01678)Ribosomal_L31 (pfam01197)→ |
| env-726 | SRS062752_LANL_scaffold_621 | + | 1742 | 1773 | **RNA**→ rpmE2 (PRK01678)Ribosomal_L31 (pfam01197)→ |
| env-727 | SRS064376_LANL_scaffold_486 | + | 2458 | 2489 | **RNA**→ rpmE2 (PRK01678)Ribosomal_L31 (pfam01197)→ |
| Lcr-7-1 | NZ_ADGR01000062.1 | + | 8752 | 8783 | **RNA**→ rpmE2 (PRK01678)Ribosomal_L31 (pfam01197)→ rpmE2 (PRK01678)Ribosomal_L31 (pfam01197 |
| env-728 | Contig2087.1 | + | 770 | 801 | **RNA**→ rpmE (PRK00019)rpmE2 (PRK01678)L31 (TIGR00105)Ribosomal_L31 (pfam01197)→ |
| env-729 | SRS024310_LANL_scaffold_284 | - | 5059 | 5028 | **RNA**→ rpmE2 (PRK01678)Ribosomal_L31 (pfam01197)→ |

| | | | | | |
|---|---|---|---|---|---|
| env-730 | SRS011355_Baylor_scaffold_238 | - | 4759 | 4728 | **RNA**→ rpmE2 (PRK01678)Ribosomal_L31 (pfam01197)→ |
| env-731 | SRS014465_C104071 | + | 8731 | 8762 | **RNA**→ rpmE2 (PRK01678)Ribosomal_L31 (pfam01197)→ |
| env-732 | SRS014466_C33055 | + | 8731 | 8762 | **RNA**→ rpmE2 (PRK01678)Ribosomal_L31 (pfam01197)→ |
| env-733 | SRS014494_C170078 | - | 239 | 208 | **RNA**→ rpmE2 (PRK01678)Ribosomal_L31 (pfam01197)→ |
| env-734 | SRS015054_WUGC_scaffold_298 | + | 8828 | 8859 | **RNA**→ rpmE2 (PRK01678)Ribosomal_L31 (pfam01197)→ |
| env-735 | SRS015072_WUGC_scaffold_197 | + | 8688 | 8719 | **RNA**→ rpmE2 (PRK01678)Ribosomal_L31 (pfam01197)→ |
| env-736 | SRS015073_C87312 | + | 548 | 579 | **RNA**→ rpmE (PRK00019)rpmE2 (PRK01678)L31 (TIGR00105)Ribosomal_L31 (pfam01197)→ |
| env-737 | SRS015168_WUGC_scaffold_217 | - | 11230 | 11199 | **RNA**→ rpmE2 (PRK01678)Ribosomal_L31 (pfam01197)→ |
| env-738 | SRS016191_WUGC_scaffold_240 | - | 4886 | 4855 | **RNA**→ rpmE2 (PRK01678)Ribosomal_L31 (pfam01197)→ |
| env-739 | SRS016516_LANL_scaffold_146 | - | 19252 | 19221 | **RNA**→ rpmE2 (PRK01678)Ribosomal_L31 (pfam01197)→ |
| env-740 | SRS016559_Baylor_scaffold_103 | + | 8762 | 8793 | **RNA**→ rpmE2 (PRK01678)Ribosomal_L31 (pfam01197)→ |
| env-741 | SRS017520_Baylor_scaffold_434 | + | 8671 | 8702 | **RNA**→ rpmE2 (PRK01678)Ribosomal_L31 (pfam01197)→ |
| env-742 | SRS017700_Baylor_scaffold_322 | + | 924 | 955 | **RNA**→ rpmE2 (PRK01678)Ribosomal_L31 (pfam01197)→ |
| env-743 | SRS018769_WUGC_scaffold_137 | + | 8731 | 8762 | **RNA**→ rpmE2 (PRK01678)Ribosomal_L31 (pfam01197)→ |
| env-744 | SRS019600_WUGC_scaffold_105 | - | 4891 | 4860 | **RNA**→ rpmE2 (PRK01678)Ribosomal_L31 (pfam01197)→ |
| env-745 | SRS019989_Baylor_scaffold_99 | - | 4890 | 4859 | **RNA**→ rpmE2 (PRK01678)Ribosomal_L31 (pfam01197)→ |
| env-746 | SRS020349_Baylor_scaffold_168 | - | 19254 | 19223 | **RNA**→ rpmE2 (PRK01678)Ribosomal_L31 (pfam01197)→ |
| env-747 | SRS021969_C38256 | + | 8828 | 8859 | **RNA**→ rpmE2 (PRK01678)Ribosomal_L31 (pfam01197)→ |
| env-748 | SRS022158_LANL_scaffold_360 | + | 8827 | 8858 | **RNA**→ rpmE2 (PRK01678)Ribosomal_L31 (pfam01197)→ |
| env-749 | SRS042858_LANL_scaffold_175 | + | 8751 | 8782 | **RNA**→ rpmE2 (PRK01678)Ribosomal_L31 (pfam01197)→ |
| env-750 | SRS049237_C38452 | + | 8826 | 8857 | **RNA**→ rpmE2 (PRK01678)Ribosomal_L31 (pfam01197)→ |
| env-751 | SRS051505_C116175 | - | 244 | 213 | **RNA**→ rpmE2 (PRK01678)Ribosomal_L31 (pfam01197)→ |
| env-752 | SRS056695_LANL_scaffold_347 | + | 8772 | 8803 | **RNA**→ rpmE2 (PRK01678)Ribosomal_L31 (pfam01197)→ |
| env-753 | SRS057807_LANL_scaffold_81 | + | 7964 | 7995 | **RNA**→ rpmE2 (PRK01678)Ribosomal_L31 (pfam01197)→ |
| env-754 | SRS065347_LANL_scaffold_257 | - | 4891 | 4860 | **RNA**→ rpmE2 (PRK01678)Ribosomal_L31 (pfam01197)→ |
| env-755 | SRS077751_C52761 | + | 8762 | 8793 | **RNA**→ rpmE2 (PRK01678)Ribosomal_L31 (pfam01197)→ |
| Lcr-9-1 | NZ_GG669835.1 | + | 9537 | 9568 | **RNA**→ rpmE2 (PRK01678)Ribosomal_L31 (pfam01197)→ |
| Lcr-10-1 | NZ_GG704612.1 | - | 70406 | 70375 | **RNA**→ rpmE2 (PRK01678)Ribosomal_L31 (pfam01197)→ rpmE2 (PRK01678)Ribosomal_L31 (pfam01197 |
| Lcr-6-1 | NZ_GG698762.1 | + | 215551 | 215582 | **RNA**→ rpmE2 (PRK01678)Ribosomal_L31 (pfam01197)→ rpmE2 (PRK01678)Ribosomal_L31 (pfam01197 |
| Lcr-2-1 | NZ_JH932290.1 | - | 67347 | 67316 | **RNA**→ rpmE2 (PRK01678)Ribosomal_L31 (pfam01197)→ |
| Lcr-3-1 | NZ_JH932272.1 | + | 214526 | 214557 | **RNA**→ rpmE2 (PRK01678)Ribosomal_L31 (pfam01197)→ |
| Lcr-4-1 | NZ_KI391980.1 | - | 180476 | 180445 | **RNA**→ rpmE2 (PRK01678)Ribosomal_L31 (pfam01197)→ |
| Lcr-8-1 | NZ_GL531743.1 | + | 68214 | 68245 | **RNA**→ rpmE2 (PRK01678)Ribosomal_L31 (pfam01197)→ rpmE2 (PRK01678)Ribosomal_L31 (pfam01197 |
| Lcr-5-1 | NZ_GL877787.1 | + | 47874 | 47905 | **RNA**→ rpmE2 (PRK01678)Ribosomal_L31 (pfam01197)→ |
| Lcr-1-1 | NZ_AXLM01000027.1 | - | 82277 | 82246 | **RNA**→ rpmE2 (PRK01678)Ribosomal_L31 (pfam01197)→ |
| Lam-1-1 | NZ_ADNY01000029.1 | - | 80869 | 80837 | **RNA**→ rpmE2 (PRK01678)Ribosomal_L31 (pfam01197)→ rpmE2 (PRK01678)Ribosomal_L31 (pfam01197 |
| Lha-1-1 | NZ_BALY01000007.1 | - | 20680 | 20648 | **RNA**→ rpmE2 (PRK01678)Ribosomal_L31 (pfam01197)→ |
| env-756 | C4681636_1_MH0065 | - | 1489 | 1459 | **RNA**→ rpmE (PRK00019)Ribosomal_L31 (pfam01197)→ ATPgrasp_ST (pfam14397)→<br>PRK05571 (PRK05571)LacAB_rpiB (pfam02502)→ |
| env-757 | AUXO014458590.1 | + | 1212 | 1242 | **RNA**→ hypo→ rpmE (PRK00019)Ribosomal_L31 (pfam01197)→<br>PRK05571 (PRK05571)LacAB_rpiB (pfam02502)→ DUF1146 (pfam06612)→<br>SpoIID (COG2385)SpoIID (pfam08486)→ |
| env-758 | AUXO016429116.1 | + | 81 | 111 | **RNA**→ rpmE (PRK00019)Ribosomal_L31 (pfam01197)→ |
| env-759 | AUXO011676208.1 | - | 513 | 483 | **RNA**→ rpmE (PRK00019)Ribosomal_L31 (pfam01197)→ PRK05571 (PRK05571)LacAB_rpiB (pfam02502 |
| env-760 | AUXO011678380.1 | + | 143 | 173 | **RNA**→ rpmE (PRK00019)Ribosomal_L31 (pfam01197)→ |
| env-761 | RUMENNODE_4330698_1 | + | 57826 | 57856 | **RNA**→ rpmE (PRK00019)Ribosomal_L31 (pfam01197)→ PRK05571 (PRK05571)LacAB_rpiB (pfam02502<br>DUF1146 (pfam06612)→ SpoIID (COG2385)SpoIID_LytB (TIGR02669)→<br>PRK11649 (PRK11649)Peptidase_M23 (pfam01551)→ hypo→ LytTR (smart00850)LytTR (pfam04397)→ |
| env-762 | AUXO017045166.1 | - | 207 | 177 | **RNA**→ rpmE (PRK00019)Ribosomal_L31 (pfam01197)→ |
| env-763 | 08_07_CONTROL_DNA_scaffold35288_1 | + | 373 | 404 | **RNA**→ rpmE (PRK00019)Ribosomal_L31 (pfam01197)→ |
| Csp-2-1 | NZ_JQNG01000002.1 | + | 962368 | 962401 | **RNA**→ rpmE2 (PRK01678)Ribosomal_L31 (pfam01197)→ |
| Csp-8-1 | NC_015391.1 | + | 305155 | 305188 | **RNA**→ rpmE2 (PRK01678)Ribosomal_L31 (pfam01197)→ |
| env-764 | AUXO010645718.1 | - | 720 | 689 | **RNA**→ rpmE (PRK00019)Ribosomal_L31 (pfam01197)→ hypo→ |

| | | | | | |
|---|---|---|---|---|---|
| env-765 | 2227219885 | + | 182 | 213 | **RNA**→ rpmE (PRK00019)Ribosomal_L31 (pfam01197)→ |
| env-766 | HCF12C_86226 | - | 167 | 134 | **RNA**→ rpmE (PRK00019)Ribosomal_L31 (pfam01197)→ |
| env-767 | HCF12C_91840 | - | 3217 | 3187 | **RNA**→ rpmE (PRK00019)Ribosomal_L31 (pfam01197)→ rpoC2 (CHL00117)CYK3 (COG5279)→ |
| | | | | | RimK (COG0189)ATPgrasp_ST (pfam14397)→ PRK05571 (PRK05571)LacAB_rpiB (pfam02502)→ |
| env-768 | JGI24728J21555_1043850 | - | 433 | 404 | **RNA**→ rpmE (PRK00019)Ribosomal_L31 (pfam01197)→ Tdk (COG1435)PRK04296 (PRK04296)→ |
| Lhe-5-1 | NZ_HG531246.1 | + | 152 | 183 | **RNA**→ rpmE2 (PRK01678)Ribosomal_L31 (pfam01197)→ |
| Lhe-9-1 | NC_021744.1 | - | 2084137 | 2084106 | **RNA**→ rpmE2 (PRK01678)Ribosomal_L31 (pfam01197)→ |
| env-769 | DOF008_scaffold29004_2 | - | 2511 | 2480 | **RNA**→ rpmE2 (PRK01678)Ribosomal_L31 (pfam01197)→ |
| Lga-1-1 | NZ_BALB01000002.1 | - | 148004 | 147973 | **RNA**→ rpmE2 (PRK01678)Ribosomal_L31 (pfam01197)→ |
| env-770 | SRS017103_C2505406 | - | 285 | 254 | **RNA**→ rpmE2 (PRK01678)Ribosomal_L31 (pfam01197)→ |
| Lac-13-1 | NC_006814.3 | + | 256007 | 256038 | **RNA**→ rpmE2 (PRK01678)Ribosomal_L31 (pfam01197)→ |
| Lac-5-1 | NZ_HG313783.1 | + | 85062 | 85093 | **RNA**→ rpmE2 (PRK01678)Ribosomal_L31 (pfam01197)→ hypo→ MurF (COG0770)Mur_ligase_M (pfam08 |
| | | | | | HELICc (smart00490)DEXDc (smart00487)SrmB (COG0513)→ AcpS (COG0736)acpS (PRK00070)→ |
| | | | | | PLPDE_III_AR (cd00430)alr (PRK00053)→ |
| Lac-3-1 | NZ_HG313752.1 | + | 48471 | 48502 | **RNA**→ rpmE2 (PRK01678)Ribosomal_L31 (pfam01197)→ hypo→ MurF (COG0770)Mur_ligase_M (pfam08 |
| | | | | | HELICc (smart00490)DEXDc (smart00487)SrmB (COG0513)→ AcpS (COG0736)acpS (PRK00070)→ |
| | | | | | PLPDE_III_AR (cd00430)alr (PRK00053)→ |
| Lac-8-1 | NZ_HG313829.1 | + | 85041 | 85072 | **RNA**→ rpmE2 (PRK01678)Ribosomal_L31 (pfam01197)→ hypo→ MurF (COG0770)Mur_ligase_M (pfam08 |
| | | | | | HELICc (smart00490)DEXDc (smart00487)SrmB (COG0513)→ AcpS (COG0736)acpS (PRK00070)→ |
| | | | | | PLPDE_III_AR (cd00430)alr (PRK00053)→ |
| Lac-9-1 | NC_021181.2 | + | 256004 | 256035 | **RNA**→ rpmE2 (PRK01678)Ribosomal_L31 (pfam01197)→ |
| Lac-1-1 | NZ_CP010432.1 | + | 256305 | 256336 | **RNA**→ rpmE2 (PRK01678)Ribosomal_L31 (pfam01197)→ |
| Lac-12-1 | NZ_GG669567.1 | + | 189473 | 189504 | **RNA**→ rpmE2 (PRK01678)Ribosomal_L31 (pfam01197)→ |
| Lac-4-1 | NZ_CBLP010000004.1 | + | 134601 | 134632 | **RNA**→ rpmE2 (PRK01678)Ribosomal_L31 (pfam01197)→ hypo→ MurF (COG0770)Mur_ligase_M (pfam08 |
| | | | | | HELICc (smart00490)DEXDc (smart00487)SrmB (COG0513)→ AcpS (COG0736)acpS (PRK00070)→ |
| | | | | | PLPDE_III_AR (cd00430)alr (PRK00053)→ |
| Lac-7-1 | NZ_CBLT010000021.1 | + | 93153 | 93184 | **RNA**→ rpmE2 (PRK01678)Ribosomal_L31 (pfam01197)→ hypo→ MurF (COG0770)Mur_ligase_M (pfam08 |
| | | | | | HELICc (smart00490)DEXDc (smart00487)SrmB (COG0513)→ AcpS (COG0736)acpS (PRK00070)→ |
| | | | | | PLPDE_III_AR (cd00430)alr (PRK00053)→ |
| Lac-2-1 | NZ_AYUB01000060.1 | + | 523 | 554 | **RNA**→ rpmE2 (PRK01678)Ribosomal_L31 (pfam01197)→ |
| Lac-6-1 | NZ_BALR01000003.1 | - | 87861 | 87830 | **RNA**→ rpmE2 (PRK01678)Ribosomal_L31 (pfam01197)→ |
| Lhe-13-1 | NZ_GG700752.1 | + | 521294 | 521325 | **RNA**→ rpmE2 (PRK01678)Ribosomal_L31 (pfam01197)→ |
| Lhe-6-1 | NZ_HG531010.1 | + | 1181 | 1212 | **RNA**→ rpmE2 (PRK01678)Ribosomal_L31 (pfam01197)→ |
| Lhe-4-1 | NZ_CBUN010000034.1 | + | 1463 | 1494 | **RNA**→ rpmE2 (PRK01678)Ribosomal_L31 (pfam01197)→ |
| Lhe-12-1 | NC_010080.1 | + | 286737 | 286768 | **RNA**→ rpmE2 (PRK01678)Ribosomal_L31 (pfam01197)→ |
| Lhe-7-1 | NZ_HG530779.1 | - | 32078 | 32047 | **RNA**→ rpmE2 (PRK01678)Ribosomal_L31 (pfam01197)→ |
| Lhe-10-1 | NZ_CP002427.1 | + | 229346 | 229377 | **RNA**→ rpmE2 (PRK01678)Ribosomal_L31 (pfam01197)→ |
| Lhe-3-1 | NZ_CP009907.1 | + | 452665 | 452696 | **RNA**→ rpmE2 (PRK01678)Ribosomal_L31 (pfam01197)→ |
| Lhe-3-2 | NZ_CP011386.1 | + | 241847 | 241878 | **RNA**→ rpmE2 (PRK01678)Ribosomal_L31 (pfam01197)→ |
| Lhe-14-1 | NC_017467.1 | + | 267989 | 268020 | **RNA**→ rpmE2 (PRK01678)Ribosomal_L31 (pfam01197)→ |
| Lhe-8-1 | NZ_CBUH010000066.1 | - | 32297 | 32266 | **RNA**→ rpmE2 (PRK01678)Ribosomal_L31 (pfam01197)→ |
| Lhe-15-1 | NC_018528.1 | + | 260860 | 260891 | **RNA**→ rpmE2 (PRK01678)Ribosomal_L31 (pfam01197)→ |
| env-771 | scaffold10515_1_V1.CD-11 | + | 1726 | 1757 | **RNA**→ rpmE2 (PRK01678)Ribosomal_L31 (pfam01197)→ |
| env-772 | DLM027_scaffold20299_1 | + | 2704 | 2735 | **RNA**→ rpmE2 (PRK01678)Ribosomal_L31 (pfam01197)→ |
| env-773 | 2205115470 | - | 638 | 607 | **RNA**→ rpmE2 (PRK01678)Ribosomal_L31 (pfam01197)→ |
| env-774 | 2210389846 | - | 188 | 157 | **RNA**→ rpmE2 (PRK01678)Ribosomal_L31 (pfam01197)→ |
| env-775 | 2157274238 | - | 188 | 157 | **RNA**→ rpmE2 (PRK01678)Ribosomal_L31 (pfam01197)→ |
| env-776 | 2157676175 | - | 638 | 607 | **RNA**→ rpmE2 (PRK01678)Ribosomal_L31 (pfam01197)→ |
| Lsp-12-1 | NZ_CDLD01000004.1 | + | 49487 | 49518 | **RNA**→ rpmE2 (PRK01678)Ribosomal_L31 (pfam01197)→ |
| Lam-3-1 | NC_017470.1 | + | 259874 | 259905 | **RNA**→ rpmE2 (PRK01678)Ribosomal_L31 (pfam01197)→ |
| Lac-11-1 | NC_015214.1 | + | 258247 | 258278 | **RNA**→ rpmE2 (PRK01678)Ribosomal_L31 (pfam01197)→ hypo→ hypo→ |
| Lam-2-1 | NC_014724.1 | + | 283638 | 283669 | **RNA**→ rpmE2 (PRK01678)Ribosomal_L31 (pfam01197)→ |
| Lki-3-1 | NZ_BALU01000004.1 | - | 89390 | 89359 | **RNA**→ rpmE2 (PRK01678)Ribosomal_L31 (pfam01197)→ |
| Lke-2-1 | NC_015602.1 | + | 2075582 | 2075613 | **RNA**→ rpmE2 (PRK01678)Ribosomal_L31 (pfam01197)→ |

| | | | | | |
|---|---|---|---|---|---|
| Lke-1-1 | NZ_BAMG01000001.1 | + | 24696 | 24727 | **RNA**→ rpmE2 (PRK01678)Ribosomal_L31 (pfam01197)→ |
| Lul-1-1 | NZ_GG693256.1 | + | 130929 | 130960 | **RNA**→ rpmE2 (PRK01678)Ribosomal_L31 (pfam01197)→ |
| env-777 | AUXO018661423.1 | + | 171 | 201 | **RNA**→ rpmE (PRK00528)rpmE (PRK00019)rpmE2 (PRK01678)Ribosomal_L31 (pfam01197)→ |
| env-778 | AUXO016587397.1 | + | 680 | 712 | **RNA**→ |
| env-779 | NLF011_scaffold650_4 | + | 511 | 541 | **RNA**→ rpmE (PRK00019)Ribosomal_L31 (pfam01197)→ |
| env-780 | scaffold17049_11_MH0065 | - | 1359 | 1329 | **RNA**→ rpmE (PRK00019)Ribosomal_L31 (pfam01197)→ PRK05571 (PRK05571)LacAB_rpiB (pfam02502 |
| | | | | | DUF1146 (pfam06612)→ SpoIID (COG2385)SpoIID (pfam08486)→ |
| Slu-3-1 | NZ_JONV01000007.1 | - | 39962 | 39933 | **RNA**→ rpmE (PRK00019)Ribosomal_L31 (pfam01197)→ PrfA (COG0216)prfA (PRK00591)→ |
| | | | | | HemK (COG2890)PRK09328 (PRK09328)→ SUA5 (COG0009)TIGR00057 (TIGR00057)SUA5 (pfam03481 |
| | | | | | LMWPc (smart00226)Wzb (COG0394)→ PRK05571 (PRK05571)LacAB_rpiB (pfam02502)→ |
| | | | | | PRK13690 (PRK13690)DUF436 (pfam04260)→ GlyA (COG0112)glyA (PRK00011)→ |
| | | | | | Upp (COG0035)upp (PRK00129)→ WecB (COG0381)wecB (TIGR00236)→ |
| Sro-1-1 | NZ_JXII01000006.1 | - | 39825 | 39796 | **RNA**→ rpmE (PRK00019)Ribosomal_L31 (pfam01197)→ PrfA (COG0216)prfA (PRK00591)→ |
| | | | | | HemK (COG2890)PRK09328 (PRK09328)→ SUA5 (COG0009)TIGR00057 (TIGR00057)SUA5 (pfam03481 |
| | | | | | LMWPc (smart00226)Wzb (COG0394)→ PRK05571 (PRK05571)LacAB_rpiB (pfam02502)→ |
| | | | | | PRK13690 (PRK13690)DUF436 (pfam04260)→ GlyA (COG0112)glyA (PRK00011)→ |
| | | | | | Upp (COG0035)upp (PRK00129)→ WecB (COG0381)wecB (TIGR00236)→ |
| env-781 | 4491423.3_NODE_115429 | + | 19212 | 19243 | **RNA**→ rpmE (PRK00019)Ribosomal_L31 (pfam01197)→ PRK05571 (PRK05571)LacAB_rpiB (pfam02502 |
| | | | | | DUF1146 (pfam06612)→ SpoIID (COG2385)SpoIID_LytB (TIGR02669)→ |
| | | | | | PRK11649 (PRK11649)Peptidase_M23 (pfam01551)→ |
| env-782 | scaffold74606_2_MH0009 | - | 3256 | 3225 | **RNA**→ rpmE (PRK00019)Ribosomal_L31 (pfam01197)→ PRK05571 (PRK05571)LacAB_rpiB (pfam02502 |
| | | | | | DUF1146 (pfam06612)→ SpoIID (COG2385)SpoIID_LytB (TIGR02669)→ |
| | | | | | PRK11649 (PRK11649)Peptidase_M23 (pfam01551)→ |
| env-783 | scaffold25277_3_MH0012 | + | 3494 | 3525 | **RNA**→ rpmE (PRK00019)Ribosomal_L31 (pfam01197)→ PRK05571 (PRK05571)LacAB_rpiB (pfam02502 |
| | | | | | DUF1146 (pfam06612)→ SpoIID (COG2385)SpoIID_LytB (TIGR02669)→ |
| | | | | | PRK11649 (PRK11649)Peptidase_M23 (pfam01551)→ |
| env-784 | scaffold97436_3_MH0042 | + | 652 | 683 | **RNA**→ rpmE (PRK00019)Ribosomal_L31 (pfam01197)→ |
| env-785 | scaffold96667_1_MH0050 | + | 19222 | 19253 | **RNA**→ rpmE (PRK00019)Ribosomal_L31 (pfam01197)→ PRK05571 (PRK05571)LacAB_rpiB (pfam02502 |
| | | | | | DUF1146 (pfam06612)→ SpoIID (COG2385)SpoIID_LytB (TIGR02669)→ |
| | | | | | PRK11649 (PRK11649)Peptidase_M23 (pfam01551)→ |
| env-786 | scaffold29407_4_MH0059 | + | 1082 | 1113 | **RNA**→ rpmE (PRK00019)Ribosomal_L31 (pfam01197)→ PRK05571 (PRK05571)LacAB_rpiB (pfam02502 |
| | | | | | DUF1146 (pfam06612)→ SpoIID (COG2385)SpoIID_LytB (TIGR02669)→ |
| | | | | | PRK11649 (PRK11649)Peptidase_M23 (pfam01551)→ hypo→ |
| | | | | | PorA (COG0674)PorB (COG1013)PorG (COG1014)NapF (COG1145)Fer4_7 (pfam12838)→ |
| env-787 | scaffold81381_2_MH0060 | - | 1784 | 1753 | **RNA**→ rpmE (PRK00019)Ribosomal_L31 (pfam01197)→ PRK05571 (PRK05571)LacAB_rpiB (pfam02502 |
| | | | | | DUF1146 (pfam06612)→ SpoIID (COG2385)SpoIID (pfam08486)→ |
| env-788 | scaffold27176_1_MH0065 | + | 124 | 155 | **RNA**→ rpmE (PRK00019)Ribosomal_L31 (pfam01197)→ PRK05571 (PRK05571)LacAB_rpiB (pfam02502 |
| | | | | | DUF1146 (pfam06612)→ SpoIID (COG2385)SpoIID_LytB (TIGR02669)→ |
| | | | | | PRK11649 (PRK11649)Peptidase_M23 (pfam01551)→ hypo→ pyruv_ox_red (TIGR02176)PorA (COG0674) |
| env-789 | scaffold62388_2_MH0077 | - | 688 | 657 | **RNA**→ rpmE (PRK00019)Ribosomal_L31 (pfam01197)→ PRK05571 (PRK05571)LacAB_rpiB (pfam02502 |
| env-790 | scaffold19282_4_MH0081 | - | 2750 | 2719 | **RNA**→ rpmE (PRK00019)Ribosomal_L31 (pfam01197)→ PRK05571 (PRK05571)LacAB_rpiB (pfam02502 |
| | | | | | DUF1146 (pfam06612)→ SpoIID (COG2385)SpoIID_LytB (TIGR02669)→ |
| | | | | | PRK11649 (PRK11649)Peptidase_M23 (pfam01551)→ |
| env-791 | scaffold70033_3_V1.UC-14 | + | 2592 | 2623 | **RNA**→ rpmE (PRK00019)Ribosomal_L31 (pfam01197)→ PRK05571 (PRK05571)LacAB_rpiB (pfam02502 |
| | | | | | DUF1146 (pfam06612)→ SpoIID (pfam08486)→ |
| env-792 | 4491477.3_NODE_7390 | + | 65032 | 65063 | **RNA**→ rpmE (PRK00019)Ribosomal_L31 (pfam01197)→ PRK05571 (PRK05571)LacAB_rpiB (pfam02502 |
| | | | | | DUF1146 (pfam06612)→ SpoIID (COG2385)SpoIID_LytB (TIGR02669)→ |
| | | | | | PRK11649 (PRK11649)Peptidase_M23 (pfam01551)→ hypo→ |
| | | | | | PorA (COG0674)PorB (COG1013)PorG (COG1014)NapF (COG1145)Fer4_7 (pfam12838)→ |
| env-793 | scaffold16126_4_MH0031 | + | 2616 | 2647 | **RNA**→ rpmE (PRK00019)Ribosomal_L31 (pfam01197)→ |
| env-794 | C2024528_1_MH0079 | + | 208 | 239 | **RNA**→ rpmE (PRK00019)Ribosomal_L31 (pfam01197)→ PRK05571 (PRK05571)LacAB_rpiB (pfam02502 |
| Aot-1-1 | NZ_JH992958.1 | + | 468018 | 468051 | **RNA**→ rpmE2 (PRK01678)Ribosomal_L31 (pfam01197)→ |
| env-795 | scaffold28040_3_MH0086 | - | 152 | 127 | **RNA**→ hypo→ |
| env-796 | UnmappedStool_Broad_scaffold_784042 | - | 503 | 478 | **RNA**→ hypo→ |
| env-797 | longitudinal_415_272 | + | 46843 | 46868 | **RNA**→ BID_2 (smart00635)YjdB (COG5492)Big_2 (pfam02368)T2SS-T3SS_pil_N (pfam13629)→ |
| Tae-1-1 | NZ_KI783301.1 | - | 3374080 | 3374047 | **RNA**→ rpmE (PRK00019)Ribosomal_L31 (pfam01197)→ |

| | | | | | |
|---|---|---|---|---|---|
| Cul-1-1 | NZ_HG764817.1 | - | 1769012 | 1768979 | **RNA**→ rpmE (PRK00019)Ribosomal_L31 (pfam01197)→ |
| env-798 | JGI25320J50211_1018643 | + | 13 | 44 | **RNA**→ rpmE (PRK00019)Ribosomal_L31 (pfam01197)→ Tdk (COG1435)PRK04296 (PRK04296)→ |
| env-799 | IMNBL1_c0022815 | - | 113 | 83 | **RNA**→ rpmE (PRK00019)Ribosomal_L31 (pfam01197)→ |
| env-800 | JGI24707J26582_10026421 | - | 2395 | 2364 | **RNA**→ rpmE2 (PRK01678)Ribosomal_L31 (pfam01197)→ |
| env-801 | JGI24708J26588_10036079 | - | 1320 | 1289 | **RNA**→ rpmE2 (PRK01678)Ribosomal_L31 (pfam01197)→ |
| env-802 | JGI24707J26582_10002967 | - | 12238 | 12207 | **RNA**→ rpmE2 (PRK01678)Ribosomal_L31 (pfam01197)→ |
| env-803 | JGI24708J26588_10002912 | + | 155 | 186 | **RNA**→ rpmE2 (PRK01678)Ribosomal_L31 (pfam01197)→ |
| Axy-1-1 | NC_018704.1 | - | 2161096 | 2161065 | **RNA**→ rpmE2 (PRK01678)Ribosomal_L31 (pfam01197)→ |
| Bsp-24-1 | NZ_JVDT01000240.1 | + | 19685 | 19715 | **RNA**→ rpmE (PRK00019)Ribosomal_L31 (pfam01197)→ |
| Bsp-71-1 | NZ_JNJJ01000072.1 | - | 7691 | 7661 | **RNA**→ rpmE (PRK00019)Ribosomal_L31 (pfam01197)→ |
| env-804 | 2222084013.11956_1 | - | 385 | 353 | **RNA**→ ←hypo |
| Lro-2-1 | NZ_AKZK01000018.1 | - | 29624 | 29592 | **RNA**→ rpmE2 (PRK01678)Ribosomal_L31 (pfam01197)→ |
| Lro-2-2 | NZ_AUAW01000007.1 | - | 29920 | 29888 | **RNA**→ rpmE2 (PRK01678)Ribosomal_L31 (pfam01197)→ |
| env-805 | RUMENNODE_2902945_304500 | + | 18085 | 18113 | **RNA**→ rpmE (PRK00019)Ribosomal_L31 (pfam01197)→ |
| env-806 | AUXO013799381.1 | - | 329 | 301 | **RNA**→ rpmE (PRK00019)Ribosomal_L31 (pfam01197)→ |
| env-807 | JGI24707J26582_10473385 | - | 185 | 151 | **RNA**→ rpmE2 (PRK01678)Ribosomal_L31 (pfam01197)→ |
| Aha-1-1 | NZ_AKIF01000052.1 | + | 15837 | 15871 | **RNA**→ rpmE2 (PRK01678)Ribosomal_L31 (pfam01197)→ |
| Hha-2-1 | NZ_KE383978.1 | + | 253591 | 253623 | **RNA**→ rpmE2 (PRK01678)Ribosomal_L31 (pfam01197)→ |
| Ssp-1-1 | NZ_AMPQ01000010.1 | - | 54400 | 54366 | **RNA**→ rpmE2 (PRK01678)Ribosomal_L31 (pfam01197)→ rpmE2 (PRK01678)Ribosomal_L31 (pfam01197 |
| Lgi-1-1 | NZ_CP012034.1 | - | 1055820 | 1055788 | **RNA**→ rpmE2 (PRK01678)Ribosomal_L31 (pfam01197)→ |
| Lve-1-1 | NZ_BACR01000034.1 | - | 75029 | 74997 | **RNA**→ rpmE2 (PRK01678)Ribosomal_L31 (pfam01197)→ rpmE2 (PRK01678)Ribosomal_L31 (pfam01197 |
| env-808 | SRS015578_WUGC_scaffold_46456 | - | 16584 | 16552 | **RNA**→ rpmE (PRK00019)Ribosomal_L31 (pfam01197)→ |
| Tsp-2-1 | NZ_ADMN01000100.1 | - | 335 | 303 | **RNA**→ rpmE (PRK00019)Ribosomal_L31 (pfam01197)→ rpmE (PRK00019)Ribosomal_L31 (pfam01197)– |
| Tsp-1-1 | NZ_AEXQ01000118.1 | - | 20412 | 20380 | **RNA**→ rpmE (PRK00019)Ribosomal_L31 (pfam01197)→ rpmE (PRK00019)Ribosomal_L31 (pfam01197)– |
| env-809 | MC6.MG3.AS1.C1503 | + | 221 | 253 | **RNA**→ rpmE (PRK00019)Ribosomal_L31 (pfam01197)→ |
| env-810 | UnmappedStool_Broad_C252689365 | - | 137 | 105 | **RNA**→ rpmE (PRK00019)Ribosomal_L31 (pfam01197)→ |
| env-811 | 2204775796 | + | 330 | 362 | **RNA**→ |
| env-812 | 2205048070 | - | 223 | 191 | **RNA**→ rpmE (PRK00019)Ribosomal_L31 (pfam01197)→ |
| env-813 | 2157595394 | - | 223 | 191 | **RNA**→ hypo→ |
| env-814 | 2157503615 | + | 330 | 363 | **RNA**→ |
| env-815 | 2033367913 | + | 113 | 152 | **RNA**→ rpmE (PRK00019)Ribosomal_L31 (pfam01197)→ |
| env-816 | AUXO013136741.1 | - | 78 | 48 | **RNA**→ |
| env-817 | RUMENNODE_3528681_1 | + | 6693 | 6724 | **RNA**→ rpmE (PRK00019)Ribosomal_L31 (pfam01197)→ |
| env-818 | JGI24503J29689_10133851 | - | 152 | 124 | **RNA**→ rpmE (PRK00019)Ribosomal_L31 (pfam01197)→ |
| env-819 | DLM022_scaffold4159_5 | - | 1098 | 1067 | **RNA**→ rpmE (PRK00019)Ribosomal_L31 (pfam01197)→ hypo→ PRK05571 (PRK05571)LacAB_rpiB (pfam02502)→ |
| env-820 | AUXO011650987.1 | + | 229 | 260 | **RNA**→ rpmE (PRK00019)Ribosomal_L31 (pfam01197)→ |
| env-821 | BMHBC_2_6286536 | + | 6 | 40 | **RNA**→ |
| env-822 | BMHBC_2_6988570 | + | 6 | 40 | **RNA**→ rpmE (PRK00019)rpmE2 (PRK01678)Ribosomal_L31 (pfam01197)→ |
| env-823 | 3300001197_1008646 | - | 7405 | 7372 | **RNA**→ rpmE2 (PRK01678)Ribosomal_L31 (pfam01197)→ |
| env-824 | 3300001196_1009145 | - | 2558 | 2525 | **RNA**→ rpmE2 (PRK01678)Ribosomal_L31 (pfam01197)→ |
| env-825 | BMHBC_1_1_newblercontig70055 | + | 6 | 39 | **RNA**→ rpmE2 (PRK01678)Ribosomal_L31 (pfam01197)→ |
| env-826 | 3300000053_013632 | - | 6432 | 6399 | **RNA**→ rpmE (PRK00019)Ribosomal_L31 (pfam01197)→ spore_YhcN_YlaJ (TIGR02898)→ |
| Bsp-84-1 | NZ_JAQV01000014.1 | + | 67184 | 67217 | **RNA**→ rpmE (PRK00019)Ribosomal_L31 (pfam01197)→ |
| env-827 | P_1C_Sed_2_UnCty_1183832 | - | 137 | 104 | **RNA**→ rpmE (PRK00019)Ribosomal_L31 (pfam01197)→ |
| env-828 | AUXO014850948.1 | - | 56 | 28 | **RNA**→ |
| env-829 | BS_KBA_SWE12_21m_c10145057 | + | 465 | 493 | **RNA**→ rpmE (PRK00019)rpmE2 (PRK01678)Ribosomal_L31 (pfam01197)→ |
| env-830 | BS_KBA_SWE02_21m_10031375 | + | 450 | 478 | **RNA**→ rpmE (PRK00019)Ribosomal_L31 (pfam01197)→ |
| env-831 | AUXO010522381.1 | + | 1127 | 1157 | **RNA**→ |
| env-832 | JGI24709J26583_10053541 | + | 1520 | 1554 | **RNA**→ |
| env-833 | JGI24709J26583_10110457 | + | 290 | 324 | **RNA**→ rpmE (PRK00019)Ribosomal_L31 (pfam01197)→ |
| env-834 | JGI24709J26583_10523343 | + | 18 | 52 | **RNA**→ rpmE (PRK00019)Ribosomal_L31 (pfam01197)→ |

| | | | | | |
|---|---|---|---|---|---|
| env-835 | JGI24710J26742_10524727 | + | 26 | 60 | **RNA**→ rpmE (PRK00019)Ribosomal_L31 (pfam01197)→ |
| env-836 | JGI24500J29687_10015483 | - | 3498 | 3464 | **RNA**→ rpmE (PRK00019)Ribosomal_L31 (pfam01197)→ |
| env-837 | JGI24503J29689_10008717 | + | 739 | 773 | **RNA**→ rpmE (PRK00019)Ribosomal_L31 (pfam01197)→ |
| env-838 | JGI24710J26742_10079609 | + | 457 | 491 | **RNA**→ rpmE (PRK00019)Ribosomal_L31 (pfam01197)→ |
| env-839 | JGI24502J29692_10016940 | + | 738 | 772 | **RNA**→ rpmE (PRK00019)Ribosomal_L31 (pfam01197)→ |
| env-840 | JGI24710J26742_10183948 | + | 545 | 579 | **RNA**→ |
| env-841 | AUXO015967212.1 | - | 302 | 272 | **RNA**→ rpmE (PRK00019)Ribosomal_L31 (pfam01197)→ |
| env-842 | JGI24023J19991_10028404 | - | 525 | 497 | **RNA**→ rpmE (PRK00019)Ribosomal_L31 (pfam01197)→ |
| env-843 | AUXO016697288.1 | - | 2054 | 2024 | **RNA**→ rpmE (PRK00019)Ribosomal_L31 (pfam01197)→ FemAB (pfam02388)→ |
| env-844 | AUXO012356518.1 | + | 680 | 712 | **RNA**→ rpmE (PRK00019)Ribosomal_L31 (pfam01197)→ RimK (COG0189)ATPgrasp_ST (pfam14397)→ PRK05571 (PRK05571)LacAB_rpiB (pfam02502)→ int_mem_ywzB (TIGR02327)DUF1146 (pfam06612)→ |
| env-845 | AUXO013477821.1 | - | 57 | 25 | **RNA**→ |
| env-846 | AUXO018606034.1 | + | 3910 | 3942 | **RNA**→ rpmE (PRK00019)Ribosomal_L31 (pfam01197)→ |
| env-847 | AUXO016358659.1 | + | 2136 | 2166 | **RNA**→ rpmE (PRK00019)Ribosomal_L31 (pfam01197)→ |
| env-848 | scaffold24133_4_MH0077 | + | 267 | 297 | **RNA**→ rpmE (PRK00019)Ribosomal_L31 (pfam01197)→ |
| env-849 | AUXO014157167.1 | + | 27 | 57 | **RNA**→ rpmE (PRK00019)Ribosomal_L31 (pfam01197)→ PRK05571 (PRK05571)LacAB_rpiB (pfam02502 |
| env-850 | SL_8KL_010_SED_10000546 | + | 24347 | 24375 | **RNA**→ rpmE (PRK00019)Ribosomal_L31 (pfam01197)→ TadA (COG0590)MafB19-deam (pfam14437)→ [illegible] hypo→ hypo→ |
| env-851 | AUXO015488917.1 | - | 184 | 154 | **RNA**→ rpmE (PRK00019)Ribosomal_L31 (pfam01197)→ |
| env-852 | 2157781566 | + | 227 | 261 | **RNA**→ ←hypo |
| env-853 | 2204752571 | + | 238 | 270 | **RNA**→ rpmE (PRK00019)Ribosomal_L31 (pfam01197)→ |
| env-854 | 2158287102 | + | 231 | 263 | **RNA**→ ←hypo |
| env-855 | SRS022137_WUGC_scaffold_49347 | + | 225 | 256 | **RNA**→ rpmE (PRK00019)Ribosomal_L31 (pfam01197)→ |
| env-856 | SRS055982_C1494925 | + | 133 | 164 | **RNA**→ rpmE (PRK00019)Ribosomal_L31 (pfam01197)→ |
| env-857 | scaffold4806_3_MH0070 | - | 1189 | 1159 | **RNA**→ rpmE (PRK00019)Ribosomal_L31 (pfam01197)→ |
| env-858 | scaffold29628_1_MH0069 | - | 3846 | 3816 | **RNA**→ rpmE (PRK00019)Ribosomal_L31 (pfam01197)→ |
| env-859 | SRS011134_C5190448 | + | 4072 | 4102 | **RNA**→ rpmE (PRK00019)Ribosomal_L31 (pfam01197)→ |
| env-860 | scaffold84902_4_MH0077 | + | 2414 | 2444 | **RNA**→ rpmE (PRK00019)Ribosomal_L31 (pfam01197)→ ATPgrasp_ST (pfam14397)→ PRK05571 (PRK05571)LacAB_rpiB (pfam02502)→ int_mem_ywzB (TIGR02327)DUF1146 (pfam06612)→ SpoIID (COG2385)SpoIID_LytB (TIGR02669)→ |
| env-861 | AUXO014546959.1 | - | 171 | 139 | **RNA**→ rpmE (PRK00019)Ribosomal_L31 (pfam01197)→ |
| env-862 | AUXO018657893.1 | - | 531 | 499 | **RNA**→ rpmE (PRK00019)Ribosomal_L31 (pfam01197)→ PRK05571 (PRK05571)LacAB_rpiB (pfam02502 |
| env-863 | 3300001592_10049945 | - | 1173 | 1142 | **RNA**→ rpmE (PRK00019)Ribosomal_L31 (pfam01197)→ |
| Sko-1-1 | NZ_KB898626.1 | - | 216304 | 216273 | **RNA**→ rpmE2 (PRK01678)Ribosomal_L31 (pfam01197)→ |
| env-864 | 2204604922 | + | 33 | 77 | **RNA**→ rpmE2 (PRK01678)Ribosomal_L31 (pfam01197)→ |
| env-865 | 2158230023 | + | 33 | 77 | **RNA**→ rpmE2 (PRK01678)Ribosomal_L31 (pfam01197)→ |
| Cma-4-1 | NZ_AMYT01000021.1 | + | 149937 | 149969 | **RNA**→ rpmE2 (PRK01678)Ribosomal_L31 (pfam01197)→ |
| env-866 | AUXO018393888.1 | - | 164 | 134 | **RNA**→ rpmE (PRK00019)Ribosomal_L31 (pfam01197)→ |
| env-867 | SL_3KL_010_SED_10124865 | + | 289 | 318 | **RNA**→ rpmE (PRK00019)Ribosomal_L31 (pfam01197)→ Tdk (COG1435)PRK04296 (PRK04296)→ TadA (COG0590)MafB19-deam (pfam14437)→ |
| env-868 | SL_3KL_010_SED_10254284 | + | 507 | 536 | **RNA**→ rpmE (PRK00019)Ribosomal_L31 (pfam01197)→ |
| Cth-1-1 | NZ_AFCE01000162.1 | - | 72996 | 72965 | **RNA**→ rpmE (PRK00019)Ribosomal_L31 (pfam01197)→ rpmE (PRK00019)Ribosomal_L31 (pfam01197)– |
| Pya-1-1 | NZ_AVBF01000041.1 | - | 31350 | 31318 | **RNA**→ rpmE2 (PRK01678)Ribosomal_L31 (pfam01197)→ |
| Lsa-1-1 | NZ_ANAG01000022.1 | - | 33056 | 33024 | **RNA**→ rpmE2 (PRK01678)Ribosomal_L31 (pfam01197)→ |
| Lsa-2-1 | NZ_AUHQ01000006.1 | + | 37298 | 37330 | **RNA**→ rpmE2 (PRK01678)Ribosomal_L31 (pfam01197)→ |
| env-869 | 3300000053_219615 | - | 205 | 173 | **RNA**→ rpmE2 (PRK01678)Ribosomal_L31 (pfam01197)→ |
| Lko-1-1 | NZ_CP012033.1 | - | 493955 | 493923 | **RNA**→ rpmE2 (PRK01678)Ribosomal_L31 (pfam01197)→ |
| Bak-1-1 | NZ_BAUV01000003.1 | + | 60587 | 60617 | **RNA**→ rpmE2 (PRK01678)Ribosomal_L31 (pfam01197)→ |
| Lsp-3-1 | NZ_JPUW01000022.1 | + | 34339 | 34372 | **RNA**→ rpmE (PRK00019)Ribosomal_L31 (pfam01197)→ Tdk (COG1435)PRK04296 (PRK04296)→ PrfA (COG0216)prfA (PRK00591)→ HemK (COG2890)PRK09328 (PRK09328)→ SUA5 (COG0009)TIGR00057 (TIGR00057)SUA5 (pfam03481)→ LMWPc (smart00226)LMWPAP (cd1634 HAMP (smart00304)MA (smart00283)Tar (COG0840)→ |
| Efa-8-1 | NZ_KE351649.1 | - | 195752 | 195720 | **RNA**→ rpmE2 (PRK01678)Ribosomal_L31 (pfam01197)→ |
| Lna-1-1 | NZ_AUEG01000014.1 | - | 9119 | 9086 | **RNA**→ rpmE2 (PRK01678)Ribosomal_L31 (pfam01197)→ |

| | | | | | |
|---|---|---|---|---|---|
| Efa-538-1 | NZ_ASAM01000214.1 | + | 9303 | 9335 | **RNA**→ rpmE2 (PRK01678)Ribosomal_L31 (pfam01197)→ |
| Bma-6-1 | NZ_HG428742.1 | - | 441263 | 441230 | **RNA**→ rpmE (PRK00019)Ribosomal_L31 (pfam01197)→ |
| Pla-3-1 | NZ_LAWY01000043.1 | - | 5907 | 5874 | **RNA**→ rpmE (PRK00019)Ribosomal_L31 (pfam01197)→ |
| Pla-2-1 | NZ_CH981517.1 | - | 2321 | 2288 | **RNA**→ rpmE (PRK00019)Ribosomal_L31 (pfam01197)→ |
| Pla-4-1 | NC_023134.1 | - | 3814299 | 3814266 | **RNA**→ rpmE (PRK00019)Ribosomal_L31 (pfam01197)→ |
| Pla-5-1 | NZ_ADFW01000001.1 | + | 2677802 | 2677835 | **RNA**→ rpmE (PRK00019)Ribosomal_L31 (pfam01197)→ |
| Pal-2-1 | NZ_AUGY01000067.1 | + | 40909 | 40942 | **RNA**→ rpmE (PRK00019)Ribosomal_L31 (pfam01197)→ DUF3501 (pfam12007)→ hypo→ hypo→ hypo→ |
| env-870 | ABMY01031648.1 | - | 61 | 28 | **RNA**→ hypo→ |
| Pch-3-1 | NZ_JUGY01000072.1 | - | 6755 | 6722 | **RNA**→ rpmE (PRK00019)Ribosomal_L31 (pfam01197)→ DUF3501 (pfam12007)→ |
| Psp-31-1 | NZ_JIAZ01000009.1 | - | 5052 | 5019 | **RNA**→ rpmE (PRK00019)Ribosomal_L31 (pfam01197)→ hypo→ |
| Lpa-2-1 | NZ_KB911377.1 | - | 1976 | 1944 | **RNA**→ rpmE2 (PRK01678)Ribosomal_L31 (pfam01197)→ |
| Cdi-1-1 | NZ_JQLO01000001.1 | - | 1825701 | 1825669 | **RNA**→ rpmE2 (PRK01678)Ribosomal_L31 (pfam01197)→ |
| Lag-1-1 | NZ_LN851172.1 | + | 41190 | 41222 | **RNA**→ rpmE2 (PRK01678)Ribosomal_L31 (pfam01197)→ |
| env-871 | JGI24707J26582_10010170 | + | 4961 | 4993 | **RNA**→ rpmE2 (PRK01678)Ribosomal_L31 (pfam01197)→ |
| env-872 | JGI24708J26588_10015525 | - | 566 | 534 | **RNA**→ rpmE2 (PRK01678)Ribosomal_L31 (pfam01197)→ |
| env-873 | JGI24711J26586_10162716 | - | 244 | 212 | **RNA**→ rpmE2 (PRK01678)Ribosomal_L31 (pfam01197)→ |
| Ece-1-1 | NZ_LDEA01000016.1 | + | 14732 | 14764 | **RNA**→ rpmE2 (PRK01678)Ribosomal_L31 (pfam01197)→ |
| Ece-1-2 | NZ_LDOX01000011.1 | + | 32174 | 32206 | **RNA**→ rpmE2 (PRK01678)Ribosomal_L31 (pfam01197)→ |
| Ece-1-3 | NZ_LDOW01000005.1 | - | 101274 | 101242 | **RNA**→ rpmE2 (PRK01678)Ribosomal_L31 (pfam01197)→ |
| Ece-1-4 | NZ_LDEB01000042.1 | - | 2043 | 2011 | **RNA**→ rpmE2 (PRK01678)Ribosomal_L31 (pfam01197)→ |
| Ece-1-5 | NZ_LDED01000027.1 | + | 14737 | 14769 | **RNA**→ rpmE2 (PRK01678)Ribosomal_L31 (pfam01197)→ |
| Ece-1-6 | NZ_LDEC01000037.1 | + | 3278 | 3310 | **RNA**→ rpmE2 (PRK01678)Ribosomal_L31 (pfam01197)→ |
| Ece-1-7 | NZ_LDDZ01000019.1 | - | 1843 | 1811 | **RNA**→ rpmE2 (PRK01678)Ribosomal_L31 (pfam01197)→ |
| Ece-2-1 | NZ_KI535313.1 | + | 35980 | 36012 | **RNA**→ rpmE2 (PRK01678)Ribosomal_L31 (pfam01197)→ |
| Ema-3-1 | NZ_CVRN01000007.1 | + | 579351 | 579383 | **RNA**→ rpmE2 (PRK01678)Ribosomal_L31 (pfam01197)→ |
| env-874 | JGI26052J44787_1053424 | + | 319 | 352 | **RNA**→ |
| Esp-8-1 | NZ_JFYW01000008.1 | - | 343971 | 343938 | **RNA**→ rpmE2 (PRK01678)Ribosomal_L31 (pfam01197)→ |
| Eun-1-1 | NZ_KK211188.1 | + | 424175 | 424208 | **RNA**→ rpmE2 (PRK01678)Ribosomal_L31 (pfam01197)→ |
| Ean-1-1 | NC_018665.1 | - | 2501740 | 2501707 | **RNA**→ rpmE2 (PRK01678)Ribosomal_L31 (pfam01197)→ rpmE2 (PRK01678)Ribosomal_L31 (pfam01197 rpmE2 (PRK01678)Ribosomal_L31 (pfam01197)→ |
| Ean-2-1 | NZ_JMKS01000003.1 | - | 1069913 | 1069880 | **RNA**→ rpmE2 (PRK01678)Ribosomal_L31 (pfam01197)→ |
| Esi-1-1 | NC_010556.1 | - | 2710763 | 2710730 | **RNA**→ rpmE2 (PRK01678)Ribosomal_L31 (pfam01197)→ |
| Esi-2-1 | NZ_KK211190.1 | + | 331383 | 331416 | **RNA**→ rpmE2 (PRK01678)Ribosomal_L31 (pfam01197)→ |
| Eun-2-1 | NZ_KK211187.1 | + | 420534 | 420567 | **RNA**→ rpmE2 (PRK01678)Ribosomal_L31 (pfam01197)→ |
| env-875 | JGI26053J44789_1022240 | - | 446 | 413 | **RNA**→ rpmE2 (PRK01678)Ribosomal_L31 (pfam01197)→ |
| env-876 | JGI26055J44792_1047538 | - | 417 | 384 | **RNA**→ rpmE2 (PRK01678)Ribosomal_L31 (pfam01197)→ |
| env-877 | JGI26057J44791_1003846 | - | 2204 | 2171 | **RNA**→ rpmE2 (PRK01678)Ribosomal_L31 (pfam01197)→ |
| env-878 | JGI26051J44788_1013255 | + | 79 | 112 | **RNA**→ rpmE2 (PRK01678)Ribosomal_L31 (pfam01197)→ |
| env-879 | JGI26052J44787_1019914 | - | 555 | 522 | **RNA**→ rpmE2 (PRK01678)Ribosomal_L31 (pfam01197)→ |
| env-880 | JGI26054J46599_1007412 | - | 248 | 215 | **RNA**→ rpmE2 (PRK01678)Ribosomal_L31 (pfam01197)→ |
| Eac-1-1 | NZ_LFQN01000004.1 | - | 62759 | 62726 | **RNA**→ rpmE2 (PRK01678)Ribosomal_L31 (pfam01197)→ |
| Eac-2-1 | NZ_JNIR01000001.1 | - | 2404781 | 2404748 | **RNA**→ rpmE2 (PRK01678)Ribosomal_L31 (pfam01197)→ |
| Esp-11-1 | NZ_JTLL01000020.1 | + | 64609 | 64642 | **RNA**→ rpmE2 (PRK01678)Ribosomal_L31 (pfam01197)→ |
| Esp-5-1 | NC_022794.1 | - | 2814663 | 2814630 | **RNA**→ rpmE2 (PRK01678)Ribosomal_L31 (pfam01197)→ |
| Esp-7-1 | NZ_BARY01000003.1 | + | 65109 | 65142 | **RNA**→ rpmE2 (PRK01678)Ribosomal_L31 (pfam01197)→ |
| env-881 | JGI26050J44790_1013333 | - | 517 | 484 | **RNA**→ rpmE2 (PRK01678)Ribosomal_L31 (pfam01197)→ |
| Eox-1-1 | NZ_JNIS01000001.1 | - | 1240167 | 1240134 | **RNA**→ rpmE2 (PRK01678)Ribosomal_L31 (pfam01197)→ |
| Lvi-2-1 | NZ_AHYZ01000045.1 | - | 4802 | 4770 | **RNA**→ rpmE2 (PRK01678)Ribosomal_L31 (pfam01197)→ rpmE2 (PRK01678)Ribosomal_L31 (pfam01197 |
| Lvi-1-1 | NZ_AHZA01000166.1 | - | 3988 | 3956 | **RNA**→ rpmE2 (PRK01678)Ribosomal_L31 (pfam01197)→ |
| Fge-1-1 | NZ_AUMP01000005.1 | + | 28726 | 28757 | **RNA**→ rpmE (PRK00019)Ribosomal_L31 (pfam01197)→ |
| Apa-1-1 | NZ_JYCD01000078.1 | - | 32801 | 32767 | **RNA**→ rpmE (PRK00019)Ribosomal_L31 (pfam01197)→ |
| Bth-46-1 | NZ_KE695653.1 | - | 188547 | 188513 | **RNA**→ rpmE (PRK00019)Ribosomal_L31 (pfam01197)→ |

| | | | | | |
|---|---|---|---|---|---|
| Bth-47-1 | NZ_JQMH01000003.1 | - | 220631 | 220597 | **RNA**→ rpmE (PRK00019)Ribosomal_L31 (pfam01197)→ |
| env-882 | 2200565360 | - | 568 | 535 | **RNA**→ rpmE (PRK00019)Ribosomal_L31 (pfam01197)→ |
| | | | | | crypto_DASH (TIGR02765)DNA_photolyase (pfam00875)→ |
| Bsp-74-1 | NZ_KI301970.1 | - | 166095 | 166062 | **RNA**→ rpmE (PRK00019)Ribosomal_L31 (pfam01197)→ |
| Bag-1-1 | NZ_JATL01000001.1 | - | 68309 | 68276 | **RNA**→ rpmE (PRK00019)Ribosomal_L31 (pfam01197)→ PRK10674 (PRK10674)FAD_binding_7 (pfam034 |
| | | | | | hypo→ Tdk (COG1435)PRK04296 (PRK04296)→ |
| Bsp-10-1 | NZ_AKKB01000126.1 | - | 33889 | 33856 | **RNA**→ rpmE (PRK00019)Ribosomal_L31 (pfam01197)→ rpmE (PRK00019)Ribosomal_L31 (pfam01197)– |
| Bag-2-1 | NZ_AOBR01000012.1 | - | 25158 | 25125 | **RNA**→ rpmE (PRK00019)Ribosomal_L31 (pfam01197)→ |
| Bpa-4-1 | NZ_KI629787.1 | + | 57528 | 57561 | **RNA**→ rpmE (PRK00019)Ribosomal_L31 (pfam01197)→ Tdk (COG1435)PRK04296 (PRK04296)→ |
| | | | | | Spore_YhcN_YlaJ (pfam09580)→ |
| Bbr-4-1 | NZ_AKYF01000027.1 | - | 190401 | 190368 | **RNA**→ rpmE (PRK00019)Ribosomal_L31 (pfam01197)→ PhrB (COG0415)photolyase_8HDF (TIGR03556) |
| | | | | | Tdk (COG1435)PRK04296 (PRK04296)→ Spore_YhcN_YlaJ (pfam09580)→ |
| Bbr-2-1 | NZ_LDZV01000001.1 | - | 5378424 | 5378391 | **RNA**→ rpmE (PRK00019)Ribosomal_L31 (pfam01197)→ PhrB (COG0415)photolyase_8HDF (TIGR03556) |
| | | | | | Tdk (COG1435)PRK04296 (PRK04296)→ Spore_YhcN_YlaJ (pfam09580)→ |
| Bsp-43-1 | NZ_LGJF01000001.1 | + | 5142336 | 5142369 | **RNA**→ rpmE (PRK00019)Ribosomal_L31 (pfam01197)→ PhrB (COG0415)PRK10674 (PRK10674)→ |
| | | | | | Tdk (COG1435)PRK04296 (PRK04296)→ hypo→ |
| Bbr-3-1 | NZ_AHKL01000034.1 | + | 80549 | 80582 | **RNA**→ rpmE (PRK00019)Ribosomal_L31 (pfam01197)→ PhrB (COG0415)PRK10674 (PRK10674)→ |
| | | | | | Tdk (COG1435)PRK04296 (PRK04296)→ hypo→ |
| Bbr-1-1 | NC_012491.1 | - | 5785760 | 5785727 | **RNA**→ rpmE (PRK00019)Ribosomal_L31 (pfam01197)→ PhrB (COG0415)PRK10674 (PRK10674)→ |
| | | | | | Tdk (COG1435)PRK04296 (PRK04296)→ hypo→ |
| Bsp-9-1 | NZ_AKIX01000074.1 | - | 36982 | 36949 | **RNA**→ rpmE (PRK00019)Ribosomal_L31 (pfam01197)→ rpmE (PRK00019)Ribosomal_L31 (pfam01197)– |
| | | | | | PhrB (COG0415)photolyase_8HDF (TIGR03556)→ PhrB (COG0415)photolyase_8HDF (TIGR03556)→ |
| | | | | | Tdk (COG1435)PRK04296 (PRK04296)→ Tdk (COG1435)PRK04296 (PRK04296)→ |
| | | | | | Spore_YhcN_YlaJ (pfam09580)→ Spore_YhcN_YlaJ (pfam09580)→ PrfA (COG0216)prfA (PRK00591)→ |
| | | | | | PrfA (COG0216)prfA (PRK00591)→ PrfA (COG0216)prfA (PRK00591)→ |
| | | | | | HemK (COG2890)PRK09328 (PRK09328)→ HemK (COG2890)PRK09328 (PRK09328)→ |
| Bfo-2-1 | NZ_LDCN01000003.1 | - | 391782 | 391749 | **RNA**→ rpmE (PRK00019)Ribosomal_L31 (pfam01197)→ PhrB (COG0415)photolyase_8HDF (TIGR03556) |
| | | | | | Tdk (COG1435)PRK04296 (PRK04296)→ hypo→ |
| Bre-1-1 | NZ_LGIQ01000009.1 | + | 2036993 | 2037026 | **RNA**→ rpmE (PRK00019)Ribosomal_L31 (pfam01197)→ hypo→ hypo→ |
| | | | | | Tdk (COG1435)PRK04296 (PRK04296)→ Tdk (COG1435)PRK04296 (PRK04296)→ |
| Bbo-4-1 | NZ_JXAV01000005.1 | - | 191334 | 191300 | **RNA**→ rpmE (PRK00019)Ribosomal_L31 (pfam01197)→ |
| Bbo-7-1 | NZ_JPRB01000009.1 | - | 190353 | 190319 | **RNA**→ rpmE (PRK00019)Ribosomal_L31 (pfam01197)→ |
| Bbo-6-1 | NZ_APBN01000004.1 | - | 192154 | 192120 | **RNA**→ rpmE (PRK00019)Ribosomal_L31 (pfam01197)→ |
| Bbo-5-1 | NZ_JAQG01000025.1 | + | 126951 | 126985 | **RNA**→ rpmE (PRK00019)Ribosomal_L31 (pfam01197)→ |
| Bla-3-1 | NZ_CAGD01000009.1 | + | 76465 | 76498 | **RNA**→ rpmE (PRK00019)Ribosomal_L31 (pfam01197)→ |
| Bla-4-1 | NZ_CP007806.1 | - | 4868784 | 4868751 | **RNA**→ rpmE (PRK00019)Ribosomal_L31 (pfam01197)→ |
| Bla-1-1 | NZ_JNFS01000001.1 | + | 2136709 | 2136742 | **RNA**→ rpmE (PRK00019)Ribosomal_L31 (pfam01197)→ |
| Bla-5-1 | NZ_AXBT01000034.1 | + | 271405 | 271438 | **RNA**→ rpmE (PRK00019)Ribosomal_L31 (pfam01197)→ |
| Bla-2-1 | NZ_KB894284.1 | + | 113737 | 113770 | **RNA**→ rpmE (PRK00019)Ribosomal_L31 (pfam01197)→ |
| Bsp-99-1 | NZ_HE978538.1 | + | 272130 | 272163 | **RNA**→ rpmE (PRK00019)Ribosomal_L31 (pfam01197)→ |
| Bsp-45-1 | NZ_LIGG01000001.1 | - | 934797 | 934765 | **RNA**→ rpmE (PRK00019)Ribosomal_L31 (pfam01197)→ |
| Bca-1-1 | NZ_LAYY01000010.1 | - | 43036 | 43004 | **RNA**→ rpmE (PRK00019)Ribosomal_L31 (pfam01197)→ |
| Aji-1-1 | NZ_JH976434.1 | - | 707904 | 707873 | **RNA**→ rpmE2 (PRK01678)Ribosomal_L31 (pfam01197)→ |
| Lmu-1-1 | NZ_JSWI01000015.1 | + | 46921 | 46954 | **RNA**→ rpmE2 (PRK01678)Ribosomal_L31 (pfam01197)→ sortase_fam (TIGR01076)Sortase_A (cd06165)– |
| env-883 | DLF014_scaffold12613_4 | + | 22418 | 22451 | **RNA**→ rpmE2 (PRK01678)Ribosomal_L31 (pfam01197)→ sortase_fam (TIGR01076)Sortase_A (cd06165)– |
| env-884 | DOF012_scaffold14126_6 | + | 21527 | 21560 | **RNA**→ rpmE2 (PRK01678)Ribosomal_L31 (pfam01197)→ sortase_fam (TIGR01076)Sortase_A (cd06165)– |
| env-885 | 2204848799 | - | 1517 | 1484 | **RNA**→ rpmE2 (PRK01678)Ribosomal_L31 (pfam01197)→ sortase_fam (TIGR01076)Sortase_A (cd06165)– |
| env-886 | 2157775506 | - | 1517 | 1484 | **RNA**→ rpmE2 (PRK01678)Ribosomal_L31 (pfam01197)→ sortase_fam (TIGR01076)Sortase_A (cd06165)– |
| Lmu-2-1 | NZ_CP011013.1 | - | 2121237 | 2121204 | **RNA**→ rpmE2 (PRK01678)Ribosomal_L31 (pfam01197)→ sortase_fam (TIGR01076)Sortase_A (cd06165)– |
| Lmu-1-2 | NZ_JROC01000031.1 | - | 24602 | 24569 | **RNA**→ rpmE2 (PRK01678)Ribosomal_L31 (pfam01197)→ hypo→ hypo→ |
| Lwa-1-1 | NZ_AWTT01000003.1 | + | 39579 | 39611 | **RNA**→ rpmE2 (PRK01678)Ribosomal_L31 (pfam01197)→ |
| Lho-1-1 | NZ_AP014680.1 | + | 411316 | 411348 | **RNA**→ rpmE2 (PRK01678)Ribosomal_L31 (pfam01197)→ |
| env-887 | SRS011111_Baylor_scaffold_3888 | - | 333 | 301 | **RNA**→ rpmE2 (PRK01678)Ribosomal_L31 (pfam01197)→ |
| Lco-3-1 | NZ_GG698805.1 | + | 187441 | 187473 | **RNA**→ rpmE2 (PRK01678)Ribosomal_L31 (pfam01197)→ rpmE2 (PRK01678)Ribosomal_L31 (pfam01197 |
| Lin-6-1 | NZ_HE997174.1 | + | 292603 | 292635 | **RNA**→ rpmE2 (PRK01678)Ribosomal_L31 (pfam01197)→ |

| | | | | | |
|---|---|---|---|---|---|
| env-888 | 2205128598 | + | 89 | 121 | **RNA**→ rpmE2 (PRK01678)Ribosomal_L31 (pfam01197)→ |
| env-889 | 2157599913 | + | 89 | 121 | **RNA**→ |
| env-890 | 2158270211 | - | 125 | 92 | **RNA**→ rpmE (PRK00019)rpmE2 (PRK01678)L31 (TIGR00105)Ribosomal_L31 (pfam01197)→ |
| Lpa-1-1 | NZ_LDPB01000046.1 | + | 40254 | 40287 | **RNA**→ rpmE2 (PRK01678)Ribosomal_L31 (pfam01197)→ |
| Lan-3-1 | NZ_GG700738.1 | + | 134448 | 134481 | **RNA**→ rpmE2 (PRK01678)Ribosomal_L31 (pfam01197)→ |
| env-891 | 2158307163 | - | 217 | 184 | **RNA**→ rpmE2 (PRK01678)Ribosomal_L31 (pfam01197)→ |
| Lor-3-1 | NZ_AEKL01000068.1 | - | 23595 | 23562 | **RNA**→ rpmE2 (PRK01678)Ribosomal_L31 (pfam01197)→ rpmE2 (PRK01678)Ribosomal_L31 (pfam01197 |
| env-892 | 2205774333 | - | 527 | 494 | **RNA**→ rpmE2 (PRK01678)Ribosomal_L31 (pfam01197)→ |
| Lor-1-1 | NZ_AFTL01000008.1 | - | 55820 | 55787 | **RNA**→ rpmE2 (PRK01678)Ribosomal_L31 (pfam01197)→ |
| Lfe-7-1 | NZ_AVAB01000001.1 | + | 249983 | 250016 | **RNA**→ rpmE2 (PRK01678)Ribosomal_L31 (pfam01197)→ |
| env-893 | scaffold63933_1_V1.CD-11 | - | 667 | 634 | **RNA**→ rpmE2 (PRK01678)Ribosomal_L31 (pfam01197)→ |
| Lfe-1-1 | NZ_LAIK01000005.1 | - | 289790 | 289757 | **RNA**→ rpmE2 (PRK01678)Ribosomal_L31 (pfam01197)→ |
| Lfe-1-2 | NZ_LBDH01000005.1 | + | 42885 | 42918 | **RNA**→ rpmE2 (PRK01678)Ribosomal_L31 (pfam01197)→ |
| Lfe-1-3 | NZ_LBDG01000019.1 | - | 2690 | 2657 | **RNA**→ rpmE2 (PRK01678)Ribosomal_L31 (pfam01197)→ |
| Lfe-5-1 | NZ_CBUR010000031.1 | - | 23728 | 23695 | **RNA**→ rpmE2 (PRK01678)Ribosomal_L31 (pfam01197)→ |
| Lfe-1-4 | NZ_JUTH01000116.1 | + | 4117 | 4150 | **RNA**→ rpmE2 (PRK01678)Ribosomal_L31 (pfam01197)→ |
| Lfe-11-1 | NC_017465.1 | + | 250243 | 250276 | **RNA**→ rpmE2 (PRK01678)Ribosomal_L31 (pfam01197)→ rpmE2 (PRK01678)Ribosomal_L31 (pfam01197 rpmE2 (PRK01678)Ribosomal_L31 (pfam01197)→ |
| Lfe-12-1 | NC_010610.1 | + | 249776 | 249809 | **RNA**→ rpmE2 (PRK01678)Ribosomal_L31 (pfam01197)→ |
| Lfe-9-1 | NZ_GG704707.1 | + | 45973 | 46006 | **RNA**→ rpmE2 (PRK01678)Ribosomal_L31 (pfam01197)→ rpmE2 (PRK01678)Ribosomal_L31 (pfam01197 |
| Lfe-10-1 | NZ_GG669907.1 | + | 47108 | 47141 | **RNA**→ rpmE2 (PRK01678)Ribosomal_L31 (pfam01197)→ |
| Lfe-3-1 | NC_021235.1 | + | 246048 | 246081 | **RNA**→ rpmE2 (PRK01678)Ribosomal_L31 (pfam01197)→ |
| Lfe-4-1 | NZ_KE356737.1 | + | 45908 | 45941 | **RNA**→ rpmE2 (PRK01678)Ribosomal_L31 (pfam01197)→ |
| Lfe-8-1 | NZ_KI546217.1 | + | 36406 | 36439 | **RNA**→ rpmE2 (PRK01678)Ribosomal_L31 (pfam01197)→ |
| Lfe-2-1 | NZ_CP011536.1 | + | 270394 | 270427 | **RNA**→ rpmE2 (PRK01678)Ribosomal_L31 (pfam01197)→ |
| Lfe-6-1 | NZ_AWXS01000029.1 | + | 34177 | 34210 | **RNA**→ rpmE2 (PRK01678)Ribosomal_L31 (pfam01197)→ |
| env-894 | scaffold29246_2_V1.CD-11 | - | 3405 | 3372 | **RNA**→ rpmE2 (PRK01678)Ribosomal_L31 (pfam01197)→ |
| env-895 | SRS014343_WUGC_scaffold_819 | - | 226 | 193 | **RNA**→ rpmE2 (PRK01678)Ribosomal_L31 (pfam01197)→ |
| env-896 | DLM027_scaffold17_2 | - | 19027 | 18994 | **RNA**→ rpmE2 (PRK01678)Ribosomal_L31 (pfam01197)→ |
| env-897 | 2157436120 | - | 110 | 77 | **RNA**→ hypo→ |
| Lre-12-1 | NZ_ACGX02000004.1 | + | 87043 | 87076 | **RNA**→ rpmE2 (PRK01678)Ribosomal_L31 (pfam01197)→ rpmE2 (PRK01678)Ribosomal_L31 (pfam01197 |
| Lre-9-1 | NC_009513.1 | + | 269553 | 269586 | **RNA**→ rpmE2 (PRK01678)Ribosomal_L31 (pfam01197)→ |
| Lre-5-1 | NZ_AEAX01000117.1 | + | 5106 | 5139 | **RNA**→ rpmE2 (PRK01678)Ribosomal_L31 (pfam01197)→ |
| Lre-10-1 | NC_010609.1 | + | 269373 | 269406 | **RNA**→ rpmE2 (PRK01678)Ribosomal_L31 (pfam01197)→ |
| Lre-11-1 | NZ_GG693761.1 | + | 83277 | 83310 | **RNA**→ rpmE2 (PRK01678)Ribosomal_L31 (pfam01197)→ |
| Lre-4-1 | NC_021872.1 | + | 288927 | 288960 | **RNA**→ rpmE2 (PRK01678)Ribosomal_L31 (pfam01197)→ |
| Lre-1-1 | NZ_CP011024.1 | + | 269552 | 269585 | **RNA**→ rpmE2 (PRK01678)Ribosomal_L31 (pfam01197)→ |
| env-898 | Contig9771.1 | + | 339 | 372 | **RNA**→ |
| env-899 | JGI24707J26582_10423514 | - | 118 | 85 | **RNA**→ |
| Lre-13-1 | NC_015697.1 | + | 1423149 | 1423182 | **RNA**→ rpmE2 (PRK01678)Ribosomal_L31 (pfam01197)→ |
| Lre-8-1 | NZ_GG693671.1 | - | 45330 | 45297 | **RNA**→ rpmE2 (PRK01678)Ribosomal_L31 (pfam01197)→ |
| Lva-2-1 | NZ_GG693416.1 | + | 80452 | 80485 | **RNA**→ rpmE2 (PRK01678)Ribosomal_L31 (pfam01197)→ |
| env-900 | 2157771745 | + | 128 | 161 | **RNA**→ rpmE (PRK00019)rpmE2 (PRK01678)L31 (TIGR00105)Ribosomal_L31 (pfam01197)→ |
| env-901 | CTBMPR_c23 | + | 46289 | 46321 | **RNA**→ rpmE (PRK00019)Ribosomal_L31 (pfam01197)→ Tdk (COG1435)PRK04296 (PRK04296)→ |
| Ate-2-1 | NZ_AUMJ01000018.1 | - | 70832 | 70800 | **RNA**→ rpmE (PRK00019)Ribosomal_L31 (pfam01197)→ |
| Bth-45-1 | NZ_AODI01000040.1 | - | 38625 | 38596 | **RNA**→ rpmE2 (PRK01678)Ribosomal_L31 (pfam01197)→ |
| Bth-45-2 | NZ_JHZM01000007.1 | - | 126566 | 126537 | **RNA**→ rpmE2 (PRK01678)Ribosomal_L31 (pfam01197)→ |
| env-902 | JGI26050J44790_1004361 | + | 1747 | 1778 | **RNA**→ rpmE (PRK00019)Ribosomal_L31 (pfam01197)→ |
| Bth-44-1 | NZ_JXLU01000136.1 | + | 78859 | 78890 | **RNA**→ rpmE (PRK00019)Ribosomal_L31 (pfam01197)→ |
| Bth-44-2 | NZ_JXLR01000022.1 | + | 37521 | 37552 | **RNA**→ rpmE (PRK00019)Ribosomal_L31 (pfam01197)→ |
| Bth-44-3 | NZ_JXLS01000066.1 | + | 66077 | 66108 | **RNA**→ rpmE (PRK00019)Ribosomal_L31 (pfam01197)→ |
| Bth-44-4 | NZ_JXLT01000090.1 | - | 39361 | 39330 | **RNA**→ rpmE (PRK00019)Ribosomal_L31 (pfam01197)→ |

| | | | | | |
|---|---|---|---|---|---|
| Bth-44-5 | NZ_CCRF01000101.1 | - | 36758 | 36727 | **RNA**→ rpmE (PRK00019)Ribosomal_L31 (pfam01197)→ |
| env-903 | JGI26050J44790_1000142 | - | 20101 | 20070 | **RNA**→ rpmE (PRK00019)Ribosomal_L31 (pfam01197)→ |
| Bca-2-1 | NZ_AODH01000022.1 | - | 112821 | 112792 | **RNA**→ rpmE2 (PRK01678)Ribosomal_L31 (pfam01197)→ |
| Pda-2-1 | NZ_JANK01000056.1 | + | 12835 | 12867 | **RNA**→ rpmE2 (PRK01678)Ribosomal_L31 (pfam01197)→ |
| env-904 | 2205852404 | + | 261 | 293 | **RNA**→ rpmE2 (PRK01678)Ribosomal_L31 (pfam01197)→ |
| env-905 | 2157565684 | + | 721 | 753 | **RNA**→ rpmE2 (PRK01678)Ribosomal_L31 (pfam01197)→ |
| Lsu-2-1 | NZ_BACO01000082.1 | - | 343 | 311 | **RNA**→ rpmE2 (PRK01678)Ribosomal_L31 (pfam01197)→ rpmE2 (PRK01678)Ribosomal_L31 (pfam01197 |
| Lmu-3-1 | NZ_KB822405.1 | - | 369578 | 369546 | **RNA**→ rpmE2 (PRK01678)Ribosomal_L31 (pfam01197)→ |
| env-906 | 2204772568 | - | 71 | 39 | **RNA**→ |
| env-907 | 2158052713 | - | 71 | 39 | **RNA**→ |
| Lan-2-1 | NZ_GL573157.1 | - | 796104 | 796072 | **RNA**→ rpmE2 (PRK01678)Ribosomal_L31 (pfam01197)→ |
| Lan-1-1 | NZ_JMHU01000004.1 | - | 157949 | 157917 | **RNA**→ rpmE2 (PRK01678)Ribosomal_L31 (pfam01197)→ |
| env-908 | 2205292183 | - | 156 | 124 | **RNA**→ rpmE2 (PRK01678)Ribosomal_L31 (pfam01197)→ |
| env-909 | 2204671542 | - | 405 | 373 | **RNA**→ rpmE2 (PRK01678)Ribosomal_L31 (pfam01197)→ |
| env-910 | 4491487.3_NODE_48029 | - | 135 | 103 | **RNA**→ rpmE2 (PRK01678)Ribosomal_L31 (pfam01197)→ |
| env-911 | DLM009_scaffold15_5 | - | 432 | 400 | **RNA**→ rpmE2 (PRK01678)Ribosomal_L31 (pfam01197)→ |
| env-912 | SRS020869_C3352487 | + | 513 | 545 | **RNA**→ rpmE2 (PRK01678)Ribosomal_L31 (pfam01197)→ |
| Lru-3-1 | NZ_AWYA01000122.1 | - | 13216 | 13184 | **RNA**→ rpmE2 (PRK01678)Ribosomal_L31 (pfam01197)→ |
| Lru-1-1 | NZ_JHAB01000034.1 | - | 29783 | 29751 | **RNA**→ rpmE2 (PRK01678)Ribosomal_L31 (pfam01197)→ |
| Lru-4-1 | NZ_AWYB01000385.1 | + | 1155 | 1187 | **RNA**→ rpmE2 (PRK01678)Ribosomal_L31 (pfam01197)→ |
| Lru-6-1 | NZ_AFOJ01000007.1 | - | 259420 | 259388 | **RNA**→ rpmE2 (PRK01678)Ribosomal_L31 (pfam01197)→ rpmE2 (PRK01678)Ribosomal_L31 (pfam01197 hypo→ hypo→ |
| Lru-2-1 | NZ_GL833115.1 | + | 94026 | 94058 | **RNA**→ rpmE2 (PRK01678)Ribosomal_L31 (pfam01197)→ rpmE2 (PRK01678)Ribosomal_L31 (pfam01197 hypo→ |
| Lru-1-2 | NZ_JHAJ01000117.1 | + | 20548 | 20580 | **RNA**→ rpmE2 (PRK01678)Ribosomal_L31 (pfam01197)→ |
| Lru-5-1 | NC_015975.1 | + | 524423 | 524455 | **RNA**→ rpmE2 (PRK01678)Ribosomal_L31 (pfam01197)→ |
| env-913 | JXWR01003518.1 | - | 29634 | 29602 | **RNA**→ rpmE2 (PRK01678)Ribosomal_L31 (pfam01197)→ |
| Ppe-6-1 | NC_008525.1 | - | 1537440 | 1537408 | **RNA**→ rpmE2 (PRK01678)Ribosomal_L31 (pfam01197)→ |
| Ppe-5-1 | NC_022780.1 | - | 1501615 | 1501583 | **RNA**→ rpmE2 (PRK01678)Ribosomal_L31 (pfam01197)→ |
| Ppe-3-1 | NZ_JDVW01000008.1 | - | 330588 | 330556 | **RNA**→ rpmE2 (PRK01678)Ribosomal_L31 (pfam01197)→ |
| Ppe-4-1 | NZ_CAHU01000002.1 | + | 140218 | 140250 | **RNA**→ rpmE2 (PRK01678)Ribosomal_L31 (pfam01197)→ |
| Pac-2-1 | NZ_AGKB01000010.1 | - | 350604 | 350572 | **RNA**→ rpmE2 (PRK01678)Ribosomal_L31 (pfam01197)→ |
| Pac-5-1 | NZ_GL397067.1 | - | 1011122 | 1011090 | **RNA**→ rpmE2 (PRK01678)Ribosomal_L31 (pfam01197)→ |
| Pac-4-1 | NZ_GG730083.1 | + | 429682 | 429714 | **RNA**→ rpmE2 (PRK01678)Ribosomal_L31 (pfam01197)→ rpmE2 (PRK01678)Ribosomal_L31 (pfam01197 |
| Pac-1-1 | NZ_KB889550.1 | + | 1414333 | 1414365 | **RNA**→ rpmE2 (PRK01678)Ribosomal_L31 (pfam01197)→ |
| Pac-3-1 | NZ_JAGU01000001.1 | + | 1340990 | 1341022 | **RNA**→ rpmE2 (PRK01678)Ribosomal_L31 (pfam01197)→ |
| Plo-1-1 | NZ_BANK01000009.1 | + | 51598 | 51630 | **RNA**→ rpmE2 (PRK01678)Ribosomal_L31 (pfam01197)→ |
| Lco-2-1 | NZ_BAMK01000007.1 | + | 99264 | 99296 | **RNA**→ rpmE2 (PRK01678)Ribosomal_L31 (pfam01197)→ |
| Lce-1-1 | NZ_AUHP01000014.1 | - | 58515 | 58483 | **RNA**→ rpmE2 (PRK01678)Ribosomal_L31 (pfam01197)→ |
| Esa-2-1 | NZ_KE136389.1 | + | 614122 | 614154 | **RNA**→ rpmE2 (PRK01678)Ribosomal_L31 (pfam01197)→ |
| Esa-2-2 | NZ_KE136523.1 | + | 816131 | 816163 | **RNA**→ rpmE2 (PRK01678)Ribosomal_L31 (pfam01197)→ |
| Eph-1-1 | NZ_KB946326.1 | + | 83226 | 83258 | **RNA**→ rpmE2 (PRK01678)Ribosomal_L31 (pfam01197)→ |
| Mpl-1-1 | NZ_JSAX01000004.1 | - | 863 | 831 | **RNA**→ rpmE2 (PRK01678)Ribosomal_L31 (pfam01197)→ |
| Mpl-1-2 | NZ_JSAY01000004.1 | - | 166842 | 166810 | **RNA**→ rpmE2 (PRK01678)Ribosomal_L31 (pfam01197)→ |
| Mpl-1-3 | NZ_JSAV01000005.1 | - | 866 | 834 | **RNA**→ rpmE2 (PRK01678)Ribosomal_L31 (pfam01197)→ |
| Mpl-1-4 | NZ_JSBC01000004.1 | - | 866 | 834 | **RNA**→ rpmE2 (PRK01678)Ribosomal_L31 (pfam01197)→ |
| Mpl-1-5 | NZ_JSBA01000004.1 | - | 597 | 565 | **RNA**→ rpmE2 (PRK01678)Ribosomal_L31 (pfam01197)→ |
| Mpl-1-6 | NZ_JSBF01000004.1 | - | 866 | 834 | **RNA**→ rpmE2 (PRK01678)Ribosomal_L31 (pfam01197)→ |
| Mpl-1-7 | NZ_JSBD01000006.1 | - | 866 | 834 | **RNA**→ rpmE2 (PRK01678)Ribosomal_L31 (pfam01197)→ |
| Mpl-1-8 | NZ_JSAW01000002.1 | - | 1025447 | 1025415 | **RNA**→ rpmE2 (PRK01678)Ribosomal_L31 (pfam01197)→ |
| Mpl-1-9 | NZ_JSBE01000004.1 | - | 857 | 825 | **RNA**→ rpmE2 (PRK01678)Ribosomal_L31 (pfam01197)→ |
| Mpl-2-1 | NC_015516.1 | - | 1279834 | 1279802 | **RNA**→ rpmE2 (PRK01678)Ribosomal_L31 (pfam01197)→ |
| Mpl-3-1 | NC_016938.1 | + | 824026 | 824058 | **RNA**→ rpmE2 (PRK01678)Ribosomal_L31 (pfam01197)→ |

| | | | | | |
|---|---|---|---|---|---|
| Mpl-1-10 | NZ_JSAZ01000004.1 | - | 1471 | 1439 | **RNA**→ rpmE2 (PRK01678)Ribosomal_L31 (pfam01197)→ |
| Mpl-1-11 | NZ_JSBB01000006.1 | - | 626 | 594 | **RNA**→ rpmE2 (PRK01678)Ribosomal_L31 (pfam01197)→ |
| Lsu-1-1 | NZ_BALC01000008.1 | + | 59522 | 59554 | **RNA**→ rpmE2 (PRK01678)Ribosomal_L31 (pfam01197)→ |
| env-914 | 2205830766 | - | 165 | 133 | **RNA**→ ←hypo |
| env-915 | 2157581444 | - | 165 | 133 | **RNA**→ ←hypo |
| Lfa-2-1 | NZ_KK036465.1 | + | 21547 | 21579 | **RNA**→ rpmE2 (PRK01678)Ribosomal_L31 (pfam01197)→ |
| Lfa-1-1 | NZ_AYGX01000076.1 | - | 1369 | 1337 | **RNA**→ rpmE2 (PRK01678)Ribosomal_L31 (pfam01197)→ |
| Lor-2-1 | NZ_BBJM01000005.1 | - | 31903 | 31871 | **RNA**→ rpmE2 (PRK01678)Ribosomal_L31 (pfam01197)→ |
| Lma-4-1 | NZ_BACN01000090.1 | - | 11120 | 11088 | **RNA**→ rpmE2 (PRK01678)Ribosomal_L31 (pfam01197)→ rpmE2 (PRK01678)Ribosomal_L31 (pfam01197 |
| Lhe-11-1 | NZ_LFEE01000004.1 | + | 26276 | 26308 | **RNA**→ rpmE2 (PRK01678)Ribosomal_L31 (pfam01197)→ |
| env-916 | 2205834061 | + | 655 | 687 | **RNA**→ rpmE2 (PRK01678)Ribosomal_L31 (pfam01197)→ |
| env-917 | JXWR01000016.1 | + | 2483 | 2515 | **RNA**→ rpmE2 (PRK01678)Ribosomal_L31 (pfam01197)→ |
| env-918 | 2157560113 | + | 655 | 687 | **RNA**→ rpmE2 (PRK01678)Ribosomal_L31 (pfam01197)→ |
| Lpl-9-1 | NZ_ASJE01000081.1 | + | 29802 | 29834 | **RNA**→ rpmE2 (PRK01678)Ribosomal_L31 (pfam01197)→ |
| Lpl-10-1 | NZ_AXDQ01000018.1 | + | 2888 | 2920 | **RNA**→ rpmE2 (PRK01678)Ribosomal_L31 (pfam01197)→ |
| Lpl-12-1 | NZ_AUTE01000054.1 | - | 8536 | 8504 | **RNA**→ rpmE2 (PRK01678)Ribosomal_L31 (pfam01197)→ |
| Lpl-16-1 | NZ_BALV01000016.1 | + | 25883 | 25915 | **RNA**→ rpmE2 (PRK01678)Ribosomal_L31 (pfam01197)→ |
| Lpl-17-1 | NZ_AGRI01000003.1 | + | 112355 | 112387 | **RNA**→ rpmE2 (PRK01678)Ribosomal_L31 (pfam01197)→ |
| Lpl-3-1 | NZ_AWTS01000023.1 | + | 15075 | 15107 | **RNA**→ rpmE2 (PRK01678)Ribosomal_L31 (pfam01197)→ |
| Lpl-4-1 | NZ_AVFI01000125.1 | - | 215 | 183 | **RNA**→ rpmE2 (PRK01678)Ribosomal_L31 (pfam01197)→ |
| Lpl-20-1 | NC_004567.2 | + | 458540 | 458572 | **RNA**→ rpmE2 (PRK01678)Ribosomal_L31 (pfam01197)→ |
| Lpl-5-1 | NZ_AYTU01000005.1 | - | 6825 | 6793 | **RNA**→ rpmE2 (PRK01678)Ribosomal_L31 (pfam01197)→ |
| Lpl-21-1 | NC_014554.1 | + | 434946 | 434978 | **RNA**→ rpmE2 (PRK01678)Ribosomal_L31 (pfam01197)→ |
| Lpl-2-1 | NC_021514.1 | + | 466087 | 466119 | **RNA**→ rpmE2 (PRK01678)Ribosomal_L31 (pfam01197)→ |
| Lpl-19-1 | NC_012984.1 | + | 438813 | 438845 | **RNA**→ rpmE2 (PRK01678)Ribosomal_L31 (pfam01197)→ |
| Lpl-15-1 | NZ_GL379763.1 | + | 506009 | 506041 | **RNA**→ rpmE2 (PRK01678)Ribosomal_L31 (pfam01197)→ |
| Lpl-11-1 | NZ_KB644964.1 | + | 98775 | 98807 | **RNA**→ rpmE2 (PRK01678)Ribosomal_L31 (pfam01197)→ |
| Lpl-18-1 | NC_021224.2 | + | 416447 | 416479 | **RNA**→ rpmE2 (PRK01678)Ribosomal_L31 (pfam01197)→ |
| Lpl-1-1 | NZ_CP010528.1 | + | 455988 | 456020 | **RNA**→ rpmE2 (PRK01678)Ribosomal_L31 (pfam01197)→ |
| Lpl-7-1 | NZ_CM002918.1 | + | 447842 | 447874 | **RNA**→ rpmE2 (PRK01678)Ribosomal_L31 (pfam01197)→ |
| Lpl-14-1 | NZ_CM003439.1 | + | 3104335 | 3104367 | **RNA**→ rpmE2 (PRK01678)Ribosomal_L31 (pfam01197)→ |
| Lpl-8-1 | NZ_AVAQ01000014.1 | + | 3252 | 3284 | **RNA**→ rpmE2 (PRK01678)Ribosomal_L31 (pfam01197)→ |
| Lpl-6-1 | NZ_AVAI01000131.1 | - | 14495 | 14463 | **RNA**→ rpmE2 (PRK01678)Ribosomal_L31 (pfam01197)→ hypo→ |
| Lpl-13-1 | NC_020229.1 | + | 683468 | 683500 | **RNA**→ rpmE2 (PRK01678)Ribosomal_L31 (pfam01197)→ |
| Lpe-2-1 | NZ_CM001538.1 | + | 477939 | 477971 | **RNA**→ rpmE2 (PRK01678)Ribosomal_L31 (pfam01197)→ |
| Lpe-1-1 | NZ_LFLY01000001.1 | - | 100060 | 100028 | **RNA**→ rpmE2 (PRK01678)Ribosomal_L31 (pfam01197)→ |
| Lpa-41-1 | NZ_KN173718.1 | - | 48885 | 48853 | **RNA**→ rpmE2 (PRK01678)Ribosomal_L31 (pfam01197)→ |
| Eit-1-1 | NZ_GL622241.1 | - | 1826854 | 1826822 | **RNA**→ rpmE2 (PRK01678)Ribosomal_L31 (pfam01197)→ |
| env-919 | 2210741799 | + | 170 | 204 | **RNA**→ |
| Vlu-1-1 | NZ_AYSH01000010.1 | + | 32998 | 33030 | **RNA**→ rpmE2 (PRK01678)Ribosomal_L31 (pfam01197)→ |
| env-920 | 2158154984 | + | 1083 | 1115 | **RNA**→ rpmE (PRK00019)rpmE2 (PRK01678)Ribosomal_L31 (pfam01197)→ |
| env-921 | 2205359904 | + | 1083 | 1115 | **RNA**→ rpmE2 (PRK01678)Ribosomal_L31 (pfam01197)→ |
| Tha-1-1 | NC_016052.1 | + | 990867 | 990899 | **RNA**→ rpmE2 (PRK01678)Ribosomal_L31 (pfam01197)→ |
| Tmu-2-1 | NZ_KE384092.1 | - | 4460 | 4428 | **RNA**→ rpmE2 (PRK01678)Ribosomal_L31 (pfam01197)→ |
| Tmu-1-1 | NZ_JPVT01000082.1 | + | 2641 | 2673 | **RNA**→ rpmE2 (PRK01678)Ribosomal_L31 (pfam01197)→ |
| Tmu-3-1 | NZ_JPVU01000102.1 | + | 5752 | 5784 | **RNA**→ rpmE2 (PRK01678)Ribosomal_L31 (pfam01197)→ |
| env-922 | 4491401.3_NODE_92843 | + | 4662 | 4694 | **RNA**→ rpmE2 (PRK01678)Ribosomal_L31 (pfam01197)→ |
| Lsa-11-1 | NC_017481.1 | + | 378680 | 378712 | **RNA**→ rpmE2 (PRK01678)Ribosomal_L31 (pfam01197)→ rpmE2 (PRK01678)Ribosomal_L31 (pfam01197 rpmE2 (PRK01678)Ribosomal_L31 (pfam01197)→ hypo→ |
| Lsa-9-1 | NZ_AEBA01000017.1 | + | 18818 | 18850 | **RNA**→ rpmE2 (PRK01678)Ribosomal_L31 (pfam01197)→ rpmE2 (PRK01678)Ribosomal_L31 (pfam01197 hypo→ hypo→ |
| Lsa-5-1 | NZ_AFMN01000001.1 | - | 7360 | 7328 | **RNA**→ rpmE2 (PRK01678)Ribosomal_L31 (pfam01197)→ hypo→ |
| Lsa-10-1 | NZ_GG693222.1 | - | 64281 | 64249 | **RNA**→ rpmE2 (PRK01678)Ribosomal_L31 (pfam01197)→ hypo→ |

| | | | | | |
|---|---|---|---|---|---|
| Lsa-6-1 | NZ_AICL01000001.1 | - | 7340 | 7308 | **RNA**→ rpmE2 (PRK01678)Ribosomal_L31 (pfam01197)→ hypo→ |
| Lsa-8-1 | NZ_CP011403.1 | + | 377103 | 377135 | **RNA**→ rpmE2 (PRK01678)Ribosomal_L31 (pfam01197)→ hypo→ |
| env-923 | NLM027_scaffold41938_1 | + | 38289 | 38321 | **RNA**→ rpmE2 (PRK01678)Ribosomal_L31 (pfam01197)→ hypo→ |
| Lsa-4-1 | NZ_AFOI01000007.1 | - | 10694 | 10662 | **RNA**→ rpmE2 (PRK01678)Ribosomal_L31 (pfam01197)→ hypo→ |
| Lsa-3-1 | NZ_CP007646.1 | + | 359733 | 359765 | **RNA**→ rpmE2 (PRK01678)Ribosomal_L31 (pfam01197)→ hypo→ hypo→ hypo→ hypo→ |
| Lsa-7-1 | NZ_CBVR010000053.1 | - | 1525 | 1493 | **RNA**→ rpmE2 (PRK01678)Ribosomal_L31 (pfam01197)→ hypo→ |
| Pcl-1-1 | NC_016605.1 | + | 219415 | 219447 | **RNA**→ rpmE2 (PRK01678)Ribosomal_L31 (pfam01197)→ |
| Bfo-1-1 | NZ_KB894703.1 | + | 46890 | 46924 | **RNA**→ rpmE (PRK00019)Ribosomal_L31 (pfam01197)→ |
| Gic-1-1 | NZ_JPYA01000113.1 | - | 36992 | 36961 | **RNA**→ rpmE (PRK00019)Ribosomal_L31 (pfam01197)→ |
| Gsu-1-1 | NZ_JQMN01000001.1 | + | 536242 | 536273 | **RNA**→ rpmE (PRK00019)Ribosomal_L31 (pfam01197)→ |
| Gth-6-1 | NZ_JFHZ01000051.1 | - | 48767 | 48736 | **RNA**→ rpmE (PRK00019)Ribosomal_L31 (pfam01197)→ |
| Gsp-16-1 | NZ_LDNZ01000071.1 | + | 7084 | 7115 | **RNA**→ rpmE (PRK00019)Ribosomal_L31 (pfam01197)→ |
| Gst-1-1 | NZ_LDNS01000203.1 | - | 1968 | 1937 | **RNA**→ rpmE (PRK00019)Ribosomal_L31 (pfam01197)→ |
| Gst-1-2 | NZ_LDNT01000178.1 | - | 1956 | 1925 | **RNA**→ rpmE (PRK00019)Ribosomal_L31 (pfam01197)→ |
| Gth-1-1 | NC_009328.1 | - | 3431917 | 3431886 | **RNA**→ rpmE (PRK00019)Ribosomal_L31 (pfam01197)→ |
| Gsp-2-1 | NZ_ABVH01000007.1 | - | 54857 | 54826 | **RNA**→ rpmE (PRK00019)Ribosomal_L31 (pfam01197)→ rpmE (PRK00019)Ribosomal_L31 (pfam01197)→ |
| Age-1-1 | NZ_JYCG01000286.1 | - | 19333 | 19302 | **RNA**→ rpmE (PRK00019)Ribosomal_L31 (pfam01197)→ |
| Gst-1-3 | NZ_JQCS01000028.1 | - | 13433 | 13402 | **RNA**→ rpmE (PRK00019)Ribosomal_L31 (pfam01197)→ |
| Gst-1-4 | NZ_JPYV01000032.1 | - | 54116 | 54085 | **RNA**→ rpmE (PRK00019)Ribosomal_L31 (pfam01197)→ |
| Gsp-14-1 | NZ_CP008903.1 | - | 1776671 | 1776640 | **RNA**→ rpmE (PRK00019)Ribosomal_L31 (pfam01197)→ |
| Gsp-8-1 | NZ_CP011832.1 | - | 3291606 | 3291575 | **RNA**→ rpmE (PRK00019)Ribosomal_L31 (pfam01197)→ |
| Gth-4-1 | NZ_AYKT01000007.1 | + | 122211 | 122242 | **RNA**→ rpmE (PRK00019)Ribosomal_L31 (pfam01197)→ |
| Gst-1-5 | NZ_LDNU01000010.1 | + | 23005 | 23036 | **RNA**→ rpmE (PRK00019)Ribosomal_L31 (pfam01197)→ |
| Gst-3-1 | NZ_JALS01000096.1 | - | 12601 | 12570 | **RNA**→ rpmE (PRK00019)Ribosomal_L31 (pfam01197)→ |
| Gka-3-1 | NZ_JYCF01000134.1 | - | 47171 | 47140 | **RNA**→ rpmE (PRK00019)Ribosomal_L31 (pfam01197)→ |
| Gka-3-2 | NZ_JYBP01000003.1 | - | 3309423 | 3309392 | **RNA**→ rpmE (PRK00019)Ribosomal_L31 (pfam01197)→ |
| Gvu-1-1 | NZ_JPOI01000001.1 | + | 1071989 | 1072020 | **RNA**→ rpmE (PRK00019)Ribosomal_L31 (pfam01197)→ |
| Gsp-9-1 | NZ_AUXP01000009.1 | + | 97898 | 97929 | **RNA**→ rpmE (PRK00019)Ribosomal_L31 (pfam01197)→ |
| Gsp-7-1 | NC_013411.1 | - | 3495641 | 3495610 | **RNA**→ rpmE (PRK00019)Ribosomal_L31 (pfam01197)→ |
| Gsp-6-1 | NC_014915.1 | - | 3501607 | 3501576 | **RNA**→ rpmE (PRK00019)Ribosomal_L31 (pfam01197)→ |
| Gsp-1-1 | NC_014206.1 | - | 3537049 | 3537018 | **RNA**→ rpmE (PRK00019)Ribosomal_L31 (pfam01197)→ |
| Gst-2-1 | NZ_CP008934.1 | - | 1637831 | 1637800 | **RNA**→ rpmE (PRK00019)Ribosomal_L31 (pfam01197)→ |
| Gka-4-1 | NZ_BBJV01000020.1 | - | 289 | 258 | **RNA**→ rpmE (PRK00019)Ribosomal_L31 (pfam01197)→ |
| Gsp-18-1 | NZ_LDPD01000028.1 | - | 53895 | 53864 | **RNA**→ rpmE (PRK00019)Ribosomal_L31 (pfam01197)→ |
| Gka-2-1 | NZ_BASG01000006.1 | + | 56589 | 56620 | **RNA**→ rpmE (PRK00019)Ribosomal_L31 (pfam01197)→ |
| Gsp-15-1 | NZ_AYSF01000001.1 | - | 65897 | 65866 | **RNA**→ rpmE (PRK00019)Ribosomal_L31 (pfam01197)→ |
| Gsp-11-1 | NZ_JGCJ01000027.1 | + | 42199 | 42230 | **RNA**→ rpmE (PRK00019)Ribosomal_L31 (pfam01197)→ |
| Gsp-10-1 | NZ_JHUS01000002.1 | - | 21989 | 21958 | **RNA**→ rpmE (PRK00019)Ribosomal_L31 (pfam01197)→ |
| Gka-1-1 | NC_006510.1 | - | 3419800 | 3419769 | **RNA**→ rpmE (PRK00019)Ribosomal_L31 (pfam01197)→ |
| Gth-3-1 | NC_016593.1 | - | 3486231 | 3486200 | **RNA**→ rpmE (PRK00019)Ribosomal_L31 (pfam01197)→ |
| Gsp-12-1 | NC_020210.1 | - | 3468659 | 3468628 | **RNA**→ rpmE (PRK00019)Ribosomal_L31 (pfam01197)→ |
| Gth-7-1 | NZ_BATY01000060.1 | + | 41926 | 41957 | **RNA**→ rpmE (PRK00019)Ribosomal_L31 (pfam01197)→ |
| Gsp-17-1 | NZ_ATCO01000040.1 | + | 14050 | 14081 | **RNA**→ rpmE (PRK00019)Ribosomal_L31 (pfam01197)→ rpmE (PRK00019)Ribosomal_L31 (pfam01197)→ |
| Bin-1-1 | NZ_JNVC02000013.1 | + | 220889 | 220922 | **RNA**→ rpmE (PRK00019)Ribosomal_L31 (pfam01197)→ |
| Bin-2-1 | NZ_JGVU02000005.1 | + | 287101 | 287134 | **RNA**→ rpmE (PRK00019)Ribosomal_L31 (pfam01197)→ |
| env-924 | JGI25317J35169_1191885 | + | 62 | 94 | **RNA**→ rpmE (PRK00019)Ribosomal_L31 (pfam01197)→ |
| env-925 | JGI25319J35699_1005535 | + | 969 | 1001 | **RNA**→ rpmE (PRK00019)Ribosomal_L31 (pfam01197)→ Tdk (COG1435)PRK04296 (PRK04296)→ spore_YhcN_YlaJ (TIGR02898)→ |
| env-926 | JGI25318J35512_1000658 | - | 8238 | 8206 | **RNA**→ rpmE (PRK00019)Ribosomal_L31 (pfam01197)→ Tdk (COG1435)PRK04296 (PRK04296)→ spore_YhcN_YlaJ (TIGR02898)→ |
| env-927 | JGI25321J39296_1018370 | + | 355 | 387 | **RNA**→ rpmE (PRK00019)Ribosomal_L31 (pfam01197)→ Tdk (COG1435)PRK04296 (PRK04296)→ hypo→ |
| env-928 | JGI25320J50211_1070748 | - | 116 | 84 | **RNA**→ hypo→ |

| | | | | | |
|---|---|---|---|---|---|
| env-929 | JGI25321J50212_10003866 | + | 2270 | 2302 | RNA→ rpmE (PRK00019)Ribosomal_L31 (pfam01197)→ Tdk (COG1435)PRK04296 (PRK04296)→ spore_YhcN_YlaJ (TIGR02898)→ |
| Aan-1-1 | NZ_KE952903.1 | - | 93698 | 93667 | RNA→ rpmE (PRK00019)Ribosomal_L31 (pfam01197)→ rpmE (PRK00019)Ribosomal_L31 (pfam01197)→ |
| Ami-1-1 | NZ_JYBN01000089.1 | - | 268857 | 268826 | RNA→ rpmE (PRK00019)Ribosomal_L31 (pfam01197)→ |
| Ami-1-2 | NZ_JYBO01000103.1 | + | 23763 | 23794 | RNA→ rpmE (PRK00019)Ribosomal_L31 (pfam01197)→ |
| Ami-1-3 | NZ_LGUG01000004.1 | + | 127332 | 127363 | RNA→ rpmE (PRK00019)Ribosomal_L31 (pfam01197)→ |
| Aty-1-1 | NZ_BBWZ01000002.1 | + | 104374 | 104405 | RNA→ rpmE (PRK00019)Ribosomal_L31 (pfam01197)→ |
| env-930 | 2211283201 | + | 84 | 115 | RNA→ |
| env-931 | 2156330801 | + | 84 | 115 | RNA→ |
| Bma-7-1 | NZ_HE978512.1 | - | 417836 | 417805 | RNA→ rpmE (PRK00019)Ribosomal_L31 (pfam01197)→ |
| Jal-1-1 | NZ_JXRQ01000028.1 | - | 38185 | 38155 | RNA→ rpmE (PRK00019)Ribosomal_L31 (pfam01197)→ |
| Jma-1-1 | NZ_CP009416.1 | - | 2865774 | 2865744 | RNA→ rpmE (PRK00019)Ribosomal_L31 (pfam01197)→ |
| Jso-1-1 | NZ_JXRP01000009.1 | + | 567308 | 567338 | RNA→ rpmE (PRK00019)Ribosomal_L31 (pfam01197)→ |
| Jca-1-1 | NZ_JXRR01000001.1 | - | 1685 | 1655 | RNA→ rpmE (PRK00019)Ribosomal_L31 (pfam01197)→ |
| env-932 | JGI24024J18818_10362070 | - | 302 | 268 | RNA→ rpmE (PRK00019)Ribosomal_L31 (pfam01197)→ |
| Bsp-8-1 | NZ_ACPC01000050.1 | - | 203195 | 203164 | RNA→ rpmE2 (PRK01678)Ribosomal_L31 (pfam01197)→ rpmE2 (PRK01678)Ribosomal_L31 (pfam01197) |
| Bsp-66-1 | NZ_LFEL01000002.1 | - | 551243 | 551212 | RNA→ rpmE2 (PRK01678)Ribosomal_L31 (pfam01197)→ |
| env-933 | JGI25151J46595_10000061 | - | 4986 | 4953 | RNA→ rpmE (PRK00019)Ribosomal_L31 (pfam01197)→ Tdk (COG1435)PRK04296 (PRK04296)→ |
| Bsp-95-1 | NZ_CP010075.1 | - | 3326407 | 3326374 | RNA→ rpmE (PRK00019)Ribosomal_L31 (pfam01197)→ Tdk (COG1435)PRK04296 (PRK04296)→ |
| Bst-1-1 | NZ_LDWL01000001.1 | - | 56445 | 56412 | RNA→ rpmE (PRK00019)Ribosomal_L31 (pfam01197)→ Tdk (COG1435)PRK04296 (PRK04296)→ |
| Bst-1-2 | NZ_LDWU01000019.1 | - | 72041 | 72008 | RNA→ rpmE (PRK00019)Ribosomal_L31 (pfam01197)→ Tdk (COG1435)PRK04296 (PRK04296)→ |
| Bst-1-3 | NZ_LDUH01000009.1 | + | 243731 | 243764 | RNA→ rpmE (PRK00019)Ribosomal_L31 (pfam01197)→ Tdk (COG1435)PRK04296 (PRK04296)→ |
| Bsp-63-1 | NZ_LDUQ01000001.1 | + | 281801 | 281834 | RNA→ rpmE (PRK00019)Ribosomal_L31 (pfam01197)→ Tdk (COG1435)PRK04296 (PRK04296)→ |
| Bsp-4-1 | NZ_AMSH01000002.1 | - | 87481 | 87448 | RNA→ rpmE (PRK00019)Ribosomal_L31 (pfam01197)→ rpmE (PRK00019)Ribosomal_L31 (pfam01197)→ Tdk (COG1435)PRK04296 (PRK04296)→ Tdk (COG1435)PRK04296 (PRK04296)→ |
| Bst-1-4 | NZ_LDWS01000018.1 | - | 87668 | 87635 | RNA→ rpmE (PRK00019)Ribosomal_L31 (pfam01197)→ Tdk (COG1435)PRK04296 (PRK04296)→ |
| Bst-1-5 | NZ_LDWI01000005.1 | - | 421554 | 421521 | RNA→ rpmE (PRK00019)Ribosomal_L31 (pfam01197)→ Tdk (COG1435)PRK04296 (PRK04296)→ |
| Bst-1-6 | NZ_LFMM01000007.1 | - | 76273 | 76240 | RNA→ rpmE (PRK00019)Ribosomal_L31 (pfam01197)→ Tdk (COG1435)PRK04296 (PRK04296)→ |
| Bsp-68-1 | NZ_LAVH01000010.1 | - | 169347 | 169314 | RNA→ rpmE (PRK00019)Ribosomal_L31 (pfam01197)→ Tdk (COG1435)PRK04296 (PRK04296)→ |
| Bpu-3-1 | NZ_CP009108.1 | - | 589531 | 589498 | RNA→ rpmE (PRK00019)Ribosomal_L31 (pfam01197)→ Tdk (COG1435)PRK04296 (PRK04296)→ |
| Bpu-3-2 | NZ_CP007436.1 | - | 3366927 | 3366894 | RNA→ rpmE (PRK00019)Ribosomal_L31 (pfam01197)→ Tdk (COG1435)PRK04296 (PRK04296)→ |
| Bpu-3-3 | NZ_CP011150.1 | - | 3353307 | 3353274 | RNA→ rpmE (PRK00019)Ribosomal_L31 (pfam01197)→ Tdk (COG1435)PRK04296 (PRK04296)→ |
| Bso-1-1 | NZ_AOFM01000009.1 | - | 593767 | 593734 | RNA→ rpmE (PRK00019)Ribosomal_L31 (pfam01197)→ Tdk (COG1435)PRK04296 (PRK04296)→ |
| Bso-2-1 | NZ_AYTN01000054.1 | - | 411813 | 411780 | RNA→ rpmE (PRK00019)Ribosomal_L31 (pfam01197)→ Tdk (COG1435)PRK04296 (PRK04296)→ |
| Bli-2-1 | NZ_JYBQ01000001.1 | - | 358665 | 358632 | RNA→ rpmE (PRK00019)Ribosomal_L31 (pfam01197)→ Tdk (COG1435)PRK04296 (PRK04296)→ |
| Bsp-33-1 | NZ_GL635727.1 | - | 941278 | 941245 | RNA→ rpmE (PRK00019)Ribosomal_L31 (pfam01197)→ Tdk (COG1435)PRK04296 (PRK04296)→ |
| Bli-11-1 | NZ_CP012110.1 | - | 3865351 | 3865318 | RNA→ rpmE (PRK00019)Ribosomal_L31 (pfam01197)→ Tdk (COG1435)PRK04296 (PRK04296)→ |
| Bli-3-1 | NZ_AJLV01000020.1 | + | 429445 | 429478 | RNA→ rpmE (PRK00019)Ribosomal_L31 (pfam01197)→ Tdk (COG1435)PRK04296 (PRK04296)→ |
| Bli-4-1 | NZ_AJLW01000015.1 | - | 100805 | 100772 | RNA→ rpmE (PRK00019)Ribosomal_L31 (pfam01197)→ Tdk (COG1435)PRK04296 (PRK04296)→ |
| Bli-1-1 | NC_006270.3 | - | 3780071 | 3780038 | RNA→ rpmE (PRK00019)Ribosomal_L31 (pfam01197)→ Tdk (COG1435)PRK04296 (PRK04296)→ |
| Bli-1-2 | NC_006322.1 | - | 3779923 | 3779890 | RNA→ rpmE (PRK00019)Ribosomal_L31 (pfam01197)→ rpmE (PRK00019)Ribosomal_L31 (pfam01197)→ Tdk (COG1435)PRK04296 (PRK04296)→ Tdk (COG1435)PRK04296 (PRK04296)→ |
| Bli-7-1 | NZ_AMWQ01000015.1 | - | 37002 | 36969 | RNA→ rpmE (PRK00019)Ribosomal_L31 (pfam01197)→ Tdk (COG1435)PRK04296 (PRK04296)→ |
| Bli-9-1 | NZ_AZSM01000001.1 | - | 309421 | 309388 | RNA→ rpmE (PRK00019)Ribosomal_L31 (pfam01197)→ Tdk (COG1435)PRK04296 (PRK04296)→ |
| Bli-8-1 | NZ_AZSL01000025.1 | + | 7611 | 7644 | RNA→ rpmE (PRK00019)Ribosomal_L31 (pfam01197)→ Tdk (COG1435)PRK04296 (PRK04296)→ |
| Bli-2-2 | NZ_JFYM01000008.1 | + | 428524 | 428557 | RNA→ rpmE (PRK00019)Ribosomal_L31 (pfam01197)→ Tdk (COG1435)PRK04296 (PRK04296)→ |
| Bli-6-1 | NZ_AVEZ01000039.1 | - | 520363 | 520330 | RNA→ rpmE (PRK00019)Ribosomal_L31 (pfam01197)→ Tdk (COG1435)PRK04296 (PRK04296)→ |
| Bsp-85-1 | NZ_JZBT01000024.1 | + | 30280 | 30313 | RNA→ rpmE (PRK00019)Ribosomal_L31 (pfam01197)→ Tdk (COG1435)PRK04296 (PRK04296)→ |
| Bgl-1-1 | NZ_LECV01000100.1 | + | 7406 | 7439 | RNA→ rpmE (PRK00019)Ribosomal_L31 (pfam01197)→ Tdk (COG1435)PRK04296 (PRK04296)→ |
| Bsp-48-1 | NZ_LECW01000089.1 | + | 25275 | 25308 | RNA→ rpmE (PRK00019)Ribosomal_L31 (pfam01197)→ Tdk (COG1435)PRK04296 (PRK04296)→ |
| Bli-2-3 | NZ_LFIM01000002.1 | - | 153023 | 152990 | RNA→ rpmE (PRK00019)Ribosomal_L31 (pfam01197)→ Tdk (COG1435)PRK04296 (PRK04296)→ |
| Bli-5-1 | NC_021362.1 | - | 3959595 | 3959562 | RNA→ rpmE (PRK00019)Ribosomal_L31 (pfam01197)→ Tdk (COG1435)PRK04296 (PRK04296)→ |
| Bsp-73-1 | NZ_JXAP01000005.1 | - | 592676 | 592643 | RNA→ rpmE (PRK00019)Ribosomal_L31 (pfam01197)→ Tdk (COG1435)PRK04296 (PRK04296)→ |

| | | | | | |
|---|---|---|---|---|---|
| Bli-10-1 | NZ_AZYP01000003.1 | - | 89779 | 89746 | **RNA**→ rpmE (PRK00019)Ribosomal_L31 (pfam01197)→ Tdk (COG1435)PRK04296 (PRK04296)→ |
| Bsp-35-1 | NZ_ANNR02000031.1 | + | 423248 | 423281 | **RNA**→ rpmE (PRK00019)Ribosomal_L31 (pfam01197)→ Tdk (COG1435)PRK04296 (PRK04296)→ |
| Bsp-82-1 | NZ_KE819960.1 | - | 635927 | 635894 | **RNA**→ rpmE (PRK00019)Ribosomal_L31 (pfam01197)→ Tdk (COG1435)PRK04296 (PRK04296)→ |
| Bpa-2-1 | NZ_CP010524.1 | - | 3979133 | 3979100 | **RNA**→ rpmE (PRK00019)Ribosomal_L31 (pfam01197)→ Tdk (COG1435)PRK04296 (PRK04296)→ |
| Bsp-60-1 | NZ_LBMN01000080.1 | + | 48203 | 48236 | **RNA**→ rpmE (PRK00019)Ribosomal_L31 (pfam01197)→ Tdk (COG1435)PRK04296 (PRK04296)→ |
| Bpa-3-1 | NZ_AZSK01000014.1 | + | 416910 | 416943 | **RNA**→ rpmE (PRK00019)Ribosomal_L31 (pfam01197)→ Tdk (COG1435)PRK04296 (PRK04296)→ |
| Bsp-76-1 | NZ_KE819965.1 | - | 716299 | 716266 | **RNA**→ rpmE (PRK00019)Ribosomal_L31 (pfam01197)→ Tdk (COG1435)PRK04296 (PRK04296)→ |
| Bsp-98-1 | NZ_KB223527.1 | - | 1669518 | 1669487 | **RNA**→ rpmE (PRK00019)Ribosomal_L31 (pfam01197)→ |
| Ama-1-1 | NZ_LELK01000004.1 | + | 263335 | 263366 | **RNA**→ rpmE (PRK00019)Ribosomal_L31 (pfam01197)→ |
| Baz-1-1 | NZ_AJLR01000149.1 | - | 16199 | 16168 | **RNA**→ rpmE (PRK00019)Ribosomal_L31 (pfam01197)→ rpmE (PRK00019)Ribosomal_L31 (pfam01197) |
| Baz-2-1 | NZ_JJRY01000009.1 | + | 91459 | 91490 | **RNA**→ rpmE (PRK00019)Ribosomal_L31 (pfam01197)→ |
| Bsp-7-1 | NZ_CH672356.1 | + | 28004 | 28034 | **RNA**→ rpmE (PRK00019)Ribosomal_L31 (pfam01197)→ |
| Bin-3-1 | NC_022524.1 | - | 4499937 | 4499907 | **RNA**→ rpmE (PRK00019)Ribosomal_L31 (pfam01197)→ |
| Bse-1-1 | NC_014219.1 | - | 3427416 | 3427386 | **RNA**→ rpmE2 (PRK01678)Ribosomal_L31 (pfam01197)→ |
| Bsp-13-1 | NZ_HE610986.1 | - | 937568 | 937535 | **RNA**→ rpmE (PRK00019)Ribosomal_L31 (pfam01197)→ |
| Bch-1-1 | NZ_KE386944.1 | + | 305910 | 305940 | **RNA**→ rpmE2 (PRK01678)Ribosomal_L31 (pfam01197)→ |
| env-934 | SR_TP_S2_1011259 | - | 203 | 173 | **RNA**→ rpmE2 (PRK01678)Ribosomal_L31 (pfam01197)→ |
| Bsp-58-1 | NZ_BAWC01000008.1 | + | 52801 | 52832 | **RNA**→ rpmE2 (PRK01678)Ribosomal_L31 (pfam01197)→ Tdk (COG1435)PRK04296 (PRK04296)→ hypo |
| Ble-1-1 | NZ_CP003923.1 | - | 3605244 | 3605213 | **RNA**→ rpmE2 (PRK01678)Ribosomal_L31 (pfam01197)→ Tdk (COG1435)PRK04296 (PRK04296)→ hypo |
| Bsp-86-1 | NZ_BAWL01000044.1 | - | 10324 | 10293 | **RNA**→ rpmE2 (PRK01678)Ribosomal_L31 (pfam01197)→ |
| Bcl-1-1 | NC_006582.1 | - | 4034646 | 4034615 | **RNA**→ rpmE2 (PRK01678)Ribosomal_L31 (pfam01197)→ Tdk (COG1435)PRK04296 (PRK04296)→ |
| Bcl-2-1 | NZ_JMCA01000031.1 | + | 272380 | 272411 | **RNA**→ rpmE2 (PRK01678)Ribosomal_L31 (pfam01197)→ Tdk (COG1435)PRK04296 (PRK04296)→ |
| Bal-1-1 | NZ_ALPT02000041.1 | - | 32350 | 32319 | **RNA**→ rpmE2 (PRK01678)Ribosomal_L31 (pfam01197)→ |
| Bwa-1-1 | NZ_BAUT01000070.1 | + | 16215 | 16246 | **RNA**→ rpmE2 (PRK01678)Ribosomal_L31 (pfam01197)→ |
| Bps-3-1 | NZ_LFJO01000001.1 | + | 460473 | 460504 | **RNA**→ rpmE2 (PRK01678)Ribosomal_L31 (pfam01197)→ |
| env-935 | JGI25318J35512_1127720 | - | 223 | 192 | **RNA**→ rpmE2 (PRK01678)Ribosomal_L31 (pfam01197)→ |
| Bhe-1-1 | NZ_BAUU01000018.1 | + | 23923 | 23954 | **RNA**→ rpmE2 (PRK01678)Ribosomal_L31 (pfam01197)→ |
| Bok-1-1 | NZ_JRJU01000042.1 | - | 26874 | 26843 | **RNA**→ rpmE2 (PRK01678)Ribosomal_L31 (pfam01197)→ |
| Bsp-56-1 | NZ_BAWA01000009.1 | + | 100911 | 100942 | **RNA**→ rpmE2 (PRK01678)Ribosomal_L31 (pfam01197)→ Tdk (COG1435)PRK04296 (PRK04296)→ hypo |
| Bsp-57-1 | NZ_BAWB01000007.1 | - | 27806 | 27775 | **RNA**→ rpmE2 (PRK01678)Ribosomal_L31 (pfam01197)→ Tdk (COG1435)PRK04296 (PRK04296)→ hypo |
| Bco-3-1 | NZ_ABFU01000064.1 | + | 21206 | 21236 | **RNA**→ rpmE (PRK00019)Ribosomal_L31 (pfam01197)→ rpmE (PRK00019)Ribosomal_L31 (pfam01197) |
| Din-1-1 | NZ_LAJB01000003.1 | - | 562219 | 562188 | **RNA**→ rpmE (PRK00019)Ribosomal_L31 (pfam01197)→ |
| Dro-1-1 | NZ_LAHL01000040.1 | - | 107117 | 107086 | **RNA**→ rpmE (PRK00019)Ribosomal_L31 (pfam01197)→ |
| Den-1-1 | NZ_LAHK01000087.1 | - | 2617 | 2586 | **RNA**→ rpmE (PRK00019)Ribosomal_L31 (pfam01197)→ Tdk (COG1435)PRK04296 (PRK04296)→ |
| Dtu-1-1 | NZ_LAJA01000003.1 | - | 28076 | 28045 | **RNA**→ rpmE (PRK00019)Ribosomal_L31 (pfam01197)→ Tdk (COG1435)PRK04296 (PRK04296)→<br>PrfA (COG0216)prfA (PRK00591)→ HemK (COG2890)PRK09328 (PRK09328)→<br>HemK (COG2890)RF_mod_PrmC (TIGR03534)→ Spore_II_R (pfam09551)→<br>SUA5 (COG0009)TIGR00057 (TIGR00057)SUA5 (pfam03481)→ |
| Bsp-22-1 | NZ_KB889752.1 | - | 2982398 | 2982367 | **RNA**→ rpmE (PRK00019)Ribosomal_L31 (pfam01197)→ Tdk (COG1435)PRK04296 (PRK04296)→<br>PrfA (COG0216)prfA (PRK00591)→ HemK (COG2890)PRK09328 (PRK09328)→ Spore_II_R (pfam09551) |
| Bsp-25-1 | NZ_JQMI01000018.1 | - | 92785 | 92754 | **RNA**→ rpmE (PRK00019)Ribosomal_L31 (pfam01197)→ Tdk (COG1435)PRK04296 (PRK04296)→<br>PrfA (COG0216)prfA (PRK00591)→ HemK (COG2890)PRK09328 (PRK09328)→ Spore_II_R (pfam09551) |
| Baq-1-1 | NZ_CCMN01000031.1 | - | 310685 | 310655 | **RNA**→ hypo→ |
| Bma-4-1 | NZ_LDWO01000012.1 | - | 159466 | 159436 | **RNA**→ rpmE (PRK00019)Ribosomal_L31 (pfam01197)→ |
| Bma-4-2 | NZ_LDWH01000003.1 | + | 320886 | 320916 | **RNA**→ rpmE (PRK00019)Ribosomal_L31 (pfam01197)→ |
| Bma-4-3 | NZ_LDWJ01000024.1 | + | 11492 | 11522 | **RNA**→ rpmE (PRK00019)Ribosomal_L31 (pfam01197)→ |
| Bma-4-4 | NZ_LGUE01000008.1 | - | 276011 | 275981 | **RNA**→ rpmE (PRK00019)Ribosomal_L31 (pfam01197)→ |
| Baq-1-2 | NZ_JXZC01000015.1 | - | 12969 | 12939 | **RNA**→ rpmE (PRK00019)Ribosomal_L31 (pfam01197)→ |
| Bvi-1-1 | NZ_CCDN010000007.1 | + | 193142 | 193172 | **RNA**→ rpmE (PRK00019)Ribosomal_L31 (pfam01197)→ |
| Lfl-3-1 | NZ_AODF01000010.1 | - | 26924 | 26895 | **RNA**→ rpmE2 (PRK01678)Ribosomal_L31 (pfam01197)→ |
| Lse-2-1 | NZ_JYOM01000013.1 | + | 11559 | 11588 | **RNA**→ rpmE2 (PRK01678)Ribosomal_L31 (pfam01197)→ |
| Lse-3-1 | NZ_CM001051.1 | - | 2676151 | 2676122 | **RNA**→ rpmE2 (PRK01678)Ribosomal_L31 (pfam01197)→ |
| Lin-4-1 | NZ_CM001049.1 | - | 2550007 | 2549978 | **RNA**→ rpmE2 (PRK01678)Ribosomal_L31 (pfam01197)→ |

| | | | | | | |
|---|---|---|---|---|---|---|
| Liv-4-1 | NZ_CP009576.1 | - | 2778969 | 2778940 | **RNA**→ | rpmE2 (PRK01678)Ribosomal_L31 (pfam01197)→ |
| Liv-4-2 | NZ_CP009575.1 | - | 2712836 | 2712807 | **RNA**→ | rpmE2 (PRK01678)Ribosomal_L31 (pfam01197)→ |
| Lmo-60-1 | NZ_APID01000006.1 | - | 151683 | 151654 | **RNA**→ | rpmE2 (PRK01678)Ribosomal_L31 (pfam01197)→ |
| Lmo-48-1 | NZ_KK073984.1 | + | 11327 | 11356 | **RNA**→ | rpmE2 (PRK01678)Ribosomal_L31 (pfam01197)→ |
| Lmo-53-1 | NZ_KK074072.1 | - | 99569 | 99540 | **RNA**→ | rpmE2 (PRK01678)Ribosomal_L31 (pfam01197)→ |
| Lmo-2-1 | NC_013768.1 | - | 2704320 | 2704291 | **RNA**→ | rpmE2 (PRK01678)Ribosomal_L31 (pfam01197)→ |
| Lmo-13-1 | NC_017547.1 | - | 2563087 | 2563058 | **RNA**→ | rpmE2 (PRK01678)Ribosomal_L31 (pfam01197)→ |
| Lmo-33-1 | NC_021823.1 | - | 2117345 | 2117316 | **RNA**→ | rpmE2 (PRK01678)Ribosomal_L31 (pfam01197)→ |
| Lmo-57-1 | NZ_HG813247.1 | - | 2626503 | 2626474 | **RNA**→ | rpmE2 (PRK01678)Ribosomal_L31 (pfam01197)→ |
| Lmo-34-1 | NC_013766.2 | - | 2737889 | 2737860 | **RNA**→ | rpmE2 (PRK01678)Ribosomal_L31 (pfam01197)→ |
| Lmo-45-1 | NZ_KK211194.1 | - | 179779 | 179750 | **RNA**→ | rpmE2 (PRK01678)Ribosomal_L31 (pfam01197)→ |
| Lmo-58-1 | NZ_APIB01000006.1 | + | 47839 | 47868 | **RNA**→ | rpmE2 (PRK01678)Ribosomal_L31 (pfam01197)→ |
| Liv-1-2 | NZ_CVPH01000006.1 | - | 842440 | 842411 | **RNA**→ | rpmE2 (PRK01678)Ribosomal_L31 (pfam01197)→ |
| Liv-1-1 | NC_016011.1 | - | 2607332 | 2607303 | **RNA**→ | rpmE2 (PRK01678)Ribosomal_L31 (pfam01197)→ |
| Liv-2-1 | NZ_CP007172.1 | - | 2653509 | 2653480 | **RNA**→ | rpmE2 (PRK01678)Ribosomal_L31 (pfam01197)→ |
| Liv-3-1 | NZ_CP009577.1 | - | 2653506 | 2653477 | **RNA**→ | rpmE2 (PRK01678)Ribosomal_L31 (pfam01197)→ |
| Lwe-1-1 | NC_008555.1 | - | 2535354 | 2535325 | **RNA**→ | rpmE2 (PRK01678)Ribosomal_L31 (pfam01197)→ |
| Laq-1-1 | NZ_AOCG01000011.1 | - | 164581 | 164552 | **RNA**→ | rpmE2 (PRK01678)Ribosomal_L31 (pfam01197)→ |
| Lmo-69-1 | NZ_APIM01000007.1 | - | 151862 | 151833 | **RNA**→ | rpmE2 (PRK01678)Ribosomal_L31 (pfam01197)→ |
| Lmo-67-1 | NZ_APIK01000004.1 | + | 47841 | 47870 | **RNA**→ | rpmE2 (PRK01678)Ribosomal_L31 (pfam01197)→ |
| Lmo-62-1 | NZ_APIF01000007.1 | - | 153381 | 153352 | **RNA**→ | rpmE2 (PRK01678)Ribosomal_L31 (pfam01197)→ |
| Lmo-65-1 | NZ_APII01000006.1 | + | 47845 | 47874 | **RNA**→ | rpmE2 (PRK01678)Ribosomal_L31 (pfam01197)→ |
| Lmo-63-1 | NZ_APIG01000008.1 | - | 113788 | 113759 | **RNA**→ | rpmE2 (PRK01678)Ribosomal_L31 (pfam01197)→ |
| Lmo-64-1 | NZ_APIH01000007.1 | - | 153278 | 153249 | **RNA**→ | rpmE2 (PRK01678)Ribosomal_L31 (pfam01197)→ |
| Lmo-66-1 | NZ_APIJ01000011.1 | - | 84673 | 84644 | **RNA**→ | rpmE2 (PRK01678)Ribosomal_L31 (pfam01197)→ |
| Lmo-59-1 | NZ_APIC01000008.1 | - | 153115 | 153086 | **RNA**→ | rpmE2 (PRK01678)Ribosomal_L31 (pfam01197)→ |
| Lmo-68-1 | NZ_APIL01000007.1 | - | 113787 | 113758 | **RNA**→ | rpmE2 (PRK01678)Ribosomal_L31 (pfam01197)→ |
| Lmo-17-1 | NZ_AARX02000004.1 | - | 163846 | 163817 | **RNA**→ | rpmE2 (PRK01678)Ribosomal_L31 (pfam01197)→ rpmE2 (PRK01678)Ribosomal_L31 (pfam01197 |
| Lmo-11-1 | NZ_AARR02000015.1 | + | 11297 | 11326 | **RNA**→ | rpmE2 (PRK01678)Ribosomal_L31 (pfam01197)→ rpmE2 (PRK01678)Ribosomal_L31 (pfam01197 |
| Lmo-7-1 | NZ_AARU02000004.1 | - | 163847 | 163818 | **RNA**→ | rpmE2 (PRK01678)Ribosomal_L31 (pfam01197)→ rpmE2 (PRK01678)Ribosomal_L31 (pfam01197 |
| Lmo-9-1 | NZ_AARP04000011.1 | + | 2287 | 2316 | **RNA**→ | rpmE2 (PRK01678)Ribosomal_L31 (pfam01197)→ rpmE2 (PRK01678)Ribosomal_L31 (pfam01197 |
| Lmo-46-1 | NZ_JHZR01000009.1 | - | 28003 | 27974 | **RNA**→ | rpmE2 (PRK01678)Ribosomal_L31 (pfam01197)→ |
| Lmo-10-1 | NZ_AARQ02000005.1 | - | 151823 | 151794 | **RNA**→ | rpmE2 (PRK01678)Ribosomal_L31 (pfam01197)→ rpmE2 (PRK01678)Ribosomal_L31 (pfam01197 |
| Lmo-30-1 | NZ_AADQ01000100.1 | - | 57 | 28 | **RNA**→ | |
| Lmo-56-1 | NZ_CBVW010000004.1 | - | 151863 | 151834 | **RNA**→ | rpmE2 (PRK01678)Ribosomal_L31 (pfam01197)→ |
| Lmo-8-1 | NZ_AARJ02000005.1 | + | 11105 | 11134 | **RNA**→ | rpmE2 (PRK01678)Ribosomal_L31 (pfam01197)→ rpmE2 (PRK01678)Ribosomal_L31 (pfam01197 |
| Lmo-55-1 | NZ_CBVZ010000006.1 | - | 152406 | 152377 | **RNA**→ | rpmE2 (PRK01678)Ribosomal_L31 (pfam01197)→ |
| Lmo-47-1 | NZ_JHUY01000011.1 | - | 182393 | 182364 | **RNA**→ | rpmE2 (PRK01678)Ribosomal_L31 (pfam01197)→ |
| Lmo-72-1 | NZ_APJT01000001.1 | - | 2489181 | 2489152 | **RNA**→ | rpmE2 (PRK01678)Ribosomal_L31 (pfam01197)→ |
| Lmo-6-1 | NC_003210.1 | - | 2624918 | 2624889 | **RNA**→ | rpmE2 (PRK01678)Ribosomal_L31 (pfam01197)→ |
| Lmo-50-1 | NZ_KK074038.1 | - | 152233 | 152204 | **RNA**→ | rpmE2 (PRK01678)Ribosomal_L31 (pfam01197)→ |
| Lmo-52-1 | NZ_KK074046.1 | + | 41826 | 41855 | **RNA**→ | rpmE2 (PRK01678)Ribosomal_L31 (pfam01197)→ |
| Lmo-26-1 | NC_018587.1 | - | 2636237 | 2636208 | **RNA**→ | rpmE2 (PRK01678)Ribosomal_L31 (pfam01197)→ |
| Lmo-39-1 | NZ_KN050647.1 | - | 713680 | 713651 | **RNA**→ | rpmE2 (PRK01678)Ribosomal_L31 (pfam01197)→ |
| Lmo-16-1 | NC_017545.1 | - | 2634195 | 2634166 | **RNA**→ | rpmE2 (PRK01678)Ribosomal_L31 (pfam01197)→ |
| Lmo-3-1 | NC_017544.1 | - | 2579403 | 2579374 | **RNA**→ | rpmE2 (PRK01678)Ribosomal_L31 (pfam01197)→ |
| Lmo-12-1 | NC_017546.1 | - | 2643374 | 2643345 | **RNA**→ | rpmE2 (PRK01678)Ribosomal_L31 (pfam01197)→ |
| Lmo-28-1 | NC_018593.1 | - | 2525570 | 2525541 | **RNA**→ | rpmE2 (PRK01678)Ribosomal_L31 (pfam01197)→ |
| Lmo-27-1 | NC_018592.1 | - | 2583378 | 2583349 | **RNA**→ | rpmE2 (PRK01678)Ribosomal_L31 (pfam01197)→ |
| Lmo-24-1 | NC_018589.1 | - | 2642056 | 2642027 | **RNA**→ | rpmE2 (PRK01678)Ribosomal_L31 (pfam01197)→ |
| Lmo-21-1 | NC_018588.1 | - | 2642444 | 2642415 | **RNA**→ | rpmE2 (PRK01678)Ribosomal_L31 (pfam01197)→ |
| Lmo-25-1 | NC_018586.1 | - | 2643186 | 2643157 | **RNA**→ | rpmE2 (PRK01678)Ribosomal_L31 (pfam01197)→ |

| | | | | | |
|---|---|---|---|---|---|
| Lmo-33-2 | NC_021838.1 | - | 242759 | 242730 | RNA→ rpmE2 (PRK01678)Ribosomal_L31 (pfam01197)→ |
| Lmo-33-3 | NC_021824.1 | - | 186543 | 186514 | RNA→ rpmE2 (PRK01678)Ribosomal_L31 (pfam01197)→ |
| Lmo-33-4 | NC_021837.1 | + | 850722 | 850751 | RNA→ rpmE2 (PRK01678)Ribosomal_L31 (pfam01197)→ |
| Lmo-33-5 | NC_021826.1 | - | 1589345 | 1589316 | RNA→ rpmE2 (PRK01678)Ribosomal_L31 (pfam01197)→ |
| Lmo-38-1 | NC_022568.1 | - | 2583826 | 2583797 | RNA→ rpmE2 (PRK01678)Ribosomal_L31 (pfam01197)→ |
| Lmo-70-1 | NZ_CP007160.1 | - | 1351849 | 1351820 | RNA→ rpmE2 (PRK01678)Ribosomal_L31 (pfam01197)→ |
| Lmo-29-1 | NC_018591.1 | - | 2606775 | 2606746 | RNA→ rpmE2 (PRK01678)Ribosomal_L31 (pfam01197)→ rpmE2 (PRK01678)Ribosomal_L31 (pfam01197 |
| Lmo-44-1 | NZ_KK211033.1 | + | 43201 | 43230 | RNA→ rpmE2 (PRK01678)Ribosomal_L31 (pfam01197)→ |
| Lmo-61-1 | NZ_APIE01000007.1 | - | 153398 | 153369 | RNA→ rpmE2 (PRK01678)Ribosomal_L31 (pfam01197)→ |
| Lin-2-1 | NZ_JZCS01000009.1 | + | 49208 | 49237 | RNA→ rpmE2 (PRK01678)Ribosomal_L31 (pfam01197)→ |
| Lin-1-1 | NC_003212.1 | - | 2709363 | 2709334 | RNA→ rpmE2 (PRK01678)Ribosomal_L31 (pfam01197)→ |
| Lin-2-2 | NZ_JZCU01000001.1 | - | 177547 | 177518 | RNA→ rpmE2 (PRK01678)Ribosomal_L31 (pfam01197)→ |
| Lin-2-3 | NZ_JRYX01000006.1 | - | 193559 | 193530 | RNA→ rpmE2 (PRK01678)Ribosomal_L31 (pfam01197)→ |
| Lmo-14-1 | NC_011660.1 | + | 46092 | 46121 | RNA→ rpmE2 (PRK01678)Ribosomal_L31 (pfam01197)→ |
| Lmo-20-1 | NC_017537.1 | - | 2607884 | 2607855 | RNA→ rpmE2 (PRK01678)Ribosomal_L31 (pfam01197)→ |
| Lmo-19-1 | NC_017529.1 | - | 2609930 | 2609901 | RNA→ rpmE2 (PRK01678)Ribosomal_L31 (pfam01197)→ |
| Lin-3-1 | NZ_JH556648.1 | + | 28282 | 28311 | RNA→ rpmE2 (PRK01678)Ribosomal_L31 (pfam01197)→ |
| Lmo-22-1 | NC_018590.1 | - | 2520256 | 2520227 | RNA→ rpmE2 (PRK01678)Ribosomal_L31 (pfam01197)→ |
| Lba-1-1 | NZ_AGUG01000004.1 | - | 22582 | 22553 | RNA→ rpmE2 (PRK01678)Ribosomal_L31 (pfam01197)→ rpmE2 (PRK01678)Ribosomal_L31 (pfam01197 |
| Lfl-2-1 | NZ_ALWW01000045.1 | + | 21645 | 21674 | RNA→ rpmE2 (PRK01678)Ribosomal_L31 (pfam01197)→ |
| Lfl-1-1 | NZ_AODM01000033.1 | + | 275 | 304 | RNA→ rpmE2 (PRK01678)Ribosomal_L31 (pfam01197)→ |
| Lse-1-1 | NC_013891.1 | - | 2523334 | 2523305 | RNA→ rpmE2 (PRK01678)Ribosomal_L31 (pfam01197)→ |
| Lmo-40-1 | NZ_CM001469.1 | - | 2232849 | 2232820 | RNA→ rpmE2 (PRK01678)Ribosomal_L31 (pfam01197)→ |
| Lmo-32-1 | NZ_AADR01000056.1 | - | 5830 | 5801 | RNA→ rpmE2 (PRK01678)Ribosomal_L31 (pfam01197)→ rpmE2 (PRK01678)Ribosomal_L31 (pfam01197 |
| Lmo-42-1 | NC_021829.1 | + | 1457392 | 1457421 | RNA→ rpmE2 (PRK01678)Ribosomal_L31 (pfam01197)→ |
| Lmo-33-6 | NC_021840.1 | + | 2164270 | 2164299 | RNA→ rpmE2 (PRK01678)Ribosomal_L31 (pfam01197)→ |
| Lmo-33-7 | NC_021839.1 | - | 2383018 | 2382989 | RNA→ rpmE2 (PRK01678)Ribosomal_L31 (pfam01197)→ |
| Lmo-33-8 | NC_021827.1 | + | 563127 | 563156 | RNA→ rpmE2 (PRK01678)Ribosomal_L31 (pfam01197)→ |
| Lgr-2-1 | NZ_GL538353.1 | + | 812725 | 812754 | RNA→ rpmE2 (PRK01678)Ribosomal_L31 (pfam01197)→ |
| Lgr-3-1 | NZ_AODG01000008.1 | + | 5832 | 5861 | RNA→ rpmE2 (PRK01678)Ribosomal_L31 (pfam01197)→ |
| Lco-1-1 | NZ_AODE01000004.1 | + | 8681 | 8710 | RNA→ rpmE2 (PRK01678)Ribosomal_L31 (pfam01197)→ |
| Lro-1-1 | NZ_AODK01000010.1 | - | 13603 | 13574 | RNA→ rpmE2 (PRK01678)Ribosomal_L31 (pfam01197)→ |
| Lri-1-1 | NZ_AODL01000006.1 | - | 88179 | 88150 | RNA→ rpmE2 (PRK01678)Ribosomal_L31 (pfam01197)→ |
| Lgr-1-1 | NZ_AODD01000005.1 | + | 122902 | 122931 | RNA→ rpmE2 (PRK01678)Ribosomal_L31 (pfam01197)→ |
| Lba-2-1 | NZ_JNEZ01000001.1 | + | 49115 | 49144 | RNA→ rpmE2 (PRK01678)Ribosomal_L31 (pfam01197)→ |
| Lne-1-1 | NZ_JNFB01000007.1 | + | 9799 | 9828 | RNA→ rpmE2 (PRK01678)Ribosomal_L31 (pfam01197)→ |
| Lbo-2-1 | NZ_JNFA01000028.1 | + | 27486 | 27515 | RNA→ rpmE2 (PRK01678)Ribosomal_L31 (pfam01197)→ |
| Bsp-94-1 | NZ_KE384493.1 | - | 53262 | 53231 | RNA→ rpmE (PRK00019)Ribosomal_L31 (pfam01197)→ |
| Bsm-2-1 | NZ_JH414746.1 | - | 53667 | 53635 | RNA→ rpmE (PRK00019)Ribosomal_L31 (pfam01197)→ hypo→ Tdk (COG1435)PRK04296 (PRK04296)- hypo→ |
| Bsm-1-1 | NZ_CP012024.1 | - | 3221214 | 3221182 | RNA→ rpmE (PRK00019)Ribosomal_L31 (pfam01197)→ |
| Blo-1-1 | NZ_LFZW01000001.1 | - | 4911598 | 4911567 | RNA→ rpmE (PRK00019)Ribosomal_L31 (pfam01197)→ |
| Bci-1-1 | NZ_KI632498.1 | + | 78172 | 78203 | RNA→ rpmE (PRK00019)Ribosomal_L31 (pfam01197)→ |
| Pth-2-1 | NZ_BAWP01000010.1 | - | 111204 | 111173 | RNA→ rpmE (PRK00019)Ribosomal_L31 (pfam01197)→ |
| Pth-1-1 | NZ_LAKX01000004.1 | - | 74606 | 74575 | RNA→ rpmE (PRK00019)Ribosomal_L31 (pfam01197)→ |
| Gsp-4-1 | NC_014650.1 | - | 3717664 | 3717633 | RNA→ rpmE (PRK00019)Ribosomal_L31 (pfam01197)→ |
| Gth-2-1 | NC_015660.1 | - | 3764572 | 3764541 | RNA→ rpmE (PRK00019)Ribosomal_L31 (pfam01197)→ |
| Gth-5-1 | NZ_CM001483.1 | - | 3617423 | 3617392 | RNA→ rpmE (PRK00019)Ribosomal_L31 (pfam01197)→ |
| Bps-1-1 | NC_013791.2 | - | 3665301 | 3665270 | RNA→ rpmE2 (PRK01678)Ribosomal_L31 (pfam01197)→ |
| Bma-5-1 | NZ_ATAE01000042.1 | - | 128216 | 128185 | RNA→ rpmE2 (PRK01678)Ribosomal_L31 (pfam01197)→ |
| Bsp-66-2 | NZ_LFEL01000017.1 | - | 3671 | 3640 | RNA→ rpmE2 (PRK01678)Ribosomal_L31 (pfam01197)→ |
| Bsp-92-1 | NZ_AXVA01000010.1 | + | 75959 | 75989 | RNA→ rpmE (PRK00019)Ribosomal_L31 (pfam01197)→ |
| env-936 | SL_3KL_010_SED_10258389 | - | 407 | 377 | RNA→ rpmE2 (PRK01678)Ribosomal_L31 (pfam01197)→ |

| | | | | | |
|---|---|---|---|---|---|
| Pge-1-1 | NZ_CM002692.1 | - | 3407512 | 3407481 | **RNA**→ rpmE (PRK00019) Ribosomal_L31 (pfam01197)→ |
| Pca-1-1 | NZ_BAWO01000004.1 | - | 18666 | 18635 | **RNA**→ rpmE (PRK00019) Ribosomal_L31 (pfam01197)→ |
| Gca-1-1 | NZ_AMRO01000071.1 | - | 76191 | 76160 | **RNA**→ rpmE (PRK00019) Ribosomal_L31 (pfam01197)→ |
| Bsp-79-1 | NZ_JXAB01000020.1 | - | 297461 | 297431 | **RNA**→ rpmE (PRK00019) Ribosomal_L31 (pfam01197)→ |
| Ben-1-1 | NZ_CP011974.1 | - | 4727388 | 4727358 | **RNA**→ rpmE (PRK00019) Ribosomal_L31 (pfam01197)→ |
| Ben-2-1 | NZ_ALIM01000035.1 | + | 148874 | 148904 | **RNA**→ rpmE (PRK00019) Ribosomal_L31 (pfam01197)→ |
| env-937 | C917852_1 | + | 160 | 193 | **RNA**→ rpmE (PRK00019) Ribosomal_L31 (pfam01197)→ |
| Bsp-54-1 | NZ_KI912515.1 | + | 222431 | 222461 | **RNA**→ rpmE (PRK00019) Ribosomal_L31 (pfam01197)→ |
| Bsp-30-1 | NZ_CCXR01000001.1 | - | 4062665 | 4062634 | **RNA**→ rpmE (PRK00019) Ribosomal_L31 (pfam01197)→ |
| Bba-2-1 | NZ_JXLO01000001.1 | - | 609426 | 609395 | **RNA**→ rpmE (PRK00019) Ribosomal_L31 (pfam01197)→ |
| Bba-2-2 | NZ_JXLP01000009.1 | - | 431011 | 430980 | **RNA**→ rpmE (PRK00019) Ribosomal_L31 (pfam01197)→ |
| Bkr-1-1 | NZ_KE387244.1 | + | 195969 | 196000 | **RNA**→ rpmE (PRK00019) Ribosomal_L31 (pfam01197)→ |
| Bsp-17-1 | NZ_KI547188.1 | + | 304476 | 304508 | **RNA**→ rpmE (PRK00019) Ribosomal_L31 (pfam01197)→ hypo→ |
| Bbo-3-1 | NZ_BAUW01000047.1 | - | 11315 | 11283 | **RNA**→ rpmE (PRK00019) Ribosomal_L31 (pfam01197)→ |
| Bse-2-1 | NZ_BASE01000017.1 | - | 87101 | 87069 | **RNA**→ rpmE (PRK00019) Ribosomal_L31 (pfam01197)→ |
| Bsu-11-1 | NZ_JXIQ01000068.1 | - | 277 | 245 | **RNA**→ rpmE (PRK00019) Ribosomal_L31 (pfam01197)→ |
| Bsp-37-1 | NZ_HG964497.1 | - | 510465 | 510434 | **RNA**→ rpmE (PRK00019) Ribosomal_L31 (pfam01197)→ |
| Bru-1-1 | NZ_CCFE01000013.1 | - | 20014 | 19983 | **RNA**→ rpmE (PRK00019) Ribosomal_L31 (pfam01197)→ |
| Bba-3-1 | NZ_JWIQ02000001.1 | + | 37168 | 37199 | **RNA**→ rpmE (PRK00019) Ribosomal_L31 (pfam01197)→ |
| Qth-1-1 | NZ_JWIR02000025.1 | - | 127899 | 127868 | **RNA**→ rpmE (PRK00019) Ribosomal_L31 (pfam01197)→ |
| Qth-1-2 | NZ_JWJE02000018.1 | + | 61399 | 61430 | **RNA**→ rpmE (PRK00019) Ribosomal_L31 (pfam01197)→ |
| Bsp-40-1 | NZ_AYSE01000005.1 | + | 137089 | 137120 | **RNA**→ rpmE (PRK00019) Ribosomal_L31 (pfam01197)→ |
| Bsp-42-1 | NZ_LFZU01000001.1 | - | 3871902 | 3871871 | **RNA**→ rpmE (PRK00019) Ribosomal_L31 (pfam01197)→ |
| Bsp-96-1 | NZ_CP008855.1 | - | 1758854 | 1758824 | **RNA**→ rpmE (PRK00019) Ribosomal_L31 (pfam01197)→ |
| Bfl-3-1 | NZ_JANV01000077.1 | + | 41386 | 41417 | **RNA**→ rpmE (PRK00019) Ribosomal_L31 (pfam01197)→ |
| Bsp-81-1 | NZ_LATZ01000128.1 | + | 256709 | 256739 | **RNA**→ rpmE (PRK00019) Ribosomal_L31 (pfam01197)→ |
| Bme-1-1 | NC_014103.1 | - | 4971544 | 4971513 | **RNA**→ rpmE (PRK00019) Ribosomal_L31 (pfam01197)→ hypo→ |
| Bfl-1-1 | NZ_LFQJ01000038.1 | + | 103356 | 103387 | **RNA**→ rpmE (PRK00019) Ribosomal_L31 (pfam01197)→ |
| Bsp-16-1 | NZ_AZVK01000006.1 | - | 237288 | 237257 | **RNA**→ rpmE (PRK00019) Ribosomal_L31 (pfam01197)→ |
| env-938 | JGI25151J46595_10060291 | - | 778 | 747 | **RNA**→ rpmE (PRK00019) Ribosomal_L31 (pfam01197)→ |
| Bar-1-1 | NZ_JPIE01000004.1 | + | 106798 | 106829 | **RNA**→ rpmE (PRK00019) Ribosomal_L31 (pfam01197)→ |
| Bar-1-2 | NZ_JXRC01000013.1 | - | 240051 | 240020 | **RNA**→ rpmE (PRK00019) Ribosomal_L31 (pfam01197)→ |
| Bar-2-1 | NZ_JYOO01000022.1 | - | 64775 | 64744 | **RNA**→ rpmE (PRK00019) Ribosomal_L31 (pfam01197)→ |
| Bar-1-3 | NZ_JMTL01000003.1 | + | 275950 | 275981 | **RNA**→ rpmE (PRK00019) Ribosomal_L31 (pfam01197)→ |
| Bar-1-4 | NZ_LDWN01000019.1 | + | 134934 | 134965 | **RNA**→ rpmE (PRK00019) Ribosomal_L31 (pfam01197)→ |
| Bar-1-5 | NZ_LDUV01000024.1 | + | 134934 | 134965 | **RNA**→ rpmE (PRK00019) Ribosomal_L31 (pfam01197)→ |
| Bme-2-1 | NC_014019.1 | - | 4974394 | 4974363 | **RNA**→ rpmE (PRK00019) Ribosomal_L31 (pfam01197)→ hypo→ |
| Bme-3-1 | NC_017138.1 | + | 117060 | 117091 | **RNA**→ rpmE (PRK00019) Ribosomal_L31 (pfam01197)→ hypo→ |
| Bme-7-1 | NZ_CP009920.1 | - | 1998706 | 1998675 | **RNA**→ rpmE (PRK00019) Ribosomal_L31 (pfam01197)→ |
| Bme-5-1 | NZ_KN125581.1 | + | 150847 | 150878 | **RNA**→ rpmE (PRK00019) Ribosomal_L31 (pfam01197)→ |
| Bme-9-1 | NZ_CP010586.1 | - | 5000760 | 5000729 | **RNA**→ rpmE (PRK00019) Ribosomal_L31 (pfam01197)→ |
| Bme-8-1 | NZ_AHTF01000099.1 | - | 240086 | 240055 | **RNA**→ rpmE (PRK00019) Ribosomal_L31 (pfam01197)→ |
| Bsp-29-1 | NZ_ANNS01000002.1 | + | 6424 | 6455 | **RNA**→ rpmE (PRK00019) Ribosomal_L31 (pfam01197)→ |
| Bsp-80-1 | NZ_AXZS01000001.1 | - | 241083 | 241052 | **RNA**→ rpmE (PRK00019) Ribosomal_L31 (pfam01197)→ |
| Bsp-20-1 | NZ_AZVI01000006.1 | + | 135180 | 135211 | **RNA**→ rpmE (PRK00019) Ribosomal_L31 (pfam01197)→ |
| Bfl-2-1 | NZ_AZUJ01000010.1 | + | 127695 | 127726 | **RNA**→ rpmE (PRK00019) Ribosomal_L31 (pfam01197)→ |
| Bpa-1-1 | NZ_AUMR01000038.1 | + | 35351 | 35383 | **RNA**→ rpmE2 (PRK01678) Ribosomal_L31 (pfam01197)→ |
| Ban-4-1 | NZ_ABKF01000017.1 | - | 739 | 707 | **RNA**→ rpmE2 (PRK01678) Ribosomal_L31 (pfam01197)→ rpmE2 (PRK01678) Ribosomal_L31 (pfam01197 |
| Ban-3-1 | NZ_ABLT01000007.1 | - | 8251 | 8219 | **RNA**→ rpmE2 (PRK01678) Ribosomal_L31 (pfam01197)→ rpmE2 (PRK01678) Ribosomal_L31 (pfam01197 |
| Ban-9-1 | NZ_ABJC01000012.1 | - | 8278 | 8246 | **RNA**→ rpmE2 (PRK01678) Ribosomal_L31 (pfam01197)→ rpmE2 (PRK01678) Ribosomal_L31 (pfam01197 |
| Ban-8-1 | NZ_ABLH01000015.1 | - | 7672 | 7640 | **RNA**→ rpmE2 (PRK01678) Ribosomal_L31 (pfam01197)→ rpmE2 (PRK01678) Ribosomal_L31 (pfam01197 |
| Ban-7-1 | NZ_ABKG01000040.1 | - | 8162 | 8130 | **RNA**→ rpmE2 (PRK01678) Ribosomal_L31 (pfam01197)→ rpmE2 (PRK01678) Ribosomal_L31 (pfam01197 |

| | | | | | |
|---|---|---|---|---|---|
| Ban-6-1 | NZ_ABLB01000054.1 | - | 8201 | 8169 | **RNA**→ rpmE2 (PRK01678)Ribosomal_L31 (pfam01197)→ rpmE2 (PRK01678)Ribosomal_L31 (pfam01197 |
| Ban-33-1 | NZ_AGQP01000037.1 | - | 62679 | 62647 | **RNA**→ rpmE2 (PRK01678)Ribosomal_L31 (pfam01197)→ |
| Ban-10-1 | NZ_AAEO01000027.3 | + | 266 | 298 | **RNA**→ rpmE2 (PRK01678)Ribosomal_L31 (pfam01197)→ rpmE2 (PRK01678)Ribosomal_L31 (pfam01197 |
| Ban-18-1 | NZ_AAEP01000040.1 | - | 8079 | 8047 | **RNA**→ rpmE2 (PRK01678)Ribosomal_L31 (pfam01197)→ rpmE2 (PRK01678)Ribosomal_L31 (pfam01197 |
| Bce-32-1 | NZ_ABDK02000015.1 | + | 132650 | 132682 | **RNA**→ hypo→ rpmE2 (PRK01678)Ribosomal_L31 (pfam01197)→ |
| Ban-14-1 | NZ_AAEN01000018.1 | + | 262 | 294 | **RNA**→ rpmE2 (PRK01678)Ribosomal_L31 (pfam01197)→ rpmE2 (PRK01678)Ribosomal_L31 (pfam01197 |
| Ban-1-1 | NZ_ABDN02000023.1 | + | 47027 | 47059 | **RNA**→ rpmE2 (PRK01678)Ribosomal_L31 (pfam01197)→ rpmE2 (PRK01678)Ribosomal_L31 (pfam01197 |
| Ban-26-1 | NZ_ANFF01000003.1 | - | 1790045 | 1790013 | **RNA**→ rpmE2 (PRK01678)Ribosomal_L31 (pfam01197)→ |
| Ban-27-1 | NZ_ANFG01000003.1 | - | 4312050 | 4312018 | **RNA**→ rpmE2 (PRK01678)Ribosomal_L31 (pfam01197)→ |
| Ban-39-1 | NZ_AZQO01000036.1 | - | 61859 | 61827 | **RNA**→ rpmE2 (PRK01678)Ribosomal_L31 (pfam01197)→ |
| Ban-11-1 | NC_003997.3 | - | 5058754 | 5058722 | **RNA**→ rpmE2 (PRK01678)Ribosomal_L31 (pfam01197)→ |
| Ban-17-1 | NC_005945.1 | - | 5060089 | 5060057 | **RNA**→ rpmE2 (PRK01678)Ribosomal_L31 (pfam01197)→ |
| Bce-2-1 | NC_012472.1 | - | 5099695 | 5099663 | **RNA**→ rpmE2 (PRK01678)Ribosomal_L31 (pfam01197)→ |
| Ban-5-1 | NC_012659.1 | - | 5058780 | 5058748 | **RNA**→ rpmE2 (PRK01678)Ribosomal_L31 (pfam01197)→ |
| Ban-13-1 | NC_012581.1 | - | 5061412 | 5061380 | **RNA**→ rpmE2 (PRK01678)Ribosomal_L31 (pfam01197)→ |
| Bce-44-1 | NC_014335.1 | - | 5012311 | 5012279 | **RNA**→ rpmE2 (PRK01678)Ribosomal_L31 (pfam01197)→ |
| Ban-15-1 | NC_017729.1 | - | 5050417 | 5050385 | **RNA**→ rpmE2 (PRK01678)Ribosomal_L31 (pfam01197)→ rpmE2 (PRK01678)Ribosomal_L31 (pfam01197 |
| Bce-26-1 | NC_016779.1 | - | 5052968 | 5052936 | **RNA**→ rpmE2 (PRK01678)Ribosomal_L31 (pfam01197)→ |
| Ban-34-1 | NZ_CP006742.1 | - | 5042149 | 5042117 | **RNA**→ rpmE2 (PRK01678)Ribosomal_L31 (pfam01197)→ |
| Ban-21-1 | NZ_CP008752.1 | + | 436721 | 436753 | **RNA**→ rpmE2 (PRK01678)Ribosomal_L31 (pfam01197)→ |
| Ban-28-1 | NZ_CP001970.1 | - | 5059354 | 5059322 | **RNA**→ rpmE2 (PRK01678)Ribosomal_L31 (pfam01197)→ |
| Ban-29-1 | NZ_CP001974.1 | - | 5059103 | 5059071 | **RNA**→ rpmE2 (PRK01678)Ribosomal_L31 (pfam01197)→ |
| Ban-21-2 | NZ_CP007618.1 | + | 4193405 | 4193437 | **RNA**→ rpmE2 (PRK01678)Ribosomal_L31 (pfam01197)→ |
| Ban-18-2 | NZ_CP007666.1 | - | 3933034 | 3933002 | **RNA**→ rpmE2 (PRK01678)Ribosomal_L31 (pfam01197)→ |
| Ban-21-3 | NZ_CP007704.1 | - | 3393140 | 3393108 | **RNA**→ rpmE2 (PRK01678)Ribosomal_L31 (pfam01197)→ |
| Ban-21-4 | NZ_CP008846.1 | - | 5044915 | 5044883 | **RNA**→ rpmE2 (PRK01678)Ribosomal_L31 (pfam01197)→ |
| Ban-25-1 | NZ_BAVT01000022.1 | - | 402375 | 402343 | **RNA**→ rpmE2 (PRK01678)Ribosomal_L31 (pfam01197)→ |
| Ban-16-1 | NZ_AAEQ01000042.3 | - | 254390 | 254358 | **RNA**→ rpmE2 (PRK01678)Ribosomal_L31 (pfam01197)→ rpmE2 (PRK01678)Ribosomal_L31 (pfam01197 |
| Ban-19-1 | NZ_AAER01000027.3 | - | 85456 | 85424 | **RNA**→ rpmE2 (PRK01678)Ribosomal_L31 (pfam01197)→ rpmE2 (PRK01678)Ribosomal_L31 (pfam01197 |
| Ban-12-1 | NZ_AAES01000044.3 | - | 7983 | 7951 | **RNA**→ rpmE2 (PRK01678)Ribosomal_L31 (pfam01197)→ rpmE2 (PRK01678)Ribosomal_L31 (pfam01197 |
| Bce-25-1 | NZ_CM000736.1 | - | 4877394 | 4877362 | **RNA**→ rpmE2 (PRK01678)Ribosomal_L31 (pfam01197)→ rpmE2 (PRK01678)Ribosomal_L31 (pfam01197 |
| Bth-17-1 | NZ_CM000757.1 | + | 4954695 | 4954727 | **RNA**→ rpmE2 (PRK01678)Ribosomal_L31 (pfam01197)→ rpmE2 (PRK01678)Ribosomal_L31 (pfam01197 |
| Ban-31-1 | NZ_KL649663.1 | - | 404429 | 404397 | **RNA**→ rpmE2 (PRK01678)Ribosomal_L31 (pfam01197)→ |
| Bce-104-1 | NZ_JH792132.1 | + | 255191 | 255223 | **RNA**→ rpmE2 (PRK01678)Ribosomal_L31 (pfam01197)→ |
| Ban-24-1 | NZ_CM002398.1 | - | 5063652 | 5063620 | **RNA**→ rpmE2 (PRK01678)Ribosomal_L31 (pfam01197)→ |
| Ban-21-5 | NZ_CP010342.1 | - | 5058896 | 5058864 | **RNA**→ rpmE2 (PRK01678)Ribosomal_L31 (pfam01197)→ |
| Ban-2-1 | NC_007530.2 | - | 5058880 | 5058848 | **RNA**→ rpmE2 (PRK01678)Ribosomal_L31 (pfam01197)→ |
| Bth-7-1 | NZ_CM000754.1 | - | 4909086 | 4909054 | **RNA**→ rpmE2 (PRK01678)Ribosomal_L31 (pfam01197)→ rpmE2 (PRK01678)Ribosomal_L31 (pfam01197 |
| Ban-21-6 | NZ_CP009325.1 | + | 150594 | 150626 | **RNA**→ rpmE2 (PRK01678)Ribosomal_L31 (pfam01197)→ |
| Ban-21-7 | NZ_CP009328.1 | - | 3853585 | 3853553 | **RNA**→ rpmE2 (PRK01678)Ribosomal_L31 (pfam01197)→ |
| Ban-21-8 | NZ_CP009331.1 | + | 1617335 | 1617367 | **RNA**→ rpmE2 (PRK01678)Ribosomal_L31 (pfam01197)→ |
| Ban-21-9 | NZ_CP010792.1 | - | 5058855 | 5058823 | **RNA**→ rpmE2 (PRK01678)Ribosomal_L31 (pfam01197)→ |
| Ban-21-10 | NZ_CP009341.1 | - | 2358106 | 2358074 | **RNA**→ rpmE2 (PRK01678)Ribosomal_L31 (pfam01197)→ |
| Ban-21-11 | NZ_CP010813.1 | - | 5058858 | 5058826 | **RNA**→ rpmE2 (PRK01678)Ribosomal_L31 (pfam01197)→ |
| Ban-21-12 | NZ_CP009464.1 | + | 1265383 | 1265415 | **RNA**→ rpmE2 (PRK01678)Ribosomal_L31 (pfam01197)→ |
| Ban-21-13 | NZ_CP009544.1 | - | 5118540 | 5118508 | **RNA**→ rpmE2 (PRK01678)Ribosomal_L31 (pfam01197)→ |
| Ban-37-1 | NZ_CP009315.1 | - | 1636902 | 1636870 | **RNA**→ rpmE2 (PRK01678)Ribosomal_L31 (pfam01197)→ |
| Ban-21-14 | NZ_CP009700.1 | + | 2531860 | 2531892 | **RNA**→ rpmE2 (PRK01678)Ribosomal_L31 (pfam01197)→ |
| Bce-85-1 | NZ_CP009300.1 | + | 4697208 | 4697240 | **RNA**→ rpmE2 (PRK01678)Ribosomal_L31 (pfam01197)→ |
| Bce-2-2 | NZ_CP009318.1 | - | 5169359 | 5169327 | **RNA**→ rpmE2 (PRK01678)Ribosomal_L31 (pfam01197)→ |
| Ban-21-15 | NZ_CP009476.1 | + | 2929400 | 2929432 | **RNA**→ rpmE2 (PRK01678)Ribosomal_L31 (pfam01197)→ |
| Ban-21-16 | NZ_CP009902.1 | + | 2026472 | 2026504 | **RNA**→ rpmE2 (PRK01678)Ribosomal_L31 (pfam01197)→ |

| | | | | | |
|---|---|---|---|---|---|
| Ban-21-17 | NZ_CP009697.1 | - | 3050717 | 3050685 | **RNA**→ rpmE2 (PRK01678)Ribosomal_L31 (pfam01197)→ |
| Ban-21-18 | NZ_CP010322.1 | - | 2776009 | 2775977 | **RNA**→ rpmE2 (PRK01678)Ribosomal_L31 (pfam01197)→ |
| Ban-17-2 | NZ_CP009541.1 | + | 2794066 | 2794098 | **RNA**→ rpmE2 (PRK01678)Ribosomal_L31 (pfam01197)→ |
| Ban-38-1 | NZ_CP009598.1 | + | 5033290 | 5033322 | **RNA**→ rpmE2 (PRK01678)Ribosomal_L31 (pfam01197)→ |
| Ban-21-19 | NZ_CP009981.1 | + | 4660357 | 4660389 | **RNA**→ rpmE2 (PRK01678)Ribosomal_L31 (pfam01197)→ |
| Ban-21-20 | NZ_CP010852.1 | - | 113594 | 113562 | **RNA**→ rpmE2 (PRK01678)Ribosomal_L31 (pfam01197)→ |
| Ban-23-1 | NZ_CM002401.1 | - | 5059774 | 5059742 | **RNA**→ rpmE2 (PRK01678)Ribosomal_L31 (pfam01197)→ |
| Ban-22-1 | NZ_CM002395.1 | - | 5059426 | 5059394 | **RNA**→ rpmE2 (PRK01678)Ribosomal_L31 (pfam01197)→ |
| Bce-23-1 | NZ_CM000716.1 | - | 4974285 | 4974253 | **RNA**→ rpmE2 (PRK01678)Ribosomal_L31 (pfam01197)→ rpmE2 (PRK01678)Ribosomal_L31 (pfam01197 |
| Ban-32-1 | NZ_CAVE010000038.1 | - | 61746 | 61714 | **RNA**→ rpmE2 (PRK01678)Ribosomal_L31 (pfam01197)→ |
| Ban-35-1 | NZ_CAVC010000043.1 | - | 61881 | 61849 | **RNA**→ rpmE2 (PRK01678)Ribosomal_L31 (pfam01197)→ |
| Ban-36-1 | NZ_CAVD010000049.1 | - | 61891 | 61859 | **RNA**→ rpmE2 (PRK01678)Ribosomal_L31 (pfam01197)→ |
| Ban-30-1 | NZ_AMDT01000019.1 | - | 27183 | 27151 | **RNA**→ rpmE2 (PRK01678)Ribosomal_L31 (pfam01197)→ |
| Bce-63-1 | NZ_JH792376.1 | + | 195540 | 195572 | **RNA**→ rpmE2 (PRK01678)Ribosomal_L31 (pfam01197)→ |
| env-939 | JGI25159J45721_1011137 | + | 238 | 270 | **RNA**→ rpmE2 (PRK01678)Ribosomal_L31 (pfam01197)→ |
| env-940 | JGI25151J46595_10014267 | - | 737 | 705 | **RNA**→ rpmE2 (PRK01678)Ribosomal_L31 (pfam01197)→ |
| Bps-2-1 | NZ_CM000745.1 | - | 4565126 | 4565094 | **RNA**→ rpmE2 (PRK01678)Ribosomal_L31 (pfam01197)→ rpmE2 (PRK01678)Ribosomal_L31 (pfam01197 |
| Bmy-2-1 | NZ_CM000743.1 | + | 4269323 | 4269355 | **RNA**→ rpmE2 (PRK01678)Ribosomal_L31 (pfam01197)→ |
| Bmy-3-1 | NZ_CM000744.1 | - | 4468592 | 4468560 | **RNA**→ rpmE2 (PRK01678)Ribosomal_L31 (pfam01197)→ |
| Bsp-26-1 | NZ_KB895693.1 | + | 28519 | 28551 | **RNA**→ rpmE2 (PRK01678)Ribosomal_L31 (pfam01197)→ |
| Bsp-14-1 | NZ_KB895568.1 | + | 173140 | 173172 | **RNA**→ rpmE2 (PRK01678)Ribosomal_L31 (pfam01197)→ |
| Bce-142-1 | NZ_KB976905.1 | - | 423230 | 423198 | **RNA**→ rpmE2 (PRK01678)Ribosomal_L31 (pfam01197)→ |
| Bce-127-1 | NZ_KB976717.1 | - | 420881 | 420849 | **RNA**→ rpmE2 (PRK01678)Ribosomal_L31 (pfam01197)→ |
| Bce-140-1 | NZ_KB976864.1 | - | 344767 | 344735 | **RNA**→ rpmE2 (PRK01678)Ribosomal_L31 (pfam01197)→ |
| Bmy-4-1 | NZ_CP009651.1 | + | 2534010 | 2534042 | **RNA**→ rpmE2 (PRK01678)Ribosomal_L31 (pfam01197)→ |
| Bmy-4-2 | NZ_CP007626.1 | - | 2305106 | 2305074 | **RNA**→ rpmE2 (PRK01678)Ribosomal_L31 (pfam01197)→ |
| Bma-2-1 | NZ_JOTN01000001.1 | - | 308890 | 308858 | **RNA**→ rpmE2 (PRK01678)Ribosomal_L31 (pfam01197)→ |
| Bcy-2-1 | NZ_JYPG01000024.1 | - | 242719 | 242687 | **RNA**→ rpmE2 (PRK01678)Ribosomal_L31 (pfam01197)→ |
| Bcy-1-1 | NC_009674.1 | - | 3909710 | 3909678 | **RNA**→ rpmE2 (PRK01678)Ribosomal_L31 (pfam01197)→ rpmE2 (PRK01678)Ribosomal_L31 (pfam01197 |
| Bsp-39-1 | NZ_JH921519.1 | - | 365021 | 364989 | **RNA**→ rpmE2 (PRK01678)Ribosomal_L31 (pfam01197)→ |
| Bdi-1-1 | NZ_HG942082.1 | - | 119037 | 119005 | **RNA**→ rpmE (PRK00019)Ribosomal_L31 (pfam01197)→ |
| Bfi-1-1 | NZ_LDWQ01000106.1 | - | 1219 | 1189 | **RNA**→ rpmE (PRK00019)Ribosomal_L31 (pfam01197)→ |
| Bba-1-1 | NZ_AJLS01000139.1 | - | 4099 | 4069 | **RNA**→ rpmE (PRK00019)Ribosomal_L31 (pfam01197)→ rpmE (PRK00019)Ribosomal_L31 (pfam01197)– |
| | | | | | Tdk (COG1435)PRK04296 (PRK04296)→ Tdk (COG1435)PRK04296 (PRK04296)→ |
| Bni-1-1 | NZ_BAWM01000007.1 | - | 1879 | 1849 | **RNA**→ rpmE (PRK00019)Ribosomal_L31 (pfam01197)→ |
| Bsp-53-1 | NZ_JAEL01000147.1 | - | 1653 | 1623 | **RNA**→ rpmE (PRK00019)Ribosomal_L31 (pfam01197)→ |
| Age-1-2 | NZ_JYCG01000013.1 | + | 44653 | 44684 | **RNA**→ rpmE (PRK00019)Ribosomal_L31 (pfam01197)→ |
| Ate-1-1 | NZ_JHVN01000001.1 | - | 392015 | 391984 | **RNA**→ rpmE (PRK00019)Ribosomal_L31 (pfam01197)→ |
| Bce-6-1 | NZ_CM000739.1 | + | 4869077 | 4869109 | **RNA**→ rpmE2 (PRK01678)Ribosomal_L31 (pfam01197)→ rpmE2 (PRK01678)Ribosomal_L31 (pfam01197 |
| Bce-106-1 | NZ_KE504110.1 | + | 260069 | 260101 | **RNA**→ rpmE2 (PRK01678)Ribosomal_L31 (pfam01197)→ |
| Bce-108-1 | NZ_KE504107.1 | + | 259819 | 259851 | **RNA**→ rpmE2 (PRK01678)Ribosomal_L31 (pfam01197)→ |
| Bce-107-1 | NZ_JH636004.1 | + | 259670 | 259702 | **RNA**→ rpmE2 (PRK01678)Ribosomal_L31 (pfam01197)→ |
| env-941 | JGI25151J46595_10017439 | + | 2866 | 2898 | **RNA**→ rpmE2 (PRK01678)Ribosomal_L31 (pfam01197)→ |
| Bce-40-1 | NZ_CM000733.1 | + | 4212588 | 4212620 | **RNA**→ rpmE2 (PRK01678)Ribosomal_L31 (pfam01197)→ rpmE2 (PRK01678)Ribosomal_L31 (pfam01197 |
| Bsp-91-1 | NZ_AXVB01000011.1 | + | 29512 | 29544 | **RNA**→ rpmE2 (PRK01678)Ribosomal_L31 (pfam01197)→ |
| env-942 | JGI25151J46595_10061557 | - | 473 | 441 | **RNA**→ rpmE2 (PRK01678)Ribosomal_L31 (pfam01197)→ |
| Bsp-89-1 | NZ_JQKQ01000011.1 | - | 200394 | 200362 | **RNA**→ rpmE2 (PRK01678)Ribosomal_L31 (pfam01197)→ |
| Bsp-15-1 | NZ_KB910926.1 | + | 143065 | 143097 | **RNA**→ rpmE2 (PRK01678)Ribosomal_L31 (pfam01197)→ |
| Bga-1-1 | NZ_JOTM01000002.1 | - | 410823 | 410791 | **RNA**→ rpmE2 (PRK01678)Ribosomal_L31 (pfam01197)→ |
| env-943 | JGI12273J12029_10707276 | + | 98 | 128 | **RNA**→ rpmE (PRK00019)Ribosomal_L31 (pfam01197)→ |
| env-944 | JGI25159J45721_1056023 | - | 243 | 211 | **RNA**→ rpmE2 (PRK01678)Ribosomal_L31 (pfam01197)→ |
| Bth-31-1 | NZ_AMXT02000005.1 | + | 2115 | 2147 | **RNA**→ rpmE2 (PRK01678)Ribosomal_L31 (pfam01197)→ |

| | | | | | |
|---|---|---|---|---|---|
| Bth-29-1 | NZ_ARXZ02000005.1 | - | 247882 | 247850 | **RNA**→ rpmE2 (PRK01678)Ribosomal_L31 (pfam01197)→ |
| Bsp-70-1 | NZ_JXIT01000003.1 | + | 4627 | 4659 | **RNA**→ rpmE2 (PRK01678)Ribosomal_L31 (pfam01197)→ |
| Bsp-49-1 | NZ_CAVI010000079.1 | + | 24082 | 24114 | **RNA**→ rpmE2 (PRK01678)Ribosomal_L31 (pfam01197)→ |
| Bsp-69-1 | NZ_JXIS01000025.1 | - | 246925 | 246893 | **RNA**→ rpmE2 (PRK01678)Ribosomal_L31 (pfam01197)→ |
| Bsp-87-1 | NZ_JSFD01000003.1 | + | 203781 | 203813 | **RNA**→ rpmE2 (PRK01678)Ribosomal_L31 (pfam01197)→ |
| Bth-40-1 | NZ_JWJY01000024.1 | - | 60949 | 60917 | **RNA**→ rpmE2 (PRK01678)Ribosomal_L31 (pfam01197)→ |
| Spn-1-1 | NZ_CVOG01000306.1 | - | 327 | 295 | **RNA**→ rpmE2 (PRK01678)Ribosomal_L31 (pfam01197)→ |
| Bce-5-1 | NZ_ABDA02000004.1 | - | 370284 | 370252 | **RNA**→ rpmE2 (PRK01678)Ribosomal_L31 (pfam01197)→ rpmE2 (PRK01678)Ribosomal_L31 (pfam01197 |
| Bce-28-1 | NZ_AAEK01000096.1 | + | 280 | 312 | **RNA**→ rpmE2 (PRK01678)Ribosomal_L31 (pfam01197)→ rpmE2 (PRK01678)Ribosomal_L31 (pfam01197 |
| Bce-43-1 | NZ_ABCZ02000035.1 | + | 6718 | 6750 | **RNA**→ rpmE2 (PRK01678)Ribosomal_L31 (pfam01197)→ rpmE2 (PRK01678)Ribosomal_L31 (pfam01197 |
| Bsp-51-1 | NZ_AYMM01000025.1 | + | 100909 | 100941 | **RNA**→ rpmE2 (PRK01678)Ribosomal_L31 (pfam01197)→ |
| Bth-27-1 | NZ_AYPV01000046.1 | - | 235676 | 235644 | **RNA**→ rpmE2 (PRK01678)Ribosomal_L31 (pfam01197)→ |
| Spn-1-2 | NZ_CKVP01000064.1 | - | 2654 | 2622 | **RNA**→ rpmE2 (PRK01678)Ribosomal_L31 (pfam01197)→ |
| Bth-26-1 | NZ_AYSM01000032.1 | - | 413082 | 413050 | **RNA**→ rpmE2 (PRK01678)Ribosomal_L31 (pfam01197)→ |
| Spn-1-3 | NZ_CLRF01000045.1 | - | 5931 | 5899 | **RNA**→ rpmE2 (PRK01678)Ribosomal_L31 (pfam01197)→ |
| Bmy-6-1 | NZ_ASPZ01000051.1 | + | 183225 | 183257 | **RNA**→ rpmE2 (PRK01678)Ribosomal_L31 (pfam01197)→ |
| Bth-32-1 | NZ_AMXS02000116.1 | + | 31044 | 31076 | **RNA**→ rpmE2 (PRK01678)Ribosomal_L31 (pfam01197)→ |
| Bsu-13-1 | NZ_AZNI01000006.1 | + | 152247 | 152279 | **RNA**→ rpmE2 (PRK01678)Ribosomal_L31 (pfam01197)→ |
| Spn-1-4 | NZ_CLBL01000054.1 | + | 4140 | 4172 | **RNA**→ rpmE2 (PRK01678)Ribosomal_L31 (pfam01197)→ |
| Spn-1-5 | NZ_CKVY01000040.1 | + | 5832 | 5864 | **RNA**→ rpmE2 (PRK01678)Ribosomal_L31 (pfam01197)→ |
| Bth-43-1 | NZ_AYXQ01000009.1 | - | 85575 | 85543 | **RNA**→ rpmE2 (PRK01678)Ribosomal_L31 (pfam01197)→ |
| Spn-1-6 | NZ_CMDP01000044.1 | + | 4474 | 4506 | **RNA**→ rpmE2 (PRK01678)Ribosomal_L31 (pfam01197)→ |
| Bth-35-1 | NZ_JEOC01000013.1 | - | 7003 | 6971 | **RNA**→ rpmE2 (PRK01678)Ribosomal_L31 (pfam01197)→ |
| Bth-12-1 | NC_005957.1 | - | 5064430 | 5064398 | **RNA**→ rpmE2 (PRK01678)Ribosomal_L31 (pfam01197)→ |
| Bce-16-1 | NC_004722.1 | - | 5237467 | 5237435 | **RNA**→ rpmE2 (PRK01678)Ribosomal_L31 (pfam01197)→ rpmE2 (PRK01678)Ribosomal_L31 (pfam01197 |
| Bce-24-1 | NC_006274.1 | - | 5128915 | 5128883 | **RNA**→ rpmE2 (PRK01678)Ribosomal_L31 (pfam01197)→ |
| Bce-29-1 | NC_011772.1 | - | 5199166 | 5199134 | **RNA**→ rpmE2 (PRK01678)Ribosomal_L31 (pfam01197)→ |
| Bce-33-1 | NC_011969.1 | - | 5036818 | 5036786 | **RNA**→ rpmE2 (PRK01678)Ribosomal_L31 (pfam01197)→ rpmE2 (PRK01678)Ribosomal_L31 (pfam01197 rpmE2 (PRK01678)Ribosomal_L31 (pfam01197)→ |
| Bce-13-1 | NC_011773.1 | - | 5118565 | 5118533 | **RNA**→ rpmE2 (PRK01678)Ribosomal_L31 (pfam01197)→ |
| Bce-15-1 | NC_003909.8 | - | 5038240 | 5038208 | **RNA**→ rpmE2 (PRK01678)Ribosomal_L31 (pfam01197)→ |
| Bce-9-1 | NC_011658.1 | - | 5092666 | 5092634 | **RNA**→ rpmE2 (PRK01678)Ribosomal_L31 (pfam01197)→ |
| Bth-10-1 | NC_017200.1 | - | 5177466 | 5177434 | **RNA**→ rpmE2 (PRK01678)Ribosomal_L31 (pfam01197)→ |
| Bth-9-1 | NC_017208.1 | - | 5312273 | 5312241 | **RNA**→ rpmE2 (PRK01678)Ribosomal_L31 (pfam01197)→ |
| Bce-31-1 | NC_016771.1 | - | 5044989 | 5044957 | **RNA**→ rpmE2 (PRK01678)Ribosomal_L31 (pfam01197)→ |
| Bth-2-1 | NC_018877.1 | - | 5325945 | 5325913 | **RNA**→ rpmE2 (PRK01678)Ribosomal_L31 (pfam01197)→ |
| Bth-24-1 | NC_018508.1 | - | 4822974 | 4822942 | **RNA**→ rpmE2 (PRK01678)Ribosomal_L31 (pfam01197)→ |
| Bth-38-1 | NC_020238.1 | - | 5466196 | 5466164 | **RNA**→ rpmE2 (PRK01678)Ribosomal_L31 (pfam01197)→ |
| Bth-3-1 | NC_018500.1 | + | 4315030 | 4315062 | **RNA**→ rpmE2 (PRK01678)Ribosomal_L31 (pfam01197)→ |
| Bth-42-1 | NC_020376.1 | - | 5315959 | 5315927 | **RNA**→ rpmE2 (PRK01678)Ribosomal_L31 (pfam01197)→ |
| Bth-6-1 | NC_018693.1 | - | 4426098 | 4426066 | **RNA**→ rpmE2 (PRK01678)Ribosomal_L31 (pfam01197)→ rpmE2 (PRK01678)Ribosomal_L31 (pfam01197 rpmE2 (PRK01678)Ribosomal_L31 (pfam01197)→ |
| Bce-27-1 | NC_018491.1 | + | 1766387 | 1766419 | **RNA**→ rpmE2 (PRK01678)Ribosomal_L31 (pfam01197)→ |
| Bto-1-1 | NC_022781.1 | - | 2454301 | 2454269 | **RNA**→ rpmE2 (PRK01678)Ribosomal_L31 (pfam01197)→ rpmE2 (PRK01678)Ribosomal_L31 (pfam01197 |
| Bth-39-1 | NZ_CP004858.1 | - | 5422066 | 5422034 | **RNA**→ rpmE2 (PRK01678)Ribosomal_L31 (pfam01197)→ |
| Bth-39-2 | NZ_CP007607.1 | + | 517856 | 517888 | **RNA**→ rpmE2 (PRK01678)Ribosomal_L31 (pfam01197)→ |
| Bth-37-1 | NZ_CP004870.1 | - | 5435621 | 5435589 | **RNA**→ rpmE2 (PRK01678)Ribosomal_L31 (pfam01197)→ |
| Bbo-2-1 | NZ_CP007512.1 | - | 4973045 | 4973013 | **RNA**→ rpmE2 (PRK01678)Ribosomal_L31 (pfam01197)→ |
| Bce-36-1 | NZ_CM000728.1 | - | 5006615 | 5006583 | **RNA**→ rpmE2 (PRK01678)Ribosomal_L31 (pfam01197)→ rpmE2 (PRK01678)Ribosomal_L31 (pfam01197 |
| Bce-12-1 | NZ_CM000738.1 | - | 4797858 | 4797826 | **RNA**→ rpmE2 (PRK01678)Ribosomal_L31 (pfam01197)→ rpmE2 (PRK01678)Ribosomal_L31 (pfam01197 |
| Bce-17-1 | NZ_CM000721.1 | + | 5030069 | 5030101 | **RNA**→ rpmE2 (PRK01678)Ribosomal_L31 (pfam01197)→ rpmE2 (PRK01678)Ribosomal_L31 (pfam01197 |
| Bth-11-1 | NZ_CM000756.1 | + | 5063643 | 5063675 | **RNA**→ rpmE2 (PRK01678)Ribosomal_L31 (pfam01197)→ rpmE2 (PRK01678)Ribosomal_L31 (pfam01197 |
| Bth-8-1 | NZ_CM000753.1 | + | 4929359 | 4929391 | **RNA**→ rpmE2 (PRK01678)Ribosomal_L31 (pfam01197)→ rpmE2 (PRK01678)Ribosomal_L31 (pfam01197 |

| | | | | | |
|---|---|---|---|---|---|
| Bce-37-1 | NZ_CM000730.1 | - | 4893712 | 4893680 | **RNA**→ rpmE2 (PRK01678)Ribosomal_L31 (pfam01197)→ rpmE2 (PRK01678)Ribosomal_L31 (pfam01197 |
| Bth-14-1 | NZ_CM000752.1 | - | 5032858 | 5032826 | **RNA**→ rpmE2 (PRK01678)Ribosomal_L31 (pfam01197)→ rpmE2 (PRK01678)Ribosomal_L31 (pfam01197 |
| Bce-41-1 | NZ_CM000735.1 | + | 4903796 | 4903828 | **RNA**→ rpmE2 (PRK01678)Ribosomal_L31 (pfam01197)→ rpmE2 (PRK01678)Ribosomal_L31 (pfam01197 |
| Bce-3-1 | NZ_CM000717.1 | - | 4853496 | 4853464 | **RNA**→ rpmE2 (PRK01678)Ribosomal_L31 (pfam01197)→ rpmE2 (PRK01678)Ribosomal_L31 (pfam01197 |
| Bce-46-1 | NZ_CM000722.1 | - | 4959910 | 4959878 | **RNA**→ rpmE2 (PRK01678)Ribosomal_L31 (pfam01197)→ rpmE2 (PRK01678)Ribosomal_L31 (pfam01197 |
| Bce-34-1 | NZ_CM000720.1 | + | 4913556 | 4913588 | **RNA**→ rpmE2 (PRK01678)Ribosomal_L31 (pfam01197)→ rpmE2 (PRK01678)Ribosomal_L31 (pfam01197 |
| Bce-14-1 | NZ_CM000715.1 | - | 5133278 | 5133246 | **RNA**→ rpmE2 (PRK01678)Ribosomal_L31 (pfam01197)→ rpmE2 (PRK01678)Ribosomal_L31 (pfam01197 |
| Bth-16-1 | NZ_CM000755.1 | + | 4886738 | 4886770 | **RNA**→ rpmE2 (PRK01678)Ribosomal_L31 (pfam01197)→ rpmE2 (PRK01678)Ribosomal_L31 (pfam01197 |
| Bth-4-1 | NZ_CM000758.1 | + | 5153478 | 5153510 | **RNA**→ rpmE2 (PRK01678)Ribosomal_L31 (pfam01197)→ rpmE2 (PRK01678)Ribosomal_L31 (pfam01197 |
| Bth-18-1 | NZ_CM000748.1 | + | 4896324 | 4896356 | **RNA**→ rpmE2 (PRK01678)Ribosomal_L31 (pfam01197)→ rpmE2 (PRK01678)Ribosomal_L31 (pfam01197 |
| Bce-38-1 | NZ_CM000731.1 | - | 4919707 | 4919675 | **RNA**→ rpmE2 (PRK01678)Ribosomal_L31 (pfam01197)→ rpmE2 (PRK01678)Ribosomal_L31 (pfam01197 |
| Bce-21-1 | NZ_CM000723.1 | + | 5024748 | 5024780 | **RNA**→ rpmE2 (PRK01678)Ribosomal_L31 (pfam01197)→ rpmE2 (PRK01678)Ribosomal_L31 (pfam01197 |
| Bth-15-1 | NZ_CM000750.1 | - | 4887859 | 4887827 | **RNA**→ rpmE2 (PRK01678)Ribosomal_L31 (pfam01197)→ rpmE2 (PRK01678)Ribosomal_L31 (pfam01197 |
| Bce-42-1 | NZ_CM000734.1 | - | 5035278 | 5035246 | **RNA**→ rpmE2 (PRK01678)Ribosomal_L31 (pfam01197)→ rpmE2 (PRK01678)Ribosomal_L31 (pfam01197 |
| Bce-35-1 | NZ_CM000729.1 | + | 4857814 | 4857846 | **RNA**→ rpmE2 (PRK01678)Ribosomal_L31 (pfam01197)→ rpmE2 (PRK01678)Ribosomal_L31 (pfam01197 |
| Bth-21-1 | NZ_KK088334.1 | - | 242518 | 242486 | **RNA**→ rpmE2 (PRK01678)Ribosomal_L31 (pfam01197)→ |
| Bce-14-2 | NZ_KN050654.1 | - | 491347 | 491315 | **RNA**→ rpmE2 (PRK01678)Ribosomal_L31 (pfam01197)→ |
| Bce-45-1 | NZ_CM000714.1 | - | 4850950 | 4850918 | **RNA**→ rpmE2 (PRK01678)Ribosomal_L31 (pfam01197)→ rpmE2 (PRK01678)Ribosomal_L31 (pfam01197 |
| Bsp-11-1 | NZ_JH414711.1 | + | 182856 | 182888 | **RNA**→ rpmE2 (PRK01678)Ribosomal_L31 (pfam01197)→ |
| Bce-69-1 | NZ_JH791968.1 | - | 5087536 | 5087504 | **RNA**→ rpmE2 (PRK01678)Ribosomal_L31 (pfam01197)→ |
| Bce-131-1 | NZ_JH792156.1 | + | 248131 | 248163 | **RNA**→ rpmE2 (PRK01678)Ribosomal_L31 (pfam01197)→ |
| Bce-55-1 | NZ_JH792223.1 | - | 409655 | 409623 | **RNA**→ rpmE2 (PRK01678)Ribosomal_L31 (pfam01197)→ |
| Bce-49-1 | NZ_JH792265.1 | + | 266039 | 266071 | **RNA**→ rpmE2 (PRK01678)Ribosomal_L31 (pfam01197)→ |
| Bce-134-1 | NZ_JH791855.1 | - | 412679 | 412647 | **RNA**→ rpmE2 (PRK01678)Ribosomal_L31 (pfam01197)→ |
| Bce-115-1 | NZ_JH792025.1 | - | 5004746 | 5004714 | **RNA**→ rpmE2 (PRK01678)Ribosomal_L31 (pfam01197)→ |
| Bce-67-1 | NZ_JH791899.1 | + | 254004 | 254036 | **RNA**→ rpmE2 (PRK01678)Ribosomal_L31 (pfam01197)→ |
| Bce-87-1 | NZ_JH804653.1 | + | 246240 | 246272 | **RNA**→ rpmE2 (PRK01678)Ribosomal_L31 (pfam01197)→ |
| Bce-121-1 | NZ_JH792240.1 | + | 257458 | 257490 | **RNA**→ rpmE2 (PRK01678)Ribosomal_L31 (pfam01197)→ |
| Bce-71-1 | NZ_JH791951.1 | - | 4892243 | 4892211 | **RNA**→ rpmE2 (PRK01678)Ribosomal_L31 (pfam01197)→ |
| Bce-52-1 | NZ_JH792321.1 | + | 243223 | 243255 | **RNA**→ rpmE2 (PRK01678)Ribosomal_L31 (pfam01197)→ |
| Bce-111-1 | NZ_JH792105.1 | + | 243201 | 243233 | **RNA**→ rpmE2 (PRK01678)Ribosomal_L31 (pfam01197)→ |
| Bce-138-1 | NZ_JH791819.1 | - | 5219376 | 5219344 | **RNA**→ rpmE2 (PRK01678)Ribosomal_L31 (pfam01197)→ |
| Bce-77-1 | NZ_JH804624.1 | + | 259696 | 259728 | **RNA**→ rpmE2 (PRK01678)Ribosomal_L31 (pfam01197)→ |
| Bce-96-1 | NZ_JH804690.1 | + | 254845 | 254877 | **RNA**→ rpmE2 (PRK01678)Ribosomal_L31 (pfam01197)→ |
| Bce-73-1 | NZ_JH791945.1 | - | 4359897 | 4359865 | **RNA**→ rpmE2 (PRK01678)Ribosomal_L31 (pfam01197)→ |
| Bce-97-1 | NZ_JH792136.1 | - | 384123 | 384091 | **RNA**→ rpmE2 (PRK01678)Ribosomal_L31 (pfam01197)→ |
| Bce-72-1 | NZ_JH804617.1 | + | 315988 | 316020 | **RNA**→ rpmE2 (PRK01678)Ribosomal_L31 (pfam01197)→ |
| Bce-113-1 | NZ_JH792018.1 | + | 281199 | 281231 | **RNA**→ rpmE2 (PRK01678)Ribosomal_L31 (pfam01197)→ |
| Bce-99-1 | NZ_JH792120.1 | - | 5021047 | 5021015 | **RNA**→ rpmE2 (PRK01678)Ribosomal_L31 (pfam01197)→ |
| Bce-133-1 | NZ_JH791865.1 | + | 265832 | 265864 | **RNA**→ rpmE2 (PRK01678)Ribosomal_L31 (pfam01197)→ |
| Bce-117-1 | NZ_JH792282.1 | + | 186404 | 186436 | **RNA**→ rpmE2 (PRK01678)Ribosomal_L31 (pfam01197)→ |
| Bce-123-1 | NZ_JH792166.1 | + | 257716 | 257748 | **RNA**→ rpmE2 (PRK01678)Ribosomal_L31 (pfam01197)→ |
| Bce-132-1 | NZ_JH791884.1 | - | 152349 | 152317 | **RNA**→ rpmE2 (PRK01678)Ribosomal_L31 (pfam01197)→ |
| Bce-135-1 | NZ_JH791833.1 | - | 4979784 | 4979752 | **RNA**→ rpmE2 (PRK01678)Ribosomal_L31 (pfam01197)→ |
| Bce-118-1 | NZ_JH792081.1 | - | 393135 | 393103 | **RNA**→ rpmE2 (PRK01678)Ribosomal_L31 (pfam01197)→ |
| Bce-70-1 | NZ_JH791942.1 | - | 5132776 | 5132744 | **RNA**→ rpmE2 (PRK01678)Ribosomal_L31 (pfam01197)→ |
| Bce-94-1 | NZ_JH804683.1 | + | 440203 | 440235 | **RNA**→ rpmE2 (PRK01678)Ribosomal_L31 (pfam01197)→ |
| Bce-112-1 | NZ_JH792039.1 | + | 21430 | 21462 | **RNA**→ rpmE2 (PRK01678)Ribosomal_L31 (pfam01197)→ |
| Bce-86-1 | NZ_CM001787.1 | - | 5054594 | 5054562 | **RNA**→ rpmE2 (PRK01678)Ribosomal_L31 (pfam01197)→ |
| Bth-22-1 | NZ_CM001804.1 | - | 5584093 | 5584061 | **RNA**→ rpmE2 (PRK01678)Ribosomal_L31 (pfam01197)→ rpmE2 (PRK01678)Ribosomal_L31 (pfam01197 |
| Bth-22-2 | NZ_CM001804.1 | - | 5582913 | 5582881 | **RNA**→ rpmE2 (PRK01678)Ribosomal_L31 (pfam01197)→ rpmE2 (PRK01678)Ribosomal_L31 (pfam01197 |
| Bce-95-1 | NZ_KB976627.1 | - | 250768 | 250736 | **RNA**→ rpmE2 (PRK01678)Ribosomal_L31 (pfam01197)→ |

| | | | | | |
|---|---|---|---|---|---|
| Bce-60-1 | NZ_KB976517.1 | - | 5040982 | 5040950 | **RNA**→ rpmE2 (PRK01678)Ribosomal_L31 (pfam01197)→ |
| Bce-81-1 | NZ_KB976593.1 | + | 263858 | 263890 | **RNA**→ rpmE2 (PRK01678)Ribosomal_L31 (pfam01197)→ |
| Bce-61-1 | NZ_KE332507.1 | - | 5030046 | 5030014 | **RNA**→ rpmE2 (PRK01678)Ribosomal_L31 (pfam01197)→ |
| Bce-114-1 | NZ_KB976039.1 | + | 263486 | 263518 | **RNA**→ rpmE2 (PRK01678)Ribosomal_L31 (pfam01197)→ |
| Bce-126-1 | NZ_KB976184.1 | - | 868527 | 868495 | **RNA**→ rpmE2 (PRK01678)Ribosomal_L31 (pfam01197)→ |
| Bce-98-1 | NZ_KB976545.1 | - | 220766 | 220734 | **RNA**→ rpmE2 (PRK01678)Ribosomal_L31 (pfam01197)→ |
| Bce-58-1 | NZ_KB976220.1 | + | 193877 | 193909 | **RNA**→ rpmE2 (PRK01678)Ribosomal_L31 (pfam01197)→ |
| Bth-23-1 | NZ_HG810020.1 | - | 443455 | 443423 | **RNA**→ rpmE2 (PRK01678)Ribosomal_L31 (pfam01197)→ |
| Bce-89-1 | NZ_KB976661.1 | - | 379081 | 379049 | **RNA**→ rpmE2 (PRK01678)Ribosomal_L31 (pfam01197)→ |
| Bce-102-1 | NZ_KB976311.1 | - | 1309599 | 1309567 | **RNA**→ rpmE2 (PRK01678)Ribosomal_L31 (pfam01197)→ |
| Bce-128-1 | NZ_KB976808.1 | - | 405722 | 405690 | **RNA**→ rpmE2 (PRK01678)Ribosomal_L31 (pfam01197)→ |
| Bce-147-1 | NZ_KB976117.1 | - | 411767 | 411735 | **RNA**→ rpmE2 (PRK01678)Ribosomal_L31 (pfam01197)→ |
| Bce-125-1 | NZ_KB976650.1 | - | 386069 | 386037 | **RNA**→ rpmE2 (PRK01678)Ribosomal_L31 (pfam01197)→ |
| Bce-57-1 | NZ_KB976207.1 | - | 5024371 | 5024339 | **RNA**→ rpmE2 (PRK01678)Ribosomal_L31 (pfam01197)→ |
| Bce-103-1 | NZ_KB976769.1 | - | 440019 | 439987 | **RNA**→ rpmE2 (PRK01678)Ribosomal_L31 (pfam01197)→ |
| Bce-54-1 | NZ_KB976160.1 | + | 255650 | 255682 | **RNA**→ rpmE2 (PRK01678)Ribosomal_L31 (pfam01197)→ |
| Bce-75-1 | NZ_KB976554.1 | - | 419271 | 419239 | **RNA**→ rpmE2 (PRK01678)Ribosomal_L31 (pfam01197)→ |
| Bce-137-1 | NZ_KB976266.1 | - | 422107 | 422075 | **RNA**→ rpmE2 (PRK01678)Ribosomal_L31 (pfam01197)→ |
| Bce-53-1 | NZ_KE332537.1 | - | 411040 | 411008 | **RNA**→ rpmE2 (PRK01678)Ribosomal_L31 (pfam01197)→ |
| Bce-59-1 | NZ_KB976237.1 | - | 4422472 | 4422440 | **RNA**→ rpmE2 (PRK01678)Ribosomal_L31 (pfam01197)→ |
| Bce-136-1 | NZ_KB976843.1 | - | 70671 | 70639 | **RNA**→ rpmE2 (PRK01678)Ribosomal_L31 (pfam01197)→ |
| Bth-13-1 | NZ_CM000751.1 | + | 4809697 | 4809729 | **RNA**→ rpmE2 (PRK01678)Ribosomal_L31 (pfam01197)→ rpmE2 (PRK01678)Ribosomal_L31 (pfam01197 |
| Bth-5-1 | NZ_CM000759.1 | + | 4781622 | 4781654 | **RNA**→ rpmE2 (PRK01678)Ribosomal_L31 (pfam01197)→ rpmE2 (PRK01678)Ribosomal_L31 (pfam01197 |
| Bth-34-1 | NZ_CP010106.1 | - | 5260251 | 5260219 | **RNA**→ rpmE2 (PRK01678)Ribosomal_L31 (pfam01197)→ |
| Bce-47-1 | NZ_CP009941.1 | + | 100239 | 100271 | **RNA**→ rpmE2 (PRK01678)Ribosomal_L31 (pfam01197)→ |
| Bth-33-1 | NZ_CP010089.1 | - | 5435610 | 5435578 | **RNA**→ rpmE2 (PRK01678)Ribosomal_L31 (pfam01197)→ |
| Bth-20-1 | NC_008600.1 | - | 5087437 | 5087405 | **RNA**→ rpmE2 (PRK01678)Ribosomal_L31 (pfam01197)→ rpmE2 (PRK01678)Ribosomal_L31 (pfam01197 rpmE2 (PRK01678)Ribosomal_L31 (pfam01197)→ |
| Bth-1-1 | NC_014171.1 | - | 5149570 | 5149538 | **RNA**→ rpmE2 (PRK01678)Ribosomal_L31 (pfam01197)→ rpmE2 (PRK01678)Ribosomal_L31 (pfam01197 rpmE2 (PRK01678)Ribosomal_L31 (pfam01197)→ |
| Bce-139-1 | NZ_KB976695.1 | - | 4408716 | 4408684 | **RNA**→ rpmE2 (PRK01678)Ribosomal_L31 (pfam01197)→ |
| Bce-105-1 | NZ_KB976300.1 | + | 263669 | 263701 | **RNA**→ rpmE2 (PRK01678)Ribosomal_L31 (pfam01197)→ |
| Bce-101-1 | NZ_KB976325.1 | - | 1320401 | 1320369 | **RNA**→ rpmE2 (PRK01678)Ribosomal_L31 (pfam01197)→ |
| Bth-30-1 | NC_022873.1 | - | 5825716 | 5825684 | **RNA**→ rpmE2 (PRK01678)Ribosomal_L31 (pfam01197)→ |
| Bce-28-2 | NZ_CP009590.1 | + | 2646537 | 2646569 | **RNA**→ rpmE2 (PRK01678)Ribosomal_L31 (pfam01197)→ |
| Bce-17-2 | NZ_CP009628.1 | + | 2800811 | 2800843 | **RNA**→ rpmE2 (PRK01678)Ribosomal_L31 (pfam01197)→ |
| Bce-47-2 | NZ_CP009596.1 | + | 3811663 | 3811695 | **RNA**→ rpmE2 (PRK01678)Ribosomal_L31 (pfam01197)→ |
| Bce-48-1 | NZ_CP009641.1 | - | 3825469 | 3825437 | **RNA**→ rpmE2 (PRK01678)Ribosomal_L31 (pfam01197)→ |
| Bce-47-3 | NZ_CP009369.1 | + | 5167894 | 5167926 | **RNA**→ rpmE2 (PRK01678)Ribosomal_L31 (pfam01197)→ |
| Bce-24-2 | NZ_CP009968.1 | + | 333073 | 333105 | **RNA**→ rpmE2 (PRK01678)Ribosomal_L31 (pfam01197)→ |
| Bth-25-1 | NZ_CP009351.1 | + | 4083244 | 4083276 | **RNA**→ rpmE2 (PRK01678)Ribosomal_L31 (pfam01197)→ |
| Bce-47-4 | NZ_CP009605.1 | + | 4356486 | 4356518 | **RNA**→ rpmE2 (PRK01678)Ribosomal_L31 (pfam01197)→ |
| Bth-36-1 | NZ_CP010005.1 | + | 361032 | 361064 | **RNA**→ rpmE2 (PRK01678)Ribosomal_L31 (pfam01197)→ |
| Bth-41-1 | NZ_CP010577.1 | - | 5464481 | 5464449 | **RNA**→ rpmE2 (PRK01678)Ribosomal_L31 (pfam01197)→ |
| Bth-28-1 | NZ_KN849185.1 | - | 117278 | 117246 | **RNA**→ rpmE2 (PRK01678)Ribosomal_L31 (pfam01197)→ |
| Bce-47-5 | NZ_CP009686.1 | - | 5177931 | 5177899 | **RNA**→ rpmE2 (PRK01678)Ribosomal_L31 (pfam01197)→ |
| Bth-19-1 | NZ_CM000746.1 | + | 5020833 | 5020865 | **RNA**→ rpmE2 (PRK01678)Ribosomal_L31 (pfam01197)→ rpmE2 (PRK01678)Ribosomal_L31 (pfam01197 |
| Bce-19-1 | NZ_CM000726.1 | - | 4943444 | 4943412 | **RNA**→ rpmE2 (PRK01678)Ribosomal_L31 (pfam01197)→ rpmE2 (PRK01678)Ribosomal_L31 (pfam01197 |
| Bce-22-1 | NZ_CM000724.1 | + | 4921172 | 4921204 | **RNA**→ rpmE2 (PRK01678)Ribosomal_L31 (pfam01197)→ rpmE2 (PRK01678)Ribosomal_L31 (pfam01197 |
| Bce-39-1 | NZ_CM000732.1 | + | 4928815 | 4928847 | **RNA**→ rpmE2 (PRK01678)Ribosomal_L31 (pfam01197)→ rpmE2 (PRK01678)Ribosomal_L31 (pfam01197 |
| Bce-4-1 | NZ_CM000727.1 | - | 4964402 | 4964370 | **RNA**→ rpmE2 (PRK01678)Ribosomal_L31 (pfam01197)→ rpmE2 (PRK01678)Ribosomal_L31 (pfam01197 |
| Bce-100-1 | NZ_KE150020.1 | + | 186887 | 186919 | **RNA**→ rpmE2 (PRK01678)Ribosomal_L31 (pfam01197)→ |
| Bce-18-1 | NC_011725.1 | - | 5245447 | 5245415 | **RNA**→ rpmE2 (PRK01678)Ribosomal_L31 (pfam01197)→ |

| | | | | | |
|---|---|---|---|---|---|
| Spn-1-7 | NZ_CRRT01000061.1 | + | 4487 | 4519 | **RNA**→ rpmE2 (PRK01678)Ribosomal_L31 (pfam01197)→ |
| env-945 | 3300000053_144823 | - | 80 | 50 | **RNA**→ hypo→ |
| Bsp-21-1 | NZ_GL635751.1 | - | 651664 | 651634 | **RNA**→ rpmE (PRK00019)Ribosomal_L31 (pfam01197)→ |
| Boc-1-1 | NZ_ALEG01000028.1 | + | 150357 | 150387 | **RNA**→ rpmE (PRK00019)Ribosomal_L31 (pfam01197)→ |
| Bfi-2-1 | NZ_APVL01000003.1 | - | 169647 | 169617 | **RNA**→ rpmE (PRK00019)Ribosomal_L31 (pfam01197)→ |
| Sgl-1-1 | NZ_LGUF01000007.1 | + | 808393 | 808423 | **RNA**→ rpmE (PRK00019)Ribosomal_L31 (pfam01197)→ |
| Bsp-44-1 | NZ_LIGI01000001.1 | - | 4212263 | 4212232 | **RNA**→ rpmE (PRK00019)Ribosomal_L31 (pfam01197)→ |
| env-946 | JGI25159J45721_1006073 | + | 2550 | 2582 | **RNA**→ rpmE2 (PRK01678)Ribosomal_L31 (pfam01197)→ |
| env-947 | JGI25151J46595_10041049 | + | 116 | 148 | **RNA**→ rpmE2 (PRK01678)Ribosomal_L31 (pfam01197)→ |
| Bsp-64-1 | NZ_LDUK01000040.1 | - | 27734 | 27702 | **RNA**→ rpmE2 (PRK01678)Ribosomal_L31 (pfam01197)→ |
| Bsp-50-1 | NZ_AYMH01000001.1 | + | 178427 | 178459 | **RNA**→ rpmE2 (PRK01678)Ribosomal_L31 (pfam01197)→ |
| Bwe-2-1 | NZ_JNLY01000008.1 | + | 176304 | 176336 | **RNA**→ rpmE2 (PRK01678)Ribosomal_L31 (pfam01197)→ |
| Bmy-5-1 | NZ_ASPY01000107.1 | - | 7718 | 7686 | **RNA**→ rpmE2 (PRK01678)Ribosomal_L31 (pfam01197)→ |
| Bwe-1-1 | NC_010184.1 | - | 5092651 | 5092619 | **RNA**→ rpmE2 (PRK01678)Ribosomal_L31 (pfam01197)→ rpmE2 (PRK01678)Ribosomal_L31 (pfam01197 |
| Bmy-7-1 | NZ_BAUY01000041.1 | - | 41558 | 41526 | **RNA**→ rpmE2 (PRK01678)Ribosomal_L31 (pfam01197)→ |
| Bce-91-1 | NZ_KB976154.1 | - | 405933 | 405901 | **RNA**→ rpmE2 (PRK01678)Ribosomal_L31 (pfam01197)→ |
| Bce-10-1 | NZ_CM000737.1 | + | 4934442 | 4934474 | **RNA**→ rpmE2 (PRK01678)Ribosomal_L31 (pfam01197)→ rpmE2 (PRK01678)Ribosomal_L31 (pfam01197 |
| Bce-11-1 | NZ_CM000719.1 | - | 4835424 | 4835392 | **RNA**→ rpmE2 (PRK01678)Ribosomal_L31 (pfam01197)→ |
| Bce-20-1 | NZ_CM000725.1 | + | 4915535 | 4915567 | **RNA**→ rpmE2 (PRK01678)Ribosomal_L31 (pfam01197)→ rpmE2 (PRK01678)Ribosomal_L31 (pfam01197 |
| Bce-30-1 | NZ_CM000718.1 | + | 5004963 | 5004995 | **RNA**→ rpmE2 (PRK01678)Ribosomal_L31 (pfam01197)→ rpmE2 (PRK01678)Ribosomal_L31 (pfam01197 |
| Bce-7-1 | NZ_CM000740.1 | - | 4811048 | 4811016 | **RNA**→ rpmE2 (PRK01678)Ribosomal_L31 (pfam01197)→ rpmE2 (PRK01678)Ribosomal_L31 (pfam01197 |
| Bmy-1-1 | NZ_CM000742.1 | + | 4692039 | 4692071 | **RNA**→ rpmE2 (PRK01678)Ribosomal_L31 (pfam01197)→ rpmE2 (PRK01678)Ribosomal_L31 (pfam01197 |
| Bce-8-1 | NZ_CM000741.1 | + | 4823193 | 4823225 | **RNA**→ rpmE2 (PRK01678)Ribosomal_L31 (pfam01197)→ rpmE2 (PRK01678)Ribosomal_L31 (pfam01197 |
| Bce-83-1 | NZ_JH792011.1 | - | 5058984 | 5058952 | **RNA**→ rpmE2 (PRK01678)Ribosomal_L31 (pfam01197)→ |
| Bce-144-1 | NZ_JH791809.1 | - | 4517571 | 4517539 | **RNA**→ rpmE2 (PRK01678)Ribosomal_L31 (pfam01197)→ |
| Bce-110-1 | NZ_JH792113.1 | + | 2442694 | 2442726 | **RNA**→ rpmE2 (PRK01678)Ribosomal_L31 (pfam01197)→ |
| Bce-84-1 | NZ_JH791991.1 | + | 263206 | 263238 | **RNA**→ rpmE2 (PRK01678)Ribosomal_L31 (pfam01197)→ |
| Bce-93-1 | NZ_JH792148.1 | + | 254398 | 254430 | **RNA**→ rpmE2 (PRK01678)Ribosomal_L31 (pfam01197)→ |
| Bce-119-1 | NZ_JH792311.1 | + | 243877 | 243909 | **RNA**→ rpmE2 (PRK01678)Ribosomal_L31 (pfam01197)→ |
| Bce-90-1 | NZ_JH792097.1 | - | 5038004 | 5037972 | **RNA**→ rpmE2 (PRK01678)Ribosomal_L31 (pfam01197)→ |
| Bce-120-1 | NZ_JH792251.1 | + | 232233 | 232265 | **RNA**→ rpmE2 (PRK01678)Ribosomal_L31 (pfam01197)→ |
| Bce-143-1 | NZ_JH791837.1 | + | 232980 | 233012 | **RNA**→ rpmE2 (PRK01678)Ribosomal_L31 (pfam01197)→ |
| Bce-82-1 | NZ_JH804642.1 | - | 5078164 | 5078132 | **RNA**→ rpmE2 (PRK01678)Ribosomal_L31 (pfam01197)→ |
| Bce-146-1 | NZ_JH791802.1 | + | 495409 | 495441 | **RNA**→ rpmE2 (PRK01678)Ribosomal_L31 (pfam01197)→ |
| Bce-88-1 | NZ_JH804673.1 | + | 262212 | 262244 | **RNA**→ rpmE2 (PRK01678)Ribosomal_L31 (pfam01197)→ |
| Bce-80-1 | NZ_JH791985.1 | + | 199165 | 199197 | **RNA**→ rpmE2 (PRK01678)Ribosomal_L31 (pfam01197)→ |
| Bce-122-1 | NZ_JH792233.1 | - | 403322 | 403290 | **RNA**→ rpmE2 (PRK01678)Ribosomal_L31 (pfam01197)→ |
| Bce-64-1 | NZ_JH791910.1 | + | 261506 | 261538 | **RNA**→ rpmE2 (PRK01678)Ribosomal_L31 (pfam01197)→ |
| Bce-76-1 | NZ_JH791976.1 | + | 253248 | 253280 | **RNA**→ rpmE2 (PRK01678)Ribosomal_L31 (pfam01197)→ |
| Bce-79-1 | NZ_JH804637.1 | - | 4782587 | 4782555 | **RNA**→ rpmE2 (PRK01678)Ribosomal_L31 (pfam01197)→ |
| Bce-65-1 | NZ_JH791914.1 | + | 33644 | 33676 | **RNA**→ rpmE2 (PRK01678)Ribosomal_L31 (pfam01197)→ |
| Bce-68-1 | NZ_JH791923.1 | + | 273803 | 273835 | **RNA**→ rpmE2 (PRK01678)Ribosomal_L31 (pfam01197)→ |
| Bce-74-1 | NZ_JH791996.1 | + | 184561 | 184593 | **RNA**→ rpmE2 (PRK01678)Ribosomal_L31 (pfam01197)→ |
| Bce-78-1 | NZ_JH804628.1 | - | 2665814 | 2665782 | **RNA**→ rpmE2 (PRK01678)Ribosomal_L31 (pfam01197)→ |
| Bce-56-1 | NZ_JH792182.1 | + | 234573 | 234605 | **RNA**→ rpmE2 (PRK01678)Ribosomal_L31 (pfam01197)→ |
| Bce-66-1 | NZ_KB976982.1 | - | 397475 | 397443 | **RNA**→ rpmE2 (PRK01678)Ribosomal_L31 (pfam01197)→ |
| Bce-124-1 | NZ_KB976800.1 | - | 1744786 | 1744754 | **RNA**→ rpmE2 (PRK01678)Ribosomal_L31 (pfam01197)→ |
| Bce-92-1 | NZ_KB976147.1 | + | 190193 | 190225 | **RNA**→ rpmE2 (PRK01678)Ribosomal_L31 (pfam01197)→ |
| Bce-116-1 | NZ_KB976281.1 | - | 389591 | 389559 | **RNA**→ rpmE2 (PRK01678)Ribosomal_L31 (pfam01197)→ |
| Bce-109-1 | NZ_KB976394.1 | + | 45840 | 45872 | **RNA**→ rpmE2 (PRK01678)Ribosomal_L31 (pfam01197)→ |
| Bce-50-1 | NZ_KB976960.1 | - | 380250 | 380218 | **RNA**→ rpmE2 (PRK01678)Ribosomal_L31 (pfam01197)→ |
| Bce-130-1 | NZ_KB976678.1 | - | 402912 | 402880 | **RNA**→ rpmE2 (PRK01678)Ribosomal_L31 (pfam01197)→ |

| | | | | | |
|---|---|---|---|---|---|
| Bce-141-1 | NZ_KB976199.1 | + | 251413 | 251445 | **RNA**→ rpmE2 (PRK01678)Ribosomal_L31 (pfam01197)→ |
| Bce-145-1 | NZ_KB976714.1 | - | 422602 | 422570 | **RNA**→ rpmE2 (PRK01678)Ribosomal_L31 (pfam01197)→ |
| Bce-51-1 | NZ_KB976291.1 | - | 415314 | 415282 | **RNA**→ rpmE2 (PRK01678)Ribosomal_L31 (pfam01197)→ |
| Bce-129-1 | NZ_KE150047.1 | + | 205009 | 205041 | **RNA**→ rpmE2 (PRK01678)Ribosomal_L31 (pfam01197)→ |
| Bmy-4-3 | NZ_CP009746.1 | + | 4223781 | 4223813 | **RNA**→ rpmE2 (PRK01678)Ribosomal_L31 (pfam01197)→ |
| Bce-62-1 | NZ_KB976530.1 | + | 248857 | 248889 | **RNA**→ rpmE2 (PRK01678)Ribosomal_L31 (pfam01197)→ |
| Bmy-4-4 | NZ_CP009692.1 | + | 659279 | 659311 | **RNA**→ rpmE2 (PRK01678)Ribosomal_L31 (pfam01197)→ |
| Afl-1-1 | NC_011567.1 | - | 2710818 | 2710787 | **RNA**→ rpmE (PRK00019)Ribosomal_L31 (pfam01197)→ |
| Afl-3-1 | NZ_APCD01000007.1 | - | 45792 | 45761 | **RNA**→ rpmE (PRK00019)Ribosomal_L31 (pfam01197)→ Tdk (COG1435)PRK04296 (PRK04296)→ |
| | | | | | PrfA (COG0216)prfA (PRK00591)→ HemK (COG2890)PRK09328 (PRK09328)→ Spore_II_R (pfam09551) |
| Ath-1-1 | NZ_JXTH01000088.1 | + | 6006 | 6037 | **RNA**→ rpmE (PRK00019)Ribosomal_L31 (pfam01197)→ Tdk (COG1435)PRK04296 (PRK04296)→ |
| Afl-6-1 | NZ_KN127223.1 | - | 319655 | 319624 | **RNA**→ rpmE (PRK00019)Ribosomal_L31 (pfam01197)→ Tdk (COG1435)PRK04296 (PRK04296)→ |
| | | | | | PrfA (COG0216)prfA (PRK00591)→ HemK (COG2890)RF_mod_PrmC (TIGR03534)→ Spore_II_R (pfam0 |
| Afl-4-1 | NZ_BARH01000011.1 | - | 11777 | 11746 | **RNA**→ rpmE (PRK00019)Ribosomal_L31 (pfam01197)→ Tdk (COG1435)PRK04296 (PRK04296)→ |
| | | | | | PrfA (COG0216)prfA (PRK00591)→ HemK (COG2890)RF_mod_PrmC (TIGR03534)→ Spore_II_R (pfam0 |
| Afl-5-1 | NZ_KB205935.1 | - | 2540628 | 2540597 | **RNA**→ rpmE (PRK00019)Ribosomal_L31 (pfam01197)→ Tdk (COG1435)PRK04296 (PRK04296)→ |
| Asp-3-1 | NZ_JPZN01000073.1 | + | 26746 | 26777 | **RNA**→ rpmE (PRK00019)Ribosomal_L31 (pfam01197)→ Tdk (COG1435)PRK04296 (PRK04296)→ |
| | | | | | PrfA (COG0216)prfA (PRK00591)→ HemK (COG2890)RF_mod_PrmC (TIGR03534)→ Spore_II_R (pfam0 |
| Ago-1-1 | NZ_CP012152.1 | - | 2661090 | 2661059 | **RNA**→ rpmE (PRK00019)Ribosomal_L31 (pfam01197)→ Tdk (COG1435)PRK04296 (PRK04296)→ |
| | | | | | PrfA (COG0216)prfA (PRK00591)→ HemK (COG2890)RF_mod_PrmC (TIGR03534)→ Spore_II_R (pfam0 |
| Aka-1-1 | NZ_ALJT01000059.1 | - | 42386 | 42355 | **RNA**→ rpmE (PRK00019)Ribosomal_L31 (pfam01197)→ Tdk (COG1435)PRK04296 (PRK04296)→ |
| | | | | | PrfA (COG0216)prfA (PRK00591)→ HemK (COG2890)RF_mod_PrmC (TIGR03534)→ Spore_II_R (pfam0 |
| Asp-2-1 | NZ_ANMT01000010.1 | - | 18982 | 18951 | **RNA**→ rpmE (PRK00019)Ribosomal_L31 (pfam01197)→ Tdk (COG1435)PRK04296 (PRK04296)→ |
| | | | | | PrfA (COG0216)prfA (PRK00591)→ HemK (COG2890)RF_mod_PrmC (TIGR03534)→ Spore_II_R (pfam0 |
| Afl-2-1 | NZ_JPZO01000009.1 | + | 56873 | 56904 | **RNA**→ rpmE (PRK00019)Ribosomal_L31 (pfam01197)→ hypo→ |
| Aay-1-1 | NZ_JXTG01000005.1 | + | 108572 | 108603 | **RNA**→ rpmE (PRK00019)Ribosomal_L31 (pfam01197)→ Tdk (COG1435)PRK04296 (PRK04296)→ |
| Asp-4-1 | NZ_ANOC01000013.1 | - | 1020 | 989 | **RNA**→ rpmE (PRK00019)Ribosomal_L31 (pfam01197)→ Tdk (COG1435)PRK04296 (PRK04296)→ |
| Asp-1-1 | NZ_JRLC01000005.1 | + | 88913 | 88944 | **RNA**→ rpmE (PRK00019)Ribosomal_L31 (pfam01197)→ Tdk (COG1435)PRK04296 (PRK04296)→ |
| Bsp-18-1 | NC_021171.1 | - | 4666274 | 4666244 | **RNA**→ rpmE (PRK00019)Ribosomal_L31 (pfam01197)→ |
| Bje-1-1 | NZ_CCAS010000008.1 | - | 57067 | 57037 | **RNA**→ rpmE (PRK00019)Ribosomal_L31 (pfam01197)→ |
| Bme-4-1 | NZ_AFEU01000002.1 | + | 535147 | 535177 | **RNA**→ rpmE (PRK00019)Ribosomal_L31 (pfam01197)→ rpmE (PRK00019)Ribosomal_L31 (pfam01197)– |
| | | | | | hypo→ Tdk (COG1435)PRK04296 (PRK04296)→ Tdk (COG1435)PRK04296 (PRK04296)→ |
| Bvi-2-1 | NZ_LDNB01000013.1 | - | 34089 | 34059 | **RNA**→ rpmE (PRK00019)Ribosomal_L31 (pfam01197)→ |
| Bvi-3-1 | NZ_ALAN01000227.1 | - | 990 | 960 | **RNA**→ rpmE (PRK00019)Ribosomal_L31 (pfam01197)→ |
| env-948 | JGI12467J12023_1238085 | + | 6 | 36 | **RNA**→ rpmE (PRK00019)Ribosomal_L31 (pfam01197)→ |
| Bsp-55-1 | NZ_CCYL01000041.1 | + | 25429 | 25459 | **RNA**→ rpmE (PRK00019)Ribosomal_L31 (pfam01197)→ |
| Bme-10-1 | NZ_CP007739.1 | - | 3209895 | 3209865 | **RNA**→ rpmE (PRK00019)Ribosomal_L31 (pfam01197)→ |
| Bsp-90-1 | NZ_KK366020.1 | + | 182519 | 182549 | **RNA**→ rpmE (PRK00019)Ribosomal_L31 (pfam01197)→ |
| Bal-4-1 | NZ_JYCE01000071.1 | + | 107377 | 107408 | **RNA**→ rpmE (PRK00019)Ribosomal_L31 (pfam01197)→ |
| Bce-1-1 | NC_014829.1 | - | 4418863 | 4418833 | **RNA**→ rpmE2 (PRK01678)Ribosomal_L31 (pfam01197)→ |
| env-949 | JGI12273J12029_10233837 | + | 19 | 49 | **RNA**→ rpmE (PRK00019)Ribosomal_L31 (pfam01197)→ |
| env-950 | JGI12273J12029_10168107 | + | 25 | 55 | **RNA**→ rpmE (PRK00019)Ribosomal_L31 (pfam01197)→ |
| Bgi-2-1 | NZ_JAGM01000007.1 | + | 56131 | 56162 | **RNA**→ rpmE (PRK00019)Ribosomal_L31 (pfam01197)→ |
| Bgi-1-1 | NZ_JRUN01000002.1 | - | 45090 | 45059 | **RNA**→ rpmE (PRK00019)Ribosomal_L31 (pfam01197)→ |
| env-951 | JGI26050J44790_1003763 | - | 2351 | 2320 | **RNA**→ rpmE (PRK00019)Ribosomal_L31 (pfam01197)→ |
| Bco-9-1 | NZ_GL995240.1 | - | 74763 | 74732 | **RNA**→ rpmE (PRK00019)Ribosomal_L31 (pfam01197)→ |
| env-952 | JGI26053J44789_1114936 | - | 54 | 23 | **RNA**→ ←hypo |
| env-953 | JGI26052J44787_1127615 | - | 54 | 23 | **RNA**→ ←hypo |
| Bco-1-1 | NC_015634.1 | - | 2895571 | 2895540 | **RNA**→ rpmE (PRK00019)Ribosomal_L31 (pfam01197)→ |
| Bco-6-1 | NZ_CP009709.1 | - | 2007760 | 2007729 | **RNA**→ rpmE (PRK00019)Ribosomal_L31 (pfam01197)→ |
| Bco-7-1 | NZ_ANAQ01000305.1 | - | 8529 | 8498 | **RNA**→ rpmE (PRK00019)Ribosomal_L31 (pfam01197)→ |
| Bco-5-1 | NZ_AXVW01000015.1 | - | 43301 | 43270 | **RNA**→ rpmE (PRK00019)Ribosomal_L31 (pfam01197)→ |
| env-954 | JGI26050J44790_1001615 | + | 5122 | 5153 | **RNA**→ rpmE (PRK00019)Ribosomal_L31 (pfam01197)→ |
| env-955 | JGI26051J44788_1000227 | - | 27870 | 27839 | **RNA**→ rpmE (PRK00019)Ribosomal_L31 (pfam01197)→ |

| | | | | | |
|---|---|---|---|---|---|
| env-956 | JGI26053J44789_1000222 | + | 6061 | 6092 | **RNA**→ rpmE (PRK00019)Ribosomal_L31 (pfam01197)→ |
| env-957 | JGI26052J44787_1000106 | + | 3807 | 3838 | **RNA**→ rpmE (PRK00019)Ribosomal_L31 (pfam01197)→ |
| Bco-4-1 | NZ_CP010525.1 | + | 190910 | 190941 | **RNA**→ rpmE (PRK00019)Ribosomal_L31 (pfam01197)→ |
| Bco-4-2 | NZ_CP011939.1 | - | 899232 | 899201 | **RNA**→ rpmE (PRK00019)Ribosomal_L31 (pfam01197)→ |
| Bco-8-1 | NZ_JSVI01000029.1 | - | 42813 | 42782 | **RNA**→ rpmE (PRK00019)Ribosomal_L31 (pfam01197)→ |
| Bal-4-2 | NZ_JYCE01000129.1 | + | 44156 | 44188 | **RNA**→ rpmE (PRK00019)Ribosomal_L31 (pfam01197)→ |
| Bps-4-1 | NZ_AJTN02000012.1 | + | 1662 | 1692 | **RNA**→ rpmE (PRK00019)Ribosomal_L31 (pfam01197)→ |
| Bac-1-1 | NZ_KB893980.1 | - | 42221 | 42189 | **RNA**→ rpmE (PRK00019)Ribosomal_L31 (pfam01197)→ Tdk (COG1435)PRK04296 (PRK04296)→ hypo |
| Bbu-1-1 | NZ_LGYA01000005.1 | + | 190988 | 191018 | **RNA**→ rpmE (PRK00019)Ribosomal_L31 (pfam01197)→ |
| env-958 | JGI25151J46595_10101225 | - | 365 | 335 | **RNA**→ rpmE (PRK00019)Ribosomal_L31 (pfam01197)→ |
| Bsi-3-1 | NZ_KN360957.1 | - | 114752 | 114722 | **RNA**→ rpmE (PRK00019)Ribosomal_L31 (pfam01197)→ |
| Bsp-19-1 | NZ_JVPL01000166.1 | - | 5209 | 5179 | **RNA**→ rpmE (PRK00019)Ribosomal_L31 (pfam01197)→ |
| Bsp-23-1 | NZ_JVGU01000213.1 | + | 50658 | 50688 | **RNA**→ rpmE (PRK00019)Ribosomal_L31 (pfam01197)→ |
| Lwe-2-1 | NZ_AODJ01000048.1 | + | 3854 | 3883 | **RNA**→ rpmE2 (PRK01678)Ribosomal_L31 (pfam01197)→ |
| Bsp-47-1 | NZ_LFZV01000001.1 | + | 123654 | 123684 | **RNA**→ rpmE (PRK00019)Ribosomal_L31 (pfam01197)→ |
| Bko-1-1 | NZ_LILC01000037.1 | - | 253710 | 253680 | **RNA**→ rpmE (PRK00019)Ribosomal_L31 (pfam01197)→ |
| Bsp-28-1 | NZ_CAPG01000064.1 | - | 31592 | 31559 | **RNA**→ rpmE (PRK00019)Ribosomal_L31 (pfam01197)→ |
| Bbo-1-1 | NZ_JHYI01000015.1 | + | 60410 | 60441 | **RNA**→ rpmE2 (PRK01678)Ribosomal_L31 (pfam01197)→ |
| Bsp-62-1 | NZ_ANNK01000010.1 | + | 7189 | 7219 | **RNA**→ rpmE2 (PRK01678)Ribosomal_L31 (pfam01197)→ |
| Ssp-6-1 | NZ_LFKO01000001.1 | - | 486806 | 486777 | **RNA**→ rpmE2 (PRK01678)Ribosomal_L31 (pfam01197)→ |
| Sxy-3-1 | NZ_ANMR01000002.1 | + | 49266 | 49295 | **RNA**→ rpmE2 (PRK01678)Ribosomal_L31 (pfam01197)→ |
| Sau-1739-1 | NZ_JBBJ01000003.1 | - | 216439 | 216410 | **RNA**→ rpmE2 (PRK01678)Ribosomal_L31 (pfam01197)→ |
| Sau-1769-1 | NZ_KI992435.1 | - | 25185 | 25156 | **RNA**→ rpmE2 (PRK01678)Ribosomal_L31 (pfam01197)→ |
| Sau-1679-1 | NZ_KI967511.1 | - | 112495 | 112466 | **RNA**→ rpmE2 (PRK01678)Ribosomal_L31 (pfam01197)→ |
| Sau-1491-1 | NZ_KI971326.1 | - | 56746 | 56717 | **RNA**→ rpmE2 (PRK01678)Ribosomal_L31 (pfam01197)→ |
| Sau-1693-1 | NZ_KI968014.1 | - | 35351 | 35322 | **RNA**→ rpmE2 (PRK01678)Ribosomal_L31 (pfam01197)→ |
| Sau-1542-1 | NZ_KI971399.1 | - | 61596 | 61567 | **RNA**→ rpmE2 (PRK01678)Ribosomal_L31 (pfam01197)→ |
| Sau-1546-1 | NZ_KK013510.1 | - | 216438 | 216409 | **RNA**→ rpmE2 (PRK01678)Ribosomal_L31 (pfam01197)→ |
| Sau-1575-1 | NZ_KK014639.1 | - | 35451 | 35422 | **RNA**→ rpmE2 (PRK01678)Ribosomal_L31 (pfam01197)→ |
| Sau-1532-1 | NZ_KK012782.1 | - | 216438 | 216409 | **RNA**→ rpmE2 (PRK01678)Ribosomal_L31 (pfam01197)→ |
| Sau-1880-1 | NZ_KI985727.1 | - | 216439 | 216410 | **RNA**→ rpmE2 (PRK01678)Ribosomal_L31 (pfam01197)→ |
| Sau-1592-1 | NZ_KK015414.1 | - | 216633 | 216604 | **RNA**→ rpmE2 (PRK01678)Ribosomal_L31 (pfam01197)→ |
| Sau-1618-1 | NZ_KK016761.1 | - | 216617 | 216588 | **RNA**→ rpmE2 (PRK01678)Ribosomal_L31 (pfam01197)→ |
| Sau-1602-1 | NZ_KK015955.1 | - | 64567 | 64538 | **RNA**→ rpmE2 (PRK01678)Ribosomal_L31 (pfam01197)→ |
| Sau-1573-1 | NZ_KK014537.1 | - | 58791 | 58762 | **RNA**→ rpmE2 (PRK01678)Ribosomal_L31 (pfam01197)→ |
| Sau-1639-1 | NZ_KK017820.1 | - | 217330 | 217301 | **RNA**→ rpmE2 (PRK01678)Ribosomal_L31 (pfam01197)→ |
| Sau-1595-1 | NZ_KK015575.1 | - | 35621 | 35592 | **RNA**→ rpmE2 (PRK01678)Ribosomal_L31 (pfam01197)→ |
| Sau-305-1 | NZ_AZJQ01000024.1 | - | 16337 | 16308 | **RNA**→ rpmE2 (PRK01678)Ribosomal_L31 (pfam01197)→ |
| env-959 | JGI25842J50178_100820 | - | 933 | 904 | **RNA**→ hypo→ rpmE2 (PRK01678)Ribosomal_L31 (pfam01197)→ hypo→ |
| Sma-1-1 | NZ_JH815597.1 | + | 44577 | 44606 | **RNA**→ rpmE2 (PRK01678)Ribosomal_L31 (pfam01197)→ |
| Sma-2-1 | NZ_AMSQ01000012.1 | - | 44261 | 44232 | **RNA**→ rpmE2 (PRK01678)Ribosomal_L31 (pfam01197)→ |
| Smi-1-1 | NZ_JXWY01000016.1 | + | 12700 | 12729 | **RNA**→ rpmE2 (PRK01678)Ribosomal_L31 (pfam01197)→ |
| Sco-3-1 | NZ_LAQN01000018.1 | + | 45418 | 45447 | **RNA**→ rpmE2 (PRK01678)Ribosomal_L31 (pfam01197)→ |
| Sca-8-1 | NZ_LAIU01000009.1 | + | 42687 | 42716 | **RNA**→ rpmE2 (PRK01678)Ribosomal_L31 (pfam01197)→ |
| Sca-3-1 | NC_012121.1 | - | 1639890 | 1639861 | **RNA**→ rpmE2 (PRK01678)Ribosomal_L31 (pfam01197)→ |
| env-960 | SRS024482_C750297 | + | 80 | 109 | **RNA**→ hypo→ rpmE2 (PRK01678)Ribosomal_L31 (pfam01197)→ |
| Sca-7-1 | NZ_JXXP01000003.1 | - | 44879 | 44850 | **RNA**→ rpmE2 (PRK01678)Ribosomal_L31 (pfam01197)→ |
| Swa-2-1 | NZ_JVOH01000089.1 | + | 43702 | 43731 | **RNA**→ rpmE2 (PRK01678)Ribosomal_L31 (pfam01197)→ |
| Sep-3-1 | NZ_GG696776.1 | - | 44864 | 44835 | **RNA**→ rpmE2 (PRK01678)Ribosomal_L31 (pfam01197)→ |
| env-961 | 2204530485 | - | 260 | 231 | **RNA**→ rpmE2 (PRK01678)Ribosomal_L31 (pfam01197)→ |
| Sca-5-1 | NZ_JVVH01000157.1 | + | 44024 | 44053 | **RNA**→ rpmE2 (PRK01678)Ribosomal_L31 (pfam01197)→ |
| Sca-5-2 | NZ_JWAZ01000042.1 | - | 19537 | 19508 | **RNA**→ rpmE2 (PRK01678)Ribosomal_L31 (pfam01197)→ |

| | | | | | |
|---|---|---|---|---|---|
| Sca-5-3 | NZ_JVAJ01000041.1 | + | 44028 | 44057 | **RNA**→ rpmE2 (PRK01678)Ribosomal_L31 (pfam01197)→ |
| Sca-5-4 | NZ_JVOP01000070.1 | + | 37095 | 37124 | **RNA**→ rpmE2 (PRK01678)Ribosomal_L31 (pfam01197)→ |
| Sca-5-5 | NZ_JVCF01000017.1 | + | 44034 | 44063 | **RNA**→ rpmE2 (PRK01678)Ribosomal_L31 (pfam01197)→ |
| Sca-5-6 | NZ_JVEM01000037.1 | + | 44014 | 44043 | **RNA**→ rpmE2 (PRK01678)Ribosomal_L31 (pfam01197)→ |
| Sca-5-7 | NZ_JUXY01000026.1 | - | 18262 | 18233 | **RNA**→ rpmE2 (PRK01678)Ribosomal_L31 (pfam01197)→ |
| Sca-5-8 | NZ_JVGZ01000010.1 | + | 43166 | 43195 | **RNA**→ rpmE2 (PRK01678)Ribosomal_L31 (pfam01197)→ |
| Sca-5-9 | NZ_JWFD01000012.1 | - | 18272 | 18243 | **RNA**→ rpmE2 (PRK01678)Ribosomal_L31 (pfam01197)→ |
| Sca-5-10 | NZ_JWDH01000031.1 | - | 44279 | 44250 | **RNA**→ rpmE2 (PRK01678)Ribosomal_L31 (pfam01197)→ |
| Sca-5-11 | NZ_JWED01000063.1 | + | 44037 | 44066 | **RNA**→ rpmE2 (PRK01678)Ribosomal_L31 (pfam01197)→ |
| Sca-5-12 | NZ_JVSX01000017.1 | + | 44023 | 44052 | **RNA**→ rpmE2 (PRK01678)Ribosomal_L31 (pfam01197)→ |
| Sca-5-13 | NZ_JUZV01000013.1 | - | 18262 | 18233 | **RNA**→ rpmE2 (PRK01678)Ribosomal_L31 (pfam01197)→ |
| Swa-2-2 | NZ_JVCA01000015.1 | + | 43256 | 43285 | **RNA**→ rpmE2 (PRK01678)Ribosomal_L31 (pfam01197)→ |
| Sca-5-14 | NZ_JUZS01000030.1 | - | 18266 | 18237 | **RNA**→ rpmE2 (PRK01678)Ribosomal_L31 (pfam01197)→ |
| Sca-5-15 | NZ_JUYG01000060.1 | - | 18397 | 18368 | **RNA**→ rpmE2 (PRK01678)Ribosomal_L31 (pfam01197)→ |
| Sca-5-16 | NZ_JUXJ01000067.1 | - | 18197 | 18168 | **RNA**→ rpmE2 (PRK01678)Ribosomal_L31 (pfam01197)→ |
| Sca-5-17 | NZ_JUYU01000011.1 | + | 44037 | 44066 | **RNA**→ rpmE2 (PRK01678)Ribosomal_L31 (pfam01197)→ |
| Sca-5-18 | NZ_JUPO01000096.1 | - | 18246 | 18217 | **RNA**→ rpmE2 (PRK01678)Ribosomal_L31 (pfam01197)→ |
| Sca-5-19 | NZ_JUVR01000034.1 | + | 25145 | 25174 | **RNA**→ rpmE2 (PRK01678)Ribosomal_L31 (pfam01197)→ |
| Sca-5-20 | NZ_CVUF01000004.1 | - | 44302 | 44273 | **RNA**→ rpmE2 (PRK01678)Ribosomal_L31 (pfam01197)→ |
| Sca-1-1 | NZ_ACFR01000009.1 | - | 44309 | 44280 | **RNA**→ rpmE2 (PRK01678)Ribosomal_L31 (pfam01197)→ rpmE2 (PRK01678)Ribosomal_L31 (pfam01197 |
| Sca-11-1 | NZ_AFTX01000024.1 | - | 44300 | 44271 | **RNA**→ rpmE2 (PRK01678)Ribosomal_L31 (pfam01197)→ |
| Sca-2-1 | NZ_GL545276.1 | - | 45468 | 45439 | **RNA**→ rpmE2 (PRK01678)Ribosomal_L31 (pfam01197)→ rpmE2 (PRK01678)Ribosomal_L31 (pfam01197 |
| Sca-10-1 | NZ_AJTH01000006.1 | + | 43719 | 43748 | **RNA**→ rpmE2 (PRK01678)Ribosomal_L31 (pfam01197)→ |
| Sca-9-1 | NZ_HG737333.1 | + | 976669 | 976698 | **RNA**→ rpmE2 (PRK01678)Ribosomal_L31 (pfam01197)→ |
| Sca-6-1 | NZ_CP007601.1 | - | 1335238 | 1335209 | **RNA**→ rpmE2 (PRK01678)Ribosomal_L31 (pfam01197)→ |
| Ssp-8-1 | NZ_JMGB01000004.1 | - | 44292 | 44263 | **RNA**→ rpmE2 (PRK01678)Ribosomal_L31 (pfam01197)→ |
| Sca-5-21 | NZ_JGYJ01000035.1 | + | 44028 | 44057 | **RNA**→ rpmE2 (PRK01678)Ribosomal_L31 (pfam01197)→ |
| Sca-4-1 | NZ_ANAM01000008.1 | - | 746220 | 746191 | **RNA**→ rpmE (PRK00019)Ribosomal_L31 (pfam01197)→ |
| Sse-1-1 | NZ_LAYZ01000002.1 | + | 453097 | 453126 | **RNA**→ rpmE (PRK00019)Ribosomal_L31 (pfam01197)→ |
| Sha-3-1 | NZ_CP011366.1 | - | 2293755 | 2293726 | **RNA**→ rpmE (PRK00019)Ribosomal_L31 (pfam01197)→ |
| Sal-1-1 | NZ_ARQJ01000026.1 | - | 43414 | 43385 | **RNA**→ rpmE (PRK00019)Ribosomal_L31 (pfam01197)→ |
| Sep-24-1 | NZ_ATDF02000020.1 | + | 416334 | 416363 | **RNA**→ rpmE2 (PRK01678)Ribosomal_L31 (pfam01197)→ |
| Sep-25-1 | NZ_ATDE02000029.1 | - | 44595 | 44566 | **RNA**→ rpmE2 (PRK01678)Ribosomal_L31 (pfam01197)→ |
| Sep-32-1 | NZ_ATCX02000207.1 | - | 29244 | 29215 | **RNA**→ rpmE2 (PRK01678)Ribosomal_L31 (pfam01197)→ |
| Sep-87-1 | NZ_ATDG02000039.1 | + | 247203 | 247232 | **RNA**→ rpmE2 (PRK01678)Ribosomal_L31 (pfam01197)→ |
| Sep-88-1 | NZ_ATDH02000068.1 | - | 1665536 | 1665507 | **RNA**→ rpmE2 (PRK01678)Ribosomal_L31 (pfam01197)→ |
| Sep-34-1 | NZ_ATCZ02000376.1 | - | 26212 | 26183 | **RNA**→ rpmE2 (PRK01678)Ribosomal_L31 (pfam01197)→ |
| Sep-33-1 | NZ_ATCY02000151.1 | + | 36943 | 36972 | **RNA**→ rpmE2 (PRK01678)Ribosomal_L31 (pfam01197)→ |
| Sep-26-1 | NZ_ATCW02000120.1 | + | 37472 | 37501 | **RNA**→ rpmE2 (PRK01678)Ribosomal_L31 (pfam01197)→ |
| Sep-20-1 | NZ_ATCU02000163.1 | + | 37472 | 37501 | **RNA**→ rpmE2 (PRK01678)Ribosomal_L31 (pfam01197)→ |
| Sep-64-1 | NZ_ATDA02000118.1 | - | 202474 | 202445 | **RNA**→ rpmE2 (PRK01678)Ribosomal_L31 (pfam01197)→ |
| Sep-52-1 | NZ_AKGX01000053.1 | - | 46122 | 46093 | **RNA**→ rpmE2 (PRK01678)Ribosomal_L31 (pfam01197)→ |
| Sep-78-1 | NZ_AHLB01000045.1 | - | 45382 | 45353 | **RNA**→ rpmE2 (PRK01678)Ribosomal_L31 (pfam01197)→ |
| Sep-28-1 | NZ_AENR01000048.1 | - | 45384 | 45355 | **RNA**→ rpmE2 (PRK01678)Ribosomal_L31 (pfam01197)→ |
| Sep-65-1 | NZ_KE367254.1 | + | 1492 | 1521 | **RNA**→ rpmE2 (PRK01678)Ribosomal_L31 (pfam01197)→ |
| Sep-21-1 | NZ_KE365561.1 | + | 6822 | 6851 | **RNA**→ rpmE2 (PRK01678)Ribosomal_L31 (pfam01197)→ |
| Sep-8-1 | NZ_CP009046.1 | - | 687865 | 687836 | **RNA**→ rpmE2 (PRK01678)Ribosomal_L31 (pfam01197)→ |
| env-962 | JGI26050J44790_1048163 | + | 124 | 153 | **RNA**→ rpmE2 (PRK01678)Ribosomal_L31 (pfam01197)→ |
| env-963 | JGI26053J44789_1028520 | + | 99 | 128 | **RNA**→ rpmE2 (PRK01678)Ribosomal_L31 (pfam01197)→ |
| env-964 | JGI26055J44792_1004441 | + | 1972 | 2001 | **RNA**→ rpmE2 (PRK01678)Ribosomal_L31 (pfam01197)→ |
| env-965 | JGI26057J44791_1004975 | + | 2013 | 2042 | **RNA**→ rpmE2 (PRK01678)Ribosomal_L31 (pfam01197)→ |
| env-966 | JGI26054J46599_1003011 | + | 3005 | 3034 | **RNA**→ rpmE2 (PRK01678)Ribosomal_L31 (pfam01197)→ |

| | | | | | |
|---|---|---|---|---|---|
| env-967 | JGI26054J46599_1013284 | + | 556 | 585 | **RNA**→ rpmE2 (PRK01678)Ribosomal_L31 (pfam01197)→ |
| Ssc-3-1 | NZ_LEOS01000012.1 | - | 44440 | 44411 | **RNA**→ rpmE2 (PRK01678)Ribosomal_L31 (pfam01197)→ |
| Ssc-4-1 | NZ_JANE01000008.1 | + | 23593 | 23622 | **RNA**→ rpmE2 (PRK01678)Ribosomal_L31 (pfam01197)→ |
| Sco-1-1 | NZ_AYOS02000031.1 | + | 49441 | 49470 | **RNA**→ rpmE2 (PRK01678)Ribosomal_L31 (pfam01197)→ |
| Sco-2-1 | NZ_LAKJ01000007.1 | - | 19353 | 19324 | **RNA**→ rpmE2 (PRK01678)Ribosomal_L31 (pfam01197)→ |
| Ssp-11-1 | NZ_ALOZ01000051.1 | + | 21417 | 21446 | **RNA**→ rpmE2 (PRK01678)Ribosomal_L31 (pfam01197)→ |
| Sho-3-1 | NZ_JWBV01000049.1 | - | 19534 | 19505 | **RNA**→ rpmE2 (PRK01678)Ribosomal_L31 (pfam01197)→ |
| Sho-1-1 | NZ_ACLP01000009.1 | + | 36054 | 36083 | **RNA**→ rpmE2 (PRK01678)Ribosomal_L31 (pfam01197)→ rpmE2 (PRK01678)Ribosomal_L31 (pfam01197 |
| Sho-2-1 | NZ_GL545257.1 | + | 36461 | 36490 | **RNA**→ rpmE2 (PRK01678)Ribosomal_L31 (pfam01197)→ rpmE2 (PRK01678)Ribosomal_L31 (pfam01197 |
| Sau-255-1 | NZ_KK222755.1 | - | 45892 | 45863 | **RNA**→ rpmE2 (PRK01678)Ribosomal_L31 (pfam01197)→ |
| env-968 | SRS015381_WUGC_scaffold_18506 | - | 443 | 414 | **RNA**→ rpmE2 (PRK01678)Ribosomal_L31 (pfam01197)→ |
| env-969 | SRS015937_C169098 | + | 201 | 230 | **RNA**→ rpmE2 (PRK01678)Ribosomal_L31 (pfam01197)→ |
| env-970 | SRS016752_C178754 | + | 488 | 517 | **RNA**→ rpmE2 (PRK01678)Ribosomal_L31 (pfam01197)→ |
| env-971 | SRS017851_Baylor_scaffold_8802 | - | 406 | 377 | **RNA**→ rpmE2 (PRK01678)Ribosomal_L31 (pfam01197)→ |
| env-972 | SRS018978_WUGC_scaffold_15620 | - | 7277 | 7248 | **RNA**→ rpmE2 (PRK01678)Ribosomal_L31 (pfam01197)→ |
| env-973 | SRS019015_WUGC_scaffold_1731 | + | 3050 | 3079 | **RNA**→ rpmE2 (PRK01678)Ribosomal_L31 (pfam01197)→ |
| env-974 | SRS019033_C268848 | + | 114 | 143 | **RNA**→ rpmE2 (PRK01678)Ribosomal_L31 (pfam01197)→ |
| env-975 | SRS019064_WUGC_scaffold_6269 | + | 5563 | 5592 | **RNA**→ rpmE2 (PRK01678)Ribosomal_L31 (pfam01197)→ |
| env-976 | SRS019081_WUGC_scaffold_29513 | + | 2508 | 2537 | **RNA**→ rpmE2 (PRK01678)Ribosomal_L31 (pfam01197)→ |
| env-977 | SRS020263_C2700470 | + | 162 | 191 | **RNA**→ rpmE2 (PRK01678)Ribosomal_L31 (pfam01197)→ |
| env-978 | SRS020386_C348936 | + | 3349 | 3378 | **RNA**→ rpmE2 (PRK01678)Ribosomal_L31 (pfam01197)→ |
| env-979 | SRS022006_C151996 | - | 510 | 481 | **RNA**→ rpmE2 (PRK01678)Ribosomal_L31 (pfam01197)→ |
| env-980 | SRS023847_Baylor_scaffold_786 | - | 3438 | 3409 | **RNA**→ rpmE2 (PRK01678)Ribosomal_L31 (pfam01197)→ |
| env-981 | SRS024424_C245768 | + | 366 | 395 | **RNA**→ rpmE2 (PRK01678)Ribosomal_L31 (pfam01197)→ |
| env-982 | SRS024482_C760912 | + | 1367 | 1396 | **RNA**→ rpmE2 (PRK01678)Ribosomal_L31 (pfam01197)→ |
| env-983 | SRS024598_C1602584 | - | 406 | 377 | **RNA**→ rpmE2 (PRK01678)Ribosomal_L31 (pfam01197)→ |
| env-984 | SRS024620_C112548 | + | 227 | 256 | **RNA**→ rpmE2 (PRK01678)Ribosomal_L31 (pfam01197)→ |
| env-985 | SRS024655_LANL_scaffold_2040 | + | 198 | 227 | **RNA**→ rpmE2 (PRK01678)Ribosomal_L31 (pfam01197)→ |
| env-986 | SRS047225_C228287 | - | 184 | 155 | **RNA**→ rpmE2 (PRK01678)Ribosomal_L31 (pfam01197)→ |
| env-987 | SRS058221_C352952 | - | 181 | 152 | **RNA**→ rpmE2 (PRK01678)Ribosomal_L31 (pfam01197)→ |
| env-988 | LWFCAn2_GMINUVW02I5FNM | + | 286 | 315 | **RNA**→ RpmE (COG0254)rpmE (PRK00019)rpmE2 (PRK01678)L31 (TIGR00105)→ |
| env-989 | GMINUVW02I5FNM | + | 286 | 315 | **RNA**→ ←hypo |
| Sep-73-1 | NZ_AGUB01000039.1 | - | 45436 | 45407 | **RNA**→ rpmE2 (PRK01678)Ribosomal_L31 (pfam01197)→ |
| Sep-79-1 | NZ_AHLC01000046.1 | - | 45370 | 45341 | **RNA**→ rpmE2 (PRK01678)Ribosomal_L31 (pfam01197)→ |
| Sep-27-1 | NZ_AURC01000030.1 | - | 45614 | 45585 | **RNA**→ rpmE2 (PRK01678)Ribosomal_L31 (pfam01197)→ |
| Sep-68-1 | NZ_AFEH01000030.1 | - | 45376 | 45347 | **RNA**→ rpmE2 (PRK01678)Ribosomal_L31 (pfam01197)→ |
| Sep-8-2 | NZ_JWEE01001238.1 | + | 36923 | 36952 | **RNA**→ rpmE2 (PRK01678)Ribosomal_L31 (pfam01197)→ |
| Ecl-1-1 | NZ_JUOF01000855.1 | - | 2934 | 2905 | **RNA**→ rpmE2 (PRK01678)Ribosomal_L31 (pfam01197)→ |
| Sep-67-1 | NZ_ATDD02000322.1 | - | 45352 | 45323 | **RNA**→ rpmE2 (PRK01678)Ribosomal_L31 (pfam01197)→ |
| Sep-66-1 | NZ_ATDC02000013.1 | + | 455134 | 455163 | **RNA**→ rpmE2 (PRK01678)Ribosomal_L31 (pfam01197)→ |
| Sep-12-1 | NZ_ARWU01000119.1 | + | 37127 | 37156 | **RNA**→ rpmE2 (PRK01678)Ribosomal_L31 (pfam01197)→ |
| Sep-18-1 | NZ_AMSJ01000011.1 | - | 45378 | 45349 | **RNA**→ rpmE2 (PRK01678)Ribosomal_L31 (pfam01197)→ |
| Sep-54-1 | NZ_AKGS01000031.1 | - | 45430 | 45401 | **RNA**→ rpmE2 (PRK01678)Ribosomal_L31 (pfam01197)→ |
| Sep-44-1 | NZ_AKHC01000051.1 | - | 45770 | 45741 | **RNA**→ rpmE2 (PRK01678)Ribosomal_L31 (pfam01197)→ |
| Sep-71-1 | NZ_AFEI01000056.1 | - | 45365 | 45336 | **RNA**→ rpmE2 (PRK01678)Ribosomal_L31 (pfam01197)→ |
| Sep-42-1 | NZ_AKHH01000041.1 | - | 45857 | 45828 | **RNA**→ rpmE2 (PRK01678)Ribosomal_L31 (pfam01197)→ |
| Sep-83-1 | NZ_AHLH01000072.1 | + | 37358 | 37387 | **RNA**→ rpmE2 (PRK01678)Ribosomal_L31 (pfam01197)→ |
| Sep-30-1 | NZ_AJJS01000025.1 | - | 45803 | 45774 | **RNA**→ rpmE2 (PRK01678)Ribosomal_L31 (pfam01197)→ |
| Sep-60-1 | NZ_AKGM01000033.1 | - | 45003 | 44974 | **RNA**→ rpmE2 (PRK01678)Ribosomal_L31 (pfam01197)→ |
| Sep-50-1 | NZ_AKGV01000047.1 | - | 45770 | 45741 | **RNA**→ rpmE2 (PRK01678)Ribosomal_L31 (pfam01197)→ |
| Sep-55-1 | NZ_AKGR01000047.1 | - | 45779 | 45750 | **RNA**→ rpmE2 (PRK01678)Ribosomal_L31 (pfam01197)→ |
| Sep-56-1 | NZ_AKGQ01000033.1 | - | 45918 | 45889 | **RNA**→ rpmE2 (PRK01678)Ribosomal_L31 (pfam01197)→ |

| | | | | | | | |
|---|---|---|---|---|---|---|---|
| Sep-49-1 | NZ_AKGW01000040.1 | - | 42734 | 42705 | **RNA**→ | rpmE2 (PRK01678)Ribosomal_L31 (pfam01197)→ | |
| Sep-43-1 | NZ_AKHG01000037.1 | + | 37775 | 37804 | **RNA**→ | rpmE2 (PRK01678)Ribosomal_L31 (pfam01197)→ | |
| Sep-45-1 | NZ_AKHB01000044.1 | - | 45769 | 45740 | **RNA**→ | rpmE2 (PRK01678)Ribosomal_L31 (pfam01197)→ | |
| Sep-40-1 | NZ_AKHL01000049.1 | - | 45769 | 45740 | **RNA**→ | rpmE2 (PRK01678)Ribosomal_L31 (pfam01197)→ | |
| Sep-75-1 | NZ_AFTZ01000029.1 | - | 45374 | 45345 | **RNA**→ | rpmE2 (PRK01678)Ribosomal_L31 (pfam01197)→ | |
| Sep-86-1 | NZ_AFED01000053.1 | + | 37355 | 37384 | **RNA**→ | rpmE2 (PRK01678)Ribosomal_L31 (pfam01197)→ | |
| Sep-46-1 | NZ_AKHA01000034.1 | - | 45852 | 45823 | **RNA**→ | rpmE2 (PRK01678)Ribosomal_L31 (pfam01197)→ | |
| Sep-37-1 | NZ_AKHE01000040.1 | - | 45851 | 45822 | **RNA**→ | rpmE2 (PRK01678)Ribosomal_L31 (pfam01197)→ | |
| Sep-35-1 | NZ_AKHJ01000042.1 | - | 45795 | 45766 | **RNA**→ | rpmE2 (PRK01678)Ribosomal_L31 (pfam01197)→ | |
| Sep-9-1 | NZ_JNLI01000004.1 | - | 2344918 | 2344889 | **RNA**→ | rpmE2 (PRK01678)Ribosomal_L31 (pfam01197)→ | |
| Sep-41-1 | NZ_AKHK01000049.1 | - | 45785 | 45756 | **RNA**→ | rpmE2 (PRK01678)Ribosomal_L31 (pfam01197)→ | |
| Sep-14-1 | NZ_JHQB01000048.1 | - | 45371 | 45342 | **RNA**→ | rpmE2 (PRK01678)Ribosomal_L31 (pfam01197)→ | |
| Sep-17-1 | NZ_JHUB01000024.1 | + | 37356 | 37385 | **RNA**→ | rpmE2 (PRK01678)Ribosomal_L31 (pfam01197)→ | |
| Sep-15-1 | NZ_JHUA01000050.1 | + | 37358 | 37387 | **RNA**→ | rpmE2 (PRK01678)Ribosomal_L31 (pfam01197)→ | |
| Sep-62-1 | NZ_AKGJ01000034.1 | - | 44971 | 44942 | **RNA**→ | rpmE2 (PRK01678)Ribosomal_L31 (pfam01197)→ | |
| Sep-74-1 | NZ_AHLU01000006.1 | - | 45373 | 45344 | **RNA**→ | rpmE2 (PRK01678)Ribosomal_L31 (pfam01197)→ | |
| Sep-48-1 | NZ_AKGY01000040.1 | - | 45875 | 45846 | **RNA**→ | rpmE2 (PRK01678)Ribosomal_L31 (pfam01197)→ | |
| Sep-39-1 | NZ_AKHD01000041.1 | - | 45762 | 45733 | **RNA**→ | rpmE2 (PRK01678)Ribosomal_L31 (pfam01197)→ | |
| Sep-82-1 | NZ_AHLG01000074.1 | + | 37361 | 37390 | **RNA**→ | rpmE2 (PRK01678)Ribosomal_L31 (pfam01197)→ | |
| Sep-38-1 | NZ_AKHI01000037.1 | - | 20728 | 20699 | **RNA**→ | rpmE2 (PRK01678)Ribosomal_L31 (pfam01197)→ | |
| Sep-81-1 | NZ_AHLF01000023.1 | + | 3788 | 3817 | **RNA**→ | rpmE2 (PRK01678)Ribosomal_L31 (pfam01197)→ | |
| Sep-29-1 | NZ_AJJR01000021.1 | - | 45750 | 45721 | **RNA**→ | rpmE2 (PRK01678)Ribosomal_L31 (pfam01197)→ | |
| Sep-80-1 | NZ_AHLE01000046.1 | + | 33561 | 33590 | **RNA**→ | rpmE2 (PRK01678)Ribosomal_L31 (pfam01197)→ | |
| Sep-16-1 | NZ_JHQC01000010.1 | - | 45434 | 45405 | **RNA**→ | rpmE2 (PRK01678)Ribosomal_L31 (pfam01197)→ | |
| Sep-13-1 | NZ_JHTZ01000129.1 | - | 45369 | 45340 | **RNA**→ | rpmE2 (PRK01678)Ribosomal_L31 (pfam01197)→ | |
| Sep-72-1 | NZ_AHKZ01000035.1 | - | 45377 | 45348 | **RNA**→ | rpmE2 (PRK01678)Ribosomal_L31 (pfam01197)→ | |
| Sep-70-1 | NZ_AHKX01000019.1 | - | 45378 | 45349 | **RNA**→ | rpmE2 (PRK01678)Ribosomal_L31 (pfam01197)→ | |
| Sep-69-1 | NZ_AFTY01000016.1 | - | 45374 | 45345 | **RNA**→ | rpmE2 (PRK01678)Ribosomal_L31 (pfam01197)→ | |
| Sep-77-1 | NZ_AHLA01000084.1 | + | 37357 | 37386 | **RNA**→ | rpmE2 (PRK01678)Ribosomal_L31 (pfam01197)→ | |
| Sep-6-1 | NZ_ADEY01000022.1 | + | 42387 | 42416 | **RNA**→ | rpmE2 (PRK01678)Ribosomal_L31 (pfam01197)→ | rpmE2 (PRK01678)Ribosomal_L31 (pfam01197 |
| Sep-76-1 | NZ_AFUA01000029.1 | - | 45373 | 45344 | **RNA**→ | rpmE2 (PRK01678)Ribosomal_L31 (pfam01197)→ | |
| Sep-1-1 | NC_004461.1 | - | 1769319 | 1769290 | **RNA**→ | rpmE2 (PRK01678)Ribosomal_L31 (pfam01197)→ | |
| Sep-4-1 | NZ_GG749263.1 | - | 45373 | 45344 | **RNA**→ | rpmE2 (PRK01678)Ribosomal_L31 (pfam01197)→ | |
| Sep-23-1 | NZ_KB373339.1 | + | 37355 | 37384 | **RNA**→ | rpmE2 (PRK01678)Ribosomal_L31 (pfam01197)→ | |
| Sep-2-1 | NZ_GG696732.1 | + | 38912 | 38941 | **RNA**→ | rpmE2 (PRK01678)Ribosomal_L31 (pfam01197)→ | |
| Sep-10-1 | NZ_KI999596.1 | - | 46485 | 46456 | **RNA**→ | rpmE2 (PRK01678)Ribosomal_L31 (pfam01197)→ | |
| Sep-31-1 | NZ_KB821725.1 | - | 47636 | 47607 | **RNA**→ | rpmE2 (PRK01678)Ribosomal_L31 (pfam01197)→ | |
| Sep-11-1 | NZ_HG813242.1 | - | 1766133 | 1766104 | **RNA**→ | rpmE2 (PRK01678)Ribosomal_L31 (pfam01197)→ | |
| Sep-5-1 | NC_002976.3 | - | 1764344 | 1764315 | **RNA**→ | rpmE2 (PRK01678)Ribosomal_L31 (pfam01197)→ | |
| Sep-63-1 | NZ_AKGI01000029.1 | - | 45860 | 45831 | **RNA**→ | rpmE2 (PRK01678)Ribosomal_L31 (pfam01197)→ | |
| Sep-36-1 | NZ_AKHF01000108.1 | - | 45856 | 45827 | **RNA**→ | rpmE2 (PRK01678)Ribosomal_L31 (pfam01197)→ | |
| Sep-61-1 | NZ_AKGK01000061.1 | + | 37967 | 37996 | **RNA**→ | rpmE2 (PRK01678)Ribosomal_L31 (pfam01197)→ | |
| Sep-8-3 | NZ_JTAY01000002.1 | - | 2023535 | 2023506 | **RNA**→ | rpmE2 (PRK01678)Ribosomal_L31 (pfam01197)→ | |
| Sep-22-1 | NZ_AMCS01000045.1 | - | 10592 | 10563 | **RNA**→ | rpmE2 (PRK01678)Ribosomal_L31 (pfam01197)→ | |
| Sho-3-2 | NZ_LFKQ01000006.1 | - | 87471 | 87442 | **RNA**→ | rpmE2 (PRK01678)Ribosomal_L31 (pfam01197)→ | |
| Sho-3-3 | NZ_LFKR01000016.1 | - | 44376 | 44347 | **RNA**→ | rpmE2 (PRK01678)Ribosomal_L31 (pfam01197)→ | |
| Sho-3-4 | NZ_JWCO01000048.1 | - | 44146 | 44117 | **RNA**→ | rpmE2 (PRK01678)Ribosomal_L31 (pfam01197)→ | |
| Sho-3-5 | NZ_JVOX01000029.1 | + | 15211 | 15240 | **RNA**→ | rpmE2 (PRK01678)Ribosomal_L31 (pfam01197)→ | |
| Sho-3-6 | NZ_JVEB01000043.1 | + | 25236 | 25265 | **RNA**→ | rpmE2 (PRK01678)Ribosomal_L31 (pfam01197)→ | |
| Sho-3-7 | NZ_JVHC01000034.1 | + | 17068 | 17097 | **RNA**→ | rpmE2 (PRK01678)Ribosomal_L31 (pfam01197)→ | |
| Sho-3-8 | NZ_JVHB01000032.1 | + | 17072 | 17101 | **RNA**→ | rpmE2 (PRK01678)Ribosomal_L31 (pfam01197)→ | |
| Sho-3-9 | NZ_JVED01000048.1 | - | 45867 | 45838 | **RNA**→ | rpmE2 (PRK01678)Ribosomal_L31 (pfam01197)→ | |

| | | | | | |
|---|---|---|---|---|---|
| Sho-3-10 | NZ_JUWW01000005.1 | - | 44144 | 44115 | **RNA**→ rpmE2 (PRK01678)Ribosomal_L31 (pfam01197)→ |
| Sho-3-11 | NZ_JUYH01000010.1 | - | 44135 | 44106 | **RNA**→ rpmE2 (PRK01678)Ribosomal_L31 (pfam01197)→ |
| Sho-3-12 | NZ_JVQD01000023.1 | + | 17103 | 17132 | **RNA**→ rpmE2 (PRK01678)Ribosomal_L31 (pfam01197)→ |
| Sho-3-13 | NZ_JUQI01000082.1 | + | 6927 | 6956 | **RNA**→ rpmE2 (PRK01678)Ribosomal_L31 (pfam01197)→ |
| Sho-3-14 | NZ_JUSP01000026.1 | + | 17103 | 17132 | **RNA**→ rpmE2 (PRK01678)Ribosomal_L31 (pfam01197)→ |
| Ssp-7-1 | NZ_KK013383.1 | - | 44200 | 44171 | **RNA**→ rpmE2 (PRK01678)Ribosomal_L31 (pfam01197)→ |
| Sho-5-1 | NZ_AKGC01000005.1 | + | 15228 | 15257 | **RNA**→ rpmE2 (PRK01678)Ribosomal_L31 (pfam01197)→ |
| Sho-3-15 | NZ_JVLC01000025.1 | - | 44260 | 44231 | **RNA**→ rpmE2 (PRK01678)Ribosomal_L31 (pfam01197)→ |
| Sho-3-16 | NZ_JVVJ01000020.1 | + | 35612 | 35641 | **RNA**→ rpmE2 (PRK01678)Ribosomal_L31 (pfam01197)→ |
| Sho-3-17 | NZ_JUSR01000027.1 | + | 35703 | 35732 | **RNA**→ rpmE2 (PRK01678)Ribosomal_L31 (pfam01197)→ |
| Sho-4-1 | NZ_AHLD01000027.1 | - | 44211 | 44182 | **RNA**→ rpmE2 (PRK01678)Ribosomal_L31 (pfam01197)→ |
| Sar-1-1 | NZ_ALWK01000038.1 | - | 15430 | 15401 | **RNA**→ rpmE2 (PRK01678)Ribosomal_L31 (pfam01197)→ rpmE2 (PRK01678)Ribosomal_L31 (pfam01197 |
| Ssp-13-1 | NZ_AVOQ01000005.1 | - | 15464 | 15435 | **RNA**→ rpmE2 (PRK01678)Ribosomal_L31 (pfam01197)→ |
| env-990 | SRS013261_Baylor_scaffold_8914 | - | 2516 | 2487 | **RNA**→ rpmE2 (PRK01678)Ribosomal_L31 (pfam01197)→ |
| env-991 | SRS017849_Baylor_scaffold_7958 | + | 46 | 75 | **RNA**→ hypo→ rpmE2 (PRK01678)Ribosomal_L31 (pfam01197)→ |
| env-992 | SRS019016_C462472 | - | 361 | 332 | **RNA**→ hypo→ rpmE2 (PRK01678)Ribosomal_L31 (pfam01197)→ |
| env-993 | SRS019986_C240789 | - | 293 | 264 | **RNA**→ hypo→ rpmE2 (PRK01678)Ribosomal_L31 (pfam01197)→ |
| env-994 | SRS020261_C1261455 | + | 44 | 73 | **RNA**→ hypo→ rpmE2 (PRK01678)Ribosomal_L31 (pfam01197)→ |
| env-995 | SRS024596_C1351232 | + | 18 | 47 | **RNA**→ rpmE2 (PRK01678)Ribosomal_L31 (pfam01197)→ |
| env-996 | SRS044474_C210137 | - | 285 | 256 | **RNA**→ hypo→ rpmE2 (PRK01678)Ribosomal_L31 (pfam01197)→ |
| Sep-59-1 | NZ_AKGN01000019.1 | - | 46028 | 45999 | **RNA**→ rpmE2 (PRK01678)Ribosomal_L31 (pfam01197)→ |
| Sep-84-1 | NZ_AHLI01000052.1 | - | 45571 | 45542 | **RNA**→ rpmE2 (PRK01678)Ribosomal_L31 (pfam01197)→ |
| Sep-51-1 | NZ_AKGU01000068.1 | - | 46036 | 46007 | **RNA**→ rpmE2 (PRK01678)Ribosomal_L31 (pfam01197)→ |
| Sep-47-1 | NZ_AKGZ01000025.1 | - | 45840 | 45811 | **RNA**→ rpmE2 (PRK01678)Ribosomal_L31 (pfam01197)→ |
| Sep-85-1 | NZ_AHLJ01000016.1 | + | 37388 | 37417 | **RNA**→ rpmE2 (PRK01678)Ribosomal_L31 (pfam01197)→ |
| Sep-53-1 | NZ_AKGT01000051.1 | - | 45819 | 45790 | **RNA**→ rpmE2 (PRK01678)Ribosomal_L31 (pfam01197)→ |
| Sep-19-1 | NZ_AGUC01000054.1 | - | 26493 | 26464 | **RNA**→ rpmE2 (PRK01678)Ribosomal_L31 (pfam01197)→ |
| Sep-7-1 | NZ_GG696805.1 | - | 45383 | 45354 | **RNA**→ rpmE2 (PRK01678)Ribosomal_L31 (pfam01197)→ |
| Sep-57-1 | NZ_AKGP01000102.1 | - | 46107 | 46078 | **RNA**→ rpmE2 (PRK01678)Ribosomal_L31 (pfam01197)→ |
| Sep-58-1 | NZ_AKGO01000080.1 | - | 46140 | 46111 | **RNA**→ rpmE2 (PRK01678)Ribosomal_L31 (pfam01197)→ |
| env-997 | SRS058213_C232285 | - | 939 | 910 | **RNA**→ hypo→ rpmE2 (PRK01678)Ribosomal_L31 (pfam01197)→ |
| Sco-2-2 | NZ_LATV01000002.1 | + | 49828 | 49857 | **RNA**→ rpmE2 (PRK01678)Ribosomal_L31 (pfam01197)→ |
| Sco-2-3 | NZ_LATU01000008.1 | + | 50130 | 50159 | **RNA**→ rpmE2 (PRK01678)Ribosomal_L31 (pfam01197)→ |
| Swa-2-3 | NZ_LAKH01000008.1 | + | 42884 | 42913 | **RNA**→ rpmE2 (PRK01678)Ribosomal_L31 (pfam01197)→ |
| Swa-2-4 | NZ_JUWX01000004.1 | - | 44955 | 44926 | **RNA**→ rpmE2 (PRK01678)Ribosomal_L31 (pfam01197)→ |
| Swa-7-1 | NZ_AFEC01000100.1 | - | 45052 | 45023 | **RNA**→ rpmE2 (PRK01678)Ribosomal_L31 (pfam01197)→ |
| Swa-6-1 | NC_020164.1 | + | 811839 | 811868 | **RNA**→ rpmE2 (PRK01678)Ribosomal_L31 (pfam01197)→ |
| Spn-1-8 | NZ_CRBR01000009.1 | - | 45357 | 45328 | **RNA**→ rpmE2 (PRK01678)Ribosomal_L31 (pfam01197)→ |
| Swa-5-1 | NZ_CANQ01000031.1 | - | 44983 | 44954 | **RNA**→ rpmE2 (PRK01678)Ribosomal_L31 (pfam01197)→ |
| Swa-4-1 | NZ_JOPV01000022.1 | + | 43084 | 43113 | **RNA**→ rpmE2 (PRK01678)Ribosomal_L31 (pfam01197)→ |
| Swa-3-1 | NZ_JOPU01000025.1 | + | 43225 | 43254 | **RNA**→ rpmE2 (PRK01678)Ribosomal_L31 (pfam01197)→ |
| Swa-2-5 | NZ_JPOW01000004.1 | + | 45929 | 45958 | **RNA**→ rpmE2 (PRK01678)Ribosomal_L31 (pfam01197)→ |
| Spa-2-1 | NC_022737.1 | - | 619055 | 619026 | **RNA**→ rpmE2 (PRK01678)Ribosomal_L31 (pfam01197)→ |
| Swa-2-6 | NZ_JUVB01000015.1 | - | 26153 | 26124 | **RNA**→ rpmE2 (PRK01678)Ribosomal_L31 (pfam01197)→ |
| Spa-1-2 | NZ_LAKF01000005.1 | + | 42862 | 42891 | **RNA**→ rpmE2 (PRK01678)Ribosomal_L31 (pfam01197)→ |
| Ssp-12-1 | NZ_ANBY01000017.1 | - | 33706 | 33677 | **RNA**→ rpmE2 (PRK01678)Ribosomal_L31 (pfam01197)→ |
| Ssp-9-1 | NZ_JHVS01000011.1 | + | 42798 | 42827 | **RNA**→ rpmE2 (PRK01678)Ribosomal_L31 (pfam01197)→ |
| Ssu-1-1 | NZ_LCSH01000018.1 | - | 44953 | 44924 | **RNA**→ rpmE2 (PRK01678)Ribosomal_L31 (pfam01197)→ |
| Mab-1-1 | NZ_CSZP01001570.1 | - | 287 | 258 | **RNA**→ rpmE2 (PRK01678)Ribosomal_L31 (pfam01197)→ |
| Swa-1-1 | NZ_ACPZ01000047.1 | - | 44428 | 44399 | **RNA**→ rpmE2 (PRK01678)Ribosomal_L31 (pfam01197)→ rpmE2 (PRK01678)Ribosomal_L31 (pfam01197 |
| Ssa-2-1 | NZ_JUTO01000006.1 | + | 44286 | 44315 | **RNA**→ rpmE2 (PRK01678)Ribosomal_L31 (pfam01197)→ |
| Sga-1-1 | NZ_JXCF01000018.1 | + | 48546 | 48575 | **RNA**→ rpmE2 (PRK01678)Ribosomal_L31 (pfam01197)→ |

| | | | | | |
|---|---|---|---|---|---|
| Sps-2-1 | NC_014925.1 | - | 1856805 | 1856776 | **RNA**→ rpmE2 (PRK01678)Ribosomal_L31 (pfam01197)→ |
| Sps-1-1 | NC_017568.1 | + | 768136 | 768165 | **RNA**→ rpmE2 (PRK01678)Ribosomal_L31 (pfam01197)→ |
| Sps-3-1 | NZ_ANOI01000001.1 | + | 901751 | 901780 | **RNA**→ rpmE2 (PRK01678)Ribosomal_L31 (pfam01197)→ |
| Sin-2-1 | NZ_CAIB01000243.1 | + | 11672 | 11701 | **RNA**→ rpmE2 (PRK01678)Ribosomal_L31 (pfam01197)→ |
| Sde-1-1 | NZ_CAIA01000073.1 | + | 28519 | 28548 | **RNA**→ rpmE2 (PRK01678)Ribosomal_L31 (pfam01197)→ |
| Mca-1-1 | NC_011999.1 | - | 1910962 | 1910933 | **RNA**→ rpmE2 (PRK01678)Ribosomal_L31 (pfam01197)→ |
| Sch-1-1 | NZ_JMJF01000010.1 | + | 39072 | 39101 | **RNA**→ rpmE2 (PRK01678)Ribosomal_L31 (pfam01197)→ |
| Sha-4-1 | NZ_JQHA01000012.1 | - | 44057 | 44028 | **RNA**→ rpmE2 (PRK01678)Ribosomal_L31 (pfam01197)→ |
| Sha-4-2 | NZ_CUGE01000009.1 | - | 19556 | 19527 | **RNA**→ rpmE2 (PRK01678)Ribosomal_L31 (pfam01197)→ |
| Sha-4-3 | NZ_CUHF01000018.1 | - | 17451 | 17422 | **RNA**→ rpmE2 (PRK01678)Ribosomal_L31 (pfam01197)→ |
| Sha-4-4 | NZ_JQHB01000033.1 | - | 44057 | 44028 | **RNA**→ rpmE2 (PRK01678)Ribosomal_L31 (pfam01197)→ |
| Ssc-1-1 | NZ_CP010309.1 | + | 858128 | 858157 | **RNA**→ rpmE2 (PRK01678)Ribosomal_L31 (pfam01197)→ |
| Ssc-1-2 | NZ_CP009762.1 | + | 793853 | 793882 | **RNA**→ rpmE2 (PRK01678)Ribosomal_L31 (pfam01197)→ |
| Ssc-1-3 | NZ_CP009676.1 | + | 818914 | 818943 | **RNA**→ rpmE2 (PRK01678)Ribosomal_L31 (pfam01197)→ |
| Ssc-1-4 | NZ_CP009470.1 | + | 790202 | 790231 | **RNA**→ rpmE2 (PRK01678)Ribosomal_L31 (pfam01197)→ |
| Ssp-14-1 | NZ_KE150418.1 | + | 949481 | 949510 | **RNA**→ rpmE2 (PRK01678)Ribosomal_L31 (pfam01197)→ |
| Sha-4-5 | NZ_LAKG01000019.1 | - | 24111 | 24082 | **RNA**→ rpmE2 (PRK01678)Ribosomal_L31 (pfam01197)→ |
| Sha-4-6 | NZ_CUEY01000017.1 | + | 28755 | 28784 | **RNA**→ rpmE2 (PRK01678)Ribosomal_L31 (pfam01197)→ |
| Sha-4-7 | NZ_CUEJ01000017.1 | + | 28692 | 28721 | **RNA**→ rpmE2 (PRK01678)Ribosomal_L31 (pfam01197)→ |
| Sha-4-8 | NZ_CUEF01000018.1 | + | 15555 | 15584 | **RNA**→ rpmE2 (PRK01678)Ribosomal_L31 (pfam01197)→ |
| Sha-4-9 | NZ_CUHK01000019.1 | + | 28699 | 28728 | **RNA**→ rpmE2 (PRK01678)Ribosomal_L31 (pfam01197)→ |
| Sha-4-10 | NZ_CUES01000010.1 | - | 44033 | 44004 | **RNA**→ rpmE2 (PRK01678)Ribosomal_L31 (pfam01197)→ |
| Sha-4-11 | NZ_CUEH01000020.1 | + | 28721 | 28750 | **RNA**→ rpmE2 (PRK01678)Ribosomal_L31 (pfam01197)→ |
| Sha-4-12 | NZ_CUHH01000016.1 | + | 28696 | 28725 | **RNA**→ rpmE2 (PRK01678)Ribosomal_L31 (pfam01197)→ |
| Sha-4-13 | NZ_CUHA01000012.1 | + | 33827 | 33856 | **RNA**→ rpmE2 (PRK01678)Ribosomal_L31 (pfam01197)→ |
| Sha-4-14 | NZ_CUEZ01000009.1 | - | 43979 | 43950 | **RNA**→ rpmE2 (PRK01678)Ribosomal_L31 (pfam01197)→ |
| Sha-4-15 | NZ_CUCM01000013.1 | - | 23830 | 23801 | **RNA**→ rpmE2 (PRK01678)Ribosomal_L31 (pfam01197)→ |
| Sha-4-16 | NZ_CUGK01000016.1 | - | 23911 | 23882 | **RNA**→ rpmE2 (PRK01678)Ribosomal_L31 (pfam01197)→ |
| Sha-4-17 | NZ_CUFM01000019.1 | + | 28709 | 28738 | **RNA**→ rpmE2 (PRK01678)Ribosomal_L31 (pfam01197)→ |
| Sha-4-18 | NZ_CUFQ01000015.1 | + | 28720 | 28749 | **RNA**→ rpmE2 (PRK01678)Ribosomal_L31 (pfam01197)→ |
| Sha-4-19 | NZ_CUFJ01000015.1 | + | 28722 | 28751 | **RNA**→ rpmE2 (PRK01678)Ribosomal_L31 (pfam01197)→ |
| Sha-4-20 | NZ_CUHB01000013.1 | - | 23912 | 23883 | **RNA**→ rpmE2 (PRK01678)Ribosomal_L31 (pfam01197)→ |
| Sha-4-21 | NZ_CUGI01000021.1 | + | 15611 | 15640 | **RNA**→ rpmE2 (PRK01678)Ribosomal_L31 (pfam01197)→ |
| Sha-4-22 | NZ_CUFR01000016.1 | + | 28709 | 28738 | **RNA**→ rpmE2 (PRK01678)Ribosomal_L31 (pfam01197)→ |
| Sha-4-23 | NZ_CUDZ01000003.1 | - | 23912 | 23883 | **RNA**→ rpmE2 (PRK01678)Ribosomal_L31 (pfam01197)→ |
| Sha-4-24 | NZ_CUGW01000017.1 | + | 28731 | 28760 | **RNA**→ rpmE2 (PRK01678)Ribosomal_L31 (pfam01197)→ |
| Sha-4-25 | NZ_CUGZ01000014.1 | - | 23911 | 23882 | **RNA**→ rpmE2 (PRK01678)Ribosomal_L31 (pfam01197)→ |
| Sha-4-26 | NZ_CUDY01000028.1 | - | 23835 | 23806 | **RNA**→ rpmE2 (PRK01678)Ribosomal_L31 (pfam01197)→ |
| Sha-4-27 | NZ_CUGN01000014.1 | + | 15549 | 15578 | **RNA**→ rpmE2 (PRK01678)Ribosomal_L31 (pfam01197)→ |
| Sha-4-28 | NZ_CUDQ01000009.1 | + | 28732 | 28761 | **RNA**→ rpmE2 (PRK01678)Ribosomal_L31 (pfam01197)→ |
| Sha-4-29 | NZ_CUEI01000018.1 | + | 15557 | 15586 | **RNA**→ rpmE2 (PRK01678)Ribosomal_L31 (pfam01197)→ |
| Sha-4-30 | NZ_CUEQ01000020.1 | + | 15546 | 15575 | **RNA**→ rpmE2 (PRK01678)Ribosomal_L31 (pfam01197)→ |
| Sha-4-31 | NZ_CUFS01000017.1 | + | 28821 | 28850 | **RNA**→ rpmE2 (PRK01678)Ribosomal_L31 (pfam01197)→ |
| Sha-4-32 | NZ_CUFE01000014.1 | - | 23893 | 23864 | **RNA**→ rpmE2 (PRK01678)Ribosomal_L31 (pfam01197)→ |
| Sha-4-33 | NZ_CUEX01000014.1 | - | 23899 | 23870 | **RNA**→ rpmE2 (PRK01678)Ribosomal_L31 (pfam01197)→ |
| Sha-4-34 | NZ_CUFY01000018.1 | + | 28706 | 28735 | **RNA**→ rpmE2 (PRK01678)Ribosomal_L31 (pfam01197)→ |
| Sha-4-35 | NZ_CUGP01000016.1 | + | 28690 | 28719 | **RNA**→ rpmE2 (PRK01678)Ribosomal_L31 (pfam01197)→ |
| Sha-4-36 | NZ_CUFX01000017.1 | + | 28755 | 28784 | **RNA**→ rpmE2 (PRK01678)Ribosomal_L31 (pfam01197)→ |
| Sha-4-37 | NZ_CUFL01000026.1 | + | 15613 | 15642 | **RNA**→ rpmE2 (PRK01678)Ribosomal_L31 (pfam01197)→ |
| Sha-4-38 | NZ_CUDS01000018.1 | + | 28713 | 28742 | **RNA**→ rpmE2 (PRK01678)Ribosomal_L31 (pfam01197)→ |
| Sha-4-39 | NZ_CUDX01000015.1 | - | 18076 | 18047 | **RNA**→ rpmE2 (PRK01678)Ribosomal_L31 (pfam01197)→ |
| Sha-4-40 | NZ_CUDU01000005.1 | - | 23912 | 23883 | **RNA**→ rpmE2 (PRK01678)Ribosomal_L31 (pfam01197)→ |

| | | | | | |
|---|---|---|---|---|---|
| Sha-4-41 | NZ_CUHQ01000018.1 | + | 28694 | 28723 | **RNA**→ rpmE2 (PRK01678)Ribosomal_L31 (pfam01197)→ |
| Sha-4-42 | NZ_CUCK01000012.1 | + | 28720 | 28749 | **RNA**→ rpmE2 (PRK01678)Ribosomal_L31 (pfam01197)→ |
| Sha-4-43 | NZ_CUGA01000014.1 | + | 28691 | 28720 | **RNA**→ rpmE2 (PRK01678)Ribosomal_L31 (pfam01197)→ |
| Sha-4-44 | NZ_CUGC01000018.1 | + | 15618 | 15647 | **RNA**→ rpmE2 (PRK01678)Ribosomal_L31 (pfam01197)→ |
| Sha-4-45 | NZ_CUDI01000019.1 | + | 28702 | 28731 | **RNA**→ rpmE2 (PRK01678)Ribosomal_L31 (pfam01197)→ |
| Sha-4-46 | NZ_CUER01000015.1 | - | 23834 | 23805 | **RNA**→ rpmE2 (PRK01678)Ribosomal_L31 (pfam01197)→ |
| Sha-4-47 | NZ_CUDO01000015.1 | - | 23830 | 23801 | **RNA**→ rpmE2 (PRK01678)Ribosomal_L31 (pfam01197)→ |
| Sha-4-48 | NZ_CUDW01000027.1 | - | 23825 | 23796 | **RNA**→ rpmE2 (PRK01678)Ribosomal_L31 (pfam01197)→ |
| Sha-4-49 | NZ_CUCL01000019.1 | + | 30377 | 30406 | **RNA**→ rpmE2 (PRK01678)Ribosomal_L31 (pfam01197)→ |
| Sha-4-50 | NZ_CUEV01000014.1 | + | 28711 | 28740 | **RNA**→ rpmE2 (PRK01678)Ribosomal_L31 (pfam01197)→ |
| Sha-4-51 | NZ_CUFD01000015.1 | + | 28739 | 28768 | **RNA**→ rpmE2 (PRK01678)Ribosomal_L31 (pfam01197)→ |
| Sha-4-52 | NZ_CUGQ01000018.1 | + | 28691 | 28720 | **RNA**→ rpmE2 (PRK01678)Ribosomal_L31 (pfam01197)→ |
| Sha-4-53 | NZ_CUDF01000019.1 | + | 28709 | 28738 | **RNA**→ rpmE2 (PRK01678)Ribosomal_L31 (pfam01197)→ |
| Sha-4-119 | NZ_CUED01000013.1 | - | 23912 | 23883 | **RNA**→ rpmE2 (PRK01678)Ribosomal_L31 (pfam01197)→ |
| Sha-4-54 | NZ_CUGU01000016.1 | + | 28755 | 28784 | **RNA**→ rpmE2 (PRK01678)Ribosomal_L31 (pfam01197)→ |
| Sha-4-55 | NZ_CUDC01000014.1 | - | 33897 | 33868 | **RNA**→ rpmE2 (PRK01678)Ribosomal_L31 (pfam01197)→ |
| Sha-4-56 | NZ_CUGT01000014.1 | - | 23922 | 23893 | **RNA**→ rpmE2 (PRK01678)Ribosomal_L31 (pfam01197)→ |
| Sha-4-57 | NZ_CUFN01000017.1 | - | 23836 | 23807 | **RNA**→ rpmE2 (PRK01678)Ribosomal_L31 (pfam01197)→ |
| Sha-4-58 | NZ_CUDG01000021.1 | + | 21475 | 21504 | **RNA**→ rpmE2 (PRK01678)Ribosomal_L31 (pfam01197)→ |
| Sha-4-59 | NZ_CUEK01000015.1 | - | 23830 | 23801 | **RNA**→ rpmE2 (PRK01678)Ribosomal_L31 (pfam01197)→ |
| Sha-4-60 | NZ_CUCS01000032.1 | - | 23842 | 23813 | **RNA**→ rpmE2 (PRK01678)Ribosomal_L31 (pfam01197)→ |
| Sha-4-61 | NZ_CUCI01000016.1 | - | 23842 | 23813 | **RNA**→ rpmE2 (PRK01678)Ribosomal_L31 (pfam01197)→ |
| Sha-4-62 | NZ_CUCR01000015.1 | + | 28672 | 28701 | **RNA**→ rpmE2 (PRK01678)Ribosomal_L31 (pfam01197)→ |
| Sha-4-63 | NZ_CUFW01000024.1 | - | 23791 | 23762 | **RNA**→ rpmE2 (PRK01678)Ribosomal_L31 (pfam01197)→ |
| Sha-4-64 | NZ_CUGD01000018.1 | + | 28719 | 28748 | **RNA**→ rpmE2 (PRK01678)Ribosomal_L31 (pfam01197)→ |
| Sha-4-65 | NZ_CUEL01000014.1 | - | 23842 | 23813 | **RNA**→ rpmE2 (PRK01678)Ribosomal_L31 (pfam01197)→ |
| Sha-4-66 | NZ_CUHI01000017.1 | + | 22937 | 22966 | **RNA**→ rpmE2 (PRK01678)Ribosomal_L31 (pfam01197)→ |
| Sha-4-67 | NZ_CUEN01000018.1 | + | 28728 | 28757 | **RNA**→ rpmE2 (PRK01678)Ribosomal_L31 (pfam01197)→ |
| Sha-4-68 | NZ_CUDM01000018.1 | + | 28755 | 28784 | **RNA**→ rpmE2 (PRK01678)Ribosomal_L31 (pfam01197)→ |
| Sha-4-69 | NZ_CUDL01000012.1 | - | 23830 | 23801 | **RNA**→ rpmE2 (PRK01678)Ribosomal_L31 (pfam01197)→ |
| Sha-4-70 | NZ_CUGR01000015.1 | - | 23922 | 23893 | **RNA**→ rpmE2 (PRK01678)Ribosomal_L31 (pfam01197)→ |
| Sha-4-71 | NZ_CUFI01000017.1 | + | 26650 | 26679 | **RNA**→ rpmE2 (PRK01678)Ribosomal_L31 (pfam01197)→ |
| Sha-4-72 | NZ_CUHE01000019.1 | + | 21344 | 21373 | **RNA**→ rpmE2 (PRK01678)Ribosomal_L31 (pfam01197)→ |
| Sha-4-73 | NZ_CUGY01000021.1 | + | 15570 | 15599 | **RNA**→ rpmE2 (PRK01678)Ribosomal_L31 (pfam01197)→ |
| Sha-4-74 | NZ_CUGO01000018.1 | - | 24793 | 24764 | **RNA**→ rpmE2 (PRK01678)Ribosomal_L31 (pfam01197)→ |
| Sha-4-75 | NZ_CUCY01000018.1 | - | 23911 | 23882 | **RNA**→ rpmE2 (PRK01678)Ribosomal_L31 (pfam01197)→ |
| Sha-4-76 | NZ_CUHC01000010.1 | - | 23911 | 23882 | **RNA**→ rpmE2 (PRK01678)Ribosomal_L31 (pfam01197)→ |
| Sha-4-77 | NZ_CUFH01000015.1 | - | 23912 | 23883 | **RNA**→ rpmE2 (PRK01678)Ribosomal_L31 (pfam01197)→ |
| Sha-4-78 | NZ_CUFA01000016.1 | + | 28691 | 28720 | **RNA**→ rpmE2 (PRK01678)Ribosomal_L31 (pfam01197)→ |
| Sha-4-79 | NZ_CUCP01000012.1 | - | 23830 | 23801 | **RNA**→ rpmE2 (PRK01678)Ribosomal_L31 (pfam01197)→ |
| Sha-4-80 | NZ_CUEB01000022.1 | + | 15564 | 15593 | **RNA**→ rpmE2 (PRK01678)Ribosomal_L31 (pfam01197)→ |
| Sha-4-81 | NZ_CUDV01000014.1 | - | 44049 | 44020 | **RNA**→ rpmE2 (PRK01678)Ribosomal_L31 (pfam01197)→ |
| Sha-4-82 | NZ_CUHG01000015.1 | + | 23286 | 23315 | **RNA**→ rpmE2 (PRK01678)Ribosomal_L31 (pfam01197)→ |
| Sha-4-83 | NZ_CUCW01000018.1 | + | 28694 | 28723 | **RNA**→ rpmE2 (PRK01678)Ribosomal_L31 (pfam01197)→ |
| Sha-4-84 | NZ_CUFT01000020.1 | + | 20131 | 20160 | **RNA**→ rpmE2 (PRK01678)Ribosomal_L31 (pfam01197)→ |
| Sha-4-85 | NZ_CUFG01000017.1 | + | 22976 | 23005 | **RNA**→ rpmE2 (PRK01678)Ribosomal_L31 (pfam01197)→ |
| Sha-4-86 | NZ_CUCO01000013.1 | - | 23842 | 23813 | **RNA**→ rpmE2 (PRK01678)Ribosomal_L31 (pfam01197)→ |
| Sha-4-87 | NZ_CUCU01000018.1 | - | 23911 | 23882 | **RNA**→ rpmE2 (PRK01678)Ribosomal_L31 (pfam01197)→ |
| Sha-4-88 | NZ_CUFK01000001.1 | + | 248538 | 248567 | **RNA**→ rpmE2 (PRK01678)Ribosomal_L31 (pfam01197)→ |
| Sha-4-89 | NZ_CUEW01000014.1 | - | 23922 | 23893 | **RNA**→ rpmE2 (PRK01678)Ribosomal_L31 (pfam01197)→ |
| Sha-4-90 | NZ_CUEC01000012.1 | + | 11448 | 11477 | **RNA**→ rpmE2 (PRK01678)Ribosomal_L31 (pfam01197)→ |
| Sha-4-91 | NZ_CUCN01000006.1 | + | 59873 | 59902 | **RNA**→ rpmE2 (PRK01678)Ribosomal_L31 (pfam01197)→ |

| | | | | | |
|---|---|---|---|---|---|
| Sha-4-92 | NZ_CUGF01000017.1 | + | 28713 | 28742 | **RNA**→ rpmE2 (PRK01678)Ribosomal_L31 (pfam01197)→ |
| Sha-4-93 | NZ_CUEA01000021.1 | + | 5363 | 5392 | **RNA**→ rpmE2 (PRK01678)Ribosomal_L31 (pfam01197)→ |
| Sha-4-94 | NZ_CUCQ01000013.1 | - | 23830 | 23801 | **RNA**→ rpmE2 (PRK01678)Ribosomal_L31 (pfam01197)→ |
| Sha-4-95 | NZ_CUET01000018.1 | + | 28738 | 28767 | **RNA**→ rpmE2 (PRK01678)Ribosomal_L31 (pfam01197)→ |
| Sha-4-96 | NZ_CUFU01000016.1 | + | 28711 | 28740 | **RNA**→ rpmE2 (PRK01678)Ribosomal_L31 (pfam01197)→ |
| Sha-4-97 | NZ_CUFV01000018.1 | + | 28699 | 28728 | **RNA**→ rpmE2 (PRK01678)Ribosomal_L31 (pfam01197)→ |
| Sha-4-98 | NZ_CUDR01000018.1 | + | 28709 | 28738 | **RNA**→ rpmE2 (PRK01678)Ribosomal_L31 (pfam01197)→ |
| Sha-4-99 | NZ_CUHL01000016.1 | + | 28755 | 28784 | **RNA**→ rpmE2 (PRK01678)Ribosomal_L31 (pfam01197)→ |
| Sha-4-100 | NZ_CUHD01000018.1 | + | 28731 | 28760 | **RNA**→ rpmE2 (PRK01678)Ribosomal_L31 (pfam01197)→ |
| Sha-4-101 | NZ_CUFF01000017.1 | + | 28705 | 28734 | **RNA**→ rpmE2 (PRK01678)Ribosomal_L31 (pfam01197)→ |
| Sha-4-102 | NZ_CUEU01000016.1 | - | 23911 | 23882 | **RNA**→ rpmE2 (PRK01678)Ribosomal_L31 (pfam01197)→ |
| Sha-4-103 | NZ_CUHM01000013.1 | + | 28722 | 28751 | **RNA**→ rpmE2 (PRK01678)Ribosomal_L31 (pfam01197)→ |
| Sha-4-104 | NZ_CUWC01000015.1 | + | 28730 | 28759 | **RNA**→ rpmE2 (PRK01678)Ribosomal_L31 (pfam01197)→ |
| Sha-4-105 | NZ_CUFB01000016.1 | - | 24282 | 24253 | **RNA**→ rpmE2 (PRK01678)Ribosomal_L31 (pfam01197)→ |
| Sha-4-106 | NZ_CUDT01000017.1 | + | 28729 | 28758 | **RNA**→ rpmE2 (PRK01678)Ribosomal_L31 (pfam01197)→ |
| Sha-4-107 | NZ_CUEO01000007.1 | - | 43987 | 43958 | **RNA**→ rpmE2 (PRK01678)Ribosomal_L31 (pfam01197)→ |
| Sha-4-108 | NZ_CUDN01000016.1 | - | 23911 | 23882 | **RNA**→ rpmE2 (PRK01678)Ribosomal_L31 (pfam01197)→ |
| Sha-4-109 | NZ_CVRV01000016.1 | + | 28691 | 28720 | **RNA**→ rpmE2 (PRK01678)Ribosomal_L31 (pfam01197)→ |
| Sha-4-110 | NZ_JVTM01000047.1 | + | 28698 | 28727 | **RNA**→ rpmE2 (PRK01678)Ribosomal_L31 (pfam01197)→ |
| Sha-4-111 | NZ_JVVE01000004.1 | - | 23752 | 23723 | **RNA**→ rpmE2 (PRK01678)Ribosomal_L31 (pfam01197)→ |
| Sha-4-112 | NZ_JVRQ01000084.1 | + | 15522 | 15551 | **RNA**→ rpmE2 (PRK01678)Ribosomal_L31 (pfam01197)→ |
| Sha-4-113 | NZ_JVMX01000090.1 | + | 28692 | 28721 | **RNA**→ rpmE2 (PRK01678)Ribosomal_L31 (pfam01197)→ |
| Sha-4-114 | NZ_JVPA01000099.1 | - | 23890 | 23861 | **RNA**→ rpmE2 (PRK01678)Ribosomal_L31 (pfam01197)→ |
| Sha-1-1 | NC_007168.1 | + | 923690 | 923719 | **RNA**→ rpmE2 (PRK01678)Ribosomal_L31 (pfam01197)→ |
| Sha-6-1 | NZ_AJVA01000015.1 | - | 23726 | 23697 | **RNA**→ rpmE2 (PRK01678)Ribosomal_L31 (pfam01197)→ |
| Sau-4078-1 | NZ_JHTU01000062.1 | + | 15552 | 15581 | **RNA**→ rpmE2 (PRK01678)Ribosomal_L31 (pfam01197)→ |
| Sha-4-115 | NZ_JPRW01000011.1 | - | 23862 | 23833 | **RNA**→ rpmE2 (PRK01678)Ribosomal_L31 (pfam01197)→ |
| Sha-4-116 | NZ_JFOJ01000025.1 | - | 23956 | 23927 | **RNA**→ rpmE2 (PRK01678)Ribosomal_L31 (pfam01197)→ |
| Sha-4-117 | NZ_CUFO01000010.1 | - | 43975 | 43946 | **RNA**→ rpmE2 (PRK01678)Ribosomal_L31 (pfam01197)→ |
| Sha-5-1 | NZ_JRNK01000025.1 | + | 36628 | 36657 | **RNA**→ rpmE2 (PRK01678)Ribosomal_L31 (pfam01197)→ |
| Sha-4-118 | NZ_CUEE01000012.1 | + | 20689 | 20718 | **RNA**→ rpmE2 (PRK01678)Ribosomal_L31 (pfam01197)→ |
| Spe-1-1 | NZ_JVAY01000023.1 | + | 42057 | 42086 | **RNA**→ rpmE2 (PRK01678)Ribosomal_L31 (pfam01197)→ |
| Spe-2-1 | NZ_AGUA01000050.1 | - | 22376 | 22347 | **RNA**→ rpmE2 (PRK01678)Ribosomal_L31 (pfam01197)→ |
| Ssi-2-1 | NZ_AEUN01000494.1 | + | 1616 | 1645 | **RNA**→ rpmE2 (PRK01678)Ribosomal_L31 (pfam01197)→ rpmE2 (PRK01678)Ribosomal_L31 (pfam01197 |
| Ssp-10-1 | NZ_JTLN01000023.1 | - | 19694 | 19665 | **RNA**→ rpmE2 (PRK01678)Ribosomal_L31 (pfam01197)→ |
| Ssa-2-2 | NZ_JXBG01000018.1 | - | 45141 | 45112 | **RNA**→ rpmE2 (PRK01678)Ribosomal_L31 (pfam01197)→ |
| Mab-1-2 | NZ_CSZG01001468.1 | - | 266 | 237 | **RNA**→ rpmE2 (PRK01678)Ribosomal_L31 (pfam01197)→ |
| Ssa-2-3 | NZ_JUUE01000025.1 | + | 26596 | 26625 | **RNA**→ rpmE2 (PRK01678)Ribosomal_L31 (pfam01197)→ |
| Ssa-1-1 | NC_007350.1 | + | 796466 | 796495 | **RNA**→ rpmE2 (PRK01678)Ribosomal_L31 (pfam01197)→ |
| Ssa-3-1 | NZ_AHKB01000013.1 | + | 47770 | 47799 | **RNA**→ rpmE2 (PRK01678)Ribosomal_L31 (pfam01197)→ |
| Shy-1-1 | NZ_CP008747.1 | + | 799781 | 799810 | **RNA**→ rpmE2 (PRK01678)Ribosomal_L31 (pfam01197)→ |
| Sag-1-1 | NZ_JPRT01000013.1 | + | 14025 | 14054 | **RNA**→ rpmE2 (PRK01678)Ribosomal_L31 (pfam01197)→ |
| Ssi-3-1 | NZ_KB373330.1 | - | 50416 | 50387 | **RNA**→ rpmE2 (PRK01678)Ribosomal_L31 (pfam01197)→ |
| Ssi-4-1 | NZ_AXDY01000013.1 | - | 48926 | 48897 | **RNA**→ rpmE2 (PRK01678)Ribosomal_L31 (pfam01197)→ |
| env-998 | Anterior_nares_Baylor_scaffold_40137 | + | 669 | 698 | **RNA**→ rpmE2 (PRK01678)Ribosomal_L31 (pfam01197)→ |
| env-999 | SRS011105_WUGC_scaffold_1835 | + | 18318 | 18347 | **RNA**→ rpmE2 (PRK01678)Ribosomal_L31 (pfam01197)→ |
| env-1000 | SRS024424_C251219 | - | 635 | 606 | **RNA**→ rpmE2 (PRK01678)Ribosomal_L31 (pfam01197)→ |
| Sau-4004-1 | NZ_AGRN01000012.1 | - | 54847 | 54818 | **RNA**→ rpmE2 (PRK01678)Ribosomal_L31 (pfam01197)→ |
| Sau-4084-1 | NZ_AICH01000086.1 | - | 47226 | 47197 | **RNA**→ rpmE2 (PRK01678)Ribosomal_L31 (pfam01197)→ |
| Sau-4117-1 | NZ_AHBQ01000042.1 | - | 25505 | 25476 | **RNA**→ rpmE2 (PRK01678)Ribosomal_L31 (pfam01197)→ |
| Sau-4095-1 | NZ_AHZL01000017.1 | + | 18318 | 18347 | **RNA**→ rpmE2 (PRK01678)Ribosomal_L31 (pfam01197)→ |
| Sau-3020-1 | NZ_JBMO01000011.1 | - | 56759 | 56730 | **RNA**→ rpmE2 (PRK01678)Ribosomal_L31 (pfam01197)→ |

| | | | | | |
|---|---|---|---|---|---|
| Sau-808-1 | NZ_JEUZ01000006.1 | - | 56894 | 56865 | **RNA**→ rpmE2 (PRK01678)Ribosomal_L31 (pfam01197)→ |
| Sau-306-1 | NZ_AZMZ01000011.1 | + | 54437 | 54466 | **RNA**→ rpmE2 (PRK01678)Ribosomal_L31 (pfam01197)→ |
| Sau-1117-1 | NZ_AVPR01000004.1 | - | 48328 | 48299 | **RNA**→ rpmE2 (PRK01678)Ribosomal_L31 (pfam01197)→ |
| Sau-2628-1 | NZ_JANQ01000006.1 | - | 31901 | 31872 | **RNA**→ rpmE2 (PRK01678)Ribosomal_L31 (pfam01197)→ |
| Sau-1164-1 | NZ_JANM01000001.1 | - | 103797 | 103768 | **RNA**→ rpmE2 (PRK01678)Ribosomal_L31 (pfam01197)→ |
| Sau-2630-1 | NZ_LBIJ01000052.1 | - | 15717 | 15688 | **RNA**→ rpmE2 (PRK01678)Ribosomal_L31 (pfam01197)→ |
| Sau-3721-1 | NZ_LAKI01000016.1 | + | 20378 | 20407 | **RNA**→ rpmE2 (PRK01678)Ribosomal_L31 (pfam01197)→ |
| Sau-1163-1 | NZ_JANO01000008.1 | + | 54412 | 54441 | **RNA**→ rpmE2 (PRK01678)Ribosomal_L31 (pfam01197)→ |
| Sau-4074-1 | NZ_AJKD01000046.1 | - | 54814 | 54785 | **RNA**→ rpmE2 (PRK01678)Ribosomal_L31 (pfam01197)→ |
| Sau-59-2 | NZ_CTYB01000015.1 | - | 25255 | 25226 | **RNA**→ rpmE2 (PRK01678)Ribosomal_L31 (pfam01197)→ |
| Sau-2631-1 | NZ_LBII01000034.1 | - | 54843 | 54814 | **RNA**→ rpmE2 (PRK01678)Ribosomal_L31 (pfam01197)→ |
| Sau-2629-1 | NZ_LBIF01000025.1 | - | 54839 | 54810 | **RNA**→ rpmE2 (PRK01678)Ribosomal_L31 (pfam01197)→ |
| Sau-3759-1 | NZ_AOFU01000007.1 | - | 54626 | 54597 | **RNA**→ rpmE2 (PRK01678)Ribosomal_L31 (pfam01197)→ |
| Sau-3984-1 | NZ_BAFI01000013.1 | - | 34888 | 34859 | **RNA**→ rpmE2 (PRK01678)Ribosomal_L31 (pfam01197)→ |
| Sau-14-1 | NZ_ACKI01000009.1 | + | 51796 | 51825 | **RNA**→ rpmE2 (PRK01678)Ribosomal_L31 (pfam01197)→ rpmE2 (PRK01678)Ribosomal_L31 (pfam01197 |
| Sau-3768-1 | NZ_AIVJ01000015.1 | - | 56838 | 56809 | **RNA**→ rpmE2 (PRK01678)Ribosomal_L31 (pfam01197)→ |
| Sau-6-1 | NZ_ACKD01000061.1 | + | 18428 | 18457 | **RNA**→ rpmE2 (PRK01678)Ribosomal_L31 (pfam01197)→ rpmE2 (PRK01678)Ribosomal_L31 (pfam01197 |
| Sau-3779-1 | NZ_AOCQ01000048.1 | + | 53422 | 53451 | **RNA**→ rpmE2 (PRK01678)Ribosomal_L31 (pfam01197)→ |
| Sau-3985-1 | NZ_AEUQ01000018.1 | + | 18711 | 18740 | **RNA**→ rpmE2 (PRK01678)Ribosomal_L31 (pfam01197)→ |
| Sau-3758-1 | NZ_BABZ01000029.1 | + | 54326 | 54355 | **RNA**→ rpmE2 (PRK01678)Ribosomal_L31 (pfam01197)→ |
| Sau-12-1 | NZ_ADJK01000010.1 | - | 35339 | 35310 | **RNA**→ rpmE2 (PRK01678)Ribosomal_L31 (pfam01197)→ rpmE2 (PRK01678)Ribosomal_L31 (pfam01197 |
| Sau-4-1 | NZ_ACSO01000022.1 | - | 25134 | 25105 | **RNA**→ rpmE2 (PRK01678)Ribosomal_L31 (pfam01197)→ rpmE2 (PRK01678)Ribosomal_L31 (pfam01197 |
| Sau-19-1 | NZ_ACKL01000025.1 | - | 34879 | 34850 | **RNA**→ rpmE2 (PRK01678)Ribosomal_L31 (pfam01197)→ rpmE2 (PRK01678)Ribosomal_L31 (pfam01197 |
| Sau-20-1 | NZ_ABFB01000013.1 | + | 18336 | 18365 | **RNA**→ rpmE2 (PRK01678)Ribosomal_L31 (pfam01197)→ rpmE2 (PRK01678)Ribosomal_L31 (pfam01197 |
| Sau-9-1 | NZ_ACKG01000001.1 | + | 18295 | 18324 | **RNA**→ rpmE2 (PRK01678)Ribosomal_L31 (pfam01197)→ rpmE2 (PRK01678)Ribosomal_L31 (pfam01197 |
| Sau-3780-1 | NZ_AOFV01000008.1 | + | 53462 | 53491 | **RNA**→ rpmE2 (PRK01678)Ribosomal_L31 (pfam01197)→ |
| Sau-16-1 | NZ_ADJI01000011.1 | - | 25055 | 25026 | **RNA**→ rpmE2 (PRK01678)Ribosomal_L31 (pfam01197)→ rpmE2 (PRK01678)Ribosomal_L31 (pfam01197 |
| Sau-3778-1 | NZ_AOCP01000035.1 | - | 54850 | 54821 | **RNA**→ rpmE2 (PRK01678)Ribosomal_L31 (pfam01197)→ |
| Sau-17-1 | NZ_ACKK01000004.1 | - | 34778 | 34749 | **RNA**→ rpmE2 (PRK01678)Ribosomal_L31 (pfam01197)→ rpmE2 (PRK01678)Ribosomal_L31 (pfam01197 |
| Sau-8-1 | NZ_ACKF01000025.1 | + | 18300 | 18329 | **RNA**→ rpmE2 (PRK01678)Ribosomal_L31 (pfam01197)→ rpmE2 (PRK01678)Ribosomal_L31 (pfam01197 |
| Sau-5-1 | NZ_ACKC01000004.1 | - | 34879 | 34850 | **RNA**→ rpmE2 (PRK01678)Ribosomal_L31 (pfam01197)→ rpmE2 (PRK01678)Ribosomal_L31 (pfam01197 |
| Sau-3781-1 | NZ_AOFW01000023.1 | - | 55027 | 54998 | **RNA**→ rpmE2 (PRK01678)Ribosomal_L31 (pfam01197)→ |
| Sau-10-1 | NZ_ACYO01000010.1 | - | 25192 | 25163 | **RNA**→ rpmE2 (PRK01678)Ribosomal_L31 (pfam01197)→ rpmE2 (PRK01678)Ribosomal_L31 (pfam01197 |
| Sau-11-1 | NZ_ADJJ01000011.1 | - | 25197 | 25168 | **RNA**→ rpmE2 (PRK01678)Ribosomal_L31 (pfam01197)→ rpmE2 (PRK01678)Ribosomal_L31 (pfam01197 |
| Sau-18-1 | NZ_ACSN01000014.1 | - | 54849 | 54820 | **RNA**→ rpmE2 (PRK01678)Ribosomal_L31 (pfam01197)→ rpmE2 (PRK01678)Ribosomal_L31 (pfam01197 |
| Sau-3766-1 | NZ_ANCJ01000020.1 | - | 54772 | 54743 | **RNA**→ rpmE2 (PRK01678)Ribosomal_L31 (pfam01197)→ |
| Sau-15-1 | NZ_ACKJ01000053.1 | - | 34879 | 34850 | **RNA**→ rpmE2 (PRK01678)Ribosomal_L31 (pfam01197)→ rpmE2 (PRK01678)Ribosomal_L31 (pfam01197 |
| Sau-13-1 | NZ_ACKH01000023.1 | + | 18294 | 18323 | **RNA**→ rpmE2 (PRK01678)Ribosomal_L31 (pfam01197)→ rpmE2 (PRK01678)Ribosomal_L31 (pfam01197 |
| Sau-7-1 | NZ_ACKE01000044.1 | + | 18294 | 18323 | **RNA**→ rpmE2 (PRK01678)Ribosomal_L31 (pfam01197)→ rpmE2 (PRK01678)Ribosomal_L31 (pfam01197 |
| Sau-3983-1 | NZ_AMTC01000034.1 | - | 15812 | 15783 | **RNA**→ rpmE2 (PRK01678)Ribosomal_L31 (pfam01197)→ |
| Sau-4108-1 | NZ_AHBK01000071.1 | - | 35128 | 35099 | **RNA**→ rpmE2 (PRK01678)Ribosomal_L31 (pfam01197)→ |
| Sau-4057-1 | NZ_AHKE01000014.1 | - | 59009 | 58980 | **RNA**→ rpmE2 (PRK01678)Ribosomal_L31 (pfam01197)→ |
| Sau-4109-1 | NZ_AHBT01000029.1 | - | 25283 | 25254 | **RNA**→ rpmE2 (PRK01678)Ribosomal_L31 (pfam01197)→ |
| Sau-77-1 | NZ_ABSA01000077.1 | - | 54759 | 54730 | **RNA**→ rpmE2 (PRK01678)Ribosomal_L31 (pfam01197)→ rpmE2 (PRK01678)Ribosomal_L31 (pfam01197 |
| Sau-4107-1 | NZ_ALWG01000007.1 | - | 54626 | 54597 | **RNA**→ rpmE2 (PRK01678)Ribosomal_L31 (pfam01197)→ |
| Sau-4070-1 | NZ_AHVT01000022.1 | - | 59124 | 59095 | **RNA**→ rpmE2 (PRK01678)Ribosomal_L31 (pfam01197)→ |
| Sau-4066-1 | NZ_AHVY01000011.1 | - | 20421 | 20392 | **RNA**→ rpmE2 (PRK01678)Ribosomal_L31 (pfam01197)→ |
| Sau-4100-1 | NZ_AFGU01000104.1 | + | 55716 | 55745 | **RNA**→ rpmE2 (PRK01678)Ribosomal_L31 (pfam01197)→ |
| Sau-4069-1 | NZ_AHVO01000012.1 | - | 58156 | 58127 | **RNA**→ rpmE2 (PRK01678)Ribosomal_L31 (pfam01197)→ |
| Sau-61-1 | NZ_BABM01000001.1 | - | 2248261 | 2248232 | **RNA**→ rpmE2 (PRK01678)Ribosomal_L31 (pfam01197)→ rpmE2 (PRK01678)Ribosomal_L31 (pfam01197 |
| Sau-4110-1 | NZ_AHBU01000033.1 | - | 35144 | 35115 | **RNA**→ rpmE2 (PRK01678)Ribosomal_L31 (pfam01197)→ |
| Sau-4116-1 | NZ_AHBP01000045.1 | - | 35049 | 35020 | **RNA**→ rpmE2 (PRK01678)Ribosomal_L31 (pfam01197)→ |

| | | | | | |
|---|---|---|---|---|---|
| Sau-4080-1 | NZ_AJLX01000024.1 | - | 54759 | 54730 | **RNA**→ rpmE2 (PRK01678)Ribosomal_L31 (pfam01197)→ |
| Sau-4112-1 | NZ_AHBL01000051.1 | - | 35238 | 35209 | **RNA**→ rpmE2 (PRK01678)Ribosomal_L31 (pfam01197)→ |
| Sau-3744-1 | NZ_ANHW01000011.1 | + | 18294 | 18323 | **RNA**→ rpmE2 (PRK01678)Ribosomal_L31 (pfam01197)→ |
| Sau-4064-1 | NZ_AIES01000019.1 | - | 57510 | 57481 | **RNA**→ rpmE2 (PRK01678)Ribosomal_L31 (pfam01197)→ |
| Sau-4068-1 | NZ_AHVL01000023.1 | - | 58578 | 58549 | **RNA**→ rpmE2 (PRK01678)Ribosomal_L31 (pfam01197)→ |
| Sau-4065-1 | NZ_AHVQ01000012.1 | - | 57736 | 57707 | **RNA**→ rpmE2 (PRK01678)Ribosomal_L31 (pfam01197)→ |
| Sau-4079-1 | NZ_AIDT01000006.1 | - | 56349 | 56320 | **RNA**→ rpmE2 (PRK01678)Ribosomal_L31 (pfam01197)→ |
| Sau-4060-1 | NZ_AHVG01000019.1 | - | 59351 | 59322 | **RNA**→ rpmE2 (PRK01678)Ribosomal_L31 (pfam01197)→ |
| Sau-4087-1 | NZ_AHLL01000033.1 | - | 35267 | 35238 | **RNA**→ rpmE2 (PRK01678)Ribosomal_L31 (pfam01197)→ |
| Sau-4111-1 | NZ_AHBV01000033.1 | - | 35172 | 35143 | **RNA**→ rpmE2 (PRK01678)Ribosomal_L31 (pfam01197)→ |
| Sau-4062-1 | NZ_AIEQ01000010.1 | - | 58114 | 58085 | **RNA**→ rpmE2 (PRK01678)Ribosomal_L31 (pfam01197)→ |
| Sau-4059-1 | NZ_AHVX01000043.1 | - | 60169 | 60140 | **RNA**→ rpmE2 (PRK01678)Ribosomal_L31 (pfam01197)→ |
| Sau-4071-1 | NZ_AHVD01000025.1 | - | 48130 | 48101 | **RNA**→ rpmE2 (PRK01678)Ribosomal_L31 (pfam01197)→ |
| Sau-4121-1 | NZ_AKYW01000010.1 | + | 28884 | 28913 | **RNA**→ rpmE2 (PRK01678)Ribosomal_L31 (pfam01197)→ |
| Sau-4115-1 | NZ_AHBO01000053.1 | - | 25113 | 25084 | **RNA**→ rpmE2 (PRK01678)Ribosomal_L31 (pfam01197)→ |
| Sau-4083-1 | NZ_AHLT01000090.1 | + | 53400 | 53429 | **RNA**→ rpmE2 (PRK01678)Ribosomal_L31 (pfam01197)→ |
| Sau-54-1 | NZ_ACZQ01000016.1 | + | 18318 | 18347 | **RNA**→ rpmE2 (PRK01678)Ribosomal_L31 (pfam01197)→ rpmE2 (PRK01678)Ribosomal_L31 (pfam01197 |
| Sau-4088-1 | NZ_AHLO01000010.1 | - | 41581 | 41552 | **RNA**→ rpmE2 (PRK01678)Ribosomal_L31 (pfam01197)→ |
| Sau-3746-1 | NZ_ANIB01000005.1 | + | 55076 | 55105 | **RNA**→ rpmE2 (PRK01678)Ribosomal_L31 (pfam01197)→ |
| Sau-4067-1 | NZ_AHVR01000009.1 | - | 58247 | 58218 | **RNA**→ rpmE2 (PRK01678)Ribosomal_L31 (pfam01197)→ |
| Sau-3743-2 | NZ_AMYL01000009.1 | - | 20386 | 20357 | **RNA**→ rpmE2 (PRK01678)Ribosomal_L31 (pfam01197)→ |
| Sau-4098-1 | NZ_BAEA01000005.1 | - | 54851 | 54822 | **RNA**→ rpmE2 (PRK01678)Ribosomal_L31 (pfam01197)→ |
| Sau-4119-1 | NZ_AHBS01000048.1 | - | 35093 | 35064 | **RNA**→ rpmE2 (PRK01678)Ribosomal_L31 (pfam01197)→ |
| Sau-4114-1 | NZ_AHBN01000052.1 | - | 25427 | 25398 | **RNA**→ rpmE2 (PRK01678)Ribosomal_L31 (pfam01197)→ |
| Sau-4113-1 | NZ_AHBM01000041.1 | - | 25182 | 25153 | **RNA**→ rpmE2 (PRK01678)Ribosomal_L31 (pfam01197)→ |
| Sau-4118-1 | NZ_AHBR01000039.1 | - | 35181 | 35152 | **RNA**→ rpmE2 (PRK01678)Ribosomal_L31 (pfam01197)→ |
| Sau-4009-1 | NZ_ANAL01000031.1 | - | 18161 | 18132 | **RNA**→ rpmE2 (PRK01678)Ribosomal_L31 (pfam01197)→ |
| Sau-3995-1 | NZ_AOFX01000036.1 | - | 20485 | 20456 | **RNA**→ rpmE2 (PRK01678)Ribosomal_L31 (pfam01197)→ |
| Sau-4028-1 | NZ_AGTV01000033.1 | + | 42513 | 42542 | **RNA**→ rpmE2 (PRK01678)Ribosomal_L31 (pfam01197)→ |
| Sau-4014-1 | NZ_AGTY01000039.1 | - | 54848 | 54819 | **RNA**→ rpmE2 (PRK01678)Ribosomal_L31 (pfam01197)→ |
| Sau-4051-1 | NZ_AHVU01000014.1 | - | 15829 | 15800 | **RNA**→ rpmE2 (PRK01678)Ribosomal_L31 (pfam01197)→ |
| Sau-4042-1 | NZ_ABWY01000017.1 | - | 34879 | 34850 | **RNA**→ rpmE2 (PRK01678)Ribosomal_L31 (pfam01197)→ |
| Sau-3999-1 | NZ_AHZK01000003.1 | + | 18314 | 18343 | **RNA**→ rpmE2 (PRK01678)Ribosomal_L31 (pfam01197)→ |
| Sau-4036-1 | NZ_ALWH01000007.1 | - | 54626 | 54597 | **RNA**→ rpmE2 (PRK01678)Ribosomal_L31 (pfam01197)→ |
| Sau-4054-1 | NZ_AHVP01000029.1 | - | 55599 | 55570 | **RNA**→ rpmE2 (PRK01678)Ribosomal_L31 (pfam01197)→ |
| Sau-4019-1 | NZ_AHJW01000027.1 | - | 55010 | 54981 | **RNA**→ rpmE2 (PRK01678)Ribosomal_L31 (pfam01197)→ |
| Sau-4048-1 | NZ_AHVJ01000017.1 | - | 20743 | 20714 | **RNA**→ rpmE2 (PRK01678)Ribosomal_L31 (pfam01197)→ |
| Sau-4047-1 | NZ_AHVE01000012.1 | - | 36420 | 36391 | **RNA**→ rpmE2 (PRK01678)Ribosomal_L31 (pfam01197)→ |
| Sau-4031-1 | NZ_AGTX01000020.1 | - | 54805 | 54776 | **RNA**→ rpmE2 (PRK01678)Ribosomal_L31 (pfam01197)→ |
| Sau-4000-1 | NZ_AJGE01000006.1 | + | 18324 | 18353 | **RNA**→ rpmE2 (PRK01678)Ribosomal_L31 (pfam01197)→ |
| Sau-4030-1 | NZ_AGTW01000035.1 | - | 54768 | 54739 | **RNA**→ rpmE2 (PRK01678)Ribosomal_L31 (pfam01197)→ |
| Sau-3997-1 | NZ_AHZH01000007.1 | - | 22857 | 22828 | **RNA**→ rpmE2 (PRK01678)Ribosomal_L31 (pfam01197)→ |
| Sau-4055-1 | NZ_AIEP01000019.1 | - | 22405 | 22376 | **RNA**→ rpmE2 (PRK01678)Ribosomal_L31 (pfam01197)→ |
| Sau-4008-1 | NZ_AFTM01000022.1 | - | 28890 | 28861 | **RNA**→ rpmE2 (PRK01678)Ribosomal_L31 (pfam01197)→ |
| Sau-4027-1 | NZ_AFNP01000048.1 | - | 54627 | 54598 | **RNA**→ rpmE2 (PRK01678)Ribosomal_L31 (pfam01197)→ |
| Sau-4001-1 | NZ_ALWE01000045.1 | - | 54609 | 54580 | **RNA**→ rpmE2 (PRK01678)Ribosomal_L31 (pfam01197)→ |
| Sau-4032-1 | NZ_AHKU01000089.1 | - | 20842 | 20813 | **RNA**→ rpmE2 (PRK01678)Ribosomal_L31 (pfam01197)→ |
| Sau-4043-1 | NZ_AHVK01000015.1 | - | 56807 | 56778 | **RNA**→ rpmE2 (PRK01678)Ribosomal_L31 (pfam01197)→ |
| Sau-3996-1 | NZ_ARXY01000030.1 | - | 54931 | 54902 | **RNA**→ rpmE2 (PRK01678)Ribosomal_L31 (pfam01197)→ |
| Sau-4049-1 | NZ_AHVF01000014.1 | - | 56620 | 56591 | **RNA**→ rpmE2 (PRK01678)Ribosomal_L31 (pfam01197)→ |
| Sau-4003-1 | NZ_AFEF01000009.1 | - | 34890 | 34861 | **RNA**→ rpmE2 (PRK01678)Ribosomal_L31 (pfam01197)→ |
| Sau-4039-1 | NZ_ASHS01000035.1 | - | 34898 | 34869 | **RNA**→ rpmE2 (PRK01678)Ribosomal_L31 (pfam01197)→ |

| | | | | | |
|---|---|---|---|---|---|
| Sau-4002-1 | NZ_AHZJ01000009.1 | + | 18316 | 18345 | **RNA**→ rpmE2 (PRK01678)Ribosomal_L31 (pfam01197)→ |
| Sau-4010-1 | NZ_AFTN01000024.1 | - | 54601 | 54572 | **RNA**→ rpmE2 (PRK01678)Ribosomal_L31 (pfam01197)→ |
| Sau-4016-1 | NZ_ANAK01000023.1 | - | 25281 | 25252 | **RNA**→ rpmE2 (PRK01678)Ribosomal_L31 (pfam01197)→ |
| Sau-3987-1 | NZ_BAFA01000003.1 | + | 54805 | 54834 | **RNA**→ rpmE2 (PRK01678)Ribosomal_L31 (pfam01197)→ |
| Sau-4035-1 | NZ_ALWF01000054.1 | - | 54609 | 54580 | **RNA**→ rpmE2 (PRK01678)Ribosomal_L31 (pfam01197)→ |
| Sau-3998-1 | NZ_AHZI01000003.1 | - | 54609 | 54580 | **RNA**→ rpmE2 (PRK01678)Ribosomal_L31 (pfam01197)→ |
| Sau-4053-1 | NZ_AHVS01000012.1 | - | 29787 | 29758 | **RNA**→ rpmE2 (PRK01678)Ribosomal_L31 (pfam01197)→ |
| Sau-3994-1 | NZ_ASTH01000013.1 | + | 53397 | 53426 | **RNA**→ rpmE2 (PRK01678)Ribosomal_L31 (pfam01197)→ |
| Sau-4023-1 | NZ_AHJY01000021.1 | + | 53399 | 53428 | **RNA**→ rpmE2 (PRK01678)Ribosomal_L31 (pfam01197)→ |
| Sau-4056-1 | NZ_AHVI01000012.1 | - | 58959 | 58930 | **RNA**→ rpmE2 (PRK01678)Ribosomal_L31 (pfam01197)→ |
| Sau-4041-1 | NZ_ABWT01000015.1 | + | 18336 | 18365 | **RNA**→ rpmE2 (PRK01678)Ribosomal_L31 (pfam01197)→ |
| Sau-4017-1 | NZ_AHJV01000045.1 | - | 54800 | 54771 | **RNA**→ rpmE2 (PRK01678)Ribosomal_L31 (pfam01197)→ |
| Sau-3986-1 | NZ_AEUR01000019.1 | - | 55149 | 55120 | **RNA**→ rpmE2 (PRK01678)Ribosomal_L31 (pfam01197)→ |
| Sau-4029-1 | NZ_AHKA01000008.1 | - | 54672 | 54643 | **RNA**→ rpmE2 (PRK01678)Ribosomal_L31 (pfam01197)→ |
| Sau-4025-1 | NZ_AHJZ01000033.1 | - | 15521 | 15492 | **RNA**→ rpmE2 (PRK01678)Ribosomal_L31 (pfam01197)→ |
| Sau-4033-1 | NZ_AHKV01000021.1 | + | 18430 | 18459 | **RNA**→ rpmE2 (PRK01678)Ribosomal_L31 (pfam01197)→ |
| Sau-4034-1 | NZ_AHKW01000042.1 | + | 28774 | 28803 | **RNA**→ rpmE2 (PRK01678)Ribosomal_L31 (pfam01197)→ |
| Sau-4050-1 | NZ_AHVM01000014.1 | - | 59158 | 59129 | **RNA**→ rpmE2 (PRK01678)Ribosomal_L31 (pfam01197)→ |
| Sau-4046-1 | NZ_AHVH01000038.1 | - | 35168 | 35139 | **RNA**→ rpmE2 (PRK01678)Ribosomal_L31 (pfam01197)→ |
| Sau-4005-1 | NZ_AFEE01000059.1 | - | 55023 | 54994 | **RNA**→ rpmE2 (PRK01678)Ribosomal_L31 (pfam01197)→ |
| Sau-23-1 | NZ_ACOT01000040.1 | - | 54845 | 54816 | **RNA**→ rpmE2 (PRK01678)Ribosomal_L31 (pfam01197)→ rpmE2 (PRK01678)Ribosomal_L31 (pfam01197 |
| Sau-4012-1 | NZ_AGRO01000055.1 | - | 15808 | 15779 | **RNA**→ rpmE2 (PRK01678)Ribosomal_L31 (pfam01197)→ |
| Sau-4020-1 | NZ_AHJX01000009.1 | - | 20656 | 20627 | **RNA**→ rpmE2 (PRK01678)Ribosomal_L31 (pfam01197)→ |
| Sau-4044-1 | NZ_AIER01000007.1 | - | 61496 | 61467 | **RNA**→ rpmE2 (PRK01678)Ribosomal_L31 (pfam01197)→ |
| Sau-2704-1 | NZ_CBWW010000060.1 | + | 7143 | 7172 | **RNA**→ rpmE2 (PRK01678)Ribosomal_L31 (pfam01197)→ |
| Sau-500-1 | NZ_JFOW01000018.1 | - | 36391 | 36362 | **RNA**→ rpmE2 (PRK01678)Ribosomal_L31 (pfam01197)→ |
| Sau-505-1 | NZ_JFPB01000020.1 | - | 36391 | 36362 | **RNA**→ rpmE2 (PRK01678)Ribosomal_L31 (pfam01197)→ |
| Sau-510-1 | NZ_JFPG01000019.1 | - | 35659 | 35630 | **RNA**→ rpmE2 (PRK01678)Ribosomal_L31 (pfam01197)→ |
| Sau-537-1 | NZ_JFQF01000015.1 | - | 36391 | 36362 | **RNA**→ rpmE2 (PRK01678)Ribosomal_L31 (pfam01197)→ |
| Sau-538-1 | NZ_JFQG01000016.1 | - | 36391 | 36362 | **RNA**→ rpmE2 (PRK01678)Ribosomal_L31 (pfam01197)→ |
| Sau-409-1 | NZ_JFRP01000016.1 | - | 36265 | 36236 | **RNA**→ rpmE2 (PRK01678)Ribosomal_L31 (pfam01197)→ |
| Sau-501-1 | NZ_JFOX01000017.1 | - | 36391 | 36362 | **RNA**→ rpmE2 (PRK01678)Ribosomal_L31 (pfam01197)→ |
| Sau-506-1 | NZ_JFPC01000017.1 | - | 35086 | 35057 | **RNA**→ rpmE2 (PRK01678)Ribosomal_L31 (pfam01197)→ |
| Sau-507-1 | NZ_JFPD01000019.1 | - | 36391 | 36362 | **RNA**→ rpmE2 (PRK01678)Ribosomal_L31 (pfam01197)→ |
| Sau-511-1 | NZ_JFPH01000018.1 | - | 36391 | 36362 | **RNA**→ rpmE2 (PRK01678)Ribosomal_L31 (pfam01197)→ |
| Sau-512-1 | NZ_JFPI01000018.1 | - | 36391 | 36362 | **RNA**→ rpmE2 (PRK01678)Ribosomal_L31 (pfam01197)→ |
| Sau-534-1 | NZ_JFQC01000018.1 | - | 36391 | 36362 | **RNA**→ rpmE2 (PRK01678)Ribosomal_L31 (pfam01197)→ |
| Sau-539-1 | NZ_JFQH01000018.1 | - | 36391 | 36362 | **RNA**→ rpmE2 (PRK01678)Ribosomal_L31 (pfam01197)→ |
| Sau-413-1 | NZ_JFRL01000015.1 | - | 36391 | 36362 | **RNA**→ rpmE2 (PRK01678)Ribosomal_L31 (pfam01197)→ |
| Sau-412-1 | NZ_JFRM01000016.1 | - | 35284 | 35255 | **RNA**→ rpmE2 (PRK01678)Ribosomal_L31 (pfam01197)→ |
| Sau-408-1 | NZ_JFRQ01000020.1 | - | 25129 | 25100 | **RNA**→ rpmE2 (PRK01678)Ribosomal_L31 (pfam01197)→ |
| Sau-457-1 | NZ_JGNN01000006.1 | - | 56762 | 56733 | **RNA**→ rpmE2 (PRK01678)Ribosomal_L31 (pfam01197)→ |
| Sau-499-1 | NZ_JFOV01000016.1 | - | 36391 | 36362 | **RNA**→ rpmE2 (PRK01678)Ribosomal_L31 (pfam01197)→ |
| Sau-504-1 | NZ_JFPA01000017.1 | - | 36391 | 36362 | **RNA**→ rpmE2 (PRK01678)Ribosomal_L31 (pfam01197)→ |
| Sau-509-1 | NZ_JFPF01000019.1 | - | 35659 | 35630 | **RNA**→ rpmE2 (PRK01678)Ribosomal_L31 (pfam01197)→ |
| Sau-536-1 | NZ_JFQE01000017.1 | - | 36391 | 36362 | **RNA**→ rpmE2 (PRK01678)Ribosomal_L31 (pfam01197)→ |
| Sau-410-1 | NZ_JFRO01000016.1 | - | 36390 | 36361 | **RNA**→ rpmE2 (PRK01678)Ribosomal_L31 (pfam01197)→ |
| Sau-497-1 | NZ_JFOT01000022.1 | - | 25361 | 25332 | **RNA**→ rpmE2 (PRK01678)Ribosomal_L31 (pfam01197)→ |
| Sau-498-1 | NZ_JFOU01000016.1 | - | 36391 | 36362 | **RNA**→ rpmE2 (PRK01678)Ribosomal_L31 (pfam01197)→ |
| Sau-502-1 | NZ_JFOY01000020.1 | - | 36391 | 36362 | **RNA**→ rpmE2 (PRK01678)Ribosomal_L31 (pfam01197)→ |
| Sau-503-1 | NZ_JFOZ01000016.1 | - | 36391 | 36362 | **RNA**→ rpmE2 (PRK01678)Ribosomal_L31 (pfam01197)→ |
| Sau-508-1 | NZ_JFPE01000019.1 | - | 36391 | 36362 | **RNA**→ rpmE2 (PRK01678)Ribosomal_L31 (pfam01197)→ |

| | | | | | |
|---|---|---|---|---|---|
| Sau-535-1 | NZ_JFQD01000018.1 | - | 36391 | 36362 | **RNA**→ rpmE2 (PRK01678)Ribosomal_L31 (pfam01197)→ |
| Sau-540-1 | NZ_JFQI01000018.1 | - | 36391 | 36362 | **RNA**→ rpmE2 (PRK01678)Ribosomal_L31 (pfam01197)→ |
| Sau-411-1 | NZ_JFRN01000019.1 | - | 36149 | 36120 | **RNA**→ rpmE2 (PRK01678)Ribosomal_L31 (pfam01197)→ |
| Sau-488-1 | NZ_JGNO01000001.1 | - | 1681337 | 1681308 | **RNA**→ rpmE2 (PRK01678)Ribosomal_L31 (pfam01197)→ |
| Sau-529-1 | NZ_JFPX01000018.1 | - | 36391 | 36362 | **RNA**→ rpmE2 (PRK01678)Ribosomal_L31 (pfam01197)→ |
| Sau-527-1 | NZ_JFPV01000016.1 | - | 36492 | 36463 | **RNA**→ rpmE2 (PRK01678)Ribosomal_L31 (pfam01197)→ |
| Sau-528-1 | NZ_JFPW01000019.1 | - | 34974 | 34945 | **RNA**→ rpmE2 (PRK01678)Ribosomal_L31 (pfam01197)→ |
| Sau-532-1 | NZ_JFQA01000016.1 | - | 35261 | 35232 | **RNA**→ rpmE2 (PRK01678)Ribosomal_L31 (pfam01197)→ |
| Sau-533-1 | NZ_JFQB01000018.1 | - | 36391 | 36362 | **RNA**→ rpmE2 (PRK01678)Ribosomal_L31 (pfam01197)→ |
| Sau-515-1 | NZ_JFPL01000018.1 | - | 36391 | 36362 | **RNA**→ rpmE2 (PRK01678)Ribosomal_L31 (pfam01197)→ |
| Sau-521-1 | NZ_JFPQ01000016.1 | - | 36391 | 36362 | **RNA**→ rpmE2 (PRK01678)Ribosomal_L31 (pfam01197)→ |
| Sau-523-1 | NZ_JFPR01000018.1 | - | 36391 | 36362 | **RNA**→ rpmE2 (PRK01678)Ribosomal_L31 (pfam01197)→ |
| Sau-516-1 | NZ_JFPM01000019.1 | - | 36391 | 36362 | **RNA**→ rpmE2 (PRK01678)Ribosomal_L31 (pfam01197)→ |
| Sau-517-1 | NZ_JFPN01000018.1 | - | 36391 | 36362 | **RNA**→ rpmE2 (PRK01678)Ribosomal_L31 (pfam01197)→ |
| Sau-524-1 | NZ_JFPS01000015.1 | - | 35659 | 35630 | **RNA**→ rpmE2 (PRK01678)Ribosomal_L31 (pfam01197)→ |
| Sau-526-1 | NZ_JFPU01000020.1 | - | 36391 | 36362 | **RNA**→ rpmE2 (PRK01678)Ribosomal_L31 (pfam01197)→ |
| Sau-531-1 | NZ_JFPZ01000015.1 | - | 36392 | 36363 | **RNA**→ rpmE2 (PRK01678)Ribosomal_L31 (pfam01197)→ |
| Sau-514-1 | NZ_JFPK01000019.1 | - | 35659 | 35630 | **RNA**→ rpmE2 (PRK01678)Ribosomal_L31 (pfam01197)→ |
| Sau-520-1 | NZ_JFPP01000018.1 | - | 36391 | 36362 | **RNA**→ rpmE2 (PRK01678)Ribosomal_L31 (pfam01197)→ |
| Sau-525-1 | NZ_JFPT01000019.1 | - | 36391 | 36362 | **RNA**→ rpmE2 (PRK01678)Ribosomal_L31 (pfam01197)→ |
| Sau-530-1 | NZ_JFPY01000018.1 | - | 36391 | 36362 | **RNA**→ rpmE2 (PRK01678)Ribosomal_L31 (pfam01197)→ |
| Sau-513-1 | NZ_JFPJ01000016.1 | - | 36391 | 36362 | **RNA**→ rpmE2 (PRK01678)Ribosomal_L31 (pfam01197)→ |
| Sau-519-1 | NZ_JFPO01000018.1 | - | 36391 | 36362 | **RNA**→ rpmE2 (PRK01678)Ribosomal_L31 (pfam01197)→ |
| Sau-1746-1 | NZ_JBBD01000006.1 | - | 49963 | 49934 | **RNA**→ rpmE2 (PRK01678)Ribosomal_L31 (pfam01197)→ |
| Sau-1817-1 | NZ_JDCJ01000001.1 | - | 49963 | 49934 | **RNA**→ rpmE2 (PRK01678)Ribosomal_L31 (pfam01197)→ |
| Sau-1847-1 | NZ_JDCO01000002.1 | - | 24749 | 24720 | **RNA**→ rpmE2 (PRK01678)Ribosomal_L31 (pfam01197)→ |
| Sau-1865-1 | NZ_JDCT01000008.1 | - | 91437 | 91408 | **RNA**→ rpmE2 (PRK01678)Ribosomal_L31 (pfam01197)→ |
| Sau-1908-1 | NZ_JDCY01000012.1 | - | 55496 | 55467 | **RNA**→ rpmE2 (PRK01678)Ribosomal_L31 (pfam01197)→ |
| Sau-1949-1 | NZ_JDDD01000016.1 | - | 21401 | 21372 | **RNA**→ rpmE2 (PRK01678)Ribosomal_L31 (pfam01197)→ |
| Sau-1829-1 | NZ_JDCK01000012.1 | - | 24749 | 24720 | **RNA**→ rpmE2 (PRK01678)Ribosomal_L31 (pfam01197)→ |
| Sau-1850-1 | NZ_JDCP01000011.1 | - | 36391 | 36362 | **RNA**→ rpmE2 (PRK01678)Ribosomal_L31 (pfam01197)→ |
| Sau-1879-1 | NZ_JDCU01000001.1 | - | 49963 | 49934 | **RNA**→ rpmE2 (PRK01678)Ribosomal_L31 (pfam01197)→ |
| Sau-1915-1 | NZ_JDCZ01000010.1 | - | 35154 | 35125 | **RNA**→ rpmE2 (PRK01678)Ribosomal_L31 (pfam01197)→ |
| Sau-1950-1 | NZ_JDDE01000006.1 | - | 133588 | 133559 | **RNA**→ rpmE2 (PRK01678)Ribosomal_L31 (pfam01197)→ |
| Sau-1952-1 | NZ_JDDF01000002.1 | - | 238695 | 238666 | **RNA**→ rpmE2 (PRK01678)Ribosomal_L31 (pfam01197)→ |
| Sau-1758-1 | NZ_JDDZ01000011.1 | - | 36491 | 36462 | **RNA**→ rpmE2 (PRK01678)Ribosomal_L31 (pfam01197)→ |
| Sau-1735-1 | NZ_JBBN01000017.1 | - | 35603 | 35574 | **RNA**→ rpmE2 (PRK01678)Ribosomal_L31 (pfam01197)→ |
| Sau-1830-1 | NZ_JDCL01000012.1 | - | 35918 | 35889 | **RNA**→ rpmE2 (PRK01678)Ribosomal_L31 (pfam01197)→ |
| Sau-1836-1 | NZ_JDCM01000007.1 | - | 62156 | 62127 | **RNA**→ rpmE2 (PRK01678)Ribosomal_L31 (pfam01197)→ |
| Sau-1852-1 | NZ_JDCQ01000009.1 | - | 62354 | 62325 | **RNA**→ rpmE2 (PRK01678)Ribosomal_L31 (pfam01197)→ |
| Sau-1859-1 | NZ_JDCR01000003.1 | - | 24749 | 24720 | **RNA**→ rpmE2 (PRK01678)Ribosomal_L31 (pfam01197)→ |
| Sau-1895-1 | NZ_JDCW01000003.1 | - | 24749 | 24720 | **RNA**→ rpmE2 (PRK01678)Ribosomal_L31 (pfam01197)→ |
| Sau-1922-1 | NZ_JDDA01000006.1 | - | 24749 | 24720 | **RNA**→ rpmE2 (PRK01678)Ribosomal_L31 (pfam01197)→ |
| Sau-1939-1 | NZ_JDDB01000001.1 | - | 238862 | 238833 | **RNA**→ rpmE2 (PRK01678)Ribosomal_L31 (pfam01197)→ |
| Sau-1767-1 | NZ_JDEA01000006.1 | - | 24749 | 24720 | **RNA**→ rpmE2 (PRK01678)Ribosomal_L31 (pfam01197)→ |
| Sau-1798-1 | NZ_JDFE01000010.1 | - | 61261 | 61232 | **RNA**→ rpmE2 (PRK01678)Ribosomal_L31 (pfam01197)→ |
| Sau-1681-1 | NZ_JBAY01000005.1 | - | 57586 | 57557 | **RNA**→ rpmE2 (PRK01678)Ribosomal_L31 (pfam01197)→ |
| Sau-1732-1 | NZ_JBBP01000013.1 | - | 61797 | 61768 | **RNA**→ rpmE2 (PRK01678)Ribosomal_L31 (pfam01197)→ |
| Sau-1748-1 | NZ_JDCI01000012.1 | - | 35617 | 35588 | **RNA**→ rpmE2 (PRK01678)Ribosomal_L31 (pfam01197)→ |
| Sau-1838-1 | NZ_JDCN01000012.1 | - | 36391 | 36362 | **RNA**→ rpmE2 (PRK01678)Ribosomal_L31 (pfam01197)→ |
| Sau-1863-1 | NZ_JDCS01000002.1 | - | 238606 | 238577 | **RNA**→ rpmE2 (PRK01678)Ribosomal_L31 (pfam01197)→ |
| Sau-1897-1 | NZ_JDCX01000012.1 | - | 35605 | 35576 | **RNA**→ rpmE2 (PRK01678)Ribosomal_L31 (pfam01197)→ |

| | | | | | |
|---|---|---|---|---|---|
| Sau-1940-1 | NZ_JDDC01000013.1 | - | 35656 | 35627 | **RNA**→ rpmE2 (PRK01678)Ribosomal_L31 (pfam01197)→ |
| Sau-1684-1 | NZ_JBDH01000007.1 | - | 111776 | 111747 | **RNA**→ rpmE2 (PRK01678)Ribosomal_L31 (pfam01197)→ |
| Sau-1680-1 | NZ_JBDK01000007.1 | - | 109916 | 109887 | **RNA**→ rpmE2 (PRK01678)Ribosomal_L31 (pfam01197)→ |
| Sau-1715-1 | NZ_JBCF01000011.1 | - | 36100 | 36071 | **RNA**→ rpmE2 (PRK01678)Ribosomal_L31 (pfam01197)→ |
| Sau-1666-1 | NZ_JBDX01000010.1 | - | 58580 | 58551 | **RNA**→ rpmE2 (PRK01678)Ribosomal_L31 (pfam01197)→ |
| Sau-1977-1 | NZ_JDDK01000001.1 | - | 58248 | 58219 | **RNA**→ rpmE2 (PRK01678)Ribosomal_L31 (pfam01197)→ |
| Sau-1720-1 | NZ_JBCA01000017.1 | - | 3620 | 3591 | **RNA**→ rpmE2 (PRK01678)Ribosomal_L31 (pfam01197)→ |
| Sau-1664-1 | NZ_JBDZ01000012.1 | - | 19785 | 19756 | **RNA**→ rpmE2 (PRK01678)Ribosomal_L31 (pfam01197)→ |
| Sau-1972-1 | NZ_JDDI01000010.1 | - | 21997 | 21968 | **RNA**→ rpmE2 (PRK01678)Ribosomal_L31 (pfam01197)→ |
| Sau-1992-1 | NZ_JDDN01000007.1 | - | 62153 | 62124 | **RNA**→ rpmE2 (PRK01678)Ribosomal_L31 (pfam01197)→ |
| Sau-1730-1 | NZ_JBBR01000006.1 | - | 112235 | 112206 | **RNA**→ rpmE2 (PRK01678)Ribosomal_L31 (pfam01197)→ |
| Sau-1653-1 | NZ_JBEJ01000011.1 | - | 32842 | 32813 | **RNA**→ rpmE2 (PRK01678)Ribosomal_L31 (pfam01197)→ |
| Sau-1966-1 | NZ_JDDH01000008.1 | - | 24749 | 24720 | **RNA**→ rpmE2 (PRK01678)Ribosomal_L31 (pfam01197)→ |
| Sau-1988-1 | NZ_JDDM01000012.1 | - | 36292 | 36263 | **RNA**→ rpmE2 (PRK01678)Ribosomal_L31 (pfam01197)→ |
| Sau-1707-1 | NZ_JBCM01000016.1 | - | 35136 | 35107 | **RNA**→ rpmE2 (PRK01678)Ribosomal_L31 (pfam01197)→ |
| Sau-1705-1 | NZ_JBCO01000009.1 | - | 24749 | 24720 | **RNA**→ rpmE2 (PRK01678)Ribosomal_L31 (pfam01197)→ |
| Sau-1671-1 | NZ_JBDT01000006.1 | - | 112159 | 112130 | **RNA**→ rpmE2 (PRK01678)Ribosomal_L31 (pfam01197)→ |
| Sau-1675-1 | NZ_JBDP01000014.1 | - | 36391 | 36362 | **RNA**→ rpmE2 (PRK01678)Ribosomal_L31 (pfam01197)→ |
| Sau-1959-1 | NZ_JDDG01000010.1 | - | 34416 | 34387 | **RNA**→ rpmE2 (PRK01678)Ribosomal_L31 (pfam01197)→ |
| Sau-1982-1 | NZ_JDDL01000012.1 | - | 36391 | 36362 | **RNA**→ rpmE2 (PRK01678)Ribosomal_L31 (pfam01197)→ |
| Sau-1142-1 | NZ_JBFF01000002.1 | - | 56894 | 56865 | **RNA**→ rpmE2 (PRK01678)Ribosomal_L31 (pfam01197)→ |
| Sau-1149-1 | NZ_JBRF01000002.1 | - | 49802 | 49773 | **RNA**→ rpmE2 (PRK01678)Ribosomal_L31 (pfam01197)→ |
| Sau-1151-1 | NZ_JBNB01000002.1 | - | 56893 | 56864 | **RNA**→ rpmE2 (PRK01678)Ribosomal_L31 (pfam01197)→ |
| Sau-1007-1 | NZ_JDZL01000002.1 | - | 54924 | 54895 | **RNA**→ rpmE2 (PRK01678)Ribosomal_L31 (pfam01197)→ |
| Sau-996-1 | NZ_JDZM01000003.1 | - | 56894 | 56865 | **RNA**→ rpmE2 (PRK01678)Ribosomal_L31 (pfam01197)→ |
| Sau-1051-1 | NZ_JFSI01000003.1 | - | 54887 | 54858 | **RNA**→ rpmE2 (PRK01678)Ribosomal_L31 (pfam01197)→ |
| Sau-1012-1 | NZ_JGFD01000019.1 | - | 34972 | 34943 | **RNA**→ rpmE2 (PRK01678)Ribosomal_L31 (pfam01197)→ |
| Sau-979-1 | NZ_JGIB01000018.1 | - | 34873 | 34844 | **RNA**→ rpmE2 (PRK01678)Ribosomal_L31 (pfam01197)→ |
| Sau-1116-1 | NZ_AVPA01000008.1 | + | 55340 | 55369 | **RNA**→ rpmE2 (PRK01678)Ribosomal_L31 (pfam01197)→ |
| Sau-952-1 | NZ_JDGS01000010.1 | - | 56694 | 56665 | **RNA**→ rpmE2 (PRK01678)Ribosomal_L31 (pfam01197)→ |
| Sau-1044-1 | NZ_JFSJ01000001.1 | - | 56794 | 56765 | **RNA**→ rpmE2 (PRK01678)Ribosomal_L31 (pfam01197)→ |
| Sau-951-1 | NZ_JEUI01000002.1 | - | 56994 | 56965 | **RNA**→ rpmE2 (PRK01678)Ribosomal_L31 (pfam01197)→ |
| Sau-1013-1 | NZ_JGFE01000018.1 | - | 35013 | 34984 | **RNA**→ rpmE2 (PRK01678)Ribosomal_L31 (pfam01197)→ |
| Sau-2351-1 | NZ_JDEI01000006.1 | - | 24749 | 24720 | **RNA**→ rpmE2 (PRK01678)Ribosomal_L31 (pfam01197)→ |
| Sau-2256-1 | NZ_JDEN01000007.1 | - | 49944 | 49915 | **RNA**→ rpmE2 (PRK01678)Ribosomal_L31 (pfam01197)→ |
| Sau-2278-1 | NZ_JDEE01000003.1 | - | 215968 | 215939 | **RNA**→ rpmE2 (PRK01678)Ribosomal_L31 (pfam01197)→ |
| Sau-2356-1 | NZ_JDEJ01000004.1 | - | 24749 | 24720 | **RNA**→ rpmE2 (PRK01678)Ribosomal_L31 (pfam01197)→ |
| Sau-2259-1 | NZ_JDEO01000013.1 | - | 35331 | 35302 | **RNA**→ rpmE2 (PRK01678)Ribosomal_L31 (pfam01197)→ |
| Sau-2166-1 | NZ_JEFF01000010.1 | - | 55960 | 55931 | **RNA**→ rpmE2 (PRK01678)Ribosomal_L31 (pfam01197)→ |
| Sau-2120-1 | NZ_JEFK01000013.1 | - | 35321 | 35292 | **RNA**→ rpmE2 (PRK01678)Ribosomal_L31 (pfam01197)→ |
| Sau-2217-1 | NZ_JEFC01000001.1 | - | 217173 | 217144 | **RNA**→ rpmE2 (PRK01678)Ribosomal_L31 (pfam01197)→ |
| Sau-2160-1 | NZ_JEFG01000011.1 | - | 72453 | 72424 | **RNA**→ rpmE2 (PRK01678)Ribosomal_L31 (pfam01197)→ |
| Sau-2156-1 | NZ_JEFH01000015.1 | - | 35809 | 35780 | **RNA**→ rpmE2 (PRK01678)Ribosomal_L31 (pfam01197)→ |
| Sau-2117-1 | NZ_JEFL01000001.1 | - | 56498 | 56469 | **RNA**→ rpmE2 (PRK01678)Ribosomal_L31 (pfam01197)→ |
| Sau-2230-1 | NZ_JEFB01000013.1 | - | 36391 | 36362 | **RNA**→ rpmE2 (PRK01678)Ribosomal_L31 (pfam01197)→ |
| Sau-2281-1 | NZ_JDEF01000001.1 | - | 238570 | 238541 | **RNA**→ rpmE2 (PRK01678)Ribosomal_L31 (pfam01197)→ |
| Sau-2237-1 | NZ_JDEK01000015.1 | - | 35331 | 35302 | **RNA**→ rpmE2 (PRK01678)Ribosomal_L31 (pfam01197)→ |
| Sau-2263-1 | NZ_JDEB01000008.1 | - | 61334 | 61305 | **RNA**→ rpmE2 (PRK01678)Ribosomal_L31 (pfam01197)→ |
| Sau-2295-1 | NZ_JDEG01000006.1 | - | 112279 | 112250 | **RNA**→ rpmE2 (PRK01678)Ribosomal_L31 (pfam01197)→ |
| Sau-2307-1 | NZ_JDEH01000002.1 | - | 236160 | 236131 | **RNA**→ rpmE2 (PRK01678)Ribosomal_L31 (pfam01197)→ |
| Sau-2240-1 | NZ_JDEL01000012.1 | - | 15617 | 15588 | **RNA**→ rpmE2 (PRK01678)Ribosomal_L31 (pfam01197)→ |
| Sau-2255-1 | NZ_JDEM01000010.1 | - | 15617 | 15588 | **RNA**→ rpmE2 (PRK01678)Ribosomal_L31 (pfam01197)→ |

| | | | | | |
|---|---|---|---|---|---|
| Sau-2183-1 | NZ_JEFE01000002.1 | - | 109185 | 109156 | **RNA**→ rpmE2 (PRK01678)Ribosomal_L31 (pfam01197)→ |
| Sau-2125-1 | NZ_JEFJ01000009.1 | - | 36391 | 36362 | **RNA**→ rpmE2 (PRK01678)Ribosomal_L31 (pfam01197)→ |
| Sau-2206-1 | NZ_JEFD01000004.1 | - | 217229 | 217200 | **RNA**→ rpmE2 (PRK01678)Ribosomal_L31 (pfam01197)→ |
| Sau-2135-1 | NZ_JEFI01000012.1 | - | 35661 | 35632 | **RNA**→ rpmE2 (PRK01678)Ribosomal_L31 (pfam01197)→ |
| Sau-2026-1 | NZ_JDDU01000006.1 | - | 58331 | 58302 | **RNA**→ rpmE2 (PRK01678)Ribosomal_L31 (pfam01197)→ |
| Sau-2008-1 | NZ_JDDS01000012.1 | - | 17025 | 16996 | **RNA**→ rpmE2 (PRK01678)Ribosomal_L31 (pfam01197)→ |
| Sau-2010-1 | NZ_JDDT01000010.1 | - | 36391 | 36362 | **RNA**→ rpmE2 (PRK01678)Ribosomal_L31 (pfam01197)→ |
| Sau-2065-1 | NZ_JEFV01000001.1 | - | 24749 | 24720 | **RNA**→ rpmE2 (PRK01678)Ribosomal_L31 (pfam01197)→ |
| Sau-2034-1 | NZ_JEFZ01000009.1 | - | 111533 | 111504 | **RNA**→ rpmE2 (PRK01678)Ribosomal_L31 (pfam01197)→ |
| Sau-2033-1 | NZ_JEGA01000001.1 | - | 117350 | 117321 | **RNA**→ rpmE2 (PRK01678)Ribosomal_L31 (pfam01197)→ |
| Sau-2006-1 | NZ_JDDR01000012.1 | - | 35129 | 35100 | **RNA**→ rpmE2 (PRK01678)Ribosomal_L31 (pfam01197)→ |
| Sau-2029-1 | NZ_JDDW01000011.1 | - | 43765 | 43736 | **RNA**→ rpmE2 (PRK01678)Ribosomal_L31 (pfam01197)→ |
| Sau-2048-1 | NZ_JEFW01000009.1 | - | 115262 | 115233 | **RNA**→ rpmE2 (PRK01678)Ribosomal_L31 (pfam01197)→ |
| Sau-995-1 | NZ_JGHN01000005.1 | - | 56893 | 56864 | **RNA**→ rpmE2 (PRK01678)Ribosomal_L31 (pfam01197)→ |
| Sau-1027-1 | NZ_JGIC01000015.1 | - | 34817 | 34788 | **RNA**→ rpmE2 (PRK01678)Ribosomal_L31 (pfam01197)→ |
| Sau-1090-1 | NZ_JGID01000011.1 | - | 34971 | 34942 | **RNA**→ rpmE2 (PRK01678)Ribosomal_L31 (pfam01197)→ |
| Sau-1137-1 | NZ_JBEM01000001.1 | - | 228947 | 228918 | **RNA**→ rpmE2 (PRK01678)Ribosomal_L31 (pfam01197)→ |
| Sau-1110-1 | NZ_JBMR01000004.1 | - | 56790 | 56761 | **RNA**→ rpmE2 (PRK01678)Ribosomal_L31 (pfam01197)→ |
| Sau-1143-1 | NZ_JBFE01000003.1 | - | 56893 | 56864 | **RNA**→ rpmE2 (PRK01678)Ribosomal_L31 (pfam01197)→ |
| Sau-1093-1 | NZ_JDGA01000018.1 | - | 34873 | 34844 | **RNA**→ rpmE2 (PRK01678)Ribosomal_L31 (pfam01197)→ |
| Sau-1010-1 | NZ_JDZK01000005.1 | - | 56892 | 56863 | **RNA**→ rpmE2 (PRK01678)Ribosomal_L31 (pfam01197)→ |
| Sau-1076-1 | NZ_JFSG01000002.1 | - | 56794 | 56765 | **RNA**→ rpmE2 (PRK01678)Ribosomal_L31 (pfam01197)→ |
| Sau-1058-1 | NZ_JFSH01000001.1 | - | 56894 | 56865 | **RNA**→ rpmE2 (PRK01678)Ribosomal_L31 (pfam01197)→ |
| Sau-938-1 | NZ_JFSL01000006.1 | - | 56794 | 56765 | **RNA**→ rpmE2 (PRK01678)Ribosomal_L31 (pfam01197)→ |
| Sau-942-1 | NZ_JEUF01000007.1 | - | 56894 | 56865 | **RNA**→ rpmE2 (PRK01678)Ribosomal_L31 (pfam01197)→ |
| Sau-1047-1 | NZ_JEVJ01000006.1 | - | 56789 | 56760 | **RNA**→ rpmE2 (PRK01678)Ribosomal_L31 (pfam01197)→ |
| Sau-1049-1 | NZ_JEVK01000016.1 | - | 34913 | 34884 | **RNA**→ rpmE2 (PRK01678)Ribosomal_L31 (pfam01197)→ |
| Sau-1077-1 | NZ_JFSF01000001.1 | - | 56794 | 56765 | **RNA**→ rpmE2 (PRK01678)Ribosomal_L31 (pfam01197)→ |
| Sau-1026-1 | NZ_JFSK01000009.1 | - | 55117 | 55088 | **RNA**→ rpmE2 (PRK01678)Ribosomal_L31 (pfam01197)→ |
| Sau-1025-1 | NZ_JEVI01000008.1 | - | 56894 | 56865 | **RNA**→ rpmE2 (PRK01678)Ribosomal_L31 (pfam01197)→ |
| Sau-1015-1 | NZ_JGFF01000005.1 | - | 56894 | 56865 | **RNA**→ rpmE2 (PRK01678)Ribosomal_L31 (pfam01197)→ |
| Sau-1120-1 | NZ_AZTI01000010.1 | + | 7310 | 7339 | **RNA**→ rpmE2 (PRK01678)Ribosomal_L31 (pfam01197)→ |
| Sau-1105-1 | NZ_JBPF01000002.1 | - | 238686 | 238657 | **RNA**→ rpmE2 (PRK01678)Ribosomal_L31 (pfam01197)→ |
| Sau-1118-1 | NZ_AZTF01000013.1 | - | 54734 | 54705 | **RNA**→ rpmE2 (PRK01678)Ribosomal_L31 (pfam01197)→ |
| Sau-1082-1 | NZ_JDYS01000002.1 | - | 56893 | 56864 | **RNA**→ rpmE2 (PRK01678)Ribosomal_L31 (pfam01197)→ |
| Sau-1068-1 | NZ_JDYT01000002.1 | - | 56894 | 56865 | **RNA**→ rpmE2 (PRK01678)Ribosomal_L31 (pfam01197)→ |
| Sau-1115-1 | NZ_JBKG01000002.1 | - | 56926 | 56897 | **RNA**→ rpmE2 (PRK01678)Ribosomal_L31 (pfam01197)→ |
| Sau-1119-1 | NZ_AZTH01000013.1 | + | 18429 | 18458 | **RNA**→ rpmE2 (PRK01678)Ribosomal_L31 (pfam01197)→ |
| Sau-1104-1 | NZ_JBPX01000004.1 | - | 238686 | 238657 | **RNA**→ rpmE2 (PRK01678)Ribosomal_L31 (pfam01197)→ |
| Sau-1121-1 | NZ_AZTG01000009.1 | - | 54919 | 54890 | **RNA**→ rpmE2 (PRK01678)Ribosomal_L31 (pfam01197)→ |
| Sau-961-1 | NZ_JEUQ01000005.1 | - | 55117 | 55088 | **RNA**→ rpmE2 (PRK01678)Ribosomal_L31 (pfam01197)→ |
| Sau-963-1 | NZ_JEUR01000001.1 | - | 56994 | 56965 | **RNA**→ rpmE2 (PRK01678)Ribosomal_L31 (pfam01197)→ |
| Sau-980-1 | NZ_JEUV01000003.1 | - | 56894 | 56865 | **RNA**→ rpmE2 (PRK01678)Ribosomal_L31 (pfam01197)→ |
| Sau-983-1 | NZ_JEUW01000011.1 | - | 34849 | 34820 | **RNA**→ rpmE2 (PRK01678)Ribosomal_L31 (pfam01197)→ |
| Sau-978-1 | NZ_JGGS01000006.1 | - | 55136 | 55107 | **RNA**→ rpmE2 (PRK01678)Ribosomal_L31 (pfam01197)→ |
| Sau-1069-1 | NZ_JGGT01000012.1 | - | 34913 | 34884 | **RNA**→ rpmE2 (PRK01678)Ribosomal_L31 (pfam01197)→ |
| Sau-1066-1 | NZ_JDYV01000004.1 | - | 56894 | 56865 | **RNA**→ rpmE2 (PRK01678)Ribosomal_L31 (pfam01197)→ |
| Sau-969-1 | NZ_JGGR01000001.1 | - | 56894 | 56865 | **RNA**→ rpmE2 (PRK01678)Ribosomal_L31 (pfam01197)→ |
| Sau-1067-1 | NZ_JDYU01000002.1 | - | 56893 | 56864 | **RNA**→ rpmE2 (PRK01678)Ribosomal_L31 (pfam01197)→ |
| Sau-972-1 | NZ_JEUT01000006.1 | - | 25226 | 25197 | **RNA**→ rpmE2 (PRK01678)Ribosomal_L31 (pfam01197)→ |
| Sau-289-1 | NZ_JBLW01000002.1 | - | 56894 | 56865 | **RNA**→ rpmE2 (PRK01678)Ribosomal_L31 (pfam01197)→ |
| Sau-287-1 | NZ_JBNU01000003.1 | - | 56894 | 56865 | **RNA**→ rpmE2 (PRK01678)Ribosomal_L31 (pfam01197)→ |

| | | | | | |
|---|---|---|---|---|---|
| Sau-282-1 | NZ_JBMG01000007.1 | - | 56893 | 56864 | **RNA**→ rpmE2 (PRK01678)Ribosomal_L31 (pfam01197)→ |
| Sau-237-1 | NZ_JBOA01000007.1 | - | 188315 | 188286 | **RNA**→ rpmE2 (PRK01678)Ribosomal_L31 (pfam01197)→ |
| Sau-310-1 | NZ_JFOR01000024.1 | - | 25129 | 25100 | **RNA**→ rpmE2 (PRK01678)Ribosomal_L31 (pfam01197)→ |
| Sau-249-1 | NZ_JDXI01000001.1 | - | 1674371 | 1674342 | **RNA**→ rpmE2 (PRK01678)Ribosomal_L31 (pfam01197)→ |
| Sau-256-1 | NZ_JIZR01000010.1 | - | 56310 | 56281 | **RNA**→ rpmE2 (PRK01678)Ribosomal_L31 (pfam01197)→ |
| Sau-147-1 | NZ_JOQA01000063.1 | - | 55594 | 55565 | **RNA**→ rpmE2 (PRK01678)Ribosomal_L31 (pfam01197)→ |
| Sau-101-1 | NZ_JOPR01000046.1 | - | 55273 | 55244 | **RNA**→ rpmE2 (PRK01678)Ribosomal_L31 (pfam01197)→ |
| Sau-148-1 | NZ_JOPT01000040.1 | - | 56454 | 56425 | **RNA**→ rpmE2 (PRK01678)Ribosomal_L31 (pfam01197)→ |
| Sau-385-1 | NZ_JFOS01000021.1 | - | 36291 | 36262 | **RNA**→ rpmE2 (PRK01678)Ribosomal_L31 (pfam01197)→ |
| Sau-252-1 | NZ_JIZV01000007.1 | - | 52502 | 52473 | **RNA**→ rpmE2 (PRK01678)Ribosomal_L31 (pfam01197)→ |
| Sau-81-1 | NZ_JJEY01000022.1 | - | 20463 | 20434 | **RNA**→ rpmE2 (PRK01678)Ribosomal_L31 (pfam01197)→ |
| Sau-100-1 | NZ_JOPY01000062.1 | - | 55189 | 55160 | **RNA**→ rpmE2 (PRK01678)Ribosomal_L31 (pfam01197)→ |
| Sau-102-1 | NZ_JOPZ01000029.1 | - | 55106 | 55077 | **RNA**→ rpmE2 (PRK01678)Ribosomal_L31 (pfam01197)→ |
| Sau-285-1 | NZ_JBQY01000001.1 | - | 120363 | 120334 | **RNA**→ rpmE2 (PRK01678)Ribosomal_L31 (pfam01197)→ |
| Sau-244-1 | NZ_JBLJ01000007.1 | - | 61838 | 61809 | **RNA**→ rpmE2 (PRK01678)Ribosomal_L31 (pfam01197)→ |
| Sau-303-1 | NZ_AZMY01000008.1 | - | 54950 | 54921 | **RNA**→ rpmE2 (PRK01678)Ribosomal_L31 (pfam01197)→ |
| Sau-301-1 | NZ_JBFS01000008.1 | - | 62157 | 62128 | **RNA**→ rpmE2 (PRK01678)Ribosomal_L31 (pfam01197)→ |
| Sau-298-1 | NZ_JBHF01000004.1 | - | 56894 | 56865 | **RNA**→ rpmE2 (PRK01678)Ribosomal_L31 (pfam01197)→ |
| Sau-284-1 | NZ_JBRA01000002.1 | - | 56893 | 56864 | **RNA**→ rpmE2 (PRK01678)Ribosomal_L31 (pfam01197)→ |
| Sau-320-1 | NZ_JFOQ01000016.1 | - | 15539 | 15510 | **RNA**→ rpmE2 (PRK01678)Ribosomal_L31 (pfam01197)→ |
| Sau-381-1 | NZ_JGNP01000003.1 | - | 462809 | 462780 | **RNA**→ rpmE2 (PRK01678)Ribosomal_L31 (pfam01197)→ |
| Sau-268-1 | NZ_JIZF01000012.1 | - | 26024 | 25995 | **RNA**→ rpmE2 (PRK01678)Ribosomal_L31 (pfam01197)→ |
| Sau-99-1 | NZ_JOPQ01000036.1 | - | 55216 | 55187 | **RNA**→ rpmE2 (PRK01678)Ribosomal_L31 (pfam01197)→ |
| Sau-338-1 | NZ_JFOO01000014.1 | - | 36224 | 36195 | **RNA**→ rpmE2 (PRK01678)Ribosomal_L31 (pfam01197)→ |
| Sau-327-1 | NZ_JFOP01000014.1 | - | 36391 | 36362 | **RNA**→ rpmE2 (PRK01678)Ribosomal_L31 (pfam01197)→ |
| Sau-210-1 | NZ_JJAH01000015.1 | - | 26024 | 25995 | **RNA**→ rpmE2 (PRK01678)Ribosomal_L31 (pfam01197)→ |
| Sau-149-1 | NZ_JOQB01000066.1 | - | 55222 | 55193 | **RNA**→ rpmE2 (PRK01678)Ribosomal_L31 (pfam01197)→ |
| Sau-223-1 | NZ_JBQV01000005.1 | - | 58248 | 58219 | **RNA**→ rpmE2 (PRK01678)Ribosomal_L31 (pfam01197)→ |
| Sau-224-1 | NZ_JBQD01000005.1 | - | 56994 | 56965 | **RNA**→ rpmE2 (PRK01678)Ribosomal_L31 (pfam01197)→ |
| Sau-93-1 | NZ_JJEM01000015.1 | - | 26024 | 25995 | **RNA**→ rpmE2 (PRK01678)Ribosomal_L31 (pfam01197)→ |
| Sau-95-1 | NZ_JJEK01000015.1 | - | 26024 | 25995 | **RNA**→ rpmE2 (PRK01678)Ribosomal_L31 (pfam01197)→ |
| Sau-82-1 | NZ_JJEX01000015.1 | - | 26024 | 25995 | **RNA**→ rpmE2 (PRK01678)Ribosomal_L31 (pfam01197)→ |
| Sau-192-1 | NZ_JJAZ01000013.1 | - | 26024 | 25995 | **RNA**→ rpmE2 (PRK01678)Ribosomal_L31 (pfam01197)→ |
| Sau-138-1 | NZ_JJCZ01000013.1 | - | 54858 | 54829 | **RNA**→ rpmE2 (PRK01678)Ribosomal_L31 (pfam01197)→ |
| Sau-304-1 | NZ_AZMX01000004.1 | - | 54950 | 54921 | **RNA**→ rpmE2 (PRK01678)Ribosomal_L31 (pfam01197)→ |
| Sau-183-1 | NZ_JJBI01000015.1 | - | 26024 | 25995 | **RNA**→ rpmE2 (PRK01678)Ribosomal_L31 (pfam01197)→ |
| Sau-128-1 | NZ_JJDH01000013.1 | - | 54814 | 54785 | **RNA**→ rpmE2 (PRK01678)Ribosomal_L31 (pfam01197)→ |
| Sau-96-1 | NZ_JJEJ01000010.1 | - | 26024 | 25995 | **RNA**→ rpmE2 (PRK01678)Ribosomal_L31 (pfam01197)→ |
| Sau-204-1 | NZ_JJAN01000014.1 | - | 26024 | 25995 | **RNA**→ rpmE2 (PRK01678)Ribosomal_L31 (pfam01197)→ |
| Sau-195-1 | NZ_JJAW01000011.1 | - | 26024 | 25995 | **RNA**→ rpmE2 (PRK01678)Ribosomal_L31 (pfam01197)→ |
| Sau-143-1 | NZ_JJCS01000014.1 | - | 26024 | 25995 | **RNA**→ rpmE2 (PRK01678)Ribosomal_L31 (pfam01197)→ |
| Sau-97-1 | NZ_JJEI01000012.1 | - | 25790 | 25761 | **RNA**→ rpmE2 (PRK01678)Ribosomal_L31 (pfam01197)→ |
| Sau-86-1 | NZ_JJET01000013.1 | - | 26024 | 25995 | **RNA**→ rpmE2 (PRK01678)Ribosomal_L31 (pfam01197)→ |
| Sau-83-1 | NZ_JJEW01000012.1 | - | 26024 | 25995 | **RNA**→ rpmE2 (PRK01678)Ribosomal_L31 (pfam01197)→ |
| Sau-168-1 | NZ_JJBX01000016.1 | - | 26024 | 25995 | **RNA**→ rpmE2 (PRK01678)Ribosomal_L31 (pfam01197)→ |
| Sau-167-1 | NZ_JJBY01000016.1 | - | 26024 | 25995 | **RNA**→ rpmE2 (PRK01678)Ribosomal_L31 (pfam01197)→ |
| Sau-115-1 | NZ_JJDU01000014.1 | - | 26024 | 25995 | **RNA**→ rpmE2 (PRK01678)Ribosomal_L31 (pfam01197)→ |
| Sau-3592-1 | NZ_JEUH01000005.1 | - | 56549 | 56520 | **RNA**→ rpmE2 (PRK01678)Ribosomal_L31 (pfam01197)→ |
| Sau-3598-1 | NZ_JEUL01000008.1 | - | 56993 | 56964 | **RNA**→ rpmE2 (PRK01678)Ribosomal_L31 (pfam01197)→ |
| Sau-3623-1 | NZ_JGGC01000005.1 | - | 34873 | 34844 | **RNA**→ rpmE2 (PRK01678)Ribosomal_L31 (pfam01197)→ |
| Sau-3639-1 | NZ_JGGD01000007.1 | - | 56894 | 56865 | **RNA**→ rpmE2 (PRK01678)Ribosomal_L31 (pfam01197)→ |
| Sau-3636-1 | NZ_JGHW01000008.1 | - | 55000 | 54971 | **RNA**→ rpmE2 (PRK01678)Ribosomal_L31 (pfam01197)→ |

| | | | | | |
|---|---|---|---|---|---|
| Sau-3565-1 | NZ_JGIG01000013.1 | - | 35013 | 34984 | **RNA**→ rpmE2 (PRK01678)Ribosomal_L31 (pfam01197)→ |
| Sau-3473-1 | NZ_JIFS01000014.1 | - | 26024 | 25995 | **RNA**→ rpmE2 (PRK01678)Ribosomal_L31 (pfam01197)→ |
| Sau-3457-1 | NZ_JIGI01000014.1 | - | 25997 | 25968 | **RNA**→ rpmE2 (PRK01678)Ribosomal_L31 (pfam01197)→ |
| Sau-3455-1 | NZ_JIGK01000015.1 | - | 25968 | 25939 | **RNA**→ rpmE2 (PRK01678)Ribosomal_L31 (pfam01197)→ |
| Sau-3419-1 | NZ_JIHU01000011.1 | - | 26024 | 25995 | **RNA**→ rpmE2 (PRK01678)Ribosomal_L31 (pfam01197)→ |
| Sau-3414-1 | NZ_JIHZ01000011.1 | - | 26024 | 25995 | **RNA**→ rpmE2 (PRK01678)Ribosomal_L31 (pfam01197)→ |
| Sau-3621-1 | NZ_JDZC01000002.1 | - | 56894 | 56865 | **RNA**→ rpmE2 (PRK01678)Ribosomal_L31 (pfam01197)→ |
| Sau-3600-1 | NZ_JDZD01000004.1 | - | 56894 | 56865 | **RNA**→ rpmE2 (PRK01678)Ribosomal_L31 (pfam01197)→ |
| Sau-3564-1 | NZ_JFRU01000011.1 | - | 34913 | 34884 | **RNA**→ rpmE2 (PRK01678)Ribosomal_L31 (pfam01197)→ |
| Sau-3615-1 | NZ_JEVR01000008.1 | - | 55017 | 54988 | **RNA**→ rpmE2 (PRK01678)Ribosomal_L31 (pfam01197)→ |
| Sau-3529-1 | NZ_JGFO01000005.1 | - | 34873 | 34844 | **RNA**→ rpmE2 (PRK01678)Ribosomal_L31 (pfam01197)→ |
| Sau-3563-1 | NZ_JGFP01000010.1 | - | 27857 | 27828 | **RNA**→ rpmE2 (PRK01678)Ribosomal_L31 (pfam01197)→ |
| Sau-3637-1 | NZ_JGHX01000008.1 | - | 56894 | 56865 | **RNA**→ rpmE2 (PRK01678)Ribosomal_L31 (pfam01197)→ |
| Sau-3471-1 | NZ_JIFU01000012.1 | - | 26024 | 25995 | **RNA**→ rpmE2 (PRK01678)Ribosomal_L31 (pfam01197)→ |
| Sau-3446-1 | NZ_JIGT01000015.1 | - | 26024 | 25995 | **RNA**→ rpmE2 (PRK01678)Ribosomal_L31 (pfam01197)→ |
| Sau-3400-1 | NZ_JIIN01000014.1 | - | 26024 | 25995 | **RNA**→ rpmE2 (PRK01678)Ribosomal_L31 (pfam01197)→ |
| Sau-3479-1 | NZ_JDGG01000002.1 | - | 56894 | 56865 | **RNA**→ rpmE2 (PRK01678)Ribosomal_L31 (pfam01197)→ |
| Sau-3483-1 | NZ_JFRW01000003.1 | - | 54900 | 54871 | **RNA**→ rpmE2 (PRK01678)Ribosomal_L31 (pfam01197)→ |
| Sau-3590-1 | NZ_JEUK01000012.1 | - | 56894 | 56865 | **RNA**→ rpmE2 (PRK01678)Ribosomal_L31 (pfam01197)→ |
| Sau-3480-1 | NZ_JGFM01000002.1 | - | 56994 | 56965 | **RNA**→ rpmE2 (PRK01678)Ribosomal_L31 (pfam01197)→ |
| Sau-3622-1 | NZ_JGGB01000014.1 | - | 56893 | 56864 | **RNA**→ rpmE2 (PRK01678)Ribosomal_L31 (pfam01197)→ |
| Sau-3634-1 | NZ_JGHV01000012.1 | - | 34913 | 34884 | **RNA**→ rpmE2 (PRK01678)Ribosomal_L31 (pfam01197)→ |
| Sau-3517-1 | NZ_JGIF01000026.1 | - | 18838 | 18809 | **RNA**→ rpmE2 (PRK01678)Ribosomal_L31 (pfam01197)→ |
| Sau-3485-1 | NZ_JGMY01000009.1 | - | 56790 | 56761 | **RNA**→ rpmE2 (PRK01678)Ribosomal_L31 (pfam01197)→ |
| Sau-3463-1 | NZ_JIGC01000014.1 | - | 26024 | 25995 | **RNA**→ rpmE2 (PRK01678)Ribosomal_L31 (pfam01197)→ |
| Sau-3433-1 | NZ_JIHG01000011.1 | - | 26024 | 25995 | **RNA**→ rpmE2 (PRK01678)Ribosomal_L31 (pfam01197)→ |
| Sau-3528-1 | NZ_JDFZ01000009.1 | - | 56694 | 56665 | **RNA**→ rpmE2 (PRK01678)Ribosomal_L31 (pfam01197)→ |
| Sau-3501-1 | NZ_JFRV01000012.1 | - | 34962 | 34933 | **RNA**→ rpmE2 (PRK01678)Ribosomal_L31 (pfam01197)→ |
| Sau-3633-1 | NZ_JEVS01000016.1 | - | 34913 | 34884 | **RNA**→ rpmE2 (PRK01678)Ribosomal_L31 (pfam01197)→ |
| Sau-3512-1 | NZ_JGHT01000010.1 | - | 55117 | 55088 | **RNA**→ rpmE2 (PRK01678)Ribosomal_L31 (pfam01197)→ |
| Sau-3550-1 | NZ_JGHU01000010.1 | - | 56790 | 56761 | **RNA**→ rpmE2 (PRK01678)Ribosomal_L31 (pfam01197)→ |
| Sau-3466-1 | NZ_JIFZ01000014.1 | - | 26024 | 25995 | **RNA**→ rpmE2 (PRK01678)Ribosomal_L31 (pfam01197)→ |
| Sau-3441-1 | NZ_JIGY01000015.1 | - | 26024 | 25995 | **RNA**→ rpmE2 (PRK01678)Ribosomal_L31 (pfam01197)→ |
| Sau-3553-1 | NZ_JGHD01000011.1 | - | 34872 | 34843 | **RNA**→ rpmE2 (PRK01678)Ribosomal_L31 (pfam01197)→ |
| Sau-3541-1 | NZ_JGHC01000005.1 | - | 56894 | 56865 | **RNA**→ rpmE2 (PRK01678)Ribosomal_L31 (pfam01197)→ |
| Sau-3523-1 | NZ_JDYM01000011.1 | - | 56793 | 56764 | **RNA**→ rpmE2 (PRK01678)Ribosomal_L31 (pfam01197)→ |
| Sau-3498-1 | NZ_JGHB01000006.1 | - | 55218 | 55189 | **RNA**→ rpmE2 (PRK01678)Ribosomal_L31 (pfam01197)→ |
| Sau-3580-1 | NZ_JGHF01000009.1 | - | 55116 | 55087 | **RNA**→ rpmE2 (PRK01678)Ribosomal_L31 (pfam01197)→ |
| Sau-3547-1 | NZ_JDYL01000002.1 | - | 56893 | 56864 | **RNA**→ rpmE2 (PRK01678)Ribosomal_L31 (pfam01197)→ |
| Sau-3603-1 | NZ_JEUP01000007.1 | - | 56894 | 56865 | **RNA**→ rpmE2 (PRK01678)Ribosomal_L31 (pfam01197)→ |
| Sau-3611-1 | NZ_JEUU01000002.1 | - | 56894 | 56865 | **RNA**→ rpmE2 (PRK01678)Ribosomal_L31 (pfam01197)→ |
| Sau-3394-1 | NZ_JIIT01000011.1 | - | 26024 | 25995 | **RNA**→ rpmE2 (PRK01678)Ribosomal_L31 (pfam01197)→ |
| Sau-3392-1 | NZ_JIIV01000015.1 | - | 54582 | 54553 | **RNA**→ rpmE2 (PRK01678)Ribosomal_L31 (pfam01197)→ |
| Sau-3555-1 | NZ_JGHE01000006.1 | - | 56893 | 56864 | **RNA**→ rpmE2 (PRK01678)Ribosomal_L31 (pfam01197)→ |
| Sau-3395-1 | NZ_JIIS01000014.1 | - | 26024 | 25995 | **RNA**→ rpmE2 (PRK01678)Ribosomal_L31 (pfam01197)→ |
| Sau-612-1 | NZ_JDGR01000007.1 | - | 56894 | 56865 | **RNA**→ rpmE2 (PRK01678)Ribosomal_L31 (pfam01197)→ |
| Sau-637-1 | NZ_JDZQ01000002.1 | - | 56894 | 56865 | **RNA**→ rpmE2 (PRK01678)Ribosomal_L31 (pfam01197)→ |
| Sau-632-1 | NZ_JDZR01000002.1 | - | 56993 | 56964 | **RNA**→ rpmE2 (PRK01678)Ribosomal_L31 (pfam01197)→ |
| Sau-543-1 | NZ_JFQL01000019.1 | - | 36391 | 36362 | **RNA**→ rpmE2 (PRK01678)Ribosomal_L31 (pfam01197)→ |
| Sau-548-1 | NZ_JFQQ01000018.1 | - | 36391 | 36362 | **RNA**→ rpmE2 (PRK01678)Ribosomal_L31 (pfam01197)→ |
| Sau-555-1 | NZ_JFQV01000017.1 | - | 36392 | 36363 | **RNA**→ rpmE2 (PRK01678)Ribosomal_L31 (pfam01197)→ |
| Sau-561-1 | NZ_JFRA01000020.1 | - | 36391 | 36362 | **RNA**→ rpmE2 (PRK01678)Ribosomal_L31 (pfam01197)→ |

| | | | | | |
|---|---|---|---|---|---|
| Sau-568-1 | NZ_JFRF01000018.1 | - | 36391 | 36362 | **RNA**→ rpmE2 (PRK01678)Ribosomal_L31 (pfam01197)→ |
| Sau-574-1 | NZ_JFRK01000024.1 | - | 28808 | 28779 | **RNA**→ rpmE2 (PRK01678)Ribosomal_L31 (pfam01197)→ |
| Sau-620-1 | NZ_JGEY01000017.1 | - | 34972 | 34943 | **RNA**→ rpmE2 (PRK01678)Ribosomal_L31 (pfam01197)→ |
| Sau-631-1 | NZ_JGFS01000010.1 | - | 34973 | 34944 | **RNA**→ rpmE2 (PRK01678)Ribosomal_L31 (pfam01197)→ |
| Sau-665-1 | NZ_JGGH01000012.1 | - | 34971 | 34942 | **RNA**→ rpmE2 (PRK01678)Ribosomal_L31 (pfam01197)→ |
| Sau-684-1 | NZ_JGGI01000011.1 | - | 34817 | 34788 | **RNA**→ rpmE2 (PRK01678)Ribosomal_L31 (pfam01197)→ |
| Sau-666-1 | NZ_JGHH01000012.1 | - | 34872 | 34843 | **RNA**→ rpmE2 (PRK01678)Ribosomal_L31 (pfam01197)→ |
| Sau-654-1 | NZ_JDGN01000011.1 | - | 56894 | 56865 | **RNA**→ rpmE2 (PRK01678)Ribosomal_L31 (pfam01197)→ |
| Sau-669-1 | NZ_JFSO01000007.1 | - | 56894 | 56865 | **RNA**→ rpmE2 (PRK01678)Ribosomal_L31 (pfam01197)→ |
| Sau-544-1 | NZ_JFQM01000014.1 | - | 36385 | 36356 | **RNA**→ rpmE2 (PRK01678)Ribosomal_L31 (pfam01197)→ |
| Sau-550-1 | NZ_JFQR01000016.1 | - | 36391 | 36362 | **RNA**→ rpmE2 (PRK01678)Ribosomal_L31 (pfam01197)→ |
| Sau-556-1 | NZ_JFQW01000017.1 | - | 36280 | 36251 | **RNA**→ rpmE2 (PRK01678)Ribosomal_L31 (pfam01197)→ |
| Sau-563-1 | NZ_JFRB01000017.1 | - | 36391 | 36362 | **RNA**→ rpmE2 (PRK01678)Ribosomal_L31 (pfam01197)→ |
| Sau-569-1 | NZ_JFRG01000017.1 | - | 36391 | 36362 | **RNA**→ rpmE2 (PRK01678)Ribosomal_L31 (pfam01197)→ |
| Sau-623-1 | NZ_JGEZ01000001.1 | - | 56894 | 56865 | **RNA**→ rpmE2 (PRK01678)Ribosomal_L31 (pfam01197)→ |
| Sau-638-1 | NZ_JDZP01000004.1 | - | 235328 | 235299 | **RNA**→ rpmE2 (PRK01678)Ribosomal_L31 (pfam01197)→ |
| Sau-541-1 | NZ_JFQJ01000018.1 | - | 36391 | 36362 | **RNA**→ rpmE2 (PRK01678)Ribosomal_L31 (pfam01197)→ |
| Sau-542-1 | NZ_JFQK01000019.1 | - | 36391 | 36362 | **RNA**→ rpmE2 (PRK01678)Ribosomal_L31 (pfam01197)→ |
| Sau-546-1 | NZ_JFQO01000019.1 | - | 36391 | 36362 | **RNA**→ rpmE2 (PRK01678)Ribosomal_L31 (pfam01197)→ |
| Sau-547-1 | NZ_JFQP01000017.1 | - | 35397 | 35368 | **RNA**→ rpmE2 (PRK01678)Ribosomal_L31 (pfam01197)→ |
| Sau-553-1 | NZ_JFQT01000018.1 | - | 35225 | 35196 | **RNA**→ rpmE2 (PRK01678)Ribosomal_L31 (pfam01197)→ |
| Sau-554-1 | NZ_JFQU01000017.1 | - | 36391 | 36362 | **RNA**→ rpmE2 (PRK01678)Ribosomal_L31 (pfam01197)→ |
| Sau-560-1 | NZ_JFQZ01000016.1 | - | 36391 | 36362 | **RNA**→ rpmE2 (PRK01678)Ribosomal_L31 (pfam01197)→ |
| Sau-566-1 | NZ_JFRE01000022.1 | - | 18347 | 18318 | **RNA**→ rpmE2 (PRK01678)Ribosomal_L31 (pfam01197)→ |
| Sau-572-1 | NZ_JFRJ01000015.1 | - | 36391 | 36362 | **RNA**→ rpmE2 (PRK01678)Ribosomal_L31 (pfam01197)→ |
| Sau-616-1 | NZ_JGEX01000037.1 | - | 4832 | 4803 | **RNA**→ rpmE2 (PRK01678)Ribosomal_L31 (pfam01197)→ |
| Sau-657-1 | NZ_JGFC01000011.1 | - | 32964 | 32935 | **RNA**→ rpmE2 (PRK01678)Ribosomal_L31 (pfam01197)→ |
| Sau-611-1 | NZ_JGFR01000013.1 | - | 34817 | 34788 | **RNA**→ rpmE2 (PRK01678)Ribosomal_L31 (pfam01197)→ |
| Sau-655-1 | NZ_JDGO01000005.1 | - | 56893 | 56864 | **RNA**→ rpmE2 (PRK01678)Ribosomal_L31 (pfam01197)→ |
| Sau-650-1 | NZ_JDZO01000001.1 | - | 56894 | 56865 | **RNA**→ rpmE2 (PRK01678)Ribosomal_L31 (pfam01197)→ |
| Sau-545-1 | NZ_JFQN01000020.1 | - | 36391 | 36362 | **RNA**→ rpmE2 (PRK01678)Ribosomal_L31 (pfam01197)→ |
| Sau-552-1 | NZ_JFQS01000032.1 | - | 15721 | 15692 | **RNA**→ rpmE2 (PRK01678)Ribosomal_L31 (pfam01197)→ |
| Sau-557-1 | NZ_JFQX01000016.1 | - | 36391 | 36362 | **RNA**→ rpmE2 (PRK01678)Ribosomal_L31 (pfam01197)→ |
| Sau-558-1 | NZ_JFQY01000018.1 | - | 36391 | 36362 | **RNA**→ rpmE2 (PRK01678)Ribosomal_L31 (pfam01197)→ |
| Sau-564-1 | NZ_JFRC01000016.1 | - | 36391 | 36362 | **RNA**→ rpmE2 (PRK01678)Ribosomal_L31 (pfam01197)→ |
| Sau-565-1 | NZ_JFRD01000015.1 | - | 36391 | 36362 | **RNA**→ rpmE2 (PRK01678)Ribosomal_L31 (pfam01197)→ |
| Sau-570-1 | NZ_JFRH01000018.1 | - | 36391 | 36362 | **RNA**→ rpmE2 (PRK01678)Ribosomal_L31 (pfam01197)→ |
| Sau-571-1 | NZ_JFRI01000019.1 | - | 36391 | 36362 | **RNA**→ rpmE2 (PRK01678)Ribosomal_L31 (pfam01197)→ |
| Sau-629-1 | NZ_JGFA01000016.1 | - | 34872 | 34843 | **RNA**→ rpmE2 (PRK01678)Ribosomal_L31 (pfam01197)→ |
| Sau-630-1 | NZ_JGFB01000026.1 | - | 19182 | 19153 | **RNA**→ rpmE2 (PRK01678)Ribosomal_L31 (pfam01197)→ |
| Sau-610-1 | NZ_JGFQ01000023.1 | - | 34972 | 34943 | **RNA**→ rpmE2 (PRK01678)Ribosomal_L31 (pfam01197)→ |
| Sau-672-1 | NZ_JDYY01000005.1 | - | 56894 | 56865 | **RNA**→ rpmE2 (PRK01678)Ribosomal_L31 (pfam01197)→ |
| Sau-662-1 | NZ_JGHG01000005.1 | - | 56894 | 56865 | **RNA**→ rpmE2 (PRK01678)Ribosomal_L31 (pfam01197)→ |
| Sau-2272-1 | NZ_JDEC01000003.1 | - | 24747 | 24718 | **RNA**→ rpmE2 (PRK01678)Ribosomal_L31 (pfam01197)→ |
| Sau-2277-1 | NZ_JDED01000012.1 | - | 55865 | 55836 | **RNA**→ rpmE2 (PRK01678)Ribosomal_L31 (pfam01197)→ |
| Sau-1996-1 | NZ_JDDO01000011.1 | - | 36390 | 36361 | **RNA**→ rpmE2 (PRK01678)Ribosomal_L31 (pfam01197)→ |
| Sau-1998-1 | NZ_JDDP01000008.1 | - | 24749 | 24720 | **RNA**→ rpmE2 (PRK01678)Ribosomal_L31 (pfam01197)→ |
| Sau-2001-1 | NZ_JDDQ01000007.1 | - | 24749 | 24720 | **RNA**→ rpmE2 (PRK01678)Ribosomal_L31 (pfam01197)→ |
| Sau-2028-1 | NZ_JDDV01000012.1 | - | 55495 | 55466 | **RNA**→ rpmE2 (PRK01678)Ribosomal_L31 (pfam01197)→ |
| Sau-2097-1 | NZ_JEFP01000012.1 | - | 14444 | 14415 | **RNA**→ rpmE2 (PRK01678)Ribosomal_L31 (pfam01197)→ |
| Sau-2083-1 | NZ_JEFQ01000002.1 | - | 217273 | 217244 | **RNA**→ rpmE2 (PRK01678)Ribosomal_L31 (pfam01197)→ |
| Sau-2067-1 | NZ_JEFU01000008.1 | - | 113736 | 113707 | **RNA**→ rpmE2 (PRK01678)Ribosomal_L31 (pfam01197)→ |

| | | | | | |
|---|---|---|---|---|---|
| Sau-2204-1 | NZ_JDYR01000002.1 | - | 56893 | 56864 | **RNA**→ rpmE2 (PRK01678)Ribosomal_L31 (pfam01197)→ |
| Sau-2109-1 | NZ_JEFM01000011.1 | - | 36391 | 36362 | **RNA**→ rpmE2 (PRK01678)Ribosomal_L31 (pfam01197)→ |
| Sau-2079-1 | NZ_JEFR01000002.1 | - | 139075 | 139046 | **RNA**→ rpmE2 (PRK01678)Ribosomal_L31 (pfam01197)→ |
| Sau-2038-1 | NZ_JEFY01000008.1 | - | 61496 | 61467 | **RNA**→ rpmE2 (PRK01678)Ribosomal_L31 (pfam01197)→ |
| Sau-2098-1 | NZ_JEFO01000010.1 | - | 43610 | 43581 | **RNA**→ rpmE2 (PRK01678)Ribosomal_L31 (pfam01197)→ |
| Sau-2072-1 | NZ_JEFT01000011.1 | - | 56148 | 56119 | **RNA**→ rpmE2 (PRK01678)Ribosomal_L31 (pfam01197)→ |
| Sau-2042-1 | NZ_JEFX01000012.1 | - | 36390 | 36361 | **RNA**→ rpmE2 (PRK01678)Ribosomal_L31 (pfam01197)→ |
| Sau-2108-1 | NZ_JEFN01000005.1 | - | 59118 | 59089 | **RNA**→ rpmE2 (PRK01678)Ribosomal_L31 (pfam01197)→ |
| Sau-2076-1 | NZ_JEFS01000016.1 | - | 25227 | 25198 | **RNA**→ rpmE2 (PRK01678)Ribosomal_L31 (pfam01197)→ |
| Sau-3351-1 | NZ_JIKK01000013.1 | - | 26024 | 25995 | **RNA**→ rpmE2 (PRK01678)Ribosomal_L31 (pfam01197)→ |
| Sau-3326-1 | NZ_JILJ01000008.1 | - | 56390 | 56361 | **RNA**→ rpmE2 (PRK01678)Ribosomal_L31 (pfam01197)→ |
| Sau-3299-1 | NZ_JIMK01000012.1 | - | 26024 | 25995 | **RNA**→ rpmE2 (PRK01678)Ribosomal_L31 (pfam01197)→ |
| Sau-1388-1 | NZ_JBUH01000001.1 | - | 231344 | 231315 | **RNA**→ rpmE2 (PRK01678)Ribosomal_L31 (pfam01197)→ |
| Sau-3271-1 | NZ_JINM01000015.1 | - | 26024 | 25995 | **RNA**→ rpmE2 (PRK01678)Ribosomal_L31 (pfam01197)→ |
| Sau-1423-1 | NZ_JBUM01000011.1 | - | 81129 | 81100 | **RNA**→ rpmE2 (PRK01678)Ribosomal_L31 (pfam01197)→ |
| Sau-3237-1 | NZ_JIOU01000001.1 | - | 56224 | 56195 | **RNA**→ rpmE2 (PRK01678)Ribosomal_L31 (pfam01197)→ |
| Sau-1465-1 | NZ_JBUR01000008.1 | - | 61690 | 61661 | **RNA**→ rpmE2 (PRK01678)Ribosomal_L31 (pfam01197)→ |
| Sau-3193-1 | NZ_JIQM01000010.1 | - | 26024 | 25995 | **RNA**→ rpmE2 (PRK01678)Ribosomal_L31 (pfam01197)→ |
| Sau-1508-1 | NZ_JBUW01000015.1 | - | 35370 | 35341 | **RNA**→ rpmE2 (PRK01678)Ribosomal_L31 (pfam01197)→ |
| Sau-3367-1 | NZ_JIJU01000014.1 | - | 26024 | 25995 | **RNA**→ rpmE2 (PRK01678)Ribosomal_L31 (pfam01197)→ |
| Sau-1555-1 | NZ_JBVB01000007.1 | - | 111475 | 111446 | **RNA**→ rpmE2 (PRK01678)Ribosomal_L31 (pfam01197)→ |
| Sau-3350-1 | NZ_JIKL01000010.1 | - | 56354 | 56325 | **RNA**→ rpmE2 (PRK01678)Ribosomal_L31 (pfam01197)→ |
| Sau-1393-1 | NZ_JBUI01000001.1 | - | 25185 | 25156 | **RNA**→ rpmE2 (PRK01678)Ribosomal_L31 (pfam01197)→ |
| Sau-3318-1 | NZ_JILR01000013.1 | - | 56439 | 56410 | **RNA**→ rpmE2 (PRK01678)Ribosomal_L31 (pfam01197)→ |
| Sau-3283-1 | NZ_JINA01000007.1 | - | 18519 | 18490 | **RNA**→ rpmE2 (PRK01678)Ribosomal_L31 (pfam01197)→ |
| Sau-1431-1 | NZ_JBUN01000009.1 | - | 7477 | 7448 | **RNA**→ rpmE2 (PRK01678)Ribosomal_L31 (pfam01197)→ |
| Sau-3270-1 | NZ_JINN01000013.1 | - | 26024 | 25995 | **RNA**→ rpmE2 (PRK01678)Ribosomal_L31 (pfam01197)→ |
| Sau-1480-1 | NZ_JBUS01000012.1 | - | 36391 | 36362 | **RNA**→ rpmE2 (PRK01678)Ribosomal_L31 (pfam01197)→ |
| Sau-3234-1 | NZ_JIOX01000011.1 | - | 26024 | 25995 | **RNA**→ rpmE2 (PRK01678)Ribosomal_L31 (pfam01197)→ |
| Sau-1513-1 | NZ_JBUX01000008.1 | - | 109368 | 109339 | **RNA**→ rpmE2 (PRK01678)Ribosomal_L31 (pfam01197)→ |
| Sau-3229-1 | NZ_JIPC01000010.1 | - | 26024 | 25995 | **RNA**→ rpmE2 (PRK01678)Ribosomal_L31 (pfam01197)→ |
| Sau-1558-1 | NZ_JBVC01000008.1 | - | 25029 | 25000 | **RNA**→ rpmE2 (PRK01678)Ribosomal_L31 (pfam01197)→ |
| Sau-3192-1 | NZ_JIQN01000013.1 | - | 23148 | 23119 | **RNA**→ rpmE2 (PRK01678)Ribosomal_L31 (pfam01197)→ |
| Sau-3190-1 | NZ_JIQP01000009.1 | - | 26024 | 25995 | **RNA**→ rpmE2 (PRK01678)Ribosomal_L31 (pfam01197)→ |
| Sau-1158-1 | NZ_JBIB01000002.1 | - | 54889 | 54860 | **RNA**→ rpmE2 (PRK01678)Ribosomal_L31 (pfam01197)→ |
| Sau-3358-1 | NZ_JIKD01000012.1 | - | 26024 | 25995 | **RNA**→ rpmE2 (PRK01678)Ribosomal_L31 (pfam01197)→ |
| Sau-3352-1 | NZ_JIKJ01000007.1 | - | 56354 | 56325 | **RNA**→ rpmE2 (PRK01678)Ribosomal_L31 (pfam01197)→ |
| Sau-1172-1 | NZ_JOPW01000064.1 | - | 55306 | 55277 | **RNA**→ rpmE2 (PRK01678)Ribosomal_L31 (pfam01197)→ |
| Sau-3346-1 | NZ_JIKP01000012.1 | - | 26024 | 25995 | **RNA**→ rpmE2 (PRK01678)Ribosomal_L31 (pfam01197)→ |
| Sau-1176-1 | NZ_JOPN01000049.1 | - | 55239 | 55210 | **RNA**→ rpmE2 (PRK01678)Ribosomal_L31 (pfam01197)→ |
| Sau-3327-1 | NZ_JILI01000015.1 | - | 26024 | 25995 | **RNA**→ rpmE2 (PRK01678)Ribosomal_L31 (pfam01197)→ |
| Sau-1175-1 | NZ_JOPM01000079.1 | - | 48479 | 48450 | **RNA**→ rpmE2 (PRK01678)Ribosomal_L31 (pfam01197)→ |
| Sau-3309-1 | NZ_JIMA01000009.1 | - | 26024 | 25995 | **RNA**→ rpmE2 (PRK01678)Ribosomal_L31 (pfam01197)→ |
| Sau-3273-1 | NZ_JINK01000012.1 | - | 26024 | 25995 | **RNA**→ rpmE2 (PRK01678)Ribosomal_L31 (pfam01197)→ |
| Sau-1403-1 | NZ_JBUJ01000008.1 | - | 58248 | 58219 | **RNA**→ rpmE2 (PRK01678)Ribosomal_L31 (pfam01197)→ |
| Sau-3242-1 | NZ_JIOP01000015.1 | - | 26024 | 25995 | **RNA**→ rpmE2 (PRK01678)Ribosomal_L31 (pfam01197)→ |
| Sau-1452-1 | NZ_JBUO01000008.1 | - | 111533 | 111504 | **RNA**→ rpmE2 (PRK01678)Ribosomal_L31 (pfam01197)→ |
| Sau-3198-1 | NZ_JIQH01000007.1 | - | 56439 | 56410 | **RNA**→ rpmE2 (PRK01678)Ribosomal_L31 (pfam01197)→ |
| Sau-1482-1 | NZ_JBUT01000007.1 | - | 49861 | 49832 | **RNA**→ rpmE2 (PRK01678)Ribosomal_L31 (pfam01197)→ |
| Sau-3361-1 | NZ_JIKA01000011.1 | - | 26024 | 25995 | **RNA**→ rpmE2 (PRK01678)Ribosomal_L31 (pfam01197)→ |
| Sau-1521-1 | NZ_JBUY01000010.1 | - | 62303 | 62274 | **RNA**→ rpmE2 (PRK01678)Ribosomal_L31 (pfam01197)→ |
| Sau-3348-1 | NZ_JIKN01000014.1 | - | 25790 | 25761 | **RNA**→ rpmE2 (PRK01678)Ribosomal_L31 (pfam01197)→ |

| | | | | | |
|---|---|---|---|---|---|
| Sau-3316-1 | NZ_JILT01000015.1 | - | 26023 | 25994 | **RNA**→ rpmE2 (PRK01678)Ribosomal_L31 (pfam01197)→ |
| Sau-1533-1 | NZ_JBUZ01000007.1 | - | 111873 | 111844 | **RNA**→ rpmE2 (PRK01678)Ribosomal_L31 (pfam01197)→ |
| Sau-3314-1 | NZ_JILV01000006.1 | - | 56352 | 56323 | **RNA**→ rpmE2 (PRK01678)Ribosomal_L31 (pfam01197)→ |
| Sau-1560-1 | NZ_JBVD01000009.1 | - | 62053 | 62024 | **RNA**→ rpmE2 (PRK01678)Ribosomal_L31 (pfam01197)→ |
| Sau-3279-1 | NZ_JINE01000013.1 | - | 26024 | 25995 | **RNA**→ rpmE2 (PRK01678)Ribosomal_L31 (pfam01197)→ |
| Sau-1583-1 | NZ_JBVE01000001.1 | - | 49963 | 49934 | **RNA**→ rpmE2 (PRK01678)Ribosomal_L31 (pfam01197)→ |
| Sau-3276-1 | NZ_JINH01000013.1 | - | 26024 | 25995 | **RNA**→ rpmE2 (PRK01678)Ribosomal_L31 (pfam01197)→ |
| Sau-1409-1 | NZ_JBUK01000012.1 | - | 25998 | 25969 | **RNA**→ rpmE2 (PRK01678)Ribosomal_L31 (pfam01197)→ |
| Sau-3261-1 | NZ_JINW01000011.1 | - | 26024 | 25995 | **RNA**→ rpmE2 (PRK01678)Ribosomal_L31 (pfam01197)→ |
| Sau-1415-1 | NZ_JBUL01000002.1 | - | 217273 | 217244 | **RNA**→ rpmE2 (PRK01678)Ribosomal_L31 (pfam01197)→ |
| Sau-3253-1 | NZ_JIOE01000008.1 | - | 56354 | 56325 | **RNA**→ rpmE2 (PRK01678)Ribosomal_L31 (pfam01197)→ |
| Sau-3201-1 | NZ_JIQE01000003.1 | - | 56305 | 56276 | **RNA**→ rpmE2 (PRK01678)Ribosomal_L31 (pfam01197)→ |
| Sau-1456-1 | NZ_JBUP01000010.1 | - | 56229 | 56200 | **RNA**→ rpmE2 (PRK01678)Ribosomal_L31 (pfam01197)→ |
| Sau-1462-1 | NZ_JBUQ01000009.1 | - | 110386 | 110357 | **RNA**→ rpmE2 (PRK01678)Ribosomal_L31 (pfam01197)→ |
| Sau-3126-1 | NZ_JITB01000012.1 | - | 26024 | 25995 | **RNA**→ rpmE2 (PRK01678)Ribosomal_L31 (pfam01197)→ |
| Sau-1484-1 | NZ_JBUU01000002.1 | - | 61690 | 61661 | **RNA**→ rpmE2 (PRK01678)Ribosomal_L31 (pfam01197)→ |
| Sau-3138-1 | NZ_JISP01000015.1 | - | 26024 | 25995 | **RNA**→ rpmE2 (PRK01678)Ribosomal_L31 (pfam01197)→ |
| Sau-1493-1 | NZ_JBUV01000006.1 | - | 117314 | 117285 | **RNA**→ rpmE2 (PRK01678)Ribosomal_L31 (pfam01197)→ |
| Sau-3129-1 | NZ_JISY01000012.1 | - | 56390 | 56361 | **RNA**→ rpmE2 (PRK01678)Ribosomal_L31 (pfam01197)→ |
| Sau-1551-1 | NZ_JBVA01000011.1 | - | 56828 | 56799 | **RNA**→ rpmE2 (PRK01678)Ribosomal_L31 (pfam01197)→ |
| Sau-3391-1 | NZ_JIIW01000010.1 | - | 54536 | 54507 | **RNA**→ rpmE2 (PRK01678)Ribosomal_L31 (pfam01197)→ |
| Sau-1604-1 | NZ_JBVF01000006.1 | - | 112234 | 112205 | **RNA**→ rpmE2 (PRK01678)Ribosomal_L31 (pfam01197)→ |
| Sau-3173-1 | NZ_JIRG01000010.1 | - | 26024 | 25995 | **RNA**→ rpmE2 (PRK01678)Ribosomal_L31 (pfam01197)→ |
| Sau-3149-1 | NZ_JISE01000012.1 | - | 26024 | 25995 | **RNA**→ rpmE2 (PRK01678)Ribosomal_L31 (pfam01197)→ |
| Sau-3386-1 | NZ_JIJB01000012.1 | - | 26024 | 25995 | **RNA**→ rpmE2 (PRK01678)Ribosomal_L31 (pfam01197)→ |
| Sau-3168-1 | NZ_JIRL01000019.1 | - | 26024 | 25995 | **RNA**→ rpmE2 (PRK01678)Ribosomal_L31 (pfam01197)→ |
| Sau-1173-1 | NZ_JOPP01000047.1 | - | 55266 | 55237 | **RNA**→ rpmE2 (PRK01678)Ribosomal_L31 (pfam01197)→ |
| Sau-3164-1 | NZ_JIRP01000014.1 | - | 26024 | 25995 | **RNA**→ rpmE2 (PRK01678)Ribosomal_L31 (pfam01197)→ |
| Sau-1171-1 | NZ_JOPO01000040.1 | - | 55193 | 55164 | **RNA**→ rpmE2 (PRK01678)Ribosomal_L31 (pfam01197)→ |
| Sau-3148-1 | NZ_JISF01000014.1 | - | 26023 | 25994 | **RNA**→ rpmE2 (PRK01678)Ribosomal_L31 (pfam01197)→ |
| Sau-1234-1 | NZ_JBTY01000001.1 | - | 36391 | 36362 | **RNA**→ rpmE2 (PRK01678)Ribosomal_L31 (pfam01197)→ |
| Sau-3145-1 | NZ_JISI01000011.1 | - | 26024 | 25995 | **RNA**→ rpmE2 (PRK01678)Ribosomal_L31 (pfam01197)→ |
| Sau-1322-1 | NZ_JBUD01000004.1 | - | 212475 | 212446 | **RNA**→ rpmE2 (PRK01678)Ribosomal_L31 (pfam01197)→ |
| Sau-3140-1 | NZ_JISN01000008.1 | - | 2541 | 2512 | **RNA**→ rpmE2 (PRK01678)Ribosomal_L31 (pfam01197)→ |
| Sau-3176-1 | NZ_JIRD01000012.1 | - | 26024 | 25995 | **RNA**→ rpmE2 (PRK01678)Ribosomal_L31 (pfam01197)→ |
| Sau-1283-1 | NZ_JBUB01000010.1 | - | 62005 | 61976 | **RNA**→ rpmE2 (PRK01678)Ribosomal_L31 (pfam01197)→ |
| Sau-3150-1 | NZ_JISD01000013.1 | - | 26024 | 25995 | **RNA**→ rpmE2 (PRK01678)Ribosomal_L31 (pfam01197)→ |
| Sau-1303-1 | NZ_JBUC01000009.1 | - | 62354 | 62325 | **RNA**→ rpmE2 (PRK01678)Ribosomal_L31 (pfam01197)→ |
| Sau-3142-1 | NZ_JISL01000013.1 | - | 26024 | 25995 | **RNA**→ rpmE2 (PRK01678)Ribosomal_L31 (pfam01197)→ |
| Sau-1386-1 | NZ_JBUG01000011.1 | - | 81189 | 81160 | **RNA**→ rpmE2 (PRK01678)Ribosomal_L31 (pfam01197)→ |
| Sau-3370-1 | NZ_JIJR01000008.1 | - | 56439 | 56410 | **RNA**→ rpmE2 (PRK01678)Ribosomal_L31 (pfam01197)→ |
| Sau-1277-1 | NZ_JBUA01000006.1 | - | 94681 | 94652 | **RNA**→ rpmE2 (PRK01678)Ribosomal_L31 (pfam01197)→ |
| Sau-3184-1 | NZ_JIQV01000013.1 | - | 26024 | 25995 | **RNA**→ rpmE2 (PRK01678)Ribosomal_L31 (pfam01197)→ |
| Sau-1381-1 | NZ_JBUF01000011.1 | - | 36390 | 36361 | **RNA**→ rpmE2 (PRK01678)Ribosomal_L31 (pfam01197)→ |
| Sau-3163-1 | NZ_JIRQ01000012.1 | - | 26024 | 25995 | **RNA**→ rpmE2 (PRK01678)Ribosomal_L31 (pfam01197)→ |
| Sau-2442-1 | NZ_JDDY01000011.1 | - | 36391 | 36362 | **RNA**→ rpmE2 (PRK01678)Ribosomal_L31 (pfam01197)→ |
| Sau-1216-1 | NZ_JBTT01000019.1 | - | 1860 | 1831 | **RNA**→ rpmE2 (PRK01678)Ribosomal_L31 (pfam01197)→ |
| Sau-1191-1 | NZ_JBTR01000006.1 | - | 61373 | 61344 | **RNA**→ rpmE2 (PRK01678)Ribosomal_L31 (pfam01197)→ |
| Sau-2389-1 | NZ_JDES01000013.1 | - | 35318 | 35289 | **RNA**→ rpmE2 (PRK01678)Ribosomal_L31 (pfam01197)→ |
| Sau-1206-1 | NZ_JBTS01000015.1 | - | 36413 | 36384 | **RNA**→ rpmE2 (PRK01678)Ribosomal_L31 (pfam01197)→ |
| Sau-2414-1 | NZ_JDEX01000004.1 | - | 24749 | 24720 | **RNA**→ rpmE2 (PRK01678)Ribosomal_L31 (pfam01197)→ |
| Sau-1229-1 | NZ_JBTW01000008.1 | - | 62105 | 62076 | **RNA**→ rpmE2 (PRK01678)Ribosomal_L31 (pfam01197)→ |

| | | | | | |
|---|---|---|---|---|---|
| Sau-1230-1 | NZ_JBTX01000001.1 | - | 238687 | 238658 | **RNA**→ rpmE2 (PRK01678)Ribosomal_L31 (pfam01197)→ |
| Sau-2424-1 | NZ_JDFC01000011.1 | - | 62331 | 62302 | **RNA**→ rpmE2 (PRK01678)Ribosomal_L31 (pfam01197)→ |
| Sau-1155-1 | NZ_JBJQ01000004.1 | - | 56894 | 56865 | **RNA**→ rpmE2 (PRK01678)Ribosomal_L31 (pfam01197)→ |
| Sau-2398-1 | NZ_JDET01000011.1 | - | 35318 | 35289 | **RNA**→ rpmE2 (PRK01678)Ribosomal_L31 (pfam01197)→ |
| Sau-1265-1 | NZ_JBTZ01000013.1 | - | 36391 | 36362 | **RNA**→ rpmE2 (PRK01678)Ribosomal_L31 (pfam01197)→ |
| Sau-2416-1 | NZ_JDEY01000008.1 | - | 24714 | 24685 | **RNA**→ rpmE2 (PRK01678)Ribosomal_L31 (pfam01197)→ |
| Sau-1373-1 | NZ_JBUE01000006.1 | - | 108030 | 108001 | **RNA**→ rpmE2 (PRK01678)Ribosomal_L31 (pfam01197)→ |
| Sau-2426-1 | NZ_JDFD01000013.1 | - | 35847 | 35818 | **RNA**→ rpmE2 (PRK01678)Ribosomal_L31 (pfam01197)→ |
| Sau-1174-1 | NZ_JOPX01000065.1 | - | 55322 | 55293 | **RNA**→ rpmE2 (PRK01678)Ribosomal_L31 (pfam01197)→ |
| Sau-2602-1 | NZ_JBMQ01000004.1 | - | 56893 | 56864 | **RNA**→ rpmE2 (PRK01678)Ribosomal_L31 (pfam01197)→ |
| Sau-1225-1 | NZ_JBTV01000007.1 | - | 61112 | 61083 | **RNA**→ rpmE2 (PRK01678)Ribosomal_L31 (pfam01197)→ |
| Sau-2606-1 | NZ_JBNI01000001.1 | - | 25229 | 25200 | **RNA**→ rpmE2 (PRK01678)Ribosomal_L31 (pfam01197)→ |
| Sau-1220-1 | NZ_JBTU01000016.1 | - | 35881 | 35852 | **RNA**→ rpmE2 (PRK01678)Ribosomal_L31 (pfam01197)→ |
| Sau-2580-1 | NZ_JDGH01000008.1 | - | 56939 | 56910 | **RNA**→ rpmE2 (PRK01678)Ribosomal_L31 (pfam01197)→ |
| Sau-2566-1 | NZ_JDZG01000011.1 | - | 34913 | 34884 | **RNA**→ rpmE2 (PRK01678)Ribosomal_L31 (pfam01197)→ |
| Sau-2520-1 | NZ_JDZH01000007.1 | - | 121803 | 121774 | **RNA**→ rpmE2 (PRK01678)Ribosomal_L31 (pfam01197)→ |
| Sau-2479-1 | NZ_JEUG01000001.1 | - | 233210 | 233181 | **RNA**→ rpmE2 (PRK01678)Ribosomal_L31 (pfam01197)→ |
| Sau-2489-1 | NZ_JEUM01000013.1 | - | 34873 | 34844 | **RNA**→ rpmE2 (PRK01678)Ribosomal_L31 (pfam01197)→ |
| Sau-2518-1 | NZ_JEVL01000007.1 | - | 56894 | 56865 | **RNA**→ rpmE2 (PRK01678)Ribosomal_L31 (pfam01197)→ |
| Sau-2559-1 | NZ_JGFX01000009.1 | - | 34871 | 34842 | **RNA**→ rpmE2 (PRK01678)Ribosomal_L31 (pfam01197)→ |
| Sau-2573-1 | NZ_JGHR01000013.1 | - | 34913 | 34884 | **RNA**→ rpmE2 (PRK01678)Ribosomal_L31 (pfam01197)→ |
| Sau-2633-1 | NZ_JIXP01000012.1 | - | 26024 | 25995 | **RNA**→ rpmE2 (PRK01678)Ribosomal_L31 (pfam01197)→ |
| Sau-2446-1 | NZ_JDGD01000014.1 | - | 15438 | 15409 | **RNA**→ rpmE2 (PRK01678)Ribosomal_L31 (pfam01197)→ |
| Sau-2515-1 | NZ_JDZI01000002.1 | - | 56994 | 56965 | **RNA**→ rpmE2 (PRK01678)Ribosomal_L31 (pfam01197)→ |
| Sau-2578-1 | NZ_JFSE01000001.1 | - | 56894 | 56865 | **RNA**→ rpmE2 (PRK01678)Ribosomal_L31 (pfam01197)→ |
| Sau-2491-1 | NZ_JEUN01000013.1 | - | 34872 | 34843 | **RNA**→ rpmE2 (PRK01678)Ribosomal_L31 (pfam01197)→ |
| Sau-2632-1 | NZ_JIXQ01000009.1 | - | 56348 | 56319 | **RNA**→ rpmE2 (PRK01678)Ribosomal_L31 (pfam01197)→ |
| Sau-2370-1 | NZ_JDEP01000015.1 | - | 35216 | 35187 | **RNA**→ rpmE2 (PRK01678)Ribosomal_L31 (pfam01197)→ |
| Sau-2402-1 | NZ_JDEU01000003.1 | - | 39489 | 39460 | **RNA**→ rpmE2 (PRK01678)Ribosomal_L31 (pfam01197)→ |
| Sau-2409-1 | NZ_JDEV01000007.1 | - | 24749 | 24720 | **RNA**→ rpmE2 (PRK01678)Ribosomal_L31 (pfam01197)→ |
| Sau-2417-1 | NZ_JDEZ01000006.1 | - | 25059 | 25030 | **RNA**→ rpmE2 (PRK01678)Ribosomal_L31 (pfam01197)→ |
| Sau-2418-1 | NZ_JDFA01000008.1 | - | 61335 | 61306 | **RNA**→ rpmE2 (PRK01678)Ribosomal_L31 (pfam01197)→ |
| Sau-2647-1 | NZ_JBLP01000001.1 | - | 215968 | 215939 | **RNA**→ rpmE2 (PRK01678)Ribosomal_L31 (pfam01197)→ |
| Sau-2376-1 | NZ_JDEQ01000012.1 | - | 35318 | 35289 | **RNA**→ rpmE2 (PRK01678)Ribosomal_L31 (pfam01197)→ |
| Sau-2383-1 | NZ_JDER01000017.1 | - | 35042 | 35013 | **RNA**→ rpmE2 (PRK01678)Ribosomal_L31 (pfam01197)→ |
| Sau-2411-1 | NZ_JDEW01000013.1 | - | 36055 | 36026 | **RNA**→ rpmE2 (PRK01678)Ribosomal_L31 (pfam01197)→ |
| Sau-2423-1 | NZ_JDFB01000001.1 | - | 217173 | 217144 | **RNA**→ rpmE2 (PRK01678)Ribosomal_L31 (pfam01197)→ |
| Sau-2503-1 | NZ_JDGQ01000002.1 | - | 56893 | 56864 | **RNA**→ rpmE2 (PRK01678)Ribosomal_L31 (pfam01197)→ |
| Sau-2540-1 | NZ_JGFH01000002.1 | - | 56894 | 56865 | **RNA**→ rpmE2 (PRK01678)Ribosomal_L31 (pfam01197)→ |
| Sau-2558-1 | NZ_JGFW01000015.1 | - | 56894 | 56865 | **RNA**→ rpmE2 (PRK01678)Ribosomal_L31 (pfam01197)→ |
| Sau-2634-1 | NZ_JIXO01000015.1 | - | 25829 | 25800 | **RNA**→ rpmE2 (PRK01678)Ribosomal_L31 (pfam01197)→ |
| Sau-2508-1 | NZ_JDZJ01000012.1 | - | 34873 | 34844 | **RNA**→ rpmE2 (PRK01678)Ribosomal_L31 (pfam01197)→ |
| Sau-2533-1 | NZ_JGFG01000001.1 | - | 4918 | 4889 | **RNA**→ rpmE2 (PRK01678)Ribosomal_L31 (pfam01197)→ |
| Sau-2538-1 | NZ_JGHO01000011.1 | - | 34817 | 34788 | **RNA**→ rpmE2 (PRK01678)Ribosomal_L31 (pfam01197)→ |
| Sau-2544-1 | NZ_JGHP01000003.1 | - | 56794 | 56765 | **RNA**→ rpmE2 (PRK01678)Ribosomal_L31 (pfam01197)→ |
| Sau-2429-1 | NZ_JDDX01000014.1 | - | 36107 | 36078 | **RNA**→ rpmE2 (PRK01678)Ribosomal_L31 (pfam01197)→ |
| Sau-2593-1 | NZ_JGGY01000001.1 | - | 15617 | 15588 | **RNA**→ rpmE2 (PRK01678)Ribosomal_L31 (pfam01197)→ |
| Sau-2575-1 | NZ_JGGX01000007.1 | - | 56894 | 56865 | **RNA**→ rpmE2 (PRK01678)Ribosomal_L31 (pfam01197)→ |
| Sau-2625-1 | NZ_JMBT01000062.1 | - | 15983 | 15954 | **RNA**→ rpmE2 (PRK01678)Ribosomal_L31 (pfam01197)→ |
| Sau-2449-1 | NZ_JDYQ01000003.1 | - | 56894 | 56865 | **RNA**→ rpmE2 (PRK01678)Ribosomal_L31 (pfam01197)→ |
| Sau-2495-1 | NZ_JGGV01000002.1 | - | 55197 | 55168 | **RNA**→ rpmE2 (PRK01678)Ribosomal_L31 (pfam01197)→ |
| Sau-2526-1 | NZ_JGGW01000003.1 | - | 55216 | 55187 | **RNA**→ rpmE2 (PRK01678)Ribosomal_L31 (pfam01197)→ |

| | | | | | |
|---|---|---|---|---|---|
| Sau-2467-1 | NZ_JDGT01000002.1 | - | 56893 | 56864 | **RNA**→ rpmE2 (PRK01678)Ribosomal_L31 (pfam01197)→ |
| Sau-2583-1 | NZ_JDYP01000017.1 | - | 34873 | 34844 | **RNA**→ rpmE2 (PRK01678)Ribosomal_L31 (pfam01197)→ |
| Sau-2458-1 | NZ_JGGU01000008.1 | - | 56173 | 56144 | **RNA**→ rpmE2 (PRK01678)Ribosomal_L31 (pfam01197)→ |
| Sau-857-1 | NZ_JBHL01000002.1 | - | 56894 | 56865 | **RNA**→ rpmE2 (PRK01678)Ribosomal_L31 (pfam01197)→ |
| Sau-864-1 | NZ_JBFO01000002.1 | - | 56895 | 56866 | **RNA**→ rpmE2 (PRK01678)Ribosomal_L31 (pfam01197)→ |
| Sau-844-1 | NZ_JBSM01000001.1 | - | 35014 | 34985 | **RNA**→ rpmE2 (PRK01678)Ribosomal_L31 (pfam01197)→ |
| Sau-812-1 | NZ_JDGM01000009.1 | - | 56895 | 56866 | **RNA**→ rpmE2 (PRK01678)Ribosomal_L31 (pfam01197)→ |
| Sau-697-1 | NZ_JFSN01000008.1 | - | 55000 | 54971 | **RNA**→ rpmE2 (PRK01678)Ribosomal_L31 (pfam01197)→ |
| Sau-695-1 | NZ_JGFN01000035.1 | - | 14467 | 14438 | **RNA**→ rpmE2 (PRK01678)Ribosomal_L31 (pfam01197)→ |
| Sau-772-1 | NZ_JGGM01000002.1 | - | 55117 | 55088 | **RNA**→ rpmE2 (PRK01678)Ribosomal_L31 (pfam01197)→ |
| Sau-901-1 | NZ_JGGN01000002.1 | - | 34873 | 34844 | **RNA**→ rpmE2 (PRK01678)Ribosomal_L31 (pfam01197)→ |
| Sau-914-1 | NZ_JGHM01000008.1 | - | 55315 | 55286 | **RNA**→ rpmE2 (PRK01678)Ribosomal_L31 (pfam01197)→ |
| Sau-698-1 | NZ_JDGI01000012.1 | - | 56940 | 56911 | **RNA**→ rpmE2 (PRK01678)Ribosomal_L31 (pfam01197)→ |
| Sau-784-1 | NZ_JDZN01000010.1 | - | 56502 | 56473 | **RNA**→ rpmE2 (PRK01678)Ribosomal_L31 (pfam01197)→ |
| Sau-813-1 | NZ_JGFT01000009.1 | - | 34482 | 34453 | **RNA**→ rpmE2 (PRK01678)Ribosomal_L31 (pfam01197)→ |
| Sau-827-1 | NZ_JGFU01000011.1 | - | 56894 | 56865 | **RNA**→ rpmE2 (PRK01678)Ribosomal_L31 (pfam01197)→ |
| Sau-710-1 | NZ_JGGJ01000004.1 | - | 55164 | 55135 | **RNA**→ rpmE2 (PRK01678)Ribosomal_L31 (pfam01197)→ |
| Sau-707-1 | NZ_JGHI01000005.1 | - | 55117 | 55088 | **RNA**→ rpmE2 (PRK01678)Ribosomal_L31 (pfam01197)→ |
| Sau-751-1 | NZ_JGHY01000012.1 | - | 34873 | 34844 | **RNA**→ rpmE2 (PRK01678)Ribosomal_L31 (pfam01197)→ |
| Sau-883-1 | NZ_JBOF01000005.1 | - | 56994 | 56965 | **RNA**→ rpmE2 (PRK01678)Ribosomal_L31 (pfam01197)→ |
| Sau-893-1 | NZ_JBEQ01000002.1 | - | 56894 | 56865 | **RNA**→ rpmE2 (PRK01678)Ribosomal_L31 (pfam01197)→ |
| Sau-896-1 | NZ_JDGB01000010.1 | - | 56939 | 56910 | **RNA**→ rpmE2 (PRK01678)Ribosomal_L31 (pfam01197)→ |
| Sau-727-1 | NZ_JDGK01000010.1 | - | 56894 | 56865 | **RNA**→ rpmE2 (PRK01678)Ribosomal_L31 (pfam01197)→ |
| Sau-754-1 | NZ_JFSM01000005.1 | - | 56993 | 56964 | **RNA**→ rpmE2 (PRK01678)Ribosomal_L31 (pfam01197)→ |
| Sau-763-1 | NZ_JGGL01000008.1 | - | 56894 | 56865 | **RNA**→ rpmE2 (PRK01678)Ribosomal_L31 (pfam01197)→ |
| Sau-897-1 | NZ_JGHL01000002.1 | - | 54839 | 54810 | **RNA**→ rpmE2 (PRK01678)Ribosomal_L31 (pfam01197)→ |
| Sau-829-1 | NZ_JGIA01000006.1 | - | 56894 | 56865 | **RNA**→ rpmE2 (PRK01678)Ribosomal_L31 (pfam01197)→ |
| Sau-877-1 | NZ_JIYB01000010.1 | - | 26024 | 25995 | **RNA**→ rpmE2 (PRK01678)Ribosomal_L31 (pfam01197)→ |
| Sau-741-1 | NZ_JDGJ01000011.1 | - | 34817 | 34788 | **RNA**→ rpmE2 (PRK01678)Ribosomal_L31 (pfam01197)→ |
| Sau-745-1 | NZ_JETZ01000011.1 | - | 34913 | 34884 | **RNA**→ rpmE2 (PRK01678)Ribosomal_L31 (pfam01197)→ |
| Sau-747-1 | NZ_JEUE01000003.1 | - | 55117 | 55088 | **RNA**→ rpmE2 (PRK01678)Ribosomal_L31 (pfam01197)→ |
| Sau-760-1 | NZ_JEUJ01000021.1 | - | 32991 | 32962 | **RNA**→ rpmE2 (PRK01678)Ribosomal_L31 (pfam01197)→ |
| Sau-828-1 | NZ_JGFV01000005.1 | - | 56894 | 56865 | **RNA**→ rpmE2 (PRK01678)Ribosomal_L31 (pfam01197)→ |
| Sau-715-1 | NZ_JGGK01000013.1 | - | 34968 | 34939 | **RNA**→ rpmE2 (PRK01678)Ribosomal_L31 (pfam01197)→ |
| Sau-708-1 | NZ_JGHJ01000001.1 | - | 56896 | 56867 | **RNA**→ rpmE2 (PRK01678)Ribosomal_L31 (pfam01197)→ |
| Sau-764-1 | NZ_JGHZ01000008.1 | - | 55117 | 55088 | **RNA**→ rpmE2 (PRK01678)Ribosomal_L31 (pfam01197)→ |
| Sau-879-1 | NZ_JIXZ01000015.1 | - | 26023 | 25994 | **RNA**→ rpmE2 (PRK01678)Ribosomal_L31 (pfam01197)→ |
| Sau-711-1 | NZ_JDYX01000002.1 | - | 56894 | 56865 | **RNA**→ rpmE2 (PRK01678)Ribosomal_L31 (pfam01197)→ |
| Sau-821-1 | NZ_JEVC01000011.1 | - | 55117 | 55088 | **RNA**→ rpmE2 (PRK01678)Ribosomal_L31 (pfam01197)→ |
| Sau-838-1 | NZ_JEVH01000002.1 | - | 56894 | 56865 | **RNA**→ rpmE2 (PRK01678)Ribosomal_L31 (pfam01197)→ |
| Sau-714-1 | NZ_JDYW01000002.1 | - | 56893 | 56864 | **RNA**→ rpmE2 (PRK01678)Ribosomal_L31 (pfam01197)→ |
| Sau-820-1 | NZ_JEVB01000002.1 | - | 56894 | 56865 | **RNA**→ rpmE2 (PRK01678)Ribosomal_L31 (pfam01197)→ |
| Sau-837-1 | NZ_JEVG01000001.1 | - | 56894 | 56865 | **RNA**→ rpmE2 (PRK01678)Ribosomal_L31 (pfam01197)→ |
| Sau-774-1 | NZ_JEUS01000002.1 | - | 56893 | 56864 | **RNA**→ rpmE2 (PRK01678)Ribosomal_L31 (pfam01197)→ |
| Sau-909-1 | NZ_JGGO01000004.1 | - | 56894 | 56865 | **RNA**→ rpmE2 (PRK01678)Ribosomal_L31 (pfam01197)→ |
| Sau-818-1 | NZ_JEVA01000007.1 | - | 56894 | 56865 | **RNA**→ rpmE2 (PRK01678)Ribosomal_L31 (pfam01197)→ |
| Sau-833-1 | NZ_JEVE01000001.1 | - | 56794 | 56765 | **RNA**→ rpmE2 (PRK01678)Ribosomal_L31 (pfam01197)→ |
| Sau-836-1 | NZ_JEVF01000008.1 | - | 56894 | 56865 | **RNA**→ rpmE2 (PRK01678)Ribosomal_L31 (pfam01197)→ |
| Sau-911-1 | NZ_JGGQ01000003.1 | - | 56894 | 56865 | **RNA**→ rpmE2 (PRK01678)Ribosomal_L31 (pfam01197)→ |
| Sau-831-1 | NZ_JEVD01000006.1 | - | 56893 | 56864 | **RNA**→ rpmE2 (PRK01678)Ribosomal_L31 (pfam01197)→ |
| Sau-806-1 | NZ_JEUY01000001.1 | - | 56890 | 56861 | **RNA**→ rpmE2 (PRK01678)Ribosomal_L31 (pfam01197)→ |
| Sau-910-1 | NZ_JGGP01000013.1 | - | 34873 | 34844 | **RNA**→ rpmE2 (PRK01678)Ribosomal_L31 (pfam01197)→ |

| | | | | | |
|---|---|---|---|---|---|
| Sau-1881-1 | NZ_JDCV01000012.1 | - | 58694 | 58665 | **RNA**→ rpmE2 (PRK01678)Ribosomal_L31 (pfam01197)→ |
| Sau-1161-1 | NZ_JBGI01000001.1 | - | 115280 | 115251 | **RNA**→ rpmE2 (PRK01678)Ribosomal_L31 (pfam01197)→ |
| Sau-1976-1 | NZ_JDDJ01000009.1 | - | 61461 | 61432 | **RNA**→ rpmE2 (PRK01678)Ribosomal_L31 (pfam01197)→ |
| Sau-2702-1 | NZ_CBWV010000285.1 | + | 3236 | 3265 | **RNA**→ rpmE2 (PRK01678)Ribosomal_L31 (pfam01197)→ |
| Sau-3050-1 | NZ_JBGU01000002.1 | - | 56894 | 56865 | **RNA**→ rpmE2 (PRK01678)Ribosomal_L31 (pfam01197)→ |
| Sau-2935-1 | NZ_JDGC01000008.1 | - | 56894 | 56865 | **RNA**→ rpmE2 (PRK01678)Ribosomal_L31 (pfam01197)→ |
| Sau-2996-1 | NZ_JFRX01000004.1 | - | 56894 | 56865 | **RNA**→ rpmE2 (PRK01678)Ribosomal_L31 (pfam01197)→ |
| Sau-2894-1 | NZ_JFRY01000002.1 | - | 56894 | 56865 | **RNA**→ rpmE2 (PRK01678)Ribosomal_L31 (pfam01197)→ |
| Sau-2973-1 | NZ_JEUB01000002.1 | - | 56894 | 56865 | **RNA**→ rpmE2 (PRK01678)Ribosomal_L31 (pfam01197)→ |
| Sau-2892-1 | NZ_JEVQ01000001.1 | - | 56894 | 56865 | **RNA**→ rpmE2 (PRK01678)Ribosomal_L31 (pfam01197)→ |
| Sau-2862-1 | NZ_JGFI01000005.1 | - | 56894 | 56865 | **RNA**→ rpmE2 (PRK01678)Ribosomal_L31 (pfam01197)→ |
| Sau-3013-1 | NZ_JIVX01000016.1 | - | 26024 | 25995 | **RNA**→ rpmE2 (PRK01678)Ribosomal_L31 (pfam01197)→ |
| Sau-2876-1 | NZ_JFRZ01000002.1 | - | 55051 | 55022 | **RNA**→ rpmE2 (PRK01678)Ribosomal_L31 (pfam01197)→ |
| Sau-2961-1 | NZ_JETY01000002.1 | - | 56894 | 56865 | **RNA**→ rpmE2 (PRK01678)Ribosomal_L31 (pfam01197)→ |
| Sau-2970-1 | NZ_JEUC01000005.1 | - | 56894 | 56865 | **RNA**→ rpmE2 (PRK01678)Ribosomal_L31 (pfam01197)→ |
| Sau-2972-1 | NZ_JEUD01000005.1 | - | 56790 | 56761 | **RNA**→ rpmE2 (PRK01678)Ribosomal_L31 (pfam01197)→ |
| Sau-3026-1 | NZ_JDXY01000001.1 | - | 1675824 | 1675795 | **RNA**→ rpmE2 (PRK01678)Ribosomal_L31 (pfam01197)→ |
| Sau-2883-1 | NZ_JGFJ01000014.1 | - | 34971 | 34942 | **RNA**→ rpmE2 (PRK01678)Ribosomal_L31 (pfam01197)→ |
| Sau-2857-1 | NZ_JGHS01000010.1 | - | 54968 | 54939 | **RNA**→ rpmE2 (PRK01678)Ribosomal_L31 (pfam01197)→ |
| Sau-3092-1 | NZ_JIUJ01000012.1 | - | 26024 | 25995 | **RNA**→ rpmE2 (PRK01678)Ribosomal_L31 (pfam01197)→ |
| Sau-3069-1 | NZ_JIVG01000012.1 | - | 26024 | 25995 | **RNA**→ rpmE2 (PRK01678)Ribosomal_L31 (pfam01197)→ |
| Sau-3054-1 | NZ_JBEZ01000002.1 | - | 56894 | 56865 | **RNA**→ rpmE2 (PRK01678)Ribosomal_L31 (pfam01197)→ |
| Sau-3042-1 | NZ_JBIW01000007.1 | - | 56994 | 56965 | **RNA**→ rpmE2 (PRK01678)Ribosomal_L31 (pfam01197)→ |
| Sau-3006-1 | NZ_JBMK01000003.1 | - | 56894 | 56865 | **RNA**→ rpmE2 (PRK01678)Ribosomal_L31 (pfam01197)→ |
| Sau-3051-1 | NZ_JBGO01000002.1 | - | 56894 | 56865 | **RNA**→ rpmE2 (PRK01678)Ribosomal_L31 (pfam01197)→ |
| Sau-3022-1 | NZ_JBMM01000002.1 | - | 56894 | 56865 | **RNA**→ rpmE2 (PRK01678)Ribosomal_L31 (pfam01197)→ |
| Sau-2951-1 | NZ_JDGF01000011.1 | - | 56894 | 56865 | **RNA**→ rpmE2 (PRK01678)Ribosomal_L31 (pfam01197)→ |
| Sau-2974-1 | NZ_JDGL01000007.1 | - | 56940 | 56911 | **RNA**→ rpmE2 (PRK01678)Ribosomal_L31 (pfam01197)→ |
| Sau-2872-1 | NZ_JDZF01000002.1 | - | 56894 | 56865 | **RNA**→ rpmE2 (PRK01678)Ribosomal_L31 (pfam01197)→ |
| Sau-2969-1 | NZ_JEUA01000013.1 | - | 34872 | 34843 | **RNA**→ rpmE2 (PRK01678)Ribosomal_L31 (pfam01197)→ |
| Sau-2889-1 | NZ_JEVP01000011.1 | - | 56993 | 56964 | **RNA**→ rpmE2 (PRK01678)Ribosomal_L31 (pfam01197)→ |
| Sau-3062-1 | NZ_JIVO01000013.1 | - | 26162 | 26133 | **RNA**→ rpmE2 (PRK01678)Ribosomal_L31 (pfam01197)→ |
| Sau-3014-1 | NZ_JIVW01000016.1 | - | 26024 | 25995 | **RNA**→ rpmE2 (PRK01678)Ribosomal_L31 (pfam01197)→ |
| Sau-2948-1 | NZ_JDGE01000007.1 | - | 56894 | 56865 | **RNA**→ rpmE2 (PRK01678)Ribosomal_L31 (pfam01197)→ |
| Sau-2965-1 | NZ_JDGP01000009.1 | - | 56794 | 56765 | **RNA**→ rpmE2 (PRK01678)Ribosomal_L31 (pfam01197)→ |
| Sau-2891-1 | NZ_JDZE01000003.1 | - | 56894 | 56865 | **RNA**→ rpmE2 (PRK01678)Ribosomal_L31 (pfam01197)→ |
| Sau-2861-1 | NZ_JFSA01000002.1 | - | 54854 | 54825 | **RNA**→ rpmE2 (PRK01678)Ribosomal_L31 (pfam01197)→ |
| Sau-2987-1 | NZ_JEUO01000013.1 | - | 34874 | 34845 | **RNA**→ rpmE2 (PRK01678)Ribosomal_L31 (pfam01197)→ |
| Sau-2866-1 | NZ_JEVO01000016.1 | - | 34913 | 34884 | **RNA**→ rpmE2 (PRK01678)Ribosomal_L31 (pfam01197)→ |
| Sau-2884-1 | NZ_JGFK01000007.1 | - | 56894 | 56865 | **RNA**→ rpmE2 (PRK01678)Ribosomal_L31 (pfam01197)→ |
| Sau-2885-1 | NZ_JGFL01000019.1 | - | 34913 | 34884 | **RNA**→ rpmE2 (PRK01678)Ribosomal_L31 (pfam01197)→ |
| Sau-2895-1 | NZ_JGIE01000009.1 | - | 34972 | 34943 | **RNA**→ rpmE2 (PRK01678)Ribosomal_L31 (pfam01197)→ |
| Sau-3076-1 | NZ_JIUZ01000012.1 | - | 26024 | 25995 | **RNA**→ rpmE2 (PRK01678)Ribosomal_L31 (pfam01197)→ |
| Sau-3067-1 | NZ_JIVI01000014.1 | - | 26024 | 25995 | **RNA**→ rpmE2 (PRK01678)Ribosomal_L31 (pfam01197)→ |
| Sau-3031-1 | NZ_JBPA01000003.1 | - | 56894 | 56865 | **RNA**→ rpmE2 (PRK01678)Ribosomal_L31 (pfam01197)→ |
| Sau-3112-1 | NZ_JITP01000016.1 | - | 26024 | 25995 | **RNA**→ rpmE2 (PRK01678)Ribosomal_L31 (pfam01197)→ |
| Sau-3094-1 | NZ_JIUH01000008.1 | - | 53212 | 53183 | **RNA**→ rpmE2 (PRK01678)Ribosomal_L31 (pfam01197)→ |
| Sau-2938-1 | NZ_JDYN01000002.1 | - | 56894 | 56865 | **RNA**→ rpmE2 (PRK01678)Ribosomal_L31 (pfam01197)→ |
| Sau-2913-1 | NZ_JGHA01000007.1 | - | 56894 | 56865 | **RNA**→ rpmE2 (PRK01678)Ribosomal_L31 (pfam01197)→ |
| Sau-3118-1 | NZ_JITJ01000011.1 | - | 26024 | 25995 | **RNA**→ rpmE2 (PRK01678)Ribosomal_L31 (pfam01197)→ |
| Sau-2901-1 | NZ_JGGZ01000007.1 | - | 55217 | 55188 | **RNA**→ rpmE2 (PRK01678)Ribosomal_L31 (pfam01197)→ |
| Sau-3120-1 | NZ_JITH01000016.1 | - | 26024 | 25995 | **RNA**→ rpmE2 (PRK01678)Ribosomal_L31 (pfam01197)→ |

| | | | | | |
|---|---|---|---|---|---|
| Sau-2859-1 | NZ_JDGU01000008.1 | - | 56894 | 56865 | **RNA**→ rpmE2 (PRK01678)Ribosomal_L31 (pfam01197)→ |
| Sau-2924-1 | NZ_JDYO01000002.1 | - | 56993 | 56964 | **RNA**→ rpmE2 (PRK01678)Ribosomal_L31 (pfam01197)→ |
| Sau-2738-1 | NZ_JBET01000002.1 | - | 25029 | 25000 | **RNA**→ rpmE2 (PRK01678)Ribosomal_L31 (pfam01197)→ |
| Sau-2671-1 | NZ_JBGE01000011.1 | - | 110326 | 110297 | **RNA**→ rpmE2 (PRK01678)Ribosomal_L31 (pfam01197)→ |
| Sau-2686-1 | NZ_JBEU01000002.1 | - | 56893 | 56864 | **RNA**→ rpmE2 (PRK01678)Ribosomal_L31 (pfam01197)→ |
| Sau-2668-1 | NZ_JBGL01000001.1 | - | 35179 | 35150 | **RNA**→ rpmE2 (PRK01678)Ribosomal_L31 (pfam01197)→ |
| Sau-2664-1 | NZ_JBGW01000001.1 | - | 626825 | 626796 | **RNA**→ rpmE2 (PRK01678)Ribosomal_L31 (pfam01197)→ |
| Sau-2756-1 | NZ_JBOD01000002.1 | - | 56894 | 56865 | **RNA**→ rpmE2 (PRK01678)Ribosomal_L31 (pfam01197)→ |
| Sau-2834-1 | NZ_JFSD01000003.1 | - | 55116 | 55087 | **RNA**→ rpmE2 (PRK01678)Ribosomal_L31 (pfam01197)→ |
| Sau-2851-1 | NZ_JEVM01000005.1 | - | 56894 | 56865 | **RNA**→ rpmE2 (PRK01678)Ribosomal_L31 (pfam01197)→ |
| Sau-2843-1 | NZ_JGFY01000026.1 | - | 15832 | 15803 | **RNA**→ rpmE2 (PRK01678)Ribosomal_L31 (pfam01197)→ |
| Sau-2844-1 | NZ_JGFZ01000004.1 | - | 27750 | 27721 | **RNA**→ rpmE2 (PRK01678)Ribosomal_L31 (pfam01197)→ |
| Sau-2713-1 | NZ_JIWP01000013.1 | - | 26024 | 25995 | **RNA**→ rpmE2 (PRK01678)Ribosomal_L31 (pfam01197)→ |
| Sau-2681-1 | NZ_JBMH01000008.1 | - | 113883 | 113854 | **RNA**→ rpmE2 (PRK01678)Ribosomal_L31 (pfam01197)→ |
| Sau-2812-1 | NZ_JBMW01000001.1 | - | 238687 | 238658 | **RNA**→ rpmE2 (PRK01678)Ribosomal_L31 (pfam01197)→ |
| Sau-2785-1 | NZ_JBNL01000003.1 | - | 56894 | 56865 | **RNA**→ rpmE2 (PRK01678)Ribosomal_L31 (pfam01197)→ |
| Sau-2726-1 | NZ_JHEA01000010.1 | - | 54919 | 54890 | **RNA**→ rpmE2 (PRK01678)Ribosomal_L31 (pfam01197)→ |
| Sau-3993-1 | NZ_ASZO01000074.1 | - | 61790 | 61761 | **RNA**→ rpmE2 (PRK01678)Ribosomal_L31 (pfam01197)→ |
| Sau-3992-1 | NZ_AUPW01000016.1 | + | 65946 | 65975 | **RNA**→ rpmE2 (PRK01678)Ribosomal_L31 (pfam01197)→ |
| Sau-3988-1 | NZ_AUPS01000015.1 | + | 42609 | 42638 | **RNA**→ rpmE2 (PRK01678)Ribosomal_L31 (pfam01197)→ |
| Sau-2718-1 | NZ_JHEB01000009.1 | - | 56494 | 56465 | **RNA**→ rpmE2 (PRK01678)Ribosomal_L31 (pfam01197)→ |
| Sau-2808-1 | NZ_JBRE01000003.1 | - | 25227 | 25198 | **RNA**→ rpmE2 (PRK01678)Ribosomal_L31 (pfam01197)→ |
| Sau-2743-1 | NZ_JBMX01000003.1 | - | 56994 | 56965 | **RNA**→ rpmE2 (PRK01678)Ribosomal_L31 (pfam01197)→ |
| Sau-2803-1 | NZ_JBNS01000002.1 | - | 56894 | 56865 | **RNA**→ rpmE2 (PRK01678)Ribosomal_L31 (pfam01197)→ |
| Sau-2753-1 | NZ_JBOI01000004.1 | - | 56894 | 56865 | **RNA**→ rpmE2 (PRK01678)Ribosomal_L31 (pfam01197)→ |
| Sau-2836-1 | NZ_JFSB01000009.1 | - | 56993 | 56964 | **RNA**→ rpmE2 (PRK01678)Ribosomal_L31 (pfam01197)→ |
| Sau-2835-1 | NZ_JFSC01000005.1 | - | 56893 | 56864 | **RNA**→ rpmE2 (PRK01678)Ribosomal_L31 (pfam01197)→ |
| Sau-2694-1 | NZ_JIXA01000010.1 | - | 56395 | 56366 | **RNA**→ rpmE2 (PRK01678)Ribosomal_L31 (pfam01197)→ |
| Sau-2852-1 | NZ_JEVN01000007.1 | - | 56893 | 56864 | **RNA**→ rpmE2 (PRK01678)Ribosomal_L31 (pfam01197)→ |
| Sau-2845-1 | NZ_JGGA01000010.1 | - | 34872 | 34843 | **RNA**→ rpmE2 (PRK01678)Ribosomal_L31 (pfam01197)→ |
| Sau-2712-1 | NZ_JIWQ01000014.1 | - | 26024 | 25995 | **RNA**→ rpmE2 (PRK01678)Ribosomal_L31 (pfam01197)→ |
| Sau-2720-1 | NZ_JHED01000008.1 | - | 54920 | 54891 | **RNA**→ rpmE2 (PRK01678)Ribosomal_L31 (pfam01197)→ |
| Sau-3989-1 | NZ_AUPV01000011.1 | + | 18390 | 18419 | **RNA**→ rpmE2 (PRK01678)Ribosomal_L31 (pfam01197)→ |
| Sau-2719-1 | NZ_JHEC01000014.1 | - | 28996 | 28967 | **RNA**→ rpmE2 (PRK01678)Ribosomal_L31 (pfam01197)→ |
| Sau-2778-1 | NZ_JBPS01000005.1 | - | 56894 | 56865 | **RNA**→ rpmE2 (PRK01678)Ribosomal_L31 (pfam01197)→ |
| Sau-2682-1 | NZ_JBPD01000001.1 | - | 57499 | 57470 | **RNA**→ rpmE2 (PRK01678)Ribosomal_L31 (pfam01197)→ |
| Sau-2730-1 | NZ_JBQF01000003.1 | - | 56893 | 56864 | **RNA**→ rpmE2 (PRK01678)Ribosomal_L31 (pfam01197)→ |
| Sau-2799-1 | NZ_JBJV01000008.1 | - | 57499 | 57470 | **RNA**→ rpmE2 (PRK01678)Ribosomal_L31 (pfam01197)→ |
| Sau-2816-1 | NZ_JBLA01000002.1 | - | 56994 | 56965 | **RNA**→ rpmE2 (PRK01678)Ribosomal_L31 (pfam01197)→ |
| Sau-2793-1 | NZ_JBKV01000001.1 | - | 238854 | 238825 | **RNA**→ rpmE2 (PRK01678)Ribosomal_L31 (pfam01197)→ |
| Sau-2725-1 | NZ_JHDZ01000008.1 | + | 52043 | 52072 | **RNA**→ rpmE2 (PRK01678)Ribosomal_L31 (pfam01197)→ |
| Sau-2724-1 | NZ_JHDY01000010.1 | + | 55746 | 55775 | **RNA**→ rpmE2 (PRK01678)Ribosomal_L31 (pfam01197)→ |
| Sau-2831-1 | NZ_JEUX01000013.1 | - | 15617 | 15588 | **RNA**→ rpmE2 (PRK01678)Ribosomal_L31 (pfam01197)→ |
| Sau-2721-1 | NZ_JHDV01000014.1 | - | 54961 | 54932 | **RNA**→ rpmE2 (PRK01678)Ribosomal_L31 (pfam01197)→ |
| Sau-3990-1 | NZ_AUPU01000015.1 | + | 65947 | 65976 | **RNA**→ rpmE2 (PRK01678)Ribosomal_L31 (pfam01197)→ |
| Sau-2770-1 | NZ_JBJF01000002.1 | - | 56894 | 56865 | **RNA**→ rpmE2 (PRK01678)Ribosomal_L31 (pfam01197)→ |
| Sau-2652-1 | NZ_JBKF01000004.1 | - | 56894 | 56865 | **RNA**→ rpmE2 (PRK01678)Ribosomal_L31 (pfam01197)→ |
| Sau-2723-1 | NZ_JHDX01000009.1 | + | 55746 | 55775 | **RNA**→ rpmE2 (PRK01678)Ribosomal_L31 (pfam01197)→ |
| Sau-3991-1 | NZ_AUPT01000016.1 | - | 25833 | 25804 | **RNA**→ rpmE2 (PRK01678)Ribosomal_L31 (pfam01197)→ |
| Sau-2798-1 | NZ_JBJX01000002.1 | - | 57001 | 56972 | **RNA**→ rpmE2 (PRK01678)Ribosomal_L31 (pfam01197)→ |
| Sau-2764-1 | NZ_JBLC01000001.1 | - | 110684 | 110655 | **RNA**→ rpmE2 (PRK01678)Ribosomal_L31 (pfam01197)→ |
| Sau-2722-1 | NZ_JHDW01000009.1 | + | 55746 | 55775 | **RNA**→ rpmE2 (PRK01678)Ribosomal_L31 (pfam01197)→ |

| | | | | | |
|---|---|---|---|---|---|
| Sau-3683-1 | NZ_JBGQ01000002.1 | - | 56894 | 56865 | **RNA**→ rpmE2 (PRK01678)Ribosomal_L31 (pfam01197)→ |
| Sau-3697-1 | NZ_JBHG01000002.1 | - | 56894 | 56865 | **RNA**→ rpmE2 (PRK01678)Ribosomal_L31 (pfam01197)→ |
| Sau-3680-1 | NZ_JBEW01000006.1 | - | 56894 | 56865 | **RNA**→ rpmE2 (PRK01678)Ribosomal_L31 (pfam01197)→ |
| Sau-3695-1 | NZ_JBFM01000003.1 | - | 56894 | 56865 | **RNA**→ rpmE2 (PRK01678)Ribosomal_L31 (pfam01197)→ |
| Sau-3725-1 | NZ_JHPT01000009.1 | - | 54839 | 54810 | **RNA**→ rpmE2 (PRK01678)Ribosomal_L31 (pfam01197)→ |
| Sau-3731-1 | NZ_JHPY01000018.1 | - | 54601 | 54572 | **RNA**→ rpmE2 (PRK01678)Ribosomal_L31 (pfam01197)→ |
| Sau-3711-1 | NZ_JIFL01000015.1 | - | 26024 | 25995 | **RNA**→ rpmE2 (PRK01678)Ribosomal_L31 (pfam01197)→ |
| Sau-3733-1 | NZ_JHTY01000067.1 | + | 18293 | 18322 | **RNA**→ rpmE2 (PRK01678)Ribosomal_L31 (pfam01197)→ |
| Sau-3652-1 | NZ_JGGE01000007.1 | - | 55082 | 55053 | **RNA**→ rpmE2 (PRK01678)Ribosomal_L31 (pfam01197)→ |
| Sau-3641-1 | NZ_JGIH01000019.1 | - | 34817 | 34788 | **RNA**→ rpmE2 (PRK01678)Ribosomal_L31 (pfam01197)→ |
| Sau-3739-1 | NZ_JHQD01000034.1 | - | 1801 | 1772 | **RNA**→ rpmE2 (PRK01678)Ribosomal_L31 (pfam01197)→ |
| Sau-3742-1 | NZ_JHTV01000022.1 | - | 25152 | 25123 | **RNA**→ rpmE2 (PRK01678)Ribosomal_L31 (pfam01197)→ |
| Sau-3732-1 | NZ_JHTX01000003.1 | - | 54710 | 54681 | **RNA**→ rpmE2 (PRK01678)Ribosomal_L31 (pfam01197)→ |
| Sau-3692-1 | NZ_JBGM01000002.1 | - | 56994 | 56965 | **RNA**→ rpmE2 (PRK01678)Ribosomal_L31 (pfam01197)→ |
| Sau-3690-1 | NZ_JBGZ01000002.1 | - | 56790 | 56761 | **RNA**→ rpmE2 (PRK01678)Ribosomal_L31 (pfam01197)→ |
| Sau-3660-1 | NZ_JEVU01000005.1 | - | 56890 | 56861 | **RNA**→ rpmE2 (PRK01678)Ribosomal_L31 (pfam01197)→ |
| Sau-3663-1 | NZ_JGGG01000011.1 | - | 34867 | 34838 | **RNA**→ rpmE2 (PRK01678)Ribosomal_L31 (pfam01197)→ |
| Sau-3727-1 | NZ_JHPU01000026.1 | - | 54858 | 54829 | **RNA**→ rpmE2 (PRK01678)Ribosomal_L31 (pfam01197)→ |
| Sau-3741-1 | NZ_JHQF01000031.1 | - | 55187 | 55158 | **RNA**→ rpmE2 (PRK01678)Ribosomal_L31 (pfam01197)→ |
| Sau-3735-1 | NZ_JHQA01000009.1 | - | 54489 | 54460 | **RNA**→ rpmE2 (PRK01678)Ribosomal_L31 (pfam01197)→ |
| Sau-3713-1 | NZ_JIFJ01000017.1 | - | 26024 | 25995 | **RNA**→ rpmE2 (PRK01678)Ribosomal_L31 (pfam01197)→ |
| Sau-3726-1 | NZ_JHTW01000035.1 | - | 54627 | 54598 | **RNA**→ rpmE2 (PRK01678)Ribosomal_L31 (pfam01197)→ |
| Sau-3738-1 | NZ_AJKC01000001.1 | + | 18638 | 18667 | **RNA**→ rpmE2 (PRK01678)Ribosomal_L31 (pfam01197)→ |
| Sau-4105-1 | NZ_AVBR01000007.1 | - | 54901 | 54872 | **RNA**→ rpmE2 (PRK01678)Ribosomal_L31 (pfam01197)→ |
| Sau-3645-1 | NZ_JEVT01000001.1 | - | 56893 | 56864 | **RNA**→ rpmE2 (PRK01678)Ribosomal_L31 (pfam01197)→ |
| Sau-3655-1 | NZ_JGGF01000011.1 | - | 56894 | 56865 | **RNA**→ rpmE2 (PRK01678)Ribosomal_L31 (pfam01197)→ |
| Sau-3740-1 | NZ_JHQE01000027.1 | - | 55261 | 55232 | **RNA**→ rpmE2 (PRK01678)Ribosomal_L31 (pfam01197)→ |
| Sau-3729-1 | NZ_JHPW01000011.1 | - | 54780 | 54751 | **RNA**→ rpmE2 (PRK01678)Ribosomal_L31 (pfam01197)→ |
| Sau-3720-1 | NZ_JIFC01000012.1 | - | 56390 | 56361 | **RNA**→ rpmE2 (PRK01678)Ribosomal_L31 (pfam01197)→ |
| Sau-3745-1 | NZ_ANPO01000022.1 | - | 54581 | 54552 | **RNA**→ rpmE2 (PRK01678)Ribosomal_L31 (pfam01197)→ |
| Sau-3668-1 | NZ_JBQW01000006.1 | - | 117087 | 117058 | **RNA**→ rpmE2 (PRK01678)Ribosomal_L31 (pfam01197)→ |
| Sau-3649-1 | NZ_JDZB01000004.1 | - | 56894 | 56865 | **RNA**→ rpmE2 (PRK01678)Ribosomal_L31 (pfam01197)→ |
| Sau-3736-1 | NZ_JASL01000006.1 | - | 54744 | 54715 | **RNA**→ rpmE2 (PRK01678)Ribosomal_L31 (pfam01197)→ |
| Sau-3723-1 | NZ_JHUH01000009.1 | - | 18886 | 18857 | **RNA**→ rpmE2 (PRK01678)Ribosomal_L31 (pfam01197)→ |
| Sau-3665-1 | NZ_JDZA01000002.1 | - | 56894 | 56865 | **RNA**→ rpmE2 (PRK01678)Ribosomal_L31 (pfam01197)→ |
| Sau-3737-1 | NZ_JASK01000008.1 | + | 52532 | 52561 | **RNA**→ rpmE2 (PRK01678)Ribosomal_L31 (pfam01197)→ |
| Sau-3666-1 | NZ_JDYZ01000006.1 | - | 56894 | 56865 | **RNA**→ rpmE2 (PRK01678)Ribosomal_L31 (pfam01197)→ |
| Sau-3-1 | NZ_ABFA01000032.1 | - | 15628 | 15599 | **RNA**→ rpmE2 (PRK01678)Ribosomal_L31 (pfam01197)→ rpmE2 (PRK01678)Ribosomal_L31 (pfam01197 |
| Sau-63-1 | NC_007795.1 | - | 2182895 | 2182866 | **RNA**→ rpmE2 (PRK01678)Ribosomal_L31 (pfam01197)→ |
| Sau-60-1 | NC_002758.2 | - | 2247063 | 2247034 | **RNA**→ rpmE2 (PRK01678)Ribosomal_L31 (pfam01197)→ |
| Sau-55-1 | NC_002952.2 | - | 2274990 | 2274961 | **RNA**→ rpmE2 (PRK01678)Ribosomal_L31 (pfam01197)→ |
| Sau-59-1 | NC_009782.1 | - | 2248701 | 2248672 | **RNA**→ rpmE2 (PRK01678)Ribosomal_L31 (pfam01197)→ |
| Sau-58-1 | NC_003923.1 | - | 2197274 | 2197245 | **RNA**→ rpmE2 (PRK01678)Ribosomal_L31 (pfam01197)→ |
| Sau-21-1 | NC_007622.1 | - | 2134777 | 2134748 | **RNA**→ rpmE2 (PRK01678)Ribosomal_L31 (pfam01197)→ |
| Sau-35-1 | NC_002951.2 | - | 2172171 | 2172142 | **RNA**→ rpmE2 (PRK01678)Ribosomal_L31 (pfam01197)→ |
| Sau-57-1 | NC_002953.3 | - | 2176379 | 2176350 | **RNA**→ rpmE2 (PRK01678)Ribosomal_L31 (pfam01197)→ |
| Sau-45-1 | NC_009487.1 | - | 2277078 | 2277049 | **RNA**→ rpmE2 (PRK01678)Ribosomal_L31 (pfam01197)→ |
| Sau-78-1 | NC_009641.1 | - | 2240554 | 2240525 | **RNA**→ rpmE2 (PRK01678)Ribosomal_L31 (pfam01197)→ |
| Sau-62-1 | NC_002745.2 | - | 2170631 | 2170602 | **RNA**→ rpmE2 (PRK01678)Ribosomal_L31 (pfam01197)→ |
| Sau-44-1 | NC_009632.1 | - | 2276953 | 2276924 | **RNA**→ rpmE2 (PRK01678)Ribosomal_L31 (pfam01197)→ |
| Sau-40-1 | NC_013450.1 | - | 2193468 | 2193439 | **RNA**→ rpmE2 (PRK01678)Ribosomal_L31 (pfam01197)→ |
| Sau-76-1 | NC_017341.1 | - | 2266296 | 2266267 | **RNA**→ rpmE2 (PRK01678)Ribosomal_L31 (pfam01197)→ |

| | | | | | |
|---|---|---|---|---|---|
| Sau-67-1 | NC_017342.1 | + | 1146277 | 1146306 | **RNA**→ rpmE2 (PRK01678)Ribosomal_L31 (pfam01197)→ |
| Sau-4024-1 | NZ_AMPF01000068.1 | - | 15521 | 15492 | **RNA**→ rpmE2 (PRK01678)Ribosomal_L31 (pfam01197)→ |
| Sau-47-1 | NC_017349.1 | - | 2132497 | 2132468 | **RNA**→ rpmE2 (PRK01678)Ribosomal_L31 (pfam01197)→ |
| Sau-48-1 | NC_016928.1 | - | 2140391 | 2140362 | **RNA**→ rpmE2 (PRK01678)Ribosomal_L31 (pfam01197)→ |
| Sau-43-1 | NC_017763.1 | - | 2198775 | 2198746 | **RNA**→ rpmE2 (PRK01678)Ribosomal_L31 (pfam01197)→ |
| Sau-4038-1 | NC_022222.1 | - | 2087639 | 2087610 | **RNA**→ rpmE2 (PRK01678)Ribosomal_L31 (pfam01197)→ |
| Sau-73-1 | NZ_GG730215.1 | - | 55940 | 55911 | **RNA**→ rpmE2 (PRK01678)Ribosomal_L31 (pfam01197)→ rpmE2 (PRK01678)Ribosomal_L31 (pfam01197 |
| Sau-74-1 | NZ_GG730248.1 | - | 57166 | 57137 | **RNA**→ rpmE2 (PRK01678)Ribosomal_L31 (pfam01197)→ rpmE2 (PRK01678)Ribosomal_L31 (pfam01197 |
| Sau-32-1 | NZ_GG730136.1 | - | 56479 | 56450 | **RNA**→ rpmE2 (PRK01678)Ribosomal_L31 (pfam01197)→ rpmE2 (PRK01678)Ribosomal_L31 (pfam01197 |
| Sau-31-1 | NZ_GG730266.1 | - | 56408 | 56379 | **RNA**→ rpmE2 (PRK01678)Ribosomal_L31 (pfam01197)→ rpmE2 (PRK01678)Ribosomal_L31 (pfam01197 |
| Sau-25-1 | NZ_GG700568.1 | - | 56291 | 56262 | **RNA**→ rpmE2 (PRK01678)Ribosomal_L31 (pfam01197)→ rpmE2 (PRK01678)Ribosomal_L31 (pfam01197 |
| Sau-4096-1 | NZ_GL638179.1 | + | 18326 | 18355 | **RNA**→ rpmE2 (PRK01678)Ribosomal_L31 (pfam01197)→ |
| Sau-50-1 | NZ_GG749318.1 | - | 55764 | 55735 | **RNA**→ rpmE2 (PRK01678)Ribosomal_L31 (pfam01197)→ rpmE2 (PRK01678)Ribosomal_L31 (pfam01197 |
| Sau-41-1 | NZ_GG770528.1 | - | 56131 | 56102 | **RNA**→ rpmE2 (PRK01678)Ribosomal_L31 (pfam01197)→ rpmE2 (PRK01678)Ribosomal_L31 (pfam01197 |
| Sau-33-1 | NZ_GG730293.1 | - | 56576 | 56547 | **RNA**→ rpmE2 (PRK01678)Ribosomal_L31 (pfam01197)→ rpmE2 (PRK01678)Ribosomal_L31 (pfam01197 |
| Sau-52-1 | NZ_GG730203.1 | - | 55609 | 55580 | **RNA**→ rpmE2 (PRK01678)Ribosomal_L31 (pfam01197)→ rpmE2 (PRK01678)Ribosomal_L31 (pfam01197 |
| Sau-37-1 | NZ_GG700611.1 | - | 55640 | 55611 | **RNA**→ rpmE2 (PRK01678)Ribosomal_L31 (pfam01197)→ rpmE2 (PRK01678)Ribosomal_L31 (pfam01197 |
| Sau-29-1 | NZ_GG774481.1 | + | 1087427 | 1087456 | **RNA**→ rpmE2 (PRK01678)Ribosomal_L31 (pfam01197)→ |
| Sau-28-1 | NZ_GG731509.1 | - | 56401 | 56372 | **RNA**→ rpmE2 (PRK01678)Ribosomal_L31 (pfam01197)→ rpmE2 (PRK01678)Ribosomal_L31 (pfam01197 |
| Sau-4097-1 | NZ_GL638212.1 | + | 18325 | 18354 | **RNA**→ rpmE2 (PRK01678)Ribosomal_L31 (pfam01197)→ |
| Sau-24-1 | NZ_GG749070.1 | - | 56614 | 56585 | **RNA**→ rpmE2 (PRK01678)Ribosomal_L31 (pfam01197)→ rpmE2 (PRK01678)Ribosomal_L31 (pfam01197 |
| Sau-51-1 | NZ_GG700626.1 | - | 56171 | 56142 | **RNA**→ rpmE2 (PRK01678)Ribosomal_L31 (pfam01197)→ rpmE2 (PRK01678)Ribosomal_L31 (pfam01197 |
| Sau-49-1 | NZ_GG749011.1 | - | 56128 | 56099 | **RNA**→ rpmE2 (PRK01678)Ribosomal_L31 (pfam01197)→ rpmE2 (PRK01678)Ribosomal_L31 (pfam01197 |
| Sau-66-1 | NZ_GG698162.1 | + | 52322 | 52351 | **RNA**→ rpmE2 (PRK01678)Ribosomal_L31 (pfam01197)→ |
| Sau-26-1 | NZ_GG700588.1 | - | 56037 | 56008 | **RNA**→ rpmE2 (PRK01678)Ribosomal_L31 (pfam01197)→ rpmE2 (PRK01678)Ribosomal_L31 (pfam01197 |
| Sau-4104-1 | NZ_GL538356.1 | + | 1145007 | 1145036 | **RNA**→ rpmE2 (PRK01678)Ribosomal_L31 (pfam01197)→ |
| Sau-3769-1 | NZ_KB820900.1 | - | 56894 | 56865 | **RNA**→ rpmE2 (PRK01678)Ribosomal_L31 (pfam01197)→ |
| Sau-3803-1 | NZ_KB821084.1 | - | 129644 | 129615 | **RNA**→ rpmE2 (PRK01678)Ribosomal_L31 (pfam01197)→ |
| Sau-3808-1 | NZ_KB821121.1 | - | 234536 | 234507 | **RNA**→ rpmE2 (PRK01678)Ribosomal_L31 (pfam01197)→ |
| Sau-3770-1 | NZ_KB820903.1 | - | 2188304 | 2188275 | **RNA**→ rpmE2 (PRK01678)Ribosomal_L31 (pfam01197)→ |
| Sau-3783-1 | NZ_KB820952.1 | - | 170256 | 170227 | **RNA**→ rpmE2 (PRK01678)Ribosomal_L31 (pfam01197)→ |
| Sau-3776-1 | NZ_KB820925.1 | - | 1613986 | 1613957 | **RNA**→ rpmE2 (PRK01678)Ribosomal_L31 (pfam01197)→ |
| Sau-3795-1 | NZ_KB821037.1 | - | 472750 | 472721 | **RNA**→ rpmE2 (PRK01678)Ribosomal_L31 (pfam01197)→ |
| Sau-3773-1 | NZ_KB820909.1 | - | 2209641 | 2209612 | **RNA**→ rpmE2 (PRK01678)Ribosomal_L31 (pfam01197)→ |
| Sau-3817-1 | NZ_KB821167.1 | - | 239480 | 239451 | **RNA**→ rpmE2 (PRK01678)Ribosomal_L31 (pfam01197)→ |
| Sau-3807-1 | NZ_KB821115.1 | - | 229502 | 229473 | **RNA**→ rpmE2 (PRK01678)Ribosomal_L31 (pfam01197)→ |
| Sau-3812-1 | NZ_KB821128.1 | - | 1585082 | 1585053 | **RNA**→ rpmE2 (PRK01678)Ribosomal_L31 (pfam01197)→ |
| Sau-3820-1 | NZ_KB821195.1 | - | 180341 | 180312 | **RNA**→ rpmE2 (PRK01678)Ribosomal_L31 (pfam01197)→ |
| Sau-3767-1 | NZ_KB820894.1 | - | 56894 | 56865 | **RNA**→ rpmE2 (PRK01678)Ribosomal_L31 (pfam01197)→ |
| Sau-3824-1 | NZ_KB821208.1 | - | 56783 | 56754 | **RNA**→ rpmE2 (PRK01678)Ribosomal_L31 (pfam01197)→ |
| Sau-3801-1 | NZ_KB821081.1 | - | 126216 | 126187 | **RNA**→ rpmE2 (PRK01678)Ribosomal_L31 (pfam01197)→ |
| Sau-3802-1 | NZ_KB821076.1 | - | 1638120 | 1638091 | **RNA**→ rpmE2 (PRK01678)Ribosomal_L31 (pfam01197)→ |
| Sau-3819-1 | NZ_KB821188.1 | - | 183209 | 183180 | **RNA**→ rpmE2 (PRK01678)Ribosomal_L31 (pfam01197)→ |
| Sau-3799-1 | NZ_KB821067.1 | - | 181606 | 181577 | **RNA**→ rpmE2 (PRK01678)Ribosomal_L31 (pfam01197)→ |
| Sau-3788-1 | NZ_KB820982.1 | - | 114751 | 114722 | **RNA**→ rpmE2 (PRK01678)Ribosomal_L31 (pfam01197)→ |
| Sau-3814-1 | NZ_KB821141.1 | - | 359642 | 359613 | **RNA**→ rpmE2 (PRK01678)Ribosomal_L31 (pfam01197)→ |
| Sau-3805-1 | NZ_KB821105.1 | - | 1111629 | 1111600 | **RNA**→ rpmE2 (PRK01678)Ribosomal_L31 (pfam01197)→ |
| Sau-3822-1 | NZ_KB821198.1 | - | 2190463 | 2190434 | **RNA**→ rpmE2 (PRK01678)Ribosomal_L31 (pfam01197)→ |
| Sau-3813-1 | NZ_KB821144.1 | - | 57288 | 57259 | **RNA**→ rpmE2 (PRK01678)Ribosomal_L31 (pfam01197)→ |
| Sau-3818-1 | NZ_KB821175.1 | - | 1680280 | 1680251 | **RNA**→ rpmE2 (PRK01678)Ribosomal_L31 (pfam01197)→ |
| Sau-3811-1 | NZ_KB821137.1 | - | 114925 | 114896 | **RNA**→ rpmE2 (PRK01678)Ribosomal_L31 (pfam01197)→ |
| Sau-3775-1 | NZ_KB820921.1 | - | 1705142 | 1705113 | **RNA**→ rpmE2 (PRK01678)Ribosomal_L31 (pfam01197)→ |

| | | | | | | |
|---|---|---|---|---|---|---|
| Sau-3784-1 | NZ_KB820963.1 | - | 1638154 | 1638125 | **RNA**→ | rpmE2 (PRK01678)Ribosomal_L31 (pfam01197)→ |
| Sau-3800-1 | NZ_KB821070.1 | - | 253442 | 253413 | **RNA**→ | rpmE2 (PRK01678)Ribosomal_L31 (pfam01197)→ |
| Sau-3774-1 | NZ_KB820916.1 | - | 1896448 | 1896419 | **RNA**→ | rpmE2 (PRK01678)Ribosomal_L31 (pfam01197)→ |
| Sau-3809-1 | NZ_KB821124.1 | - | 192056 | 192027 | **RNA**→ | rpmE2 (PRK01678)Ribosomal_L31 (pfam01197)→ |
| Sau-3771-1 | NZ_KB820912.1 | - | 1676207 | 1676178 | **RNA**→ | rpmE2 (PRK01678)Ribosomal_L31 (pfam01197)→ |
| Sau-3794-1 | NZ_KB821023.1 | - | 115012 | 114983 | **RNA**→ | rpmE2 (PRK01678)Ribosomal_L31 (pfam01197)→ |
| Sau-3793-1 | NZ_KB821021.1 | - | 59768 | 59739 | **RNA**→ | rpmE2 (PRK01678)Ribosomal_L31 (pfam01197)→ |
| Sau-3816-1 | NZ_KB821155.1 | - | 56763 | 56734 | **RNA**→ | rpmE2 (PRK01678)Ribosomal_L31 (pfam01197)→ |
| Sau-3806-1 | NZ_KB821113.1 | - | 435020 | 434991 | **RNA**→ | rpmE2 (PRK01678)Ribosomal_L31 (pfam01197)→ |
| Sau-3782-1 | NZ_KB820938.1 | - | 56763 | 56734 | **RNA**→ | rpmE2 (PRK01678)Ribosomal_L31 (pfam01197)→ |
| Sau-3786-1 | NZ_KB820971.1 | - | 459385 | 459356 | **RNA**→ | rpmE2 (PRK01678)Ribosomal_L31 (pfam01197)→ |
| Sau-3840-1 | NZ_KB821307.1 | - | 468982 | 468953 | **RNA**→ | rpmE2 (PRK01678)Ribosomal_L31 (pfam01197)→ |
| Sau-3848-1 | NZ_KB821355.1 | - | 1644452 | 1644423 | **RNA**→ | rpmE2 (PRK01678)Ribosomal_L31 (pfam01197)→ |
| Sau-3837-1 | NZ_KB821284.1 | - | 124072 | 124043 | **RNA**→ | rpmE2 (PRK01678)Ribosomal_L31 (pfam01197)→ |
| Sau-3804-1 | NZ_KB821093.1 | - | 61324 | 61295 | **RNA**→ | rpmE2 (PRK01678)Ribosomal_L31 (pfam01197)→ |
| Sau-3849-1 | NZ_KB821366.1 | - | 230978 | 230949 | **RNA**→ | rpmE2 (PRK01678)Ribosomal_L31 (pfam01197)→ |
| Sau-3828-1 | NZ_KB821234.1 | - | 184607 | 184578 | **RNA**→ | rpmE2 (PRK01678)Ribosomal_L31 (pfam01197)→ |
| Sau-3797-1 | NZ_KB821056.1 | - | 198382 | 198353 | **RNA**→ | rpmE2 (PRK01678)Ribosomal_L31 (pfam01197)→ |
| Sau-3791-1 | NZ_KB821003.1 | - | 200561 | 200532 | **RNA**→ | rpmE2 (PRK01678)Ribosomal_L31 (pfam01197)→ |
| Sau-3850-1 | NZ_KB821373.1 | - | 58248 | 58219 | **RNA**→ | rpmE2 (PRK01678)Ribosomal_L31 (pfam01197)→ |
| Sau-3798-1 | NZ_KB821045.1 | - | 56967 | 56938 | **RNA**→ | rpmE2 (PRK01678)Ribosomal_L31 (pfam01197)→ |
| Sau-3785-1 | NZ_KB820966.1 | - | 1684535 | 1684506 | **RNA**→ | rpmE2 (PRK01678)Ribosomal_L31 (pfam01197)→ |
| Sau-3906-1 | NZ_KB821822.1 | - | 230922 | 230893 | **RNA**→ | rpmE2 (PRK01678)Ribosomal_L31 (pfam01197)→ |
| Sau-3877-1 | NZ_KB821568.1 | - | 232229 | 232200 | **RNA**→ | rpmE2 (PRK01678)Ribosomal_L31 (pfam01197)→ |
| Sau-3868-1 | NZ_KB821491.1 | - | 468075 | 468046 | **RNA**→ | rpmE2 (PRK01678)Ribosomal_L31 (pfam01197)→ |
| Sau-3930-1 | NZ_KB822033.1 | - | 56588 | 56559 | **RNA**→ | rpmE2 (PRK01678)Ribosomal_L31 (pfam01197)→ |
| Sau-3825-1 | NZ_KB821214.1 | - | 470766 | 470737 | **RNA**→ | rpmE2 (PRK01678)Ribosomal_L31 (pfam01197)→ |
| Sau-3938-1 | NZ_KB822062.1 | - | 468208 | 468179 | **RNA**→ | rpmE2 (PRK01678)Ribosomal_L31 (pfam01197)→ |
| Sau-3951-1 | NZ_KB822134.1 | - | 1633116 | 1633087 | **RNA**→ | rpmE2 (PRK01678)Ribosomal_L31 (pfam01197)→ |
| Sau-3929-1 | NZ_KB822014.1 | - | 49965 | 49936 | **RNA**→ | rpmE2 (PRK01678)Ribosomal_L31 (pfam01197)→ |
| Sau-3889-1 | NZ_KB821698.1 | - | 217486 | 217457 | **RNA**→ | rpmE2 (PRK01678)Ribosomal_L31 (pfam01197)→ |
| Sau-3911-1 | NZ_KB821846.1 | - | 231900 | 231871 | **RNA**→ | rpmE2 (PRK01678)Ribosomal_L31 (pfam01197)→ |
| Sau-3879-1 | NZ_KB821583.1 | - | 49963 | 49934 | **RNA**→ | rpmE2 (PRK01678)Ribosomal_L31 (pfam01197)→ |
| Sau-3815-1 | NZ_KB821139.1 | - | 417187 | 417158 | **RNA**→ | rpmE2 (PRK01678)Ribosomal_L31 (pfam01197)→ |
| Sau-3940-1 | NZ_KB822075.1 | - | 1587356 | 1587327 | **RNA**→ | rpmE2 (PRK01678)Ribosomal_L31 (pfam01197)→ |
| Sau-3905-1 | NZ_KB821817.1 | - | 60935 | 60906 | **RNA**→ | rpmE2 (PRK01678)Ribosomal_L31 (pfam01197)→ |
| Sau-3870-1 | NZ_KB821499.1 | - | 57430 | 57401 | **RNA**→ | rpmE2 (PRK01678)Ribosomal_L31 (pfam01197)→ |
| Sau-3937-1 | NZ_KB822053.1 | - | 231944 | 231915 | **RNA**→ | rpmE2 (PRK01678)Ribosomal_L31 (pfam01197)→ |
| Sau-3838-1 | NZ_KB821316.1 | - | 175381 | 175352 | **RNA**→ | rpmE2 (PRK01678)Ribosomal_L31 (pfam01197)→ |
| Sau-3942-1 | NZ_KB822082.1 | - | 925424 | 925395 | **RNA**→ | rpmE2 (PRK01678)Ribosomal_L31 (pfam01197)→ |
| Sau-3885-1 | NZ_KB821667.1 | - | 232434 | 232405 | **RNA**→ | rpmE2 (PRK01678)Ribosomal_L31 (pfam01197)→ |
| Sau-3854-1 | NZ_KB821402.1 | - | 222935 | 222906 | **RNA**→ | rpmE2 (PRK01678)Ribosomal_L31 (pfam01197)→ |
| Sau-3931-1 | NZ_KB822005.1 | - | 102504 | 102475 | **RNA**→ | rpmE2 (PRK01678)Ribosomal_L31 (pfam01197)→ |
| Sau-3887-1 | NZ_KB821650.1 | - | 172485 | 172456 | **RNA**→ | rpmE2 (PRK01678)Ribosomal_L31 (pfam01197)→ |
| Sau-3860-1 | NZ_KB821438.1 | - | 1666799 | 1666770 | **RNA**→ | rpmE2 (PRK01678)Ribosomal_L31 (pfam01197)→ |
| Sau-3892-1 | NZ_KB821706.1 | - | 100662 | 100633 | **RNA**→ | rpmE2 (PRK01678)Ribosomal_L31 (pfam01197)→ |
| Sau-3875-1 | NZ_KB821519.1 | - | 56762 | 56733 | **RNA**→ | rpmE2 (PRK01678)Ribosomal_L31 (pfam01197)→ |
| Sau-3943-1 | NZ_KB822376.1 | - | 54872 | 54843 | **RNA**→ | rpmE2 (PRK01678)Ribosomal_L31 (pfam01197)→ |
| Sau-3882-1 | NZ_KB821615.1 | - | 57538 | 57509 | **RNA**→ | rpmE2 (PRK01678)Ribosomal_L31 (pfam01197)→ |
| Sau-3982-1 | NZ_KB822322.1 | - | 233801 | 233772 | **RNA**→ | rpmE2 (PRK01678)Ribosomal_L31 (pfam01197)→ |
| Sau-3853-1 | NZ_KB821387.1 | - | 109205 | 109176 | **RNA**→ | rpmE2 (PRK01678)Ribosomal_L31 (pfam01197)→ |
| Sau-3865-1 | NZ_KB821484.1 | - | 219814 | 219785 | **RNA**→ | rpmE2 (PRK01678)Ribosomal_L31 (pfam01197)→ |

| | | | | | |
|---|---|---|---|---|---|
| Sau-3950-1 | NZ_KB822128.1 | - | 401009 | 400980 | **RNA**→ rpmE2 (PRK01678)Ribosomal_L31 (pfam01197)→ |
| Sau-3863-1 | NZ_KB821442.1 | - | 167231 | 167202 | **RNA**→ rpmE2 (PRK01678)Ribosomal_L31 (pfam01197)→ |
| Sau-3923-1 | NZ_KB822366.1 | - | 61547 | 61518 | **RNA**→ rpmE2 (PRK01678)Ribosomal_L31 (pfam01197)→ |
| Sau-3894-1 | NZ_KB821734.1 | - | 57586 | 57557 | **RNA**→ rpmE2 (PRK01678)Ribosomal_L31 (pfam01197)→ |
| Sau-3946-1 | NZ_KB822105.1 | - | 105409 | 105380 | **RNA**→ rpmE2 (PRK01678)Ribosomal_L31 (pfam01197)→ |
| Sau-3904-1 | NZ_KB821811.1 | - | 507764 | 507735 | **RNA**→ rpmE2 (PRK01678)Ribosomal_L31 (pfam01197)→ |
| Sau-3924-1 | NZ_KB821972.1 | - | 25249 | 25220 | **RNA**→ rpmE2 (PRK01678)Ribosomal_L31 (pfam01197)→ |
| Sau-3953-1 | NZ_KB822142.1 | - | 106748 | 106719 | **RNA**→ rpmE2 (PRK01678)Ribosomal_L31 (pfam01197)→ |
| Sau-3921-1 | NZ_KB821932.1 | - | 231611 | 231582 | **RNA**→ rpmE2 (PRK01678)Ribosomal_L31 (pfam01197)→ |
| Sau-3971-1 | NZ_KB822385.1 | - | 111190 | 111161 | **RNA**→ rpmE2 (PRK01678)Ribosomal_L31 (pfam01197)→ |
| Sau-3878-1 | NZ_KB821575.1 | - | 229624 | 229595 | **RNA**→ rpmE2 (PRK01678)Ribosomal_L31 (pfam01197)→ |
| Sau-3925-1 | NZ_KB821957.1 | - | 61593 | 61564 | **RNA**→ rpmE2 (PRK01678)Ribosomal_L31 (pfam01197)→ |
| Sau-3909-1 | NZ_KB821826.1 | - | 233560 | 233531 | **RNA**→ rpmE2 (PRK01678)Ribosomal_L31 (pfam01197)→ |
| Sau-3845-1 | NZ_KB821335.1 | - | 2130755 | 2130726 | **RNA**→ rpmE2 (PRK01678)Ribosomal_L31 (pfam01197)→ |
| Sau-3952-1 | NZ_KB822138.1 | - | 229498 | 229469 | **RNA**→ rpmE2 (PRK01678)Ribosomal_L31 (pfam01197)→ |
| Sau-3834-1 | NZ_KB821269.1 | - | 56603 | 56574 | **RNA**→ rpmE2 (PRK01678)Ribosomal_L31 (pfam01197)→ |
| Sau-3900-1 | NZ_KB821789.1 | + | 61055 | 61084 | **RNA**→ rpmE2 (PRK01678)Ribosomal_L31 (pfam01197)→ |
| Sau-3917-1 | NZ_KB821900.1 | - | 469540 | 469511 | **RNA**→ rpmE2 (PRK01678)Ribosomal_L31 (pfam01197)→ |
| Sau-3941-1 | NZ_KB822074.1 | - | 55982 | 55953 | **RNA**→ rpmE2 (PRK01678)Ribosomal_L31 (pfam01197)→ |
| Sau-3893-1 | NZ_KB821716.1 | + | 53720 | 53749 | **RNA**→ rpmE2 (PRK01678)Ribosomal_L31 (pfam01197)→ |
| Sau-3957-1 | NZ_KB822184.1 | - | 58538 | 58509 | **RNA**→ rpmE2 (PRK01678)Ribosomal_L31 (pfam01197)→ |
| Sau-3839-1 | NZ_KB821299.1 | - | 1190418 | 1190389 | **RNA**→ rpmE2 (PRK01678)Ribosomal_L31 (pfam01197)→ |
| Sau-3872-1 | NZ_KB821506.1 | - | 1691566 | 1691537 | **RNA**→ rpmE2 (PRK01678)Ribosomal_L31 (pfam01197)→ |
| Sau-3796-1 | NZ_KB821038.1 | - | 178180 | 178151 | **RNA**→ rpmE2 (PRK01678)Ribosomal_L31 (pfam01197)→ |
| Sau-3862-1 | NZ_KB821450.1 | - | 176222 | 176193 | **RNA**→ rpmE2 (PRK01678)Ribosomal_L31 (pfam01197)→ |
| Sau-3884-1 | NZ_KB821638.1 | - | 458897 | 458868 | **RNA**→ rpmE2 (PRK01678)Ribosomal_L31 (pfam01197)→ |
| Sau-3949-1 | NZ_KB822114.1 | + | 1120890 | 1120919 | **RNA**→ rpmE2 (PRK01678)Ribosomal_L31 (pfam01197)→ |
| Sau-3855-1 | NZ_KB821405.1 | - | 1639677 | 1639648 | **RNA**→ rpmE2 (PRK01678)Ribosomal_L31 (pfam01197)→ |
| Sau-3831-1 | NZ_KB821253.1 | - | 194612 | 194583 | **RNA**→ rpmE2 (PRK01678)Ribosomal_L31 (pfam01197)→ |
| Sau-3830-1 | NZ_KB821248.1 | - | 1686649 | 1686620 | **RNA**→ rpmE2 (PRK01678)Ribosomal_L31 (pfam01197)→ |
| Sau-3843-1 | NZ_KB821317.1 | - | 56762 | 56733 | **RNA**→ rpmE2 (PRK01678)Ribosomal_L31 (pfam01197)→ |
| Sau-3842-1 | NZ_KB821326.1 | - | 56154 | 56125 | **RNA**→ rpmE2 (PRK01678)Ribosomal_L31 (pfam01197)→ |
| Sau-3832-1 | NZ_KB821262.1 | - | 231637 | 231608 | **RNA**→ rpmE2 (PRK01678)Ribosomal_L31 (pfam01197)→ |
| Sau-3835-1 | NZ_KB821276.1 | - | 225444 | 225415 | **RNA**→ rpmE2 (PRK01678)Ribosomal_L31 (pfam01197)→ |
| Sau-3897-1 | NZ_KB821760.1 | - | 464567 | 464538 | **RNA**→ rpmE2 (PRK01678)Ribosomal_L31 (pfam01197)→ |
| Sau-3912-1 | NZ_KB821848.1 | + | 952842 | 952871 | **RNA**→ rpmE2 (PRK01678)Ribosomal_L31 (pfam01197)→ |
| Sau-1858-1 | NZ_KI985158.1 | - | 238686 | 238657 | **RNA**→ rpmE2 (PRK01678)Ribosomal_L31 (pfam01197)→ |
| Sau-1866-1 | NZ_KI985258.1 | - | 217786 | 217757 | **RNA**→ rpmE2 (PRK01678)Ribosomal_L31 (pfam01197)→ |
| Sau-1867-1 | NZ_KI985276.1 | - | 36026 | 35997 | **RNA**→ rpmE2 (PRK01678)Ribosomal_L31 (pfam01197)→ |
| Sau-1980-1 | NZ_KI989431.1 | - | 24749 | 24720 | **RNA**→ rpmE2 (PRK01678)Ribosomal_L31 (pfam01197)→ |
| Sau-1873-1 | NZ_KI985536.1 | - | 61531 | 61502 | **RNA**→ rpmE2 (PRK01678)Ribosomal_L31 (pfam01197)→ |
| Sau-2017-1 | NZ_KI990677.1 | - | 113811 | 113782 | **RNA**→ rpmE2 (PRK01678)Ribosomal_L31 (pfam01197)→ |
| Sau-1878-1 | NZ_KI985707.1 | - | 143557 | 143528 | **RNA**→ rpmE2 (PRK01678)Ribosomal_L31 (pfam01197)→ |
| Sau-2023-1 | NZ_KI990898.1 | - | 35180 | 35151 | **RNA**→ rpmE2 (PRK01678)Ribosomal_L31 (pfam01197)→ |
| Sau-1886-1 | NZ_KI985949.1 | - | 133925 | 133896 | **RNA**→ rpmE2 (PRK01678)Ribosomal_L31 (pfam01197)→ |
| Sau-1762-1 | NZ_KI992135.1 | - | 55326 | 55297 | **RNA**→ rpmE2 (PRK01678)Ribosomal_L31 (pfam01197)→ |
| Sau-1768-1 | NZ_KI992364.1 | - | 55372 | 55343 | **RNA**→ rpmE2 (PRK01678)Ribosomal_L31 (pfam01197)→ |
| Sau-2271-1 | NZ_KI992891.1 | - | 36491 | 36462 | **RNA**→ rpmE2 (PRK01678)Ribosomal_L31 (pfam01197)→ |
| Sau-279-1 | NZ_KI978876.1 | - | 56894 | 56865 | **RNA**→ rpmE2 (PRK01678)Ribosomal_L31 (pfam01197)→ |
| Sau-2280-1 | NZ_KI993053.1 | - | 20693 | 20664 | **RNA**→ rpmE2 (PRK01678)Ribosomal_L31 (pfam01197)→ |
| Sau-2287-1 | NZ_KI993232.1 | - | 123205 | 123176 | **RNA**→ rpmE2 (PRK01678)Ribosomal_L31 (pfam01197)→ |
| Sau-281-1 | NZ_KI978885.1 | - | 194479 | 194450 | **RNA**→ rpmE2 (PRK01678)Ribosomal_L31 (pfam01197)→ |

| | | | | | |
|---|---|---|---|---|---|
| Sau-2293-1 | NZ_KI993390.1 | - | 43602 | 43573 | **RNA**→ rpmE2 (PRK01678)Ribosomal_L31 (pfam01197)→ |
| Sau-1141-1 | NZ_KI979009.1 | - | 239541 | 239512 | **RNA**→ rpmE2 (PRK01678)Ribosomal_L31 (pfam01197)→ |
| Sau-2299-1 | NZ_KI993707.1 | - | 113749 | 113720 | **RNA**→ rpmE2 (PRK01678)Ribosomal_L31 (pfam01197)→ |
| Sau-2304-1 | NZ_KI993857.1 | - | 79464 | 79435 | **RNA**→ rpmE2 (PRK01678)Ribosomal_L31 (pfam01197)→ |
| Sau-859-1 | NZ_KI979238.1 | - | 56894 | 56865 | **RNA**→ rpmE2 (PRK01678)Ribosomal_L31 (pfam01197)→ |
| Sau-1969-1 | NZ_KI988913.1 | - | 118854 | 118825 | **RNA**→ rpmE2 (PRK01678)Ribosomal_L31 (pfam01197)→ |
| Sau-1755-1 | NZ_KI981742.1 | - | 61399 | 61370 | **RNA**→ rpmE2 (PRK01678)Ribosomal_L31 (pfam01197)→ |
| Sau-2009-1 | NZ_KI990354.1 | - | 28766 | 28737 | **RNA**→ rpmE2 (PRK01678)Ribosomal_L31 (pfam01197)→ |
| Sau-1764-1 | NZ_KI992236.1 | - | 113065 | 113036 | **RNA**→ rpmE2 (PRK01678)Ribosomal_L31 (pfam01197)→ |
| Sau-1777-1 | NZ_KI981979.1 | - | 54757 | 54728 | **RNA**→ rpmE2 (PRK01678)Ribosomal_L31 (pfam01197)→ |
| Sau-2276-1 | NZ_KI992998.1 | - | 78935 | 78906 | **RNA**→ rpmE2 (PRK01678)Ribosomal_L31 (pfam01197)→ |
| Sau-294-1 | NZ_KI976868.1 | - | 56894 | 56865 | **RNA**→ rpmE2 (PRK01678)Ribosomal_L31 (pfam01197)→ |
| Sau-2285-1 | NZ_KI993140.1 | - | 116825 | 116796 | **RNA**→ rpmE2 (PRK01678)Ribosomal_L31 (pfam01197)→ |
| Sau-1134-1 | NZ_KI977602.1 | - | 112534 | 112505 | **RNA**→ rpmE2 (PRK01678)Ribosomal_L31 (pfam01197)→ |
| Sau-2291-1 | NZ_KI993346.1 | - | 55372 | 55343 | **RNA**→ rpmE2 (PRK01678)Ribosomal_L31 (pfam01197)→ |
| Sau-852-1 | NZ_KI977780.1 | - | 56893 | 56864 | **RNA**→ rpmE2 (PRK01678)Ribosomal_L31 (pfam01197)→ |
| Sau-2297-1 | NZ_KI993607.1 | - | 55327 | 55298 | **RNA**→ rpmE2 (PRK01678)Ribosomal_L31 (pfam01197)→ |
| Sau-855-1 | NZ_KI977949.1 | - | 56894 | 56865 | **RNA**→ rpmE2 (PRK01678)Ribosomal_L31 (pfam01197)→ |
| Sau-2302-1 | NZ_KI993828.1 | - | 62154 | 62125 | **RNA**→ rpmE2 (PRK01678)Ribosomal_L31 (pfam01197)→ |
| Sau-2614-1 | NZ_KI978573.1 | - | 110501 | 110472 | **RNA**→ rpmE2 (PRK01678)Ribosomal_L31 (pfam01197)→ |
| Sau-2308-1 | NZ_KI993984.1 | - | 35334 | 35305 | **RNA**→ rpmE2 (PRK01678)Ribosomal_L31 (pfam01197)→ |
| Sau-888-1 | NZ_KI978673.1 | - | 56894 | 56865 | **RNA**→ rpmE2 (PRK01678)Ribosomal_L31 (pfam01197)→ |
| Sau-2242-1 | NZ_KI996121.1 | - | 25129 | 25100 | **RNA**→ rpmE2 (PRK01678)Ribosomal_L31 (pfam01197)→ |
| Sau-2273-1 | NZ_KI992923.1 | - | 25227 | 25198 | **RNA**→ rpmE2 (PRK01678)Ribosomal_L31 (pfam01197)→ |
| Sau-2283-1 | NZ_KI993098.1 | - | 15617 | 15588 | **RNA**→ rpmE2 (PRK01678)Ribosomal_L31 (pfam01197)→ |
| Sau-2294-1 | NZ_KI993454.1 | - | 29085 | 29056 | **RNA**→ rpmE2 (PRK01678)Ribosomal_L31 (pfam01197)→ |
| Sau-2296-1 | NZ_KI993543.1 | - | 56763 | 56734 | **RNA**→ rpmE2 (PRK01678)Ribosomal_L31 (pfam01197)→ |
| Sau-2305-1 | NZ_KI993943.1 | - | 24749 | 24720 | **RNA**→ rpmE2 (PRK01678)Ribosomal_L31 (pfam01197)→ |
| Sau-863-1 | NZ_KI978799.1 | - | 56894 | 56865 | **RNA**→ rpmE2 (PRK01678)Ribosomal_L31 (pfam01197)→ |
| Sau-1760-1 | NZ_KI992041.1 | - | 70324 | 70295 | **RNA**→ rpmE2 (PRK01678)Ribosomal_L31 (pfam01197)→ |
| Sau-2270-1 | NZ_KI992811.1 | - | 36041 | 36012 | **RNA**→ rpmE2 (PRK01678)Ribosomal_L31 (pfam01197)→ |
| Sau-1098-1 | NZ_KI978949.1 | - | 111685 | 111656 | **RNA**→ rpmE2 (PRK01678)Ribosomal_L31 (pfam01197)→ |
| Sau-2279-1 | NZ_KI993025.1 | - | 35767 | 35738 | **RNA**→ rpmE2 (PRK01678)Ribosomal_L31 (pfam01197)→ |
| Sau-1138-1 | NZ_KI979264.1 | - | 110974 | 110945 | **RNA**→ rpmE2 (PRK01678)Ribosomal_L31 (pfam01197)→ |
| Sau-2298-1 | NZ_KI993612.1 | - | 25185 | 25156 | **RNA**→ rpmE2 (PRK01678)Ribosomal_L31 (pfam01197)→ |
| Sau-1754-1 | NZ_KI981682.1 | - | 54757 | 54728 | **RNA**→ rpmE2 (PRK01678)Ribosomal_L31 (pfam01197)→ |
| Sau-2303-1 | NZ_KI993830.1 | - | 131178 | 131149 | **RNA**→ rpmE2 (PRK01678)Ribosomal_L31 (pfam01197)→ |
| Sau-1775-1 | NZ_KI981936.1 | - | 56120 | 56091 | **RNA**→ rpmE2 (PRK01678)Ribosomal_L31 (pfam01197)→ |
| Sau-2309-1 | NZ_KI994027.1 | - | 35968 | 35939 | **RNA**→ rpmE2 (PRK01678)Ribosomal_L31 (pfam01197)→ |
| Sau-2264-1 | NZ_KI992684.1 | - | 35290 | 35261 | **RNA**→ rpmE2 (PRK01678)Ribosomal_L31 (pfam01197)→ |
| Sau-2251-1 | NZ_KI996449.1 | - | 35284 | 35255 | **RNA**→ rpmE2 (PRK01678)Ribosomal_L31 (pfam01197)→ |
| Sau-2260-1 | NZ_KI996704.1 | - | 291284 | 291255 | **RNA**→ rpmE2 (PRK01678)Ribosomal_L31 (pfam01197)→ |
| Sau-2640-1 | NZ_KI975748.1 | - | 111952 | 111923 | **RNA**→ rpmE2 (PRK01678)Ribosomal_L31 (pfam01197)→ |
| Sau-1124-1 | NZ_KI977461.1 | - | 235312 | 235283 | **RNA**→ rpmE2 (PRK01678)Ribosomal_L31 (pfam01197)→ |
| Sau-1135-1 | NZ_KI977932.1 | - | 238639 | 238610 | **RNA**→ rpmE2 (PRK01678)Ribosomal_L31 (pfam01197)→ |
| Sau-3827-1 | NZ_KB821221.1 | - | 62154 | 62125 | **RNA**→ rpmE2 (PRK01678)Ribosomal_L31 (pfam01197)→ |
| Sau-3836-1 | NZ_KB821280.1 | - | 58197 | 58168 | **RNA**→ rpmE2 (PRK01678)Ribosomal_L31 (pfam01197)→ |
| Sau-3826-1 | NZ_KB821233.1 | - | 57586 | 57557 | **RNA**→ rpmE2 (PRK01678)Ribosomal_L31 (pfam01197)→ |
| Sau-3852-1 | NZ_KB821382.1 | - | 61675 | 61646 | **RNA**→ rpmE2 (PRK01678)Ribosomal_L31 (pfam01197)→ |
| Sau-3895-1 | NZ_KB821746.1 | - | 231536 | 231507 | **RNA**→ rpmE2 (PRK01678)Ribosomal_L31 (pfam01197)→ |
| Sau-3810-1 | NZ_KB821130.1 | - | 2768476 | 2768447 | **RNA**→ rpmE2 (PRK01678)Ribosomal_L31 (pfam01197)→ |
| Sau-3888-1 | NZ_KB821686.1 | - | 233138 | 233109 | **RNA**→ rpmE2 (PRK01678)Ribosomal_L31 (pfam01197)→ |

| | | | | | |
|---|---|---|---|---|---|
| Sau-3886-1 | NZ_KB821678.1 | - | 1113791 | 1113762 | **RNA**→ rpmE2 (PRK01678)Ribosomal_L31 (pfam01197)→ |
| Sau-3857-1 | NZ_KB821413.1 | + | 2436466 | 2436495 | **RNA**→ rpmE2 (PRK01678)Ribosomal_L31 (pfam01197)→ |
| Sau-3915-1 | NZ_KB821872.1 | - | 61469 | 61440 | **RNA**→ rpmE2 (PRK01678)Ribosomal_L31 (pfam01197)→ |
| Sau-3874-1 | NZ_KB821512.1 | - | 459630 | 459601 | **RNA**→ rpmE2 (PRK01678)Ribosomal_L31 (pfam01197)→ |
| Sau-3876-1 | NZ_KB821535.1 | - | 179512 | 179483 | **RNA**→ rpmE2 (PRK01678)Ribosomal_L31 (pfam01197)→ |
| Sau-3866-1 | NZ_KB821477.1 | - | 459748 | 459719 | **RNA**→ rpmE2 (PRK01678)Ribosomal_L31 (pfam01197)→ |
| Sau-3856-1 | NZ_KB821410.1 | - | 414274 | 414245 | **RNA**→ rpmE2 (PRK01678)Ribosomal_L31 (pfam01197)→ |
| Sau-3864-1 | NZ_KB821473.1 | - | 101203 | 101174 | **RNA**→ rpmE2 (PRK01678)Ribosomal_L31 (pfam01197)→ |
| Sau-3833-1 | NZ_KB821272.1 | - | 2101380 | 2101351 | **RNA**→ rpmE2 (PRK01678)Ribosomal_L31 (pfam01197)→ |
| Sau-3792-1 | NZ_KB821011.1 | - | 57688 | 57659 | **RNA**→ rpmE2 (PRK01678)Ribosomal_L31 (pfam01197)→ |
| Sau-3869-1 | NZ_KB821500.1 | - | 101923 | 101894 | **RNA**→ rpmE2 (PRK01678)Ribosomal_L31 (pfam01197)→ |
| Sau-3880-1 | NZ_KB821595.1 | - | 116823 | 116794 | **RNA**→ rpmE2 (PRK01678)Ribosomal_L31 (pfam01197)→ |
| Sau-3871-1 | NZ_KB821504.1 | - | 1313322 | 1313293 | **RNA**→ rpmE2 (PRK01678)Ribosomal_L31 (pfam01197)→ |
| Sau-3861-1 | NZ_KB821468.1 | - | 1493399 | 1493370 | **RNA**→ rpmE2 (PRK01678)Ribosomal_L31 (pfam01197)→ |
| Sau-3859-1 | NZ_KB821431.1 | - | 123349 | 123320 | **RNA**→ rpmE2 (PRK01678)Ribosomal_L31 (pfam01197)→ |
| Sau-3847-1 | NZ_KB821345.1 | - | 1062611 | 1062582 | **RNA**→ rpmE2 (PRK01678)Ribosomal_L31 (pfam01197)→ |
| Sau-3974-1 | NZ_KB822266.1 | - | 2762750 | 2762721 | **RNA**→ rpmE2 (PRK01678)Ribosomal_L31 (pfam01197)→ |
| Sau-3962-1 | NZ_KB822216.1 | - | 190880 | 190851 | **RNA**→ rpmE2 (PRK01678)Ribosomal_L31 (pfam01197)→ |
| Sau-3789-1 | NZ_KB820997.1 | - | 61740 | 61711 | **RNA**→ rpmE2 (PRK01678)Ribosomal_L31 (pfam01197)→ |
| Sau-3947-1 | NZ_KB822112.1 | - | 237858 | 237829 | **RNA**→ rpmE2 (PRK01678)Ribosomal_L31 (pfam01197)→ |
| Sau-3790-1 | NZ_KB822350.1 | - | 229728 | 229699 | **RNA**→ rpmE2 (PRK01678)Ribosomal_L31 (pfam01197)→ |
| Sau-3965-1 | NZ_KB822547.1 | - | 102904 | 102875 | **RNA**→ rpmE2 (PRK01678)Ribosomal_L31 (pfam01197)→ |
| Sau-3955-1 | NZ_KB822161.1 | - | 59333 | 59304 | **RNA**→ rpmE2 (PRK01678)Ribosomal_L31 (pfam01197)→ |
| Sau-3821-1 | NZ_KB822352.1 | - | 195129 | 195100 | **RNA**→ rpmE2 (PRK01678)Ribosomal_L31 (pfam01197)→ |
| Sau-3964-1 | NZ_KB822228.1 | - | 231908 | 231879 | **RNA**→ rpmE2 (PRK01678)Ribosomal_L31 (pfam01197)→ |
| Sau-3944-1 | NZ_KB822110.1 | - | 61121 | 61092 | **RNA**→ rpmE2 (PRK01678)Ribosomal_L31 (pfam01197)→ |
| Sau-3981-1 | NZ_KB822299.1 | - | 421723 | 421694 | **RNA**→ rpmE2 (PRK01678)Ribosomal_L31 (pfam01197)→ |
| Sau-3970-1 | NZ_KB822563.1 | - | 61444 | 61415 | **RNA**→ rpmE2 (PRK01678)Ribosomal_L31 (pfam01197)→ |
| Sau-3969-1 | NZ_KB822552.1 | - | 1670678 | 1670649 | **RNA**→ rpmE2 (PRK01678)Ribosomal_L31 (pfam01197)→ |
| Sau-3980-1 | NZ_KB822308.1 | - | 325379 | 325350 | **RNA**→ rpmE2 (PRK01678)Ribosomal_L31 (pfam01197)→ |
| Sau-3967-1 | NZ_KB822238.1 | - | 229453 | 229424 | **RNA**→ rpmE2 (PRK01678)Ribosomal_L31 (pfam01197)→ |
| Sau-34-1 | NZ_GG730155.1 | - | 56187 | 56158 | **RNA**→ rpmE2 (PRK01678)Ribosomal_L31 (pfam01197)→ rpmE2 (PRK01678)Ribosomal_L31 (pfam01197 |
| Sau-3956-1 | NZ_KB822177.1 | + | 1184966 | 1184995 | **RNA**→ rpmE2 (PRK01678)Ribosomal_L31 (pfam01197)→ |
| Sau-3858-1 | NZ_KB821416.1 | - | 458073 | 458044 | **RNA**→ rpmE2 (PRK01678)Ribosomal_L31 (pfam01197)→ |
| Sau-3934-1 | NZ_KB822067.1 | - | 238429 | 238400 | **RNA**→ rpmE2 (PRK01678)Ribosomal_L31 (pfam01197)→ |
| Sau-3975-1 | NZ_KB822273.1 | - | 466868 | 466839 | **RNA**→ rpmE2 (PRK01678)Ribosomal_L31 (pfam01197)→ |
| Sau-3898-1 | NZ_KB821769.1 | - | 57927 | 57898 | **RNA**→ rpmE2 (PRK01678)Ribosomal_L31 (pfam01197)→ |
| Sau-3960-1 | NZ_KB822204.1 | - | 230327 | 230298 | **RNA**→ rpmE2 (PRK01678)Ribosomal_L31 (pfam01197)→ |
| Sau-4072-1 | NZ_JH806525.1 | + | 18328 | 18357 | **RNA**→ rpmE2 (PRK01678)Ribosomal_L31 (pfam01197)→ |
| Sau-3926-1 | NZ_KB821950.1 | - | 354313 | 354284 | **RNA**→ rpmE2 (PRK01678)Ribosomal_L31 (pfam01197)→ |
| Sau-3903-1 | NZ_KB821777.1 | - | 1099170 | 1099141 | **RNA**→ rpmE2 (PRK01678)Ribosomal_L31 (pfam01197)→ |
| Sau-3977-1 | NZ_KB822288.1 | - | 61009 | 60980 | **RNA**→ rpmE2 (PRK01678)Ribosomal_L31 (pfam01197)→ |
| Sau-3945-1 | NZ_KB822087.1 | - | 215839 | 215810 | **RNA**→ rpmE2 (PRK01678)Ribosomal_L31 (pfam01197)→ |
| Sau-3918-1 | NZ_KB821918.1 | - | 233981 | 233952 | **RNA**→ rpmE2 (PRK01678)Ribosomal_L31 (pfam01197)→ |
| Sau-3896-1 | NZ_KB821749.1 | - | 57585 | 57556 | **RNA**→ rpmE2 (PRK01678)Ribosomal_L31 (pfam01197)→ |
| Sau-3958-1 | NZ_KB822187.1 | - | 60783 | 60754 | **RNA**→ rpmE2 (PRK01678)Ribosomal_L31 (pfam01197)→ |
| Sau-3973-1 | NZ_KB822261.1 | - | 61691 | 61662 | **RNA**→ rpmE2 (PRK01678)Ribosomal_L31 (pfam01197)→ |
| Sau-3963-1 | NZ_KB822226.1 | - | 56762 | 56733 | **RNA**→ rpmE2 (PRK01678)Ribosomal_L31 (pfam01197)→ |
| Sau-3961-1 | NZ_KB822208.1 | - | 1698775 | 1698746 | **RNA**→ rpmE2 (PRK01678)Ribosomal_L31 (pfam01197)→ |
| Sau-3883-1 | NZ_KB821635.1 | - | 234212 | 234183 | **RNA**→ rpmE2 (PRK01678)Ribosomal_L31 (pfam01197)→ |
| Sau-3772-1 | NZ_KB820906.1 | - | 1674159 | 1674130 | **RNA**→ rpmE2 (PRK01678)Ribosomal_L31 (pfam01197)→ |
| Sau-3959-1 | NZ_KB822201.1 | - | 1063697 | 1063668 | **RNA**→ rpmE2 (PRK01678)Ribosomal_L31 (pfam01197)→ |

| | | | | | |
|---|---|---|---|---|---|
| Sau-3891-1 | NZ_KB821711.1 | - | 60935 | 60906 | RNA→ rpmE2 (PRK01678)Ribosomal_L31 (pfam01197)→ |
| Sau-3902-1 | NZ_KB821806.1 | - | 424380 | 424351 | RNA→ rpmE2 (PRK01678)Ribosomal_L31 (pfam01197)→ |
| Sau-3881-1 | NZ_KB821605.1 | - | 115520 | 115491 | RNA→ rpmE2 (PRK01678)Ribosomal_L31 (pfam01197)→ |
| Sau-3913-1 | NZ_KB821869.1 | - | 56763 | 56734 | RNA→ rpmE2 (PRK01678)Ribosomal_L31 (pfam01197)→ |
| Sau-3761-1 | NZ_KB822326.1 | - | 232181 | 232152 | RNA→ rpmE2 (PRK01678)Ribosomal_L31 (pfam01197)→ |
| Sau-3919-1 | NZ_KB821906.1 | - | 114248 | 114219 | RNA→ rpmE2 (PRK01678)Ribosomal_L31 (pfam01197)→ |
| Sau-3916-1 | NZ_KB822368.1 | - | 109574 | 109545 | RNA→ rpmE2 (PRK01678)Ribosomal_L31 (pfam01197)→ |
| Sau-3968-1 | NZ_KB822245.1 | - | 230562 | 230533 | RNA→ rpmE2 (PRK01678)Ribosomal_L31 (pfam01197)→ |
| Sau-3867-1 | NZ_KB821486.1 | - | 1313769 | 1313740 | RNA→ rpmE2 (PRK01678)Ribosomal_L31 (pfam01197)→ |
| Sau-3762-1 | NZ_KB822339.1 | - | 232013 | 231984 | RNA→ rpmE2 (PRK01678)Ribosomal_L31 (pfam01197)→ |
| Sau-3901-1 | NZ_KB821798.1 | - | 57585 | 57556 | RNA→ rpmE2 (PRK01678)Ribosomal_L31 (pfam01197)→ |
| Sau-3972-1 | NZ_KB822251.1 | - | 237694 | 237665 | RNA→ rpmE2 (PRK01678)Ribosomal_L31 (pfam01197)→ |
| Sau-3933-1 | NZ_KB822049.1 | - | 102188 | 102159 | RNA→ rpmE2 (PRK01678)Ribosomal_L31 (pfam01197)→ |
| Sau-3927-1 | NZ_KB821985.1 | - | 60324 | 60295 | RNA→ rpmE2 (PRK01678)Ribosomal_L31 (pfam01197)→ |
| Sau-3976-1 | NZ_KB822264.1 | - | 1610982 | 1610953 | RNA→ rpmE2 (PRK01678)Ribosomal_L31 (pfam01197)→ |
| Sau-3966-1 | NZ_KB822379.1 | - | 104248 | 104219 | RNA→ rpmE2 (PRK01678)Ribosomal_L31 (pfam01197)→ |
| Sau-3910-1 | NZ_KB821839.1 | - | 57581 | 57552 | RNA→ rpmE2 (PRK01678)Ribosomal_L31 (pfam01197)→ |
| Sau-3936-1 | NZ_KB822373.1 | - | 230489 | 230460 | RNA→ rpmE2 (PRK01678)Ribosomal_L31 (pfam01197)→ |
| Sau-3922-1 | NZ_KB821953.1 | - | 105045 | 105016 | RNA→ rpmE2 (PRK01678)Ribosomal_L31 (pfam01197)→ |
| Sau-3928-1 | NZ_KB822034.1 | - | 182165 | 182136 | RNA→ rpmE2 (PRK01678)Ribosomal_L31 (pfam01197)→ |
| Sau-3914-1 | NZ_KB821925.1 | - | 230555 | 230526 | RNA→ rpmE2 (PRK01678)Ribosomal_L31 (pfam01197)→ |
| Sau-4093-1 | NZ_KI669385.1 | - | 192582 | 192553 | RNA→ rpmE2 (PRK01678)Ribosomal_L31 (pfam01197)→ |
| Sau-3979-1 | NZ_KB822296.1 | - | 230666 | 230637 | RNA→ rpmE2 (PRK01678)Ribosomal_L31 (pfam01197)→ |
| Sau-3935-1 | NZ_KB822060.1 | - | 56193 | 56164 | RNA→ rpmE2 (PRK01678)Ribosomal_L31 (pfam01197)→ |
| Sau-4092-1 | NZ_KI669391.1 | + | 1112462 | 1112491 | RNA→ rpmE2 (PRK01678)Ribosomal_L31 (pfam01197)→ |
| Sau-3763-1 | NZ_KB822345.1 | - | 235630 | 235601 | RNA→ rpmE2 (PRK01678)Ribosomal_L31 (pfam01197)→ |
| Sau-3954-1 | NZ_KB822171.1 | - | 545143 | 545114 | RNA→ rpmE2 (PRK01678)Ribosomal_L31 (pfam01197)→ |
| Sau-3932-1 | NZ_KB821989.1 | - | 61902 | 61873 | RNA→ rpmE2 (PRK01678)Ribosomal_L31 (pfam01197)→ |
| Sau-3760-1 | NZ_KB822317.1 | - | 56642 | 56613 | RNA→ rpmE2 (PRK01678)Ribosomal_L31 (pfam01197)→ |
| Sau-331-1 | NZ_KI966042.1 | - | 1673431 | 1673402 | RNA→ rpmE2 (PRK01678)Ribosomal_L31 (pfam01197)→ |
| Sau-1682-1 | NZ_KI967573.1 | - | 118186 | 118157 | RNA→ rpmE2 (PRK01678)Ribosomal_L31 (pfam01197)→ |
| Sau-339-1 | NZ_KI966058.1 | - | 57585 | 57556 | RNA→ rpmE2 (PRK01678)Ribosomal_L31 (pfam01197)→ |
| Sau-344-1 | NZ_KI966065.1 | - | 107924 | 107895 | RNA→ rpmE2 (PRK01678)Ribosomal_L31 (pfam01197)→ |
| Sau-598-1 | NZ_KI966196.1 | - | 60037 | 60008 | RNA→ rpmE2 (PRK01678)Ribosomal_L31 (pfam01197)→ |
| Sau-399-1 | NZ_KI966087.1 | - | 56209 | 56180 | RNA→ rpmE2 (PRK01678)Ribosomal_L31 (pfam01197)→ |
| Sau-1676-1 | NZ_KI967272.1 | - | 62156 | 62127 | RNA→ rpmE2 (PRK01678)Ribosomal_L31 (pfam01197)→ |
| Sau-1683-1 | NZ_KI967631.1 | - | 112163 | 112134 | RNA→ rpmE2 (PRK01678)Ribosomal_L31 (pfam01197)→ |
| Sau-1689-1 | NZ_KI967913.1 | - | 62153 | 62124 | RNA→ rpmE2 (PRK01678)Ribosomal_L31 (pfam01197)→ |
| Sau-1695-1 | NZ_KI968141.1 | - | 16918 | 16889 | RNA→ rpmE2 (PRK01678)Ribosomal_L31 (pfam01197)→ |
| Sau-1709-1 | NZ_KI969434.1 | - | 112994 | 112965 | RNA→ rpmE2 (PRK01678)Ribosomal_L31 (pfam01197)→ |
| Sau-1716-1 | NZ_KI969660.1 | - | 49963 | 49934 | RNA→ rpmE2 (PRK01678)Ribosomal_L31 (pfam01197)→ |
| Sau-1717-1 | NZ_KI969744.1 | - | 62203 | 62174 | RNA→ rpmE2 (PRK01678)Ribosomal_L31 (pfam01197)→ |
| Sau-398-1 | NZ_KI966076.1 | - | 57163 | 57134 | RNA→ rpmE2 (PRK01678)Ribosomal_L31 (pfam01197)→ |
| Sau-1724-1 | NZ_KI970048.1 | - | 54695 | 54666 | RNA→ rpmE2 (PRK01678)Ribosomal_L31 (pfam01197)→ |
| Sau-602-1 | NZ_KI966138.1 | - | 59482 | 59453 | RNA→ rpmE2 (PRK01678)Ribosomal_L31 (pfam01197)→ |
| Sau-1702-1 | NZ_KI971027.1 | - | 402695 | 402666 | RNA→ rpmE2 (PRK01678)Ribosomal_L31 (pfam01197)→ |
| Sau-603-1 | NZ_KI966131.1 | - | 59482 | 59453 | RNA→ rpmE2 (PRK01678)Ribosomal_L31 (pfam01197)→ |
| Sau-1925-1 | NZ_KI972225.1 | - | 61649 | 61620 | RNA→ rpmE2 (PRK01678)Ribosomal_L31 (pfam01197)→ |
| Sau-588-1 | NZ_KI966224.1 | - | 59482 | 59453 | RNA→ rpmE2 (PRK01678)Ribosomal_L31 (pfam01197)→ |
| Sau-2111-1 | NZ_KI972814.1 | - | 230550 | 230521 | RNA→ rpmE2 (PRK01678)Ribosomal_L31 (pfam01197)→ |
| Sau-589-1 | NZ_KI966239.1 | - | 59692 | 59663 | RNA→ rpmE2 (PRK01678)Ribosomal_L31 (pfam01197)→ |
| Sau-549-1 | NZ_KI966323.1 | - | 64187 | 64158 | RNA→ rpmE2 (PRK01678)Ribosomal_L31 (pfam01197)→ |

| | | | | | | |
|---|---|---|---|---|---|---|
| Sau-1654-1 | NZ_KI966474.1 | - | 12988 | 12959 | **RNA**→ | rpmE2 (PRK01678)Ribosomal_L31 (pfam01197)→ |
| Sau-1668-1 | NZ_KI967046.1 | - | 113920 | 113891 | **RNA**→ | rpmE2 (PRK01678)Ribosomal_L31 (pfam01197)→ |
| Sau-2268-1 | NZ_KI972451.1 | - | 55454 | 55425 | **RNA**→ | rpmE2 (PRK01678)Ribosomal_L31 (pfam01197)→ |
| Sau-1289-1 | NZ_KI971633.1 | - | 1595760 | 1595731 | **RNA**→ | rpmE2 (PRK01678)Ribosomal_L31 (pfam01197)→ |
| Sau-1194-1 | NZ_KI971786.1 | - | 465526 | 465497 | **RNA**→ | rpmE2 (PRK01678)Ribosomal_L31 (pfam01197)→ |
| Sau-1677-1 | NZ_KI967404.1 | - | 61724 | 61695 | **RNA**→ | rpmE2 (PRK01678)Ribosomal_L31 (pfam01197)→ |
| Sau-2147-1 | NZ_KI972833.1 | - | 235217 | 235188 | **RNA**→ | rpmE2 (PRK01678)Ribosomal_L31 (pfam01197)→ |
| Sau-1697-1 | NZ_KI968271.1 | - | 21252 | 21223 | **RNA**→ | rpmE2 (PRK01678)Ribosomal_L31 (pfam01197)→ |
| Sau-1883-1 | NZ_KI972146.1 | - | 234345 | 234316 | **RNA**→ | rpmE2 (PRK01678)Ribosomal_L31 (pfam01197)→ |
| Sau-2363-1 | NZ_KI973005.1 | - | 56593 | 56564 | **RNA**→ | rpmE2 (PRK01678)Ribosomal_L31 (pfam01197)→ |
| Sau-1984-1 | NZ_KI972323.1 | - | 57587 | 57558 | **RNA**→ | rpmE2 (PRK01678)Ribosomal_L31 (pfam01197)→ |
| Sau-2095-1 | NZ_KI972772.1 | - | 230296 | 230267 | **RNA**→ | rpmE2 (PRK01678)Ribosomal_L31 (pfam01197)→ |
| Sau-1719-1 | NZ_KI969805.1 | - | 54994 | 54965 | **RNA**→ | rpmE2 (PRK01678)Ribosomal_L31 (pfam01197)→ |
| Sau-1726-1 | NZ_KI970096.1 | - | 43785 | 43756 | **RNA**→ | rpmE2 (PRK01678)Ribosomal_L31 (pfam01197)→ |
| Sau-1741-1 | NZ_KI970705.1 | - | 61571 | 61542 | **RNA**→ | rpmE2 (PRK01678)Ribosomal_L31 (pfam01197)→ |
| Sau-1815-1 | NZ_KI972023.1 | - | 231732 | 231703 | **RNA**→ | rpmE2 (PRK01678)Ribosomal_L31 (pfam01197)→ |
| Sau-2368-1 | NZ_KI973034.1 | - | 58943 | 58914 | **RNA**→ | rpmE2 (PRK01678)Ribosomal_L31 (pfam01197)→ |
| Sau-1831-1 | NZ_KI972040.1 | - | 233784 | 233755 | **RNA**→ | rpmE2 (PRK01678)Ribosomal_L31 (pfam01197)→ |
| Sau-605-1 | NZ_KI966013.1 | - | 1657877 | 1657848 | **RNA**→ | rpmE2 (PRK01678)Ribosomal_L31 (pfam01197)→ |
| Sau-518-1 | NZ_KI966340.1 | - | 60905 | 60876 | **RNA**→ | rpmE2 (PRK01678)Ribosomal_L31 (pfam01197)→ |
| Sau-1656-1 | NZ_KI966575.1 | - | 63214 | 63185 | **RNA**→ | rpmE2 (PRK01678)Ribosomal_L31 (pfam01197)→ |
| Sau-1686-1 | NZ_KI967724.1 | - | 35475 | 35446 | **RNA**→ | rpmE2 (PRK01678)Ribosomal_L31 (pfam01197)→ |
| Sau-1662-1 | NZ_KI966829.1 | - | 36390 | 36361 | **RNA**→ | rpmE2 (PRK01678)Ribosomal_L31 (pfam01197)→ |
| Sau-1557-1 | NZ_KI971102.1 | - | 56763 | 56734 | **RNA**→ | rpmE2 (PRK01678)Ribosomal_L31 (pfam01197)→ |
| Sau-1669-1 | NZ_KI967058.1 | - | 36295 | 36266 | **RNA**→ | rpmE2 (PRK01678)Ribosomal_L31 (pfam01197)→ |
| Sau-1554-1 | NZ_KI971422.1 | - | 56762 | 56733 | **RNA**→ | rpmE2 (PRK01678)Ribosomal_L31 (pfam01197)→ |
| Sau-1596-1 | NZ_KI971192.1 | - | 49963 | 49934 | **RNA**→ | rpmE2 (PRK01678)Ribosomal_L31 (pfam01197)→ |
| Sau-1312-1 | NZ_KI971667.1 | - | 398213 | 398184 | **RNA**→ | rpmE2 (PRK01678)Ribosomal_L31 (pfam01197)→ |
| Sau-1461-1 | NZ_KI971570.1 | - | 402113 | 402084 | **RNA**→ | rpmE2 (PRK01678)Ribosomal_L31 (pfam01197)→ |
| Sau-1335-1 | NZ_KI971720.1 | - | 61106 | 61077 | **RNA**→ | rpmE2 (PRK01678)Ribosomal_L31 (pfam01197)→ |
| Sau-1237-1 | NZ_KI971878.1 | - | 56852 | 56823 | **RNA**→ | rpmE2 (PRK01678)Ribosomal_L31 (pfam01197)→ |
| Sau-1704-1 | NZ_KI968528.1 | - | 55116 | 55087 | **RNA**→ | rpmE2 (PRK01678)Ribosomal_L31 (pfam01197)→ |
| Sau-1750-1 | NZ_KI971930.1 | - | 272475 | 272446 | **RNA**→ | rpmE2 (PRK01678)Ribosomal_L31 (pfam01197)→ |
| Sau-1714-1 | NZ_KI969563.1 | - | 5387 | 5358 | **RNA**→ | rpmE2 (PRK01678)Ribosomal_L31 (pfam01197)→ |
| Sau-1907-1 | NZ_KI972180.1 | - | 116961 | 116932 | **RNA**→ | rpmE2 (PRK01678)Ribosomal_L31 (pfam01197)→ |
| Sau-1743-1 | NZ_KI970708.1 | - | 27279 | 27250 | **RNA**→ | rpmE2 (PRK01678)Ribosomal_L31 (pfam01197)→ |
| Sau-601-1 | NZ_KI966158.1 | - | 59483 | 59454 | **RNA**→ | rpmE2 (PRK01678)Ribosomal_L31 (pfam01197)→ |
| Sau-1744-1 | NZ_KI970818.1 | - | 62202 | 62173 | **RNA**→ | rpmE2 (PRK01678)Ribosomal_L31 (pfam01197)→ |
| Sau-1776-1 | NZ_KI971952.1 | - | 58283 | 58254 | **RNA**→ | rpmE2 (PRK01678)Ribosomal_L31 (pfam01197)→ |
| Sau-1657-1 | NZ_KI966643.1 | - | 35225 | 35196 | **RNA**→ | rpmE2 (PRK01678)Ribosomal_L31 (pfam01197)→ |
| Sau-2187-1 | NZ_KI972860.1 | - | 828427 | 828398 | **RNA**→ | rpmE2 (PRK01678)Ribosomal_L31 (pfam01197)→ |
| Sau-1663-1 | NZ_KI966880.1 | - | 34180 | 34151 | **RNA**→ | rpmE2 (PRK01678)Ribosomal_L31 (pfam01197)→ |
| Sau-1672-1 | NZ_KI967105.1 | - | 35862 | 35833 | **RNA**→ | rpmE2 (PRK01678)Ribosomal_L31 (pfam01197)→ |
| Sau-1628-1 | NZ_KI971233.1 | - | 494957 | 494928 | **RNA**→ | rpmE2 (PRK01678)Ribosomal_L31 (pfam01197)→ |
| Sau-1530-1 | NZ_KI971392.1 | - | 113065 | 113036 | **RNA**→ | rpmE2 (PRK01678)Ribosomal_L31 (pfam01197)→ |
| Sau-600-1 | NZ_KI966173.1 | - | 66056 | 66027 | **RNA**→ | rpmE2 (PRK01678)Ribosomal_L31 (pfam01197)→ |
| Sau-567-1 | NZ_KI966294.1 | - | 59482 | 59453 | **RNA**→ | rpmE2 (PRK01678)Ribosomal_L31 (pfam01197)→ |
| Sau-551-1 | NZ_KI966306.1 | - | 59679 | 59650 | **RNA**→ | rpmE2 (PRK01678)Ribosomal_L31 (pfam01197)→ |
| Sau-1658-1 | NZ_KI966704.1 | - | 62196 | 62167 | **RNA**→ | rpmE2 (PRK01678)Ribosomal_L31 (pfam01197)→ |
| Sau-1734-1 | NZ_KI970382.1 | - | 99800 | 99771 | **RNA**→ | rpmE2 (PRK01678)Ribosomal_L31 (pfam01197)→ |
| Sau-1665-1 | NZ_KI966922.1 | - | 58301 | 58272 | **RNA**→ | rpmE2 (PRK01678)Ribosomal_L31 (pfam01197)→ |
| Sau-1673-1 | NZ_KI967166.1 | - | 36412 | 36383 | **RNA**→ | rpmE2 (PRK01678)Ribosomal_L31 (pfam01197)→ |

| | | | | | |
|---|---|---|---|---|---|
| Sau-1733-1 | NZ_KI971070.1 | - | 173252 | 173223 | RNA→ rpmE2 (PRK01678)Ribosomal_L31 (pfam01197)→ |
| Sau-1674-1 | NZ_KI967226.1 | - | 81102 | 81073 | RNA→ rpmE2 (PRK01678)Ribosomal_L31 (pfam01197)→ |
| Sau-1670-1 | NZ_KI970968.1 | - | 58348 | 58319 | RNA→ rpmE2 (PRK01678)Ribosomal_L31 (pfam01197)→ |
| Sau-1280-1 | NZ_KI971631.1 | - | 56763 | 56734 | RNA→ rpmE2 (PRK01678)Ribosomal_L31 (pfam01197)→ |
| Sau-1861-1 | NZ_KI972120.1 | - | 58043 | 58014 | RNA→ rpmE2 (PRK01678)Ribosomal_L31 (pfam01197)→ |
| Sau-1740-1 | NZ_KI970586.1 | - | 61333 | 61304 | RNA→ rpmE2 (PRK01678)Ribosomal_L31 (pfam01197)→ |
| Sau-1444-1 | NZ_KI971546.1 | - | 56493 | 56464 | RNA→ rpmE2 (PRK01678)Ribosomal_L31 (pfam01197)→ |
| Sau-1483-1 | NZ_KI971311.1 | - | 183428 | 183399 | RNA→ rpmE2 (PRK01678)Ribosomal_L31 (pfam01197)→ |
| Sau-1511-1 | NZ_KI971375.1 | - | 56763 | 56734 | RNA→ rpmE2 (PRK01678)Ribosomal_L31 (pfam01197)→ |
| Sau-1696-1 | NZ_KI968191.1 | - | 36391 | 36362 | RNA→ rpmE2 (PRK01678)Ribosomal_L31 (pfam01197)→ |
| Sau-1889-1 | NZ_KI972165.1 | - | 253911 | 253882 | RNA→ rpmE2 (PRK01678)Ribosomal_L31 (pfam01197)→ |
| Sau-2199-1 | NZ_KI972918.1 | - | 464236 | 464207 | RNA→ rpmE2 (PRK01678)Ribosomal_L31 (pfam01197)→ |
| Sau-1710-1 | NZ_KI969464.1 | - | 2446 | 2417 | RNA→ rpmE2 (PRK01678)Ribosomal_L31 (pfam01197)→ |
| Sau-1711-1 | NZ_KI969482.1 | - | 35467 | 35438 | RNA→ rpmE2 (PRK01678)Ribosomal_L31 (pfam01197)→ |
| Sau-1718-1 | NZ_KI969749.1 | - | 35697 | 35668 | RNA→ rpmE2 (PRK01678)Ribosomal_L31 (pfam01197)→ |
| Sau-1298-1 | NZ_KI971655.1 | - | 56843 | 56814 | RNA→ rpmE2 (PRK01678)Ribosomal_L31 (pfam01197)→ |
| Sau-1721-1 | NZ_KI969874.1 | - | 59617 | 59588 | RNA→ rpmE2 (PRK01678)Ribosomal_L31 (pfam01197)→ |
| Sau-1727-1 | NZ_KI970143.1 | - | 35100 | 35071 | RNA→ rpmE2 (PRK01678)Ribosomal_L31 (pfam01197)→ |
| Sau-1916-1 | NZ_KI972193.1 | - | 57081 | 57052 | RNA→ rpmE2 (PRK01678)Ribosomal_L31 (pfam01197)→ |
| Sau-2374-1 | NZ_KI973037.1 | - | 213705 | 213676 | RNA→ rpmE2 (PRK01678)Ribosomal_L31 (pfam01197)→ |
| Sau-1655-1 | NZ_KI966501.1 | - | 55094 | 55065 | RNA→ rpmE2 (PRK01678)Ribosomal_L31 (pfam01197)→ |
| Sau-1723-1 | NZ_KI971047.1 | - | 494175 | 494146 | RNA→ rpmE2 (PRK01678)Ribosomal_L31 (pfam01197)→ |
| Sau-1648-1 | NZ_KI971263.1 | - | 118819 | 118790 | RNA→ rpmE2 (PRK01678)Ribosomal_L31 (pfam01197)→ |
| Sau-1319-1 | NZ_KI971674.1 | - | 173915 | 173886 | RNA→ rpmE2 (PRK01678)Ribosomal_L31 (pfam01197)→ |
| Sau-1712-1 | NZ_KI969558.1 | - | 61890 | 61861 | RNA→ rpmE2 (PRK01678)Ribosomal_L31 (pfam01197)→ |
| Sau-1706-1 | NZ_KI968629.1 | - | 36512 | 36483 | RNA→ rpmE2 (PRK01678)Ribosomal_L31 (pfam01197)→ |
| Sau-1844-1 | NZ_KI972088.1 | - | 227707 | 227678 | RNA→ rpmE2 (PRK01678)Ribosomal_L31 (pfam01197)→ |
| Sau-1703-1 | NZ_KI968479.1 | - | 35162 | 35133 | RNA→ rpmE2 (PRK01678)Ribosomal_L31 (pfam01197)→ |
| Sau-2367-1 | NZ_KI973007.1 | - | 56367 | 56338 | RNA→ rpmE2 (PRK01678)Ribosomal_L31 (pfam01197)→ |
| Sau-1728-1 | NZ_KI970221.1 | - | 32991 | 32962 | RNA→ rpmE2 (PRK01678)Ribosomal_L31 (pfam01197)→ |
| Sau-1694-1 | NZ_KI968123.1 | - | 62146 | 62117 | RNA→ rpmE2 (PRK01678)Ribosomal_L31 (pfam01197)→ |
| Sau-1745-1 | NZ_KI970848.1 | - | 36005 | 35976 | RNA→ rpmE2 (PRK01678)Ribosomal_L31 (pfam01197)→ |
| Sau-1701-1 | NZ_KI968449.1 | - | 35643 | 35614 | RNA→ rpmE2 (PRK01678)Ribosomal_L31 (pfam01197)→ |
| Sau-2094-1 | NZ_KI972762.1 | - | 230995 | 230966 | RNA→ rpmE2 (PRK01678)Ribosomal_L31 (pfam01197)→ |
| Sau-1659-1 | NZ_KI970964.1 | - | 61955 | 61926 | RNA→ rpmE2 (PRK01678)Ribosomal_L31 (pfam01197)→ |
| Sau-1637-1 | NZ_KI971244.1 | - | 58248 | 58219 | RNA→ rpmE2 (PRK01678)Ribosomal_L31 (pfam01197)→ |
| Sau-2362-1 | NZ_KI972975.1 | - | 2198240 | 2198211 | RNA→ rpmE2 (PRK01678)Ribosomal_L31 (pfam01197)→ |
| Sau-1520-1 | NZ_KI971383.1 | - | 56459 | 56430 | RNA→ rpmE2 (PRK01678)Ribosomal_L31 (pfam01197)→ |
| Sau-1201-1 | NZ_KI971799.1 | - | 180429 | 180400 | RNA→ rpmE2 (PRK01678)Ribosomal_L31 (pfam01197)→ |
| Sau-1840-1 | NZ_KI972070.1 | - | 61630 | 61601 | RNA→ rpmE2 (PRK01678)Ribosomal_L31 (pfam01197)→ |
| Sau-1466-1 | NZ_KI971272.1 | - | 56763 | 56734 | RNA→ rpmE2 (PRK01678)Ribosomal_L31 (pfam01197)→ |
| Sau-1747-1 | NZ_KI970852.1 | - | 35561 | 35532 | RNA→ rpmE2 (PRK01678)Ribosomal_L31 (pfam01197)→ |
| Sau-1585-1 | NZ_KI971155.1 | - | 56782 | 56753 | RNA→ rpmE2 (PRK01678)Ribosomal_L31 (pfam01197)→ |
| Sau-1272-1 | NZ_KI971596.1 | - | 57586 | 57557 | RNA→ rpmE2 (PRK01678)Ribosomal_L31 (pfam01197)→ |
| Sau-2269-1 | NZ_KI972465.1 | - | 58992 | 58963 | RNA→ rpmE2 (PRK01678)Ribosomal_L31 (pfam01197)→ |
| Sau-2358-1 | NZ_KI972951.1 | - | 464468 | 464439 | RNA→ rpmE2 (PRK01678)Ribosomal_L31 (pfam01197)→ |
| Sau-1612-1 | NZ_KI971209.1 | - | 465008 | 464979 | RNA→ rpmE2 (PRK01678)Ribosomal_L31 (pfam01197)→ |
| Sau-1708-1 | NZ_KI968648.1 | - | 26789 | 26760 | RNA→ rpmE2 (PRK01678)Ribosomal_L31 (pfam01197)→ |
| Sau-1652-1 | NZ_KI970947.1 | - | 56763 | 56734 | RNA→ rpmE2 (PRK01678)Ribosomal_L31 (pfam01197)→ |
| Sau-1258-1 | NZ_KI971893.1 | - | 463233 | 463204 | RNA→ rpmE2 (PRK01678)Ribosomal_L31 (pfam01197)→ |
| Sau-1872-1 | NZ_KI972138.1 | - | 54695 | 54666 | RNA→ rpmE2 (PRK01678)Ribosomal_L31 (pfam01197)→ |
| Sau-2239-1 | NZ_KI972925.1 | - | 239945 | 239916 | RNA→ rpmE2 (PRK01678)Ribosomal_L31 (pfam01197)→ |

| | | | | | | |
|---|---|---|---|---|---|---|
| Sau-2382-1 | NZ_KI973077.1 | - | 25085 | 25056 | **RNA**→ | rpmE2 (PRK01678)Ribosomal_L31 (pfam01197)→ |
| Sau-2198-1 | NZ_KI972897.1 | - | 198468 | 198439 | **RNA**→ | rpmE2 (PRK01678)Ribosomal_L31 (pfam01197)→ |
| Sau-1700-1 | NZ_KI968406.1 | - | 61630 | 61601 | **RNA**→ | rpmE2 (PRK01678)Ribosomal_L31 (pfam01197)→ |
| Sau-1722-1 | NZ_KI969940.1 | - | 35400 | 35371 | **RNA**→ | rpmE2 (PRK01678)Ribosomal_L31 (pfam01197)→ |
| Sau-1731-1 | NZ_KI970334.1 | - | 62156 | 62127 | **RNA**→ | rpmE2 (PRK01678)Ribosomal_L31 (pfam01197)→ |
| Sau-1691-1 | NZ_KI971018.1 | - | 57585 | 57556 | **RNA**→ | rpmE2 (PRK01678)Ribosomal_L31 (pfam01197)→ |
| Sau-1742-1 | NZ_KI971082.1 | - | 405094 | 405065 | **RNA**→ | rpmE2 (PRK01678)Ribosomal_L31 (pfam01197)→ |
| Sau-280-1 | NZ_KI973576.1 | - | 34872 | 34843 | **RNA**→ | rpmE2 (PRK01678)Ribosomal_L31 (pfam01197)→ |
| Sau-283-1 | NZ_KI974226.1 | - | 120491 | 120462 | **RNA**→ | rpmE2 (PRK01678)Ribosomal_L31 (pfam01197)→ |
| Sau-1699-1 | NZ_KI968398.1 | - | 54833 | 54804 | **RNA**→ | rpmE2 (PRK01678)Ribosomal_L31 (pfam01197)→ |
| Sau-1729-1 | NZ_KI970276.1 | - | 56493 | 56464 | **RNA**→ | rpmE2 (PRK01678)Ribosomal_L31 (pfam01197)→ |
| Sau-1472-1 | NZ_KI971305.1 | - | 56763 | 56734 | **RNA**→ | rpmE2 (PRK01678)Ribosomal_L31 (pfam01197)→ |
| Sau-288-1 | NZ_KI974232.1 | - | 56893 | 56864 | **RNA**→ | rpmE2 (PRK01678)Ribosomal_L31 (pfam01197)→ |
| Sau-2274-1 | NZ_KI972485.1 | - | 57586 | 57557 | **RNA**→ | rpmE2 (PRK01678)Ribosomal_L31 (pfam01197)→ |
| Sau-242-1 | NZ_KI974540.1 | - | 56894 | 56865 | **RNA**→ | rpmE2 (PRK01678)Ribosomal_L31 (pfam01197)→ |
| Sau-2379-1 | NZ_KI973048.1 | - | 231806 | 231777 | **RNA**→ | rpmE2 (PRK01678)Ribosomal_L31 (pfam01197)→ |
| Sau-227-1 | NZ_KI974962.1 | - | 55407 | 55378 | **RNA**→ | rpmE2 (PRK01678)Ribosomal_L31 (pfam01197)→ |
| Sau-2782-1 | NZ_KI973820.1 | - | 55262 | 55233 | **RNA**→ | rpmE2 (PRK01678)Ribosomal_L31 (pfam01197)→ |
| Sau-396-1 | NZ_KI966105.1 | - | 55022 | 54993 | **RNA**→ | rpmE2 (PRK01678)Ribosomal_L31 (pfam01197)→ |
| Sau-2032-1 | NZ_KI972434.1 | - | 56417 | 56388 | **RNA**→ | rpmE2 (PRK01678)Ribosomal_L31 (pfam01197)→ |
| Sau-2754-1 | NZ_KI973379.1 | - | 34913 | 34884 | **RNA**→ | rpmE2 (PRK01678)Ribosomal_L31 (pfam01197)→ |
| Sau-2742-1 | NZ_KI973985.1 | - | 56894 | 56865 | **RNA**→ | rpmE2 (PRK01678)Ribosomal_L31 (pfam01197)→ |
| Sau-238-1 | NZ_KI973531.1 | - | 56894 | 56865 | **RNA**→ | rpmE2 (PRK01678)Ribosomal_L31 (pfam01197)→ |
| Sau-2650-1 | NZ_KI974790.1 | - | 110356 | 110327 | **RNA**→ | rpmE2 (PRK01678)Ribosomal_L31 (pfam01197)→ |
| Sau-239-1 | NZ_KI973724.1 | - | 57687 | 57658 | **RNA**→ | rpmE2 (PRK01678)Ribosomal_L31 (pfam01197)→ |
| Sau-1101-1 | NZ_KI974926.1 | - | 216089 | 216060 | **RNA**→ | rpmE2 (PRK01678)Ribosomal_L31 (pfam01197)→ |
| Sau-2752-1 | NZ_KI973351.1 | - | 56894 | 56865 | **RNA**→ | rpmE2 (PRK01678)Ribosomal_L31 (pfam01197)→ |
| Sau-2757-1 | NZ_KI973508.1 | - | 56508 | 56479 | **RNA**→ | rpmE2 (PRK01678)Ribosomal_L31 (pfam01197)→ |
| Sau-2680-1 | NZ_KI974183.1 | - | 56894 | 56865 | **RNA**→ | rpmE2 (PRK01678)Ribosomal_L31 (pfam01197)→ |
| Sau-2762-1 | NZ_KI974398.1 | - | 56000 | 55971 | **RNA**→ | rpmE2 (PRK01678)Ribosomal_L31 (pfam01197)→ |
| Sau-1807-1 | NZ_KI972001.1 | - | 476505 | 476476 | **RNA**→ | rpmE2 (PRK01678)Ribosomal_L31 (pfam01197)→ |
| Sau-2679-1 | NZ_KI973605.1 | - | 172385 | 172356 | **RNA**→ | rpmE2 (PRK01678)Ribosomal_L31 (pfam01197)→ |
| Sau-2760-1 | NZ_KI974240.1 | - | 117788 | 117759 | **RNA**→ | rpmE2 (PRK01678)Ribosomal_L31 (pfam01197)→ |
| Sau-2646-1 | NZ_KI974468.1 | - | 114326 | 114297 | **RNA**→ | rpmE2 (PRK01678)Ribosomal_L31 (pfam01197)→ |
| Sau-2746-1 | NZ_KI974843.1 | - | 111732 | 111703 | **RNA**→ | rpmE2 (PRK01678)Ribosomal_L31 (pfam01197)→ |
| Sau-2745-1 | NZ_KI974880.1 | - | 56894 | 56865 | **RNA**→ | rpmE2 (PRK01678)Ribosomal_L31 (pfam01197)→ |
| Sau-2748-1 | NZ_KI974888.1 | - | 34967 | 34938 | **RNA**→ | rpmE2 (PRK01678)Ribosomal_L31 (pfam01197)→ |
| Sau-2777-1 | NZ_KI976193.1 | - | 110423 | 110394 | **RNA**→ | rpmE2 (PRK01678)Ribosomal_L31 (pfam01197)→ |
| Sau-2732-1 | NZ_KI976327.1 | - | 56894 | 56865 | **RNA**→ | rpmE2 (PRK01678)Ribosomal_L31 (pfam01197)→ |
| Sau-2651-1 | NZ_KI976449.1 | - | 56643 | 56614 | **RNA**→ | rpmE2 (PRK01678)Ribosomal_L31 (pfam01197)→ |
| Sau-2769-1 | NZ_KI976741.1 | - | 163294 | 163265 | **RNA**→ | rpmE2 (PRK01678)Ribosomal_L31 (pfam01197)→ |
| Sau-2784-1 | NZ_KI973792.1 | - | 56894 | 56865 | **RNA**→ | rpmE2 (PRK01678)Ribosomal_L31 (pfam01197)→ |
| Sau-2649-1 | NZ_KI974493.1 | - | 112597 | 112568 | **RNA**→ | rpmE2 (PRK01678)Ribosomal_L31 (pfam01197)→ |
| Sau-2776-1 | NZ_KI975309.1 | - | 55262 | 55233 | **RNA**→ | rpmE2 (PRK01678)Ribosomal_L31 (pfam01197)→ |
| Sau-2731-1 | NZ_KI976242.1 | - | 111777 | 111748 | **RNA**→ | rpmE2 (PRK01678)Ribosomal_L31 (pfam01197)→ |
| Sau-2763-1 | NZ_KI976564.1 | - | 57686 | 57657 | **RNA**→ | rpmE2 (PRK01678)Ribosomal_L31 (pfam01197)→ |
| Sau-2759-1 | NZ_KI973929.1 | - | 56894 | 56865 | **RNA**→ | rpmE2 (PRK01678)Ribosomal_L31 (pfam01197)→ |
| Sau-2774-1 | NZ_KI975249.1 | - | 118410 | 118381 | **RNA**→ | rpmE2 (PRK01678)Ribosomal_L31 (pfam01197)→ |
| Sau-2773-1 | NZ_KI975250.1 | - | 113458 | 113429 | **RNA**→ | rpmE2 (PRK01678)Ribosomal_L31 (pfam01197)→ |
| Sau-2740-1 | NZ_KI975832.1 | - | 238778 | 238749 | **RNA**→ | rpmE2 (PRK01678)Ribosomal_L31 (pfam01197)→ |
| Sau-2779-1 | NZ_KI976188.1 | - | 238384 | 238355 | **RNA**→ | rpmE2 (PRK01678)Ribosomal_L31 (pfam01197)→ |
| Sau-2765-1 | NZ_KI976626.1 | - | 237053 | 237024 | **RNA**→ | rpmE2 (PRK01678)Ribosomal_L31 (pfam01197)→ |

| | | | | | |
|---|---|---|---|---|---|
| Sau-2747-1 | NZ_KI974841.1 | - | 62196 | 62167 | **RNA**→ rpmE2 (PRK01678)Ribosomal_L31 (pfam01197)→ |
| Sau-2749-1 | NZ_KI975069.1 | - | 25249 | 25220 | **RNA**→ rpmE2 (PRK01678)Ribosomal_L31 (pfam01197)→ |
| Sau-2729-1 | NZ_KI975199.1 | - | 119992 | 119963 | **RNA**→ rpmE2 (PRK01678)Ribosomal_L31 (pfam01197)→ |
| Sau-2750-1 | NZ_KI975679.1 | - | 111359 | 111330 | **RNA**→ rpmE2 (PRK01678)Ribosomal_L31 (pfam01197)→ |
| Sau-1099-1 | NZ_KI973769.1 | - | 56488 | 56459 | **RNA**→ rpmE2 (PRK01678)Ribosomal_L31 (pfam01197)→ |
| Sau-2751-1 | NZ_KI975684.1 | - | 35284 | 35255 | **RNA**→ rpmE2 (PRK01678)Ribosomal_L31 (pfam01197)→ |
| Sau-2780-1 | NZ_KI976171.1 | - | 56894 | 56865 | **RNA**→ rpmE2 (PRK01678)Ribosomal_L31 (pfam01197)→ |
| Sau-240-1 | NZ_KI974041.1 | - | 61027 | 60998 | **RNA**→ rpmE2 (PRK01678)Ribosomal_L31 (pfam01197)→ |
| Sau-2767-1 | NZ_KI976602.1 | - | 35015 | 34986 | **RNA**→ rpmE2 (PRK01678)Ribosomal_L31 (pfam01197)→ |
| Sau-245-1 | NZ_KI974590.1 | - | 112127 | 112098 | **RNA**→ rpmE2 (PRK01678)Ribosomal_L31 (pfam01197)→ |
| Sau-2653-1 | NZ_KI977062.1 | - | 19182 | 19153 | **RNA**→ rpmE2 (PRK01678)Ribosomal_L31 (pfam01197)→ |
| Sau-1605-1 | NZ_KI971201.1 | - | 122322 | 122293 | **RNA**→ rpmE2 (PRK01678)Ribosomal_L31 (pfam01197)→ |
| Sau-1188-1 | NZ_KI971784.1 | - | 56663 | 56634 | **RNA**→ rpmE2 (PRK01678)Ribosomal_L31 (pfam01197)→ |
| Sau-1661-1 | NZ_KI966824.1 | - | 62005 | 61976 | **RNA**→ rpmE2 (PRK01678)Ribosomal_L31 (pfam01197)→ |
| Sau-1738-1 | NZ_KI970529.1 | - | 58829 | 58800 | **RNA**→ rpmE2 (PRK01678)Ribosomal_L31 (pfam01197)→ |
| Sau-1368-1 | NZ_KI971436.1 | - | 57585 | 57556 | **RNA**→ rpmE2 (PRK01678)Ribosomal_L31 (pfam01197)→ |
| Sau-236-1 | NZ_KI973490.1 | - | 56894 | 56865 | **RNA**→ rpmE2 (PRK01678)Ribosomal_L31 (pfam01197)→ |
| Sau-2328-1 | NZ_KI972569.1 | - | 118224 | 118195 | **RNA**→ rpmE2 (PRK01678)Ribosomal_L31 (pfam01197)→ |
| Sau-225-1 | NZ_KI974722.1 | - | 119048 | 119019 | **RNA**→ rpmE2 (PRK01678)Ribosomal_L31 (pfam01197)→ |
| Sau-1420-1 | NZ_KI971523.1 | - | 118776 | 118747 | **RNA**→ rpmE2 (PRK01678)Ribosomal_L31 (pfam01197)→ |
| Sau-1713-1 | NZ_KI971041.1 | - | 467476 | 467447 | **RNA**→ rpmE2 (PRK01678)Ribosomal_L31 (pfam01197)→ |
| Sau-1500-1 | NZ_KI971347.1 | - | 56521 | 56492 | **RNA**→ rpmE2 (PRK01678)Ribosomal_L31 (pfam01197)→ |
| Sau-2644-1 | NZ_KI973974.1 | - | 56894 | 56865 | **RNA**→ rpmE2 (PRK01678)Ribosomal_L31 (pfam01197)→ |
| Sau-229-1 | NZ_KI974994.1 | - | 34872 | 34843 | **RNA**→ rpmE2 (PRK01678)Ribosomal_L31 (pfam01197)→ |
| Sau-231-1 | NZ_KI975047.1 | - | 56894 | 56865 | **RNA**→ rpmE2 (PRK01678)Ribosomal_L31 (pfam01197)→ |
| Sau-233-1 | NZ_KI975535.1 | - | 469077 | 469048 | **RNA**→ rpmE2 (PRK01678)Ribosomal_L31 (pfam01197)→ |
| Sau-1103-1 | NZ_KI975861.1 | - | 107623 | 107594 | **RNA**→ rpmE2 (PRK01678)Ribosomal_L31 (pfam01197)→ |
| Sau-1154-1 | NZ_KI974747.1 | - | 56828 | 56799 | **RNA**→ rpmE2 (PRK01678)Ribosomal_L31 (pfam01197)→ |
| Sau-884-1 | NZ_KI973539.1 | - | 100527 | 100498 | **RNA**→ rpmE2 (PRK01678)Ribosomal_L31 (pfam01197)→ |
| Sau-241-1 | NZ_KI974535.1 | - | 56894 | 56865 | **RNA**→ rpmE2 (PRK01678)Ribosomal_L31 (pfam01197)→ |
| Sau-246-1 | NZ_KI974603.1 | - | 56894 | 56865 | **RNA**→ rpmE2 (PRK01678)Ribosomal_L31 (pfam01197)→ |
| Sau-230-1 | NZ_KI975061.1 | - | 56994 | 56965 | **RNA**→ rpmE2 (PRK01678)Ribosomal_L31 (pfam01197)→ |
| Sau-2605-1 | NZ_KI973836.1 | - | 239089 | 239060 | **RNA**→ rpmE2 (PRK01678)Ribosomal_L31 (pfam01197)→ |
| Sau-881-1 | NZ_KI975875.1 | - | 113460 | 113431 | **RNA**→ rpmE2 (PRK01678)Ribosomal_L31 (pfam01197)→ |
| Sau-2603-1 | NZ_KI973853.1 | - | 61889 | 61860 | **RNA**→ rpmE2 (PRK01678)Ribosomal_L31 (pfam01197)→ |
| Sau-1146-1 | NZ_KI975396.1 | - | 238621 | 238592 | **RNA**→ rpmE2 (PRK01678)Ribosomal_L31 (pfam01197)→ |
| Sau-291-1 | NZ_KI975911.1 | - | 56994 | 56965 | **RNA**→ rpmE2 (PRK01678)Ribosomal_L31 (pfam01197)→ |
| Sau-302-1 | NZ_KI976160.1 | - | 56994 | 56965 | **RNA**→ rpmE2 (PRK01678)Ribosomal_L31 (pfam01197)→ |
| Sau-2611-1 | NZ_KI975631.1 | - | 57506 | 57477 | **RNA**→ rpmE2 (PRK01678)Ribosomal_L31 (pfam01197)→ |
| Sau-293-1 | NZ_KI976683.1 | - | 239227 | 239198 | **RNA**→ rpmE2 (PRK01678)Ribosomal_L31 (pfam01197)→ |
| Sau-243-1 | NZ_KI974552.1 | - | 56894 | 56865 | **RNA**→ rpmE2 (PRK01678)Ribosomal_L31 (pfam01197)→ |
| Sau-2641-1 | NZ_KI975976.1 | - | 107609 | 107580 | **RNA**→ rpmE2 (PRK01678)Ribosomal_L31 (pfam01197)→ |
| Sau-1102-1 | NZ_KI975389.1 | - | 56894 | 56865 | **RNA**→ rpmE2 (PRK01678)Ribosomal_L31 (pfam01197)→ |
| Sau-1128-1 | NZ_KI976091.1 | - | 35048 | 35019 | **RNA**→ rpmE2 (PRK01678)Ribosomal_L31 (pfam01197)→ |
| Sau-2620-1 | NZ_KI976472.1 | - | 56994 | 56965 | **RNA**→ rpmE2 (PRK01678)Ribosomal_L31 (pfam01197)→ |
| Sau-235-1 | NZ_KI975491.1 | - | 56894 | 56865 | **RNA**→ rpmE2 (PRK01678)Ribosomal_L31 (pfam01197)→ |
| Sau-1113-1 | NZ_KI976543.1 | - | 107627 | 107598 | **RNA**→ rpmE2 (PRK01678)Ribosomal_L31 (pfam01197)→ |
| Sau-234-1 | NZ_KI975519.1 | - | 233482 | 233453 | **RNA**→ rpmE2 (PRK01678)Ribosomal_L31 (pfam01197)→ |
| Sau-1114-1 | NZ_KI976959.1 | - | 628831 | 628802 | **RNA**→ rpmE2 (PRK01678)Ribosomal_L31 (pfam01197)→ |
| Sau-1131-1 | NZ_KI977383.1 | - | 117531 | 117502 | **RNA**→ rpmE2 (PRK01678)Ribosomal_L31 (pfam01197)→ |
| Sau-1150-1 | NZ_KI973423.1 | - | 54893 | 54864 | **RNA**→ rpmE2 (PRK01678)Ribosomal_L31 (pfam01197)→ |
| Sau-1108-1 | NZ_KI973888.1 | - | 118746 | 118717 | **RNA**→ rpmE2 (PRK01678)Ribosomal_L31 (pfam01197)→ |

| | | | | | |
|---|---|---|---|---|---|
| Sau-2618-1 | NZ_KI974606.1 | - | 1344431 | 1344402 | RNA→ rpmE2 (PRK01678)Ribosomal_L31 (pfam01197)→ |
| Sau-2610-1 | NZ_KI975651.1 | - | 58425 | 58396 | RNA→ rpmE2 (PRK01678)Ribosomal_L31 (pfam01197)→ |
| Sau-1125-1 | NZ_KI975789.1 | - | 109375 | 109346 | RNA→ rpmE2 (PRK01678)Ribosomal_L31 (pfam01197)→ |
| Sau-1111-1 | NZ_KI974177.1 | - | 57506 | 57477 | RNA→ rpmE2 (PRK01678)Ribosomal_L31 (pfam01197)→ |
| Sau-286-1 | NZ_KI976422.1 | - | 56267 | 56238 | RNA→ rpmE2 (PRK01678)Ribosomal_L31 (pfam01197)→ |
| Sau-1145-1 | NZ_KI975105.1 | - | 56894 | 56865 | RNA→ rpmE2 (PRK01678)Ribosomal_L31 (pfam01197)→ |
| Sau-846-1 | NZ_KI976517.1 | - | 61690 | 61661 | RNA→ rpmE2 (PRK01678)Ribosomal_L31 (pfam01197)→ |
| Sau-292-1 | NZ_KI976620.1 | - | 56894 | 56865 | RNA→ rpmE2 (PRK01678)Ribosomal_L31 (pfam01197)→ |
| Sau-2638-1 | NZ_KI975338.1 | - | 59010 | 58981 | RNA→ rpmE2 (PRK01678)Ribosomal_L31 (pfam01197)→ |
| Sau-247-1 | NZ_KI977331.1 | - | 61890 | 61861 | RNA→ rpmE2 (PRK01678)Ribosomal_L31 (pfam01197)→ |
| Sau-1165-1 | NZ_KI975607.1 | - | 56894 | 56865 | RNA→ rpmE2 (PRK01678)Ribosomal_L31 (pfam01197)→ |
| Sau-2608-1 | NZ_KI976071.1 | - | 56894 | 56865 | RNA→ rpmE2 (PRK01678)Ribosomal_L31 (pfam01197)→ |
| Sau-1095-1 | NZ_KI975565.1 | - | 56894 | 56865 | RNA→ rpmE2 (PRK01678)Ribosomal_L31 (pfam01197)→ |
| Sau-2612-1 | NZ_KI977119.1 | - | 115884 | 115855 | RNA→ rpmE2 (PRK01678)Ribosomal_L31 (pfam01197)→ |
| Sau-3021-1 | NZ_KI974138.1 | - | 56894 | 56865 | RNA→ rpmE2 (PRK01678)Ribosomal_L31 (pfam01197)→ |
| Sau-232-1 | NZ_KI975573.1 | - | 239942 | 239913 | RNA→ rpmE2 (PRK01678)Ribosomal_L31 (pfam01197)→ |
| Sau-2817-1 | NZ_KI974715.1 | - | 238688 | 238659 | RNA→ rpmE2 (PRK01678)Ribosomal_L31 (pfam01197)→ |
| Sau-295-1 | NZ_KI976942.1 | - | 56894 | 56865 | RNA→ rpmE2 (PRK01678)Ribosomal_L31 (pfam01197)→ |
| Sau-1109-1 | NZ_KI973890.1 | - | 213618 | 213589 | RNA→ rpmE2 (PRK01678)Ribosomal_L31 (pfam01197)→ |
| Sau-1170-1 | NZ_KI974428.1 | - | 56894 | 56865 | RNA→ rpmE2 (PRK01678)Ribosomal_L31 (pfam01197)→ |
| Sau-2645-1 | NZ_KI974438.1 | - | 57225 | 57196 | RNA→ rpmE2 (PRK01678)Ribosomal_L31 (pfam01197)→ |
| Sau-1152-1 | NZ_KI974611.1 | - | 118606 | 118577 | RNA→ rpmE2 (PRK01678)Ribosomal_L31 (pfam01197)→ |
| Sau-2639-1 | NZ_KI975328.1 | - | 424201 | 424172 | RNA→ rpmE2 (PRK01678)Ribosomal_L31 (pfam01197)→ |
| Sau-1148-1 | NZ_KI975476.1 | - | 56994 | 56965 | RNA→ rpmE2 (PRK01678)Ribosomal_L31 (pfam01197)→ |
| Sau-3019-1 | NZ_KI974078.1 | - | 34873 | 34844 | RNA→ rpmE2 (PRK01678)Ribosomal_L31 (pfam01197)→ |
| Sau-1126-1 | NZ_KI975802.1 | - | 35013 | 34984 | RNA→ rpmE2 (PRK01678)Ribosomal_L31 (pfam01197)→ |
| Sau-2789-1 | NZ_KI974201.1 | - | 56893 | 56864 | RNA→ rpmE2 (PRK01678)Ribosomal_L31 (pfam01197)→ |
| Sau-2607-1 | NZ_KI976384.1 | - | 56258 | 56229 | RNA→ rpmE2 (PRK01678)Ribosomal_L31 (pfam01197)→ |
| Sau-2813-1 | NZ_KI974294.1 | - | 25249 | 25220 | RNA→ rpmE2 (PRK01678)Ribosomal_L31 (pfam01197)→ |
| Sau-1129-1 | NZ_KI976498.1 | - | 110548 | 110519 | RNA→ rpmE2 (PRK01678)Ribosomal_L31 (pfam01197)→ |
| Sau-3699-1 | NZ_KI973666.1 | - | 56588 | 56559 | RNA→ rpmE2 (PRK01678)Ribosomal_L31 (pfam01197)→ |
| Sau-2790-1 | NZ_KI974271.1 | - | 56894 | 56865 | RNA→ rpmE2 (PRK01678)Ribosomal_L31 (pfam01197)→ |
| Sau-2809-1 | NZ_KI975113.1 | - | 110250 | 110221 | RNA→ rpmE2 (PRK01678)Ribosomal_L31 (pfam01197)→ |
| Sau-3689-1 | NZ_KI973743.1 | - | 56893 | 56864 | RNA→ rpmE2 (PRK01678)Ribosomal_L31 (pfam01197)→ |
| Sau-3688-1 | NZ_KI974031.1 | - | 237053 | 237024 | RNA→ rpmE2 (PRK01678)Ribosomal_L31 (pfam01197)→ |
| Sau-3701-1 | NZ_KI974146.1 | + | 48532 | 48561 | RNA→ rpmE2 (PRK01678)Ribosomal_L31 (pfam01197)→ |
| Sau-3703-1 | NZ_KI974376.1 | - | 56894 | 56865 | RNA→ rpmE2 (PRK01678)Ribosomal_L31 (pfam01197)→ |
| Sau-3028-1 | NZ_KI974655.1 | - | 113124 | 113095 | RNA→ rpmE2 (PRK01678)Ribosomal_L31 (pfam01197)→ |
| Sau-2810-1 | NZ_KI975172.1 | - | 57695 | 57666 | RNA→ rpmE2 (PRK01678)Ribosomal_L31 (pfam01197)→ |
| Sau-3018-1 | NZ_KI975754.1 | - | 56894 | 56865 | RNA→ rpmE2 (PRK01678)Ribosomal_L31 (pfam01197)→ |
| Sau-2804-1 | NZ_KI976335.1 | - | 35013 | 34984 | RNA→ rpmE2 (PRK01678)Ribosomal_L31 (pfam01197)→ |
| Sau-3032-1 | NZ_KI977132.1 | - | 55117 | 55088 | RNA→ rpmE2 (PRK01678)Ribosomal_L31 (pfam01197)→ |
| Sau-3702-1 | NZ_KI974426.1 | - | 58231 | 58202 | RNA→ rpmE2 (PRK01678)Ribosomal_L31 (pfam01197)→ |
| Sau-3029-1 | NZ_KI974659.1 | - | 56588 | 56559 | RNA→ rpmE2 (PRK01678)Ribosomal_L31 (pfam01197)→ |
| Sau-2811-1 | NZ_KI975174.1 | - | 216298 | 216269 | RNA→ rpmE2 (PRK01678)Ribosomal_L31 (pfam01197)→ |
| Sau-3025-1 | NZ_KI975987.1 | - | 34872 | 34843 | RNA→ rpmE2 (PRK01678)Ribosomal_L31 (pfam01197)→ |
| Sau-3673-1 | NZ_KI976036.1 | - | 239284 | 239255 | RNA→ rpmE2 (PRK01678)Ribosomal_L31 (pfam01197)→ |
| Sau-3023-1 | NZ_KI976376.1 | - | 119659 | 119630 | RNA→ rpmE2 (PRK01678)Ribosomal_L31 (pfam01197)→ |
| Sau-2792-1 | NZ_KI976774.1 | - | 35284 | 35255 | RNA→ rpmE2 (PRK01678)Ribosomal_L31 (pfam01197)→ |
| Sau-3670-1 | NZ_KI974020.1 | - | 56894 | 56865 | RNA→ rpmE2 (PRK01678)Ribosomal_L31 (pfam01197)→ |
| Sau-3027-1 | NZ_KI975953.1 | - | 121790 | 121761 | RNA→ rpmE2 (PRK01678)Ribosomal_L31 (pfam01197)→ |
| Sau-2797-1 | NZ_KI976725.1 | - | 121803 | 121774 | RNA→ rpmE2 (PRK01678)Ribosomal_L31 (pfam01197)→ |

| | | | | | | |
|---|---|---|---|---|---|---|
| Sau-3705-1 | NZ_KI976952.1 | - | 56893 | 56864 | **RNA**→ | rpmE2 (PRK01678)Ribosomal_L31 (pfam01197)→ |
| Sau-2802-1 | NZ_KI973735.1 | - | 56890 | 56861 | **RNA**→ | rpmE2 (PRK01678)Ribosomal_L31 (pfam01197)→ |
| Sau-2787-1 | NZ_KI974371.1 | - | 110422 | 110393 | **RNA**→ | rpmE2 (PRK01678)Ribosomal_L31 (pfam01197)→ |
| Sau-2807-1 | NZ_KI976065.1 | - | 25258 | 25229 | **RNA**→ | rpmE2 (PRK01678)Ribosomal_L31 (pfam01197)→ |
| Sau-2794-1 | NZ_KI976690.1 | - | 58248 | 58219 | **RNA**→ | rpmE2 (PRK01678)Ribosomal_L31 (pfam01197)→ |
| Sau-1471-1 | NZ_KK010128.1 | - | 36391 | 36362 | **RNA**→ | rpmE2 (PRK01678)Ribosomal_L31 (pfam01197)→ |
| Sau-1524-1 | NZ_KK012312.1 | - | 35094 | 35065 | **RNA**→ | rpmE2 (PRK01678)Ribosomal_L31 (pfam01197)→ |
| Sau-1541-1 | NZ_KK013214.1 | - | 35398 | 35369 | **RNA**→ | rpmE2 (PRK01678)Ribosomal_L31 (pfam01197)→ |
| Sau-2350-1 | NZ_KI995645.1 | - | 36089 | 36060 | **RNA**→ | rpmE2 (PRK01678)Ribosomal_L31 (pfam01197)→ |
| Sau-2387-1 | NZ_KI997549.1 | - | 61851 | 61822 | **RNA**→ | rpmE2 (PRK01678)Ribosomal_L31 (pfam01197)→ |
| Sau-2397-1 | NZ_KI997918.1 | - | 213793 | 213764 | **RNA**→ | rpmE2 (PRK01678)Ribosomal_L31 (pfam01197)→ |
| Sau-1568-1 | NZ_KK014253.1 | - | 35305 | 35276 | **RNA**→ | rpmE2 (PRK01678)Ribosomal_L31 (pfam01197)→ |
| Sau-2399-1 | NZ_KI997937.1 | - | 118626 | 118597 | **RNA**→ | rpmE2 (PRK01678)Ribosomal_L31 (pfam01197)→ |
| Sau-1516-1 | NZ_KK012056.1 | - | 32527 | 32498 | **RNA**→ | rpmE2 (PRK01678)Ribosomal_L31 (pfam01197)→ |
| Sau-2404-1 | NZ_KI998095.1 | - | 110981 | 110952 | **RNA**→ | rpmE2 (PRK01678)Ribosomal_L31 (pfam01197)→ |
| Sau-2421-1 | NZ_KI998596.1 | - | 24749 | 24720 | **RNA**→ | rpmE2 (PRK01678)Ribosomal_L31 (pfam01197)→ |
| Sau-1527-1 | NZ_KK012536.1 | - | 110244 | 110215 | **RNA**→ | rpmE2 (PRK01678)Ribosomal_L31 (pfam01197)→ |
| Sau-2247-1 | NZ_KI996273.1 | - | 122904 | 122875 | **RNA**→ | rpmE2 (PRK01678)Ribosomal_L31 (pfam01197)→ |
| Sau-1540-1 | NZ_KK013208.1 | - | 25227 | 25198 | **RNA**→ | rpmE2 (PRK01678)Ribosomal_L31 (pfam01197)→ |
| Sau-2359-1 | NZ_KI996835.1 | - | 62199 | 62170 | **RNA**→ | rpmE2 (PRK01678)Ribosomal_L31 (pfam01197)→ |
| Sau-1494-1 | NZ_KK011060.1 | - | 35151 | 35122 | **RNA**→ | rpmE2 (PRK01678)Ribosomal_L31 (pfam01197)→ |
| Sau-2366-1 | NZ_KI997029.1 | - | 62104 | 62075 | **RNA**→ | rpmE2 (PRK01678)Ribosomal_L31 (pfam01197)→ |
| Sau-1917-1 | NZ_KI987117.1 | - | 110394 | 110365 | **RNA**→ | rpmE2 (PRK01678)Ribosomal_L31 (pfam01197)→ |
| Sau-1499-1 | NZ_KK011331.1 | - | 35760 | 35731 | **RNA**→ | rpmE2 (PRK01678)Ribosomal_L31 (pfam01197)→ |
| Sau-2375-1 | NZ_KI997218.1 | - | 24749 | 24720 | **RNA**→ | rpmE2 (PRK01678)Ribosomal_L31 (pfam01197)→ |
| Sau-2384-1 | NZ_KI997424.1 | - | 35349 | 35320 | **RNA**→ | rpmE2 (PRK01678)Ribosomal_L31 (pfam01197)→ |
| Sau-1548-1 | NZ_KK013557.1 | - | 36391 | 36362 | **RNA**→ | rpmE2 (PRK01678)Ribosomal_L31 (pfam01197)→ |
| Sau-2395-1 | NZ_KI997821.1 | - | 98284 | 98255 | **RNA**→ | rpmE2 (PRK01678)Ribosomal_L31 (pfam01197)→ |
| Sau-1556-1 | NZ_KK013787.1 | - | 35610 | 35581 | **RNA**→ | rpmE2 (PRK01678)Ribosomal_L31 (pfam01197)→ |
| Sau-1965-1 | NZ_KI988768.1 | - | 55131 | 55102 | **RNA**→ | rpmE2 (PRK01678)Ribosomal_L31 (pfam01197)→ |
| Sau-2407-1 | NZ_KI998233.1 | - | 35955 | 35926 | **RNA**→ | rpmE2 (PRK01678)Ribosomal_L31 (pfam01197)→ |
| Sau-1564-1 | NZ_KK014054.1 | - | 36391 | 36362 | **RNA**→ | rpmE2 (PRK01678)Ribosomal_L31 (pfam01197)→ |
| Sau-2415-1 | NZ_KI998474.1 | - | 21021 | 20992 | **RNA**→ | rpmE2 (PRK01678)Ribosomal_L31 (pfam01197)→ |
| Sau-2425-1 | NZ_KI998629.1 | - | 52040 | 52011 | **RNA**→ | rpmE2 (PRK01678)Ribosomal_L31 (pfam01197)→ |
| Sau-1987-1 | NZ_KI989651.1 | - | 56762 | 56733 | **RNA**→ | rpmE2 (PRK01678)Ribosomal_L31 (pfam01197)→ |
| Sau-1460-1 | NZ_KK009730.1 | - | 36391 | 36362 | **RNA**→ | rpmE2 (PRK01678)Ribosomal_L31 (pfam01197)→ |
| Sau-2322-1 | NZ_KI994432.1 | - | 24749 | 24720 | **RNA**→ | rpmE2 (PRK01678)Ribosomal_L31 (pfam01197)→ |
| Sau-2003-1 | NZ_KI990116.1 | - | 24749 | 24720 | **RNA**→ | rpmE2 (PRK01678)Ribosomal_L31 (pfam01197)→ |
| Sau-2235-1 | NZ_KI995970.1 | - | 35290 | 35261 | **RNA**→ | rpmE2 (PRK01678)Ribosomal_L31 (pfam01197)→ |
| Sau-1514-1 | NZ_KK011889.1 | - | 77885 | 77856 | **RNA**→ | rpmE2 (PRK01678)Ribosomal_L31 (pfam01197)→ |
| Sau-2012-1 | NZ_KI990411.1 | - | 83970 | 83941 | **RNA**→ | rpmE2 (PRK01678)Ribosomal_L31 (pfam01197)→ |
| Sau-1519-1 | NZ_KK012183.1 | - | 36391 | 36362 | **RNA**→ | rpmE2 (PRK01678)Ribosomal_L31 (pfam01197)→ |
| Sau-2246-1 | NZ_KI996258.1 | - | 150172 | 150143 | **RNA**→ | rpmE2 (PRK01678)Ribosomal_L31 (pfam01197)→ |
| Sau-2434-1 | NZ_KI991416.1 | - | 35701 | 35672 | **RNA**→ | rpmE2 (PRK01678)Ribosomal_L31 (pfam01197)→ |
| Sau-1526-1 | NZ_KK012427.1 | - | 36391 | 36362 | **RNA**→ | rpmE2 (PRK01678)Ribosomal_L31 (pfam01197)→ |
| Sau-2365-1 | NZ_KI997025.1 | - | 24749 | 24720 | **RNA**→ | rpmE2 (PRK01678)Ribosomal_L31 (pfam01197)→ |
| Sau-2388-1 | NZ_KI997568.1 | - | 35672 | 35643 | **RNA**→ | rpmE2 (PRK01678)Ribosomal_L31 (pfam01197)→ |
| Sau-2435-1 | NZ_KI991448.1 | - | 36391 | 36362 | **RNA**→ | rpmE2 (PRK01678)Ribosomal_L31 (pfam01197)→ |
| Sau-2394-1 | NZ_KI997810.1 | - | 514560 | 514531 | **RNA**→ | rpmE2 (PRK01678)Ribosomal_L31 (pfam01197)→ |
| Sau-1756-1 | NZ_KI991889.1 | - | 24749 | 24720 | **RNA**→ | rpmE2 (PRK01678)Ribosomal_L31 (pfam01197)→ |
| Sau-2400-1 | NZ_KI997959.1 | - | 36391 | 36362 | **RNA**→ | rpmE2 (PRK01678)Ribosomal_L31 (pfam01197)→ |
| Sau-1539-1 | NZ_KK013085.1 | - | 26900 | 26871 | **RNA**→ | rpmE2 (PRK01678)Ribosomal_L31 (pfam01197)→ |

| | | | | | | |
|---|---|---|---|---|---|---|
| Sau-2310-1 | NZ_KI994065.1 | - | 62157 | 62128 | **RNA**→ | rpmE2 (PRK01678)Ribosomal_L31 (pfam01197)→ |
| Sau-2406-1 | NZ_KI998165.1 | - | 35156 | 35127 | **RNA**→ | rpmE2 (PRK01678)Ribosomal_L31 (pfam01197)→ |
| Sau-1559-1 | NZ_KK013862.1 | - | 114870 | 114841 | **RNA**→ | rpmE2 (PRK01678)Ribosomal_L31 (pfam01197)→ |
| Sau-2422-1 | NZ_KI998604.1 | - | 24749 | 24720 | **RNA**→ | rpmE2 (PRK01678)Ribosomal_L31 (pfam01197)→ |
| Sau-1956-1 | NZ_KI988450.1 | - | 76879 | 76850 | **RNA**→ | rpmE2 (PRK01678)Ribosomal_L31 (pfam01197)→ |
| Sau-1565-1 | NZ_KK014174.1 | - | 113266 | 113237 | **RNA**→ | rpmE2 (PRK01678)Ribosomal_L31 (pfam01197)→ |
| Sau-2354-1 | NZ_KI995816.1 | - | 35969 | 35940 | **RNA**→ | rpmE2 (PRK01678)Ribosomal_L31 (pfam01197)→ |
| Sau-1983-1 | NZ_KI989506.1 | - | 3698 | 3669 | **RNA**→ | rpmE2 (PRK01678)Ribosomal_L31 (pfam01197)→ |
| Sau-2385-1 | NZ_KI997468.1 | - | 35156 | 35127 | **RNA**→ | rpmE2 (PRK01678)Ribosomal_L31 (pfam01197)→ |
| Sau-2403-1 | NZ_KI998058.1 | - | 103696 | 103667 | **RNA**→ | rpmE2 (PRK01678)Ribosomal_L31 (pfam01197)→ |
| Sau-1999-1 | NZ_KI990012.1 | - | 125991 | 125962 | **RNA**→ | rpmE2 (PRK01678)Ribosomal_L31 (pfam01197)→ |
| Sau-2408-1 | NZ_KI998319.1 | - | 61384 | 61355 | **RNA**→ | rpmE2 (PRK01678)Ribosomal_L31 (pfam01197)→ |
| Sau-2410-1 | NZ_KI998325.1 | - | 36340 | 36311 | **RNA**→ | rpmE2 (PRK01678)Ribosomal_L31 (pfam01197)→ |
| Sau-2021-1 | NZ_KI990840.1 | - | 62157 | 62128 | **RNA**→ | rpmE2 (PRK01678)Ribosomal_L31 (pfam01197)→ |
| Sau-2419-1 | NZ_KI998493.1 | - | 35170 | 35141 | **RNA**→ | rpmE2 (PRK01678)Ribosomal_L31 (pfam01197)→ |
| Sau-1180-1 | NZ_KI998844.1 | - | 55131 | 55102 | **RNA**→ | rpmE2 (PRK01678)Ribosomal_L31 (pfam01197)→ |
| Sau-2432-1 | NZ_KI991367.1 | - | 36413 | 36384 | **RNA**→ | rpmE2 (PRK01678)Ribosomal_L31 (pfam01197)→ |
| Sau-2378-1 | NZ_KI997278.1 | - | 36491 | 36462 | **RNA**→ | rpmE2 (PRK01678)Ribosomal_L31 (pfam01197)→ |
| Sau-1770-1 | NZ_KI992468.1 | - | 70571 | 70542 | **RNA**→ | rpmE2 (PRK01678)Ribosomal_L31 (pfam01197)→ |
| Sau-2412-1 | NZ_KI998382.1 | - | 35390 | 35361 | **RNA**→ | rpmE2 (PRK01678)Ribosomal_L31 (pfam01197)→ |
| Sau-2266-1 | NZ_KI992708.1 | - | 35936 | 35907 | **RNA**→ | rpmE2 (PRK01678)Ribosomal_L31 (pfam01197)→ |
| Sau-2249-1 | NZ_KI996364.1 | - | 498623 | 498594 | **RNA**→ | rpmE2 (PRK01678)Ribosomal_L31 (pfam01197)→ |
| Sau-1203-1 | NZ_KK000026.1 | - | 27865 | 27836 | **RNA**→ | rpmE2 (PRK01678)Ribosomal_L31 (pfam01197)→ |
| Sau-2257-1 | NZ_KI996580.1 | - | 3545 | 3516 | **RNA**→ | rpmE2 (PRK01678)Ribosomal_L31 (pfam01197)→ |
| Sau-2275-1 | NZ_KI992964.1 | - | 27699 | 27670 | **RNA**→ | rpmE2 (PRK01678)Ribosomal_L31 (pfam01197)→ |
| Sau-1215-1 | NZ_KK000626.1 | - | 35264 | 35235 | **RNA**→ | rpmE2 (PRK01678)Ribosomal_L31 (pfam01197)→ |
| Sau-2396-1 | NZ_KI997868.1 | - | 24749 | 24720 | **RNA**→ | rpmE2 (PRK01678)Ribosomal_L31 (pfam01197)→ |
| Sau-2284-1 | NZ_KI993105.1 | - | 35349 | 35320 | **RNA**→ | rpmE2 (PRK01678)Ribosomal_L31 (pfam01197)→ |
| Sau-1178-1 | NZ_KI998708.1 | - | 36172 | 36143 | **RNA**→ | rpmE2 (PRK01678)Ribosomal_L31 (pfam01197)→ |
| Sau-2427-1 | NZ_KI998671.1 | - | 24749 | 24720 | **RNA**→ | rpmE2 (PRK01678)Ribosomal_L31 (pfam01197)→ |
| Sau-1182-1 | NZ_KI998925.1 | - | 36391 | 36362 | **RNA**→ | rpmE2 (PRK01678)Ribosomal_L31 (pfam01197)→ |
| Sau-2289-1 | NZ_KI993273.1 | - | 216283 | 216254 | **RNA**→ | rpmE2 (PRK01678)Ribosomal_L31 (pfam01197)→ |
| Sau-1321-1 | NZ_KK004815.1 | - | 110935 | 110906 | **RNA**→ | rpmE2 (PRK01678)Ribosomal_L31 (pfam01197)→ |
| Sau-2320-1 | NZ_KI994356.1 | - | 23949 | 23920 | **RNA**→ | rpmE2 (PRK01678)Ribosomal_L31 (pfam01197)→ |
| Sau-2413-1 | NZ_KI998446.1 | - | 24749 | 24720 | **RNA**→ | rpmE2 (PRK01678)Ribosomal_L31 (pfam01197)→ |
| Sau-1235-1 | NZ_KK001260.1 | - | 109425 | 109396 | **RNA**→ | rpmE2 (PRK01678)Ribosomal_L31 (pfam01197)→ |
| Sau-1933-1 | NZ_KI987796.1 | - | 36390 | 36361 | **RNA**→ | rpmE2 (PRK01678)Ribosomal_L31 (pfam01197)→ |
| Sau-1307-1 | NZ_KK004030.1 | - | 9107 | 9078 | **RNA**→ | rpmE2 (PRK01678)Ribosomal_L31 (pfam01197)→ |
| Sau-1247-1 | NZ_KK001698.1 | - | 238687 | 238658 | **RNA**→ | rpmE2 (PRK01678)Ribosomal_L31 (pfam01197)→ |
| Sau-2401-1 | NZ_KI998009.1 | - | 24749 | 24720 | **RNA**→ | rpmE2 (PRK01678)Ribosomal_L31 (pfam01197)→ |
| Sau-2002-1 | NZ_KI990083.1 | - | 114771 | 114742 | **RNA**→ | rpmE2 (PRK01678)Ribosomal_L31 (pfam01197)→ |
| Sau-1271-1 | NZ_KK002585.1 | - | 62105 | 62076 | **RNA**→ | rpmE2 (PRK01678)Ribosomal_L31 (pfam01197)→ |
| Sau-2011-1 | NZ_KI990359.1 | - | 61128 | 61099 | **RNA**→ | rpmE2 (PRK01678)Ribosomal_L31 (pfam01197)→ |
| Sau-1186-1 | NZ_KI999175.1 | - | 2421 | 2392 | **RNA**→ | rpmE2 (PRK01678)Ribosomal_L31 (pfam01197)→ |
| Sau-2016-1 | NZ_KI990652.1 | - | 56605 | 56576 | **RNA**→ | rpmE2 (PRK01678)Ribosomal_L31 (pfam01197)→ |
| Sau-1195-1 | NZ_KI999511.1 | - | 1611 | 1582 | **RNA**→ | rpmE2 (PRK01678)Ribosomal_L31 (pfam01197)→ |
| Sau-2031-1 | NZ_KI991141.1 | - | 56154 | 56125 | **RNA**→ | rpmE2 (PRK01678)Ribosomal_L31 (pfam01197)→ |
| Sau-1205-1 | NZ_KK000098.1 | - | 2518 | 2489 | **RNA**→ | rpmE2 (PRK01678)Ribosomal_L31 (pfam01197)→ |
| Sau-2433-1 | NZ_KI991414.1 | - | 81526 | 81497 | **RNA**→ | rpmE2 (PRK01678)Ribosomal_L31 (pfam01197)→ |
| Sau-1752-1 | NZ_KI991788.1 | - | 35329 | 35300 | **RNA**→ | rpmE2 (PRK01678)Ribosomal_L31 (pfam01197)→ |
| Sau-1389-1 | NZ_KK006830.1 | - | 24749 | 24720 | **RNA**→ | rpmE2 (PRK01678)Ribosomal_L31 (pfam01197)→ |
| Sau-1211-1 | NZ_KK000497.1 | - | 25129 | 25100 | **RNA**→ | rpmE2 (PRK01678)Ribosomal_L31 (pfam01197)→ |

| | | | | | |
|---|---|---|---|---|---|
| Sau-2380-1 | NZ_KI997336.1 | - | 62256 | 62227 | **RNA**→ rpmE2 (PRK01678)Ribosomal_L31 (pfam01197)→ |
| Sau-1251-1 | NZ_KK001764.1 | - | 110415 | 110386 | **RNA**→ rpmE2 (PRK01678)Ribosomal_L31 (pfam01197)→ |
| Sau-1761-1 | NZ_KI992078.1 | - | 24749 | 24720 | **RNA**→ rpmE2 (PRK01678)Ribosomal_L31 (pfam01197)→ |
| Sau-1270-1 | NZ_KK002552.1 | - | 100428 | 100399 | **RNA**→ rpmE2 (PRK01678)Ribosomal_L31 (pfam01197)→ |
| Sau-1404-1 | NZ_KK007400.1 | - | 52215 | 52186 | **RNA**→ rpmE2 (PRK01678)Ribosomal_L31 (pfam01197)→ |
| Sau-1765-1 | NZ_KI992241.1 | - | 36491 | 36462 | **RNA**→ rpmE2 (PRK01678)Ribosomal_L31 (pfam01197)→ |
| Sau-1179-1 | NZ_KI998742.1 | - | 36390 | 36361 | **RNA**→ rpmE2 (PRK01678)Ribosomal_L31 (pfam01197)→ |
| Sau-1399-1 | NZ_KK007204.1 | - | 119705 | 119676 | **RNA**→ rpmE2 (PRK01678)Ribosomal_L31 (pfam01197)→ |
| Sau-1766-1 | NZ_KI992307.1 | - | 24749 | 24720 | **RNA**→ rpmE2 (PRK01678)Ribosomal_L31 (pfam01197)→ |
| Sau-1184-1 | NZ_KI999066.1 | - | 36391 | 36362 | **RNA**→ rpmE2 (PRK01678)Ribosomal_L31 (pfam01197)→ |
| Sau-1405-1 | NZ_KK007455.1 | - | 25264 | 25235 | **RNA**→ rpmE2 (PRK01678)Ribosomal_L31 (pfam01197)→ |
| Sau-1771-1 | NZ_KI992525.1 | - | 55426 | 55397 | **RNA**→ rpmE2 (PRK01678)Ribosomal_L31 (pfam01197)→ |
| Sau-1412-1 | NZ_KK007629.1 | - | 105070 | 105041 | **RNA**→ rpmE2 (PRK01678)Ribosomal_L31 (pfam01197)→ |
| Sau-1192-1 | NZ_KI999469.1 | - | 107905 | 107876 | **RNA**→ rpmE2 (PRK01678)Ribosomal_L31 (pfam01197)→ |
| Sau-1418-1 | NZ_KK007813.1 | - | 62057 | 62028 | **RNA**→ rpmE2 (PRK01678)Ribosomal_L31 (pfam01197)→ |
| Sau-1209-1 | NZ_KK000279.1 | - | 2421 | 2392 | **RNA**→ rpmE2 (PRK01678)Ribosomal_L31 (pfam01197)→ |
| Sau-2262-1 | NZ_KI992643.1 | - | 61990 | 61961 | **RNA**→ rpmE2 (PRK01678)Ribosomal_L31 (pfam01197)→ |
| Sau-1419-1 | NZ_KK007920.1 | - | 62157 | 62128 | **RNA**→ rpmE2 (PRK01678)Ribosomal_L31 (pfam01197)→ |
| Sau-2286-1 | NZ_KI993169.1 | - | 35990 | 35961 | **RNA**→ rpmE2 (PRK01678)Ribosomal_L31 (pfam01197)→ |
| Sau-1426-1 | NZ_KK008138.1 | - | 35323 | 35294 | **RNA**→ rpmE2 (PRK01678)Ribosomal_L31 (pfam01197)→ |
| Sau-1217-1 | NZ_KK000731.1 | - | 25029 | 25000 | **RNA**→ rpmE2 (PRK01678)Ribosomal_L31 (pfam01197)→ |
| Sau-1438-1 | NZ_KK008729.1 | - | 36391 | 36362 | **RNA**→ rpmE2 (PRK01678)Ribosomal_L31 (pfam01197)→ |
| Sau-1255-1 | NZ_KK001985.1 | - | 36391 | 36362 | **RNA**→ rpmE2 (PRK01678)Ribosomal_L31 (pfam01197)→ |
| Sau-1443-1 | NZ_KK009013.1 | - | 34375 | 34346 | **RNA**→ rpmE2 (PRK01678)Ribosomal_L31 (pfam01197)→ |
| Sau-1261-1 | NZ_KK002301.1 | - | 113854 | 113825 | **RNA**→ rpmE2 (PRK01678)Ribosomal_L31 (pfam01197)→ |
| Sau-1449-1 | NZ_KK009222.1 | - | 56828 | 56799 | **RNA**→ rpmE2 (PRK01678)Ribosomal_L31 (pfam01197)→ |
| Sau-1268-1 | NZ_KK002505.1 | - | 63488 | 63459 | **RNA**→ rpmE2 (PRK01678)Ribosomal_L31 (pfam01197)→ |
| Sau-1486-1 | NZ_KK010739.1 | - | 56205 | 56176 | **RNA**→ rpmE2 (PRK01678)Ribosomal_L31 (pfam01197)→ |
| Sau-1299-1 | NZ_KK003662.1 | - | 194176 | 194147 | **RNA**→ rpmE2 (PRK01678)Ribosomal_L31 (pfam01197)→ |
| Sau-1398-1 | NZ_KK007165.1 | - | 26132 | 26103 | **RNA**→ rpmE2 (PRK01678)Ribosomal_L31 (pfam01197)→ |
| Sau-1208-1 | NZ_KK000225.1 | - | 36390 | 36361 | **RNA**→ rpmE2 (PRK01678)Ribosomal_L31 (pfam01197)→ |
| Sau-1411-1 | NZ_KK007527.1 | - | 36177 | 36148 | **RNA**→ rpmE2 (PRK01678)Ribosomal_L31 (pfam01197)→ |
| Sau-1214-1 | NZ_KK000556.1 | - | 2418 | 2389 | **RNA**→ rpmE2 (PRK01678)Ribosomal_L31 (pfam01197)→ |
| Sau-1429-1 | NZ_KK008343.1 | - | 61334 | 61305 | **RNA**→ rpmE2 (PRK01678)Ribosomal_L31 (pfam01197)→ |
| Sau-1221-1 | NZ_KK000817.1 | - | 36279 | 36250 | **RNA**→ rpmE2 (PRK01678)Ribosomal_L31 (pfam01197)→ |
| Sau-1442-1 | NZ_KK008967.1 | - | 35730 | 35701 | **RNA**→ rpmE2 (PRK01678)Ribosomal_L31 (pfam01197)→ |
| Sau-1236-1 | NZ_KK001262.1 | - | 217172 | 217143 | **RNA**→ rpmE2 (PRK01678)Ribosomal_L31 (pfam01197)→ |
| Sau-1455-1 | NZ_KK009463.1 | - | 35974 | 35945 | **RNA**→ rpmE2 (PRK01678)Ribosomal_L31 (pfam01197)→ |
| Sau-1463-1 | NZ_KK009790.1 | - | 36145 | 36116 | **RNA**→ rpmE2 (PRK01678)Ribosomal_L31 (pfam01197)→ |
| Sau-1238-1 | NZ_KK001281.1 | - | 217172 | 217143 | **RNA**→ rpmE2 (PRK01678)Ribosomal_L31 (pfam01197)→ |
| Sau-1242-1 | NZ_KK001486.1 | - | 62157 | 62128 | **RNA**→ rpmE2 (PRK01678)Ribosomal_L31 (pfam01197)→ |
| Sau-1394-1 | NZ_KK006995.1 | - | 192301 | 192272 | **RNA**→ rpmE2 (PRK01678)Ribosomal_L31 (pfam01197)→ |
| Sau-1395-1 | NZ_KK007015.1 | - | 36391 | 36362 | **RNA**→ rpmE2 (PRK01678)Ribosomal_L31 (pfam01197)→ |
| Sau-1243-1 | NZ_KK001565.1 | - | 111533 | 111504 | **RNA**→ rpmE2 (PRK01678)Ribosomal_L31 (pfam01197)→ |
| Sau-1400-1 | NZ_KK007261.1 | - | 67891 | 67862 | **RNA**→ rpmE2 (PRK01678)Ribosomal_L31 (pfam01197)→ |
| Sau-1249-1 | NZ_KK001741.1 | - | 58247 | 58218 | **RNA**→ rpmE2 (PRK01678)Ribosomal_L31 (pfam01197)→ |
| Sau-1401-1 | NZ_KK007264.1 | - | 50765 | 50736 | **RNA**→ rpmE2 (PRK01678)Ribosomal_L31 (pfam01197)→ |
| Sau-1254-1 | NZ_KK001938.1 | - | 61872 | 61843 | **RNA**→ rpmE2 (PRK01678)Ribosomal_L31 (pfam01197)→ |
| Sau-1406-1 | NZ_KK007468.1 | - | 56418 | 56389 | **RNA**→ rpmE2 (PRK01678)Ribosomal_L31 (pfam01197)→ |
| Sau-1267-1 | NZ_KK002428.1 | - | 118636 | 118607 | **RNA**→ rpmE2 (PRK01678)Ribosomal_L31 (pfam01197)→ |
| Sau-1407-1 | NZ_KK007482.1 | - | 36225 | 36196 | **RNA**→ rpmE2 (PRK01678)Ribosomal_L31 (pfam01197)→ |
| Sau-1304-1 | NZ_KK003871.1 | - | 36491 | 36462 | **RNA**→ rpmE2 (PRK01678)Ribosomal_L31 (pfam01197)→ |

| | | | | | | |
|---|---|---|---|---|---|---|
| Sau-1413-1 | NZ_KK007634.1 | - | 36391 | 36362 | **RNA**→ | rpmE2 (PRK01678)Ribosomal_L31 (pfam01197)→ |
| Sau-1414-1 | NZ_KK007734.1 | - | 114172 | 114143 | **RNA**→ | rpmE2 (PRK01678)Ribosomal_L31 (pfam01197)→ |
| Sau-1198-1 | NZ_KI999796.1 | - | 28749 | 28720 | **RNA**→ | rpmE2 (PRK01678)Ribosomal_L31 (pfam01197)→ |
| Sau-1427-1 | NZ_KK008172.1 | - | 35163 | 35134 | **RNA**→ | rpmE2 (PRK01678)Ribosomal_L31 (pfam01197)→ |
| Sau-1204-1 | NZ_KK000032.1 | - | 55130 | 55101 | **RNA**→ | rpmE2 (PRK01678)Ribosomal_L31 (pfam01197)→ |
| Sau-1467-1 | NZ_KK009893.1 | - | 30144 | 30115 | **RNA**→ | rpmE2 (PRK01678)Ribosomal_L31 (pfam01197)→ |
| Sau-1218-1 | NZ_KK000734.1 | - | 35215 | 35186 | **RNA**→ | rpmE2 (PRK01678)Ribosomal_L31 (pfam01197)→ |
| Sau-1479-1 | NZ_KK010576.1 | - | 61380 | 61351 | **RNA**→ | rpmE2 (PRK01678)Ribosomal_L31 (pfam01197)→ |
| Sau-1390-1 | NZ_KK006869.1 | - | 25227 | 25198 | **RNA**→ | rpmE2 (PRK01678)Ribosomal_L31 (pfam01197)→ |
| Sau-1226-1 | NZ_KK000948.1 | - | 36490 | 36461 | **RNA**→ | rpmE2 (PRK01678)Ribosomal_L31 (pfam01197)→ |
| Sau-1396-1 | NZ_KK007071.1 | - | 36390 | 36361 | **RNA**→ | rpmE2 (PRK01678)Ribosomal_L31 (pfam01197)→ |
| Sau-1233-1 | NZ_KK001167.1 | - | 56086 | 56057 | **RNA**→ | rpmE2 (PRK01678)Ribosomal_L31 (pfam01197)→ |
| Sau-1402-1 | NZ_KK007347.1 | - | 62157 | 62128 | **RNA**→ | rpmE2 (PRK01678)Ribosomal_L31 (pfam01197)→ |
| Sau-1240-1 | NZ_KK001353.1 | - | 36491 | 36462 | **RNA**→ | rpmE2 (PRK01678)Ribosomal_L31 (pfam01197)→ |
| Sau-1422-1 | NZ_KK007990.1 | - | 6175 | 6146 | **RNA**→ | rpmE2 (PRK01678)Ribosomal_L31 (pfam01197)→ |
| Sau-1428-1 | NZ_KK008239.1 | - | 35607 | 35578 | **RNA**→ | rpmE2 (PRK01678)Ribosomal_L31 (pfam01197)→ |
| Sau-1245-1 | NZ_KK001682.1 | - | 61334 | 61305 | **RNA**→ | rpmE2 (PRK01678)Ribosomal_L31 (pfam01197)→ |
| Sau-1446-1 | NZ_KK009114.1 | - | 113893 | 113864 | **RNA**→ | rpmE2 (PRK01678)Ribosomal_L31 (pfam01197)→ |
| Sau-1269-1 | NZ_KK002539.1 | - | 36391 | 36362 | **RNA**→ | rpmE2 (PRK01678)Ribosomal_L31 (pfam01197)→ |
| Sau-1453-1 | NZ_KK009347.1 | - | 36413 | 36384 | **RNA**→ | rpmE2 (PRK01678)Ribosomal_L31 (pfam01197)→ |
| Sau-1282-1 | NZ_KK002976.1 | - | 36391 | 36362 | **RNA**→ | rpmE2 (PRK01678)Ribosomal_L31 (pfam01197)→ |
| Sau-1183-1 | NZ_KI999003.1 | - | 2467 | 2438 | **RNA**→ | rpmE2 (PRK01678)Ribosomal_L31 (pfam01197)→ |
| Sau-1468-1 | NZ_KK010017.1 | - | 61805 | 61776 | **RNA**→ | rpmE2 (PRK01678)Ribosomal_L31 (pfam01197)→ |
| Sau-1469-1 | NZ_KK010031.1 | - | 8424 | 8395 | **RNA**→ | rpmE2 (PRK01678)Ribosomal_L31 (pfam01197)→ |
| Sau-1187-1 | NZ_KI999294.1 | - | 28749 | 28720 | **RNA**→ | rpmE2 (PRK01678)Ribosomal_L31 (pfam01197)→ |
| Sau-1189-1 | NZ_KI999318.1 | - | 56493 | 56464 | **RNA**→ | rpmE2 (PRK01678)Ribosomal_L31 (pfam01197)→ |
| Sau-1857-1 | NZ_KI985139.1 | - | 215969 | 215940 | **RNA**→ | rpmE2 (PRK01678)Ribosomal_L31 (pfam01197)→ |
| Sau-1219-1 | NZ_KK000810.1 | - | 107780 | 107751 | **RNA**→ | rpmE2 (PRK01678)Ribosomal_L31 (pfam01197)→ |
| Sau-1946-1 | NZ_KI988207.1 | - | 112019 | 111990 | **RNA**→ | rpmE2 (PRK01678)Ribosomal_L31 (pfam01197)→ |
| Sau-1263-1 | NZ_KK002379.1 | - | 119174 | 119145 | **RNA**→ | rpmE2 (PRK01678)Ribosomal_L31 (pfam01197)→ |
| Sau-1974-1 | NZ_KI989131.1 | - | 36695 | 36666 | **RNA**→ | rpmE2 (PRK01678)Ribosomal_L31 (pfam01197)→ |
| Sau-1266-1 | NZ_KK002416.1 | - | 110123 | 110094 | **RNA**→ | rpmE2 (PRK01678)Ribosomal_L31 (pfam01197)→ |
| Sau-1302-1 | NZ_KK003802.1 | - | 36391 | 36362 | **RNA**→ | rpmE2 (PRK01678)Ribosomal_L31 (pfam01197)→ |
| Sau-1981-1 | NZ_KI989453.1 | - | 26580 | 26551 | **RNA**→ | rpmE2 (PRK01678)Ribosomal_L31 (pfam01197)→ |
| Sau-2013-1 | NZ_KI990459.1 | - | 36490 | 36461 | **RNA**→ | rpmE2 (PRK01678)Ribosomal_L31 (pfam01197)→ |
| Sau-2019-1 | NZ_KI990696.1 | - | 35503 | 35474 | **RNA**→ | rpmE2 (PRK01678)Ribosomal_L31 (pfam01197)→ |
| Sau-2025-1 | NZ_KI991022.1 | - | 36391 | 36362 | **RNA**→ | rpmE2 (PRK01678)Ribosomal_L31 (pfam01197)→ |
| Sau-2431-1 | NZ_KI991290.1 | - | 24739 | 24710 | **RNA**→ | rpmE2 (PRK01678)Ribosomal_L31 (pfam01197)→ |
| Sau-2436-1 | NZ_KI991526.1 | - | 61378 | 61349 | **RNA**→ | rpmE2 (PRK01678)Ribosomal_L31 (pfam01197)→ |
| Sau-2673-1 | NZ_KI978813.1 | - | 56893 | 56864 | **RNA**→ | rpmE2 (PRK01678)Ribosomal_L31 (pfam01197)→ |
| Sau-1757-1 | NZ_KI991896.1 | - | 36391 | 36362 | **RNA**→ | rpmE2 (PRK01678)Ribosomal_L31 (pfam01197)→ |
| Sau-2744-1 | NZ_KI976555.1 | - | 56894 | 56865 | **RNA**→ | rpmE2 (PRK01678)Ribosomal_L31 (pfam01197)→ |
| Sau-1763-1 | NZ_KI992197.1 | - | 54757 | 54728 | **RNA**→ | rpmE2 (PRK01678)Ribosomal_L31 (pfam01197)→ |
| Sau-2265-1 | NZ_KI992696.1 | - | 113715 | 113686 | **RNA**→ | rpmE2 (PRK01678)Ribosomal_L31 (pfam01197)→ |
| Sau-2288-1 | NZ_KI993242.1 | - | 35381 | 35352 | **RNA**→ | rpmE2 (PRK01678)Ribosomal_L31 (pfam01197)→ |
| Sau-2300-1 | NZ_KI993733.1 | - | 62256 | 62227 | **RNA**→ | rpmE2 (PRK01678)Ribosomal_L31 (pfam01197)→ |
| Sau-2306-1 | NZ_KI993956.1 | - | 107880 | 107851 | **RNA**→ | rpmE2 (PRK01678)Ribosomal_L31 (pfam01197)→ |
| Sau-2768-1 | NZ_KI976650.1 | - | 25095 | 25066 | **RNA**→ | rpmE2 (PRK01678)Ribosomal_L31 (pfam01197)→ |
| Sau-2800-1 | NZ_KI977253.1 | - | 25249 | 25220 | **RNA**→ | rpmE2 (PRK01678)Ribosomal_L31 (pfam01197)→ |
| Sau-2818-1 | NZ_KI977480.1 | - | 60074 | 60045 | **RNA**→ | rpmE2 (PRK01678)Ribosomal_L31 (pfam01197)→ |
| Sau-2440-1 | NZ_KI991672.1 | - | 36491 | 36462 | **RNA**→ | rpmE2 (PRK01678)Ribosomal_L31 (pfam01197)→ |
| Sau-2428-1 | NZ_KI991190.1 | - | 36391 | 36362 | **RNA**→ | rpmE2 (PRK01678)Ribosomal_L31 (pfam01197)→ |

| | | | | | |
|---|---|---|---|---|---|
| Sau-2437-1 | NZ_KI991531.1 | - | 35701 | 35672 | **RNA**→ rpmE2 (PRK01678)Ribosomal_L31 (pfam01197)→ |
| Sau-2317-1 | NZ_KI994282.1 | - | 120140 | 120111 | **RNA**→ rpmE2 (PRK01678)Ribosomal_L31 (pfam01197)→ |
| Sau-2323-1 | NZ_KI994449.1 | - | 35159 | 35130 | **RNA**→ rpmE2 (PRK01678)Ribosomal_L31 (pfam01197)→ |
| Sau-2443-1 | NZ_KI991753.1 | - | 36391 | 36362 | **RNA**→ rpmE2 (PRK01678)Ribosomal_L31 (pfam01197)→ |
| Sau-2441-1 | NZ_KI991745.1 | - | 25124 | 25095 | **RNA**→ rpmE2 (PRK01678)Ribosomal_L31 (pfam01197)→ |
| Sau-2331-1 | NZ_KI994746.1 | - | 56149 | 56120 | **RNA**→ rpmE2 (PRK01678)Ribosomal_L31 (pfam01197)→ |
| Sau-2336-1 | NZ_KI994986.1 | - | 35090 | 35061 | **RNA**→ rpmE2 (PRK01678)Ribosomal_L31 (pfam01197)→ |
| Sau-2352-1 | NZ_KI995731.1 | - | 24749 | 24720 | **RNA**→ rpmE2 (PRK01678)Ribosomal_L31 (pfam01197)→ |
| Sau-2313-1 | NZ_KI994181.1 | - | 37036 | 37007 | **RNA**→ rpmE2 (PRK01678)Ribosomal_L31 (pfam01197)→ |
| Sau-2325-1 | NZ_KI994561.1 | - | 111473 | 111444 | **RNA**→ rpmE2 (PRK01678)Ribosomal_L31 (pfam01197)→ |
| Sau-2333-1 | NZ_KI994804.1 | - | 36390 | 36361 | **RNA**→ rpmE2 (PRK01678)Ribosomal_L31 (pfam01197)→ |
| Sau-2338-1 | NZ_KI995065.1 | - | 35148 | 35119 | **RNA**→ rpmE2 (PRK01678)Ribosomal_L31 (pfam01197)→ |
| Sau-2405-1 | NZ_KI998115.1 | - | 24749 | 24720 | **RNA**→ rpmE2 (PRK01678)Ribosomal_L31 (pfam01197)→ |
| Sau-2349-1 | NZ_KI995640.1 | - | 61396 | 61367 | **RNA**→ rpmE2 (PRK01678)Ribosomal_L31 (pfam01197)→ |
| Sau-1197-1 | NZ_KI999666.1 | - | 36413 | 36384 | **RNA**→ rpmE2 (PRK01678)Ribosomal_L31 (pfam01197)→ |
| Sau-2430-1 | NZ_KI991285.1 | - | 61214 | 61185 | **RNA**→ rpmE2 (PRK01678)Ribosomal_L31 (pfam01197)→ |
| Sau-1210-1 | NZ_KK000352.1 | - | 26067 | 26038 | **RNA**→ rpmE2 (PRK01678)Ribosomal_L31 (pfam01197)→ |
| Sau-2319-1 | NZ_KI994335.1 | - | 36391 | 36362 | **RNA**→ rpmE2 (PRK01678)Ribosomal_L31 (pfam01197)→ |
| Sau-1222-1 | NZ_KK000868.1 | - | 35170 | 35141 | **RNA**→ rpmE2 (PRK01678)Ribosomal_L31 (pfam01197)→ |
| Sau-2337-1 | NZ_KI995029.1 | - | 35284 | 35255 | **RNA**→ rpmE2 (PRK01678)Ribosomal_L31 (pfam01197)→ |
| Sau-2342-1 | NZ_KI995318.1 | - | 26613 | 26584 | **RNA**→ rpmE2 (PRK01678)Ribosomal_L31 (pfam01197)→ |
| Sau-1224-1 | NZ_KK000942.1 | - | 117871 | 117842 | **RNA**→ rpmE2 (PRK01678)Ribosomal_L31 (pfam01197)→ |
| Sau-1231-1 | NZ_KK001077.1 | - | 119874 | 119845 | **RNA**→ rpmE2 (PRK01678)Ribosomal_L31 (pfam01197)→ |
| Sau-2438-1 | NZ_KI991621.1 | - | 94599 | 94570 | **RNA**→ rpmE2 (PRK01678)Ribosomal_L31 (pfam01197)→ |
| Sau-1232-1 | NZ_KK001101.1 | - | 36391 | 36362 | **RNA**→ rpmE2 (PRK01678)Ribosomal_L31 (pfam01197)→ |
| Sau-1244-1 | NZ_KK001577.1 | - | 36391 | 36362 | **RNA**→ rpmE2 (PRK01678)Ribosomal_L31 (pfam01197)→ |
| Sau-2316-1 | NZ_KI994268.1 | - | 26559 | 26530 | **RNA**→ rpmE2 (PRK01678)Ribosomal_L31 (pfam01197)→ |
| Sau-2321-1 | NZ_KI994414.1 | - | 24749 | 24720 | **RNA**→ rpmE2 (PRK01678)Ribosomal_L31 (pfam01197)→ |
| Sau-1250-1 | NZ_KK001743.1 | - | 215988 | 215959 | **RNA**→ rpmE2 (PRK01678)Ribosomal_L31 (pfam01197)→ |
| Sau-2326-1 | NZ_KI994570.1 | - | 36391 | 36362 | **RNA**→ rpmE2 (PRK01678)Ribosomal_L31 (pfam01197)→ |
| Sau-1293-1 | NZ_KK003417.1 | - | 36390 | 36361 | **RNA**→ rpmE2 (PRK01678)Ribosomal_L31 (pfam01197)→ |
| Sau-2329-1 | NZ_KI994621.1 | - | 22790 | 22761 | **RNA**→ rpmE2 (PRK01678)Ribosomal_L31 (pfam01197)→ |
| Sau-1300-1 | NZ_KK003735.1 | - | 61791 | 61762 | **RNA**→ rpmE2 (PRK01678)Ribosomal_L31 (pfam01197)→ |
| Sau-1305-1 | NZ_KK003928.1 | - | 36491 | 36462 | **RNA**→ rpmE2 (PRK01678)Ribosomal_L31 (pfam01197)→ |
| Sau-2335-1 | NZ_KI994913.1 | - | 54634 | 54605 | **RNA**→ rpmE2 (PRK01678)Ribosomal_L31 (pfam01197)→ |
| Sau-1317-1 | NZ_KK004598.1 | - | 111224 | 111195 | **RNA**→ rpmE2 (PRK01678)Ribosomal_L31 (pfam01197)→ |
| Sau-1324-1 | NZ_KK004885.1 | - | 60436 | 60407 | **RNA**→ rpmE2 (PRK01678)Ribosomal_L31 (pfam01197)→ |
| Sau-1329-1 | NZ_KK005157.1 | - | 61801 | 61772 | **RNA**→ rpmE2 (PRK01678)Ribosomal_L31 (pfam01197)→ |
| Sau-2311-1 | NZ_KI994155.1 | - | 190898 | 190869 | **RNA**→ rpmE2 (PRK01678)Ribosomal_L31 (pfam01197)→ |
| Sau-1358-1 | NZ_KK006237.1 | - | 37057 | 37028 | **RNA**→ rpmE2 (PRK01678)Ribosomal_L31 (pfam01197)→ |
| Sau-2318-1 | NZ_KI994328.1 | - | 62150 | 62121 | **RNA**→ rpmE2 (PRK01678)Ribosomal_L31 (pfam01197)→ |
| Sau-2341-1 | NZ_KI995272.1 | - | 35284 | 35255 | **RNA**→ rpmE2 (PRK01678)Ribosomal_L31 (pfam01197)→ |
| Sau-2390-1 | NZ_KI997593.1 | - | 35188 | 35159 | **RNA**→ rpmE2 (PRK01678)Ribosomal_L31 (pfam01197)→ |
| Sau-2346-1 | NZ_KI995457.1 | - | 24749 | 24720 | **RNA**→ rpmE2 (PRK01678)Ribosomal_L31 (pfam01197)→ |
| Sau-1200-1 | NZ_KI999903.1 | - | 151572 | 151543 | **RNA**→ rpmE2 (PRK01678)Ribosomal_L31 (pfam01197)→ |
| Sau-2364-1 | NZ_KI996937.1 | - | 35318 | 35289 | **RNA**→ rpmE2 (PRK01678)Ribosomal_L31 (pfam01197)→ |
| Sau-1212-1 | NZ_KK000534.1 | - | 65122 | 65093 | **RNA**→ rpmE2 (PRK01678)Ribosomal_L31 (pfam01197)→ |
| Sau-1252-1 | NZ_KK001778.1 | - | 119887 | 119858 | **RNA**→ rpmE2 (PRK01678)Ribosomal_L31 (pfam01197)→ |
| Sau-2372-1 | NZ_KI997187.1 | - | 24749 | 24720 | **RNA**→ rpmE2 (PRK01678)Ribosomal_L31 (pfam01197)→ |
| Sau-1278-1 | NZ_KK002782.1 | - | 37084 | 37055 | **RNA**→ rpmE2 (PRK01678)Ribosomal_L31 (pfam01197)→ |
| Sau-2373-1 | NZ_KI997194.1 | - | 213232 | 213203 | **RNA**→ rpmE2 (PRK01678)Ribosomal_L31 (pfam01197)→ |
| Sau-1291-1 | NZ_KK003367.1 | - | 115636 | 115607 | **RNA**→ rpmE2 (PRK01678)Ribosomal_L31 (pfam01197)→ |

| | | | | | | |
|---|---|---|---|---|---|---|
| Sau-2381-1 | NZ_KI997418.1 | - | 58459 | 58430 | **RNA**→ | rpmE2 (PRK01678)Ribosomal_L31 (pfam01197)→ |
| Sau-1296-1 | NZ_KK003598.1 | - | 151040 | 151011 | **RNA**→ | rpmE2 (PRK01678)Ribosomal_L31 (pfam01197)→ |
| Sau-2343-1 | NZ_KI995366.1 | - | 36391 | 36362 | **RNA**→ | rpmE2 (PRK01678)Ribosomal_L31 (pfam01197)→ |
| Sau-1308-1 | NZ_KK004075.1 | - | 58187 | 58158 | **RNA**→ | rpmE2 (PRK01678)Ribosomal_L31 (pfam01197)→ |
| Sau-2355-1 | NZ_KI995873.1 | - | 36226 | 36197 | **RNA**→ | rpmE2 (PRK01678)Ribosomal_L31 (pfam01197)→ |
| Sau-1314-1 | NZ_KK004401.1 | - | 1461 | 1432 | **RNA**→ | rpmE2 (PRK01678)Ribosomal_L31 (pfam01197)→ |
| Sau-2360-1 | NZ_KI996842.1 | - | 34585 | 34556 | **RNA**→ | rpmE2 (PRK01678)Ribosomal_L31 (pfam01197)→ |
| Sau-1326-1 | NZ_KK004974.1 | - | 116790 | 116761 | **RNA**→ | rpmE2 (PRK01678)Ribosomal_L31 (pfam01197)→ |
| Sau-1327-1 | NZ_KK004981.1 | - | 37010 | 36981 | **RNA**→ | rpmE2 (PRK01678)Ribosomal_L31 (pfam01197)→ |
| Sau-1338-1 | NZ_KK005486.1 | - | 116912 | 116883 | **RNA**→ | rpmE2 (PRK01678)Ribosomal_L31 (pfam01197)→ |
| Sau-2369-1 | NZ_KI997125.1 | - | 60974 | 60945 | **RNA**→ | rpmE2 (PRK01678)Ribosomal_L31 (pfam01197)→ |
| Sau-1339-1 | NZ_KK005523.1 | - | 171217 | 171188 | **RNA**→ | rpmE2 (PRK01678)Ribosomal_L31 (pfam01197)→ |
| Sau-2377-1 | NZ_KI997228.1 | - | 35318 | 35289 | **RNA**→ | rpmE2 (PRK01678)Ribosomal_L31 (pfam01197)→ |
| Sau-1193-1 | NZ_KI999496.1 | - | 116493 | 116464 | **RNA**→ | rpmE2 (PRK01678)Ribosomal_L31 (pfam01197)→ |
| Sau-2391-1 | NZ_KI997674.1 | - | 109253 | 109224 | **RNA**→ | rpmE2 (PRK01678)Ribosomal_L31 (pfam01197)→ |
| Sau-1275-1 | NZ_KK002702.1 | - | 117105 | 117076 | **RNA**→ | rpmE2 (PRK01678)Ribosomal_L31 (pfam01197)→ |
| Sau-1290-1 | NZ_KK003277.1 | - | 35572 | 35543 | **RNA**→ | rpmE2 (PRK01678)Ribosomal_L31 (pfam01197)→ |
| Sau-2420-1 | NZ_KI998546.1 | - | 35801 | 35772 | **RNA**→ | rpmE2 (PRK01678)Ribosomal_L31 (pfam01197)→ |
| Sau-1348-1 | NZ_KK005851.1 | - | 130981 | 130952 | **RNA**→ | rpmE2 (PRK01678)Ribosomal_L31 (pfam01197)→ |
| Sau-2334-1 | NZ_KI994905.1 | - | 61334 | 61305 | **RNA**→ | rpmE2 (PRK01678)Ribosomal_L31 (pfam01197)→ |
| Sau-1202-1 | NZ_KI999924.1 | - | 36491 | 36462 | **RNA**→ | rpmE2 (PRK01678)Ribosomal_L31 (pfam01197)→ |
| Sau-2339-1 | NZ_KI995125.1 | - | 55231 | 55202 | **RNA**→ | rpmE2 (PRK01678)Ribosomal_L31 (pfam01197)→ |
| Sau-1228-1 | NZ_KK001064.1 | - | 109928 | 109899 | **RNA**→ | rpmE2 (PRK01678)Ribosomal_L31 (pfam01197)→ |
| Sau-1248-1 | NZ_KK001705.1 | - | 232070 | 232041 | **RNA**→ | rpmE2 (PRK01678)Ribosomal_L31 (pfam01197)→ |
| Sau-2340-1 | NZ_KI995267.1 | - | 24749 | 24720 | **RNA**→ | rpmE2 (PRK01678)Ribosomal_L31 (pfam01197)→ |
| Sau-1273-1 | NZ_KK002613.1 | - | 108221 | 108192 | **RNA**→ | rpmE2 (PRK01678)Ribosomal_L31 (pfam01197)→ |
| Sau-2344-1 | NZ_KI995407.1 | - | 35284 | 35255 | **RNA**→ | rpmE2 (PRK01678)Ribosomal_L31 (pfam01197)→ |
| Sau-1279-1 | NZ_KK002853.1 | - | 2466 | 2437 | **RNA**→ | rpmE2 (PRK01678)Ribosomal_L31 (pfam01197)→ |
| Sau-2345-1 | NZ_KI995448.1 | - | 215968 | 215939 | **RNA**→ | rpmE2 (PRK01678)Ribosomal_L31 (pfam01197)→ |
| Sau-1286-1 | NZ_KK003159.1 | - | 36393 | 36364 | **RNA**→ | rpmE2 (PRK01678)Ribosomal_L31 (pfam01197)→ |
| Sau-2357-1 | NZ_KI995960.1 | - | 62204 | 62175 | **RNA**→ | rpmE2 (PRK01678)Ribosomal_L31 (pfam01197)→ |
| Sau-1292-1 | NZ_KK003413.1 | - | 25129 | 25100 | **RNA**→ | rpmE2 (PRK01678)Ribosomal_L31 (pfam01197)→ |
| Sau-2361-1 | NZ_KI996884.1 | - | 25291 | 25262 | **RNA**→ | rpmE2 (PRK01678)Ribosomal_L31 (pfam01197)→ |
| Sau-1297-1 | NZ_KK003607.1 | - | 36492 | 36463 | **RNA**→ | rpmE2 (PRK01678)Ribosomal_L31 (pfam01197)→ |
| Sau-2371-1 | NZ_KI997144.1 | - | 12936 | 12907 | **RNA**→ | rpmE2 (PRK01678)Ribosomal_L31 (pfam01197)→ |
| Sau-1309-1 | NZ_KK004207.1 | - | 108037 | 108008 | **RNA**→ | rpmE2 (PRK01678)Ribosomal_L31 (pfam01197)→ |
| Sau-1323-1 | NZ_KK004825.1 | - | 36391 | 36362 | **RNA**→ | rpmE2 (PRK01678)Ribosomal_L31 (pfam01197)→ |
| Sau-2392-1 | NZ_KI997700.1 | - | 35990 | 35961 | **RNA**→ | rpmE2 (PRK01678)Ribosomal_L31 (pfam01197)→ |
| Sau-1334-1 | NZ_KK005353.1 | - | 118726 | 118697 | **RNA**→ | rpmE2 (PRK01678)Ribosomal_L31 (pfam01197)→ |
| Sau-2393-1 | NZ_KI997749.1 | - | 35711 | 35682 | **RNA**→ | rpmE2 (PRK01678)Ribosomal_L31 (pfam01197)→ |
| Sau-1351-1 | NZ_KK005962.1 | - | 36391 | 36362 | **RNA**→ | rpmE2 (PRK01678)Ribosomal_L31 (pfam01197)→ |
| Sau-2324-1 | NZ_KI994478.1 | - | 56277 | 56248 | **RNA**→ | rpmE2 (PRK01678)Ribosomal_L31 (pfam01197)→ |
| Sau-2332-1 | NZ_KI994753.1 | - | 36391 | 36362 | **RNA**→ | rpmE2 (PRK01678)Ribosomal_L31 (pfam01197)→ |
| Sau-2347-1 | NZ_KI995510.1 | - | 36391 | 36362 | **RNA**→ | rpmE2 (PRK01678)Ribosomal_L31 (pfam01197)→ |
| Sau-2353-1 | NZ_KI995810.1 | - | 36413 | 36384 | **RNA**→ | rpmE2 (PRK01678)Ribosomal_L31 (pfam01197)→ |
| Sau-1534-1 | NZ_KK012823.1 | - | 122391 | 122362 | **RNA**→ | rpmE2 (PRK01678)Ribosomal_L31 (pfam01197)→ |
| Sau-1518-1 | NZ_KK012119.1 | - | 35384 | 35355 | **RNA**→ | rpmE2 (PRK01678)Ribosomal_L31 (pfam01197)→ |
| Sau-1888-1 | NZ_KI986039.1 | - | 24746 | 24717 | **RNA**→ | rpmE2 (PRK01678)Ribosomal_L31 (pfam01197)→ |
| Sau-1547-1 | NZ_KK013532.1 | - | 21758 | 21729 | **RNA**→ | rpmE2 (PRK01678)Ribosomal_L31 (pfam01197)→ |
| Sau-1890-1 | NZ_KI986058.1 | - | 54570 | 54541 | **RNA**→ | rpmE2 (PRK01678)Ribosomal_L31 (pfam01197)→ |
| Sau-1522-1 | NZ_KK012212.1 | - | 36391 | 36362 | **RNA**→ | rpmE2 (PRK01678)Ribosomal_L31 (pfam01197)→ |
| Sau-1894-1 | NZ_KI986254.1 | - | 35104 | 35075 | **RNA**→ | rpmE2 (PRK01678)Ribosomal_L31 (pfam01197)→ |

| | | | | | | |
|---|---|---|---|---|---|---|
| Sau-1567-1 | NZ_KK014247.1 | - | 61380 | 61351 | RNA→ | rpmE2 (PRK01678)Ribosomal_L31 (pfam01197)→ |
| Sau-1896-1 | NZ_KI986312.1 | - | 56289 | 56260 | RNA→ | rpmE2 (PRK01678)Ribosomal_L31 (pfam01197)→ |
| Sau-1906-1 | NZ_KI986770.1 | - | 54695 | 54666 | RNA→ | rpmE2 (PRK01678)Ribosomal_L31 (pfam01197)→ |
| Sau-1549-1 | NZ_KK013599.1 | - | 55101 | 55072 | RNA→ | rpmE2 (PRK01678)Ribosomal_L31 (pfam01197)→ |
| Sau-1909-1 | NZ_KI986816.1 | - | 36119 | 36090 | RNA→ | rpmE2 (PRK01678)Ribosomal_L31 (pfam01197)→ |
| Sau-1921-1 | NZ_KI987324.1 | - | 43760 | 43731 | RNA→ | rpmE2 (PRK01678)Ribosomal_L31 (pfam01197)→ |
| Sau-1898-1 | NZ_KI986353.1 | - | 36391 | 36362 | RNA→ | rpmE2 (PRK01678)Ribosomal_L31 (pfam01197)→ |
| Sau-1811-1 | NZ_KI983623.1 | - | 54651 | 54622 | RNA→ | rpmE2 (PRK01678)Ribosomal_L31 (pfam01197)→ |
| Sau-1910-1 | NZ_KI986850.1 | - | 35287 | 35258 | RNA→ | rpmE2 (PRK01678)Ribosomal_L31 (pfam01197)→ |
| Sau-1812-1 | NZ_KI983692.1 | - | 54757 | 54728 | RNA→ | rpmE2 (PRK01678)Ribosomal_L31 (pfam01197)→ |
| Sau-1935-1 | NZ_KI987851.1 | - | 61891 | 61862 | RNA→ | rpmE2 (PRK01678)Ribosomal_L31 (pfam01197)→ |
| Sau-1887-1 | NZ_KI985955.1 | - | 61433 | 61404 | RNA→ | rpmE2 (PRK01678)Ribosomal_L31 (pfam01197)→ |
| Sau-1819-1 | NZ_KI983880.1 | - | 61430 | 61401 | RNA→ | rpmE2 (PRK01678)Ribosomal_L31 (pfam01197)→ |
| Sau-1893-1 | NZ_KI986224.1 | - | 24749 | 24720 | RNA→ | rpmE2 (PRK01678)Ribosomal_L31 (pfam01197)→ |
| Sau-1905-1 | NZ_KI986766.1 | - | 116758 | 116729 | RNA→ | rpmE2 (PRK01678)Ribosomal_L31 (pfam01197)→ |
| Sau-1823-1 | NZ_KI984111.1 | - | 24749 | 24720 | RNA→ | rpmE2 (PRK01678)Ribosomal_L31 (pfam01197)→ |
| Sau-1913-1 | NZ_KI987032.1 | - | 36391 | 36362 | RNA→ | rpmE2 (PRK01678)Ribosomal_L31 (pfam01197)→ |
| Sau-1824-1 | NZ_KI984172.1 | - | 36391 | 36362 | RNA→ | rpmE2 (PRK01678)Ribosomal_L31 (pfam01197)→ |
| Sau-1892-1 | NZ_KI986135.1 | - | 35377 | 35348 | RNA→ | rpmE2 (PRK01678)Ribosomal_L31 (pfam01197)→ |
| Sau-1832-1 | NZ_KI984424.1 | - | 61797 | 61768 | RNA→ | rpmE2 (PRK01678)Ribosomal_L31 (pfam01197)→ |
| Sau-1924-1 | NZ_KI987398.1 | - | 24749 | 24720 | RNA→ | rpmE2 (PRK01678)Ribosomal_L31 (pfam01197)→ |
| Sau-1901-1 | NZ_KI986519.1 | - | 35292 | 35263 | RNA→ | rpmE2 (PRK01678)Ribosomal_L31 (pfam01197)→ |
| Sau-1839-1 | NZ_KI984673.1 | - | 54994 | 54965 | RNA→ | rpmE2 (PRK01678)Ribosomal_L31 (pfam01197)→ |
| Sau-1902-1 | NZ_KI986631.1 | - | 111669 | 111640 | RNA→ | rpmE2 (PRK01678)Ribosomal_L31 (pfam01197)→ |
| Sau-1846-1 | NZ_KI984939.1 | - | 54651 | 54622 | RNA→ | rpmE2 (PRK01678)Ribosomal_L31 (pfam01197)→ |
| Sau-1914-1 | NZ_KI987041.1 | - | 111532 | 111503 | RNA→ | rpmE2 (PRK01678)Ribosomal_L31 (pfam01197)→ |
| Sau-1855-1 | NZ_KI985103.1 | - | 61792 | 61763 | RNA→ | rpmE2 (PRK01678)Ribosomal_L31 (pfam01197)→ |
| Sau-1928-1 | NZ_KI987591.1 | - | 24749 | 24720 | RNA→ | rpmE2 (PRK01678)Ribosomal_L31 (pfam01197)→ |
| Sau-1862-1 | NZ_KI985187.1 | - | 49963 | 49934 | RNA→ | rpmE2 (PRK01678)Ribosomal_L31 (pfam01197)→ |
| Sau-1942-1 | NZ_KI988047.1 | - | 24103 | 24074 | RNA→ | rpmE2 (PRK01678)Ribosomal_L31 (pfam01197)→ |
| Sau-1875-1 | NZ_KI985644.1 | - | 56762 | 56733 | RNA→ | rpmE2 (PRK01678)Ribosomal_L31 (pfam01197)→ |
| Sau-1948-1 | NZ_KI988232.1 | - | 36390 | 36361 | RNA→ | rpmE2 (PRK01678)Ribosomal_L31 (pfam01197)→ |
| Sau-1882-1 | NZ_KI985801.1 | - | 111898 | 111869 | RNA→ | rpmE2 (PRK01678)Ribosomal_L31 (pfam01197)→ |
| Sau-1957-1 | NZ_KI988497.1 | - | 21173 | 21144 | RNA→ | rpmE2 (PRK01678)Ribosomal_L31 (pfam01197)→ |
| Sau-1813-1 | NZ_KI983712.1 | - | 24749 | 24720 | RNA→ | rpmE2 (PRK01678)Ribosomal_L31 (pfam01197)→ |
| Sau-1820-1 | NZ_KI983954.1 | - | 36391 | 36362 | RNA→ | rpmE2 (PRK01678)Ribosomal_L31 (pfam01197)→ |
| Sau-1912-1 | NZ_KI986932.1 | - | 55136 | 55107 | RNA→ | rpmE2 (PRK01678)Ribosomal_L31 (pfam01197)→ |
| Sau-1825-1 | NZ_KI984221.1 | - | 36391 | 36362 | RNA→ | rpmE2 (PRK01678)Ribosomal_L31 (pfam01197)→ |
| Sau-1919-1 | NZ_KI987227.1 | - | 111530 | 111501 | RNA→ | rpmE2 (PRK01678)Ribosomal_L31 (pfam01197)→ |
| Sau-1833-1 | NZ_KI984431.1 | - | 35144 | 35115 | RNA→ | rpmE2 (PRK01678)Ribosomal_L31 (pfam01197)→ |
| Sau-1920-1 | NZ_KI987293.1 | - | 61749 | 61720 | RNA→ | rpmE2 (PRK01678)Ribosomal_L31 (pfam01197)→ |
| Sau-1926-1 | NZ_KI987485.1 | - | 126282 | 126253 | RNA→ | rpmE2 (PRK01678)Ribosomal_L31 (pfam01197)→ |
| Sau-1841-1 | NZ_KI984804.1 | - | 26803 | 26774 | RNA→ | rpmE2 (PRK01678)Ribosomal_L31 (pfam01197)→ |
| Sau-1927-1 | NZ_KI987518.1 | - | 203212 | 203183 | RNA→ | rpmE2 (PRK01678)Ribosomal_L31 (pfam01197)→ |
| Sau-1848-1 | NZ_KI984951.1 | - | 111102 | 111073 | RNA→ | rpmE2 (PRK01678)Ribosomal_L31 (pfam01197)→ |
| Sau-1856-1 | NZ_KI985109.1 | - | 113831 | 113802 | RNA→ | rpmE2 (PRK01678)Ribosomal_L31 (pfam01197)→ |
| Sau-1932-1 | NZ_KI987777.1 | - | 108267 | 108238 | RNA→ | rpmE2 (PRK01678)Ribosomal_L31 (pfam01197)→ |
| Sau-1864-1 | NZ_KI985225.1 | - | 62105 | 62076 | RNA→ | rpmE2 (PRK01678)Ribosomal_L31 (pfam01197)→ |
| Sau-1941-1 | NZ_KI987988.1 | - | 25168 | 25139 | RNA→ | rpmE2 (PRK01678)Ribosomal_L31 (pfam01197)→ |
| Sau-1947-1 | NZ_KI988220.1 | - | 24749 | 24720 | RNA→ | rpmE2 (PRK01678)Ribosomal_L31 (pfam01197)→ |
| Sau-1871-1 | NZ_KI985527.1 | - | 61434 | 61405 | RNA→ | rpmE2 (PRK01678)Ribosomal_L31 (pfam01197)→ |
| Sau-1899-1 | NZ_KI986431.1 | - | 212475 | 212446 | RNA→ | rpmE2 (PRK01678)Ribosomal_L31 (pfam01197)→ |

| | | | | | |
|---|---|---|---|---|---|
| Sau-1876-1 | NZ_KI985685.1 | - | 177827 | 177798 | RNA→ rpmE2 (PRK01678)Ribosomal_L31 (pfam01197)→ |
| Sau-1904-1 | NZ_KI986675.1 | - | 36390 | 36361 | RNA→ rpmE2 (PRK01678)Ribosomal_L31 (pfam01197)→ |
| Sau-1877-1 | NZ_KI985700.1 | - | 24749 | 24720 | RNA→ rpmE2 (PRK01678)Ribosomal_L31 (pfam01197)→ |
| Sau-1884-1 | NZ_KI985807.1 | - | 36391 | 36362 | RNA→ rpmE2 (PRK01678)Ribosomal_L31 (pfam01197)→ |
| Sau-1911-1 | NZ_KI986876.1 | - | 36391 | 36362 | RNA→ rpmE2 (PRK01678)Ribosomal_L31 (pfam01197)→ |
| Sau-1810-1 | NZ_KI983559.1 | - | 36391 | 36362 | RNA→ rpmE2 (PRK01678)Ribosomal_L31 (pfam01197)→ |
| Sau-1918-1 | NZ_KI987165.1 | - | 24749 | 24720 | RNA→ rpmE2 (PRK01678)Ribosomal_L31 (pfam01197)→ |
| Sau-1930-1 | NZ_KI987653.1 | - | 81097 | 81068 | RNA→ rpmE2 (PRK01678)Ribosomal_L31 (pfam01197)→ |
| Sau-1816-1 | NZ_KI983765.1 | - | 36391 | 36362 | RNA→ rpmE2 (PRK01678)Ribosomal_L31 (pfam01197)→ |
| Sau-1822-1 | NZ_KI984061.1 | - | 36391 | 36362 | RNA→ rpmE2 (PRK01678)Ribosomal_L31 (pfam01197)→ |
| Sau-1931-1 | NZ_KI987730.1 | - | 32891 | 32862 | RNA→ rpmE2 (PRK01678)Ribosomal_L31 (pfam01197)→ |
| Sau-1936-1 | NZ_KI987962.1 | - | 113737 | 113708 | RNA→ rpmE2 (PRK01678)Ribosomal_L31 (pfam01197)→ |
| Sau-1827-1 | NZ_KI984308.1 | - | 36490 | 36461 | RNA→ rpmE2 (PRK01678)Ribosomal_L31 (pfam01197)→ |
| Sau-1938-1 | NZ_KI987963.1 | - | 239096 | 239067 | RNA→ rpmE2 (PRK01678)Ribosomal_L31 (pfam01197)→ |
| Sau-1828-1 | NZ_KI984334.1 | - | 61334 | 61305 | RNA→ rpmE2 (PRK01678)Ribosomal_L31 (pfam01197)→ |
| Sau-1944-1 | NZ_KI988094.1 | - | 25058 | 25029 | RNA→ rpmE2 (PRK01678)Ribosomal_L31 (pfam01197)→ |
| Sau-1837-1 | NZ_KI984667.1 | - | 56763 | 56734 | RNA→ rpmE2 (PRK01678)Ribosomal_L31 (pfam01197)→ |
| Sau-1954-1 | NZ_KI988428.1 | - | 26791 | 26762 | RNA→ rpmE2 (PRK01678)Ribosomal_L31 (pfam01197)→ |
| Sau-1843-1 | NZ_KI984882.1 | - | 62156 | 62127 | RNA→ rpmE2 (PRK01678)Ribosomal_L31 (pfam01197)→ |
| Sau-1961-1 | NZ_KI988618.1 | - | 212692 | 212663 | RNA→ rpmE2 (PRK01678)Ribosomal_L31 (pfam01197)→ |
| Sau-1845-1 | NZ_KI984898.1 | - | 113835 | 113806 | RNA→ rpmE2 (PRK01678)Ribosomal_L31 (pfam01197)→ |
| Sau-1891-1 | NZ_KI986108.1 | - | 49963 | 49934 | RNA→ rpmE2 (PRK01678)Ribosomal_L31 (pfam01197)→ |
| Sau-1854-1 | NZ_KI985077.1 | - | 61542 | 61513 | RNA→ rpmE2 (PRK01678)Ribosomal_L31 (pfam01197)→ |
| Sau-1923-1 | NZ_KI987331.1 | - | 61334 | 61305 | RNA→ rpmE2 (PRK01678)Ribosomal_L31 (pfam01197)→ |
| Sau-1860-1 | NZ_KI985163.1 | - | 110439 | 110410 | RNA→ rpmE2 (PRK01678)Ribosomal_L31 (pfam01197)→ |
| Sau-1929-1 | NZ_KI987601.1 | - | 35801 | 35772 | RNA→ rpmE2 (PRK01678)Ribosomal_L31 (pfam01197)→ |
| Sau-1868-1 | NZ_KI985296.1 | - | 151137 | 151108 | RNA→ rpmE2 (PRK01678)Ribosomal_L31 (pfam01197)→ |
| Sau-1943-1 | NZ_KI988056.1 | - | 36103 | 36074 | RNA→ rpmE2 (PRK01678)Ribosomal_L31 (pfam01197)→ |
| Sau-1874-1 | NZ_KI985617.1 | - | 212631 | 212602 | RNA→ rpmE2 (PRK01678)Ribosomal_L31 (pfam01197)→ |
| Sau-1951-1 | NZ_KI988269.1 | - | 116669 | 116640 | RNA→ rpmE2 (PRK01678)Ribosomal_L31 (pfam01197)→ |
| Sau-1953-1 | NZ_KI988356.1 | - | 112019 | 111990 | RNA→ rpmE2 (PRK01678)Ribosomal_L31 (pfam01197)→ |
| Sau-1814-1 | NZ_KI983758.1 | - | 64944 | 64915 | RNA→ rpmE2 (PRK01678)Ribosomal_L31 (pfam01197)→ |
| Sau-1821-1 | NZ_KI984014.1 | - | 58248 | 58219 | RNA→ rpmE2 (PRK01678)Ribosomal_L31 (pfam01197)→ |
| Sau-1958-1 | NZ_KI988547.1 | - | 111812 | 111783 | RNA→ rpmE2 (PRK01678)Ribosomal_L31 (pfam01197)→ |
| Sau-1826-1 | NZ_KI984299.1 | - | 35013 | 34984 | RNA→ rpmE2 (PRK01678)Ribosomal_L31 (pfam01197)→ |
| Sau-1960-1 | NZ_KI988571.1 | - | 212691 | 212662 | RNA→ rpmE2 (PRK01678)Ribosomal_L31 (pfam01197)→ |
| Sau-1834-1 | NZ_KI984496.1 | - | 36390 | 36361 | RNA→ rpmE2 (PRK01678)Ribosomal_L31 (pfam01197)→ |
| Sau-1842-1 | NZ_KI984850.1 | - | 215908 | 215879 | RNA→ rpmE2 (PRK01678)Ribosomal_L31 (pfam01197)→ |
| Sau-1849-1 | NZ_KI984965.1 | - | 49963 | 49934 | RNA→ rpmE2 (PRK01678)Ribosomal_L31 (pfam01197)→ |
| Sau-1851-1 | NZ_KI984994.1 | - | 49963 | 49934 | RNA→ rpmE2 (PRK01678)Ribosomal_L31 (pfam01197)→ |
| Sau-2244-1 | NZ_KI996151.1 | - | 56764 | 56735 | RNA→ rpmE2 (PRK01678)Ribosomal_L31 (pfam01197)→ |
| Sau-885-1 | NZ_KI978043.1 | - | 56894 | 56865 | RNA→ rpmE2 (PRK01678)Ribosomal_L31 (pfam01197)→ |
| Sau-2267-1 | NZ_KI992801.1 | - | 62204 | 62175 | RNA→ rpmE2 (PRK01678)Ribosomal_L31 (pfam01197)→ |
| Sau-2292-1 | NZ_KI993348.1 | - | 61238 | 61209 | RNA→ rpmE2 (PRK01678)Ribosomal_L31 (pfam01197)→ |
| Sau-226-1 | NZ_KI978154.1 | - | 111076 | 111047 | RNA→ rpmE2 (PRK01678)Ribosomal_L31 (pfam01197)→ |
| Sau-2245-1 | NZ_KI996196.1 | - | 54989 | 54960 | RNA→ rpmE2 (PRK01678)Ribosomal_L31 (pfam01197)→ |
| Sau-853-1 | NZ_KI978238.1 | - | 220164 | 220135 | RNA→ rpmE2 (PRK01678)Ribosomal_L31 (pfam01197)→ |
| Sau-2250-1 | NZ_KI996381.1 | - | 35318 | 35289 | RNA→ rpmE2 (PRK01678)Ribosomal_L31 (pfam01197)→ |
| Sau-886-1 | NZ_KI978243.1 | - | 239301 | 239272 | RNA→ rpmE2 (PRK01678)Ribosomal_L31 (pfam01197)→ |
| Sau-2258-1 | NZ_KI996669.1 | - | 5943 | 5914 | RNA→ rpmE2 (PRK01678)Ribosomal_L31 (pfam01197)→ |
| Sau-1159-1 | NZ_KI978422.1 | - | 20728 | 20699 | RNA→ rpmE2 (PRK01678)Ribosomal_L31 (pfam01197)→ |
| Sau-887-1 | NZ_KI978548.1 | - | 35013 | 34984 | RNA→ rpmE2 (PRK01678)Ribosomal_L31 (pfam01197)→ |

| | | | | | | |
|---|---|---|---|---|---|---|
| Sau-861-1 | NZ_KI978733.1 | - | 56403 | 56374 | **RNA**→ | rpmE2 (PRK01678)Ribosomal_L31 (pfam01197)→ |
| Sau-2301-1 | NZ_KI993776.1 | - | 119058 | 119029 | **RNA**→ | rpmE2 (PRK01678)Ribosomal_L31 (pfam01197)→ |
| Sau-2241-1 | NZ_KI996065.1 | - | 36110 | 36081 | **RNA**→ | rpmE2 (PRK01678)Ribosomal_L31 (pfam01197)→ |
| Sau-2248-1 | NZ_KI996301.1 | - | 55372 | 55343 | **RNA**→ | rpmE2 (PRK01678)Ribosomal_L31 (pfam01197)→ |
| Sau-2252-1 | NZ_KI996482.1 | - | 35349 | 35320 | **RNA**→ | rpmE2 (PRK01678)Ribosomal_L31 (pfam01197)→ |
| Sau-2254-1 | NZ_KI996531.1 | - | 35898 | 35869 | **RNA**→ | rpmE2 (PRK01678)Ribosomal_L31 (pfam01197)→ |
| Sau-2261-1 | NZ_KI996725.1 | - | 35672 | 35643 | **RNA**→ | rpmE2 (PRK01678)Ribosomal_L31 (pfam01197)→ |
| Sau-890-1 | NZ_KI979155.1 | - | 34872 | 34843 | **RNA**→ | rpmE2 (PRK01678)Ribosomal_L31 (pfam01197)→ |
| Sau-1749-1 | NZ_KI981588.1 | - | 36412 | 36383 | **RNA**→ | rpmE2 (PRK01678)Ribosomal_L31 (pfam01197)→ |
| Sau-296-1 | NZ_KI976905.1 | - | 107173 | 107144 | **RNA**→ | rpmE2 (PRK01678)Ribosomal_L31 (pfam01197)→ |
| Sau-850-1 | NZ_KI977560.1 | - | 107669 | 107640 | **RNA**→ | rpmE2 (PRK01678)Ribosomal_L31 (pfam01197)→ |
| Sau-250-1 | NZ_KI977699.1 | - | 34817 | 34788 | **RNA**→ | rpmE2 (PRK01678)Ribosomal_L31 (pfam01197)→ |
| Sau-2613-1 | NZ_KI977844.1 | - | 56852 | 56823 | **RNA**→ | rpmE2 (PRK01678)Ribosomal_L31 (pfam01197)→ |
| Sau-2615-1 | NZ_KI978633.1 | - | 61789 | 61760 | **RNA**→ | rpmE2 (PRK01678)Ribosomal_L31 (pfam01197)→ |
| Sau-299-1 | NZ_KI978780.1 | - | 56894 | 56865 | **RNA**→ | rpmE2 (PRK01678)Ribosomal_L31 (pfam01197)→ |
| Sau-1097-1 | NZ_KI978896.1 | - | 56994 | 56965 | **RNA**→ | rpmE2 (PRK01678)Ribosomal_L31 (pfam01197)→ |
| Sau-1772-1 | NZ_KI981781.1 | - | 115444 | 115415 | **RNA**→ | rpmE2 (PRK01678)Ribosomal_L31 (pfam01197)→ |
| Sau-2734-1 | NZ_KI977873.1 | - | 56880 | 56851 | **RNA**→ | rpmE2 (PRK01678)Ribosomal_L31 (pfam01197)→ |
| Sau-1778-1 | NZ_KI982017.1 | - | 143557 | 143528 | **RNA**→ | rpmE2 (PRK01678)Ribosomal_L31 (pfam01197)→ |
| Sau-3043-1 | NZ_KI977944.1 | - | 56645 | 56616 | **RNA**→ | rpmE2 (PRK01678)Ribosomal_L31 (pfam01197)→ |
| Sau-891-1 | NZ_KI976989.1 | - | 56894 | 56865 | **RNA**→ | rpmE2 (PRK01678)Ribosomal_L31 (pfam01197)→ |
| Sau-3045-1 | NZ_KI978159.1 | - | 90758 | 90729 | **RNA**→ | rpmE2 (PRK01678)Ribosomal_L31 (pfam01197)→ |
| Sau-3672-1 | NZ_KI978274.1 | - | 118921 | 118892 | **RNA**→ | rpmE2 (PRK01678)Ribosomal_L31 (pfam01197)→ |
| Sau-1133-1 | NZ_KI977398.1 | - | 111528 | 111499 | **RNA**→ | rpmE2 (PRK01678)Ribosomal_L31 (pfam01197)→ |
| Sau-2678-1 | NZ_KI976313.1 | - | 56894 | 56865 | **RNA**→ | rpmE2 (PRK01678)Ribosomal_L31 (pfam01197)→ |
| Sau-1123-1 | NZ_KI977413.1 | - | 240524 | 240495 | **RNA**→ | rpmE2 (PRK01678)Ribosomal_L31 (pfam01197)→ |
| Sau-2814-1 | NZ_KI976836.1 | - | 238123 | 238094 | **RNA**→ | rpmE2 (PRK01678)Ribosomal_L31 (pfam01197)→ |
| Sau-248-1 | NZ_KI977505.1 | - | 34816 | 34787 | **RNA**→ | rpmE2 (PRK01678)Ribosomal_L31 (pfam01197)→ |
| Sau-3035-1 | NZ_KI977215.1 | - | 59010 | 58981 | **RNA**→ | rpmE2 (PRK01678)Ribosomal_L31 (pfam01197)→ |
| Sau-1156-1 | NZ_KI977697.1 | - | 49965 | 49936 | **RNA**→ | rpmE2 (PRK01678)Ribosomal_L31 (pfam01197)→ |
| Sau-3671-1 | NZ_KI977580.1 | - | 117552 | 117523 | **RNA**→ | rpmE2 (PRK01678)Ribosomal_L31 (pfam01197)→ |
| Sau-2657-1 | NZ_KI977740.1 | - | 34872 | 34843 | **RNA**→ | rpmE2 (PRK01678)Ribosomal_L31 (pfam01197)→ |
| Sau-856-1 | NZ_KI977962.1 | - | 56894 | 56865 | **RNA**→ | rpmE2 (PRK01678)Ribosomal_L31 (pfam01197)→ |
| Sau-2819-1 | NZ_KI977852.1 | - | 238406 | 238377 | **RNA**→ | rpmE2 (PRK01678)Ribosomal_L31 (pfam01197)→ |
| Sau-2616-1 | NZ_KI978193.1 | - | 106543 | 106514 | **RNA**→ | rpmE2 (PRK01678)Ribosomal_L31 (pfam01197)→ |
| Sau-2659-1 | NZ_KI977995.1 | - | 19102 | 19073 | **RNA**→ | rpmE2 (PRK01678)Ribosomal_L31 (pfam01197)→ |
| Sau-3044-1 | NZ_KI978127.1 | - | 25029 | 25000 | **RNA**→ | rpmE2 (PRK01678)Ribosomal_L31 (pfam01197)→ |
| Sau-2737-1 | NZ_KI978649.1 | - | 115251 | 115222 | **RNA**→ | rpmE2 (PRK01678)Ribosomal_L31 (pfam01197)→ |
| Sau-2739-1 | NZ_KI975825.1 | - | 56894 | 56865 | **RNA**→ | rpmE2 (PRK01678)Ribosomal_L31 (pfam01197)→ |
| Sau-1538-1 | NZ_KK013038.1 | - | 36391 | 36362 | **RNA**→ | rpmE2 (PRK01678)Ribosomal_L31 (pfam01197)→ |
| Sau-2781-1 | NZ_KI976300.1 | - | 114787 | 114758 | **RNA**→ | rpmE2 (PRK01678)Ribosomal_L31 (pfam01197)→ |
| Sau-2795-1 | NZ_KI976795.1 | - | 117314 | 117285 | **RNA**→ | rpmE2 (PRK01678)Ribosomal_L31 (pfam01197)→ |
| Sau-1544-1 | NZ_KK013332.1 | - | 110285 | 110256 | **RNA**→ | rpmE2 (PRK01678)Ribosomal_L31 (pfam01197)→ |
| Sau-2654-1 | NZ_KI977000.1 | - | 19182 | 19153 | **RNA**→ | rpmE2 (PRK01678)Ribosomal_L31 (pfam01197)→ |
| Sau-1613-1 | NZ_KK016426.1 | - | 56093 | 56064 | **RNA**→ | rpmE2 (PRK01678)Ribosomal_L31 (pfam01197)→ |
| Sau-3034-1 | NZ_KI977152.1 | - | 107193 | 107164 | **RNA**→ | rpmE2 (PRK01678)Ribosomal_L31 (pfam01197)→ |
| Sau-2771-1 | NZ_KI977188.1 | - | 35071 | 35042 | **RNA**→ | rpmE2 (PRK01678)Ribosomal_L31 (pfam01197)→ |
| Sau-1619-1 | NZ_KK016776.1 | - | 36492 | 36463 | **RNA**→ | rpmE2 (PRK01678)Ribosomal_L31 (pfam01197)→ |
| Sau-3041-1 | NZ_KI977433.1 | - | 59010 | 58981 | **RNA**→ | rpmE2 (PRK01678)Ribosomal_L31 (pfam01197)→ |
| Sau-2735-1 | NZ_KI977911.1 | - | 35284 | 35255 | **RNA**→ | rpmE2 (PRK01678)Ribosomal_L31 (pfam01197)→ |
| Sau-1630-1 | NZ_KK017358.1 | - | 35246 | 35217 | **RNA**→ | rpmE2 (PRK01678)Ribosomal_L31 (pfam01197)→ |
| Sau-2660-1 | NZ_KI977970.1 | - | 238655 | 238626 | **RNA**→ | rpmE2 (PRK01678)Ribosomal_L31 (pfam01197)→ |

| | | | | | |
|---|---|---|---|---|---|
| Sau-3046-1 | NZ_KI978198.1 | - | 118242 | 118213 | **RNA**→ rpmE2 (PRK01678)Ribosomal_L31 (pfam01197)→ |
| Sau-1635-1 | NZ_KK017643.1 | - | 16230 | 16201 | **RNA**→ rpmE2 (PRK01678)Ribosomal_L31 (pfam01197)→ |
| Sau-3047-1 | NZ_KI978290.1 | - | 962431 | 962402 | **RNA**→ rpmE2 (PRK01678)Ribosomal_L31 (pfam01197)→ |
| Sau-3049-1 | NZ_KI978514.1 | - | 149156 | 149127 | **RNA**→ rpmE2 (PRK01678)Ribosomal_L31 (pfam01197)→ |
| Sau-1370-1 | NZ_KK018920.1 | - | 111870 | 111841 | **RNA**→ rpmE2 (PRK01678)Ribosomal_L31 (pfam01197)→ |
| Sau-2667-1 | NZ_KI978541.1 | - | 111456 | 111427 | **RNA**→ rpmE2 (PRK01678)Ribosomal_L31 (pfam01197)→ |
| Sau-2683-1 | NZ_KI978725.1 | - | 56894 | 56865 | **RNA**→ rpmE2 (PRK01678)Ribosomal_L31 (pfam01197)→ |
| Sau-2674-1 | NZ_KI978826.1 | - | 56993 | 56964 | **RNA**→ rpmE2 (PRK01678)Ribosomal_L31 (pfam01197)→ |
| Sau-3676-1 | NZ_KI978841.1 | - | 56894 | 56865 | **RNA**→ rpmE2 (PRK01678)Ribosomal_L31 (pfam01197)→ |
| Sau-2588-1 | NZ_KK021149.1 | - | 56940 | 56911 | **RNA**→ rpmE2 (PRK01678)Ribosomal_L31 (pfam01197)→ |
| Sau-3053-1 | NZ_KI979026.1 | - | 56894 | 56865 | **RNA**→ rpmE2 (PRK01678)Ribosomal_L31 (pfam01197)→ |
| Sau-724-1 | NZ_KK022333.1 | - | 34817 | 34788 | **RNA**→ rpmE2 (PRK01678)Ribosomal_L31 (pfam01197)→ |
| Sau-2672-1 | NZ_KI979249.1 | - | 56894 | 56865 | **RNA**→ rpmE2 (PRK01678)Ribosomal_L31 (pfam01197)→ |
| Sau-3030-1 | NZ_KI976484.1 | - | 25119 | 25090 | **RNA**→ rpmE2 (PRK01678)Ribosomal_L31 (pfam01197)→ |
| Sau-1492-1 | NZ_KK011005.1 | - | 37315 | 37286 | **RNA**→ rpmE2 (PRK01678)Ribosomal_L31 (pfam01197)→ |
| Sau-2815-1 | NZ_KI976882.1 | - | 34912 | 34883 | **RNA**→ rpmE2 (PRK01678)Ribosomal_L31 (pfam01197)→ |
| Sau-3033-1 | NZ_KI977135.1 | - | 237869 | 237840 | **RNA**→ rpmE2 (PRK01678)Ribosomal_L31 (pfam01197)→ |
| Sau-1512-1 | NZ_KK011855.1 | - | 36391 | 36362 | **RNA**→ rpmE2 (PRK01678)Ribosomal_L31 (pfam01197)→ |
| Sau-2801-1 | NZ_KI977270.1 | - | 118641 | 118612 | **RNA**→ rpmE2 (PRK01678)Ribosomal_L31 (pfam01197)→ |
| Sau-1525-1 | NZ_KK012360.1 | - | 37317 | 37288 | **RNA**→ rpmE2 (PRK01678)Ribosomal_L31 (pfam01197)→ |
| Sau-2656-1 | NZ_KI977550.1 | - | 238684 | 238655 | **RNA**→ rpmE2 (PRK01678)Ribosomal_L31 (pfam01197)→ |
| Sau-3040-1 | NZ_KI977638.1 | - | 58910 | 58881 | **RNA**→ rpmE2 (PRK01678)Ribosomal_L31 (pfam01197)→ |
| Sau-1543-1 | NZ_KK013328.1 | - | 111936 | 111907 | **RNA**→ rpmE2 (PRK01678)Ribosomal_L31 (pfam01197)→ |
| Sau-2658-1 | NZ_KI977831.1 | - | 61890 | 61861 | **RNA**→ rpmE2 (PRK01678)Ribosomal_L31 (pfam01197)→ |
| Sau-1553-1 | NZ_KK013748.1 | - | 36491 | 36462 | **RNA**→ rpmE2 (PRK01678)Ribosomal_L31 (pfam01197)→ |
| Sau-3007-1 | NZ_KI977892.1 | - | 56894 | 56865 | **RNA**→ rpmE2 (PRK01678)Ribosomal_L31 (pfam01197)→ |
| Sau-1563-1 | NZ_KK014046.1 | - | 109373 | 109344 | **RNA**→ rpmE2 (PRK01678)Ribosomal_L31 (pfam01197)→ |
| Sau-2736-1 | NZ_KI978070.1 | - | 56894 | 56865 | **RNA**→ rpmE2 (PRK01678)Ribosomal_L31 (pfam01197)→ |
| Sau-2661-1 | NZ_KI978282.1 | - | 56894 | 56865 | **RNA**→ rpmE2 (PRK01678)Ribosomal_L31 (pfam01197)→ |
| Sau-1569-1 | NZ_KK014352.1 | - | 111576 | 111547 | **RNA**→ rpmE2 (PRK01678)Ribosomal_L31 (pfam01197)→ |
| Sau-2665-1 | NZ_KI978489.1 | - | 60162 | 60133 | **RNA**→ rpmE2 (PRK01678)Ribosomal_L31 (pfam01197)→ |
| Sau-1611-1 | NZ_KK016386.1 | - | 36390 | 36361 | **RNA**→ rpmE2 (PRK01678)Ribosomal_L31 (pfam01197)→ |
| Sau-1369-1 | NZ_KK018861.1 | - | 217144 | 217115 | **RNA**→ rpmE2 (PRK01678)Ribosomal_L31 (pfam01197)→ |
| Sau-2669-1 | NZ_KI978611.1 | - | 58910 | 58881 | **RNA**→ rpmE2 (PRK01678)Ribosomal_L31 (pfam01197)→ |
| Sau-2670-1 | NZ_KI978696.1 | - | 56892 | 56863 | **RNA**→ rpmE2 (PRK01678)Ribosomal_L31 (pfam01197)→ |
| Sau-1375-1 | NZ_KK019079.1 | - | 35364 | 35335 | **RNA**→ rpmE2 (PRK01678)Ribosomal_L31 (pfam01197)→ |
| Sau-682-1 | NZ_KK021052.1 | - | 35169 | 35140 | **RNA**→ rpmE2 (PRK01678)Ribosomal_L31 (pfam01197)→ |
| Sau-2684-1 | NZ_KI978769.1 | - | 211074 | 211045 | **RNA**→ rpmE2 (PRK01678)Ribosomal_L31 (pfam01197)→ |
| Sau-2517-1 | NZ_KK022591.1 | - | 2239603 | 2239574 | **RNA**→ rpmE2 (PRK01678)Ribosomal_L31 (pfam01197)→ |
| Sau-2504-1 | NZ_KK022875.1 | - | 56994 | 56965 | **RNA**→ rpmE2 (PRK01678)Ribosomal_L31 (pfam01197)→ |
| Sau-2536-1 | NZ_KK023228.1 | - | 2240385 | 2240356 | **RNA**→ rpmE2 (PRK01678)Ribosomal_L31 (pfam01197)→ |
| Sau-1509-1 | NZ_KK011811.1 | - | 115367 | 115338 | **RNA**→ rpmE2 (PRK01678)Ribosomal_L31 (pfam01197)→ |
| Sau-1536-1 | NZ_KK012906.1 | - | 35687 | 35658 | **RNA**→ rpmE2 (PRK01678)Ribosomal_L31 (pfam01197)→ |
| Sau-1552-1 | NZ_KK013743.1 | - | 123305 | 123276 | **RNA**→ rpmE2 (PRK01678)Ribosomal_L31 (pfam01197)→ |
| Sau-1574-1 | NZ_KK014582.1 | - | 28890 | 28861 | **RNA**→ rpmE2 (PRK01678)Ribosomal_L31 (pfam01197)→ |
| Sau-1587-1 | NZ_KK015127.1 | - | 29142 | 29113 | **RNA**→ rpmE2 (PRK01678)Ribosomal_L31 (pfam01197)→ |
| Sau-1598-1 | NZ_KK015734.1 | - | 62157 | 62128 | **RNA**→ rpmE2 (PRK01678)Ribosomal_L31 (pfam01197)→ |
| Sau-1621-1 | NZ_KK016884.1 | - | 49963 | 49934 | **RNA**→ rpmE2 (PRK01678)Ribosomal_L31 (pfam01197)→ |
| Sau-1632-1 | NZ_KK017470.1 | - | 61337 | 61308 | **RNA**→ rpmE2 (PRK01678)Ribosomal_L31 (pfam01197)→ |
| Sau-1644-1 | NZ_KK018094.1 | - | 62090 | 62061 | **RNA**→ rpmE2 (PRK01678)Ribosomal_L31 (pfam01197)→ |
| Sau-1072-1 | NZ_KK021514.1 | - | 27192 | 27163 | **RNA**→ rpmE2 (PRK01678)Ribosomal_L31 (pfam01197)→ |
| Sau-803-1 | NZ_KK021567.1 | - | 34817 | 34788 | **RNA**→ rpmE2 (PRK01678)Ribosomal_L31 (pfam01197)→ |

| | | | | | |
|---|---|---|---|---|---|
| Sau-1523-1 | NZ_KK012278.1 | - | 60738 | 60709 | **RNA**→ rpmE2 (PRK01678)Ribosomal_L31 (pfam01197)→ |
| Sau-1535-1 | NZ_KK012841.1 | - | 107715 | 107686 | **RNA**→ rpmE2 (PRK01678)Ribosomal_L31 (pfam01197)→ |
| Sau-2685-1 | NZ_KI979084.1 | - | 153683 | 153654 | **RNA**→ rpmE2 (PRK01678)Ribosomal_L31 (pfam01197)→ |
| Sau-1545-1 | NZ_KK013431.1 | - | 27285 | 27256 | **RNA**→ rpmE2 (PRK01678)Ribosomal_L31 (pfam01197)→ |
| Sau-1550-1 | NZ_KK013704.1 | - | 111824 | 111795 | **RNA**→ rpmE2 (PRK01678)Ribosomal_L31 (pfam01197)→ |
| Sau-1642-1 | NZ_KK017973.1 | - | 35605 | 35576 | **RNA**→ rpmE2 (PRK01678)Ribosomal_L31 (pfam01197)→ |
| Sau-679-1 | NZ_KK020982.1 | - | 56941 | 56912 | **RNA**→ rpmE2 (PRK01678)Ribosomal_L31 (pfam01197)→ |
| Sau-2238-1 | NZ_KK021925.1 | - | 55904 | 55875 | **RNA**→ rpmE2 (PRK01678)Ribosomal_L31 (pfam01197)→ |
| Sau-1016-1 | NZ_KK022619.1 | - | 2237211 | 2237182 | **RNA**→ rpmE2 (PRK01678)Ribosomal_L31 (pfam01197)→ |
| Sau-1591-1 | NZ_KK015352.1 | - | 35169 | 35140 | **RNA**→ rpmE2 (PRK01678)Ribosomal_L31 (pfam01197)→ |
| Sau-1372-1 | NZ_KK018965.1 | - | 63907 | 63878 | **RNA**→ rpmE2 (PRK01678)Ribosomal_L31 (pfam01197)→ |
| Sau-2706-1 | NZ_KK021460.1 | - | 34873 | 34844 | **RNA**→ rpmE2 (PRK01678)Ribosomal_L31 (pfam01197)→ |
| Sau-1577-1 | NZ_KK014667.1 | - | 36391 | 36362 | **RNA**→ rpmE2 (PRK01678)Ribosomal_L31 (pfam01197)→ |
| Sau-1589-1 | NZ_KK015302.1 | - | 111429 | 111400 | **RNA**→ rpmE2 (PRK01678)Ribosomal_L31 (pfam01197)→ |
| Sau-694-1 | NZ_KK021452.1 | - | 56940 | 56911 | **RNA**→ rpmE2 (PRK01678)Ribosomal_L31 (pfam01197)→ |
| Sau-152-1 | NZ_KK220206.1 | - | 34754 | 34725 | **RNA**→ rpmE2 (PRK01678)Ribosomal_L31 (pfam01197)→ |
| Sau-673-1 | NZ_KK039814.1 | - | 34873 | 34844 | **RNA**→ rpmE2 (PRK01678)Ribosomal_L31 (pfam01197)→ |
| Sau-419-1 | NZ_KK097822.1 | - | 222887 | 222858 | **RNA**→ rpmE2 (PRK01678)Ribosomal_L31 (pfam01197)→ |
| Sau-2509-1 | NZ_KK041010.1 | - | 34912 | 34883 | **RNA**→ rpmE2 (PRK01678)Ribosomal_L31 (pfam01197)→ |
| Sau-337-1 | NZ_KK099176.1 | - | 464015 | 463986 | **RNA**→ rpmE2 (PRK01678)Ribosomal_L31 (pfam01197)→ |
| Sau-3071-1 | NZ_KK226598.1 | - | 26024 | 25995 | **RNA**→ rpmE2 (PRK01678)Ribosomal_L31 (pfam01197)→ |
| Sau-1085-1 | NZ_KK039088.1 | - | 35209 | 35180 | **RNA**→ rpmE2 (PRK01678)Ribosomal_L31 (pfam01197)→ |
| Sau-151-1 | NZ_KK220145.1 | - | 56395 | 56366 | **RNA**→ rpmE2 (PRK01678)Ribosomal_L31 (pfam01197)→ |
| Sau-1036-1 | NZ_KK041081.1 | - | 34817 | 34788 | **RNA**→ rpmE2 (PRK01678)Ribosomal_L31 (pfam01197)→ |
| Sau-185-1 | NZ_KK221386.1 | - | 26024 | 25995 | **RNA**→ rpmE2 (PRK01678)Ribosomal_L31 (pfam01197)→ |
| Sau-2207-1 | NZ_KK039044.1 | - | 34862 | 34833 | **RNA**→ rpmE2 (PRK01678)Ribosomal_L31 (pfam01197)→ |
| Sau-2900-1 | NZ_KK095025.1 | - | 56894 | 56865 | **RNA**→ rpmE2 (PRK01678)Ribosomal_L31 (pfam01197)→ |
| Sau-313-1 | NZ_KK099030.1 | - | 180328 | 180299 | **RNA**→ rpmE2 (PRK01678)Ribosomal_L31 (pfam01197)→ |
| Sau-1035-1 | NZ_KK041118.1 | - | 34862 | 34833 | **RNA**→ rpmE2 (PRK01678)Ribosomal_L31 (pfam01197)→ |
| Sau-139-1 | NZ_KK219769.1 | - | 25612 | 25583 | **RNA**→ rpmE2 (PRK01678)Ribosomal_L31 (pfam01197)→ |
| Sau-2710-1 | NZ_KK225426.1 | - | 26024 | 25995 | **RNA**→ rpmE2 (PRK01678)Ribosomal_L31 (pfam01197)→ |
| Sau-731-1 | NZ_KK039411.1 | - | 34872 | 34843 | **RNA**→ rpmE2 (PRK01678)Ribosomal_L31 (pfam01197)→ |
| Sau-251-1 | NZ_KK222651.1 | - | 26024 | 25995 | **RNA**→ rpmE2 (PRK01678)Ribosomal_L31 (pfam01197)→ |
| Sau-2700-1 | NZ_KK225337.1 | - | 53513 | 53484 | **RNA**→ rpmE2 (PRK01678)Ribosomal_L31 (pfam01197)→ |
| Sau-1022-1 | NZ_KK041284.1 | - | 55804 | 55775 | **RNA**→ rpmE2 (PRK01678)Ribosomal_L31 (pfam01197)→ |
| Sau-3057-1 | NZ_KK226422.1 | - | 56441 | 56412 | **RNA**→ rpmE2 (PRK01678)Ribosomal_L31 (pfam01197)→ |
| Sau-416-1 | NZ_KK097804.1 | - | 56588 | 56559 | **RNA**→ rpmE2 (PRK01678)Ribosomal_L31 (pfam01197)→ |
| Sau-1021-1 | NZ_KK041296.1 | - | 34873 | 34844 | **RNA**→ rpmE2 (PRK01678)Ribosomal_L31 (pfam01197)→ |
| Sau-2822-1 | NZ_KK225732.1 | - | 26024 | 25995 | **RNA**→ rpmE2 (PRK01678)Ribosomal_L31 (pfam01197)→ |
| Sau-145-1 | NZ_KK220079.1 | - | 56310 | 56281 | **RNA**→ rpmE2 (PRK01678)Ribosomal_L31 (pfam01197)→ |
| Sau-804-1 | NZ_KK041645.1 | - | 34873 | 34844 | **RNA**→ rpmE2 (PRK01678)Ribosomal_L31 (pfam01197)→ |
| Sau-3061-1 | NZ_KK226326.1 | - | 56210 | 56181 | **RNA**→ rpmE2 (PRK01678)Ribosomal_L31 (pfam01197)→ |
| Sau-270-1 | NZ_KK223267.1 | - | 26024 | 25995 | **RNA**→ rpmE2 (PRK01678)Ribosomal_L31 (pfam01197)→ |
| Sau-898-1 | NZ_KK039334.1 | - | 34872 | 34843 | **RNA**→ rpmE2 (PRK01678)Ribosomal_L31 (pfam01197)→ |
| Sau-2922-1 | NZ_KK095176.1 | - | 56894 | 56865 | **RNA**→ rpmE2 (PRK01678)Ribosomal_L31 (pfam01197)→ |
| Sau-94-1 | NZ_KK218433.1 | - | 26024 | 25995 | **RNA**→ rpmE2 (PRK01678)Ribosomal_L31 (pfam01197)→ |
| Sau-1034-1 | NZ_KK041147.1 | - | 34873 | 34844 | **RNA**→ rpmE2 (PRK01678)Ribosomal_L31 (pfam01197)→ |
| Sau-3081-1 | NZ_KK226999.1 | - | 26023 | 25994 | **RNA**→ rpmE2 (PRK01678)Ribosomal_L31 (pfam01197)→ |
| Sau-3086-1 | NZ_KK227206.1 | - | 26024 | 25995 | **RNA**→ rpmE2 (PRK01678)Ribosomal_L31 (pfam01197)→ |
| Sau-202-1 | NZ_KK221958.1 | - | 26024 | 25995 | **RNA**→ rpmE2 (PRK01678)Ribosomal_L31 (pfam01197)→ |
| Sau-1008-1 | NZ_KK041409.1 | - | 56894 | 56865 | **RNA**→ rpmE2 (PRK01678)Ribosomal_L31 (pfam01197)→ |
| Sau-815-1 | NZ_KK041546.1 | - | 34872 | 34843 | **RNA**→ rpmE2 (PRK01678)Ribosomal_L31 (pfam01197)→ |

| | | | | | |
|---|---|---|---|---|---|
| Sau-2927-1 | NZ_KK095216.1 | - | 29170 | 29141 | **RNA**→ rpmE2 (PRK01678)Ribosomal_L31 (pfam01197)→ |
| Sau-2675-1 | NZ_KK224694.1 | - | 26024 | 25995 | **RNA**→ rpmE2 (PRK01678)Ribosomal_L31 (pfam01197)→ |
| Sau-174-1 | NZ_KK221018.1 | - | 26024 | 25995 | **RNA**→ rpmE2 (PRK01678)Ribosomal_L31 (pfam01197)→ |
| Sau-2823-1 | NZ_KK225762.1 | - | 26024 | 25995 | **RNA**→ rpmE2 (PRK01678)Ribosomal_L31 (pfam01197)→ |
| Sau-190-1 | NZ_KK221571.1 | - | 56395 | 56366 | **RNA**→ rpmE2 (PRK01678)Ribosomal_L31 (pfam01197)→ |
| Sau-2828-1 | NZ_KK225934.1 | - | 26024 | 25995 | **RNA**→ rpmE2 (PRK01678)Ribosomal_L31 (pfam01197)→ |
| Sau-114-1 | NZ_KK218941.1 | - | 55009 | 54980 | **RNA**→ rpmE2 (PRK01678)Ribosomal_L31 (pfam01197)→ |
| Sau-3079-1 | NZ_KK226919.1 | - | 26024 | 25995 | **RNA**→ rpmE2 (PRK01678)Ribosomal_L31 (pfam01197)→ |
| Sau-3089-1 | NZ_KK227338.1 | - | 26024 | 25995 | **RNA**→ rpmE2 (PRK01678)Ribosomal_L31 (pfam01197)→ |
| Sau-275-1 | NZ_KK223430.1 | - | 26024 | 25995 | **RNA**→ rpmE2 (PRK01678)Ribosomal_L31 (pfam01197)→ |
| Sau-3009-1 | NZ_KK226022.1 | - | 26024 | 25995 | **RNA**→ rpmE2 (PRK01678)Ribosomal_L31 (pfam01197)→ |
| Sau-2827-1 | NZ_KK225900.1 | - | 24379 | 24350 | **RNA**→ rpmE2 (PRK01678)Ribosomal_L31 (pfam01197)→ |
| Sau-3075-1 | NZ_KK226773.1 | - | 53698 | 53669 | **RNA**→ rpmE2 (PRK01678)Ribosomal_L31 (pfam01197)→ |
| Sau-3084-1 | NZ_KK227106.1 | - | 56390 | 56361 | **RNA**→ rpmE2 (PRK01678)Ribosomal_L31 (pfam01197)→ |
| Sau-2577-1 | NZ_KK094808.1 | - | 35013 | 34984 | **RNA**→ rpmE2 (PRK01678)Ribosomal_L31 (pfam01197)→ |
| Sau-2691-1 | NZ_KK225002.1 | - | 26024 | 25995 | **RNA**→ rpmE2 (PRK01678)Ribosomal_L31 (pfam01197)→ |
| Sau-2825-1 | NZ_KK225836.1 | - | 24806 | 24777 | **RNA**→ rpmE2 (PRK01678)Ribosomal_L31 (pfam01197)→ |
| Sau-2688-1 | NZ_KK224861.1 | - | 26024 | 25995 | **RNA**→ rpmE2 (PRK01678)Ribosomal_L31 (pfam01197)→ |
| Sau-2693-1 | NZ_KK225109.1 | - | 56438 | 56409 | **RNA**→ rpmE2 (PRK01678)Ribosomal_L31 (pfam01197)→ |
| Sau-2907-1 | NZ_KK038479.1 | - | 92275 | 92246 | **RNA**→ rpmE2 (PRK01678)Ribosomal_L31 (pfam01197)→ |
| Sau-2567-1 | NZ_KK038957.1 | - | 56894 | 56865 | **RNA**→ rpmE2 (PRK01678)Ribosomal_L31 (pfam01197)→ |
| Sau-2904-1 | NZ_KK038550.1 | - | 34872 | 34843 | **RNA**→ rpmE2 (PRK01678)Ribosomal_L31 (pfam01197)→ |
| Sau-2898-1 | NZ_KK038628.1 | - | 34872 | 34843 | **RNA**→ rpmE2 (PRK01678)Ribosomal_L31 (pfam01197)→ |
| Sau-2903-1 | NZ_KK038592.1 | - | 29243 | 29214 | **RNA**→ rpmE2 (PRK01678)Ribosomal_L31 (pfam01197)→ |
| Sau-2568-1 | NZ_KK038929.1 | - | 56894 | 56865 | **RNA**→ rpmE2 (PRK01678)Ribosomal_L31 (pfam01197)→ |
| Sau-2854-1 | NZ_KK040538.1 | - | 54999 | 54970 | **RNA**→ rpmE2 (PRK01678)Ribosomal_L31 (pfam01197)→ |
| Sau-2847-1 | NZ_KK040724.1 | - | 54739 | 54710 | **RNA**→ rpmE2 (PRK01678)Ribosomal_L31 (pfam01197)→ |
| Sau-2555-1 | NZ_KK040779.1 | - | 34817 | 34788 | **RNA**→ rpmE2 (PRK01678)Ribosomal_L31 (pfam01197)→ |
| Sau-2525-1 | NZ_KK040917.1 | - | 54858 | 54829 | **RNA**→ rpmE2 (PRK01678)Ribosomal_L31 (pfam01197)→ |
| Sau-3004-1 | NZ_KK040206.1 | - | 34817 | 34788 | **RNA**→ rpmE2 (PRK01678)Ribosomal_L31 (pfam01197)→ |
| Sau-2709-1 | NZ_KK040748.1 | - | 54695 | 54666 | **RNA**→ rpmE2 (PRK01678)Ribosomal_L31 (pfam01197)→ |
| Sau-2563-1 | NZ_KK040762.1 | - | 56894 | 56865 | **RNA**→ rpmE2 (PRK01678)Ribosomal_L31 (pfam01197)→ |
| Sau-2541-1 | NZ_KK040861.1 | - | 54905 | 54876 | **RNA**→ rpmE2 (PRK01678)Ribosomal_L31 (pfam01197)→ |
| Sau-2511-1 | NZ_KK040948.1 | - | 56895 | 56866 | **RNA**→ rpmE2 (PRK01678)Ribosomal_L31 (pfam01197)→ |
| Sau-3005-1 | NZ_KK040197.1 | - | 57094 | 57065 | **RNA**→ rpmE2 (PRK01678)Ribosomal_L31 (pfam01197)→ |
| Sau-2853-1 | NZ_KK040540.1 | - | 34872 | 34843 | **RNA**→ rpmE2 (PRK01678)Ribosomal_L31 (pfam01197)→ |
| Sau-2846-1 | NZ_KK040730.1 | - | 56888 | 56859 | **RNA**→ rpmE2 (PRK01678)Ribosomal_L31 (pfam01197)→ |
| Sau-3011-1 | NZ_KK226077.1 | - | 56336 | 56307 | **RNA**→ rpmE2 (PRK01678)Ribosomal_L31 (pfam01197)→ |
| Sau-3002-1 | NZ_KK040256.1 | - | 34817 | 34788 | **RNA**→ rpmE2 (PRK01678)Ribosomal_L31 (pfam01197)→ |
| Sau-3001-1 | NZ_KK040296.1 | - | 34872 | 34843 | **RNA**→ rpmE2 (PRK01678)Ribosomal_L31 (pfam01197)→ |
| Sau-2871-1 | NZ_KK040488.1 | - | 56894 | 56865 | **RNA**→ rpmE2 (PRK01678)Ribosomal_L31 (pfam01197)→ |
| Sau-2860-1 | NZ_KK040503.1 | - | 35442 | 35413 | **RNA**→ rpmE2 (PRK01678)Ribosomal_L31 (pfam01197)→ |
| Sau-2848-1 | NZ_KK040659.1 | - | 34817 | 34788 | **RNA**→ rpmE2 (PRK01678)Ribosomal_L31 (pfam01197)→ |
| Sau-2570-1 | NZ_KK038869.1 | - | 34873 | 34844 | **RNA**→ rpmE2 (PRK01678)Ribosomal_L31 (pfam01197)→ |
| Sau-817-1 | NZ_KK041442.1 | - | 34817 | 34788 | **RNA**→ rpmE2 (PRK01678)Ribosomal_L31 (pfam01197)→ |
| Sau-796-1 | NZ_KK041680.1 | - | 34872 | 34843 | **RNA**→ rpmE2 (PRK01678)Ribosomal_L31 (pfam01197)→ |
| Sau-814-1 | NZ_KK041583.1 | - | 34817 | 34788 | **RNA**→ rpmE2 (PRK01678)Ribosomal_L31 (pfam01197)→ |
| Sau-3136-1 | NZ_KK228932.1 | - | 26023 | 25994 | **RNA**→ rpmE2 (PRK01678)Ribosomal_L31 (pfam01197)→ |
| Sau-2880-1 | NZ_KK040429.1 | - | 56894 | 56865 | **RNA**→ rpmE2 (PRK01678)Ribosomal_L31 (pfam01197)→ |
| Sau-2554-1 | NZ_KK040817.1 | - | 56894 | 56865 | **RNA**→ rpmE2 (PRK01678)Ribosomal_L31 (pfam01197)→ |
| Sau-3495-1 | NZ_KK070557.1 | - | 34913 | 34884 | **RNA**→ rpmE2 (PRK01678)Ribosomal_L31 (pfam01197)→ |
| Sau-2485-1 | NZ_KK071187.1 | - | 35013 | 34984 | **RNA**→ rpmE2 (PRK01678)Ribosomal_L31 (pfam01197)→ |

| | | | | | |
|---|---|---|---|---|---|
| Sau-2474-1 | NZ_KK070701.1 | - | 56894 | 56865 | **RNA**→ rpmE2 (PRK01678)Ribosomal_L31 (pfam01197)→ |
| Sau-1043-1 | NZ_KK041058.1 | - | 56994 | 56965 | **RNA**→ rpmE2 (PRK01678)Ribosomal_L31 (pfam01197)→ |
| Sau-816-1 | NZ_KK041539.1 | - | 56894 | 56865 | **RNA**→ rpmE2 (PRK01678)Ribosomal_L31 (pfam01197)→ |
| Sau-2064-1 | NZ_KK068774.1 | - | 36390 | 36361 | **RNA**→ rpmE2 (PRK01678)Ribosomal_L31 (pfam01197)→ |
| Sau-316-1 | NZ_KK099049.1 | - | 1657695 | 1657666 | **RNA**→ rpmE2 (PRK01678)Ribosomal_L31 (pfam01197)→ |
| Sau-427-1 | NZ_KK097890.1 | - | 61690 | 61661 | **RNA**→ rpmE2 (PRK01678)Ribosomal_L31 (pfam01197)→ |
| Sau-176-1 | NZ_KK221090.1 | - | 26024 | 25995 | **RNA**→ rpmE2 (PRK01678)Ribosomal_L31 (pfam01197)→ |
| Sau-187-1 | NZ_KK221466.1 | - | 56305 | 56276 | **RNA**→ rpmE2 (PRK01678)Ribosomal_L31 (pfam01197)→ |
| Sau-205-1 | NZ_KK222053.1 | - | 56438 | 56409 | **RNA**→ rpmE2 (PRK01678)Ribosomal_L31 (pfam01197)→ |
| Sau-2099-1 | NZ_KK224356.1 | - | 56390 | 56361 | **RNA**→ rpmE2 (PRK01678)Ribosomal_L31 (pfam01197)→ |
| Sau-2626-1 | NZ_KK224503.1 | - | 56353 | 56324 | **RNA**→ rpmE2 (PRK01678)Ribosomal_L31 (pfam01197)→ |
| Sau-3497-1 | NZ_KK095369.1 | - | 34913 | 34884 | **RNA**→ rpmE2 (PRK01678)Ribosomal_L31 (pfam01197)→ |
| Sau-2984-1 | NZ_KK095312.1 | - | 2231827 | 2231798 | **RNA**→ rpmE2 (PRK01678)Ribosomal_L31 (pfam01197)→ |
| Sau-3544-1 | NZ_KK095555.1 | - | 34972 | 34943 | **RNA**→ rpmE2 (PRK01678)Ribosomal_L31 (pfam01197)→ |
| Sau-2890-1 | NZ_KK040372.1 | - | 34873 | 34844 | **RNA**→ rpmE2 (PRK01678)Ribosomal_L31 (pfam01197)→ |
| Sau-1053-1 | NZ_KK041050.1 | - | 328859 | 328830 | **RNA**→ rpmE2 (PRK01678)Ribosomal_L31 (pfam01197)→ |
| Sau-2562-1 | NZ_KK040771.1 | - | 235119 | 235090 | **RNA**→ rpmE2 (PRK01678)Ribosomal_L31 (pfam01197)→ |
| Sau-2532-1 | NZ_KK040873.1 | - | 28693 | 28664 | **RNA**→ rpmE2 (PRK01678)Ribosomal_L31 (pfam01197)→ |
| Sau-2514-1 | NZ_KK073114.1 | - | 55192 | 55163 | **RNA**→ rpmE2 (PRK01678)Ribosomal_L31 (pfam01197)→ |
| Sau-2582-1 | NZ_KK038808.1 | - | 34817 | 34788 | **RNA**→ rpmE2 (PRK01678)Ribosomal_L31 (pfam01197)→ |
| Sau-1024-1 | NZ_KK073057.1 | - | 56894 | 56865 | **RNA**→ rpmE2 (PRK01678)Ribosomal_L31 (pfam01197)→ |
| Sau-2455-1 | NZ_KK038978.1 | - | 56893 | 56864 | **RNA**→ rpmE2 (PRK01678)Ribosomal_L31 (pfam01197)→ |
| Sau-3489-1 | NZ_KK074863.1 | - | 34873 | 34844 | **RNA**→ rpmE2 (PRK01678)Ribosomal_L31 (pfam01197)→ |
| Sau-2542-1 | NZ_KK040825.1 | - | 34913 | 34884 | **RNA**→ rpmE2 (PRK01678)Ribosomal_L31 (pfam01197)→ |
| Sau-962-1 | NZ_KK071614.1 | - | 54903 | 54874 | **RNA**→ rpmE2 (PRK01678)Ribosomal_L31 (pfam01197)→ |
| Sau-1435-1 | NZ_KI971533.1 | - | 116484 | 116455 | **RNA**→ rpmE2 (PRK01678)Ribosomal_L31 (pfam01197)→ |
| Sau-810-1 | NZ_KK072686.1 | - | 34913 | 34884 | **RNA**→ rpmE2 (PRK01678)Ribosomal_L31 (pfam01197)→ |
| Sau-2591-1 | NZ_KK038780.1 | - | 34913 | 34884 | **RNA**→ rpmE2 (PRK01678)Ribosomal_L31 (pfam01197)→ |
| Sau-704-1 | NZ_KK096038.1 | - | 56789 | 56760 | **RNA**→ rpmE2 (PRK01678)Ribosomal_L31 (pfam01197)→ |
| Sau-3619-1 | NZ_KK040006.1 | - | 34873 | 34844 | **RNA**→ rpmE2 (PRK01678)Ribosomal_L31 (pfam01197)→ |
| Sau-1000-1 | NZ_KK073013.1 | - | 56794 | 56765 | **RNA**→ rpmE2 (PRK01678)Ribosomal_L31 (pfam01197)→ |
| Sau-1042-1 | NZ_KK041070.1 | - | 56894 | 56865 | **RNA**→ rpmE2 (PRK01678)Ribosomal_L31 (pfam01197)→ |
| Sau-2893-1 | NZ_KK073491.1 | - | 56994 | 56965 | **RNA**→ rpmE2 (PRK01678)Ribosomal_L31 (pfam01197)→ |
| Sau-1006-1 | NZ_KK041418.1 | - | 56893 | 56864 | **RNA**→ rpmE2 (PRK01678)Ribosomal_L31 (pfam01197)→ |
| Sau-3538-1 | NZ_KK097292.1 | - | 34913 | 34884 | **RNA**→ rpmE2 (PRK01678)Ribosomal_L31 (pfam01197)→ |
| Sau-1331-1 | NZ_KI971688.1 | - | 61711 | 61682 | **RNA**→ rpmE2 (PRK01678)Ribosomal_L31 (pfam01197)→ |
| Sau-2939-1 | NZ_KK038351.1 | - | 25254 | 25225 | **RNA**→ rpmE2 (PRK01678)Ribosomal_L31 (pfam01197)→ |
| Sau-2920-1 | NZ_KK038460.1 | - | 56894 | 56865 | **RNA**→ rpmE2 (PRK01678)Ribosomal_L31 (pfam01197)→ |
| Sau-2513-1 | NZ_KK072349.1 | - | 2229291 | 2229262 | **RNA**→ rpmE2 (PRK01678)Ribosomal_L31 (pfam01197)→ |
| Sau-2944-1 | NZ_KK075050.1 | - | 56645 | 56616 | **RNA**→ rpmE2 (PRK01678)Ribosomal_L31 (pfam01197)→ |
| Sau-2572-1 | NZ_KK075358.1 | - | 56988 | 56959 | **RNA**→ rpmE2 (PRK01678)Ribosomal_L31 (pfam01197)→ |
| Sau-2055-1 | NZ_KI972673.1 | - | 57557 | 57528 | **RNA**→ rpmE2 (PRK01678)Ribosomal_L31 (pfam01197)→ |
| Sau-2545-1 | NZ_KK075468.1 | - | 54839 | 54810 | **RNA**→ rpmE2 (PRK01678)Ribosomal_L31 (pfam01197)→ |
| Sau-3511-1 | NZ_KK038216.1 | - | 56894 | 56865 | **RNA**→ rpmE2 (PRK01678)Ribosomal_L31 (pfam01197)→ |
| Sau-2519-1 | NZ_KK040927.1 | - | 232923 | 232894 | **RNA**→ rpmE2 (PRK01678)Ribosomal_L31 (pfam01197)→ |
| Sau-834-1 | NZ_KK072896.1 | - | 56892 | 56863 | **RNA**→ rpmE2 (PRK01678)Ribosomal_L31 (pfam01197)→ |
| Sau-809-1 | NZ_KK072626.1 | - | 34913 | 34884 | **RNA**→ rpmE2 (PRK01678)Ribosomal_L31 (pfam01197)→ |
| Sau-647-1 | NZ_KK076222.1 | - | 54999 | 54970 | **RNA**→ rpmE2 (PRK01678)Ribosomal_L31 (pfam01197)→ |
| Sau-2118-1 | NZ_KK066962.1 | - | 36213 | 36184 | **RNA**→ rpmE2 (PRK01678)Ribosomal_L31 (pfam01197)→ |
| Sau-2497-1 | NZ_KK071854.1 | - | 34913 | 34884 | **RNA**→ rpmE2 (PRK01678)Ribosomal_L31 (pfam01197)→ |
| Sau-2887-1 | NZ_KK075168.1 | - | 56741 | 56712 | **RNA**→ rpmE2 (PRK01678)Ribosomal_L31 (pfam01197)→ |
| Sau-1092-1 | NZ_KK096426.1 | - | 34874 | 34845 | **RNA**→ rpmE2 (PRK01678)Ribosomal_L31 (pfam01197)→ |

| | | | | | |
|---|---|---|---|---|---|
| Sau-2553-1 | NZ_KK075434.1 | - | 56893 | 56864 | **RNA**→ rpmE2 (PRK01678)Ribosomal_L31 (pfam01197)→ |
| Sau-805-1 | NZ_KK041613.1 | - | 34873 | 34844 | **RNA**→ rpmE2 (PRK01678)Ribosomal_L31 (pfam01197)→ |
| Sau-1089-1 | NZ_KK075606.1 | - | 34873 | 34844 | **RNA**→ rpmE2 (PRK01678)Ribosomal_L31 (pfam01197)→ |
| Sau-3642-1 | NZ_KK074550.1 | - | 34913 | 34884 | **RNA**→ rpmE2 (PRK01678)Ribosomal_L31 (pfam01197)→ |
| Sau-1032-1 | NZ_KK075777.1 | - | 55218 | 55189 | **RNA**→ rpmE2 (PRK01678)Ribosomal_L31 (pfam01197)→ |
| Sau-3482-1 | NZ_KK075000.1 | - | 56794 | 56765 | **RNA**→ rpmE2 (PRK01678)Ribosomal_L31 (pfam01197)→ |
| Sau-646-1 | NZ_KK076258.1 | - | 56894 | 56865 | **RNA**→ rpmE2 (PRK01678)Ribosomal_L31 (pfam01197)→ |
| Sau-761-1 | NZ_KK071031.1 | - | 34817 | 34788 | **RNA**→ rpmE2 (PRK01678)Ribosomal_L31 (pfam01197)→ |
| Sau-3531-1 | NZ_KK038182.1 | - | 56091 | 56062 | **RNA**→ rpmE2 (PRK01678)Ribosomal_L31 (pfam01197)→ |
| Sau-2579-1 | NZ_KK075298.1 | - | 56894 | 56865 | **RNA**→ rpmE2 (PRK01678)Ribosomal_L31 (pfam01197)→ |
| Sau-990-1 | NZ_KK072916.1 | - | 34913 | 34884 | **RNA**→ rpmE2 (PRK01678)Ribosomal_L31 (pfam01197)→ |
| Sau-671-1 | NZ_KK076023.1 | - | 34970 | 34941 | **RNA**→ rpmE2 (PRK01678)Ribosomal_L31 (pfam01197)→ |
| Sau-1037-1 | NZ_KK075763.1 | - | 55117 | 55088 | **RNA**→ rpmE2 (PRK01678)Ribosomal_L31 (pfam01197)→ |
| Sau-1028-1 | NZ_KK075856.1 | - | 34872 | 34843 | **RNA**→ rpmE2 (PRK01678)Ribosomal_L31 (pfam01197)→ |
| Sau-2594-1 | NZ_KK075244.1 | - | 55129 | 55100 | **RNA**→ rpmE2 (PRK01678)Ribosomal_L31 (pfam01197)→ |
| Sau-1055-1 | NZ_KK075684.1 | - | 55117 | 55088 | **RNA**→ rpmE2 (PRK01678)Ribosomal_L31 (pfam01197)→ |
| Sau-752-1 | NZ_KK075969.1 | - | 34873 | 34844 | **RNA**→ rpmE2 (PRK01678)Ribosomal_L31 (pfam01197)→ |
| Sau-991-1 | NZ_KK072423.1 | - | 34913 | 34884 | **RNA**→ rpmE2 (PRK01678)Ribosomal_L31 (pfam01197)→ |
| Sau-2527-1 | NZ_KK094731.1 | - | 25249 | 25220 | **RNA**→ rpmE2 (PRK01678)Ribosomal_L31 (pfam01197)→ |
| Sau-2906-1 | NZ_KK038504.1 | - | 34913 | 34884 | **RNA**→ rpmE2 (PRK01678)Ribosomal_L31 (pfam01197)→ |
| Sau-2855-1 | NZ_KK073344.1 | - | 34913 | 34884 | **RNA**→ rpmE2 (PRK01678)Ribosomal_L31 (pfam01197)→ |
| Sau-2468-1 | NZ_KK096641.1 | - | 15617 | 15588 | **RNA**→ rpmE2 (PRK01678)Ribosomal_L31 (pfam01197)→ |
| Sau-3499-1 | NZ_KK074762.1 | - | 34962 | 34933 | **RNA**→ rpmE2 (PRK01678)Ribosomal_L31 (pfam01197)→ |
| Sau-2481-1 | NZ_KK224412.1 | - | 56439 | 56410 | **RNA**→ rpmE2 (PRK01678)Ribosomal_L31 (pfam01197)→ |
| Sau-2699-1 | NZ_KK225314.1 | - | 26024 | 25995 | **RNA**→ rpmE2 (PRK01678)Ribosomal_L31 (pfam01197)→ |
| Sau-3058-1 | NZ_KK226264.1 | - | 26024 | 25995 | **RNA**→ rpmE2 (PRK01678)Ribosomal_L31 (pfam01197)→ |
| Sau-2695-1 | NZ_KK225169.1 | - | 26024 | 25995 | **RNA**→ rpmE2 (PRK01678)Ribosomal_L31 (pfam01197)→ |
| Sau-2727-1 | NZ_KK225600.1 | - | 26024 | 25995 | **RNA**→ rpmE2 (PRK01678)Ribosomal_L31 (pfam01197)→ |
| Sau-436-1 | NZ_KK097984.1 | - | 56858 | 56829 | **RNA**→ rpmE2 (PRK01678)Ribosomal_L31 (pfam01197)→ |
| Sau-363-1 | NZ_KK099356.1 | - | 465519 | 465490 | **RNA**→ rpmE2 (PRK01678)Ribosomal_L31 (pfam01197)→ |
| Sau-2946-1 | NZ_KK097098.1 | - | 19182 | 19153 | **RNA**→ rpmE2 (PRK01678)Ribosomal_L31 (pfam01197)→ |
| Sau-645-1 | NZ_KK072543.1 | - | 55176 | 55147 | **RNA**→ rpmE2 (PRK01678)Ribosomal_L31 (pfam01197)→ |
| Sau-197-1 | NZ_KK221759.1 | - | 25790 | 25761 | **RNA**→ rpmE2 (PRK01678)Ribosomal_L31 (pfam01197)→ |
| Sau-477-1 | NZ_KK098514.1 | - | 1677866 | 1677837 | **RNA**→ rpmE2 (PRK01678)Ribosomal_L31 (pfam01197)→ |
| Sau-372-1 | NZ_KK099440.1 | - | 57593 | 57564 | **RNA**→ rpmE2 (PRK01678)Ribosomal_L31 (pfam01197)→ |
| Sau-357-1 | NZ_KK099307.1 | - | 2100797 | 2100768 | **RNA**→ rpmE2 (PRK01678)Ribosomal_L31 (pfam01197)→ |
| Sau-142-1 | NZ_KK219997.1 | - | 56395 | 56366 | **RNA**→ rpmE2 (PRK01678)Ribosomal_L31 (pfam01197)→ |
| Sau-216-1 | NZ_KK222467.1 | - | 26024 | 25995 | **RNA**→ rpmE2 (PRK01678)Ribosomal_L31 (pfam01197)→ |
| Sau-173-1 | NZ_KK220987.1 | - | 55471 | 55442 | **RNA**→ rpmE2 (PRK01678)Ribosomal_L31 (pfam01197)→ |
| Sau-429-1 | NZ_KK097926.1 | - | 57675 | 57646 | **RNA**→ rpmE2 (PRK01678)Ribosomal_L31 (pfam01197)→ |
| Sau-472-1 | NZ_KK098444.1 | - | 61574 | 61545 | **RNA**→ rpmE2 (PRK01678)Ribosomal_L31 (pfam01197)→ |
| Sau-264-1 | NZ_KK223042.1 | - | 56390 | 56361 | **RNA**→ rpmE2 (PRK01678)Ribosomal_L31 (pfam01197)→ |
| Sau-162-1 | NZ_KK220613.1 | - | 55707 | 55678 | **RNA**→ rpmE2 (PRK01678)Ribosomal_L31 (pfam01197)→ |
| Sau-2587-1 | NZ_KK094871.1 | - | 34972 | 34943 | **RNA**→ rpmE2 (PRK01678)Ribosomal_L31 (pfam01197)→ |
| Sau-211-1 | NZ_KK222290.1 | - | 1768 | 1739 | **RNA**→ rpmE2 (PRK01678)Ribosomal_L31 (pfam01197)→ |
| Sau-442-1 | NZ_KK098091.1 | - | 409713 | 409684 | **RNA**→ rpmE2 (PRK01678)Ribosomal_L31 (pfam01197)→ |
| Sau-621-1 | NZ_KK072459.1 | - | 34873 | 34844 | **RNA**→ rpmE2 (PRK01678)Ribosomal_L31 (pfam01197)→ |
| Sau-390-1 | NZ_KK099582.1 | - | 59481 | 59452 | **RNA**→ rpmE2 (PRK01678)Ribosomal_L31 (pfam01197)→ |
| Sau-522-1 | NZ_KK074253.1 | - | 35659 | 35630 | **RNA**→ rpmE2 (PRK01678)Ribosomal_L31 (pfam01197)→ |
| Sau-184-1 | NZ_KK221358.1 | - | 25790 | 25761 | **RNA**→ rpmE2 (PRK01678)Ribosomal_L31 (pfam01197)→ |
| Sau-189-1 | NZ_KK221549.1 | - | 56395 | 56366 | **RNA**→ rpmE2 (PRK01678)Ribosomal_L31 (pfam01197)→ |
| Sau-324-1 | NZ_KK099086.1 | - | 1640651 | 1640622 | **RNA**→ rpmE2 (PRK01678)Ribosomal_L31 (pfam01197)→ |

| | | | | | | |
|---|---|---|---|---|---|---|
| Sau-360-1 | NZ_KK099334.1 | - | 61691 | 61662 | **RNA**→ | rpmE2 (PRK01678)Ribosomal_L31 (pfam01197)→ |
| Sau-137-1 | NZ_KK219743.1 | - | 56439 | 56410 | **RNA**→ | rpmE2 (PRK01678)Ribosomal_L31 (pfam01197)→ |
| Sau-122-1 | NZ_KK219261.1 | - | 3263 | 3234 | **RNA**→ | rpmE2 (PRK01678)Ribosomal_L31 (pfam01197)→ |
| Sau-562-1 | NZ_KK074340.1 | - | 36224 | 36195 | **RNA**→ | rpmE2 (PRK01678)Ribosomal_L31 (pfam01197)→ |
| Sau-2599-1 | NZ_KK096816.1 | - | 25255 | 25226 | **RNA**→ | rpmE2 (PRK01678)Ribosomal_L31 (pfam01197)→ |
| Sau-171-1 | NZ_KK220922.1 | - | 26024 | 25995 | **RNA**→ | rpmE2 (PRK01678)Ribosomal_L31 (pfam01197)→ |
| Sau-876-1 | NZ_KK224237.1 | - | 56390 | 56361 | **RNA**→ | rpmE2 (PRK01678)Ribosomal_L31 (pfam01197)→ |
| Sau-462-1 | NZ_KK098326.1 | - | 226837 | 226808 | **RNA**→ | rpmE2 (PRK01678)Ribosomal_L31 (pfam01197)→ |
| Sau-2833-1 | NZ_KK094970.1 | - | 34913 | 34884 | **RNA**→ | rpmE2 (PRK01678)Ribosomal_L31 (pfam01197)→ |
| Sau-2921-1 | NZ_KK095162.1 | - | 55084 | 55055 | **RNA**→ | rpmE2 (PRK01678)Ribosomal_L31 (pfam01197)→ |
| Sau-874-1 | NZ_KK224189.1 | - | 56441 | 56412 | **RNA**→ | rpmE2 (PRK01678)Ribosomal_L31 (pfam01197)→ |
| Sau-2915-1 | NZ_KK095113.1 | - | 35168 | 35139 | **RNA**→ | rpmE2 (PRK01678)Ribosomal_L31 (pfam01197)→ |
| Sau-2516-1 | NZ_KK224471.1 | - | 26024 | 25995 | **RNA**→ | rpmE2 (PRK01678)Ribosomal_L31 (pfam01197)→ |
| Sau-2997-1 | NZ_KK095339.1 | - | 56894 | 56865 | **RNA**→ | rpmE2 (PRK01678)Ribosomal_L31 (pfam01197)→ |
| Sau-595-1 | NZ_KK099762.1 | - | 59750 | 59721 | **RNA**→ | rpmE2 (PRK01678)Ribosomal_L31 (pfam01197)→ |
| Sau-1651-1 | NZ_KK018430.1 | - | 217332 | 217303 | **RNA**→ | rpmE2 (PRK01678)Ribosomal_L31 (pfam01197)→ |
| Sau-1963-1 | NZ_KI988686.1 | - | 113409 | 113380 | **RNA**→ | rpmE2 (PRK01678)Ribosomal_L31 (pfam01197)→ |
| Sau-1970-1 | NZ_KI988920.1 | - | 35742 | 35713 | **RNA**→ | rpmE2 (PRK01678)Ribosomal_L31 (pfam01197)→ |
| Sau-1978-1 | NZ_KI989249.1 | - | 36391 | 36362 | **RNA**→ | rpmE2 (PRK01678)Ribosomal_L31 (pfam01197)→ |
| Sau-1979-1 | NZ_KI989299.1 | - | 24749 | 24720 | **RNA**→ | rpmE2 (PRK01678)Ribosomal_L31 (pfam01197)→ |
| Sau-1985-1 | NZ_KI989585.1 | - | 61571 | 61542 | **RNA**→ | rpmE2 (PRK01678)Ribosomal_L31 (pfam01197)→ |
| Sau-1986-1 | NZ_KI989593.1 | - | 25029 | 25000 | **RNA**→ | rpmE2 (PRK01678)Ribosomal_L31 (pfam01197)→ |
| Sau-1991-1 | NZ_KI989778.1 | - | 62206 | 62177 | **RNA**→ | rpmE2 (PRK01678)Ribosomal_L31 (pfam01197)→ |
| Sau-1994-1 | NZ_KI989785.1 | - | 36391 | 36362 | **RNA**→ | rpmE2 (PRK01678)Ribosomal_L31 (pfam01197)→ |
| Sau-2022-1 | NZ_KI990875.1 | - | 175788 | 175759 | **RNA**→ | rpmE2 (PRK01678)Ribosomal_L31 (pfam01197)→ |
| Sau-1962-1 | NZ_KI988673.1 | - | 35180 | 35151 | **RNA**→ | rpmE2 (PRK01678)Ribosomal_L31 (pfam01197)→ |
| Sau-1975-1 | NZ_KI989201.1 | - | 56091 | 56062 | **RNA**→ | rpmE2 (PRK01678)Ribosomal_L31 (pfam01197)→ |
| Sau-1990-1 | NZ_KI989707.1 | - | 217268 | 217239 | **RNA**→ | rpmE2 (PRK01678)Ribosomal_L31 (pfam01197)→ |
| Sau-2000-1 | NZ_KI990071.1 | - | 61380 | 61351 | **RNA**→ | rpmE2 (PRK01678)Ribosomal_L31 (pfam01197)→ |
| Sau-2005-1 | NZ_KI990204.1 | - | 35421 | 35392 | **RNA**→ | rpmE2 (PRK01678)Ribosomal_L31 (pfam01197)→ |
| Sau-1967-1 | NZ_KI988891.1 | - | 62202 | 62173 | **RNA**→ | rpmE2 (PRK01678)Ribosomal_L31 (pfam01197)→ |
| Sau-1989-1 | NZ_KI989655.1 | - | 36391 | 36362 | **RNA**→ | rpmE2 (PRK01678)Ribosomal_L31 (pfam01197)→ |
| Sau-2004-1 | NZ_KI990131.1 | - | 35829 | 35800 | **RNA**→ | rpmE2 (PRK01678)Ribosomal_L31 (pfam01197)→ |
| Sau-2020-1 | NZ_KI990737.1 | - | 62257 | 62228 | **RNA**→ | rpmE2 (PRK01678)Ribosomal_L31 (pfam01197)→ |
| Sau-3706-1 | NZ_KI977880.1 | - | 56997 | 56968 | **RNA**→ | rpmE2 (PRK01678)Ribosomal_L31 (pfam01197)→ |
| Sau-2024-1 | NZ_KI990951.1 | - | 54994 | 54965 | **RNA**→ | rpmE2 (PRK01678)Ribosomal_L31 (pfam01197)→ |
| Sau-3677-1 | NZ_KI978852.1 | - | 239539 | 239510 | **RNA**→ | rpmE2 (PRK01678)Ribosomal_L31 (pfam01197)→ |
| Sau-3681-1 | NZ_KI979204.1 | - | 58513 | 58484 | **RNA**→ | rpmE2 (PRK01678)Ribosomal_L31 (pfam01197)→ |
| Sau-3678-1 | NZ_KI978909.1 | - | 175828 | 175799 | **RNA**→ | rpmE2 (PRK01678)Ribosomal_L31 (pfam01197)→ |
| Sau-3687-1 | NZ_KI977321.1 | - | 56894 | 56865 | **RNA**→ | rpmE2 (PRK01678)Ribosomal_L31 (pfam01197)→ |
| Sau-3707-1 | NZ_KI978114.1 | - | 57605 | 57576 | **RNA**→ | rpmE2 (PRK01678)Ribosomal_L31 (pfam01197)→ |
| Sau-1964-1 | NZ_KI988761.1 | - | 62331 | 62302 | **RNA**→ | rpmE2 (PRK01678)Ribosomal_L31 (pfam01197)→ |
| Sau-3679-1 | NZ_KI979142.1 | - | 56893 | 56864 | **RNA**→ | rpmE2 (PRK01678)Ribosomal_L31 (pfam01197)→ |
| Sau-3698-1 | NZ_KI975542.1 | - | 60192 | 60163 | **RNA**→ | rpmE2 (PRK01678)Ribosomal_L31 (pfam01197)→ |
| Sau-1973-1 | NZ_KI989113.1 | - | 62156 | 62127 | **RNA**→ | rpmE2 (PRK01678)Ribosomal_L31 (pfam01197)→ |
| Sau-3686-1 | NZ_KI977286.1 | - | 34913 | 34884 | **RNA**→ | rpmE2 (PRK01678)Ribosomal_L31 (pfam01197)→ |
| Sau-1995-1 | NZ_KI989832.1 | - | 35563 | 35534 | **RNA**→ | rpmE2 (PRK01678)Ribosomal_L31 (pfam01197)→ |
| Sau-1356-1 | NZ_KK006186.1 | - | 58196 | 58167 | **RNA**→ | rpmE2 (PRK01678)Ribosomal_L31 (pfam01197)→ |
| Sau-1384-1 | NZ_KK006658.1 | - | 36391 | 36362 | **RNA**→ | rpmE2 (PRK01678)Ribosomal_L31 (pfam01197)→ |
| Sau-1392-1 | NZ_KK006921.1 | - | 36491 | 36462 | **RNA**→ | rpmE2 (PRK01678)Ribosomal_L31 (pfam01197)→ |
| Sau-1430-1 | NZ_KK008404.1 | - | 107989 | 107960 | **RNA**→ | rpmE2 (PRK01678)Ribosomal_L31 (pfam01197)→ |
| Sau-1432-1 | NZ_KK008462.1 | - | 62050 | 62021 | **RNA**→ | rpmE2 (PRK01678)Ribosomal_L31 (pfam01197)→ |

| | | | | | |
|---|---|---|---|---|---|
| Sau-1457-1 | NZ_KK009531.1 | - | 28985 | 28956 | **RNA**→ rpmE2 (PRK01678)Ribosomal_L31 (pfam01197)→ |
| Sau-1464-1 | NZ_KK009890.1 | - | 24262 | 24233 | **RNA**→ rpmE2 (PRK01678)Ribosomal_L31 (pfam01197)→ |
| Sau-1477-1 | NZ_KK010450.1 | - | 61373 | 61344 | **RNA**→ rpmE2 (PRK01678)Ribosomal_L31 (pfam01197)→ |
| Sau-3841-1 | NZ_KK010958.1 | - | 35474 | 35445 | **RNA**→ rpmE2 (PRK01678)Ribosomal_L31 (pfam01197)→ |
| Sau-1497-1 | NZ_KK011269.1 | - | 58247 | 58218 | **RNA**→ rpmE2 (PRK01678)Ribosomal_L31 (pfam01197)→ |
| Sau-1498-1 | NZ_KK011277.1 | - | 36188 | 36159 | **RNA**→ rpmE2 (PRK01678)Ribosomal_L31 (pfam01197)→ |
| Sau-1391-1 | NZ_KK006918.1 | - | 111834 | 111805 | **RNA**→ rpmE2 (PRK01678)Ribosomal_L31 (pfam01197)→ |
| Sau-1417-1 | NZ_KK007810.1 | - | 26012 | 25983 | **RNA**→ rpmE2 (PRK01678)Ribosomal_L31 (pfam01197)→ |
| Sau-1424-1 | NZ_KK008053.1 | - | 108168 | 108139 | **RNA**→ rpmE2 (PRK01678)Ribosomal_L31 (pfam01197)→ |
| Sau-3038-1 | NZ_KK019246.1 | - | 56418 | 56389 | **RNA**→ rpmE2 (PRK01678)Ribosomal_L31 (pfam01197)→ |
| Sau-1436-1 | NZ_KK008599.1 | - | 36391 | 36362 | **RNA**→ rpmE2 (PRK01678)Ribosomal_L31 (pfam01197)→ |
| Sau-1437-1 | NZ_KK008721.1 | - | 61334 | 61305 | **RNA**→ rpmE2 (PRK01678)Ribosomal_L31 (pfam01197)→ |
| Sau-1441-1 | NZ_KK008960.1 | - | 114097 | 114068 | **RNA**→ rpmE2 (PRK01678)Ribosomal_L31 (pfam01197)→ |
| Sau-1447-1 | NZ_KK009126.1 | - | 36392 | 36363 | **RNA**→ rpmE2 (PRK01678)Ribosomal_L31 (pfam01197)→ |
| Sau-3648-1 | NZ_KK022617.1 | - | 56894 | 56865 | **RNA**→ rpmE2 (PRK01678)Ribosomal_L31 (pfam01197)→ |
| Sau-1448-1 | NZ_KK009212.1 | - | 114343 | 114314 | **RNA**→ rpmE2 (PRK01678)Ribosomal_L31 (pfam01197)→ |
| Sau-3037-1 | NZ_KK019319.1 | - | 35193 | 35164 | **RNA**→ rpmE2 (PRK01678)Ribosomal_L31 (pfam01197)→ |
| Sau-1470-1 | NZ_KK010081.1 | - | 2000 | 1971 | **RNA**→ rpmE2 (PRK01678)Ribosomal_L31 (pfam01197)→ |
| Sau-3527-1 | NZ_KK021203.1 | - | 34872 | 34843 | **RNA**→ rpmE2 (PRK01678)Ribosomal_L31 (pfam01197)→ |
| Sau-1485-1 | NZ_KK010686.1 | - | 119529 | 119500 | **RNA**→ rpmE2 (PRK01678)Ribosomal_L31 (pfam01197)→ |
| Sau-1490-1 | NZ_KK010894.1 | - | 37315 | 37286 | **RNA**→ rpmE2 (PRK01678)Ribosomal_L31 (pfam01197)→ |
| Sau-3514-1 | NZ_KK020708.1 | - | 95189 | 95160 | **RNA**→ rpmE2 (PRK01678)Ribosomal_L31 (pfam01197)→ |
| Sau-1496-1 | NZ_KK011178.1 | - | 55103 | 55074 | **RNA**→ rpmE2 (PRK01678)Ribosomal_L31 (pfam01197)→ |
| Sau-2950-1 | NZ_KK022846.1 | - | 56894 | 56865 | **RNA**→ rpmE2 (PRK01678)Ribosomal_L31 (pfam01197)→ |
| Sau-1502-1 | NZ_KK011440.1 | - | 61555 | 61526 | **RNA**→ rpmE2 (PRK01678)Ribosomal_L31 (pfam01197)→ |
| Sau-1408-1 | NZ_KK007517.1 | - | 195989 | 195960 | **RNA**→ rpmE2 (PRK01678)Ribosomal_L31 (pfam01197)→ |
| Sau-1416-1 | NZ_KK007738.1 | - | 113597 | 113568 | **RNA**→ rpmE2 (PRK01678)Ribosomal_L31 (pfam01197)→ |
| Sau-1434-1 | NZ_KK008532.1 | - | 36391 | 36362 | **RNA**→ rpmE2 (PRK01678)Ribosomal_L31 (pfam01197)→ |
| Sau-1440-1 | NZ_KK008864.1 | - | 61213 | 61184 | **RNA**→ rpmE2 (PRK01678)Ribosomal_L31 (pfam01197)→ |
| Sau-1454-1 | NZ_KK009420.1 | - | 36491 | 36462 | **RNA**→ rpmE2 (PRK01678)Ribosomal_L31 (pfam01197)→ |
| Sau-1459-1 | NZ_KK009661.1 | - | 36391 | 36362 | **RNA**→ rpmE2 (PRK01678)Ribosomal_L31 (pfam01197)→ |
| Sau-1185-1 | NZ_KI999129.1 | - | 36391 | 36362 | **RNA**→ rpmE2 (PRK01678)Ribosomal_L31 (pfam01197)→ |
| Sau-1475-1 | NZ_KK010309.1 | - | 111814 | 111785 | **RNA**→ rpmE2 (PRK01678)Ribosomal_L31 (pfam01197)→ |
| Sau-1481-1 | NZ_KK010625.1 | - | 111577 | 111548 | **RNA**→ rpmE2 (PRK01678)Ribosomal_L31 (pfam01197)→ |
| Sau-1281-1 | NZ_KK002970.1 | - | 62354 | 62325 | **RNA**→ rpmE2 (PRK01678)Ribosomal_L31 (pfam01197)→ |
| Sau-1489-1 | NZ_KK010871.1 | - | 62005 | 61976 | **RNA**→ rpmE2 (PRK01678)Ribosomal_L31 (pfam01197)→ |
| Sau-1495-1 | NZ_KK011169.1 | - | 61380 | 61351 | **RNA**→ rpmE2 (PRK01678)Ribosomal_L31 (pfam01197)→ |
| Sau-1287-1 | NZ_KK003215.1 | - | 126484 | 126455 | **RNA**→ rpmE2 (PRK01678)Ribosomal_L31 (pfam01197)→ |
| Sau-1421-1 | NZ_KK007928.1 | - | 36391 | 36362 | **RNA**→ rpmE2 (PRK01678)Ribosomal_L31 (pfam01197)→ |
| Sau-1288-1 | NZ_KK003234.1 | - | 43829 | 43800 | **RNA**→ rpmE2 (PRK01678)Ribosomal_L31 (pfam01197)→ |
| Sau-1445-1 | NZ_KK009087.1 | - | 145173 | 145144 | **RNA**→ rpmE2 (PRK01678)Ribosomal_L31 (pfam01197)→ |
| Sau-1294-1 | NZ_KK003529.1 | - | 61480 | 61451 | **RNA**→ rpmE2 (PRK01678)Ribosomal_L31 (pfam01197)→ |
| Sau-1458-1 | NZ_KK009644.1 | - | 108047 | 108018 | **RNA**→ rpmE2 (PRK01678)Ribosomal_L31 (pfam01197)→ |
| Sau-1306-1 | NZ_KK003976.1 | - | 36861 | 36832 | **RNA**→ rpmE2 (PRK01678)Ribosomal_L31 (pfam01197)→ |
| Sau-1473-1 | NZ_KK010162.1 | - | 82709 | 82680 | **RNA**→ rpmE2 (PRK01678)Ribosomal_L31 (pfam01197)→ |
| Sau-1311-1 | NZ_KK004318.1 | - | 114419 | 114390 | **RNA**→ rpmE2 (PRK01678)Ribosomal_L31 (pfam01197)→ |
| Sau-1474-1 | NZ_KK010214.1 | - | 25255 | 25226 | **RNA**→ rpmE2 (PRK01678)Ribosomal_L31 (pfam01197)→ |
| Sau-1336-1 | NZ_KK005381.1 | - | 15850 | 15821 | **RNA**→ rpmE2 (PRK01678)Ribosomal_L31 (pfam01197)→ |
| Sau-1478-1 | NZ_KK010455.1 | - | 36021 | 35992 | **RNA**→ rpmE2 (PRK01678)Ribosomal_L31 (pfam01197)→ |
| Sau-1341-1 | NZ_KK005590.1 | - | 62005 | 61976 | **RNA**→ rpmE2 (PRK01678)Ribosomal_L31 (pfam01197)→ |
| Sau-1487-1 | NZ_KK010743.1 | - | 35170 | 35141 | **RNA**→ rpmE2 (PRK01678)Ribosomal_L31 (pfam01197)→ |
| Sau-1505-1 | NZ_KK011613.1 | - | 35397 | 35368 | **RNA**→ rpmE2 (PRK01678)Ribosomal_L31 (pfam01197)→ |

| | | | | | | |
|---|---|---|---|---|---|---|
| Sau-1352-1 | NZ_KK006057.1 | - | 114518 | 114489 | **RNA**→ | rpmE2 (PRK01678)Ribosomal_L31 (pfam01197)→ |
| Sau-1366-1 | NZ_KK006305.1 | - | 35908 | 35879 | **RNA**→ | rpmE2 (PRK01678)Ribosomal_L31 (pfam01197)→ |
| Sau-1488-1 | NZ_KK010792.1 | - | 178313 | 178284 | **RNA**→ | rpmE2 (PRK01678)Ribosomal_L31 (pfam01197)→ |
| Sau-1476-1 | NZ_KK010315.1 | - | 36391 | 36362 | **RNA**→ | rpmE2 (PRK01678)Ribosomal_L31 (pfam01197)→ |
| Sau-1380-1 | NZ_KK006558.1 | - | 117031 | 117002 | **RNA**→ | rpmE2 (PRK01678)Ribosomal_L31 (pfam01197)→ |
| Sau-1507-1 | NZ_KK011728.1 | - | 35371 | 35342 | **RNA**→ | rpmE2 (PRK01678)Ribosomal_L31 (pfam01197)→ |
| Sau-1382-1 | NZ_KK006578.1 | - | 116022 | 115993 | **RNA**→ | rpmE2 (PRK01678)Ribosomal_L31 (pfam01197)→ |
| Sau-1503-1 | NZ_KK011501.1 | - | 36513 | 36484 | **RNA**→ | rpmE2 (PRK01678)Ribosomal_L31 (pfam01197)→ |
| Sau-1199-1 | NZ_KI999874.1 | - | 28749 | 28720 | **RNA**→ | rpmE2 (PRK01678)Ribosomal_L31 (pfam01197)→ |
| Sau-1504-1 | NZ_KK011582.1 | - | 178484 | 178455 | **RNA**→ | rpmE2 (PRK01678)Ribosomal_L31 (pfam01197)→ |
| Sau-1284-1 | NZ_KK003030.1 | - | 65411 | 65382 | **RNA**→ | rpmE2 (PRK01678)Ribosomal_L31 (pfam01197)→ |
| Sau-1510-1 | NZ_KK011817.1 | - | 56781 | 56752 | **RNA**→ | rpmE2 (PRK01678)Ribosomal_L31 (pfam01197)→ |
| Sau-1295-1 | NZ_KK003538.1 | - | 36220 | 36191 | **RNA**→ | rpmE2 (PRK01678)Ribosomal_L31 (pfam01197)→ |
| Sau-1517-1 | NZ_KK012068.1 | - | 35253 | 35224 | **RNA**→ | rpmE2 (PRK01678)Ribosomal_L31 (pfam01197)→ |
| Sau-1301-1 | NZ_KK003741.1 | - | 36391 | 36362 | **RNA**→ | rpmE2 (PRK01678)Ribosomal_L31 (pfam01197)→ |
| Sau-1501-1 | NZ_KK011400.1 | - | 2500 | 2471 | **RNA**→ | rpmE2 (PRK01678)Ribosomal_L31 (pfam01197)→ |
| Sau-1313-1 | NZ_KK004322.1 | - | 36491 | 36462 | **RNA**→ | rpmE2 (PRK01678)Ribosomal_L31 (pfam01197)→ |
| Sau-1318-1 | NZ_KK004611.1 | - | 1824 | 1795 | **RNA**→ | rpmE2 (PRK01678)Ribosomal_L31 (pfam01197)→ |
| Sau-1325-1 | NZ_KK004904.1 | - | 2235 | 2206 | **RNA**→ | rpmE2 (PRK01678)Ribosomal_L31 (pfam01197)→ |
| Sau-1330-1 | NZ_KK005165.1 | - | 36491 | 36462 | **RNA**→ | rpmE2 (PRK01678)Ribosomal_L31 (pfam01197)→ |
| Sau-1344-1 | NZ_KK005683.1 | - | 62156 | 62127 | **RNA**→ | rpmE2 (PRK01678)Ribosomal_L31 (pfam01197)→ |
| Sau-1349-1 | NZ_KK005857.1 | - | 36390 | 36361 | **RNA**→ | rpmE2 (PRK01678)Ribosomal_L31 (pfam01197)→ |
| Sau-1354-1 | NZ_KK006093.1 | - | 114114 | 114085 | **RNA**→ | rpmE2 (PRK01678)Ribosomal_L31 (pfam01197)→ |
| Sau-1383-1 | NZ_KK006590.1 | - | 22238 | 22209 | **RNA**→ | rpmE2 (PRK01678)Ribosomal_L31 (pfam01197)→ |
| Sau-1310-1 | NZ_KK004215.1 | - | 36391 | 36362 | **RNA**→ | rpmE2 (PRK01678)Ribosomal_L31 (pfam01197)→ |
| Sau-1315-1 | NZ_KK004456.1 | - | 35757 | 35728 | **RNA**→ | rpmE2 (PRK01678)Ribosomal_L31 (pfam01197)→ |
| Sau-573-1 | NZ_KI966278.1 | - | 59905 | 59876 | **RNA**→ | rpmE2 (PRK01678)Ribosomal_L31 (pfam01197)→ |
| Sau-1316-1 | NZ_KK004546.1 | - | 109745 | 109716 | **RNA**→ | rpmE2 (PRK01678)Ribosomal_L31 (pfam01197)→ |
| Sau-1346-1 | NZ_KK005738.1 | - | 35986 | 35957 | **RNA**→ | rpmE2 (PRK01678)Ribosomal_L31 (pfam01197)→ |
| Sau-1687-1 | NZ_KI967761.1 | - | 36281 | 36252 | **RNA**→ | rpmE2 (PRK01678)Ribosomal_L31 (pfam01197)→ |
| Sau-1357-1 | NZ_KK006191.1 | - | 56146 | 56117 | **RNA**→ | rpmE2 (PRK01678)Ribosomal_L31 (pfam01197)→ |
| Sau-1692-1 | NZ_KI967978.1 | - | 36579 | 36550 | **RNA**→ | rpmE2 (PRK01678)Ribosomal_L31 (pfam01197)→ |
| Sau-1385-1 | NZ_KK006727.1 | - | 36016 | 35987 | **RNA**→ | rpmE2 (PRK01678)Ribosomal_L31 (pfam01197)→ |
| Sau-1698-1 | NZ_KI968309.1 | - | 36391 | 36362 | **RNA**→ | rpmE2 (PRK01678)Ribosomal_L31 (pfam01197)→ |
| Sau-1196-1 | NZ_KI999558.1 | - | 36632 | 36603 | **RNA**→ | rpmE2 (PRK01678)Ribosomal_L31 (pfam01197)→ |
| Sau-1207-1 | NZ_KK000219.1 | - | 28749 | 28720 | **RNA**→ | rpmE2 (PRK01678)Ribosomal_L31 (pfam01197)→ |
| Sau-1227-1 | NZ_KK000992.1 | - | 7425 | 7396 | **RNA**→ | rpmE2 (PRK01678)Ribosomal_L31 (pfam01197)→ |
| Sau-1320-1 | NZ_KK004674.1 | - | 36391 | 36362 | **RNA**→ | rpmE2 (PRK01678)Ribosomal_L31 (pfam01197)→ |
| Sau-1332-1 | NZ_KK005270.1 | - | 114270 | 114241 | **RNA**→ | rpmE2 (PRK01678)Ribosomal_L31 (pfam01197)→ |
| Sau-1333-1 | NZ_KK005279.1 | - | 36391 | 36362 | **RNA**→ | rpmE2 (PRK01678)Ribosomal_L31 (pfam01197)→ |
| Sau-1345-1 | NZ_KK005685.1 | - | 36391 | 36362 | **RNA**→ | rpmE2 (PRK01678)Ribosomal_L31 (pfam01197)→ |
| Sau-1350-1 | NZ_KK005907.1 | - | 32991 | 32962 | **RNA**→ | rpmE2 (PRK01678)Ribosomal_L31 (pfam01197)→ |
| Sau-1241-1 | NZ_KK001441.1 | - | 25227 | 25198 | **RNA**→ | rpmE2 (PRK01678)Ribosomal_L31 (pfam01197)→ |
| Sau-1285-1 | NZ_KK003118.1 | - | 25185 | 25156 | **RNA**→ | rpmE2 (PRK01678)Ribosomal_L31 (pfam01197)→ |
| Sau-1378-1 | NZ_KK006456.1 | - | 94742 | 94713 | **RNA**→ | rpmE2 (PRK01678)Ribosomal_L31 (pfam01197)→ |
| Sau-1337-1 | NZ_KK005431.1 | - | 36391 | 36362 | **RNA**→ | rpmE2 (PRK01678)Ribosomal_L31 (pfam01197)→ |
| Sau-1782-1 | NZ_KI982238.1 | - | 113825 | 113796 | **RNA**→ | rpmE2 (PRK01678)Ribosomal_L31 (pfam01197)→ |
| Sau-1343-1 | NZ_KK005632.1 | - | 130992 | 130963 | **RNA**→ | rpmE2 (PRK01678)Ribosomal_L31 (pfam01197)→ |
| Sau-1788-1 | NZ_KI982535.1 | - | 61379 | 61350 | **RNA**→ | rpmE2 (PRK01678)Ribosomal_L31 (pfam01197)→ |
| Sau-1355-1 | NZ_KK006099.1 | - | 81276 | 81247 | **RNA**→ | rpmE2 (PRK01678)Ribosomal_L31 (pfam01197)→ |
| Sau-1800-1 | NZ_KI983038.1 | - | 36391 | 36362 | **RNA**→ | rpmE2 (PRK01678)Ribosomal_L31 (pfam01197)→ |
| Sau-1805-1 | NZ_KI983336.1 | - | 36390 | 36361 | **RNA**→ | rpmE2 (PRK01678)Ribosomal_L31 (pfam01197)→ |

| | | | | | |
|---|---|---|---|---|---|
| Sau-1377-1 | NZ_KK006414.1 | - | 113836 | 113807 | **RNA**→ rpmE2 (PRK01678)Ribosomal_L31 (pfam01197)→ |
| Sau-1190-1 | NZ_KI999369.1 | - | 35872 | 35843 | **RNA**→ rpmE2 (PRK01678)Ribosomal_L31 (pfam01197)→ |
| Sau-1340-1 | NZ_KK005535.1 | - | 9413 | 9384 | **RNA**→ rpmE2 (PRK01678)Ribosomal_L31 (pfam01197)→ |
| Sau-1379-1 | NZ_KK006465.1 | - | 36491 | 36462 | **RNA**→ rpmE2 (PRK01678)Ribosomal_L31 (pfam01197)→ |
| Sau-1783-1 | NZ_KI982243.1 | - | 35387 | 35358 | **RNA**→ rpmE2 (PRK01678)Ribosomal_L31 (pfam01197)→ |
| Sau-1347-1 | NZ_KK005768.1 | - | 35179 | 35150 | **RNA**→ rpmE2 (PRK01678)Ribosomal_L31 (pfam01197)→ |
| Sau-1789-1 | NZ_KI982539.1 | - | 47628 | 47599 | **RNA**→ rpmE2 (PRK01678)Ribosomal_L31 (pfam01197)→ |
| Sau-1790-1 | NZ_KI982579.1 | - | 55496 | 55467 | **RNA**→ rpmE2 (PRK01678)Ribosomal_L31 (pfam01197)→ |
| Sau-1795-1 | NZ_KI982920.1 | - | 107814 | 107785 | **RNA**→ rpmE2 (PRK01678)Ribosomal_L31 (pfam01197)→ |
| Sau-1801-1 | NZ_KI983087.1 | - | 11532 | 11503 | **RNA**→ rpmE2 (PRK01678)Ribosomal_L31 (pfam01197)→ |
| Sau-1802-1 | NZ_KI983209.1 | - | 61379 | 61350 | **RNA**→ rpmE2 (PRK01678)Ribosomal_L31 (pfam01197)→ |
| Sau-1806-1 | NZ_KI983445.1 | - | 25184 | 25155 | **RNA**→ rpmE2 (PRK01678)Ribosomal_L31 (pfam01197)→ |
| Sau-583-1 | NZ_KI966250.1 | - | 110430 | 110401 | **RNA**→ rpmE2 (PRK01678)Ribosomal_L31 (pfam01197)→ |
| Sau-1690-1 | NZ_KI967971.1 | - | 26596 | 26567 | **RNA**→ rpmE2 (PRK01678)Ribosomal_L31 (pfam01197)→ |
| Sau-1725-1 | NZ_KI970090.1 | - | 25227 | 25198 | **RNA**→ rpmE2 (PRK01678)Ribosomal_L31 (pfam01197)→ |
| Sau-1624-1 | NZ_KK017043.1 | - | 15812 | 15783 | **RNA**→ rpmE2 (PRK01678)Ribosomal_L31 (pfam01197)→ |
| Sau-1626-1 | NZ_KK017149.1 | - | 11533 | 11504 | **RNA**→ rpmE2 (PRK01678)Ribosomal_L31 (pfam01197)→ |
| Sau-892-1 | NZ_KI979231.1 | - | 113578 | 113549 | **RNA**→ rpmE2 (PRK01678)Ribosomal_L31 (pfam01197)→ |
| Sau-1751-1 | NZ_KI981591.1 | - | 81045 | 81016 | **RNA**→ rpmE2 (PRK01678)Ribosomal_L31 (pfam01197)→ |
| Sau-1650-1 | NZ_KK018407.1 | - | 62157 | 62128 | **RNA**→ rpmE2 (PRK01678)Ribosomal_L31 (pfam01197)→ |
| Sau-1781-1 | NZ_KI982144.1 | - | 36391 | 36362 | **RNA**→ rpmE2 (PRK01678)Ribosomal_L31 (pfam01197)→ |
| Sau-1593-1 | NZ_KK015493.1 | - | 62156 | 62127 | **RNA**→ rpmE2 (PRK01678)Ribosomal_L31 (pfam01197)→ |
| Sau-1787-1 | NZ_KI982431.1 | - | 54695 | 54666 | **RNA**→ rpmE2 (PRK01678)Ribosomal_L31 (pfam01197)→ |
| Sau-1617-1 | NZ_KK016736.1 | - | 96856 | 96827 | **RNA**→ rpmE2 (PRK01678)Ribosomal_L31 (pfam01197)→ |
| Sau-1633-1 | NZ_KK017532.1 | - | 35482 | 35453 | **RNA**→ rpmE2 (PRK01678)Ribosomal_L31 (pfam01197)→ |
| Sau-1792-1 | NZ_KI982725.1 | - | 56593 | 56564 | **RNA**→ rpmE2 (PRK01678)Ribosomal_L31 (pfam01197)→ |
| Sau-1799-1 | NZ_KI982988.1 | - | 24749 | 24720 | **RNA**→ rpmE2 (PRK01678)Ribosomal_L31 (pfam01197)→ |
| Sau-1804-1 | NZ_KI983259.1 | - | 25185 | 25156 | **RNA**→ rpmE2 (PRK01678)Ribosomal_L31 (pfam01197)→ |
| Sau-604-1 | NZ_KI966123.1 | - | 63210 | 63181 | **RNA**→ rpmE2 (PRK01678)Ribosomal_L31 (pfam01197)→ |
| Sau-599-1 | NZ_KI966182.1 | - | 61249 | 61220 | **RNA**→ rpmE2 (PRK01678)Ribosomal_L31 (pfam01197)→ |
| Sau-1597-1 | NZ_KK015613.1 | - | 28883 | 28854 | **RNA**→ rpmE2 (PRK01678)Ribosomal_L31 (pfam01197)→ |
| Sau-300-1 | NZ_KI978791.1 | - | 56893 | 56864 | **RNA**→ rpmE2 (PRK01678)Ribosomal_L31 (pfam01197)→ |
| Sau-889-1 | NZ_KI978849.1 | - | 56894 | 56865 | **RNA**→ rpmE2 (PRK01678)Ribosomal_L31 (pfam01197)→ |
| Sau-278-1 | NZ_KI979258.1 | - | 56995 | 56966 | **RNA**→ rpmE2 (PRK01678)Ribosomal_L31 (pfam01197)→ |
| Sau-1773-1 | NZ_KI981785.1 | - | 70324 | 70295 | **RNA**→ rpmE2 (PRK01678)Ribosomal_L31 (pfam01197)→ |
| Sau-1342-1 | NZ_KK018458.1 | - | 36391 | 36362 | **RNA**→ rpmE2 (PRK01678)Ribosomal_L31 (pfam01197)→ |
| Sau-1774-1 | NZ_KI981898.1 | - | 61468 | 61439 | **RNA**→ rpmE2 (PRK01678)Ribosomal_L31 (pfam01197)→ |
| Sau-1779-1 | NZ_KI982023.1 | - | 61894 | 61865 | **RNA**→ rpmE2 (PRK01678)Ribosomal_L31 (pfam01197)→ |
| Sau-1371-1 | NZ_KK018935.1 | - | 35170 | 35141 | **RNA**→ rpmE2 (PRK01678)Ribosomal_L31 (pfam01197)→ |
| Sau-1780-1 | NZ_KI982082.1 | - | 36320 | 36291 | **RNA**→ rpmE2 (PRK01678)Ribosomal_L31 (pfam01197)→ |
| Sau-619-1 | NZ_KK022635.1 | - | 34873 | 34844 | **RNA**→ rpmE2 (PRK01678)Ribosomal_L31 (pfam01197)→ |
| Sau-1785-1 | NZ_KI982294.1 | - | 24749 | 24720 | **RNA**→ rpmE2 (PRK01678)Ribosomal_L31 (pfam01197)→ |
| Sau-1786-1 | NZ_KI982353.1 | - | 36491 | 36462 | **RNA**→ rpmE2 (PRK01678)Ribosomal_L31 (pfam01197)→ |
| Sau-1791-1 | NZ_KI982675.1 | - | 25129 | 25100 | **RNA**→ rpmE2 (PRK01678)Ribosomal_L31 (pfam01197)→ |
| Sau-1579-1 | NZ_KK014783.1 | - | 36139 | 36110 | **RNA**→ rpmE2 (PRK01678)Ribosomal_L31 (pfam01197)→ |
| Sau-1797-1 | NZ_KI982953.1 | - | 49963 | 49934 | **RNA**→ rpmE2 (PRK01678)Ribosomal_L31 (pfam01197)→ |
| Sau-1638-1 | NZ_KK017770.1 | - | 36391 | 36362 | **RNA**→ rpmE2 (PRK01678)Ribosomal_L31 (pfam01197)→ |
| Sau-1803-1 | NZ_KI983216.1 | - | 35204 | 35175 | **RNA**→ rpmE2 (PRK01678)Ribosomal_L31 (pfam01197)→ |
| Sau-1809-1 | NZ_KI983518.1 | - | 50063 | 50034 | **RNA**→ rpmE2 (PRK01678)Ribosomal_L31 (pfam01197)→ |
| Sau-1649-1 | NZ_KK018347.1 | - | 62154 | 62125 | **RNA**→ rpmE2 (PRK01678)Ribosomal_L31 (pfam01197)→ |
| Sau-1374-1 | NZ_KK019025.1 | - | 16360 | 16331 | **RNA**→ rpmE2 (PRK01678)Ribosomal_L31 (pfam01197)→ |
| Sau-1759-1 | NZ_KI991973.1 | - | 36391 | 36362 | **RNA**→ rpmE2 (PRK01678)Ribosomal_L31 (pfam01197)→ |

| | | | | | |
|---|---|---|---|---|---|
| Sau-1753-1 | NZ_KI991857.1 | - | 61333 | 61304 | **RNA**→ rpmE2 (PRK01678)Ribosomal_L31 (pfam01197)→ |
| Sau-3525-1 | NZ_KK021290.1 | - | 34873 | 34844 | **RNA**→ rpmE2 (PRK01678)Ribosomal_L31 (pfam01197)→ |
| Sau-895-1 | NZ_KK022550.1 | - | 34874 | 34845 | **RNA**→ rpmE2 (PRK01678)Ribosomal_L31 (pfam01197)→ |
| Sau-1599-1 | NZ_KK015745.1 | - | 36750 | 36721 | **RNA**→ rpmE2 (PRK01678)Ribosomal_L31 (pfam01197)→ |
| Sau-1623-1 | NZ_KK017004.1 | - | 107853 | 107824 | **RNA**→ rpmE2 (PRK01678)Ribosomal_L31 (pfam01197)→ |
| Sau-1629-1 | NZ_KK017303.1 | - | 35334 | 35305 | **RNA**→ rpmE2 (PRK01678)Ribosomal_L31 (pfam01197)→ |
| Sau-1645-1 | NZ_KK018155.1 | - | 62156 | 62127 | **RNA**→ rpmE2 (PRK01678)Ribosomal_L31 (pfam01197)→ |
| Sau-678-1 | NZ_KK021099.1 | - | 34874 | 34845 | **RNA**→ rpmE2 (PRK01678)Ribosomal_L31 (pfam01197)→ |
| Sau-3551-1 | NZ_KK021976.1 | - | 25255 | 25226 | **RNA**→ rpmE2 (PRK01678)Ribosomal_L31 (pfam01197)→ |
| Sau-1600-1 | NZ_KK015795.1 | - | 35719 | 35690 | **RNA**→ rpmE2 (PRK01678)Ribosomal_L31 (pfam01197)→ |
| Sau-2457-1 | NZ_KK022793.1 | - | 123801 | 123772 | **RNA**→ rpmE2 (PRK01678)Ribosomal_L31 (pfam01197)→ |
| Sau-2949-1 | NZ_KK022818.1 | - | 56940 | 56911 | **RNA**→ rpmE2 (PRK01678)Ribosomal_L31 (pfam01197)→ |
| Sau-2121-1 | NZ_KK066900.1 | - | 62105 | 62076 | **RNA**→ rpmE2 (PRK01678)Ribosomal_L31 (pfam01197)→ |
| Sau-2115-1 | NZ_KK067042.1 | - | 111347 | 111318 | **RNA**→ rpmE2 (PRK01678)Ribosomal_L31 (pfam01197)→ |
| Sau-2106-1 | NZ_KK067270.1 | - | 36390 | 36361 | **RNA**→ rpmE2 (PRK01678)Ribosomal_L31 (pfam01197)→ |
| Sau-2146-1 | NZ_KK065789.1 | - | 36748 | 36719 | **RNA**→ rpmE2 (PRK01678)Ribosomal_L31 (pfam01197)→ |
| Sau-2124-1 | NZ_KK066757.1 | - | 35379 | 35350 | **RNA**→ rpmE2 (PRK01678)Ribosomal_L31 (pfam01197)→ |
| Sau-2110-1 | NZ_KK067172.1 | - | 35461 | 35432 | **RNA**→ rpmE2 (PRK01678)Ribosomal_L31 (pfam01197)→ |
| Sau-2154-1 | NZ_KK065505.1 | - | 61121 | 61092 | **RNA**→ rpmE2 (PRK01678)Ribosomal_L31 (pfam01197)→ |
| Sau-2149-1 | NZ_KK065701.1 | - | 54674 | 54645 | **RNA**→ rpmE2 (PRK01678)Ribosomal_L31 (pfam01197)→ |
| Sau-2143-1 | NZ_KK065925.1 | - | 36181 | 36152 | **RNA**→ rpmE2 (PRK01678)Ribosomal_L31 (pfam01197)→ |
| Sau-2093-1 | NZ_KK067569.1 | - | 24083 | 24054 | **RNA**→ rpmE2 (PRK01678)Ribosomal_L31 (pfam01197)→ |
| Sau-2128-1 | NZ_KK066638.1 | - | 60124 | 60095 | **RNA**→ rpmE2 (PRK01678)Ribosomal_L31 (pfam01197)→ |
| Sau-681-1 | NZ_KK020994.1 | - | 34816 | 34787 | **RNA**→ rpmE2 (PRK01678)Ribosomal_L31 (pfam01197)→ |
| Sau-699-1 | NZ_KK023877.1 | - | 33076 | 33047 | **RNA**→ rpmE2 (PRK01678)Ribosomal_L31 (pfam01197)→ |
| Sau-800-1 | NZ_KK027143.1 | - | 34869 | 34840 | **RNA**→ rpmE2 (PRK01678)Ribosomal_L31 (pfam01197)→ |
| Sau-824-1 | NZ_KK027509.1 | - | 34873 | 34844 | **RNA**→ rpmE2 (PRK01678)Ribosomal_L31 (pfam01197)→ |
| Sau-664-1 | NZ_KK029115.1 | - | 34873 | 34844 | **RNA**→ rpmE2 (PRK01678)Ribosomal_L31 (pfam01197)→ |
| Sau-667-1 | NZ_KK029215.1 | - | 34863 | 34834 | **RNA**→ rpmE2 (PRK01678)Ribosomal_L31 (pfam01197)→ |
| Sau-757-1 | NZ_KK029350.1 | - | 34872 | 34843 | **RNA**→ rpmE2 (PRK01678)Ribosomal_L31 (pfam01197)→ |
| Sau-785-1 | NZ_KK041807.1 | - | 34873 | 34844 | **RNA**→ rpmE2 (PRK01678)Ribosomal_L31 (pfam01197)→ |
| Sau-705-1 | NZ_KK021611.1 | - | 34873 | 34844 | **RNA**→ rpmE2 (PRK01678)Ribosomal_L31 (pfam01197)→ |
| Sau-693-1 | NZ_KK023840.1 | - | 2239382 | 2239353 | **RNA**→ rpmE2 (PRK01678)Ribosomal_L31 (pfam01197)→ |
| Sau-755-1 | NZ_KK025445.1 | - | 34873 | 34844 | **RNA**→ rpmE2 (PRK01678)Ribosomal_L31 (pfam01197)→ |
| Sau-762-1 | NZ_KK025890.1 | + | 18543 | 18572 | **RNA**→ rpmE2 (PRK01678)Ribosomal_L31 (pfam01197)→ |
| Sau-770-1 | NZ_KK026102.1 | - | 56893 | 56864 | **RNA**→ rpmE2 (PRK01678)Ribosomal_L31 (pfam01197)→ |
| Sau-790-1 | NZ_KK026722.1 | - | 34872 | 34843 | **RNA**→ rpmE2 (PRK01678)Ribosomal_L31 (pfam01197)→ |
| Sau-801-1 | NZ_KK027198.1 | - | 32356 | 32327 | **RNA**→ rpmE2 (PRK01678)Ribosomal_L31 (pfam01197)→ |
| Sau-825-1 | NZ_KK027558.1 | - | 34817 | 34788 | **RNA**→ rpmE2 (PRK01678)Ribosomal_L31 (pfam01197)→ |
| Sau-840-1 | NZ_KK027859.1 | - | 56502 | 56473 | **RNA**→ rpmE2 (PRK01678)Ribosomal_L31 (pfam01197)→ |
| Sau-783-1 | NZ_KK029395.1 | - | 54878 | 54849 | **RNA**→ rpmE2 (PRK01678)Ribosomal_L31 (pfam01197)→ |
| Sau-689-1 | NZ_KK039752.1 | - | 34817 | 34788 | **RNA**→ rpmE2 (PRK01678)Ribosomal_L31 (pfam01197)→ |
| Sau-782-1 | NZ_KK041853.1 | - | 34864 | 34835 | **RNA**→ rpmE2 (PRK01678)Ribosomal_L31 (pfam01197)→ |
| Sau-732-1 | NZ_KK024821.1 | - | 57040 | 57011 | **RNA**→ rpmE2 (PRK01678)Ribosomal_L31 (pfam01197)→ |
| Sau-739-1 | NZ_KK024990.1 | - | 34872 | 34843 | **RNA**→ rpmE2 (PRK01678)Ribosomal_L31 (pfam01197)→ |
| Sau-743-1 | NZ_KK029255.1 | - | 34730 | 34701 | **RNA**→ rpmE2 (PRK01678)Ribosomal_L31 (pfam01197)→ |
| Sau-2934-1 | NZ_KK021933.1 | - | 54878 | 54849 | **RNA**→ rpmE2 (PRK01678)Ribosomal_L31 (pfam01197)→ |
| Sau-2174-1 | NZ_KK024197.1 | - | 34873 | 34844 | **RNA**→ rpmE2 (PRK01678)Ribosomal_L31 (pfam01197)→ |
| Sau-2869-1 | NZ_KK023742.1 | - | 34872 | 34843 | **RNA**→ rpmE2 (PRK01678)Ribosomal_L31 (pfam01197)→ |
| Sau-2460-1 | NZ_KK025154.1 | - | 34872 | 34843 | **RNA**→ rpmE2 (PRK01678)Ribosomal_L31 (pfam01197)→ |
| Sau-2942-1 | NZ_KK024911.1 | - | 34864 | 34835 | **RNA**→ rpmE2 (PRK01678)Ribosomal_L31 (pfam01197)→ |
| Sau-2989-1 | NZ_KK026829.1 | - | 34872 | 34843 | **RNA**→ rpmE2 (PRK01678)Ribosomal_L31 (pfam01197)→ |

| | | | | | | |
|---|---|---|---|---|---|---|
| Sau-2988-1 | NZ_KK026878.1 | - | 34872 | 34843 | **RNA**→ | rpmE2 (PRK01678)Ribosomal_L31 (pfam01197)→ |
| Sau-2231-1 | NZ_KK062836.1 | - | 101112 | 101083 | **RNA**→ | rpmE2 (PRK01678)Ribosomal_L31 (pfam01197)→ |
| Sau-2966-1 | NZ_KK028360.1 | - | 34873 | 34844 | **RNA**→ | rpmE2 (PRK01678)Ribosomal_L31 (pfam01197)→ |
| Sau-2224-1 | NZ_KK063022.1 | - | 15595 | 15566 | **RNA**→ | rpmE2 (PRK01678)Ribosomal_L31 (pfam01197)→ |
| Sau-2219-1 | NZ_KK063290.1 | - | 115797 | 115768 | **RNA**→ | rpmE2 (PRK01678)Ribosomal_L31 (pfam01197)→ |
| Sau-2878-1 | NZ_KK031157.1 | - | 56894 | 56865 | **RNA**→ | rpmE2 (PRK01678)Ribosomal_L31 (pfam01197)→ |
| Sau-2956-1 | NZ_KK031325.1 | - | 30759 | 30730 | **RNA**→ | rpmE2 (PRK01678)Ribosomal_L31 (pfam01197)→ |
| Sau-2218-1 | NZ_KK063294.1 | - | 36281 | 36252 | **RNA**→ | rpmE2 (PRK01678)Ribosomal_L31 (pfam01197)→ |
| Sau-907-1 | NZ_KK024505.1 | - | 34817 | 34788 | **RNA**→ | rpmE2 (PRK01678)Ribosomal_L31 (pfam01197)→ |
| Sau-2968-1 | NZ_KK031580.1 | - | 35073 | 35044 | **RNA**→ | rpmE2 (PRK01678)Ribosomal_L31 (pfam01197)→ |
| Sau-931-1 | NZ_KK025407.1 | - | 34874 | 34845 | **RNA**→ | rpmE2 (PRK01678)Ribosomal_L31 (pfam01197)→ |
| Sau-2212-1 | NZ_KK063495.1 | - | 49963 | 49934 | **RNA**→ | rpmE2 (PRK01678)Ribosomal_L31 (pfam01197)→ |
| Sau-2991-1 | NZ_KK031859.1 | - | 30812 | 30783 | **RNA**→ | rpmE2 (PRK01678)Ribosomal_L31 (pfam01197)→ |
| Sau-947-1 | NZ_KK025832.1 | - | 25029 | 25000 | **RNA**→ | rpmE2 (PRK01678)Ribosomal_L31 (pfam01197)→ |
| Sau-3492-1 | NZ_KK032127.1 | - | 34872 | 34843 | **RNA**→ | rpmE2 (PRK01678)Ribosomal_L31 (pfam01197)→ |
| Sau-936-1 | NZ_KK026316.1 | - | 34872 | 34843 | **RNA**→ | rpmE2 (PRK01678)Ribosomal_L31 (pfam01197)→ |
| Sau-2201-1 | NZ_KK063757.1 | - | 61749 | 61720 | **RNA**→ | rpmE2 (PRK01678)Ribosomal_L31 (pfam01197)→ |
| Sau-2940-1 | NZ_KK022765.1 | - | 121855 | 121826 | **RNA**→ | rpmE2 (PRK01678)Ribosomal_L31 (pfam01197)→ |
| Sau-2194-1 | NZ_KK063959.1 | - | 36391 | 36362 | **RNA**→ | rpmE2 (PRK01678)Ribosomal_L31 (pfam01197)→ |
| Sau-2963-1 | NZ_KK025704.1 | - | 56894 | 56865 | **RNA**→ | rpmE2 (PRK01678)Ribosomal_L31 (pfam01197)→ |
| Sau-966-1 | NZ_KK026630.1 | - | 56502 | 56473 | **RNA**→ | rpmE2 (PRK01678)Ribosomal_L31 (pfam01197)→ |
| Sau-2990-1 | NZ_KK026950.1 | - | 34872 | 34843 | **RNA**→ | rpmE2 (PRK01678)Ribosomal_L31 (pfam01197)→ |
| Sau-976-1 | NZ_KK027088.1 | - | 56976 | 56947 | **RNA**→ | rpmE2 (PRK01678)Ribosomal_L31 (pfam01197)→ |
| Sau-2188-1 | NZ_KK064220.1 | - | 35462 | 35433 | **RNA**→ | rpmE2 (PRK01678)Ribosomal_L31 (pfam01197)→ |
| Sau-988-1 | NZ_KK027269.1 | - | 25256 | 25227 | **RNA**→ | rpmE2 (PRK01678)Ribosomal_L31 (pfam01197)→ |
| Sau-2896-1 | NZ_KK031188.1 | - | 57033 | 57004 | **RNA**→ | rpmE2 (PRK01678)Ribosomal_L31 (pfam01197)→ |
| Sau-932-1 | NZ_KK029535.1 | - | 34872 | 34843 | **RNA**→ | rpmE2 (PRK01678)Ribosomal_L31 (pfam01197)→ |
| Sau-2897-1 | NZ_KK031197.1 | - | 34872 | 34843 | **RNA**→ | rpmE2 (PRK01678)Ribosomal_L31 (pfam01197)→ |
| Sau-2181-1 | NZ_KK064488.1 | - | 108622 | 108593 | **RNA**→ | rpmE2 (PRK01678)Ribosomal_L31 (pfam01197)→ |
| Sau-950-1 | NZ_KK029765.1 | - | 34913 | 34884 | **RNA**→ | rpmE2 (PRK01678)Ribosomal_L31 (pfam01197)→ |
| Sau-2957-1 | NZ_KK031395.1 | - | 34872 | 34843 | **RNA**→ | rpmE2 (PRK01678)Ribosomal_L31 (pfam01197)→ |
| Sau-1005-1 | NZ_KK030022.1 | - | 34817 | 34788 | **RNA**→ | rpmE2 (PRK01678)Ribosomal_L31 (pfam01197)→ |
| Sau-2975-1 | NZ_KK031628.1 | - | 34817 | 34788 | **RNA**→ | rpmE2 (PRK01678)Ribosomal_L31 (pfam01197)→ |
| Sau-2176-1 | NZ_KK064706.1 | - | 54695 | 54666 | **RNA**→ | rpmE2 (PRK01678)Ribosomal_L31 (pfam01197)→ |
| Sau-2976-1 | NZ_KK031665.1 | - | 34873 | 34844 | **RNA**→ | rpmE2 (PRK01678)Ribosomal_L31 (pfam01197)→ |
| Sau-858-1 | NZ_KK047717.1 | - | 1678825 | 1678796 | **RNA**→ | rpmE2 (PRK01678)Ribosomal_L31 (pfam01197)→ |
| Sau-900-1 | NZ_KK023995.1 | - | 34872 | 34843 | **RNA**→ | rpmE2 (PRK01678)Ribosomal_L31 (pfam01197)→ |
| Sau-2992-1 | NZ_KK031903.1 | - | 34772 | 34743 | **RNA**→ | rpmE2 (PRK01678)Ribosomal_L31 (pfam01197)→ |
| Sau-2167-1 | NZ_KK064958.1 | - | 36391 | 36362 | **RNA**→ | rpmE2 (PRK01678)Ribosomal_L31 (pfam01197)→ |
| Sau-3493-1 | NZ_KK032159.1 | - | 34817 | 34788 | **RNA**→ | rpmE2 (PRK01678)Ribosomal_L31 (pfam01197)→ |
| Sau-905-1 | NZ_KK024562.1 | - | 34873 | 34844 | **RNA**→ | rpmE2 (PRK01678)Ribosomal_L31 (pfam01197)→ |
| Sau-1622-1 | NZ_KK016918.1 | - | 36391 | 36362 | **RNA**→ | rpmE2 (PRK01678)Ribosomal_L31 (pfam01197)→ |
| Sau-2914-1 | NZ_KK023853.1 | - | 56893 | 56864 | **RNA**→ | rpmE2 (PRK01678)Ribosomal_L31 (pfam01197)→ |
| Sau-854-1 | NZ_KK047726.1 | - | 56893 | 56864 | **RNA**→ | rpmE2 (PRK01678)Ribosomal_L31 (pfam01197)→ |
| Sau-2964-1 | NZ_KK025714.1 | - | 34872 | 34843 | **RNA**→ | rpmE2 (PRK01678)Ribosomal_L31 (pfam01197)→ |
| Sau-860-1 | NZ_KK019362.1 | - | 34872 | 34843 | **RNA**→ | rpmE2 (PRK01678)Ribosomal_L31 (pfam01197)→ |
| Sau-1634-1 | NZ_KK017640.1 | - | 36390 | 36361 | **RNA**→ | rpmE2 (PRK01678)Ribosomal_L31 (pfam01197)→ |
| Sau-2918-1 | NZ_KK028274.1 | - | 34961 | 34932 | **RNA**→ | rpmE2 (PRK01678)Ribosomal_L31 (pfam01197)→ |
| Sau-929-1 | NZ_KK025290.1 | - | 34873 | 34844 | **RNA**→ | rpmE2 (PRK01678)Ribosomal_L31 (pfam01197)→ |
| Sau-2902-1 | NZ_KK031248.1 | - | 34846 | 34817 | **RNA**→ | rpmE2 (PRK01678)Ribosomal_L31 (pfam01197)→ |
| Sau-948-1 | NZ_KK025784.1 | - | 56893 | 56864 | **RNA**→ | rpmE2 (PRK01678)Ribosomal_L31 (pfam01197)→ |
| Sau-2234-1 | NZ_KK047930.1 | - | 43687 | 43658 | **RNA**→ | rpmE2 (PRK01678)Ribosomal_L31 (pfam01197)→ |

| | | | | | |
|---|---|---|---|---|---|
| Sau-2959-1 | NZ_KK031455.1 | - | 34872 | 34843 | **RNA**→ rpmE2 (PRK01678)Ribosomal_L31 (pfam01197)→ |
| Sau-2977-1 | NZ_KK031711.1 | - | 56894 | 56865 | **RNA**→ rpmE2 (PRK01678)Ribosomal_L31 (pfam01197)→ |
| Sau-2228-1 | NZ_KK062861.1 | - | 35170 | 35141 | **RNA**→ rpmE2 (PRK01678)Ribosomal_L31 (pfam01197)→ |
| Sau-985-1 | NZ_KK028757.1 | - | 19182 | 19153 | **RNA**→ rpmE2 (PRK01678)Ribosomal_L31 (pfam01197)→ |
| Sau-2994-1 | NZ_KK032014.1 | - | 34873 | 34844 | **RNA**→ rpmE2 (PRK01678)Ribosomal_L31 (pfam01197)→ |
| Sau-940-1 | NZ_KK029729.1 | - | 34773 | 34744 | **RNA**→ rpmE2 (PRK01678)Ribosomal_L31 (pfam01197)→ |
| Sau-2227-1 | NZ_KK062912.1 | - | 36391 | 36362 | **RNA**→ rpmE2 (PRK01678)Ribosomal_L31 (pfam01197)→ |
| Sau-2917-1 | NZ_KK021166.1 | - | 34773 | 34744 | **RNA**→ rpmE2 (PRK01678)Ribosomal_L31 (pfam01197)→ |
| Sau-1018-1 | NZ_KK030137.1 | - | 34872 | 34843 | **RNA**→ rpmE2 (PRK01678)Ribosomal_L31 (pfam01197)→ |
| Sau-2223-1 | NZ_KK063053.1 | - | 36390 | 36361 | **RNA**→ rpmE2 (PRK01678)Ribosomal_L31 (pfam01197)→ |
| Sau-848-1 | NZ_KK019147.1 | - | 59010 | 58981 | **RNA**→ rpmE2 (PRK01678)Ribosomal_L31 (pfam01197)→ |
| Sau-2882-1 | NZ_KK023322.1 | - | 56894 | 56865 | **RNA**→ rpmE2 (PRK01678)Ribosomal_L31 (pfam01197)→ |
| Sau-2943-1 | NZ_KK024836.1 | - | 34872 | 34843 | **RNA**→ rpmE2 (PRK01678)Ribosomal_L31 (pfam01197)→ |
| Sau-2216-1 | NZ_KK063340.1 | - | 64842 | 64813 | **RNA**→ rpmE2 (PRK01678)Ribosomal_L31 (pfam01197)→ |
| Sau-906-1 | NZ_KK024727.1 | - | 34874 | 34845 | **RNA**→ rpmE2 (PRK01678)Ribosomal_L31 (pfam01197)→ |
| Sau-2999-1 | NZ_KK027403.1 | - | 34872 | 34843 | **RNA**→ rpmE2 (PRK01678)Ribosomal_L31 (pfam01197)→ |
| Sau-949-1 | NZ_KK025817.1 | - | 56941 | 56912 | **RNA**→ rpmE2 (PRK01678)Ribosomal_L31 (pfam01197)→ |
| Sau-2445-1 | NZ_KK030612.1 | - | 34862 | 34833 | **RNA**→ rpmE2 (PRK01678)Ribosomal_L31 (pfam01197)→ |
| Sau-2877-1 | NZ_KK031078.1 | - | 34873 | 34844 | **RNA**→ rpmE2 (PRK01678)Ribosomal_L31 (pfam01197)→ |
| Sau-935-1 | NZ_KK026006.1 | - | 34863 | 34834 | **RNA**→ rpmE2 (PRK01678)Ribosomal_L31 (pfam01197)→ |
| Sau-2941-1 | NZ_KK031298.1 | - | 56894 | 56865 | **RNA**→ rpmE2 (PRK01678)Ribosomal_L31 (pfam01197)→ |
| Sau-2233-1 | NZ_KK047970.1 | - | 36391 | 36362 | **RNA**→ rpmE2 (PRK01678)Ribosomal_L31 (pfam01197)→ |
| Sau-965-1 | NZ_KK026604.1 | - | 56894 | 56865 | **RNA**→ rpmE2 (PRK01678)Ribosomal_L31 (pfam01197)→ |
| Sau-2981-1 | NZ_KK031750.1 | - | 34873 | 34844 | **RNA**→ rpmE2 (PRK01678)Ribosomal_L31 (pfam01197)→ |
| Sau-923-1 | NZ_KK028350.1 | - | 57199 | 57170 | **RNA**→ rpmE2 (PRK01678)Ribosomal_L31 (pfam01197)→ |
| Sau-2226-1 | NZ_KK062953.1 | - | 96895 | 96866 | **RNA**→ rpmE2 (PRK01678)Ribosomal_L31 (pfam01197)→ |
| Sau-986-1 | NZ_KK028808.1 | - | 19182 | 19153 | **RNA**→ rpmE2 (PRK01678)Ribosomal_L31 (pfam01197)→ |
| Sau-3039-1 | NZ_KK019290.1 | - | 54764 | 54735 | **RNA**→ rpmE2 (PRK01678)Ribosomal_L31 (pfam01197)→ |
| Sau-2221-1 | NZ_KK063145.1 | - | 49861 | 49832 | **RNA**→ rpmE2 (PRK01678)Ribosomal_L31 (pfam01197)→ |
| Sau-925-1 | NZ_KK029504.1 | - | 25062 | 25033 | **RNA**→ rpmE2 (PRK01678)Ribosomal_L31 (pfam01197)→ |
| Sau-2930-1 | NZ_KK021792.1 | - | 56894 | 56865 | **RNA**→ rpmE2 (PRK01678)Ribosomal_L31 (pfam01197)→ |
| Sau-1003-1 | NZ_KK029896.1 | - | 34874 | 34845 | **RNA**→ rpmE2 (PRK01678)Ribosomal_L31 (pfam01197)→ |
| Sau-2215-1 | NZ_KK063417.1 | - | 111834 | 111805 | **RNA**→ rpmE2 (PRK01678)Ribosomal_L31 (pfam01197)→ |
| Sau-2962-1 | NZ_KK025626.1 | - | 56894 | 56865 | **RNA**→ rpmE2 (PRK01678)Ribosomal_L31 (pfam01197)→ |
| Sau-1019-1 | NZ_KK030205.1 | - | 56894 | 56865 | **RNA**→ rpmE2 (PRK01678)Ribosomal_L31 (pfam01197)→ |
| Sau-2205-1 | NZ_KK063588.1 | - | 36391 | 36362 | **RNA**→ rpmE2 (PRK01678)Ribosomal_L31 (pfam01197)→ |
| Sau-2983-1 | NZ_KK031848.1 | - | 34817 | 34788 | **RNA**→ rpmE2 (PRK01678)Ribosomal_L31 (pfam01197)→ |
| Sau-851-1 | NZ_KK047742.1 | - | 1679220 | 1679191 | **RNA**→ rpmE2 (PRK01678)Ribosomal_L31 (pfam01197)→ |
| Sau-2993-1 | NZ_KK031941.1 | - | 34872 | 34843 | **RNA**→ rpmE2 (PRK01678)Ribosomal_L31 (pfam01197)→ |
| Sau-997-1 | NZ_KK072945.1 | - | 34874 | 34845 | **RNA**→ rpmE2 (PRK01678)Ribosomal_L31 (pfam01197)→ |
| Sau-2185-1 | NZ_KK064346.1 | + | 117742 | 117771 | **RNA**→ rpmE2 (PRK01678)Ribosomal_L31 (pfam01197)→ |
| Sau-3491-1 | NZ_KK032093.1 | - | 2229537 | 2229508 | **RNA**→ rpmE2 (PRK01678)Ribosomal_L31 (pfam01197)→ |
| Sau-918-1 | NZ_KK024778.1 | - | 2236157 | 2236128 | **RNA**→ rpmE2 (PRK01678)Ribosomal_L31 (pfam01197)→ |
| Sau-2958-1 | NZ_KK031432.1 | - | 56894 | 56865 | **RNA**→ rpmE2 (PRK01678)Ribosomal_L31 (pfam01197)→ |
| Sau-958-1 | NZ_KK029827.1 | - | 34817 | 34788 | **RNA**→ rpmE2 (PRK01678)Ribosomal_L31 (pfam01197)→ |
| Sau-2179-1 | NZ_KK064605.1 | - | 59118 | 59089 | **RNA**→ rpmE2 (PRK01678)Ribosomal_L31 (pfam01197)→ |
| Sau-3036-1 | NZ_KK019241.1 | - | 56894 | 56865 | **RNA**→ rpmE2 (PRK01678)Ribosomal_L31 (pfam01197)→ |
| Sau-1039-1 | NZ_KK030281.1 | - | 2238815 | 2238786 | **RNA**→ rpmE2 (PRK01678)Ribosomal_L31 (pfam01197)→ |
| Sau-1625-1 | NZ_KK017092.1 | - | 36490 | 36461 | **RNA**→ rpmE2 (PRK01678)Ribosomal_L31 (pfam01197)→ |
| Sau-2881-1 | NZ_KK023296.1 | - | 56894 | 56865 | **RNA**→ rpmE2 (PRK01678)Ribosomal_L31 (pfam01197)→ |
| Sau-917-1 | NZ_KK024797.1 | - | 56893 | 56864 | **RNA**→ rpmE2 (PRK01678)Ribosomal_L31 (pfam01197)→ |
| Sau-2982-1 | NZ_KK027319.1 | - | 34872 | 34843 | **RNA**→ rpmE2 (PRK01678)Ribosomal_L31 (pfam01197)→ |

| | | | | | |
|---|---|---|---|---|---|
| Sau-987-1 | NZ_KK027315.1 | - | 56939 | 56910 | **RNA**→ rpmE2 (PRK01678)Ribosomal_L31 (pfam01197)→ |
| Sau-1631-1 | NZ_KK017466.1 | - | 62257 | 62228 | **RNA**→ rpmE2 (PRK01678)Ribosomal_L31 (pfam01197)→ |
| Sau-2899-1 | NZ_KK023472.1 | - | 34872 | 34843 | **RNA**→ rpmE2 (PRK01678)Ribosomal_L31 (pfam01197)→ |
| Sau-959-1 | NZ_KK029882.1 | - | 2252120 | 2252091 | **RNA**→ rpmE2 (PRK01678)Ribosomal_L31 (pfam01197)→ |
| Sau-75-1 | NZ_CABA01000062.1 | - | 16773 | 16744 | **RNA**→ rpmE2 (PRK01678)Ribosomal_L31 (pfam01197)→ rpmE2 (PRK01678)Ribosomal_L31 (pfam01197 |
| Sau-2998-1 | NZ_KK027252.1 | - | 2235915 | 2235886 | **RNA**→ rpmE2 (PRK01678)Ribosomal_L31 (pfam01197)→ |
| Sau-847-1 | NZ_KK019119.1 | - | 56794 | 56765 | **RNA**→ rpmE2 (PRK01678)Ribosomal_L31 (pfam01197)→ |
| Sau-2444-1 | NZ_KK024715.1 | - | 34873 | 34844 | **RNA**→ rpmE2 (PRK01678)Ribosomal_L31 (pfam01197)→ |
| Sau-3000-1 | NZ_KK027444.1 | - | 34872 | 34843 | **RNA**→ rpmE2 (PRK01678)Ribosomal_L31 (pfam01197)→ |
| Sau-1001-1 | NZ_KK022887.1 | - | 34872 | 34843 | **RNA**→ rpmE2 (PRK01678)Ribosomal_L31 (pfam01197)→ |
| Sau-2459-1 | NZ_KK025101.1 | - | 34817 | 34788 | **RNA**→ rpmE2 (PRK01678)Ribosomal_L31 (pfam01197)→ |
| Sau-2967-1 | NZ_KK031544.1 | - | 35471 | 35442 | **RNA**→ rpmE2 (PRK01678)Ribosomal_L31 (pfam01197)→ |
| Sau-922-1 | NZ_KK028309.1 | - | 56894 | 56865 | **RNA**→ rpmE2 (PRK01678)Ribosomal_L31 (pfam01197)→ |
| Sau-3484-1 | NZ_KK032086.1 | - | 56894 | 56865 | **RNA**→ rpmE2 (PRK01678)Ribosomal_L31 (pfam01197)→ |
| Sau-2470-1 | NZ_KK025595.1 | - | 34872 | 34843 | **RNA**→ rpmE2 (PRK01678)Ribosomal_L31 (pfam01197)→ |
| Sau-924-1 | NZ_KK029438.1 | - | 25255 | 25226 | **RNA**→ rpmE2 (PRK01678)Ribosomal_L31 (pfam01197)→ |
| Sau-960-1 | NZ_KK029884.1 | - | 2237296 | 2237267 | **RNA**→ rpmE2 (PRK01678)Ribosomal_L31 (pfam01197)→ |
| Sau-2232-1 | NZ_KK048054.1 | - | 101158 | 101129 | **RNA**→ rpmE2 (PRK01678)Ribosomal_L31 (pfam01197)→ |
| Sau-849-1 | NZ_KK019137.1 | - | 36391 | 36362 | **RNA**→ rpmE2 (PRK01678)Ribosomal_L31 (pfam01197)→ |
| Sau-913-1 | NZ_KK022144.1 | - | 55982 | 55953 | **RNA**→ rpmE2 (PRK01678)Ribosomal_L31 (pfam01197)→ |
| Sau-2225-1 | NZ_KK063004.1 | - | 28766 | 28737 | **RNA**→ rpmE2 (PRK01678)Ribosomal_L31 (pfam01197)→ |
| Sau-904-1 | NZ_KK024427.1 | - | 34817 | 34788 | **RNA**→ rpmE2 (PRK01678)Ribosomal_L31 (pfam01197)→ |
| Sau-2214-1 | NZ_KK063464.1 | - | 26628 | 26599 | **RNA**→ rpmE2 (PRK01678)Ribosomal_L31 (pfam01197)→ |
| Sau-977-1 | NZ_KK027082.1 | - | 56894 | 56865 | **RNA**→ rpmE2 (PRK01678)Ribosomal_L31 (pfam01197)→ |
| Sau-998-1 | NZ_KK027716.1 | - | 34873 | 34844 | **RNA**→ rpmE2 (PRK01678)Ribosomal_L31 (pfam01197)→ |
| Sau-2052-1 | NZ_KK069221.1 | - | 36391 | 36362 | **RNA**→ rpmE2 (PRK01678)Ribosomal_L31 (pfam01197)→ |
| Sau-941-1 | NZ_KK029756.1 | - | 2237277 | 2237248 | **RNA**→ rpmE2 (PRK01678)Ribosomal_L31 (pfam01197)→ |
| Sau-1004-1 | NZ_KK029957.1 | - | 34817 | 34788 | **RNA**→ rpmE2 (PRK01678)Ribosomal_L31 (pfam01197)→ |
| Sau-1616-1 | NZ_KK016624.1 | - | 36391 | 36362 | **RNA**→ rpmE2 (PRK01678)Ribosomal_L31 (pfam01197)→ |
| Sau-2175-1 | NZ_KK024257.1 | - | 34872 | 34843 | **RNA**→ rpmE2 (PRK01678)Ribosomal_L31 (pfam01197)→ |
| Sau-2569-1 | NZ_KK038922.1 | - | 56895 | 56866 | **RNA**→ rpmE2 (PRK01678)Ribosomal_L31 (pfam01197)→ |
| Sau-1640-1 | NZ_KK017841.1 | - | 28307 | 28278 | **RNA**→ rpmE2 (PRK01678)Ribosomal_L31 (pfam01197)→ |
| Sau-1641-1 | NZ_KK017902.1 | - | 2421 | 2392 | **RNA**→ rpmE2 (PRK01678)Ribosomal_L31 (pfam01197)→ |
| Sau-2584-1 | NZ_KK020717.1 | - | 34869 | 34840 | **RNA**→ rpmE2 (PRK01678)Ribosomal_L31 (pfam01197)→ |
| Sau-1646-1 | NZ_KK018231.1 | - | 2421 | 2392 | **RNA**→ rpmE2 (PRK01678)Ribosomal_L31 (pfam01197)→ |
| Sau-2863-1 | NZ_KK023001.1 | - | 34817 | 34788 | **RNA**→ rpmE2 (PRK01678)Ribosomal_L31 (pfam01197)→ |
| Sau-1580-1 | NZ_KK014893.1 | - | 62256 | 62227 | **RNA**→ rpmE2 (PRK01678)Ribosomal_L31 (pfam01197)→ |
| Sau-2868-1 | NZ_KK023547.1 | - | 34872 | 34843 | **RNA**→ rpmE2 (PRK01678)Ribosomal_L31 (pfam01197)→ |
| Sau-2463-1 | NZ_KK025514.1 | - | 56894 | 56865 | **RNA**→ rpmE2 (PRK01678)Ribosomal_L31 (pfam01197)→ |
| Sau-2581-1 | NZ_KK023799.1 | - | 34817 | 34788 | **RNA**→ rpmE2 (PRK01678)Ribosomal_L31 (pfam01197)→ |
| Sau-1586-1 | NZ_KK015113.1 | - | 61277 | 61248 | **RNA**→ rpmE2 (PRK01678)Ribosomal_L31 (pfam01197)→ |
| Sau-2462-1 | NZ_KK025523.1 | - | 34873 | 34844 | **RNA**→ rpmE2 (PRK01678)Ribosomal_L31 (pfam01197)→ |
| Sau-2841-1 | NZ_KK028984.1 | - | 34862 | 34833 | **RNA**→ rpmE2 (PRK01678)Ribosomal_L31 (pfam01197)→ |
| Sau-1603-1 | NZ_KK016033.1 | - | 61228 | 61199 | **RNA**→ rpmE2 (PRK01678)Ribosomal_L31 (pfam01197)→ |
| Sau-1620-1 | NZ_KK016818.1 | - | 36391 | 36362 | **RNA**→ rpmE2 (PRK01678)Ribosomal_L31 (pfam01197)→ |
| Sau-2492-1 | NZ_KK030749.1 | - | 34873 | 34844 | **RNA**→ rpmE2 (PRK01678)Ribosomal_L31 (pfam01197)→ |
| Sau-1610-1 | NZ_KK016378.1 | - | 61369 | 61340 | **RNA**→ rpmE2 (PRK01678)Ribosomal_L31 (pfam01197)→ |
| Sau-2597-1 | NZ_KK038693.1 | - | 34913 | 34884 | **RNA**→ rpmE2 (PRK01678)Ribosomal_L31 (pfam01197)→ |
| Sau-1359-1 | NZ_KK018504.1 | - | 36392 | 36363 | **RNA**→ rpmE2 (PRK01678)Ribosomal_L31 (pfam01197)→ |
| Sau-2622-1 | NZ_KK047755.1 | - | 57585 | 57556 | **RNA**→ rpmE2 (PRK01678)Ribosomal_L31 (pfam01197)→ |
| Sau-1636-1 | NZ_KK017766.1 | - | 107508 | 107479 | **RNA**→ rpmE2 (PRK01678)Ribosomal_L31 (pfam01197)→ |
| Sau-1130-1 | NZ_KK019217.1 | - | 56763 | 56734 | **RNA**→ rpmE2 (PRK01678)Ribosomal_L31 (pfam01197)→ |

| | | | | | |
|---|---|---|---|---|---|
| Sau-2788-1 | NZ_KK047909.1 | - | 1673122 | 1673093 | **RNA**→ rpmE2 (PRK01678)Ribosomal_L31 (pfam01197)→ |
| Sau-1647-1 | NZ_KK018303.1 | - | 117245 | 117216 | **RNA**→ rpmE2 (PRK01678)Ribosomal_L31 (pfam01197)→ |
| Sau-1060-1 | NZ_KK030519.1 | - | 34873 | 34844 | **RNA**→ rpmE2 (PRK01678)Ribosomal_L31 (pfam01197)→ |
| Sau-2556-1 | NZ_KK023258.1 | - | 56894 | 56865 | **RNA**→ rpmE2 (PRK01678)Ribosomal_L31 (pfam01197)→ |
| Sau-2236-1 | NZ_KK021870.1 | - | 57040 | 57011 | **RNA**→ rpmE2 (PRK01678)Ribosomal_L31 (pfam01197)→ |
| Sau-2564-1 | NZ_KK023376.1 | - | 34817 | 34788 | **RNA**→ rpmE2 (PRK01678)Ribosomal_L31 (pfam01197)→ |
| Sau-1107-1 | NZ_KK047803.1 | - | 1683613 | 1683584 | **RNA**→ rpmE2 (PRK01678)Ribosomal_L31 (pfam01197)→ |
| Sau-2829-1 | NZ_KK028877.1 | - | 56940 | 56911 | **RNA**→ rpmE2 (PRK01678)Ribosomal_L31 (pfam01197)→ |
| Sau-1581-1 | NZ_KK014900.1 | - | 36391 | 36362 | **RNA**→ rpmE2 (PRK01678)Ribosomal_L31 (pfam01197)→ |
| Sau-2171-1 | NZ_KK024130.1 | - | 34872 | 34843 | **RNA**→ rpmE2 (PRK01678)Ribosomal_L31 (pfam01197)→ |
| Sau-2666-1 | NZ_KK047697.1 | - | 1708894 | 1708865 | **RNA**→ rpmE2 (PRK01678)Ribosomal_L31 (pfam01197)→ |
| Sau-1073-1 | NZ_KK023603.1 | - | 34817 | 34788 | **RNA**→ rpmE2 (PRK01678)Ribosomal_L31 (pfam01197)→ |
| Sau-2806-1 | NZ_KK047849.1 | - | 2267822 | 2267793 | **RNA**→ rpmE2 (PRK01678)Ribosomal_L31 (pfam01197)→ |
| Sau-2617-1 | NZ_KK047918.1 | + | 620873 | 620902 | **RNA**→ rpmE2 (PRK01678)Ribosomal_L31 (pfam01197)→ |
| Sau-1084-1 | NZ_KK024366.1 | - | 34913 | 34884 | **RNA**→ rpmE2 (PRK01678)Ribosomal_L31 (pfam01197)→ |
| Sau-2600-1 | NZ_KK021676.1 | - | 34873 | 34844 | **RNA**→ rpmE2 (PRK01678)Ribosomal_L31 (pfam01197)→ |
| Sau-1040-1 | NZ_KK030295.1 | - | 34870 | 34841 | **RNA**→ rpmE2 (PRK01678)Ribosomal_L31 (pfam01197)→ |
| Sau-2535-1 | NZ_KK023119.1 | - | 56940 | 56911 | **RNA**→ rpmE2 (PRK01678)Ribosomal_L31 (pfam01197)→ |
| Sau-2832-1 | NZ_KK027642.1 | - | 56894 | 56865 | **RNA**→ rpmE2 (PRK01678)Ribosomal_L31 (pfam01197)→ |
| Sau-1061-1 | NZ_KK030574.1 | - | 34773 | 34744 | **RNA**→ rpmE2 (PRK01678)Ribosomal_L31 (pfam01197)→ |
| Sau-2703-1 | NZ_KK027868.1 | - | 34863 | 34834 | **RNA**→ rpmE2 (PRK01678)Ribosomal_L31 (pfam01197)→ |
| Sau-1132-1 | NZ_KK047765.1 | - | 61246 | 61217 | **RNA**→ rpmE2 (PRK01678)Ribosomal_L31 (pfam01197)→ |
| Sau-2842-1 | NZ_KK027964.1 | - | 34817 | 34788 | **RNA**→ rpmE2 (PRK01678)Ribosomal_L31 (pfam01197)→ |
| Sau-2537-1 | NZ_KK030874.1 | - | 2244515 | 2244486 | **RNA**→ rpmE2 (PRK01678)Ribosomal_L31 (pfam01197)→ |
| Sau-1106-1 | NZ_KK047819.1 | - | 252534 | 252505 | **RNA**→ rpmE2 (PRK01678)Ribosomal_L31 (pfam01197)→ |
| Sau-2548-1 | NZ_KK030899.1 | - | 56893 | 56864 | **RNA**→ rpmE2 (PRK01678)Ribosomal_L31 (pfam01197)→ |
| Sau-1608-1 | NZ_KK016202.1 | - | 36491 | 36462 | **RNA**→ rpmE2 (PRK01678)Ribosomal_L31 (pfam01197)→ |
| Sau-2592-1 | NZ_KK038742.1 | - | 34871 | 34842 | **RNA**→ rpmE2 (PRK01678)Ribosomal_L31 (pfam01197)→ |
| Sau-2663-1 | NZ_KK047703.1 | - | 240492 | 240463 | **RNA**→ rpmE2 (PRK01678)Ribosomal_L31 (pfam01197)→ |
| Sau-2624-1 | NZ_KK047734.1 | - | 1731888 | 1731859 | **RNA**→ rpmE2 (PRK01678)Ribosomal_L31 (pfam01197)→ |
| Sau-2772-1 | NZ_KK047773.1 | - | 56894 | 56865 | **RNA**→ rpmE2 (PRK01678)Ribosomal_L31 (pfam01197)→ |
| Sau-1167-1 | NZ_KK047866.1 | - | 55137 | 55108 | **RNA**→ rpmE2 (PRK01678)Ribosomal_L31 (pfam01197)→ |
| Sau-2805-1 | NZ_KK047827.1 | - | 56893 | 56864 | **RNA**→ rpmE2 (PRK01678)Ribosomal_L31 (pfam01197)→ |
| Sau-2733-1 | NZ_KK047830.1 | - | 2196141 | 2196112 | **RNA**→ rpmE2 (PRK01678)Ribosomal_L31 (pfam01197)→ |
| Sau-1168-1 | NZ_KK047922.1 | - | 1709647 | 1709618 | **RNA**→ rpmE2 (PRK01678)Ribosomal_L31 (pfam01197)→ |
| Sau-2609-1 | NZ_KK047855.1 | - | 1123865 | 1123836 | **RNA**→ rpmE2 (PRK01678)Ribosomal_L31 (pfam01197)→ |
| Sau-1572-1 | NZ_KK014503.1 | - | 115765 | 115736 | **RNA**→ rpmE2 (PRK01678)Ribosomal_L31 (pfam01197)→ |
| Sau-2623-1 | NZ_KK019284.1 | - | 25062 | 25033 | **RNA**→ rpmE2 (PRK01678)Ribosomal_L31 (pfam01197)→ |
| Sau-2493-1 | NZ_KK022189.1 | - | 34873 | 34844 | **RNA**→ rpmE2 (PRK01678)Ribosomal_L31 (pfam01197)→ |
| Sau-1590-1 | NZ_KK015343.1 | - | 108273 | 108244 | **RNA**→ rpmE2 (PRK01678)Ribosomal_L31 (pfam01197)→ |
| Sau-2475-1 | NZ_KK030727.1 | - | 56993 | 56964 | **RNA**→ rpmE2 (PRK01678)Ribosomal_L31 (pfam01197)→ |
| Sau-1363-1 | NZ_KK018751.1 | - | 194096 | 194067 | **RNA**→ rpmE2 (PRK01678)Ribosomal_L31 (pfam01197)→ |
| Sau-2549-1 | NZ_KK030912.1 | - | 34872 | 34843 | **RNA**→ rpmE2 (PRK01678)Ribosomal_L31 (pfam01197)→ |
| Sau-1059-1 | NZ_KK030508.1 | - | 25166 | 25137 | **RNA**→ rpmE2 (PRK01678)Ribosomal_L31 (pfam01197)→ |
| Sau-2662-1 | NZ_KK047710.1 | - | 56776 | 56747 | **RNA**→ rpmE2 (PRK01678)Ribosomal_L31 (pfam01197)→ |
| Sau-2621-1 | NZ_KK047748.1 | - | 1679024 | 1678995 | **RNA**→ rpmE2 (PRK01678)Ribosomal_L31 (pfam01197)→ |
| Sau-1094-1 | NZ_KK039071.1 | - | 56892 | 56863 | **RNA**→ rpmE2 (PRK01678)Ribosomal_L31 (pfam01197)→ |
| Sau-2791-1 | NZ_KK047784.1 | - | 55958 | 55929 | **RNA**→ rpmE2 (PRK01678)Ribosomal_L31 (pfam01197)→ |
| Sau-1080-1 | NZ_KK039235.1 | - | 34873 | 34844 | **RNA**→ rpmE2 (PRK01678)Ribosomal_L31 (pfam01197)→ |
| Sau-2766-1 | NZ_KK047798.1 | - | 230868 | 230839 | **RNA**→ rpmE2 (PRK01678)Ribosomal_L31 (pfam01197)→ |
| Sau-2642-1 | NZ_KK047838.1 | - | 56762 | 56733 | **RNA**→ rpmE2 (PRK01678)Ribosomal_L31 (pfam01197)→ |
| Sau-1166-1 | NZ_KK047870.1 | - | 2236752 | 2236723 | **RNA**→ rpmE2 (PRK01678)Ribosomal_L31 (pfam01197)→ |

| | | | | | |
|---|---|---|---|---|---|
| Sau-2741-1 | NZ_KK047902.1 | - | 1723084 | 1723055 | RNA→ rpmE2 (PRK01678)Ribosomal_L31 (pfam01197)→ |
| Sau-2557-1 | NZ_KK031009.1 | - | 56893 | 56864 | RNA→ rpmE2 (PRK01678)Ribosomal_L31 (pfam01197)→ |
| Sau-1562-1 | NZ_KK013946.1 | - | 36391 | 36362 | RNA→ rpmE2 (PRK01678)Ribosomal_L31 (pfam01197)→ |
| Sau-2531-1 | NZ_KK030789.1 | - | 34872 | 34843 | RNA→ rpmE2 (PRK01678)Ribosomal_L31 (pfam01197)→ |
| Sau-1360-1 | NZ_KK018605.1 | - | 36413 | 36384 | RNA→ rpmE2 (PRK01678)Ribosomal_L31 (pfam01197)→ |
| Sau-2715-1 | NZ_KK024671.1 | - | 34807 | 34778 | RNA→ rpmE2 (PRK01678)Ribosomal_L31 (pfam01197)→ |
| Sau-2487-1 | NZ_KK026516.1 | - | 34826 | 34797 | RNA→ rpmE2 (PRK01678)Ribosomal_L31 (pfam01197)→ |
| Sau-1367-1 | NZ_KK018811.1 | - | 2420 | 2391 | RNA→ rpmE2 (PRK01678)Ribosomal_L31 (pfam01197)→ |
| Sau-1054-1 | NZ_KK023207.1 | - | 56940 | 56911 | RNA→ rpmE2 (PRK01678)Ribosomal_L31 (pfam01197)→ |
| Sau-2830-1 | NZ_KK028950.1 | - | 56894 | 56865 | RNA→ rpmE2 (PRK01678)Ribosomal_L31 (pfam01197)→ |
| Sau-2471-1 | NZ_KK030648.1 | - | 34817 | 34788 | RNA→ rpmE2 (PRK01678)Ribosomal_L31 (pfam01197)→ |
| Sau-2864-1 | NZ_KK031045.1 | - | 2239804 | 2239775 | RNA→ rpmE2 (PRK01678)Ribosomal_L31 (pfam01197)→ |
| Sau-2586-1 | NZ_KK020798.1 | - | 34872 | 34843 | RNA→ rpmE2 (PRK01678)Ribosomal_L31 (pfam01197)→ |
| Sau-2601-1 | NZ_KK021735.1 | - | 30842 | 30813 | RNA→ rpmE2 (PRK01678)Ribosomal_L31 (pfam01197)→ |
| Sau-1433-1 | NZ_KK008467.1 | - | 36391 | 36362 | RNA→ rpmE2 (PRK01678)Ribosomal_L31 (pfam01197)→ |
| Sau-2529-1 | NZ_KK073242.1 | - | 34871 | 34842 | RNA→ rpmE2 (PRK01678)Ribosomal_L31 (pfam01197)→ |
| Sau-1570-1 | NZ_KK014417.1 | - | 36391 | 36362 | RNA→ rpmE2 (PRK01678)Ribosomal_L31 (pfam01197)→ |
| Sau-1361-1 | NZ_KK018612.1 | - | 36391 | 36362 | RNA→ rpmE2 (PRK01678)Ribosomal_L31 (pfam01197)→ |
| Sau-1074-1 | NZ_KK023648.1 | - | 2239930 | 2239901 | RNA→ rpmE2 (PRK01678)Ribosomal_L31 (pfam01197)→ |
| Sau-3539-1 | NZ_KK021748.1 | - | 34872 | 34843 | RNA→ rpmE2 (PRK01678)Ribosomal_L31 (pfam01197)→ |
| Sau-1571-1 | NZ_KK014495.1 | - | 94568 | 94539 | RNA→ rpmE2 (PRK01678)Ribosomal_L31 (pfam01197)→ |
| Sau-1582-1 | NZ_KK014951.1 | - | 118923 | 118894 | RNA→ rpmE2 (PRK01678)Ribosomal_L31 (pfam01197)→ |
| Sau-1614-1 | NZ_KK016472.1 | - | 35286 | 35257 | RNA→ rpmE2 (PRK01678)Ribosomal_L31 (pfam01197)→ |
| Sau-1362-1 | NZ_KK018658.1 | - | 1772 | 1743 | RNA→ rpmE2 (PRK01678)Ribosomal_L31 (pfam01197)→ |
| Sau-3595-1 | NZ_KK026049.1 | - | 56652 | 56623 | RNA→ rpmE2 (PRK01678)Ribosomal_L31 (pfam01197)→ |
| Sau-1065-1 | NZ_KK023463.1 | - | 2236651 | 2236622 | RNA→ rpmE2 (PRK01678)Ribosomal_L31 (pfam01197)→ |
| Sau-1075-1 | NZ_KK023651.1 | - | 55804 | 55775 | RNA→ rpmE2 (PRK01678)Ribosomal_L31 (pfam01197)→ |
| Sau-3709-1 | NZ_KK047684.1 | - | 56894 | 56865 | RNA→ rpmE2 (PRK01678)Ribosomal_L31 (pfam01197)→ |
| Sau-1041-1 | NZ_KK030362.1 | - | 34873 | 34844 | RNA→ rpmE2 (PRK01678)Ribosomal_L31 (pfam01197)→ |
| Sau-3708-1 | NZ_KK047688.1 | - | 1704238 | 1704209 | RNA→ rpmE2 (PRK01678)Ribosomal_L31 (pfam01197)→ |
| Sau-3674-1 | NZ_KK047843.1 | - | 1673624 | 1673595 | RNA→ rpmE2 (PRK01678)Ribosomal_L31 (pfam01197)→ |
| Sau-64-1 | NC_017333.1 | - | 2238912 | 2238883 | RNA→ rpmE2 (PRK01678)Ribosomal_L31 (pfam01197)→ |
| Sau-1561-1 | NZ_KK013917.1 | - | 62156 | 62127 | RNA→ rpmE2 (PRK01678)Ribosomal_L31 (pfam01197)→ |
| Sau-1578-1 | NZ_KK014718.1 | - | 55189 | 55160 | RNA→ rpmE2 (PRK01678)Ribosomal_L31 (pfam01197)→ |
| Sau-3593-1 | NZ_KK025962.1 | - | 33076 | 33047 | RNA→ rpmE2 (PRK01678)Ribosomal_L31 (pfam01197)→ |
| Sau-1584-1 | NZ_KK014983.1 | - | 36390 | 36361 | RNA→ rpmE2 (PRK01678)Ribosomal_L31 (pfam01197)→ |
| Sau-3586-1 | NZ_KK028408.1 | - | 46630 | 46601 | RNA→ rpmE2 (PRK01678)Ribosomal_L31 (pfam01197)→ |
| Sau-1601-1 | NZ_KK015846.1 | - | 36391 | 36362 | RNA→ rpmE2 (PRK01678)Ribosomal_L31 (pfam01197)→ |
| Sau-3577-1 | NZ_KK032474.1 | - | 34872 | 34843 | RNA→ rpmE2 (PRK01678)Ribosomal_L31 (pfam01197)→ |
| Sau-1052-1 | NZ_KK030433.1 | - | 34872 | 34843 | RNA→ rpmE2 (PRK01678)Ribosomal_L31 (pfam01197)→ |
| Sau-3591-1 | NZ_KK032745.1 | - | 34873 | 34844 | RNA→ rpmE2 (PRK01678)Ribosomal_L31 (pfam01197)→ |
| Sau-3704-1 | NZ_KK047880.1 | - | 1658923 | 1658894 | RNA→ rpmE2 (PRK01678)Ribosomal_L31 (pfam01197)→ |
| Sau-3594-1 | NZ_KK025972.1 | - | 34872 | 34843 | RNA→ rpmE2 (PRK01678)Ribosomal_L31 (pfam01197)→ |
| Sau-3587-1 | NZ_KK028466.1 | - | 46746 | 46717 | RNA→ rpmE2 (PRK01678)Ribosomal_L31 (pfam01197)→ |
| Sau-3588-1 | NZ_KK028533.1 | - | 15176 | 15147 | RNA→ rpmE2 (PRK01678)Ribosomal_L31 (pfam01197)→ |
| Sau-3578-1 | NZ_KK032489.1 | - | 34872 | 34843 | RNA→ rpmE2 (PRK01678)Ribosomal_L31 (pfam01197)→ |
| Sau-3601-1 | NZ_KK032800.1 | - | 34817 | 34788 | RNA→ rpmE2 (PRK01678)Ribosomal_L31 (pfam01197)→ |
| Sau-3658-1 | NZ_KK033002.1 | - | 34817 | 34788 | RNA→ rpmE2 (PRK01678)Ribosomal_L31 (pfam01197)→ |
| Sau-3696-1 | NZ_KK047674.1 | - | 56228 | 56199 | RNA→ rpmE2 (PRK01678)Ribosomal_L31 (pfam01197)→ |
| Sau-3684-1 | NZ_KK047888.1 | - | 237730 | 237701 | RNA→ rpmE2 (PRK01678)Ribosomal_L31 (pfam01197)→ |
| Sau-3496-1 | NZ_KK023149.1 | - | 34817 | 34788 | RNA→ rpmE2 (PRK01678)Ribosomal_L31 (pfam01197)→ |
| Sau-3631-1 | NZ_KK027664.1 | - | 34873 | 34844 | RNA→ rpmE2 (PRK01678)Ribosomal_L31 (pfam01197)→ |

| | | | | | | |
|---|---|---|---|---|---|---|
| Sau-3507-1 | NZ_KK032260.1 | - | 54968 | 54939 | RNA→ | rpmE2 (PRK01678)Ribosomal_L31 (pfam01197)→ |
| Sau-3664-1 | NZ_KK033064.1 | - | 34873 | 34844 | RNA→ | rpmE2 (PRK01678)Ribosomal_L31 (pfam01197)→ |
| Sau-3675-1 | NZ_KK047678.1 | - | 2241127 | 2241098 | RNA→ | rpmE2 (PRK01678)Ribosomal_L31 (pfam01197)→ |
| Sau-3691-1 | NZ_KK047705.1 | - | 1675481 | 1675452 | RNA→ | rpmE2 (PRK01678)Ribosomal_L31 (pfam01197)→ |
| Sau-3568-1 | NZ_KK022849.1 | - | 2229307 | 2229278 | RNA→ | rpmE2 (PRK01678)Ribosomal_L31 (pfam01197)→ |
| Sau-3504-1 | NZ_KK023326.1 | - | 33076 | 33047 | RNA→ | rpmE2 (PRK01678)Ribosomal_L31 (pfam01197)→ |
| Sau-3548-1 | NZ_KK023919.1 | - | 34873 | 34844 | RNA→ | rpmE2 (PRK01678)Ribosomal_L31 (pfam01197)→ |
| Sau-3508-1 | NZ_KK032308.1 | - | 34841 | 34812 | RNA→ | rpmE2 (PRK01678)Ribosomal_L31 (pfam01197)→ |
| Sau-3576-1 | NZ_KK032381.1 | - | 34873 | 34844 | RNA→ | rpmE2 (PRK01678)Ribosomal_L31 (pfam01197)→ |
| Sau-3584-1 | NZ_KK032670.1 | - | 34872 | 34843 | RNA→ | rpmE2 (PRK01678)Ribosomal_L31 (pfam01197)→ |
| Sau-3650-1 | NZ_KK032923.1 | - | 55477 | 55448 | RNA→ | rpmE2 (PRK01678)Ribosomal_L31 (pfam01197)→ |
| Sau-795-1 | NZ_KK041736.1 | - | 56993 | 56964 | RNA→ | rpmE2 (PRK01678)Ribosomal_L31 (pfam01197)→ |
| Sau-3526-1 | NZ_KK021419.1 | - | 34817 | 34788 | RNA→ | rpmE2 (PRK01678)Ribosomal_L31 (pfam01197)→ |
| Sau-787-1 | NZ_KK041753.1 | - | 35209 | 35180 | RNA→ | rpmE2 (PRK01678)Ribosomal_L31 (pfam01197)→ |
| Sau-3545-1 | NZ_KK021833.1 | - | 56790 | 56761 | RNA→ | rpmE2 (PRK01678)Ribosomal_L31 (pfam01197)→ |
| Sau-781-1 | NZ_KK041899.1 | - | 19182 | 19153 | RNA→ | rpmE2 (PRK01678)Ribosomal_L31 (pfam01197)→ |
| Sau-3607-1 | NZ_KK032863.1 | - | 34872 | 34843 | RNA→ | rpmE2 (PRK01678)Ribosomal_L31 (pfam01197)→ |
| Sau-661-1 | NZ_KK041939.1 | - | 35066 | 35037 | RNA→ | rpmE2 (PRK01678)Ribosomal_L31 (pfam01197)→ |
| Sau-3651-1 | NZ_KK032992.1 | - | 2236249 | 2236220 | RNA→ | rpmE2 (PRK01678)Ribosomal_L31 (pfam01197)→ |
| Sau-725-1 | NZ_KK022396.1 | - | 30868 | 30839 | RNA→ | rpmE2 (PRK01678)Ribosomal_L31 (pfam01197)→ |
| Sau-3685-1 | NZ_KK019345.1 | - | 56994 | 56965 | RNA→ | rpmE2 (PRK01678)Ribosomal_L31 (pfam01197)→ |
| Sau-740-1 | NZ_KK025058.1 | - | 34873 | 34844 | RNA→ | rpmE2 (PRK01678)Ribosomal_L31 (pfam01197)→ |
| Sau-791-1 | NZ_KK026781.1 | - | 34817 | 34788 | RNA→ | rpmE2 (PRK01678)Ribosomal_L31 (pfam01197)→ |
| Sau-3558-1 | NZ_KK024371.1 | - | 57040 | 57011 | RNA→ | rpmE2 (PRK01678)Ribosomal_L31 (pfam01197)→ |
| Sau-756-1 | NZ_KK029293.1 | - | 34872 | 34843 | RNA→ | rpmE2 (PRK01678)Ribosomal_L31 (pfam01197)→ |
| Sau-3575-1 | NZ_KK025280.1 | - | 56994 | 56965 | RNA→ | rpmE2 (PRK01678)Ribosomal_L31 (pfam01197)→ |
| Sau-786-1 | NZ_KK041769.1 | - | 34817 | 34788 | RNA→ | rpmE2 (PRK01678)Ribosomal_L31 (pfam01197)→ |
| Sau-3503-1 | NZ_KK028175.1 | - | 34872 | 34843 | RNA→ | rpmE2 (PRK01678)Ribosomal_L31 (pfam01197)→ |
| Sau-660-1 | NZ_KK041973.1 | - | 34872 | 34843 | RNA→ | rpmE2 (PRK01678)Ribosomal_L31 (pfam01197)→ |
| Sau-3506-1 | NZ_KK032215.1 | - | 19182 | 19153 | RNA→ | rpmE2 (PRK01678)Ribosomal_L31 (pfam01197)→ |
| Sau-722-1 | NZ_KK022244.1 | - | 34873 | 34844 | RNA→ | rpmE2 (PRK01678)Ribosomal_L31 (pfam01197)→ |
| Sau-3534-1 | NZ_KK023872.1 | - | 2238473 | 2238444 | RNA→ | rpmE2 (PRK01678)Ribosomal_L31 (pfam01197)→ |
| Sau-788-1 | NZ_KK026645.1 | - | 34872 | 34843 | RNA→ | rpmE2 (PRK01678)Ribosomal_L31 (pfam01197)→ |
| Sau-3559-1 | NZ_KK024423.1 | - | 56940 | 56911 | RNA→ | rpmE2 (PRK01678)Ribosomal_L31 (pfam01197)→ |
| Sau-839-1 | NZ_KK027771.1 | - | 34772 | 34743 | RNA→ | rpmE2 (PRK01678)Ribosomal_L31 (pfam01197)→ |
| Sau-3569-1 | NZ_KK024897.1 | - | 56894 | 56865 | RNA→ | rpmE2 (PRK01678)Ribosomal_L31 (pfam01197)→ |
| Sau-771-1 | NZ_KK029389.1 | - | 2236533 | 2236504 | RNA→ | rpmE2 (PRK01678)Ribosomal_L31 (pfam01197)→ |
| Sau-3612-1 | NZ_KK027253.1 | - | 2240851 | 2240822 | RNA→ | rpmE2 (PRK01678)Ribosomal_L31 (pfam01197)→ |
| Sau-723-1 | NZ_KK022328.1 | - | 56894 | 56865 | RNA→ | rpmE2 (PRK01678)Ribosomal_L31 (pfam01197)→ |
| Sau-759-1 | NZ_KK026434.1 | - | 34873 | 34844 | RNA→ | rpmE2 (PRK01678)Ribosomal_L31 (pfam01197)→ |
| Sau-3583-1 | NZ_KK032605.1 | - | 34873 | 34844 | RNA→ | rpmE2 (PRK01678)Ribosomal_L31 (pfam01197)→ |
| Sau-794-1 | NZ_KK028873.1 | - | 2236659 | 2236630 | RNA→ | rpmE2 (PRK01678)Ribosomal_L31 (pfam01197)→ |
| Sau-3659-1 | NZ_KK033045.1 | - | 56894 | 56865 | RNA→ | rpmE2 (PRK01678)Ribosomal_L31 (pfam01197)→ |
| Sau-742-1 | NZ_KK029243.1 | - | 56891 | 56862 | RNA→ | rpmE2 (PRK01678)Ribosomal_L31 (pfam01197)→ |
| Sau-826-1 | NZ_KK022877.1 | - | 2236712 | 2236683 | RNA→ | rpmE2 (PRK01678)Ribosomal_L31 (pfam01197)→ |
| Sau-777-1 | NZ_KK026178.1 | - | 34913 | 34884 | RNA→ | rpmE2 (PRK01678)Ribosomal_L31 (pfam01197)→ |
| Sau-2105-1 | NZ_KK067350.1 | - | 36490 | 36461 | RNA→ | rpmE2 (PRK01678)Ribosomal_L31 (pfam01197)→ |
| Sau-792-1 | NZ_KK026776.1 | - | 56894 | 56865 | RNA→ | rpmE2 (PRK01678)Ribosomal_L31 (pfam01197)→ |
| Sau-2092-1 | NZ_KK067612.1 | - | 236801 | 236772 | RNA→ | rpmE2 (PRK01678)Ribosomal_L31 (pfam01197)→ |
| Sau-802-1 | NZ_KK027250.1 | - | 56894 | 56865 | RNA→ | rpmE2 (PRK01678)Ribosomal_L31 (pfam01197)→ |
| Sau-712-1 | NZ_KK070316.1 | - | 2245370 | 2245341 | RNA→ | rpmE2 (PRK01678)Ribosomal_L31 (pfam01197)→ |
| Sau-3599-1 | NZ_KK071292.1 | - | 56894 | 56865 | RNA→ | rpmE2 (PRK01678)Ribosomal_L31 (pfam01197)→ |

| | | | | | | |
|---|---|---|---|---|---|---|
| Sau-726-1 | NZ_KK022489.1 | - | 34873 | 34844 | **RNA**→ | rpmE2 (PRK01678)Ribosomal_L31 (pfam01197)→ |
| Sau-2152-1 | NZ_KK065541.1 | - | 36806 | 36777 | **RNA**→ | rpmE2 (PRK01678)Ribosomal_L31 (pfam01197)→ |
| Sau-677-1 | NZ_KK023680.1 | - | 35161 | 35132 | **RNA**→ | rpmE2 (PRK01678)Ribosomal_L31 (pfam01197)→ |
| Sau-778-1 | NZ_KK026259.1 | - | 19182 | 19153 | **RNA**→ | rpmE2 (PRK01678)Ribosomal_L31 (pfam01197)→ |
| Sau-2130-1 | NZ_KK066572.1 | - | 55458 | 55429 | **RNA**→ | rpmE2 (PRK01678)Ribosomal_L31 (pfam01197)→ |
| Sau-767-1 | NZ_KK028585.1 | - | 34873 | 34844 | **RNA**→ | rpmE2 (PRK01678)Ribosomal_L31 (pfam01197)→ |
| Sau-2090-1 | NZ_KK067652.1 | - | 56044 | 56015 | **RNA**→ | rpmE2 (PRK01678)Ribosomal_L31 (pfam01197)→ |
| Sau-3571-1 | NZ_KK070199.1 | - | 1094703 | 1094674 | **RNA**→ | rpmE2 (PRK01678)Ribosomal_L31 (pfam01197)→ |
| Sau-2978-1 | NZ_KK070975.1 | - | 34870 | 34841 | **RNA**→ | rpmE2 (PRK01678)Ribosomal_L31 (pfam01197)→ |
| Sau-3606-1 | NZ_KK071564.1 | - | 56894 | 56865 | **RNA**→ | rpmE2 (PRK01678)Ribosomal_L31 (pfam01197)→ |
| Sau-2502-1 | NZ_KK070240.1 | - | 236894 | 236865 | **RNA**→ | rpmE2 (PRK01678)Ribosomal_L31 (pfam01197)→ |
| Sau-2447-1 | NZ_KK070432.1 | - | 58560 | 58531 | **RNA**→ | rpmE2 (PRK01678)Ribosomal_L31 (pfam01197)→ |
| Sau-2506-1 | NZ_KK071935.1 | - | 56894 | 56865 | **RNA**→ | rpmE2 (PRK01678)Ribosomal_L31 (pfam01197)→ |
| Sau-3610-1 | NZ_KK071957.1 | - | 34913 | 34884 | **RNA**→ | rpmE2 (PRK01678)Ribosomal_L31 (pfam01197)→ |
| Sau-2598-1 | NZ_KK038666.1 | - | 34913 | 34884 | **RNA**→ | rpmE2 (PRK01678)Ribosomal_L31 (pfam01197)→ |
| Sau-2161-1 | NZ_KK065316.1 | - | 54697 | 54668 | **RNA**→ | rpmE2 (PRK01678)Ribosomal_L31 (pfam01197)→ |
| Sau-2483-1 | NZ_KK071147.1 | - | 34954 | 34925 | **RNA**→ | rpmE2 (PRK01678)Ribosomal_L31 (pfam01197)→ |
| Sau-2158-1 | NZ_KK065367.1 | - | 35190 | 35161 | **RNA**→ | rpmE2 (PRK01678)Ribosomal_L31 (pfam01197)→ |
| Sau-2101-1 | NZ_KK067484.1 | - | 35296 | 35267 | **RNA**→ | rpmE2 (PRK01678)Ribosomal_L31 (pfam01197)→ |
| Sau-2498-1 | NZ_KK071690.1 | - | 34973 | 34944 | **RNA**→ | rpmE2 (PRK01678)Ribosomal_L31 (pfam01197)→ |
| Sau-2122-1 | NZ_KK066887.1 | - | 62203 | 62174 | **RNA**→ | rpmE2 (PRK01678)Ribosomal_L31 (pfam01197)→ |
| Sau-2114-1 | NZ_KK067114.1 | - | 190679 | 190650 | **RNA**→ | rpmE2 (PRK01678)Ribosomal_L31 (pfam01197)→ |
| Sau-2059-1 | NZ_KK069017.1 | - | 36579 | 36550 | **RNA**→ | rpmE2 (PRK01678)Ribosomal_L31 (pfam01197)→ |
| Sau-2084-1 | NZ_KK067969.1 | - | 61690 | 61661 | **RNA**→ | rpmE2 (PRK01678)Ribosomal_L31 (pfam01197)→ |
| Sau-1387-1 | NZ_KI971464.1 | - | 56763 | 56734 | **RNA**→ | rpmE2 (PRK01678)Ribosomal_L31 (pfam01197)→ |
| Sau-2087-1 | NZ_KK067883.1 | - | 61546 | 61517 | **RNA**→ | rpmE2 (PRK01678)Ribosomal_L31 (pfam01197)→ |
| Sau-2071-1 | NZ_KK068467.1 | - | 25128 | 25099 | **RNA**→ | rpmE2 (PRK01678)Ribosomal_L31 (pfam01197)→ |
| Sau-2051-1 | NZ_KK069266.1 | - | 25029 | 25000 | **RNA**→ | rpmE2 (PRK01678)Ribosomal_L31 (pfam01197)→ |
| Sau-2044-1 | NZ_KK069594.1 | - | 24749 | 24720 | **RNA**→ | rpmE2 (PRK01678)Ribosomal_L31 (pfam01197)→ |
| Sau-2062-1 | NZ_KK068890.1 | - | 36390 | 36361 | **RNA**→ | rpmE2 (PRK01678)Ribosomal_L31 (pfam01197)→ |
| Sau-1062-1 | NZ_KK033122.1 | - | 56644 | 56615 | **RNA**→ | rpmE2 (PRK01678)Ribosomal_L31 (pfam01197)→ |
| Sau-2081-1 | NZ_KK068115.1 | - | 8576 | 8547 | **RNA**→ | rpmE2 (PRK01678)Ribosomal_L31 (pfam01197)→ |
| Sau-717-1 | NZ_KK070061.1 | - | 1690011 | 1689982 | **RNA**→ | rpmE2 (PRK01678)Ribosomal_L31 (pfam01197)→ |
| Sau-769-1 | NZ_KK070254.1 | - | 1658420 | 1658391 | **RNA**→ | rpmE2 (PRK01678)Ribosomal_L31 (pfam01197)→ |
| Sau-2069-1 | NZ_KK068572.1 | - | 35669 | 35640 | **RNA**→ | rpmE2 (PRK01678)Ribosomal_L31 (pfam01197)→ |
| Sau-2080-1 | NZ_KK068152.1 | - | 32991 | 32962 | **RNA**→ | rpmE2 (PRK01678)Ribosomal_L31 (pfam01197)→ |
| Sau-2073-1 | NZ_KK068424.1 | - | 54564 | 54535 | **RNA**→ | rpmE2 (PRK01678)Ribosomal_L31 (pfam01197)→ |
| Sau-2066-1 | NZ_KK068711.1 | - | 2421 | 2392 | **RNA**→ | rpmE2 (PRK01678)Ribosomal_L31 (pfam01197)→ |
| Sau-2060-1 | NZ_KK068997.1 | - | 62153 | 62124 | **RNA**→ | rpmE2 (PRK01678)Ribosomal_L31 (pfam01197)→ |
| Sau-2047-1 | NZ_KK069374.1 | - | 24749 | 24720 | **RNA**→ | rpmE2 (PRK01678)Ribosomal_L31 (pfam01197)→ |
| Sau-2037-1 | NZ_KK069794.1 | - | 111530 | 111501 | **RNA**→ | rpmE2 (PRK01678)Ribosomal_L31 (pfam01197)→ |
| Sau-957-1 | NZ_KK071339.1 | - | 34894 | 34865 | **RNA**→ | rpmE2 (PRK01678)Ribosomal_L31 (pfam01197)→ |
| Sau-659-1 | NZ_KK042057.1 | - | 34874 | 34845 | **RNA**→ | rpmE2 (PRK01678)Ribosomal_L31 (pfam01197)→ |
| Sau-633-1 | NZ_KK042196.1 | - | 54968 | 54939 | **RNA**→ | rpmE2 (PRK01678)Ribosomal_L31 (pfam01197)→ |
| Sau-635-1 | NZ_KK029047.1 | - | 29278 | 29249 | **RNA**→ | rpmE2 (PRK01678)Ribosomal_L31 (pfam01197)→ |
| Sau-651-1 | NZ_KK042077.1 | - | 56894 | 56865 | **RNA**→ | rpmE2 (PRK01678)Ribosomal_L31 (pfam01197)→ |
| Sau-625-1 | NZ_KK042274.1 | - | 25255 | 25226 | **RNA**→ | rpmE2 (PRK01678)Ribosomal_L31 (pfam01197)→ |
| Sau-624-1 | NZ_KK042282.1 | - | 56894 | 56865 | **RNA**→ | rpmE2 (PRK01678)Ribosomal_L31 (pfam01197)→ |
| Sau-290-1 | NZ_KK047913.1 | - | 2192944 | 2192915 | **RNA**→ | rpmE2 (PRK01678)Ribosomal_L31 (pfam01197)→ |
| Sau-636-1 | NZ_KK029097.1 | - | 121888 | 121859 | **RNA**→ | rpmE2 (PRK01678)Ribosomal_L31 (pfam01197)→ |
| Sau-649-1 | NZ_KK042095.1 | - | 33080 | 33051 | **RNA**→ | rpmE2 (PRK01678)Ribosomal_L31 (pfam01197)→ |
| Sau-640-1 | NZ_KK042104.1 | - | 34913 | 34884 | **RNA**→ | rpmE2 (PRK01678)Ribosomal_L31 (pfam01197)→ |

| | | | | | | |
|---|---|---|---|---|---|---|
| Sau-618-1 | NZ_KK042299.1 | - | 34873 | 34844 | **RNA**→ | rpmE2 (PRK01678)Ribosomal_L31 (pfam01197)→ |
| Sau-955-1 | NZ_KK071439.1 | - | 34873 | 34844 | **RNA**→ | rpmE2 (PRK01678)Ribosomal_L31 (pfam01197)→ |
| Sau-613-1 | NZ_KK022975.1 | - | 34872 | 34843 | **RNA**→ | rpmE2 (PRK01678)Ribosomal_L31 (pfam01197)→ |
| Sau-639-1 | NZ_KK042131.1 | - | 25255 | 25226 | **RNA**→ | rpmE2 (PRK01678)Ribosomal_L31 (pfam01197)→ |
| Sau-617-1 | NZ_KK042394.1 | - | 34817 | 34788 | **RNA**→ | rpmE2 (PRK01678)Ribosomal_L31 (pfam01197)→ |
| Sau-765-1 | NZ_KK071370.1 | - | 34873 | 34844 | **RNA**→ | rpmE2 (PRK01678)Ribosomal_L31 (pfam01197)→ |
| Sau-2040-1 | NZ_KK069665.1 | - | 35499 | 35470 | **RNA**→ | rpmE2 (PRK01678)Ribosomal_L31 (pfam01197)→ |
| Sau-744-1 | NZ_KK070618.1 | - | 56894 | 56865 | **RNA**→ | rpmE2 (PRK01678)Ribosomal_L31 (pfam01197)→ |
| Sau-799-1 | NZ_KK072266.1 | - | 15832 | 15803 | **RNA**→ | rpmE2 (PRK01678)Ribosomal_L31 (pfam01197)→ |
| Sau-1376-1 | NZ_KI971452.1 | - | 56763 | 56734 | **RNA**→ | rpmE2 (PRK01678)Ribosomal_L31 (pfam01197)→ |
| Sau-973-1 | NZ_KK071835.1 | - | 56794 | 56765 | **RNA**→ | rpmE2 (PRK01678)Ribosomal_L31 (pfam01197)→ |
| Sau-944-1 | NZ_KK070909.1 | - | 56894 | 56865 | **RNA**→ | rpmE2 (PRK01678)Ribosomal_L31 (pfam01197)→ |
| Sau-797-1 | NZ_KK072197.1 | - | 56883 | 56854 | **RNA**→ | rpmE2 (PRK01678)Ribosomal_L31 (pfam01197)→ |
| Sau-1410-1 | NZ_KI971503.1 | - | 58300 | 58271 | **RNA**→ | rpmE2 (PRK01678)Ribosomal_L31 (pfam01197)→ |
| Sau-2088-1 | NZ_KK067776.1 | - | 35384 | 35355 | **RNA**→ | rpmE2 (PRK01678)Ribosomal_L31 (pfam01197)→ |
| Sau-819-1 | NZ_KK072733.1 | - | 56894 | 56865 | **RNA**→ | rpmE2 (PRK01678)Ribosomal_L31 (pfam01197)→ |
| Sau-981-1 | NZ_KK071993.1 | - | 34865 | 34836 | **RNA**→ | rpmE2 (PRK01678)Ribosomal_L31 (pfam01197)→ |
| Sau-1064-1 | NZ_KK072299.1 | - | 31125 | 31096 | **RNA**→ | rpmE2 (PRK01678)Ribosomal_L31 (pfam01197)→ |
| Sau-749-1 | NZ_KK070937.1 | - | 56894 | 56865 | **RNA**→ | rpmE2 (PRK01678)Ribosomal_L31 (pfam01197)→ |
| Sau-2127-1 | NZ_KK066657.1 | - | 35451 | 35422 | **RNA**→ | rpmE2 (PRK01678)Ribosomal_L31 (pfam01197)→ |
| Sau-644-1 | NZ_KK022681.1 | - | 34817 | 34788 | **RNA**→ | rpmE2 (PRK01678)Ribosomal_L31 (pfam01197)→ |
| Sau-648-1 | NZ_KK029104.1 | - | 2228793 | 2228764 | **RNA**→ | rpmE2 (PRK01678)Ribosomal_L31 (pfam01197)→ |
| Sau-720-1 | NZ_KK070340.1 | - | 58240 | 58211 | **RNA**→ | rpmE2 (PRK01678)Ribosomal_L31 (pfam01197)→ |
| Sau-974-1 | NZ_KK070262.1 | - | 1639617 | 1639588 | **RNA**→ | rpmE2 (PRK01678)Ribosomal_L31 (pfam01197)→ |
| Sau-823-1 | NZ_KK072797.1 | - | 56714 | 56685 | **RNA**→ | rpmE2 (PRK01678)Ribosomal_L31 (pfam01197)→ |
| Sau-701-1 | NZ_KK070279.1 | - | 56894 | 56865 | **RNA**→ | rpmE2 (PRK01678)Ribosomal_L31 (pfam01197)→ |
| Sau-2136-1 | NZ_KK066349.1 | - | 91457 | 91428 | **RNA**→ | rpmE2 (PRK01678)Ribosomal_L31 (pfam01197)→ |
| Sau-2141-1 | NZ_KK065990.1 | - | 35404 | 35375 | **RNA**→ | rpmE2 (PRK01678)Ribosomal_L31 (pfam01197)→ |
| Sau-2488-1 | NZ_KK070113.1 | - | 54795 | 54766 | **RNA**→ | rpmE2 (PRK01678)Ribosomal_L31 (pfam01197)→ |
| Sau-2971-1 | NZ_KK070751.1 | - | 35013 | 34984 | **RNA**→ | rpmE2 (PRK01678)Ribosomal_L31 (pfam01197)→ |
| Sau-2954-1 | NZ_KK070449.1 | - | 1725436 | 1725407 | **RNA**→ | rpmE2 (PRK01678)Ribosomal_L31 (pfam01197)→ |
| Sau-3574-1 | NZ_KK070474.1 | - | 224376 | 224347 | **RNA**→ | rpmE2 (PRK01678)Ribosomal_L31 (pfam01197)→ |
| Sau-2138-1 | NZ_KK066185.1 | - | 35169 | 35140 | **RNA**→ | rpmE2 (PRK01678)Ribosomal_L31 (pfam01197)→ |
| Sau-2132-1 | NZ_KK066446.1 | - | 36391 | 36362 | **RNA**→ | rpmE2 (PRK01678)Ribosomal_L31 (pfam01197)→ |
| Sau-3604-1 | NZ_KK071651.1 | - | 25255 | 25226 | **RNA**→ | rpmE2 (PRK01678)Ribosomal_L31 (pfam01197)→ |
| Sau-2496-1 | NZ_KK071775.1 | - | 34913 | 34884 | **RNA**→ | rpmE2 (PRK01678)Ribosomal_L31 (pfam01197)→ |
| Sau-2448-1 | NZ_KK038990.1 | - | 34817 | 34788 | **RNA**→ | rpmE2 (PRK01678)Ribosomal_L31 (pfam01197)→ |
| Sau-2986-1 | NZ_KK071519.1 | - | 34872 | 34843 | **RNA**→ | rpmE2 (PRK01678)Ribosomal_L31 (pfam01197)→ |
| Sau-2550-1 | NZ_KK073273.1 | - | 55214 | 55185 | **RNA**→ | rpmE2 (PRK01678)Ribosomal_L31 (pfam01197)→ |
| Sau-2499-1 | NZ_KK072095.1 | - | 109235 | 109206 | **RNA**→ | rpmE2 (PRK01678)Ribosomal_L31 (pfam01197)→ |
| Sau-1353-1 | NZ_KI971738.1 | - | 56763 | 56734 | **RNA**→ | rpmE2 (PRK01678)Ribosomal_L31 (pfam01197)→ |
| Sau-2500-1 | NZ_KK072188.1 | - | 34967 | 34938 | **RNA**→ | rpmE2 (PRK01678)Ribosomal_L31 (pfam01197)→ |
| Sau-1934-1 | NZ_KI972241.1 | - | 57675 | 57646 | **RNA**→ | rpmE2 (PRK01678)Ribosomal_L31 (pfam01197)→ |
| Sau-2523-1 | NZ_KK073134.1 | - | 34862 | 34833 | **RNA**→ | rpmE2 (PRK01678)Ribosomal_L31 (pfam01197)→ |
| Sau-1784-1 | NZ_KI971981.1 | - | 56762 | 56733 | **RNA**→ | rpmE2 (PRK01678)Ribosomal_L31 (pfam01197)→ |
| Sau-1945-1 | NZ_KI972277.1 | - | 141138 | 141109 | **RNA**→ | rpmE2 (PRK01678)Ribosomal_L31 (pfam01197)→ |
| Sau-3613-1 | NZ_KK072381.1 | - | 34913 | 34884 | **RNA**→ | rpmE2 (PRK01678)Ribosomal_L31 (pfam01197)→ |
| Sau-1365-1 | NZ_KI971747.1 | - | 56763 | 56734 | **RNA**→ | rpmE2 (PRK01678)Ribosomal_L31 (pfam01197)→ |
| Sau-1796-1 | NZ_KI971983.1 | - | 234880 | 234851 | **RNA**→ | rpmE2 (PRK01678)Ribosomal_L31 (pfam01197)→ |
| Sau-1937-1 | NZ_KI972261.1 | - | 61715 | 61686 | **RNA**→ | rpmE2 (PRK01678)Ribosomal_L31 (pfam01197)→ |
| Sau-1955-1 | NZ_KI972284.1 | - | 880152 | 880123 | **RNA**→ | rpmE2 (PRK01678)Ribosomal_L31 (pfam01197)→ |
| Sau-2007-1 | NZ_KI972388.1 | - | 59768 | 59739 | **RNA**→ | rpmE2 (PRK01678)Ribosomal_L31 (pfam01197)→ |

| | | | | | | |
|---|---|---|---|---|---|---|
| Sau-2314-1 | NZ_KI972526.1 | - | 228594 | 228565 | **RNA**→ | rpmE2 (PRK01678)Ribosomal_L31 (pfam01197)→ |
| Sau-2327-1 | NZ_KI972558.1 | - | 115301 | 115272 | **RNA**→ | rpmE2 (PRK01678)Ribosomal_L31 (pfam01197)→ |
| Sau-615-1 | NZ_KK092943.1 | - | 34972 | 34943 | **RNA**→ | rpmE2 (PRK01678)Ribosomal_L31 (pfam01197)→ |
| Sau-2867-1 | NZ_KK093446.1 | - | 55983 | 55954 | **RNA**→ | rpmE2 (PRK01678)Ribosomal_L31 (pfam01197)→ |
| Sau-2085-1 | NZ_KI972742.1 | - | 234815 | 234786 | **RNA**→ | rpmE2 (PRK01678)Ribosomal_L31 (pfam01197)→ |
| Sau-1968-1 | NZ_KI972321.1 | - | 119786 | 119757 | **RNA**→ | rpmE2 (PRK01678)Ribosomal_L31 (pfam01197)→ |
| Sau-2018-1 | NZ_KI972401.1 | - | 56763 | 56734 | **RNA**→ | rpmE2 (PRK01678)Ribosomal_L31 (pfam01197)→ |
| Sau-1002-1 | NZ_KK028024.1 | - | 34872 | 34843 | **RNA**→ | rpmE2 (PRK01678)Ribosomal_L31 (pfam01197)→ |
| Sau-2282-1 | NZ_KI972493.1 | - | 401361 | 401332 | **RNA**→ | rpmE2 (PRK01678)Ribosomal_L31 (pfam01197)→ |
| Sau-2315-1 | NZ_KI972541.1 | - | 55259 | 55230 | **RNA**→ | rpmE2 (PRK01678)Ribosomal_L31 (pfam01197)→ |
| Sau-2330-1 | NZ_KI972603.1 | - | 474378 | 474349 | **RNA**→ | rpmE2 (PRK01678)Ribosomal_L31 (pfam01197)→ |
| Sau-3532-1 | NZ_KK038133.1 | - | 56894 | 56865 | **RNA**→ | rpmE2 (PRK01678)Ribosomal_L31 (pfam01197)→ |
| Sau-686-1 | NZ_KK093676.1 | - | 34872 | 34843 | **RNA**→ | rpmE2 (PRK01678)Ribosomal_L31 (pfam01197)→ |
| Sau-2476-1 | NZ_KK092439.1 | - | 35556 | 35527 | **RNA**→ | rpmE2 (PRK01678)Ribosomal_L31 (pfam01197)→ |
| Sau-2077-1 | NZ_KI972732.1 | - | 56763 | 56734 | **RNA**→ | rpmE2 (PRK01678)Ribosomal_L31 (pfam01197)→ |
| Sau-1853-1 | NZ_KI972093.1 | - | 56763 | 56734 | **RNA**→ | rpmE2 (PRK01678)Ribosomal_L31 (pfam01197)→ |
| Sau-3552-1 | NZ_KK022059.1 | - | 34872 | 34843 | **RNA**→ | rpmE2 (PRK01678)Ribosomal_L31 (pfam01197)→ |
| Sau-2041-1 | NZ_KI972629.1 | - | 468563 | 468534 | **RNA**→ | rpmE2 (PRK01678)Ribosomal_L31 (pfam01197)→ |
| Sau-3543-1 | NZ_KK038080.1 | - | 56894 | 56865 | **RNA**→ | rpmE2 (PRK01678)Ribosomal_L31 (pfam01197)→ |
| Sau-3520-1 | NZ_KK020928.1 | - | 56940 | 56911 | **RNA**→ | rpmE2 (PRK01678)Ribosomal_L31 (pfam01197)→ |
| Sau-2484-1 | NZ_KK092708.1 | - | 34913 | 34884 | **RNA**→ | rpmE2 (PRK01678)Ribosomal_L31 (pfam01197)→ |
| Sau-3567-1 | NZ_KK038067.1 | - | 2160060 | 2160031 | **RNA**→ | rpmE2 (PRK01678)Ribosomal_L31 (pfam01197)→ |
| Sau-2050-1 | NZ_KI972665.1 | - | 61740 | 61711 | **RNA**→ | rpmE2 (PRK01678)Ribosomal_L31 (pfam01197)→ |
| Sau-2058-1 | NZ_KI972713.1 | - | 56763 | 56734 | **RNA**→ | rpmE2 (PRK01678)Ribosomal_L31 (pfam01197)→ |
| Sau-2057-1 | NZ_KI972702.1 | - | 216427 | 216398 | **RNA**→ | rpmE2 (PRK01678)Ribosomal_L31 (pfam01197)→ |
| Sau-3540-1 | NZ_KK022054.1 | - | 57040 | 57011 | **RNA**→ | rpmE2 (PRK01678)Ribosomal_L31 (pfam01197)→ |
| Sau-3627-1 | NZ_KK028083.1 | - | 34874 | 34845 | **RNA**→ | rpmE2 (PRK01678)Ribosomal_L31 (pfam01197)→ |
| Sau-683-1 | NZ_KK093607.1 | - | 29170 | 29141 | **RNA**→ | rpmE2 (PRK01678)Ribosomal_L31 (pfam01197)→ |
| Sau-3542-1 | NZ_KK038087.1 | - | 2155644 | 2155615 | **RNA**→ | rpmE2 (PRK01678)Ribosomal_L31 (pfam01197)→ |
| Sau-3533-1 | NZ_KK038099.1 | - | 56940 | 56911 | **RNA**→ | rpmE2 (PRK01678)Ribosomal_L31 (pfam01197)→ |
| Sau-2585-1 | NZ_KK020778.1 | - | 56893 | 56864 | **RNA**→ | rpmE2 (PRK01678)Ribosomal_L31 (pfam01197)→ |
| Sau-3573-1 | NZ_KK038019.1 | - | 56877 | 56848 | **RNA**→ | rpmE2 (PRK01678)Ribosomal_L31 (pfam01197)→ |
| Sau-3510-1 | NZ_KK038233.1 | - | 34872 | 34843 | **RNA**→ | rpmE2 (PRK01678)Ribosomal_L31 (pfam01197)→ |
| Sau-2933-1 | NZ_KK038394.1 | - | 56894 | 56865 | **RNA**→ | rpmE2 (PRK01678)Ribosomal_L31 (pfam01197)→ |
| Sau-2955-1 | NZ_KK038274.1 | - | 34817 | 34788 | **RNA**→ | rpmE2 (PRK01678)Ribosomal_L31 (pfam01197)→ |
| Sau-2925-1 | NZ_KK038441.1 | - | 56894 | 56865 | **RNA**→ | rpmE2 (PRK01678)Ribosomal_L31 (pfam01197)→ |
| Sau-3509-1 | NZ_KK038264.1 | - | 56794 | 56765 | **RNA**→ | rpmE2 (PRK01678)Ribosomal_L31 (pfam01197)→ |
| Sau-2931-1 | NZ_KK038399.1 | - | 34873 | 34844 | **RNA**→ | rpmE2 (PRK01678)Ribosomal_L31 (pfam01197)→ |
| Sau-696-1 | NZ_KK093807.1 | - | 34913 | 34884 | **RNA**→ | rpmE2 (PRK01678)Ribosomal_L31 (pfam01197)→ |
| Sau-2947-1 | NZ_KK038331.1 | - | 121725 | 121696 | **RNA**→ | rpmE2 (PRK01678)Ribosomal_L31 (pfam01197)→ |
| Sau-2923-1 | NZ_KK038451.1 | - | 56894 | 56865 | **RNA**→ | rpmE2 (PRK01678)Ribosomal_L31 (pfam01197)→ |
| Sau-2919-1 | NZ_KK038477.1 | - | 56789 | 56760 | **RNA**→ | rpmE2 (PRK01678)Ribosomal_L31 (pfam01197)→ |
| Sau-3572-1 | NZ_KK038059.1 | - | 56894 | 56865 | **RNA**→ | rpmE2 (PRK01678)Ribosomal_L31 (pfam01197)→ |
| Sau-688-1 | NZ_KK093755.1 | - | 34872 | 34843 | **RNA**→ | rpmE2 (PRK01678)Ribosomal_L31 (pfam01197)→ |
| Sau-793-1 | NZ_KK093938.1 | - | 55100 | 55071 | **RNA**→ | rpmE2 (PRK01678)Ribosomal_L31 (pfam01197)→ |
| Sau-920-1 | NZ_KK094093.1 | - | 54889 | 54860 | **RNA**→ | rpmE2 (PRK01678)Ribosomal_L31 (pfam01197)→ |
| Sau-908-1 | NZ_KK094033.1 | - | 34872 | 34843 | **RNA**→ | rpmE2 (PRK01678)Ribosomal_L31 (pfam01197)→ |
| Sau-1048-1 | NZ_KK093310.1 | - | 56894 | 56865 | **RNA**→ | rpmE2 (PRK01678)Ribosomal_L31 (pfam01197)→ |
| Sau-894-1 | NZ_KK093961.1 | - | 56889 | 56860 | **RNA**→ | rpmE2 (PRK01678)Ribosomal_L31 (pfam01197)→ |
| Sau-967-1 | NZ_KK094117.1 | - | 55199 | 55170 | **RNA**→ | rpmE2 (PRK01678)Ribosomal_L31 (pfam01197)→ |
| Sau-446-1 | NZ_KK098159.1 | - | 400765 | 400736 | **RNA**→ | rpmE2 (PRK01678)Ribosomal_L31 (pfam01197)→ |
| Sau-395-1 | NZ_KK099647.1 | - | 56709 | 56680 | **RNA**→ | rpmE2 (PRK01678)Ribosomal_L31 (pfam01197)→ |

| | | | | | | |
|---|---|---|---|---|---|---|
| Sau-424-1 | NZ_KK097860.1 | - | 470236 | 470207 | **RNA**→ | rpmE2 (PRK01678)Ribosomal_L31 (pfam01197)→ |
| Sau-444-1 | NZ_KK098125.1 | - | 56419 | 56390 | **RNA**→ | rpmE2 (PRK01678)Ribosomal_L31 (pfam01197)→ |
| Sau-465-1 | NZ_KK098357.1 | - | 121861 | 121832 | **RNA**→ | rpmE2 (PRK01678)Ribosomal_L31 (pfam01197)→ |
| Sau-2480-1 | NZ_KK094627.1 | - | 34973 | 34944 | **RNA**→ | rpmE2 (PRK01678)Ribosomal_L31 (pfam01197)→ |
| Sau-3579-1 | NZ_KK095731.1 | - | 55116 | 55087 | **RNA**→ | rpmE2 (PRK01678)Ribosomal_L31 (pfam01197)→ |
| Sau-111-1 | NZ_KK218822.1 | - | 56439 | 56410 | **RNA**→ | rpmE2 (PRK01678)Ribosomal_L31 (pfam01197)→ |
| Sau-2453-1 | NZ_KK096504.1 | - | 34972 | 34943 | **RNA**→ | rpmE2 (PRK01678)Ribosomal_L31 (pfam01197)→ |
| Sau-493-1 | NZ_KK098720.1 | - | 36391 | 36362 | **RNA**→ | rpmE2 (PRK01678)Ribosomal_L31 (pfam01197)→ |
| Sau-468-1 | NZ_KK098407.1 | - | 58283 | 58254 | **RNA**→ | rpmE2 (PRK01678)Ribosomal_L31 (pfam01197)→ |
| Sau-365-1 | NZ_KK099370.1 | - | 410097 | 410068 | **RNA**→ | rpmE2 (PRK01678)Ribosomal_L31 (pfam01197)→ |
| Sau-3536-1 | NZ_KK097202.1 | - | 34871 | 34842 | **RNA**→ | rpmE2 (PRK01678)Ribosomal_L31 (pfam01197)→ |
| Sau-422-1 | NZ_KK097842.1 | - | 908056 | 908027 | **RNA**→ | rpmE2 (PRK01678)Ribosomal_L31 (pfam01197)→ |
| Sau-329-1 | NZ_KK099108.1 | - | 466349 | 466320 | **RNA**→ | rpmE2 (PRK01678)Ribosomal_L31 (pfam01197)→ |
| Sau-2451-1 | NZ_KK094499.1 | - | 19182 | 19153 | **RNA**→ | rpmE2 (PRK01678)Ribosomal_L31 (pfam01197)→ |
| Sau-335-1 | NZ_KK099161.1 | - | 1644242 | 1644213 | **RNA**→ | rpmE2 (PRK01678)Ribosomal_L31 (pfam01197)→ |
| Sau-3597-1 | NZ_KK095813.1 | - | 41430 | 41401 | **RNA**→ | rpmE2 (PRK01678)Ribosomal_L31 (pfam01197)→ |
| Sau-117-1 | NZ_KK219060.1 | - | 1684 | 1655 | **RNA**→ | rpmE2 (PRK01678)Ribosomal_L31 (pfam01197)→ |
| Sau-915-1 | NZ_KK096211.1 | - | 35013 | 34984 | **RNA**→ | rpmE2 (PRK01678)Ribosomal_L31 (pfam01197)→ |
| Sau-414-1 | NZ_KK097788.1 | - | 56497 | 56468 | **RNA**→ | rpmE2 (PRK01678)Ribosomal_L31 (pfam01197)→ |
| Sau-355-1 | NZ_KK099294.1 | - | 59967 | 59938 | **RNA**→ | rpmE2 (PRK01678)Ribosomal_L31 (pfam01197)→ |
| Sau-641-1 | NZ_KK095880.1 | - | 34873 | 34844 | **RNA**→ | rpmE2 (PRK01678)Ribosomal_L31 (pfam01197)→ |
| Sau-2937-1 | NZ_KK097015.1 | - | 55117 | 55088 | **RNA**→ | rpmE2 (PRK01678)Ribosomal_L31 (pfam01197)→ |
| Sau-406-1 | NZ_KK099723.1 | - | 57585 | 57556 | **RNA**→ | rpmE2 (PRK01678)Ribosomal_L31 (pfam01197)→ |
| Sau-775-1 | NZ_KK096138.1 | - | 55250 | 55221 | **RNA**→ | rpmE2 (PRK01678)Ribosomal_L31 (pfam01197)→ |
| Sau-272-1 | NZ_KK223341.1 | - | 26024 | 25995 | **RNA**→ | rpmE2 (PRK01678)Ribosomal_L31 (pfam01197)→ |
| Sau-591-1 | NZ_KK099825.1 | - | 107062 | 107033 | **RNA**→ | rpmE2 (PRK01678)Ribosomal_L31 (pfam01197)→ |
| Sau-426-1 | NZ_KK097878.1 | - | 56763 | 56734 | **RNA**→ | rpmE2 (PRK01678)Ribosomal_L31 (pfam01197)→ |
| Sau-358-1 | NZ_KK099312.1 | - | 173069 | 173040 | **RNA**→ | rpmE2 (PRK01678)Ribosomal_L31 (pfam01197)→ |
| Sau-386-1 | NZ_KK099532.1 | - | 59566 | 59537 | **RNA**→ | rpmE2 (PRK01678)Ribosomal_L31 (pfam01197)→ |
| Sau-2637-1 | NZ_KK224655.1 | - | 26024 | 25995 | **RNA**→ | rpmE2 (PRK01678)Ribosomal_L31 (pfam01197)→ |
| Sau-154-1 | NZ_KK220279.1 | - | 56395 | 56366 | **RNA**→ | rpmE2 (PRK01678)Ribosomal_L31 (pfam01197)→ |
| Sau-970-1 | NZ_KK094165.1 | - | 34940 | 34911 | **RNA**→ | rpmE2 (PRK01678)Ribosomal_L31 (pfam01197)→ |
| Sau-458-1 | NZ_KK098272.1 | - | 61690 | 61661 | **RNA**→ | rpmE2 (PRK01678)Ribosomal_L31 (pfam01197)→ |
| Sau-125-1 | NZ_KK219385.1 | - | 54858 | 54829 | **RNA**→ | rpmE2 (PRK01678)Ribosomal_L31 (pfam01197)→ |
| Sau-2507-1 | NZ_KK094656.1 | - | 34872 | 34843 | **RNA**→ | rpmE2 (PRK01678)Ribosomal_L31 (pfam01197)→ |
| Sau-362-1 | NZ_KK099351.1 | - | 233436 | 233407 | **RNA**→ | rpmE2 (PRK01678)Ribosomal_L31 (pfam01197)→ |
| Sau-449-1 | NZ_KK098183.1 | - | 1089082 | 1089053 | **RNA**→ | rpmE2 (PRK01678)Ribosomal_L31 (pfam01197)→ |
| Sau-367-1 | NZ_KK099388.1 | - | 57279 | 57250 | **RNA**→ | rpmE2 (PRK01678)Ribosomal_L31 (pfam01197)→ |
| Sau-3070-1 | NZ_KK226556.1 | - | 56354 | 56325 | **RNA**→ | rpmE2 (PRK01678)Ribosomal_L31 (pfam01197)→ |
| Sau-309-1 | NZ_KK223593.1 | - | 29633 | 29604 | **RNA**→ | rpmE2 (PRK01678)Ribosomal_L31 (pfam01197)→ |
| Sau-2209-1 | NZ_KK094375.1 | - | 34873 | 34844 | **RNA**→ | rpmE2 (PRK01678)Ribosomal_L31 (pfam01197)→ |
| Sau-383-1 | NZ_KK099521.1 | - | 462571 | 462542 | **RNA**→ | rpmE2 (PRK01678)Ribosomal_L31 (pfam01197)→ |
| Sau-473-1 | NZ_KK098455.1 | - | 57586 | 57557 | **RNA**→ | rpmE2 (PRK01678)Ribosomal_L31 (pfam01197)→ |
| Sau-133-1 | NZ_KK219580.1 | - | 56395 | 56366 | **RNA**→ | rpmE2 (PRK01678)Ribosomal_L31 (pfam01197)→ |
| Sau-496-1 | NZ_KK098769.1 | - | 108030 | 108001 | **RNA**→ | rpmE2 (PRK01678)Ribosomal_L31 (pfam01197)→ |
| Sau-196-1 | NZ_KK221721.1 | - | 26024 | 25995 | **RNA**→ | rpmE2 (PRK01678)Ribosomal_L31 (pfam01197)→ |
| Sau-843-1 | NZ_KK223787.1 | - | 54494 | 54465 | **RNA**→ | rpmE2 (PRK01678)Ribosomal_L31 (pfam01197)→ |
| Sau-219-1 | NZ_KK222532.1 | - | 26024 | 25995 | **RNA**→ | rpmE2 (PRK01678)Ribosomal_L31 (pfam01197)→ |
| Sau-609-1 | NZ_KK223686.1 | - | 56439 | 56410 | **RNA**→ | rpmE2 (PRK01678)Ribosomal_L31 (pfam01197)→ |
| Sau-404-1 | NZ_KK099699.1 | - | 118463 | 118434 | **RNA**→ | rpmE2 (PRK01678)Ribosomal_L31 (pfam01197)→ |
| Sau-2714-1 | NZ_KK225547.1 | - | 26024 | 25995 | **RNA**→ | rpmE2 (PRK01678)Ribosomal_L31 (pfam01197)→ |
| Sau-375-1 | NZ_KK099461.1 | - | 1319543 | 1319514 | **RNA**→ | rpmE2 (PRK01678)Ribosomal_L31 (pfam01197)→ |

| | | | | | |
|---|---|---|---|---|---|
| Sau-3554-1 | NZ_KK095573.1 | - | 56893 | 56864 | **RNA**→ rpmE2 (PRK01678)Ribosomal_L31 (pfam01197)→ |
| Sau-3624-1 | NZ_KK097427.1 | - | 34913 | 34884 | **RNA**→ rpmE2 (PRK01678)Ribosomal_L31 (pfam01197)→ |
| Sau-2689-1 | NZ_KK224908.1 | - | 26024 | 25995 | **RNA**→ rpmE2 (PRK01678)Ribosomal_L31 (pfam01197)→ |
| Sau-3059-1 | NZ_KK226275.1 | - | 56395 | 56366 | **RNA**→ rpmE2 (PRK01678)Ribosomal_L31 (pfam01197)→ |
| Sau-869-1 | NZ_KK223977.1 | - | 56305 | 56276 | **RNA**→ rpmE2 (PRK01678)Ribosomal_L31 (pfam01197)→ |
| Sau-2952-1 | NZ_KK095283.1 | - | 55307 | 55278 | **RNA**→ rpmE2 (PRK01678)Ribosomal_L31 (pfam01197)→ |
| Sau-161-1 | NZ_KK220557.1 | - | 56395 | 56366 | **RNA**→ rpmE2 (PRK01678)Ribosomal_L31 (pfam01197)→ |
| Sau-593-1 | NZ_KK099779.1 | - | 56232 | 56203 | **RNA**→ rpmE2 (PRK01678)Ribosomal_L31 (pfam01197)→ |
| Sau-676-1 | NZ_KK095985.1 | - | 55117 | 55088 | **RNA**→ rpmE2 (PRK01678)Ribosomal_L31 (pfam01197)→ |
| Sau-169-1 | NZ_KK220854.1 | - | 2211 | 2182 | **RNA**→ rpmE2 (PRK01678)Ribosomal_L31 (pfam01197)→ |
| Sau-368-1 | NZ_KK099392.1 | - | 459748 | 459719 | **RNA**→ rpmE2 (PRK01678)Ribosomal_L31 (pfam01197)→ |
| Sau-590-1 | NZ_KK099840.1 | - | 59481 | 59452 | **RNA**→ rpmE2 (PRK01678)Ribosomal_L31 (pfam01197)→ |
| Sau-208-1 | NZ_KK222133.1 | - | 26024 | 25995 | **RNA**→ rpmE2 (PRK01678)Ribosomal_L31 (pfam01197)→ |
| Sau-480-1 | NZ_KK098543.1 | - | 56762 | 56733 | **RNA**→ rpmE2 (PRK01678)Ribosomal_L31 (pfam01197)→ |
| Sau-88-1 | NZ_KK218170.1 | - | 54759 | 54730 | **RNA**→ rpmE2 (PRK01678)Ribosomal_L31 (pfam01197)→ |
| Sau-451-1 | NZ_KK098208.1 | - | 56763 | 56734 | **RNA**→ rpmE2 (PRK01678)Ribosomal_L31 (pfam01197)→ |
| Sau-456-1 | NZ_KK098255.1 | - | 467917 | 467888 | **RNA**→ rpmE2 (PRK01678)Ribosomal_L31 (pfam01197)→ |
| Sau-350-1 | NZ_KK099258.1 | - | 459805 | 459776 | **RNA**→ rpmE2 (PRK01678)Ribosomal_L31 (pfam01197)→ |
| Sau-2953-1 | NZ_KK095290.1 | - | 55137 | 55108 | **RNA**→ rpmE2 (PRK01678)Ribosomal_L31 (pfam01197)→ |
| Sau-370-1 | NZ_KK099415.1 | - | 57012 | 56983 | **RNA**→ rpmE2 (PRK01678)Ribosomal_L31 (pfam01197)→ |
| Sau-460-1 | NZ_KK098302.1 | - | 434310 | 434281 | **RNA**→ rpmE2 (PRK01678)Ribosomal_L31 (pfam01197)→ |
| Sau-261-1 | NZ_KK222954.1 | - | 26024 | 25995 | **RNA**→ rpmE2 (PRK01678)Ribosomal_L31 (pfam01197)→ |
| Sau-441-1 | NZ_KK098077.1 | - | 238988 | 238959 | **RNA**→ rpmE2 (PRK01678)Ribosomal_L31 (pfam01197)→ |
| Sau-377-1 | NZ_KK099474.1 | - | 57059 | 57030 | **RNA**→ rpmE2 (PRK01678)Ribosomal_L31 (pfam01197)→ |
| Sau-91-1 | NZ_KK218289.1 | - | 26024 | 25995 | **RNA**→ rpmE2 (PRK01678)Ribosomal_L31 (pfam01197)→ |
| Sau-193-1 | NZ_KK221644.1 | - | 26024 | 25995 | **RNA**→ rpmE2 (PRK01678)Ribosomal_L31 (pfam01197)→ |
| Sau-157-1 | NZ_KK220407.1 | - | 56439 | 56410 | **RNA**→ rpmE2 (PRK01678)Ribosomal_L31 (pfam01197)→ |
| Sau-421-1 | NZ_KK097837.1 | - | 246307 | 246278 | **RNA**→ rpmE2 (PRK01678)Ribosomal_L31 (pfam01197)→ |
| Sau-127-1 | NZ_KK219466.1 | - | 54814 | 54785 | **RNA**→ rpmE2 (PRK01678)Ribosomal_L31 (pfam01197)→ |
| Sau-434-1 | NZ_KK097963.1 | - | 399430 | 399401 | **RNA**→ rpmE2 (PRK01678)Ribosomal_L31 (pfam01197)→ |
| Sau-393-1 | NZ_KK099625.1 | - | 1650562 | 1650533 | **RNA**→ rpmE2 (PRK01678)Ribosomal_L31 (pfam01197)→ |
| Sau-311-1 | NZ_KK099010.1 | + | 55670 | 55699 | **RNA**→ rpmE2 (PRK01678)Ribosomal_L31 (pfam01197)→ |
| Sau-342-1 | NZ_KK099199.1 | - | 59941 | 59912 | **RNA**→ rpmE2 (PRK01678)Ribosomal_L31 (pfam01197)→ |
| Sau-388-1 | NZ_KK099558.1 | - | 63060 | 63031 | **RNA**→ rpmE2 (PRK01678)Ribosomal_L31 (pfam01197)→ |
| Sau-179-1 | NZ_KK221204.1 | - | 1732 | 1703 | **RNA**→ rpmE2 (PRK01678)Ribosomal_L31 (pfam01197)→ |
| Sau-373-1 | NZ_KK099448.1 | - | 1640441 | 1640412 | **RNA**→ rpmE2 (PRK01678)Ribosomal_L31 (pfam01197)→ |
| Sau-2465-1 | NZ_KK094548.1 | - | 56894 | 56865 | **RNA**→ rpmE2 (PRK01678)Ribosomal_L31 (pfam01197)→ |
| Sau-104-1 | NZ_KK218547.1 | - | 56439 | 56410 | **RNA**→ rpmE2 (PRK01678)Ribosomal_L31 (pfam01197)→ |
| Sau-109-1 | NZ_KK218732.1 | - | 26024 | 25995 | **RNA**→ rpmE2 (PRK01678)Ribosomal_L31 (pfam01197)→ |
| Sau-482-1 | NZ_KK098582.1 | - | 56588 | 56559 | **RNA**→ rpmE2 (PRK01678)Ribosomal_L31 (pfam01197)→ |
| Sau-432-1 | NZ_KK097945.1 | - | 372981 | 372952 | **RNA**→ rpmE2 (PRK01678)Ribosomal_L31 (pfam01197)→ |
| Sau-470-1 | NZ_KK098421.1 | - | 109456 | 109427 | **RNA**→ rpmE2 (PRK01678)Ribosomal_L31 (pfam01197)→ |
| Sau-378-1 | NZ_KK099485.1 | - | 466934 | 466905 | **RNA**→ rpmE2 (PRK01678)Ribosomal_L31 (pfam01197)→ |
| Sau-397-1 | NZ_KK099653.1 | - | 1568744 | 1568715 | **RNA**→ rpmE2 (PRK01678)Ribosomal_L31 (pfam01197)→ |
| Sau-580-1 | NZ_KK098940.1 | - | 59584 | 59555 | **RNA**→ rpmE2 (PRK01678)Ribosomal_L31 (pfam01197)→ |
| Sau-596-1 | NZ_KK099745.1 | - | 59482 | 59453 | **RNA**→ rpmE2 (PRK01678)Ribosomal_L31 (pfam01197)→ |
| Sau-135-1 | NZ_KK219653.1 | - | 54858 | 54829 | **RNA**→ rpmE2 (PRK01678)Ribosomal_L31 (pfam01197)→ |
| Sau-452-1 | NZ_KK098220.1 | + | 681871 | 681900 | **RNA**→ rpmE2 (PRK01678)Ribosomal_L31 (pfam01197)→ |
| Sau-3108-1 | NZ_KK227993.1 | - | 26024 | 25995 | **RNA**→ rpmE2 (PRK01678)Ribosomal_L31 (pfam01197)→ |
| Sau-463-1 | NZ_KK098335.1 | - | 58446 | 58417 | **RNA**→ rpmE2 (PRK01678)Ribosomal_L31 (pfam01197)→ |
| Sau-131-1 | NZ_KK219851.1 | - | 54858 | 54829 | **RNA**→ rpmE2 (PRK01678)Ribosomal_L31 (pfam01197)→ |
| Sau-1087-1 | NZ_KK094299.1 | - | 34913 | 34884 | **RNA**→ rpmE2 (PRK01678)Ribosomal_L31 (pfam01197)→ |

| | | | | | |
|---|---|---|---|---|---|
| Sau-3096-1 | NZ_KK227523.1 | - | 26024 | 25995 | **RNA**→ rpmE2 (PRK01678)Ribosomal_L31 (pfam01197)→ |
| Sau-1086-1 | NZ_KK094251.1 | - | 34873 | 34844 | **RNA**→ rpmE2 (PRK01678)Ribosomal_L31 (pfam01197)→ |
| Sau-2450-1 | NZ_KK094446.1 | - | 34871 | 34842 | **RNA**→ rpmE2 (PRK01678)Ribosomal_L31 (pfam01197)→ |
| Sau-431-1 | NZ_KK097940.1 | - | 61789 | 61760 | **RNA**→ rpmE2 (PRK01678)Ribosomal_L31 (pfam01197)→ |
| Sau-3661-1 | NZ_KK097614.1 | - | 34872 | 34843 | **RNA**→ rpmE2 (PRK01678)Ribosomal_L31 (pfam01197)→ |
| Sau-3091-1 | NZ_KK227403.1 | - | 56305 | 56276 | **RNA**→ rpmE2 (PRK01678)Ribosomal_L31 (pfam01197)→ |
| Sau-164-1 | NZ_KK220717.1 | - | 26024 | 25995 | **RNA**→ rpmE2 (PRK01678)Ribosomal_L31 (pfam01197)→ |
| Sau-3098-1 | NZ_KK227605.1 | - | 26024 | 25995 | **RNA**→ rpmE2 (PRK01678)Ribosomal_L31 (pfam01197)→ |
| Sau-181-1 | NZ_KK221262.1 | - | 26024 | 25995 | **RNA**→ rpmE2 (PRK01678)Ribosomal_L31 (pfam01197)→ |
| Sau-866-1 | NZ_KK223867.1 | - | 26024 | 25995 | **RNA**→ rpmE2 (PRK01678)Ribosomal_L31 (pfam01197)→ |
| Sau-3103-1 | NZ_KK227783.1 | - | 26024 | 25995 | **RNA**→ rpmE2 (PRK01678)Ribosomal_L31 (pfam01197)→ |
| Sau-199-1 | NZ_KK221848.1 | - | 56395 | 56366 | **RNA**→ rpmE2 (PRK01678)Ribosomal_L31 (pfam01197)→ |
| Sau-871-1 | NZ_KK224053.1 | - | 15457 | 15428 | **RNA**→ rpmE2 (PRK01678)Ribosomal_L31 (pfam01197)→ |
| Sau-3114-1 | NZ_KK228187.1 | - | 26024 | 25995 | **RNA**→ rpmE2 (PRK01678)Ribosomal_L31 (pfam01197)→ |
| Sau-487-1 | NZ_KK098646.1 | - | 404926 | 404897 | **RNA**→ rpmE2 (PRK01678)Ribosomal_L31 (pfam01197)→ |
| Sau-3106-1 | NZ_KK227907.1 | - | 56439 | 56410 | **RNA**→ rpmE2 (PRK01678)Ribosomal_L31 (pfam01197)→ |
| Sau-439-1 | NZ_KK098013.1 | - | 56745 | 56716 | **RNA**→ rpmE2 (PRK01678)Ribosomal_L31 (pfam01197)→ |
| Sau-3117-1 | NZ_KK228324.1 | - | 26024 | 25995 | **RNA**→ rpmE2 (PRK01678)Ribosomal_L31 (pfam01197)→ |
| Sau-3131-1 | NZ_KK228748.1 | - | 26024 | 25995 | **RNA**→ rpmE2 (PRK01678)Ribosomal_L31 (pfam01197)→ |
| Sau-467-1 | NZ_KK098392.1 | - | 56588 | 56559 | **RNA**→ rpmE2 (PRK01678)Ribosomal_L31 (pfam01197)→ |
| Sau-475-1 | NZ_KK098491.1 | - | 58997 | 58968 | **RNA**→ rpmE2 (PRK01678)Ribosomal_L31 (pfam01197)→ |
| Sau-3518-1 | NZ_KK095445.1 | - | 34913 | 34884 | **RNA**→ rpmE2 (PRK01678)Ribosomal_L31 (pfam01197)→ |
| Sau-332-1 | NZ_KK099134.1 | - | 56588 | 56559 | **RNA**→ rpmE2 (PRK01678)Ribosomal_L31 (pfam01197)→ |
| Sau-485-1 | NZ_KK098623.1 | - | 58248 | 58219 | **RNA**→ rpmE2 (PRK01678)Ribosomal_L31 (pfam01197)→ |
| Sau-166-1 | NZ_KK220803.1 | - | 26185 | 26156 | **RNA**→ rpmE2 (PRK01678)Ribosomal_L31 (pfam01197)→ |
| Sau-579-1 | NZ_KK099886.1 | - | 237855 | 237826 | **RNA**→ rpmE2 (PRK01678)Ribosomal_L31 (pfam01197)→ |
| Sau-417-1 | NZ_KK097808.1 | - | 1414448 | 1414419 | **RNA**→ rpmE2 (PRK01678)Ribosomal_L31 (pfam01197)→ |
| Sau-577-1 | NZ_KK099906.1 | - | 58726 | 58697 | **RNA**→ rpmE2 (PRK01678)Ribosomal_L31 (pfam01197)→ |
| Sau-582-1 | NZ_KK098916.1 | - | 337935 | 337906 | **RNA**→ rpmE2 (PRK01678)Ribosomal_L31 (pfam01197)→ |
| Sau-348-1 | NZ_KK099241.1 | - | 119478 | 119449 | **RNA**→ rpmE2 (PRK01678)Ribosomal_L31 (pfam01197)→ |
| Sau-2456-1 | NZ_KK038959.1 | - | 233095 | 233066 | **RNA**→ rpmE2 (PRK01678)Ribosomal_L31 (pfam01197)→ |
| Sau-3139-1 | NZ_KK229019.1 | - | 53234 | 53205 | **RNA**→ rpmE2 (PRK01678)Ribosomal_L31 (pfam01197)→ |
| Sau-106-1 | NZ_KK218624.1 | - | 56439 | 56410 | **RNA**→ rpmE2 (PRK01678)Ribosomal_L31 (pfam01197)→ |
| Sau-3101-1 | NZ_KK227710.1 | - | 56439 | 56410 | **RNA**→ rpmE2 (PRK01678)Ribosomal_L31 (pfam01197)→ |
| Sau-2208-1 | NZ_KK039024.1 | - | 36293 | 36264 | **RNA**→ rpmE2 (PRK01678)Ribosomal_L31 (pfam01197)→ |
| Sau-254-1 | NZ_KK222726.1 | - | 26024 | 25995 | **RNA**→ rpmE2 (PRK01678)Ribosomal_L31 (pfam01197)→ |
| Sau-3124-1 | NZ_KK228551.1 | - | 26024 | 25995 | **RNA**→ rpmE2 (PRK01678)Ribosomal_L31 (pfam01197)→ |
| Sau-1079-1 | NZ_KK039281.1 | - | 34793 | 34764 | **RNA**→ rpmE2 (PRK01678)Ribosomal_L31 (pfam01197)→ |
| Sau-318-1 | NZ_KK099065.1 | - | 56568 | 56539 | **RNA**→ rpmE2 (PRK01678)Ribosomal_L31 (pfam01197)→ |
| Sau-3144-1 | NZ_KK229110.1 | - | 56390 | 56361 | **RNA**→ rpmE2 (PRK01678)Ribosomal_L31 (pfam01197)→ |
| Sau-735-1 | NZ_KK039382.1 | - | 34970 | 34941 | **RNA**→ rpmE2 (PRK01678)Ribosomal_L31 (pfam01197)→ |
| Sau-120-1 | NZ_KK219160.1 | - | 26024 | 25995 | **RNA**→ rpmE2 (PRK01678)Ribosomal_L31 (pfam01197)→ |
| Sau-899-1 | NZ_KK039331.1 | - | 56994 | 56965 | **RNA**→ rpmE2 (PRK01678)Ribosomal_L31 (pfam01197)→ |
| Sau-3620-1 | NZ_KK039998.1 | - | 70352 | 70323 | **RNA**→ rpmE2 (PRK01678)Ribosomal_L31 (pfam01197)→ |
| Sau-352-1 | NZ_KK099276.1 | - | 57586 | 57557 | **RNA**→ rpmE2 (PRK01678)Ribosomal_L31 (pfam01197)→ |
| Sau-692-1 | NZ_KK039589.1 | - | 34817 | 34788 | **RNA**→ rpmE2 (PRK01678)Ribosomal_L31 (pfam01197)→ |
| Sau-3502-1 | NZ_KK040133.1 | - | 34817 | 34788 | **RNA**→ rpmE2 (PRK01678)Ribosomal_L31 (pfam01197)→ |
| Sau-691-1 | NZ_KK039640.1 | - | 34873 | 34844 | **RNA**→ rpmE2 (PRK01678)Ribosomal_L31 (pfam01197)→ |
| Sau-3626-1 | NZ_KK039943.1 | - | 34873 | 34844 | **RNA**→ rpmE2 (PRK01678)Ribosomal_L31 (pfam01197)→ |
| Sau-345-1 | NZ_KK099219.1 | - | 461736 | 461707 | **RNA**→ rpmE2 (PRK01678)Ribosomal_L31 (pfam01197)→ |
| Sau-3494-1 | NZ_KK040167.1 | - | 34872 | 34843 | **RNA**→ rpmE2 (PRK01678)Ribosomal_L31 (pfam01197)→ |
| Sau-2290-1 | NZ_KI972513.1 | - | 476429 | 476400 | **RNA**→ rpmE2 (PRK01678)Ribosomal_L31 (pfam01197)→ |

| | | | | | |
|---|---|---|---|---|---|
| Sau-401-1 | NZ_KK099682.1 | - | 1641421 | 1641392 | **RNA**→ rpmE2 (PRK01678)Ribosomal_L31 (pfam01197)→ |
| Sau-3121-1 | NZ_KK228395.1 | - | 56354 | 56325 | **RNA**→ rpmE2 (PRK01678)Ribosomal_L31 (pfam01197)→ |
| Sau-1083-1 | NZ_KK039143.1 | - | 34873 | 34844 | **RNA**→ rpmE2 (PRK01678)Ribosomal_L31 (pfam01197)→ |
| Sau-178-1 | NZ_KK221159.1 | - | 56395 | 56366 | **RNA**→ rpmE2 (PRK01678)Ribosomal_L31 (pfam01197)→ |
| Sau-3152-1 | NZ_KK229299.1 | - | 26024 | 25995 | **RNA**→ rpmE2 (PRK01678)Ribosomal_L31 (pfam01197)→ |
| Sau-729-1 | NZ_KK039514.1 | - | 34913 | 34884 | **RNA**→ rpmE2 (PRK01678)Ribosomal_L31 (pfam01197)→ |
| Sau-3111-1 | NZ_KK228117.1 | - | 26024 | 25995 | **RNA**→ rpmE2 (PRK01678)Ribosomal_L31 (pfam01197)→ |
| Sau-266-1 | NZ_KK223125.1 | - | 56439 | 56410 | **RNA**→ rpmE2 (PRK01678)Ribosomal_L31 (pfam01197)→ |
| Sau-1081-1 | NZ_KK039198.1 | - | 34873 | 34844 | **RNA**→ rpmE2 (PRK01678)Ribosomal_L31 (pfam01197)→ |
| Sau-3638-1 | NZ_KK039930.1 | - | 56893 | 56864 | **RNA**→ rpmE2 (PRK01678)Ribosomal_L31 (pfam01197)→ |
| Sau-201-1 | NZ_KK221926.1 | - | 26024 | 25995 | **RNA**→ rpmE2 (PRK01678)Ribosomal_L31 (pfam01197)→ |
| Sau-675-1 | NZ_KK039788.1 | - | 56894 | 56865 | **RNA**→ rpmE2 (PRK01678)Ribosomal_L31 (pfam01197)→ |
| Sau-3602-1 | NZ_KK040125.1 | - | 56894 | 56865 | **RNA**→ rpmE2 (PRK01678)Ribosomal_L31 (pfam01197)→ |
| Sau-928-1 | NZ_KK039302.1 | - | 34817 | 34788 | **RNA**→ rpmE2 (PRK01678)Ribosomal_L31 (pfam01197)→ |
| Sau-213-1 | NZ_KK222349.1 | - | 56439 | 56410 | **RNA**→ rpmE2 (PRK01678)Ribosomal_L31 (pfam01197)→ |
| Sau-690-1 | NZ_KK039683.1 | - | 34817 | 34788 | **RNA**→ rpmE2 (PRK01678)Ribosomal_L31 (pfam01197)→ |
| Sau-85-1 | NZ_KK218112.1 | - | 26024 | 25995 | **RNA**→ rpmE2 (PRK01678)Ribosomal_L31 (pfam01197)→ |
| Sau-2510-1 | NZ_KK040956.1 | - | 34913 | 34884 | **RNA**→ rpmE2 (PRK01678)Ribosomal_L31 (pfam01197)→ |
| Sau-4040-1 | NZ_ABWS01000010.1 | - | 56850 | 56821 | **RNA**→ rpmE2 (PRK01678)Ribosomal_L31 (pfam01197)→ |
| Sau-4021-1 | NZ_AFTT01000003.1 | - | 54788 | 54759 | **RNA**→ rpmE2 (PRK01678)Ribosomal_L31 (pfam01197)→ |
| Sau-4026-1 | NZ_AFNO01000007.1 | - | 54488 | 54459 | **RNA**→ rpmE2 (PRK01678)Ribosomal_L31 (pfam01197)→ |
| Sau-4011-1 | NZ_AFTO01000031.1 | - | 25119 | 25090 | **RNA**→ rpmE2 (PRK01678)Ribosomal_L31 (pfam01197)→ |
| Sau-4018-1 | NZ_AFTS01000008.1 | - | 54482 | 54453 | **RNA**→ rpmE2 (PRK01678)Ribosomal_L31 (pfam01197)→ |
| Sau-4007-1 | NZ_AGTU01000208.1 | + | 6464 | 6493 | **RNA**→ rpmE2 (PRK01678)Ribosomal_L31 (pfam01197)→ |
| Sau-4022-1 | NZ_AFTU01000056.1 | - | 54711 | 54682 | **RNA**→ rpmE2 (PRK01678)Ribosomal_L31 (pfam01197)→ |
| Sau-3073-1 | NZ_KK226674.1 | - | 26024 | 25995 | **RNA**→ rpmE2 (PRK01678)Ribosomal_L31 (pfam01197)→ |
| Sau-2697-1 | NZ_KK225236.1 | - | 25968 | 25939 | **RNA**→ rpmE2 (PRK01678)Ribosomal_L31 (pfam01197)→ |
| Sau-454-1 | NZ_KK098243.1 | - | 56418 | 56389 | **RNA**→ rpmE2 (PRK01678)Ribosomal_L31 (pfam01197)→ |
| Sau-3012-1 | NZ_KK226129.1 | - | 26024 | 25995 | **RNA**→ rpmE2 (PRK01678)Ribosomal_L31 (pfam01197)→ |
| Sau-353-1 | NZ_KK099285.1 | - | 232864 | 232835 | **RNA**→ rpmE2 (PRK01678)Ribosomal_L31 (pfam01197)→ |
| Sau-2677-1 | NZ_KK224759.1 | - | 56390 | 56361 | **RNA**→ rpmE2 (PRK01678)Ribosomal_L31 (pfam01197)→ |
| Sau-437-1 | NZ_KK097990.1 | - | 467385 | 467356 | **RNA**→ rpmE2 (PRK01678)Ribosomal_L31 (pfam01197)→ |
| Sau-159-1 | NZ_KK220472.1 | - | 18748 | 18719 | **RNA**→ rpmE2 (PRK01678)Ribosomal_L31 (pfam01197)→ |
| Sau-491-1 | NZ_KK098694.1 | - | 1600347 | 1600318 | **RNA**→ rpmE2 (PRK01678)Ribosomal_L31 (pfam01197)→ |
| Sau-403-1 | NZ_KK099696.1 | - | 57586 | 57557 | **RNA**→ rpmE2 (PRK01678)Ribosomal_L31 (pfam01197)→ |
| Sau-259-1 | NZ_KK222863.1 | - | 56935 | 56906 | **RNA**→ rpmE2 (PRK01678)Ribosomal_L31 (pfam01197)→ |
| Sau-3016-1 | NZ_KK226203.1 | - | 55550 | 55521 | **RNA**→ rpmE2 (PRK01678)Ribosomal_L31 (pfam01197)→ |
| Sau-277-1 | NZ_KK223489.1 | - | 26024 | 25995 | **RNA**→ rpmE2 (PRK01678)Ribosomal_L31 (pfam01197)→ |
| Sau-3064-1 | NZ_KK226405.1 | - | 26024 | 25995 | **RNA**→ rpmE2 (PRK01678)Ribosomal_L31 (pfam01197)→ |
| Sau-141-1 | NZ_KK219948.1 | - | 54814 | 54785 | **RNA**→ rpmE2 (PRK01678)Ribosomal_L31 (pfam01197)→ |
| Sau-993-1 | NZ_KK096312.1 | - | 56894 | 56865 | **RNA**→ rpmE2 (PRK01678)Ribosomal_L31 (pfam01197)→ |
| Sau-1014-1 | NZ_KK096345.1 | - | 56790 | 56761 | **RNA**→ rpmE2 (PRK01678)Ribosomal_L31 (pfam01197)→ |
| Sau-3066-1 | NZ_KK226495.1 | - | 56354 | 56325 | **RNA**→ rpmE2 (PRK01678)Ribosomal_L31 (pfam01197)→ |
| Sau-322-1 | NZ_KK099076.1 | - | 1682860 | 1682831 | **RNA**→ rpmE2 (PRK01678)Ribosomal_L31 (pfam01197)→ |
| Sau-2820-1 | NZ_KK225657.1 | - | 56395 | 56366 | **RNA**→ rpmE2 (PRK01678)Ribosomal_L31 (pfam01197)→ |
| Sau-391-1 | NZ_KK099601.1 | - | 59482 | 59453 | **RNA**→ rpmE2 (PRK01678)Ribosomal_L31 (pfam01197)→ |
| Sau-30-1 | NZ_GL397103.1 | - | 275482 | 275453 | **RNA**→ rpmE2 (PRK01678)Ribosomal_L31 (pfam01197)→ |
| Sau-2936-1 | NZ_KK095262.1 | - | 34873 | 34844 | **RNA**→ rpmE2 (PRK01678)Ribosomal_L31 (pfam01197)→ |
| Sau-2985-1 | NZ_KK095314.1 | - | 56790 | 56761 | **RNA**→ rpmE2 (PRK01678)Ribosomal_L31 (pfam01197)→ |
| Sau-3530-1 | NZ_KK095505.1 | - | 25255 | 25226 | **RNA**→ rpmE2 (PRK01678)Ribosomal_L31 (pfam01197)→ |
| Sau-3656-1 | NZ_KK093561.1 | - | 15617 | 15588 | **RNA**→ rpmE2 (PRK01678)Ribosomal_L31 (pfam01197)→ |
| Sau-3581-1 | NZ_KK092334.1 | - | 25129 | 25100 | **RNA**→ rpmE2 (PRK01678)Ribosomal_L31 (pfam01197)→ |

| | | | | | | |
|---|---|---|---|---|---|---|
| Sau-3561-1 | NZ_KK095655.1 | - | 34913 | 34884 | **RNA**→ | rpmE2 (PRK01678)Ribosomal_L31 (pfam01197)→ |
| Sau-3519-1 | NZ_KK097131.1 | - | 56994 | 56965 | **RNA**→ | rpmE2 (PRK01678)Ribosomal_L31 (pfam01197)→ |
| Sau-3549-1 | NZ_KK097344.1 | - | 35013 | 34984 | **RNA**→ | rpmE2 (PRK01678)Ribosomal_L31 (pfam01197)→ |
| Sau-3635-1 | NZ_KK097502.1 | - | 25255 | 25226 | **RNA**→ | rpmE2 (PRK01678)Ribosomal_L31 (pfam01197)→ |
| Sau-3585-1 | NZ_KK095753.1 | - | 56992 | 56963 | **RNA**→ | rpmE2 (PRK01678)Ribosomal_L31 (pfam01197)→ |
| Sau-3562-1 | NZ_KK095700.1 | - | 56894 | 56865 | **RNA**→ | rpmE2 (PRK01678)Ribosomal_L31 (pfam01197)→ |
| Sau-3640-1 | NZ_KK097564.1 | - | 34873 | 34844 | **RNA**→ | rpmE2 (PRK01678)Ribosomal_L31 (pfam01197)→ |
| Sau-2926-1 | NZ_KK095190.1 | - | 56893 | 56864 | **RNA**→ | rpmE2 (PRK01678)Ribosomal_L31 (pfam01197)→ |
| Sau-2960-1 | NZ_KK095306.1 | - | 55166 | 55137 | **RNA**→ | rpmE2 (PRK01678)Ribosomal_L31 (pfam01197)→ |
| Sau-3556-1 | NZ_KK095597.1 | - | 55082 | 55053 | **RNA**→ | rpmE2 (PRK01678)Ribosomal_L31 (pfam01197)→ |
| Sau-3596-1 | NZ_KK095778.1 | - | 34913 | 34884 | **RNA**→ | rpmE2 (PRK01678)Ribosomal_L31 (pfam01197)→ |
| Sau-2945-1 | NZ_KK097035.1 | - | 19182 | 19153 | **RNA**→ | rpmE2 (PRK01678)Ribosomal_L31 (pfam01197)→ |
| Sau-3537-1 | NZ_KK097238.1 | - | 34873 | 34844 | **RNA**→ | rpmE2 (PRK01678)Ribosomal_L31 (pfam01197)→ |
| Sau-3625-1 | NZ_KK097452.1 | - | 56894 | 56865 | **RNA**→ | rpmE2 (PRK01678)Ribosomal_L31 (pfam01197)→ |
| Sau-626-1 | NZ_KK095848.1 | - | 55116 | 55087 | **RNA**→ | rpmE2 (PRK01678)Ribosomal_L31 (pfam01197)→ |
| Sau-447-1 | NZ_KK098166.1 | - | 120192 | 120163 | **RNA**→ | rpmE2 (PRK01678)Ribosomal_L31 (pfam01197)→ |
| Sau-628-1 | NZ_KK093018.1 | - | 34972 | 34943 | **RNA**→ | rpmE2 (PRK01678)Ribosomal_L31 (pfam01197)→ |
| Sau-687-1 | NZ_KK093705.1 | - | 34872 | 34843 | **RNA**→ | rpmE2 (PRK01678)Ribosomal_L31 (pfam01197)→ |
| Sau-652-1 | NZ_KK095905.1 | - | 56894 | 56865 | **RNA**→ | rpmE2 (PRK01678)Ribosomal_L31 (pfam01197)→ |
| Sau-706-1 | NZ_KK096064.1 | - | 56894 | 56865 | **RNA**→ | rpmE2 (PRK01678)Ribosomal_L31 (pfam01197)→ |
| Sau-448-1 | NZ_KK098177.1 | - | 36791 | 36762 | **RNA**→ | rpmE2 (PRK01678)Ribosomal_L31 (pfam01197)→ |
| Sau-474-1 | NZ_KK098474.1 | - | 471096 | 471067 | **RNA**→ | rpmE2 (PRK01678)Ribosomal_L31 (pfam01197)→ |
| Sau-614-1 | NZ_KK092835.1 | - | 34972 | 34943 | **RNA**→ | rpmE2 (PRK01678)Ribosomal_L31 (pfam01197)→ |
| Sau-685-1 | NZ_KK093635.1 | - | 34756 | 34727 | **RNA**→ | rpmE2 (PRK01678)Ribosomal_L31 (pfam01197)→ |
| Sau-709-1 | NZ_KK093823.1 | - | 56893 | 56864 | **RNA**→ | rpmE2 (PRK01678)Ribosomal_L31 (pfam01197)→ |
| Sau-658-1 | NZ_KK093149.1 | - | 55082 | 55053 | **RNA**→ | rpmE2 (PRK01678)Ribosomal_L31 (pfam01197)→ |
| Sau-430-1 | NZ_KK097929.1 | - | 1652335 | 1652306 | **RNA**→ | rpmE2 (PRK01678)Ribosomal_L31 (pfam01197)→ |
| Sau-435-1 | NZ_KK097974.1 | - | 61771 | 61742 | **RNA**→ | rpmE2 (PRK01678)Ribosomal_L31 (pfam01197)→ |
| Sau-440-1 | NZ_KK098041.1 | - | 56763 | 56734 | **RNA**→ | rpmE2 (PRK01678)Ribosomal_L31 (pfam01197)→ |
| Sau-445-1 | NZ_KK098149.1 | - | 57586 | 57557 | **RNA**→ | rpmE2 (PRK01678)Ribosomal_L31 (pfam01197)→ |
| Sau-450-1 | NZ_KK098195.1 | - | 57586 | 57557 | **RNA**→ | rpmE2 (PRK01678)Ribosomal_L31 (pfam01197)→ |
| Sau-606-1 | NZ_KK098961.1 | - | 62155 | 62126 | **RNA**→ | rpmE2 (PRK01678)Ribosomal_L31 (pfam01197)→ |
| Sau-438-1 | NZ_KK097995.1 | - | 466008 | 465979 | **RNA**→ | rpmE2 (PRK01678)Ribosomal_L31 (pfam01197)→ |
| Sau-425-1 | NZ_KK097867.1 | - | 231594 | 231565 | **RNA**→ | rpmE2 (PRK01678)Ribosomal_L31 (pfam01197)→ |
| Sau-443-1 | NZ_KK098104.1 | - | 469570 | 469541 | **RNA**→ | rpmE2 (PRK01678)Ribosomal_L31 (pfam01197)→ |
| Sau-453-1 | NZ_KK098229.1 | - | 254240 | 254211 | **RNA**→ | rpmE2 (PRK01678)Ribosomal_L31 (pfam01197)→ |
| Sau-459-1 | NZ_KK098290.1 | - | 56763 | 56734 | **RNA**→ | rpmE2 (PRK01678)Ribosomal_L31 (pfam01197)→ |
| Sau-469-1 | NZ_KK098416.1 | - | 56419 | 56390 | **RNA**→ | rpmE2 (PRK01678)Ribosomal_L31 (pfam01197)→ |
| Sau-479-1 | NZ_KK098532.1 | - | 1650917 | 1650888 | **RNA**→ | rpmE2 (PRK01678)Ribosomal_L31 (pfam01197)→ |
| Sau-575-1 | NZ_KK099949.1 | - | 59485 | 59456 | **RNA**→ | rpmE2 (PRK01678)Ribosomal_L31 (pfam01197)→ |
| Sau-703-1 | NZ_KK096018.1 | - | 54999 | 54970 | **RNA**→ | rpmE2 (PRK01678)Ribosomal_L31 (pfam01197)→ |
| Sau-478-1 | NZ_KK098526.1 | - | 784642 | 784613 | **RNA**→ | rpmE2 (PRK01678)Ribosomal_L31 (pfam01197)→ |
| Sau-483-1 | NZ_KK098587.1 | - | 826998 | 826969 | **RNA**→ | rpmE2 (PRK01678)Ribosomal_L31 (pfam01197)→ |
| Sau-484-1 | NZ_KK098605.1 | - | 56763 | 56734 | **RNA**→ | rpmE2 (PRK01678)Ribosomal_L31 (pfam01197)→ |
| Sau-490-1 | NZ_KK098680.1 | - | 56763 | 56734 | **RNA**→ | rpmE2 (PRK01678)Ribosomal_L31 (pfam01197)→ |
| Sau-607-1 | NZ_KK099001.1 | - | 976089 | 976060 | **RNA**→ | rpmE2 (PRK01678)Ribosomal_L31 (pfam01197)→ |
| Sau-495-1 | NZ_KK098762.1 | - | 61689 | 61660 | **RNA**→ | rpmE2 (PRK01678)Ribosomal_L31 (pfam01197)→ |
| Sau-585-1 | NZ_KK098836.1 | - | 60916 | 60887 | **RNA**→ | rpmE2 (PRK01678)Ribosomal_L31 (pfam01197)→ |
| Sau-728-1 | NZ_KK096097.1 | - | 34913 | 34884 | **RNA**→ | rpmE2 (PRK01678)Ribosomal_L31 (pfam01197)→ |
| Sau-455-1 | NZ_KK098247.1 | - | 57585 | 57556 | **RNA**→ | rpmE2 (PRK01678)Ribosomal_L31 (pfam01197)→ |
| Sau-461-1 | NZ_KK098312.1 | - | 1097381 | 1097352 | **RNA**→ | rpmE2 (PRK01678)Ribosomal_L31 (pfam01197)→ |
| Sau-466-1 | NZ_KK098379.1 | - | 56763 | 56734 | **RNA**→ | rpmE2 (PRK01678)Ribosomal_L31 (pfam01197)→ |

| | | | | | | |
|---|---|---|---|---|---|---|
| Sau-471-1 | NZ_KK098433.1 | - | 1674196 | 1674167 | **RNA**→ | rpmE2 (PRK01678)Ribosomal_L31 (pfam01197)→ |
| Sau-481-1 | NZ_KK098572.1 | - | 1944061 | 1944032 | **RNA**→ | rpmE2 (PRK01678)Ribosomal_L31 (pfam01197)→ |
| Sau-486-1 | NZ_KK098636.1 | - | 488835 | 488806 | **RNA**→ | rpmE2 (PRK01678)Ribosomal_L31 (pfam01197)→ |
| Sau-492-1 | NZ_KK098700.1 | - | 56416 | 56387 | **RNA**→ | rpmE2 (PRK01678)Ribosomal_L31 (pfam01197)→ |
| Sau-592-1 | NZ_KK099811.1 | - | 58064 | 58035 | **RNA**→ | rpmE2 (PRK01678)Ribosomal_L31 (pfam01197)→ |
| Sau-586-1 | NZ_KK098796.1 | - | 59482 | 59453 | **RNA**→ | rpmE2 (PRK01678)Ribosomal_L31 (pfam01197)→ |
| Sau-581-1 | NZ_KK098920.1 | - | 59482 | 59453 | **RNA**→ | rpmE2 (PRK01678)Ribosomal_L31 (pfam01197)→ |
| Sau-733-1 | NZ_KK093853.1 | - | 34873 | 34844 | **RNA**→ | rpmE2 (PRK01678)Ribosomal_L31 (pfam01197)→ |
| Sau-2211-1 | NZ_KK094430.1 | - | 56434 | 56405 | **RNA**→ | rpmE2 (PRK01678)Ribosomal_L31 (pfam01197)→ |
| Sau-2837-1 | NZ_KK095014.1 | - | 56793 | 56764 | **RNA**→ | rpmE2 (PRK01678)Ribosomal_L31 (pfam01197)→ |
| Sau-2916-1 | NZ_KK095141.1 | - | 34872 | 34843 | **RNA**→ | rpmE2 (PRK01678)Ribosomal_L31 (pfam01197)→ |
| Sau-1091-1 | NZ_KK096393.1 | - | 35074 | 35045 | **RNA**→ | rpmE2 (PRK01678)Ribosomal_L31 (pfam01197)→ |
| Sau-415-1 | NZ_KK097796.1 | - | 1631360 | 1631331 | **RNA**→ | rpmE2 (PRK01678)Ribosomal_L31 (pfam01197)→ |
| Sau-2560-1 | NZ_KK096785.1 | - | 55181 | 55152 | **RNA**→ | rpmE2 (PRK01678)Ribosomal_L31 (pfam01197)→ |
| Sau-314-1 | NZ_KK099035.1 | - | 60257 | 60228 | **RNA**→ | rpmE2 (PRK01678)Ribosomal_L31 (pfam01197)→ |
| Sau-1070-1 | NZ_KK094214.1 | - | 34874 | 34845 | **RNA**→ | rpmE2 (PRK01678)Ribosomal_L31 (pfam01197)→ |
| Sau-319-1 | NZ_KK099068.1 | - | 1269188 | 1269159 | **RNA**→ | rpmE2 (PRK01678)Ribosomal_L31 (pfam01197)→ |
| Sau-340-1 | NZ_KK099188.1 | - | 57320 | 57291 | **RNA**→ | rpmE2 (PRK01678)Ribosomal_L31 (pfam01197)→ |
| Sau-351-1 | NZ_KK099273.1 | - | 1590185 | 1590156 | **RNA**→ | rpmE2 (PRK01678)Ribosomal_L31 (pfam01197)→ |
| Sau-356-1 | NZ_KK099300.1 | - | 1636052 | 1636023 | **RNA**→ | rpmE2 (PRK01678)Ribosomal_L31 (pfam01197)→ |
| Sau-361-1 | NZ_KK099337.1 | - | 61553 | 61524 | **RNA**→ | rpmE2 (PRK01678)Ribosomal_L31 (pfam01197)→ |
| Sau-371-1 | NZ_KK099427.1 | - | 1594352 | 1594323 | **RNA**→ | rpmE2 (PRK01678)Ribosomal_L31 (pfam01197)→ |
| Sau-746-1 | NZ_KK092605.1 | - | 602 | 573 | **RNA**→ | rpmE2 (PRK01678)Ribosomal_L31 (pfam01197)→ |
| Sau-428-1 | NZ_KK097904.1 | - | 114869 | 114840 | **RNA**→ | rpmE2 (PRK01678)Ribosomal_L31 (pfam01197)→ |
| Sau-349-1 | NZ_KK099251.1 | - | 459747 | 459718 | **RNA**→ | rpmE2 (PRK01678)Ribosomal_L31 (pfam01197)→ |
| Sau-359-1 | NZ_KK099321.1 | - | 1593274 | 1593245 | **RNA**→ | rpmE2 (PRK01678)Ribosomal_L31 (pfam01197)→ |
| Sau-364-1 | NZ_KK099361.1 | - | 255990 | 255961 | **RNA**→ | rpmE2 (PRK01678)Ribosomal_L31 (pfam01197)→ |
| Sau-369-1 | NZ_KK099403.1 | - | 460546 | 460517 | **RNA**→ | rpmE2 (PRK01678)Ribosomal_L31 (pfam01197)→ |
| Sau-374-1 | NZ_KK099456.1 | - | 60524 | 60495 | **RNA**→ | rpmE2 (PRK01678)Ribosomal_L31 (pfam01197)→ |
| Sau-405-1 | NZ_KK099718.1 | - | 56726 | 56697 | **RNA**→ | rpmE2 (PRK01678)Ribosomal_L31 (pfam01197)→ |
| Sau-1088-1 | NZ_KK094350.1 | - | 56794 | 56765 | **RNA**→ | rpmE2 (PRK01678)Ribosomal_L31 (pfam01197)→ |
| Sau-2464-1 | NZ_KK094533.1 | - | 56894 | 56865 | **RNA**→ | rpmE2 (PRK01678)Ribosomal_L31 (pfam01197)→ |
| Sau-2521-1 | NZ_KK094681.1 | - | 15617 | 15588 | **RNA**→ | rpmE2 (PRK01678)Ribosomal_L31 (pfam01197)→ |
| Sau-2595-1 | NZ_KK094924.1 | - | 34913 | 34884 | **RNA**→ | rpmE2 (PRK01678)Ribosomal_L31 (pfam01197)→ |
| Sau-2546-1 | NZ_KK096768.1 | - | 56894 | 56865 | **RNA**→ | rpmE2 (PRK01678)Ribosomal_L31 (pfam01197)→ |
| Sau-2840-1 | NZ_KK096936.1 | - | 34913 | 34884 | **RNA**→ | rpmE2 (PRK01678)Ribosomal_L31 (pfam01197)→ |
| Sau-2858-1 | NZ_KK096966.1 | - | 54878 | 54849 | **RNA**→ | rpmE2 (PRK01678)Ribosomal_L31 (pfam01197)→ |
| Sau-902-1 | NZ_KK093986.1 | - | 34913 | 34884 | **RNA**→ | rpmE2 (PRK01678)Ribosomal_L31 (pfam01197)→ |
| Sau-968-1 | NZ_KK094143.1 | - | 34874 | 34845 | **RNA**→ | rpmE2 (PRK01678)Ribosomal_L31 (pfam01197)→ |
| Sau-418-1 | NZ_KK097815.1 | - | 2121940 | 2121911 | **RNA**→ | rpmE2 (PRK01678)Ribosomal_L31 (pfam01197)→ |
| Sau-423-1 | NZ_KK097848.1 | - | 1643406 | 1643377 | **RNA**→ | rpmE2 (PRK01678)Ribosomal_L31 (pfam01197)→ |
| Sau-2909-1 | NZ_KK095085.1 | - | 57030 | 57001 | **RNA**→ | rpmE2 (PRK01678)Ribosomal_L31 (pfam01197)→ |
| Sau-392-1 | NZ_KK099615.1 | - | 244736 | 244707 | **RNA**→ | rpmE2 (PRK01678)Ribosomal_L31 (pfam01197)→ |
| Sau-945-1 | NZ_KK096297.1 | - | 56894 | 56865 | **RNA**→ | rpmE2 (PRK01678)Ribosomal_L31 (pfam01197)→ |
| Sau-2534-1 | NZ_KK096682.1 | - | 19182 | 19153 | **RNA**→ | rpmE2 (PRK01678)Ribosomal_L31 (pfam01197)→ |
| Sau-2886-1 | NZ_KK093486.1 | - | 56794 | 56765 | **RNA**→ | rpmE2 (PRK01678)Ribosomal_L31 (pfam01197)→ |
| Sau-315-1 | NZ_KK099043.1 | - | 56467 | 56438 | **RNA**→ | rpmE2 (PRK01678)Ribosomal_L31 (pfam01197)→ |
| Sau-321-1 | NZ_KK099074.1 | - | 238688 | 238659 | **RNA**→ | rpmE2 (PRK01678)Ribosomal_L31 (pfam01197)→ |
| Sau-734-1 | NZ_KK093901.1 | - | 34873 | 34844 | **RNA**→ | rpmE2 (PRK01678)Ribosomal_L31 (pfam01197)→ |
| Sau-334-1 | NZ_KK099151.1 | - | 1670304 | 1670275 | **RNA**→ | rpmE2 (PRK01678)Ribosomal_L31 (pfam01197)→ |
| Sau-341-1 | NZ_KK099197.1 | - | 122934 | 122905 | **RNA**→ | rpmE2 (PRK01678)Ribosomal_L31 (pfam01197)→ |
| Sau-916-1 | NZ_KK096255.1 | - | 34913 | 34884 | **RNA**→ | rpmE2 (PRK01678)Ribosomal_L31 (pfam01197)→ |

| | | | | | |
|---|---|---|---|---|---|
| Sau-347-1 | NZ_KK099239.1 | - | 299829 | 299800 | RNA→ rpmE2 (PRK01678)Ribosomal_L31 (pfam01197)→ |
| Sau-1020-1 | NZ_KK096369.1 | - | 55328 | 55299 | RNA→ rpmE2 (PRK01678)Ribosomal_L31 (pfam01197)→ |
| Sau-2466-1 | NZ_KK094572.1 | - | 34871 | 34842 | RNA→ rpmE2 (PRK01678)Ribosomal_L31 (pfam01197)→ |
| Sau-2574-1 | NZ_KK094758.1 | - | 35013 | 34984 | RNA→ rpmE2 (PRK01678)Ribosomal_L31 (pfam01197)→ |
| Sau-835-1 | NZ_KK096188.1 | - | 56987 | 56958 | RNA→ rpmE2 (PRK01678)Ribosomal_L31 (pfam01197)→ |
| Sau-2873-1 | NZ_KK097003.1 | - | 55000 | 54971 | RNA→ rpmE2 (PRK01678)Ribosomal_L31 (pfam01197)→ |
| Sau-2253-1 | NZ_KK096457.1 | - | 34913 | 34884 | RNA→ rpmE2 (PRK01678)Ribosomal_L31 (pfam01197)→ |
| Sau-2839-1 | NZ_KK096872.1 | - | 34913 | 34884 | RNA→ rpmE2 (PRK01678)Ribosomal_L31 (pfam01197)→ |
| Sau-312-1 | NZ_KK099020.1 | - | 1063131 | 1063102 | RNA→ rpmE2 (PRK01678)Ribosomal_L31 (pfam01197)→ |
| Sau-317-1 | NZ_KK099057.1 | - | 1584677 | 1584648 | RNA→ rpmE2 (PRK01678)Ribosomal_L31 (pfam01197)→ |
| Sau-330-1 | NZ_KK099121.1 | - | 267595 | 267566 | RNA→ rpmE2 (PRK01678)Ribosomal_L31 (pfam01197)→ |
| Sau-336-1 | NZ_KK099170.1 | - | 1586288 | 1586259 | RNA→ rpmE2 (PRK01678)Ribosomal_L31 (pfam01197)→ |
| Sau-402-1 | NZ_KK099688.1 | - | 1609421 | 1609392 | RNA→ rpmE2 (PRK01678)Ribosomal_L31 (pfam01197)→ |
| Sau-2454-1 | NZ_KK096555.1 | - | 34872 | 34843 | RNA→ rpmE2 (PRK01678)Ribosomal_L31 (pfam01197)→ |
| Sau-346-1 | NZ_KK099233.1 | - | 234593 | 234564 | RNA→ rpmE2 (PRK01678)Ribosomal_L31 (pfam01197)→ |
| Sau-3662-1 | NZ_KK097659.1 | - | 34872 | 34843 | RNA→ rpmE2 (PRK01678)Ribosomal_L31 (pfam01197)→ |
| Sau-464-1 | NZ_KK098341.1 | - | 468366 | 468337 | RNA→ rpmE2 (PRK01678)Ribosomal_L31 (pfam01197)→ |
| Sau-394-1 | NZ_KK099633.1 | - | 133140 | 133111 | RNA→ rpmE2 (PRK01678)Ribosomal_L31 (pfam01197)→ |
| Sau-343-1 | NZ_KK099211.1 | - | 56663 | 56634 | RNA→ rpmE2 (PRK01678)Ribosomal_L31 (pfam01197)→ |
| Sau-354-1 | NZ_KK099287.1 | - | 465984 | 465955 | RNA→ rpmE2 (PRK01678)Ribosomal_L31 (pfam01197)→ |
| Sau-379-1 | NZ_KK099491.1 | - | 231563 | 231534 | RNA→ rpmE2 (PRK01678)Ribosomal_L31 (pfam01197)→ |
| Sau-400-1 | NZ_KK099667.1 | - | 59483 | 59454 | RNA→ rpmE2 (PRK01678)Ribosomal_L31 (pfam01197)→ |
| Sau-594-1 | NZ_KK099766.1 | - | 130624 | 130595 | RNA→ rpmE2 (PRK01678)Ribosomal_L31 (pfam01197)→ |
| Sau-587-1 | NZ_KK099859.1 | - | 2340216 | 2340187 | RNA→ rpmE2 (PRK01678)Ribosomal_L31 (pfam01197)→ |
| Sau-387-1 | NZ_KK099544.1 | - | 61697 | 61668 | RNA→ rpmE2 (PRK01678)Ribosomal_L31 (pfam01197)→ |
| Sau-3515-1 | NZ_KK095401.1 | - | 34867 | 34838 | RNA→ rpmE2 (PRK01678)Ribosomal_L31 (pfam01197)→ |
| Sau-380-1 | NZ_KK099504.1 | - | 61507 | 61478 | RNA→ rpmE2 (PRK01678)Ribosomal_L31 (pfam01197)→ |
| Sau-407-1 | NZ_KK099728.1 | - | 60837 | 60808 | RNA→ rpmE2 (PRK01678)Ribosomal_L31 (pfam01197)→ |
| Sau-494-1 | NZ_KK098743.1 | - | 122322 | 122293 | RNA→ rpmE2 (PRK01678)Ribosomal_L31 (pfam01197)→ |
| Sau-576-1 | NZ_KK099921.1 | - | 61323 | 61294 | RNA→ rpmE2 (PRK01678)Ribosomal_L31 (pfam01197)→ |
| Sau-919-1 | NZ_KK094076.1 | - | 25255 | 25226 | RNA→ rpmE2 (PRK01678)Ribosomal_L31 (pfam01197)→ |
| Sau-584-1 | NZ_KK098856.1 | - | 56240 | 56211 | RNA→ rpmE2 (PRK01678)Ribosomal_L31 (pfam01197)→ |
| Sau-656-1 | NZ_KK095933.1 | - | 34972 | 34943 | RNA→ rpmE2 (PRK01678)Ribosomal_L31 (pfam01197)→ |
| Sau-112-1 | NZ_KK218847.1 | - | 55733 | 55704 | RNA→ rpmE2 (PRK01678)Ribosomal_L31 (pfam01197)→ |
| Sau-123-1 | NZ_KK219305.1 | - | 56310 | 56281 | RNA→ rpmE2 (PRK01678)Ribosomal_L31 (pfam01197)→ |
| Sau-150-1 | NZ_KK220114.1 | - | 31722 | 31693 | RNA→ rpmE2 (PRK01678)Ribosomal_L31 (pfam01197)→ |
| Sau-92-1 | NZ_KK218349.1 | - | 56395 | 56366 | RNA→ rpmE2 (PRK01678)Ribosomal_L31 (pfam01197)→ |
| Sau-130-1 | NZ_KK219549.1 | - | 54860 | 54831 | RNA→ rpmE2 (PRK01678)Ribosomal_L31 (pfam01197)→ |
| Sau-98-1 | NZ_KK218470.1 | - | 55289 | 55260 | RNA→ rpmE2 (PRK01678)Ribosomal_L31 (pfam01197)→ |
| Sau-107-1 | NZ_KK218658.1 | - | 26024 | 25995 | RNA→ rpmE2 (PRK01678)Ribosomal_L31 (pfam01197)→ |
| Sau-129-1 | NZ_KK219518.1 | - | 26024 | 25995 | RNA→ rpmE2 (PRK01678)Ribosomal_L31 (pfam01197)→ |
| Sau-132-1 | NZ_KK219900.1 | - | 26024 | 25995 | RNA→ rpmE2 (PRK01678)Ribosomal_L31 (pfam01197)→ |
| Sau-155-1 | NZ_KK220312.1 | - | 26024 | 25995 | RNA→ rpmE2 (PRK01678)Ribosomal_L31 (pfam01197)→ |
| Sau-160-1 | NZ_KK220514.1 | - | 26024 | 25995 | RNA→ rpmE2 (PRK01678)Ribosomal_L31 (pfam01197)→ |
| Sau-165-1 | NZ_KK220763.1 | - | 26024 | 25995 | RNA→ rpmE2 (PRK01678)Ribosomal_L31 (pfam01197)→ |
| Sau-172-1 | NZ_KK220954.1 | - | 55963 | 55934 | RNA→ rpmE2 (PRK01678)Ribosomal_L31 (pfam01197)→ |
| Sau-177-1 | NZ_KK221128.1 | - | 26024 | 25995 | RNA→ rpmE2 (PRK01678)Ribosomal_L31 (pfam01197)→ |
| Sau-260-1 | NZ_KK222914.1 | - | 26024 | 25995 | RNA→ rpmE2 (PRK01678)Ribosomal_L31 (pfam01197)→ |
| Sau-3080-1 | NZ_KK226958.1 | - | 53265 | 53236 | RNA→ rpmE2 (PRK01678)Ribosomal_L31 (pfam01197)→ |
| Sau-271-1 | NZ_KK223311.1 | - | 26024 | 25995 | RNA→ rpmE2 (PRK01678)Ribosomal_L31 (pfam01197)→ |
| Sau-3085-1 | NZ_KK227159.1 | - | 25965 | 25936 | RNA→ rpmE2 (PRK01678)Ribosomal_L31 (pfam01197)→ |
| Sau-276-1 | NZ_KK223462.1 | - | 30762 | 30733 | RNA→ rpmE2 (PRK01678)Ribosomal_L31 (pfam01197)→ |

| | | | | | |
|---|---|---|---|---|---|
| Sau-3099-1 | NZ_KK227638.1 | - | 26024 | 25995 | **RNA**→ rpmE2 (PRK01678)Ribosomal_L31 (pfam01197)→ |
| Sau-608-1 | NZ_KK223626.1 | - | 56394 | 56365 | **RNA**→ rpmE2 (PRK01678)Ribosomal_L31 (pfam01197)→ |
| Sau-3143-1 | NZ_KK229081.1 | - | 26024 | 25995 | **RNA**→ rpmE2 (PRK01678)Ribosomal_L31 (pfam01197)→ |
| Sau-865-1 | NZ_KK223829.1 | - | 56395 | 56366 | **RNA**→ rpmE2 (PRK01678)Ribosomal_L31 (pfam01197)→ |
| Sau-188-1 | NZ_KK221506.1 | - | 26024 | 25995 | **RNA**→ rpmE2 (PRK01678)Ribosomal_L31 (pfam01197)→ |
| Sau-194-1 | NZ_KK221684.1 | - | 26024 | 25995 | **RNA**→ rpmE2 (PRK01678)Ribosomal_L31 (pfam01197)→ |
| Sau-136-1 | NZ_KK219698.1 | - | 46806 | 46777 | **RNA**→ rpmE2 (PRK01678)Ribosomal_L31 (pfam01197)→ |
| Sau-3063-1 | NZ_KK226363.1 | - | 26024 | 25995 | **RNA**→ rpmE2 (PRK01678)Ribosomal_L31 (pfam01197)→ |
| Sau-206-1 | NZ_KK222184.1 | - | 20478 | 20449 | **RNA**→ rpmE2 (PRK01678)Ribosomal_L31 (pfam01197)→ |
| Sau-3068-1 | NZ_KK226529.1 | - | 26024 | 25995 | **RNA**→ rpmE2 (PRK01678)Ribosomal_L31 (pfam01197)→ |
| Sau-214-1 | NZ_KK222381.1 | - | 18988 | 18959 | **RNA**→ rpmE2 (PRK01678)Ribosomal_L31 (pfam01197)→ |
| Sau-3166-1 | NZ_KK229800.1 | - | 56439 | 56410 | **RNA**→ rpmE2 (PRK01678)Ribosomal_L31 (pfam01197)→ |
| Sau-267-1 | NZ_KK223163.1 | - | 26024 | 25995 | **RNA**→ rpmE2 (PRK01678)Ribosomal_L31 (pfam01197)→ |
| Sau-273-1 | NZ_KK223372.1 | - | 56439 | 56410 | **RNA**→ rpmE2 (PRK01678)Ribosomal_L31 (pfam01197)→ |
| Sau-307-1 | NZ_KK223517.1 | - | 26024 | 25995 | **RNA**→ rpmE2 (PRK01678)Ribosomal_L31 (pfam01197)→ |
| Sau-872-1 | NZ_KK224086.1 | - | 56439 | 56410 | **RNA**→ rpmE2 (PRK01678)Ribosomal_L31 (pfam01197)→ |
| Sau-878-1 | NZ_KK224279.1 | - | 26024 | 25995 | **RNA**→ rpmE2 (PRK01678)Ribosomal_L31 (pfam01197)→ |
| Sau-2692-1 | NZ_KK225056.1 | - | 26024 | 25995 | **RNA**→ rpmE2 (PRK01678)Ribosomal_L31 (pfam01197)→ |
| Sau-3090-1 | NZ_KK227375.1 | - | 56354 | 56325 | **RNA**→ rpmE2 (PRK01678)Ribosomal_L31 (pfam01197)→ |
| Sau-175-1 | NZ_KK221058.1 | - | 26024 | 25995 | **RNA**→ rpmE2 (PRK01678)Ribosomal_L31 (pfam01197)→ |
| Sau-2821-1 | NZ_KK225696.1 | - | 56395 | 56366 | **RNA**→ rpmE2 (PRK01678)Ribosomal_L31 (pfam01197)→ |
| Sau-2505-1 | NZ_KK224434.1 | - | 56439 | 56410 | **RNA**→ rpmE2 (PRK01678)Ribosomal_L31 (pfam01197)→ |
| Sau-257-1 | NZ_KK222794.1 | - | 56395 | 56366 | **RNA**→ rpmE2 (PRK01678)Ribosomal_L31 (pfam01197)→ |
| Sau-126-1 | NZ_KK219434.1 | - | 56261 | 56232 | **RNA**→ rpmE2 (PRK01678)Ribosomal_L31 (pfam01197)→ |
| Sau-269-1 | NZ_KK223216.1 | - | 52550 | 52521 | **RNA**→ rpmE2 (PRK01678)Ribosomal_L31 (pfam01197)→ |
| Sau-134-1 | NZ_KK219611.1 | - | 26024 | 25995 | **RNA**→ rpmE2 (PRK01678)Ribosomal_L31 (pfam01197)→ |
| Sau-274-1 | NZ_KK223399.1 | - | 26024 | 25995 | **RNA**→ rpmE2 (PRK01678)Ribosomal_L31 (pfam01197)→ |
| Sau-200-1 | NZ_KK221878.1 | - | 54642 | 54613 | **RNA**→ rpmE2 (PRK01678)Ribosomal_L31 (pfam01197)→ |
| Sau-873-1 | NZ_KK224144.1 | - | 56395 | 56366 | **RNA**→ rpmE2 (PRK01678)Ribosomal_L31 (pfam01197)→ |
| Sau-207-1 | NZ_KK222088.1 | - | 26024 | 25995 | **RNA**→ rpmE2 (PRK01678)Ribosomal_L31 (pfam01197)→ |
| Sau-3082-1 | NZ_KK227042.1 | - | 26024 | 25995 | **RNA**→ rpmE2 (PRK01678)Ribosomal_L31 (pfam01197)→ |
| Sau-217-1 | NZ_KK222503.1 | - | 26024 | 25995 | **RNA**→ rpmE2 (PRK01678)Ribosomal_L31 (pfam01197)→ |
| Sau-3087-1 | NZ_KK227240.1 | - | 26024 | 25995 | **RNA**→ rpmE2 (PRK01678)Ribosomal_L31 (pfam01197)→ |
| Sau-253-1 | NZ_KK222701.1 | - | 18846 | 18817 | **RNA**→ rpmE2 (PRK01678)Ribosomal_L31 (pfam01197)→ |
| Sau-3095-1 | NZ_KK227491.1 | - | 26024 | 25995 | **RNA**→ rpmE2 (PRK01678)Ribosomal_L31 (pfam01197)→ |
| Sau-875-1 | NZ_KK224222.1 | - | 26024 | 25995 | **RNA**→ rpmE2 (PRK01678)Ribosomal_L31 (pfam01197)→ |
| Sau-2676-1 | NZ_KK224731.1 | - | 26024 | 25995 | **RNA**→ rpmE2 (PRK01678)Ribosomal_L31 (pfam01197)→ |
| Sau-2701-1 | NZ_KK225372.1 | - | 56390 | 56361 | **RNA**→ rpmE2 (PRK01678)Ribosomal_L31 (pfam01197)→ |
| Sau-144-1 | NZ_KK220038.1 | - | 26024 | 25995 | **RNA**→ rpmE2 (PRK01678)Ribosomal_L31 (pfam01197)→ |
| Sau-153-1 | NZ_KK220240.1 | - | 55477 | 55448 | **RNA**→ rpmE2 (PRK01678)Ribosomal_L31 (pfam01197)→ |
| Sau-158-1 | NZ_KK220436.1 | - | 25968 | 25939 | **RNA**→ rpmE2 (PRK01678)Ribosomal_L31 (pfam01197)→ |
| Sau-163-1 | NZ_KK220641.1 | - | 56390 | 56361 | **RNA**→ rpmE2 (PRK01678)Ribosomal_L31 (pfam01197)→ |
| Sau-170-1 | NZ_KK220886.1 | - | 31818 | 31789 | **RNA**→ rpmE2 (PRK01678)Ribosomal_L31 (pfam01197)→ |
| Sau-180-1 | NZ_KK221232.1 | - | 26024 | 25995 | **RNA**→ rpmE2 (PRK01678)Ribosomal_L31 (pfam01197)→ |
| Sau-186-1 | NZ_KK221416.1 | - | 56310 | 56281 | **RNA**→ rpmE2 (PRK01678)Ribosomal_L31 (pfam01197)→ |
| Sau-209-1 | NZ_KK222240.1 | - | 26024 | 25995 | **RNA**→ rpmE2 (PRK01678)Ribosomal_L31 (pfam01197)→ |
| Sau-220-1 | NZ_KK222577.1 | - | 26024 | 25995 | **RNA**→ rpmE2 (PRK01678)Ribosomal_L31 (pfam01197)→ |
| Sau-215-1 | NZ_KK222425.1 | - | 56395 | 56366 | **RNA**→ rpmE2 (PRK01678)Ribosomal_L31 (pfam01197)→ |
| Sau-841-1 | NZ_KK223720.1 | - | 26024 | 25995 | **RNA**→ rpmE2 (PRK01678)Ribosomal_L31 (pfam01197)→ |
| Sau-222-1 | NZ_KK222619.1 | - | 7609 | 7580 | **RNA**→ rpmE2 (PRK01678)Ribosomal_L31 (pfam01197)→ |
| Sau-2698-1 | NZ_KK225257.1 | - | 56390 | 56361 | **RNA**→ rpmE2 (PRK01678)Ribosomal_L31 (pfam01197)→ |
| Sau-258-1 | NZ_KK222826.1 | - | 26024 | 25995 | **RNA**→ rpmE2 (PRK01678)Ribosomal_L31 (pfam01197)→ |

| | | | | | |
|---|---|---|---|---|---|
| Sau-2711-1 | NZ_KK225491.1 | - | 26024 | 25995 | **RNA**→ rpmE2 (PRK01678)Ribosomal_L31 (pfam01197)→ |
| Sau-263-1 | NZ_KK223018.1 | - | 25790 | 25761 | **RNA**→ rpmE2 (PRK01678)Ribosomal_L31 (pfam01197)→ |
| Sau-308-1 | NZ_KK223557.1 | - | 26024 | 25995 | **RNA**→ rpmE2 (PRK01678)Ribosomal_L31 (pfam01197)→ |
| Sau-2826-1 | NZ_KK225869.1 | - | 26024 | 25995 | **RNA**→ rpmE2 (PRK01678)Ribosomal_L31 (pfam01197)→ |
| Sau-842-1 | NZ_KK223754.1 | - | 30994 | 30965 | **RNA**→ rpmE2 (PRK01678)Ribosomal_L31 (pfam01197)→ |
| Sau-3115-1 | NZ_KK228218.1 | - | 26024 | 25995 | **RNA**→ rpmE2 (PRK01678)Ribosomal_L31 (pfam01197)→ |
| Sau-868-1 | NZ_KK223946.1 | - | 56390 | 56361 | **RNA**→ rpmE2 (PRK01678)Ribosomal_L31 (pfam01197)→ |
| Sau-3160-1 | NZ_KK229644.1 | - | 56439 | 56410 | **RNA**→ rpmE2 (PRK01678)Ribosomal_L31 (pfam01197)→ |
| Sau-191-1 | NZ_KK221597.1 | - | 26024 | 25995 | **RNA**→ rpmE2 (PRK01678)Ribosomal_L31 (pfam01197)→ |
| Sau-2636-1 | NZ_KK224621.1 | - | 26024 | 25995 | **RNA**→ rpmE2 (PRK01678)Ribosomal_L31 (pfam01197)→ |
| Sau-198-1 | NZ_KK221811.1 | - | 56165 | 56136 | **RNA**→ rpmE2 (PRK01678)Ribosomal_L31 (pfam01197)→ |
| Sau-212-1 | NZ_KK222314.1 | - | 26024 | 25995 | **RNA**→ rpmE2 (PRK01678)Ribosomal_L31 (pfam01197)→ |
| Sau-2690-1 | NZ_KK224950.1 | - | 56395 | 56366 | **RNA**→ rpmE2 (PRK01678)Ribosomal_L31 (pfam01197)→ |
| Sau-3065-1 | NZ_KK226453.1 | - | 26024 | 25995 | **RNA**→ rpmE2 (PRK01678)Ribosomal_L31 (pfam01197)→ |
| Sau-2469-1 | NZ_KK224385.1 | - | 55969 | 55940 | **RNA**→ rpmE2 (PRK01678)Ribosomal_L31 (pfam01197)→ |
| Sau-2627-1 | NZ_KK224533.1 | - | 26024 | 25995 | **RNA**→ rpmE2 (PRK01678)Ribosomal_L31 (pfam01197)→ |
| Sau-118-1 | NZ_KK219075.1 | - | 56395 | 56366 | **RNA**→ rpmE2 (PRK01678)Ribosomal_L31 (pfam01197)→ |
| Sau-87-1 | NZ_KK218139.1 | - | 55553 | 55524 | **RNA**→ rpmE2 (PRK01678)Ribosomal_L31 (pfam01197)→ |
| Sau-262-1 | NZ_KK222984.1 | - | 56438 | 56409 | **RNA**→ rpmE2 (PRK01678)Ribosomal_L31 (pfam01197)→ |
| Sau-3109-1 | NZ_KK228029.1 | - | 26024 | 25995 | **RNA**→ rpmE2 (PRK01678)Ribosomal_L31 (pfam01197)→ |
| Sau-3122-1 | NZ_KK228451.1 | - | 53737 | 53708 | **RNA**→ rpmE2 (PRK01678)Ribosomal_L31 (pfam01197)→ |
| Sau-80-1 | NZ_KK218041.1 | - | 54768 | 54739 | **RNA**→ rpmE2 (PRK01678)Ribosomal_L31 (pfam01197)→ |
| Sau-3175-1 | NZ_KK230124.1 | - | 56354 | 56325 | **RNA**→ rpmE2 (PRK01678)Ribosomal_L31 (pfam01197)→ |
| Sau-84-1 | NZ_KK218077.1 | - | 56261 | 56232 | **RNA**→ rpmE2 (PRK01678)Ribosomal_L31 (pfam01197)→ |
| Sau-880-1 | NZ_KK224318.1 | - | 56390 | 56361 | **RNA**→ rpmE2 (PRK01678)Ribosomal_L31 (pfam01197)→ |
| Sau-89-1 | NZ_KK218219.1 | - | 26024 | 25995 | **RNA**→ rpmE2 (PRK01678)Ribosomal_L31 (pfam01197)→ |
| Sau-3077-1 | NZ_KK226834.1 | - | 26024 | 25995 | **RNA**→ rpmE2 (PRK01678)Ribosomal_L31 (pfam01197)→ |
| Sau-90-1 | NZ_KK218244.1 | - | 26024 | 25995 | **RNA**→ rpmE2 (PRK01678)Ribosomal_L31 (pfam01197)→ |
| Sau-3083-1 | NZ_KK227076.1 | - | 26024 | 25995 | **RNA**→ rpmE2 (PRK01678)Ribosomal_L31 (pfam01197)→ |
| Sau-113-1 | NZ_KK218892.1 | - | 54808 | 54779 | **RNA**→ rpmE2 (PRK01678)Ribosomal_L31 (pfam01197)→ |
| Sau-3088-1 | NZ_KK227288.1 | - | 26024 | 25995 | **RNA**→ rpmE2 (PRK01678)Ribosomal_L31 (pfam01197)→ |
| Sau-119-1 | NZ_KK219116.1 | - | 26024 | 25995 | **RNA**→ rpmE2 (PRK01678)Ribosomal_L31 (pfam01197)→ |
| Sau-3116-1 | NZ_KK228289.1 | - | 18618 | 18589 | **RNA**→ rpmE2 (PRK01678)Ribosomal_L31 (pfam01197)→ |
| Sau-124-1 | NZ_KK219338.1 | - | 56310 | 56281 | **RNA**→ rpmE2 (PRK01678)Ribosomal_L31 (pfam01197)→ |
| Sau-3123-1 | NZ_KK228505.1 | - | 26024 | 25995 | **RNA**→ rpmE2 (PRK01678)Ribosomal_L31 (pfam01197)→ |
| Sau-3130-1 | NZ_KK228701.1 | - | 26024 | 25995 | **RNA**→ rpmE2 (PRK01678)Ribosomal_L31 (pfam01197)→ |
| Sau-3157-1 | NZ_KK229539.1 | - | 26024 | 25995 | **RNA**→ rpmE2 (PRK01678)Ribosomal_L31 (pfam01197)→ |
| Sau-3162-1 | NZ_KK229729.1 | - | 26024 | 25995 | **RNA**→ rpmE2 (PRK01678)Ribosomal_L31 (pfam01197)→ |
| Sau-3093-1 | NZ_KK227448.1 | - | 56353 | 56324 | **RNA**→ rpmE2 (PRK01678)Ribosomal_L31 (pfam01197)→ |
| Sau-140-1 | NZ_KK219813.1 | - | 26024 | 25995 | **RNA**→ rpmE2 (PRK01678)Ribosomal_L31 (pfam01197)→ |
| Sau-2824-1 | NZ_KK225802.1 | - | 56165 | 56136 | **RNA**→ rpmE2 (PRK01678)Ribosomal_L31 (pfam01197)→ |
| Sau-3060-1 | NZ_KK226308.1 | - | 26024 | 25995 | **RNA**→ rpmE2 (PRK01678)Ribosomal_L31 (pfam01197)→ |
| Sau-3153-1 | NZ_KK229332.1 | - | 26024 | 25995 | **RNA**→ rpmE2 (PRK01678)Ribosomal_L31 (pfam01197)→ |
| Sau-3154-1 | NZ_KK229377.1 | - | 26024 | 25995 | **RNA**→ rpmE2 (PRK01678)Ribosomal_L31 (pfam01197)→ |
| Sau-3165-1 | NZ_KK229754.1 | - | 56395 | 56366 | **RNA**→ rpmE2 (PRK01678)Ribosomal_L31 (pfam01197)→ |
| Sau-3183-1 | NZ_KK230455.1 | - | 26024 | 25995 | **RNA**→ rpmE2 (PRK01678)Ribosomal_L31 (pfam01197)→ |
| Sau-182-1 | NZ_KK221300.1 | - | 56439 | 56410 | **RNA**→ rpmE2 (PRK01678)Ribosomal_L31 (pfam01197)→ |
| Sau-265-1 | NZ_KK223082.1 | - | 56395 | 56366 | **RNA**→ rpmE2 (PRK01678)Ribosomal_L31 (pfam01197)→ |
| Sau-870-1 | NZ_KK224019.1 | - | 26024 | 25995 | **RNA**→ rpmE2 (PRK01678)Ribosomal_L31 (pfam01197)→ |
| Sau-103-1 | NZ_KK218519.1 | - | 26024 | 25995 | **RNA**→ rpmE2 (PRK01678)Ribosomal_L31 (pfam01197)→ |
| Sau-3010-1 | NZ_KK226052.1 | - | 56395 | 56366 | **RNA**→ rpmE2 (PRK01678)Ribosomal_L31 (pfam01197)→ |
| Sau-108-1 | NZ_KK218685.1 | - | 26024 | 25995 | **RNA**→ rpmE2 (PRK01678)Ribosomal_L31 (pfam01197)→ |

| | | | | | | |
|---|---|---|---|---|---|---|
| Sau-3017-1 | NZ_KK226235.1 | - | 26024 | 25995 | **RNA**→ | rpmE2 (PRK01678)Ribosomal_L31 (pfam01197)→ |
| Sau-3119-1 | NZ_KK228363.1 | - | 26024 | 25995 | **RNA**→ | rpmE2 (PRK01678)Ribosomal_L31 (pfam01197)→ |
| Sau-3132-1 | NZ_KK228781.1 | - | 57286 | 57257 | **RNA**→ | rpmE2 (PRK01678)Ribosomal_L31 (pfam01197)→ |
| Sau-3147-1 | NZ_KK229226.1 | - | 56354 | 56325 | **RNA**→ | rpmE2 (PRK01678)Ribosomal_L31 (pfam01197)→ |
| Sau-3179-1 | NZ_KK230245.1 | - | 26023 | 25994 | **RNA**→ | rpmE2 (PRK01678)Ribosomal_L31 (pfam01197)→ |
| Sau-3097-1 | NZ_KK227565.1 | - | 26024 | 25995 | **RNA**→ | rpmE2 (PRK01678)Ribosomal_L31 (pfam01197)→ |
| Sau-110-1 | NZ_KK218762.1 | - | 54882 | 54853 | **RNA**→ | rpmE2 (PRK01678)Ribosomal_L31 (pfam01197)→ |
| Sau-116-1 | NZ_KK219000.1 | - | 54756 | 54727 | **RNA**→ | rpmE2 (PRK01678)Ribosomal_L31 (pfam01197)→ |
| Sau-121-1 | NZ_KK219204.1 | - | 43868 | 43839 | **RNA**→ | rpmE2 (PRK01678)Ribosomal_L31 (pfam01197)→ |
| Sau-105-1 | NZ_KK218599.1 | - | 26024 | 25995 | **RNA**→ | rpmE2 (PRK01678)Ribosomal_L31 (pfam01197)→ |
| Sau-3312-1 | NZ_KK235067.1 | - | 26024 | 25995 | **RNA**→ | rpmE2 (PRK01678)Ribosomal_L31 (pfam01197)→ |
| Sau-3320-1 | NZ_KK235290.1 | - | 26024 | 25995 | **RNA**→ | rpmE2 (PRK01678)Ribosomal_L31 (pfam01197)→ |
| Sau-3321-1 | NZ_KK235319.1 | - | 56439 | 56410 | **RNA**→ | rpmE2 (PRK01678)Ribosomal_L31 (pfam01197)→ |
| Sau-3328-1 | NZ_KK235526.1 | - | 26024 | 25995 | **RNA**→ | rpmE2 (PRK01678)Ribosomal_L31 (pfam01197)→ |
| Sau-3332-1 | NZ_KK235680.1 | - | 26024 | 25995 | **RNA**→ | rpmE2 (PRK01678)Ribosomal_L31 (pfam01197)→ |
| Sau-3333-1 | NZ_KK235727.1 | - | 26024 | 25995 | **RNA**→ | rpmE2 (PRK01678)Ribosomal_L31 (pfam01197)→ |
| Sau-3338-1 | NZ_KK235952.1 | - | 26024 | 25995 | **RNA**→ | rpmE2 (PRK01678)Ribosomal_L31 (pfam01197)→ |
| Sau-3379-1 | NZ_KK237259.1 | - | 54480 | 54451 | **RNA**→ | rpmE2 (PRK01678)Ribosomal_L31 (pfam01197)→ |
| Sau-3384-1 | NZ_KK237467.1 | - | 56317 | 56288 | **RNA**→ | rpmE2 (PRK01678)Ribosomal_L31 (pfam01197)→ |
| Sau-3390-1 | NZ_KK237651.1 | - | 54281 | 54252 | **RNA**→ | rpmE2 (PRK01678)Ribosomal_L31 (pfam01197)→ |
| Sau-3399-1 | NZ_KK237828.1 | - | 26024 | 25995 | **RNA**→ | rpmE2 (PRK01678)Ribosomal_L31 (pfam01197)→ |
| Sau-3405-1 | NZ_KK238039.1 | - | 53570 | 53541 | **RNA**→ | rpmE2 (PRK01678)Ribosomal_L31 (pfam01197)→ |
| Sau-3410-1 | NZ_KK238232.1 | - | 56439 | 56410 | **RNA**→ | rpmE2 (PRK01678)Ribosomal_L31 (pfam01197)→ |
| Sau-3417-1 | NZ_KK238513.1 | - | 26024 | 25995 | **RNA**→ | rpmE2 (PRK01678)Ribosomal_L31 (pfam01197)→ |
| Sau-3422-1 | NZ_KK238712.1 | - | 26024 | 25995 | **RNA**→ | rpmE2 (PRK01678)Ribosomal_L31 (pfam01197)→ |
| Sau-3423-1 | NZ_KK238762.1 | - | 26024 | 25995 | **RNA**→ | rpmE2 (PRK01678)Ribosomal_L31 (pfam01197)→ |
| Sau-3427-1 | NZ_KK238898.1 | - | 26024 | 25995 | **RNA**→ | rpmE2 (PRK01678)Ribosomal_L31 (pfam01197)→ |
| Sau-3428-1 | NZ_KK238957.1 | - | 26024 | 25995 | **RNA**→ | rpmE2 (PRK01678)Ribosomal_L31 (pfam01197)→ |
| Sau-3372-1 | NZ_KK236944.1 | - | 26024 | 25995 | **RNA**→ | rpmE2 (PRK01678)Ribosomal_L31 (pfam01197)→ |
| Sau-3373-1 | NZ_KK236999.1 | - | 29618 | 29589 | **RNA**→ | rpmE2 (PRK01678)Ribosomal_L31 (pfam01197)→ |
| Sau-3378-1 | NZ_KK237210.1 | - | 53941 | 53912 | **RNA**→ | rpmE2 (PRK01678)Ribosomal_L31 (pfam01197)→ |
| Sau-3383-1 | NZ_KK237441.1 | - | 56279 | 56250 | **RNA**→ | rpmE2 (PRK01678)Ribosomal_L31 (pfam01197)→ |
| Sau-3389-1 | NZ_KK237594.1 | - | 56353 | 56324 | **RNA**→ | rpmE2 (PRK01678)Ribosomal_L31 (pfam01197)→ |
| Sau-3398-1 | NZ_KK237796.1 | - | 26332 | 26303 | **RNA**→ | rpmE2 (PRK01678)Ribosomal_L31 (pfam01197)→ |
| Sau-3404-1 | NZ_KK237995.1 | - | 54477 | 54448 | **RNA**→ | rpmE2 (PRK01678)Ribosomal_L31 (pfam01197)→ |
| Sau-3409-1 | NZ_KK238213.1 | - | 26024 | 25995 | **RNA**→ | rpmE2 (PRK01678)Ribosomal_L31 (pfam01197)→ |
| Sau-3415-1 | NZ_KK238417.1 | - | 26024 | 25995 | **RNA**→ | rpmE2 (PRK01678)Ribosomal_L31 (pfam01197)→ |
| Sau-3431-1 | NZ_KK239111.1 | - | 26024 | 25995 | **RNA**→ | rpmE2 (PRK01678)Ribosomal_L31 (pfam01197)→ |
| Sau-3432-1 | NZ_KK239165.1 | - | 56395 | 56366 | **RNA**→ | rpmE2 (PRK01678)Ribosomal_L31 (pfam01197)→ |
| Sau-3421-1 | NZ_KK238654.1 | - | 26024 | 25995 | **RNA**→ | rpmE2 (PRK01678)Ribosomal_L31 (pfam01197)→ |
| Sau-3426-1 | NZ_KK238862.1 | - | 26024 | 25995 | **RNA**→ | rpmE2 (PRK01678)Ribosomal_L31 (pfam01197)→ |
| Sau-3376-1 | NZ_KK237117.1 | - | 54536 | 54507 | **RNA**→ | rpmE2 (PRK01678)Ribosomal_L31 (pfam01197)→ |
| Sau-3377-1 | NZ_KK237165.1 | - | 54481 | 54452 | **RNA**→ | rpmE2 (PRK01678)Ribosomal_L31 (pfam01197)→ |
| Sau-3382-1 | NZ_KK237395.1 | - | 56390 | 56361 | **RNA**→ | rpmE2 (PRK01678)Ribosomal_L31 (pfam01197)→ |
| Sau-3387-1 | NZ_KK237537.1 | - | 26024 | 25995 | **RNA**→ | rpmE2 (PRK01678)Ribosomal_L31 (pfam01197)→ |
| Sau-3388-1 | NZ_KK237569.1 | - | 26024 | 25995 | **RNA**→ | rpmE2 (PRK01678)Ribosomal_L31 (pfam01197)→ |
| Sau-3396-1 | NZ_KK237741.1 | - | 2607 | 2578 | **RNA**→ | rpmE2 (PRK01678)Ribosomal_L31 (pfam01197)→ |
| Sau-3397-1 | NZ_KK237767.1 | - | 56395 | 56366 | **RNA**→ | rpmE2 (PRK01678)Ribosomal_L31 (pfam01197)→ |
| Sau-3403-1 | NZ_KK237967.1 | - | 26024 | 25995 | **RNA**→ | rpmE2 (PRK01678)Ribosomal_L31 (pfam01197)→ |
| Sau-3408-1 | NZ_KK238161.1 | - | 56394 | 56365 | **RNA**→ | rpmE2 (PRK01678)Ribosomal_L31 (pfam01197)→ |
| Sau-3413-1 | NZ_KK238364.1 | - | 56395 | 56366 | **RNA**→ | rpmE2 (PRK01678)Ribosomal_L31 (pfam01197)→ |
| Sau-3430-1 | NZ_KK239060.1 | - | 26024 | 25995 | **RNA**→ | rpmE2 (PRK01678)Ribosomal_L31 (pfam01197)→ |

| | | | | | |
|---|---|---|---|---|---|
| Sau-3420-1 | NZ_KK238600.1 | - | 55512 | 55483 | **RNA**→ rpmE2 (PRK01678)Ribosomal_L31 (pfam01197)→ |
| Sau-3425-1 | NZ_KK238827.1 | - | 26025 | 25996 | **RNA**→ rpmE2 (PRK01678)Ribosomal_L31 (pfam01197)→ |
| Sau-3393-1 | NZ_KK237699.1 | - | 26024 | 25995 | **RNA**→ rpmE2 (PRK01678)Ribosomal_L31 (pfam01197)→ |
| Sau-3401-1 | NZ_KK237884.1 | - | 19182 | 19153 | **RNA**→ rpmE2 (PRK01678)Ribosomal_L31 (pfam01197)→ |
| Sau-3402-1 | NZ_KK237920.1 | - | 56438 | 56409 | **RNA**→ rpmE2 (PRK01678)Ribosomal_L31 (pfam01197)→ |
| Sau-3406-1 | NZ_KK238084.1 | - | 26024 | 25995 | **RNA**→ rpmE2 (PRK01678)Ribosomal_L31 (pfam01197)→ |
| Sau-3407-1 | NZ_KK238117.1 | - | 26024 | 25995 | **RNA**→ rpmE2 (PRK01678)Ribosomal_L31 (pfam01197)→ |
| Sau-3411-1 | NZ_KK238270.1 | - | 25968 | 25939 | **RNA**→ rpmE2 (PRK01678)Ribosomal_L31 (pfam01197)→ |
| Sau-3412-1 | NZ_KK238306.1 | - | 56390 | 56361 | **RNA**→ rpmE2 (PRK01678)Ribosomal_L31 (pfam01197)→ |
| Sau-3418-1 | NZ_KK238556.1 | - | 17345 | 17316 | **RNA**→ rpmE2 (PRK01678)Ribosomal_L31 (pfam01197)→ |
| Sau-3424-1 | NZ_KK238790.1 | - | 26025 | 25996 | **RNA**→ rpmE2 (PRK01678)Ribosomal_L31 (pfam01197)→ |
| Sau-3429-1 | NZ_KK239008.1 | - | 26024 | 25995 | **RNA**→ rpmE2 (PRK01678)Ribosomal_L31 (pfam01197)→ |
| Sau-3337-1 | NZ_KK235896.1 | - | 26024 | 25995 | **RNA**→ rpmE2 (PRK01678)Ribosomal_L31 (pfam01197)→ |
| Sau-3341-1 | NZ_KK236088.1 | - | 26024 | 25995 | **RNA**→ rpmE2 (PRK01678)Ribosomal_L31 (pfam01197)→ |
| Sau-3362-1 | NZ_KK236638.1 | - | 54582 | 54553 | **RNA**→ rpmE2 (PRK01678)Ribosomal_L31 (pfam01197)→ |
| Sau-3434-1 | NZ_KK239222.1 | - | 26024 | 25995 | **RNA**→ rpmE2 (PRK01678)Ribosomal_L31 (pfam01197)→ |
| Sau-3439-1 | NZ_KK239416.1 | - | 26024 | 25995 | **RNA**→ rpmE2 (PRK01678)Ribosomal_L31 (pfam01197)→ |
| Sau-3445-1 | NZ_KK239632.1 | - | 26024 | 25995 | **RNA**→ rpmE2 (PRK01678)Ribosomal_L31 (pfam01197)→ |
| Sau-3458-1 | NZ_KK240009.1 | - | 25997 | 25968 | **RNA**→ rpmE2 (PRK01678)Ribosomal_L31 (pfam01197)→ |
| Sau-3464-1 | NZ_KK240210.1 | - | 26024 | 25995 | **RNA**→ rpmE2 (PRK01678)Ribosomal_L31 (pfam01197)→ |
| Sau-3284-1 | NZ_KK233856.1 | - | 26024 | 25995 | **RNA**→ rpmE2 (PRK01678)Ribosomal_L31 (pfam01197)→ |
| Sau-3470-1 | NZ_KK240383.1 | - | 56384 | 56355 | **RNA**→ rpmE2 (PRK01678)Ribosomal_L31 (pfam01197)→ |
| Sau-3477-1 | NZ_KK240551.1 | - | 56439 | 56410 | **RNA**→ rpmE2 (PRK01678)Ribosomal_L31 (pfam01197)→ |
| Sau-3715-1 | NZ_KK240724.1 | - | 56439 | 56410 | **RNA**→ rpmE2 (PRK01678)Ribosomal_L31 (pfam01197)→ |
| Sau-3437-1 | NZ_KK239359.1 | - | 26027 | 25998 | **RNA**→ rpmE2 (PRK01678)Ribosomal_L31 (pfam01197)→ |
| Sau-3438-1 | NZ_KK239396.1 | - | 26024 | 25995 | **RNA**→ rpmE2 (PRK01678)Ribosomal_L31 (pfam01197)→ |
| Sau-3443-1 | NZ_KK239562.1 | - | 5833 | 5804 | **RNA**→ rpmE2 (PRK01678)Ribosomal_L31 (pfam01197)→ |
| Sau-3444-1 | NZ_KK239596.1 | - | 26024 | 25995 | **RNA**→ rpmE2 (PRK01678)Ribosomal_L31 (pfam01197)→ |
| Sau-3450-1 | NZ_KK239796.1 | - | 26024 | 25995 | **RNA**→ rpmE2 (PRK01678)Ribosomal_L31 (pfam01197)→ |
| Sau-3456-1 | NZ_KK239979.1 | - | 25389 | 25360 | **RNA**→ rpmE2 (PRK01678)Ribosomal_L31 (pfam01197)→ |
| Sau-3225-1 | NZ_KK231905.1 | - | 56309 | 56280 | **RNA**→ rpmE2 (PRK01678)Ribosomal_L31 (pfam01197)→ |
| Sau-3469-1 | NZ_KK240355.1 | - | 56384 | 56355 | **RNA**→ rpmE2 (PRK01678)Ribosomal_L31 (pfam01197)→ |
| Sau-3266-1 | NZ_KK233355.1 | - | 26024 | 25995 | **RNA**→ rpmE2 (PRK01678)Ribosomal_L31 (pfam01197)→ |
| Sau-3476-1 | NZ_KK240520.1 | - | 26332 | 26303 | **RNA**→ rpmE2 (PRK01678)Ribosomal_L31 (pfam01197)→ |
| Sau-3323-1 | NZ_KK235408.1 | - | 26024 | 25995 | **RNA**→ rpmE2 (PRK01678)Ribosomal_L31 (pfam01197)→ |
| Sau-3714-1 | NZ_KK240688.1 | - | 56441 | 56412 | **RNA**→ rpmE2 (PRK01678)Ribosomal_L31 (pfam01197)→ |
| Sau-3719-1 | NZ_KK240874.1 | - | 26024 | 25995 | **RNA**→ rpmE2 (PRK01678)Ribosomal_L31 (pfam01197)→ |
| Sau-3330-1 | NZ_KK235606.1 | - | 25733 | 25704 | **RNA**→ rpmE2 (PRK01678)Ribosomal_L31 (pfam01197)→ |
| Sau-3353-1 | NZ_KK236370.1 | - | 26024 | 25995 | **RNA**→ rpmE2 (PRK01678)Ribosomal_L31 (pfam01197)→ |
| Sau-3365-1 | NZ_KK236756.1 | - | 54478 | 54449 | **RNA**→ rpmE2 (PRK01678)Ribosomal_L31 (pfam01197)→ |
| Sau-3366-1 | NZ_KK236796.1 | - | 54582 | 54553 | **RNA**→ rpmE2 (PRK01678)Ribosomal_L31 (pfam01197)→ |
| Sau-3436-1 | NZ_KK239317.1 | - | 26024 | 25995 | **RNA**→ rpmE2 (PRK01678)Ribosomal_L31 (pfam01197)→ |
| Sau-3442-1 | NZ_KK239489.1 | - | 26024 | 25995 | **RNA**→ rpmE2 (PRK01678)Ribosomal_L31 (pfam01197)→ |
| Sau-3448-1 | NZ_KK239707.1 | - | 26024 | 25995 | **RNA**→ rpmE2 (PRK01678)Ribosomal_L31 (pfam01197)→ |
| Sau-597-1 | NZ_KI966209.1 | - | 59482 | 59453 | **RNA**→ rpmE2 (PRK01678)Ribosomal_L31 (pfam01197)→ |
| Sau-3449-1 | NZ_KK239758.1 | - | 26024 | 25995 | **RNA**→ rpmE2 (PRK01678)Ribosomal_L31 (pfam01197)→ |
| Sau-3453-1 | NZ_KK239901.1 | - | 26024 | 25995 | **RNA**→ rpmE2 (PRK01678)Ribosomal_L31 (pfam01197)→ |
| Sau-3288-1 | NZ_KK234034.1 | - | 26024 | 25995 | **RNA**→ rpmE2 (PRK01678)Ribosomal_L31 (pfam01197)→ |
| Sau-3454-1 | NZ_KK239942.1 | - | 26024 | 25995 | **RNA**→ rpmE2 (PRK01678)Ribosomal_L31 (pfam01197)→ |
| Sau-3460-1 | NZ_KK240086.1 | - | 26024 | 25995 | **RNA**→ rpmE2 (PRK01678)Ribosomal_L31 (pfam01197)→ |
| Sau-3461-1 | NZ_KK240125.1 | - | 26024 | 25995 | **RNA**→ rpmE2 (PRK01678)Ribosomal_L31 (pfam01197)→ |
| Sau-3467-1 | NZ_KK240284.1 | - | 56403 | 56374 | **RNA**→ rpmE2 (PRK01678)Ribosomal_L31 (pfam01197)→ |

| | | | | | |
|---|---|---|---|---|---|
| Sau-3468-1 | NZ_KK240323.1 | - | 26024 | 25995 | **RNA**→ rpmE2 (PRK01678)Ribosomal_L31 (pfam01197)→ |
| Sau-3475-1 | NZ_KK240494.1 | - | 26024 | 25995 | **RNA**→ rpmE2 (PRK01678)Ribosomal_L31 (pfam01197)→ |
| Sau-3712-1 | NZ_KK240655.1 | - | 56395 | 56366 | **RNA**→ rpmE2 (PRK01678)Ribosomal_L31 (pfam01197)→ |
| Sau-3718-1 | NZ_KK240836.1 | - | 26024 | 25995 | **RNA**→ rpmE2 (PRK01678)Ribosomal_L31 (pfam01197)→ |
| Sau-3440-1 | NZ_KK239446.1 | - | 26024 | 25995 | **RNA**→ rpmE2 (PRK01678)Ribosomal_L31 (pfam01197)→ |
| Sau-3447-1 | NZ_KK239680.1 | - | 26024 | 25995 | **RNA**→ rpmE2 (PRK01678)Ribosomal_L31 (pfam01197)→ |
| Sau-3223-1 | NZ_KK231839.1 | - | 55801 | 55772 | **RNA**→ rpmE2 (PRK01678)Ribosomal_L31 (pfam01197)→ |
| Sau-3452-1 | NZ_KK239862.1 | - | 56310 | 56281 | **RNA**→ rpmE2 (PRK01678)Ribosomal_L31 (pfam01197)→ |
| Sau-3291-1 | NZ_KK234207.1 | - | 33041 | 33012 | **RNA**→ rpmE2 (PRK01678)Ribosomal_L31 (pfam01197)→ |
| Sau-3459-1 | NZ_KK240053.1 | - | 26024 | 25995 | **RNA**→ rpmE2 (PRK01678)Ribosomal_L31 (pfam01197)→ |
| Sau-3472-1 | NZ_KK240411.1 | - | 26024 | 25995 | **RNA**→ rpmE2 (PRK01678)Ribosomal_L31 (pfam01197)→ |
| Sau-3296-1 | NZ_KK234444.1 | - | 56354 | 56325 | **RNA**→ rpmE2 (PRK01678)Ribosomal_L31 (pfam01197)→ |
| Sau-3474-1 | NZ_KK240451.1 | - | 26059 | 26030 | **RNA**→ rpmE2 (PRK01678)Ribosomal_L31 (pfam01197)→ |
| Sau-3313-1 | NZ_KK235112.1 | - | 26024 | 25995 | **RNA**→ rpmE2 (PRK01678)Ribosomal_L31 (pfam01197)→ |
| Sau-3478-1 | NZ_KK240580.1 | - | 26024 | 25995 | **RNA**→ rpmE2 (PRK01678)Ribosomal_L31 (pfam01197)→ |
| Sau-3322-1 | NZ_KK235373.1 | - | 26024 | 25995 | **RNA**→ rpmE2 (PRK01678)Ribosomal_L31 (pfam01197)→ |
| Sau-3246-1 | NZ_KK232545.1 | - | 56439 | 56410 | **RNA**→ rpmE2 (PRK01678)Ribosomal_L31 (pfam01197)→ |
| Sau-3344-1 | NZ_KK236227.1 | - | 324 | 295 | **RNA**→ rpmE2 (PRK01678)Ribosomal_L31 (pfam01197)→ |
| Sau-3710-1 | NZ_KK240619.1 | - | 26024 | 25995 | **RNA**→ rpmE2 (PRK01678)Ribosomal_L31 (pfam01197)→ |
| Sau-3349-1 | NZ_KK236311.1 | - | 56305 | 56276 | **RNA**→ rpmE2 (PRK01678)Ribosomal_L31 (pfam01197)→ |
| Sau-3247-1 | NZ_KK232579.1 | - | 26024 | 25995 | **RNA**→ rpmE2 (PRK01678)Ribosomal_L31 (pfam01197)→ |
| Sau-3716-1 | NZ_KK240757.1 | - | 26024 | 25995 | **RNA**→ rpmE2 (PRK01678)Ribosomal_L31 (pfam01197)→ |
| Sau-3717-1 | NZ_KK240801.1 | - | 26024 | 25995 | **RNA**→ rpmE2 (PRK01678)Ribosomal_L31 (pfam01197)→ |
| Sau-3357-1 | NZ_KK236523.1 | - | 19385 | 19356 | **RNA**→ rpmE2 (PRK01678)Ribosomal_L31 (pfam01197)→ |
| Sau-3251-1 | NZ_KK232759.1 | - | 25789 | 25760 | **RNA**→ rpmE2 (PRK01678)Ribosomal_L31 (pfam01197)→ |
| Sau-3364-1 | NZ_KK236725.1 | - | 2562 | 2533 | **RNA**→ rpmE2 (PRK01678)Ribosomal_L31 (pfam01197)→ |
| Sau-3252-1 | NZ_KK232802.1 | - | 26023 | 25994 | **RNA**→ rpmE2 (PRK01678)Ribosomal_L31 (pfam01197)→ |
| Sau-3257-1 | NZ_KK233004.1 | - | 26024 | 25995 | **RNA**→ rpmE2 (PRK01678)Ribosomal_L31 (pfam01197)→ |
| Sau-3258-1 | NZ_KK233060.1 | - | 26024 | 25995 | **RNA**→ rpmE2 (PRK01678)Ribosomal_L31 (pfam01197)→ |
| Sau-3264-1 | NZ_KK233293.1 | - | 56395 | 56366 | **RNA**→ rpmE2 (PRK01678)Ribosomal_L31 (pfam01197)→ |
| Sau-3269-1 | NZ_KK233491.1 | - | 26024 | 25995 | **RNA**→ rpmE2 (PRK01678)Ribosomal_L31 (pfam01197)→ |
| Sau-3301-1 | NZ_KK234667.1 | - | 26024 | 25995 | **RNA**→ rpmE2 (PRK01678)Ribosomal_L31 (pfam01197)→ |
| Sau-3278-1 | NZ_KK233668.1 | - | 26024 | 25995 | **RNA**→ rpmE2 (PRK01678)Ribosomal_L31 (pfam01197)→ |
| Sau-3347-1 | NZ_KK236266.1 | - | 26024 | 25995 | **RNA**→ rpmE2 (PRK01678)Ribosomal_L31 (pfam01197)→ |
| Sau-3285-1 | NZ_KK233916.1 | - | 26024 | 25995 | **RNA**→ rpmE2 (PRK01678)Ribosomal_L31 (pfam01197)→ |
| Sau-3356-1 | NZ_KK236493.1 | - | 56439 | 56410 | **RNA**→ rpmE2 (PRK01678)Ribosomal_L31 (pfam01197)→ |
| Sau-3243-1 | NZ_KK232425.1 | - | 56395 | 56366 | **RNA**→ rpmE2 (PRK01678)Ribosomal_L31 (pfam01197)→ |
| Sau-3369-1 | NZ_KK236870.1 | - | 18726 | 18697 | **RNA**→ rpmE2 (PRK01678)Ribosomal_L31 (pfam01197)→ |
| Sau-3248-1 | NZ_KK232626.1 | - | 56390 | 56361 | **RNA**→ rpmE2 (PRK01678)Ribosomal_L31 (pfam01197)→ |
| Sau-3375-1 | NZ_KK237079.1 | - | 56389 | 56360 | **RNA**→ rpmE2 (PRK01678)Ribosomal_L31 (pfam01197)→ |
| Sau-3259-1 | NZ_KK233125.1 | - | 3349 | 3320 | **RNA**→ rpmE2 (PRK01678)Ribosomal_L31 (pfam01197)→ |
| Sau-3380-1 | NZ_KK237308.1 | - | 26024 | 25995 | **RNA**→ rpmE2 (PRK01678)Ribosomal_L31 (pfam01197)→ |
| Sau-3265-1 | NZ_KK233320.1 | - | 26024 | 25995 | **RNA**→ rpmE2 (PRK01678)Ribosomal_L31 (pfam01197)→ |
| Sau-3385-1 | NZ_KK237500.1 | - | 56354 | 56325 | **RNA**→ rpmE2 (PRK01678)Ribosomal_L31 (pfam01197)→ |
| Sau-3272-1 | NZ_KK233526.1 | - | 26024 | 25995 | **RNA**→ rpmE2 (PRK01678)Ribosomal_L31 (pfam01197)→ |
| Sau-3315-1 | NZ_KK235151.1 | - | 26023 | 25994 | **RNA**→ rpmE2 (PRK01678)Ribosomal_L31 (pfam01197)→ |
| Sau-3302-1 | NZ_KK234707.1 | - | 26024 | 25995 | **RNA**→ rpmE2 (PRK01678)Ribosomal_L31 (pfam01197)→ |
| Sau-3329-1 | NZ_KK235571.1 | - | 26024 | 25995 | **RNA**→ rpmE2 (PRK01678)Ribosomal_L31 (pfam01197)→ |
| Sau-3303-1 | NZ_KK234760.1 | - | 26024 | 25995 | **RNA**→ rpmE2 (PRK01678)Ribosomal_L31 (pfam01197)→ |
| Sau-3334-1 | NZ_KK235765.1 | - | 56310 | 56281 | **RNA**→ rpmE2 (PRK01678)Ribosomal_L31 (pfam01197)→ |
| Sau-3307-1 | NZ_KK234915.1 | - | 26024 | 25995 | **RNA**→ rpmE2 (PRK01678)Ribosomal_L31 (pfam01197)→ |
| Sau-3339-1 | NZ_KK235994.1 | - | 26024 | 25995 | **RNA**→ rpmE2 (PRK01678)Ribosomal_L31 (pfam01197)→ |

| | | | | | | |
|---|---|---|---|---|---|---|
| Sau-3308-1 | NZ_KK234944.1 | - | 26024 | 25995 | **RNA**→ | rpmE2 (PRK01678)Ribosomal_L31 (pfam01197)→ |
| Sau-3371-1 | NZ_KK236898.1 | - | 2481 | 2452 | **RNA**→ | rpmE2 (PRK01678)Ribosomal_L31 (pfam01197)→ |
| Sau-3290-1 | NZ_KK234154.1 | - | 26024 | 25995 | **RNA**→ | rpmE2 (PRK01678)Ribosomal_L31 (pfam01197)→ |
| Sau-3280-1 | NZ_KK233714.1 | - | 26024 | 25995 | **RNA**→ | rpmE2 (PRK01678)Ribosomal_L31 (pfam01197)→ |
| Sau-3381-1 | NZ_KK237351.1 | - | 26024 | 25995 | **RNA**→ | rpmE2 (PRK01678)Ribosomal_L31 (pfam01197)→ |
| Sau-3295-1 | NZ_KK234384.1 | - | 26024 | 25995 | **RNA**→ | rpmE2 (PRK01678)Ribosomal_L31 (pfam01197)→ |
| Sau-3286-1 | NZ_KK233953.1 | - | 56439 | 56410 | **RNA**→ | rpmE2 (PRK01678)Ribosomal_L31 (pfam01197)→ |
| Sau-3306-1 | NZ_KK234874.1 | - | 56440 | 56411 | **RNA**→ | rpmE2 (PRK01678)Ribosomal_L31 (pfam01197)→ |
| Sau-3245-1 | NZ_KK232506.1 | - | 54852 | 54823 | **RNA**→ | rpmE2 (PRK01678)Ribosomal_L31 (pfam01197)→ |
| Sau-3319-1 | NZ_KK235253.1 | - | 56439 | 56410 | **RNA**→ | rpmE2 (PRK01678)Ribosomal_L31 (pfam01197)→ |
| Sau-3325-1 | NZ_KK235486.1 | - | 26024 | 25995 | **RNA**→ | rpmE2 (PRK01678)Ribosomal_L31 (pfam01197)→ |
| Sau-3250-1 | NZ_KK232698.1 | - | 26024 | 25995 | **RNA**→ | rpmE2 (PRK01678)Ribosomal_L31 (pfam01197)→ |
| Sau-3324-1 | NZ_KK235447.1 | - | 25968 | 25939 | **RNA**→ | rpmE2 (PRK01678)Ribosomal_L31 (pfam01197)→ |
| Sau-3343-1 | NZ_KK236171.1 | - | 25968 | 25939 | **RNA**→ | rpmE2 (PRK01678)Ribosomal_L31 (pfam01197)→ |
| Sau-3331-1 | NZ_KK235649.1 | - | 26024 | 25995 | **RNA**→ | rpmE2 (PRK01678)Ribosomal_L31 (pfam01197)→ |
| Sau-3256-1 | NZ_KK232949.1 | - | 26024 | 25995 | **RNA**→ | rpmE2 (PRK01678)Ribosomal_L31 (pfam01197)→ |
| Sau-3336-1 | NZ_KK235845.1 | - | 26024 | 25995 | **RNA**→ | rpmE2 (PRK01678)Ribosomal_L31 (pfam01197)→ |
| Sau-3363-1 | NZ_KK236694.1 | - | 26024 | 25995 | **RNA**→ | rpmE2 (PRK01678)Ribosomal_L31 (pfam01197)→ |
| Sau-3262-1 | NZ_KK233186.1 | - | 56439 | 56410 | **RNA**→ | rpmE2 (PRK01678)Ribosomal_L31 (pfam01197)→ |
| Sau-3342-1 | NZ_KK236132.1 | - | 26024 | 25995 | **RNA**→ | rpmE2 (PRK01678)Ribosomal_L31 (pfam01197)→ |
| Sau-3345-1 | NZ_KK328125.1 | - | 25997 | 25968 | **RNA**→ | rpmE2 (PRK01678)Ribosomal_L31 (pfam01197)→ |
| Sau-3263-1 | NZ_KK233245.1 | - | 26024 | 25995 | **RNA**→ | rpmE2 (PRK01678)Ribosomal_L31 (pfam01197)→ |
| Sau-3354-1 | NZ_KK236396.1 | - | 56390 | 56361 | **RNA**→ | rpmE2 (PRK01678)Ribosomal_L31 (pfam01197)→ |
| Sau-3267-1 | NZ_KK233392.1 | - | 26024 | 25995 | **RNA**→ | rpmE2 (PRK01678)Ribosomal_L31 (pfam01197)→ |
| Sau-3355-1 | NZ_KK236455.1 | - | 56395 | 56366 | **RNA**→ | rpmE2 (PRK01678)Ribosomal_L31 (pfam01197)→ |
| Sau-3268-1 | NZ_KK233439.1 | - | 26024 | 25995 | **RNA**→ | rpmE2 (PRK01678)Ribosomal_L31 (pfam01197)→ |
| Sau-3368-1 | NZ_KK236834.1 | - | 56279 | 56250 | **RNA**→ | rpmE2 (PRK01678)Ribosomal_L31 (pfam01197)→ |
| Sau-3275-1 | NZ_KK233602.1 | - | 26024 | 25995 | **RNA**→ | rpmE2 (PRK01678)Ribosomal_L31 (pfam01197)→ |
| Sau-3374-1 | NZ_KK237046.1 | - | 26024 | 25995 | **RNA**→ | rpmE2 (PRK01678)Ribosomal_L31 (pfam01197)→ |
| Sau-3277-1 | NZ_KK233637.1 | - | 26024 | 25995 | **RNA**→ | rpmE2 (PRK01678)Ribosomal_L31 (pfam01197)→ |
| Sau-3317-1 | NZ_KK235201.1 | - | 26024 | 25995 | **RNA**→ | rpmE2 (PRK01678)Ribosomal_L31 (pfam01197)→ |
| Sau-3294-1 | NZ_KK234337.1 | - | 26024 | 25995 | **RNA**→ | rpmE2 (PRK01678)Ribosomal_L31 (pfam01197)→ |
| Sau-3300-1 | NZ_KK234614.1 | - | 53444 | 53415 | **RNA**→ | rpmE2 (PRK01678)Ribosomal_L31 (pfam01197)→ |
| Sau-3340-1 | NZ_KK236040.1 | - | 26024 | 25995 | **RNA**→ | rpmE2 (PRK01678)Ribosomal_L31 (pfam01197)→ |
| Sau-3305-1 | NZ_KK234843.1 | - | 26024 | 25995 | **RNA**→ | rpmE2 (PRK01678)Ribosomal_L31 (pfam01197)→ |
| Sau-3311-1 | NZ_KK235031.1 | - | 26024 | 25995 | **RNA**→ | rpmE2 (PRK01678)Ribosomal_L31 (pfam01197)→ |
| Sau-3359-1 | NZ_KK236567.1 | - | 26024 | 25995 | **RNA**→ | rpmE2 (PRK01678)Ribosomal_L31 (pfam01197)→ |
| Sau-3282-1 | NZ_KK233806.1 | - | 26024 | 25995 | **RNA**→ | rpmE2 (PRK01678)Ribosomal_L31 (pfam01197)→ |
| Sau-3360-1 | NZ_KK236596.1 | - | 26024 | 25995 | **RNA**→ | rpmE2 (PRK01678)Ribosomal_L31 (pfam01197)→ |
| Sau-3289-1 | NZ_KK234091.1 | - | 56439 | 56410 | **RNA**→ | rpmE2 (PRK01678)Ribosomal_L31 (pfam01197)→ |
| Sau-3244-1 | NZ_KK232479.1 | - | 26023 | 25994 | **RNA**→ | rpmE2 (PRK01678)Ribosomal_L31 (pfam01197)→ |
| Sau-3249-1 | NZ_KK232671.1 | - | 56439 | 56410 | **RNA**→ | rpmE2 (PRK01678)Ribosomal_L31 (pfam01197)→ |
| Sau-3255-1 | NZ_KK232899.1 | - | 26024 | 25995 | **RNA**→ | rpmE2 (PRK01678)Ribosomal_L31 (pfam01197)→ |
| Sau-3260-1 | NZ_KK233161.1 | - | 26024 | 25995 | **RNA**→ | rpmE2 (PRK01678)Ribosomal_L31 (pfam01197)→ |
| Sau-3274-1 | NZ_KK233565.1 | - | 26024 | 25995 | **RNA**→ | rpmE2 (PRK01678)Ribosomal_L31 (pfam01197)→ |
| Sau-3292-1 | NZ_KK234244.1 | - | 26024 | 25995 | **RNA**→ | rpmE2 (PRK01678)Ribosomal_L31 (pfam01197)→ |
| Sau-3293-1 | NZ_KK234279.1 | - | 26024 | 25995 | **RNA**→ | rpmE2 (PRK01678)Ribosomal_L31 (pfam01197)→ |
| Sau-3297-1 | NZ_KK234517.1 | - | 26024 | 25995 | **RNA**→ | rpmE2 (PRK01678)Ribosomal_L31 (pfam01197)→ |
| Sau-3298-1 | NZ_KK234558.1 | - | 26024 | 25995 | **RNA**→ | rpmE2 (PRK01678)Ribosomal_L31 (pfam01197)→ |
| Sau-3304-1 | NZ_KK234801.1 | - | 26024 | 25995 | **RNA**→ | rpmE2 (PRK01678)Ribosomal_L31 (pfam01197)→ |
| Sau-3310-1 | NZ_KK234978.1 | - | 26024 | 25995 | **RNA**→ | rpmE2 (PRK01678)Ribosomal_L31 (pfam01197)→ |
| Sau-3281-1 | NZ_KK233762.1 | - | 26024 | 25995 | **RNA**→ | rpmE2 (PRK01678)Ribosomal_L31 (pfam01197)→ |

| | | | | | | |
|---|---|---|---|---|---|---|
| Sau-3287-1 | NZ_KK233998.1 | - | 26024 | 25995 | **RNA**→ | rpmE2 (PRK01678)Ribosomal_L31 (pfam01197)→ |
| Sau-3180-1 | NZ_KK230294.1 | - | 26024 | 25995 | **RNA**→ | rpmE2 (PRK01678)Ribosomal_L31 (pfam01197)→ |
| Sau-3181-1 | NZ_KK230346.1 | - | 29589 | 29560 | **RNA**→ | rpmE2 (PRK01678)Ribosomal_L31 (pfam01197)→ |
| Sau-3202-1 | NZ_KK230956.1 | - | 56439 | 56410 | **RNA**→ | rpmE2 (PRK01678)Ribosomal_L31 (pfam01197)→ |
| Sau-867-1 | NZ_KK223898.1 | - | 56351 | 56322 | **RNA**→ | rpmE2 (PRK01678)Ribosomal_L31 (pfam01197)→ |
| Sau-3207-1 | NZ_KK231188.1 | - | 26024 | 25995 | **RNA**→ | rpmE2 (PRK01678)Ribosomal_L31 (pfam01197)→ |
| Sau-3212-1 | NZ_KK231386.1 | - | 26024 | 25995 | **RNA**→ | rpmE2 (PRK01678)Ribosomal_L31 (pfam01197)→ |
| Sau-2687-1 | NZ_KK224793.1 | - | 56395 | 56366 | **RNA**→ | rpmE2 (PRK01678)Ribosomal_L31 (pfam01197)→ |
| Sau-4013-1 | NZ_AGRP01000086.1 | - | 54853 | 54824 | **RNA**→ | rpmE2 (PRK01678)Ribosomal_L31 (pfam01197)→ |
| Sau-4015-1 | NZ_AFTQ01000078.1 | - | 55162 | 55133 | **RNA**→ | rpmE2 (PRK01678)Ribosomal_L31 (pfam01197)→ |
| Sau-3217-1 | NZ_KK231605.1 | - | 26024 | 25995 | **RNA**→ | rpmE2 (PRK01678)Ribosomal_L31 (pfam01197)→ |
| Sau-3222-1 | NZ_KK231797.1 | - | 26024 | 25995 | **RNA**→ | rpmE2 (PRK01678)Ribosomal_L31 (pfam01197)→ |
| Sau-3104-1 | NZ_KK227826.1 | - | 26023 | 25994 | **RNA**→ | rpmE2 (PRK01678)Ribosomal_L31 (pfam01197)→ |
| Sau-3227-1 | NZ_KK231957.1 | - | 56309 | 56280 | **RNA**→ | rpmE2 (PRK01678)Ribosomal_L31 (pfam01197)→ |
| Sau-3128-1 | NZ_KK228659.1 | - | 26024 | 25995 | **RNA**→ | rpmE2 (PRK01678)Ribosomal_L31 (pfam01197)→ |
| Sau-3233-1 | NZ_KK232123.1 | - | 26024 | 25995 | **RNA**→ | rpmE2 (PRK01678)Ribosomal_L31 (pfam01197)→ |
| Sau-3240-1 | NZ_KK232321.1 | - | 53699 | 53670 | **RNA**→ | rpmE2 (PRK01678)Ribosomal_L31 (pfam01197)→ |
| Sau-3134-1 | NZ_KK228852.1 | - | 26024 | 25995 | **RNA**→ | rpmE2 (PRK01678)Ribosomal_L31 (pfam01197)→ |
| Sau-3186-1 | NZ_KK230550.1 | - | 56420 | 56391 | **RNA**→ | rpmE2 (PRK01678)Ribosomal_L31 (pfam01197)→ |
| Sau-3141-1 | NZ_KK229051.1 | - | 56395 | 56366 | **RNA**→ | rpmE2 (PRK01678)Ribosomal_L31 (pfam01197)→ |
| Sau-3187-1 | NZ_KK230593.1 | - | 56395 | 56366 | **RNA**→ | rpmE2 (PRK01678)Ribosomal_L31 (pfam01197)→ |
| Sau-3195-1 | NZ_KK230781.1 | - | 56304 | 56275 | **RNA**→ | rpmE2 (PRK01678)Ribosomal_L31 (pfam01197)→ |
| Sau-3151-1 | NZ_KK229266.1 | - | 26024 | 25995 | **RNA**→ | rpmE2 (PRK01678)Ribosomal_L31 (pfam01197)→ |
| Sau-3182-1 | NZ_KK230400.1 | - | 56395 | 56366 | **RNA**→ | rpmE2 (PRK01678)Ribosomal_L31 (pfam01197)→ |
| Sau-3203-1 | NZ_KK231012.1 | - | 26024 | 25995 | **RNA**→ | rpmE2 (PRK01678)Ribosomal_L31 (pfam01197)→ |
| Sau-3155-1 | NZ_KK229434.1 | - | 56395 | 56366 | **RNA**→ | rpmE2 (PRK01678)Ribosomal_L31 (pfam01197)→ |
| Sau-3208-1 | NZ_KK231221.1 | - | 56439 | 56410 | **RNA**→ | rpmE2 (PRK01678)Ribosomal_L31 (pfam01197)→ |
| Sau-3213-1 | NZ_KK231435.1 | - | 26024 | 25995 | **RNA**→ | rpmE2 (PRK01678)Ribosomal_L31 (pfam01197)→ |
| Sau-3167-1 | NZ_KK229842.1 | - | 56395 | 56366 | **RNA**→ | rpmE2 (PRK01678)Ribosomal_L31 (pfam01197)→ |
| Sau-3218-1 | NZ_KK231633.1 | - | 26024 | 25995 | **RNA**→ | rpmE2 (PRK01678)Ribosomal_L31 (pfam01197)→ |
| Sau-3174-1 | NZ_KK230081.1 | - | 26024 | 25995 | **RNA**→ | rpmE2 (PRK01678)Ribosomal_L31 (pfam01197)→ |
| Sau-3228-1 | NZ_KK231982.1 | - | 26024 | 25995 | **RNA**→ | rpmE2 (PRK01678)Ribosomal_L31 (pfam01197)→ |
| Sau-3230-1 | NZ_KK232006.1 | - | 56353 | 56324 | **RNA**→ | rpmE2 (PRK01678)Ribosomal_L31 (pfam01197)→ |
| Sau-3235-1 | NZ_KK232151.1 | - | 26024 | 25995 | **RNA**→ | rpmE2 (PRK01678)Ribosomal_L31 (pfam01197)→ |
| Sau-3236-1 | NZ_KK232180.1 | - | 26025 | 25996 | **RNA**→ | rpmE2 (PRK01678)Ribosomal_L31 (pfam01197)→ |
| Sau-3241-1 | NZ_KK232371.1 | - | 56394 | 56365 | **RNA**→ | rpmE2 (PRK01678)Ribosomal_L31 (pfam01197)→ |
| Sau-203-1 | NZ_KK222014.1 | - | 34754 | 34725 | **RNA**→ | rpmE2 (PRK01678)Ribosomal_L31 (pfam01197)→ |
| Sau-3188-1 | NZ_KK230635.1 | - | 26024 | 25995 | **RNA**→ | rpmE2 (PRK01678)Ribosomal_L31 (pfam01197)→ |
| Sau-3196-1 | NZ_KK230828.1 | - | 26024 | 25995 | **RNA**→ | rpmE2 (PRK01678)Ribosomal_L31 (pfam01197)→ |
| Sau-3105-1 | NZ_KK227870.1 | - | 26023 | 25994 | **RNA**→ | rpmE2 (PRK01678)Ribosomal_L31 (pfam01197)→ |
| Sau-3185-1 | NZ_KK230491.1 | - | 56439 | 56410 | **RNA**→ | rpmE2 (PRK01678)Ribosomal_L31 (pfam01197)→ |
| Sau-3110-1 | NZ_KK228070.1 | - | 26024 | 25995 | **RNA**→ | rpmE2 (PRK01678)Ribosomal_L31 (pfam01197)→ |
| Sau-3200-1 | NZ_KK230930.1 | - | 53254 | 53225 | **RNA**→ | rpmE2 (PRK01678)Ribosomal_L31 (pfam01197)→ |
| Sau-3205-1 | NZ_KK231074.1 | - | 26287 | 26258 | **RNA**→ | rpmE2 (PRK01678)Ribosomal_L31 (pfam01197)→ |
| Sau-3135-1 | NZ_KK228890.1 | - | 26024 | 25995 | **RNA**→ | rpmE2 (PRK01678)Ribosomal_L31 (pfam01197)→ |
| Sau-3206-1 | NZ_KK231155.1 | - | 56394 | 56365 | **RNA**→ | rpmE2 (PRK01678)Ribosomal_L31 (pfam01197)→ |
| Sau-3210-1 | NZ_KK231317.1 | - | 26024 | 25995 | **RNA**→ | rpmE2 (PRK01678)Ribosomal_L31 (pfam01197)→ |
| Sau-3156-1 | NZ_KK229493.1 | - | 26024 | 25995 | **RNA**→ | rpmE2 (PRK01678)Ribosomal_L31 (pfam01197)→ |
| Sau-3211-1 | NZ_KK231348.1 | - | 25968 | 25939 | **RNA**→ | rpmE2 (PRK01678)Ribosomal_L31 (pfam01197)→ |
| Sau-3216-1 | NZ_KK231552.1 | - | 9739 | 9710 | **RNA**→ | rpmE2 (PRK01678)Ribosomal_L31 (pfam01197)→ |
| Sau-3161-1 | NZ_KK229686.1 | - | 26024 | 25995 | **RNA**→ | rpmE2 (PRK01678)Ribosomal_L31 (pfam01197)→ |
| Sau-3221-1 | NZ_KK231752.1 | - | 26024 | 25995 | **RNA**→ | rpmE2 (PRK01678)Ribosomal_L31 (pfam01197)→ |

| | | | | | |
|---|---|---|---|---|---|
| Sau-3226-1 | NZ_KK231934.1 | - | 54426 | 54397 | **RNA**→ rpmE2 (PRK01678)Ribosomal_L31 (pfam01197)→ |
| Sau-3169-1 | NZ_KK229903.1 | - | 56339 | 56310 | **RNA**→ rpmE2 (PRK01678)Ribosomal_L31 (pfam01197)→ |
| Sau-3232-1 | NZ_KK232090.1 | - | 324 | 295 | **RNA**→ rpmE2 (PRK01678)Ribosomal_L31 (pfam01197)→ |
| Sau-3239-1 | NZ_KK232282.1 | - | 56310 | 56281 | **RNA**→ rpmE2 (PRK01678)Ribosomal_L31 (pfam01197)→ |
| Sau-3170-1 | NZ_KK229956.1 | - | 26024 | 25995 | **RNA**→ rpmE2 (PRK01678)Ribosomal_L31 (pfam01197)→ |
| Sau-3191-1 | NZ_KK230709.1 | - | 56439 | 56410 | **RNA**→ rpmE2 (PRK01678)Ribosomal_L31 (pfam01197)→ |
| Sau-3194-1 | NZ_KK230757.1 | - | 56310 | 56281 | **RNA**→ rpmE2 (PRK01678)Ribosomal_L31 (pfam01197)→ |
| Sau-3177-1 | NZ_KK230171.1 | - | 26023 | 25994 | **RNA**→ rpmE2 (PRK01678)Ribosomal_L31 (pfam01197)→ |
| Sau-3199-1 | NZ_KK230896.1 | - | 26024 | 25995 | **RNA**→ rpmE2 (PRK01678)Ribosomal_L31 (pfam01197)→ |
| Sau-3204-1 | NZ_KK231053.1 | - | 26024 | 25995 | **RNA**→ rpmE2 (PRK01678)Ribosomal_L31 (pfam01197)→ |
| Sau-3209-1 | NZ_KK231265.1 | - | 26024 | 25995 | **RNA**→ rpmE2 (PRK01678)Ribosomal_L31 (pfam01197)→ |
| Sau-3214-1 | NZ_KK231470.1 | - | 26149 | 26120 | **RNA**→ rpmE2 (PRK01678)Ribosomal_L31 (pfam01197)→ |
| Sau-3215-1 | NZ_KK231503.1 | - | 26024 | 25995 | **RNA**→ rpmE2 (PRK01678)Ribosomal_L31 (pfam01197)→ |
| Sau-3219-1 | NZ_KK231663.1 | - | 53737 | 53708 | **RNA**→ rpmE2 (PRK01678)Ribosomal_L31 (pfam01197)→ |
| Sau-3220-1 | NZ_KK231710.1 | - | 26024 | 25995 | **RNA**→ rpmE2 (PRK01678)Ribosomal_L31 (pfam01197)→ |
| Sau-3224-1 | NZ_KK231877.1 | - | 26024 | 25995 | **RNA**→ rpmE2 (PRK01678)Ribosomal_L31 (pfam01197)→ |
| Sau-3231-1 | NZ_KK232044.1 | - | 26024 | 25995 | **RNA**→ rpmE2 (PRK01678)Ribosomal_L31 (pfam01197)→ |
| Sau-3238-1 | NZ_KK232221.1 | - | 26024 | 25995 | **RNA**→ rpmE2 (PRK01678)Ribosomal_L31 (pfam01197)→ |
| Sau-3100-1 | NZ_KK227661.1 | - | 57233 | 57204 | **RNA**→ rpmE2 (PRK01678)Ribosomal_L31 (pfam01197)→ |
| Sau-3189-1 | NZ_KK230662.1 | - | 56439 | 56410 | **RNA**→ rpmE2 (PRK01678)Ribosomal_L31 (pfam01197)→ |
| Sau-3197-1 | NZ_KK230870.1 | - | 26024 | 25995 | **RNA**→ rpmE2 (PRK01678)Ribosomal_L31 (pfam01197)→ |
| Sau-3074-1 | NZ_KK226711.1 | - | 26024 | 25995 | **RNA**→ rpmE2 (PRK01678)Ribosomal_L31 (pfam01197)→ |
| Sau-3107-1 | NZ_KK227954.1 | - | 26024 | 25995 | **RNA**→ rpmE2 (PRK01678)Ribosomal_L31 (pfam01197)→ |
| Sau-3113-1 | NZ_KK228153.1 | - | 26024 | 25995 | **RNA**→ rpmE2 (PRK01678)Ribosomal_L31 (pfam01197)→ |
| Sau-3125-1 | NZ_KK228588.1 | - | 26024 | 25995 | **RNA**→ rpmE2 (PRK01678)Ribosomal_L31 (pfam01197)→ |
| Sau-3127-1 | NZ_KK228618.1 | - | 26024 | 25995 | **RNA**→ rpmE2 (PRK01678)Ribosomal_L31 (pfam01197)→ |
| Sau-3133-1 | NZ_KK228823.1 | - | 56380 | 56351 | **RNA**→ rpmE2 (PRK01678)Ribosomal_L31 (pfam01197)→ |
| Sau-3159-1 | NZ_KK229612.1 | - | 26024 | 25995 | **RNA**→ rpmE2 (PRK01678)Ribosomal_L31 (pfam01197)→ |
| Sau-3172-1 | NZ_KK230044.1 | - | 26024 | 25995 | **RNA**→ rpmE2 (PRK01678)Ribosomal_L31 (pfam01197)→ |
| Sau-2635-1 | NZ_KK224580.1 | - | 31457 | 31428 | **RNA**→ rpmE2 (PRK01678)Ribosomal_L31 (pfam01197)→ |
| Sau-2696-1 | NZ_KK225209.1 | - | 26024 | 25995 | **RNA**→ rpmE2 (PRK01678)Ribosomal_L31 (pfam01197)→ |
| Sau-2728-1 | NZ_KK225630.1 | - | 26024 | 25995 | **RNA**→ rpmE2 (PRK01678)Ribosomal_L31 (pfam01197)→ |
| Sau-3008-1 | NZ_KK225971.1 | - | 26024 | 25995 | **RNA**→ rpmE2 (PRK01678)Ribosomal_L31 (pfam01197)→ |
| Sau-3015-1 | NZ_KK226169.1 | - | 26024 | 25995 | **RNA**→ rpmE2 (PRK01678)Ribosomal_L31 (pfam01197)→ |
| Sau-3072-1 | NZ_KK226639.1 | - | 26024 | 25995 | **RNA**→ rpmE2 (PRK01678)Ribosomal_L31 (pfam01197)→ |
| Sau-3078-1 | NZ_KK226872.1 | - | 56439 | 56410 | **RNA**→ rpmE2 (PRK01678)Ribosomal_L31 (pfam01197)→ |
| Sau-3137-1 | NZ_KK228988.1 | - | 56439 | 56410 | **RNA**→ rpmE2 (PRK01678)Ribosomal_L31 (pfam01197)→ |
| Sau-3146-1 | NZ_KK229172.1 | - | 26024 | 25995 | **RNA**→ rpmE2 (PRK01678)Ribosomal_L31 (pfam01197)→ |
| Sau-3158-1 | NZ_KK229579.1 | - | 26024 | 25995 | **RNA**→ rpmE2 (PRK01678)Ribosomal_L31 (pfam01197)→ |
| Sau-3171-1 | NZ_KK229998.1 | - | 26024 | 25995 | **RNA**→ rpmE2 (PRK01678)Ribosomal_L31 (pfam01197)→ |
| Sau-3178-1 | NZ_KK230206.1 | - | 56440 | 56411 | **RNA**→ rpmE2 (PRK01678)Ribosomal_L31 (pfam01197)→ |
| Sau-3102-1 | NZ_KK227736.1 | - | 56395 | 56366 | **RNA**→ rpmE2 (PRK01678)Ribosomal_L31 (pfam01197)→ |
| Sau-3416-1 | NZ_KK238465.1 | - | 26024 | 25995 | **RNA**→ rpmE2 (PRK01678)Ribosomal_L31 (pfam01197)→ |
| Sau-3451-1 | NZ_KK239825.1 | - | 56352 | 56323 | **RNA**→ rpmE2 (PRK01678)Ribosomal_L31 (pfam01197)→ |
| Sau-366-1 | NZ_KK099382.1 | - | 57558 | 57529 | **RNA**→ rpmE2 (PRK01678)Ribosomal_L31 (pfam01197)→ |
| Sau-384-1 | NZ_KK098824.1 | - | 58224 | 58195 | **RNA**→ rpmE2 (PRK01678)Ribosomal_L31 (pfam01197)→ |
| Sau-1071-1 | NZ_KK097754.1 | - | 34817 | 34788 | **RNA**→ rpmE2 (PRK01678)Ribosomal_L31 (pfam01197)→ |
| Sau-994-1 | NZ_KK096328.1 | - | 56794 | 56765 | **RNA**→ rpmE2 (PRK01678)Ribosomal_L31 (pfam01197)→ |
| Sau-3435-1 | NZ_KK239279.1 | - | 56439 | 56410 | **RNA**→ rpmE2 (PRK01678)Ribosomal_L31 (pfam01197)→ |
| Sau-3465-1 | NZ_KK240248.1 | - | 26024 | 25995 | **RNA**→ rpmE2 (PRK01678)Ribosomal_L31 (pfam01197)→ |
| Sau-2705-1 | NZ_KK093341.1 | - | 25162 | 25133 | **RNA**→ rpmE2 (PRK01678)Ribosomal_L31 (pfam01197)→ |
| Sau-3254-1 | NZ_KK232846.1 | - | 26024 | 25995 | **RNA**→ rpmE2 (PRK01678)Ribosomal_L31 (pfam01197)→ |

| | | | | | |
|---|---|---|---|---|---|
| Sau-3560-1 | NZ_KK097381.1 | - | 56894 | 56865 | **RNA**→ rpmE2 (PRK01678)Ribosomal_L31 (pfam01197)→ |
| Sau-3335-1 | NZ_KK235791.1 | - | 26024 | 25995 | **RNA**→ rpmE2 (PRK01678)Ribosomal_L31 (pfam01197)→ |
| Sau-323-1 | NZ_KK099083.1 | - | 423928 | 423899 | **RNA**→ rpmE2 (PRK01678)Ribosomal_L31 (pfam01197)→ |
| Sau-3462-1 | NZ_KK240163.1 | - | 26024 | 25995 | **RNA**→ rpmE2 (PRK01678)Ribosomal_L31 (pfam01197)→ |
| Sau-2908-1 | NZ_KK095057.1 | - | 34913 | 34884 | **RNA**→ rpmE2 (PRK01678)Ribosomal_L31 (pfam01197)→ |
| Sau-433-1 | NZ_KK097952.1 | - | 402019 | 401990 | **RNA**→ rpmE2 (PRK01678)Ribosomal_L31 (pfam01197)→ |
| Sau-476-1 | NZ_KK098504.1 | - | 56762 | 56733 | **RNA**→ rpmE2 (PRK01678)Ribosomal_L31 (pfam01197)→ |
| Sau-578-1 | NZ_KK099895.1 | - | 64875 | 64846 | **RNA**→ rpmE2 (PRK01678)Ribosomal_L31 (pfam01197)→ |
| Sau-3535-1 | NZ_KK097153.1 | - | 34913 | 34884 | **RNA**→ rpmE2 (PRK01678)Ribosomal_L31 (pfam01197)→ |
| Sau-382-1 | NZ_KK099509.1 | - | 119448 | 119419 | **RNA**→ rpmE2 (PRK01678)Ribosomal_L31 (pfam01197)→ |
| Sau-420-1 | NZ_KK097832.1 | - | 238441 | 238412 | **RNA**→ rpmE2 (PRK01678)Ribosomal_L31 (pfam01197)→ |
| Sau-389-1 | NZ_KK099568.1 | - | 59482 | 59453 | **RNA**→ rpmE2 (PRK01678)Ribosomal_L31 (pfam01197)→ |
| Sau-489-1 | NZ_KK098660.1 | - | 247385 | 247356 | **RNA**→ rpmE2 (PRK01678)Ribosomal_L31 (pfam01197)→ |
| Sau-326-1 | NZ_KK099092.1 | - | 56763 | 56734 | **RNA**→ rpmE2 (PRK01678)Ribosomal_L31 (pfam01197)→ |
| Sau-325-1 | NZ_KI966037.1 | - | 60735 | 60706 | **RNA**→ rpmE2 (PRK01678)Ribosomal_L31 (pfam01197)→ |
| Sau-1993-1 | NZ_KI972366.1 | - | 56763 | 56734 | **RNA**→ rpmE2 (PRK01678)Ribosomal_L31 (pfam01197)→ |
| Sau-3052-1 | NZ_KI979037.1 | - | 56994 | 56965 | **RNA**→ rpmE2 (PRK01678)Ribosomal_L31 (pfam01197)→ |
| Sau-3048-1 | NZ_KI978310.1 | + | 30719 | 30748 | **RNA**→ rpmE2 (PRK01678)Ribosomal_L31 (pfam01197)→ |
| Sau-2386-1 | NZ_KI997506.1 | - | 94282 | 94253 | **RNA**→ rpmE2 (PRK01678)Ribosomal_L31 (pfam01197)→ |
| Sau-3643-1 | NZ_KK027977.1 | - | 34874 | 34845 | **RNA**→ rpmE2 (PRK01678)Ribosomal_L31 (pfam01197)→ |
| Sau-2786-1 | NZ_KI974346.1 | - | 238794 | 238765 | **RNA**→ rpmE2 (PRK01678)Ribosomal_L31 (pfam01197)→ |
| Sau-2796-1 | NZ_KI976735.1 | - | 230191 | 230162 | **RNA**→ rpmE2 (PRK01678)Ribosomal_L31 (pfam01197)→ |
| Sau-627-1 | NZ_KK029040.1 | - | 56891 | 56862 | **RNA**→ rpmE2 (PRK01678)Ribosomal_L31 (pfam01197)→ |
| Sau-2708-1 | NZ_KK028975.1 | - | 56893 | 56864 | **RNA**→ rpmE2 (PRK01678)Ribosomal_L31 (pfam01197)→ |
| Sau-297-1 | NZ_KI978058.1 | - | 56894 | 56865 | **RNA**→ rpmE2 (PRK01678)Ribosomal_L31 (pfam01197)→ |
| Sau-2229-1 | NZ_KK021866.1 | - | 92275 | 92246 | **RNA**→ rpmE2 (PRK01678)Ribosomal_L31 (pfam01197)→ |
| Sau-228-1 | NZ_KI974986.1 | - | 56894 | 56865 | **RNA**→ rpmE2 (PRK01678)Ribosomal_L31 (pfam01197)→ |
| Sau-2870-1 | NZ_KK031065.1 | - | 56894 | 56865 | **RNA**→ rpmE2 (PRK01678)Ribosomal_L31 (pfam01197)→ |
| Sau-2312-1 | NZ_KI994172.1 | - | 218046 | 218017 | **RNA**→ rpmE2 (PRK01678)Ribosomal_L31 (pfam01197)→ |
| Sau-2707-1 | NZ_KK028867.1 | - | 56893 | 56864 | **RNA**→ rpmE2 (PRK01678)Ribosomal_L31 (pfam01197)→ |
| Sau-3700-1 | NZ_KI974015.1 | - | 108552 | 108523 | **RNA**→ rpmE2 (PRK01678)Ribosomal_L31 (pfam01197)→ |
| Sau-3693-1 | NZ_KI978641.1 | - | 55562 | 55533 | **RNA**→ rpmE2 (PRK01678)Ribosomal_L31 (pfam01197)→ |
| Sau-2439-1 | NZ_KI991670.1 | - | 36391 | 36362 | **RNA**→ rpmE2 (PRK01678)Ribosomal_L31 (pfam01197)→ |
| Sau-2565-1 | NZ_KK031039.1 | - | 56891 | 56862 | **RNA**→ rpmE2 (PRK01678)Ribosomal_L31 (pfam01197)→ |
| Sau-933-1 | NZ_KK029598.1 | - | 34817 | 34788 | **RNA**→ rpmE2 (PRK01678)Ribosomal_L31 (pfam01197)→ |
| Sau-2461-1 | NZ_KK025203.1 | - | 34913 | 34884 | **RNA**→ rpmE2 (PRK01678)Ribosomal_L31 (pfam01197)→ |
| Sau-716-1 | NZ_KK024072.1 | - | 36850 | 36821 | **RNA**→ rpmE2 (PRK01678)Ribosomal_L31 (pfam01197)→ |
| Sau-934-1 | NZ_KK029661.1 | - | 34873 | 34844 | **RNA**→ rpmE2 (PRK01678)Ribosomal_L31 (pfam01197)→ |
| Sau-2643-1 | NZ_KI973965.1 | - | 113174 | 113145 | **RNA**→ rpmE2 (PRK01678)Ribosomal_L31 (pfam01197)→ |
| Sau-2980-1 | NZ_KK026111.1 | - | 34913 | 34884 | **RNA**→ rpmE2 (PRK01678)Ribosomal_L31 (pfam01197)→ |
| Sau-3582-1 | NZ_KK032539.1 | - | 34872 | 34843 | **RNA**→ rpmE2 (PRK01678)Ribosomal_L31 (pfam01197)→ |
| Sau-2543-1 | NZ_KK023131.1 | - | 122766 | 122737 | **RNA**→ rpmE2 (PRK01678)Ribosomal_L31 (pfam01197)→ |
| Sau-780-1 | NZ_KK028731.1 | - | 56894 | 56865 | **RNA**→ rpmE2 (PRK01678)Ribosomal_L31 (pfam01197)→ |
| Sau-2596-1 | NZ_KK038732.1 | - | 56894 | 56865 | **RNA**→ rpmE2 (PRK01678)Ribosomal_L31 (pfam01197)→ |
| Sau-2210-1 | NZ_KK063536.1 | - | 36340 | 36311 | **RNA**→ rpmE2 (PRK01678)Ribosomal_L31 (pfam01197)→ |
| Sau-2203-1 | NZ_KK063687.1 | - | 55327 | 55298 | **RNA**→ rpmE2 (PRK01678)Ribosomal_L31 (pfam01197)→ |
| Sau-882-1 | NZ_KK047892.1 | - | 1743431 | 1743402 | **RNA**→ rpmE2 (PRK01678)Ribosomal_L31 (pfam01197)→ |
| Sau-2879-1 | NZ_KK040438.1 | - | 34873 | 34844 | **RNA**→ rpmE2 (PRK01678)Ribosomal_L31 (pfam01197)→ |
| Sau-2213-1 | NZ_KK063477.1 | - | 61690 | 61661 | **RNA**→ rpmE2 (PRK01678)Ribosomal_L31 (pfam01197)→ |
| Sau-2202-1 | NZ_KK063693.1 | - | 89866 | 89837 | **RNA**→ rpmE2 (PRK01678)Ribosomal_L31 (pfam01197)→ |
| Sau-2202-2 | NZ_KK063701.1 | - | 729 | 700 | **RNA**→ rpmE2 (PRK01678)Ribosomal_L31 (pfam01197)→ |
| Sau-2200-1 | NZ_KK063759.1 | - | 62102 | 62073 | **RNA**→ rpmE2 (PRK01678)Ribosomal_L31 (pfam01197)→ |

| | | | | | |
|---|---|---|---|---|---|
| Sau-3524-1 | NZ_KK038189.1 | - | 34817 | 34788 | **RNA**→ rpmE2 (PRK01678)Ribosomal_L31 (pfam01197)→ |
| Sau-3570-1 | NZ_KK070196.1 | - | 56763 | 56734 | **RNA**→ rpmE2 (PRK01678)Ribosomal_L31 (pfam01197)→ |
| Sau-2197-1 | NZ_KK063823.1 | - | 49861 | 49832 | **RNA**→ rpmE2 (PRK01678)Ribosomal_L31 (pfam01197)→ |
| Sau-1038-1 | NZ_KK023042.1 | - | 34817 | 34788 | **RNA**→ rpmE2 (PRK01678)Ribosomal_L31 (pfam01197)→ |
| Sau-1594-1 | NZ_KK015498.1 | - | 36391 | 36362 | **RNA**→ rpmE2 (PRK01678)Ribosomal_L31 (pfam01197)→ |
| Sau-1017-1 | NZ_KK030121.1 | - | 34874 | 34845 | **RNA**→ rpmE2 (PRK01678)Ribosomal_L31 (pfam01197)→ |
| Sau-1808-1 | NZ_KI983480.1 | - | 159047 | 159018 | **RNA**→ rpmE2 (PRK01678)Ribosomal_L31 (pfam01197)→ |
| Sau-2911-1 | NZ_KK028256.1 | - | 56893 | 56864 | **RNA**→ rpmE2 (PRK01678)Ribosomal_L31 (pfam01197)→ |
| Sau-1033-1 | NZ_KK030215.1 | - | 34817 | 34788 | **RNA**→ rpmE2 (PRK01678)Ribosomal_L31 (pfam01197)→ |
| Sau-2910-1 | NZ_KK070146.1 | - | 2279611 | 2279582 | **RNA**→ rpmE2 (PRK01678)Ribosomal_L31 (pfam01197)→ |
| Sau-1364-1 | NZ_KK018792.1 | - | 54650 | 54621 | **RNA**→ rpmE2 (PRK01678)Ribosomal_L31 (pfam01197)→ |
| Sau-3667-1 | NZ_KK039880.1 | - | 34723 | 34694 | **RNA**→ rpmE2 (PRK01678)Ribosomal_L31 (pfam01197)→ |
| Sau-1136-1 | NZ_KI978371.1 | - | 114662 | 114633 | **RNA**→ rpmE2 (PRK01678)Ribosomal_L31 (pfam01197)→ |
| Sau-1140-1 | NZ_KI978955.1 | + | 28862 | 28891 | **RNA**→ rpmE2 (PRK01678)Ribosomal_L31 (pfam01197)→ |
| Sau-2849-1 | NZ_KK040624.1 | - | 34913 | 34884 | **RNA**→ rpmE2 (PRK01678)Ribosomal_L31 (pfam01197)→ |
| Sau-975-1 | NZ_KK027004.1 | - | 35252 | 35223 | **RNA**→ rpmE2 (PRK01678)Ribosomal_L31 (pfam01197)→ |
| Sau-3644-1 | NZ_KK070141.1 | - | 58499 | 58470 | **RNA**→ rpmE2 (PRK01678)Ribosomal_L31 (pfam01197)→ |
| Sau-2850-1 | NZ_KK040575.1 | - | 34872 | 34843 | **RNA**→ rpmE2 (PRK01678)Ribosomal_L31 (pfam01197)→ |
| Sau-1045-1 | NZ_KK028163.1 | - | 56839 | 56810 | **RNA**→ rpmE2 (PRK01678)Ribosomal_L31 (pfam01197)→ |
| Sau-3618-1 | NZ_KK040065.1 | - | 34913 | 34884 | **RNA**→ rpmE2 (PRK01678)Ribosomal_L31 (pfam01197)→ |
| Sau-1157-1 | NZ_KI977819.1 | - | 56711 | 56682 | **RNA**→ rpmE2 (PRK01678)Ribosomal_L31 (pfam01197)→ |
| Sau-3546-1 | NZ_KK070463.1 | - | 233759 | 233730 | **RNA**→ rpmE2 (PRK01678)Ribosomal_L31 (pfam01197)→ |
| Sau-1900-1 | NZ_KI986469.1 | - | 36391 | 36362 | **RNA**→ rpmE2 (PRK01678)Ribosomal_L31 (pfam01197)→ |
| Sau-3566-1 | NZ_KK070522.1 | - | 56894 | 56865 | **RNA**→ rpmE2 (PRK01678)Ribosomal_L31 (pfam01197)→ |
| Sau-1046-1 | NZ_KK028120.1 | - | 56840 | 56811 | **RNA**→ rpmE2 (PRK01678)Ribosomal_L31 (pfam01197)→ |
| Sau-2932-1 | NZ_KK070434.1 | - | 1317508 | 1317479 | **RNA**→ rpmE2 (PRK01678)Ribosomal_L31 (pfam01197)→ |
| Sau-1506-1 | NZ_KK011654.1 | - | 29343 | 29314 | **RNA**→ rpmE2 (PRK01678)Ribosomal_L31 (pfam01197)→ |
| Sau-3589-1 | NZ_KK070796.1 | - | 34817 | 34788 | **RNA**→ rpmE2 (PRK01678)Ribosomal_L31 (pfam01197)→ |
| Sau-1144-1 | NZ_KI979048.1 | + | 28862 | 28891 | **RNA**→ rpmE2 (PRK01678)Ribosomal_L31 (pfam01197)→ |
| Sau-1425-1 | NZ_KK008095.1 | - | 16412 | 16383 | **RNA**→ rpmE2 (PRK01678)Ribosomal_L31 (pfam01197)→ |
| Sau-2089-1 | NZ_KK067715.1 | - | 35432 | 35403 | **RNA**→ rpmE2 (PRK01678)Ribosomal_L31 (pfam01197)→ |
| Sau-1903-1 | NZ_KI986668.1 | - | 61890 | 61861 | **RNA**→ rpmE2 (PRK01678)Ribosomal_L31 (pfam01197)→ |
| Sau-1515-1 | NZ_KK011935.1 | - | 28890 | 28861 | **RNA**→ rpmE2 (PRK01678)Ribosomal_L31 (pfam01197)→ |
| Sau-2070-1 | NZ_KK068525.1 | - | 25129 | 25100 | **RNA**→ rpmE2 (PRK01678)Ribosomal_L31 (pfam01197)→ |
| Sau-2137-1 | NZ_KK066290.1 | - | 100337 | 100308 | **RNA**→ rpmE2 (PRK01678)Ribosomal_L31 (pfam01197)→ |
| Sau-1971-1 | NZ_KI988985.1 | - | 35965 | 35936 | **RNA**→ rpmE2 (PRK01678)Ribosomal_L31 (pfam01197)→ |
| Sau-2133-1 | NZ_KK066410.1 | - | 36391 | 36362 | **RNA**→ rpmE2 (PRK01678)Ribosomal_L31 (pfam01197)→ |
| Sau-2222-1 | NZ_KK063117.1 | - | 113837 | 113808 | **RNA**→ rpmE2 (PRK01678)Ribosomal_L31 (pfam01197)→ |
| Sau-2014-1 | NZ_KI990584.1 | - | 97672 | 97643 | **RNA**→ rpmE2 (PRK01678)Ribosomal_L31 (pfam01197)→ |
| Sau-2015-1 | NZ_KI990591.1 | - | 35811 | 35782 | **RNA**→ rpmE2 (PRK01678)Ribosomal_L31 (pfam01197)→ |
| Sau-2184-1 | NZ_KK064370.1 | - | 238606 | 238577 | **RNA**→ rpmE2 (PRK01678)Ribosomal_L31 (pfam01197)→ |
| Sau-954-1 | NZ_KK028703.1 | - | 56794 | 56765 | **RNA**→ rpmE2 (PRK01678)Ribosomal_L31 (pfam01197)→ |
| Sau-779-1 | NZ_KK070244.1 | - | 1717691 | 1717662 | **RNA**→ rpmE2 (PRK01678)Ribosomal_L31 (pfam01197)→ |
| Sau-953-1 | NZ_KK028665.1 | - | 56794 | 56765 | **RNA**→ rpmE2 (PRK01678)Ribosomal_L31 (pfam01197)→ |
| Sau-2178-1 | NZ_KK064644.1 | - | 39446 | 39417 | **RNA**→ rpmE2 (PRK01678)Ribosomal_L31 (pfam01197)→ |
| Sau-1537-1 | NZ_KK012963.1 | - | 28970 | 28941 | **RNA**→ rpmE2 (PRK01678)Ribosomal_L31 (pfam01197)→ |
| Sau-2116-1 | NZ_KK067026.1 | - | 190797 | 190768 | **RNA**→ rpmE2 (PRK01678)Ribosomal_L31 (pfam01197)→ |
| Sau-2096-1 | NZ_KI972794.1 | - | 132173 | 132144 | **RNA**→ rpmE2 (PRK01678)Ribosomal_L31 (pfam01197)→ |
| Sau-946-1 | NZ_KK070928.1 | - | 56894 | 56865 | **RNA**→ rpmE2 (PRK01678)Ribosomal_L31 (pfam01197)→ |
| Sau-2113-1 | NZ_KI972829.1 | - | 1690075 | 1690046 | **RNA**→ rpmE2 (PRK01678)Ribosomal_L31 (pfam01197)→ |
| Sau-912-1 | NZ_KK070406.1 | - | 60273 | 60244 | **RNA**→ rpmE2 (PRK01678)Ribosomal_L31 (pfam01197)→ |
| Sau-2148-1 | NZ_KK065750.1 | - | 14631 | 14602 | **RNA**→ rpmE2 (PRK01678)Ribosomal_L31 (pfam01197)→ |

| | | | | | |
|---|---|---|---|---|---|
| Sau-964-1 | NZ_KK070258.1 | - | 2212058 | 2212029 | **RNA**→ rpmE2 (PRK01678)Ribosomal_L31 (pfam01197)→ |
| Sau-1276-1 | NZ_KK002718.1 | - | 2467 | 2438 | **RNA**→ rpmE2 (PRK01678)Ribosomal_L31 (pfam01197)→ |
| Sau-2153-1 | NZ_KK065535.1 | - | 35318 | 35289 | **RNA**→ rpmE2 (PRK01678)Ribosomal_L31 (pfam01197)→ |
| Sau-2177-1 | NZ_KK064649.1 | + | 92353 | 92382 | **RNA**→ rpmE2 (PRK01678)Ribosomal_L31 (pfam01197)→ |
| Sau-2046-1 | NZ_KK069412.1 | - | 36391 | 36362 | **RNA**→ rpmE2 (PRK01678)Ribosomal_L31 (pfam01197)→ |
| Sau-2168-1 | NZ_KK064917.1 | - | 35170 | 35141 | **RNA**→ rpmE2 (PRK01678)Ribosomal_L31 (pfam01197)→ |
| Sau-2195-1 | NZ_KK063955.1 | - | 65032 | 65003 | **RNA**→ rpmE2 (PRK01678)Ribosomal_L31 (pfam01197)→ |
| Sau-927-1 | NZ_KK070415.1 | - | 56894 | 56865 | **RNA**→ rpmE2 (PRK01678)Ribosomal_L31 (pfam01197)→ |
| Sau-2155-1 | NZ_KK065442.1 | - | 36391 | 36362 | **RNA**→ rpmE2 (PRK01678)Ribosomal_L31 (pfam01197)→ |
| Sau-2180-1 | NZ_KK064502.1 | - | 54695 | 54666 | **RNA**→ rpmE2 (PRK01678)Ribosomal_L31 (pfam01197)→ |
| Sau-2104-1 | NZ_KI972810.1 | - | 272641 | 272612 | **RNA**→ rpmE2 (PRK01678)Ribosomal_L31 (pfam01197)→ |
| Sau-2173-1 | NZ_KK064758.1 | - | 35655 | 35626 | **RNA**→ rpmE2 (PRK01678)Ribosomal_L31 (pfam01197)→ |
| Sau-2182-1 | NZ_KK064385.1 | - | 36390 | 36361 | **RNA**→ rpmE2 (PRK01678)Ribosomal_L31 (pfam01197)→ |
| Sau-2142-1 | NZ_KK065987.1 | - | 36390 | 36361 | **RNA**→ rpmE2 (PRK01678)Ribosomal_L31 (pfam01197)→ |
| Sau-2165-1 | NZ_KK065079.1 | - | 56863 | 56834 | **RNA**→ rpmE2 (PRK01678)Ribosomal_L31 (pfam01197)→ |
| Sau-2054-1 | NZ_KK069179.1 | - | 111617 | 111588 | **RNA**→ rpmE2 (PRK01678)Ribosomal_L31 (pfam01197)→ |
| Sau-2053-1 | NZ_KK069187.1 | - | 62157 | 62128 | **RNA**→ rpmE2 (PRK01678)Ribosomal_L31 (pfam01197)→ |
| Sau-2068-1 | NZ_KK068648.1 | - | 29575 | 29546 | **RNA**→ rpmE2 (PRK01678)Ribosomal_L31 (pfam01197)→ |
| Sau-2027-1 | NZ_KI972421.1 | - | 58040 | 58011 | **RNA**→ rpmE2 (PRK01678)Ribosomal_L31 (pfam01197)→ |
| Sau-2473-1 | NZ_KK070688.1 | - | 56894 | 56865 | **RNA**→ rpmE2 (PRK01678)Ribosomal_L31 (pfam01197)→ |
| Sau-2102-1 | NZ_KK067436.1 | - | 440 | 411 | **RNA**→ rpmE2 (PRK01678)Ribosomal_L31 (pfam01197)→ |
| Sau-2129-1 | NZ_KK066608.1 | - | 215967 | 215938 | **RNA**→ rpmE2 (PRK01678)Ribosomal_L31 (pfam01197)→ |
| Sau-2131-1 | NZ_KK066564.1 | - | 111834 | 111805 | **RNA**→ rpmE2 (PRK01678)Ribosomal_L31 (pfam01197)→ |
| Sau-2126-1 | NZ_KK066755.1 | - | 54695 | 54666 | **RNA**→ rpmE2 (PRK01678)Ribosomal_L31 (pfam01197)→ |
| Sau-2119-1 | NZ_KK066925.1 | - | 36391 | 36362 | **RNA**→ rpmE2 (PRK01678)Ribosomal_L31 (pfam01197)→ |
| Sau-1009-1 | NZ_KK041351.1 | - | 34912 | 34883 | **RNA**→ rpmE2 (PRK01678)Ribosomal_L31 (pfam01197)→ |
| Sau-2186-1 | NZ_KK064262.1 | - | 36340 | 36311 | **RNA**→ rpmE2 (PRK01678)Ribosomal_L31 (pfam01197)→ |
| Sau-2091-1 | NZ_KK067619.1 | - | 43771 | 43742 | **RNA**→ rpmE2 (PRK01678)Ribosomal_L31 (pfam01197)→ |
| Sau-2074-1 | NZ_KK068353.1 | - | 36391 | 36362 | **RNA**→ rpmE2 (PRK01678)Ribosomal_L31 (pfam01197)→ |
| Sau-2086-1 | NZ_KK067890.1 | - | 36391 | 36362 | **RNA**→ rpmE2 (PRK01678)Ribosomal_L31 (pfam01197)→ |
| Sau-2192-1 | NZ_KK064013.1 | - | 35704 | 35675 | **RNA**→ rpmE2 (PRK01678)Ribosomal_L31 (pfam01197)→ |
| Sau-2063-1 | NZ_KK068829.1 | - | 35670 | 35641 | **RNA**→ rpmE2 (PRK01678)Ribosomal_L31 (pfam01197)→ |
| Sau-713-1 | NZ_KK070332.1 | - | 60061 | 60032 | **RNA**→ rpmE2 (PRK01678)Ribosomal_L31 (pfam01197)→ |
| Sau-2145-1 | NZ_KK065895.1 | - | 62202 | 62173 | **RNA**→ rpmE2 (PRK01678)Ribosomal_L31 (pfam01197)→ |
| Sau-2035-1 | NZ_KK069868.1 | - | 35148 | 35119 | **RNA**→ rpmE2 (PRK01678)Ribosomal_L31 (pfam01197)→ |
| Sau-2150-1 | NZ_KK065628.1 | - | 61380 | 61351 | **RNA**→ rpmE2 (PRK01678)Ribosomal_L31 (pfam01197)→ |
| Sau-943-1 | NZ_KK070851.1 | - | 34873 | 34844 | **RNA**→ rpmE2 (PRK01678)Ribosomal_L31 (pfam01197)→ |
| Sau-2112-1 | NZ_KK067126.1 | - | 36391 | 36362 | **RNA**→ rpmE2 (PRK01678)Ribosomal_L31 (pfam01197)→ |
| Sau-2043-1 | NZ_KK069655.1 | - | 110269 | 110240 | **RNA**→ rpmE2 (PRK01678)Ribosomal_L31 (pfam01197)→ |
| Sau-2107-1 | NZ_KK067213.1 | - | 36391 | 36362 | **RNA**→ rpmE2 (PRK01678)Ribosomal_L31 (pfam01197)→ |
| Sau-1023-1 | NZ_KK041205.1 | - | 34913 | 34884 | **RNA**→ rpmE2 (PRK01678)Ribosomal_L31 (pfam01197)→ |
| Sau-984-1 | NZ_KK041433.1 | - | 56986 | 56957 | **RNA**→ rpmE2 (PRK01678)Ribosomal_L31 (pfam01197)→ |
| Sau-2189-1 | NZ_KK064216.1 | - | 101158 | 101129 | **RNA**→ rpmE2 (PRK01678)Ribosomal_L31 (pfam01197)→ |
| Sau-2162-1 | NZ_KK065224.1 | - | 36491 | 36462 | **RNA**→ rpmE2 (PRK01678)Ribosomal_L31 (pfam01197)→ |
| Sau-2144-1 | NZ_KK065899.1 | - | 54697 | 54668 | **RNA**→ rpmE2 (PRK01678)Ribosomal_L31 (pfam01197)→ |
| Sau-2139-1 | NZ_KK066119.1 | - | 35307 | 35278 | **RNA**→ rpmE2 (PRK01678)Ribosomal_L31 (pfam01197)→ |
| Sau-1063-1 | NZ_KK070162.1 | - | 61782 | 61753 | **RNA**→ rpmE2 (PRK01678)Ribosomal_L31 (pfam01197)→ |
| Sau-903-1 | NZ_KK070387.1 | - | 1675784 | 1675755 | **RNA**→ rpmE2 (PRK01678)Ribosomal_L31 (pfam01197)→ |
| Sau-2157-1 | NZ_KK065402.1 | - | 35959 | 35930 | **RNA**→ rpmE2 (PRK01678)Ribosomal_L31 (pfam01197)→ |
| Sau-2082-1 | NZ_KK068024.1 | - | 62005 | 61976 | **RNA**→ rpmE2 (PRK01678)Ribosomal_L31 (pfam01197)→ |
| Sau-2151-1 | NZ_KK065582.1 | - | 35786 | 35757 | **RNA**→ rpmE2 (PRK01678)Ribosomal_L31 (pfam01197)→ |
| Sau-700-1 | NZ_KK070274.1 | - | 1613796 | 1613767 | **RNA**→ rpmE2 (PRK01678)Ribosomal_L31 (pfam01197)→ |

| | | | | | | |
|---|---|---|---|---|---|---|
| Sau-2045-1 | NZ_KK069459.1 | - | 35844 | 35815 | **RNA**→ | rpmE2 (PRK01678)Ribosomal_L31 (pfam01197)→ |
| Sau-2169-1 | NZ_KK064864.1 | - | 36391 | 36362 | **RNA**→ | rpmE2 (PRK01678)Ribosomal_L31 (pfam01197)→ |
| Sau-2140-1 | NZ_KK066058.1 | - | 10628 | 10599 | **RNA**→ | rpmE2 (PRK01678)Ribosomal_L31 (pfam01197)→ |
| Sau-680-1 | NZ_KK070046.1 | - | 57065 | 57036 | **RNA**→ | rpmE2 (PRK01678)Ribosomal_L31 (pfam01197)→ |
| Sau-2100-1 | NZ_KK067537.1 | - | 11745 | 11716 | **RNA**→ | rpmE2 (PRK01678)Ribosomal_L31 (pfam01197)→ |
| Sau-736-1 | NZ_KK070488.1 | - | 34988 | 34959 | **RNA**→ | rpmE2 (PRK01678)Ribosomal_L31 (pfam01197)→ |
| Sau-2172-1 | NZ_KK070074.1 | - | 62003 | 61974 | **RNA**→ | rpmE2 (PRK01678)Ribosomal_L31 (pfam01197)→ |
| Sau-702-1 | NZ_KK070315.1 | - | 60739 | 60710 | **RNA**→ | rpmE2 (PRK01678)Ribosomal_L31 (pfam01197)→ |
| Sau-2036-1 | NZ_KK069801.1 | - | 36390 | 36361 | **RNA**→ | rpmE2 (PRK01678)Ribosomal_L31 (pfam01197)→ |
| Sau-738-1 | NZ_KK070550.1 | - | 56893 | 56864 | **RNA**→ | rpmE2 (PRK01678)Ribosomal_L31 (pfam01197)→ |
| Sau-2159-1 | NZ_KK065320.1 | - | 36391 | 36362 | **RNA**→ | rpmE2 (PRK01678)Ribosomal_L31 (pfam01197)→ |
| Sau-2061-1 | NZ_KK068950.1 | - | 36391 | 36362 | **RNA**→ | rpmE2 (PRK01678)Ribosomal_L31 (pfam01197)→ |
| Sau-768-1 | NZ_KK070248.1 | - | 1659107 | 1659078 | **RNA**→ | rpmE2 (PRK01678)Ribosomal_L31 (pfam01197)→ |
| Sau-730-1 | NZ_KK039465.1 | - | 34872 | 34843 | **RNA**→ | rpmE2 (PRK01678)Ribosomal_L31 (pfam01197)→ |
| Sau-758-1 | NZ_KK070953.1 | - | 56991 | 56962 | **RNA**→ | rpmE2 (PRK01678)Ribosomal_L31 (pfam01197)→ |
| Sau-721-1 | NZ_KK070384.1 | - | 61506 | 61477 | **RNA**→ | rpmE2 (PRK01678)Ribosomal_L31 (pfam01197)→ |
| Sau-718-1 | NZ_KK039541.1 | - | 36294 | 36265 | **RNA**→ | rpmE2 (PRK01678)Ribosomal_L31 (pfam01197)→ |
| Sau-663-1 | NZ_KK070269.1 | - | 1624827 | 1624798 | **RNA**→ | rpmE2 (PRK01678)Ribosomal_L31 (pfam01197)→ |
| Sau-2482-1 | NZ_KK071089.1 | - | 34873 | 34844 | **RNA**→ | rpmE2 (PRK01678)Ribosomal_L31 (pfam01197)→ |
| Sau-2039-1 | NZ_KK069750.1 | - | 31193 | 31164 | **RNA**→ | rpmE2 (PRK01678)Ribosomal_L31 (pfam01197)→ |
| Sau-2191-1 | NZ_KK064074.1 | - | 54917 | 54888 | **RNA**→ | rpmE2 (PRK01678)Ribosomal_L31 (pfam01197)→ |
| Sau-2190-1 | NZ_KK064116.1 | - | 59845 | 59816 | **RNA**→ | rpmE2 (PRK01678)Ribosomal_L31 (pfam01197)→ |
| Sau-2472-1 | NZ_KK070711.1 | - | 56892 | 56863 | **RNA**→ | rpmE2 (PRK01678)Ribosomal_L31 (pfam01197)→ |
| Sau-2979-1 | NZ_KK071027.1 | - | 56993 | 56964 | **RNA**→ | rpmE2 (PRK01678)Ribosomal_L31 (pfam01197)→ |
| Sau-748-1 | NZ_KK070598.1 | - | 56994 | 56965 | **RNA**→ | rpmE2 (PRK01678)Ribosomal_L31 (pfam01197)→ |
| Sau-737-1 | NZ_KK070534.1 | - | 56790 | 56761 | **RNA**→ | rpmE2 (PRK01678)Ribosomal_L31 (pfam01197)→ |
| Sau-2078-1 | NZ_KK068249.1 | - | 62156 | 62127 | **RNA**→ | rpmE2 (PRK01678)Ribosomal_L31 (pfam01197)→ |
| Sau-830-1 | NZ_KK072816.1 | - | 56894 | 56865 | **RNA**→ | rpmE2 (PRK01678)Ribosomal_L31 (pfam01197)→ |
| Sau-2123-1 | NZ_KK066844.1 | - | 52141 | 52112 | **RNA**→ | rpmE2 (PRK01678)Ribosomal_L31 (pfam01197)→ |
| Sau-2486-1 | NZ_KK071243.1 | - | 34872 | 34843 | **RNA**→ | rpmE2 (PRK01678)Ribosomal_L31 (pfam01197)→ |
| Sau-3003-1 | NZ_KK071918.1 | - | 55194 | 55165 | **RNA**→ | rpmE2 (PRK01678)Ribosomal_L31 (pfam01197)→ |
| Sau-2501-1 | NZ_KK070232.1 | - | 1675361 | 1675332 | **RNA**→ | rpmE2 (PRK01678)Ribosomal_L31 (pfam01197)→ |
| Sau-956-1 | NZ_KK071309.1 | - | 34873 | 34844 | **RNA**→ | rpmE2 (PRK01678)Ribosomal_L31 (pfam01197)→ |
| Sau-3609-1 | NZ_KK071940.1 | - | 56893 | 56864 | **RNA**→ | rpmE2 (PRK01678)Ribosomal_L31 (pfam01197)→ |
| Sau-798-1 | NZ_KK072206.1 | - | 34913 | 34884 | **RNA**→ | rpmE2 (PRK01678)Ribosomal_L31 (pfam01197)→ |
| Sau-2838-1 | NZ_KK073322.1 | - | 34973 | 34944 | **RNA**→ | rpmE2 (PRK01678)Ribosomal_L31 (pfam01197)→ |
| Sau-2478-1 | NZ_KK070966.1 | - | 56894 | 56865 | **RNA**→ | rpmE2 (PRK01678)Ribosomal_L31 (pfam01197)→ |
| Sau-971-1 | NZ_KK071827.1 | - | 56994 | 56965 | **RNA**→ | rpmE2 (PRK01678)Ribosomal_L31 (pfam01197)→ |
| Sau-2490-1 | NZ_KK071516.1 | - | 55217 | 55188 | **RNA**→ | rpmE2 (PRK01678)Ribosomal_L31 (pfam01197)→ |
| Sau-2856-1 | NZ_KK073413.1 | - | 54968 | 54939 | **RNA**→ | rpmE2 (PRK01678)Ribosomal_L31 (pfam01197)→ |
| Sau-807-1 | NZ_KK072576.1 | - | 34872 | 34843 | **RNA**→ | rpmE2 (PRK01678)Ribosomal_L31 (pfam01197)→ |
| Sau-3505-1 | NZ_KK070053.1 | - | 1681745 | 1681716 | **RNA**→ | rpmE2 (PRK01678)Ribosomal_L31 (pfam01197)→ |
| Sau-811-1 | NZ_KK072713.1 | - | 56894 | 56865 | **RNA**→ | rpmE2 (PRK01678)Ribosomal_L31 (pfam01197)→ |
| Sau-992-1 | NZ_KK072419.1 | - | 54895 | 54866 | **RNA**→ | rpmE2 (PRK01678)Ribosomal_L31 (pfam01197)→ |
| Sau-1011-1 | NZ_KK073025.1 | - | 58371 | 58342 | **RNA**→ | rpmE2 (PRK01678)Ribosomal_L31 (pfam01197)→ |
| Sau-3617-1 | NZ_KK073538.1 | - | 19183 | 19154 | **RNA**→ | rpmE2 (PRK01678)Ribosomal_L31 (pfam01197)→ |
| Sau-1056-1 | NZ_KK073074.1 | - | 56894 | 56865 | **RNA**→ | rpmE2 (PRK01678)Ribosomal_L31 (pfam01197)→ |
| Sau-2524-1 | NZ_KK073170.1 | - | 56893 | 56864 | **RNA**→ | rpmE2 (PRK01678)Ribosomal_L31 (pfam01197)→ |
| Sau-3605-1 | NZ_KK071587.1 | - | 56790 | 56761 | **RNA**→ | rpmE2 (PRK01678)Ribosomal_L31 (pfam01197)→ |
| Sau-999-1 | NZ_KK072994.1 | - | 55000 | 54971 | **RNA**→ | rpmE2 (PRK01678)Ribosomal_L31 (pfam01197)→ |
| Sau-2528-1 | NZ_KK073191.1 | - | 55116 | 55087 | **RNA**→ | rpmE2 (PRK01678)Ribosomal_L31 (pfam01197)→ |
| Sau-2134-1 | NZ_KK066353.1 | - | 54695 | 54666 | **RNA**→ | rpmE2 (PRK01678)Ribosomal_L31 (pfam01197)→ |

| | | | | | |
|---|---|---|---|---|---|
| Sau-989-1 | NZ_KK072905.1 | - | 56645 | 56616 | **RNA**→ rpmE2 (PRK01678)Ribosomal_L31 (pfam01197)→ |
| Sau-3646-1 | NZ_KK073630.1 | - | 34873 | 34844 | **RNA**→ rpmE2 (PRK01678)Ribosomal_L31 (pfam01197)→ |
| Sau-982-1 | NZ_KK072140.1 | - | 34872 | 34843 | **RNA**→ rpmE2 (PRK01678)Ribosomal_L31 (pfam01197)→ |
| Sau-3616-1 | NZ_KK072352.1 | - | 54968 | 54939 | **RNA**→ rpmE2 (PRK01678)Ribosomal_L31 (pfam01197)→ |
| Sau-2995-1 | NZ_KK071742.1 | - | 29186 | 29157 | **RNA**→ rpmE2 (PRK01678)Ribosomal_L31 (pfam01197)→ |
| Sau-750-1 | NZ_KK070624.1 | - | 34913 | 34884 | **RNA**→ rpmE2 (PRK01678)Ribosomal_L31 (pfam01197)→ |
| Sau-3608-1 | NZ_KK072054.1 | - | 34873 | 34844 | **RNA**→ rpmE2 (PRK01678)Ribosomal_L31 (pfam01197)→ |
| Sau-2512-1 | NZ_KK072288.1 | - | 56894 | 56865 | **RNA**→ rpmE2 (PRK01678)Ribosomal_L31 (pfam01197)→ |
| Sau-3628-1 | NZ_KK073585.1 | - | 56894 | 56865 | **RNA**→ rpmE2 (PRK01678)Ribosomal_L31 (pfam01197)→ |
| Sau-2530-1 | NZ_KK073263.1 | - | 55117 | 55088 | **RNA**→ rpmE2 (PRK01678)Ribosomal_L31 (pfam01197)→ |
| Sau-3614-1 | NZ_KK073511.1 | - | 34913 | 34884 | **RNA**→ rpmE2 (PRK01678)Ribosomal_L31 (pfam01197)→ |
| Sau-822-1 | NZ_KK072745.1 | - | 34913 | 34884 | **RNA**→ rpmE2 (PRK01678)Ribosomal_L31 (pfam01197)→ |
| Sau-2888-1 | NZ_KK073475.1 | - | 2229157 | 2229128 | **RNA**→ rpmE2 (PRK01678)Ribosomal_L31 (pfam01197)→ |
| Sau-3632-1 | NZ_KK073602.1 | - | 56994 | 56965 | **RNA**→ rpmE2 (PRK01678)Ribosomal_L31 (pfam01197)→ |
| Sau-2874-1 | NZ_KK073433.1 | - | 35013 | 34984 | **RNA**→ rpmE2 (PRK01678)Ribosomal_L31 (pfam01197)→ |
| Sau-2522-1 | NZ_KK073121.1 | - | 55150 | 55121 | **RNA**→ rpmE2 (PRK01678)Ribosomal_L31 (pfam01197)→ |
| Sau-3657-1 | NZ_KK073725.1 | - | 35266 | 35237 | **RNA**→ rpmE2 (PRK01678)Ribosomal_L31 (pfam01197)→ |
| Sau-766-1 | NZ_KK071427.1 | - | 56894 | 56865 | **RNA**→ rpmE2 (PRK01678)Ribosomal_L31 (pfam01197)→ |
| Sau-832-1 | NZ_KK072828.1 | - | 34913 | 34884 | **RNA**→ rpmE2 (PRK01678)Ribosomal_L31 (pfam01197)→ |
| Sau-1057-1 | NZ_KK073082.1 | - | 56894 | 56865 | **RNA**→ rpmE2 (PRK01678)Ribosomal_L31 (pfam01197)→ |
| Sau-2865-1 | NZ_KK073426.1 | - | 56994 | 56965 | **RNA**→ rpmE2 (PRK01678)Ribosomal_L31 (pfam01197)→ |
| Sau-3486-1 | NZ_KK073499.1 | - | 55180 | 55151 | **RNA**→ rpmE2 (PRK01678)Ribosomal_L31 (pfam01197)→ |
| Sau-2551-1 | NZ_KK073287.1 | - | 25259 | 25230 | **RNA**→ rpmE2 (PRK01678)Ribosomal_L31 (pfam01197)→ |
| Sau-773-1 | NZ_KK071626.1 | - | 34913 | 34884 | **RNA**→ rpmE2 (PRK01678)Ribosomal_L31 (pfam01197)→ |
| Sau-3647-1 | NZ_KK073682.1 | - | 34873 | 34844 | **RNA**→ rpmE2 (PRK01678)Ribosomal_L31 (pfam01197)→ |
| Sau-2452-1 | NZ_KK075555.1 | - | 34913 | 34884 | **RNA**→ rpmE2 (PRK01678)Ribosomal_L31 (pfam01197)→ |
| Sau-622-1 | NZ_KK072501.1 | - | 34972 | 34943 | **RNA**→ rpmE2 (PRK01678)Ribosomal_L31 (pfam01197)→ |
| Sau-3557-1 | NZ_KK074598.1 | - | 54895 | 54866 | **RNA**→ rpmE2 (PRK01678)Ribosomal_L31 (pfam01197)→ |
| Sau-2875-1 | NZ_KK075183.1 | - | 56794 | 56765 | **RNA**→ rpmE2 (PRK01678)Ribosomal_L31 (pfam01197)→ |
| Sau-1078-1 | NZ_KK075649.1 | - | 56545 | 56516 | **RNA**→ rpmE2 (PRK01678)Ribosomal_L31 (pfam01197)→ |
| Sau-3654-1 | NZ_KK074429.1 | - | 55082 | 55053 | **RNA**→ rpmE2 (PRK01678)Ribosomal_L31 (pfam01197)→ |
| Sau-3653-1 | NZ_KK074489.1 | - | 54955 | 54926 | **RNA**→ rpmE2 (PRK01678)Ribosomal_L31 (pfam01197)→ |
| Sau-2576-1 | NZ_KK075313.1 | - | 34817 | 34788 | **RNA**→ rpmE2 (PRK01678)Ribosomal_L31 (pfam01197)→ |
| Sau-1050-1 | NZ_KK075688.1 | - | 34817 | 34788 | **RNA**→ rpmE2 (PRK01678)Ribosomal_L31 (pfam01197)→ |
| Sau-2589-1 | NZ_KK075278.1 | - | 56893 | 56864 | **RNA**→ rpmE2 (PRK01678)Ribosomal_L31 (pfam01197)→ |
| Sau-2571-1 | NZ_KK075413.1 | - | 56893 | 56864 | **RNA**→ rpmE2 (PRK01678)Ribosomal_L31 (pfam01197)→ |
| Sau-2552-1 | NZ_KK075464.1 | - | 56893 | 56864 | **RNA**→ rpmE2 (PRK01678)Ribosomal_L31 (pfam01197)→ |
| Sau-3490-1 | NZ_KK074798.1 | - | 34817 | 34788 | **RNA**→ rpmE2 (PRK01678)Ribosomal_L31 (pfam01197)→ |
| Sau-3488-1 | NZ_KK074916.1 | - | 34873 | 34844 | **RNA**→ rpmE2 (PRK01678)Ribosomal_L31 (pfam01197)→ |
| Sau-3521-1 | NZ_KK074673.1 | - | 55184 | 55155 | **RNA**→ rpmE2 (PRK01678)Ribosomal_L31 (pfam01197)→ |
| Sau-1029-1 | NZ_KK075843.1 | - | 56993 | 56964 | **RNA**→ rpmE2 (PRK01678)Ribosomal_L31 (pfam01197)→ |
| Sau-1031-1 | NZ_KK075811.1 | - | 56892 | 56863 | **RNA**→ rpmE2 (PRK01678)Ribosomal_L31 (pfam01197)→ |
| Sau-1030-1 | NZ_KK075827.1 | - | 55165 | 55136 | **RNA**→ rpmE2 (PRK01678)Ribosomal_L31 (pfam01197)→ |
| Sau-3481-1 | NZ_KK075042.1 | - | 56893 | 56864 | **RNA**→ rpmE2 (PRK01678)Ribosomal_L31 (pfam01197)→ |
| Sau-3487-1 | NZ_KK074958.1 | - | 34873 | 34844 | **RNA**→ rpmE2 (PRK01678)Ribosomal_L31 (pfam01197)→ |
| Sau-2928-1 | NZ_KK075123.1 | - | 34913 | 34884 | **RNA**→ rpmE2 (PRK01678)Ribosomal_L31 (pfam01197)→ |
| Sau-2590-1 | NZ_KK075264.1 | - | 56893 | 56864 | **RNA**→ rpmE2 (PRK01678)Ribosomal_L31 (pfam01197)→ |
| Sau-937-1 | NZ_KK075909.1 | - | 56794 | 56765 | **RNA**→ rpmE2 (PRK01678)Ribosomal_L31 (pfam01197)→ |
| Sau-939-1 | NZ_KK075889.1 | - | 56794 | 56765 | **RNA**→ rpmE2 (PRK01678)Ribosomal_L31 (pfam01197)→ |
| Sau-668-1 | NZ_KK076136.1 | - | 56894 | 56865 | **RNA**→ rpmE2 (PRK01678)Ribosomal_L31 (pfam01197)→ |
| Sau-670-1 | NZ_KK076076.1 | - | 34873 | 34844 | **RNA**→ rpmE2 (PRK01678)Ribosomal_L31 (pfam01197)→ |
| Sau-634-1 | NZ_KK076386.1 | - | 34817 | 34788 | **RNA**→ rpmE2 (PRK01678)Ribosomal_L31 (pfam01197)→ |

| | | | | | | | |
|---|---|---|---|---|---|---|---|
| Sau-642-1 | NZ_KK076331.1 | - | 34873 | 34844 | **RNA**→ | rpmE2 (PRK01678)Ribosomal_L31 (pfam01197)→ | |
| Sau-921-1 | NZ_KK075946.1 | - | 56894 | 56865 | **RNA**→ | rpmE2 (PRK01678)Ribosomal_L31 (pfam01197)→ | |
| Sau-559-1 | NZ_KK074307.1 | - | 35225 | 35196 | **RNA**→ | rpmE2 (PRK01678)Ribosomal_L31 (pfam01197)→ | |
| Sau-3500-1 | NZ_KK074734.1 | - | 34862 | 34833 | **RNA**→ | rpmE2 (PRK01678)Ribosomal_L31 (pfam01197)→ | |
| Sau-2561-1 | NZ_KK075418.1 | - | 55143 | 55114 | **RNA**→ | rpmE2 (PRK01678)Ribosomal_L31 (pfam01197)→ | |
| Sau-2929-1 | NZ_KK075070.1 | - | 34913 | 34884 | **RNA**→ | rpmE2 (PRK01678)Ribosomal_L31 (pfam01197)→ | |
| Sau-2494-1 | NZ_KK075544.1 | - | 55129 | 55100 | **RNA**→ | rpmE2 (PRK01678)Ribosomal_L31 (pfam01197)→ | |
| Sau-653-1 | NZ_KK076162.1 | - | 55117 | 55088 | **RNA**→ | rpmE2 (PRK01678)Ribosomal_L31 (pfam01197)→ | |
| Sau-3516-1 | NZ_KK074684.1 | - | 18844 | 18815 | **RNA**→ | rpmE2 (PRK01678)Ribosomal_L31 (pfam01197)→ | |
| Sau-3948-1 | NZ_KB822121.1 | - | 61693 | 61664 | **RNA**→ | rpmE2 (PRK01678)Ribosomal_L31 (pfam01197)→ | |
| Sau-3846-1 | NZ_KB821353.1 | - | 423199 | 423170 | **RNA**→ | rpmE2 (PRK01678)Ribosomal_L31 (pfam01197)→ | |
| Sau-3978-1 | NZ_KB822284.1 | - | 61557 | 61528 | **RNA**→ | rpmE2 (PRK01678)Ribosomal_L31 (pfam01197)→ | |
| Sau-46-1 | NC_017338.1 | - | 2183672 | 2183643 | **RNA**→ | rpmE2 (PRK01678)Ribosomal_L31 (pfam01197)→ | |
| Sau-42-1 | NZ_GG730345.1 | - | 54966 | 54937 | **RNA**→ | rpmE2 (PRK01678)Ribosomal_L31 (pfam01197)→ | rpmE2 (PRK01678)Ribosomal_L31 (pfam01197 |
| Sau-753-1 | NZ_KK075956.1 | - | 56988 | 56959 | **RNA**→ | rpmE2 (PRK01678)Ribosomal_L31 (pfam01197)→ | |
| Sau-53-1 | NZ_CM000952.1 | + | 1580336 | 1580365 | **RNA**→ | rpmE2 (PRK01678)Ribosomal_L31 (pfam01197)→ | |
| Sau-36-1 | NZ_GG730180.1 | - | 55039 | 55010 | **RNA**→ | rpmE2 (PRK01678)Ribosomal_L31 (pfam01197)→ | rpmE2 (PRK01678)Ribosomal_L31 (pfam01197 |
| Sau-65-1 | NC_017347.1 | - | 2231645 | 2231616 | **RNA**→ | rpmE2 (PRK01678)Ribosomal_L31 (pfam01197)→ | |
| Sau-39-1 | NC_017337.1 | - | 2207066 | 2207037 | **RNA**→ | rpmE2 (PRK01678)Ribosomal_L31 (pfam01197)→ | |
| Sau-22-1 | NC_017351.1 | - | 2218864 | 2218835 | **RNA**→ | rpmE2 (PRK01678)Ribosomal_L31 (pfam01197)→ | |
| Sau-72-1 | NC_016912.1 | - | 2054189 | 2054160 | **RNA**→ | rpmE2 (PRK01678)Ribosomal_L31 (pfam01197)→ | |
| Sau-3722-5 | NZ_CP012120.1 | + | 1585676 | 1585705 | **RNA**→ | rpmE2 (PRK01678)Ribosomal_L31 (pfam01197)→ | |
| Sau-27-1 | NC_017673.1 | - | 2081692 | 2081663 | **RNA**→ | rpmE2 (PRK01678)Ribosomal_L31 (pfam01197)→ | |
| Sau-2-1 | NC_018608.1 | - | 2149095 | 2149066 | **RNA**→ | rpmE2 (PRK01678)Ribosomal_L31 (pfam01197)→ | |
| Sau-3844-1 | NZ_KB821340.1 | - | 240089 | 240060 | **RNA**→ | rpmE2 (PRK01678)Ribosomal_L31 (pfam01197)→ | |
| Sau-3823-1 | NZ_KB821215.1 | - | 1776670 | 1776641 | **RNA**→ | rpmE2 (PRK01678)Ribosomal_L31 (pfam01197)→ | |
| Sau-3851-1 | NZ_KB821359.1 | - | 1730139 | 1730110 | **RNA**→ | rpmE2 (PRK01678)Ribosomal_L31 (pfam01197)→ | |
| Sau-3920-1 | NZ_KB821944.1 | - | 59999 | 59970 | **RNA**→ | rpmE2 (PRK01678)Ribosomal_L31 (pfam01197)→ | |
| Sau-3787-1 | NZ_KB820976.1 | - | 2184946 | 2184917 | **RNA**→ | rpmE2 (PRK01678)Ribosomal_L31 (pfam01197)→ | |
| Sau-3873-1 | NZ_KB821509.1 | - | 242380 | 242351 | **RNA**→ | rpmE2 (PRK01678)Ribosomal_L31 (pfam01197)→ | |
| Sau-3908-1 | NZ_KB821832.1 | - | 56781 | 56752 | **RNA**→ | rpmE2 (PRK01678)Ribosomal_L31 (pfam01197)→ | |
| Sau-3829-1 | NZ_KB821242.1 | - | 56893 | 56864 | **RNA**→ | rpmE2 (PRK01678)Ribosomal_L31 (pfam01197)→ | |
| Sau-3890-1 | NZ_KB821688.1 | - | 1733325 | 1733296 | **RNA**→ | rpmE2 (PRK01678)Ribosomal_L31 (pfam01197)→ | |
| Sau-3899-1 | NZ_KB821782.1 | - | 56963 | 56934 | **RNA**→ | rpmE2 (PRK01678)Ribosomal_L31 (pfam01197)→ | |
| Sau-3907-1 | NC_021059.1 | - | 2228162 | 2228133 | **RNA**→ | rpmE2 (PRK01678)Ribosomal_L31 (pfam01197)→ | |
| Sau-4103-1 | NC_020566.1 | - | 2120659 | 2120630 | **RNA**→ | rpmE2 (PRK01678)Ribosomal_L31 (pfam01197)→ | rpmE2 (PRK01678)Ribosomal_L31 (pfam01197 |
| Sau-4103-2 | NC_020533.1 | - | 2121038 | 2121009 | **RNA**→ | rpmE2 (PRK01678)Ribosomal_L31 (pfam01197)→ | rpmE2 (PRK01678)Ribosomal_L31 (pfam01197 |
| Sau-4103-3 | NC_020568.1 | - | 2120524 | 2120495 | **RNA**→ | rpmE2 (PRK01678)Ribosomal_L31 (pfam01197)→ | rpmE2 (PRK01678)Ribosomal_L31 (pfam01197 |
| Sau-38-1 | NC_017343.1 | - | 2100233 | 2100204 | **RNA**→ | rpmE2 (PRK01678)Ribosomal_L31 (pfam01197)→ | |
| Sau-3764-1 | NC_021670.1 | - | 2323309 | 2323280 | **RNA**→ | rpmE2 (PRK01678)Ribosomal_L31 (pfam01197)→ | |
| Sau-3765-1 | NC_021554.1 | - | 2228321 | 2228292 | **RNA**→ | rpmE2 (PRK01678)Ribosomal_L31 (pfam01197)→ | |
| Sau-4037-1 | NC_022113.1 | - | 2130139 | 2130110 | **RNA**→ | rpmE2 (PRK01678)Ribosomal_L31 (pfam01197)→ | |
| Sau-4073-1 | NC_022226.1 | - | 2123384 | 2123355 | **RNA**→ | rpmE2 (PRK01678)Ribosomal_L31 (pfam01197)→ | |
| Sau-4101-1 | NC_022443.1 | - | 2079264 | 2079235 | **RNA**→ | rpmE2 (PRK01678)Ribosomal_L31 (pfam01197)→ | |
| Sau-4102-1 | NC_022442.1 | - | 2140502 | 2140473 | **RNA**→ | rpmE2 (PRK01678)Ribosomal_L31 (pfam01197)→ | |
| Sau-4099-1 | NZ_KI542650.1 | - | 54850 | 54821 | **RNA**→ | rpmE2 (PRK01678)Ribosomal_L31 (pfam01197)→ | |
| Sau-3722-1 | NZ_CP009681.1 | - | 2385668 | 2385639 | **RNA**→ | rpmE2 (PRK01678)Ribosomal_L31 (pfam01197)→ | |
| Sau-2348-1 | NZ_KI995558.1 | - | 94306 | 94277 | **RNA**→ | rpmE2 (PRK01678)Ribosomal_L31 (pfam01197)→ | |
| Sau-221-1 | NZ_KN716005.1 | - | 83047 | 83018 | **RNA**→ | rpmE2 (PRK01678)Ribosomal_L31 (pfam01197)→ | |
| Sau-69-1 | NC_007793.1 | - | 2235973 | 2235944 | **RNA**→ | rpmE2 (PRK01678)Ribosomal_L31 (pfam01197)→ | |
| Sau-70-1 | NC_010079.1 | - | 2236367 | 2236338 | **RNA**→ | rpmE2 (PRK01678)Ribosomal_L31 (pfam01197)→ | |
| Sau-68-1 | NC_017331.1 | - | 2388545 | 2388516 | **RNA**→ | rpmE2 (PRK01678)Ribosomal_L31 (pfam01197)→ | |

| | | | | | | |
|---|---|---|---|---|---|---|
| Sau-71-1 | NZ_GG697988.1 | + | 54821 | 54850 | **RNA**→ | rpmE2 (PRK01678)Ribosomal_L31 (pfam01197)→ |
| Sau-4120-1 | NC_022604.1 | - | 2332120 | 2332091 | **RNA**→ | rpmE2 (PRK01678)Ribosomal_L31 (pfam01197)→ |
| Sau-719-1 | NZ_KK070335.1 | - | 2250080 | 2250051 | **RNA**→ | rpmE2 (PRK01678)Ribosomal_L31 (pfam01197)→ |
| Sau-3682-1 | NZ_KI978453.1 | - | 54812 | 54783 | **RNA**→ | rpmE2 (PRK01678)Ribosomal_L31 (pfam01197)→ |
| Sau-674-1 | NZ_KK039802.1 | - | 54900 | 54871 | **RNA**→ | rpmE2 (PRK01678)Ribosomal_L31 (pfam01197)→ |
| Sau-3055-1 | NZ_KI979115.1 | - | 56894 | 56865 | **RNA**→ | rpmE2 (PRK01678)Ribosomal_L31 (pfam01197)→ |
| Sau-2758-1 | NZ_KI973663.1 | - | 56625 | 56596 | **RNA**→ | rpmE2 (PRK01678)Ribosomal_L31 (pfam01197)→ |
| Sau-845-1 | NZ_KI974683.1 | - | 56626 | 56597 | **RNA**→ | rpmE2 (PRK01678)Ribosomal_L31 (pfam01197)→ |
| Sau-1147-1 | NZ_KI975430.1 | - | 56660 | 56631 | **RNA**→ | rpmE2 (PRK01678)Ribosomal_L31 (pfam01197)→ |
| Sau-3024-1 | NZ_KI976149.1 | - | 60297 | 60268 | **RNA**→ | rpmE2 (PRK01678)Ribosomal_L31 (pfam01197)→ |
| Sau-2755-1 | NZ_KI973439.1 | - | 60262 | 60233 | **RNA**→ | rpmE2 (PRK01678)Ribosomal_L31 (pfam01197)→ |
| Sau-1439-1 | NZ_KK008820.1 | - | 54837 | 54808 | **RNA**→ | rpmE2 (PRK01678)Ribosomal_L31 (pfam01197)→ |
| Sau-1139-1 | NZ_KI979287.1 | - | 56523 | 56494 | **RNA**→ | rpmE2 (PRK01678)Ribosomal_L31 (pfam01197)→ |
| Sau-3722-2 | NZ_CP009554.1 | - | 2260619 | 2260590 | **RNA**→ | rpmE2 (PRK01678)Ribosomal_L31 (pfam01197)→ |
| Sau-1667-1 | NZ_KI966968.1 | - | 54889 | 54860 | **RNA**→ | rpmE2 (PRK01678)Ribosomal_L31 (pfam01197)→ |
| Sau-1688-1 | NZ_KI967791.1 | - | 55129 | 55100 | **RNA**→ | rpmE2 (PRK01678)Ribosomal_L31 (pfam01197)→ |
| Sau-2243-1 | NZ_KI972937.1 | - | 1063329 | 1063300 | **RNA**→ | rpmE2 (PRK01678)Ribosomal_L31 (pfam01197)→ |
| Sau-1264-1 | NZ_KI971916.1 | - | 237016 | 236987 | **RNA**→ | rpmE2 (PRK01678)Ribosomal_L31 (pfam01197)→ |
| Sau-1450-1 | NZ_KI971554.1 | - | 1106971 | 1106942 | **RNA**→ | rpmE2 (PRK01678)Ribosomal_L31 (pfam01197)→ |
| Sau-1685-1 | NZ_KI967653.1 | - | 55327 | 55298 | **RNA**→ | rpmE2 (PRK01678)Ribosomal_L31 (pfam01197)→ |
| Sau-1177-1 | NZ_KI971760.1 | - | 57191 | 57162 | **RNA**→ | rpmE2 (PRK01678)Ribosomal_L31 (pfam01197)→ |
| Sau-2193-1 | NZ_KI972880.1 | - | 56529 | 56500 | **RNA**→ | rpmE2 (PRK01678)Ribosomal_L31 (pfam01197)→ |
| Sau-1660-1 | NZ_KI966712.1 | - | 55956 | 55927 | **RNA**→ | rpmE2 (PRK01678)Ribosomal_L31 (pfam01197)→ |
| Sau-1566-1 | NZ_KI971107.1 | - | 1157441 | 1157412 | **RNA**→ | rpmE2 (PRK01678)Ribosomal_L31 (pfam01197)→ |
| Sau-2905-1 | NZ_KK038542.1 | - | 56893 | 56864 | **RNA**→ | rpmE2 (PRK01678)Ribosomal_L31 (pfam01197)→ |
| Sau-1576-1 | NZ_KI971132.1 | - | 56529 | 56500 | **RNA**→ | rpmE2 (PRK01678)Ribosomal_L31 (pfam01197)→ |
| Sau-1223-1 | NZ_KI971849.1 | - | 60286 | 60257 | **RNA**→ | rpmE2 (PRK01678)Ribosomal_L31 (pfam01197)→ |
| Sau-1678-1 | NZ_KI967412.1 | - | 56005 | 55976 | **RNA**→ | rpmE2 (PRK01678)Ribosomal_L31 (pfam01197)→ |
| Sau-862-1 | NZ_KI978748.1 | - | 792665 | 792636 | **RNA**→ | rpmE2 (PRK01678)Ribosomal_L31 (pfam01197)→ |
| Sau-1246-1 | NZ_KI971881.1 | - | 1655839 | 1655810 | **RNA**→ | rpmE2 (PRK01678)Ribosomal_L31 (pfam01197)→ |
| Sau-2163-1 | NZ_KK065158.1 | - | 55513 | 55484 | **RNA**→ | rpmE2 (PRK01678)Ribosomal_L31 (pfam01197)→ |
| Sau-1153-1 | NZ_KI974632.1 | - | 56894 | 56865 | **RNA**→ | rpmE2 (PRK01678)Ribosomal_L31 (pfam01197)→ |
| Sau-1736-1 | NZ_KI970390.1 | - | 60042 | 60013 | **RNA**→ | rpmE2 (PRK01678)Ribosomal_L31 (pfam01197)→ |
| Sau-1737-1 | NZ_KI970462.1 | - | 55043 | 55014 | **RNA**→ | rpmE2 (PRK01678)Ribosomal_L31 (pfam01197)→ |
| Sau-3629-1 | NZ_KK073589.1 | - | 56894 | 56865 | **RNA**→ | rpmE2 (PRK01678)Ribosomal_L31 (pfam01197)→ |
| Sau-3669-1 | NZ_KI973764.1 | - | 56993 | 56964 | **RNA**→ | rpmE2 (PRK01678)Ribosomal_L31 (pfam01197)→ |
| Sau-2648-1 | NZ_KI974486.1 | - | 56892 | 56863 | **RNA**→ | rpmE2 (PRK01678)Ribosomal_L31 (pfam01197)→ |
| Sau-1100-1 | NZ_KK047877.1 | - | 55212 | 55183 | **RNA**→ | rpmE2 (PRK01678)Ribosomal_L31 (pfam01197)→ |
| Sau-2103-1 | NZ_KI972799.1 | - | 160062 | 160033 | **RNA**→ | rpmE2 (PRK01678)Ribosomal_L31 (pfam01197)→ |
| Sau-1835-1 | NZ_KI984554.1 | - | 55326 | 55297 | **RNA**→ | rpmE2 (PRK01678)Ribosomal_L31 (pfam01197)→ |
| Sau-1213-1 | NZ_KI971833.1 | - | 56529 | 56500 | **RNA**→ | rpmE2 (PRK01678)Ribosomal_L31 (pfam01197)→ |
| Sau-1529-1 | NZ_KK012683.1 | - | 55280 | 55251 | **RNA**→ | rpmE2 (PRK01678)Ribosomal_L31 (pfam01197)→ |
| Sau-789-1 | NZ_KK070096.1 | - | 237242 | 237213 | **RNA**→ | rpmE2 (PRK01678)Ribosomal_L31 (pfam01197)→ |
| Sau-1793-1 | NZ_KI982734.1 | - | 55094 | 55065 | **RNA**→ | rpmE2 (PRK01678)Ribosomal_L31 (pfam01197)→ |
| Sau-1607-1 | NZ_KK016117.1 | - | 54989 | 54960 | **RNA**→ | rpmE2 (PRK01678)Ribosomal_L31 (pfam01197)→ |
| Sau-3513-1 | NZ_KK023462.1 | - | 56940 | 56911 | **RNA**→ | rpmE2 (PRK01678)Ribosomal_L31 (pfam01197)→ |
| Sau-1169-1 | NZ_KI976235.1 | - | 56894 | 56865 | **RNA**→ | rpmE2 (PRK01678)Ribosomal_L31 (pfam01197)→ |
| Sau-1160-1 | NZ_KI978692.1 | - | 56894 | 56865 | **RNA**→ | rpmE2 (PRK01678)Ribosomal_L31 (pfam01197)→ |
| Sau-1397-1 | NZ_KI971474.1 | - | 56724 | 56695 | **RNA**→ | rpmE2 (PRK01678)Ribosomal_L31 (pfam01197)→ |
| Sau-926-1 | NZ_KK024977.1 | - | 56894 | 56865 | **RNA**→ | rpmE2 (PRK01678)Ribosomal_L31 (pfam01197)→ |
| Sau-1112-1 | NZ_KI974788.1 | - | 56994 | 56965 | **RNA**→ | rpmE2 (PRK01678)Ribosomal_L31 (pfam01197)→ |
| Sau-1096-1 | NZ_KK047666.1 | - | 56405 | 56376 | **RNA**→ | rpmE2 (PRK01678)Ribosomal_L31 (pfam01197)→ |

| | | | | | |
|---|---|---|---|---|---|
| Sau-2030-1 | NZ_KI991135.1 | - | 54894 | 54865 | **RNA**→ rpmE2 (PRK01678)Ribosomal_L31 (pfam01197)→ |
| Sau-3056-1 | NZ_CP007176.1 | - | 2285144 | 2285115 | **RNA**→ rpmE2 (PRK01678)Ribosomal_L31 (pfam01197)→ |
| Sau-3757-1 | NZ_CP010526.1 | - | 2229272 | 2229243 | **RNA**→ rpmE2 (PRK01678)Ribosomal_L31 (pfam01197)→ |
| Sau-2619-1 | NZ_KI976828.1 | - | 56994 | 56965 | **RNA**→ rpmE2 (PRK01678)Ribosomal_L31 (pfam01197)→ |
| Sau-1260-1 | NZ_KK002230.1 | - | 55140 | 55111 | **RNA**→ rpmE2 (PRK01678)Ribosomal_L31 (pfam01197)→ |
| Sau-1588-1 | NZ_KK015179.1 | - | 55189 | 55160 | **RNA**→ rpmE2 (PRK01678)Ribosomal_L31 (pfam01197)→ |
| Sau-1615-1 | NZ_KK016620.1 | - | 54989 | 54960 | **RNA**→ rpmE2 (PRK01678)Ribosomal_L31 (pfam01197)→ |
| Sau-2220-1 | NZ_KK063185.1 | - | 55259 | 55230 | **RNA**→ rpmE2 (PRK01678)Ribosomal_L31 (pfam01197)→ |
| Sau-1870-1 | NZ_KI985398.1 | - | 55233 | 55204 | **RNA**→ rpmE2 (PRK01678)Ribosomal_L31 (pfam01197)→ |
| Sau-1997-1 | NZ_KI989969.1 | - | 54889 | 54860 | **RNA**→ rpmE2 (PRK01678)Ribosomal_L31 (pfam01197)→ |
| Sau-2056-1 | NZ_KK069096.1 | - | 55426 | 55397 | **RNA**→ rpmE2 (PRK01678)Ribosomal_L31 (pfam01197)→ |
| Sau-3522-1 | NZ_KK070456.1 | - | 279710 | 279681 | **RNA**→ rpmE2 (PRK01678)Ribosomal_L31 (pfam01197)→ |
| Sau-2196-1 | NZ_KK063830.1 | - | 55189 | 55160 | **RNA**→ rpmE2 (PRK01678)Ribosomal_L31 (pfam01197)→ |
| Sau-1451-1 | NZ_KK009269.1 | - | 54889 | 54860 | **RNA**→ rpmE2 (PRK01678)Ribosomal_L31 (pfam01197)→ |
| Sau-2655-1 | NZ_KK019219.1 | - | 792664 | 792635 | **RNA**→ rpmE2 (PRK01678)Ribosomal_L31 (pfam01197)→ |
| Sau-1257-1 | NZ_KK002069.1 | - | 55131 | 55102 | **RNA**→ rpmE2 (PRK01678)Ribosomal_L31 (pfam01197)→ |
| Sau-2912-1 | NZ_KK020889.1 | - | 41304 | 41275 | **RNA**→ rpmE2 (PRK01678)Ribosomal_L31 (pfam01197)→ |
| Sau-1127-1 | NZ_KI975919.1 | - | 56894 | 56865 | **RNA**→ rpmE2 (PRK01678)Ribosomal_L31 (pfam01197)→ |
| Sau-1239-1 | NZ_KK001294.1 | - | 55148 | 55119 | **RNA**→ rpmE2 (PRK01678)Ribosomal_L31 (pfam01197)→ |
| Sau-1274-1 | NZ_KK002623.1 | - | 55426 | 55397 | **RNA**→ rpmE2 (PRK01678)Ribosomal_L31 (pfam01197)→ |
| Sau-1794-1 | NZ_KI982815.1 | - | 55130 | 55101 | **RNA**→ rpmE2 (PRK01678)Ribosomal_L31 (pfam01197)→ |
| Sau-1885-1 | NZ_KI985844.1 | - | 55424 | 55395 | **RNA**→ rpmE2 (PRK01678)Ribosomal_L31 (pfam01197)→ |
| Sau-1262-1 | NZ_KK002316.1 | - | 55407 | 55378 | **RNA**→ rpmE2 (PRK01678)Ribosomal_L31 (pfam01197)→ |
| Sau-1528-1 | NZ_KK012556.1 | - | 55135 | 55106 | **RNA**→ rpmE2 (PRK01678)Ribosomal_L31 (pfam01197)→ |
| Sau-1609-1 | NZ_KK016307.1 | - | 55043 | 55014 | **RNA**→ rpmE2 (PRK01678)Ribosomal_L31 (pfam01197)→ |
| Sau-776-1 | NZ_KK026432.1 | - | 29424 | 29395 | **RNA**→ rpmE2 (PRK01678)Ribosomal_L31 (pfam01197)→ |
| Sau-2604-1 | NZ_KI973827.1 | - | 56894 | 56865 | **RNA**→ rpmE2 (PRK01678)Ribosomal_L31 (pfam01197)→ |
| Sau-1328-1 | NZ_KK005051.1 | - | 55231 | 55202 | **RNA**→ rpmE2 (PRK01678)Ribosomal_L31 (pfam01197)→ |
| Sau-1531-1 | NZ_KK012699.1 | - | 55135 | 55106 | **RNA**→ rpmE2 (PRK01678)Ribosomal_L31 (pfam01197)→ |
| Sau-1181-1 | NZ_KI998921.1 | - | 54889 | 54860 | **RNA**→ rpmE2 (PRK01678)Ribosomal_L31 (pfam01197)→ |
| Sau-1627-1 | NZ_KK017224.1 | - | 54989 | 54960 | **RNA**→ rpmE2 (PRK01678)Ribosomal_L31 (pfam01197)→ |
| Sau-930-1 | NZ_KK025367.1 | - | 56894 | 56865 | **RNA**→ rpmE2 (PRK01678)Ribosomal_L31 (pfam01197)→ |
| Sau-2075-1 | NZ_KK068270.1 | - | 55231 | 55202 | **RNA**→ rpmE2 (PRK01678)Ribosomal_L31 (pfam01197)→ |
| Sau-2164-1 | NZ_KK065099.1 | - | 25205 | 25176 | **RNA**→ rpmE2 (PRK01678)Ribosomal_L31 (pfam01197)→ |
| Sau-2775-1 | NZ_KI975292.1 | - | 56894 | 56865 | **RNA**→ rpmE2 (PRK01678)Ribosomal_L31 (pfam01197)→ |
| Sau-1869-1 | NZ_KI985316.1 | - | 55331 | 55302 | **RNA**→ rpmE2 (PRK01678)Ribosomal_L31 (pfam01197)→ |
| Sau-1259-1 | NZ_KK002154.1 | - | 55231 | 55202 | **RNA**→ rpmE2 (PRK01678)Ribosomal_L31 (pfam01197)→ |
| Sau-1643-1 | NZ_KK018033.1 | - | 60065 | 60036 | **RNA**→ rpmE2 (PRK01678)Ribosomal_L31 (pfam01197)→ |
| Sau-1606-1 | NZ_KK016041.1 | - | 55331 | 55302 | **RNA**→ rpmE2 (PRK01678)Ribosomal_L31 (pfam01197)→ |
| Sau-1818-1 | NZ_KI983816.1 | - | 27093 | 27064 | **RNA**→ rpmE2 (PRK01678)Ribosomal_L31 (pfam01197)→ |
| Sau-1256-1 | NZ_KK001992.1 | - | 55372 | 55343 | **RNA**→ rpmE2 (PRK01678)Ribosomal_L31 (pfam01197)→ |
| Sau-2170-1 | NZ_KK064806.1 | - | 55454 | 55425 | **RNA**→ rpmE2 (PRK01678)Ribosomal_L31 (pfam01197)→ |
| Sau-1253-1 | NZ_KK001790.1 | - | 55131 | 55102 | **RNA**→ rpmE2 (PRK01678)Ribosomal_L31 (pfam01197)→ |
| Sau-2049-1 | NZ_KK069306.1 | - | 54994 | 54965 | **RNA**→ rpmE2 (PRK01678)Ribosomal_L31 (pfam01197)→ |
| Sau-2539-1 | NZ_KK096734.1 | - | 34913 | 34884 | **RNA**→ rpmE2 (PRK01678)Ribosomal_L31 (pfam01197)→ |
| Sau-2477-1 | NZ_KK092636.1 | - | 56894 | 56865 | **RNA**→ rpmE2 (PRK01678)Ribosomal_L31 (pfam01197)→ |
| Sau-3630-1 | NZ_KK097480.1 | - | 57198 | 57169 | **RNA**→ rpmE2 (PRK01678)Ribosomal_L31 (pfam01197)→ |
| Sau-643-1 | NZ_KK076265.1 | - | 34873 | 34844 | **RNA**→ rpmE2 (PRK01678)Ribosomal_L31 (pfam01197)→ |
| Sau-328-1 | NZ_KK099102.1 | - | 348428 | 348399 | **RNA**→ rpmE2 (PRK01678)Ribosomal_L31 (pfam01197)→ |
| Sau-333-1 | NZ_KK099145.1 | + | 57760 | 57789 | **RNA**→ rpmE2 (PRK01678)Ribosomal_L31 (pfam01197)→ |
| Sau-376-1 | NZ_KK099464.1 | - | 350084 | 350055 | **RNA**→ rpmE2 (PRK01678)Ribosomal_L31 (pfam01197)→ |
| Sau-156-1 | NZ_KK220355.1 | - | 26024 | 25995 | **RNA**→ rpmE2 (PRK01678)Ribosomal_L31 (pfam01197)→ |

| | | | | | |
|---|---|---|---|---|---|
| Sau-3722-3 | NZ_CP007659.1 | - | 2212795 | 2212766 | **RNA**→ rpmE2 (PRK01678)Ribosomal_L31 (pfam01197)→ |
| Sau-3755-1 | NZ_CP006630.1 | - | 2169570 | 2169541 | **RNA**→ rpmE2 (PRK01678)Ribosomal_L31 (pfam01197)→ |
| Sau-3756-1 | NZ_KL662263.1 | - | 54767 | 54738 | **RNA**→ rpmE2 (PRK01678)Ribosomal_L31 (pfam01197)→ |
| Sau-3722-4 | NZ_CP009361.1 | - | 2152383 | 2152354 | **RNA**→ rpmE2 (PRK01678)Ribosomal_L31 (pfam01197)→ |
| Sau-79-1 | NZ_LN626917.1 | - | 2210527 | 2210498 | **RNA**→ rpmE2 (PRK01678)Ribosomal_L31 (pfam01197)→ |
| Sau-4103-4 | NC_020529.1 | - | 2120722 | 2120693 | **RNA**→ rpmE2 (PRK01678)Ribosomal_L31 (pfam01197)→ rpmE2 (PRK01678)Ribosomal_L31 (pfam01197 |
| Sau-4103-5 | NC_020564.1 | - | 2120709 | 2120680 | **RNA**→ rpmE2 (PRK01678)Ribosomal_L31 (pfam01197)→ rpmE2 (PRK01678)Ribosomal_L31 (pfam01197 |
| Sau-4103-6 | NC_020536.1 | - | 2120708 | 2120679 | **RNA**→ rpmE2 (PRK01678)Ribosomal_L31 (pfam01197)→ rpmE2 (PRK01678)Ribosomal_L31 (pfam01197 |
| Sau-4103-7 | NC_020537.1 | - | 2120471 | 2120442 | **RNA**→ rpmE2 (PRK01678)Ribosomal_L31 (pfam01197)→ rpmE2 (PRK01678)Ribosomal_L31 (pfam01197 |
| Sau-4103-8 | NC_020532.1 | - | 2121018 | 2120989 | **RNA**→ rpmE2 (PRK01678)Ribosomal_L31 (pfam01197)→ rpmE2 (PRK01678)Ribosomal_L31 (pfam01197 |
| Sau-218-1 | NZ_KK240911.1 | + | 56430 | 56459 | **RNA**→ rpmE2 (PRK01678)Ribosomal_L31 (pfam01197)→ |
| Sau-3939-1 | NZ_KB822069.1 | - | 1303682 | 1303653 | **RNA**→ rpmE2 (PRK01678)Ribosomal_L31 (pfam01197)→ |
| Sau-79-2 | NZ_CP007539.1 | - | 2184540 | 2184511 | **RNA**→ rpmE2 (PRK01678)Ribosomal_L31 (pfam01197)→ |
| Sau-1-1 | NC_017340.1 | - | 2190574 | 2190545 | **RNA**→ rpmE2 (PRK01678)Ribosomal_L31 (pfam01197)→ |
| Sau-3722-6 | NZ_CP012119.1 | + | 2562238 | 2562267 | **RNA**→ rpmE2 (PRK01678)Ribosomal_L31 (pfam01197)→ |
| Sau-3743-1 | NZ_CP011526.1 | - | 2117394 | 2117365 | **RNA**→ rpmE2 (PRK01678)Ribosomal_L31 (pfam01197)→ |
| Sau-4094-1 | NZ_JH691956.1 | - | 1623707 | 1623678 | **RNA**→ rpmE2 (PRK01678)Ribosomal_L31 (pfam01197)→ |
| Sau-1162-1 | NZ_KE503211.1 | + | 1132666 | 1132695 | **RNA**→ rpmE2 (PRK01678)Ribosomal_L31 (pfam01197)→ |
| Sau-4075-1 | NZ_AJKF01000031.1 | - | 3066 | 3037 | **RNA**→ rpmE2 (PRK01678)Ribosomal_L31 (pfam01197)→ |
| Sau-4106-1 | NZ_AGTZ01000005.1 | + | 28862 | 28891 | **RNA**→ rpmE2 (PRK01678)Ribosomal_L31 (pfam01197)→ |
| Sau-4085-1 | NZ_AICI01000200.1 | - | 15991 | 15962 | **RNA**→ rpmE2 (PRK01678)Ribosomal_L31 (pfam01197)→ |
| Sau-4089-1 | NZ_AHLP01000002.1 | - | 55170 | 55141 | **RNA**→ rpmE2 (PRK01678)Ribosomal_L31 (pfam01197)→ |
| Sau-4061-1 | NZ_AHVN01000011.1 | - | 57147 | 57118 | **RNA**→ rpmE2 (PRK01678)Ribosomal_L31 (pfam01197)→ |
| Sau-4090-1 | NZ_AHLQ01000001.1 | - | 25176 | 25147 | **RNA**→ rpmE2 (PRK01678)Ribosomal_L31 (pfam01197)→ |
| Sau-4063-1 | NZ_AHVV01000014.1 | - | 59240 | 59211 | **RNA**→ rpmE2 (PRK01678)Ribosomal_L31 (pfam01197)→ |
| Sau-4081-1 | NZ_AHLR01000057.1 | - | 54627 | 54598 | **RNA**→ rpmE2 (PRK01678)Ribosomal_L31 (pfam01197)→ |
| Sau-4082-1 | NZ_AHLS01000123.1 | - | 9926 | 9897 | **RNA**→ rpmE2 (PRK01678)Ribosomal_L31 (pfam01197)→ |
| Sau-4045-1 | NZ_AHKD01000013.1 | - | 59992 | 59963 | **RNA**→ rpmE2 (PRK01678)Ribosomal_L31 (pfam01197)→ |
| Sau-4086-1 | NZ_AHLM01000010.1 | - | 55240 | 55211 | **RNA**→ rpmE2 (PRK01678)Ribosomal_L31 (pfam01197)→ |
| Sau-4006-1 | NZ_AFEG01000024.1 | - | 54491 | 54462 | **RNA**→ rpmE2 (PRK01678)Ribosomal_L31 (pfam01197)→ |
| Sau-4091-1 | NZ_AJLD01000001.1 | - | 35341 | 35312 | **RNA**→ rpmE2 (PRK01678)Ribosomal_L31 (pfam01197)→ |
| Sau-3749-1 | NZ_CAWB010000325.1 | + | 6522 | 6551 | **RNA**→ rpmE2 (PRK01678)Ribosomal_L31 (pfam01197)→ |
| Sau-3694-1 | NZ_JBFR01000002.1 | - | 56894 | 56865 | **RNA**→ rpmE2 (PRK01678)Ribosomal_L31 (pfam01197)→ |
| Sau-3747-1 | NZ_CAVH010000032.1 | - | 25734 | 25705 | **RNA**→ rpmE2 (PRK01678)Ribosomal_L31 (pfam01197)→ |
| Sau-1122-1 | NZ_JBJK01000006.1 | - | 56894 | 56865 | **RNA**→ rpmE2 (PRK01678)Ribosomal_L31 (pfam01197)→ |
| Sau-2783-1 | NZ_JBNJ01000001.1 | - | 1322998 | 1322969 | **RNA**→ rpmE2 (PRK01678)Ribosomal_L31 (pfam01197)→ |
| Sau-3752-1 | NZ_CAWA010000012.1 | + | 48028 | 48057 | **RNA**→ rpmE2 (PRK01678)Ribosomal_L31 (pfam01197)→ |
| Sau-2761-1 | NZ_JBLV01000002.1 | - | 56894 | 56865 | **RNA**→ rpmE2 (PRK01678)Ribosomal_L31 (pfam01197)→ |
| Sau-3754-1 | NZ_CAVW010000042.1 | + | 18308 | 18337 | **RNA**→ rpmE2 (PRK01678)Ribosomal_L31 (pfam01197)→ |
| Sau-3753-1 | NZ_CAVV010000049.1 | - | 15697 | 15668 | **RNA**→ rpmE2 (PRK01678)Ribosomal_L31 (pfam01197)→ |
| Sau-3748-1 | NZ_CAVX010000008.1 | - | 25736 | 25707 | **RNA**→ rpmE2 (PRK01678)Ribosomal_L31 (pfam01197)→ |
| Sau-3750-1 | NZ_CAVY010000101.1 | - | 20464 | 20435 | **RNA**→ rpmE2 (PRK01678)Ribosomal_L31 (pfam01197)→ |
| Sau-3751-1 | NZ_CAVU010000011.1 | + | 48082 | 48111 | **RNA**→ rpmE2 (PRK01678)Ribosomal_L31 (pfam01197)→ |
| Sau-2547-1 | NZ_JGHQ01000002.1 | - | 56894 | 56865 | **RNA**→ rpmE2 (PRK01678)Ribosomal_L31 (pfam01197)→ |
| Sau-2716-1 | NZ_JEMM01000011.1 | - | 54849 | 54820 | **RNA**→ rpmE2 (PRK01678)Ribosomal_L31 (pfam01197)→ |
| Sau-3734-1 | NZ_JHPZ01000040.1 | - | 54712 | 54683 | **RNA**→ rpmE2 (PRK01678)Ribosomal_L31 (pfam01197)→ |
| Sau-3730-1 | NZ_JHPX01000001.1 | - | 54958 | 54929 | **RNA**→ rpmE2 (PRK01678)Ribosomal_L31 (pfam01197)→ |
| Sau-3724-1 | NZ_JHPS01000026.1 | - | 29607 | 29578 | **RNA**→ rpmE2 (PRK01678)Ribosomal_L31 (pfam01197)→ |
| Sau-4076-1 | NZ_JHQG01000095.1 | - | 503 | 474 | **RNA**→ rpmE2 (PRK01678)Ribosomal_L31 (pfam01197)→ |
| Sau-3728-1 | NZ_JHPV01000038.1 | - | 56085 | 56056 | **RNA**→ rpmE2 (PRK01678)Ribosomal_L31 (pfam01197)→ |
| Sau-4077-1 | NZ_JHTT01000096.1 | - | 55218 | 55189 | **RNA**→ rpmE2 (PRK01678)Ribosomal_L31 (pfam01197)→ |
| Sau-2717-1 | NZ_JDRT01000006.1 | + | 46152 | 46181 | **RNA**→ rpmE2 (PRK01678)Ribosomal_L31 (pfam01197)→ |

| | | | | | |
|---|---|---|---|---|---|
| Sau-146-1 | NZ_JOPS01000038.1 | - | 55248 | 55219 | RNA→ rpmE2 (PRK01678)Ribosomal_L31 (pfam01197)→ |
| Sau-3777-1 | NZ_APFH01000011.1 | - | 159007 | 158978 | RNA→ rpmE2 (PRK01678)Ribosomal_L31 (pfam01197)→ |
| Sau-4052-1 | NZ_AIEO01000052.1 | - | 61224 | 61195 | RNA→ rpmE2 (PRK01678)Ribosomal_L31 (pfam01197)→ |
| Slu-1-1 | NC_013893.1 | + | 967452 | 967481 | RNA→ rpmE2 (PRK01678)Ribosomal_L31 (pfam01197)→ |
| Slu-8-1 | NZ_GL622353.1 | - | 44362 | 44333 | RNA→ rpmE2 (PRK01678)Ribosomal_L31 (pfam01197)→ |
| Slu-2-1 | NC_017353.1 | + | 998627 | 998656 | RNA→ rpmE2 (PRK01678)Ribosomal_L31 (pfam01197)→ |
| Slu-7-1 | NZ_KB373322.1 | + | 1720229 | 1720258 | RNA→ rpmE2 (PRK01678)Ribosomal_L31 (pfam01197)→ |
| Slu-4-1 | NZ_KI978380.1 | - | 16661 | 16632 | RNA→ rpmE2 (PRK01678)Ribosomal_L31 (pfam01197)→ |
| Slu-9-1 | NZ_AHLK01000035.1 | - | 16664 | 16635 | RNA→ rpmE2 (PRK01678)Ribosomal_L31 (pfam01197)→ |
| Slu-6-1 | NZ_JIBS01000015.1 | - | 44361 | 44332 | RNA→ rpmE2 (PRK01678)Ribosomal_L31 (pfam01197)→ |
| Slu-5-1 | NZ_JIBR01000008.1 | + | 39788 | 39817 | RNA→ rpmE2 (PRK01678)Ribosomal_L31 (pfam01197)→ |
| Ssc-2-1 | NZ_CCEG01000010.1 | - | 50595 | 50566 | RNA→ rpmE2 (PRK01678)Ribosomal_L31 (pfam01197)→ |
| Ssc-2-2 | NZ_CCEQ01000008.1 | - | 28157 | 28128 | RNA→ rpmE2 (PRK01678)Ribosomal_L31 (pfam01197)→ |
| Ssc-2-3 | NZ_CCEO01000012.1 | - | 55074 | 55045 | RNA→ rpmE2 (PRK01678)Ribosomal_L31 (pfam01197)→ |
| Spe-1-2 | NZ_JVVL01000017.1 | + | 42051 | 42080 | RNA→ rpmE2 (PRK01678)Ribosomal_L31 (pfam01197)→ |
| Svi-2-1 | NZ_AJTR01000004.1 | - | 43878 | 43849 | RNA→ rpmE2 (PRK01678)Ribosomal_L31 (pfam01197)→ |
| Sau-4058-1 | NZ_AHVW01000007.1 | - | 59446 | 59417 | RNA→ rpmE2 (PRK01678)Ribosomal_L31 (pfam01197)→ |
| Seq-2-1 | NZ_LAKE01000007.1 | - | 45187 | 45158 | RNA→ rpmE2 (PRK01678)Ribosomal_L31 (pfam01197)→ |
| Seq-1-1 | NZ_CAJL01000004.1 | - | 45027 | 44998 | RNA→ rpmE2 (PRK01678)Ribosomal_L31 (pfam01197)→ rpmE2 (PRK01678)Ribosomal_L31 (pfam01197 |
| Seq-3-1 | NZ_AVBD01000013.1 | - | 17682 | 17653 | RNA→ rpmE2 (PRK01678)Ribosomal_L31 (pfam01197)→ |
| Ssp-2-1 | NZ_ALPU01000004.1 | + | 52056 | 52085 | RNA→ rpmE2 (PRK01678)Ribosomal_L31 (pfam01197)→ rpmE2 (PRK01678)Ribosomal_L31 (pfam01197 |
| Jma-3-1 | NZ_AUEE01000004.1 | - | 25421 | 25392 | RNA→ rpmE2 (PRK01678)Ribosomal_L31 (pfam01197)→ |
| env-1001 | JGI24707J26582_10439517 | - | 249 | 220 | RNA→ rpmE2 (PRK01678)Ribosomal_L31 (pfam01197)→ |
| env-1002 | JGI24708J26588_10298792 | + | 134 | 163 | RNA→ rpmE2 (PRK01678)Ribosomal_L31 (pfam01197)→ |
| Jsa-1-1 | NZ_CCSE01000001.1 | + | 1953030 | 1953059 | RNA→ rpmE2 (PRK01678)Ribosomal_L31 (pfam01197)→ |
| Jps-1-1 | NZ_AUEF01000015.1 | - | 44498 | 44469 | RNA→ rpmE2 (PRK01678)Ribosomal_L31 (pfam01197)→ |
| env-1003 | JGI24707J26582_10106987 | + | 208 | 237 | RNA→ rpmE2 (PRK01678)Ribosomal_L31 (pfam01197)→ |
| env-1004 | 2204520978 | - | 38187 | 38158 | RNA→ rpmE2 (PRK01678)Ribosomal_L31 (pfam01197)→ PrfA (COG0216)prfA (PRK00591)→<br>HemK (COG2890)PRK09328 (PRK09328)→ SUA5 (COG0009)TIGR00057 (TIGR00057)SUA5 (pfam03481<br>LMWPc (smart00226)Wzb (COG0394)→ PRK05571 (PRK05571)LacAB_rpiB (pfam02502)→<br>PRK13690 (PRK13690)DUF436 (pfam04260)→ GlyA (COG0112)glyA (PRK00011)→<br>Upp (COG0035)upp (PRK00129)→ WecB (COG0381)wecB (TIGR00236)→ |
| env-1005 | 2157730857 | - | 1915 | 1886 | RNA→ rpmE2 (PRK01678)Ribosomal_L31 (pfam01197)→ PrfA (COG0216)prfA (PRK00591)→<br>HemK (COG2890)Methyltransf_25 (pfam13649)→ |
| Gsp-13-1 | NC_022080.4 | - | 3302086 | 3302054 | RNA→ rpmE (PRK00019)Ribosomal_L31 (pfam01197)→ |
| Bma-1-1 | NZ_AKKV01000043.1 | - | 43743 | 43712 | RNA→ rpmE2 (PRK01678)Ribosomal_L31 (pfam01197)→ rpmE2 (PRK01678)Ribosomal_L31 (pfam01197 |
| Gla-1-1 | NZ_KB898662.1 | - | 511085 | 511054 | RNA→ rpmE2 (PRK01678)Ribosomal_L31 (pfam01197)→ |
| Gbo-1-1 | NZ_BAVS01000005.1 | - | 89857 | 89826 | RNA→ rpmE2 (PRK01678)Ribosomal_L31 (pfam01197)→ |
| Pha-1-1 | NZ_AULI01000001.1 | - | 227155 | 227123 | RNA→ rpmE2 (PRK01678)Ribosomal_L31 (pfam01197)→ |
| Pha-1-2 | NZ_AVPE01000001.1 | + | 286167 | 286199 | RNA→ rpmE2 (PRK01678)Ribosomal_L31 (pfam01197)→ |
| Pma-1-1 | NZ_KE384330.1 | - | 164379 | 164347 | RNA→ rpmE2 (PRK01678)Ribosomal_L31 (pfam01197)→ |
| env-1006 | LV_Brine_h2_0102_1152597 | - | 205 | 173 | RNA→ rpmE2 (PRK01678)Ribosomal_L31 (pfam01197)→ |
| Tai-1-1 | NZ_CP008876.1 | + | 2935582 | 2935614 | RNA→ rpmE2 (PRK01678)Ribosomal_L31 (pfam01197)→ |
| Osc-1-1 | NZ_AEWH01000071.1 | - | 116507 | 116476 | RNA→ rpmE2 (PRK01678)Ribosomal_L31 (pfam01197)→ rpmE2 (PRK01678)Ribosomal_L31 (pfam01197 |
| Vse-1-1 | NZ_CCXU01000020.1 | + | 89039 | 89070 | RNA→ rpmE2 (PRK01678)Ribosomal_L31 (pfam01197)→ |
| Bsp-75-1 | NZ_KI530538.1 | - | 102853 | 102822 | RNA→ rpmE2 (PRK01678)Ribosomal_L31 (pfam01197)→ |
| Sha-2-1 | NZ_KI543242.1 | - | 205703 | 205672 | RNA→ rpmE2 (PRK01678)Ribosomal_L31 (pfam01197)→ |
| Vha-1-1 | NZ_ALEF01000051.1 | - | 39336 | 39304 | RNA→ rpmE2 (PRK01678)Ribosomal_L31 (pfam01197)→ |
| Vsp-2-1 | NZ_CP007161.1 | - | 3356021 | 3355989 | RNA→ rpmE2 (PRK01678)Ribosomal_L31 (pfam01197)→ hypo→ |
| Oje-1-1 | NZ_CCDM010000001.1 | + | 767568 | 767600 | RNA→ rpmE2 (PRK01678)Ribosomal_L31 (pfam01197)→ |
| Oon-1-1 | NZ_CDGG01000001.1 | + | 590892 | 590924 | RNA→ rpmE2 (PRK01678)Ribosomal_L31 (pfam01197)→ |
| Hku-1-1 | NZ_AUDY01000009.1 | + | 56142 | 56174 | RNA→ rpmE2 (PRK01678)Ribosomal_L31 (pfam01197)→ |
| Hha-1-1 | NC_017668.1 | - | 3554991 | 3554959 | RNA→ rpmE2 (PRK01678)Ribosomal_L31 (pfam01197)→ hypo→ |

| | | | | | |
|---|---|---|---|---|---|
| Oki-1-1 | NZ_CM001792.1 | - | 3174116 | 3174084 | **RNA**→ rpmE2 (PRK01678)Ribosomal_L31 (pfam01197)→ |
| Oih-1-1 | NC_004193.1 | - | 3108848 | 3108816 | **RNA**→ rpmE2 (PRK01678)Ribosomal_L31 (pfam01197)→ |
| Pgl-1-1 | NZ_KI519435.1 | - | 42469 | 42438 | **RNA**→ rpmE2 (PRK01678)Ribosomal_L31 (pfam01197)→ |
| Osp-1-1 | NZ_HE610976.1 | - | 738196 | 738164 | **RNA**→ rpmE2 (PRK01678)Ribosomal_L31 (pfam01197)→ |
| Oma-1-1 | NZ_CCDL010000001.1 | + | 930544 | 930575 | **RNA**→ rpmE2 (PRK01678)Ribosomal_L31 (pfam01197)→ |
| Bba-4-1 | NZ_LAZH01000073.1 | + | 72597 | 72628 | **RNA**→ rpmE2 (PRK01678)Ribosomal_L31 (pfam01197)→ |
| Sai-1-1 | NZ_KE007196.1 | - | 146719 | 146687 | **RNA**→ rpmE2 (PRK01678)Ribosomal_L31 (pfam01197)→ |
| Pch-1-1 | NZ_AVBG01000002.1 | - | 81446 | 81414 | **RNA**→ rpmE2 (PRK01678)Ribosomal_L31 (pfam01197)→ |
| Bme-6-1 | NZ_KI518593.1 | - | 65186 | 65154 | **RNA**→ rpmE2 (PRK01678)Ribosomal_L31 (pfam01197)→ |
| Pli-1-1 | NZ_AVPG01000013.1 | - | 37362 | 37330 | **RNA**→ rpmE2 (PRK01678)Ribosomal_L31 (pfam01197)→ |
| Opi-1-1 | NZ_CCAX010000004.1 | - | 131537 | 131506 | **RNA**→ rpmE2 (PRK01678)Ribosomal_L31 (pfam01197)→ |
| Bsp-83-1 | NZ_KI270940.1 | + | 507592 | 507624 | **RNA**→ rpmE2 (PRK01678)Ribosomal_L31 (pfam01197)→ |
| Hsp-1-1 | NZ_ANPF01000001.1 | + | 62661 | 62693 | **RNA**→ rpmE2 (PRK01678)Ribosomal_L31 (pfam01197)→ |
| Hsp-2-1 | NZ_JRNX01000108.1 | - | 3575 | 3543 | **RNA**→ rpmE2 (PRK01678)Ribosomal_L31 (pfam01197)→ |
| Tde-1-1 | NZ_KI543237.1 | - | 735753 | 735721 | **RNA**→ rpmE2 (PRK01678)Ribosomal_L31 (pfam01197)→ |
| Htr-1-1 | NZ_CCDJ010000001.1 | - | 775405 | 775373 | **RNA**→ rpmE2 (PRK01678)Ribosomal_L31 (pfam01197)→ |
| Hda-1-1 | NZ_CCDH010000001.1 | + | 1071371 | 1071403 | **RNA**→ rpmE2 (PRK01678)Ribosomal_L31 (pfam01197)→ |
| Hka-1-1 | NZ_CCDI010000001.1 | - | 1280886 | 1280854 | **RNA**→ rpmE2 (PRK01678)Ribosomal_L31 (pfam01197)→ |
| Vsp-1-1 | NZ_AUQA01000002.1 | - | 1068398 | 1068366 | **RNA**→ rpmE2 (PRK01678)Ribosomal_L31 (pfam01197)→ |
| Vma-1-1 | NZ_CCDP010000001.1 | + | 560592 | 560624 | **RNA**→ rpmE2 (PRK01678)Ribosomal_L31 (pfam01197)→ |
| Vpa-1-1 | NZ_LGTO01000005.1 | + | 626906 | 626938 | **RNA**→ rpmE2 (PRK01678)Ribosomal_L31 (pfam01197)→ |
| Oco-1-1 | NZ_LDPV02000007.1 | - | 9420 | 9389 | **RNA**→ rpmE2 (PRK01678)Ribosomal_L31 (pfam01197)→ |
| Psp-6-1 | NZ_CBYO010000007.1 | - | 292529 | 292498 | **RNA**→ rpmE2 (PRK01678)Ribosomal_L31 (pfam01197)→ |
| Oca-1-1 | NZ_LDUE01000021.1 | - | 69650 | 69619 | **RNA**→ rpmE2 (PRK01678)Ribosomal_L31 (pfam01197)→ |
| Val-1-1 | NZ_JFBD01000064.1 | - | 10605 | 10572 | **RNA**→ rpmE2 (PRK01678)Ribosomal_L31 (pfam01197)→ |
| env-1007 | TV14_100008 | - | 68254 | 68222 | **RNA**→ rpmE2 (PRK01678)Ribosomal_L31 (pfam01197)→ |
| Gha-2-1 | NZ_APML01000077.1 | - | 3139 | 3107 | **RNA**→ rpmE2 (PRK01678)Ribosomal_L31 (pfam01197)→ |
| Pgl-2-1 | NZ_LGAF01000018.1 | - | 43190 | 43163 | **RNA**→ rpmE (PRK00019)Ribosomal_L31 (pfam01197)→ |
| Pgl-3-1 | NZ_AUYR01000095.1 | + | 35852 | 35879 | **RNA**→ rpmE (PRK00019)Ribosomal_L31 (pfam01197)→ |
| Pma-5-1 | NZ_CCXS01000001.1 | + | 811716 | 811743 | **RNA**→ rpmE (PRK00019)Ribosomal_L31 (pfam01197)→ |
| env-1008 | 3300001196_1064895 | - | 266 | 236 | **RNA**→ rpmE (PRK00019)Ribosomal_L31 (pfam01197)→ |
| env-1009 | 3300001197_1032424 | - | 97 | 67 | **RNA**→ hypo→ |
| Sur-1-1 | NZ_AUDQ01000005.1 | - | 34401 | 34371 | **RNA**→ rpmE (PRK00019)Ribosomal_L31 (pfam01197)→ |
| Var-2-1 | NZ_LILB01000005.1 | - | 1317698 | 1317668 | **RNA**→ rpmE (PRK00019)Ribosomal_L31 (pfam01197)→ |
| Var-1-1 | NZ_ASQA01000009.1 | - | 79603 | 79573 | **RNA**→ rpmE (PRK00019)Ribosomal_L31 (pfam01197)→ |
| Bce-148-1 | NZ_AOFT01000001.1 | + | 212893 | 212923 | **RNA**→ rpmE (PRK00019)Ribosomal_L31 (pfam01197)→ Tdk (COG1435)PRK04296 (PRK04296)→ Tdk (COG1435)PRK04296 (PRK04296)→ |
| env-1010 | topACOD_FV90NF401DKND5 | - | 503 | 472 | **RNA**→ rpmE (PRK00019)Ribosomal_L31 (pfam01197)→ Tdk (COG1435)PRK04296 (PRK04296)→ |
| env-1011 | 2204573945 | - | 257 | 227 | **RNA**→ rpmE (PRK00019)Ribosomal_L31 (pfam01197)→ |
| env-1012 | 2157810416 | - | 257 | 227 | **RNA**→ rpmE (PRK00019)Ribosomal_L31 (pfam01197)→ |
| env-1013 | MA40A_contig22098 | + | 215 | 245 | **RNA**→ rpmE (PRK00019)Ribosomal_L31 (pfam01197)→ |
| Bsp-77-1 | NZ_AP013294.1 | + | 1455730 | 1455760 | **RNA**→ rpmE (PRK00019)Ribosomal_L31 (pfam01197)→ |
| Sko-2-1 | NZ_CDGU01000008.1 | - | 326072 | 326042 | **RNA**→ rpmE (PRK00019)Ribosomal_L31 (pfam01197)→ |
| Sne-1-1 | NZ_GL982997.1 | - | 1321838 | 1321808 | **RNA**→ rpmE (PRK00019)Ribosomal_L31 (pfam01197)→ |
| env-1014 | 3300001196_1006164 | - | 2809 | 2779 | **RNA**→ rpmE (PRK00019)Ribosomal_L31 (pfam01197)→ |
| env-1015 | JGI24024J18818_10026976 | + | 2203 | 2233 | **RNA**→ ←hypo |
| env-1016 | JGI24024J18818_10198670 | + | 28 | 58 | **RNA**→ rpmE (PRK00019)Ribosomal_L31 (pfam01197)→ |
| env-1017 | JGI24024J18818_10035368 | - | 167 | 137 | **RNA**→ rpmE (PRK00019)Ribosomal_L31 (pfam01197)→ |
| Bce-149-1 | NZ_CDGP01000041.1 | + | 2627 | 2657 | **RNA**→ rpmE (PRK00019)Ribosomal_L31 (pfam01197)→ |
| Bce-149-2 | NZ_CDGP01000042.1 | + | 2627 | 2657 | **RNA**→ rpmE (PRK00019)Ribosomal_L31 (pfam01197)→ |
| Psp-37-1 | NZ_JRGN01000034.1 | - | 1132 | 1102 | **RNA**→ rpmE (PRK00019)Ribosomal_L31 (pfam01197)→ |
| Pan-1-1 | NZ_AJYB01000017.1 | - | 68220 | 68190 | **RNA**→ rpmE (PRK00019)Ribosomal_L31 (pfam01197)→ rpmE (PRK00019)Ribosomal_L31 (pfam01197)– |

| | | | | | |
|---|---|---|---|---|---|
| Psp-38-1 | NZ_JHDT01000001.1 | - | 6165 | 6135 | **RNA**→ rpmE (PRK00019)Ribosomal_L31 (pfam01197)→ |
| Pha-3-1 | NZ_ANBV01000010.1 | + | 146734 | 146764 | **RNA**→ rpmE (PRK00019)Ribosomal_L31 (pfam01197)→ |
| Psp-39-1 | NZ_CP009129.1 | - | 2482118 | 2482088 | **RNA**→ rpmE (PRK00019)Ribosomal_L31 (pfam01197)→ |
| Pdo-1-1 | NZ_AEPB01000064.1 | + | 9346 | 9376 | **RNA**→ rpmE (PRK00019)Ribosomal_L31 (pfam01197)→ rpmE (PRK00019)Ribosomal_L31 (pfam01197)– |
| env-1018 | KGI_S2_ANT04_2345m_c1001284 | - | 9677 | 9647 | **RNA**→ rpmE (PRK00019)Ribosomal_L31 (pfam01197)→ hypo→ |
| env-1019 | KGI_S2_ANT04_2345m_c1014326 | + | 879 | 909 | **RNA**→ rpmE (PRK00019)Ribosomal_L31 (pfam01197)→ |
| env-1020 | KGI_S2_ANT05_2345m_c1009878 | - | 1148 | 1118 | **RNA**→ rpmE (PRK00019)Ribosomal_L31 (pfam01197)→ hypo→ |
| env-1021 | KGI_S1_ANT02_95m_c10007926 | - | 333 | 303 | **RNA**→ rpmE (PRK00019)Ribosomal_L31 (pfam01197)→ |
| Uth-1-1 | NZ_AJIK01000003.1 | - | 25693 | 25662 | **RNA**→ rpmE (PRK00019)Ribosomal_L31 (pfam01197)→ Tdk (COG1435)PRK04296 (PRK04296)→ |
| env-1022 | JGI24505J29691_1082918 | + | 38 | 68 | **RNA**→ rpmE (PRK00019)Ribosomal_L31 (pfam01197)→ |
| Bnd-1-1 | NZ_HG964403.1 | + | 361882 | 361912 | **RNA**→ rpmE (PRK00019)Ribosomal_L31 (pfam01197)→ Tdk (COG1435)PRK04296 (PRK04296)→ |
| | | | | | PrfA (COG0216)prfA (PRK00591)→ HemK (COG2890)RF_mod_PrmC (TIGR03534)→ Spore_II_R (pfam0 |
| Lbo-1-1 | NZ_JPVR01000063.1 | + | 12881 | 12911 | **RNA**→ rpmE (PRK00019)Ribosomal_L31 (pfam01197)→ |
| env-1023 | JGI25151J46595_10149639 | + | 515 | 545 | **RNA**→ |
| Lsp-8-1 | NZ_KB933398.1 | + | 1093231 | 1093261 | **RNA**→ rpmE (PRK00019)Ribosomal_L31 (pfam01197)→ |
| Lsp-5-1 | NZ_JPDM01000191.1 | - | 6944 | 6914 | **RNA**→ rpmE (PRK00019)Ribosomal_L31 (pfam01197)→ |
| env-1024 | JGI25151J46595_10018623 | - | 794 | 764 | **RNA**→ rpmE (PRK00019)Ribosomal_L31 (pfam01197)→ |
| Bsp-3-1 | NZ_AAXV01000038.1 | - | 28491 | 28461 | **RNA**→ rpmE (PRK00019)Ribosomal_L31 (pfam01197)→ rpmE (PRK00019)Ribosomal_L31 (pfam01197)– |
| Lsp-2-1 | NZ_JSZM01000017.1 | + | 118321 | 118351 | **RNA**→ rpmE (PRK00019)Ribosomal_L31 (pfam01197)→ |
| Lfu-3-1 | NZ_AYMK01000007.1 | - | 69241 | 69211 | **RNA**→ rpmE (PRK00019)Ribosomal_L31 (pfam01197)→ |
| Lsp-5-2 | NZ_JMMU01000037.1 | + | 118169 | 118199 | **RNA**→ rpmE (PRK00019)Ribosomal_L31 (pfam01197)→ |
| Lfu-2-1 | NZ_JRBA01000001.1 | + | 118968 | 118998 | **RNA**→ rpmE (PRK00019)Ribosomal_L31 (pfam01197)→ |
| Lfu-1-1 | NZ_ADJR01000052.1 | + | 46794 | 46824 | **RNA**→ rpmE (PRK00019)Ribosomal_L31 (pfam01197)→ rpmE (PRK00019)Ribosomal_L31 (pfam01197)– |
| Lfu-4-1 | NZ_AMQZ01000018.1 | - | 70817 | 70787 | **RNA**→ rpmE (PRK00019)Ribosomal_L31 (pfam01197)→ |
| Lva-1-1 | NZ_CP006837.1 | + | 564657 | 564687 | **RNA**→ rpmE (PRK00019)Ribosomal_L31 (pfam01197)→ |
| Lsp-7-1 | NZ_AUOZ01000018.1 | + | 205124 | 205154 | **RNA**→ rpmE (PRK00019)Ribosomal_L31 (pfam01197)→ |
| env-1025 | JGI25151J46595_10124138 | - | 251 | 221 | **RNA**→ rpmE (PRK00019)Ribosomal_L31 (pfam01197)→ |
| Lxy-1-1 | NZ_LFXJ01000010.1 | + | 171030 | 171060 | **RNA**→ rpmE (PRK00019)Ribosomal_L31 (pfam01197)→ |
| Lsp-6-1 | NZ_KK037167.1 | + | 1006983 | 1007013 | **RNA**→ rpmE (PRK00019)Ribosomal_L31 (pfam01197)→ |
| Lsp-5-3 | NZ_JPDJ01000203.1 | + | 13735 | 13765 | **RNA**→ rpmE (PRK00019)Ribosomal_L31 (pfam01197)→ |
| Lsp-5-4 | NZ_JPDK01000128.1 | + | 109979 | 110009 | **RNA**→ rpmE (PRK00019)Ribosomal_L31 (pfam01197)→ |
| Lsp-5-5 | NZ_JPDL01000110.1 | + | 13742 | 13772 | **RNA**→ rpmE (PRK00019)Ribosomal_L31 (pfam01197)→ |
| Lma-2-1 | NZ_JPVQ01000001.1 | + | 254351 | 254381 | **RNA**→ rpmE (PRK00019)Ribosomal_L31 (pfam01197)→ |
| Lsp-4-1 | NZ_LDUJ01000035.1 | + | 13772 | 13802 | **RNA**→ rpmE (PRK00019)Ribosomal_L31 (pfam01197)→ |
| Lod-1-1 | NZ_JPVP01000053.1 | - | 366 | 336 | **RNA**→ rpmE (PRK00019)Ribosomal_L31 (pfam01197)→ |
| Ssi-1-1 | NC_018065.1 | - | 3510116 | 3510086 | **RNA**→ rpmE (PRK00019)Ribosomal_L31 (pfam01197)→ |
| Bis-1-1 | NZ_AMCK01000003.1 | - | 71858 | 71828 | **RNA**→ rpmE (PRK00019)Ribosomal_L31 (pfam01197)→ rpmE (PRK00019)Ribosomal_L31 (pfam01197)– |
| Lma-1-1 | NZ_JPVN01000002.1 | - | 107895 | 107865 | **RNA**→ rpmE (PRK00019)Ribosomal_L31 (pfam01197)→ |
| Khu-1-1 | NZ_AYTB01000002.1 | + | 218270 | 218300 | **RNA**→ rpmE (PRK00019)Ribosomal_L31 (pfam01197)→ |
| Kma-1-1 | NZ_HE610996.1 | - | 681420 | 681390 | **RNA**→ rpmE (PRK00019)Ribosomal_L31 (pfam01197)→ |
| Ksp-1-1 | NZ_HE578956.1 | + | 151508 | 151538 | **RNA**→ rpmE (PRK00019)Ribosomal_L31 (pfam01197)→ |
| env-1026 | GBANfinal_FHNL2OP03R1YNQ | - | 115 | 85 | **RNA**→ rpmE (PRK00019)rpmE2 (PRK01678)Ribosomal_L31 (pfam01197)→ |
| Lsi-1-1 | NZ_JPVO01000048.1 | + | 100905 | 100935 | **RNA**→ rpmE (PRK00019)Ribosomal_L31 (pfam01197)→ |
| Ksp-2-1 | NZ_HE611064.1 | - | 867109 | 867079 | **RNA**→ rpmE (PRK00019)Ribosomal_L31 (pfam01197)→ |
| env-1027 | JGI24024J18818_10110073 | + | 370 | 400 | **RNA**→ rpmE (PRK00019)Ribosomal_L31 (pfam01197)→ |
| env-1028 | JGI24024J18818_10018649 | - | 700 | 670 | **RNA**→ rpmE (PRK00019)Ribosomal_L31 (pfam01197)→ |
| env-1029 | JGI24024J18818_10129236 | - | 495 | 465 | **RNA**→ rpmE (PRK00019)Ribosomal_L31 (pfam01197)→ |
| env-1030 | JGI24024J18818_10222139 | + | 165 | 195 | **RNA**→ rpmE (PRK00019)Ribosomal_L31 (pfam01197)→ |
| Psp-34-1 | NZ_KE150421.1 | - | 1801271 | 1801241 | **RNA**→ rpmE (PRK00019)Ribosomal_L31 (pfam01197)→ |
| Psp-35-1 | NZ_AMGD01000100.1 | + | 11731 | 11761 | **RNA**→ rpmE (PRK00019)Ribosomal_L31 (pfam01197)→ |
| Psp-36-1 | NZ_ALJG01000332.1 | + | 26178 | 26208 | **RNA**→ rpmE (PRK00019)Ribosomal_L31 (pfam01197)→ hypo→ |
| env-1031 | JGI24024J18818_10369468 | - | 280 | 250 | **RNA**→ rpmE (PRK00019)Ribosomal_L31 (pfam01197)→ |

| | | | | | |
|---|---|---|---|---|---|
| env-1032 | JGI24024J18818_10113359 | - | 224 | 194 | **RNA**→ rpmE (PRK00019)Ribosomal_L31 (pfam01197)→ |
| Ssp-4-1 | NZ_AXVE01000011.1 | - | 120561 | 120531 | **RNA**→ rpmE (PRK00019)Ribosomal_L31 (pfam01197)→ |
| env-1033 | JGI24024J18818_10141399 | + | 497 | 527 | **RNA**→ rpmE (PRK00019)Ribosomal_L31 (pfam01197)→ |
| env-1034 | 2204743222 | - | 7004 | 6972 | **RNA**→ rpmE2 (PRK01678)Ribosomal_L31 (pfam01197)→ |
| env-1035 | 2157908317 | - | 7005 | 6973 | **RNA**→ rpmE2 (PRK01678)Ribosomal_L31 (pfam01197)→ |
| Lpo-1-1 | NZ_APCP01000064.1 | - | 31463 | 31431 | **RNA**→ rpmE2 (PRK01678)Ribosomal_L31 (pfam01197)→ |
| Lac-10-1 | NZ_BACS01000323.1 | + | 3193 | 3225 | **RNA**→ rpmE2 (PRK01678)Ribosomal_L31 (pfam01197)→ rpmE2 (PRK01678)Ribosomal_L31 (pfam01197 |
| env-1036 | 2204529503 | + | 210 | 242 | **RNA**→ rpmE2 (PRK01678)Ribosomal_L31 (pfam01197)→ |
| env-1037 | 2157700330 | - | 115 | 83 | **RNA**→ hypo→ |
| Lha-2-1 | NZ_AUEH01000024.1 | + | 33012 | 33044 | **RNA**→ rpmE2 (PRK01678)Ribosomal_L31 (pfam01197)→ |
| Lsh-1-1 | NZ_KI271609.1 | + | 4007 | 4039 | **RNA**→ rpmE2 (PRK01678)Ribosomal_L31 (pfam01197)→ |
| env-1038 | JGI25151J46595_10001160 | + | 404 | 435 | **RNA**→ rpmE2 (PRK01678)Ribosomal_L31 (pfam01197)→ |
| env-1039 | JGI25151J46595_10040961 | - | 97 | 66 | **RNA**→ rpmE (PRK00019)rpmE2 (PRK01678)Ribosomal_L31 (pfam01197)→ |
| env-1040 | JGI25151J46595_10160259 | + | 129 | 160 | **RNA**→ rpmE2 (PRK01678)Ribosomal_L31 (pfam01197)→ |
| Bsp-36-1 | NZ_CVPA01000001.1 | - | 2715811 | 2715778 | **RNA**→ rpmE (PRK00019)Ribosomal_L31 (pfam01197)→ Tdk (COG1435)PRK04296 (PRK04296)→ |
| Bam-14-1 | NZ_AWQY01000001.1 | + | 376504 | 376537 | **RNA**→ rpmE (PRK00019)Ribosomal_L31 (pfam01197)→ Tdk (COG1435)PRK04296 (PRK04296)→ |
| Bsp-65-1 | NZ_LDUN01000006.1 | - | 82154 | 82121 | **RNA**→ rpmE (PRK00019)Ribosomal_L31 (pfam01197)→ Tdk (COG1435)PRK04296 (PRK04296)→ |
| Bam-19-1 | NZ_KB206087.1 | + | 429369 | 429402 | **RNA**→ rpmE (PRK00019)Ribosomal_L31 (pfam01197)→ Tdk (COG1435)PRK04296 (PRK04296)→ |
| Bam-7-1 | NC_016784.1 | - | 3652345 | 3652312 | **RNA**→ rpmE (PRK00019)Ribosomal_L31 (pfam01197)→ Tdk (COG1435)PRK04296 (PRK04296)→ |
| Bam-12-1 | NC_020272.1 | + | 365167 | 365200 | **RNA**→ rpmE (PRK00019)Ribosomal_L31 (pfam01197)→ Tdk (COG1435)PRK04296 (PRK04296)→ |
| Bam-13-1 | NC_023073.1 | - | 3586155 | 3586122 | **RNA**→ rpmE (PRK00019)Ribosomal_L31 (pfam01197)→ Tdk (COG1435)PRK04296 (PRK04296)→ |
| Bsp-78-1 | NZ_CP010406.1 | + | 2434203 | 2434236 | **RNA**→ rpmE (PRK00019)Ribosomal_L31 (pfam01197)→ Tdk (COG1435)PRK04296 (PRK04296)→ |
| Bam-9-2 | NZ_CP010556.1 | - | 3527608 | 3527575 | **RNA**→ rpmE (PRK00019)Ribosomal_L31 (pfam01197)→ Tdk (COG1435)PRK04296 (PRK04296)→ |
| Bve-4-1 | NZ_CP006890.1 | - | 3733812 | 3733779 | **RNA**→ rpmE (PRK00019)Ribosomal_L31 (pfam01197)→ Tdk (COG1435)PRK04296 (PRK04296)→ |
| Bve-2-1 | NZ_CP007165.1 | - | 3688006 | 3687973 | **RNA**→ rpmE (PRK00019)Ribosomal_L31 (pfam01197)→ Tdk (COG1435)PRK04296 (PRK04296)→ |
| Bam-9-3 | NZ_CP011278.1 | - | 3524652 | 3524619 | **RNA**→ rpmE (PRK00019)Ribosomal_L31 (pfam01197)→ Tdk (COG1435)PRK04296 (PRK04296)→ |
| Bve-1-1 | NZ_CP011346.1 | - | 3740648 | 3740615 | **RNA**→ rpmE (PRK00019)Ribosomal_L31 (pfam01197)→ Tdk (COG1435)PRK04296 (PRK04296)→ |
| Bam-21-1 | NZ_AMPK01000004.1 | + | 201753 | 201786 | **RNA**→ rpmE (PRK00019)Ribosomal_L31 (pfam01197)→ Tdk (COG1435)PRK04296 (PRK04296)→ |
| Bsp-34-1 | NZ_LAKC01000010.1 | - | 628356 | 628323 | **RNA**→ rpmE (PRK00019)Ribosomal_L31 (pfam01197)→ Tdk (COG1435)PRK04296 (PRK04296)→ |
| Bsu-18-1 | NZ_AMCA01000303.1 | + | 4300 | 4333 | **RNA**→ rpmE (PRK00019)Ribosomal_L31 (pfam01197)→ Tdk (COG1435)PRK04296 (PRK04296)→ |
| Bmo-2-1 | NZ_ASJT01000105.1 | - | 9779 | 9746 | **RNA**→ rpmE (PRK00019)Ribosomal_L31 (pfam01197)→ Tdk (COG1435)PRK04296 (PRK04296)→ |
| Sai-1-2 | NZ_KE007193.1 | - | 641613 | 641580 | **RNA**→ rpmE (PRK00019)Ribosomal_L31 (pfam01197)→ Tdk (COG1435)PRK04296 (PRK04296)→ |
| Bam-4-1 | NC_017188.1 | - | 3565005 | 3564972 | **RNA**→ rpmE (PRK00019)Ribosomal_L31 (pfam01197)→ Tdk (COG1435)PRK04296 (PRK04296)→ |
| Bam-3-1 | NC_017190.1 | - | 3611825 | 3611792 | **RNA**→ rpmE (PRK00019)Ribosomal_L31 (pfam01197)→ Tdk (COG1435)PRK04296 (PRK04296)→ |
| Bam-5-1 | NC_017191.1 | - | 3566668 | 3566635 | **RNA**→ rpmE (PRK00019)Ribosomal_L31 (pfam01197)→ Tdk (COG1435)PRK04296 (PRK04296)→ |
| Bam-1-1 | NC_014551.1 | - | 3595990 | 3595957 | **RNA**→ rpmE (PRK00019)Ribosomal_L31 (pfam01197)→ Tdk (COG1435)PRK04296 (PRK04296)→ |
| Bsu-23-1 | NZ_AWXG01000034.1 | + | 431293 | 431326 | **RNA**→ rpmE (PRK00019)Ribosomal_L31 (pfam01197)→ Tdk (COG1435)PRK04296 (PRK04296)→ |
| Bsu-47-1 | NZ_CP007409.1 | - | 3624807 | 3624774 | **RNA**→ rpmE (PRK00019)Ribosomal_L31 (pfam01197)→ Tdk (COG1435)PRK04296 (PRK04296)→ |
| Bsp-27-1 | NZ_JXAJ01000001.1 | + | 580404 | 580437 | **RNA**→ rpmE (PRK00019)Ribosomal_L31 (pfam01197)→ Tdk (COG1435)PRK04296 (PRK04296)→ |
| Bsp-5-1 | NC_017743.1 | - | 3700469 | 3700436 | **RNA**→ rpmE (PRK00019)Ribosomal_L31 (pfam01197)→ Tdk (COG1435)PRK04296 (PRK04296)→ |
| Bva-1-1 | NZ_JH600237.1 | - | 355087 | 355054 | **RNA**→ rpmE (PRK00019)Ribosomal_L31 (pfam01197)→ Tdk (COG1435)PRK04296 (PRK04296)→ |
| Bsu-35-1 | NZ_AJHM01000011.1 | - | 146308 | 146275 | **RNA**→ rpmE (PRK00019)Ribosomal_L31 (pfam01197)→ Tdk (COG1435)PRK04296 (PRK04296)→ |
| Bsu-34-1 | NZ_AJHL01000019.1 | - | 135154 | 135121 | **RNA**→ rpmE (PRK00019)Ribosomal_L31 (pfam01197)→ Tdk (COG1435)PRK04296 (PRK04296)→ |
| Bsu-37-1 | NZ_AJHO01000010.1 | - | 629627 | 629594 | **RNA**→ rpmE (PRK00019)Ribosomal_L31 (pfam01197)→ Tdk (COG1435)PRK04296 (PRK04296)→ |
| Bsu-36-1 | NZ_AJHN01000013.1 | - | 368376 | 368343 | **RNA**→ rpmE (PRK00019)Ribosomal_L31 (pfam01197)→ Tdk (COG1435)PRK04296 (PRK04296)→ |
| Bsu-29-1 | NZ_AMXN01000001.1 | - | 386697 | 386664 | **RNA**→ rpmE (PRK00019)Ribosomal_L31 (pfam01197)→ Tdk (COG1435)PRK04296 (PRK04296)→ |
| Bsu-33-1 | NZ_AFSG01000020.1 | - | 365508 | 365475 | **RNA**→ rpmE (PRK00019)Ribosomal_L31 (pfam01197)→ Tdk (COG1435)PRK04296 (PRK04296)→ |
| Bsu-3-1 | NZ_ADGS01000016.1 | - | 106516 | 106483 | **RNA**→ rpmE (PRK00019)Ribosomal_L31 (pfam01197)→ rpmE (PRK00019)Ribosomal_L31 (pfam01197)– Tdk (COG1435)PRK04296 (PRK04296)→ Tdk (COG1435)PRK04296 (PRK04296)→ |
| Jma-2-1 | NZ_LILA01000013.1 | - | 203741 | 203708 | **RNA**→ rpmE (PRK00019)Ribosomal_L31 (pfam01197)→ Tdk (COG1435)PRK04296 (PRK04296)→ |
| Bsu-28-1 | NZ_AEHM01000061.1 | - | 125708 | 125675 | **RNA**→ rpmE (PRK00019)Ribosomal_L31 (pfam01197)→ Tdk (COG1435)PRK04296 (PRK04296)→ |
| Bsu-5-1 | NC_014479.1 | - | 3613005 | 3612972 | **RNA**→ rpmE (PRK00019)Ribosomal_L31 (pfam01197)→ Tdk (COG1435)PRK04296 (PRK04296)→ |

| | | | | | |
|---|---|---|---|---|---|
| Bsu-32-1 | NZ_CP010434.1 | - | 3612998 | 3612965 | **RNA**→ rpmE (PRK00019)Ribosomal_L31 (pfam01197)→ Tdk (COG1435)PRK04296 (PRK04296)→ |
| Bsu-4-1 | NC_016047.1 | - | 3790206 | 3790173 | **RNA**→ rpmE (PRK00019)Ribosomal_L31 (pfam01197)→ Tdk (COG1435)PRK04296 (PRK04296)→ |
| Bsp-32-1 | NZ_JPWY01000014.1 | + | 425314 | 425347 | **RNA**→ rpmE (PRK00019)Ribosomal_L31 (pfam01197)→ Tdk (COG1435)PRK04296 (PRK04296)→ |
| env-1041 | JGI11877J14254_119565 | - | 68 | 35 | **RNA**→ |
| env-1042 | GBANfinal_FHNL2OP03QJMDD | - | 298 | 265 | **RNA**→ rpmE (PRK00019)Ribosomal_L31 (pfam01197)→ |
| Bsp-72-1 | NZ_JXAQ01000010.1 | + | 225785 | 225818 | **RNA**→ rpmE (PRK00019)Ribosomal_L31 (pfam01197)→ Tdk (COG1435)PRK04296 (PRK04296)→ |
| Bsp-38-1 | NZ_AVPM01000060.1 | - | 644910 | 644877 | **RNA**→ rpmE (PRK00019)Ribosomal_L31 (pfam01197)→ Tdk (COG1435)PRK04296 (PRK04296)→ hypo |
| Bsp-46-1 | NZ_LFZT01000003.1 | + | 506089 | 506122 | **RNA**→ rpmE (PRK00019)Ribosomal_L31 (pfam01197)→ Tdk (COG1435)PRK04296 (PRK04296)→ |
| Bsu-48-1 | NZ_AGFW01000001.1 | + | 494266 | 494299 | **RNA**→ rpmE (PRK00019)Ribosomal_L31 (pfam01197)→ Tdk (COG1435)PRK04296 (PRK04296)→ |
| Bsu-15-1 | NZ_CAUC01000015.1 | - | 644806 | 644773 | **RNA**→ rpmE (PRK00019)Ribosomal_L31 (pfam01197)→ Tdk (COG1435)PRK04296 (PRK04296)→ |
| Bmu-1-1 | NZ_LGUH01000009.1 | - | 624647 | 624614 | **RNA**→ rpmE (PRK00019)Ribosomal_L31 (pfam01197)→ Tdk (COG1435)PRK04296 (PRK04296)→ |
| Bsu-6-1 | NC_000964.3 | - | 3803324 | 3803291 | **RNA**→ rpmE (PRK00019)Ribosomal_L31 (pfam01197)→ Tdk (COG1435)PRK04296 (PRK04296)→ |
| Bsu-30-1 | NC_017196.2 | - | 3675052 | 3675019 | **RNA**→ rpmE (PRK00019)Ribosomal_L31 (pfam01197)→ Tdk (COG1435)PRK04296 (PRK04296)→ |
| Bsu-10-1 | NZ_CM000490.1 | - | 3802359 | 3802326 | **RNA**→ rpmE (PRK00019)Ribosomal_L31 (pfam01197)→ Tdk (COG1435)PRK04296 (PRK04296)→ |
| Bsu-7-1 | NZ_CM000489.1 | - | 3775336 | 3775303 | **RNA**→ rpmE (PRK00019)Ribosomal_L31 (pfam01197)→ Tdk (COG1435)PRK04296 (PRK04296)→ |
| Bsu-41-1 | NZ_JH600164.1 | + | 17381 | 17414 | **RNA**→ rpmE (PRK00019)Ribosomal_L31 (pfam01197)→ Tdk (COG1435)PRK04296 (PRK04296)→ |
| Bsu-1-1 | NC_014976.1 | - | 1804697 | 1804664 | **RNA**→ rpmE (PRK00019)Ribosomal_L31 (pfam01197)→ Tdk (COG1435)PRK04296 (PRK04296)→ |
| Bsu-9-1 | NC_017195.1 | - | 3606494 | 3606461 | **RNA**→ rpmE (PRK00019)Ribosomal_L31 (pfam01197)→ Tdk (COG1435)PRK04296 (PRK04296)→ |
| Bsu-2-1 | NC_018520.1 | - | 3734766 | 3734733 | **RNA**→ rpmE (PRK00019)Ribosomal_L31 (pfam01197)→ rpmE (PRK00019)Ribosomal_L31 (pfam01197)– Tdk (COG1435)PRK04296 (PRK04296)→ |
| Bsu-43-1 | NC_019896.1 | + | 403519 | 403552 | **RNA**→ rpmE (PRK00019)Ribosomal_L31 (pfam01197)→ Tdk (COG1435)PRK04296 (PRK04296)→ |
| Bsu-27-1 | NC_020244.1 | - | 3642206 | 3642173 | **RNA**→ rpmE (PRK00019)Ribosomal_L31 (pfam01197)→ Tdk (COG1435)PRK04296 (PRK04296)→ |
| Bsu-42-1 | NC_020832.1 | - | 3605096 | 3605063 | **RNA**→ rpmE (PRK00019)Ribosomal_L31 (pfam01197)→ Tdk (COG1435)PRK04296 (PRK04296)→ |
| Bsu-39-1 | NC_020507.1 | - | 3803329 | 3803296 | **RNA**→ rpmE (PRK00019)Ribosomal_L31 (pfam01197)→ Tdk (COG1435)PRK04296 (PRK04296)→ |
| Bsu-24-1 | NC_022898.1 | - | 3621179 | 3621146 | **RNA**→ rpmE (PRK00019)Ribosomal_L31 (pfam01197)→ Tdk (COG1435)PRK04296 (PRK04296)→ hypo |
| Bsu-14-1 | NZ_AP012496.1 | - | 3630878 | 3630845 | **RNA**→ rpmE (PRK00019)Ribosomal_L31 (pfam01197)→ Tdk (COG1435)PRK04296 (PRK04296)→ |
| Bsu-6-2 | NZ_CP010052.1 | - | 3803337 | 3803304 | **RNA**→ rpmE (PRK00019)Ribosomal_L31 (pfam01197)→ Tdk (COG1435)PRK04296 (PRK04296)→ |
| Bsu-38-1 | NZ_CP010314.1 | - | 3782820 | 3782787 | **RNA**→ rpmE (PRK00019)Ribosomal_L31 (pfam01197)→ Tdk (COG1435)PRK04296 (PRK04296)→ |
| Bsp-97-1 | NZ_CP010014.1 | - | 3439669 | 3439636 | **RNA**→ rpmE (PRK00019)Ribosomal_L31 (pfam01197)→ Tdk (COG1435)PRK04296 (PRK04296)→ |
| Bsu-44-1 | NZ_CP007800.1 | - | 3776092 | 3776059 | **RNA**→ rpmE (PRK00019)Ribosomal_L31 (pfam01197)→ Tdk (COG1435)PRK04296 (PRK04296)→ |
| Bsu-40-1 | NZ_CP008698.1 | - | 3781363 | 3781330 | **RNA**→ rpmE (PRK00019)Ribosomal_L31 (pfam01197)→ Tdk (COG1435)PRK04296 (PRK04296)→ |
| Bsp-12-1 | NZ_LN680001.1 | - | 3821333 | 3821300 | **RNA**→ rpmE (PRK00019)Ribosomal_L31 (pfam01197)→ Tdk (COG1435)PRK04296 (PRK04296)→ |
| Bsu-12-1 | NZ_LN649259.1 | - | 3839370 | 3839337 | **RNA**→ rpmE (PRK00019)Ribosomal_L31 (pfam01197)→ Tdk (COG1435)PRK04296 (PRK04296)→ |
| Bsu-17-1 | NZ_CP007173.1 | - | 3594029 | 3593996 | **RNA**→ rpmE (PRK00019)Ribosomal_L31 (pfam01197)→ Tdk (COG1435)PRK04296 (PRK04296)→ |
| Bsp-67-1 | NZ_CP011101.1 | - | 3656783 | 3656750 | **RNA**→ rpmE (PRK00019)Ribosomal_L31 (pfam01197)→ Tdk (COG1435)PRK04296 (PRK04296)→ |
| Bsu-19-1 | NZ_CP011115.1 | - | 3803352 | 3803319 | **RNA**→ rpmE (PRK00019)Ribosomal_L31 (pfam01197)→ Tdk (COG1435)PRK04296 (PRK04296)→ |
| Bsu-8-1 | NZ_CM000488.1 | - | 3802318 | 3802285 | **RNA**→ rpmE (PRK00019)Ribosomal_L31 (pfam01197)→ Tdk (COG1435)PRK04296 (PRK04296)→ |
| Bsu-26-1 | NZ_ANIP01000016.1 | + | 8075 | 8108 | **RNA**→ rpmE (PRK00019)Ribosomal_L31 (pfam01197)→ Tdk (COG1435)PRK04296 (PRK04296)→ |
| Bsu-22-1 | NZ_AQGR01000122.1 | + | 80257 | 80290 | **RNA**→ rpmE (PRK00019)Ribosomal_L31 (pfam01197)→ Tdk (COG1435)PRK04296 (PRK04296)→ |
| Psp-40-1 | NZ_AVOF01000755.1 | + | 314 | 347 | **RNA**→ rpmE (PRK00019)Ribosomal_L31 (pfam01197)→ rpmE (PRK00019)Ribosomal_L31 (pfam01197)– rpmE (PRK00019)Ribosomal_L31 (pfam01197)→ hypo→ |
| Bsu-46-1 | NZ_APMW01000035.1 | + | 8558 | 8591 | **RNA**→ rpmE (PRK00019)Ribosomal_L31 (pfam01197)→ Tdk (COG1435)PRK04296 (PRK04296)→ |
| Bsu-45-1 | NZ_APMX01000221.1 | + | 12037 | 12070 | **RNA**→ rpmE (PRK00019)Ribosomal_L31 (pfam01197)→ Tdk (COG1435)PRK04296 (PRK04296)→ |
| Bsu-25-1 | NZ_AZQS01000001.1 | + | 419264 | 419297 | **RNA**→ rpmE (PRK00019)Ribosomal_L31 (pfam01197)→ Tdk (COG1435)PRK04296 (PRK04296)→ |
| Bsu-20-1 | NZ_AOTY01000001.1 | - | 624810 | 624777 | **RNA**→ rpmE (PRK00019)Ribosomal_L31 (pfam01197)→ Tdk (COG1435)PRK04296 (PRK04296)→ hypo |
| Bsu-21-1 | NZ_BALZ01000225.1 | - | 36042 | 36009 | **RNA**→ rpmE (PRK00019)Ribosomal_L31 (pfam01197)→ Tdk (COG1435)PRK04296 (PRK04296)→ |
| Bte-1-1 | NZ_AYTO01000009.1 | - | 97581 | 97548 | **RNA**→ rpmE (PRK00019)Ribosomal_L31 (pfam01197)→ Tdk (COG1435)PRK04296 (PRK04296)→ |
| Bsi-1-1 | NZ_LAGT01000025.1 | - | 10366 | 10333 | **RNA**→ rpmE (PRK00019)Ribosomal_L31 (pfam01197)→ Tdk (COG1435)PRK04296 (PRK04296)→ |
| Bsi-2-1 | NZ_AJVF01000039.1 | + | 357272 | 357305 | **RNA**→ rpmE (PRK00019)Ribosomal_L31 (pfam01197)→ Tdk (COG1435)PRK04296 (PRK04296)→ |
| Bsp-59-1 | NZ_KQ236689.1 | - | 1238300 | 1238267 | **RNA**→ rpmE (PRK00019)Ribosomal_L31 (pfam01197)→ Tdk (COG1435)PRK04296 (PRK04296)→ |
| Bat-8-1 | NZ_AEFU01000001.1 | + | 154389 | 154422 | **RNA**→ rpmE (PRK00019)Ribosomal_L31 (pfam01197)→ Tdk (COG1435)PRK04296 (PRK04296)→ |
| Bat-7-1 | NZ_AEFM01000002.1 | + | 154389 | 154422 | **RNA**→ rpmE (PRK00019)Ribosomal_L31 (pfam01197)→ Tdk (COG1435)PRK04296 (PRK04296)→ |
| Bat-6-1 | NZ_AEFW01000008.1 | + | 154400 | 154433 | **RNA**→ rpmE (PRK00019)Ribosomal_L31 (pfam01197)→ Tdk (COG1435)PRK04296 (PRK04296)→ |

| | | | | | |
|---|---|---|---|---|---|
| Bat-5-1 | NZ_AEFV01000008.1 | - | 321837 | 321804 | RNA→ rpmE (PRK00019)Ribosomal_L31 (pfam01197)→ Tdk (COG1435)PRK04296 (PRK04296)→ |
| Bat-1-1 | NC_014639.1 | - | 3225185 | 3225152 | RNA→ rpmE (PRK00019)Ribosomal_L31 (pfam01197)→ Tdk (COG1435)PRK04296 (PRK04296)→ |
| Bat-14-1 | NZ_AEFR01000017.1 | - | 96226 | 96193 | RNA→ rpmE (PRK00019)Ribosomal_L31 (pfam01197)→ Tdk (COG1435)PRK04296 (PRK04296)→ |
| Bat-3-1 | NZ_AEFS01000014.1 | + | 412950 | 412983 | RNA→ rpmE (PRK00019)Ribosomal_L31 (pfam01197)→ Tdk (COG1435)PRK04296 (PRK04296)→ |
| Bat-4-1 | NZ_AEFT01000005.1 | + | 154391 | 154424 | RNA→ rpmE (PRK00019)Ribosomal_L31 (pfam01197)→ Tdk (COG1435)PRK04296 (PRK04296)→ |
| Bat-9-1 | NZ_AEFX01000018.1 | - | 116510 | 116477 | RNA→ rpmE (PRK00019)Ribosomal_L31 (pfam01197)→ Tdk (COG1435)PRK04296 (PRK04296)→ |
| Bat-13-1 | NZ_AEFQ01000015.1 | - | 96075 | 96042 | RNA→ rpmE (PRK00019)Ribosomal_L31 (pfam01197)→ Tdk (COG1435)PRK04296 (PRK04296)→ |
| Bat-12-1 | NZ_AEFP01000005.1 | + | 154382 | 154415 | RNA→ rpmE (PRK00019)Ribosomal_L31 (pfam01197)→ Tdk (COG1435)PRK04296 (PRK04296)→ |
| Bat-10-1 | NZ_AEFY01000001.1 | - | 7160 | 7127 | RNA→ rpmE (PRK00019)Ribosomal_L31 (pfam01197)→ Tdk (COG1435)PRK04296 (PRK04296)→ |
| Bat-16-1 | NZ_AEFO01000028.1 | - | 116543 | 116510 | RNA→ rpmE (PRK00019)Ribosomal_L31 (pfam01197)→ Tdk (COG1435)PRK04296 (PRK04296)→ |
| Bat-2-1 | NZ_CP010778.1 | - | 3150625 | 3150592 | RNA→ rpmE (PRK00019)Ribosomal_L31 (pfam01197)→ Tdk (COG1435)PRK04296 (PRK04296)→ |
| Bat-17-1 | NZ_CP007640.1 | - | 3150622 | 3150589 | RNA→ rpmE (PRK00019)Ribosomal_L31 (pfam01197)→ Tdk (COG1435)PRK04296 (PRK04296)→ |
| Bsu-31-1 | NZ_KN049968.1 | + | 491770 | 491803 | RNA→ rpmE (PRK00019)Ribosomal_L31 (pfam01197)→ Tdk (COG1435)PRK04296 (PRK04296)→ |
| Bat-11-1 | NZ_AJRJ01000028.1 | + | 60488 | 60521 | RNA→ rpmE (PRK00019)Ribosomal_L31 (pfam01197)→ Tdk (COG1435)PRK04296 (PRK04296)→ |
| Bat-15-1 | NZ_CP011802.1 | - | 3582239 | 3582206 | RNA→ rpmE (PRK00019)Ribosomal_L31 (pfam01197)→ Tdk (COG1435)PRK04296 (PRK04296)→ |
| Gsp-3-1 | NC_012793.1 | - | 3352755 | 3352724 | RNA→ rpmE (PRK00019)Ribosomal_L31 (pfam01197)→ |
| Pmu-2-1 | NC_015690.1 | - | 8332068 | 8332036 | RNA→ rpmE (PRK00019)Ribosomal_L31 (pfam01197)→ rpmE (PRK00019)Ribosomal_L31 (pfam01197)→ rpmE (PRK00019)Ribosomal_L31 (pfam01197)→ hypo→ |
| Pmu-1-1 | NC_016935.1 | - | 8401920 | 8401888 | RNA→ rpmE (PRK00019)Ribosomal_L31 (pfam01197)→ hypo→ |
| Pmu-3-1 | NC_017672.3 | - | 8491650 | 8491618 | RNA→ rpmE (PRK00019)Ribosomal_L31 (pfam01197)→ hypo→ |
| Pty-1-1 | NZ_JNVM01000021.1 | + | 226105 | 226137 | RNA→ rpmE (PRK00019)Ribosomal_L31 (pfam01197)→ hypo→ hypo→ hypo→ hypo→ hypo→ |
| Pel-1-1 | NZ_AFHW01000134.1 | + | 17241 | 17273 | RNA→ rpmE (PRK00019)Ribosomal_L31 (pfam01197)→ rpmE (PRK00019)Ribosomal_L31 (pfam01197)→ hypo→ hypo→ hypo→ hypo→ hypo→ hypo→ hypo→ hypo→ hypo→ |
| Peh-1-1 | NZ_JFHX01000117.1 | - | 6087 | 6055 | RNA→ rpmE (PRK00019)Ribosomal_L31 (pfam01197)→ hypo→ |
| Pgi-1-1 | NZ_KB905580.1 | + | 47463 | 47495 | RNA→ rpmE (PRK00019)Ribosomal_L31 (pfam01197)→ hypo→ hypo→ hypo→ hypo→ hypo→ hypo→ |
| Psp-29-1 | NZ_JMLS01000043.1 | - | 6126 | 6094 | RNA→ rpmE (PRK00019)Ribosomal_L31 (pfam01197)→ |
| Nma-1-1 | NZ_CAVG010000025.1 | - | 41899 | 41870 | RNA→ rpmE (PRK00019)Ribosomal_L31 (pfam01197)→ |
| env-1043 | JGI24731J21663_1114416 | + | 271 | 291 | RNA→ yfiA (TIGR00741)→ |
| Bte-2-1 | NZ_LN851186.1 | - | 3875235 | 3875202 | RNA→ rpmE (PRK00019)Ribosomal_L31 (pfam01197)→ |
| Lga-13-1 | NZ_AICN01000021.1 | - | 56274 | 56241 | RNA→ rpmE2 (PRK01678)Ribosomal_L31 (pfam01197)→ rpmE2 (PRK01678)Ribosomal_L31 (pfam01197)→ sortase_fam (TIGR01076)Sortase_A (cd06165)→ sortase_fam (TIGR01076)Sortase_A (cd06165)→ |
| env-1044 | Subgingival_plaque_LANL_C11222194 | - | 533 | 502 | RNA→ rpmE (PRK00019)Ribosomal_L31 (pfam01197)→ PRK05571 (PRK05571)LacAB_rpiB (pfam02502)→ |
| env-1045 | Supragingival_plaque_LANL_scaffold_212735 | - | 387 | 356 | RNA→ rpmE (PRK00019)Ribosomal_L31 (pfam01197)→ PRK05571 (PRK05571)LacAB_rpiB (pfam02502)→ |
| env-1046 | SRS015215_WUGC_scaffold_60585 | + | 10090 | 10121 | RNA→ rpmE (PRK00019)Ribosomal_L31 (pfam01197)→ PRK05571 (PRK05571)LacAB_rpiB (pfam02502)→ GT_WecA_like (cd06853)mraY (PRK00108)→ |
| env-1047 | SRS058808_LANL_scaffold_75639 | - | 12573 | 12542 | RNA→ rpmE (PRK00019)Ribosomal_L31 (pfam01197)→ PRK05571 (PRK05571)LacAB_rpiB (pfam02502)→ GT_WecA_like (cd06853)mraY (PRK00108)→ |
| env-1048 | SRS014691_C2747958 | - | 181 | 150 | RNA→ rpmE (PRK00019)Ribosomal_L31 (pfam01197)→ |
| env-1049 | SRS016331_C3935637 | + | 51 | 82 | RNA→ rpmE (PRK00019)Ribosomal_L31 (pfam01197)→ |
| env-1050 | AUXO010054219.1 | + | 1093 | 1119 | RNA→ rpmE (PRK00019)Ribosomal_L31 (pfam01197)→ |
| env-1051 | AUXO011270623.1 | - | 268 | 242 | RNA→ rpmE (PRK00019)Ribosomal_L31 (pfam01197)→ |
| env-1052 | AUXO013500089.1 | - | 2645 | 2619 | RNA→ rpmE (PRK00019)Ribosomal_L31 (pfam01197)→ PRK05571 (PRK05571)LacAB_rpiB (pfam02502)→ hypo→ PRK11649 (PRK11649)Peptidase_M23 (pfam01551)→ PLN02662 (PLN02662)→ MreB_like (cd10225)PRK13930 (PRK13930)→ |
| env-1053 | AUXO016543257.1 | + | 22 | 48 | RNA→ rpmE (PRK00019)Ribosomal_L31 (pfam01197)→ |
| env-1054 | RUMENNODE_4189451_1 | - | 803 | 777 | RNA→ rpmE (PRK00019)Ribosomal_L31 (pfam01197)→ PRK05571 (PRK05571)LacAB_rpiB (pfam02502)→ |
| env-1055 | AUXO015505626.1 | - | 271 | 245 | RNA→ rpmE (PRK00019)Ribosomal_L31 (pfam01197)→ |
| env-1056 | AUXO010144317.1 | - | 519 | 493 | RNA→ rpmE (PRK00019)Ribosomal_L31 (pfam01197)→ PRK05571 (PRK05571)LacAB_rpiB (pfam02502)→ |
| env-1057 | AUXO011567691.1 | + | 201 | 227 | RNA→ rpmE (PRK00019)Ribosomal_L31 (pfam01197)→ PRK05571 (PRK05571)LacAB_rpiB (pfam02502)→ |
| env-1058 | AUXO017972327.1 | - | 2127 | 2101 | RNA→ rpmE (PRK00019)Ribosomal_L31 (pfam01197)→ PRK05571 (PRK05571)LacAB_rpiB (pfam02502)→ |
| env-1059 | AUXO014943907.1 | + | 26 | 52 | RNA→ rpmE (PRK00019)Ribosomal_L31 (pfam01197)→ PRK05571 (PRK05571)LacAB_rpiB (pfam02502)→ |
| Esu-1-1 | NZ_KE136394.1 | - | 693694 | 693662 | RNA→ rpmE2 (PRK01678)Ribosomal_L31 (pfam01197)→ |
| Lno-1-1 | NZ_BAMN01000015.1 | + | 13011 | 13043 | RNA→ rpmE2 (PRK01678)Ribosomal_L31 (pfam01197)→ |
| env-1060 | 2157768806 | - | 3632 | 3600 | RNA→ rpmE2 (PRK01678)Ribosomal_L31 (pfam01197)→ |

| | | | | | |
|---|---|---|---|---|---|
| env-1061 | 2204671115 | - | 175 | 143 | **RNA**→ rpmE2 (PRK01678)Ribosomal_L31 (pfam01197)→ |
| env-1062 | 2204576607 | - | 346 | 314 | **RNA**→ rpmE2 (PRK01678)Ribosomal_L31 (pfam01197)→ |
| env-1063 | 2158343614 | + | 370 | 402 | **RNA**→ rpmE2 (PRK01678)Ribosomal_L31 (pfam01197)→ |
| Lhe-1-1 | NZ_JXRB02000018.1 | + | 19659 | 19691 | **RNA**→ rpmE2 (PRK01678)Ribosomal_L31 (pfam01197)→ |
| env-1064 | 2205034447 | + | 3 | 35 | **RNA**→ rpmE2 (PRK01678)Ribosomal_L31 (pfam01197)→ |
| env-1065 | 2157868829 | + | 3 | 35 | **RNA**→ rpmE2 (PRK01678)Ribosomal_L31 (pfam01197)→ |
| env-1066 | 2157842730 | + | 168 | 200 | **RNA**→ rpmE2 (PRK01678)Ribosomal_L31 (pfam01197)→ |
| Lfa-3-1 | NZ_GL575016.1 | + | 462497 | 462529 | **RNA**→ rpmE2 (PRK01678)Ribosomal_L31 (pfam01197)→ |
| env-1067 | 2205326546 | - | 113 | 81 | **RNA**→ RpmE (COG0254)rpmE (PRK00019)rpmE2 (PRK01678)→ |
| env-1068 | 2205768956 | - | 49 | 17 | **RNA**→ hypo→ |
| env-1069 | 2157958570 | - | 130 | 98 | **RNA**→ rpmE2 (PRK01678)Ribosomal_L31 (pfam01197)→ |
| env-1070 | 2158360337 | - | 1026 | 994 | **RNA**→ rpmE2 (PRK01678)Ribosomal_L31 (pfam01197)→ |
| env-1071 | 2158215248 | + | 1955 | 1987 | **RNA**→ rpmE2 (PRK01678)Ribosomal_L31 (pfam01197)→ |
| Lpa-40-1 | NZ_BAMH01000001.1 | + | 33535 | 33567 | **RNA**→ rpmE2 (PRK01678)Ribosomal_L31 (pfam01197)→ |
| env-1072 | 2204592145 | - | 130 | 98 | **RNA**→ rpmE2 (PRK01678)Ribosomal_L31 (pfam01197)→ |
| env-1073 | 2204759623 | - | 130 | 98 | **RNA**→ rpmE2 (PRK01678)Ribosomal_L31 (pfam01197)→ |
| env-1074 | 2204902668 | - | 219 | 185 | **RNA**→ rpmE2 (PRK01678)Ribosomal_L31 (pfam01197)→ |
| env-1075 | 2205863775 | - | 892 | 860 | **RNA**→ rpmE2 (PRK01678)Ribosomal_L31 (pfam01197)→ |
| env-1076 | 2205378811 | + | 115 | 147 | **RNA**→ |
| env-1077 | AUXO011544979.1 | + | 300 | 333 | **RNA**→ rpmE (PRK00019)Ribosomal_L31 (pfam01197)→ |
| env-1078 | HCE12Call500_c0130983 | - | 383 | 350 | **RNA**→ rpmE (PRK00019)Ribosomal_L31 (pfam01197)→ |
| env-1079 | AUXO016321813.1 | + | 31 | 59 | **RNA**→ rpmE (PRK00019)Ribosomal_L31 (pfam01197)→ PRK05571 (PRK05571)LacAB_rpiB (pfam02502)→ |
| env-1080 | AUXO010547256.1 | - | 1503 | 1473 | **RNA**→ rpmE (PRK00019)Ribosomal_L31 (pfam01197)→ hypo→ PRK05571 (PRK05571)LacAB_rpiB (pfam02502)→ |
| env-1081 | 3300000558_10011235 | - | 8705 | 8678 | **RNA**→ rpmE (PRK00019)Ribosomal_L31 (pfam01197)→ Tdk (COG1435)PRK04296 (PRK04296)→ TadA (COG0590)MafB19-deam (pfam14437)→ hypo→ hypo→ TktA (COG0021)PRK05899 (PRK05899)→ MnmG (COG0445)PRK05192 (PRK05192)→ RsmG (COG0357)gidB (PRK00107)→ BcsQ (COG1192)AAA_31 (pfam13614)→ ParB (smart00470)Spo0J (COG1475)→ DUF951 (pfam06107)→ |
| env-1082 | 3300000558_10025197 | + | 11494 | 11521 | **RNA**→ rpmE (PRK00019)Ribosomal_L31 (pfam01197)→ Tdk (COG1435)PRK04296 (PRK04296)→ |
| env-1083 | JGI24732J26686_1021304 | - | 562 | 535 | **RNA**→ rpmE (PRK00019)Ribosomal_L31 (pfam01197)→ Tdk (COG1435)PRK04296 (PRK04296)→ |
| env-1084 | 3300000558_11733789 | + | 20189 | 20216 | **RNA**→ rpmE (PRK00019)Ribosomal_L31 (pfam01197)→ Tdk (COG1435)PRK04296 (PRK04296)→ TadA (COG0590)MafB19-deam (pfam14437)→ hypo→ |
| env-1085 | 3300000568_10173220 | - | 1263 | 1236 | **RNA**→ rpmE (PRK00019)Ribosomal_L31 (pfam01197)→ Tdk (COG1435)PRK04296 (PRK04296)→ TadA (COG0590)MafB19-deam (pfam14437)→ |
| env-1086 | 3300001592_10282939 | + | 145 | 172 | **RNA**→ rpmE (PRK00019)Ribosomal_L31 (pfam01197)→ Tdk (COG1435)PRK04296 (PRK04296)→ |
| env-1087 | 3300001605_10004551 | - | 1637 | 1610 | **RNA**→ rpmE (PRK00019)Ribosomal_L31 (pfam01197)→ Tdk (COG1435)PRK04296 (PRK04296)→ TadA (COG0590)MafB19-deam (pfam14437)→ hypo→ hypo→ |
| env-1088 | C687J13896_1000025 | - | 8927 | 8900 | **RNA**→ rpmE (PRK00019)Ribosomal_L31 (pfam01197)→ Tdk (COG1435)PRK04296 (PRK04296)→ TadA (COG0590)MafB19-deam (pfam14437)→ hypo→ hypo→ TktA (COG0021)PRK05899 (PRK05899)→ MnmG (COG0445)PRK05192 (PRK05192)→ RsmG (COG0357)gidB (PRK00107)→ BcsQ (COG1192)AAA_31 (pfam13614)→ ParB (smart00470)Spo0J (COG1475)→ DUF951 (pfam06107)→ |
| env-1089 | C687J26661_1000263 | + | 11891 | 11918 | **RNA**→ rpmE (PRK00019)Ribosomal_L31 (pfam01197)→ Tdk (COG1435)PRK04296 (PRK04296)→ TadA (COG0590)MafB19-deam (pfam14437)→ hypo→ hypo→ TktA (COG0021)PRK05899 (PRK05899)→ MnmG (COG0445)PRK05192 (PRK05192)→ RsmG (COG0357)gidB (PRK00107)→ |
| env-1090 | C687J26609_1000080 | + | 30209 | 30236 | **RNA**→ rpmE (PRK00019)Ribosomal_L31 (pfam01197)→ Tdk (COG1435)PRK04296 (PRK04296)→ TadA (COG0590)MafB19-deam (pfam14437)→ hypo→ hypo→ TktA (COG0021)PRK05899 (PRK05899)→ MnmG (COG0445)PRK05192 (PRK05192)→ RsmG (COG0357)gidB (PRK00107)→ BcsQ (COG1192)AAA_31 (pfam13614)→ ParB (smart00470)Spo0J (COG1475)→ DUF951 (pfam06107)→ |
| env-1091 | AUXO014680939.1 | - | 290 | 261 | **RNA**→ rpmE (PRK00019)Ribosomal_L31 (pfam01197)→ |
| env-1092 | AUXO018499996.1 | + | 1229 | 1256 | **RNA**→ hypo→ |
| env-1093 | AUXO015654006.1 | - | 159 | 129 | **RNA**→ rpmE (PRK00019)rpmE2 (PRK01678)Ribosomal_L31 (pfam01197)→ |
| env-1094 | AUXO015665870.1 | - | 168 | 139 | **RNA**→ rpmE (PRK00019)Ribosomal_L31 (pfam01197)→ |
| env-1095 | scaffold116237_1_MH0011 | + | 11558 | 11589 | **RNA**→ rpmE (PRK00019)Ribosomal_L31 (pfam01197)→ |
| env-1096 | DLM006_scaffold1604_48 | - | 400 | 369 | **RNA**→ rpmE (PRK00019)Ribosomal_L31 (pfam01197)→ |

| | | | | | |
|---|---|---|---|---|---|
| env-1097 | DLF005_scaffold873_28 | - | 1819 | 1788 | **RNA**→ rpmE (PRK00019)Ribosomal_L31 (pfam01197)→ |
| env-1098 | scaffold3783_3_MH0012 | - | 1230 | 1199 | **RNA**→ rpmE (PRK00019)Ribosomal_L31 (pfam01197)→ |
| env-1099 | C8408919_1_O2.UC-16 | + | 1665 | 1696 | **RNA**→ rpmE (PRK00019)→ |
| env-1100 | scaffold3759_1_V1.UC-14 | + | 4949 | 4980 | **RNA**→ rpmE (PRK00019)Ribosomal_L31 (pfam01197)→ |
| env-1101 | AUXO018500014.1 | + | 475 | 506 | **RNA**→ rpmE (PRK00019)Ribosomal_L31 (pfam01197)→ |
| env-1102 | AUXO011648946.1 | - | 105 | 74 | **RNA**→ rpmE (PRK00019)Ribosomal_L31 (pfam01197)→ |
| env-1103 | SL_7KL_010_BRINE_10014548 | + | 2019 | 2047 | **RNA**→ rpmE (PRK00019)Ribosomal_L31 (pfam01197)→ Tdk (COG1435)PRK04296 (PRK04296)→ |
| | | | | | TadA (COG0590)MafB19-deam (pfam14437)→ hypo→ |
| env-1104 | SL_8KL_010_SED_10014681 | - | 2941 | 2913 | **RNA**→ rpmE (PRK00019)Ribosomal_L31 (pfam01197)→ Tdk (COG1435)PRK04296 (PRK04296)→ |
| | | | | | TadA (COG0590)MafB19-deam (pfam14437)→ hypo→ hypo→ |
| env-1105 | SL_8KL_010_SED_10058133 | + | 931 | 959 | **RNA**→ rpmE (PRK00019)Ribosomal_L31 (pfam01197)→ Tdk (COG1435)PRK04296 (PRK04296)→ |
| env-1106 | SL_5KL_010_BRINE_10000476 | - | 7556 | 7527 | **RNA**→ rpmE (PRK00019)Ribosomal_L31 (pfam01197)→ Tdk (COG1435)PRK04296 (PRK04296)→ |
| | | | | | TadA (COG0590)MafB19-deam (pfam14437)→ hypo→ hypo→ Cof (COG0561)Hydrolase_3 (pfam08282)→ |
| | | | | | MnmG (COG0445)PRK05192 (PRK05192)→ RsmG (COG0357)gidB (PRK00107)→ TehA (COG1275)→ hypo |
| env-1107 | scaffold18398_5_MH0031 | - | 1738 | 1709 | **RNA**→ rpmE (PRK00019)Ribosomal_L31 (pfam01197)→ PRK05571 (PRK05571)LacAB_rpiB (pfam02502)→ |
| | | | | | DUF1146 (pfam06612)→ PRK11649 (PRK11649)Peptidase_M23 (pfam01551)→ |
| env-1108 | AUXO015774907.1 | + | 144 | 170 | **RNA**→ rpmE (PRK00019)Ribosomal_L31 (pfam01197)→ |
| env-1109 | AUXO017297246.1 | + | 839 | 867 | **RNA**→ rpmE (PRK00019)Ribosomal_L31 (pfam01197)→ pssA (TIGR00473)CDP-OH_P_transf (pfam01066)→ |
| | | | | | PRK00723 (PRK00723)PS_Dcarbxylase (pfam02666)→ |
| | | | | | EpsL (COG4632)NAGPA (pfam09992)DUF5011 (pfam16403)→ |
| | | | | | EpsL (COG4632)NAGPA (pfam09992)DUF5011 (pfam16403)→ |
| env-1110 | SRS020869_Baylor_scaffold_16775 | - | 3733 | 3695 | **RNA**→ rpmE (PRK00019)Ribosomal_L31 (pfam01197)→ |
| env-1111 | SRS052697_LANL_scaffold_55776 | + | 28 | 66 | **RNA**→ rpmE (PRK00019)Ribosomal_L31 (pfam01197)→ |
| Agr-1-1 | NZ_JAFR01000008.1 | + | 156062 | 156094 | **RNA**→ rpmE2 (PRK01678)Ribosomal_L31 (pfam01197)→ Tdk (COG1435)PRK04296 (PRK04296)→ |
| | | | | | TadA (COG0590)MafB19-deam (pfam14437)→ hypo→ hypo→ ND5 (MTH00151)DUF1510 (pfam07423)→ |
| | | | | | TktA (COG0021)PRK05899 (PRK05899)→ MnmG (COG0445)PRK05192 (PRK05192)→ |
| | | | | | RsmG (COG0357)gidB (PRK00107)→ DUF951 (pfam06107)→ |
| env-1112 | AUXO012476243.1 | + | 239 | 265 | **RNA**→ rpmE (PRK00019)Ribosomal_L31 (pfam01197)→ |
| env-1113 | AUXO016581159.1 | + | 430 | 458 | **RNA**→ rpmE (PRK00019)Ribosomal_L31 (pfam01197)→ TadA (COG0590)MafB19-deam (pfam14437)→ |
| env-1114 | AUXO016692971.1 | + | 59 | 85 | **RNA**→ rpmE (PRK00019)Ribosomal_L31 (pfam01197)→ Tdk (COG1435)PRK04296 (PRK04296)→ |
| | | | | | TadA (COG0590)MafB19-deam (pfam14437)→ hypo→ |
| env-1115 | AUXO018402175.1 | + | 2437 | 2463 | **RNA**→ rpmE (PRK00019)Ribosomal_L31 (pfam01197)→ |
| env-1116 | AUXO016106023.1 | - | 236 | 204 | **RNA**→ rpmE (PRK00019)Ribosomal_L31 (pfam01197)→ |
| env-1117 | AUXO014725215.1 | + | 295 | 325 | **RNA**→ rpmE (PRK00019)Ribosomal_L31 (pfam01197)→ PRK05571 (PRK05571)LacAB_rpiB (pfam02502)→ |
| | | | | | rim_protein (TIGR01257)DUF1146 (pfam06612)→ PRK11649 (PRK11649)Peptidase_M23 (pfam01551)→ |
| env-1118 | SRS011586_WUGC_scaffold_12590 | - | 3042 | 3012 | **RNA**→ rpmE (PRK00019)Ribosomal_L31 (pfam01197)→ |
| env-1119 | AUXO015284327.1 | + | 113 | 143 | **RNA**→ rpmE (PRK00019)Ribosomal_L31 (pfam01197)→ |
| env-1120 | AUXO014723598.1 | - | 601 | 571 | **RNA**→ rpmE (PRK00019)Ribosomal_L31 (pfam01197)→ |
| env-1121 | 4491414.3_NODE_610 | + | 253 | 284 | **RNA**→ rpmE (PRK00019)Ribosomal_L31 (pfam01197)→ |
| env-1122 | NLF005_scaffold40809_5 | + | 841 | 872 | **RNA**→ rpmE (PRK00019)Ribosomal_L31 (pfam01197)→ |
| env-1123 | NOM007_scaffold19881_17 | - | 705 | 674 | **RNA**→ rpmE (PRK00019)Ribosomal_L31 (pfam01197)→ |
| env-1124 | SRS019267_WUGC_scaffold_19133 | + | 141 | 171 | **RNA**→ rpmE (PRK00019)Ribosomal_L31 (pfam01197)→ PRK05571 (PRK05571)LacAB_rpiB (pfam02502)→ |
| env-1125 | scaffold41612_1_MH0035 | - | 275 | 245 | **RNA**→ rpmE (PRK00019)Ribosomal_L31 (pfam01197)→ |
| env-1126 | scaffold10444_1_O2.UC-4 | - | 2270 | 2240 | **RNA**→ rpmE (PRK00019)Ribosomal_L31 (pfam01197)→ |
| env-1127 | UnmappedStool_Broad_scaffold_1068660 | - | 502 | 472 | **RNA**→ rpmE (PRK00019)Ribosomal_L31 (pfam01197)→ hypo→ |
| env-1128 | SRS016335_WUGC_scaffold_10606 | + | 18701 | 18731 | **RNA**→ rpmE (PRK00019)Ribosomal_L31 (pfam01197)→ hypo→ |
| | | | | | PRK05571 (PRK05571)LacAB_rpiB (pfam02502)→ DUF1146 (pfam06612)→ |
| | | | | | PRK11649 (PRK11649)Peptidase_M23 (pfam01551)→ |
| env-1129 | 4491401.3_NODE_20020 | - | 5413 | 5376 | **RNA**→ rpmE (PRK00019)Ribosomal_L31 (pfam01197)→ |
| env-1130 | 4491412.3_NODE_76939 | + | 15872 | 15909 | **RNA**→ rpmE (PRK00019)Ribosomal_L31 (pfam01197)→ |
| env-1131 | SRS049995_LANL_scaffold_7413 | + | 31138 | 31175 | **RNA**→ rpmE (PRK00019)Ribosomal_L31 (pfam01197)→ |
| env-1132 | AUXO016739573.1 | - | 186 | 155 | **RNA**→ rpmE (PRK00019)Ribosomal_L31 (pfam01197)→ |
| env-1133 | AUXO011888757.1 | - | 275 | 242 | **RNA**→ rpmE (PRK00019)Ribosomal_L31 (pfam01197)→ |
| env-1134 | JGI994J12354_10205797 | - | 625 | 593 | **RNA**→ rpmE (PRK00019)Ribosomal_L31 (pfam01197)→ PRK05571 (PRK05571)LacAB_rpiB (pfam02502)→ |

| | | | | | |
|---|---|---|---|---|---|
| env-1135 | 07_22_PECTIN_DNA_scaffold7405_9 | + | 1633 | 1665 | **RNA**→ rpmE (PRK00019)Ribosomal_L31 (pfam01197)→ |
| env-1136 | O8_19_POLYDEXT_DNA_scaffold8118_8 | + | 655 | 687 | **RNA**→ rpmE (PRK00019)Ribosomal_L31 (pfam01197)→ |
| env-1137 | scaffold70386_2_MH0028 | + | 1501 | 1538 | **RNA**→ rpmE (PRK00019)Ribosomal_L31 (pfam01197)→ |
| env-1138 | scaffold177066_1_MH0012 | - | 3152 | 3115 | **RNA**→ rpmE (PRK00019)Ribosomal_L31 (pfam01197)→ |
| env-1139 | scaffold63114_3_MH0038 | - | 266 | 229 | **RNA**→ rpmE (PRK00019)Ribosomal_L31 (pfam01197)→ |
| env-1140 | scaffold111032_6_MH0043 | - | 25214 | 25177 | **RNA**→ rpmE (PRK00019)Ribosomal_L31 (pfam01197)→ |
| env-1141 | 4491410.3_NODE_3015 | + | 6163 | 6200 | **RNA**→ rpmE (PRK00019)Ribosomal_L31 (pfam01197)→ |
| env-1142 | 4491417.3_NODE_443 | - | 12219 | 12182 | **RNA**→ rpmE (PRK00019)Ribosomal_L31 (pfam01197)→ |
| env-1143 | SRS015782_C3503917 | + | 101 | 138 | **RNA**→ rpmE (PRK00019)Ribosomal_L31 (pfam01197)→ |
| env-1144 | SRS016335_C5149495 | + | 101 | 138 | **RNA**→ rpmE (PRK00019)Ribosomal_L31 (pfam01197)→ |
| env-1145 | SRS048870_WUGC_scaffold_10582 | + | 47080 | 47117 | **RNA**→ rpmE (PRK00019)Ribosomal_L31 (pfam01197)→ |
| env-1146 | scaffold78818_1_MH0009 | + | 44 | 81 | **RNA**→ rpmE (PRK00019)Ribosomal_L31 (pfam01197)→ |
| env-1147 | scaffold73433_1_MH0066 | - | 11981 | 11944 | **RNA**→ rpmE (PRK00019)Ribosomal_L31 (pfam01197)→ |
| env-1148 | scaffold75138_2_MH0075 | - | 4600 | 4563 | **RNA**→ rpmE (PRK00019)Ribosomal_L31 (pfam01197)→ |
| env-1149 | scaffold77905_1_V1.CD-9 | + | 6094 | 6131 | **RNA**→ rpmE (PRK00019)Ribosomal_L31 (pfam01197)→ |
| env-1150 | 4491402.3_NODE_210 | + | 11945 | 11982 | **RNA**→ rpmE (PRK00019)Ribosomal_L31 (pfam01197)→ |
| env-1151 | 4491407.3_NODE_396 | - | 93635 | 93598 | **RNA**→ rpmE (PRK00019)Ribosomal_L31 (pfam01197)→ |
| env-1152 | 4491488.3_NODE_1475 | + | 52307 | 52344 | **RNA**→ rpmE (PRK00019)Ribosomal_L31 (pfam01197)→ |
| env-1153 | 4491490.3_NODE_508 | + | 123464 | 123501 | **RNA**→ rpmE (PRK00019)Ribosomal_L31 (pfam01197)→ |
| env-1154 | SRS016541_C1560402 | - | 510 | 473 | **RNA**→ rpmE (PRK00019)Ribosomal_L31 (pfam01197)→ |
| env-1155 | SRS011239_Baylor_scaffold_7795 | - | 1685 | 1648 | **RNA**→ rpmE (PRK00019)Ribosomal_L31 (pfam01197)→ |
| env-1156 | SRS013476_C3124393 | + | 101 | 138 | **RNA**→ rpmE (PRK00019)Ribosomal_L31 (pfam01197)→ |
| env-1157 | SRS013687_C2512595 | - | 510 | 473 | **RNA**→ rpmE (PRK00019)Ribosomal_L31 (pfam01197)→ |
| env-1158 | SRS014235_WUGC_scaffold_15445 | + | 6150 | 6187 | **RNA**→ rpmE (PRK00019)Ribosomal_L31 (pfam01197)→ |
| env-1159 | SRS016203_C1523113 | - | 510 | 473 | **RNA**→ rpmE (PRK00019)Ribosomal_L31 (pfam01197)→ |
| env-1160 | SRS019685_WUGC_scaffold_52356 | - | 2456 | 2419 | **RNA**→ rpmE (PRK00019)Ribosomal_L31 (pfam01197)→ |
| env-1161 | SRS022071_C3064241 | - | 4175 | 4138 | **RNA**→ rpmE (PRK00019)Ribosomal_L31 (pfam01197)→ |
| env-1162 | SRS024009_Baylor_scaffold_14978 | - | 11946 | 11909 | **RNA**→ rpmE (PRK00019)Ribosomal_L31 (pfam01197)→ |
| env-1163 | SRS065504_C3190398 | - | 326 | 289 | **RNA**→ rpmE (PRK00019)Ribosomal_L31 (pfam01197)→ |
| env-1164 | SRS014979_WUGC_scaffold_36099 | + | 2031 | 2068 | **RNA**→ rpmE (PRK00019)Ribosomal_L31 (pfam01197)→ |
| env-1165 | AUXO014604482.1 | - | 186 | 156 | **RNA**→ rpmE (PRK00019)Ribosomal_L31 (pfam01197)→ |
| env-1166 | SS_3KL_010_SOIL_10002557 | - | 6348 | 6319 | **RNA**→ rpmE (PRK00019)Ribosomal_L31 (pfam01197)→ Tdk (COG1435)PRK04296 (PRK04296)→ |
| | | | | | TadA (COG0590)MafB19-deam (pfam14437)→ hypo→ hypo→ Hydrolase_3 (pfam08282)HAD_Pase (cd07516)– |
| | | | | | MnmG (COG0445)PRK05192 (PRK05192)→ RsmG (COG0357)gidB (PRK00107)→ TrkG (COG0168)→ hypo |
| env-1167 | SL_3KL_010_SED_10143958 | + | 940 | 969 | **RNA**→ rpmE (PRK00019)Ribosomal_L31 (pfam01197)→ |
| env-1168 | AUXO013330472.1 | + | 709 | 739 | **RNA**→ rpmE (PRK00019)Ribosomal_L31 (pfam01197)→ |
| env-1169 | C687J13239_1000702 | + | 13944 | 13972 | **RNA**→ rpmE (PRK00019)Ribosomal_L31 (pfam01197)→ Tdk (COG1435)PRK04296 (PRK04296)→ |
| | | | | | TadA (COG0590)MafB19-deam (pfam14437)→ hypo→ hypo→ |
| env-1170 | SakLake120mDRAFT_c0000643 | - | 13148 | 13120 | **RNA**→ rpmE (PRK00019)Ribosomal_L31 (pfam01197)→ Tdk (COG1435)PRK04296 (PRK04296)→ |
| | | | | | TadA (COG0590)MafB19-deam (pfam14437)→ hypo→ hypo→ hypo→ TktA (COG0021)PRK05899 (PRK0589 |
| | | | | | MnmG (COG0445)PRK05192 (PRK05192)→ RsmG (COG0357)gidB (PRK00107)→ |
| | | | | | BcsQ (COG1192)AAA_31 (pfam13614)→ ParB (smart00470)Spo0J (COG1475)→ DUF951 (pfam06107)→ |
| env-1171 | BS_KBA_SWE12_21m_c10169119 | + | 346 | 374 | **RNA**→ rpmE (PRK00019)Ribosomal_L31 (pfam01197)→ |
| env-1172 | BS_KBA_SWE12_21m_c10534462 | + | 125 | 153 | **RNA**→ rpmE (PRK00019)Ribosomal_L31 (pfam01197)→ |
| env-1173 | BS_KBA_SWE12_21m_c10227724 | - | 46 | 18 | **RNA**→ |
| env-1174 | BS_KBA_SWE02_21m_10520892 | + | 149 | 177 | **RNA**→ rpmE (PRK00019)Ribosomal_L31 (pfam01197)→ |
| env-1175 | 3300000558_10171620 | - | 3674 | 3646 | **RNA**→ rpmE (PRK00019)Ribosomal_L31 (pfam01197)→ Tdk (COG1435)PRK04296 (PRK04296)→ |
| | | | | | TadA (COG0590)MafB19-deam (pfam14437)→ hypo→ RseC_MucC (pfam04246)→ |
| | | | | | TktA (COG0021)PRK05899 (PRK05899)→ |
| env-1176 | 3300001605_10461389 | - | 587 | 559 | **RNA**→ rpmE (PRK00019)Ribosomal_L31 (pfam01197)→ Tdk (COG1435)PRK04296 (PRK04296)→ |
| env-1177 | AUXO018660770.1 | + | 201 | 234 | **RNA**→ |
| env-1178 | DLM022_scaffold47491_2 | - | 2757 | 2724 | **RNA**→ rpmE (PRK00019)Ribosomal_L31 (pfam01197)→ |
| env-1179 | 4491423.3_NODE_55225 | + | 274 | 307 | **RNA**→ hypo→ |

| | | | | | |
|---|---|---|---|---|---|
| env-1180 | UnmappedStool_Broad_C252973095 | + | 238 | 271 | **RNA**→ rpmE (PRK00019)Ribosomal_L31 (pfam01197)→ |
| env-1181 | DOF002_scaffold67521_1 | - | 8868 | 8835 | **RNA**→ rpmE (PRK00019)Ribosomal_L31 (pfam01197)→ |
| env-1182 | scaffold4818_2_MH0033 | - | 6655 | 6624 | **RNA**→ rpmE (PRK00019)Ribosomal_L31 (pfam01197)→ |
| env-1183 | 4491487.3_NODE_36255 | - | 194 | 163 | **RNA**→ rpmE (PRK00019)Ribosomal_L31 (pfam01197)→ |
| env-1184 | UnmappedStool_Broad_C253824745 | + | 1657 | 1688 | **RNA**→ rpmE (PRK00019)Ribosomal_L31 (pfam01197)→ |
| env-1185 | SRS049995_LANL_scaffold_41479 | - | 17405 | 17374 | **RNA**→ rpmE (PRK00019)Ribosomal_L31 (pfam01197)→ |
| env-1186 | DLF005_scaffold22529_20 | - | 1910 | 1879 | **RNA**→ rpmE (PRK00019)Ribosomal_L31 (pfam01197)→ |
| env-1187 | scaffold55864_1_MH0024 | + | 736 | 767 | **RNA**→ rpmE (PRK00019)Ribosomal_L31 (pfam01197)→ |
| env-1188 | SL_3KL_010_SED_10386470 | - | 84 | 54 | **RNA**→ |
| env-1189 | AUXO016893802.1 | - | 348 | 317 | **RNA**→ rpmE (PRK00019)Ribosomal_L31 (pfam01197)→ |
| env-1190 | AUXO013205533.1 | - | 927 | 896 | **RNA**→ rpmE (PRK00019)Ribosomal_L31 (pfam01197)→ PRK05571 (PRK05571)LacAB_rpiB (pfam02502)→ DUF1146 (pfam06612)→ |
| env-1191 | scaffold110634_1_MH0033 | + | 575 | 607 | **RNA**→ rpmE (PRK00019)Ribosomal_L31 (pfam01197)→ PRK05571 (PRK05571)LacAB_rpiB (pfam02502)→ SpoIID (COG2385)SpoIID_LytB (TIGR02669)→ PRK11649 (PRK11649)Peptidase_M23 (pfam01551)→ |
| env-1192 | scaffold73146_1_MH0035 | - | 2482 | 2450 | **RNA**→ rpmE (PRK00019)Ribosomal_L31 (pfam01197)→ PRK05571 (PRK05571)LacAB_rpiB (pfam02502)→ SpoIID (COG2385)SpoIID_LytB (TIGR02669)→ PRK11649 (PRK11649)Peptidase_M23 (pfam01551)→ |
| env-1193 | UnmappedStool_Broad_C253558353 | - | 162 | 130 | **RNA**→ rpmE (PRK00019)Ribosomal_L31 (pfam01197)→ |
| env-1194 | DOM014_scaffold175_1 | + | 11760 | 11792 | **RNA**→ rpmE (PRK00019)Ribosomal_L31 (pfam01197)→ PRK05571 (PRK05571)LacAB_rpiB (pfam02502)→ |
| env-1195 | SRS049995_LANL_scaffold_25862 | - | 46739 | 46707 | **RNA**→ rpmE (PRK00019)Ribosomal_L31 (pfam01197)→ PRK05571 (PRK05571)LacAB_rpiB (pfam02502)→ SpoIID (COG2385)SpoIID_LytB (TIGR02669)→ PRK11649 (PRK11649)Peptidase_M23 (pfam01551)→ Tra8 (COG2826)SpoIIID (pfam12116)→ PRK13930 (PRK13930)MreB_Mbl (pfam06723)→ ECA_wecA (TIGR02380)GT_WecA_like (cd06853)→ CheY (COG0784)→ HATPase_c (smart00387)HATPase_c (pfam02518)→ |
| env-1196 | SRS078176_C2331252 | + | 1688 | 1720 | **RNA**→ rpmE (PRK00019)Ribosomal_L31 (pfam01197)→ PRK05571 (PRK05571)LacAB_rpiB (pfam02502)→ SpoIID (COG2385)SpoIID (pfam08486)→ |
| env-1197 | 3300000558_10030915 | - | 447 | 420 | **RNA**→ rpmE (PRK00019)Ribosomal_L31 (pfam01197)→ Tdk (COG1435)PRK04296 (PRK04296)→ |
| env-1198 | AUXO010231042.1 | + | 5787 | 5817 | **RNA**→ rpmE (PRK00019)Ribosomal_L31 (pfam01197)→ PRK05571 (PRK05571)LacAB_rpiB (pfam02502)→ hypo→ PRK11649 (PRK11649)Peptidase_M23 (pfam01551)→ MreB_like (cd10225)PRK13930 (PRK13930)→ ECA_wecA (TIGR02380)GT_WecA_like (cd06853)→ LytTR (smart00850)LytTR (pfam04397)→ CSP (smart00357)CspC (COG1278)→ yfiA (TIGR00741)RaiA (COG1544)Ribosom_S30AE_C (pfam16321)→ SecA (COG0653)PRK12904 (PRK12904)→ |
| env-1199 | AUXO016989467.1 | - | 1131 | 1100 | **RNA**→ rpmE (PRK00019)Ribosomal_L31 (pfam01197)→ PRK05571 (PRK05571)LacAB_rpiB (pfam02502)→ SpoIID (COG2385)SpoIID (pfam08486)→ |
| env-1200 | AUXO014231841.1 | + | 3562 | 3595 | **RNA**→ rpmE (PRK00019)Ribosomal_L31 (pfam01197)→ |
| env-1201 | AUXO017411865.1 | + | 2453 | 2484 | **RNA**→ rpmE (PRK00019)Ribosomal_L31 (pfam01197)→ PRK05571 (PRK05571)LacAB_rpiB (pfam02502)→ hypo→ ECA_wecA (TIGR02380)GT_WecA_like (cd06853)→ yfiA (TIGR00741)Ribosomal_S30AE (pfam02482)- |
| env-1202 | AUXO013834857.1 | - | 8878 | 8847 | **RNA**→ rpmE (PRK00019)Ribosomal_L31 (pfam01197)→ |
| env-1203 | 2227602434 | - | 1896 | 1866 | **RNA**→ rpmE (PRK00019)Ribosomal_L31 (pfam01197)→ PRK05571 (PRK05571)LacAB_rpiB (pfam02502)→ hypo→ Peptidase_M23 (pfam01551)→ SpoIIID (pfam12116)→ |
| env-1204 | JGI994J12354_10000780 | + | 359 | 389 | **RNA**→ rpmE (PRK00019)Ribosomal_L31 (pfam01197)→ PRK05571 (PRK05571)LacAB_rpiB (pfam02502)→ hypo→ Peptidase_M23 (pfam01551)→ SpoIIID (pfam12116)→ MreB_like (cd10225)PRK13930 (PRK13930)→ |
| env-1205 | IMNBL1_c0133606 | - | 538 | 508 | **RNA**→ rpmE (PRK00019)Ribosomal_L31 (pfam01197)→ PRK05571 (PRK05571)LacAB_rpiB (pfam02502)→ |
| env-1206 | AUXO015478559.1 | + | 273 | 302 | **RNA**→ rpmE (PRK00019)Ribosomal_L31 (pfam01197)→ |
| env-1207 | AUXO013760466.1 | - | 331 | 301 | **RNA**→ rpmE (PRK00019)Ribosomal_L31 (pfam01197)→ |
| env-1208 | 4491477.3_NODE_1076 | - | 3022 | 2991 | **RNA**→ rpmE (PRK00019)Ribosomal_L31 (pfam01197)→ |
| env-1209 | 3300001592_10522345 | - | 327 | 300 | **RNA**→ rpmE (PRK00019)Ribosomal_L31 (pfam01197)→ |
| env-1210 | AUXO013710643.1 | - | 2677 | 2647 | **RNA**→ rpmE (PRK00019)Ribosomal_L31 (pfam01197)→ PRK05571 (PRK05571)LacAB_rpiB (pfam02502)→ hypo→ SpoIID (COG2385)SpoIID_LytB (TIGR02669)→ PRK11649 (PRK11649)Peptidase_M23 (pfam01551)→ |
| env-1211 | AUXO013726310.1 | - | 252 | 222 | **RNA**→ rpmE (PRK00019)Ribosomal_L31 (pfam01197)→ |
| env-1212 | AUXO012175234.1 | + | 1448 | 1480 | **RNA**→ rpmE (PRK00019)Ribosomal_L31 (pfam01197)→ |
| env-1213 | AUXO013960360.1 | - | 83 | 52 | **RNA**→ rpmE (PRK00019)Ribosomal_L31 (pfam01197)→ |
| env-1214 | AUXO013939667.1 | + | 1121 | 1152 | **RNA**→ rpmE (PRK00019)Ribosomal_L31 (pfam01197)→ |
| env-1215 | RUMENNODE_3847397_1 | + | 357 | 387 | **RNA**→ rpmE (PRK00019)Ribosomal_L31 (pfam01197)→ |
| env-1216 | AUXO012691611.1 | + | 29 | 59 | **RNA**→ rpmE (PRK00019)Ribosomal_L31 (pfam01197)→ YcaP (COG2323)→ PRK10776 (PRK10776)Nudix_Hydrolase_1 (cd03674)→ hypo→ |

| | | | | | |
|---|---|---|---|---|---|
| env-1217 | AUXO011512029.1 | + | 159 | 189 | **RNA**→ hypo→ rpmE (PRK00019)Ribosomal_L31 (pfam01197)→ |
| env-1218 | O8_19_POLYDEXT_DNA_scaffold41145_1 | + | 5890 | 5919 | **RNA**→ rpmE (PRK00019)Ribosomal_L31 (pfam01197)→ YcaP (COG2323)→ Nudix_Hydrolase_1 (cd03674)IPP_isom_1 (TIGR02150)→ hypo→ |
| env-1219 | AUXO011585839.1 | - | 183 | 153 | **RNA**→ rpmE (PRK00019)Ribosomal_L31 (pfam01197)→ |
| env-1220 | AUXO012730652.1 | + | 843 | 873 | **RNA**→ rpmE (PRK00019)Ribosomal_L31 (pfam01197)→ |
| env-1221 | RUMENNODE_4178079_1 | - | 13408 | 13378 | **RNA**→ rpmE (PRK00019)Ribosomal_L31 (pfam01197)→ YcaP (COG2323)→ |
| env-1222 | SRS011134_C5183664 | - | 3140 | 3109 | **RNA**→ ←COG3415 (COG3415)HTH_28 (pfam13518) |
| env-1223 | SRS049959_WUGC_scaffold_60445 | + | 13512 | 13543 | **RNA**→ rpmE (PRK00019)Ribosomal_L31 (pfam01197)→ |
| env-1224 | AUXO015487162.1 | - | 138 | 107 | **RNA**→ rpmE (PRK00019)Ribosomal_L31 (pfam01197)→ |
| env-1225 | SRS049164_WUGC_scaffold_33709 | + | 1599 | 1630 | **RNA**→ rpmE (PRK00019)Ribosomal_L31 (pfam01197)→ |
| env-1226 | SRS016335_WUGC_scaffold_45869 | - | 8316 | 8285 | **RNA**→ rpmE (PRK00019)Ribosomal_L31 (pfam01197)→ |
| env-1227 | SRS047014_WUGC_scaffold_63392 | - | 774 | 743 | **RNA**→ rpmE (PRK00019)Ribosomal_L31 (pfam01197)→ |
| env-1228 | 08_07_CONTROL_DNA_scaffold13296_18 | + | 817 | 848 | **RNA**→ rpmE (PRK00019)Ribosomal_L31 (pfam01197)→ |
| env-1229 | 08_07_XOS_DNA_scaffold66974_1 | + | 19539 | 19570 | **RNA**→ rpmE (PRK00019)Ribosomal_L31 (pfam01197)→ |
| env-1230 | AUXO014710346.1 | - | 273 | 242 | **RNA**→ rpmE (PRK00019)Ribosomal_L31 (pfam01197)→ |
| env-1231 | AUXO017881809.1 | + | 781 | 812 | **RNA**→ |
| env-1232 | AUXO012078207.1 | - | 183 | 153 | **RNA**→ rpmE (PRK00019)Ribosomal_L31 (pfam01197)→ |
| env-1233 | SRS015663_WUGC_scaffold_1238 | - | 20838 | 20806 | **RNA**→ rpmE (PRK00019)Ribosomal_L31 (pfam01197)→ |
| env-1234 | AUXO013159041.1 | + | 29 | 59 | **RNA**→ rpmE (PRK00019)Ribosomal_L31 (pfam01197)→ YcaP (COG2323)→ |
| env-1235 | 07_22_CONTROL_DNA_scaffold85199_3 | + | 352 | 378 | **RNA**→ rpmE (PRK00019)Ribosomal_L31 (pfam01197)→ |
| env-1236 | AUXO012376566.1 | + | 1531 | 1559 | **RNA**→ rpmE (PRK00019)Ribosomal_L31 (pfam01197)→ Lactamase_B (smart00849)metallo-hydrolase-like_MBL-fold (cd07743)→ |
| env-1237 | AUXO015435427.1 | - | 1944 | 1915 | **RNA**→ rpmE (PRK00019)Ribosomal_L31 (pfam01197)→ PRK05571 (PRK05571)LacAB_rpiB (pfam02502)→ int_mem_ywzB (TIGR02327)DUF1146 (pfam06612)→ SpoIID (COG2385)SpoIID_LytB (TIGR02669)→ |
| env-1238 | JGI24708J26588_10236894 | - | 59 | 34 | **RNA**→ |
| env-1239 | SR_TTP_S3_1000105 | + | 34613 | 34642 | **RNA**→ rpmE (PRK00019)Ribosomal_L31 (pfam01197)→ Tdk (COG1435)PRK04296 (PRK04296)→ TadA (COG0590)MafB19-deam (pfam14437)→ hypo→ hypo→ hypo→ TktA (COG0021)PRK05899 (PRK0589 MnmG (COG0445)PRK05192 (PRK05192)→ RsmG (COG0357)gidB (PRK00107)→ BcsQ (COG1192)AAA_31 (pfam13614)→ ParB (smart00470)Spo0J (COG1475)→ DUF951 (pfam06107)→ |
| env-1240 | AUXO016960809.1 | - | 487 | 455 | **RNA**→ rpmE (PRK00019)Ribosomal_L31 (pfam01197)→ Pacs-1 (pfam10254)→ |
| env-1241 | AUXO016034446.1 | - | 1236 | 1204 | **RNA**→ rpmE (PRK00019)Ribosomal_L31 (pfam01197)→ Pacs-1 (pfam10254)→ HTH_XRE (smart00530)XRE (COG1476)→ hypo→ |
| env-1242 | AUXO011570238.1 | + | 4931 | 4963 | **RNA**→ |
| env-1243 | AUXO012487438.1 | + | 2243 | 2275 | **RNA**→ |
| env-1244 | AUXO010195109.1 | + | 4196 | 4228 | **RNA**→ rpmE (PRK00019)Ribosomal_L31 (pfam01197)→ |
| env-1245 | AUXO014081958.1 | + | 1464 | 1496 | **RNA**→ rpmE (PRK00019)Ribosomal_L31 (pfam01197)→ |
| env-1246 | NLF005_scaffold13642_7 | + | 92 | 122 | **RNA**→ rpmE (PRK00019)Ribosomal_L31 (pfam01197)→ PRK05571 (PRK05571)LacAB_rpiB (pfam02502)→ DUF1146 (pfam06612)→ SpoIID (COG2385)SpoIID_LytB (TIGR02669)→ PRK11649 (PRK11649)Peptidase_M23 (pfam01551)→ SpoIIID (pfam12116)→ MreB_like (cd10225)PRK13930 (PRK13930)→ |
| env-1247 | scaffold87272_2_MH0081 | - | 271 | 241 | **RNA**→ rpmE (PRK00019)Ribosomal_L31 (pfam01197)→ |
| env-1248 | AUXO018221316.1 | - | 119 | 89 | **RNA**→ rpmE (PRK00019)rpmE2 (PRK01678)Ribosomal_L31 (pfam01197)→ |
| env-1249 | scaffold1635_5_MH0082 | + | 249 | 278 | **RNA**→ rpmE (PRK00019)Ribosomal_L31 (pfam01197)→ |
| env-1250 | SR_TP_S2_1000079 | + | 14131 | 14157 | **RNA**→ rpmE (PRK00019)Ribosomal_L31 (pfam01197)→ |
| env-1251 | 3300001605_10172276 | + | 1019 | 1045 | **RNA**→ rpmE (PRK00019)Ribosomal_L31 (pfam01197)→ |
| env-1252 | AUXO018756927.1 | + | 1687 | 1717 | **RNA**→ rpmE (PRK00019)Ribosomal_L31 (pfam01197)→ |
| env-1253 | JGI24710J26742_10162176 | + | 368 | 399 | **RNA**→ rpmE (PRK00019)Ribosomal_L31 (pfam01197)→ |
| env-1254 | AUXO016786490.1 | + | 2740 | 2766 | **RNA**→ rpmE (PRK00019)Ribosomal_L31 (pfam01197)→ PRK05571 (PRK05571)LacAB_rpiB (pfam02502)→ |
| env-1255 | AUXO011123210.1 | - | 174 | 143 | **RNA**→ rpmE (PRK00019)Ribosomal_L31 (pfam01197)→ |
| env-1256 | AUXO015180417.1 | + | 171 | 201 | **RNA**→ rpmE (PRK00019)Ribosomal_L31 (pfam01197)→ YcaP (COG2323)→ |
| env-1257 | O8_19_POLYDEXT_DNA_scaffold85068_3 | + | 626 | 656 | **RNA**→ rpmE (PRK00019)Ribosomal_L31 (pfam01197)→ |
| env-1258 | scaffold4959_2_MH0065 | + | 497 | 523 | **RNA**→ rpmE (PRK00019)Ribosomal_L31 (pfam01197)→ |
| env-1259 | JGI24713J26584_10183766 | + | 158 | 188 | **RNA**→ rpmE (PRK00019)Ribosomal_L31 (pfam01197)→ |
| env-1260 | AUXO013227359.1 | - | 65 | 34 | **RNA**→ |

| | | | | | |
|---|---|---|---|---|---|
| env-1261 | AUXO010399693.1 | - | 1188 | 1160 | **RNA**→ rpmE (PRK00019)Ribosomal_L31 (pfam01197)→ PRK05571 (PRK05571)LacAB_rpiB (pfam02502)→ DUF1146 (pfam06612)→ hypo→ |
| env-1262 | AUXO012301092.1 | - | 266 | 236 | **RNA**→ rpmE (PRK00019)Ribosomal_L31 (pfam01197)→ |
| env-1263 | AUXO014076405.1 | + | 105 | 135 | **RNA**→ rpmE (PRK00019)Ribosomal_L31 (pfam01197)→ PRK05571 (PRK05571)LacAB_rpiB (pfam02502)→ rim_protein (TIGR01257)DUF1146 (pfam06612)→ PRK11649 (PRK11649)Peptidase_M23 (pfam01551)→ MreB_like (cd10225)PRK13930 (PRK13930)→ ECA_wecA (TIGR02380)GT_WecA_like (cd06853)→ LytTR (smart00850)LytTR (pfam04397)→ |
| env-1264 | SRMUA_GNVGR9Q01AT41D | + | 16 | 47 | **RNA**→ rpmE (PRK00019)Ribosomal_L31 (pfam01197)→ |
| env-1265 | AUXO010283157.1 | + | 10331 | 10362 | **RNA**→ rpmE (PRK00019)Ribosomal_L31 (pfam01197)→ PRK05571 (PRK05571)LacAB_rpiB (pfam02502)→ DUF1146 (pfam06612)→ PRK11649 (PRK11649)Peptidase_M23 (pfam01551)→ MreB_like (cd10225)PRK13930 (PRK13930)→ ECA_wecA (TIGR02380)GT_WecA_like (cd06853)→ LytTR (smart00850)LytTR (pfam04397)→ |
| env-1266 | AUXO014310563.1 | + | 270 | 301 | **RNA**→ rpmE (PRK00019)Ribosomal_L31 (pfam01197)→ PRK05571 (PRK05571)LacAB_rpiB (pfam02502)→ rim_protein (TIGR01257)DUF1146 (pfam06612)→ nlpD (PRK10871)Peptidase_M23 (pfam01551)→ MreB_like (cd10225)PRK13930 (PRK13930)→ ECA_wecA (TIGR02380)GT_WecA_like (cd06853)→ hypo→ |
| env-1267 | AUXO011380099.1 | + | 513 | 544 | **RNA**→ |
| env-1268 | AUXO014310801.1 | - | 4992 | 4964 | **RNA**→ rpmE (PRK00019)Ribosomal_L31 (pfam01197)→ PRK05571 (PRK05571)LacAB_rpiB (pfam02502)→ DUF1146 (pfam06612)→ PRK11649 (PRK11649)Peptidase_M23 (pfam01551)→ MreB_like (cd10225)PRK13930 (PRK13930)→ ECA_wecA (TIGR02380)GT_WecA_like (cd06853)→ yfiA (TIGR00741)RaiA (COG1544)Ribosom_S30AE_C (pfam16321)→ SecA (COG0653)PRK12904 (PRK12904 |
| env-1269 | AUXO017364526.1 | - | 66 | 35 | **RNA**→ |
| env-1270 | AUXO011848655.1 | + | 2629 | 2660 | **RNA**→ rpmE (PRK00019)Ribosomal_L31 (pfam01197)→ PRK05571 (PRK05571)LacAB_rpiB (pfam02502)→ DUF1146 (pfam06612)→ MreB_like (cd10225)PRK13930 (PRK13930)→ ECA_wecA (TIGR02380)GT_WecA_like (cd06853)→ |
| env-1271 | AUXO014937547.1 | - | 5403 | 5374 | **RNA**→ rpmE (PRK00019)Ribosomal_L31 (pfam01197)→ ycf1 (CHL00204)Amidoligase_2 (pfam12224)→ PRK05571 (PRK05571)LacAB_rpiB (pfam02502)→ DUF1146 (pfam06612)→ PRK11649 (PRK11649)Peptidase_M23 (pfam01551)→ MreB_like (cd10225)PRK13930 (PRK13930)→ ECA_wecA (TIGR02380)GT_WecA_like (cd06853)→ CheY (COG0784)→ |
| env-1272 | AUXO015447990.1 | - | 61 | 32 | **RNA**→ |
| env-1273 | AUXO011771154.1 | - | 1416 | 1383 | **RNA**→ rpmE (PRK00019)Ribosomal_L31 (pfam01197)→ Acetyltransf_1 (pfam00583)Acetyltransf_7 (pfam1350 hypo→ PRK05571 (PRK05571)LacAB_rpiB (pfam02502)→ |
| env-1274 | SRS013951_C2824150 | - | 311 | 281 | **RNA**→ rpmE (PRK00019)Ribosomal_L31 (pfam01197)→ |
| env-1275 | AUXO016081430.1 | + | 434 | 465 | **RNA**→ |
| env-1276 | AUXO016398220.1 | - | 197 | 166 | **RNA**→ rpmE (PRK00019)Ribosomal_L31 (pfam01197)→ |
| env-1277 | AUXO016483557.1 | + | 979 | 1009 | **RNA**→ rpmE (PRK00019)SGNH_hydrolase (cd00229)Ribosomal_L31 (pfam01197)Lipase_GDSL_2 (pfam13472 PRK05571 (PRK05571)LacAB_rpiB (pfam02502)→ hypo→ PRK11649 (PRK11649)Peptidase_M23 (pfam01551 |
| env-1278 | AUXO016479613.1 | + | 343 | 374 | **RNA**→ |
| env-1279 | AUXO013409902.1 | + | 482 | 514 | **RNA**→ rpmE (PRK00019)Ribosomal_L31 (pfam01197)→ |
| env-1280 | AUXO017875554.1 | + | 7952 | 7983 | **RNA**→ SGNH_hydrolase (cd00229)→ rpmE (PRK00019)Ribosomal_L31 (pfam01197)→ PRK05571 (PRK05571)LacAB_rpiB (pfam02502)→ DUF1146 (pfam06612)→ PRK11649 (PRK11649)Peptidase_M23 (pfam01551)→ |
| env-1281 | AUXO012311456.1 | - | 82 | 52 | **RNA**→ rpmE (PRK00528)Ribosomal_L31 (pfam01197)→ |
| env-1282 | AUXO013805451.1 | - | 82 | 52 | **RNA**→ rpmE (PRK00528)Ribosomal_L31 (pfam01197)→ |
| env-1283 | AUXO013112282.1 | - | 284 | 252 | **RNA**→ rpmE (PRK00019)Ribosomal_L31 (pfam01197)→ |
| env-1284 | AUXO014919177.1 | - | 130 | 100 | **RNA**→ rpmE (PRK00019)Ribosomal_L31 (pfam01197)→ |
| env-1285 | AUXO012946948.1 | - | 173 | 142 | **RNA**→ rpmE (PRK00019)Ribosomal_L31 (pfam01197)→ |
| env-1286 | AUXO013252499.1 | + | 233 | 263 | **RNA**→ rpmE (PRK00528)rpmE (PRK00019)Ribosomal_L31 (pfam01197)→ |
| env-1287 | SRS019582_C2881833 | - | 282 | 252 | **RNA**→ rpmE (PRK00019)Ribosomal_L31 (pfam01197)→ |
| env-1288 | SRS012273_C3602910 | - | 368 | 337 | **RNA**→ rpmE (PRK00019)Ribosomal_L31 (pfam01197)→ |
| env-1289 | SRS022071_C2951214 | + | 292 | 323 | **RNA**→ rpmE (PRK00019)Ribosomal_L31 (pfam01197)→ |
| env-1290 | AUXO012038790.1 | + | 252 | 281 | **RNA**→ rpmE (PRK00019)Ribosomal_L31 (pfam01197)→ hypo→ thioredoxin (TIGR01068)TRX_family (cd02947)→ bPH_3 (pfam14470)→ hypo→ |
| env-1291 | AUXO011416548.1 | - | 68 | 41 | **RNA**→ |
| env-1292 | AUXO017181277.1 | - | 182 | 152 | **RNA**→ rpmE (PRK00019)Ribosomal_L31 (pfam01197)→ |

| | | | | | |
|---|---|---|---|---|---|
| env-1293 | AUXO016134728.1 | + | 1040 | 1070 | **RNA**→ rpmE (PRK00019)Ribosomal_L31 (pfam01197)→ PRK05571 (PRK05571)LacAB_rpiB (pfam02502)→ DUF1146 (pfam06612)→ |
| env-1294 | AUXO011423476.1 | + | 2455 | 2485 | **RNA**→ |
| env-1295 | AUXO011517276.1 | + | 28 | 58 | **RNA**→ rpmE (PRK00019)Ribosomal_L31 (pfam01197)→ |
| env-1296 | JGI994J12354_10524623 | + | 105 | 134 | **RNA**→ rpmE (PRK00019)Ribosomal_L31 (pfam01197)→ |
| env-1297 | AUXO012504192.1 | + | 415 | 446 | **RNA**→ rpmE (PRK00019)Ribosomal_L31 (pfam01197)→ |
| env-1298 | AUXO014624430.1 | + | 363 | 389 | **RNA**→ rpmE (PRK00019)→ |
| env-1299 | AUXO012372966.1 | + | 82 | 112 | **RNA**→ rpmE (PRK00019)Ribosomal_L31 (pfam01197)→ |
| env-1300 | SRMUA_GNVGR9Q01DNPCS | + | 60 | 87 | **RNA**→ rpmE (PRK00019)Ribosomal_L31 (pfam01197)→ |
| env-1301 | AUXO016542428.1 | - | 1502 | 1471 | **RNA**→ rpmE (PRK00019)Ribosomal_L31 (pfam01197)→ |
| env-1302 | AUXO014856272.1 | - | 1536 | 1505 | **RNA**→ rpmE (PRK00019)Ribosomal_L31 (pfam01197)→ |
| env-1303 | AUXO012441124.1 | - | 149 | 121 | **RNA**→ rpmE (PRK00019)Ribosomal_L31 (pfam01197)→ |
| env-1304 | 3300000558_10084062 | + | 2958 | 2985 | **RNA**→ rpmE (PRK00019)Ribosomal_L31 (pfam01197)→ Tdk (COG1435)PRK04296 (PRK04296)→ |
| env-1305 | AUXO012809103.1 | - | 1236 | 1207 | **RNA**→ rpmE (PRK00019)Ribosomal_L31 (pfam01197)→ PRK05571 (PRK05571)LacAB_rpiB (pfam02502)→ DUF1146 (pfam06612)→ hypo→ |
| env-1306 | AUXO010815353.1 | + | 82 | 112 | **RNA**→ rpmE (PRK00019)Ribosomal_L31 (pfam01197)→ YcaP (COG2323)→ |
| env-1307 | AUXO015985898.1 | - | 109 | 79 | **RNA**→ rpmE (PRK00019)rpmE2 (PRK01678)Ribosomal_L31 (pfam01197)→ |
| env-1308 | RUMENNODE_4377773_84001 | + | 25108 | 25136 | **RNA**→ rpmE (PRK00019)Ribosomal_L31 (pfam01197)→ PRK13759 (PRK13759)LTA_synthase (cd16015)→ PRK11361 (PRK11361)LytTR (smart00850)LytT (COG3279)→ CitA (COG3290)WcaJ_sugtrans (TIGR03023)HATPase_c_5 (pfam14501)→ |
| env-1309 | AUXO011141601.1 | - | 618 | 590 | **RNA**→ rpmE (PRK00019)Ribosomal_L31 (pfam01197)→ hypo→ |
| env-1310 | AUXO010916397.1 | + | 5920 | 5948 | **RNA**→ rpmE (PRK00019)Ribosomal_L31 (pfam01197)→ MdoB (COG1368)PRK13759 (PRK13759)→ PRK11361 (PRK11361)LytTR (smart00850)LytT (COG3279)→ CitA (COG3290)PRK11086 (PRK11086)→ |
| env-1311 | AUXO017784779.1 | - | 2057 | 2027 | **RNA**→ rpmE (PRK00019)Ribosomal_L31 (pfam01197)→ PRK05571 (PRK05571)LacAB_rpiB (pfam02502)→ DUF1146 (pfam06612)→ PRK11649 (PRK11649)Peptidase_M23 (pfam01551)→ |
| env-1312 | scaffold30364_4_MH0066 | - | 520 | 493 | **RNA**→ rpmE (PRK00019)Ribosomal_L31 (pfam01197)→ |
| env-1313 | C4277000_1_V1.UC-14 | - | 882 | 855 | **RNA**→ rpmE (PRK00019)Ribosomal_L31 (pfam01197)→ |
| env-1314 | scaffold4347_1_V1.UC-19 | + | 160 | 187 | **RNA**→ rpmE (PRK00019)Ribosomal_L31 (pfam01197)→ |
| env-1315 | SRS049164_WUGC_scaffold_87487 | - | 15418 | 15391 | **RNA**→ rpmE (PRK00019)Ribosomal_L31 (pfam01197)→ |
| env-1316 | DOF002_scaffold60922_2 | + | 3588 | 3615 | **RNA**→ rpmE (PRK00019)Ribosomal_L31 (pfam01197)→ |
| env-1317 | DOM013_scaffold66250_2 | + | 114 | 141 | **RNA**→ rpmE (PRK00019)Ribosomal_L31 (pfam01197)→ |
| env-1318 | DOM014_scaffold2248_7_2 | - | 1915 | 1888 | **RNA**→ rpmE (PRK00019)Ribosomal_L31 (pfam01197)→ |
| env-1319 | NOF001_scaffold58173_1 | + | 4467 | 4494 | **RNA**→ rpmE (PRK00019)Ribosomal_L31 (pfam01197)→ |
| env-1320 | NOF014_scaffold69538_9 | - | 2379 | 2352 | **RNA**→ rpmE (PRK00019)Ribosomal_L31 (pfam01197)→ |
| env-1321 | NOM013_scaffold55288_1 | - | 18149 | 18122 | **RNA**→ rpmE (PRK00019)Ribosomal_L31 (pfam01197)→ |
| env-1322 | NOM019_scaffold49208_1 | + | 249 | 276 | **RNA**→ rpmE (PRK00019)Ribosomal_L31 (pfam01197)→ |
| env-1323 | SRS011271_WUGC_scaffold_4131 | - | 16966 | 16939 | **RNA**→ rpmE (PRK00019)Ribosomal_L31 (pfam01197)→ |
| env-1324 | SRS011586_WUGC_scaffold_25822 | - | 10363 | 10336 | **RNA**→ rpmE (PRK00019)Ribosomal_L31 (pfam01197)→ |
| env-1325 | SRS015578_C2525326 | - | 256 | 229 | **RNA**→ rpmE (PRK00019)Ribosomal_L31 (pfam01197)→ |
| env-1326 | SRS016335_WUGC_scaffold_45818 | - | 9930 | 9903 | **RNA**→ rpmE (PRK00019)Ribosomal_L31 (pfam01197)→ |
| env-1327 | SRS020233_C4068279 | + | 109 | 136 | **RNA**→ rpmE (PRK00019)Ribosomal_L31 (pfam01197)→ |
| env-1328 | SRS020869_C3081547 | - | 165 | 138 | **RNA**→ rpmE (PRK00019)Ribosomal_L31 (pfam01197)→ |
| env-1329 | SRS021948_C2732547 | - | 488 | 461 | **RNA**→ rpmE (PRK00019)Ribosomal_L31 (pfam01197)→ |
| env-1330 | SRS022071_C3057294 | - | 2824 | 2797 | **RNA**→ rpmE (PRK00019)Ribosomal_L31 (pfam01197)→ |
| env-1331 | SRS023526_C2997413 | - | 272 | 245 | **RNA**→ rpmE (PRK00019)Ribosomal_L31 (pfam01197)→ |
| env-1332 | SRS047014_C3660042 | + | 154 | 181 | **RNA**→ rpmE (PRK00019)Ribosomal_L31 (pfam01197)→ |
| env-1333 | SRS048870_WUGC_scaffold_52503 | + | 97 | 124 | **RNA**→ rpmE (PRK00019)Ribosomal_L31 (pfam01197)→ |
| env-1334 | SRS049959_C3987040 | - | 11396 | 11369 | **RNA**→ rpmE (PRK00019)Ribosomal_L31 (pfam01197)→ |
| env-1335 | SRS049995_LANL_scaffold_33915 | - | 4520 | 4493 | **RNA**→ rpmE (PRK00019)Ribosomal_L31 (pfam01197)→ |
| env-1336 | SRS078176_LANL_scaffold_951 | - | 4520 | 4493 | **RNA**→ rpmE (PRK00019)Ribosomal_L31 (pfam01197)→ |
| env-1337 | scaffold129929_1_MH0012 | + | 17 | 44 | **RNA**→ rpmE (PRK00019)Ribosomal_L31 (pfam01197)→ |
| env-1338 | SRS012273_C3781305 | + | 19 | 46 | **RNA**→ rpmE (PRK00019)Ribosomal_L31 (pfam01197)→ |
| env-1339 | SRS014979_C3210300 | + | 19 | 46 | **RNA**→ rpmE (PRK00019)Ribosomal_L31 (pfam01197)→ |

| | | | | | |
|---|---|---|---|---|---|
| env-1340 | SRS024435_C3024763 | + | 19 | 46 | RNA→ rpmE (PRK00019)Ribosomal_L31 (pfam01197)→ |
| env-1341 | SRS045645_C2053142 | - | 2027 | 2000 | RNA→ rpmE (PRK00019)Ribosomal_L31 (pfam01197)→ |
| env-1342 | scaffold4802_1_MH0014 | - | 1015 | 989 | RNA→ rpmE (PRK00019)Ribosomal_L31 (pfam01197)→ hypo→ |
| env-1343 | C4062675_1_MH0016 | - | 447 | 421 | RNA→ rpmE (PRK00019)Ribosomal_L31 (pfam01197)→ hypo→ |
| env-1344 | scaffold207_2_MH0026 | - | 473 | 447 | RNA→ rpmE (PRK00019)Ribosomal_L31 (pfam01197)→ hypo→ |
| env-1345 | scaffold29535_1_MH0035 | + | 8024 | 8050 | RNA→ rpmE (PRK00019)Ribosomal_L31 (pfam01197)→ PRK13759 (PRK13759)LTA_synthase (cd16015)→ PRK11361 (PRK11361)LytTR (smart00850)LytT (COG3279)→ CitA (COG3290)HATPase_c_5 (pfam14501)→ |
| env-1346 | scaffold78402_1_MH0064 | - | 449 | 423 | RNA→ rpmE (PRK00019)Ribosomal_L31 (pfam01197)→ hypo→ |
| env-1347 | scaffold87074_1_MH0075 | + | 319 | 345 | RNA→ rpmE (PRK00019)Ribosomal_L31 (pfam01197)→ |
| env-1348 | scaffold104927_2_V1.CD-14 | - | 230 | 204 | RNA→ rpmE (PRK00019)Ribosomal_L31 (pfam01197)→ |
| env-1349 | 4491482.3_NODE_16610 | - | 635 | 609 | RNA→ rpmE (PRK00019)Ribosomal_L31 (pfam01197)→ hypo→ |
| env-1350 | 4491487.3_NODE_42294 | - | 2870 | 2844 | RNA→ rpmE (PRK00019)SGNH_hydrolase (cd00229)Ribosomal_L31 (pfam01197)→ PRK13759 (PRK13759)LTA_synthase (cd16015)→ PRK11361 (PRK11361)REC (cd00156)→ |
| env-1351 | NLM025_scaffold35660_1 | - | 351 | 325 | RNA→ rpmE (PRK00019)Lipase_GDSL (pfam00657)Ribosomal_L31 (pfam01197)→ |
| env-1352 | NLM029_scaffold44136_3 | - | 436 | 410 | RNA→ rpmE (PRK00019)Ribosomal_L31 (pfam01197)→ hypo→ |
| env-1353 | SRS011134_C5157013 | - | 291 | 265 | RNA→ rpmE (PRK00019)Ribosomal_L31 (pfam01197)→ |
| env-1354 | SRS011239_Baylor_scaffold_29786 | + | 725 | 751 | RNA→ rpmE (PRK00019)Ribosomal_L31 (pfam01197)→ hypo→ |
| env-1355 | SRS012273_Baylor_scaffold_10464 | + | 11441 | 11467 | RNA→ rpmE (PRK00019)Ribosomal_L31 (pfam01197)→ PRK13759 (PRK13759)LTA_synthase (cd16015)→ PRK11361 (PRK11361)LytTR (smart00850)LytT (COG3279)→ HATPase_c (pfam02518)HATPase_c_5 (pfam14501)→ |
| env-1356 | SRS017701_Baylor_scaffold_18247 | + | 358 | 384 | RNA→ rpmE (PRK00019)Ribosomal_L31 (pfam01197)→ hypo→ |
| env-1357 | SRS018656_WUGC_scaffold_5423 | - | 1241 | 1215 | RNA→ rpmE (PRK00019)Ribosomal_L31 (pfam01197)→ LTA_synthase (cd16015)→ |
| env-1358 | SRS022071_C2877461 | + | 104 | 130 | RNA→ rpmE (PRK00019)Ribosomal_L31 (pfam01197)→ |
| env-1359 | SRS050752_LANL_scaffold_27273 | + | 33215 | 33241 | RNA→ rpmE (PRK00019)Ribosomal_L31 (pfam01197)→ PRK13759 (PRK13759)LTA_synthase (cd16015)→ PRK11361 (PRK11361)LytTR (smart00850)LytT (COG3279)→ CitA (COG3290)HATPase_c_5 (pfam14501)→ |
| env-1360 | 4491410.3_NODE_39918 | + | 398 | 424 | RNA→ rpmE (PRK00019)Ribosomal_L31 (pfam01197)→ |
| env-1361 | SRS011302_Baylor_scaffold_316 | + | 5248 | 5274 | RNA→ rpmE (PRK00019)Ribosomal_L31 (pfam01197)→ PRK13759 (PRK13759)LTA_synthase (cd16015)→ PRK11361 (PRK11361)LytTR (smart00850)LytT (COG3279)→ |
| env-1362 | scaffold70250_1_MH0012 | + | 907 | 933 | RNA→ rpmE (PRK00019)Ribosomal_L31 (pfam01197)→ MdoB (COG1368)→ |
| env-1363 | scaffold70665_1_MH0028 | + | 5114 | 5140 | RNA→ rpmE (PRK00019)Ribosomal_L31 (pfam01197)→ PRK13759 (PRK13759)LTA_synthase (cd16015)→ PRK11361 (PRK11361)LytTR (smart00850)LytT (COG3279)→ CitA (COG3290)HATPase_c_5 (pfam14501)→ |
| env-1364 | scaffold109810_1_MH0033 | - | 175 | 149 | RNA→ rpmE (PRK00019)Ribosomal_L31 (pfam01197)→ |
| env-1365 | scaffold62702_1_MH0066 | + | 253 | 279 | RNA→ rpmE (PRK00019)Ribosomal_L31 (pfam01197)→ hypo→ |
| env-1366 | scaffold25252_2_MH0081 | + | 5634 | 5660 | RNA→ rpmE (PRK00019)Ribosomal_L31 (pfam01197)→ LTA_synthase (cd16015)→ |
| env-1367 | 4491423.3_NODE_40207 | - | 509 | 483 | RNA→ rpmE (PRK00019)Ribosomal_L31 (pfam01197)→ hypo→ |
| env-1368 | DLF003_scaffold13796_5 | - | 219 | 193 | RNA→ rpmE (PRK00019)Ribosomal_L31 (pfam01197)→ |
| env-1369 | DOM012_scaffold215_3 | + | 9962 | 9988 | RNA→ rpmE (PRK00019)Lipase_GDSL (pfam00657)Ribosomal_L31 (pfam01197)→ PRK13759 (PRK13759)LTA_synthase (cd16015)→ PRK11361 (PRK11361)LytTR (smart00850)LytT (COG327 CitA (COG3290)HATPase_c_5 (pfam14501)→ |
| env-1370 | NLM015_scaffold4789_7 | + | 420 | 446 | RNA→ rpmE (PRK00019)Ribosomal_L31 (pfam01197)→ |
| env-1371 | SRS011529_WUGC_scaffold_2327 | + | 32692 | 32718 | RNA→ rpmE (PRK00019)Ribosomal_L31 (pfam01197)→ PRK13759 (PRK13759)LTA_synthase (cd16015)→ PRK11361 (PRK11361)LytTR (smart00850)LytT (COG3279)→ CitA (COG3290)HATPase_c_5 (pfam14501)→ |
| env-1372 | SRS015782_WUGC_scaffold_16117 | - | 5795 | 5769 | RNA→ rpmE (PRK00019)Ribosomal_L31 (pfam01197)→ PRK13759 (PRK13759)LTA_synthase (cd16015)→ PRK11361 (PRK11361)LytTR (smart00850)LytT (COG3279)→ CitA (COG3290)HATPase_c_5 (pfam14501)→ |
| env-1373 | SRS018817_C1984407 | - | 174 | 148 | RNA→ rpmE (PRK00019)Ribosomal_L31 (pfam01197)→ |
| env-1374 | SRS020233_C4276649 | + | 1616 | 1642 | RNA→ rpmE (PRK00019)Ribosomal_L31 (pfam01197)→ PRK13759 (PRK13759)LTA_synthase (cd16015)→ PRK11361 (PRK11361)LytTR (smart00850)LytT (COG3279)→ CitA (COG3290)HATPase_c_5 (pfam14501)→ |
| env-1375 | SRS021948_C2860438 | + | 1458 | 1484 | RNA→ rpmE (PRK00019)Ribosomal_L31 (pfam01197)→ hypo→ |
| env-1376 | SRS047044_WUGC_scaffold_15413 | + | 11469 | 11495 | RNA→ rpmE (PRK00019)Ribosomal_L31 (pfam01197)→ PRK13759 (PRK13759)LTA_synthase (cd16015)→ PRK11361 (PRK11361)LytTR (smart00850)LytT (COG3279)→ hypo→ |
| env-1377 | SRS052697_C3372869 | - | 2793 | 2767 | RNA→ rpmE (PRK00019)Ribosomal_L31 (pfam01197)→ PRK13759 (PRK13759)LTA_synthase (cd16015)→ PRK11361 (PRK11361)REC (cd00156)→ |
| env-1378 | SRS065504_LANL_scaffold_33624 | - | 18639 | 18613 | RNA→ rpmE (PRK00019)Ribosomal_L31 (pfam01197)→ PRK13759 (PRK13759)LTA_synthase (cd16015)→ PRK11361 (PRK11361)LytTR (smart00850)LytT (COG3279)→ CitA (COG3290)HATPase_c_5 (pfam14501)→ |
| env-1379 | C2077074_1_MH0079 | + | 2299 | 2326 | RNA→ rpmE (PRK00019)Ribosomal_L31 (pfam01197)→ |

| | | | | | |
|---|---|---|---|---|---|
| env-1380 | DLF003_scaffold7759_1 | - | 143 | 116 | RNA→ rpmE (PRK00019)Ribosomal_L31 (pfam01197)→ |
| env-1381 | SRS011271_WUGC_scaffold_24401 | - | 16266 | 16240 | RNA→ rpmE (PRK00019)Ribosomal_L31 (pfam01197)→ PRK13759 (PRK13759)LTA_synthase (cd16015)→ hypo→ |
| env-1382 | SRS023526_Baylor_scaffold_35082 | + | 302 | 328 | RNA→ rpmE (PRK00019)Ribosomal_L31 (pfam01197)→ PRK13759 (PRK13759)LTA_synthase (cd16015)→ PRK11361 (PRK11361)LytTR (smart00850)LytT (COG3279)→ CitA (COG3290)HATPase_c_5 (pfam14501)→ |
| env-1383 | AUXO018649474.1 | + | 79 | 105 | RNA→ rpmE (PRK00019)Ribosomal_L31 (pfam01197)→ |
| env-1384 | SRS011134_C4857448 | + | 310 | 337 | RNA→ Ribosomal_L31 (pfam01197)→ |
| env-1385 | DLF004_scaffold6617_4 | - | 1944 | 1917 | RNA→ rpmE (PRK00019)SGNH_hydrolase (cd00229)Ribosomal_L31 (pfam01197)Lipase_GDSL_2 (pfam13472 PRK12363 (PRK12363)LTA_synthase (cd16015)→ |
| env-1386 | DLF010_scaffold22656_1 | - | 107791 | 107764 | RNA→ rpmE (PRK00019)Ribosomal_L31 (pfam01197)→ PRK12363 (PRK12363)MdoB (COG1368)→ cztR_silR_copR (TIGR01387)LytTR (smart00850)LytT (COG3279)→ CitA (COG3290)HATPase_c_5 (pfam14501)→ |
| env-1387 | 4491414.3_NODE_12936 | + | 3149 | 3176 | RNA→ SGNH_hydrolase (cd00229)Lipase_GDSL_2 (pfam13472)→ rpmE (PRK00019)Ribosomal_L31 (pfam01197)→ PRK12363 (PRK12363)LTA_synthase (cd16015)→ cztR_silR_copR (TIGR01387)LytTR (smart00850)LytT (COG3279)→ CitA (COG3290)HATPase_c_5 (pfam14501)→ |
| env-1388 | AUXO017324133.1 | - | 24508 | 24481 | RNA→ SGNH_hydrolase (cd00229)Lipase_GDSL_2 (pfam13472)→ rpmE (PRK00019)Ribosomal_L31 (pfam01197)→ MdoB (COG1368)PRK13759 (PRK13759)→ PRK11361 (PRK11361)LytTR (smart00850)LytT (COG3279)→ PRK11086 (PRK11086)HATPase_c (pfam0251 |
| env-1389 | AUXO015658258.1 | - | 64 | 35 | RNA→ |
| env-1390 | AUXO014149459.1 | - | 2076 | 2045 | RNA→ rpmE (PRK00019)Ribosomal_L31 (pfam01197)→ CYK3 (COG5279)→ |
| env-1391 | scaffold84569_2_MH0038 | - | 567 | 538 | RNA→ rpmE (PRK00019)Ribosomal_L31 (pfam01197)→ |
| env-1392 | SRS011586_WUGC_scaffold_19197 | - | 7279 | 7250 | RNA→ rpmE (PRK00019)Ribosomal_L31 (pfam01197)→ |
| env-1393 | AUXO015167000.1 | + | 837 | 867 | RNA→ rpmE (PRK00019)Ribosomal_L31 (pfam01197)→ |
| env-1394 | AUXO017088504.1 | + | 293 | 323 | RNA→ rpmE (PRK00019)Ribosomal_L31 (pfam01197)→ |
| env-1395 | UnmappedStool_Broad_C253180942 | - | 644 | 614 | RNA→ rpmE (PRK00019)Ribosomal_L31 (pfam01197)→ |
| env-1396 | NLM026_scaffold61948_3 | + | 70 | 100 | RNA→ rpmE (PRK00019)Ribosomal_L31 (pfam01197)→ |
| env-1397 | AUXO016873222.1 | + | 8429 | 8460 | RNA→ rpmE (PRK00019)Ribosomal_L31 (pfam01197)→ |
| env-1398 | AUXO017217394.1 | + | 2799 | 2830 | RNA→ rpmE (PRK00019)Ribosomal_L31 (pfam01197)→ PRK05571 (PRK05571)LacAB_rpiB (pfam02502)→ DUF1146 (pfam06612)→ MreB_like (cd10225)PRK13930 (PRK13930)→ ECA_wecA (TIGR02380)GT_WecA_like (cd06853)→ |
| env-1399 | C1643059_1_MH0013 | + | 4131 | 4161 | RNA→ rpmE (PRK00019)Ribosomal_L31 (pfam01197)→ PRK05571 (PRK05571)LacAB_rpiB (pfam02502)→ DUF1146 (pfam06612)→ hypo→ |
| env-1400 | scaffold22971_1_MH0030 | + | 23528 | 23558 | RNA→ rpmE (PRK00019)Ribosomal_L31 (pfam01197)→ PRK05571 (PRK05571)LacAB_rpiB (pfam02502)→ |
| env-1401 | SRS011134_C5179588 | + | 1191 | 1221 | RNA→ rpmE (PRK00019)Ribosomal_L31 (pfam01197)→ PRK05571 (PRK05571)LacAB_rpiB (pfam02502)→ DUF1146 (pfam06612)→ PRK11649 (PRK11649)Peptidase_M23 (pfam01551)→ SpoIIID (pfam12116)→ |
| env-1402 | 2227080773 | - | 117461 | 117431 | RNA→ rpmE (PRK00019)Ribosomal_L31 (pfam01197)→ YcaP (COG2323)→ hypo→ |
| env-1403 | JGI996J12353_1033205 | + | 84 | 114 | RNA→ rpmE (PRK00019)Ribosomal_L31 (pfam01197)→ hypo→ |
| env-1404 | AUXO017836399.1 | - | 2207 | 2180 | RNA→ rpmE (PRK00019)Ribosomal_L31 (pfam01197)→ |
| env-1405 | AUXO014543416.1 | + | 2000 | 2027 | RNA→ |
| env-1406 | AUXO014166827.1 | + | 585 | 612 | RNA→ |
| env-1407 | AUXO013533263.1 | - | 71 | 44 | RNA→ |
| env-1408 | AUXO015166946.1 | + | 1763 | 1790 | RNA→ rpmE (PRK00019)Ribosomal_L31 (pfam01197)→ |
| env-1409 | SRS053917_LANL_scaffold_40441 | - | 12785 | 12754 | RNA→ rpmE (PRK00019)Ribosomal_L31 (pfam01197)→ PRK05571 (PRK05571)LacAB_rpiB (pfam02502)→ GT_WecA_like (cd06853)mraY (PRK00108)→ hypo→ yfiA (TIGR00741)Ribosomal_S30AE (pfam02482)Ribosom_S30AE_C (pfam16321)→ SecA (COG0653)PRK12904 (PRK12904)SEC-C (pfam02810)→ |
| env-1410 | SRS018157_C4833607 | + | 933 | 964 | RNA→ rpmE (PRK00019)Ribosomal_L31 (pfam01197)→ |
| env-1411 | SRS022725_LANL_scaffold_39231 | - | 12149 | 12118 | RNA→ rpmE (PRK00019)Ribosomal_L31 (pfam01197)→ PRK05571 (PRK05571)LacAB_rpiB (pfam02502)→ GT_WecA_like (cd06853)mraY (PRK00108)→ hypo→ yfiA (TIGR00741)Ribosomal_S30AE (pfam02482)Ribosom_S30AE_C (pfam16321)→ SecA (COG0653)PRK12904 (PRK12904)SEC-C (pfam02810)→ |
| env-1412 | SRS047113_LANL_scaffold_83317 | + | 5415 | 5446 | RNA→ rpmE (PRK00019)Ribosomal_L31 (pfam01197)→ PRK05571 (PRK05571)LacAB_rpiB (pfam02502)→ ECA_wecA (TIGR02380)GT_WecA_like (cd06853)→ hypo→ |
| env-1413 | SRS018394_C3985949 | - | 346 | 315 | RNA→ rpmE (PRK00019)Ribosomal_L31 (pfam01197)→ |

| | | | | | |
|---|---|---|---|---|---|
| env-1414 | SRS018739_C3705271 | + | 412 | 443 | RNA→ rpmE (PRK00019)Ribosomal_L31 (pfam01197)→ PRK05571 (PRK05571)LacAB_rpiB (pfam02502)→ GT_WecA_like (cd06853)mraY (PRK00108)→ |
| env-1415 | SRS019028_WUGC_scaffold_67797 | - | 1216 | 1185 | RNA→ rpmE (PRK00019)Ribosomal_L31 (pfam01197)→ PRK05571 (PRK05571)LacAB_rpiB (pfam02502)→ ECA_wecA (TIGR02380)GT_WecA_like (cd06853)→ |
| env-1416 | SRS013705_Baylor_scaffold_29455 | - | 274 | 244 | RNA→ rpmE (PRK00019)Ribosomal_L31 (pfam01197)→ |
| env-1417 | SRS015434_C5843902 | + | 15 | 45 | RNA→ rpmE (PRK00019)Ribosomal_L31 (pfam01197)→ PRK05571 (PRK05571)LacAB_rpiB (pfam02502)→ |
| env-1418 | SRS049389_C4729994 | - | 1752 | 1722 | RNA→ rpmE (PRK00019)Ribosomal_L31 (pfam01197)→ PRK05571 (PRK05571)LacAB_rpiB (pfam02502)→ GT_WecA_like (cd06853)mraY (PRK00108)→ |
| env-1419 | SRS017209_C2431801 | + | 21 | 51 | RNA→ rpmE (PRK00019)Ribosomal_L31 (pfam01197)→ |
| env-1420 | SRS022530_LANL_scaffold_38958 | + | 323 | 353 | RNA→ rpmE (PRK00019)Ribosomal_L31 (pfam01197)→ |
| env-1421 | SRS021496_C2799659 | - | 346 | 316 | RNA→ rpmE (PRK00019)Ribosomal_L31 (pfam01197)→ |
| env-1422 | SRS057692_C2577720 | - | 245 | 215 | RNA→ rpmE (PRK00019)Ribosomal_L31 (pfam01197)→ |
| env-1423 | SRS015762_C3673319 | - | 5711 | 5680 | RNA→ rpmE (PRK00019)Ribosomal_L31 (pfam01197)→ PRK05571 (PRK05571)LacAB_rpiB (pfam02502)→ GT_WecA_like (cd06853)mraY (PRK00108)→ hypo→ yfiA (TIGR00741)RaiA (cd00552)Ribosom_S30AE_C (pfam16321)→ |
| env-1424 | SRS064329_LANL_scaffold_43584 | - | 16350 | 16319 | RNA→ rpmE (PRK00019)Ribosomal_L31 (pfam01197)→ PRK05571 (PRK05571)LacAB_rpiB (pfam02502)→ GT_WecA_like (cd06853)mraY (PRK00108)→ hypo→ yfiA (TIGR00741)RaiA (cd00552)Ribosom_S30AE_C (pfam16321)→ SecA (COG0653)PRK12904 (PRK12904)SEC-C (pfam02810)→ |
| env-1425 | SRS014271_C2962237 | - | 3039 | 3008 | RNA→ rpmE (PRK00019)Ribosomal_L31 (pfam01197)→ PRK05571 (PRK05571)LacAB_rpiB (pfam02502)→ GT_WecA_like (cd06853)mraY (PRK00108)→ hypo→ |
| env-1426 | SRS016002_C2638962 | + | 1640 | 1671 | RNA→ rpmE (PRK00019)Ribosomal_L31 (pfam01197)→ PRK05571 (PRK05571)LacAB_rpiB (pfam02502)→ Rfe (COG0472)→ |
| env-1427 | SRS016319_WUGC_scaffold_23354 | + | 464 | 495 | RNA→ rpmE (PRK00019)Ribosomal_L31 (pfam01197)→ PRK05571 (PRK05571)LacAB_rpiB (pfam02502)→ GT_WecA_like (cd06853)mraY (PRK00108)→ hypo→ yfiA (TIGR00741)RaiA (cd00552)Ribosom_S30AE_C (pfam16321)→ SecA (COG0653)PRK12904 (PRK12904)SEC-C (pfam02810)→ |
| env-1428 | SRS016569_C2512869 | + | 593 | 624 | RNA→ rpmE (PRK00019)Ribosomal_L31 (pfam01197)→ PRK05571 (PRK05571)LacAB_rpiB (pfam02502)→ GT_WecA_like (cd06853)mraY (PRK00108)→ hypo→ yfiA (TIGR00741)RaiA (cd00552)Ribosom_S30AE_C (pfam16321)→ |
| env-1429 | SRS017209_C2525129 | + | 21 | 52 | RNA→ rpmE (PRK00019)Ribosomal_L31 (pfam01197)→ PRK05571 (PRK05571)LacAB_rpiB (pfam02502)→ |
| env-1430 | SRS018300_C3361889 | - | 1806 | 1775 | RNA→ rpmE (PRK00019)Ribosomal_L31 (pfam01197)→ PRK05571 (PRK05571)LacAB_rpiB (pfam02502)→ GT_WecA_like (cd06853)mraY (PRK00108)→ |
| env-1431 | SRS021496_C2821855 | + | 460 | 491 | RNA→ rpmE (PRK00019)Ribosomal_L31 (pfam01197)→ PRK05571 (PRK05571)LacAB_rpiB (pfam02502)→ |
| env-1432 | SRS024081_LANL_scaffold_71404 | - | 16350 | 16319 | RNA→ rpmE (PRK00019)Ribosomal_L31 (pfam01197)→ PRK05571 (PRK05571)LacAB_rpiB (pfam02502)→ GT_WecA_like (cd06853)mraY (PRK00108)→ hypo→ yfiA (TIGR00741)RaiA (cd00552)Ribosom_S30AE_C (pfam16321)→ SecA (COG0653)PRK12904 (PRK12904)SEC-C (pfam02810)→ |
| env-1433 | SRS042643_WUGC_scaffold_65941 | - | 5725 | 5694 | RNA→ rpmE (PRK00019)Ribosomal_L31 (pfam01197)→ PRK05571 (PRK05571)LacAB_rpiB (pfam02502)→ GT_WecA_like (cd06853)mraY (PRK00108)→ hypo→ |
| env-1434 | SRS049389_WUGC_scaffold_9485 | + | 15 | 46 | RNA→ rpmE (PRK00019)Ribosomal_L31 (pfam01197)→ PRK05571 (PRK05571)LacAB_rpiB (pfam02502)→ GT_WecA_like (cd06853)mraY (PRK00108)→ hypo→ |
| env-1435 | SRS050244_LANL_scaffold_89207 | + | 322 | 353 | RNA→ rpmE (PRK00019)Ribosomal_L31 (pfam01197)→ |
| env-1436 | SRS057205_C3408538 | - | 470 | 439 | RNA→ rpmE (PRK00019)Ribosomal_L31 (pfam01197)→ PRK05571 (PRK05571)LacAB_rpiB (pfam02502)→ |
| env-1437 | SRS062544_LANL_scaffold_15860 | + | 10967 | 10998 | RNA→ rpmE (PRK00019)Ribosomal_L31 (pfam01197)→ PRK05571 (PRK05571)LacAB_rpiB (pfam02502)→ GT_WecA_like (cd06853)mraY (PRK00108)→ hypo→ yfiA (TIGR00741)RaiA (cd00552)Ribosom_S30AE_C (pfam16321)→ SecA (COG0653)PRK12904 (PRK12904)SEC-C (pfam02810)→ |
| env-1438 | SRS015434_C5843986 | - | 559 | 528 | RNA→ rpmE (PRK00019)Ribosomal_L31 (pfam01197)→ PRK05571 (PRK05571)LacAB_rpiB (pfam02502)→ |
| env-1439 | JGI24707J26582_10016376 | + | 3058 | 3088 | RNA→ rpmE (PRK00019)Ribosomal_L31 (pfam01197)→ |
| env-1440 | JGI24708J26588_10079158 | + | 766 | 796 | RNA→ rpmE (PRK00019)Ribosomal_L31 (pfam01197)→ |
| env-1441 | JGI24709J26583_10002472 | - | 2550 | 2520 | RNA→ rpmE (PRK00019)Ribosomal_L31 (pfam01197)→ |
| env-1442 | JGI24710J26742_10002773 | - | 2550 | 2520 | RNA→ rpmE (PRK00019)Ribosomal_L31 (pfam01197)→ |
| env-1443 | AUXO011514504.1 | - | 134 | 105 | RNA→ rpmE (PRK00019)Ribosomal_L31 (pfam01197)→ |

| | | | | | |
|---|---|---|---|---|---|
| Aoc-1-1 | NZ_LK028559.1 | - | 1499011 | 1498981 | **RNA**→ rpmE2 (PRK01678)Ribosomal_L31 (pfam01197)→ Tdk (COG1435)PRK04296 (PRK04296)→ |
| | | | | | TadA (COG0590)MafB19-deam (pfam14437)→ hypo→ hypo→ hypo→ TktA (COG0021)PRK05899 (PRK0589[illegible] |
| | | | | | MnmG (COG0445)PRK05192 (PRK05192)→ RsmG (COG0357)gidB (PRK00107)→ DUF951 (pfam06107)→ |
| env-1444 | AUXO011017412.1 | - | 412 | 381 | **RNA**→ rpmE (PRK00019)Ribosomal_L31 (pfam01197)→ |
| env-1445 | SL_3KL_010_SED_10003070 | + | 5652 | 5682 | **RNA**→ rpmE (PRK00019)Ribosomal_L31 (pfam01197)→ Tdk (COG1435)PRK04296 (PRK04296)→ |
| | | | | | TadA (COG0590)MafB19-deam (pfam14437)→ hypo→ hypo→ |
| env-1446 | BGMT1_NODE_617975 | - | 5130 | 5099 | **RNA**→ rpmE2 (PRK01678)Ribosomal_L31 (pfam01197)→ |
| env-1447 | HBC_scaff_10321092 | + | 809 | 840 | **RNA**→ rpmE2 (PRK01678)Ribosomal_L31 (pfam01197)→ |
| env-1448 | HBC_ctgs_1025513 | + | 809 | 840 | **RNA**→ rpmE2 (PRK01678)Ribosomal_L31 (pfam01197)→ |
| Lap-1-1 | NZ_KQ034000.1 | - | 943418 | 943387 | **RNA**→ rpmE2 (PRK01678)Ribosomal_L31 (pfam01197)→ |
| env-1449 | BGMT1_NODE_652147 | - | 6023 | 5992 | **RNA**→ rpmE2 (PRK01678)Ribosomal_L31 (pfam01197)→ |
| env-1450 | HBC_scaff_10316565 | - | 378 | 346 | **RNA**→ rpmE2 (PRK01678)Ribosomal_L31 (pfam01197)→ |
| Lsp-13-1 | NZ_JRJB01000014.1 | - | 24691 | 24659 | **RNA**→ rpmE2 (PRK01678)Ribosomal_L31 (pfam01197)→ |
| env-1451 | BGMT1_NODE_398163 | + | 8659 | 8691 | **RNA**→ rpmE2 (PRK01678)Ribosomal_L31 (pfam01197)→ |
| Lsp-14-1 | NZ_CP009531.1 | - | 1466989 | 1466957 | **RNA**→ rpmE2 (PRK01678)Ribosomal_L31 (pfam01197)→ |
| Lhe-2-1 | NZ_KQ034047.1 | - | 141369 | 141337 | **RNA**→ rpmE2 (PRK01678)Ribosomal_L31 (pfam01197)→ |
| Lku-1-1 | NZ_KQ033871.1 | - | 1476483 | 1476451 | **RNA**→ rpmE2 (PRK01678)Ribosomal_L31 (pfam01197)→ |
| Lki-1-1 | NZ_KQ034007.1 | - | 1482946 | 1482914 | **RNA**→ rpmE2 (PRK01678)Ribosomal_L31 (pfam01197)→ |
| env-1452 | BGMT1_NODE_274826 | - | 167 | 135 | **RNA**→ rpmE2 (PRK01678)Ribosomal_L31 (pfam01197)→ |
| Lme-1-1 | NZ_KQ034022.1 | - | 150781 | 150749 | **RNA**→ rpmE2 (PRK01678)Ribosomal_L31 (pfam01197)→ |
| env-1453 | HBC_scaff_10010740 | + | 12482 | 12514 | **RNA**→ |
| env-1454 | HBC_ctgs_1000528 | + | 8470 | 8502 | **RNA**→ |
| env-1455 | HBC_scaff_10029449 | - | 222 | 190 | **RNA**→ rpmE2 (PRK01678)Ribosomal_L31 (pfam01197)→ |
| env-1456 | HBC_scaff_10290669 | + | 700 | 732 | **RNA**→ |
| env-1457 | HBC_ctgs_1243392 | - | 222 | 190 | **RNA**→ rpmE2 (PRK01678)Ribosomal_L31 (pfam01197)→ |
| env-1458 | HBC_ctgs_1078860 | + | 700 | 732 | **RNA**→ |
| env-1459 | AUXO017032561.1 | - | 631 | 600 | **RNA**→ rpmE (PRK00019)Ribosomal_L31 (pfam01197)→ PRK05571 (PRK05571)LacAB_rpiB (pfam02502)→ |
| env-1460 | AUXO011120786.1 | + | 459 | 485 | **RNA**→ rpmE (PRK00019)Ribosomal_L31 (pfam01197)→ PRK05571 (PRK05571)LacAB_rpiB (pfam02502)→ |
| env-1461 | 4491477.3_NODE_17856 | - | 7482 | 7445 | **RNA**→ rpmE (PRK00019)Ribosomal_L31 (pfam01197)→ |
| env-1462 | scaffold72512_1_MH0030 | + | 4464 | 4501 | **RNA**→ rpmE (PRK00019)Ribosomal_L31 (pfam01197)→ |
| env-1463 | scaffold36821_2_MH0063 | + | 24014 | 24051 | **RNA**→ rpmE (PRK00019)Ribosomal_L31 (pfam01197)→ |
| env-1464 | C3728251_1_MH0081 | - | 290 | 253 | **RNA**→ rpmE (PRK00019)Ribosomal_L31 (pfam01197)→ |
| env-1465 | 4491403.3_NODE_270 | - | 47741 | 47704 | **RNA**→ rpmE (PRK00019)Ribosomal_L31 (pfam01197)→ |
| env-1466 | SRS011134_Baylor_scaffold_23202 | - | 157 | 120 | **RNA**→ rpmE (PRK00019)Ribosomal_L31 (pfam01197)→ |
| env-1467 | SRS011271_WUGC_scaffold_22780 | - | 25462 | 25425 | **RNA**→ rpmE (PRK00019)Ribosomal_L31 (pfam01197)→ |
| env-1468 | SRS017307_Baylor_scaffold_2382 | - | 311 | 274 | **RNA**→ rpmE (PRK00019)Ribosomal_L31 (pfam01197)→ |
| env-1469 | SRS022609_C3091801 | - | 312 | 275 | **RNA**→ rpmE (PRK00019)Ribosomal_L31 (pfam01197)→ |
| env-1470 | SRS023526_C2985960 | + | 101 | 138 | **RNA**→ rpmE (PRK00019)Ribosomal_L31 (pfam01197)→ |
| env-1471 | scaffold16809_2_MH0022 | - | 760 | 723 | **RNA**→ rpmE (PRK00019)Ribosomal_L31 (pfam01197)→ |
| env-1472 | C4139562_1_MH0024 | + | 184 | 221 | **RNA**→ rpmE (PRK00019)Ribosomal_L31 (pfam01197)→ |
| env-1473 | scaffold76898_1_MH0054 | - | 1475 | 1438 | **RNA**→ rpmE (PRK00019)Ribosomal_L31 (pfam01197)→ |
| env-1474 | 4491421.3_NODE_66616 | + | 1471 | 1508 | **RNA**→ rpmE (PRK00019)Ribosomal_L31 (pfam01197)→ |
| env-1475 | DLM019_scaffold25402_3 | + | 922 | 959 | **RNA**→ rpmE (PRK00019)Ribosomal_L31 (pfam01197)→ |
| env-1476 | AUXO017642701.1 | - | 963 | 933 | **RNA**→ rpmE (PRK00019)Ribosomal_L31 (pfam01197)→ |
| env-1477 | AUXO014022239.1 | + | 67 | 93 | **RNA**→ rpmE (PRK00019)Ribosomal_L31 (pfam01197)→ |
| env-1478 | RUMENNODE_3890134_54500 | + | 4623 | 4649 | **RNA**→ rpmE (PRK00019)Ribosomal_L31 (pfam01197)→ Tdk (COG1435)PRK04296 (PRK04296)→ |
| | | | | | PRK10860 (PRK10860)TadA (COG0590)→ |
| Efa-353-1 | NZ_KB029550.1 | - | 223357 | 223325 | **RNA**→ rpmE2 (PRK01678)Ribosomal_L31 (pfam01197)→ |
| Efa-526-1 | NZ_KE351936.1 | + | 9306 | 9338 | **RNA**→ rpmE2 (PRK01678)Ribosomal_L31 (pfam01197)→ |
| Efa-537-1 | NZ_AWWM01000112.1 | + | 6178 | 6210 | **RNA**→ rpmE2 (PRK01678)Ribosomal_L31 (pfam01197)→ |
| env-1479 | DLM012_scaffold178_4 | - | 1663 | 1631 | **RNA**→ rpmE2 (PRK01678)Ribosomal_L31 (pfam01197)→ |
| env-1480 | longitudinal_178_386 | + | 58976 | 59008 | **RNA**→ rpmE2 (PRK01678)Ribosomal_L31 (pfam01197)→ |
| Efa-544-1 | NZ_JAAM01000007.1 | - | 69039 | 69007 | **RNA**→ rpmE2 (PRK01678)Ribosomal_L31 (pfam01197)→ |

| | | | | | | | |
|---|---|---|---|---|---|---|---|
| Efa-335-1 | NZ_AUWX01000117.1 | + | 5605 | 5637 | **RNA**→ | rpmE2 (PRK01678)Ribosomal_L31 (pfam01197)→ | |
| Efa-505-1 | NZ_LBIN01000026.1 | + | 8852 | 8884 | **RNA**→ | rpmE2 (PRK01678)Ribosomal_L31 (pfam01197)→ | |
| Efa-506-1 | NZ_LBIM01000105.1 | + | 9159 | 9191 | **RNA**→ | rpmE2 (PRK01678)Ribosomal_L31 (pfam01197)→ | |
| Efa-425-1 | NZ_AITP01000010.1 | + | 63892 | 63924 | **RNA**→ | rpmE2 (PRK01678)Ribosomal_L31 (pfam01197)→ | |
| Efa-334-1 | NZ_AUWW01000110.1 | - | 3516 | 3484 | **RNA**→ | rpmE2 (PRK01678)Ribosomal_L31 (pfam01197)→ | |
| Efa-340-1 | NZ_AUWV01000070.1 | - | 7538 | 7506 | **RNA**→ | rpmE2 (PRK01678)Ribosomal_L31 (pfam01197)→ | |
| Efa-535-1 | NZ_AWWP01000021.1 | - | 7580 | 7548 | **RNA**→ | rpmE2 (PRK01678)Ribosomal_L31 (pfam01197)→ | |
| Efa-539-1 | NZ_AWWN01000088.1 | + | 41487 | 41519 | **RNA**→ | rpmE2 (PRK01678)Ribosomal_L31 (pfam01197)→ | |
| Efa-540-1 | NZ_AWWO01000049.1 | - | 7504 | 7472 | **RNA**→ | rpmE2 (PRK01678)Ribosomal_L31 (pfam01197)→ | |
| Efa-536-1 | NZ_ASAL01000014.1 | + | 40701 | 40733 | **RNA**→ | rpmE2 (PRK01678)Ribosomal_L31 (pfam01197)→ | |
| Efa-587-1 | NZ_ABQJ01000011.1 | - | 45637 | 45605 | **RNA**→ | rpmE2 (PRK01678)Ribosomal_L31 (pfam01197)→ | rpmE2 (PRK01678)Ribosomal_L31 (pfam01197)→ |
| Efa-588-1 | NZ_ABRY01000165.1 | + | 23876 | 23908 | **RNA**→ | rpmE2 (PRK01678)Ribosomal_L31 (pfam01197)→ | rpmE2 (PRK01678)Ribosomal_L31 (pfam01197)→ |
| Esp-20-1 | NZ_AJQZ01000052.1 | - | 7648 | 7616 | **RNA**→ | rpmE2 (PRK01678)Ribosomal_L31 (pfam01197)→ | rpmE2 (PRK01678)Ribosomal_L31 (pfam01197)→ |
| Efa-545-1 | NZ_JAAJ01000007.1 | - | 74320 | 74288 | **RNA**→ | rpmE2 (PRK01678)Ribosomal_L31 (pfam01197)→ | |
| Efa-338-1 | NZ_AEOU01000253.1 | + | 6244 | 6276 | **RNA**→ | rpmE2 (PRK01678)Ribosomal_L31 (pfam01197)→ | |
| Efa-339-1 | NZ_AEDZ01000214.1 | + | 6288 | 6320 | **RNA**→ | rpmE2 (PRK01678)Ribosomal_L31 (pfam01197)→ | |
| Esp-16-1 | NZ_AJRF02000013.1 | + | 59040 | 59072 | **RNA**→ | rpmE2 (PRK01678)Ribosomal_L31 (pfam01197)→ | rpmE2 (PRK01678)Ribosomal_L31 (pfam01197)→ |
| Efa-542-1 | NZ_JAAK01000063.1 | - | 6286 | 6254 | **RNA**→ | rpmE2 (PRK01678)Ribosomal_L31 (pfam01197)→ | |
| Efa-586-1 | NZ_ABQI01000014.1 | - | 45654 | 45622 | **RNA**→ | rpmE2 (PRK01678)Ribosomal_L31 (pfam01197)→ | rpmE2 (PRK01678)Ribosomal_L31 (pfam01197)→ |
| Efa-543-1 | NZ_JAAL01000009.1 | - | 75132 | 75100 | **RNA**→ | rpmE2 (PRK01678)Ribosomal_L31 (pfam01197)→ | |
| Efa-589-1 | NZ_ABSC01000180.1 | - | 3586 | 3554 | **RNA**→ | rpmE2 (PRK01678)Ribosomal_L31 (pfam01197)→ | rpmE2 (PRK01678)Ribosomal_L31 (pfam01197)→ |
| Efa-330-1 | NZ_AJTW01000196.1 | + | 6625 | 6657 | **RNA**→ | rpmE2 (PRK01678)Ribosomal_L31 (pfam01197)→ | |
| Efa-585-1 | NZ_ACOS01000122.1 | + | 43812 | 43844 | **RNA**→ | rpmE2 (PRK01678)Ribosomal_L31 (pfam01197)→ | rpmE2 (PRK01678)Ribosomal_L31 (pfam01197)→ |
| Efa-591-1 | NZ_ADMM01000075.1 | - | 28289 | 28257 | **RNA**→ | rpmE2 (PRK01678)Ribosomal_L31 (pfam01197)→ | rpmE2 (PRK01678)Ribosomal_L31 (pfam01197)→ |
| Efa-583-1 | NZ_ACZZ01000032.1 | + | 23875 | 23907 | **RNA**→ | rpmE2 (PRK01678)Ribosomal_L31 (pfam01197)→ | rpmE2 (PRK01678)Ribosomal_L31 (pfam01197)→ |
| Esp-21-1 | NZ_AJRD01000052.1 | + | 27318 | 27350 | **RNA**→ | rpmE2 (PRK01678)Ribosomal_L31 (pfam01197)→ | rpmE2 (PRK01678)Ribosomal_L31 (pfam01197)→ |
| Esp-19-1 | NZ_AJQX01000101.1 | + | 6288 | 6320 | **RNA**→ | rpmE2 (PRK01678)Ribosomal_L31 (pfam01197)→ | rpmE2 (PRK01678)Ribosomal_L31 (pfam01197)→ |
| Efa-333-1 | NZ_ACJQ02000001.1 | + | 927330 | 927362 | **RNA**→ | rpmE2 (PRK01678)Ribosomal_L31 (pfam01197)→ | |
| Efa-584-1 | NC_017960.1 | + | 728948 | 728980 | **RNA**→ | rpmE2 (PRK01678)Ribosomal_L31 (pfam01197)→ | |
| Esp-15-1 | NZ_AJRB03000048.1 | + | 20083 | 20115 | **RNA**→ | rpmE2 (PRK01678)Ribosomal_L31 (pfam01197)→ | |
| Efa-580-1 | NC_017022.1 | + | 732071 | 732103 | **RNA**→ | rpmE2 (PRK01678)Ribosomal_L31 (pfam01197)→ | |
| Efa-508-1 | NC_020207.1 | - | 2017175 | 2017143 | **RNA**→ | rpmE2 (PRK01678)Ribosomal_L31 (pfam01197)→ | |
| Efa-590-1 | NZ_ABQA01000008.1 | - | 7649 | 7617 | **RNA**→ | rpmE2 (PRK01678)Ribosomal_L31 (pfam01197)→ | rpmE2 (PRK01678)Ribosomal_L31 (pfam01197)→ |
| Efa-594-1 | NZ_ABSW01000073.1 | + | 6353 | 6385 | **RNA**→ | rpmE2 (PRK01678)Ribosomal_L31 (pfam01197)→ | rpmE2 (PRK01678)Ribosomal_L31 (pfam01197)→ |
| Efa-592-1 | NZ_GG703586.1 | + | 23876 | 23908 | **RNA**→ | rpmE2 (PRK01678)Ribosomal_L31 (pfam01197)→ | rpmE2 (PRK01678)Ribosomal_L31 (pfam01197)→ |
| Efa-581-1 | NZ_GG670309.1 | - | 219486 | 219454 | **RNA**→ | rpmE2 (PRK01678)Ribosomal_L31 (pfam01197)→ | rpmE2 (PRK01678)Ribosomal_L31 (pfam01197)→ |
| Efa-574-1 | NZ_GG688464.1 | + | 40627 | 40659 | **RNA**→ | rpmE2 (PRK01678)Ribosomal_L31 (pfam01197)→ | rpmE2 (PRK01678)Ribosomal_L31 (pfam01197)→ |
| Efa-577-1 | NZ_GG692469.1 | + | 248894 | 248926 | **RNA**→ | rpmE2 (PRK01678)Ribosomal_L31 (pfam01197)→ | rpmE2 (PRK01678)Ribosomal_L31 (pfam01197)→ |
| Efa-576-1 | NZ_GG688550.1 | + | 52869 | 52901 | **RNA**→ | rpmE2 (PRK01678)Ribosomal_L31 (pfam01197)→ | rpmE2 (PRK01678)Ribosomal_L31 (pfam01197)→ |
| Efa-575-1 | NZ_GG692539.1 | + | 253964 | 253996 | **RNA**→ | rpmE2 (PRK01678)Ribosomal_L31 (pfam01197)→ | rpmE2 (PRK01678)Ribosomal_L31 (pfam01197)→ |
| Efa-579-1 | NZ_GG688488.1 | - | 158427 | 158395 | **RNA**→ | rpmE2 (PRK01678)Ribosomal_L31 (pfam01197)→ | rpmE2 (PRK01678)Ribosomal_L31 (pfam01197)→ |
| Efa-593-1 | NZ_GG669062.1 | + | 41557 | 41589 | **RNA**→ | rpmE2 (PRK01678)Ribosomal_L31 (pfam01197)→ | |
| Efa-578-1 | NZ_GG688437.1 | + | 44705 | 44737 | **RNA**→ | rpmE2 (PRK01678)Ribosomal_L31 (pfam01197)→ | rpmE2 (PRK01678)Ribosomal_L31 (pfam01197)→ |
| Efa-582-1 | NZ_GG670326.1 | - | 265655 | 265623 | **RNA**→ | rpmE2 (PRK01678)Ribosomal_L31 (pfam01197)→ | rpmE2 (PRK01678)Ribosomal_L31 (pfam01197)→ |
| Efa-379-1 | NZ_KB029743.1 | - | 212272 | 212240 | **RNA**→ | rpmE2 (PRK01678)Ribosomal_L31 (pfam01197)→ | |
| Efa-429-1 | NZ_KN150989.1 | - | 1835052 | 1835020 | **RNA**→ | rpmE2 (PRK01678)Ribosomal_L31 (pfam01197)→ | |
| Efa-533-1 | NZ_GL454927.1 | - | 99217 | 99185 | **RNA**→ | rpmE2 (PRK01678)Ribosomal_L31 (pfam01197)→ | |
| Efa-528-1 | NZ_GL455899.1 | + | 2806 | 2838 | **RNA**→ | rpmE2 (PRK01678)Ribosomal_L31 (pfam01197)→ | |
| Efa-530-1 | NZ_GL456960.1 | - | 7536 | 7504 | **RNA**→ | rpmE2 (PRK01678)Ribosomal_L31 (pfam01197)→ | |
| Efa-529-1 | NZ_GL456579.1 | + | 8753 | 8785 | **RNA**→ | rpmE2 (PRK01678)Ribosomal_L31 (pfam01197)→ | |
| Efa-532-1 | NZ_GL476228.1 | - | 58872 | 58840 | **RNA**→ | rpmE2 (PRK01678)Ribosomal_L31 (pfam01197)→ | |
| Efa-520-1 | NZ_JH807512.1 | - | 73032 | 73000 | **RNA**→ | rpmE2 (PRK01678)Ribosomal_L31 (pfam01197)→ | |

| | | | | | |
|---|---|---|---|---|---|
| Efa-517-1 | NZ_JH807894.1 | - | 7534 | 7502 | **RNA**→ rpmE2 (PRK01678)Ribosomal_L31 (pfam01197)→ |
| Efa-534-1 | NZ_JH804714.1 | + | 41265 | 41297 | **RNA**→ rpmE2 (PRK01678)Ribosomal_L31 (pfam01197)→ |
| Efa-521-1 | NZ_JH807180.1 | - | 7534 | 7502 | **RNA**→ rpmE2 (PRK01678)Ribosomal_L31 (pfam01197)→ |
| Efa-511-1 | NZ_JH810149.1 | - | 7536 | 7504 | **RNA**→ rpmE2 (PRK01678)Ribosomal_L31 (pfam01197)→ |
| Efa-329-1 | NZ_JH811665.1 | + | 6276 | 6308 | **RNA**→ rpmE2 (PRK01678)Ribosomal_L31 (pfam01197)→ |
| Efa-331-1 | NZ_JH812054.1 | - | 7536 | 7504 | **RNA**→ rpmE2 (PRK01678)Ribosomal_L31 (pfam01197)→ |
| Efa-317-1 | NZ_JH811476.1 | - | 75604 | 75572 | **RNA**→ rpmE2 (PRK01678)Ribosomal_L31 (pfam01197)→ |
| Efa-349-1 | NZ_JH810334.1 | - | 7536 | 7504 | **RNA**→ rpmE2 (PRK01678)Ribosomal_L31 (pfam01197)→ |
| Efa-541-1 | NZ_JH806654.1 | + | 6276 | 6308 | **RNA**→ rpmE2 (PRK01678)Ribosomal_L31 (pfam01197)→ |
| Efa-513-1 | NZ_JH808985.1 | + | 6274 | 6306 | **RNA**→ rpmE2 (PRK01678)Ribosomal_L31 (pfam01197)→ |
| Efa-346-1 | NZ_JH809915.1 | + | 6276 | 6308 | **RNA**→ rpmE2 (PRK01678)Ribosomal_L31 (pfam01197)→ |
| Efa-316-1 | NZ_JH809460.1 | + | 6276 | 6308 | **RNA**→ rpmE2 (PRK01678)Ribosomal_L31 (pfam01197)→ |
| Efa-347-1 | NZ_JH811408.1 | + | 2744 | 2776 | **RNA**→ rpmE2 (PRK01678)Ribosomal_L31 (pfam01197)→ |
| Efa-519-1 | NZ_JH807528.1 | - | 7534 | 7502 | **RNA**→ rpmE2 (PRK01678)Ribosomal_L31 (pfam01197)→ |
| Efa-344-1 | NZ_JH810983.1 | + | 6276 | 6308 | **RNA**→ rpmE2 (PRK01678)Ribosomal_L31 (pfam01197)→ |
| Efa-325-1 | NZ_JH812576.1 | + | 6274 | 6306 | **RNA**→ rpmE2 (PRK01678)Ribosomal_L31 (pfam01197)→ |
| Efa-320-1 | NZ_JH814080.1 | + | 6276 | 6308 | **RNA**→ rpmE2 (PRK01678)Ribosomal_L31 (pfam01197)→ |
| Efa-516-1 | NZ_JH808111.1 | - | 7536 | 7504 | **RNA**→ rpmE2 (PRK01678)Ribosomal_L31 (pfam01197)→ |
| Efa-522-1 | NZ_JH806976.1 | + | 6276 | 6308 | **RNA**→ rpmE2 (PRK01678)Ribosomal_L31 (pfam01197)→ |
| Efa-514-1 | NZ_JH808870.1 | + | 6276 | 6308 | **RNA**→ rpmE2 (PRK01678)Ribosomal_L31 (pfam01197)→ |
| Efa-319-1 | NZ_JH813837.1 | - | 7536 | 7504 | **RNA**→ rpmE2 (PRK01678)Ribosomal_L31 (pfam01197)→ |
| Efa-512-1 | NZ_JH809227.1 | - | 7535 | 7503 | **RNA**→ rpmE2 (PRK01678)Ribosomal_L31 (pfam01197)→ |
| Efa-342-1 | NZ_JH811828.1 | + | 6272 | 6304 | **RNA**→ rpmE2 (PRK01678)Ribosomal_L31 (pfam01197)→ |
| Efa-341-1 | NZ_JH813598.1 | + | 6276 | 6308 | **RNA**→ rpmE2 (PRK01678)Ribosomal_L31 (pfam01197)→ |
| Efa-322-1 | NZ_JH812725.1 | + | 6276 | 6308 | **RNA**→ rpmE2 (PRK01678)Ribosomal_L31 (pfam01197)→ |
| Efa-510-1 | NZ_JH810528.1 | + | 6251 | 6283 | **RNA**→ rpmE2 (PRK01678)Ribosomal_L31 (pfam01197)→ |
| Efa-345-1 | NZ_JH809769.1 | + | 6276 | 6308 | **RNA**→ rpmE2 (PRK01678)Ribosomal_L31 (pfam01197)→ |
| Efa-332-1 | NZ_JH813224.1 | + | 6276 | 6308 | **RNA**→ rpmE2 (PRK01678)Ribosomal_L31 (pfam01197)→ |
| Efa-324-1 | NZ_JH812225.1 | + | 6276 | 6308 | **RNA**→ rpmE2 (PRK01678)Ribosomal_L31 (pfam01197)→ |
| Efa-323-1 | NZ_JH812970.1 | + | 6276 | 6308 | **RNA**→ rpmE2 (PRK01678)Ribosomal_L31 (pfam01197)→ |
| Efa-518-1 | NZ_JH808617.1 | + | 6276 | 6308 | **RNA**→ rpmE2 (PRK01678)Ribosomal_L31 (pfam01197)→ |
| Efa-336-1 | NZ_JH813478.1 | - | 7536 | 7504 | **RNA**→ rpmE2 (PRK01678)Ribosomal_L31 (pfam01197)→ |
| Efa-337-1 | NZ_JH814258.1 | + | 6276 | 6308 | **RNA**→ rpmE2 (PRK01678)Ribosomal_L31 (pfam01197)→ |
| Edu-3-1 | NZ_JH992913.1 | - | 613071 | 613039 | **RNA**→ rpmE2 (PRK01678)Ribosomal_L31 (pfam01197)→ |
| Efa-402-1 | NZ_KB030118.1 | + | 255203 | 255235 | **RNA**→ rpmE2 (PRK01678)Ribosomal_L31 (pfam01197)→ |
| Efa-393-1 | NZ_KB030094.1 | + | 695384 | 695416 | **RNA**→ rpmE2 (PRK01678)Ribosomal_L31 (pfam01197)→ |
| Efa-394-1 | NZ_KB029982.1 | + | 44399 | 44431 | **RNA**→ rpmE2 (PRK01678)Ribosomal_L31 (pfam01197)→ |
| Efa-368-1 | NZ_KB029483.1 | - | 213888 | 213856 | **RNA**→ rpmE2 (PRK01678)Ribosomal_L31 (pfam01197)→ |
| Efa-361-1 | NZ_KB029403.1 | + | 60851 | 60883 | **RNA**→ rpmE2 (PRK01678)Ribosomal_L31 (pfam01197)→ |
| Efa-356-1 | NZ_KB029682.1 | - | 1934997 | 1934965 | **RNA**→ rpmE2 (PRK01678)Ribosomal_L31 (pfam01197)→ |
| Efa-386-1 | NZ_KB029828.1 | + | 57930 | 57962 | **RNA**→ rpmE2 (PRK01678)Ribosomal_L31 (pfam01197)→ |
| Efa-400-1 | NZ_KB030037.1 | + | 67862 | 67894 | **RNA**→ rpmE2 (PRK01678)Ribosomal_L31 (pfam01197)→ |
| Efa-366-1 | NZ_KB029503.1 | - | 253720 | 253688 | **RNA**→ rpmE2 (PRK01678)Ribosomal_L31 (pfam01197)→ |
| Efa-354-1 | NZ_KB029387.1 | - | 40753 | 40721 | **RNA**→ rpmE2 (PRK01678)Ribosomal_L31 (pfam01197)→ |
| Efa-369-1 | NZ_KB029676.1 | + | 67496 | 67528 | **RNA**→ rpmE2 (PRK01678)Ribosomal_L31 (pfam01197)→ |
| Efa-380-1 | NZ_KB029722.1 | - | 211836 | 211804 | **RNA**→ rpmE2 (PRK01678)Ribosomal_L31 (pfam01197)→ |
| Efa-392-1 | NZ_KB030159.1 | + | 265610 | 265642 | **RNA**→ rpmE2 (PRK01678)Ribosomal_L31 (pfam01197)→ |
| Efa-397-1 | NZ_KB030080.1 | - | 108961 | 108929 | **RNA**→ rpmE2 (PRK01678)Ribosomal_L31 (pfam01197)→ |
| Efa-395-1 | NZ_KB030014.1 | - | 219917 | 219885 | **RNA**→ rpmE2 (PRK01678)Ribosomal_L31 (pfam01197)→ |
| Efa-399-1 | NZ_KB029993.1 | + | 62268 | 62300 | **RNA**→ rpmE2 (PRK01678)Ribosomal_L31 (pfam01197)→ |
| Efa-363-1 | NZ_KB029436.1 | - | 960038 | 960006 | **RNA**→ rpmE2 (PRK01678)Ribosomal_L31 (pfam01197)→ |
| Efa-357-1 | NZ_KB029490.1 | - | 459393 | 459361 | **RNA**→ rpmE2 (PRK01678)Ribosomal_L31 (pfam01197)→ |

| | | | | | | |
|---|---|---|---|---|---|---|
| Efa-358-1 | NZ_KB029598.1 | + | 46434 | 46466 | **RNA**→ | rpmE2 (PRK01678)Ribosomal_L31 (pfam01197)→ |
| Efa-391-1 | NZ_KB029933.1 | - | 53494 | 53462 | **RNA**→ | rpmE2 (PRK01678)Ribosomal_L31 (pfam01197)→ |
| Efa-398-1 | NZ_KB030041.1 | + | 682995 | 683027 | **RNA**→ | rpmE2 (PRK01678)Ribosomal_L31 (pfam01197)→ |
| Efa-401-1 | NZ_KB029969.1 | + | 44072 | 44104 | **RNA**→ | rpmE2 (PRK01678)Ribosomal_L31 (pfam01197)→ |
| Efa-365-1 | NZ_KB029577.1 | + | 250792 | 250824 | **RNA**→ | rpmE2 (PRK01678)Ribosomal_L31 (pfam01197)→ |
| Efa-370-1 | NZ_KB029606.1 | + | 45552 | 45584 | **RNA**→ | rpmE2 (PRK01678)Ribosomal_L31 (pfam01197)→ |
| Efa-359-1 | NZ_KB029431.1 | + | 44918 | 44950 | **RNA**→ | rpmE2 (PRK01678)Ribosomal_L31 (pfam01197)→ |
| Efa-367-1 | NZ_KB029659.1 | - | 211055 | 211023 | **RNA**→ | rpmE2 (PRK01678)Ribosomal_L31 (pfam01197)→ |
| Efa-378-1 | NZ_KB029972.1 | + | 432907 | 432939 | **RNA**→ | rpmE2 (PRK01678)Ribosomal_L31 (pfam01197)→ |
| Efa-375-1 | NZ_KB029716.1 | + | 48115 | 48147 | **RNA**→ | rpmE2 (PRK01678)Ribosomal_L31 (pfam01197)→ |
| Efa-382-1 | NZ_KB029943.1 | - | 105802 | 105770 | **RNA**→ | rpmE2 (PRK01678)Ribosomal_L31 (pfam01197)→ |
| Efa-377-1 | NZ_KB029704.1 | - | 1594555 | 1594523 | **RNA**→ | rpmE2 (PRK01678)Ribosomal_L31 (pfam01197)→ |
| Efa-351-1 | NZ_KB029533.1 | + | 233100 | 233132 | **RNA**→ | rpmE2 (PRK01678)Ribosomal_L31 (pfam01197)→ |
| Efa-381-1 | NZ_KB029905.1 | - | 1598264 | 1598232 | **RNA**→ | rpmE2 (PRK01678)Ribosomal_L31 (pfam01197)→ |
| Efa-389-1 | NZ_KB029866.1 | + | 45982 | 46014 | **RNA**→ | rpmE2 (PRK01678)Ribosomal_L31 (pfam01197)→ |
| Efa-360-1 | NZ_KB029516.1 | - | 210212 | 210180 | **RNA**→ | rpmE2 (PRK01678)Ribosomal_L31 (pfam01197)→ |
| Efa-350-1 | NZ_KB029609.1 | - | 213120 | 213088 | **RNA**→ | rpmE2 (PRK01678)Ribosomal_L31 (pfam01197)→ |
| Efa-373-1 | NZ_KB029748.1 | + | 68700 | 68732 | **RNA**→ | rpmE2 (PRK01678)Ribosomal_L31 (pfam01197)→ |
| Efa-396-1 | NZ_KB030126.1 | + | 888746 | 888778 | **RNA**→ | rpmE2 (PRK01678)Ribosomal_L31 (pfam01197)→ |
| Efa-500-1 | NZ_KB206111.1 | - | 78995 | 78963 | **RNA**→ | rpmE2 (PRK01678)Ribosomal_L31 (pfam01197)→ |
| Efa-362-1 | NZ_KB029455.1 | + | 253636 | 253668 | **RNA**→ | rpmE2 (PRK01678)Ribosomal_L31 (pfam01197)→ |
| Efa-352-1 | NZ_KB029696.1 | + | 44617 | 44649 | **RNA**→ | rpmE2 (PRK01678)Ribosomal_L31 (pfam01197)→ |
| Efa-384-1 | NZ_KB029840.1 | + | 281406 | 281438 | **RNA**→ | rpmE2 (PRK01678)Ribosomal_L31 (pfam01197)→ |
| Efa-385-1 | NZ_KB029962.1 | + | 326959 | 326991 | **RNA**→ | rpmE2 (PRK01678)Ribosomal_L31 (pfam01197)→ |
| Efa-376-1 | NZ_KB029699.1 | + | 411594 | 411626 | **RNA**→ | rpmE2 (PRK01678)Ribosomal_L31 (pfam01197)→ |
| Efa-501-1 | NZ_KB206122.1 | - | 78218 | 78186 | **RNA**→ | rpmE2 (PRK01678)Ribosomal_L31 (pfam01197)→ |
| Efa-372-1 | NZ_KB029904.1 | + | 77834 | 77866 | **RNA**→ | rpmE2 (PRK01678)Ribosomal_L31 (pfam01197)→ |
| Efa-314-1 | NZ_KI518313.1 | + | 70480 | 70512 | **RNA**→ | rpmE2 (PRK01678)Ribosomal_L31 (pfam01197)→ |
| Efa-507-1 | NZ_KI518270.1 | + | 253442 | 253474 | **RNA**→ | rpmE2 (PRK01678)Ribosomal_L31 (pfam01197)→ |
| Efa-328-1 | NZ_KI518258.1 | + | 257242 | 257274 | **RNA**→ | rpmE2 (PRK01678)Ribosomal_L31 (pfam01197)→ |
| Efa-498-1 | NZ_KE352418.1 | + | 6276 | 6308 | **RNA**→ | rpmE2 (PRK01678)Ribosomal_L31 (pfam01197)→ |
| Efa-403-1 | NZ_KB932617.1 | - | 925358 | 925326 | **RNA**→ | rpmE2 (PRK01678)Ribosomal_L31 (pfam01197)→ |
| Efa-408-1 | NZ_KB932649.1 | + | 43998 | 44030 | **RNA**→ | rpmE2 (PRK01678)Ribosomal_L31 (pfam01197)→ |
| Efa-417-1 | NZ_KB932697.1 | + | 45823 | 45855 | **RNA**→ | rpmE2 (PRK01678)Ribosomal_L31 (pfam01197)→ |
| Efa-423-1 | NZ_KB932732.1 | - | 232272 | 232240 | **RNA**→ | rpmE2 (PRK01678)Ribosomal_L31 (pfam01197)→ |
| Efa-388-1 | NZ_KB932310.1 | + | 486449 | 486481 | **RNA**→ | rpmE2 (PRK01678)Ribosomal_L31 (pfam01197)→ |
| Efa-416-1 | NZ_KB932687.1 | + | 40891 | 40923 | **RNA**→ | rpmE2 (PRK01678)Ribosomal_L31 (pfam01197)→ |
| Efa-494-1 | NZ_KB944883.1 | - | 208506 | 208474 | **RNA**→ | rpmE2 (PRK01678)Ribosomal_L31 (pfam01197)→ |
| Efa-326-1 | NZ_KB946243.1 | - | 119734 | 119702 | **RNA**→ | rpmE2 (PRK01678)Ribosomal_L31 (pfam01197)→ |
| Efa-191-1 | NZ_KB946795.1 | + | 45506 | 45538 | **RNA**→ | rpmE2 (PRK01678)Ribosomal_L31 (pfam01197)→ |
| Efa-422-1 | NZ_KB932724.1 | + | 45780 | 45812 | **RNA**→ | rpmE2 (PRK01678)Ribosomal_L31 (pfam01197)→ |
| Efa-493-1 | NZ_KB944903.1 | - | 207618 | 207586 | **RNA**→ | rpmE2 (PRK01678)Ribosomal_L31 (pfam01197)→ |
| Efa-413-1 | NZ_KB932656.1 | + | 329826 | 329858 | **RNA**→ | rpmE2 (PRK01678)Ribosomal_L31 (pfam01197)→ |
| Efa-355-1 | NZ_KB932305.1 | - | 1992059 | 1992027 | **RNA**→ | rpmE2 (PRK01678)Ribosomal_L31 (pfam01197)→ |
| Efa-424-1 | NZ_KB945952.1 | + | 60323 | 60355 | **RNA**→ | rpmE2 (PRK01678)Ribosomal_L31 (pfam01197)→ |
| Efa-475-1 | NZ_KB946010.1 | - | 253687 | 253655 | **RNA**→ | rpmE2 (PRK01678)Ribosomal_L31 (pfam01197)→ |
| Efa-480-1 | NZ_KB946685.1 | + | 64486 | 64518 | **RNA**→ | rpmE2 (PRK01678)Ribosomal_L31 (pfam01197)→ |
| Efa-409-1 | NZ_KB932665.1 | - | 232386 | 232354 | **RNA**→ | rpmE2 (PRK01678)Ribosomal_L31 (pfam01197)→ |
| Efa-404-1 | NZ_KB932624.1 | + | 43998 | 44030 | **RNA**→ | rpmE2 (PRK01678)Ribosomal_L31 (pfam01197)→ |
| Efa-485-1 | NZ_KB946908.1 | + | 771396 | 771428 | **RNA**→ | rpmE2 (PRK01678)Ribosomal_L31 (pfam01197)→ |
| Efa-411-1 | NZ_KB945924.1 | + | 45600 | 45632 | **RNA**→ | rpmE2 (PRK01678)Ribosomal_L31 (pfam01197)→ |
| Efa-418-1 | NZ_KB932703.1 | - | 1430349 | 1430317 | **RNA**→ | rpmE2 (PRK01678)Ribosomal_L31 (pfam01197)→ |

| | | | | | |
|---|---|---|---|---|---|
| Efa-421-1 | NZ_KB932718.1 | + | 45649 | 45681 | **RNA**→ rpmE2 (PRK01678)Ribosomal_L31 (pfam01197)→ |
| Efa-426-1 | NZ_KB932745.1 | + | 43998 | 44030 | **RNA**→ rpmE2 (PRK01678)Ribosomal_L31 (pfam01197)→ |
| Efa-405-1 | NZ_KB932639.1 | - | 230561 | 230529 | **RNA**→ rpmE2 (PRK01678)Ribosomal_L31 (pfam01197)→ |
| Efa-492-1 | NZ_KB947018.1 | - | 253035 | 253003 | **RNA**→ rpmE2 (PRK01678)Ribosomal_L31 (pfam01197)→ |
| Efa-407-1 | NZ_KB932671.1 | - | 230654 | 230622 | **RNA**→ rpmE2 (PRK01678)Ribosomal_L31 (pfam01197)→ |
| Efa-420-1 | NZ_KB945940.1 | + | 43998 | 44030 | **RNA**→ rpmE2 (PRK01678)Ribosomal_L31 (pfam01197)→ |
| Efa-428-1 | NZ_KB932766.1 | - | 210553 | 210521 | **RNA**→ rpmE2 (PRK01678)Ribosomal_L31 (pfam01197)→ |
| Efa-478-1 | NZ_KB946134.1 | + | 255837 | 255869 | **RNA**→ rpmE2 (PRK01678)Ribosomal_L31 (pfam01197)→ |
| Efa-444-1 | NZ_KB947903.1 | - | 206421 | 206389 | **RNA**→ rpmE2 (PRK01678)Ribosomal_L31 (pfam01197)→ |
| Efa-490-1 | NZ_KB946990.1 | - | 151958 | 151926 | **RNA**→ rpmE2 (PRK01678)Ribosomal_L31 (pfam01197)→ |
| Efa-482-1 | NZ_KB946712.1 | + | 70071 | 70103 | **RNA**→ rpmE2 (PRK01678)Ribosomal_L31 (pfam01197)→ |
| Efa-483-1 | NZ_KB946903.1 | - | 1377984 | 1377952 | **RNA**→ rpmE2 (PRK01678)Ribosomal_L31 (pfam01197)→ |
| Efa-487-1 | NZ_KB946953.1 | + | 258359 | 258391 | **RNA**→ rpmE2 (PRK01678)Ribosomal_L31 (pfam01197)→ |
| Efa-433-1 | NZ_KB947584.1 | - | 1000879 | 1000847 | **RNA**→ rpmE2 (PRK01678)Ribosomal_L31 (pfam01197)→ |
| Efa-439-1 | NZ_KB947721.1 | + | 66082 | 66114 | **RNA**→ rpmE2 (PRK01678)Ribosomal_L31 (pfam01197)→ |
| Efa-449-1 | NZ_KB948008.1 | + | 60821 | 60853 | **RNA**→ rpmE2 (PRK01678)Ribosomal_L31 (pfam01197)→ |
| Efa-491-1 | NZ_KB946830.1 | - | 1257899 | 1257867 | **RNA**→ rpmE2 (PRK01678)Ribosomal_L31 (pfam01197)→ |
| Efa-458-1 | NZ_KB949403.1 | + | 44027 | 44059 | **RNA**→ rpmE2 (PRK01678)Ribosomal_L31 (pfam01197)→ |
| Efa-489-1 | NZ_KB946824.1 | + | 264128 | 264160 | **RNA**→ rpmE2 (PRK01678)Ribosomal_L31 (pfam01197)→ |
| Efa-414-1 | NZ_KB932642.1 | + | 43998 | 44030 | **RNA**→ rpmE2 (PRK01678)Ribosomal_L31 (pfam01197)→ |
| Efa-479-1 | NZ_KB946149.1 | - | 76166 | 76134 | **RNA**→ rpmE2 (PRK01678)Ribosomal_L31 (pfam01197)→ |
| Efa-445-1 | NZ_KB947916.1 | - | 208388 | 208356 | **RNA**→ rpmE2 (PRK01678)Ribosomal_L31 (pfam01197)→ |
| Efa-419-1 | NZ_KB932708.1 | + | 43998 | 44030 | **RNA**→ rpmE2 (PRK01678)Ribosomal_L31 (pfam01197)→ |
| Efa-476-1 | NZ_KB946063.1 | + | 254275 | 254307 | **RNA**→ rpmE2 (PRK01678)Ribosomal_L31 (pfam01197)→ |
| Efa-481-1 | NZ_KB947049.1 | - | 255740 | 255708 | **RNA**→ rpmE2 (PRK01678)Ribosomal_L31 (pfam01197)→ |
| Efa-436-1 | NZ_KB947641.1 | + | 252182 | 252214 | **RNA**→ rpmE2 (PRK01678)Ribosomal_L31 (pfam01197)→ |
| Efa-446-1 | NZ_KB947936.1 | - | 247560 | 247528 | **RNA**→ rpmE2 (PRK01678)Ribosomal_L31 (pfam01197)→ |
| Efa-451-1 | NZ_KB948052.1 | - | 225651 | 225619 | **RNA**→ rpmE2 (PRK01678)Ribosomal_L31 (pfam01197)→ |
| Efa-456-1 | NZ_KB948239.1 | + | 8756 | 8788 | **RNA**→ rpmE2 (PRK01678)Ribosomal_L31 (pfam01197)→ |
| Efa-461-1 | NZ_KB949440.1 | + | 43998 | 44030 | **RNA**→ rpmE2 (PRK01678)Ribosomal_L31 (pfam01197)→ |
| Efa-440-1 | NZ_KB947747.1 | + | 98352 | 98384 | **RNA**→ rpmE2 (PRK01678)Ribosomal_L31 (pfam01197)→ |
| Efa-450-1 | NZ_KB948035.1 | + | 269630 | 269662 | **RNA**→ rpmE2 (PRK01678)Ribosomal_L31 (pfam01197)→ |
| Efa-459-1 | NZ_KB949415.1 | - | 904667 | 904635 | **RNA**→ rpmE2 (PRK01678)Ribosomal_L31 (pfam01197)→ |
| Efa-460-1 | NZ_KB949423.1 | + | 43998 | 44030 | **RNA**→ rpmE2 (PRK01678)Ribosomal_L31 (pfam01197)→ |
| Efa-469-1 | NZ_KB949696.1 | - | 260133 | 260101 | **RNA**→ rpmE2 (PRK01678)Ribosomal_L31 (pfam01197)→ |
| Efa-412-1 | NZ_KB945908.1 | + | 40855 | 40887 | **RNA**→ rpmE2 (PRK01678)Ribosomal_L31 (pfam01197)→ |
| Efa-443-1 | NZ_KB947850.1 | + | 252543 | 252575 | **RNA**→ rpmE2 (PRK01678)Ribosomal_L31 (pfam01197)→ |
| Efa-448-1 | NZ_KB947954.1 | + | 66239 | 66271 | **RNA**→ rpmE2 (PRK01678)Ribosomal_L31 (pfam01197)→ |
| Efa-454-1 | NZ_KB948170.1 | - | 254051 | 254019 | **RNA**→ rpmE2 (PRK01678)Ribosomal_L31 (pfam01197)→ |
| Efa-463-1 | NZ_KB949473.1 | + | 44026 | 44058 | **RNA**→ rpmE2 (PRK01678)Ribosomal_L31 (pfam01197)→ |
| Efa-464-1 | NZ_KB949489.1 | + | 65425 | 65457 | **RNA**→ rpmE2 (PRK01678)Ribosomal_L31 (pfam01197)→ |
| Efa-472-1 | NZ_KB949806.1 | + | 255072 | 255104 | **RNA**→ rpmE2 (PRK01678)Ribosomal_L31 (pfam01197)→ |
| Efa-441-1 | NZ_KB947813.1 | + | 267269 | 267301 | **RNA**→ rpmE2 (PRK01678)Ribosomal_L31 (pfam01197)→ |
| Efa-466-1 | NZ_KB949541.1 | + | 252270 | 252302 | **RNA**→ rpmE2 (PRK01678)Ribosomal_L31 (pfam01197)→ |
| Efa-488-1 | NZ_KB946960.1 | + | 771616 | 771648 | **RNA**→ rpmE2 (PRK01678)Ribosomal_L31 (pfam01197)→ |
| Efa-437-1 | NZ_KB947662.1 | - | 210708 | 210676 | **RNA**→ rpmE2 (PRK01678)Ribosomal_L31 (pfam01197)→ |
| Efa-442-1 | NZ_KB947831.1 | + | 264751 | 264783 | **RNA**→ rpmE2 (PRK01678)Ribosomal_L31 (pfam01197)→ |
| Efa-447-1 | NZ_KB947952.1 | + | 67274 | 67306 | **RNA**→ rpmE2 (PRK01678)Ribosomal_L31 (pfam01197)→ |
| Efa-452-1 | NZ_KB948120.1 | - | 309201 | 309169 | **RNA**→ rpmE2 (PRK01678)Ribosomal_L31 (pfam01197)→ |
| Efa-453-1 | NZ_KB948121.1 | + | 719806 | 719838 | **RNA**→ rpmE2 (PRK01678)Ribosomal_L31 (pfam01197)→ |
| Efa-457-1 | NZ_KB948267.1 | + | 336995 | 337027 | **RNA**→ rpmE2 (PRK01678)Ribosomal_L31 (pfam01197)→ |
| Efa-462-1 | NZ_KB949451.1 | + | 44122 | 44154 | **RNA**→ rpmE2 (PRK01678)Ribosomal_L31 (pfam01197)→ |

| | | | | | |
|---|---|---|---|---|---|
| Efa-471-1 | NZ_KB949770.1 | - | 251932 | 251900 | **RNA**→ rpmE2 (PRK01678)Ribosomal_L31 (pfam01197)→ |
| Efa-430-1 | NZ_KB947513.1 | + | 43998 | 44030 | **RNA**→ rpmE2 (PRK01678)Ribosomal_L31 (pfam01197)→ |
| Efa-465-1 | NZ_KB949525.1 | + | 67931 | 67963 | **RNA**→ rpmE2 (PRK01678)Ribosomal_L31 (pfam01197)→ |
| Efa-473-1 | NZ_KB949826.1 | - | 237195 | 237163 | **RNA**→ rpmE2 (PRK01678)Ribosomal_L31 (pfam01197)→ |
| Efa-470-1 | NZ_KB949740.1 | + | 704139 | 704171 | **RNA**→ rpmE2 (PRK01678)Ribosomal_L31 (pfam01197)→ |
| Efa-497-1 | NZ_KE136460.1 | - | 106574 | 106542 | **RNA**→ rpmE2 (PRK01678)Ribosomal_L31 (pfam01197)→ |
| Efa-188-1 | NZ_KB946782.1 | - | 205906 | 205874 | **RNA**→ rpmE2 (PRK01678)Ribosomal_L31 (pfam01197)→ |
| Efa-474-1 | NZ_KB949842.1 | + | 63724 | 63756 | **RNA**→ rpmE2 (PRK01678)Ribosomal_L31 (pfam01197)→ |
| Efa-468-1 | NZ_KB949632.1 | - | 176857 | 176825 | **RNA**→ rpmE2 (PRK01678)Ribosomal_L31 (pfam01197)→ |
| Efa-496-1 | NZ_KE136534.1 | + | 253712 | 253744 | **RNA**→ rpmE2 (PRK01678)Ribosomal_L31 (pfam01197)→ |
| Efa-497-2 | NZ_KE136554.1 | - | 152402 | 152370 | **RNA**→ rpmE2 (PRK01678)Ribosomal_L31 (pfam01197)→ |
| Efa-495-1 | NZ_KE136411.1 | - | 134559 | 134527 | **RNA**→ rpmE2 (PRK01678)Ribosomal_L31 (pfam01197)→ |
| Efa-496-2 | NZ_KE136427.1 | + | 256865 | 256897 | **RNA**→ rpmE2 (PRK01678)Ribosomal_L31 (pfam01197)→ |
| Efa-503-1 | NZ_KE150233.1 | + | 502299 | 502331 | **RNA**→ rpmE2 (PRK01678)Ribosomal_L31 (pfam01197)→ |
| Efa-524-1 | NZ_KE352068.1 | - | 7536 | 7504 | **RNA**→ rpmE2 (PRK01678)Ribosomal_L31 (pfam01197)→ |
| Efa-502-1 | NZ_KE150065.1 | + | 429121 | 429153 | **RNA**→ rpmE2 (PRK01678)Ribosomal_L31 (pfam01197)→ |
| Efa-509-1 | NZ_KE352399.1 | + | 52532 | 52564 | **RNA**→ rpmE2 (PRK01678)Ribosomal_L31 (pfam01197)→ |
| Efa-525-1 | NZ_KE351986.1 | - | 34152 | 34120 | **RNA**→ rpmE2 (PRK01678)Ribosomal_L31 (pfam01197)→ |
| Efa-531-1 | NZ_GL455151.1 | - | 99216 | 99184 | **RNA**→ rpmE2 (PRK01678)Ribosomal_L31 (pfam01197)→ |
| Efa-427-1 | NZ_KB932760.1 | - | 105635 | 105603 | **RNA**→ rpmE2 (PRK01678)Ribosomal_L31 (pfam01197)→ |
| Efa-343-1 | NZ_JH811220.1 | - | 379 | 347 | **RNA**→ rpmE2 (PRK01678)Ribosomal_L31 (pfam01197)→ |
| Efa-327-1 | NC_021994.1 | + | 687115 | 687147 | **RNA**→ rpmE2 (PRK01678)Ribosomal_L31 (pfam01197)→ |
| Efa-515-1 | NZ_JH808280.1 | - | 7536 | 7504 | **RNA**→ rpmE2 (PRK01678)Ribosomal_L31 (pfam01197)→ |
| Efa-321-1 | NZ_JH814453.1 | - | 7536 | 7504 | **RNA**→ rpmE2 (PRK01678)Ribosomal_L31 (pfam01197)→ |
| Efa-348-1 | NZ_JH810749.1 | - | 7536 | 7504 | **RNA**→ rpmE2 (PRK01678)Ribosomal_L31 (pfam01197)→ |
| Efa-313-1 | NZ_KN985989.1 | - | 2662842 | 2662810 | **RNA**→ rpmE2 (PRK01678)Ribosomal_L31 (pfam01197)→ |
| Efa-504-1 | NZ_JH636590.1 | - | 73897 | 73865 | **RNA**→ rpmE2 (PRK01678)Ribosomal_L31 (pfam01197)→ |
| Efa-499-1 | NZ_JH691958.1 | - | 1923172 | 1923140 | **RNA**→ rpmE2 (PRK01678)Ribosomal_L31 (pfam01197)→ |
| Emu-4-1 | NZ_AFWZ01000014.1 | - | 14870 | 14838 | **RNA**→ rpmE2 (PRK01678)Ribosomal_L31 (pfam01197)→ |
| Emu-1-1 | NZ_KB946218.1 | - | 1566975 | 1566943 | **RNA**→ rpmE2 (PRK01678)Ribosomal_L31 (pfam01197)→ |
| Emu-3-1 | NC_022878.1 | + | 768682 | 768714 | **RNA**→ rpmE2 (PRK01678)Ribosomal_L31 (pfam01197)→ |
| Emu-2-1 | NZ_JDFT01000010.1 | - | 62363 | 62331 | **RNA**→ rpmE2 (PRK01678)Ribosomal_L31 (pfam01197)→ |
| Cga-1-1 | NZ_JQLU01000005.1 | + | 2105866 | 2105898 | **RNA**→ rpmE2 (PRK01678)Ribosomal_L31 (pfam01197)→ |
| env-1481 | JXWR01031192.1 | - | 596 | 564 | **RNA**→ rpmE2 (PRK01678)Ribosomal_L31 (pfam01197)→ |
| Lbr-1-1 | NZ_JOKA01000068.1 | + | 76768 | 76800 | **RNA**→ rpmE2 (PRK01678)Ribosomal_L31 (pfam01197)→ |
| Lbr-1-2 | NZ_BBOW01000057.1 | + | 73685 | 73717 | **RNA**→ rpmE2 (PRK01678)Ribosomal_L31 (pfam01197)→ |
| Lbr-1-3 | NZ_JXUE01000015.1 | - | 385418 | 385386 | **RNA**→ rpmE2 (PRK01678)Ribosomal_L31 (pfam01197)→ |
| Lbr-1-4 | NZ_JXCD01000008.1 | - | 3214 | 3182 | **RNA**→ rpmE2 (PRK01678)Ribosomal_L31 (pfam01197)→ |
| Lbr-1-5 | NZ_LBHR01000024.1 | - | 33108 | 33076 | **RNA**→ rpmE2 (PRK01678)Ribosomal_L31 (pfam01197)→ |
| Lbr-5-1 | NC_020819.1 | + | 528420 | 528452 | **RNA**→ rpmE2 (PRK01678)Ribosomal_L31 (pfam01197)→ hypo→ sortase_fam (TIGR01076)Sortase_A (cd0616 |
| Lbr-6-1 | NC_008497.1 | + | 531348 | 531380 | **RNA**→ rpmE2 (PRK01678)Ribosomal_L31 (pfam01197)→ |
| Lbr-3-1 | NZ_KI271225.1 | + | 5864 | 5896 | **RNA**→ rpmE2 (PRK01678)Ribosomal_L31 (pfam01197)→ hypo→ sortase_fam (TIGR01076)Sortase_A (cd0616 |
| Lbr-2-1 | NZ_JAGR01000001.1 | - | 737769 | 737737 | **RNA**→ rpmE2 (PRK01678)Ribosomal_L31 (pfam01197)→ |
| Lbr-1-6 | NZ_JXUG01000016.1 | - | 41734 | 41702 | **RNA**→ rpmE2 (PRK01678)Ribosomal_L31 (pfam01197)→ |
| Lbr-4-1 | NZ_AUTD01000009.1 | - | 11717 | 11685 | **RNA**→ rpmE2 (PRK01678)Ribosomal_L31 (pfam01197)→ |
| Lna-2-1 | NZ_BAOT01000056.1 | - | 26005 | 25972 | **RNA**→ rpmE2 (PRK01678)Ribosomal_L31 (pfam01197)→ |
| env-1482 | 2204712971 | - | 160 | 127 | **RNA**→ rpmE2 (PRK01678)Ribosomal_L31 (pfam01197)→ |
| env-1483 | 2157898307 | + | 162 | 195 | **RNA**→ rpmE2 (PRK01678)Ribosomal_L31 (pfam01197)→ |
| Lsp-15-1 | NZ_JZCR01000012.1 | - | 34731 | 34698 | **RNA**→ rpmE2 (PRK01678)Ribosomal_L31 (pfam01197)→ |
| Lma-3-2 | NZ_BACP01000068.1 | + | 42656 | 42688 | **RNA**→ rpmE2 (PRK01678)Ribosomal_L31 (pfam01197)→ rpmE2 (PRK01678)Ribosomal_L31 (pfam01197)→ |
| Lma-3-1 | NZ_AKKT01000010.1 | + | 8614 | 8646 | **RNA**→ rpmE2 (PRK01678)Ribosomal_L31 (pfam01197)→ |
| Cfu-1-1 | NZ_JQLL01000001.1 | - | 1384361 | 1384329 | **RNA**→ rpmE2 (PRK01678)Ribosomal_L31 (pfam01197)→ |

| | | | | | |
|---|---|---|---|---|---|
| Cma-5-1 | NZ_AGNS01000026.1 | + | 96248 | 96281 | **RNA**→ rpmE2 (PRK01678)Ribosomal_L31 (pfam01197)→ rpmE2 (PRK01678)Ribosomal_L31 (pfam01197)→ |
| Cma-3-1 | NC_019425.2 | - | 2842409 | 2842376 | **RNA**→ rpmE2 (PRK01678)Ribosomal_L31 (pfam01197)→ |
| Cma-2-1 | NZ_JQMX01000001.1 | + | 3115749 | 3115782 | **RNA**→ rpmE2 (PRK01678)Ribosomal_L31 (pfam01197)→ |
| Ema-2-1 | NZ_KB946249.1 | - | 604276 | 604244 | **RNA**→ rpmE2 (PRK01678)Ribosomal_L31 (pfam01197)→ |
| Eav-1-1 | NZ_JUZB01000046.1 | + | 16390 | 16422 | **RNA**→ rpmE2 (PRK01678)Ribosomal_L31 (pfam01197)→ |
| Era-1-1 | NZ_KB946257.1 | + | 170182 | 170214 | **RNA**→ rpmE2 (PRK01678)Ribosomal_L31 (pfam01197)→ |
| Ehi-1-1 | NZ_JUPQ01000184.1 | + | 9015 | 9047 | **RNA**→ rpmE2 (PRK01678)Ribosomal_L31 (pfam01197)→ |
| Eav-2-1 | NZ_KE136363.1 | + | 39237 | 39269 | **RNA**→ rpmE2 (PRK01678)Ribosomal_L31 (pfam01197)→ |
| Eav-2-2 | NZ_KE136509.1 | - | 615844 | 615812 | **RNA**→ rpmE2 (PRK01678)Ribosomal_L31 (pfam01197)→ |
| env-1484 | 2205839497 | - | 267 | 235 | **RNA**→ rpmE2 (PRK01678)Ribosomal_L31 (pfam01197)→ |
| env-1485 | 2158422988 | - | 267 | 235 | **RNA**→ rpmE2 (PRK01678)Ribosomal_L31 (pfam01197)→ |
| Ehi-2-1 | NZ_KB946231.1 | - | 1167763 | 1167731 | **RNA**→ rpmE2 (PRK01678)Ribosomal_L31 (pfam01197)→ |
| Efa-406-1 | NZ_KB932610.1 | - | 340657 | 340625 | **RNA**→ rpmE2 (PRK01678)Ribosomal_L31 (pfam01197)→ |
| Ehi-2-2 | NC_018081.1 | + | 1570676 | 1570708 | **RNA**→ rpmE2 (PRK01678)Ribosomal_L31 (pfam01197)→ |
| Ehi-1-2 | NZ_JMIG01000005.1 | + | 48625 | 48657 | **RNA**→ rpmE2 (PRK01678)Ribosomal_L31 (pfam01197)→ |
| Edu-2-1 | NZ_KE136383.1 | + | 49835 | 49867 | **RNA**→ rpmE2 (PRK01678)Ribosomal_L31 (pfam01197)→ |
| Edu-2-2 | NZ_KE136516.1 | + | 45291 | 45323 | **RNA**→ rpmE2 (PRK01678)Ribosomal_L31 (pfam01197)→ |
| Edu-1-1 | NZ_CP012384.1 | - | 2493264 | 2493232 | **RNA**→ rpmE2 (PRK01678)Ribosomal_L31 (pfam01197)→ |
| Edu-1-2 | NZ_CP012366.1 | - | 697844 | 697812 | **RNA**→ rpmE2 (PRK01678)Ribosomal_L31 (pfam01197)→ |
| Edu-4-1 | NZ_AOSM01000013.1 | + | 23749 | 23781 | **RNA**→ rpmE2 (PRK01678)Ribosomal_L31 (pfam01197)→ |
| env-1486 | ACOFG987_F36MELC02HSZ0T | + | 32 | 64 | **RNA**→ hypo→ rpmE2 (PRK01678)Ribosomal_L31 (pfam01197)→ |
| Evi-1-1 | NZ_KB946287.1 | - | 791418 | 791386 | **RNA**→ rpmE2 (PRK01678)Ribosomal_L31 (pfam01197)→ |
| Efa-172-1 | NZ_AJET01000006.1 | + | 629973 | 630005 | **RNA**→ rpmE2 (PRK01678)Ribosomal_L31 (pfam01197)→ |
| Esp-12-1 | NZ_JWBQ01000066.1 | + | 72115 | 72147 | **RNA**→ rpmE2 (PRK01678)Ribosomal_L31 (pfam01197)→ |
| Efa-1-1 | NZ_JVQP01000016.1 | + | 72248 | 72280 | **RNA**→ rpmE2 (PRK01678)Ribosomal_L31 (pfam01197)→ |
| Efa-1-2 | NZ_JVOA01000123.1 | + | 50507 | 50539 | **RNA**→ rpmE2 (PRK01678)Ribosomal_L31 (pfam01197)→ |
| Efa-1-3 | NZ_JVOF01000024.1 | - | 136699 | 136667 | **RNA**→ rpmE2 (PRK01678)Ribosomal_L31 (pfam01197)→ |
| Efa-1-4 | NZ_JVZS01000075.1 | - | 80355 | 80323 | **RNA**→ rpmE2 (PRK01678)Ribosomal_L31 (pfam01197)→ |
| Efa-1-5 | NZ_JVQT01000017.1 | - | 155630 | 155598 | **RNA**→ rpmE2 (PRK01678)Ribosomal_L31 (pfam01197)→ |
| Efa-1-6 | NZ_JVQS01000014.1 | + | 72248 | 72280 | **RNA**→ rpmE2 (PRK01678)Ribosomal_L31 (pfam01197)→ |
| Efa-1-7 | NZ_JVIK01000038.1 | - | 75811 | 75779 | **RNA**→ rpmE2 (PRK01678)Ribosomal_L31 (pfam01197)→ |
| Efa-1-8 | NZ_JVBG01000102.1 | - | 20244 | 20212 | **RNA**→ rpmE2 (PRK01678)Ribosomal_L31 (pfam01197)→ |
| Efa-1-9 | NZ_JVPG01000107.1 | - | 20252 | 20220 | **RNA**→ rpmE2 (PRK01678)Ribosomal_L31 (pfam01197)→ |
| Efa-1-10 | NZ_JVIY01000046.1 | - | 20244 | 20212 | **RNA**→ rpmE2 (PRK01678)Ribosomal_L31 (pfam01197)→ |
| Esp-13-1 | NZ_JVOE01000070.1 | + | 50458 | 50490 | **RNA**→ rpmE2 (PRK01678)Ribosomal_L31 (pfam01197)→ |
| Efa-1-11 | NZ_JVQQ01000033.1 | - | 155729 | 155697 | **RNA**→ rpmE2 (PRK01678)Ribosomal_L31 (pfam01197)→ |
| Efa-1-12 | NZ_JVQF01000009.1 | + | 50546 | 50578 | **RNA**→ rpmE2 (PRK01678)Ribosomal_L31 (pfam01197)→ |
| Efa-1-13 | NZ_JVBD01000057.1 | + | 50551 | 50583 | **RNA**→ rpmE2 (PRK01678)Ribosomal_L31 (pfam01197)→ |
| Efa-1-14 | NZ_JVPS01000092.1 | + | 50507 | 50539 | **RNA**→ rpmE2 (PRK01678)Ribosomal_L31 (pfam01197)→ |
| Efa-1-15 | NZ_JUZT01000074.1 | + | 50507 | 50539 | **RNA**→ rpmE2 (PRK01678)Ribosomal_L31 (pfam01197)→ |
| Efa-1-16 | NZ_JVOB01000025.1 | + | 50497 | 50529 | **RNA**→ rpmE2 (PRK01678)Ribosomal_L31 (pfam01197)→ |
| Efa-1-17 | NZ_JVOC01000134.1 | - | 75987 | 75955 | **RNA**→ rpmE2 (PRK01678)Ribosomal_L31 (pfam01197)→ |
| Efa-1-18 | NZ_JVAM01000128.1 | - | 78901 | 78869 | **RNA**→ rpmE2 (PRK01678)Ribosomal_L31 (pfam01197)→ |
| Efa-1-19 | NZ_JUWK01000014.1 | + | 66857 | 66889 | **RNA**→ rpmE2 (PRK01678)Ribosomal_L31 (pfam01197)→ |
| Efa-1-20 | NZ_JVJB01000035.1 | - | 75865 | 75833 | **RNA**→ rpmE2 (PRK01678)Ribosomal_L31 (pfam01197)→ |
| Efa-1-21 | NZ_JUMK01000040.1 | + | 72186 | 72218 | **RNA**→ rpmE2 (PRK01678)Ribosomal_L31 (pfam01197)→ |
| Efa-1-22 | NZ_JUOO01000146.1 | - | 20246 | 20214 | **RNA**→ rpmE2 (PRK01678)Ribosomal_L31 (pfam01197)→ |
| Efa-1-23 | NZ_JUVP01000129.1 | - | 75833 | 75801 | **RNA**→ rpmE2 (PRK01678)Ribosomal_L31 (pfam01197)→ |
| Efa-567-1 | NZ_ACOX02000001.1 | - | 165398 | 165366 | **RNA**→ rpmE2 (PRK01678)Ribosomal_L31 (pfam01197)→ rpmE2 (PRK01678)Ribosomal_L31 (pfam01197)→ |
| Efa-559-1 | NZ_ADKN01000064.1 | + | 5576 | 5608 | **RNA**→ rpmE2 (PRK01678)Ribosomal_L31 (pfam01197)→ rpmE2 (PRK01678)Ribosomal_L31 (pfam01197)→ |
| Efa-278-1 | NZ_AYKU01000027.1 | - | 144500 | 144468 | **RNA**→ rpmE2 (PRK01678)Ribosomal_L31 (pfam01197)→ |
| Efa-22-1 | NZ_AYLU01000042.1 | + | 97278 | 97310 | **RNA**→ rpmE2 (PRK01678)Ribosomal_L31 (pfam01197)→ |

| | | | | | |
|---|---|---|---|---|---|
| Efa-1-24 | NZ_JPTY01000024.1 | + | 98124 | 98156 | **RNA**→ rpmE2 (PRK01678)Ribosomal_L31 (pfam01197)→ |
| Efa-549-1 | NC_018221.1 | + | 1022628 | 1022660 | **RNA**→ rpmE2 (PRK01678)Ribosomal_L31 (pfam01197)→ |
| Efa-312-1 | NC_019770.1 | + | 1028524 | 1028556 | **RNA**→ rpmE2 (PRK01678)Ribosomal_L31 (pfam01197)→ |
| Efa-19-1 | NZ_CP008816.1 | + | 231325 | 231357 | **RNA**→ rpmE2 (PRK01678)Ribosomal_L31 (pfam01197)→ |
| Efa-560-1 | NZ_GG739846.1 | - | 421383 | 421351 | **RNA**→ rpmE2 (PRK01678)Ribosomal_L31 (pfam01197)→ |
| Efa-562-1 | NZ_GG670358.1 | - | 1399663 | 1399631 | **RNA**→ rpmE2 (PRK01678)Ribosomal_L31 (pfam01197)→ rpmE2 (PRK01678)Ribosomal_L31 (pfam01197)→ |
| Efa-551-1 | NZ_GG692891.1 | + | 121192 | 121224 | **RNA**→ rpmE2 (PRK01678)Ribosomal_L31 (pfam01197)→ rpmE2 (PRK01678)Ribosomal_L31 (pfam01197)→ |
| Efa-18-2 | NZ_GG668755.1 | - | 1351023 | 1350991 | **RNA**→ rpmE2 (PRK01678)Ribosomal_L31 (pfam01197)→ |
| Efa-557-1 | NZ_GG692929.1 | - | 1368435 | 1368403 | **RNA**→ rpmE2 (PRK01678)Ribosomal_L31 (pfam01197)→ rpmE2 (PRK01678)Ribosomal_L31 (pfam01197)→ |
| Efa-573-1 | NZ_GG688434.1 | - | 1376486 | 1376454 | **RNA**→ rpmE2 (PRK01678)Ribosomal_L31 (pfam01197)→ rpmE2 (PRK01678)Ribosomal_L31 (pfam01197)→ |
| Efa-553-1 | NZ_GG692670.1 | - | 1446803 | 1446771 | **RNA**→ rpmE2 (PRK01678)Ribosomal_L31 (pfam01197)→ rpmE2 (PRK01678)Ribosomal_L31 (pfam01197)→ |
| Efa-556-1 | NZ_GG692696.1 | - | 38543 | 38511 | **RNA**→ rpmE2 (PRK01678)Ribosomal_L31 (pfam01197)→ rpmE2 (PRK01678)Ribosomal_L31 (pfam01197)→ |
| Efa-552-1 | NZ_GG692683.1 | - | 1370149 | 1370117 | **RNA**→ rpmE2 (PRK01678)Ribosomal_L31 (pfam01197)→ rpmE2 (PRK01678)Ribosomal_L31 (pfam01197)→ |
| Efa-289-1 | NZ_GL454506.1 | - | 3215 | 3183 | **RNA**→ rpmE2 (PRK01678)Ribosomal_L31 (pfam01197)→ hypo→ DUF916 (pfam06030)DUF3324 (pfam11797 |
| | | | | | hypo→ hypo→ NagC (COG1940)ROK_glcA_fam (TIGR00744)→ CsbD (pfam05532)→ |
| Efa-300-1 | NZ_GL454537.1 | + | 136485 | 136517 | **RNA**→ rpmE2 (PRK01678)Ribosomal_L31 (pfam01197)→ |
| Efa-28-1 | NZ_GL455718.1 | + | 15340 | 15372 | **RNA**→ rpmE2 (PRK01678)Ribosomal_L31 (pfam01197)→ |
| Efa-285-1 | NZ_GL456486.1 | + | 15327 | 15359 | **RNA**→ rpmE2 (PRK01678)Ribosomal_L31 (pfam01197)→ |
| Efa-27-1 | NZ_GL455808.1 | + | 15343 | 15375 | **RNA**→ rpmE2 (PRK01678)Ribosomal_L31 (pfam01197)→ |
| Efa-294-1 | NZ_GL456782.1 | - | 70795 | 70763 | **RNA**→ rpmE2 (PRK01678)Ribosomal_L31 (pfam01197)→ |
| Efa-290-1 | NZ_GL455121.1 | + | 15383 | 15415 | **RNA**→ rpmE2 (PRK01678)Ribosomal_L31 (pfam01197)→ |
| Efa-572-1 | NZ_GL454430.1 | - | 79041 | 79009 | **RNA**→ rpmE2 (PRK01678)Ribosomal_L31 (pfam01197)→ hypo→ DUF916 (pfam06030)DUF3324 (pfam11797 |
| | | | | | hypo→ hypo→ NagC (COG1940)ROK_glcA_fam (TIGR00744)→ CsbD (pfam05532)YtxH (pfam12732)→ |
| Efa-571-1 | NZ_GL455597.1 | + | 15307 | 15339 | **RNA**→ rpmE2 (PRK01678)Ribosomal_L31 (pfam01197)→ |
| Efa-303-1 | NZ_GL454593.1 | + | 75214 | 75246 | **RNA**→ rpmE2 (PRK01678)Ribosomal_L31 (pfam01197)→ |
| Efa-288-1 | NZ_GL454737.1 | - | 70823 | 70791 | **RNA**→ rpmE2 (PRK01678)Ribosomal_L31 (pfam01197)→ |
| Efa-291-1 | NZ_GL455457.1 | - | 99932 | 99900 | **RNA**→ rpmE2 (PRK01678)Ribosomal_L31 (pfam01197)→ |
| Efa-286-1 | NZ_GL454210.1 | + | 100704 | 100736 | **RNA**→ rpmE2 (PRK01678)Ribosomal_L31 (pfam01197)→ |
| Efa-287-1 | NZ_GL454392.1 | - | 79063 | 79031 | **RNA**→ rpmE2 (PRK01678)Ribosomal_L31 (pfam01197)→ |
| Efa-252-1 | NZ_KI913024.1 | + | 162848 | 162880 | **RNA**→ rpmE2 (PRK01678)Ribosomal_L31 (pfam01197)→ |
| Efa-245-1 | NZ_KI912990.1 | + | 249634 | 249666 | **RNA**→ rpmE2 (PRK01678)Ribosomal_L31 (pfam01197)→ |
| Efa-258-1 | NZ_KI913061.1 | + | 883273 | 883305 | **RNA**→ rpmE2 (PRK01678)Ribosomal_L31 (pfam01197)→ |
| Efa-241-1 | NZ_KI912966.1 | + | 125732 | 125764 | **RNA**→ rpmE2 (PRK01678)Ribosomal_L31 (pfam01197)→ |
| Efa-254-1 | NZ_KI913040.1 | + | 118788 | 118820 | **RNA**→ rpmE2 (PRK01678)Ribosomal_L31 (pfam01197)→ |
| Efa-243-1 | NZ_KI912979.1 | + | 120525 | 120557 | **RNA**→ rpmE2 (PRK01678)Ribosomal_L31 (pfam01197)→ |
| Efa-240-1 | NZ_KI912960.1 | + | 882120 | 882152 | **RNA**→ rpmE2 (PRK01678)Ribosomal_L31 (pfam01197)→ |
| Efa-263-1 | NZ_KI913093.1 | + | 1162405 | 1162437 | **RNA**→ rpmE2 (PRK01678)Ribosomal_L31 (pfam01197)→ |
| Efa-239-1 | NZ_KI912956.1 | + | 166079 | 166111 | **RNA**→ rpmE2 (PRK01678)Ribosomal_L31 (pfam01197)→ |
| Efa-250-1 | NZ_KI913015.1 | + | 134778 | 134810 | **RNA**→ rpmE2 (PRK01678)Ribosomal_L31 (pfam01197)→ |
| Efa-262-1 | NZ_KI913086.1 | + | 243014 | 243046 | **RNA**→ rpmE2 (PRK01678)Ribosomal_L31 (pfam01197)→ |
| Efa-249-1 | NZ_KI913011.1 | + | 1118035 | 1118067 | **RNA**→ rpmE2 (PRK01678)Ribosomal_L31 (pfam01197)→ |
| Efa-238-1 | NZ_KI912939.1 | + | 893011 | 893043 | **RNA**→ rpmE2 (PRK01678)Ribosomal_L31 (pfam01197)→ |
| Efa-242-1 | NZ_KI912974.1 | + | 120793 | 120825 | **RNA**→ rpmE2 (PRK01678)Ribosomal_L31 (pfam01197)→ |
| Efa-255-1 | NZ_KI913051.1 | + | 1065840 | 1065872 | **RNA**→ rpmE2 (PRK01678)Ribosomal_L31 (pfam01197)→ |
| Efa-260-1 | NZ_KI913083.1 | - | 1336694 | 1336662 | **RNA**→ rpmE2 (PRK01678)Ribosomal_L31 (pfam01197)→ |
| Efa-259-1 | NZ_KI913069.1 | + | 815731 | 815763 | **RNA**→ rpmE2 (PRK01678)Ribosomal_L31 (pfam01197)→ |
| Efa-228-1 | NZ_KB944632.1 | + | 125914 | 125946 | **RNA**→ rpmE2 (PRK01678)Ribosomal_L31 (pfam01197)→ |
| Efa-217-1 | NZ_KB944671.1 | + | 124502 | 124534 | **RNA**→ rpmE2 (PRK01678)Ribosomal_L31 (pfam01197)→ |
| Efa-214-1 | NZ_KB944586.1 | + | 118308 | 118340 | **RNA**→ rpmE2 (PRK01678)Ribosomal_L31 (pfam01197)→ |
| Efa-215-1 | NZ_KB944615.1 | + | 121371 | 121403 | **RNA**→ rpmE2 (PRK01678)Ribosomal_L31 (pfam01197)→ |
| Efa-222-1 | NZ_KB944766.1 | + | 142847 | 142879 | **RNA**→ rpmE2 (PRK01678)Ribosomal_L31 (pfam01197)→ |
| Efa-68-1 | NZ_KB932378.1 | + | 125977 | 126009 | **RNA**→ rpmE2 (PRK01678)Ribosomal_L31 (pfam01197)→ |
| Efa-20-1 | NZ_KB944642.1 | + | 125027 | 125059 | **RNA**→ rpmE2 (PRK01678)Ribosomal_L31 (pfam01197)→ |

| | | | | | |
|---|---|---|---|---|---|
| Efa-195-1 | NZ_KB945025.1 | + | 125183 | 125215 | **RNA**→ rpmE2 (PRK01678)Ribosomal_L31 (pfam01197)→ |
| Efa-203-1 | NZ_KB945029.1 | + | 168632 | 168664 | **RNA**→ rpmE2 (PRK01678)Ribosomal_L31 (pfam01197)→ |
| Efa-216-1 | NZ_KB944663.1 | + | 122100 | 122132 | **RNA**→ rpmE2 (PRK01678)Ribosomal_L31 (pfam01197)→ |
| Efa-14-1 | NZ_KB944873.1 | + | 123793 | 123825 | **RNA**→ rpmE2 (PRK01678)Ribosomal_L31 (pfam01197)→ |
| Efa-200-1 | NZ_KB944979.1 | + | 120581 | 120613 | **RNA**→ rpmE2 (PRK01678)Ribosomal_L31 (pfam01197)→ |
| Efa-198-1 | NZ_KB945018.1 | + | 120521 | 120553 | **RNA**→ rpmE2 (PRK01678)Ribosomal_L31 (pfam01197)→ |
| Efa-221-1 | NZ_KB944759.1 | + | 143579 | 143611 | **RNA**→ rpmE2 (PRK01678)Ribosomal_L31 (pfam01197)→ hypo→ DUF916 (pfam06030)DUF3324 (pfam11797 hypo→ |
| Efa-196-1 | NZ_KB944971.1 | + | 149512 | 149544 | **RNA**→ rpmE2 (PRK01678)Ribosomal_L31 (pfam01197)→ |
| Efa-16-1 | NZ_KB947429.1 | + | 157881 | 157913 | **RNA**→ rpmE2 (PRK01678)Ribosomal_L31 (pfam01197)→ |
| Efa-67-1 | NZ_KB932366.1 | + | 139890 | 139922 | **RNA**→ rpmE2 (PRK01678)Ribosomal_L31 (pfam01197)→ |
| Efa-162-1 | NZ_KB946671.1 | + | 158011 | 158043 | **RNA**→ rpmE2 (PRK01678)Ribosomal_L31 (pfam01197)→ |
| Efa-152-1 | NZ_KB946409.1 | + | 160128 | 160160 | **RNA**→ rpmE2 (PRK01678)Ribosomal_L31 (pfam01197)→ |
| Efa-218-1 | NZ_KB944674.1 | + | 123199 | 123231 | **RNA**→ rpmE2 (PRK01678)Ribosomal_L31 (pfam01197)→ |
| Efa-165-1 | NZ_KB947327.1 | + | 97950 | 97982 | **RNA**→ rpmE2 (PRK01678)Ribosomal_L31 (pfam01197)→ |
| Efa-179-1 | NZ_KB947400.1 | + | 122702 | 122734 | **RNA**→ rpmE2 (PRK01678)Ribosomal_L31 (pfam01197)→ |
| Efa-174-1 | NZ_KB947439.1 | + | 149395 | 149427 | **RNA**→ rpmE2 (PRK01678)Ribosomal_L31 (pfam01197)→ |
| Efa-176-1 | NZ_KB947300.1 | + | 865457 | 865489 | **RNA**→ rpmE2 (PRK01678)Ribosomal_L31 (pfam01197)→ |
| Efa-167-1 | NZ_KB947338.1 | + | 123113 | 123145 | **RNA**→ rpmE2 (PRK01678)Ribosomal_L31 (pfam01197)→ |
| Efa-177-1 | NZ_KB947421.1 | + | 121290 | 121322 | **RNA**→ rpmE2 (PRK01678)Ribosomal_L31 (pfam01197)→ |
| Efa-175-1 | NZ_KB947485.1 | + | 846302 | 846334 | **RNA**→ rpmE2 (PRK01678)Ribosomal_L31 (pfam01197)→ |
| Efa-17-1 | NZ_KB947435.1 | + | 136478 | 136510 | **RNA**→ rpmE2 (PRK01678)Ribosomal_L31 (pfam01197)→ |
| Efa-187-1 | NZ_KB947334.1 | + | 785472 | 785504 | **RNA**→ rpmE2 (PRK01678)Ribosomal_L31 (pfam01197)→ |
| Efa-171-1 | NZ_KB947466.1 | + | 150408 | 150440 | **RNA**→ rpmE2 (PRK01678)Ribosomal_L31 (pfam01197)→ |
| Efa-168-1 | NZ_KB947308.1 | + | 122445 | 122477 | **RNA**→ rpmE2 (PRK01678)Ribosomal_L31 (pfam01197)→ |
| Efa-18-1 | NZ_KB947320.1 | + | 123023 | 123055 | **RNA**→ rpmE2 (PRK01678)Ribosomal_L31 (pfam01197)→ |
| Efa-169-1 | NZ_KB947506.1 | + | 123365 | 123397 | **RNA**→ rpmE2 (PRK01678)Ribosomal_L31 (pfam01197)→ |
| Efa-467-1 | NZ_KB949610.1 | + | 123805 | 123837 | **RNA**→ rpmE2 (PRK01678)Ribosomal_L31 (pfam01197)→ |
| Efa-145-1 | NZ_KB949669.1 | + | 125345 | 125377 | **RNA**→ rpmE2 (PRK01678)Ribosomal_L31 (pfam01197)→ |
| Efa-23-1 | NZ_KE351093.1 | + | 66098 | 66130 | **RNA**→ rpmE2 (PRK01678)Ribosomal_L31 (pfam01197)→ |
| Efa-10-1 | NZ_KE351720.1 | + | 133898 | 133930 | **RNA**→ rpmE2 (PRK01678)Ribosomal_L31 (pfam01197)→ |
| Efa-9-1 | NZ_KE350863.1 | + | 133929 | 133961 | **RNA**→ rpmE2 (PRK01678)Ribosomal_L31 (pfam01197)→ |
| Efa-311-1 | NZ_KE352562.1 | - | 571410 | 571378 | **RNA**→ rpmE2 (PRK01678)Ribosomal_L31 (pfam01197)→ |
| Efa-523-1 | NZ_KE352305.1 | + | 15319 | 15351 | **RNA**→ rpmE2 (PRK01678)Ribosomal_L31 (pfam01197)→ |
| Efa-308-1 | NZ_KE352699.1 | + | 133744 | 133776 | **RNA**→ rpmE2 (PRK01678)Ribosomal_L31 (pfam01197)→ |
| Efa-3-1 | NZ_KE350832.1 | + | 97516 | 97548 | **RNA**→ rpmE2 (PRK01678)Ribosomal_L31 (pfam01197)→ |
| Efa-11-1 | NZ_KE351747.1 | + | 101809 | 101841 | **RNA**→ rpmE2 (PRK01678)Ribosomal_L31 (pfam01197)→ |
| Efa-561-1 | NZ_GG739749.1 | + | 72311 | 72343 | **RNA**→ rpmE2 (PRK01678)Ribosomal_L31 (pfam01197)→ |
| Efa-104-1 | NZ_KB932565.1 | + | 120769 | 120801 | **RNA**→ rpmE2 (PRK01678)Ribosomal_L31 (pfam01197)→ |
| Efa-247-1 | NZ_KI913001.1 | + | 251023 | 251055 | **RNA**→ rpmE2 (PRK01678)Ribosomal_L31 (pfam01197)→ |
| Efa-268-1 | NZ_AYLV01000014.1 | + | 15346 | 15378 | **RNA**→ rpmE2 (PRK01678)Ribosomal_L31 (pfam01197)→ |
| Efa-1-25 | NZ_JTKW01000031.1 | - | 79072 | 79040 | **RNA**→ rpmE2 (PRK01678)Ribosomal_L31 (pfam01197)→ |
| Efa-1-26 | NZ_JTKT01000016.1 | - | 79072 | 79040 | **RNA**→ rpmE2 (PRK01678)Ribosomal_L31 (pfam01197)→ |
| Efa-1-27 | NZ_JUPP01000126.1 | - | 23368 | 23336 | **RNA**→ rpmE2 (PRK01678)Ribosomal_L31 (pfam01197)→ |
| env-1487 | AEQT01000844.1 | + | 5466 | 5498 | **RNA**→ rpmE2 (PRK01678)Ribosomal_L31 (pfam01197)→ |
| Efa-1-28 | NZ_JPTZ01000018.1 | + | 98763 | 98795 | **RNA**→ rpmE2 (PRK01678)Ribosomal_L31 (pfam01197)→ |
| Efa-30-1 | NZ_AWPI01000053.1 | - | 23340 | 23308 | **RNA**→ rpmE2 (PRK01678)Ribosomal_L31 (pfam01197)→ |
| Efa-193-1 | NZ_AOPW01000055.1 | - | 6414 | 6382 | **RNA**→ rpmE2 (PRK01678)Ribosomal_L31 (pfam01197)→ |
| Efa-1-29 | NZ_LAEB01000001.1 | + | 120172 | 120204 | **RNA**→ rpmE2 (PRK01678)Ribosomal_L31 (pfam01197)→ |
| Efa-281-1 | NZ_AFHH01000009.1 | + | 5589 | 5621 | **RNA**→ rpmE2 (PRK01678)Ribosomal_L31 (pfam01197)→ |
| Efa-273-1 | NZ_AGVN01000048.1 | - | 71733 | 71701 | **RNA**→ rpmE2 (PRK01678)Ribosomal_L31 (pfam01197)→ |
| Efa-1-30 | NZ_JWAW01000015.1 | - | 23381 | 23349 | **RNA**→ rpmE2 (PRK01678)Ribosomal_L31 (pfam01197)→ |
| Efa-1-31 | NZ_JWBU01000140.1 | - | 3760 | 3728 | **RNA**→ rpmE2 (PRK01678)Ribosomal_L31 (pfam01197)→ |

| | | | | | |
|---|---|---|---|---|---|
| Efa-1-32 | NZ_JVYW01000023.1 | - | 39746 | 39714 | **RNA**→ rpmE2 (PRK01678)Ribosomal_L31 (pfam01197)→ |
| Efa-1-33 | NZ_JVTX01000024.1 | - | 139814 | 139782 | **RNA**→ rpmE2 (PRK01678)Ribosomal_L31 (pfam01197)→ |
| Efa-1-34 | NZ_JVVS01000038.1 | + | 50558 | 50590 | **RNA**→ rpmE2 (PRK01678)Ribosomal_L31 (pfam01197)→ |
| Efa-1-35 | NZ_JVSW01000024.1 | + | 67849 | 67881 | **RNA**→ rpmE2 (PRK01678)Ribosomal_L31 (pfam01197)→ |
| Efa-1-36 | NZ_JVTK01000025.1 | - | 139814 | 139782 | **RNA**→ rpmE2 (PRK01678)Ribosomal_L31 (pfam01197)→ |
| Efa-1-37 | NZ_JVZM01000052.1 | + | 50559 | 50591 | **RNA**→ rpmE2 (PRK01678)Ribosomal_L31 (pfam01197)→ |
| Efa-1-38 | NZ_JVTP01000163.1 | - | 139757 | 139725 | **RNA**→ rpmE2 (PRK01678)Ribosomal_L31 (pfam01197)→ |
| Efa-1-39 | NZ_JVXY01000154.1 | + | 72669 | 72701 | **RNA**→ rpmE2 (PRK01678)Ribosomal_L31 (pfam01197)→ |
| Efa-1-40 | NZ_JVUK01000009.1 | + | 50550 | 50582 | **RNA**→ rpmE2 (PRK01678)Ribosomal_L31 (pfam01197)→ |
| Efa-1-41 | NZ_JVCH01000106.1 | + | 50558 | 50590 | **RNA**→ rpmE2 (PRK01678)Ribosomal_L31 (pfam01197)→ |
| Efa-1-42 | NZ_JVSV01000004.1 | - | 139638 | 139606 | **RNA**→ rpmE2 (PRK01678)Ribosomal_L31 (pfam01197)→ |
| Efa-1-43 | NZ_JVTG01000037.1 | + | 100342 | 100374 | **RNA**→ rpmE2 (PRK01678)Ribosomal_L31 (pfam01197)→ |
| Efa-1-44 | NZ_JVKC01000009.1 | + | 66961 | 66993 | **RNA**→ rpmE2 (PRK01678)Ribosomal_L31 (pfam01197)→ |
| Efa-1-45 | NZ_JVGB01000067.1 | - | 41207 | 41175 | **RNA**→ rpmE2 (PRK01678)Ribosomal_L31 (pfam01197)→ |
| Efa-1-46 | NZ_JVAI01000107.1 | + | 72273 | 72305 | **RNA**→ rpmE2 (PRK01678)Ribosomal_L31 (pfam01197)→ |
| Efa-1-47 | NZ_JVPV01000250.1 | + | 3341 | 3373 | **RNA**→ rpmE2 (PRK01678)Ribosomal_L31 (pfam01197)→ |
| Efa-1-48 | NZ_JVXC01000110.1 | + | 72810 | 72842 | **RNA**→ rpmE2 (PRK01678)Ribosomal_L31 (pfam01197)→ |
| Efa-1-49 | NZ_JVQY01000105.1 | - | 40275 | 40243 | **RNA**→ rpmE2 (PRK01678)Ribosomal_L31 (pfam01197)→ |
| Efa-1-50 | NZ_JVPJ01000031.1 | - | 150815 | 150783 | **RNA**→ rpmE2 (PRK01678)Ribosomal_L31 (pfam01197)→ |
| Efa-1-51 | NZ_JVBW01000037.1 | + | 66956 | 66988 | **RNA**→ rpmE2 (PRK01678)Ribosomal_L31 (pfam01197)→ |
| Efa-1-52 | NZ_JVBV01000042.1 | + | 72347 | 72379 | **RNA**→ rpmE2 (PRK01678)Ribosomal_L31 (pfam01197)→ |
| Efa-1-53 | NZ_JVHL01000037.1 | - | 260046 | 260014 | **RNA**→ rpmE2 (PRK01678)Ribosomal_L31 (pfam01197)→ |
| Efa-1-54 | NZ_JVOQ01000069.1 | - | 77612 | 77580 | **RNA**→ rpmE2 (PRK01678)Ribosomal_L31 (pfam01197)→ |
| Efa-1-55 | NZ_JVOG01000063.1 | - | 40276 | 40244 | **RNA**→ rpmE2 (PRK01678)Ribosomal_L31 (pfam01197)→ |
| Efa-1-56 | NZ_JVNY01000006.1 | + | 50558 | 50590 | **RNA**→ rpmE2 (PRK01678)Ribosomal_L31 (pfam01197)→ |
| Efa-1-57 | NZ_JVID01000037.1 | + | 89220 | 89252 | **RNA**→ rpmE2 (PRK01678)Ribosomal_L31 (pfam01197)→ |
| Efa-1-58 | NZ_JUUJ01000069.1 | - | 3396 | 3364 | **RNA**→ rpmE2 (PRK01678)Ribosomal_L31 (pfam01197)→ |
| Efa-1-59 | NZ_JUXT01000064.1 | - | 3510 | 3478 | **RNA**→ rpmE2 (PRK01678)Ribosomal_L31 (pfam01197)→ |
| Efa-1-60 | NZ_JUQC01000149.1 | - | 23381 | 23349 | **RNA**→ rpmE2 (PRK01678)Ribosomal_L31 (pfam01197)→ |
| Efa-1-61 | NZ_JULA01000068.1 | - | 39749 | 39717 | **RNA**→ rpmE2 (PRK01678)Ribosomal_L31 (pfam01197)→ |
| Efa-1-62 | NZ_JUNN01000112.1 | + | 8486 | 8518 | **RNA**→ rpmE2 (PRK01678)Ribosomal_L31 (pfam01197)→ |
| Efa-1-63 | NZ_JUPR01000265.1 | - | 45791 | 45759 | **RNA**→ rpmE2 (PRK01678)Ribosomal_L31 (pfam01197)→ |
| Efa-1-64 | NZ_JUUM01000251.1 | + | 72669 | 72701 | **RNA**→ rpmE2 (PRK01678)Ribosomal_L31 (pfam01197)→ |
| Efa-1-65 | NZ_JUXC01000167.1 | - | 17001 | 16969 | **RNA**→ rpmE2 (PRK01678)Ribosomal_L31 (pfam01197)→ |
| Efa-1-66 | NZ_JUOP01000166.1 | + | 50540 | 50572 | **RNA**→ rpmE2 (PRK01678)Ribosomal_L31 (pfam01197)→ |
| Efa-1-67 | NZ_JUVA01000209.1 | + | 4910 | 4942 | **RNA**→ rpmE2 (PRK01678)Ribosomal_L31 (pfam01197)→ |
| Efa-1-68 | NZ_JUMJ01000040.1 | + | 50550 | 50582 | **RNA**→ rpmE2 (PRK01678)Ribosomal_L31 (pfam01197)→ |
| Efa-1-69 | NZ_JUVH01000090.1 | + | 50547 | 50579 | **RNA**→ rpmE2 (PRK01678)Ribosomal_L31 (pfam01197)→ |
| Efa-1-70 | NZ_JUWE01000037.1 | - | 23379 | 23347 | **RNA**→ rpmE2 (PRK01678)Ribosomal_L31 (pfam01197)→ |
| Efa-1-71 | NZ_JVAN01000023.1 | - | 19069 | 19037 | **RNA**→ rpmE2 (PRK01678)Ribosomal_L31 (pfam01197)→ |
| Efa-1-72 | NZ_JUNQ01000033.1 | - | 139778 | 139746 | **RNA**→ rpmE2 (PRK01678)Ribosomal_L31 (pfam01197)→ |
| Efa-1-73 | NZ_JUNL01000200.1 | + | 1215 | 1247 | **RNA**→ rpmE2 (PRK01678)Ribosomal_L31 (pfam01197)→ |
| Efa-7-1 | NZ_ASWX01000016.1 | - | 74824 | 74792 | **RNA**→ rpmE2 (PRK01678)Ribosomal_L31 (pfam01197)→ |
| Efa-267-1 | NZ_AYKL01000013.1 | - | 139637 | 139605 | **RNA**→ rpmE2 (PRK01678)Ribosomal_L31 (pfam01197)→ |
| Efa-1-74 | NZ_JPWN01000003.1 | - | 424614 | 424582 | **RNA**→ rpmE2 (PRK01678)Ribosomal_L31 (pfam01197)→ |
| Efa-271-1 | NZ_AYND01000026.1 | - | 83291 | 83259 | **RNA**→ rpmE2 (PRK01678)Ribosomal_L31 (pfam01197)→ |
| Efa-275-1 | NZ_AYLN01000020.1 | - | 38399 | 38367 | **RNA**→ rpmE2 (PRK01678)Ribosomal_L31 (pfam01197)→ |
| Efa-279-1 | NZ_AYOK01000014.1 | - | 39604 | 39572 | **RNA**→ rpmE2 (PRK01678)Ribosomal_L31 (pfam01197)→ |
| Efa-280-1 | NZ_AYOL01000011.1 | + | 15280 | 15312 | **RNA**→ rpmE2 (PRK01678)Ribosomal_L31 (pfam01197)→ |
| Efa-266-1 | NZ_AYKK01000037.1 | + | 15346 | 15378 | **RNA**→ rpmE2 (PRK01678)Ribosomal_L31 (pfam01197)→ |
| Efa-276-1 | NZ_AYKM01000017.1 | + | 50490 | 50522 | **RNA**→ rpmE2 (PRK01678)Ribosomal_L31 (pfam01197)→ |
| Efa-274-1 | NZ_ANMP01000017.1 | + | 15298 | 15330 | **RNA**→ rpmE2 (PRK01678)Ribosomal_L31 (pfam01197)→ |

| | | | | | | | |
|---|---|---|---|---|---|---|---|
| Efa-309-3 | NC_004668.1 | + | 1142020 | 1142052 | **RNA**→ | rpmE2 (PRK01678)Ribosomal_L31 (pfam01197)→ | |
| Efa-558-1 | NC_017316.1 | + | 990713 | 990745 | **RNA**→ | rpmE2 (PRK01678)Ribosomal_L31 (pfam01197)→ | |
| Efa-29-1 | NZ_CP004081.1 | + | 1156388 | 1156420 | **RNA**→ | rpmE2 (PRK01678)Ribosomal_L31 (pfam01197)→ | |
| Efa-550-1 | NZ_GG688629.1 | + | 120120 | 120152 | **RNA**→ | rpmE2 (PRK01678)Ribosomal_L31 (pfam01197)→ | rpmE2 (PRK01678)Ribosomal_L31 (pfam01197)→ |
| Efa-547-1 | NZ_GG670383.1 | + | 188194 | 188226 | **RNA**→ | rpmE2 (PRK01678)Ribosomal_L31 (pfam01197)→ | rpmE2 (PRK01678)Ribosomal_L31 (pfam01197)→ |
| Efa-565-1 | NZ_GG670369.1 | + | 117194 | 117226 | **RNA**→ | rpmE2 (PRK01678)Ribosomal_L31 (pfam01197)→ | rpmE2 (PRK01678)Ribosomal_L31 (pfam01197)→ |
| Efa-555-1 | NZ_GG692656.1 | - | 743973 | 743941 | **RNA**→ | rpmE2 (PRK01678)Ribosomal_L31 (pfam01197)→ | rpmE2 (PRK01678)Ribosomal_L31 (pfam01197)→ |
| Efa-570-1 | NZ_GG669017.1 | - | 1360589 | 1360557 | **RNA**→ | rpmE2 (PRK01678)Ribosomal_L31 (pfam01197)→ | |
| Efa-564-1 | NZ_GG692851.1 | - | 1436735 | 1436703 | **RNA**→ | rpmE2 (PRK01678)Ribosomal_L31 (pfam01197)→ | rpmE2 (PRK01678)Ribosomal_L31 (pfam01197)→ |
| Efa-554-1 | NZ_GG668819.1 | + | 119410 | 119442 | **RNA**→ | rpmE2 (PRK01678)Ribosomal_L31 (pfam01197)→ | |
| Efa-546-1 | NZ_GG692906.1 | + | 759872 | 759904 | **RNA**→ | rpmE2 (PRK01678)Ribosomal_L31 (pfam01197)→ | rpmE2 (PRK01678)Ribosomal_L31 (pfam01197)→ |
| Efa-563-1 | NZ_GG688647.1 | + | 124828 | 124860 | **RNA**→ | rpmE2 (PRK01678)Ribosomal_L31 (pfam01197)→ | rpmE2 (PRK01678)Ribosomal_L31 (pfam01197)→ |
| Efa-566-1 | NZ_GG698878.1 | + | 119116 | 119148 | **RNA**→ | rpmE2 (PRK01678)Ribosomal_L31 (pfam01197)→ | rpmE2 (PRK01678)Ribosomal_L31 (pfam01197)→ |
| Efa-548-1 | NZ_GG688666.1 | + | 121000 | 121032 | **RNA**→ | rpmE2 (PRK01678)Ribosomal_L31 (pfam01197)→ | rpmE2 (PRK01678)Ribosomal_L31 (pfam01197)→ |
| Efa-13-1 | NZ_KK640475.1 | + | 209811 | 209843 | **RNA**→ | rpmE2 (PRK01678)Ribosomal_L31 (pfam01197)→ | |
| Efa-307-1 | NZ_GL454691.1 | - | 3308 | 3276 | **RNA**→ | rpmE2 (PRK01678)Ribosomal_L31 (pfam01197)→ | |
| Efa-302-1 | NZ_GL454579.1 | + | 99322 | 99354 | **RNA**→ | rpmE2 (PRK01678)Ribosomal_L31 (pfam01197)→ | |
| Efa-569-1 | NZ_GL455684.1 | + | 72259 | 72291 | **RNA**→ | rpmE2 (PRK01678)Ribosomal_L31 (pfam01197)→ | |
| Efa-306-1 | NZ_GL476280.1 | - | 39719 | 39687 | **RNA**→ | rpmE2 (PRK01678)Ribosomal_L31 (pfam01197)→ | |
| Efa-305-1 | NZ_GL457030.1 | + | 98133 | 98165 | **RNA**→ | rpmE2 (PRK01678)Ribosomal_L31 (pfam01197)→ | |
| Efa-296-1 | NZ_GL456521.1 | + | 98921 | 98953 | **RNA**→ | rpmE2 (PRK01678)Ribosomal_L31 (pfam01197)→ | |
| Efa-301-1 | NZ_GL457177.1 | + | 98525 | 98557 | **RNA**→ | rpmE2 (PRK01678)Ribosomal_L31 (pfam01197)→ | |
| Efa-292-1 | NZ_GL454300.1 | - | 6431 | 6399 | **RNA**→ | rpmE2 (PRK01678)Ribosomal_L31 (pfam01197)→ | |
| Efa-293-1 | NZ_GL454272.1 | + | 99142 | 99174 | **RNA**→ | rpmE2 (PRK01678)Ribosomal_L31 (pfam01197)→ | |
| Efa-298-1 | NZ_GL457134.1 | - | 6431 | 6399 | **RNA**→ | rpmE2 (PRK01678)Ribosomal_L31 (pfam01197)→ | |
| Efa-304-1 | NZ_GL454064.1 | - | 6435 | 6403 | **RNA**→ | rpmE2 (PRK01678)Ribosomal_L31 (pfam01197)→ | |
| Efa-299-1 | NZ_GL455510.1 | - | 6363 | 6331 | **RNA**→ | rpmE2 (PRK01678)Ribosomal_L31 (pfam01197)→ | |
| Efa-295-1 | NZ_GL456440.1 | + | 97150 | 97182 | **RNA**→ | rpmE2 (PRK01678)Ribosomal_L31 (pfam01197)→ | |
| Efa-297-1 | NZ_GL454840.1 | + | 73694 | 73726 | **RNA**→ | rpmE2 (PRK01678)Ribosomal_L31 (pfam01197)→ | |
| Efa-12-1 | NZ_JH805581.1 | - | 73970 | 73938 | **RNA**→ | rpmE2 (PRK01678)Ribosomal_L31 (pfam01197)→ | |
| Efa-282-1 | NZ_JH804753.1 | - | 81850 | 81818 | **RNA**→ | rpmE2 (PRK01678)Ribosomal_L31 (pfam01197)→ | |
| Efa-269-1 | NZ_KI518257.1 | + | 122898 | 122930 | **RNA**→ | rpmE2 (PRK01678)Ribosomal_L31 (pfam01197)→ | |
| Efa-24-1 | NZ_KI518253.1 | + | 125179 | 125211 | **RNA**→ | rpmE2 (PRK01678)Ribosomal_L31 (pfam01197)→ | |
| Efa-25-1 | NZ_KI518248.1 | + | 118972 | 119004 | **RNA**→ | rpmE2 (PRK01678)Ribosomal_L31 (pfam01197)→ | |
| Efa-257-1 | NZ_KI913060.1 | + | 999492 | 999524 | **RNA**→ | rpmE2 (PRK01678)Ribosomal_L31 (pfam01197)→ | |
| Efa-261-1 | NZ_KI913085.1 | + | 715722 | 715754 | **RNA**→ | rpmE2 (PRK01678)Ribosomal_L31 (pfam01197)→ | |
| Efa-251-1 | NZ_KI913020.1 | + | 730724 | 730756 | **RNA**→ | rpmE2 (PRK01678)Ribosomal_L31 (pfam01197)→ | |
| Efa-244-1 | NZ_KI912988.1 | + | 1001606 | 1001638 | **RNA**→ | rpmE2 (PRK01678)Ribosomal_L31 (pfam01197)→ | |
| Efa-248-1 | NZ_KI913005.1 | + | 642798 | 642830 | **RNA**→ | rpmE2 (PRK01678)Ribosomal_L31 (pfam01197)→ | |
| Efa-253-1 | NZ_KI913031.1 | + | 861627 | 861659 | **RNA**→ | rpmE2 (PRK01678)Ribosomal_L31 (pfam01197)→ | |
| Efa-237-1 | NZ_KI912926.1 | + | 1097807 | 1097839 | **RNA**→ | rpmE2 (PRK01678)Ribosomal_L31 (pfam01197)→ | |
| Efa-256-1 | NZ_KI913055.1 | + | 119066 | 119098 | **RNA**→ | rpmE2 (PRK01678)Ribosomal_L31 (pfam01197)→ | |
| Efa-246-1 | NZ_KI912997.1 | + | 996595 | 996627 | **RNA**→ | rpmE2 (PRK01678)Ribosomal_L31 (pfam01197)→ | |
| Efa-264-1 | NZ_KI913096.1 | + | 119261 | 119293 | **RNA**→ | rpmE2 (PRK01678)Ribosomal_L31 (pfam01197)→ | |
| Efa-50-1 | NZ_KB932381.1 | + | 121925 | 121957 | **RNA**→ | rpmE2 (PRK01678)Ribosomal_L31 (pfam01197)→ | |
| Efa-213-1 | NZ_KB944581.1 | + | 247558 | 247590 | **RNA**→ | rpmE2 (PRK01678)Ribosomal_L31 (pfam01197)→ | |
| Efa-232-1 | NZ_KB944838.1 | + | 119466 | 119498 | **RNA**→ | rpmE2 (PRK01678)Ribosomal_L31 (pfam01197)→ | |
| Efa-56-1 | NZ_KB932331.1 | + | 160762 | 160794 | **RNA**→ | rpmE2 (PRK01678)Ribosomal_L31 (pfam01197)→ | |
| Efa-15-1 | NZ_KB944590.1 | + | 122729 | 122761 | **RNA**→ | rpmE2 (PRK01678)Ribosomal_L31 (pfam01197)→ | |
| Efa-206-1 | NZ_KB944678.1 | + | 122933 | 122965 | **RNA**→ | rpmE2 (PRK01678)Ribosomal_L31 (pfam01197)→ | |
| Efa-231-1 | NZ_KB944545.1 | + | 233956 | 233988 | **RNA**→ | rpmE2 (PRK01678)Ribosomal_L31 (pfam01197)→ | |
| Efa-49-1 | NZ_KB932337.1 | + | 157468 | 157500 | **RNA**→ | rpmE2 (PRK01678)Ribosomal_L31 (pfam01197)→ | |

| | | | | | | |
|---|---|---|---|---|---|---|
| Efa-63-1 | NZ_KB932363.1 | + | 104909 | 104941 | **RNA**→ | rpmE2 (PRK01678)Ribosomal_L31 (pfam01197)→ |
| Efa-52-1 | NZ_KB932317.1 | - | 1903453 | 1903421 | **RNA**→ | rpmE2 (PRK01678)Ribosomal_L31 (pfam01197)→ |
| Efa-229-1 | NZ_KB944658.1 | + | 124147 | 124179 | **RNA**→ | rpmE2 (PRK01678)Ribosomal_L31 (pfam01197)→ |
| Efa-66-1 | NZ_KB932344.1 | + | 120296 | 120328 | **RNA**→ | rpmE2 (PRK01678)Ribosomal_L31 (pfam01197)→ |
| Efa-233-1 | NZ_KB944807.1 | + | 190819 | 190851 | **RNA**→ | rpmE2 (PRK01678)Ribosomal_L31 (pfam01197)→ |
| Efa-224-1 | NZ_KB944819.1 | + | 123718 | 123750 | **RNA**→ | rpmE2 (PRK01678)Ribosomal_L31 (pfam01197)→ |
| Efa-202-1 | NZ_KB945035.1 | + | 122741 | 122773 | **RNA**→ | rpmE2 (PRK01678)Ribosomal_L31 (pfam01197)→ |
| Efa-201-1 | NZ_KB933473.1 | + | 116902 | 116934 | **RNA**→ | rpmE2 (PRK01678)Ribosomal_L31 (pfam01197)→ |
| Efa-73-1 | NZ_KB932398.1 | - | 1315435 | 1315403 | **RNA**→ | rpmE2 (PRK01678)Ribosomal_L31 (pfam01197)→ |
| Efa-235-1 | NZ_KB944800.1 | + | 125536 | 125568 | **RNA**→ | rpmE2 (PRK01678)Ribosomal_L31 (pfam01197)→ |
| Efa-207-1 | NZ_KB944930.1 | + | 854300 | 854332 | **RNA**→ | rpmE2 (PRK01678)Ribosomal_L31 (pfam01197)→ |
| Efa-70-1 | NZ_KB932385.1 | + | 120253 | 120285 | **RNA**→ | rpmE2 (PRK01678)Ribosomal_L31 (pfam01197)→ |
| Efa-210-1 | NZ_KB944537.1 | + | 641238 | 641270 | **RNA**→ | rpmE2 (PRK01678)Ribosomal_L31 (pfam01197)→ |
| Efa-62-1 | NZ_KB932405.1 | - | 1393944 | 1393912 | **RNA**→ | rpmE2 (PRK01678)Ribosomal_L31 (pfam01197)→ |
| Efa-227-1 | NZ_KB944601.1 | + | 125725 | 125757 | **RNA**→ | rpmE2 (PRK01678)Ribosomal_L31 (pfam01197)→ |
| Efa-95-1 | NZ_KB932427.1 | + | 123804 | 123836 | **RNA**→ | rpmE2 (PRK01678)Ribosomal_L31 (pfam01197)→ |
| Efa-205-1 | NZ_KB944666.1 | + | 1303327 | 1303359 | **RNA**→ | rpmE2 (PRK01678)Ribosomal_L31 (pfam01197)→ |
| Efa-92-1 | NZ_KB932581.1 | + | 230841 | 230873 | **RNA**→ | rpmE2 (PRK01678)Ribosomal_L31 (pfam01197)→ |
| Efa-225-1 | NZ_KB944826.1 | + | 121386 | 121418 | **RNA**→ | rpmE2 (PRK01678)Ribosomal_L31 (pfam01197)→ |
| Efa-54-1 | NZ_KB932324.1 | - | 1596841 | 1596809 | **RNA**→ | rpmE2 (PRK01678)Ribosomal_L31 (pfam01197)→ |
| Efa-197-1 | NZ_KB945039.1 | + | 117789 | 117821 | **RNA**→ | rpmE2 (PRK01678)Ribosomal_L31 (pfam01197)→ |
| Efa-65-1 | NZ_KB945185.1 | + | 122889 | 122921 | **RNA**→ | rpmE2 (PRK01678)Ribosomal_L31 (pfam01197)→ |
| Efa-219-1 | NZ_KB944718.1 | + | 119546 | 119578 | **RNA**→ | rpmE2 (PRK01678)Ribosomal_L31 (pfam01197)→ |
| Efa-71-1 | NZ_KB932389.1 | + | 120253 | 120285 | **RNA**→ | rpmE2 (PRK01678)Ribosomal_L31 (pfam01197)→ |
| Efa-61-1 | NZ_KB945241.1 | + | 125221 | 125253 | **RNA**→ | rpmE2 (PRK01678)Ribosomal_L31 (pfam01197)→ |
| Efa-209-1 | NZ_KB944833.1 | + | 125453 | 125485 | **RNA**→ | rpmE2 (PRK01678)Ribosomal_L31 (pfam01197)→ |
| Efa-57-1 | NZ_KB932410.1 | + | 121919 | 121951 | **RNA**→ | rpmE2 (PRK01678)Ribosomal_L31 (pfam01197)→ |
| Efa-77-1 | NZ_KB932431.1 | + | 683611 | 683643 | **RNA**→ | rpmE2 (PRK01678)Ribosomal_L31 (pfam01197)→ |
| Efa-184-1 | NZ_KB946767.1 | + | 124364 | 124396 | **RNA**→ | rpmE2 (PRK01678)Ribosomal_L31 (pfam01197)→ |
| Efa-99-1 | NZ_KB932453.1 | + | 124004 | 124036 | **RNA**→ | rpmE2 (PRK01678)Ribosomal_L31 (pfam01197)→ |
| Efa-234-1 | NZ_KB944754.1 | + | 837249 | 837281 | **RNA**→ | rpmE2 (PRK01678)Ribosomal_L31 (pfam01197)→ |
| Efa-86-1 | NZ_KB932513.1 | + | 681664 | 681696 | **RNA**→ | rpmE2 (PRK01678)Ribosomal_L31 (pfam01197)→ |
| Efa-93-1 | NZ_KB932585.1 | + | 229409 | 229441 | **RNA**→ | rpmE2 (PRK01678)Ribosomal_L31 (pfam01197)→ |
| Efa-75-1 | NZ_KB932562.1 | + | 663647 | 663679 | **RNA**→ | rpmE2 (PRK01678)Ribosomal_L31 (pfam01197)→ |
| Efa-51-1 | NZ_KB932320.1 | - | 1554080 | 1554048 | **RNA**→ | rpmE2 (PRK01678)Ribosomal_L31 (pfam01197)→ |
| Efa-108-1 | NZ_KB932479.1 | + | 121580 | 121612 | **RNA**→ | rpmE2 (PRK01678)Ribosomal_L31 (pfam01197)→ |
| Efa-64-1 | NZ_KB932347.1 | - | 696492 | 696460 | **RNA**→ | rpmE2 (PRK01678)Ribosomal_L31 (pfam01197)→ |
| Efa-103-1 | NZ_KB932539.1 | + | 121716 | 121748 | **RNA**→ | rpmE2 (PRK01678)Ribosomal_L31 (pfam01197)→ |
| Efa-69-1 | NZ_KB932372.1 | - | 1782054 | 1782022 | **RNA**→ | rpmE2 (PRK01678)Ribosomal_L31 (pfam01197)→ |
| Efa-156-1 | NZ_KB946490.1 | + | 123693 | 123725 | **RNA**→ | rpmE2 (PRK01678)Ribosomal_L31 (pfam01197)→ |
| Efa-59-1 | NZ_KB932417.1 | + | 122720 | 122752 | **RNA**→ | rpmE2 (PRK01678)Ribosomal_L31 (pfam01197)→ |
| Efa-78-1 | NZ_KB932437.1 | + | 2252823 | 2252855 | **RNA**→ | rpmE2 (PRK01678)Ribosomal_L31 (pfam01197)→ |
| Efa-181-1 | NZ_KB947238.1 | + | 125685 | 125717 | **RNA**→ | rpmE2 (PRK01678)Ribosomal_L31 (pfam01197)→ |
| Efa-83-1 | NZ_KB932490.1 | + | 123557 | 123589 | **RNA**→ | rpmE2 (PRK01678)Ribosomal_L31 (pfam01197)→ |
| Efa-90-1 | NZ_KB932574.1 | + | 123230 | 123262 | **RNA**→ | rpmE2 (PRK01678)Ribosomal_L31 (pfam01197)→ |
| Efa-48-1 | NZ_KB932356.1 | + | 156551 | 156583 | **RNA**→ | rpmE2 (PRK01678)Ribosomal_L31 (pfam01197)→ |
| Efa-230-1 | NZ_KB944688.1 | + | 122685 | 122717 | **RNA**→ | rpmE2 (PRK01678)Ribosomal_L31 (pfam01197)→ |
| Efa-100-1 | NZ_KB932473.1 | + | 698382 | 698414 | **RNA**→ | rpmE2 (PRK01678)Ribosomal_L31 (pfam01197)→ |
| Efa-211-1 | NZ_KB944694.1 | + | 122838 | 122870 | **RNA**→ | rpmE2 (PRK01678)Ribosomal_L31 (pfam01197)→ |
| Efa-105-1 | NZ_KB932593.1 | + | 185323 | 185355 | **RNA**→ | rpmE2 (PRK01678)Ribosomal_L31 (pfam01197)→ |
| Efa-85-1 | NZ_KB932506.1 | + | 121940 | 121972 | **RNA**→ | rpmE2 (PRK01678)Ribosomal_L31 (pfam01197)→ |
| Efa-153-1 | NZ_KB946414.1 | + | 120628 | 120660 | **RNA**→ | rpmE2 (PRK01678)Ribosomal_L31 (pfam01197)→ |

| | | | | | |
|---|---|---|---|---|---|
| Efa-97-1 | NZ_KB932559.1 | + | 123243 | 123275 | **RNA**→ rpmE2 (PRK01678)Ribosomal_L31 (pfam01197)→ |
| Efa-158-1 | NZ_KB946508.1 | + | 125411 | 125443 | **RNA**→ rpmE2 (PRK01678)Ribosomal_L31 (pfam01197)→ |
| Efa-55-1 | NZ_KB945106.1 | + | 124044 | 124076 | **RNA**→ rpmE2 (PRK01678)Ribosomal_L31 (pfam01197)→ |
| Efa-182-1 | NZ_KB946719.1 | + | 944180 | 944212 | **RNA**→ rpmE2 (PRK01678)Ribosomal_L31 (pfam01197)→ |
| Efa-212-1 | NZ_KB944552.1 | + | 121578 | 121610 | **RNA**→ rpmE2 (PRK01678)Ribosomal_L31 (pfam01197)→ |
| Efa-72-1 | NZ_KB932393.1 | - | 1371754 | 1371722 | **RNA**→ rpmE2 (PRK01678)Ribosomal_L31 (pfam01197)→ |
| Efa-58-1 | NZ_KB932415.1 | + | 909832 | 909864 | **RNA**→ rpmE2 (PRK01678)Ribosomal_L31 (pfam01197)→ |
| Efa-185-1 | NZ_KB947273.1 | + | 125433 | 125465 | **RNA**→ rpmE2 (PRK01678)Ribosomal_L31 (pfam01197)→ |
| Efa-60-1 | NZ_KB932422.1 | - | 1981852 | 1981820 | **RNA**→ rpmE2 (PRK01678)Ribosomal_L31 (pfam01197)→ |
| Efa-226-1 | NZ_KB944566.1 | + | 121785 | 121817 | **RNA**→ rpmE2 (PRK01678)Ribosomal_L31 (pfam01197)→ |
| Efa-199-1 | NZ_KB945052.1 | + | 119630 | 119662 | **RNA**→ rpmE2 (PRK01678)Ribosomal_L31 (pfam01197)→ |
| Efa-79-1 | NZ_KB932442.1 | + | 121860 | 121892 | **RNA**→ rpmE2 (PRK01678)Ribosomal_L31 (pfam01197)→ |
| Efa-102-1 | NZ_KB932532.1 | + | 118858 | 118890 | **RNA**→ rpmE2 (PRK01678)Ribosomal_L31 (pfam01197)→ |
| Efa-80-1 | NZ_KB932457.1 | + | 123784 | 123816 | **RNA**→ rpmE2 (PRK01678)Ribosomal_L31 (pfam01197)→ |
| Efa-74-1 | NZ_KB932469.1 | + | 697454 | 697486 | **RNA**→ rpmE2 (PRK01678)Ribosomal_L31 (pfam01197)→ |
| Efa-84-1 | NZ_KB932497.1 | + | 124103 | 124135 | **RNA**→ rpmE2 (PRK01678)Ribosomal_L31 (pfam01197)→ |
| Efa-101-1 | NZ_KB932518.1 | + | 685526 | 685558 | **RNA**→ rpmE2 (PRK01678)Ribosomal_L31 (pfam01197)→ |
| Efa-180-1 | NZ_KB947112.1 | + | 654286 | 654318 | **RNA**→ rpmE2 (PRK01678)Ribosomal_L31 (pfam01197)→ |
| Efa-88-1 | NZ_KB932527.1 | + | 119530 | 119562 | **RNA**→ rpmE2 (PRK01678)Ribosomal_L31 (pfam01197)→ |
| Efa-220-1 | NZ_KB944728.1 | + | 122601 | 122633 | **RNA**→ rpmE2 (PRK01678)Ribosomal_L31 (pfam01197)→ |
| Efa-91-1 | NZ_KB932577.1 | + | 229782 | 229814 | **RNA**→ rpmE2 (PRK01678)Ribosomal_L31 (pfam01197)→ |
| Efa-154-1 | NZ_KB946433.1 | + | 119307 | 119339 | **RNA**→ rpmE2 (PRK01678)Ribosomal_L31 (pfam01197)→ |
| Efa-161-1 | NZ_KB946666.1 | + | 115624 | 115656 | **RNA**→ rpmE2 (PRK01678)Ribosomal_L31 (pfam01197)→ |
| Efa-163-1 | NZ_KB946616.1 | + | 125518 | 125550 | **RNA**→ rpmE2 (PRK01678)Ribosomal_L31 (pfam01197)→ |
| Efa-53-1 | NZ_KB932336.1 | + | 887741 | 887773 | **RNA**→ rpmE2 (PRK01678)Ribosomal_L31 (pfam01197)→ |
| Efa-98-1 | NZ_KB932447.1 | + | 118980 | 119012 | **RNA**→ rpmE2 (PRK01678)Ribosomal_L31 (pfam01197)→ |
| Efa-107-1 | NZ_KB932601.1 | + | 230184 | 230216 | **RNA**→ rpmE2 (PRK01678)Ribosomal_L31 (pfam01197)→ |
| Efa-157-1 | NZ_KB946499.1 | + | 125503 | 125535 | **RNA**→ rpmE2 (PRK01678)Ribosomal_L31 (pfam01197)→ |
| Efa-160-1 | NZ_KB946551.1 | + | 122102 | 122134 | **RNA**→ rpmE2 (PRK01678)Ribosomal_L31 (pfam01197)→ |
| Efa-94-1 | NZ_KB932589.1 | + | 223552 | 223584 | **RNA**→ rpmE2 (PRK01678)Ribosomal_L31 (pfam01197)→ |
| Efa-186-1 | NZ_KB947205.1 | + | 126658 | 126690 | **RNA**→ rpmE2 (PRK01678)Ribosomal_L31 (pfam01197)→ |
| Efa-123-1 | NZ_KB948613.1 | + | 692248 | 692280 | **RNA**→ rpmE2 (PRK01678)Ribosomal_L31 (pfam01197)→ |
| Efa-109-1 | NZ_KB932435.1 | + | 903029 | 903061 | **RNA**→ rpmE2 (PRK01678)Ribosomal_L31 (pfam01197)→ |
| Efa-89-1 | NZ_KB932545.1 | + | 118833 | 118865 | **RNA**→ rpmE2 (PRK01678)Ribosomal_L31 (pfam01197)→ |
| Efa-76-1 | NZ_KB932553.1 | + | 688587 | 688619 | **RNA**→ rpmE2 (PRK01678)Ribosomal_L31 (pfam01197)→ |
| Efa-106-1 | NZ_KB932597.1 | + | 227160 | 227192 | **RNA**→ rpmE2 (PRK01678)Ribosomal_L31 (pfam01197)→ |
| Efa-178-1 | NZ_KB947149.1 | + | 1211418 | 1211450 | **RNA**→ rpmE2 (PRK01678)Ribosomal_L31 (pfam01197)→ |
| Efa-170-1 | NZ_KB947405.1 | + | 113537 | 113569 | **RNA**→ rpmE2 (PRK01678)Ribosomal_L31 (pfam01197)→ |
| Efa-21-1 | NZ_KB944862.1 | + | 135281 | 135313 | **RNA**→ rpmE2 (PRK01678)Ribosomal_L31 (pfam01197)→ |
| Efa-208-1 | NZ_KB944938.1 | + | 119215 | 119247 | **RNA**→ rpmE2 (PRK01678)Ribosomal_L31 (pfam01197)→ |
| Efa-151-1 | NZ_KB946400.1 | + | 121933 | 121965 | **RNA**→ rpmE2 (PRK01678)Ribosomal_L31 (pfam01197)→ |
| Efa-144-1 | NZ_KB946638.1 | + | 122019 | 122051 | **RNA**→ rpmE2 (PRK01678)Ribosomal_L31 (pfam01197)→ |
| Efa-166-1 | NZ_KB947183.1 | + | 852047 | 852079 | **RNA**→ rpmE2 (PRK01678)Ribosomal_L31 (pfam01197)→ |
| Efa-173-1 | NZ_KB947341.1 | + | 861817 | 861849 | **RNA**→ rpmE2 (PRK01678)Ribosomal_L31 (pfam01197)→ |
| Efa-118-1 | NZ_KB948491.1 | + | 118961 | 118993 | **RNA**→ rpmE2 (PRK01678)Ribosomal_L31 (pfam01197)→ |
| Efa-81-1 | NZ_KB932465.1 | + | 122091 | 122123 | **RNA**→ rpmE2 (PRK01678)Ribosomal_L31 (pfam01197)→ |
| Efa-87-1 | NZ_KB932522.1 | + | 707656 | 707688 | **RNA**→ rpmE2 (PRK01678)Ribosomal_L31 (pfam01197)→ |
| Efa-96-1 | NZ_KB932550.1 | + | 119174 | 119206 | **RNA**→ rpmE2 (PRK01678)Ribosomal_L31 (pfam01197)→ |
| Efa-147-1 | NZ_KB946069.1 | + | 834258 | 834290 | **RNA**→ rpmE2 (PRK01678)Ribosomal_L31 (pfam01197)→ |
| Efa-148-1 | NZ_KB946181.1 | + | 121938 | 121970 | **RNA**→ rpmE2 (PRK01678)Ribosomal_L31 (pfam01197)→ |
| Efa-192-1 | NZ_KB947132.1 | + | 903557 | 903589 | **RNA**→ rpmE2 (PRK01678)Ribosomal_L31 (pfam01197)→ |
| Efa-164-1 | NZ_KB946584.1 | + | 846944 | 846976 | **RNA**→ rpmE2 (PRK01678)Ribosomal_L31 (pfam01197)→ |

| | | | | | |
|---|---|---|---|---|---|
| Efa-183-1 | NZ_KB947346.1 | + | 912443 | 912475 | **RNA**→ rpmE2 (PRK01678)Ribosomal_L31 (pfam01197)→ |
| Efa-117-1 | NZ_KB948452.1 | + | 121982 | 122014 | **RNA**→ rpmE2 (PRK01678)Ribosomal_L31 (pfam01197)→ |
| Efa-121-1 | NZ_KB948631.1 | + | 118986 | 119018 | **RNA**→ rpmE2 (PRK01678)Ribosomal_L31 (pfam01197)→ |
| Efa-139-1 | NZ_KB949292.1 | + | 118895 | 118927 | **RNA**→ rpmE2 (PRK01678)Ribosomal_L31 (pfam01197)→ |
| Efa-159-1 | NZ_KB946519.1 | + | 746880 | 746912 | **RNA**→ rpmE2 (PRK01678)Ribosomal_L31 (pfam01197)→ |
| Efa-111-1 | NZ_KB948306.1 | + | 692749 | 692781 | **RNA**→ rpmE2 (PRK01678)Ribosomal_L31 (pfam01197)→ |
| Efa-113-1 | NZ_KB948374.1 | + | 121434 | 121466 | **RNA**→ rpmE2 (PRK01678)Ribosomal_L31 (pfam01197)→ |
| Efa-134-1 | NZ_KB949098.1 | + | 679266 | 679298 | **RNA**→ rpmE2 (PRK01678)Ribosomal_L31 (pfam01197)→ |
| Efa-115-1 | NZ_KB948411.1 | + | 124207 | 124239 | **RNA**→ rpmE2 (PRK01678)Ribosomal_L31 (pfam01197)→ |
| Efa-128-1 | NZ_KB948910.1 | + | 125113 | 125145 | **RNA**→ rpmE2 (PRK01678)Ribosomal_L31 (pfam01197)→ |
| Efa-142-1 | NZ_KB949380.1 | + | 121687 | 121719 | **RNA**→ rpmE2 (PRK01678)Ribosomal_L31 (pfam01197)→ |
| Efa-194-1 | NZ_KB947266.1 | + | 125568 | 125600 | **RNA**→ rpmE2 (PRK01678)Ribosomal_L31 (pfam01197)→ |
| Efa-116-1 | NZ_KB948432.1 | + | 119775 | 119807 | **RNA**→ rpmE2 (PRK01678)Ribosomal_L31 (pfam01197)→ |
| Efa-124-1 | NZ_KB948773.1 | + | 122070 | 122102 | **RNA**→ rpmE2 (PRK01678)Ribosomal_L31 (pfam01197)→ |
| Efa-129-1 | NZ_KB948924.1 | + | 681265 | 681297 | **RNA**→ rpmE2 (PRK01678)Ribosomal_L31 (pfam01197)→ |
| Efa-189-1 | NZ_KB947291.1 | + | 120577 | 120609 | **RNA**→ rpmE2 (PRK01678)Ribosomal_L31 (pfam01197)→ |
| Efa-146-1 | NZ_KB949772.1 | + | 1111167 | 1111199 | **RNA**→ rpmE2 (PRK01678)Ribosomal_L31 (pfam01197)→ |
| Efa-110-1 | NZ_KB948281.1 | + | 119213 | 119245 | **RNA**→ rpmE2 (PRK01678)Ribosomal_L31 (pfam01197)→ |
| Efa-120-1 | NZ_KB948590.1 | + | 120205 | 120237 | **RNA**→ rpmE2 (PRK01678)Ribosomal_L31 (pfam01197)→ |
| Efa-190-1 | NZ_KB947168.1 | + | 124005 | 124037 | **RNA**→ rpmE2 (PRK01678)Ribosomal_L31 (pfam01197)→ |
| Efa-204-1 | NZ_KB947484.1 | + | 121690 | 121722 | **RNA**→ rpmE2 (PRK01678)Ribosomal_L31 (pfam01197)→ |
| Efa-137-1 | NZ_KB949231.1 | + | 119523 | 119555 | **RNA**→ rpmE2 (PRK01678)Ribosomal_L31 (pfam01197)→ |
| Efa-125-1 | NZ_KB948803.1 | + | 121909 | 121941 | **RNA**→ rpmE2 (PRK01678)Ribosomal_L31 (pfam01197)→ |
| Efa-133-1 | NZ_KB949059.1 | + | 676773 | 676805 | **RNA**→ rpmE2 (PRK01678)Ribosomal_L31 (pfam01197)→ |
| Efa-130-1 | NZ_KB948962.1 | + | 121896 | 121928 | **RNA**→ rpmE2 (PRK01678)Ribosomal_L31 (pfam01197)→ |
| Efa-135-1 | NZ_KB949160.1 | + | 121821 | 121853 | **RNA**→ rpmE2 (PRK01678)Ribosomal_L31 (pfam01197)→ |
| Efa-119-1 | NZ_KB948517.1 | + | 121976 | 122008 | **RNA**→ rpmE2 (PRK01678)Ribosomal_L31 (pfam01197)→ |
| Efa-143-1 | NZ_KB948681.1 | + | 121876 | 121908 | **RNA**→ rpmE2 (PRK01678)Ribosomal_L31 (pfam01197)→ |
| Efa-122-1 | NZ_KB948722.1 | + | 687418 | 687450 | **RNA**→ rpmE2 (PRK01678)Ribosomal_L31 (pfam01197)→ |
| Efa-126-1 | NZ_KB948846.1 | + | 119055 | 119087 | **RNA**→ rpmE2 (PRK01678)Ribosomal_L31 (pfam01197)→ |
| Efa-127-1 | NZ_KB948878.1 | + | 118805 | 118837 | **RNA**→ rpmE2 (PRK01678)Ribosomal_L31 (pfam01197)→ |
| Efa-141-1 | NZ_KB949363.1 | + | 123019 | 123051 | **RNA**→ rpmE2 (PRK01678)Ribosomal_L31 (pfam01197)→ |
| Efa-131-1 | NZ_KB948990.1 | + | 122095 | 122127 | **RNA**→ rpmE2 (PRK01678)Ribosomal_L31 (pfam01197)→ |
| Efa-132-1 | NZ_KB949021.1 | + | 123650 | 123682 | **RNA**→ rpmE2 (PRK01678)Ribosomal_L31 (pfam01197)→ |
| Efa-136-1 | NZ_KB949171.1 | + | 652090 | 652122 | **RNA**→ rpmE2 (PRK01678)Ribosomal_L31 (pfam01197)→ |
| Efa-112-1 | NZ_KB948356.1 | + | 692356 | 692388 | **RNA**→ rpmE2 (PRK01678)Ribosomal_L31 (pfam01197)→ |
| Efa-140-1 | NZ_KB949324.1 | + | 694227 | 694259 | **RNA**→ rpmE2 (PRK01678)Ribosomal_L31 (pfam01197)→ |
| Efa-236-1 | NZ_KB948740.1 | + | 118896 | 118928 | **RNA**→ rpmE2 (PRK01678)Ribosomal_L31 (pfam01197)→ |
| Efa-138-1 | NZ_KB949265.1 | + | 686455 | 686487 | **RNA**→ rpmE2 (PRK01678)Ribosomal_L31 (pfam01197)→ |
| Efa-309-1 | NZ_KE136528.1 | + | 119156 | 119188 | **RNA**→ rpmE2 (PRK01678)Ribosomal_L31 (pfam01197)→ |
| Efa-309-2 | NZ_KE136402.1 | - | 2324068 | 2324036 | **RNA**→ rpmE2 (PRK01678)Ribosomal_L31 (pfam01197)→ |
| Efa-283-1 | NZ_KE351799.1 | - | 6437 | 6405 | **RNA**→ rpmE2 (PRK01678)Ribosomal_L31 (pfam01197)→ |
| Efa-26-1 | NZ_KE351073.1 | + | 72178 | 72210 | **RNA**→ rpmE2 (PRK01678)Ribosomal_L31 (pfam01197)→ |
| Efa-5-1 | NZ_KE351325.1 | + | 7087 | 7119 | **RNA**→ rpmE2 (PRK01678)Ribosomal_L31 (pfam01197)→ |
| Efa-2-1 | NZ_KE350972.1 | + | 65344 | 65376 | **RNA**→ rpmE2 (PRK01678)Ribosomal_L31 (pfam01197)→ hypo→ DUF916 (pfam06030)DUF3324 (pfam11797 hypo→ |
| Efa-310-1 | NZ_KE352642.1 | + | 15337 | 15369 | **RNA**→ rpmE2 (PRK01678)Ribosomal_L31 (pfam01197)→ |
| Efa-284-1 | NZ_KE352762.1 | + | 15329 | 15361 | **RNA**→ rpmE2 (PRK01678)Ribosomal_L31 (pfam01197)→ |
| Efa-265-1 | NZ_KE351588.1 | - | 236352 | 236320 | **RNA**→ rpmE2 (PRK01678)Ribosomal_L31 (pfam01197)→ |
| Efa-6-1 | NZ_KE351218.1 | + | 7091 | 7123 | **RNA**→ rpmE2 (PRK01678)Ribosomal_L31 (pfam01197)→ |
| Efa-270-1 | NZ_KE350323.1 | - | 128689 | 128657 | **RNA**→ rpmE2 (PRK01678)Ribosomal_L31 (pfam01197)→ |
| Efa-272-1 | NZ_KE352792.1 | + | 15345 | 15377 | **RNA**→ rpmE2 (PRK01678)Ribosomal_L31 (pfam01197)→ |
| Efa-47-1 | NZ_KL584693.1 | - | 424542 | 424510 | **RNA**→ rpmE2 (PRK01678)Ribosomal_L31 (pfam01197)→ |

| | | | | | |
|---|---|---|---|---|---|
| Efa-277-1 | NZ_AYOJ01000045.1 | + | 50557 | 50589 | **RNA**→ rpmE2 (PRK01678)Ribosomal_L31 (pfam01197)→ |
| Efa-1-75 | NZ_JSES01000004.1 | - | 229328 | 229296 | **RNA**→ rpmE2 (PRK01678)Ribosomal_L31 (pfam01197)→ |
| Efa-1-76 | NZ_JTKS01000008.1 | + | 98901 | 98933 | **RNA**→ rpmE2 (PRK01678)Ribosomal_L31 (pfam01197)→ |
| Efa-1-77 | NZ_JTKU01000015.1 | - | 557832 | 557800 | **RNA**→ rpmE2 (PRK01678)Ribosomal_L31 (pfam01197)→ |
| Efa-1-78 | NZ_JTKX01000056.1 | + | 101015 | 101047 | **RNA**→ rpmE2 (PRK01678)Ribosomal_L31 (pfam01197)→ |
| Efa-1-79 | NZ_JTKV01000020.1 | - | 229146 | 229114 | **RNA**→ rpmE2 (PRK01678)Ribosomal_L31 (pfam01197)→ |
| Efa-1-80 | NZ_JQHD01000026.1 | - | 3289 | 3257 | **RNA**→ rpmE2 (PRK01678)Ribosomal_L31 (pfam01197)→ |
| Efa-1-81 | NZ_JWAR01000070.1 | - | 87014 | 86982 | **RNA**→ rpmE2 (PRK01678)Ribosomal_L31 (pfam01197)→ |
| Efa-1-82 | NZ_JVAD01000032.1 | - | 86836 | 86804 | **RNA**→ rpmE2 (PRK01678)Ribosomal_L31 (pfam01197)→ |
| Efa-1-83 | NZ_JVDH01000056.1 | - | 120626 | 120594 | **RNA**→ rpmE2 (PRK01678)Ribosomal_L31 (pfam01197)→ |
| Efa-1-84 | NZ_JUXV01000036.1 | - | 86836 | 86804 | **RNA**→ rpmE2 (PRK01678)Ribosomal_L31 (pfam01197)→ |
| Efa-1-85 | NZ_JUXZ01000087.1 | + | 50554 | 50586 | **RNA**→ rpmE2 (PRK01678)Ribosomal_L31 (pfam01197)→ |
| Efa-1-86 | NZ_JUXL01000154.1 | + | 5045 | 5077 | **RNA**→ rpmE2 (PRK01678)Ribosomal_L31 (pfam01197)→ |
| Efa-1-87 | NZ_JUYS01000097.1 | + | 15659 | 15691 | **RNA**→ rpmE2 (PRK01678)Ribosomal_L31 (pfam01197)→ |
| Efa-568-1 | NZ_GG668922.1 | + | 118017 | 118049 | **RNA**→ rpmE2 (PRK01678)Ribosomal_L31 (pfam01197)→ |
| Efa-38-1 | NZ_JH805817.1 | + | 98302 | 98334 | **RNA**→ rpmE2 (PRK01678)Ribosomal_L31 (pfam01197)→ |
| Efa-33-1 | NZ_JH806227.1 | + | 15338 | 15370 | **RNA**→ rpmE2 (PRK01678)Ribosomal_L31 (pfam01197)→ |
| Efa-32-1 | NZ_JH806069.1 | + | 98267 | 98299 | **RNA**→ rpmE2 (PRK01678)Ribosomal_L31 (pfam01197)→ |
| Efa-35-1 | NZ_JH806464.1 | - | 198218 | 198186 | **RNA**→ rpmE2 (PRK01678)Ribosomal_L31 (pfam01197)→ |
| Efa-34-1 | NZ_JH806329.1 | + | 72149 | 72181 | **RNA**→ rpmE2 (PRK01678)Ribosomal_L31 (pfam01197)→ |
| Efa-37-1 | NZ_JH805939.1 | - | 198203 | 198171 | **RNA**→ rpmE2 (PRK01678)Ribosomal_L31 (pfam01197)→ |
| Efa-42-1 | NZ_JH805527.1 | + | 75536 | 75568 | **RNA**→ rpmE2 (PRK01678)Ribosomal_L31 (pfam01197)→ |
| Efa-31-1 | NZ_JH805886.1 | + | 72171 | 72203 | **RNA**→ rpmE2 (PRK01678)Ribosomal_L31 (pfam01197)→ |
| Efa-36-1 | NZ_JH806144.1 | - | 198240 | 198208 | **RNA**→ rpmE2 (PRK01678)Ribosomal_L31 (pfam01197)→ |
| Efa-43-1 | NZ_JH805255.1 | - | 198654 | 198622 | **RNA**→ rpmE2 (PRK01678)Ribosomal_L31 (pfam01197)→ |
| Efa-46-1 | NZ_JH804975.1 | + | 15342 | 15374 | **RNA**→ rpmE2 (PRK01678)Ribosomal_L31 (pfam01197)→ |
| Efa-41-1 | NZ_JH805339.1 | + | 72173 | 72205 | **RNA**→ rpmE2 (PRK01678)Ribosomal_L31 (pfam01197)→ |
| Efa-44-1 | NZ_JH805118.1 | + | 15343 | 15375 | **RNA**→ rpmE2 (PRK01678)Ribosomal_L31 (pfam01197)→ |
| Efa-39-1 | NZ_JH805694.1 | - | 198161 | 198129 | **RNA**→ rpmE2 (PRK01678)Ribosomal_L31 (pfam01197)→ |
| Efa-45-1 | NZ_JH804884.1 | - | 198224 | 198192 | **RNA**→ rpmE2 (PRK01678)Ribosomal_L31 (pfam01197)→ |
| Efa-223-1 | NZ_KB944788.1 | + | 121130 | 121162 | **RNA**→ rpmE2 (PRK01678)Ribosomal_L31 (pfam01197)→ |
| Efa-150-1 | NZ_KB946369.1 | + | 121423 | 121455 | **RNA**→ rpmE2 (PRK01678)Ribosomal_L31 (pfam01197)→ |
| Efa-149-1 | NZ_KB946341.1 | + | 120776 | 120808 | **RNA**→ rpmE2 (PRK01678)Ribosomal_L31 (pfam01197)→ |
| Efa-114-1 | NZ_KB948391.1 | + | 123030 | 123062 | **RNA**→ rpmE2 (PRK01678)Ribosomal_L31 (pfam01197)→ |
| Efa-155-1 | NZ_KB946441.1 | + | 121214 | 121246 | **RNA**→ rpmE2 (PRK01678)Ribosomal_L31 (pfam01197)→ |
| Efa-40-1 | NZ_JH805440.1 | - | 198081 | 198049 | **RNA**→ rpmE2 (PRK01678)Ribosomal_L31 (pfam01197)→ |
| Efa-82-1 | NZ_KB932483.1 | + | 121271 | 121303 | **RNA**→ rpmE2 (PRK01678)Ribosomal_L31 (pfam01197)→ |
| Csp-3-1 | NZ_JTLM01000039.1 | + | 16014 | 16047 | **RNA**→ rpmE2 (PRK01678)Ribosomal_L31 (pfam01197)→ |
| Eca-1-1 | NZ_KB946333.1 | + | 919506 | 919538 | **RNA**→ rpmE2 (PRK01678)Ribosomal_L31 (pfam01197)→ |
| Emo-1-1 | NZ_KB946318.1 | + | 836233 | 836265 | **RNA**→ rpmE2 (PRK01678)Ribosomal_L31 (pfam01197)→ |
| Eha-1-1 | NZ_KB946315.1 | + | 715011 | 715043 | **RNA**→ rpmE2 (PRK01678)Ribosomal_L31 (pfam01197)→ |
| Esp-17-1 | NZ_AYPQ01000003.1 | - | 139381 | 139349 | **RNA**→ rpmE2 (PRK01678)Ribosomal_L31 (pfam01197)→ |
| Edi-1-1 | NZ_KE136354.1 | - | 1524939 | 1524907 | **RNA**→ rpmE2 (PRK01678)Ribosomal_L31 (pfam01197)→ rpmE2 (PRK01678)Ribosomal_L31 (pfam01197)→ |
| Lap-2-1 | NZ_BAMM01000001.1 | + | 55628 | 55660 | **RNA**→ rpmE2 (PRK01678)Ribosomal_L31 (pfam01197)→ |
| env-1488 | AglaG_GBVDVLI01EDQX8 | + | 207 | 239 | **RNA**→ |
| Ega-1-1 | NZ_JMGP01000013.1 | + | 62327 | 62359 | **RNA**→ rpmE2 (PRK01678)Ribosomal_L31 (pfam01197)→ |
| Ega-2-1 | NZ_GG670288.1 | + | 67876 | 67908 | **RNA**→ rpmE2 (PRK01678)Ribosomal_L31 (pfam01197)→ hypo→ |
| Esa-1-1 | NZ_JH376940.1 | + | 68614 | 68646 | **RNA**→ rpmE2 (PRK01678)Ribosomal_L31 (pfam01197)→ |
| Ega-1-2 | NZ_JNLR01000003.1 | - | 2516883 | 2516851 | **RNA**→ rpmE2 (PRK01678)Ribosomal_L31 (pfam01197)→ |
| Eco-1-1 | NZ_KB890286.1 | + | 27222 | 27254 | **RNA**→ rpmE2 (PRK01678)Ribosomal_L31 (pfam01197)→ |
| Lha-3-1 | NZ_BAML01000008.1 | - | 24487 | 24455 | **RNA**→ rpmE2 (PRK01678)Ribosomal_L31 (pfam01197)→ hypo→ |
| Leq-1-1 | NZ_AWWH01000135.1 | + | 28867 | 28900 | **RNA**→ rpmE2 (PRK01678)Ribosomal_L31 (pfam01197)→ |

| | | | | | |
|---|---|---|---|---|---|
| Leq-2-1 | NZ_BAMI01000008.1 | - | 21982 | 21949 | **RNA**→ rpmE2 (PRK01678)Ribosomal_L31 (pfam01197)→ |
| env-1489 | 2204882122 | - | 2442 | 2410 | **RNA**→ rpmE2 (PRK01678)Ribosomal_L31 (pfam01197)→ |
| env-1490 | 2158238682 | + | 11762 | 11794 | **RNA**→ rpmE2 (PRK01678)Ribosomal_L31 (pfam01197)→ |
| Bok-2-1 | NZ_LILD01000001.1 | + | 243048 | 243079 | **RNA**→ rpmE2 (PRK01678)Ribosomal_L31 (pfam01197)→ |
| Bha-1-1 | NC_002570.2 | - | 3899644 | 3899613 | **RNA**→ rpmE2 (PRK01678)Ribosomal_L31 (pfam01197)→ |
| Wko-2-1 | NC_015759.1 | - | 274266 | 274233 | **RNA**→ rpmE2 (PRK01678)Ribosomal_L31 (pfam01197)→ |
| Wko-1-1 | NZ_AKGG01000014.1 | - | 93148 | 93115 | **RNA**→ rpmE2 (PRK01678)Ribosomal_L31 (pfam01197)→ |
| Bma-8-1 | NZ_CDPP01000002.1 | - | 333009 | 332973 | **RNA**→ rpmE (PRK00019)Ribosomal_L31 (pfam01197)→ Lactamase_B (smart00849)PhnP (COG1235)→ PRK00103 (PRK00103)SPOUT_MTase (pfam02590)→ |
| env-1491 | AKYZ01003788.1 | - | 1483 | 1450 | **RNA**→ rpmE (PRK00019)Ribosomal_L31 (pfam01197)→ |
| env-1492 | AKZA01003178.1 | + | 2474 | 2507 | **RNA**→ rpmE (PRK00019)Ribosomal_L31 (pfam01197)→ |
| env-1493 | SRS011586_C3401226 | - | 110 | 77 | **RNA**→ rpmE (PRK00019)rpmE2 (PRK01678)Ribosomal_L31 (pfam01197)→ |
| env-1494 | SRS017521_C2775917 | + | 102 | 135 | **RNA**→ rpmE (PRK00019)Ribosomal_L31 (pfam01197)→ |
| Csp-7-1 | NZ_AKCB01000001.1 | - | 1308492 | 1308459 | **RNA**→ rpmE (PRK00019)Ribosomal_L31 (pfam01197)→ |
| Csp-5-1 | NZ_GL636577.1 | + | 412437 | 412470 | **RNA**→ rpmE (PRK00019)Ribosomal_L31 (pfam01197)→ |
| env-1495 | BAAV01014627.1 | - | 242 | 209 | **RNA**→ rpmE (PRK00019)Ribosomal_L31 (pfam01197)→ |
| env-1496 | scaffold83146_5_MH0053 | - | 142 | 109 | **RNA**→ rpmE (PRK00019)Ribosomal_L31 (pfam01197)→ |
| env-1497 | scaffold10346_4_MH0082 | - | 929 | 896 | **RNA**→ rpmE (PRK00019)Ribosomal_L31 (pfam01197)→ |
| env-1498 | scaffold21466_3_MH0083 | - | 331 | 298 | **RNA**→ rpmE (PRK00019)Ribosomal_L31 (pfam01197)→ |
| env-1499 | scaffold30922_4_O2.UC-12 | + | 780 | 813 | **RNA**→ rpmE (PRK00019)Ribosomal_L31 (pfam01197)→ |
| env-1500 | scaffold47149_2_V1.CD-6 | + | 2716 | 2749 | **RNA**→ rpmE (PRK00019)Ribosomal_L31 (pfam01197)→ |
| env-1501 | scaffold42088_2_V1.UC-14 | - | 149 | 116 | **RNA**→ rpmE (PRK00019)Ribosomal_L31 (pfam01197)→ |
| env-1502 | 4491405.3_NODE_38711 | - | 355 | 322 | **RNA**→ rpmE (PRK00019)Ribosomal_L31 (pfam01197)→ |
| env-1503 | NLM004_scaffold5032_16 | + | 1211 | 1244 | **RNA**→ rpmE (PRK00019)Ribosomal_L31 (pfam01197)→ |
| env-1504 | SRS011134_Baylor_scaffold_20481 | + | 1045 | 1078 | **RNA**→ |
| env-1505 | SRS014683_WUGC_scaffold_50673 | + | 1869 | 1902 | **RNA**→ rpmE (PRK00019)Ribosomal_L31 (pfam01197)→ |
| env-1506 | SRS019582_C3007814 | + | 421 | 454 | **RNA**→ rpmE (PRK00019)Ribosomal_L31 (pfam01197)→ |
| env-1507 | SRS049712_C15539391 | - | 84 | 51 | **RNA**→ |
| env-1508 | 4448807.3_Contig4303 | + | 447 | 480 | **RNA**→ rpmE (PRK00019)Ribosomal_L31 (pfam01197)→ |
| env-1509 | scaffold112155_1_MH0012 | - | 1337 | 1304 | **RNA**→ rpmE (PRK00019)Ribosomal_L31 (pfam01197)→ |
| Cmi-1-1 | NZ_ACCK01000449.1 | - | 23456 | 23423 | **RNA**→ rpmE (PRK00019)Ribosomal_L31 (pfam01197)→ rpmE (PRK00019)Ribosomal_L31 (pfam01197)→ |
| env-1510 | 4448819.3_Contig3697 | - | 316 | 283 | **RNA**→ rpmE (PRK00019)Ribosomal_L31 (pfam01197)→ |
| CSt-1-1 | NZ_HG005284.1 | - | 229433 | 229400 | **RNA**→ rpmE (PRK00019)Ribosomal_L31 (pfam01197)→ |
| env-1511 | SRS019601_C2033027 | + | 1675 | 1708 | **RNA**→ rpmE (PRK00019)Ribosomal_L31 (pfam01197)→ |
| env-1512 | 2211711892 | + | 393 | 426 | **RNA**→ |
| env-1513 | 2156523182 | + | 393 | 426 | **RNA**→ hypo→ |
| env-1514 | TB_AS07_7DRAFT_10000101 | - | 48063 | 48032 | **RNA**→ rpmE (PRK00019)Ribosomal_L31 (pfam01197)→ Tdk (COG1435)PRK04296 (PRK04296)→ spore_YhcN_YlaJ (TIGR02898)→ |
| Bsp-100-1 | NZ_JHXX01000001.1 | - | 753957 | 753936 | **RNA**→ Acetyltransf_1 (pfam00583)Acetyltransf_7 (pfam13508)→ CAT (smart01059)CAT (pfam00302)→ |
| Lmo-37-1 | NZ_LHRX01000005.1 | + | 41811 | 41840 | **RNA**→ rpmE2 (PRK01678)Ribosomal_L31 (pfam01197)→ |
| Lmo-74-1 | NZ_AVQQ01000001.1 | + | 10648 | 10677 | **RNA**→ rpmE2 (PRK01678)Ribosomal_L31 (pfam01197)→ |
| Lmo-15-1 | NZ_AATL02000007.1 | + | 41815 | 41844 | **RNA**→ rpmE2 (PRK01678)Ribosomal_L31 (pfam01197)→ rpmE2 (PRK01678)Ribosomal_L31 (pfam01197)→ |
| Lmo-75-1 | NZ_AVQM01000001.1 | - | 155185 | 155156 | **RNA**→ rpmE2 (PRK01678)Ribosomal_L31 (pfam01197)→ |
| Lmo-76-1 | NZ_AVQN01000027.1 | + | 41830 | 41859 | **RNA**→ rpmE2 (PRK01678)Ribosomal_L31 (pfam01197)→ |
| Lmo-77-1 | NZ_AVQO01000004.1 | + | 11588 | 11617 | **RNA**→ rpmE2 (PRK01678)Ribosomal_L31 (pfam01197)→ |
| Lmo-78-1 | NZ_AVQP01000004.1 | + | 41830 | 41859 | **RNA**→ rpmE2 (PRK01678)Ribosomal_L31 (pfam01197)→ |
| Lmo-73-1 | NC_019556.1 | - | 2579159 | 2579130 | **RNA**→ rpmE2 (PRK01678)Ribosomal_L31 (pfam01197)→ |
| Lmo-79-1 | NZ_CM001159.1 | - | 2710091 | 2710062 | **RNA**→ rpmE2 (PRK01678)Ribosomal_L31 (pfam01197)→ |
| Lmo-49-1 | NZ_KK074009.1 | - | 101216 | 101187 | **RNA**→ rpmE2 (PRK01678)Ribosomal_L31 (pfam01197)→ |
| Lmo-5-1 | NC_012488.1 | - | 2597658 | 2597629 | **RNA**→ rpmE2 (PRK01678)Ribosomal_L31 (pfam01197)→ |
| Lmo-1-1 | NC_017728.1 | - | 2576506 | 2576477 | **RNA**→ rpmE2 (PRK01678)Ribosomal_L31 (pfam01197)→ |
| Lmo-18-1 | NC_018642.1 | - | 2597808 | 2597779 | **RNA**→ rpmE2 (PRK01678)Ribosomal_L31 (pfam01197)→ |
| Lmo-4-1 | NC_018584.1 | - | 2640176 | 2640147 | **RNA**→ rpmE2 (PRK01678)Ribosomal_L31 (pfam01197)→ |

| | | | | | |
|---|---|---|---|---|---|
| Lmo-23-1 | NC_018585.1 | - | 2616367 | 2616338 | **RNA**→ rpmE2 (PRK01678)Ribosomal_L31 (pfam01197)→ |
| Lmo-33-9 | NC_021825.1 | + | 457670 | 457699 | **RNA**→ rpmE2 (PRK01678)Ribosomal_L31 (pfam01197)→ |
| Lmo-41-1 | NC_021830.1 | - | 2435601 | 2435572 | **RNA**→ rpmE2 (PRK01678)Ribosomal_L31 (pfam01197)→ |
| Lmo-31-1 | NC_002973.6 | - | 2579689 | 2579660 | **RNA**→ rpmE2 (PRK01678)Ribosomal_L31 (pfam01197)→ |
| Lmo-71-1 | NZ_CP007210.1 | - | 2025111 | 2025082 | **RNA**→ rpmE2 (PRK01678)Ribosomal_L31 (pfam01197)→ |
| Lmo-43-1 | NZ_AYPT01000018.1 | - | 101218 | 101189 | **RNA**→ rpmE2 (PRK01678)Ribosomal_L31 (pfam01197)→ |
| Bsa-1-1 | NZ_JYKW01000002.1 | - | 36424 | 36391 | **RNA**→ rpmE (PRK00019)Ribosomal_L31 (pfam01197)→ Tdk (COG1435)PRK04296 (PRK04296)→ |
| Bsa-1-2 | NZ_LATH01000002.1 | + | 378778 | 378811 | **RNA**→ rpmE (PRK00019)Ribosomal_L31 (pfam01197)→ Tdk (COG1435)PRK04296 (PRK04296)→ |
| Bpu-5-1 | NZ_AUYP01000013.1 | - | 87433 | 87400 | **RNA**→ rpmE (PRK00019)Ribosomal_L31 (pfam01197)→ Tdk (COG1435)PRK04296 (PRK04296)→ |
| Bsa-3-1 | NZ_AUPF01000002.1 | - | 584661 | 584628 | **RNA**→ rpmE (PRK00019)Ribosomal_L31 (pfam01197)→ Tdk (COG1435)PRK04296 (PRK04296)→ |
| Bsa-2-1 | NZ_ASJD01000002.1 | - | 402680 | 402647 | **RNA**→ rpmE (PRK00019)Ribosomal_L31 (pfam01197)→ Tdk (COG1435)PRK04296 (PRK04296)→ |
| Bsa-1-3 | NZ_JNBO01000025.1 | + | 19403 | 19436 | **RNA**→ rpmE (PRK00019)Ribosomal_L31 (pfam01197)→ Tdk (COG1435)PRK04296 (PRK04296)→ |
| Bae-1-1 | NZ_JXRO01000016.1 | - | 570625 | 570592 | **RNA**→ rpmE (PRK00019)Ribosomal_L31 (pfam01197)→ Tdk (COG1435)PRK04296 (PRK04296)→ |
| Bal-2-1 | NZ_JXAI01000001.1 | + | 295974 | 296007 | **RNA**→ rpmE (PRK00019)Ribosomal_L31 (pfam01197)→ Tdk (COG1435)PRK04296 (PRK04296)→ |
| Bal-2-2 | NZ_LDPI01000008.1 | - | 79586 | 79553 | **RNA**→ rpmE (PRK00019)Ribosomal_L31 (pfam01197)→ Tdk (COG1435)PRK04296 (PRK04296)→ |
| Bpu-7-1 | NZ_JH137675.1 | + | 97459 | 97492 | **RNA**→ rpmE (PRK00019)Ribosomal_L31 (pfam01197)→ Tdk (COG1435)PRK04296 (PRK04296)→ |
| Bpu-4-1 | NZ_AMDH01000015.1 | - | 232565 | 232532 | **RNA**→ rpmE (PRK00019)Ribosomal_L31 (pfam01197)→ Tdk (COG1435)PRK04296 (PRK04296)→ |
| Bst-2-1 | NZ_APAS01000013.1 | - | 90146 | 90113 | **RNA**→ rpmE (PRK00019)Ribosomal_L31 (pfam01197)→ rpmE (PRK00019)Ribosomal_L31 (pfam01197)→ Tdk (COG1435)PRK04296 (PRK04296)→ |
| Bpu-6-1 | NZ_AYTK01000014.1 | - | 90600 | 90567 | **RNA**→ rpmE (PRK00019)Ribosomal_L31 (pfam01197)→ Tdk (COG1435)PRK04296 (PRK04296)→ |
| Bal-3-1 | NZ_ASJC01000004.1 | - | 121394 | 121361 | **RNA**→ rpmE (PRK00019)Ribosomal_L31 (pfam01197)→ Tdk (COG1435)PRK04296 (PRK04296)→ |
| Bsp-88-1 | NZ_JNIG01000009.1 | - | 635721 | 635688 | **RNA**→ rpmE (PRK00019)Ribosomal_L31 (pfam01197)→ Tdk (COG1435)PRK04296 (PRK04296)→ |
| Bzh-1-1 | NZ_JOTP01000003.1 | + | 53546 | 53579 | **RNA**→ rpmE (PRK00019)Ribosomal_L31 (pfam01197)→ Tdk (COG1435)PRK04296 (PRK04296)→ |
| Bsp-6-1 | NZ_AJWW01000012.1 | - | 126382 | 126349 | **RNA**→ rpmE (PRK00019)Ribosomal_L31 (pfam01197)→ rpmE (PRK00019)Ribosomal_L31 (pfam01197)→ Tdk (COG1435)PRK04296 (PRK04296)→ Tdk (COG1435)PRK04296 (PRK04296)→ |
| Bal-2-3 | NZ_JOVS01000001.1 | + | 321014 | 321047 | **RNA**→ rpmE (PRK00019)Ribosomal_L31 (pfam01197)→ Tdk (COG1435)PRK04296 (PRK04296)→ |
| Bpu-2-1 | NC_009848.1 | - | 3327937 | 3327904 | **RNA**→ rpmE (PRK00019)Ribosomal_L31 (pfam01197)→ Tdk (COG1435)PRK04296 (PRK04296)→ |
| Bpu-1-1 | NZ_ABRX01000004.1 | - | 80659 | 80626 | **RNA**→ rpmE (PRK00019)Ribosomal_L31 (pfam01197)→ rpmE (PRK00019)Ribosomal_L31 (pfam01197)→ Tdk (COG1435)PRK04296 (PRK04296)→ Tdk (COG1435)PRK04296 (PRK04296)→ |
| Efa-374-1 | NZ_KB029918.1 | - | 250318 | 250286 | **RNA**→ rpmE2 (PRK01678)Ribosomal_L31 (pfam01197)→ |
| Efa-364-1 | NZ_KB029496.1 | + | 333459 | 333491 | **RNA**→ rpmE2 (PRK01678)Ribosomal_L31 (pfam01197)→ |
| Bam-9-1 | NZ_LGYP01000016.1 | - | 457684 | 457651 | **RNA**→ rpmE (PRK00019)Ribosomal_L31 (pfam01197)→ Tdk (COG1435)PRK04296 (PRK04296)→ |
| Bam-18-1 | NZ_AUNG01000003.1 | + | 391952 | 391985 | **RNA**→ rpmE (PRK00019)Ribosomal_L31 (pfam01197)→ Tdk (COG1435)PRK04296 (PRK04296)→ |
| Bsp-93-1 | NZ_JQKM01000003.1 | + | 344087 | 344120 | **RNA**→ rpmE (PRK00019)Ribosomal_L31 (pfam01197)→ Tdk (COG1435)PRK04296 (PRK04296)→ |
| Bam-16-1 | NZ_AUWK01000002.1 | + | 389923 | 389956 | **RNA**→ rpmE (PRK00019)Ribosomal_L31 (pfam01197)→ Tdk (COG1435)PRK04296 (PRK04296)→ |
| Bsu-16-1 | NZ_AYTJ01000005.1 | - | 83015 | 82982 | **RNA**→ rpmE (PRK00019)Ribosomal_L31 (pfam01197)→ Tdk (COG1435)PRK04296 (PRK04296)→ |
| Bsp-2-1 | NZ_AFSU01000009.1 | - | 134410 | 134377 | **RNA**→ rpmE (PRK00019)Ribosomal_L31 (pfam01197)→ rpmE (PRK00019)Ribosomal_L31 (pfam01197)→ Tdk (COG1435)PRK04296 (PRK04296)→ Tdk (COG1435)PRK04296 (PRK04296)→ |
| Bve-3-1 | NZ_AOFO01000001.1 | - | 406706 | 406673 | **RNA**→ rpmE (PRK00019)Ribosomal_L31 (pfam01197)→ Tdk (COG1435)PRK04296 (PRK04296)→ |
| Bam-15-1 | NZ_JCOC01000009.1 | + | 36110 | 36143 | **RNA**→ rpmE (PRK00019)Ribosomal_L31 (pfam01197)→ Tdk (COG1435)PRK04296 (PRK04296)→ |
| Bam-2-1 | NC_009725.1 | - | 3531629 | 3531596 | **RNA**→ rpmE (PRK00019)Ribosomal_L31 (pfam01197)→ Tdk (COG1435)PRK04296 (PRK04296)→ |
| Bam-6-1 | NC_017912.1 | - | 3841226 | 3841193 | **RNA**→ rpmE (PRK00019)Ribosomal_L31 (pfam01197)→ rpmE (PRK00019)Ribosomal_L31 (pfam01197)→ rpmE (PRK00019)Ribosomal_L31 (pfam01197)→ Tdk (COG1435)PRK04296 (PRK04296)→ |
| Bam-8-1 | NC_017061.1 | - | 3845102 | 3845069 | **RNA**→ rpmE (PRK00019)Ribosomal_L31 (pfam01197)→ Tdk (COG1435)PRK04296 (PRK04296)→ |
| Bam-20-1 | NC_019842.1 | - | 3566773 | 3566740 | **RNA**→ rpmE (PRK00019)Ribosomal_L31 (pfam01197)→ Tdk (COG1435)PRK04296 (PRK04296)→ |
| Bam-24-1 | NC_020410.1 | - | 3525764 | 3525731 | **RNA**→ rpmE (PRK00019)Ribosomal_L31 (pfam01197)→ Tdk (COG1435)PRK04296 (PRK04296)→ |
| Bam-25-1 | NC_022081.1 | - | 3517148 | 3517115 | **RNA**→ rpmE (PRK00019)Ribosomal_L31 (pfam01197)→ Tdk (COG1435)PRK04296 (PRK04296)→ |
| Bam-23-1 | NC_022075.1 | - | 3685897 | 3685864 | **RNA**→ rpmE (PRK00019)Ribosomal_L31 (pfam01197)→ Tdk (COG1435)PRK04296 (PRK04296)→ |
| Bam-10-1 | NC_022653.1 | - | 3529867 | 3529834 | **RNA**→ rpmE (PRK00019)Ribosomal_L31 (pfam01197)→ rpmE (PRK00019)Ribosomal_L31 (pfam01197)→ Tdk (COG1435)PRK04296 (PRK04296)→ |
| Bam-22-1 | NC_022530.1 | - | 3806043 | 3806010 | **RNA**→ rpmE (PRK00019)Ribosomal_L31 (pfam01197)→ Tdk (COG1435)PRK04296 (PRK04296)→ |
| Bve-1-2 | NZ_CP009679.1 | - | 3615874 | 3615841 | **RNA**→ rpmE (PRK00019)Ribosomal_L31 (pfam01197)→ Tdk (COG1435)PRK04296 (PRK04296)→ |
| Bve-5-1 | NZ_CP007244.1 | - | 3565547 | 3565514 | **RNA**→ rpmE (PRK00019)Ribosomal_L31 (pfam01197)→ Tdk (COG1435)PRK04296 (PRK04296)→ |
| Bsp-31-1 | NZ_CP009938.1 | - | 151413 | 151380 | **RNA**→ rpmE (PRK00019)Ribosomal_L31 (pfam01197)→ Tdk (COG1435)PRK04296 (PRK04296)→ |

| | | | | | |
|---|---|---|---|---|---|
| Bam-17-1 | NZ_CP007242.1 | - | 3385240 | 3385207 | **RNA**→ rpmE (PRK00019)Ribosomal_L31 (pfam01197)→ Tdk (COG1435)PRK04296 (PRK04296)→ |
| Bve-1-3 | NZ_CP011347.1 | + | 385791 | 385824 | **RNA**→ rpmE (PRK00019)Ribosomal_L31 (pfam01197)→ Tdk (COG1435)PRK04296 (PRK04296)→ |
| Bve-1-4 | NZ_CP011686.1 | - | 3621002 | 3620969 | **RNA**→ rpmE (PRK00019)Ribosomal_L31 (pfam01197)→ Tdk (COG1435)PRK04296 (PRK04296)→ |
| Bsp-1-1 | NZ_AJST01000001.1 | - | 3334151 | 3334118 | **RNA**→ rpmE (PRK00019)Ribosomal_L31 (pfam01197)→ rpmE (PRK00019)Ribosomal_L31 (pfam01197)→ Tdk (COG1435)PRK04296 (PRK04296)→ Tdk (COG1435)PRK04296 (PRK04296)→ |
| Bam-11-1 | NZ_AVQH01000023.1 | + | 411156 | 411189 | **RNA**→ rpmE (PRK00019)Ribosomal_L31 (pfam01197)→ Tdk (COG1435)PRK04296 (PRK04296)→ |
| Ban-20-1 | NZ_LN868937.1 | + | 267111 | 267142 | **RNA**→ rpmE (PRK00019)Ribosomal_L31 (pfam01197)→ |
| Bsp-61-1 | NZ_CCNM01000041.1 | + | 187611 | 187642 | **RNA**→ rpmE (PRK00019)Ribosomal_L31 (pfam01197)→ |
| Efa-527-1 | NZ_CP006030.1 | - | 2037832 | 2037800 | **RNA**→ rpmE2 (PRK01678)Ribosomal_L31 (pfam01197)→ |
| Efa-318-1 | NZ_JH813136.1 | - | 88872 | 88840 | **RNA**→ rpmE2 (PRK01678)Ribosomal_L31 (pfam01197)→ |
| Efa-477-1 | NZ_KB946089.1 | + | 43740 | 43772 | **RNA**→ rpmE2 (PRK01678)Ribosomal_L31 (pfam01197)→ |
| Bmo-1-1 | NZ_JH600279.1 | - | 659896 | 659863 | **RNA**→ rpmE (PRK00019)Ribosomal_L31 (pfam01197)→ Tdk (COG1435)PRK04296 (PRK04296)→ |
| Egi-1-1 | NZ_KB946874.1 | - | 1793866 | 1793834 | **RNA**→ rpmE2 (PRK01678)Ribosomal_L31 (pfam01197)→ |
| Bco-2-1 | NC_016023.1 | + | 1562815 | 1562846 | **RNA**→ rpmE (PRK00019)Ribosomal_L31 (pfam01197)→ |
| Bco-10-1 | NZ_ANAP01000032.1 | - | 35355 | 35324 | **RNA**→ rpmE (PRK00019)Ribosomal_L31 (pfam01197)→ |
| Efa-371-1 | NZ_KB029417.1 | - | 209198 | 209166 | **RNA**→ rpmE2 (PRK01678)Ribosomal_L31 (pfam01197)→ |
| Efa-387-1 | NZ_KB029838.1 | + | 68007 | 68039 | **RNA**→ rpmE2 (PRK01678)Ribosomal_L31 (pfam01197)→ |
| Efa-390-1 | NZ_KB029937.1 | + | 330992 | 331024 | **RNA**→ rpmE2 (PRK01678)Ribosomal_L31 (pfam01197)→ |
| Efa-383-1 | NZ_KB029808.1 | - | 211176 | 211144 | **RNA**→ rpmE2 (PRK01678)Ribosomal_L31 (pfam01197)→ |
| Efa-415-1 | NZ_KB945936.1 | + | 53148 | 53180 | **RNA**→ rpmE2 (PRK01678)Ribosomal_L31 (pfam01197)→ |
| Efa-484-1 | NZ_KB946737.1 | + | 59491 | 59523 | **RNA**→ rpmE2 (PRK01678)Ribosomal_L31 (pfam01197)→ |
| Efa-435-1 | NZ_KB947613.1 | - | 208560 | 208528 | **RNA**→ rpmE2 (PRK01678)Ribosomal_L31 (pfam01197)→ |
| Efa-486-1 | NZ_KB946762.1 | - | 207793 | 207761 | **RNA**→ rpmE2 (PRK01678)Ribosomal_L31 (pfam01197)→ |
| Efa-434-1 | NZ_KB947589.1 | - | 209551 | 209519 | **RNA**→ rpmE2 (PRK01678)Ribosomal_L31 (pfam01197)→ |
| Efa-438-1 | NZ_KB947681.1 | + | 53471 | 53503 | **RNA**→ rpmE2 (PRK01678)Ribosomal_L31 (pfam01197)→ |
| Efa-431-1 | NZ_KB947540.1 | + | 60225 | 60257 | **RNA**→ rpmE2 (PRK01678)Ribosomal_L31 (pfam01197)→ |
| Efa-432-1 | NZ_KB947550.1 | - | 209167 | 209135 | **RNA**→ rpmE2 (PRK01678)Ribosomal_L31 (pfam01197)→ |
| Efa-455-1 | NZ_KB948198.1 | - | 210063 | 210031 | **RNA**→ rpmE2 (PRK01678)Ribosomal_L31 (pfam01197)→ |
| Efa-410-1 | NZ_KB932608.1 | - | 210349 | 210317 | **RNA**→ rpmE2 (PRK01678)Ribosomal_L31 (pfam01197)→ |
| Sar-2-1 | NZ_KK096598.1 | - | 56203 | 56174 | **RNA**→ rpmE2 (PRK01678)Ribosomal_L31 (pfam01197)→ |
| Sau-56-1 | NC_016941.1 | - | 2116146 | 2116117 | **RNA**→ rpmE2 (PRK01678)Ribosomal_L31 (pfam01197)→ |
| Lre-7-1 | NZ_AAPZ02000001.1 | + | 1432470 | 1432503 | **RNA**→ rpmE2 (PRK01678)Ribosomal_L31 (pfam01197)→ rpmE2 (PRK01678)Ribosomal_L31 (pfam01197)→ |
| Lre-6-1 | NZ_AEAW01000058.1 | + | 71250 | 71283 | **RNA**→ rpmE2 (PRK01678)Ribosomal_L31 (pfam01197)→ |
| Lre-2-1 | NZ_FR854362.1 | + | 95442 | 95475 | **RNA**→ rpmE2 (PRK01678)Ribosomal_L31 (pfam01197)→ |
| Lre-3-1 | NC_021494.1 | - | 1774307 | 1774274 | **RNA**→ rpmE2 (PRK01678)Ribosomal_L31 (pfam01197)→ |
| Lsa-12-1 | NC_007929.1 | + | 378133 | 378165 | **RNA**→ rpmE2 (PRK01678)Ribosomal_L31 (pfam01197)→ hypo→ |
| Lmo-35-1 | NZ_CBXR010000002.1 | - | 221773 | 221744 | **RNA**→ rpmE2 (PRK01678)Ribosomal_L31 (pfam01197)→ |
| Lmo-54-1 | NZ_AXDU01000020.1 | + | 11271 | 11300 | **RNA**→ rpmE2 (PRK01678)Ribosomal_L31 (pfam01197)→ |
| Lmo-51-1 | NZ_KK074120.1 | - | 120845 | 120816 | **RNA**→ rpmE2 (PRK01678)Ribosomal_L31 (pfam01197)→ |
| Lmo-36-1 | NZ_HG813249.1 | + | 1613739 | 1613768 | **RNA**→ rpmE2 (PRK01678)Ribosomal_L31 (pfam01197)→ |
| Sxy-1-1 | NZ_LN554884.1 | + | 870932 | 870961 | **RNA**→ rpmE2 (PRK01678)Ribosomal_L31 (pfam01197)→ |
| Sxy-2-1 | NZ_AURW01000004.1 | - | 45188 | 45159 | **RNA**→ rpmE2 (PRK01678)Ribosomal_L31 (pfam01197)→ |

#### 11.4 Conserved domains

Conserved domains found in protein-coding genes listed in Section 11.3 are shown below, with the first sentence in their description from the Conserved Domain Database (if any). Conserved domains associated with more than one L31-Firmicutes RNA are assigned a color, while others are shown in gray. The symbols 'd' and 'u' (if any) to the left of the colored domain name indicate whether the domain occurs upstream or downstream of the RNA motif. Domains marked with lower-case 'd' occur (at least once) present downstream of some L31-Firmicutes RNA in the opposite orientation. Domains marked with capital 'D' occur downstream and in the same orientation (i.e., a potential *cis*-regulatory arrangement). Domains marked with a 'u' occur upstream of the RNA in either orientation. If the 'd'/'u' symbols are missing, then the domain is downstream and in the same strand. (This mode is used for motifs predicted as *cis*-regulatory.) The number in parentheses after the colored domain name is the number of occurrences in Section 11.3.

cd00156 (2) Signal receiver domain; originally thought to be unique to bacteria (CheY, OmpR, NtrC, and PhoB), now recently identified in eukaroytes ETR1 Arabidopsis thaliana; this domain receives the signal from the sensor partner in a two-component systems; contains a phosphoacceptor site that is phosphorylated by histidine kinase homologs; usually found N-terminal to a DNA binding effector domain; forms homodimers
cd00198 (3) Von Willebrand factor type A (vWA) domain was originally found in the blood coagulation protein von Willebrand factor (vWF).
cd00229 (6) SGNH_hydrolase, or GDSL_hydrolase, is a diverse family of lipases and esterases.
cd00293 (2) Usp: Universal stress protein family.
cd00383 (1) DNA-binding effector domain of two-component system response regulators.
cd00430 (5) Type III Pyridoxal 5-phosphate (PLP)-Dependent Enzyme Alanine Racemase.
cd00552 (6) RaiA ("ribosome-associated inhibitor A", also known as Protein Y (PY), YfiA, and SpotY, is a stress-response protein that binds the ribosomal subunit interface and arrests translation by interfering with aminoacyl-tRNA binding to the ribosomal A site.
cd00616 (1) 3-amino-5-hydroxybenzoic acid synthase family (AHBA_syn).
cd02947 (1) TRX family; composed of two groups: Group I, which includes proteins that exclusively encode a TRX domain; and Group II, which are composed of fusion proteins of TRX and additional domains.
cd03674 (2) Members of the Nudix hydrolase superfamily catalyze the hydrolysis of NUcleoside DIphosphates linked to other moieties, X.
cd04250 (1) AAK_NAGK-C: N-Acetyl-L-glutamate kinase - cyclic (NAGK-C) catalyzes the phosphorylation of the gamma-COOH group of N-acetyl-L-glutamate (NAG) by ATP in the second step of arginine biosynthesis found in some bacteria and photosynthetic organisms using the non-acetylated, cyclic route of ornithine biosynthesis.
cd06165 (10) Sortase domain found in class A sortases.
cd06262 (4) mainly hydrolytic enzymes and related proteins which carry out various biological functions; MBL-fold metallohydrolase domain.
cd06850 (1) The biotinyl-domain or biotin carboxyl carrier protein (BCCP) domain is present in all biotin-dependent enzymes, such as acetyl-CoA carboxylase, pyruvate carboxylase, propionyl-CoA carboxylase, methylcrotonyl-CoA carboxylase, geranyl-CoA carboxylase, oxaloacetate decarboxylase, methylmalonyl-CoA decarboxylase, transcarboxylase and urea amidolyase.
cd06853 (29) This subfamily contains Escherichia coli WecA, Bacillus subtilis TagO and related proteins.
cd07516 (1) phosphatase, similar to Escherichia coli Cof and Thermotoga maritima TM0651; belongs to the haloacid dehalogenase-like superfamily.
cd07557 (4) Trimeric dUTP diphosphatases.
cd07733 (3) uncharacterized subgroup which includes Bacillus subtilis YycJ and related proteins; MBL-fold metallo hydrolase domain.
cd07743 (1) uncharacterized subgroup of the MBL-fold_metallo-hydrolase superfamily; MBL-fold metallo hydrolase domain.
cd07937 (1) Pyruvate carboxylase and Transcarboxylase 5S, carboxyltransferase domain.
cd10225 (12) MreB and similar proteins.
cd12114 (1) The adenylation domain of nonribosomal peptide synthetases (NRPS), including Streptoalloteichus tallysomycin biosynthesis genes.
cd16015 (20) Lipoteichoic acid synthase like.
cd16344 (1) low molecular weight protein arginine phosphatase.
CHL00117 (1) RNA polymerase beta" subunit; Reviewed
CHL00204 (1) Ycf1; Provisional
COG0002 (2) N-acetyl-gamma-glutamylphosphate reductase [Amino acid transport and metabolism]
COG0009 (5) tRNA A37 threonylcarbamoyladenosine synthetase subunit TsaC/SUA5/YrdC [Translation, ribosomal structure and biogenesis]
COG0021 (12) Transketolase [Carbohydrate transport and metabolism]
COG0025 (6) NhaP-type Na+/H+ or K+/H+ antiporter [Inorganic ion transport and metabolism]
COG0035 (3) Uracil phosphoribosyltransferase [Nucleotide transport and metabolism]
COG0053 (1) Divalent metal cation (Fe/Co/Zn/Cd) transporter [Inorganic ion transport and metabolism]
COG0078 (1) Ornithine carbamoyltransferase [Amino acid transport and metabolism]
COG0112 (3) Glycine/serine hydroxymethyltransferase [Amino acid transport and metabolism]
COG0143 (3) Methionyl-tRNA synthetase [Translation, ribosomal structure and biogenesis]
COG0150 (1) Phosphoribosylaminoimidazole (AIR) synthetase [Nucleotide transport and metabolism]
COG0168 (1) Trk-type K+ transport system, membrane component [Inorganic ion transport and metabolism]
COG0189 (2) Glutathione synthase/RimK-type ligase, ATP-grasp superfamily [Coenzyme transport and metabolism, Translation, ribosomal structure and biogenesis]
COG0216 (90) Protein chain release factor A [Translation, ribosomal structure and biogenesis]
COG0254 (3) Ribosomal protein L31 [Translation, ribosomal structure and biogenesis]
COG0357 (12) 16S rRNA G527 N7-methylase RsmG (former glucose-inhibited division protein B) [Translation, ribosomal structure and biogenesis]
COG0381 (3) UDP-N-acetylglucosamine 2-epimerase [Cell wall/membrane/envelope biogenesis]
COG0394 (3) Protein-tyrosine-phosphatase [Signal transduction mechanisms]
COG0415 (8) Deoxyribodipyrimidine photolyase [Replication, recombination and repair]
COG0436 (1) Aspartate/methionine/tyrosine aminotransferase [Amino acid transport and metabolism]
COG0439 (1) Biotin carboxylase [Lipid transport and metabolism]
COG0445 (13) tRNA U34 5-carboxymethylaminomethyl modifying enzyme MnmG/GidA [Translation, ribosomal structure and biogenesis]
COG0456 (1) Ribosomal protein S18 acetylase RimI and related acetyltransferases [Translation, ribosomal structure and biogenesis]
COG0464 (3) AAA+-type ATPase, SpoVK/Ycf46/Vps4 family [Cell wall/membrane/envelope biogenesis, Cell cycle control, cell division, chromosome partitioning, Signal transduction mechanisms]
COG0472 (1) UDP-N-acetylmuramyl pentapeptide phosphotransferase/UDP-N-acetylglucosamine-1-phosphate transferase [Cell wall/membrane/envelope biogenesis]
COG0513 (5) Superfamily II DNA and RNA helicase [Replication, recombination and repair]
COG0561 (1) Hydroxymethylpyrimidine pyrophosphatase and other HAD family phosphatases [Coenzyme transport and metabolism, General function prediction only]
COG0590 (43) tRNA(Arg) A34 adenosine deaminase TadA [Translation, ribosomal structure

and biogenesis]
COG0602 (21) Organic radical activating enzyme [General function prediction only]
COG0653 (8) Preprotein translocase subunit SecA (ATPase, RNA helicase) [Intracellular trafficking, secretion, and vesicular transport]
COG0657 (1) Acetyl esterase/lipase [Lipid transport and metabolism]
COG0674 (3) Pyruvate:ferredoxin oxidoreductase or related 2-oxoacid:ferredoxin oxidoreductase, alpha subunit [Energy production and conversion]
COG0697 (2) Permease of the drug/metabolite transporter (DMT) superfamily [Carbohydrate transport and metabolism, Amino acid transport and metabolism, General function prediction only]
COG0736 (5) Phosphopantetheinyl transferase (holo-ACP synthase) [Lipid transport and metabolism]
COG0745 (1) DNA-binding response regulator, OmpR family, contains REC and winged-helix (wHTH) domain [Signal transduction mechanisms, Transcription]
COG0756 (13) dUTPase [Nucleotide transport and metabolism, Defense mechanisms]
COG0770 (5) UDP-N-acetylmuramyl pentapeptide synthase [Cell wall/membrane/envelope biogenesis]
COG0784 (2) CheY chemotaxis protein or a CheY-like REC (receiver) domain [Signal transduction mechanisms]
COG0840 (1) Methyl-accepting chemotaxis protein [Cell motility, Signal transduction mechanisms]
COG1013 (2) Pyruvate:ferredoxin oxidoreductase or related 2-oxoacid:ferredoxin oxidoreductase, beta subunit [Energy production and conversion]
COG1014 (2) Pyruvate:ferredoxin oxidoreductase or related 2-oxoacid:ferredoxin oxidoreductase, gamma subunit [Energy production and conversion]
COG1145 (2) Ferredoxin [Energy production and conversion]
COG1192 (5) Cellulose biosynthesis protein BcsQ [Cell motility]
COG1235 (12) Phosphoribosyl 1,2-cyclic phosphodiesterase [Inorganic ion transport and metabolism]
COG1275 (1) Tellurite resistance protein TehA and related permeases [Defense mechanisms]
COG1278 (1) Cold shock protein, CspA family [Transcription]
COG1368 (4) Phosphoglycerol transferase MdoB or a related enzyme of AlkP superfamily [Cell wall/membrane/envelope biogenesis]
COG1404 (1) Serine protease, subtilisin family [Posttranslational modification, protein turnover, chaperones]
COG1435 (423) Thymidine kinase [Nucleotide transport and metabolism]
COG1475 (6) Chromosome segregation protein Spo0J, contains ParB-like nuclease domain [Cell cycle control, cell division, chromosome partitioning]
COG1476 (1) DNA-binding transcriptional regulator, XRE-family HTH domain [Transcription]
COG1544 (2) Ribosome-associated translation inhibitor RaiA [Translation, ribosomal structure and biogenesis]
COG1737 (4) DNA-binding transcriptional regulator, MurR/RpiR family, contains HTH and SIS domains [Transcription]
COG1940 (2) Sugar kinase of the NBD/HSP70 family, may contain an N-terminal HTH domain [Transcription, Carbohydrate transport and metabolism]
COG2235 (4) Arginine deiminase [Amino acid transport and metabolism]
COG2323 (7) Uncharacterized membrane protein YcaP, DUF421 family [Function unknown]
COG2385 (21) Peptidoglycan hydrolase (amidase) enhancer domain [Cell wall/membrane/envelope biogenesis]
COG2812 (2) DNA polymerase III, gamma/tau subunits [Replication, recombination and repair]
COG2826 (1) Transposase and inactivated derivatives, IS30 family [Mobilome: prophages, transposons]
COG2890 (62) Methylase of polypeptide chain release factors [Translation, ribosomal structure and biogenesis]
COG3279 (17) DNA-binding response regulator, LytR/AlgR family [Transcription, Signal transduction mechanisms]
COG3290 (13) Sensor histidine kinase regulating citrate/malate metabolism [Signal transduction mechanisms]
COG3415 (1) Transposase [Mobilome: prophages, transposons]
COG4603 (2) ABC-type uncharacterized transport system, permease component [General function prediction only]
COG4632 (2) Exopolysaccharide biosynthesis protein related to N-acetylglucosamine-1-phosphodiester alpha-N-acety. [Carbohydrate transport and metabolism]
COG5279 (2) Cytokinesis protein 3, contains TGc (transglutaminase/protease-like) domain [Cell cycle control, cell division, chromosome partitioning]
COG5421 (1) Transposase [Mobilome: prophages, transposons]
COG5492 (1) Uncharacterized conserved protein YjdB, contains Ig-like domain [General function prediction only]
MTH00151 (1) NADH dehydrogenase subunit 5; Provisional
pfam00301 (1) Rubredoxin.
pfam00302 (1) Chloramphenicol acetyltransferase.
pfam00334 (2) Nucleoside diphosphate kinase.
pfam00472 (1) RF-1 domain.
pfam00486 (1) Transcriptional regulatory protein, C terminal.
pfam00583 (2) Acetyltransferase (GNAT) family.
pfam00657 (2) GDSL-like Lipase/Acylhydrolase.
pfam00875 (1) DNA photolyase.
pfam00892 (2) EamA-like transporter family.
pfam01066 (1) CDP-alcohol phosphatidyltransferase.
pfam01197 (8437) Ribosomal protein L31.
pfam01551 (31) Peptidase family M23.
pfam01648 (4) 4'-phosphopantetheinyl transferase superfamily.
pfam01960 (1) ArgJ family.
pfam01996 (1) F420-0:Gamma-glutamyl ligase.
pfam02368 (1) Bacterial Ig-like domain (group 2).
pfam02388 (1) FemAB family.
pfam02436 (1) Conserved carboxylase domain.
pfam02482 (3) Sigma 54 modulation protein / S30EA ribosomal protein.
pfam02502 (104) Ribose/Galactose Isomerase.
pfam02518 (3) Histidine kinase-, DNA gyrase B-, and HSP90-like ATPase.
pfam02575 (1) YbaB/EbfC DNA-binding family.
pfam02590 (12) Predicted SPOUT methyltransferase.
pfam02645 (1) Uncharacterized protein, DegV family COG1307.
pfam02666 (1) Phosphatidylserine decarboxylase.
pfam02810 (6) SEC-C motif. [HC]
pfam02897 (3) Prolyl oligopeptidase, N-terminal beta-propeller domain.
pfam03441 (1) FAD binding domain of DNA photolyase.

pfam03462 (3) PCRF domain.
pfam03481 (5) Putative GTP-binding controlling metal-binding.
pfam03840 (1) Preprotein translocase SecG subunit.
pfam03947 (1) Ribosomal Proteins L2, C-terminal domain.
pfam04011 (1) LemA family.
pfam04246 (1) Positive regulator of sigma(E), RseC/MucC.
pfam04260 (3) Protein of unknown function (DUF436).
pfam04313 (9) Type I restriction enzyme R protein N terminus (HSDR_N).
pfam04397 (5) LytTr DNA-binding domain.
pfam05532 (2) CsbD-like.
pfam05932 (1) Tir chaperone protein (CesT) family.
pfam06030 (4) Bacterial protein of unknown function (DUF916).
pfam06107 (8) Bacterial protein of unknown function (DUF951).
pfam06612 (38) Protein of unknown function (DUF1146).
pfam06723 (1) MreB/Mbl protein.
pfam07423 (1) Protein of unknown function (DUF1510).
pfam07586 (1) Protein of unknown function (DUF1552).
pfam07859 (1) alpha/beta hydrolase fold.
pfam08245 (5) Mur ligase middle domain.
pfam08282 (2) haloacid dehalogenase-like hydrolase.
pfam08297 (1) U3 snoRNA associated.
pfam08345 (1) Flagellar M-ring protein C-terminal.
pfam08486 (6) Stage II sporulation protein.
pfam09551 (11) Stage II sporulation protein R (spore_II_R).
pfam09580 (6) Sporulation lipoprotein YhcN/YlaJ (Spore_YhcN_YlaJ).
pfam09992 (2) Phosphodiester glycosidase.
pfam10254 (2) PACS-1 cytosolic sorting protein.
pfam10722 (1) Putative bacterial sensory transduction regulator.
pfam11797 (4) Protein of unknown function C-terminal (DUF3324).
pfam12007 (2) Protein of unknown function (DUF3501).
pfam12116 (5) Stage III sporulation protein D.
pfam12146 (1) Serine aminopeptidase, S33.
pfam12224 (1) Putative amidoligase enzyme.
pfam12732 (1) YtxH-like protein.
pfam12838 (2) 4Fe-4S dicluster domain.
pfam13353 (21) 4Fe-4S single cluster domain.
pfam13472 (4) GDSL-like Lipase/Acylhydrolase family.
pfam13508 (3) Acetyltransferase (GNAT) domain.
pfam13518 (1) Helix-turn-helix domain.
pfam13588 (9) Type I restriction enzyme R protein N terminus (HSDR_N).
pfam13614 (5) AAA domain.
pfam13629 (1) Pilus formation protein N terminal region.
pfam13649 (3) Methyltransferase domain.
pfam14397 (4) Sugar-transfer associated ATP-grasp.
pfam14437 (40) MafB19-like deaminase.
pfam14450 (1) Cell division protein FtsA.
pfam14470 (1) Bacterial PH domain.
pfam14501 (13) GHKL domain.
pfam16321 (10) Sigma 54 modulation/S30EA ribosomal protein C terminus.
pfam16339 (1) Domain of unknown function (DUF4969).
pfam16403 (2) Domain of unknown function (DUF5011).
pfam16916 (1) Dimerisation domain of Zinc Transporter.

PHA02601 (1) integrase; Provisional
PLN02662 (1) cinnamyl-alcohol dehydrogenase family protein
PRK00011 (3) serine hydroxymethyltransferase; Reviewed
PRK00019 (1362) 50S ribosomal protein L31; Reviewed
PRK00053 (5) alanine racemase; Reviewed
PRK00070 (5) 4'-phosphopantetheinyl transferase; Provisional
PRK00103 (12) rRNA large subunit methyltransferase; Provisional
PRK00107 (12) 16S rRNA methyltransferase GidB; Reviewed
PRK00108 (16) phospho-N-acetylmuramoyl-pentapeptide-transferase; Provisional
PRK00129 (3) uracil phosphoribosyltransferase; Reviewed
PRK00153 (1) hypothetical protein; Validated
PRK00436 (2) N-acetyl-gamma-glutamyl-phosphate reductase; Validated
PRK00528 (5) 50S ribosomal protein L31; Reviewed
PRK00591 (83) peptide chain release factor 1; Validated
PRK00601 (18) deoxyuridine 5'-triphosphate nucleotidohydrolase; Provisional
PRK00723 (1) phosphatidylserine decarboxylase; Provisional
PRK00942 (1) acetylglutamate kinase; Provisional
PRK01388 (4) arginine deiminase; Provisional
PRK01678 (7101) 50S ribosomal protein L31 type B; Reviewed
PRK02102 (1) ornithine carbamoyltransferase; Validated
PRK04296 (423) thymidine kinase; Provisional
PRK05192 (13) tRNA uridine 5-carboxymethylaminomethyl modification enzyme GidA; Validated
PRK05326 (6) potassium/proton antiporter; Reviewed
PRK05385 (1) phosphoribosylaminoimidazole synthetase; Provisional
PRK05388 (1) bifunctional ornithine acetyltransferase/N-acetylglutamate synthase protein; Validated
PRK05452 (1) anaerobic nitric oxide reductase flavorubredoxin; Provisional
PRK05563 (2) DNA polymerase III subunits gamma and tau; Validated
PRK05571 (104) ribose-5-phosphate isomerase B; Provisional
PRK05667 (2) DNA primase; Validated
PRK05899 (12) transketolase; Reviewed
PRK07681 (1) aspartate aminotransferase; Provisional
PRK09328 (51) N5-glutamine S-adenosyl-L-methionine-dependent methyltransferase; Provisional
PRK09374 (1) 50S ribosomal protein L2; Validated
PRK10351 (3) holo-(acyl carrier protein) synthase 2; Provisional
PRK10674 (4) deoxyribodipyrimidine photolyase; Provisional
PRK10776 (1) nucleoside triphosphate pyrophosphohydrolase; Provisional
PRK10860 (3) tRNA-specific adenosine deaminase; Provisional
PRK10871 (1) lipoprotein NlpD; Provisional
PRK11086 (2) sensory histidine kinase DcuS; Provisional
PRK11361 (17) acetoacetate metabolism regulatory protein AtoC; Provisional
PRK11649 (28) putative peptidase; Provisional
PRK11893 (1) methionyl-tRNA synthetase; Reviewed
PRK12267 (2) methionyl-tRNA synthetase; Reviewed
PRK12363 (3) phosphoglycerol transferase I; Provisional
PRK12904 (8) preprotein translocase subunit SecA; Reviewed
PRK12999 (1) pyruvate carboxylase; Reviewed
PRK13690 (3) hypothetical protein; Provisional
PRK13759 (18) arylsulfatase; Provisional
PRK13930 (13) rod shape-determining protein MreB; Provisional

PRK15482 (4) transcriptional regulator MurR; Provisional
smart00226 (4) Low molecular weight phosphatase family.
smart00283 (1) Methyl-accepting chemotaxis-like domains (chemotaxis sensory transducer).
smart00304 (1) HAMP (Histidine kinases, Adenylyl cyclases, Methyl binding proteins, Phosphatases) domain.
smart00357 (1) Cold shock protein domain.
smart00382 (3) ATPases associated with a variety of cellular activities.
smart00387 (1) Histidine kinase-like ATPases.
smart00448 (1) cheY-homologous receiver domain.
smart00470 (6) ParB-like nuclease domain.
smart00487 (5) DEAD-like helicases superfamily.
smart00490 (5) helicase superfamily c-terminal domain.
smart00530 (1) Helix-turn-helix XRE-family like proteins.
smart00562 (2) Enzymes that catalyze nonsubstrate specific conversions of nucleoside diphosphates to nucleoside triphosphates.
smart00635 (1) Bacterial Ig-like domain 2.
smart00849 (17) Metallo-beta-lactamase superfamily.
smart00850 (22) LytTr DNA-binding domain.
smart00862 (2) Transcriptional regulatory protein, C terminal.
smart00937 (10) This domain is found in peptide chain release factors.
smart01059 (1) Chloramphenicol acetyltransferase.
TIGR00057 (5) tRNA threonylcarbamoyl adenosine modification protein, Sua5/YciO/YrdC/YwlC family. [Protein synthesis, tRNA and rRNA base modification]
TIGR00105 (6) ribosomal protein L31. [Protein synthesis, Ribosomal proteins: synthesis and modification]
TIGR00236 (3) UDP-N-acetylglucosamine 2-epimerase. [Cell envelope, Biosynthesis and degradation of surface polysaccharides and lipopolysaccharides]
TIGR00473 (1) CDP-diacylglycerol–serine O-phosphatidyltransferase. [Fatty acid and phospholipid metabolism, Biosynthesis]
TIGR00556 (1) phosphopantetheine–protein transferase domain. [Protein fate, Protein modification and repair]
TIGR00741 (12) ribosomal subunit interface protein. [Protein synthesis, Translation factors]
TIGR00744 (2) ROK family protein (putative glucokinase). [Unknown function, General]
TIGR00762 (1) EDD domain protein, DegV family. [Unknown function, General]
TIGR00810 (1) protein translocase, SecG subunit. [Protein fate, Protein and peptide secretion and trafficking]
TIGR00950 (1) Carboxylate/Amino Acid/Amine Transporter. [Transport and binding proteins, Amino acids, peptides and amines]
TIGR01068 (1) thioredoxin. [Energy metabolism, Electron transport]
TIGR01076 (10) LPXTG-site transpeptidase (sortase) family protein. [Cell envelope, Other, Protein fate, Protein and peptide secretion and trafficking]
TIGR01243 (3) AAA family ATPase, CDC48 subfamily.
TIGR01257 (3) retinal-specific rim ABC transporter. [Transport and binding proteins, Other]
TIGR01387 (2) heavy metal response regulator. [Regulatory functions, DNA interactions]
TIGR02150 (1) isopentenyl-diphosphate delta-isomerase, type 1. [Biosynthesis of cofactors, prosthetic groups, and carriers, Other]
TIGR02176 (1) pyruvate:ferredoxin (flavodoxin) oxidoreductase, homodimeric.
TIGR02327 (5) conserved hypothetical integral membrane protein.
TIGR02380 (13) undecaprenyl-phosphate alpha-N-acetylglucosaminyl 1-phosphatetransferase. [Cell envelope, Biosynthesis and degradation of surface polysaccharides and lipopolysaccharides]
TIGR02669 (16) SpoIID/LytB domain. [Unknown function, General]
TIGR02765 (1) cryptochrome, DASH family. [Cellular processes, Adaptations to atypical conditions]
TIGR02898 (7) sporulation lipoprotein, YhcN/YlaJ family. [Cellular processes, Sporulation and germination]
TIGR03023 (1) Undecaprenyl-phosphate glucose phosphotransferase.
TIGR03534 (8) protein-(glutamine-N5) methyltransferase, release factor-specific. [LIV]PRx[DE]TE (in Escherichia coli, IPRPDTE) confers specificity for the release factors rather than for ribosomal protein L3. [Protein fate, Protein modification and repair]
TIGR03556 (5) deoxyribodipyrimidine photo-lyase, 8-HDF type. [DNA metabolism, DNA replication, recombination, and repair]
TIGR03921 (1) type VII secretion-associated serine protease mycosin. [Protein fate, Protein and peptide secretion and trafficking, Protein fate, Protein modification and repair]
TIGR04181 (1) aminotransferase, LLPSF_NHT_00031 family.

#### 11.5 Multiple-sequence alignment

Each L31-Firmicutes RNA is denoted by an abbreviation (like “Eco-1-1”) that refers to a taxonomy given in Section 11.1. (Superscript numbers refer to annotations listed in Section 11.2) The alignment may include sequences containing the RNA motif, as well as flanking sequence. The L31-Firmicutes RNA itself is denoted by the line underneath marked 5′ and 3′ on either end. Nucleotides in flanking sequences (i.e., not part of the motif) are written in gray letters. Stems of predicted rho-independent transcription terminators, if any, are shaded yellow. (Note: terminator predictions have not been analyzed manually, and many are likely to be false positives. Terminator predictions are those of the RNIE software.) Nucleotides predicted to function (as DNA) as transcription-factor binding sites, if any, are shaded in green. (But note: these predictions are manually annotated, so they might be under-predicted.) Annotated start codons, if any, are shaded green. (Note: start codons are frequently misannotated, especially in environmental samples.) Nucleotides proposed to basepair as part of the consensus structure are shaded in color when they comprise Watson-Crick or G-U pairs. Otherwise they are shaded gray. Conserved stems are also indicated at the bottom of the alignment by angle brackets, where matching < and > denote base-paired columns. Below these angle brackets, the symbol “2” denotes base pairs exhibiting covariation according to the statistically well-founded R-scape method. “1” denotes base pairs exhibiting covariation according to R2R’s simplistic method. “0” denotes base pairs that are not observed to mutate and “?” denotes base pairs that have a significant frequency of non-canonical nucleotides for Watson-Crick or G-U pairs (> 5%). Below these base pair annotation is the consensus sequence: "R" = "A" or "G", "Y" = "C" or "U", red nucleotides: nucleotide identity conserved more than 97% of the time, black nucleotides: 90%, gray nucleotides: 75%, red circle (•): nucleotide is present 97% of the time, black circle (•): 90%, gray circle (•): 75%, white circle (◦): 50%. All percentages of sequences just described (e.g. 97% conserved) assume that sequences have been weighted by the GSC algorithm implemented by the Infernal software package.

Duplicate sequences: the following putative homologs are not shown in the alignment because their sequences are identical to a homolog already shown: Aka-1-1, Ami-1-3, Asp-1-1, Asp-2-1, Asp-4-1, Axy-1-1, Bag-1-1, Bag-2-1, Bal-2-1, Bal-2-2, Bal-2-3, Bal-3-1, Bam-1-1, Bam-10-1, Bam-11-1, Bam-12-1, Bam-13-1, Bam-14-1, Bam-15-1, Bam-16-1, Bam-17-1, Bam-18-1, Bam-19-1, Bam-2-1, Bam-20-1, Bam-21-1, Bam-22-1, Bam-23-1, Bam-24-1, Bam-25-1, Bam-3-1, Bam-5-1, Bam-6-1, Bam-7-1, Bam-8-1, Bam-9-2, Bam-9-3, Ban-1-1, Ban-10-1, Ban-11-1, Ban-12-1, Ban-13-1, Ban-14-1, Ban-15-1, Ban-16-1, Ban-17-1, Ban-17-2, Ban-18-1, Ban-18-2, Ban-19-1, Ban-2-1, Ban-21-1, Ban-21-10, Ban-21-11, Ban-21-12, Ban-21-13, Ban-21-14, Ban-21-15, Ban-21-16, Ban-21-17, Ban-21-18, Ban-21-19, Ban-21-2, Ban-21-20, Ban-21-3, Ban-21-4, Ban-21-5, Ban-21-6, Ban-21-7, Ban-21-8, Ban-21-9, Ban-22-1, Ban-23-1, Ban-24-1, Ban-25-1, Ban-26-1, Ban-27-1, Ban-28-1, Ban-29-1, Ban-3-1, Ban-30-1, Ban-31-1, Ban-32-1, Ban-33-1, Ban-34-1, Ban-35-1, Ban-36-1, Ban-37-1, Ban-38-1, Ban-39-1, Ban-5-1, Ban-6-1, Ban-7-1, Ban-8-1, Ban-9-1, Bar-1-1, Bar-1-2, Bar-1-3, Bar-1-4, Bar-1-5, Bar-2-1, Bat-1-1, Bat-10-1, Bat-11-1, Bat-12-1, Bat-13-1, Bat-14-1, Bat-16-1, Bat-17-1, Bat-2-1, Bat-3-1, Bat-4-1, Bat-5-1, Bat-6-1, Bat-7-1, Bat-9-1, Baz-2-1, Bba-2-2, Bbo-2-1, Bbo-3-1, Bbo-6-1, Bbo-7-1, Bbr-3-1, Bce-10-1, Bce-100-1, Bce-101-1, Bce-102-1, Bce-103-1, Bce-104-1, Bce-105-1, Bce-107-1, Bce-108-1, Bce-109-1, Bce-11-1, Bce-110-1, Bce-111-1, Bce-112-1, Bce-113-1, Bce-114-1, Bce-115-1, Bce-116-1, Bce-117-1, Bce-118-1, Bce-119-1, Bce-12-1, Bce-120-1, Bce-121-1, Bce-122-1, Bce-123-1, Bce-124-1, Bce-125-1, Bce-126-1, Bce-127-1, Bce-128-1, Bce-129-1, Bce-13-1, Bce-130-1, Bce-131-1, Bce-132-1, Bce-133-1, Bce-134-1, Bce-135-1, Bce-136-1, Bce-137-1, Bce-138-1, Bce-139-1, Bce-14-1, Bce-14-2, Bce-140-1, Bce-141-1, Bce-142-1, Bce-143-1, Bce-144-1, Bce-145-1, Bce-146-1, Bce-147-1, Bce-149-2, Bce-15-1, Bce-16-1, Bce-17-1, Bce-17-2, Bce-18-1, Bce-19-1, Bce-2-1, Bce-2-2, Bce-20-1, Bce-21-1, Bce-22-1, Bce-23-1, Bce-24-1, Bce-24-2, Bce-25-1, Bce-26-1, Bce-27-1, Bce-28-1, Bce-28-2, Bce-29-1, Bce-3-1, Bce-30-1, Bce-31-1, Bce-32-1, Bce-33-1, Bce-34-1, Bce-35-1, Bce-36-1, Bce-37-1, Bce-38-1, Bce-39-1, Bce-4-1, Bce-40-1, Bce-41-1, Bce-42-1, Bce-43-1, Bce-44-1, Bce-45-1, Bce-46-1, Bce-47-1, Bce-47-2, Bce-47-3, Bce-47-4, Bce-47-5, Bce-48-1, Bce-49-1, Bce-5-1, Bce-50-1, Bce-51-1, Bce-52-1, Bce-53-1, Bce-54-1, Bce-55-1, Bce-56-1, Bce-57-1, Bce-58-1, Bce-59-1, Bce-60-1, Bce-61-1, Bce-62-1, Bce-64-1, Bce-65-1, Bce-66-1, Bce-67-1, Bce-68-1, Bce-69-1, Bce-7-1, Bce-70-1, Bce-71-1, Bce-72-1, Bce-73-1, Bce-74-1, Bce-75-1, Bce-76-1, Bce-77-1, Bce-78-1, Bce-79-1, Bce-8-1, Bce-80-1, Bce-81-1, Bce-82-1, Bce-83-1, Bce-84-1, Bce-85-1, Bce-86-1, Bce-87-1, Bce-88-1, Bce-89-1, Bce-9-1, Bce-90-1, Bce-91-1, Bce-92-1, Bce-93-1, Bce-94-1, Bce-95-1, Bce-96-1, Bce-97-1, Bce-98-1, Bce-99-1, Bcl-2-1, Bco-1-1, Bco-10-1, Bco-4-1, Bco-4-2, Bco-6-1, Bco-7-1, Bco-8-1, Bcy-1-1, Ben-1-1, Ben-2-1, Bfl-2-1, Bgi-1-1, Bgl-1-1, Bha-1-1, Bin-2-1, Bin-3-1, Bis-1-1, Bla-1-1, Bla-4-1, Bla-5-1, Bli-1-1, Bli-1-2, Bli-10-1, Bli-11-1, Bli-2-2, Bli-3-1, Bli-4-1, Bli-5-1, Bli-6-1, Bli-7-1, Bli-8-1, Bli-9-1, Bma-4-2, Bma-4-3, Bme-10-1, Bme-2-1, Bme-3-1, Bme-5-1, Bme-7-1, Bme-8-1, Bme-9-1, Bmu-1-1, Bmy-1-1, Bmy-2-1, Bmy-3-1, Bmy-4-1, Bmy-4-2, Bmy-4-3, Bmy-4-4, Bmy-5-1, Bmy-6-1, Bmy-7-1, Boc-1-1, Bpa-2-1, Bpa-3-1, Bps-2-1, Bpu-1-1, Bpu-3-1, Bpu-3-2, Bpu-3-3, Bpu-4-1, Bpu-5-1, Bpu-6-1, Bpu-7-1, Bsa-1-2, Bsa-1-3, Bsa-2-1, Bsa-3-1, Bsi-2-1, Bsi-3-1, Bsm-1-1, Bso-2-1, Bsp-1-1, Bsp-10-1, Bsp-11-1, Bsp-12-1, Bsp-14-1, Bsp-16-1, Bsp-2-1, Bsp-20-1, Bsp-23-1, Bsp-25-1, Bsp-26-1, Bsp-29-1, Bsp-31-1, Bsp-32-1, Bsp-33-1, Bsp-35-1, Bsp-38-1, Bsp-46-1, Bsp-48-1, Bsp-49-1, Bsp-5-1, Bsp-50-1, Bsp-51-1, Bsp-55-1, Bsp-57-1, Bsp-59-1, Bsp-6-1, Bsp-61-1, Bsp-63-1, Bsp-64-1, Bsp-65-1, Bsp-67-1, Bsp-68-1, Bsp-69-1, Bsp-70-1, Bsp-71-1, Bsp-72-1, Bsp-78-1, Bsp-80-1, Bsp-82-1, Bsp-87-1, Bsp-88-1, Bsp-89-1, Bsp-91-1, Bsp-93-1, Bsp-95-1, Bsp-97-1, Bst-1-2, Bst-1-3, Bst-1-5, Bst-2-1, Bsu-1-1, Bsu-10-1, Bsu-12-1, Bsu-13-1, Bsu-14-1, Bsu-15-1, Bsu-16-1, Bsu-17-1, Bsu-18-1, Bsu-19-1, Bsu-2-1, Bsu-20-1, Bsu-21-1, Bsu-22-1, Bsu-24-1, Bsu-25-1, Bsu-26-1, Bsu-27-1, Bsu-28-1, Bsu-29-1, Bsu-3-1, Bsu-30-1, Bsu-31-1, Bsu-32-1, Bsu-33-1, Bsu-34-1, Bsu-36-1, Bsu-37-1, Bsu-38-1, Bsu-39-1, Bsu-4-1, Bsu-40-1, Bsu-41-1, Bsu-42-1, Bsu-43-1, Bsu-44-1, Bsu-45-1, Bsu-46-1, Bsu-47-1, Bsu-48-1, Bsu-5-1, Bsu-6-1, Bsu-6-2, Bsu-7-1, Bsu-8-1, Bsu-9-1, Bth-1-1, Bth-10-1, Bth-11-1, Bth-12-1, Bth-13-1, Bth-14-1, Bth-15-1, Bth-16-1, Bth-17-1, Bth-18-1, Bth-19-1, Bth-2-1, Bth-20-1, Bth-21-1, Bth-22-1, Bth-22-2, Bth-23-1, Bth-24-1, Bth-25-1, Bth-26-1, Bth-27-1, Bth-28-1, Bth-29-1, Bth-3-1, Bth-30-1, Bth-31-1, Bth-32-1, Bth-33-1, Bth-34-1, Bth-35-1, Bth-36-1, Bth-37-1, Bth-38-1, Bth-39-1, Bth-39-2, Bth-4-1, Bth-40-1, Bth-41-1, Bth-42-1, Bth-43-1, Bth-44-1, Bth-44-2, Bth-44-3, Bth-44-5, Bth-45-2, Bth-46-1, Bth-47-1, Bth-5-1, Bth-6-1, Bth-7-1, Bth-8-1, Bth-9-1, Bto-1-1, Bva-1-1, Bve-1-1, Bve-1-2, Bve-1-3, Bve-1-4, Bve-2-1, Bve-3-1, Bve-4-1, Bve-5-1, Bvi-3-1, Bwe-1-1, Bwe-2-1, Bzh-1-1, Cde-1-1, Cin-1-1, Cje-1-1, Cje-2-1, Cma-1-1, Cma-2-1, Cmi-1-1, Csp-4-1, Csp-5-1, Csp-6-1, Csp-7-1, Ean-2-1, Eav-2-1, Eav-2-2, Eba-1-1, Eca-2-1, Eca-4-1, Eca-6-1, Eca-7-1, Ece-1-1, Ece-1-2, Ece-1-3, Ece-1-4, Ece-1-5, Ece-1-6, Ece-1-7, Ece-2-1, Ecl-1-1, Edo-1-1, Edu-1-1, Edu-1-2, Edu-2-2, Edu-3-1, Edu-4-1, Efa-1-1, Efa-1-10, Efa-1-11, Efa-1-12, Efa-1-13, Efa-1-14, Efa-1-15, Efa-1-16, Efa-1-17, Efa-1-18, Efa-1-19, Efa-1-2, Efa-1-20, Efa-1-21, Efa-1-22, Efa-1-23, Efa-1-24, Efa-1-25, Efa-1-26, Efa-1-28, Efa-1-29, Efa-1-3, Efa-1-30, Efa-1-31, Efa-1-32, Efa-1-33, Efa-1-34, Efa-1-35, Efa-1-36, Efa-1-37, Efa-1-38, Efa-1-39, Efa-1-4, Efa-1-40, Efa-1-41, Efa-1-42, Efa-1-43, Efa-1-44, Efa-1-45, Efa-1-46, Efa-1-47, Efa-1-48, Efa-1-49, Efa-1-5, Efa-1-50, Efa-1-51, Efa-1-52, Efa-1-53, Efa-1-54, Efa-1-55, Efa-1-56, Efa-1-57, Efa-1-58, Efa-1-59, Efa-1-6, Efa-1-60, Efa-1-61, Efa-1-62, Efa-1-63, Efa-1-64, Efa-1-65, Efa-1-66, Efa-1-67, Efa-1-68, Efa-1-69, Efa-1-7, Efa-1-70, Efa-1-71, Efa-1-72, Efa-1-73, Efa-1-74, Efa-1-75, Efa-1-76, Efa-1-77, Efa-1-78, Efa-1-79, Efa-1-8, Efa-1-80, Efa-1-82, Efa-1-83, Efa-1-84, Efa-1-85, Efa-1-86, Efa-1-87, Efa-1-9, Efa-10-1, Efa-100-1, Efa-101-1,

Efa-102-1, Efa-103-1, Efa-104-1, Efa-105-1, Efa-106-1, Efa-107-1, Efa-108-1, Efa-109-1, Efa-11-1, Efa-110-1, Efa-111-1, Efa-112-1, Efa-113-1, Efa-114-1, Efa-115-1, Efa-116-1, Efa-117-1, Efa-118-1, Efa-119-1, Efa-12-1, Efa-120-1, Efa-121-1, Efa-122-1, Efa-123-1, Efa-124-1, Efa-125-1, Efa-126-1, Efa-127-1, Efa-128-1, Efa-129-1, Efa-13-1, Efa-130-1, Efa-131-1, Efa-132-1, Efa-133-1, Efa-134-1, Efa-135-1, Efa-136-1, Efa-137-1, Efa-138-1, Efa-139-1, Efa-14-1, Efa-140-1, Efa-141-1, Efa-142-1, Efa-143-1, Efa-144-1, Efa-145-1, Efa-146-1, Efa-147-1, Efa-148-1, Efa-149-1, Efa-15-1, Efa-150-1, Efa-151-1, Efa-152-1, Efa-153-1, Efa-154-1, Efa-155-1, Efa-156-1, Efa-157-1, Efa-158-1, Efa-159-1, Efa-16-1, Efa-160-1, Efa-161-1, Efa-162-1, Efa-163-1, Efa-164-1, Efa-165-1, Efa-166-1, Efa-167-1, Efa-168-1, Efa-169-1, Efa-17-1, Efa-170-1, Efa-171-1, Efa-173-1, Efa-174-1, Efa-175-1, Efa-176-1, Efa-177-1, Efa-178-1, Efa-179-1, Efa-18-1, Efa-18-2, Efa-180-1, Efa-181-1, Efa-182-1, Efa-183-1, Efa-184-1, Efa-185-1, Efa-186-1, Efa-187-1, Efa-188-1, Efa-189-1, Efa-19-1, Efa-190-1, Efa-191-1, Efa-192-1, Efa-193-1, Efa-194-1, Efa-195-1, Efa-196-1, Efa-197-1, Efa-198-1, Efa-199-1, Efa-2-1, Efa-20-1, Efa-200-1, Efa-201-1, Efa-202-1, Efa-203-1, Efa-204-1, Efa-205-1, Efa-206-1, Efa-207-1, Efa-208-1, Efa-209-1, Efa-21-1, Efa-210-1, Efa-211-1, Efa-212-1, Efa-213-1, Efa-214-1, Efa-215-1, Efa-216-1, Efa-217-1, Efa-218-1, Efa-219-1, Efa-22-1, Efa-220-1, Efa-221-1, Efa-222-1, Efa-223-1, Efa-224-1, Efa-225-1, Efa-226-1, Efa-227-1, Efa-228-1, Efa-229-1, Efa-23-1, Efa-230-1, Efa-231-1, Efa-232-1, Efa-233-1, Efa-234-1, Efa-235-1, Efa-236-1, Efa-237-1, Efa-238-1, Efa-239-1, Efa-24-1, Efa-240-1, Efa-241-1, Efa-242-1, Efa-243-1, Efa-244-1, Efa-245-1, Efa-246-1, Efa-247-1, Efa-248-1, Efa-249-1, Efa-25-1, Efa-250-1, Efa-251-1, Efa-252-1, Efa-253-1, Efa-254-1, Efa-255-1, Efa-256-1, Efa-257-1, Efa-258-1, Efa-259-1, Efa-26-1, Efa-260-1, Efa-261-1, Efa-262-1, Efa-263-1, Efa-264-1, Efa-265-1, Efa-266-1, Efa-267-1, Efa-268-1, Efa-269-1, Efa-27-1, Efa-270-1, Efa-271-1, Efa-272-1, Efa-273-1, Efa-274-1, Efa-275-1, Efa-276-1, Efa-277-1, Efa-278-1, Efa-279-1, Efa-28-1, Efa-280-1, Efa-281-1, Efa-282-1, Efa-283-1, Efa-284-1, Efa-285-1, Efa-286-1, Efa-287-1, Efa-288-1, Efa-289-1, Efa-29-1, Efa-290-1, Efa-291-1, Efa-292-1, Efa-293-1, Efa-294-1, Efa-295-1, Efa-296-1, Efa-297-1, Efa-298-1, Efa-299-1, Efa-3-1, Efa-30-1, Efa-300-1, Efa-301-1, Efa-302-1, Efa-303-1, Efa-304-1, Efa-305-1, Efa-306-1, Efa-307-1, Efa-308-1, Efa-309-1, Efa-309-2, Efa-309-3, Efa-31-1, Efa-310-1, Efa-311-1, Efa-312-1, Efa-313-1, Efa-314-1, Efa-315-1, Efa-316-1, Efa-317-1, Efa-318-1, Efa-319-1, Efa-32-1, Efa-320-1, Efa-321-1, Efa-322-1, Efa-323-1, Efa-324-1, Efa-325-1, Efa-326-1, Efa-327-1, Efa-328-1, Efa-329-1, Efa-33-1, Efa-330-1, Efa-331-1, Efa-332-1, Efa-333-1, Efa-334-1, Efa-335-1, Efa-336-1, Efa-337-1, Efa-338-1, Efa-339-1, Efa-34-1, Efa-340-1, Efa-341-1, Efa-342-1, Efa-343-1, Efa-344-1, Efa-345-1, Efa-346-1, Efa-347-1, Efa-348-1, Efa-349-1, Efa-35-1, Efa-350-1, Efa-351-1, Efa-352-1, Efa-354-1, Efa-355-1, Efa-356-1, Efa-357-1, Efa-358-1, Efa-359-1, Efa-36-1, Efa-360-1, Efa-361-1, Efa-362-1, Efa-363-1, Efa-364-1, Efa-365-1, Efa-366-1, Efa-367-1, Efa-368-1, Efa-369-1, Efa-37-1, Efa-370-1, Efa-372-1, Efa-373-1, Efa-375-1, Efa-376-1, Efa-377-1, Efa-378-1, Efa-379-1, Efa-38-1, Efa-380-1, Efa-381-1, Efa-382-1, Efa-383-1, Efa-384-1, Efa-385-1, Efa-386-1, Efa-387-1, Efa-388-1, Efa-389-1, Efa-39-1, Efa-390-1, Efa-391-1, Efa-392-1, Efa-393-1, Efa-394-1, Efa-395-1, Efa-396-1, Efa-397-1, Efa-398-1, Efa-399-1, Efa-4-1, Efa-40-1, Efa-400-1, Efa-401-1, Efa-402-1, Efa-403-1, Efa-404-1, Efa-405-1, Efa-406-1, Efa-407-1, Efa-408-1, Efa-409-1, Efa-41-1, Efa-410-1, Efa-411-1, Efa-412-1, Efa-413-1, Efa-414-1, Efa-415-1, Efa-416-1, Efa-417-1, Efa-418-1, Efa-419-1, Efa-42-1, Efa-420-1, Efa-421-1, Efa-422-1, Efa-423-1, Efa-424-1, Efa-425-1, Efa-426-1, Efa-427-1, Efa-428-1, Efa-429-1, Efa-43-1, Efa-430-1, Efa-431-1, Efa-432-1, Efa-433-1, Efa-434-1, Efa-435-1, Efa-436-1, Efa-437-1, Efa-438-1, Efa-439-1, Efa-44-1, Efa-440-1, Efa-441-1, Efa-442-1, Efa-443-1, Efa-444-1, Efa-445-1, Efa-446-1, Efa-447-1, Efa-448-1, Efa-449-1, Efa-45-1, Efa-450-1, Efa-451-1, Efa-452-1, Efa-453-1, Efa-454-1, Efa-455-1, Efa-456-1, Efa-457-1, Efa-458-1, Efa-459-1, Efa-46-1, Efa-460-1, Efa-461-1, Efa-462-1, Efa-463-1, Efa-464-1, Efa-465-1, Efa-466-1, Efa-467-1, Efa-468-1, Efa-469-1, Efa-47-1, Efa-470-1, Efa-471-1, Efa-472-1, Efa-473-1, Efa-474-1, Efa-475-1, Efa-476-1, Efa-477-1, Efa-478-1, Efa-479-1, Efa-48-1, Efa-480-1, Efa-481-1, Efa-482-1, Efa-483-1, Efa-484-1, Efa-485-1, Efa-486-1, Efa-487-1, Efa-488-1, Efa-489-1, Efa-49-1, Efa-490-1, Efa-491-1, Efa-492-1, Efa-493-1, Efa-494-1, Efa-495-1, Efa-496-1, Efa-496-2, Efa-497-1, Efa-497-2, Efa-498-1, Efa-499-1, Efa-5-1, Efa-50-1, Efa-500-1, Efa-501-1, Efa-502-1, Efa-503-1, Efa-504-1, Efa-505-1, Efa-506-1, Efa-507-1, Efa-508-1, Efa-509-1, Efa-51-1, Efa-510-1, Efa-511-1, Efa-512-1, Efa-513-1, Efa-514-1, Efa-515-1, Efa-516-1, Efa-517-1, Efa-518-1, Efa-519-1, Efa-52-1, Efa-520-1, Efa-521-1, Efa-522-1, Efa-523-1, Efa-524-1, Efa-525-1, Efa-526-1, Efa-528-1, Efa-529-1, Efa-53-1, Efa-530-1, Efa-531-1, Efa-532-1, Efa-533-1, Efa-534-1, Efa-535-1, Efa-536-1, Efa-539-1, Efa-54-1, Efa-540-1, Efa-541-1, Efa-542-1, Efa-543-1, Efa-544-1, Efa-545-1, Efa-546-1, Efa-547-1, Efa-548-1, Efa-549-1, Efa-55-1, Efa-550-1, Efa-551-1, Efa-552-1, Efa-553-1, Efa-554-1, Efa-555-1, Efa-556-1, Efa-557-1, Efa-558-1, Efa-559-1, Efa-56-1, Efa-560-1, Efa-561-1, Efa-562-1, Efa-563-1, Efa-564-1, Efa-565-1, Efa-566-1, Efa-567-1, Efa-568-1, Efa-569-1, Efa-57-1, Efa-570-1, Efa-571-1, Efa-572-1, Efa-573-1, Efa-574-1, Efa-575-1, Efa-576-1, Efa-577-1, Efa-578-1, Efa-579-1, Efa-58-1, Efa-580-1, Efa-581-1, Efa-582-1, Efa-583-1, Efa-584-1, Efa-585-1, Efa-586-1, Efa-587-1, Efa-588-1, Efa-589-1, Efa-59-1, Efa-590-1, Efa-591-1, Efa-592-1, Efa-593-1, Efa-594-1, Efa-6-1, Efa-60-1, Efa-61-1, Efa-62-1, Efa-63-1, Efa-64-1, Efa-65-1, Efa-66-1, Efa-67-1, Efa-68-1, Efa-69-1, Efa-7-1, Efa-70-1, Efa-71-1, Efa-72-1, Efa-73-1, Efa-74-1, Efa-75-1, Efa-76-1, Efa-77-1, Efa-78-1, Efa-79-1, Efa-80-1, Efa-81-1, Efa-82-1, Efa-83-1, Efa-84-1, Efa-85-1, Efa-86-1, Efa-87-1, Efa-88-1, Efa-89-1, Efa-9-1, Efa-90-1, Efa-91-1, Efa-92-1, Efa-93-1, Efa-94-1, Efa-95-1, Efa-96-1, Efa-97-1, Efa-98-1, Efa-99-1, Efl-1-1, Ega-1-2, Ega-2-1, Ehi-1-2, Ehi-2-1, Ehi-2-2, Emu-1-1, Emu-2-1, Emu-3-1, Epa-1-1, Esa-1-1, Esa-2-2, Esp-1-1, Esp-10-1, Esp-11-1, Esp-12-1, Esp-13-1, Esp-14-1, Esp-15-1, Esp-16-1, Esp-18-1, Esp-19-1, Esp-20-1, Esp-21-1, Esp-6-1, Esp-7-1, Gad-1-1, Gca-1-1, Gha-1-1, Gha-3-1, Gka-1-1, Gka-2-1, Gka-3-2, Gmo-1-1, Gsa-1-1, Gsa-2-1, Gsp-10-1, Gsp-11-1, Gsp-12-1, Gsp-14-1, Gsp-15-1, Gsp-18-1, Gsp-19-1, Gsp-2-1, Gsp-20-1, Gsp-4-1, Gsp-6-1, Gsp-8-1, Gst-1-2, Gst-1-3, Gst-1-4, Gst-2-1, Gst-3-1, Gth-2-1, Gth-3-1, Gth-5-1, Gth-7-1, Hda-1-1, Hka-1-1, Hsp-1-1, Jma-2-1, Kvi-1-1, Kvi-2-1, Kvi-3-1, Lac-1-1, Lac-11-1, Lac-12-1, Lac-13-1, Lac-2-1, Lac-3-1, Lac-4-1, Lac-5-1, Lac-6-1, Lac-7-1, Lac-8-1, Lac-9-1, Lam-2-1, Lam-3-1, Lan-1-1, Lan-2-1, Lbr-1-1, Lbr-1-2, Lbr-1-3, Lbr-1-4, Lbr-1-5, Lbr-2-1, Lbr-3-1, Lbr-5-1, Lbr-6-1, Lca-1-1, Lca-1-2, Lca-1-3, Lca-1-4, Lca-1-5, Lca-10-1, Lca-11-1,

Lca-12-1, Lca-13-1, Lca-14-1, Lca-15-1, Lca-16-1, Lca-17-1, Lca-17-2, Lca-18-1, Lca-19-1, Lca-2-1, Lca-20-1, Lca-21-1, Lca-22-1, Lca-23-1, Lca-24-1, Lca-3-1, Lca-4-1, Lca-5-1, Lca-6-2, Lca-7-1, Lca-8-1, Lca-9-1, Lci-1-1, Lci-2-1, Lci-3-1, Lci-4-1, Lci-4-2, Lci-4-3, Lci-5-1, Lcr-10-1, Lcr-2-1, Lcr-3-1, Lcr-4-1, Lcr-5-1, Lcr-6-1, Lcr-7-1, Lcr-8-1, Lcr-9-1, Lfa-1-1, Lfe-1-1, Lfe-1-2, Lfe-1-3, Lfe-1-4, Lfe-10-1, Lfe-11-1, Lfe-12-1, Lfe-2-1, Lfe-3-1, Lfe-4-1, Lfe-5-1, Lfe-6-1, Lfe-8-1, Lfe-9-1, Lfl-1-1, Lfl-2-1, Lfu-2-1, Lfu-3-1, Lfu-4-1, Lga-1-1, Lga-10-1, Lga-11-1, Lga-12-1, Lga-2-2, Lga-2-3, Lga-2-4, Lga-2-5, Lga-3-1, Lga-4-1, Lga-5-1, Lga-6-1, Lga-7-1, Lga-8-1, Lga-9-1, Lhe-10-1, Lhe-2-1, Lhe-3-1, Lhe-3-2, Lhe-4-1, Lhe-6-1, Lhe-7-1, Lhe-8-1, Lhe-9-1, Lhi-1-1, Lin-1-1, Lin-11-1, Lin-12-1, Lin-14-1, Lin-15-1, Lin-16-1, Lin-17-1, Lin-18-1, Lin-19-1, Lin-2-3, Lin-20-1, Lin-3-1, Lin-5-1, Lin-7-1, Lin-8-1, Lin-9-1, Liv-1-1, Liv-2-1, Liv-3-1, Liv-4-2, Ljo-2-1, Ljo-7-1, Lke-1-1, Lki-3-1, Lku-2-1, Lku-2-2, Lma-3-1, Lme-1-1, Lmo-1-1, Lmo-10-1, Lmo-11-1, Lmo-12-1, Lmo-13-1, Lmo-14-1, Lmo-15-1, Lmo-16-1, Lmo-17-1, Lmo-18-1, Lmo-19-1, Lmo-2-1, Lmo-20-1, Lmo-21-1, Lmo-22-1, Lmo-23-1, Lmo-24-1, Lmo-25-1, Lmo-26-1, Lmo-27-1, Lmo-28-1, Lmo-29-1, Lmo-3-1, Lmo-30-1, Lmo-31-1, Lmo-33-1, Lmo-33-2, Lmo-33-3, Lmo-33-4, Lmo-33-5, Lmo-33-6, Lmo-33-7, Lmo-33-8, Lmo-33-9, Lmo-34-1, Lmo-36-1, Lmo-38-1, Lmo-39-1, Lmo-4-1, Lmo-41-1, Lmo-42-1, Lmo-43-1, Lmo-44-1, Lmo-45-1, Lmo-46-1, Lmo-47-1, Lmo-48-1, Lmo-49-1, Lmo-5-1, Lmo-50-1, Lmo-51-1, Lmo-52-1, Lmo-53-1, Lmo-54-1, Lmo-55-1, Lmo-56-1, Lmo-57-1, Lmo-58-1, Lmo-59-1, Lmo-6-1, Lmo-61-1, Lmo-62-1, Lmo-63-1, Lmo-64-1, Lmo-65-1, Lmo-66-1, Lmo-67-1, Lmo-68-1, Lmo-7-1, Lmo-70-1, Lmo-71-1, Lmo-72-1, Lmo-73-1, Lmo-74-1, Lmo-75-1, Lmo-76-1, Lmo-77-1, Lmo-78-1, Lmo-79-1, Lmo-8-1, Lmo-9-1, Lmu-1-2, Lmu-2-1, Lne-1-1, Lpa-1-1, Lpa-10-1, Lpa-11-1, Lpa-12-1, Lpa-13-1, Lpa-14-1, Lpa-15-1, Lpa-16-1, Lpa-17-1, Lpa-18-1, Lpa-19-1, Lpa-20-1, Lpa-21-1, Lpa-22-1, Lpa-23-1, Lpa-24-1, Lpa-25-1, Lpa-26-1, Lpa-27-1, Lpa-28-1, Lpa-29-1, Lpa-3-1, Lpa-3-2, Lpa-30-1, Lpa-31-1, Lpa-32-1, Lpa-33-1, Lpa-34-1, Lpa-35-1, Lpa-36-1, Lpa-37-1, Lpa-38-1, Lpa-4-1, Lpa-40-1, Lpa-42-1, Lpa-5-1, Lpa-6-1, Lpa-7-1, Lpa-8-1, Lpa-9-1, Lpl-1-1, Lpl-10-1, Lpl-11-1, Lpl-12-1, Lpl-14-1, Lpl-15-1, Lpl-16-1, Lpl-17-1, Lpl-18-1, Lpl-19-1, Lpl-2-1, Lpl-20-1, Lpl-21-1, Lpl-3-1, Lpl-4-1, Lpl-5-1, Lpl-7-1, Lpl-9-1, Lre-1-1, Lre-10-1, Lre-11-1, Lre-12-1, Lre-13-1, Lre-2-1, Lre-3-1, Lre-4-1, Lre-5-1, Lre-6-1, Lre-8-1, Lre-9-1, Lrh-1-1, Lrh-1-11, Lrh-1-12, Lrh-1-13, Lrh-1-14, Lrh-1-15, Lrh-1-16, Lrh-1-17, Lrh-1-18, Lrh-1-19, Lrh-1-2, Lrh-1-20, Lrh-1-21, Lrh-1-22, Lrh-1-23, Lrh-1-24, Lrh-1-25, Lrh-1-26, Lrh-1-27, Lrh-1-28, Lrh-1-3, Lrh-1-4, Lrh-1-5, Lrh-1-6, Lrh-1-7, Lrh-1-8, Lrh-1-9, Lrh-11-1, Lrh-12-1, Lrh-13-1, Lrh-15-1, Lrh-16-1, Lrh-17-1, Lrh-18-1, Lrh-19-1, Lrh-2-1, Lrh-20-1, Lrh-20-2, Lrh-21-1, Lrh-22-1, Lrh-23-1, Lrh-3-1, Lrh-4-1, Lrh-5-1, Lrh-6-1, Lrh-7-1, Lrh-8-1, Lrh-9-1, Lro-2-2, Lru-1-1, Lru-2-1, Lru-3-1, Lru-4-1, Lru-5-1, Lru-6-1, Lsa-10-1, Lsa-11-1, Lsa-2-1, Lsa-4-1, Lsa-5-1, Lsa-6-1, Lsa-7-1, Lsa-8-1, Lsa-9-1, Lse-3-1, Lsp-11-1, Lsp-12-1, Lsp-14-1, Lsp-15-1, Lsp-16-1, Lsp-2-1, Lsp-5-2, Lsp-5-3, Lsp-5-4, Lsp-5-5, Lsp-7-1, Lsp-9-1, Lvi-1-1, Mab-1-2, Mpl-1-11, Mpl-1-2, Mpl-1-3, Mpl-1-4, Mpl-1-5, Mpl-1-6, Mpl-1-7, Mpl-1-8, Mpl-1-9, Mpl-2-1, Mpl-3-1, Pac-3-1, Pac-4-1, Pac-5-1, Pal-4-1, Pda-1-2, Pda-1-3, Pgl-3-1, Pha-1-2, Pla-2-1, Pla-4-1, Pla-5-1, Plo-1-1, Pma-4-1, Pmu-1-1, Pmu-3-1, Ppe-2-1, Ppe-3-1, Ppe-5-1, Ppe-6-1, Ppo-2-1, Ppo-2-2, Ppo-5-1, Ppo-7-1, Ppo-8-1, Ppo-9-1, Psp-13-1, Psp-14-1, Psp-15-1, Psp-16-1, Psp-17-1, Psp-18-1, Psp-22-1, Psp-28-1, Psp-40-1, Psp-9-1, Pte-1-1, Pth-1-1, Qth-1-1, Qth-1-2, Sai-1-2, Sau-1-1, Sau-10-1, Sau-100-1, Sau-1000-1, Sau-1001-1, Sau-1002-1, Sau-1003-1, Sau-1004-1, Sau-1005-1, Sau-1006-1, Sau-1007-1, Sau-1008-1, Sau-1009-1, Sau-101-1, Sau-1010-1, Sau-1011-1, Sau-1012-1, Sau-1013-1, Sau-1014-1, Sau-1015-1, Sau-1016-1, Sau-1017-1, Sau-1018-1, Sau-1019-1, Sau-102-1, Sau-1020-1, Sau-1021-1, Sau-1022-1, Sau-1023-1, Sau-1024-1, Sau-1025-1, Sau-1026-1, Sau-1027-1, Sau-1028-1, Sau-1029-1, Sau-103-1, Sau-1030-1, Sau-1031-1, Sau-1032-1, Sau-1033-1, Sau-1034-1, Sau-1035-1, Sau-1036-1, Sau-1037-1, Sau-1038-1, Sau-1039-1, Sau-104-1, Sau-1040-1, Sau-1041-1, Sau-1042-1, Sau-1043-1, Sau-1044-1, Sau-1045-1, Sau-1046-1, Sau-1047-1, Sau-1048-1, Sau-1049-1, Sau-105-1, Sau-1050-1, Sau-1051-1, Sau-1052-1, Sau-1053-1, Sau-1054-1, Sau-1055-1, Sau-1056-1, Sau-1057-1, Sau-1058-1, Sau-1059-1, Sau-106-1, Sau-1060-1, Sau-1061-1, Sau-1062-1, Sau-1063-1, Sau-1064-1, Sau-1065-1, Sau-1066-1, Sau-1067-1, Sau-1068-1, Sau-1069-1, Sau-107-1, Sau-1070-1, Sau-1071-1, Sau-1072-1, Sau-1073-1, Sau-1074-1, Sau-1075-1, Sau-1076-1, Sau-1077-1, Sau-1078-1, Sau-1079-1, Sau-108-1, Sau-1080-1, Sau-1081-1, Sau-1082-1, Sau-1083-1, Sau-1084-1, Sau-1085-1, Sau-1086-1, Sau-1087-1, Sau-1088-1, Sau-1089-1, Sau-109-1, Sau-1090-1, Sau-1091-1, Sau-1092-1, Sau-1093-1, Sau-1094-1, Sau-1095-1, Sau-1096-1, Sau-1097-1, Sau-1098-1, Sau-1099-1, Sau-11-1, Sau-110-1, Sau-1100-1, Sau-1101-1, Sau-1102-1, Sau-1103-1, Sau-1104-1, Sau-1105-1, Sau-1106-1, Sau-1107-1, Sau-1108-1, Sau-1109-1, Sau-111-1, Sau-1110-1, Sau-1111-1, Sau-1112-1, Sau-1113-1, Sau-1114-1, Sau-1115-1, Sau-1116-1, Sau-1117-1, Sau-1118-1, Sau-1119-1, Sau-112-1, Sau-1120-1, Sau-1121-1, Sau-1122-1, Sau-1123-1, Sau-1124-1, Sau-1125-1, Sau-1126-1, Sau-1127-1, Sau-1128-1, Sau-1129-1, Sau-113-1, Sau-1130-1, Sau-1131-1, Sau-1132-1, Sau-1133-1, Sau-1134-1, Sau-1135-1, Sau-1136-1, Sau-1137-1, Sau-1138-1, Sau-1139-1, Sau-114-1, Sau-1140-1, Sau-1141-1, Sau-1142-1, Sau-1143-1, Sau-1144-1, Sau-1145-1, Sau-1146-1, Sau-1147-1, Sau-1148-1, Sau-1149-1, Sau-115-1, Sau-1150-1, Sau-1151-1, Sau-1152-1, Sau-1153-1, Sau-1154-1, Sau-1155-1, Sau-1156-1, Sau-1157-1, Sau-1158-1, Sau-1159-1, Sau-116-1, Sau-1160-1, Sau-1161-1, Sau-1162-1, Sau-1163-1, Sau-1164-1, Sau-1165-1, Sau-1166-1, Sau-1167-1, Sau-1168-1, Sau-1169-1, Sau-117-1, Sau-1170-1, Sau-1171-1, Sau-1172-1, Sau-1173-1, Sau-1174-1, Sau-1175-1, Sau-1176-1, Sau-1177-1, Sau-1178-1, Sau-1179-1, Sau-118-1, Sau-1180-1, Sau-1181-1, Sau-1182-1, Sau-1183-1, Sau-1184-1, Sau-1185-1, Sau-1186-1, Sau-1187-1, Sau-1188-1, Sau-1189-1, Sau-119-1, Sau-1190-1, Sau-1191-1, Sau-1192-1, Sau-1193-1, Sau-1194-1, Sau-1195-1, Sau-1196-1, Sau-1197-1, Sau-1198-1, Sau-1199-1, Sau-12-1, Sau-120-1, Sau-1200-1, Sau-1201-1, Sau-1202-1, Sau-1203-1, Sau-1204-1, Sau-1205-1, Sau-1206-1, Sau-1207-1, Sau-1208-1, Sau-1209-1, Sau-121-1, Sau-1210-1, Sau-1211-1, Sau-1212-1, Sau-1213-1, Sau-1214-1, Sau-1215-1, Sau-1216-1, Sau-1217-1, Sau-1218-1, Sau-1219-1, Sau-122-1, Sau-1220-1, Sau-1221-1, Sau-1222-1, Sau-1223-1, Sau-1224-1, Sau-1225-1, Sau-1226-1, Sau-1227-1, Sau-1228-1, Sau-1229-1, Sau-123-1, Sau-1230-1, Sau-1231-1, Sau-1232-1, Sau-1233-1, Sau-1234-1, Sau-1235-1, Sau-1236-1, Sau-1237-1, Sau-1238-1, Sau-1239-1, Sau-124-1, Sau-1240-1, Sau-1241-1, Sau-1242-1, Sau-1243-1, Sau-1244-1, Sau-1245-1, Sau-1246-1, Sau-1247-1, Sau-1248-1, Sau-1249-1, Sau-125-1, Sau-1250-1, Sau-1251-1, Sau-1252-1, Sau-1253-1, Sau-1254-1, Sau-1255-1,

Sau-1256-1, Sau-1257-1, Sau-1258-1, Sau-1259-1, Sau-126-1, Sau-1260-1, Sau-1261-1, Sau-1262-1, Sau-1263-1, Sau-1264-1, Sau-1265-1, Sau-1266-1, Sau-1267-1, Sau-1268-1, Sau-1269-1, Sau-127-1, Sau-1270-1, Sau-1271-1, Sau-1272-1, Sau-1273-1, Sau-1274-1, Sau-1275-1, Sau-1276-1, Sau-1277-1, Sau-1278-1, Sau-1279-1, Sau-128-1, Sau-1280-1, Sau-1281-1, Sau-1282-1, Sau-1283-1, Sau-1284-1, Sau-1285-1, Sau-1286-1, Sau-1287-1, Sau-1288-1, Sau-1289-1, Sau-129-1, Sau-1290-1, Sau-1291-1, Sau-1292-1, Sau-1293-1, Sau-1294-1, Sau-1295-1, Sau-1296-1, Sau-1297-1, Sau-1298-1, Sau-1299-1, Sau-13-1, Sau-130-1, Sau-1300-1, Sau-1301-1, Sau-1302-1, Sau-1303-1, Sau-1304-1, Sau-1305-1, Sau-1306-1, Sau-1307-1, Sau-1308-1, Sau-1309-1, Sau-131-1, Sau-1310-1, Sau-1311-1, Sau-1312-1, Sau-1313-1, Sau-1314-1, Sau-1315-1, Sau-1316-1, Sau-1317-1, Sau-1318-1, Sau-1319-1, Sau-132-1, Sau-1320-1, Sau-1321-1, Sau-1322-1, Sau-1323-1, Sau-1324-1, Sau-1325-1, Sau-1326-1, Sau-1327-1, Sau-1328-1, Sau-1329-1, Sau-133-1, Sau-1330-1, Sau-1331-1, Sau-1332-1, Sau-1333-1, Sau-1334-1, Sau-1335-1, Sau-1336-1, Sau-1337-1, Sau-1338-1, Sau-1339-1, Sau-134-1, Sau-1340-1, Sau-1341-1, Sau-1342-1, Sau-1343-1, Sau-1344-1, Sau-1345-1, Sau-1346-1, Sau-1347-1, Sau-1348-1, Sau-1349-1, Sau-135-1, Sau-1350-1, Sau-1351-1, Sau-1352-1, Sau-1353-1, Sau-1354-1, Sau-1355-1, Sau-1356-1, Sau-1357-1, Sau-1358-1, Sau-1359-1, Sau-136-1, Sau-1360-1, Sau-1361-1, Sau-1362-1, Sau-1363-1, Sau-1364-1, Sau-1365-1, Sau-1366-1, Sau-1367-1, Sau-1368-1, Sau-1369-1, Sau-137-1, Sau-1370-1, Sau-1371-1, Sau-1372-1, Sau-1373-1, Sau-1374-1, Sau-1375-1, Sau-1376-1, Sau-1377-1, Sau-1378-1, Sau-1379-1, Sau-138-1, Sau-1380-1, Sau-1381-1, Sau-1382-1, Sau-1383-1, Sau-1384-1, Sau-1385-1, Sau-1386-1, Sau-1387-1, Sau-1388-1, Sau-1389-1, Sau-139-1, Sau-1390-1, Sau-1391-1, Sau-1392-1, Sau-1393-1, Sau-1394-1, Sau-1395-1, Sau-1396-1, Sau-1397-1, Sau-1398-1, Sau-1399-1, Sau-14-1, Sau-140-1, Sau-1400-1, Sau-1401-1, Sau-1402-1, Sau-1403-1, Sau-1404-1, Sau-1405-1, Sau-1406-1, Sau-1407-1, Sau-1408-1, Sau-1409-1, Sau-141-1, Sau-1410-1, Sau-1411-1, Sau-1412-1, Sau-1413-1, Sau-1414-1, Sau-1415-1, Sau-1416-1, Sau-1417-1, Sau-1418-1, Sau-1419-1, Sau-142-1, Sau-1420-1, Sau-1421-1, Sau-1422-1, Sau-1423-1, Sau-1424-1, Sau-1425-1, Sau-1426-1, Sau-1427-1, Sau-1428-1, Sau-1429-1, Sau-143-1, Sau-1430-1, Sau-1431-1, Sau-1432-1, Sau-1433-1, Sau-1434-1, Sau-1435-1, Sau-1436-1, Sau-1437-1, Sau-1438-1, Sau-1439-1, Sau-144-1, Sau-1440-1, Sau-1441-1, Sau-1442-1, Sau-1443-1, Sau-1444-1, Sau-1445-1, Sau-1446-1, Sau-1447-1, Sau-1448-1, Sau-1449-1, Sau-145-1, Sau-1450-1, Sau-1451-1, Sau-1452-1, Sau-1453-1, Sau-1454-1, Sau-1455-1, Sau-1456-1, Sau-1457-1, Sau-1458-1, Sau-1459-1, Sau-146-1, Sau-1460-1, Sau-1461-1, Sau-1462-1, Sau-1463-1, Sau-1464-1, Sau-1465-1, Sau-1466-1, Sau-1467-1, Sau-1468-1, Sau-1469-1, Sau-147-1, Sau-1470-1, Sau-1471-1, Sau-1472-1, Sau-1473-1, Sau-1474-1, Sau-1475-1, Sau-1476-1, Sau-1477-1, Sau-1478-1, Sau-1479-1, Sau-148-1, Sau-1480-1, Sau-1481-1, Sau-1482-1, Sau-1483-1, Sau-1484-1, Sau-1485-1, Sau-1486-1, Sau-1487-1, Sau-1488-1, Sau-1489-1, Sau-149-1, Sau-1490-1, Sau-1491-1, Sau-1492-1, Sau-1493-1, Sau-1494-1, Sau-1495-1, Sau-1496-1, Sau-1497-1, Sau-1498-1, Sau-1499-1, Sau-15-1, Sau-150-1, Sau-1500-1, Sau-1501-1, Sau-1502-1, Sau-1503-1, Sau-1504-1, Sau-1505-1, Sau-1506-1, Sau-1507-1, Sau-1508-1, Sau-1509-1, Sau-151-1, Sau-1510-1, Sau-1511-1, Sau-1512-1, Sau-1513-1, Sau-1514-1, Sau-1515-1, Sau-1516-1, Sau-1517-1, Sau-1518-1, Sau-1519-1, Sau-152-1, Sau-1520-1, Sau-1521-1, Sau-1522-1, Sau-1523-1, Sau-1524-1, Sau-1525-1, Sau-1526-1, Sau-1527-1, Sau-1528-1, Sau-1529-1, Sau-153-1, Sau-1530-1, Sau-1531-1, Sau-1532-1, Sau-1533-1, Sau-1534-1, Sau-1535-1, Sau-1536-1, Sau-1537-1, Sau-1538-1, Sau-1539-1, Sau-154-1, Sau-1540-1, Sau-1541-1, Sau-1542-1, Sau-1543-1, Sau-1544-1, Sau-1545-1, Sau-1546-1, Sau-1547-1, Sau-1548-1, Sau-1549-1, Sau-155-1, Sau-1550-1, Sau-1551-1, Sau-1552-1, Sau-1553-1, Sau-1554-1, Sau-1555-1, Sau-1556-1, Sau-1557-1, Sau-1558-1, Sau-1559-1, Sau-156-1, Sau-1560-1, Sau-1561-1, Sau-1562-1, Sau-1563-1, Sau-1564-1, Sau-1565-1, Sau-1566-1, Sau-1567-1, Sau-1568-1, Sau-1569-1, Sau-157-1, Sau-1570-1, Sau-1571-1, Sau-1572-1, Sau-1573-1, Sau-1574-1, Sau-1575-1, Sau-1576-1, Sau-1577-1, Sau-1578-1, Sau-1579-1, Sau-158-1, Sau-1580-1, Sau-1581-1, Sau-1582-1, Sau-1583-1, Sau-1584-1, Sau-1585-1, Sau-1586-1, Sau-1587-1, Sau-1588-1, Sau-1589-1, Sau-159-1, Sau-1590-1, Sau-1591-1, Sau-1592-1, Sau-1593-1, Sau-1594-1, Sau-1595-1, Sau-1596-1, Sau-1597-1, Sau-1598-1, Sau-1599-1, Sau-16-1, Sau-160-1, Sau-1600-1, Sau-1601-1, Sau-1602-1, Sau-1603-1, Sau-1604-1, Sau-1605-1, Sau-1606-1, Sau-1607-1, Sau-1608-1, Sau-1609-1, Sau-161-1, Sau-1610-1, Sau-1611-1, Sau-1612-1, Sau-1613-1, Sau-1614-1, Sau-1615-1, Sau-1616-1, Sau-1617-1, Sau-1618-1, Sau-1619-1, Sau-162-1, Sau-1620-1, Sau-1621-1, Sau-1622-1, Sau-1623-1, Sau-1624-1, Sau-1625-1, Sau-1626-1, Sau-1627-1, Sau-1628-1, Sau-1629-1, Sau-163-1, Sau-1630-1, Sau-1631-1, Sau-1632-1, Sau-1633-1, Sau-1634-1, Sau-1635-1, Sau-1636-1, Sau-1637-1, Sau-1638-1, Sau-1639-1, Sau-164-1, Sau-1640-1, Sau-1641-1, Sau-1642-1, Sau-1643-1, Sau-1644-1, Sau-1645-1, Sau-1646-1, Sau-1647-1, Sau-1648-1, Sau-1649-1, Sau-165-1, Sau-1650-1, Sau-1651-1, Sau-1652-1, Sau-1653-1, Sau-1654-1, Sau-1655-1, Sau-1656-1, Sau-1657-1, Sau-1658-1, Sau-1659-1, Sau-166-1, Sau-1660-1, Sau-1661-1, Sau-1662-1, Sau-1663-1, Sau-1664-1, Sau-1665-1, Sau-1666-1, Sau-1667-1, Sau-1668-1, Sau-1669-1, Sau-167-1, Sau-1670-1, Sau-1671-1, Sau-1672-1, Sau-1673-1, Sau-1674-1, Sau-1675-1, Sau-1676-1, Sau-1677-1, Sau-1678-1, Sau-1679-1, Sau-168-1, Sau-1680-1, Sau-1681-1, Sau-1682-1, Sau-1683-1, Sau-1684-1, Sau-1685-1, Sau-1686-1, Sau-1687-1, Sau-1688-1, Sau-1689-1, Sau-169-1, Sau-1690-1, Sau-1691-1, Sau-1692-1, Sau-1693-1, Sau-1694-1, Sau-1695-1, Sau-1696-1, Sau-1697-1, Sau-1698-1, Sau-1699-1, Sau-17-1, Sau-170-1, Sau-1700-1, Sau-1701-1, Sau-1702-1, Sau-1703-1, Sau-1704-1, Sau-1705-1, Sau-1706-1, Sau-1707-1, Sau-1708-1, Sau-1709-1, Sau-171-1, Sau-1710-1, Sau-1711-1, Sau-1712-1, Sau-1713-1, Sau-1714-1, Sau-1715-1, Sau-1716-1, Sau-1717-1, Sau-1718-1, Sau-1719-1, Sau-172-1, Sau-1720-1, Sau-1721-1, Sau-1722-1, Sau-1723-1, Sau-1724-1, Sau-1725-1, Sau-1726-1, Sau-1727-1, Sau-1728-1, Sau-1729-1, Sau-173-1, Sau-1730-1, Sau-1731-1, Sau-1732-1, Sau-1733-1, Sau-1734-1, Sau-1735-1, Sau-1736-1, Sau-1737-1, Sau-1738-1, Sau-174-1, Sau-1740-1, Sau-1741-1, Sau-1742-1, Sau-1743-1, Sau-1744-1, Sau-1745-1, Sau-1746-1, Sau-1747-1, Sau-1748-1, Sau-1749-1, Sau-175-1, Sau-1750-1, Sau-1751-1, Sau-1752-1, Sau-1753-1, Sau-1754-1, Sau-1755-1, Sau-1756-1, Sau-1757-1, Sau-1758-1, Sau-1759-1, Sau-176-1, Sau-1760-1, Sau-1761-1, Sau-1762-1, Sau-1763-1, Sau-1764-1, Sau-1765-1, Sau-1766-1, Sau-1767-1, Sau-1768-1, Sau-1769-1, Sau-177-1, Sau-1770-1, Sau-1771-1, Sau-1772-1, Sau-1773-1, Sau-1774-1, Sau-1775-1, Sau-1776-1, Sau-1777-1, Sau-1778-1, Sau-1779-1, Sau-178-1, Sau-1780-1, Sau-1781-1, Sau-1782-1, Sau-1783-1, Sau-1784-1, Sau-1785-1, Sau-1786-1, Sau-1787-1, Sau-1788-1, Sau-1789-1, Sau-179-1, Sau-1790-1, Sau-1791-1, Sau-

1792-1, Sau-1793-1, Sau-1794-1, Sau-1795-1, Sau-1796-1, Sau-1797-1, Sau-1798-1, Sau-1799-1, Sau-18-1, Sau-180-1, Sau-1800-1, Sau-1801-1, Sau-1802-1, Sau-1803-1, Sau-1804-1, Sau-1805-1, Sau-1806-1, Sau-1807-1, Sau-1808-1, Sau-1809-1, Sau-181-1, Sau-1810-1, Sau-1811-1, Sau-1812-1, Sau-1813-1, Sau-1814-1, Sau-1815-1, Sau-1816-1, Sau-1817-1, Sau-1818-1, Sau-1819-1, Sau-182-1, Sau-1820-1, Sau-1821-1, Sau-1822-1, Sau-1823-1, Sau-1824-1, Sau-1825-1, Sau-1826-1, Sau-1827-1, Sau-1828-1, Sau-1829-1, Sau-183-1, Sau-1830-1, Sau-1831-1, Sau-1832-1, Sau-1833-1, Sau-1834-1, Sau-1835-1, Sau-1836-1, Sau-1837-1, Sau-1838-1, Sau-1839-1, Sau-184-1, Sau-1840-1, Sau-1841-1, Sau-1842-1, Sau-1843-1, Sau-1844-1, Sau-1845-1, Sau-1846-1, Sau-1847-1, Sau-1848-1, Sau-1849-1, Sau-185-1, Sau-1850-1, Sau-1851-1, Sau-1852-1, Sau-1853-1, Sau-1854-1, Sau-1855-1, Sau-1856-1, Sau-1857-1, Sau-1858-1, Sau-1859-1, Sau-186-1, Sau-1860-1, Sau-1861-1, Sau-1862-1, Sau-1863-1, Sau-1864-1, Sau-1865-1, Sau-1866-1, Sau-1867-1, Sau-1868-1, Sau-1869-1, Sau-187-1, Sau-1870-1, Sau-1871-1, Sau-1872-1, Sau-1873-1, Sau-1874-1, Sau-1875-1, Sau-1876-1, Sau-1877-1, Sau-1878-1, Sau-1879-1, Sau-188-1, Sau-1880-1, Sau-1881-1, Sau-1882-1, Sau-1883-1, Sau-1884-1, Sau-1885-1, Sau-1886-1, Sau-1887-1, Sau-1888-1, Sau-1889-1, Sau-189-1, Sau-1890-1, Sau-1891-1, Sau-1892-1, Sau-1893-1, Sau-1894-1, Sau-1895-1, Sau-1896-1, Sau-1897-1, Sau-1898-1, Sau-1899-1, Sau-19-1, Sau-190-1, Sau-1900-1, Sau-1901-1, Sau-1902-1, Sau-1903-1, Sau-1904-1, Sau-1905-1, Sau-1906-1, Sau-1907-1, Sau-1908-1, Sau-1909-1, Sau-191-1, Sau-1910-1, Sau-1911-1, Sau-1912-1, Sau-1913-1, Sau-1914-1, Sau-1915-1, Sau-1916-1, Sau-1917-1, Sau-1918-1, Sau-1919-1, Sau-192-1, Sau-1920-1, Sau-1921-1, Sau-1922-1, Sau-1923-1, Sau-1924-1, Sau-1925-1, Sau-1926-1, Sau-1927-1, Sau-1928-1, Sau-1929-1, Sau-193-1, Sau-1930-1, Sau-1931-1, Sau-1932-1, Sau-1933-1, Sau-1934-1, Sau-1935-1, Sau-1936-1, Sau-1937-1, Sau-1938-1, Sau-1939-1, Sau-194-1, Sau-1940-1, Sau-1941-1, Sau-1942-1, Sau-1943-1, Sau-1944-1, Sau-1945-1, Sau-1946-1, Sau-1947-1, Sau-1948-1, Sau-1949-1, Sau-195-1, Sau-1950-1, Sau-1951-1, Sau-1952-1, Sau-1953-1, Sau-1954-1, Sau-1955-1, Sau-1956-1, Sau-1957-1, Sau-1958-1, Sau-1959-1, Sau-196-1, Sau-1960-1, Sau-1961-1, Sau-1962-1, Sau-1963-1, Sau-1964-1, Sau-1965-1, Sau-1966-1, Sau-1967-1, Sau-1968-1, Sau-1969-1, Sau-197-1, Sau-1970-1, Sau-1971-1, Sau-1972-1, Sau-1973-1, Sau-1974-1, Sau-1975-1, Sau-1976-1, Sau-1977-1, Sau-1978-1, Sau-1979-1, Sau-198-1, Sau-1980-1, Sau-1981-1, Sau-1982-1, Sau-1983-1, Sau-1984-1, Sau-1985-1, Sau-1986-1, Sau-1987-1, Sau-1988-1, Sau-1989-1, Sau-199-1, Sau-1990-1, Sau-1991-1, Sau-1992-1, Sau-1993-1, Sau-1994-1, Sau-1995-1, Sau-1996-1, Sau-1997-1, Sau-1998-1, Sau-1999-1, Sau-2-1, Sau-20-1, Sau-200-1, Sau-2000-1, Sau-2001-1, Sau-2002-1, Sau-2003-1, Sau-2004-1, Sau-2005-1, Sau-2006-1, Sau-2007-1, Sau-2008-1, Sau-2009-1, Sau-201-1, Sau-2010-1, Sau-2011-1, Sau-2012-1, Sau-2013-1, Sau-2014-1, Sau-2015-1, Sau-2016-1, Sau-2017-1, Sau-2018-1, Sau-2019-1, Sau-202-1, Sau-2020-1, Sau-2021-1, Sau-2022-1, Sau-2023-1, Sau-2024-1, Sau-2025-1, Sau-2026-1, Sau-2027-1, Sau-2028-1, Sau-2029-1, Sau-203-1, Sau-2030-1, Sau-2031-1, Sau-2032-1, Sau-2033-1, Sau-2034-1, Sau-2035-1, Sau-2036-1, Sau-2037-1, Sau-2038-1, Sau-2039-1, Sau-204-1, Sau-2040-1, Sau-2041-1, Sau-2042-1, Sau-2043-1, Sau-2044-1, Sau-2045-1, Sau-2046-1, Sau-2047-1, Sau-2048-1, Sau-2049-1, Sau-205-1, Sau-2050-1, Sau-2051-1, Sau-2052-1, Sau-2053-1, Sau-2054-1, Sau-2055-1, Sau-2056-1, Sau-2057-1, Sau-2058-1, Sau-2059-1, Sau-206-1, Sau-2060-1, Sau-2061-1, Sau-2062-1, Sau-2063-1, Sau-2064-1, Sau-2065-1, Sau-2066-1, Sau-2067-1, Sau-2068-1, Sau-2069-1, Sau-207-1, Sau-2070-1, Sau-2071-1, Sau-2072-1, Sau-2073-1, Sau-2074-1, Sau-2075-1, Sau-2076-1, Sau-2077-1, Sau-2078-1, Sau-2079-1, Sau-208-1, Sau-2080-1, Sau-2081-1, Sau-2082-1, Sau-2083-1, Sau-2084-1, Sau-2085-1, Sau-2086-1, Sau-2087-1, Sau-2088-1, Sau-2089-1, Sau-209-1, Sau-2090-1, Sau-2091-1, Sau-2092-1, Sau-2093-1, Sau-2094-1, Sau-2095-1, Sau-2096-1, Sau-2097-1, Sau-2098-1, Sau-2099-1, Sau-21-1, Sau-210-1, Sau-2100-1, Sau-2101-1, Sau-2102-1, Sau-2103-1, Sau-2104-1, Sau-2105-1, Sau-2106-1, Sau-2107-1, Sau-2108-1, Sau-2109-1, Sau-211-1, Sau-2110-1, Sau-2111-1, Sau-2112-1, Sau-2113-1, Sau-2114-1, Sau-2115-1, Sau-2116-1, Sau-2117-1, Sau-2118-1, Sau-2119-1, Sau-212-1, Sau-2120-1, Sau-2121-1, Sau-2122-1, Sau-2123-1, Sau-2124-1, Sau-2125-1, Sau-2126-1, Sau-2127-1, Sau-2128-1, Sau-2129-1, Sau-213-1, Sau-2130-1, Sau-2131-1, Sau-2132-1, Sau-2133-1, Sau-2134-1, Sau-2135-1, Sau-2136-1, Sau-2137-1, Sau-2138-1, Sau-2139-1, Sau-214-1, Sau-2140-1, Sau-2141-1, Sau-2142-1, Sau-2143-1, Sau-2144-1, Sau-2145-1, Sau-2146-1, Sau-2147-1, Sau-2148-1, Sau-2149-1, Sau-215-1, Sau-2150-1, Sau-2151-1, Sau-2152-1, Sau-2153-1, Sau-2154-1, Sau-2155-1, Sau-2156-1, Sau-2157-1, Sau-2158-1, Sau-2159-1, Sau-216-1, Sau-2160-1, Sau-2161-1, Sau-2162-1, Sau-2163-1, Sau-2164-1, Sau-2165-1, Sau-2166-1, Sau-2167-1, Sau-2168-1, Sau-2169-1, Sau-217-1, Sau-2170-1, Sau-2171-1, Sau-2172-1, Sau-2173-1, Sau-2174-1, Sau-2175-1, Sau-2176-1, Sau-2177-1, Sau-2178-1, Sau-2179-1, Sau-218-1, Sau-2180-1, Sau-2181-1, Sau-2182-1, Sau-2183-1, Sau-2184-1, Sau-2185-1, Sau-2186-1, Sau-2187-1, Sau-2188-1, Sau-2189-1, Sau-219-1, Sau-2190-1, Sau-2191-1, Sau-2192-1, Sau-2193-1, Sau-2194-1, Sau-2195-1, Sau-2196-1, Sau-2197-1, Sau-2198-1, Sau-2199-1, Sau-22-1, Sau-220-1, Sau-2200-1, Sau-2201-1, Sau-2202-1, Sau-2202-2, Sau-2203-1, Sau-2204-1, Sau-2205-1, Sau-2206-1, Sau-2207-1, Sau-2208-1, Sau-2209-1, Sau-221-1, Sau-2210-1, Sau-2211-1, Sau-2212-1, Sau-2213-1, Sau-2214-1, Sau-2215-1, Sau-2216-1, Sau-2217-1, Sau-2218-1, Sau-2219-1, Sau-222-1, Sau-2220-1, Sau-2221-1, Sau-2222-1, Sau-2223-1, Sau-2224-1, Sau-2225-1, Sau-2226-1, Sau-2227-1, Sau-2228-1, Sau-2229-1, Sau-223-1, Sau-2230-1, Sau-2231-1, Sau-2232-1, Sau-2233-1, Sau-2234-1, Sau-2235-1, Sau-2236-1, Sau-2237-1, Sau-2238-1, Sau-2239-1, Sau-224-1, Sau-2240-1, Sau-2241-1, Sau-2242-1, Sau-2243-1, Sau-2244-1, Sau-2245-1, Sau-2246-1, Sau-2247-1, Sau-2248-1, Sau-2249-1, Sau-225-1, Sau-2250-1, Sau-2251-1, Sau-2252-1, Sau-2253-1, Sau-2254-1, Sau-2255-1, Sau-2256-1, Sau-2257-1, Sau-2258-1, Sau-2259-1, Sau-226-1, Sau-2260-1, Sau-2261-1, Sau-2262-1, Sau-2263-1, Sau-2264-1, Sau-2265-1, Sau-2266-1, Sau-2267-1, Sau-2268-1, Sau-2269-1, Sau-227-1, Sau-2270-1, Sau-2271-1, Sau-2272-1, Sau-2273-1, Sau-2274-1, Sau-2275-1, Sau-2276-1, Sau-2277-1, Sau-2278-1, Sau-2279-1, Sau-228-1, Sau-2280-1, Sau-2281-1, Sau-2282-1, Sau-2283-1, Sau-2284-1, Sau-2285-1, Sau-2286-1, Sau-2287-1, Sau-2288-1, Sau-2289-1, Sau-229-1, Sau-2290-1, Sau-2291-1, Sau-2292-1, Sau-2293-1, Sau-2294-1, Sau-2295-1, Sau-2296-1, Sau-2297-1, Sau-2298-1, Sau-2299-1, Sau-23-1, Sau-230-1, Sau-2300-1, Sau-2301-1, Sau-2302-1, Sau-2303-1, Sau-2304-1, Sau-2305-1, Sau-2306-1, Sau-2307-1, Sau-2308-1, Sau-2309-1, Sau-231-1, Sau-2310-1, Sau-2311-1, Sau-2312-1, Sau-2313-1, Sau-2314-1, Sau-2315-1, Sau-2316-1, Sau-2317-1, Sau-2318-1, Sau-2319-1, Sau-232-1, Sau-2320-1, Sau-2321-1, Sau-2322-1, Sau-2323-1, Sau-2324-1, Sau-2325-1,

Sau-2326-1, Sau-2327-1, Sau-2328-1, Sau-2329-1, Sau-233-1, Sau-2330-1, Sau-2331-1, Sau-2332-1, Sau-2333-1, Sau-2334-1, Sau-2335-1, Sau-2336-1, Sau-2337-1, Sau-2338-1, Sau-2339-1, Sau-234-1, Sau-2340-1, Sau-2341-1, Sau-2342-1, Sau-2343-1, Sau-2344-1, Sau-2345-1, Sau-2346-1, Sau-2347-1, Sau-2348-1, Sau-2349-1, Sau-235-1, Sau-2350-1, Sau-2351-1, Sau-2352-1, Sau-2353-1, Sau-2354-1, Sau-2355-1, Sau-2356-1, Sau-2357-1, Sau-2358-1, Sau-2359-1, Sau-236-1, Sau-2360-1, Sau-2361-1, Sau-2362-1, Sau-2363-1, Sau-2364-1, Sau-2365-1, Sau-2366-1, Sau-2367-1, Sau-2368-1, Sau-2369-1, Sau-237-1, Sau-2370-1, Sau-2371-1, Sau-2372-1, Sau-2373-1, Sau-2374-1, Sau-2375-1, Sau-2376-1, Sau-2377-1, Sau-2378-1, Sau-2379-1, Sau-238-1, Sau-2380-1, Sau-2381-1, Sau-2382-1, Sau-2383-1, Sau-2384-1, Sau-2385-1, Sau-2386-1, Sau-2387-1, Sau-2388-1, Sau-2389-1, Sau-239-1, Sau-2390-1, Sau-2391-1, Sau-2392-1, Sau-2393-1, Sau-2394-1, Sau-2395-1, Sau-2396-1, Sau-2397-1, Sau-2398-1, Sau-2399-1, Sau-24-1, Sau-240-1, Sau-2400-1, Sau-2401-1, Sau-2402-1, Sau-2403-1, Sau-2404-1, Sau-2405-1, Sau-2406-1, Sau-2407-1, Sau-2408-1, Sau-2409-1, Sau-241-1, Sau-2410-1, Sau-2411-1, Sau-2412-1, Sau-2413-1, Sau-2414-1, Sau-2415-1, Sau-2416-1, Sau-2417-1, Sau-2418-1, Sau-2419-1, Sau-242-1, Sau-2420-1, Sau-2421-1, Sau-2422-1, Sau-2423-1, Sau-2424-1, Sau-2425-1, Sau-2426-1, Sau-2427-1, Sau-2428-1, Sau-2429-1, Sau-243-1, Sau-2430-1, Sau-2431-1, Sau-2432-1, Sau-2433-1, Sau-2434-1, Sau-2435-1, Sau-2436-1, Sau-2437-1, Sau-2438-1, Sau-2439-1, Sau-244-1, Sau-2440-1, Sau-2441-1, Sau-2442-1, Sau-2443-1, Sau-2444-1, Sau-2445-1, Sau-2446-1, Sau-2447-1, Sau-2448-1, Sau-2449-1, Sau-245-1, Sau-2450-1, Sau-2451-1, Sau-2452-1, Sau-2453-1, Sau-2454-1, Sau-2455-1, Sau-2456-1, Sau-2457-1, Sau-2458-1, Sau-2459-1, Sau-246-1, Sau-2460-1, Sau-2461-1, Sau-2462-1, Sau-2463-1, Sau-2464-1, Sau-2465-1, Sau-2466-1, Sau-2467-1, Sau-2468-1, Sau-2469-1, Sau-247-1, Sau-2470-1, Sau-2471-1, Sau-2472-1, Sau-2473-1, Sau-2474-1, Sau-2475-1, Sau-2476-1, Sau-2477-1, Sau-2478-1, Sau-2479-1, Sau-248-1, Sau-2480-1, Sau-2481-1, Sau-2482-1, Sau-2483-1, Sau-2484-1, Sau-2485-1, Sau-2486-1, Sau-2487-1, Sau-2488-1, Sau-2489-1, Sau-249-1, Sau-2490-1, Sau-2491-1, Sau-2492-1, Sau-2493-1, Sau-2494-1, Sau-2495-1, Sau-2496-1, Sau-2497-1, Sau-2498-1, Sau-2499-1, Sau-25-1, Sau-250-1, Sau-2500-1, Sau-2501-1, Sau-2502-1, Sau-2503-1, Sau-2504-1, Sau-2505-1, Sau-2506-1, Sau-2507-1, Sau-2508-1, Sau-2509-1, Sau-251-1, Sau-2510-1, Sau-2511-1, Sau-2512-1, Sau-2513-1, Sau-2514-1, Sau-2515-1, Sau-2516-1, Sau-2517-1, Sau-2518-1, Sau-2519-1, Sau-252-1, Sau-2520-1, Sau-2521-1, Sau-2522-1, Sau-2523-1, Sau-2524-1, Sau-2525-1, Sau-2526-1, Sau-2527-1, Sau-2528-1, Sau-2529-1, Sau-253-1, Sau-2530-1, Sau-2531-1, Sau-2532-1, Sau-2533-1, Sau-2534-1, Sau-2535-1, Sau-2536-1, Sau-2537-1, Sau-2538-1, Sau-2539-1, Sau-254-1, Sau-2540-1, Sau-2541-1, Sau-2542-1, Sau-2543-1, Sau-2544-1, Sau-2545-1, Sau-2546-1, Sau-2547-1, Sau-2548-1, Sau-2549-1, Sau-2550-1, Sau-2551-1, Sau-2552-1, Sau-2553-1, Sau-2554-1, Sau-2555-1, Sau-2556-1, Sau-2557-1, Sau-2558-1, Sau-2559-1, Sau-256-1, Sau-2560-1, Sau-2561-1, Sau-2562-1, Sau-2563-1, Sau-2564-1, Sau-2565-1, Sau-2566-1, Sau-2567-1, Sau-2568-1, Sau-2569-1, Sau-257-1, Sau-2570-1, Sau-2571-1, Sau-2572-1, Sau-2573-1, Sau-2574-1, Sau-2575-1, Sau-2576-1, Sau-2577-1, Sau-2578-1, Sau-2579-1, Sau-258-1, Sau-2580-1, Sau-2581-1, Sau-2582-1, Sau-2583-1, Sau-2584-1, Sau-2585-1, Sau-2586-1, Sau-2587-1, Sau-2588-1, Sau-2589-1, Sau-259-1, Sau-2590-1, Sau-2591-1, Sau-2592-1, Sau-2593-1, Sau-2594-1, Sau-2595-1, Sau-2596-1, Sau-2597-1, Sau-2598-1, Sau-2599-1, Sau-26-1, Sau-260-1, Sau-2600-1, Sau-2601-1, Sau-2602-1, Sau-2603-1, Sau-2604-1, Sau-2605-1, Sau-2606-1, Sau-2607-1, Sau-2608-1, Sau-2609-1, Sau-261-1, Sau-2610-1, Sau-2611-1, Sau-2612-1, Sau-2613-1, Sau-2614-1, Sau-2615-1, Sau-2616-1, Sau-2617-1, Sau-2618-1, Sau-2619-1, Sau-262-1, Sau-2620-1, Sau-2621-1, Sau-2622-1, Sau-2623-1, Sau-2624-1, Sau-2625-1, Sau-2626-1, Sau-2627-1, Sau-2628-1, Sau-2629-1, Sau-263-1, Sau-2630-1, Sau-2631-1, Sau-2632-1, Sau-2633-1, Sau-2634-1, Sau-2635-1, Sau-2636-1, Sau-2637-1, Sau-2638-1, Sau-2639-1, Sau-264-1, Sau-2640-1, Sau-2641-1, Sau-2642-1, Sau-2643-1, Sau-2644-1, Sau-2645-1, Sau-2646-1, Sau-2647-1, Sau-2648-1, Sau-2649-1, Sau-265-1, Sau-2650-1, Sau-2651-1, Sau-2652-1, Sau-2653-1, Sau-2654-1, Sau-2655-1, Sau-2656-1, Sau-2657-1, Sau-2658-1, Sau-2659-1, Sau-266-1, Sau-2660-1, Sau-2661-1, Sau-2662-1, Sau-2663-1, Sau-2664-1, Sau-2665-1, Sau-2666-1, Sau-2667-1, Sau-2668-1, Sau-2669-1, Sau-267-1, Sau-2670-1, Sau-2671-1, Sau-2672-1, Sau-2673-1, Sau-2674-1, Sau-2675-1, Sau-2676-1, Sau-2677-1, Sau-2678-1, Sau-2679-1, Sau-268-1, Sau-2680-1, Sau-2681-1, Sau-2682-1, Sau-2683-1, Sau-2684-1, Sau-2685-1, Sau-2686-1, Sau-2687-1, Sau-2688-1, Sau-2689-1, Sau-269-1, Sau-2690-1, Sau-2691-1, Sau-2692-1, Sau-2693-1, Sau-2694-1, Sau-2695-1, Sau-2696-1, Sau-2697-1, Sau-2698-1, Sau-2699-1, Sau-27-1, Sau-270-1, Sau-2700-1, Sau-2701-1, Sau-2702-1, Sau-2703-1, Sau-2704-1, Sau-2705-1, Sau-2706-1, Sau-2707-1, Sau-2708-1, Sau-2709-1, Sau-271-1, Sau-2710-1, Sau-2711-1, Sau-2712-1, Sau-2713-1, Sau-2714-1, Sau-2715-1, Sau-2716-1, Sau-2717-1, Sau-2718-1, Sau-2719-1, Sau-272-1, Sau-2720-1, Sau-2721-1, Sau-2722-1, Sau-2723-1, Sau-2724-1, Sau-2725-1, Sau-2726-1, Sau-2727-1, Sau-2728-1, Sau-2729-1, Sau-273-1, Sau-2730-1, Sau-2731-1, Sau-2732-1, Sau-2733-1, Sau-2734-1, Sau-2735-1, Sau-2736-1, Sau-2737-1, Sau-2738-1, Sau-2739-1, Sau-274-1, Sau-2740-1, Sau-2741-1, Sau-2742-1, Sau-2743-1, Sau-2744-1, Sau-2745-1, Sau-2746-1, Sau-2747-1, Sau-2748-1, Sau-2749-1, Sau-275-1, Sau-2750-1, Sau-2751-1, Sau-2752-1, Sau-2753-1, Sau-2754-1, Sau-2755-1, Sau-2756-1, Sau-2757-1, Sau-2758-1, Sau-2759-1, Sau-276-1, Sau-2760-1, Sau-2761-1, Sau-2762-1, Sau-2763-1, Sau-2764-1, Sau-2765-1, Sau-2766-1, Sau-2767-1, Sau-2768-1, Sau-2769-1, Sau-277-1, Sau-2770-1, Sau-2771-1, Sau-2772-1, Sau-2773-1, Sau-2774-1, Sau-2775-1, Sau-2776-1, Sau-2777-1, Sau-2778-1, Sau-2779-1, Sau-278-1, Sau-2780-1, Sau-2781-1, Sau-2782-1, Sau-2783-1, Sau-2784-1, Sau-2785-1, Sau-2786-1, Sau-2787-1, Sau-2788-1, Sau-2789-1, Sau-279-1, Sau-2790-1, Sau-2791-1, Sau-2792-1, Sau-2793-1, Sau-2794-1, Sau-2795-1, Sau-2796-1, Sau-2797-1, Sau-2798-1, Sau-2799-1, Sau-28-1, Sau-280-1, Sau-2800-1, Sau-2801-1, Sau-2802-1, Sau-2803-1, Sau-2804-1, Sau-2805-1, Sau-2806-1, Sau-2807-1, Sau-2808-1, Sau-2809-1, Sau-281-1, Sau-2810-1, Sau-2811-1, Sau-2812-1, Sau-2813-1, Sau-2814-1, Sau-2815-1, Sau-2816-1, Sau-2817-1, Sau-2818-1, Sau-2819-1, Sau-282-1, Sau-2820-1, Sau-2821-1, Sau-2822-1, Sau-2823-1, Sau-2824-1, Sau-2825-1, Sau-2826-1, Sau-2827-1, Sau-2828-1, Sau-2829-1, Sau-283-1, Sau-2830-1, Sau-2831-1, Sau-2832-1, Sau-2833-1, Sau-2834-1, Sau-2835-1, Sau-2836-1, Sau-2837-1, Sau-2838-1, Sau-2839-1, Sau-284-1, Sau-2840-1, Sau-2841-1, Sau-2842-1, Sau-2843-1, Sau-2844-1, Sau-2845-1, Sau-2846-1, Sau-2847-1, Sau-2848-1, Sau-2849-1, Sau-285-1, Sau-2850-1, Sau-2851-1, Sau-2852-1, Sau-2853-1, Sau-2854-1, Sau-2855-1, Sau-2856-1, Sau-2857-1, Sau-2858-1, Sau-2859-1, Sau-286-1, Sau-2860-1, Sau-2861-1, Sau-

2862-1, Sau-2863-1, Sau-2864-1, Sau-2865-1, Sau-2866-1, Sau-2867-1, Sau-2868-1, Sau-2869-1, Sau-287-1, Sau-2870-1, Sau-2871-1, Sau-2872-1, Sau-2873-1, Sau-2874-1, Sau-2875-1, Sau-2876-1, Sau-2877-1, Sau-2878-1, Sau-2879-1, Sau-288-1, Sau-2880-1, Sau-2881-1, Sau-2882-1, Sau-2883-1, Sau-2884-1, Sau-2885-1, Sau-2886-1, Sau-2887-1, Sau-2888-1, Sau-2889-1, Sau-289-1, Sau-2890-1, Sau-2891-1, Sau-2892-1, Sau-2893-1, Sau-2894-1, Sau-2895-1, Sau-2896-1, Sau-2897-1, Sau-2898-1, Sau-2899-1, Sau-29-1, Sau-290-1, Sau-2900-1, Sau-2901-1, Sau-2902-1, Sau-2903-1, Sau-2904-1, Sau-2905-1, Sau-2906-1, Sau-2907-1, Sau-2908-1, Sau-2909-1, Sau-291-1, Sau-2910-1, Sau-2911-1, Sau-2912-1, Sau-2913-1, Sau-2914-1, Sau-2915-1, Sau-2916-1, Sau-2917-1, Sau-2918-1, Sau-2919-1, Sau-292-1, Sau-2920-1, Sau-2921-1, Sau-2922-1, Sau-2923-1, Sau-2924-1, Sau-2925-1, Sau-2926-1, Sau-2927-1, Sau-2928-1, Sau-2929-1, Sau-293-1, Sau-2930-1, Sau-2931-1, Sau-2932-1, Sau-2933-1, Sau-2934-1, Sau-2935-1, Sau-2936-1, Sau-2937-1, Sau-2938-1, Sau-2939-1, Sau-294-1, Sau-2940-1, Sau-2941-1, Sau-2942-1, Sau-2943-1, Sau-2944-1, Sau-2945-1, Sau-2946-1, Sau-2947-1, Sau-2948-1, Sau-2949-1, Sau-295-1, Sau-2950-1, Sau-2951-1, Sau-2952-1, Sau-2953-1, Sau-2954-1, Sau-2955-1, Sau-2956-1, Sau-2957-1, Sau-2958-1, Sau-2959-1, Sau-296-1, Sau-2960-1, Sau-2961-1, Sau-2962-1, Sau-2963-1, Sau-2964-1, Sau-2965-1, Sau-2966-1, Sau-2967-1, Sau-2968-1, Sau-2969-1, Sau-297-1, Sau-2970-1, Sau-2971-1, Sau-2972-1, Sau-2973-1, Sau-2974-1, Sau-2975-1, Sau-2976-1, Sau-2977-1, Sau-2978-1, Sau-2979-1, Sau-298-1, Sau-2980-1, Sau-2981-1, Sau-2982-1, Sau-2983-1, Sau-2984-1, Sau-2985-1, Sau-2986-1, Sau-2987-1, Sau-2988-1, Sau-2989-1, Sau-299-1, Sau-2990-1, Sau-2991-1, Sau-2992-1, Sau-2993-1, Sau-2994-1, Sau-2995-1, Sau-2996-1, Sau-2997-1, Sau-2998-1, Sau-2999-1, Sau-3-1, Sau-30-1, Sau-300-1, Sau-3000-1, Sau-3001-1, Sau-3002-1, Sau-3003-1, Sau-3004-1, Sau-3005-1, Sau-3006-1, Sau-3007-1, Sau-3008-1, Sau-3009-1, Sau-301-1, Sau-3010-1, Sau-3011-1, Sau-3012-1, Sau-3013-1, Sau-3014-1, Sau-3015-1, Sau-3016-1, Sau-3017-1, Sau-3018-1, Sau-3019-1, Sau-302-1, Sau-3020-1, Sau-3021-1, Sau-3022-1, Sau-3023-1, Sau-3024-1, Sau-3025-1, Sau-3026-1, Sau-3027-1, Sau-3028-1, Sau-3029-1, Sau-303-1, Sau-3030-1, Sau-3031-1, Sau-3032-1, Sau-3033-1, Sau-3034-1, Sau-3035-1, Sau-3036-1, Sau-3037-1, Sau-3038-1, Sau-3039-1, Sau-304-1, Sau-3040-1, Sau-3041-1, Sau-3042-1, Sau-3043-1, Sau-3044-1, Sau-3045-1, Sau-3046-1, Sau-3047-1, Sau-3048-1, Sau-3049-1, Sau-3050-1, Sau-3051-1, Sau-3052-1, Sau-3053-1, Sau-3054-1, Sau-3055-1, Sau-3056-1, Sau-3057-1, Sau-3058-1, Sau-3059-1, Sau-306-1, Sau-3060-1, Sau-3061-1, Sau-3062-1, Sau-3063-1, Sau-3064-1, Sau-3065-1, Sau-3066-1, Sau-3067-1, Sau-3068-1, Sau-3069-1, Sau-307-1, Sau-3070-1, Sau-3071-1, Sau-3072-1, Sau-3073-1, Sau-3074-1, Sau-3075-1, Sau-3076-1, Sau-3077-1, Sau-3078-1, Sau-3079-1, Sau-308-1, Sau-3080-1, Sau-3081-1, Sau-3082-1, Sau-3083-1, Sau-3084-1, Sau-3085-1, Sau-3086-1, Sau-3087-1, Sau-3088-1, Sau-3089-1, Sau-309-1, Sau-3090-1, Sau-3091-1, Sau-3092-1, Sau-3093-1, Sau-3094-1, Sau-3095-1, Sau-3096-1, Sau-3097-1, Sau-3098-1, Sau-3099-1, Sau-31-1, Sau-310-1, Sau-3100-1, Sau-3101-1, Sau-3102-1, Sau-3103-1, Sau-3104-1, Sau-3105-1, Sau-3106-1, Sau-3107-1, Sau-3108-1, Sau-3109-1, Sau-311-1, Sau-3110-1, Sau-3111-1, Sau-3112-1, Sau-3113-1, Sau-3114-1, Sau-3115-1, Sau-3116-1, Sau-3117-1, Sau-3118-1, Sau-3119-1, Sau-312-1, Sau-3120-1, Sau-3121-1, Sau-3122-1, Sau-3123-1, Sau-3124-1, Sau-3125-1, Sau-3126-1, Sau-3127-1, Sau-3128-1, Sau-3129-1, Sau-313-1, Sau-3130-1, Sau-3131-1, Sau-3132-1, Sau-3133-1, Sau-3134-1, Sau-3135-1, Sau-3136-1, Sau-3137-1, Sau-3138-1, Sau-3139-1, Sau-314-1, Sau-3140-1, Sau-3141-1, Sau-3142-1, Sau-3143-1, Sau-3144-1, Sau-3145-1, Sau-3146-1, Sau-3147-1, Sau-3148-1, Sau-3149-1, Sau-315-1, Sau-3150-1, Sau-3151-1, Sau-3152-1, Sau-3153-1, Sau-3154-1, Sau-3155-1, Sau-3156-1, Sau-3157-1, Sau-3158-1, Sau-3159-1, Sau-316-1, Sau-3160-1, Sau-3161-1, Sau-3162-1, Sau-3163-1, Sau-3164-1, Sau-3165-1, Sau-3166-1, Sau-3167-1, Sau-3168-1, Sau-3169-1, Sau-317-1, Sau-3170-1, Sau-3171-1, Sau-3172-1, Sau-3173-1, Sau-3174-1, Sau-3175-1, Sau-3176-1, Sau-3177-1, Sau-3178-1, Sau-3179-1, Sau-318-1, Sau-3180-1, Sau-3181-1, Sau-3182-1, Sau-3183-1, Sau-3184-1, Sau-3185-1, Sau-3186-1, Sau-3187-1, Sau-3188-1, Sau-3189-1, Sau-319-1, Sau-3190-1, Sau-3191-1, Sau-3192-1, Sau-3193-1, Sau-3194-1, Sau-3195-1, Sau-3196-1, Sau-3197-1, Sau-3198-1, Sau-3199-1, Sau-32-1, Sau-320-1, Sau-3200-1, Sau-3201-1, Sau-3202-1, Sau-3203-1, Sau-3204-1, Sau-3205-1, Sau-3206-1, Sau-3207-1, Sau-3208-1, Sau-3209-1, Sau-321-1, Sau-3210-1, Sau-3211-1, Sau-3212-1, Sau-3213-1, Sau-3214-1, Sau-3215-1, Sau-3216-1, Sau-3217-1, Sau-3218-1, Sau-3219-1, Sau-322-1, Sau-3220-1, Sau-3221-1, Sau-3222-1, Sau-3223-1, Sau-3224-1, Sau-3225-1, Sau-3226-1, Sau-3227-1, Sau-3228-1, Sau-3229-1, Sau-323-1, Sau-3230-1, Sau-3231-1, Sau-3232-1, Sau-3233-1, Sau-3234-1, Sau-3235-1, Sau-3236-1, Sau-3237-1, Sau-3238-1, Sau-3239-1, Sau-324-1, Sau-3240-1, Sau-3241-1, Sau-3242-1, Sau-3243-1, Sau-3244-1, Sau-3245-1, Sau-3246-1, Sau-3247-1, Sau-3248-1, Sau-3249-1, Sau-325-1, Sau-3250-1, Sau-3251-1, Sau-3252-1, Sau-3253-1, Sau-3254-1, Sau-3255-1, Sau-3256-1, Sau-3257-1, Sau-3258-1, Sau-3259-1, Sau-326-1, Sau-3260-1, Sau-3261-1, Sau-3262-1, Sau-3263-1, Sau-3264-1, Sau-3265-1, Sau-3266-1, Sau-3267-1, Sau-3268-1, Sau-3269-1, Sau-327-1, Sau-3270-1, Sau-3271-1, Sau-3272-1, Sau-3273-1, Sau-3274-1, Sau-3275-1, Sau-3276-1, Sau-3277-1, Sau-3278-1, Sau-3279-1, Sau-328-1, Sau-3280-1, Sau-3281-1, Sau-3282-1, Sau-3283-1, Sau-3284-1, Sau-3285-1, Sau-3286-1, Sau-3287-1, Sau-3288-1, Sau-3289-1, Sau-329-1, Sau-3290-1, Sau-3291-1, Sau-3292-1, Sau-3293-1, Sau-3294-1, Sau-3295-1, Sau-3296-1, Sau-3297-1, Sau-3298-1, Sau-3299-1, Sau-33-1, Sau-330-1, Sau-3300-1, Sau-3301-1, Sau-3302-1, Sau-3303-1, Sau-3304-1, Sau-3305-1, Sau-3306-1, Sau-3307-1, Sau-3308-1, Sau-3309-1, Sau-331-1, Sau-3310-1, Sau-3311-1, Sau-3312-1, Sau-3313-1, Sau-3314-1, Sau-3315-1, Sau-3316-1, Sau-3317-1, Sau-3318-1, Sau-3319-1, Sau-332-1, Sau-3320-1, Sau-3321-1, Sau-3322-1, Sau-3323-1, Sau-3324-1, Sau-3325-1, Sau-3326-1, Sau-3327-1, Sau-3328-1, Sau-3329-1, Sau-333-1, Sau-3330-1, Sau-3331-1, Sau-3332-1, Sau-3333-1, Sau-3334-1, Sau-3335-1, Sau-3336-1, Sau-3337-1, Sau-3338-1, Sau-3339-1, Sau-334-1, Sau-3340-1, Sau-3341-1, Sau-3342-1, Sau-3343-1, Sau-3344-1, Sau-3345-1, Sau-3346-1, Sau-3347-1, Sau-3348-1, Sau-3349-1, Sau-335-1, Sau-3350-1, Sau-3351-1, Sau-3352-1, Sau-3353-1, Sau-3354-1, Sau-3355-1, Sau-3356-1, Sau-3357-1, Sau-3358-1, Sau-3359-1, Sau-336-1, Sau-3360-1, Sau-3361-1, Sau-3362-1, Sau-3363-1, Sau-3364-1, Sau-3365-1, Sau-3366-1, Sau-3367-1, Sau-3368-1, Sau-3369-1, Sau-337-1, Sau-3370-1, Sau-3371-1, Sau-3372-1, Sau-3373-1, Sau-3374-1, Sau-3375-1, Sau-3376-1, Sau-3377-1, Sau-3378-1, Sau-3379-1, Sau-338-1, Sau-3380-1, Sau-3381-1, Sau-3382-1, Sau-3383-1, Sau-3384-1, Sau-3385-1, Sau-3386-1, Sau-3387-1, Sau-3388-1, Sau-3389-1, Sau-339-1, Sau-3390-1, Sau-3391-1, Sau-3392-1, Sau-3393-1, Sau-3394-1, Sau-3395-1, Sau-3396-1, Sau-3397-1, Sau-

3398-1, Sau-3399-1, Sau-34-1, Sau-340-1, Sau-3400-1, Sau-3401-1, Sau-3402-1, Sau-3403-1, Sau-3404-1, Sau-3405-1, Sau-3406-1, Sau-3407-1, Sau-3408-1, Sau-3409-1, Sau-341-1, Sau-3410-1, Sau-3411-1, Sau-3412-1, Sau-3413-1, Sau-3414-1, Sau-3415-1, Sau-3416-1, Sau-3417-1, Sau-3418-1, Sau-3419-1, Sau-342-1, Sau-3420-1, Sau-3421-1, Sau-3422-1, Sau-3423-1, Sau-3424-1, Sau-3425-1, Sau-3426-1, Sau-3427-1, Sau-3428-1, Sau-3429-1, Sau-343-1, Sau-3430-1, Sau-3431-1, Sau-3432-1, Sau-3433-1, Sau-3434-1, Sau-3435-1, Sau-3436-1, Sau-3437-1, Sau-3438-1, Sau-3439-1, Sau-344-1, Sau-3440-1, Sau-3441-1, Sau-3442-1, Sau-3443-1, Sau-3444-1, Sau-3445-1, Sau-3446-1, Sau-3447-1, Sau-3448-1, Sau-3449-1, Sau-345-1, Sau-3450-1, Sau-3451-1, Sau-3452-1, Sau-3453-1, Sau-3454-1, Sau-3455-1, Sau-3456-1, Sau-3457-1, Sau-3458-1, Sau-3459-1, Sau-346-1, Sau-3460-1, Sau-3461-1, Sau-3462-1, Sau-3463-1, Sau-3464-1, Sau-3465-1, Sau-3466-1, Sau-3467-1, Sau-3468-1, Sau-3469-1, Sau-347-1, Sau-3470-1, Sau-3471-1, Sau-3472-1, Sau-3473-1, Sau-3474-1, Sau-3475-1, Sau-3476-1, Sau-3477-1, Sau-3478-1, Sau-3479-1, Sau-348-1, Sau-3480-1, Sau-3481-1, Sau-3482-1, Sau-3483-1, Sau-3484-1, Sau-3485-1, Sau-3486-1, Sau-3487-1, Sau-3488-1, Sau-3489-1, Sau-349-1, Sau-3490-1, Sau-3491-1, Sau-3492-1, Sau-3493-1, Sau-3494-1, Sau-3495-1, Sau-3496-1, Sau-3497-1, Sau-3498-1, Sau-3499-1, Sau-35-1, Sau-350-1, Sau-3500-1, Sau-3501-1, Sau-3502-1, Sau-3503-1, Sau-3504-1, Sau-3505-1, Sau-3506-1, Sau-3507-1, Sau-3508-1, Sau-3509-1, Sau-351-1, Sau-3510-1, Sau-3511-1, Sau-3512-1, Sau-3513-1, Sau-3514-1, Sau-3515-1, Sau-3516-1, Sau-3517-1, Sau-3518-1, Sau-3519-1, Sau-352-1, Sau-3520-1, Sau-3521-1, Sau-3522-1, Sau-3523-1, Sau-3524-1, Sau-3525-1, Sau-3526-1, Sau-3527-1, Sau-3528-1, Sau-3529-1, Sau-353-1, Sau-3530-1, Sau-3531-1, Sau-3532-1, Sau-3533-1, Sau-3534-1, Sau-3535-1, Sau-3536-1, Sau-3537-1, Sau-3538-1, Sau-3539-1, Sau-354-1, Sau-3540-1, Sau-3541-1, Sau-3542-1, Sau-3543-1, Sau-3544-1, Sau-3545-1, Sau-3546-1, Sau-3547-1, Sau-3548-1, Sau-3549-1, Sau-355-1, Sau-3550-1, Sau-3551-1, Sau-3552-1, Sau-3553-1, Sau-3554-1, Sau-3555-1, Sau-3556-1, Sau-3557-1, Sau-3558-1, Sau-3559-1, Sau-356-1, Sau-3560-1, Sau-3561-1, Sau-3562-1, Sau-3563-1, Sau-3564-1, Sau-3565-1, Sau-3566-1, Sau-3567-1, Sau-3568-1, Sau-3569-1, Sau-357-1, Sau-3570-1, Sau-3571-1, Sau-3572-1, Sau-3573-1, Sau-3574-1, Sau-3575-1, Sau-3576-1, Sau-3577-1, Sau-3578-1, Sau-3579-1, Sau-358-1, Sau-3580-1, Sau-3581-1, Sau-3582-1, Sau-3583-1, Sau-3584-1, Sau-3585-1, Sau-3586-1, Sau-3587-1, Sau-3588-1, Sau-3589-1, Sau-359-1, Sau-3590-1, Sau-3591-1, Sau-3592-1, Sau-3593-1, Sau-3594-1, Sau-3595-1, Sau-3596-1, Sau-3597-1, Sau-3598-1, Sau-3599-1, Sau-36-1, Sau-360-1, Sau-3600-1, Sau-3601-1, Sau-3602-1, Sau-3603-1, Sau-3604-1, Sau-3605-1, Sau-3606-1, Sau-3607-1, Sau-3608-1, Sau-3609-1, Sau-361-1, Sau-3610-1, Sau-3611-1, Sau-3612-1, Sau-3613-1, Sau-3614-1, Sau-3615-1, Sau-3616-1, Sau-3617-1, Sau-3618-1, Sau-3619-1, Sau-362-1, Sau-3620-1, Sau-3621-1, Sau-3622-1, Sau-3623-1, Sau-3624-1, Sau-3625-1, Sau-3626-1, Sau-3627-1, Sau-3628-1, Sau-3629-1, Sau-363-1, Sau-3630-1, Sau-3631-1, Sau-3632-1, Sau-3633-1, Sau-3634-1, Sau-3635-1, Sau-3636-1, Sau-3637-1, Sau-3638-1, Sau-3639-1, Sau-364-1, Sau-3640-1, Sau-3641-1, Sau-3642-1, Sau-3643-1, Sau-3644-1, Sau-3645-1, Sau-3646-1, Sau-3647-1, Sau-3648-1, Sau-3649-1, Sau-365-1, Sau-3650-1, Sau-3651-1, Sau-3652-1, Sau-3653-1, Sau-3654-1, Sau-3655-1, Sau-3656-1, Sau-3657-1, Sau-3658-1, Sau-3659-1, Sau-366-1, Sau-3660-1, Sau-3661-1, Sau-3662-1, Sau-3663-1, Sau-3664-1, Sau-3665-1, Sau-3666-1, Sau-3667-1, Sau-3668-1, Sau-3669-1, Sau-367-1, Sau-3670-1, Sau-3671-1, Sau-3672-1, Sau-3673-1, Sau-3674-1, Sau-3675-1, Sau-3676-1, Sau-3677-1, Sau-3678-1, Sau-3679-1, Sau-368-1, Sau-3680-1, Sau-3681-1, Sau-3682-1, Sau-3683-1, Sau-3684-1, Sau-3685-1, Sau-3686-1, Sau-3687-1, Sau-3688-1, Sau-3689-1, Sau-369-1, Sau-3690-1, Sau-3691-1, Sau-3692-1, Sau-3693-1, Sau-3694-1, Sau-3695-1, Sau-3696-1, Sau-3697-1, Sau-3698-1, Sau-3699-1, Sau-37-1, Sau-370-1, Sau-3700-1, Sau-3701-1, Sau-3702-1, Sau-3703-1, Sau-3704-1, Sau-3705-1, Sau-3706-1, Sau-3707-1, Sau-3708-1, Sau-3709-1, Sau-371-1, Sau-3710-1, Sau-3711-1, Sau-3712-1, Sau-3713-1, Sau-3714-1, Sau-3715-1, Sau-3716-1, Sau-3717-1, Sau-3718-1, Sau-3719-1, Sau-372-1, Sau-3720-1, Sau-3721-1, Sau-3722-1, Sau-3722-2, Sau-3722-3, Sau-3722-4, Sau-3722-5, Sau-3722-6, Sau-3723-1, Sau-3724-1, Sau-3725-1, Sau-3726-1, Sau-3727-1, Sau-3728-1, Sau-3729-1, Sau-373-1, Sau-3730-1, Sau-3731-1, Sau-3732-1, Sau-3733-1, Sau-3734-1, Sau-3735-1, Sau-3736-1, Sau-3737-1, Sau-3738-1, Sau-3739-1, Sau-374-1, Sau-3740-1, Sau-3741-1, Sau-3742-1, Sau-3743-1, Sau-3743-2, Sau-3744-1, Sau-3745-1, Sau-3746-1, Sau-3747-1, Sau-3748-1, Sau-3749-1, Sau-375-1, Sau-3750-1, Sau-3751-1, Sau-3752-1, Sau-3753-1, Sau-3754-1, Sau-3755-1, Sau-3756-1, Sau-3757-1, Sau-3758-1, Sau-3759-1, Sau-376-1, Sau-3760-1, Sau-3761-1, Sau-3762-1, Sau-3763-1, Sau-3764-1, Sau-3765-1, Sau-3766-1, Sau-3767-1, Sau-3768-1, Sau-3769-1, Sau-377-1, Sau-3770-1, Sau-3771-1, Sau-3772-1, Sau-3773-1, Sau-3774-1, Sau-3775-1, Sau-3776-1, Sau-3777-1, Sau-3778-1, Sau-3779-1, Sau-378-1, Sau-3780-1, Sau-3781-1, Sau-3782-1, Sau-3783-1, Sau-3784-1, Sau-3785-1, Sau-3786-1, Sau-3787-1, Sau-3788-1, Sau-3789-1, Sau-379-1, Sau-3790-1, Sau-3791-1, Sau-3792-1, Sau-3793-1, Sau-3794-1, Sau-3795-1, Sau-3796-1, Sau-3797-1, Sau-3798-1, Sau-3799-1, Sau-38-1, Sau-380-1, Sau-3800-1, Sau-3801-1, Sau-3802-1, Sau-3803-1, Sau-3804-1, Sau-3805-1, Sau-3806-1, Sau-3807-1, Sau-3808-1, Sau-3809-1, Sau-381-1, Sau-3810-1, Sau-3811-1, Sau-3812-1, Sau-3813-1, Sau-3814-1, Sau-3815-1, Sau-3816-1, Sau-3817-1, Sau-3818-1, Sau-3819-1, Sau-382-1, Sau-3820-1, Sau-3821-1, Sau-3822-1, Sau-3823-1, Sau-3824-1, Sau-3825-1, Sau-3826-1, Sau-3827-1, Sau-3828-1, Sau-3829-1, Sau-383-1, Sau-3830-1, Sau-3831-1, Sau-3832-1, Sau-3833-1, Sau-3834-1, Sau-3835-1, Sau-3836-1, Sau-3837-1, Sau-3838-1, Sau-3839-1, Sau-384-1, Sau-3840-1, Sau-3841-1, Sau-3842-1, Sau-3843-1, Sau-3844-1, Sau-3845-1, Sau-3846-1, Sau-3847-1, Sau-3848-1, Sau-3849-1, Sau-385-1, Sau-3850-1, Sau-3851-1, Sau-3852-1, Sau-3853-1, Sau-3854-1, Sau-3855-1, Sau-3856-1, Sau-3857-1, Sau-3858-1, Sau-3859-1, Sau-386-1, Sau-3860-1, Sau-3861-1, Sau-3862-1, Sau-3863-1, Sau-3864-1, Sau-3865-1, Sau-3866-1, Sau-3867-1, Sau-3868-1, Sau-3869-1, Sau-387-1, Sau-3870-1, Sau-3871-1, Sau-3872-1, Sau-3873-1, Sau-3874-1, Sau-3875-1, Sau-3876-1, Sau-3877-1, Sau-3878-1, Sau-3879-1, Sau-388-1, Sau-3880-1, Sau-3881-1, Sau-3882-1, Sau-3883-1, Sau-3884-1, Sau-3885-1, Sau-3886-1, Sau-3887-1, Sau-3888-1, Sau-3889-1, Sau-389-1, Sau-3890-1, Sau-3891-1, Sau-3892-1, Sau-3893-1, Sau-3894-1, Sau-3895-1, Sau-3896-1, Sau-3897-1, Sau-3898-1, Sau-3899-1, Sau-39-1, Sau-390-1, Sau-3900-1, Sau-3901-1, Sau-3902-1, Sau-3903-1, Sau-3904-1, Sau-3905-1, Sau-3906-1, Sau-3907-1, Sau-3908-1, Sau-3909-1, Sau-391-1, Sau-3910-1, Sau-3911-1, Sau-3912-1, Sau-3913-1, Sau-3914-1, Sau-3915-1, Sau-3916-1, Sau-3917-1, Sau-3918-1, Sau-3919-1, Sau-392-1, Sau-3920-1, Sau-3921-1, Sau-3922-1, Sau-3923-1, Sau-3924-1, Sau-3925-1, Sau-3926-1, Sau-

3927-1, Sau-3928-1, Sau-3929-1, Sau-393-1, Sau-3930-1, Sau-3931-1, Sau-3932-1, Sau-3933-1, Sau-3934-1, Sau-3935-1, Sau-3936-1, Sau-3937-1, Sau-3938-1, Sau-3939-1, Sau-394-1, Sau-3940-1, Sau-3941-1, Sau-3942-1, Sau-3943-1, Sau-3944-1, Sau-3945-1, Sau-3946-1, Sau-3947-1, Sau-3948-1, Sau-3949-1, Sau-395-1, Sau-3950-1, Sau-3951-1, Sau-3952-1, Sau-3953-1, Sau-3954-1, Sau-3955-1, Sau-3956-1, Sau-3957-1, Sau-3958-1, Sau-3959-1, Sau-396-1, Sau-3960-1, Sau-3961-1, Sau-3962-1, Sau-3963-1, Sau-3964-1, Sau-3965-1, Sau-3966-1, Sau-3967-1, Sau-3968-1, Sau-3969-1, Sau-397-1, Sau-3970-1, Sau-3971-1, Sau-3972-1, Sau-3973-1, Sau-3974-1, Sau-3975-1, Sau-3976-1, Sau-3977-1, Sau-3978-1, Sau-3979-1, Sau-398-1, Sau-3980-1, Sau-3981-1, Sau-3982-1, Sau-3983-1, Sau-3984-1, Sau-3985-1, Sau-3986-1, Sau-3987-1, Sau-3988-1, Sau-3989-1, Sau-399-1, Sau-3990-1, Sau-3991-1, Sau-3992-1, Sau-3993-1, Sau-3994-1, Sau-3995-1, Sau-3996-1, Sau-3997-1, Sau-3998-1, Sau-3999-1, Sau-4-1, Sau-40-1, Sau-400-1, Sau-4000-1, Sau-4001-1, Sau-4002-1, Sau-4003-1, Sau-4004-1, Sau-4005-1, Sau-4006-1, Sau-4007-1, Sau-4008-1, Sau-4009-1, Sau-401-1, Sau-4010-1, Sau-4011-1, Sau-4012-1, Sau-4013-1, Sau-4014-1, Sau-4015-1, Sau-4016-1, Sau-4017-1, Sau-4018-1, Sau-4019-1, Sau-402-1, Sau-4020-1, Sau-4021-1, Sau-4022-1, Sau-4023-1, Sau-4024-1, Sau-4025-1, Sau-4026-1, Sau-4027-1, Sau-4028-1, Sau-4029-1, Sau-403-1, Sau-4030-1, Sau-4031-1, Sau-4032-1, Sau-4033-1, Sau-4034-1, Sau-4035-1, Sau-4036-1, Sau-4037-1, Sau-4038-1, Sau-4039-1, Sau-404-1, Sau-4040-1, Sau-4041-1, Sau-4042-1, Sau-4043-1, Sau-4044-1, Sau-4045-1, Sau-4046-1, Sau-4047-1, Sau-4048-1, Sau-4049-1, Sau-405-1, Sau-4050-1, Sau-4051-1, Sau-4052-1, Sau-4053-1, Sau-4054-1, Sau-4055-1, Sau-4056-1, Sau-4057-1, Sau-4059-1, Sau-406-1, Sau-4060-1, Sau-4061-1, Sau-4062-1, Sau-4063-1, Sau-4064-1, Sau-4065-1, Sau-4066-1, Sau-4067-1, Sau-4068-1, Sau-4069-1, Sau-407-1, Sau-4070-1, Sau-4071-1, Sau-4072-1, Sau-4073-1, Sau-4074-1, Sau-4075-1, Sau-4076-1, Sau-4077-1, Sau-4078-1, Sau-4079-1, Sau-408-1, Sau-4080-1, Sau-4081-1, Sau-4082-1, Sau-4083-1, Sau-4084-1, Sau-4085-1, Sau-4086-1, Sau-4087-1, Sau-4088-1, Sau-4089-1, Sau-409-1, Sau-4090-1, Sau-4091-1, Sau-4092-1, Sau-4093-1, Sau-4094-1, Sau-4095-1, Sau-4096-1, Sau-4097-1, Sau-4098-1, Sau-4099-1, Sau-41-1, Sau-410-1, Sau-4100-1, Sau-4101-1, Sau-4102-1, Sau-4103-1, Sau-4103-2, Sau-4103-3, Sau-4103-4, Sau-4103-5, Sau-4103-6, Sau-4103-7, Sau-4103-8, Sau-4104-1, Sau-4105-1, Sau-4106-1, Sau-4107-1, Sau-4108-1, Sau-4109-1, Sau-411-1, Sau-4110-1, Sau-4111-1, Sau-4112-1, Sau-4113-1, Sau-4114-1, Sau-4115-1, Sau-4116-1, Sau-4117-1, Sau-4118-1, Sau-4119-1, Sau-412-1, Sau-4120-1, Sau-4121-1, Sau-413-1, Sau-414-1, Sau-415-1, Sau-416-1, Sau-417-1, Sau-418-1, Sau-419-1, Sau-42-1, Sau-420-1, Sau-421-1, Sau-422-1, Sau-423-1, Sau-424-1, Sau-425-1, Sau-426-1, Sau-427-1, Sau-428-1, Sau-429-1, Sau-43-1, Sau-430-1, Sau-431-1, Sau-432-1, Sau-433-1, Sau-434-1, Sau-435-1, Sau-436-1, Sau-437-1, Sau-438-1, Sau-439-1, Sau-44-1, Sau-440-1, Sau-441-1, Sau-442-1, Sau-443-1, Sau-444-1, Sau-445-1, Sau-446-1, Sau-447-1, Sau-448-1, Sau-449-1, Sau-45-1, Sau-450-1, Sau-451-1, Sau-452-1, Sau-453-1, Sau-454-1, Sau-455-1, Sau-456-1, Sau-457-1, Sau-458-1, Sau-459-1, Sau-46-1, Sau-460-1, Sau-461-1, Sau-462-1, Sau-463-1, Sau-464-1, Sau-465-1, Sau-466-1, Sau-467-1, Sau-468-1, Sau-469-1, Sau-47-1, Sau-470-1, Sau-471-1, Sau-472-1, Sau-473-1, Sau-474-1, Sau-475-1, Sau-476-1, Sau-477-1, Sau-478-1, Sau-479-1, Sau-48-1, Sau-480-1, Sau-481-1, Sau-482-1, Sau-483-1, Sau-484-1, Sau-485-1, Sau-486-1, Sau-487-1, Sau-488-1, Sau-489-1, Sau-49-1, Sau-490-1, Sau-491-1, Sau-492-1, Sau-493-1, Sau-494-1, Sau-495-1, Sau-496-1, Sau-497-1, Sau-498-1, Sau-499-1, Sau-5-1, Sau-50-1, Sau-500-1, Sau-501-1, Sau-502-1, Sau-503-1, Sau-504-1, Sau-505-1, Sau-506-1, Sau-507-1, Sau-508-1, Sau-509-1, Sau-51-1, Sau-510-1, Sau-511-1, Sau-512-1, Sau-513-1, Sau-514-1, Sau-515-1, Sau-516-1, Sau-517-1, Sau-518-1, Sau-519-1, Sau-52-1, Sau-520-1, Sau-521-1, Sau-522-1, Sau-523-1, Sau-524-1, Sau-525-1, Sau-526-1, Sau-527-1, Sau-528-1, Sau-529-1, Sau-53-1, Sau-530-1, Sau-531-1, Sau-532-1, Sau-533-1, Sau-534-1, Sau-535-1, Sau-536-1, Sau-537-1, Sau-538-1, Sau-539-1, Sau-54-1, Sau-540-1, Sau-541-1, Sau-542-1, Sau-543-1, Sau-544-1, Sau-545-1, Sau-546-1, Sau-547-1, Sau-548-1, Sau-549-1, Sau-55-1, Sau-550-1, Sau-551-1, Sau-552-1, Sau-553-1, Sau-554-1, Sau-555-1, Sau-556-1, Sau-557-1, Sau-558-1, Sau-559-1, Sau-56-1, Sau-560-1, Sau-561-1, Sau-562-1, Sau-563-1, Sau-564-1, Sau-565-1, Sau-566-1, Sau-567-1, Sau-568-1, Sau-569-1, Sau-57-1, Sau-570-1, Sau-571-1, Sau-572-1, Sau-573-1, Sau-574-1, Sau-575-1, Sau-576-1, Sau-577-1, Sau-578-1, Sau-579-1, Sau-58-1, Sau-580-1, Sau-581-1, Sau-582-1, Sau-583-1, Sau-584-1, Sau-585-1, Sau-586-1, Sau-587-1, Sau-588-1, Sau-589-1, Sau-59-1, Sau-59-2, Sau-590-1, Sau-591-1, Sau-592-1, Sau-593-1, Sau-594-1, Sau-595-1, Sau-596-1, Sau-597-1, Sau-598-1, Sau-599-1, Sau-6-1, Sau-60-1, Sau-600-1, Sau-601-1, Sau-602-1, Sau-603-1, Sau-604-1, Sau-605-1, Sau-606-1, Sau-607-1, Sau-608-1, Sau-609-1, Sau-61-1, Sau-610-1, Sau-611-1, Sau-612-1, Sau-613-1, Sau-614-1, Sau-615-1, Sau-616-1, Sau-617-1, Sau-618-1, Sau-619-1, Sau-62-1, Sau-620-1, Sau-621-1, Sau-622-1, Sau-623-1, Sau-624-1, Sau-625-1, Sau-626-1, Sau-627-1, Sau-628-1, Sau-629-1, Sau-63-1, Sau-630-1, Sau-631-1, Sau-632-1, Sau-633-1, Sau-634-1, Sau-635-1, Sau-636-1, Sau-637-1, Sau-638-1, Sau-639-1, Sau-64-1, Sau-640-1, Sau-641-1, Sau-642-1, Sau-643-1, Sau-644-1, Sau-645-1, Sau-646-1, Sau-647-1, Sau-648-1, Sau-649-1, Sau-65-1, Sau-650-1, Sau-651-1, Sau-652-1, Sau-653-1, Sau-654-1, Sau-655-1, Sau-656-1, Sau-657-1, Sau-658-1, Sau-659-1, Sau-66-1, Sau-660-1, Sau-661-1, Sau-662-1, Sau-663-1, Sau-664-1, Sau-665-1, Sau-666-1, Sau-667-1, Sau-668-1, Sau-669-1, Sau-67-1, Sau-670-1, Sau-671-1, Sau-672-1, Sau-673-1, Sau-674-1, Sau-675-1, Sau-676-1, Sau-677-1, Sau-678-1, Sau-679-1, Sau-68-1, Sau-680-1, Sau-681-1, Sau-682-1, Sau-683-1, Sau-684-1, Sau-685-1, Sau-686-1, Sau-687-1, Sau-688-1, Sau-689-1, Sau-69-1, Sau-690-1, Sau-691-1, Sau-692-1, Sau-693-1, Sau-694-1, Sau-695-1, Sau-696-1, Sau-697-1, Sau-698-1, Sau-699-1, Sau-7-1, Sau-70-1, Sau-700-1, Sau-701-1, Sau-702-1, Sau-703-1, Sau-704-1, Sau-705-1, Sau-706-1, Sau-707-1, Sau-708-1, Sau-709-1, Sau-71-1, Sau-710-1, Sau-711-1, Sau-712-1, Sau-713-1, Sau-714-1, Sau-715-1, Sau-716-1, Sau-717-1, Sau-718-1, Sau-719-1, Sau-72-1, Sau-720-1, Sau-721-1, Sau-722-1, Sau-723-1, Sau-724-1, Sau-725-1, Sau-726-1, Sau-727-1, Sau-728-1, Sau-729-1, Sau-73-1, Sau-730-1, Sau-731-1, Sau-732-1, Sau-733-1, Sau-734-1, Sau-735-1, Sau-736-1, Sau-737-1, Sau-738-1, Sau-739-1, Sau-74-1, Sau-740-1, Sau-741-1, Sau-742-1, Sau-743-1, Sau-744-1, Sau-745-1, Sau-746-1, Sau-747-1, Sau-748-1, Sau-749-1, Sau-75-1, Sau-750-1, Sau-751-1, Sau-752-1, Sau-753-1, Sau-754-1, Sau-755-1, Sau-756-1, Sau-757-1, Sau-758-1, Sau-759-1, Sau-76-1, Sau-760-1, Sau-761-1, Sau-762-1, Sau-763-1, Sau-764-1, Sau-765-1, Sau-766-1, Sau-767-1, Sau-768-1, Sau-769-1, Sau-77-1, Sau-770-1, Sau-771-1, Sau-772-1, Sau-773-1, Sau-774-1, Sau-775-1, Sau-776-1, Sau-777-1, Sau-778-1, Sau-779-1, Sau-78-1, Sau-780-1, Sau-781-1, Sau-782-1, Sau-783-1,

Sau-784-1, Sau-785-1, Sau-786-1, Sau-787-1, Sau-788-1, Sau-789-1, Sau-79-1, Sau-79-2, Sau-790-1, Sau-791-1, Sau-792-1, Sau-793-1, Sau-794-1, Sau-795-1, Sau-796-1, Sau-797-1, Sau-798-1, Sau-799-1, Sau-8-1, Sau-80-1, Sau-800-1, Sau-801-1, Sau-802-1, Sau-803-1, Sau-804-1, Sau-805-1, Sau-806-1, Sau-807-1, Sau-808-1, Sau-809-1, Sau-81-1, Sau-810-1, Sau-811-1, Sau-812-1, Sau-813-1, Sau-814-1, Sau-815-1, Sau-816-1, Sau-817-1, Sau-818-1, Sau-819-1, Sau-82-1, Sau-820-1, Sau-821-1, Sau-822-1, Sau-823-1, Sau-824-1, Sau-825-1, Sau-826-1, Sau-827-1, Sau-828-1, Sau-829-1, Sau-83-1, Sau-830-1, Sau-831-1, Sau-832-1, Sau-833-1, Sau-834-1, Sau-835-1, Sau-836-1, Sau-837-1, Sau-838-1, Sau-839-1, Sau-84-1, Sau-840-1, Sau-841-1, Sau-842-1, Sau-843-1, Sau-844-1, Sau-845-1, Sau-846-1, Sau-847-1, Sau-848-1, Sau-849-1, Sau-85-1, Sau-850-1, Sau-851-1, Sau-852-1, Sau-853-1, Sau-854-1, Sau-855-1, Sau-856-1, Sau-857-1, Sau-858-1, Sau-859-1, Sau-86-1, Sau-860-1, Sau-861-1, Sau-862-1, Sau-863-1, Sau-864-1, Sau-865-1, Sau-866-1, Sau-867-1, Sau-868-1, Sau-869-1, Sau-87-1, Sau-870-1, Sau-871-1, Sau-872-1, Sau-873-1, Sau-874-1, Sau-875-1, Sau-876-1, Sau-877-1, Sau-878-1, Sau-879-1, Sau-88-1, Sau-880-1, Sau-881-1, Sau-882-1, Sau-883-1, Sau-884-1, Sau-885-1, Sau-886-1, Sau-887-1, Sau-888-1, Sau-889-1, Sau-89-1, Sau-890-1, Sau-891-1, Sau-892-1, Sau-893-1, Sau-894-1, Sau-895-1, Sau-896-1, Sau-897-1, Sau-898-1, Sau-899-1, Sau-9-1, Sau-90-1, Sau-900-1, Sau-901-1, Sau-902-1, Sau-903-1, Sau-904-1, Sau-905-1, Sau-906-1, Sau-907-1, Sau-908-1, Sau-909-1, Sau-91-1, Sau-910-1, Sau-911-1, Sau-912-1, Sau-913-1, Sau-914-1, Sau-915-1, Sau-916-1, Sau-917-1, Sau-918-1, Sau-919-1, Sau-92-1, Sau-920-1, Sau-921-1, Sau-922-1, Sau-923-1, Sau-924-1, Sau-925-1, Sau-926-1, Sau-927-1, Sau-928-1, Sau-929-1, Sau-93-1, Sau-930-1, Sau-931-1, Sau-932-1, Sau-933-1, Sau-934-1, Sau-935-1, Sau-936-1, Sau-937-1, Sau-938-1, Sau-939-1, Sau-94-1, Sau-940-1, Sau-941-1, Sau-942-1, Sau-943-1, Sau-944-1, Sau-945-1, Sau-946-1, Sau-947-1, Sau-948-1, Sau-949-1, Sau-95-1, Sau-950-1, Sau-951-1, Sau-952-1, Sau-953-1, Sau-954-1, Sau-955-1, Sau-956-1, Sau-957-1, Sau-958-1, Sau-959-1, Sau-96-1, Sau-960-1, Sau-961-1, Sau-962-1, Sau-963-1, Sau-964-1, Sau-965-1, Sau-966-1, Sau-967-1, Sau-968-1, Sau-969-1, Sau-97-1, Sau-970-1, Sau-971-1, Sau-972-1, Sau-973-1, Sau-974-1, Sau-975-1, Sau-976-1, Sau-977-1, Sau-978-1, Sau-979-1, Sau-98-1, Sau-980-1, Sau-981-1, Sau-982-1, Sau-983-1, Sau-984-1, Sau-985-1, Sau-986-1, Sau-987-1, Sau-988-1, Sau-989-1, Sau-99-1, Sau-990-1, Sau-991-1, Sau-992-1, Sau-993-1, Sau-994-1, Sau-995-1, Sau-996-1, Sau-997-1, Sau-998-1, Sau-999-1, Sca-1-1, Sca-10-1, Sca-11-1, Sca-2-1, Sca-3-1, Sca-5-10, Sca-5-11, Sca-5-12, Sca-5-13, Sca-5-14, Sca-5-15, Sca-5-16, Sca-5-17, Sca-5-18, Sca-5-19, Sca-5-2, Sca-5-20, Sca-5-21, Sca-5-3, Sca-5-4, Sca-5-5, Sca-5-6, Sca-5-7, Sca-5-8, Sca-5-9, Sca-6-1, Sca-9-1, Sco-2-1, Sco-2-3, Sep-1-1, Sep-10-1, Sep-11-1, Sep-12-1, Sep-13-1, Sep-14-1, Sep-15-1, Sep-16-1, Sep-17-1, Sep-18-1, Sep-19-1, Sep-2-1, Sep-20-1, Sep-21-1, Sep-22-1, Sep-23-1, Sep-25-1, Sep-26-1, Sep-27-1, Sep-28-1, Sep-29-1, Sep-3-1, Sep-30-1, Sep-31-1, Sep-32-1, Sep-33-1, Sep-34-1, Sep-35-1, Sep-36-1, Sep-37-1, Sep-38-1, Sep-39-1, Sep-4-1, Sep-40-1, Sep-41-1, Sep-42-1, Sep-43-1, Sep-44-1, Sep-45-1, Sep-46-1, Sep-47-1, Sep-48-1, Sep-49-1, Sep-5-1, Sep-50-1, Sep-51-1, Sep-52-1, Sep-53-1, Sep-54-1, Sep-55-1, Sep-56-1, Sep-57-1, Sep-58-1, Sep-59-1, Sep-6-1, Sep-60-1, Sep-61-1, Sep-62-1, Sep-63-1, Sep-64-1, Sep-65-1, Sep-66-1, Sep-67-1, Sep-68-1, Sep-69-1, Sep-7-1, Sep-70-1, Sep-71-1, Sep-72-1, Sep-73-1, Sep-74-1, Sep-75-1, Sep-76-1, Sep-77-1, Sep-78-1, Sep-79-1, Sep-8-2, Sep-8-3, Sep-80-1, Sep-81-1, Sep-82-1, Sep-83-1, Sep-84-1, Sep-85-1, Sep-86-1, Sep-87-1, Sep-88-1, Sep-9-1, Seq-1-1, Seq-3-1, Sha-1-1, Sha-2-1, Sha-4-10, Sha-4-100, Sha-4-101, Sha-4-102, Sha-4-103, Sha-4-104, Sha-4-105, Sha-4-106, Sha-4-107, Sha-4-108, Sha-4-109, Sha-4-11, Sha-4-110, Sha-4-111, Sha-4-112, Sha-4-113, Sha-4-114, Sha-4-115, Sha-4-116, Sha-4-119, Sha-4-12, Sha-4-13, Sha-4-14, Sha-4-15, Sha-4-16, Sha-4-17, Sha-4-18, Sha-4-19, Sha-4-2, Sha-4-20, Sha-4-21, Sha-4-22, Sha-4-23, Sha-4-24, Sha-4-25, Sha-4-26, Sha-4-27, Sha-4-28, Sha-4-29, Sha-4-3, Sha-4-30, Sha-4-31, Sha-4-32, Sha-4-33, Sha-4-34, Sha-4-35, Sha-4-36, Sha-4-37, Sha-4-38, Sha-4-39, Sha-4-4, Sha-4-40, Sha-4-41, Sha-4-42, Sha-4-43, Sha-4-44, Sha-4-45, Sha-4-46, Sha-4-47, Sha-4-48, Sha-4-49, Sha-4-50, Sha-4-51, Sha-4-52, Sha-4-53, Sha-4-54, Sha-4-55, Sha-4-56, Sha-4-57, Sha-4-58, Sha-4-59, Sha-4-6, Sha-4-60, Sha-4-61, Sha-4-62, Sha-4-63, Sha-4-64, Sha-4-65, Sha-4-66, Sha-4-67, Sha-4-68, Sha-4-69, Sha-4-7, Sha-4-70, Sha-4-71, Sha-4-72, Sha-4-73, Sha-4-74, Sha-4-75, Sha-4-76, Sha-4-77, Sha-4-78, Sha-4-79, Sha-4-8, Sha-4-80, Sha-4-81, Sha-4-82, Sha-4-83, Sha-4-84, Sha-4-85, Sha-4-86, Sha-4-87, Sha-4-88, Sha-4-89, Sha-4-9, Sha-4-90, Sha-4-91, Sha-4-92, Sha-4-93, Sha-4-94, Sha-4-95, Sha-4-96, Sha-4-97, Sha-4-98, Sha-4-99, Sha-5-1, Sha-6-1, Sho-1-1, Sho-2-1, Sho-3-10, Sho-3-11, Sho-3-12, Sho-3-13, Sho-3-14, Sho-3-16, Sho-3-17, Sho-3-3, Sho-3-4, Sho-3-5, Sho-3-6, Sho-3-7, Sho-3-8, Sho-3-9, Sho-4-1, Sho-5-1, Slu-2-1, Slu-4-1, Slu-5-1, Slu-6-1, Slu-7-1, Slu-8-1, Slu-9-1, Sma-2-1, Spe-2-1, Spn-1-1, Spn-1-2, Spn-1-3, Spn-1-4, Spn-1-5, Spn-1-6, Spn-1-7, Spn-1-8, Sps-1-1, Sps-3-1, Ssa-1-1, Ssa-2-2, Ssa-2-3, Ssa-3-1, Ssc-1-2, Ssc-1-3, Ssc-1-4, Ssc-2-2, Ssc-2-3, Ssc-3-1, Ssc-4-1, Ssi-4-1, Ssp-11-1, Ssp-12-1, Ssp-13-1, Ssp-15-1, Ssp-2-1, Ssp-4-1, Ssp-7-1, Ssp-8-1, Ssp-9-1, Ste-1-1, Svi-1-1, Swa-2-1, Swa-2-2, Swa-2-4, Swa-2-5, Swa-3-1, Swa-4-1, Swa-5-1, Swa-6-1, Swa-7-1, Sxy-2-1, Tmu-1-1, Tmu-3-1, Tsp-1-1, Tsp-2-1, Vma-1-1, Wci-1-2, Wci-1-3, Wci-2-1, Wko-1-1, Xca-1-1, Xca-2-1, Xca-3-1, Xca-4-1, Xca-5-1, Xva-1-1, Xva-2-1, bLF-1-1, env-100, env-1000, env-101, env-1012, env-1016, env-1019, env-102, env-1020, env-1021, env-103, env-1030, env-1035, env-1037, env-104, env-1042, env-1046, env-1047, env-105, env-106, env-1063, env-1065, env-1068, env-1069, env-107, env-1070, env-1071, env-108, env-1083, env-1085, env-1086, env-1087, env-1089, env-109, env-1090, env-1096, env-1099, env-11, env-110, env-1100, env-111, env-1111, env-112, env-1123, env-1126, env-1127, env-1128, env-113, env-1130, env-1131, env-1136, env-114, env-1140, env-1141, env-1142, env-1143, env-1144, env-1145, env-1147, env-1148, env-1149, env-115, env-1150, env-1151, env-1152, env-1153, env-1154, env-1155, env-1156, env-1157, env-1158, env-1159, env-116, env-1160, env-1161, env-1162, env-1163, env-1167, env-117, env-1173, env-1174, env-1176, env-118, env-1180, env-1181, env-1183, env-1185, env-119, env-1192, env-1193, env-1194, env-1195, env-1196, env-12, env-120, env-121, env-122, env-1223, env-1226, env-1227, env-1229, env-123, env-124, env-125, env-126, env-127, env-128, env-1289, env-129, env-130, env-131, env-1313, env-1314, env-1315, env-1316, env-1317, env-1318, env-1319, env-132, env-1320, env-1321, env-1322, env-1323, env-1324, env-1325, env-1326, env-1327, env-1328, env-1329, env-133, env-1330, env-1331, env-1332, env-1333, env-1334, env-1335, env-1336, env-1338, env-1339, env-134, env-1340, env-1341, env-1343, env-1344, env-1345, env-1346, env-1347, env-1348, env-1349, env-135, env-1350, env-1351, env-1352, env-1353, env-1354, env-1355, env-1356, env-1357, env-1358, env-1359, env-136, env-1363, env-1364, env-1365, env-1366, env-1367, env-1368, env-1369, env-137, env-1370, env-1371, env-1372, env-1373, env-1374, env-1375, env-1376, env-1377, env-1378, env-138, env-1380, env-1382, env-1386, env-139, env-1392, env-1394, env-1396, env-14, env-140, env-1400, env-1401, env-1403, env-141, env-1410, env-1411, env-1412, env-1417, env-1418, env-142, env-1422, env-1425, env-1426, env-1427, env-1428, env-1429, env-143, env-1430, env-1431, env-1432, env-1433, env-1434, env-1435, env-1436, env-1437, env-144, env-1440, env-1441, env-1442, env-1447, env-1448, env-145, env-1454, env-1456, env-1457, env-1458, env-146, env-1463, env-1464, env-1465, env-1466, env-1467, env-1468, env-1469, env-147, env-1470, env-1473, env-1474, env-1475, env-148, env-1480, env-1483, env-1485, env-149, env-1490, env-1492, env-1494, env-1496, env-1497, env-1498, env-1499, env-150, env-1500, env-1501, env-1502, env-1503, env-1504, env-1505, env-1506, env-1507, env-151, env-1513, env-152, env-153, env-154, env-155, env-156, env-157, env-158, env-159, env-160, env-161, env-162, env-163, env-164, env-165, env-166, env-167, env-168, env-169, env-170, env-171, env-172, env-173, env-174, env-175, env-176, env-177, env-178, env-179, env-180, env-181, env-182, env-183, env-184, env-187, env-188, env-189, env-190, env-191, env-192, env-193, env-194, env-195, env-196, env-197, env-198, env-199, env-200, env-201, env-202, env-203, env-204, env-205, env-206, env-207, env-208, env-21, env-210, env-211, env-212, env-217, env-22, env-221, env-222, env-223, env-231, env-232, env-235, env-236, env-24, env-243, env-248, env-252, env-255, env-256, env-257, env-259, env-26, env-260, env-261, env-263, env-264, env-265, env-266, env-27, env-271, env-272, env-273, env-274, env-275, env-276, env-277, env-278, env-279, env-281, env-282, env-283, env-285, env-286, env-29, env-295, env-296, env-298, env-30, env-300, env-304, env-305, env-306, env-307, env-308, env-309, env-31, env-310, env-311, env-312, env-313, env-314, env-315, env-316, env-317, env-318, env-319, env-32, env-320, env-321, env-322, env-323, env-324, env-325, env-326, env-327, env-328, env-329, env-33, env-330, env-331, env-332, env-333, env-334, env-335, env-336, env-337, env-338, env-339, env-34, env-340, env-341, env-342, env-343, env-344, env-345, env-346, env-347, env-348, env-349, env-35, env-350, env-351, env-352, env-353, env-354, env-355, env-356, env-357, env-358, env-359, env-36, env-360, env-361, env-362, env-363, env-364, env-365, env-366, env-367, env-368, env-369, env-37, env-370, env-371, env-372, env-373, env-374, env-375, env-376, env-377, env-378, env-379, env-38, env-380, env-381, env-382, env-383, env-384, env-385, env-386, env-387, env-388, env-389, env-39, env-390, env-391, env-392, env-393, env-394, env-396, env-397, env-398, env-399, env-40, env-400, env-401, env-402, env-403, env-404, env-405, env-406, env-407, env-408, env-409, env-41, env-410, env-411, env-412, env-413, env-414, env-415, env-417, env-418, env-419, env-42, env-420, env-421, env-422, env-423, env-424, env-425, env-426, env-427, env-428, env-429, env-43, env-430, env-431, env-432, env-433, env-434, env-435, env-436, env-437, env-438, env-439, env-44, env-440, env-441, env-442, env-443, env-444, env-445, env-446, env-447, env-448, env-449, env-45, env-450, env-451, env-452, env-453, env-454, env-455, env-456, env-457, env-458, env-459, env-46, env-460, env-461, env-462, env-463, env-464, env-465, env-466, env-468, env-469, env-47, env-470, env-471, env-472, env-473, env-474, env-475, env-476, env-477, env-478, env-48, env-484, env-485, env-486, env-488, env-489, env-49, env-493, env-496, env-497, env-498, env-50, env-500, env-503, env-504, env-505, env-509, env-51, env-510, env-511, env-512, env-52, env-521, env-528, env-53, env-530, env-531, env-534, env-537, env-539, env-54, env-540, env-541, env-542, env-543, env-544, env-545, env-546, env-547, env-55, env-56, env-560, env-561, env-562, env-563, env-564, env-565, env-567, env-568, env-569, env-57, env-570, env-571, env-572, env-573, env-574, env-575, env-58, env-583, env-585, env-586, env-587, env-589, env-59, env-590, env-591, env-592, env-593, env-594, env-595, env-60, env-602, env-603, env-61, env-62, env-621, env-622, env-627, env-629, env-63, env-637, env-638, env-639, env-64, env-640, env-641, env-642, env-643, env-644, env-645, env-646, env-647, env-649, env-65, env-651, env-652, env-653, env-654, env-658, env-66, env-661, env-666, env-667, env-668, env-669, env-67, env-670, env-671, env-672, env-673, env-674, env-675, env-677, env-68, env-687, env-69, env-690, env-693, env-694, env-695, env-696, env-697, env-698, env-699, env-70, env-700, env-701, env-702, env-703, env-705, env-706, env-708, env-709, env-71, env-711, env-713, env-714, env-715, env-716, env-717, env-718, env-719, env-72, env-724, env-725, env-726, env-727, env-729, env-730, env-731, env-732, env-733, env-734, env-735, env-736, env-737, env-738, env-739, env-74, env-740, env-741, env-742, env-743, env-744, env-745, env-746, env-747, env-748, env-749, env-75, env-750, env-751, env-752, env-753, env-754, env-755, env-76, env-77, env-772, env-773, env-774, env-775, env-776, env-78, env-783, env-784, env-785, env-786, env-787, env-788, env-789, env-79, env-790, env-791, env-792, env-796, env-797, env-80, env-801, env-803, env-81, env-810, env-813, env-82, env-822, env-825, env-83, env-830, env-833, env-834, env-835, env-836, env-837, env-84, env-85, env-854, env-856, env-859, env-86, env-865, env-868, env-87, env-872, env-873, env-877, env-88, env-884, env-885, env-886, env-889, env-89, env-895, env-896, env-897, env-90, env-905, env-907, env-91, env-911, env-912, env-915, env-917, env-918, env-92, env-926, env-927, env-928, env-929, env-93, env-931, env-94, env-940, env-947, env-95, env-953, env-956, env-957, env-96, env-963, env-964, env-965, env-966, env-967, env-969, env-97, env-970, env-971, env-972, env-973, env-974, env-975, env-976, env-977, env-978, env-979, env-98, env-980, env-981, env-982, env-983, env-984, env-985, env-986, env-987, env-988, env-989, env-99, env-991, env-992, env-993, env-994, env-995, env-996, env-999

alignment positions 1 · · · 175

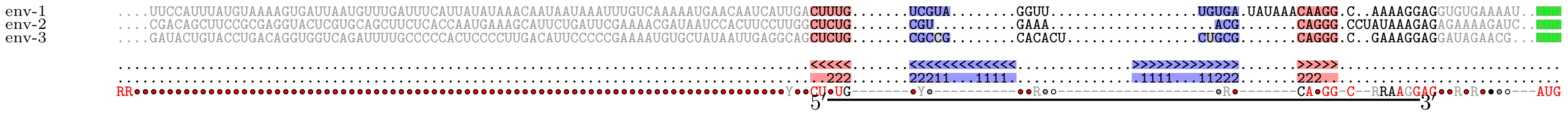

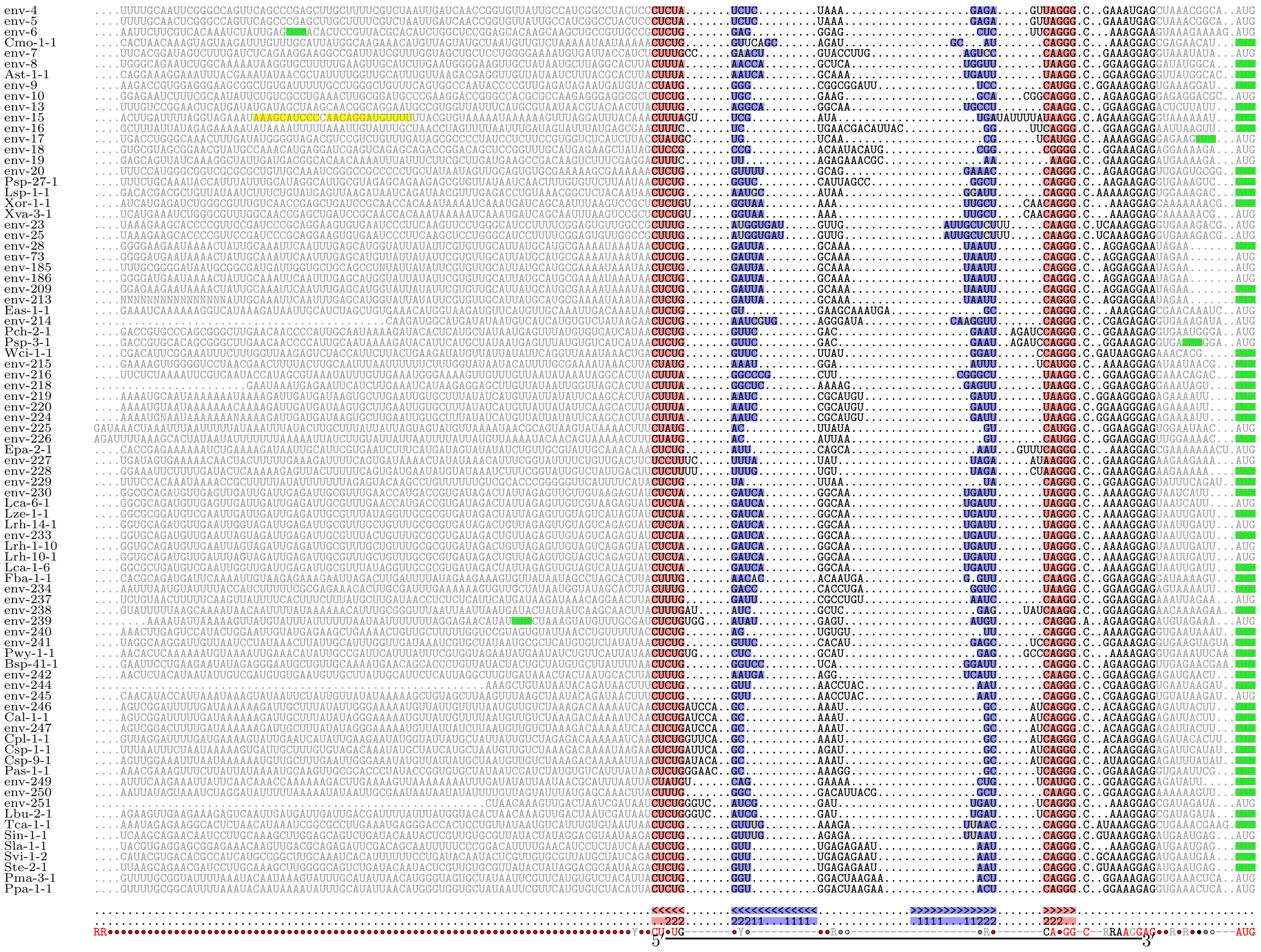
env-4
env-5
env-6
Cmo-1-1
env-7
env-8
Ast-1-1
env-9
env-10
env-13
env-15
env-16
env-17
env-18
env-19
env-20
Psp-27-1
Lsp-1-1
Xor-1-1
Xva-3-1
env-23
env-25
env-28
env-73
env-185
env-186
env-209
env-213
Eas-1-1
env-214
Pch-2-1
Psp-3-1
Wci-1-1
env-215
env-216
env-218
env-219
env-220
env-224
env-225
env-226
Epa-2-1
env-227
env-228
env-229
env-230
Lca-6-1
Lze-1-1
Lrh-14-1
env-233
Lrh-1-10
Lrh-10-1
Lca-1-6
Fba-1-1
env-234
env-237
env-238
env-239
env-240
env-241
Pwy-1-1
Bsp-41-1
env-242
env-244
env-245
env-246
Cal-1-1
env-247
Cpl-1-1
Csp-1-1
Csp-9-1
Pas-1-1
env-249
env-250
env-251
Lbu-2-1
Tca-1-1
Sin-1-1
Sla-1-1
Svi-1-2
Ste-2-1
Pma-3-1
Ppa-1-1
5′
3′

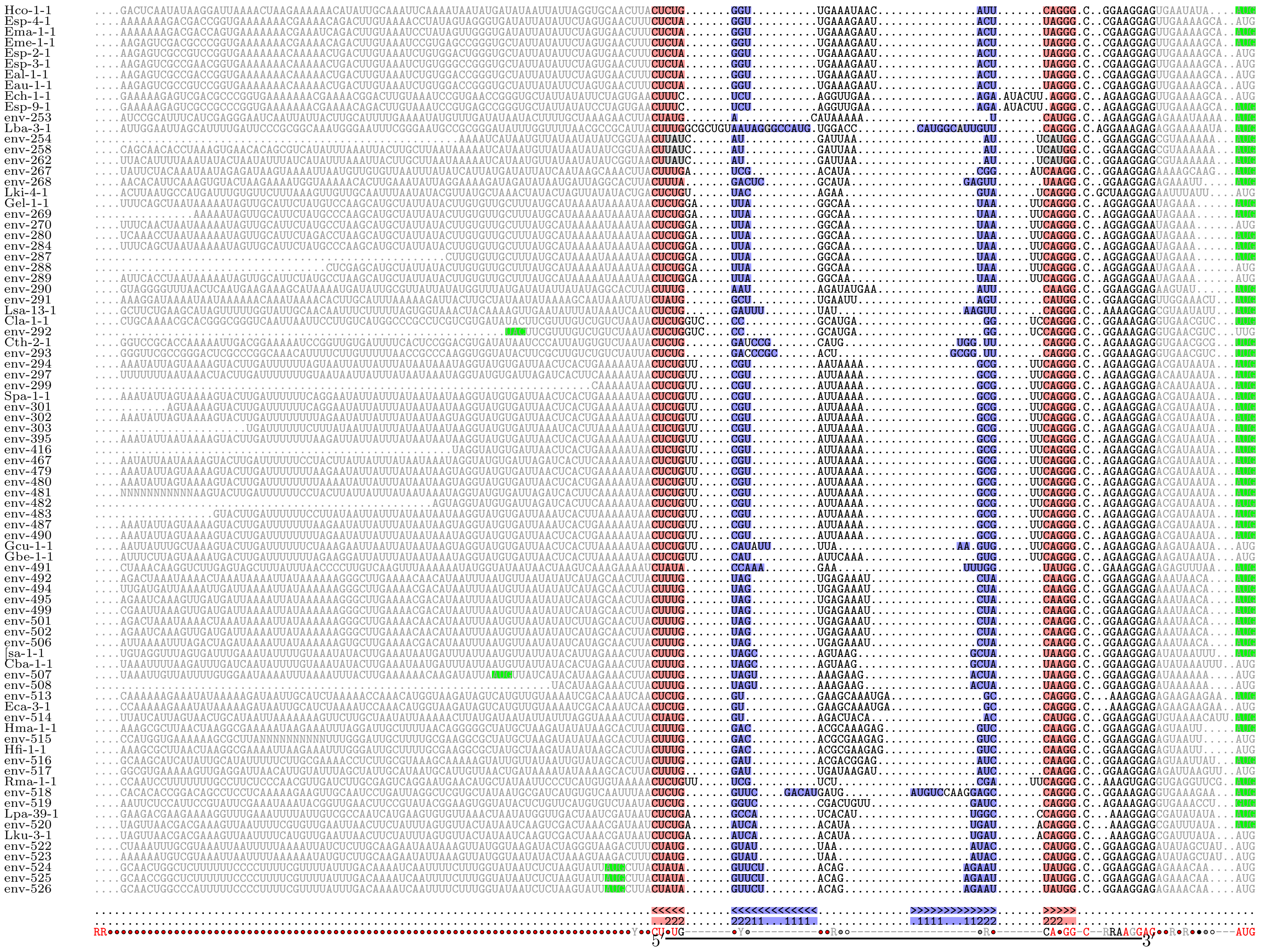

Hco-1-1
Esp-4-1
Ema-1-1
Eme-1-1
Esp-2-1
Esp-3-1
Eal-1-1
Eau-1-1
Ech-1-1
Esp-9-1
env-253
Lba-3-1
env-254
env-258
env-262
env-267
env-268
Lki-4-1
Gel-1-1
env-269
env-270
env-280
env-284
env-287
env-288
env-289
env-290
env-291
Lsa-13-1
Cla-1-1
env-292
Cth-2-1
env-293
env-294
env-297
env-299
Spa-1-1
env-301
env-302
env-303
env-395
env-416
env-467
env-479
env-480
env-481
env-482
env-483
env-487
env-490
Gcu-1-1
Gbe-1-1
env-491
env-492
env-494
env-495
env-499
env-501
env-502
env-506
Isa-1-1
Cba-1-1
env-507
env-508
env-513
Eca-3-1
env-514
Hma-1-1
env-515
Hfi-1-1
env-516
env-517
Rma-1-1
env-518
env-519
Lpa-39-1
env-520
Lku-3-1
env-522
env-523
env-524
env-525
env-526
5′
3′

```
env-527    ....ACCUGUCCCUGCGGUUCCGUGGCUCGGAGAACCAGCGGCUCAUUGACGUUCCAGCUUCUCUGGCUGCGGGAAAAACCAUUCUCUG......GCA.........AGAC..................UGG.......CAGGG.A..GAAAAGAGAGGAAUGAA...AUG
env-529    ....ACCUGUCCCUGCGGUUCCGUGGCUCGGAGAACCAGCGGCUCAUUGACGUUCGGCGUCUCUGGCCGCGGGAAAAACCAUUCUCUG......GCA.........GGAC..................UGG.......CAGGG.A..GAAAAGAGAGGAAUGAA...AUG
env-532    ....ACCUGUCCCUGCGGUUCCGUGGCUCGGAGAACCAGCGGCUCAUUGACGUUCGGCUUCUCUGGCCGCGGGAAAAACUAUUCUCUG......GCA.........GGAC..................UGG.......CAGGG.A..GAAAAGAGAGGAAUGAA...AUG
env-533    ....AUCUGUCCCUGCGGUUCCGUGGCUCGGAGAACCAGCGGCUCAUUGACGUUCGGCUUCUCUGGCCGCGGGAAAAACCAUUCUCUG......GCA.........GGAC..................UGG.......CAGGG.A..GAAAAGAGAGGAAUGAA...AUG
env-535    ....ACCUGUCCCUGCGGUUCCGUGGCUCGGAGAACCAGCGGCUCAUUGACGUUCGGCGUCUCUGGCCGAGGGAAAAACCAUUCUCUG......GCA.........GGAC..................UGG.......CAGGG.A..GAAAAGAGAGGAAUGAA...AUG
env-536    ....ACCUGUCCCUGCGGUUCGUGGCUCGGAGAACCAGCGGCUCAUUGACGUUCGGCGUCUCUGGCCGCGGGAAAAACCAUUCUCUG......GCA.........GGAC..................UGG.......CAGGG.A..GAAAAGAGAGGAAUGAA...AUG
env-538    ....ACCUGUCCCUGCGGUUCCGUGGCUCGGAGAACCAGCGGCUCAUUGACGUUCGGCUUCUCUGGCCGCGGGAAAAACCAUUCUCUG......GCA.........GGAC..................UGG.......CAGGG.A..GAAAAGAGAGGAAUGAA...AUG
Fso-1-1    ....CCUGAAAUUCAUCGAUAAAGUCUUUGCAUUCUCUAACGAGUCAUGGUAUUAUUCUAUUGUUGCAUUUCAUGCAUGAAUAACUGUGUU....CCA.........GCCUA.................UGG.....UUCACGG.C.AGAAAGGAAGUAUAAAU...AUG
env-548    ....GGGCUAAGAUAAAUUUAAAAGAAAUUUAGUAACAUACUUGAAAAUUAAUUAAUUUUUUGAUAAUAUAUUCUGGUAUUUAUCUAUG......GACU........UAUUUU................GGUU.......CAUGG.C...AAAGGAGGGUAAAAUU...AUG
env-549    ....GAACUAAGAUAAAUUUAAAAGAAAUUUAGUAACAUACUUGAAAAUUAAUUAAUUUUUUGAUAAUAUAUUCUGGUAUUUAUCUAUG......GACU........UAUUUU................GGUU.......CAUGG.C...AAAGGAGGGUAAAAUU...AUG
env-550    ..............CCUACAAUUUUUUUCUUGCUUUUUAAAAUGCUAUUUGAUAUAAUACCUUUGCGAUAUAGUAUGAACUUACUAUGA.....UU..........GAAAUGA..................AA.....AUCAUGG.C..AGAAGGAGAAAGAU......AUG
Psp-21-1   ....AGUUCACCCAGUAAAUUCUGCAAGAGAUCCGCUUGCUAUGCCAAUAUAAUCGUGAUAUAAUUCGUCCAGUAUGCAUAUAACUCUG......GUCU........GCGGGCUUAAG............GGAU.......CAGGG.C..GGAAAGAGGUGAAAGAUU..AUG
env-551    ....CUUUGUCUACUCAUACGUGUGUAUAAAAAAGUAUUUGAUAAACAUAAUUGUAUUAUGGUAAUAUCUAUCUCGCAAACUUUCUUUG......GAU.........AAAGUAAC..............AUC.......CAAGG.C..GGAAGGAGAAACAAA......AUG
env-552    ....UUUGGGGUUUCUUCAUAUGCGUGUAAAAAGUAUUUGAUAAACAUAAUUGUAUUAUGAUAUAUCUAUCUCGCAAACUUUCUUUG......GAU.........AAAGUAAC..............AUC.......CAAGG.C..GGAAGGAGAAAAAU......AUG
env-553    ....UUAUAAAUAAAUAAGUUUUUUCAAAUAAAUUAAUCUUUACUAUUAAUAAAGAUUUUUGUUAUUAUCAUUAAGCAAACUUUCUUUG......GGCG........AAGUAC...............UGCU.......CAAGG.C..GGAAGGAGAAACAAAAUUU.AUG
env-554    ....UUUAAAACCUAUCUUUACAAUUAGCCUUACUUUUUACUUGCGAAAGCAAGUUCCUCGCGUAUGAUUAGACCCGAACUCUCUUUG......GCA.........AAAGUAAG..............UGC.......CAAGG.C..GAAAGGAGUAAAGAA......AUG
env-555    ....CUCAAUACUUAUCUUUACAAUUAGCCUUACUUUUUGCUUGCGAAAGCAAGUUCCUCGCGUAUGAUUAGACGCGAACUCUCUUUG......GCA.........AAAGUAAG..............UGC.......CAAGG.C..GAAAGGAGUAAAAAA......AUG
Ssp-3-1    ....AUUCAUAUACGACUGACAUACACGGGUCGUGUAAGGCAUCUGUCUUUUACUGGGAAGGCGAAAGCCAACCAGAAUAAAAACUCUGU.....ACC.........AGAU..................GGU.....UCAGGG.C..GAGAGGAGAGAGACCG.....AUG
Ssp-5-1    ....AUUCAUAUACGACUGACAUACACGGGUCGUGUAAGGCAUCUGUCUUUUACUGGGAAGGCGCAAGCCAACCAGAAUAAAAACUCUGU.....ACC.........AGAC..................GGU.....UCAGGG.C..GAGAGGAGAGAGACCG.....AUG
env-556    ....AUAAGUCGGAACAAUACCAUAUUCUGUGACAUAUGCGACUGGCAUUGACGGGUCGUGUAAGGUGACAGUUAAAUAACCAACUCUGU.....GCC.........AGAU..................GGU.....UCAGGG.C..GAGAGGAGAUUUAAG......AUG
env-557    ....UAUAAGUCGGUAGAUUUACCAUAUCUGUGACAUAUUCGACUGAAUUGAACGGGUCGUGUAAGGUGACAGUCUAAUAACCACUCUGU.....ACC.........AGAU..................GGU.....CCAGGG.C..GAGAGGAGAUUUAAG......AUG
env-558    ....GCGUGAGCAGGUCGACUGAAAAAGGCCUCAAACACAGCUGCUUGCAUUGACAUUUUGCCUAUGGUACUAUAGUAAGGCACUUACUUUGGA....UC..........GAUCGGA.................GA.....CCCAAGG.C..GGAAGGAGAAAAAAGAA..AUG
env-559    ....CGACAUUCGGAAAAUUCUUUGGUUAAGAGUCUACCAUUCUUACUGAAGAUAUGUUAUUAUAUUCAGGUUAAAUAAACUGACUCUGGU....UCU.........UAU..................AGA.....UCCAGGG.C.GAUAAGGAGAAACACG.....AUG
Whe-1-1    ....UAAAAUAAAAUGAUAGCUUUUAGUGAAAAACUAUCAUUUUAUAAAAUUUUAUGUUAAUAUAGUUUAGUUAAAUAAACUGACUCUGAU....UUU.........UAU..................AAA.....UUCAGGG.C.GAUAAGGAGAAAAUA......AUG
1 env-566  ...........................................................AUAUUAAACACGCACUUACUUUG......AAUU........GAUCAGA.................AAUU.......CAAGG.C..GGAAGGAGAAAAUGUU...AUG
env-576    ....CUGACGGUGAUCUCCUGGAGCCGAUCGAGGUCGCGCAGGAUAUCGGCCCAUUCCGGCGCGAGCCCGGCGCGGCGGAGCUUCUCUGUCGAGGGCCUG........AUG..................CAGG.....AUCAGGG.CGAGGAAGGAGCAGAAGAC....AUG
env-577    ....UAAUUUAAUGCCUGGAACCAUUGAUAAAAUAUGUUGCAACGUAAAUCAUGAUAUGAUAAACUAUACUAGUUAUAUACUGACUCUGUU....AC..........AGAA..................GU.....UUCAGGG.C.GCUAAGGAGAAAUAGUU....AUG
Lca-25-1   ....UUGAACUCGGUAAUCAAACUUUUGAUUAAAAACUUGUCAUGCAAUAUAUUGUAUGUCAAAAUAUACUAGUUAUAAUCUGACUCUGUU....AC..........AGAA..................GU.....UUCAGGG.C.GCUAAGGAGAAAUUAUU....AUG
env-578    ....UGAACUCGGUAAUCAAACUUUUGNUUAAAAACUUAGUCAUGCAAUAUAUUGUAUGUCAAAAUAUACUAGUUAUAAUCUGACUCUGUU....AC..........AGAA..................GU.....UUCAGGG.C.GCUAAGGAGAAAUUAUU....AUG
env-579    ..................AAUUGACAGCUUUUUUUGUCUUGAUUAUAUUGUAUUUCUGUGUUAAAGUUAGUAAGCACUUACUUUG......GAUU........CGCAUGA.................AAUC.......CAAGG.C..GGAAGGAGAAAAAC......AUG
Psa-1-1    ....UCGGGUCAAGGUUCGACUAAAAUACAAGCGAUUGCGUCAAGAGAUGGGCCGUGCUAUAAUUCGUUAGUGUAUCAUUAUAACUCUGU.....GCAC........GAUUGA.................G.GC.....UCAGGG.C..GGAAAGAGGUGAAAGUG....AUG
env-580    ....AUUUUAAAGUAAUACUAGAAAUCUACUAAUUCAUUGUUGAUUUAAACUACGUUUUAUGUAUACUAAUUAAGCAAGCUUUCUUUG......AACUA.......UAGG..................UAGUU......CAAGG.C..GGAAGGAGUUAAAUUU....AUG
env-581    ....CAUACGCCGGAUAGCCUUCGUAGAAAGCAGUUGCAAUCGAAAUAAAGCUCAUGAUAUAAUGCCUUCAUGUGUCCAUAUAACUCUG......GUUC...GACG.UAA..............UGUCAAUGGGAC......CAGGG.C..GGAAAGAGGUGAAAGAU...AUG
env-582    ....GAAGAAAAAUAAAUAAAAAUUGUAAAAUUAACUUGCAUAAAAGUAAAAUCUAUGGUAACAUAAAUCGGUAAAAAACUUACUAUA......GUU.........ACCUAC.................AAC.......UAUGG.C..GGAAGGAGAUUAAAGACU..AUG
env-584    ....AAAAAGAAGAAAUAAAUAAAUUUGUAAAUAAACUUGCACAAAAAUAAAAACUAUGGUAUUAUAAAUCGGUAAAAACUUACUAUA......GUU.........ACCUAC.................AAC.......UAUGG.C..GGAAGGAGAUUAAAGACU..AUG
env-588    ....AAAAAGAAAAAAUAAAUAAAAUUUGUAAAAUAAACUUGCACAAAAAUAAAAACUAUGGUAUUAUAAAUCGGUAAAAACUUACUAUA......GUU.........ACCUAC.................AAC.......UAUGG.C..GGAAGGAGAUUAAAGACU..AUG
Cko-1-1    ....ACGAAAAAAAUAAUCGGGUUCAGACACCAUUGUUGCUAUAACGCCCUGCCGCGUGAUAUAAUCCGUUUAUGUGUCCAUUACUCUG......AAUCCG......CCUG................UUGAUU......CAGGG.C..AGAAAGAGGUGAACCCU...AUG
env-596    ....GGUAAAGUUUAGCGGCCUUGCCGGAGGCAUUGCUAUUCCGGCAUGCGGCGUGAUAUAAUCCGUUUAUGUGCGUAUUUUAACUCUG......GGCU.CG.....CAA................CGUAGCC......CAGGG.C..GGAAAGAGGUGAACGAG...AUG
env-597    ....GGGCCGCGAAGCGGGCUGUUGCGAAGACUAUUGCGCUUUCGAUUGCAUCCGUGAUAUAAUCCGCCUAUGUGUCGCAAUUUAACUCUG......CGUC........CGCAUGC...............GAUG.......CAGGG.C..AGAAAGAGGUGAACGAG...AUG
env-598    ....................................GGCACGGGAUGAAGCCCGUGUUAUAAUCGCUUUAUGUGUCAGUCUAUACUCUG......GAUU........CGAAGGC...............AAUC.......CAGGG.C..GGAAAGAGGUGAAUCUC...AUG
Pta-1-1    ....UUCUAGUGCUUAUGGAUGAAUGCUAAUAUGACUUGCGCCACGAGGCAAGCUUGUGUUAUACUGCACAUUGUUCCAUAUAACUCUG......GGCAC.......GCAAA..............GGGCU.......CAGGG.C..CGAAAGAGGUGAAUUCG...AUG
Psp-11-1   ....GCGGACAAGAAUAUGGUUAUAACAUGAGAUUGCGAACUAGAUGCAAACAUGCUAUAAUCCAUUCAUGUGUUUUUAUAUACUCUG......GGCAC.......GCAAA..............GGGCU.......CAGGG.C..GGAAAGAGGUGAAUUCGA..GUG
env-599    ..............AUAUGGUUAUAACAUGAGAUUGCGAACUAGAUGCAAACAUGCUAUAAUCCAUUCAUGUGUUUUUAUAUACUCUG......GGCAC.......GCAAA..............GGGCU.......CAGGG.C..GGAAAGAGGUGAAUUCGA..AUG
Pal-1-1    ....GCGGACAAGAAUAUGGUUAUAACAUGAGAUUGCGAACUAGAUGCAAACAUGCUAUAAUCCAUUCAUGCGUUUUUAUAUACUCUG......GGCAC.......GCAAA..............GGGCU.......CAGGG.C..GGAAAGAGGUGAAUUCA...AUG
Pal-5-1    ....GCGGACAAGCAUAUGGUUAUAACAUGAGAUUGCGAACUAGAUGCAAACAUGCUAUAAUCCAUUCAUGUGUUUUUAUAUACUCUG......GGCAC.......GCAAA..............GGGCU.......CAGGG.C..GGAAAGAGGUGAAUUCGA..GUG
Pte-3-1    ....UAUAAGGACGAUGAAGCCGUUUCUUUUUGUAUUGCAGUCCAUGUGACAACAUGCUAUAAUGCAUCUAUUGUGUCAAUAUACUCUG......GGGUC.......GCAAA..............GAGCC.......CAGGG.C..GGAAAGAGGUGAAUUCG...AUG
env-600    ....UAUAAGGACGAUGAAGCCGUUUCAUUUUGUAUUGCAGUCCAUGUGACAACAUGCUAUAAUGCAUCUAUUGUGUCAAUAUACUCUG......GGGUC.......GCAAA..............GAGCC.......CAGGG.C..GGAAAGAGGUGAAUUCG...AUG
Psp-32-1   ....GUUCCAACUCGCUAAGGUGCAGGCUAUGCUAUUGCGCCAGGCGACGCGUUCGUGUUAUAAUCGAACUCGUGUCCAUUGAACUCUG......GAUUC.......GCAAA..............GAGUC.......CAGGG.C..GGAAAGAGGUGAACGAA...AUG
env-601    ....GGAUGACGGCCGGAAAAAUCGCGGCACGGGCGGGUUGGCACGGAGCUUCCUUCGUGCUAUAAUCGUCAAGUGUCCACAAUACUCUG......GAUUC.......GCAGG..............GAAUC.......CAGGG.C..GGAAAGAGGUGAAGGGCC..AUG
env-604    ....UUCGGUAACAUUUUAGCUGAGUGACUGCGGCGGGUUGGCACGGAGCUUCCUUCGUGCUAUAAUCGUUCAGUGUCCAUAAUACUCUG......GAUUC.......GCGGG..............GAAUC.......CAGGG.C..GGAAAGAGGUGAACGGCC..AUG
env-605    ....CGAUCCGCGUCUGGUCGGGUAACCGCGACGGGUUGGCACGGGGCUUCCUUCGUGCUAUAAUCGUUCAGUGUCCAUAAUACUCUG......GAUUC.......GCGUG..............GAAUC.......CAGGG.C..GGAAAGAGGUGAACGGCC..AUG
env-606    ....CUUCGCCGCUUGCAAUGCGGAUAAACGAUGGUUGCAUCGGGGGUUGGUGCGUGCUAUAAUGUCUUCAUGUGUCCAAUUUUACUCUG......GGCGC.......AAG...............GCGCU.......CAGGG.C..GGAAAGAGGUGAACGAUCG.AUG
env-607    ....CGGAAAUGAAGUGCCAAACAAAUAGAAACAUUGCGCCAAUCGGGCAGACUGUGCUAUAAUCCGCUUAUGUGUCUAGAUAACUCUG......GGCCG.......CAGAUCGUCA...........UGGCU.......CAGGG.C..GGAAAGAGGUGAUGAGGCUU.AUG
env-608    ..........................................................CGUUUUACUGUGUCAAUUUACUCUGGGCU....GCC.........AACAAA.................GGC.....UCAGGG.C..GGAAAGAGGUGAAAGAC...AUG
Pan-2-1    ....CGAAGGAUGCAAAAUUUCCUGAGAUAGACAAUUGCGCAGGGAAUAUCGCCGUGCUAUAAUCAUUUUAUGUGUCAUGAUAACUCUGU.....GCC.C.......GCAU..............GAGGC......CCAGGG.C..GGAAAGAGGUGAAAGUC...AUG
env-609    ....GAAGGCCGUCCGGGCAGCGGCAAUUUGAUGUUGCCAGCGGAUCGGCCGACGUGAUAUAAUGCUUUAUGUGUCCAUAAAACUCUG......GCCCG.......ACGGCGAC...........UGGGC.......CAGGG.C..GGAAAGAGGUGAAAGGG...AUG
Pte-2-1    ....UUACGCCAUUUUUGGAUACACAAAAGAGUGUUGAAACGCUCAUAUUCAGGUGCUAUAAUGUGAACGUGUGUUCAUAUAACUCUG......GGC.........CGAUAGGAC.............GCU.......CAGGG.C..GGAAAGAGGUGAAAGGUUCA.AUG
Psp-7-1    ....UACACCAGUUUUAAGAUGCGCAAAACAGGUAUUGAAACACUUAUAUUCAAGUGCUAUAAUGUGAACAUGUGUUCAUACAACUCUG......GAC.........UGAUAGGAC.............GUU.......CAGGG.C..GGAAAGAGGUGAAAGUUA..AUG
Psp-1-1    ....UUCACGUCAGUUUGAGUGCGCAAAACAGGUAUUGAAACACUUAUAUUCAAGUGCUAUAAUUUGAACAUGUGUUCAUACAACUCUG......GAC.........UGAUAGGAC.............GUU.......CAGGG.C..GGAAAGAGGUGAAAGUUA..AUG
Ppe-1-1    ....UUUACGUCGGUUUGGGUGCGCAAAACAGGUAUUGAAACACUUAUAUUCAAGUGCUAUAAUUUGAACAUGUGUUCAUACAACUCUG......GAC.........UGAUAGGAC.............GUU.......CAGGG.C..GGAAAGAGGUGAAAGUUA..AUG
Ppo-3-1    ....AUGUUACACUGUUUUGUGCACUAAAAAGAUGUUGAAACACUUAUAUUCAAGUGCUAUAAUUUGAACAUGUGUUCAUACAACUCUG......GGC.........UGAUAGGAC.............GUU.......CAGGG.C..GGAAAGAGGUGAAAGUUA..AUG
Ppo-4-1    ....AUCUUACACUGUUUUGUGUGCUAAAAAGAUGUUGAAACACUUAUAUUCAAGUGCUAUAAUUUGAACAUGUGUUCAUACAACUCUG......GGC.........UGAUAGGAC.............GUU.......CAGGG.C..GGAAAGAGGUGAAAGUUA..AUG
Ppo-1-1    ....AUCUUACACUGUUUUGUGUACUAAAAAGAUGUUGAAACACUUAUAUUCAAGUGCUAUAAUUUGAACAUGUGUUCAUACAACUCUG......GGC.........UGAUAGGAC.............GUU.......CAGGG.C..GGAAAGAGGUGAAAGUUA..AUG
Psp-8-1    ....UUACGCCAUUUUUGGAUACACAAAAGAGUGUUGAAACGCUCAUAUCCAGGUGCUAUAAUGUGAACGUGUGUUCAUACAACUCUG......GGC.........UGAUAGGAC.............GUU.......CAGGG.C..GGAAAGAGGUGAAAGUUCA.AUG
Psp-30-1   ....UUAUGCCAUCUUUGAAUGCAGAAAAGAGUGUUGAAACGCUCAUAUUCAAGUGCUAUAAUGUGAACAUGUGUUCAUACAACUCUG......GAC.........UGAUAGGAC.............GUU.......CAGGG.C..GGAAAGAGGUGAAAGUUCA.AUG
Ppo-6-1    ....UUACGCUAUCUUUGAAUGCGGAAAAGAGGUGUUGAAACGCUCAUAUUCAAGUGCUAUAAUGUGAACAUGUGUUCAUACAACUCUG......GAC.........UGAUAGGAC.............GUU.......CAGGG.C..GGAAAGAGGUGAAAGUUCA.AUG
Ppo-10-1   ....UUACGCCACCUUUGAAUGCAGAAAAGAGUGUUGAAACGCUCAUAUUCAAGUGCUAUAAUGUGAACAUGUGUUCAUACAACUCUG......GAC.........UGAUAGGAC.............GUU.......CAGGG.C..GGAAAGAGGUGAAAGUUCA.AUG
env-610    ........................................................GAUGAUCAGAGCGCGGGCUCUAUG......GU..........CAG...................GU.......CAUGG.C...GAAAGAGGGGAAAGU....AUG
Ppi-1-1    ....AGUCUUUUAAGAAAGAACAAAUUCAAUAAUGAUUGAUUAUUAUAUUUGUCCCUGUUAUAAUUAGAAGUGUAUCGAUAUAACUCUG......GGC.........ACGCAUGAG.............GUU.......CAGGG.C..AAAAAGAGGUGAAAGUC...AUG
Psp-24-1   ....GGCGCGGGACUAACAGAUUCUAACAUUUCGUUGAUUAUUGCGUUUGUCCCUGUUAUAAUAUAAGUAUGUGUCAUUAUAACUCUG......UGU.........ACGCAUGAG.............ACU.......CAGGG.C..AAAAAGAGGUGAAAGAC...AUG
Psp-19-1   ....AUUAUUACCAUAUAAACAAUAAUGGAUUAUGAUUGCUUAUUGAGAAUCAUCCUGCUAUAAUCGUGUUAUGUCCAAUUAACUCUG......UGU.........ACACAUGAG.............ACA.......CAGGG.C..GGAAAGAGGUGAAAGUC...AUG
env-611    ....CCGCGUCAAAGUGCGCGGAUAAAACAGGCAAUUGCGUGAACGGAUGGCCGUGCUAUAAUGCGAUUGUGUGUCAUUAUAACUCUG......GGC.........ACGACUGAG.............GCU.......CAGGG.C..GGAAAGAGAUGGAAAGUG...AUG
env-612    ....AAAACGGCUUGAUUACCUCUUAACAAUAAUGGAUUGCAACGGGCACAAGGCCGUGUUAUAAUCUCAGAGUUAUGCCAUUUACUCUG......GGUC........CGCUUGA..............GGCU.......CAGGG.C..GGAAAGAGGUGAAAGGUA..AUG
env-613    ....AAACGUCUGACCACCUCAUGACAAUAAUGGAUUGCAACCGGCAUUAGGCCGUGUUAUAAUCUUCGAUGUUGUGCCAUUUACUCUG......GGUC........CGCUUGA..............GGCU.......CAGGG.C..GGAAAGAGGUGAAUGGUA..AUG
Psp-33-1   ....AAAACGGUUGAUUACCUCUUAACAAUAAUGGAUUGCAACAGGCGCAAGGUCGUGUUAUAAUCUCAGGGUUAUGCCAUUUACUCUG......GGUC........CGCUUGA..............GGCU.......CAGGG.C..GGAAAGAGGUGAAAGGUA..AUG
Psp-26-1   ....AAAACGGUUGAUUACCUCUUGAUAAUAAUGGAUUGCAACCGGCAUUAGGCCGUGUUAUAAUUUCUGAGUUAUGCCAUUUACUCUG......GGUC........CGCUUGA..............GGCU.......CAGGG.C..GGAAAGAGGUGAAAGGUA..AUG
Psp-23-1   ....AAACGCCUGAUCACUUCUUGACAAUAAUGGAUUGCAACCGGCAUUAGGCCGUGUUAUAAUCUUCGAUGUUGUGCCAUUUACUCUG......GGUC........CGCUUGA..............GGCU.......CAGGG.C..GGAAAGAGGUGAAAGGUA..AUG
Psp-2-1    ....CGGACUUGUCUGAUGAUUUGUGUAUGGACUAUUGCGCAUUCAAUAUGCUCGUGCUAUAUCUCUUUUAUGUGUCCAUAUAACUCUG......GGUC........CGCAUGA..............GGCU.......CAGGG.C..GGAAAGAGGUGAAAGCUACAAUG
Pvo-1-1    ....UGAACUUGUCUGAUGUUUUGUGUAUGGGCUAUUGCGCAAUCAAUAUGCUCGUGAUAUACUCGGUUUAUGUGUCCAUAUAACUCUG......GGUC........CGCAUGA..............GGCU.......CAGGG.C..GGAAAGAGGUGAAAGCUACAAUG
Gsp-5-1    ....CGGACUUGUCUGAUGAUUUGUGUAUGGACUAUUGCGCAUUCAAUAUGCUCGUGCUAUACUCCUUUUAUGUGUCCAUAUAACUCUG......GGUC........CGCAUGA..............GGCU.......CAGGG.C..GGAAAGAGGUGAAAGCUACAAUG
Psp-10-1   ....GGGCGUCCCUUGUUGUUUCCUCAUAGGCUAUUGCACAAUUAAAAAUGCCGUGAUAUACUUUUUUUAUGUGUCCAUAUAACUCUG......AGUCC.......GCAUGA..............GG.CU.......CAGGG.C..GGAAAGAGGUGAAAGCUA..AUG
Bsp-52-1   ....UGGGCUCGGCUGAGUAUUUGUGUGCGGGCUAUUGCGCAAUGAAAAAUGCCGUGAUAUACUGAUUUUAUGUGUCCAUAUAACUCUG......GGUCC.......GCAUGA..............GG.CU.......CAGGG.C..GGAAAGAGGUGAAAGGUAUAAUG
Pla-1-1    ....UGGGCUUGGCUGAGUAUUUGUGUGCGGGCUAUUGCGCAAUGAAAAAUGCCGUGAUAUACUGAUUUUAUGUGUCCAUAUAACUCUG......GGUCC.......GCAUGA..............GG.CU.......CAGGG.C..GGAAAGAGGUGAAAGGUAUAAUG

           .....................................................................................&lt;&lt;&lt;&lt;&lt;......&lt;&lt;&lt;&lt;&lt;&lt;&lt;&lt;&lt;&lt;&lt;&lt;&lt;.............&gt;&gt;&gt;&gt;&gt;&gt;&gt;&gt;&gt;&gt;&gt;&gt;&gt;.......&gt;&gt;&gt;&gt;&gt;..............................
           ......................................................................................222.......22211...1111..............1111...11222.......222...............................
           RR••••••••••••••••••••••••••••••••••••••••••••••••••••••••••••••••••••••••••••••Y••CU•UG-------•Y•-----------••R∘∘------------------∘R•-------CA•GG-C--RRAAGGAG••R•R••∘∘---AUG
                                                                                             5'                                                                                  3'
```

```
env-614   ..............................................................................UCAUUACUCUG.......GGUCC........GCUUGA................GG.CU.......CAGGG.C..GGAAAGAGGUGAAAGGAAA.AUG
Pst-1-1   ....ACGGCGGGGCUGAUUUGUCAAGGAAAAGUAUUGCCUCAGUCCAAGGUUCGUGAUAAACUACCUCAUGUGUCUUAUAUUAACUCUG.......GGUCC........GCAUGA................GG.CU.......CAGGG.C...AAAAGAGGUGAAAUCAAA.AUG
Psp-20-1  ....CUCGGGUUAAGGUACGAGGAUAAAACAGGCAAUUGCGCGAAAGGUUGCCCCAUGCUAUAAUGCGUUUGUGUGUCAUAUAACUCUG.......GGCAC........GACUGA................G.GCU.......CAGGG.C..GGAAAGAGGUGAAAGAAG..AUG
env-615   ......................................................CAUGCUAUAAUGCGUUUGUGUGUCAUAUAACUCUG.......GGCAC........GACUGA................G.GCU.......CAGGG.C..GGAAAGAGGUGAAAGAAG..AUG
Pba-1-1   ....CCCGCGUCAAGGUGCGAGGAUAAAACAGGCAAUUGCGCGAAAGGUUGCCCCGUGCUAUAAUGCGUUUGUGUGUCAUAUAACUCUG.......GGCAC........GACUGA................G.GCU.......CAGGG.C..GGAAAGAGGUGAAAGAAG..AUG
Psp-5-1   ....CCGUGUCAAAGUGGGCGGAUAAAACAGGCAAUUGCGUGAACGAGGGGCCCGUGCUAUAAUGCGAUUAUGUGUCAUUAUAACUCUG.......GGCAC........GACUGA................G.GCU.......CAGGG.C..GGAAAGAGGUGAAAGAAG..AUG
Pma-2-1   ....AAAGCCAAGGUGCAUGGAUAAAACAGGUAAUUGCGCUGGCGGUAUUAGCCGUGCUAUAAUGCUUUUAUGUGUCAUUAUAACUCUG.......GGCAC........GAUCUGA...............G.GCU.......CAGGG.C..GGAAAGAGGUGAAAGUG...AUG
Pbe-1-1   ....UAAGGGAGAACAGCAGGCGACGAAUUCAUAUUGCAACAGAGUGCGGCGCGUGCUAUACUUGAUCUACAUGUGUAUUAUAACUCUG.......GGUC.........CGCCUGUU...............GACU.......CAGGG.C..AGAAAGAGGUGAACGAG...AUG
Pha-2-1   ....AUUCAAAACGAUCGAUAUGCAUGAUUAUUCAUUGCAACGGGAAAGAGGUUCAUGCUAUAAUCCGUAGAGUGUGAUUUUAACUCUG.......GGUC.........CGCAUGC................GAUU.......CAGGG.C..AGAAAGAGGUGAACGUG...AUG
Ppi-2-1   ....GUAAGACCCUUUUUAACGGCGUGCAAAAAGCAUUGCAACUGCUUGCCUUCCUAUGUUAUAAUCCCCUAUGUUAGAUUGAACUCUG.......GGUC.........CAUCAGUC...............GGCU.......CAGGG.C..AGAAAGAGGUGAACGAG...AUG
env-616   ....GAGGCGUUCCGGCGCCGGAAAUGAAUUGACCGUUGCGCGCUUUCGCGCACUCGUGCUAUAAUCUCAUCAUGUGUCAUUUACUCUG.......GGCC.........GGCGCUUCAUG............GGCC.......CAGGG.C.GUGAAAGAGGUGAACGAG...AUG
env-617   ....UAUCGGGCGCCGCGUCCGUCCGCAUGAAGCCGUUGCGUUCAGACCGGCCGGCGUGAUAUAAUCGUUUCAUGUGUCAAUAACUCUG.......GCCCG........GCGUUCA...............UGGGC......ACAGGG.C.GUGAAAGAGGUGAACGAG...AUG
env-618   ....CGGGUGCCGCGCCGUUCCGAAUGAAGCCGUUGCACCGGGGCAAGCCGGCGUGUUAUAAUCACUUCAUGUGUGUCAAUAAACUCUG.......GCCCG........CAUUUCA...............UGGGC......ACAGGG.C.GCGAAAGAGGUGAACGAG...AUG
Psp-12-1  ....ACAUCUAAUAUUGACAUAUGAGCGAUUAUUCAUUGCAACUGCAAAACAGUUCAUGCUAUAAUCCGAGAUGUGUGAUUUAACUCUG.......GGUC.........CGCAUGC................GAUU.......CAGGG.C..AGAAAGAGGUGAACGAG...AUG
env-619   ....AUAGAGGACCGGAAAUUGACGGAACAGACCGUUGCGCGAUUUUGCGCAUUCGUGCUAUAAUCCUUUCAUGUGUCAUUUAACUCUG.......GGCCU........GCUGCAA...............GGGCU.......CAGGG.C.GUGAAAGAGGUGAACGAG...AUG
env-620   ....GCCGGGAAGGAAUGACGGACUGGAAAGACCAUUGCGCGAGUCUGACCAUCCGUGCUAUAAUCCUCUCAUGUGUCAUUUAACUCUG.......GGCCUG.......CAUUA................UGGGCU.......CAGGG.C.GUGAAAGAGGUGAACGAG...AUG
Tco-1-1   ....GCCGGCCGGGAUUGACGUGCCGGAAAGACCGUUGCGCGAGUUUGGCUAUCCGUGCUAUAAUCCUCUCAUGUGUCAUUUAACUCUG.......GGCCU........GCUGUCUAA.............GGGCU.......CAGGG.C.GUGAAAGAGGUGAACGAG...AUG
env-623   ..................................GUUGCGCGAUUUUGCGCGUUCGUGCUAUAAUCCUUUCAUGUGUCAUUUAACUCUG.......GGCCU........GCCAUCCAA.............GGGCU.......CAGGG.C.GUGAAAGAGGUGAACGAG...AUG
Pal-3-1   ....UUCCGGUAUGGGACAGCGCCUUCUUUUGUGUUUUACAACAAGGACUUGGCUAUGCUAUAAUGCAGAGAUGUAUGACUUAAUUCUG.......GGCC.C.......GCCU.................GUGGCU.......CAGGG.C..AGAAAGAGGUGAAAGAG...AUG
Pgo-1-1   ....AGAUAGUUGGCGGAAUUGGAAAUGACAAUGGCAUUGCAAGGCGGAUAGUGUCAUGCUAUAAUCCGUAUGUGUAAUUUUAACUCUG.......GGUC.C.......GCAU.................GUGGCU.......CAGGG.C..AGAAAGAGGUGAACGAG...AUG
Psp-4-1   ....CUGCACGCAAGUAGCUGGCAAAACAUCGGCAUUGCAUGCUUGUAGUCGUCGUGCUAUAAUCCGUGUUGUGUGAUUUAUAACUCUG.......GACC.C.......GCAU.................GUGGUU.......CAGGG.C..AGAAAGAGGUGAACGAG...AUG
env-624   ....CUGCAUUUAAGUAGCUGGCAAAAACAUCGGCAUUGCAUGCUUGUAAUCGUCGUGCUAUAAUCCGUGUUGUGUGAUUUUAACUCUG.......GACC.C.......GCAU.................GUGGUU.......CAGGG.C..AGAAAGAGGUGAACGAG...AUG
Pda-1-1   ....GCGCCGGUUCGCAGAAUGUCAUAAUUAGGAAUUGCACCAUACGGGGUUGAGUGCUAUAAUCCGAUUAUGUGUCAUUUUAACUCUG.......GGCC.........CGCCUGUA...............GGCU.......CAGGG.C..AGAAAGAGGUGAACGAG...AUG
Ppa-2-1   ....UGGCCCGCCGCCGCUGAGAGCUGCCGAAGGGCAUUGCGCGGUACGGGCAUGCAUGAUAUAGUAUUCAAGUAUUGCGAUAACUCUG.......GGCU.C.......GCAU.................GAAGCU.......CAGGG.C...AAAAGAGGUGAAAGACG..AUG
Psp-25-1  ....UGGACGUCCUGCGCCGGCGGACCGCAGCAAUCCUGUUGCCUAAGGCACUUGUCCGUGAUAUACUAUUCAGGUAUGCAUAACUCUG.......GGCU.C.......GCUU.................GAAGCU.......CAGGG.C..AGAAAGAGGUGAAAGACG..AUG
env-625   ....GCGGUUCGGUGACCGGCCCGAAAAACAGACCAUUGCGCAAGUUUGGCGGGUCGUGCUAUAAUCACUUCAUGUGUCAUUUACUCUG.......GGUC.........GGUGUCAAAAG............GAUC.......CAGGG.C.GUGAAAGAGGUGAACGAG...AUG
Gma-1-1   ....GCGGGAUUUGGAAAGUGCAGUUCUUUAUUCUUGCAAUAAGUUAUUUUCCUGUGAUAUAAUCCCGUUAUGUGUCCAGACAACUCUG.......GGUC.........GCUAACAAA..............GAUC.......CAGGG.C...AAAAGAGGUGAAAGAA...AUG
env-626   ....GUUGAGAGAUCUGUCGAAUGGAUUUUUUAUAUUGCGGAGGAACGGGGAGUUUGGUAUAAUCAUUUAGUGCGUCCAAUUUACUCUG.......GAAC.........CGAAUGUU...............GUUC.......CAGGG.C..GGAAAGAGGUGAACGAAGCCAUG
env-628   ....UAAGUAAUCGAAAUGAACGGAAAUUGGGAAUUGUACAACGUAUAUGGAUAUGAUAUAAUGCAAUCAUGUGUCAAAUUGAACUCUGU......UCCC.........AUG....................GGGA......UCAGGG.C..GGAAAGAGGUGAAACAG...AUG
Bma-3-1   ....UGCCUUGAUGAACAUUGCAGAGGAAAGUUAUUGCAAGUCUAAGACCGAUCAUGAUAUAAUCUUUUUAUGUGUUUAGUUAACUCUGUU.....UCU..........UCA.....................AGA....UUCAGGG.C..GGAAGGAGUGAACCAAG...AUG
env-630   ....CCGGCGGAUGUCAGUCAUUGCAUUUUCUGUACAGCCUUGAUAUAAUGAAGAAUGGUGUCUUUUAUGAACCAAUUGAAAAACUCUG.......CAUCC........CCG...................GGAUG.......CAGGG.C..AGAAGGAGAUGAACACAGG.AUG
Bau-1-1   ....UGCAAACCGGACUGCUAAUUGAUAUAAUUUCGUAUGUAUGAAAAACUCGUGUGCGCAUGUCUAUGUGUACUUGAAUUUAACUCUG.......AUUC.........GAAA...................GUUU.......CAGGG.C..GGAAGGAGAUGAAAAGA...AUG
Bne-1-1   ....UGUUGCAAAACAAGAAUGUCCUUGUUAUAAUAUAAUCAAUGUGUUUUAACAAUUACAGUGCUUAUGGCAUUGUUUUAUAACUCUG.......UUUC.........GAAA...................GUUG.......CAGGG.C..GGAAGGAGAUGAAAAGGA..AUG
Bci-2-1   ....AGUGUUGCAAAACAAGAAUGUACUUGUUAUAAUAUAAUCAAUGUGUUUUACAAUUACAGUGUCUAAGCACUGUUUUAGAACUCUG.......UUUC.........GAAA...................GUUG.......CAGGG.C..GGAAGGAGAUGAAAAGGA..AUG
env-631   ....UAGUUUUUUCUUGCUGUUUGGAAACAAAAGUGUUGAAAGUAUUGUAGUUUUGUUGCUAUAAUACAAAAGAUGUUUAAAAACUAUA.......UCAG.........AAGAAA.................UUGA.......UAUGG.C..GGAAGGAGAAAAGAA....AUG
env-632   ...........................................................................GUUUAAAAACUAUA.......UCAG.........AAGUAA.................UUGA.......UAUGG.C..GGAAGGAGAGAAAUUAA...AUG
Eca-5-1   ....AUAUGGUAGAAGACUGAUAAAAUAAUUUUAAAAACUAUUGUAUUAUUAAAUUAUUAGUGCUAUUAUACUAGAGCAACUUACUUUG.......GAGUA........GCAUA.................UAUUU.......CAAGG.C..GGAAGGAGAGACAACU...AUG
CSt-2-1   ....AUAUGAAAUAGAAGAUUAAAGAAAUAUAAAAAAAGUGUUGUAUUUAUGGAUAAGAAAUGCUAUUAUAAUUGAGCAACUUACUUUG.......AAGUA........GCAGA.................UAUUU.......CAAGG.C..GGAAGGAGAAGAGAAA...AUG
env-633   ....UCGAAAAAAAUUGAUGAAGAAAAUUUUUAAAAAACUAUUGUAAUUCCCAUUUGAUAAUGGUAUUAUGUGUUAGCAACUUACUUUG.......AAGUA........GCAUA.................UAUUU.......CAAGG.C..GGAAGGAGAUUAUAAA...AUG
env-634   ....UUUAUUUAAAACUGUAUUUAUUUAGCUUGAAUUUUAGUUGUAAAAAUGGUAUUAUUAGUUGGCUUAAAGAUAAUAACUUUCUUUG.......AAGUA........GUGAA.................UAUUU.......CAAGG.C..GGAAGGAGAAAAUGAAA..AUG
env-635   ....GUAAAAUUAUAUUUAUAGUAAAAUUAUAAAAAAGGGCUUGAAAACGACAUAAUUUAAUGUUAAUAUAUCAUAGCAACUUACUUUG.......UAG..........UGAGAAAC................CUA.......CAAGG.C..GGAAGGAGAAAUAAAA...AUG
env-636   ....AGAAUCAAAGUUGAUGAUUAAAAUUAUAAAAAAGGGCUUGAAAACGACAUAAUUUAAUGUUAAUAUAUCAUAGCAACUUACUUUG.......UAG..........UGAGAAAU................CUA.......CAAGG.C..GGAAGGAGAAAUUAAAA..AUG
env-648   ....GACACGGUAUUUUGAAAGCCAUUAAAAUCGCUUGCAUGUUAUGGGUGAUUUUGUUAUAAUUGACACGCUGAGAAAACUUACUAUG.......AU...........GAAAA....................AU.......CAUGG.C..GGAAGGAGCAGUAGCA...AUG
env-650   ....AGAAGAAAAAAAAGUUUAGCAUUAGUUAAAAAUUAUUGACUAAUGUUUCACAUGUGAUAUAAUUUUUAAGUGGAAACUUACUAUG.......GAC..........GUGA....................GUU.......CAUGG.C..GGAAGGAGAAGGAA.....AUG
env-655   ....UUAUUAUAGAGAAAAAUUAAAAUUUUUAAAAUGUAUUUGAUAUACUACUUUUUAUUUGUUAGUAUUUAUGAGCAAACUUUCUCUG.......AAC..........GACAUUACA...............GUU.......CAGGG.C..GGAAGGAGAAUUAAAGCU..AUG
env-656   ....CAUUAUAGACUAAAAACUAUAUUUGUAAAAAUAUGUUUGCGAACAUUUUUAUAAUUUGCUACUAUUUAUGGGCAAACUUUCUUUG.......GACG.........ACAUUUAC...............UGUC.......CAAGG.C..GGAAGGAGAAUUAAAGCU..AUG
env-657   ....CAUUAUAAUCAUAAAAAAAUAUUUUUUAAAUUCUAAUUGCUAACCUAGUUUUAAUUUGUUAGUAUUUAUGAGCAAACUUUCUUUG.......AACG.........ACAUUUAC...............UGUU.......CAAGG.C..GGAAGGAGAAUUAAAGUC..AUG
env-659   ....CAUUAUAAACAUUAAAGGAUAUUUUUUAAAAUAUAAUUGCCAAUAGUAUUUUAUUUUGUUAGUAUUAGUGAGCAAACUUUCUUUG.......AGC..........GACAUUACG...............GCU.......CAAGG.C..GGAAGGAGAAAAAAGUU...AUG
env-660   ....AAUUAUAGUAAAUCUAGGAUAUUUUUAAAAAUAUAAUUGCGAAUAAUAAUGUAUUUUGUUAGUAUUUAAGGCAAACUUACUUUG.......GGC..........GACAUUACG...............GCU.......CAAGG.C..GGAAGGAGAAAAAGUU....AUG
env-662   ....UUUUACAAAAAUAAACAAAUUAUUUUUAAAAUCUAUUUGCUAACCUAAUUUUUAUUUGAUAGUAUUUAUGAGCAAUCUUUCUUUG.......AAC..........GACAUUACG...............GUU.......CAAGG.C..GGAAGGAGAAUUAAGUU...AUG
env-663   ....UAAUUAUAGUCAAAAAGGAUAUUUUUUAAAAUAUAUUUGCCAAUAGAGGCAUUAUUUGUUAGUAUUUAUGAGCAAACUUACUUUG.......AAC..........GACAUUACA...............GUU.......CAAGG.C..GGAAGGAGAAAAAAGUU...AUG
env-664   ....AUUUAUAAAAUUAACCUCUAUUAAUUAAAAAAAGAUUGAAUAAAAUUCAAGGCUUUGAUAUAUUCUUUAGUGCAAACUUUCUUUG.......AACG.........ACAUUAC................UGUU.......CAAGG.C..GGAAGGAGAAAAAACGAA..AUG
env-665   ....AAGAAGAUGAAACACAAAAAUAAUUAAAUUAAACGCUUGUAAUUUAUAUUCAAUGUGUUAUAAUAAAUCGUGCGAACUUACUAUG.......AACU.........AUAGA..................AGUU.......CAUGG.C..GGAAGGAGUAGAGAGU....AUG
env-676   ....UUUAAAUUUAAAAACCAAAAAAGGCGGGAAAAAACAUUUGACAAGAUUGGUCUUUAUGUUAUAAUAAAUGAGCGUACUUACUAUG.......AAC..........GUAAA...................GUU.......CAUGG.C..GGAAGGAGAAAAGGUU...AUG
env-678   ....AAAAAAAGUGUAAAUUUAAAGUGAUUUUUAGUUGAUUAAUUAAAUUCUUAUGAUAUAAUACAUAAGACGAAAGAAAAAUACUAUG.......UUAU.........GGAU...................AUAA.......CAUGG.C..GAAAGGAGAAUUAUUA....AUG
env-679   ....AAAGAAAAUGUUAAGUAUAAGGUAGAUUUAUGCUUGAAUAAGCAUAAAAGUAUGAUAUAAUACCUAUGCGAAUGAAAAAACUAUG.......UUA..........UGGAUU..................UAA.......CAUGG.C..GAAAGGAGAAUAUCUAG...AUG
env-680   ....UCAUCAAUAAUUCCUAUAAAUUAAGAAAAAGUAAUUGAUUUAUUUAAAACUUAUGAUAUAAUAUCUAAGUCAAUCAAUUACUAUG.......UUUU.........UAUAAC.................AAAA.......CAUGG.C..GGAAGGAGCGUAAUUU....AUG
env-681   ....GAUAUUUAUGAUAGAGAUUAAAAAAAAUAAAAAAGUAUUUGCUAUUGAAAAAUAAAUAUGUUACAAUGUCAAAGUAACUUACUAUG.......GUUA.........UAAUAU.................UUAC.......CAUGG.C..GGAAGGAGUGAAUUAGC...AUG
Lga-2-1   ....ACUAGGAUUUGAAGUCGUAACUAUAAAAUAGCUUGAAUUAUGGUUAGAAAGGUGCUAUUCUUAACUAGUCGUGAAAUAAUCUAUG.......AUCUG........AAAUA.................UUGGU.......CAUGG.C...AAAGGAGUAUUAAAUA...AUG
Lho-2-1   ....GCAAGUUUAACCGGGAAGAUAAUAUAAAAACUCCUUGAAAUUAGUGACUGAAGGUGCUAUUCUUAAUUAGUCAUAUUAAUCUAUG.......AUCUG........AAAGA.................CUGAU.......CAUGG.C...AAAGGAGAUUAAAAUU...AUG
env-682   .....................................................................CCUCUCUUUUUAGCGCUCUG.......GCGCUGGACAA..AGAGC...........UUGAUCA...C.......CAGGG.C...AAAGGAGCUGAAAUAA...AUG
env-683   ....AGGGGGAUCGCGAACAUCAUGGUCUGCUGGAACAGGAAGUAAAAAAUACUCAUUUGCUUAAGCCCCCUCUCUUUUUAGCGCUCUG.......GCGCUGGACAA..AGAGC...........UUGAUCA...C.......CAGGG.C...AAAGGAGCUGAAAUAA...AUG
env-684   ....................CGUGGUAUUAUUUCAAUGUUAUGGGCAUUGACCCAGAUAUUUGUGUUGCAAUUGUUUUGAAUAACUCUG.......GUUUA........GGAAA.................UAAAU.......CAGGG.CA.UAAAGGAGAGAGAAGAAGA.AUG
Bse-3-1   ....UUUGCUAGCAAGAUGAAAUAGACUUGCAUUGGCUUUUUUCUCAUGUUAUUAUGACUCUUGUUGAUUAAUAUUAAGAGGAACUCUA.......GUUUA........GCAGA.................UAAAU.......UAGGG.C...AAAGGAGAGAACAGACG..AUG
Aph-1-1   ....AAUAAGUCUCAAAAGUAUUGCAAAAUCAGUCUAGCUUUGCUAAUAUUAACCGUGUUGAUUAAUUUUAUUAUUAUGUAAAUCUCUA.......GUUUA........GCAAA.................UAGAC.......UAGGG.C...AAAGGAGAGAAUACAA...AUG
env-685   ....AAGCAGAACAAAUUCGUAGAAGGCACCAUCUGCUACACUGGCGAGAUCACCGAUACUUCUCGUAACUACACCUUGAAGUACUAUG....................UUGA..............................CAUGG.C...AAAGGAACUGAAAAAG...AUG
env-686   ....UAGAAUCCUGGACAGAAAAUAAUAAGUAAAUAUUCGUUGACUAAAAGAGAAUAUUUUGAUAUAAUACCAACGUAAACUUACUAUG.......GUUA.........AGGU...................UAAC.......UAUGG.C..GGAAGGAGAGAAGUAAA...AUG
Lbu-1-1   ....GAAGAUGAAGAAAGAGUUUAAAAGAUGAUUGAUUGGUAUCAAUAUUUGUGGUAAACUAACAAGGUUGACUAAUCGAUAAUCUCUG.......GGCCA........UCGUAU................UGGUU.......CAGGG.C...AAAGGAGCGAUAUAUU...AUG
Lbr-7-1   ....GAAGAUGAAGAAAGAGUUUAAAAGAUGAUUGAUUGGUAUAAAUAAUUGUGGUAAACUAACAAGGUUGACUAAUCGAUAAUCUCUG.......GGCCA........UCGUAU................UGGUU.......CAGGG.C...AAAGGAGCGAUAUAUU...AUG
Lfa-4-1   ....GGUGACGAAGAAAGAGUUUAAAAGAUUAUUGAUUGGACUUGUUAAUUGUGUUAAACUAACAAGGUUGACUAAUCGAUAAUCUCUG.......GGCCA........UCAUAU................UGGCU.......CAGGG.C...AAAGGAGCGAUAUAAUU..AUG
Lcu-1-1   ....GCGACUAAUUAUUUCUGUAAUUUUCAUAUUGAACCAAGUUUUAAAUUAUGUUAAACUAACAAAGUUGACUAAUCGAUAAUCUCUG.......GGUCA........UCACAU................UGGUU.......CAGGG.C...AAAGGAGCGAUAUAUU...AUG
Lki-2-1   ....UCGAAGAAAAAGUUUAGCUAAAAACGUGUUGAAUGCCUAAUUAUUUUAUGUUAAACUUACAAGGUUGACUAAUCGAUAAUCUCUG.......GGUCA........UCGAAU................UGAUU.......CAGGG.C...AAAGGAGCGAUAUAUU...AUG
Lot-1-1   ....GGAAGUUGAAGAAAGAGUAAAUUGAUUAUUGAUUGCCAUUUUCAUUUAUGGUACACUAACAAAGUUGACUAAUCGAUAAUCUCUG.......GGUCA........UCAGAU................UGAUU.......CAGGG.C...AAAGGAGCGAUAUAUA...AUG
env-688   ....UGGAAAUUGAGAAAGAGUAAAUUGAUUAUUGAUUGCCAUUUUUAUUUAUGGUACACUUACAAAGUUGACUAAUCGAUAAUCUCUG.......GGUCA........UCAGAU................UGAUU.......CAGGG.C...AAAGGAGCGAUAUAUA...AUG
env-689   ....GUAAAUUGAAGAAAGAGUAAAUUGAUUAUUGAUUGCCAUUUUUAUUUAUGGUACACUUACAAAGUUGACUAAUCGAUAAUCUCUG.......GGUCA........UCAGAU................UGAUU.......CAGGG.C...AAAGGAGCGAUAUAUA...AUG
Lbu-3-1   ....AGAAGUUGAAGAAAGAGUCAAUUGAUGAUUGAUUGACGAUUUUAUUUAUGGUACACUAACAAAGUUGACUAAUCGAUAAUCUCUG.......GGUCA........UCGGAU................UGAUU.......CAGGG.C...AAAGGAGCGAUAUAUA...AUG
Ata-1-1   ....UUCAUUUCCAAAAAUGAAAAAGUAUUGCGCUUGAAAAAAGUGUAUGGUAUUAUAUCUGAUGUUGCCUAAGGGUAAAAUAACUCUG.......GUACA........AAAA..................UGGAC.......CAGGG.C..AUAAGGAGUGAAAAUGAAC.AUG
env-691   ....CAGGCGGUUAACACAGGAUAUACAUGUCAUUGCGCAGUUUGAAUGACACGUGCUAUAAUCACCGUUGUAUGUGCUAUUUACUCUG.......GAUU.........CGCAUGA................AGUC.......CAGGG.C..GGAAAGAGGUGAACGAG...AUG
Pcu-1-1   ....UUUUGGCAUUAGCUCAGGCGACGGAUUGCAUUGCAUAUUUUGAAUGGACCGUGCUAUAAUCACGUUGUCUGUGCAAUUUACUCUG.......GAUU.........CGCAUGU................AAUC.......CAGGG.C..GGAAAGAGGUGAACGCG...AUG
env-692   ....AAUCAGAAUAAUUUGGAGCAAAAAUUAUUUUUAUCUUGAAUUAGUUAGUUACUGAUGAUAUAAUUUAUUAGUCAUGUUAAUCUAUG.......GUC..........UGAGAAAU................GGC.......UAUGG.C...AAAGGAGAUAAUAAUU..AUG
Lin-10-1  ....AAUAAGAAUAAUUUGGAGCAAAAAUUAUUUUGUCUUGAAUUAGUUAGUUACUGAUGAUAUAAUUUAUUAGUCAUGUUAAUCUAUG.......GUC..........UGAGAAAU................GGC.......UAUGG.C...AAAGGAGAUAAUAAUU..AUG
env-704   ....AAAUCAGAAUAAUUGGAGCAAAAAUUAUUUUUAUCUUGAAUUAGUUAGUUACUGAUGAUAUAAUUUAUUAGUCAUGUUAAUCUAUG.......GUC..........UGAGAAAU................GGC.......UAUGG.C...AAAGGAGAUAAUAAUU..AUG
env-707   ....AAUAAGAAUAAUUUGGAGCAAAAAUUAUUUUUAUCUUGAAUUAGUUAGUUACUGAUGAUAUAAUUUAUUAGUCAUGUUAAUCUAUG.......GUC..........UGAGAAAU................GGC.......UAUGG.C...AAAGGAGAUAAUAAUU..AUG

          ....................................................................................<<<<<.......<<<<<<<<<<<<<..............>>>>>>>>>>>>>.......>>>>>.......................
          ......................................................................................222.......22211...1111...............1111...11222.......222.........................
          RR•••••••••••••••••••••••••••••••••••••••••••••••••••••••••••••••••••••••••••••Y••CU•UG-------•Y○-----------••R○○------------------○R•-------CA•GG-C--RRAAGGAG••R•R•••○---AUG
                                                                                            5′                                                                      3′
```

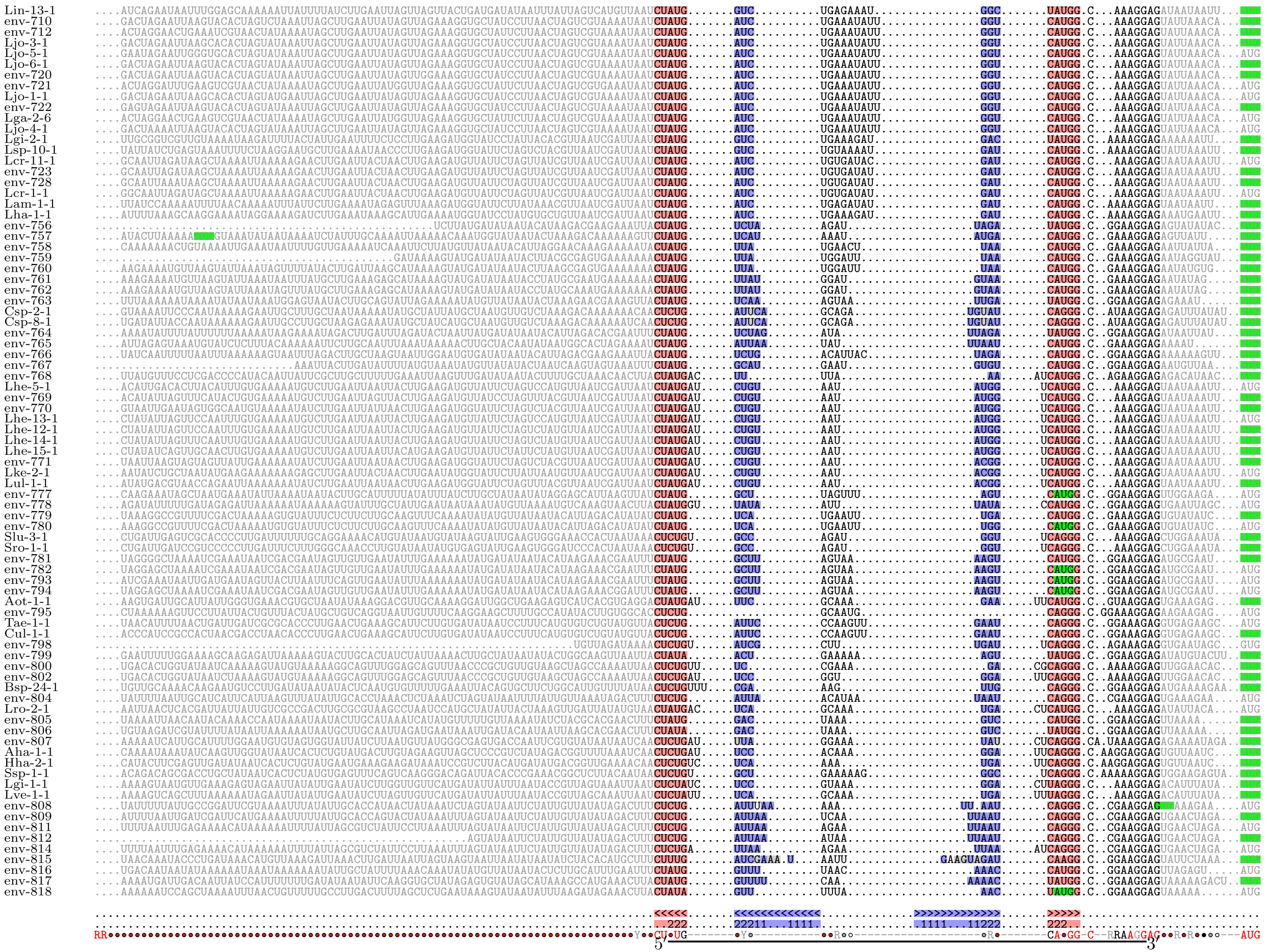
Lin-13-1
env-710
env-712
Ljo-3-1
Ljo-5-1
Ljo-6-1
env-720
env-721
Ljo-1-1
env-722
Lga-2-6
Ljo-4-1
Lgi-2-1
Lsp-10-1
Lcr-11-1
env-723
env-728
Lcr-1-1
Lam-1-1
Lha-1-1
env-756
env-757
env-758
env-759
env-760
env-761
env-762
env-763
Csp-2-1
Csp-8-1
env-764
env-765
env-766
env-767
env-768
Lhe-5-1
env-769
env-770
Lhe-13-1
Lhe-12-1
Lhe-14-1
Lhe-15-1
env-771
Lke-2-1
Lul-1-1
env-777
env-778
env-779
env-780
Slu-3-1
Sro-1-1
env-781
env-782
env-793
env-794
Aot-1-1
env-795
Tae-1-1
Cul-1-1
env-798
env-799
env-800
env-802
Bsp-24-1
env-804
Lro-2-1
env-805
env-806
env-807
Aha-1-1
Hha-2-1
Ssp-1-1
Lgi-1-1
Lve-1-1
env-808
env-809
env-811
env-812
env-814
env-815
env-816
env-817
env-818

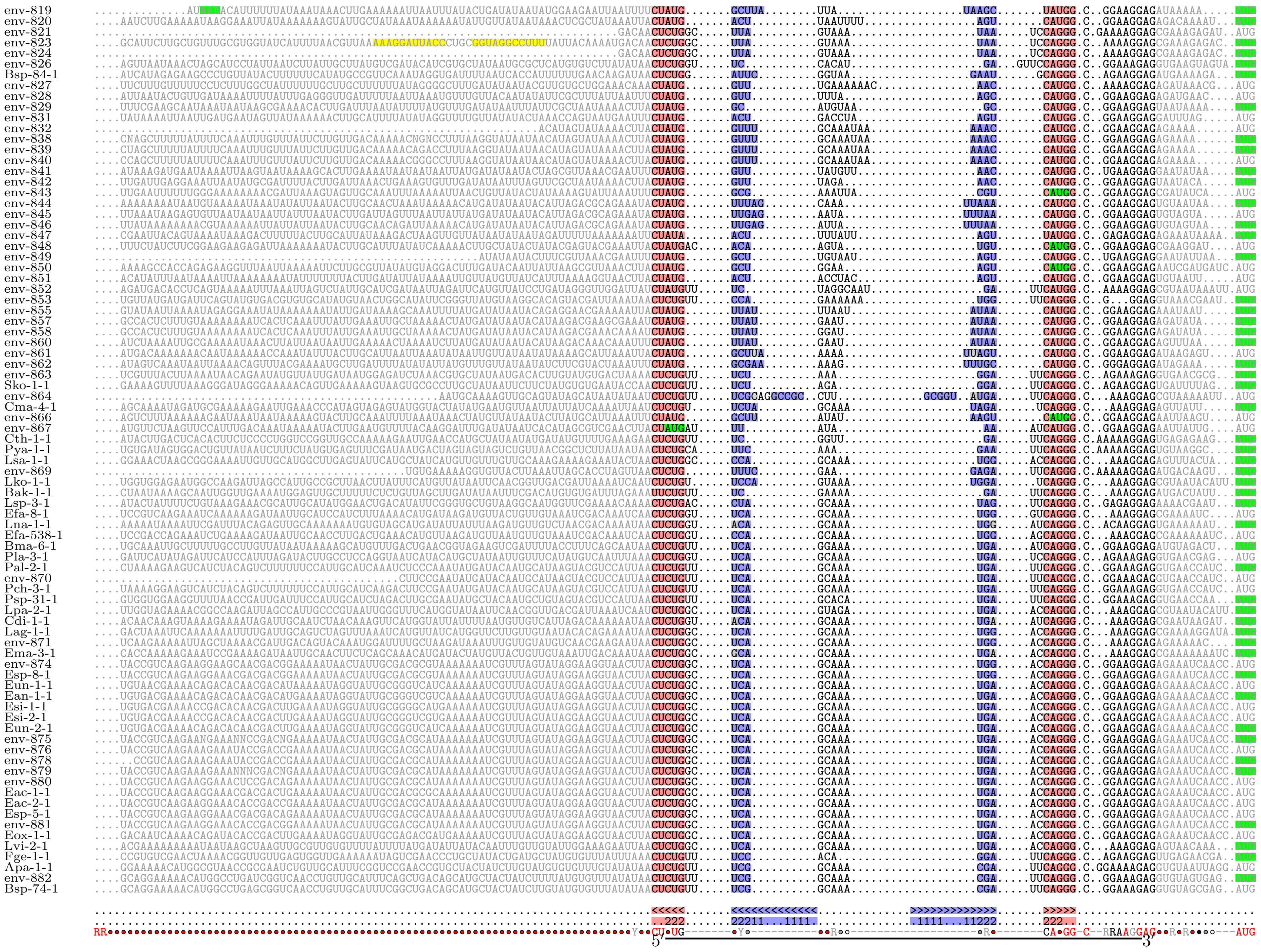
env-819
env-820
env-821
env-823
env-824
env-826
Bsp-84-1
env-827
env-828
env-829
env-831
env-832
env-838
env-839
env-840
env-841
env-842
env-843
env-844
env-845
env-846
env-847
env-848
env-849
env-850
env-851
env-852
env-853
env-855
env-857
env-858
env-860
env-861
env-862
env-863
Sko-1-1
env-864
Cma-4-1
env-866
env-867
Cth-1-1
Pya-1-1
Lsa-1-1
env-869
Lko-1-1
Bak-1-1
Lsp-3-1
Efa-8-1
Lna-1-1
Efa-538-1
Bma-6-1
Pla-3-1
Pal-2-1
env-870
Pch-3-1
Psp-31-1
Lpa-2-1
Cdi-1-1
Lag-1-1
env-871
Ema-3-1
env-874
Esp-8-1
Eun-1-1
Ean-1-1
Esi-1-1
Esi-2-1
Eun-2-1
env-875
env-876
env-878
env-879
env-880
Eac-1-1
Eac-2-1
Esp-5-1
env-881
Eox-1-1
Lvi-2-1
Fge-1-1
Apa-1-1
env-882
Bsp-74-1
5′
3′

```
Bpa-4-1    ....CGCAGGAAAAGCAUGGAAUAACACGGCAGAUUGUUGCAAAUCGCCGCUAGGCAUGCUAAUCUGUAAGGUGUGUUUAAUAACUCUGUU.....UCG..........GCAAA...................CGA.....UUCAGGG.C..AAAAAGAGGUGUAGCGAG..AUG
Bbr-4-1    ....CAGGAAAAACAUGGCGUAAUCGGCCAACCAGUUGCAUUUCGGCAGUGUUUCAUGCUACUAUCUGAUAUGUGUUUAUAUAACUCUGUU.....UCG..........GCAAA...................CGA.....UUCAGGG.C..GGAAAGAGGUGUAGAAGAG.AUG
Bbr-2-1    ....UAGGAAAAACAUGGCGUAAUCGGCCAACCAGUUGCAUUUUGGCAGUGUUUCAUGCUACUAUCUGAUAUGUGUUUAUAUAACUCUGUU.....UCG..........GCAAA...................CGA.....UUCAGGG.C..GGAAAGAGGUGUAGAAGAG.AUG
Bsp-43-1   ....UAGGAAAAACAUGGCGUAAUCGGCCAACCAGUUGCAUUUCGACAGUGUUUCAUGCUACUAUCUUGUAUGUGUUUAUAUAACUCUGUU.....UCG..........GCAAA...................CGA.....UUCAGGG.C..GGAAAGAGGUGUAGAAGAG.AUG
Bbr-1-1    ....UAGGAAAAACAUGGCGUAAUCGGUCAACCAGUUGCAUUUCGACAGUGUUUCAUGCUACUAUCUGAUAUGUGUUUAUAUAACUCUGUU.....UCG..........GCAAA...................CGA.....UUCAGGG.C..GGAAAGAGGUGUAGAAGAG.AUG
Bsp-9-1    ....UAGGAAAAACAUGGCGUAAUCGGCCAACCAGUUGCAUUUCGGCAGUGAUUCAUGCUACUAUCUGAUAUGUGUUUAUAUAACUCUGUU.....UCG..........GCAAA...................CGA.....UUCAGGG.C..GGAAAGAGGUGUAGAAGAG.AUG
Bfo-2-1    ....CAGGAAAAACAUGGCGUAAUCGGCCAACCAGUUGCAUUUCGGCAGUGUUUCAUGCUACUAUCUGUUAUGUGUUUAUAUAACUCUGUU.....UCG..........GCAAA...................CGA.....UUCAGGG.C..GGAAAGAGGUGUAGAUAAG.AUG
Bre-1-1    ....GCAGGAAAAACAUGGCUUGAUCGGUCAACCUGUUGCAUUUCUGCUGACAGCGUGCUACUAUCUGAUAUGUGUUGAUAUAACUCUGUU.....UCG..........GCAAA...................CGA.....UUCAGGG.C..GGAAAGAGGUGUAGAUGAG.AUG
Bbo-4-1    ....AAAAGGCUGGAUAAGAAGUAAAUUCAGUUGCUUUUCGAUGCUGGCCAUGCUACUAUUUUCUAUGUGUAUUUUCUAUAUAACUCUGUU.....UCG..........GCAAA...................CGA.....UUCAGGG.C.GGAAAAGAGGUGUAUGUGAG.AUG
Bbo-5-1    ....AAAAGGCUGGAUAAGAAGUAAAUUCAGUUGCUUUUCGAUACUGGCCAUGCUACUAUUUUCUAUGUGUAUUUUCUAUAUAACUCUGUU.....UCG..........GCAAA...................CGA.....UUCAGGG.C.GGAAAAGAGGUGUAUGUGAG.AUG
Bla-3-1    ....UAGACUUUUCCUUCCAAUCAAGUUAGGAUGGUUGCAAUAGAAACAUAGACGUGAUACUAUUCACUUAUGUGUUUAUAUAACUCUGAU.....UCG..........GCAAA...................CGA.....UUCAGGG.C..GGAAAGAGGUGCAUAGAG..AUG
Bla-2-1    ....AGACUUUUGUUUCCAAUCAAGUCAGUAUGGUUGCAAUAGACACAUAGACGUGAUACUAUUCAUUUAUGUGUUUUAUAUAACUCUGAU.....UCG..........GCAAA...................CGA.....UUCAGGG.C..GGAAAGAGGUGCAUAGAG..AUG
Bsp-99-1   ....GAAAAAUAUGUAAACCACCUUCCUGCUAUUGCAAAUCGGCACCAGUCGUGCUACUAUCUUUUUAUGUCUGUUUUAUAUAACUCUGUU.....UCG..........GUAUA...................CGA.....UUCAGGG.C..GGAAAGAGGUGGAUAAC...AUG
Bsp-45-1   ....AUUAUUUGAAAUAGGCUUGUUAUAAUAAUUCCAGUGUGCUUUUAAUGAUUGCGGAGGCCAAUUAGCCCCGUACAAUAUAACUCUGUC.....UCG..........CAUA....................UGA.....UUCAGGG.C..AAAAGGAGAUGAAAGGGA..AUG
Bca-1-1    ....AUUAUUUGAAAUAGGCUUGUUAUAAUAAAUCCAGUGUGUUUUUAAUGAUUGCGGAGGUCUUUAGGCCCCGUACAAUAUAACUCUGUC.....UCG..........CAUA....................UGA.....UUCAGGG.C..AAAAGGAGAUGAAAGGGA..AUG
Aji-1-1    ....UGACACUGUUAUAAUUUUAAAGUGUGCAAAAAGGUAGCUUGAAUUCGUUGCUAGAAUGGUUUUAAGCAUACUUAAAUUAACUCUGUU.....UCC..........AAG.....................GGA.....UUCAGGG.C..AGAAGGAGUUGUAAAAAC..AUG
Lmu-1-1    ....GUUAGAAUUGAAGAAGAAUAGAAAAUGACAGAGUAGUUGCAAGUUGAACAAAGAUAUGUUAAUCUGUGAAGGUUGAUUAUCUAUGUU.....UCA..........GGCAA...................UGA.....UUCAUGG.C..AAAAGGAGCGUAAAAUUU..AUG
env-883    ....GUUAGGAUUGAAGAAGAAUAGAAAAUGACAGAGUAGUUGCAAGUUGAACAAAGAUAUGUUAAUCUGUGAAGGUUGAUUAUCUAUGUU.....UCA..........GGCAA...................UGA.....UUCAUGG.C..AAAAGGAGCGUAAAAUUU..AUG
Lwa-1-1    ....AAUCUGGCGUCGAAGUUACAGAAUAAUGUGUUGCGAAAAAUAGCGAUUCAUGGUAAUCUAUUGUGGUUAACGAUAAAUAACUAUGUU.....UCA..........GUCAA...................UGA.....UUCAUGG.C...AAAGGAGCGUAUUAAACU.AUG
Lho-1-1    ....AAUCUGGUGUGGAAGUAAUGGAAUAAUGUGUUGCUAAAAAUGUAAAUUCAUGGUAAUCUAUUGUGGUUAACGAUAAAUAACUAUGUC.....UCA..........GUCAA...................UGA.....UUCAUGG.C...AAAGGAGCGUAUUAAACU.AUG
env-887    ....GAUGUUACAAUUGAAGAUGAUGAUUAAUUAGUUUGCGUUCAGCUUCAAAUUAUGUUAUCCUAUCGUAGUUGUAAAAAUAACUAUGUU.....UCA..........GGCAA...................UGA.....UUCAUGG.C...AAAGGAGCGUAUAAAUU..AUG
Lco-3-1    ....GACGUUACAAUUGAAGACGACGAUUAAUUAGUUUGCGUUCAGCUUCAAAUUAUGUUAUUCUAUCGUAGUUGUAAAAAUAACUAUGUU.....UCA..........GGCAA...................UGA.....UUCAUGG.C...AAAGGAGCGUAUAAAUU..AUG
Lin-6-1    ....CCUUUUUGGCGGUUGGCCCGGGUUCGUUGGUUGCUAAUUGGAGCAAACUGUGUUAUUCUGUCUAGGUUAUUGAAAAUAAUCUAUGUU.....UCA..........GGCAA...................UGA.....UUCAUGG.C...AAAGGAGCGUAUAAAUU..AUG
env-888    ....GGACGUUACAAUUGAAGAUGACGAUUAAUUAGUUUGCGUUCAGCUUCAAAUUAUGUUAUUCUAUCGUAGUUGUAAAAAUAACUAUGUU.....UCA..........GGCAA...................UGA.....UUCAUGG.C...AAAGGAGCGUAUAAAUU..AUG
env-890    ....UUGAGGACGAUUCAUCAGUAAAAAUUAGCUAAACUGUUGCGUUGAUAAUUAAUUCAUGUUAUUCUGUUAGGGUUGAUUAUCUAUGUU.....UCA..........GGCAA...................UGA.....UUCAUGG.C..AAAAGGAGCGUAUUAAGUU.AUG
Lan-3-1    ....UUGAAGACGAUUCGUCGGUAAAAAUUGGUUAGGCGGUUGCGUUGCUAACUGAUUCAUGAUAUUCUAUUAGGGUUGAUUAUCUAUGUU.....UCA..........GGCAA...................UGA.....UUCAUGG.C..AAAAGGAGCGUAUAAAGUU.AUG
env-891    ....CGAAGAACGAUACAUCAGUAAAAAUUAGCUAGGCUGUUGCGUUACUAACUGAUUCAUGAUAUUCUAUUAGGGUUGAUUAUCUAUGUU.....UCA..........GGCAA...................UGA.....UUCAUGG.C..AAAAGGAGCGUAUAAAGUU.AUG
Lor-3-1    ....UCGAAGACGAUACAUCAGUAAAAAUUAGCUAGGCUGUUGCGUUACUAACUGAUUUAUGAUAUUCUAUUAGGGUUGAUUAUCUAUGUU.....UCA..........GGCAA...................UGA.....UUCAUGG.C..AAAAGGAGCGUAUAAAGUU.AUG
env-892    ....UUGAGGACGAUUCAUCAGUAAAAAUUAGCUAAACUGUUGCGUUGAUAAUUAAUUCAUGUUAUUCUGUUAGGGUUGAUUAUCUAUGUU.....UCA..........GGCAA...................UGA.....UUCAUGG.C..AAAAGGAGCGUAUAAAGUU.AUG
Lor-1-1    ....UCGAAGACGAUACAUCAGUAAAAAUUAGCUAGGCUGUUGCGUUACUAACUGAUUCAUGAUAUUCUAUUAGGGUUGAUUAUCUAUGUU.....UCA..........GGCAA...................UGA.....UUCAUGG.C..AAAAGGAGCGUAUAAAGUU.AUG
Lfe-7-1    ....ACGUCCGGAUUGAAGACGAUGAGCAAGACUAGGUUGCUUUUUGACUGAGCACGUGUUACUCUAUUAGGGUUAUUGAAAUACUAUGUU.....UCA..........GGCAA...................UGA.....UUCAUGG.C..AAAAGGAGCGUAUUAAAUU.AUG
env-893    ....CGUCCGGAUUGAAGACGAUGAGCAAGACUAGGUUGCUUUUUGACUGAGCACGUGUUACUCUAUUAGGGUUAUUGAAAUAACUAUGUU.....UCA..........GGCAA...................UGA.....UUCAUGG.C..AAAAGGAGCGUAUUAAAUU.AUG
env-894    ...............................................................................UAUCUAUGUU.....UCA..........GGCAA...................UGA.....UUCAUGG.C..AAAAGGAGCGUAAUAAAUU.AUG
env-898    ....UCGAAGAUGACAGCUCAGUAAAAAUCUAAUUGUCUAUUGCAUCGAUAAUGGAUCCAUGUUAUUCUGUUAGGGUUGAUUAUCUAUGUU.....UCA..........GGCAA...................UGA.....UUCAUGG.C..AAAAGGAGCGUAAUAAAUU.AUG
env-899    ....AUUGAGGACGAUUCUUCAGUAAAAAUUGGUUAGAAGUUGCGUACUUCUUUAAAUCGUGGUAUCCUGUUAGAGUUGAUUAUCUAUGUU.....UCA..........GGCAA...................UGA.....UUCAUGG.C..AAAAGGAGCGUAAUAAAUU.AUG
Lva-2-1    ....CGAAGACGAUAGCUCAGUAAAAAUUUAAUCAGUCUAUUGCAUCGAUAAUUGAUUCGUGUUAUUCUGUUAGGGUUGAUUAUCUAUGUU.....UCA..........GGCAA...................UGA.....UUCAUGG.C..AAAAGGAGCGUAAUAAAUU.AUG
env-900    ....UGAGGACGAUUCUUCAGUAAAAAUUGGUUAGAAGUUGCGUACUUCUUUAAAUCGUGGUAUCCUGUUUAGAGUUUGAUUAUCUAUGUU.....UCA..........GGCAA...................UGA.....UUCAUGG.C..AAAAGGAGCGUAAUAAAUU.AUG
env-901    ....GUAAACCCUUGUUAUAAUUUUUCAUGUGUGUUUAAGAAUGUUGAAGCGUGAAAUAAUUGCACGUUUCCGAAUAGAUAAACUCUGUU.....UCG..........AAA.....................UGA.....UUCAGGG.C.GUAAAGGAGUUGAAAACU...AUG
Ate-2-1    ....AAGCAUAAGGAUUUAAAAAUGGAAACGAGUCCUUGCAAUACUGUAAAUACCGUGCUAUACUCACAAUAUUGUGAUUGAAACUCUGUU.....UC...........GUAAA....................GA.....UUCAGGG.C.AGAAAGGGUGAAAUAA.....AUG
Bth-45-1   ....UAUAAUUUCAGCUUUGGUUUGUUUGCAUAAUUGACUUUUGCGUAGUAUAAUGUUUUUAUGUGCCAAUAGGGCGUAAAUAACUCUGUU.....UC...........AGAU.....................GA.....UUCAGGG.C...AAAGGAGUGAAAUAAUC..AUG
env-902    ....ACGAUUAUUGCUUUUUAGGAGGCAACUUGCUAUAAUAGUAAUAGUGUUUUUUUAACGAUACCUUUAUGGAUGAGAAAUAACUCUGUU.....UCA..........GAA.....................UGA.....UUCAGGG.C..AAAAGGAGUGAAAUAGG...AUG
Bth-44-4   ....ACGAUUAUUGCUUUUUAGGAGGCAACUUGCUAUAAUAGUAAUAGUGUUUUUUUAGCGAUACCUUUAUGGAUGAGAAAUAACUCUGUU.....UCA..........GAA.....................UGA.....UUCAGGG.C..AAAAGGAGUGAAAUAGG...AUG
env-903    ....ACGGUUAUUGCUUUUUAGGGAUUGCCUUGCUAUAAUAAUAACAGUGUUUUUAAAUGAUACCUUUUAUGGUGAGAAAUAACUCUGUU.....UCA..........GAA.....................UGA.....UUCAGGG.C..AAAAGGAGUGAAAUAGG...AUG
Bca-2-1    ....UUGUAGAAGUGAAACGUUUAUUUGCAAAUUUGACGUGCGCAUAGUAUAAUCAUAUUAUGUGCCAAUUGGCCGUAAAAUAACUCUGUU.....UC...........AGAU.....................GA.....UUCAGGG.C...AAAGGAGUGAAAACAAUC.AUG
Pda-2-1    ....CGCCAAUCUCGGUAAAAUUGGUUUGCGUAUAACAGAUAUUCAUGAUAUUAUAAUGAACGUUGAUUAAUUAUGAAAAAUAACUAUGUC.....UCA..........GUAAA...................UGA.....CUCAUGG.C...AAAGGAGUGUACUAACA..AUG
env-904    ....AUGAAUCUGGUGUUGAAGUCAUUGAAUAACCUUGCAGUCUAAAAAAAAUAUGUUAGUCUGUUAGAGUUAAUGAUAAAUAACUAUGUC.....UCA..........GCAAA...................UGA.....CUCAUGG.C...AAAGGAGCGUAUAAACU..AUG
Lsu-2-1    ....UGAGUCUGGUGUCGAAGUGAUUGAAUAGCCUUGCAGUUUUAAAUAGAUUAUGUUAAUCUGUUAGGGUUAAUGAUAAAUAACUAUGUC.....UCA..........GCAAA...................UGA.....CUCAUGG.C...AAAGGAGCGUAUAAAC...AUG
Lmu-3-1    ....GAUGAUAAAAAUCGUAAAAGCUAAUUGCAACAAAUUUCGAUUUGUGUUAUCAUGUUAUUUGUUGUAUAACACGAAAAUAACUCUGAC.....CCA..........GCAAA...................UGG.....AUCAGGG.C...AAAGGAGCGAACUUAGA..AUG
env-906    ....UCGACACAAAAUCGCAAAAGAUAAUUGCAACGAAGUAAAAUUCAUGGUAUCAUGUUGUUUGUUGUAUAACACGAAAAUAACUCUGAC.....CCA..........GCAAA...................UGG.....AUCAGGG.C...AAAGGAGCGAAAUUAGA..AUG
env-908    ..........................................GAUUCAUGUUAUUAUGUUAUUUGUUGUAUAACACGAAAAUAACUCUGAC.....CCA..........GCAAA...................UGG.....AUCAGGG.C...AAAGGAGCGAAAUUAGA..AUG
env-909    ............................................................................CGAAAUAACUCUGAC.....UCA..........GCAAA...................UGG.....AUCAGGG.C...AAAGGAGCGAAAUUAGUA.AUG
env-910    ....GGCAAAAAAAUCGAAAAAAGAUAAUUGCAUGACUUGAAGUUUCAUGUUAUUAUGAAGUAUGUUGCGAUAGCACAGAAUAACUCUGAU.....CCA..........GCAAA...................UGG.....UUCAGGG.C...AAAGGAGCGAAAUUAUA..AUG
Lru-1-2    ....AGCAAAAAAAUCGGAAAAAGAUAAUUGCAUGACUUGAAGUUUCAUGUUAUUAUGAAGUAUGUUGCGAUAGCACAGAAUAACUCUGAU.....CCA..........GCAAA...................UGG.....UUCAGGG.C...AAAGGAGCGAAAUUAUA..AUG
env-913    ....UGGUAGUAAAAAAUCUUAAUCGACCUUGUUUAAGUUGUUUUUCUAUGCUAUUAUUGGGGACGUUGAUUUAUAAUAAAUAACUAUGAC.....UCA..........GCAAA...................UGA.....CUCAUGG.C...AAAGGAGCGUAAUAUC...AUG
Ppe-4-1    ....UGGUAGUAAAAAAUCUUAAUCGACCUUGUUUAAGUUGUUUUUCUAUGCUAUUAUUGAGGACGUUGAUUUAUAAUAAAUAACUAUGAC.....UCA..........GCAAA...................UGA.....CUCAUGG.C...AAAGGAGCGUAAUAUC...AUG
Pac-2-1    ....UAAUUUCGGCAAAAAGUCCUAAAUGAGGUUGUUUGAAAUAAUAAUUUAUGCUAUGAUUGUGGUCGUUGGUUUUAUAAUAACUAUGAU.....UCA..........GCAAA...................UGA.....UUCAUGG.C...AAAGGAGCGUAAUAUC...AUG
Pac-1-1    ....UAAUUUCGGCAAAAAGUCCUAAAUGAGGUUGUUUGAAAUGAUAAUUUAUGCUAUGAUUGUGGUCGUUGGUUUUAUAAUAACUAUGAU.....UCA..........GCAAA...................UGA.....UUCAUGG.C...AAAGGAGCGUAAUAUC...AUG
Lco-2-1    ....CCUGGGAAAUAUUUAAAAAGUCAUUGUUAUUCAAUUAAAUCCAUGCUAUCAUACUAUGAUGUGCAUGAAGGAUAUUAUAUCUCUAUU.....UCA..........GCAAA...................UGA.....UUUAGGG.C...AAAGGAGCGAAAACAUU..AUG
Lce-1-1    ....UUUGUUUUUAAAAACUAAAAGAUGAUUGCACCAAAAUAAAAUUCAUGCUAUUAUGAUGAUUGUUAUAUUACAAAGAAUAACUCUAUU.....UCA..........GCAAA...................UGA.....AUUAGGG.C...AAAGGAGCGAAAUUAAUA.AUG
Esa-2-1    ....AAAGAGUCAAAAGAAAUGUGAAAAGAUAAUUGCAAAUGAUUUUAACUCAUGUUAAGAUAGUAACGUUGUAAACAAAUCAACUCUGAC.....UCA..........GCAAA...................UGA.....UUCAGGG.C...AAAGGAGAGAAAGCAAA..AUG
Eph-1-1    ....ACCAAGAAGAAAUCAAAAAAGAUAAUUGCAUAUUGAAACAAAUCAUGGUACGAUAUUACUGUUGUAAAUCGACAAAUCAACUCUGAC.....UCA..........GCAAA...................UGA.....UUCAGGG.C...AAAGGAGCGAACGAAAG..AUG
Mpl-1-1    ....AUUAAAAGAAAUUAUAAAAGAUAAUUGUAACUUGAUUCAAUUUAUGUUAAAAUAAAUGUAAUUGUUUAAUGAAAAAACAACUCUGAC.....UCA..........GCAAA...................UGA.....UUCAGGG.C...AAAGGAGCGAAUGAGAA..AUG
Mpl-1-10   ....AUUAAAAGAAAUUAUAAAAGAUAAUUGUAACUUGAUUCAAUUUAUGUUAAAUUAAAUGUAAUUGUUUAAUGAAAAAACAACUCUGAC.....UCA..........GCAAA...................UGA.....UUCAGGG.C...AAAGGAGCGAAUGAGAA..AUG
Lsu-1-1    ....GCGAGGAGAUAACAAAAAAAGAUAAUUGCGUGCAUUGUUUUUUCGUGAUACAAUAUCUUGAGUUGUGGUACUGAGAAUAACUCUGAU.....UCA..........GCAAA...................UGA.....UUCAGGG.C...AAAGGAGAGUAAUAUA...AUG
env-914    ....AAGGCCUGAACAGUAAUUCAGCUUGACGAUGCUAACGGGUUCAUGAUAAGAUGUUACUCGUUGAUUAAAACUAUGAUAAUCUCUGUU.....UCA..........GCAAA...................UGA.....UUCAGGG.C...AAAGGAGCGUAAUAUU...AUG
Lfa-2-1    ....GCAAUGCUCGUCCCUAACCCGGCUUGACGGACGCUCUGACACAUGAUAAACUACAAAGGUUGAUUUAAAACAAUGAUAAUCUCUGUU.....UCA..........GCAAA...................UGA.....UUCAGGG.C...AAAGGAGCGUAAUAAUU..AUG
Lor-2-1    ....CGAGUCCUAAUUUUCUUAAUUUAGUGCAUUGCCAGAAAAAACGUUUCGUGUUAUAGUACAAAGGUUGGUUAUUCAUGAAUCUCUGUU.....UCA..........GCAAA...................UGA.....UUCAGGG.C...AAAGGAGCGAAGAAAUU..AUG
Lma-4-1    ....UGUCGAAUCAUAAAAAUUAAUUAGUCCAUUGAUUGAAGUUUAAAUUCAUGUUAUAGUACAAAGGUUAGUUAUUUAUGAAUCUCUGUC.....UCA..........GCAAA...................UGA.....UUCAGGG.C...AAAGGAGCGAAGAAAAUU.AUG
Lhe-11-1   ....UACGGUUAGGGCAUUGGACAGGUUGACGGAACAACGUCAUUCAUGAUAAUAUGUGUAACGUUGAUUAAAACUAUGAUAAUCUCUGUU.....UCA..........GCAAA...................UGA.....UUCAGGG.C...AAAGGAGCGUUAUACA...AUG
env-916    ....GGCAGGCAGGCUACUUUGUAGGUUGACGGGAAGCUGUGAUUCGUGAUAACAUGUUUAUCGUUGGUUAAAACUAUGAUAAUCUCUGUU.....UCA..........GCAAA...................UGA.....UUCAGGG.C...AAAGGAGCGAAAUACA...AUG
Lpl-8-1    ....GCUAGUCCCCUGGUUUUAGAGGUUGACGGGGCCUAGUCAUUCGUGAUAACAUGUUUAUCGUUGGUUAAAACUAUGAUAAUCUCUGUU.....UCA..........GCAAA...................UGA.....UUCAGGG.C...AAAGGAGCGAAAUACA...AUG
Lpl-6-1    ....GACAGUUAGGCUACUUUGUAGGUUGACGAGAAGCUGUGAUUCGUGAUAACAUGUUUAUCGUUGGUUAAAACUAUGAUAAUCUCUGUU.....UCA..........GCAAA...................UGA.....UUCAGGG.C...AAAGGAGCGAAAUACA...AUG
Lpl-13-1   ....GGCAGACGGGCUACUUUGUAGGUUGACGGGAAGCUGUGAUUCGUGAUAACAUGUUUAUCGUUGGUUAAAACUAUGAUAAUCUCUGUU.....UCA..........GCAAA...................UGA.....UUCAGGG.C...AAAGGAGCGAAAUACA...AUG
Lpe-2-1    ....GUUAGUCCCCUGGUUUUAGAGGUUGACGGGGCCUAGUCAUUCGUGAUAACAUGUUUAUCGUUGGUUAAAACUAUGAUAAUCUCUGUU.....UCA..........GCAAA...................UGA.....UUCAGGG.C...AAAGGAGCGAAAUACA...AUG
Lpe-1-1    ....GCUAGUCCAGUCCUUUUAGUGGUUGACGGGGCCUAGUCAUUCGUGAUAACAUGUUUAUCGUUGGUUAAAACUAUGAUAAUCUCUGUU.....UCA..........GCAAA...................UGA.....UUCAGGG.C...AAAGGAGCGAAAUACA...AUG
Lpa-41-1   ....GACAGUUAGGCUACUUUGUAGGUUGACGGGAAGCUGUGAUUCGUGAUAACAUGUUUAUCGUUGGUUAAAACUAUGAUAAUCUCUGUU.....UCA..........GCAAA...................UGA.....UUCAGGG.C...AAAGGAGCGAAAUACA...AUG
Eit-1-1    ....AAAUGCAAAAAAUCGCAAAAGAUAAUUGCCAAGAAGACUGAAAGAUGAUAGUAUAUUCUUGUUGCAUUAUAGACAAAUAACUCUGUU.....UCA..........GCAAA...................UGA.....CUCAGGG.C...AAAGGAGAGAAAAACAC..AUG
env-919    ....GGUAAUCAGGCUACUUUGUAGGUUGACGGGAAGCUGUGAUUCGUGAUAACAUGUUUAUCGUUGGUUAAAACUAUGAUAAUCUCUGUU.....UCA..........GCAAA...................UGA.....UUCAGGGGC..AAAAGGAGCGAAAAUACA..AUG
Vlu-1-1    ....UUUCUUUAUCGAACAAAAGGUAGUUGUCUUUUAUAGUGAAACGUGCUAUCAUAGUUCUGUUAUAAAAACUUAAGAAGAAACUCUGUU.....UCA..........GCAAA...................UGA.....UUCAGGG.C...AAAGGAGAGAAAAUC....AUG
env-920    ....AUCGGGUAGAAGAAUUAGUGAGUGACUUGAAUUAGGACAAUUCUUAUGAUAUUCUAUUUAGGUUGACGAUAACAAUCAAUCUCUGUU.....UCA..........GUAGA...................UGA.....UUCAGGG.C...AAAGGAGCGAAACAAUU..AUG
env-921    ....AUCGGUAGAAGAAUUAUAAGAGUGACUUGAAUUAGACCAAUUCUUAUGAUAUUCUAUUUAGGUUGACGAUAACAAUCAAUCUCUGUU.....UCA..........GUACA...................UGA.....UUCAGGG.C...AAAGGAGCGAAACAAUU..AUG
Tha-1-1    ....GUAGAGUUUCAAAUAAAUUAAAGCCCCUGAUUGCAUUAAGUUCUAAAAAAUGUUAAUAUGUUACGUGUUGUAAAGAAACACUCUGAU.....UUA..........GCAAA...................UAA.....UUCAGGG.C...AAAGGAGAGUAAUAAU...AUG
Tmu-2-1    ....UUUUCAGUAGAAAAAGCUUAUAAGAAUGAAUUGCAUCCCAUAAUAAAAAAUGUUAAUAUGUUCAUGUUACAAAGAAAUAACUCUGAU.....UUA..........GUAAA...................UAA.....CUCAGGG.C...AAAGGAGAGUAGUCAUU..AUG

           ....................................................................................<<<<<.......<<<<<<<<<<<<<...............>>>>>>>>>>>>>.......>>>>>.................................
           ....................................................................................222.......22211...1111...............1111...11222.......222....................................
           RR••••••••••••••••••••••••••••••••••••••••••••••••••••••••••••••••••••••••••••••••Y••CU•UG-------•Y•----------••R••------------------•R•-------CA•GG-C--RRAAGGAG••R•R••••---AUG
                                                                                               5′                                                                                     3′
```

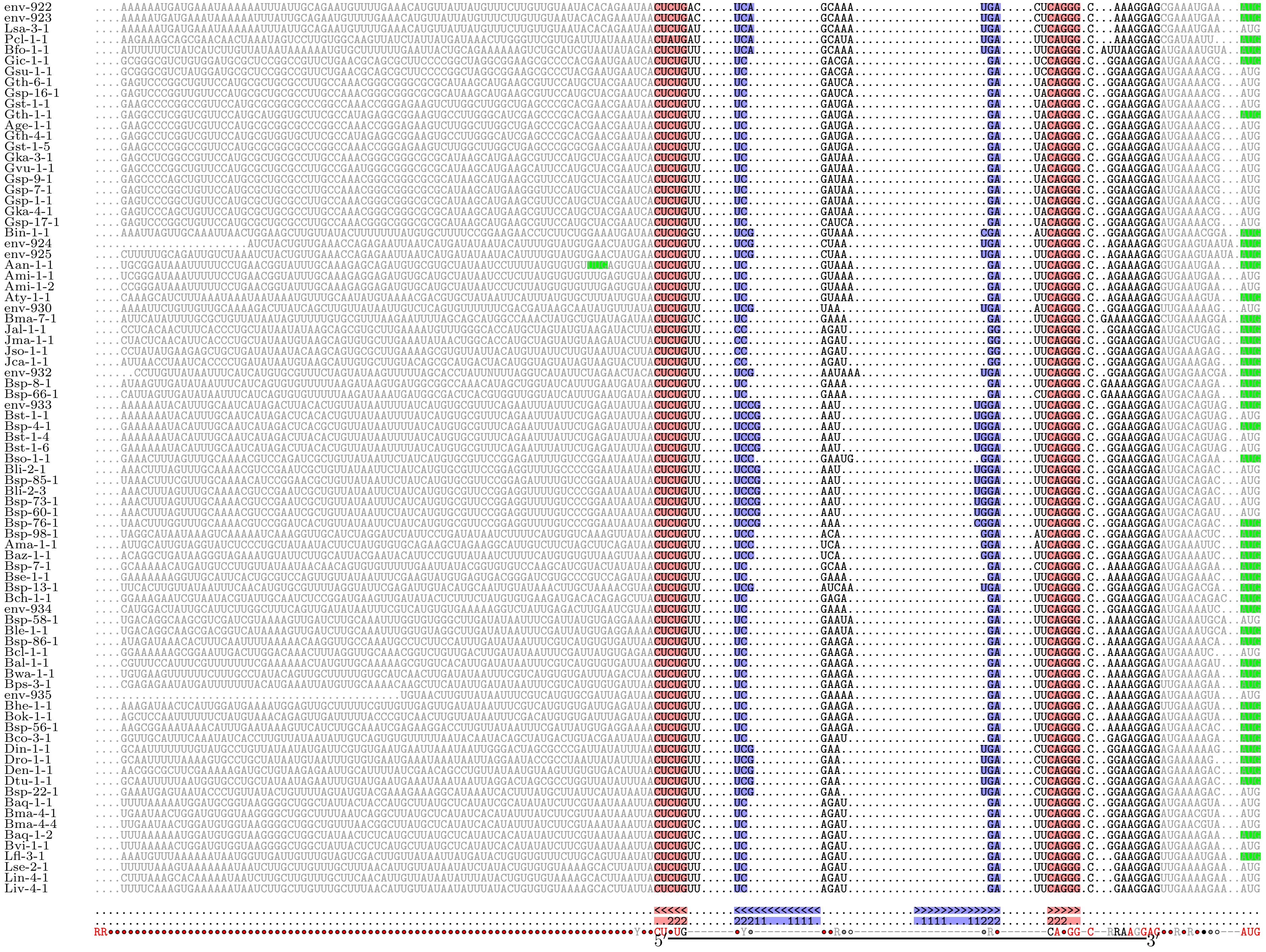
env-922
env-923
Lsa-3-1
Pcl-1-1
Bfo-1-1
Gic-1-1
Gsu-1-1
Gth-6-1
Gsp-16-1
Gst-1-1
Gth-1-1
Age-1-1
Gth-4-1
Gst-1-5
Gka-3-1
Gvu-1-1
Gsp-9-1
Gsp-7-1
Gsp-1-1
Gka-4-1
Gsp-17-1
Bin-1-1
env-924
env-925
Aan-1-1
Ami-1-1
Ami-1-2
Aty-1-1
env-930
Bma-7-1
Jal-1-1
Jma-1-1
Jso-1-1
Jca-1-1
env-932
Bsp-8-1
Bsp-66-1
env-933
Bst-1-1
Bsp-4-1
Bst-1-4
Bst-1-6
Bso-1-1
Bli-2-1
Bsp-85-1
Bli-2-3
Bsp-73-1
Bsp-60-1
Bsp-76-1
Bsp-98-1
Ama-1-1
Baz-1-1
Bsp-7-1
Bse-1-1
Bsp-13-1
Bch-1-1
env-934
Bsp-58-1
Ble-1-1
Bsp-86-1
Bcl-1-1
Bal-1-1
Bwa-1-1
Bps-3-1
env-935
Bhe-1-1
Bok-1-1
Bsp-56-1
Bco-3-1
Din-1-1
Dro-1-1
Den-1-1
Dtu-1-1
Bsp-22-1
Baq-1-1
Bma-4-1
Bma-4-4
Baq-1-2
Bvi-1-1
Lfl-3-1
Lse-2-1
Lin-4-1
Liv-4-1
5′
3′

```
Lmo-60-1   ....UUUCAAAGUGAAAAAAUAAUCUUGCUUGUUUGCUUCAACAUUGUUAUAAUAUCUAUACUGUGUGUAAAAACACUUAAUUACUCUGUU.....UC...........AGAU.....................GA.....UUCAGGG.C...GAAGGAGUUGAAAAGAA..AUG
Liv-1-2    ....UUUUCAAAGUAAAAAAAAUAAUCUUGCUUGUUUGCUUUAACAUUGUUAUAAUAUCUAUACUGUGUGUAAAAGCACUUAUUACUCUGUU.....UC...........AGAU.....................GA.....UUCAGGG.C...GAAGGAGUUGAAAAGAA..AUG
Lwe-1-1    ....CUUCAAAGUGAAAAAAUAAUCUUGCUUGUUUGCUAUAACAUUGUUAUAAUACUUAUACUGUGUGUAAAAGCACUUAAUUACUCUGUU.....UC...........AGAU.....................GA.....UUCAGGG.C...GAAGGAGUUGAAAAGAA..AUG
Laq-1-1    ....AAAUGUUUAAAAAAUAACUAUUGCUUGUUUGCGCAUCAGUUGUUAUAAUUAUCAUACUGUGUGUUUCUUGCAGUGAAUUUCUCUGUU.....UC...........AGAU.....................GA.....UUCAGGG.C...GAAGGAGUUGAAAAGAA..AUG
Lmo-69-1   ....UUUCAAAGUGAAAAAAUAAUCUUGCUUGUUUGCUUCAACAUUGUUAUAAUAUCUAUACUGUGUGUAAAAGCACUUAAUUACUCUGUU.....UC...........AGAU.....................GA.....UUCAGGG.C...GAAGGAGUUGAAAAGAA..AUG
Lin-2-1    ....AUCUAAAGCACAAAAAUAAUCUUGCUUGUUUGCUUCAACAUUGUUAUAAUAUUUAUACUGUGUGUAAAAGCACUUAAUUACUCUGUU.....UC...........AGAU.....................GA.....UUCAGGG.C...GAAGGAGUUGAAAAGAA..AUG
Lin-2-2    ....CUUUAAAGUGCAAAAAUAAUCUUGCUUGUUUGCUUCAACAUUGUUAUAAUAUUUAUACUGUGUGUAAAAGCACUUAAUUACUCUGUU.....UC...........AGAU.....................GA.....UUCAGGG.C...GAAGGAGUUGAAAAGAA..AUG
Lba-1-1    ....CAAAUGUUUAAAAAAUAACAGUUGCUUGUUUUCGUCUCACUUGUUAUAAUUAUGGUACUGUGUUUUCGUGCAGUUAAUUACUCUGUU.....UC...........AGAU.....................GA.....UUCAGGG.C...GAAGGAGUUGAAAAGAA..AUG
Lse-1-1    ....UCUUCAAAGUGAAAAAAUAAUCUUGCUUGUUUGCUUUAACAUUGUUAUAAUAUCUAUACUGUGUGUAAAAGCACUUAUUACUCUGUU.....UC...........AGAU.....................GA.....UUCAGGG.C...GAAGGAGUUGAAAAGAA..AUG
Lmo-40-1   ....CUUUAAAGUACAAAAAUAAUCUUGCUUGUUUGCUUCAACAUUGUUAUAAUAUUUAUACUGUGUGUAAAAGCACUUAAUUACUCUGUU.....UC...........AGAU.....................GA.....UUCAGGG.C...GAAGGAGUUGAAAAGAA..AUG
Lmo-32-1   ....CUUUAAAGUGAAAAAAUAAUCUUGCUUGUUUGCUUCAACAUUGUUAUAAUAUCUAUACUGUGUGUAAAAGCACUUAAUUACUCUGUU.....UC...........AGAU.....................GA.....UUCAGGG.C...GAAGGAGUUGAAAAGAA..AUG
Lgr-2-1    ....GAAAUGUUUAAAAAAUAACCUUGCUUGUUUGCUUUACGCUUGUUAUAAUGAAGAUACUGUGCAAGUAAUGCAGUUAAUAACUCUGUU.....UC...........AGAU.....................GA.....UUCAGGG.C...GAAGGAGUUGAAAAUA...AUG
Lgr-3-1    ....AAAUGUUUAAAAAAAUAACCUUGCUUGUUUGCUUUACGCUUGUUAUAAUGAAGAUACUGUGCAAGUAAUGCAGUUAAUAACUCUGUU.....UC...........AGAU.....................GA.....UUCAGGG.C...GAAGGAGUUGAAAAUA...AUG
Lco-1-1    ....AAAUGUUUAAAAAAUAAGCCUUGCUAAAAUAGAAAAGCAUUGGUAUAAUGAGUUUACUGUGUGCAAUUGUGCAGAAAUUACUCUGUU.....UC...........AGAU.....................GA.....UUCAGGG.C...GAAGGAGUUGAAAACA...AUG
Lro-1-1    ....AAAUGUUUAAAAAAUAAGCCUUGCUAAAAUAGAAAAGCAUUGUUAUAAUGGUUUUACUGUGUGCAAUUGUGCAGAAAUUACUCUGUU.....UC...........AGAU.....................GA.....UUCAGGG.C...GAAGGAGUUGAAAACA...AUG
Lri-1-1    ....AAAUGUUUAAAAAAUAAGCCUUGCUAAAAGUGAAAAGCAUUGUUAUAAUAAAUUUACUGUGUGCAAUUGUGCAGAAAUUACUCUGUU.....UC...........AGAU.....................GA.....UUCAGGG.C...GAAGGAGUUGAAAACA...AUG
Lgr-1-1    ....AAAUGUUUAAAAAAUAAGCCUUGCUAAAAUAGAGAAGCAUUGUUAUAAUGAGUUUACUGUGUGCAAUUGUGCAGAAAUUACUCUGUU.....UC...........AGAU.....................GA.....UUCAGGG.C...GAAGGAGUUGAAAACA...AUG
Lba-2-1    ....AAAUGUUUAAAAAAUAAGCCUUGCUAAAUUAGAAAAGCAUUGUUAUAAUGAGUUUACUGUGUGCAAUUGUGCAGAAAUUACUCUGUU.....UC...........AGAU.....................GA.....UUCAGGG.C...GAAGGAGUUGAAAACA...AUG
Lbo-2-1    ....AAAUGUUUAAAAAAUAAGCCUUGCUAAAAAUGAAAAGCAUUGUUAUAAUAAAUUUACUGUGUGCAAUUGUGCAGAAAUUACUCUGUU.....UC...........AGAU.....................GA.....UUCAGGG.C...GAAGGAGUUGAAAACA...AUG
Bsp-94-1   ....ACUAGGAUCCCUUGCUAUAAUGAUCAUAGUGUGUUUUUAGCUUAUUAAUGGUUGUUAAAAAGACAAGCAUUAAUUUAUAACUCUGUU.....UCA..........GAA.....................UGA.....CUCAGGG.C..AGAAGGAGAUGAAAAGA...AUG
Bsm-2-1    ....AUGAAAUACAUUUUCUGCUAUAAUCCUACUAGUGUUUUUUGAAAAAAGAAGUAUGCGGUUGUUUCCGCACUUUCAUAGAACUCUGUU.....UC...........AGAU.....................GA.....CUCAGGG.CAUAAAAGGAGAUGAAAAGA...AUG
Blo-1-1    ....CUCGACUACAGUUGAUAUAAUAAGUGCAUGUAUUUUUACAAAUUAUGUGAGGGUCACUUAUCUUCGCAUUGAAGUUAUAACUCUGUU.....UCA..........GAA.....................UGA.....UUCAGGG.C..AGAAGGAGCUGAAAAGA...AUG
Bci-1-1    ....CUCGACUACAGUUGUUAUAAUAAGUGCAUGUGUUUUUACAAUUUAUGUGAGGGUUACUUCUCUUCGCAUUGAAGUUAUAACUCUGUU.....UCA..........GAA.....................UGA.....UUCAGGG.C..AGAAGGAGCUGAAAAGA...AUG
Pth-2-1    ....GCAAAAGGAAGACAAAGUUGUUAUAAUUUUAACAUGUGUGUUUUGGUAAAUGCCAAUAGAUGAACAUAACGAUAUAUAAACUCUGUU.....UCG..........AAA.....................UGA.....UACAGGG.C..GGAAGGAGAUGAGAAUA...AUG
Bps-1-1    ....AACAACGUAUAAAAUCAUCGAAAAUACAGUUGCAAAACGAAGCUUGGCUUGAUAUAAUUUCGUCAUGUGUGAUUAGAUAACUCUGUU.....UC...........GAAGA....................GA.....UUCAGGG.C..GGAAGGAGAUGAAAUCA...AUG
Bma-5-1    ....AACAACGUAUAAAAUCAUCGAAAAUACAGUUGCAAAACAAGGCUUGGCUUGAUAUAAUUUCGUCAUGUGUGAUUAGAUAACUCUGUU.....UC...........GAAGA....................GA.....UUCAGGG.C..GGAAGGAGAUGAAAUCA...AUG
Bsp-66-2   ....CAUACCUUUUAAAUAGAUGUAAAAUACAGUUGCAAAAGGAAGCUUAGCUUGUUAUAAUUUCGUCAUGUGUGAUUAGAUAACUCUGUU.....UC...........GAAAA....................GA.....UUCAGGG.C..GGAAGGAGAUGAAAGCA...AUG
Bsp-92-1   ....AAAGUAAACACUCGUUGUUAUAAUGGGUACAGUGUGUUUUAAGUAUUUCAGUGCCUGUUUAAGACAAGCACUUAUGAUAACUCUGUU.....UC...........GAAA.....................GA.....UUCAGGG.C..GGAAGGAGCUGAAAAAA...AUG
env-936    ....GACGUCAUAAUCUUUUAUUGCAUUCUCCUUCCCGCCUUGUUAUACUUUUUUUAUGUGUGAAAGAGUUCAACACAGACUUACUCUGUU.....UC...........GAAA.....................GA.....UUCAGGG.C..GGAAGGAGCUGAAAAGAC..AUG
Pge-1-1    ....GUUGCAAAGAAACAGGAAAGUUGUUAUAAUGUUAGCAUGUGUGUUUUGAUAAACGCCAAUUCAUUUAAUUGAUAUAUCAACUCUGUU.....UCG..........AAA.....................UGA.....UACAGGG.C..GGAAGGAGAUGAAAAGA...AUG
Pca-1-1    ....GUUGCAAAGGAACAGGAAAGUUGUUAUAAUCUUAAUAUGUGUGUUUUGGUAAACGCCAAUUCAUUUUAUUGAUAUAUCAACUCUGUU.....UCG..........AAA.....................UGA.....UACAGGG.C..GGAAGGAGAUGAAAAGA...AUG
Bsp-79-1   ....AAAACAAUGCUAUUGCAAAAGAAGUAAACCCUUGUUAUAAUUCAAAGAUGUGCGACAUCACGCAAUUAGCGUGGAUAUAACUCUGUU.....UC...........AGAU.....................GA.....UUCAGGG.C..GGAAGGAGAUGAAAAAC...AUG
env-937    ....UUGCAAAUCACGGAUUAGCUUGCUAUAAUAAAAAGCAUGUUUGAUUAAAGAUGGGUGAAGUUUUACCCUUCAUUAUAUAACUCUGUU.....UCA..........GGCAA...................UGA.....UUCAGGG.C..GGAAGGAGAUGAAAAGA...AUG
Bsp-54-1   ....GCACUUGUUAUAAUUUCAUCAUGUGCUUUUCUAGGAUAUUAGUUCGUCAAUUGGACGAGUUGACCUACCUAGAUAUAUAACUCUGUC.....UC...........GAAU.....................GA.....UUCAGGG.C..GGAAGGAGAUGAAAAGA...AUG
Bsp-30-1   ....CCCAUCUUGCAAAACAGCGAUGCGCUUGUUAUAAUGAUAACAGUGUUUUUUACCGAGGGCUGCUAUGGCCCAGGCUAUAACUCUGUU.....UCG..........AAA.....................UGA.....CUCAGGG.C..GGAAGGAGAUGACAAGA...AUG
Bba-2-1    ....UUGGACAAACCCUUGUUAUAAUAAUACAGUGUGCUUAGCCUAUAAGUACGGGGUAAUCUUGCCGUACUUCGUAUAGACAACUCUGUU.....UCG..........AAA.....................UGA.....UUCAGGG.C..GGAAGGAGAUGACAAGA...AUG
Bkr-1-1    ....AAUGUAAGAUUGUUGCUAUAAUAAAGUUAAUGUGUUUGAAUGUUUUAUGCGGAAUAACACAGCCGCAUUUAAGGUAUAAACUCUGUU.....UCG..........GUA.....................UGA.....CUCAGGG.C..GGAAGGAGAUGAAAAGA...AUG
Bsp-17-1   ....AAAUAUAUAGAUGGACUUGUUAUAAUGAUUGCAGUGUGUUUUUCCAAUAACGGCAGGAUUGCUAUAGCCGUUGAUUAUAACUCUGUU.....UCG..........GAUA....................UGA.....UUCAGGG.C..GGAAGGAGAUGAAAAGA...AUG
Bse-2-1    ....CUAACUUGAAAUAUACAGUUGGACUUGUUAUAAUGAGUGCAGUGUGUUUUUCCAAUAACGGCCUAAGCCGUUGAUUAUAACUCUGUU.....UCG..........GAUA....................UGA.....UUCAGGG.C..GGAAGGAGAUGAAAAGA...AUG
Bsu-11-1   ....UUUGGUUUGUAAAUUGAGCUUGAAAUAUAAAGAUAGACUUGGUAUAAUAGUGUCAGUGUGUUUUUUCAAUAACGGCAUAACUCUGUU.....UCG..........GAUA....................UGA.....UUCAGGG.C..GGAAGGAGAUGAAAAGA...AUG
Bsp-37-1   ....CUUGCAAAAAGGACACGUCCCUGAUAUAAUGAUACCAGUGUGUUUUUCCACCUCGGGCUUAUAGGUGCCCGAUUCUAUAACUCUGUU.....UCG..........GAA.....................UGA.....CUCAGGG.C..GGAAGGAGAUGAAAAGA...AUG
Bru-1-1    ....GAUAGGGCGUUGGUAUAAUGUGUACAGUGUGUUUUUAGUAAUAUAAGUGCUGUUUUUAGACAAGCACUUUUGAAUGAUAACUCUGUU.....UCG..........GAA.....................UGA.....UUCAGGG.C..GGAAGGAGAUGAAAAGA...AUG
Bba-3-1    ....UAGCAGAACACUUGCUAUAAUAAUGUCAGUGUGCUUAGGACAUAAGUGCAGAGCAAGAAGUCUGCGCUUCGUAUAGACAACUCUGUU.....UCG..........GAA.....................UGA.....UUCAGGG.C..GGAAGGAGAUGAAAAGA...AUG
Bsp-40-1   ....UCUUCAUGUGUGUUUAGUAUACGAUGUCGAAAUGAGCAUAUCGAAUGCUGAUGUUUCGUCUAUCGUAUUGGUUUAUAUAACUCUGUU.....UCA..........GAA.....................UGA.....UUCAGGG.C..GGAAGGAGAUGAAAAGC...AUG
Bsp-42-1   ....UGGACAAACACUUGUUAUAAUAAUACCAGUGUGUUCAGCCUAUAAGUACGGAGUAAUAACACCGUACUUCGUAUAGACAACUCUGUU.....UCA..........GAA.....................UGA.....UUCAGGG.C..GGAAGGAGAUGACAAGA...AUG
Bsp-96-1   ....UAGGGGAGUCCCCUGUUAUAAUGGGGACAGUGUGUUUUUAGUCAUUGAGUGUAGUCUUAAAGACAAGCACUUCAUUUUAACUCUGUU.....UC...........AGAU.....................GA.....UUCAGGG.C..GGAAGGAGAUGAAAAGA...AUG
Bfl-3-1    ....CCCUUGUUAUAAUUUCAUCAUGUGUGUUGAUAAGAUACACGAUGGUGCGAUUAUGCUCAUCAUUUUUUCUUAUCAAUAAACUCUGUU.....UCA..........GAA.....................UGA.....UUCAGGG.C..GGAAGGAGAUGAAAAGA...AUG
Bsp-81-1   ....CUCUUAGCACUUGUUAUAAUUUCAUCAUGUGCUUUUCUAGAAAUGAUAUCGUCAAUCGGACGAGUCACCUAGAUAUUUAACUCUGUU.....UC...........AGAU.....................GA.....UUCAGGG.C..GGAAGGAGAUGAAAAGA...AUG
Bme-1-1    ....CCUUGUUAUAAUUUCAUCAUGUGUGUUGAUAAAGUACACAGUGAAAUAGUUAUUAUUUCAUCACUAUCUUUAUCAAUAAACUCUGUU.....UCA..........GAA.....................UGA.....UUCAGGG.C..GGAAGGAGAUGAAAUGA...AUG
Bfl-1-1    ....CCUUGUUAUAAUUUCAUCAUGUGUGUUGAUAAAGUACACAGUGAAAUAGUUACUAUUUCAUCACUAUCUUUAUCAAUAAACUCUGUU.....UCA..........GAA.....................UGA.....UUCAGGG.C..GGAAGGAGAUGAAACGA...AUG
env-938    ....CCUUGUUAUAAUUUCAUCAUGUGUGUUGAUAAAGUACACAGUGAAAUAGUUAUUAUUUCAUCACUAUCUUUAUCAAUAAACUCUGUU.....UCA..........GAA.....................UGA.....UUCAGGG.C..GGAAGGAGAUGAAACGA...AUG
Bpa-1-1    ....CAAAAUGCAUCCUACCUUGUUAUAAUUUCAUCAUGUGUGUUUCAUCGUUAGACUAUGCUGCUAUAGCAUUUGAUGAAAAACUCUGUU.....UC...........UAAAAU...................GA.....UUCAGGG.C..GGAAGGAGAUGAAAAGA...AUG
Ban-4-1    ....GCAAAAUGAAGUUAGCCUUGUUAUAAUUUCAUCAUAUGUGUUUCAUCAAUAUAGUUGUGUAUGCAGCUAAAGAUGAAAAACUCUGUU.....UCG..........AAAA....................UGA.....UUCAGGG.C..GGAAGGAGAUGAAAAGA...AUG
Bce-63-1   ....GCAAAAUGGAGUUAGCCUUGUUAUAAUUUCAUCAUAUGUGUUUCAUCAAUAUAGUUGUGCAAGCAGCUGAAGAUGAAAAACUCUGUU.....UCG..........AAAA....................UGA.....UUCAGGG.C..GGAAGGAGAUGAAAAGA...AUG
env-939    ....GCAAAAUGAAGUUAGCCUUGUUAUAAUUUCAUCAUAUGUGUUUCAUCAUUAUAGUUGUAUUCGCAGCUAAAGAUGAAAAACUCUGUU.....UCG..........AAAA....................UGA.....UUCAGGG.C..GGAAGGAGAUGAAAAGA...AUG
Bma-2-1    ....CAAAAUGAGGAUAGCCUUGUUAUAAUUUCCUCAUGUGUGUUUCAUCAUUAUAGUUGUAUACGCAGCUAAAGAUGAAAAAACUCUGUU.....UCG..........AAAA....................UGA.....UUCAGGG.C..GGAAGGAGAUGAAAAGA...AUG
Bcy-2-1    ....GCAAAAUGGUGUGAGCCUUGUUAUAAUUUCAUCAUAUGUGCUUCAUCAAUAUAGUUGUGAAUGCAGCUAGAGAUGAAAAACUCUGUU.....UCG..........AAAA....................UGA.....UUCAGGG.C..GGAAGGAGAUGAAAAGA...AUG
Bsp-39-1   ....GCAAAAUGAAGUUAGCCUUGUUAUAAUUUCAUCAUAUGUGUUUCAUCAUUAUAGUUGUGAUUGCAGCUAAAGAUGAAAAACUCUGUU.....UCG..........AAAA....................UGA.....UUCAGGG.C..GGAAGGAGAUGAAAAGA...AUG
Bdi-1-1    ....CCUGUUAUAAUGGGGUCAGUGUGUUUUUAGUUUUUUUGAUACCAAGUGUUUGUUCUAUCCAGACAAGCACUUUACGAUAACUCUGUU.....UCG..........AACA....................UGA.....UUCAGGG.C..GGAAGGAGAUGAAAAGA...AUG
Bfi-1-1    ....AAAAAAUGUAUGGACUUGUUAUAAUGAUGACAGUGUGUUUUUGAAGUUUACGGCAAUUCAUGAUUGCCCGUAAUAUAUAACUCUGUU.....UC...........GAAU.....................GA.....UUCAGGG.C..GGAAGGAGAUGAAAAGA...AUG
Bba-1-1    ....AAAUAAGUUGACCUUGUUAUAAUGUUAGCAGUGUGUUUUAAGUAUAUUACUGCUUGGUUUUUGACAAGCAUUUAUGAUAACUCUGUU.....UC...........GAAU.....................GA.....UUCAGGG.C..GGAAGGAGAUGAAAAGA...AUG
Bni-1-1    ....AAAACAGGUUCACCUUGUUAUAAUAGUUGCAGUGUGUUUUUAGUAUUUGGUGCUUGUUUUUAAGACAGGCAUUUAUAUAACUCUGUU.....UC...........GAAU.....................GA.....UUCAGGG.C..GGAAGGAGAUGAAAAGA...AUG
Bsp-53-1   ....AAAUUAUACUUGUACUUGUUAUAAUGAUCACAGUGUGUUUUUGAAGUUUACGGCAAUCGCUGGUUGCCCGUAAUAUAUAACUCUGUU.....UC...........GAAU.....................GA.....UUCAGGG.C..GGAAGGAGAUGAAAAGA...AUG
Age-1-2    ....AAAACCAUAACACCCUUGCUAUAAUUUUAUCAUGUGUGUUUUGUCAAACAAUGAGAAAUGACUCACAACGGAUAGAGAAACUCUGUU.....UCG..........AAA.....................UGA.....UUCAGGG.C..GGAAGGAGAUGAAAAGAA..AUG
Ate-1-1    ....CAAAACAAAGGCACCCUUGCUAUAAUUUUGUCAUGUGUGUUUUGUUAAACGGUGAGAAAUGACUCACAACGAUAGAGAAACUCUGUU.....UCG..........AAA.....................UGA.....UUCAGGG.C..GGAAGGAGAUGAAAAGAA..AUG
Bce-6-1    ....GCAAAAUAGAGUUAGCCUUGUUAUAAUUUCAUCAUAUGUGUUUCAUCAAUAUAGUUGUGCAAGCAGCUGAAGAUGAAAAACUCUGUU.....UCG..........AAAA....................UGA.....UUCAGGG.C..GGAAGGAGAUGAAAAGA...AUG
Bce-106-1  ....GCAAAAUGAAGUUAGCCUUGUUAUAAUUUCAUCAUAUGUGUUUCAUCAAUAUAGUUGUGUACGCAGCUGAAGAUGAAAAACUCUGUU.....UCG..........AAAA....................UGA.....UUCAGGG.C..GGAAGGAGAUGAAAAGA...AUG
env-941    ....GCAAAAUGAAGUUAGCCUUGUUAUAAUUUCAUCAUAUGUGUUUCAUCAAUAUAGUUGUGAUUGCAGCUAAAGAUGAAAAACUCUGUU.....UCG..........AAAA....................UGA.....UUCAGGG.C..GGAAGGAGAUGAAAAGA...AUG
env-942    ....AAAAUGUAGAUAGCCUUGUUAUAAUUUCAUCAUGUGCGUUUCAUCAACUAUAGUUGUGAUUGCAGCUAACGAUGAAAAAACUCUGUU.....UCG..........AAAA....................UGA.....UUCAGGG.C..GGAAGGAGAUGAAAAGA...AUG
Bsp-15-1   ....CAAAAUGCAGAUAGCCUUGUUAUAAUUUCAUCAUGUGCGUUUCAUCACUACAGUUGUGAUAACAGCUAAUGAUGAAAAAACUCUGUU.....UCG..........AAAA....................UGA.....UUCAGGG.C..GGAAGGAGAUGAAAAGA...AUG
Bga-1-1    ....GCAAAAUGAAGUUAGCCUUGUUAUAAUUUCAUCAUAUGUGUUUCAUCAUUAUAGUUGUAUUUGCAGCUAAAGAUGAAAAACUCUGUU.....UCG..........AAAA....................UGA.....UUCAGGG.C..GGAAGGAGAUGAAAAGA...AUG
env-943    ....ACAUGUUUCCCUUGUUAUAAUGGAUGCAGUGUGUUUUUAGUAUUUCAGUGUUUGUUUUAGACAAGCACUUCAUAUGAUAACUCUGUU.....UC...........GAAU.....................GA.....UUCAGGG.C..GGAAGGAGAUGAAAAGA...AUG
env-944    ....GCAAAAUGAAGUUAGCCUUGUUAUAAUUUCAUCAUAUGUGUUUCAUCAAUAUAGUUGUGUAUGCAGCUGAAGAUGAAAAACUCUGUU.....UCG..........AAAA....................UGA.....UUCAGGG.C..GGAAGGAGAUGAAAAGA...AUG
env-945    ....AAAUACAAUUGGACUUGUUAUAAUGAGGACAGUGUGUUUUUCCAAUAACGGCUGGUAUGAACAAUAGCCGUUGAUUAUAACUCUGUU.....UC...........GAAU.....................GA.....UUCAGGG.C..GGAAGGAGAUGAAAAGA...AUG
Bsp-21-1   ....AAAAUAUACAUGGACUUGUUAUAAUGAUGACAGUGUGUUUUUGAAGUUUACGGCAAUUCAUGAUUGCCCGUAAUAUAUAACUCUGUU.....UC...........GAAU.....................GA.....UUCAGGG.C..GGAAGGAGAUGAAAAGA...AUG
Bfi-2-1    ....AAAAAACACAUGGACUUGUUAUAAUGAUGACAGUGUGUUUUUGAAGUUUACGGCAAUUCAUGAUUGCCCGUAAUAUAUAACUCUGUU.....UC...........GAAU.....................GA.....UUCAGGG.C..GGAAGGAGAUGAAAAGA...AUG
Sgl-1-1    ....AAAAAACAAUUGGACUUGUUAUAAUGAUGACAGUGUGUUUUUGAAGUUUACGGCAAUUCAUGAUUGCCCGUAAUAUAUAACUCUGUU.....UC...........GAAU.....................GA.....UUCAGGG.C..GGAAGGAGAUGAAAAGA...AUG
Bsp-44-1   ....UGCAAAAGCAGAACAGGCUUGUUAUAAUAAUAACAGUGUUUUUUUACCACUUCGGGGCUGAAAUGCACCCGGGUCUAUAACUCUGUU.....UCG..........AAA.....................UGA.....UUCAGGG.C..GGAAGGAGAUGAAAAGA...AUG
env-946    ....GCAAAAUGGAGUUAGCCUUGUUAUAAUUUCAUCAUAUGUGUUUCAUCAAUAUAGUUGUGUAAGCAGCUGAAGAUGAAAAACUCUGUU.....UCG..........AAAA....................UGA.....UUCAGGG.C..GGAAGGAGAUGAAAAGA...AUG
Afl-1-1    ....AAACGGUUAUAAAUGGAUAGUUGCAAAACAGUGAUGCGGUUGUUAUAAUUUUGUCAUGUGUGUUUUUCAUUGCAUAUAAACUCUGUU.....UC...........GAAAA....................GA.....UUCAGGG.C..GGAAGGAGAUGAAAUGAG..AUG
Afl-3-1    ....AAACGGUUAUAAAUGGAUAGUUGCAAAACAUCGGGACGGUUGUUAUAAUUUUAUCAUGUGUGUUUUUCAUUGCAUAUAAACUCUGUU.....UC...........GAAAA....................GA.....UUCAGGG.C..GGAAGGAGAUGAAAUGAG..AUG
Ath-1-1    ....AAACGGUUAUAAAUGGAUAGUUGCAAAACAGUGAUACGGUUGUUAUAAUUUUGUCAUGUGUGUUUUUUAUUGCAUAUAAACUCUGUU.....UC...........GAAAA....................GA.....UUCAGGG.C..GGAAGGAGAUGAAAUGAG..AUG
Afl-6-1    ....AAACGGUUAUAAAUGGAUAGUUGCAAAACAGCGGGACGGUUGUUAUAAUUUUAUCAUGUGUGUUUUUGAUUGCAUAUAAACUCUGUU.....UC...........GAAAA....................GA.....UUCAGGG.C..GGAAGGAGAUGAAAUGAG..AUG

           ........................................................................................<<<<<.......<<<<<<<<<<<<<..............>>>>>>>>>>>>>.......>>>>>...........................
           ..........................................................................................222.......22211...1111...............1111...11222.......222.........................
           RR••••••••••••••••••••••••••••••••••••••••••••••••••••••••••••••••••••••••••••••••••••Y••CU•UG------•Y○----------••R○○-------------------○R•-------CA•GG-C--RRAAGGAG••R•R••○○---AUG
                                                                                                 5′                                                                          3′
```

```
Afl-4-1     ....AAACGGUUAUAAAUGGAUAGUUGCAAAACAGCGAGGCGGUUGUUAUAAUUUUAUCAUGUGUGUUUUUCAUUGCAUAUAAACUCUGUU.....UC...........GAAAA....................GA.....UUCAGGG.C..GGAAGGAGAUGAAAUGAG..AUG
Afl-5-1     ....AAACGGUUAUAAAUGGAUAGUUGCAAAACAACGGAACGGUUGUUAUAAUUUUAUCAUGUGUGUUUUUCAUUGCAUAUAAACUCUGUU.....UC...........GAAAA....................GA.....UUCAGGG.C..GGAAGGAGAUGAAAUGAG..AUG
Asp-3-1     ....AAACGGUUAUAAAUGGAUAGUUGCAAAACAACGGGACGAUUGUUAUAAUUUUAUCAUGUGUGUUUUUCAUUGCAUAUAAACUCUGUU.....UC...........GAAAA....................GA.....UUCAGGG.C..GGAAGGAGAUGAAAUGAG..AUG
Ago-1-1     ....AAACGGUUAUAAAUGGAUAGUUGCAAAACAGCGAUACGGUUGUUAUAAUUUUAUCAUGUGUGUUUUUCAUUGCAUAUAAACUCUGUU.....UC...........GAAAA....................GA.....UUCAGGG.C..GGAAGGAGAUGAAAUGAG..AUG
Afl-2-1     ....AAAACGGUUAUAAAUGGAUAGUUGCAAAACAGUGAUACGAUUGUUAUAAUUUUAUCAUGUGUGUUUUCAUUGCAUAUAAACUCUGUU.....UC...........GAAAA....................GA.....UUCAGGG.C..GGAAGGAGAUGAAAUAAG..AUG
Aay-1-1     ....AAACGGUUAUAAAUGGAUAGUUGCAAAACAGUGAUACGAUUGUUAUAAUUUUAUCAUGUGUGUUUUUCAUUGCAUAUAAACUCUGUU.....UC...........GAAAA....................GA.....UUCAGGG.C..GGAAGGAGAUGAAAUAAG..AUG
Bsp-18-1    ....GCAACAUGCUUGUUAUAAUAAUUUCAGUGUGUUUUUGAAUUUUACGGAGUUAGGUAUUUGUUCUAACACGUAACAUAUAACUCUGUU.....UC...........GAAA.....................GA.....UUCAGGG.C..GGAAGGAGAUGAAAAGA...AUG
Bje-1-1     ....AAAUAAGGUUCCCCUUGUUAUAAUGGGUGCAGUGUGUUUGAAGUAUUUCAUGCUUGUUUUUACGACAGGCAUUUUUAUAACUCUGUU.....UC...........GAAA.....................GA.....UUCAGGG.C..GGAAGGAGAUGAAAAGA...AUG
Bme-4-1     ....UGCAAAAUGAAGGUCUGGUUGUUAUAAUAGGUUCAGUGUUUUUUUGAAGAUUGCGGUUUAUGAAGCCGUAACAUAUAUAACUCUGUU.....UC...........GAAA.....................GA.....UUCAGGG.C..GGAAGGAGAUGAAAAGA...AUG
Bvi-2-1     ....AAACAGAUAUCCCUUGUUAUAAUUGUUGCAGUGUGCUAUAAGUAUUUCAGUGCUCGUCGACAGACAAGCACUUUAGCUAACUCUGUU.....UC...........GAAA.....................GA.....UUCAGGG.C..GGAAGGAGAUGAAAAGA...AUG
env-948     ...............................................................................UAUAACUCUGUU.....UC...........GAAA.....................GA.....UUCAGGG.C..GGAAGGAGAUGAAAAGA...AUG
Bsp-90-1    ....AAAUAGGUUGCCCUUGUUAUAAUGGGUGCAGUGUGUUUUUAGUAUUUCAGUGCUUGUUUUUGGACAGGUGCUUUAUAUAACUCUGUU.....UC...........GAAA.....................GA.....UUCAGGG.C..GGAAGGAGAUGAAAAGA...AUG
Bal-4-1     ....AAAGACGGCUUGUUAUAAUUUUAUCAUGUGUGUUUUGCUAAAACGUGAGAAGAUGCUCUUACUCACAACUUAAUAUAUAACUCUGUU.....UC...........GAAAA....................GA.....AUCAGGG.C..GGAAGGAGAUGAAAAGA...AUG
Bce-1-1     ....UGCAAAUUAAAUGCACACUUGAUAUAAUUUCACAGUAUGCAAAAAUGUACGAUGCAGGGCGUAUGCUACUGCUCAUUAAACUCUGUU.....UC...........GAAA.....................GA.....CUCAGGG.C..GGAAGGAGAUGAAAAGA...AUG
env-949     ..................................................................AGUCUGUAUCAUAGAUAACUCUGUU.....UC...........GAAA.....................GA.....CUCAGGG.C..GGAAGGAGAUGAAAAGA...AUG
env-950     ............................................................ACUAGCUAUAGUGCUAUAAUAUAACUCUGUU.....UC...........GAAA.....................GA.....UCCAGGG.C..GGAAGGAGAUGAAAAGA...AUG
Bgi-2-1     ....CAUUUCACAGUGGCACUUGUUAUAAUAAUCACAGUGUGUUUCUGGAUAUUUUACUGGGUUAAGCCCAUAAGAAUAUAUAACUCUGUU.....UCG..........CAA.....................UGA.....UUCAGGG.C..UGAAGGAGAUGAAAAGA...AUG
env-951     .....................................AGUGUGUUUAUGAAAUUUUACAGGGCGUGUCCCGUAAAACAAUAGAACUCUGUU.....UCG..........CAA.....................UGA.....CUCAGGG.C..UGAAGGAGAUGAAAAGC...AUG
Bco-9-1     ....GCAUUUCCACUCCGGACUUGUUAUAAUAAUGACAGUGUGUUUAUGAAAUUUUACAGGGCGUGUCCUGUAAAACAAUAGAACUCUGUU.....UCG..........CAA.....................UGA.....CUCAGGG.C..UGAAGGAGAUGAAAAGC...AUG
env-952     ....GCAUUUCCACUCCGGACUUGUUAUAAUAAUGACAGUGUGUUUAUGAAAUUUUACAGGGCGUGUCCCGUAAAACAAUAGAACUCUGUU.....UCG..........CAA.....................UGA.....CUCAGGG.C..UGAAGGAGAUGAAAAGC...AUG
Bco-5-1     ....GCAUUUCCACUCCGGACUUGCUAUAAUAAUGACAGUGUGUUUAUGAAAUUUUACAGGGCGUGUCCCGUAAAACAAUAGAACUCUGUU.....UCG..........CAA.....................UGA.....CUCAGGG.C..UGAAGGAGAUGAAAAGC...AUG
env-954     ....GCAUUUCCACUUCGGACUUGCUAUACUAAGCAUAGUGUGUUUAUGAAAUUUUACAGGNCGUGUCCCGUAAAACAAUAGAACUCUGUU.....UCG..........CAA.....................UGA.....CUCAGGG.C..UGAAGGAGAUGAAAAGC...AUG
env-955     ....GCAUUUCCACUUCGGACUUGCUAUACUAAGCAUAGUGUGUUUAUGAAAUUUUACAGGGCGUGUCCCGUAAAACAAUAGAACUCUGUU.....UCG..........CAA.....................UGA.....CUCAGGG.C..UGAAGGAGAUGAAAAGC...AUG
Bal-4-2     ....CGAAAAGGCUUGAUAUAAUUUCAUCAUGUGUAUUUCAGACAUUGAAGCGUGAUUUUCGAUCACGUUUCUAUAAUAGAUAACUCUGUU.....UCG..........AAA.....................UGA.....UUCAGGG.C.GAAAAGGAGAUGAAUAGA...AUG
Bps-4-1     ....GAGUUUUGUUGCUAUAAUAAAGGCAUGUGUUUUUACAAAAUAUGUGAGGGGUACUACUUACCUCACCUGAAAGUAUAUAACUCUGUU.....UC...........GAAU.....................GA.....UUCAGGG.C..AUAAGGAGAUGAAAAAGA..AUG
Bac-1-1     ....CAUUUGGGGACUUGUUAUAAUAAUCACAGUGUGUUUUAGAAAUUUUACUGGGCAGGACUGACACCCGUAAAAAUAUACAACUCUGUU.....UCG..........AAA.....................UGA.....CUCAGGG.C.AUUAAGGAGAUGAAAAGA...AUG
Bbu-1-1     ....UAGUCUUAAAUUGCUAUAAUGAGUUCAUGUGUUUUUACAAUAAAUGUGAGGGUACUUUUCUCUCACCUUGAAGUAUAUAACUCUGUU.....UC...........GAAU.....................GA.....UUCAGGG.C..AGAAGGAGCUGAAAAGA...AUG
env-958     ....UAGUCUUAAGUUGUUAUAAUGAGUUCAUGUGUUUUUACAAUAAAUGUGAGGGUGCUUUUCCCUCAACUUGAAGUAUAUAACUCUGUU.....UC...........GAAU.....................GA.....UUCAGGG.C..AUAAGGAGAUGAAAAGA...AUG
Bsp-19-1    ....UAGUUUUAAGUUGUUAUAAUGAGUUCAUGUGUUUUUACAAUAAAUGUGAGGGUGCUUUUCCCUCAACUUGAAGUAUAUAACUCUGUU.....UC...........GAAU.....................GA.....UUCAGGG.C..AUAAGGAGCUGAAAAGA...AUG
Lwe-2-1     ....AAAUGUUUAAAAAAUAAGCCUUGCUAAAUUGAAAAAGCAUUGGUAUAAUGAGUUUACUGUGUGAAAUUGUGCAGAAAUUACUCUGUU.....UC...........AGAU.....................GA.....UUCAGGG.C...CAAGGAGUUGAAAGCA...AUG
Bsp-47-1    ....AGGCGAUUUGUUGAUUUAUAAAAGUAUGUCUGUUAUACUGUUUUAAGUGCACUUCACUUGUGUGAAAAGGCACCUUAUAACUCUGUU.....UC...........GAAA.....................GA.....CUCAGGG.C..GGAAGGAGUGGAAAAG....AUG
Bko-1-1     ....AAUUUCAUCAUGUGUGUAUUAGCUAAGUGUAUGUGAAGGUGAAAAGAUCGUUGUAGUUUUCAUAUAUGCUUUCAAUAUAACUCUGUU.....UC...........AAAA.....................GA.....UUCAGGG.C..GGAAGGAGAUGAAAUC....AUG
Bsp-28-1    ....CAAAUCCAUUUAUCGCUUGAUAUAAUAAAAAGCAUGUUUGAUUAUGAAGCUGAGAGGUUCUUUUACCUUUCAACAUAUAACUCUGAU.....UCA..........GGAAA...................UGA.....UUCAGGG.C..GGAAGGAGCUGAAAAGA...AUG
Bbo-1-1     ....GGCCAAAGCGAAAAUAAAUUCGAUUUCCAGUUGCAAAUUCAUUCUCACCUUGGUAUAAUUUCGUCAUGUGUGUUAGAUAACUCUGUU.....UC...........GAAUA....................GA.....UUCAGGG.C..GGAAGGAGUUGAAAAGCA..AUG
Bsp-62-1    ....AAAAUAAGCUGGUUUCGUUAAAACAGCAGUUGCAAAAACGGAUAGGGCUUGUUAUAAUUUCGUCAUGUGUGAUUAGAUAACUCUGUU.....UC...........GAAA.....................GA.....AUCAGGG.C..GGAAGGAGCUGAAAACG...AUG
Ssp-6-1     ....AAAAUAAUUUGGAAAUGGUUGAUUUUUGAAGUGCUACAAUGUAUAAUACACAAGUAUUGUGUGGCAAAACCACAAAAUAACUCUGUU.....CC...........AGAU.....................GG.....UUCAGGG.C...AAGGGAGCUGAAUAUC...AUG
Sxy-3-1     ....AAGCAAUAUUUGAUUUGGGGUUGCGUUUUAUAUUUAAAGCCAUUAUAAUUAUCAAGUAUUGGGUAAAAACUCACAAAAAACUCUGUG.....CC...........AGAU.....................GG.....UUCAGGG.C...AAAGGAGCUGAAAAGUA..AUG
Sau-1739-1  ....AAUAAACAUUAUAUAGGGGCUUGCGUUUUGAAUUAAUUACCUUUAUAAUUACACAGUAUUGGGUAAAAACUCACAAAUAACUCUAUU.....CC...........AGAU.....................GG.....UUCAGGG.C...AAAGGAGCUGAAAAUA...AUG
Sau-305-1   ....AAUAAACAUUAUAUAGGGGCUUGCGUUUUGAAUUAAUUACCUUUAUAAUUACACAGUAUUGGGUAAAAACUCACAAAUAACUCUGUU.....CC...........AGAC.....................GG.....UUCAGGG.C...AAAGGAGCUGAAAAUA...AUG
env-959     ....CCAAUAUUUGAUUUGGGGUUGCGUUUUAGAAAUAAAGCCAUUAUAAUUAUCAAGUAUUGGGUAAAAAAACUCAGAAAUAACUCUGUU.....CC...........AGAU.....................GG.....UCCAGGG.C...AAAGGAGCUGAAAAUA...AUG
Sma-1-1     ....AAAGGAUUUCCAUAUGGGGUUGCUUUUUAAAAGGUAAACCGAUAUAAUACCUAAGUAUUGGGUAAAAAAACUCACAAAUAACUCUGUU.....CC...........AGAU.....................GG.....UUCAGGG.C...AAAGGAGCUGAAUAGAA..AUG
Smi-1-1     ....AUAAAUAAUCGUGUAUAGGGUUGCUUUUUGAAUACACACCCUUUAUAAUAACUAAGUAUUGGGUAAAAACUCACAAAUAACUCUGUU.....CC...........AGAU.....................GG.....UUCAGGG.C...AAAGGAGCUGAAAAGA...AUG
Sco-3-1     ....UAGAAGAAUCUUAGAAUGGGUUGCAAUUUAUACUUCUACCAAUUAUAAUUACUAAGUAUUGGGUAAAAACUCACAAAUAACUCUGUU.....CC...........AGAU.....................GG.....UUCAGGG.C...AAAGGAGCUGAAAUAAUA.AUG
Sca-8-1     ....UAGAAGAAUUUGAGAAUGGGUUGCAAUUUAUACUGCUACCAAUUAUAAUUACUAAGUAUUGGGUAAAAACUCACAAAUAACUCUGUU.....CC...........AGAU.....................GG.....UUCAGGG.C...AAAGGAGCUGAAAUAAUA.AUG
env-960     .....GUUAUCUUUGAUAAAGGGUUGCGUUUUGAUUUGAUAACUAUUAUAAUUACACAGUAUUGGGUAAAAAACUCACAAAUAACUCUGUU.....CC...........AGAU.....................GG.....UUCAGGG.C...AAAGGAGCUGAAAGAA...AUG
Sca-7-1     ....AACGAAUAUUGAUAAAGGGUUGCGUUUUAAAUUGAUUCCUAUUAUAAUUACACAGUAUUGGGUAAAAAACUCACAAAUAACUCUGUU.....CC...........AGAU.....................GG.....UUCAGGG.C...AAAGGAGCUGAAAGAA...AUG
env-961     ....AUCAUAUAAAAUUUAAGGUUGCAAUUUAAUUACUCAACAUAUAUAAUAUGUAAGUAUUGAGUGAAUAAGUCACAAAUAAACUCUGUU.....CC...........AGAU.....................GG.....UUCAGGG.C...AAAGGAGCUGAAAGAA...AUG
Sca-5-1     ....AGUUAUCUUUGAUAAAGGGUUGCGUUUUGAUUUGAUAACUAUUAUAAUUACACAGUAUUGGGUAAAAAACUCACAAAUAACUCUGUU.....CC...........AGAU.....................GG.....UUCAGGG.C...AAAGGAGCUGAAAGAA...AUG
Sca-4-1     ....AAGAAAAAUGCAUCCCUUGAUUUUUUACACUGCAGGGUGUAUAAUAUAUUAAUGUCGAAGUGGCGGCUGCCACUAAUCAACUCUGUU.....CC...........AGAU.....................GG.....UUCAGGG.C...AAAGGAGCUGAAAGUA...AUG
Sse-1-1     ....AAAUGAAAUGCAUCUCUUGAUUUUUUACACUGCAGGGUGUAUAAUGUAUUAAUGUCGAAGUGGCGGAUGCCACUAAUCAACUCUGUU.....CC...........AGAU.....................GG.....UUCAGGG.C...AAAGGAGCUGAAAGUA...AUG
Sha-3-1     ....AACUGAAACACGCCUCUUGAUUUUUUACACCGCGGGGUGUAUAAUAUAUUAAUGUCGAAGUGGUAAAAGCCACUAAUAAACUCUGUU.....CC...........AGAU.....................GG.....UUCAGGG.C...AAAGGAGCUGAAAGUA...AUG
Sal-1-1     ....UAAGUUGAAACAGAACUAUUGAUUUUAAGCGAUACUCAGUGUAUAAUGUAUAAGUACUGAAGUGGAAACCCACUAUUAACUCUGUU.....CC...........AGAU.....................GG.....UUCAGGG.C...AAAGGAGCUGAAUGUA...AUG
Sep-24-1    ....UAAACUCGUAGAUUAAGGGUUGCGUUUUCCAUAUAUUACUAUUAUAAUUACACAGUAUUGGGUAAAAACUCACAAAUAACUCUGUU.....CC...........AGAU.....................GG.....UUCAGGG.C...AAAGGAGCUGAAAAAA...AUG
Sep-8-1     ....UAAACUCGUAGAUUAAGGGUUGCGUUUUCCAAAUAUUACUAUUAUAAUUACACAGUAUUGGGUAAAAAACUCACAAAUAACUCUGUU.....CC...........AGAU.....................GG.....UUCAGGG.C...AAAGGAGCUGAAAAA....AUG
env-962     ....GAUUAUAUAAAUUUAAGGUUGCAUUUUAAUUCUUCAACAUAUAUAAUAUGUAAGUAUUGAGUGAAUAAGUCACAAAUAAACUCUGUU.....CC...........AGAU.....................GG.....UUCAGGG.C...AAAGGAGCUGAAAAAA...AUG
Sco-1-1     ....UUAAAAUGCGAUUUAGGGUUGCGUUUUAGUUUUAUAACAAUUAUAAUUAUCAAGUAUUGGGUAAAAAAACUCACAAACAACUCUGUU.....CC...........AGAU.....................GG.....UUCAGGG.C...AAAGGAGCUGAAAAAA...AUG
Sho-3-1     ....CUAAAAUAUGUGAUUGAGGGUUGCGUUUUCCUUCAAUAACUAUUAUAAUUACACAGUAUUGGGUAAAAACUUACAAAUAACUCUGUU.....CC...........AGAU.....................GG.....UUCAGGG.C...AAAGGAGCUGAAAAAA...AUG
Sau-255-1   ....AAUUAUAUAAAUUUAAGGUUGCAUUUUAAUUCUUCAACAUAUAUAAUAUGUAAGUAUUGAGUGAAUAAGUCACAAAUAAACUCUGUU.....CC...........AGAU.....................GG.....UUCAGGG.C...AAAGGAGCUGAAAAAA...AUG
env-968     ....UAAACUCGUAGAUUAAGGGUUGCGUUUUCCAAAUAUUACUAUUAUAAUUACACAGUAUUGGGUAAAAAACUCACAAAUAACUCUGUU.....CC...........AGAU.....................GG.....UUCAGGG.C...AAAGGAGCUGAAAAAA...AUG
Sho-3-2     ....CUAAAAUAUAUGAUUGAGGGUUGCGUUUUCCUUCAAUAACUAUUAUAAUUACACAGUAUUGGGUAAAAACUCACAAAUAACUCUGUU.....CC...........AGAU.....................GG.....UUCAGGG.C...AAAGGAGCUGAAAAAA...AUG
Sho-3-15    ....CUAAAAUAUGUGAUUGAGGGUUGCGUUUUCCUUCAAUAACUAUUAUAAUUACACAGUAUUGGGUAAAAACUCACAAAUAACUCUGUU.....CC...........AGAU.....................GG.....UUCAGGG.C...AAAGGAGCUGAAAAAA...AUG
Sar-1-1     ....UGAAACUUUAGAUAAAGGGUUGUGUUUUUCAAUAAUAGCAAUUAUAAUUAUUUAGUAUUGGGUAAAAAACUCACAAAUAACUCUGUU.....CC...........AGAU.....................GG.....UUCAGGG.C...AAAGGAGCUGAAAAAA...AUG
env-990     ...................................................................AAAAAACUCACAAAUAACUCUGUU.....CC...........AGAU.....................GG.....UUCAGGG.C...AAAGGAGCUGAAAAAA...AUG
env-997     ..........GUAGAUUAAGGGUUGCGUUUUCCAUAUAUUACUAUUAUAAUUACACAGUAUUGGGUAAAAAACUCACAAAUAACUCUGUU.....CC...........AGAU.....................GG.....UUCAGGG.C...AAAGGAGCUGAAAAAA...AUG
Sco-2-2     ....UAAAUAUGUGAUUUAGGGUUGCGUUUUAGUUUUAUAACAAUUAUAAUUAUCAAGUAUUGGGUAAAAAAACUCACAAACAACUCUGUU.....CC...........AGAU.....................GG.....UUCAGGG.C...AAAGGAGCUGAAAAAA...AUG
Swa-2-3     ....UUAUUUAAUUUUUCAAAGGGUUGCGUUUUACAUUAAUUACUAUUAUAAUUACCAAGUAUUGGGUAAAAACUCACAAAUAACUCUGUU.....CC...........AGAU.....................GG.....UUCAGGG.C...AAAGGAGCUGAAAACA...AUG
Spa-2-1     ....UUAUUUAAUUUUACAAAGGGUUGCGUUUUACAUUAAUUACUAUUAUAAUUACCAAGUAUUGGGUAAAAACUCACAAAUAACUCUGUU.....CC...........AGAU.....................GG.....UUCAGGG.C...AAAGGAGCUGAAAACA...AUG
Swa-2-6     ....UUAUUUAAUUUUACAAAGGGUUGCGUUUUACAUUAAUUACUAUUAUAAUUAUCAAGUAUUGGGUAAAAACUCACAAAUAACUCUGUU.....CC...........AGAU.....................GG.....UUCAGGG.C...AAAGGAGCUGAAAACA...AUG
Spa-1-2     ....UAUUUAAUGUUACAAAGGGUUGCGUUUUACAUUAAAUACUAUUAUAAUUACACAGUAUUGGGUAAAAAACUCACAAAUAACUCUGUU.....CC...........AGAU.....................GG.....UUCAGGG.C...AAAGGAGCUGAAAACA...AUG
Ssu-1-1     ....UUAAUAUCGGAUAAAGGGUUGCGUUUUAUAAUUAAAGCAAUUAUAAUUAUCAAGUAUUGGGUAAAAAAACUCACAAAUAACUCUGUU.....CC...........AGAU.....................GG.....UUCAGGG.C...AAAGGAGCUGAAAACA...AUG
Mab-1-1     ..................AAAGGGUUGCGUUUUACAUUAAUUACUAUUAUAAUUACCAAGUAUUGGGUAAAAACUCACAAAUAACUCUGUU.....CC...........AGAU.....................GG.....UUCAGGG.C...AAAGGAGCUGAAAACA...AUG
Swa-1-1     ....UUAUUUAAUUUUACAAAGGGUUGCGUUUUACCUUAAUUACUAUUAUAAUUACCAAGUAUUGGGUAAAAACUCACAAAUAACUCUGUU.....CC...........AGAU.....................GG.....UUCAGGG.C...AAAGGAGCUGAAAACA...AUG
Ssa-2-1     ....CGAAACUUUAUAAAGAGGGUUGCGUUUUAUAUGUAUAACAAUUAUAAUGACUUAGUAUUGGGUAAAAAACUCACAAACAACUCUGUU.....CC...........GGAU.....................GG.....UUCAGGG.C...AAAGGAGCUGAAAACA...AUG
Sga-1-1     ....UGUAAUGUUAGAUUUAGGGUUGCGUUUUACAUGUAUAGCUAUUAUAAUUACGUAGUAUUGGGUAAAAAACUCACAAACAACUCUGUU.....CC...........GGAU.....................GG.....UUCAGGG.C...AAAGGAGCUGAAAGUA...AUG
Sps-2-1     ....GAAUAAUUUUAGAUGUAGGGUUGCAAUUUCAUCAUGCGGCCUAUAUAAUGUUUAAGUAUUGGGUAAAAACUCAGAAAUAACUCUGUU.....CC...........GGAU.....................GG.....UUCAGGG.C...AAAGGAGCUGAAAAUU...AUG
Sin-2-1     ....UAAUUAUUUCAGAUGUAGGGUUGCAAUUUCAUCAUCCGGCCUAUAUAAUGUUUAAGUAUUGGGUAAAAACUCACAAAUAACUCUGUU.....CC...........GGAU.....................GG.....UUCAGGG.C...AAAGGAGCUGAAAAUU...AUG
Sde-1-1     ....GAAUAAUUUCAGAUGUAGGGUUGCAAUUUCAUCAUGCGGCCUAUAUAAUGUUUAAGUAUUGGGUAAAAACUCAGAAAUAACUCUGUU.....CC...........GGAU.....................GG.....UUCAGGG.C...AAAGGAGCUGAAAAUU...AUG
Mca-1-1     ....CAUAUAAAUUUAUAGUUGCAUUCUACACACAAUCCAUAUAUAAUAUGUAAGUAUUGACGUGAAAAUUAUUCACGAAUAAACUCUGUU.....CC...........GGAU.....................GG.....UUCAGGG.C...AAAGGAGCUGAAAAAUA..AUG
Sch-1-1     ....UGAAUAAAUGUUAAAGUAGGGUUGCAUUUUAAAAAUGCAACCUUUAUAAUGUUUAAGUAUUGGUGAAAACUCACAAAUAACUCUGUU.....CC...........GGAU.....................GG.....UUCAGGG.C...AAAGGAGCUGAAAAUA...AUG
Sha-4-1     ....ACGAAUGUCUGUUUGAGGGUUGCGUUUUAAUCUAAUAACUAUUAUAAUUACCAAGUAUUGGGUAAAAACUCACAAAUAACUCUGUU.....CC...........AGAU.....................GG.....UUCAGGG.C...AAAGGAGCUGAAAAUUA..AUG
Ssc-1-1     ....CGAAUAAUUAAAAAGUAGGGUUGCAUUUUCGGGAUACGCCAUUUAUAAUGUUUAAGUAUUGGGUAAAAACUCACAAAUAACUCUGUU.....CC...........AGAU.....................GG.....UUCAGGG.C...AAAGGAGCUGAAAAUU...AUG
Ssp-14-1    ....CGAAUAAUUAAAAAGUAGGGUUGCAUUUUCGUGAUACGCCAUUUAUAAUGUUUAAGUAUUGGGUAAAAACUCACAAAUAACUCUGUU.....CC...........AGAU.....................GG.....UUCAGGG.C...AAAGGAGCUGAAAAUU...AUG
Sha-4-5     ....ACGAAUGUCAGUUUGAGGGUUGCGUUUUAAUCUAAUAACUAUUAUAAUUACCAAGUAUUGGGUAAAAACUCACAAAUAACUCUGUU.....CC...........AGAU.....................GG.....UUCAGGG.C...AAAGGAGCUGAAAAUUA..AUG

            ......................................................................................<<<<<.......<<<<<<<<<<<<<..............>>>>>>>>>>>>>.......>>>>>...........................
            ........................................................................................222.......22211...1111...............1111...11222.......222.............................
            RR••••••••••••••••••••••••••••••••••••••••••••••••••••••••••••••••••••••••••••••••••Y••CU•UG-------•Y∘----------••R∘∘------------------∘R•-------CA•GG-C--RRAAGGAG••R•R••∘∘---AUG
                                                                                                  5′                                                                         3′
```

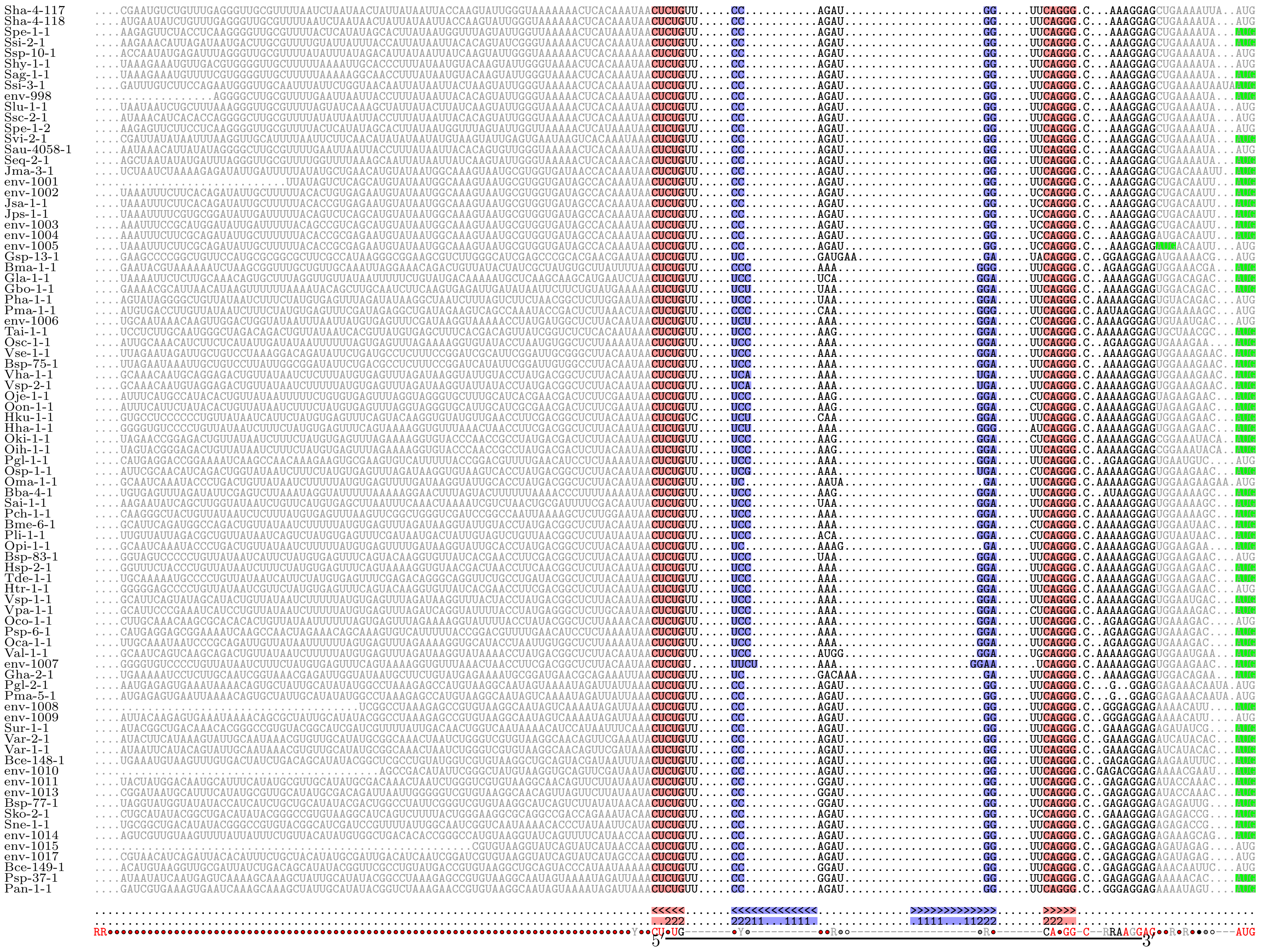
Sha-4-117
Sha-4-118
Spe-1-1
Ssi-2-1
Ssp-10-1
Shy-1-1
Sag-1-1
Ssi-3-1
env-998
Slu-1-1
Ssc-2-1
Spe-1-2
Svi-2-1
Sau-4058-1
Seq-2-1
Jma-3-1
env-1001
env-1002
Jsa-1-1
Jps-1-1
env-1003
env-1004
env-1005
Gsp-13-1
Bma-1-1
Gla-1-1
Gbo-1-1
Pha-1-1
Pma-1-1
env-1006
Tai-1-1
Osc-1-1
Vse-1-1
Bsp-75-1
Vha-1-1
Vsp-2-1
Oje-1-1
Oon-1-1
Hku-1-1
Hha-1-1
Oki-1-1
Oih-1-1
Pgl-1-1
Osp-1-1
Oma-1-1
Bba-4-1
Sai-1-1
Pch-1-1
Bme-6-1
Pli-1-1
Opi-1-1
Bsp-83-1
Hsp-2-1
Tde-1-1
Htr-1-1
Vsp-1-1
Vpa-1-1
Oco-1-1
Psp-6-1
Oca-1-1
Val-1-1
env-1007
Gha-2-1
Pgl-2-1
Pma-5-1
env-1008
env-1009
Sur-1-1
Var-2-1
Var-1-1
Bce-148-1
env-1010
env-1011
env-1013
Bsp-77-1
Sko-2-1
Sne-1-1
env-1014
env-1015
env-1017
Bce-149-1
Psp-37-1
Pan-1-1
5′
3′

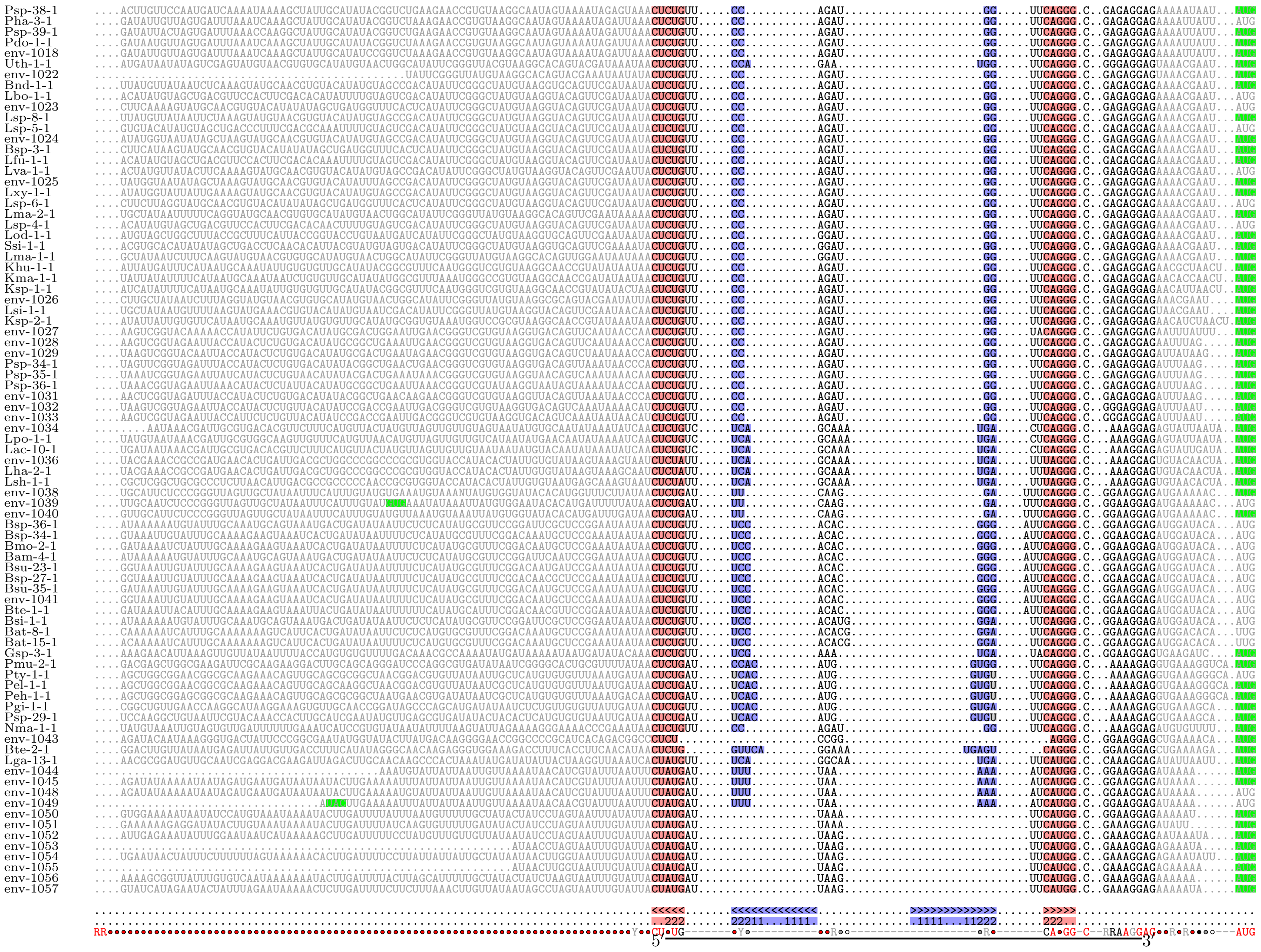
Psp-38-1
Pha-3-1
Psp-39-1
Pdo-1-1
env-1018
Uth-1-1
env-1022
Bnd-1-1
Lbo-1-1
env-1023
Lsp-8-1
Lsp-5-1
env-1024
Bsp-3-1
Lfu-1-1
Lva-1-1
env-1025
Lxy-1-1
Lsp-6-1
Lma-2-1
Lsp-4-1
Lod-1-1
Ssi-1-1
Lma-1-1
Khu-1-1
Kma-1-1
Ksp-1-1
env-1026
Lsi-1-1
Ksp-2-1
env-1027
env-1028
env-1029
Psp-34-1
Psp-35-1
Psp-36-1
env-1031
env-1032
env-1033
env-1034
Lpo-1-1
Lac-10-1
env-1036
Lha-2-1
Lsh-1-1
env-1038
env-1039
env-1040
Bsp-36-1
Bsp-34-1
Bmo-2-1
Bam-4-1
Bsu-23-1
Bsp-27-1
Bsu-35-1
env-1041
Bte-1-1
Bsi-1-1
Bat-8-1
Bat-15-1
Gsp-3-1
Pmu-2-1
Pty-1-1
Pel-1-1
Peh-1-1
Pgi-1-1
Psp-29-1
Nma-1-1
env-1043
Bte-2-1
Lga-13-1
env-1044
env-1045
env-1048
env-1049
env-1050
env-1051
env-1052
env-1053
env-1054
env-1055
env-1056
env-1057
5′
3′

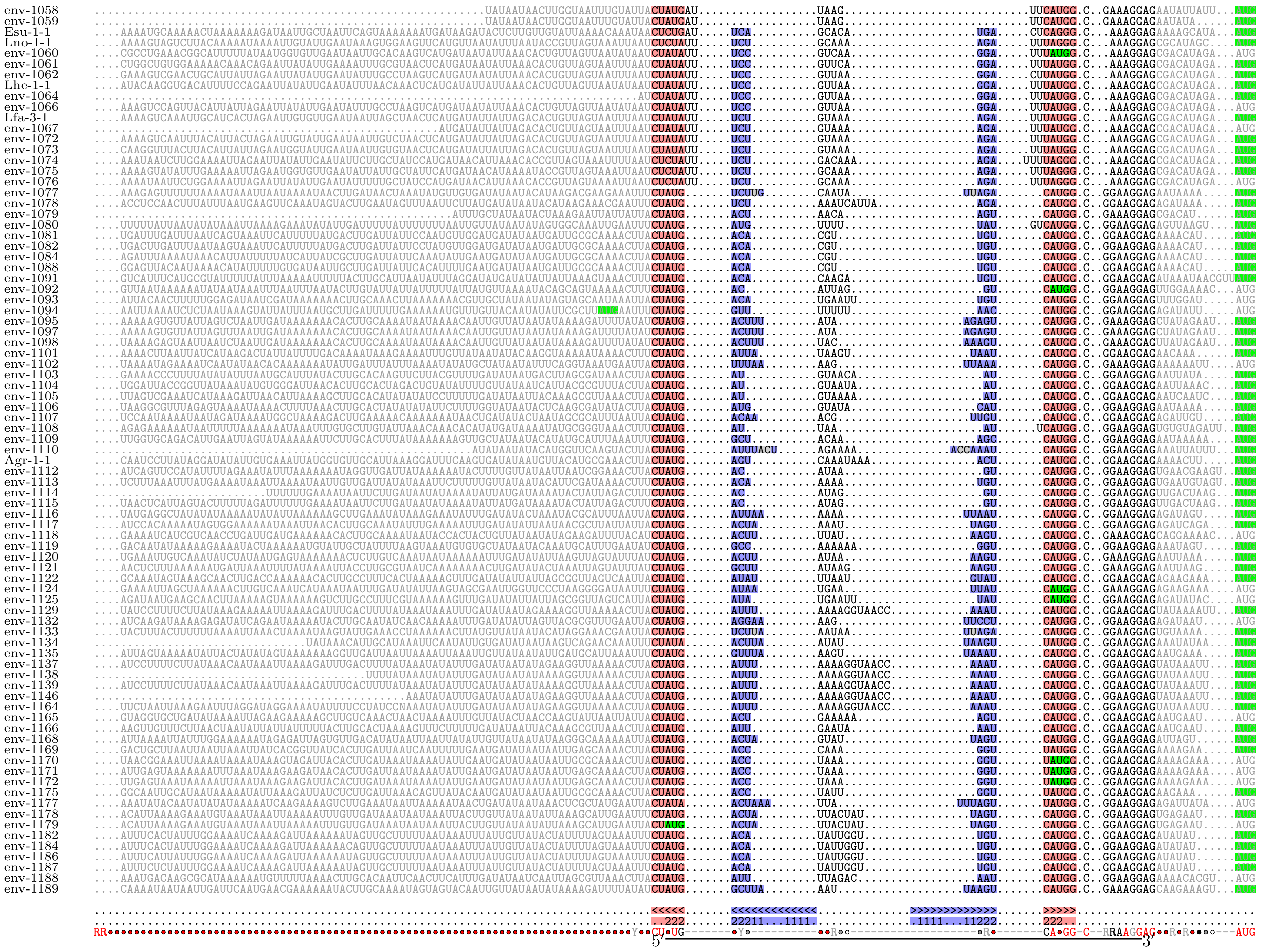
env-1058
env-1059
Esu-1-1
Lno-1-1
env-1060
env-1061
env-1062
Lhe-1-1
env-1064
env-1066
Lfa-3-1
env-1067
env-1072
env-1073
env-1074
env-1075
env-1076
env-1077
env-1078
env-1079
env-1080
env-1081
env-1082
env-1084
env-1088
env-1091
env-1092
env-1093
env-1094
env-1095
env-1097
env-1098
env-1101
env-1102
env-1103
env-1104
env-1105
env-1106
env-1107
env-1108
env-1109
env-1110
Agr-1-1
env-1112
env-1113
env-1114
env-1115
env-1116
env-1117
env-1118
env-1119
env-1120
env-1121
env-1122
env-1124
env-1125
env-1129
env-1132
env-1133
env-1134
env-1135
env-1137
env-1138
env-1139
env-1146
env-1164
env-1165
env-1166
env-1168
env-1169
env-1170
env-1171
env-1172
env-1175
env-1177
env-1178
env-1179
env-1182
env-1184
env-1186
env-1187
env-1188
env-1189
5′
3′

```
env-1190 ....CAAAUGAAAUACUAAAAAAAUUAUAAAAUAUAUUGAUAAAUAUUAAAAUAGUUGUUAUAAUAUGAAAGACUUUGAAAUUACUAUG......ACUAA........UUA..................UUAGU.......CAUGG.C..GGAAGGAGAGUGUAUU....AUG
env-1191 ....AUAAAUUAUAUCGUAAAAUUGAUCAAUUAUACUUGCAUAAACAACGCUAAAUUGUUAUAAUAUUAAAGAUUUUAGUAAUUCUAUG......ACUA.........AAAAAA...................UAGU.......UAUGG.C..GGAAGGAGAAAUAAAA....AUG
env-1197 ....AGCANUAGGAUAUAAAUAAUAUUUUAUUUAACCCUUGAUAAUUUAAUUUUACAAUGAUAUAAUAAUUGAGCGAAACUUACUAUG......AC...........ACAUC....................GU.......CAUGG.C..GGAAGGAGAAGAAAAAAA...AUG
env-1198 ....AAAGCAAUAGCACAAAAAAUAAUUGAAAAAACACUUGAAAAAAAACAAAAUAUUUGAUAUAGUAACAAUGCAUUUAAUUACUAUG......ACAA.........GUAU...................UUGU.......CAUGG.C..GGAAGGAGAGAAUAAU....AUG
env-1199 ....ACAAAAUAUAUAGUAAAAUUGCUAAAUUAUACUUGCAUAAACAACUGUAAAUUGUUAUAAUAGUAAAGAUUUUAGUAAUUCUAUA......ACUAA........AAA..................UUAGU.......UAUGG.C..GGAAGGAGAUAUGAAAAU...AUG
env-1200 ....AUAUAAUUAAUAUAUCUAUAAAAAUAAAGAAAAGUCUUGAAAUAAUUAAAAAUUAUUGAUAUAAUAAACUCGCUAUGAAUUACUAUA......ACUAA........AAUUA................UUAGU.......UAUGG.C..GGAAGGAGAGAUAAAA....AUG
env-1201 ....AAAAGAAGAAGAAAGUACUAAAACUCGAUAAAAACUUGCAUUUUGUAAAAAUUGUUUGUUAUAUUAUAAAAGCAAUAUUUUCUAUG......AUUAA........UAU..................UUAAU.......CAUGG.C..GGAAGGAGUGUCGAAAU...AUG
env-1202 ....AGCUUAUUAAAACAUACGAAAAAAUAAAAAAAUACUUGCAAAAAUAUUAAUUAUUUGAUAUAUUAUAAUACGCUUAUUAUUACUAUG......AUUUA........UAU..................UAAAU.......CAUGG.C..GGAAGGAGAGAUAUAA....AUG
env-1203 ....UAAUAUCAAAAUUACCAAUAUUAAACAAAUUAGUGUUGCAAAAGUAAGUAUUGAUUGCUAUAAUAUACAAGAAUUAAUUACUAUG......AUUA.........AUAU...................UAAU.......CAUGG.C..GGAAGGAGAUAAAU......AUG
env-1204 ....UAAUAUCAAAAUUACCAAUAUUAAACAAAUUAGUGUUGCAAAAACAAAUAUAGAUUGCUAUAAUAUACAAGAAUUAAUUACUAUG......AUUA.........ACAU...................UAAU.......CAUGG.C..GGAAGGAGAUAAAU......AUG
env-1205 ....UAGCAUCAAAAGUACCAAUAUUAAACAGAUUAUAUUGCCAAAACAAAUAUAGACUGAUAUAAUAUACAAGAAUUAAUUACUAUG......AUUA.........AUAU...................UAAU.......CAUGG.C..GGAAGGAGUUAAU.......AUG
env-1206 ....UAAGUAAAAUUCCUACUUUUAUAAAAAAAACACUUGCAAAUAUUAAAAUAAUUUGAUAAACUAUGUAAGUAGUUAAAUUUCUAUG......GCA..........GGAAA...................UGU.......CAUGG.C..GGAAGGAGAUAAGU......AUG
env-1207 ....AUGUAAACAUUUAUAAAAAAAUUAAGAAAAAACUAUUGUUUUAACCUUAUUUUAUGUUAUAUAUGUAUGCAUUAAAUUUCUAUG......GCA..........UGAAUU..................UGU.......CAUGG.C..GGAAGGAGAGUAGUUU....AUG
env-1208 ....UCAUAAUUUAAUGAACAAAGUAUAAUUUGUCUCUUGCAUUUUUCAAAUAUAUGUAUACUAUCGAAGUAAAAACGAAUUUCUAUG......AUUUA........AUU..................UAAAU.......CAUGG.C..GGAAGGAGAAUAAAAG....AUG
env-1209 ..........GAAUAAAAAAAGAUUUUAGCAUUUACGCUUGAUUAUUCAAAUAUUGAAUGAUAUAAUGAUUGCGCAAAACUUACUAUG......AC...........UUAUU....................GU.......CAUGG.C..GGAAGGAGAUAGACAA....AUG
env-1210 ....UUUAGAAUAAUUGGUAAAAGUGCUAGAAAUAACUUGAAAAAAAAUAUUAUUAUGUUAUAAUAUAGAACGUAUUUAUUAUCUAUG......GC...........UUUAUUUU.................GU.......CAUGG.C..GAAGGAGAGAAAAAGUA..AUG
env-1211 ....ACAAAUUAUAUAGUAAAAUUGAUUGUUUGUACUUGCAUAAACGUAAAUUGUUAUAAUAUUGAUAUAAUAUUAGGAUUUUAUAAUCUAUA......ACU..........UAAUUU..................AGU.......UAUGG.C..GGAAGGAGAAAUGUAAAA..AUG
env-1212 ....AACUUAUAAGAUUAUUUGGAAAAAGCAAAGAUUAGUCUUGCUUUUUUUCUAAAACUUGUUAUACUAAACAAGUAAAUUUCUAUG......GCUUG........AAUU..................UAAGU.......CAUGG.C..GGAAGGAGAGAACU......AUG
env-1213 ....GUAGGAGCAAAAAUAUCAUUAAUAGAAGUUUAAACUUGCAUUUAUUAAAGAAAUUUGUUAUAUUAUAAUAGGCAUUUAUAUCUAUG......ACUUA........UAU..................UUAGU.......CAUGG.C..GAGAGGAGGAAAGUAUU...AUG
env-1214 ....AUAAAGAUAAAAGAGGUUUAAGUGUAAAAAACACUUGAAAACAAAGAAAAUAUUUGGUAUAUUAUAAUACGCGUCAAAUUUCUAUG......ACA..........AGAUUGC................UGU.......CAUGG.C..GGAAGGAGAGAAGUUU....AUG
env-1215 ....GGCAAAUAAUUGUAAUUAAAUUAAGAAAAAACAGUUGAUUUUAGUAAUAAAGCUUGUUAUAAUAUAGCAGUAAUAAAUUUCUAUG......ACU..........UGAAUU..................AGU.......CAUGG.C..GGAAGGAGACACAAGAA...AUG
env-1216 ..................................................................UUUGUAUAAAAUAUGAACGUAUUAAAUUUCUAUG......ACU..........UGAAUU..................AGU.......CAUGG.C..GGAAGGAGAGAGAAC.....AUG
env-1217 ....AAUGAUGGAAUUAACCAAAUUCAUCAAAUAAUUGUUGAAAAAUUAAAAAAAUACUGAUAUAAUAUAAAAAGUAUUUAUUACUAUG......ACA..........UUUAUU..................UGU.......CAUGG.C..GAAAGGAGAAAUAAU.....AUG
env-1218 ....AUGAUUAAAAUGUAAUUUGCUUUACAUUUUUUUCUUGCAAUUAUUAUAUAAUAUUUGUUAGAAUAUGUAUGAAUUAAAUUUCUAUG......ACU..........UAUUU...................AGU.......CAUGG.C..GGAAGGAGCGUAGAAU....AUG
env-1219 ....AAAUCUAAAAGUCGGAGAAAAAGUAUUAAAAUACUUGCAUUUAAAGAUAUAACUUGCUAUAUAUAGGAGCAAUAAAAUUUCUAUG......GCA..........UGAAUU..................UGU.......CAUGG.C..GGAAGGAGUGCUAAAUU...AUG
env-1220 ....UUUAUUUGAUAAAUAAAAUGUAAGAAAUAUAUUGAUUUUUGUUAUAUUACUUGUUAUAAUAUAAUGGCAAAUUGAAUUUCUAUG......ACU..........UGUAUU..................AGU.......CAUGG.C..GGAAGGAGAGUAUAGU....AUG
env-1221 ....UAUUGGAAAACAAAAAAUUAGUAUAAAGUACUUGCAAAUAUCAAAAUAUUUGAUAAACUAUGUAAGUAAUAAAUUUCUAUG......ACU..........UGAAUU..................AGU.......CAUGG.C..GGGAGGAGAGUAUAU.....AUG
env-1222 ....AUGCCGGAAAACAAGAAAUAAUUAAAUUAAUAGUUGAAAAUAUUAAAAAAAUAAUUGAUAUAAUAUAAAACGAUAUUUAUUACUAUG......ACUUA........AUU..................UUAGU.......CAUGG.C..GAAAGGAGAAUAAAG.....AUG
env-1224 ....UACUUAGAAUUUUUGGAAAAGAUAAUUAAAAUACUUGCAAAUAAAAAAAAAGCCUGCUAUAUAUAGGAGCAAUAAAAUUUCUAUG......ACUUA........AAU..................UUAGU.......CAUGG.C..GGAAGGAGAAUAAAU.....AUG
env-1225 ....UUUGAAAAUAUUAAAGCAAAUGUCGAAAAUAGCUUGAAAAAUAAUAAAAUAGUUGAUAUACUAAUAACGCAUUUAAUUACUAUG......ACCUA........AAU..................UGAGU.......CAUGG.C..GGAAGGAGAGAUAAUU....AUG
env-1228 ....ACACAAUCAAAAUAGGUAAAAUAUAAAAAAAAGCUUGAUAUAUCUAAAAAUACUUGUUAUAAUAAAUUCGCUAUUAAUUACUAUG......ACUUA........AAU..................UGAGU.......CAUGG.C..GGAAGGAGAGACAGAA....AUG
env-1230 ....ACGCAGUAUAAAUAAGUAAAAAUACAAAAAAAAGCUUGCUAUAUCUGAAAAUACUUGUUAUAAUAAAUUCGCUAUUAAUUACUAUG......ACUUA........AAU..................UAAGU.......CAUGG.C..GGAAGGAGAGACAGAA....AUG
env-1231 ....AUAUAUAGAAAGUAAGUAAAAAAAAGAAAAAGUAUUGCCAUAAAUCAAAAUUACUGUUAUAAUAAACUCGCUAUUAAUUACUAUG......ACUUA........AAU..................UAAGU.......CAUGG.C..GGAAGGAGAGACAAUA....AUG
env-1232 ..............AAGAUAAAAAACAAAAAAAGCUUGAUAUAUCUAAUAAAUACUUGCUAUAUAAUAAAUUCGCUAUUAAUUACUAUG......ACU..........UGAAUU..................AGU.......CAUGG.C..GGAAGGAGAGACAAAA....AUG
env-1233 ....GAUAUUAUCAAUAAAAAAUGAAUUAAUAAAAUACUUGCCUUUUAUAAAUUACUUGUUAUAAUAUAGAAGCAUUAAAUUUCUAUG......ACUU.........AAAUGA.................AAGU.......CAUGG.C..GGAAGGAGAGUAUCAA....AUG
env-1234 ..........................................................UUUUAUAAAAUAUGAACGUAUUAAAUUUCUAUG......ACU..........UGAAUU..................AGU.......CAUGG.C..GGAAGGAGCGAUAAAA....AUG
env-1235 ....UGCAAGAUUUUCUAUCCAAUAAAAAAAUAAAAAUCAGUUGCAAUCUCUUUCUUAGUAUGGUAAAAUACUAUGGAAACUUACUAUG......AC...........UUUA....................GU.......CAUGG.C..GGAAGGAGUGAAAUGUAGUUAUG
env-1236 ....AUUAAAACAUAAUUAUACAAAUUAAAAAAAAGUAGUUGAUUAUUAAAACAACUAUGAUAGAAUACUUUCGUAAAACUUUCUAUG......AC...........UUAAGU..................GU.......CAUGG.C..GGAAGGAGUGUAAUGUAGU.AUG
env-1237 ....UUGACAAUAAUAUAAAAAAUAAUAAAAACUUGUUGCUAUUUAAAUAAAACUGUGUUAUAAUACUUUUGCAUUGAAUUUCUAUG......GCA..........UAUUG...................UGU.......CAUGG.C..GGAAGGAGUAAUAAAU....AUG
env-1238 ....CAGAGAUCAUUAGAAACAUAAUGAAAUAAAUGUGUUGAUAAAUAUAAAUACAAGUGUUAUAAUACGUUGUGCGAAACUUACUAUG......AC...........UAU.....................GU.......CAUGG.C..GGAAGGAGAAAGAAA.....AUG
env-1239 ....AGAUUGCUUGAUUUGAUGAAAUUUGAUUUUAAUACUUGAUUAAACAUUUUUCAAAUGAUAUAAUAAUUGAGCGAAACUUACUAUG......ACUU.........AAA..................GAGU.......CAUGG.C..GGAAGGAGAAAAAAACUA..AUG
env-1240 ..........................................................AAAACUUGCUAUACUAAACAAGUAAAUUUCUAUG......ACUU.........GAAAUA.................AAGU.......CAUGG.C..GGAAGGAGAGAAUU......AUG
env-1241 ....AACUAUUAAGUUUAUUUGGAAAAAACAGAGAUUAGACUUGAUUUUUAUAAAAAAAGUGCUAUACUAAACAAGUAAAUUUCUAUG......ACUU.........GAAUUA.................AAGU.......CAUGG.C..GGAAGGAGAGAAUU......AUG
env-1242 ....AACUUUAAAAUUAUUUGGAAAAAGCAAAGAUUAGACUUGCUUUUUAUUAUAAAACUUGUUAUACUAUACGAGUAAAUUUCUAUG......ACUU.........GAAAAA.................AAGU.......CAUGG.C..GGAAGGAGAGACUAAA....AUG
env-1243 ....ACUAAUAAGAUUAUUUGGAAAAAGCAAAGAUUAAUACUUGCUUUUUUUUUGAAAACUUGUUAUACUAUACAAGUAAAUUUCUAUG......ACUU.........GAAAUA.................AAGU.......CAUGG.C..GGAAGGAGAGACUAAA....AUG
env-1244 ....AACUUUUAUAAGAUUAUUUGGAAAAAAGCAAAGAUUAGAUCUUGCUUUUUUUUUAAAACUUGCUAUACUAAACAAGUAAAUUUCUAUG......GCUU.........GAAAUA.................AAGU.......CAUGG.C..GGAAGGAGAGAACU......AUG
env-1245 ....AACUUCUAAGAUUAUUUGGAAAAAGCAAAGAUUAGUCUUGCUUUUUUUUAAAAUAGCUGGUAUACUAAACAAGUAAAUUUCUAUG......GCUU.........GAAUUA.................AAGU.......CAUGG.C..GGAAGGAGAGAAACU.....AUG
env-1246 ....AAAUAAUAAAAUUAUAGAAAAUUUAAAAAAUACCUUGCUUUUUAAUAGAAUAUUUGAUAUAAUAUAAAGGAUUAAAUUUCUAUG......ACU..........UAUUUU..................AGU.......CAUGG.C..GGAAGGAGAUAUGU......AUG
env-1247 ....UUAAAGAAAGUUUGGAUUCUUUUUAAUAUUUUAGUUGAUUUUUCUUUUUUUAUUGUUAUACUAGUAACGAAUUAAAUUUCUAUG......ACU..........UAUUUU..................AGU.......CAUGG.C..GGAAGGAGAUAUUU......AUG
env-1248 ....UUGACAUGUUUCGUAUAAAAUAGUAAAAAAUGCUUGAUUUUUUUAAUAAUUGUUUGCUAUAAUAUAGAAGCAUUAAAUUUCUAUG......ACUA.........AUUU..................UAGU.......CAUGG.C..GGAAGGAGAUAAAGU.....AUG
env-1249 ....UACAAGGAUAUUACAAAUAUAUGAAAAAAUUACUUGAUAAAAUUUUAAAAAAACUGAUAUAAUACUACCUAGUUCAAGUUACUAUG......ACUA.........AUU..................UAGU.......CAUGG.C..GGAAGGAGAGUAAAC.....AUG
env-1250 ....AAGAUAAAAUUUUAAUUUCAUGAUAAAUCACUUGCACAAAAGAGUCCUGUUUGUUAUAAUUGUAUCGCUUAUAAAACUUACUAUG......AC...........AUGA....................GU.......CAUGG.C..GGAAGGAGAGAAAGACCU..AUG
env-1251 ....CGAAAGUUGAUUCUUCAUUUUUCAAUCGAUUGGGCUUGAUUAACCCGAUUUGUAAUGAUAUAAUCAUUGUGCAAACUUACUAUG......AC...........UCUU....................GU.......CAUGG.C..GGAAGGAGAAAAAGAA....AUG
env-1252 ....UUGACAUGAUUAAUGUAAAAUAAUAAAAAAUGCUUGAUUUUUUGCUUUCUAAUUUGCUAUAAUAUAAAAGCAUUAAAUUUCUAUG......ACUU.........UAUU..................AAGU.......CAUGG.C..GGAAGGAGAAAAGU......AUG
env-1253 ....AUAAAAAAAAUUAAUCAAACUCUACUAAAGUAGUUGCGAAGAUACGAAAAAGUUGCUAUAAUAUAGAGGCAUUAAAGUUACUAUG......ACUAA........UUU..................UUAGU.......CAUGG.C..GGAAGGAGAGAAUAAA....AUG
env-1254 ....CAAAUAUUAAAAUAUAGAUGAUAAAAAAUAGUUGAUUUUAUAUAAAAAAAUUGUUAUAUAAACAAGAAUUUAUUUAUUACUAUG......AC...........UUUA....................GU.......CAUGG.C..GGAAGGAGAAAAUU......AUG
env-1255 ....AUGAUGCUAUAUCAAAACCAAUACAAAAAACAUCUUGCAUAAUAUUUAAAUAUUUGCUACAAUAGGUAGGAAUUGAAUUUCUAUG......ACUAA........UAU..................UUAGU.......CAUGG.C..GGAAGGAGCGAAGAAA....AUG
env-1256 ....UAAUAGUUAUCUGACUAUUUUUAUUAUAAACACUUGCAUUUUUAUUAAAUAAUUUGAUAAACUAUGUAAGUAAUUAAAUUUCUAUG......ACA..........UAUAUU..................UGU.......CAUGG.C..GGGAGGAGAGAAAAU.....AUG
env-1257 ....UUAAUAAAAAAAUAUGAAAUUUCUUUAAAUAUAUUGCAAAUUUAUAAAUAUUUGAUAAACUAUGUAAGUAAUUAAAUUUCUAUG......ACA..........UAUAUU..................UGU.......CAUGG.C..GGGAGGAGAGUAAAAA....AUG
env-1258 ....AUUUUUGUUUUAAAAGUUAUUAAAAAAGAGUUGUCAAAUCGUUUUUUUAUUGAUAUAAUAUAUAGAGCGUAUUUUAUAUUACUAUG......AC...........UUUU....................GU.......CAUGG.C..GAAGGGAGAGAAAAAAAU..AUG
env-1259 ....AUUAAAAAAAUCAACUCUACACUUCUAAAAAUAACAGUUGCGCAUUUCACUAAUUACUGCUAUAAUAUAUUAAAGAAAAUUACUAUG......ACUU.........AAAA..................AAGU.......CAUGG.C..GGAAGGAGAGAAAAAAC...AUG
env-1260 ....GAUUCUUUAAUAAAAAAAUAUAAUUCUAAAAUAUCUUGCAAAAAAUAAAACAUUUGAUAUACUAAUAACGCUUAUUAUUACUAUG......ACUUA........AUU..................UUAGU.......CAUGG.C..GGAAGGAGAGAUAUAA....AUG
env-1261 ....UAUCAAAUAAAAUAUACAAAAUGAUAGAAAAAGGUUGAAAAAUAAUAAAAAUAAUGAUAUAAUUAUAAUACGCAUUUAAUUACUAUG......AC...........UUUUUU..................GU.......CAUGG.C..GGAAGGAGCGAUAGAA....AUG
env-1262 ....UAUAGAAAAAUUGCGGAAAAUAUAAUAAAAAGCCUUGAAAAUAAAUAAAUUAUUUGAUAUAUUAAUAACGCAUUAAAUUACUAUG......GCA..........GAAAAA..................UGU.......CAUGG.C..GGAAGGAGAGAUAGA.....AUG
env-1263 ....UAUAGAAAAAUUGCGGAAAAUAUAAUAAAAAGCCUUGAAAAUAAAUAAAUUAUUUGGUAUAUUAAUAACGCAUUAAAUUACUAUG......GCA..........AAAAAA..................UGU.......CAUGG.C..GGAAGGAGAGAUAGA.....AUG
env-1264 ............................................................................UGAGCUAAUGAAUUUCUAUG......GCAAA........AAA..................UUUGU.......CAUGG.C..GGAAGGAGAGAUAGA.....AUG
env-1265 ....AAUCAAGUAAUAUCAAAGAAAAUAUAUAAAAGUACUUGAAAAUAAUAAAAAUAUUUGAUAUAUUAAUAAAGCAAUUUAUAUCUAUG......GCAAG........UCU..................UUUGU.......CAUGG.C..GGAAGGAGAGAUAAU.....AUG
env-1266 ....CAAGCAAUUGCAAAGAAAAUAUAUUAGAAAAACACUUGAAAAAUGAAAAAAUAUUUGAUAUAUUAAGUACGCACUUAAUUACUAUG......GCAA.........CUAUU..................UUGU.......CAUGG.C..GGAAGGAGAGAUAAU.....AUG
env-1267 ....CAAAAGAUAGCACAAGAAAUAAUUACAAAAACACUUGAAAAAGAAGAAAUAAUUUGAUAUAGUAUAUGGCAUUAAAUUACUAUG......ACAAG........UAA..................CUUGU.......CAUGG.C..GGAAGGAGAGAUAGU.....AUG
env-1268 ....AAACAAAAUAUACAAAAAGUAUACAAAAAUGCUUGAAAAAUAAGUAAAACCUUGCUAUACUAAUAACGCGAUUUGAAUUACUAUG......AC...........UUAAUU..................GU.......CAUGG.C..GGAAGGAGAGAUAGU.....AUG
env-1269 ....UAACUCUAAAAGACGUAUAAUAGCAAGAAAUACUUGCAAAAUAAAGAAAUAUUUGAUAUAUUAAAAAAGCAUUUGAAUUACUAUG......ACAU.........AAAUU..................GUGU.......CAUGG.C..GGAAGGAGAGAUAAU.....AUG
env-1270 ....UGCUAAAAUAAAACUAAAAGAUGUAUAAAACACUUGCAAAAUAAAGAAAUAGUUGUUAUACUAAUAACGCAUUUGAAUUACUAUG......ACAU.........AAAUU..................GUGU.......CAUGG.C..GGAAGGAGAGAUAGU.....AUG
env-1271 ....AGAAAAAUAUCGGAGAAAAUAUUAUAAUAAACACUUGAAAAAUAAAGAAAAGUUUGAUAUAUUAAUAACGCAUUUGAAUUACUAUG......ACAA.........AAU..................UUGU.......CAUGG.C..GGAAGGAGAGAUAAU.....AUG
env-1272 ....AUAGGAAAAUAUAUGAAGAAAUCAAAAAAACACUUGAUAAAAUAAGAAAAAAUUUGAUAUAUUAAUAACGCGAUUGAAUUACUAUG......ACAA.........AAU..................UUGU.......CAUGG.C..GGAAGGAGAGAUAAU.....AUG
env-1273 ....AAAGAGAUUUAAAAUCUCUUUUUCGUUGAAAAUACUUGCUAAAAUGCAAAAACUUUGAUAUAUUAAUAACGCAUUUAAUUACUAUG......ACGAG........AAAUA.................UUUGU.......CAUGG.C..GGAAGGAGAGAUAAU.....AUG
env-1274 ....UAAUUGGGGCAAUAAAUAAAACAUUGUUAAACUCUUGAUUUUUAUAGUUUUAUCUGUUACAAUAUAAAGGCAUUGAAUUUCUAUG......GCAA.........AAUU..................UUGU.......CAUGG.C..GGAAGGAGAGAUACU.....AUG
env-1275 ....UUUAUUUCUUUUUACUAAAAUGCUAAAAAAUGCUUGAAAAUACAACAAAUAUUUGUUAUAUUAAUAACGCAUUUGAAUUACUAUG......ACAAA........UAU..................UUUGU.......CAUGG.C..GGAAGGAGAGAUAAU.....AUG
env-1276 ....AAUACUUAUAUUUCUUUAUUUUUAAAGAAAUAACUUGAAAAAAUAUAAAAAUAUUUGAUAUAUUAAUAACGCAUUUGAAUUACUAUG......ACAAA........UAU..................UUUGU.......CAUGG.C..GGAAGGAGAGAUAAU.....AUG
env-1277 ....UAAAGAGAUAUCAAAGAAAAUAAUAUACUAAAAACACUUGAAAAAAAAGAAAAUAUUUGAUAUAUUAAUAACGCAUUUAAUUACUAUG......ACAA.........AUUU..................UUGU.......CAUGG.C..GGAAGGAGAGAUAGU.....AUG
env-1278 ....CAAGCUAUUGCUAAGAAAAUAUUUGAAAAAACACUUGAAAAACAAGAAAUAUUUGAUAUAUUAAGUACGCAUUUGAAUUACUAUG......ACAAA........UAG..................UUUGU.......CAUGG.C..GGAAGGAGAGAUAAU.....AUG
env-1279 ....UCAAAGAAUAGCAACUAAAAUAAUUGAAAAAACUCUUGAAAAAUCAUAAAAAGUUUGAUAUAUUAAGUACGCAUUUAAUUACUAUG......ACAAA........UUAU..................UUUGU.......CAUGG.C..GGAAGGAGAGAUAGU.....AUG
env-1280 ....CAUCAAGUCAUAUCCAAAAAACUUAUAAAAACACUUGAAAAUAAUAAAAAUAUUUGAUAUAUUAAUAAGGCAAUUUAUAUCUAUG......GCAAA........UUA..................UUUGU.......CAUGG.C..GGAAGGAGAGAUAAU.....AUG
env-1281 ....ACACAGAAAGAUGUUAAUAAUUAAAAAACACUUGAUAAAUAAUGAAAUAUUUGAUAUAUUAAUAACGCGAAUUGAAUUACUAUG......AC...........UAAUUAUU................GU.......CAUGG.C..GGAAGGAGAGAUAAU.....AUG

...........................................................................................&lt;&lt;&lt;&lt;&lt;......&lt;&lt;&lt;&lt;&lt;&lt;&lt;&lt;&lt;&lt;&lt;&lt;&lt;..............&gt;&gt;&gt;&gt;&gt;&gt;&gt;&gt;&gt;&gt;&gt;&gt;&gt;.......&gt;&gt;&gt;&gt;&gt;.............................
...........................................................................................222......22211...1111...............1111...11222.......222..................................
RR.........................................................................................Y..CU.UG------.Y.-----------..R..------------------.R.-------CA.GG-C--RRAAGGAG..R.R......---AUG
                                                                                            5'                                                                         3'
```

```
env-1282  ....ACAGUGAAAGAACUUAAUAUUUAAAAAAACACUUGAUAAAUAAUGAAAUAUUUGAUAUAUUAAUAACGCAAAUUGAAUUACUAUG......AC..........UAAUUAUU................GU.......CAUGG.C..GGAAGGAGAGAUAAU.....AUG
env-1283  ..................................................UUAAAUGUUAUAUAUAUGGUUGCUAUAAAUUUCUAUG......GCAAG........AAAA..................UUUGU.......CAUGG.C..GGAAGGAGAGAGAUU.....AUG
env-1284  ....AAUUGAUUGCAAAUGAAAUAAUAGAAAAAACACUUGAAAAAUAGAACUAAAUUUGAUAUAUUAAUAACGCGAUUUAAUUACUAUG......ACAA.........AAAA..................UUGU.......CAUGG.C..GGAAGGAGAGAUAUU.....AUG
env-1285  ....UUUUUCAAGAAAUUAUUAAACAAACAAAAAAGACUUGAAAAGUAAAAAAAAAUUUGAUAUAUUAAUAACGCAUUUAAUUACUAUG......UCAG.........AAAAA.................CUGU.......CAUGG.C..GGAAGGAGAGAUACU.....AUG
env-1286  ....UUUUUCAAGAAAUAAUAUAUCAAACAAAAAAAGCUUGAAAAGUUUAAAAAAAUUUGAUAUAUUAAUAACGCAUUUAAUUACUAUG......UCAG.........GUAA..................CUGU.......CAUGG.C..GGAAGGAGAGAUAUU.....AUG
env-1287  ....AUUUUUGGAAAUAGUAGAGAUUAAAUAAAAAUACUUGCCAAAUAAUUAGAAAUUUGAUAUAUUAUAUAAGUAUUUAUUAUCUAUG......ACA..........UGAAUU.................UGU.......CAUGG.C..GGAAGGAGAGAUUACAGU..AUG
env-1288  ....GCAAAUAGUAAAGCAACUUGACCAAAAAAUGCUUGCCUUUCGCUAAAAAGUUUGAUAUAUUAUUAGCGGUUAGUCAAUUACUAUG......AUAU.........UUAAU.................GUAU.......CAUGG.C..GGAAGGAGAGAAAAA.....AUG
env-1290  ....GUUGGAGCUGACAUUGAAUUAAUUUAAAAAAAAGUUGCAAUCUUAAAAAAAAGUUGUUAUAAUACAAAUGCAUUGAAUUUCUAUG......ACU..........UAACU.................AGU.......CAUGG.C..GGAAGGAGAGUUAAA.....AUG
env-1291  ....AUUUUAAAGCAUUAUAAAAAAAUAUAUUAUUUUUCUUGUAUUAUUGUUUUUAAUAUGUUAAAAUGCAAUAGUAAAACUUUCUAUG......AC...........UAAAU..................GU.......CAUGG.C..GGAAGGAGUUGGAAAC....AUG
env-1292  ....UGCUAAUAAAAUAAUAAAAAACAGUAAAAAUACUUGCAAUUAAAAUAAUUACUUGAUAAUAUAAAAGCAAUAAAUUACUAUG......ACU..........UGAAUU................AGU.......CAUGG.C..GGAAGGAGUUGGAAAU....AUG
env-1293  ....AUUAAGACUUUUUGGCGAUAAUAAUUAAAAAUACUUGCAAACUUCUAAAAACGUUGCUAUAAUAUAGGAGCAAUAAAUUACUAUG......ACU..........UGAAUU................AGU.......CAUGG.C..GGAAGGAGAAAUAAU.....AUG
env-1294  ....AACAUAGGAUAUCGUCAAAUUUAUAAAUUAAUAGUUGAAAAAUUAAAAAAAACUUGAUAUAAUAUCAAAUGUGUUUAUUACUAUG......ACU..........UGAAUU................AGU.......CAUGG.C..GAAAGGAGAAAUAAA.....AUG
env-1295  ......................................................UUAAAUAUAUAUCAAAUGUAUUUAUUACUAUG......ACU..........UGUAUU................AGU.......CAUGG.C..GAAAGGAGAAAUAAA.....AUG
env-1296  ....AUAAGUAGAUGAACUUAUCAAAUUAAUAAAAUAAUGUUGCAAAACAAUUAAAAUUGUUAUAUAAAUUAGCAAUUAAUUACUAUG......ACU..........UGUAU.................AGU.......CAUGG.C..GGAAGGAGAGCUAUAU....AUG
env-1297  ....AUUGAGUAAUAUACAAUAUUUUUUAUGAAAAGUAUUGCCAGAAUUAAAAAUAAUGUUAUAUAAUAACUCGCUAUAAAUUACUAUG......AC...........UUAUAUUUU..............GU.......CAUGG.C..GGAAGGAGAGAUAAAA....AUG
env-1298  ....UGCUGGUUAUGAGAAAAUUUUUCAAAUAAUAGUUGAUAAAAUAUAAAAUUAUUGUUAUAAUAGUGAACGUAAUUUUAUUACUAUG......AC...........UUUG..................GU.......CAUGG.C..GGAAGGAGAGAUAAAA....AUG
env-1299  ....UAUGCAGAUUUCUAACAAAGUAUUAUUUGAUACUUGAUUUUAAAAAAUGUUUGAUAUAUUCGCUUGUGAAUUUACUAUG......ACUU.........UUUU..................AAGU.......CAUGG.C..GGAAGGAGAGAUAGU.....AUG
env-1300  ..........................AUUAAAAAUACUUGAAAUAUUUAAAAAUACUUGUUAUAAUAUUUGUCGUGUUUUAUUACUAUG......ACU..........UUU...................GGU.......CAUGG.C..GGAAGGAGAGAUAU.....AUG
env-1301  ....GAUUAAAAUUUCGGACAAAGUAUUAUUUAGUGCUUGAUUUUUGAAAAAAUGUUUGCUACAAUAUACUCGCUUAUGAAUUUCUAUG......ACUU.........UAUUU.................AGGU.......CAUGG.C..GGAAGGAGAGAUAU.....AUG
env-1302  ....UAUUAGUGCUUCUGAUAAAAUUUUGUUUGAUACUUGAUUUUUAAAAAAAUGUUUGCUACAAUAUACUCGCUUAUGAAUUUCUAUG......GCUUA........UUU...................UGAGU.......CAUGG.C..GGAAGGAGAGAUAU.....AUG
env-1303  ....AGAAGGAUACGAAAAAAUUUCACAAAUAAUUAUUGAAAAUUCAAAAAUACCUGAUAUAAUAUACUUCGUAAUUUUAUUACUAUG......AC...........UUAAUU................GU.......CAUGG.C..GGAAGGAGAGAAAAAAAU..AUG
env-1304  ....UUUGAAAUUAAAAACAAAAAUACGUUGUUAUCUCUUGAUUAAUAAAAUAUUGAAUGAUAUAAUGAAUGCGCUAAAUCUUACUAUG......AC...........UUUUU.................GU.......CAUGG.C..GGAAGGAGAAAGAAA.....AUG
env-1305  ....AUCUGAGUCUAUAUAUAAAACUUAUGAAAAAAGCUUGAAAAAUAAUAAAUUAAUGGUAUAUUAAUAACGCAAUUUAAUUUCUAUG......ACAA.........AUU...................UUGU.......CAUGG.C..GGAAGGAGAAGAAGAAA.....AUG
env-1306  ....AAUAGUCAUAUGACUAUUUUUUCUAUAAAAUACUUGCAUUUUUUGCAAAUAGUUUGAUAAACUAUGUAAGUAUUUAUUACUAUG......ACA..........UUAUUU................UGU.......CAUGG.C..GGAAGGAGAGUAAAAA....AUG
env-1307  ....AGAUAACGCUAUUGAAAAACCUAUAGAAAAAAGGUUGCAAUUACUUUAAAUAUUUGUUAAAAUAGGAACGUAUUAAAUUACUAUG......ACU..........UGAAUU................AGU.......CAUGG.C..GGAAGGAGAGAAAAAA....AUG
env-1308  ....UAAUGGUUAUGAAAAAAUUUCACAAUUAAUUAUUGAAAAUUUUAAAAAUAACUGAUAUAAUAUACUUCGUAAUUUUAUUACUAUG......AC...........UUAAUU................GU.......CAUGG.C..GGAAGGAGAGAAAAA.....AUG
env-1309  ....AAGCGGUUAUGAAAAAAUUUCGCAAAUAAUUAUUGAAAAUUUUAAAAAUAACUGAUAUAAUAUACUCGUAAUUUUAUUACUAUG......AC...........UUAAUU................GU.......CAUGG.C..GGAAGGAGAGAAAAAA....AUG
env-1310  ....AGAGGGUUAUGAAAAAAUUUCGCAAAUAAUUAUUGAAAAAAUUAAAAAUAACUGAUAUAAUAUCCGUAAUUUUAUUACUAUG......AC...........UUAAUU................GU.......CAUGG.C..GGAAGGAGAGAAAAC.....AUG
env-1311  ....AUAUUAGAAAAGUACAAUUAAAUACUAAAAAACUUGAAAAAAUCAAAAAAUAAUGUUACACUAAUAACGCGAUUUAAUAUCUAUG......GCU..........UGAAUU................AGU.......CAUGG.C..GGAAGGAGAGAUAAA.....AUG
env-1312  ....UAAAGGUUAUCAAAAAAUUUCACAAAUAAUAGUUGAAAAUUUAAAAAAUACUGAUAUAAUAUACCAUGUAAUUUUAUUACUAUG......AC...........UUUUU.................GU.......CAUGG.C..GGAAGGAGAGAGAAUA....AUG
env-1337  ...........................................................................CCAUGUAAUUUUAUUACUAUG......AC...........UUUUU.................GU.......CAUGG.C..GGAAGGAGAGAGAAUA....AUG
env-1342  ....AGAAGGAUAUGAAAAAAUUUCUCAAAUAAUAGUUGAAAAAAUACAAAAUAACUGAUAUAAUAUACCAUGUAAUUUUAUUACUAUG......AC...........AUUU..................GU.......CAUGG.C..GGAAGGAGAGAAAAAA....AUG
env-1360  ....NNNNNNNNNNNNNNNNNNNNNNNNNNNNNNNNNNNNNNNAAAUGCAAAAUAACUGAUAUAAUAUACCAUGUAAUUUUAUUACUAUG......AC...........AUUU..................GU.......CAUGG.C..GGAAGGAGAGAAAAAA....AUG
env-1361  ....AGAAGGAUAUGAAAAAAUUUCUCAAAUAAUAGUUGAAAAAAUACAAAAUAACUGAUAUAAUAUACCAUGUAAUUUUAUUACUAUG......AC...........AUUU..................GU.......CAUGG.C..GGAAGGAGAGAAAAA.....AUG
env-1362  ....AGAAGGAUAUGAAAAAAUUUCUCAAAUAAUAGUUGAAAAAAUGCAAAAUAACUGAUAUAAUAUACCAUGUAAUUUUAUUACUAUG......AC...........AUUU..................GU.......CAUGG.C..GGAAGGAGAGAAAAAA....AUG
env-1379  ....GAAGAAGGAAAUCAAUUAAUUAGUAAAUUAAUAGUUGAAAAGUUUAAAAAUAGUUGAUAUAAUAUACCAUGUAUUUUAUUACUAUG......ACA..........UUU...................UGU.......CAUGG.C..GGAAGGAGAGAAAAAA....AUG
env-1381  ....AGAAGGAUAUGAAAAAAUUUCUCAAAUAAUAGUUGAAAAAAUACAAAAUAACUGAUAUAAUAUACCAUGUAAUUUUAUUACUAUG......AC...........GUUU..................GU.......CAUGG.C..GGAAGGAGAGAAAAAA....AUG
env-1383  ......AAGGUUAUCAAAAAAUUUCACAAAUAAUAGUUGAAAAUUUAGAAAAUAACUGAUAUAAUAUAUACACGUAGUUUUAUUACUAUG......AC...........UUUU..................GU.......CAUGG.C..GGAAGGAGAGAAAAAAGU...AUG
env-1384  ....CAAGGGAUAUGAAAAAAUUUUUCACAAAUAAUAGUUGAAAAUUUAGAAAAUAACUGAUAUAAUAUACCAUGUAAUUUUAUUACUAUG......AC...........UUUUU.................GU.......CAUGG.C..GGAAGGAGAGAAAAA.....AUG
env-1385  ....CAAAGGAUAUGAAAAAAUUUCACAAAUAAUAGUUGAAAAAAUAAAAAAUAACUGAUAUAAUAUACCAUGUAAUUUUAUUACUAUG......AC...........UUUUU.................GU.......CAUGG.C..GGAAGGAGAGAAAAU.....AUG
env-1387  ....CAAAGGAUAUGAAAAAAUUUCACAAAUAAUAGUUGAAAAAAUAAAAAAUAACUGAUAUAAUAUACCAUGUAAUUUUAUUACUAUG......AC...........UUUUU.................GU.......CAUGG.C..GGAAGGAGAGAAAAAU....AUG
env-1388  ....UAGCGGUUAUGAAAAAAUUUCACAAAUAAUUAUUGAAAAAAUAAAAAAUACCUGAUAUAAUAUACUUCGUAAUUUUAUUACUAUG......AC...........UUUUU.................GU.......CAUGG.C..GGAAGGAGAGAAAA......AUG
env-1389  ....UAAAAUUAUUUGGUAAAGAUAAUUAAAAAACACUUGCAUACAUAAGAAAACAUUGCUAUAAUAUAAAAGCAAAGAAAUUACUAUG......ACAU.........UUU...................AUGU.......CAUGG.C..GGAAGGAGAUUAAAAA....AUG
env-1390  ....UAUAAAUAGAAUGAAUAAAGUUUUAUUUAGGCUUGAUUUUAAAAAAUGUUUGCUACAAUAUAUCGCUUGUGAAUUUACUAUG......ACUU.........UAAGU.................AAGU.......CAUGG.C..GGAAGGAGAGAUAU.....AUG
env-1391  ....UCAAGACAAAUAGUCUUAAAAAACAUCAAAAAAACAUGAAAAGUCUAGAAAUAACUGAUAUACUAAUAACGCAUUUAAUUACUAUG......AC...........UAAUGUU...............GU.......CAUGG.C..GGAAGGAGAAGAAGAAU....AUG
env-1393  ....UUACUUAGAAUUUUUGCAAAUCGUGAUUAAAUACUUGCAAAGUAUAAAAACUUGCUAUAAUAGAGGGCAAAGAAAUUACUAUG......ACU..........UGAAUU................AGU.......CAUGG.C..GGAAGGAGAAUGAAA.....AUG
env-1395  .............AAAUAAAAUAUUAAGUUAAUAGCUUGCAUUUGUAUAAAAAUACUUGCUAUAAUAUAAAAGCAAUAAAUUACUAUG......ACU..........UGAAUU................AGU.......CAUGG.C..GGAAGGAGAAUGAAG.....AUG
env-1397  ....ACACCUAUUACAAAUAUUUGGUGAUAGAAACUAGUUGAUUUUAAAUAUAAAUCUGUUAUACUAUAAAGGCAAUUUAAUUACUAUG......ACU..........UGAAUUU...............AGU.......CAUGG.C..GGAAGGAGAGUUACAU....AUG
env-1398  ....UAUCAUUAAAAGAUGUUUAAACUAACAAAACACUUGCAAAAUAAGACAAUUCUUGCUAUACUAUAAACGAUUUGAAUUACUAUG......ACUU.........AUAGG.................AAGU.......CAUGG.C..GGAAGGAGAGAUAGU.....AUG
env-1399  ....AAUAUAGUAAAAAAUAGCAUAAAUUAAAAAAAUGUGCAAAACUAAAAAUAUCUUGAUAUACUAUAUUAGCAAUUAAUUACUAUG......ACU..........UGAAUU................AGU.......CAUGG.C..GGGAGGAGAGAUUAGA....AUG
env-1402  ....AAAUGUAUAAUCAAAAAUAUUAGGUUAUUUUUGGUGCAUUUAUAAAAAUAUUUUGCUACAAUAUGGUGGUAUUGAAUUACUAUG......AC...........UUAGGUUU..............GU.......CAUGG.C..GGAAGGAGAGUAAAAA....AUG
env-1404  ....UUUUUCUUUAUUUAUAAAUAAAAAUAAAAAUGUUAUUGCAAAUAUCAUAAUUCUAUGAUAAAAUAACGCACGUAAGACUUUCUAUG......AC...........UUAUU.................GU.......CAUGG.C..GGAAGGAGUGAAUUGUAGU.AUG
env-1405  ....AUCAGUUCCACUAUUUAAAAAAUUAUAUAAUUUUUUGUUGCAAAUAUAAAAAUGUAUGCUAAAAUAUCACCCGUAGACUUUCUAUG......AC...........UAUUU.................GU.......CAUGG.C..GGAAGGAGUGAAUUGUAGU.AUG
env-1406  ....CAGUUCCACUAUUUAAAAAAUUUAUAUUUUUUAUAUUGCAAAUAGUAAAAUAGUAUGUUAAAAUAUCACCCGUAAACUUUCUAUG......ACU..........AGA...................AGU.......CAUGG.C..GGAAGGAGUGAAUUGUAGU.AUG
env-1407  ....UCAGUACCACUAUUUAAAAAAUUAUAAUUUGUUGUUGCAAACAAUAAAAUAGUAUGCUAAAAUAUCACCCGUAAGACUUUCUAUG......ACU..........UAA...................GGU.......CAUGG.C..GGAAGGAGUGAAUUGUAGU.AUG
env-1408  ....CAGUUCCACUAUUUAAAAAUUUAUAUUUUAUAUCUUGCAAAUGAUAUAAUAGUAUGUUAAAAUAUCACCCGUAAACUUUCUAUG......AC...........AUUAG.................GU.......CAUGG.C..GGAAGGAGUGAAUUGUAGU.AUG
env-1409  ....AGAUUUAAAAUAAUAGAUGAAUGAUAAUAAUACUUGAAAAAAUUUAUUAUUAAUGUUAAAAUAACAUCGUAUUUAAUUUCUAUG......AUUU.........UUAAU.................AAAU.......CAUGG.C..GGAAGGAGAUAAAA......AUG
env-1413  ....AGAUAUAAAAAUAAUAGAUGAAUGAUAAUAAUACUUGAAAAAUUUAUUAUUAAUGUUAAAAUAACAUCGUAUUUAAUUUCUAUG......AUUU.........UUAAU.................AAAU.......CAUGG.C..GGAAGGAGAUAAAA......AUG
env-1414  ....AGAUAUAAAAAUAAUAGAUGAAUGAUAAUAAUACUUGAAAAAUGUAUUAUUAAUGUUAAAAUAACAUCGUAUUUAAUUUCUAUG......AUUUU........AAA...................AAAAU.......CAUGG.C..GGAAGGAGAUAAAA......AUG
env-1415  ....AGAUAUAAAAAUAAUAGAUGAAUGAUAAUAAUACUUGAAAAAUGUAUUAUUAAUGUUAAAAUAACAUCGUAUUUAAUUUCUAUG......AUUUU........UAA...................AAAGU.......CAUGG.C..GGAAGGAGAUAAAA......AUG
env-1416  ............................................................................AUCGUAUUUAAUUUCUAUG......AUUU.........UUUU..................AAAU.......CAUGG.C..GGAAGGAGAUAAA.......AUG
env-1419  ...................................................................AAUAACAUCGUAUUUAAUUUCUAUG......GUUU.........UUUU..................AAAU.......CAUGG.C..GGAAGGAGAUAAA.......AUG
env-1420  ....NNNNNNNNNNNNNNNNNNNNNNNNNNNNNNNNNNNNNNNNNNNGUUUUAAUUACUUGUUAAAAUAACAUCGUAUUUAAUUUCUAUG......GUUU.........UUUU..................AAAU.......CAUGG.C..GGAAGGAGAUAAA.......AUG
env-1421  ......AUAUAAAAAUAUUAGAUGAAUAGUAAUAAUACUUGAAAAGUUUUUAAUUACUUGUUAAAAUAACAUCGUAUUUAAUUUCUAUG......GUUU.........UUUU..................AAAU.......CAUGG.C..GGAAGGAGAUAAA.......AUG
env-1423  ....CAGAUAUAAAAAUAUUAGAUGAUAAUAAUAAUACUUGAAAAGUUUUUGAUUACUUGUUAAAAUAACAUCGUAUUUAAUUUCUAUG......AUUU.........UUAAU.................AAAU.......CAUGG.C..GGAAGGAGAUAAA.......AUG
env-1424  ...................................................................................UUUCUAUG......AUUU.........UUAAU.................AAAU.......CAUGG.C..GGAAGGAGAUAAA.......AUG
env-1438  ............................................................................AUCGUAUUUAAUUACUAUG......AUUU.........UUAAU.................AAAU.......CAUGG.C..GGAAGGAGAUAAA.......AUG
env-1439  ....AUUAAUGAAACAUUACUAAGAAAUUAAUAAAAAAGUGCUUAAUUCAAUAAUAAUGUUAUAUAAUUAACGAAUUUAAUUACUAUG......AUUU.........UAAA..................AAAU.......CAUGG.C..GGAAGGAGAAUUAAAC....AUG
env-1443  ....UGUCGGAGCAUCAAUAAAAUUAGAAGAAAUCCCUUGCCAAAACGAUUGAAAUUUGUUAUACUACUAAAGAAUUAUUAUUACUAUG......ACU..........AAAAA.................AGU.......CAUGG.C..GGAAGGAGAAAAC......AUG
Aoc-1-1   ....UCUUAAUCUUCAAAUGAAAUUGUGAAAUUAUGGAGUUGCAUAAAAGAUAUGUGAAUGAUAUAAUAAUAGGCGAAACUUACUAUG......AGU..........AGAAAC................ACU.......CAUGG.C..GGAAGGAGAAAAUU.....AUG
env-1444  ....AUUAACAAAACUUUAUUAUAUAAUAAAAUAAUUGUUGCAUUUAUUAAAUAUUUGUUGAUAUAUAAUAAACGCAUUGAAUUUCUAUG......GCA..........UAUAUUU...............UGU.......CAUGG.C..GGAAGGAGAGUUAAGU....AUG
env-1445  ....UGUAGAAACGAAUUUAAAAAAUGACUUUUAAACUUGCACAAAUGAUGUUCAUUUGAUAUAAUCAAUUAGCGUAAACUUACUAUG......AUU..........AUAAGA................AAU.......CAUGG.C..GGAAGGAGAAAACACGU...AUG
env-1446  ....AUUAAUAAAAAGCUAGUUUAGUAAUAUAUCAUGAAAUUGACUCAAAAAAAAUGUUUAUUGUAAUAAACGUUAAUCGAUUAAUCUAUG......GCUU.........GAAAGA................GAGU.......CAUGG.C...AAAGGAGUAUAAAAC....AUG
Lap-1-1   ....AUUAAUAAAAAGCUAGUUUAGUAAUAUAUCUUGAAAUUGACUCGGAAAAAUGUUUAUUGUAAUAAACGUUAAUCGAUUAAUCUAUG......GCUU.........GAAAGA................GAGU.......CAUGG.C...AAAGGAGUAUAAAAC....AUG
env-1449  ....AAUUGAUGCAGCUGCUAUAAUAAAUAAGUCUUGAAUUUUAUUAUCAAAAAUGAUGAUAAGGUAUUAAGCGUUAAUCGAUUAAUCUAUG......ACUU.........GAAUAG................GAGU.......CAUGG.C...AAAGGAGUAUAAAAUU...AUG
env-1450  ....AAUUGAUGCAGCUGCUAUAAUAAAUAAGUCUUGAAUUUUUGCCUUUAAAGAUGAUAAGGUAUUAAACGUUAAUCGAUUAAUCUAUG......ACUU.........GCAAUAG...............GAGU.......CAUGG.C...AAAGGAGUAUAAAAU....AUG
Lsp-13-1  ....AAUUGAUGCAGCUGCUAUAAUAAAUAAGUCUUGAAUUUUAUUAUCAAAAAUGAUGAUAAGGUAUUAAACGUUAAUCGAUUAAUCUAUG......ACUU.........GAAAUAG...............GAGU.......CAUGG.C...AAAGGAGUAUAAUU....AUG
env-1451  ....AAUUAAUCCAGCUGCUAUAAUAAAUAAGUCUUGAAAUUUUACUAUUAAAAAUGUUAAGGUAUUAAACGUUAAUCGAUUAAUCUAUG......ACUU.........GCAAUAG...............GAGU.......CAUGG.C...AAAGGAGUAUAAAAUU...AUG
Lku-1-1   ....AAUUGAUGCAGCUGCUAUAAUAAAUAAGUCUUGAAUUUUUGCCUUUAAAGAUGAUAAGGUAGUAAACGUUAAUCGAUUAAUCUAUG......ACUU.........GAAAUAG...............GAGU.......CAUGG.C...AAAGGAGUAUAAAAU....AUG
Lki-1-1   ....AAUUGAUGCAGCUGCUAUAAUAAAUAAGUCUUGAAUUUUGCCUUUAAAGAUGAUAAGGUAUUAAACGUUAAUCGAUUAAUCUAUG......ACUU.........GAAAUAG...............GAGU.......CAUGG.C...AAAGGAGUAUAAAAU....AUG
env-1452  ....AAUUAACUCUGCUGCUAUAAUAAAUAAGUCUUGAAUUUUACUAUUAAAAAUGUUAAGGUAUUAAGCGUUAAUCGAUUAAUCUAUG......ACUU.........GAAAUAG...............GAGU.......CAUGG.C...AAAGGAGUAUAAAAU....AUG
env-1453  ....AAUUAACUCUGUUGCUAUAAUAAAUAAGUCUUGAAUUUUACUUUUAAAAAUGUUAAGGUAUUAGACGUUAAUCGAUUAAUCUAUG......ACUU.........GAAAUAG...............GAGU.......CAUGG.C...AAAGGAGUAUAAAAU....AUG
env-1455  ....AAUUAACUCUGCUGCUAUAAUAAAUAAGUCUUGAAUUUUACUAUUAAAAAUGUUAAGGUAUUAAACGUUAAUCGAUUAAUCUAUG......ACUU.........GAAAUAG...............GAGU.......CAUGG.C...AAAGGAGUAUAAAAU....AUG

          ..................................................................................<<<<<......<<<<<<<<<<<<<..............>>>>>>>>>>>>>.......>>>>>.........................
          .................................................................................. .222......22211...1111...............1111...11222.......222............................
          RR••••••••••••••••••••••••••••••••••••••••••••••••••••••••••••••••••••••••••••••••Y••CU•UG------•Y•-----------••R•∘---------------------∘R•-------CA•GG-C--RRAAGGAG••R•R•••∘∘---AUG
                                                                                          5'                                                                              3'
```

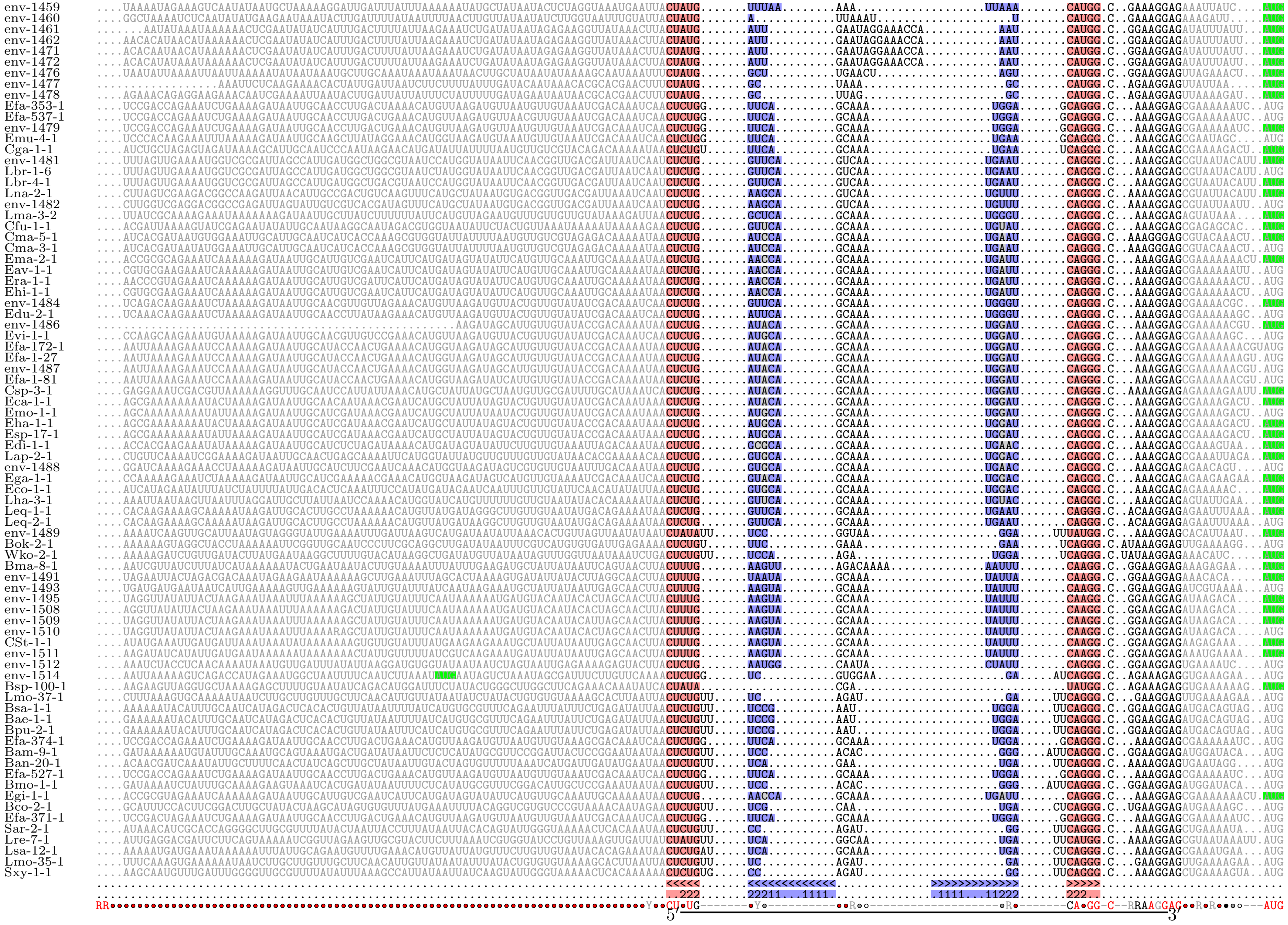
env-1459
env-1460
env-1461
env-1462
env-1471
env-1472
env-1476
env-1477
env-1478
Efa-353-1
Efa-537-1
env-1479
Emu-4-1
Cga-1-1
env-1481
Lbr-1-6
Lbr-4-1
Lna-2-1
env-1482
Lma-3-2
Cfu-1-1
Cma-5-1
Cma-3-1
Ema-2-1
Eav-1-1
Era-1-1
Ehi-1-1
env-1484
Edu-2-1
env-1486
Evi-1-1
Efa-172-1
Efa-1-27
env-1487
Efa-1-81
Csp-3-1
Eca-1-1
Emo-1-1
Eha-1-1
Esp-17-1
Edi-1-1
Lap-2-1
env-1488
Ega-1-1
Eco-1-1
Lha-3-1
Leq-1-1
Leq-2-1
env-1489
Bok-2-1
Wko-2-1
Bma-8-1
env-1491
env-1493
env-1495
env-1508
env-1509
env-1510
CSt-1-1
env-1511
env-1512
env-1514
Bsp-100-1
Lmo-37-1
Bsa-1-1
Bae-1-1
Bpu-2-1
Efa-374-1
Bam-9-1
Ban-20-1
Efa-527-1
Bmo-1-1
Egi-1-1
Bco-2-1
Efa-371-1
Sar-2-1
Lre-7-1
Lsa-12-1
Lmo-35-1
Sxy-1-1
5′
3′

### 12 L31-Gammaproteobacteria

#### 12.1 Taxa

The taxonomy of each organism containing a putative L31-Gammaproteobacteria RNA is listed, with abbreviations identifying each hit (e.g., "Eco-1-1" and "Eco-1-2" might hypothetically represent two distinct RNAs in *E. coli*). The abbreviations will be used to identify each individual L31-Gammaproteobacteria RNA in Sections 12.2 and 12.4:.

| abbrev. of hits | taxonomy of species |
|---|---|
| Sen-1-1 | AKU_12601Bacteria Proteobacteria Gammaproteobacteria Enterobacteriales Enterobacteriaceae *Salmonella enterica subsp. enterica serovar* Paratyphi A str |
| Ban-1-1 | Bacteria Actinobacteria Actinobacteridae Bifidobacteriales Bifidobacteriaceae *Bifidobacterium animalis subsp. lactis* HN019 |
| bsy-1-1 | Bacteria bacteria symbiont BFo1 of Frankliniella occidentalis |
| Cmi-1-1 | Bacteria Proteobacteria Betaproteobacteria Burkholderiales Burkholderiaceae *Caballeronia mineralivorans* PML1(12) |
| Aal-1-1 to Aal-1-2 | Bacteria Proteobacteria Gammaproteobacteria Aeromonadales Aeromonadaceae *Aeromonas allosaccharophila* |
| Aaq-1-1 | Bacteria Proteobacteria Gammaproteobacteria Aeromonadales Aeromonadaceae *Aeromonas aquariorum* AAK1 |
| Aaq-2-1 | Bacteria Proteobacteria Gammaproteobacteria Aeromonadales Aeromonadaceae *Aeromonas aquatica* |
| Aau-1-1 | Bacteria Proteobacteria Gammaproteobacteria Aeromonadales Aeromonadaceae *Aeromonas australiensis* |
| Abe-1-1 | Bacteria Proteobacteria Gammaproteobacteria Aeromonadales Aeromonadaceae *Aeromonas bestiarum* |
| Abi-1-1 | Bacteria Proteobacteria Gammaproteobacteria Aeromonadales Aeromonadaceae *Aeromonas bivalvium* |
| Aca-1-1 | Bacteria Proteobacteria Gammaproteobacteria Aeromonadales Aeromonadaceae *Aeromonas caviae* |
| Aca-2-1 | Bacteria Proteobacteria Gammaproteobacteria Aeromonadales Aeromonadaceae *Aeromonas caviae* Ae398 |
| Adh-1-1 to Adh-1-3 | Bacteria Proteobacteria Gammaproteobacteria Aeromonadales Aeromonadaceae *Aeromonas dhakensis* |
| Adi-1-1 | Bacteria Proteobacteria Gammaproteobacteria Aeromonadales Aeromonadaceae *Aeromonas diversa* 2478-85 |
| Adi-2-1 | Bacteria Proteobacteria Gammaproteobacteria Aeromonadales Aeromonadaceae *Aeromonas diversa* CDC 2478-85 |
| Aen-1-1 | Bacteria Proteobacteria Gammaproteobacteria Aeromonadales Aeromonadaceae *Aeromonas encheleia* |
| Aen-2-1 to Aen-2-3 | Bacteria Proteobacteria Gammaproteobacteria Aeromonadales Aeromonadaceae *Aeromonas enteropelogenes* |
| Aeu-1-1 | Bacteria Proteobacteria Gammaproteobacteria Aeromonadales Aeromonadaceae *Aeromonas eucrenophila* |
| Afi-1-1 | Bacteria Proteobacteria Gammaproteobacteria Aeromonadales Aeromonadaceae *Aeromonas finlandiensis* |
| Afl-1-1 | Bacteria Proteobacteria Gammaproteobacteria Aeromonadales Aeromonadaceae *Aeromonas fluvialis* |
| Ahy-1-1 to Ahy-1-2 | Bacteria Proteobacteria Gammaproteobacteria Aeromonadales Aeromonadaceae *Aeromonas hydrophila* |
| Ahy-2-1 | Bacteria Proteobacteria Gammaproteobacteria Aeromonadales Aeromonadaceae *Aeromonas hydrophila* 113 |
| Ahy-3-1 | Bacteria Proteobacteria Gammaproteobacteria Aeromonadales Aeromonadaceae *Aeromonas hydrophila* 14 |
| Ahy-4-1 | Bacteria Proteobacteria Gammaproteobacteria Aeromonadales Aeromonadaceae *Aeromonas hydrophila* 145 |
| Ahy-5-1 | Bacteria Proteobacteria Gammaproteobacteria Aeromonadales Aeromonadaceae *Aeromonas hydrophila* 173 |
| Ahy-6-1 | Bacteria Proteobacteria Gammaproteobacteria Aeromonadales Aeromonadaceae *Aeromonas hydrophila* 187 |
| Ahy-7-1 | Bacteria Proteobacteria Gammaproteobacteria Aeromonadales Aeromonadaceae *Aeromonas hydrophila* 226 |
| Ahy-8-1 | Bacteria Proteobacteria Gammaproteobacteria Aeromonadales Aeromonadaceae *Aeromonas hydrophila* 259 |
| Ahy-9-1 | Bacteria Proteobacteria Gammaproteobacteria Aeromonadales Aeromonadaceae *Aeromonas hydrophila* 277 |
| Ahy-10-1 | Bacteria Proteobacteria Gammaproteobacteria Aeromonadales Aeromonadaceae *Aeromonas hydrophila* 4AK4 |
| Ahy-11-1 | Bacteria Proteobacteria Gammaproteobacteria Aeromonadales Aeromonadaceae *Aeromonas hydrophila* AD9 |
| Ahy-12-1 | Bacteria Proteobacteria Gammaproteobacteria Aeromonadales Aeromonadaceae *Aeromonas hydrophila* AL09-71 |
| Ahy-13-1 | Bacteria Proteobacteria Gammaproteobacteria Aeromonadales Aeromonadaceae *Aeromonas hydrophila* J-1 |
| Ahy-14-1 | Bacteria Proteobacteria Gammaproteobacteria Aeromonadales Aeromonadaceae *Aeromonas hydrophila* ML09-119 |
| Ahy-15-1 | Bacteria Proteobacteria Gammaproteobacteria Aeromonadales Aeromonadaceae *Aeromonas hydrophila* NJ-35 |
| Ahy-19-1 | Bacteria Proteobacteria Gammaproteobacteria Aeromonadales Aeromonadaceae *Aeromonas hydrophila pc*104A |
| Ahy-16-1 | Bacteria Proteobacteria Gammaproteobacteria Aeromonadales Aeromonadaceae *Aeromonas hydrophila* SNUFPC-A8 |
| Ahy-17-1 | Bacteria Proteobacteria Gammaproteobacteria Aeromonadales Aeromonadaceae *Aeromonas hydrophila* SSU |
| Ahy-20-1 | Bacteria Proteobacteria Gammaproteobacteria Aeromonadales Aeromonadaceae *Aeromonas hydrophila subsp. hydrophila* ATCC 7966 |
| Ahy-21-1 | Bacteria Proteobacteria Gammaproteobacteria Aeromonadales Aeromonadaceae *Aeromonas hydrophila subsp. ranae* |
| Ahy-18-1 | Bacteria Proteobacteria Gammaproteobacteria Aeromonadales Aeromonadaceae *Aeromonas hydrophila* YL17 |
| Aja-1-1 to Aja-1-2 | Bacteria Proteobacteria Gammaproteobacteria Aeromonadales Aeromonadaceae *Aeromonas jandaei* |
| Aja-2-1 | Bacteria Proteobacteria Gammaproteobacteria Aeromonadales Aeromonadaceae *Aeromonas jandaei* Riv2 |
| Ala-1-1 | Bacteria Proteobacteria Gammaproteobacteria Aeromonadales Aeromonadaceae *Aeromonas lacus* |
| Ame-1-1 to Ame-1-3 | Bacteria Proteobacteria Gammaproteobacteria Aeromonadales Aeromonadaceae *Aeromonas media* |
| Ame-2-1 | Bacteria Proteobacteria Gammaproteobacteria Aeromonadales Aeromonadaceae *Aeromonas media* WS |
| Amo-1-1 | Bacteria Proteobacteria Gammaproteobacteria Aeromonadales Aeromonadaceae *Aeromonas molluscorum* 848 |
| Api-1-1 | Bacteria Proteobacteria Gammaproteobacteria Aeromonadales Aeromonadaceae *Aeromonas piscicola* |
| Apo-1-1 | Bacteria Proteobacteria Gammaproteobacteria Aeromonadales Aeromonadaceae *Aeromonas popoffii* |

| | |
|---|---|
| Ari-1-1 | Bacteria Proteobacteria Gammaproteobacteria Aeromonadales Aeromonadaceae *Aeromonas rivuli* |
| Asa-1-1 | Bacteria Proteobacteria Gammaproteobacteria Aeromonadales Aeromonadaceae *Aeromonas salmonicida* |
| Asa-2-1 | Bacteria Proteobacteria Gammaproteobacteria Aeromonadales Aeromonadaceae *Aeromonas salmonicida subsp. achromogenes* AS03 |
| Asa-3-1 | Bacteria Proteobacteria Gammaproteobacteria Aeromonadales Aeromonadaceae *Aeromonas salmonicida subsp. masoucida* NBRC 13784 |
| Asa-4-1 | Bacteria Proteobacteria Gammaproteobacteria Aeromonadales Aeromonadaceae *Aeromonas salmonicida subsp. pectinolytica* 34mel |
| Asa-5-1 to Asa-5-4 | Bacteria Proteobacteria Gammaproteobacteria Aeromonadales Aeromonadaceae *Aeromonas salmonicida subsp. salmonicida* |
| Asa-6-1 | Bacteria Proteobacteria Gammaproteobacteria Aeromonadales Aeromonadaceae *Aeromonas salmonicida subsp. salmonicida* 01-B526 |
| Asa-7-1 | Bacteria Proteobacteria Gammaproteobacteria Aeromonadales Aeromonadaceae *Aeromonas salmonicida subsp. salmonicida* A449 |
| Asa-8-1 | Bacteria Proteobacteria Gammaproteobacteria Aeromonadales Aeromonadaceae *Aeromonas sanarellii* |
| Asc-1-1 | Bacteria Proteobacteria Gammaproteobacteria Aeromonadales Aeromonadaceae *Aeromonas schubertii* |
| Asi-1-1 | Bacteria Proteobacteria Gammaproteobacteria Aeromonadales Aeromonadaceae *Aeromonas simiae* |
| Aso-1-1 | Bacteria Proteobacteria Gammaproteobacteria Aeromonadales Aeromonadaceae *Aeromonas sobria* |
| Asp-1-1 | Bacteria Proteobacteria Gammaproteobacteria Aeromonadales Aeromonadaceae *Aeromonas sp.* HZM |
| Asp-2-1 | Bacteria Proteobacteria Gammaproteobacteria Aeromonadales Aeromonadaceae *Aeromonas sp.* L_1B5_3 |
| Asp-3-1 | Bacteria Proteobacteria Gammaproteobacteria Aeromonadales Aeromonadaceae *Aeromonas sp.* ZOR0001 |
| Asp-4-1 | Bacteria Proteobacteria Gammaproteobacteria Aeromonadales Aeromonadaceae *Aeromonas sp.* ZOR0002 |
| Ata-1-1 to Ata-1-2 | Bacteria Proteobacteria Gammaproteobacteria Aeromonadales Aeromonadaceae *Aeromonas taiwanensis* |
| Ate-1-1 | Bacteria Proteobacteria Gammaproteobacteria Aeromonadales Aeromonadaceae *Aeromonas tecta* |
| Ave-1-1 to Ave-1-3 | Bacteria Proteobacteria Gammaproteobacteria Aeromonadales Aeromonadaceae *Aeromonas veronii* |
| Ave-2-1 | Bacteria Proteobacteria Gammaproteobacteria Aeromonadales Aeromonadaceae *Aeromonas veronii* AER39 |
| Ave-3-1 | Bacteria Proteobacteria Gammaproteobacteria Aeromonadales Aeromonadaceae *Aeromonas veronii* AER397 |
| Ave-4-1 | Bacteria Proteobacteria Gammaproteobacteria Aeromonadales Aeromonadaceae *Aeromonas veronii* AMC34 |
| Ave-5-1 | Bacteria Proteobacteria Gammaproteobacteria Aeromonadales Aeromonadaceae *Aeromonas veronii* AMC35 |
| Ave-6-1 | Bacteria Proteobacteria Gammaproteobacteria Aeromonadales Aeromonadaceae *Aeromonas veronii* B565 |
| Ave-8-1 | Bacteria Proteobacteria Gammaproteobacteria Aeromonadales Aeromonadaceae *Aeromonas veronii bv. sobria* |
| Ave-9-1 | Bacteria Proteobacteria Gammaproteobacteria Aeromonadales Aeromonadaceae *Aeromonas veronii bv. veronii* |
| Ave-7-1 | Bacteria Proteobacteria Gammaproteobacteria Aeromonadales Aeromonadaceae *Aeromonas veronii* Hm21 |
| Osm-1-1 | Bacteria Proteobacteria Gammaproteobacteria Aeromonadales Aeromonadaceae *Oceanimonas smirnovii* ATCC BAA-899 |
| Osp-1-1 | Bacteria Proteobacteria Gammaproteobacteria Aeromonadales Aeromonadaceae *Oceanimonas sp.* GK1 |
| Tau-1-1 | Bacteria Proteobacteria Gammaproteobacteria Aeromonadales Aeromonadaceae *Tolumonas auensis* DSM 9187 |
| Tli-1-1 | Bacteria Proteobacteria Gammaproteobacteria Aeromonadales Aeromonadaceae *Tolumonas lignilytica* |
| Sam-1-1 | Bacteria Proteobacteria Gammaproteobacteria Aeromonadales Succinivibrionaceae *Succinimonas amylolytica* DSM 2873 |
| Asa-9-1 | Bacteria Proteobacteria Gammaproteobacteria Alteromonadales Alteromonadaceae *Aestuariibacter salexigens* DSM 15300 |
| Aal-2-1 | Bacteria Proteobacteria Gammaproteobacteria Alteromonadales Alteromonadaceae *Agarivorans albus* MKT 106 |
| Ama-1-1 | Bacteria Proteobacteria Gammaproteobacteria Alteromonadales Alteromonadaceae *Aliagarivorans marinus* DSM 23064 |
| Ata-2-1 | Bacteria Proteobacteria Gammaproteobacteria Alteromonadales Alteromonadaceae *Aliagarivorans taiwanensis* DSM 22990 |
| Aae-1-1 | Bacteria Proteobacteria Gammaproteobacteria Alteromonadales Alteromonadaceae *Alishewanella aestuarii* B11 |
| Aag-1-1 | Bacteria Proteobacteria Gammaproteobacteria Alteromonadales Alteromonadaceae *Alishewanella agri* BL06 |
| Aje-1-1 | Bacteria Proteobacteria Gammaproteobacteria Alteromonadales Alteromonadaceae *Alishewanella jeotgali* KCTC 22429 |
| Aau-2-1 to Aau-2-2 | Bacteria Proteobacteria Gammaproteobacteria Alteromonadales Alteromonadaceae *Alteromonas australica* |
| Ama-2-1 to Ama-2-3 | Bacteria Proteobacteria Gammaproteobacteria Alteromonadales Alteromonadaceae *Alteromonas macleodii* |
| Ama-4-1 | Bacteria Proteobacteria Gammaproteobacteria Alteromonadales Alteromonadaceae *Alteromonas macleodii* AltDE1 |
| Ama-3-1 | Bacteria Proteobacteria Gammaproteobacteria Alteromonadales Alteromonadaceae *Alteromonas macleodii* ATCC 27126 |
| Ama-5-1 | Bacteria Proteobacteria Gammaproteobacteria Alteromonadales Alteromonadaceae *Alteromonas macleodii str.* 'Aegean Sea MED64' |
| Ama-6-1 | Bacteria Proteobacteria Gammaproteobacteria Alteromonadales Alteromonadaceae *Alteromonas macleodii str.* 'Balearic Sea AD45' |
| Ama-7-1 | Bacteria Proteobacteria Gammaproteobacteria Alteromonadales Alteromonadaceae *Alteromonas macleodii str.* 'Black Sea 11' |
| Ama-8-1 | Bacteria Proteobacteria Gammaproteobacteria Alteromonadales Alteromonadaceae *Alteromonas macleodii str.* 'Deep ecotype' |
| Ama-9-1 | Bacteria Proteobacteria Gammaproteobacteria Alteromonadales Alteromonadaceae *Alteromonas macleodii str.* 'English Channel 673' |
| Ama-10-1 | Bacteria Proteobacteria Gammaproteobacteria Alteromonadales Alteromonadaceae *Alteromonas macleodii str.* 'Ionian Sea U7' |
| Ama-11-1 | Bacteria Proteobacteria Gammaproteobacteria Alteromonadales Alteromonadaceae *Alteromonas macleodii str.* 'Ionian Sea U8' |
| Ama-12-1 | Bacteria Proteobacteria Gammaproteobacteria Alteromonadales Alteromonadaceae *Alteromonas macleodii str.* 'Ionian Sea UM4b' |
| Ama-13-1 | Bacteria Proteobacteria Gammaproteobacteria Alteromonadales Alteromonadaceae *Alteromonas macleodii str.* 'Ionian Sea UM7' |
| Ama-14-1 | Bacteria Proteobacteria Gammaproteobacteria Alteromonadales Alteromonadaceae *Alteromonas marina* |
| Asp-5-1 | Bacteria Proteobacteria Gammaproteobacteria Alteromonadales Alteromonadaceae *Alteromonas sp.* ALT199 |
| Asp-6-1 | Bacteria Proteobacteria Gammaproteobacteria Alteromonadales Alteromonadaceae *Alteromonas sp.* LOR |
| Asp-7-1 | Bacteria Proteobacteria Gammaproteobacteria Alteromonadales Alteromonadaceae *Alteromonas sp.* LTR |
| Asp-8-1 | Bacteria Proteobacteria Gammaproteobacteria Alteromonadales Alteromonadaceae *Alteromonas sp.* SN2 |
| Gag-1-1 | Bacteria Proteobacteria Gammaproteobacteria Alteromonadales Alteromonadaceae *Glaciecola agarilytica* NO2 |
| Gar-1-1 | Bacteria Proteobacteria Gammaproteobacteria Alteromonadales Alteromonadaceae *Glaciecola arctica* BSs20135 |
| Gch-1-1 | Bacteria Proteobacteria Gammaproteobacteria Alteromonadales Alteromonadaceae *Glaciecola chathamensis* S18K6 |
| Gli-1-1 | Bacteria Proteobacteria Gammaproteobacteria Alteromonadales Alteromonadaceae *Glaciecola lipolytica* E3 |
| Gme-1-1 | Bacteria Proteobacteria Gammaproteobacteria Alteromonadales Alteromonadaceae *Glaciecola mesophila* KMM 241 |
| Gni-1-1 | Bacteria Proteobacteria Gammaproteobacteria Alteromonadales Alteromonadaceae *Glaciecola nitratireducens* FR1064 |
| Gpa-1-1 | Bacteria Proteobacteria Gammaproteobacteria Alteromonadales Alteromonadaceae *Glaciecola pallidula* DSM 14239 = ACAM 615 |
| Gpo-1-1 | Bacteria Proteobacteria Gammaproteobacteria Alteromonadales Alteromonadaceae *Glaciecola polaris* LMG 21857 |

| | |
|---|---|
| Gps-1-1 | Bacteria Proteobacteria Gammaproteobacteria Alteromonadales Alteromonadaceae *Glaciecola psychrophila* 170 |
| Gpu-1-1 | Bacteria Proteobacteria Gammaproteobacteria Alteromonadales Alteromonadaceae *Glaciecola punicea* DSM 14233 = ACAM 611 |
| Gsp-1-1 | Bacteria Proteobacteria Gammaproteobacteria Alteromonadales Alteromonadaceae *Glaciecola sp.* 4H-3-7+YE-5 |
| Gsp-2-1 | Bacteria Proteobacteria Gammaproteobacteria Alteromonadales Alteromonadaceae *Glaciecola sp.* HTCC2999 |
| Maq-1-1 | Bacteria Proteobacteria Gammaproteobacteria Alteromonadales Alteromonadaceae *Marinobacter aquaeolei* VT8 |
| Mhy-1-1 | Bacteria Proteobacteria Gammaproteobacteria Alteromonadales Alteromonadaceae *Marinobacter hydrocarbonoclasticus* ATCC 49840 |
| Msp-1-1 | Bacteria Proteobacteria Gammaproteobacteria Alteromonadales Alteromonadaceae *Marinobacter sp.* C1S70 |
| Msp-2-1 | Bacteria Proteobacteria Gammaproteobacteria Alteromonadales Alteromonadaceae *Marinobacter sp.* EN3 |
| Msp-3-1 | Bacteria Proteobacteria Gammaproteobacteria Alteromonadales Alteromonadaceae *Marinobacter sp.* EVN1 |
| Msu-1-1 | Bacteria Proteobacteria Gammaproteobacteria Alteromonadales Alteromonadaceae *Marinobacter subterrani* |
| Sch-1-1 | Bacteria Proteobacteria Gammaproteobacteria Alteromonadales Alteromonadaceae *Salinimonas chungwhensis* DSM 16280 |
| Aba-1-1 | Bacteria Proteobacteria Gammaproteobacteria *Alteromonadales bacterium* TW-7 |
| Csp-1-1 | Bacteria Proteobacteria Gammaproteobacteria Alteromonadales Colwelliaceae *Colwellia sp.* 8_GOM-1096m |
| Tac-1-1 | Bacteria Proteobacteria Gammaproteobacteria Alteromonadales Colwelliaceae *Thalassomonas actiniarum* |
| Tvi-1-1 | Bacteria Proteobacteria Gammaproteobacteria Alteromonadales Colwelliaceae *Thalassomonas viridans* |
| Tsp-1-1 | Bacteria Proteobacteria Gammaproteobacteria Alteromonadales Colwelliaceae *Thalassotalea sp.* ND16A |
| Fba-1-1 | Bacteria Proteobacteria Gammaproteobacteria Alteromonadales Ferrimonadaceae *Ferrimonas balearica* DSM 9799 |
| Ffu-1-1 | Bacteria Proteobacteria Gammaproteobacteria Alteromonadales Ferrimonadaceae *Ferrimonas futtsuensis* DSM 18154 |
| Fky-1-1 | Bacteria Proteobacteria Gammaproteobacteria Alteromonadales Ferrimonadaceae *Ferrimonas kyonanensis* DSM 18153 |
| Fse-1-1 | Bacteria Proteobacteria Gammaproteobacteria Alteromonadales Ferrimonadaceae *Ferrimonas senticii* DSM 18821 |
| Iat-1-1 | Bacteria Proteobacteria Gammaproteobacteria Alteromonadales Idiomarinaceae *Idiomarina atlantica* |
| Iba-1-1 | Bacteria Proteobacteria Gammaproteobacteria Alteromonadales Idiomarinaceae *Idiomarina baltica* OS145 |
| Ilo-1-1 | Bacteria Proteobacteria Gammaproteobacteria Alteromonadales Idiomarinaceae *Idiomarina loihiensis* GSL 199 |
| Ilo-2-1 | Bacteria Proteobacteria Gammaproteobacteria Alteromonadales Idiomarinaceae *Idiomarina loihiensis* L2TR |
| Isa-1-1 | Bacteria Proteobacteria Gammaproteobacteria Alteromonadales Idiomarinaceae *Idiomarina salinarum* |
| Ise-1-1 | Bacteria Proteobacteria Gammaproteobacteria Alteromonadales Idiomarinaceae *Idiomarina sediminum* DSM 21906 |
| Isp-1-1 | Bacteria Proteobacteria Gammaproteobacteria Alteromonadales Idiomarinaceae *Idiomarina sp.* 28-8 |
| Isp-2-1 | Bacteria Proteobacteria Gammaproteobacteria Alteromonadales Idiomarinaceae *Idiomarina sp.* A28L |
| Ixi-1-1 | Bacteria Proteobacteria Gammaproteobacteria Alteromonadales Idiomarinaceae *Idiomarina xiamenensis* 10-D-4 |
| Mma-1-1 | Bacteria Proteobacteria Gammaproteobacteria Alteromonadales Moritellaceae *Moritella marina* ATCC 15381 |
| Asa-10-1 | Bacteria Proteobacteria Gammaproteobacteria Alteromonadales Pseudoalteromonadaceae *Algicola sagamiensis* DSM 14643 |
| Pag-1-1 | Bacteria Proteobacteria Gammaproteobacteria Alteromonadales Pseudoalteromonadaceae *Pseudoalteromonas agarivorans* S816 |
| Par-1-1 | Bacteria Proteobacteria Gammaproteobacteria Alteromonadales Pseudoalteromonadaceae *Pseudoalteromonas arctica* A 37-1-2 |
| Pat-1-1 | Bacteria Proteobacteria Gammaproteobacteria Alteromonadales Pseudoalteromonadaceae *Pseudoalteromonas atlantica* T6c |
| Pci-1-1 | Bacteria Proteobacteria Gammaproteobacteria Alteromonadales Pseudoalteromonadaceae *Pseudoalteromonas citrea* NCIMB 1889 |
| Pdi-1-1 | Bacteria Proteobacteria Gammaproteobacteria Alteromonadales Pseudoalteromonadaceae *Pseudoalteromonas distincta* |
| Pel-1-1 | Bacteria Proteobacteria Gammaproteobacteria Alteromonadales Pseudoalteromonadaceae *Pseudoalteromonas elyakovii* |
| Pfl-1-1 | Bacteria Proteobacteria Gammaproteobacteria Alteromonadales Pseudoalteromonadaceae *Pseudoalteromonas flavipulchra* JG1 |
| Pfl-2-1 | Bacteria Proteobacteria Gammaproteobacteria Alteromonadales Pseudoalteromonadaceae *Pseudoalteromonas flavipulchra* NCIMB 2033 = ATCC BAA-314 |
| Pfu-1-1 | Bacteria Proteobacteria Gammaproteobacteria Alteromonadales Pseudoalteromonadaceae *Pseudoalteromonas fuliginea* |
| Pha-1-1 | Bacteria Proteobacteria Gammaproteobacteria Alteromonadales Pseudoalteromonadaceae *Pseudoalteromonas haloplanktis* ANT/505 |
| Pha-2-1 | Bacteria Proteobacteria Gammaproteobacteria Alteromonadales Pseudoalteromonadaceae *Pseudoalteromonas haloplanktis* ATCC 14393 |
| Pha-3-1 | Bacteria Proteobacteria Gammaproteobacteria Alteromonadales Pseudoalteromonadaceae *Pseudoalteromonas haloplanktis* TAC125 |
| Pli-1-1 | Bacteria Proteobacteria Gammaproteobacteria Alteromonadales Pseudoalteromonadaceae *Pseudoalteromonas lipolytica* SCSIO 04301 |
| Plu-1-1 | Bacteria Proteobacteria Gammaproteobacteria Alteromonadales Pseudoalteromonadaceae *Pseudoalteromonas luteoviolacea* |
| Plu-2-1 | Bacteria Proteobacteria Gammaproteobacteria Alteromonadales Pseudoalteromonadaceae *Pseudoalteromonas luteoviolacea* 2ta16 |
| Plu-3-1 | Bacteria Proteobacteria Gammaproteobacteria Alteromonadales Pseudoalteromonadaceae *Pseudoalteromonas luteoviolacea* B = ATCC 29581 |
| Plu-4-1 | Bacteria Proteobacteria Gammaproteobacteria Alteromonadales Pseudoalteromonadaceae *Pseudoalteromonas luteoviolacea* S4054 |
| Pma-1-1 | Bacteria Proteobacteria Gammaproteobacteria Alteromonadales Pseudoalteromonadaceae *Pseudoalteromonas marina mano*4 |
| Ppi-1-1 | Bacteria Proteobacteria Gammaproteobacteria Alteromonadales Pseudoalteromonadaceae *Pseudoalteromonas piratica* |
| Ppi-2-1 to Ppi-2-2 | Bacteria Proteobacteria Gammaproteobacteria Alteromonadales Pseudoalteromonadaceae *Pseudoalteromonas piscicida* |
| Ppi-3-1 | Bacteria Proteobacteria Gammaproteobacteria Alteromonadales Pseudoalteromonadaceae *Pseudoalteromonas piscicida* ATCC 15057 |
| Ppi-4-1 | Bacteria Proteobacteria Gammaproteobacteria Alteromonadales Pseudoalteromonadaceae *Pseudoalteromonas piscicida* JCM 20779 |
| Pru-1-1 to Pru-1-2 | Bacteria Proteobacteria Gammaproteobacteria Alteromonadales Pseudoalteromonadaceae *Pseudoalteromonas rubra* |
| Pru-2-1 | Bacteria Proteobacteria Gammaproteobacteria Alteromonadales Pseudoalteromonadaceae *Pseudoalteromonas rubra* ATCC 29570 |
| Pru-3-1 to Pru-3-2 | Bacteria Proteobacteria Gammaproteobacteria Alteromonadales Pseudoalteromonadaceae *Pseudoalteromonas ruthenica* |
| Pru-4-1 | Bacteria Proteobacteria Gammaproteobacteria Alteromonadales Pseudoalteromonadaceae *Pseudoalteromonas ruthenica* CP76 |
| Psp-1-1 | Bacteria Proteobacteria Gammaproteobacteria Alteromonadales Pseudoalteromonadaceae *Pseudoalteromonas sp.* '520P1 No. 412' |
| Psp-2-1 | Bacteria Proteobacteria Gammaproteobacteria Alteromonadales Pseudoalteromonadaceae *Pseudoalteromonas sp.* 23_GOM-1509m |
| Psp-3-1 | Bacteria Proteobacteria Gammaproteobacteria Alteromonadales Pseudoalteromonadaceae *Pseudoalteromonas sp.* 6BO_GOM-1096m |
| Psp-4-1 | Bacteria Proteobacteria Gammaproteobacteria Alteromonadales Pseudoalteromonadaceae *Pseudoalteromonas sp.* A2 |
| Psp-5-1 | Bacteria Proteobacteria Gammaproteobacteria Alteromonadales Pseudoalteromonadaceae *Pseudoalteromonas sp.* AC163 |
| Psp-6-1 | Bacteria Proteobacteria Gammaproteobacteria Alteromonadales Pseudoalteromonadaceae *Pseudoalteromonas sp.* BSi20311 |
| Psp-7-1 | Bacteria Proteobacteria Gammaproteobacteria Alteromonadales Pseudoalteromonadaceae *Pseudoalteromonas sp.* BSi20429 |
| Psp-8-1 | Bacteria Proteobacteria Gammaproteobacteria Alteromonadales Pseudoalteromonadaceae *Pseudoalteromonas sp.* BSi20439 |

| | |
|---|---|
| Psp-9-1 | Bacteria Proteobacteria Gammaproteobacteria Alteromonadales Pseudoalteromonadaceae *Pseudoalteromonas sp.* BSi20480 |
| Psp-10-1 | Bacteria Proteobacteria Gammaproteobacteria Alteromonadales Pseudoalteromonadaceae *Pseudoalteromonas sp.* BSi20495 |
| Psp-11-1 | Bacteria Proteobacteria Gammaproteobacteria Alteromonadales Pseudoalteromonadaceae *Pseudoalteromonas sp.* BSi20652 |
| Psp-12-1 | Bacteria Proteobacteria Gammaproteobacteria Alteromonadales Pseudoalteromonadaceae *Pseudoalteromonas sp.* Bsw20308 |
| Psp-13-1 | Bacteria Proteobacteria Gammaproteobacteria Alteromonadales Pseudoalteromonadaceae *Pseudoalteromonas sp.* ECSMB14103 |
| Psp-14-1 | Bacteria Proteobacteria Gammaproteobacteria Alteromonadales Pseudoalteromonadaceae *Pseudoalteromonas sp.* ND6B |
| Psp-15-1 | Bacteria Proteobacteria Gammaproteobacteria Alteromonadales Pseudoalteromonadaceae *Pseudoalteromonas sp.* NJ631 |
| Psp-16-1 | Bacteria Proteobacteria Gammaproteobacteria Alteromonadales Pseudoalteromonadaceae *Pseudoalteromonas sp.* NW 4327 |
| Psp-17-1 | Bacteria Proteobacteria Gammaproteobacteria Alteromonadales Pseudoalteromonadaceae *Pseudoalteromonas sp.* PAMC 22718 |
| Psp-18-1 | Bacteria Proteobacteria Gammaproteobacteria Alteromonadales Pseudoalteromonadaceae *Pseudoalteromonas sp.* PLSV |
| Psp-19-1 | Bacteria Proteobacteria Gammaproteobacteria Alteromonadales Pseudoalteromonadaceae *Pseudoalteromonas sp.* S2292 |
| Psp-20-1 | Bacteria Proteobacteria Gammaproteobacteria Alteromonadales Pseudoalteromonadaceae *Pseudoalteromonas sp.* S3431 |
| Psp-21-1 | Bacteria Proteobacteria Gammaproteobacteria Alteromonadales Pseudoalteromonadaceae *Pseudoalteromonas sp.* S8-38 |
| Psp-22-1 | Bacteria Proteobacteria Gammaproteobacteria Alteromonadales Pseudoalteromonadaceae *Pseudoalteromonas sp.* S8-8 |
| Psp-23-1 | Bacteria Proteobacteria Gammaproteobacteria Alteromonadales Pseudoalteromonadaceae *Pseudoalteromonas sp.* SCSIO_11900 |
| Psp-24-1 | Bacteria Proteobacteria Gammaproteobacteria Alteromonadales Pseudoalteromonadaceae *Pseudoalteromonas sp.* SM9913 |
| Psp-25-1 | Bacteria Proteobacteria Gammaproteobacteria Alteromonadales Pseudoalteromonadaceae *Pseudoalteromonas sp.* TAB23 |
| Psp-26-1 | Bacteria Proteobacteria Gammaproteobacteria Alteromonadales Pseudoalteromonadaceae *Pseudoalteromonas sp.* TAE56 |
| Psp-27-1 | Bacteria Proteobacteria Gammaproteobacteria Alteromonadales Pseudoalteromonadaceae *Pseudoalteromonas sp.* TAE79 |
| Psp-28-1 | Bacteria Proteobacteria Gammaproteobacteria Alteromonadales Pseudoalteromonadaceae *Pseudoalteromonas sp.* TAE80 |
| Psp-29-1 | Bacteria Proteobacteria Gammaproteobacteria Alteromonadales Pseudoalteromonadaceae *Pseudoalteromonas sp.* TB13 |
| Psp-30-1 | Bacteria Proteobacteria Gammaproteobacteria Alteromonadales Pseudoalteromonadaceae *Pseudoalteromonas sp.* TB25 |
| Psp-31-1 | Bacteria Proteobacteria Gammaproteobacteria Alteromonadales Pseudoalteromonadaceae *Pseudoalteromonas sp.* TB41 |
| Psp-32-1 | Bacteria Proteobacteria Gammaproteobacteria Alteromonadales Pseudoalteromonadaceae *Pseudoalteromonas sp.* TB64 |
| Psp-33-1 | Bacteria Proteobacteria Gammaproteobacteria Alteromonadales Pseudoalteromonadaceae *Pseudoalteromonas spongiae* UST010723-006 |
| Ptu-1-1 | Bacteria Proteobacteria Gammaproteobacteria Alteromonadales Pseudoalteromonadaceae *Pseudoalteromonas tunicata* D2 |
| Pun-1-1 | Bacteria Proteobacteria Gammaproteobacteria Alteromonadales Pseudoalteromonadaceae *Pseudoalteromonas undina* NCIMB 2128 |
| Paq-1-1 | Bacteria Proteobacteria Gammaproteobacteria Alteromonadales Psychromonadaceae *Psychromonas aquimarina* ATCC BAA-1526 |
| Par-2-1 | Bacteria Proteobacteria Gammaproteobacteria Alteromonadales Psychromonadaceae *Psychromonas arctica* DSM 14288 |
| Pha-4-1 | Bacteria Proteobacteria Gammaproteobacteria Alteromonadales Psychromonadaceae *Psychromonas hadalis* ATCC BAA-638 |
| Pin-1-1 | Bacteria Proteobacteria Gammaproteobacteria Alteromonadales Psychromonadaceae *Psychromonas ingrahamii* 37 |
| Pos-1-1 | Bacteria Proteobacteria Gammaproteobacteria Alteromonadales Psychromonadaceae *Psychromonas ossibalaenae* ATCC BAA-1528 |
| Psp-34-1 | Bacteria Proteobacteria Gammaproteobacteria Alteromonadales Psychromonadaceae *Psychromonas sp.* CNPT3 |
| Psp-35-1 | Bacteria Proteobacteria Gammaproteobacteria Alteromonadales Psychromonadaceae *Psychromonas sp.* SP041 |
| Sal-1-1 | Bacteria Proteobacteria Gammaproteobacteria Alteromonadales Shewanellaceae *Shewanella algae* |
| Sal-2-1 | Bacteria Proteobacteria Gammaproteobacteria Alteromonadales Shewanellaceae *Shewanella algae* JCM 21037 = NBRC 103173 |
| Sam-2-1 | Bacteria Proteobacteria Gammaproteobacteria Alteromonadales Shewanellaceae *Shewanella amazonensis* SB2B |
| Sba-1-1 | Bacteria Proteobacteria Gammaproteobacteria Alteromonadales Shewanellaceae *Shewanella baltica* BA175 |
| Sba-2-1 | Bacteria Proteobacteria Gammaproteobacteria Alteromonadales Shewanellaceae *Shewanella baltica* OS117 |
| Sba-3-1 | Bacteria Proteobacteria Gammaproteobacteria Alteromonadales Shewanellaceae *Shewanella baltica* OS155 |
| Sba-4-1 | Bacteria Proteobacteria Gammaproteobacteria Alteromonadales Shewanellaceae *Shewanella baltica* OS183 |
| Sba-5-1 | Bacteria Proteobacteria Gammaproteobacteria Alteromonadales Shewanellaceae *Shewanella baltica* OS185 |
| Sba-6-1 | Bacteria Proteobacteria Gammaproteobacteria Alteromonadales Shewanellaceae *Shewanella baltica* OS195 |
| Sba-7-1 | Bacteria Proteobacteria Gammaproteobacteria Alteromonadales Shewanellaceae *Shewanella baltica* OS223 |
| Sba-8-1 | Bacteria Proteobacteria Gammaproteobacteria Alteromonadales Shewanellaceae *Shewanella baltica* OS625 |
| Sba-9-1 | Bacteria Proteobacteria Gammaproteobacteria Alteromonadales Shewanellaceae *Shewanella baltica* OS678 |
| Sbe-1-1 | Bacteria Proteobacteria Gammaproteobacteria Alteromonadales Shewanellaceae *Shewanella benthica* KT99 |
| Sco-1-1 | Bacteria Proteobacteria Gammaproteobacteria Alteromonadales Shewanellaceae *Shewanella colwelliana* ATCC 39565 |
| Sde-1-1 | Bacteria Proteobacteria Gammaproteobacteria Alteromonadales Shewanellaceae *Shewanella decolorationis* S12 |
| Sde-2-1 | Bacteria Proteobacteria Gammaproteobacteria Alteromonadales Shewanellaceae *Shewanella denitrificans* OS217 |
| Sfi-1-1 | Bacteria Proteobacteria Gammaproteobacteria Alteromonadales Shewanellaceae *Shewanella fidelis* ATCC BAA-318 |
| Sfr-1-1 | Bacteria Proteobacteria Gammaproteobacteria Alteromonadales Shewanellaceae *Shewanella frigidimarina* NCIMB 400 |
| Sha-1-1 | Bacteria Proteobacteria Gammaproteobacteria Alteromonadales Shewanellaceae *Shewanella halifaxensis* HAW-EB4 |
| Sha-2-1 | Bacteria Proteobacteria Gammaproteobacteria Alteromonadales Shewanellaceae *Shewanella haliotis* JCM 14758 |
| Slo-1-1 | Bacteria Proteobacteria Gammaproteobacteria Alteromonadales Shewanellaceae *Shewanella loihica* PV-4 |
| Sma-1-1 | Bacteria Proteobacteria Gammaproteobacteria Alteromonadales Shewanellaceae *Shewanella mangrovi* |
| Sma-2-1 | Bacteria Proteobacteria Gammaproteobacteria Alteromonadales Shewanellaceae *Shewanella marina* JCM 15074 |
| Son-1-1 | Bacteria Proteobacteria Gammaproteobacteria Alteromonadales Shewanellaceae *Shewanella oneidensis* MR-1 |
| Spe-1-1 | Bacteria Proteobacteria Gammaproteobacteria Alteromonadales Shewanellaceae *Shewanella pealeana* ATCC 700345 |
| Spi-1-1 | Bacteria Proteobacteria Gammaproteobacteria Alteromonadales Shewanellaceae *Shewanella piezotolerans* WP3 |
| Spu-1-1 | Bacteria Proteobacteria Gammaproteobacteria Alteromonadales Shewanellaceae *Shewanella putrefaciens* 200 |
| Spu-2-1 | Bacteria Proteobacteria Gammaproteobacteria Alteromonadales Shewanellaceae *Shewanella putrefaciens* CN-32 |
| Sse-1-1 | Bacteria Proteobacteria Gammaproteobacteria Alteromonadales Shewanellaceae *Shewanella sediminis* HAW-EB3 |
| Ssp-1-1 | Bacteria Proteobacteria Gammaproteobacteria Alteromonadales Shewanellaceae *Shewanella sp.* 38A_GOM-205m |
| Ssp-2-1 | Bacteria Proteobacteria Gammaproteobacteria Alteromonadales Shewanellaceae *Shewanella sp.* ANA-3 |

| | |
|---|---|
| Ssp-11-1 | Bacteria Proteobacteria Gammaproteobacteria Alteromonadales Shewanellaceae *Shewanella sp. cp*20 |
| Ssp-3-1 | Bacteria Proteobacteria Gammaproteobacteria Alteromonadales Shewanellaceae *Shewanella sp.* ECSMB14101 |
| Ssp-4-1 | Bacteria Proteobacteria Gammaproteobacteria Alteromonadales Shewanellaceae *Shewanella sp.* ECSMB14102 |
| Ssp-5-1 | Bacteria Proteobacteria Gammaproteobacteria Alteromonadales Shewanellaceae *Shewanella sp.* HN-41 |
| Ssp-6-1 | Bacteria Proteobacteria Gammaproteobacteria Alteromonadales Shewanellaceae *Shewanella sp.* MR-4 |
| Ssp-7-1 | Bacteria Proteobacteria Gammaproteobacteria Alteromonadales Shewanellaceae *Shewanella sp.* MR-7 |
| Ssp-8-1 | Bacteria Proteobacteria Gammaproteobacteria Alteromonadales Shewanellaceae *Shewanella sp.* POL2 |
| Ssp-9-1 | Bacteria Proteobacteria Gammaproteobacteria Alteromonadales Shewanellaceae *Shewanella sp.* W3-18-1 |
| Ssp-10-1 | Bacteria Proteobacteria Gammaproteobacteria Alteromonadales Shewanellaceae *Shewanella sp.* ZOR0012 |
| Svi-1-1 | Bacteria Proteobacteria Gammaproteobacteria Alteromonadales Shewanellaceae *Shewanella violacea* DSS12 |
| Swa-1-1 | Bacteria Proteobacteria Gammaproteobacteria Alteromonadales Shewanellaceae *Shewanella waksmanii* ATCC BAA-643 |
| Swo-1-1 | Bacteria Proteobacteria Gammaproteobacteria Alteromonadales Shewanellaceae *Shewanella woodyi* ATCC 51908 |
| Sxi-1-1 | Bacteria Proteobacteria Gammaproteobacteria Alteromonadales Shewanellaceae *Shewanella xiamenensis* |
| CBa-1-1 | Bacteria Proteobacteria Gammaproteobacteria CandidatusBaumannia *Candidatus* Baumannia cicadellinicola |
| Aik-1-1 | Bacteria Proteobacteria Gammaproteobacteria Chromatiales Chromatiaceae *Arsukibacterium ikkense* |
| Asp-9-1 | Bacteria Proteobacteria Gammaproteobacteria Chromatiales Chromatiaceae *Arsukibacterium sp.* MJ3 |
| Pme-1-1 | Bacteria Proteobacteria Gammaproteobacteria Chromatiales Chromatiaceae *Pararheinheimera mesophila* |
| Pte-1-1 | Bacteria Proteobacteria Gammaproteobacteria Chromatiales Chromatiaceae *Pararheinheimera texasensis* DSM 17496 |
| Rba-1-1 | Bacteria Proteobacteria Gammaproteobacteria Chromatiales Chromatiaceae *Rheinheimera baltica* DSM 14885 |
| Rna-1-1 | Bacteria Proteobacteria Gammaproteobacteria Chromatiales Chromatiaceae *Rheinheimera nanhaiensis* E407-8 |
| Rpe-1-1 | Bacteria Proteobacteria Gammaproteobacteria Chromatiales Chromatiaceae *Rheinheimera perlucida* DSM 18276 |
| Rsp-1-1 | Bacteria Proteobacteria Gammaproteobacteria Chromatiales Chromatiaceae *Rheinheimera sp.* A13L |
| Rsp-2-1 | Bacteria Proteobacteria Gammaproteobacteria Chromatiales Chromatiaceae *Rheinheimera sp.* KL1 |
| Baq-1-1 | Bacteria Proteobacteria Gammaproteobacteria Enterobacterales Budviciaceae *Budvicia aquatica* DSM 5075 = ATCC 35567 |
| Lgr-1-1 to Lgr-1-2 | Bacteria Proteobacteria Gammaproteobacteria Enterobacterales Budviciaceae *Leminorella grimontii* ATCC 33999 = DSM 5078 |
| Pfo-1-1 | Bacteria Proteobacteria Gammaproteobacteria Enterobacterales Budviciaceae *Pragia fontium* |
| Bag-1-1 | Bacteria Proteobacteria Gammaproteobacteria Enterobacterales Enterobacteriaceae *Buttiauxella agrestis* |
| Bag-2-1 | Bacteria Proteobacteria Gammaproteobacteria Enterobacterales Enterobacteriaceae *Buttiauxella agrestis* ATCC 33320 |
| Cne-1-1 to Cne-1-3 | Bacteria Proteobacteria Gammaproteobacteria Enterobacterales Enterobacteriaceae *Cedecea neteri* |
| Cam-1-1 to Cam-1-3 | Bacteria Proteobacteria Gammaproteobacteria Enterobacterales Enterobacteriaceae *Citrobacter amalonaticus* |
| Cam-2-1 | Bacteria Proteobacteria Gammaproteobacteria Enterobacterales Enterobacteriaceae *Citrobacter amalonaticus* Y19 |
| Cfa-1-1 | Bacteria Proteobacteria Gammaproteobacteria Enterobacterales Enterobacteriaceae *Citrobacter farmeri* GTC 1319 |
| Cbr-1-1 to Cbr-1-7 | Bacteria Proteobacteria Gammaproteobacteria Enterobacterales Enterobacteriaceae Citrobacter Citrobacter freundii complex *Citrobacter braakii* |
| Cfr-1-1 to Cfr-1-4 | Bacteria Proteobacteria Gammaproteobacteria Enterobacterales Enterobacteriaceae Citrobacter Citrobacter freundii complex *Citrobacter freundii* |
| Cfr-2-1 to Cfr-2-2 | Bacteria Proteobacteria Gammaproteobacteria Enterobacterales Enterobacteriaceae Citrobacter Citrobacter freundii complex *Citrobacter freundii* ATCC 8090 = MTCC 1658 |
| Cfr-3-1 | Bacteria Proteobacteria Gammaproteobacteria Enterobacterales Enterobacteriaceae Citrobacter Citrobacter freundii complex *Citrobacter freundii* CFNIH1 |
| Cfr-4-1 | Bacteria Proteobacteria Gammaproteobacteria Enterobacterales Enterobacteriaceae Citrobacter Citrobacter freundii complex *Citrobacter freundii* MGH 56 |
| Cfr-5-1 | Bacteria Proteobacteria Gammaproteobacteria Enterobacterales Enterobacteriaceae Citrobacter Citrobacter freundii complex *Citrobacter freundii* NBRC 12681 |
| Cfr-6-1 | Bacteria Proteobacteria Gammaproteobacteria Enterobacterales Enterobacteriaceae Citrobacter Citrobacter freundii complex *Citrobacter freundii* RLS1 |
| Cfr-7-1 | Bacteria Proteobacteria Gammaproteobacteria Enterobacterales Enterobacteriaceae Citrobacter Citrobacter freundii complex *Citrobacter freundii* UCI 31 |
| Cfr-8-1 | Bacteria Proteobacteria Gammaproteobacteria Enterobacterales Enterobacteriaceae Citrobacter Citrobacter freundii complex *Citrobacter freundii* UCI 32 |
| Cse-1-1 | Bacteria Proteobacteria Gammaproteobacteria Enterobacterales Enterobacteriaceae Citrobacter Citrobacter freundii complex *Citrobacter sedlakii* NBRC 105722 |
| Csp-2-1 | Bacteria Proteobacteria Gammaproteobacteria Enterobacterales Enterobacteriaceae Citrobacter Citrobacter freundii complex *Citrobacter sp.* BIDMC107 |
| Csp-3-1 | Bacteria Proteobacteria Gammaproteobacteria Enterobacterales Enterobacteriaceae Citrobacter Citrobacter freundii complex *Citrobacter sp.* BIDMC108 |
| Csp-4-1 | Bacteria Proteobacteria Gammaproteobacteria Enterobacterales Enterobacteriaceae Citrobacter Citrobacter freundii complex *Citrobacter sp.* MGH100 |
| Csp-5-1 | Bacteria Proteobacteria Gammaproteobacteria Enterobacterales Enterobacteriaceae Citrobacter Citrobacter freundii complex *Citrobacter sp.* MGH103 |
| Csp-6-1 | Bacteria Proteobacteria Gammaproteobacteria Enterobacterales Enterobacteriaceae Citrobacter Citrobacter freundii complex *Citrobacter sp.* MGH104 |
| Csp-7-1 | Bacteria Proteobacteria Gammaproteobacteria Enterobacterales Enterobacteriaceae Citrobacter Citrobacter freundii complex *Citrobacter sp.* MGH105 |
| Csp-8-1 | Bacteria Proteobacteria Gammaproteobacteria Enterobacterales Enterobacteriaceae Citrobacter Citrobacter freundii complex *Citrobacter sp.* MGH106 |
| Csp-9-1 | Bacteria Proteobacteria Gammaproteobacteria Enterobacterales Enterobacteriaceae Citrobacter Citrobacter freundii complex *Citrobacter sp.* MGH109 |
| Csp-10-1 | Bacteria Proteobacteria Gammaproteobacteria Enterobacterales Enterobacteriaceae Citrobacter Citrobacter freundii complex *Citrobacter sp.* MGH110 |
| Csp-11-1 | Bacteria Proteobacteria Gammaproteobacteria Enterobacterales Enterobacteriaceae Citrobacter Citrobacter freundii complex *Citrobacter sp.* MGH99 |
| Cwe-1-1 | Bacteria Proteobacteria Gammaproteobacteria Enterobacterales Enterobacteriaceae Citrobacter Citrobacter freundii complex *Citrobacter werkmanii* NBRC 105721 |
| Cko-1-1 to Cko-1-2 | Bacteria Proteobacteria Gammaproteobacteria Enterobacterales Enterobacteriaceae *Citrobacter koseri* |
| Cpa-1-1 | Bacteria Proteobacteria Gammaproteobacteria Enterobacterales Enterobacteriaceae *Citrobacter pasteurii* |
| Cro-1-1 | Bacteria Proteobacteria Gammaproteobacteria Enterobacterales Enterobacteriaceae *Citrobacter rodentium* |
| Cro-2-1 | Bacteria Proteobacteria Gammaproteobacteria Enterobacterales Enterobacteriaceae *Citrobacter rodentium* NBRC 105723 |
| Csp-12-1 | Bacteria Proteobacteria Gammaproteobacteria Enterobacterales Enterobacteriaceae *Citrobacter sp.* JT3 |
| Csp-13-1 | Bacteria Proteobacteria Gammaproteobacteria Enterobacterales Enterobacteriaceae *Citrobacter sp.* MGH 55 |

| | |
|---|---|
| Csp-14-1 | Bacteria Proteobacteria Gammaproteobacteria Enterobacterales Enterobacteriaceae *Citrobacter sp.* S-77 |
| Cdu-1-1 | Bacteria Proteobacteria Gammaproteobacteria Enterobacterales Enterobacteriaceae *Cronobacter dublinensis subsp. dublinensis* LMG 23823 |
| Cdu-2-1 | Bacteria Proteobacteria Gammaproteobacteria Enterobacterales Enterobacteriaceae *Cronobacter dublinensis subsp. lactaridi* LMG 23825 |
| Cdu-3-1 | Bacteria Proteobacteria Gammaproteobacteria Enterobacterales Enterobacteriaceae *Cronobacter dublinensis subsp. lausannensis* LMG 23824 |
| Cma-1-1 to Cma-1-5 | Bacteria Proteobacteria Gammaproteobacteria Enterobacterales Enterobacteriaceae *Cronobacter malonaticus* |
| Cma-2-1 | Bacteria Proteobacteria Gammaproteobacteria Enterobacterales Enterobacteriaceae *Cronobacter malonaticus* ENBT0334 |
| Cma-3-1 | Bacteria Proteobacteria Gammaproteobacteria Enterobacterales Enterobacteriaceae *Cronobacter malonaticus* LMG 23826 |
| Cmu-1-1 | Bacteria Proteobacteria Gammaproteobacteria Enterobacterales Enterobacteriaceae *Cronobacter muytjensii* ATCC 51329 |
| Csa-1-1 | Bacteria Proteobacteria Gammaproteobacteria Enterobacterales Enterobacteriaceae *Cronobacter sakazakii* |
| Csa-2-1 | Bacteria Proteobacteria Gammaproteobacteria Enterobacterales Enterobacteriaceae *Cronobacter sakazakii* 2151 |
| Csa-3-1 | Bacteria Proteobacteria Gammaproteobacteria Enterobacterales Enterobacteriaceae *Cronobacter sakazakii* 8399 |
| Csa-4-1 | Bacteria Proteobacteria Gammaproteobacteria Enterobacterales Enterobacteriaceae *Cronobacter sakazakii* E764 |
| Csa-5-1 | Bacteria Proteobacteria Gammaproteobacteria Enterobacterales Enterobacteriaceae *Cronobacter sakazakii* ES35 |
| Csa-6-1 | Bacteria Proteobacteria Gammaproteobacteria Enterobacterales Enterobacteriaceae *Cronobacter sakazakii* ES713 |
| Csa-7-1 | Bacteria Proteobacteria Gammaproteobacteria Enterobacterales Enterobacteriaceae *Cronobacter sakazakii* NBRC 102416 |
| Csa-8-1 | Bacteria Proteobacteria Gammaproteobacteria Enterobacterales Enterobacteriaceae *Cronobacter sakazakii* NCIMB 8272 |
| Ctu-1-1 | Bacteria Proteobacteria Gammaproteobacteria Enterobacterales Enterobacteriaceae *Cronobacter turicensis z*508 |
| Ctu-2-1 | Bacteria Proteobacteria Gammaproteobacteria Enterobacterales Enterobacteriaceae *Cronobacter turicensis z*610 |
| Cun-1-1 to Cun-1-2 | Bacteria Proteobacteria Gammaproteobacteria Enterobacterales Enterobacteriaceae *Cronobacter universalis* NCTC 9529 |
| Eas-1-1 to Eas-1-2 | Bacteria Proteobacteria Gammaproteobacteria Enterobacterales Enterobacteriaceae Enterobacter Enterobacter cloacae complex *Enterobacter asburiae* |
| Eas-2-1 | Bacteria Proteobacteria Gammaproteobacteria Enterobacterales Enterobacteriaceae Enterobacter Enterobacter cloacae complex *Enterobacter asburiae* C1 |
| Eas-3-1 | Bacteria Proteobacteria Gammaproteobacteria Enterobacterales Enterobacteriaceae Enterobacter Enterobacter cloacae complex *Enterobacter asburiae* L1 |
| Eca-1-1 | Bacteria Proteobacteria Gammaproteobacteria Enterobacterales Enterobacteriaceae Enterobacter Enterobacter cloacae complex *Enterobacter cancerogenus* |
| Eca-2-1 | Bacteria Proteobacteria Gammaproteobacteria Enterobacterales Enterobacteriaceae Enterobacter Enterobacter cloacae complex *Enterobacter cancerogenus* YZ1 |
| Ecl-1-1 to Ecl-1-12 | Bacteria Proteobacteria Gammaproteobacteria Enterobacterales Enterobacteriaceae Enterobacter Enterobacter cloacae complex *Enterobacter cloacae* |
| Ecl-2-1 | Bacteria Proteobacteria Gammaproteobacteria Enterobacterales Enterobacteriaceae Enterobacter Enterobacter cloacae complex *Enterobacter cloacae* 'Hoffmann cluster III' MGH 13 |
| Ecl-3-1 | Bacteria Proteobacteria Gammaproteobacteria Enterobacterales Enterobacteriaceae Enterobacter Enterobacter cloacae complex *Enterobacter cloacae* BIDMC 33A |
| Ecl-4-1 | Bacteria Proteobacteria Gammaproteobacteria Enterobacterales Enterobacteriaceae Enterobacter Enterobacter cloacae complex *Enterobacter cloacae* BIDMC 66 |
| Ecl-5-1 | Bacteria Proteobacteria Gammaproteobacteria Enterobacterales Enterobacteriaceae Enterobacter Enterobacter cloacae complex *Enterobacter cloacae* BIDMC 67 |
| Ecl-6-1 | Bacteria Proteobacteria Gammaproteobacteria Enterobacterales Enterobacteriaceae Enterobacter Enterobacter cloacae complex *Enterobacter cloacae* BIDMC 8 |
| Ecl-7-1 | Bacteria Proteobacteria Gammaproteobacteria Enterobacterales Enterobacteriaceae Enterobacter Enterobacter cloacae complex *Enterobacter cloacae* BWH 43 |
| Ecl-8-1 | Bacteria Proteobacteria Gammaproteobacteria Enterobacterales Enterobacteriaceae Enterobacter Enterobacter cloacae complex *Enterobacter cloacae* CHS 79 |
| Ecl-27-1 | Bacteria Proteobacteria Gammaproteobacteria Enterobacterales Enterobacteriaceae Enterobacter *Enterobacter cloacae complex* 'Hoffmann cluster III' |
| Ecl-28-1 to Ecl-28-6 | Bacteria Proteobacteria Gammaproteobacteria Enterobacterales Enterobacteriaceae Enterobacter *Enterobacter cloacae complex* 'Hoffmann cluster IV' |
| Ecl-29-1 | Bacteria Proteobacteria Gammaproteobacteria Enterobacterales Enterobacteriaceae Enterobacter *Enterobacter cloacae complex sp.* 20432 |
| Ecl-30-1 | Bacteria Proteobacteria Gammaproteobacteria Enterobacterales Enterobacteriaceae Enterobacter *Enterobacter cloacae complex sp.* 35009 |
| Ecl-31-1 | Bacteria Proteobacteria Gammaproteobacteria Enterobacterales Enterobacteriaceae Enterobacter *Enterobacter cloacae complex sp.* 35618 |
| Ecl-32-1 | Bacteria Proteobacteria Gammaproteobacteria Enterobacterales Enterobacteriaceae Enterobacter *Enterobacter cloacae complex sp.* 35651 |
| Ecl-33-1 | Bacteria Proteobacteria Gammaproteobacteria Enterobacterales Enterobacteriaceae Enterobacter *Enterobacter cloacae complex sp.* 35669 |
| Ecl-34-1 | Bacteria Proteobacteria Gammaproteobacteria Enterobacterales Enterobacteriaceae Enterobacter *Enterobacter cloacae complex sp.* 35699 |
| Ecl-35-1 | Bacteria Proteobacteria Gammaproteobacteria Enterobacterales Enterobacteriaceae Enterobacter *Enterobacter cloacae complex sp.* 35731 |
| Ecl-36-1 | Bacteria Proteobacteria Gammaproteobacteria Enterobacterales Enterobacteriaceae Enterobacter *Enterobacter cloacae complex sp.* 35734 |
| Ecl-37-1 | Bacteria Proteobacteria Gammaproteobacteria Enterobacterales Enterobacteriaceae Enterobacter *Enterobacter cloacae complex sp.* 42192 |
| Ecl-38-1 | Bacteria Proteobacteria Gammaproteobacteria Enterobacterales Enterobacteriaceae Enterobacter *Enterobacter cloacae complex sp.* 43444 |
| Ecl-39-1 | Bacteria Proteobacteria Gammaproteobacteria Enterobacterales Enterobacteriaceae Enterobacter *Enterobacter cloacae complex sp.* 44242 |
| Ecl-40-1 | Bacteria Proteobacteria Gammaproteobacteria Enterobacterales Enterobacteriaceae Enterobacter *Enterobacter cloacae complex sp.* 44246 |
| Ecl-41-1 | Bacteria Proteobacteria Gammaproteobacteria Enterobacterales Enterobacteriaceae Enterobacter *Enterobacter cloacae complex sp.* CIDEIMsCOL1 |
| Ecl-42-1 | Bacteria Proteobacteria Gammaproteobacteria Enterobacterales Enterobacteriaceae Enterobacter *Enterobacter cloacae complex sp.* CIDEIMsCOL9 |
| Ecl-43-1 | Bacteria Proteobacteria Gammaproteobacteria Enterobacterales Enterobacteriaceae Enterobacter *Enterobacter cloacae complex sp.* GN02141 |
| Ecl-44-1 | Bacteria Proteobacteria Gammaproteobacteria Enterobacterales Enterobacteriaceae Enterobacter *Enterobacter cloacae complex sp.* GN02208 |
| Ecl-45-1 | Bacteria Proteobacteria Gammaproteobacteria Enterobacterales Enterobacteriaceae Enterobacter *Enterobacter cloacae complex sp.* GN02283 |
| Ecl-46-1 | Bacteria Proteobacteria Gammaproteobacteria Enterobacterales Enterobacteriaceae Enterobacter *Enterobacter cloacae complex sp.* GN02440 |
| Ecl-47-1 | Bacteria Proteobacteria Gammaproteobacteria Enterobacterales Enterobacteriaceae Enterobacter *Enterobacter cloacae complex sp.* GN02548 |

| | |
|---|---|
| Ecl-48-1 | Bacteria Proteobacteria Gammaproteobacteria Enterobacterales Enterobacteriaceae Enterobacter *Enterobacter cloacae complex sp.* GN02570 |
| Ecl-49-1 | Bacteria Proteobacteria Gammaproteobacteria Enterobacterales Enterobacteriaceae Enterobacter *Enterobacter cloacae complex sp.* GN02587 |
| Ecl-50-1 | Bacteria Proteobacteria Gammaproteobacteria Enterobacterales Enterobacteriaceae Enterobacter *Enterobacter cloacae complex sp.* GN02692 |
| Ecl-51-1 | Bacteria Proteobacteria Gammaproteobacteria Enterobacterales Enterobacteriaceae Enterobacter *Enterobacter cloacae complex sp.* GN03164 |
| Ecl-9-1 | Bacteria Proteobacteria Gammaproteobacteria Enterobacterales Enterobacteriaceae Enterobacter Enterobacter cloacae complex *Enterobacter cloacae* EC-NIH2 |
| Ecl-10-1 | Bacteria Proteobacteria Gammaproteobacteria Enterobacterales Enterobacteriaceae Enterobacter Enterobacter cloacae complex *Enterobacter cloacae* EC-NIH3 |
| Ecl-11-1 | Bacteria Proteobacteria Gammaproteobacteria Enterobacterales Enterobacteriaceae Enterobacter Enterobacter cloacae complex *Enterobacter cloacae* ECR091 |
| Ecl-12-1 | Bacteria Proteobacteria Gammaproteobacteria Enterobacterales Enterobacteriaceae Enterobacter Enterobacter cloacae complex *Enterobacter cloacae* IIT-BT 08 |
| Ecl-13-1 | Bacteria Proteobacteria Gammaproteobacteria Enterobacterales Enterobacteriaceae Enterobacter Enterobacter cloacae complex *Enterobacter cloacae* JD6301 |
| Ecl-14-1 | Bacteria Proteobacteria Gammaproteobacteria Enterobacterales Enterobacteriaceae Enterobacter Enterobacter cloacae complex *Enterobacter cloacae* JD8715 |
| Ecl-15-1 | Bacteria Proteobacteria Gammaproteobacteria Enterobacterales Enterobacteriaceae Enterobacter Enterobacter cloacae complex *Enterobacter cloacae* MGH 53 |
| Ecl-16-1 | Bacteria Proteobacteria Gammaproteobacteria Enterobacterales Enterobacteriaceae Enterobacter Enterobacter cloacae complex *Enterobacter cloacae* MGH 54 |
| Ecl-17-1 | Bacteria Proteobacteria Gammaproteobacteria Enterobacterales Enterobacteriaceae Enterobacter Enterobacter cloacae complex *Enterobacter cloacae* P101 |
| Ecl-52-1 to Ecl-52-5 | Bacteria Proteobacteria Gammaproteobacteria Enterobacterales Enterobacteriaceae Enterobacter Enterobacter cloacae complex *Enterobacter cloacae subsp. cloacae* |
| Ecl-53-1 | Bacteria Proteobacteria Gammaproteobacteria Enterobacterales Enterobacteriaceae Enterobacter Enterobacter cloacae complex *Enterobacter cloacae subsp. cloacae* SY-70 |
| Ecl-54-1 | Bacteria Proteobacteria Gammaproteobacteria Enterobacterales Enterobacteriaceae Enterobacter Enterobacter cloacae complex *Enterobacter cloacae subsp. dissolvens* |
| Ecl-18-1 | Bacteria Proteobacteria Gammaproteobacteria Enterobacterales Enterobacteriaceae Enterobacter Enterobacter cloacae complex *Enterobacter cloacae* UCI 23 |
| Ecl-19-1 | Bacteria Proteobacteria Gammaproteobacteria Enterobacterales Enterobacteriaceae Enterobacter Enterobacter cloacae complex *Enterobacter cloacae* UCI 24 |
| Ecl-20-1 | Bacteria Proteobacteria Gammaproteobacteria Enterobacterales Enterobacteriaceae Enterobacter Enterobacter cloacae complex *Enterobacter cloacae* UCI 29 |
| Ecl-21-1 | Bacteria Proteobacteria Gammaproteobacteria Enterobacterales Enterobacteriaceae Enterobacter Enterobacter cloacae complex *Enterobacter cloacae* UCI 30 |
| Ecl-22-1 | Bacteria Proteobacteria Gammaproteobacteria Enterobacterales Enterobacteriaceae Enterobacter Enterobacter cloacae complex *Enterobacter cloacae* UCI 35 |
| Ecl-23-1 | Bacteria Proteobacteria Gammaproteobacteria Enterobacterales Enterobacteriaceae Enterobacter Enterobacter cloacae complex *Enterobacter cloacae* UCI 36 |
| Ecl-24-1 | Bacteria Proteobacteria Gammaproteobacteria Enterobacterales Enterobacteriaceae Enterobacter Enterobacter cloacae complex *Enterobacter cloacae* UCI 39 |
| Ecl-25-1 | Bacteria Proteobacteria Gammaproteobacteria Enterobacterales Enterobacteriaceae Enterobacter Enterobacter cloacae complex *Enterobacter cloacae* UCI 49 |
| Ecl-26-1 | Bacteria Proteobacteria Gammaproteobacteria Enterobacterales Enterobacteriaceae Enterobacter Enterobacter cloacae complex *Enterobacter cloacae* UCI 50 |
| Eho-1-1 to Eho-1-9 | Bacteria Proteobacteria Gammaproteobacteria Enterobacterales Enterobacteriaceae Enterobacter Enterobacter cloacae complex *Enterobacter hormaechei* |
| Eho-2-1 | Bacteria Proteobacteria Gammaproteobacteria Enterobacterales Enterobacteriaceae Enterobacter Enterobacter cloacae complex *Enterobacter hormaechei subsp. hormaechei* |
| Eho-3-1 to Eho-3-2 | Bacteria Proteobacteria Gammaproteobacteria Enterobacterales Enterobacteriaceae Enterobacter Enterobacter cloacae complex *Enterobacter hormaechei subsp. oharae* |
| Eho-4-1 to Eho-4-2 | Bacteria Proteobacteria Gammaproteobacteria Enterobacterales Enterobacteriaceae Enterobacter Enterobacter cloacae complex *Enterobacter hormaechei subsp. steigerwaltii* |
| Eko-1-1 to Eko-1-11 | Bacteria Proteobacteria Gammaproteobacteria Enterobacterales Enterobacteriaceae Enterobacter Enterobacter cloacae complex *Enterobacter kobei* |
| Elu-1-1 to Elu-1-5 | Bacteria Proteobacteria Gammaproteobacteria Enterobacterales Enterobacteriaceae Enterobacter Enterobacter cloacae complex *Enterobacter ludwigii* |
| Esp-1-1 | Bacteria Proteobacteria Gammaproteobacteria Enterobacterales Enterobacteriaceae Enterobacter Enterobacter cloacae complex *Enterobacter sp.* 5-4 |

| | |
|---|---|
| Esp-2-1 | Bacteria Proteobacteria Gammaproteobacteria Enterobacterales Enterobacteriaceae Enterobacter Enterobacter cloacae complex *Enterobacter sp.* BIDMC 26 |
| Esp-3-1 | Bacteria Proteobacteria Gammaproteobacteria Enterobacterales Enterobacteriaceae Enterobacter Enterobacter cloacae complex *Enterobacter sp.* BIDMC 27 |
| Esp-4-1 | Bacteria Proteobacteria Gammaproteobacteria Enterobacterales Enterobacteriaceae Enterobacter Enterobacter cloacae complex *Enterobacter sp.* BIDMC 28 |
| Esp-5-1 | Bacteria Proteobacteria Gammaproteobacteria Enterobacterales Enterobacteriaceae Enterobacter Enterobacter cloacae complex *Enterobacter sp.* BIDMC 29 |
| Esp-6-1 | Bacteria Proteobacteria Gammaproteobacteria Enterobacterales Enterobacteriaceae Enterobacter Enterobacter cloacae complex *Enterobacter sp.* BIDMC 30 |
| Esp-7-1 | Bacteria Proteobacteria Gammaproteobacteria Enterobacterales Enterobacteriaceae Enterobacter Enterobacter cloacae complex *Enterobacter sp.* BIDMC100 |
| Esp-8-1 | Bacteria Proteobacteria Gammaproteobacteria Enterobacterales Enterobacteriaceae Enterobacter Enterobacter cloacae complex *Enterobacter sp.* BIDMC109 |
| Esp-9-1 | Bacteria Proteobacteria Gammaproteobacteria Enterobacterales Enterobacteriaceae Enterobacter Enterobacter cloacae complex *Enterobacter sp.* BIDMC87 |
| Esp-10-1 | Bacteria Proteobacteria Gammaproteobacteria Enterobacterales Enterobacteriaceae Enterobacter Enterobacter cloacae complex *Enterobacter sp.* BIDMC93 |
| Esp-11-1 | Bacteria Proteobacteria Gammaproteobacteria Enterobacterales Enterobacteriaceae Enterobacter Enterobacter cloacae complex *Enterobacter sp.* BIDMC94 |
| Esp-12-1 | Bacteria Proteobacteria Gammaproteobacteria Enterobacterales Enterobacteriaceae Enterobacter Enterobacter cloacae complex *Enterobacter sp.* BIDMC99 |
| Esp-13-1 | Bacteria Proteobacteria Gammaproteobacteria Enterobacterales Enterobacteriaceae Enterobacter Enterobacter cloacae complex *Enterobacter sp.* BWH 27 |
| Esp-14-1 | Bacteria Proteobacteria Gammaproteobacteria Enterobacterales Enterobacteriaceae Enterobacter Enterobacter cloacae complex *Enterobacter sp.* BWH 37 |
| Esp-15-1 | Bacteria Proteobacteria Gammaproteobacteria Enterobacterales Enterobacteriaceae Enterobacter Enterobacter cloacae complex *Enterobacter sp.* BWH 39 |
| Esp-16-1 | Bacteria Proteobacteria Gammaproteobacteria Enterobacterales Enterobacteriaceae Enterobacter Enterobacter cloacae complex *Enterobacter sp.* BWH52 |
| Esp-17-1 | Bacteria Proteobacteria Gammaproteobacteria Enterobacterales Enterobacteriaceae Enterobacter Enterobacter cloacae complex *Enterobacter sp.* BWH63 |
| Esp-18-1 | Bacteria Proteobacteria Gammaproteobacteria Enterobacterales Enterobacteriaceae Enterobacter Enterobacter cloacae complex *Enterobacter sp.* BWH64 |
| Esp-19-1 | Bacteria Proteobacteria Gammaproteobacteria Enterobacterales Enterobacteriaceae Enterobacter Enterobacter cloacae complex *Enterobacter sp.* FB |
| Esp-20-1 | Bacteria Proteobacteria Gammaproteobacteria Enterobacterales Enterobacteriaceae Enterobacter Enterobacter cloacae complex *Enterobacter sp.* MGH 1 |
| Esp-21-1 | Bacteria Proteobacteria Gammaproteobacteria Enterobacterales Enterobacteriaceae Enterobacter Enterobacter cloacae complex *Enterobacter sp.* MGH 10 |
| Esp-22-1 | Bacteria Proteobacteria Gammaproteobacteria Enterobacterales Enterobacteriaceae Enterobacter Enterobacter cloacae complex *Enterobacter sp.* MGH 11 |
| Esp-23-1 | Bacteria Proteobacteria Gammaproteobacteria Enterobacterales Enterobacteriaceae Enterobacter Enterobacter cloacae complex *Enterobacter sp.* MGH 12 |
| Esp-24-1 | Bacteria Proteobacteria Gammaproteobacteria Enterobacterales Enterobacteriaceae Enterobacter Enterobacter cloacae complex *Enterobacter sp.* MGH 15 |
| Esp-25-1 | Bacteria Proteobacteria Gammaproteobacteria Enterobacterales Enterobacteriaceae Enterobacter Enterobacter cloacae complex *Enterobacter sp.* MGH 2 |
| Esp-26-1 | Bacteria Proteobacteria Gammaproteobacteria Enterobacterales Enterobacteriaceae Enterobacter Enterobacter cloacae complex *Enterobacter sp.* MGH 3 |
| Esp-27-1 | Bacteria Proteobacteria Gammaproteobacteria Enterobacterales Enterobacteriaceae Enterobacter Enterobacter cloacae complex *Enterobacter sp.* MGH 33 |
| Esp-28-1 | Bacteria Proteobacteria Gammaproteobacteria Enterobacterales Enterobacteriaceae Enterobacter Enterobacter cloacae complex *Enterobacter sp.* MGH 37 |
| Esp-29-1 | Bacteria Proteobacteria Gammaproteobacteria Enterobacterales Enterobacteriaceae Enterobacter Enterobacter cloacae complex *Enterobacter sp.* MGH 4 |
| Esp-30-1 | Bacteria Proteobacteria Gammaproteobacteria Enterobacterales Enterobacteriaceae Enterobacter Enterobacter cloacae complex *Enterobacter sp.* MGH 5 |
| Esp-31-1 | Bacteria Proteobacteria Gammaproteobacteria Enterobacterales Enterobacteriaceae Enterobacter Enterobacter cloacae complex *Enterobacter sp.* MGH 6 |
| Esp-32-1 | Bacteria Proteobacteria Gammaproteobacteria Enterobacterales Enterobacteriaceae Enterobacter Enterobacter cloacae complex *Enterobacter sp.* MGH 7 |
| Esp-33-1 | Bacteria Proteobacteria Gammaproteobacteria Enterobacterales Enterobacteriaceae Enterobacter Enterobacter cloacae complex *Enterobacter sp.* MGH 9 |
| Esp-34-1 | Bacteria Proteobacteria Gammaproteobacteria Enterobacterales Enterobacteriaceae Enterobacter Enterobacter cloacae complex *Enterobacter sp.* MGH119 |
| Esp-35-1 | Bacteria Proteobacteria Gammaproteobacteria Enterobacterales Enterobacteriaceae Enterobacter Enterobacter cloacae complex *Enterobacter sp.* MGH120 |
| Esp-36-1 | Bacteria Proteobacteria Gammaproteobacteria Enterobacterales Enterobacteriaceae Enterobacter Enterobacter cloacae complex *Enterobacter sp.* MGH127 |
| Esp-37-1 | Bacteria Proteobacteria Gammaproteobacteria Enterobacterales Enterobacteriaceae Enterobacter Enterobacter cloacae complex *Enterobacter sp.* MGH128 |
| Esp-38-1 | Bacteria Proteobacteria Gammaproteobacteria Enterobacterales Enterobacteriaceae Enterobacter Enterobacter cloacae complex *Enterobacter sp.* MGH85 |
| Esp-39-1 | Bacteria Proteobacteria Gammaproteobacteria Enterobacterales Enterobacteriaceae Enterobacter Enterobacter cloacae complex *Enterobacter sp.* MGH86 |
| Esp-40-1 | Bacteria Proteobacteria Gammaproteobacteria Enterobacterales Enterobacteriaceae *Enterobacter sp.* 940_PEND |
| Esp-41-1 | Bacteria Proteobacteria Gammaproteobacteria Enterobacterales Enterobacteriaceae *Enterobacter sp.* BIDMC92 |
| Esp-42-1 | Bacteria Proteobacteria Gammaproteobacteria Enterobacterales Enterobacteriaceae *Enterobacter sp.* Bisph1 |
| Esp-43-1 | Bacteria Proteobacteria Gammaproteobacteria Enterobacterales Enterobacteriaceae *Enterobacter sp.* Bisph2 |
| Esp-44-1 | Bacteria Proteobacteria Gammaproteobacteria Enterobacterales Enterobacteriaceae *Enterobacter sp.* DC1 |
| Esp-45-1 | Bacteria Proteobacteria Gammaproteobacteria Enterobacterales Enterobacteriaceae *Enterobacter sp.* DC3 |
| Esp-46-1 | Bacteria Proteobacteria Gammaproteobacteria Enterobacterales Enterobacteriaceae *Enterobacter sp.* DC4 |
| Esp-47-1 | Bacteria Proteobacteria Gammaproteobacteria Enterobacterales Enterobacteriaceae *Enterobacter sp.* EGD-HP1 |
| Esp-48-1 | Bacteria Proteobacteria Gammaproteobacteria Enterobacterales Enterobacteriaceae *Enterobacter sp.* GN02600 |
| Esp-49-1 | Bacteria Proteobacteria Gammaproteobacteria Enterobacterales Enterobacteriaceae *Enterobacter sp.* T1-1 |
| Esp-50-1 | Bacteria Proteobacteria Gammaproteobacteria Enterobacterales Enterobacteriaceae *Enterobacter sp.* UCD-UG_FMILLET |
| Esp-51-1 | Bacteria Proteobacteria Gammaproteobacteria Enterobacterales Enterobacteriaceae *Enterobacter sp.* ZOR0014 |
| Eba-1-1 | Bacteria Proteobacteria Gammaproteobacteria Enterobacterales *Enterobacteriaceae bacterium* ATCC 29904 |
| Eba-2-1 | Bacteria Proteobacteria Gammaproteobacteria Enterobacterales *Enterobacteriaceae bacterium* B14 |
| Eal-1-1 | Bacteria Proteobacteria Gammaproteobacteria Enterobacterales Enterobacteriaceae *Escherichia albertii* KF1 |
| Eal-2-1 | Bacteria Proteobacteria Gammaproteobacteria Enterobacterales Enterobacteriaceae *Escherichia albertii* NBRC 107761 |
| Eco-1-1 to Eco-1-4 | Bacteria Proteobacteria Gammaproteobacteria Enterobacterales Enterobacteriaceae *Escherichia coli* |
| Eco-2-1 | Bacteria Proteobacteria Gammaproteobacteria Enterobacterales Enterobacteriaceae *Escherichia coli* 1-110-08_S1_C1 |
| Eco-3-1 | Bacteria Proteobacteria Gammaproteobacteria Enterobacterales Enterobacteriaceae *Escherichia coli* 1-110-08_S1_C2 |
| Eco-4-1 | Bacteria Proteobacteria Gammaproteobacteria Enterobacterales Enterobacteriaceae *Escherichia coli* 1-110-08_S1_C3 |

| | |
|---|---|
| Eco-5-1 | Bacteria Proteobacteria Gammaproteobacteria Enterobacterales Enterobacteriaceae *Escherichia coli* 1-110-08_S3_C1 |
| Eco-6-1 | Bacteria Proteobacteria Gammaproteobacteria Enterobacterales Enterobacteriaceae *Escherichia coli* 1-110-08_S3_C2 |
| Eco-7-1 | Bacteria Proteobacteria Gammaproteobacteria Enterobacterales Enterobacteriaceae *Escherichia coli* 1-110-08_S3_C3 |
| Eco-8-1 | Bacteria Proteobacteria Gammaproteobacteria Enterobacterales Enterobacteriaceae *Escherichia coli* 1-110-08_S4_C1 |
| Eco-9-1 | Bacteria Proteobacteria Gammaproteobacteria Enterobacterales Enterobacteriaceae *Escherichia coli* 1-110-08_S4_C2 |
| Eco-10-1 | Bacteria Proteobacteria Gammaproteobacteria Enterobacterales Enterobacteriaceae *Escherichia coli* 1-110-08_S4_C3 |
| Eco-11-1 | Bacteria Proteobacteria Gammaproteobacteria Enterobacterales Enterobacteriaceae *Escherichia coli* 1-176-05_S1_C1 |
| Eco-12-1 | Bacteria Proteobacteria Gammaproteobacteria Enterobacterales Enterobacteriaceae *Escherichia coli* 1-176-05_S1_C2 |
| Eco-13-1 | Bacteria Proteobacteria Gammaproteobacteria Enterobacterales Enterobacteriaceae *Escherichia coli* 1-176-05_S1_C3 |
| Eco-14-1 | Bacteria Proteobacteria Gammaproteobacteria Enterobacterales Enterobacteriaceae *Escherichia coli* 1-176-05_S3_C1 |
| Eco-15-1 | Bacteria Proteobacteria Gammaproteobacteria Enterobacterales Enterobacteriaceae *Escherichia coli* 1-176-05_S3_C2 |
| Eco-16-1 | Bacteria Proteobacteria Gammaproteobacteria Enterobacterales Enterobacteriaceae *Escherichia coli* 1-176-05_S4_C1 |
| Eco-17-1 | Bacteria Proteobacteria Gammaproteobacteria Enterobacterales Enterobacteriaceae *Escherichia coli* 1-176-05_S4_C2 |
| Eco-18-1 | Bacteria Proteobacteria Gammaproteobacteria Enterobacterales Enterobacteriaceae *Escherichia coli* 1-176-05_S4_C3 |
| Eco-19-1 | Bacteria Proteobacteria Gammaproteobacteria Enterobacterales Enterobacteriaceae *Escherichia coli* 1-182-04_S1_C1 |
| Eco-20-1 | Bacteria Proteobacteria Gammaproteobacteria Enterobacterales Enterobacteriaceae *Escherichia coli* 1-182-04_S1_C2 |
| Eco-21-1 | Bacteria Proteobacteria Gammaproteobacteria Enterobacterales Enterobacteriaceae *Escherichia coli* 1-182-04_S1_C3 |
| Eco-22-1 | Bacteria Proteobacteria Gammaproteobacteria Enterobacterales Enterobacteriaceae *Escherichia coli* 1-182-04_S3_C1 |
| Eco-23-1 | Bacteria Proteobacteria Gammaproteobacteria Enterobacterales Enterobacteriaceae *Escherichia coli* 1-182-04_S3_C2 |
| Eco-24-1 | Bacteria Proteobacteria Gammaproteobacteria Enterobacterales Enterobacteriaceae *Escherichia coli* 1-182-04_S3_C3 |
| Eco-25-1 | Bacteria Proteobacteria Gammaproteobacteria Enterobacterales Enterobacteriaceae *Escherichia coli* 1-182-04_S4_C1 |
| Eco-26-1 | Bacteria Proteobacteria Gammaproteobacteria Enterobacterales Enterobacteriaceae *Escherichia coli* 1-182-04_S4_C2 |
| Eco-27-1 | Bacteria Proteobacteria Gammaproteobacteria Enterobacterales Enterobacteriaceae *Escherichia coli* 1-182-04_S4_C3 |
| Eco-28-1 | Bacteria Proteobacteria Gammaproteobacteria Enterobacterales Enterobacteriaceae *Escherichia coli* 1-250-04_S1_C1 |
| Eco-29-1 | Bacteria Proteobacteria Gammaproteobacteria Enterobacterales Enterobacteriaceae *Escherichia coli* 1-250-04_S1_C2 |
| Eco-30-1 | Bacteria Proteobacteria Gammaproteobacteria Enterobacterales Enterobacteriaceae *Escherichia coli* 1-250-04_S1_C3 |
| Eco-31-1 | Bacteria Proteobacteria Gammaproteobacteria Enterobacterales Enterobacteriaceae *Escherichia coli* 1-250-04_S3_C1 |
| Eco-32-1 | Bacteria Proteobacteria Gammaproteobacteria Enterobacterales Enterobacteriaceae *Escherichia coli* 1-250-04_S3_C2 |
| Eco-33-1 | Bacteria Proteobacteria Gammaproteobacteria Enterobacterales Enterobacteriaceae *Escherichia coli* 1-250-04_S4_C1 |
| Eco-34-1 | Bacteria Proteobacteria Gammaproteobacteria Enterobacterales Enterobacteriaceae *Escherichia coli* 1-250-04_S4_C2 |
| Eco-35-1 | Bacteria Proteobacteria Gammaproteobacteria Enterobacterales Enterobacteriaceae *Escherichia coli* 1-392-07_S1_C1 |
| Eco-36-1 | Bacteria Proteobacteria Gammaproteobacteria Enterobacterales Enterobacteriaceae *Escherichia coli* 1-392-07_S1_C2 |
| Eco-37-1 | Bacteria Proteobacteria Gammaproteobacteria Enterobacterales Enterobacteriaceae *Escherichia coli* 1-392-07_S3_C1 |
| Eco-38-1 | Bacteria Proteobacteria Gammaproteobacteria Enterobacterales Enterobacteriaceae *Escherichia coli* 1-392-07_S3_C2 |
| Eco-39-1 | Bacteria Proteobacteria Gammaproteobacteria Enterobacterales Enterobacteriaceae *Escherichia coli* 1-392-07_S3_C3 |
| Eco-40-1 | Bacteria Proteobacteria Gammaproteobacteria Enterobacterales Enterobacteriaceae *Escherichia coli* 1-392-07_S4_C1 |
| Eco-41-1 | Bacteria Proteobacteria Gammaproteobacteria Enterobacterales Enterobacteriaceae *Escherichia coli* 1-392-07_S4_C2 |
| Eco-42-1 | Bacteria Proteobacteria Gammaproteobacteria Enterobacterales Enterobacteriaceae *Escherichia coli* 1-392-07_S4_C3 |
| Eco-43-1 | Bacteria Proteobacteria Gammaproteobacteria Enterobacterales Enterobacteriaceae *Escherichia coli* 100329 |
| Eco-44-1 | Bacteria Proteobacteria Gammaproteobacteria Enterobacterales Enterobacteriaceae *Escherichia coli* 103385 |
| Eco-45-1 | Bacteria Proteobacteria Gammaproteobacteria Enterobacterales Enterobacteriaceae *Escherichia coli* 103573 |
| Eco-46-1 | Bacteria Proteobacteria Gammaproteobacteria Enterobacterales Enterobacteriaceae *Escherichia coli* 1047 |
| Eco-47-1 | Bacteria Proteobacteria Gammaproteobacteria Enterobacterales Enterobacteriaceae *Escherichia coli* 10810 |
| Eco-48-1 | Bacteria Proteobacteria Gammaproteobacteria Enterobacterales Enterobacteriaceae *Escherichia coli* 11117 |
| Eco-49-1 | Bacteria Proteobacteria Gammaproteobacteria Enterobacterales Enterobacteriaceae *Escherichia coli* 112469215-isolate1 |
| Eco-50-1 | Bacteria Proteobacteria Gammaproteobacteria Enterobacterales Enterobacteriaceae *Escherichia coli* 112469218-isolate1 |
| Eco-51-1 | Bacteria Proteobacteria Gammaproteobacteria Enterobacterales Enterobacteriaceae *Escherichia coli* 1240 |
| Eco-52-1 | Bacteria Proteobacteria Gammaproteobacteria Enterobacterales Enterobacteriaceae *Escherichia coli* 1303 |
| Eco-53-1 | Bacteria Proteobacteria Gammaproteobacteria Enterobacterales Enterobacteriaceae *Escherichia coli* 1350 |
| Eco-54-1 | Bacteria Proteobacteria Gammaproteobacteria Enterobacterales Enterobacteriaceae *Escherichia coli* 1365 |
| Eco-55-1 | Bacteria Proteobacteria Gammaproteobacteria Enterobacterales Enterobacteriaceae *Escherichia coli* 148 |
| Eco-56-1 | Bacteria Proteobacteria Gammaproteobacteria Enterobacterales Enterobacteriaceae *Escherichia coli* 1A |
| Eco-57-1 | Bacteria Proteobacteria Gammaproteobacteria Enterobacterales Enterobacteriaceae *Escherichia coli* 2-005-03_S1_C1 |
| Eco-58-1 | Bacteria Proteobacteria Gammaproteobacteria Enterobacterales Enterobacteriaceae *Escherichia coli* 2-005-03_S1_C2 |
| Eco-59-1 | Bacteria Proteobacteria Gammaproteobacteria Enterobacterales Enterobacteriaceae *Escherichia coli* 2-005-03_S1_C3 |
| Eco-60-1 | Bacteria Proteobacteria Gammaproteobacteria Enterobacterales Enterobacteriaceae *Escherichia coli* 2-005-03_S3_C1 |
| Eco-61-1 | Bacteria Proteobacteria Gammaproteobacteria Enterobacterales Enterobacteriaceae *Escherichia coli* 2-005-03_S3_C2 |
| Eco-62-1 | Bacteria Proteobacteria Gammaproteobacteria Enterobacterales Enterobacteriaceae *Escherichia coli* 2-005-03_S3_C3 |
| Eco-63-1 | Bacteria Proteobacteria Gammaproteobacteria Enterobacterales Enterobacteriaceae *Escherichia coli* 2-005-03_S4_C1 |
| Eco-64-1 | Bacteria Proteobacteria Gammaproteobacteria Enterobacterales Enterobacteriaceae *Escherichia coli* 2-005-03_S4_C2 |
| Eco-65-1 | Bacteria Proteobacteria Gammaproteobacteria Enterobacterales Enterobacteriaceae *Escherichia coli* 2-005-03_S4_C3 |
| Eco-66-1 | Bacteria Proteobacteria Gammaproteobacteria Enterobacterales Enterobacteriaceae *Escherichia coli* 2-011-08_S1_C1 |
| Eco-67-1 | Bacteria Proteobacteria Gammaproteobacteria Enterobacterales Enterobacteriaceae *Escherichia coli* 2-011-08_S1_C2 |
| Eco-68-1 | Bacteria Proteobacteria Gammaproteobacteria Enterobacterales Enterobacteriaceae *Escherichia coli* 2-011-08_S1_C3 |
| Eco-69-1 | Bacteria Proteobacteria Gammaproteobacteria Enterobacterales Enterobacteriaceae *Escherichia coli* 2-011-08_S3_C1 |

| | |
|---|---|
| Eco-70-1 | Bacteria Proteobacteria Gammaproteobacteria Enterobacterales Enterobacteriaceae *Escherichia coli* 2-011-08_S3_C2 |
| Eco-71-1 | Bacteria Proteobacteria Gammaproteobacteria Enterobacterales Enterobacteriaceae *Escherichia coli* 2-011-08_S3_C3 |
| Eco-72-1 | Bacteria Proteobacteria Gammaproteobacteria Enterobacterales Enterobacteriaceae *Escherichia coli* 2-011-08_S4_C1 |
| Eco-73-1 | Bacteria Proteobacteria Gammaproteobacteria Enterobacterales Enterobacteriaceae *Escherichia coli* 2-011-08_S4_C3 |
| Eco-74-1 | Bacteria Proteobacteria Gammaproteobacteria Enterobacterales Enterobacteriaceae *Escherichia coli* 2-052-05_S1_C1 |
| Eco-75-1 | Bacteria Proteobacteria Gammaproteobacteria Enterobacterales Enterobacteriaceae *Escherichia coli* 2-052-05_S1_C3 |
| Eco-76-1 | Bacteria Proteobacteria Gammaproteobacteria Enterobacterales Enterobacteriaceae *Escherichia coli* 2-052-05_S3_C1 |
| Eco-77-1 | Bacteria Proteobacteria Gammaproteobacteria Enterobacterales Enterobacteriaceae *Escherichia coli* 2-052-05_S3_C2 |
| Eco-78-1 | Bacteria Proteobacteria Gammaproteobacteria Enterobacterales Enterobacteriaceae *Escherichia coli* 2-052-05_S3_C3 |
| Eco-79-1 | Bacteria Proteobacteria Gammaproteobacteria Enterobacterales Enterobacteriaceae *Escherichia coli* 2-052-05_S4_C1 |
| Eco-80-1 | Bacteria Proteobacteria Gammaproteobacteria Enterobacterales Enterobacteriaceae *Escherichia coli* 2-052-05_S4_C2 |
| Eco-81-1 | Bacteria Proteobacteria Gammaproteobacteria Enterobacterales Enterobacteriaceae *Escherichia coli* 2-052-05_S4_C3 |
| Eco-82-1 | Bacteria Proteobacteria Gammaproteobacteria Enterobacterales Enterobacteriaceae *Escherichia coli* 2-156-04_S1_C3 |
| Eco-83-1 | Bacteria Proteobacteria Gammaproteobacteria Enterobacterales Enterobacteriaceae *Escherichia coli* 2-156-04_S3_C1 |
| Eco-84-1 | Bacteria Proteobacteria Gammaproteobacteria Enterobacterales Enterobacteriaceae *Escherichia coli* 2-156-04_S3_C2 |
| Eco-85-1 | Bacteria Proteobacteria Gammaproteobacteria Enterobacterales Enterobacteriaceae *Escherichia coli* 2-156-04_S3_C3 |
| Eco-86-1 | Bacteria Proteobacteria Gammaproteobacteria Enterobacterales Enterobacteriaceae *Escherichia coli* 2-156-04_S4_C1 |
| Eco-87-1 | Bacteria Proteobacteria Gammaproteobacteria Enterobacterales Enterobacteriaceae *Escherichia coli* 2-156-04_S4_C2 |
| Eco-88-1 | Bacteria Proteobacteria Gammaproteobacteria Enterobacterales Enterobacteriaceae *Escherichia coli* 2-156-04_S4_C3 |
| Eco-89-1 | Bacteria Proteobacteria Gammaproteobacteria Enterobacterales Enterobacteriaceae *Escherichia coli* 2-177-06_S1_C1 |
| Eco-90-1 | Bacteria Proteobacteria Gammaproteobacteria Enterobacterales Enterobacteriaceae *Escherichia coli* 2-177-06_S1_C2 |
| Eco-91-1 | Bacteria Proteobacteria Gammaproteobacteria Enterobacterales Enterobacteriaceae *Escherichia coli* 2-177-06_S1_C3 |
| Eco-92-1 | Bacteria Proteobacteria Gammaproteobacteria Enterobacterales Enterobacteriaceae *Escherichia coli* 2-177-06_S3_C1 |
| Eco-93-1 | Bacteria Proteobacteria Gammaproteobacteria Enterobacterales Enterobacteriaceae *Escherichia coli* 2-177-06_S3_C2 |
| Eco-94-1 | Bacteria Proteobacteria Gammaproteobacteria Enterobacterales Enterobacteriaceae *Escherichia coli* 2-177-06_S3_C3 |
| Eco-95-1 | Bacteria Proteobacteria Gammaproteobacteria Enterobacterales Enterobacteriaceae *Escherichia coli* 2-177-06_S4_C1 |
| Eco-96-1 | Bacteria Proteobacteria Gammaproteobacteria Enterobacterales Enterobacteriaceae *Escherichia coli* 2-177-06_S4_C2 |
| Eco-97-1 | Bacteria Proteobacteria Gammaproteobacteria Enterobacterales Enterobacteriaceae *Escherichia coli* 2-177-06_S4_C3 |
| Eco-98-1 | Bacteria Proteobacteria Gammaproteobacteria Enterobacterales Enterobacteriaceae *Escherichia coli* 2-210-07_S1_C2 |
| Eco-99-1 | Bacteria Proteobacteria Gammaproteobacteria Enterobacterales Enterobacteriaceae *Escherichia coli* 2-210-07_S1_C3 |
| Eco-100-1 | Bacteria Proteobacteria Gammaproteobacteria Enterobacterales Enterobacteriaceae *Escherichia coli* 2-210-07_S3_C1 |
| Eco-101-1 | Bacteria Proteobacteria Gammaproteobacteria Enterobacterales Enterobacteriaceae *Escherichia coli* 2-210-07_S3_C2 |
| Eco-102-1 | Bacteria Proteobacteria Gammaproteobacteria Enterobacterales Enterobacteriaceae *Escherichia coli* 2-210-07_S3_C3 |
| Eco-103-1 | Bacteria Proteobacteria Gammaproteobacteria Enterobacterales Enterobacteriaceae *Escherichia coli* 2-210-07_S4_C1 |
| Eco-104-1 | Bacteria Proteobacteria Gammaproteobacteria Enterobacterales Enterobacteriaceae *Escherichia coli* 2-210-07_S4_C2 |
| Eco-105-1 | Bacteria Proteobacteria Gammaproteobacteria Enterobacterales Enterobacteriaceae *Escherichia coli* 2-210-07_S4_C3 |
| Eco-106-1 | Bacteria Proteobacteria Gammaproteobacteria Enterobacterales Enterobacteriaceae *Escherichia coli* 2-222-05_S1_C1 |
| Eco-107-1 | Bacteria Proteobacteria Gammaproteobacteria Enterobacterales Enterobacteriaceae *Escherichia coli* 2-222-05_S1_C2 |
| Eco-108-1 | Bacteria Proteobacteria Gammaproteobacteria Enterobacterales Enterobacteriaceae *Escherichia coli* 2-222-05_S1_C3 |
| Eco-109-1 | Bacteria Proteobacteria Gammaproteobacteria Enterobacterales Enterobacteriaceae *Escherichia coli* 2-222-05_S3_C1 |
| Eco-110-1 | Bacteria Proteobacteria Gammaproteobacteria Enterobacterales Enterobacteriaceae *Escherichia coli* 2-222-05_S3_C2 |
| Eco-111-1 | Bacteria Proteobacteria Gammaproteobacteria Enterobacterales Enterobacteriaceae *Escherichia coli* 2-222-05_S3_C3 |
| Eco-112-1 | Bacteria Proteobacteria Gammaproteobacteria Enterobacterales Enterobacteriaceae *Escherichia coli* 2-222-05_S4_C1 |
| Eco-113-1 | Bacteria Proteobacteria Gammaproteobacteria Enterobacterales Enterobacteriaceae *Escherichia coli* 2-222-05_S4_C2 |
| Eco-114-1 | Bacteria Proteobacteria Gammaproteobacteria Enterobacterales Enterobacteriaceae *Escherichia coli* 2-222-05_S4_C3 |
| Eco-115-1 | Bacteria Proteobacteria Gammaproteobacteria Enterobacterales Enterobacteriaceae *Escherichia coli* 2-316-03_S1_C1 |
| Eco-116-1 | Bacteria Proteobacteria Gammaproteobacteria Enterobacterales Enterobacteriaceae *Escherichia coli* 2-316-03_S1_C2 |
| Eco-117-1 | Bacteria Proteobacteria Gammaproteobacteria Enterobacterales Enterobacteriaceae *Escherichia coli* 2-316-03_S3_C1 |
| Eco-118-1 | Bacteria Proteobacteria Gammaproteobacteria Enterobacterales Enterobacteriaceae *Escherichia coli* 2-316-03_S3_C2 |
| Eco-119-1 | Bacteria Proteobacteria Gammaproteobacteria Enterobacterales Enterobacteriaceae *Escherichia coli* 2-316-03_S3_C3 |
| Eco-120-1 | Bacteria Proteobacteria Gammaproteobacteria Enterobacterales Enterobacteriaceae *Escherichia coli* 2-316-03_S4_C1 |
| Eco-121-1 | Bacteria Proteobacteria Gammaproteobacteria Enterobacterales Enterobacteriaceae *Escherichia coli* 2-316-03_S4_C2 |
| Eco-122-1 | Bacteria Proteobacteria Gammaproteobacteria Enterobacterales Enterobacteriaceae *Escherichia coli* 2-316-03_S4_C3 |
| Eco-123-1 | Bacteria Proteobacteria Gammaproteobacteria Enterobacterales Enterobacteriaceae *Escherichia coli* 2-427-07_S1_C1 |
| Eco-124-1 | Bacteria Proteobacteria Gammaproteobacteria Enterobacterales Enterobacteriaceae *Escherichia coli* 2-427-07_S1_C2 |
| Eco-125-1 | Bacteria Proteobacteria Gammaproteobacteria Enterobacterales Enterobacteriaceae *Escherichia coli* 2-427-07_S1_C3 |
| Eco-126-1 | Bacteria Proteobacteria Gammaproteobacteria Enterobacterales Enterobacteriaceae *Escherichia coli* 2-427-07_S3_C1 |
| Eco-127-1 | Bacteria Proteobacteria Gammaproteobacteria Enterobacterales Enterobacteriaceae *Escherichia coli* 2-427-07_S3_C3 |
| Eco-128-1 | Bacteria Proteobacteria Gammaproteobacteria Enterobacterales Enterobacteriaceae *Escherichia coli* 2-427-07_S4_C1 |
| Eco-129-1 | Bacteria Proteobacteria Gammaproteobacteria Enterobacterales Enterobacteriaceae *Escherichia coli* 2-427-07_S4_C2 |
| Eco-130-1 | Bacteria Proteobacteria Gammaproteobacteria Enterobacterales Enterobacteriaceae *Escherichia coli* 2-427-07_S4_C3 |
| Eco-131-1 | Bacteria Proteobacteria Gammaproteobacteria Enterobacterales Enterobacteriaceae *Escherichia coli* 2-460-02_S1_C1 |
| Eco-132-1 | Bacteria Proteobacteria Gammaproteobacteria Enterobacterales Enterobacteriaceae *Escherichia coli* 2-460-02_S1_C2 |
| Eco-133-1 | Bacteria Proteobacteria Gammaproteobacteria Enterobacterales Enterobacteriaceae *Escherichia coli* 2-460-02_S1_C3 |
| Eco-134-1 | Bacteria Proteobacteria Gammaproteobacteria Enterobacterales Enterobacteriaceae *Escherichia coli* 2-460-02_S3_C1 |

| | |
|---|---|
| Eco-135-1 | Bacteria Proteobacteria Gammaproteobacteria Enterobacterales Enterobacteriaceae *Escherichia coli* 2-460-02_S3_C2 |
| Eco-136-1 | Bacteria Proteobacteria Gammaproteobacteria Enterobacterales Enterobacteriaceae *Escherichia coli* 2-460-02_S3_C3 |
| Eco-137-1 | Bacteria Proteobacteria Gammaproteobacteria Enterobacterales Enterobacteriaceae *Escherichia coli* 2-460-02_S4_C1 |
| Eco-138-1 | Bacteria Proteobacteria Gammaproteobacteria Enterobacterales Enterobacteriaceae *Escherichia coli* 2-460-02_S4_C2 |
| Eco-139-1 | Bacteria Proteobacteria Gammaproteobacteria Enterobacterales Enterobacteriaceae *Escherichia coli* 2-460-02_S4_C3 |
| Eco-140-1 | Bacteria Proteobacteria Gammaproteobacteria Enterobacterales Enterobacteriaceae *Escherichia coli* 2-474-04_S1_C1 |
| Eco-141-1 | Bacteria Proteobacteria Gammaproteobacteria Enterobacterales Enterobacteriaceae *Escherichia coli* 2-474-04_S1_C2 |
| Eco-142-1 | Bacteria Proteobacteria Gammaproteobacteria Enterobacterales Enterobacteriaceae *Escherichia coli* 2-474-04_S3_C1 |
| Eco-143-1 | Bacteria Proteobacteria Gammaproteobacteria Enterobacterales Enterobacteriaceae *Escherichia coli* 2-474-04_S3_C2 |
| Eco-144-1 | Bacteria Proteobacteria Gammaproteobacteria Enterobacterales Enterobacteriaceae *Escherichia coli* 2-474-04_S3_C3 |
| Eco-145-1 | Bacteria Proteobacteria Gammaproteobacteria Enterobacterales Enterobacteriaceae *Escherichia coli* 2-474-04_S4_C1 |
| Eco-146-1 | Bacteria Proteobacteria Gammaproteobacteria Enterobacterales Enterobacteriaceae *Escherichia coli* 2-474-04_S4_C2 |
| Eco-147-1 | Bacteria Proteobacteria Gammaproteobacteria Enterobacterales Enterobacteriaceae *Escherichia coli* 2-474-04_S4_C3 |
| Eco-148-1 | Bacteria Proteobacteria Gammaproteobacteria Enterobacterales Enterobacteriaceae *Escherichia coli* 27A |
| Eco-149-1 | Bacteria Proteobacteria Gammaproteobacteria Enterobacterales Enterobacteriaceae *Escherichia coli* 3-020-07_S1_C1 |
| Eco-150-1 | Bacteria Proteobacteria Gammaproteobacteria Enterobacterales Enterobacteriaceae *Escherichia coli* 3-020-07_S1_C2 |
| Eco-151-1 | Bacteria Proteobacteria Gammaproteobacteria Enterobacterales Enterobacteriaceae *Escherichia coli* 3-020-07_S1_C3 |
| Eco-152-1 | Bacteria Proteobacteria Gammaproteobacteria Enterobacterales Enterobacteriaceae *Escherichia coli* 3-020-07_S3_C1 |
| Eco-153-1 | Bacteria Proteobacteria Gammaproteobacteria Enterobacterales Enterobacteriaceae *Escherichia coli* 3-020-07_S3_C2 |
| Eco-154-1 | Bacteria Proteobacteria Gammaproteobacteria Enterobacterales Enterobacteriaceae *Escherichia coli* 3-020-07_S4_C1 |
| Eco-155-1 | Bacteria Proteobacteria Gammaproteobacteria Enterobacterales Enterobacteriaceae *Escherichia coli* 3-020-07_S4_C2 |
| Eco-156-1 | Bacteria Proteobacteria Gammaproteobacteria Enterobacterales Enterobacteriaceae *Escherichia coli* 3-020-07_S4_C3 |
| Eco-157-1 | Bacteria Proteobacteria Gammaproteobacteria Enterobacterales Enterobacteriaceae *Escherichia coli* 3-073-06_S1_C1 |
| Eco-158-1 | Bacteria Proteobacteria Gammaproteobacteria Enterobacterales Enterobacteriaceae *Escherichia coli* 3-073-06_S1_C2 |
| Eco-159-1 | Bacteria Proteobacteria Gammaproteobacteria Enterobacterales Enterobacteriaceae *Escherichia coli* 3-073-06_S3_C1 |
| Eco-160-1 | Bacteria Proteobacteria Gammaproteobacteria Enterobacterales Enterobacteriaceae *Escherichia coli* 3-073-06_S3_C2 |
| Eco-161-1 | Bacteria Proteobacteria Gammaproteobacteria Enterobacterales Enterobacteriaceae *Escherichia coli* 3-073-06_S4_C1 |
| Eco-162-1 | Bacteria Proteobacteria Gammaproteobacteria Enterobacterales Enterobacteriaceae *Escherichia coli* 3-073-06_S4_C2 |
| Eco-163-1 | Bacteria Proteobacteria Gammaproteobacteria Enterobacterales Enterobacteriaceae *Escherichia coli* 3-073-06_S4_C3 |
| Eco-164-1 | Bacteria Proteobacteria Gammaproteobacteria Enterobacterales Enterobacteriaceae *Escherichia coli* 3-105-05_S1_C1 |
| Eco-165-1 | Bacteria Proteobacteria Gammaproteobacteria Enterobacterales Enterobacteriaceae *Escherichia coli* 3-105-05_S1_C2 |
| Eco-166-1 | Bacteria Proteobacteria Gammaproteobacteria Enterobacterales Enterobacteriaceae *Escherichia coli* 3-105-05_S1_C3 |
| Eco-167-1 | Bacteria Proteobacteria Gammaproteobacteria Enterobacterales Enterobacteriaceae *Escherichia coli* 3-105-05_S3_C1 |
| Eco-168-1 | Bacteria Proteobacteria Gammaproteobacteria Enterobacterales Enterobacteriaceae *Escherichia coli* 3-105-05_S3_C2 |
| Eco-169-1 | Bacteria Proteobacteria Gammaproteobacteria Enterobacterales Enterobacteriaceae *Escherichia coli* 3-105-05_S3_C3 |
| Eco-170-1 | Bacteria Proteobacteria Gammaproteobacteria Enterobacterales Enterobacteriaceae *Escherichia coli* 3-105-05_S4_C1 |
| Eco-171-1 | Bacteria Proteobacteria Gammaproteobacteria Enterobacterales Enterobacteriaceae *Escherichia coli* 3-105-05_S4_C2 |
| Eco-172-1 | Bacteria Proteobacteria Gammaproteobacteria Enterobacterales Enterobacteriaceae *Escherichia coli* 3-105-05_S4_C3 |
| Eco-173-1 | Bacteria Proteobacteria Gammaproteobacteria Enterobacterales Enterobacteriaceae *Escherichia coli* 3-267-03_S1_C1 |
| Eco-174-1 | Bacteria Proteobacteria Gammaproteobacteria Enterobacterales Enterobacteriaceae *Escherichia coli* 3-267-03_S1_C2 |
| Eco-175-1 | Bacteria Proteobacteria Gammaproteobacteria Enterobacterales Enterobacteriaceae *Escherichia coli* 3-267-03_S1_C3 |
| Eco-176-1 | Bacteria Proteobacteria Gammaproteobacteria Enterobacterales Enterobacteriaceae *Escherichia coli* 3-267-03_S3_C1 |
| Eco-177-1 | Bacteria Proteobacteria Gammaproteobacteria Enterobacterales Enterobacteriaceae *Escherichia coli* 3-267-03_S3_C2 |
| Eco-178-1 | Bacteria Proteobacteria Gammaproteobacteria Enterobacterales Enterobacteriaceae *Escherichia coli* 3-267-03_S4_C1 |
| Eco-179-1 | Bacteria Proteobacteria Gammaproteobacteria Enterobacterales Enterobacteriaceae *Escherichia coli* 3-267-03_S4_C2 |
| Eco-180-1 | Bacteria Proteobacteria Gammaproteobacteria Enterobacterales Enterobacteriaceae *Escherichia coli* 3-373-03_S1_C1 |
| Eco-181-1 | Bacteria Proteobacteria Gammaproteobacteria Enterobacterales Enterobacteriaceae *Escherichia coli* 3-373-03_S1_C2 |
| Eco-182-1 | Bacteria Proteobacteria Gammaproteobacteria Enterobacterales Enterobacteriaceae *Escherichia coli* 3-373-03_S1_C3 |
| Eco-183-1 | Bacteria Proteobacteria Gammaproteobacteria Enterobacterales Enterobacteriaceae *Escherichia coli* 3-373-03_S3_C1 |
| Eco-184-1 | Bacteria Proteobacteria Gammaproteobacteria Enterobacterales Enterobacteriaceae *Escherichia coli* 3-373-03_S3_C2 |
| Eco-185-1 | Bacteria Proteobacteria Gammaproteobacteria Enterobacterales Enterobacteriaceae *Escherichia coli* 3-373-03_S3_C3 |
| Eco-186-1 | Bacteria Proteobacteria Gammaproteobacteria Enterobacterales Enterobacteriaceae *Escherichia coli* 3-373-03_S4_C1 |
| Eco-187-1 | Bacteria Proteobacteria Gammaproteobacteria Enterobacterales Enterobacteriaceae *Escherichia coli* 3-373-03_S4_C2 |
| Eco-188-1 | Bacteria Proteobacteria Gammaproteobacteria Enterobacterales Enterobacteriaceae *Escherichia coli* 3-373-03_S4_C3 |
| Eco-189-1 | Bacteria Proteobacteria Gammaproteobacteria Enterobacterales Enterobacteriaceae *Escherichia coli* 3-475-03_S1_C1 |
| Eco-190-1 | Bacteria Proteobacteria Gammaproteobacteria Enterobacterales Enterobacteriaceae *Escherichia coli* 3-475-03_S1_C2 |
| Eco-191-1 | Bacteria Proteobacteria Gammaproteobacteria Enterobacterales Enterobacteriaceae *Escherichia coli* 3-475-03_S3_C1 |
| Eco-192-1 | Bacteria Proteobacteria Gammaproteobacteria Enterobacterales Enterobacteriaceae *Escherichia coli* 3-475-03_S3_C2 |
| Eco-193-1 | Bacteria Proteobacteria Gammaproteobacteria Enterobacterales Enterobacteriaceae *Escherichia coli* 3-475-03_S4_C1 |
| Eco-194-1 | Bacteria Proteobacteria Gammaproteobacteria Enterobacterales Enterobacteriaceae *Escherichia coli* 3-475-03_S4_C2 |
| Eco-195-1 | Bacteria Proteobacteria Gammaproteobacteria Enterobacterales Enterobacteriaceae *Escherichia coli* 300059 |
| Eco-196-1 | Bacteria Proteobacteria Gammaproteobacteria Enterobacterales Enterobacteriaceae *Escherichia coli* 302053 |
| Eco-197-1 | Bacteria Proteobacteria Gammaproteobacteria Enterobacterales Enterobacteriaceae *Escherichia coli* 320 |
| Eco-198-1 | Bacteria Proteobacteria Gammaproteobacteria Enterobacterales Enterobacteriaceae *Escherichia coli* 328 |
| Eco-199-1 | Bacteria Proteobacteria Gammaproteobacteria Enterobacterales Enterobacteriaceae *Escherichia coli* 38.16 |

| | |
|---|---|
| Eco-200-1 | Bacteria Proteobacteria Gammaproteobacteria Enterobacterales Enterobacteriaceae *Escherichia coli* 38.27 |
| Eco-201-1 | Bacteria Proteobacteria Gammaproteobacteria Enterobacterales Enterobacteriaceae *Escherichia coli* 38.34 |
| Eco-202-1 | Bacteria Proteobacteria Gammaproteobacteria Enterobacterales Enterobacteriaceae *Escherichia coli* 38.52 |
| Eco-203-1 | Bacteria Proteobacteria Gammaproteobacteria Enterobacterales Enterobacteriaceae *Escherichia coli* 4-203-08_S1_C1 |
| Eco-204-1 | Bacteria Proteobacteria Gammaproteobacteria Enterobacterales Enterobacteriaceae *Escherichia coli* 4-203-08_S1_C2 |
| Eco-205-1 | Bacteria Proteobacteria Gammaproteobacteria Enterobacterales Enterobacteriaceae *Escherichia coli* 4-203-08_S1_C3 |
| Eco-206-1 | Bacteria Proteobacteria Gammaproteobacteria Enterobacterales Enterobacteriaceae *Escherichia coli* 4-203-08_S3_C1 |
| Eco-207-1 | Bacteria Proteobacteria Gammaproteobacteria Enterobacterales Enterobacteriaceae *Escherichia coli* 4-203-08_S3_C2 |
| Eco-208-1 | Bacteria Proteobacteria Gammaproteobacteria Enterobacterales Enterobacteriaceae *Escherichia coli* 4-203-08_S3_C3 |
| Eco-209-1 | Bacteria Proteobacteria Gammaproteobacteria Enterobacterales Enterobacteriaceae *Escherichia coli* 4-203-08_S4_C2 |
| Eco-210-1 | Bacteria Proteobacteria Gammaproteobacteria Enterobacterales Enterobacteriaceae *Escherichia coli* 4-203-08_S4_C3 |
| Eco-211-1 | Bacteria Proteobacteria Gammaproteobacteria Enterobacterales Enterobacteriaceae *Escherichia coli* 401140 |
| Eco-212-1 | Bacteria Proteobacteria Gammaproteobacteria Enterobacterales Enterobacteriaceae *Escherichia coli* 402290 |
| Eco-213-1 | Bacteria Proteobacteria Gammaproteobacteria Enterobacterales Enterobacteriaceae *Escherichia coli* 435 |
| Eco-214-1 | Bacteria Proteobacteria Gammaproteobacteria Enterobacterales Enterobacteriaceae *Escherichia coli* 4541-1 |
| Eco-215-1 to Eco-215-2 | Bacteria Proteobacteria Gammaproteobacteria Enterobacterales Enterobacteriaceae *Escherichia coli* 4552-1 |
| Eco-216-1 | Bacteria Proteobacteria Gammaproteobacteria Enterobacterales Enterobacteriaceae *Escherichia coli* 5-172-05_S1_C3 |
| Eco-217-1 | Bacteria Proteobacteria Gammaproteobacteria Enterobacterales Enterobacteriaceae *Escherichia coli* 5-172-05_S3_C1 |
| Eco-218-1 | Bacteria Proteobacteria Gammaproteobacteria Enterobacterales Enterobacteriaceae *Escherichia coli* 5-172-05_S3_C3 |
| Eco-219-1 | Bacteria Proteobacteria Gammaproteobacteria Enterobacterales Enterobacteriaceae *Escherichia coli* 5-172-05_S4_C1 |
| Eco-220-1 | Bacteria Proteobacteria Gammaproteobacteria Enterobacterales Enterobacteriaceae *Escherichia coli* 5-172-05_S4_C2 |
| Eco-221-1 | Bacteria Proteobacteria Gammaproteobacteria Enterobacterales Enterobacteriaceae *Escherichia coli* 5-172-05_S4_C3 |
| Eco-222-1 | Bacteria Proteobacteria Gammaproteobacteria Enterobacterales Enterobacteriaceae *Escherichia coli* 5-366-08_S1_C1 |
| Eco-223-1 | Bacteria Proteobacteria Gammaproteobacteria Enterobacterales Enterobacteriaceae *Escherichia coli* 5-366-08_S1_C3 |
| Eco-224-1 | Bacteria Proteobacteria Gammaproteobacteria Enterobacterales Enterobacteriaceae *Escherichia coli* 5-366-08_S3_C1 |
| Eco-225-1 | Bacteria Proteobacteria Gammaproteobacteria Enterobacterales Enterobacteriaceae *Escherichia coli* 5-366-08_S3_C2 |
| Eco-226-1 | Bacteria Proteobacteria Gammaproteobacteria Enterobacterales Enterobacteriaceae *Escherichia coli* 5-366-08_S3_C3 |
| Eco-227-1 | Bacteria Proteobacteria Gammaproteobacteria Enterobacterales Enterobacteriaceae *Escherichia coli* 5-366-08_S4_C1 |
| Eco-228-1 | Bacteria Proteobacteria Gammaproteobacteria Enterobacterales Enterobacteriaceae *Escherichia coli* 5-366-08_S4_C2 |
| Eco-229-1 | Bacteria Proteobacteria Gammaproteobacteria Enterobacterales Enterobacteriaceae *Escherichia coli* 53A |
| Eco-230-1 to Eco-230-2 | Bacteria Proteobacteria Gammaproteobacteria Enterobacterales Enterobacteriaceae *Escherichia coli* 53C |
| Eco-231-1 | Bacteria Proteobacteria Gammaproteobacteria Enterobacterales Enterobacteriaceae *Escherichia coli* 597 |
| Eco-232-1 | Bacteria Proteobacteria Gammaproteobacteria Enterobacterales Enterobacteriaceae *Escherichia coli* 6-175-07_S1_C1 |
| Eco-233-1 | Bacteria Proteobacteria Gammaproteobacteria Enterobacterales Enterobacteriaceae *Escherichia coli* 6-175-07_S1_C2 |
| Eco-234-1 | Bacteria Proteobacteria Gammaproteobacteria Enterobacterales Enterobacteriaceae *Escherichia coli* 6-175-07_S1_C3 |
| Eco-235-1 | Bacteria Proteobacteria Gammaproteobacteria Enterobacterales Enterobacteriaceae *Escherichia coli* 6-175-07_S3_C1 |
| Eco-236-1 | Bacteria Proteobacteria Gammaproteobacteria Enterobacterales Enterobacteriaceae *Escherichia coli* 6-175-07_S3_C2 |
| Eco-237-1 | Bacteria Proteobacteria Gammaproteobacteria Enterobacterales Enterobacteriaceae *Escherichia coli* 6-175-07_S3_C3 |
| Eco-238-1 | Bacteria Proteobacteria Gammaproteobacteria Enterobacterales Enterobacteriaceae *Escherichia coli* 6-175-07_S4_C1 |
| Eco-239-1 | Bacteria Proteobacteria Gammaproteobacteria Enterobacterales Enterobacteriaceae *Escherichia coli* 6-175-07_S4_C2 |
| Eco-240-1 | Bacteria Proteobacteria Gammaproteobacteria Enterobacterales Enterobacteriaceae *Escherichia coli* 6-175-07_S4_C3 |
| Eco-241-1 | Bacteria Proteobacteria Gammaproteobacteria Enterobacterales Enterobacteriaceae *Escherichia coli* 6-319-05_S1_C1 |
| Eco-242-1 | Bacteria Proteobacteria Gammaproteobacteria Enterobacterales Enterobacteriaceae *Escherichia coli* 6-319-05_S1_C2 |
| Eco-243-1 | Bacteria Proteobacteria Gammaproteobacteria Enterobacterales Enterobacteriaceae *Escherichia coli* 6-319-05_S1_C3 |
| Eco-244-1 | Bacteria Proteobacteria Gammaproteobacteria Enterobacterales Enterobacteriaceae *Escherichia coli* 6-319-05_S3_C1 |
| Eco-245-1 | Bacteria Proteobacteria Gammaproteobacteria Enterobacterales Enterobacteriaceae *Escherichia coli* 6-319-05_S3_C2 |
| Eco-246-1 | Bacteria Proteobacteria Gammaproteobacteria Enterobacterales Enterobacteriaceae *Escherichia coli* 6-319-05_S3_C3 |
| Eco-247-1 | Bacteria Proteobacteria Gammaproteobacteria Enterobacterales Enterobacteriaceae *Escherichia coli* 6-319-05_S4_C2 |
| Eco-248-1 | Bacteria Proteobacteria Gammaproteobacteria Enterobacterales Enterobacteriaceae *Escherichia coli* 6-319-05_S4_C3 |
| Eco-249-1 | Bacteria Proteobacteria Gammaproteobacteria Enterobacterales Enterobacteriaceae *Escherichia coli* 6-537-08_S1_C1 |
| Eco-250-1 | Bacteria Proteobacteria Gammaproteobacteria Enterobacterales Enterobacteriaceae *Escherichia coli* 6-537-08_S1_C2 |
| Eco-251-1 | Bacteria Proteobacteria Gammaproteobacteria Enterobacterales Enterobacteriaceae *Escherichia coli* 6-537-08_S1_C3 |
| Eco-252-1 | Bacteria Proteobacteria Gammaproteobacteria Enterobacterales Enterobacteriaceae *Escherichia coli* 6-537-08_S3_C1 |
| Eco-253-1 | Bacteria Proteobacteria Gammaproteobacteria Enterobacterales Enterobacteriaceae *Escherichia coli* 6-537-08_S3_C2 |
| Eco-254-1 | Bacteria Proteobacteria Gammaproteobacteria Enterobacterales Enterobacteriaceae *Escherichia coli* 6-537-08_S3_C3 |
| Eco-255-1 | Bacteria Proteobacteria Gammaproteobacteria Enterobacterales Enterobacteriaceae *Escherichia coli* 6-537-08_S4_C1 |
| Eco-256-1 | Bacteria Proteobacteria Gammaproteobacteria Enterobacterales Enterobacteriaceae *Escherichia coli* 6-537-08_S4_C2 |
| Eco-257-1 | Bacteria Proteobacteria Gammaproteobacteria Enterobacterales Enterobacteriaceae *Escherichia coli* 606 |
| Eco-258-1 | Bacteria Proteobacteria Gammaproteobacteria Enterobacterales Enterobacteriaceae *Escherichia coli* 668 |
| Eco-259-1 | Bacteria Proteobacteria Gammaproteobacteria Enterobacterales Enterobacteriaceae *Escherichia coli* 681 |
| Eco-260-1 | Bacteria Proteobacteria Gammaproteobacteria Enterobacterales Enterobacteriaceae *Escherichia coli* 7-233-03_S1_C2 |
| Eco-261-1 | Bacteria Proteobacteria Gammaproteobacteria Enterobacterales Enterobacteriaceae *Escherichia coli* 7-233-03_S1_C3 |
| Eco-262-1 | Bacteria Proteobacteria Gammaproteobacteria Enterobacterales Enterobacteriaceae *Escherichia coli* 7-233-03_S3_C1 |
| Eco-263-1 | Bacteria Proteobacteria Gammaproteobacteria Enterobacterales Enterobacteriaceae *Escherichia coli* 7-233-03_S3_C2 |
| Eco-264-1 | Bacteria Proteobacteria Gammaproteobacteria Enterobacterales Enterobacteriaceae *Escherichia coli* 7-233-03_S3_C3 |

| | |
|---|---|
| Eco-265-1 | Bacteria Proteobacteria Gammaproteobacteria Enterobacterales Enterobacteriaceae *Escherichia coli* 7-233-03_S4_C1 |
| Eco-266-1 | Bacteria Proteobacteria Gammaproteobacteria Enterobacterales Enterobacteriaceae *Escherichia coli* 7-233-03_S4_C2 |
| Eco-267-1 | Bacteria Proteobacteria Gammaproteobacteria Enterobacterales Enterobacteriaceae *Escherichia coli* 7-233-03_S4_C3 |
| Eco-268-1 | Bacteria Proteobacteria Gammaproteobacteria Enterobacterales Enterobacteriaceae *Escherichia coli* 7996-1 |
| Eco-269-1 | Bacteria Proteobacteria Gammaproteobacteria Enterobacterales Enterobacteriaceae *Escherichia coli* 8-415-05_S1_C1 |
| Eco-270-1 | Bacteria Proteobacteria Gammaproteobacteria Enterobacterales Enterobacteriaceae *Escherichia coli* 8-415-05_S1_C2 |
| Eco-271-1 | Bacteria Proteobacteria Gammaproteobacteria Enterobacterales Enterobacteriaceae *Escherichia coli* 8-415-05_S3_C1 |
| Eco-272-1 | Bacteria Proteobacteria Gammaproteobacteria Enterobacterales Enterobacteriaceae *Escherichia coli* 8-415-05_S3_C2 |
| Eco-273-1 | Bacteria Proteobacteria Gammaproteobacteria Enterobacterales Enterobacteriaceae *Escherichia coli* 8-415-05_S3_C3 |
| Eco-274-1 | Bacteria Proteobacteria Gammaproteobacteria Enterobacterales Enterobacteriaceae *Escherichia coli* 8-415-05_S4_C1 |
| Eco-275-1 | Bacteria Proteobacteria Gammaproteobacteria Enterobacterales Enterobacteriaceae *Escherichia coli* 8-415-05_S4_C2 |
| Eco-276-1 | Bacteria Proteobacteria Gammaproteobacteria Enterobacterales Enterobacteriaceae *Escherichia coli* 8-415-05_S4_C3 |
| Eco-277-1 | Bacteria Proteobacteria Gammaproteobacteria Enterobacterales Enterobacteriaceae *Escherichia coli* 85B |
| Eco-278-1 | Bacteria Proteobacteria Gammaproteobacteria Enterobacterales Enterobacteriaceae *Escherichia coli* 8624 |
| Eco-279-1 | Bacteria Proteobacteria Gammaproteobacteria Enterobacterales Enterobacteriaceae *Escherichia coli* 87A |
| Eco-280-1 | Bacteria Proteobacteria Gammaproteobacteria Enterobacterales Enterobacteriaceae *Escherichia coli* 897 |
| Eco-281-1 | Bacteria Proteobacteria Gammaproteobacteria Enterobacterales Enterobacteriaceae *Escherichia coli* 9.1649 |
| Eco-282-1 | Bacteria Proteobacteria Gammaproteobacteria Enterobacterales Enterobacteriaceae *Escherichia coli* 97.0264 |
| Eco-283-1 | Bacteria Proteobacteria Gammaproteobacteria Enterobacterales Enterobacteriaceae *Escherichia coli* 9B |
| Eco-284-1 | Bacteria Proteobacteria Gammaproteobacteria Enterobacterales Enterobacteriaceae *Escherichia coli* AB42410445-isolate1 |
| Eco-285-1 | Bacteria Proteobacteria Gammaproteobacteria Enterobacterales Enterobacteriaceae *Escherichia coli* AB42554418-isolate1 |
| Eco-286-1 | Bacteria Proteobacteria Gammaproteobacteria Enterobacterales Enterobacteriaceae *Escherichia coli* AB42602061-isolate1 |
| Eco-287-1 | Bacteria Proteobacteria Gammaproteobacteria Enterobacterales Enterobacteriaceae *Escherichia coli* AB43739056-isolate1 |
| Eco-288-1 | Bacteria Proteobacteria Gammaproteobacteria Enterobacterales Enterobacteriaceae *Escherichia coli* ACN001 |
| Eco-289-1 to Eco-289-2 | Bacteria Proteobacteria Gammaproteobacteria Enterobacterales Enterobacteriaceae *Escherichia coli* AD30 |
| Eco-290-1 | Bacteria Proteobacteria Gammaproteobacteria Enterobacterales Enterobacteriaceae *Escherichia coli* APEC IMT5155 |
| Eco-291-1 to Eco-291-2 | Bacteria Proteobacteria Gammaproteobacteria Enterobacterales Enterobacteriaceae *Escherichia coli* ATCC 25922 |
| Eco-292-1 | Bacteria Proteobacteria Gammaproteobacteria Enterobacterales Enterobacteriaceae *Escherichia coli* ATCC 35150 |
| Eco-293-1 | Bacteria Proteobacteria Gammaproteobacteria Enterobacterales Enterobacteriaceae *Escherichia coli* ATCC BAA-2192 |
| Eco-294-1 | Bacteria Proteobacteria Gammaproteobacteria Enterobacterales Enterobacteriaceae *Escherichia coli* ATCC BAA-2193 |
| Eco-295-1 | Bacteria Proteobacteria Gammaproteobacteria Enterobacterales Enterobacteriaceae *Escherichia coli* BIDMC 15 |
| Eco-296-1 | Bacteria Proteobacteria Gammaproteobacteria Enterobacterales Enterobacteriaceae *Escherichia coli* BIDMC 17A |
| Eco-297-1 | Bacteria Proteobacteria Gammaproteobacteria Enterobacterales Enterobacteriaceae *Escherichia coli* BIDMC 17B |
| Eco-298-1 | Bacteria Proteobacteria Gammaproteobacteria Enterobacterales Enterobacteriaceae *Escherichia coli* BIDMC 19A |
| Eco-299-1 | Bacteria Proteobacteria Gammaproteobacteria Enterobacterales Enterobacteriaceae *Escherichia coli* BIDMC 19B |
| Eco-300-1 | Bacteria Proteobacteria Gammaproteobacteria Enterobacterales Enterobacteriaceae *Escherichia coli* BIDMC 20A |
| Eco-301-1 | Bacteria Proteobacteria Gammaproteobacteria Enterobacterales Enterobacteriaceae *Escherichia coli* BIDMC 20B |
| Eco-302-1 | Bacteria Proteobacteria Gammaproteobacteria Enterobacterales Enterobacteriaceae *Escherichia coli* BIDMC 2B |
| Eco-303-1 | Bacteria Proteobacteria Gammaproteobacteria Enterobacterales Enterobacteriaceae *Escherichia coli* BIDMC 3 |
| Eco-304-1 | Bacteria Proteobacteria Gammaproteobacteria Enterobacterales Enterobacteriaceae *Escherichia coli* BIDMC 43a |
| Eco-305-1 | Bacteria Proteobacteria Gammaproteobacteria Enterobacterales Enterobacteriaceae *Escherichia coli* BIDMC 43b |
| Eco-306-1 | Bacteria Proteobacteria Gammaproteobacteria Enterobacterales Enterobacteriaceae *Escherichia coli* BIDMC 49a |
| Eco-307-1 | Bacteria Proteobacteria Gammaproteobacteria Enterobacterales Enterobacteriaceae *Escherichia coli* BIDMC 49b |
| Eco-308-1 | Bacteria Proteobacteria Gammaproteobacteria Enterobacterales Enterobacteriaceae *Escherichia coli* BIDMC 58 |
| Eco-309-1 | Bacteria Proteobacteria Gammaproteobacteria Enterobacterales Enterobacteriaceae *Escherichia coli* BIDMC 59 |
| Eco-310-1 | Bacteria Proteobacteria Gammaproteobacteria Enterobacterales Enterobacteriaceae *Escherichia coli* BIDMC 6 |
| Eco-311-1 | Bacteria Proteobacteria Gammaproteobacteria Enterobacterales Enterobacteriaceae *Escherichia coli* BIDMC 62 |
| Eco-312-1 | Bacteria Proteobacteria Gammaproteobacteria Enterobacterales Enterobacteriaceae *Escherichia coli* BIDMC 63 |
| Eco-313-1 | Bacteria Proteobacteria Gammaproteobacteria Enterobacterales Enterobacteriaceae *Escherichia coli* BIDMC 64 |
| Eco-314-1 | Bacteria Proteobacteria Gammaproteobacteria Enterobacterales Enterobacteriaceae *Escherichia coli* BIDMC 65 |
| Eco-315-1 | Bacteria Proteobacteria Gammaproteobacteria Enterobacterales Enterobacteriaceae *Escherichia coli* BIDMC 70 |
| Eco-316-1 | Bacteria Proteobacteria Gammaproteobacteria Enterobacterales Enterobacteriaceae *Escherichia coli* BIDMC 71 |
| Eco-317-1 | Bacteria Proteobacteria Gammaproteobacteria Enterobacterales Enterobacteriaceae *Escherichia coli* BIDMC 72 |
| Eco-318-1 | Bacteria Proteobacteria Gammaproteobacteria Enterobacterales Enterobacteriaceae *Escherichia coli* BIDMC 73 |
| Eco-319-1 | Bacteria Proteobacteria Gammaproteobacteria Enterobacterales Enterobacteriaceae *Escherichia coli* BIDMC 74 |
| Eco-320-1 | Bacteria Proteobacteria Gammaproteobacteria Enterobacterales Enterobacteriaceae *Escherichia coli* BIDMC 75 |
| Eco-321-1 | Bacteria Proteobacteria Gammaproteobacteria Enterobacterales Enterobacteriaceae *Escherichia coli* BIDMC 76 |
| Eco-322-1 | Bacteria Proteobacteria Gammaproteobacteria Enterobacterales Enterobacteriaceae *Escherichia coli* BIDMC 77 |
| Eco-323-1 | Bacteria Proteobacteria Gammaproteobacteria Enterobacterales Enterobacteriaceae *Escherichia coli* BIDMC 78 |
| Eco-324-1 | Bacteria Proteobacteria Gammaproteobacteria Enterobacterales Enterobacteriaceae *Escherichia coli* BIDMC 79 |
| Eco-325-1 | Bacteria Proteobacteria Gammaproteobacteria Enterobacterales Enterobacteriaceae *Escherichia coli* BIDMC 82 |
| Eco-326-1 | Bacteria Proteobacteria Gammaproteobacteria Enterobacterales Enterobacteriaceae *Escherichia coli* BIDMC 83 |
| Eco-327-1 | Bacteria Proteobacteria Gammaproteobacteria Enterobacterales Enterobacteriaceae *Escherichia coli* BIDMC 9 |
| Eco-328-1 | Bacteria Proteobacteria Gammaproteobacteria Enterobacterales Enterobacteriaceae *Escherichia coli* BW25113 |
| Eco-329-1 | Bacteria Proteobacteria Gammaproteobacteria Enterobacterales Enterobacteriaceae *Escherichia coli* BWH 34 |

| | |
|---|---|
| Eco-330-1 | Bacteria Proteobacteria Gammaproteobacteria Enterobacterales Enterobacteriaceae *Escherichia coli* BWH 40 |
| Eco-331-1 | Bacteria Proteobacteria Gammaproteobacteria Enterobacterales Enterobacteriaceae *Escherichia coli* C1214_90 |
| Eco-332-1 | Bacteria Proteobacteria Gammaproteobacteria Enterobacterales Enterobacteriaceae *Escherichia coli* C1244_91 |
| Eco-333-1 | Bacteria Proteobacteria Gammaproteobacteria Enterobacterales Enterobacteriaceae *Escherichia coli* C12_92 |
| Eco-334-1 | Bacteria Proteobacteria Gammaproteobacteria Enterobacterales Enterobacteriaceae *Escherichia coli* C154_11 |
| Eco-335-1 | Bacteria Proteobacteria Gammaproteobacteria Enterobacterales Enterobacteriaceae *Escherichia coli* C155_11 |
| Eco-336-1 | Bacteria Proteobacteria Gammaproteobacteria Enterobacterales Enterobacteriaceae *Escherichia coli* C157_11 |
| Eco-337-1 | Bacteria Proteobacteria Gammaproteobacteria Enterobacterales Enterobacteriaceae *Escherichia coli* C161_11 |
| Eco-338-1 | Bacteria Proteobacteria Gammaproteobacteria Enterobacterales Enterobacteriaceae *Escherichia coli* C166_11 |
| Eco-339-1 | Bacteria Proteobacteria Gammaproteobacteria Enterobacterales Enterobacteriaceae *Escherichia coli* C170_11 |
| Eco-340-1 | Bacteria Proteobacteria Gammaproteobacteria Enterobacterales Enterobacteriaceae *Escherichia coli* C2139_99 |
| Eco-341-1 | Bacteria Proteobacteria Gammaproteobacteria Enterobacterales Enterobacteriaceae *Escherichia coli* C213_10 |
| Eco-342-1 | Bacteria Proteobacteria Gammaproteobacteria Enterobacterales Enterobacteriaceae *Escherichia coli* C238_91 |
| Eco-343-1 | Bacteria Proteobacteria Gammaproteobacteria Enterobacterales Enterobacteriaceae *Escherichia coli* C260_92 |
| Eco-344-1 | Bacteria Proteobacteria Gammaproteobacteria Enterobacterales Enterobacteriaceae *Escherichia coli* C262_10 |
| Eco-345-1 | Bacteria Proteobacteria Gammaproteobacteria Enterobacterales Enterobacteriaceae *Escherichia coli* C283_09 |
| Eco-346-1 | Bacteria Proteobacteria Gammaproteobacteria Enterobacterales Enterobacteriaceae *Escherichia coli* C295_10 |
| Eco-347-1 | Bacteria Proteobacteria Gammaproteobacteria Enterobacterales Enterobacteriaceae *Escherichia coli* C341_10 |
| Eco-348-1 | Bacteria Proteobacteria Gammaproteobacteria Enterobacterales Enterobacteriaceae *Escherichia coli* C343_08 |
| Eco-349-1 | Bacteria Proteobacteria Gammaproteobacteria Enterobacterales Enterobacteriaceae *Escherichia coli* C347_93 |
| Eco-350-1 | Bacteria Proteobacteria Gammaproteobacteria Enterobacterales Enterobacteriaceae *Escherichia coli* C353_09 |
| Eco-351-1 | Bacteria Proteobacteria Gammaproteobacteria Enterobacterales Enterobacteriaceae *Escherichia coli* C354_03B |
| Eco-352-1 | Bacteria Proteobacteria Gammaproteobacteria Enterobacterales Enterobacteriaceae *Escherichia coli* C40_11 |
| Eco-353-1 | Bacteria Proteobacteria Gammaproteobacteria Enterobacterales Enterobacteriaceae *Escherichia coli* C418_89 |
| Eco-354-1 | Bacteria Proteobacteria Gammaproteobacteria Enterobacterales Enterobacteriaceae *Escherichia coli* C458_10 |
| Eco-355-1 | Bacteria Proteobacteria Gammaproteobacteria Enterobacterales Enterobacteriaceae *Escherichia coli* C488_07 |
| Eco-356-1 | Bacteria Proteobacteria Gammaproteobacteria Enterobacterales Enterobacteriaceae *Escherichia coli* C496_10 |
| Eco-357-1 | Bacteria Proteobacteria Gammaproteobacteria Enterobacterales Enterobacteriaceae *Escherichia coli* C497_10 |
| Eco-358-1 | Bacteria Proteobacteria Gammaproteobacteria Enterobacterales Enterobacteriaceae *Escherichia coli* C527_94 |
| Eco-359-1 | Bacteria Proteobacteria Gammaproteobacteria Enterobacterales Enterobacteriaceae *Escherichia coli* C581_05 |
| Eco-360-1 | Bacteria Proteobacteria Gammaproteobacteria Enterobacterales Enterobacteriaceae *Escherichia coli* C586_05 |
| Eco-361-1 | Bacteria Proteobacteria Gammaproteobacteria Enterobacterales Enterobacteriaceae *Escherichia coli* C58_11 |
| Eco-362-1 | Bacteria Proteobacteria Gammaproteobacteria Enterobacterales Enterobacteriaceae *Escherichia coli* C639_08 |
| Eco-363-1 | Bacteria Proteobacteria Gammaproteobacteria Enterobacterales Enterobacteriaceae *Escherichia coli* C652_10 |
| Eco-364-1 | Bacteria Proteobacteria Gammaproteobacteria Enterobacterales Enterobacteriaceae *Escherichia coli* C654_09 |
| Eco-365-1 | Bacteria Proteobacteria Gammaproteobacteria Enterobacterales Enterobacteriaceae *Escherichia coli* C691-71 (14b) |
| Eco-366-1 | Bacteria Proteobacteria Gammaproteobacteria Enterobacterales Enterobacteriaceae *Escherichia coli* C717_10 |
| Eco-367-1 | Bacteria Proteobacteria Gammaproteobacteria Enterobacterales Enterobacteriaceae *Escherichia coli* C725_88 |
| Eco-368-1 | Bacteria Proteobacteria Gammaproteobacteria Enterobacterales Enterobacteriaceae *Escherichia coli* C732_98 |
| Eco-369-1 | Bacteria Proteobacteria Gammaproteobacteria Enterobacterales Enterobacteriaceae *Escherichia coli* C743_03 |
| Eco-370-1 | Bacteria Proteobacteria Gammaproteobacteria Enterobacterales Enterobacteriaceae *Escherichia coli* C751_03 |
| Eco-371-1 | Bacteria Proteobacteria Gammaproteobacteria Enterobacterales Enterobacteriaceae *Escherichia coli* C78_09C |
| Eco-372-1 | Bacteria Proteobacteria Gammaproteobacteria Enterobacterales Enterobacteriaceae *Escherichia coli* C792_92 |
| Eco-373-1 | Bacteria Proteobacteria Gammaproteobacteria Enterobacterales Enterobacteriaceae *Escherichia coli* C796_10 |
| Eco-374-1 | Bacteria Proteobacteria Gammaproteobacteria Enterobacterales Enterobacteriaceae *Escherichia coli* C799_92 |
| Eco-375-1 | Bacteria Proteobacteria Gammaproteobacteria Enterobacterales Enterobacteriaceae *Escherichia coli* C79_08 |
| Eco-376-1 | Bacteria Proteobacteria Gammaproteobacteria Enterobacterales Enterobacteriaceae *Escherichia coli* C807_09 |
| Eco-377-1 | Bacteria Proteobacteria Gammaproteobacteria Enterobacterales Enterobacteriaceae *Escherichia coli* C80_08 |
| Eco-378-1 | Bacteria Proteobacteria Gammaproteobacteria Enterobacterales Enterobacteriaceae *Escherichia coli* C824_10 |
| Eco-379-1 | Bacteria Proteobacteria Gammaproteobacteria Enterobacterales Enterobacteriaceae *Escherichia coli* C82_11 |
| Eco-380-1 | Bacteria Proteobacteria Gammaproteobacteria Enterobacterales Enterobacteriaceae *Escherichia coli* C842_97 |
| Eco-381-1 | Bacteria Proteobacteria Gammaproteobacteria Enterobacterales Enterobacteriaceae *Escherichia coli* C844_97 |
| Eco-382-1 | Bacteria Proteobacteria Gammaproteobacteria Enterobacterales Enterobacteriaceae *Escherichia coli* C87_11 |
| Eco-383-1 | Bacteria Proteobacteria Gammaproteobacteria Enterobacterales Enterobacteriaceae *Escherichia coli* C887_10 |
| Eco-384-1 | Bacteria Proteobacteria Gammaproteobacteria Enterobacterales Enterobacteriaceae *Escherichia coli* C900_01 |
| Eco-385-1 | Bacteria Proteobacteria Gammaproteobacteria Enterobacterales Enterobacteriaceae *Escherichia coli* C93_11 |
| Eco-386-1 | Bacteria Proteobacteria Gammaproteobacteria Enterobacterales Enterobacteriaceae *Escherichia coli* C9_92 |
| Eco-387-1 | Bacteria Proteobacteria Gammaproteobacteria Enterobacterales Enterobacteriaceae *Escherichia coli* CHS 68 |
| Eco-388-1 | Bacteria Proteobacteria Gammaproteobacteria Enterobacterales Enterobacteriaceae *Escherichia coli* CHS 69 |
| Eco-389-1 | Bacteria Proteobacteria Gammaproteobacteria Enterobacterales Enterobacteriaceae *Escherichia coli* CHS 77 |
| Eco-390-1 | Bacteria Proteobacteria Gammaproteobacteria Enterobacterales Enterobacteriaceae *Escherichia coli* D6-113.11 |
| Eco-391-1 | Bacteria Proteobacteria Gammaproteobacteria Enterobacterales Enterobacteriaceae *Escherichia coli* D6-117.29 |
| Eco-392-1 | Bacteria Proteobacteria Gammaproteobacteria Enterobacterales Enterobacteriaceae *Escherichia coli* DH5[alpha] |
| Eco-393-1 | Bacteria Proteobacteria Gammaproteobacteria Enterobacterales Enterobacteriaceae *Escherichia coli* DSM 30083 = JCM 1649 = ATCC 11775 |
| Eco-394-1 | Bacteria Proteobacteria Gammaproteobacteria Enterobacterales Enterobacteriaceae *Escherichia coli* E1140 |

| | |
|---|---|
| Eco-395-1 | Bacteria Proteobacteria Gammaproteobacteria Enterobacterales Enterobacteriaceae *Escherichia coli* E1728 |
| Eco-396-1 | Bacteria Proteobacteria Gammaproteobacteria Enterobacterales Enterobacteriaceae *Escherichia coli* E1777 |
| Eco-397-1 | Bacteria Proteobacteria Gammaproteobacteria Enterobacterales Enterobacteriaceae *Escherichia coli* EC096/10 |
| Eco-398-1 | Bacteria Proteobacteria Gammaproteobacteria Enterobacterales Enterobacteriaceae *Escherichia coli* ECC-1470 |
| Eco-399-1 | Bacteria Proteobacteria Gammaproteobacteria Enterobacterales Enterobacteriaceae *Escherichia coli* ER2796 |
| Eco-400-1 | Bacteria Proteobacteria Gammaproteobacteria Enterobacterales Enterobacteriaceae *Escherichia coli* FAH1 |
| Eco-401-1 | Bacteria Proteobacteria Gammaproteobacteria Enterobacterales Enterobacteriaceae *Escherichia coli* FAH2 |
| Eco-402-1 | Bacteria Proteobacteria Gammaproteobacteria Enterobacterales Enterobacteriaceae *Escherichia coli* FAP1 |
| Eco-403-1 | Bacteria Proteobacteria Gammaproteobacteria Enterobacterales Enterobacteriaceae *Escherichia coli* FAP2 |
| Eco-404-1 | Bacteria Proteobacteria Gammaproteobacteria Enterobacterales Enterobacteriaceae *Escherichia coli* FBH1 |
| Eco-405-1 | Bacteria Proteobacteria Gammaproteobacteria Enterobacterales Enterobacteriaceae *Escherichia coli* FBP1 |
| Eco-406-1 | Bacteria Proteobacteria Gammaproteobacteria Enterobacterales Enterobacteriaceae *Escherichia coli* FCH1 |
| Eco-407-1 | Bacteria Proteobacteria Gammaproteobacteria Enterobacterales Enterobacteriaceae *Escherichia coli* FCP1 |
| Eco-408-1 | Bacteria Proteobacteria Gammaproteobacteria Enterobacterales Enterobacteriaceae *Escherichia coli* G3/10 |
| Eco-409-1 | Bacteria Proteobacteria Gammaproteobacteria Enterobacterales Enterobacteriaceae *Escherichia coli* GSK2022 |
| Eco-410-1 | Bacteria Proteobacteria Gammaproteobacteria Enterobacterales Enterobacteriaceae *Escherichia coli* GSK2023 |
| Eco-411-1 | Bacteria Proteobacteria Gammaproteobacteria Enterobacterales Enterobacteriaceae *Escherichia coli* GSK2024 |
| Eco-412-1 | Bacteria Proteobacteria Gammaproteobacteria Enterobacterales Enterobacteriaceae *Escherichia coli* GSK202B |
| Eco-413-1 | Bacteria Proteobacteria Gammaproteobacteria Enterobacterales Enterobacteriaceae *Escherichia coli* GSK202U |
| Eco-414-1 | Bacteria Proteobacteria Gammaproteobacteria Enterobacterales Enterobacteriaceae *Escherichia coli* IMT8073 |
| Eco-415-1 | Bacteria Proteobacteria Gammaproteobacteria Enterobacterales Enterobacteriaceae *Escherichia coli* IS1 |
| Eco-416-1 | Bacteria Proteobacteria Gammaproteobacteria Enterobacterales Enterobacteriaceae *Escherichia coli* IS25 |
| Eco-417-1 | Bacteria Proteobacteria Gammaproteobacteria Enterobacterales Enterobacteriaceae *Escherichia coli* IS9 |
| Eco-418-1 | Bacteria Proteobacteria Gammaproteobacteria Enterobacterales Enterobacteriaceae *Escherichia coli* JT5 |
| Eco-419-1 | Bacteria Proteobacteria Gammaproteobacteria Enterobacterales Enterobacteriaceae *Escherichia coli* K-12 |
| Eco-420-1 | Bacteria Proteobacteria Gammaproteobacteria Enterobacterales Enterobacteriaceae *Escherichia coli* K02 |
| Eco-421-1 to Eco-421-4 | Bacteria Proteobacteria Gammaproteobacteria Enterobacterales Enterobacteriaceae *Escherichia coli* K1 |
| Eco-422-1 | Bacteria Proteobacteria Gammaproteobacteria Enterobacterales Enterobacteriaceae *Escherichia coli* K2 |
| Eco-423-1 | Bacteria Proteobacteria Gammaproteobacteria Enterobacterales Enterobacteriaceae *Escherichia coli* KLY |
| Eco-424-1 | Bacteria Proteobacteria Gammaproteobacteria Enterobacterales Enterobacteriaceae *Escherichia coli* LAU-EC2 |
| Eco-425-1 | Bacteria Proteobacteria Gammaproteobacteria Enterobacterales Enterobacteriaceae *Escherichia coli* LAU-EC3 |
| Eco-426-1 | Bacteria Proteobacteria Gammaproteobacteria Enterobacterales Enterobacteriaceae *Escherichia coli* LAU-EC4 |
| Eco-427-1 | Bacteria Proteobacteria Gammaproteobacteria Enterobacterales Enterobacteriaceae *Escherichia coli* LAU-EC5 |
| Eco-428-1 | Bacteria Proteobacteria Gammaproteobacteria Enterobacterales Enterobacteriaceae *Escherichia coli* LR09 |
| Eco-429-1 | Bacteria Proteobacteria Gammaproteobacteria Enterobacterales Enterobacteriaceae *Escherichia coli* M1 |
| Eco-430-1 | Bacteria Proteobacteria Gammaproteobacteria Enterobacterales Enterobacteriaceae *Escherichia coli* M10 |
| Eco-431-1 | Bacteria Proteobacteria Gammaproteobacteria Enterobacterales Enterobacteriaceae *Escherichia coli* M11 |
| Eco-432-1 | Bacteria Proteobacteria Gammaproteobacteria Enterobacterales Enterobacteriaceae *Escherichia coli* M12 |
| Eco-433-1 | Bacteria Proteobacteria Gammaproteobacteria Enterobacterales Enterobacteriaceae *Escherichia coli* M13 |
| Eco-434-1 | Bacteria Proteobacteria Gammaproteobacteria Enterobacterales Enterobacteriaceae *Escherichia coli* M14 |
| Eco-435-1 | Bacteria Proteobacteria Gammaproteobacteria Enterobacterales Enterobacteriaceae *Escherichia coli* M15 |
| Eco-436-1 | Bacteria Proteobacteria Gammaproteobacteria Enterobacterales Enterobacteriaceae *Escherichia coli* M16 |
| Eco-437-1 | Bacteria Proteobacteria Gammaproteobacteria Enterobacterales Enterobacteriaceae *Escherichia coli* M17 |
| Eco-438-1 | Bacteria Proteobacteria Gammaproteobacteria Enterobacterales Enterobacteriaceae *Escherichia coli* M18 |
| Eco-439-1 | Bacteria Proteobacteria Gammaproteobacteria Enterobacterales Enterobacteriaceae *Escherichia coli* M19 |
| Eco-440-1 | Bacteria Proteobacteria Gammaproteobacteria Enterobacterales Enterobacteriaceae *Escherichia coli* M2 |
| Eco-441-1 | Bacteria Proteobacteria Gammaproteobacteria Enterobacterales Enterobacteriaceae *Escherichia coli* M20 |
| Eco-442-1 | Bacteria Proteobacteria Gammaproteobacteria Enterobacterales Enterobacteriaceae *Escherichia coli* M21 |
| Eco-443-1 | Bacteria Proteobacteria Gammaproteobacteria Enterobacterales Enterobacteriaceae *Escherichia coli* M22 |
| Eco-444-1 | Bacteria Proteobacteria Gammaproteobacteria Enterobacterales Enterobacteriaceae *Escherichia coli* M23 |
| Eco-445-1 | Bacteria Proteobacteria Gammaproteobacteria Enterobacterales Enterobacteriaceae *Escherichia coli* M3 |
| Eco-446-1 | Bacteria Proteobacteria Gammaproteobacteria Enterobacterales Enterobacteriaceae *Escherichia coli* M4 |
| Eco-447-1 | Bacteria Proteobacteria Gammaproteobacteria Enterobacterales Enterobacteriaceae *Escherichia coli* M5 |
| Eco-448-1 | Bacteria Proteobacteria Gammaproteobacteria Enterobacterales Enterobacteriaceae *Escherichia coli* M6 |
| Eco-449-1 | Bacteria Proteobacteria Gammaproteobacteria Enterobacterales Enterobacteriaceae *Escherichia coli* M7 |
| Eco-450-1 | Bacteria Proteobacteria Gammaproteobacteria Enterobacterales Enterobacteriaceae *Escherichia coli* M8 |
| Eco-451-1 | Bacteria Proteobacteria Gammaproteobacteria Enterobacterales Enterobacteriaceae *Escherichia coli* M9 |
| Eco-452-1 | Bacteria Proteobacteria Gammaproteobacteria Enterobacterales Enterobacteriaceae *Escherichia coli* MC19 |
| Eco-453-1 | Bacteria Proteobacteria Gammaproteobacteria Enterobacterales Enterobacteriaceae *Escherichia coli* MC21 |
| Eco-454-1 | Bacteria Proteobacteria Gammaproteobacteria Enterobacterales Enterobacteriaceae *Escherichia coli* MC23 |
| Eco-455-1 | Bacteria Proteobacteria Gammaproteobacteria Enterobacterales Enterobacteriaceae *Escherichia coli* MC6002 |
| Eco-456-1 | Bacteria Proteobacteria Gammaproteobacteria Enterobacterales Enterobacteriaceae *Escherichia coli* MC6003 |
| Eco-457-1 | Bacteria Proteobacteria Gammaproteobacteria Enterobacterales Enterobacteriaceae *Escherichia coli* MGH 57 |
| Eco-458-1 | Bacteria Proteobacteria Gammaproteobacteria Enterobacterales Enterobacteriaceae *Escherichia coli* MGH 58 |
| Eco-459-1 | Bacteria Proteobacteria Gammaproteobacteria Enterobacterales Enterobacteriaceae *Escherichia coli* MP1 |

| | |
|---|---|
| Eco-460-1 | Bacteria Proteobacteria Gammaproteobacteria Enterobacterales Enterobacteriaceae *Escherichia coli* MRSN 10204 |
| Eco-461-1 | Bacteria Proteobacteria Gammaproteobacteria Enterobacterales Enterobacteriaceae *Escherichia coli* MS499 |
| Eco-462-1 | Bacteria Proteobacteria Gammaproteobacteria Enterobacterales Enterobacteriaceae *Escherichia coli* NCTC 50110 |
| Eco-463-1 to Eco-463-4 | Bacteria Proteobacteria Gammaproteobacteria Enterobacterales Enterobacteriaceae *Escherichia coli* Nissle 1917 |
| Eco-464-1 | Bacteria Proteobacteria Gammaproteobacteria Enterobacterales Enterobacteriaceae *Escherichia coli* O103:H11 str. 04-3023 |
| Eco-465-1 | Bacteria Proteobacteria Gammaproteobacteria Enterobacterales Enterobacteriaceae *Escherichia coli* O103:H11 str. 2010C-3214 |
| Eco-466-1 | Bacteria Proteobacteria Gammaproteobacteria Enterobacterales Enterobacteriaceae *Escherichia coli* O103:H2 str. 2009C-3279 |
| Eco-467-1 | Bacteria Proteobacteria Gammaproteobacteria Enterobacterales Enterobacteriaceae *Escherichia coli* O103:H2 str. 2010C-4433 |
| Eco-468-1 | Bacteria Proteobacteria Gammaproteobacteria Enterobacterales Enterobacteriaceae *Escherichia coli* O103:H2 str. 2011C-3750 |
| Eco-469-1 | Bacteria Proteobacteria Gammaproteobacteria Enterobacterales Enterobacteriaceae *Escherichia coli* O103:H25 str. 2010C-4529 |
| Eco-470-1 | Bacteria Proteobacteria Gammaproteobacteria Enterobacterales Enterobacteriaceae *Escherichia coli* O104:H21 str. 94-3025 |
| Eco-471-1 | Bacteria Proteobacteria Gammaproteobacteria Enterobacterales Enterobacteriaceae *Escherichia coli* O104:H4 str. 2011EL-1675A |
| Eco-472-1 to Eco-472-2 | Bacteria Proteobacteria Gammaproteobacteria Enterobacterales Enterobacteriaceae *Escherichia coli* O104:H4 str. C227-11 |
| Eco-473-1 | Bacteria Proteobacteria Gammaproteobacteria Enterobacterales Enterobacteriaceae *Escherichia coli* O111:H8 str. 2009C-4126 |
| Eco-474-1 | Bacteria Proteobacteria Gammaproteobacteria Enterobacterales Enterobacteriaceae *Escherichia coli* O111:H8 str. 2009EL-2169 |
| Eco-475-1 | Bacteria Proteobacteria Gammaproteobacteria Enterobacterales Enterobacteriaceae *Escherichia coli* O111:H8 str. 2011C-3453 |
| Eco-476-1 | Bacteria Proteobacteria Gammaproteobacteria Enterobacterales Enterobacteriaceae *Escherichia coli* O111:H8 str. F6627 |
| Eco-477-1 | Bacteria Proteobacteria Gammaproteobacteria Enterobacterales Enterobacteriaceae *Escherichia coli* O111:NM str. 01-3076 |
| Eco-478-1 | Bacteria Proteobacteria Gammaproteobacteria Enterobacterales Enterobacteriaceae *Escherichia coli* O111:NM str. 03-3484 |
| Eco-479-1 | Bacteria Proteobacteria Gammaproteobacteria Enterobacterales Enterobacteriaceae *Escherichia coli* O111:NM str. 04-3211 |
| Eco-480-1 | Bacteria Proteobacteria Gammaproteobacteria Enterobacterales Enterobacteriaceae *Escherichia coli* O111:NM str. 08-4487 |
| Eco-481-1 | Bacteria Proteobacteria Gammaproteobacteria Enterobacterales Enterobacteriaceae *Escherichia coli* O111:NM str. 2009C-4006 |
| Eco-482-1 | Bacteria Proteobacteria Gammaproteobacteria Enterobacterales Enterobacteriaceae *Escherichia coli* O111:NM str. 2009C-4052 |
| Eco-483-1 | Bacteria Proteobacteria Gammaproteobacteria Enterobacterales Enterobacteriaceae *Escherichia coli* O111:NM str. 2010C-3053 |
| Eco-484-1 | Bacteria Proteobacteria Gammaproteobacteria Enterobacterales Enterobacteriaceae *Escherichia coli* O111:NM str. 2010C-3977 |
| Eco-485-1 | Bacteria Proteobacteria Gammaproteobacteria Enterobacterales Enterobacteriaceae *Escherichia coli* O111:NM str. 2010C-4086 |
| Eco-486-1 | Bacteria Proteobacteria Gammaproteobacteria Enterobacterales Enterobacteriaceae *Escherichia coli* O111:NM str. 2010C-4221 |
| Eco-487-1 | Bacteria Proteobacteria Gammaproteobacteria Enterobacterales Enterobacteriaceae *Escherichia coli* O111:NM str. 2010C-4592 |
| Eco-488-1 | Bacteria Proteobacteria Gammaproteobacteria Enterobacterales Enterobacteriaceae *Escherichia coli* O111:NM str. 2010C-4622 |
| Eco-489-1 | Bacteria Proteobacteria Gammaproteobacteria Enterobacterales Enterobacteriaceae *Escherichia coli* O111:NM str. 2010C-4715 |
| Eco-490-1 | Bacteria Proteobacteria Gammaproteobacteria Enterobacterales Enterobacteriaceae *Escherichia coli* O111:NM str. 2010C-4735 |
| Eco-491-1 | Bacteria Proteobacteria Gammaproteobacteria Enterobacterales Enterobacteriaceae *Escherichia coli* O111:NM str. 2010C-4746 |
| Eco-492-1 | Bacteria Proteobacteria Gammaproteobacteria Enterobacterales Enterobacteriaceae *Escherichia coli* O111:NM str. 2010C-4799 |
| Eco-493-1 | Bacteria Proteobacteria Gammaproteobacteria Enterobacterales Enterobacteriaceae *Escherichia coli* O111:NM str. 2010C-4818 |
| Eco-494-1 | Bacteria Proteobacteria Gammaproteobacteria Enterobacterales Enterobacteriaceae *Escherichia coli* O111:NM str. 2011C-3170 |
| Eco-495-1 | Bacteria Proteobacteria Gammaproteobacteria Enterobacterales Enterobacteriaceae *Escherichia coli* O111:NM str. 2011C-3362 |
| Eco-496-1 | Bacteria Proteobacteria Gammaproteobacteria Enterobacterales Enterobacteriaceae *Escherichia coli* O111:NM str. 2011C-3573 |
| Eco-497-1 | Bacteria Proteobacteria Gammaproteobacteria Enterobacterales Enterobacteriaceae *Escherichia coli* O111:NM str. 2011C-3632 |
| Eco-498-1 | Bacteria Proteobacteria Gammaproteobacteria Enterobacterales Enterobacteriaceae *Escherichia coli* O111:NM str. 2011C-3679 |
| Eco-499-1 | Bacteria Proteobacteria Gammaproteobacteria Enterobacterales Enterobacteriaceae *Escherichia coli* O111:NM str. K6722 |
| Eco-500-1 | Bacteria Proteobacteria Gammaproteobacteria Enterobacterales Enterobacteriaceae *Escherichia coli* O111:NM str. K6723 |
| Eco-501-1 | Bacteria Proteobacteria Gammaproteobacteria Enterobacterales Enterobacteriaceae *Escherichia coli* O111:NM str. K6728 |
| Eco-502-1 | Bacteria Proteobacteria Gammaproteobacteria Enterobacterales Enterobacteriaceae *Escherichia coli* O111:NM str. K6890 |
| Eco-503-1 | Bacteria Proteobacteria Gammaproteobacteria Enterobacterales Enterobacteriaceae *Escherichia coli* O111:NM str. K6895 |
| Eco-504-1 | Bacteria Proteobacteria Gammaproteobacteria Enterobacterales Enterobacteriaceae *Escherichia coli* O111:NM str. K6897 |
| Eco-505-1 | Bacteria Proteobacteria Gammaproteobacteria Enterobacterales Enterobacteriaceae *Escherichia coli* O111:NM str. K6898 |
| Eco-506-1 | Bacteria Proteobacteria Gammaproteobacteria Enterobacterales Enterobacteriaceae *Escherichia coli* O111:NM str. K6904 |
| Eco-507-1 | Bacteria Proteobacteria Gammaproteobacteria Enterobacterales Enterobacteriaceae *Escherichia coli* O111:NM str. K6908 |
| Eco-508-1 | Bacteria Proteobacteria Gammaproteobacteria Enterobacterales Enterobacteriaceae *Escherichia coli* O111:NM str. K6915 |
| Eco-509-1 | Bacteria Proteobacteria Gammaproteobacteria Enterobacterales Enterobacteriaceae *Escherichia coli* O118:H16 str. 06-3256 |
| Eco-510-1 | Bacteria Proteobacteria Gammaproteobacteria Enterobacterales Enterobacteriaceae *Escherichia coli* O118:H16 str. 06-3612 |
| Eco-511-1 | Bacteria Proteobacteria Gammaproteobacteria Enterobacterales Enterobacteriaceae *Escherichia coli* O118:H16 str. 07-4255 |
| Eco-512-1 | Bacteria Proteobacteria Gammaproteobacteria Enterobacterales Enterobacteriaceae *Escherichia coli* O118:H16 str. 08-3651 |
| Eco-513-1 | Bacteria Proteobacteria Gammaproteobacteria Enterobacterales Enterobacteriaceae *Escherichia coli* O118:H16 str. 2009C-4446 |
| Eco-514-1 | Bacteria Proteobacteria Gammaproteobacteria Enterobacterales Enterobacteriaceae *Escherichia coli* O119:H4 str. 03-3458 |
| Eco-515-1 | Bacteria Proteobacteria Gammaproteobacteria Enterobacterales Enterobacteriaceae *Escherichia coli* O121:H19 str. 03-3227 |
| Eco-516-1 | Bacteria Proteobacteria Gammaproteobacteria Enterobacterales Enterobacteriaceae *Escherichia coli* O121:H19 str. 06-3003 |
| Eco-517-1 | Bacteria Proteobacteria Gammaproteobacteria Enterobacterales Enterobacteriaceae *Escherichia coli* O121:H19 str. 06-3822 |
| Eco-518-1 | Bacteria Proteobacteria Gammaproteobacteria Enterobacterales Enterobacteriaceae *Escherichia coli* O121:H19 str. 2009C-4050 |
| Eco-519-1 | Bacteria Proteobacteria Gammaproteobacteria Enterobacterales Enterobacteriaceae *Escherichia coli* O121:H19 str. 2009C-4659 |
| Eco-520-1 | Bacteria Proteobacteria Gammaproteobacteria Enterobacterales Enterobacteriaceae *Escherichia coli* O121:H19 str. 2009C-4750 |
| Eco-521-1 | Bacteria Proteobacteria Gammaproteobacteria Enterobacterales Enterobacteriaceae *Escherichia coli* O121:H19 str. 2009EL1302 |
| Eco-522-1 | Bacteria Proteobacteria Gammaproteobacteria Enterobacterales Enterobacteriaceae *Escherichia coli* O121:H19 str. 2009EL1412 |
| Eco-523-1 | Bacteria Proteobacteria Gammaproteobacteria Enterobacterales Enterobacteriaceae *Escherichia coli* O121:H19 str. 2010C-3609 |
| Eco-524-1 | Bacteria Proteobacteria Gammaproteobacteria Enterobacterales Enterobacteriaceae *Escherichia coli* O121:H19 str. 2010C-3794 |

| | |
|---|---|
| Eco-525-1 | Bacteria Proteobacteria Gammaproteobacteria Enterobacterales Enterobacteriaceae *Escherichia coli* O121:H19 str. 2010C-3840 |
| Eco-526-1 | Bacteria Proteobacteria Gammaproteobacteria Enterobacterales Enterobacteriaceae *Escherichia coli* O121:H19 str. 2010C-4254 |
| Eco-527-1 | Bacteria Proteobacteria Gammaproteobacteria Enterobacterales Enterobacteriaceae *Escherichia coli* O121:H19 str. 2010C-4732 |
| Eco-528-1 | Bacteria Proteobacteria Gammaproteobacteria Enterobacterales Enterobacteriaceae *Escherichia coli* O121:H19 str. 2010C-4824 |
| Eco-529-1 | Bacteria Proteobacteria Gammaproteobacteria Enterobacterales Enterobacteriaceae *Escherichia coli* O121:H19 str. 2010C-4966 |
| Eco-530-1 | Bacteria Proteobacteria Gammaproteobacteria Enterobacterales Enterobacteriaceae *Escherichia coli* O121:H19 str. 2010C-4989 |
| Eco-531-1 | Bacteria Proteobacteria Gammaproteobacteria Enterobacterales Enterobacteriaceae *Escherichia coli* O121:H19 str. 2010EL1058 |
| Eco-532-1 | Bacteria Proteobacteria Gammaproteobacteria Enterobacterales Enterobacteriaceae *Escherichia coli* O121:H19 str. 2011C-3072 |
| Eco-533-1 | Bacteria Proteobacteria Gammaproteobacteria Enterobacterales Enterobacteriaceae *Escherichia coli* O121:H19 str. 2011C-3108 |
| Eco-534-1 | Bacteria Proteobacteria Gammaproteobacteria Enterobacterales Enterobacteriaceae *Escherichia coli* O121:H19 str. 2011C-3216 |
| Eco-535-1 | Bacteria Proteobacteria Gammaproteobacteria Enterobacterales Enterobacteriaceae *Escherichia coli* O121:H19 str. 2011C-3500 |
| Eco-536-1 | Bacteria Proteobacteria Gammaproteobacteria Enterobacterales Enterobacteriaceae *Escherichia coli* O121:H19 str. 2011C-3537 |
| Eco-537-1 | Bacteria Proteobacteria Gammaproteobacteria Enterobacterales Enterobacteriaceae *Escherichia coli* O121:H19 str. 2011C-3609 |
| Eco-538-1 | Bacteria Proteobacteria Gammaproteobacteria Enterobacterales Enterobacteriaceae *Escherichia coli* O121:H19 str. F6714 |
| Eco-539-1 | Bacteria Proteobacteria Gammaproteobacteria Enterobacterales Enterobacteriaceae *Escherichia coli* O121:H19 str. K5198 |
| Eco-540-1 | Bacteria Proteobacteria Gammaproteobacteria Enterobacterales Enterobacteriaceae *Escherichia coli* O121:H19 str. K5269 |
| Eco-541-1 | Bacteria Proteobacteria Gammaproteobacteria Enterobacterales Enterobacteriaceae *Escherichia coli* O121:H7 str. 2009C-3299 |
| Eco-542-1 | Bacteria Proteobacteria Gammaproteobacteria Enterobacterales Enterobacteriaceae *Escherichia coli* O123:H11 str. 2009C-3307 |
| Eco-543-1 | Bacteria Proteobacteria Gammaproteobacteria Enterobacterales Enterobacteriaceae *Escherichia coli* O127:H6 str. E2348/69 substr. CVDNalr |
| Eco-544-1 | Bacteria Proteobacteria Gammaproteobacteria Enterobacterales Enterobacteriaceae *Escherichia coli* O127:H6 str. E2348/69 substr. UMD753 |
| Eco-545-1 | Bacteria Proteobacteria Gammaproteobacteria Enterobacterales Enterobacteriaceae *Escherichia coli* O128:H2 str. 2011C-3317 |
| Eco-546-1 | Bacteria Proteobacteria Gammaproteobacteria Enterobacterales Enterobacteriaceae *Escherichia coli* O145:H25 str. 07-3858 |
| Eco-547-1 | Bacteria Proteobacteria Gammaproteobacteria Enterobacterales Enterobacteriaceae *Escherichia coli* O145:H28 str. 2009C-3292 |
| Eco-548-1 to Eco-548-2 | Bacteria Proteobacteria Gammaproteobacteria Enterobacterales Enterobacteriaceae *Escherichia coli* O145:H28 str. 4865/96 |
| Eco-549-1 | Bacteria Proteobacteria Gammaproteobacteria Enterobacterales Enterobacteriaceae *Escherichia coli* O145:H28 str. RM12581 |
| Eco-550-1 | Bacteria Proteobacteria Gammaproteobacteria Enterobacterales Enterobacteriaceae *Escherichia coli* O145:H28 str. RM12761 |
| Eco-551-1 | Bacteria Proteobacteria Gammaproteobacteria Enterobacterales Enterobacteriaceae *Escherichia coli* O145:H28 str. RM13514 |
| Eco-552-1 | Bacteria Proteobacteria Gammaproteobacteria Enterobacterales Enterobacteriaceae *Escherichia coli* O145:H28 str. RM13516 |
| Eco-553-1 | Bacteria Proteobacteria Gammaproteobacteria Enterobacterales Enterobacteriaceae *Escherichia coli* O145:NM str. 06-3484 |
| Eco-554-1 | Bacteria Proteobacteria Gammaproteobacteria Enterobacterales Enterobacteriaceae *Escherichia coli* O145:NM str. 08-4270 |
| Eco-555-1 | Bacteria Proteobacteria Gammaproteobacteria Enterobacterales Enterobacteriaceae *Escherichia coli* O145:NM str. 2010C-3507 |
| Eco-556-1 | Bacteria Proteobacteria Gammaproteobacteria Enterobacterales Enterobacteriaceae *Escherichia coli* O145:NM str. 2010C-3508 |
| Eco-557-1 | Bacteria Proteobacteria Gammaproteobacteria Enterobacterales Enterobacteriaceae *Escherichia coli* O145:NM str. 2010C-3509 |
| Eco-558-1 | Bacteria Proteobacteria Gammaproteobacteria Enterobacterales Enterobacteriaceae *Escherichia coli* O145:NM str. 2010C-3510 |
| Eco-559-1 | Bacteria Proteobacteria Gammaproteobacteria Enterobacterales Enterobacteriaceae *Escherichia coli* O145:NM str. 2010C-3511 |
| Eco-560-1 | Bacteria Proteobacteria Gammaproteobacteria Enterobacterales Enterobacteriaceae *Escherichia coli* O145:NM str. 2010C-3516 |
| Eco-561-1 | Bacteria Proteobacteria Gammaproteobacteria Enterobacterales Enterobacteriaceae *Escherichia coli* O145:NM str. 2010C-3517 |
| Eco-562-1 | Bacteria Proteobacteria Gammaproteobacteria Enterobacterales Enterobacteriaceae *Escherichia coli* O145:NM str. 2010C-3518 |
| Eco-563-1 | Bacteria Proteobacteria Gammaproteobacteria Enterobacterales Enterobacteriaceae *Escherichia coli* O145:NM str. 2010C-3521 |
| Eco-564-1 | Bacteria Proteobacteria Gammaproteobacteria Enterobacterales Enterobacteriaceae *Escherichia coli* O145:NM str. 2010C-3526 |
| Eco-565-1 | Bacteria Proteobacteria Gammaproteobacteria Enterobacterales Enterobacteriaceae *Escherichia coli* O145:NM str. 2010C-4557C2 |
| Eco-566-1 | Bacteria Proteobacteria Gammaproteobacteria Enterobacterales Enterobacteriaceae *Escherichia coli* O146:H21 str. 2010C-3325 |
| Eco-567-1 | Bacteria Proteobacteria Gammaproteobacteria Enterobacterales Enterobacteriaceae *Escherichia coli* O153:H2 str. 2010C-5034 |
| Eco-568-1 | Bacteria Proteobacteria Gammaproteobacteria Enterobacterales Enterobacteriaceae *Escherichia coli* O156:H25 str. 2011C-3602 |
| Eco-569-1 | Bacteria Proteobacteria Gammaproteobacteria Enterobacterales Enterobacteriaceae *Escherichia coli* O157 str. NCCP15739 |
| Eco-570-1 | Bacteria Proteobacteria Gammaproteobacteria Enterobacterales Enterobacteriaceae *Escherichia coli* O157: str. 2010EL-2044 |
| Eco-571-1 | Bacteria Proteobacteria Gammaproteobacteria Enterobacterales Enterobacteriaceae *Escherichia coli* O157: str. 2010EL-2045 |
| Eco-572-1 | Bacteria Proteobacteria Gammaproteobacteria Enterobacterales Enterobacteriaceae *Escherichia coli* O157:H16 |
| Eco-573-1 | Bacteria Proteobacteria Gammaproteobacteria Enterobacterales Enterobacteriaceae *Escherichia coli* O157:H16 str. 98-3133 |
| Eco-574-1 to Eco-574-3 | Bacteria Proteobacteria Gammaproteobacteria Enterobacterales Enterobacteriaceae *Escherichia coli* O157:H7 |
| Eco-575-1 | Bacteria Proteobacteria Gammaproteobacteria Enterobacterales Enterobacteriaceae *Escherichia coli* O157:H7 str. 06-3745 |
| Eco-576-1 | Bacteria Proteobacteria Gammaproteobacteria Enterobacterales Enterobacteriaceae *Escherichia coli* O157:H7 str. 06-4039 |
| Eco-577-1 | Bacteria Proteobacteria Gammaproteobacteria Enterobacterales Enterobacteriaceae *Escherichia coli* O157:H7 str. 07-3091 |
| Eco-578-1 | Bacteria Proteobacteria Gammaproteobacteria Enterobacterales Enterobacteriaceae *Escherichia coli* O157:H7 str. 07-3391 |
| Eco-579-1 | Bacteria Proteobacteria Gammaproteobacteria Enterobacterales Enterobacteriaceae *Escherichia coli* O157:H7 str. 08-3037 |
| Eco-580-1 | Bacteria Proteobacteria Gammaproteobacteria Enterobacterales Enterobacteriaceae *Escherichia coli* O157:H7 str. 08-3527 |
| Eco-581-1 | Bacteria Proteobacteria Gammaproteobacteria Enterobacterales Enterobacteriaceae *Escherichia coli* O157:H7 str. 08-4169 |
| Eco-582-1 | Bacteria Proteobacteria Gammaproteobacteria Enterobacterales Enterobacteriaceae *Escherichia coli* O157:H7 str. 08-4529 |
| Eco-583-1 | Bacteria Proteobacteria Gammaproteobacteria Enterobacterales Enterobacteriaceae *Escherichia coli* O157:H7 str. 2009C-4258 |
| Eco-584-1 | Bacteria Proteobacteria Gammaproteobacteria Enterobacterales Enterobacteriaceae *Escherichia coli* O157:H7 str. 2009EL1449 |
| Eco-585-1 | Bacteria Proteobacteria Gammaproteobacteria Enterobacterales Enterobacteriaceae *Escherichia coli* O157:H7 str. 2009EL1705 |
| Eco-586-1 | Bacteria Proteobacteria Gammaproteobacteria Enterobacterales Enterobacteriaceae *Escherichia coli* O157:H7 str. 2009EL1913 |
| Eco-587-1 | Bacteria Proteobacteria Gammaproteobacteria Enterobacterales Enterobacteriaceae *Escherichia coli* O157:H7 str. 2009EL2109 |
| Eco-588-1 | Bacteria Proteobacteria Gammaproteobacteria Enterobacterales Enterobacteriaceae *Escherichia coli* O157:H7 str. 2010C-4979C1 |
| Eco-589-1 | Bacteria Proteobacteria Gammaproteobacteria Enterobacterales Enterobacteriaceae *Escherichia coli* O157:H7 str. 2011EL-1107 |

| | |
|---|---|
| Eco-590-1 | Bacteria Proteobacteria Gammaproteobacteria Enterobacterales Enterobacteriaceae *Escherichia coli* O157:H7 str. 2011EL-2090 |
| Eco-591-1 | Bacteria Proteobacteria Gammaproteobacteria Enterobacterales Enterobacteriaceae *Escherichia coli* O157:H7 str. 2011EL-2091 |
| Eco-592-1 | Bacteria Proteobacteria Gammaproteobacteria Enterobacterales Enterobacteriaceae *Escherichia coli* O157:H7 str. 2011EL-2092 |
| Eco-593-1 | Bacteria Proteobacteria Gammaproteobacteria Enterobacterales Enterobacteriaceae *Escherichia coli* O157:H7 str. 2011EL-2093 |
| Eco-594-1 | Bacteria Proteobacteria Gammaproteobacteria Enterobacterales Enterobacteriaceae *Escherichia coli* O157:H7 str. 2011EL-2094 |
| Eco-595-1 | Bacteria Proteobacteria Gammaproteobacteria Enterobacterales Enterobacteriaceae *Escherichia coli* O157:H7 str. 2011EL-2096 |
| Eco-596-1 | Bacteria Proteobacteria Gammaproteobacteria Enterobacterales Enterobacteriaceae *Escherichia coli* O157:H7 str. 2011EL-2097 |
| Eco-597-1 | Bacteria Proteobacteria Gammaproteobacteria Enterobacterales Enterobacteriaceae *Escherichia coli* O157:H7 str. 2011EL-2098 |
| Eco-598-1 | Bacteria Proteobacteria Gammaproteobacteria Enterobacterales Enterobacteriaceae *Escherichia coli* O157:H7 str. 2011EL-2099 |
| Eco-599-1 | Bacteria Proteobacteria Gammaproteobacteria Enterobacterales Enterobacteriaceae *Escherichia coli* O157:H7 str. 2011EL-2101 |
| Eco-600-1 | Bacteria Proteobacteria Gammaproteobacteria Enterobacterales Enterobacteriaceae *Escherichia coli* O157:H7 str. 2011EL-2103 |
| Eco-601-1 | Bacteria Proteobacteria Gammaproteobacteria Enterobacterales Enterobacteriaceae *Escherichia coli* O157:H7 str. 2011EL-2104 |
| Eco-602-1 | Bacteria Proteobacteria Gammaproteobacteria Enterobacterales Enterobacteriaceae *Escherichia coli* O157:H7 str. 2011EL-2105 |
| Eco-603-1 | Bacteria Proteobacteria Gammaproteobacteria Enterobacterales Enterobacteriaceae *Escherichia coli* O157:H7 str. 2011EL-2106 |
| Eco-604-1 | Bacteria Proteobacteria Gammaproteobacteria Enterobacterales Enterobacteriaceae *Escherichia coli* O157:H7 str. 2011EL-2107 |
| Eco-605-1 | Bacteria Proteobacteria Gammaproteobacteria Enterobacterales Enterobacteriaceae *Escherichia coli* O157:H7 str. 2011EL-2108 |
| Eco-606-1 | Bacteria Proteobacteria Gammaproteobacteria Enterobacterales Enterobacteriaceae *Escherichia coli* O157:H7 str. 2011EL-2109 |
| Eco-607-1 | Bacteria Proteobacteria Gammaproteobacteria Enterobacterales Enterobacteriaceae *Escherichia coli* O157:H7 str. 2011EL-2111 |
| Eco-608-1 | Bacteria Proteobacteria Gammaproteobacteria Enterobacterales Enterobacteriaceae *Escherichia coli* O157:H7 str. 2011EL-2112 |
| Eco-609-1 | Bacteria Proteobacteria Gammaproteobacteria Enterobacterales Enterobacteriaceae *Escherichia coli* O157:H7 str. 2011EL-2113 |
| Eco-610-1 | Bacteria Proteobacteria Gammaproteobacteria Enterobacterales Enterobacteriaceae *Escherichia coli* O157:H7 str. 2011EL-2114 |
| Eco-611-1 | Bacteria Proteobacteria Gammaproteobacteria Enterobacterales Enterobacteriaceae *Escherichia coli* O157:H7 str. 2011EL-2286 |
| Eco-612-1 | Bacteria Proteobacteria Gammaproteobacteria Enterobacterales Enterobacteriaceae *Escherichia coli* O157:H7 str. 2011EL-2287 |
| Eco-613-1 | Bacteria Proteobacteria Gammaproteobacteria Enterobacterales Enterobacteriaceae *Escherichia coli* O157:H7 str. 2011EL-2288 |
| Eco-614-1 | Bacteria Proteobacteria Gammaproteobacteria Enterobacterales Enterobacteriaceae *Escherichia coli* O157:H7 str. 2011EL-2289 |
| Eco-615-1 | Bacteria Proteobacteria Gammaproteobacteria Enterobacterales Enterobacteriaceae *Escherichia coli* O157:H7 str. 2011EL-2290 |
| Eco-616-1 | Bacteria Proteobacteria Gammaproteobacteria Enterobacterales Enterobacteriaceae *Escherichia coli* O157:H7 str. 2011EL-2312 |
| Eco-617-1 | Bacteria Proteobacteria Gammaproteobacteria Enterobacterales Enterobacteriaceae *Escherichia coli* O157:H7 str. 2011EL-2313 |
| Eco-618-1 | Bacteria Proteobacteria Gammaproteobacteria Enterobacterales Enterobacteriaceae *Escherichia coli* O157:H7 str. EC4127 |
| Eco-619-1 | Bacteria Proteobacteria Gammaproteobacteria Enterobacterales Enterobacteriaceae *Escherichia coli* O157:H7 str. EC4192 |
| Eco-620-1 | Bacteria Proteobacteria Gammaproteobacteria Enterobacterales Enterobacteriaceae *Escherichia coli* O157:H7 str. EC536 |
| Eco-621-1 | Bacteria Proteobacteria Gammaproteobacteria Enterobacterales Enterobacteriaceae *Escherichia coli* O157:H7 str. EDL933 |
| Eco-622-1 | Bacteria Proteobacteria Gammaproteobacteria Enterobacterales Enterobacteriaceae *Escherichia coli* O157:H7 str. F6142 |
| Eco-623-1 | Bacteria Proteobacteria Gammaproteobacteria Enterobacterales Enterobacteriaceae *Escherichia coli* O157:H7 str. F6749 |
| Eco-624-1 | Bacteria Proteobacteria Gammaproteobacteria Enterobacterales Enterobacteriaceae *Escherichia coli* O157:H7 str. F6750 |
| Eco-625-1 | Bacteria Proteobacteria Gammaproteobacteria Enterobacterales Enterobacteriaceae *Escherichia coli* O157:H7 str. F6751 |
| Eco-626-1 | Bacteria Proteobacteria Gammaproteobacteria Enterobacterales Enterobacteriaceae *Escherichia coli* O157:H7 str. F7350 |
| Eco-627-1 | Bacteria Proteobacteria Gammaproteobacteria Enterobacterales Enterobacteriaceae *Escherichia coli* O157:H7 str. F7377 |
| Eco-628-1 | Bacteria Proteobacteria Gammaproteobacteria Enterobacterales Enterobacteriaceae *Escherichia coli* O157:H7 str. F7384 |
| Eco-629-1 | Bacteria Proteobacteria Gammaproteobacteria Enterobacterales Enterobacteriaceae *Escherichia coli* O157:H7 str. F7410 |
| Eco-630-1 | Bacteria Proteobacteria Gammaproteobacteria Enterobacterales Enterobacteriaceae *Escherichia coli* O157:H7 str. F8092B |
| Eco-631-1 | Bacteria Proteobacteria Gammaproteobacteria Enterobacterales Enterobacteriaceae *Escherichia coli* O157:H7 str. G5303 |
| Eco-632-1 | Bacteria Proteobacteria Gammaproteobacteria Enterobacterales Enterobacteriaceae *Escherichia coli* O157:H7 str. H093800014 |
| Eco-633-1 | Bacteria Proteobacteria Gammaproteobacteria Enterobacterales Enterobacteriaceae *Escherichia coli* O157:H7 str. H2495 |
| Eco-634-1 | Bacteria Proteobacteria Gammaproteobacteria Enterobacterales Enterobacteriaceae *Escherichia coli* O157:H7 str. H2498 |
| Eco-635-1 | Bacteria Proteobacteria Gammaproteobacteria Enterobacterales Enterobacteriaceae *Escherichia coli* O157:H7 str. K1420 |
| Eco-636-1 | Bacteria Proteobacteria Gammaproteobacteria Enterobacterales Enterobacteriaceae *Escherichia coli* O157:H7 str. K1792 |
| Eco-637-1 | Bacteria Proteobacteria Gammaproteobacteria Enterobacterales Enterobacteriaceae *Escherichia coli* O157:H7 str. K1793 |
| Eco-638-1 | Bacteria Proteobacteria Gammaproteobacteria Enterobacterales Enterobacteriaceae *Escherichia coli* O157:H7 str. K1795 |
| Eco-639-1 | Bacteria Proteobacteria Gammaproteobacteria Enterobacterales Enterobacteriaceae *Escherichia coli* O157:H7 str. K1796 |
| Eco-640-1 | Bacteria Proteobacteria Gammaproteobacteria Enterobacterales Enterobacteriaceae *Escherichia coli* O157:H7 str. K1845 |
| Eco-641-1 | Bacteria Proteobacteria Gammaproteobacteria Enterobacterales Enterobacteriaceae *Escherichia coli* O157:H7 str. K1921 |
| Eco-642-1 | Bacteria Proteobacteria Gammaproteobacteria Enterobacterales Enterobacteriaceae *Escherichia coli* O157:H7 str. K1927 |
| Eco-643-1 | Bacteria Proteobacteria Gammaproteobacteria Enterobacterales Enterobacteriaceae *Escherichia coli* O157:H7 str. K2188 |
| Eco-644-1 | Bacteria Proteobacteria Gammaproteobacteria Enterobacterales Enterobacteriaceae *Escherichia coli* O157:H7 str. K2191 |
| Eco-645-1 | Bacteria Proteobacteria Gammaproteobacteria Enterobacterales Enterobacteriaceae *Escherichia coli* O157:H7 str. K2192 |
| Eco-646-1 | Bacteria Proteobacteria Gammaproteobacteria Enterobacterales Enterobacteriaceae *Escherichia coli* O157:H7 str. K2324 |
| Eco-647-1 | Bacteria Proteobacteria Gammaproteobacteria Enterobacterales Enterobacteriaceae *Escherichia coli* O157:H7 str. K2581 |
| Eco-648-1 | Bacteria Proteobacteria Gammaproteobacteria Enterobacterales Enterobacteriaceae *Escherichia coli* O157:H7 str. K2622 |
| Eco-649-1 | Bacteria Proteobacteria Gammaproteobacteria Enterobacterales Enterobacteriaceae *Escherichia coli* O157:H7 str. K2845 |
| Eco-650-1 | Bacteria Proteobacteria Gammaproteobacteria Enterobacterales Enterobacteriaceae *Escherichia coli* O157:H7 str. K2854 |
| Eco-651-1 | Bacteria Proteobacteria Gammaproteobacteria Enterobacterales Enterobacteriaceae *Escherichia coli* O157:H7 str. K4396 |
| Eco-652-1 | Bacteria Proteobacteria Gammaproteobacteria Enterobacterales Enterobacteriaceae *Escherichia coli* O157:H7 str. K4405 |
| Eco-653-1 | Bacteria Proteobacteria Gammaproteobacteria Enterobacterales Enterobacteriaceae *Escherichia coli* O157:H7 str. K4406 |
| Eco-654-1 | Bacteria Proteobacteria Gammaproteobacteria Enterobacterales Enterobacteriaceae *Escherichia coli* O157:H7 str. K4527 |

| | |
|---|---|
| Eco-655-1 | Bacteria Proteobacteria Gammaproteobacteria Enterobacterales Enterobacteriaceae *Escherichia coli* O157:H7 str. K5418 |
| Eco-656-1 | Bacteria Proteobacteria Gammaproteobacteria Enterobacterales Enterobacteriaceae *Escherichia coli* O157:H7 str. K5448 |
| Eco-657-1 | Bacteria Proteobacteria Gammaproteobacteria Enterobacterales Enterobacteriaceae *Escherichia coli* O157:H7 str. K5449 |
| Eco-658-1 | Bacteria Proteobacteria Gammaproteobacteria Enterobacterales Enterobacteriaceae *Escherichia coli* O157:H7 str. K5453 |
| Eco-659-1 | Bacteria Proteobacteria Gammaproteobacteria Enterobacterales Enterobacteriaceae *Escherichia coli* O157:H7 str. K5460 |
| Eco-660-1 | Bacteria Proteobacteria Gammaproteobacteria Enterobacterales Enterobacteriaceae *Escherichia coli* O157:H7 str. K5467 |
| Eco-661-1 | Bacteria Proteobacteria Gammaproteobacteria Enterobacterales Enterobacteriaceae *Escherichia coli* O157:H7 str. K5602 |
| Eco-662-1 | Bacteria Proteobacteria Gammaproteobacteria Enterobacterales Enterobacteriaceae *Escherichia coli* O157:H7 str. K5607 |
| Eco-663-1 | Bacteria Proteobacteria Gammaproteobacteria Enterobacterales Enterobacteriaceae *Escherichia coli* O157:H7 str. K5609 |
| Eco-664-1 | Bacteria Proteobacteria Gammaproteobacteria Enterobacterales Enterobacteriaceae *Escherichia coli* O157:H7 str. K5806 |
| Eco-665-1 | Bacteria Proteobacteria Gammaproteobacteria Enterobacterales Enterobacteriaceae *Escherichia coli* O157:H7 str. K5852 |
| Eco-666-1 | Bacteria Proteobacteria Gammaproteobacteria Enterobacterales Enterobacteriaceae *Escherichia coli* O157:H7 str. K6590 |
| Eco-667-1 | Bacteria Proteobacteria Gammaproteobacteria Enterobacterales Enterobacteriaceae *Escherichia coli* O157:H7 str. K6676 |
| Eco-668-1 | Bacteria Proteobacteria Gammaproteobacteria Enterobacterales Enterobacteriaceae *Escherichia coli* O157:H7 str. K6687 |
| Eco-669-1 | Bacteria Proteobacteria Gammaproteobacteria Enterobacterales Enterobacteriaceae *Escherichia coli* O157:H7 str. K7140 |
| Eco-670-1 | Bacteria Proteobacteria Gammaproteobacteria Enterobacterales Enterobacteriaceae *Escherichia coli* O157:H7 str. SS17 |
| Eco-671-1 | Bacteria Proteobacteria Gammaproteobacteria Enterobacterales Enterobacteriaceae *Escherichia coli* O157:H7 str. SS52 |
| Eco-672-1 | Bacteria Proteobacteria Gammaproteobacteria Enterobacterales Enterobacteriaceae *Escherichia coli* O157:H7 str. T1543_06 |
| Eco-673-1 | Bacteria Proteobacteria Gammaproteobacteria Enterobacterales Enterobacteriaceae *Escherichia coli* O157:NM str. 08-4540 |
| Eco-674-1 | Bacteria Proteobacteria Gammaproteobacteria Enterobacterales Enterobacteriaceae *Escherichia coli* O15:H18 str. K1516 |
| Eco-675-1 | Bacteria Proteobacteria Gammaproteobacteria Enterobacterales Enterobacteriaceae *Escherichia coli* O165:H25 str. 2010C-4874 |
| Eco-676-1 | Bacteria Proteobacteria Gammaproteobacteria Enterobacterales Enterobacteriaceae *Escherichia coli* O169:H41 str. F9792 |
| Eco-677-1 | Bacteria Proteobacteria Gammaproteobacteria Enterobacterales Enterobacteriaceae *Escherichia coli* O174:H21 |
| Eco-678-1 | Bacteria Proteobacteria Gammaproteobacteria Enterobacterales Enterobacteriaceae *Escherichia coli* O174:H21 str. 03-3269 |
| Eco-679-1 | Bacteria Proteobacteria Gammaproteobacteria Enterobacterales Enterobacteriaceae *Escherichia coli* O174:H8 str. 04-3038 |
| Eco-680-1 | Bacteria Proteobacteria Gammaproteobacteria Enterobacterales Enterobacteriaceae *Escherichia coli* O177:NM str. 2010C-4558 |
| Eco-681-1 | Bacteria Proteobacteria Gammaproteobacteria Enterobacterales Enterobacteriaceae *Escherichia coli* O25:NM str. E2539C1 |
| Eco-682-1 | Bacteria Proteobacteria Gammaproteobacteria Enterobacterales Enterobacteriaceae *Escherichia coli* O25b:H4-ST131 |
| Eco-683-1 | Bacteria Proteobacteria Gammaproteobacteria Enterobacterales Enterobacteriaceae *Escherichia coli* O26:H1 str. 2009C-4747 |
| Eco-684-1 | Bacteria Proteobacteria Gammaproteobacteria Enterobacterales Enterobacteriaceae *Escherichia coli* O26:H11 str. 03-3500 |
| Eco-685-1 | Bacteria Proteobacteria Gammaproteobacteria Enterobacterales Enterobacteriaceae *Escherichia coli* O26:H11 str. 05-3646 |
| Eco-686-1 | Bacteria Proteobacteria Gammaproteobacteria Enterobacterales Enterobacteriaceae *Escherichia coli* O26:H11 str. 06-3464 |
| Eco-687-1 | Bacteria Proteobacteria Gammaproteobacteria Enterobacterales Enterobacteriaceae *Escherichia coli* O26:H11 str. 2009C-3612 |
| Eco-688-1 | Bacteria Proteobacteria Gammaproteobacteria Enterobacterales Enterobacteriaceae *Escherichia coli* O26:H11 str. 2009C-3689 |
| Eco-689-1 | Bacteria Proteobacteria Gammaproteobacteria Enterobacterales Enterobacteriaceae *Escherichia coli* O26:H11 str. 2009C-3996 |
| Eco-690-1 | Bacteria Proteobacteria Gammaproteobacteria Enterobacterales Enterobacteriaceae *Escherichia coli* O26:H11 str. 2009C-4760 |
| Eco-691-1 | Bacteria Proteobacteria Gammaproteobacteria Enterobacterales Enterobacteriaceae *Escherichia coli* O26:H11 str. 2009C-4826 |
| Eco-692-1 | Bacteria Proteobacteria Gammaproteobacteria Enterobacterales Enterobacteriaceae *Escherichia coli* O26:H11 str. 2010C-3051 |
| Eco-693-1 | Bacteria Proteobacteria Gammaproteobacteria Enterobacterales Enterobacteriaceae *Escherichia coli* O26:H11 str. 2010C-3472 |
| Eco-694-1 | Bacteria Proteobacteria Gammaproteobacteria Enterobacterales Enterobacteriaceae *Escherichia coli* O26:H11 str. 2010C-3871 |
| Eco-695-1 | Bacteria Proteobacteria Gammaproteobacteria Enterobacterales Enterobacteriaceae *Escherichia coli* O26:H11 str. 2010C-3902 |
| Eco-696-1 | Bacteria Proteobacteria Gammaproteobacteria Enterobacterales Enterobacteriaceae *Escherichia coli* O26:H11 str. 2010C-4244 |
| Eco-697-1 | Bacteria Proteobacteria Gammaproteobacteria Enterobacterales Enterobacteriaceae *Escherichia coli* O26:H11 str. 2010C-4430 |
| Eco-698-1 | Bacteria Proteobacteria Gammaproteobacteria Enterobacterales Enterobacteriaceae *Escherichia coli* O26:H11 str. 2010C-4819 |
| Eco-699-1 | Bacteria Proteobacteria Gammaproteobacteria Enterobacterales Enterobacteriaceae *Escherichia coli* O26:H11 str. 2010C-4834 |
| Eco-700-1 | Bacteria Proteobacteria Gammaproteobacteria Enterobacterales Enterobacteriaceae *Escherichia coli* O26:H11 str. 2010C-5028 |
| Eco-701-1 | Bacteria Proteobacteria Gammaproteobacteria Enterobacterales Enterobacteriaceae *Escherichia coli* O26:H11 str. 2010EL-1699 |
| Eco-702-1 | Bacteria Proteobacteria Gammaproteobacteria Enterobacterales Enterobacteriaceae *Escherichia coli* O26:H11 str. 2011C-3270 |
| Eco-703-1 | Bacteria Proteobacteria Gammaproteobacteria Enterobacterales Enterobacteriaceae *Escherichia coli* O26:H11 str. 2011C-3282 |
| Eco-704-1 | Bacteria Proteobacteria Gammaproteobacteria Enterobacterales Enterobacteriaceae *Escherichia coli* O26:H11 str. 2011C-3387 |
| Eco-705-1 | Bacteria Proteobacteria Gammaproteobacteria Enterobacterales Enterobacteriaceae *Escherichia coli* O26:H11 str. 2011C-3506 |
| Eco-706-1 | Bacteria Proteobacteria Gammaproteobacteria Enterobacterales Enterobacteriaceae *Escherichia coli* O26:H11 str. 2011C-3655 |
| Eco-707-1 | Bacteria Proteobacteria Gammaproteobacteria Enterobacterales Enterobacteriaceae *Escherichia coli* O26:NM str. 2010C-4347 |
| Eco-708-1 | Bacteria Proteobacteria Gammaproteobacteria Enterobacterales Enterobacteriaceae *Escherichia coli* O26:NM str. 2010C-4788 |
| Eco-709-1 | Bacteria Proteobacteria Gammaproteobacteria Enterobacterales Enterobacteriaceae *Escherichia coli* O28ac:NM str. 02-3404 |
| Eco-710-1 | Bacteria Proteobacteria Gammaproteobacteria Enterobacterales Enterobacteriaceae *Escherichia coli* O39:NM str. F8704-2 |
| Eco-711-1 | Bacteria Proteobacteria Gammaproteobacteria Enterobacterales Enterobacteriaceae *Escherichia coli* O45:H2 str. 01-3147 |
| Eco-712-1 | Bacteria Proteobacteria Gammaproteobacteria Enterobacterales Enterobacteriaceae *Escherichia coli* O45:H2 str. 2009C-3686 |
| Eco-713-1 | Bacteria Proteobacteria Gammaproteobacteria Enterobacterales Enterobacteriaceae *Escherichia coli* O45:H2 str. 2009C-4780 |
| Eco-714-1 | Bacteria Proteobacteria Gammaproteobacteria Enterobacterales Enterobacteriaceae *Escherichia coli* O45:H2 str. 2010C-3876 |
| Eco-715-1 | Bacteria Proteobacteria Gammaproteobacteria Enterobacterales Enterobacteriaceae *Escherichia coli* O45:H2 str. 2010C-4211 |
| Eco-716-1 | Bacteria Proteobacteria Gammaproteobacteria Enterobacterales Enterobacteriaceae *Escherichia coli* O69:H11 str. 06-3325 |
| Eco-717-1 | Bacteria Proteobacteria Gammaproteobacteria Enterobacterales Enterobacteriaceae *Escherichia coli* O69:H11 str. 07-3763 |
| Eco-718-1 | Bacteria Proteobacteria Gammaproteobacteria Enterobacterales Enterobacteriaceae *Escherichia coli* O69:H11 str. 07-4281 |
| Eco-719-1 | Bacteria Proteobacteria Gammaproteobacteria Enterobacterales Enterobacteriaceae *Escherichia coli* O69:H11 str. 08-4661 |

| | |
|---|---|
| Eco-720-1 | Bacteria Proteobacteria Gammaproteobacteria Enterobacterales Enterobacteriaceae *Escherichia coli* O69:H11 str. 2009C-3601 |
| Eco-721-1 | Bacteria Proteobacteria Gammaproteobacteria Enterobacterales Enterobacteriaceae *Escherichia coli* O6:H16 str. 99-3165 |
| Eco-722-1 | Bacteria Proteobacteria Gammaproteobacteria Enterobacterales Enterobacteriaceae *Escherichia coli* O6:H16 str. F5656C1 |
| Eco-723-1 | Bacteria Proteobacteria Gammaproteobacteria Enterobacterales Enterobacteriaceae *Escherichia coli* O6:H16:CFA/II str. B2C |
| Eco-724-1 | Bacteria Proteobacteria Gammaproteobacteria Enterobacterales Enterobacteriaceae *Escherichia coli* O78:H12 str. 00-3279 |
| Eco-725-1 | Bacteria Proteobacteria Gammaproteobacteria Enterobacterales Enterobacteriaceae *Escherichia coli* O79:H7 str. 06-3501 |
| Eco-726-1 | Bacteria Proteobacteria Gammaproteobacteria Enterobacterales Enterobacteriaceae *Escherichia coli* O81:NM str. 02-3012 |
| Eco-727-1 | Bacteria Proteobacteria Gammaproteobacteria Enterobacterales Enterobacteriaceae *Escherichia coli* O86:H34 str. 99-3124 |
| Eco-728-1 to Eco-728-2 | Bacteria Proteobacteria Gammaproteobacteria Enterobacterales Enterobacteriaceae *Escherichia coli* O91 |
| Eco-729-1 | Bacteria Proteobacteria Gammaproteobacteria Enterobacterales Enterobacteriaceae *Escherichia coli* O91:H14 str. 06-3691 |
| Eco-730-1 | Bacteria Proteobacteria Gammaproteobacteria Enterobacterales Enterobacteriaceae *Escherichia coli* O91:H14 str. 2009C-3227 |
| Eco-731-1 | Bacteria Proteobacteria Gammaproteobacteria Enterobacterales Enterobacteriaceae *Escherichia coli* O91:H21 str. 2009C-3740 |
| Eco-732-1 | Bacteria Proteobacteria Gammaproteobacteria Enterobacterales Enterobacteriaceae *Escherichia coli* O91:H21 str. 2009C-4646 |
| Eco-733-1 | Bacteria Proteobacteria Gammaproteobacteria Enterobacterales Enterobacteriaceae *Escherichia coli* O91:NM str. 2009C-3745 |
| Eco-734-1 | Bacteria Proteobacteria Gammaproteobacteria Enterobacterales Enterobacteriaceae *Escherichia coli* PCN033 |
| Eco-735-1 | Bacteria Proteobacteria Gammaproteobacteria Enterobacterales Enterobacteriaceae *Escherichia coli* PCN061 |
| Eco-736-1 | Bacteria Proteobacteria Gammaproteobacteria Enterobacterales Enterobacteriaceae *Escherichia coli* RS218 |
| Eco-737-1 | Bacteria Proteobacteria Gammaproteobacteria Enterobacterales Enterobacteriaceae *Escherichia coli* STEC O174:H2 str. 02-04446 |
| Eco-738-1 | Bacteria Proteobacteria Gammaproteobacteria Enterobacterales Enterobacteriaceae *Escherichia coli* STEC O174:H46 str. I-151 |
| Eco-739-1 | Bacteria Proteobacteria Gammaproteobacteria Enterobacterales Enterobacteriaceae *Escherichia coli* STEC O174:H8 str. 02-07607 |
| Eco-751-1 | Bacteria Proteobacteria Gammaproteobacteria Enterobacterales Enterobacteriaceae *Escherichia coli str.* K-12 substr. MC4100 |
| Eco-752-1 to Eco-752-2 | Bacteria Proteobacteria Gammaproteobacteria Enterobacterales Enterobacteriaceae *Escherichia coli str.* K-12 substr. MG1655 |
| Eco-740-1 | Bacteria Proteobacteria Gammaproteobacteria Enterobacterales Enterobacteriaceae *Escherichia coli* UCI 51 |
| Eco-741-1 | Bacteria Proteobacteria Gammaproteobacteria Enterobacterales Enterobacteriaceae *Escherichia coli* UCI 53 |
| Eco-742-1 | Bacteria Proteobacteria Gammaproteobacteria Enterobacterales Enterobacteriaceae *Escherichia coli* UCI 57 |
| Eco-743-1 | Bacteria Proteobacteria Gammaproteobacteria Enterobacterales Enterobacteriaceae *Escherichia coli* UCI 58 |
| Eco-744-1 | Bacteria Proteobacteria Gammaproteobacteria Enterobacterales Enterobacteriaceae *Escherichia coli* UCI 65 |
| Eco-745-1 | Bacteria Proteobacteria Gammaproteobacteria Enterobacterales Enterobacteriaceae *Escherichia coli* UCI 66 |
| Eco-746-1 | Bacteria Proteobacteria Gammaproteobacteria Enterobacterales Enterobacteriaceae *Escherichia coli* VR50 |
| Eco-749-1 | Bacteria Proteobacteria Gammaproteobacteria Enterobacterales Enterobacteriaceae *Escherichia coli* Wa1 |
| Eco-750-1 | Bacteria Proteobacteria Gammaproteobacteria Enterobacterales Enterobacteriaceae *Escherichia coli* Wa2 |
| Eco-747-1 | Bacteria Proteobacteria Gammaproteobacteria Enterobacterales Enterobacteriaceae *Escherichia coli* WC1 |
| Eco-748-1 | Bacteria Proteobacteria Gammaproteobacteria Enterobacterales Enterobacteriaceae *Escherichia coli* WC2 |
| Ema-1-1 | Bacteria Proteobacteria Gammaproteobacteria Enterobacterales Enterobacteriaceae *Escherichia marmotae* |
| Esp-52-1 | Bacteria Proteobacteria Gammaproteobacteria Enterobacterales Enterobacteriaceae *Escherichia sp.* 3_2_53FAA |
| Esp-53-1 | Bacteria Proteobacteria Gammaproteobacteria Enterobacterales Enterobacteriaceae *Escherichia sp.* TW14182 |
| Esp-54-1 | Bacteria Proteobacteria Gammaproteobacteria Enterobacterales Enterobacteriaceae *Escherichia sp.* TW15838 |
| Fhe-1-1 | Bacteria Proteobacteria Gammaproteobacteria Enterobacterales Enterobacteriaceae *Franconibacter helveticus* |
| Fhe-2-1 | Bacteria Proteobacteria Gammaproteobacteria Enterobacterales Enterobacteriaceae *Franconibacter helveticus* 1159 |
| Fhe-3-1 | Bacteria Proteobacteria Gammaproteobacteria Enterobacterales Enterobacteriaceae *Franconibacter helveticus* 513 |
| Fhe-4-1 | Bacteria Proteobacteria Gammaproteobacteria Enterobacterales Enterobacteriaceae *Franconibacter helveticus* LMG 23732 |
| Fpu-1-1 | Bacteria Proteobacteria Gammaproteobacteria Enterobacterales Enterobacteriaceae *Franconibacter pulveris* |
| Fpu-2-1 | Bacteria Proteobacteria Gammaproteobacteria Enterobacterales Enterobacteriaceae *Franconibacter pulveris* 1160 |
| Fpu-3-1 | Bacteria Proteobacteria Gammaproteobacteria Enterobacterales Enterobacteriaceae *Franconibacter pulveris* 601 |
| Fpu-4-1 | Bacteria Proteobacteria Gammaproteobacteria Enterobacterales Enterobacteriaceae *Franconibacter pulveris* DSM 19144 |
| Fpu-5-1 | Bacteria Proteobacteria Gammaproteobacteria Enterobacterales Enterobacteriaceae *Franconibacter pulveris* LMG 24059 |
| Kae-1-1 | Bacteria Proteobacteria Gammaproteobacteria Enterobacterales Enterobacteriaceae *Klebsiella aerogenes* |
| Kae-2-1 | Bacteria Proteobacteria Gammaproteobacteria Enterobacterales Enterobacteriaceae *Klebsiella aerogenes* MGH 61 |
| Kae-3-1 | Bacteria Proteobacteria Gammaproteobacteria Enterobacterales Enterobacteriaceae *Klebsiella aerogenes* MGH 62 |
| Kae-4-1 | Bacteria Proteobacteria Gammaproteobacteria Enterobacterales Enterobacteriaceae *Klebsiella aerogenes* MGH 77 |
| Kae-5-1 | Bacteria Proteobacteria Gammaproteobacteria Enterobacterales Enterobacteriaceae *Klebsiella aerogenes* MGH 78 |
| Kae-6-1 | Bacteria Proteobacteria Gammaproteobacteria Enterobacterales Enterobacteriaceae *Klebsiella aerogenes* UCI 15 |
| Kae-7-1 | Bacteria Proteobacteria Gammaproteobacteria Enterobacterales Enterobacteriaceae *Klebsiella aerogenes* UCI 16 |
| Kae-8-1 | Bacteria Proteobacteria Gammaproteobacteria Enterobacterales Enterobacteriaceae *Klebsiella aerogenes* UCI 27 |
| Kae-9-1 | Bacteria Proteobacteria Gammaproteobacteria Enterobacterales Enterobacteriaceae *Klebsiella aerogenes* UCI 28 |
| Kae-10-1 | Bacteria Proteobacteria Gammaproteobacteria Enterobacterales Enterobacteriaceae *Klebsiella aerogenes* UCI 45 |
| Kae-11-1 | Bacteria Proteobacteria Gammaproteobacteria Enterobacterales Enterobacteriaceae *Klebsiella aerogenes* UCI 46 |
| Kae-12-1 | Bacteria Proteobacteria Gammaproteobacteria Enterobacterales Enterobacteriaceae *Klebsiella aerogenes* UCI 47 |
| Kae-13-1 | Bacteria Proteobacteria Gammaproteobacteria Enterobacterales Enterobacteriaceae *Klebsiella aerogenes* UCI 48 |
| Kcf-1-1 | Bacteria Proteobacteria Gammaproteobacteria Enterobacterales Enterobacteriaceae *Klebsiella cf. planticola* B43 |
| Kmi-1-1 to Kmi-1-2 | Bacteria Proteobacteria Gammaproteobacteria Enterobacterales Enterobacteriaceae *Klebsiella michiganensis* |
| Kmi-2-1 | Bacteria Proteobacteria Gammaproteobacteria Enterobacterales Enterobacteriaceae *Klebsiella michiganensis* H1g |
| Kmi-3-1 | Bacteria Proteobacteria Gammaproteobacteria Enterobacterales Enterobacteriaceae *Klebsiella michiganensis* HKOPL1 |
| Kox-1-1 to Kox-1-4 | Bacteria Proteobacteria Gammaproteobacteria Enterobacterales Enterobacteriaceae *Klebsiella oxytoca* |
| Kox-2-1 | Bacteria Proteobacteria Gammaproteobacteria Enterobacterales Enterobacteriaceae *Klebsiella oxytoca* 09-7231 |

| | |
|---|---|
| Kox-3-1 | Bacteria Proteobacteria Gammaproteobacteria Enterobacterales Enterobacteriaceae *Klebsiella oxytoca* 10-5244 |
| Kox-4-1 to Kox-4-2 | Bacteria Proteobacteria Gammaproteobacteria Enterobacterales Enterobacteriaceae *Klebsiella oxytoca* 10-5248 |
| Kox-5-1 | Bacteria Proteobacteria Gammaproteobacteria Enterobacterales Enterobacteriaceae *Klebsiella oxytoca* 10-5249 |
| Kox-6-1 | Bacteria Proteobacteria Gammaproteobacteria Enterobacterales Enterobacteriaceae *Klebsiella oxytoca* BRL6-2 |
| Kox-7-1 | Bacteria Proteobacteria Gammaproteobacteria Enterobacterales Enterobacteriaceae *Klebsiella oxytoca* G54 |
| Kox-8-1 | Bacteria Proteobacteria Gammaproteobacteria Enterobacterales Enterobacteriaceae *Klebsiella oxytoca* KA-2 |
| Kox-9-1 | Bacteria Proteobacteria Gammaproteobacteria Enterobacterales Enterobacteriaceae *Klebsiella oxytoca* KONIH1 |
| Kox-10-1 | Bacteria Proteobacteria Gammaproteobacteria Enterobacterales Enterobacteriaceae *Klebsiella oxytoca* OK-1 |
| Kpn-1-1 to Kpn-1-9 | Bacteria Proteobacteria Gammaproteobacteria Enterobacterales Enterobacteriaceae *Klebsiella pneumoniae* |
| Kpn-2-1 | Bacteria Proteobacteria Gammaproteobacteria Enterobacterales Enterobacteriaceae *Klebsiella pneumoniae* 11227-1 |
| Kpn-3-1 to Kpn-3-2 | Bacteria Proteobacteria Gammaproteobacteria Enterobacterales Enterobacteriaceae *Klebsiella pneumoniae* 160_1080 |
| Kpn-4-1 | Bacteria Proteobacteria Gammaproteobacteria Enterobacterales Enterobacteriaceae *Klebsiella pneumoniae* 30660/NJST258_1 |
| Kpn-5-1 | Bacteria Proteobacteria Gammaproteobacteria Enterobacterales Enterobacteriaceae *Klebsiella pneumoniae* 30684/NJST258_2 |
| Kpn-6-1 | Bacteria Proteobacteria Gammaproteobacteria Enterobacterales Enterobacteriaceae *Klebsiella pneumoniae* 4006 |
| Kpn-7-1 | Bacteria Proteobacteria Gammaproteobacteria Enterobacterales Enterobacteriaceae *Klebsiella pneumoniae* 4541-2 |
| Kpn-8-1 | Bacteria Proteobacteria Gammaproteobacteria Enterobacterales Enterobacteriaceae *Klebsiella pneumoniae* 500_1420 |
| Kpn-9-1 | Bacteria Proteobacteria Gammaproteobacteria Enterobacterales Enterobacteriaceae *Klebsiella pneumoniae* 7699 |
| Kpn-10-1 | Bacteria Proteobacteria Gammaproteobacteria Enterobacterales Enterobacteriaceae *Klebsiella pneumoniae* ATCC 25955 |
| Kpn-11-1 | Bacteria Proteobacteria Gammaproteobacteria Enterobacterales Enterobacteriaceae *Klebsiella pneumoniae* BIDMC 1 |
| Kpn-12-1 | Bacteria Proteobacteria Gammaproteobacteria Enterobacterales Enterobacteriaceae *Klebsiella pneumoniae* BIDMC 10 |
| Kpn-13-1 | Bacteria Proteobacteria Gammaproteobacteria Enterobacterales Enterobacteriaceae *Klebsiella pneumoniae* BIDMC 11 |
| Kpn-14-1 | Bacteria Proteobacteria Gammaproteobacteria Enterobacterales Enterobacteriaceae *Klebsiella pneumoniae* BIDMC 12A |
| Kpn-15-1 | Bacteria Proteobacteria Gammaproteobacteria Enterobacterales Enterobacteriaceae *Klebsiella pneumoniae* BIDMC 12B |
| Kpn-16-1 | Bacteria Proteobacteria Gammaproteobacteria Enterobacterales Enterobacteriaceae *Klebsiella pneumoniae* BIDMC 13 |
| Kpn-17-1 | Bacteria Proteobacteria Gammaproteobacteria Enterobacterales Enterobacteriaceae *Klebsiella pneumoniae* BIDMC 14 |
| Kpn-18-1 | Bacteria Proteobacteria Gammaproteobacteria Enterobacterales Enterobacteriaceae *Klebsiella pneumoniae* BIDMC 18A |
| Kpn-19-1 | Bacteria Proteobacteria Gammaproteobacteria Enterobacterales Enterobacteriaceae *Klebsiella pneumoniae* BIDMC 18B |
| Kpn-20-1 | Bacteria Proteobacteria Gammaproteobacteria Enterobacterales Enterobacteriaceae *Klebsiella pneumoniae* BIDMC 18D |
| Kpn-21-1 | Bacteria Proteobacteria Gammaproteobacteria Enterobacterales Enterobacteriaceae *Klebsiella pneumoniae* BIDMC 2A |
| Kpn-22-1 | Bacteria Proteobacteria Gammaproteobacteria Enterobacterales Enterobacteriaceae *Klebsiella pneumoniae* BIDMC 31 |
| Kpn-23-1 | Bacteria Proteobacteria Gammaproteobacteria Enterobacterales Enterobacteriaceae *Klebsiella pneumoniae* BIDMC 32 |
| Kpn-24-1 | Bacteria Proteobacteria Gammaproteobacteria Enterobacterales Enterobacteriaceae *Klebsiella pneumoniae* BIDMC 33B |
| Kpn-25-1 | Bacteria Proteobacteria Gammaproteobacteria Enterobacterales Enterobacteriaceae *Klebsiella pneumoniae* BIDMC 34 |
| Kpn-26-1 | Bacteria Proteobacteria Gammaproteobacteria Enterobacterales Enterobacteriaceae *Klebsiella pneumoniae* BIDMC 4 |
| Kpn-27-1 | Bacteria Proteobacteria Gammaproteobacteria Enterobacterales Enterobacteriaceae *Klebsiella pneumoniae* BIDMC 42a |
| Kpn-28-1 | Bacteria Proteobacteria Gammaproteobacteria Enterobacterales Enterobacteriaceae *Klebsiella pneumoniae* BIDMC 42b |
| Kpn-29-1 | Bacteria Proteobacteria Gammaproteobacteria Enterobacterales Enterobacteriaceae *Klebsiella pneumoniae* BIDMC 45 |
| Kpn-30-1 | Bacteria Proteobacteria Gammaproteobacteria Enterobacterales Enterobacteriaceae *Klebsiella pneumoniae* BIDMC 46a |
| Kpn-31-1 | Bacteria Proteobacteria Gammaproteobacteria Enterobacterales Enterobacteriaceae *Klebsiella pneumoniae* BIDMC 46b |
| Kpn-32-1 | Bacteria Proteobacteria Gammaproteobacteria Enterobacterales Enterobacteriaceae *Klebsiella pneumoniae* BIDMC 47 |
| Kpn-33-1 | Bacteria Proteobacteria Gammaproteobacteria Enterobacterales Enterobacteriaceae *Klebsiella pneumoniae* BIDMC 48 |
| Kpn-34-1 | Bacteria Proteobacteria Gammaproteobacteria Enterobacterales Enterobacteriaceae *Klebsiella pneumoniae* BIDMC 5 |
| Kpn-35-1 | Bacteria Proteobacteria Gammaproteobacteria Enterobacterales Enterobacteriaceae *Klebsiella pneumoniae* BIDMC 51 |
| Kpn-36-1 | Bacteria Proteobacteria Gammaproteobacteria Enterobacterales Enterobacteriaceae *Klebsiella pneumoniae* BIDMC 52 |
| Kpn-37-1 | Bacteria Proteobacteria Gammaproteobacteria Enterobacterales Enterobacteriaceae *Klebsiella pneumoniae* BIDMC 53 |
| Kpn-38-1 | Bacteria Proteobacteria Gammaproteobacteria Enterobacterales Enterobacteriaceae *Klebsiella pneumoniae* BIDMC 54 |
| Kpn-39-1 | Bacteria Proteobacteria Gammaproteobacteria Enterobacterales Enterobacteriaceae *Klebsiella pneumoniae* BIDMC 55 |
| Kpn-40-1 | Bacteria Proteobacteria Gammaproteobacteria Enterobacterales Enterobacteriaceae *Klebsiella pneumoniae* BIDMC 60 |
| Kpn-41-1 | Bacteria Proteobacteria Gammaproteobacteria Enterobacterales Enterobacteriaceae *Klebsiella pneumoniae* BIDMC 68 |
| Kpn-42-1 | Bacteria Proteobacteria Gammaproteobacteria Enterobacterales Enterobacteriaceae *Klebsiella pneumoniae* BIDMC 69 |
| Kpn-43-1 | Bacteria Proteobacteria Gammaproteobacteria Enterobacterales Enterobacteriaceae *Klebsiella pneumoniae* BIDMC 7A |
| Kpn-44-1 | Bacteria Proteobacteria Gammaproteobacteria Enterobacterales Enterobacteriaceae *Klebsiella pneumoniae* BIDMC 7B |
| Kpn-45-1 | Bacteria Proteobacteria Gammaproteobacteria Enterobacterales Enterobacteriaceae *Klebsiella pneumoniae* BWH 15 |
| Kpn-46-1 | Bacteria Proteobacteria Gammaproteobacteria Enterobacterales Enterobacteriaceae *Klebsiella pneumoniae* BWH 2 |
| Kpn-47-1 | Bacteria Proteobacteria Gammaproteobacteria Enterobacterales Enterobacteriaceae *Klebsiella pneumoniae* BWH 22 |
| Kpn-48-1 | Bacteria Proteobacteria Gammaproteobacteria Enterobacterales Enterobacteriaceae *Klebsiella pneumoniae* BWH 36 |
| Kpn-49-1 | Bacteria Proteobacteria Gammaproteobacteria Enterobacterales Enterobacteriaceae *Klebsiella pneumoniae* BWH 41 |
| Kpn-50-1 | Bacteria Proteobacteria Gammaproteobacteria Enterobacterales Enterobacteriaceae *Klebsiella pneumoniae* BWH 45 |
| Kpn-51-1 | Bacteria Proteobacteria Gammaproteobacteria Enterobacterales Enterobacteriaceae *Klebsiella pneumoniae* BWH 46 |
| Kpn-52-1 | Bacteria Proteobacteria Gammaproteobacteria Enterobacterales Enterobacteriaceae *Klebsiella pneumoniae* BWH 47 |
| Kpn-53-1 | Bacteria Proteobacteria Gammaproteobacteria Enterobacterales Enterobacteriaceae *Klebsiella pneumoniae* BWH 48 |
| Kpn-54-1 | Bacteria Proteobacteria Gammaproteobacteria Enterobacterales Enterobacteriaceae *Klebsiella pneumoniae* CHS 02 |
| Kpn-55-1 | Bacteria Proteobacteria Gammaproteobacteria Enterobacterales Enterobacteriaceae *Klebsiella pneumoniae* CHS 03 |
| Kpn-56-1 | Bacteria Proteobacteria Gammaproteobacteria Enterobacterales Enterobacteriaceae *Klebsiella pneumoniae* CHS 05 |
| Kpn-57-1 | Bacteria Proteobacteria Gammaproteobacteria Enterobacterales Enterobacteriaceae *Klebsiella pneumoniae* CHS 06 |

| | |
|---|---|
| Kpn-58-1 | Bacteria Proteobacteria Gammaproteobacteria Enterobacterales Enterobacteriaceae *Klebsiella pneumoniae* CHS 07 |
| Kpn-59-1 | Bacteria Proteobacteria Gammaproteobacteria Enterobacterales Enterobacteriaceae *Klebsiella pneumoniae* CHS 08 |
| Kpn-60-1 | Bacteria Proteobacteria Gammaproteobacteria Enterobacterales Enterobacteriaceae *Klebsiella pneumoniae* CHS 09 |
| Kpn-61-1 | Bacteria Proteobacteria Gammaproteobacteria Enterobacterales Enterobacteriaceae *Klebsiella pneumoniae* CHS 11 |
| Kpn-62-1 | Bacteria Proteobacteria Gammaproteobacteria Enterobacterales Enterobacteriaceae *Klebsiella pneumoniae* CHS 12 |
| Kpn-63-1 | Bacteria Proteobacteria Gammaproteobacteria Enterobacterales Enterobacteriaceae *Klebsiella pneumoniae* CHS 13 |
| Kpn-64-1 | Bacteria Proteobacteria Gammaproteobacteria Enterobacterales Enterobacteriaceae *Klebsiella pneumoniae* CHS 14 |
| Kpn-65-1 | Bacteria Proteobacteria Gammaproteobacteria Enterobacterales Enterobacteriaceae *Klebsiella pneumoniae* CHS 16 |
| Kpn-66-1 | Bacteria Proteobacteria Gammaproteobacteria Enterobacterales Enterobacteriaceae *Klebsiella pneumoniae* CHS 17 |
| Kpn-67-1 | Bacteria Proteobacteria Gammaproteobacteria Enterobacterales Enterobacteriaceae *Klebsiella pneumoniae* CHS 18 |
| Kpn-68-1 | Bacteria Proteobacteria Gammaproteobacteria Enterobacterales Enterobacteriaceae *Klebsiella pneumoniae* CHS 19 |
| Kpn-69-1 | Bacteria Proteobacteria Gammaproteobacteria Enterobacterales Enterobacteriaceae *Klebsiella pneumoniae* CHS 20 |
| Kpn-70-1 | Bacteria Proteobacteria Gammaproteobacteria Enterobacterales Enterobacteriaceae *Klebsiella pneumoniae* CHS 21 |
| Kpn-71-1 | Bacteria Proteobacteria Gammaproteobacteria Enterobacterales Enterobacteriaceae *Klebsiella pneumoniae* CHS 22 |
| Kpn-72-1 | Bacteria Proteobacteria Gammaproteobacteria Enterobacterales Enterobacteriaceae *Klebsiella pneumoniae* CHS 23 |
| Kpn-73-1 | Bacteria Proteobacteria Gammaproteobacteria Enterobacterales Enterobacteriaceae *Klebsiella pneumoniae* CHS 24 |
| Kpn-74-1 | Bacteria Proteobacteria Gammaproteobacteria Enterobacterales Enterobacteriaceae *Klebsiella pneumoniae* CHS 25 |
| Kpn-75-1 | Bacteria Proteobacteria Gammaproteobacteria Enterobacterales Enterobacteriaceae *Klebsiella pneumoniae* CHS 26 |
| Kpn-76-1 | Bacteria Proteobacteria Gammaproteobacteria Enterobacterales Enterobacteriaceae *Klebsiella pneumoniae* CHS 27 |
| Kpn-77-1 | Bacteria Proteobacteria Gammaproteobacteria Enterobacterales Enterobacteriaceae *Klebsiella pneumoniae* CHS 28 |
| Kpn-78-1 | Bacteria Proteobacteria Gammaproteobacteria Enterobacterales Enterobacteriaceae *Klebsiella pneumoniae* CHS 29 |
| Kpn-79-1 | Bacteria Proteobacteria Gammaproteobacteria Enterobacterales Enterobacteriaceae *Klebsiella pneumoniae* CHS 30 |
| Kpn-80-1 | Bacteria Proteobacteria Gammaproteobacteria Enterobacterales Enterobacteriaceae *Klebsiella pneumoniae* CHS 31 |
| Kpn-81-1 | Bacteria Proteobacteria Gammaproteobacteria Enterobacterales Enterobacteriaceae *Klebsiella pneumoniae* CHS 32 |
| Kpn-82-1 | Bacteria Proteobacteria Gammaproteobacteria Enterobacterales Enterobacteriaceae *Klebsiella pneumoniae* CHS 33 |
| Kpn-83-1 | Bacteria Proteobacteria Gammaproteobacteria Enterobacterales Enterobacteriaceae *Klebsiella pneumoniae* CHS 34 |
| Kpn-84-1 | Bacteria Proteobacteria Gammaproteobacteria Enterobacterales Enterobacteriaceae *Klebsiella pneumoniae* CHS 35 |
| Kpn-85-1 | Bacteria Proteobacteria Gammaproteobacteria Enterobacterales Enterobacteriaceae *Klebsiella pneumoniae* CHS 36 |
| Kpn-86-1 | Bacteria Proteobacteria Gammaproteobacteria Enterobacterales Enterobacteriaceae *Klebsiella pneumoniae* CHS 37 |
| Kpn-87-1 | Bacteria Proteobacteria Gammaproteobacteria Enterobacterales Enterobacteriaceae *Klebsiella pneumoniae* CHS 38 |
| Kpn-88-1 | Bacteria Proteobacteria Gammaproteobacteria Enterobacterales Enterobacteriaceae *Klebsiella pneumoniae* CHS 39 |
| Kpn-89-1 | Bacteria Proteobacteria Gammaproteobacteria Enterobacterales Enterobacteriaceae *Klebsiella pneumoniae* CHS 40 |
| Kpn-90-1 | Bacteria Proteobacteria Gammaproteobacteria Enterobacterales Enterobacteriaceae *Klebsiella pneumoniae* CHS 41 |
| Kpn-91-1 | Bacteria Proteobacteria Gammaproteobacteria Enterobacterales Enterobacteriaceae *Klebsiella pneumoniae* CHS 42 |
| Kpn-92-1 | Bacteria Proteobacteria Gammaproteobacteria Enterobacterales Enterobacteriaceae *Klebsiella pneumoniae* CHS 43 |
| Kpn-93-1 | Bacteria Proteobacteria Gammaproteobacteria Enterobacterales Enterobacteriaceae *Klebsiella pneumoniae* CHS 44 |
| Kpn-94-1 | Bacteria Proteobacteria Gammaproteobacteria Enterobacterales Enterobacteriaceae *Klebsiella pneumoniae* CHS 45 |
| Kpn-95-1 | Bacteria Proteobacteria Gammaproteobacteria Enterobacterales Enterobacteriaceae *Klebsiella pneumoniae* CHS 46 |
| Kpn-96-1 | Bacteria Proteobacteria Gammaproteobacteria Enterobacterales Enterobacteriaceae *Klebsiella pneumoniae* CHS 47 |
| Kpn-97-1 | Bacteria Proteobacteria Gammaproteobacteria Enterobacterales Enterobacteriaceae *Klebsiella pneumoniae* CHS 48 |
| Kpn-98-1 | Bacteria Proteobacteria Gammaproteobacteria Enterobacterales Enterobacteriaceae *Klebsiella pneumoniae* CHS 49 |
| Kpn-99-1 | Bacteria Proteobacteria Gammaproteobacteria Enterobacterales Enterobacteriaceae *Klebsiella pneumoniae* CHS 50 |
| Kpn-100-1 | Bacteria Proteobacteria Gammaproteobacteria Enterobacterales Enterobacteriaceae *Klebsiella pneumoniae* CHS 51 |
| Kpn-101-1 | Bacteria Proteobacteria Gammaproteobacteria Enterobacterales Enterobacteriaceae *Klebsiella pneumoniae* CHS 52 |
| Kpn-102-1 | Bacteria Proteobacteria Gammaproteobacteria Enterobacterales Enterobacteriaceae *Klebsiella pneumoniae* CHS 53 |
| Kpn-103-1 | Bacteria Proteobacteria Gammaproteobacteria Enterobacterales Enterobacteriaceae *Klebsiella pneumoniae* CHS 54 |
| Kpn-104-1 | Bacteria Proteobacteria Gammaproteobacteria Enterobacterales Enterobacteriaceae *Klebsiella pneumoniae* CHS 55 |
| Kpn-105-1 | Bacteria Proteobacteria Gammaproteobacteria Enterobacterales Enterobacteriaceae *Klebsiella pneumoniae* CHS 56 |
| Kpn-106-1 | Bacteria Proteobacteria Gammaproteobacteria Enterobacterales Enterobacteriaceae *Klebsiella pneumoniae* CHS 57 |
| Kpn-107-1 | Bacteria Proteobacteria Gammaproteobacteria Enterobacterales Enterobacteriaceae *Klebsiella pneumoniae* CHS 58 |
| Kpn-108-1 | Bacteria Proteobacteria Gammaproteobacteria Enterobacterales Enterobacteriaceae *Klebsiella pneumoniae* CHS 59 |
| Kpn-109-1 | Bacteria Proteobacteria Gammaproteobacteria Enterobacterales Enterobacteriaceae *Klebsiella pneumoniae* CHS 60 |
| Kpn-110-1 | Bacteria Proteobacteria Gammaproteobacteria Enterobacterales Enterobacteriaceae *Klebsiella pneumoniae* CHS 61 |
| Kpn-111-1 | Bacteria Proteobacteria Gammaproteobacteria Enterobacterales Enterobacteriaceae *Klebsiella pneumoniae* CHS 62 |
| Kpn-112-1 | Bacteria Proteobacteria Gammaproteobacteria Enterobacterales Enterobacteriaceae *Klebsiella pneumoniae* CHS 63 |
| Kpn-113-1 | Bacteria Proteobacteria Gammaproteobacteria Enterobacterales Enterobacteriaceae *Klebsiella pneumoniae* CHS 64 |
| Kpn-114-1 | Bacteria Proteobacteria Gammaproteobacteria Enterobacterales Enterobacteriaceae *Klebsiella pneumoniae* CHS 65 |
| Kpn-115-1 | Bacteria Proteobacteria Gammaproteobacteria Enterobacterales Enterobacteriaceae *Klebsiella pneumoniae* CHS 66 |
| Kpn-116-1 | Bacteria Proteobacteria Gammaproteobacteria Enterobacterales Enterobacteriaceae *Klebsiella pneumoniae* CHS 67 |
| Kpn-117-1 | Bacteria Proteobacteria Gammaproteobacteria Enterobacterales Enterobacteriaceae *Klebsiella pneumoniae* CHS 70 |
| Kpn-118-1 | Bacteria Proteobacteria Gammaproteobacteria Enterobacterales Enterobacteriaceae *Klebsiella pneumoniae* CHS 71 |
| Kpn-119-1 | Bacteria Proteobacteria Gammaproteobacteria Enterobacterales Enterobacteriaceae *Klebsiella pneumoniae* CHS 72 |
| Kpn-120-1 | Bacteria Proteobacteria Gammaproteobacteria Enterobacterales Enterobacteriaceae *Klebsiella pneumoniae* CHS 73 |
| Kpn-121-1 | Bacteria Proteobacteria Gammaproteobacteria Enterobacterales Enterobacteriaceae *Klebsiella pneumoniae* CHS 74 |
| Kpn-122-1 | Bacteria Proteobacteria Gammaproteobacteria Enterobacterales Enterobacteriaceae *Klebsiella pneumoniae* CHS 75 |

| | |
|---|---|
| Kpn-123-1 | Bacteria Proteobacteria Gammaproteobacteria Enterobacterales Enterobacteriaceae *Klebsiella pneumoniae* CHS 76 |
| Kpn-124-1 | Bacteria Proteobacteria Gammaproteobacteria Enterobacterales Enterobacteriaceae *Klebsiella pneumoniae* CHS 80 |
| Kpn-125-1 | Bacteria Proteobacteria Gammaproteobacteria Enterobacterales Enterobacteriaceae *Klebsiella pneumoniae* DMC1097 |
| Kpn-126-1 | Bacteria Proteobacteria Gammaproteobacteria Enterobacterales Enterobacteriaceae *Klebsiella pneumoniae* EGD-HP19-C |
| Kpn-127-1 | Bacteria Proteobacteria Gammaproteobacteria Enterobacterales Enterobacteriaceae *Klebsiella pneumoniae* G5-2 |
| Kpn-128-1 | Bacteria Proteobacteria Gammaproteobacteria Enterobacterales Enterobacteriaceae *Klebsiella pneumoniae* HE12 |
| Kpn-129-1 | Bacteria Proteobacteria Gammaproteobacteria Enterobacterales Enterobacteriaceae *Klebsiella pneumoniae* HK787 |
| Kpn-130-1 | Bacteria Proteobacteria Gammaproteobacteria Enterobacterales Enterobacteriaceae *Klebsiella pneumoniae* JT4 |
| Kpn-136-1 | Bacteria Proteobacteria Gammaproteobacteria Enterobacterales Enterobacteriaceae *Klebsiella pneumoniae* Kb140 |
| Kpn-137-1 | Bacteria Proteobacteria Gammaproteobacteria Enterobacterales Enterobacteriaceae *Klebsiella pneumoniae* Kb677 |
| Kpn-131-1 | Bacteria Proteobacteria Gammaproteobacteria Enterobacterales Enterobacteriaceae *Klebsiella pneumoniae* KP1-I |
| Kpn-132-1 | Bacteria Proteobacteria Gammaproteobacteria Enterobacterales Enterobacteriaceae *Klebsiella pneumoniae* KP2-R |
| Kpn-133-1 | Bacteria Proteobacteria Gammaproteobacteria Enterobacterales Enterobacteriaceae *Klebsiella pneumoniae* KP3-S |
| Kpn-134-1 | Bacteria Proteobacteria Gammaproteobacteria Enterobacterales Enterobacteriaceae *Klebsiella pneumoniae* KP4-R |
| Kpn-135-1 | Bacteria Proteobacteria Gammaproteobacteria Enterobacterales Enterobacteriaceae *Klebsiella pneumoniae* KP5-R |
| Kpn-138-1 | Bacteria Proteobacteria Gammaproteobacteria Enterobacterales Enterobacteriaceae *Klebsiella pneumoniae* LAU-KP1 |
| Kpn-139-1 | Bacteria Proteobacteria Gammaproteobacteria Enterobacterales Enterobacteriaceae *Klebsiella pneumoniae* LCT-KP182 |
| Kpn-140-1 | Bacteria Proteobacteria Gammaproteobacteria Enterobacterales Enterobacteriaceae *Klebsiella pneumoniae* LCT-KP289 |
| Kpn-141-1 | Bacteria Proteobacteria Gammaproteobacteria Enterobacterales Enterobacteriaceae *Klebsiella pneumoniae* MGH 29 |
| Kpn-142-1 | Bacteria Proteobacteria Gammaproteobacteria Enterobacterales Enterobacteriaceae *Klebsiella pneumoniae* MGH 35 |
| Kpn-143-1 | Bacteria Proteobacteria Gammaproteobacteria Enterobacterales Enterobacteriaceae *Klebsiella pneumoniae* MGH 39 |
| Kpn-144-1 | Bacteria Proteobacteria Gammaproteobacteria Enterobacterales Enterobacteriaceae *Klebsiella pneumoniae* MGH 43 |
| Kpn-145-1 | Bacteria Proteobacteria Gammaproteobacteria Enterobacterales Enterobacteriaceae *Klebsiella pneumoniae* MGH 45 |
| Kpn-146-1 | Bacteria Proteobacteria Gammaproteobacteria Enterobacterales Enterobacteriaceae *Klebsiella pneumoniae* MGH 47 |
| Kpn-147-1 | Bacteria Proteobacteria Gammaproteobacteria Enterobacterales Enterobacteriaceae *Klebsiella pneumoniae* MGH 51 |
| Kpn-148-1 | Bacteria Proteobacteria Gammaproteobacteria Enterobacterales Enterobacteriaceae *Klebsiella pneumoniae* MGH 52 |
| Kpn-149-1 | Bacteria Proteobacteria Gammaproteobacteria Enterobacterales Enterobacteriaceae *Klebsiella pneumoniae* MGH 60 |
| Kpn-150-1 | Bacteria Proteobacteria Gammaproteobacteria Enterobacterales Enterobacteriaceae *Klebsiella pneumoniae* MGH 63 |
| Kpn-151-1 | Bacteria Proteobacteria Gammaproteobacteria Enterobacterales Enterobacteriaceae *Klebsiella pneumoniae* MGH 64 |
| Kpn-152-1 | Bacteria Proteobacteria Gammaproteobacteria Enterobacterales Enterobacteriaceae *Klebsiella pneumoniae* MGH 65 |
| Kpn-153-1 | Bacteria Proteobacteria Gammaproteobacteria Enterobacterales Enterobacteriaceae *Klebsiella pneumoniae* MGH 66 |
| Kpn-154-1 | Bacteria Proteobacteria Gammaproteobacteria Enterobacterales Enterobacteriaceae *Klebsiella pneumoniae* MGH 67 |
| Kpn-155-1 | Bacteria Proteobacteria Gammaproteobacteria Enterobacterales Enterobacteriaceae *Klebsiella pneumoniae* MGH 69 |
| Kpn-156-1 | Bacteria Proteobacteria Gammaproteobacteria Enterobacterales Enterobacteriaceae *Klebsiella pneumoniae* MGH 70 |
| Kpn-157-1 | Bacteria Proteobacteria Gammaproteobacteria Enterobacterales Enterobacteriaceae *Klebsiella pneumoniae* MGH 71 |
| Kpn-158-1 | Bacteria Proteobacteria Gammaproteobacteria Enterobacterales Enterobacteriaceae *Klebsiella pneumoniae* MGH 72 |
| Kpn-159-1 | Bacteria Proteobacteria Gammaproteobacteria Enterobacterales Enterobacteriaceae *Klebsiella pneumoniae* MGH 73 |
| Kpn-160-1 | Bacteria Proteobacteria Gammaproteobacteria Enterobacterales Enterobacteriaceae *Klebsiella pneumoniae* MGH 74 |
| Kpn-161-1 | Bacteria Proteobacteria Gammaproteobacteria Enterobacterales Enterobacteriaceae *Klebsiella pneumoniae* MGH 75 |
| Kpn-162-1 | Bacteria Proteobacteria Gammaproteobacteria Enterobacterales Enterobacteriaceae *Klebsiella pneumoniae* MGH 79 |
| Kpn-163-1 | Bacteria Proteobacteria Gammaproteobacteria Enterobacterales Enterobacteriaceae *Klebsiella pneumoniae* MRSN 1319 |
| Kpn-164-1 | Bacteria Proteobacteria Gammaproteobacteria Enterobacterales Enterobacteriaceae *Klebsiella pneumoniae* MRSN 2404 |
| Kpn-165-1 | Bacteria Proteobacteria Gammaproteobacteria Enterobacterales Enterobacteriaceae *Klebsiella pneumoniae* MRSN 3562 |
| Kpn-166-1 | Bacteria Proteobacteria Gammaproteobacteria Enterobacterales Enterobacteriaceae *Klebsiella pneumoniae* MRSN 3852 |
| Kpn-167-1 | Bacteria Proteobacteria Gammaproteobacteria Enterobacterales Enterobacteriaceae *Klebsiella pneumoniae* MRSN 6902 |
| Kpn-168-1 | Bacteria Proteobacteria Gammaproteobacteria Enterobacterales Enterobacteriaceae *Klebsiella pneumoniae* NB60 |
| Kpn-169-1 | Bacteria Proteobacteria Gammaproteobacteria Enterobacterales Enterobacteriaceae *Klebsiella pneumoniae* PR04 |
| Kpn-205-1 to Kpn-205-4 | Bacteria Proteobacteria Gammaproteobacteria Enterobacterales Enterobacteriaceae *Klebsiella pneumoniae subsp. ozaenae* |
| Kpn-206-1 to Kpn-206-7 | Bacteria Proteobacteria Gammaproteobacteria Enterobacterales Enterobacteriaceae *Klebsiella pneumoniae subsp. pneumoniae* |
| Kpn-207-1 | Bacteria Proteobacteria Gammaproteobacteria Enterobacterales Enterobacteriaceae *Klebsiella pneumoniae subsp. pneumoniae* 1158 |
| Kpn-208-1 | Bacteria Proteobacteria Gammaproteobacteria Enterobacterales Enterobacteriaceae *Klebsiella pneumoniae subsp. pneumoniae* BJ1-GA |
| Kpn-209-1 | Bacteria Proteobacteria Gammaproteobacteria Enterobacterales Enterobacteriaceae *Klebsiella pneumoniae subsp. pneumoniae* KKBO-1 |
| Kpn-210-1 | Bacteria Proteobacteria Gammaproteobacteria Enterobacterales Enterobacteriaceae *Klebsiella pneumoniae subsp. pneumoniae* KKBO-4 |
| Kpn-218-1 | Bacteria Proteobacteria Gammaproteobacteria Enterobacterales Enterobacteriaceae *Klebsiella pneumoniae subsp. pneumoniae* Kp13 |
| Kpn-211-1 | Bacteria Proteobacteria Gammaproteobacteria Enterobacterales Enterobacteriaceae *Klebsiella pneumoniae subsp. pneumoniae* KPB-1 |
| Kpn-212-1 | Bacteria Proteobacteria Gammaproteobacteria Enterobacterales Enterobacteriaceae *Klebsiella pneumoniae subsp. pneumoniae* KPB-2 |
| Kpn-213-1 | Bacteria Proteobacteria Gammaproteobacteria Enterobacterales Enterobacteriaceae *Klebsiella pneumoniae subsp. pneumoniae* KPNIH1 |
| Kpn-214-1 | Bacteria Proteobacteria Gammaproteobacteria Enterobacterales Enterobacteriaceae *Klebsiella pneumoniae subsp. pneumoniae* KPNIH10 |
| Kpn-215-1 | Bacteria Proteobacteria Gammaproteobacteria Enterobacterales Enterobacteriaceae *Klebsiella pneumoniae subsp. pneumoniae* KPNIH24 |
| Kpn-216-1 | Bacteria Proteobacteria Gammaproteobacteria Enterobacterales Enterobacteriaceae *Klebsiella pneumoniae subsp. pneumoniae* KPNIH27 |
| Kpn-219-1 | Bacteria Proteobacteria Gammaproteobacteria Enterobacterales Enterobacteriaceae *Klebsiella pneumoniae subsp. pneumoniae* KpQ15 |
| Kpn-220-1 | Bacteria Proteobacteria Gammaproteobacteria Enterobacterales Enterobacteriaceae *Klebsiella pneumoniae subsp. pneumoniae* KpQ24 |
| Kpn-217-1 | Bacteria Proteobacteria Gammaproteobacteria Enterobacterales Enterobacteriaceae *Klebsiella pneumoniae subsp. pneumoniae* KPR0928 |
| Kpn-221-1 | Bacteria Proteobacteria Gammaproteobacteria Enterobacterales Enterobacteriaceae *Klebsiella pneumoniae subsp. pneumoniae* LCT-KP214 |
| Kpn-222-1 | Bacteria Proteobacteria Gammaproteobacteria Enterobacterales Enterobacteriaceae *Klebsiella pneumoniae subsp. pneumoniae* PittNDM01 |

| | |
|---|---|
| Kpn-223-1 | Bacteria Proteobacteria Gammaproteobacteria Enterobacterales Enterobacteriaceae *Klebsiella pneumoniae subsp. pneumoniae* SA1 |
| Kpn-224-1 | Bacteria Proteobacteria Gammaproteobacteria Enterobacterales Enterobacteriaceae *Klebsiella pneumoniae subsp. pneumoniae* T69 |
| Kpn-225-1 | Bacteria Proteobacteria Gammaproteobacteria Enterobacterales Enterobacteriaceae *Klebsiella pneumoniae subsp. rhinoscleromatis* SB3432 |
| Kpn-170-1 | Bacteria Proteobacteria Gammaproteobacteria Enterobacterales Enterobacteriaceae *Klebsiella pneumoniae* T2-1-1 |
| Kpn-171-1 | Bacteria Proteobacteria Gammaproteobacteria Enterobacterales Enterobacteriaceae *Klebsiella pneumoniae* T2-1-2 |
| Kpn-172-1 | Bacteria Proteobacteria Gammaproteobacteria Enterobacterales Enterobacteriaceae *Klebsiella pneumoniae* UCI 17 |
| Kpn-173-1 | Bacteria Proteobacteria Gammaproteobacteria Enterobacterales Enterobacteriaceae *Klebsiella pneumoniae* UCI 19 |
| Kpn-174-1 | Bacteria Proteobacteria Gammaproteobacteria Enterobacterales Enterobacteriaceae *Klebsiella pneumoniae* UCI 20 |
| Kpn-175-1 | Bacteria Proteobacteria Gammaproteobacteria Enterobacterales Enterobacteriaceae *Klebsiella pneumoniae* UCI 21 |
| Kpn-176-1 | Bacteria Proteobacteria Gammaproteobacteria Enterobacterales Enterobacteriaceae *Klebsiella pneumoniae* UCI 22 |
| Kpn-177-1 | Bacteria Proteobacteria Gammaproteobacteria Enterobacterales Enterobacteriaceae *Klebsiella pneumoniae* UCI 25 |
| Kpn-178-1 | Bacteria Proteobacteria Gammaproteobacteria Enterobacterales Enterobacteriaceae *Klebsiella pneumoniae* UCI 26 |
| Kpn-179-1 | Bacteria Proteobacteria Gammaproteobacteria Enterobacterales Enterobacteriaceae *Klebsiella pneumoniae* UCI 33 |
| Kpn-180-1 | Bacteria Proteobacteria Gammaproteobacteria Enterobacterales Enterobacteriaceae *Klebsiella pneumoniae* UCI 34 |
| Kpn-181-1 | Bacteria Proteobacteria Gammaproteobacteria Enterobacterales Enterobacteriaceae *Klebsiella pneumoniae* UCI 37 |
| Kpn-182-1 | Bacteria Proteobacteria Gammaproteobacteria Enterobacterales Enterobacteriaceae *Klebsiella pneumoniae* UCI 38 |
| Kpn-183-1 | Bacteria Proteobacteria Gammaproteobacteria Enterobacterales Enterobacteriaceae *Klebsiella pneumoniae* UCI 41 |
| Kpn-184-1 | Bacteria Proteobacteria Gammaproteobacteria Enterobacterales Enterobacteriaceae *Klebsiella pneumoniae* UCI 42 |
| Kpn-185-1 | Bacteria Proteobacteria Gammaproteobacteria Enterobacterales Enterobacteriaceae *Klebsiella pneumoniae* UCI 43 |
| Kpn-186-1 | Bacteria Proteobacteria Gammaproteobacteria Enterobacterales Enterobacteriaceae *Klebsiella pneumoniae* UCI 44 |
| Kpn-187-1 | Bacteria Proteobacteria Gammaproteobacteria Enterobacterales Enterobacteriaceae *Klebsiella pneumoniae* UCI 55 |
| Kpn-188-1 | Bacteria Proteobacteria Gammaproteobacteria Enterobacterales Enterobacteriaceae *Klebsiella pneumoniae* UCI 56 |
| Kpn-189-1 | Bacteria Proteobacteria Gammaproteobacteria Enterobacterales Enterobacteriaceae *Klebsiella pneumoniae* UCI 59 |
| Kpn-190-1 | Bacteria Proteobacteria Gammaproteobacteria Enterobacterales Enterobacteriaceae *Klebsiella pneumoniae* UCI 60 |
| Kpn-191-1 | Bacteria Proteobacteria Gammaproteobacteria Enterobacterales Enterobacteriaceae *Klebsiella pneumoniae* UCI 61 |
| Kpn-192-1 | Bacteria Proteobacteria Gammaproteobacteria Enterobacterales Enterobacteriaceae *Klebsiella pneumoniae* UCI 62 |
| Kpn-193-1 | Bacteria Proteobacteria Gammaproteobacteria Enterobacterales Enterobacteriaceae *Klebsiella pneumoniae* UCI 63 |
| Kpn-194-1 | Bacteria Proteobacteria Gammaproteobacteria Enterobacterales Enterobacteriaceae *Klebsiella pneumoniae* UCI 64 |
| Kpn-195-1 | Bacteria Proteobacteria Gammaproteobacteria Enterobacterales Enterobacteriaceae *Klebsiella pneumoniae* UCI 67 |
| Kpn-196-1 | Bacteria Proteobacteria Gammaproteobacteria Enterobacterales Enterobacteriaceae *Klebsiella pneumoniae* UCI 68 |
| Kpn-197-1 | Bacteria Proteobacteria Gammaproteobacteria Enterobacterales Enterobacteriaceae *Klebsiella pneumoniae* UCICRE 1 |
| Kpn-198-1 | Bacteria Proteobacteria Gammaproteobacteria Enterobacterales Enterobacteriaceae *Klebsiella pneumoniae* UCICRE 13 |
| Kpn-199-1 to Kpn-199-2 | Bacteria Proteobacteria Gammaproteobacteria Enterobacterales Enterobacteriaceae *Klebsiella pneumoniae* UHKPC 52 |
| Kpn-200-1 to Kpn-200-2 | Bacteria Proteobacteria Gammaproteobacteria Enterobacterales Enterobacteriaceae *Klebsiella pneumoniae* UHKPC05 |
| Kpn-201-1 | Bacteria Proteobacteria Gammaproteobacteria Enterobacterales Enterobacteriaceae *Klebsiella pneumoniae* UHKPC07 |
| Kpn-202-1 | Bacteria Proteobacteria Gammaproteobacteria Enterobacterales Enterobacteriaceae *Klebsiella pneumoniae* UHKPC33 |
| Kpn-203-1 to Kpn-203-2 | Bacteria Proteobacteria Gammaproteobacteria Enterobacterales Enterobacteriaceae *Klebsiella pneumoniae* UHKPC45 |
| Kpn-204-1 to Kpn-204-2 | Bacteria Proteobacteria Gammaproteobacteria Enterobacterales Enterobacteriaceae *Klebsiella pneumoniae* VAKPC278 |
| Kqu-1-1 | Bacteria Proteobacteria Gammaproteobacteria Enterobacterales Enterobacteriaceae *Klebsiella quasipneumoniae subsp. quasipneumoniae* |
| Kqu-2-1 to Kqu-2-4 | Bacteria Proteobacteria Gammaproteobacteria Enterobacterales Enterobacteriaceae *Klebsiella quasipneumoniae subsp. similipneumoniae* |
| Ksp-1-1 | Bacteria Proteobacteria Gammaproteobacteria Enterobacterales Enterobacteriaceae *Klebsiella sp.* 10982 |
| Ksp-2-1 | Bacteria Proteobacteria Gammaproteobacteria Enterobacterales Enterobacteriaceae *Klebsiella sp.* AS10 |
| Ksp-3-1 | Bacteria Proteobacteria Gammaproteobacteria Enterobacterales Enterobacteriaceae *Klebsiella sp.* RIT-PI-d |
| Kva-1-1 to Kva-1-7 | Bacteria Proteobacteria Gammaproteobacteria Enterobacterales Enterobacteriaceae *Klebsiella variicola* |
| Kas-1-1 | Bacteria Proteobacteria Gammaproteobacteria Enterobacterales Enterobacteriaceae *Kluyvera ascorbata* ATCC 33433 |
| Kcr-1-1 | Bacteria Proteobacteria Gammaproteobacteria Enterobacterales Enterobacteriaceae *Kluyvera cryocrescens* |
| Kor-1-1 | Bacteria Proteobacteria Gammaproteobacteria Enterobacterales Enterobacteriaceae *Kosakonia oryzae* |
| Kra-1-1 | Bacteria Proteobacteria Gammaproteobacteria Enterobacterales Enterobacteriaceae *Kosakonia radicincitans* UMEnt01/12 |
| Kra-2-1 | Bacteria Proteobacteria Gammaproteobacteria Enterobacterales Enterobacteriaceae *Kosakonia radicincitans* YD4 |
| Lad-1-1 to Lad-1-2 | Bacteria Proteobacteria Gammaproteobacteria Enterobacterales Enterobacteriaceae *Leclercia adecarboxylata* |
| Lad-2-1 | Bacteria Proteobacteria Gammaproteobacteria Enterobacterales Enterobacteriaceae *Leclercia adecarboxylata* ATCC 23216 = NBRC 102595 |
| Lsp-1-1 | Bacteria Proteobacteria Gammaproteobacteria Enterobacterales Enterobacteriaceae *Leclercia sp.* LK8 |
| Lam-1-1 | Bacteria Proteobacteria Gammaproteobacteria Enterobacterales Enterobacteriaceae *Lelliottia amnigena* CHS 78 |
| Msp-4-1 | Bacteria Proteobacteria Gammaproteobacteria Enterobacterales Enterobacteriaceae *Mangrovibacter sp.* MFB070 |
| Mma-2-1 | Bacteria Proteobacteria Gammaproteobacteria Enterobacterales Enterobacteriaceae *Metakosakonia massiliensis* JC163 |
| Pge-1-1 | Bacteria Proteobacteria Gammaproteobacteria Enterobacterales Enterobacteriaceae *Pluralibacter gergoviae* |
| Pvu-1-1 | Bacteria Proteobacteria Gammaproteobacteria Enterobacterales Enterobacteriaceae *Pseudescherichia vulneris* NBRC 102420 |
| Ror-1-1 | Bacteria Proteobacteria Gammaproteobacteria Enterobacterales Enterobacteriaceae *Raoultella ornithinolytica* |
| Ror-2-1 | Bacteria Proteobacteria Gammaproteobacteria Enterobacterales Enterobacteriaceae *Raoultella ornithinolytica* 2-156-04_S1_C1 |
| Ror-3-1 | Bacteria Proteobacteria Gammaproteobacteria Enterobacterales Enterobacteriaceae *Raoultella ornithinolytica* 2-156-04_S1_C2 |
| Ror-4-1 to Ror-4-2 | Bacteria Proteobacteria Gammaproteobacteria Enterobacterales Enterobacteriaceae *Raoultella ornithinolytica* B6 |
| Rpl-1-1 to Rpl-1-3 | Bacteria Proteobacteria Gammaproteobacteria Enterobacterales Enterobacteriaceae *Raoultella planticola* |
| Rpl-2-1 | Bacteria Proteobacteria Gammaproteobacteria Enterobacterales Enterobacteriaceae *Raoultella planticola* ATCC 33531 |
| Rte-1-1 | Bacteria Proteobacteria Gammaproteobacteria Enterobacterales Enterobacteriaceae *Raoultella terrigena* |
| Sbo-1-1 | Bacteria Proteobacteria Gammaproteobacteria Enterobacterales Enterobacteriaceae *Salmonella bongori serovar* 48:z41:– str. RKS3044 |

| | |
|---|---|
| Sen-2-1 to Sen-2-2 | Bacteria Proteobacteria Gammaproteobacteria Enterobacterales Enterobacteriaceae *Salmonella enterica* |
| Sen-3-1 | Bacteria Proteobacteria Gammaproteobacteria Enterobacterales Enterobacteriaceae *Salmonella enterica subsp. arizonae serovar* 62:z36:- str. RKS2983 |
| Sen-4-1 | Bacteria Proteobacteria Gammaproteobacteria Enterobacterales Enterobacteriaceae *Salmonella enterica subsp. enterica* |
| Sen-5-1 to Sen-5-6 | Bacteria Proteobacteria Gammaproteobacteria Enterobacterales Enterobacteriaceae *Salmonella enterica subsp. enterica serovar* 4,12:i:- |
| Sen-6-1 to Sen-6-5 | Bacteria Proteobacteria Gammaproteobacteria Enterobacterales Enterobacteriaceae *Salmonella enterica subsp. enterica serovar* 4,5,12:i- |
| Sen-7-1 to Sen-7-3 | Bacteria Proteobacteria Gammaproteobacteria Enterobacterales Enterobacteriaceae *Salmonella enterica subsp. enterica serovar* 4,[5],12:i:- |
| Sen-8-1 | Bacteria Proteobacteria Gammaproteobacteria Enterobacterales Enterobacteriaceae *Salmonella enterica subsp. enterica serovar* 8,(20):-:z6 |
| Sen-9-1 | Bacteria Proteobacteria Gammaproteobacteria Enterobacterales Enterobacteriaceae *Salmonella enterica subsp. enterica serovar* 9,12:l,v:- str. 94293 |
| Sen-10-1 | Bacteria Proteobacteria Gammaproteobacteria Enterobacterales Enterobacteriaceae *Salmonella enterica subsp. enterica serovar* Abaetetuba str. ATCC 35640 |
| Sen-11-1 | Bacteria Proteobacteria Gammaproteobacteria Enterobacterales Enterobacteriaceae *Salmonella enterica subsp. enterica serovar* Abony str. 0014 |
| Sen-12-1 to Sen-12-3 | Bacteria Proteobacteria Gammaproteobacteria Enterobacterales Enterobacteriaceae *Salmonella enterica subsp. enterica serovar* Agona |
| Sen-13-1 | Bacteria Proteobacteria Gammaproteobacteria Enterobacterales Enterobacteriaceae *Salmonella enterica subsp. enterica serovar* Agona str. 460004 2-1 |
| Sen-14-1 to Sen-14-4 | Bacteria Proteobacteria Gammaproteobacteria Enterobacterales Enterobacteriaceae *Salmonella enterica subsp. enterica serovar* Albany |
| Sen-15-1 to Sen-15-3 | Bacteria Proteobacteria Gammaproteobacteria Enterobacterales Enterobacteriaceae *Salmonella enterica subsp. enterica serovar* Anatum |
| Sen-16-1 | Bacteria Proteobacteria Gammaproteobacteria Enterobacterales Enterobacteriaceae *Salmonella enterica subsp. enterica serovar* Anatum str. 06-0532 |
| Sen-17-1 | Bacteria Proteobacteria Gammaproteobacteria Enterobacterales Enterobacteriaceae *Salmonella enterica subsp. enterica serovar* Anatum str. ATCC BAA-1592 |
| Sen-18-1 | Bacteria Proteobacteria Gammaproteobacteria Enterobacterales Enterobacteriaceae *Salmonella enterica subsp. enterica serovar* Aqua str. NVSL2001 |
| Sen-19-1 | Bacteria Proteobacteria Gammaproteobacteria Enterobacterales Enterobacteriaceae *Salmonella enterica subsp. enterica serovar* Bareilly str. 07-0873 |
| Sen-20-1 | Bacteria Proteobacteria Gammaproteobacteria Enterobacterales Enterobacteriaceae *Salmonella enterica subsp. enterica serovar* Bareilly str. CFSAN000178 |
| Sen-21-1 | Bacteria Proteobacteria Gammaproteobacteria Enterobacterales Enterobacteriaceae *Salmonella enterica subsp. enterica serovar* Bareilly str. CFSAN000179 |
| Sen-22-1 | Bacteria Proteobacteria Gammaproteobacteria Enterobacterales Enterobacteriaceae *Salmonella enterica subsp. enterica serovar* Bareilly str. CFSAN000180 |
| Sen-23-1 | Bacteria Proteobacteria Gammaproteobacteria Enterobacterales Enterobacteriaceae *Salmonella enterica subsp. enterica serovar* Bareilly str. CFSAN000181 |
| Sen-24-1 | Bacteria Proteobacteria Gammaproteobacteria Enterobacterales Enterobacteriaceae *Salmonella enterica subsp. enterica serovar* Bareilly str. CFSAN000182 |
| Sen-25-1 | Bacteria Proteobacteria Gammaproteobacteria Enterobacterales Enterobacteriaceae *Salmonella enterica subsp. enterica serovar* Bareilly str. CFSAN000184 |
| Sen-26-1 | Bacteria Proteobacteria Gammaproteobacteria Enterobacterales Enterobacteriaceae *Salmonella enterica subsp. enterica serovar* Bareilly str. CFSAN000185 |
| Sen-27-1 | Bacteria Proteobacteria Gammaproteobacteria Enterobacterales Enterobacteriaceae *Salmonella enterica subsp. enterica serovar* Bareilly str. CFSAN000186 |
| Sen-28-1 | Bacteria Proteobacteria Gammaproteobacteria Enterobacterales Enterobacteriaceae *Salmonella enterica subsp. enterica serovar* Bareilly str. CFSAN000187 |
| Sen-29-1 | Bacteria Proteobacteria Gammaproteobacteria Enterobacterales Enterobacteriaceae *Salmonella enterica subsp. enterica serovar* Bareilly str. CFSAN000188 |
| Sen-30-1 | Bacteria Proteobacteria Gammaproteobacteria Enterobacterales Enterobacteriaceae *Salmonella enterica subsp. enterica serovar* Bareilly str. CFSAN000190 |
| Sen-31-1 | Bacteria Proteobacteria Gammaproteobacteria Enterobacterales Enterobacteriaceae *Salmonella enterica subsp. enterica serovar* Bareilly str. CFSAN000191 |
| Sen-32-1 | Bacteria Proteobacteria Gammaproteobacteria Enterobacterales Enterobacteriaceae *Salmonella enterica subsp. enterica serovar* Bareilly str. CFSAN000192 |
| Sen-33-1 | Bacteria Proteobacteria Gammaproteobacteria Enterobacterales Enterobacteriaceae *Salmonella enterica subsp. enterica serovar* Bareilly str. CFSAN000193 |
| Sen-34-1 | Bacteria Proteobacteria Gammaproteobacteria Enterobacterales Enterobacteriaceae *Salmonella enterica subsp. enterica serovar* Bareilly str. CFSAN000194 |
| Sen-35-1 | Bacteria Proteobacteria Gammaproteobacteria Enterobacterales Enterobacteriaceae *Salmonella enterica subsp. enterica serovar* Bareilly str. CFSAN000195 |
| Sen-36-1 | Bacteria Proteobacteria Gammaproteobacteria Enterobacterales Enterobacteriaceae *Salmonella enterica subsp. enterica serovar* Bareilly str. CFSAN000196 |
| Sen-37-1 | Bacteria Proteobacteria Gammaproteobacteria Enterobacterales Enterobacteriaceae *Salmonella enterica subsp. enterica serovar* Bareilly str. CFSAN000198 |
| Sen-38-1 | Bacteria Proteobacteria Gammaproteobacteria Enterobacterales Enterobacteriaceae *Salmonella enterica subsp. enterica serovar* Bareilly str. CFSAN000199 |
| Sen-39-1 | Bacteria Proteobacteria Gammaproteobacteria Enterobacterales Enterobacteriaceae *Salmonella enterica subsp. enterica serovar* Bareilly str. CFSAN000201 |
| Sen-40-1 | Bacteria Proteobacteria Gammaproteobacteria Enterobacterales Enterobacteriaceae *Salmonella enterica subsp. enterica serovar* Bareilly str. CFSAN000202 |
| Sen-41-1 | Bacteria Proteobacteria Gammaproteobacteria Enterobacterales Enterobacteriaceae *Salmonella enterica subsp. enterica serovar* Bareilly str. CFSAN000203 |
| Sen-42-1 | Bacteria Proteobacteria Gammaproteobacteria Enterobacterales Enterobacteriaceae *Salmonella enterica subsp. enterica serovar* Bareilly str. CFSAN000204 |
| Sen-43-1 | Bacteria Proteobacteria Gammaproteobacteria Enterobacterales Enterobacteriaceae *Salmonella enterica subsp. enterica serovar* Bareilly str. CFSAN000205 |
| Sen-44-1 | Bacteria Proteobacteria Gammaproteobacteria Enterobacterales Enterobacteriaceae *Salmonella enterica subsp. enterica serovar* Bareilly str. CFSAN000206 |
| Sen-45-1 | Bacteria Proteobacteria Gammaproteobacteria Enterobacterales Enterobacteriaceae *Salmonella enterica subsp. enterica serovar* Bareilly str. CFSAN000207 |
| Sen-46-1 | Bacteria Proteobacteria Gammaproteobacteria Enterobacterales Enterobacteriaceae *Salmonella enterica subsp. enterica serovar* Bareilly str. CFSAN000208 |
| Sen-47-1 | Bacteria Proteobacteria Gammaproteobacteria Enterobacterales Enterobacteriaceae *Salmonella enterica subsp. enterica serovar* Bareilly str. CFSAN000209 |
| Sen-48-1 | Bacteria Proteobacteria Gammaproteobacteria Enterobacterales Enterobacteriaceae *Salmonella enterica subsp. enterica serovar* Bareilly str. CFSAN000210 |
| Sen-49-1 | Bacteria Proteobacteria Gammaproteobacteria Enterobacterales Enterobacteriaceae *Salmonella enterica subsp. enterica serovar* Bareilly str. CFSAN000211 |
| Sen-50-1 | Bacteria Proteobacteria Gammaproteobacteria Enterobacterales Enterobacteriaceae *Salmonella enterica subsp. enterica serovar* Bareilly str. CFSAN000212 |
| Sen-51-1 | Bacteria Proteobacteria Gammaproteobacteria Enterobacterales Enterobacteriaceae *Salmonella enterica subsp. enterica serovar* Bareilly str. CFSAN000213 |
| Sen-52-1 | Bacteria Proteobacteria Gammaproteobacteria Enterobacterales Enterobacteriaceae *Salmonella enterica subsp. enterica serovar* Bareilly str. CFSAN000214 |
| Sen-53-1 | Bacteria Proteobacteria Gammaproteobacteria Enterobacterales Enterobacteriaceae *Salmonella enterica subsp. enterica serovar* Bareilly str. CFSAN000215 |
| Sen-54-1 | Bacteria Proteobacteria Gammaproteobacteria Enterobacterales Enterobacteriaceae *Salmonella enterica subsp. enterica serovar* Bareilly str. CFSAN000216 |
| Sen-55-1 | Bacteria Proteobacteria Gammaproteobacteria Enterobacterales Enterobacteriaceae *Salmonella enterica subsp. enterica serovar* Bareilly str. CFSAN000217 |
| Sen-56-1 | Bacteria Proteobacteria Gammaproteobacteria Enterobacterales Enterobacteriaceae *Salmonella enterica subsp. enterica serovar* Bareilly str. CFSAN000218 |
| Sen-57-1 | Bacteria Proteobacteria Gammaproteobacteria Enterobacterales Enterobacteriaceae *Salmonella enterica subsp. enterica serovar* Bareilly str. CFSAN000219 |
| Sen-58-1 | Bacteria Proteobacteria Gammaproteobacteria Enterobacterales Enterobacteriaceae *Salmonella enterica subsp. enterica serovar* Bareilly str. CFSAN000220 |
| Sen-59-1 | Bacteria Proteobacteria Gammaproteobacteria Enterobacterales Enterobacteriaceae *Salmonella enterica subsp. enterica serovar* Bareilly str. CFSAN000221 |
| Sen-60-1 | Bacteria Proteobacteria Gammaproteobacteria Enterobacterales Enterobacteriaceae *Salmonella enterica subsp. enterica serovar* Bareilly str. CFSAN000222 |
| Sen-61-1 | Bacteria Proteobacteria Gammaproteobacteria Enterobacterales Enterobacteriaceae *Salmonella enterica subsp. enterica serovar* Bareilly str. CFSAN000223 |
| Sen-62-1 | Bacteria Proteobacteria Gammaproteobacteria Enterobacterales Enterobacteriaceae *Salmonella enterica subsp. enterica serovar* Bareilly str. CFSAN000224 |
| Sen-63-1 | Bacteria Proteobacteria Gammaproteobacteria Enterobacterales Enterobacteriaceae *Salmonella enterica subsp. enterica serovar* Bareilly str. CFSAN000225 |

| | |
|---|---|
| Sen-64-1 | Bacteria Proteobacteria Gammaproteobacteria Enterobacterales Enterobacteriaceae *Salmonella enterica subsp. enterica serovar* Bareilly str. CFSAN000227 |
| Sen-65-1 | Bacteria Proteobacteria Gammaproteobacteria Enterobacterales Enterobacteriaceae *Salmonella enterica subsp. enterica serovar* Bareilly str. CFSAN000228 |
| Sen-66-1 | Bacteria Proteobacteria Gammaproteobacteria Enterobacterales Enterobacteriaceae *Salmonella enterica subsp. enterica serovar* Bareilly str. CFSAN000231 |
| Sen-67-1 | Bacteria Proteobacteria Gammaproteobacteria Enterobacterales Enterobacteriaceae *Salmonella enterica subsp. enterica serovar* Bareilly str. CFSAN000235 |
| Sen-68-1 | Bacteria Proteobacteria Gammaproteobacteria Enterobacterales Enterobacteriaceae *Salmonella enterica subsp. enterica serovar* Bareilly str. CFSAN000236 |
| Sen-69-1 | Bacteria Proteobacteria Gammaproteobacteria Enterobacterales Enterobacteriaceae *Salmonella enterica subsp. enterica serovar* Bareilly str. CFSAN000661 |
| Sen-70-1 | Bacteria Proteobacteria Gammaproteobacteria Enterobacterales Enterobacteriaceae *Salmonella enterica subsp. enterica serovar* Bareilly str. CFSAN000662 |
| Sen-71-1 | Bacteria Proteobacteria Gammaproteobacteria Enterobacterales Enterobacteriaceae *Salmonella enterica subsp. enterica serovar* Bareilly str. CFSAN000669 |
| Sen-72-1 | Bacteria Proteobacteria Gammaproteobacteria Enterobacterales Enterobacteriaceae *Salmonella enterica subsp. enterica serovar* Bareilly str. CFSAN000680 |
| Sen-73-1 | Bacteria Proteobacteria Gammaproteobacteria Enterobacterales Enterobacteriaceae *Salmonella enterica subsp. enterica serovar* Bareilly str. CFSAN000700 |
| Sen-74-1 | Bacteria Proteobacteria Gammaproteobacteria Enterobacterales Enterobacteriaceae *Salmonella enterica subsp. enterica serovar* Bareilly str. CFSAN000752 |
| Sen-75-1 | Bacteria Proteobacteria Gammaproteobacteria Enterobacterales Enterobacteriaceae *Salmonella enterica subsp. enterica serovar* Bareilly str. CFSAN000753 |
| Sen-76-1 | Bacteria Proteobacteria Gammaproteobacteria Enterobacterales Enterobacteriaceae *Salmonella enterica subsp. enterica serovar* Bareilly str. CFSAN000754 |
| Sen-77-1 | Bacteria Proteobacteria Gammaproteobacteria Enterobacterales Enterobacteriaceae *Salmonella enterica subsp. enterica serovar* Bareilly str. CFSAN000755 |
| Sen-78-1 | Bacteria Proteobacteria Gammaproteobacteria Enterobacterales Enterobacteriaceae *Salmonella enterica subsp. enterica serovar* Bareilly str. CFSAN000951 |
| Sen-79-1 | Bacteria Proteobacteria Gammaproteobacteria Enterobacterales Enterobacteriaceae *Salmonella enterica subsp. enterica serovar* Bareilly str. CFSAN000952 |
| Sen-80-1 | Bacteria Proteobacteria Gammaproteobacteria Enterobacterales Enterobacteriaceae *Salmonella enterica subsp. enterica serovar* Bareilly str. CFSAN000953 |
| Sen-81-1 | Bacteria Proteobacteria Gammaproteobacteria Enterobacterales Enterobacteriaceae *Salmonella enterica subsp. enterica serovar* Bareilly str. CFSAN000954 |
| Sen-82-1 | Bacteria Proteobacteria Gammaproteobacteria Enterobacterales Enterobacteriaceae *Salmonella enterica subsp. enterica serovar* Bareilly str. CFSAN000955 |
| Sen-83-1 | Bacteria Proteobacteria Gammaproteobacteria Enterobacterales Enterobacteriaceae *Salmonella enterica subsp. enterica serovar* Bareilly str. CFSAN000956 |
| Sen-84-1 | Bacteria Proteobacteria Gammaproteobacteria Enterobacterales Enterobacteriaceae *Salmonella enterica subsp. enterica serovar* Bareilly str. CFSAN000957 |
| Sen-85-1 | Bacteria Proteobacteria Gammaproteobacteria Enterobacterales Enterobacteriaceae *Salmonella enterica subsp. enterica serovar* Bareilly str. CFSAN000958 |
| Sen-86-1 | Bacteria Proteobacteria Gammaproteobacteria Enterobacterales Enterobacteriaceae *Salmonella enterica subsp. enterica serovar* Bareilly str. CFSAN000959 |
| Sen-87-1 | Bacteria Proteobacteria Gammaproteobacteria Enterobacterales Enterobacteriaceae *Salmonella enterica subsp. enterica serovar* Bareilly str. CFSAN000960 |
| Sen-88-1 | Bacteria Proteobacteria Gammaproteobacteria Enterobacterales Enterobacteriaceae *Salmonella enterica subsp. enterica serovar* Bareilly str. CFSAN000961 |
| Sen-89-1 | Bacteria Proteobacteria Gammaproteobacteria Enterobacterales Enterobacteriaceae *Salmonella enterica subsp. enterica serovar* Bareilly str. CFSAN000962 |
| Sen-90-1 | Bacteria Proteobacteria Gammaproteobacteria Enterobacterales Enterobacteriaceae *Salmonella enterica subsp. enterica serovar* Bareilly str. CFSAN000963 |
| Sen-91-1 | Bacteria Proteobacteria Gammaproteobacteria Enterobacterales Enterobacteriaceae *Salmonella enterica subsp. enterica serovar* Bareilly str. CFSAN000964 |
| Sen-92-1 | Bacteria Proteobacteria Gammaproteobacteria Enterobacterales Enterobacteriaceae *Salmonella enterica subsp. enterica serovar* Bareilly str. CFSAN000965 |
| Sen-93-1 | Bacteria Proteobacteria Gammaproteobacteria Enterobacterales Enterobacteriaceae *Salmonella enterica subsp. enterica serovar* Bareilly str. CFSAN000966 |
| Sen-94-1 | Bacteria Proteobacteria Gammaproteobacteria Enterobacterales Enterobacteriaceae *Salmonella enterica subsp. enterica serovar* Bareilly str. CFSAN000967 |
| Sen-95-1 | Bacteria Proteobacteria Gammaproteobacteria Enterobacterales Enterobacteriaceae *Salmonella enterica subsp. enterica serovar* Bareilly str. CFSAN000968 |
| Sen-96-1 | Bacteria Proteobacteria Gammaproteobacteria Enterobacterales Enterobacteriaceae *Salmonella enterica subsp. enterica serovar* Bareilly str. CFSAN000969 |
| Sen-97-1 | Bacteria Proteobacteria Gammaproteobacteria Enterobacterales Enterobacteriaceae *Salmonella enterica subsp. enterica serovar* Bareilly str. CFSAN000970 |
| Sen-98-1 | Bacteria Proteobacteria Gammaproteobacteria Enterobacterales Enterobacteriaceae *Salmonella enterica subsp. enterica serovar* Bareilly str. CFSAN001087 |
| Sen-99-1 | Bacteria Proteobacteria Gammaproteobacteria Enterobacterales Enterobacteriaceae *Salmonella enterica subsp. enterica serovar* Bareilly str. CFSAN001088 |
| Sen-100-1 | Bacteria Proteobacteria Gammaproteobacteria Enterobacterales Enterobacteriaceae *Salmonella enterica subsp. enterica serovar* Bareilly str. CFSAN001089 |
| Sen-101-1 | Bacteria Proteobacteria Gammaproteobacteria Enterobacterales Enterobacteriaceae *Salmonella enterica subsp. enterica serovar* Bareilly str. CFSAN001090 |
| Sen-102-1 | Bacteria Proteobacteria Gammaproteobacteria Enterobacterales Enterobacteriaceae *Salmonella enterica subsp. enterica serovar* Bareilly str. CFSAN001099 |
| Sen-103-1 | Bacteria Proteobacteria Gammaproteobacteria Enterobacterales Enterobacteriaceae *Salmonella enterica subsp. enterica serovar* Bareilly str. CFSAN001102 |
| Sen-104-1 | Bacteria Proteobacteria Gammaproteobacteria Enterobacterales Enterobacteriaceae *Salmonella enterica subsp. enterica serovar* Bareilly str. CFSAN001105 |
| Sen-105-1 | Bacteria Proteobacteria Gammaproteobacteria Enterobacterales Enterobacteriaceae *Salmonella enterica subsp. enterica serovar* Bareilly str. CFSAN001108 |
| Sen-106-1 | Bacteria Proteobacteria Gammaproteobacteria Enterobacterales Enterobacteriaceae *Salmonella enterica subsp. enterica serovar* Bareilly str. CFSAN001109 |
| Sen-107-1 | Bacteria Proteobacteria Gammaproteobacteria Enterobacterales Enterobacteriaceae *Salmonella enterica subsp. enterica serovar* Bareilly str. CFSAN001112 |
| Sen-108-1 | Bacteria Proteobacteria Gammaproteobacteria Enterobacterales Enterobacteriaceae *Salmonella enterica subsp. enterica serovar* Bareilly str. CFSAN001115 |
| Sen-109-1 | Bacteria Proteobacteria Gammaproteobacteria Enterobacterales Enterobacteriaceae *Salmonella enterica subsp. enterica serovar* Bareilly str. CFSAN001118 |
| Sen-110-1 | Bacteria Proteobacteria Gammaproteobacteria Enterobacterales Enterobacteriaceae *Salmonella enterica subsp. enterica serovar* Bareilly str. CFSAN001140 |
| Sen-111-1 | Bacteria Proteobacteria Gammaproteobacteria Enterobacterales Enterobacteriaceae *Salmonella enterica subsp. enterica serovar* Bareilly str. CFSAN001591 |
| Sen-112-1 to Sen-112-3 | Bacteria Proteobacteria Gammaproteobacteria Enterobacterales Enterobacteriaceae *Salmonella enterica subsp. enterica serovar* Berta |
| Sen-113-1 to Sen-113-2 | Bacteria Proteobacteria Gammaproteobacteria Enterobacterales Enterobacteriaceae *Salmonella enterica subsp. enterica serovar* Berta str. ATCC 8392 |
| Sen-114-1 | Bacteria Proteobacteria Gammaproteobacteria Enterobacterales Enterobacteriaceae *Salmonella enterica subsp. enterica serovar* Bovismorbificans str. Sal185 |
| Sen-115-1 | Bacteria Proteobacteria Gammaproteobacteria Enterobacterales Enterobacteriaceae *Salmonella enterica subsp. enterica serovar* Bovismorbificans str. Sal609 |
| Sen-116-1 | Bacteria Proteobacteria Gammaproteobacteria Enterobacterales Enterobacteriaceae *Salmonella enterica subsp. enterica serovar* Bovismorbificans str. Sal610 |
| Sen-117-1 | Bacteria Proteobacteria Gammaproteobacteria Enterobacterales Enterobacteriaceae *Salmonella enterica subsp. enterica serovar* Bovismorbificans str. Sal615 |
| Sen-118-1 | Bacteria Proteobacteria Gammaproteobacteria Enterobacterales Enterobacteriaceae *Salmonella enterica subsp. enterica serovar* Bovismorbificans str. Sal616 |
| Sen-119-1 | Bacteria Proteobacteria Gammaproteobacteria Enterobacterales Enterobacteriaceae *Salmonella enterica subsp. enterica serovar* Bovismorbificans str. Sal617 |
| Sen-120-1 | Bacteria Proteobacteria Gammaproteobacteria Enterobacterales Enterobacteriaceae *Salmonella enterica subsp. enterica serovar* Bovismorbificans str. Sal644 |
| Sen-121-1 | Bacteria Proteobacteria Gammaproteobacteria Enterobacterales Enterobacteriaceae *Salmonella enterica subsp. enterica serovar* Bovismorbificans str. Sal676 |
| Sen-122-1 | Bacteria Proteobacteria Gammaproteobacteria Enterobacterales Enterobacteriaceae *Salmonella enterica subsp. enterica serovar* Bovismorbificans str. Sal677 |
| Sen-123-1 | Bacteria Proteobacteria Gammaproteobacteria Enterobacterales Enterobacteriaceae *Salmonella enterica subsp. enterica serovar* Bovismorbificans str. Sal678 |
| Sen-124-1 | Bacteria Proteobacteria Gammaproteobacteria Enterobacterales Enterobacteriaceae *Salmonella enterica subsp. enterica serovar* Bovismorbificans str. Sal679 |
| Sen-125-1 | Bacteria Proteobacteria Gammaproteobacteria Enterobacterales Enterobacteriaceae *Salmonella enterica subsp. enterica serovar* Bovismorbificans str. Sal680 |
| Sen-126-1 | Bacteria Proteobacteria Gammaproteobacteria Enterobacterales Enterobacteriaceae *Salmonella enterica subsp. enterica serovar* Bovismorbificans str. Sal681 |
| Sen-127-1 | Bacteria Proteobacteria Gammaproteobacteria Enterobacterales Enterobacteriaceae *Salmonella enterica subsp. enterica serovar* Bovismorbificans str. Sal682 |
| Sen-128-1 | Bacteria Proteobacteria Gammaproteobacteria Enterobacterales Enterobacteriaceae *Salmonella enterica subsp. enterica serovar* Bovismorbificans str. Sal683 |

| | |
|---|---|
| Sen-129-1 to Sen-129-2 | Bacteria Proteobacteria Gammaproteobacteria Enterobacterales Enterobacteriaceae *Salmonella enterica subsp. enterica serovar* Braenderup |
| Sen-130-1 to Sen-130-3 | Bacteria Proteobacteria Gammaproteobacteria Enterobacterales Enterobacteriaceae *Salmonella enterica subsp. enterica serovar* Brandenburg |
| Sen-131-1 | Bacteria Proteobacteria Gammaproteobacteria Enterobacterales Enterobacteriaceae *Salmonella enterica subsp. enterica serovar* Bredeney |
| Sen-132-1 | Bacteria Proteobacteria Gammaproteobacteria Enterobacterales Enterobacteriaceae *Salmonella enterica subsp. enterica serovar* Bredeney str. CF-SAN001080 |
| Sen-133-1 | Bacteria Proteobacteria Gammaproteobacteria Enterobacterales Enterobacteriaceae *Salmonella enterica subsp. enterica serovar* Cerro |
| Sen-134-1 | Bacteria Proteobacteria Gammaproteobacteria Enterobacterales Enterobacteriaceae *Salmonella enterica subsp. enterica serovar* Choleraesuis |
| Sen-135-1 | Bacteria Proteobacteria Gammaproteobacteria Enterobacterales Enterobacteriaceae *Salmonella enterica subsp. enterica serovar* Choleraesuis str. ATCC 10708 |
| Sen-136-1 to Sen-136-2 | Bacteria Proteobacteria Gammaproteobacteria Enterobacterales Enterobacteriaceae *Salmonella enterica subsp. enterica serovar* Cubana str. 76814 |
| Sen-137-1 to Sen-137-6 | Bacteria Proteobacteria Gammaproteobacteria Enterobacterales Enterobacteriaceae *Salmonella enterica subsp. enterica serovar* Derby |
| Sen-138-1 to Sen-138-5 | Bacteria Proteobacteria Gammaproteobacteria Enterobacterales Enterobacteriaceae *Salmonella enterica subsp. enterica serovar* Dublin |
| Sen-139-1 to Sen-139-15 | Bacteria Proteobacteria Gammaproteobacteria Enterobacterales Enterobacteriaceae *Salmonella enterica subsp. enterica serovar* Enteritidis |
| Sen-140-1 | Bacteria Proteobacteria Gammaproteobacteria Enterobacterales Enterobacteriaceae *Salmonella enterica subsp. enterica serovar* Enteritidis str. 18569 |
| Sen-141-1 | Bacteria Proteobacteria Gammaproteobacteria Enterobacterales Enterobacteriaceae *Salmonella enterica subsp. enterica serovar* Enteritidis str. 77-1427 |
| Sen-142-1 | Bacteria Proteobacteria Gammaproteobacteria Enterobacterales Enterobacteriaceae *Salmonella enterica subsp. enterica serovar* Enteritidis str. CDC_2010K_0968 |
| Sen-143-1 | Bacteria Proteobacteria Gammaproteobacteria Enterobacterales Enterobacteriaceae *Salmonella enterica subsp. enterica serovar* Enteritidis str. EC20090641 |
| Sen-144-1 | Bacteria Proteobacteria Gammaproteobacteria Enterobacterales Enterobacteriaceae *Salmonella enterica subsp. enterica serovar* Enteritidis str. EC20090698 |
| Sen-145-1 | Bacteria Proteobacteria Gammaproteobacteria Enterobacterales Enterobacteriaceae *Salmonella enterica subsp. enterica serovar* Enteritidis str. EC20100101 |
| Sen-146-1 | Bacteria Proteobacteria Gammaproteobacteria Enterobacterales Enterobacteriaceae *Salmonella enterica subsp. enterica serovar* Enteritidis str. EC20110221 |
| Sen-147-1 | Bacteria Proteobacteria Gammaproteobacteria Enterobacterales Enterobacteriaceae *Salmonella enterica subsp. enterica serovar* Enteritidis str. EC20110354 |
| Sen-148-1 | Bacteria Proteobacteria Gammaproteobacteria Enterobacterales Enterobacteriaceae *Salmonella enterica subsp. enterica serovar* Enteritidis str. EC20110355 |
| Sen-149-1 | Bacteria Proteobacteria Gammaproteobacteria Enterobacterales Enterobacteriaceae *Salmonella enterica subsp. enterica serovar* Enteritidis str. EC20110356 |
| Sen-150-1 | Bacteria Proteobacteria Gammaproteobacteria Enterobacterales Enterobacteriaceae *Salmonella enterica subsp. enterica serovar* Enteritidis str. EC20110357 |
| Sen-151-1 | Bacteria Proteobacteria Gammaproteobacteria Enterobacterales Enterobacteriaceae *Salmonella enterica subsp. enterica serovar* Enteritidis str. EC20110358 |
| Sen-152-1 | Bacteria Proteobacteria Gammaproteobacteria Enterobacterales Enterobacteriaceae *Salmonella enterica subsp. enterica serovar* Enteritidis str. EC20110359 |
| Sen-153-1 | Bacteria Proteobacteria Gammaproteobacteria Enterobacterales Enterobacteriaceae *Salmonella enterica subsp. enterica serovar* Enteritidis str. EC20110360 |
| Sen-154-1 | Bacteria Proteobacteria Gammaproteobacteria Enterobacterales Enterobacteriaceae *Salmonella enterica subsp. enterica serovar* Enteritidis str. EC20110361 |
| Sen-155-1 | Bacteria Proteobacteria Gammaproteobacteria Enterobacterales Enterobacteriaceae *Salmonella enterica subsp. enterica serovar* Enteritidis str. EC20111095 |
| Sen-156-1 | Bacteria Proteobacteria Gammaproteobacteria Enterobacterales Enterobacteriaceae *Salmonella enterica subsp. enterica serovar* Enteritidis str. EC20111174 |
| Sen-157-1 | Bacteria Proteobacteria Gammaproteobacteria Enterobacterales Enterobacteriaceae *Salmonella enterica subsp. enterica serovar* Enteritidis str. EC20111175 |
| Sen-158-1 | Bacteria Proteobacteria Gammaproteobacteria Enterobacterales Enterobacteriaceae *Salmonella enterica subsp. enterica serovar* Enteritidis str. EC20120002 |
| Sen-159-1 | Bacteria Proteobacteria Gammaproteobacteria Enterobacterales Enterobacteriaceae *Salmonella enterica subsp. enterica serovar* Enteritidis str. EC20120005 |
| Sen-160-1 | Bacteria Proteobacteria Gammaproteobacteria Enterobacterales Enterobacteriaceae *Salmonella enterica subsp. enterica serovar* Enteritidis str. EC20120008 |
| Sen-161-1 | Bacteria Proteobacteria Gammaproteobacteria Enterobacterales Enterobacteriaceae *Salmonella enterica subsp. enterica serovar* Enteritidis str. EC20120916 |
| Sen-162-1 | Bacteria Proteobacteria Gammaproteobacteria Enterobacterales Enterobacteriaceae *Salmonella enterica subsp. enterica serovar* Enteritidis str. EC20121175 |
| Sen-163-1 | Bacteria Proteobacteria Gammaproteobacteria Enterobacterales Enterobacteriaceae *Salmonella enterica subsp. enterica serovar* Enteritidis str. LA5 |
| Sen-164-1 | Bacteria Proteobacteria Gammaproteobacteria Enterobacterales Enterobacteriaceae *Salmonella enterica subsp. enterica serovar* Enteritidis str. SA19940857 |
| Sen-165-1 | Bacteria Proteobacteria Gammaproteobacteria Enterobacterales Enterobacteriaceae *Salmonella enterica subsp. enterica serovar* Gallinarum/pullorum str. FCAV198 |
| Sen-166-1 | Bacteria Proteobacteria Gammaproteobacteria Enterobacterales Enterobacteriaceae *Salmonella enterica subsp. enterica serovar* Give |
| Sen-167-1 to Sen-167-18 | Bacteria Proteobacteria Gammaproteobacteria Enterobacterales Enterobacteriaceae *Salmonella enterica subsp. enterica serovar* Hadar |
| Sen-168-1 | Bacteria Proteobacteria Gammaproteobacteria Enterobacterales Enterobacteriaceae *Salmonella enterica subsp. enterica serovar* Hadar str. 2012K-0272 |
| Sen-169-1 | Bacteria Proteobacteria Gammaproteobacteria Enterobacterales Enterobacteriaceae *Salmonella enterica subsp. enterica serovar* Hartford str. 2012K-0272 |
| Sen-170-1 to Sen-170-52 | Bacteria Proteobacteria Gammaproteobacteria Enterobacterales Enterobacteriaceae *Salmonella enterica subsp. enterica serovar* Heidelberg |
| Sen-171-1 | Bacteria Proteobacteria Gammaproteobacteria Enterobacterales Enterobacteriaceae *Salmonella enterica subsp. enterica serovar* Heidelberg str. 21381 |
| Sen-172-1 | Bacteria Proteobacteria Gammaproteobacteria Enterobacterales Enterobacteriaceae *Salmonella enterica subsp. enterica serovar* Heidelberg str. 29169 |
| Sen-173-1 | Bacteria Proteobacteria Gammaproteobacteria Enterobacterales Enterobacteriaceae *Salmonella enterica subsp. enterica serovar* Heidelberg str. 32507 |
| Sen-174-1 | Bacteria Proteobacteria Gammaproteobacteria Enterobacterales Enterobacteriaceae *Salmonella enterica subsp. enterica serovar* Heidelberg str. 41567 |
| Sen-175-1 | Bacteria Proteobacteria Gammaproteobacteria Enterobacterales Enterobacteriaceae *Salmonella enterica subsp. enterica serovar* Heidelberg str. 41584 |
| Sen-176-1 | Bacteria Proteobacteria Gammaproteobacteria Enterobacterales Enterobacteriaceae *Salmonella enterica subsp. enterica serovar* Heidelberg str. 579083-10 |
| Sen-177-1 | Bacteria Proteobacteria Gammaproteobacteria Enterobacterales Enterobacteriaceae *Salmonella enterica subsp. enterica serovar* Heidelberg str. 607309-34 |
| Sen-178-1 | Bacteria Proteobacteria Gammaproteobacteria Enterobacterales Enterobacteriaceae *Salmonella enterica subsp. enterica serovar* Heidelberg str. 607309-5 |
| Sen-179-1 | Bacteria Proteobacteria Gammaproteobacteria Enterobacterales Enterobacteriaceae *Salmonella enterica subsp. enterica serovar* Heidelberg str. 622737-21 |
| Sen-180-1 | Bacteria Proteobacteria Gammaproteobacteria Enterobacterales Enterobacteriaceae *Salmonella enterica subsp. enterica serovar* Heidelberg str. 632675-24 |
| Sen-181-1 | Bacteria Proteobacteria Gammaproteobacteria Enterobacterales Enterobacteriaceae *Salmonella enterica subsp. enterica serovar* Heidelberg str. 640151-11 |
| Sen-182-1 | Bacteria Proteobacteria Gammaproteobacteria Enterobacterales Enterobacteriaceae *Salmonella enterica subsp. enterica serovar* Heidelberg str. 670102-3 |
| Sen-183-1 | Bacteria Proteobacteria Gammaproteobacteria Enterobacterales Enterobacteriaceae *Salmonella enterica subsp. enterica serovar* Heidelberg str. 670102-5 |
| Sen-184-1 | Bacteria Proteobacteria Gammaproteobacteria Enterobacterales Enterobacteriaceae *Salmonella enterica subsp. enterica serovar* Heidelberg str. 82-1617 |
| Sen-185-1 | Bacteria Proteobacteria Gammaproteobacteria Enterobacterales Enterobacteriaceae *Salmonella enterica subsp. enterica serovar* Heidelberg str. 84-1004 |
| Sen-186-1 | Bacteria Proteobacteria Gammaproteobacteria Enterobacterales Enterobacteriaceae *Salmonella enterica subsp. enterica serovar* Heidelberg str. 85-0486 |
| Sen-187-1 | Bacteria Proteobacteria Gammaproteobacteria Enterobacterales Enterobacteriaceae *Salmonella enterica subsp. enterica serovar* Heidelberg str. 86-0255 |

| | |
|---|---|
| Sen-188-1 | Bacteria Proteobacteria Gammaproteobacteria Enterobacterales Enterobacteriaceae *Salmonella enterica subsp. enterica serovar* Heidelberg str. 87-0208 |
| Sen-189-1 | Bacteria Proteobacteria Gammaproteobacteria Enterobacterales Enterobacteriaceae *Salmonella enterica subsp. enterica serovar* Heidelberg str. 88-0312 |
| Sen-190-1 | Bacteria Proteobacteria Gammaproteobacteria Enterobacterales Enterobacteriaceae *Salmonella enterica subsp. enterica serovar* Heidelberg str. 89-0213 |
| Sen-191-1 | Bacteria Proteobacteria Gammaproteobacteria Enterobacterales Enterobacteriaceae *Salmonella enterica subsp. enterica serovar* Heidelberg str. 92-0138 |
| Sen-192-1 | Bacteria Proteobacteria Gammaproteobacteria Enterobacterales Enterobacteriaceae *Salmonella enterica subsp. enterica serovar* Heidelberg str. 92-0144 |
| Sen-193-1 | Bacteria Proteobacteria Gammaproteobacteria Enterobacterales Enterobacteriaceae *Salmonella enterica subsp. enterica serovar* Heidelberg str. CF-SAN002064 |
| Sen-194-1 | Bacteria Proteobacteria Gammaproteobacteria Enterobacterales Enterobacteriaceae *Salmonella enterica subsp. enterica serovar* Heidelberg str. CF-SAN00323 |
| Sen-195-1 | Bacteria Proteobacteria Gammaproteobacteria Enterobacterales Enterobacteriaceae *Salmonella enterica subsp. enterica serovar* Heidelberg str. CF-SAN00324 |
| Sen-196-1 | Bacteria Proteobacteria Gammaproteobacteria Enterobacterales Enterobacteriaceae *Salmonella enterica subsp. enterica serovar* Heidelberg str. CF-SAN00327 |
| Sen-197-1 | Bacteria Proteobacteria Gammaproteobacteria Enterobacterales Enterobacteriaceae *Salmonella enterica subsp. enterica serovar* Heidelberg str. CVM20752 |
| Sen-198-1 | Bacteria Proteobacteria Gammaproteobacteria Enterobacterales Enterobacteriaceae *Salmonella enterica subsp. enterica serovar* Heidelberg str. CVM24359 |
| Sen-199-1 | Bacteria Proteobacteria Gammaproteobacteria Enterobacterales Enterobacteriaceae *Salmonella enterica subsp. enterica serovar* Heidelberg str. CVM24388 |
| Sen-200-1 | Bacteria Proteobacteria Gammaproteobacteria Enterobacterales Enterobacteriaceae *Salmonella enterica subsp. enterica serovar* Heidelberg str. CVM24391 |
| Sen-201-1 | Bacteria Proteobacteria Gammaproteobacteria Enterobacterales Enterobacteriaceae *Salmonella enterica subsp. enterica serovar* Heidelberg str. N1536 |
| Sen-202-1 | Bacteria Proteobacteria Gammaproteobacteria Enterobacterales Enterobacteriaceae *Salmonella enterica subsp. enterica serovar* Heidelberg str. N15757 |
| Sen-203-1 | Bacteria Proteobacteria Gammaproteobacteria Enterobacterales Enterobacteriaceae *Salmonella enterica subsp. enterica serovar* Heidelberg str. N18393 |
| Sen-204-1 | Bacteria Proteobacteria Gammaproteobacteria Enterobacterales Enterobacteriaceae *Salmonella enterica subsp. enterica serovar* Heidelberg str. N18413 |
| Sen-205-1 | Bacteria Proteobacteria Gammaproteobacteria Enterobacterales Enterobacteriaceae *Salmonella enterica subsp. enterica serovar* Heidelberg str. N18440 |
| Sen-206-1 | Bacteria Proteobacteria Gammaproteobacteria Enterobacterales Enterobacteriaceae *Salmonella enterica subsp. enterica serovar* Heidelberg str. N18453 |
| Sen-207-1 | Bacteria Proteobacteria Gammaproteobacteria Enterobacterales Enterobacteriaceae *Salmonella enterica subsp. enterica serovar* Heidelberg str. N19871 |
| Sen-208-1 | Bacteria Proteobacteria Gammaproteobacteria Enterobacterales Enterobacteriaceae *Salmonella enterica subsp. enterica serovar* Heidelberg str. N19992 |
| Sen-209-1 | Bacteria Proteobacteria Gammaproteobacteria Enterobacterales Enterobacteriaceae *Salmonella enterica subsp. enterica serovar* Heidelberg str. N20134 |
| Sen-210-1 | Bacteria Proteobacteria Gammaproteobacteria Enterobacterales Enterobacteriaceae *Salmonella enterica subsp. enterica serovar* Heidelberg str. N26457 |
| Sen-211-1 | Bacteria Proteobacteria Gammaproteobacteria Enterobacterales Enterobacteriaceae *Salmonella enterica subsp. enterica serovar* Heidelberg str. N29341 |
| Sen-212-1 | Bacteria Proteobacteria Gammaproteobacteria Enterobacterales Enterobacteriaceae *Salmonella enterica subsp. enterica serovar* Heidelberg str. N30678 |
| Sen-213-1 | Bacteria Proteobacteria Gammaproteobacteria Enterobacterales Enterobacteriaceae *Salmonella enterica subsp. enterica serovar* Heidelberg str. N418 |
| Sen-214-1 | Bacteria Proteobacteria Gammaproteobacteria Enterobacterales Enterobacteriaceae *Salmonella enterica subsp. enterica serovar* Heidelberg str. N4630 |
| Sen-215-1 | Bacteria Proteobacteria Gammaproteobacteria Enterobacterales Enterobacteriaceae *Salmonella enterica subsp. enterica serovar* Heidelberg str. N653 |
| Sen-216-1 | Bacteria Proteobacteria Gammaproteobacteria Enterobacterales Enterobacteriaceae *Salmonella enterica subsp. enterica serovar* Heidelberg str. RI-11-013193 |
| Sen-217-1 | Bacteria Proteobacteria Gammaproteobacteria Enterobacterales Enterobacteriaceae *Salmonella enterica subsp. enterica serovar* Heidelberg str. RI-11-014319 |
| Sen-218-1 | Bacteria Proteobacteria Gammaproteobacteria Enterobacterales Enterobacteriaceae *Salmonella enterica subsp. enterica serovar* Heidelberg str. SARA31 |
| Sen-219-1 | Bacteria Proteobacteria Gammaproteobacteria Enterobacterales Enterobacteriaceae *Salmonella enterica subsp. enterica serovar* Heidelberg str. SARA32 |
| Sen-220-1 | Bacteria Proteobacteria Gammaproteobacteria Enterobacterales Enterobacteriaceae *Salmonella enterica subsp. enterica serovar* Heidelberg str. SARA36 |
| Sen-221-1 | Bacteria Proteobacteria Gammaproteobacteria Enterobacterales Enterobacteriaceae *Salmonella enterica subsp. enterica serovar* Heidelberg str. SARA37 |
| Sen-222-1 | Bacteria Proteobacteria Gammaproteobacteria Enterobacterales Enterobacteriaceae *Salmonella enterica subsp. enterica serovar* Heidelberg str. SARA40 |
| Sen-223-1 | Bacteria Proteobacteria Gammaproteobacteria Enterobacterales Enterobacteriaceae *Salmonella enterica subsp. enterica serovar* Infantis |
| Sen-224-1 to Sen-224-3 | Bacteria Proteobacteria Gammaproteobacteria Enterobacterales Enterobacteriaceae *Salmonella enterica subsp. enterica serovar* Johannesburg |
| Sen-225-1 to Sen-225-52 | Bacteria Proteobacteria Gammaproteobacteria Enterobacterales Enterobacteriaceae *Salmonella enterica subsp. enterica serovar* Kentucky |
| Sen-226-1 to Sen-226-4 | Bacteria Proteobacteria Gammaproteobacteria Enterobacterales Enterobacteriaceae *Salmonella enterica subsp. enterica serovar* Litchfield |
| Sen-227-1 to Sen-227-2 | Bacteria Proteobacteria Gammaproteobacteria Enterobacterales Enterobacteriaceae *Salmonella enterica subsp. enterica serovar* Liverpool |
| Sen-228-1 to Sen-228-2 | Bacteria Proteobacteria Gammaproteobacteria Enterobacterales Enterobacteriaceae *Salmonella enterica subsp. enterica serovar* Livingstone |
| Sen-229-1 | Bacteria Proteobacteria Gammaproteobacteria Enterobacterales Enterobacteriaceae *Salmonella enterica subsp. enterica serovar* Manhattan str. 111113 |
| Sen-230-1 to Sen-230-4 | Bacteria Proteobacteria Gammaproteobacteria Enterobacterales Enterobacteriaceae *Salmonella enterica subsp. enterica serovar* Mbandaka |
| Sen-231-1 | Bacteria Proteobacteria Gammaproteobacteria Enterobacterales Enterobacteriaceae *Salmonella enterica subsp. enterica serovar* Mbandaka str. 2012K-0273 |
| Sen-232-1 | Bacteria Proteobacteria Gammaproteobacteria Enterobacterales Enterobacteriaceae *Salmonella enterica subsp. enterica serovar* Mishmarhaemek |
| Sen-233-1 to Sen-233-5 | Bacteria Proteobacteria Gammaproteobacteria Enterobacterales Enterobacteriaceae *Salmonella enterica subsp. enterica serovar* Montevideo |
| Sen-234-1 | Bacteria Proteobacteria Gammaproteobacteria Enterobacterales Enterobacteriaceae *Salmonella enterica subsp. enterica serovar* Montevideo str. 316111868 |
| Sen-235-1 | Bacteria Proteobacteria Gammaproteobacteria Enterobacterales Enterobacteriaceae *Salmonella enterica subsp. enterica serovar* Montevideo str. 507440-20 |
| Sen-236-1 | Bacteria Proteobacteria Gammaproteobacteria Enterobacterales Enterobacteriaceae *Salmonella enterica subsp. enterica serovar* Montevideo str. USDA-ARS-USMARC-1903 |
| Sen-237-1 to Sen-237-16 | Bacteria Proteobacteria Gammaproteobacteria Enterobacterales Enterobacteriaceae *Salmonella enterica subsp. enterica serovar* Muenchen |
| Sen-238-1 | Bacteria Proteobacteria Gammaproteobacteria Enterobacterales Enterobacteriaceae *Salmonella enterica subsp. enterica serovar* Muenchen str. 08-1125 |
| Sen-239-1 | Bacteria Proteobacteria Gammaproteobacteria Enterobacterales Enterobacteriaceae *Salmonella enterica subsp. enterica serovar* Muenchen str. 2009K-0951 |
| Sen-240-1 | Bacteria Proteobacteria Gammaproteobacteria Enterobacterales Enterobacteriaceae *Salmonella enterica subsp. enterica serovar* Muenster |
| Sen-241-1 | Bacteria Proteobacteria Gammaproteobacteria Enterobacterales Enterobacteriaceae *Salmonella enterica subsp. enterica serovar* Namur str. 05-2929 |
| Sen-242-1 | Bacteria Proteobacteria Gammaproteobacteria Enterobacterales Enterobacteriaceae *Salmonella enterica subsp. enterica serovar* Napoli |

| | |
|---|---|
| Sen-243-1 to Sen-243-2 | Bacteria Proteobacteria Gammaproteobacteria Enterobacterales Enterobacteriaceae *Salmonella enterica subsp. enterica serovar* Newport |
| Sen-244-1 | Bacteria Proteobacteria Gammaproteobacteria Enterobacterales Enterobacteriaceae *Salmonella enterica subsp. enterica serovar* Newport str. 36799 |
| Sen-245-1 | Bacteria Proteobacteria Gammaproteobacteria Enterobacterales Enterobacteriaceae *Salmonella enterica subsp. enterica serovar* Newport str. 36801 |
| Sen-246-1 | Bacteria Proteobacteria Gammaproteobacteria Enterobacterales Enterobacteriaceae *Salmonella enterica subsp. enterica serovar* Newport str. 36802 |
| Sen-247-1 | Bacteria Proteobacteria Gammaproteobacteria Enterobacterales Enterobacteriaceae *Salmonella enterica subsp. enterica serovar* Newport str. 36803 |
| Sen-248-1 | Bacteria Proteobacteria Gammaproteobacteria Enterobacterales Enterobacteriaceae *Salmonella enterica subsp. enterica serovar* Newport str. 36804 |
| Sen-249-1 | Bacteria Proteobacteria Gammaproteobacteria Enterobacterales Enterobacteriaceae *Salmonella enterica subsp. enterica serovar* Newport str. 36805 |
| Sen-250-1 | Bacteria Proteobacteria Gammaproteobacteria Enterobacterales Enterobacteriaceae *Salmonella enterica subsp. enterica serovar* Newport str. 36807 |
| Sen-251-1 | Bacteria Proteobacteria Gammaproteobacteria Enterobacterales Enterobacteriaceae *Salmonella enterica subsp. enterica serovar* Newport str. A182RVB |
| Sen-252-1 | Bacteria Proteobacteria Gammaproteobacteria Enterobacterales Enterobacteriaceae *Salmonella enterica subsp. enterica serovar* Newport str. CVM 21538 |
| Sen-253-1 | Bacteria Proteobacteria Gammaproteobacteria Enterobacterales Enterobacteriaceae *Salmonella enterica subsp. enterica serovar* Newport str. CVM 21550 |
| Sen-254-1 | Bacteria Proteobacteria Gammaproteobacteria Enterobacterales Enterobacteriaceae *Salmonella enterica subsp. enterica serovar* Newport str. CVM 22425 |
| Sen-255-1 | Bacteria Proteobacteria Gammaproteobacteria Enterobacterales Enterobacteriaceae *Salmonella enterica subsp. enterica serovar* Newport str. CVM 22462 |
| Sen-256-1 | Bacteria Proteobacteria Gammaproteobacteria Enterobacterales Enterobacteriaceae *Salmonella enterica subsp. enterica serovar* Newport str. CVM 22513 |
| Sen-257-1 | Bacteria Proteobacteria Gammaproteobacteria Enterobacterales Enterobacteriaceae *Salmonella enterica subsp. enterica serovar* Newport str. CVM N1543 |
| Sen-258-1 | Bacteria Proteobacteria Gammaproteobacteria Enterobacterales Enterobacteriaceae *Salmonella enterica subsp. enterica serovar* Newport str. CVM N18486 |
| Sen-259-1 | Bacteria Proteobacteria Gammaproteobacteria Enterobacterales Enterobacteriaceae *Salmonella enterica subsp. enterica serovar* Newport str. DC_10-446 |
| Sen-260-1 | Bacteria Proteobacteria Gammaproteobacteria Enterobacterales Enterobacteriaceae *Salmonella enterica subsp. enterica serovar* Newport str. DC_10-449 |
| Sen-261-1 | Bacteria Proteobacteria Gammaproteobacteria Enterobacterales Enterobacteriaceae *Salmonella enterica subsp. enterica serovar* Newport str. DC_10-450 |
| Sen-262-1 | Bacteria Proteobacteria Gammaproteobacteria Enterobacterales Enterobacteriaceae *Salmonella enterica subsp. enterica serovar* Newport str. MA_10EN1469 |
| Sen-264-1 | Bacteria Proteobacteria Gammaproteobacteria Enterobacterales Enterobacteriaceae *Salmonella enterica subsp. enterica serovar* Newport str. Pond080-2TTA |
| Sen-263-1 | Bacteria Proteobacteria Gammaproteobacteria Enterobacterales Enterobacteriaceae *Salmonella enterica subsp. enterica serovar* Newport str. PRS_2010_0624 |
| Sen-265-1 | Bacteria Proteobacteria Gammaproteobacteria Enterobacterales Enterobacteriaceae *Salmonella enterica subsp. enterica serovar* Newport str. S103RVX |
| Sen-266-1 to Sen-266-2 | Bacteria Proteobacteria Gammaproteobacteria Enterobacterales Enterobacteriaceae *Salmonella enterica subsp. enterica serovar* Newport str. SL254 |
| Sen-267-1 | Bacteria Proteobacteria Gammaproteobacteria Enterobacterales Enterobacteriaceae *Salmonella enterica subsp. enterica serovar* Newport str. USDA-ARS-USMARC-1927 |
| Sen-268-1 | Bacteria Proteobacteria Gammaproteobacteria Enterobacterales Enterobacteriaceae *Salmonella enterica subsp. enterica serovar* Newport str. VA_R100804798 |
| Sen-269-1 | Bacteria Proteobacteria Gammaproteobacteria Enterobacterales Enterobacteriaceae *Salmonella enterica subsp. enterica serovar* Newport str. VA_R100808502 |
| Sen-270-1 | Bacteria Proteobacteria Gammaproteobacteria Enterobacterales Enterobacteriaceae *Salmonella enterica subsp. enterica serovar* Ohio |
| Sen-271-1 | Bacteria Proteobacteria Gammaproteobacteria Enterobacterales Enterobacteriaceae *Salmonella enterica subsp. enterica serovar* Ouakam |
| Sen-272-1 | Bacteria Proteobacteria Gammaproteobacteria Enterobacterales Enterobacteriaceae *Salmonella enterica subsp. enterica serovar* Panama str. ATCC 7378 |
| Sen-273-1 to Sen-273-3 | Bacteria Proteobacteria Gammaproteobacteria Enterobacterales Enterobacteriaceae *Salmonella enterica subsp. enterica serovar* Paratyphi A |
| Sen-274-1 | Bacteria Proteobacteria Gammaproteobacteria Enterobacterales Enterobacteriaceae *Salmonella enterica subsp. enterica serovar* Poona str. 2010K-2244 |
| Sen-275-1 | Bacteria Proteobacteria Gammaproteobacteria Enterobacterales Enterobacteriaceae *Salmonella enterica subsp. enterica serovar* Pullorum |
| Sen-276-1 | Bacteria Proteobacteria Gammaproteobacteria Enterobacterales Enterobacteriaceae *Salmonella enterica subsp. enterica serovar* Pullorum str. ATCC 9120 |
| Sen-277-1 to Sen-277-5 | Bacteria Proteobacteria Gammaproteobacteria Enterobacterales Enterobacteriaceae *Salmonella enterica subsp. enterica serovar* Reading |
| Sen-278-1 to Sen-278-22 | Bacteria Proteobacteria Gammaproteobacteria Enterobacterales Enterobacteriaceae *Salmonella enterica subsp. enterica serovar* Saintpaul |
| Sen-279-1 to Sen-279-14 | Bacteria Proteobacteria Gammaproteobacteria Enterobacterales Enterobacteriaceae *Salmonella enterica subsp. enterica serovar* Schwarzengrund |
| Sen-280-1 to Sen-280-3 | Bacteria Proteobacteria Gammaproteobacteria Enterobacterales Enterobacteriaceae *Salmonella enterica subsp. enterica serovar* Senftenberg |
| Sen-281-1 | Bacteria Proteobacteria Gammaproteobacteria Enterobacterales Enterobacteriaceae *Salmonella enterica subsp. enterica serovar* Sloterdijk str. ATCC 15791 |
| Sen-282-1 | Bacteria Proteobacteria Gammaproteobacteria Enterobacterales Enterobacteriaceae *Salmonella enterica subsp. enterica serovar* Stanley str. 06-0538 |
| Sen-283-1 | Bacteria Proteobacteria Gammaproteobacteria Enterobacterales Enterobacteriaceae *Salmonella enterica subsp. enterica serovar* Tennessee str. 4535 |
| Sen-284-1 | Bacteria Proteobacteria Gammaproteobacteria Enterobacterales Enterobacteriaceae *Salmonella enterica subsp. enterica serovar* Tennessee str. CDC07-0191 |
| Sen-285-1 | Bacteria Proteobacteria Gammaproteobacteria Enterobacterales Enterobacteriaceae *Salmonella enterica subsp. enterica serovar* Tennessee str. TXSC_TXSC08-19 |
| Sen-286-1 to Sen-286-5 | Bacteria Proteobacteria Gammaproteobacteria Enterobacterales Enterobacteriaceae *Salmonella enterica subsp. enterica serovar* Thompson |
| Sen-287-1 | Bacteria Proteobacteria Gammaproteobacteria Enterobacterales Enterobacteriaceae *Salmonella enterica subsp. enterica serovar* Thompson str. ATCC 8391 |
| Sen-288-1 to Sen-288-1125 | Bacteria Proteobacteria Gammaproteobacteria Enterobacterales Enterobacteriaceae *Salmonella enterica subsp. enterica serovar* Typhi |
| Sen-289-1 | Bacteria Proteobacteria Gammaproteobacteria Enterobacterales Enterobacteriaceae *Salmonella enterica subsp. enterica serovar* Typhi str. CR0063 |
| Sen-290-1 | Bacteria Proteobacteria Gammaproteobacteria Enterobacterales Enterobacteriaceae *Salmonella enterica subsp. enterica serovar* Typhi str. STH2370 |
| Sen-291-1 to Sen-291-7 | Bacteria Proteobacteria Gammaproteobacteria Enterobacterales Enterobacteriaceae *Salmonella enterica subsp. enterica serovar* Typhimurium |
| Sen-292-1 | Bacteria Proteobacteria Gammaproteobacteria Enterobacterales Enterobacteriaceae *Salmonella enterica subsp. enterica serovar* Typhimurium str. 104772 |
| Sen-293-1 | Bacteria Proteobacteria Gammaproteobacteria Enterobacterales Enterobacteriaceae *Salmonella enterica subsp. enterica serovar* Typhimurium str. 108402 |
| Sen-294-1 | Bacteria Proteobacteria Gammaproteobacteria Enterobacterales Enterobacteriaceae *Salmonella enterica subsp. enterica serovar* Typhimurium str. 116045 |
| Sen-295-1 | Bacteria Proteobacteria Gammaproteobacteria Enterobacterales Enterobacteriaceae *Salmonella enterica subsp. enterica serovar* Typhimurium str. 86-0368 |
| Sen-296-1 | Bacteria Proteobacteria Gammaproteobacteria Enterobacterales Enterobacteriaceae *Salmonella enterica subsp. enterica serovar* Typhimurium str. 95799 |
| Sen-297-1 | Bacteria Proteobacteria Gammaproteobacteria Enterobacterales Enterobacteriaceae *Salmonella enterica subsp. enterica serovar* Typhimurium str. 98346 |

| | |
|---|---|
| Sen-298-1 | Bacteria Proteobacteria Gammaproteobacteria Enterobacterales Enterobacteriaceae *Salmonella enterica subsp. enterica serovar* Typhimurium str. CDC 2011K-0870 |
| Sen-299-1 | Bacteria Proteobacteria Gammaproteobacteria Enterobacterales Enterobacteriaceae *Salmonella enterica subsp. enterica serovar* Typhimurium str. CDC_2009K1153 |
| Sen-300-1 to Sen-300-330 | Bacteria Proteobacteria Gammaproteobacteria Enterobacterales Enterobacteriaceae *Salmonella enterica subsp. enterica serovar* Typhimurium str. DT104 |
| Sen-301-1 | Bacteria Proteobacteria Gammaproteobacteria Enterobacterales Enterobacteriaceae *Salmonella enterica subsp. enterica serovar* Typhimurium str. L-3553 |
| Sen-302-1 | Bacteria Proteobacteria Gammaproteobacteria Enterobacterales Enterobacteriaceae *Salmonella enterica subsp. enterica serovar* Typhimurium str. LT2-4 |
| Sen-303-1 | Bacteria Proteobacteria Gammaproteobacteria Enterobacterales Enterobacteriaceae *Salmonella enterica subsp. enterica serovar* Typhimurium str. LT2-4_delta.ramA::kan |
| Sen-304-1 | Bacteria Proteobacteria Gammaproteobacteria Enterobacterales Enterobacteriaceae *Salmonella enterica subsp. enterica serovar* Typhimurium str. ST1489 |
| Sen-305-1 | Bacteria Proteobacteria Gammaproteobacteria Enterobacterales Enterobacteriaceae *Salmonella enterica subsp. enterica serovar* Typhimurium str. ST4848 |
| Sen-306-1 to Sen-306-63 | Bacteria Proteobacteria Gammaproteobacteria Enterobacterales Enterobacteriaceae *Salmonella enterica subsp. enterica serovar* Typhimurium var. 5- |
| Sen-307-1 | Bacteria Proteobacteria Gammaproteobacteria Enterobacterales Enterobacteriaceae *Salmonella enterica subsp. enterica serovar* Virchow str. SVQ1 |
| Sen-308-1 to Sen-308-4 | Bacteria Proteobacteria Gammaproteobacteria Enterobacterales Enterobacteriaceae *Salmonella enterica subsp. enterica serovar* Worthington |
| Sen-309-1 | Bacteria Proteobacteria Gammaproteobacteria Enterobacterales Enterobacteriaceae *Salmonella enterica subsp. salamae* |
| Sen-310-1 | Bacteria Proteobacteria Gammaproteobacteria Enterobacterales Enterobacteriaceae *Salmonella enterica subsp. salamae str.* DMA-1 |
| Sbo-2-1 to Sbo-2-10 | Bacteria Proteobacteria Gammaproteobacteria Enterobacterales Enterobacteriaceae *Shigella boydii* |
| Sdy-1-1 | Bacteria Proteobacteria Gammaproteobacteria Enterobacterales Enterobacteriaceae *Shigella dysenteriae* S6205 |
| Sdy-2-1 | Bacteria Proteobacteria Gammaproteobacteria Enterobacterales Enterobacteriaceae *Shigella dysenteriae* SD1D |
| Sfl-1-1 to Sfl-1-2 | Bacteria Proteobacteria Gammaproteobacteria Enterobacterales Enterobacteriaceae *Shigella flexneri* |
| Sfl-2-1 | Bacteria Proteobacteria Gammaproteobacteria Enterobacterales Enterobacteriaceae *Shigella flexneri* 1997005 |
| Sfl-3-1 | Bacteria Proteobacteria Gammaproteobacteria Enterobacterales Enterobacteriaceae *Shigella flexneri* 2000019 |
| Sfl-4-1 | Bacteria Proteobacteria Gammaproteobacteria Enterobacterales Enterobacteriaceae *Shigella flexneri* 2001004 |
| Sfl-5-1 | Bacteria Proteobacteria Gammaproteobacteria Enterobacterales Enterobacteriaceae *Shigella flexneri* 2001025 |
| Sfl-6-1 | Bacteria Proteobacteria Gammaproteobacteria Enterobacterales Enterobacteriaceae *Shigella flexneri* 2001027 |
| Sfl-7-1 | Bacteria Proteobacteria Gammaproteobacteria Enterobacterales Enterobacteriaceae *Shigella flexneri* 2001042 |
| Sfl-8-1 | Bacteria Proteobacteria Gammaproteobacteria Enterobacterales Enterobacteriaceae *Shigella flexneri* 2001044 |
| Sfl-9-1 | Bacteria Proteobacteria Gammaproteobacteria Enterobacterales Enterobacteriaceae *Shigella flexneri* 2001048 |
| Sfl-10-1 | Bacteria Proteobacteria Gammaproteobacteria Enterobacterales Enterobacteriaceae *Shigella flexneri* 2002007 |
| Sfl-11-1 | Bacteria Proteobacteria Gammaproteobacteria Enterobacterales Enterobacteriaceae *Shigella flexneri* 2002021 |
| Sfl-12-1 | Bacteria Proteobacteria Gammaproteobacteria Enterobacterales Enterobacteriaceae *Shigella flexneri* 2002035 |
| Sfl-13-1 | Bacteria Proteobacteria Gammaproteobacteria Enterobacterales Enterobacteriaceae *Shigella flexneri* 2002069 |
| Sfl-14-1 | Bacteria Proteobacteria Gammaproteobacteria Enterobacterales Enterobacteriaceae *Shigella flexneri* 2002091 |
| Sfl-15-1 | Bacteria Proteobacteria Gammaproteobacteria Enterobacterales Enterobacteriaceae *Shigella flexneri* 2002103 |
| Sfl-16-1 | Bacteria Proteobacteria Gammaproteobacteria Enterobacterales Enterobacteriaceae *Shigella flexneri* 2002110 |
| Sfl-17-1 | Bacteria Proteobacteria Gammaproteobacteria Enterobacterales Enterobacteriaceae *Shigella flexneri* 2002127 |
| Sfl-18-1 | Bacteria Proteobacteria Gammaproteobacteria Enterobacterales Enterobacteriaceae *Shigella flexneri* 2002140 |
| Sfl-19-1 | Bacteria Proteobacteria Gammaproteobacteria Enterobacterales Enterobacteriaceae *Shigella flexneri* 2002141 |
| Sfl-20-1 | Bacteria Proteobacteria Gammaproteobacteria Enterobacterales Enterobacteriaceae *Shigella flexneri* 2002142 |
| Sfl-21-1 | Bacteria Proteobacteria Gammaproteobacteria Enterobacterales Enterobacteriaceae *Shigella flexneri* 2003055 |
| Sfl-22-1 | Bacteria Proteobacteria Gammaproteobacteria Enterobacterales Enterobacteriaceae *Shigella flexneri* 2005051 |
| Sfl-23-1 | Bacteria Proteobacteria Gammaproteobacteria Enterobacterales Enterobacteriaceae *Shigella flexneri* 2005AH264 |
| Sfl-24-1 | Bacteria Proteobacteria Gammaproteobacteria Enterobacterales Enterobacteriaceae *Shigella flexneri* 2005GS061 |
| Sfl-25-1 to Sfl-25-14 | Bacteria Proteobacteria Gammaproteobacteria Enterobacterales Enterobacteriaceae *Shigella flexneri* 2a |
| Sfl-26-1 | Bacteria Proteobacteria Gammaproteobacteria Enterobacterales Enterobacteriaceae *Shigella flexneri* 51575 |
| Sfl-27-1 | Bacteria Proteobacteria Gammaproteobacteria Enterobacterales Enterobacteriaceae *Shigella flexneri* 51576 |
| Sfl-28-1 | Bacteria Proteobacteria Gammaproteobacteria Enterobacterales Enterobacteriaceae *Shigella flexneri* 51577 |
| Sfl-29-1 | Bacteria Proteobacteria Gammaproteobacteria Enterobacterales Enterobacteriaceae *Shigella flexneri* Shi05SX04 |
| Sfl-30-1 | Bacteria Proteobacteria Gammaproteobacteria Enterobacterales Enterobacteriaceae *Shigella flexneri* Shi06AH028 |
| Sfl-31-1 | Bacteria Proteobacteria Gammaproteobacteria Enterobacterales Enterobacteriaceae *Shigella flexneri* Shi06AH091 |
| Sfl-32-1 | Bacteria Proteobacteria Gammaproteobacteria Enterobacterales Enterobacteriaceae *Shigella flexneri* Shi06AH116 |
| Sfl-33-1 | Bacteria Proteobacteria Gammaproteobacteria Enterobacterales Enterobacteriaceae *Shigella flexneri* Shi06AH130 |
| Sfl-34-1 | Bacteria Proteobacteria Gammaproteobacteria Enterobacterales Enterobacteriaceae *Shigella flexneri* Shi06AH135 |
| Sfl-35-1 | Bacteria Proteobacteria Gammaproteobacteria Enterobacterales Enterobacteriaceae *Shigella flexneri* Shi06AH66 |
| Sfl-36-1 | Bacteria Proteobacteria Gammaproteobacteria Enterobacterales Enterobacteriaceae *Shigella flexneri* Shi06GS07 |
| Sfl-37-1 | Bacteria Proteobacteria Gammaproteobacteria Enterobacterales Enterobacteriaceae *Shigella flexneri* Shi06GS37 |
| Sfl-38-1 | Bacteria Proteobacteria Gammaproteobacteria Enterobacterales Enterobacteriaceae *Shigella flexneri* Shi06GS43 |
| Sfl-39-1 | Bacteria Proteobacteria Gammaproteobacteria Enterobacterales Enterobacteriaceae *Shigella flexneri* Shi06GS48 |
| Sfl-40-1 | Bacteria Proteobacteria Gammaproteobacteria Enterobacterales Enterobacteriaceae *Shigella flexneri* Shi06GS55 |
| Sfl-41-1 | Bacteria Proteobacteria Gammaproteobacteria Enterobacterales Enterobacteriaceae *Shigella flexneri* Shi06HN016 |
| Sfl-42-1 | Bacteria Proteobacteria Gammaproteobacteria Enterobacterales Enterobacteriaceae *Shigella flexneri* Shi06HN023 |
| Sfl-43-1 | Bacteria Proteobacteria Gammaproteobacteria Enterobacterales Enterobacteriaceae *Shigella flexneri* Shi06HN091 |
| Sfl-44-1 | Bacteria Proteobacteria Gammaproteobacteria Enterobacterales Enterobacteriaceae *Shigella flexneri* Shi06HN118 |
| Sfl-45-1 | Bacteria Proteobacteria Gammaproteobacteria Enterobacterales Enterobacteriaceae *Shigella flexneri* Shi06HN159 |

| | |
|---|---|
| Sfl-46-1 | Bacteria Proteobacteria Gammaproteobacteria Enterobacterales Enterobacteriaceae *Shigella flexneri* Shi06HN244 |
| Sfl-47-1 | Bacteria Proteobacteria Gammaproteobacteria Enterobacterales Enterobacteriaceae *Shigella flexneri* Shi06HN250 |
| Sfl-48-1 | Bacteria Proteobacteria Gammaproteobacteria Enterobacterales Enterobacteriaceae *Shigella flexneri* Shi06HN378 |
| Sfl-49-1 | Bacteria Proteobacteria Gammaproteobacteria Enterobacterales Enterobacteriaceae *Shigella flexneri* Shi06SX36 |
| Sfl-50-1 | Bacteria Proteobacteria Gammaproteobacteria Enterobacterales Enterobacteriaceae *Shigella flexneri* Shi06SX53 |
| Sfl-51-1 to Sfl-51-2 | Bacteria Proteobacteria Gammaproteobacteria Enterobacterales Enterobacteriaceae *Shigella flexneri* Y |
| Sso-1-1 to Sso-1-547 | Bacteria Proteobacteria Gammaproteobacteria Enterobacterales Enterobacteriaceae *Shigella sonnei* |
| Sso-2-1 | Bacteria Proteobacteria Gammaproteobacteria Enterobacterales Enterobacteriaceae *Shigella sonnei* 1DT-1 |
| Sco-2-1 | Bacteria Proteobacteria Gammaproteobacteria Enterobacterales Enterobacteriaceae *Siccibacter colletis* |
| Stu-1-1 | Bacteria Proteobacteria Gammaproteobacteria Enterobacterales Enterobacteriaceae *Siccibacter turicensis* LMG 23730 |
| Tgu-1-1 | Bacteria Proteobacteria Gammaproteobacteria Enterobacterales Enterobacteriaceae *Trabulsiella guamensis* ATCC 49490 |
| Tod-1-1 to Tod-1-2 | Bacteria Proteobacteria Gammaproteobacteria Enterobacterales Enterobacteriaceae *Trabulsiella odontotermitis* |
| Eam-1-1 | Bacteria Proteobacteria Gammaproteobacteria Enterobacterales Erwiniaceae *Erwinia amylovora* NBRC 12687 |
| Ebi-1-1 | Bacteria Proteobacteria Gammaproteobacteria Enterobacterales Erwiniaceae *Erwinia billingiae* |
| Ein-1-1 to Ein-1-2 | Bacteria Proteobacteria Gammaproteobacteria Enterobacterales Erwiniaceae *Erwinia iniecta* |
| Ema-2-1 | Bacteria Proteobacteria Gammaproteobacteria Enterobacterales Erwiniaceae *Erwinia mallotivora* |
| Eol-1-1 | Bacteria Proteobacteria Gammaproteobacteria Enterobacterales Erwiniaceae *Erwinia oleae* |
| Esp-55-1 | Bacteria Proteobacteria Gammaproteobacteria Enterobacterales Erwiniaceae *Erwinia sp.* 9145 |
| Etr-1-1 | Bacteria Proteobacteria Gammaproteobacteria Enterobacterales Erwiniaceae *Erwinia tracheiphila* |
| Ety-1-1 | Bacteria Proteobacteria Gammaproteobacteria Enterobacterales Erwiniaceae *Erwinia typographi* |
| Pag-2-1 to Pag-2-6 | Bacteria Proteobacteria Gammaproteobacteria Enterobacterales Erwiniaceae Pantoea Pantoea agglomerans group *Pantoea agglomerans* |
| Pag-3-1 | Bacteria Proteobacteria Gammaproteobacteria Enterobacterales Erwiniaceae Pantoea Pantoea agglomerans group *Pantoea agglomerans* Eh318 |
| Pan-1-1 to Pan-1-5 | Bacteria Proteobacteria Gammaproteobacteria Enterobacterales Erwiniaceae *Pantoea ananatis* |
| Pan-2-1 | Bacteria Proteobacteria Gammaproteobacteria Enterobacterales Erwiniaceae *Pantoea ananatis* BD442 |
| Pan-3-1 to Pan-3-2 | Bacteria Proteobacteria Gammaproteobacteria Enterobacterales Erwiniaceae *Pantoea ananatis* LMG 2665 |
| Pan-4-1 | Bacteria Proteobacteria Gammaproteobacteria Enterobacterales Erwiniaceae *Pantoea ananatis* Sd-1 |
| Pan-5-1 | Bacteria Proteobacteria Gammaproteobacteria Enterobacterales Erwiniaceae *Pantoea anthophila* |
| Pro-1-1 | Bacteria Proteobacteria Gammaproteobacteria Enterobacterales Erwiniaceae *Pantoea rodasii* |
| Prw-1-1 | Bacteria Proteobacteria Gammaproteobacteria Enterobacterales Erwiniaceae *Pantoea rwandensis* |
| Pse-1-1 | Bacteria Proteobacteria Gammaproteobacteria Enterobacterales Erwiniaceae *Pantoea septica* |
| Psp-36-1 | Bacteria Proteobacteria Gammaproteobacteria Enterobacterales Erwiniaceae *Pantoea sp.* 3.5.1 |
| Psp-37-1 | Bacteria Proteobacteria Gammaproteobacteria Enterobacterales Erwiniaceae *Pantoea sp.* 9140 |
| Psp-38-1 | Bacteria Proteobacteria Gammaproteobacteria Enterobacterales Erwiniaceae *Pantoea sp.* BL1 |
| Psp-39-1 | Bacteria Proteobacteria Gammaproteobacteria Enterobacterales Erwiniaceae *Pantoea sp.* CFSAN033090 |
| Psp-40-1 | Bacteria Proteobacteria Gammaproteobacteria Enterobacterales Erwiniaceae *Pantoea sp.* IMH |
| Psp-41-1 | Bacteria Proteobacteria Gammaproteobacteria Enterobacterales Erwiniaceae *Pantoea sp.* MBLJ3 |
| Psp-42-1 | Bacteria Proteobacteria Gammaproteobacteria Enterobacterales Erwiniaceae *Pantoea sp.* NGS-ED-1003 |
| Psp-43-1 | Bacteria Proteobacteria Gammaproteobacteria Enterobacterales Erwiniaceae *Pantoea sp.* PSNIH1 |
| Psp-44-1 | Bacteria Proteobacteria Gammaproteobacteria Enterobacterales Erwiniaceae *Pantoea sp.* PSNIH2 |
| Psp-45-1 | Bacteria Proteobacteria Gammaproteobacteria Enterobacterales Erwiniaceae *Pantoea sp.* RIT-PI-b |
| Psp-46-1 | Bacteria Proteobacteria Gammaproteobacteria Enterobacterales Erwiniaceae *Pantoea sp.* SM3 |
| Pst-1-1 to Pst-1-2 | Bacteria Proteobacteria Gammaproteobacteria Enterobacterales Erwiniaceae *Pantoea stewartii* |
| Pva-1-1 to Pva-1-3 | Bacteria Proteobacteria Gammaproteobacteria Enterobacterales Erwiniaceae *Pantoea vagans* |
| Pfl-3-1 | Bacteria Proteobacteria Gammaproteobacteria Enterobacterales Erwiniaceae *Phaseolibacter flectens* ATCC 12775 |
| Tmo-1-1 | Bacteria Proteobacteria Gammaproteobacteria Enterobacterales Erwiniaceae *Tatumella morbirosei* |
| Tpt-1-1 to Tpt-1-2 | Bacteria Proteobacteria Gammaproteobacteria Enterobacterales Erwiniaceae *Tatumella ptyseos* ATCC 33301 |
| Tsa-1-1 | Bacteria Proteobacteria Gammaproteobacteria Enterobacterales Erwiniaceae *Tatumella saanichensis* |
| Tsp-2-1 | Bacteria Proteobacteria Gammaproteobacteria Enterobacterales Erwiniaceae *Tatumella sp.* UCD-D_suzukii |
| Ean-1-1 | Bacteria Proteobacteria Gammaproteobacteria Enterobacterales Hafniaceae *Edwardsiella anguillarum* ET070829 |
| Ean-2-1 | Bacteria Proteobacteria Gammaproteobacteria Enterobacterales Hafniaceae *Edwardsiella anguillarum* ET080729 |
| Ean-3-1 | Bacteria Proteobacteria Gammaproteobacteria Enterobacterales Hafniaceae *Edwardsiella anguillarum* ET080813 |
| Ean-4-1 | Bacteria Proteobacteria Gammaproteobacteria Enterobacterales Hafniaceae *Edwardsiella anguillarum* ET081126R |
| Eho-5-1 | Bacteria Proteobacteria Gammaproteobacteria Enterobacterales Hafniaceae *Edwardsiella hoshinae* NBRC 105699 = ATCC 33379 |
| Eic-1-1 | Bacteria Proteobacteria Gammaproteobacteria Enterobacterales Hafniaceae *Edwardsiella ictaluri* ATCC 33202 |
| Epi-1-1 to Epi-1-2 | Bacteria Proteobacteria Gammaproteobacteria Enterobacterales Hafniaceae *Edwardsiella piscicida* |
| Esp-56-1 | Bacteria Proteobacteria Gammaproteobacteria Enterobacterales Hafniaceae *Edwardsiella sp.* EA181011 |
| Eta-1-1 | Bacteria Proteobacteria Gammaproteobacteria Enterobacterales Hafniaceae *Edwardsiella tarda* |
| Eta-2-1 | Bacteria Proteobacteria Gammaproteobacteria Enterobacterales Hafniaceae *Edwardsiella tarda* ATCC 15947 = NBRC 105688 |
| Eta-3-1 | Bacteria Proteobacteria Gammaproteobacteria Enterobacterales Hafniaceae *Edwardsiella tarda* DT |
| Hal-1-1 to Hal-1-4 | Bacteria Proteobacteria Gammaproteobacteria Enterobacterales Hafniaceae *Hafnia alvei* |
| Hal-2-1 | Bacteria Proteobacteria Gammaproteobacteria Enterobacterales Hafniaceae *Hafnia alvei* ATCC 13337 |
| Hal-3-1 | Bacteria Proteobacteria Gammaproteobacteria Enterobacterales Hafniaceae *Hafnia alvei* FB1 |
| Hpa-1-1 to Hpa-1-2 | Bacteria Proteobacteria Gammaproteobacteria Enterobacterales Hafniaceae *Hafnia paralvei* |
| Aen-3-1 | Bacteria Proteobacteria Gammaproteobacteria Enterobacterales Morganellaceae *Arsenophonus endosymbiont str.* Hangzhou of Nilaparvata lugens |
| Ana-1-1 | Bacteria Proteobacteria Gammaproteobacteria Enterobacterales Morganellaceae *Arsenophonus nasoniae* DSM 15247 |

| | |
|---|---|
| Mwi-1-1 | Bacteria Proteobacteria Gammaproteobacteria Enterobacterales Morganellaceae *Moellerella wisconsensis* |
| Mmo-1-1 to Mmo-1-8 | Bacteria Proteobacteria Gammaproteobacteria Enterobacterales Morganellaceae *Morganella morganii* |
| Mmo-2-1 | Bacteria Proteobacteria Gammaproteobacteria Enterobacterales Morganellaceae *Morganella morganii* F675 |
| Mmo-3-1 | Bacteria Proteobacteria Gammaproteobacteria Enterobacterales Morganellaceae *Morganella morganii* H1r |
| Mmo-4-1 | Bacteria Proteobacteria Gammaproteobacteria Enterobacterales Morganellaceae *Morganella morganii* IS15 |
| Msp-5-1 | Bacteria Proteobacteria Gammaproteobacteria Enterobacterales Morganellaceae *Morganella sp.* EGD-HP17 |
| Pas-1-1 | Bacteria Proteobacteria Gammaproteobacteria Enterobacterales Morganellaceae *Photorhabdus asymbiotica subsp. australis* DSM 17609 |
| Plu-5-1 to Plu-5-2 | Bacteria Proteobacteria Gammaproteobacteria Enterobacterales Morganellaceae *Photorhabdus luminescens* |
| Plu-6-1 | Bacteria Proteobacteria Gammaproteobacteria Enterobacterales Morganellaceae *Photorhabdus luminescens* BA1 |
| Plu-7-1 | Bacteria Proteobacteria Gammaproteobacteria Enterobacterales Morganellaceae *Photorhabdus luminescens* NBAII H75HRPL105 |
| Plu-8-1 | Bacteria Proteobacteria Gammaproteobacteria Enterobacterales Morganellaceae *Photorhabdus luminescens* NBAII HiPL101 |
| Plu-9-1 | Bacteria Proteobacteria Gammaproteobacteria Enterobacterales Morganellaceae *Photorhabdus luminescens subsp. luminescens* |
| Pte-2-1 | Bacteria Proteobacteria Gammaproteobacteria Enterobacterales Morganellaceae *Photorhabdus temperata subsp. khanii* NC19 |
| Pte-3-1 | Bacteria Proteobacteria Gammaproteobacteria Enterobacterales Morganellaceae *Photorhabdus temperata subsp. temperata* Meg1 |
| Pte-4-1 | Bacteria Proteobacteria Gammaproteobacteria Enterobacterales Morganellaceae *Photorhabdus temperata subsp. thracensis* |
| Pmi-1-1 to Pmi-1-6 | Bacteria Proteobacteria Gammaproteobacteria Enterobacterales Morganellaceae *Proteus mirabilis* |
| Pvu-2-1 | Bacteria Proteobacteria Gammaproteobacteria Enterobacterales Morganellaceae *Proteus vulgaris* |
| Pal-1-1 | Bacteria Proteobacteria Gammaproteobacteria Enterobacterales Morganellaceae *Providencia alcalifaciens* 205/92 |
| Pal-2-1 | Bacteria Proteobacteria Gammaproteobacteria Enterobacterales Morganellaceae *Providencia alcalifaciens* F90-2004 |
| Pal-3-1 | Bacteria Proteobacteria Gammaproteobacteria Enterobacterales Morganellaceae *Providencia alcalifaciens* PAL-1 |
| Pal-4-1 | Bacteria Proteobacteria Gammaproteobacteria Enterobacterales Morganellaceae *Providencia alcalifaciens* PAL-2 |
| Pal-5-1 | Bacteria Proteobacteria Gammaproteobacteria Enterobacterales Morganellaceae *Providencia alcalifaciens* PAL-3 |
| Pal-6-1 | Bacteria Proteobacteria Gammaproteobacteria Enterobacterales Morganellaceae *Providencia alcalifaciens* R90-1475 |
| Pal-7-1 | Bacteria Proteobacteria Gammaproteobacteria Enterobacterales Morganellaceae *Providencia alcalifaciens* RIMD 1656011 |
| Pre-1-1 to Pre-1-2 | Bacteria Proteobacteria Gammaproteobacteria Enterobacterales Morganellaceae *Providencia rettgeri* |
| Pst-2-1 | Bacteria Proteobacteria Gammaproteobacteria Enterobacterales Morganellaceae *Providencia stuartii* |
| Xbo-1-1 | Bacteria Proteobacteria Gammaproteobacteria Enterobacterales Morganellaceae *Xenorhabdus bovienii* |
| Xbo-4-1 | Bacteria Proteobacteria Gammaproteobacteria Enterobacterales Morganellaceae *Xenorhabdus bovienii str. feltiae* Florida |
| Xbo-5-1 | Bacteria Proteobacteria Gammaproteobacteria Enterobacterales Morganellaceae *Xenorhabdus bovienii str. feltiae* France |
| Xbo-6-1 | Bacteria Proteobacteria Gammaproteobacteria Enterobacterales Morganellaceae *Xenorhabdus bovienii str. feltiae* Moldova |
| Xbo-2-1 | Bacteria Proteobacteria Gammaproteobacteria Enterobacterales Morganellaceae *Xenorhabdus bovienii str.* Intermedium |
| Xbo-3-1 | Bacteria Proteobacteria Gammaproteobacteria Enterobacterales Morganellaceae *Xenorhabdus bovienii str.* Jollieti |
| Xbo-7-1 | Bacteria Proteobacteria Gammaproteobacteria Enterobacterales Morganellaceae *Xenorhabdus bovienii str. kraussei* Becker Underwood |
| Xbo-8-1 | Bacteria Proteobacteria Gammaproteobacteria Enterobacterales Morganellaceae *Xenorhabdus bovienii str. kraussei* Quebec |
| Xbo-9-1 | Bacteria Proteobacteria Gammaproteobacteria Enterobacterales Morganellaceae *Xenorhabdus bovienii str. oregonense* |
| Xbo-10-1 | Bacteria Proteobacteria Gammaproteobacteria Enterobacterales Morganellaceae *Xenorhabdus bovienii str. puntauvense* |
| Xca-1-1 | Bacteria Proteobacteria Gammaproteobacteria Enterobacterales Morganellaceae *Xenorhabdus cabanillasii* JM26 |
| Xdo-1-1 | Bacteria Proteobacteria Gammaproteobacteria Enterobacterales Morganellaceae *Xenorhabdus doucetiae* |
| Xgr-1-1 | Bacteria Proteobacteria Gammaproteobacteria Enterobacterales Morganellaceae *Xenorhabdus griffiniae* |
| Xkh-1-1 | Bacteria Proteobacteria Gammaproteobacteria Enterobacterales Morganellaceae *Xenorhabdus khoisanae* |
| Xne-1-1 | Bacteria Proteobacteria Gammaproteobacteria Enterobacterales Morganellaceae *Xenorhabdus nematophila* |
| Xne-2-1 | Bacteria Proteobacteria Gammaproteobacteria Enterobacterales Morganellaceae *Xenorhabdus nematophila* AN6/1 |
| Xpo-1-1 | Bacteria Proteobacteria Gammaproteobacteria Enterobacterales Morganellaceae *Xenorhabdus poinarii* G6 |
| Xsp-1-1 | Bacteria Proteobacteria Gammaproteobacteria Enterobacterales Morganellaceae *Xenorhabdus sp.* NBAII XenSa04 |
| Xsz-1-1 | Bacteria Proteobacteria Gammaproteobacteria Enterobacterales Morganellaceae *Xenorhabdus szentirmaii* DSM 16338 |
| Bgo-1-1 | Bacteria Proteobacteria Gammaproteobacteria Enterobacterales Pectobacteriaceae *Brenneria goodwinii* |
| Dch-1-1 to Dch-1-2 | Bacteria Proteobacteria Gammaproteobacteria Enterobacterales Pectobacteriaceae *Dickeya chrysanthemi* |
| Dch-2-1 | Bacteria Proteobacteria Gammaproteobacteria Enterobacterales Pectobacteriaceae *Dickeya chrysanthemi* NCPPB 3533 |
| Dch-3-1 | Bacteria Proteobacteria Gammaproteobacteria Enterobacterales Pectobacteriaceae *Dickeya chrysanthemi* NCPPB 402 |
| Dch-4-1 | Bacteria Proteobacteria Gammaproteobacteria Enterobacterales Pectobacteriaceae *Dickeya chrysanthemi* NCPPB 516 |
| Dda-1-1 | Bacteria Proteobacteria Gammaproteobacteria Enterobacterales Pectobacteriaceae *Dickeya dadantii* NCPPB 3537 |
| Dda-2-1 | Bacteria Proteobacteria Gammaproteobacteria Enterobacterales Pectobacteriaceae *Dickeya dadantii* NCPPB 898 |
| Dda-3-1 | Bacteria Proteobacteria Gammaproteobacteria Enterobacterales Pectobacteriaceae *Dickeya dadantii subsp. dieffenbachiae* NCPPB 2976 |
| Ddi-1-1 | Bacteria Proteobacteria Gammaproteobacteria Enterobacterales Pectobacteriaceae *Dickeya dianthicola* IPO 980 |
| Ddi-2-1 | Bacteria Proteobacteria Gammaproteobacteria Enterobacterales Pectobacteriaceae *Dickeya dianthicola* RNS04.9 |
| Dso-1-1 | Bacteria Proteobacteria Gammaproteobacteria Enterobacterales Pectobacteriaceae *Dickeya solani* |
| Dsp-1-1 | Bacteria Proteobacteria Gammaproteobacteria Enterobacterales Pectobacteriaceae *Dickeya sp.* 2B12 |
| Dsp-2-1 | Bacteria Proteobacteria Gammaproteobacteria Enterobacterales Pectobacteriaceae *Dickeya sp.* B16 |
| Dsp-3-1 | Bacteria Proteobacteria Gammaproteobacteria Enterobacterales Pectobacteriaceae *Dickeya sp.* DW 0440 |
| Dsp-4-1 | Bacteria Proteobacteria Gammaproteobacteria Enterobacterales Pectobacteriaceae *Dickeya sp.* MK7 |
| Dsp-5-1 | Bacteria Proteobacteria Gammaproteobacteria Enterobacterales Pectobacteriaceae *Dickeya sp.* NCPPB 3274 |
| Dsp-6-1 | Bacteria Proteobacteria Gammaproteobacteria Enterobacterales Pectobacteriaceae *Dickeya sp.* S1 |
| Dze-1-1 | Bacteria Proteobacteria Gammaproteobacteria Enterobacterales Pectobacteriaceae *Dickeya zeae* CSL RW192 |
| Dze-2-1 | Bacteria Proteobacteria Gammaproteobacteria Enterobacterales Pectobacteriaceae *Dickeya zeae* DZ2Q |
| Dze-3-1 | Bacteria Proteobacteria Gammaproteobacteria Enterobacterales Pectobacteriaceae *Dickeya zeae* EC1 |

| | |
|---|---|
| Dze-4-1 | Bacteria Proteobacteria Gammaproteobacteria Enterobacterales Pectobacteriaceae *Dickeya zeae* MK19 |
| Dze-5-1 | Bacteria Proteobacteria Gammaproteobacteria Enterobacterales Pectobacteriaceae *Dickeya zeae* NCPPB 2538 |
| Dze-6-1 | Bacteria Proteobacteria Gammaproteobacteria Enterobacterales Pectobacteriaceae *Dickeya zeae* NCPPB 3531 |
| Lqu-1-1 to Lqu-1-3 | Bacteria Proteobacteria Gammaproteobacteria Enterobacterales Pectobacteriaceae *Lonsdalea quercina subsp. quercina* |
| Pat-2-1 to Pat-2-2 | Bacteria Proteobacteria Gammaproteobacteria Enterobacterales Pectobacteriaceae *Pectobacterium atrosepticum* |
| Pat-3-1 | Bacteria Proteobacteria Gammaproteobacteria Enterobacterales Pectobacteriaceae *Pectobacterium atrosepticum* CFBP 6276 |
| Pbe-1-1 to Pbe-1-2 | Bacteria Proteobacteria Gammaproteobacteria Enterobacterales Pectobacteriaceae *Pectobacterium betavasculorum* |
| Pca-1-1 | Bacteria Proteobacteria Gammaproteobacteria Enterobacterales Pectobacteriaceae *Pectobacterium carotovorum* |
| Pca-2-1 | Bacteria Proteobacteria Gammaproteobacteria Enterobacterales Pectobacteriaceae *Pectobacterium carotovorum subsp. actinidiae* |
| Pca-3-1 to Pca-3-15 | Bacteria Proteobacteria Gammaproteobacteria Enterobacterales Pectobacteriaceae *Pectobacterium carotovorum subsp. brasiliense* |
| Pca-4-1 | Bacteria Proteobacteria Gammaproteobacteria Enterobacterales Pectobacteriaceae *Pectobacterium carotovorum subsp. brasiliensis* ICMP 19477 |
| Pca-5-1 to Pca-5-11 | Bacteria Proteobacteria Gammaproteobacteria Enterobacterales Pectobacteriaceae *Pectobacterium carotovorum subsp. carotovorum* |
| Pca-6-1 | Bacteria Proteobacteria Gammaproteobacteria Enterobacterales Pectobacteriaceae *Pectobacterium carotovorum subsp. carotovorum* ICMP 5702 |
| Pca-7-1 | Bacteria Proteobacteria Gammaproteobacteria Enterobacterales Pectobacteriaceae *Pectobacterium carotovorum subsp. carotovorum* UGC32 |
| Pca-8-1 to Pca-8-2 | Bacteria Proteobacteria Gammaproteobacteria Enterobacterales Pectobacteriaceae *Pectobacterium carotovorum subsp. odoriferum* |
| Ppa-1-1 | Bacteria Proteobacteria Gammaproteobacteria Enterobacterales Pectobacteriaceae *Pectobacterium parmentieri* |
| Ppa-2-1 | Bacteria Proteobacteria Gammaproteobacteria Enterobacterales Pectobacteriaceae *Pectobacterium parmentieri* CFIA1002 |
| Ppo-1-1 | Bacteria Proteobacteria Gammaproteobacteria Enterobacterales Pectobacteriaceae *Pectobacterium polaris* |
| Pwa-1-1 to Pwa-1-2 | Bacteria Proteobacteria Gammaproteobacteria Enterobacterales Pectobacteriaceae *Pectobacterium wasabiae* |
| CSo-1-1 | Bacteria Proteobacteria Gammaproteobacteria Enterobacterales Pectobacteriaceae Sodalis *Candidatus* Sodalis pierantonius str. SOPE |
| Spr-1-1 | Bacteria Proteobacteria Gammaproteobacteria Enterobacterales Pectobacteriaceae *Sodalis praecaptivus* |
| Pur-1-1 | Bacteria Proteobacteria Gammaproteobacteria Enterobacterales unclassified Enterobacterales *Phytobacter ursingii* |
| Psh-1-1 | Bacteria Proteobacteria Gammaproteobacteria Enterobacterales unclassified Enterobacterales *Plesiomonas shigelloides* |
| Psp-47-1 | Bacteria Proteobacteria Gammaproteobacteria Enterobacterales unclassified Enterobacterales *Plesiomonas sp.* ZOR0011 |
| Cmu-2-1 | Bacteria Proteobacteria Gammaproteobacteria Enterobacterales Yersiniaceae *Chania multitudinisentens* RB-25 |
| Eam-2-1 | Bacteria Proteobacteria Gammaproteobacteria Enterobacterales Yersiniaceae *Ewingella americana* ATCC 33852 |
| Raq-1-1 to Raq-1-2 | Bacteria Proteobacteria Gammaproteobacteria Enterobacterales Yersiniaceae *Rahnella aquatilis* |
| Raq-2-1 to Raq-2-2 | Bacteria Proteobacteria Gammaproteobacteria Enterobacterales Yersiniaceae *Rahnella aquatilis* CIP 78.65 = ATCC 33071 |
| Rsp-3-1 | Bacteria Proteobacteria Gammaproteobacteria Enterobacterales Yersiniaceae *Rahnella sp.* WP5 |
| Rch-1-1 | Bacteria Proteobacteria Gammaproteobacteria Enterobacterales Yersiniaceae *Rouxiella chamberiensis* |
| Sfo-1-1 | Bacteria Proteobacteria Gammaproteobacteria Enterobacterales Yersiniaceae *Serratia fonticola* |
| Sfo-2-1 | Bacteria Proteobacteria Gammaproteobacteria Enterobacterales Yersiniaceae *Serratia fonticola* LMG 7882 |
| Sfo-3-1 | Bacteria Proteobacteria Gammaproteobacteria Enterobacterales Yersiniaceae *Serratia fonticola* UTAD54 |
| Sgr-1-1 | Bacteria Proteobacteria Gammaproteobacteria Enterobacterales Yersiniaceae *Serratia grimesii* |
| Sli-1-1 | Bacteria Proteobacteria Gammaproteobacteria Enterobacterales Yersiniaceae *Serratia liquefaciens* |
| Sli-2-1 | Bacteria Proteobacteria Gammaproteobacteria Enterobacterales Yersiniaceae *Serratia liquefaciens* FK01 |
| Sma-3-1 to Sma-3-4 | Bacteria Proteobacteria Gammaproteobacteria Enterobacterales Yersiniaceae *Serratia marcescens* |
| Sma-4-1 | Bacteria Proteobacteria Gammaproteobacteria Enterobacterales Yersiniaceae *Serratia marcescens* AB42556419-isolate1 |
| Sma-5-1 | Bacteria Proteobacteria Gammaproteobacteria Enterobacterales Yersiniaceae *Serratia marcescens* BIDMC 44 |
| Sma-6-1 | Bacteria Proteobacteria Gammaproteobacteria Enterobacterales Yersiniaceae *Serratia marcescens* BIDMC 50 |
| Sma-7-1 | Bacteria Proteobacteria Gammaproteobacteria Enterobacterales Yersiniaceae *Serratia marcescens* BIDMC 80 |
| Sma-8-1 | Bacteria Proteobacteria Gammaproteobacteria Enterobacterales Yersiniaceae *Serratia marcescens* BIDMC 81 |
| Sma-9-1 | Bacteria Proteobacteria Gammaproteobacteria Enterobacterales Yersiniaceae *Serratia marcescens* H1q |
| Sma-10-1 | Bacteria Proteobacteria Gammaproteobacteria Enterobacterales Yersiniaceae *Serratia marcescens* LCT-SM166 |
| Sma-11-1 | Bacteria Proteobacteria Gammaproteobacteria Enterobacterales Yersiniaceae *Serratia marcescens* LCT-SM262 |
| Sma-12-1 | Bacteria Proteobacteria Gammaproteobacteria Enterobacterales Yersiniaceae *Serratia marcescens* MC458 |
| Sma-13-1 | Bacteria Proteobacteria Gammaproteobacteria Enterobacterales Yersiniaceae *Serratia marcescens* MC459 |
| Sma-14-1 | Bacteria Proteobacteria Gammaproteobacteria Enterobacterales Yersiniaceae *Serratia marcescens* MC460 |
| Sma-15-1 | Bacteria Proteobacteria Gammaproteobacteria Enterobacterales Yersiniaceae *Serratia marcescens* MC6000 |
| Sma-16-1 | Bacteria Proteobacteria Gammaproteobacteria Enterobacterales Yersiniaceae *Serratia marcescens* MC6001 |
| Sma-17-1 | Bacteria Proteobacteria Gammaproteobacteria Enterobacterales Yersiniaceae *Serratia marcescens* MC620 |
| Sma-18-1 | Bacteria Proteobacteria Gammaproteobacteria Enterobacterales Yersiniaceae *Serratia marcescens* PH1a |
| Sma-19-1 | Bacteria Proteobacteria Gammaproteobacteria Enterobacterales Yersiniaceae *Serratia marcescens* SM39 |
| Sma-20-1 to Sma-20-2 | Bacteria Proteobacteria Gammaproteobacteria Enterobacterales Yersiniaceae *Serratia marcescens subsp. marcescens* |
| Sma-21-1 | Bacteria Proteobacteria Gammaproteobacteria Enterobacterales Yersiniaceae *Serratia marcescens subsp. marcescens* ATCC 13880 |
| Sma-22-1 | Bacteria Proteobacteria Gammaproteobacteria Enterobacterales Yersiniaceae *Serratia marcescens subsp. marcescens* Db11 |
| Sne-1-1 | Bacteria Proteobacteria Gammaproteobacteria Enterobacterales Yersiniaceae *Serratia nematodiphila* DZ0503SBS1 |
| Spl-1-1 | Bacteria Proteobacteria Gammaproteobacteria Enterobacterales Yersiniaceae *Serratia plymuthica* RVH1 |
| Ssp-12-1 | Bacteria Proteobacteria Gammaproteobacteria Enterobacterales Yersiniaceae *Serratia sp.* 506_PEND |
| Ssp-13-1 | Bacteria Proteobacteria Gammaproteobacteria Enterobacterales Yersiniaceae *Serratia sp.* Ag1 |
| Ssp-14-1 | Bacteria Proteobacteria Gammaproteobacteria Enterobacterales Yersiniaceae *Serratia sp.* Ag2 |
| Ssp-15-1 | Bacteria Proteobacteria Gammaproteobacteria Enterobacterales Yersiniaceae *Serratia sp.* DD3 |
| Ssp-16-1 | Bacteria Proteobacteria Gammaproteobacteria Enterobacterales Yersiniaceae *Serratia sp.* FS14 |
| Ssp-17-1 | Bacteria Proteobacteria Gammaproteobacteria Enterobacterales Yersiniaceae *Serratia sp.* H1n |
| Ssp-18-1 | Bacteria Proteobacteria Gammaproteobacteria Enterobacterales Yersiniaceae *Serratia sp.* H1w |

| | |
|---|---|
| Ssp-19-1 | Bacteria Proteobacteria Gammaproteobacteria Enterobacterales Yersiniaceae *Serratia sp.* SCBI |
| Ssp-20-1 to Ssp-20-2 | Bacteria Proteobacteria Gammaproteobacteria Enterobacterales Yersiniaceae *Serratia sp.* TEL |
| Ssy-1-1 | Bacteria Proteobacteria Gammaproteobacteria Enterobacterales Yersiniaceae *Serratia symbiotica* |
| Sur-1-1 | Bacteria Proteobacteria Gammaproteobacteria Enterobacterales Yersiniaceae *Serratia ureilytica* |
| Yal-1-1 to Yal-1-4 | Bacteria Proteobacteria Gammaproteobacteria Enterobacterales Yersiniaceae *Yersinia aldovae* |
| Yal-2-1 | Bacteria Proteobacteria Gammaproteobacteria Enterobacterales Yersiniaceae *Yersinia aldovae* 670-83 |
| Yal-3-1 | Bacteria Proteobacteria Gammaproteobacteria Enterobacterales Yersiniaceae *Yersinia aleksiciae* |
| Ybe-1-1 | Bacteria Proteobacteria Gammaproteobacteria Enterobacterales Yersiniaceae *Yersinia bercovieri* |
| Yen-1-1 to Yen-1-4 | Bacteria Proteobacteria Gammaproteobacteria Enterobacterales Yersiniaceae *Yersinia enterocolitica* |
| Yen-2-1 | Bacteria Proteobacteria Gammaproteobacteria Enterobacterales Yersiniaceae *Yersinia enterocolitica (type* O:2) str. YE3094/96 |
| Yen-3-1 | Bacteria Proteobacteria Gammaproteobacteria Enterobacterales Yersiniaceae *Yersinia enterocolitica (type* O:5) str. YE53/03 |
| Yen-4-1 | Bacteria Proteobacteria Gammaproteobacteria Enterobacterales Yersiniaceae *Yersinia enterocolitica (type* O:9) str. YE56/03 |
| Yen-5-1 | Bacteria Proteobacteria Gammaproteobacteria Enterobacterales Yersiniaceae *Yersinia enterocolitica subsp. palearctica* 556/8265 |
| Yen-6-1 | Bacteria Proteobacteria Gammaproteobacteria Enterobacterales Yersiniaceae *Yersinia enterocolitica subsp. palearctica* 647/5307 |
| Yen-7-1 | Bacteria Proteobacteria Gammaproteobacteria Enterobacterales Yersiniaceae *Yersinia enterocolitica subsp. palearctica* YO527 |
| Yfr-1-1 to Yfr-1-17 | Bacteria Proteobacteria Gammaproteobacteria Enterobacterales Yersiniaceae *Yersinia frederiksenii* |
| Yfr-2-1 to Yfr-2-2 | Bacteria Proteobacteria Gammaproteobacteria Enterobacterales Yersiniaceae *Yersinia frederiksenii* ATCC 33641 |
| Yfr-3-1 | Bacteria Proteobacteria Gammaproteobacteria Enterobacterales Yersiniaceae *Yersinia frederiksenii* Y225 |
| Yin-1-1 | Bacteria Proteobacteria Gammaproteobacteria Enterobacterales Yersiniaceae *Yersinia intermedia* |
| Ykr-1-1 to Ykr-1-2 | Bacteria Proteobacteria Gammaproteobacteria Enterobacterales Yersiniaceae *Yersinia kristensenii* |
| Yma-1-1 | Bacteria Proteobacteria Gammaproteobacteria Enterobacterales Yersiniaceae *Yersinia massiliensis* |
| Yma-2-1 | Bacteria Proteobacteria Gammaproteobacteria Enterobacterales Yersiniaceae *Yersinia massiliensis* CCUG 53443 |
| Ymo-1-1 to Ymo-1-7 | Bacteria Proteobacteria Gammaproteobacteria Enterobacterales Yersiniaceae *Yersinia mollaretii* |
| Ynu-1-1 | Bacteria Proteobacteria Gammaproteobacteria Enterobacterales Yersiniaceae *Yersinia nurmii* |
| Ype-1-1 to Ype-1-2 | Bacteria Proteobacteria Gammaproteobacteria Enterobacterales Yersiniaceae *Yersinia pekkanenii* |
| Ype-2-1 to Ype-2-9 | Bacteria Proteobacteria Gammaproteobacteria Enterobacterales Yersiniaceae *Yersinia pestis* |
| Ype-3-1 | Bacteria Proteobacteria Gammaproteobacteria Enterobacterales Yersiniaceae *Yersinia pestis* 1045 |
| Ype-4-1 | Bacteria Proteobacteria Gammaproteobacteria Enterobacterales Yersiniaceae *Yersinia pestis* 12 |
| Ype-5-1 | Bacteria Proteobacteria Gammaproteobacteria Enterobacterales Yersiniaceae *Yersinia pestis* 1412 |
| Ype-6-1 | Bacteria Proteobacteria Gammaproteobacteria Enterobacterales Yersiniaceae *Yersinia pestis* 1413 |
| Ype-7-1 | Bacteria Proteobacteria Gammaproteobacteria Enterobacterales Yersiniaceae *Yersinia pestis* 14735 |
| Ype-8-1 | Bacteria Proteobacteria Gammaproteobacteria Enterobacterales Yersiniaceae *Yersinia pestis* 1522 |
| Ype-9-1 | Bacteria Proteobacteria Gammaproteobacteria Enterobacterales Yersiniaceae *Yersinia pestis* 164 |
| Ype-10-1 | Bacteria Proteobacteria Gammaproteobacteria Enterobacterales Yersiniaceae *Yersinia pestis* 1670 |
| Ype-11-1 | Bacteria Proteobacteria Gammaproteobacteria Enterobacterales Yersiniaceae *Yersinia pestis* 2 |
| Ype-12-1 | Bacteria Proteobacteria Gammaproteobacteria Enterobacterales Yersiniaceae *Yersinia pestis* 2944 |
| Ype-13-1 | Bacteria Proteobacteria Gammaproteobacteria Enterobacterales Yersiniaceae *Yersinia pestis* 3067 |
| Ype-14-1 | Bacteria Proteobacteria Gammaproteobacteria Enterobacterales Yersiniaceae *Yersinia pestis* 34008 |
| Ype-15-1 | Bacteria Proteobacteria Gammaproteobacteria Enterobacterales Yersiniaceae *Yersinia pestis* 34202 |
| Ype-16-1 | Bacteria Proteobacteria Gammaproteobacteria Enterobacterales Yersiniaceae *Yersinia pestis* 351001 |
| Ype-17-1 | Bacteria Proteobacteria Gammaproteobacteria Enterobacterales Yersiniaceae *Yersinia pestis* 3770 |
| Ype-18-1 | Bacteria Proteobacteria Gammaproteobacteria Enterobacterales Yersiniaceae *Yersinia pestis* 42095 |
| Ype-19-1 | Bacteria Proteobacteria Gammaproteobacteria Enterobacterales Yersiniaceae *Yersinia pestis* 5 |
| Ype-20-1 | Bacteria Proteobacteria Gammaproteobacteria Enterobacterales Yersiniaceae *Yersinia pestis* 620024 |
| Ype-21-1 | Bacteria Proteobacteria Gammaproteobacteria Enterobacterales Yersiniaceae *Yersinia pestis* 71021 |
| Ype-22-1 | Bacteria Proteobacteria Gammaproteobacteria Enterobacterales Yersiniaceae *Yersinia pestis* 710317 |
| Ype-23-1 | Bacteria Proteobacteria Gammaproteobacteria Enterobacterales Yersiniaceae *Yersinia pestis* 7338 |
| Ype-24-1 | Bacteria Proteobacteria Gammaproteobacteria Enterobacterales Yersiniaceae *Yersinia pestis* 790 |
| Ype-25-1 | Bacteria Proteobacteria Gammaproteobacteria Enterobacterales Yersiniaceae *Yersinia pestis* 8787 |
| Ype-26-1 | Bacteria Proteobacteria Gammaproteobacteria Enterobacterales Yersiniaceae *Yersinia pestis* 9 |
| Ype-27-1 | Bacteria Proteobacteria Gammaproteobacteria Enterobacterales Yersiniaceae *Yersinia pestis* 945 |
| Ype-28-1 | Bacteria Proteobacteria Gammaproteobacteria Enterobacterales Yersiniaceae *Yersinia pestis* 970754 |
| Ype-29-1 to Ype-29-2 | Bacteria Proteobacteria Gammaproteobacteria Enterobacterales Yersiniaceae *Yersinia pestis* A1122 |
| Ype-30-1 | Bacteria Proteobacteria Gammaproteobacteria Enterobacterales Yersiniaceae *Yersinia pestis* A1973001 |
| Ype-31-1 to Ype-31-2 | Bacteria Proteobacteria Gammaproteobacteria Enterobacterales Yersiniaceae *Yersinia pestis* Angola |
| Ype-32-1 to Ype-32-2 | Bacteria Proteobacteria Gammaproteobacteria Enterobacterales Yersiniaceae *Yersinia pestis* Antiqua |
| Ype-104-1 | Bacteria Proteobacteria Gammaproteobacteria Enterobacterales Yersiniaceae *Yersinia pestis biovar* Orientalis str. AS200901156 |
| Ype-105-1 | Bacteria Proteobacteria Gammaproteobacteria Enterobacterales Yersiniaceae *Yersinia pestis biovar* Orientalis str. AS200901434 |
| Ype-106-1 | Bacteria Proteobacteria Gammaproteobacteria Enterobacterales Yersiniaceae *Yersinia pestis biovar* Orientalis str. AS200901509 |
| Ype-107-1 | Bacteria Proteobacteria Gammaproteobacteria Enterobacterales Yersiniaceae *Yersinia pestis biovar* Orientalis str. AS200901539 |
| Ype-108-1 | Bacteria Proteobacteria Gammaproteobacteria Enterobacterales Yersiniaceae *Yersinia pestis biovar* Orientalis str. AS200902147 |
| Ype-109-1 | Bacteria Proteobacteria Gammaproteobacteria Enterobacterales Yersiniaceae *Yersinia pestis biovar* Orientalis str. BA200901703 |
| Ype-110-1 | Bacteria Proteobacteria Gammaproteobacteria Enterobacterales Yersiniaceae *Yersinia pestis biovar* Orientalis str. BA200901799 |
| Ype-111-1 | Bacteria Proteobacteria Gammaproteobacteria Enterobacterales Yersiniaceae *Yersinia pestis biovar* Orientalis str. BA200901990 |
| Ype-112-1 | Bacteria Proteobacteria Gammaproteobacteria Enterobacterales Yersiniaceae *Yersinia pestis biovar* Orientalis str. BA200902009 |

| | |
|---|---|
| Ype-33-1 | Bacteria Proteobacteria Gammaproteobacteria Enterobacterales Yersiniaceae *Yersinia pestis* C1975003 |
| Ype-34-1 | Bacteria Proteobacteria Gammaproteobacteria Enterobacterales Yersiniaceae *Yersinia pestis* CMCC02041 |
| Ype-35-1 | Bacteria Proteobacteria Gammaproteobacteria Enterobacterales Yersiniaceae *Yersinia pestis* CMCC03001 |
| Ype-36-1 | Bacteria Proteobacteria Gammaproteobacteria Enterobacterales Yersiniaceae *Yersinia pestis* CMCC05009 |
| Ype-37-1 | Bacteria Proteobacteria Gammaproteobacteria Enterobacterales Yersiniaceae *Yersinia pestis* CMCC05013 |
| Ype-38-1 | Bacteria Proteobacteria Gammaproteobacteria Enterobacterales Yersiniaceae *Yersinia pestis* CMCC10012 |
| Ype-39-1 | Bacteria Proteobacteria Gammaproteobacteria Enterobacterales Yersiniaceae *Yersinia pestis* CMCC104003 |
| Ype-40-1 | Bacteria Proteobacteria Gammaproteobacteria Enterobacterales Yersiniaceae *Yersinia pestis* CMCC106002 |
| Ype-41-1 | Bacteria Proteobacteria Gammaproteobacteria Enterobacterales Yersiniaceae *Yersinia pestis* CMCC11001 |
| Ype-42-1 | Bacteria Proteobacteria Gammaproteobacteria Enterobacterales Yersiniaceae *Yersinia pestis* CMCC114001 |
| Ype-43-1 | Bacteria Proteobacteria Gammaproteobacteria Enterobacterales Yersiniaceae *Yersinia pestis* CMCC12003 |
| Ype-44-1 | Bacteria Proteobacteria Gammaproteobacteria Enterobacterales Yersiniaceae *Yersinia pestis* CMCC18019 |
| Ype-45-1 | Bacteria Proteobacteria Gammaproteobacteria Enterobacterales Yersiniaceae *Yersinia pestis* CMCC21106 |
| Ype-46-1 | Bacteria Proteobacteria Gammaproteobacteria Enterobacterales Yersiniaceae *Yersinia pestis* CMCC27002 |
| Ype-47-1 | Bacteria Proteobacteria Gammaproteobacteria Enterobacterales Yersiniaceae *Yersinia pestis* CMCC31004 |
| Ype-48-1 | Bacteria Proteobacteria Gammaproteobacteria Enterobacterales Yersiniaceae *Yersinia pestis* CMCC347001 |
| Ype-49-1 | Bacteria Proteobacteria Gammaproteobacteria Enterobacterales Yersiniaceae *Yersinia pestis* CMCC38001 |
| Ype-50-1 | Bacteria Proteobacteria Gammaproteobacteria Enterobacterales Yersiniaceae *Yersinia pestis* CMCC42007 |
| Ype-51-1 | Bacteria Proteobacteria Gammaproteobacteria Enterobacterales Yersiniaceae *Yersinia pestis* CMCC43032 |
| Ype-52-1 | Bacteria Proteobacteria Gammaproteobacteria Enterobacterales Yersiniaceae *Yersinia pestis* CMCC51020 |
| Ype-53-1 | Bacteria Proteobacteria Gammaproteobacteria Enterobacterales Yersiniaceae *Yersinia pestis* CMCC640047 |
| Ype-54-1 | Bacteria Proteobacteria Gammaproteobacteria Enterobacterales Yersiniaceae *Yersinia pestis* CMCC67001 |
| Ype-55-1 | Bacteria Proteobacteria Gammaproteobacteria Enterobacterales Yersiniaceae *Yersinia pestis* CMCC71001 |
| Ype-56-1 | Bacteria Proteobacteria Gammaproteobacteria Enterobacterales Yersiniaceae *Yersinia pestis* CMCC84033 |
| Ype-57-1 | Bacteria Proteobacteria Gammaproteobacteria Enterobacterales Yersiniaceae *Yersinia pestis* CMCC84046 |
| Ype-58-1 | Bacteria Proteobacteria Gammaproteobacteria Enterobacterales Yersiniaceae *Yersinia pestis* CMCC90027 |
| Ype-59-1 | Bacteria Proteobacteria Gammaproteobacteria Enterobacterales Yersiniaceae *Yersinia pestis* CMCC92004 |
| Ype-60-1 | Bacteria Proteobacteria Gammaproteobacteria Enterobacterales Yersiniaceae *Yersinia pestis* CMCC92010 |
| Ype-61-1 | Bacteria Proteobacteria Gammaproteobacteria Enterobacterales Yersiniaceae *Yersinia pestis* CMCC95001 |
| Ype-62-1 | Bacteria Proteobacteria Gammaproteobacteria Enterobacterales Yersiniaceae *Yersinia pestis* CMCC96007 |
| Ype-63-1 | Bacteria Proteobacteria Gammaproteobacteria Enterobacterales Yersiniaceae *Yersinia pestis* CMCC99103 |
| Ype-64-1 | Bacteria Proteobacteria Gammaproteobacteria Enterobacterales Yersiniaceae *Yersinia pestis* CMCCK100001 |
| Ype-65-1 | Bacteria Proteobacteria Gammaproteobacteria Enterobacterales Yersiniaceae *Yersinia pestis* CMCCK110001 |
| Ype-66-1 | Bacteria Proteobacteria Gammaproteobacteria Enterobacterales Yersiniaceae *Yersinia pestis* CMCCN010025 |
| Ype-67-1 | Bacteria Proteobacteria Gammaproteobacteria Enterobacterales Yersiniaceae *Yersinia pestis* D1964001 |
| Ype-68-1 | Bacteria Proteobacteria Gammaproteobacteria Enterobacterales Yersiniaceae *Yersinia pestis* D1964002 |
| Ype-69-1 | Bacteria Proteobacteria Gammaproteobacteria Enterobacterales Yersiniaceae *Yersinia pestis* D1991004 |
| Ype-70-1 | Bacteria Proteobacteria Gammaproteobacteria Enterobacterales Yersiniaceae *Yersinia pestis* E1977001 |
| Ype-71-1 | Bacteria Proteobacteria Gammaproteobacteria Enterobacterales Yersiniaceae *Yersinia pestis* EBD10-058 |
| Ype-72-1 | Bacteria Proteobacteria Gammaproteobacteria Enterobacterales Yersiniaceae *Yersinia pestis* EV76-CN |
| Ype-73-1 | Bacteria Proteobacteria Gammaproteobacteria Enterobacterales Yersiniaceae *Yersinia pestis* F1946001 |
| Ype-74-1 | Bacteria Proteobacteria Gammaproteobacteria Enterobacterales Yersiniaceae *Yersinia pestis* F1954001 |
| Ype-75-1 | Bacteria Proteobacteria Gammaproteobacteria Enterobacterales Yersiniaceae *Yersinia pestis* F1984001 |
| Ype-76-1 | Bacteria Proteobacteria Gammaproteobacteria Enterobacterales Yersiniaceae *Yersinia pestis* G1670A |
| Ype-77-1 | Bacteria Proteobacteria Gammaproteobacteria Enterobacterales Yersiniaceae *Yersinia pestis* G1670E |
| Ype-78-1 | Bacteria Proteobacteria Gammaproteobacteria Enterobacterales Yersiniaceae *Yersinia pestis* H1958004 |
| Ype-79-1 | Bacteria Proteobacteria Gammaproteobacteria Enterobacterales Yersiniaceae *Yersinia pestis* I160001 |
| Ype-80-1 | Bacteria Proteobacteria Gammaproteobacteria Enterobacterales Yersiniaceae *Yersinia pestis* I1969003 |
| Ype-81-1 | Bacteria Proteobacteria Gammaproteobacteria Enterobacterales Yersiniaceae *Yersinia pestis* I1970005 |
| Ype-82-1 | Bacteria Proteobacteria Gammaproteobacteria Enterobacterales Yersiniaceae *Yersinia pestis* I1991001 |
| Ype-83-1 | Bacteria Proteobacteria Gammaproteobacteria Enterobacterales Yersiniaceae *Yersinia pestis* I1994006 |
| Ype-84-1 | Bacteria Proteobacteria Gammaproteobacteria Enterobacterales Yersiniaceae *Yersinia pestis* I2001001 |
| Ype-85-1 | Bacteria Proteobacteria Gammaproteobacteria Enterobacterales Yersiniaceae *Yersinia pestis* J1963002 |
| Ype-86-1 | Bacteria Proteobacteria Gammaproteobacteria Enterobacterales Yersiniaceae *Yersinia pestis* J1978002 |
| Ype-87-1 | Bacteria Proteobacteria Gammaproteobacteria Enterobacterales Yersiniaceae *Yersinia pestis* K21985002 |
| Ype-88-1 | Bacteria Proteobacteria Gammaproteobacteria Enterobacterales Yersiniaceae *Yersinia pestis* M0000002 |
| Ype-89-1 | Bacteria Proteobacteria Gammaproteobacteria Enterobacterales Yersiniaceae *Yersinia pestis* MGJZ12 |
| Ype-90-1 | Bacteria Proteobacteria Gammaproteobacteria Enterobacterales Yersiniaceae *Yersinia pestis* MGJZ3 |
| Ype-91-1 | Bacteria Proteobacteria Gammaproteobacteria Enterobacterales Yersiniaceae *Yersinia pestis* MGJZ6 |
| Ype-92-1 | Bacteria Proteobacteria Gammaproteobacteria Enterobacterales Yersiniaceae *Yersinia pestis* MGJZ7 |
| Ype-93-1 | Bacteria Proteobacteria Gammaproteobacteria Enterobacterales Yersiniaceae *Yersinia pestis* MGJZ9 |
| Ype-94-1 to Ype-94-2 | Bacteria Proteobacteria Gammaproteobacteria Enterobacterales Yersiniaceae *Yersinia pestis* Pestoides F |
| Ype-95-1 | Bacteria Proteobacteria Gammaproteobacteria Enterobacterales Yersiniaceae *Yersinia pestis* Pestoides G |
| Ype-96-1 | Bacteria Proteobacteria Gammaproteobacteria Enterobacterales Yersiniaceae *Yersinia pestis* SHAN11 |
| Ype-97-1 | Bacteria Proteobacteria Gammaproteobacteria Enterobacterales Yersiniaceae *Yersinia pestis* SHAN12 |

| | |
|---|---|
| Ype-113-1 | Bacteria Proteobacteria Gammaproteobacteria Enterobacterales Yersiniaceae *Yersinia pestis str.* Pestoides B |
| Ype-114-1 to Ype-114-2 | Bacteria Proteobacteria Gammaproteobacteria Enterobacterales Yersiniaceae *Yersinia pestis subsp. microtus bv.* Ulegeica |
| Ype-115-1 | Bacteria Proteobacteria Gammaproteobacteria Enterobacterales Yersiniaceae *Yersinia pestis subsp. pestis bv.* Orientalis |
| Ype-98-1 | Bacteria Proteobacteria Gammaproteobacteria Enterobacterales Yersiniaceae *Yersinia pestis* YN1065 |
| Ype-99-1 | Bacteria Proteobacteria Gammaproteobacteria Enterobacterales Yersiniaceae *Yersinia pestis* YN1683 |
| Ype-100-1 | Bacteria Proteobacteria Gammaproteobacteria Enterobacterales Yersiniaceae *Yersinia pestis* YN2179 |
| Ype-101-1 | Bacteria Proteobacteria Gammaproteobacteria Enterobacterales Yersiniaceae *Yersinia pestis* YN2588 |
| Ype-102-1 | Bacteria Proteobacteria Gammaproteobacteria Enterobacterales Yersiniaceae *Yersinia pestis* YN472 |
| Ype-103-1 | Bacteria Proteobacteria Gammaproteobacteria Enterobacterales Yersiniaceae *Yersinia pestis* YN663 |
| Yps-1-1 to Yps-1-4 | Bacteria Proteobacteria Gammaproteobacteria Enterobacterales Yersiniaceae *Yersinia pseudotuberculosis* |
| Yps-2-1 | Bacteria Proteobacteria Gammaproteobacteria Enterobacterales Yersiniaceae *Yersinia pseudotuberculosis* B-6796 |
| Yps-3-1 | Bacteria Proteobacteria Gammaproteobacteria Enterobacterales Yersiniaceae *Yersinia pseudotuberculosis* B-6862 |
| Yps-4-1 | Bacteria Proteobacteria Gammaproteobacteria Enterobacterales Yersiniaceae *Yersinia pseudotuberculosis* B-6863 |
| Yps-5-1 | Bacteria Proteobacteria Gammaproteobacteria Enterobacterales Yersiniaceae *Yersinia pseudotuberculosis* B-6864 |
| Yps-6-1 | Bacteria Proteobacteria Gammaproteobacteria Enterobacterales Yersiniaceae *Yersinia pseudotuberculosis* B-6865 |
| Yps-7-1 | Bacteria Proteobacteria Gammaproteobacteria Enterobacterales Yersiniaceae *Yersinia pseudotuberculosis* B-6866 |
| Yps-8-1 | Bacteria Proteobacteria Gammaproteobacteria Enterobacterales Yersiniaceae *Yersinia pseudotuberculosis* B-7194 |
| Yps-9-1 | Bacteria Proteobacteria Gammaproteobacteria Enterobacterales Yersiniaceae *Yersinia pseudotuberculosis* B-7195 |
| Yps-10-1 to Yps-10-2 | Bacteria Proteobacteria Gammaproteobacteria Enterobacterales Yersiniaceae *Yersinia pseudotuberculosis* IP 32953 |
| Yps-11-1 to Yps-11-2 | Bacteria Proteobacteria Gammaproteobacteria Enterobacterales Yersiniaceae *Yersinia pseudotuberculosis* PB1/+ |
| Yps-13-1 | Bacteria Proteobacteria Gammaproteobacteria Enterobacterales Yersiniaceae *Yersinia pseudotuberculosis str.* PA3606 |
| Yps-12-1 to Yps-12-2 | Bacteria Proteobacteria Gammaproteobacteria Enterobacterales Yersiniaceae *Yersinia pseudotuberculosis* YPIII |
| Yro-1-1 | Bacteria Proteobacteria Gammaproteobacteria Enterobacterales Yersiniaceae *Yersinia rohdei* |
| Yru-1-1 to Yru-1-3 | Bacteria Proteobacteria Gammaproteobacteria Enterobacterales Yersiniaceae *Yersinia ruckeri* |
| Yru-2-1 to Yru-2-2 | Bacteria Proteobacteria Gammaproteobacteria Enterobacterales Yersiniaceae *Yersinia ruckeri* ATCC 29473 |
| Ysi-1-1 | Bacteria Proteobacteria Gammaproteobacteria Enterobacterales Yersiniaceae *Yersinia similis* |
| Ywa-1-1 to Ywa-1-2 | Bacteria Proteobacteria Gammaproteobacteria Enterobacterales Yersiniaceae *Yersinia wautersii* |
| CRe-1-1 | Bacteria Proteobacteria Gammaproteobacteria Enterobacteriales Enterobacteriaceae aphid secondary symbionts *Candidatus* Regiella insecticola LSR1 |
| Bsp-1-1 | Bacteria Proteobacteria Gammaproteobacteria Enterobacteriales Enterobacteriaceae *Brenneria sp.* EniD312 |
| Cda-1-1 | Bacteria Proteobacteria Gammaproteobacteria Enterobacteriales Enterobacteriaceae *Cedecea davisae* DSM 4568 |
| Cfr-1-1 to Cfr-1-4 | Bacteria Proteobacteria Gammaproteobacteria Enterobacteriales Enterobacteriaceae Citrobacter Citrobacter freundii complex *Citrobacter freundii* |
| Cfr-9-1 | Bacteria Proteobacteria Gammaproteobacteria Enterobacteriales Enterobacteriaceae Citrobacter Citrobacter freundii complex *Citrobacter freundii* 4_7_47CFAA |
| Cfr-2-1 to Cfr-2-2 | Bacteria Proteobacteria Gammaproteobacteria Enterobacteriales Enterobacteriaceae Citrobacter Citrobacter freundii complex *Citrobacter freundii* ATCC 8090 = MTCC 1658 |
| Cfr-10-1 | Bacteria Proteobacteria Gammaproteobacteria Enterobacteriales Enterobacteriaceae Citrobacter Citrobacter freundii complex *Citrobacter freundii* GTC 09479 |
| Cfr-11-1 | Bacteria Proteobacteria Gammaproteobacteria Enterobacteriales Enterobacteriaceae Citrobacter Citrobacter freundii complex *Citrobacter freundii* GTC 09629 |
| Cko-1-1 to Cko-1-2 | Bacteria Proteobacteria Gammaproteobacteria Enterobacteriales Enterobacteriaceae *Citrobacter koseri* |
| Cko-2-1 | Bacteria Proteobacteria Gammaproteobacteria Enterobacteriales Enterobacteriaceae *Citrobacter koseri* ATCC BAA-895 |
| Cro-3-1 | Bacteria Proteobacteria Gammaproteobacteria Enterobacteriales Enterobacteriaceae *Citrobacter rodentium* ICC168 |
| Csp-15-1 | Bacteria Proteobacteria Gammaproteobacteria Enterobacteriales Enterobacteriaceae *Citrobacter sp.* 30_2 |
| Csp-16-1 | Bacteria Proteobacteria Gammaproteobacteria Enterobacteriales Enterobacteriaceae *Citrobacter sp.* A1 |
| Csp-17-1 | Bacteria Proteobacteria Gammaproteobacteria Enterobacteriales Enterobacteriaceae *Citrobacter sp.* KTE151 |
| Csp-18-1 | Bacteria Proteobacteria Gammaproteobacteria Enterobacteriales Enterobacteriaceae *Citrobacter sp.* KTE30 |
| Csp-19-1 | Bacteria Proteobacteria Gammaproteobacteria Enterobacteriales Enterobacteriaceae *Citrobacter sp.* KTE32 |
| Csp-20-1 | Bacteria Proteobacteria Gammaproteobacteria Enterobacteriales Enterobacteriaceae *Citrobacter sp.* L17 |
| Cyo-1-1 | Bacteria Proteobacteria Gammaproteobacteria Enterobacteriales Enterobacteriaceae *Citrobacter youngae* ATCC 29220 |
| Cco-1-1 | Bacteria Proteobacteria Gammaproteobacteria Enterobacteriales Enterobacteriaceae *Cronobacter condimenti* 1330 |
| Cdu-4-1 | Bacteria Proteobacteria Gammaproteobacteria Enterobacteriales Enterobacteriaceae *Cronobacter dublinensis* 1210 |
| Cdu-5-1 | Bacteria Proteobacteria Gammaproteobacteria Enterobacteriales Enterobacteriaceae *Cronobacter dublinensis* 582 |
| Cma-4-1 | Bacteria Proteobacteria Gammaproteobacteria Enterobacteriales Enterobacteriaceae *Cronobacter malonaticus* 507 |
| Cma-5-1 | Bacteria Proteobacteria Gammaproteobacteria Enterobacteriales Enterobacteriaceae *Cronobacter malonaticus* 681 |
| Csa-9-1 | Bacteria Proteobacteria Gammaproteobacteria Enterobacteriales Enterobacteriaceae *Cronobacter sakazakii* 680 |
| Csa-10-1 | Bacteria Proteobacteria Gammaproteobacteria Enterobacteriales Enterobacteriaceae *Cronobacter sakazakii* 696 |
| Csa-11-1 | Bacteria Proteobacteria Gammaproteobacteria Enterobacteriales Enterobacteriaceae *Cronobacter sakazakii* 701 |
| Csa-12-1 | Bacteria Proteobacteria Gammaproteobacteria Enterobacteriales Enterobacteriaceae *Cronobacter sakazakii* ATCC BAA-894 |
| Csa-13-1 | Bacteria Proteobacteria Gammaproteobacteria Enterobacteriales Enterobacteriaceae *Cronobacter sakazakii* CMCC 45402 |
| Csa-14-1 | Bacteria Proteobacteria Gammaproteobacteria Enterobacteriales Enterobacteriaceae *Cronobacter sakazakii* E899 |
| Csa-15-1 | Bacteria Proteobacteria Gammaproteobacteria Enterobacteriales Enterobacteriaceae *Cronobacter sakazakii* ES15 |
| Csa-16-1 | Bacteria Proteobacteria Gammaproteobacteria Enterobacteriales Enterobacteriaceae *Cronobacter sakazakii* SP291 |
| Ctu-3-1 | Bacteria Proteobacteria Gammaproteobacteria Enterobacteriales Enterobacteriaceae *Cronobacter turicensis* 564 |

| | |
|---|---|
| Ctu-4-1 | Bacteria Proteobacteria Gammaproteobacteria Enterobacteriales Enterobacteriaceae *Cronobacter turicensis* z3032 |
| Cun-1-1 to Cun-1-2 | Bacteria Proteobacteria Gammaproteobacteria Enterobacteriales Enterobacteriaceae *Cronobacter universalis* NCTC 9529 |
| Dda-4-1 | Bacteria Proteobacteria Gammaproteobacteria Enterobacteriales Enterobacteriaceae *Dickeya dadantii* 3937 |
| Dda-5-1 | Bacteria Proteobacteria Gammaproteobacteria Enterobacteriales Enterobacteriaceae *Dickeya dadantii* Ech586 |
| Ddi-3-1 | Bacteria Proteobacteria Gammaproteobacteria Enterobacteriales Enterobacteriaceae *Dickeya dianthicola* GBBC 2039 |
| Ddi-4-1 | Bacteria Proteobacteria Gammaproteobacteria Enterobacteriales Enterobacteriaceae *Dickeya dianthicola* NCPPB 3534 |
| Ddi-5-1 | Bacteria Proteobacteria Gammaproteobacteria Enterobacteriales Enterobacteriaceae *Dickeya dianthicola* NCPPB 453 |
| Dpa-1-1 | Bacteria Proteobacteria Gammaproteobacteria Enterobacteriales Enterobacteriaceae *Dickeya paradisiaca* NCPPB 2511 |
| Dso-2-1 | Bacteria Proteobacteria Gammaproteobacteria Enterobacteriales Enterobacteriaceae *Dickeya solani* GBBC 2040 |
| Dso-3-1 | Bacteria Proteobacteria Gammaproteobacteria Enterobacteriales Enterobacteriaceae *Dickeya solani* IPO 2222 |
| Dso-4-1 | Bacteria Proteobacteria Gammaproteobacteria Enterobacteriales Enterobacteriaceae *Dickeya solani* MK10 |
| Dso-5-1 | Bacteria Proteobacteria Gammaproteobacteria Enterobacteriales Enterobacteriaceae *Dickeya solani* MK16 |
| Dso-6-1 | Bacteria Proteobacteria Gammaproteobacteria Enterobacteriales Enterobacteriaceae *Dickeya solani* RNS 08.23.3.1.A |
| Dsp-7-1 | Bacteria Proteobacteria Gammaproteobacteria Enterobacteriales Enterobacteriaceae *Dickeya sp.* CSL RW240 |
| Dsp-8-1 | Bacteria Proteobacteria Gammaproteobacteria Enterobacteriales Enterobacteriaceae *Dickeya sp.* D s0432-1 |
| Dsp-9-1 | Bacteria Proteobacteria Gammaproteobacteria Enterobacteriales Enterobacteriaceae *Dickeya sp.* NCPPB 569 |
| Dze-7-1 | Bacteria Proteobacteria Gammaproteobacteria Enterobacteriales Enterobacteriaceae *Dickeya zeae* MS1 |
| Dze-8-1 | Bacteria Proteobacteria Gammaproteobacteria Enterobacteriales Enterobacteriaceae *Dickeya zeae* NCPPB 3532 |
| Dze-9-1 | Bacteria Proteobacteria Gammaproteobacteria Enterobacteriales Enterobacteriaceae *Dickeya zeae* ZJU1202 |
| Eic-2-1 | Bacteria Proteobacteria Gammaproteobacteria Enterobacteriales Enterobacteriaceae *Edwardsiella ictaluri* 93-146 |
| Eta-4-1 | Bacteria Proteobacteria Gammaproteobacteria Enterobacteriales Enterobacteriaceae *Edwardsiella tarda* ATCC 23685 |
| Eta-5-1 | Bacteria Proteobacteria Gammaproteobacteria Enterobacteriales Enterobacteriaceae *Edwardsiella tarda* C07-087 |
| Eta-6-1 | Bacteria Proteobacteria Gammaproteobacteria Enterobacteriales Enterobacteriaceae *Edwardsiella tarda* EIB202 |
| Eta-7-1 | Bacteria Proteobacteria Gammaproteobacteria Enterobacteriales Enterobacteriaceae *Edwardsiella tarda* FL6-60 |
| Eta-8-1 | Bacteria Proteobacteria Gammaproteobacteria Enterobacteriales Enterobacteriaceae *Edwardsiella tarda* NBRC 105688 |
| Eae-1-1 | Bacteria Proteobacteria Gammaproteobacteria Enterobacteriales Enterobacteriaceae *Enterobacter aerogenes* EA1509E |
| Eae-2-1 | Bacteria Proteobacteria Gammaproteobacteria Enterobacteriales Enterobacteriaceae *Enterobacter aerogenes* FGI35 |
| Eae-3-1 | Bacteria Proteobacteria Gammaproteobacteria Enterobacteriales Enterobacteriaceae *Enterobacter aerogenes* KCTC 2190 |
| Eca-3-1 | Bacteria Proteobacteria Gammaproteobacteria Enterobacteriales Enterobacteriaceae *Enterobacter cancerogenus* ATCC 35316 |
| Eas-1-1 to Eas-1-2 | Bacteria Proteobacteria Gammaproteobacteria Enterobacteriales Enterobacteriaceae Enterobacter Enterobacter cloacae complex *Enterobacter asburiae* |
| Eas-4-1 | Bacteria Proteobacteria Gammaproteobacteria Enterobacteriales Enterobacteriaceae Enterobacter Enterobacter cloacae complex *Enterobacter asburiae* LF7a |
| Ecl-1-1 to Ecl-1-12 | Bacteria Proteobacteria Gammaproteobacteria Enterobacteriales Enterobacteriaceae Enterobacter Enterobacter cloacae complex *Enterobacter cloacae* |
| Ecl-55-1 | Bacteria Proteobacteria Gammaproteobacteria Enterobacteriales Enterobacteriaceae Enterobacter Enterobacter cloacae complex *Enterobacter cloacae* BWH 29 |
| Ecl-56-1 | Bacteria Proteobacteria Gammaproteobacteria Enterobacteriales Enterobacteriaceae Enterobacter Enterobacter cloacae complex *Enterobacter cloacae* BWH 31 |
| Ecl-57-1 | Bacteria Proteobacteria Gammaproteobacteria Enterobacteriales Enterobacteriaceae Enterobacter Enterobacter cloacae complex *Enterobacter cloacae* EC_38VIM1 |
| Ecl-58-1 | Bacteria Proteobacteria Gammaproteobacteria Enterobacteriales Enterobacteriaceae Enterobacter Enterobacter cloacae complex *Enterobacter cloacae* EcWSU1 |
| Ecl-59-1 | Bacteria Proteobacteria Gammaproteobacteria Enterobacteriales Enterobacteriaceae Enterobacter Enterobacter cloacae complex *Enterobacter cloacae* S611 |
| Ecl-60-1 | Bacteria Proteobacteria Gammaproteobacteria Enterobacteriales Enterobacteriaceae Enterobacter Enterobacter cloacae complex *Enterobacter cloacae* SCF1 |
| Ecl-66-1 | Bacteria Proteobacteria Gammaproteobacteria Enterobacteriales Enterobacteriaceae Enterobacter Enterobacter cloacae complex *Enterobacter cloacae str.* Hanford |
| Ecl-52-1 to Ecl-52-5 | Bacteria Proteobacteria Gammaproteobacteria Enterobacteriales Enterobacteriaceae Enterobacter Enterobacter cloacae complex *Enterobacter cloacae subsp. cloacae* |
| Ecl-67-1 | Bacteria Proteobacteria Gammaproteobacteria Enterobacteriales Enterobacteriaceae Enterobacter Enterobacter cloacae complex *Enterobacter cloacae subsp. cloacae* 08XA1 |
| Ecl-68-1 | Bacteria Proteobacteria Gammaproteobacteria Enterobacteriales Enterobacteriaceae Enterobacter Enterobacter cloacae complex *Enterobacter cloacae subsp. cloacae* ATCC 13047 |
| Ecl-69-1 | Bacteria Proteobacteria Gammaproteobacteria Enterobacteriales Enterobacteriaceae Enterobacter Enterobacter cloacae complex *Enterobacter cloacae subsp. cloacae* ENHKU01 |
| Ecl-70-1 | Bacteria Proteobacteria Gammaproteobacteria Enterobacteriales Enterobacteriaceae Enterobacter Enterobacter cloacae complex *Enterobacter cloacae subsp. dissolvens* SDM |
| Ecl-61-1 | Bacteria Proteobacteria Gammaproteobacteria Enterobacteriales Enterobacteriaceae Enterobacter Enterobacter cloacae complex *Enterobacter cloacae* UCI-CRE 11 |
| Ecl-62-1 | Bacteria Proteobacteria Gammaproteobacteria Enterobacteriales Enterobacteriaceae Enterobacter Enterobacter cloacae complex *Enterobacter cloacae* UCI-CRE 12 |

| | |
|---|---|
| Ecl-63-1 | Bacteria Proteobacteria Gammaproteobacteria Enterobacteriales Enterobacteriaceae Enterobacter Enterobacter cloacae complex *Enterobacter cloacae* UCI-CRE 3 |
| Ecl-64-1 | Bacteria Proteobacteria Gammaproteobacteria Enterobacteriales Enterobacteriaceae Enterobacter Enterobacter cloacae complex *Enterobacter cloacae* UCI-CRE 5 |
| Ecl-65-1 | Bacteria Proteobacteria Gammaproteobacteria Enterobacteriales Enterobacteriaceae Enterobacter Enterobacter cloacae complex *Enterobacter cloacae* UCI-CRE 9 |
| Eho-6-1 | Bacteria Proteobacteria Gammaproteobacteria Enterobacteriales Enterobacteriaceae Enterobacter Enterobacter cloacae complex *Enterobacter hormaechei* ATCC 49162 |
| Eho-7-1 | Bacteria Proteobacteria Gammaproteobacteria Enterobacteriales Enterobacteriaceae Enterobacter Enterobacter cloacae complex *Enterobacter hormaechei* YT2 |
| Eho-8-1 | Bacteria Proteobacteria Gammaproteobacteria Enterobacteriales Enterobacteriaceae Enterobacter Enterobacter cloacae complex *Enterobacter hormaechei* YT3 |
| Esp-57-1 | Bacteria Proteobacteria Gammaproteobacteria Enterobacteriales Enterobacteriaceae Enterobacter Enterobacter cloacae complex *Enterobacter sp.* MGH 14 |
| Esp-58-1 | Bacteria Proteobacteria Gammaproteobacteria Enterobacteriales Enterobacteriaceae Enterobacter Enterobacter cloacae complex *Enterobacter sp.* MGH 16 |
| Esp-59-1 | Bacteria Proteobacteria Gammaproteobacteria Enterobacteriales Enterobacteriaceae Enterobacter Enterobacter cloacae complex *Enterobacter sp.* MGH 22 |
| Esp-60-1 | Bacteria Proteobacteria Gammaproteobacteria Enterobacteriales Enterobacteriaceae Enterobacter Enterobacter cloacae complex *Enterobacter sp.* MGH 23 |
| Esp-61-1 | Bacteria Proteobacteria Gammaproteobacteria Enterobacteriales Enterobacteriaceae Enterobacter Enterobacter cloacae complex *Enterobacter sp.* MGH 24 |
| Esp-62-1 | Bacteria Proteobacteria Gammaproteobacteria Enterobacteriales Enterobacteriaceae Enterobacter Enterobacter cloacae complex *Enterobacter sp.* MGH 25 |
| Esp-63-1 | Bacteria Proteobacteria Gammaproteobacteria Enterobacteriales Enterobacteriaceae Enterobacter Enterobacter cloacae complex *Enterobacter sp.* MGH 26 |
| Esp-64-1 | Bacteria Proteobacteria Gammaproteobacteria Enterobacteriales Enterobacteriaceae Enterobacter Enterobacter cloacae complex *Enterobacter sp.* MGH 34 |
| Esp-65-1 | Bacteria Proteobacteria Gammaproteobacteria Enterobacteriales Enterobacteriaceae Enterobacter Enterobacter cloacae complex *Enterobacter sp.* MGH 38 |
| Esp-66-1 | Bacteria Proteobacteria Gammaproteobacteria Enterobacteriales Enterobacteriaceae Enterobacter Enterobacter cloacae complex *Enterobacter sp.* MGH 8 |
| Emo-1-1 | Bacteria Proteobacteria Gammaproteobacteria Enterobacteriales Enterobacteriaceae *Enterobacter mori* LMG 25706 |
| Esp-67-1 | Bacteria Proteobacteria Gammaproteobacteria Enterobacteriales Enterobacteriaceae *Enterobacter sp.* 638 |
| Esp-68-1 | Bacteria Proteobacteria Gammaproteobacteria Enterobacteriales Enterobacteriaceae *Enterobacter sp.* Ag1 |
| Esp-69-1 | Bacteria Proteobacteria Gammaproteobacteria Enterobacteriales Enterobacteriaceae *Enterobacter sp.* E20 |
| Esp-70-1 | Bacteria Proteobacteria Gammaproteobacteria Enterobacteriales Enterobacteriaceae *Enterobacter sp.* GN02454 |
| Esp-71-1 | Bacteria Proteobacteria Gammaproteobacteria Enterobacteriales Enterobacteriaceae *Enterobacter sp.* GN02457 |
| Esp-72-1 | Bacteria Proteobacteria Gammaproteobacteria Enterobacteriales Enterobacteriaceae *Enterobacter sp.* R4-368 |
| Esp-73-1 | Bacteria Proteobacteria Gammaproteobacteria Enterobacteriales Enterobacteriaceae *Enterobacter sp.* SST3 |
| Eba-3-1 | Bacteria Proteobacteria Gammaproteobacteria Enterobacteriales *Enterobacteriaceae bacterium* 9_2_54FAA |
| Eba-4-1 | Bacteria Proteobacteria Gammaproteobacteria Enterobacteriales *Enterobacteriaceae bacterium* LSJC7 |
| Eba-5-1 | Bacteria Proteobacteria Gammaproteobacteria Enterobacteriales *Enterobacteriaceae bacterium strain* FGI 57 |
| Eam-3-1 | Bacteria Proteobacteria Gammaproteobacteria Enterobacteriales Enterobacteriaceae *Erwinia amylovora* 01SFR-BO |
| Eam-4-1 | Bacteria Proteobacteria Gammaproteobacteria Enterobacteriales Enterobacteriaceae *Erwinia amylovora* ACW56400 |
| Eam-5-1 | Bacteria Proteobacteria Gammaproteobacteria Enterobacteriales Enterobacteriaceae *Erwinia amylovora* ATCC 49946 |
| Eam-6-1 | Bacteria Proteobacteria Gammaproteobacteria Enterobacteriales Enterobacteriaceae *Erwinia amylovora* CFBP 1232 |
| Eam-7-1 | Bacteria Proteobacteria Gammaproteobacteria Enterobacteriales Enterobacteriaceae *Erwinia amylovora* CFBP 2585 |
| Eam-8-1 | Bacteria Proteobacteria Gammaproteobacteria Enterobacteriales Enterobacteriaceae *Erwinia amylovora* CFBP1430 |
| Eam-9-1 | Bacteria Proteobacteria Gammaproteobacteria Enterobacteriales Enterobacteriaceae *Erwinia amylovora* Ea266 |
| Eam-10-1 | Bacteria Proteobacteria Gammaproteobacteria Enterobacteriales Enterobacteriaceae *Erwinia amylovora* Ea356 |
| Eam-11-1 | Bacteria Proteobacteria Gammaproteobacteria Enterobacteriales Enterobacteriaceae *Erwinia amylovora* Ea644 |
| Eam-12-1 | Bacteria Proteobacteria Gammaproteobacteria Enterobacteriales Enterobacteriaceae *Erwinia amylovora* LA635 |
| Eam-13-1 | Bacteria Proteobacteria Gammaproteobacteria Enterobacteriales Enterobacteriaceae *Erwinia amylovora* LA636 |
| Eam-14-1 | Bacteria Proteobacteria Gammaproteobacteria Enterobacteriales Enterobacteriaceae *Erwinia amylovora* LA637 |
| Eam-15-1 | Bacteria Proteobacteria Gammaproteobacteria Enterobacteriales Enterobacteriaceae *Erwinia amylovora* UPN527 |
| Ebi-2-1 | Bacteria Proteobacteria Gammaproteobacteria Enterobacteriales Enterobacteriaceae *Erwinia billingiae* Eb661 |
| Epi-2-1 | Bacteria Proteobacteria Gammaproteobacteria Enterobacteriales Enterobacteriaceae *Erwinia piriflorinigrans* CFBP 5888 |
| Epy-1-1 | Bacteria Proteobacteria Gammaproteobacteria Enterobacteriales Enterobacteriaceae *Erwinia pyrifoliae* DSM 12163 |
| Epy-2-1 | Bacteria Proteobacteria Gammaproteobacteria Enterobacteriales Enterobacteriaceae *Erwinia pyrifoliae* Ep1/96 |
| Esp-74-1 | Bacteria Proteobacteria Gammaproteobacteria Enterobacteriales Enterobacteriaceae *Erwinia sp.* Ejp617 |
| Eta-9-1 | Bacteria Proteobacteria Gammaproteobacteria Enterobacteriales Enterobacteriaceae *Erwinia tasmaniensis* Et1/99 |
| Eto-1-1 | Bacteria Proteobacteria Gammaproteobacteria Enterobacteriales Enterobacteriaceae *Erwinia toletana* DAPP-PG 735 |
| Etr-2-1 | Bacteria Proteobacteria Gammaproteobacteria Enterobacteriales Enterobacteriaceae *Erwinia tracheiphila* PSU-1 |
| Eal-3-1 | Bacteria Proteobacteria Gammaproteobacteria Enterobacteriales Enterobacteriaceae *Escherichia albertii* TW07627 |
| Eal-4-1 | Bacteria Proteobacteria Gammaproteobacteria Enterobacteriales Enterobacteriaceae *Escherichia albertii* TW08933 |
| Eal-5-1 | Bacteria Proteobacteria Gammaproteobacteria Enterobacteriales Enterobacteriaceae *Escherichia albertii* TW15818 |
| Ebl-1-1 | Bacteria Proteobacteria Gammaproteobacteria Enterobacteriales Enterobacteriaceae *Escherichia blattae* DSM 4481 |
| Eco-1-1 to Eco-1-4 | Bacteria Proteobacteria Gammaproteobacteria Enterobacteriales Enterobacteriaceae *Escherichia coli* |
| Eco-753-1 | Bacteria Proteobacteria Gammaproteobacteria Enterobacteriales Enterobacteriaceae *Escherichia coli* 0.1288 |
| Eco-754-1 | Bacteria Proteobacteria Gammaproteobacteria Enterobacteriales Enterobacteriaceae *Escherichia coli* 0.1304 |
| Eco-755-1 | Bacteria Proteobacteria Gammaproteobacteria Enterobacteriales Enterobacteriaceae *Escherichia coli* 042 |

| | |
|---|---|
| Eco-756-1 | Bacteria Proteobacteria Gammaproteobacteria Enterobacteriales Enterobacteriaceae *Escherichia coli* 07798 |
| Eco-757-1 | Bacteria Proteobacteria Gammaproteobacteria Enterobacteriales Enterobacteriaceae *Escherichia coli* 08BKT055439 |
| Eco-758-1 | Bacteria Proteobacteria Gammaproteobacteria Enterobacteriales Enterobacteriaceae *Escherichia coli* 08BKT77219 |
| Eco-759-1 | Bacteria Proteobacteria Gammaproteobacteria Enterobacteriales Enterobacteriaceae *Escherichia coli* 09BKT024447 |
| Eco-760-1 | Bacteria Proteobacteria Gammaproteobacteria Enterobacteriales Enterobacteriaceae *Escherichia coli* 09BKT076207 |
| Eco-761-1 | Bacteria Proteobacteria Gammaproteobacteria Enterobacteriales Enterobacteriaceae *Escherichia coli* 09BKT078844 |
| Eco-762-1 | Bacteria Proteobacteria Gammaproteobacteria Enterobacteriales Enterobacteriaceae *Escherichia coli* 1.2264 |
| Eco-763-1 | Bacteria Proteobacteria Gammaproteobacteria Enterobacteriales Enterobacteriaceae *Escherichia coli* 1.2741 |
| Eco-764-1 | Bacteria Proteobacteria Gammaproteobacteria Enterobacteriales Enterobacteriaceae *Escherichia coli* 10.0821 |
| Eco-765-1 | Bacteria Proteobacteria Gammaproteobacteria Enterobacteriales Enterobacteriaceae *Escherichia coli* 10.0833 |
| Eco-766-1 | Bacteria Proteobacteria Gammaproteobacteria Enterobacteriales Enterobacteriaceae *Escherichia coli* 10.0869 |
| Eco-767-1 | Bacteria Proteobacteria Gammaproteobacteria Enterobacteriales Enterobacteriaceae *Escherichia coli* 101-1 |
| Eco-768-1 | Bacteria Proteobacteria Gammaproteobacteria Enterobacteriales Enterobacteriaceae *Escherichia coli* 110957 |
| Eco-769-1 | Bacteria Proteobacteria Gammaproteobacteria Enterobacteriales Enterobacteriaceae *Escherichia coli* 113290 |
| Eco-770-1 | Bacteria Proteobacteria Gammaproteobacteria Enterobacteriales Enterobacteriaceae *Escherichia coli* 113302 |
| Eco-771-1 | Bacteria Proteobacteria Gammaproteobacteria Enterobacteriales Enterobacteriaceae *Escherichia coli* 113303 |
| Eco-772-1 | Bacteria Proteobacteria Gammaproteobacteria Enterobacteriales Enterobacteriaceae *Escherichia coli* 14A |
| Eco-773-1 | Bacteria Proteobacteria Gammaproteobacteria Enterobacteriales Enterobacteriaceae *Escherichia coli* 174750 |
| Eco-774-1 | Bacteria Proteobacteria Gammaproteobacteria Enterobacteriales Enterobacteriaceae *Escherichia coli* 174900 |
| Eco-775-1 | Bacteria Proteobacteria Gammaproteobacteria Enterobacteriales Enterobacteriaceae *Escherichia coli* 178200 |
| Eco-776-1 | Bacteria Proteobacteria Gammaproteobacteria Enterobacteriales Enterobacteriaceae *Escherichia coli* 178850 |
| Eco-777-1 | Bacteria Proteobacteria Gammaproteobacteria Enterobacteriales Enterobacteriaceae *Escherichia coli* 178900 |
| Eco-778-1 | Bacteria Proteobacteria Gammaproteobacteria Enterobacteriales Enterobacteriaceae *Escherichia coli* 179100 |
| Eco-779-1 | Bacteria Proteobacteria Gammaproteobacteria Enterobacteriales Enterobacteriaceae *Escherichia coli* 179550 |
| Eco-780-1 | Bacteria Proteobacteria Gammaproteobacteria Enterobacteriales Enterobacteriaceae *Escherichia coli* 180050 |
| Eco-781-1 | Bacteria Proteobacteria Gammaproteobacteria Enterobacteriales Enterobacteriaceae *Escherichia coli* 180200 |
| Eco-782-1 | Bacteria Proteobacteria Gammaproteobacteria Enterobacteriales Enterobacteriaceae *Escherichia coli* 180600 |
| Eco-783-1 | Bacteria Proteobacteria Gammaproteobacteria Enterobacteriales Enterobacteriaceae *Escherichia coli* 199900.1 |
| Eco-784-1 | Bacteria Proteobacteria Gammaproteobacteria Enterobacteriales Enterobacteriaceae *Escherichia coli* 2.3916 |
| Eco-785-1 | Bacteria Proteobacteria Gammaproteobacteria Enterobacteriales Enterobacteriaceae *Escherichia coli* 2.4168 |
| Eco-786-1 | Bacteria Proteobacteria Gammaproteobacteria Enterobacteriales Enterobacteriaceae *Escherichia coli* 201600.1 |
| Eco-787-1 | Bacteria Proteobacteria Gammaproteobacteria Enterobacteriales Enterobacteriaceae *Escherichia coli* 2362-75 |
| Eco-788-1 | Bacteria Proteobacteria Gammaproteobacteria Enterobacteriales Enterobacteriaceae *Escherichia coli* 2534-86 |
| Eco-789-1 | Bacteria Proteobacteria Gammaproteobacteria Enterobacteriales Enterobacteriaceae *Escherichia coli* 2719100 |
| Eco-790-1 | Bacteria Proteobacteria Gammaproteobacteria Enterobacteriales Enterobacteriaceae *Escherichia coli* 2720900 |
| Eco-791-1 | Bacteria Proteobacteria Gammaproteobacteria Enterobacteriales Enterobacteriaceae *Escherichia coli* 2722950 |
| Eco-792-1 | Bacteria Proteobacteria Gammaproteobacteria Enterobacteriales Enterobacteriaceae *Escherichia coli* 2726800 |
| Eco-793-1 | Bacteria Proteobacteria Gammaproteobacteria Enterobacteriales Enterobacteriaceae *Escherichia coli* 2726950 |
| Eco-794-1 | Bacteria Proteobacteria Gammaproteobacteria Enterobacteriales Enterobacteriaceae *Escherichia coli* 2729250 |
| Eco-795-1 | Bacteria Proteobacteria Gammaproteobacteria Enterobacteriales Enterobacteriaceae *Escherichia coli* 2730350 |
| Eco-796-1 | Bacteria Proteobacteria Gammaproteobacteria Enterobacteriales Enterobacteriaceae *Escherichia coli* 2730450 |
| Eco-797-1 | Bacteria Proteobacteria Gammaproteobacteria Enterobacteriales Enterobacteriaceae *Escherichia coli* 2731150 |
| Eco-798-1 | Bacteria Proteobacteria Gammaproteobacteria Enterobacteriales Enterobacteriaceae *Escherichia coli* 2733950 |
| Eco-799-1 | Bacteria Proteobacteria Gammaproteobacteria Enterobacteriales Enterobacteriaceae *Escherichia coli* 2735000 |
| Eco-800-1 | Bacteria Proteobacteria Gammaproteobacteria Enterobacteriales Enterobacteriaceae *Escherichia coli* 2741950 |
| Eco-801-1 | Bacteria Proteobacteria Gammaproteobacteria Enterobacteriales Enterobacteriaceae *Escherichia coli* 2747800 |
| Eco-802-1 | Bacteria Proteobacteria Gammaproteobacteria Enterobacteriales Enterobacteriaceae *Escherichia coli* 2749250 |
| Eco-803-1 | Bacteria Proteobacteria Gammaproteobacteria Enterobacteriales Enterobacteriaceae *Escherichia coli* 2756500 |
| Eco-804-1 | Bacteria Proteobacteria Gammaproteobacteria Enterobacteriales Enterobacteriaceae *Escherichia coli* 2762100 |
| Eco-805-1 | Bacteria Proteobacteria Gammaproteobacteria Enterobacteriales Enterobacteriaceae *Escherichia coli* 2770900 |
| Eco-806-1 | Bacteria Proteobacteria Gammaproteobacteria Enterobacteriales Enterobacteriaceae *Escherichia coli* 2780750 |
| Eco-807-1 | Bacteria Proteobacteria Gammaproteobacteria Enterobacteriales Enterobacteriaceae *Escherichia coli* 2785200 |
| Eco-808-1 | Bacteria Proteobacteria Gammaproteobacteria Enterobacteriales Enterobacteriaceae *Escherichia coli* 2788150 |
| Eco-809-1 | Bacteria Proteobacteria Gammaproteobacteria Enterobacteriales Enterobacteriaceae *Escherichia coli* 2845350 |
| Eco-810-1 | Bacteria Proteobacteria Gammaproteobacteria Enterobacteriales Enterobacteriaceae *Escherichia coli* 2845650 |
| Eco-811-1 | Bacteria Proteobacteria Gammaproteobacteria Enterobacteriales Enterobacteriaceae *Escherichia coli* 2846750 |
| Eco-812-1 | Bacteria Proteobacteria Gammaproteobacteria Enterobacteriales Enterobacteriaceae *Escherichia coli* 2848050 |
| Eco-813-1 | Bacteria Proteobacteria Gammaproteobacteria Enterobacteriales Enterobacteriaceae *Escherichia coli* 2850400 |
| Eco-814-1 | Bacteria Proteobacteria Gammaproteobacteria Enterobacteriales Enterobacteriaceae *Escherichia coli* 2850750 |
| Eco-815-1 | Bacteria Proteobacteria Gammaproteobacteria Enterobacteriales Enterobacteriaceae *Escherichia coli* 2851500 |
| Eco-816-1 | Bacteria Proteobacteria Gammaproteobacteria Enterobacteriales Enterobacteriaceae *Escherichia coli* 2853500 |
| Eco-817-1 | Bacteria Proteobacteria Gammaproteobacteria Enterobacteriales Enterobacteriaceae *Escherichia coli* 2854350 |
| Eco-818-1 | Bacteria Proteobacteria Gammaproteobacteria Enterobacteriales Enterobacteriaceae *Escherichia coli* 2860050 |
| Eco-819-1 | Bacteria Proteobacteria Gammaproteobacteria Enterobacteriales Enterobacteriaceae *Escherichia coli* 2860650 |
| Eco-820-1 | Bacteria Proteobacteria Gammaproteobacteria Enterobacteriales Enterobacteriaceae *Escherichia coli* 2861200 |

| | |
|---|---|
| Eco-821-1 | Bacteria Proteobacteria Gammaproteobacteria Enterobacteriales Enterobacteriaceae *Escherichia coli* 2862600 |
| Eco-822-1 | Bacteria Proteobacteria Gammaproteobacteria Enterobacteriales Enterobacteriaceae *Escherichia coli* 2864350 |
| Eco-823-1 | Bacteria Proteobacteria Gammaproteobacteria Enterobacteriales Enterobacteriaceae *Escherichia coli* 2865200 |
| Eco-824-1 | Bacteria Proteobacteria Gammaproteobacteria Enterobacteriales Enterobacteriaceae *Escherichia coli* 2866350 |
| Eco-825-1 | Bacteria Proteobacteria Gammaproteobacteria Enterobacteriales Enterobacteriaceae *Escherichia coli* 2866450 |
| Eco-826-1 | Bacteria Proteobacteria Gammaproteobacteria Enterobacteriales Enterobacteriaceae *Escherichia coli* 2866550 |
| Eco-827-1 | Bacteria Proteobacteria Gammaproteobacteria Enterobacteriales Enterobacteriaceae *Escherichia coli* 2866750 |
| Eco-828-1 | Bacteria Proteobacteria Gammaproteobacteria Enterobacteriales Enterobacteriaceae *Escherichia coli* 2867750 |
| Eco-829-1 | Bacteria Proteobacteria Gammaproteobacteria Enterobacteriales Enterobacteriaceae *Escherichia coli* 2871950 |
| Eco-830-1 | Bacteria Proteobacteria Gammaproteobacteria Enterobacteriales Enterobacteriaceae *Escherichia coli* 2872000 |
| Eco-831-1 | Bacteria Proteobacteria Gammaproteobacteria Enterobacteriales Enterobacteriaceae *Escherichia coli* 2872800 |
| Eco-832-1 | Bacteria Proteobacteria Gammaproteobacteria Enterobacteriales Enterobacteriaceae *Escherichia coli* 2875000 |
| Eco-833-1 | Bacteria Proteobacteria Gammaproteobacteria Enterobacteriales Enterobacteriaceae *Escherichia coli* 2875150 |
| Eco-834-1 | Bacteria Proteobacteria Gammaproteobacteria Enterobacteriales Enterobacteriaceae *Escherichia coli* 2886-75 |
| Eco-835-1 | Bacteria Proteobacteria Gammaproteobacteria Enterobacteriales Enterobacteriaceae *Escherichia coli* 3.2303 |
| Eco-836-1 | Bacteria Proteobacteria Gammaproteobacteria Enterobacteriales Enterobacteriaceae *Escherichia coli* 3.2608 |
| Eco-837-1 | Bacteria Proteobacteria Gammaproteobacteria Enterobacteriales Enterobacteriaceae *Escherichia coli* 3.3884 |
| Eco-838-1 | Bacteria Proteobacteria Gammaproteobacteria Enterobacteriales Enterobacteriaceae *Escherichia coli* 3.4870 |
| Eco-839-1 | Bacteria Proteobacteria Gammaproteobacteria Enterobacteriales Enterobacteriaceae *Escherichia coli* 3.4880 |
| Eco-840-1 | Bacteria Proteobacteria Gammaproteobacteria Enterobacteriales Enterobacteriaceae *Escherichia coli* 3003 |
| Eco-841-1 | Bacteria Proteobacteria Gammaproteobacteria Enterobacteriales Enterobacteriaceae *Escherichia coli* 3006 |
| Eco-842-1 | Bacteria Proteobacteria Gammaproteobacteria Enterobacteriales Enterobacteriaceae *Escherichia coli* 3431 |
| Eco-843-1 | Bacteria Proteobacteria Gammaproteobacteria Enterobacteriales Enterobacteriaceae *Escherichia coli* 4.0522 |
| Eco-844-1 | Bacteria Proteobacteria Gammaproteobacteria Enterobacteriales Enterobacteriaceae *Escherichia coli* 4.0967 |
| Eco-845-1 | Bacteria Proteobacteria Gammaproteobacteria Enterobacteriales Enterobacteriaceae *Escherichia coli* 5.0588 |
| Eco-846-1 | Bacteria Proteobacteria Gammaproteobacteria Enterobacteriales Enterobacteriaceae *Escherichia coli* 5.0959 |
| Eco-847-1 | Bacteria Proteobacteria Gammaproteobacteria Enterobacteriales Enterobacteriaceae *Escherichia coli* 5.2239 |
| Eco-848-1 | Bacteria Proteobacteria Gammaproteobacteria Enterobacteriales Enterobacteriaceae *Escherichia coli* 536 |
| Eco-849-1 | Bacteria Proteobacteria Gammaproteobacteria Enterobacteriales Enterobacteriaceae *Escherichia coli* 53638 |
| Eco-850-1 | Bacteria Proteobacteria Gammaproteobacteria Enterobacteriales Enterobacteriaceae *Escherichia coli* 541-1 |
| Eco-851-1 | Bacteria Proteobacteria Gammaproteobacteria Enterobacteriales Enterobacteriaceae *Escherichia coli* 541-15 |
| Eco-852-1 | Bacteria Proteobacteria Gammaproteobacteria Enterobacteriales Enterobacteriaceae *Escherichia coli* 5412 |
| Eco-853-1 | Bacteria Proteobacteria Gammaproteobacteria Enterobacteriales Enterobacteriaceae *Escherichia coli* 55989 |
| Eco-854-1 | Bacteria Proteobacteria Gammaproteobacteria Enterobacteriales Enterobacteriaceae *Escherichia coli* 576-1 |
| Eco-855-1 | Bacteria Proteobacteria Gammaproteobacteria Enterobacteriales Enterobacteriaceae *Escherichia coli* 5905 |
| Eco-856-1 | Bacteria Proteobacteria Gammaproteobacteria Enterobacteriales Enterobacteriaceae *Escherichia coli* 6.0172 |
| Eco-857-1 | Bacteria Proteobacteria Gammaproteobacteria Enterobacteriales Enterobacteriaceae *Escherichia coli* 7.1982 |
| Eco-858-1 | Bacteria Proteobacteria Gammaproteobacteria Enterobacteriales Enterobacteriaceae *Escherichia coli* 75 |
| Eco-859-1 | Bacteria Proteobacteria Gammaproteobacteria Enterobacteriales Enterobacteriaceae *Escherichia coli* 8.0416 |
| Eco-860-1 | Bacteria Proteobacteria Gammaproteobacteria Enterobacteriales Enterobacteriaceae *Escherichia coli* 8.0566 |
| Eco-861-1 | Bacteria Proteobacteria Gammaproteobacteria Enterobacteriales Enterobacteriaceae *Escherichia coli* 8.0569 |
| Eco-862-1 | Bacteria Proteobacteria Gammaproteobacteria Enterobacteriales Enterobacteriaceae *Escherichia coli* 8.0586 |
| Eco-863-1 | Bacteria Proteobacteria Gammaproteobacteria Enterobacteriales Enterobacteriaceae *Escherichia coli* 8.2524 |
| Eco-864-1 | Bacteria Proteobacteria Gammaproteobacteria Enterobacteriales Enterobacteriaceae *Escherichia coli* 83972 |
| Eco-865-1 | Bacteria Proteobacteria Gammaproteobacteria Enterobacteriales Enterobacteriaceae *Escherichia coli* 88.0221 |
| Eco-866-1 | Bacteria Proteobacteria Gammaproteobacteria Enterobacteriales Enterobacteriaceae *Escherichia coli* 88.1042 |
| Eco-867-1 | Bacteria Proteobacteria Gammaproteobacteria Enterobacteriales Enterobacteriaceae *Escherichia coli* 88.1467 |
| Eco-868-1 | Bacteria Proteobacteria Gammaproteobacteria Enterobacteriales Enterobacteriaceae *Escherichia coli* 89.0511 |
| Eco-869-1 | Bacteria Proteobacteria Gammaproteobacteria Enterobacteriales Enterobacteriaceae *Escherichia coli* 9.0111 |
| Eco-870-1 | Bacteria Proteobacteria Gammaproteobacteria Enterobacteriales Enterobacteriaceae *Escherichia coli* 90.0039 |
| Eco-871-1 | Bacteria Proteobacteria Gammaproteobacteria Enterobacteriales Enterobacteriaceae *Escherichia coli* 90.0091 |
| Eco-872-1 | Bacteria Proteobacteria Gammaproteobacteria Enterobacteriales Enterobacteriaceae *Escherichia coli* 90.2281 |
| Eco-873-1 | Bacteria Proteobacteria Gammaproteobacteria Enterobacteriales Enterobacteriaceae *Escherichia coli* 900105 (10e) |
| Eco-874-1 | Bacteria Proteobacteria Gammaproteobacteria Enterobacteriales Enterobacteriaceae *Escherichia coli* 907357 |
| Eco-875-1 | Bacteria Proteobacteria Gammaproteobacteria Enterobacteriales Enterobacteriaceae *Escherichia coli* 907391 |
| Eco-876-1 | Bacteria Proteobacteria Gammaproteobacteria Enterobacteriales Enterobacteriaceae *Escherichia coli* 907446 |
| Eco-877-1 | Bacteria Proteobacteria Gammaproteobacteria Enterobacteriales Enterobacteriaceae *Escherichia coli* 907672 |
| Eco-878-1 | Bacteria Proteobacteria Gammaproteobacteria Enterobacteriales Enterobacteriaceae *Escherichia coli* 907700 |
| Eco-879-1 | Bacteria Proteobacteria Gammaproteobacteria Enterobacteriales Enterobacteriaceae *Escherichia coli* 907701 |
| Eco-880-1 | Bacteria Proteobacteria Gammaproteobacteria Enterobacteriales Enterobacteriaceae *Escherichia coli* 907710 |
| Eco-881-1 | Bacteria Proteobacteria Gammaproteobacteria Enterobacteriales Enterobacteriaceae *Escherichia coli* 907713 |
| Eco-882-1 | Bacteria Proteobacteria Gammaproteobacteria Enterobacteriales Enterobacteriaceae *Escherichia coli* 907715 |
| Eco-883-1 | Bacteria Proteobacteria Gammaproteobacteria Enterobacteriales Enterobacteriaceae *Escherichia coli* 907779 |
| Eco-884-1 | Bacteria Proteobacteria Gammaproteobacteria Enterobacteriales Enterobacteriaceae *Escherichia coli* 907889 |
| Eco-885-1 | Bacteria Proteobacteria Gammaproteobacteria Enterobacteriales Enterobacteriaceae *Escherichia coli* 907892 |

| | |
|---|---|
| Eco-886-1 | Bacteria Proteobacteria Gammaproteobacteria Enterobacteriales Enterobacteriaceae *Escherichia coli* 908519 |
| Eco-887-1 | Bacteria Proteobacteria Gammaproteobacteria Enterobacteriales Enterobacteriaceae *Escherichia coli* 908521 |
| Eco-888-1 | Bacteria Proteobacteria Gammaproteobacteria Enterobacteriales Enterobacteriaceae *Escherichia coli* 908522 |
| Eco-889-1 | Bacteria Proteobacteria Gammaproteobacteria Enterobacteriales Enterobacteriaceae *Escherichia coli* 908524 |
| Eco-890-1 | Bacteria Proteobacteria Gammaproteobacteria Enterobacteriales Enterobacteriaceae *Escherichia coli* 908525 |
| Eco-891-1 | Bacteria Proteobacteria Gammaproteobacteria Enterobacteriales Enterobacteriaceae *Escherichia coli* 908541 |
| Eco-892-1 | Bacteria Proteobacteria Gammaproteobacteria Enterobacteriales Enterobacteriaceae *Escherichia coli* 908555 |
| Eco-893-1 | Bacteria Proteobacteria Gammaproteobacteria Enterobacteriales Enterobacteriaceae *Escherichia coli* 908573 |
| Eco-894-1 | Bacteria Proteobacteria Gammaproteobacteria Enterobacteriales Enterobacteriaceae *Escherichia coli* 908585 |
| Eco-895-1 | Bacteria Proteobacteria Gammaproteobacteria Enterobacteriales Enterobacteriaceae *Escherichia coli* 908616 |
| Eco-896-1 | Bacteria Proteobacteria Gammaproteobacteria Enterobacteriales Enterobacteriaceae *Escherichia coli* 908624 |
| Eco-897-1 | Bacteria Proteobacteria Gammaproteobacteria Enterobacteriales Enterobacteriaceae *Escherichia coli* 908632 |
| Eco-898-1 | Bacteria Proteobacteria Gammaproteobacteria Enterobacteriales Enterobacteriaceae *Escherichia coli* 908658 |
| Eco-899-1 | Bacteria Proteobacteria Gammaproteobacteria Enterobacteriales Enterobacteriaceae *Escherichia coli* 908675 |
| Eco-900-1 | Bacteria Proteobacteria Gammaproteobacteria Enterobacteriales Enterobacteriaceae *Escherichia coli* 908691 |
| Eco-901-1 | Bacteria Proteobacteria Gammaproteobacteria Enterobacteriales Enterobacteriaceae *Escherichia coli* 909945-2 |
| Eco-902-1 | Bacteria Proteobacteria Gammaproteobacteria Enterobacteriales Enterobacteriaceae *Escherichia coli* 909957 |
| Eco-903-1 | Bacteria Proteobacteria Gammaproteobacteria Enterobacteriales Enterobacteriaceae *Escherichia coli* 910096-2 |
| Eco-904-1 | Bacteria Proteobacteria Gammaproteobacteria Enterobacteriales Enterobacteriaceae *Escherichia coli* 93-001 |
| Eco-905-1 | Bacteria Proteobacteria Gammaproteobacteria Enterobacteriales Enterobacteriaceae *Escherichia coli* 93.0055 |
| Eco-906-1 | Bacteria Proteobacteria Gammaproteobacteria Enterobacteriales Enterobacteriaceae *Escherichia coli* 93.0056 |
| Eco-907-1 | Bacteria Proteobacteria Gammaproteobacteria Enterobacteriales Enterobacteriaceae *Escherichia coli* 93.0624 |
| Eco-908-1 | Bacteria Proteobacteria Gammaproteobacteria Enterobacteriales Enterobacteriaceae *Escherichia coli* 94.0618 |
| Eco-909-1 | Bacteria Proteobacteria Gammaproteobacteria Enterobacteriales Enterobacteriaceae *Escherichia coli* 95.0083 |
| Eco-910-1 | Bacteria Proteobacteria Gammaproteobacteria Enterobacteriales Enterobacteriaceae *Escherichia coli* 95.0183 |
| Eco-911-1 | Bacteria Proteobacteria Gammaproteobacteria Enterobacteriales Enterobacteriaceae *Escherichia coli* 95.0941 |
| Eco-912-1 | Bacteria Proteobacteria Gammaproteobacteria Enterobacteriales Enterobacteriaceae *Escherichia coli* 95.0943 |
| Eco-913-1 | Bacteria Proteobacteria Gammaproteobacteria Enterobacteriales Enterobacteriaceae *Escherichia coli* 95.1288 |
| Eco-914-1 | Bacteria Proteobacteria Gammaproteobacteria Enterobacteriales Enterobacteriaceae *Escherichia coli* 95JB1 |
| Eco-915-1 | Bacteria Proteobacteria Gammaproteobacteria Enterobacteriales Enterobacteriaceae *Escherichia coli* 95NR1 |
| Eco-916-1 | Bacteria Proteobacteria Gammaproteobacteria Enterobacteriales Enterobacteriaceae *Escherichia coli* 96.0107 |
| Eco-917-1 | Bacteria Proteobacteria Gammaproteobacteria Enterobacteriales Enterobacteriaceae *Escherichia coli* 96.0109 |
| Eco-918-1 | Bacteria Proteobacteria Gammaproteobacteria Enterobacteriales Enterobacteriaceae *Escherichia coli* 96.0427 |
| Eco-919-1 | Bacteria Proteobacteria Gammaproteobacteria Enterobacteriales Enterobacteriaceae *Escherichia coli* 96.0428 |
| Eco-920-1 | Bacteria Proteobacteria Gammaproteobacteria Enterobacteriales Enterobacteriaceae *Escherichia coli* 96.0497 |
| Eco-921-1 | Bacteria Proteobacteria Gammaproteobacteria Enterobacteriales Enterobacteriaceae *Escherichia coli* 96.0932 |
| Eco-922-1 | Bacteria Proteobacteria Gammaproteobacteria Enterobacteriales Enterobacteriaceae *Escherichia coli* 96.0939 |
| Eco-923-1 | Bacteria Proteobacteria Gammaproteobacteria Enterobacteriales Enterobacteriaceae *Escherichia coli* 96.154 |
| Eco-924-1 | Bacteria Proteobacteria Gammaproteobacteria Enterobacteriales Enterobacteriaceae *Escherichia coli* 97.0003 |
| Eco-925-1 | Bacteria Proteobacteria Gammaproteobacteria Enterobacteriales Enterobacteriaceae *Escherichia coli* 97.0007 |
| Eco-926-1 | Bacteria Proteobacteria Gammaproteobacteria Enterobacteriales Enterobacteriaceae *Escherichia coli* 97.0010 |
| Eco-927-1 | Bacteria Proteobacteria Gammaproteobacteria Enterobacteriales Enterobacteriaceae *Escherichia coli* 97.0246 |
| Eco-928-1 | Bacteria Proteobacteria Gammaproteobacteria Enterobacteriales Enterobacteriaceae *Escherichia coli* 97.0259 |
| Eco-929-1 | Bacteria Proteobacteria Gammaproteobacteria Enterobacteriales Enterobacteriaceae *Escherichia coli* 97.1742 |
| Eco-930-1 | Bacteria Proteobacteria Gammaproteobacteria Enterobacteriales Enterobacteriaceae *Escherichia coli* 99.0670 |
| Eco-931-1 | Bacteria Proteobacteria Gammaproteobacteria Enterobacteriales Enterobacteriaceae *Escherichia coli* 99.0672 |
| Eco-932-1 | Bacteria Proteobacteria Gammaproteobacteria Enterobacteriales Enterobacteriaceae *Escherichia coli* 99.0678 |
| Eco-933-1 | Bacteria Proteobacteria Gammaproteobacteria Enterobacteriales Enterobacteriaceae *Escherichia coli* 99.0713 |
| Eco-934-1 | Bacteria Proteobacteria Gammaproteobacteria Enterobacteriales Enterobacteriaceae *Escherichia coli* 99.0741 |
| Eco-935-1 | Bacteria Proteobacteria Gammaproteobacteria Enterobacteriales Enterobacteriaceae *Escherichia coli* 99.0814 |
| Eco-936-1 | Bacteria Proteobacteria Gammaproteobacteria Enterobacteriales Enterobacteriaceae *Escherichia coli* 99.0815 |
| Eco-937-1 | Bacteria Proteobacteria Gammaproteobacteria Enterobacteriales Enterobacteriaceae *Escherichia coli* 99.0816 |
| Eco-938-1 | Bacteria Proteobacteria Gammaproteobacteria Enterobacteriales Enterobacteriaceae *Escherichia coli* 99.0839 |
| Eco-939-1 | Bacteria Proteobacteria Gammaproteobacteria Enterobacteriales Enterobacteriaceae *Escherichia coli* 99.0848 |
| Eco-940-1 | Bacteria Proteobacteria Gammaproteobacteria Enterobacteriales Enterobacteriaceae *Escherichia coli* 99.1753 |
| Eco-941-1 | Bacteria Proteobacteria Gammaproteobacteria Enterobacteriales Enterobacteriaceae *Escherichia coli* 99.1762 |
| Eco-942-1 | Bacteria Proteobacteria Gammaproteobacteria Enterobacteriales Enterobacteriaceae *Escherichia coli* 99.1775 |
| Eco-943-1 | Bacteria Proteobacteria Gammaproteobacteria Enterobacteriales Enterobacteriaceae *Escherichia coli* 99.1781 |
| Eco-944-1 | Bacteria Proteobacteria Gammaproteobacteria Enterobacteriales Enterobacteriaceae *Escherichia coli* 99.1793 |
| Eco-945-1 | Bacteria Proteobacteria Gammaproteobacteria Enterobacteriales Enterobacteriaceae *Escherichia coli* A25922R |
| Eco-946-1 | Bacteria Proteobacteria Gammaproteobacteria Enterobacteriales Enterobacteriaceae *Escherichia coli* A35218R |
| Eco-947-1 | Bacteria Proteobacteria Gammaproteobacteria Enterobacteriales Enterobacteriaceae *Escherichia coli* AA86 |
| Eco-948-1 | Bacteria Proteobacteria Gammaproteobacteria Enterobacteriales Enterobacteriaceae *Escherichia coli* ABU 83972 |
| Eco-289-1 to Eco-289-2 | Bacteria Proteobacteria Gammaproteobacteria Enterobacteriales Enterobacteriaceae *Escherichia coli* AD30 |
| Eco-949-1 | Bacteria Proteobacteria Gammaproteobacteria Enterobacteriales Enterobacteriaceae *Escherichia coli* AI27 |

| | |
|---|---|
| Eco-950-1 | Bacteria Proteobacteria Gammaproteobacteria Enterobacteriales Enterobacteriaceae *Escherichia coli* APEC O1 |
| Eco-951-1 | Bacteria Proteobacteria Gammaproteobacteria Enterobacteriales Enterobacteriaceae *Escherichia coli* APEC O78 |
| Eco-952-1 | Bacteria Proteobacteria Gammaproteobacteria Enterobacteriales Enterobacteriaceae *Escherichia coli* ARS4.2123 |
| Eco-291-1 to Eco-291-2 | Bacteria Proteobacteria Gammaproteobacteria Enterobacteriales Enterobacteriaceae *Escherichia coli* ATCC 25922 |
| Eco-953-1 | Bacteria Proteobacteria Gammaproteobacteria Enterobacteriales Enterobacteriaceae *Escherichia coli* ATCC 700728 |
| Eco-954-1 | Bacteria Proteobacteria Gammaproteobacteria Enterobacteriales Enterobacteriaceae *Escherichia coli* ATCC 8739 |
| Eco-955-1 | Bacteria Proteobacteria Gammaproteobacteria Enterobacteriales Enterobacteriaceae *Escherichia coli* ATCC BAA-2196 |
| Eco-956-1 | Bacteria Proteobacteria Gammaproteobacteria Enterobacteriales Enterobacteriaceae *Escherichia coli* ATCC BAA-2209 |
| Eco-957-1 | Bacteria Proteobacteria Gammaproteobacteria Enterobacteriales Enterobacteriaceae *Escherichia coli* ATCC BAA-2215 |
| Eco-958-1 | Bacteria Proteobacteria Gammaproteobacteria Enterobacteriales Enterobacteriaceae *Escherichia coli* ATCC BAA-2219 |
| Eco-959-1 | Bacteria Proteobacteria Gammaproteobacteria Enterobacteriales Enterobacteriaceae *Escherichia coli* B str. REL606 |
| Eco-960-1 | Bacteria Proteobacteria Gammaproteobacteria Enterobacteriales Enterobacteriaceae *Escherichia coli* B088 |
| Eco-961-1 | Bacteria Proteobacteria Gammaproteobacteria Enterobacteriales Enterobacteriaceae *Escherichia coli* B093 |
| Eco-962-1 | Bacteria Proteobacteria Gammaproteobacteria Enterobacteriales Enterobacteriaceae *Escherichia coli* B102 |
| Eco-963-1 | Bacteria Proteobacteria Gammaproteobacteria Enterobacteriales Enterobacteriaceae *Escherichia coli* B103 |
| Eco-964-1 | Bacteria Proteobacteria Gammaproteobacteria Enterobacteriales Enterobacteriaceae *Escherichia coli* B104 |
| Eco-965-1 | Bacteria Proteobacteria Gammaproteobacteria Enterobacteriales Enterobacteriaceae *Escherichia coli* B105 |
| Eco-966-1 | Bacteria Proteobacteria Gammaproteobacteria Enterobacteriales Enterobacteriaceae *Escherichia coli* B106 |
| Eco-967-1 | Bacteria Proteobacteria Gammaproteobacteria Enterobacteriales Enterobacteriaceae *Escherichia coli* B107 |
| Eco-968-1 | Bacteria Proteobacteria Gammaproteobacteria Enterobacteriales Enterobacteriaceae *Escherichia coli* B108 |
| Eco-969-1 | Bacteria Proteobacteria Gammaproteobacteria Enterobacteriales Enterobacteriaceae *Escherichia coli* B109 |
| Eco-970-1 | Bacteria Proteobacteria Gammaproteobacteria Enterobacteriales Enterobacteriaceae *Escherichia coli* B112 |
| Eco-971-1 | Bacteria Proteobacteria Gammaproteobacteria Enterobacteriales Enterobacteriaceae *Escherichia coli* B113 |
| Eco-972-1 | Bacteria Proteobacteria Gammaproteobacteria Enterobacteriales Enterobacteriaceae *Escherichia coli* B114 |
| Eco-973-1 | Bacteria Proteobacteria Gammaproteobacteria Enterobacteriales Enterobacteriaceae *Escherichia coli* B15 |
| Eco-974-1 | Bacteria Proteobacteria Gammaproteobacteria Enterobacteriales Enterobacteriaceae *Escherichia coli* B17 |
| Eco-975-1 | Bacteria Proteobacteria Gammaproteobacteria Enterobacteriales Enterobacteriaceae *Escherichia coli* B171 |
| Eco-976-1 | Bacteria Proteobacteria Gammaproteobacteria Enterobacteriales Enterobacteriaceae *Escherichia coli* B185 |
| Eco-977-1 | Bacteria Proteobacteria Gammaproteobacteria Enterobacteriales Enterobacteriaceae *Escherichia coli* B26-1 |
| Eco-978-1 | Bacteria Proteobacteria Gammaproteobacteria Enterobacteriales Enterobacteriaceae *Escherichia coli* B26-2 |
| Eco-979-1 | Bacteria Proteobacteria Gammaproteobacteria Enterobacteriales Enterobacteriaceae *Escherichia coli* B28-1 |
| Eco-980-1 | Bacteria Proteobacteria Gammaproteobacteria Enterobacteriales Enterobacteriaceae *Escherichia coli* B28-2 |
| Eco-981-1 | Bacteria Proteobacteria Gammaproteobacteria Enterobacteriales Enterobacteriaceae *Escherichia coli* B29-1 |
| Eco-982-1 | Bacteria Proteobacteria Gammaproteobacteria Enterobacteriales Enterobacteriaceae *Escherichia coli* B29-2 |
| Eco-983-1 | Bacteria Proteobacteria Gammaproteobacteria Enterobacteriales Enterobacteriaceae *Escherichia coli* B354 |
| Eco-984-1 | Bacteria Proteobacteria Gammaproteobacteria Enterobacteriales Enterobacteriaceae *Escherichia coli* B36-1 |
| Eco-985-1 | Bacteria Proteobacteria Gammaproteobacteria Enterobacteriales Enterobacteriaceae *Escherichia coli* B36-2 |
| Eco-986-1 | Bacteria Proteobacteria Gammaproteobacteria Enterobacteriales Enterobacteriaceae *Escherichia coli* B40-1 |
| Eco-987-1 | Bacteria Proteobacteria Gammaproteobacteria Enterobacteriales Enterobacteriaceae *Escherichia coli* B40-2 |
| Eco-988-1 | Bacteria Proteobacteria Gammaproteobacteria Enterobacteriales Enterobacteriaceae *Escherichia coli* B41 |
| Eco-989-1 | Bacteria Proteobacteria Gammaproteobacteria Enterobacteriales Enterobacteriaceae *Escherichia coli* B49-2 |
| Eco-990-1 | Bacteria Proteobacteria Gammaproteobacteria Enterobacteriales Enterobacteriaceae *Escherichia coli* B5-2 |
| Eco-991-1 | Bacteria Proteobacteria Gammaproteobacteria Enterobacteriales Enterobacteriaceae *Escherichia coli* B7-1 |
| Eco-992-1 | Bacteria Proteobacteria Gammaproteobacteria Enterobacteriales Enterobacteriaceae *Escherichia coli* B7-2 |
| Eco-993-1 | Bacteria Proteobacteria Gammaproteobacteria Enterobacteriales Enterobacteriaceae *Escherichia coli* B799 |
| Eco-994-1 | Bacteria Proteobacteria Gammaproteobacteria Enterobacteriales Enterobacteriaceae *Escherichia coli* B7A |
| Eco-995-1 | Bacteria Proteobacteria Gammaproteobacteria Enterobacteriales Enterobacteriaceae *Escherichia coli* B83 |
| Eco-996-1 | Bacteria Proteobacteria Gammaproteobacteria Enterobacteriales Enterobacteriaceae *Escherichia coli* B84 |
| Eco-997-1 | Bacteria Proteobacteria Gammaproteobacteria Enterobacteriales Enterobacteriaceae *Escherichia coli* B85 |
| Eco-998-1 | Bacteria Proteobacteria Gammaproteobacteria Enterobacteriales Enterobacteriaceae *Escherichia coli* B86 |
| Eco-999-1 | Bacteria Proteobacteria Gammaproteobacteria Enterobacteriales Enterobacteriaceae *Escherichia coli* B89 |
| Eco-1000-1 | Bacteria Proteobacteria Gammaproteobacteria Enterobacteriales Enterobacteriaceae *Escherichia coli* B90 |
| Eco-1001-1 | Bacteria Proteobacteria Gammaproteobacteria Enterobacteriales Enterobacteriaceae *Escherichia coli* B93 |
| Eco-1002-1 | Bacteria Proteobacteria Gammaproteobacteria Enterobacteriales Enterobacteriaceae *Escherichia coli* B94 |
| Eco-1003-1 | Bacteria Proteobacteria Gammaproteobacteria Enterobacteriales Enterobacteriaceae *Escherichia coli* B95 |
| Eco-1004-1 | Bacteria Proteobacteria Gammaproteobacteria Enterobacteriales Enterobacteriaceae *Escherichia coli* BCE001_MS16 |
| Eco-1005-1 | Bacteria Proteobacteria Gammaproteobacteria Enterobacteriales Enterobacteriaceae *Escherichia coli* BCE002_MS12 |
| Eco-1006-1 | Bacteria Proteobacteria Gammaproteobacteria Enterobacteriales Enterobacteriaceae *Escherichia coli* BCE006_MS-23 |
| Eco-1007-1 | Bacteria Proteobacteria Gammaproteobacteria Enterobacteriales Enterobacteriaceae *Escherichia coli* BCE007_MS-11 |
| Eco-1008-1 | Bacteria Proteobacteria Gammaproteobacteria Enterobacteriales Enterobacteriaceae *Escherichia coli* BCE008_MS-01 |
| Eco-1009-1 | Bacteria Proteobacteria Gammaproteobacteria Enterobacteriales Enterobacteriaceae *Escherichia coli* BCE008_MS-13 |
| Eco-1010-1 | Bacteria Proteobacteria Gammaproteobacteria Enterobacteriales Enterobacteriaceae *Escherichia coli* BCE011_MS-01 |
| Eco-1011-1 | Bacteria Proteobacteria Gammaproteobacteria Enterobacteriales Enterobacteriaceae *Escherichia coli* BCE019_MS-13 |
| Eco-1012-1 | Bacteria Proteobacteria Gammaproteobacteria Enterobacteriales Enterobacteriaceae *Escherichia coli* BCE030_MS-09 |
| Eco-1013-1 | Bacteria Proteobacteria Gammaproteobacteria Enterobacteriales Enterobacteriaceae *Escherichia coli* BCE032_MS-12 |

| | |
|---|---|
| Eco-1014-1 | Bacteria Proteobacteria Gammaproteobacteria Enterobacteriales Enterobacteriaceae *Escherichia coli* BCE034_MS-14 |
| Eco-1023-1 | Bacteria Proteobacteria Gammaproteobacteria Enterobacteriales Enterobacteriaceae *Escherichia coli* Bd5610_99 |
| Eco-1015-1 | Bacteria Proteobacteria Gammaproteobacteria Enterobacteriales Enterobacteriaceae *Escherichia coli* BIDMC 19C |
| Eco-1016-1 | Bacteria Proteobacteria Gammaproteobacteria Enterobacteriales Enterobacteriaceae *Escherichia coli* BIDMC 37 |
| Eco-1017-1 | Bacteria Proteobacteria Gammaproteobacteria Enterobacteriales Enterobacteriaceae *Escherichia coli* BIDMC 38 |
| Eco-1018-1 | Bacteria Proteobacteria Gammaproteobacteria Enterobacteriales Enterobacteriaceae *Escherichia coli* BIDMC 39 |
| Eco-1019-1 to Eco-1019-3 | Bacteria Proteobacteria Gammaproteobacteria Enterobacteriales Enterobacteriaceae *Escherichia coli* BL21(DE3) |
| Eco-1020-1 | Bacteria Proteobacteria Gammaproteobacteria Enterobacteriales Enterobacteriaceae *Escherichia coli* BW2952 |
| Eco-1021-1 | Bacteria Proteobacteria Gammaproteobacteria Enterobacteriales Enterobacteriaceae *Escherichia coli* BWH 24 |
| Eco-1022-1 | Bacteria Proteobacteria Gammaproteobacteria Enterobacteriales Enterobacteriaceae *Escherichia coli* BWH 32 |
| Eco-1024-1 | Bacteria Proteobacteria Gammaproteobacteria Enterobacteriales Enterobacteriaceae *Escherichia coli* C-34666 |
| Eco-1025-1 | Bacteria Proteobacteria Gammaproteobacteria Enterobacteriales Enterobacteriaceae *Escherichia coli* CB7326 |
| Eco-1026-1 | Bacteria Proteobacteria Gammaproteobacteria Enterobacteriales Enterobacteriaceae *Escherichia coli* CE418 |
| Eco-1027-1 | Bacteria Proteobacteria Gammaproteobacteria Enterobacteriales Enterobacteriaceae *Escherichia coli* CE516 |
| Eco-1028-1 | Bacteria Proteobacteria Gammaproteobacteria Enterobacteriales Enterobacteriaceae *Escherichia coli* CE549 |
| Eco-1029-1 | Bacteria Proteobacteria Gammaproteobacteria Enterobacteriales Enterobacteriaceae *Escherichia coli* CFT073 |
| Eco-2163-1 | Bacteria Proteobacteria Gammaproteobacteria Enterobacteriales Enterobacteriaceae *Escherichia coli* *chi*7122 |
| Eco-2164-1 | Bacteria Proteobacteria Gammaproteobacteria Enterobacteriales Enterobacteriaceae *Escherichia coli* *clone*A_i1 |
| Eco-1030-1 | Bacteria Proteobacteria Gammaproteobacteria Enterobacteriales Enterobacteriaceae *Escherichia coli* CUMT8 |
| Eco-1031-1 | Bacteria Proteobacteria Gammaproteobacteria Enterobacteriales Enterobacteriaceae *Escherichia coli* DEC10E |
| Eco-1032-1 | Bacteria Proteobacteria Gammaproteobacteria Enterobacteriales Enterobacteriaceae *Escherichia coli* DEC10F |
| Eco-1033-1 | Bacteria Proteobacteria Gammaproteobacteria Enterobacteriales Enterobacteriaceae *Escherichia coli* DEC11A |
| Eco-1034-1 | Bacteria Proteobacteria Gammaproteobacteria Enterobacteriales Enterobacteriaceae *Escherichia coli* DEC11B |
| Eco-1035-1 | Bacteria Proteobacteria Gammaproteobacteria Enterobacteriales Enterobacteriaceae *Escherichia coli* DEC11C |
| Eco-1036-1 | Bacteria Proteobacteria Gammaproteobacteria Enterobacteriales Enterobacteriaceae *Escherichia coli* DEC11D |
| Eco-1037-1 | Bacteria Proteobacteria Gammaproteobacteria Enterobacteriales Enterobacteriaceae *Escherichia coli* DEC11E |
| Eco-1038-1 | Bacteria Proteobacteria Gammaproteobacteria Enterobacteriales Enterobacteriaceae *Escherichia coli* DEC12A |
| Eco-1039-1 | Bacteria Proteobacteria Gammaproteobacteria Enterobacteriales Enterobacteriaceae *Escherichia coli* DEC12B |
| Eco-1040-1 | Bacteria Proteobacteria Gammaproteobacteria Enterobacteriales Enterobacteriaceae *Escherichia coli* DEC12C |
| Eco-1041-1 | Bacteria Proteobacteria Gammaproteobacteria Enterobacteriales Enterobacteriaceae *Escherichia coli* DEC12D |
| Eco-1042-1 | Bacteria Proteobacteria Gammaproteobacteria Enterobacteriales Enterobacteriaceae *Escherichia coli* DEC12E |
| Eco-1043-1 | Bacteria Proteobacteria Gammaproteobacteria Enterobacteriales Enterobacteriaceae *Escherichia coli* DEC13A |
| Eco-1044-1 | Bacteria Proteobacteria Gammaproteobacteria Enterobacteriales Enterobacteriaceae *Escherichia coli* DEC13B |
| Eco-1045-1 | Bacteria Proteobacteria Gammaproteobacteria Enterobacteriales Enterobacteriaceae *Escherichia coli* DEC13C |
| Eco-1046-1 | Bacteria Proteobacteria Gammaproteobacteria Enterobacteriales Enterobacteriaceae *Escherichia coli* DEC13D |
| Eco-1047-1 | Bacteria Proteobacteria Gammaproteobacteria Enterobacteriales Enterobacteriaceae *Escherichia coli* DEC13E |
| Eco-1048-1 | Bacteria Proteobacteria Gammaproteobacteria Enterobacteriales Enterobacteriaceae *Escherichia coli* DEC14A |
| Eco-1049-1 | Bacteria Proteobacteria Gammaproteobacteria Enterobacteriales Enterobacteriaceae *Escherichia coli* DEC14B |
| Eco-1050-1 | Bacteria Proteobacteria Gammaproteobacteria Enterobacteriales Enterobacteriaceae *Escherichia coli* DEC14C |
| Eco-1051-1 | Bacteria Proteobacteria Gammaproteobacteria Enterobacteriales Enterobacteriaceae *Escherichia coli* DEC14D |
| Eco-1052-1 | Bacteria Proteobacteria Gammaproteobacteria Enterobacteriales Enterobacteriaceae *Escherichia coli* DEC15A |
| Eco-1053-1 | Bacteria Proteobacteria Gammaproteobacteria Enterobacteriales Enterobacteriaceae *Escherichia coli* DEC15B |
| Eco-1054-1 | Bacteria Proteobacteria Gammaproteobacteria Enterobacteriales Enterobacteriaceae *Escherichia coli* DEC15C |
| Eco-1055-1 | Bacteria Proteobacteria Gammaproteobacteria Enterobacteriales Enterobacteriaceae *Escherichia coli* DEC15D |
| Eco-1056-1 | Bacteria Proteobacteria Gammaproteobacteria Enterobacteriales Enterobacteriaceae *Escherichia coli* DEC15E |
| Eco-1057-1 | Bacteria Proteobacteria Gammaproteobacteria Enterobacteriales Enterobacteriaceae *Escherichia coli* DEC1A |
| Eco-1058-1 | Bacteria Proteobacteria Gammaproteobacteria Enterobacteriales Enterobacteriaceae *Escherichia coli* DEC1B |
| Eco-1059-1 | Bacteria Proteobacteria Gammaproteobacteria Enterobacteriales Enterobacteriaceae *Escherichia coli* DEC1C |
| Eco-1060-1 | Bacteria Proteobacteria Gammaproteobacteria Enterobacteriales Enterobacteriaceae *Escherichia coli* DEC1D |
| Eco-1061-1 | Bacteria Proteobacteria Gammaproteobacteria Enterobacteriales Enterobacteriaceae *Escherichia coli* DEC1E |
| Eco-1062-1 | Bacteria Proteobacteria Gammaproteobacteria Enterobacteriales Enterobacteriaceae *Escherichia coli* DEC2B |
| Eco-1063-1 | Bacteria Proteobacteria Gammaproteobacteria Enterobacteriales Enterobacteriaceae *Escherichia coli* DEC2C |
| Eco-1064-1 | Bacteria Proteobacteria Gammaproteobacteria Enterobacteriales Enterobacteriaceae *Escherichia coli* DEC2D |
| Eco-1065-1 | Bacteria Proteobacteria Gammaproteobacteria Enterobacteriales Enterobacteriaceae *Escherichia coli* DEC2E |
| Eco-1066-1 | Bacteria Proteobacteria Gammaproteobacteria Enterobacteriales Enterobacteriaceae *Escherichia coli* DEC3A |
| Eco-1067-1 | Bacteria Proteobacteria Gammaproteobacteria Enterobacteriales Enterobacteriaceae *Escherichia coli* DEC3B |
| Eco-1068-1 | Bacteria Proteobacteria Gammaproteobacteria Enterobacteriales Enterobacteriaceae *Escherichia coli* DEC3C |
| Eco-1069-1 | Bacteria Proteobacteria Gammaproteobacteria Enterobacteriales Enterobacteriaceae *Escherichia coli* DEC3D |
| Eco-1070-1 | Bacteria Proteobacteria Gammaproteobacteria Enterobacteriales Enterobacteriaceae *Escherichia coli* DEC3E |
| Eco-1071-1 | Bacteria Proteobacteria Gammaproteobacteria Enterobacteriales Enterobacteriaceae *Escherichia coli* DEC3F |
| Eco-1072-1 | Bacteria Proteobacteria Gammaproteobacteria Enterobacteriales Enterobacteriaceae *Escherichia coli* DEC4A |
| Eco-1073-1 | Bacteria Proteobacteria Gammaproteobacteria Enterobacteriales Enterobacteriaceae *Escherichia coli* DEC4B |
| Eco-1074-1 | Bacteria Proteobacteria Gammaproteobacteria Enterobacteriales Enterobacteriaceae *Escherichia coli* DEC4C |
| Eco-1075-1 | Bacteria Proteobacteria Gammaproteobacteria Enterobacteriales Enterobacteriaceae *Escherichia coli* DEC4D |
| Eco-1076-1 | Bacteria Proteobacteria Gammaproteobacteria Enterobacteriales Enterobacteriaceae *Escherichia coli* DEC4E |

| | |
|---|---|
| Eco-1077-1 | Bacteria Proteobacteria Gammaproteobacteria Enterobacteriales Enterobacteriaceae *Escherichia coli* DEC4F |
| Eco-1078-1 | Bacteria Proteobacteria Gammaproteobacteria Enterobacteriales Enterobacteriaceae *Escherichia coli* DEC5A |
| Eco-1079-1 | Bacteria Proteobacteria Gammaproteobacteria Enterobacteriales Enterobacteriaceae *Escherichia coli* DEC5B |
| Eco-1080-1 | Bacteria Proteobacteria Gammaproteobacteria Enterobacteriales Enterobacteriaceae *Escherichia coli* DEC5C |
| Eco-1081-1 | Bacteria Proteobacteria Gammaproteobacteria Enterobacteriales Enterobacteriaceae *Escherichia coli* DEC5D |
| Eco-1082-1 | Bacteria Proteobacteria Gammaproteobacteria Enterobacteriales Enterobacteriaceae *Escherichia coli* DEC5E |
| Eco-1083-1 | Bacteria Proteobacteria Gammaproteobacteria Enterobacteriales Enterobacteriaceae *Escherichia coli* DEC6A |
| Eco-1084-1 | Bacteria Proteobacteria Gammaproteobacteria Enterobacteriales Enterobacteriaceae *Escherichia coli* DEC6B |
| Eco-1085-1 | Bacteria Proteobacteria Gammaproteobacteria Enterobacteriales Enterobacteriaceae *Escherichia coli* DEC6E |
| Eco-1086-1 | Bacteria Proteobacteria Gammaproteobacteria Enterobacteriales Enterobacteriaceae *Escherichia coli* DEC7A |
| Eco-1087-1 | Bacteria Proteobacteria Gammaproteobacteria Enterobacteriales Enterobacteriaceae *Escherichia coli* DEC7B |
| Eco-1088-1 | Bacteria Proteobacteria Gammaproteobacteria Enterobacteriales Enterobacteriaceae *Escherichia coli* DEC7C |
| Eco-1089-1 | Bacteria Proteobacteria Gammaproteobacteria Enterobacteriales Enterobacteriaceae *Escherichia coli* DEC7D |
| Eco-1090-1 | Bacteria Proteobacteria Gammaproteobacteria Enterobacteriales Enterobacteriaceae *Escherichia coli* DEC7E |
| Eco-1091-1 | Bacteria Proteobacteria Gammaproteobacteria Enterobacteriales Enterobacteriaceae *Escherichia coli* DEC8A |
| Eco-1092-1 | Bacteria Proteobacteria Gammaproteobacteria Enterobacteriales Enterobacteriaceae *Escherichia coli* DEC8B |
| Eco-1093-1 | Bacteria Proteobacteria Gammaproteobacteria Enterobacteriales Enterobacteriaceae *Escherichia coli* DEC8C |
| Eco-1094-1 | Bacteria Proteobacteria Gammaproteobacteria Enterobacteriales Enterobacteriaceae *Escherichia coli* DEC8D |
| Eco-1095-1 | Bacteria Proteobacteria Gammaproteobacteria Enterobacteriales Enterobacteriaceae *Escherichia coli* DEC8E |
| Eco-1096-1 | Bacteria Proteobacteria Gammaproteobacteria Enterobacteriales Enterobacteriaceae *Escherichia coli* DEC9A |
| Eco-1097-1 | Bacteria Proteobacteria Gammaproteobacteria Enterobacteriales Enterobacteriaceae *Escherichia coli* DEC9B |
| Eco-1098-1 | Bacteria Proteobacteria Gammaproteobacteria Enterobacteriales Enterobacteriaceae *Escherichia coli* DEC9C |
| Eco-1099-1 | Bacteria Proteobacteria Gammaproteobacteria Enterobacteriales Enterobacteriaceae *Escherichia coli* DEC9D |
| Eco-1100-1 | Bacteria Proteobacteria Gammaproteobacteria Enterobacteriales Enterobacteriaceae *Escherichia coli* DEC9E |
| Eco-1101-1 to Eco-1101-2 | Bacteria Proteobacteria Gammaproteobacteria Enterobacteriales Enterobacteriaceae *Escherichia coli* DH1 |
| Eco-1102-1 | Bacteria Proteobacteria Gammaproteobacteria Enterobacteriales Enterobacteriaceae *Escherichia coli* E101 |
| Eco-1103-1 | Bacteria Proteobacteria Gammaproteobacteria Enterobacteriales Enterobacteriaceae *Escherichia coli* E110019 |
| Eco-1104-1 | Bacteria Proteobacteria Gammaproteobacteria Enterobacteriales Enterobacteriaceae *Escherichia coli* E1167 |
| Eco-1105-1 | Bacteria Proteobacteria Gammaproteobacteria Enterobacteriales Enterobacteriaceae *Escherichia coli* E128010 |
| Eco-1106-1 | Bacteria Proteobacteria Gammaproteobacteria Enterobacteriales Enterobacteriaceae *Escherichia coli* E1520 |
| Eco-1107-1 | Bacteria Proteobacteria Gammaproteobacteria Enterobacteriales Enterobacteriaceae *Escherichia coli* E22 |
| Eco-1108-1 | Bacteria Proteobacteria Gammaproteobacteria Enterobacteriales Enterobacteriaceae *Escherichia coli* E2265 |
| Eco-1109-1 | Bacteria Proteobacteria Gammaproteobacteria Enterobacteriales Enterobacteriaceae *Escherichia coli* E24377A |
| Eco-1110-1 | Bacteria Proteobacteria Gammaproteobacteria Enterobacteriales Enterobacteriaceae *Escherichia coli* E482 |
| Eco-1111-1 | Bacteria Proteobacteria Gammaproteobacteria Enterobacteriales Enterobacteriaceae *Escherichia coli* EC1734 |
| Eco-1112-1 | Bacteria Proteobacteria Gammaproteobacteria Enterobacteriales Enterobacteriaceae *Escherichia coli* EC1735 |
| Eco-1113-1 | Bacteria Proteobacteria Gammaproteobacteria Enterobacteriales Enterobacteriaceae *Escherichia coli* EC1736 |
| Eco-1114-1 | Bacteria Proteobacteria Gammaproteobacteria Enterobacteriales Enterobacteriaceae *Escherichia coli* EC1737 |
| Eco-1115-1 | Bacteria Proteobacteria Gammaproteobacteria Enterobacteriales Enterobacteriaceae *Escherichia coli* EC1738 |
| Eco-1116-1 | Bacteria Proteobacteria Gammaproteobacteria Enterobacteriales Enterobacteriaceae *Escherichia coli* EC1845 |
| Eco-1117-1 | Bacteria Proteobacteria Gammaproteobacteria Enterobacteriales Enterobacteriaceae *Escherichia coli* EC1846 |
| Eco-1118-1 | Bacteria Proteobacteria Gammaproteobacteria Enterobacteriales Enterobacteriaceae *Escherichia coli* EC1847 |
| Eco-1119-1 | Bacteria Proteobacteria Gammaproteobacteria Enterobacteriales Enterobacteriaceae *Escherichia coli* EC1848 |
| Eco-1120-1 | Bacteria Proteobacteria Gammaproteobacteria Enterobacteriales Enterobacteriaceae *Escherichia coli* EC1849 |
| Eco-1121-1 | Bacteria Proteobacteria Gammaproteobacteria Enterobacteriales Enterobacteriaceae *Escherichia coli* EC1850 |
| Eco-1122-1 | Bacteria Proteobacteria Gammaproteobacteria Enterobacteriales Enterobacteriaceae *Escherichia coli* EC1856 |
| Eco-1123-1 | Bacteria Proteobacteria Gammaproteobacteria Enterobacteriales Enterobacteriaceae *Escherichia coli* EC1862 |
| Eco-1124-1 | Bacteria Proteobacteria Gammaproteobacteria Enterobacteriales Enterobacteriaceae *Escherichia coli* EC1863 |
| Eco-1125-1 | Bacteria Proteobacteria Gammaproteobacteria Enterobacteriales Enterobacteriaceae *Escherichia coli* EC1864 |
| Eco-1126-1 | Bacteria Proteobacteria Gammaproteobacteria Enterobacteriales Enterobacteriaceae *Escherichia coli* EC1865 |
| Eco-1127-1 | Bacteria Proteobacteria Gammaproteobacteria Enterobacteriales Enterobacteriaceae *Escherichia coli* EC1866 |
| Eco-1128-1 | Bacteria Proteobacteria Gammaproteobacteria Enterobacteriales Enterobacteriaceae *Escherichia coli* EC1868 |
| Eco-1129-1 | Bacteria Proteobacteria Gammaproteobacteria Enterobacteriales Enterobacteriaceae *Escherichia coli* EC1869 |
| Eco-1130-1 | Bacteria Proteobacteria Gammaproteobacteria Enterobacteriales Enterobacteriaceae *Escherichia coli* EC1870 |
| Eco-1131-1 | Bacteria Proteobacteria Gammaproteobacteria Enterobacteriales Enterobacteriaceae *Escherichia coli* EC302/04 |
| Eco-1132-1 | Bacteria Proteobacteria Gammaproteobacteria Enterobacteriales Enterobacteriaceae *Escherichia coli* EC4013 |
| Eco-1133-1 | Bacteria Proteobacteria Gammaproteobacteria Enterobacteriales Enterobacteriaceae *Escherichia coli* EC4100B |
| Eco-1134-1 | Bacteria Proteobacteria Gammaproteobacteria Enterobacteriales Enterobacteriaceae *Escherichia coli* EC4196 |
| Eco-1135-1 | Bacteria Proteobacteria Gammaproteobacteria Enterobacteriales Enterobacteriaceae *Escherichia coli* EC4203 |
| Eco-1136-1 | Bacteria Proteobacteria Gammaproteobacteria Enterobacteriales Enterobacteriaceae *Escherichia coli* EC4402 |
| Eco-1137-1 | Bacteria Proteobacteria Gammaproteobacteria Enterobacteriales Enterobacteriaceae *Escherichia coli* EC4421 |
| Eco-1138-1 | Bacteria Proteobacteria Gammaproteobacteria Enterobacteriales Enterobacteriaceae *Escherichia coli* EC4422 |
| Eco-1139-1 | Bacteria Proteobacteria Gammaproteobacteria Enterobacteriales Enterobacteriaceae *Escherichia coli* EC4436 |
| Eco-1140-1 | Bacteria Proteobacteria Gammaproteobacteria Enterobacteriales Enterobacteriaceae *Escherichia coli* EC4437 |
| Eco-1141-1 | Bacteria Proteobacteria Gammaproteobacteria Enterobacteriales Enterobacteriaceae *Escherichia coli* EC4439 |

| | |
|---|---|
| Eco-1142-1 | Bacteria Proteobacteria Gammaproteobacteria Enterobacteriales Enterobacteriaceae *Escherichia coli* EC4448 |
| Eco-1143-1 | Bacteria Proteobacteria Gammaproteobacteria Enterobacteriales Enterobacteriaceae *Escherichia coli* EC96038 |
| Eco-1144-1 | Bacteria Proteobacteria Gammaproteobacteria Enterobacteriales Enterobacteriaceae *Escherichia coli* ECC-Z |
| Eco-1145-1 | Bacteria Proteobacteria Gammaproteobacteria Enterobacteriales Enterobacteriaceae *Escherichia coli* ED1a |
| Eco-1150-1 | Bacteria Proteobacteria Gammaproteobacteria Enterobacteriales Enterobacteriaceae *Escherichia coli* Envira 10/1 |
| Eco-1151-1 | Bacteria Proteobacteria Gammaproteobacteria Enterobacteriales Enterobacteriaceae *Escherichia coli* Envira 8/11 |
| Eco-1146-1 | Bacteria Proteobacteria Gammaproteobacteria Enterobacteriales Enterobacteriaceae *Escherichia coli* EPEC C342-62 |
| Eco-1147-1 | Bacteria Proteobacteria Gammaproteobacteria Enterobacteriales Enterobacteriaceae *Escherichia coli* EPECa12 |
| Eco-1148-1 | Bacteria Proteobacteria Gammaproteobacteria Enterobacteriales Enterobacteriaceae *Escherichia coli* EPECa14 |
| Eco-1149-1 | Bacteria Proteobacteria Gammaproteobacteria Enterobacteriales Enterobacteriaceae *Escherichia coli* ETEC H10407 |
| Eco-1152-1 | Bacteria Proteobacteria Gammaproteobacteria Enterobacteriales Enterobacteriaceae *Escherichia coli* F11 |
| Eco-1153-1 | Bacteria Proteobacteria Gammaproteobacteria Enterobacteriales Enterobacteriaceae *Escherichia coli* FDA504 |
| Eco-1154-1 | Bacteria Proteobacteria Gammaproteobacteria Enterobacteriales Enterobacteriaceae *Escherichia coli* FDA505 |
| Eco-1155-1 | Bacteria Proteobacteria Gammaproteobacteria Enterobacteriales Enterobacteriaceae *Escherichia coli* FDA506 |
| Eco-1156-1 | Bacteria Proteobacteria Gammaproteobacteria Enterobacteriales Enterobacteriaceae *Escherichia coli* FDA507 |
| Eco-1157-1 | Bacteria Proteobacteria Gammaproteobacteria Enterobacteriales Enterobacteriaceae *Escherichia coli* FDA517 |
| Eco-1158-1 | Bacteria Proteobacteria Gammaproteobacteria Enterobacteriales Enterobacteriaceae *Escherichia coli* FRIK1985 |
| Eco-1159-1 | Bacteria Proteobacteria Gammaproteobacteria Enterobacteriales Enterobacteriaceae *Escherichia coli* FRIK1990 |
| Eco-1160-1 | Bacteria Proteobacteria Gammaproteobacteria Enterobacteriales Enterobacteriaceae *Escherichia coli* FRIK1996 |
| Eco-1161-1 | Bacteria Proteobacteria Gammaproteobacteria Enterobacteriales Enterobacteriaceae *Escherichia coli* FRIK1997 |
| Eco-1162-1 | Bacteria Proteobacteria Gammaproteobacteria Enterobacteriales Enterobacteriaceae *Escherichia coli* FRIK1999 |
| Eco-1163-1 | Bacteria Proteobacteria Gammaproteobacteria Enterobacteriales Enterobacteriaceae *Escherichia coli* FRIK2001 |
| Eco-1164-1 | Bacteria Proteobacteria Gammaproteobacteria Enterobacteriales Enterobacteriaceae *Escherichia coli* FRIK523 |
| Eco-1165-1 | Bacteria Proteobacteria Gammaproteobacteria Enterobacteriales Enterobacteriaceae *Escherichia coli* FRIK920 |
| Eco-1166-1 | Bacteria Proteobacteria Gammaproteobacteria Enterobacteriales Enterobacteriaceae *Escherichia coli* FVEC1302 |
| Eco-1167-1 | Bacteria Proteobacteria Gammaproteobacteria Enterobacteriales Enterobacteriaceae *Escherichia coli* FVEC1412 |
| Eco-1168-1 | Bacteria Proteobacteria Gammaproteobacteria Enterobacteriales Enterobacteriaceae *Escherichia coli* G58-1 |
| Eco-1169-1 to Eco-1169-2 | Bacteria Proteobacteria Gammaproteobacteria Enterobacteriales Enterobacteriaceae *Escherichia coli* GM4792 |
| Eco-1170-1 | Bacteria Proteobacteria Gammaproteobacteria Enterobacteriales Enterobacteriaceae *Escherichia coli* H120 |
| Eco-1171-1 | Bacteria Proteobacteria Gammaproteobacteria Enterobacteriales Enterobacteriaceae *Escherichia coli* H252 |
| Eco-1172-1 | Bacteria Proteobacteria Gammaproteobacteria Enterobacteriales Enterobacteriaceae *Escherichia coli* H263 |
| Eco-1173-1 | Bacteria Proteobacteria Gammaproteobacteria Enterobacteriales Enterobacteriaceae *Escherichia coli* H299 |
| Eco-1174-1 | Bacteria Proteobacteria Gammaproteobacteria Enterobacteriales Enterobacteriaceae *Escherichia coli* H30 |
| Eco-1175-1 | Bacteria Proteobacteria Gammaproteobacteria Enterobacteriales Enterobacteriaceae *Escherichia coli* H397 |
| Eco-1176-1 | Bacteria Proteobacteria Gammaproteobacteria Enterobacteriales Enterobacteriaceae *Escherichia coli* H489 |
| Eco-1177-1 | Bacteria Proteobacteria Gammaproteobacteria Enterobacteriales Enterobacteriaceae *Escherichia coli* H494 |
| Eco-1178-1 | Bacteria Proteobacteria Gammaproteobacteria Enterobacteriales Enterobacteriaceae *Escherichia coli* H591 |
| Eco-1179-1 | Bacteria Proteobacteria Gammaproteobacteria Enterobacteriales Enterobacteriaceae *Escherichia coli* H730 |
| Eco-1180-1 | Bacteria Proteobacteria Gammaproteobacteria Enterobacteriales Enterobacteriaceae *Escherichia coli* H736 |
| Eco-1181-1 | Bacteria Proteobacteria Gammaproteobacteria Enterobacteriales Enterobacteriaceae *Escherichia coli* HM26 |
| Eco-1182-1 | Bacteria Proteobacteria Gammaproteobacteria Enterobacteriales Enterobacteriaceae *Escherichia coli* HM27 |
| Eco-1183-1 | Bacteria Proteobacteria Gammaproteobacteria Enterobacteriales Enterobacteriaceae *Escherichia coli* HM46 |
| Eco-1184-1 | Bacteria Proteobacteria Gammaproteobacteria Enterobacteriales Enterobacteriaceae *Escherichia coli* HM605 |
| Eco-1185-1 | Bacteria Proteobacteria Gammaproteobacteria Enterobacteriales Enterobacteriaceae *Escherichia coli* HM65 |
| Eco-1186-1 | Bacteria Proteobacteria Gammaproteobacteria Enterobacteriales Enterobacteriaceae *Escherichia coli* HM69 |
| Eco-1187-1 | Bacteria Proteobacteria Gammaproteobacteria Enterobacteriales Enterobacteriaceae *Escherichia coli* HS |
| Eco-1188-1 | Bacteria Proteobacteria Gammaproteobacteria Enterobacteriales Enterobacteriaceae *Escherichia coli* HVH 1 (4-6876161) |
| Eco-1189-1 | Bacteria Proteobacteria Gammaproteobacteria Enterobacteriales Enterobacteriaceae *Escherichia coli* HVH 10 (4-6832164) |
| Eco-1190-1 | Bacteria Proteobacteria Gammaproteobacteria Enterobacteriales Enterobacteriaceae *Escherichia coli* HVH 100 (4-2850729) |
| Eco-1191-1 | Bacteria Proteobacteria Gammaproteobacteria Enterobacteriales Enterobacteriaceae *Escherichia coli* HVH 102 (4-6906788) |
| Eco-1192-1 | Bacteria Proteobacteria Gammaproteobacteria Enterobacteriales Enterobacteriaceae *Escherichia coli* HVH 103 (4-5904188) |
| Eco-1193-1 | Bacteria Proteobacteria Gammaproteobacteria Enterobacteriales Enterobacteriaceae *Escherichia coli* HVH 104 (4-6977960) |
| Eco-1194-1 | Bacteria Proteobacteria Gammaproteobacteria Enterobacteriales Enterobacteriaceae *Escherichia coli* HVH 106 (4-6881831) |
| Eco-1195-1 | Bacteria Proteobacteria Gammaproteobacteria Enterobacteriales Enterobacteriaceae *Escherichia coli* HVH 107 (4-5860571) |
| Eco-1196-1 | Bacteria Proteobacteria Gammaproteobacteria Enterobacteriales Enterobacteriaceae *Escherichia coli* HVH 108 (4-6924867) |
| Eco-1197-1 | Bacteria Proteobacteria Gammaproteobacteria Enterobacteriales Enterobacteriaceae *Escherichia coli* HVH 109 (4-6977162) |
| Eco-1198-1 | Bacteria Proteobacteria Gammaproteobacteria Enterobacteriales Enterobacteriaceae *Escherichia coli* HVH 110 (4-6978754) |
| Eco-1199-1 | Bacteria Proteobacteria Gammaproteobacteria Enterobacteriales Enterobacteriaceae *Escherichia coli* HVH 111 (4-7039018) |
| Eco-1200-1 | Bacteria Proteobacteria Gammaproteobacteria Enterobacteriales Enterobacteriaceae *Escherichia coli* HVH 112 (4-5987253) |
| Eco-1201-1 | Bacteria Proteobacteria Gammaproteobacteria Enterobacteriales Enterobacteriaceae *Escherichia coli* HVH 113 (4-7535473) |
| Eco-1202-1 | Bacteria Proteobacteria Gammaproteobacteria Enterobacteriales Enterobacteriaceae *Escherichia coli* HVH 114 (4-7037740) |
| Eco-1203-1 | Bacteria Proteobacteria Gammaproteobacteria Enterobacteriales Enterobacteriaceae *Escherichia coli* HVH 115 (4-4465989) |
| Eco-1204-1 | Bacteria Proteobacteria Gammaproteobacteria Enterobacteriales Enterobacteriaceae *Escherichia coli* HVH 115 (4-4465997) |
| Eco-1205-1 | Bacteria Proteobacteria Gammaproteobacteria Enterobacteriales Enterobacteriaceae *Escherichia coli* HVH 116 (4-6879942) |
| Eco-1206-1 | Bacteria Proteobacteria Gammaproteobacteria Enterobacteriales Enterobacteriaceae *Escherichia coli* HVH 117 (4-6857191) |

| | |
|---|---|
| Eco-1207-1 | Bacteria Proteobacteria Gammaproteobacteria Enterobacteriales Enterobacteriaceae *Escherichia coli* HVH 118 (4-7345399) |
| Eco-1208-1 | Bacteria Proteobacteria Gammaproteobacteria Enterobacteriales Enterobacteriaceae *Escherichia coli* HVH 119 (4-6879578) |
| Eco-1209-1 | Bacteria Proteobacteria Gammaproteobacteria Enterobacteriales Enterobacteriaceae *Escherichia coli* HVH 12 (4-7653042) |
| Eco-1210-1 | Bacteria Proteobacteria Gammaproteobacteria Enterobacteriales Enterobacteriaceae *Escherichia coli* HVH 120 (4-6978681) |
| Eco-1211-1 | Bacteria Proteobacteria Gammaproteobacteria Enterobacteriales Enterobacteriaceae *Escherichia coli* HVH 121 (4-6877826) |
| Eco-1212-1 | Bacteria Proteobacteria Gammaproteobacteria Enterobacteriales Enterobacteriaceae *Escherichia coli* HVH 122 (4-6851606) |
| Eco-1213-1 | Bacteria Proteobacteria Gammaproteobacteria Enterobacteriales Enterobacteriaceae *Escherichia coli* HVH 125 (4-2634716) |
| Eco-1214-1 | Bacteria Proteobacteria Gammaproteobacteria Enterobacteriales Enterobacteriaceae *Escherichia coli* HVH 126 (4-6034225) |
| Eco-1215-1 | Bacteria Proteobacteria Gammaproteobacteria Enterobacteriales Enterobacteriaceae *Escherichia coli* HVH 127 (4-7303629) |
| Eco-1216-1 | Bacteria Proteobacteria Gammaproteobacteria Enterobacteriales Enterobacteriaceae *Escherichia coli* HVH 128 (4-7030436) |
| Eco-1217-1 | Bacteria Proteobacteria Gammaproteobacteria Enterobacteriales Enterobacteriaceae *Escherichia coli* HVH 13 (4-7634056) |
| Eco-1218-1 | Bacteria Proteobacteria Gammaproteobacteria Enterobacteriales Enterobacteriaceae *Escherichia coli* HVH 130 (4-7036876) |
| Eco-1219-1 | Bacteria Proteobacteria Gammaproteobacteria Enterobacteriales Enterobacteriaceae *Escherichia coli* HVH 132 (4-6876862) |
| Eco-1220-1 | Bacteria Proteobacteria Gammaproteobacteria Enterobacteriales Enterobacteriaceae *Escherichia coli* HVH 133 (4-4466519) |
| Eco-1221-1 | Bacteria Proteobacteria Gammaproteobacteria Enterobacteriales Enterobacteriaceae *Escherichia coli* HVH 134 (4-6073441) |
| Eco-1222-1 | Bacteria Proteobacteria Gammaproteobacteria Enterobacteriales Enterobacteriaceae *Escherichia coli* HVH 135 (4-4449320) |
| Eco-1223-1 | Bacteria Proteobacteria Gammaproteobacteria Enterobacteriales Enterobacteriaceae *Escherichia coli* HVH 136 (4-5970458) |
| Eco-1224-1 | Bacteria Proteobacteria Gammaproteobacteria Enterobacteriales Enterobacteriaceae *Escherichia coli* HVH 137 (4-2124971) |
| Eco-1225-1 | Bacteria Proteobacteria Gammaproteobacteria Enterobacteriales Enterobacteriaceae *Escherichia coli* HVH 138 (4-6066704) |
| Eco-1226-1 | Bacteria Proteobacteria Gammaproteobacteria Enterobacteriales Enterobacteriaceae *Escherichia coli* HVH 139 (4-3192644) |
| Eco-1227-1 | Bacteria Proteobacteria Gammaproteobacteria Enterobacteriales Enterobacteriaceae *Escherichia coli* HVH 140 (4-5894387) |
| Eco-1228-1 | Bacteria Proteobacteria Gammaproteobacteria Enterobacteriales Enterobacteriaceae *Escherichia coli* HVH 141 (4-5995973) |
| Eco-1229-1 | Bacteria Proteobacteria Gammaproteobacteria Enterobacteriales Enterobacteriaceae *Escherichia coli* HVH 142 (4-5627451) |
| Eco-1230-1 | Bacteria Proteobacteria Gammaproteobacteria Enterobacteriales Enterobacteriaceae *Escherichia coli* HVH 143 (4-5674999) |
| Eco-1231-1 | Bacteria Proteobacteria Gammaproteobacteria Enterobacteriales Enterobacteriaceae *Escherichia coli* HVH 144 (4-4451937) |
| Eco-1232-1 | Bacteria Proteobacteria Gammaproteobacteria Enterobacteriales Enterobacteriaceae *Escherichia coli* HVH 145 (4-5672112) |
| Eco-1233-1 | Bacteria Proteobacteria Gammaproteobacteria Enterobacteriales Enterobacteriaceae *Escherichia coli* HVH 146 (4-3189767) |
| Eco-1234-1 | Bacteria Proteobacteria Gammaproteobacteria Enterobacteriales Enterobacteriaceae *Escherichia coli* HVH 147 (4-5893887) |
| Eco-1235-1 | Bacteria Proteobacteria Gammaproteobacteria Enterobacteriales Enterobacteriaceae *Escherichia coli* HVH 148 (4-3192490) |
| Eco-1236-1 | Bacteria Proteobacteria Gammaproteobacteria Enterobacteriales Enterobacteriaceae *Escherichia coli* HVH 149 (4-4451880) |
| Eco-1237-1 | Bacteria Proteobacteria Gammaproteobacteria Enterobacteriales Enterobacteriaceae *Escherichia coli* HVH 150 (4-3258106) |
| Eco-1238-1 | Bacteria Proteobacteria Gammaproteobacteria Enterobacteriales Enterobacteriaceae *Escherichia coli* HVH 151 (4-5755573) |
| Eco-1239-1 | Bacteria Proteobacteria Gammaproteobacteria Enterobacteriales Enterobacteriaceae *Escherichia coli* HVH 152 (4-3447545) |
| Eco-1240-1 | Bacteria Proteobacteria Gammaproteobacteria Enterobacteriales Enterobacteriaceae *Escherichia coli* HVH 153 (3-9344314) |
| Eco-1241-1 | Bacteria Proteobacteria Gammaproteobacteria Enterobacteriales Enterobacteriaceae *Escherichia coli* HVH 154 (4-5636698) |
| Eco-1242-1 | Bacteria Proteobacteria Gammaproteobacteria Enterobacteriales Enterobacteriaceae *Escherichia coli* HVH 155 (4-4509048) |
| Eco-1243-1 | Bacteria Proteobacteria Gammaproteobacteria Enterobacteriales Enterobacteriaceae *Escherichia coli* HVH 156 (4-3206505) |
| Eco-1244-1 | Bacteria Proteobacteria Gammaproteobacteria Enterobacteriales Enterobacteriaceae *Escherichia coli* HVH 157 (4-3406229) |
| Eco-1245-1 | Bacteria Proteobacteria Gammaproteobacteria Enterobacteriales Enterobacteriaceae *Escherichia coli* HVH 158 (4-3224287) |
| Eco-1246-1 | Bacteria Proteobacteria Gammaproteobacteria Enterobacteriales Enterobacteriaceae *Escherichia coli* HVH 159 (4-5818141) |
| Eco-1247-1 | Bacteria Proteobacteria Gammaproteobacteria Enterobacteriales Enterobacteriaceae *Escherichia coli* HVH 16 (4-7649002) |
| Eco-1248-1 | Bacteria Proteobacteria Gammaproteobacteria Enterobacteriales Enterobacteriaceae *Escherichia coli* HVH 160 (4-5695937) |
| Eco-1249-1 | Bacteria Proteobacteria Gammaproteobacteria Enterobacteriales Enterobacteriaceae *Escherichia coli* HVH 161 (4-3119890) |
| Eco-1250-1 | Bacteria Proteobacteria Gammaproteobacteria Enterobacteriales Enterobacteriaceae *Escherichia coli* HVH 162 (4-5627982) |
| Eco-1251-1 | Bacteria Proteobacteria Gammaproteobacteria Enterobacteriales Enterobacteriaceae *Escherichia coli* HVH 163 (4-4697553) |
| Eco-1252-1 | Bacteria Proteobacteria Gammaproteobacteria Enterobacteriales Enterobacteriaceae *Escherichia coli* HVH 164 (4-5953081) |
| Eco-1253-1 | Bacteria Proteobacteria Gammaproteobacteria Enterobacteriales Enterobacteriaceae *Escherichia coli* HVH 167 (4-6073565) |
| Eco-1254-1 | Bacteria Proteobacteria Gammaproteobacteria Enterobacteriales Enterobacteriaceae *Escherichia coli* HVH 169 (4-1075578) |
| Eco-1255-1 | Bacteria Proteobacteria Gammaproteobacteria Enterobacteriales Enterobacteriaceae *Escherichia coli* HVH 17 (4-7473087) |
| Eco-1256-1 | Bacteria Proteobacteria Gammaproteobacteria Enterobacteriales Enterobacteriaceae *Escherichia coli* HVH 170 (4-3026949) |
| Eco-1257-1 | Bacteria Proteobacteria Gammaproteobacteria Enterobacteriales Enterobacteriaceae *Escherichia coli* HVH 171 (4-3191958) |
| Eco-1258-1 | Bacteria Proteobacteria Gammaproteobacteria Enterobacteriales Enterobacteriaceae *Escherichia coli* HVH 172 (4-3248542) |
| Eco-1259-1 | Bacteria Proteobacteria Gammaproteobacteria Enterobacteriales Enterobacteriaceae *Escherichia coli* HVH 173 (3-9175482) |
| Eco-1260-1 | Bacteria Proteobacteria Gammaproteobacteria Enterobacteriales Enterobacteriaceae *Escherichia coli* HVH 175 (4-3405184) |
| Eco-1261-1 | Bacteria Proteobacteria Gammaproteobacteria Enterobacteriales Enterobacteriaceae *Escherichia coli* HVH 176 (4-3428664) |
| Eco-1262-1 | Bacteria Proteobacteria Gammaproteobacteria Enterobacteriales Enterobacteriaceae *Escherichia coli* HVH 177 (4-2876612) |
| Eco-1263-1 | Bacteria Proteobacteria Gammaproteobacteria Enterobacteriales Enterobacteriaceae *Escherichia coli* HVH 178 (4-3189163) |
| Eco-1264-1 | Bacteria Proteobacteria Gammaproteobacteria Enterobacteriales Enterobacteriaceae *Escherichia coli* HVH 18 (4-8589585) |
| Eco-1265-1 | Bacteria Proteobacteria Gammaproteobacteria Enterobacteriales Enterobacteriaceae *Escherichia coli* HVH 180 (4-3051617) |
| Eco-1266-1 | Bacteria Proteobacteria Gammaproteobacteria Enterobacteriales Enterobacteriaceae *Escherichia coli* HVH 182 (4-0985554) |
| Eco-1267-1 | Bacteria Proteobacteria Gammaproteobacteria Enterobacteriales Enterobacteriaceae *Escherichia coli* HVH 183 (4-3205932) |
| Eco-1268-1 | Bacteria Proteobacteria Gammaproteobacteria Enterobacteriales Enterobacteriaceae *Escherichia coli* HVH 184 (4-3343286) |
| Eco-1269-1 | Bacteria Proteobacteria Gammaproteobacteria Enterobacteriales Enterobacteriaceae *Escherichia coli* HVH 185 (4-2876639) |
| Eco-1270-1 | Bacteria Proteobacteria Gammaproteobacteria Enterobacteriales Enterobacteriaceae *Escherichia coli* HVH 186 (4-3405044) |
| Eco-1271-1 | Bacteria Proteobacteria Gammaproteobacteria Enterobacteriales Enterobacteriaceae *Escherichia coli* HVH 187 (4-4471660) |

| | |
|---|---|
| Eco-1272-1 | Bacteria Proteobacteria Gammaproteobacteria Enterobacteriales Enterobacteriaceae *Escherichia coli* HVH 188 (4-2356988) |
| Eco-1273-1 | Bacteria Proteobacteria Gammaproteobacteria Enterobacteriales Enterobacteriaceae *Escherichia coli* HVH 189 (4-3220125) |
| Eco-1274-1 | Bacteria Proteobacteria Gammaproteobacteria Enterobacteriales Enterobacteriaceae *Escherichia coli* HVH 19 (4-7154984) |
| Eco-1275-1 | Bacteria Proteobacteria Gammaproteobacteria Enterobacteriales Enterobacteriaceae *Escherichia coli* HVH 190 (4-3255514) |
| Eco-1276-1 | Bacteria Proteobacteria Gammaproteobacteria Enterobacteriales Enterobacteriaceae *Escherichia coli* HVH 191 (3-9341900) |
| Eco-1277-1 | Bacteria Proteobacteria Gammaproteobacteria Enterobacteriales Enterobacteriaceae *Escherichia coli* HVH 192 (4-3054470) |
| Eco-1278-1 | Bacteria Proteobacteria Gammaproteobacteria Enterobacteriales Enterobacteriaceae *Escherichia coli* HVH 193 (4-3331423) |
| Eco-1279-1 | Bacteria Proteobacteria Gammaproteobacteria Enterobacteriales Enterobacteriaceae *Escherichia coli* HVH 194 (4-2356805) |
| Eco-1280-1 | Bacteria Proteobacteria Gammaproteobacteria Enterobacteriales Enterobacteriaceae *Escherichia coli* HVH 195 (3-7155360) |
| Eco-1281-1 | Bacteria Proteobacteria Gammaproteobacteria Enterobacteriales Enterobacteriaceae *Escherichia coli* HVH 196 (4-4530470) |
| Eco-1282-1 | Bacteria Proteobacteria Gammaproteobacteria Enterobacteriales Enterobacteriaceae *Escherichia coli* HVH 197 (4-4466217) |
| Eco-1283-1 | Bacteria Proteobacteria Gammaproteobacteria Enterobacteriales Enterobacteriaceae *Escherichia coli* HVH 198 (4-3206106) |
| Eco-1284-1 | Bacteria Proteobacteria Gammaproteobacteria Enterobacteriales Enterobacteriaceae *Escherichia coli* HVH 199 (4-5670322) |
| Eco-1285-1 | Bacteria Proteobacteria Gammaproteobacteria Enterobacteriales Enterobacteriaceae *Escherichia coli* HVH 2 (4-6943160) |
| Eco-1286-1 | Bacteria Proteobacteria Gammaproteobacteria Enterobacteriales Enterobacteriaceae *Escherichia coli* HVH 20 (4-5865042) |
| Eco-1287-1 | Bacteria Proteobacteria Gammaproteobacteria Enterobacteriales Enterobacteriaceae *Escherichia coli* HVH 200 (4-4449924) |
| Eco-1288-1 | Bacteria Proteobacteria Gammaproteobacteria Enterobacteriales Enterobacteriaceae *Escherichia coli* HVH 201 (4-4459431) |
| Eco-1289-1 | Bacteria Proteobacteria Gammaproteobacteria Enterobacteriales Enterobacteriaceae *Escherichia coli* HVH 202 (4-3163997) |
| Eco-1290-1 | Bacteria Proteobacteria Gammaproteobacteria Enterobacteriales Enterobacteriaceae *Escherichia coli* HVH 203 (4-3126218) |
| Eco-1291-1 | Bacteria Proteobacteria Gammaproteobacteria Enterobacteriales Enterobacteriaceae *Escherichia coli* HVH 204 (4-3112802) |
| Eco-1292-1 | Bacteria Proteobacteria Gammaproteobacteria Enterobacteriales Enterobacteriaceae *Escherichia coli* HVH 205 (4-3094677) |
| Eco-1293-1 | Bacteria Proteobacteria Gammaproteobacteria Enterobacteriales Enterobacteriaceae *Escherichia coli* HVH 206 (4-3128229) |
| Eco-1294-1 | Bacteria Proteobacteria Gammaproteobacteria Enterobacteriales Enterobacteriaceae *Escherichia coli* HVH 207 (4-3113221) |
| Eco-1295-1 | Bacteria Proteobacteria Gammaproteobacteria Enterobacteriales Enterobacteriaceae *Escherichia coli* HVH 208 (4-3112292) |
| Eco-1296-1 | Bacteria Proteobacteria Gammaproteobacteria Enterobacteriales Enterobacteriaceae *Escherichia coli* HVH 209 (4-3062651) |
| Eco-1297-1 | Bacteria Proteobacteria Gammaproteobacteria Enterobacteriales Enterobacteriaceae *Escherichia coli* HVH 21 (4-4517873) |
| Eco-1298-1 | Bacteria Proteobacteria Gammaproteobacteria Enterobacteriales Enterobacteriaceae *Escherichia coli* HVH 210 (4-3042480) |
| Eco-1299-1 | Bacteria Proteobacteria Gammaproteobacteria Enterobacteriales Enterobacteriaceae *Escherichia coli* HVH 211 (4-3041891) |
| Eco-1300-1 | Bacteria Proteobacteria Gammaproteobacteria Enterobacteriales Enterobacteriaceae *Escherichia coli* HVH 212 (3-9305343) |
| Eco-1301-1 | Bacteria Proteobacteria Gammaproteobacteria Enterobacteriales Enterobacteriaceae *Escherichia coli* HVH 213 (4-3042928) |
| Eco-1302-1 | Bacteria Proteobacteria Gammaproteobacteria Enterobacteriales Enterobacteriaceae *Escherichia coli* HVH 214 (4-3062198) |
| Eco-1303-1 | Bacteria Proteobacteria Gammaproteobacteria Enterobacteriales Enterobacteriaceae *Escherichia coli* HVH 215 (4-3008371) |
| Eco-1304-1 | Bacteria Proteobacteria Gammaproteobacteria Enterobacteriales Enterobacteriaceae *Escherichia coli* HVH 216 (4-3042952) |
| Eco-1305-1 | Bacteria Proteobacteria Gammaproteobacteria Enterobacteriales Enterobacteriaceae *Escherichia coli* HVH 217 (4-1022806) |
| Eco-1306-1 | Bacteria Proteobacteria Gammaproteobacteria Enterobacteriales Enterobacteriaceae *Escherichia coli* HVH 218 (4-4500903) |
| Eco-1307-1 | Bacteria Proteobacteria Gammaproteobacteria Enterobacteriales Enterobacteriaceae *Escherichia coli* HVH 22 (4-2258986) |
| Eco-1308-1 | Bacteria Proteobacteria Gammaproteobacteria Enterobacteriales Enterobacteriaceae *Escherichia coli* HVH 220 (4-5876842) |
| Eco-1309-1 | Bacteria Proteobacteria Gammaproteobacteria Enterobacteriales Enterobacteriaceae *Escherichia coli* HVH 221 (4-3136817) |
| Eco-1310-1 | Bacteria Proteobacteria Gammaproteobacteria Enterobacteriales Enterobacteriaceae *Escherichia coli* HVH 222 (4-2977443) |
| Eco-1311-1 | Bacteria Proteobacteria Gammaproteobacteria Enterobacteriales Enterobacteriaceae *Escherichia coli* HVH 223 (4-2976528) |
| Eco-1312-1 | Bacteria Proteobacteria Gammaproteobacteria Enterobacteriales Enterobacteriaceae *Escherichia coli* HVH 225 (4-1273116) |
| Eco-1313-1 | Bacteria Proteobacteria Gammaproteobacteria Enterobacteriales Enterobacteriaceae *Escherichia coli* HVH 227 (4-2277670) |
| Eco-1314-1 | Bacteria Proteobacteria Gammaproteobacteria Enterobacteriales Enterobacteriaceae *Escherichia coli* HVH 228 (4-7787030) |
| Eco-1315-1 | Bacteria Proteobacteria Gammaproteobacteria Enterobacteriales Enterobacteriaceae *Escherichia coli* HVH 23 (4-6066488) |
| Eco-1316-1 | Bacteria Proteobacteria Gammaproteobacteria Enterobacteriales Enterobacteriaceae *Escherichia coli* HVH 24 (4-5985145) |
| Eco-1317-1 | Bacteria Proteobacteria Gammaproteobacteria Enterobacteriales Enterobacteriaceae *Escherichia coli* HVH 25 (4-5851939) |
| Eco-1318-1 | Bacteria Proteobacteria Gammaproteobacteria Enterobacteriales Enterobacteriaceae *Escherichia coli* HVH 26 (4-5703913) |
| Eco-1319-1 | Bacteria Proteobacteria Gammaproteobacteria Enterobacteriales Enterobacteriaceae *Escherichia coli* HVH 27 (4-7449267) |
| Eco-1320-1 | Bacteria Proteobacteria Gammaproteobacteria Enterobacteriales Enterobacteriaceae *Escherichia coli* HVH 28 (4-0907367) |
| Eco-1321-1 | Bacteria Proteobacteria Gammaproteobacteria Enterobacteriales Enterobacteriaceae *Escherichia coli* HVH 29 (4-3418073) |
| Eco-1322-1 | Bacteria Proteobacteria Gammaproteobacteria Enterobacteriales Enterobacteriaceae *Escherichia coli* HVH 3 (4-7276001) |
| Eco-1323-1 | Bacteria Proteobacteria Gammaproteobacteria Enterobacteriales Enterobacteriaceae *Escherichia coli* HVH 30 (4-2661829) |
| Eco-1324-1 | Bacteria Proteobacteria Gammaproteobacteria Enterobacteriales Enterobacteriaceae *Escherichia coli* HVH 31 (4-2602156) |
| Eco-1325-1 | Bacteria Proteobacteria Gammaproteobacteria Enterobacteriales Enterobacteriaceae *Escherichia coli* HVH 32 (4-3773988) |
| Eco-1326-1 | Bacteria Proteobacteria Gammaproteobacteria Enterobacteriales Enterobacteriaceae *Escherichia coli* HVH 33 (4-2174936) |
| Eco-1327-1 | Bacteria Proteobacteria Gammaproteobacteria Enterobacteriales Enterobacteriaceae *Escherichia coli* HVH 35 (4-2962667) |
| Eco-1328-1 | Bacteria Proteobacteria Gammaproteobacteria Enterobacteriales Enterobacteriaceae *Escherichia coli* HVH 36 (4-5675286) |
| Eco-1329-1 | Bacteria Proteobacteria Gammaproteobacteria Enterobacteriales Enterobacteriaceae *Escherichia coli* HVH 37 (4-2773848) |
| Eco-1330-1 | Bacteria Proteobacteria Gammaproteobacteria Enterobacteriales Enterobacteriaceae *Escherichia coli* HVH 38 (4-2774682) |
| Eco-1331-1 | Bacteria Proteobacteria Gammaproteobacteria Enterobacteriales Enterobacteriaceae *Escherichia coli* HVH 39 (4-2679949) |
| Eco-1332-1 | Bacteria Proteobacteria Gammaproteobacteria Enterobacteriales Enterobacteriaceae *Escherichia coli* HVH 4 (4-7276109) |
| Eco-1333-1 | Bacteria Proteobacteria Gammaproteobacteria Enterobacteriales Enterobacteriaceae *Escherichia coli* HVH 40 (4-1219782) |
| Eco-1334-1 | Bacteria Proteobacteria Gammaproteobacteria Enterobacteriales Enterobacteriaceae *Escherichia coli* HVH 41 (4-2677849) |
| Eco-1335-1 | Bacteria Proteobacteria Gammaproteobacteria Enterobacteriales Enterobacteriaceae *Escherichia coli* HVH 42 (4-2100061) |
| Eco-1336-1 | Bacteria Proteobacteria Gammaproteobacteria Enterobacteriales Enterobacteriaceae *Escherichia coli* HVH 43 (4-2173468) |

| | |
|---|---|
| Eco-1337-1 | Bacteria Proteobacteria Gammaproteobacteria Enterobacteriales Enterobacteriaceae *Escherichia coli* HVH 44 (4-2298570) |
| Eco-1338-1 | Bacteria Proteobacteria Gammaproteobacteria Enterobacteriales Enterobacteriaceae *Escherichia coli* HVH 45 (4-3129918) |
| Eco-1339-1 | Bacteria Proteobacteria Gammaproteobacteria Enterobacteriales Enterobacteriaceae *Escherichia coli* HVH 46 (4-2758776) |
| Eco-1340-1 | Bacteria Proteobacteria Gammaproteobacteria Enterobacteriales Enterobacteriaceae *Escherichia coli* HVH 48 (4-2658593) |
| Eco-1341-1 | Bacteria Proteobacteria Gammaproteobacteria Enterobacteriales Enterobacteriaceae *Escherichia coli* HVH 5 (4-7148410) |
| Eco-1342-1 | Bacteria Proteobacteria Gammaproteobacteria Enterobacteriales Enterobacteriaceae *Escherichia coli* HVH 50 (4-2593475) |
| Eco-1343-1 | Bacteria Proteobacteria Gammaproteobacteria Enterobacteriales Enterobacteriaceae *Escherichia coli* HVH 51 (4-2172526) |
| Eco-1344-1 | Bacteria Proteobacteria Gammaproteobacteria Enterobacteriales Enterobacteriaceae *Escherichia coli* HVH 53 (4-0631051) |
| Eco-1345-1 | Bacteria Proteobacteria Gammaproteobacteria Enterobacteriales Enterobacteriaceae *Escherichia coli* HVH 55 (4-2646161) |
| Eco-1346-1 | Bacteria Proteobacteria Gammaproteobacteria Enterobacteriales Enterobacteriaceae *Escherichia coli* HVH 56 (4-2153033) |
| Eco-1347-1 | Bacteria Proteobacteria Gammaproteobacteria Enterobacteriales Enterobacteriaceae *Escherichia coli* HVH 58 (4-2839709) |
| Eco-1348-1 | Bacteria Proteobacteria Gammaproteobacteria Enterobacteriales Enterobacteriaceae *Escherichia coli* HVH 59 (4-1119338) |
| Eco-1349-1 | Bacteria Proteobacteria Gammaproteobacteria Enterobacteriales Enterobacteriaceae *Escherichia coli* HVH 6 (3-8296502) |
| Eco-1350-1 | Bacteria Proteobacteria Gammaproteobacteria Enterobacteriales Enterobacteriaceae *Escherichia coli* HVH 61 (4-2736020) |
| Eco-1351-1 | Bacteria Proteobacteria Gammaproteobacteria Enterobacteriales Enterobacteriaceae *Escherichia coli* HVH 63 (4-2542528) |
| Eco-1352-1 | Bacteria Proteobacteria Gammaproteobacteria Enterobacteriales Enterobacteriaceae *Escherichia coli* HVH 65 (4-2262045) |
| Eco-1353-1 | Bacteria Proteobacteria Gammaproteobacteria Enterobacteriales Enterobacteriaceae *Escherichia coli* HVH 68 (4-0888028) |
| Eco-1354-1 | Bacteria Proteobacteria Gammaproteobacteria Enterobacteriales Enterobacteriaceae *Escherichia coli* HVH 69 (4-2837072) |
| Eco-1355-1 | Bacteria Proteobacteria Gammaproteobacteria Enterobacteriales Enterobacteriaceae *Escherichia coli* HVH 7 (4-7315031) |
| Eco-1356-1 | Bacteria Proteobacteria Gammaproteobacteria Enterobacteriales Enterobacteriaceae *Escherichia coli* HVH 70 (4-2963531) |
| Eco-1357-1 | Bacteria Proteobacteria Gammaproteobacteria Enterobacteriales Enterobacteriaceae *Escherichia coli* HVH 73 (4-2393174) |
| Eco-1358-1 | Bacteria Proteobacteria Gammaproteobacteria Enterobacteriales Enterobacteriaceae *Escherichia coli* HVH 74 (4-1034782) |
| Eco-1359-1 | Bacteria Proteobacteria Gammaproteobacteria Enterobacteriales Enterobacteriaceae *Escherichia coli* HVH 76 (4-2538717) |
| Eco-1360-1 | Bacteria Proteobacteria Gammaproteobacteria Enterobacteriales Enterobacteriaceae *Escherichia coli* HVH 77 (4-2605759) |
| Eco-1361-1 | Bacteria Proteobacteria Gammaproteobacteria Enterobacteriales Enterobacteriaceae *Escherichia coli* HVH 78 (4-2735946) |
| Eco-1362-1 | Bacteria Proteobacteria Gammaproteobacteria Enterobacteriales Enterobacteriaceae *Escherichia coli* HVH 79 (4-2512823) |
| Eco-1363-1 | Bacteria Proteobacteria Gammaproteobacteria Enterobacteriales Enterobacteriaceae *Escherichia coli* HVH 80 (4-2428830) |
| Eco-1364-1 | Bacteria Proteobacteria Gammaproteobacteria Enterobacteriales Enterobacteriaceae *Escherichia coli* HVH 82 (4-2209276) |
| Eco-1365-1 | Bacteria Proteobacteria Gammaproteobacteria Enterobacteriales Enterobacteriaceae *Escherichia coli* HVH 83 (4-2051087) |
| Eco-1366-1 | Bacteria Proteobacteria Gammaproteobacteria Enterobacteriales Enterobacteriaceae *Escherichia coli* HVH 84 (4-1021478) |
| Eco-1367-1 | Bacteria Proteobacteria Gammaproteobacteria Enterobacteriales Enterobacteriaceae *Escherichia coli* HVH 85 (4-0792144) |
| Eco-1368-1 | Bacteria Proteobacteria Gammaproteobacteria Enterobacteriales Enterobacteriaceae *Escherichia coli* HVH 86 (4-7026218) |
| Eco-1369-1 | Bacteria Proteobacteria Gammaproteobacteria Enterobacteriales Enterobacteriaceae *Escherichia coli* HVH 87 (4-5977630) |
| Eco-1370-1 | Bacteria Proteobacteria Gammaproteobacteria Enterobacteriales Enterobacteriaceae *Escherichia coli* HVH 88 (4-5854636) |
| Eco-1371-1 | Bacteria Proteobacteria Gammaproteobacteria Enterobacteriales Enterobacteriaceae *Escherichia coli* HVH 89 (4-5885604) |
| Eco-1372-1 | Bacteria Proteobacteria Gammaproteobacteria Enterobacteriales Enterobacteriaceae *Escherichia coli* HVH 9 (4-6942539) |
| Eco-1373-1 | Bacteria Proteobacteria Gammaproteobacteria Enterobacteriales Enterobacteriaceae *Escherichia coli* HVH 90 (4-3191362) |
| Eco-1374-1 | Bacteria Proteobacteria Gammaproteobacteria Enterobacteriales Enterobacteriaceae *Escherichia coli* HVH 91 (4-4638751) |
| Eco-1375-1 | Bacteria Proteobacteria Gammaproteobacteria Enterobacteriales Enterobacteriaceae *Escherichia coli* HVH 92 (4-5930790) |
| Eco-1376-1 | Bacteria Proteobacteria Gammaproteobacteria Enterobacteriales Enterobacteriaceae *Escherichia coli* HVH 95 (4-6074464) |
| Eco-1377-1 | Bacteria Proteobacteria Gammaproteobacteria Enterobacteriales Enterobacteriaceae *Escherichia coli* HVH 96 (4-5934869) |
| Eco-1378-1 | Bacteria Proteobacteria Gammaproteobacteria Enterobacteriales Enterobacteriaceae *Escherichia coli* HVH 98 (4-5799287) |
| Eco-1379-1 | Bacteria Proteobacteria Gammaproteobacteria Enterobacteriales Enterobacteriaceae *Escherichia coli* IAI1 |
| Eco-1380-1 | Bacteria Proteobacteria Gammaproteobacteria Enterobacteriales Enterobacteriaceae *Escherichia coli* IAI39 |
| Eco-1381-1 | Bacteria Proteobacteria Gammaproteobacteria Enterobacteriales Enterobacteriaceae *Escherichia coli* IHE3034 |
| Eco-1382-1 | Bacteria Proteobacteria Gammaproteobacteria Enterobacteriales Enterobacteriaceae *Escherichia coli* IS29 |
| Eco-1383-1 | Bacteria Proteobacteria Gammaproteobacteria Enterobacteriales Enterobacteriaceae *Escherichia coli* IS5 |
| Eco-1384-1 | Bacteria Proteobacteria Gammaproteobacteria Enterobacteriales Enterobacteriaceae *Escherichia coli* J53 |
| Eco-1385-1 | Bacteria Proteobacteria Gammaproteobacteria Enterobacteriales Enterobacteriaceae *Escherichia coli* J96 |
| Eco-1386-1 | Bacteria Proteobacteria Gammaproteobacteria Enterobacteriales Enterobacteriaceae *Escherichia coli* JB1-95 |
| Eco-1387-1 | Bacteria Proteobacteria Gammaproteobacteria Enterobacteriales Enterobacteriaceae *Escherichia coli* JJ1886 |
| Eco-1388-1 | Bacteria Proteobacteria Gammaproteobacteria Enterobacteriales Enterobacteriaceae *Escherichia coli* Jurua 18/11 |
| Eco-1389-1 | Bacteria Proteobacteria Gammaproteobacteria Enterobacteriales Enterobacteriaceae *Escherichia coli* Jurua 20/10 |
| Eco-1390-1 | Bacteria Proteobacteria Gammaproteobacteria Enterobacteriales Enterobacteriaceae *Escherichia coli* KD1 |
| Eco-1391-1 | Bacteria Proteobacteria Gammaproteobacteria Enterobacteriales Enterobacteriaceae *Escherichia coli* KD2 |
| Eco-1392-1 to Eco-1392-2 | Bacteria Proteobacteria Gammaproteobacteria Enterobacteriales Enterobacteriaceae *Escherichia coli* KO11FL |
| Eco-1393-1 | Bacteria Proteobacteria Gammaproteobacteria Enterobacteriales Enterobacteriaceae *Escherichia coli* KOEGE 10 (25a) |
| Eco-1394-1 | Bacteria Proteobacteria Gammaproteobacteria Enterobacteriales Enterobacteriaceae *Escherichia coli* KOEGE 118 (317a) |
| Eco-1395-1 | Bacteria Proteobacteria Gammaproteobacteria Enterobacteriales Enterobacteriaceae *Escherichia coli* KOEGE 131 (358a) |
| Eco-1396-1 | Bacteria Proteobacteria Gammaproteobacteria Enterobacteriales Enterobacteriaceae *Escherichia coli* KOEGE 3 (4a) |
| Eco-1397-1 | Bacteria Proteobacteria Gammaproteobacteria Enterobacteriales Enterobacteriaceae *Escherichia coli* KOEGE 30 (63a) |
| Eco-1398-1 | Bacteria Proteobacteria Gammaproteobacteria Enterobacteriales Enterobacteriaceae *Escherichia coli* KOEGE 32 (66a) |
| Eco-1399-1 | Bacteria Proteobacteria Gammaproteobacteria Enterobacteriales Enterobacteriaceae *Escherichia coli* KOEGE 33 (68a) |
| Eco-1400-1 | Bacteria Proteobacteria Gammaproteobacteria Enterobacteriales Enterobacteriaceae *Escherichia coli* KOEGE 40 (102a) |
| Eco-1401-1 | Bacteria Proteobacteria Gammaproteobacteria Enterobacteriales Enterobacteriaceae *Escherichia coli* KOEGE 43 (105a) |

| | |
|---|---|
| Eco-1402-1 | Bacteria Proteobacteria Gammaproteobacteria Enterobacteriales Enterobacteriaceae *Escherichia coli* KOEGE 44 (106a) |
| Eco-1403-1 | Bacteria Proteobacteria Gammaproteobacteria Enterobacteriales Enterobacteriaceae *Escherichia coli* KOEGE 56 (169a) |
| Eco-1404-1 | Bacteria Proteobacteria Gammaproteobacteria Enterobacteriales Enterobacteriaceae *Escherichia coli* KOEGE 58 (171a) |
| Eco-1405-1 | Bacteria Proteobacteria Gammaproteobacteria Enterobacteriales Enterobacteriaceae *Escherichia coli* KOEGE 61 (174a) |
| Eco-1406-1 | Bacteria Proteobacteria Gammaproteobacteria Enterobacteriales Enterobacteriaceae *Escherichia coli* KOEGE 62 (175a) |
| Eco-1407-1 | Bacteria Proteobacteria Gammaproteobacteria Enterobacteriales Enterobacteriaceae *Escherichia coli* KOEGE 68 (182a) |
| Eco-1408-1 | Bacteria Proteobacteria Gammaproteobacteria Enterobacteriales Enterobacteriaceae *Escherichia coli* KOEGE 7 (16a) |
| Eco-1409-1 | Bacteria Proteobacteria Gammaproteobacteria Enterobacteriales Enterobacteriaceae *Escherichia coli* KOEGE 70 (185a) |
| Eco-1410-1 | Bacteria Proteobacteria Gammaproteobacteria Enterobacteriales Enterobacteriaceae *Escherichia coli* KOEGE 71 (186a) |
| Eco-1411-1 | Bacteria Proteobacteria Gammaproteobacteria Enterobacteriales Enterobacteriaceae *Escherichia coli* KOEGE 73 (195a) |
| Eco-1412-1 | Bacteria Proteobacteria Gammaproteobacteria Enterobacteriales Enterobacteriaceae *Escherichia coli* KOEGE 77 (202a) |
| Eco-1413-1 | Bacteria Proteobacteria Gammaproteobacteria Enterobacteriales Enterobacteriaceae *Escherichia coli* KTE1 |
| Eco-1414-1 | Bacteria Proteobacteria Gammaproteobacteria Enterobacteriales Enterobacteriaceae *Escherichia coli* KTE10 |
| Eco-1415-1 | Bacteria Proteobacteria Gammaproteobacteria Enterobacteriales Enterobacteriaceae *Escherichia coli* KTE100 |
| Eco-1416-1 | Bacteria Proteobacteria Gammaproteobacteria Enterobacteriales Enterobacteriaceae *Escherichia coli* KTE101 |
| Eco-1417-1 | Bacteria Proteobacteria Gammaproteobacteria Enterobacteriales Enterobacteriaceae *Escherichia coli* KTE102 |
| Eco-1418-1 | Bacteria Proteobacteria Gammaproteobacteria Enterobacteriales Enterobacteriaceae *Escherichia coli* KTE103 |
| Eco-1419-1 | Bacteria Proteobacteria Gammaproteobacteria Enterobacteriales Enterobacteriaceae *Escherichia coli* KTE104 |
| Eco-1420-1 | Bacteria Proteobacteria Gammaproteobacteria Enterobacteriales Enterobacteriaceae *Escherichia coli* KTE105 |
| Eco-1421-1 | Bacteria Proteobacteria Gammaproteobacteria Enterobacteriales Enterobacteriaceae *Escherichia coli* KTE106 |
| Eco-1422-1 | Bacteria Proteobacteria Gammaproteobacteria Enterobacteriales Enterobacteriaceae *Escherichia coli* KTE107 |
| Eco-1423-1 | Bacteria Proteobacteria Gammaproteobacteria Enterobacteriales Enterobacteriaceae *Escherichia coli* KTE108 |
| Eco-1424-1 | Bacteria Proteobacteria Gammaproteobacteria Enterobacteriales Enterobacteriaceae *Escherichia coli* KTE109 |
| Eco-1425-1 | Bacteria Proteobacteria Gammaproteobacteria Enterobacteriales Enterobacteriaceae *Escherichia coli* KTE11 |
| Eco-1426-1 | Bacteria Proteobacteria Gammaproteobacteria Enterobacteriales Enterobacteriaceae *Escherichia coli* KTE111 |
| Eco-1427-1 | Bacteria Proteobacteria Gammaproteobacteria Enterobacteriales Enterobacteriaceae *Escherichia coli* KTE112 |
| Eco-1428-1 | Bacteria Proteobacteria Gammaproteobacteria Enterobacteriales Enterobacteriaceae *Escherichia coli* KTE113 |
| Eco-1429-1 | Bacteria Proteobacteria Gammaproteobacteria Enterobacteriales Enterobacteriaceae *Escherichia coli* KTE114 |
| Eco-1430-1 | Bacteria Proteobacteria Gammaproteobacteria Enterobacteriales Enterobacteriaceae *Escherichia coli* KTE115 |
| Eco-1431-1 | Bacteria Proteobacteria Gammaproteobacteria Enterobacteriales Enterobacteriaceae *Escherichia coli* KTE116 |
| Eco-1432-1 | Bacteria Proteobacteria Gammaproteobacteria Enterobacteriales Enterobacteriaceae *Escherichia coli* KTE117 |
| Eco-1433-1 | Bacteria Proteobacteria Gammaproteobacteria Enterobacteriales Enterobacteriaceae *Escherichia coli* KTE118 |
| Eco-1434-1 | Bacteria Proteobacteria Gammaproteobacteria Enterobacteriales Enterobacteriaceae *Escherichia coli* KTE119 |
| Eco-1435-1 | Bacteria Proteobacteria Gammaproteobacteria Enterobacteriales Enterobacteriaceae *Escherichia coli* KTE12 |
| Eco-1436-1 | Bacteria Proteobacteria Gammaproteobacteria Enterobacteriales Enterobacteriaceae *Escherichia coli* KTE120 |
| Eco-1437-1 | Bacteria Proteobacteria Gammaproteobacteria Enterobacteriales Enterobacteriaceae *Escherichia coli* KTE121 |
| Eco-1438-1 | Bacteria Proteobacteria Gammaproteobacteria Enterobacteriales Enterobacteriaceae *Escherichia coli* KTE122 |
| Eco-1439-1 | Bacteria Proteobacteria Gammaproteobacteria Enterobacteriales Enterobacteriaceae *Escherichia coli* KTE123 |
| Eco-1440-1 | Bacteria Proteobacteria Gammaproteobacteria Enterobacteriales Enterobacteriaceae *Escherichia coli* KTE124 |
| Eco-1441-1 | Bacteria Proteobacteria Gammaproteobacteria Enterobacteriales Enterobacteriaceae *Escherichia coli* KTE125 |
| Eco-1442-1 | Bacteria Proteobacteria Gammaproteobacteria Enterobacteriales Enterobacteriaceae *Escherichia coli* KTE126 |
| Eco-1443-1 | Bacteria Proteobacteria Gammaproteobacteria Enterobacteriales Enterobacteriaceae *Escherichia coli* KTE127 |
| Eco-1444-1 | Bacteria Proteobacteria Gammaproteobacteria Enterobacteriales Enterobacteriaceae *Escherichia coli* KTE128 |
| Eco-1445-1 | Bacteria Proteobacteria Gammaproteobacteria Enterobacteriales Enterobacteriaceae *Escherichia coli* KTE129 |
| Eco-1446-1 | Bacteria Proteobacteria Gammaproteobacteria Enterobacteriales Enterobacteriaceae *Escherichia coli* KTE13 |
| Eco-1447-1 | Bacteria Proteobacteria Gammaproteobacteria Enterobacteriales Enterobacteriaceae *Escherichia coli* KTE130 |
| Eco-1448-1 | Bacteria Proteobacteria Gammaproteobacteria Enterobacteriales Enterobacteriaceae *Escherichia coli* KTE131 |
| Eco-1449-1 | Bacteria Proteobacteria Gammaproteobacteria Enterobacteriales Enterobacteriaceae *Escherichia coli* KTE132 |
| Eco-1450-1 | Bacteria Proteobacteria Gammaproteobacteria Enterobacteriales Enterobacteriaceae *Escherichia coli* KTE133 |
| Eco-1451-1 | Bacteria Proteobacteria Gammaproteobacteria Enterobacteriales Enterobacteriaceae *Escherichia coli* KTE134 |
| Eco-1452-1 | Bacteria Proteobacteria Gammaproteobacteria Enterobacteriales Enterobacteriaceae *Escherichia coli* KTE135 |
| Eco-1453-1 | Bacteria Proteobacteria Gammaproteobacteria Enterobacteriales Enterobacteriaceae *Escherichia coli* KTE136 |
| Eco-1454-1 | Bacteria Proteobacteria Gammaproteobacteria Enterobacteriales Enterobacteriaceae *Escherichia coli* KTE137 |
| Eco-1455-1 | Bacteria Proteobacteria Gammaproteobacteria Enterobacteriales Enterobacteriaceae *Escherichia coli* KTE138 |
| Eco-1456-1 | Bacteria Proteobacteria Gammaproteobacteria Enterobacteriales Enterobacteriaceae *Escherichia coli* KTE139 |
| Eco-1457-1 | Bacteria Proteobacteria Gammaproteobacteria Enterobacteriales Enterobacteriaceae *Escherichia coli* KTE14 |
| Eco-1458-1 | Bacteria Proteobacteria Gammaproteobacteria Enterobacteriales Enterobacteriaceae *Escherichia coli* KTE140 |
| Eco-1459-1 | Bacteria Proteobacteria Gammaproteobacteria Enterobacteriales Enterobacteriaceae *Escherichia coli* KTE141 |
| Eco-1460-1 | Bacteria Proteobacteria Gammaproteobacteria Enterobacteriales Enterobacteriaceae *Escherichia coli* KTE142 |
| Eco-1461-1 | Bacteria Proteobacteria Gammaproteobacteria Enterobacteriales Enterobacteriaceae *Escherichia coli* KTE143 |
| Eco-1462-1 | Bacteria Proteobacteria Gammaproteobacteria Enterobacteriales Enterobacteriaceae *Escherichia coli* KTE144 |
| Eco-1463-1 | Bacteria Proteobacteria Gammaproteobacteria Enterobacteriales Enterobacteriaceae *Escherichia coli* KTE145 |
| Eco-1464-1 | Bacteria Proteobacteria Gammaproteobacteria Enterobacteriales Enterobacteriaceae *Escherichia coli* KTE146 |
| Eco-1465-1 | Bacteria Proteobacteria Gammaproteobacteria Enterobacteriales Enterobacteriaceae *Escherichia coli* KTE147 |
| Eco-1466-1 | Bacteria Proteobacteria Gammaproteobacteria Enterobacteriales Enterobacteriaceae *Escherichia coli* KTE148 |

| | |
|---|---|
| Eco-1467-1 | Bacteria Proteobacteria Gammaproteobacteria Enterobacteriales Enterobacteriaceae *Escherichia coli* KTE15 |
| Eco-1468-1 | Bacteria Proteobacteria Gammaproteobacteria Enterobacteriales Enterobacteriaceae *Escherichia coli* KTE150 |
| Eco-1469-1 | Bacteria Proteobacteria Gammaproteobacteria Enterobacteriales Enterobacteriaceae *Escherichia coli* KTE153 |
| Eco-1470-1 | Bacteria Proteobacteria Gammaproteobacteria Enterobacteriales Enterobacteriaceae *Escherichia coli* KTE154 |
| Eco-1471-1 | Bacteria Proteobacteria Gammaproteobacteria Enterobacteriales Enterobacteriaceae *Escherichia coli* KTE155 |
| Eco-1472-1 | Bacteria Proteobacteria Gammaproteobacteria Enterobacteriales Enterobacteriaceae *Escherichia coli* KTE156 |
| Eco-1473-1 | Bacteria Proteobacteria Gammaproteobacteria Enterobacteriales Enterobacteriaceae *Escherichia coli* KTE157 |
| Eco-1474-1 | Bacteria Proteobacteria Gammaproteobacteria Enterobacteriales Enterobacteriaceae *Escherichia coli* KTE158 |
| Eco-1475-1 | Bacteria Proteobacteria Gammaproteobacteria Enterobacteriales Enterobacteriaceae *Escherichia coli* KTE159 |
| Eco-1476-1 | Bacteria Proteobacteria Gammaproteobacteria Enterobacteriales Enterobacteriaceae *Escherichia coli* KTE16 |
| Eco-1477-1 | Bacteria Proteobacteria Gammaproteobacteria Enterobacteriales Enterobacteriaceae *Escherichia coli* KTE160 |
| Eco-1478-1 | Bacteria Proteobacteria Gammaproteobacteria Enterobacteriales Enterobacteriaceae *Escherichia coli* KTE161 |
| Eco-1479-1 | Bacteria Proteobacteria Gammaproteobacteria Enterobacteriales Enterobacteriaceae *Escherichia coli* KTE162 |
| Eco-1480-1 | Bacteria Proteobacteria Gammaproteobacteria Enterobacteriales Enterobacteriaceae *Escherichia coli* KTE163 |
| Eco-1481-1 | Bacteria Proteobacteria Gammaproteobacteria Enterobacteriales Enterobacteriaceae *Escherichia coli* KTE165 |
| Eco-1482-1 | Bacteria Proteobacteria Gammaproteobacteria Enterobacteriales Enterobacteriaceae *Escherichia coli* KTE166 |
| Eco-1483-1 | Bacteria Proteobacteria Gammaproteobacteria Enterobacteriales Enterobacteriaceae *Escherichia coli* KTE167 |
| Eco-1484-1 | Bacteria Proteobacteria Gammaproteobacteria Enterobacteriales Enterobacteriaceae *Escherichia coli* KTE168 |
| Eco-1485-1 | Bacteria Proteobacteria Gammaproteobacteria Enterobacteriales Enterobacteriaceae *Escherichia coli* KTE169 |
| Eco-1486-1 | Bacteria Proteobacteria Gammaproteobacteria Enterobacteriales Enterobacteriaceae *Escherichia coli* KTE17 |
| Eco-1487-1 | Bacteria Proteobacteria Gammaproteobacteria Enterobacteriales Enterobacteriaceae *Escherichia coli* KTE170 |
| Eco-1488-1 | Bacteria Proteobacteria Gammaproteobacteria Enterobacteriales Enterobacteriaceae *Escherichia coli* KTE171 |
| Eco-1489-1 | Bacteria Proteobacteria Gammaproteobacteria Enterobacteriales Enterobacteriaceae *Escherichia coli* KTE172 |
| Eco-1490-1 | Bacteria Proteobacteria Gammaproteobacteria Enterobacteriales Enterobacteriaceae *Escherichia coli* KTE173 |
| Eco-1491-1 | Bacteria Proteobacteria Gammaproteobacteria Enterobacteriales Enterobacteriaceae *Escherichia coli* KTE174 |
| Eco-1492-1 | Bacteria Proteobacteria Gammaproteobacteria Enterobacteriales Enterobacteriaceae *Escherichia coli* KTE175 |
| Eco-1493-1 | Bacteria Proteobacteria Gammaproteobacteria Enterobacteriales Enterobacteriaceae *Escherichia coli* KTE176 |
| Eco-1494-1 | Bacteria Proteobacteria Gammaproteobacteria Enterobacteriales Enterobacteriaceae *Escherichia coli* KTE177 |
| Eco-1495-1 | Bacteria Proteobacteria Gammaproteobacteria Enterobacteriales Enterobacteriaceae *Escherichia coli* KTE178 |
| Eco-1496-1 | Bacteria Proteobacteria Gammaproteobacteria Enterobacteriales Enterobacteriaceae *Escherichia coli* KTE179 |
| Eco-1497-1 | Bacteria Proteobacteria Gammaproteobacteria Enterobacteriales Enterobacteriaceae *Escherichia coli* KTE18 |
| Eco-1498-1 | Bacteria Proteobacteria Gammaproteobacteria Enterobacteriales Enterobacteriaceae *Escherichia coli* KTE180 |
| Eco-1499-1 | Bacteria Proteobacteria Gammaproteobacteria Enterobacteriales Enterobacteriaceae *Escherichia coli* KTE181 |
| Eco-1500-1 | Bacteria Proteobacteria Gammaproteobacteria Enterobacteriales Enterobacteriaceae *Escherichia coli* KTE182 |
| Eco-1501-1 | Bacteria Proteobacteria Gammaproteobacteria Enterobacteriales Enterobacteriaceae *Escherichia coli* KTE183 |
| Eco-1502-1 | Bacteria Proteobacteria Gammaproteobacteria Enterobacteriales Enterobacteriaceae *Escherichia coli* KTE184 |
| Eco-1503-1 | Bacteria Proteobacteria Gammaproteobacteria Enterobacteriales Enterobacteriaceae *Escherichia coli* KTE185 |
| Eco-1504-1 | Bacteria Proteobacteria Gammaproteobacteria Enterobacteriales Enterobacteriaceae *Escherichia coli* KTE186 |
| Eco-1505-1 | Bacteria Proteobacteria Gammaproteobacteria Enterobacteriales Enterobacteriaceae *Escherichia coli* KTE187 |
| Eco-1506-1 | Bacteria Proteobacteria Gammaproteobacteria Enterobacteriales Enterobacteriaceae *Escherichia coli* KTE188 |
| Eco-1507-1 | Bacteria Proteobacteria Gammaproteobacteria Enterobacteriales Enterobacteriaceae *Escherichia coli* KTE189 |
| Eco-1508-1 | Bacteria Proteobacteria Gammaproteobacteria Enterobacteriales Enterobacteriaceae *Escherichia coli* KTE19 |
| Eco-1509-1 | Bacteria Proteobacteria Gammaproteobacteria Enterobacteriales Enterobacteriaceae *Escherichia coli* KTE190 |
| Eco-1510-1 | Bacteria Proteobacteria Gammaproteobacteria Enterobacteriales Enterobacteriaceae *Escherichia coli* KTE191 |
| Eco-1511-1 | Bacteria Proteobacteria Gammaproteobacteria Enterobacteriales Enterobacteriaceae *Escherichia coli* KTE192 |
| Eco-1512-1 | Bacteria Proteobacteria Gammaproteobacteria Enterobacteriales Enterobacteriaceae *Escherichia coli* KTE193 |
| Eco-1513-1 | Bacteria Proteobacteria Gammaproteobacteria Enterobacteriales Enterobacteriaceae *Escherichia coli* KTE194 |
| Eco-1514-1 | Bacteria Proteobacteria Gammaproteobacteria Enterobacteriales Enterobacteriaceae *Escherichia coli* KTE195 |
| Eco-1515-1 | Bacteria Proteobacteria Gammaproteobacteria Enterobacteriales Enterobacteriaceae *Escherichia coli* KTE196 |
| Eco-1516-1 | Bacteria Proteobacteria Gammaproteobacteria Enterobacteriales Enterobacteriaceae *Escherichia coli* KTE197 |
| Eco-1517-1 | Bacteria Proteobacteria Gammaproteobacteria Enterobacteriales Enterobacteriaceae *Escherichia coli* KTE198 |
| Eco-1518-1 | Bacteria Proteobacteria Gammaproteobacteria Enterobacteriales Enterobacteriaceae *Escherichia coli* KTE199 |
| Eco-1519-1 | Bacteria Proteobacteria Gammaproteobacteria Enterobacteriales Enterobacteriaceae *Escherichia coli* KTE2 |
| Eco-1520-1 | Bacteria Proteobacteria Gammaproteobacteria Enterobacteriales Enterobacteriaceae *Escherichia coli* KTE20 |
| Eco-1521-1 | Bacteria Proteobacteria Gammaproteobacteria Enterobacteriales Enterobacteriaceae *Escherichia coli* KTE200 |
| Eco-1522-1 | Bacteria Proteobacteria Gammaproteobacteria Enterobacteriales Enterobacteriaceae *Escherichia coli* KTE201 |
| Eco-1523-1 | Bacteria Proteobacteria Gammaproteobacteria Enterobacteriales Enterobacteriaceae *Escherichia coli* KTE202 |
| Eco-1524-1 | Bacteria Proteobacteria Gammaproteobacteria Enterobacteriales Enterobacteriaceae *Escherichia coli* KTE203 |
| Eco-1525-1 | Bacteria Proteobacteria Gammaproteobacteria Enterobacteriales Enterobacteriaceae *Escherichia coli* KTE204 |
| Eco-1526-1 | Bacteria Proteobacteria Gammaproteobacteria Enterobacteriales Enterobacteriaceae *Escherichia coli* KTE205 |
| Eco-1527-1 | Bacteria Proteobacteria Gammaproteobacteria Enterobacteriales Enterobacteriaceae *Escherichia coli* KTE206 |
| Eco-1528-1 | Bacteria Proteobacteria Gammaproteobacteria Enterobacteriales Enterobacteriaceae *Escherichia coli* KTE207 |
| Eco-1529-1 | Bacteria Proteobacteria Gammaproteobacteria Enterobacteriales Enterobacteriaceae *Escherichia coli* KTE208 |
| Eco-1530-1 | Bacteria Proteobacteria Gammaproteobacteria Enterobacteriales Enterobacteriaceae *Escherichia coli* KTE209 |
| Eco-1531-1 | Bacteria Proteobacteria Gammaproteobacteria Enterobacteriales Enterobacteriaceae *Escherichia coli* KTE21 |

| | |
|---|---|
| Eco-1532-1 | Bacteria Proteobacteria Gammaproteobacteria Enterobacteriales Enterobacteriaceae *Escherichia coli* KTE210 |
| Eco-1533-1 | Bacteria Proteobacteria Gammaproteobacteria Enterobacteriales Enterobacteriaceae *Escherichia coli* KTE211 |
| Eco-1534-1 | Bacteria Proteobacteria Gammaproteobacteria Enterobacteriales Enterobacteriaceae *Escherichia coli* KTE212 |
| Eco-1535-1 | Bacteria Proteobacteria Gammaproteobacteria Enterobacteriales Enterobacteriaceae *Escherichia coli* KTE213 |
| Eco-1536-1 | Bacteria Proteobacteria Gammaproteobacteria Enterobacteriales Enterobacteriaceae *Escherichia coli* KTE214 |
| Eco-1537-1 | Bacteria Proteobacteria Gammaproteobacteria Enterobacteriales Enterobacteriaceae *Escherichia coli* KTE215 |
| Eco-1538-1 | Bacteria Proteobacteria Gammaproteobacteria Enterobacteriales Enterobacteriaceae *Escherichia coli* KTE216 |
| Eco-1539-1 | Bacteria Proteobacteria Gammaproteobacteria Enterobacteriales Enterobacteriaceae *Escherichia coli* KTE217 |
| Eco-1540-1 | Bacteria Proteobacteria Gammaproteobacteria Enterobacteriales Enterobacteriaceae *Escherichia coli* KTE218 |
| Eco-1541-1 | Bacteria Proteobacteria Gammaproteobacteria Enterobacteriales Enterobacteriaceae *Escherichia coli* KTE219 |
| Eco-1542-1 | Bacteria Proteobacteria Gammaproteobacteria Enterobacteriales Enterobacteriaceae *Escherichia coli* KTE22 |
| Eco-1543-1 | Bacteria Proteobacteria Gammaproteobacteria Enterobacteriales Enterobacteriaceae *Escherichia coli* KTE220 |
| Eco-1544-1 | Bacteria Proteobacteria Gammaproteobacteria Enterobacteriales Enterobacteriaceae *Escherichia coli* KTE221 |
| Eco-1545-1 | Bacteria Proteobacteria Gammaproteobacteria Enterobacteriales Enterobacteriaceae *Escherichia coli* KTE222 |
| Eco-1546-1 | Bacteria Proteobacteria Gammaproteobacteria Enterobacteriales Enterobacteriaceae *Escherichia coli* KTE223 |
| Eco-1547-1 | Bacteria Proteobacteria Gammaproteobacteria Enterobacteriales Enterobacteriaceae *Escherichia coli* KTE224 |
| Eco-1548-1 | Bacteria Proteobacteria Gammaproteobacteria Enterobacteriales Enterobacteriaceae *Escherichia coli* KTE225 |
| Eco-1549-1 | Bacteria Proteobacteria Gammaproteobacteria Enterobacteriales Enterobacteriaceae *Escherichia coli* KTE226 |
| Eco-1550-1 | Bacteria Proteobacteria Gammaproteobacteria Enterobacteriales Enterobacteriaceae *Escherichia coli* KTE227 |
| Eco-1551-1 | Bacteria Proteobacteria Gammaproteobacteria Enterobacteriales Enterobacteriaceae *Escherichia coli* KTE228 |
| Eco-1552-1 | Bacteria Proteobacteria Gammaproteobacteria Enterobacteriales Enterobacteriaceae *Escherichia coli* KTE229 |
| Eco-1553-1 | Bacteria Proteobacteria Gammaproteobacteria Enterobacteriales Enterobacteriaceae *Escherichia coli* KTE23 |
| Eco-1554-1 | Bacteria Proteobacteria Gammaproteobacteria Enterobacteriales Enterobacteriaceae *Escherichia coli* KTE230 |
| Eco-1555-1 | Bacteria Proteobacteria Gammaproteobacteria Enterobacteriales Enterobacteriaceae *Escherichia coli* KTE231 |
| Eco-1556-1 | Bacteria Proteobacteria Gammaproteobacteria Enterobacteriales Enterobacteriaceae *Escherichia coli* KTE232 |
| Eco-1557-1 | Bacteria Proteobacteria Gammaproteobacteria Enterobacteriales Enterobacteriaceae *Escherichia coli* KTE233 |
| Eco-1558-1 | Bacteria Proteobacteria Gammaproteobacteria Enterobacteriales Enterobacteriaceae *Escherichia coli* KTE234 |
| Eco-1559-1 | Bacteria Proteobacteria Gammaproteobacteria Enterobacteriales Enterobacteriaceae *Escherichia coli* KTE235 |
| Eco-1560-1 | Bacteria Proteobacteria Gammaproteobacteria Enterobacteriales Enterobacteriaceae *Escherichia coli* KTE236 |
| Eco-1561-1 | Bacteria Proteobacteria Gammaproteobacteria Enterobacteriales Enterobacteriaceae *Escherichia coli* KTE237 |
| Eco-1562-1 | Bacteria Proteobacteria Gammaproteobacteria Enterobacteriales Enterobacteriaceae *Escherichia coli* KTE24 |
| Eco-1563-1 | Bacteria Proteobacteria Gammaproteobacteria Enterobacteriales Enterobacteriaceae *Escherichia coli* KTE240 |
| Eco-1564-1 | Bacteria Proteobacteria Gammaproteobacteria Enterobacteriales Enterobacteriaceae *Escherichia coli* KTE25 |
| Eco-1565-1 | Bacteria Proteobacteria Gammaproteobacteria Enterobacteriales Enterobacteriaceae *Escherichia coli* KTE26 |
| Eco-1566-1 | Bacteria Proteobacteria Gammaproteobacteria Enterobacteriales Enterobacteriaceae *Escherichia coli* KTE27 |
| Eco-1567-1 | Bacteria Proteobacteria Gammaproteobacteria Enterobacteriales Enterobacteriaceae *Escherichia coli* KTE28 |
| Eco-1568-1 | Bacteria Proteobacteria Gammaproteobacteria Enterobacteriales Enterobacteriaceae *Escherichia coli* KTE29 |
| Eco-1569-1 | Bacteria Proteobacteria Gammaproteobacteria Enterobacteriales Enterobacteriaceae *Escherichia coli* KTE3 |
| Eco-1570-1 | Bacteria Proteobacteria Gammaproteobacteria Enterobacteriales Enterobacteriaceae *Escherichia coli* KTE31 |
| Eco-1571-1 | Bacteria Proteobacteria Gammaproteobacteria Enterobacteriales Enterobacteriaceae *Escherichia coli* KTE33 |
| Eco-1572-1 | Bacteria Proteobacteria Gammaproteobacteria Enterobacteriales Enterobacteriaceae *Escherichia coli* KTE34 |
| Eco-1573-1 | Bacteria Proteobacteria Gammaproteobacteria Enterobacteriales Enterobacteriaceae *Escherichia coli* KTE35 |
| Eco-1574-1 | Bacteria Proteobacteria Gammaproteobacteria Enterobacteriales Enterobacteriaceae *Escherichia coli* KTE36 |
| Eco-1575-1 | Bacteria Proteobacteria Gammaproteobacteria Enterobacteriales Enterobacteriaceae *Escherichia coli* KTE37 |
| Eco-1576-1 | Bacteria Proteobacteria Gammaproteobacteria Enterobacteriales Enterobacteriaceae *Escherichia coli* KTE38 |
| Eco-1577-1 | Bacteria Proteobacteria Gammaproteobacteria Enterobacteriales Enterobacteriaceae *Escherichia coli* KTE39 |
| Eco-1578-1 | Bacteria Proteobacteria Gammaproteobacteria Enterobacteriales Enterobacteriaceae *Escherichia coli* KTE4 |
| Eco-1579-1 | Bacteria Proteobacteria Gammaproteobacteria Enterobacteriales Enterobacteriaceae *Escherichia coli* KTE40 |
| Eco-1580-1 | Bacteria Proteobacteria Gammaproteobacteria Enterobacteriales Enterobacteriaceae *Escherichia coli* KTE41 |
| Eco-1581-1 | Bacteria Proteobacteria Gammaproteobacteria Enterobacteriales Enterobacteriaceae *Escherichia coli* KTE42 |
| Eco-1582-1 | Bacteria Proteobacteria Gammaproteobacteria Enterobacteriales Enterobacteriaceae *Escherichia coli* KTE43 |
| Eco-1583-1 | Bacteria Proteobacteria Gammaproteobacteria Enterobacteriales Enterobacteriaceae *Escherichia coli* KTE44 |
| Eco-1584-1 | Bacteria Proteobacteria Gammaproteobacteria Enterobacteriales Enterobacteriaceae *Escherichia coli* KTE45 |
| Eco-1585-1 | Bacteria Proteobacteria Gammaproteobacteria Enterobacteriales Enterobacteriaceae *Escherichia coli* KTE46 |
| Eco-1586-1 | Bacteria Proteobacteria Gammaproteobacteria Enterobacteriales Enterobacteriaceae *Escherichia coli* KTE47 |
| Eco-1587-1 | Bacteria Proteobacteria Gammaproteobacteria Enterobacteriales Enterobacteriaceae *Escherichia coli* KTE48 |
| Eco-1588-1 | Bacteria Proteobacteria Gammaproteobacteria Enterobacteriales Enterobacteriaceae *Escherichia coli* KTE49 |
| Eco-1589-1 | Bacteria Proteobacteria Gammaproteobacteria Enterobacteriales Enterobacteriaceae *Escherichia coli* KTE5 |
| Eco-1590-1 | Bacteria Proteobacteria Gammaproteobacteria Enterobacteriales Enterobacteriaceae *Escherichia coli* KTE50 |
| Eco-1591-1 | Bacteria Proteobacteria Gammaproteobacteria Enterobacteriales Enterobacteriaceae *Escherichia coli* KTE51 |
| Eco-1592-1 | Bacteria Proteobacteria Gammaproteobacteria Enterobacteriales Enterobacteriaceae *Escherichia coli* KTE52 |
| Eco-1593-1 | Bacteria Proteobacteria Gammaproteobacteria Enterobacteriales Enterobacteriaceae *Escherichia coli* KTE53 |
| Eco-1594-1 | Bacteria Proteobacteria Gammaproteobacteria Enterobacteriales Enterobacteriaceae *Escherichia coli* KTE54 |
| Eco-1595-1 | Bacteria Proteobacteria Gammaproteobacteria Enterobacteriales Enterobacteriaceae *Escherichia coli* KTE55 |
| Eco-1596-1 | Bacteria Proteobacteria Gammaproteobacteria Enterobacteriales Enterobacteriaceae *Escherichia coli* KTE56 |

| | |
|---|---|
| Eco-1597-1 | Bacteria Proteobacteria Gammaproteobacteria Enterobacteriales Enterobacteriaceae *Escherichia coli* KTE57 |
| Eco-1598-1 | Bacteria Proteobacteria Gammaproteobacteria Enterobacteriales Enterobacteriaceae *Escherichia coli* KTE58 |
| Eco-1599-1 | Bacteria Proteobacteria Gammaproteobacteria Enterobacteriales Enterobacteriaceae *Escherichia coli* KTE59 |
| Eco-1600-1 | Bacteria Proteobacteria Gammaproteobacteria Enterobacteriales Enterobacteriaceae *Escherichia coli* KTE6 |
| Eco-1601-1 | Bacteria Proteobacteria Gammaproteobacteria Enterobacteriales Enterobacteriaceae *Escherichia coli* KTE60 |
| Eco-1602-1 | Bacteria Proteobacteria Gammaproteobacteria Enterobacteriales Enterobacteriaceae *Escherichia coli* KTE61 |
| Eco-1603-1 | Bacteria Proteobacteria Gammaproteobacteria Enterobacteriales Enterobacteriaceae *Escherichia coli* KTE62 |
| Eco-1604-1 | Bacteria Proteobacteria Gammaproteobacteria Enterobacteriales Enterobacteriaceae *Escherichia coli* KTE63 |
| Eco-1605-1 | Bacteria Proteobacteria Gammaproteobacteria Enterobacteriales Enterobacteriaceae *Escherichia coli* KTE64 |
| Eco-1606-1 | Bacteria Proteobacteria Gammaproteobacteria Enterobacteriales Enterobacteriaceae *Escherichia coli* KTE65 |
| Eco-1607-1 | Bacteria Proteobacteria Gammaproteobacteria Enterobacteriales Enterobacteriaceae *Escherichia coli* KTE66 |
| Eco-1608-1 | Bacteria Proteobacteria Gammaproteobacteria Enterobacteriales Enterobacteriaceae *Escherichia coli* KTE67 |
| Eco-1609-1 | Bacteria Proteobacteria Gammaproteobacteria Enterobacteriales Enterobacteriaceae *Escherichia coli* KTE68 |
| Eco-1610-1 | Bacteria Proteobacteria Gammaproteobacteria Enterobacteriales Enterobacteriaceae *Escherichia coli* KTE69 |
| Eco-1611-1 | Bacteria Proteobacteria Gammaproteobacteria Enterobacteriales Enterobacteriaceae *Escherichia coli* KTE7 |
| Eco-1612-1 | Bacteria Proteobacteria Gammaproteobacteria Enterobacteriales Enterobacteriaceae *Escherichia coli* KTE70 |
| Eco-1613-1 | Bacteria Proteobacteria Gammaproteobacteria Enterobacteriales Enterobacteriaceae *Escherichia coli* KTE71 |
| Eco-1614-1 | Bacteria Proteobacteria Gammaproteobacteria Enterobacteriales Enterobacteriaceae *Escherichia coli* KTE72 |
| Eco-1615-1 | Bacteria Proteobacteria Gammaproteobacteria Enterobacteriales Enterobacteriaceae *Escherichia coli* KTE73 |
| Eco-1616-1 | Bacteria Proteobacteria Gammaproteobacteria Enterobacteriales Enterobacteriaceae *Escherichia coli* KTE74 |
| Eco-1617-1 | Bacteria Proteobacteria Gammaproteobacteria Enterobacteriales Enterobacteriaceae *Escherichia coli* KTE75 |
| Eco-1618-1 | Bacteria Proteobacteria Gammaproteobacteria Enterobacteriales Enterobacteriaceae *Escherichia coli* KTE76 |
| Eco-1619-1 | Bacteria Proteobacteria Gammaproteobacteria Enterobacteriales Enterobacteriaceae *Escherichia coli* KTE77 |
| Eco-1620-1 | Bacteria Proteobacteria Gammaproteobacteria Enterobacteriales Enterobacteriaceae *Escherichia coli* KTE78 |
| Eco-1621-1 | Bacteria Proteobacteria Gammaproteobacteria Enterobacteriales Enterobacteriaceae *Escherichia coli* KTE79 |
| Eco-1622-1 | Bacteria Proteobacteria Gammaproteobacteria Enterobacteriales Enterobacteriaceae *Escherichia coli* KTE8 |
| Eco-1623-1 | Bacteria Proteobacteria Gammaproteobacteria Enterobacteriales Enterobacteriaceae *Escherichia coli* KTE80 |
| Eco-1624-1 | Bacteria Proteobacteria Gammaproteobacteria Enterobacteriales Enterobacteriaceae *Escherichia coli* KTE81 |
| Eco-1625-1 | Bacteria Proteobacteria Gammaproteobacteria Enterobacteriales Enterobacteriaceae *Escherichia coli* KTE82 |
| Eco-1626-1 | Bacteria Proteobacteria Gammaproteobacteria Enterobacteriales Enterobacteriaceae *Escherichia coli* KTE83 |
| Eco-1627-1 | Bacteria Proteobacteria Gammaproteobacteria Enterobacteriales Enterobacteriaceae *Escherichia coli* KTE84 |
| Eco-1628-1 | Bacteria Proteobacteria Gammaproteobacteria Enterobacteriales Enterobacteriaceae *Escherichia coli* KTE85 |
| Eco-1629-1 | Bacteria Proteobacteria Gammaproteobacteria Enterobacteriales Enterobacteriaceae *Escherichia coli* KTE86 |
| Eco-1630-1 | Bacteria Proteobacteria Gammaproteobacteria Enterobacteriales Enterobacteriaceae *Escherichia coli* KTE87 |
| Eco-1631-1 | Bacteria Proteobacteria Gammaproteobacteria Enterobacteriales Enterobacteriaceae *Escherichia coli* KTE88 |
| Eco-1632-1 | Bacteria Proteobacteria Gammaproteobacteria Enterobacteriales Enterobacteriaceae *Escherichia coli* KTE89 |
| Eco-1633-1 | Bacteria Proteobacteria Gammaproteobacteria Enterobacteriales Enterobacteriaceae *Escherichia coli* KTE9 |
| Eco-1634-1 | Bacteria Proteobacteria Gammaproteobacteria Enterobacteriales Enterobacteriaceae *Escherichia coli* KTE90 |
| Eco-1635-1 | Bacteria Proteobacteria Gammaproteobacteria Enterobacteriales Enterobacteriaceae *Escherichia coli* KTE91 |
| Eco-1636-1 | Bacteria Proteobacteria Gammaproteobacteria Enterobacteriales Enterobacteriaceae *Escherichia coli* KTE93 |
| Eco-1637-1 | Bacteria Proteobacteria Gammaproteobacteria Enterobacteriales Enterobacteriaceae *Escherichia coli* KTE94 |
| Eco-1638-1 | Bacteria Proteobacteria Gammaproteobacteria Enterobacteriales Enterobacteriaceae *Escherichia coli* KTE95 |
| Eco-1639-1 | Bacteria Proteobacteria Gammaproteobacteria Enterobacteriales Enterobacteriaceae *Escherichia coli* KTE96 |
| Eco-1640-1 | Bacteria Proteobacteria Gammaproteobacteria Enterobacteriales Enterobacteriaceae *Escherichia coli* KTE97 |
| Eco-1641-1 | Bacteria Proteobacteria Gammaproteobacteria Enterobacteriales Enterobacteriaceae *Escherichia coli* KTE98 |
| Eco-1642-1 | Bacteria Proteobacteria Gammaproteobacteria Enterobacteriales Enterobacteriaceae *Escherichia coli* KTE99 |
| Eco-1643-1 | Bacteria Proteobacteria Gammaproteobacteria Enterobacteriales Enterobacteriaceae *Escherichia coli* LAU-EC10 |
| Eco-1644-1 | Bacteria Proteobacteria Gammaproteobacteria Enterobacteriales Enterobacteriaceae *Escherichia coli* LAU-EC6 |
| Eco-1645-1 | Bacteria Proteobacteria Gammaproteobacteria Enterobacteriales Enterobacteriaceae *Escherichia coli* LAU-EC7 |
| Eco-1646-1 | Bacteria Proteobacteria Gammaproteobacteria Enterobacteriales Enterobacteriaceae *Escherichia coli* LAU-EC8 |
| Eco-1647-1 | Bacteria Proteobacteria Gammaproteobacteria Enterobacteriales Enterobacteriaceae *Escherichia coli* LAU-EC9 |
| Eco-1648-1 | Bacteria Proteobacteria Gammaproteobacteria Enterobacteriales Enterobacteriaceae *Escherichia coli* LCT-EC106 |
| Eco-1649-1 | Bacteria Proteobacteria Gammaproteobacteria Enterobacteriales Enterobacteriaceae *Escherichia coli* LCT-EC52 |
| Eco-1650-1 | Bacteria Proteobacteria Gammaproteobacteria Enterobacteriales Enterobacteriaceae *Escherichia coli* LCT-EC59 |
| Eco-1651-1 | Bacteria Proteobacteria Gammaproteobacteria Enterobacteriales Enterobacteriaceae *Escherichia coli* LF82 |
| Eco-1652-1 | Bacteria Proteobacteria Gammaproteobacteria Enterobacteriales Enterobacteriaceae *Escherichia coli* LY180 |
| Eco-1653-1 | Bacteria Proteobacteria Gammaproteobacteria Enterobacteriales Enterobacteriaceae *Escherichia coli* M605 |
| Eco-1654-1 | Bacteria Proteobacteria Gammaproteobacteria Enterobacteriales Enterobacteriaceae *Escherichia coli* M718 |
| Eco-1655-1 | Bacteria Proteobacteria Gammaproteobacteria Enterobacteriales Enterobacteriaceae *Escherichia coli* M863 |
| Eco-1656-1 | Bacteria Proteobacteria Gammaproteobacteria Enterobacteriales Enterobacteriaceae *Escherichia coli* M919 |
| Eco-1657-1 | Bacteria Proteobacteria Gammaproteobacteria Enterobacteriales Enterobacteriaceae *Escherichia coli* MA6 |
| Eco-1658-1 | Bacteria Proteobacteria Gammaproteobacteria Enterobacteriales Enterobacteriaceae *Escherichia coli* MP020940.1 |
| Eco-1659-1 | Bacteria Proteobacteria Gammaproteobacteria Enterobacteriales Enterobacteriaceae *Escherichia coli* MP020980.1 |
| Eco-1660-1 | Bacteria Proteobacteria Gammaproteobacteria Enterobacteriales Enterobacteriaceae *Escherichia coli* MP020980.2 |
| Eco-1661-1 | Bacteria Proteobacteria Gammaproteobacteria Enterobacteriales Enterobacteriaceae *Escherichia coli* MP021017.1 |

| | |
|---|---|
| Eco-1662-1 | Bacteria Proteobacteria Gammaproteobacteria Enterobacteriales Enterobacteriaceae *Escherichia coli* MP021017.10 |
| Eco-1663-1 | Bacteria Proteobacteria Gammaproteobacteria Enterobacteriales Enterobacteriaceae *Escherichia coli* MP021017.11 |
| Eco-1664-1 | Bacteria Proteobacteria Gammaproteobacteria Enterobacteriales Enterobacteriaceae *Escherichia coli* MP021017.12 |
| Eco-1665-1 | Bacteria Proteobacteria Gammaproteobacteria Enterobacteriales Enterobacteriaceae *Escherichia coli* MP021017.2 |
| Eco-1666-1 | Bacteria Proteobacteria Gammaproteobacteria Enterobacteriales Enterobacteriaceae *Escherichia coli* MP021017.3 |
| Eco-1667-1 | Bacteria Proteobacteria Gammaproteobacteria Enterobacteriales Enterobacteriaceae *Escherichia coli* MP021017.4 |
| Eco-1668-1 | Bacteria Proteobacteria Gammaproteobacteria Enterobacteriales Enterobacteriaceae *Escherichia coli* MP021017.5 |
| Eco-1669-1 | Bacteria Proteobacteria Gammaproteobacteria Enterobacteriales Enterobacteriaceae *Escherichia coli* MP021017.6 |
| Eco-1670-1 | Bacteria Proteobacteria Gammaproteobacteria Enterobacteriales Enterobacteriaceae *Escherichia coli* MP021017.9 |
| Eco-1671-1 | Bacteria Proteobacteria Gammaproteobacteria Enterobacteriales Enterobacteriaceae *Escherichia coli* MP021552.11 |
| Eco-1672-1 | Bacteria Proteobacteria Gammaproteobacteria Enterobacteriales Enterobacteriaceae *Escherichia coli* MP021552.12 |
| Eco-1673-1 | Bacteria Proteobacteria Gammaproteobacteria Enterobacteriales Enterobacteriaceae *Escherichia coli* MP021552.7 |
| Eco-1674-1 | Bacteria Proteobacteria Gammaproteobacteria Enterobacteriales Enterobacteriaceae *Escherichia coli* MP021552.8 |
| Eco-1675-1 | Bacteria Proteobacteria Gammaproteobacteria Enterobacteriales Enterobacteriaceae *Escherichia coli* MP021561.2 |
| Eco-1676-1 | Bacteria Proteobacteria Gammaproteobacteria Enterobacteriales Enterobacteriaceae *Escherichia coli* MP021561.3 |
| Eco-1677-1 | Bacteria Proteobacteria Gammaproteobacteria Enterobacteriales Enterobacteriaceae *Escherichia coli* MP021566.1 |
| Eco-1678-1 | Bacteria Proteobacteria Gammaproteobacteria Enterobacteriales Enterobacteriaceae *Escherichia coli* MS 107-1 |
| Eco-1679-1 | Bacteria Proteobacteria Gammaproteobacteria Enterobacteriales Enterobacteriaceae *Escherichia coli* MS 110-3 |
| Eco-1680-1 | Bacteria Proteobacteria Gammaproteobacteria Enterobacteriales Enterobacteriaceae *Escherichia coli* MS 115-1 |
| Eco-1681-1 | Bacteria Proteobacteria Gammaproteobacteria Enterobacteriales Enterobacteriaceae *Escherichia coli* MS 116-1 |
| Eco-1682-1 | Bacteria Proteobacteria Gammaproteobacteria Enterobacteriales Enterobacteriaceae *Escherichia coli* MS 117-3 |
| Eco-1683-1 | Bacteria Proteobacteria Gammaproteobacteria Enterobacteriales Enterobacteriaceae *Escherichia coli* MS 119-7 |
| Eco-1684-1 | Bacteria Proteobacteria Gammaproteobacteria Enterobacteriales Enterobacteriaceae *Escherichia coli* MS 124-1 |
| Eco-1685-1 | Bacteria Proteobacteria Gammaproteobacteria Enterobacteriales Enterobacteriaceae *Escherichia coli* MS 145-7 |
| Eco-1686-1 | Bacteria Proteobacteria Gammaproteobacteria Enterobacteriales Enterobacteriaceae *Escherichia coli* MS 146-1 |
| Eco-1687-1 | Bacteria Proteobacteria Gammaproteobacteria Enterobacteriales Enterobacteriaceae *Escherichia coli* MS 153-1 |
| Eco-1688-1 | Bacteria Proteobacteria Gammaproteobacteria Enterobacteriales Enterobacteriaceae *Escherichia coli* MS 16-3 |
| Eco-1689-1 | Bacteria Proteobacteria Gammaproteobacteria Enterobacteriales Enterobacteriaceae *Escherichia coli* MS 175-1 |
| Eco-1690-1 | Bacteria Proteobacteria Gammaproteobacteria Enterobacteriales Enterobacteriaceae *Escherichia coli* MS 182-1 |
| Eco-1691-1 | Bacteria Proteobacteria Gammaproteobacteria Enterobacteriales Enterobacteriaceae *Escherichia coli* MS 185-1 |
| Eco-1692-1 | Bacteria Proteobacteria Gammaproteobacteria Enterobacteriales Enterobacteriaceae *Escherichia coli* MS 187-1 |
| Eco-1693-1 | Bacteria Proteobacteria Gammaproteobacteria Enterobacteriales Enterobacteriaceae *Escherichia coli* MS 196-1 |
| Eco-1694-1 | Bacteria Proteobacteria Gammaproteobacteria Enterobacteriales Enterobacteriaceae *Escherichia coli* MS 198-1 |
| Eco-1695-1 | Bacteria Proteobacteria Gammaproteobacteria Enterobacteriales Enterobacteriaceae *Escherichia coli* MS 200-1 |
| Eco-1696-1 | Bacteria Proteobacteria Gammaproteobacteria Enterobacteriales Enterobacteriaceae *Escherichia coli* MS 21-1 |
| Eco-1697-1 | Bacteria Proteobacteria Gammaproteobacteria Enterobacteriales Enterobacteriaceae *Escherichia coli* MS 45-1 |
| Eco-1698-1 | Bacteria Proteobacteria Gammaproteobacteria Enterobacteriales Enterobacteriaceae *Escherichia coli* MS 57-2 |
| Eco-1699-1 | Bacteria Proteobacteria Gammaproteobacteria Enterobacteriales Enterobacteriaceae *Escherichia coli* MS 60-1 |
| Eco-1700-1 | Bacteria Proteobacteria Gammaproteobacteria Enterobacteriales Enterobacteriaceae *Escherichia coli* MS 69-1 |
| Eco-1701-1 | Bacteria Proteobacteria Gammaproteobacteria Enterobacteriales Enterobacteriaceae *Escherichia coli* MS 78-1 |
| Eco-1702-1 | Bacteria Proteobacteria Gammaproteobacteria Enterobacteriales Enterobacteriaceae *Escherichia coli* MS 79-10 |
| Eco-1703-1 | Bacteria Proteobacteria Gammaproteobacteria Enterobacteriales Enterobacteriaceae *Escherichia coli* MS 84-1 |
| Eco-1704-1 | Bacteria Proteobacteria Gammaproteobacteria Enterobacteriales Enterobacteriaceae *Escherichia coli* MS 85-1 |
| Eco-1705-1 | Bacteria Proteobacteria Gammaproteobacteria Enterobacteriales Enterobacteriaceae *Escherichia coli* N1 |
| Eco-1706-1 | Bacteria Proteobacteria Gammaproteobacteria Enterobacteriales Enterobacteriaceae *Escherichia coli* NA114 |
| Eco-1707-1 | Bacteria Proteobacteria Gammaproteobacteria Enterobacteriales Enterobacteriaceae *Escherichia coli* NC101 |
| Eco-1708-1 | Bacteria Proteobacteria Gammaproteobacteria Enterobacteriales Enterobacteriaceae *Escherichia coli* NCCP15647 |
| Eco-1709-1 | Bacteria Proteobacteria Gammaproteobacteria Enterobacteriales Enterobacteriaceae *Escherichia coli* NCCP15657 |
| Eco-1710-1 | Bacteria Proteobacteria Gammaproteobacteria Enterobacteriales Enterobacteriaceae *Escherichia coli* NCCP15658 |
| Eco-1711-1 | Bacteria Proteobacteria Gammaproteobacteria Enterobacteriales Enterobacteriaceae *Escherichia coli* NE037 |
| Eco-1712-1 | Bacteria Proteobacteria Gammaproteobacteria Enterobacteriales Enterobacteriaceae *Escherichia coli* NE098 |
| Eco-1713-1 | Bacteria Proteobacteria Gammaproteobacteria Enterobacteriales Enterobacteriaceae *Escherichia coli* NE1487 |
| Eco-463-1 to Eco-463-4 | Bacteria Proteobacteria Gammaproteobacteria Enterobacteriales Enterobacteriaceae *Escherichia coli* Nissle 1917 |
| Eco-1714-1 | Bacteria Proteobacteria Gammaproteobacteria Enterobacteriales Enterobacteriaceae *Escherichia coli* O08 |
| Eco-1715-1 | Bacteria Proteobacteria Gammaproteobacteria Enterobacteriales Enterobacteriaceae *Escherichia coli* O103:H2 str. 12009 |
| Eco-1716-1 | Bacteria Proteobacteria Gammaproteobacteria Enterobacteriales Enterobacteriaceae *Escherichia coli* O103:H2 str. CVM9450 |
| Eco-1717-1 | Bacteria Proteobacteria Gammaproteobacteria Enterobacteriales Enterobacteriaceae *Escherichia coli* O103:H25 str. CVM9340 |
| Eco-1718-1 | Bacteria Proteobacteria Gammaproteobacteria Enterobacteriales Enterobacteriaceae *Escherichia coli* O103:H25 str. NIPH-11060424 |
| Eco-1719-1 | Bacteria Proteobacteria Gammaproteobacteria Enterobacteriales Enterobacteriaceae *Escherichia coli* O104:H4 str. 04-8351 |
| Eco-1720-1 | Bacteria Proteobacteria Gammaproteobacteria Enterobacteriales Enterobacteriaceae *Escherichia coli* O104:H4 str. 09-7901 |
| Eco-1721-1 | Bacteria Proteobacteria Gammaproteobacteria Enterobacteriales Enterobacteriaceae *Escherichia coli* O104:H4 str. 11-02030 |
| Eco-1722-1 | Bacteria Proteobacteria Gammaproteobacteria Enterobacteriales Enterobacteriaceae *Escherichia coli* O104:H4 str. 11-02033-1 |
| Eco-1723-1 | Bacteria Proteobacteria Gammaproteobacteria Enterobacteriales Enterobacteriaceae *Escherichia coli* O104:H4 str. 11-02092 |
| Eco-1724-1 | Bacteria Proteobacteria Gammaproteobacteria Enterobacteriales Enterobacteriaceae *Escherichia coli* O104:H4 str. 11-02093 |
| Eco-1725-1 | Bacteria Proteobacteria Gammaproteobacteria Enterobacteriales Enterobacteriaceae *Escherichia coli* O104:H4 str. 11-02281 |

| | |
|---|---|
| Eco-1726-1 | Bacteria Proteobacteria Gammaproteobacteria Enterobacteriales Enterobacteriaceae *Escherichia coli* O104:H4 str. 11-02318 |
| Eco-1727-1 | Bacteria Proteobacteria Gammaproteobacteria Enterobacteriales Enterobacteriaceae *Escherichia coli* O104:H4 str. 11-02913 |
| Eco-1728-1 | Bacteria Proteobacteria Gammaproteobacteria Enterobacteriales Enterobacteriaceae *Escherichia coli* O104:H4 str. 11-03439 |
| Eco-1729-1 | Bacteria Proteobacteria Gammaproteobacteria Enterobacteriales Enterobacteriaceae *Escherichia coli* O104:H4 str. 11-03943 |
| Eco-1730-1 | Bacteria Proteobacteria Gammaproteobacteria Enterobacteriales Enterobacteriaceae *Escherichia coli* O104:H4 str. 11-04080 |
| Eco-1731-1 | Bacteria Proteobacteria Gammaproteobacteria Enterobacteriales Enterobacteriaceae *Escherichia coli* O104:H4 str. 11-3677 |
| Eco-1732-1 | Bacteria Proteobacteria Gammaproteobacteria Enterobacteriales Enterobacteriaceae *Escherichia coli* O104:H4 str. 11-4404 |
| Eco-1733-1 | Bacteria Proteobacteria Gammaproteobacteria Enterobacteriales Enterobacteriaceae *Escherichia coli* O104:H4 str. 11-4522 |
| Eco-1734-1 | Bacteria Proteobacteria Gammaproteobacteria Enterobacteriales Enterobacteriaceae *Escherichia coli* O104:H4 str. 11-4623 |
| Eco-1735-1 | Bacteria Proteobacteria Gammaproteobacteria Enterobacteriales Enterobacteriaceae *Escherichia coli* O104:H4 str. 11-4632 C1 |
| Eco-1736-1 | Bacteria Proteobacteria Gammaproteobacteria Enterobacteriales Enterobacteriaceae *Escherichia coli* O104:H4 str. 11-4632 C2 |
| Eco-1737-1 | Bacteria Proteobacteria Gammaproteobacteria Enterobacteriales Enterobacteriaceae *Escherichia coli* O104:H4 str. 11-4632 C3 |
| Eco-1738-1 | Bacteria Proteobacteria Gammaproteobacteria Enterobacteriales Enterobacteriaceae *Escherichia coli* O104:H4 str. 11-4632 C4 |
| Eco-1739-1 | Bacteria Proteobacteria Gammaproteobacteria Enterobacteriales Enterobacteriaceae *Escherichia coli* O104:H4 str. 11-4632 C5 |
| Eco-1740-1 | Bacteria Proteobacteria Gammaproteobacteria Enterobacteriales Enterobacteriaceae *Escherichia coli* O104:H4 str. 2009EL-2050 |
| Eco-1741-1 | Bacteria Proteobacteria Gammaproteobacteria Enterobacteriales Enterobacteriaceae *Escherichia coli* O104:H4 str. 2009EL-2071 |
| Eco-1742-1 | Bacteria Proteobacteria Gammaproteobacteria Enterobacteriales Enterobacteriaceae *Escherichia coli* O104:H4 str. 2011C-3493 |
| Eco-472-1 to Eco-472-2 | Bacteria Proteobacteria Gammaproteobacteria Enterobacteriales Enterobacteriaceae *Escherichia coli* O104:H4 str. C227-11 |
| Eco-1743-1 | Bacteria Proteobacteria Gammaproteobacteria Enterobacteriales Enterobacteriaceae *Escherichia coli* O104:H4 str. C236-11 |
| Eco-1744-1 | Bacteria Proteobacteria Gammaproteobacteria Enterobacteriales Enterobacteriaceae *Escherichia coli* O104:H4 str. E112/10 |
| Eco-1745-1 | Bacteria Proteobacteria Gammaproteobacteria Enterobacteriales Enterobacteriaceae *Escherichia coli* O104:H4 str. E92/11 |
| Eco-1746-1 | Bacteria Proteobacteria Gammaproteobacteria Enterobacteriales Enterobacteriaceae *Escherichia coli* O104:H4 str. Ec11-4984 |
| Eco-1747-1 | Bacteria Proteobacteria Gammaproteobacteria Enterobacteriales Enterobacteriaceae *Escherichia coli* O104:H4 str. Ec11-4986 |
| Eco-1748-1 | Bacteria Proteobacteria Gammaproteobacteria Enterobacteriales Enterobacteriaceae *Escherichia coli* O104:H4 str. Ec11-4987 |
| Eco-1749-1 | Bacteria Proteobacteria Gammaproteobacteria Enterobacteriales Enterobacteriaceae *Escherichia coli* O104:H4 str. Ec11-4988 |
| Eco-1750-1 | Bacteria Proteobacteria Gammaproteobacteria Enterobacteriales Enterobacteriaceae *Escherichia coli* O104:H4 str. Ec11-5603 |
| Eco-1751-1 | Bacteria Proteobacteria Gammaproteobacteria Enterobacteriales Enterobacteriaceae *Escherichia coli* O104:H4 str. Ec11-5604 |
| Eco-1752-1 | Bacteria Proteobacteria Gammaproteobacteria Enterobacteriales Enterobacteriaceae *Escherichia coli* O104:H4 str. Ec11-6006 |
| Eco-1753-1 | Bacteria Proteobacteria Gammaproteobacteria Enterobacteriales Enterobacteriaceae *Escherichia coli* O104:H4 str. Ec11-9450 |
| Eco-1754-1 | Bacteria Proteobacteria Gammaproteobacteria Enterobacteriales Enterobacteriaceae *Escherichia coli* O104:H4 str. Ec11-9941 |
| Eco-1755-1 | Bacteria Proteobacteria Gammaproteobacteria Enterobacteriales Enterobacteriaceae *Escherichia coli* O104:H4 str. Ec11-9990 |
| Eco-1756-1 | Bacteria Proteobacteria Gammaproteobacteria Enterobacteriales Enterobacteriaceae *Escherichia coli* O104:H4 str. Ec12-0465 |
| Eco-1757-1 | Bacteria Proteobacteria Gammaproteobacteria Enterobacteriales Enterobacteriaceae *Escherichia coli* O104:H4 str. Ec12-0466 |
| Eco-1758-1 | Bacteria Proteobacteria Gammaproteobacteria Enterobacteriales Enterobacteriaceae *Escherichia coli* O104:H4 str. GOS1 |
| Eco-1759-1 | Bacteria Proteobacteria Gammaproteobacteria Enterobacteriales Enterobacteriaceae *Escherichia coli* O104:H4 str. GOS2 |
| Eco-1760-1 | Bacteria Proteobacteria Gammaproteobacteria Enterobacteriales Enterobacteriaceae *Escherichia coli* O104:H4 str. H112180280 |
| Eco-1761-1 | Bacteria Proteobacteria Gammaproteobacteria Enterobacteriales Enterobacteriaceae *Escherichia coli* O104:H4 str. H112180282 |
| Eco-1762-1 | Bacteria Proteobacteria Gammaproteobacteria Enterobacteriales Enterobacteriaceae *Escherichia coli* O104:H4 str. ON2010 |
| Eco-1763-1 | Bacteria Proteobacteria Gammaproteobacteria Enterobacteriales Enterobacteriaceae *Escherichia coli* O104:H4 str. ON2011 |
| Eco-1764-1 | Bacteria Proteobacteria Gammaproteobacteria Enterobacteriales Enterobacteriaceae *Escherichia coli* O104:H4 str. TY-2482 |
| Eco-1765-1 | Bacteria Proteobacteria Gammaproteobacteria Enterobacteriales Enterobacteriaceae *Escherichia coli* O10:K5(L):H4 str. ATCC 23506 |
| Eco-1766-1 | Bacteria Proteobacteria Gammaproteobacteria Enterobacteriales Enterobacteriaceae *Escherichia coli* O111:H- str. 11128 |
| Eco-1767-1 | Bacteria Proteobacteria Gammaproteobacteria Enterobacteriales Enterobacteriaceae *Escherichia coli* O111:H11 str. CFSAN001630 |
| Eco-1768-1 | Bacteria Proteobacteria Gammaproteobacteria Enterobacteriales Enterobacteriaceae *Escherichia coli* O111:H11 str. CVM9455 |
| Eco-1769-1 | Bacteria Proteobacteria Gammaproteobacteria Enterobacteriales Enterobacteriaceae *Escherichia coli* O111:H11 str. CVM9534 |
| Eco-1770-1 | Bacteria Proteobacteria Gammaproteobacteria Enterobacteriales Enterobacteriaceae *Escherichia coli* O111:H11 str. CVM9545 |
| Eco-1771-1 | Bacteria Proteobacteria Gammaproteobacteria Enterobacteriales Enterobacteriaceae *Escherichia coli* O111:H11 str. CVM9553 |
| Eco-1772-1 | Bacteria Proteobacteria Gammaproteobacteria Enterobacteriales Enterobacteriaceae *Escherichia coli* O111:H8 str. CFSAN001632 |
| Eco-1773-1 | Bacteria Proteobacteria Gammaproteobacteria Enterobacteriales Enterobacteriaceae *Escherichia coli* O111:H8 str. CVM9570 |
| Eco-1774-1 | Bacteria Proteobacteria Gammaproteobacteria Enterobacteriales Enterobacteriaceae *Escherichia coli* O111:H8 str. CVM9574 |
| Eco-1775-1 | Bacteria Proteobacteria Gammaproteobacteria Enterobacteriales Enterobacteriaceae *Escherichia coli* O111:H8 str. CVM9602 |
| Eco-1776-1 | Bacteria Proteobacteria Gammaproteobacteria Enterobacteriales Enterobacteriaceae *Escherichia coli* O111:H8 str. CVM9634 |
| Eco-1777-1 | Bacteria Proteobacteria Gammaproteobacteria Enterobacteriales Enterobacteriaceae *Escherichia coli* O113:H21 str. CL-3 |
| Eco-1778-1 | Bacteria Proteobacteria Gammaproteobacteria Enterobacteriales Enterobacteriaceae *Escherichia coli* O121:H19 str. MT#2 |
| Eco-1779-1 | Bacteria Proteobacteria Gammaproteobacteria Enterobacteriales Enterobacteriaceae *Escherichia coli* O127:H27 str. C43/90 |
| Eco-1780-1 | Bacteria Proteobacteria Gammaproteobacteria Enterobacteriales Enterobacteriaceae *Escherichia coli* O127:H6 str. E2348/69 |
| Eco-548-1 to Eco-548-2 | Bacteria Proteobacteria Gammaproteobacteria Enterobacteriales Enterobacteriaceae *Escherichia coli* O145:H28 str. 4865/96 |
| Eco-1781-1 | Bacteria Proteobacteria Gammaproteobacteria Enterobacteriales Enterobacteriaceae *Escherichia coli* O157 str. NCCP15738 |
| Eco-1782-1 to Eco-1782-2 | Bacteria Proteobacteria Gammaproteobacteria Enterobacteriales Enterobacteriaceae *Escherichia coli* O157:H- str. 493-89 |
| Eco-1783-1 | Bacteria Proteobacteria Gammaproteobacteria Enterobacteriales Enterobacteriaceae *Escherichia coli* O157:H- str. H 2687 |
| Eco-1784-1 | Bacteria Proteobacteria Gammaproteobacteria Enterobacteriales Enterobacteriaceae *Escherichia coli* O157:H43 str. T22 |
| Eco-1785-1 | Bacteria Proteobacteria Gammaproteobacteria Enterobacteriales Enterobacteriaceae *Escherichia coli* O157:H7 EDL933 |
| Eco-1786-1 | Bacteria Proteobacteria Gammaproteobacteria Enterobacteriales Enterobacteriaceae *Escherichia coli* O157:H7 str. 08BKT061141 |
| Eco-1787-1 | Bacteria Proteobacteria Gammaproteobacteria Enterobacteriales Enterobacteriaceae *Escherichia coli* O157:H7 str. 09BKT048303 |
| Eco-1788-1 | Bacteria Proteobacteria Gammaproteobacteria Enterobacteriales Enterobacteriaceae *Escherichia coli* O157:H7 str. 1044 |

| | |
|---|---|
| Eco-1789-1 | Bacteria Proteobacteria Gammaproteobacteria Enterobacteriales Enterobacteriaceae *Escherichia coli* O157:H7 str. 1125 |
| Eco-1790-1 | Bacteria Proteobacteria Gammaproteobacteria Enterobacteriales Enterobacteriaceae *Escherichia coli* O157:H7 str. EC1212 |
| Eco-1791-1 | Bacteria Proteobacteria Gammaproteobacteria Enterobacteriales Enterobacteriaceae *Escherichia coli* O157:H7 str. EC4009 |
| Eco-1792-1 | Bacteria Proteobacteria Gammaproteobacteria Enterobacteriales Enterobacteriaceae *Escherichia coli* O157:H7 str. EC4042 |
| Eco-1793-1 | Bacteria Proteobacteria Gammaproteobacteria Enterobacteriales Enterobacteriaceae *Escherichia coli* O157:H7 str. EC4045 |
| Eco-1794-1 | Bacteria Proteobacteria Gammaproteobacteria Enterobacteriales Enterobacteriaceae *Escherichia coli* O157:H7 str. EC4076 |
| Eco-1795-1 | Bacteria Proteobacteria Gammaproteobacteria Enterobacteriales Enterobacteriaceae *Escherichia coli* O157:H7 str. EC4113 |
| Eco-1796-1 | Bacteria Proteobacteria Gammaproteobacteria Enterobacteriales Enterobacteriaceae *Escherichia coli* O157:H7 str. EC4115 |
| Eco-1797-1 | Bacteria Proteobacteria Gammaproteobacteria Enterobacteriales Enterobacteriaceae *Escherichia coli* O157:H7 str. EC4191 |
| Eco-1798-1 | Bacteria Proteobacteria Gammaproteobacteria Enterobacteriales Enterobacteriaceae *Escherichia coli* O157:H7 str. EC4196 |
| Eco-1799-1 | Bacteria Proteobacteria Gammaproteobacteria Enterobacteriales Enterobacteriaceae *Escherichia coli* O157:H7 str. EC4206 |
| Eco-1800-1 | Bacteria Proteobacteria Gammaproteobacteria Enterobacteriales Enterobacteriaceae *Escherichia coli* O157:H7 str. EC4401 |
| Eco-1801-1 | Bacteria Proteobacteria Gammaproteobacteria Enterobacteriales Enterobacteriaceae *Escherichia coli* O157:H7 str. EC4486 |
| Eco-1802-1 | Bacteria Proteobacteria Gammaproteobacteria Enterobacteriales Enterobacteriaceae *Escherichia coli* O157:H7 str. EC4501 |
| Eco-1803-1 | Bacteria Proteobacteria Gammaproteobacteria Enterobacteriales Enterobacteriaceae *Escherichia coli* O157:H7 str. EC508 |
| Eco-1804-1 | Bacteria Proteobacteria Gammaproteobacteria Enterobacteriales Enterobacteriaceae *Escherichia coli* O157:H7 str. EC869 |
| Eco-1805-1 | Bacteria Proteobacteria Gammaproteobacteria Enterobacteriales Enterobacteriaceae *Escherichia coli* O157:H7 str. FRIK2000 |
| Eco-1806-1 | Bacteria Proteobacteria Gammaproteobacteria Enterobacteriales Enterobacteriaceae *Escherichia coli* O157:H7 str. FRIK966 |
| Eco-1807-1 | Bacteria Proteobacteria Gammaproteobacteria Enterobacteriales Enterobacteriaceae *Escherichia coli* O157:H7 str. G5101 |
| Eco-1808-1 | Bacteria Proteobacteria Gammaproteobacteria Enterobacteriales Enterobacteriaceae *Escherichia coli* O157:H7 str. LSU-61 |
| Eco-1809-1 | Bacteria Proteobacteria Gammaproteobacteria Enterobacteriales Enterobacteriaceae *Escherichia coli* O157:H7 str. Sakai |
| Eco-1810-1 | Bacteria Proteobacteria Gammaproteobacteria Enterobacteriales Enterobacteriaceae *Escherichia coli* O157:H7 str. TW14359 |
| Eco-1811-1 | Bacteria Proteobacteria Gammaproteobacteria Enterobacteriales Enterobacteriaceae *Escherichia coli* O157:H7 str. TW14588 |
| Eco-1812-1 | Bacteria Proteobacteria Gammaproteobacteria Enterobacteriales Enterobacteriaceae *Escherichia coli* O26:H11 str. 11368 |
| Eco-1813-1 | Bacteria Proteobacteria Gammaproteobacteria Enterobacteriales Enterobacteriaceae *Escherichia coli* O26:H11 str. 2011C-3274 |
| Eco-1814-1 | Bacteria Proteobacteria Gammaproteobacteria Enterobacteriales Enterobacteriaceae *Escherichia coli* O26:H11 str. CFSAN001629 |
| Eco-1815-1 | Bacteria Proteobacteria Gammaproteobacteria Enterobacteriales Enterobacteriaceae *Escherichia coli* O26:H11 str. CVM10021 |
| Eco-1816-1 | Bacteria Proteobacteria Gammaproteobacteria Enterobacteriales Enterobacteriaceae *Escherichia coli* O26:H11 str. CVM10026 |
| Eco-1817-1 | Bacteria Proteobacteria Gammaproteobacteria Enterobacteriales Enterobacteriaceae *Escherichia coli* O26:H11 str. CVM10030 |
| Eco-1818-1 | Bacteria Proteobacteria Gammaproteobacteria Enterobacteriales Enterobacteriaceae *Escherichia coli* O26:H11 str. CVM10224 |
| Eco-1819-1 | Bacteria Proteobacteria Gammaproteobacteria Enterobacteriales Enterobacteriaceae *Escherichia coli* O26:H11 str. CVM9942 |
| Eco-1820-1 | Bacteria Proteobacteria Gammaproteobacteria Enterobacteriales Enterobacteriaceae *Escherichia coli* O26:H11 str. CVM9952 |
| Eco-1821-1 | Bacteria Proteobacteria Gammaproteobacteria Enterobacteriales Enterobacteriaceae *Escherichia coli* O32:H37 str. P4 |
| Eco-1822-1 | Bacteria Proteobacteria Gammaproteobacteria Enterobacteriales Enterobacteriaceae *Escherichia coli* O45:H2 str. 03-EN-705 |
| Eco-1823-1 | Bacteria Proteobacteria Gammaproteobacteria Enterobacteriales Enterobacteriaceae *Escherichia coli* O55:H7 str. 3256-97 |
| Eco-1824-1 | Bacteria Proteobacteria Gammaproteobacteria Enterobacteriales Enterobacteriaceae *Escherichia coli* O55:H7 str. CB9615 |
| Eco-1825-1 | Bacteria Proteobacteria Gammaproteobacteria Enterobacteriales Enterobacteriaceae *Escherichia coli* O55:H7 str. RM12579 |
| Eco-1826-1 | Bacteria Proteobacteria Gammaproteobacteria Enterobacteriales Enterobacteriaceae *Escherichia coli* O55:H7 str. USDA 5905 |
| Eco-1827-1 | Bacteria Proteobacteria Gammaproteobacteria Enterobacteriales Enterobacteriaceae *Escherichia coli* O5:K4(L):H4 str. ATCC 23502 |
| Eco-1828-1 | Bacteria Proteobacteria Gammaproteobacteria Enterobacteriales Enterobacteriaceae *Escherichia coli* O7:K1 str. CE10 |
| Eco-1829-1 | Bacteria Proteobacteria Gammaproteobacteria Enterobacteriales Enterobacteriaceae *Escherichia coli* O83:H1 str. NRG 857C |
| Eco-1830-1 | Bacteria Proteobacteria Gammaproteobacteria Enterobacteriales Enterobacteriaceae *Escherichia coli* O91:H21 str. B2F1 |
| Eco-1831-1 | Bacteria Proteobacteria Gammaproteobacteria Enterobacteriales Enterobacteriaceae *Escherichia coli* OK1114 |
| Eco-1832-1 | Bacteria Proteobacteria Gammaproteobacteria Enterobacteriales Enterobacteriaceae *Escherichia coli* OK1180 |
| Eco-1833-1 | Bacteria Proteobacteria Gammaproteobacteria Enterobacteriales Enterobacteriaceae *Escherichia coli* OK1357 |
| Eco-1834-1 | Bacteria Proteobacteria Gammaproteobacteria Enterobacteriales Enterobacteriaceae *Escherichia coli* ONT:H33 str. C48/93 |
| Eco-1835-1 | Bacteria Proteobacteria Gammaproteobacteria Enterobacteriales Enterobacteriaceae *Escherichia coli* P0298942.1 |
| Eco-1836-1 | Bacteria Proteobacteria Gammaproteobacteria Enterobacteriales Enterobacteriaceae *Escherichia coli* P0298942.10 |
| Eco-1837-1 | Bacteria Proteobacteria Gammaproteobacteria Enterobacteriales Enterobacteriaceae *Escherichia coli* P0298942.11 |
| Eco-1838-1 | Bacteria Proteobacteria Gammaproteobacteria Enterobacteriales Enterobacteriaceae *Escherichia coli* P0298942.12 |
| Eco-1839-1 | Bacteria Proteobacteria Gammaproteobacteria Enterobacteriales Enterobacteriaceae *Escherichia coli* P0298942.14 |
| Eco-1840-1 | Bacteria Proteobacteria Gammaproteobacteria Enterobacteriales Enterobacteriaceae *Escherichia coli* P0298942.15 |
| Eco-1841-1 | Bacteria Proteobacteria Gammaproteobacteria Enterobacteriales Enterobacteriaceae *Escherichia coli* P0298942.2 |
| Eco-1842-1 | Bacteria Proteobacteria Gammaproteobacteria Enterobacteriales Enterobacteriaceae *Escherichia coli* P0298942.3 |
| Eco-1843-1 | Bacteria Proteobacteria Gammaproteobacteria Enterobacteriales Enterobacteriaceae *Escherichia coli* P0298942.4 |
| Eco-1844-1 | Bacteria Proteobacteria Gammaproteobacteria Enterobacteriales Enterobacteriaceae *Escherichia coli* P0298942.6 |
| Eco-1845-1 | Bacteria Proteobacteria Gammaproteobacteria Enterobacteriales Enterobacteriaceae *Escherichia coli* P0298942.7 |
| Eco-1846-1 | Bacteria Proteobacteria Gammaproteobacteria Enterobacteriales Enterobacteriaceae *Escherichia coli* P0298942.8 |
| Eco-1847-1 | Bacteria Proteobacteria Gammaproteobacteria Enterobacteriales Enterobacteriaceae *Escherichia coli* P0298942.9 |
| Eco-1848-1 | Bacteria Proteobacteria Gammaproteobacteria Enterobacteriales Enterobacteriaceae *Escherichia coli* P0299438.10 |
| Eco-1849-1 | Bacteria Proteobacteria Gammaproteobacteria Enterobacteriales Enterobacteriaceae *Escherichia coli* P0299438.11 |
| Eco-1850-1 | Bacteria Proteobacteria Gammaproteobacteria Enterobacteriales Enterobacteriaceae *Escherichia coli* P0299438.2 |
| Eco-1851-1 | Bacteria Proteobacteria Gammaproteobacteria Enterobacteriales Enterobacteriaceae *Escherichia coli* P0299438.3 |
| Eco-1852-1 | Bacteria Proteobacteria Gammaproteobacteria Enterobacteriales Enterobacteriaceae *Escherichia coli* P0299438.4 |
| Eco-1853-1 | Bacteria Proteobacteria Gammaproteobacteria Enterobacteriales Enterobacteriaceae *Escherichia coli* P0299438.5 |

| | |
|---|---|
| Eco-1854-1 | Bacteria Proteobacteria Gammaproteobacteria Enterobacteriales Enterobacteriaceae *Escherichia coli* P0299438.6 |
| Eco-1855-1 | Bacteria Proteobacteria Gammaproteobacteria Enterobacteriales Enterobacteriaceae *Escherichia coli* P0299438.7 |
| Eco-1856-1 | Bacteria Proteobacteria Gammaproteobacteria Enterobacteriales Enterobacteriaceae *Escherichia coli* P0299438.8 |
| Eco-1857-1 | Bacteria Proteobacteria Gammaproteobacteria Enterobacteriales Enterobacteriaceae *Escherichia coli* P0299438.9 |
| Eco-1858-1 | Bacteria Proteobacteria Gammaproteobacteria Enterobacteriales Enterobacteriaceae *Escherichia coli* P0299483.1 |
| Eco-1859-1 | Bacteria Proteobacteria Gammaproteobacteria Enterobacteriales Enterobacteriaceae *Escherichia coli* P0299483.2 |
| Eco-1860-1 | Bacteria Proteobacteria Gammaproteobacteria Enterobacteriales Enterobacteriaceae *Escherichia coli* P0299483.3 |
| Eco-1861-1 | Bacteria Proteobacteria Gammaproteobacteria Enterobacteriales Enterobacteriaceae *Escherichia coli* P02997067.6 |
| Eco-1862-1 | Bacteria Proteobacteria Gammaproteobacteria Enterobacteriales Enterobacteriaceae *Escherichia coli* P0299917.1 |
| Eco-1863-1 | Bacteria Proteobacteria Gammaproteobacteria Enterobacteriales Enterobacteriaceae *Escherichia coli* P0299917.10 |
| Eco-1864-1 | Bacteria Proteobacteria Gammaproteobacteria Enterobacteriales Enterobacteriaceae *Escherichia coli* P0299917.2 |
| Eco-1865-1 | Bacteria Proteobacteria Gammaproteobacteria Enterobacteriales Enterobacteriaceae *Escherichia coli* P0299917.3 |
| Eco-1866-1 | Bacteria Proteobacteria Gammaproteobacteria Enterobacteriales Enterobacteriaceae *Escherichia coli* P0299917.4 |
| Eco-1867-1 | Bacteria Proteobacteria Gammaproteobacteria Enterobacteriales Enterobacteriaceae *Escherichia coli* P0299917.5 |
| Eco-1868-1 | Bacteria Proteobacteria Gammaproteobacteria Enterobacteriales Enterobacteriaceae *Escherichia coli* P0299917.6 |
| Eco-1869-1 | Bacteria Proteobacteria Gammaproteobacteria Enterobacteriales Enterobacteriaceae *Escherichia coli* P0299917.7 |
| Eco-1870-1 | Bacteria Proteobacteria Gammaproteobacteria Enterobacteriales Enterobacteriaceae *Escherichia coli* P0299917.8 |
| Eco-1871-1 | Bacteria Proteobacteria Gammaproteobacteria Enterobacteriales Enterobacteriaceae *Escherichia coli* P0299917.9 |
| Eco-1872-1 | Bacteria Proteobacteria Gammaproteobacteria Enterobacteriales Enterobacteriaceae *Escherichia coli* P0301867.1 |
| Eco-1873-1 | Bacteria Proteobacteria Gammaproteobacteria Enterobacteriales Enterobacteriaceae *Escherichia coli* P0301867.11 |
| Eco-1874-1 | Bacteria Proteobacteria Gammaproteobacteria Enterobacteriales Enterobacteriaceae *Escherichia coli* P0301867.13 |
| Eco-1875-1 | Bacteria Proteobacteria Gammaproteobacteria Enterobacteriales Enterobacteriaceae *Escherichia coli* P0301867.2 |
| Eco-1876-1 | Bacteria Proteobacteria Gammaproteobacteria Enterobacteriales Enterobacteriaceae *Escherichia coli* P0301867.3 |
| Eco-1877-1 | Bacteria Proteobacteria Gammaproteobacteria Enterobacteriales Enterobacteriaceae *Escherichia coli* P0301867.4 |
| Eco-1878-1 | Bacteria Proteobacteria Gammaproteobacteria Enterobacteriales Enterobacteriaceae *Escherichia coli* P0301867.5 |
| Eco-1879-1 | Bacteria Proteobacteria Gammaproteobacteria Enterobacteriales Enterobacteriaceae *Escherichia coli* P0301867.7 |
| Eco-1880-1 | Bacteria Proteobacteria Gammaproteobacteria Enterobacteriales Enterobacteriaceae *Escherichia coli* P0301867.8 |
| Eco-1881-1 | Bacteria Proteobacteria Gammaproteobacteria Enterobacteriales Enterobacteriaceae *Escherichia coli* P0301904.3 |
| Eco-1882-1 | Bacteria Proteobacteria Gammaproteobacteria Enterobacteriales Enterobacteriaceae *Escherichia coli* P0302293.10 |
| Eco-1883-1 | Bacteria Proteobacteria Gammaproteobacteria Enterobacteriales Enterobacteriaceae *Escherichia coli* P0302293.2 |
| Eco-1884-1 | Bacteria Proteobacteria Gammaproteobacteria Enterobacteriales Enterobacteriaceae *Escherichia coli* P0302293.3 |
| Eco-1885-1 | Bacteria Proteobacteria Gammaproteobacteria Enterobacteriales Enterobacteriaceae *Escherichia coli* P0302293.4 |
| Eco-1886-1 | Bacteria Proteobacteria Gammaproteobacteria Enterobacteriales Enterobacteriaceae *Escherichia coli* P0302293.6 |
| Eco-1887-1 | Bacteria Proteobacteria Gammaproteobacteria Enterobacteriales Enterobacteriaceae *Escherichia coli* P0302293.7 |
| Eco-1888-1 | Bacteria Proteobacteria Gammaproteobacteria Enterobacteriales Enterobacteriaceae *Escherichia coli* P0302293.8 |
| Eco-1889-1 | Bacteria Proteobacteria Gammaproteobacteria Enterobacteriales Enterobacteriaceae *Escherichia coli* P0302293.9 |
| Eco-1890-1 | Bacteria Proteobacteria Gammaproteobacteria Enterobacteriales Enterobacteriaceae *Escherichia coli* P0302308.1 |
| Eco-1891-1 | Bacteria Proteobacteria Gammaproteobacteria Enterobacteriales Enterobacteriaceae *Escherichia coli* P0302308.10 |
| Eco-1892-1 | Bacteria Proteobacteria Gammaproteobacteria Enterobacteriales Enterobacteriaceae *Escherichia coli* P0302308.11 |
| Eco-1893-1 | Bacteria Proteobacteria Gammaproteobacteria Enterobacteriales Enterobacteriaceae *Escherichia coli* P0302308.12 |
| Eco-1894-1 | Bacteria Proteobacteria Gammaproteobacteria Enterobacteriales Enterobacteriaceae *Escherichia coli* P0302308.13 |
| Eco-1895-1 | Bacteria Proteobacteria Gammaproteobacteria Enterobacteriales Enterobacteriaceae *Escherichia coli* P0302308.14 |
| Eco-1896-1 | Bacteria Proteobacteria Gammaproteobacteria Enterobacteriales Enterobacteriaceae *Escherichia coli* P0302308.2 |
| Eco-1897-1 | Bacteria Proteobacteria Gammaproteobacteria Enterobacteriales Enterobacteriaceae *Escherichia coli* P0302308.3 |
| Eco-1898-1 | Bacteria Proteobacteria Gammaproteobacteria Enterobacteriales Enterobacteriaceae *Escherichia coli* P0302308.4 |
| Eco-1899-1 | Bacteria Proteobacteria Gammaproteobacteria Enterobacteriales Enterobacteriaceae *Escherichia coli* P0302308.5 |
| Eco-1900-1 | Bacteria Proteobacteria Gammaproteobacteria Enterobacteriales Enterobacteriaceae *Escherichia coli* P0304777.1 |
| Eco-1901-1 | Bacteria Proteobacteria Gammaproteobacteria Enterobacteriales Enterobacteriaceae *Escherichia coli* P0304777.10 |
| Eco-1902-1 | Bacteria Proteobacteria Gammaproteobacteria Enterobacteriales Enterobacteriaceae *Escherichia coli* P0304777.11 |
| Eco-1903-1 | Bacteria Proteobacteria Gammaproteobacteria Enterobacteriales Enterobacteriaceae *Escherichia coli* P0304777.12 |
| Eco-1904-1 | Bacteria Proteobacteria Gammaproteobacteria Enterobacteriales Enterobacteriaceae *Escherichia coli* P0304777.13 |
| Eco-1905-1 | Bacteria Proteobacteria Gammaproteobacteria Enterobacteriales Enterobacteriaceae *Escherichia coli* P0304777.14 |
| Eco-1906-1 | Bacteria Proteobacteria Gammaproteobacteria Enterobacteriales Enterobacteriaceae *Escherichia coli* P0304777.15 |
| Eco-1907-1 | Bacteria Proteobacteria Gammaproteobacteria Enterobacteriales Enterobacteriaceae *Escherichia coli* P0304777.2 |
| Eco-1908-1 | Bacteria Proteobacteria Gammaproteobacteria Enterobacteriales Enterobacteriaceae *Escherichia coli* P0304777.3 |
| Eco-1909-1 to Eco-1909-2 | Bacteria Proteobacteria Gammaproteobacteria Enterobacteriales Enterobacteriaceae *Escherichia coli* P0304777.4 |
| Eco-1910-1 | Bacteria Proteobacteria Gammaproteobacteria Enterobacteriales Enterobacteriaceae *Escherichia coli* P0304777.5 |
| Eco-1911-1 | Bacteria Proteobacteria Gammaproteobacteria Enterobacteriales Enterobacteriaceae *Escherichia coli* P0304777.7 |
| Eco-1912-1 | Bacteria Proteobacteria Gammaproteobacteria Enterobacteriales Enterobacteriaceae *Escherichia coli* P0304777.8 |
| Eco-1913-1 | Bacteria Proteobacteria Gammaproteobacteria Enterobacteriales Enterobacteriaceae *Escherichia coli* P0304777.9 |
| Eco-1914-1 | Bacteria Proteobacteria Gammaproteobacteria Enterobacteriales Enterobacteriaceae *Escherichia coli* P0304799.3 |
| Eco-1915-1 | Bacteria Proteobacteria Gammaproteobacteria Enterobacteriales Enterobacteriaceae *Escherichia coli* P0304816.1 |
| Eco-1916-1 | Bacteria Proteobacteria Gammaproteobacteria Enterobacteriales Enterobacteriaceae *Escherichia coli* P0304816.10 |
| Eco-1917-1 | Bacteria Proteobacteria Gammaproteobacteria Enterobacteriales Enterobacteriaceae *Escherichia coli* P0304816.11 |
| Eco-1918-1 | Bacteria Proteobacteria Gammaproteobacteria Enterobacteriales Enterobacteriaceae *Escherichia coli* P0304816.12 |

| | |
|---|---|
| Eco-1919-1 | Bacteria Proteobacteria Gammaproteobacteria Enterobacteriales Enterobacteriaceae *Escherichia coli* P0304816.13 |
| Eco-1920-1 | Bacteria Proteobacteria Gammaproteobacteria Enterobacteriales Enterobacteriaceae *Escherichia coli* P0304816.14 |
| Eco-1921-1 | Bacteria Proteobacteria Gammaproteobacteria Enterobacteriales Enterobacteriaceae *Escherichia coli* P0304816.15 |
| Eco-1922-1 | Bacteria Proteobacteria Gammaproteobacteria Enterobacteriales Enterobacteriaceae *Escherichia coli* P0304816.2 |
| Eco-1923-1 | Bacteria Proteobacteria Gammaproteobacteria Enterobacteriales Enterobacteriaceae *Escherichia coli* P0304816.3 |
| Eco-1924-1 | Bacteria Proteobacteria Gammaproteobacteria Enterobacteriales Enterobacteriaceae *Escherichia coli* P0304816.4 |
| Eco-1925-1 | Bacteria Proteobacteria Gammaproteobacteria Enterobacteriales Enterobacteriaceae *Escherichia coli* P0304816.5 |
| Eco-1926-1 | Bacteria Proteobacteria Gammaproteobacteria Enterobacteriales Enterobacteriaceae *Escherichia coli* P0304816.6 |
| Eco-1927-1 | Bacteria Proteobacteria Gammaproteobacteria Enterobacteriales Enterobacteriaceae *Escherichia coli* P0304816.7 |
| Eco-1928-1 | Bacteria Proteobacteria Gammaproteobacteria Enterobacteriales Enterobacteriaceae *Escherichia coli* P0304816.8 |
| Eco-1929-1 | Bacteria Proteobacteria Gammaproteobacteria Enterobacteriales Enterobacteriaceae *Escherichia coli* P0304816.9 |
| Eco-1930-1 | Bacteria Proteobacteria Gammaproteobacteria Enterobacteriales Enterobacteriaceae *Escherichia coli* P0305260.1 |
| Eco-1931-1 | Bacteria Proteobacteria Gammaproteobacteria Enterobacteriales Enterobacteriaceae *Escherichia coli* P0305260.10 |
| Eco-1932-1 | Bacteria Proteobacteria Gammaproteobacteria Enterobacteriales Enterobacteriaceae *Escherichia coli* P0305260.11 |
| Eco-1933-1 | Bacteria Proteobacteria Gammaproteobacteria Enterobacteriales Enterobacteriaceae *Escherichia coli* P0305260.12 |
| Eco-1934-1 | Bacteria Proteobacteria Gammaproteobacteria Enterobacteriales Enterobacteriaceae *Escherichia coli* P0305260.13 |
| Eco-1935-1 | Bacteria Proteobacteria Gammaproteobacteria Enterobacteriales Enterobacteriaceae *Escherichia coli* P0305260.15 |
| Eco-1936-1 | Bacteria Proteobacteria Gammaproteobacteria Enterobacteriales Enterobacteriaceae *Escherichia coli* P0305260.2 |
| Eco-1937-1 | Bacteria Proteobacteria Gammaproteobacteria Enterobacteriales Enterobacteriaceae *Escherichia coli* P0305260.3 |
| Eco-1938-1 | Bacteria Proteobacteria Gammaproteobacteria Enterobacteriales Enterobacteriaceae *Escherichia coli* P0305260.4 |
| Eco-1939-1 | Bacteria Proteobacteria Gammaproteobacteria Enterobacteriales Enterobacteriaceae *Escherichia coli* P0305260.5 |
| Eco-1940-1 | Bacteria Proteobacteria Gammaproteobacteria Enterobacteriales Enterobacteriaceae *Escherichia coli* P0305260.6 |
| Eco-1941-1 | Bacteria Proteobacteria Gammaproteobacteria Enterobacteriales Enterobacteriaceae *Escherichia coli* P0305260.7 |
| Eco-1942-1 | Bacteria Proteobacteria Gammaproteobacteria Enterobacteriales Enterobacteriaceae *Escherichia coli* P0305260.8 |
| Eco-1943-1 | Bacteria Proteobacteria Gammaproteobacteria Enterobacteriales Enterobacteriaceae *Escherichia coli* P0305260.9 |
| Eco-2165-1 | Bacteria Proteobacteria Gammaproteobacteria Enterobacteriales Enterobacteriaceae *Escherichia coli p*0305293.1 |
| Eco-2166-1 | Bacteria Proteobacteria Gammaproteobacteria Enterobacteriales Enterobacteriaceae *Escherichia coli p*0305293.10 |
| Eco-2167-1 | Bacteria Proteobacteria Gammaproteobacteria Enterobacteriales Enterobacteriaceae *Escherichia coli p*0305293.11 |
| Eco-2168-1 | Bacteria Proteobacteria Gammaproteobacteria Enterobacteriales Enterobacteriaceae *Escherichia coli p*0305293.12 |
| Eco-2169-1 | Bacteria Proteobacteria Gammaproteobacteria Enterobacteriales Enterobacteriaceae *Escherichia coli p*0305293.13 |
| Eco-2170-1 | Bacteria Proteobacteria Gammaproteobacteria Enterobacteriales Enterobacteriaceae *Escherichia coli p*0305293.14 |
| Eco-2171-1 | Bacteria Proteobacteria Gammaproteobacteria Enterobacteriales Enterobacteriaceae *Escherichia coli p*0305293.15 |
| Eco-2172-1 | Bacteria Proteobacteria Gammaproteobacteria Enterobacteriales Enterobacteriaceae *Escherichia coli p*0305293.2 |
| Eco-2173-1 | Bacteria Proteobacteria Gammaproteobacteria Enterobacteriales Enterobacteriaceae *Escherichia coli p*0305293.3 |
| Eco-2174-1 | Bacteria Proteobacteria Gammaproteobacteria Enterobacteriales Enterobacteriaceae *Escherichia coli p*0305293.4 |
| Eco-2175-1 | Bacteria Proteobacteria Gammaproteobacteria Enterobacteriales Enterobacteriaceae *Escherichia coli p*0305293.5 |
| Eco-2176-1 | Bacteria Proteobacteria Gammaproteobacteria Enterobacteriales Enterobacteriaceae *Escherichia coli p*0305293.6 |
| Eco-2177-1 | Bacteria Proteobacteria Gammaproteobacteria Enterobacteriales Enterobacteriaceae *Escherichia coli p*0305293.7 |
| Eco-2178-1 | Bacteria Proteobacteria Gammaproteobacteria Enterobacteriales Enterobacteriaceae *Escherichia coli p*0305293.8 |
| Eco-2179-1 | Bacteria Proteobacteria Gammaproteobacteria Enterobacteriales Enterobacteriaceae *Escherichia coli p*0305293.9 |
| Eco-1944-1 | Bacteria Proteobacteria Gammaproteobacteria Enterobacteriales Enterobacteriaceae *Escherichia coli* P12b |
| Eco-1945-1 | Bacteria Proteobacteria Gammaproteobacteria Enterobacteriales Enterobacteriaceae *Escherichia coli* P4-96 |
| Eco-1946-1 | Bacteria Proteobacteria Gammaproteobacteria Enterobacteriales Enterobacteriaceae *Escherichia coli* P4-NR |
| Eco-1947-1 | Bacteria Proteobacteria Gammaproteobacteria Enterobacteriales Enterobacteriaceae *Escherichia coli* PA10 |
| Eco-1948-1 | Bacteria Proteobacteria Gammaproteobacteria Enterobacteriales Enterobacteriaceae *Escherichia coli* PA11 |
| Eco-1949-1 | Bacteria Proteobacteria Gammaproteobacteria Enterobacteriales Enterobacteriaceae *Escherichia coli* PA13 |
| Eco-1950-1 | Bacteria Proteobacteria Gammaproteobacteria Enterobacteriales Enterobacteriaceae *Escherichia coli* PA14 |
| Eco-1951-1 | Bacteria Proteobacteria Gammaproteobacteria Enterobacteriales Enterobacteriaceae *Escherichia coli* PA15 |
| Eco-1952-1 | Bacteria Proteobacteria Gammaproteobacteria Enterobacteriales Enterobacteriaceae *Escherichia coli* PA19 |
| Eco-1953-1 | Bacteria Proteobacteria Gammaproteobacteria Enterobacteriales Enterobacteriaceae *Escherichia coli* PA2 |
| Eco-1954-1 | Bacteria Proteobacteria Gammaproteobacteria Enterobacteriales Enterobacteriaceae *Escherichia coli* PA22 |
| Eco-1955-1 | Bacteria Proteobacteria Gammaproteobacteria Enterobacteriales Enterobacteriaceae *Escherichia coli* PA23 |
| Eco-1956-1 | Bacteria Proteobacteria Gammaproteobacteria Enterobacteriales Enterobacteriaceae *Escherichia coli* PA24 |
| Eco-1957-1 | Bacteria Proteobacteria Gammaproteobacteria Enterobacteriales Enterobacteriaceae *Escherichia coli* PA25 |
| Eco-1958-1 | Bacteria Proteobacteria Gammaproteobacteria Enterobacteriales Enterobacteriaceae *Escherichia coli* PA28 |
| Eco-1959-1 | Bacteria Proteobacteria Gammaproteobacteria Enterobacteriales Enterobacteriaceae *Escherichia coli* PA3 |
| Eco-1960-1 | Bacteria Proteobacteria Gammaproteobacteria Enterobacteriales Enterobacteriaceae *Escherichia coli* PA31 |
| Eco-1961-1 | Bacteria Proteobacteria Gammaproteobacteria Enterobacteriales Enterobacteriaceae *Escherichia coli* PA32 |
| Eco-1962-1 | Bacteria Proteobacteria Gammaproteobacteria Enterobacteriales Enterobacteriaceae *Escherichia coli* PA33 |
| Eco-1963-1 | Bacteria Proteobacteria Gammaproteobacteria Enterobacteriales Enterobacteriaceae *Escherichia coli* PA34 |
| Eco-1964-1 | Bacteria Proteobacteria Gammaproteobacteria Enterobacteriales Enterobacteriaceae *Escherichia coli* PA35 |
| Eco-1965-1 | Bacteria Proteobacteria Gammaproteobacteria Enterobacteriales Enterobacteriaceae *Escherichia coli* PA38 |
| Eco-1966-1 | Bacteria Proteobacteria Gammaproteobacteria Enterobacteriales Enterobacteriaceae *Escherichia coli* PA39 |
| Eco-1967-1 | Bacteria Proteobacteria Gammaproteobacteria Enterobacteriales Enterobacteriaceae *Escherichia coli* PA4 |
| Eco-1968-1 | Bacteria Proteobacteria Gammaproteobacteria Enterobacteriales Enterobacteriaceae *Escherichia coli* PA40 |

| | |
|---|---|
| Eco-1969-1 | Bacteria Proteobacteria Gammaproteobacteria Enterobacteriales Enterobacteriaceae *Escherichia coli* PA41 |
| Eco-1970-1 | Bacteria Proteobacteria Gammaproteobacteria Enterobacteriales Enterobacteriaceae *Escherichia coli* PA42 |
| Eco-1971-1 | Bacteria Proteobacteria Gammaproteobacteria Enterobacteriales Enterobacteriaceae *Escherichia coli* PA45 |
| Eco-1972-1 | Bacteria Proteobacteria Gammaproteobacteria Enterobacteriales Enterobacteriaceae *Escherichia coli* PA47 |
| Eco-1973-1 | Bacteria Proteobacteria Gammaproteobacteria Enterobacteriales Enterobacteriaceae *Escherichia coli* PA48 |
| Eco-1974-1 | Bacteria Proteobacteria Gammaproteobacteria Enterobacteriales Enterobacteriaceae *Escherichia coli* PA49 |
| Eco-1975-1 | Bacteria Proteobacteria Gammaproteobacteria Enterobacteriales Enterobacteriaceae *Escherichia coli* PA5 |
| Eco-1976-1 | Bacteria Proteobacteria Gammaproteobacteria Enterobacteriales Enterobacteriaceae *Escherichia coli* PA7 |
| Eco-1977-1 | Bacteria Proteobacteria Gammaproteobacteria Enterobacteriales Enterobacteriaceae *Escherichia coli* PA8 |
| Eco-1978-1 | Bacteria Proteobacteria Gammaproteobacteria Enterobacteriales Enterobacteriaceae *Escherichia coli* PA9 |
| Eco-1979-1 | Bacteria Proteobacteria Gammaproteobacteria Enterobacteriales Enterobacteriaceae *Escherichia coli* PMV-1 |
| Eco-1980-1 | Bacteria Proteobacteria Gammaproteobacteria Enterobacteriales Enterobacteriaceae *Escherichia coli* RN587/1 |
| Eco-1981-1 | Bacteria Proteobacteria Gammaproteobacteria Enterobacteriales Enterobacteriaceae *Escherichia coli* S17 |
| Eco-1982-1 | Bacteria Proteobacteria Gammaproteobacteria Enterobacteriales Enterobacteriaceae *Escherichia coli* SCD1 |
| Eco-1983-1 | Bacteria Proteobacteria Gammaproteobacteria Enterobacteriales Enterobacteriaceae *Escherichia coli* SCD2 |
| Eco-1984-1 | Bacteria Proteobacteria Gammaproteobacteria Enterobacteriales Enterobacteriaceae *Escherichia coli* SCI-07 |
| Eco-1985-1 | Bacteria Proteobacteria Gammaproteobacteria Enterobacteriales Enterobacteriaceae *Escherichia coli* SE11 |
| Eco-1986-1 | Bacteria Proteobacteria Gammaproteobacteria Enterobacteriales Enterobacteriaceae *Escherichia coli* SE15 |
| Eco-1987-1 | Bacteria Proteobacteria Gammaproteobacteria Enterobacteriales Enterobacteriaceae *Escherichia coli* SEPT362 |
| Eco-1988-1 | Bacteria Proteobacteria Gammaproteobacteria Enterobacteriales Enterobacteriaceae *Escherichia coli* SMS-3-5 |
| Eco-1989-1 | Bacteria Proteobacteria Gammaproteobacteria Enterobacteriales Enterobacteriaceae *Escherichia coli* STEC_7v |
| Eco-1990-1 | Bacteria Proteobacteria Gammaproteobacteria Enterobacteriales Enterobacteriaceae *Escherichia coli* STEC_B2F1 |
| Eco-1991-1 | Bacteria Proteobacteria Gammaproteobacteria Enterobacteriales Enterobacteriaceae *Escherichia coli* STEC_C165-02 |
| Eco-1992-1 | Bacteria Proteobacteria Gammaproteobacteria Enterobacteriales Enterobacteriaceae *Escherichia coli* STEC_EH250 |
| Eco-1993-1 | Bacteria Proteobacteria Gammaproteobacteria Enterobacteriales Enterobacteriaceae *Escherichia coli* STEC_MHI813 |
| Eco-1994-1 | Bacteria Proteobacteria Gammaproteobacteria Enterobacteriales Enterobacteriaceae *Escherichia coli* STEC_O31 |
| Eco-1995-1 | Bacteria Proteobacteria Gammaproteobacteria Enterobacteriales Enterobacteriaceae *Escherichia coli* STEC_S1191 |
| Eco-2180-1 | Bacteria Proteobacteria Gammaproteobacteria Enterobacteriales Enterobacteriaceae *Escherichia coli str. 'clone* D i14' |
| Eco-2181-1 | Bacteria Proteobacteria Gammaproteobacteria Enterobacteriales Enterobacteriaceae *Escherichia coli str. 'clone* D i2' |
| Eco-2182-1 | Bacteria Proteobacteria Gammaproteobacteria Enterobacteriales Enterobacteriaceae *Escherichia coli str.* K-12 substr. DH10B |
| Eco-2183-1 | Bacteria Proteobacteria Gammaproteobacteria Enterobacteriales Enterobacteriaceae *Escherichia coli str.* K-12 substr. MDS42 |
| Eco-752-1 to Eco-752-2 | Bacteria Proteobacteria Gammaproteobacteria Enterobacteriales Enterobacteriaceae *Escherichia coli str.* K-12 substr. MG1655 |
| Eco-2184-1 | Bacteria Proteobacteria Gammaproteobacteria Enterobacteriales Enterobacteriaceae *Escherichia coli str.* K-12 substr. MG1655star |
| Eco-2185-1 | Bacteria Proteobacteria Gammaproteobacteria Enterobacteriales Enterobacteriaceae *Escherichia coli str.* K-12 substr. W3110 |
| Eco-1996-1 | Bacteria Proteobacteria Gammaproteobacteria Enterobacteriales Enterobacteriaceae *Escherichia coli* SWW33 |
| Eco-1997-1 | Bacteria Proteobacteria Gammaproteobacteria Enterobacteriales Enterobacteriaceae *Escherichia coli* T1282_01 |
| Eco-1998-1 | Bacteria Proteobacteria Gammaproteobacteria Enterobacteriales Enterobacteriaceae *Escherichia coli* T1840_97 |
| Eco-1999-1 | Bacteria Proteobacteria Gammaproteobacteria Enterobacteriales Enterobacteriaceae *Escherichia coli* T234_00 |
| Eco-2000-1 | Bacteria Proteobacteria Gammaproteobacteria Enterobacteriales Enterobacteriaceae *Escherichia coli* T924_01 |
| Eco-2001-1 | Bacteria Proteobacteria Gammaproteobacteria Enterobacteriales Enterobacteriaceae *Escherichia coli* TA124 |
| Eco-2002-1 | Bacteria Proteobacteria Gammaproteobacteria Enterobacteriales Enterobacteriaceae *Escherichia coli* TA143 |
| Eco-2003-1 | Bacteria Proteobacteria Gammaproteobacteria Enterobacteriales Enterobacteriaceae *Escherichia coli* TA206 |
| Eco-2004-1 | Bacteria Proteobacteria Gammaproteobacteria Enterobacteriales Enterobacteriaceae *Escherichia coli* TA271 |
| Eco-2043-1 | Bacteria Proteobacteria Gammaproteobacteria Enterobacteriales Enterobacteriaceae *Escherichia coli* ThroopD |
| Eco-2005-1 to Eco-2005-2 | Bacteria Proteobacteria Gammaproteobacteria Enterobacteriales Enterobacteriaceae *Escherichia coli* TOP2386 |
| Eco-2006-1 | Bacteria Proteobacteria Gammaproteobacteria Enterobacteriales Enterobacteriaceae *Escherichia coli* TOP2396-1 |
| Eco-2007-1 | Bacteria Proteobacteria Gammaproteobacteria Enterobacteriales Enterobacteriaceae *Escherichia coli* TOP2396-2 |
| Eco-2008-1 | Bacteria Proteobacteria Gammaproteobacteria Enterobacteriales Enterobacteriaceae *Escherichia coli* TOP2396-3 |
| Eco-2009-1 | Bacteria Proteobacteria Gammaproteobacteria Enterobacteriales Enterobacteriaceae *Escherichia coli* TOP2522-1 |
| Eco-2010-1 to Eco-2010-2 | Bacteria Proteobacteria Gammaproteobacteria Enterobacteriales Enterobacteriaceae *Escherichia coli* TOP2652 |
| Eco-2011-1 | Bacteria Proteobacteria Gammaproteobacteria Enterobacteriales Enterobacteriaceae *Escherichia coli* TOP2662-1 |
| Eco-2012-1 | Bacteria Proteobacteria Gammaproteobacteria Enterobacteriales Enterobacteriaceae *Escherichia coli* TOP2662-2 |
| Eco-2013-1 | Bacteria Proteobacteria Gammaproteobacteria Enterobacteriales Enterobacteriaceae *Escherichia coli* TOP2662-3 |
| Eco-2014-1 | Bacteria Proteobacteria Gammaproteobacteria Enterobacteriales Enterobacteriaceae *Escherichia coli* TOP2662-4 |
| Eco-2015-1 | Bacteria Proteobacteria Gammaproteobacteria Enterobacteriales Enterobacteriaceae *Escherichia coli* TOP382-1 |
| Eco-2016-1 | Bacteria Proteobacteria Gammaproteobacteria Enterobacteriales Enterobacteriaceae *Escherichia coli* TOP382-2 |
| Eco-2017-1 | Bacteria Proteobacteria Gammaproteobacteria Enterobacteriales Enterobacteriaceae *Escherichia coli* TOP382-3 |
| Eco-2018-1 | Bacteria Proteobacteria Gammaproteobacteria Enterobacteriales Enterobacteriaceae *Escherichia coli* TOP550-2 |
| Eco-2019-1 | Bacteria Proteobacteria Gammaproteobacteria Enterobacteriales Enterobacteriaceae *Escherichia coli* TOP550-3 |
| Eco-2020-1 | Bacteria Proteobacteria Gammaproteobacteria Enterobacteriales Enterobacteriaceae *Escherichia coli* TOP550-4 |
| Eco-2021-1 | Bacteria Proteobacteria Gammaproteobacteria Enterobacteriales Enterobacteriaceae *Escherichia coli* TT12B |
| Eco-2022-1 | Bacteria Proteobacteria Gammaproteobacteria Enterobacteriales Enterobacteriaceae *Escherichia coli* TW00353 |
| Eco-2023-1 | Bacteria Proteobacteria Gammaproteobacteria Enterobacteriales Enterobacteriaceae *Escherichia coli* TW06591 |
| Eco-2024-1 | Bacteria Proteobacteria Gammaproteobacteria Enterobacteriales Enterobacteriaceae *Escherichia coli* TW07509 |
| Eco-2025-1 | Bacteria Proteobacteria Gammaproteobacteria Enterobacteriales Enterobacteriaceae *Escherichia coli* TW07793 |

| | |
|---|---|
| Eco-2026-1 | Bacteria Proteobacteria Gammaproteobacteria Enterobacteriales Enterobacteriaceae *Escherichia coli* TW07945 |
| Eco-2027-1 | Bacteria Proteobacteria Gammaproteobacteria Enterobacteriales Enterobacteriaceae *Escherichia coli* TW09098 |
| Eco-2028-1 | Bacteria Proteobacteria Gammaproteobacteria Enterobacteriales Enterobacteriaceae *Escherichia coli* TW09109 |
| Eco-2029-1 | Bacteria Proteobacteria Gammaproteobacteria Enterobacteriales Enterobacteriaceae *Escherichia coli* TW09195 |
| Eco-2030-1 | Bacteria Proteobacteria Gammaproteobacteria Enterobacteriales Enterobacteriaceae *Escherichia coli* TW10119 |
| Eco-2031-1 | Bacteria Proteobacteria Gammaproteobacteria Enterobacteriales Enterobacteriaceae *Escherichia coli* TW10246 |
| Eco-2032-1 | Bacteria Proteobacteria Gammaproteobacteria Enterobacteriales Enterobacteriaceae *Escherichia coli* TW10509 |
| Eco-2033-1 | Bacteria Proteobacteria Gammaproteobacteria Enterobacteriales Enterobacteriaceae *Escherichia coli* TW10598 |
| Eco-2034-1 | Bacteria Proteobacteria Gammaproteobacteria Enterobacteriales Enterobacteriaceae *Escherichia coli* TW10722 |
| Eco-2035-1 | Bacteria Proteobacteria Gammaproteobacteria Enterobacteriales Enterobacteriaceae *Escherichia coli* TW10828 |
| Eco-2036-1 | Bacteria Proteobacteria Gammaproteobacteria Enterobacteriales Enterobacteriaceae *Escherichia coli* TW11039 |
| Eco-2037-1 | Bacteria Proteobacteria Gammaproteobacteria Enterobacteriales Enterobacteriaceae *Escherichia coli* TW11681 |
| Eco-2038-1 | Bacteria Proteobacteria Gammaproteobacteria Enterobacteriales Enterobacteriaceae *Escherichia coli* TW14301 |
| Eco-2039-1 | Bacteria Proteobacteria Gammaproteobacteria Enterobacteriales Enterobacteriaceae *Escherichia coli* TW14313 |
| Eco-2040-1 | Bacteria Proteobacteria Gammaproteobacteria Enterobacteriales Enterobacteriaceae *Escherichia coli* TW14425 |
| Eco-2041-1 | Bacteria Proteobacteria Gammaproteobacteria Enterobacteriales Enterobacteriaceae *Escherichia coli* TW15901 |
| Eco-2044-1 | Bacteria Proteobacteria Gammaproteobacteria Enterobacteriales Enterobacteriaceae *Escherichia coli* Tx1686 |
| Eco-2042-1 | Bacteria Proteobacteria Gammaproteobacteria Enterobacteriales Enterobacteriaceae *Escherichia coli* TX1999 |
| Eco-2045-1 | Bacteria Proteobacteria Gammaproteobacteria Enterobacteriales Enterobacteriaceae *Escherichia coli* Tx3800 |
| Eco-2046-1 | Bacteria Proteobacteria Gammaproteobacteria Enterobacteriales Enterobacteriaceae *Escherichia coli* UM146 |
| Eco-2047-1 | Bacteria Proteobacteria Gammaproteobacteria Enterobacteriales Enterobacteriaceae *Escherichia coli* UMEA 3014-1 |
| Eco-2048-1 | Bacteria Proteobacteria Gammaproteobacteria Enterobacteriales Enterobacteriaceae *Escherichia coli* UMEA 3022-1 |
| Eco-2049-1 | Bacteria Proteobacteria Gammaproteobacteria Enterobacteriales Enterobacteriaceae *Escherichia coli* UMEA 3033-1 |
| Eco-2050-1 | Bacteria Proteobacteria Gammaproteobacteria Enterobacteriales Enterobacteriaceae *Escherichia coli* UMEA 3041-1 |
| Eco-2051-1 | Bacteria Proteobacteria Gammaproteobacteria Enterobacteriales Enterobacteriaceae *Escherichia coli* UMEA 3052-1 |
| Eco-2052-1 | Bacteria Proteobacteria Gammaproteobacteria Enterobacteriales Enterobacteriaceae *Escherichia coli* UMEA 3053-1 |
| Eco-2053-1 | Bacteria Proteobacteria Gammaproteobacteria Enterobacteriales Enterobacteriaceae *Escherichia coli* UMEA 3065-1 |
| Eco-2054-1 | Bacteria Proteobacteria Gammaproteobacteria Enterobacteriales Enterobacteriaceae *Escherichia coli* UMEA 3087-1 |
| Eco-2055-1 | Bacteria Proteobacteria Gammaproteobacteria Enterobacteriales Enterobacteriaceae *Escherichia coli* UMEA 3088-1 |
| Eco-2056-1 | Bacteria Proteobacteria Gammaproteobacteria Enterobacteriales Enterobacteriaceae *Escherichia coli* UMEA 3097-1 |
| Eco-2057-1 | Bacteria Proteobacteria Gammaproteobacteria Enterobacteriales Enterobacteriaceae *Escherichia coli* UMEA 3108-1 |
| Eco-2058-1 | Bacteria Proteobacteria Gammaproteobacteria Enterobacteriales Enterobacteriaceae *Escherichia coli* UMEA 3113-1 |
| Eco-2059-1 | Bacteria Proteobacteria Gammaproteobacteria Enterobacteriales Enterobacteriaceae *Escherichia coli* UMEA 3117-1 |
| Eco-2060-1 | Bacteria Proteobacteria Gammaproteobacteria Enterobacteriales Enterobacteriaceae *Escherichia coli* UMEA 3121-1 |
| Eco-2061-1 | Bacteria Proteobacteria Gammaproteobacteria Enterobacteriales Enterobacteriaceae *Escherichia coli* UMEA 3122-1 |
| Eco-2062-1 | Bacteria Proteobacteria Gammaproteobacteria Enterobacteriales Enterobacteriaceae *Escherichia coli* UMEA 3124-1 |
| Eco-2063-1 | Bacteria Proteobacteria Gammaproteobacteria Enterobacteriales Enterobacteriaceae *Escherichia coli* UMEA 3139-1 |
| Eco-2064-1 | Bacteria Proteobacteria Gammaproteobacteria Enterobacteriales Enterobacteriaceae *Escherichia coli* UMEA 3140-1 |
| Eco-2065-1 | Bacteria Proteobacteria Gammaproteobacteria Enterobacteriales Enterobacteriaceae *Escherichia coli* UMEA 3144-1 |
| Eco-2066-1 | Bacteria Proteobacteria Gammaproteobacteria Enterobacteriales Enterobacteriaceae *Escherichia coli* UMEA 3148-1 |
| Eco-2067-1 | Bacteria Proteobacteria Gammaproteobacteria Enterobacteriales Enterobacteriaceae *Escherichia coli* UMEA 3150-1 |
| Eco-2068-1 | Bacteria Proteobacteria Gammaproteobacteria Enterobacteriales Enterobacteriaceae *Escherichia coli* UMEA 3151-1 |
| Eco-2069-1 | Bacteria Proteobacteria Gammaproteobacteria Enterobacteriales Enterobacteriaceae *Escherichia coli* UMEA 3152-1 |
| Eco-2070-1 | Bacteria Proteobacteria Gammaproteobacteria Enterobacteriales Enterobacteriaceae *Escherichia coli* UMEA 3155-1 |
| Eco-2071-1 | Bacteria Proteobacteria Gammaproteobacteria Enterobacteriales Enterobacteriaceae *Escherichia coli* UMEA 3159-1 |
| Eco-2072-1 | Bacteria Proteobacteria Gammaproteobacteria Enterobacteriales Enterobacteriaceae *Escherichia coli* UMEA 3160-1 |
| Eco-2073-1 | Bacteria Proteobacteria Gammaproteobacteria Enterobacteriales Enterobacteriaceae *Escherichia coli* UMEA 3161-1 |
| Eco-2074-1 | Bacteria Proteobacteria Gammaproteobacteria Enterobacteriales Enterobacteriaceae *Escherichia coli* UMEA 3162-1 |
| Eco-2075-1 | Bacteria Proteobacteria Gammaproteobacteria Enterobacteriales Enterobacteriaceae *Escherichia coli* UMEA 3163-1 |
| Eco-2076-1 | Bacteria Proteobacteria Gammaproteobacteria Enterobacteriales Enterobacteriaceae *Escherichia coli* UMEA 3172-1 |
| Eco-2077-1 | Bacteria Proteobacteria Gammaproteobacteria Enterobacteriales Enterobacteriaceae *Escherichia coli* UMEA 3173-1 |
| Eco-2078-1 | Bacteria Proteobacteria Gammaproteobacteria Enterobacteriales Enterobacteriaceae *Escherichia coli* UMEA 3174-1 |
| Eco-2079-1 | Bacteria Proteobacteria Gammaproteobacteria Enterobacteriales Enterobacteriaceae *Escherichia coli* UMEA 3175-1 |
| Eco-2080-1 | Bacteria Proteobacteria Gammaproteobacteria Enterobacteriales Enterobacteriaceae *Escherichia coli* UMEA 3176-1 |
| Eco-2081-1 | Bacteria Proteobacteria Gammaproteobacteria Enterobacteriales Enterobacteriaceae *Escherichia coli* UMEA 3178-1 |
| Eco-2082-1 | Bacteria Proteobacteria Gammaproteobacteria Enterobacteriales Enterobacteriaceae *Escherichia coli* UMEA 3180-1 |
| Eco-2083-1 | Bacteria Proteobacteria Gammaproteobacteria Enterobacteriales Enterobacteriaceae *Escherichia coli* UMEA 3185-1 |
| Eco-2084-1 | Bacteria Proteobacteria Gammaproteobacteria Enterobacteriales Enterobacteriaceae *Escherichia coli* UMEA 3190-1 |
| Eco-2085-1 | Bacteria Proteobacteria Gammaproteobacteria Enterobacteriales Enterobacteriaceae *Escherichia coli* UMEA 3193-1 |
| Eco-2086-1 | Bacteria Proteobacteria Gammaproteobacteria Enterobacteriales Enterobacteriaceae *Escherichia coli* UMEA 3199-1 |
| Eco-2087-1 | Bacteria Proteobacteria Gammaproteobacteria Enterobacteriales Enterobacteriaceae *Escherichia coli* UMEA 3200-1 |
| Eco-2088-1 | Bacteria Proteobacteria Gammaproteobacteria Enterobacteriales Enterobacteriaceae *Escherichia coli* UMEA 3201-1 |
| Eco-2089-1 | Bacteria Proteobacteria Gammaproteobacteria Enterobacteriales Enterobacteriaceae *Escherichia coli* UMEA 3203-1 |
| Eco-2090-1 | Bacteria Proteobacteria Gammaproteobacteria Enterobacteriales Enterobacteriaceae *Escherichia coli* UMEA 3206-1 |
| Eco-2091-1 | Bacteria Proteobacteria Gammaproteobacteria Enterobacteriales Enterobacteriaceae *Escherichia coli* UMEA 3208-1 |

| | |
|---|---|
| Eco-2092-1 | Bacteria Proteobacteria Gammaproteobacteria Enterobacteriales Enterobacteriaceae *Escherichia coli* UMEA 3212-1 |
| Eco-2093-1 | Bacteria Proteobacteria Gammaproteobacteria Enterobacteriales Enterobacteriaceae *Escherichia coli* UMEA 3215-1 |
| Eco-2094-1 | Bacteria Proteobacteria Gammaproteobacteria Enterobacteriales Enterobacteriaceae *Escherichia coli* UMEA 3216-1 |
| Eco-2095-1 | Bacteria Proteobacteria Gammaproteobacteria Enterobacteriales Enterobacteriaceae *Escherichia coli* UMEA 3217-1 |
| Eco-2096-1 | Bacteria Proteobacteria Gammaproteobacteria Enterobacteriales Enterobacteriaceae *Escherichia coli* UMEA 3220-1 |
| Eco-2097-1 | Bacteria Proteobacteria Gammaproteobacteria Enterobacteriales Enterobacteriaceae *Escherichia coli* UMEA 3221-1 |
| Eco-2098-1 | Bacteria Proteobacteria Gammaproteobacteria Enterobacteriales Enterobacteriaceae *Escherichia coli* UMEA 3222-1 |
| Eco-2099-1 | Bacteria Proteobacteria Gammaproteobacteria Enterobacteriales Enterobacteriaceae *Escherichia coli* UMEA 3230-1 |
| Eco-2100-1 | Bacteria Proteobacteria Gammaproteobacteria Enterobacteriales Enterobacteriaceae *Escherichia coli* UMEA 3233-1 |
| Eco-2101-1 | Bacteria Proteobacteria Gammaproteobacteria Enterobacteriales Enterobacteriaceae *Escherichia coli* UMEA 3240-1 |
| Eco-2102-1 | Bacteria Proteobacteria Gammaproteobacteria Enterobacteriales Enterobacteriaceae *Escherichia coli* UMEA 3244-1 |
| Eco-2103-1 | Bacteria Proteobacteria Gammaproteobacteria Enterobacteriales Enterobacteriaceae *Escherichia coli* UMEA 3257-1 |
| Eco-2104-1 | Bacteria Proteobacteria Gammaproteobacteria Enterobacteriales Enterobacteriaceae *Escherichia coli* UMEA 3264-1 |
| Eco-2105-1 | Bacteria Proteobacteria Gammaproteobacteria Enterobacteriales Enterobacteriaceae *Escherichia coli* UMEA 3268-1 |
| Eco-2106-1 | Bacteria Proteobacteria Gammaproteobacteria Enterobacteriales Enterobacteriaceae *Escherichia coli* UMEA 3271-1 |
| Eco-2107-1 | Bacteria Proteobacteria Gammaproteobacteria Enterobacteriales Enterobacteriaceae *Escherichia coli* UMEA 3290-1 |
| Eco-2108-1 | Bacteria Proteobacteria Gammaproteobacteria Enterobacteriales Enterobacteriaceae *Escherichia coli* UMEA 3292-1 |
| Eco-2109-1 | Bacteria Proteobacteria Gammaproteobacteria Enterobacteriales Enterobacteriaceae *Escherichia coli* UMEA 3298-1 |
| Eco-2110-1 | Bacteria Proteobacteria Gammaproteobacteria Enterobacteriales Enterobacteriaceae *Escherichia coli* UMEA 3304-1 |
| Eco-2111-1 | Bacteria Proteobacteria Gammaproteobacteria Enterobacteriales Enterobacteriaceae *Escherichia coli* UMEA 3314-1 |
| Eco-2112-1 | Bacteria Proteobacteria Gammaproteobacteria Enterobacteriales Enterobacteriaceae *Escherichia coli* UMEA 3317-1 |
| Eco-2113-1 | Bacteria Proteobacteria Gammaproteobacteria Enterobacteriales Enterobacteriaceae *Escherichia coli* UMEA 3318-1 |
| Eco-2114-1 | Bacteria Proteobacteria Gammaproteobacteria Enterobacteriales Enterobacteriaceae *Escherichia coli* UMEA 3323-1 |
| Eco-2115-1 | Bacteria Proteobacteria Gammaproteobacteria Enterobacteriales Enterobacteriaceae *Escherichia coli* UMEA 3329-1 |
| Eco-2116-1 | Bacteria Proteobacteria Gammaproteobacteria Enterobacteriales Enterobacteriaceae *Escherichia coli* UMEA 3336-1 |
| Eco-2117-1 | Bacteria Proteobacteria Gammaproteobacteria Enterobacteriales Enterobacteriaceae *Escherichia coli* UMEA 3337-1 |
| Eco-2118-1 | Bacteria Proteobacteria Gammaproteobacteria Enterobacteriales Enterobacteriaceae *Escherichia coli* UMEA 3341-1 |
| Eco-2119-1 | Bacteria Proteobacteria Gammaproteobacteria Enterobacteriales Enterobacteriaceae *Escherichia coli* UMEA 3342-1 |
| Eco-2120-1 | Bacteria Proteobacteria Gammaproteobacteria Enterobacteriales Enterobacteriaceae *Escherichia coli* UMEA 3355-1 |
| Eco-2121-1 | Bacteria Proteobacteria Gammaproteobacteria Enterobacteriales Enterobacteriaceae *Escherichia coli* UMEA 3391-1 |
| Eco-2122-1 | Bacteria Proteobacteria Gammaproteobacteria Enterobacteriales Enterobacteriaceae *Escherichia coli* UMEA 3426-1 |
| Eco-2123-1 | Bacteria Proteobacteria Gammaproteobacteria Enterobacteriales Enterobacteriaceae *Escherichia coli* UMEA 3489-1 |
| Eco-2124-1 | Bacteria Proteobacteria Gammaproteobacteria Enterobacteriales Enterobacteriaceae *Escherichia coli* UMEA 3490-1 |
| Eco-2125-1 | Bacteria Proteobacteria Gammaproteobacteria Enterobacteriales Enterobacteriaceae *Escherichia coli* UMEA 3585-1 |
| Eco-2126-1 | Bacteria Proteobacteria Gammaproteobacteria Enterobacteriales Enterobacteriaceae *Escherichia coli* UMEA 3592-1 |
| Eco-2127-1 | Bacteria Proteobacteria Gammaproteobacteria Enterobacteriales Enterobacteriaceae *Escherichia coli* UMEA 3609-1 |
| Eco-2128-1 | Bacteria Proteobacteria Gammaproteobacteria Enterobacteriales Enterobacteriaceae *Escherichia coli* UMEA 3617-1 |
| Eco-2129-1 | Bacteria Proteobacteria Gammaproteobacteria Enterobacteriales Enterobacteriaceae *Escherichia coli* UMEA 3632-1 |
| Eco-2130-1 | Bacteria Proteobacteria Gammaproteobacteria Enterobacteriales Enterobacteriaceae *Escherichia coli* UMEA 3652-1 |
| Eco-2131-1 | Bacteria Proteobacteria Gammaproteobacteria Enterobacteriales Enterobacteriaceae *Escherichia coli* UMEA 3656-1 |
| Eco-2132-1 | Bacteria Proteobacteria Gammaproteobacteria Enterobacteriales Enterobacteriaceae *Escherichia coli* UMEA 3662-1 |
| Eco-2133-1 | Bacteria Proteobacteria Gammaproteobacteria Enterobacteriales Enterobacteriaceae *Escherichia coli* UMEA 3671-1 |
| Eco-2134-1 | Bacteria Proteobacteria Gammaproteobacteria Enterobacteriales Enterobacteriaceae *Escherichia coli* UMEA 3682-1 |
| Eco-2135-1 | Bacteria Proteobacteria Gammaproteobacteria Enterobacteriales Enterobacteriaceae *Escherichia coli* UMEA 3687-1 |
| Eco-2136-1 | Bacteria Proteobacteria Gammaproteobacteria Enterobacteriales Enterobacteriaceae *Escherichia coli* UMEA 3693-1 |
| Eco-2137-1 | Bacteria Proteobacteria Gammaproteobacteria Enterobacteriales Enterobacteriaceae *Escherichia coli* UMEA 3694-1 |
| Eco-2138-1 | Bacteria Proteobacteria Gammaproteobacteria Enterobacteriales Enterobacteriaceae *Escherichia coli* UMEA 3702-1 |
| Eco-2139-1 | Bacteria Proteobacteria Gammaproteobacteria Enterobacteriales Enterobacteriaceae *Escherichia coli* UMEA 3703-1 |
| Eco-2140-1 | Bacteria Proteobacteria Gammaproteobacteria Enterobacteriales Enterobacteriaceae *Escherichia coli* UMEA 3705-1 |
| Eco-2141-1 | Bacteria Proteobacteria Gammaproteobacteria Enterobacteriales Enterobacteriaceae *Escherichia coli* UMEA 3707-1 |
| Eco-2142-1 | Bacteria Proteobacteria Gammaproteobacteria Enterobacteriales Enterobacteriaceae *Escherichia coli* UMEA 3718-1 |
| Eco-2143-1 | Bacteria Proteobacteria Gammaproteobacteria Enterobacteriales Enterobacteriaceae *Escherichia coli* UMEA 3805-1 |
| Eco-2144-1 | Bacteria Proteobacteria Gammaproteobacteria Enterobacteriales Enterobacteriaceae *Escherichia coli* UMEA 3821-1 |
| Eco-2145-1 | Bacteria Proteobacteria Gammaproteobacteria Enterobacteriales Enterobacteriaceae *Escherichia coli* UMEA 3834-1 |
| Eco-2146-1 | Bacteria Proteobacteria Gammaproteobacteria Enterobacteriales Enterobacteriaceae *Escherichia coli* UMEA 3889-1 |
| Eco-2147-1 | Bacteria Proteobacteria Gammaproteobacteria Enterobacteriales Enterobacteriaceae *Escherichia coli* UMEA 3893-1 |
| Eco-2148-1 | Bacteria Proteobacteria Gammaproteobacteria Enterobacteriales Enterobacteriaceae *Escherichia coli* UMEA 3899-1 |
| Eco-2149-1 | Bacteria Proteobacteria Gammaproteobacteria Enterobacteriales Enterobacteriaceae *Escherichia coli* UMEA 3955-1 |
| Eco-2150-1 | Bacteria Proteobacteria Gammaproteobacteria Enterobacteriales Enterobacteriaceae *Escherichia coli* UMEA 4075-1 |
| Eco-2151-1 | Bacteria Proteobacteria Gammaproteobacteria Enterobacteriales Enterobacteriaceae *Escherichia coli* UMEA 4076-1 |
| Eco-2152-1 | Bacteria Proteobacteria Gammaproteobacteria Enterobacteriales Enterobacteriaceae *Escherichia coli* UMEA 4207-1 |
| Eco-2153-1 | Bacteria Proteobacteria Gammaproteobacteria Enterobacteriales Enterobacteriaceae *Escherichia coli* UMN026 |
| Eco-2154-1 | Bacteria Proteobacteria Gammaproteobacteria Enterobacteriales Enterobacteriaceae *Escherichia coli* UMNF18 |
| Eco-2155-1 | Bacteria Proteobacteria Gammaproteobacteria Enterobacteriales Enterobacteriaceae *Escherichia coli* UMNK88 |
| Eco-2156-1 | Bacteria Proteobacteria Gammaproteobacteria Enterobacteriales Enterobacteriaceae *Escherichia coli* UTI89 |

| | |
|---|---|
| Eco-2157-1 to Eco-2157-2 | Bacteria Proteobacteria Gammaproteobacteria Enterobacteriales Enterobacteriaceae *Escherichia coli* W |
| Eco-2158-1 | Bacteria Proteobacteria Gammaproteobacteria Enterobacteriales Enterobacteriaceae *Escherichia coli* W26 |
| Eco-2159-1 | Bacteria Proteobacteria Gammaproteobacteria Enterobacteriales Enterobacteriaceae *Escherichia coli* WV_060327 |
| Eco-2160-1 | Bacteria Proteobacteria Gammaproteobacteria Enterobacteriales Enterobacteriaceae *Escherichia coli* XH001 |
| Eco-2161-1 | Bacteria Proteobacteria Gammaproteobacteria Enterobacteriales Enterobacteriaceae *Escherichia coli* XH140A |
| Eco-2162-1 | Bacteria Proteobacteria Gammaproteobacteria Enterobacteriales Enterobacteriaceae *Escherichia coli* Xuzhou21 |
| Efe-1-1 to Efe-1-3 | Bacteria Proteobacteria Gammaproteobacteria Enterobacteriales Enterobacteriaceae *Escherichia fergusonii* |
| Efe-2-1 | Bacteria Proteobacteria Gammaproteobacteria Enterobacteriales Enterobacteriaceae *Escherichia fergusonii* ATCC 35469 |
| Efe-3-1 | Bacteria Proteobacteria Gammaproteobacteria Enterobacteriales Enterobacteriaceae *Escherichia fergusonii* B253 |
| Efe-4-1 | Bacteria Proteobacteria Gammaproteobacteria Enterobacteriales Enterobacteriaceae *Escherichia fergusonii* ECD227 |
| Ehe-1-1 | Bacteria Proteobacteria Gammaproteobacteria Enterobacteriales Enterobacteriaceae *Escherichia hermannii* NBRC 105704 |
| Esp-75-1 | Bacteria Proteobacteria Gammaproteobacteria Enterobacteriales Enterobacteriaceae *Escherichia sp.* 1_1_43 |
| Esp-76-1 | Bacteria Proteobacteria Gammaproteobacteria Enterobacteriales Enterobacteriaceae *Escherichia sp.* 4_1_40B |
| Esp-77-1 | Bacteria Proteobacteria Gammaproteobacteria Enterobacteriales Enterobacteriaceae *Escherichia sp.* TW09231 |
| Esp-78-1 | Bacteria Proteobacteria Gammaproteobacteria Enterobacteriales Enterobacteriaceae *Escherichia sp.* TW09276 |
| Esp-79-1 | Bacteria Proteobacteria Gammaproteobacteria Enterobacteriales Enterobacteriaceae *Escherichia sp.* TW09308 |
| Esp-80-1 | Bacteria Proteobacteria Gammaproteobacteria Enterobacteriales Enterobacteriaceae *Escherichia sp.* TW10509 |
| Hal-4-1 | Bacteria Proteobacteria Gammaproteobacteria Enterobacteriales Enterobacteriaceae *Hafnia alvei* ATCC 51873 |
| Kox-1-1 to Kox-1-4 | Bacteria Proteobacteria Gammaproteobacteria Enterobacteriales Enterobacteriaceae *Klebsiella oxytoca* |
| Kox-11-1 | Bacteria Proteobacteria Gammaproteobacteria Enterobacteriales Enterobacteriaceae *Klebsiella oxytoca* 10-5242 |
| Kox-12-1 | Bacteria Proteobacteria Gammaproteobacteria Enterobacteriales Enterobacteriaceae *Klebsiella oxytoca* 10-5243 |
| Kox-13-1 | Bacteria Proteobacteria Gammaproteobacteria Enterobacteriales Enterobacteriaceae *Klebsiella oxytoca* 10-5245 |
| Kox-14-1 | Bacteria Proteobacteria Gammaproteobacteria Enterobacteriales Enterobacteriaceae *Klebsiella oxytoca* 10-5246 |
| Kox-4-1 to Kox-4-2 | Bacteria Proteobacteria Gammaproteobacteria Enterobacteriales Enterobacteriaceae *Klebsiella oxytoca* 10-5248 |
| Kox-15-1 | Bacteria Proteobacteria Gammaproteobacteria Enterobacteriales Enterobacteriaceae *Klebsiella oxytoca* 10-5250 |
| Kox-16-1 | Bacteria Proteobacteria Gammaproteobacteria Enterobacteriales Enterobacteriaceae *Klebsiella oxytoca* 11492-1 |
| Kox-17-1 | Bacteria Proteobacteria Gammaproteobacteria Enterobacteriales Enterobacteriaceae *Klebsiella oxytoca* E718 |
| Kox-18-1 | Bacteria Proteobacteria Gammaproteobacteria Enterobacteriales Enterobacteriaceae *Klebsiella oxytoca* KCTC 1686 |
| Kox-19-1 | Bacteria Proteobacteria Gammaproteobacteria Enterobacteriales Enterobacteriaceae *Klebsiella oxytoca* MGH 28 |
| Kox-20-1 | Bacteria Proteobacteria Gammaproteobacteria Enterobacteriales Enterobacteriaceae *Klebsiella oxytoca* MGH 42 |
| Kpn-1-1 to Kpn-1-9 | Bacteria Proteobacteria Gammaproteobacteria Enterobacteriales Enterobacteriaceae *Klebsiella pneumoniae* |
| Kpn-226-1 | Bacteria Proteobacteria Gammaproteobacteria Enterobacteriales Enterobacteriaceae *Klebsiella pneumoniae* 1162281 |
| Kpn-227-1 | Bacteria Proteobacteria Gammaproteobacteria Enterobacteriales Enterobacteriaceae *Klebsiella pneumoniae* 1191100241 |
| Kpn-228-1 | Bacteria Proteobacteria Gammaproteobacteria Enterobacteriales Enterobacteriaceae *Klebsiella pneumoniae* 120_1020 |
| Kpn-229-1 | Bacteria Proteobacteria Gammaproteobacteria Enterobacteriales Enterobacteriaceae *Klebsiella pneumoniae* 140_1040 |
| Kpn-3-1 to Kpn-3-2 | Bacteria Proteobacteria Gammaproteobacteria Enterobacteriales Enterobacteriaceae *Klebsiella pneumoniae* 160_1080 |
| Kpn-230-1 | Bacteria Proteobacteria Gammaproteobacteria Enterobacteriales Enterobacteriaceae *Klebsiella pneumoniae* 280_1220 |
| Kpn-231-1 | Bacteria Proteobacteria Gammaproteobacteria Enterobacteriales Enterobacteriaceae *Klebsiella pneumoniae* 303K |
| Kpn-232-1 | Bacteria Proteobacteria Gammaproteobacteria Enterobacteriales Enterobacteriaceae *Klebsiella pneumoniae* 342 |
| Kpn-233-1 | Bacteria Proteobacteria Gammaproteobacteria Enterobacteriales Enterobacteriaceae *Klebsiella pneumoniae* 361_1301 |
| Kpn-234-1 | Bacteria Proteobacteria Gammaproteobacteria Enterobacteriales Enterobacteriaceae *Klebsiella pneumoniae* 440_1540 |
| Kpn-235-1 | Bacteria Proteobacteria Gammaproteobacteria Enterobacteriales Enterobacteriaceae *Klebsiella pneumoniae* 540_1460 |
| Kpn-236-1 | Bacteria Proteobacteria Gammaproteobacteria Enterobacteriales Enterobacteriaceae *Klebsiella pneumoniae* 646_1568 |
| Kpn-237-1 | Bacteria Proteobacteria Gammaproteobacteria Enterobacteriales Enterobacteriaceae *Klebsiella pneumoniae* 700603 |
| Kpn-238-1 | Bacteria Proteobacteria Gammaproteobacteria Enterobacteriales Enterobacteriaceae *Klebsiella pneumoniae* ATCC BAA-1705 |
| Kpn-239-1 | Bacteria Proteobacteria Gammaproteobacteria Enterobacteriales Enterobacteriaceae *Klebsiella pneumoniae* ATCC BAA-2146 |
| Kpn-240-1 | Bacteria Proteobacteria Gammaproteobacteria Enterobacteriales Enterobacteriaceae *Klebsiella pneumoniae* BIDMC 12C |
| Kpn-241-1 | Bacteria Proteobacteria Gammaproteobacteria Enterobacteriales Enterobacteriaceae *Klebsiella pneumoniae* BIDMC 16 |
| Kpn-242-1 | Bacteria Proteobacteria Gammaproteobacteria Enterobacteriales Enterobacteriaceae *Klebsiella pneumoniae* BIDMC 18C |
| Kpn-243-1 | Bacteria Proteobacteria Gammaproteobacteria Enterobacteriales Enterobacteriaceae *Klebsiella pneumoniae* BIDMC 21 |
| Kpn-244-1 | Bacteria Proteobacteria Gammaproteobacteria Enterobacteriales Enterobacteriaceae *Klebsiella pneumoniae* BIDMC 22 |
| Kpn-245-1 | Bacteria Proteobacteria Gammaproteobacteria Enterobacteriales Enterobacteriaceae *Klebsiella pneumoniae* BIDMC 23 |
| Kpn-246-1 | Bacteria Proteobacteria Gammaproteobacteria Enterobacteriales Enterobacteriaceae *Klebsiella pneumoniae* BIDMC 24 |
| Kpn-247-1 | Bacteria Proteobacteria Gammaproteobacteria Enterobacteriales Enterobacteriaceae *Klebsiella pneumoniae* BIDMC 25 |
| Kpn-248-1 | Bacteria Proteobacteria Gammaproteobacteria Enterobacteriales Enterobacteriaceae *Klebsiella pneumoniae* BIDMC 35 |
| Kpn-249-1 | Bacteria Proteobacteria Gammaproteobacteria Enterobacteriales Enterobacteriaceae *Klebsiella pneumoniae* BIDMC 36 |
| Kpn-250-1 | Bacteria Proteobacteria Gammaproteobacteria Enterobacteriales Enterobacteriaceae *Klebsiella pneumoniae* BIDMC 40 |
| Kpn-251-1 | Bacteria Proteobacteria Gammaproteobacteria Enterobacteriales Enterobacteriaceae *Klebsiella pneumoniae* BIDMC 41 |
| Kpn-252-1 | Bacteria Proteobacteria Gammaproteobacteria Enterobacteriales Enterobacteriaceae *Klebsiella pneumoniae* BWH 28 |
| Kpn-253-1 | Bacteria Proteobacteria Gammaproteobacteria Enterobacteriales Enterobacteriaceae *Klebsiella pneumoniae* BWH 30 |
| Kpn-254-1 | Bacteria Proteobacteria Gammaproteobacteria Enterobacteriales Enterobacteriaceae *Klebsiella pneumoniae* CG43 |
| Kpn-255-1 | Bacteria Proteobacteria Gammaproteobacteria Enterobacteriales Enterobacteriaceae *Klebsiella pneumoniae* DMC0526 |
| Kpn-256-1 | Bacteria Proteobacteria Gammaproteobacteria Enterobacteriales Enterobacteriaceae *Klebsiella pneumoniae* DMC0799 |
| Kpn-257-1 | Bacteria Proteobacteria Gammaproteobacteria Enterobacteriales Enterobacteriaceae *Klebsiella pneumoniae* DMC1316 |
| Kpn-324-1 | Bacteria Proteobacteria Gammaproteobacteria Enterobacteriales Enterobacteriaceae *Klebsiella pneumoniae hv*KP1 |

| | |
|---|---|
| Kpn-258-1 | Bacteria Proteobacteria Gammaproteobacteria Enterobacteriales Enterobacteriaceae *Klebsiella pneumoniae* JH1 |
| Kpn-259-1 | Bacteria Proteobacteria Gammaproteobacteria Enterobacteriales Enterobacteriaceae *Klebsiella pneumoniae* JHCK1 |
| Kpn-260-1 | Bacteria Proteobacteria Gammaproteobacteria Enterobacteriales Enterobacteriaceae *Klebsiella pneumoniae* JM45 |
| Kpn-261-1 | Bacteria Proteobacteria Gammaproteobacteria Enterobacteriales Enterobacteriaceae *Klebsiella pneumoniae* KCTC 2242 |
| Kpn-262-1 | Bacteria Proteobacteria Gammaproteobacteria Enterobacteriales Enterobacteriaceae *Klebsiella pneumoniae* KP-1 |
| Kpn-263-1 | Bacteria Proteobacteria Gammaproteobacteria Enterobacteriales Enterobacteriaceae *Klebsiella pneumoniae* KP-11 |
| Kpn-264-1 | Bacteria Proteobacteria Gammaproteobacteria Enterobacteriales Enterobacteriaceae *Klebsiella pneumoniae* KP-7 |
| Kpn-265-1 | Bacteria Proteobacteria Gammaproteobacteria Enterobacteriales Enterobacteriaceae *Klebsiella pneumoniae* MGH 17 |
| Kpn-266-1 | Bacteria Proteobacteria Gammaproteobacteria Enterobacteriales Enterobacteriaceae *Klebsiella pneumoniae* MGH 18 |
| Kpn-267-1 | Bacteria Proteobacteria Gammaproteobacteria Enterobacteriales Enterobacteriaceae *Klebsiella pneumoniae* MGH 19 |
| Kpn-268-1 | Bacteria Proteobacteria Gammaproteobacteria Enterobacteriales Enterobacteriaceae *Klebsiella pneumoniae* MGH 20 |
| Kpn-269-1 | Bacteria Proteobacteria Gammaproteobacteria Enterobacteriales Enterobacteriaceae *Klebsiella pneumoniae* MGH 21 |
| Kpn-270-1 | Bacteria Proteobacteria Gammaproteobacteria Enterobacteriales Enterobacteriaceae *Klebsiella pneumoniae* MGH 30 |
| Kpn-271-1 | Bacteria Proteobacteria Gammaproteobacteria Enterobacteriales Enterobacteriaceae *Klebsiella pneumoniae* MGH 31 |
| Kpn-272-1 | Bacteria Proteobacteria Gammaproteobacteria Enterobacteriales Enterobacteriaceae *Klebsiella pneumoniae* MGH 32 |
| Kpn-273-1 | Bacteria Proteobacteria Gammaproteobacteria Enterobacteriales Enterobacteriaceae *Klebsiella pneumoniae* MGH 36 |
| Kpn-274-1 | Bacteria Proteobacteria Gammaproteobacteria Enterobacteriales Enterobacteriaceae *Klebsiella pneumoniae* MGH 40 |
| Kpn-275-1 | Bacteria Proteobacteria Gammaproteobacteria Enterobacteriales Enterobacteriaceae *Klebsiella pneumoniae* MGH 44 |
| Kpn-276-1 | Bacteria Proteobacteria Gammaproteobacteria Enterobacteriales Enterobacteriaceae *Klebsiella pneumoniae* MGH 46 |
| Kpn-277-1 | Bacteria Proteobacteria Gammaproteobacteria Enterobacteriales Enterobacteriaceae *Klebsiella pneumoniae* MGH 48 |
| Kpn-278-1 | Bacteria Proteobacteria Gammaproteobacteria Enterobacteriales Enterobacteriaceae *Klebsiella pneumoniae* MGH 59 |
| Kpn-279-1 | Bacteria Proteobacteria Gammaproteobacteria Enterobacteriales Enterobacteriaceae *Klebsiella pneumoniae* NTUH-K2044 |
| Kpn-205-1 to Kpn-205-4 | Bacteria Proteobacteria Gammaproteobacteria Enterobacteriales Enterobacteriaceae *Klebsiella pneumoniae subsp. ozaenae* |
| Kpn-325-1 | Bacteria Proteobacteria Gammaproteobacteria Enterobacteriales Enterobacteriaceae *Klebsiella pneumoniae subsp. pneumoniae* 1084 |
| Kpn-326-1 | Bacteria Proteobacteria Gammaproteobacteria Enterobacteriales Enterobacteriaceae *Klebsiella pneumoniae subsp. pneumoniae* 12-3578 |
| Kpn-327-1 | Bacteria Proteobacteria Gammaproteobacteria Enterobacteriales Enterobacteriaceae *Klebsiella pneumoniae subsp. pneumoniae* B5055 |
| Kpn-328-1 | Bacteria Proteobacteria Gammaproteobacteria Enterobacteriales Enterobacteriaceae *Klebsiella pneumoniae subsp. pneumoniae* DSM 30104 |
| Kpn-329-1 | Bacteria Proteobacteria Gammaproteobacteria Enterobacteriales Enterobacteriaceae *Klebsiella pneumoniae subsp. pneumoniae* Ecl8 |
| Kpn-330-1 | Bacteria Proteobacteria Gammaproteobacteria Enterobacteriales Enterobacteriaceae *Klebsiella pneumoniae subsp. pneumoniae* HS11286 |
| Kpn-331-1 | Bacteria Proteobacteria Gammaproteobacteria Enterobacteriales Enterobacteriaceae *Klebsiella pneumoniae subsp. pneumoniae* KPNIH11 |
| Kpn-332-1 | Bacteria Proteobacteria Gammaproteobacteria Enterobacteriales Enterobacteriaceae *Klebsiella pneumoniae subsp. pneumoniae* KPNIH12 |
| Kpn-333-1 | Bacteria Proteobacteria Gammaproteobacteria Enterobacteriales Enterobacteriaceae *Klebsiella pneumoniae subsp. pneumoniae* KPNIH14 |
| Kpn-334-1 | Bacteria Proteobacteria Gammaproteobacteria Enterobacteriales Enterobacteriaceae *Klebsiella pneumoniae subsp. pneumoniae* KPNIH16 |
| Kpn-335-1 | Bacteria Proteobacteria Gammaproteobacteria Enterobacteriales Enterobacteriaceae *Klebsiella pneumoniae subsp. pneumoniae* KPNIH17 |
| Kpn-336-1 | Bacteria Proteobacteria Gammaproteobacteria Enterobacteriales Enterobacteriaceae *Klebsiella pneumoniae subsp. pneumoniae* KPNIH18 |
| Kpn-337-1 | Bacteria Proteobacteria Gammaproteobacteria Enterobacteriales Enterobacteriaceae *Klebsiella pneumoniae subsp. pneumoniae* KPNIH19 |
| Kpn-338-1 | Bacteria Proteobacteria Gammaproteobacteria Enterobacteriales Enterobacteriaceae *Klebsiella pneumoniae subsp. pneumoniae* KPNIH2 |
| Kpn-339-1 | Bacteria Proteobacteria Gammaproteobacteria Enterobacteriales Enterobacteriaceae *Klebsiella pneumoniae subsp. pneumoniae* KPNIH20 |
| Kpn-340-1 | Bacteria Proteobacteria Gammaproteobacteria Enterobacteriales Enterobacteriaceae *Klebsiella pneumoniae subsp. pneumoniae* KPNIH21 |
| Kpn-341-1 | Bacteria Proteobacteria Gammaproteobacteria Enterobacteriales Enterobacteriaceae *Klebsiella pneumoniae subsp. pneumoniae* KPNIH22 |
| Kpn-342-1 | Bacteria Proteobacteria Gammaproteobacteria Enterobacteriales Enterobacteriaceae *Klebsiella pneumoniae subsp. pneumoniae* KPNIH23 |
| Kpn-343-1 | Bacteria Proteobacteria Gammaproteobacteria Enterobacteriales Enterobacteriaceae *Klebsiella pneumoniae subsp. pneumoniae* KPNIH4 |
| Kpn-344-1 | Bacteria Proteobacteria Gammaproteobacteria Enterobacteriales Enterobacteriaceae *Klebsiella pneumoniae subsp. pneumoniae* KPNIH5 |
| Kpn-345-1 | Bacteria Proteobacteria Gammaproteobacteria Enterobacteriales Enterobacteriaceae *Klebsiella pneumoniae subsp. pneumoniae* KPNIH6 |
| Kpn-346-1 | Bacteria Proteobacteria Gammaproteobacteria Enterobacteriales Enterobacteriaceae *Klebsiella pneumoniae subsp. pneumoniae* KPNIH7 |
| Kpn-347-1 | Bacteria Proteobacteria Gammaproteobacteria Enterobacteriales Enterobacteriaceae *Klebsiella pneumoniae subsp. pneumoniae* KPNIH8 |
| Kpn-348-1 | Bacteria Proteobacteria Gammaproteobacteria Enterobacteriales Enterobacteriaceae *Klebsiella pneumoniae subsp. pneumoniae* KPNIH9 |
| Kpn-349-1 | Bacteria Proteobacteria Gammaproteobacteria Enterobacteriales Enterobacteriaceae *Klebsiella pneumoniae subsp. pneumoniae* KpQ3 |
| Kpn-350-1 | Bacteria Proteobacteria Gammaproteobacteria Enterobacteriales Enterobacteriaceae *Klebsiella pneumoniae subsp. pneumoniae* LZ |
| Kpn-351-1 | Bacteria Proteobacteria Gammaproteobacteria Enterobacteriales Enterobacteriaceae *Klebsiella pneumoniae subsp. pneumoniae* MGH 78578 |
| Kpn-352-1 | Bacteria Proteobacteria Gammaproteobacteria Enterobacteriales Enterobacteriaceae *Klebsiella pneumoniae subsp. pneumoniae* MP14 |
| Kpn-353-1 | Bacteria Proteobacteria Gammaproteobacteria Enterobacteriales Enterobacteriaceae *Klebsiella pneumoniae subsp. pneumoniae* ST258-490 |
| Kpn-354-1 | Bacteria Proteobacteria Gammaproteobacteria Enterobacteriales Enterobacteriaceae *Klebsiella pneumoniae subsp. pneumoniae* ST258-K26BO |
| Kpn-355-1 | Bacteria Proteobacteria Gammaproteobacteria Enterobacteriales Enterobacteriaceae *Klebsiella pneumoniae subsp. pneumoniae* ST258-K28BO |
| Kpn-356-1 | Bacteria Proteobacteria Gammaproteobacteria Enterobacteriales Enterobacteriaceae *Klebsiella pneumoniae subsp. pneumoniae* ST512-K30BO |
| Kpn-357-1 | Bacteria Proteobacteria Gammaproteobacteria Enterobacteriales Enterobacteriaceae *Klebsiella pneumoniae subsp. pneumoniae* UKKV901664 |
| Kpn-358-1 | Bacteria Proteobacteria Gammaproteobacteria Enterobacteriales Enterobacteriaceae *Klebsiella pneumoniae subsp. pneumoniae* WGLW1 |
| Kpn-359-1 | Bacteria Proteobacteria Gammaproteobacteria Enterobacteriales Enterobacteriaceae *Klebsiella pneumoniae subsp. pneumoniae* WGLW2 |
| Kpn-360-1 | Bacteria Proteobacteria Gammaproteobacteria Enterobacteriales Enterobacteriaceae *Klebsiella pneumoniae subsp. pneumoniae* WGLW3 |
| Kpn-361-1 | Bacteria Proteobacteria Gammaproteobacteria Enterobacteriales Enterobacteriaceae *Klebsiella pneumoniae subsp. pneumoniae* WGLW5 |
| Kpn-362-1 | Bacteria Proteobacteria Gammaproteobacteria Enterobacteriales Enterobacteriaceae *Klebsiella pneumoniae subsp. rhinoscleromatis* ATCC 13884 |
| Kpn-280-1 | Bacteria Proteobacteria Gammaproteobacteria Enterobacteriales Enterobacteriaceae *Klebsiella pneumoniae* UCICRE 10 |
| Kpn-281-1 | Bacteria Proteobacteria Gammaproteobacteria Enterobacteriales Enterobacteriaceae *Klebsiella pneumoniae* UCICRE 14 |
| Kpn-282-1 | Bacteria Proteobacteria Gammaproteobacteria Enterobacteriales Enterobacteriaceae *Klebsiella pneumoniae* UCICRE 2 |
| Kpn-283-1 | Bacteria Proteobacteria Gammaproteobacteria Enterobacteriales Enterobacteriaceae *Klebsiella pneumoniae* UCICRE 4 |

| | |
|---|---|
| Kpn-284-1 | Bacteria Proteobacteria Gammaproteobacteria Enterobacteriales Enterobacteriaceae *Klebsiella pneumoniae* UCICRE 6 |
| Kpn-285-1 | Bacteria Proteobacteria Gammaproteobacteria Enterobacteriales Enterobacteriaceae *Klebsiella pneumoniae* UCICRE 7 |
| Kpn-286-1 | Bacteria Proteobacteria Gammaproteobacteria Enterobacteriales Enterobacteriaceae *Klebsiella pneumoniae* UCICRE 8 |
| Kpn-199-1 to Kpn-199-2 | Bacteria Proteobacteria Gammaproteobacteria Enterobacteriales Enterobacteriaceae *Klebsiella pneumoniae* UHKPC 52 |
| Kpn-287-1 | Bacteria Proteobacteria Gammaproteobacteria Enterobacteriales Enterobacteriaceae *Klebsiella pneumoniae* UHKPC01 |
| Kpn-288-1 | Bacteria Proteobacteria Gammaproteobacteria Enterobacteriales Enterobacteriaceae *Klebsiella pneumoniae* UHKPC02 |
| Kpn-289-1 | Bacteria Proteobacteria Gammaproteobacteria Enterobacteriales Enterobacteriaceae *Klebsiella pneumoniae* UHKPC04 |
| Kpn-200-1 to Kpn-200-2 | Bacteria Proteobacteria Gammaproteobacteria Enterobacteriales Enterobacteriaceae *Klebsiella pneumoniae* UHKPC05 |
| Kpn-290-1 | Bacteria Proteobacteria Gammaproteobacteria Enterobacteriales Enterobacteriaceae *Klebsiella pneumoniae* UHKPC06 |
| Kpn-291-1 | Bacteria Proteobacteria Gammaproteobacteria Enterobacteriales Enterobacteriaceae *Klebsiella pneumoniae* UHKPC09 |
| Kpn-292-1 | Bacteria Proteobacteria Gammaproteobacteria Enterobacteriales Enterobacteriaceae *Klebsiella pneumoniae* UHKPC17 |
| Kpn-293-1 | Bacteria Proteobacteria Gammaproteobacteria Enterobacteriales Enterobacteriaceae *Klebsiella pneumoniae* UHKPC179 |
| Kpn-294-1 | Bacteria Proteobacteria Gammaproteobacteria Enterobacteriales Enterobacteriaceae *Klebsiella pneumoniae* UHKPC18 |
| Kpn-295-1 | Bacteria Proteobacteria Gammaproteobacteria Enterobacteriales Enterobacteriaceae *Klebsiella pneumoniae* UHKPC22 |
| Kpn-296-1 | Bacteria Proteobacteria Gammaproteobacteria Enterobacteriales Enterobacteriaceae *Klebsiella pneumoniae* UHKPC23 |
| Kpn-297-1 | Bacteria Proteobacteria Gammaproteobacteria Enterobacteriales Enterobacteriaceae *Klebsiella pneumoniae* UHKPC24 |
| Kpn-298-1 | Bacteria Proteobacteria Gammaproteobacteria Enterobacteriales Enterobacteriaceae *Klebsiella pneumoniae* UHKPC26 |
| Kpn-299-1 | Bacteria Proteobacteria Gammaproteobacteria Enterobacteriales Enterobacteriaceae *Klebsiella pneumoniae* UHKPC27 |
| Kpn-300-1 | Bacteria Proteobacteria Gammaproteobacteria Enterobacteriales Enterobacteriaceae *Klebsiella pneumoniae* UHKPC28 |
| Kpn-301-1 | Bacteria Proteobacteria Gammaproteobacteria Enterobacteriales Enterobacteriaceae *Klebsiella pneumoniae* UHKPC29 |
| Kpn-302-1 | Bacteria Proteobacteria Gammaproteobacteria Enterobacteriales Enterobacteriaceae *Klebsiella pneumoniae* UHKPC31 |
| Kpn-303-1 | Bacteria Proteobacteria Gammaproteobacteria Enterobacteriales Enterobacteriaceae *Klebsiella pneumoniae* UHKPC32 |
| Kpn-304-1 | Bacteria Proteobacteria Gammaproteobacteria Enterobacteriales Enterobacteriaceae *Klebsiella pneumoniae* UHKPC40 |
| Kpn-203-1 to Kpn-203-2 | Bacteria Proteobacteria Gammaproteobacteria Enterobacteriales Enterobacteriaceae *Klebsiella pneumoniae* UHKPC45 |
| Kpn-305-1 | Bacteria Proteobacteria Gammaproteobacteria Enterobacteriales Enterobacteriaceae *Klebsiella pneumoniae* UHKPC47 |
| Kpn-306-1 | Bacteria Proteobacteria Gammaproteobacteria Enterobacteriales Enterobacteriaceae *Klebsiella pneumoniae* UHKPC48 |
| Kpn-307-1 | Bacteria Proteobacteria Gammaproteobacteria Enterobacteriales Enterobacteriaceae *Klebsiella pneumoniae* UHKPC57 |
| Kpn-308-1 | Bacteria Proteobacteria Gammaproteobacteria Enterobacteriales Enterobacteriaceae *Klebsiella pneumoniae* UHKPC59 |
| Kpn-309-1 | Bacteria Proteobacteria Gammaproteobacteria Enterobacteriales Enterobacteriaceae *Klebsiella pneumoniae* UHKPC61 |
| Kpn-310-1 | Bacteria Proteobacteria Gammaproteobacteria Enterobacteriales Enterobacteriaceae *Klebsiella pneumoniae* UHKPC67 |
| Kpn-311-1 | Bacteria Proteobacteria Gammaproteobacteria Enterobacteriales Enterobacteriaceae *Klebsiella pneumoniae* UHKPC69 |
| Kpn-312-1 | Bacteria Proteobacteria Gammaproteobacteria Enterobacteriales Enterobacteriaceae *Klebsiella pneumoniae* UHKPC77 |
| Kpn-313-1 | Bacteria Proteobacteria Gammaproteobacteria Enterobacteriales Enterobacteriaceae *Klebsiella pneumoniae* UHKPC81 |
| Kpn-314-1 | Bacteria Proteobacteria Gammaproteobacteria Enterobacteriales Enterobacteriaceae *Klebsiella pneumoniae* UHKPC96 |
| Kpn-315-1 | Bacteria Proteobacteria Gammaproteobacteria Enterobacteriales Enterobacteriaceae *Klebsiella pneumoniae* VA360 |
| Kpn-316-1 | Bacteria Proteobacteria Gammaproteobacteria Enterobacteriales Enterobacteriaceae *Klebsiella pneumoniae* VAKPC252 |
| Kpn-317-1 | Bacteria Proteobacteria Gammaproteobacteria Enterobacteriales Enterobacteriaceae *Klebsiella pneumoniae* VAKPC254 |
| Kpn-318-1 | Bacteria Proteobacteria Gammaproteobacteria Enterobacteriales Enterobacteriaceae *Klebsiella pneumoniae* VAKPC269 |
| Kpn-319-1 | Bacteria Proteobacteria Gammaproteobacteria Enterobacteriales Enterobacteriaceae *Klebsiella pneumoniae* VAKPC270 |
| Kpn-320-1 | Bacteria Proteobacteria Gammaproteobacteria Enterobacteriales Enterobacteriaceae *Klebsiella pneumoniae* VAKPC276 |
| Kpn-204-1 to Kpn-204-2 | Bacteria Proteobacteria Gammaproteobacteria Enterobacteriales Enterobacteriaceae *Klebsiella pneumoniae* VAKPC278 |
| Kpn-321-1 | Bacteria Proteobacteria Gammaproteobacteria Enterobacteriales Enterobacteriaceae *Klebsiella pneumoniae* VAKPC280 |
| Kpn-322-1 | Bacteria Proteobacteria Gammaproteobacteria Enterobacteriales Enterobacteriaceae *Klebsiella pneumoniae* VAKPC297 |
| Kpn-323-1 | Bacteria Proteobacteria Gammaproteobacteria Enterobacteriales Enterobacteriaceae *Klebsiella pneumoniae* VAKPC309 |
| Ksp-4-1 | Bacteria Proteobacteria Gammaproteobacteria Enterobacteriales Enterobacteriaceae *Klebsiella sp.* 1_1_55 |
| Ksp-5-1 | Bacteria Proteobacteria Gammaproteobacteria Enterobacteriales Enterobacteriaceae *Klebsiella sp.* 4_1_44FAA |
| Ksp-6-1 | Bacteria Proteobacteria Gammaproteobacteria Enterobacteriales Enterobacteriaceae *Klebsiella sp.* KTE92 |
| Ksp-7-1 | Bacteria Proteobacteria Gammaproteobacteria Enterobacteriales Enterobacteriaceae *Klebsiella sp.* MS 92-3 |
| Ksp-8-1 | Bacteria Proteobacteria Gammaproteobacteria Enterobacteriales Enterobacteriaceae *Klebsiella sp.* OBRC7 |
| Kva-1-1 to Kva-1-7 | Bacteria Proteobacteria Gammaproteobacteria Enterobacteriales Enterobacteriaceae *Klebsiella variicola* |
| Kva-2-1 | Bacteria Proteobacteria Gammaproteobacteria Enterobacteriales Enterobacteriaceae *Klebsiella variicola* At-22 |
| Era-1-1 | Bacteria Proteobacteria Gammaproteobacteria Enterobacteriales Enterobacteriaceae Kosakonia *Enterobacter radicincitans* DSM 16656 |
| Ksa-1-1 | Bacteria Proteobacteria Gammaproteobacteria Enterobacteriales Enterobacteriaceae *Kosakonia sacchari* SP1 |
| Mmo-1-1 to Mmo-1-8 | Bacteria Proteobacteria Gammaproteobacteria Enterobacteriales Enterobacteriaceae *Morganella morganii* |
| Mmo-5-1 | Bacteria Proteobacteria Gammaproteobacteria Enterobacteriales Enterobacteriaceae *Morganella morganii* SC01 |
| Mmo-6-1 | Bacteria Proteobacteria Gammaproteobacteria Enterobacteriales Enterobacteriaceae *Morganella morganii subsp. morganii* KT |
| Pag-4-1 | Bacteria Proteobacteria Gammaproteobacteria Enterobacteriales Enterobacteriaceae *Pantoea agglomerans* 299R |
| Pag-2-1 to Pag-2-6 | Bacteria Proteobacteria Gammaproteobacteria Enterobacteriales Enterobacteriaceae Pantoea Pantoea agglomerans group *Pantoea agglomerans* |
| Pag-5-1 | Bacteria Proteobacteria Gammaproteobacteria Enterobacteriales Enterobacteriaceae *Pantoea agglomerans* IG1 |
| Pag-6-1 | Bacteria Proteobacteria Gammaproteobacteria Enterobacteriales Enterobacteriaceae *Pantoea agglomerans* Tx10 |
| Pan-1-1 to Pan-1-5 | Bacteria Proteobacteria Gammaproteobacteria Enterobacteriales Enterobacteriaceae *Pantoea ananatis* |
| Pan-6-1 | Bacteria Proteobacteria Gammaproteobacteria Enterobacteriales Enterobacteriaceae *Pantoea ananatis* AJ13355 |
| Pan-7-1 | Bacteria Proteobacteria Gammaproteobacteria Enterobacteriales Enterobacteriaceae *Pantoea ananatis* BRT175 |
| Pan-8-1 | Bacteria Proteobacteria Gammaproteobacteria Enterobacteriales Enterobacteriaceae *Pantoea ananatis* LMG 20103 |
| Pan-9-1 | Bacteria Proteobacteria Gammaproteobacteria Enterobacteriales Enterobacteriaceae *Pantoea ananatis* LMG 5342 |

| | |
|---|---|
| Pan-10-1 | Bacteria Proteobacteria Gammaproteobacteria Enterobacteriales Enterobacteriaceae *Pantoea ananatis* PA13 |
| Pdi-2-1 | Bacteria Proteobacteria Gammaproteobacteria Enterobacteriales Enterobacteriaceae *Pantoea dispersa* EGD-AAK13 |
| Psp-48-1 | Bacteria Proteobacteria Gammaproteobacteria Enterobacteriales Enterobacteriaceae *Pantoea sp.* A4 |
| Psp-54-1 | Bacteria Proteobacteria Gammaproteobacteria Enterobacteriales Enterobacteriaceae *Pantoea sp.* aB |
| Psp-49-1 | Bacteria Proteobacteria Gammaproteobacteria Enterobacteriales Enterobacteriaceae *Pantoea sp.* At-9b |
| Psp-50-1 | Bacteria Proteobacteria Gammaproteobacteria Enterobacteriales Enterobacteriaceae *Pantoea sp.* GM01 |
| Psp-52-1 | Bacteria Proteobacteria Gammaproteobacteria Enterobacteriales Enterobacteriaceae *Pantoea sp.* Sc1 |
| Psp-51-1 | Bacteria Proteobacteria Gammaproteobacteria Enterobacteriales Enterobacteriaceae *Pantoea sp.* SL1_M5 |
| Psp-53-1 | Bacteria Proteobacteria Gammaproteobacteria Enterobacteriales Enterobacteriaceae *Pantoea sp.* YR343 |
| Pst-3-1 | Bacteria Proteobacteria Gammaproteobacteria Enterobacteriales Enterobacteriaceae *Pantoea stewartii subsp. indologenes* |
| Pst-4-1 | Bacteria Proteobacteria Gammaproteobacteria Enterobacteriales Enterobacteriaceae *Pantoea stewartii subsp. stewartii* DC283 |
| Pva-2-1 | Bacteria Proteobacteria Gammaproteobacteria Enterobacteriales Enterobacteriaceae *Pantoea vagans* C9-1 |
| Pat-4-1 | Bacteria Proteobacteria Gammaproteobacteria Enterobacteriales Enterobacteriaceae *Pectobacterium atrosepticum* SCRI1043 |
| Pca-9-1 | Bacteria Proteobacteria Gammaproteobacteria Enterobacteriales Enterobacteriaceae *Pectobacterium carotovorum subsp. brasiliensis* PBR1692 |
| Pca-5-1 to Pca-5-11 | Bacteria Proteobacteria Gammaproteobacteria Enterobacteriales Enterobacteriaceae *Pectobacterium carotovorum subsp. carotovorum* |
| Pca-10-1 | Bacteria Proteobacteria Gammaproteobacteria Enterobacteriales Enterobacteriaceae *Pectobacterium carotovorum subsp. carotovorum* PC1 |
| Pca-11-1 | Bacteria Proteobacteria Gammaproteobacteria Enterobacteriales Enterobacteriaceae *Pectobacterium carotovorum subsp. carotovorum* PCC21 |
| Pca-12-1 | Bacteria Proteobacteria Gammaproteobacteria Enterobacteriales Enterobacteriaceae *Pectobacterium carotovorum subsp. carotovorum* WPP14 |
| Psp-55-1 | Bacteria Proteobacteria Gammaproteobacteria Enterobacteriales Enterobacteriaceae *Pectobacterium sp.* SCC3193 |
| Pwa-2-1 | Bacteria Proteobacteria Gammaproteobacteria Enterobacteriales Enterobacteriaceae *Pectobacterium wasabiae* CFBP 3304 |
| Pwa-3-1 | Bacteria Proteobacteria Gammaproteobacteria Enterobacteriales Enterobacteriaceae *Pectobacterium wasabiae* WPP163 |
| Pas-2-1 | Bacteria Proteobacteria Gammaproteobacteria Enterobacteriales Enterobacteriaceae *Photorhabdus asymbiotica* |
| Plu-10-1 | Bacteria Proteobacteria Gammaproteobacteria Enterobacteriales Enterobacteriaceae *Photorhabdus luminescens subsp. laumondii* TTO1 |
| Pte-5-1 | Bacteria Proteobacteria Gammaproteobacteria Enterobacteriales Enterobacteriaceae *Photorhabdus temperata* J3 |
| Pte-6-1 | Bacteria Proteobacteria Gammaproteobacteria Enterobacteriales Enterobacteriaceae *Photorhabdus temperata subsp. temperata* M1021 |
| Pst-5-1 | Bacteria Proteobacteria Gammaproteobacteria Enterobacteriales Enterobacteriaceae *Plautia stali symbiont* |
| Psh-2-1 | Bacteria Proteobacteria Gammaproteobacteria Enterobacteriales Enterobacteriaceae *Plesiomonas shigelloides* 302-73 |
| Pha-5-1 | Bacteria Proteobacteria Gammaproteobacteria Enterobacteriales Enterobacteriaceae *Proteus hauseri* ZMd44 |
| Pmi-1-1 to Pmi-1-6 | Bacteria Proteobacteria Gammaproteobacteria Enterobacteriales Enterobacteriaceae *Proteus mirabilis* |
| Pmi-2-1 | Bacteria Proteobacteria Gammaproteobacteria Enterobacteriales Enterobacteriaceae *Proteus mirabilis* ATCC 29906 |
| Pmi-3-1 | Bacteria Proteobacteria Gammaproteobacteria Enterobacteriales Enterobacteriaceae *Proteus mirabilis* BB2000 |
| Pmi-4-1 | Bacteria Proteobacteria Gammaproteobacteria Enterobacteriales Enterobacteriaceae *Proteus mirabilis* C05028 |
| Pmi-5-1 | Bacteria Proteobacteria Gammaproteobacteria Enterobacteriales Enterobacteriaceae *Proteus mirabilis* HI4320 |
| Pmi-6-1 | Bacteria Proteobacteria Gammaproteobacteria Enterobacteriales Enterobacteriaceae *Proteus mirabilis* PR03 |
| Pmi-7-1 | Bacteria Proteobacteria Gammaproteobacteria Enterobacteriales Enterobacteriaceae *Proteus mirabilis* WGLW4 |
| Pmi-8-1 | Bacteria Proteobacteria Gammaproteobacteria Enterobacteriales Enterobacteriaceae *Proteus mirabilis* WGLW6 |
| Pal-9-1 | Bacteria Proteobacteria Gammaproteobacteria Enterobacteriales Enterobacteriaceae *Providencia alcalifaciens* Dmel2 |
| Pal-8-1 | Bacteria Proteobacteria Gammaproteobacteria Enterobacteriales Enterobacteriaceae *Providencia alcalifaciens* DSM 30120 |
| Pbu-1-1 | Bacteria Proteobacteria Gammaproteobacteria Enterobacteriales Enterobacteriaceae *Providencia burhodogranariea* DSM 19968 |
| Pre-3-1 | Bacteria Proteobacteria Gammaproteobacteria Enterobacteriales Enterobacteriaceae *Providencia rettgeri* Dmel1 |
| Pre-2-1 | Bacteria Proteobacteria Gammaproteobacteria Enterobacteriales Enterobacteriaceae *Providencia rettgeri* DSM 1131 |
| Pru-5-1 | Bacteria Proteobacteria Gammaproteobacteria Enterobacteriales Enterobacteriaceae *Providencia rustigianii* DSM 4541 |
| Psn-1-1 | Bacteria Proteobacteria Gammaproteobacteria Enterobacteriales Enterobacteriaceae *Providencia sneebia* DSM 19967 |
| Pst-6-1 | Bacteria Proteobacteria Gammaproteobacteria Enterobacteriales Enterobacteriaceae *Providencia stuartii* ATCC 25827 |
| Pst-7-1 | Bacteria Proteobacteria Gammaproteobacteria Enterobacteriales Enterobacteriaceae *Providencia stuartii* MRSN 2154 |
| Raq-2-1 to Raq-2-2 | Bacteria Proteobacteria Gammaproteobacteria Enterobacteriales Enterobacteriaceae *Rahnella aquatilis* CIP 78.65 = ATCC 33071 |
| Raq-3-1 | Bacteria Proteobacteria Gammaproteobacteria Enterobacteriales Enterobacteriaceae *Rahnella aquatilis* HX2 |
| Rsp-4-1 | Bacteria Proteobacteria Gammaproteobacteria Enterobacteriales Enterobacteriaceae *Rahnella sp.* Y9602 |
| Ror-4-1 to Ror-4-2 | Bacteria Proteobacteria Gammaproteobacteria Enterobacteriales Enterobacteriaceae *Raoultella ornithinolytica* B6 |
| Rpl-1-1 to Rpl-1-3 | Bacteria Proteobacteria Gammaproteobacteria Enterobacteriales Enterobacteriaceae *Raoultella planticola* |
| Sbo-3-1 | Bacteria Proteobacteria Gammaproteobacteria Enterobacteriales Enterobacteriaceae *Salmonella bongori* N268-08 |
| Sbo-4-1 | Bacteria Proteobacteria Gammaproteobacteria Enterobacteriales Enterobacteriaceae *Salmonella bongori* NCTC 12419 |
| Sen-2-1 to Sen-2-2 | Bacteria Proteobacteria Gammaproteobacteria Enterobacteriales Enterobacteriaceae *Salmonella enterica* |
| Sen-311-1 | Bacteria Proteobacteria Gammaproteobacteria Enterobacteriales Enterobacteriaceae *Salmonella enterica subsp. arizonae serovar* 62:z4,z23:– |
| Sen-312-1 | Bacteria Proteobacteria Gammaproteobacteria Enterobacteriales Enterobacteriaceae *Salmonella enterica subsp. diarizonae serovar* 60:r:e,n,x,z15 str. 01-0170 |
| Sen-313-1 | Bacteria Proteobacteria Gammaproteobacteria Enterobacteriales Enterobacteriaceae *Salmonella enterica subsp. enterica serovar* 4,[5],12:i:- str. 08-1700 |
| Sen-314-1 | Bacteria Proteobacteria Gammaproteobacteria Enterobacteriales Enterobacteriaceae *Salmonella enterica subsp. enterica serovar* 4,[5],12:i:- str. 08-1736 |
| Sen-315-1 | Bacteria Proteobacteria Gammaproteobacteria Enterobacteriales Enterobacteriaceae *Salmonella enterica subsp. enterica serovar* 4,[5],12:i:- str. 08-1739 |
| Sen-316-1 | Bacteria Proteobacteria Gammaproteobacteria Enterobacteriales Enterobacteriaceae *Salmonella enterica subsp. enterica serovar* Agona str. 01.O.05 |
| Sen-317-1 | Bacteria Proteobacteria Gammaproteobacteria Enterobacteriales Enterobacteriaceae *Salmonella enterica subsp. enterica serovar* Agona str. 02.O.05 |
| Sen-318-1 | Bacteria Proteobacteria Gammaproteobacteria Enterobacteriales Enterobacteriaceae *Salmonella enterica subsp. enterica serovar* Agona str. 0292 |
| Sen-319-1 | Bacteria Proteobacteria Gammaproteobacteria Enterobacteriales Enterobacteriaceae *Salmonella enterica subsp. enterica serovar* Agona str. 03.O.05 |
| Sen-320-1 | Bacteria Proteobacteria Gammaproteobacteria Enterobacteriales Enterobacteriaceae *Salmonella enterica subsp. enterica serovar* Agona str. 0322 |
| Sen-321-1 | Bacteria Proteobacteria Gammaproteobacteria Enterobacteriales Enterobacteriaceae *Salmonella enterica subsp. enterica serovar* Agona str. 04.O.05 |

| | |
|---|---|
| Sen-322-1 | Bacteria Proteobacteria Gammaproteobacteria Enterobacteriales Enterobacteriaceae *Salmonella enterica subsp. enterica serovar* Agona str. 05.O.06 |
| Sen-323-1 | Bacteria Proteobacteria Gammaproteobacteria Enterobacteriales Enterobacteriaceae *Salmonella enterica subsp. enterica serovar* Agona str. 06.O.05 |
| Sen-324-1 | Bacteria Proteobacteria Gammaproteobacteria Enterobacteriales Enterobacteriaceae *Salmonella enterica subsp. enterica serovar* Agona str. 07.O.05 |
| Sen-325-1 | Bacteria Proteobacteria Gammaproteobacteria Enterobacteriales Enterobacteriaceae *Salmonella enterica subsp. enterica serovar* Agona str. 08.A.05 |
| Sen-326-1 | Bacteria Proteobacteria Gammaproteobacteria Enterobacteriales Enterobacteriaceae *Salmonella enterica subsp. enterica serovar* Agona str. 09.F.08 |
| Sen-327-1 | Bacteria Proteobacteria Gammaproteobacteria Enterobacteriales Enterobacteriaceae *Salmonella enterica subsp. enterica serovar* Agona str. 10.A.05 |
| Sen-328-1 | Bacteria Proteobacteria Gammaproteobacteria Enterobacteriales Enterobacteriaceae *Salmonella enterica subsp. enterica serovar* Agona str. 11.A.05 |
| Sen-329-1 | Bacteria Proteobacteria Gammaproteobacteria Enterobacteriales Enterobacteriaceae *Salmonella enterica subsp. enterica serovar* Agona str. 12.A.06 |
| Sen-330-1 | Bacteria Proteobacteria Gammaproteobacteria Enterobacteriales Enterobacteriaceae *Salmonella enterica subsp. enterica serovar* Agona str. 13.E.05 |
| Sen-331-1 | Bacteria Proteobacteria Gammaproteobacteria Enterobacteriales Enterobacteriaceae *Salmonella enterica subsp. enterica serovar* Agona str. 14.E.05 |
| Sen-332-1 | Bacteria Proteobacteria Gammaproteobacteria Enterobacteriales Enterobacteriaceae *Salmonella enterica subsp. enterica serovar* Agona str. 15.H.03 |
| Sen-333-1 | Bacteria Proteobacteria Gammaproteobacteria Enterobacteriales Enterobacteriaceae *Salmonella enterica subsp. enterica serovar* Agona str. 16.H.08 |
| Sen-334-1 | Bacteria Proteobacteria Gammaproteobacteria Enterobacteriales Enterobacteriaceae *Salmonella enterica subsp. enterica serovar* Agona str. 17.H.06 |
| Sen-335-1 | Bacteria Proteobacteria Gammaproteobacteria Enterobacteriales Enterobacteriaceae *Salmonella enterica subsp. enterica serovar* Agona str. 18.H.07 |
| Sen-336-1 | Bacteria Proteobacteria Gammaproteobacteria Enterobacteriales Enterobacteriaceae *Salmonella enterica subsp. enterica serovar* Agona str. 19.F.03 |
| Sen-337-1 | Bacteria Proteobacteria Gammaproteobacteria Enterobacteriales Enterobacteriaceae *Salmonella enterica subsp. enterica serovar* Agona str. 20.H.06 |
| Sen-338-1 | Bacteria Proteobacteria Gammaproteobacteria Enterobacteriales Enterobacteriaceae *Salmonella enterica subsp. enterica serovar* Agona str. 21.H.10 |
| Sen-339-1 | Bacteria Proteobacteria Gammaproteobacteria Enterobacteriales Enterobacteriaceae *Salmonella enterica subsp. enterica serovar* Agona str. 22.H.04 |
| Sen-340-1 | Bacteria Proteobacteria Gammaproteobacteria Enterobacteriales Enterobacteriaceae *Salmonella enterica subsp. enterica serovar* Agona str. 23.F.01 |
| Sen-341-1 | Bacteria Proteobacteria Gammaproteobacteria Enterobacteriales Enterobacteriaceae *Salmonella enterica subsp. enterica serovar* Agona str. 24.H.04 |
| Sen-342-1 | Bacteria Proteobacteria Gammaproteobacteria Enterobacteriales Enterobacteriaceae *Salmonella enterica subsp. enterica serovar* Agona str. 241981 |
| Sen-343-1 | Bacteria Proteobacteria Gammaproteobacteria Enterobacteriales Enterobacteriaceae *Salmonella enterica subsp. enterica serovar* Agona str. 24249 |
| Sen-344-1 | Bacteria Proteobacteria Gammaproteobacteria Enterobacteriales Enterobacteriaceae *Salmonella enterica subsp. enterica serovar* Agona str. 246555-3 |
| Sen-345-1 | Bacteria Proteobacteria Gammaproteobacteria Enterobacteriales Enterobacteriaceae *Salmonella enterica subsp. enterica serovar* Agona str. 26.F.98 |
| Sen-346-1 | Bacteria Proteobacteria Gammaproteobacteria Enterobacteriales Enterobacteriaceae *Salmonella enterica subsp. enterica serovar* Agona str. 266757-1 |
| Sen-347-1 | Bacteria Proteobacteria Gammaproteobacteria Enterobacteriales Enterobacteriaceae *Salmonella enterica subsp. enterica serovar* Agona str. 27.O.98 |
| Sen-348-1 | Bacteria Proteobacteria Gammaproteobacteria Enterobacteriales Enterobacteriaceae *Salmonella enterica subsp. enterica serovar* Agona str. 28.O.08 |
| Sen-349-1 | Bacteria Proteobacteria Gammaproteobacteria Enterobacteriales Enterobacteriaceae *Salmonella enterica subsp. enterica serovar* Agona str. 29.O.08 |
| Sen-350-1 | Bacteria Proteobacteria Gammaproteobacteria Enterobacteriales Enterobacteriaceae *Salmonella enterica subsp. enterica serovar* Agona str. 30.H.04 |
| Sen-351-1 | Bacteria Proteobacteria Gammaproteobacteria Enterobacteriales Enterobacteriaceae *Salmonella enterica subsp. enterica serovar* Agona str. 31.H.09 |
| Sen-352-1 | Bacteria Proteobacteria Gammaproteobacteria Enterobacteriales Enterobacteriaceae *Salmonella enterica subsp. enterica serovar* Agona str. 311387-1 |
| Sen-353-1 | Bacteria Proteobacteria Gammaproteobacteria Enterobacteriales Enterobacteriaceae *Salmonella enterica subsp. enterica serovar* Agona str. 32.A.00 |
| Sen-354-1 | Bacteria Proteobacteria Gammaproteobacteria Enterobacteriales Enterobacteriaceae *Salmonella enterica subsp. enterica serovar* Agona str. 33.A.05 |
| Sen-355-1 | Bacteria Proteobacteria Gammaproteobacteria Enterobacteriales Enterobacteriaceae *Salmonella enterica subsp. enterica serovar* Agona str. 339787 |
| Sen-356-1 | Bacteria Proteobacteria Gammaproteobacteria Enterobacteriales Enterobacteriaceae *Salmonella enterica subsp. enterica serovar* Agona str. 34.H.09 |
| Sen-357-1 | Bacteria Proteobacteria Gammaproteobacteria Enterobacteriales Enterobacteriaceae *Salmonella enterica subsp. enterica serovar* Agona str. 35.H.08 |
| Sen-358-1 | Bacteria Proteobacteria Gammaproteobacteria Enterobacteriales Enterobacteriaceae *Salmonella enterica subsp. enterica serovar* Agona str. 36.H.00 |
| Sen-359-1 | Bacteria Proteobacteria Gammaproteobacteria Enterobacteriales Enterobacteriaceae *Salmonella enterica subsp. enterica serovar* Agona str. 37.F.02 |
| Sen-360-1 | Bacteria Proteobacteria Gammaproteobacteria Enterobacteriales Enterobacteriaceae *Salmonella enterica subsp. enterica serovar* Agona str. 38.O.03 |
| Sen-361-1 | Bacteria Proteobacteria Gammaproteobacteria Enterobacteriales Enterobacteriaceae *Salmonella enterica subsp. enterica serovar* Agona str. 39.O.03 |
| Sen-362-1 | Bacteria Proteobacteria Gammaproteobacteria Enterobacteriales Enterobacteriaceae *Salmonella enterica subsp. enterica serovar* Agona str. 392869-1 |
| Sen-363-1 | Bacteria Proteobacteria Gammaproteobacteria Enterobacteriales Enterobacteriaceae *Salmonella enterica subsp. enterica serovar* Agona str. 392869-2 |
| Sen-364-1 | Bacteria Proteobacteria Gammaproteobacteria Enterobacteriales Enterobacteriaceae *Salmonella enterica subsp. enterica serovar* Agona str. 40.E.08 |
| Sen-365-1 | Bacteria Proteobacteria Gammaproteobacteria Enterobacteriales Enterobacteriaceae *Salmonella enterica subsp. enterica serovar* Agona str. 400095 11 |
| Sen-366-1 | Bacteria Proteobacteria Gammaproteobacteria Enterobacteriales Enterobacteriaceae *Salmonella enterica subsp. enterica serovar* Agona str. 400095 13 |
| Sen-367-1 | Bacteria Proteobacteria Gammaproteobacteria Enterobacteriales Enterobacteriaceae *Salmonella enterica subsp. enterica serovar* Agona str. 400095 14 |
| Sen-368-1 | Bacteria Proteobacteria Gammaproteobacteria Enterobacteriales Enterobacteriaceae *Salmonella enterica subsp. enterica serovar* Agona str. 400095 17 |
| Sen-369-1 | Bacteria Proteobacteria Gammaproteobacteria Enterobacteriales Enterobacteriaceae *Salmonella enterica subsp. enterica serovar* Agona str. 400095 18 |
| Sen-370-1 | Bacteria Proteobacteria Gammaproteobacteria Enterobacteriales Enterobacteriaceae *Salmonella enterica subsp. enterica serovar* Agona str. 409753-6 |
| Sen-371-1 | Bacteria Proteobacteria Gammaproteobacteria Enterobacteriales Enterobacteriaceae *Salmonella enterica subsp. enterica serovar* Agona str. 41.E.09 |
| Sen-372-1 | Bacteria Proteobacteria Gammaproteobacteria Enterobacteriales Enterobacteriaceae *Salmonella enterica subsp. enterica serovar* Agona str. 419639 2-1 |
| Sen-373-1 | Bacteria Proteobacteria Gammaproteobacteria Enterobacteriales Enterobacteriaceae *Salmonella enterica subsp. enterica serovar* Agona str. 42.E.09 |
| Sen-374-1 | Bacteria Proteobacteria Gammaproteobacteria Enterobacteriales Enterobacteriaceae *Salmonella enterica subsp. enterica serovar* Agona str. 43.E.09 |
| Sen-375-1 | Bacteria Proteobacteria Gammaproteobacteria Enterobacteriales Enterobacteriaceae *Salmonella enterica subsp. enterica serovar* Agona str. 432613 |
| Sen-376-1 | Bacteria Proteobacteria Gammaproteobacteria Enterobacteriales Enterobacteriaceae *Salmonella enterica subsp. enterica serovar* Agona str. 44.E.09 |
| Sen-377-1 | Bacteria Proteobacteria Gammaproteobacteria Enterobacteriales Enterobacteriaceae *Salmonella enterica subsp. enterica serovar* Agona str. 442692 2-4 |
| Sen-378-1 | Bacteria Proteobacteria Gammaproteobacteria Enterobacteriales Enterobacteriaceae *Salmonella enterica subsp. enterica serovar* Agona str. 442692 2-5 |
| Sen-379-1 | Bacteria Proteobacteria Gammaproteobacteria Enterobacteriales Enterobacteriaceae *Salmonella enterica subsp. enterica serovar* Agona str. 447967 1-1 |
| Sen-380-1 | Bacteria Proteobacteria Gammaproteobacteria Enterobacteriales Enterobacteriaceae *Salmonella enterica subsp. enterica serovar* Agona str. 447967 2-1 |
| Sen-381-1 | Bacteria Proteobacteria Gammaproteobacteria Enterobacteriales Enterobacteriaceae *Salmonella enterica subsp. enterica serovar* Agona str. 45.E.09 |
| Sen-382-1 | Bacteria Proteobacteria Gammaproteobacteria Enterobacteriales Enterobacteriaceae *Salmonella enterica subsp. enterica serovar* Agona str. 46.E.09 |
| Sen-383-1 | Bacteria Proteobacteria Gammaproteobacteria Enterobacteriales Enterobacteriaceae *Salmonella enterica subsp. enterica serovar* Agona str. 460004 1-1 |
| Sen-384-1 | Bacteria Proteobacteria Gammaproteobacteria Enterobacteriales Enterobacteriaceae *Salmonella enterica subsp. enterica serovar* Agona str. 467481 |
| Sen-385-1 | Bacteria Proteobacteria Gammaproteobacteria Enterobacteriales Enterobacteriaceae *Salmonella enterica subsp. enterica serovar* Agona str. 47.E.09 |
| Sen-386-1 | Bacteria Proteobacteria Gammaproteobacteria Enterobacteriales Enterobacteriaceae *Salmonella enterica subsp. enterica serovar* Agona str. 48.E.08 |

| | |
|---|---|
| Sen-387-1 | Bacteria Proteobacteria Gammaproteobacteria Enterobacteriales Enterobacteriaceae *Salmonella enterica subsp. enterica serovar* Agona str. 49.E.09 |
| Sen-388-1 | Bacteria Proteobacteria Gammaproteobacteria Enterobacteriales Enterobacteriaceae *Salmonella enterica subsp. enterica serovar* Agona str. 50.E.08 |
| Sen-389-1 | Bacteria Proteobacteria Gammaproteobacteria Enterobacteriales Enterobacteriaceae *Salmonella enterica subsp. enterica serovar* Agona str. 51.E.09 |
| Sen-390-1 | Bacteria Proteobacteria Gammaproteobacteria Enterobacteriales Enterobacteriaceae *Salmonella enterica subsp. enterica serovar* Agona str. 52.F.08 |
| Sen-391-1 | Bacteria Proteobacteria Gammaproteobacteria Enterobacteriales Enterobacteriaceae *Salmonella enterica subsp. enterica serovar* Agona str. 53.F.08 |
| Sen-392-1 | Bacteria Proteobacteria Gammaproteobacteria Enterobacteriales Enterobacteriaceae *Salmonella enterica subsp. enterica serovar* Agona str. 54.O.08 |
| Sen-393-1 | Bacteria Proteobacteria Gammaproteobacteria Enterobacteriales Enterobacteriaceae *Salmonella enterica subsp. enterica serovar* Agona str. 55.U.08 |
| Sen-394-1 | Bacteria Proteobacteria Gammaproteobacteria Enterobacteriales Enterobacteriaceae *Salmonella enterica subsp. enterica serovar* Agona str. 557928 |
| Sen-395-1 | Bacteria Proteobacteria Gammaproteobacteria Enterobacteriales Enterobacteriaceae *Salmonella enterica subsp. enterica serovar* Agona str. 56.O.08 |
| Sen-396-1 | Bacteria Proteobacteria Gammaproteobacteria Enterobacteriales Enterobacteriaceae *Salmonella enterica subsp. enterica serovar* Agona str. 57.A.08 |
| Sen-397-1 | Bacteria Proteobacteria Gammaproteobacteria Enterobacteriales Enterobacteriaceae *Salmonella enterica subsp. enterica serovar* Agona str. 58.E.08 |
| Sen-398-1 | Bacteria Proteobacteria Gammaproteobacteria Enterobacteriales Enterobacteriaceae *Salmonella enterica subsp. enterica serovar* Agona str. 59.F.08 |
| Sen-399-1 | Bacteria Proteobacteria Gammaproteobacteria Enterobacteriales Enterobacteriaceae *Salmonella enterica subsp. enterica serovar* Agona str. 60.O.08 |
| Sen-400-1 | Bacteria Proteobacteria Gammaproteobacteria Enterobacteriales Enterobacteriaceae *Salmonella enterica subsp. enterica serovar* Agona str. 61.O.08 |
| Sen-401-1 | Bacteria Proteobacteria Gammaproteobacteria Enterobacteriales Enterobacteriaceae *Salmonella enterica subsp. enterica serovar* Agona str. 62.H.72 |
| Sen-402-1 | Bacteria Proteobacteria Gammaproteobacteria Enterobacteriales Enterobacteriaceae *Salmonella enterica subsp. enterica serovar* Agona str. 620239 |
| Sen-403-1 | Bacteria Proteobacteria Gammaproteobacteria Enterobacteriales Enterobacteriaceae *Salmonella enterica subsp. enterica serovar* Agona str. 63.H.87 |
| Sen-404-1 | Bacteria Proteobacteria Gammaproteobacteria Enterobacteriales Enterobacteriaceae *Salmonella enterica subsp. enterica serovar* Agona str. 632182-2 |
| Sen-405-1 | Bacteria Proteobacteria Gammaproteobacteria Enterobacteriales Enterobacteriaceae *Salmonella enterica subsp. enterica serovar* Agona str. 64.H.00 |
| Sen-406-1 | Bacteria Proteobacteria Gammaproteobacteria Enterobacteriales Enterobacteriaceae *Salmonella enterica subsp. enterica serovar* Agona str. 648586-1 |
| Sen-407-1 | Bacteria Proteobacteria Gammaproteobacteria Enterobacteriales Enterobacteriaceae *Salmonella enterica subsp. enterica serovar* Agona str. 65.H.72 |
| Sen-408-1 | Bacteria Proteobacteria Gammaproteobacteria Enterobacteriales Enterobacteriaceae *Salmonella enterica subsp. enterica serovar* Agona str. 66.F.99 |
| Sen-409-1 | Bacteria Proteobacteria Gammaproteobacteria Enterobacteriales Enterobacteriaceae *Salmonella enterica subsp. enterica serovar* Agona str. 67.H.09 |
| Sen-410-1 | Bacteria Proteobacteria Gammaproteobacteria Enterobacteriales Enterobacteriaceae *Salmonella enterica subsp. enterica serovar* Agona str. 68.U.05 |
| Sen-411-1 | Bacteria Proteobacteria Gammaproteobacteria Enterobacteriales Enterobacteriaceae *Salmonella enterica subsp. enterica serovar* Agona str. 69.H.06 |
| Sen-412-1 | Bacteria Proteobacteria Gammaproteobacteria Enterobacteriales Enterobacteriaceae *Salmonella enterica subsp. enterica serovar* Agona str. 70.E.05 |
| Sen-413-1 | Bacteria Proteobacteria Gammaproteobacteria Enterobacteriales Enterobacteriaceae *Salmonella enterica subsp. enterica serovar* Agona str. 71.E.05 |
| Sen-414-1 | Bacteria Proteobacteria Gammaproteobacteria Enterobacteriales Enterobacteriaceae *Salmonella enterica subsp. enterica serovar* Agona str. 72.A.52 |
| Sen-415-1 | Bacteria Proteobacteria Gammaproteobacteria Enterobacteriales Enterobacteriaceae *Salmonella enterica subsp. enterica serovar* Agona str. 73.H.09 |
| Sen-416-1 | Bacteria Proteobacteria Gammaproteobacteria Enterobacteriales Enterobacteriaceae *Salmonella enterica subsp. enterica serovar* Agona str. ATCC 51957 |
| Sen-417-1 | Bacteria Proteobacteria Gammaproteobacteria Enterobacteriales Enterobacteriaceae *Salmonella enterica subsp. enterica serovar* Agona str. ATCC BAA-707 |
| Sen-418-1 | Bacteria Proteobacteria Gammaproteobacteria Enterobacteriales Enterobacteriaceae *Salmonella enterica subsp. enterica serovar* Agona str. SA-1 |
| Sen-419-1 | Bacteria Proteobacteria Gammaproteobacteria Enterobacteriales Enterobacteriaceae *Salmonella enterica subsp. enterica serovar* Agona str. SA-2 |
| Sen-420-1 | Bacteria Proteobacteria Gammaproteobacteria Enterobacteriales Enterobacteriaceae *Salmonella enterica subsp. enterica serovar* Agona str. SA-3 |
| Sen-421-1 | Bacteria Proteobacteria Gammaproteobacteria Enterobacteriales Enterobacteriaceae *Salmonella enterica subsp. enterica serovar* Agona str. SA-4 |
| Sen-422-1 | Bacteria Proteobacteria Gammaproteobacteria Enterobacteriales Enterobacteriaceae *Salmonella enterica subsp. enterica serovar* Agona str. SA-5 |
| Sen-423-1 | Bacteria Proteobacteria Gammaproteobacteria Enterobacteriales Enterobacteriaceae *Salmonella enterica subsp. enterica serovar* Agona str. SH08SF124 |
| Sen-424-1 | Bacteria Proteobacteria Gammaproteobacteria Enterobacteriales Enterobacteriaceae *Salmonella enterica subsp. enterica serovar* Agona str. SH10GFN094 |
| Sen-425-1 | Bacteria Proteobacteria Gammaproteobacteria Enterobacteriales Enterobacteriaceae *Salmonella enterica subsp. enterica serovar* Agona str. SH11G1113 |
| Sen-426-1 | Bacteria Proteobacteria Gammaproteobacteria Enterobacteriales Enterobacteriaceae *Salmonella enterica subsp. enterica serovar* Albany str. ATCC 51960 |
| Sen-427-1 | Bacteria Proteobacteria Gammaproteobacteria Enterobacteriales Enterobacteriaceae *Salmonella enterica subsp. enterica serovar* Anatum str. CDC 06-0532 |
| Sen-428-1 | Bacteria Proteobacteria Gammaproteobacteria Enterobacteriales Enterobacteriaceae *Salmonella enterica subsp. enterica serovar* Anatum str. USDA 100 |
| Sen-429-1 | Bacteria Proteobacteria Gammaproteobacteria Enterobacteriales Enterobacteriaceae *Salmonella enterica subsp. enterica serovar* Anatum str. USDA-ARS-USMARC-1175 |
| Sen-430-1 | Bacteria Proteobacteria Gammaproteobacteria Enterobacteriales Enterobacteriaceae *Salmonella enterica subsp. enterica serovar* Anatum str. USDA-ARS-USMARC-1735 |
| Sen-431-1 | Bacteria Proteobacteria Gammaproteobacteria Enterobacteriales Enterobacteriaceae *Salmonella enterica subsp. enterica serovar* Baildon str. R6-199 |
| Sen-432-1 | Bacteria Proteobacteria Gammaproteobacteria Enterobacteriales Enterobacteriaceae *Salmonella enterica subsp. enterica serovar* Bareilly str. 06-0784 |
| Sen-433-1 | Bacteria Proteobacteria Gammaproteobacteria Enterobacteriales Enterobacteriaceae *Salmonella enterica subsp. enterica serovar* Bareilly str. 2780 |
| Sen-434-1 | Bacteria Proteobacteria Gammaproteobacteria Enterobacteriales Enterobacteriaceae *Salmonella enterica subsp. enterica serovar* Bareilly str. ATCC 9115 |
| Sen-435-1 | Bacteria Proteobacteria Gammaproteobacteria Enterobacteriales Enterobacteriaceae *Salmonella enterica subsp. enterica serovar* Bareilly str. CFSAN000183 |
| Sen-436-1 | Bacteria Proteobacteria Gammaproteobacteria Enterobacteriales Enterobacteriaceae *Salmonella enterica subsp. enterica serovar* Bareilly str. CFSAN000189 |
| Sen-437-1 | Bacteria Proteobacteria Gammaproteobacteria Enterobacteriales Enterobacteriaceae *Salmonella enterica subsp. enterica serovar* Bareilly str. CFSAN000197 |
| Sen-438-1 | Bacteria Proteobacteria Gammaproteobacteria Enterobacteriales Enterobacteriaceae *Salmonella enterica subsp. enterica serovar* Bareilly str. CFSAN000200 |
| Sen-113-1 to Sen-113-2 | Bacteria Proteobacteria Gammaproteobacteria Enterobacteriales Enterobacteriaceae *Salmonella enterica subsp. enterica serovar* Berta str. ATCC 8392 |
| Sen-439-1 to Sen-439-11 | Bacteria Proteobacteria Gammaproteobacteria Enterobacteriales Enterobacteriaceae *Salmonella enterica subsp. enterica serovar* Bovismorbificans |
| Sen-440-1 | Bacteria Proteobacteria Gammaproteobacteria Enterobacteriales Enterobacteriaceae *Salmonella enterica subsp. enterica serovar* Bovismorbificans str. 3114 |
| Sen-441-1 | Bacteria Proteobacteria Gammaproteobacteria Enterobacteriales Enterobacteriaceae *Salmonella enterica subsp. enterica serovar* Braenderup str. ATCC BAA-664 |
| Sen-442-1 | Bacteria Proteobacteria Gammaproteobacteria Enterobacteriales Enterobacteriaceae *Salmonella enterica subsp. enterica serovar* Braenderup str. CF-SAN000756 |
| Sen-443-1 | Bacteria Proteobacteria Gammaproteobacteria Enterobacteriales Enterobacteriaceae *Salmonella enterica subsp. enterica serovar* Cerro str. 5569 |
| Sen-444-1 | Bacteria Proteobacteria Gammaproteobacteria Enterobacteriales Enterobacteriaceae *Salmonella enterica subsp. enterica serovar* Cerro str. 818 |

| | |
|---|---|
| Sen-445-1 | Bacteria Proteobacteria Gammaproteobacteria Enterobacteriales Enterobacteriaceae *Salmonella enterica subsp. enterica serovar* Cerro str. CFSAN001587 |
| Sen-446-1 | Bacteria Proteobacteria Gammaproteobacteria Enterobacteriales Enterobacteriaceae *Salmonella enterica subsp. enterica serovar* Cerro str. CFSAN001588 |
| Sen-447-1 | Bacteria Proteobacteria Gammaproteobacteria Enterobacteriales Enterobacteriaceae *Salmonella enterica subsp. enterica serovar* Cerro str. CFSAN001589 |
| Sen-448-1 | Bacteria Proteobacteria Gammaproteobacteria Enterobacteriales Enterobacteriaceae *Salmonella enterica subsp. enterica serovar* Cerro str. CFSAN001590 |
| Sen-449-1 | Bacteria Proteobacteria Gammaproteobacteria Enterobacteriales Enterobacteriaceae *Salmonella enterica subsp. enterica serovar* Cerro str. CFSAN001669 |
| Sen-450-1 | Bacteria Proteobacteria Gammaproteobacteria Enterobacteriales Enterobacteriaceae *Salmonella enterica subsp. enterica serovar* Cerro str. CFSAN001670 |
| Sen-451-1 | Bacteria Proteobacteria Gammaproteobacteria Enterobacteriales Enterobacteriaceae *Salmonella enterica subsp. enterica serovar* Cerro str. CFSAN001671 |
| Sen-452-1 | Bacteria Proteobacteria Gammaproteobacteria Enterobacteriales Enterobacteriaceae *Salmonella enterica subsp. enterica serovar* Cerro str. CFSAN001673 |
| Sen-453-1 | Bacteria Proteobacteria Gammaproteobacteria Enterobacteriales Enterobacteriaceae *Salmonella enterica subsp. enterica serovar* Cerro str. CFSAN001674 |
| Sen-454-1 | Bacteria Proteobacteria Gammaproteobacteria Enterobacteriales Enterobacteriaceae *Salmonella enterica subsp. enterica serovar* Cerro str. CFSAN001679 |
| Sen-455-1 | Bacteria Proteobacteria Gammaproteobacteria Enterobacteriales Enterobacteriaceae *Salmonella enterica subsp. enterica serovar* Cerro str. CFSAN001680 |
| Sen-456-1 | Bacteria Proteobacteria Gammaproteobacteria Enterobacteriales Enterobacteriaceae *Salmonella enterica subsp. enterica serovar* Cerro str. CFSAN001681 |
| Sen-457-1 | Bacteria Proteobacteria Gammaproteobacteria Enterobacteriales Enterobacteriaceae *Salmonella enterica subsp. enterica serovar* Cerro str. CFSAN001690 |
| Sen-458-1 | Bacteria Proteobacteria Gammaproteobacteria Enterobacteriales Enterobacteriaceae *Salmonella enterica subsp. enterica serovar* Cerro str. CFSAN001691 |
| Sen-459-1 | Bacteria Proteobacteria Gammaproteobacteria Enterobacteriales Enterobacteriaceae *Salmonella enterica subsp. enterica serovar* Cerro str. CFSAN001692 |
| Sen-460-1 | Bacteria Proteobacteria Gammaproteobacteria Enterobacteriales Enterobacteriaceae *Salmonella enterica subsp. enterica serovar* Cerro str. CFSAN001697 |
| Sen-461-1 | Bacteria Proteobacteria Gammaproteobacteria Enterobacteriales Enterobacteriaceae *Salmonella enterica subsp. enterica serovar* Chester str. ATCC 11997 |
| Sen-136-1 to Sen-136-2 | Bacteria Proteobacteria Gammaproteobacteria Enterobacteriales Enterobacteriaceae *Salmonella enterica subsp. enterica serovar* Cubana str. 76814 |
| Sen-462-1 | Bacteria Proteobacteria Gammaproteobacteria Enterobacteriales Enterobacteriaceae *Salmonella enterica subsp. enterica serovar* Cubana str. CFSAN001083 |
| Sen-463-1 | Bacteria Proteobacteria Gammaproteobacteria Enterobacteriales Enterobacteriaceae *Salmonella enterica subsp. enterica serovar* Cubana str. CFSAN002050 |
| Sen-464-1 | Bacteria Proteobacteria Gammaproteobacteria Enterobacteriales Enterobacteriaceae *Salmonella enterica subsp. enterica serovar* Cubana str. CVM42234 |
| Sen-465-1 | Bacteria Proteobacteria Gammaproteobacteria Enterobacteriales Enterobacteriaceae *Salmonella enterica subsp. enterica serovar* Derby str. 626 |
| Sen-466-1 | Bacteria Proteobacteria Gammaproteobacteria Enterobacteriales Enterobacteriaceae *Salmonella enterica subsp. enterica serovar* Dublin str. DG22 |
| Sen-467-1 | Bacteria Proteobacteria Gammaproteobacteria Enterobacteriales Enterobacteriaceae *Salmonella enterica subsp. enterica serovar* Dublin str. HWS51 |
| Sen-468-1 | Bacteria Proteobacteria Gammaproteobacteria Enterobacteriales Enterobacteriaceae *Salmonella enterica subsp. enterica serovar* Dublin str. SD3246 |
| Sen-469-1 | Bacteria Proteobacteria Gammaproteobacteria Enterobacteriales Enterobacteriaceae *Salmonella enterica subsp. enterica serovar* Dublin str. SL1438 |
| Sen-470-1 | Bacteria Proteobacteria Gammaproteobacteria Enterobacteriales Enterobacteriaceae *Salmonella enterica subsp. enterica serovar* Dublin str. UC16 |
| Sen-471-1 | Bacteria Proteobacteria Gammaproteobacteria Enterobacteriales Enterobacteriaceae *Salmonella enterica subsp. enterica serovar* Eastbourne str. CFSAN001084 |
| Sen-472-1 | Bacteria Proteobacteria Gammaproteobacteria Enterobacteriales Enterobacteriaceae *Salmonella enterica subsp. enterica serovar* Enteritidis str. 08-0047 |
| Sen-473-1 | Bacteria Proteobacteria Gammaproteobacteria Enterobacteriales Enterobacteriaceae *Salmonella enterica subsp. enterica serovar* Enteritidis str. 08-0128 |
| Sen-474-1 | Bacteria Proteobacteria Gammaproteobacteria Enterobacteriales Enterobacteriaceae *Salmonella enterica subsp. enterica serovar* Enteritidis str. 08-0627 |
| Sen-475-1 | Bacteria Proteobacteria Gammaproteobacteria Enterobacteriales Enterobacteriaceae *Salmonella enterica subsp. enterica serovar* Enteritidis str. 08-1080 |
| Sen-476-1 | Bacteria Proteobacteria Gammaproteobacteria Enterobacteriales Enterobacteriaceae *Salmonella enterica subsp. enterica serovar* Enteritidis str. 10-34587 |
| Sen-477-1 | Bacteria Proteobacteria Gammaproteobacteria Enterobacteriales Enterobacteriaceae *Salmonella enterica subsp. enterica serovar* Enteritidis str. 13-1 |
| Sen-478-1 | Bacteria Proteobacteria Gammaproteobacteria Enterobacteriales Enterobacteriaceae *Salmonella enterica subsp. enterica serovar* Enteritidis str. 13183-1 |
| Sen-479-1 | Bacteria Proteobacteria Gammaproteobacteria Enterobacteriales Enterobacteriaceae *Salmonella enterica subsp. enterica serovar* Enteritidis str. 17927 |
| Sen-480-1 | Bacteria Proteobacteria Gammaproteobacteria Enterobacteriales Enterobacteriaceae *Salmonella enterica subsp. enterica serovar* Enteritidis str. 20037 |
| Sen-481-1 | Bacteria Proteobacteria Gammaproteobacteria Enterobacteriales Enterobacteriaceae *Salmonella enterica subsp. enterica serovar* Enteritidis str. 2009K0958 |
| Sen-482-1 | Bacteria Proteobacteria Gammaproteobacteria Enterobacteriales Enterobacteriaceae *Salmonella enterica subsp. enterica serovar* Enteritidis str. 2009K1651 |
| Sen-483-1 | Bacteria Proteobacteria Gammaproteobacteria Enterobacteriales Enterobacteriaceae *Salmonella enterica subsp. enterica serovar* Enteritidis str. 2009K1726 |
| Sen-484-1 | Bacteria Proteobacteria Gammaproteobacteria Enterobacteriales Enterobacteriaceae *Salmonella enterica subsp. enterica serovar* Enteritidis str. 2010-1237 |
| Sen-485-1 | Bacteria Proteobacteria Gammaproteobacteria Enterobacteriales Enterobacteriaceae *Salmonella enterica subsp. enterica serovar* Enteritidis str. 2010K-0262 |
| Sen-486-1 | Bacteria Proteobacteria Gammaproteobacteria Enterobacteriales Enterobacteriaceae *Salmonella enterica subsp. enterica serovar* Enteritidis str. 2010K-0267 |
| Sen-487-1 | Bacteria Proteobacteria Gammaproteobacteria Enterobacteriales Enterobacteriaceae *Salmonella enterica subsp. enterica serovar* Enteritidis str. 2010K-0271 |
| Sen-488-1 | Bacteria Proteobacteria Gammaproteobacteria Enterobacteriales Enterobacteriaceae *Salmonella enterica subsp. enterica serovar* Enteritidis str. 2010K-0284 |
| Sen-489-1 | Bacteria Proteobacteria Gammaproteobacteria Enterobacteriales Enterobacteriaceae *Salmonella enterica subsp. enterica serovar* Enteritidis str. 2010K-0286 |
| Sen-490-1 | Bacteria Proteobacteria Gammaproteobacteria Enterobacteriales Enterobacteriaceae *Salmonella enterica subsp. enterica serovar* Enteritidis str. 2010K-0669 |
| Sen-491-1 | Bacteria Proteobacteria Gammaproteobacteria Enterobacteriales Enterobacteriaceae *Salmonella enterica subsp. enterica serovar* Enteritidis str. 2010K-2599 |
| Sen-492-1 | Bacteria Proteobacteria Gammaproteobacteria Enterobacteriales Enterobacteriaceae *Salmonella enterica subsp. enterica serovar* Enteritidis str. 22510-1 |
| Sen-493-1 | Bacteria Proteobacteria Gammaproteobacteria Enterobacteriales Enterobacteriaceae *Salmonella enterica subsp. enterica serovar* Enteritidis str. 22704 |
| Sen-494-1 | Bacteria Proteobacteria Gammaproteobacteria Enterobacteriales Enterobacteriaceae *Salmonella enterica subsp. enterica serovar* Enteritidis str. 33944 |
| Sen-495-1 | Bacteria Proteobacteria Gammaproteobacteria Enterobacteriales Enterobacteriaceae *Salmonella enterica subsp. enterica serovar* Enteritidis str. 3402 |
| Sen-496-1 | Bacteria Proteobacteria Gammaproteobacteria Enterobacteriales Enterobacteriaceae *Salmonella enterica subsp. enterica serovar* Enteritidis str. 436 |
| Sen-497-1 | Bacteria Proteobacteria Gammaproteobacteria Enterobacteriales Enterobacteriaceae *Salmonella enterica subsp. enterica serovar* Enteritidis str. 485549-17 |
| Sen-498-1 | Bacteria Proteobacteria Gammaproteobacteria Enterobacteriales Enterobacteriaceae *Salmonella enterica subsp. enterica serovar* Enteritidis str. 50-3079 |
| Sen-499-1 | Bacteria Proteobacteria Gammaproteobacteria Enterobacteriales Enterobacteriaceae *Salmonella enterica subsp. enterica serovar* Enteritidis str. 50-5646 |
| Sen-500-1 | Bacteria Proteobacteria Gammaproteobacteria Enterobacteriales Enterobacteriaceae *Salmonella enterica subsp. enterica serovar* Enteritidis str. 53-407 |
| Sen-501-1 | Bacteria Proteobacteria Gammaproteobacteria Enterobacteriales Enterobacteriaceae *Salmonella enterica subsp. enterica serovar* Enteritidis str. 543463 22-17 |
| Sen-502-1 | Bacteria Proteobacteria Gammaproteobacteria Enterobacteriales Enterobacteriaceae *Salmonella enterica subsp. enterica serovar* Enteritidis str. 543463 40-18 |

| | |
|---|---|
| Sen-503-1 | Bacteria Proteobacteria Gammaproteobacteria Enterobacteriales Enterobacteriaceae *Salmonella enterica subsp. enterica serovar* Enteritidis str. 543463 42-20 |
| Sen-504-1 | Bacteria Proteobacteria Gammaproteobacteria Enterobacteriales Enterobacteriaceae *Salmonella enterica subsp. enterica serovar* Enteritidis str. 561362 1-1 |
| Sen-505-1 | Bacteria Proteobacteria Gammaproteobacteria Enterobacteriales Enterobacteriaceae *Salmonella enterica subsp. enterica serovar* Enteritidis str. 561362 9-7 |
| Sen-506-1 | Bacteria Proteobacteria Gammaproteobacteria Enterobacteriales Enterobacteriaceae *Salmonella enterica subsp. enterica serovar* Enteritidis str. 576709 |
| Sen-507-1 | Bacteria Proteobacteria Gammaproteobacteria Enterobacteriales Enterobacteriaceae *Salmonella enterica subsp. enterica serovar* Enteritidis str. 58-6482 |
| Sen-508-1 | Bacteria Proteobacteria Gammaproteobacteria Enterobacteriales Enterobacteriaceae *Salmonella enterica subsp. enterica serovar* Enteritidis str. 596866-22 |
| Sen-509-1 | Bacteria Proteobacteria Gammaproteobacteria Enterobacteriales Enterobacteriaceae *Salmonella enterica subsp. enterica serovar* Enteritidis str. 596866-70 |
| Sen-510-1 | Bacteria Proteobacteria Gammaproteobacteria Enterobacteriales Enterobacteriaceae *Salmonella enterica subsp. enterica serovar* Enteritidis str. 6.0562-1 |
| Sen-511-1 | Bacteria Proteobacteria Gammaproteobacteria Enterobacteriales Enterobacteriaceae *Salmonella enterica subsp. enterica serovar* Enteritidis str. 607307-2 |
| Sen-512-1 | Bacteria Proteobacteria Gammaproteobacteria Enterobacteriales Enterobacteriaceae *Salmonella enterica subsp. enterica serovar* Enteritidis str. 607307-6 |
| Sen-513-1 | Bacteria Proteobacteria Gammaproteobacteria Enterobacteriales Enterobacteriaceae *Salmonella enterica subsp. enterica serovar* Enteritidis str. 607308-16 |
| Sen-514-1 | Bacteria Proteobacteria Gammaproteobacteria Enterobacteriales Enterobacteriaceae *Salmonella enterica subsp. enterica serovar* Enteritidis str. 607308-19 |
| Sen-515-1 | Bacteria Proteobacteria Gammaproteobacteria Enterobacteriales Enterobacteriaceae *Salmonella enterica subsp. enterica serovar* Enteritidis str. 607308-9 |
| Sen-516-1 | Bacteria Proteobacteria Gammaproteobacteria Enterobacteriales Enterobacteriaceae *Salmonella enterica subsp. enterica serovar* Enteritidis str. 62-1976 |
| Sen-517-1 | Bacteria Proteobacteria Gammaproteobacteria Enterobacteriales Enterobacteriaceae *Salmonella enterica subsp. enterica serovar* Enteritidis str. 622731-39 |
| Sen-518-1 | Bacteria Proteobacteria Gammaproteobacteria Enterobacteriales Enterobacteriaceae *Salmonella enterica subsp. enterica serovar* Enteritidis str. 629163 |
| Sen-519-1 | Bacteria Proteobacteria Gammaproteobacteria Enterobacteriales Enterobacteriaceae *Salmonella enterica subsp. enterica serovar* Enteritidis str. 629164-26 |
| Sen-520-1 | Bacteria Proteobacteria Gammaproteobacteria Enterobacteriales Enterobacteriaceae *Salmonella enterica subsp. enterica serovar* Enteritidis str. 629164-37 |
| Sen-521-1 | Bacteria Proteobacteria Gammaproteobacteria Enterobacteriales Enterobacteriaceae *Salmonella enterica subsp. enterica serovar* Enteritidis str. 635290-58 |
| Sen-522-1 | Bacteria Proteobacteria Gammaproteobacteria Enterobacteriales Enterobacteriaceae *Salmonella enterica subsp. enterica serovar* Enteritidis str. 638970-15 |
| Sen-523-1 | Bacteria Proteobacteria Gammaproteobacteria Enterobacteriales Enterobacteriaceae *Salmonella enterica subsp. enterica serovar* Enteritidis str. 639016-6 |
| Sen-524-1 | Bacteria Proteobacteria Gammaproteobacteria Enterobacteriales Enterobacteriaceae *Salmonella enterica subsp. enterica serovar* Enteritidis str. 639672-46 |
| Sen-525-1 | Bacteria Proteobacteria Gammaproteobacteria Enterobacteriales Enterobacteriaceae *Salmonella enterica subsp. enterica serovar* Enteritidis str. 639672-50 |
| Sen-526-1 | Bacteria Proteobacteria Gammaproteobacteria Enterobacteriales Enterobacteriaceae *Salmonella enterica subsp. enterica serovar* Enteritidis str. 640631 |
| Sen-527-1 | Bacteria Proteobacteria Gammaproteobacteria Enterobacteriales Enterobacteriaceae *Salmonella enterica subsp. enterica serovar* Enteritidis str. 642044 4-1 |
| Sen-528-1 | Bacteria Proteobacteria Gammaproteobacteria Enterobacteriales Enterobacteriaceae *Salmonella enterica subsp. enterica serovar* Enteritidis str. 642044 8-1 |
| Sen-529-1 | Bacteria Proteobacteria Gammaproteobacteria Enterobacteriales Enterobacteriaceae *Salmonella enterica subsp. enterica serovar* Enteritidis str. 642046 4-7 |
| Sen-530-1 | Bacteria Proteobacteria Gammaproteobacteria Enterobacteriales Enterobacteriaceae *Salmonella enterica subsp. enterica serovar* Enteritidis str. 648898 4-5 |
| Sen-531-1 | Bacteria Proteobacteria Gammaproteobacteria Enterobacteriales Enterobacteriaceae *Salmonella enterica subsp. enterica serovar* Enteritidis str. 648899 3-17 |
| Sen-532-1 | Bacteria Proteobacteria Gammaproteobacteria Enterobacteriales Enterobacteriaceae *Salmonella enterica subsp. enterica serovar* Enteritidis str. 648900 1-16 |
| Sen-533-1 | Bacteria Proteobacteria Gammaproteobacteria Enterobacteriales Enterobacteriaceae *Salmonella enterica subsp. enterica serovar* Enteritidis str. 648901 1-17 |
| Sen-534-1 | Bacteria Proteobacteria Gammaproteobacteria Enterobacteriales Enterobacteriaceae *Salmonella enterica subsp. enterica serovar* Enteritidis str. 648901 16-16 |
| Sen-535-1 | Bacteria Proteobacteria Gammaproteobacteria Enterobacteriales Enterobacteriaceae *Salmonella enterica subsp. enterica serovar* Enteritidis str. 648901 39-2 |
| Sen-536-1 | Bacteria Proteobacteria Gammaproteobacteria Enterobacteriales Enterobacteriaceae *Salmonella enterica subsp. enterica serovar* Enteritidis str. 648901 6-18 |
| Sen-537-1 | Bacteria Proteobacteria Gammaproteobacteria Enterobacteriales Enterobacteriaceae *Salmonella enterica subsp. enterica serovar* Enteritidis str. 648902 6-8 |
| Sen-538-1 | Bacteria Proteobacteria Gammaproteobacteria Enterobacteriales Enterobacteriaceae *Salmonella enterica subsp. enterica serovar* Enteritidis str. 648903 1-6 |
| Sen-539-1 | Bacteria Proteobacteria Gammaproteobacteria Enterobacteriales Enterobacteriaceae *Salmonella enterica subsp. enterica serovar* Enteritidis str. 648904 3-6 |
| Sen-540-1 | Bacteria Proteobacteria Gammaproteobacteria Enterobacteriales Enterobacteriaceae *Salmonella enterica subsp. enterica serovar* Enteritidis str. 648905 5-18 |
| Sen-541-1 | Bacteria Proteobacteria Gammaproteobacteria Enterobacteriales Enterobacteriaceae *Salmonella enterica subsp. enterica serovar* Enteritidis str. 653049 13-19 |
| Sen-542-1 | Bacteria Proteobacteria Gammaproteobacteria Enterobacteriales Enterobacteriaceae *Salmonella enterica subsp. enterica serovar* Enteritidis str. 76-2651 |
| Sen-543-1 | Bacteria Proteobacteria Gammaproteobacteria Enterobacteriales Enterobacteriaceae *Salmonella enterica subsp. enterica serovar* Enteritidis str. 77-0424 |
| Sen-544-1 | Bacteria Proteobacteria Gammaproteobacteria Enterobacteriales Enterobacteriaceae *Salmonella enterica subsp. enterica serovar* Enteritidis str. 77-2659 |
| Sen-545-1 | Bacteria Proteobacteria Gammaproteobacteria Enterobacteriales Enterobacteriaceae *Salmonella enterica subsp. enterica serovar* Enteritidis str. 78-1757 |
| Sen-546-1 | Bacteria Proteobacteria Gammaproteobacteria Enterobacteriales Enterobacteriaceae *Salmonella enterica subsp. enterica serovar* Enteritidis str. 81-2625 |
| Sen-547-1 | Bacteria Proteobacteria Gammaproteobacteria Enterobacteriales Enterobacteriaceae *Salmonella enterica subsp. enterica serovar* Enteritidis str. 8b-1 |
| Sen-548-1 | Bacteria Proteobacteria Gammaproteobacteria Enterobacteriales Enterobacteriaceae *Salmonella enterica subsp. enterica serovar* Enteritidis str. CDC_2010K_0895 |
| Sen-549-1 | Bacteria Proteobacteria Gammaproteobacteria Enterobacteriales Enterobacteriaceae *Salmonella enterica subsp. enterica serovar* Enteritidis str. CDC_2010K_0899 |

| | |
|---|---|
| Sen-550-1 | Bacteria Proteobacteria Gammaproteobacteria Enterobacteriales Enterobacteriaceae *Salmonella enterica subsp. enterica serovar* Enteritidis str. CDC_2010K_0956 |
| Sen-551-1 | Bacteria Proteobacteria Gammaproteobacteria Enterobacteriales Enterobacteriaceae *Salmonella enterica subsp. enterica serovar* Enteritidis str. CDC_2010K_1010 |
| Sen-552-1 | Bacteria Proteobacteria Gammaproteobacteria Enterobacteriales Enterobacteriaceae *Salmonella enterica subsp. enterica serovar* Enteritidis str. CDC_2010K_1018 |
| Sen-553-1 | Bacteria Proteobacteria Gammaproteobacteria Enterobacteriales Enterobacteriaceae *Salmonella enterica subsp. enterica serovar* Enteritidis str. CDC_2010K_1441 |
| Sen-554-1 | Bacteria Proteobacteria Gammaproteobacteria Enterobacteriales Enterobacteriaceae *Salmonella enterica subsp. enterica serovar* Enteritidis str. CDC_2010K_1444 |
| Sen-555-1 | Bacteria Proteobacteria Gammaproteobacteria Enterobacteriales Enterobacteriaceae *Salmonella enterica subsp. enterica serovar* Enteritidis str. CDC_2010K_1445 |
| Sen-556-1 | Bacteria Proteobacteria Gammaproteobacteria Enterobacteriales Enterobacteriaceae *Salmonella enterica subsp. enterica serovar* Enteritidis str. CDC_2010K_1455 |
| Sen-557-1 | Bacteria Proteobacteria Gammaproteobacteria Enterobacteriales Enterobacteriaceae *Salmonella enterica subsp. enterica serovar* Enteritidis str. CDC_2010K_1457 |
| Sen-558-1 | Bacteria Proteobacteria Gammaproteobacteria Enterobacteriales Enterobacteriaceae *Salmonella enterica subsp. enterica serovar* Enteritidis str. CDC_2010K_1543 |
| Sen-559-1 | Bacteria Proteobacteria Gammaproteobacteria Enterobacteriales Enterobacteriaceae *Salmonella enterica subsp. enterica serovar* Enteritidis str. CDC_2010K_1558 |
| Sen-560-1 | Bacteria Proteobacteria Gammaproteobacteria Enterobacteriales Enterobacteriaceae *Salmonella enterica subsp. enterica serovar* Enteritidis str. CDC_2010K_1559 |
| Sen-561-1 | Bacteria Proteobacteria Gammaproteobacteria Enterobacteriales Enterobacteriaceae *Salmonella enterica subsp. enterica serovar* Enteritidis str. CDC_2010K_1565 |
| Sen-562-1 | Bacteria Proteobacteria Gammaproteobacteria Enterobacteriales Enterobacteriaceae *Salmonella enterica subsp. enterica serovar* Enteritidis str. CDC_2010K_1566 |
| Sen-563-1 | Bacteria Proteobacteria Gammaproteobacteria Enterobacteriales Enterobacteriaceae *Salmonella enterica subsp. enterica serovar* Enteritidis str. CDC_2010K_1575 |
| Sen-564-1 | Bacteria Proteobacteria Gammaproteobacteria Enterobacteriales Enterobacteriaceae *Salmonella enterica subsp. enterica serovar* Enteritidis str. CDC_2010K_1580 |
| Sen-565-1 | Bacteria Proteobacteria Gammaproteobacteria Enterobacteriales Enterobacteriaceae *Salmonella enterica subsp. enterica serovar* Enteritidis str. CDC_2010K_1594 |
| Sen-566-1 | Bacteria Proteobacteria Gammaproteobacteria Enterobacteriales Enterobacteriaceae *Salmonella enterica subsp. enterica serovar* Enteritidis str. CDC_2010K_1725 |
| Sen-567-1 | Bacteria Proteobacteria Gammaproteobacteria Enterobacteriales Enterobacteriaceae *Salmonella enterica subsp. enterica serovar* Enteritidis str. CDC_2010K_1729 |
| Sen-568-1 | Bacteria Proteobacteria Gammaproteobacteria Enterobacteriales Enterobacteriaceae *Salmonella enterica subsp. enterica serovar* Enteritidis str. CDC_2010K_1745 |
| Sen-569-1 | Bacteria Proteobacteria Gammaproteobacteria Enterobacteriales Enterobacteriaceae *Salmonella enterica subsp. enterica serovar* Enteritidis str. CDC_2010K_1747 |
| Sen-570-1 | Bacteria Proteobacteria Gammaproteobacteria Enterobacteriales Enterobacteriaceae *Salmonella enterica subsp. enterica serovar* Enteritidis str. CDC_2010K_1791 |
| Sen-571-1 | Bacteria Proteobacteria Gammaproteobacteria Enterobacteriales Enterobacteriaceae *Salmonella enterica subsp. enterica serovar* Enteritidis str. CDC_2010K_1795 |
| Sen-572-1 | Bacteria Proteobacteria Gammaproteobacteria Enterobacteriales Enterobacteriaceae *Salmonella enterica subsp. enterica serovar* Enteritidis str. CDC_2010K_1808 |
| Sen-573-1 | Bacteria Proteobacteria Gammaproteobacteria Enterobacteriales Enterobacteriaceae *Salmonella enterica subsp. enterica serovar* Enteritidis str. CDC_2010K_1810 |
| Sen-574-1 | Bacteria Proteobacteria Gammaproteobacteria Enterobacteriales Enterobacteriaceae *Salmonella enterica subsp. enterica serovar* Enteritidis str. CDC_2010K_1811 |
| Sen-575-1 | Bacteria Proteobacteria Gammaproteobacteria Enterobacteriales Enterobacteriaceae *Salmonella enterica subsp. enterica serovar* Enteritidis str. CDC_2010K_1882 |

| | |
|---|---|
| Sen-576-1 | Bacteria Proteobacteria Gammaproteobacteria Enterobacteriales Enterobacteriaceae *Salmonella enterica subsp. enterica serovar* Enteritidis str. CDC_2010K_1884 |
| Sen-577-1 | Bacteria Proteobacteria Gammaproteobacteria Enterobacteriales Enterobacteriaceae *Salmonella enterica subsp. enterica serovar* Enteritidis str. CHS4 |
| Sen-578-1 | Bacteria Proteobacteria Gammaproteobacteria Enterobacteriales Enterobacteriaceae *Salmonella enterica subsp. enterica serovar* Enteritidis str. CHS44 |
| Sen-579-1 | Bacteria Proteobacteria Gammaproteobacteria Enterobacteriales Enterobacteriaceae *Salmonella enterica subsp. enterica serovar* Enteritidis str. CVM_56-3991 |
| Sen-580-1 | Bacteria Proteobacteria Gammaproteobacteria Enterobacteriales Enterobacteriaceae *Salmonella enterica subsp. enterica serovar* Enteritidis str. CVM_69-4941 |
| Sen-581-1 | Bacteria Proteobacteria Gammaproteobacteria Enterobacteriales Enterobacteriaceae *Salmonella enterica subsp. enterica serovar* Enteritidis str. CVM_76-3618 |
| Sen-582-1 | Bacteria Proteobacteria Gammaproteobacteria Enterobacteriales Enterobacteriaceae *Salmonella enterica subsp. enterica serovar* Enteritidis str. CVM_81-2490 |
| Sen-583-1 | Bacteria Proteobacteria Gammaproteobacteria Enterobacteriales Enterobacteriaceae *Salmonella enterica subsp. enterica serovar* Enteritidis str. CVM_N202 |
| Sen-584-1 | Bacteria Proteobacteria Gammaproteobacteria Enterobacteriales Enterobacteriaceae *Salmonella enterica subsp. enterica serovar* Enteritidis str. EC20090135 |
| Sen-585-1 | Bacteria Proteobacteria Gammaproteobacteria Enterobacteriales Enterobacteriaceae *Salmonella enterica subsp. enterica serovar* Enteritidis str. EC20090193 |
| Sen-586-1 | Bacteria Proteobacteria Gammaproteobacteria Enterobacteriales Enterobacteriaceae *Salmonella enterica subsp. enterica serovar* Enteritidis str. EC20090195 |
| Sen-587-1 | Bacteria Proteobacteria Gammaproteobacteria Enterobacteriales Enterobacteriaceae *Salmonella enterica subsp. enterica serovar* Enteritidis str. EC20090332 |
| Sen-588-1 | Bacteria Proteobacteria Gammaproteobacteria Enterobacteriales Enterobacteriaceae *Salmonella enterica subsp. enterica serovar* Enteritidis str. EC20090530 |
| Sen-589-1 | Bacteria Proteobacteria Gammaproteobacteria Enterobacteriales Enterobacteriaceae *Salmonella enterica subsp. enterica serovar* Enteritidis str. EC20090531 |
| Sen-590-1 | Bacteria Proteobacteria Gammaproteobacteria Enterobacteriales Enterobacteriaceae *Salmonella enterica subsp. enterica serovar* Enteritidis str. EC20090884 |
| Sen-591-1 | Bacteria Proteobacteria Gammaproteobacteria Enterobacteriales Enterobacteriaceae *Salmonella enterica subsp. enterica serovar* Enteritidis str. EC20100088 |
| Sen-592-1 | Bacteria Proteobacteria Gammaproteobacteria Enterobacteriales Enterobacteriaceae *Salmonella enterica subsp. enterica serovar* Enteritidis str. EC20100089 |
| Sen-593-1 | Bacteria Proteobacteria Gammaproteobacteria Enterobacteriales Enterobacteriaceae *Salmonella enterica subsp. enterica serovar* Enteritidis str. EC20100100 |
| Sen-594-1 | Bacteria Proteobacteria Gammaproteobacteria Enterobacteriales Enterobacteriaceae *Salmonella enterica subsp. enterica serovar* Enteritidis str. EC20100103 |
| Sen-595-1 | Bacteria Proteobacteria Gammaproteobacteria Enterobacteriales Enterobacteriaceae *Salmonella enterica subsp. enterica serovar* Enteritidis str. EC20100130 |
| Sen-596-1 | Bacteria Proteobacteria Gammaproteobacteria Enterobacteriales Enterobacteriaceae *Salmonella enterica subsp. enterica serovar* Enteritidis str. EC20100131 |
| Sen-597-1 | Bacteria Proteobacteria Gammaproteobacteria Enterobacteriales Enterobacteriaceae *Salmonella enterica subsp. enterica serovar* Enteritidis str. EC20100134 |
| Sen-598-1 | Bacteria Proteobacteria Gammaproteobacteria Enterobacteriales Enterobacteriaceae *Salmonella enterica subsp. enterica serovar* Enteritidis str. EC20100325 |
| Sen-599-1 | Bacteria Proteobacteria Gammaproteobacteria Enterobacteriales Enterobacteriaceae *Salmonella enterica subsp. enterica serovar* Enteritidis str. EC20110222 |
| Sen-600-1 | Bacteria Proteobacteria Gammaproteobacteria Enterobacteriales Enterobacteriaceae *Salmonella enterica subsp. enterica serovar* Enteritidis str. EC20110223 |
| Sen-601-1 | Bacteria Proteobacteria Gammaproteobacteria Enterobacteriales Enterobacteriaceae *Salmonella enterica subsp. enterica serovar* Enteritidis str. EC20110353 |
| Sen-602-1 | Bacteria Proteobacteria Gammaproteobacteria Enterobacteriales Enterobacteriaceae *Salmonella enterica subsp. enterica serovar* Enteritidis str. EC20111510 |
| Sen-603-1 | Bacteria Proteobacteria Gammaproteobacteria Enterobacteriales Enterobacteriaceae *Salmonella enterica subsp. enterica serovar* Enteritidis str. EC20111514 |
| Sen-604-1 | Bacteria Proteobacteria Gammaproteobacteria Enterobacteriales Enterobacteriaceae *Salmonella enterica subsp. enterica serovar* Enteritidis str. EC20111515 |
| Sen-605-1 | Bacteria Proteobacteria Gammaproteobacteria Enterobacteriales Enterobacteriaceae *Salmonella enterica subsp. enterica serovar* Enteritidis str. EC20111554 |
| Sen-606-1 | Bacteria Proteobacteria Gammaproteobacteria Enterobacteriales Enterobacteriaceae *Salmonella enterica subsp. enterica serovar* Enteritidis str. EC20111561 |
| Sen-607-1 | Bacteria Proteobacteria Gammaproteobacteria Enterobacteriales Enterobacteriaceae *Salmonella enterica subsp. enterica serovar* Enteritidis str. EC20111576 |
| Sen-608-1 | Bacteria Proteobacteria Gammaproteobacteria Enterobacteriales Enterobacteriaceae *Salmonella enterica subsp. enterica serovar* Enteritidis str. EC20120003 |
| Sen-609-1 | Bacteria Proteobacteria Gammaproteobacteria Enterobacteriales Enterobacteriaceae *Salmonella enterica subsp. enterica serovar* Enteritidis str. EC20120007 |
| Sen-610-1 | Bacteria Proteobacteria Gammaproteobacteria Enterobacteriales Enterobacteriaceae *Salmonella enterica subsp. enterica serovar* Enteritidis str. EC20120051 |
| Sen-611-1 | Bacteria Proteobacteria Gammaproteobacteria Enterobacteriales Enterobacteriaceae *Salmonella enterica subsp. enterica serovar* Enteritidis str. EC20120200 |
| Sen-612-1 | Bacteria Proteobacteria Gammaproteobacteria Enterobacteriales Enterobacteriaceae *Salmonella enterica subsp. enterica serovar* Enteritidis str. EC20120213 |
| Sen-613-1 | Bacteria Proteobacteria Gammaproteobacteria Enterobacteriales Enterobacteriaceae *Salmonella enterica subsp. enterica serovar* Enteritidis str. EC20120219 |
| Sen-614-1 | Bacteria Proteobacteria Gammaproteobacteria Enterobacteriales Enterobacteriaceae *Salmonella enterica subsp. enterica serovar* Enteritidis str. EC20120229 |
| Sen-615-1 | Bacteria Proteobacteria Gammaproteobacteria Enterobacteriales Enterobacteriaceae *Salmonella enterica subsp. enterica serovar* Enteritidis str. EC20120240 |
| Sen-616-1 | Bacteria Proteobacteria Gammaproteobacteria Enterobacteriales Enterobacteriaceae *Salmonella enterica subsp. enterica serovar* Enteritidis str. EC20120356 |
| Sen-617-1 | Bacteria Proteobacteria Gammaproteobacteria Enterobacteriales Enterobacteriaceae *Salmonella enterica subsp. enterica serovar* Enteritidis str. EC20120469 |
| Sen-618-1 | Bacteria Proteobacteria Gammaproteobacteria Enterobacteriales Enterobacteriaceae *Salmonella enterica subsp. enterica serovar* Enteritidis str. EC20120496 |
| Sen-619-1 | Bacteria Proteobacteria Gammaproteobacteria Enterobacteriales Enterobacteriaceae *Salmonella enterica subsp. enterica serovar* Enteritidis str. EC20120497 |
| Sen-620-1 | Bacteria Proteobacteria Gammaproteobacteria Enterobacteriales Enterobacteriaceae *Salmonella enterica subsp. enterica serovar* Enteritidis str. EC20120498 |
| Sen-621-1 | Bacteria Proteobacteria Gammaproteobacteria Enterobacteriales Enterobacteriaceae *Salmonella enterica subsp. enterica serovar* Enteritidis str. EC20120505 |
| Sen-622-1 | Bacteria Proteobacteria Gammaproteobacteria Enterobacteriales Enterobacteriaceae *Salmonella enterica subsp. enterica serovar* Enteritidis str. EC20120528 |
| Sen-623-1 | Bacteria Proteobacteria Gammaproteobacteria Enterobacteriales Enterobacteriaceae *Salmonella enterica subsp. enterica serovar* Enteritidis str. EC20120544 |
| Sen-624-1 | Bacteria Proteobacteria Gammaproteobacteria Enterobacteriales Enterobacteriaceae *Salmonella enterica subsp. enterica serovar* Enteritidis str. EC20120548 |
| Sen-625-1 | Bacteria Proteobacteria Gammaproteobacteria Enterobacteriales Enterobacteriaceae *Salmonella enterica subsp. enterica serovar* Enteritidis str. EC20120555 |
| Sen-626-1 | Bacteria Proteobacteria Gammaproteobacteria Enterobacteriales Enterobacteriaceae *Salmonella enterica subsp. enterica serovar* Enteritidis str. EC20120580 |
| Sen-627-1 | Bacteria Proteobacteria Gammaproteobacteria Enterobacteriales Enterobacteriaceae *Salmonella enterica subsp. enterica serovar* Enteritidis str. EC20120581 |
| Sen-628-1 | Bacteria Proteobacteria Gammaproteobacteria Enterobacteriales Enterobacteriaceae *Salmonella enterica subsp. enterica serovar* Enteritidis str. EC20120590 |
| Sen-629-1 | Bacteria Proteobacteria Gammaproteobacteria Enterobacteriales Enterobacteriaceae *Salmonella enterica subsp. enterica serovar* Enteritidis str. EC20120597 |
| Sen-630-1 | Bacteria Proteobacteria Gammaproteobacteria Enterobacteriales Enterobacteriaceae *Salmonella enterica subsp. enterica serovar* Enteritidis str. EC20120686 |
| Sen-631-1 | Bacteria Proteobacteria Gammaproteobacteria Enterobacteriales Enterobacteriaceae *Salmonella enterica subsp. enterica serovar* Enteritidis str. EC20120687 |
| Sen-632-1 | Bacteria Proteobacteria Gammaproteobacteria Enterobacteriales Enterobacteriaceae *Salmonella enterica subsp. enterica serovar* Enteritidis str. EC20120722 |
| Sen-633-1 | Bacteria Proteobacteria Gammaproteobacteria Enterobacteriales Enterobacteriaceae *Salmonella enterica subsp. enterica serovar* Enteritidis str. EC20120734 |

| | |
|---|---|
| Sen-634-1 | Bacteria Proteobacteria Gammaproteobacteria Enterobacteriales Enterobacteriaceae *Salmonella enterica subsp. enterica serovar* Enteritidis str. EC20120738 |
| Sen-635-1 | Bacteria Proteobacteria Gammaproteobacteria Enterobacteriales Enterobacteriaceae *Salmonella enterica subsp. enterica serovar* Enteritidis str. EC20120765 |
| Sen-636-1 | Bacteria Proteobacteria Gammaproteobacteria Enterobacteriales Enterobacteriaceae *Salmonella enterica subsp. enterica serovar* Enteritidis str. EC20120773 |
| Sen-637-1 | Bacteria Proteobacteria Gammaproteobacteria Enterobacteriales Enterobacteriaceae *Salmonella enterica subsp. enterica serovar* Enteritidis str. EC20120774 |
| Sen-638-1 | Bacteria Proteobacteria Gammaproteobacteria Enterobacteriales Enterobacteriaceae *Salmonella enterica subsp. enterica serovar* Enteritidis str. EC20120775 |
| Sen-639-1 | Bacteria Proteobacteria Gammaproteobacteria Enterobacteriales Enterobacteriaceae *Salmonella enterica subsp. enterica serovar* Enteritidis str. EC20120776 |
| Sen-640-1 | Bacteria Proteobacteria Gammaproteobacteria Enterobacteriales Enterobacteriaceae *Salmonella enterica subsp. enterica serovar* Enteritidis str. EC20120917 |
| Sen-641-1 | Bacteria Proteobacteria Gammaproteobacteria Enterobacteriales Enterobacteriaceae *Salmonella enterica subsp. enterica serovar* Enteritidis str. EC20120918 |
| Sen-642-1 | Bacteria Proteobacteria Gammaproteobacteria Enterobacteriales Enterobacteriaceae *Salmonella enterica subsp. enterica serovar* Enteritidis str. EC20120925 |
| Sen-643-1 | Bacteria Proteobacteria Gammaproteobacteria Enterobacteriales Enterobacteriaceae *Salmonella enterica subsp. enterica serovar* Enteritidis str. EC20120927 |
| Sen-644-1 | Bacteria Proteobacteria Gammaproteobacteria Enterobacteriales Enterobacteriaceae *Salmonella enterica subsp. enterica serovar* Enteritidis str. EC20120929 |
| Sen-645-1 | Bacteria Proteobacteria Gammaproteobacteria Enterobacteriales Enterobacteriaceae *Salmonella enterica subsp. enterica serovar* Enteritidis str. EC20120963 |
| Sen-646-1 | Bacteria Proteobacteria Gammaproteobacteria Enterobacteriales Enterobacteriaceae *Salmonella enterica subsp. enterica serovar* Enteritidis str. EC20120968 |
| Sen-647-1 | Bacteria Proteobacteria Gammaproteobacteria Enterobacteriales Enterobacteriaceae *Salmonella enterica subsp. enterica serovar* Enteritidis str. EC20120969 |
| Sen-648-1 | Bacteria Proteobacteria Gammaproteobacteria Enterobacteriales Enterobacteriaceae *Salmonella enterica subsp. enterica serovar* Enteritidis str. EC20120970 |
| Sen-649-1 to Sen-649-2 | Bacteria Proteobacteria Gammaproteobacteria Enterobacteriales Enterobacteriaceae *Salmonella enterica subsp. enterica serovar* Enteritidis str. EC20121179 |
| Sen-650-1 | Bacteria Proteobacteria Gammaproteobacteria Enterobacteriales Enterobacteriaceae *Salmonella enterica subsp. enterica serovar* Enteritidis str. EC20121542 |
| Sen-651-1 | Bacteria Proteobacteria Gammaproteobacteria Enterobacteriales Enterobacteriaceae *Salmonella enterica subsp. enterica serovar* Enteritidis str. EC20121671 |
| Sen-652-1 | Bacteria Proteobacteria Gammaproteobacteria Enterobacteriales Enterobacteriaceae *Salmonella enterica subsp. enterica serovar* Enteritidis str. EC20121672 |
| Sen-653-1 | Bacteria Proteobacteria Gammaproteobacteria Enterobacteriales Enterobacteriaceae *Salmonella enterica subsp. enterica serovar* Enteritidis str. EC20121689 |
| Sen-654-1 | Bacteria Proteobacteria Gammaproteobacteria Enterobacteriales Enterobacteriaceae *Salmonella enterica subsp. enterica serovar* Enteritidis str. EC20121744 |
| Sen-655-1 | Bacteria Proteobacteria Gammaproteobacteria Enterobacteriales Enterobacteriaceae *Salmonella enterica subsp. enterica serovar* Enteritidis str. EC20121746 |
| Sen-656-1 | Bacteria Proteobacteria Gammaproteobacteria Enterobacteriales Enterobacteriaceae *Salmonella enterica subsp. enterica serovar* Enteritidis str. EC20121747 |
| Sen-657-1 | Bacteria Proteobacteria Gammaproteobacteria Enterobacteriales Enterobacteriaceae *Salmonella enterica subsp. enterica serovar* Enteritidis str. EC20121748 |
| Sen-658-1 | Bacteria Proteobacteria Gammaproteobacteria Enterobacteriales Enterobacteriaceae *Salmonella enterica subsp. enterica serovar* Enteritidis str. EC20121750 |
| Sen-659-1 | Bacteria Proteobacteria Gammaproteobacteria Enterobacteriales Enterobacteriaceae *Salmonella enterica subsp. enterica serovar* Enteritidis str. EC20121751 |
| Sen-660-1 | Bacteria Proteobacteria Gammaproteobacteria Enterobacteriales Enterobacteriaceae *Salmonella enterica subsp. enterica serovar* Enteritidis str. EC20121753 |
| Sen-661-1 | Bacteria Proteobacteria Gammaproteobacteria Enterobacteriales Enterobacteriaceae *Salmonella enterica subsp. enterica serovar* Enteritidis str. EC20121765 |
| Sen-662-1 | Bacteria Proteobacteria Gammaproteobacteria Enterobacteriales Enterobacteriaceae *Salmonella enterica subsp. enterica serovar* Enteritidis str. EC20121812 |
| Sen-663-1 | Bacteria Proteobacteria Gammaproteobacteria Enterobacteriales Enterobacteriaceae *Salmonella enterica subsp. enterica serovar* Enteritidis str. EC20121825 |
| Sen-664-1 | Bacteria Proteobacteria Gammaproteobacteria Enterobacteriales Enterobacteriaceae *Salmonella enterica subsp. enterica serovar* Enteritidis str. EC20121826 |
| Sen-665-1 | Bacteria Proteobacteria Gammaproteobacteria Enterobacteriales Enterobacteriaceae *Salmonella enterica subsp. enterica serovar* Enteritidis str. EC20121969 |
| Sen-666-1 | Bacteria Proteobacteria Gammaproteobacteria Enterobacteriales Enterobacteriaceae *Salmonella enterica subsp. enterica serovar* Enteritidis str. EC20121970 |
| Sen-667-1 | Bacteria Proteobacteria Gammaproteobacteria Enterobacteriales Enterobacteriaceae *Salmonella enterica subsp. enterica serovar* Enteritidis str. EC20121976 |
| Sen-668-1 | Bacteria Proteobacteria Gammaproteobacteria Enterobacteriales Enterobacteriaceae *Salmonella enterica subsp. enterica serovar* Enteritidis str. EC20121986 |
| Sen-669-1 | Bacteria Proteobacteria Gammaproteobacteria Enterobacteriales Enterobacteriaceae *Salmonella enterica subsp. enterica serovar* Enteritidis str. EC20121989 |
| Sen-670-1 | Bacteria Proteobacteria Gammaproteobacteria Enterobacteriales Enterobacteriaceae *Salmonella enterica subsp. enterica serovar* Enteritidis str. EC20121990 |
| Sen-671-1 | Bacteria Proteobacteria Gammaproteobacteria Enterobacteriales Enterobacteriaceae *Salmonella enterica subsp. enterica serovar* Enteritidis str. EC20122022 |
| Sen-672-1 | Bacteria Proteobacteria Gammaproteobacteria Enterobacteriales Enterobacteriaceae *Salmonella enterica subsp. enterica serovar* Enteritidis str. EC20122026 |
| Sen-673-1 | Bacteria Proteobacteria Gammaproteobacteria Enterobacteriales Enterobacteriaceae *Salmonella enterica subsp. enterica serovar* Enteritidis str. EC20122031 |
| Sen-674-1 | Bacteria Proteobacteria Gammaproteobacteria Enterobacteriales Enterobacteriaceae *Salmonella enterica subsp. enterica serovar* Enteritidis str. EC20122033 |
| Sen-675-1 | Bacteria Proteobacteria Gammaproteobacteria Enterobacteriales Enterobacteriaceae *Salmonella enterica subsp. enterica serovar* Enteritidis str. EC20122045 |
| Sen-676-1 | Bacteria Proteobacteria Gammaproteobacteria Enterobacteriales Enterobacteriaceae *Salmonella enterica subsp. enterica serovar* Enteritidis str. EC20130345 |
| Sen-677-1 | Bacteria Proteobacteria Gammaproteobacteria Enterobacteriales Enterobacteriaceae *Salmonella enterica subsp. enterica serovar* Enteritidis str. EC20130346 |
| Sen-678-1 | Bacteria Proteobacteria Gammaproteobacteria Enterobacteriales Enterobacteriaceae *Salmonella enterica subsp. enterica serovar* Enteritidis str. EC20130348 |
| Sen-679-1 | Bacteria Proteobacteria Gammaproteobacteria Enterobacteriales Enterobacteriaceae *Salmonella enterica subsp. enterica serovar* Enteritidis str. PT23 |
| Sen-680-1 | Bacteria Proteobacteria Gammaproteobacteria Enterobacteriales Enterobacteriaceae *Salmonella enterica subsp. enterica serovar* Enteritidis str. SA19930684 |
| Sen-681-1 | Bacteria Proteobacteria Gammaproteobacteria Enterobacteriales Enterobacteriaceae *Salmonella enterica subsp. enterica serovar* Enteritidis str. SA19942384 |
| Sen-682-1 | Bacteria Proteobacteria Gammaproteobacteria Enterobacteriales Enterobacteriaceae *Salmonella enterica subsp. enterica serovar* Enteritidis str. SA19943269 |
| Sen-683-1 | Bacteria Proteobacteria Gammaproteobacteria Enterobacteriales Enterobacteriaceae *Salmonella enterica subsp. enterica serovar* Enteritidis str. SA19960848 |
| Sen-684-1 | Bacteria Proteobacteria Gammaproteobacteria Enterobacteriales Enterobacteriaceae *Salmonella enterica subsp. enterica serovar* Enteritidis str. SA19961622 |
| Sen-685-1 | Bacteria Proteobacteria Gammaproteobacteria Enterobacteriales Enterobacteriaceae *Salmonella enterica subsp. enterica serovar* Enteritidis str. SA19970510 |
| Sen-686-1 | Bacteria Proteobacteria Gammaproteobacteria Enterobacteriales Enterobacteriaceae *Salmonella enterica subsp. enterica serovar* Enteritidis str. SA19970769 |
| Sen-687-1 | Bacteria Proteobacteria Gammaproteobacteria Enterobacteriales Enterobacteriaceae *Salmonella enterica subsp. enterica serovar* Enteritidis str. SA19971331 |
| Sen-688-1 | Bacteria Proteobacteria Gammaproteobacteria Enterobacteriales Enterobacteriaceae *Salmonella enterica subsp. enterica serovar* Enteritidis str. SA19980677 |
| Sen-689-1 | Bacteria Proteobacteria Gammaproteobacteria Enterobacteriales Enterobacteriaceae *Salmonella enterica subsp. enterica serovar* Enteritidis str. SA19981522 |
| Sen-690-1 | Bacteria Proteobacteria Gammaproteobacteria Enterobacteriales Enterobacteriaceae *Salmonella enterica subsp. enterica serovar* Enteritidis str. SA19981857 |
| Sen-691-1 | Bacteria Proteobacteria Gammaproteobacteria Enterobacteriales Enterobacteriaceae *Salmonella enterica subsp. enterica serovar* Enteritidis str. SA19982831 |
| Sen-692-1 | Bacteria Proteobacteria Gammaproteobacteria Enterobacteriales Enterobacteriaceae *Salmonella enterica subsp. enterica serovar* Enteritidis str. SA19983126 |
| Sen-693-1 | Bacteria Proteobacteria Gammaproteobacteria Enterobacteriales Enterobacteriaceae *Salmonella enterica subsp. enterica serovar* Enteritidis str. SA19992322 |
| Sen-694-1 | Bacteria Proteobacteria Gammaproteobacteria Enterobacteriales Enterobacteriaceae *Salmonella enterica subsp. enterica serovar* Enteritidis str. SA20083456 |
| Sen-695-1 | Bacteria Proteobacteria Gammaproteobacteria Enterobacteriales Enterobacteriaceae *Salmonella enterica subsp. enterica serovar* Enteritidis str. SA20083636 |
| Sen-696-1 | Bacteria Proteobacteria Gammaproteobacteria Enterobacteriales Enterobacteriaceae *Salmonella enterica subsp. enterica serovar* Enteritidis str. SA20084384 |
| Sen-697-1 | Bacteria Proteobacteria Gammaproteobacteria Enterobacteriales Enterobacteriaceae *Salmonella enterica subsp. enterica serovar* Enteritidis str. SA20090435 |
| Sen-698-1 | Bacteria Proteobacteria Gammaproteobacteria Enterobacteriales Enterobacteriaceae *Salmonella enterica subsp. enterica serovar* Enteritidis str. SA20090877 |

| | |
|---|---|
| Sen-699-1 | Bacteria Proteobacteria Gammaproteobacteria Enterobacteriales Enterobacteriaceae *Salmonella enterica subsp. enterica serovar* Enteritidis str. SA20091739 |
| Sen-700-1 | Bacteria Proteobacteria Gammaproteobacteria Enterobacteriales Enterobacteriaceae *Salmonella enterica subsp. enterica serovar* Enteritidis str. SA20092320 |
| Sen-701-1 | Bacteria Proteobacteria Gammaproteobacteria Enterobacteriales Enterobacteriaceae *Salmonella enterica subsp. enterica serovar* Enteritidis str. SA20093266 |
| Sen-702-1 | Bacteria Proteobacteria Gammaproteobacteria Enterobacteriales Enterobacteriaceae *Salmonella enterica subsp. enterica serovar* Enteritidis str. SA20093421 |
| Sen-703-1 | Bacteria Proteobacteria Gammaproteobacteria Enterobacteriales Enterobacteriaceae *Salmonella enterica subsp. enterica serovar* Enteritidis str. SA20093430 |
| Sen-704-1 | Bacteria Proteobacteria Gammaproteobacteria Enterobacteriales Enterobacteriaceae *Salmonella enterica subsp. enterica serovar* Enteritidis str. SA20093538 |
| Sen-705-1 | Bacteria Proteobacteria Gammaproteobacteria Enterobacteriales Enterobacteriaceae *Salmonella enterica subsp. enterica serovar* Enteritidis str. SA20093543 |
| Sen-706-1 | Bacteria Proteobacteria Gammaproteobacteria Enterobacteriales Enterobacteriaceae *Salmonella enterica subsp. enterica serovar* Enteritidis str. SA20093784 |
| Sen-707-1 | Bacteria Proteobacteria Gammaproteobacteria Enterobacteriales Enterobacteriaceae *Salmonella enterica subsp. enterica serovar* Enteritidis str. SA20093788 |
| Sen-708-1 | Bacteria Proteobacteria Gammaproteobacteria Enterobacteriales Enterobacteriaceae *Salmonella enterica subsp. enterica serovar* Enteritidis str. SA20093950 |
| Sen-709-1 | Bacteria Proteobacteria Gammaproteobacteria Enterobacteriales Enterobacteriaceae *Salmonella enterica subsp. enterica serovar* Enteritidis str. SA20093977 |
| Sen-710-1 | Bacteria Proteobacteria Gammaproteobacteria Enterobacteriales Enterobacteriaceae *Salmonella enterica subsp. enterica serovar* Enteritidis str. SA20094079 |
| Sen-711-1 | Bacteria Proteobacteria Gammaproteobacteria Enterobacteriales Enterobacteriaceae *Salmonella enterica subsp. enterica serovar* Enteritidis str. SA20094350 |
| Sen-712-1 | Bacteria Proteobacteria Gammaproteobacteria Enterobacteriales Enterobacteriaceae *Salmonella enterica subsp. enterica serovar* Enteritidis str. SA20094352 |
| Sen-713-1 | Bacteria Proteobacteria Gammaproteobacteria Enterobacteriales Enterobacteriaceae *Salmonella enterica subsp. enterica serovar* Enteritidis str. SA20094383 |
| Sen-714-1 | Bacteria Proteobacteria Gammaproteobacteria Enterobacteriales Enterobacteriaceae *Salmonella enterica subsp. enterica serovar* Enteritidis str. SA20094389 |
| Sen-715-1 | Bacteria Proteobacteria Gammaproteobacteria Enterobacteriales Enterobacteriaceae *Salmonella enterica subsp. enterica serovar* Enteritidis str. SA20094521 |
| Sen-716-1 | Bacteria Proteobacteria Gammaproteobacteria Enterobacteriales Enterobacteriaceae *Salmonella enterica subsp. enterica serovar* Enteritidis str. SA20094642 |
| Sen-717-1 | Bacteria Proteobacteria Gammaproteobacteria Enterobacteriales Enterobacteriaceae *Salmonella enterica subsp. enterica serovar* Enteritidis str. SA20094682 |
| Sen-718-1 | Bacteria Proteobacteria Gammaproteobacteria Enterobacteriales Enterobacteriaceae *Salmonella enterica subsp. enterica serovar* Enteritidis str. SA20094803 |
| Sen-719-1 | Bacteria Proteobacteria Gammaproteobacteria Enterobacteriales Enterobacteriaceae *Salmonella enterica subsp. enterica serovar* Enteritidis str. SA20095309 |
| Sen-720-1 | Bacteria Proteobacteria Gammaproteobacteria Enterobacteriales Enterobacteriaceae *Salmonella enterica subsp. enterica serovar* Enteritidis str. SA20121703 |
| Sen-721-1 | Bacteria Proteobacteria Gammaproteobacteria Enterobacteriales Enterobacteriaceae *Salmonella enterica subsp. enterica serovar* Enteritidis str. SA20123395 |
| Sen-722-1 | Bacteria Proteobacteria Gammaproteobacteria Enterobacteriales Enterobacteriaceae *Salmonella enterica subsp. enterica serovar* Enteritidis str. SARB17 |
| Sen-723-1 | Bacteria Proteobacteria Gammaproteobacteria Enterobacteriales Enterobacteriaceae *Salmonella enterica subsp. enterica serovar* Enteritidis str. SE10 |
| Sen-724-1 | Bacteria Proteobacteria Gammaproteobacteria Enterobacteriales Enterobacteriaceae *Salmonella enterica subsp. enterica serovar* Enteritidis str. SE15-1 |
| Sen-725-1 | Bacteria Proteobacteria Gammaproteobacteria Enterobacteriales Enterobacteriaceae *Salmonella enterica subsp. enterica serovar* Enteritidis str. SE30663 |
| Sen-726-1 | Bacteria Proteobacteria Gammaproteobacteria Enterobacteriales Enterobacteriaceae *Salmonella enterica subsp. enterica serovar* Enteritidis str. SE8a |
| Sen-727-1 | Bacteria Proteobacteria Gammaproteobacteria Enterobacteriales Enterobacteriaceae *Salmonella enterica subsp. enterica serovar* Enteritidis str. SL909 |
| Sen-728-1 | Bacteria Proteobacteria Gammaproteobacteria Enterobacteriales Enterobacteriaceae *Salmonella enterica subsp. enterica serovar* Enteritidis str. SL913 |
| Sen-729-1 | Bacteria Proteobacteria Gammaproteobacteria Enterobacteriales Enterobacteriaceae *Salmonella enterica subsp. enterica serovar* Gallinarum |
| Sen-730-1 | Bacteria Proteobacteria Gammaproteobacteria Enterobacteriales Enterobacteriaceae *Salmonella enterica subsp. enterica serovar* Gallinarum str. 9184 |
| Sen-731-1 | Bacteria Proteobacteria Gammaproteobacteria Enterobacteriales Enterobacteriaceae *Salmonella enterica subsp. enterica serovar* Gallinarum str. SG9 |
| Sen-732-1 | Bacteria Proteobacteria Gammaproteobacteria Enterobacteriales Enterobacteriaceae *Salmonella enterica subsp. enterica serovar* Gallinarum/pullorum str. CDC1983-67 |
| Sen-733-1 | Bacteria Proteobacteria Gammaproteobacteria Enterobacteriales Enterobacteriaceae *Salmonella enterica subsp. enterica serovar* Give str. 564 |
| Sen-734-1 | Bacteria Proteobacteria Gammaproteobacteria Enterobacteriales Enterobacteriaceae *Salmonella enterica subsp. enterica serovar* Give str. S5-487 |
| Sen-735-1 | Bacteria Proteobacteria Gammaproteobacteria Enterobacteriales Enterobacteriaceae *Salmonella enterica subsp. enterica serovar* Give var. 15 str. CFSAN004343 |
| Sen-736-1 | Bacteria Proteobacteria Gammaproteobacteria Enterobacteriales Enterobacteriaceae *Salmonella enterica subsp. enterica serovar* Hadar str. 2011K-0112 |
| Sen-737-1 | Bacteria Proteobacteria Gammaproteobacteria Enterobacteriales Enterobacteriaceae *Salmonella enterica subsp. enterica serovar* Hadar str. ATCC 51956 |
| Sen-738-1 | Bacteria Proteobacteria Gammaproteobacteria Enterobacteriales Enterobacteriaceae *Salmonella enterica subsp. enterica serovar* Hartford str. 06-0676 |
| Sen-739-1 | Bacteria Proteobacteria Gammaproteobacteria Enterobacteriales Enterobacteriaceae *Salmonella enterica subsp. enterica serovar* Hartford str. CFSAN001075 |
| Sen-740-1 | Bacteria Proteobacteria Gammaproteobacteria Enterobacteriales Enterobacteriaceae *Salmonella enterica subsp. enterica serovar* Havana str. CFSAN001082 |
| Sen-170-1 to Sen-170-52 | Bacteria Proteobacteria Gammaproteobacteria Enterobacteriales Enterobacteriaceae *Salmonella enterica subsp. enterica serovar* Heidelberg |
| Sen-741-1 | Bacteria Proteobacteria Gammaproteobacteria Enterobacteriales Enterobacteriaceae *Salmonella enterica subsp. enterica serovar* Heidelberg str. 24390 |
| Sen-742-1 | Bacteria Proteobacteria Gammaproteobacteria Enterobacteriales Enterobacteriaceae *Salmonella enterica subsp. enterica serovar* Heidelberg str. 24393 |
| Sen-743-1 | Bacteria Proteobacteria Gammaproteobacteria Enterobacteriales Enterobacteriaceae *Salmonella enterica subsp. enterica serovar* Heidelberg str. 41563 |
| Sen-744-1 | Bacteria Proteobacteria Gammaproteobacteria Enterobacteriales Enterobacteriaceae *Salmonella enterica subsp. enterica serovar* Heidelberg str. 41565 |
| Sen-745-1 | Bacteria Proteobacteria Gammaproteobacteria Enterobacteriales Enterobacteriaceae *Salmonella enterica subsp. enterica serovar* Heidelberg str. 41566 |
| Sen-746-1 | Bacteria Proteobacteria Gammaproteobacteria Enterobacteriales Enterobacteriaceae *Salmonella enterica subsp. enterica serovar* Heidelberg str. 41573 |
| Sen-747-1 | Bacteria Proteobacteria Gammaproteobacteria Enterobacteriales Enterobacteriaceae *Salmonella enterica subsp. enterica serovar* Heidelberg str. 41576 |
| Sen-748-1 | Bacteria Proteobacteria Gammaproteobacteria Enterobacteriales Enterobacteriaceae *Salmonella enterica subsp. enterica serovar* Heidelberg str. 41578 |
| Sen-749-1 | Bacteria Proteobacteria Gammaproteobacteria Enterobacteriales Enterobacteriaceae *Salmonella enterica subsp. enterica serovar* Heidelberg str. 41579 |
| Sen-750-1 | Bacteria Proteobacteria Gammaproteobacteria Enterobacteriales Enterobacteriaceae *Salmonella enterica subsp. enterica serovar* Heidelberg str. 579082-8 |
| Sen-751-1 | Bacteria Proteobacteria Gammaproteobacteria Enterobacteriales Enterobacteriaceae *Salmonella enterica subsp. enterica serovar* Heidelberg str. 579083-19 |
| Sen-752-1 | Bacteria Proteobacteria Gammaproteobacteria Enterobacteriales Enterobacteriaceae *Salmonella enterica subsp. enterica serovar* Heidelberg str. 607310-1 |
| Sen-753-1 | Bacteria Proteobacteria Gammaproteobacteria Enterobacteriales Enterobacteriaceae *Salmonella enterica subsp. enterica serovar* Heidelberg str. 622737-11 |
| Sen-754-1 | Bacteria Proteobacteria Gammaproteobacteria Enterobacteriales Enterobacteriaceae *Salmonella enterica subsp. enterica serovar* Heidelberg str. 622737-12 |
| Sen-755-1 | Bacteria Proteobacteria Gammaproteobacteria Enterobacteriales Enterobacteriaceae *Salmonella enterica subsp. enterica serovar* Heidelberg str. 622737-14 |
| Sen-756-1 | Bacteria Proteobacteria Gammaproteobacteria Enterobacteriales Enterobacteriaceae *Salmonella enterica subsp. enterica serovar* Heidelberg str. 79-0480 |
| Sen-757-1 | Bacteria Proteobacteria Gammaproteobacteria Enterobacteriales Enterobacteriaceae *Salmonella enterica subsp. enterica serovar* Heidelberg str. 82-2052 |
| Sen-758-1 | Bacteria Proteobacteria Gammaproteobacteria Enterobacteriales Enterobacteriaceae *Salmonella enterica subsp. enterica serovar* Heidelberg str. 83-1068 |

| | |
|---|---|
| Sen-759-1 | Bacteria Proteobacteria Gammaproteobacteria Enterobacteriales Enterobacteriaceae *Salmonella enterica subsp. enterica serovar* Heidelberg str. 90-0318 |
| Sen-760-1 | Bacteria Proteobacteria Gammaproteobacteria Enterobacteriales Enterobacteriaceae *Salmonella enterica subsp. enterica serovar* Heidelberg str. CF-SAN002069 |
| Sen-761-1 | Bacteria Proteobacteria Gammaproteobacteria Enterobacteriales Enterobacteriaceae *Salmonella enterica subsp. enterica serovar* Heidelberg str. CF-SAN00322 |
| Sen-762-1 | Bacteria Proteobacteria Gammaproteobacteria Enterobacteriales Enterobacteriaceae *Salmonella enterica subsp. enterica serovar* Heidelberg str. CF-SAN00325 |
| Sen-763-1 | Bacteria Proteobacteria Gammaproteobacteria Enterobacteriales Enterobacteriaceae *Salmonella enterica subsp. enterica serovar* Heidelberg str. CF-SAN00326 |
| Sen-764-1 | Bacteria Proteobacteria Gammaproteobacteria Enterobacteriales Enterobacteriaceae *Salmonella enterica subsp. enterica serovar* Heidelberg str. CF-SAN00328 |
| Sen-765-1 | Bacteria Proteobacteria Gammaproteobacteria Enterobacteriales Enterobacteriaceae *Salmonella enterica subsp. enterica serovar* Heidelberg str. N1514 |
| Sen-766-1 | Bacteria Proteobacteria Gammaproteobacteria Enterobacteriales Enterobacteriaceae *Salmonella enterica subsp. enterica serovar* Heidelberg str. N189 |
| Sen-767-1 | Bacteria Proteobacteria Gammaproteobacteria Enterobacteriales Enterobacteriaceae *Salmonella enterica subsp. enterica serovar* Heidelberg str. N4403 |
| Sen-768-1 | Bacteria Proteobacteria Gammaproteobacteria Enterobacteriales Enterobacteriaceae *Salmonella enterica subsp. enterica serovar* Heidelberg str. N4496 |
| Sen-769-1 | Bacteria Proteobacteria Gammaproteobacteria Enterobacteriales Enterobacteriaceae *Salmonella enterica subsp. enterica serovar* Heidelberg str. N4541 |
| Sen-770-1 | Bacteria Proteobacteria Gammaproteobacteria Enterobacteriales Enterobacteriaceae *Salmonella enterica subsp. enterica serovar* Heidelberg str. RI-11-013312 |
| Sen-771-1 | Bacteria Proteobacteria Gammaproteobacteria Enterobacteriales Enterobacteriaceae *Salmonella enterica subsp. enterica serovar* Heidelberg str. RI-11-013343 |
| Sen-772-1 | Bacteria Proteobacteria Gammaproteobacteria Enterobacteriales Enterobacteriaceae *Salmonella enterica subsp. enterica serovar* Heidelberg str. RI-11-013374 |
| Sen-773-1 | Bacteria Proteobacteria Gammaproteobacteria Enterobacteriales Enterobacteriaceae *Salmonella enterica subsp. enterica serovar* Heidelberg str. RI-11-013988 |
| Sen-774-1 | Bacteria Proteobacteria Gammaproteobacteria Enterobacteriales Enterobacteriaceae *Salmonella enterica subsp. enterica serovar* Heidelberg str. RI-11-014588 |
| Sen-775-1 | Bacteria Proteobacteria Gammaproteobacteria Enterobacteriales Enterobacteriaceae *Salmonella enterica subsp. enterica serovar* Heidelberg str. SARA 39 |
| Sen-776-1 | Bacteria Proteobacteria Gammaproteobacteria Enterobacteriales Enterobacteriaceae *Salmonella enterica subsp. enterica serovar* Heidelberg str. SARA35 |
| Sen-777-1 | Bacteria Proteobacteria Gammaproteobacteria Enterobacteriales Enterobacteriaceae *Salmonella enterica subsp. enterica serovar* Indiana str. ATCC 51959 |
| Sen-778-1 | Bacteria Proteobacteria Gammaproteobacteria Enterobacteriales Enterobacteriaceae *Salmonella enterica subsp. enterica serovar* Infantis str. 119944 |
| Sen-779-1 | Bacteria Proteobacteria Gammaproteobacteria Enterobacteriales Enterobacteriaceae *Salmonella enterica subsp. enterica serovar* Infantis str. 335-3 |
| Sen-780-1 | Bacteria Proteobacteria Gammaproteobacteria Enterobacteriales Enterobacteriaceae *Salmonella enterica subsp. enterica serovar* Infantis str. SARB27 |
| Sen-781-1 | Bacteria Proteobacteria Gammaproteobacteria Enterobacteriales Enterobacteriaceae *Salmonella enterica subsp. enterica serovar* Inverness str. ATCC 10720 |
| Sen-782-1 | Bacteria Proteobacteria Gammaproteobacteria Enterobacteriales Enterobacteriaceae *Salmonella enterica subsp. enterica serovar* Javiana str. 10721 |
| Sen-783-1 | Bacteria Proteobacteria Gammaproteobacteria Enterobacteriales Enterobacteriaceae *Salmonella enterica subsp. enterica serovar* Javiana str. ATCC BAA-1593 |
| Sen-784-1 | Bacteria Proteobacteria Gammaproteobacteria Enterobacteriales Enterobacteriaceae *Salmonella enterica subsp. enterica serovar* Javiana str. CFSAN001992 |
| Sen-785-1 | Bacteria Proteobacteria Gammaproteobacteria Enterobacteriales Enterobacteriaceae *Salmonella enterica subsp. enterica serovar* Javiana str. PRS_2010_0720 |
| Sen-786-1 | Bacteria Proteobacteria Gammaproteobacteria Enterobacteriales Enterobacteriaceae *Salmonella enterica subsp. enterica serovar* Kentucky str. 0253 |
| Sen-787-1 | Bacteria Proteobacteria Gammaproteobacteria Enterobacteriales Enterobacteriaceae *Salmonella enterica subsp. enterica serovar* Kentucky str. 13562 |
| Sen-788-1 | Bacteria Proteobacteria Gammaproteobacteria Enterobacteriales Enterobacteriaceae *Salmonella enterica subsp. enterica serovar* Kentucky str. 20793 |
| Sen-789-1 | Bacteria Proteobacteria Gammaproteobacteria Enterobacteriales Enterobacteriaceae *Salmonella enterica subsp. enterica serovar* Kentucky str. 22694 |
| Sen-790-1 | Bacteria Proteobacteria Gammaproteobacteria Enterobacteriales Enterobacteriaceae *Salmonella enterica subsp. enterica serovar* Kentucky str. 29166 |
| Sen-791-1 | Bacteria Proteobacteria Gammaproteobacteria Enterobacteriales Enterobacteriaceae *Salmonella enterica subsp. enterica serovar* Kentucky str. 29439 |
| Sen-792-1 | Bacteria Proteobacteria Gammaproteobacteria Enterobacteriales Enterobacteriaceae *Salmonella enterica subsp. enterica serovar* Kentucky str. 5349 |
| Sen-793-1 | Bacteria Proteobacteria Gammaproteobacteria Enterobacteriales Enterobacteriaceae *Salmonella enterica subsp. enterica serovar* Kentucky str. N312 |
| Sen-794-1 | Bacteria Proteobacteria Gammaproteobacteria Enterobacteriales Enterobacteriaceae *Salmonella enterica subsp. enterica serovar* Litchfield str. CF-SAN001076 |
| Sen-795-1 | Bacteria Proteobacteria Gammaproteobacteria Enterobacteriales Enterobacteriaceae *Salmonella enterica subsp. enterica serovar* Manhattan str. CF-SAN001078 |
| Sen-796-1 | Bacteria Proteobacteria Gammaproteobacteria Enterobacteriales Enterobacteriaceae *Salmonella enterica subsp. enterica serovar* Mbandaka str. 2009K-0807 |
| Sen-797-1 | Bacteria Proteobacteria Gammaproteobacteria Enterobacteriales Enterobacteriaceae *Salmonella enterica subsp. enterica serovar* Mbandaka str. ATCC 51958 |
| Sen-798-1 | Bacteria Proteobacteria Gammaproteobacteria Enterobacteriales Enterobacteriaceae *Salmonella enterica subsp. enterica serovar* Meleagridis str. 0047 |
| Sen-799-1 | Bacteria Proteobacteria Gammaproteobacteria Enterobacteriales Enterobacteriaceae *Salmonella enterica subsp. enterica serovar* Miami str. 1923 |
| Sen-800-1 | Bacteria Proteobacteria Gammaproteobacteria Enterobacteriales Enterobacteriaceae *Salmonella enterica subsp. enterica serovar* Minnesota str. ATCC 49284 |
| Sen-801-1 | Bacteria Proteobacteria Gammaproteobacteria Enterobacteriales Enterobacteriaceae *Salmonella enterica subsp. enterica serovar* Mississippi str. 2010K-1406 |

| | |
|---|---|
| Sen-802-1 | Bacteria Proteobacteria Gammaproteobacteria Enterobacteriales Enterobacteriaceae *Salmonella enterica subsp. enterica serovar* Mississippi str. A4-633 |
| Sen-803-1 | Bacteria Proteobacteria Gammaproteobacteria Enterobacteriales Enterobacteriaceae *Salmonella enterica subsp. enterica serovar* Montevideo str. 19N |
| Sen-804-1 | Bacteria Proteobacteria Gammaproteobacteria Enterobacteriales Enterobacteriaceae *Salmonella enterica subsp. enterica serovar* Montevideo str. 2009083312 |
| Sen-805-1 | Bacteria Proteobacteria Gammaproteobacteria Enterobacteriales Enterobacteriaceae *Salmonella enterica subsp. enterica serovar* Montevideo str. 2009085258 |
| Sen-806-1 | Bacteria Proteobacteria Gammaproteobacteria Enterobacteriales Enterobacteriaceae *Salmonella enterica subsp. enterica serovar* Montevideo str. 29N |
| Sen-807-1 | Bacteria Proteobacteria Gammaproteobacteria Enterobacteriales Enterobacteriaceae *Salmonella enterica subsp. enterica serovar* Montevideo str. 315731156 |
| Sen-808-1 | Bacteria Proteobacteria Gammaproteobacteria Enterobacteriales Enterobacteriaceae *Salmonella enterica subsp. enterica serovar* Montevideo str. 315996572 |
| Sen-809-1 | Bacteria Proteobacteria Gammaproteobacteria Enterobacteriales Enterobacteriaceae *Salmonella enterica subsp. enterica serovar* Montevideo str. 366867 |
| Sen-810-1 | Bacteria Proteobacteria Gammaproteobacteria Enterobacteriales Enterobacteriaceae *Salmonella enterica subsp. enterica serovar* Montevideo str. 413180 |
| Sen-811-1 | Bacteria Proteobacteria Gammaproteobacteria Enterobacteriales Enterobacteriaceae *Salmonella enterica subsp. enterica serovar* Montevideo str. 414877 |
| Sen-812-1 | Bacteria Proteobacteria Gammaproteobacteria Enterobacteriales Enterobacteriaceae *Salmonella enterica subsp. enterica serovar* Montevideo str. 42N |
| Sen-813-1 | Bacteria Proteobacteria Gammaproteobacteria Enterobacteriales Enterobacteriaceae *Salmonella enterica subsp. enterica serovar* Montevideo str. 4441 H |
| Sen-814-1 | Bacteria Proteobacteria Gammaproteobacteria Enterobacteriales Enterobacteriaceae *Salmonella enterica subsp. enterica serovar* Montevideo str. 446600 |
| Sen-815-1 | Bacteria Proteobacteria Gammaproteobacteria Enterobacteriales Enterobacteriaceae *Salmonella enterica subsp. enterica serovar* Montevideo str. 495297-1 |
| Sen-816-1 | Bacteria Proteobacteria Gammaproteobacteria Enterobacteriales Enterobacteriaceae *Salmonella enterica subsp. enterica serovar* Montevideo str. 495297-3 |
| Sen-817-1 | Bacteria Proteobacteria Gammaproteobacteria Enterobacteriales Enterobacteriaceae *Salmonella enterica subsp. enterica serovar* Montevideo str. 495297-4 |
| Sen-818-1 | Bacteria Proteobacteria Gammaproteobacteria Enterobacteriales Enterobacteriaceae *Salmonella enterica subsp. enterica serovar* Montevideo str. 515920-1 |
| Sen-819-1 | Bacteria Proteobacteria Gammaproteobacteria Enterobacteriales Enterobacteriaceae *Salmonella enterica subsp. enterica serovar* Montevideo str. 515920-2 |
| Sen-820-1 | Bacteria Proteobacteria Gammaproteobacteria Enterobacteriales Enterobacteriaceae *Salmonella enterica subsp. enterica serovar* Montevideo str. 531954 |
| Sen-821-1 | Bacteria Proteobacteria Gammaproteobacteria Enterobacteriales Enterobacteriaceae *Salmonella enterica subsp. enterica serovar* Montevideo str. 556150-1 |
| Sen-822-1 | Bacteria Proteobacteria Gammaproteobacteria Enterobacteriales Enterobacteriaceae *Salmonella enterica subsp. enterica serovar* Montevideo str. 556152 |
| Sen-823-1 | Bacteria Proteobacteria Gammaproteobacteria Enterobacteriales Enterobacteriaceae *Salmonella enterica subsp. enterica serovar* Montevideo str. 609458-1 |
| Sen-824-1 | Bacteria Proteobacteria Gammaproteobacteria Enterobacteriales Enterobacteriaceae *Salmonella enterica subsp. enterica serovar* Montevideo str. 609460 |
| Sen-825-1 | Bacteria Proteobacteria Gammaproteobacteria Enterobacteriales Enterobacteriaceae *Salmonella enterica subsp. enterica serovar* Montevideo str. 80959-06 |
| Sen-826-1 | Bacteria Proteobacteria Gammaproteobacteria Enterobacteriales Enterobacteriaceae *Salmonella enterica subsp. enterica serovar* Montevideo str. 81038-01 |
| Sen-827-1 | Bacteria Proteobacteria Gammaproteobacteria Enterobacteriales Enterobacteriaceae *Salmonella enterica subsp. enterica serovar* Montevideo str. 8387 |
| Sen-828-1 | Bacteria Proteobacteria Gammaproteobacteria Enterobacteriales Enterobacteriaceae *Salmonella enterica subsp. enterica serovar* Montevideo str. ATCC BAA710 |
| Sen-829-1 | Bacteria Proteobacteria Gammaproteobacteria Enterobacteriales Enterobacteriaceae *Salmonella enterica subsp. enterica serovar* Montevideo str. CASC_09SCPH15965 |
| Sen-830-1 | Bacteria Proteobacteria Gammaproteobacteria Enterobacteriales Enterobacteriaceae *Salmonella enterica subsp. enterica serovar* Montevideo str. CFSAN004346 |
| Sen-831-1 | Bacteria Proteobacteria Gammaproteobacteria Enterobacteriales Enterobacteriaceae *Salmonella enterica subsp. enterica serovar* Montevideo str. CT_02035278 |
| Sen-832-1 | Bacteria Proteobacteria Gammaproteobacteria Enterobacteriales Enterobacteriaceae *Salmonella enterica subsp. enterica serovar* Montevideo str. CT_02035318 |
| Sen-833-1 | Bacteria Proteobacteria Gammaproteobacteria Enterobacteriales Enterobacteriaceae *Salmonella enterica subsp. enterica serovar* Montevideo str. CT_02035320 |
| Sen-834-1 | Bacteria Proteobacteria Gammaproteobacteria Enterobacteriales Enterobacteriaceae *Salmonella enterica subsp. enterica serovar* Montevideo str. CT_02035321 |
| Sen-835-1 | Bacteria Proteobacteria Gammaproteobacteria Enterobacteriales Enterobacteriaceae *Salmonella enterica subsp. enterica serovar* Montevideo str. CT_02035327 |
| Sen-836-1 | Bacteria Proteobacteria Gammaproteobacteria Enterobacteriales Enterobacteriaceae *Salmonella enterica subsp. enterica serovar* Montevideo str. FVM_628064 |
| Sen-837-1 | Bacteria Proteobacteria Gammaproteobacteria Enterobacteriales Enterobacteriaceae *Salmonella enterica subsp. enterica serovar* Montevideo str. IA_2009159199 |
| Sen-838-1 | Bacteria Proteobacteria Gammaproteobacteria Enterobacteriales Enterobacteriaceae *Salmonella enterica subsp. enterica serovar* Montevideo str. IA_2010008282 |
| Sen-839-1 | Bacteria Proteobacteria Gammaproteobacteria Enterobacteriales Enterobacteriaceae *Salmonella enterica subsp. enterica serovar* Montevideo str. IA_2010008283 |
| Sen-840-1 | Bacteria Proteobacteria Gammaproteobacteria Enterobacteriales Enterobacteriaceae *Salmonella enterica subsp. enterica serovar* Montevideo str. IA_2010008284 |
| Sen-841-1 | Bacteria Proteobacteria Gammaproteobacteria Enterobacteriales Enterobacteriaceae *Salmonella enterica subsp. enterica serovar* Montevideo str. IA_2010008285 |
| Sen-842-1 | Bacteria Proteobacteria Gammaproteobacteria Enterobacteriales Enterobacteriaceae *Salmonella enterica subsp. enterica serovar* Montevideo str. IA_2010008286 |

Sen-843-1 Bacteria Proteobacteria Gammaproteobacteria Enterobacteriales Enterobacteriaceae *Salmonella enterica subsp. enterica serovar* Montevideo str. IA_2010008287
Sen-844-1 Bacteria Proteobacteria Gammaproteobacteria Enterobacteriales Enterobacteriaceae *Salmonella enterica subsp. enterica serovar* Montevideo str. LQC 10
Sen-845-1 Bacteria Proteobacteria Gammaproteobacteria Enterobacteriales Enterobacteriaceae *Salmonella enterica subsp. enterica serovar* Montevideo str. MB101509-0077
Sen-846-1 Bacteria Proteobacteria Gammaproteobacteria Enterobacteriales Enterobacteriaceae *Salmonella enterica subsp. enterica serovar* Montevideo str. MB102109-0047
Sen-847-1 Bacteria Proteobacteria Gammaproteobacteria Enterobacteriales Enterobacteriaceae *Salmonella enterica subsp. enterica serovar* Montevideo str. MB110209-0055
Sen-848-1 Bacteria Proteobacteria Gammaproteobacteria Enterobacteriales Enterobacteriaceae *Salmonella enterica subsp. enterica serovar* Montevideo str. MB111609-0052
Sen-849-1 Bacteria Proteobacteria Gammaproteobacteria Enterobacteriales Enterobacteriaceae *Salmonella enterica subsp. enterica serovar* Montevideo str. MD_MDA09249507
Sen-850-1 Bacteria Proteobacteria Gammaproteobacteria Enterobacteriales Enterobacteriaceae *Salmonella enterica subsp. enterica serovar* Montevideo str. NC_MB110209-0054
Sen-851-1 Bacteria Proteobacteria Gammaproteobacteria Enterobacteriales Enterobacteriaceae *Salmonella enterica subsp. enterica serovar* Montevideo str. OH_2009072675
Sen-852-1 Bacteria Proteobacteria Gammaproteobacteria Enterobacteriales Enterobacteriaceae *Salmonella enterica subsp. enterica serovar* Montevideo str. SARB30
Sen-853-1 Bacteria Proteobacteria Gammaproteobacteria Enterobacteriales Enterobacteriaceae *Salmonella enterica subsp. enterica serovar* Montevideo str. SARB31
Sen-854-1 Bacteria Proteobacteria Gammaproteobacteria Enterobacteriales Enterobacteriaceae *Salmonella enterica subsp. enterica serovar* Montevideo str. USDA-ARS-USMARC-1921
Sen-855-1 Bacteria Proteobacteria Gammaproteobacteria Enterobacteriales Enterobacteriaceae *Salmonella enterica subsp. enterica serovar* Muenchen str. ATCC 8388
Sen-856-1 Bacteria Proteobacteria Gammaproteobacteria Enterobacteriales Enterobacteriaceae *Salmonella enterica subsp. enterica serovar* Muenchen str. baa1594
Sen-857-1 Bacteria Proteobacteria Gammaproteobacteria Enterobacteriales Enterobacteriaceae *Salmonella enterica subsp. enterica serovar* Muenchen str. baa1674
Sen-858-1 Bacteria Proteobacteria Gammaproteobacteria Enterobacteriales Enterobacteriaceae *Salmonella enterica subsp. enterica serovar* Muenster str. 0315
Sen-859-1 Bacteria Proteobacteria Gammaproteobacteria Enterobacteriales Enterobacteriaceae *Salmonella enterica subsp. enterica serovar* Muenster str. 420
Sen-860-1 Bacteria Proteobacteria Gammaproteobacteria Enterobacteriales Enterobacteriaceae *Salmonella enterica subsp. enterica serovar* Muenster str. 660
Sen-861-1 Bacteria Proteobacteria Gammaproteobacteria Enterobacteriales Enterobacteriaceae *Salmonella enterica subsp. enterica serovar* Muenster str. CF-SAN004344
Sen-862-1 Bacteria Proteobacteria Gammaproteobacteria Enterobacteriales Enterobacteriaceae *Salmonella enterica subsp. enterica serovar* Nchanga str. CF-SAN001091
Sen-863-1 Bacteria Proteobacteria Gammaproteobacteria Enterobacteriales Enterobacteriaceae *Salmonella enterica subsp. enterica serovar* Nchanga str. CF-SAN001092
Sen-864-1 Bacteria Proteobacteria Gammaproteobacteria Enterobacteriales Enterobacteriaceae *Salmonella enterica subsp. enterica serovar* Newport str. #11-2
Sen-865-1 Bacteria Proteobacteria Gammaproteobacteria Enterobacteriales Enterobacteriaceae *Salmonella enterica subsp. enterica serovar* Newport str. #11-3
Sen-866-1 Bacteria Proteobacteria Gammaproteobacteria Enterobacteriales Enterobacteriaceae *Salmonella enterica subsp. enterica serovar* Newport str. #11-4
Sen-867-1 Bacteria Proteobacteria Gammaproteobacteria Enterobacteriales Enterobacteriaceae *Salmonella enterica subsp. enterica serovar* Newport str. 637564_17
Sen-868-1 Bacteria Proteobacteria Gammaproteobacteria Enterobacteriales Enterobacteriaceae *Salmonella enterica subsp. enterica serovar* Newport str. CDC 2010K-2159
Sen-869-1 Bacteria Proteobacteria Gammaproteobacteria Enterobacteriales Enterobacteriaceae *Salmonella enterica subsp. enterica serovar* Newport str. CVM 19443
Sen-870-1 Bacteria Proteobacteria Gammaproteobacteria Enterobacteriales Enterobacteriaceae *Salmonella enterica subsp. enterica serovar* Newport str. CVM 19447
Sen-871-1 Bacteria Proteobacteria Gammaproteobacteria Enterobacteriales Enterobacteriaceae *Salmonella enterica subsp. enterica serovar* Newport str. CVM 19449
Sen-872-1 Bacteria Proteobacteria Gammaproteobacteria Enterobacteriales Enterobacteriaceae *Salmonella enterica subsp. enterica serovar* Newport str. CVM 19470
Sen-873-1 Bacteria Proteobacteria Gammaproteobacteria Enterobacteriales Enterobacteriaceae *Salmonella enterica subsp. enterica serovar* Newport str. CVM 19536
Sen-874-1 Bacteria Proteobacteria Gammaproteobacteria Enterobacteriales Enterobacteriaceae *Salmonella enterica subsp. enterica serovar* Newport str. CVM 19567
Sen-875-1 Bacteria Proteobacteria Gammaproteobacteria Enterobacteriales Enterobacteriaceae *Salmonella enterica subsp. enterica serovar* Newport str. CVM 19593
Sen-876-1 Bacteria Proteobacteria Gammaproteobacteria Enterobacteriales Enterobacteriaceae *Salmonella enterica subsp. enterica serovar* Newport str. CVM 21539
Sen-877-1 Bacteria Proteobacteria Gammaproteobacteria Enterobacteriales Enterobacteriaceae *Salmonella enterica subsp. enterica serovar* Newport str. CVM 21559
Sen-878-1 Bacteria Proteobacteria Gammaproteobacteria Enterobacteriales Enterobacteriaceae *Salmonella enterica subsp. enterica serovar* Newport str. CVM 33953
Sen-879-1 Bacteria Proteobacteria Gammaproteobacteria Enterobacteriales Enterobacteriaceae *Salmonella enterica subsp. enterica serovar* Newport str. CVM 35185
Sen-880-1 Bacteria Proteobacteria Gammaproteobacteria Enterobacteriales Enterobacteriaceae *Salmonella enterica subsp. enterica serovar* Newport str. CVM 35188
Sen-881-1 Bacteria Proteobacteria Gammaproteobacteria Enterobacteriales Enterobacteriaceae *Salmonella enterica subsp. enterica serovar* Newport str. CVM 35199
Sen-882-1 Bacteria Proteobacteria Gammaproteobacteria Enterobacteriales Enterobacteriaceae *Salmonella enterica subsp. enterica serovar* Newport str. CVM 35202
Sen-883-1 Bacteria Proteobacteria Gammaproteobacteria Enterobacteriales Enterobacteriaceae *Salmonella enterica subsp. enterica serovar* Newport str. CVM 37978
Sen-884-1 Bacteria Proteobacteria Gammaproteobacteria Enterobacteriales Enterobacteriaceae *Salmonella enterica subsp. enterica serovar* Newport str. CVM 4176
Sen-885-1 Bacteria Proteobacteria Gammaproteobacteria Enterobacteriales Enterobacteriaceae *Salmonella enterica subsp. enterica serovar* Newport str. Henan_3
Sen-886-1 Bacteria Proteobacteria Gammaproteobacteria Enterobacteriales Enterobacteriaceae *Salmonella enterica subsp. enterica serovar* Newport str. JS09102
Sen-887-1 Bacteria Proteobacteria Gammaproteobacteria Enterobacteriales Enterobacteriaceae *Salmonella enterica subsp. enterica serovar* Newport str. Levine 1
Sen-888-1 Bacteria Proteobacteria Gammaproteobacteria Enterobacteriales Enterobacteriaceae *Salmonella enterica subsp. enterica serovar* Newport str. Levine 15

| | |
|---|---|
| Sen-889-1 | Bacteria Proteobacteria Gammaproteobacteria Enterobacteriales Enterobacteriaceae *Salmonella enterica subsp. enterica serovar* Newport str. RI_10P068 |
| Sen-890-1 | Bacteria Proteobacteria Gammaproteobacteria Enterobacteriales Enterobacteriaceae *Salmonella enterica subsp. enterica serovar* Newport str. RI_10P069 |
| Sen-891-1 | Bacteria Proteobacteria Gammaproteobacteria Enterobacteriales Enterobacteriaceae *Salmonella enterica subsp. enterica serovar* Newport str. RI_10P078 |
| Sen-892-1 | Bacteria Proteobacteria Gammaproteobacteria Enterobacteriales Enterobacteriaceae *Salmonella enterica subsp. enterica serovar* Newport str. RI_10P079 |
| Sen-893-1 | Bacteria Proteobacteria Gammaproteobacteria Enterobacteriales Enterobacteriaceae *Salmonella enterica subsp. enterica serovar* Newport str. SH111077 |
| Sen-894-1 | Bacteria Proteobacteria Gammaproteobacteria Enterobacteriales Enterobacteriaceae *Salmonella enterica subsp. enterica serovar* Newport str. Shandong_3 |
| Sen-895-1 | Bacteria Proteobacteria Gammaproteobacteria Enterobacteriales Enterobacteriaceae *Salmonella enterica subsp. enterica serovar* Newport str. USMARC-S3124.1 |
| Sen-896-1 | Bacteria Proteobacteria Gammaproteobacteria Enterobacteriales Enterobacteriaceae *Salmonella enterica subsp. enterica serovar* Newport str. VA_R100506907 |
| Sen-897-1 | Bacteria Proteobacteria Gammaproteobacteria Enterobacteriales Enterobacteriaceae *Salmonella enterica subsp. enterica serovar* Newport str. VA_R100512570 |
| Sen-898-1 | Bacteria Proteobacteria Gammaproteobacteria Enterobacteriales Enterobacteriaceae *Salmonella enterica subsp. enterica serovar* Newport str. VA_R100512572 |
| Sen-899-1 | Bacteria Proteobacteria Gammaproteobacteria Enterobacteriales Enterobacteriaceae *Salmonella enterica subsp. enterica serovar* Newport str. WA_14881 |
| Sen-900-1 | Bacteria Proteobacteria Gammaproteobacteria Enterobacteriales Enterobacteriaceae *Salmonella enterica subsp. enterica serovar* Newport str. WA_14882 |
| Sen-901-1 | Bacteria Proteobacteria Gammaproteobacteria Enterobacteriales Enterobacteriaceae *Salmonella enterica subsp. enterica serovar* Newport str. WA_14885 |
| Sen-902-1 | Bacteria Proteobacteria Gammaproteobacteria Enterobacteriales Enterobacteriaceae *Salmonella enterica subsp. enterica serovar* Newport str. WA_14900 |
| Sen-903-1 | Bacteria Proteobacteria Gammaproteobacteria Enterobacteriales Enterobacteriaceae *Salmonella enterica subsp. enterica serovar* Norwich str. CFSAN001077 |
| Sen-904-1 | Bacteria Proteobacteria Gammaproteobacteria Enterobacteriales Enterobacteriaceae *Salmonella enterica subsp. enterica serovar* Ohio str. CFSAN001079 |
| Sen-905-1 | Bacteria Proteobacteria Gammaproteobacteria Enterobacteriales Enterobacteriaceae *Salmonella enterica subsp. enterica serovar* Oranienburg str. 0250 |
| Sen-906-1 | Bacteria Proteobacteria Gammaproteobacteria Enterobacteriales Enterobacteriaceae *Salmonella enterica subsp. enterica serovar* Oranienburg str. 701 |
| Sen-907-1 | Bacteria Proteobacteria Gammaproteobacteria Enterobacteriales Enterobacteriaceae *Salmonella enterica subsp. enterica serovar* Oranienburg str. S-76 |
| Sen-908-1 | Bacteria Proteobacteria Gammaproteobacteria Enterobacteriales Enterobacteriaceae *Salmonella enterica subsp. enterica serovar* Paratyphi A str. ATCC |
| Sen-909-1 | Bacteria Proteobacteria Gammaproteobacteria Enterobacteriales Enterobacteriaceae *Salmonella enterica subsp. enterica serovar* Paratyphi A str. ATCC 11511 |
| Sen-910-1 | Bacteria Proteobacteria Gammaproteobacteria Enterobacteriales Enterobacteriaceae *Salmonella enterica subsp. enterica serovar* Paratyphi A str. GXS2268 |
| Sen-911-1 | Bacteria Proteobacteria Gammaproteobacteria Enterobacteriales Enterobacteriaceae *Salmonella enterica subsp. enterica serovar* Paratyphi A str. GZ9A00052 |
| Sen-912-1 | Bacteria Proteobacteria Gammaproteobacteria Enterobacteriales Enterobacteriaceae *Salmonella enterica subsp. enterica serovar* Paratyphi A str. JX05-19 |
| Sen-913-1 | Bacteria Proteobacteria Gammaproteobacteria Enterobacteriales Enterobacteriaceae *Salmonella enterica subsp. enterica serovar* Paratyphi A str. YN09620 |
| Sen-914-1 | Bacteria Proteobacteria Gammaproteobacteria Enterobacteriales Enterobacteriaceae *Salmonella enterica subsp. enterica serovar* Paratyphi A str. ZJ98-53 |
| Sen-915-1 | Bacteria Proteobacteria Gammaproteobacteria Enterobacteriales Enterobacteriaceae *Salmonella enterica subsp. enterica serovar* Paratyphi B str. ATCC 10719 |
| Sen-916-1 | Bacteria Proteobacteria Gammaproteobacteria Enterobacteriales Enterobacteriaceae *Salmonella enterica subsp. enterica serovar* Paratyphi B str. ATCC 19940 |
| Sen-917-1 | Bacteria Proteobacteria Gammaproteobacteria Enterobacteriales Enterobacteriaceae *Salmonella enterica subsp. enterica serovar* Paratyphi B str. ATCC 51962 |
| Sen-918-1 | Bacteria Proteobacteria Gammaproteobacteria Enterobacteriales Enterobacteriaceae *Salmonella enterica subsp. enterica serovar* Paratyphi B str. ATCC 8759 |
| Sen-919-1 | Bacteria Proteobacteria Gammaproteobacteria Enterobacteriales Enterobacteriaceae *Salmonella enterica subsp. enterica serovar* Paratyphi B str. ATCC BAA-1585 |
| Sen-920-1 | Bacteria Proteobacteria Gammaproteobacteria Enterobacteriales Enterobacteriaceae *Salmonella enterica subsp. enterica serovar* Paratyphi B str. SARA42 |
| Sen-921-1 | Bacteria Proteobacteria Gammaproteobacteria Enterobacteriales Enterobacteriaceae *Salmonella enterica subsp. enterica serovar* Paratyphi B str. SARA56 |
| Sen-922-1 | Bacteria Proteobacteria Gammaproteobacteria Enterobacteriales Enterobacteriaceae *Salmonella enterica subsp. enterica serovar* Paratyphi B str. SARA61 |
| Sen-923-1 | Bacteria Proteobacteria Gammaproteobacteria Enterobacteriales Enterobacteriaceae *Salmonella enterica subsp. enterica serovar* Paratyphi B str. SARA62 |
| Sen-924-1 | Bacteria Proteobacteria Gammaproteobacteria Enterobacteriales Enterobacteriaceae *Salmonella enterica subsp. enterica serovar* Pomona str. ATCC 10729 |
| Sen-925-1 | Bacteria Proteobacteria Gammaproteobacteria Enterobacteriales Enterobacteriaceae *Salmonella enterica subsp. enterica serovar* Poona str. ATCC BAA-1673 |
| Sen-926-1 | Bacteria Proteobacteria Gammaproteobacteria Enterobacteriales Enterobacteriaceae *Salmonella enterica subsp. enterica serovar* Pullorum str. 13036 |
| Sen-927-1 | Bacteria Proteobacteria Gammaproteobacteria Enterobacteriales Enterobacteriaceae *Salmonella enterica subsp. enterica serovar* Pullorum str. 19945 |
| Sen-928-1 | Bacteria Proteobacteria Gammaproteobacteria Enterobacteriales Enterobacteriaceae *Salmonella enterica subsp. enterica serovar* Pullorum str. S06004 |
| Sen-929-1 | Bacteria Proteobacteria Gammaproteobacteria Enterobacteriales Enterobacteriaceae *Salmonella enterica subsp. enterica serovar* Rissen str. 150 |
| Sen-930-1 | Bacteria Proteobacteria Gammaproteobacteria Enterobacteriales Enterobacteriaceae *Salmonella enterica subsp. enterica serovar* Rubislaw str. ATCC 10717 |
| Sen-931-1 | Bacteria Proteobacteria Gammaproteobacteria Enterobacteriales Enterobacteriaceae *Salmonella enterica subsp. enterica serovar* Saintpaul str. 9712 |
| Sen-932-1 | Bacteria Proteobacteria Gammaproteobacteria Enterobacteriales Enterobacteriaceae *Salmonella enterica subsp. enterica serovar* Saintpaul str. JO2008 |
| Sen-933-1 | Bacteria Proteobacteria Gammaproteobacteria Enterobacteriales Enterobacteriaceae *Salmonella enterica subsp. enterica serovar* Saintpaul str. S-70 |
| Sen-934-1 | Bacteria Proteobacteria Gammaproteobacteria Enterobacteriales Enterobacteriaceae *Salmonella enterica subsp. enterica serovar* Saintpaul str. SARA26 |
| Sen-935-1 | Bacteria Proteobacteria Gammaproteobacteria Enterobacteriales Enterobacteriaceae *Salmonella enterica subsp. enterica serovar* Senftenberg str. 316235162 |

| | |
|---|---|
| Sen-936-1 | Bacteria Proteobacteria Gammaproteobacteria Enterobacteriales Enterobacteriaceae *Salmonella enterica subsp. enterica serovar* Senftenberg str. 361154004 |
| Sen-937-1 | Bacteria Proteobacteria Gammaproteobacteria Enterobacteriales Enterobacteriaceae *Salmonella enterica subsp. enterica serovar* Senftenberg str. 423984-1 |
| Sen-938-1 | Bacteria Proteobacteria Gammaproteobacteria Enterobacteriales Enterobacteriaceae *Salmonella enterica subsp. enterica serovar* Senftenberg str. 423984-2 |
| Sen-939-1 | Bacteria Proteobacteria Gammaproteobacteria Enterobacteriales Enterobacteriaceae *Salmonella enterica subsp. enterica serovar* Senftenberg str. 604314 |
| Sen-940-1 | Bacteria Proteobacteria Gammaproteobacteria Enterobacteriales Enterobacteriaceae *Salmonella enterica subsp. enterica serovar* Senftenberg str. ATCC 43845 |
| Sen-941-1 | Bacteria Proteobacteria Gammaproteobacteria Enterobacteriales Enterobacteriaceae *Salmonella enterica subsp. enterica serovar* Senftenberg str. ATCC 8400 |
| Sen-942-1 | Bacteria Proteobacteria Gammaproteobacteria Enterobacteriales Enterobacteriaceae *Salmonella enterica subsp. enterica serovar* Senftenberg str. NC_MB012510-0038 |
| Sen-943-1 | Bacteria Proteobacteria Gammaproteobacteria Enterobacteriales Enterobacteriaceae *Salmonella enterica subsp. enterica serovar* Senftenberg str. SS209 |
| Sen-944-1 | Bacteria Proteobacteria Gammaproteobacteria Enterobacteriales Enterobacteriaceae *Salmonella enterica subsp. enterica serovar* Soerenga str. 695 |
| Sen-945-1 | Bacteria Proteobacteria Gammaproteobacteria Enterobacteriales Enterobacteriaceae *Salmonella enterica subsp. enterica serovar* Stanley str. ATCC 7308 |
| Sen-946-1 | Bacteria Proteobacteria Gammaproteobacteria Enterobacteriales Enterobacteriaceae *Salmonella enterica subsp. enterica serovar* Stanleyville str. CF-SAN000624 |
| Sen-947-1 | Bacteria Proteobacteria Gammaproteobacteria Enterobacteriales Enterobacteriaceae *Salmonella enterica subsp. enterica serovar* Tallahassee str. 0012 |
| Sen-948-1 | Bacteria Proteobacteria Gammaproteobacteria Enterobacteriales Enterobacteriaceae *Salmonella enterica subsp. enterica serovar* Tennessee str. TXSC_TXSC08-21 |
| Sen-949-1 | Bacteria Proteobacteria Gammaproteobacteria Enterobacteriales Enterobacteriaceae *Salmonella enterica subsp. enterica serovar* Thompson str. 2010K-1863 |
| Sen-950-1 | Bacteria Proteobacteria Gammaproteobacteria Enterobacteriales Enterobacteriaceae *Salmonella enterica subsp. enterica serovar* Thompson str. RM6836 |
| Sen-288-1 to Sen-288-1125 | Bacteria Proteobacteria Gammaproteobacteria Enterobacteriales Enterobacteriaceae *Salmonella enterica subsp. enterica serovar* Typhi |
| Sen-951-1 | Bacteria Proteobacteria Gammaproteobacteria Enterobacteriales Enterobacteriaceae *Salmonella enterica subsp. enterica serovar* Typhi str. BL196 |
| Sen-952-1 | Bacteria Proteobacteria Gammaproteobacteria Enterobacteriales Enterobacteriaceae *Salmonella enterica subsp. enterica serovar* Typhi str. CR0044 |
| Sen-953-1 | Bacteria Proteobacteria Gammaproteobacteria Enterobacteriales Enterobacteriaceae *Salmonella enterica subsp. enterica serovar* Typhi str. P-stx-12 |
| Sen-954-1 | Bacteria Proteobacteria Gammaproteobacteria Enterobacteriales Enterobacteriaceae *Salmonella enterica subsp. enterica serovar* Typhi str. ST0208 |
| Sen-955-1 | Bacteria Proteobacteria Gammaproteobacteria Enterobacteriales Enterobacteriaceae *Salmonella enterica subsp. enterica serovar* Typhi str. Ty21a |
| Sen-956-1 | Bacteria Proteobacteria Gammaproteobacteria Enterobacteriales Enterobacteriaceae *Salmonella enterica subsp. enterica serovar* Typhi str. UJ308A |
| Sen-957-1 | Bacteria Proteobacteria Gammaproteobacteria Enterobacteriales Enterobacteriaceae *Salmonella enterica subsp. enterica serovar* Typhi str. UJ816A |
| Sen-291-1 to Sen-291-7 | Bacteria Proteobacteria Gammaproteobacteria Enterobacteriales Enterobacteriaceae *Salmonella enterica subsp. enterica serovar* Typhimurium |
| Sen-958-1 | Bacteria Proteobacteria Gammaproteobacteria Enterobacteriales Enterobacteriaceae *Salmonella enterica subsp. enterica serovar* Typhimurium str. 14028S |
| Sen-959-1 | Bacteria Proteobacteria Gammaproteobacteria Enterobacteriales Enterobacteriaceae *Salmonella enterica subsp. enterica serovar* Typhimurium str. 34502 |
| Sen-960-1 | Bacteria Proteobacteria Gammaproteobacteria Enterobacteriales Enterobacteriaceae *Salmonella enterica subsp. enterica serovar* Typhimurium str. 35423 |
| Sen-961-1 | Bacteria Proteobacteria Gammaproteobacteria Enterobacteriales Enterobacteriaceae *Salmonella enterica subsp. enterica serovar* Typhimurium str. 36618 |
| Sen-962-1 | Bacteria Proteobacteria Gammaproteobacteria Enterobacteriales Enterobacteriaceae *Salmonella enterica subsp. enterica serovar* Typhimurium str. 798 |
| Sen-963-1 | Bacteria Proteobacteria Gammaproteobacteria Enterobacteriales Enterobacteriaceae *Salmonella enterica subsp. enterica serovar* Typhimurium str. AZ 057 |
| Sen-964-1 | Bacteria Proteobacteria Gammaproteobacteria Enterobacteriales Enterobacteriaceae *Salmonella enterica subsp. enterica serovar* Typhimurium str. CDC_2009K1158 |
| Sen-965-1 | Bacteria Proteobacteria Gammaproteobacteria Enterobacteriales Enterobacteriaceae *Salmonella enterica subsp. enterica serovar* Typhimurium str. CDC_2009K1277 |
| Sen-966-1 | Bacteria Proteobacteria Gammaproteobacteria Enterobacteriales Enterobacteriaceae *Salmonella enterica subsp. enterica serovar* Typhimurium str. CDC_2009K1283 |
| Sen-967-1 | Bacteria Proteobacteria Gammaproteobacteria Enterobacteriales Enterobacteriaceae *Salmonella enterica subsp. enterica serovar* Typhimurium str. CDC_2009K1288 |
| Sen-968-1 | Bacteria Proteobacteria Gammaproteobacteria Enterobacteriales Enterobacteriaceae *Salmonella enterica subsp. enterica serovar* Typhimurium str. D23580 |
| Sen-300-1 to Sen-300-330 | Bacteria Proteobacteria Gammaproteobacteria Enterobacteriales Enterobacteriaceae *Salmonella enterica subsp. enterica serovar* Typhimurium str. DT104 |
| Sen-969-1 | Bacteria Proteobacteria Gammaproteobacteria Enterobacteriales Enterobacteriaceae *Salmonella enterica subsp. enterica serovar* Typhimurium str. DT2 |
| Sen-970-1 | Bacteria Proteobacteria Gammaproteobacteria Enterobacteriales Enterobacteriaceae *Salmonella enterica subsp. enterica serovar* Typhimurium str. SARA13 |
| Sen-971-1 | Bacteria Proteobacteria Gammaproteobacteria Enterobacteriales Enterobacteriaceae *Salmonella enterica subsp. enterica serovar* Typhimurium str. SL1344 |
| Sen-972-1 | Bacteria Proteobacteria Gammaproteobacteria Enterobacteriales Enterobacteriaceae *Salmonella enterica subsp. enterica serovar* Typhimurium str. ST1660/06 |
| Sen-973-1 | Bacteria Proteobacteria Gammaproteobacteria Enterobacteriales Enterobacteriaceae *Salmonella enterica subsp. enterica serovar* Typhimurium str. ST4/74 |
| Sen-974-1 | Bacteria Proteobacteria Gammaproteobacteria Enterobacteriales Enterobacteriaceae *Salmonella enterica subsp. enterica serovar* Typhimurium str. ST4581 |
| Sen-975-1 | Bacteria Proteobacteria Gammaproteobacteria Enterobacteriales Enterobacteriaceae *Salmonella enterica subsp. enterica serovar* Typhimurium str. ST78896 |
| Sen-976-1 | Bacteria Proteobacteria Gammaproteobacteria Enterobacteriales Enterobacteriaceae *Salmonella enterica subsp. enterica serovar* Typhimurium str. STm1 |
| Sen-977-1 | Bacteria Proteobacteria Gammaproteobacteria Enterobacteriales Enterobacteriaceae *Salmonella enterica subsp. enterica serovar* Typhimurium str. STm10 |
| Sen-978-1 | Bacteria Proteobacteria Gammaproteobacteria Enterobacteriales Enterobacteriaceae *Salmonella enterica subsp. enterica serovar* Typhimurium str. STm11 |
| Sen-979-1 | Bacteria Proteobacteria Gammaproteobacteria Enterobacteriales Enterobacteriaceae *Salmonella enterica subsp. enterica serovar* Typhimurium str. STm12 |
| Sen-980-1 | Bacteria Proteobacteria Gammaproteobacteria Enterobacteriales Enterobacteriaceae *Salmonella enterica subsp. enterica serovar* Typhimurium str. STm2 |
| Sen-981-1 | Bacteria Proteobacteria Gammaproteobacteria Enterobacteriales Enterobacteriaceae *Salmonella enterica subsp. enterica serovar* Typhimurium str. STm3 |
| Sen-982-1 | Bacteria Proteobacteria Gammaproteobacteria Enterobacteriales Enterobacteriaceae *Salmonella enterica subsp. enterica serovar* Typhimurium str. STm4 |

| | |
|---|---|
| Sen-983-1 | Bacteria Proteobacteria Gammaproteobacteria Enterobacteriales Enterobacteriaceae *Salmonella enterica subsp. enterica serovar* Typhimurium str. STm5 |
| Sen-984-1 | Bacteria Proteobacteria Gammaproteobacteria Enterobacteriales Enterobacteriaceae *Salmonella enterica subsp. enterica serovar* Typhimurium str. STm6 |
| Sen-985-1 | Bacteria Proteobacteria Gammaproteobacteria Enterobacteriales Enterobacteriaceae *Salmonella enterica subsp. enterica serovar* Typhimurium str. STm7 |
| Sen-986-1 | Bacteria Proteobacteria Gammaproteobacteria Enterobacteriales Enterobacteriaceae *Salmonella enterica subsp. enterica serovar* Typhimurium str. STm8 |
| Sen-987-1 | Bacteria Proteobacteria Gammaproteobacteria Enterobacteriales Enterobacteriaceae *Salmonella enterica subsp. enterica serovar* Typhimurium str. STm9 |
| Sen-988-1 | Bacteria Proteobacteria Gammaproteobacteria Enterobacteriales Enterobacteriaceae *Salmonella enterica subsp. enterica serovar* Typhimurium str. T000240 |
| Sen-989-1 | Bacteria Proteobacteria Gammaproteobacteria Enterobacteriales Enterobacteriaceae *Salmonella enterica subsp. enterica serovar* Typhimurium str. TN061786 |
| Sen-990-1 | Bacteria Proteobacteria Gammaproteobacteria Enterobacteriales Enterobacteriaceae *Salmonella enterica subsp. enterica serovar* Typhimurium str. U288 |
| Sen-991-1 | Bacteria Proteobacteria Gammaproteobacteria Enterobacteriales Enterobacteriaceae *Salmonella enterica subsp. enterica serovar* Typhimurium str. UK-1 |
| Sen-992-1 | Bacteria Proteobacteria Gammaproteobacteria Enterobacteriales Enterobacteriaceae *Salmonella enterica subsp. enterica serovar* Typhimurium str. USDA-ARS-USMARC-1899 |
| Sen-993-1 | Bacteria Proteobacteria Gammaproteobacteria Enterobacteriales Enterobacteriaceae *Salmonella enterica subsp. enterica serovar* Typhimurium var. 5- str. CFSAN001921 |
| Sen-994-1 | Bacteria Proteobacteria Gammaproteobacteria Enterobacteriales Enterobacteriaceae *Salmonella enterica subsp. enterica serovar* Typhimurium var. 5- str. CFSAN004345 |
| Sen-995-1 | Bacteria Proteobacteria Gammaproteobacteria Enterobacteriales Enterobacteriaceae *Salmonella enterica subsp. enterica serovar* Typhimurium var. Copenhagen str. 0084 |
| Sen-996-1 | Bacteria Proteobacteria Gammaproteobacteria Enterobacteriales Enterobacteriaceae *Salmonella enterica subsp. enterica serovar* Uganda str. R8-3404 |
| Sen-997-1 | Bacteria Proteobacteria Gammaproteobacteria Enterobacteriales Enterobacteriaceae *Salmonella enterica subsp. enterica serovar* Urbana str. ATCC 9261 |
| Sen-998-1 | Bacteria Proteobacteria Gammaproteobacteria Enterobacteriales Enterobacteriaceae *Salmonella enterica subsp. enterica serovar* Worthington str. ATCC 9607 |
| Sen-999-1 | Bacteria Proteobacteria Gammaproteobacteria Enterobacteriales Enterobacteriaceae Salmonella Salmonella enterica subsp. entericaserovar Agona *Salmonella enterica subsp. enterica serovar* Agona str. SL483 |
| Sen-1000-1 | Bacteria Proteobacteria Gammaproteobacteria Enterobacteriales Enterobacteriaceae Salmonella Salmonella enterica subsp. entericaserovar Choleraesuis *Salmonella enterica subsp. enterica serovar* Choleraesuis str. SCSA50 |
| Sen-1001-1 | Bacteria Proteobacteria Gammaproteobacteria Enterobacteriales Enterobacteriaceae Salmonella Salmonella enterica subsp. entericaserovar Dublin *Salmonella enterica subsp. enterica serovar* Dublin str. CT_02021853 |
| Sen-1002-1 | Bacteria Proteobacteria Gammaproteobacteria Enterobacteriales Enterobacteriaceae Salmonella Salmonella enterica subsp. entericaserovar Gallinarum/pullorum *Salmonella enterica subsp. enterica serovar* Gallinarum/pullorum str. RKS5078 |
| Sen-1003-1 | Bacteria Proteobacteria Gammaproteobacteria Enterobacteriales Enterobacteriaceae Salmonella Salmonella enterica subsp. entericaserovar Gallinarum *Salmonella enterica subsp. enterica serovar* Gallinarum str. 287/91 |
| Sen-1004-1 | Bacteria Proteobacteria Gammaproteobacteria Enterobacteriales Enterobacteriaceae Salmonella Salmonella enterica subsp. entericaserovar Hadar *Salmonella enterica subsp. enterica serovar* Hadar str. RI_05P066 |
| Sen-1005-1 | Bacteria Proteobacteria Gammaproteobacteria Enterobacteriales Enterobacteriaceae Salmonella Salmonella enterica subsp. entericaserovar Heidelberg *Salmonella enterica subsp. enterica serovar* Heidelberg str. B182 |
| Sen-1006-1 | Bacteria Proteobacteria Gammaproteobacteria Enterobacteriales Enterobacteriaceae Salmonella Salmonella enterica subsp. entericaserovar Heidelberg *Salmonella enterica subsp. enterica serovar* Heidelberg str. SL476 |
| Sen-1007-1 | Bacteria Proteobacteria Gammaproteobacteria Enterobacteriales Enterobacteriaceae Salmonella Salmonella enterica subsp. entericaserovar Heidelberg *Salmonella enterica subsp. enterica serovar* Heidelberg str. SL486 |
| Sen-1008-1 | Bacteria Proteobacteria Gammaproteobacteria Enterobacteriales Enterobacteriaceae Salmonella Salmonella enterica subsp. entericaserovar Kentucky *Salmonella enterica subsp. enterica serovar* Kentucky str. CDC 191 |
| Sen-1009-1 | Bacteria Proteobacteria Gammaproteobacteria Enterobacteriales Enterobacteriaceae Salmonella Salmonella enterica subsp. entericaserovar Kentucky *Salmonella enterica subsp. enterica serovar* Kentucky str. CVM29188 |
| Sen-266-1 to Sen-266-2 | Bacteria Proteobacteria Gammaproteobacteria Enterobacteriales Enterobacteriaceae Salmonella Salmonella enterica subsp. entericaserovar Newport *Salmonella enterica subsp. enterica serovar* Newport str. SL254 |
| Sen-1010-1 | Bacteria Proteobacteria Gammaproteobacteria Enterobacteriales Enterobacteriaceae Salmonella Salmonella enterica subsp. entericaserovar Newport *Salmonella enterica subsp. enterica serovar* Newport str. SL317 |
| Sen-1011-1 | Bacteria Proteobacteria Gammaproteobacteria Enterobacteriales Enterobacteriaceae Salmonella Salmonella enterica subsp. entericaserovar Paratyphi B *Salmonella enterica subsp. enterica serovar* Paratyphi B str. SPB7 |
| Sen-1012-1 | Bacteria Proteobacteria Gammaproteobacteria Enterobacteriales Enterobacteriaceae Salmonella Salmonella enterica subsp. entericaserovar Saintpaul *Salmonella enterica subsp. enterica serovar* Saintpaul str. SARA23 |
| Sen-1013-1 | Bacteria Proteobacteria Gammaproteobacteria Enterobacteriales Enterobacteriaceae Salmonella Salmonella enterica subsp. entericaserovar Saintpaul *Salmonella enterica subsp. enterica serovar* Saintpaul str. SARA29 |

| | |
|---|---|
| Sen-1014-1 | Bacteria Proteobacteria Gammaproteobacteria Enterobacteriales Enterobacteriaceae Salmonella Salmonella enterica subsp. entericaserovar Typhi *Salmonella enterica subsp. enterica serovar* Typhi str. CT18 |
| Sen-1015-1 | Bacteria Proteobacteria Gammaproteobacteria Enterobacteriales Enterobacteriaceae Salmonella Salmonella enterica subsp. entericaserovar Typhi *Salmonella enterica subsp. enterica serovar* Typhi str. E00-7866 |
| Sen-1016-1 | Bacteria Proteobacteria Gammaproteobacteria Enterobacteriales Enterobacteriaceae Salmonella Salmonella enterica subsp. entericaserovar Typhi *Salmonella enterica subsp. enterica serovar* Typhi str. Ty2 |
| Sen-1017-1 | Bacteria Proteobacteria Gammaproteobacteria Enterobacteriales Enterobacteriaceae Salmonella Salmonella enterica subsp. entericaserovar Typhimurium *Salmonella enterica subsp. enterica serovar* Typhimurium str. LT2 |
| Sen-1018-1 | Bacteria Proteobacteria Gammaproteobacteria Enterobacteriales Enterobacteriaceae Salmonella Salmonella enterica subsp. entericaserovar Virchow *Salmonella enterica subsp. enterica serovar* Virchow str. SL491 |
| Sen-1019-1 | Bacteria Proteobacteria Gammaproteobacteria Enterobacteriales Enterobacteriaceae *Salmonella enterica subsp. houtenae* |
| Sen-1020-1 | Bacteria Proteobacteria Gammaproteobacteria Enterobacteriales Enterobacteriaceae *Salmonella enterica subsp. houtenae serovar* 16:z4,z32:– str. RKS3027 |
| Sen-1021-1 | Bacteria Proteobacteria Gammaproteobacteria Enterobacteriales Enterobacteriaceae *Salmonella enterica subsp. houtenae serovar* 50:g,z51:- str. 01-0133 |
| Sen-1022-1 | Bacteria Proteobacteria Gammaproteobacteria Enterobacteriales Enterobacteriaceae *Salmonella enterica subsp. houtenae str.* ATCC BAA-1581 |
| Sen-1023-1 | Bacteria Proteobacteria Gammaproteobacteria Enterobacteriales Enterobacteriaceae *Salmonella enterica subsp. indica serovar* 6,14,25:z10:1,(2),7 str. 1121 |
| Sen-1024-1 | Bacteria Proteobacteria Gammaproteobacteria Enterobacteriales Enterobacteriaceae *Salmonella enterica subsp. salamae serovar* 58:l,z13,z28:z6 str. 00-0163 |
| Sen-1025-1 | Bacteria Proteobacteria Gammaproteobacteria Enterobacteriales Enterobacteriaceae *Salmonella enterica subsp. salamae str.* 3588/07 |
| Sfo-4-1 | Bacteria Proteobacteria Gammaproteobacteria Enterobacteriales Enterobacteriaceae *Serratia fonticola* AU-AP2C |
| Sfo-5-1 | Bacteria Proteobacteria Gammaproteobacteria Enterobacteriales Enterobacteriaceae *Serratia fonticola* AU-P3(3) |
| Sli-3-1 | Bacteria Proteobacteria Gammaproteobacteria Enterobacteriales Enterobacteriaceae *Serratia liquefaciens* ATCC 27592 |
| Sma-3-1 to Sma-3-4 | Bacteria Proteobacteria Gammaproteobacteria Enterobacteriales Enterobacteriaceae *Serratia marcescens* |
| Sma-23-1 | Bacteria Proteobacteria Gammaproteobacteria Enterobacteriales Enterobacteriaceae *Serratia marcescens* EGD-HP20 |
| Sma-24-1 | Bacteria Proteobacteria Gammaproteobacteria Enterobacteriales Enterobacteriaceae *Serratia marcescens* FGI94 |
| Sma-25-1 | Bacteria Proteobacteria Gammaproteobacteria Enterobacteriales Enterobacteriaceae *Serratia marcescens* LCT-SM213 |
| Sma-26-1 | Bacteria Proteobacteria Gammaproteobacteria Enterobacteriales Enterobacteriaceae *Serratia marcescens* VGH107 |
| Sma-27-1 | Bacteria Proteobacteria Gammaproteobacteria Enterobacteriales Enterobacteriaceae *Serratia marcescens* W2.3 |
| Sma-28-1 | Bacteria Proteobacteria Gammaproteobacteria Enterobacteriales Enterobacteriaceae *Serratia marcescens* WW4 |
| Sod-1-1 | Bacteria Proteobacteria Gammaproteobacteria Enterobacteriales Enterobacteriaceae *Serratia odorifera* DSM 4582 |
| Spl-2-1 | Bacteria Proteobacteria Gammaproteobacteria Enterobacteriales Enterobacteriaceae *Serratia plymuthica* 4Rx13 |
| Spl-3-1 | Bacteria Proteobacteria Gammaproteobacteria Enterobacteriales Enterobacteriaceae *Serratia plymuthica* A30 |
| Spl-4-1 | Bacteria Proteobacteria Gammaproteobacteria Enterobacteriales Enterobacteriaceae *Serratia plymuthica* AS9 |
| Spl-5-1 | Bacteria Proteobacteria Gammaproteobacteria Enterobacteriales Enterobacteriaceae *Serratia plymuthica* PRI-2C |
| Spl-6-1 | Bacteria Proteobacteria Gammaproteobacteria Enterobacteriales Enterobacteriaceae *Serratia plymuthica* S13 |
| Spr-2-1 | Bacteria Proteobacteria Gammaproteobacteria Enterobacteriales Enterobacteriaceae *Serratia proteamaculans* 568 |
| Ssp-21-1 | Bacteria Proteobacteria Gammaproteobacteria Enterobacteriales Enterobacteriaceae *Serratia sp.* AS12 |
| Ssp-22-1 | Bacteria Proteobacteria Gammaproteobacteria Enterobacteriales Enterobacteriaceae *Serratia sp.* AS13 |
| Ssp-23-1 | Bacteria Proteobacteria Gammaproteobacteria Enterobacteriales Enterobacteriaceae *Serratia sp.* ATCC 39006 |
| Ssp-24-1 | Bacteria Proteobacteria Gammaproteobacteria Enterobacteriales Enterobacteriaceae *Serratia sp.* M24T3 |
| Ssp-25-1 | Bacteria Proteobacteria Gammaproteobacteria Enterobacteriales Enterobacteriaceae *Serratia sp.* S4 |
| Ssy-2-1 | Bacteria Proteobacteria Gammaproteobacteria Enterobacteriales Enterobacteriaceae Serratia *Serratia symbiotica str.* Tucson |
| Sbo-2-1 to Sbo-2-10 | Bacteria Proteobacteria Gammaproteobacteria Enterobacteriales Enterobacteriaceae *Shigella boydii* |
| Sbo-5-1 | Bacteria Proteobacteria Gammaproteobacteria Enterobacteriales Enterobacteriaceae *Shigella boydii* 08-0009 |
| Sbo-6-1 | Bacteria Proteobacteria Gammaproteobacteria Enterobacteriales Enterobacteriaceae *Shigella boydii* 08-0280 |
| Sbo-7-1 | Bacteria Proteobacteria Gammaproteobacteria Enterobacteriales Enterobacteriaceae *Shigella boydii* 08-2671 |
| Sbo-8-1 | Bacteria Proteobacteria Gammaproteobacteria Enterobacteriales Enterobacteriaceae *Shigella boydii* 08-2675 |
| Sbo-9-1 | Bacteria Proteobacteria Gammaproteobacteria Enterobacteriales Enterobacteriaceae *Shigella boydii* 08-6341 |
| Sbo-10-1 | Bacteria Proteobacteria Gammaproteobacteria Enterobacteriales Enterobacteriaceae *Shigella boydii* 09-0344 |
| Sbo-11-1 | Bacteria Proteobacteria Gammaproteobacteria Enterobacteriales Enterobacteriaceae *Shigella boydii* 248-1B |
| Sbo-12-1 | Bacteria Proteobacteria Gammaproteobacteria Enterobacteriales Enterobacteriaceae *Shigella boydii* 4444-74 |
| Sbo-13-1 | Bacteria Proteobacteria Gammaproteobacteria Enterobacteriales Enterobacteriaceae *Shigella boydii* 5216-82 |
| Sbo-14-1 | Bacteria Proteobacteria Gammaproteobacteria Enterobacteriales Enterobacteriaceae *Shigella boydii* ATCC 9905 |
| Sbo-15-1 | Bacteria Proteobacteria Gammaproteobacteria Enterobacteriales Enterobacteriaceae *Shigella boydii* CDC 3083-94 |
| Sbo-16-1 | Bacteria Proteobacteria Gammaproteobacteria Enterobacteriales Enterobacteriaceae *Shigella boydii* S6614 |
| Sbo-17-1 | Bacteria Proteobacteria Gammaproteobacteria Enterobacteriales Enterobacteriaceae *Shigella boydii* S7334 |
| Sbo-18-1 | Bacteria Proteobacteria Gammaproteobacteria Enterobacteriales Enterobacteriaceae *Shigella boydii* Sb227 |
| Sdy-3-1 | Bacteria Proteobacteria Gammaproteobacteria Enterobacteriales Enterobacteriaceae *Shigella dysenteriae* 1012 |
| Sdy-4-1 | Bacteria Proteobacteria Gammaproteobacteria Enterobacteriales Enterobacteriaceae *Shigella dysenteriae* 225-75 |
| Sdy-5-1 | Bacteria Proteobacteria Gammaproteobacteria Enterobacteriales Enterobacteriaceae *Shigella dysenteriae* 4 |
| Sdy-6-1 | Bacteria Proteobacteria Gammaproteobacteria Enterobacteriales Enterobacteriaceae *Shigella dysenteriae* CDC 74-1112 |
| Sdy-7-1 | Bacteria Proteobacteria Gammaproteobacteria Enterobacteriales Enterobacteriaceae *Shigella dysenteriae* S6554 |
| Sdy-8-1 | Bacteria Proteobacteria Gammaproteobacteria Enterobacteriales Enterobacteriaceae *Shigella dysenteriae* Sd197 |
| Sdy-9-1 | Bacteria Proteobacteria Gammaproteobacteria Enterobacteriales Enterobacteriaceae *Shigella dysenteriae* WRSd3 |
| Sdy-10-1 | Bacteria Proteobacteria Gammaproteobacteria Enterobacteriales Enterobacteriaceae *Shigella dysenteriae* WRSd5 |

| | |
|---|---|
| Sfl-1-1 to Sfl-1-2 | Bacteria Proteobacteria Gammaproteobacteria Enterobacteriales Enterobacteriaceae *Shigella flexneri* |
| Sfl-52-1 | Bacteria Proteobacteria Gammaproteobacteria Enterobacteriales Enterobacteriaceae *Shigella flexneri* 2001020 |
| Sfl-53-1 | Bacteria Proteobacteria Gammaproteobacteria Enterobacteriales Enterobacteriaceae *Shigella flexneri* 2002017 |
| Sfl-54-1 | Bacteria Proteobacteria Gammaproteobacteria Enterobacteriales Enterobacteriaceae *Shigella flexneri* 2002028 |
| Sfl-55-1 | Bacteria Proteobacteria Gammaproteobacteria Enterobacteriales Enterobacteriaceae *Shigella flexneri* 2002106 |
| Sfl-56-1 | Bacteria Proteobacteria Gammaproteobacteria Enterobacteriales Enterobacteriaceae *Shigella flexneri* 2003035 |
| Sfl-57-1 | Bacteria Proteobacteria Gammaproteobacteria Enterobacteriales Enterobacteriaceae *Shigella flexneri* 2003036 |
| Sfl-58-1 | Bacteria Proteobacteria Gammaproteobacteria Enterobacteriales Enterobacteriaceae *Shigella flexneri* 2005002 |
| Sfl-59-1 | Bacteria Proteobacteria Gammaproteobacteria Enterobacteriales Enterobacteriaceae *Shigella flexneri* 2005025 |
| Sfl-60-1 | Bacteria Proteobacteria Gammaproteobacteria Enterobacteriales Enterobacteriaceae *Shigella flexneri* 2005184 |
| Sfl-61-1 | Bacteria Proteobacteria Gammaproteobacteria Enterobacteriales Enterobacteriaceae *Shigella flexneri* 2930-71 |
| Sfl-25-1 to Sfl-25-14 | Bacteria Proteobacteria Gammaproteobacteria Enterobacteriales Enterobacteriaceae *Shigella flexneri* 2a |
| Sfl-62-1 | Bacteria Proteobacteria Gammaproteobacteria Enterobacteriales Enterobacteriaceae *Shigella flexneri* 2a str. 2457T |
| Sfl-63-1 | Bacteria Proteobacteria Gammaproteobacteria Enterobacteriales Enterobacteriaceae *Shigella flexneri* 2a str. 301 |
| Sfl-64-1 to Sfl-64-2 | Bacteria Proteobacteria Gammaproteobacteria Enterobacteriales Enterobacteriaceae *Shigella flexneri* 3a |
| Sfl-65-1 | Bacteria Proteobacteria Gammaproteobacteria Enterobacteriales Enterobacteriaceae *Shigella flexneri* 4343-70 |
| Sfl-66-1 | Bacteria Proteobacteria Gammaproteobacteria Enterobacteriales Enterobacteriaceae *Shigella flexneri* 5 str. 8401 |
| Sfl-67-1 | Bacteria Proteobacteria Gammaproteobacteria Enterobacteriales Enterobacteriaceae *Shigella flexneri* 51581 |
| Sfl-68-1 | Bacteria Proteobacteria Gammaproteobacteria Enterobacteriales Enterobacteriaceae *Shigella flexneri* 5a str. M90T |
| Sfl-69-1 | Bacteria Proteobacteria Gammaproteobacteria Enterobacteriales Enterobacteriaceae *Shigella flexneri* 6 |
| Sfl-70-1 | Bacteria Proteobacteria Gammaproteobacteria Enterobacteriales Enterobacteriaceae *Shigella flexneri* CCH060 |
| Sfl-71-1 | Bacteria Proteobacteria Gammaproteobacteria Enterobacteriales Enterobacteriaceae *Shigella flexneri* CDC 796-83 |
| Sfl-72-1 | Bacteria Proteobacteria Gammaproteobacteria Enterobacteriales Enterobacteriaceae *Shigella flexneri* G1663 |
| Sfl-73-1 | Bacteria Proteobacteria Gammaproteobacteria Enterobacteriales Enterobacteriaceae *Shigella flexneri* MT1457 |
| Sfl-74-1 | Bacteria Proteobacteria Gammaproteobacteria Enterobacteriales Enterobacteriaceae *Shigella flexneri* S5644 |
| Sfl-75-1 | Bacteria Proteobacteria Gammaproteobacteria Enterobacteriales Enterobacteriaceae *Shigella flexneri* S5717 |
| Sfl-76-1 | Bacteria Proteobacteria Gammaproteobacteria Enterobacteriales Enterobacteriaceae *Shigella flexneri* S6162 |
| Sfl-77-1 | Bacteria Proteobacteria Gammaproteobacteria Enterobacteriales Enterobacteriaceae *Shigella flexneri* S6585 |
| Sfl-78-1 | Bacteria Proteobacteria Gammaproteobacteria Enterobacteriales Enterobacteriaceae *Shigella flexneri* S6678 |
| Sfl-79-1 | Bacteria Proteobacteria Gammaproteobacteria Enterobacteriales Enterobacteriaceae *Shigella flexneri* S6764 |
| Sfl-80-1 | Bacteria Proteobacteria Gammaproteobacteria Enterobacteriales Enterobacteriaceae *Shigella flexneri* S7737 |
| Sfl-81-1 | Bacteria Proteobacteria Gammaproteobacteria Enterobacteriales Enterobacteriaceae *Shigella flexneri* Shi06GS02 |
| Sfl-82-1 | Bacteria Proteobacteria Gammaproteobacteria Enterobacteriales Enterobacteriaceae *Shigella flexneri* Shi06HN006 |
| Sfl-83-1 | Bacteria Proteobacteria Gammaproteobacteria Enterobacteriales Enterobacteriaceae *Shigella flexneri* Shi06HN344 |
| Sfl-84-1 | Bacteria Proteobacteria Gammaproteobacteria Enterobacteriales Enterobacteriaceae *Shigella flexneri* Shi06HN347 |
| Sfl-85-1 | Bacteria Proteobacteria Gammaproteobacteria Enterobacteriales Enterobacteriaceae *Shigella flexneri* VA-6 |
| Sfl-51-1 to Sfl-51-2 | Bacteria Proteobacteria Gammaproteobacteria Enterobacteriales Enterobacteriaceae *Shigella flexneri* Y |
| Sso-1-1 to Sso-1-547 | Bacteria Proteobacteria Gammaproteobacteria Enterobacteriales Enterobacteriaceae *Shigella sonnei* |
| Sso-3-1 | Bacteria Proteobacteria Gammaproteobacteria Enterobacteriales Enterobacteriaceae *Shigella sonnei* 08-7761 |
| Sso-4-1 | Bacteria Proteobacteria Gammaproteobacteria Enterobacteriales Enterobacteriaceae *Shigella sonnei* 08-7765 |
| Sso-5-1 | Bacteria Proteobacteria Gammaproteobacteria Enterobacteriales Enterobacteriaceae *Shigella sonnei* 09-1032 |
| Sso-6-1 | Bacteria Proteobacteria Gammaproteobacteria Enterobacteriales Enterobacteriaceae *Shigella sonnei* 09-2245 |
| Sso-7-1 | Bacteria Proteobacteria Gammaproteobacteria Enterobacteriales Enterobacteriaceae *Shigella sonnei* 09-4962 |
| Sso-8-1 | Bacteria Proteobacteria Gammaproteobacteria Enterobacteriales Enterobacteriaceae *Shigella sonnei* 3233-85 |
| Sso-9-1 | Bacteria Proteobacteria Gammaproteobacteria Enterobacteriales Enterobacteriaceae *Shigella sonnei* 4822-66 |
| Sso-10-1 | Bacteria Proteobacteria Gammaproteobacteria Enterobacteriales Enterobacteriaceae *Shigella sonnei* 53G |
| Sso-11-1 | Bacteria Proteobacteria Gammaproteobacteria Enterobacteriales Enterobacteriaceae *Shigella sonnei* S6513 |
| Sso-12-1 | Bacteria Proteobacteria Gammaproteobacteria Enterobacteriales Enterobacteriaceae *Shigella sonnei* Ss046 |
| Sso-13-1 | Bacteria Proteobacteria Gammaproteobacteria Enterobacteriales Enterobacteriaceae *Shigella sonnei str.* Moseley |
| Ssp-26-1 | Bacteria Proteobacteria Gammaproteobacteria Enterobacteriales Enterobacteriaceae *Shigella sp.* D9 |
| Sbl-1-1 | Bacteria Proteobacteria Gammaproteobacteria Enterobacteriales Enterobacteriaceae *Shimwellia blattae* DSM 4481 = NBRC 105725 |
| Sgl-1-1 | Bacteria Proteobacteria Gammaproteobacteria Enterobacteriales Enterobacteriaceae *Sodalis glossinidius str. 'morsitans'* |
| Xbo-11-1 | Bacteria Proteobacteria Gammaproteobacteria Enterobacteriales Enterobacteriaceae *Xenorhabdus bovienii* SS-2004 |
| Xne-3-1 | Bacteria Proteobacteria Gammaproteobacteria Enterobacteriales Enterobacteriaceae *Xenorhabdus nematophila* ATCC 19061 |
| Xne-4-1 | Bacteria Proteobacteria Gammaproteobacteria Enterobacteriales Enterobacteriaceae *Xenorhabdus nematophila* F1 |
| Xne-5-1 to Xne-5-2 | Bacteria Proteobacteria Gammaproteobacteria Enterobacteriales Enterobacteriaceae *Xenorhabdus nematophila str.* Anatoliense |
| Xne-6-1 | Bacteria Proteobacteria Gammaproteobacteria Enterobacteriales Enterobacteriaceae *Xenorhabdus nematophila str.* Websteri |
| Yal-4-1 | Bacteria Proteobacteria Gammaproteobacteria Enterobacteriales Enterobacteriaceae *Yersinia aldovae* ATCC 35236 |
| Ybe-2-1 | Bacteria Proteobacteria Gammaproteobacteria Enterobacteriales Enterobacteriaceae *Yersinia bercovieri* ATCC 43970 |
| Yen-8-1 | Bacteria Proteobacteria Gammaproteobacteria Enterobacteriales Enterobacteriaceae *Yersinia enterocolitica (type* O:3) str. YE12/03 |
| Yen-9-1 | Bacteria Proteobacteria Gammaproteobacteria Enterobacteriales Enterobacteriaceae *Yersinia enterocolitica (type* O:5,27) str. YE149/02 |
| Yen-10-1 | Bacteria Proteobacteria Gammaproteobacteria Enterobacteriales Enterobacteriaceae *Yersinia enterocolitica (type* O:9) str. YE212/02 |
| Yen-11-1 | Bacteria Proteobacteria Gammaproteobacteria Enterobacteriales Enterobacteriaceae *Yersinia enterocolitica* IP 10393 |
| Yen-12-1 | Bacteria Proteobacteria Gammaproteobacteria Enterobacteriales Enterobacteriaceae *Yersinia enterocolitica* IP2222 |
| Yen-13-1 | Bacteria Proteobacteria Gammaproteobacteria Enterobacteriales Enterobacteriaceae *Yersinia enterocolitica* NFO |

| | |
|---|---|
| Yen-14-1 | Bacteria Proteobacteria Gammaproteobacteria Enterobacteriales Enterobacteriaceae *Yersinia enterocolitica subsp. enterocolitica* 8081 |
| Yen-15-1 | Bacteria Proteobacteria Gammaproteobacteria Enterobacteriales Enterobacteriaceae *Yersinia enterocolitica subsp. enterocolitica* WA-314 |
| Yen-16-1 | Bacteria Proteobacteria Gammaproteobacteria Enterobacteriales Enterobacteriaceae *Yersinia enterocolitica subsp. palearctica* 105.5R(r) |
| Yen-17-1 | Bacteria Proteobacteria Gammaproteobacteria Enterobacteriales Enterobacteriaceae *Yersinia enterocolitica subsp. palearctica* PhRBD_Ye1 |
| Yen-18-1 | Bacteria Proteobacteria Gammaproteobacteria Enterobacteriales Enterobacteriaceae *Yersinia enterocolitica subsp. palearctica* Y11 |
| Yen-19-1 | Bacteria Proteobacteria Gammaproteobacteria Enterobacteriales Enterobacteriaceae *Yersinia enterocolitica subsp. palearctica* YE-149 |
| Yen-20-1 | Bacteria Proteobacteria Gammaproteobacteria Enterobacteriales Enterobacteriaceae *Yersinia enterocolitica subsp. palearctica* YE-150 |
| Yen-21-1 | Bacteria Proteobacteria Gammaproteobacteria Enterobacteriales Enterobacteriaceae *Yersinia enterocolitica subsp. palearctica* YE-P1 |
| Yen-22-1 | Bacteria Proteobacteria Gammaproteobacteria Enterobacteriales Enterobacteriaceae *Yersinia enterocolitica subsp. palearctica* YE-P4 |
| Yfr-2-1 to Yfr-2-2 | Bacteria Proteobacteria Gammaproteobacteria Enterobacteriales Enterobacteriaceae *Yersinia frederiksenii* ATCC 33641 |
| Yin-2-1 | Bacteria Proteobacteria Gammaproteobacteria Enterobacteriales Enterobacteriaceae *Yersinia intermedia* ATCC 29909 |
| Ykr-2-1 | Bacteria Proteobacteria Gammaproteobacteria Enterobacteriales Enterobacteriaceae *Yersinia kristensenii* ATCC 33638 |
| Ymo-2-1 | Bacteria Proteobacteria Gammaproteobacteria Enterobacteriales Enterobacteriaceae *Yersinia mollaretii* ATCC 43969 |
| Ype-116-1 | Bacteria Proteobacteria Gammaproteobacteria Enterobacteriales Enterobacteriaceae *Yersinia pestis* 113 |
| Ype-117-1 | Bacteria Proteobacteria Gammaproteobacteria Enterobacteriales Enterobacteriaceae *Yersinia pestis* 2330 |
| Ype-118-1 | Bacteria Proteobacteria Gammaproteobacteria Enterobacteriales Enterobacteriaceae *Yersinia pestis* 2501 |
| Ype-119-1 | Bacteria Proteobacteria Gammaproteobacteria Enterobacteriales Enterobacteriaceae *Yersinia pestis* 2504 |
| Ype-120-1 | Bacteria Proteobacteria Gammaproteobacteria Enterobacteriales Enterobacteriaceae *Yersinia pestis* 2506 |
| Ype-121-1 | Bacteria Proteobacteria Gammaproteobacteria Enterobacteriales Enterobacteriaceae *Yersinia pestis* 30017 |
| Ype-122-1 | Bacteria Proteobacteria Gammaproteobacteria Enterobacteriales Enterobacteriaceae *Yersinia pestis* 42013 |
| Ype-123-1 | Bacteria Proteobacteria Gammaproteobacteria Enterobacteriales Enterobacteriaceae *Yersinia pestis* 42091 |
| Ype-124-1 | Bacteria Proteobacteria Gammaproteobacteria Enterobacteriales Enterobacteriaceae *Yersinia pestis* 5761 |
| Ype-125-1 | Bacteria Proteobacteria Gammaproteobacteria Enterobacteriales Enterobacteriaceae *Yersinia pestis* 7 |
| Ype-126-1 | Bacteria Proteobacteria Gammaproteobacteria Enterobacteriales Enterobacteriaceae *Yersinia pestis* 735 |
| Ype-127-1 | Bacteria Proteobacteria Gammaproteobacteria Enterobacteriales Enterobacteriaceae *Yersinia pestis* 780441 |
| Ype-128-1 | Bacteria Proteobacteria Gammaproteobacteria Enterobacteriales Enterobacteriaceae *Yersinia pestis* 91 |
| Ype-29-1 to Ype-29-2 | Bacteria Proteobacteria Gammaproteobacteria Enterobacteriales Enterobacteriaceae *Yersinia pestis* A1122 |
| Ype-129-1 | Bacteria Proteobacteria Gammaproteobacteria Enterobacteriales Enterobacteriaceae *Yersinia pestis* A1956001 |
| Ype-31-1 to Ype-31-2 | Bacteria Proteobacteria Gammaproteobacteria Enterobacteriales Enterobacteriaceae *Yersinia pestis* Angola |
| Ype-32-1 to Ype-32-2 | Bacteria Proteobacteria Gammaproteobacteria Enterobacteriales Enterobacteriaceae *Yersinia pestis* Antiqua |
| Ype-218-1 | Bacteria Proteobacteria Gammaproteobacteria Enterobacteriales Enterobacteriaceae *Yersinia pestis biovar* Antiqua str. B42003004 |
| Ype-219-1 to Ype-219-2 | Bacteria Proteobacteria Gammaproteobacteria Enterobacteriales Enterobacteriaceae *Yersinia pestis biovar* Antiqua str. E1979001 |
| Ype-220-1 | Bacteria Proteobacteria Gammaproteobacteria Enterobacteriales Enterobacteriaceae *Yersinia pestis biovar* Antiqua str. UG05-0454 |
| Ype-221-1 | Bacteria Proteobacteria Gammaproteobacteria Enterobacteriales Enterobacteriaceae *Yersinia pestis biovar* Mediaevalis str. K1973002 |
| Ype-222-1 | Bacteria Proteobacteria Gammaproteobacteria Enterobacteriales Enterobacteriaceae *Yersinia pestis biovar* Medievalis str. Harbin 35 |
| Ype-223-1 | Bacteria Proteobacteria Gammaproteobacteria Enterobacteriales Enterobacteriaceae *Yersinia pestis biovar* Microtus str. 91001 |
| Ype-224-1 | Bacteria Proteobacteria Gammaproteobacteria Enterobacteriales Enterobacteriaceae *Yersinia pestis biovar* Orientalis str. F1991016 |
| Ype-226-1 | Bacteria Proteobacteria Gammaproteobacteria Enterobacteriales Enterobacteriaceae *Yersinia pestis biovar* Orientalis str. India 195 |
| Ype-225-1 | Bacteria Proteobacteria Gammaproteobacteria Enterobacteriales Enterobacteriaceae *Yersinia pestis biovar* Orientalis str. IP275 |
| Ype-227-1 | Bacteria Proteobacteria Gammaproteobacteria Enterobacteriales Enterobacteriaceae *Yersinia pestis biovar* Orientalis str. MG05-1020 |
| Ype-228-1 | Bacteria Proteobacteria Gammaproteobacteria Enterobacteriales Enterobacteriaceae *Yersinia pestis biovar* Orientalis str. PEXU2 |
| Ype-130-1 | Bacteria Proteobacteria Gammaproteobacteria Enterobacteriales Enterobacteriaceae *Yersinia pestis* C1976001 |
| Ype-131-1 | Bacteria Proteobacteria Gammaproteobacteria Enterobacteriales Enterobacteriaceae *Yersinia pestis* C1989001 |
| Ype-132-1 | Bacteria Proteobacteria Gammaproteobacteria Enterobacteriales Enterobacteriaceae *Yersinia pestis* CA88-4125 |
| Ype-133-1 | Bacteria Proteobacteria Gammaproteobacteria Enterobacteriales Enterobacteriaceae *Yersinia pestis* CMCC125002 |
| Ype-134-1 | Bacteria Proteobacteria Gammaproteobacteria Enterobacteriales Enterobacteriaceae *Yersinia pestis* CMCC348002 |
| Ype-135-1 | Bacteria Proteobacteria Gammaproteobacteria Enterobacteriales Enterobacteriaceae *Yersinia pestis* CMCC49003 |
| Ype-136-1 | Bacteria Proteobacteria Gammaproteobacteria Enterobacteriales Enterobacteriaceae *Yersinia pestis* CMCC64001 |
| Ype-137-1 | Bacteria Proteobacteria Gammaproteobacteria Enterobacteriales Enterobacteriaceae *Yersinia pestis* CMCC8211 |
| Ype-138-1 | Bacteria Proteobacteria Gammaproteobacteria Enterobacteriales Enterobacteriaceae *Yersinia pestis* CMCC87001 |
| Ype-139-1 | Bacteria Proteobacteria Gammaproteobacteria Enterobacteriales Enterobacteriaceae *Yersinia pestis* CMCC91090 |
| Ype-140-1 | Bacteria Proteobacteria Gammaproteobacteria Enterobacteriales Enterobacteriaceae *Yersinia pestis* CMCC93014 |
| Ype-141-1 | Bacteria Proteobacteria Gammaproteobacteria Enterobacteriales Enterobacteriaceae *Yersinia pestis* CMCC96001 |
| Ype-142-1 | Bacteria Proteobacteria Gammaproteobacteria Enterobacteriales Enterobacteriaceae *Yersinia pestis* CO92 |
| Ype-143-1 | Bacteria Proteobacteria Gammaproteobacteria Enterobacteriales Enterobacteriaceae *Yersinia pestis* D106004 |
| Ype-144-1 | Bacteria Proteobacteria Gammaproteobacteria Enterobacteriales Enterobacteriaceae *Yersinia pestis* D182038 |
| Ype-145-1 | Bacteria Proteobacteria Gammaproteobacteria Enterobacteriales Enterobacteriaceae *Yersinia pestis* D1982001 |
| Ype-146-1 | Bacteria Proteobacteria Gammaproteobacteria Enterobacteriales Enterobacteriaceae *Yersinia pestis* FV-1 |
| Ype-147-1 | Bacteria Proteobacteria Gammaproteobacteria Enterobacteriales Enterobacteriaceae *Yersinia pestis* G1996006 |
| Ype-148-1 | Bacteria Proteobacteria Gammaproteobacteria Enterobacteriales Enterobacteriaceae *Yersinia pestis* G1996010 |
| Ype-149-1 | Bacteria Proteobacteria Gammaproteobacteria Enterobacteriales Enterobacteriaceae *Yersinia pestis* G8786 |
| Ype-150-1 | Bacteria Proteobacteria Gammaproteobacteria Enterobacteriales Enterobacteriaceae *Yersinia pestis* H1959004 |
| Ype-151-1 | Bacteria Proteobacteria Gammaproteobacteria Enterobacteriales Enterobacteriaceae *Yersinia pestis* INS |
| Ype-152-1 | Bacteria Proteobacteria Gammaproteobacteria Enterobacteriales Enterobacteriaceae *Yersinia pestis* K11973002 |
| Ype-153-1 | Bacteria Proteobacteria Gammaproteobacteria Enterobacteriales Enterobacteriaceae *Yersinia pestis* KIM |

| | |
|---|---|
| Ype-154-1 | Bacteria Proteobacteria Gammaproteobacteria Enterobacteriales Enterobacteriaceae *Yersinia pestis* MGJZ11 |
| Ype-155-1 | Bacteria Proteobacteria Gammaproteobacteria Enterobacteriales Enterobacteriaceae *Yersinia pestis* Nepal516 |
| Ype-215-1 | Bacteria Proteobacteria Gammaproteobacteria Enterobacteriales Enterobacteriaceae *Yersinia pestis* Pestoides A |
| Ype-94-1 to Ype-94-2 | Bacteria Proteobacteria Gammaproteobacteria Enterobacteriales Enterobacteriaceae *Yersinia pestis* Pestoides F |
| Ype-156-1 | Bacteria Proteobacteria Gammaproteobacteria Enterobacteriales Enterobacteriaceae *Yersinia pestis* PY-01 |
| Ype-157-1 | Bacteria Proteobacteria Gammaproteobacteria Enterobacteriales Enterobacteriaceae *Yersinia pestis* PY-02 |
| Ype-158-1 | Bacteria Proteobacteria Gammaproteobacteria Enterobacteriales Enterobacteriaceae *Yersinia pestis* PY-03 |
| Ype-159-1 | Bacteria Proteobacteria Gammaproteobacteria Enterobacteriales Enterobacteriaceae *Yersinia pestis* PY-04 |
| Ype-160-1 | Bacteria Proteobacteria Gammaproteobacteria Enterobacteriales Enterobacteriaceae *Yersinia pestis* PY-05 |
| Ype-161-1 | Bacteria Proteobacteria Gammaproteobacteria Enterobacteriales Enterobacteriaceae *Yersinia pestis* PY-06 |
| Ype-162-1 | Bacteria Proteobacteria Gammaproteobacteria Enterobacteriales Enterobacteriaceae *Yersinia pestis* PY-07 |
| Ype-163-1 | Bacteria Proteobacteria Gammaproteobacteria Enterobacteriales Enterobacteriaceae *Yersinia pestis* PY-08 |
| Ype-164-1 | Bacteria Proteobacteria Gammaproteobacteria Enterobacteriales Enterobacteriaceae *Yersinia pestis* PY-09 |
| Ype-165-1 | Bacteria Proteobacteria Gammaproteobacteria Enterobacteriales Enterobacteriaceae *Yersinia pestis* PY-10 |
| Ype-166-1 | Bacteria Proteobacteria Gammaproteobacteria Enterobacteriales Enterobacteriaceae *Yersinia pestis* PY-100 |
| Ype-167-1 | Bacteria Proteobacteria Gammaproteobacteria Enterobacteriales Enterobacteriaceae *Yersinia pestis* PY-101 |
| Ype-168-1 | Bacteria Proteobacteria Gammaproteobacteria Enterobacteriales Enterobacteriaceae *Yersinia pestis* PY-102 |
| Ype-169-1 | Bacteria Proteobacteria Gammaproteobacteria Enterobacteriales Enterobacteriaceae *Yersinia pestis* PY-103 |
| Ype-170-1 | Bacteria Proteobacteria Gammaproteobacteria Enterobacteriales Enterobacteriaceae *Yersinia pestis* PY-11 |
| Ype-171-1 | Bacteria Proteobacteria Gammaproteobacteria Enterobacteriales Enterobacteriaceae *Yersinia pestis* PY-113 |
| Ype-172-1 | Bacteria Proteobacteria Gammaproteobacteria Enterobacteriales Enterobacteriaceae *Yersinia pestis* PY-12 |
| Ype-173-1 | Bacteria Proteobacteria Gammaproteobacteria Enterobacteriales Enterobacteriaceae *Yersinia pestis* PY-13 |
| Ype-174-1 | Bacteria Proteobacteria Gammaproteobacteria Enterobacteriales Enterobacteriaceae *Yersinia pestis* PY-14 |
| Ype-175-1 | Bacteria Proteobacteria Gammaproteobacteria Enterobacteriales Enterobacteriaceae *Yersinia pestis* PY-15 |
| Ype-176-1 | Bacteria Proteobacteria Gammaproteobacteria Enterobacteriales Enterobacteriaceae *Yersinia pestis* PY-16 |
| Ype-177-1 | Bacteria Proteobacteria Gammaproteobacteria Enterobacteriales Enterobacteriaceae *Yersinia pestis* PY-19 |
| Ype-178-1 | Bacteria Proteobacteria Gammaproteobacteria Enterobacteriales Enterobacteriaceae *Yersinia pestis* PY-25 |
| Ype-179-1 | Bacteria Proteobacteria Gammaproteobacteria Enterobacteriales Enterobacteriaceae *Yersinia pestis* PY-29 |
| Ype-180-1 | Bacteria Proteobacteria Gammaproteobacteria Enterobacteriales Enterobacteriaceae *Yersinia pestis* PY-32 |
| Ype-181-1 | Bacteria Proteobacteria Gammaproteobacteria Enterobacteriales Enterobacteriaceae *Yersinia pestis* PY-34 |
| Ype-182-1 | Bacteria Proteobacteria Gammaproteobacteria Enterobacteriales Enterobacteriaceae *Yersinia pestis* PY-36 |
| Ype-183-1 | Bacteria Proteobacteria Gammaproteobacteria Enterobacteriales Enterobacteriaceae *Yersinia pestis* PY-42 |
| Ype-184-1 | Bacteria Proteobacteria Gammaproteobacteria Enterobacteriales Enterobacteriaceae *Yersinia pestis* PY-45 |
| Ype-185-1 | Bacteria Proteobacteria Gammaproteobacteria Enterobacteriales Enterobacteriaceae *Yersinia pestis* PY-46 |
| Ype-186-1 | Bacteria Proteobacteria Gammaproteobacteria Enterobacteriales Enterobacteriaceae *Yersinia pestis* PY-47 |
| Ype-187-1 | Bacteria Proteobacteria Gammaproteobacteria Enterobacteriales Enterobacteriaceae *Yersinia pestis* PY-48 |
| Ype-188-1 | Bacteria Proteobacteria Gammaproteobacteria Enterobacteriales Enterobacteriaceae *Yersinia pestis* PY-52 |
| Ype-189-1 | Bacteria Proteobacteria Gammaproteobacteria Enterobacteriales Enterobacteriaceae *Yersinia pestis* PY-53 |
| Ype-190-1 | Bacteria Proteobacteria Gammaproteobacteria Enterobacteriales Enterobacteriaceae *Yersinia pestis* PY-54 |
| Ype-191-1 | Bacteria Proteobacteria Gammaproteobacteria Enterobacteriales Enterobacteriaceae *Yersinia pestis* PY-55 |
| Ype-192-1 | Bacteria Proteobacteria Gammaproteobacteria Enterobacteriales Enterobacteriaceae *Yersinia pestis* PY-56 |
| Ype-193-1 | Bacteria Proteobacteria Gammaproteobacteria Enterobacteriales Enterobacteriaceae *Yersinia pestis* PY-58 |
| Ype-194-1 | Bacteria Proteobacteria Gammaproteobacteria Enterobacteriales Enterobacteriaceae *Yersinia pestis* PY-59 |
| Ype-195-1 | Bacteria Proteobacteria Gammaproteobacteria Enterobacteriales Enterobacteriaceae *Yersinia pestis* PY-60 |
| Ype-196-1 | Bacteria Proteobacteria Gammaproteobacteria Enterobacteriales Enterobacteriaceae *Yersinia pestis* PY-61 |
| Ype-197-1 | Bacteria Proteobacteria Gammaproteobacteria Enterobacteriales Enterobacteriaceae *Yersinia pestis* PY-63 |
| Ype-198-1 | Bacteria Proteobacteria Gammaproteobacteria Enterobacteriales Enterobacteriaceae *Yersinia pestis* PY-64 |
| Ype-199-1 | Bacteria Proteobacteria Gammaproteobacteria Enterobacteriales Enterobacteriaceae *Yersinia pestis* PY-65 |
| Ype-200-1 | Bacteria Proteobacteria Gammaproteobacteria Enterobacteriales Enterobacteriaceae *Yersinia pestis* PY-66 |
| Ype-201-1 | Bacteria Proteobacteria Gammaproteobacteria Enterobacteriales Enterobacteriaceae *Yersinia pestis* PY-71 |
| Ype-202-1 | Bacteria Proteobacteria Gammaproteobacteria Enterobacteriales Enterobacteriaceae *Yersinia pestis* PY-72 |
| Ype-203-1 | Bacteria Proteobacteria Gammaproteobacteria Enterobacteriales Enterobacteriaceae *Yersinia pestis* PY-76 |
| Ype-204-1 | Bacteria Proteobacteria Gammaproteobacteria Enterobacteriales Enterobacteriaceae *Yersinia pestis* PY-88 |
| Ype-205-1 | Bacteria Proteobacteria Gammaproteobacteria Enterobacteriales Enterobacteriaceae *Yersinia pestis* PY-89 |
| Ype-206-1 | Bacteria Proteobacteria Gammaproteobacteria Enterobacteriales Enterobacteriaceae *Yersinia pestis* PY-90 |
| Ype-207-1 | Bacteria Proteobacteria Gammaproteobacteria Enterobacteriales Enterobacteriaceae *Yersinia pestis* PY-91 |
| Ype-208-1 | Bacteria Proteobacteria Gammaproteobacteria Enterobacteriales Enterobacteriaceae *Yersinia pestis* PY-92 |
| Ype-209-1 | Bacteria Proteobacteria Gammaproteobacteria Enterobacteriales Enterobacteriaceae *Yersinia pestis* PY-93 |
| Ype-210-1 | Bacteria Proteobacteria Gammaproteobacteria Enterobacteriales Enterobacteriaceae *Yersinia pestis* PY-94 |
| Ype-211-1 | Bacteria Proteobacteria Gammaproteobacteria Enterobacteriales Enterobacteriaceae *Yersinia pestis* PY-95 |
| Ype-212-1 | Bacteria Proteobacteria Gammaproteobacteria Enterobacteriales Enterobacteriaceae *Yersinia pestis* PY-96 |
| Ype-213-1 | Bacteria Proteobacteria Gammaproteobacteria Enterobacteriales Enterobacteriaceae *Yersinia pestis* PY-98 |
| Ype-214-1 | Bacteria Proteobacteria Gammaproteobacteria Enterobacteriales Enterobacteriaceae *Yersinia pestis* PY-99 |
| Ype-216-1 | Bacteria Proteobacteria Gammaproteobacteria Enterobacteriales Enterobacteriaceae *Yersinia pestis* YN2551 |
| Ype-217-1 | Bacteria Proteobacteria Gammaproteobacteria Enterobacteriales Enterobacteriaceae *Yersinia pestis* Z176003 |

| | |
|---|---|
| Yps-14-1 | Bacteria Proteobacteria Gammaproteobacteria Enterobacteriales Enterobacteriaceae *Yersinia pseudotuberculosis* IP 31758 |
| Yps-10-1 to Yps-10-2 | Bacteria Proteobacteria Gammaproteobacteria Enterobacteriales Enterobacteriaceae *Yersinia pseudotuberculosis* IP 32953 |
| Yps-15-1 | Bacteria Proteobacteria Gammaproteobacteria Enterobacteriales Enterobacteriaceae *Yersinia pseudotuberculosis* NBRC 105692 |
| Yps-11-1 to Yps-11-2 | Bacteria Proteobacteria Gammaproteobacteria Enterobacteriales Enterobacteriaceae *Yersinia pseudotuberculosis* PB1/+ |
| Yps-12-1 to Yps-12-2 | Bacteria Proteobacteria Gammaproteobacteria Enterobacteriales Enterobacteriaceae *Yersinia pseudotuberculosis* YPIII |
| Yro-2-1 | Bacteria Proteobacteria Gammaproteobacteria Enterobacteriales Enterobacteriaceae *Yersinia rohdei* ATCC 43380 |
| Yru-1-1 to Yru-1-3 | Bacteria Proteobacteria Gammaproteobacteria Enterobacteriales Enterobacteriaceae *Yersinia ruckeri* |
| Yru-2-1 to Yru-2-2 | Bacteria Proteobacteria Gammaproteobacteria Enterobacteriales Enterobacteriaceae *Yersinia ruckeri* ATCC 29473 |
| Gpe-1-1 | Bacteria Proteobacteria Gammaproteobacteria *Gallaecimonas pentaromativorans* |
| Gxi-1-1 | Bacteria Proteobacteria Gammaproteobacteria *Gallaecimonas xiamenensis* 3-C-1 |
| gpr-1-1 | Bacteria Proteobacteria Gammaproteobacteria gamma proteobacterium WG36 |
| Aca-3-1 | Bacteria Proteobacteria Gammaproteobacteria Pasteurellales Pasteurellaceae *Actinobacillus capsulatus* DSM 19761 |
| Aeq-1-1 | Bacteria Proteobacteria Gammaproteobacteria Pasteurellales Pasteurellaceae *Actinobacillus equuli subsp. equuli* |
| Ami-1-1 | Bacteria Proteobacteria Gammaproteobacteria Pasteurellales Pasteurellaceae *Actinobacillus minor* 202 |
| Ami-2-1 | Bacteria Proteobacteria Gammaproteobacteria Pasteurellales Pasteurellaceae *Actinobacillus minor* NM305 |
| Apl-1-1 to Apl-1-6 | Bacteria Proteobacteria Gammaproteobacteria Pasteurellales Pasteurellaceae *Actinobacillus pleuropneumoniae* |
| Apl-2-1 | Bacteria Proteobacteria Gammaproteobacteria Pasteurellales Pasteurellaceae *Actinobacillus pleuropneumoniae* L20 |
| Apl-3-1 | Bacteria Proteobacteria Gammaproteobacteria Pasteurellales Pasteurellaceae *Actinobacillus pleuropneumoniae* S8 |
| Apl-4-1 to Apl-4-2 | Bacteria Proteobacteria Gammaproteobacteria Pasteurellales Pasteurellaceae *Actinobacillus pleuropneumoniae serovar* 1 str. 4074 |
| Apl-5-1 | Bacteria Proteobacteria Gammaproteobacteria Pasteurellales Pasteurellaceae *Actinobacillus pleuropneumoniae serovar* 10 str. D13039 |
| Apl-6-1 | Bacteria Proteobacteria Gammaproteobacteria Pasteurellales Pasteurellaceae *Actinobacillus pleuropneumoniae serovar* 11 str. 56153 |
| Apl-7-1 | Bacteria Proteobacteria Gammaproteobacteria Pasteurellales Pasteurellaceae *Actinobacillus pleuropneumoniae serovar* 12 str. 1096 |
| Apl-8-1 | Bacteria Proteobacteria Gammaproteobacteria Pasteurellales Pasteurellaceae *Actinobacillus pleuropneumoniae serovar* 13 str. N273 |
| Apl-9-1 | Bacteria Proteobacteria Gammaproteobacteria Pasteurellales Pasteurellaceae *Actinobacillus pleuropneumoniae serovar* 2 str. 4226 |
| Apl-10-1 | Bacteria Proteobacteria Gammaproteobacteria Pasteurellales Pasteurellaceae *Actinobacillus pleuropneumoniae serovar* 2 str. S1536 |
| Apl-11-1 | Bacteria Proteobacteria Gammaproteobacteria Pasteurellales Pasteurellaceae *Actinobacillus pleuropneumoniae serovar* 3 str. JL03 |
| Apl-12-1 | Bacteria Proteobacteria Gammaproteobacteria Pasteurellales Pasteurellaceae *Actinobacillus pleuropneumoniae serovar* 4 str. M62 |
| Apl-13-1 to Apl-13-2 | Bacteria Proteobacteria Gammaproteobacteria Pasteurellales Pasteurellaceae *Actinobacillus pleuropneumoniae serovar* 6 str. Femo |
| Apl-14-1 | Bacteria Proteobacteria Gammaproteobacteria Pasteurellales Pasteurellaceae *Actinobacillus pleuropneumoniae serovar* 7 str. AP76 |
| Apl-15-1 | Bacteria Proteobacteria Gammaproteobacteria Pasteurellales Pasteurellaceae *Actinobacillus pleuropneumoniae serovar* 9 str. CVJ13261 |
| Asu-1-1 | Bacteria Proteobacteria Gammaproteobacteria Pasteurellales Pasteurellaceae *Actinobacillus succinogenes* 130Z |
| Asu-2-1 | Bacteria Proteobacteria Gammaproteobacteria Pasteurellales Pasteurellaceae *Actinobacillus suis* ATCC 33415 |
| Asu-3-1 | Bacteria Proteobacteria Gammaproteobacteria Pasteurellales Pasteurellaceae *Actinobacillus suis* H91-0380 |
| Aur-1-1 | Bacteria Proteobacteria Gammaproteobacteria Pasteurellales Pasteurellaceae *Actinobacillus ureae* ATCC 25976 |
| Aac-1-1 | Bacteria Proteobacteria Gammaproteobacteria Pasteurellales Pasteurellaceae *Aggregatibacter actinomycetemcomitans* ANH9381 |
| Aac-2-1 | Bacteria Proteobacteria Gammaproteobacteria Pasteurellales Pasteurellaceae *Aggregatibacter actinomycetemcomitans* D17P-3 |
| Aac-3-1 | Bacteria Proteobacteria Gammaproteobacteria Pasteurellales Pasteurellaceae *Aggregatibacter actinomycetemcomitans* D7S-1 |
| Aac-4-1 | Bacteria Proteobacteria Gammaproteobacteria Pasteurellales Pasteurellaceae *Aggregatibacter actinomycetemcomitans* DSM 8324 |
| Aac-5-1 to Aac-5-2 | Bacteria Proteobacteria Gammaproteobacteria Pasteurellales Pasteurellaceae *Aggregatibacter actinomycetemcomitans* HK1651 |
| Aac-6-1 to Aac-6-2 | Bacteria Proteobacteria Gammaproteobacteria Pasteurellales Pasteurellaceae *Aggregatibacter actinomycetemcomitans* RhAA1 |
| Aac-8-1 | Bacteria Proteobacteria Gammaproteobacteria Pasteurellales Pasteurellaceae *Aggregatibacter actinomycetemcomitans serotype a str.* A160 |
| Aac-9-1 | Bacteria Proteobacteria Gammaproteobacteria Pasteurellales Pasteurellaceae *Aggregatibacter actinomycetemcomitans serotype a str.* H5P1 |
| Aac-10-1 | Bacteria Proteobacteria Gammaproteobacteria Pasteurellales Pasteurellaceae *Aggregatibacter actinomycetemcomitans serotype b str.* I23C |
| Aac-11-1 | Bacteria Proteobacteria Gammaproteobacteria Pasteurellales Pasteurellaceae *Aggregatibacter actinomycetemcomitans serotype b str.* S23A |
| Aac-12-1 | Bacteria Proteobacteria Gammaproteobacteria Pasteurellales Pasteurellaceae *Aggregatibacter actinomycetemcomitans serotype b str.* SCC1398 |
| Aac-13-1 | Bacteria Proteobacteria Gammaproteobacteria Pasteurellales Pasteurellaceae *Aggregatibacter actinomycetemcomitans serotype b str.* SCC4092 |
| Aac-14-1 | Bacteria Proteobacteria Gammaproteobacteria Pasteurellales Pasteurellaceae *Aggregatibacter actinomycetemcomitans serotype c str.* AAS4A |
| Aac-15-1 | Bacteria Proteobacteria Gammaproteobacteria Pasteurellales Pasteurellaceae *Aggregatibacter actinomycetemcomitans serotype c str.* D17P-2 |
| Aac-16-1 | Bacteria Proteobacteria Gammaproteobacteria Pasteurellales Pasteurellaceae *Aggregatibacter actinomycetemcomitans serotype c str.* SCC2302 |
| Aac-17-1 | Bacteria Proteobacteria Gammaproteobacteria Pasteurellales Pasteurellaceae *Aggregatibacter actinomycetemcomitans serotype d str.* I63B |
| Aac-18-1 | Bacteria Proteobacteria Gammaproteobacteria Pasteurellales Pasteurellaceae *Aggregatibacter actinomycetemcomitans serotype e str.* SC1083 |
| Aac-19-1 | Bacteria Proteobacteria Gammaproteobacteria Pasteurellales Pasteurellaceae *Aggregatibacter actinomycetemcomitans serotype e str.* SCC393 |
| Aac-20-1 | Bacteria Proteobacteria Gammaproteobacteria Pasteurellales Pasteurellaceae *Aggregatibacter actinomycetemcomitans serotype f str.* D18P1 |
| Aac-7-1 | Bacteria Proteobacteria Gammaproteobacteria Pasteurellales Pasteurellaceae *Aggregatibacter actinomycetemcomitans* Y4 |
| Aap-1-1 | Bacteria Proteobacteria Gammaproteobacteria Pasteurellales Pasteurellaceae *Aggregatibacter aphrophilus* |
| Aap-2-1 | Bacteria Proteobacteria Gammaproteobacteria Pasteurellales Pasteurellaceae *Aggregatibacter aphrophilus* ATCC 33389 |
| Aap-3-1 | Bacteria Proteobacteria Gammaproteobacteria Pasteurellales Pasteurellaceae *Aggregatibacter aphrophilus* F0387 |
| Aap-4-1 | Bacteria Proteobacteria Gammaproteobacteria Pasteurellales Pasteurellaceae *Aggregatibacter aphrophilus* NJ8700 |
| Ase-1-1 | Bacteria Proteobacteria Gammaproteobacteria Pasteurellales Pasteurellaceae *Aggregatibacter segnis* |
| Ase-2-1 | Bacteria Proteobacteria Gammaproteobacteria Pasteurellales Pasteurellaceae *Aggregatibacter segnis* ATCC 33393 |
| Asp-10-1 | Bacteria Proteobacteria Gammaproteobacteria Pasteurellales Pasteurellaceae *Aggregatibacter sp. oral taxon* 458 str. W10330 |
| Apa-1-1 to Apa-1-2 | Bacteria Proteobacteria Gammaproteobacteria Pasteurellales Pasteurellaceae *Avibacterium paragallinarum* |
| Apa-2-1 | Bacteria Proteobacteria Gammaproteobacteria Pasteurellales Pasteurellaceae *Avibacterium paragallinarum* 221 |
| Apa-3-1 | Bacteria Proteobacteria Gammaproteobacteria Pasteurellales Pasteurellaceae *Avibacterium paragallinarum* 72 |
| Apa-4-1 | Bacteria Proteobacteria Gammaproteobacteria Pasteurellales Pasteurellaceae *Avibacterium paragallinarum* JF4211 |

| | |
|---|---|
| Cor-1-1 | Bacteria Proteobacteria Gammaproteobacteria Pasteurellales Pasteurellaceae *Chelonobacter oris* |
| Hae-1-1 | Bacteria Proteobacteria Gammaproteobacteria Pasteurellales Pasteurellaceae *Haemophilus aegyptius* ATCC 11116 |
| Hha-1-1 to Hha-1-4 | Bacteria Proteobacteria Gammaproteobacteria Pasteurellales Pasteurellaceae *Haemophilus haemolyticus* |
| Hha-2-1 | Bacteria Proteobacteria Gammaproteobacteria Pasteurellales Pasteurellaceae *Haemophilus haemolyticus* HK386 |
| Hha-3-1 | Bacteria Proteobacteria Gammaproteobacteria Pasteurellales Pasteurellaceae *Haemophilus haemolyticus* M19501 |
| Hha-4-1 | Bacteria Proteobacteria Gammaproteobacteria Pasteurellales Pasteurellaceae *Haemophilus haemolyticus* M21127 |
| Hha-5-1 | Bacteria Proteobacteria Gammaproteobacteria Pasteurellales Pasteurellaceae *Haemophilus haemolyticus* M21621 |
| Hha-6-1 | Bacteria Proteobacteria Gammaproteobacteria Pasteurellales Pasteurellaceae *Haemophilus haemolyticus* M21639 |
| Hin-1-1 to Hin-1-4 | Bacteria Proteobacteria Gammaproteobacteria Pasteurellales Pasteurellaceae *Haemophilus influenzae* |
| Hin-2-1 | Bacteria Proteobacteria Gammaproteobacteria Pasteurellales Pasteurellaceae *Haemophilus influenzae* 10810 |
| Hin-3-1 | Bacteria Proteobacteria Gammaproteobacteria Pasteurellales Pasteurellaceae *Haemophilus influenzae* 2019 |
| Hin-4-1 | Bacteria Proteobacteria Gammaproteobacteria Pasteurellales Pasteurellaceae *Haemophilus influenzae* 3655 |
| Hin-5-1 | Bacteria Proteobacteria Gammaproteobacteria Pasteurellales Pasteurellaceae *Haemophilus influenzae* 60294N1 |
| Hin-6-1 | Bacteria Proteobacteria Gammaproteobacteria Pasteurellales Pasteurellaceae *Haemophilus influenzae* 6P18H1 |
| Hin-7-1 | Bacteria Proteobacteria Gammaproteobacteria Pasteurellales Pasteurellaceae *Haemophilus influenzae* 7P49H1 |
| Hin-8-1 | Bacteria Proteobacteria Gammaproteobacteria Pasteurellales Pasteurellaceae *Haemophilus influenzae* 86-028NP |
| Hin-9-1 | Bacteria Proteobacteria Gammaproteobacteria Pasteurellales Pasteurellaceae *Haemophilus influenzae* CGSHiCZ412602 |
| Hin-10-1 | Bacteria Proteobacteria Gammaproteobacteria Pasteurellales Pasteurellaceae *Haemophilus influenzae* F3031 |
| Hin-11-1 | Bacteria Proteobacteria Gammaproteobacteria Pasteurellales Pasteurellaceae *Haemophilus influenzae* F3047 |
| Hin-12-1 | Bacteria Proteobacteria Gammaproteobacteria Pasteurellales Pasteurellaceae *Haemophilus influenzae* KR494 |
| Hin-13-1 | Bacteria Proteobacteria Gammaproteobacteria Pasteurellales Pasteurellaceae *Haemophilus influenzae* NT127 |
| Hin-14-1 | Bacteria Proteobacteria Gammaproteobacteria Pasteurellales Pasteurellaceae *Haemophilus influenzae* PittAA |
| Hin-15-1 | Bacteria Proteobacteria Gammaproteobacteria Pasteurellales Pasteurellaceae *Haemophilus influenzae* PittEE |
| Hin-16-1 | Bacteria Proteobacteria Gammaproteobacteria Pasteurellales Pasteurellaceae *Haemophilus influenzae* PittHH |
| Hin-17-1 to Hin-17-2 | Bacteria Proteobacteria Gammaproteobacteria Pasteurellales Pasteurellaceae *Haemophilus influenzae* PittII |
| Hin-18-1 to Hin-18-2 | Bacteria Proteobacteria Gammaproteobacteria Pasteurellales Pasteurellaceae *Haemophilus influenzae* R2846 |
| Hin-19-1 to Hin-19-2 | Bacteria Proteobacteria Gammaproteobacteria Pasteurellales Pasteurellaceae *Haemophilus influenzae* R2866 |
| Hin-20-1 | Bacteria Proteobacteria Gammaproteobacteria Pasteurellales Pasteurellaceae *Haemophilus influenzae* Rd KW20 |
| Hin-21-1 | Bacteria Proteobacteria Gammaproteobacteria Pasteurellales Pasteurellaceae *Haemophilus influenzae* RdAW |
| Hma-1-1 | Bacteria Proteobacteria Gammaproteobacteria Pasteurellales Pasteurellaceae *Haemophilus massiliensis* |
| Hpa-2-1 to Hpa-2-11 | Bacteria Proteobacteria Gammaproteobacteria Pasteurellales Pasteurellaceae *Haemophilus parainfluenzae* |
| Hpa-3-1 | Bacteria Proteobacteria Gammaproteobacteria Pasteurellales Pasteurellaceae *Haemophilus parainfluenzae* ATCC 33392 |
| Hpa-4-1 | Bacteria Proteobacteria Gammaproteobacteria Pasteurellales Pasteurellaceae *Haemophilus parainfluenzae* HK2019 |
| Hpa-5-1 | Bacteria Proteobacteria Gammaproteobacteria Pasteurellales Pasteurellaceae *Haemophilus parainfluenzae* HK262 |
| Hpa-6-1 | Bacteria Proteobacteria Gammaproteobacteria Pasteurellales Pasteurellaceae *Haemophilus parainfluenzae* T3T1 |
| Hpa-7-1 | Bacteria Proteobacteria Gammaproteobacteria Pasteurellales Pasteurellaceae *Haemophilus paraphrohaemolyticus* HK411 |
| Hsp-1-1 | Bacteria Proteobacteria Gammaproteobacteria Pasteurellales Pasteurellaceae *Haemophilus sp.* FF7 |
| Hsp-2-1 | Bacteria Proteobacteria Gammaproteobacteria Pasteurellales Pasteurellaceae *Haemophilus sp. oral taxon* 851 str. F0397 |
| Hsp-3-1 | Bacteria Proteobacteria Gammaproteobacteria Pasteurellales Pasteurellaceae *Haemophilus sputorum* CCUG 13788 |
| Hsp-4-1 | Bacteria Proteobacteria Gammaproteobacteria Pasteurellales Pasteurellaceae *Haemophilus sputorum* HK 2154 |
| Mgr-1-1 | Bacteria Proteobacteria Gammaproteobacteria Pasteurellales Pasteurellaceae *Mannheimia granulomatis* DSM 19156 |
| Mha-1-1 | Bacteria Proteobacteria Gammaproteobacteria Pasteurellales Pasteurellaceae *Mannheimia haemolytica serotype* A1/A6 str. PKL10 |
| Mma-3-1 | Bacteria Proteobacteria Gammaproteobacteria Pasteurellales Pasteurellaceae *Mannheimia massilioguelmaensis* |
| Msu-2-1 | Bacteria Proteobacteria Gammaproteobacteria Pasteurellales Pasteurellaceae *Mannheimia succiniciproducens* MBEL55E |
| Mmu-1-1 | Bacteria Proteobacteria Gammaproteobacteria Pasteurellales Pasteurellaceae *Muribacter muris* |
| Nro-1-1 | Bacteria Proteobacteria Gammaproteobacteria Pasteurellales Pasteurellaceae *Necropsobacter rosorum* |
| Pbe-2-1 | Bacteria Proteobacteria Gammaproteobacteria Pasteurellales Pasteurellaceae *Pasteurella bettyae* CCUG 2042 |
| Pda-1-1 | Bacteria Proteobacteria Gammaproteobacteria Pasteurellales Pasteurellaceae *Pasteurella dagmatis* ATCC 43325 |
| Pmu-1-1 | Bacteria Proteobacteria Gammaproteobacteria Pasteurellales Pasteurellaceae *Pasteurella multocida* |
| Pmu-2-1 | Bacteria Proteobacteria Gammaproteobacteria Pasteurellales Pasteurellaceae *Pasteurella multocida* 1500C |
| Pmu-3-1 | Bacteria Proteobacteria Gammaproteobacteria Pasteurellales Pasteurellaceae *Pasteurella multocida* 1500E |
| Pmu-4-1 | Bacteria Proteobacteria Gammaproteobacteria Pasteurellales Pasteurellaceae *Pasteurella multocida* 2000 |
| Pmu-5-1 | Bacteria Proteobacteria Gammaproteobacteria Pasteurellales Pasteurellaceae *Pasteurella multocida* 36950 |
| Pmu-6-1 | Bacteria Proteobacteria Gammaproteobacteria Pasteurellales Pasteurellaceae *Pasteurella multocida* 671/90 |
| Pmu-7-1 | Bacteria Proteobacteria Gammaproteobacteria Pasteurellales Pasteurellaceae *Pasteurella multocida* 93002 |
| Pmu-8-1 | Bacteria Proteobacteria Gammaproteobacteria Pasteurellales Pasteurellaceae *Pasteurella multocida* OH1905 |
| Pmu-9-1 | Bacteria Proteobacteria Gammaproteobacteria Pasteurellales Pasteurellaceae *Pasteurella multocida* P1933 |
| Pmu-10-1 | Bacteria Proteobacteria Gammaproteobacteria Pasteurellales Pasteurellaceae *Pasteurella multocida* RIIF |
| Pmu-11-1 | Bacteria Proteobacteria Gammaproteobacteria Pasteurellales Pasteurellaceae *Pasteurella multocida subsp. gallicida* P1059 |
| Pmu-12-1 | Bacteria Proteobacteria Gammaproteobacteria Pasteurellales Pasteurellaceae *Pasteurella multocida subsp. gallicida* X73 |
| Pmu-13-1 to Pmu-13-12 | Bacteria Proteobacteria Gammaproteobacteria Pasteurellales Pasteurellaceae *Pasteurella multocida subsp. multocida* |
| Pmu-14-1 | Bacteria Proteobacteria Gammaproteobacteria Pasteurellales Pasteurellaceae *Pasteurella multocida subsp. multocida* OH4807 |
| Pmu-15-1 | Bacteria Proteobacteria Gammaproteobacteria Pasteurellales Pasteurellaceae *Pasteurella multocida subsp. multocida* P1062 |
| Pmu-17-1 | Bacteria Proteobacteria Gammaproteobacteria Pasteurellales Pasteurellaceae *Pasteurella multocida subsp. multocida str.* 3480 |
| Pmu-18-1 | Bacteria Proteobacteria Gammaproteobacteria Pasteurellales Pasteurellaceae *Pasteurella multocida subsp. multocida str.* HB03 |

| | |
|---|---|
| Pmu-19-1 | Bacteria Proteobacteria Gammaproteobacteria Pasteurellales Pasteurellaceae *Pasteurella multocida subsp. multocida str.* HN06 |
| Pmu-21-1 | Bacteria Proteobacteria Gammaproteobacteria Pasteurellales Pasteurellaceae *Pasteurella multocida subsp. multocida str.* Pm70 |
| Pmu-20-1 | Bacteria Proteobacteria Gammaproteobacteria Pasteurellales Pasteurellaceae *Pasteurella multocida subsp. multocida str.* PMTB |
| Pmu-16-1 | Bacteria Proteobacteria Gammaproteobacteria Pasteurellales Pasteurellaceae *Pasteurella multocida subsp. multocida* VTCCBAA264 |
| Ppn-1-1 | Bacteria Proteobacteria Gammaproteobacteria Pasteurellales Pasteurellaceae *Pasteurella pneumotropica* DSM 21403 |
| Psp-56-1 to Psp-56-3 | Bacteria Proteobacteria Gammaproteobacteria Pasteurellales Pasteurellaceae *Pasteurella sp.* FF6 |
| Rpn-1-1 to Rpn-1-2 | Bacteria Proteobacteria Gammaproteobacteria Pasteurellales Pasteurellaceae *Rodentibacter pneumotropicus* |
| Vba-1-1 | Bacteria Proteobacteria Gammaproteobacteria Vibrionales unclassified Vibrionales *Vibrionales bacterium* SWAT-3 |
| Afi-2-1 | Bacteria Proteobacteria Gammaproteobacteria Vibrionales Vibrionaceae *Aliivibrio fischeri* |
| Afi-3-1 | Bacteria Proteobacteria Gammaproteobacteria Vibrionales Vibrionaceae *Aliivibrio fischeri* ZF-211 |
| Alo-1-1 | Bacteria Proteobacteria Gammaproteobacteria Vibrionales Vibrionaceae *Aliivibrio logei* 5S-186 |
| Alo-2-1 | Bacteria Proteobacteria Gammaproteobacteria Vibrionales Vibrionaceae *Aliivibrio logei* ATCC 35077 |
| Asa-11-1 | Bacteria Proteobacteria Gammaproteobacteria Vibrionales Vibrionaceae *Aliivibrio salmonicida* LFI1238 |
| Awo-1-1 | Bacteria Proteobacteria Gammaproteobacteria Vibrionales Vibrionaceae *Aliivibrio wodanis* |
| Vfi-1-1 | Bacteria Proteobacteria Gammaproteobacteria Vibrionales Vibrionaceae Aliivibrio *Vibrio fischeri* ES114 |
| Vfi-2-1 | Bacteria Proteobacteria Gammaproteobacteria Vibrionales Vibrionaceae Aliivibrio *Vibrio fischeri* MJ11 |
| Vfi-3-1 | Bacteria Proteobacteria Gammaproteobacteria Vibrionales Vibrionaceae Aliivibrio *Vibrio fischeri* SR5 |
| Eca-4-1 | Bacteria Proteobacteria Gammaproteobacteria Vibrionales Vibrionaceae *Enterovibrio calviensis* 1F-211 |
| Eca-5-1 | Bacteria Proteobacteria Gammaproteobacteria Vibrionales Vibrionaceae *Enterovibrio calviensis* 1F-230 |
| Eca-6-1 | Bacteria Proteobacteria Gammaproteobacteria Vibrionales Vibrionaceae *Enterovibrio calviensis* DSM 14347 |
| Eca-7-1 | Bacteria Proteobacteria Gammaproteobacteria Vibrionales Vibrionaceae *Enterovibrio calviensis* FF-85 |
| Eno-1-1 | Bacteria Proteobacteria Gammaproteobacteria Vibrionales Vibrionaceae *Enterovibrio norvegicus* FF-162 |
| Eno-2-1 | Bacteria Proteobacteria Gammaproteobacteria Vibrionales Vibrionaceae *Enterovibrio norvegicus* FF-33 |
| Eno-3-1 | Bacteria Proteobacteria Gammaproteobacteria Vibrionales Vibrionaceae *Enterovibrio norvegicus* FF-454 |
| Gho-1-1 | Bacteria Proteobacteria Gammaproteobacteria Vibrionales Vibrionaceae *Grimontia hollisae* CIP 101886 |
| Gsp-3-1 | Bacteria Proteobacteria Gammaproteobacteria Vibrionales Vibrionaceae *Grimontia sp.* AD028 |
| Gsp-4-1 | Bacteria Proteobacteria Gammaproteobacteria Vibrionales Vibrionaceae *Grimontia sp.* AK16 |
| Pan-11-1 to Pan-11-12 | Bacteria Proteobacteria Gammaproteobacteria Vibrionales Vibrionaceae *Photobacterium angustum* |
| Pan-12-1 | Bacteria Proteobacteria Gammaproteobacteria Vibrionales Vibrionaceae *Photobacterium angustum* S14 |
| Pap-1-1 | Bacteria Proteobacteria Gammaproteobacteria Vibrionales Vibrionaceae *Photobacterium aphoticum* |
| Paq-2-1 | Bacteria Proteobacteria Gammaproteobacteria Vibrionales Vibrionaceae *Photobacterium aquae* |
| Pda-2-1 | Bacteria Proteobacteria Gammaproteobacteria Vibrionales Vibrionaceae *Photobacterium damselae subsp. damselae* |
| Pda-3-1 | Bacteria Proteobacteria Gammaproteobacteria Vibrionales Vibrionaceae *Photobacterium damselae subsp. damselae* CIP 102761 |
| Pda-4-1 | Bacteria Proteobacteria Gammaproteobacteria Vibrionales Vibrionaceae *Photobacterium damselae subsp. piscicida* DI21 |
| Pga-1-1 | Bacteria Proteobacteria Gammaproteobacteria Vibrionales Vibrionaceae *Photobacterium gaetbulicola* |
| Pga-2-1 | Bacteria Proteobacteria Gammaproteobacteria Vibrionales Vibrionaceae *Photobacterium gaetbulicola* Gung47 |
| Pga-3-1 | Bacteria Proteobacteria Gammaproteobacteria Vibrionales Vibrionaceae *Photobacterium galatheae* |
| Pga-4-1 | Bacteria Proteobacteria Gammaproteobacteria Vibrionales Vibrionaceae *Photobacterium ganghwense* |
| Pha-6-1 | Bacteria Proteobacteria Gammaproteobacteria Vibrionales Vibrionaceae *Photobacterium halotolerans* |
| Pha-7-1 | Bacteria Proteobacteria Gammaproteobacteria Vibrionales Vibrionaceae *Photobacterium halotolerans* DSM 18316 |
| Pil-1-1 to Pil-1-2 | Bacteria Proteobacteria Gammaproteobacteria Vibrionales Vibrionaceae *Photobacterium iliopiscarium* |
| Pki-1-1 to Pki-1-5 | Bacteria Proteobacteria Gammaproteobacteria Vibrionales Vibrionaceae *Photobacterium kishitanii* |
| Ple-1-1 to Ple-1-2 | Bacteria Proteobacteria Gammaproteobacteria Vibrionales Vibrionaceae *Photobacterium leiognathi* |
| Ple-2-1 | Bacteria Proteobacteria Gammaproteobacteria Vibrionales Vibrionaceae *Photobacterium leiognathi lrivu.*4.1 |
| Ple-3-1 | Bacteria Proteobacteria Gammaproteobacteria Vibrionales Vibrionaceae *Photobacterium leiognathi subsp. mandapamensis svers.*1.1 |
| Pph-1-1 | Bacteria Proteobacteria Gammaproteobacteria Vibrionales Vibrionaceae *Photobacterium phosphoreum* |
| Ppr-1-1 | Bacteria Proteobacteria Gammaproteobacteria Vibrionales Vibrionaceae *Photobacterium profundum* 3TCK |
| Ppr-2-1 | Bacteria Proteobacteria Gammaproteobacteria Vibrionales Vibrionaceae *Photobacterium profundum* SS9 |
| Psa-1-1 | Bacteria Proteobacteria Gammaproteobacteria Vibrionales Vibrionaceae *Photobacterium sanctipauli* |
| Psp-57-1 | Bacteria Proteobacteria Gammaproteobacteria Vibrionales Vibrionaceae *Photobacterium sp.* AK15 |
| Psp-58-1 | Bacteria Proteobacteria Gammaproteobacteria Vibrionales Vibrionaceae *Photobacterium sp.* SKA34 |
| Psw-1-1 | Bacteria Proteobacteria Gammaproteobacteria Vibrionales Vibrionaceae *Photobacterium swingsii* |
| Sco-3-1 | Bacteria Proteobacteria Gammaproteobacteria Vibrionales Vibrionaceae *Salinivibrio costicola subsp. costicola* ATCC 33508 = LMG 11651 |
| Sso-14-1 | Bacteria Proteobacteria Gammaproteobacteria Vibrionales Vibrionaceae *Salinivibrio socompensis* S10B |
| Sso-15-1 | Bacteria Proteobacteria Gammaproteobacteria Vibrionales Vibrionaceae *Salinivibrio socompensis* S35 |
| Ssp-27-1 | Bacteria Proteobacteria Gammaproteobacteria Vibrionales Vibrionaceae *Salinivibrio sp.* KP-1 |
| Lan-1-1 | Bacteria Proteobacteria Gammaproteobacteria Vibrionales Vibrionaceae Vibrio *Listonella anguillarum* M3 |
| Val-1-1 | Bacteria Proteobacteria Gammaproteobacteria Vibrionales Vibrionaceae *Vibrio albensis* ATCC 14547 |
| Val-2-1 to Val-2-9 | Bacteria Proteobacteria Gammaproteobacteria Vibrionales Vibrionaceae *Vibrio alginolyticus* |
| Val-3-1 | Bacteria Proteobacteria Gammaproteobacteria Vibrionales Vibrionaceae *Vibrio alginolyticus* 12G01 |
| Val-4-1 | Bacteria Proteobacteria Gammaproteobacteria Vibrionales Vibrionaceae *Vibrio alginolyticus* 40B |
| Val-5-1 | Bacteria Proteobacteria Gammaproteobacteria Vibrionales Vibrionaceae *Vibrio alginolyticus* E0666 |
| Val-6-1 | Bacteria Proteobacteria Gammaproteobacteria Vibrionales Vibrionaceae *Vibrio alginolyticus* NBRC 15630 = ATCC 17749 |
| Van-1-1 | Bacteria Proteobacteria Gammaproteobacteria Vibrionales Vibrionaceae *Vibrio anguillarum* |
| Van-2-1 | Bacteria Proteobacteria Gammaproteobacteria Vibrionales Vibrionaceae *Vibrio anguillarum* 775 |

| | |
|---|---|
| Van-3-1 | Bacteria Proteobacteria Gammaproteobacteria Vibrionales Vibrionaceae *Vibrio anguillarum* 96F |
| Van-4-1 | Bacteria Proteobacteria Gammaproteobacteria Vibrionales Vibrionaceae *Vibrio anguillarum* RV22 |
| Vaz-1-1 to Vaz-1-2 | Bacteria Proteobacteria Gammaproteobacteria Vibrionales Vibrionaceae *Vibrio azureus* NBRC 104587 |
| Vbr-1-1 | Bacteria Proteobacteria Gammaproteobacteria Vibrionales Vibrionaceae *Vibrio brasiliensis* LMG 20546 |
| Vbr-2-1 | Bacteria Proteobacteria Gammaproteobacteria Vibrionales Vibrionaceae *Vibrio breoganii* 1C10 |
| Vbr-3-1 | Bacteria Proteobacteria Gammaproteobacteria Vibrionales Vibrionaceae *Vibrio breoganii* ZF-29 |
| Vbr-4-1 | Bacteria Proteobacteria Gammaproteobacteria Vibrionales Vibrionaceae *Vibrio breoganii* ZF-55 |
| Vca-1-1 to Vca-1-10 | Bacteria Proteobacteria Gammaproteobacteria Vibrionales Vibrionaceae *Vibrio campbellii* |
| Vca-2-1 | Bacteria Proteobacteria Gammaproteobacteria Vibrionales Vibrionaceae *Vibrio campbellii* 200612B |
| Vca-3-1 | Bacteria Proteobacteria Gammaproteobacteria Vibrionales Vibrionaceae *Vibrio campbellii* ATCC BAA-1116 |
| Vca-4-1 to Vca-4-2 | Bacteria Proteobacteria Gammaproteobacteria Vibrionales Vibrionaceae *Vibrio campbellii* CAIM 519 = NBRC 15631 |
| Vca-5-1 | Bacteria Proteobacteria Gammaproteobacteria Vibrionales Vibrionaceae *Vibrio campbellii* DS40M4 |
| Vca-6-1 | Bacteria Proteobacteria Gammaproteobacteria Vibrionales Vibrionaceae *Vibrio caribbeanicus* |
| Vca-7-1 | Bacteria Proteobacteria Gammaproteobacteria Vibrionales Vibrionaceae *Vibrio caribbenthicus* ATCC BAA-2122 |
| Vch-1-1 | Bacteria Proteobacteria Gammaproteobacteria Vibrionales Vibrionaceae *Vibrio cholera* CIRS 101 |
| Vch-2-1 to Vch-2-4 | Bacteria Proteobacteria Gammaproteobacteria Vibrionales Vibrionaceae *Vibrio cholerae* |
| Vch-3-1 | Bacteria Proteobacteria Gammaproteobacteria Vibrionales Vibrionaceae *Vibrio cholerae* 12129(1) |
| Vch-4-1 | Bacteria Proteobacteria Gammaproteobacteria Vibrionales Vibrionaceae *Vibrio cholerae* 1587 |
| Vch-5-1 | Bacteria Proteobacteria Gammaproteobacteria Vibrionales Vibrionaceae *Vibrio cholerae* 2011EL-301 |
| Vch-6-1 | Bacteria Proteobacteria Gammaproteobacteria Vibrionales Vibrionaceae *Vibrio cholerae* 2012EL-1759 |
| Vch-7-1 | Bacteria Proteobacteria Gammaproteobacteria Vibrionales Vibrionaceae *Vibrio cholerae* 2740-80 |
| Vch-8-1 | Bacteria Proteobacteria Gammaproteobacteria Vibrionales Vibrionaceae *Vibrio cholerae* 4260B |
| Vch-9-1 | Bacteria Proteobacteria Gammaproteobacteria Vibrionales Vibrionaceae *Vibrio cholerae* 623-39 |
| Vch-10-1 | Bacteria Proteobacteria Gammaproteobacteria Vibrionales Vibrionaceae *Vibrio cholerae* AM-19226 |
| Vch-11-1 | Bacteria Proteobacteria Gammaproteobacteria Vibrionales Vibrionaceae *Vibrio cholerae* ATCC 14035 |
| Vch-12-1 | Bacteria Proteobacteria Gammaproteobacteria Vibrionales Vibrionaceae *Vibrio cholerae* B33 |
| Vch-13-1 | Bacteria Proteobacteria Gammaproteobacteria Vibrionales Vibrionaceae *Vibrio cholerae* BJG-01 |
| Vch-197-1 | Bacteria Proteobacteria Gammaproteobacteria Vibrionales Vibrionaceae *Vibrio cholerae bv. albensis* VL426 |
| Vch-14-1 | Bacteria Proteobacteria Gammaproteobacteria Vibrionales Vibrionaceae *Vibrio cholerae* BX 330286 |
| Vch-15-1 | Bacteria Proteobacteria Gammaproteobacteria Vibrionales Vibrionaceae *Vibrio cholerae* CP1030(3) |
| Vch-16-1 | Bacteria Proteobacteria Gammaproteobacteria Vibrionales Vibrionaceae *Vibrio cholerae* CP1032(5) |
| Vch-17-1 | Bacteria Proteobacteria Gammaproteobacteria Vibrionales Vibrionaceae *Vibrio cholerae* CP1033(6) |
| Vch-18-1 | Bacteria Proteobacteria Gammaproteobacteria Vibrionales Vibrionaceae *Vibrio cholerae* CP1035(8) |
| Vch-19-1 | Bacteria Proteobacteria Gammaproteobacteria Vibrionales Vibrionaceae *Vibrio cholerae* CP1037(10) |
| Vch-20-1 | Bacteria Proteobacteria Gammaproteobacteria Vibrionales Vibrionaceae *Vibrio cholerae* CP1038(11) |
| Vch-21-1 | Bacteria Proteobacteria Gammaproteobacteria Vibrionales Vibrionaceae *Vibrio cholerae* CP1040(13) |
| Vch-22-1 | Bacteria Proteobacteria Gammaproteobacteria Vibrionales Vibrionaceae *Vibrio cholerae* CP1041(14) |
| Vch-23-1 | Bacteria Proteobacteria Gammaproteobacteria Vibrionales Vibrionaceae *Vibrio cholerae* CP1042(15) |
| Vch-24-1 | Bacteria Proteobacteria Gammaproteobacteria Vibrionales Vibrionaceae *Vibrio cholerae* CP1044(17) |
| Vch-25-1 | Bacteria Proteobacteria Gammaproteobacteria Vibrionales Vibrionaceae *Vibrio cholerae* CP1046(19) |
| Vch-26-1 | Bacteria Proteobacteria Gammaproteobacteria Vibrionales Vibrionaceae *Vibrio cholerae* CP1047(20) |
| Vch-27-1 | Bacteria Proteobacteria Gammaproteobacteria Vibrionales Vibrionaceae *Vibrio cholerae* CP1048(21) |
| Vch-28-1 | Bacteria Proteobacteria Gammaproteobacteria Vibrionales Vibrionaceae *Vibrio cholerae* CP1050(23) |
| Vch-29-1 | Bacteria Proteobacteria Gammaproteobacteria Vibrionales Vibrionaceae *Vibrio cholerae* CP1110 |
| Vch-30-1 | Bacteria Proteobacteria Gammaproteobacteria Vibrionales Vibrionaceae *Vibrio cholerae* CP1111 |
| Vch-31-1 | Bacteria Proteobacteria Gammaproteobacteria Vibrionales Vibrionaceae *Vibrio cholerae* CP1112 |
| Vch-32-1 | Bacteria Proteobacteria Gammaproteobacteria Vibrionales Vibrionaceae *Vibrio cholerae* CP1113 |
| Vch-33-1 | Bacteria Proteobacteria Gammaproteobacteria Vibrionales Vibrionaceae *Vibrio cholerae* CP1114 |
| Vch-34-1 | Bacteria Proteobacteria Gammaproteobacteria Vibrionales Vibrionaceae *Vibrio cholerae* CP1115 |
| Vch-35-1 | Bacteria Proteobacteria Gammaproteobacteria Vibrionales Vibrionaceae *Vibrio cholerae* CP1116 |
| Vch-36-1 | Bacteria Proteobacteria Gammaproteobacteria Vibrionales Vibrionaceae *Vibrio cholerae* CP1117 |
| Vch-37-1 | Bacteria Proteobacteria Gammaproteobacteria Vibrionales Vibrionaceae *Vibrio cholerae* CT 5369-93 |
| Vch-38-1 | Bacteria Proteobacteria Gammaproteobacteria Vibrionales Vibrionaceae *Vibrio cholerae* E306 |
| Vch-39-1 | Bacteria Proteobacteria Gammaproteobacteria Vibrionales Vibrionaceae *Vibrio cholerae* H1 |
| Vch-40-1 | Bacteria Proteobacteria Gammaproteobacteria Vibrionales Vibrionaceae *Vibrio cholerae* HC-02A1 |
| Vch-41-1 | Bacteria Proteobacteria Gammaproteobacteria Vibrionales Vibrionaceae *Vibrio cholerae* HC-02C1 |
| Vch-42-1 | Bacteria Proteobacteria Gammaproteobacteria Vibrionales Vibrionaceae *Vibrio cholerae* HC-06A1 |
| Vch-43-1 | Bacteria Proteobacteria Gammaproteobacteria Vibrionales Vibrionaceae *Vibrio cholerae* HC-17A1 |
| Vch-44-1 | Bacteria Proteobacteria Gammaproteobacteria Vibrionales Vibrionaceae *Vibrio cholerae* HC-17A2 |
| Vch-45-1 | Bacteria Proteobacteria Gammaproteobacteria Vibrionales Vibrionaceae *Vibrio cholerae* HC-19A1 |
| Vch-46-1 | Bacteria Proteobacteria Gammaproteobacteria Vibrionales Vibrionaceae *Vibrio cholerae* HC-1A2 |
| Vch-47-1 | Bacteria Proteobacteria Gammaproteobacteria Vibrionales Vibrionaceae *Vibrio cholerae* HC-20A2 |
| Vch-48-1 | Bacteria Proteobacteria Gammaproteobacteria Vibrionales Vibrionaceae *Vibrio cholerae* HC-21A1 |
| Vch-49-1 | Bacteria Proteobacteria Gammaproteobacteria Vibrionales Vibrionaceae *Vibrio cholerae* HC-22A1 |
| Vch-50-1 | Bacteria Proteobacteria Gammaproteobacteria Vibrionales Vibrionaceae *Vibrio cholerae* HC-23A1 |

| | |
|---|---|
| Vch-51-1 | Bacteria Proteobacteria Gammaproteobacteria Vibrionales Vibrionaceae *Vibrio cholerae* HC-28A1 |
| Vch-52-1 | Bacteria Proteobacteria Gammaproteobacteria Vibrionales Vibrionaceae *Vibrio cholerae* HC-32A1 |
| Vch-53-1 | Bacteria Proteobacteria Gammaproteobacteria Vibrionales Vibrionaceae *Vibrio cholerae* HC-33A2 |
| Vch-54-1 | Bacteria Proteobacteria Gammaproteobacteria Vibrionales Vibrionaceae *Vibrio cholerae* HC-36A1 |
| Vch-55-1 | Bacteria Proteobacteria Gammaproteobacteria Vibrionales Vibrionaceae *Vibrio cholerae* HC-37A1 |
| Vch-56-1 | Bacteria Proteobacteria Gammaproteobacteria Vibrionales Vibrionaceae *Vibrio cholerae* HC-38A1 |
| Vch-57-1 | Bacteria Proteobacteria Gammaproteobacteria Vibrionales Vibrionaceae *Vibrio cholerae* HC-39A1 |
| Vch-58-1 | Bacteria Proteobacteria Gammaproteobacteria Vibrionales Vibrionaceae *Vibrio cholerae* HC-40A1 |
| Vch-59-1 | Bacteria Proteobacteria Gammaproteobacteria Vibrionales Vibrionaceae *Vibrio cholerae* HC-41A1 |
| Vch-60-1 | Bacteria Proteobacteria Gammaproteobacteria Vibrionales Vibrionaceae *Vibrio cholerae* HC-41B1 |
| Vch-61-1 | Bacteria Proteobacteria Gammaproteobacteria Vibrionales Vibrionaceae *Vibrio cholerae* HC-42A1 |
| Vch-62-1 | Bacteria Proteobacteria Gammaproteobacteria Vibrionales Vibrionaceae *Vibrio cholerae* HC-43A1 |
| Vch-63-1 | Bacteria Proteobacteria Gammaproteobacteria Vibrionales Vibrionaceae *Vibrio cholerae* HC-43B1 |
| Vch-64-1 | Bacteria Proteobacteria Gammaproteobacteria Vibrionales Vibrionaceae *Vibrio cholerae* HC-44C1 |
| Vch-65-1 | Bacteria Proteobacteria Gammaproteobacteria Vibrionales Vibrionaceae *Vibrio cholerae* HC-46A1 |
| Vch-66-1 | Bacteria Proteobacteria Gammaproteobacteria Vibrionales Vibrionaceae *Vibrio cholerae* HC-46B1 |
| Vch-67-1 | Bacteria Proteobacteria Gammaproteobacteria Vibrionales Vibrionaceae *Vibrio cholerae* HC-47A1 |
| Vch-68-1 | Bacteria Proteobacteria Gammaproteobacteria Vibrionales Vibrionaceae *Vibrio cholerae* HC-48A1 |
| Vch-69-1 | Bacteria Proteobacteria Gammaproteobacteria Vibrionales Vibrionaceae *Vibrio cholerae* HC-48B2 |
| Vch-70-1 | Bacteria Proteobacteria Gammaproteobacteria Vibrionales Vibrionaceae *Vibrio cholerae* HC-49A2 |
| Vch-71-1 | Bacteria Proteobacteria Gammaproteobacteria Vibrionales Vibrionaceae *Vibrio cholerae* HC-50A1 |
| Vch-72-1 | Bacteria Proteobacteria Gammaproteobacteria Vibrionales Vibrionaceae *Vibrio cholerae* HC-50A2 |
| Vch-73-1 | Bacteria Proteobacteria Gammaproteobacteria Vibrionales Vibrionaceae *Vibrio cholerae* HC-51A1 |
| Vch-74-1 | Bacteria Proteobacteria Gammaproteobacteria Vibrionales Vibrionaceae *Vibrio cholerae* HC-52A1 |
| Vch-75-1 | Bacteria Proteobacteria Gammaproteobacteria Vibrionales Vibrionaceae *Vibrio cholerae* HC-55A1 |
| Vch-76-1 | Bacteria Proteobacteria Gammaproteobacteria Vibrionales Vibrionaceae *Vibrio cholerae* HC-55B2 |
| Vch-77-1 | Bacteria Proteobacteria Gammaproteobacteria Vibrionales Vibrionaceae *Vibrio cholerae* HC-55C2 |
| Vch-78-1 | Bacteria Proteobacteria Gammaproteobacteria Vibrionales Vibrionaceae *Vibrio cholerae* HC-56A1 |
| Vch-79-1 | Bacteria Proteobacteria Gammaproteobacteria Vibrionales Vibrionaceae *Vibrio cholerae* HC-56A2 |
| Vch-80-1 | Bacteria Proteobacteria Gammaproteobacteria Vibrionales Vibrionaceae *Vibrio cholerae* HC-57A1 |
| Vch-81-1 | Bacteria Proteobacteria Gammaproteobacteria Vibrionales Vibrionaceae *Vibrio cholerae* HC-57A2 |
| Vch-82-1 | Bacteria Proteobacteria Gammaproteobacteria Vibrionales Vibrionaceae *Vibrio cholerae* HC-59A1 |
| Vch-83-1 | Bacteria Proteobacteria Gammaproteobacteria Vibrionales Vibrionaceae *Vibrio cholerae* HC-59B1 |
| Vch-84-1 | Bacteria Proteobacteria Gammaproteobacteria Vibrionales Vibrionaceae *Vibrio cholerae* HC-60A1 |
| Vch-85-1 | Bacteria Proteobacteria Gammaproteobacteria Vibrionales Vibrionaceae *Vibrio cholerae* HC-61A1 |
| Vch-86-1 | Bacteria Proteobacteria Gammaproteobacteria Vibrionales Vibrionaceae *Vibrio cholerae* HC-61A2 |
| Vch-87-1 | Bacteria Proteobacteria Gammaproteobacteria Vibrionales Vibrionaceae *Vibrio cholerae* HC-62A1 |
| Vch-88-1 | Bacteria Proteobacteria Gammaproteobacteria Vibrionales Vibrionaceae *Vibrio cholerae* HC-62B1 |
| Vch-89-1 | Bacteria Proteobacteria Gammaproteobacteria Vibrionales Vibrionaceae *Vibrio cholerae* HC-64A1 |
| Vch-90-1 | Bacteria Proteobacteria Gammaproteobacteria Vibrionales Vibrionaceae *Vibrio cholerae* HC-65A1 |
| Vch-91-1 | Bacteria Proteobacteria Gammaproteobacteria Vibrionales Vibrionaceae *Vibrio cholerae* HC-67A1 |
| Vch-92-1 | Bacteria Proteobacteria Gammaproteobacteria Vibrionales Vibrionaceae *Vibrio cholerae* HC-68A1 |
| Vch-93-1 | Bacteria Proteobacteria Gammaproteobacteria Vibrionales Vibrionaceae *Vibrio cholerae* HC-69A1 |
| Vch-94-1 | Bacteria Proteobacteria Gammaproteobacteria Vibrionales Vibrionaceae *Vibrio cholerae* HC-70A1 |
| Vch-95-1 | Bacteria Proteobacteria Gammaproteobacteria Vibrionales Vibrionaceae *Vibrio cholerae* HC-71A1 |
| Vch-96-1 | Bacteria Proteobacteria Gammaproteobacteria Vibrionales Vibrionaceae *Vibrio cholerae* HC-72A2 |
| Vch-97-1 | Bacteria Proteobacteria Gammaproteobacteria Vibrionales Vibrionaceae *Vibrio cholerae* HC-77A1 |
| Vch-98-1 | Bacteria Proteobacteria Gammaproteobacteria Vibrionales Vibrionaceae *Vibrio cholerae* HC-78A1 |
| Vch-99-1 | Bacteria Proteobacteria Gammaproteobacteria Vibrionales Vibrionaceae *Vibrio cholerae* HC-7A1 |
| Vch-100-1 | Bacteria Proteobacteria Gammaproteobacteria Vibrionales Vibrionaceae *Vibrio cholerae* HC-80A1 |
| Vch-101-1 | Bacteria Proteobacteria Gammaproteobacteria Vibrionales Vibrionaceae *Vibrio cholerae* HC-81A1 |
| Vch-102-1 | Bacteria Proteobacteria Gammaproteobacteria Vibrionales Vibrionaceae *Vibrio cholerae* HC-81A2 |
| Vch-103-1 | Bacteria Proteobacteria Gammaproteobacteria Vibrionales Vibrionaceae *Vibrio cholerae* HCUF01 |
| Vch-104-1 | Bacteria Proteobacteria Gammaproteobacteria Vibrionales Vibrionaceae *Vibrio cholerae* HE-09 |
| Vch-105-1 | Bacteria Proteobacteria Gammaproteobacteria Vibrionales Vibrionaceae *Vibrio cholerae* HE-16 |
| Vch-106-1 | Bacteria Proteobacteria Gammaproteobacteria Vibrionales Vibrionaceae *Vibrio cholerae* HE-25 |
| Vch-107-1 | Bacteria Proteobacteria Gammaproteobacteria Vibrionales Vibrionaceae *Vibrio cholerae* HE-40 |
| Vch-108-1 | Bacteria Proteobacteria Gammaproteobacteria Vibrionales Vibrionaceae *Vibrio cholerae* HE-45 |
| Vch-109-1 | Bacteria Proteobacteria Gammaproteobacteria Vibrionales Vibrionaceae *Vibrio cholerae* HE-46 |
| Vch-110-1 | Bacteria Proteobacteria Gammaproteobacteria Vibrionales Vibrionaceae *Vibrio cholerae* HE39 |
| Vch-111-1 | Bacteria Proteobacteria Gammaproteobacteria Vibrionales Vibrionaceae *Vibrio cholerae* HE48 |
| Vch-112-1 | Bacteria Proteobacteria Gammaproteobacteria Vibrionales Vibrionaceae *Vibrio cholerae* HFU-02 |
| Vch-113-1 | Bacteria Proteobacteria Gammaproteobacteria Vibrionales Vibrionaceae *Vibrio cholerae* IEC224 |
| Vch-114-1 | Bacteria Proteobacteria Gammaproteobacteria Vibrionales Vibrionaceae *Vibrio cholerae* INDRE 91/1 |
| Vch-115-1 | Bacteria Proteobacteria Gammaproteobacteria Vibrionales Vibrionaceae *Vibrio cholerae* LMA3984-4 |

| | |
|---|---|
| Vch-116-1 | Bacteria Proteobacteria Gammaproteobacteria Vibrionales Vibrionaceae *Vibrio cholerae* M66-2 |
| Vch-117-1 | Bacteria Proteobacteria Gammaproteobacteria Vibrionales Vibrionaceae *Vibrio cholerae* MAK 757 |
| Vch-118-1 | Bacteria Proteobacteria Gammaproteobacteria Vibrionales Vibrionaceae *Vibrio cholerae* MJ-1236 |
| Vch-119-1 | Bacteria Proteobacteria Gammaproteobacteria Vibrionales Vibrionaceae *Vibrio cholerae* MO10 |
| Vch-120-1 | Bacteria Proteobacteria Gammaproteobacteria Vibrionales Vibrionaceae *Vibrio cholerae* MS6 |
| Vch-121-1 | Bacteria Proteobacteria Gammaproteobacteria Vibrionales Vibrionaceae *Vibrio cholerae* MZO-2 |
| Vch-122-1 | Bacteria Proteobacteria Gammaproteobacteria Vibrionales Vibrionaceae *Vibrio cholerae* NCTC 8457 |
| Vch-123-1 to Vch-123-41 | Bacteria Proteobacteria Gammaproteobacteria Vibrionales Vibrionaceae *Vibrio cholerae* O1 |
| Vch-124-1 | Bacteria Proteobacteria Gammaproteobacteria Vibrionales Vibrionaceae *Vibrio cholerae* O1 biovar El Tor |
| Vch-125-1 | Bacteria Proteobacteria Gammaproteobacteria Vibrionales Vibrionaceae *Vibrio cholerae* O1 biovar El Tor str. Inaba RND18826 |
| Vch-126-1 | Bacteria Proteobacteria Gammaproteobacteria Vibrionales Vibrionaceae *Vibrio cholerae* O1 biovar El Tor str. Inaba RND18899 |
| Vch-127-1 | Bacteria Proteobacteria Gammaproteobacteria Vibrionales Vibrionaceae *Vibrio cholerae* O1 biovar El Tor str. L-3226 |
| Vch-128-1 | Bacteria Proteobacteria Gammaproteobacteria Vibrionales Vibrionaceae *Vibrio cholerae* O1 biovar El Tor str. N16961 |
| Vch-129-1 | Bacteria Proteobacteria Gammaproteobacteria Vibrionales Vibrionaceae *Vibrio cholerae* O1 biovar El Tor str. Ogawa RND19187 |
| Vch-130-1 | Bacteria Proteobacteria Gammaproteobacteria Vibrionales Vibrionaceae *Vibrio cholerae* O1 biovar El Tor str. Ogawa RND6878 |
| Vch-131-1 | Bacteria Proteobacteria Gammaproteobacteria Vibrionales Vibrionaceae *Vibrio cholerae* O1 str. 116059 |
| Vch-132-1 | Bacteria Proteobacteria Gammaproteobacteria Vibrionales Vibrionaceae *Vibrio cholerae* O1 str. 116063 |
| Vch-133-1 | Bacteria Proteobacteria Gammaproteobacteria Vibrionales Vibrionaceae *Vibrio cholerae* O1 str. 2009V-1046 |
| Vch-134-1 | Bacteria Proteobacteria Gammaproteobacteria Vibrionales Vibrionaceae *Vibrio cholerae* O1 str. 2009V-1085 |
| Vch-135-1 | Bacteria Proteobacteria Gammaproteobacteria Vibrionales Vibrionaceae *Vibrio cholerae* O1 str. 2009V-1096 |
| Vch-136-1 | Bacteria Proteobacteria Gammaproteobacteria Vibrionales Vibrionaceae *Vibrio cholerae* O1 str. 2009V-1116 |
| Vch-137-1 | Bacteria Proteobacteria Gammaproteobacteria Vibrionales Vibrionaceae *Vibrio cholerae* O1 str. 2009V-1131 |
| Vch-138-1 | Bacteria Proteobacteria Gammaproteobacteria Vibrionales Vibrionaceae *Vibrio cholerae* O1 str. 2010EL-1749 |
| Vch-139-1 | Bacteria Proteobacteria Gammaproteobacteria Vibrionales Vibrionaceae *Vibrio cholerae* O1 str. 2010EL-1786 |
| Vch-140-1 | Bacteria Proteobacteria Gammaproteobacteria Vibrionales Vibrionaceae *Vibrio cholerae* O1 str. 2010EL-1792 |
| Vch-141-1 | Bacteria Proteobacteria Gammaproteobacteria Vibrionales Vibrionaceae *Vibrio cholerae* O1 str. 2010EL-1798 |
| Vch-142-1 | Bacteria Proteobacteria Gammaproteobacteria Vibrionales Vibrionaceae *Vibrio cholerae* O1 str. 2010EL-1961 |
| Vch-143-1 | Bacteria Proteobacteria Gammaproteobacteria Vibrionales Vibrionaceae *Vibrio cholerae* O1 str. 2010EL-2010H |
| Vch-144-1 | Bacteria Proteobacteria Gammaproteobacteria Vibrionales Vibrionaceae *Vibrio cholerae* O1 str. 2010EL-2010N |
| Vch-145-1 | Bacteria Proteobacteria Gammaproteobacteria Vibrionales Vibrionaceae *Vibrio cholerae* O1 str. 2010V-1014 |
| Vch-146-1 | Bacteria Proteobacteria Gammaproteobacteria Vibrionales Vibrionaceae *Vibrio cholerae* O1 str. 2011EL-1089 |
| Vch-147-1 | Bacteria Proteobacteria Gammaproteobacteria Vibrionales Vibrionaceae *Vibrio cholerae* O1 str. 2011EL-1137 |
| Vch-148-1 | Bacteria Proteobacteria Gammaproteobacteria Vibrionales Vibrionaceae *Vibrio cholerae* O1 str. 2011V-1021 |
| Vch-149-1 | Bacteria Proteobacteria Gammaproteobacteria Vibrionales Vibrionaceae *Vibrio cholerae* O1 str. 3500-05 |
| Vch-150-1 | Bacteria Proteobacteria Gammaproteobacteria Vibrionales Vibrionaceae *Vibrio cholerae* O1 str. 3546-06 |
| Vch-151-1 | Bacteria Proteobacteria Gammaproteobacteria Vibrionales Vibrionaceae *Vibrio cholerae* O1 str. 3554-08 |
| Vch-152-1 | Bacteria Proteobacteria Gammaproteobacteria Vibrionales Vibrionaceae *Vibrio cholerae* O1 str. 3582-05 |
| Vch-153-1 | Bacteria Proteobacteria Gammaproteobacteria Vibrionales Vibrionaceae *Vibrio cholerae* O1 str. 87395 |
| Vch-154-1 | Bacteria Proteobacteria Gammaproteobacteria Vibrionales Vibrionaceae *Vibrio cholerae* O1 str. 95412 |
| Vch-155-1 | Bacteria Proteobacteria Gammaproteobacteria Vibrionales Vibrionaceae *Vibrio cholerae* O1 str. AG-7404 |
| Vch-156-1 | Bacteria Proteobacteria Gammaproteobacteria Vibrionales Vibrionaceae *Vibrio cholerae* O1 str. AG-8040 |
| Vch-157-1 | Bacteria Proteobacteria Gammaproteobacteria Vibrionales Vibrionaceae *Vibrio cholerae* O1 str. Amazonia |
| Vch-158-1 | Bacteria Proteobacteria Gammaproteobacteria Vibrionales Vibrionaceae *Vibrio cholerae* O1 str. C6706 |
| Vch-159-1 | Bacteria Proteobacteria Gammaproteobacteria Vibrionales Vibrionaceae *Vibrio cholerae* O1 str. EC-0009 |
| Vch-160-1 | Bacteria Proteobacteria Gammaproteobacteria Vibrionales Vibrionaceae *Vibrio cholerae* O1 str. EC-0012 |
| Vch-161-1 | Bacteria Proteobacteria Gammaproteobacteria Vibrionales Vibrionaceae *Vibrio cholerae* O1 str. EC-0027 |
| Vch-162-1 | Bacteria Proteobacteria Gammaproteobacteria Vibrionales Vibrionaceae *Vibrio cholerae* O1 str. EC-0051 |
| Vch-163-1 | Bacteria Proteobacteria Gammaproteobacteria Vibrionales Vibrionaceae *Vibrio cholerae* O1 str. EDC-020 |
| Vch-164-1 | Bacteria Proteobacteria Gammaproteobacteria Vibrionales Vibrionaceae *Vibrio cholerae* O1 str. EDC-022 |
| Vch-165-1 | Bacteria Proteobacteria Gammaproteobacteria Vibrionales Vibrionaceae *Vibrio cholerae* O1 str. EM-1536 |
| Vch-166-1 | Bacteria Proteobacteria Gammaproteobacteria Vibrionales Vibrionaceae *Vibrio cholerae* O1 str. EM-1546 |
| Vch-167-1 | Bacteria Proteobacteria Gammaproteobacteria Vibrionales Vibrionaceae *Vibrio cholerae* O1 str. EM-1626 |
| Vch-168-1 | Bacteria Proteobacteria Gammaproteobacteria Vibrionales Vibrionaceae *Vibrio cholerae* O1 str. EM-1676A |
| Vch-169-1 | Bacteria Proteobacteria Gammaproteobacteria Vibrionales Vibrionaceae *Vibrio cholerae* O1 str. EM-1727 |
| Vch-170-1 | Bacteria Proteobacteria Gammaproteobacteria Vibrionales Vibrionaceae *Vibrio cholerae* O1 str. Inaba G4222 |
| Vch-175-1 | Bacteria Proteobacteria Gammaproteobacteria Vibrionales Vibrionaceae *Vibrio cholerae* O1 str. Nep-21106 |
| Vch-176-1 | Bacteria Proteobacteria Gammaproteobacteria Vibrionales Vibrionaceae *Vibrio cholerae* O1 str. Nep-21113 |
| Vch-171-1 | Bacteria Proteobacteria Gammaproteobacteria Vibrionales Vibrionaceae *Vibrio cholerae* O1 str. NHCC-004A |
| Vch-172-1 | Bacteria Proteobacteria Gammaproteobacteria Vibrionales Vibrionaceae *Vibrio cholerae* O1 str. NHCC-006C |
| Vch-173-1 | Bacteria Proteobacteria Gammaproteobacteria Vibrionales Vibrionaceae *Vibrio cholerae* O1 str. NHCC-008D |
| Vch-174-1 | Bacteria Proteobacteria Gammaproteobacteria Vibrionales Vibrionaceae *Vibrio cholerae* O1 str. NHCC-010F |
| Vch-177-1 | Bacteria Proteobacteria Gammaproteobacteria Vibrionales Vibrionaceae *Vibrio cholerae* O1 str. PCS-022 |
| Vch-178-1 | Bacteria Proteobacteria Gammaproteobacteria Vibrionales Vibrionaceae *Vibrio cholerae* O1 str. PCS-023 |
| Vch-179-1 to Vch-179-2 | Bacteria Proteobacteria Gammaproteobacteria Vibrionales Vibrionaceae *Vibrio cholerae* O395 |
| Vch-180-1 | Bacteria Proteobacteria Gammaproteobacteria Vibrionales Vibrionaceae *Vibrio cholerae* P-18785 |

| | |
|---|---|
| Vch-182-1 | Bacteria Proteobacteria Gammaproteobacteria Vibrionales Vibrionaceae *Vibrio cholerae* PhVC-311 |
| Vch-183-1 | Bacteria Proteobacteria Gammaproteobacteria Vibrionales Vibrionaceae *Vibrio cholerae* PhVC-326 |
| Vch-184-1 | Bacteria Proteobacteria Gammaproteobacteria Vibrionales Vibrionaceae *Vibrio cholerae* PhVE-5 |
| Vch-181-1 | Bacteria Proteobacteria Gammaproteobacteria Vibrionales Vibrionaceae *Vibrio cholerae* PS15 |
| Vch-185-1 | Bacteria Proteobacteria Gammaproteobacteria Vibrionales Vibrionaceae *Vibrio cholerae* RC27 |
| Vch-186-1 | Bacteria Proteobacteria Gammaproteobacteria Vibrionales Vibrionaceae *Vibrio cholerae* RC385 |
| Vch-187-1 | Bacteria Proteobacteria Gammaproteobacteria Vibrionales Vibrionaceae *Vibrio cholerae* RC9 |
| Vch-188-1 | Bacteria Proteobacteria Gammaproteobacteria Vibrionales Vibrionaceae *Vibrio cholerae* TM 11079-80 |
| Vch-189-1 | Bacteria Proteobacteria Gammaproteobacteria Vibrionales Vibrionaceae *Vibrio cholerae* TMA 21 |
| Vch-190-1 | Bacteria Proteobacteria Gammaproteobacteria Vibrionales Vibrionaceae *Vibrio cholerae* V51 |
| Vch-191-1 | Bacteria Proteobacteria Gammaproteobacteria Vibrionales Vibrionaceae *Vibrio cholerae* V52 |
| Vch-192-1 | Bacteria Proteobacteria Gammaproteobacteria Vibrionales Vibrionaceae *Vibrio cholerae* VC1761 |
| Vch-193-1 | Bacteria Proteobacteria Gammaproteobacteria Vibrionales Vibrionaceae *Vibrio cholerae* VC35 |
| Vch-194-1 | Bacteria Proteobacteria Gammaproteobacteria Vibrionales Vibrionaceae *Vibrio cholerae* VC4370 |
| Vch-195-1 | Bacteria Proteobacteria Gammaproteobacteria Vibrionales Vibrionaceae *Vibrio cholerae* VCC19 |
| Vch-196-1 | Bacteria Proteobacteria Gammaproteobacteria Vibrionales Vibrionaceae *Vibrio cholerae* ZWU0020 |
| Vco-1-1 to Vco-1-2 | Bacteria Proteobacteria Gammaproteobacteria Vibrionales Vibrionaceae *Vibrio coralliilyticus* |
| Vco-2-1 | Bacteria Proteobacteria Gammaproteobacteria Vibrionales Vibrionaceae *Vibrio coralliilyticus* ATCC BAA-450 |
| Vco-3-1 | Bacteria Proteobacteria Gammaproteobacteria Vibrionales Vibrionaceae *Vibrio coralliilyticus* P1 |
| Vcr-1-1 to Vcr-1-6 | Bacteria Proteobacteria Gammaproteobacteria Vibrionales Vibrionaceae *Vibrio crassostreae* |
| Vcr-2-1 | Bacteria Proteobacteria Gammaproteobacteria Vibrionales Vibrionaceae *Vibrio crassostreae* 9CS106 |
| Vcr-3-1 | Bacteria Proteobacteria Gammaproteobacteria Vibrionales Vibrionaceae *Vibrio crassostreae* 9ZC13 |
| Vcr-4-1 | Bacteria Proteobacteria Gammaproteobacteria Vibrionales Vibrionaceae *Vibrio crassostreae* 9ZC77 |
| Vcr-5-1 | Bacteria Proteobacteria Gammaproteobacteria Vibrionales Vibrionaceae *Vibrio crassostreae* 9ZC88 |
| Vcr-6-1 | Bacteria Proteobacteria Gammaproteobacteria Vibrionales Vibrionaceae *Vibrio crassostreae* ZF-91 |
| Vcy-1-1 | Bacteria Proteobacteria Gammaproteobacteria Vibrionales Vibrionaceae *Vibrio cyclitrophicus* 1F111 |
| Vcy-2-1 | Bacteria Proteobacteria Gammaproteobacteria Vibrionales Vibrionaceae *Vibrio cyclitrophicus* 1F175 |
| Vcy-3-1 | Bacteria Proteobacteria Gammaproteobacteria Vibrionales Vibrionaceae *Vibrio cyclitrophicus* 1F273 |
| Vcy-4-1 | Bacteria Proteobacteria Gammaproteobacteria Vibrionales Vibrionaceae *Vibrio cyclitrophicus* 1F289 |
| Vcy-5-1 | Bacteria Proteobacteria Gammaproteobacteria Vibrionales Vibrionaceae *Vibrio cyclitrophicus* 1F53 |
| Vcy-6-1 | Bacteria Proteobacteria Gammaproteobacteria Vibrionales Vibrionaceae *Vibrio cyclitrophicus* 1F97 |
| Vcy-7-1 | Bacteria Proteobacteria Gammaproteobacteria Vibrionales Vibrionaceae *Vibrio cyclitrophicus* FF160 |
| Vcy-8-1 | Bacteria Proteobacteria Gammaproteobacteria Vibrionales Vibrionaceae *Vibrio cyclitrophicus* FF274 |
| Vcy-9-1 | Bacteria Proteobacteria Gammaproteobacteria Vibrionales Vibrionaceae *Vibrio cyclitrophicus* FF75 |
| Vcy-10-1 | Bacteria Proteobacteria Gammaproteobacteria Vibrionales Vibrionaceae *Vibrio cyclitrophicus* ZF14 |
| Vcy-11-1 | Bacteria Proteobacteria Gammaproteobacteria Vibrionales Vibrionaceae *Vibrio cyclitrophicus* ZF170 |
| Vcy-12-1 | Bacteria Proteobacteria Gammaproteobacteria Vibrionales Vibrionaceae *Vibrio cyclitrophicus* ZF205 |
| Vcy-13-1 | Bacteria Proteobacteria Gammaproteobacteria Vibrionales Vibrionaceae *Vibrio cyclitrophicus* ZF207 |
| Vcy-14-1 | Bacteria Proteobacteria Gammaproteobacteria Vibrionales Vibrionaceae *Vibrio cyclitrophicus* ZF255 |
| Vcy-15-1 | Bacteria Proteobacteria Gammaproteobacteria Vibrionales Vibrionaceae *Vibrio cyclitrophicus* ZF264 |
| Vcy-16-1 | Bacteria Proteobacteria Gammaproteobacteria Vibrionales Vibrionaceae *Vibrio cyclitrophicus* ZF270 |
| Vcy-17-1 | Bacteria Proteobacteria Gammaproteobacteria Vibrionales Vibrionaceae *Vibrio cyclitrophicus* ZF28 |
| Vcy-18-1 | Bacteria Proteobacteria Gammaproteobacteria Vibrionales Vibrionaceae *Vibrio cyclitrophicus* ZF30 |
| Vcy-19-1 | Bacteria Proteobacteria Gammaproteobacteria Vibrionales Vibrionaceae *Vibrio cyclitrophicus* ZF65 |
| Vcy-20-1 | Bacteria Proteobacteria Gammaproteobacteria Vibrionales Vibrionaceae *Vibrio cyclitrophicus* ZF99 |
| Vdi-1-1 | Bacteria Proteobacteria Gammaproteobacteria Vibrionales Vibrionaceae *Vibrio diabolicus* |
| Vdi-2-1 | Bacteria Proteobacteria Gammaproteobacteria Vibrionales Vibrionaceae *Vibrio diazotrophicus* NBRC 103148 |
| Vez-1-1 | Bacteria Proteobacteria Gammaproteobacteria Vibrionales Vibrionaceae *Vibrio ezurae* NBRC 102218 |
| Vfl-1-1 | Bacteria Proteobacteria Gammaproteobacteria Vibrionales Vibrionaceae *Vibrio fluvialis* |
| Vfl-2-1 | Bacteria Proteobacteria Gammaproteobacteria Vibrionales Vibrionaceae *Vibrio fluvialis* 539 |
| Vfl-3-1 | Bacteria Proteobacteria Gammaproteobacteria Vibrionales Vibrionaceae *Vibrio fluvialis* 560 |
| Vfl-4-1 | Bacteria Proteobacteria Gammaproteobacteria Vibrionales Vibrionaceae *Vibrio fluvialis* I21563 |
| Vfl-5-1 | Bacteria Proteobacteria Gammaproteobacteria Vibrionales Vibrionaceae *Vibrio fluvialis* PG41 |
| Vfo-1-1 | Bacteria Proteobacteria Gammaproteobacteria Vibrionales Vibrionaceae *Vibrio fortis* |
| Vfu-1-1 | Bacteria Proteobacteria Gammaproteobacteria Vibrionales Vibrionaceae *Vibrio furnissii* CIP 102972 |
| Vfu-2-1 | Bacteria Proteobacteria Gammaproteobacteria Vibrionales Vibrionaceae *Vibrio furnissii* NCTC 11218 |
| Vfu-3-1 | Bacteria Proteobacteria Gammaproteobacteria Vibrionales Vibrionaceae *Vibrio furnissii* ZOR0035 |
| Vga-1-1 | Bacteria Proteobacteria Gammaproteobacteria Vibrionales Vibrionaceae *Vibrio galatheae* |
| Vge-1-1 | Bacteria Proteobacteria Gammaproteobacteria Vibrionales Vibrionaceae *Vibrio genomosp.* F10 str. 9ZB36 |
| Vge-2-1 | Bacteria Proteobacteria Gammaproteobacteria Vibrionales Vibrionaceae *Vibrio genomosp.* F10 str. 9ZC157 |
| Vge-3-1 | Bacteria Proteobacteria Gammaproteobacteria Vibrionales Vibrionaceae *Vibrio genomosp.* F10 str. 9ZD137 |
| Vge-4-1 | Bacteria Proteobacteria Gammaproteobacteria Vibrionales Vibrionaceae *Vibrio genomosp.* F10 str. ZF-129 |
| Vge-5-1 | Bacteria Proteobacteria Gammaproteobacteria Vibrionales Vibrionaceae *Vibrio genomosp.* F6 str. FF-238 |
| Vha-1-1 | Bacteria Proteobacteria Gammaproteobacteria Vibrionales Vibrionaceae *Vibrio halioticoli* NBRC 102217 |
| Vha-2-1 | Bacteria Proteobacteria Gammaproteobacteria Vibrionales Vibrionaceae *Vibrio harveyi* |

| | |
|---|---|
| Vha-3-1 | Bacteria Proteobacteria Gammaproteobacteria Vibrionales Vibrionaceae *Vibrio harveyi* 1DA3 |
| Vha-4-1 | Bacteria Proteobacteria Gammaproteobacteria Vibrionales Vibrionaceae *Vibrio harveyi* AOD131 |
| Vha-5-1 | Bacteria Proteobacteria Gammaproteobacteria Vibrionales Vibrionaceae *Vibrio harveyi* ATCC BAA-1116 |
| Vha-6-1 | Bacteria Proteobacteria Gammaproteobacteria Vibrionales Vibrionaceae *Vibrio harveyi* CAIM 1792 |
| Vha-7-1 | Bacteria Proteobacteria Gammaproteobacteria Vibrionales Vibrionaceae *Vibrio harveyi* E385 |
| Vha-8-1 | Bacteria Proteobacteria Gammaproteobacteria Vibrionales Vibrionaceae *Vibrio harveyi* HY01 |
| Vha-9-1 to Vha-9-2 | Bacteria Proteobacteria Gammaproteobacteria Vibrionales Vibrionaceae *Vibrio harveyi* NBRC 15634 = ATCC 14126 |
| Vha-10-1 | Bacteria Proteobacteria Gammaproteobacteria Vibrionales Vibrionaceae *Vibrio harveyi* VHJR4 |
| Vha-11-1 | Bacteria Proteobacteria Gammaproteobacteria Vibrionales Vibrionaceae *Vibrio harveyi* VHJR7 |
| Vha-12-1 | Bacteria Proteobacteria Gammaproteobacteria Vibrionales Vibrionaceae *Vibrio harveyi* ZJ0603 |
| Vhy-1-1 to Vhy-1-3 | Bacteria Proteobacteria Gammaproteobacteria Vibrionales Vibrionaceae *Vibrio hyugaensis* |
| Vic-1-1 | Bacteria Proteobacteria Gammaproteobacteria Vibrionales Vibrionaceae *Vibrio ichthyoenteri* ATCC 700023 |
| Vja-1-1 to Vja-1-2 | Bacteria Proteobacteria Gammaproteobacteria Vibrionales Vibrionaceae *Vibrio jasicida* |
| Vja-2-1 | Bacteria Proteobacteria Gammaproteobacteria Vibrionales Vibrionaceae *Vibrio jasicida* 090810c |
| Vja-3-1 | Bacteria Proteobacteria Gammaproteobacteria Vibrionales Vibrionaceae *Vibrio jasicida* CAIM 1864 = LMG 25398 |
| Vja-4-1 | Bacteria Proteobacteria Gammaproteobacteria Vibrionales Vibrionaceae *Vibrio jasicida* MWB 21 |
| Vka-1-1 | Bacteria Proteobacteria Gammaproteobacteria Vibrionales Vibrionaceae *Vibrio kanaloae* 5S-149 |
| Vle-1-1 | Bacteria Proteobacteria Gammaproteobacteria Vibrionales Vibrionaceae *Vibrio lentus* |
| Vli-1-1 | Bacteria Proteobacteria Gammaproteobacteria Vibrionales Vibrionaceae *Vibrio litoralis* DSM 17657 |
| Vma-1-1 to Vma-1-2 | Bacteria Proteobacteria Gammaproteobacteria Vibrionales Vibrionaceae *Vibrio maritimus* |
| Vme-1-1 | Bacteria Proteobacteria Gammaproteobacteria Vibrionales Vibrionaceae *Vibrio metschnikovii* CIP 69.14 |
| Vme-2-1 | Bacteria Proteobacteria Gammaproteobacteria Vibrionales Vibrionaceae *Vibrio mexicanus* |
| Vmi-1-1 | Bacteria Proteobacteria Gammaproteobacteria Vibrionales Vibrionaceae *Vibrio mimicus* CAIM 1883 |
| Vmi-2-1 | Bacteria Proteobacteria Gammaproteobacteria Vibrionales Vibrionaceae *Vibrio mimicus* CAIM 602 |
| Vmi-3-1 | Bacteria Proteobacteria Gammaproteobacteria Vibrionales Vibrionaceae *Vibrio mimicus* MB-451 |
| Vmi-4-1 | Bacteria Proteobacteria Gammaproteobacteria Vibrionales Vibrionaceae *Vibrio mimicus* SX-4 |
| Vmi-5-1 | Bacteria Proteobacteria Gammaproteobacteria Vibrionales Vibrionaceae *Vibrio mimicus* VM223 |
| Vmi-6-1 | Bacteria Proteobacteria Gammaproteobacteria Vibrionales Vibrionaceae *Vibrio mimicus* VM573 |
| Vmi-7-1 | Bacteria Proteobacteria Gammaproteobacteria Vibrionales Vibrionaceae *Vibrio mimicus* VM603 |
| Vmy-1-1 | Bacteria Proteobacteria Gammaproteobacteria Vibrionales Vibrionaceae *Vibrio mytili* |
| Vna-1-1 to Vna-1-2 | Bacteria Proteobacteria Gammaproteobacteria Vibrionales Vibrionaceae *Vibrio natriegens* NBRC 15636 = ATCC 14048 = DSM 759 |
| Vna-2-1 to Vna-2-4 | Bacteria Proteobacteria Gammaproteobacteria Vibrionales Vibrionaceae *Vibrio navarrensis* |
| Vne-1-1 | Bacteria Proteobacteria Gammaproteobacteria Vibrionales Vibrionaceae *Vibrio neptunius* |
| Vne-2-1 | Bacteria Proteobacteria Gammaproteobacteria Vibrionales Vibrionaceae *Vibrio nereis* |
| Vni-1-1 | Bacteria Proteobacteria Gammaproteobacteria Vibrionales Vibrionaceae *Vibrio nigripulchritudo* |
| Vni-2-1 | Bacteria Proteobacteria Gammaproteobacteria Vibrionales Vibrionaceae *Vibrio nigripulchritudo* AM115 |
| Vni-3-1 | Bacteria Proteobacteria Gammaproteobacteria Vibrionales Vibrionaceae *Vibrio nigripulchritudo* ATCC 27043 |
| Vni-4-1 | Bacteria Proteobacteria Gammaproteobacteria Vibrionales Vibrionaceae *Vibrio nigripulchritudo* BLFn1 |
| Vni-5-1 | Bacteria Proteobacteria Gammaproteobacteria Vibrionales Vibrionaceae *Vibrio nigripulchritudo* ENn2 |
| Vni-6-1 | Bacteria Proteobacteria Gammaproteobacteria Vibrionales Vibrionaceae *Vibrio nigripulchritudo* FTn2 |
| Vni-7-1 | Bacteria Proteobacteria Gammaproteobacteria Vibrionales Vibrionaceae *Vibrio nigripulchritudo* MADA3029 |
| Vni-8-1 | Bacteria Proteobacteria Gammaproteobacteria Vibrionales Vibrionaceae *Vibrio nigripulchritudo* Pon4 |
| Vni-9-1 | Bacteria Proteobacteria Gammaproteobacteria Vibrionales Vibrionaceae *Vibrio nigripulchritudo* SFn118 |
| Vni-10-1 | Bacteria Proteobacteria Gammaproteobacteria Vibrionales Vibrionaceae *Vibrio nigripulchritudo* SFn135 |
| Vni-11-1 | Bacteria Proteobacteria Gammaproteobacteria Vibrionales Vibrionaceae *Vibrio nigripulchritudo* SFn27 |
| Vni-12-1 | Bacteria Proteobacteria Gammaproteobacteria Vibrionales Vibrionaceae *Vibrio nigripulchritudo* SO65 |
| Vni-13-1 | Bacteria Proteobacteria Gammaproteobacteria Vibrionales Vibrionaceae *Vibrio nigripulchritudo* SOn1 |
| Vni-14-1 | Bacteria Proteobacteria Gammaproteobacteria Vibrionales Vibrionaceae *Vibrio nigripulchritudo* Wn13 |
| Vor-1-1 | Bacteria Proteobacteria Gammaproteobacteria Vibrionales Vibrionaceae *Vibrio ordalii* 12B09 |
| Vor-2-1 | Bacteria Proteobacteria Gammaproteobacteria Vibrionales Vibrionaceae *Vibrio ordalii* ATCC 33509 |
| Vor-3-1 | Bacteria Proteobacteria Gammaproteobacteria Vibrionales Vibrionaceae *Vibrio ordalii* FF-167 |
| Vor-4-1 | Bacteria Proteobacteria Gammaproteobacteria Vibrionales Vibrionaceae *Vibrio ordalii* FF-93 |
| Vor-5-1 | Bacteria Proteobacteria Gammaproteobacteria Vibrionales Vibrionaceae *Vibrio ordalii* FS-144 |
| Vor-6-1 | Bacteria Proteobacteria Gammaproteobacteria Vibrionales Vibrionaceae *Vibrio ordalii* FS-238 |
| Vhe-1-1 | Bacteria Proteobacteria Gammaproteobacteria Vibrionales Vibrionaceae Vibrio Vibrio oreintalis group *Vibrio hepatarius* |
| Vsi-1-1 to Vsi-1-4 | Bacteria Proteobacteria Gammaproteobacteria Vibrionales Vibrionaceae Vibrio Vibrio oreintalis group *Vibrio sinaloensis* |
| Vtu-1-1 | Bacteria Proteobacteria Gammaproteobacteria Vibrionales Vibrionaceae Vibrio Vibrio oreintalis group *Vibrio tubiashii* |
| Vtu-2-1 to Vtu-2-2 | Bacteria Proteobacteria Gammaproteobacteria Vibrionales Vibrionaceae Vibrio Vibrio oreintalis group *Vibrio tubiashii* ATCC 19109 |
| Vor-7-1 | Bacteria Proteobacteria Gammaproteobacteria Vibrionales Vibrionaceae *Vibrio orientalis* CIP 102891 |
| Vor-8-1 | Bacteria Proteobacteria Gammaproteobacteria Vibrionales Vibrionaceae *Vibrio orientalis* CIP 102891 = ATCC 33934 |
| Vow-2-1 to Vow-2-4 | Bacteria Proteobacteria Gammaproteobacteria Vibrionales Vibrionaceae *Vibrio owensii* |
| Vow-3-1 | Bacteria Proteobacteria Gammaproteobacteria Vibrionales Vibrionaceae *Vibrio owensii* 47666-1 |
| Vow-4-1 | Bacteria Proteobacteria Gammaproteobacteria Vibrionales Vibrionaceae *Vibrio owensii* ATCC 25919 |
| Vow-5-1 | Bacteria Proteobacteria Gammaproteobacteria Vibrionales Vibrionaceae *Vibrio owensii* CAIM 1854 = LMG 25443 |
| Vow-1-1 | Bacteria Proteobacteria Gammaproteobacteria Vibrionales Vibrionaceae Vibrio VIbrio owensii CAIM 1854 = LMG 25443 |

| | |
|---|---|
| Vow-6-1 | Bacteria Proteobacteria Gammaproteobacteria Vibrionales Vibrionaceae *Vibrio owensii* LMG 25430 |
| Vpa-1-1 | Bacteria Proteobacteria Gammaproteobacteria Vibrionales Vibrionaceae *Vibrio pacinii* DSM 19139 |
| Vpa-2-1 | Bacteria Proteobacteria Gammaproteobacteria Vibrionales Vibrionaceae *Vibrio parahaemolyticus* |
| Vpa-3-1 | Bacteria Proteobacteria Gammaproteobacteria Vibrionales Vibrionaceae *Vibrio parahaemolyticus* 10290 |
| Vpa-4-1 | Bacteria Proteobacteria Gammaproteobacteria Vibrionales Vibrionaceae *Vibrio parahaemolyticus* 10296 |
| Vpa-5-1 | Bacteria Proteobacteria Gammaproteobacteria Vibrionales Vibrionaceae *Vibrio parahaemolyticus* 10329 |
| Vpa-6-1 | Bacteria Proteobacteria Gammaproteobacteria Vibrionales Vibrionaceae *Vibrio parahaemolyticus* 12310 |
| Vpa-7-1 | Bacteria Proteobacteria Gammaproteobacteria Vibrionales Vibrionaceae *Vibrio parahaemolyticus* 12315 |
| Vpa-8-1 | Bacteria Proteobacteria Gammaproteobacteria Vibrionales Vibrionaceae *Vibrio parahaemolyticus* 16 |
| Vpa-9-1 | Bacteria Proteobacteria Gammaproteobacteria Vibrionales Vibrionaceae *Vibrio parahaemolyticus* 3256 |
| Vpa-10-1 | Bacteria Proteobacteria Gammaproteobacteria Vibrionales Vibrionaceae *Vibrio parahaemolyticus* 3259 |
| Vpa-11-1 | Bacteria Proteobacteria Gammaproteobacteria Vibrionales Vibrionaceae *Vibrio parahaemolyticus* 3324 |
| Vpa-12-1 | Bacteria Proteobacteria Gammaproteobacteria Vibrionales Vibrionaceae *Vibrio parahaemolyticus* 3355 |
| Vpa-13-1 | Bacteria Proteobacteria Gammaproteobacteria Vibrionales Vibrionaceae *Vibrio parahaemolyticus* 3631 |
| Vpa-14-1 | Bacteria Proteobacteria Gammaproteobacteria Vibrionales Vibrionaceae *Vibrio parahaemolyticus* 3644 |
| Vpa-15-1 | Bacteria Proteobacteria Gammaproteobacteria Vibrionales Vibrionaceae *Vibrio parahaemolyticus* 3646 |
| Vpa-16-1 | Bacteria Proteobacteria Gammaproteobacteria Vibrionales Vibrionaceae *Vibrio parahaemolyticus* 49 |
| Vpa-17-1 | Bacteria Proteobacteria Gammaproteobacteria Vibrionales Vibrionaceae *Vibrio parahaemolyticus* 50 |
| Vpa-18-1 | Bacteria Proteobacteria Gammaproteobacteria Vibrionales Vibrionaceae *Vibrio parahaemolyticus* 605 |
| Vpa-19-1 | Bacteria Proteobacteria Gammaproteobacteria Vibrionales Vibrionaceae *Vibrio parahaemolyticus* 846 |
| Vpa-20-1 | Bacteria Proteobacteria Gammaproteobacteria Vibrionales Vibrionaceae *Vibrio parahaemolyticus* 861 |
| Vpa-21-1 | Bacteria Proteobacteria Gammaproteobacteria Vibrionales Vibrionaceae *Vibrio parahaemolyticus* 863 |
| Vpa-22-1 | Bacteria Proteobacteria Gammaproteobacteria Vibrionales Vibrionaceae *Vibrio parahaemolyticus* 901128 |
| Vpa-23-1 | Bacteria Proteobacteria Gammaproteobacteria Vibrionales Vibrionaceae *Vibrio parahaemolyticus* 930 |
| Vpa-24-1 | Bacteria Proteobacteria Gammaproteobacteria Vibrionales Vibrionaceae *Vibrio parahaemolyticus* 949 |
| Vpa-25-1 | Bacteria Proteobacteria Gammaproteobacteria Vibrionales Vibrionaceae *Vibrio parahaemolyticus* 97-10290 |
| Vpa-26-1 | Bacteria Proteobacteria Gammaproteobacteria Vibrionales Vibrionaceae *Vibrio parahaemolyticus* 970107 |
| Vpa-27-1 | Bacteria Proteobacteria Gammaproteobacteria Vibrionales Vibrionaceae *Vibrio parahaemolyticus* 98-513-F52 |
| Vpa-28-1 | Bacteria Proteobacteria Gammaproteobacteria Vibrionales Vibrionaceae *Vibrio parahaemolyticus* AN-5034 |
| Vpa-29-1 | Bacteria Proteobacteria Gammaproteobacteria Vibrionales Vibrionaceae *Vibrio parahaemolyticus* AQ3810 |
| Vpa-30-1 | Bacteria Proteobacteria Gammaproteobacteria Vibrionales Vibrionaceae *Vibrio parahaemolyticus* AQ4037 |
| Vpa-31-1 | Bacteria Proteobacteria Gammaproteobacteria Vibrionales Vibrionaceae *Vibrio parahaemolyticus* B-265 |
| Vpa-32-1 | Bacteria Proteobacteria Gammaproteobacteria Vibrionales Vibrionaceae *Vibrio parahaemolyticus* BB22OP |
| Vpa-33-1 | Bacteria Proteobacteria Gammaproteobacteria Vibrionales Vibrionaceae *Vibrio parahaemolyticus* CFSAN001595 |
| Vpa-34-1 | Bacteria Proteobacteria Gammaproteobacteria Vibrionales Vibrionaceae *Vibrio parahaemolyticus* EKP-008 |
| Vpa-35-1 | Bacteria Proteobacteria Gammaproteobacteria Vibrionales Vibrionaceae *Vibrio parahaemolyticus* EKP-021 |
| Vpa-36-1 | Bacteria Proteobacteria Gammaproteobacteria Vibrionales Vibrionaceae *Vibrio parahaemolyticus* EKP-026 |
| Vpa-37-1 | Bacteria Proteobacteria Gammaproteobacteria Vibrionales Vibrionaceae *Vibrio parahaemolyticus* EKP-028 |
| Vpa-38-1 | Bacteria Proteobacteria Gammaproteobacteria Vibrionales Vibrionaceae *Vibrio parahaemolyticus* EN2910 |
| Vpa-39-1 | Bacteria Proteobacteria Gammaproteobacteria Vibrionales Vibrionaceae *Vibrio parahaemolyticus* EN9701072 |
| Vpa-40-1 | Bacteria Proteobacteria Gammaproteobacteria Vibrionales Vibrionaceae *Vibrio parahaemolyticus* EN9701121 |
| Vpa-41-1 | Bacteria Proteobacteria Gammaproteobacteria Vibrionales Vibrionaceae *Vibrio parahaemolyticus* EN9701173 |
| Vpa-42-1 | Bacteria Proteobacteria Gammaproteobacteria Vibrionales Vibrionaceae *Vibrio parahaemolyticus* EN9901310 |
| Vpa-43-1 | Bacteria Proteobacteria Gammaproteobacteria Vibrionales Vibrionaceae *Vibrio parahaemolyticus* IDH02189 |
| Vpa-44-1 | Bacteria Proteobacteria Gammaproteobacteria Vibrionales Vibrionaceae *Vibrio parahaemolyticus* IDH02640 |
| Vpa-45-1 | Bacteria Proteobacteria Gammaproteobacteria Vibrionales Vibrionaceae *Vibrio parahaemolyticus* K5030 |
| Vpa-46-1 | Bacteria Proteobacteria Gammaproteobacteria Vibrionales Vibrionaceae *Vibrio parahaemolyticus* M0605 |
| Vpa-47-1 | Bacteria Proteobacteria Gammaproteobacteria Vibrionales Vibrionaceae *Vibrio parahaemolyticus* NBRC 12711 |
| Vpa-48-1 | Bacteria Proteobacteria Gammaproteobacteria Vibrionales Vibrionaceae *Vibrio parahaemolyticus* NIHCB0603 |
| Vpa-49-1 | Bacteria Proteobacteria Gammaproteobacteria Vibrionales Vibrionaceae *Vibrio parahaemolyticus* NIHCB0757 |
| Vpa-50-1 | Bacteria Proteobacteria Gammaproteobacteria Vibrionales Vibrionaceae *Vibrio parahaemolyticus* O1:K33 str. CDC_K4557 |
| Vpa-51-1 | Bacteria Proteobacteria Gammaproteobacteria Vibrionales Vibrionaceae *Vibrio parahaemolyticus* O1:Kuk str. FDA_R31 |
| Vpa-52-1 | Bacteria Proteobacteria Gammaproteobacteria Vibrionales Vibrionaceae *Vibrio parahaemolyticus* O4:K12 str. K1203 |
| Vpa-53-1 | Bacteria Proteobacteria Gammaproteobacteria Vibrionales Vibrionaceae *Vibrio parahaemolyticus* PCV08-7 |
| Vpa-54-1 | Bacteria Proteobacteria Gammaproteobacteria Vibrionales Vibrionaceae *Vibrio parahaemolyticus* Peru-288 |
| Vpa-55-1 | Bacteria Proteobacteria Gammaproteobacteria Vibrionales Vibrionaceae *Vibrio parahaemolyticus* Peru-466 |
| Vpa-56-1 | Bacteria Proteobacteria Gammaproteobacteria Vibrionales Vibrionaceae *Vibrio parahaemolyticus* RIMD 2210633 |
| Vpa-57-1 | Bacteria Proteobacteria Gammaproteobacteria Vibrionales Vibrionaceae *Vibrio parahaemolyticus* S001 |
| Vpa-58-1 | Bacteria Proteobacteria Gammaproteobacteria Vibrionales Vibrionaceae *Vibrio parahaemolyticus* S002 |
| Vpa-59-1 | Bacteria Proteobacteria Gammaproteobacteria Vibrionales Vibrionaceae *Vibrio parahaemolyticus* S003 |
| Vpa-60-1 | Bacteria Proteobacteria Gammaproteobacteria Vibrionales Vibrionaceae *Vibrio parahaemolyticus* S004 |
| Vpa-61-1 | Bacteria Proteobacteria Gammaproteobacteria Vibrionales Vibrionaceae *Vibrio parahaemolyticus* S005 |
| Vpa-62-1 | Bacteria Proteobacteria Gammaproteobacteria Vibrionales Vibrionaceae *Vibrio parahaemolyticus* S008 |
| Vpa-63-1 | Bacteria Proteobacteria Gammaproteobacteria Vibrionales Vibrionaceae *Vibrio parahaemolyticus* S009 |
| Vpa-64-1 | Bacteria Proteobacteria Gammaproteobacteria Vibrionales Vibrionaceae *Vibrio parahaemolyticus* S011 |

| | |
|---|---|
| Vpa-65-1 | Bacteria Proteobacteria Gammaproteobacteria Vibrionales Vibrionaceae *Vibrio parahaemolyticus* S012 |
| Vpa-66-1 | Bacteria Proteobacteria Gammaproteobacteria Vibrionales Vibrionaceae *Vibrio parahaemolyticus* S013 |
| Vpa-67-1 | Bacteria Proteobacteria Gammaproteobacteria Vibrionales Vibrionaceae *Vibrio parahaemolyticus* S014 |
| Vpa-68-1 | Bacteria Proteobacteria Gammaproteobacteria Vibrionales Vibrionaceae *Vibrio parahaemolyticus* S015 |
| Vpa-69-1 | Bacteria Proteobacteria Gammaproteobacteria Vibrionales Vibrionaceae *Vibrio parahaemolyticus* S016 |
| Vpa-70-1 | Bacteria Proteobacteria Gammaproteobacteria Vibrionales Vibrionaceae *Vibrio parahaemolyticus* S017 |
| Vpa-71-1 | Bacteria Proteobacteria Gammaproteobacteria Vibrionales Vibrionaceae *Vibrio parahaemolyticus* S018 |
| Vpa-72-1 | Bacteria Proteobacteria Gammaproteobacteria Vibrionales Vibrionaceae *Vibrio parahaemolyticus* S019 |
| Vpa-73-1 | Bacteria Proteobacteria Gammaproteobacteria Vibrionales Vibrionaceae *Vibrio parahaemolyticus* S020 |
| Vpa-74-1 | Bacteria Proteobacteria Gammaproteobacteria Vibrionales Vibrionaceae *Vibrio parahaemolyticus* S021 |
| Vpa-75-1 | Bacteria Proteobacteria Gammaproteobacteria Vibrionales Vibrionaceae *Vibrio parahaemolyticus* S022 |
| Vpa-76-1 | Bacteria Proteobacteria Gammaproteobacteria Vibrionales Vibrionaceae *Vibrio parahaemolyticus* S023 |
| Vpa-77-1 | Bacteria Proteobacteria Gammaproteobacteria Vibrionales Vibrionaceae *Vibrio parahaemolyticus* S024 |
| Vpa-78-1 | Bacteria Proteobacteria Gammaproteobacteria Vibrionales Vibrionaceae *Vibrio parahaemolyticus* S025 |
| Vpa-79-1 | Bacteria Proteobacteria Gammaproteobacteria Vibrionales Vibrionaceae *Vibrio parahaemolyticus* S026 |
| Vpa-80-1 | Bacteria Proteobacteria Gammaproteobacteria Vibrionales Vibrionaceae *Vibrio parahaemolyticus* S027 |
| Vpa-81-1 | Bacteria Proteobacteria Gammaproteobacteria Vibrionales Vibrionaceae *Vibrio parahaemolyticus* S028 |
| Vpa-82-1 | Bacteria Proteobacteria Gammaproteobacteria Vibrionales Vibrionaceae *Vibrio parahaemolyticus* S029 |
| Vpa-83-1 | Bacteria Proteobacteria Gammaproteobacteria Vibrionales Vibrionaceae *Vibrio parahaemolyticus* S030 |
| Vpa-84-1 | Bacteria Proteobacteria Gammaproteobacteria Vibrionales Vibrionaceae *Vibrio parahaemolyticus* S031 |
| Vpa-85-1 | Bacteria Proteobacteria Gammaproteobacteria Vibrionales Vibrionaceae *Vibrio parahaemolyticus* S032 |
| Vpa-86-1 | Bacteria Proteobacteria Gammaproteobacteria Vibrionales Vibrionaceae *Vibrio parahaemolyticus* S033 |
| Vpa-87-1 | Bacteria Proteobacteria Gammaproteobacteria Vibrionales Vibrionaceae *Vibrio parahaemolyticus* S034 |
| Vpa-88-1 | Bacteria Proteobacteria Gammaproteobacteria Vibrionales Vibrionaceae *Vibrio parahaemolyticus* S035 |
| Vpa-89-1 | Bacteria Proteobacteria Gammaproteobacteria Vibrionales Vibrionaceae *Vibrio parahaemolyticus* S036 |
| Vpa-90-1 | Bacteria Proteobacteria Gammaproteobacteria Vibrionales Vibrionaceae *Vibrio parahaemolyticus* S037 |
| Vpa-91-1 | Bacteria Proteobacteria Gammaproteobacteria Vibrionales Vibrionaceae *Vibrio parahaemolyticus* S038 |
| Vpa-92-1 | Bacteria Proteobacteria Gammaproteobacteria Vibrionales Vibrionaceae *Vibrio parahaemolyticus* S039 |
| Vpa-93-1 | Bacteria Proteobacteria Gammaproteobacteria Vibrionales Vibrionaceae *Vibrio parahaemolyticus* S040 |
| Vpa-94-1 | Bacteria Proteobacteria Gammaproteobacteria Vibrionales Vibrionaceae *Vibrio parahaemolyticus* S041 |
| Vpa-95-1 | Bacteria Proteobacteria Gammaproteobacteria Vibrionales Vibrionaceae *Vibrio parahaemolyticus* S042 |
| Vpa-96-1 | Bacteria Proteobacteria Gammaproteobacteria Vibrionales Vibrionaceae *Vibrio parahaemolyticus* S043 |
| Vpa-97-1 | Bacteria Proteobacteria Gammaproteobacteria Vibrionales Vibrionaceae *Vibrio parahaemolyticus* S044 |
| Vpa-98-1 | Bacteria Proteobacteria Gammaproteobacteria Vibrionales Vibrionaceae *Vibrio parahaemolyticus* S045 |
| Vpa-99-1 | Bacteria Proteobacteria Gammaproteobacteria Vibrionales Vibrionaceae *Vibrio parahaemolyticus* S046 |
| Vpa-100-1 | Bacteria Proteobacteria Gammaproteobacteria Vibrionales Vibrionaceae *Vibrio parahaemolyticus* S047 |
| Vpa-101-1 | Bacteria Proteobacteria Gammaproteobacteria Vibrionales Vibrionaceae *Vibrio parahaemolyticus* S048 |
| Vpa-102-1 | Bacteria Proteobacteria Gammaproteobacteria Vibrionales Vibrionaceae *Vibrio parahaemolyticus* S049 |
| Vpa-103-1 | Bacteria Proteobacteria Gammaproteobacteria Vibrionales Vibrionaceae *Vibrio parahaemolyticus* S052 |
| Vpa-104-1 | Bacteria Proteobacteria Gammaproteobacteria Vibrionales Vibrionaceae *Vibrio parahaemolyticus* S053 |
| Vpa-105-1 | Bacteria Proteobacteria Gammaproteobacteria Vibrionales Vibrionaceae *Vibrio parahaemolyticus* S054 |
| Vpa-106-1 | Bacteria Proteobacteria Gammaproteobacteria Vibrionales Vibrionaceae *Vibrio parahaemolyticus* S055 |
| Vpa-107-1 | Bacteria Proteobacteria Gammaproteobacteria Vibrionales Vibrionaceae *Vibrio parahaemolyticus* S056 |
| Vpa-108-1 | Bacteria Proteobacteria Gammaproteobacteria Vibrionales Vibrionaceae *Vibrio parahaemolyticus* S057 |
| Vpa-109-1 | Bacteria Proteobacteria Gammaproteobacteria Vibrionales Vibrionaceae *Vibrio parahaemolyticus* S058 |
| Vpa-110-1 | Bacteria Proteobacteria Gammaproteobacteria Vibrionales Vibrionaceae *Vibrio parahaemolyticus* S060 |
| Vpa-111-1 | Bacteria Proteobacteria Gammaproteobacteria Vibrionales Vibrionaceae *Vibrio parahaemolyticus* S062 |
| Vpa-112-1 | Bacteria Proteobacteria Gammaproteobacteria Vibrionales Vibrionaceae *Vibrio parahaemolyticus* S063 |
| Vpa-113-1 | Bacteria Proteobacteria Gammaproteobacteria Vibrionales Vibrionaceae *Vibrio parahaemolyticus* S064 |
| Vpa-114-1 | Bacteria Proteobacteria Gammaproteobacteria Vibrionales Vibrionaceae *Vibrio parahaemolyticus* S065 |
| Vpa-115-1 | Bacteria Proteobacteria Gammaproteobacteria Vibrionales Vibrionaceae *Vibrio parahaemolyticus* S066 |
| Vpa-116-1 | Bacteria Proteobacteria Gammaproteobacteria Vibrionales Vibrionaceae *Vibrio parahaemolyticus* S067 |
| Vpa-117-1 | Bacteria Proteobacteria Gammaproteobacteria Vibrionales Vibrionaceae *Vibrio parahaemolyticus* S068 |
| Vpa-118-1 | Bacteria Proteobacteria Gammaproteobacteria Vibrionales Vibrionaceae *Vibrio parahaemolyticus* S069 |
| Vpa-119-1 | Bacteria Proteobacteria Gammaproteobacteria Vibrionales Vibrionaceae *Vibrio parahaemolyticus* S070 |
| Vpa-120-1 | Bacteria Proteobacteria Gammaproteobacteria Vibrionales Vibrionaceae *Vibrio parahaemolyticus* S071 |
| Vpa-121-1 | Bacteria Proteobacteria Gammaproteobacteria Vibrionales Vibrionaceae *Vibrio parahaemolyticus* S072 |
| Vpa-122-1 | Bacteria Proteobacteria Gammaproteobacteria Vibrionales Vibrionaceae *Vibrio parahaemolyticus* S073 |
| Vpa-123-1 | Bacteria Proteobacteria Gammaproteobacteria Vibrionales Vibrionaceae *Vibrio parahaemolyticus* S074 |
| Vpa-124-1 | Bacteria Proteobacteria Gammaproteobacteria Vibrionales Vibrionaceae *Vibrio parahaemolyticus* S075 |
| Vpa-125-1 | Bacteria Proteobacteria Gammaproteobacteria Vibrionales Vibrionaceae *Vibrio parahaemolyticus* S076 |
| Vpa-126-1 | Bacteria Proteobacteria Gammaproteobacteria Vibrionales Vibrionaceae *Vibrio parahaemolyticus* S077 |
| Vpa-127-1 | Bacteria Proteobacteria Gammaproteobacteria Vibrionales Vibrionaceae *Vibrio parahaemolyticus* S078 |
| Vpa-128-1 | Bacteria Proteobacteria Gammaproteobacteria Vibrionales Vibrionaceae *Vibrio parahaemolyticus* S079 |
| Vpa-129-1 | Bacteria Proteobacteria Gammaproteobacteria Vibrionales Vibrionaceae *Vibrio parahaemolyticus* S081 |

| | |
|---|---|
| Vpa-130-1 | Bacteria Proteobacteria Gammaproteobacteria Vibrionales Vibrionaceae *Vibrio parahaemolyticus* S082 |
| Vpa-131-1 | Bacteria Proteobacteria Gammaproteobacteria Vibrionales Vibrionaceae *Vibrio parahaemolyticus* S083 |
| Vpa-132-1 | Bacteria Proteobacteria Gammaproteobacteria Vibrionales Vibrionaceae *Vibrio parahaemolyticus* S086 |
| Vpa-133-1 | Bacteria Proteobacteria Gammaproteobacteria Vibrionales Vibrionaceae *Vibrio parahaemolyticus* S087 |
| Vpa-134-1 | Bacteria Proteobacteria Gammaproteobacteria Vibrionales Vibrionaceae *Vibrio parahaemolyticus* S088 |
| Vpa-135-1 | Bacteria Proteobacteria Gammaproteobacteria Vibrionales Vibrionaceae *Vibrio parahaemolyticus* S090 |
| Vpa-136-1 | Bacteria Proteobacteria Gammaproteobacteria Vibrionales Vibrionaceae *Vibrio parahaemolyticus* S091 |
| Vpa-137-1 | Bacteria Proteobacteria Gammaproteobacteria Vibrionales Vibrionaceae *Vibrio parahaemolyticus* S092 |
| Vpa-138-1 | Bacteria Proteobacteria Gammaproteobacteria Vibrionales Vibrionaceae *Vibrio parahaemolyticus* S093 |
| Vpa-139-1 | Bacteria Proteobacteria Gammaproteobacteria Vibrionales Vibrionaceae *Vibrio parahaemolyticus* S094 |
| Vpa-140-1 | Bacteria Proteobacteria Gammaproteobacteria Vibrionales Vibrionaceae *Vibrio parahaemolyticus* S095 |
| Vpa-141-1 | Bacteria Proteobacteria Gammaproteobacteria Vibrionales Vibrionaceae *Vibrio parahaemolyticus* S096 |
| Vpa-142-1 | Bacteria Proteobacteria Gammaproteobacteria Vibrionales Vibrionaceae *Vibrio parahaemolyticus* S097 |
| Vpa-143-1 | Bacteria Proteobacteria Gammaproteobacteria Vibrionales Vibrionaceae *Vibrio parahaemolyticus* S098 |
| Vpa-144-1 | Bacteria Proteobacteria Gammaproteobacteria Vibrionales Vibrionaceae *Vibrio parahaemolyticus* S100 |
| Vpa-145-1 | Bacteria Proteobacteria Gammaproteobacteria Vibrionales Vibrionaceae *Vibrio parahaemolyticus* S101 |
| Vpa-146-1 | Bacteria Proteobacteria Gammaproteobacteria Vibrionales Vibrionaceae *Vibrio parahaemolyticus* S104 |
| Vpa-147-1 | Bacteria Proteobacteria Gammaproteobacteria Vibrionales Vibrionaceae *Vibrio parahaemolyticus* S105 |
| Vpa-148-1 | Bacteria Proteobacteria Gammaproteobacteria Vibrionales Vibrionaceae *Vibrio parahaemolyticus* S106 |
| Vpa-149-1 | Bacteria Proteobacteria Gammaproteobacteria Vibrionales Vibrionaceae *Vibrio parahaemolyticus* S108 |
| Vpa-150-1 | Bacteria Proteobacteria Gammaproteobacteria Vibrionales Vibrionaceae *Vibrio parahaemolyticus* S109 |
| Vpa-151-1 | Bacteria Proteobacteria Gammaproteobacteria Vibrionales Vibrionaceae *Vibrio parahaemolyticus* S110 |
| Vpa-152-1 | Bacteria Proteobacteria Gammaproteobacteria Vibrionales Vibrionaceae *Vibrio parahaemolyticus* S111 |
| Vpa-153-1 | Bacteria Proteobacteria Gammaproteobacteria Vibrionales Vibrionaceae *Vibrio parahaemolyticus* S112 |
| Vpa-154-1 | Bacteria Proteobacteria Gammaproteobacteria Vibrionales Vibrionaceae *Vibrio parahaemolyticus* S113 |
| Vpa-155-1 | Bacteria Proteobacteria Gammaproteobacteria Vibrionales Vibrionaceae *Vibrio parahaemolyticus* S114 |
| Vpa-156-1 | Bacteria Proteobacteria Gammaproteobacteria Vibrionales Vibrionaceae *Vibrio parahaemolyticus* S115 |
| Vpa-157-1 | Bacteria Proteobacteria Gammaproteobacteria Vibrionales Vibrionaceae *Vibrio parahaemolyticus* S116 |
| Vpa-158-1 | Bacteria Proteobacteria Gammaproteobacteria Vibrionales Vibrionaceae *Vibrio parahaemolyticus* S117 |
| Vpa-159-1 | Bacteria Proteobacteria Gammaproteobacteria Vibrionales Vibrionaceae *Vibrio parahaemolyticus* S118 |
| Vpa-160-1 | Bacteria Proteobacteria Gammaproteobacteria Vibrionales Vibrionaceae *Vibrio parahaemolyticus* S119 |
| Vpa-161-1 | Bacteria Proteobacteria Gammaproteobacteria Vibrionales Vibrionaceae *Vibrio parahaemolyticus* S120 |
| Vpa-162-1 | Bacteria Proteobacteria Gammaproteobacteria Vibrionales Vibrionaceae *Vibrio parahaemolyticus* S121 |
| Vpa-163-1 | Bacteria Proteobacteria Gammaproteobacteria Vibrionales Vibrionaceae *Vibrio parahaemolyticus* S122 |
| Vpa-164-1 | Bacteria Proteobacteria Gammaproteobacteria Vibrionales Vibrionaceae *Vibrio parahaemolyticus* S123 |
| Vpa-165-1 | Bacteria Proteobacteria Gammaproteobacteria Vibrionales Vibrionaceae *Vibrio parahaemolyticus* S124 |
| Vpa-166-1 | Bacteria Proteobacteria Gammaproteobacteria Vibrionales Vibrionaceae *Vibrio parahaemolyticus* S125 |
| Vpa-167-1 | Bacteria Proteobacteria Gammaproteobacteria Vibrionales Vibrionaceae *Vibrio parahaemolyticus* S126 |
| Vpa-168-1 | Bacteria Proteobacteria Gammaproteobacteria Vibrionales Vibrionaceae *Vibrio parahaemolyticus* S128 |
| Vpa-169-1 | Bacteria Proteobacteria Gammaproteobacteria Vibrionales Vibrionaceae *Vibrio parahaemolyticus* S129 |
| Vpa-170-1 | Bacteria Proteobacteria Gammaproteobacteria Vibrionales Vibrionaceae *Vibrio parahaemolyticus* S130 |
| Vpa-171-1 | Bacteria Proteobacteria Gammaproteobacteria Vibrionales Vibrionaceae *Vibrio parahaemolyticus* S131 |
| Vpa-172-1 | Bacteria Proteobacteria Gammaproteobacteria Vibrionales Vibrionaceae *Vibrio parahaemolyticus* S132 |
| Vpa-173-1 | Bacteria Proteobacteria Gammaproteobacteria Vibrionales Vibrionaceae *Vibrio parahaemolyticus* S133 |
| Vpa-174-1 | Bacteria Proteobacteria Gammaproteobacteria Vibrionales Vibrionaceae *Vibrio parahaemolyticus* S134 |
| Vpa-175-1 | Bacteria Proteobacteria Gammaproteobacteria Vibrionales Vibrionaceae *Vibrio parahaemolyticus* S135 |
| Vpa-176-1 | Bacteria Proteobacteria Gammaproteobacteria Vibrionales Vibrionaceae *Vibrio parahaemolyticus* S136 |
| Vpa-177-1 | Bacteria Proteobacteria Gammaproteobacteria Vibrionales Vibrionaceae *Vibrio parahaemolyticus* S137 |
| Vpa-178-1 | Bacteria Proteobacteria Gammaproteobacteria Vibrionales Vibrionaceae *Vibrio parahaemolyticus* S138 |
| Vpa-179-1 | Bacteria Proteobacteria Gammaproteobacteria Vibrionales Vibrionaceae *Vibrio parahaemolyticus* S139 |
| Vpa-180-1 | Bacteria Proteobacteria Gammaproteobacteria Vibrionales Vibrionaceae *Vibrio parahaemolyticus* S140 |
| Vpa-181-1 | Bacteria Proteobacteria Gammaproteobacteria Vibrionales Vibrionaceae *Vibrio parahaemolyticus* S141 |
| Vpa-182-1 | Bacteria Proteobacteria Gammaproteobacteria Vibrionales Vibrionaceae *Vibrio parahaemolyticus* S142 |
| Vpa-183-1 | Bacteria Proteobacteria Gammaproteobacteria Vibrionales Vibrionaceae *Vibrio parahaemolyticus* S143 |
| Vpa-184-1 | Bacteria Proteobacteria Gammaproteobacteria Vibrionales Vibrionaceae *Vibrio parahaemolyticus* S144 |
| Vpa-185-1 | Bacteria Proteobacteria Gammaproteobacteria Vibrionales Vibrionaceae *Vibrio parahaemolyticus* S145 |
| Vpa-186-1 | Bacteria Proteobacteria Gammaproteobacteria Vibrionales Vibrionaceae *Vibrio parahaemolyticus* S146 |
| Vpa-187-1 | Bacteria Proteobacteria Gammaproteobacteria Vibrionales Vibrionaceae *Vibrio parahaemolyticus* S147 |
| Vpa-188-1 | Bacteria Proteobacteria Gammaproteobacteria Vibrionales Vibrionaceae *Vibrio parahaemolyticus* S148 |
| Vpa-189-1 | Bacteria Proteobacteria Gammaproteobacteria Vibrionales Vibrionaceae *Vibrio parahaemolyticus* S150 |
| Vpa-190-1 | Bacteria Proteobacteria Gammaproteobacteria Vibrionales Vibrionaceae *Vibrio parahaemolyticus* S151 |
| Vpa-191-1 | Bacteria Proteobacteria Gammaproteobacteria Vibrionales Vibrionaceae *Vibrio parahaemolyticus* S152 |
| Vpa-192-1 | Bacteria Proteobacteria Gammaproteobacteria Vibrionales Vibrionaceae *Vibrio parahaemolyticus* S153 |
| Vpa-193-1 | Bacteria Proteobacteria Gammaproteobacteria Vibrionales Vibrionaceae *Vibrio parahaemolyticus* S154 |
| Vpa-194-1 | Bacteria Proteobacteria Gammaproteobacteria Vibrionales Vibrionaceae *Vibrio parahaemolyticus* S155 |

| | |
|---|---|
| Vpa-195-1 | Bacteria Proteobacteria Gammaproteobacteria Vibrionales Vibrionaceae *Vibrio parahaemolyticus* S156 |
| Vpa-196-1 | Bacteria Proteobacteria Gammaproteobacteria Vibrionales Vibrionaceae *Vibrio parahaemolyticus* S157 |
| Vpa-197-1 | Bacteria Proteobacteria Gammaproteobacteria Vibrionales Vibrionaceae *Vibrio parahaemolyticus* S158 |
| Vpa-198-1 | Bacteria Proteobacteria Gammaproteobacteria Vibrionales Vibrionaceae *Vibrio parahaemolyticus* S159 |
| Vpa-199-1 | Bacteria Proteobacteria Gammaproteobacteria Vibrionales Vibrionaceae *Vibrio parahaemolyticus* S160 |
| Vpa-200-1 | Bacteria Proteobacteria Gammaproteobacteria Vibrionales Vibrionaceae *Vibrio parahaemolyticus* S161 |
| Vpa-201-1 | Bacteria Proteobacteria Gammaproteobacteria Vibrionales Vibrionaceae *Vibrio parahaemolyticus* S162 |
| Vpa-202-1 | Bacteria Proteobacteria Gammaproteobacteria Vibrionales Vibrionaceae *Vibrio parahaemolyticus* S163 |
| Vpa-203-1 | Bacteria Proteobacteria Gammaproteobacteria Vibrionales Vibrionaceae *Vibrio parahaemolyticus* S164 |
| Vpa-204-1 | Bacteria Proteobacteria Gammaproteobacteria Vibrionales Vibrionaceae *Vibrio parahaemolyticus* S165 |
| Vpa-205-1 | Bacteria Proteobacteria Gammaproteobacteria Vibrionales Vibrionaceae *Vibrio parahaemolyticus* S166 |
| Vpa-206-1 | Bacteria Proteobacteria Gammaproteobacteria Vibrionales Vibrionaceae *Vibrio parahaemolyticus* S167 |
| Vpa-207-1 | Bacteria Proteobacteria Gammaproteobacteria Vibrionales Vibrionaceae *Vibrio parahaemolyticus* S168 |
| Vpa-208-1 | Bacteria Proteobacteria Gammaproteobacteria Vibrionales Vibrionaceae *Vibrio parahaemolyticus* S170 |
| Vpa-209-1 | Bacteria Proteobacteria Gammaproteobacteria Vibrionales Vibrionaceae *Vibrio parahaemolyticus* S171 |
| Vpa-210-1 | Bacteria Proteobacteria Gammaproteobacteria Vibrionales Vibrionaceae *Vibrio parahaemolyticus* S172 |
| Vpa-211-1 | Bacteria Proteobacteria Gammaproteobacteria Vibrionales Vibrionaceae *Vibrio parahaemolyticus* S173 |
| Vpa-212-1 | Bacteria Proteobacteria Gammaproteobacteria Vibrionales Vibrionaceae *Vibrio parahaemolyticus* S174 |
| Vpa-213-1 | Bacteria Proteobacteria Gammaproteobacteria Vibrionales Vibrionaceae *Vibrio parahaemolyticus* SBR10290 |
| Vpa-214-1 | Bacteria Proteobacteria Gammaproteobacteria Vibrionales Vibrionaceae *Vibrio parahaemolyticus* SNUVpS-1 |
| Vpa-215-1 | Bacteria Proteobacteria Gammaproteobacteria Vibrionales Vibrionaceae *Vibrio parahaemolyticus* TUMSAT_D06_S3 |
| Vpa-216-1 | Bacteria Proteobacteria Gammaproteobacteria Vibrionales Vibrionaceae *Vibrio parahaemolyticus* TUMSAT_DE1_S1 |
| Vpa-217-1 | Bacteria Proteobacteria Gammaproteobacteria Vibrionales Vibrionaceae *Vibrio parahaemolyticus* TUMSAT_DE2_S2 |
| Vpa-218-1 | Bacteria Proteobacteria Gammaproteobacteria Vibrionales Vibrionaceae *Vibrio parahaemolyticus* TUMSAT_H01_S4 |
| Vpa-219-1 | Bacteria Proteobacteria Gammaproteobacteria Vibrionales Vibrionaceae *Vibrio parahaemolyticus* TUMSAT_H03_S5 |
| Vpa-220-1 | Bacteria Proteobacteria Gammaproteobacteria Vibrionales Vibrionaceae *Vibrio parahaemolyticus* TUMSAT_H10_S6 |
| Vpa-221-1 | Bacteria Proteobacteria Gammaproteobacteria Vibrionales Vibrionaceae *Vibrio parahaemolyticus* UCM-V493 |
| Vpa-244-1 | Bacteria Proteobacteria Gammaproteobacteria Vibrionales Vibrionaceae *Vibrio parahaemolyticus v*110 |
| Vpa-222-1 | Bacteria Proteobacteria Gammaproteobacteria Vibrionales Vibrionaceae *Vibrio parahaemolyticus* V14/01 |
| Vpa-223-1 | Bacteria Proteobacteria Gammaproteobacteria Vibrionales Vibrionaceae *Vibrio parahaemolyticus* VIP4-0219 |
| VPa-1-1 | Bacteria Proteobacteria Gammaproteobacteria Vibrionales Vibrionaceae *Vibrio* Parahaemolyticus VIP4-0395 |
| Vpa-224-1 | Bacteria Proteobacteria Gammaproteobacteria Vibrionales Vibrionaceae *Vibrio parahaemolyticus* VIP4-0407 |
| Vpa-225-1 | Bacteria Proteobacteria Gammaproteobacteria Vibrionales Vibrionaceae *Vibrio parahaemolyticus* VIP4-0430 |
| Vpa-226-1 | Bacteria Proteobacteria Gammaproteobacteria Vibrionales Vibrionaceae *Vibrio parahaemolyticus* VIP4-0434 |
| VPa-2-1 | Bacteria Proteobacteria Gammaproteobacteria Vibrionales Vibrionaceae *Vibrio* Parahaemolyticus VIP4-0439 |
| Vpa-227-1 | Bacteria Proteobacteria Gammaproteobacteria Vibrionales Vibrionaceae *Vibrio parahaemolyticus* VIP4-0443 |
| Vpa-228-1 | Bacteria Proteobacteria Gammaproteobacteria Vibrionales Vibrionaceae *Vibrio parahaemolyticus* VIP4-0444 |
| Vpa-229-1 | Bacteria Proteobacteria Gammaproteobacteria Vibrionales Vibrionaceae *Vibrio parahaemolyticus* VIP4-0445 |
| Vpa-230-1 | Bacteria Proteobacteria Gammaproteobacteria Vibrionales Vibrionaceae *Vibrio parahaemolyticus* VIP4-0447 |
| Vpa-231-1 | Bacteria Proteobacteria Gammaproteobacteria Vibrionales Vibrionaceae *Vibrio parahaemolyticus* VP-48 |
| Vpa-232-1 | Bacteria Proteobacteria Gammaproteobacteria Vibrionales Vibrionaceae *Vibrio parahaemolyticus* VP-NY4 |
| Vpa-233-1 | Bacteria Proteobacteria Gammaproteobacteria Vibrionales Vibrionaceae *Vibrio parahaemolyticus* VP2007-007 |
| Vpa-234-1 | Bacteria Proteobacteria Gammaproteobacteria Vibrionales Vibrionaceae *Vibrio parahaemolyticus* VP2007-095 |
| Vpa-235-1 | Bacteria Proteobacteria Gammaproteobacteria Vibrionales Vibrionaceae *Vibrio parahaemolyticus* VP232 |
| Vpa-236-1 | Bacteria Proteobacteria Gammaproteobacteria Vibrionales Vibrionaceae *Vibrio parahaemolyticus* VP250 |
| Vpa-237-1 | Bacteria Proteobacteria Gammaproteobacteria Vibrionales Vibrionaceae *Vibrio parahaemolyticus* VP551 |
| Vpa-238-1 | Bacteria Proteobacteria Gammaproteobacteria Vibrionales Vibrionaceae *Vibrio parahaemolyticus* VP766 |
| Vpa-239-1 | Bacteria Proteobacteria Gammaproteobacteria Vibrionales Vibrionaceae *Vibrio parahaemolyticus* VPCR-2009 |
| Vpa-240-1 | Bacteria Proteobacteria Gammaproteobacteria Vibrionales Vibrionaceae *Vibrio parahaemolyticus* VPCR-2010 |
| Vpa-241-1 | Bacteria Proteobacteria Gammaproteobacteria Vibrionales Vibrionaceae *Vibrio parahaemolyticus* VPTS-2009 |
| Vpa-242-1 | Bacteria Proteobacteria Gammaproteobacteria Vibrionales Vibrionaceae *Vibrio parahaemolyticus* VPTS-2010 |
| Vpa-243-1 | Bacteria Proteobacteria Gammaproteobacteria Vibrionales Vibrionaceae *Vibrio parahaemolyticus* VPTS-2010_2 |
| Vpr-1-1 | Bacteria Proteobacteria Gammaproteobacteria Vibrionales Vibrionaceae *Vibrio proteolyticus* NBRC 13287 |
| Vre-1-1 to Vre-1-2 | Bacteria Proteobacteria Gammaproteobacteria Vibrionales Vibrionaceae *Vibrio renipiscarius* |
| Vrh-1-1 | Bacteria Proteobacteria Gammaproteobacteria Vibrionales Vibrionaceae *Vibrio rhizosphaerae* DSM 18581 |
| Vro-1-1 | Bacteria Proteobacteria Gammaproteobacteria Vibrionales Vibrionaceae *Vibrio rotiferianus* |
| Vro-2-1 | Bacteria Proteobacteria Gammaproteobacteria Vibrionales Vibrionaceae *Vibrio rotiferianus* CAIM 577 = LMG 21460 |
| Vro-3-1 | Bacteria Proteobacteria Gammaproteobacteria Vibrionales Vibrionaceae *Vibrio rotiferianus* DAT722 |
| Vru-1-1 | Bacteria Proteobacteria Gammaproteobacteria Vibrionales Vibrionaceae *Vibrio rumoiensis* 1S-45 |
| Vsa-1-1 | Bacteria Proteobacteria Gammaproteobacteria Vibrionales Vibrionaceae *Vibrio sagamiensis* NBRC 104589 |
| Vsc-1-1 | Bacteria Proteobacteria Gammaproteobacteria Vibrionales Vibrionaceae *Vibrio scophthalmi* LMG 19158 |
| Vsh-1-1 | Bacteria Proteobacteria Gammaproteobacteria Vibrionales Vibrionaceae *Vibrio shiloi* A203 |
| Vsh-2-1 | Bacteria Proteobacteria Gammaproteobacteria Vibrionales Vibrionaceae *Vibrio shilonii* AK1 |
| Vsi-2-1 | Bacteria Proteobacteria Gammaproteobacteria Vibrionales Vibrionaceae *Vibrio sinaloensis* DSM 21326 |
| Vsp-1-1 | Bacteria Proteobacteria Gammaproteobacteria Vibrionales Vibrionaceae *Vibrio sp.* 712i1 |

| | |
|---|---|
| Vsp-2-1 | Bacteria Proteobacteria Gammaproteobacteria Vibrionales Vibrionaceae *Vibrio sp.* AND4 |
| Vsp-3-1 | Bacteria Proteobacteria Gammaproteobacteria Vibrionales Vibrionaceae *Vibrio sp.* B183 |
| Vsp-4-1 | Bacteria Proteobacteria Gammaproteobacteria Vibrionales Vibrionaceae *Vibrio sp.* ECSMB14105 |
| Vsp-5-1 | Bacteria Proteobacteria Gammaproteobacteria Vibrionales Vibrionaceae *Vibrio sp.* ECSMB14106 |
| Vsp-6-1 | Bacteria Proteobacteria Gammaproteobacteria Vibrionales Vibrionaceae *Vibrio sp.* EJY3 |
| Vsp-7-1 | Bacteria Proteobacteria Gammaproteobacteria Vibrionales Vibrionaceae *Vibrio sp.* ER1A |
| Vsp-8-1 | Bacteria Proteobacteria Gammaproteobacteria Vibrionales Vibrionaceae *Vibrio sp.* Ex25 |
| Vsp-9-1 | Bacteria Proteobacteria Gammaproteobacteria Vibrionales Vibrionaceae *Vibrio sp.* HENC-01 |
| Vsp-10-1 | Bacteria Proteobacteria Gammaproteobacteria Vibrionales Vibrionaceae *Vibrio sp.* HENC-03 |
| Vsp-11-1 | Bacteria Proteobacteria Gammaproteobacteria Vibrionales Vibrionaceae *Vibrio sp.* J2-1 |
| Vsp-12-1 | Bacteria Proteobacteria Gammaproteobacteria Vibrionales Vibrionaceae *Vibrio sp.* J2-12 |
| Vsp-13-1 | Bacteria Proteobacteria Gammaproteobacteria Vibrionales Vibrionaceae *Vibrio sp.* J2-15 |
| Vsp-14-1 | Bacteria Proteobacteria Gammaproteobacteria Vibrionales Vibrionaceae *Vibrio sp.* J2-17 |
| Vsp-15-1 | Bacteria Proteobacteria Gammaproteobacteria Vibrionales Vibrionaceae *Vibrio sp.* J2-26 |
| Vsp-16-1 | Bacteria Proteobacteria Gammaproteobacteria Vibrionales Vibrionaceae *Vibrio sp.* J2-29 |
| Vsp-17-1 | Bacteria Proteobacteria Gammaproteobacteria Vibrionales Vibrionaceae *Vibrio sp.* J2-3 |
| Vsp-18-1 | Bacteria Proteobacteria Gammaproteobacteria Vibrionales Vibrionaceae *Vibrio sp.* J2-31 |
| Vsp-19-1 | Bacteria Proteobacteria Gammaproteobacteria Vibrionales Vibrionaceae *Vibrio sp.* J2-4 |
| Vsp-20-1 | Bacteria Proteobacteria Gammaproteobacteria Vibrionales Vibrionaceae *Vibrio sp.* J2-6 |
| Vsp-21-1 | Bacteria Proteobacteria Gammaproteobacteria Vibrionales Vibrionaceae *Vibrio sp.* J2-8 |
| Vsp-22-1 | Bacteria Proteobacteria Gammaproteobacteria Vibrionales Vibrionaceae *Vibrio sp.* MED222 |
| Vsp-23-1 | Bacteria Proteobacteria Gammaproteobacteria Vibrionales Vibrionaceae *Vibrio sp.* N418 |
| Vsp-24-1 | Bacteria Proteobacteria Gammaproteobacteria Vibrionales Vibrionaceae *Vibrio sp.* OY15 |
| Vsp-25-1 | Bacteria Proteobacteria Gammaproteobacteria Vibrionales Vibrionaceae *Vibrio sp.* RC341 |
| Vsp-26-1 | Bacteria Proteobacteria Gammaproteobacteria Vibrionales Vibrionaceae *Vibrio sp.* RC586 |
| Vsp-27-1 | Bacteria Proteobacteria Gammaproteobacteria Vibrionales Vibrionaceae *Vibrio sp.* S234-5 |
| Vsp-28-1 | Bacteria Proteobacteria Gammaproteobacteria Vibrionales Vibrionaceae *Vibrio sp.* S457-15 |
| Vsp-29-1 | Bacteria Proteobacteria Gammaproteobacteria Vibrionales Vibrionaceae *Vibrio sp.* S512-13 |
| Vsp-31-1 | Bacteria Proteobacteria Gammaproteobacteria Vibrionales Vibrionaceae *Vibrio sp.* Vb278 |
| Vsp-30-1 | Bacteria Proteobacteria Gammaproteobacteria Vibrionales Vibrionaceae *Vibrio sp.* VPAP30 |
| Vsp-32-1 | Bacteria Proteobacteria Gammaproteobacteria Vibrionales Vibrionaceae *Vibrio sp.* ZOR0018 |
| Vsp-33-1 | Bacteria Proteobacteria Gammaproteobacteria Vibrionales Vibrionaceae *Vibrio splendidus* 0407ZC148 |
| Vsp-34-1 | Bacteria Proteobacteria Gammaproteobacteria Vibrionales Vibrionaceae *Vibrio splendidus* 12B01 |
| Vsp-35-1 | Bacteria Proteobacteria Gammaproteobacteria Vibrionales Vibrionaceae *Vibrio splendidus* 12E03 |
| Vsp-36-1 | Bacteria Proteobacteria Gammaproteobacteria Vibrionales Vibrionaceae *Vibrio splendidus* 12F01 |
| Vsp-37-1 | Bacteria Proteobacteria Gammaproteobacteria Vibrionales Vibrionaceae *Vibrio splendidus* 1F-157 |
| Vsp-38-1 | Bacteria Proteobacteria Gammaproteobacteria Vibrionales Vibrionaceae *Vibrio splendidus* 1S-124 |
| Vsp-39-1 | Bacteria Proteobacteria Gammaproteobacteria Vibrionales Vibrionaceae *Vibrio splendidus* 5S-101 |
| Vsp-40-1 | Bacteria Proteobacteria Gammaproteobacteria Vibrionales Vibrionaceae *Vibrio splendidus* ATCC 33789 |
| Vsp-41-1 | Bacteria Proteobacteria Gammaproteobacteria Vibrionales Vibrionaceae *Vibrio splendidus* FF-500 |
| Vsp-42-1 | Bacteria Proteobacteria Gammaproteobacteria Vibrionales Vibrionaceae *Vibrio splendidus* FF-6 |
| Vsp-43-1 | Bacteria Proteobacteria Gammaproteobacteria Vibrionales Vibrionaceae *Vibrio splendidus* LGP32 |
| Vsp-44-1 | Bacteria Proteobacteria Gammaproteobacteria Vibrionales Vibrionaceae *Vibrio splendidus* ZF-90 |
| Vsp-45-1 | Bacteria Proteobacteria Gammaproteobacteria Vibrionales Vibrionaceae *Vibrio splendidus* ZS-139 |
| Vta-1-1 | Bacteria Proteobacteria Gammaproteobacteria Vibrionales Vibrionaceae *Vibrio tasmaniensis* 1F-155 |
| Vta-2-1 | Bacteria Proteobacteria Gammaproteobacteria Vibrionales Vibrionaceae *Vibrio tasmaniensis* 1F-187 |
| Vta-3-1 | Bacteria Proteobacteria Gammaproteobacteria Vibrionales Vibrionaceae *Vibrio tasmaniensis* 1F-267 |
| Vtu-2-1 to Vtu-2-2 | Bacteria Proteobacteria Gammaproteobacteria Vibrionales Vibrionaceae *Vibrio tubiashii* ATCC 19109 |
| Vtu-3-1 | Bacteria Proteobacteria Gammaproteobacteria Vibrionales Vibrionaceae *Vibrio tubiashii* NCIMB 1337 = ATCC 19106 |
| Vva-1-1 | Bacteria Proteobacteria Gammaproteobacteria Vibrionales Vibrionaceae *Vibrio variabilis* |
| Vvu-1-1 | Bacteria Proteobacteria Gammaproteobacteria Vibrionales Vibrionaceae *Vibrio vulnificus* |
| Vvu-2-1 | Bacteria Proteobacteria Gammaproteobacteria Vibrionales Vibrionaceae *Vibrio vulnificus* B2 |
| Vvu-3-1 | Bacteria Proteobacteria Gammaproteobacteria Vibrionales Vibrionaceae *Vibrio vulnificus* BAA87 |
| Vvu-5-1 | Bacteria Proteobacteria Gammaproteobacteria Vibrionales Vibrionaceae *Vibrio vulnificus* CladeA-yb158 |
| Vvu-4-1 | Bacteria Proteobacteria Gammaproteobacteria Vibrionales Vibrionaceae *Vibrio vulnificus* CMCP6 |
| Vvu-6-1 | Bacteria Proteobacteria Gammaproteobacteria Vibrionales Vibrionaceae *Vibrio vulnificus* E64MW |
| Vvu-7-1 | Bacteria Proteobacteria Gammaproteobacteria Vibrionales Vibrionaceae *Vibrio vulnificus* JY1305 |
| Vvu-8-1 | Bacteria Proteobacteria Gammaproteobacteria Vibrionales Vibrionaceae *Vibrio vulnificus* MO6-24/O |
| Vvu-9-1 | Bacteria Proteobacteria Gammaproteobacteria Vibrionales Vibrionaceae *Vibrio vulnificus* NBRC 15645 = ATCC 27562 |
| Vvu-10-1 | Bacteria Proteobacteria Gammaproteobacteria Vibrionales Vibrionaceae *Vibrio vulnificus* VVyb1(BT3) |
| Vvu-11-1 | Bacteria Proteobacteria Gammaproteobacteria Vibrionales Vibrionaceae *Vibrio vulnificus* YJ016 |
| Sen-1026-1 | CDC07-0191Bacteria Proteobacteria Gammaproteobacteria Enterobacteriales Enterobacteriaceae Salmonella Salmonella enterica subsp. entericaserovar Tennessee *Salmonella enterica subsp. enterica serovar* Tennessee str |

| | |
|---|---|
| Sen-1027-1 to Sen-1027-2 | CVM19633Bacteria Proteobacteria Gammaproteobacteria Enterobacteriales Enterobacteriaceae *Salmonella enterica subsp. enterica serovar* Schwarzengrund str |
| Sen-1028-1 | CVM23701Bacteria Proteobacteria Gammaproteobacteria Enterobacteriales Enterobacteriaceae Salmonella Salmonella enterica subsp. entericaserovar 4,[5],12:i:- *Salmonella enterica subsp. enterica serovar* 4,[5],12:i:- str |
| Sen-1029-1 | GA_MM04042433Bacteria Proteobacteria Gammaproteobacteria Enterobacteriales Enterobacteriaceae *Salmonella enterica subsp. enterica serovar* Javiana str |
| Sen-1030-1 | HI_N05-537Bacteria Proteobacteria Gammaproteobacteria Enterobacteriales Enterobacteriaceae Salmonella Salmonella enterica subsp. entericaserovar Weltevreden *Salmonella enterica subsp. enterica serovar* Weltevreden str |
| Sen-1031-1 | P125109Bacteria Proteobacteria Gammaproteobacteria Enterobacteriales Enterobacteriaceae Salmonella Salmonella enterica subsp. entericaserovar Enteritidis *Salmonella enterica subsp. enterica serovar* Enteritidis str |
| Sen-1032-1 | RKS4594Bacteria Proteobacteria Gammaproteobacteria Enterobacteriales Enterobacteriaceae Salmonella Salmonella enterica subsp. entericaserovar Paratyphi C *Salmonella enterica subsp. enterica serovar* Paratyphi C strain |
| Sen-1033-1 | SC-B67Bacteria Proteobacteria Gammaproteobacteria Enterobacteriales Enterobacteriaceae Salmonella Salmonella enterica subsp. entericaserovar Choleraesuis *Salmonella enterica subsp. enterica serovar* Choleraesuis str |
| Sen-1027-1 to Sen-1027-2 | SL480Bacteria Proteobacteria Gammaproteobacteria Enterobacteriales Enterobacteriaceae *Salmonella enterica subsp. enterica serovar* Schwarzengrund str |
| env-1 to env-1113 | environmental samples |

#### 12.2 Gene contexts

Each L31-Gammaproteobacteria RNA (indicated by "RNA→") is listed. For each hit, the downstream genes predicted to reside in a regulated operon are listed. If the nearest downstream gene is encoding in the opposite strand (and therefore presumed to not be a part of a regulated operon), then that gene is still depicted. Some environmental sequences and some RefSeq entries lack gene annotations, and so no genes are available for such sequences. The direction of each gene is indicated with an arrow (→), and each predicted conserved domain in the gene is named. Conserved domains associated with more than one L31-Gammaproteobacteria RNA are assigned a color; other domains are gray. Information about these conserved domains is given in Section 12.3. The accession of the sequence containing each L31-Gammaproteobacteria RNA is given in the column named "Seq. accession". Accessions beginning with "NC_", "NS_", "NW_" or "NZ_" are contained in RefSeq. Other accession refer to environmental samples. Nucleotide coordinates are given for the 5′ and 3′ boundaries of each L31-Gammaproteobacteria RNA. If the 5′ coordinate is greater than the 3′ coordinate, the RNA is present on the reverse-complement strand of the containing genomic DNA sequence. Each hit is denoted by an abbreviation (like "Eco-1-1") that refers to a taxonomy given in Section 12.1.

| abbrev. | Seq. accession | | 5′ at | 3′ at | genes |
|---|---|---|---|---|---|
| CBa-1-1 | NZ_CP008985.1 | - | 204157 | 204106 | **RNA**→ rpmE (PRK00019)Ribosomal_L31 (pfam01197)→ |
| Apl-1-1 | NZ_JSVY01000012.1 | - | 77506 | 77438 | **RNA**→ rpmE (PRK00019)Ribosomal_L31 (pfam01197)→ |
| Apl-1-2 | NZ_JSVG01000014.1 | - | 77621 | 77553 | **RNA**→ rpmE (PRK00019)Ribosomal_L31 (pfam01197)→ |
| Apl-1-3 | NZ_JSVV01000016.1 | - | 77506 | 77438 | **RNA**→ rpmE (PRK00019)Ribosomal_L31 (pfam01197)→ |
| Apl-1-4 | NZ_JSVF01000014.1 | - | 77502 | 77434 | **RNA**→ rpmE (PRK00019)Ribosomal_L31 (pfam01197)→ |
| Apl-1-5 | NZ_JSVZ01000012.1 | - | 77500 | 77432 | **RNA**→ rpmE (PRK00019)Ribosomal_L31 (pfam01197)→ |
| Apl-1-6 | NZ_JSVX01000015.1 | - | 77504 | 77436 | **RNA**→ rpmE (PRK00019)Ribosomal_L31 (pfam01197)→ |
| Apl-7-1 | NZ_ADOL01000025.1 | - | 39997 | 39929 | **RNA**→ rpmE (PRK00019)Ribosomal_L31 (pfam01197)→ rpmE (PRK00019)Ribosomal_L31 (pfam01197)→ rpmE (PRK00019)Ribosomal_L31 (pfam01197)→ |
| Apl-9-1 | NZ_ADXN01000034.1 | - | 46762 | 46694 | **RNA**→ rpmE (PRK00019)Ribosomal_L31 (pfam01197)→ rpmE (PRK00019)Ribosomal_L31 (pfam01197)→ |
| Apl-12-1 | NZ_ADOF01000042.1 | - | 41201 | 41133 | **RNA**→ rpmE (PRK00019)Ribosomal_L31 (pfam01197)→ rpmE (PRK00019)Ribosomal_L31 (pfam01197)→ rpmE (PRK00019)Ribosomal_L31 (pfam01197)→ |
| Apl-10-1 | NZ_ADOE01000024.1 | - | 29041 | 28973 | **RNA**→ rpmE (PRK00019)Ribosomal_L31 (pfam01197)→ rpmE (PRK00019)Ribosomal_L31 (pfam01197)→ |
| Apl-5-1 | NZ_ADOJ01000013.1 | - | 61583 | 61515 | **RNA**→ rpmE (PRK00019)Ribosomal_L31 (pfam01197)→ rpmE (PRK00019)Ribosomal_L31 (pfam01197)→ rpmE (PRK00019)Ribosomal_L31 (pfam01197)→ |
| Apl-4-1 | NZ_AACK01000007.1 | + | 70556 | 70624 | **RNA**→ rpmE (PRK00019)Ribosomal_L31 (pfam01197)→ rpmE (PRK00019)Ribosomal_L31 (pfam01197)→ |
| Apl-2-1 | NC_009053.1 | - | 1135896 | 1135828 | **RNA**→ rpmE (PRK00019)Ribosomal_L31 (pfam01197)→ |
| Apl-11-1 | NC_010278.1 | - | 1103179 | 1103111 | **RNA**→ rpmE (PRK00019)Ribosomal_L31 (pfam01197)→ |
| Apl-6-1 | NZ_ADOK01000015.1 | - | 71511 | 71443 | **RNA**→ rpmE (PRK00019)Ribosomal_L31 (pfam01197)→ rpmE (PRK00019)Ribosomal_L31 (pfam01197)→ |

| | | | | | |
|---|---|---|---|---|---|
| Apl-15-1 | NZ_ADOI01000029.1 | - | 71436 | 71368 | **RNA**→ rpmE (PRK00019)Ribosomal_L31 (pfam01197)→ rpmE (PRK00019)Ribosomal_L31 (pfam01197)→ |
| Apl-4-2 | NZ_ADOD01000015.1 | - | 71510 | 71442 | **RNA**→ rpmE (PRK00019)Ribosomal_L31 (pfam01197)→ rpmE (PRK00019)Ribosomal_L31 (pfam01197)→ |
| | | | | | Glyco_transf_4 (pfam13439)Glyco_trans_1_4 (pfam13692)GT1_WabH_like (cd03811)→ |
| | | | | | Glyco_transf_4 (pfam13439)Glyco_trans_1_4 (pfam13692)GT1_WabH_like (cd03811)→ |
| | | | | | RfaB (COG0438)Glyco_trans_1_4 (pfam13692)→ RfaB (COG0438)Glyco_trans_1_4 (pfam13692)→ |
| | | | | | Glyco_transf_25 (pfam01755)→ Glyco_transf_25 (pfam01755)→ heptsyl_trn_III (TIGR02201)RfaF (COG0859)- |
| | | | | | heptsyl_trn_III (TIGR02201)RfaF (COG0859)→ |
| | | | | | PBP2_IlvY (cd08430)PRK11716 (PRK11716)HTH_1 (pfam00126)→ |
| | | | | | PBP2_IlvY (cd08430)PRK11716 (PRK11716)HTH_1 (pfam00126)→ |
| Apl-13-1 | NZ_ADXO01000023.1 | + | 90529 | 90597 | **RNA**→ rpmE (PRK00019)Ribosomal_L31 (pfam01197)→ rpmE (PRK00019)Ribosomal_L31 (pfam01197)→ |
| | | | | | Glyco_transf_4 (pfam13439)Glyco_trans_1_4 (pfam13692)GT1_WabH_like (cd03811)→ |
| | | | | | Glyco_transf_4 (pfam13439)Glyco_trans_1_4 (pfam13692)GT1_WabH_like (cd03811)→ |
| | | | | | RfaB (COG0438)Glyco_trans_1_4 (pfam13692)→ RfaB (COG0438)Glyco_trans_1_4 (pfam13692)→ |
| | | | | | Glyco_transf_25 (pfam01755)→ Glyco_transf_25 (pfam01755)→ heptsyl_trn_III (TIGR02201)RfaF (COG0859)- |
| | | | | | heptsyl_trn_III (TIGR02201)RfaF (COG0859)→ |
| | | | | | PBP2_IlvY (cd08430)PRK11716 (PRK11716)HTH_1 (pfam00126)→ |
| | | | | | PBP2_IlvY (cd08430)PRK11716 (PRK11716)HTH_1 (pfam00126)→ |
| Apl-13-2 | NZ_ADOG01000014.1 | - | 47926 | 47858 | **RNA**→ rpmE (PRK00019)Ribosomal_L31 (pfam01197)→ rpmE (PRK00019)Ribosomal_L31 (pfam01197)→ |
| | | | | | Glyco_transf_4 (pfam13439)Glyco_trans_1_4 (pfam13692)GT1_WabH_like (cd03811)→ |
| | | | | | Glyco_transf_4 (pfam13439)Glyco_trans_1_4 (pfam13692)GT1_WabH_like (cd03811)→ |
| | | | | | RfaB (COG0438)Glyco_trans_1_4 (pfam13692)→ RfaB (COG0438)Glyco_trans_1_4 (pfam13692)→ |
| | | | | | Glyco_transf_25 (pfam01755)→ Glyco_transf_25 (pfam01755)→ heptsyl_trn_III (TIGR02201)RfaF (COG0859)- |
| | | | | | heptsyl_trn_III (TIGR02201)RfaF (COG0859)→ |
| | | | | | PBP2_IlvY (cd08430)PRK11716 (PRK11716)HTH_1 (pfam00126)→ |
| | | | | | PBP2_IlvY (cd08430)PRK11716 (PRK11716)HTH_1 (pfam00126)→ |
| Apl-8-1 | NZ_ADOM01000021.1 | - | 33413 | 33345 | **RNA**→ rpmE (PRK00019)Ribosomal_L31 (pfam01197)→ rpmE (PRK00019)Ribosomal_L31 (pfam01197)→ |
| | | | | | rpmE (PRK00019)Ribosomal_L31 (pfam01197)→ |
| Apl-3-1 | NZ_ALYN01000041.1 | + | 3861 | 3929 | **RNA**→ rpmE (PRK00019)Ribosomal_L31 (pfam01197)→ |
| Apl-14-1 | NC_010939.1 | - | 1164688 | 1164620 | **RNA**→ rpmE (PRK00019)Ribosomal_L31 (pfam01197)→ |
| Aur-1-1 | NZ_GL831080.1 | - | 1453322 | 1453254 | **RNA**→ rpmE (PRK00019)Ribosomal_L31 (pfam01197)→ COG3306 (COG3306)→ |
| Aca-3-1 | NZ_KB893948.1 | - | 16328 | 16260 | **RNA**→ rpmE (PRK00019)Ribosomal_L31 (pfam01197)→ |
| Aeq-1-1 | NZ_CP007715.1 | + | 402750 | 402818 | **RNA**→ rpmE (PRK00019)Ribosomal_L31 (pfam01197)→ PRK11560 (PRK11560)OpgE (COG2194)→ |
| | | | | | PRK11560 (PRK11560)OpgE (COG2194)→ PBP2_IlvY (cd08430)PRK11716 (PRK11716)HTH_1 (pfam00126) |
| Asu-3-1 | NC_018690.1 | + | 390402 | 390470 | **RNA**→ rpmE (PRK00019)Ribosomal_L31 (pfam01197)→ |
| Asu-2-1 | NZ_CP009159.1 | + | 393106 | 393174 | **RNA**→ rpmE (PRK00019)Ribosomal_L31 (pfam01197)→ |
| Mgr-1-1 | NZ_KK211212.1 | + | 396749 | 396813 | **RNA**→ rpmE (PRK00019)Ribosomal_L31 (pfam01197)→ NT_Rel-Spo_like (cd05399)RelA_SpoT (smart00954) |
| | | | | | Glyco_trans_1_4 (pfam13692)GT1_WabH_like (cd03811)→ RfaB (COG0438)Glyco_trans_1_4 (pfam13692)→ |
| | | | | | COG3306 (COG3306)→ Glyco_transf_25 (pfam01755)→ hypo→ RfaF (COG0859)heptsyl_trn_I (TIGR02193)- |
| | | | | | heptsyl_trn_III (TIGR02201)RfaF (COG0859)→ |
| | | | | | PBP2_IlvY (cd08430)PRK11716 (PRK11716)HTH_1 (pfam00126)→ |
| Mha-1-1 | NZ_JANJ01000001.1 | - | 79621 | 79557 | **RNA**→ rpmE (PRK00019)Ribosomal_L31 (pfam01197)→ hypo→ hypo→ |
| | | | | | Glyco_trans_1_4 (pfam13692)GT1_WabH_like (cd03811)→ RfaB (COG0438)Glyco_trans_1_4 (pfam13692)→ |
| | | | | | Glyco_transf_25 (pfam01755)→ hypo→ RfaF (COG0859)heptsyl_trn_I (TIGR02193)→ |
| | | | | | heptsyl_trn_III (TIGR02201)RfaF (COG0859)→ |
| | | | | | PBP2_IlvY (cd08430)PRK11716 (PRK11716)HTH_1 (pfam00126)→ |
| Ami-1-1 | NZ_ACFT01000084.1 | - | 10282 | 10218 | **RNA**→ rpmE (PRK00019)Ribosomal_L31 (pfam01197)→ rpmE (PRK00019)Ribosomal_L31 (pfam01197)→ |
| env-1 | SRS015985_C1526570 | + | 287 | 362 | **RNA**→ rpmE (PRK00528)rpmE (PRK00019)rpmE2 (PRK01678)Ribosomal_L31 (pfam01197)→ |
| env-2 | SRS023930_C2176577 | + | 466 | 541 | **RNA**→ rpmE (PRK00019)Ribosomal_L31 (pfam01197)→ |
| env-3 | SRS018157_C4602690 | - | 233 | 158 | **RNA**→ rpmE (PRK00019)Ribosomal_L31 (pfam01197)→ |
| env-4 | SRS047634_C4407474 | - | 233 | 158 | **RNA**→ rpmE (PRK00019)Ribosomal_L31 (pfam01197)→ |
| env-5 | SRS054569_C1471628 | - | 107 | 32 | **RNA**→ ←hypo |
| env-6 | SRS054776_C1595061 | + | 330 | 405 | **RNA**→ rpmE (PRK00019)Ribosomal_L31 (pfam01197)→ |
| env-7 | SRS057692_C2665711 | - | 233 | 158 | **RNA**→ rpmE (PRK00019)Ribosomal_L31 (pfam01197)→ |
| Hpa-7-1 | NZ_AJMU01000049.1 | + | 103223 | 103298 | **RNA**→ rpmE (PRK00019)Ribosomal_L31 (pfam01197)→ rpmE (PRK00019)Ribosomal_L31 (pfam01197)→ |
| | | | | | GT8_A4GalT_like (cd04194)PRK15171 (PRK15171)→ GT8_A4GalT_like (cd04194)PRK15171 (PRK15171)→ |

| | | | | | |
|---|---|---|---|---|---|
| env-8 | SRS016746_Baylor_scaffold_40224 | + | 146 | 221 | **RNA**→ rpmE (PRK00019)Ribosomal_L31 (pfam01197)→ |
| env-9 | SRS054569_LANL_scaffold_25218 | - | 94 | 31 | **RNA**→ |
| env-10 | SRS063288_LANL_scaffold_2530 | + | 1047 | 1110 | **RNA**→ rpmE (PRK00019)Ribosomal_L31 (pfam01197)→ |
| Hsp-3-1 | NZ_AFNK01000053.1 | + | 171491 | 171554 | **RNA**→ rpmE (PRK00019)Ribosomal_L31 (pfam01197)→ rpmE (PRK00019)Ribosomal_L31 (pfam01197)→ |
| Hsp-4-1 | NZ_ALJP01000009.1 | - | 558292 | 558229 | **RNA**→ rpmE (PRK00019)Ribosomal_L31 (pfam01197)→ rpmE (PRK00019)Ribosomal_L31 (pfam01197)→ |
| Ami-2-1 | NZ_ACQL01000101.1 | + | 2353 | 2417 | **RNA**→ rpmE (PRK00019)Ribosomal_L31 (pfam01197)→ rpmE (PRK00019)Ribosomal_L31 (pfam01197)→ |
| | | | | | hypo→ hypo→ heptsyl_trn_III (TIGR02201)RfaF (COG0859)→ heptsyl_trn_III (TIGR02201)RfaF (COG0859) |
| | | | | | PBP2_IlvY (cd08430)PRK11716 (PRK11716)HTH_1 (pfam00126)→ |
| | | | | | PBP2_IlvY (cd08430)PRK11716 (PRK11716)HTH_1 (pfam00126)→ |
| Lgr-1-1 | NZ_AUUA01000002.1 | - | 220470 | 220418 | **RNA**→ rpmE (PRK00019)Ribosomal_L31 (pfam01197)→ |
| Lgr-1-2 | NZ_JMPN01000025.1 | + | 293044 | 293096 | **RNA**→ rpmE (PRK00019)Ribosomal_L31 (pfam01197)→ |
| Asu-1-1 | NC_009655.1 | - | 577302 | 577244 | **RNA**→ rpmE (PRK00019)Ribosomal_L31 (pfam01197)→ |
| | | | | | RfaB (COG0438)Glyco_transf_4 (pfam13439)Glyco_trans_1_4 (pfam13692)→ |
| env-11 | CXWF01061925.1 | - | 3308 | 3247 | **RNA**→ rpmE (PRK00019)Ribosomal_L31 (pfam01197)→ |
| env-12 | SS_2KS_010_SOIL_10275784 | - | 131 | 83 | **RNA**→ rpmE (PRK00019)Ribosomal_L31 (pfam01197)→ |
| env-13 | SS_3KL_010_SOIL_10001691 | - | 7887 | 7839 | **RNA**→ rpmE (PRK00019)Ribosomal_L31 (pfam01197)→ |
| env-14 | JGI24723J26617_10758127 | - | 170 | 102 | **RNA**→ ←hypo |
| env-15 | CESC01038687.1 | - | 729 | 663 | **RNA**→ rpmE (PRK00019)Ribosomal_L31 (pfam01197)→ |
| env-16 | CEQH01042505.1 | - | 385 | 318 | **RNA**→ rpmE (PRK00019)Ribosomal_L31 (pfam01197)→ |
| env-17 | CETW01364038.1 | + | 629 | 696 | **RNA**→ rpmE (PRK00019)Ribosomal_L31 (pfam01197)→ |
| env-18 | CETX01048977.1 | - | 408 | 341 | **RNA**→ rpmE (PRK00019)Ribosomal_L31 (pfam01197)→ |
| env-19 | CERF01019127.1 | - | 162 | 95 | **RNA**→ rpmE (PRK00019)Ribosomal_L31 (pfam01197)→ |
| env-20 | CEQG01074012.1 | - | 409 | 342 | **RNA**→ ←hypo |
| env-21 | CETV01128562.1 | + | 13281 | 13348 | **RNA**→ rpmE (PRK00019)Ribosomal_L31 (pfam01197)→ |
| env-22 | CEUN01426188.1 | + | 2011 | 2078 | **RNA**→ |
| Msu-1-1 | NZ_LFBU01000002.1 | + | 848270 | 848337 | **RNA**→ rpmE (PRK00019)Ribosomal_L31 (pfam01197)→ |
| env-23 | CEPO01041841.1 | + | 302 | 369 | **RNA**→ rpmE (PRK00019)Ribosomal_L31 (pfam01197)→ |
| env-24 | JCVI_SCAF_1096626511506 | + | 1121 | 1188 | **RNA**→ ←hypo |
| env-25 | CEQG01020404.1 | - | 227 | 160 | **RNA**→ rpmE (PRK00019)Ribosomal_L31 (pfam01197)→ |
| env-26 | CEQI01049741.1 | - | 226 | 159 | **RNA**→ rpmE (PRK00019)Ribosomal_L31 (pfam01197)→ |
| env-27 | CESF01207655.1 | - | 224 | 157 | **RNA**→ rpmE (PRK00019)Ribosomal_L31 (pfam01197)→ |
| env-28 | CESL01139457.1 | - | 5587 | 5520 | **RNA**→ rpmE (PRK00019)Ribosomal_L31 (pfam01197)→ |
| env-29 | CESZ01134189.1 | - | 245 | 178 | **RNA**→ rpmE (PRK00019)Ribosomal_L31 (pfam01197)→ |
| env-30 | CETE01075107.1 | - | 140 | 73 | **RNA**→ rpmE (PRK00019)Ribosomal_L31 (pfam01197)→ |
| env-31 | CETO01087919.1 | - | 2899 | 2832 | **RNA**→ rpmE (PRK00019)Ribosomal_L31 (pfam01197)→ |
| env-32 | CEUD01163365.1 | - | 87 | 20 | **RNA**→ |
| env-33 | CEUK01041327.1 | + | 883 | 950 | **RNA**→ rpmE (PRK00019)Ribosomal_L31 (pfam01197)→ |
| env-34 | CEUN01036455.1 | + | 243 | 310 | **RNA**→ rpmE (PRK00019)Ribosomal_L31 (pfam01197)→ |
| env-35 | CEUO01155056.1 | - | 244 | 177 | **RNA**→ rpmE (PRK00019)Ribosomal_L31 (pfam01197)→ |
| env-36 | CEUP01002300.1 | - | 807 | 740 | **RNA**→ rpmE (PRK00019)Ribosomal_L31 (pfam01197)→ |
| env-37 | CEUY01458679.1 | + | 707 | 774 | **RNA**→ rpmE (PRK00019)Ribosomal_L31 (pfam01197)→ |
| env-38 | CEVA01074234.1 | + | 2215 | 2282 | **RNA**→ rpmE (PRK00019)Ribosomal_L31 (pfam01197)→ |
| env-39 | CEVF01240641.1 | + | 177 | 244 | **RNA**→ rpmE (PRK00019)Ribosomal_L31 (pfam01197)→ |
| Maq-1-1 | NC_008740.1 | + | 934053 | 934120 | **RNA**→ rpmE (PRK00019)Ribosomal_L31 (pfam01197)→ |
| Mhy-1-1 | NC_017067.1 | + | 725999 | 726066 | **RNA**→ rpmE (PRK00019)Ribosomal_L31 (pfam01197)→ |
| env-40 | CEOQ01442252.1 | + | 97 | 164 | **RNA**→ rpmE (PRK00019)Ribosomal_L31 (pfam01197)→ |
| env-41 | CESC01020679.1 | + | 177 | 244 | **RNA**→ rpmE (PRK00019)Ribosomal_L31 (pfam01197)→ |
| env-42 | CESG01154223.1 | + | 103 | 170 | **RNA**→ rpmE (PRK00019)Ribosomal_L31 (pfam01197)→ |
| env-43 | CETV01059046.1 | + | 2077 | 2144 | **RNA**→ |
| env-44 | CETX01171622.1 | - | 76 | 9 | **RNA**→ |
| Msp-3-1 | NZ_AXCB01000001.1 | + | 105151 | 105218 | **RNA**→ rpmE (PRK00019)Ribosomal_L31 (pfam01197)→ |
| Msp-2-1 | NZ_AXCC01000018.1 | + | 77998 | 78065 | **RNA**→ rpmE (PRK00019)Ribosomal_L31 (pfam01197)→ |
| Msp-1-1 | NZ_AXBW01000028.1 | + | 103644 | 103711 | **RNA**→ rpmE (PRK00019)Ribosomal_L31 (pfam01197)→ |

| | | | | | |
|---|---|---|---|---|---|
| Vli-1-1 | NZ_AUFZ01000019.1 | + | 108185 | 108264 | **RNA**→ rpmE (PRK00019)Ribosomal_L31 (pfam01197)→ |
| Mma-1-1 | NZ_KB907177.1 | + | 14551 | 14600 | **RNA**→ rpmE (PRK00019)Ribosomal_L31 (pfam01197)→ |
| env-45 | SRS012285_C2873870 | - | 78 | 16 | **RNA**→ ←hypo |
| Ase-1-1 | NZ_JUNA01000018.1 | + | 111392 | 111454 | **RNA**→ rpmE (PRK00019)Ribosomal_L31 (pfam01197)→ PRK11204 (PRK11204)Glycos_transf_2 (pfam00535<br>Glyco_transf_25 (pfam01755)→ PRK10422 (PRK10422)RfaF (COG0859)heptsyl_trn_I (TIGR02193)→ |
| env-46 | SRS018665_C3795409 | + | 2089 | 2151 | **RNA**→ |
| env-47 | SRS013252_Baylor_scaffold_62500 | - | 785 | 723 | **RNA**→ rpmE (PRK00019)Ribosomal_L31 (pfam01197)→ |
| env-48 | SRS024561_LANL_scaffold_41247 | + | 465 | 527 | **RNA**→ |
| env-49 | SRS054653_C2095882 | + | 4516 | 4578 | **RNA**→ rpmE (PRK00019)Ribosomal_L31 (pfam01197)→ |
| env-50 | SRS019980_C2847671 | + | 273 | 335 | **RNA**→ |
| Asp-10-1 | NZ_KE952614.1 | + | 72182 | 72244 | **RNA**→ rpmE (PRK00019)Ribosomal_L31 (pfam01197)→ |
| env-51 | SRS020340_C2956907 | + | 757 | 819 | **RNA**→ |
| env-52 | SRS012285_Baylor_scaffold_9025 | - | 2205 | 2143 | **RNA**→ rpmE (PRK00019)Ribosomal_L31 (pfam01197)→ |
| env-53 | SRS022083_Baylor_scaffold_5040 | - | 9430 | 9368 | **RNA**→ rpmE (PRK00019)Ribosomal_L31 (pfam01197)→ |
| env-54 | SRS024021_C2036330 | + | 1818 | 1880 | **RNA**→ |
| env-55 | SRS063999_LANL_scaffold_32076 | + | 1170 | 1232 | **RNA**→ |
| env-56 | SRS015470_WUGC_scaffold_58201 | - | 80 | 18 | **RNA**→ ←hypo |
| env-57 | SRS022625_C2179389 | + | 516 | 578 | **RNA**→ |
| env-58 | SRS016360_Baylor_scaffold_18492 | + | 21485 | 21547 | **RNA**→ rpmE (PRK00019)Ribosomal_L31 (pfam01197)→ |
| env-59 | SRS015989_C2159326 | - | 78 | 16 | **RNA**→ ←hypo |
| env-60 | SRS018573_WUGC_scaffold_22344 | + | 439 | 501 | **RNA**→ rpmE (PRK00019)Ribosomal_L31 (pfam01197)→<br>Glyco_tranf_GTA_type (cd00761)PgaC_IcaA (TIGR03937)→ Glyco_transf_25 (cd06532)→<br>heptsyl_trn_III (TIGR02201)RfaF (COG0859)→ RfaF (COG0859)heptsyl_trn_I (TIGR02193)→ |
| env-61 | SRS017025_Baylor_scaffold_27584 | + | 2458 | 2520 | **RNA**→ rpmE (PRK00019)Ribosomal_L31 (pfam01197)→ Glyco_transf_25 (cd06532)→ |
| env-62 | SRS024381_LANL_scaffold_67893 | + | 1708 | 1770 | **RNA**→ |
| env-63 | SRS014894_WUGC_scaffold_36672 | - | 4155 | 4093 | **RNA**→ rpmE (PRK00019)Ribosomal_L31 (pfam01197)→ Glyco_transf_25 (cd06532)→ RfaF (COG0859)→<br>RfaF (COG0859)heptsyl_trn_I (TIGR02193)→ |
| env-64 | SRS015215_C2600664 | - | 78 | 16 | **RNA**→ ←hypo |
| env-65 | SRS015899_WUGC_scaffold_3106 | + | 46064 | 46126 | **RNA**→ rpmE (PRK00019)Ribosomal_L31 (pfam01197)→ Glyco_transf_25 (cd06532)→ RfaF (COG0859)→<br>RfaF (COG0859)heptsyl_trn_I (TIGR02193)→ |
| env-66 | SRS017227_Baylor_scaffold_61342 | + | 554 | 616 | **RNA**→ rpmE (PRK00019)Ribosomal_L31 (pfam01197)→ Glyco_transf_25 (cd06532)→ RfaF (COG0859)→<br>RfaF (COG0859)heptsyl_trn_I (TIGR02193)→ |
| env-67 | SRS017304_Baylor_scaffold_19757 | - | 78 | 16 | **RNA**→ ←hypo |
| env-68 | SRS019387_WUGC_scaffold_24141 | + | 6275 | 6337 | **RNA**→ rpmE (PRK00019)Ribosomal_L31 (pfam01197)→ Glyco_transf_25 (cd06532)→ RfaF (COG0859)→<br>RfaF (COG0859)heptsyl_trn_I (TIGR02193)→ |
| env-69 | SRS047113_LANL_scaffold_23525 | - | 321 | 259 | **RNA**→ rpmE (PRK00019)Ribosomal_L31 (pfam01197)→ |
| env-70 | SRS051930_LANL_scaffold_27228 | - | 252 | 190 | **RNA**→ rpmE (PRK00019)Ribosomal_L31 (pfam01197)→ |
| Aap-1-1 | NZ_CP012067.1 | - | 796610 | 796548 | **RNA**→ rpmE (PRK00019)Ribosomal_L31 (pfam01197)→ Glyco_transf_25 (cd06532)→ RfaF (COG0859)→<br>RfaF (COG0859)heptsyl_trn_I (TIGR02193)→ |
| env-71 | SRS016541_C1538931 | + | 423 | 485 | **RNA**→ |
| env-72 | SRS022602_Baylor_scaffold_61937 | - | 1250 | 1188 | **RNA**→ rpmE (PRK00019)Ribosomal_L31 (pfam01197)→ Glyco_transf_25 (pfam01755)→ RfaF (COG0859)→ |
| Aap-2-1 | NZ_AEWB02000018.1 | - | 85143 | 85081 | **RNA**→ rpmE (PRK00019)Ribosomal_L31 (pfam01197)→ Glyco_transf_25 (pfam01755)→<br>heptsyl_trn_III (TIGR02201)RfaF (COG0859)→ hypo→ RfaF (COG0859)heptsyl_trn_I (TIGR02193)→ hypo– |
| Aap-4-1 | NC_012913.1 | - | 2198554 | 2198492 | **RNA**→ rpmE (PRK00019)Ribosomal_L31 (pfam01197)→ Glyco_transf_25 (pfam01755)→<br>heptsyl_trn_III (TIGR02201)RfaF (COG0859)→ RfaF (COG0859)heptsyl_trn_I (TIGR02193)→ |
| Aap-3-1 | NZ_JH370360.1 | + | 401206 | 401268 | **RNA**→ rpmE (PRK00019)Ribosomal_L31 (pfam01197)→ Glyco_transf_25 (pfam01755)→<br>heptsyl_trn_III (TIGR02201)RfaF (COG0859)→ RfaF (COG0859)heptsyl_trn_I (TIGR02193)→ |
| env-73 | SRS043772_C1670834 | + | 6 | 68 | **RNA**→ rpmE (PRK00019)Ribosomal_L31 (pfam01197)→ |
| Aac-6-1 | NZ_AHGR01000004.1 | - | 184068 | 184006 | **RNA**→ rpmE (PRK00019)Ribosomal_L31 (pfam01197)→ Glyco_transf_25 (cd06532)→<br>RfaF (COG0859)heptsyl_trn_II (TIGR02195)→ RfaF (COG0859)heptsyl_trn_I (TIGR02193)→<br>RfaF (COG0859)heptsyl_trn_I (TIGR02193)→ |
| Aac-6-2 | NZ_JPZI01000004.1 | - | 184068 | 184006 | **RNA**→ rpmE (PRK00019)Ribosomal_L31 (pfam01197)→ Glyco_transf_25 (cd06532)→<br>RfaF (COG0859)heptsyl_trn_II (TIGR02195)→ RfaF (COG0859)heptsyl_trn_I (TIGR02193)→ |
| Aac-18-1 | NZ_AEJM01000001.1 | - | 87551 | 87489 | **RNA**→ rpmE (PRK00019)Ribosomal_L31 (pfam01197)→ Glyco_transf_25 (cd06532)→<br>heptsyl_trn_III (TIGR02201)RfaF (COG0859)→ RfaF (COG0859)heptsyl_trn_I (TIGR02193)→ |

| | | | | | |
|---|---|---|---|---|---|
| env-74 | SRS015440_WUGC_scaffold_6417 | - | 4077 | 4015 | **RNA**→ rpmE (PRK00019)Ribosomal_L31 (pfam01197)→ Glyco_transf_25 (pfam01755)→ RfaF (COG0859)heptsyl_trn_I (TIGR02193)→ RfaF (COG0859)heptsyl_trn_I (TIGR02193)→ |
| env-75 | SRS015470_WUGC_scaffold_59811 | + | 5208 | 5270 | **RNA**→ ←hypo |
| Aac-11-1 | NZ_AJMH02000011.1 | - | 24012 | 23950 | **RNA**→ rpmE (PRK00019)Ribosomal_L31 (pfam01197)→ Glyco_transf_25 (pfam01755)→ RfaF (COG0859)heptsyl_trn_II (TIGR02195)→ RfaF (COG0859)heptsyl_trn_I (TIGR02193)→ |
| Aac-13-1 | NZ_AJMF02000017.1 | + | 70410 | 70472 | **RNA**→ rpmE (PRK00019)Ribosomal_L31 (pfam01197)→ Glyco_transf_25 (pfam01755)→ RfaF (COG0859)heptsyl_trn_II (TIGR02195)→ RfaF (COG0859)heptsyl_trn_I (TIGR02193)→ |
| Aac-10-1 | NZ_AEJQ02000005.1 | + | 70473 | 70535 | **RNA**→ rpmE (PRK00019)Ribosomal_L31 (pfam01197)→ Glyco_transf_25 (pfam01755)→ RfaF (COG0859)heptsyl_trn_II (TIGR02195)→ RfaF (COG0859)heptsyl_trn_I (TIGR02193)→ |
| Aac-5-1 | NZ_CP008984.1 | - | 898809 | 898747 | **RNA**→ rpmE (PRK00019)Ribosomal_L31 (pfam01197)→ Glyco_transf_25 (pfam01755)→ RfaF (COG0859)heptsyl_trn_II (TIGR02195)→ RfaF (COG0859)heptsyl_trn_I (TIGR02193)→ |
| Aac-12-1 | NZ_AEJP02000020.1 | + | 70468 | 70530 | **RNA**→ rpmE (PRK00019)Ribosomal_L31 (pfam01197)→ Glyco_transf_25 (pfam01755)→ RfaF (COG0859)heptsyl_trn_II (TIGR02195)→ RfaF (COG0859)heptsyl_trn_I (TIGR02193)→ |
| Aac-5-2 | NZ_CP007502.1 | - | 898825 | 898763 | **RNA**→ rpmE (PRK00019)Ribosomal_L31 (pfam01197)→ Glyco_transf_25 (pfam01755)→ RfaF (COG0859)heptsyl_trn_II (TIGR02195)→ RfaF (COG0859)heptsyl_trn_I (TIGR02193)→ |
| env-76 | SRS014578_WUGC_scaffold_38482 | - | 78 | 16 | **RNA**→ ←hypo |
| Aac-17-1 | NZ_AEJL02000024.1 | + | 13375 | 13437 | **RNA**→ rpmE (PRK00019)Ribosomal_L31 (pfam01197)→ Glyco_transf_25 (pfam01755)→ heptsyl_trn_III (TIGR02201)RfaF (COG0859)→ RfaF (COG0859)heptsyl_trn_I (TIGR02193)→ |
| Aac-8-1 | NZ_AJME02000090.1 | + | 106908 | 106970 | **RNA**→ rpmE (PRK00019)Ribosomal_L31 (pfam01197)→ Glyco_transf_25 (pfam01755)→ heptsyl_trn_III (TIGR02201)RfaF (COG0859)→ RfaF (COG0859)heptsyl_trn_I (TIGR02193)→ |
| Aac-19-1 | NZ_AEJN02000055.1 | - | 25976 | 25914 | **RNA**→ rpmE (PRK00019)Ribosomal_L31 (pfam01197)→ Glyco_transf_25 (pfam01755)→ heptsyl_trn_III (TIGR02201)RfaF (COG0859)→ RfaF (COG0859)heptsyl_trn_I (TIGR02193)→ |
| Aac-20-1 | NZ_AEJO02000018.1 | - | 44504 | 44442 | **RNA**→ rpmE (PRK00019)Ribosomal_L31 (pfam01197)→ Glyco_transf_25 (pfam01755)→ heptsyl_trn_III (TIGR02201)RfaF (COG0859)→ RfaF (COG0859)heptsyl_trn_I (TIGR02193)→ |
| Aac-2-1 | NZ_ADOA02000047.1 | - | 30641 | 30579 | **RNA**→ rpmE (PRK00019)Ribosomal_L31 (pfam01197)→ Glyco_transf_25 (pfam01755)→ heptsyl_trn_III (TIGR02201)RfaF (COG0859)→ RfaF (COG0859)heptsyl_trn_I (TIGR02193)→ |
| Aac-7-1 | NZ_KB290891.1 | - | 30592 | 30530 | **RNA**→ rpmE (PRK00019)Ribosomal_L31 (pfam01197)→ Glyco_transf_25 (pfam01755)→ heptsyl_trn_III (TIGR02201)RfaF (COG0859)→ RfaF (COG0859)heptsyl_trn_I (TIGR02193)→ |
| Aac-3-1 | NC_017846.1 | + | 1029004 | 1029066 | **RNA**→ rpmE (PRK00019)Ribosomal_L31 (pfam01197)→ Glyco_transf_25 (pfam01755)→ heptsyl_trn_III (TIGR02201)RfaF (COG0859)→ RfaF (COG0859)heptsyl_trn_I (TIGR02193)→ |
| Aac-9-1 | NZ_AEJK02000020.1 | + | 101509 | 101571 | **RNA**→ rpmE (PRK00019)Ribosomal_L31 (pfam01197)→ Glyco_transf_25 (pfam01755)→ heptsyl_trn_III (TIGR02201)RfaF (COG0859)→ RfaF (COG0859)heptsyl_trn_I (TIGR02193)→ |
| Aac-1-1 | NC_016513.1 | + | 697940 | 698002 | **RNA**→ rpmE (PRK00019)Ribosomal_L31 (pfam01197)→ Glyco_transf_25 (pfam01755)→ RfaF (COG0859)heptsyl_trn_II (TIGR02195)→ RfaF (COG0859)→ RfaF (COG0859)heptsyl_trn_I (TIGR0219 RfaF (COG0859)heptsyl_trn_I (TIGR02193)→ |
| Aac-14-1 | NZ_AJMG02000014.1 | - | 113578 | 113516 | **RNA**→ rpmE (PRK00019)Ribosomal_L31 (pfam01197)→ Glyco_transf_25 (cd06532)→ heptsyl_trn_III (TIGR02201)RfaF (COG0859)→ RfaF (COG0859)heptsyl_trn_I (TIGR02193)→ |
| Aac-15-1 | NZ_ADOB02000039.1 | - | 112687 | 112625 | **RNA**→ rpmE (PRK00019)Ribosomal_L31 (pfam01197)→ Glyco_transf_25 (cd06532)→ heptsyl_trn_III (TIGR02201)RfaF (COG0859)→ RfaF (COG0859)heptsyl_trn_I (TIGR02193)→ |
| Aac-16-1 | NZ_AEJR02000001.1 | - | 113506 | 113444 | **RNA**→ rpmE (PRK00019)Ribosomal_L31 (pfam01197)→ Glyco_transf_25 (cd06532)→ heptsyl_trn_III (TIGR02201)RfaF (COG0859)→ RfaF (COG0859)heptsyl_trn_I (TIGR02193)→ |
| Aac-4-1 | NZ_AQUU01000002.1 | - | 112901 | 112839 | **RNA**→ rpmE (PRK00019)Ribosomal_L31 (pfam01197)→ Glyco_transf_25 (cd06532)→ heptsyl_trn_III (TIGR02201)RfaF (COG0859)→ |
| env-77 | SRS023841_C1659419 | - | 2614 | 2552 | **RNA**→ rpmE (PRK00019)Ribosomal_L31 (pfam01197)→ Glyco_transf_25 (pfam01755)→ RfaF (COG0859)heptsyl_trn_I (TIGR02193)→ |
| env-78 | SRS015755_WUGC_scaffold_22028 | + | 65 | 127 | **RNA**→ MdoB (COG1368)→ |
| env-79 | SRS018337_C2327845 | - | 78 | 16 | **RNA**→ ←hypo |
| env-80 | SRS021960_Baylor_scaffold_32245 | + | 12294 | 12356 | **RNA**→ |
| env-81 | SRS023358_LANL_scaffold_27865 | + | 1270 | 1332 | **RNA**→ |
| env-82 | SRS023964_Baylor_scaffold_14591 | - | 354 | 292 | **RNA**→ rpmE (PRK00019)Ribosomal_L31 (pfam01197)→ |
| env-83 | SRS024649_C3591928 | - | 72 | 10 | **RNA**→ ←hypo |
| env-84 | SRS043018_C3857017 | + | 957 | 1019 | **RNA**→ ←hypo |
| env-85 | SRS051378_C1863951 | + | 1282 | 1344 | **RNA**→ |
| env-86 | SRS013170_Baylor_scaffold_88653 | - | 228 | 166 | **RNA**→ rpmE (PRK00019)Ribosomal_L31 (pfam01197)→ |
| env-87 | SRS015044_WUGC_scaffold_71804 | + | 4985 | 5047 | **RNA**→ |
| env-88 | SRS017227_Baylor_scaffold_62075 | - | 78 | 16 | **RNA**→ ←hypo |

| | | | | | |
|---|---|---|---|---|---|
| env-89 | SRS042984_LANL_scaffold_10671 | + | 2469 | 2531 | **RNA**→ rpmE (PRK00019)Ribosomal_L31 (pfam01197)→ |
| | | | | | Glyco_tranf_GTA_type (cd00761)PgaC_IcaA (TIGR03937)→ Glyco_transf_25 (cd06532)→ |
| | | | | | heptsyl_trn_III (TIGR02201)RfaF (COG0859)→ RfaF (COG0859)heptsyl_trn_I (TIGR02193)→ |
| env-90 | SRS016575_Baylor_scaffold_17318 | - | 78 | 16 | **RNA**→ ←hypo |
| env-91 | SRS019028_WUGC_scaffold_5331 | + | 1625 | 1687 | **RNA**→ Glyco_trans_1_4 (pfam13692)GT1_WabH_like (cd03811)→ RfaB (COG0438)Glyco_trans_1_4 (pfam136 |
| | | | | | Glyco_transf_25 (pfam01755)→ heptsyl_trn_III (TIGR02201)RfaF (COG0859)→ |
| | | | | | Glyco_tranf_GTA_type (cd00761)PRK10073 (PRK10073)→ |
| env-92 | SRS022621_C4052712 | - | 78 | 16 | **RNA**→ ←hypo |
| Ase-2-1 | NZ_GL622200.1 | - | 217473 | 217411 | **RNA**→ rpmE (PRK00019)Ribosomal_L31 (pfam01197)→ |
| env-93 | SRS019980_Baylor_scaffold_31063 | - | 78 | 16 | **RNA**→ ←hypo |
| env-94 | SRS063999_C3424258 | - | 78 | 16 | **RNA**→ ←hypo |
| env-95 | SRS049318_LANL_scaffold_14633 | + | 1589 | 1651 | **RNA**→ |
| Pda-1-1 | NZ_GG704815.1 | - | 133177 | 133128 | **RNA**→ rpmE (PRK00019)Ribosomal_L31 (pfam01197)→ Glyco_transf_25 (pfam01755)→ |
| | | | | | heptsyl_trn_III (TIGR02201)RfaF (COG0859)→ |
| Pmu-14-1 | NZ_CP004391.1 | + | 1445051 | 1445101 | **RNA**→ rpmE (PRK00019)Ribosomal_L31 (pfam01197)→ GT8_A4GalT_like (cd04194)PRK15171 (PRK15171 |
| Pmu-2-1 | NZ_AROA01000259.1 | + | 533 | 586 | **RNA**→ rpmE (PRK00019)Ribosomal_L31 (pfam01197)→ |
| | | | | | Glyco_tranf_GTA_type (cd00761)Glyco_tranf_2_3 (pfam13641)→ hypo→ |
| Pmu-20-1 | NZ_AWTD01000005.1 | + | 291403 | 291456 | **RNA**→ rpmE (PRK00019)Ribosomal_L31 (pfam01197)→ |
| | | | | | Glyco_tranf_GTA_type (cd00761)Glyco_tranf_2_3 (pfam13641)→ hypo→ |
| | | | | | RfaF (COG0859)heptsyl_trn_I (TIGR02193)→ heptsyl_trn_III (TIGR02201)RfaF (COG0859)→ |
| Pmu-13-1 | NZ_JQAF01000008.1 | + | 291632 | 291685 | **RNA**→ rpmE (PRK00019)Ribosomal_L31 (pfam01197)→ |
| | | | | | Glyco_tranf_GTA_type (cd00761)Glyco_tranf_2_3 (pfam13641)→ hypo→ |
| | | | | | RfaF (COG0859)heptsyl_trn_I (TIGR02193)→ heptsyl_trn_III (TIGR02201)RfaF (COG0859)→ |
| Pmu-13-2 | NZ_JQAC01000008.1 | + | 291094 | 291147 | **RNA**→ rpmE (PRK00019)Ribosomal_L31 (pfam01197)→ |
| | | | | | Glyco_tranf_GTA_type (cd00761)Glyco_tranf_2_3 (pfam13641)→ hypo→ |
| | | | | | RfaF (COG0859)heptsyl_trn_I (TIGR02193)→ heptsyl_trn_III (TIGR02201)RfaF (COG0859)→ |
| Pmu-13-3 | NZ_JQAB01000008.1 | + | 291094 | 291147 | **RNA**→ rpmE (PRK00019)Ribosomal_L31 (pfam01197)→ |
| | | | | | Glyco_tranf_GTA_type (cd00761)Glyco_tranf_2_3 (pfam13641)→ hypo→ |
| | | | | | RfaF (COG0859)heptsyl_trn_I (TIGR02193)→ heptsyl_trn_III (TIGR02201)RfaF (COG0859)→ |
| Pmu-13-4 | NZ_JQAE01000008.1 | + | 292160 | 292213 | **RNA**→ rpmE (PRK00019)Ribosomal_L31 (pfam01197)→ |
| | | | | | Glyco_tranf_GTA_type (cd00761)Glyco_tranf_2_3 (pfam13641)→ hypo→ |
| | | | | | RfaF (COG0859)heptsyl_trn_I (TIGR02193)→ heptsyl_trn_III (TIGR02201)RfaF (COG0859)→ |
| Pmu-13-5 | NZ_JQAH01000009.1 | + | 223095 | 223148 | **RNA**→ rpmE (PRK00019)Ribosomal_L31 (pfam01197)→ |
| | | | | | Glyco_tranf_GTA_type (cd00761)Glyco_tranf_2_3 (pfam13641)→ hypo→ |
| | | | | | RfaF (COG0859)heptsyl_trn_I (TIGR02193)→ heptsyl_trn_III (TIGR02201)RfaF (COG0859)→ |
| Pmu-13-6 | NZ_JPHI01000008.1 | + | 291065 | 291118 | **RNA**→ rpmE (PRK00019)Ribosomal_L31 (pfam01197)→ |
| | | | | | Glyco_tranf_GTA_type (cd00761)Glyco_tranf_2_3 (pfam13641)→ hypo→ |
| | | | | | RfaF (COG0859)heptsyl_trn_I (TIGR02193)→ heptsyl_trn_III (TIGR02201)RfaF (COG0859)→ |
| Pmu-13-7 | NZ_JQAG01000008.1 | + | 291632 | 291685 | **RNA**→ rpmE (PRK00019)Ribosomal_L31 (pfam01197)→ |
| | | | | | Glyco_tranf_GTA_type (cd00761)Glyco_tranf_2_3 (pfam13641)→ hypo→ |
| | | | | | RfaF (COG0859)heptsyl_trn_I (TIGR02193)→ heptsyl_trn_III (TIGR02201)RfaF (COG0859)→ |
| Pmu-13-8 | NZ_JQEB01000010.1 | + | 291109 | 291162 | **RNA**→ rpmE (PRK00019)Ribosomal_L31 (pfam01197)→ |
| | | | | | Glyco_tranf_GTA_type (cd00761)Glyco_tranf_2_3 (pfam13641)→ hypo→ |
| | | | | | RfaF (COG0859)heptsyl_trn_I (TIGR02193)→ heptsyl_trn_III (TIGR02201)RfaF (COG0859)→ |
| Pmu-13-9 | NZ_JQEA01000012.1 | + | 290579 | 290632 | **RNA**→ rpmE (PRK00019)Ribosomal_L31 (pfam01197)→ |
| | | | | | Glyco_tranf_GTA_type (cd00761)Glyco_tranf_2_3 (pfam13641)→ hypo→ |
| | | | | | RfaF (COG0859)heptsyl_trn_I (TIGR02193)→ heptsyl_trn_III (TIGR02201)RfaF (COG0859)→ |
| Pmu-13-10 | NZ_JQAD01000008.1 | + | 291094 | 291147 | **RNA**→ rpmE (PRK00019)Ribosomal_L31 (pfam01197)→ |
| | | | | | Glyco_tranf_GTA_type (cd00761)Glyco_tranf_2_3 (pfam13641)→ hypo→ |
| | | | | | RfaF (COG0859)heptsyl_trn_I (TIGR02193)→ heptsyl_trn_III (TIGR02201)RfaF (COG0859)→ |
| Pmu-13-11 | NZ_JQAO01000009.1 | + | 223226 | 223279 | **RNA**→ rpmE (PRK00019)Ribosomal_L31 (pfam01197)→ |
| | | | | | Glyco_tranf_GTA_type (cd00761)Glyco_tranf_2_3 (pfam13641)→ hypo→ |
| | | | | | RfaF (COG0859)heptsyl_trn_I (TIGR02193)→ heptsyl_trn_III (TIGR02201)RfaF (COG0859)→ |
| Pmu-13-12 | NZ_JQAI01000008.1 | + | 291094 | 291147 | **RNA**→ rpmE (PRK00019)Ribosomal_L31 (pfam01197)→ |
| | | | | | Glyco_tranf_GTA_type (cd00761)Glyco_tranf_2_3 (pfam13641)→ hypo→ |
| | | | | | RfaF (COG0859)heptsyl_trn_I (TIGR02193)→ heptsyl_trn_III (TIGR02201)RfaF (COG0859)→ |
| Pmu-16-1 | NZ_ALYC02000068.1 | - | 827 | 774 | **RNA**→ rpmE (PRK00019)Ribosomal_L31 (pfam01197)→ hypo→ hypo→ |

| | | | | | |
|---|---|---|---|---|---|
| Pmu-8-1 | NZ_CP004392.1 | - | 1820422 | 1820372 | **RNA**→ rpmE (PRK00019)Ribosomal_L31 (pfam01197)→ Glyco_transf_25 (cd06532)→ Glyco_transf_25 (pfam01755)→ CgtA (pfam06306)→ CgtA (pfam06306)→ Glyco_transf_4 (pfam13439)Glyco_trans_1_4 (pfam13692)GT1_WabH_like (cd03811)→ |
| Pmu-12-1 | NZ_CM001580.1 | + | 1329398 | 1329448 | **RNA**→ rpmE (PRK00019)Ribosomal_L31 (pfam01197)→ |
| Pmu-3-1 | NZ_AQTL01000106.1 | - | 3016 | 2966 | **RNA**→ rpmE (PRK00019)Ribosomal_L31 (pfam01197)→ Glyco_transf_25 (pfam01755)→ heptsyl_trn_III (TIGR02201)RfaF (COG0859)→ |
| Pmu-10-1 | NZ_ARNZ01000104.1 | + | 1190 | 1240 | **RNA**→ rpmE (PRK00019)Ribosomal_L31 (pfam01197)→ Glyco_transf_25 (pfam01755)→ heptsyl_trn_III (TIGR02201)RfaF (COG0859)→ |
| Pmu-4-1 | NZ_ARNW01000018.1 | + | 4332 | 4382 | **RNA**→ rpmE (PRK00019)Ribosomal_L31 (pfam01197)→ Glyco_transf_25 (pfam01755)→ heptsyl_trn_III (TIGR02201)RfaF (COG0859)→ |
| Pmu-7-1 | NZ_ARNX01000001.1 | - | 43763 | 43713 | **RNA**→ rpmE (PRK00019)Ribosomal_L31 (pfam01197)→ Glyco_transf_25 (pfam01755)→ heptsyl_trn_III (TIGR02201)RfaF (COG0859)→ |
| Pmu-21-1 | NC_002663.1 | + | 1342547 | 1342597 | **RNA**→ rpmE (PRK00019)Ribosomal_L31 (pfam01197)→ Glyco_transf_25 (pfam01755)→ heptsyl_trn_III (TIGR02201)RfaF (COG0859)→ |
| Pmu-11-1 | NZ_CM001581.1 | + | 1417199 | 1417249 | **RNA**→ rpmE (PRK00019)Ribosomal_L31 (pfam01197)→ Glyco_transf_25 (pfam01755)→ heptsyl_trn_III (TIGR02201)RfaF (COG0859)→ heptsyl_trn_III (TIGR02201)RfaF (COG0859)→ |
| Pmu-5-1 | NC_016808.1 | - | 12183 | 12133 | **RNA**→ rpmE (PRK00019)Ribosomal_L31 (pfam01197)→ Glyco_transf_25 (pfam01755)→ heptsyl_trn_III (TIGR02201)RfaF (COG0859)→ |
| Pmu-18-1 | NZ_CP003328.1 | - | 12083 | 12033 | **RNA**→ rpmE (PRK00019)Ribosomal_L31 (pfam01197)→ Glyco_transf_25 (pfam01755)→ heptsyl_trn_III (TIGR02201)RfaF (COG0859)→ |
| Pmu-15-1 | NZ_CM002276.1 | + | 23245 | 23295 | **RNA**→ rpmE (PRK00019)Ribosomal_L31 (pfam01197)→ Glyco_transf_25 (pfam01755)→ heptsyl_trn_III (TIGR02201)RfaF (COG0859)→ |
| Pmu-1-1 | NZ_CP008918.1 | - | 823037 | 822987 | **RNA**→ rpmE (PRK00019)Ribosomal_L31 (pfam01197)→ Glyco_transf_25 (pfam01755)→ heptsyl_trn_III (TIGR02201)RfaF (COG0859)→ |
| Pmu-9-1 | NZ_ARNY01000003.1 | - | 60597 | 60547 | **RNA**→ rpmE (PRK00019)Ribosomal_L31 (pfam01197)→ Glyco_transf_25 (pfam01755)→ heptsyl_trn_III (TIGR02201)RfaF (COG0859)→ |
| Pmu-6-1 | NZ_ARWR01000002.1 | + | 265889 | 265939 | **RNA**→ rpmE (PRK00019)Ribosomal_L31 (pfam01197)→ Glyco_transf_25 (pfam01755)→ heptsyl_trn_III (TIGR02201)RfaF (COG0859)→ |
| Pmu-19-1 | NC_017027.1 | - | 15811 | 15761 | **RNA**→ rpmE (PRK00019)Ribosomal_L31 (pfam01197)→ Glyco_transf_25 (pfam01755)→ heptsyl_trn_III (TIGR02201)RfaF (COG0859)→ |
| Pmu-17-1 | NC_017764.1 | - | 15831 | 15781 | **RNA**→ rpmE (PRK00019)Ribosomal_L31 (pfam01197)→ Glyco_transf_25 (pfam01755)→ heptsyl_trn_III (TIGR02201)RfaF (COG0859)→ |
| Mma-3-1 | NZ_CDQL01000004.1 | - | 255958 | 255897 | **RNA**→ rpmE (PRK00019)Ribosomal_L31 (pfam01197)→ |
| Pbe-2-1 | NZ_AJSX01000004.1 | + | 13647 | 13708 | **RNA**→ rpmE (PRK00019)Ribosomal_L31 (pfam01197)→ rpmE (PRK00019)Ribosomal_L31 (pfam01197)→ |
| Msu-2-1 | NC_006300.1 | - | 416530 | 416470 | **RNA**→ rpmE (PRK00019)Ribosomal_L31 (pfam01197)→ GT8_A4GalT_like (cd04194)PRK15171 (PRK15171 Glyco_transf_25 (pfam01755)→ heptsyl_trn_III (TIGR02201)RfaF (COG0859)→ hypo→ |
| env-96 | SRS023958_Baylor_scaffold_5426 | - | 392 | 342 | **RNA**→ hypo→ |
| env-97 | SRS011271_C3031949 | + | 247 | 297 | **RNA**→ rpmE (PRK00019)Ribosomal_L31 (pfam01197)→ |
| env-98 | SRS018969_WUGC_scaffold_19392 | - | 1477 | 1427 | **RNA**→ rpmE (PRK00019)Ribosomal_L31 (pfam01197)→ |
| env-99 | SRS023526_C3187054 | - | 873 | 823 | **RNA**→ rpmE (PRK00019)Ribosomal_L31 (pfam01197)→ |
| env-100 | SRS024277_LANL_scaffold_49966 | + | 339 | 389 | **RNA**→ rpmE (PRK00019)Ribosomal_L31 (pfam01197)→ |
| Hpa-2-1 | NZ_JVSL01000025.1 | + | 13438 | 13488 | **RNA**→ rpmE (PRK00019)Ribosomal_L31 (pfam01197)→ |
| Hpa-2-2 | NZ_JVHG01000004.1 | - | 10892 | 10842 | **RNA**→ rpmE (PRK00019)Ribosomal_L31 (pfam01197)→ |
| env-101 | SRS016225_WUGC_scaffold_2971 | + | 183 | 233 | **RNA**→ rpmE (PRK00019)Ribosomal_L31 (pfam01197)→ |
| env-102 | SRS019071_WUGC_scaffold_37000 | - | 305 | 255 | **RNA**→ rpmE (PRK00019)Ribosomal_L31 (pfam01197)→ |
| env-103 | SRS024087_C3536836 | + | 1002 | 1052 | **RNA**→ |
| env-104 | SRS063932_LANL_scaffold_93582 | - | 66 | 16 | **RNA**→ |
| env-105 | SRS024021_Baylor_scaffold_3567 | + | 655 | 705 | **RNA**→ |
| env-106 | SRS014473_WUGC_scaffold_5102 | + | 2802 | 2852 | **RNA**→ rpmE (PRK00019)Ribosomal_L31 (pfam01197)→ |
| env-107 | SRS015060_WUGC_scaffold_6035 | - | 267 | 217 | **RNA**→ rpmE (PRK00019)Ribosomal_L31 (pfam01197)→ |
| env-108 | SRS023534_C140704 | + | 184 | 234 | **RNA**→ rpmE (PRK00019)Ribosomal_L31 (pfam01197)→ |
| env-109 | scaffold9440_1_MH0072 | + | 805 | 855 | **RNA**→ rpmE (PRK00019)Ribosomal_L31 (pfam01197)→ |
| env-110 | SRS014468_C479594 | + | 50 | 100 | **RNA**→ rpmE (PRK00019)Ribosomal_L31 (pfam01197)→ |
| env-111 | SRS012902_C495709 | + | 47 | 97 | **RNA**→ rpmE (PRK00019)Ribosomal_L31 (pfam01197)→ |
| env-112 | SRS014107_C1310462 | + | 78 | 128 | **RNA**→ rpmE (PRK00019)Ribosomal_L31 (pfam01197)→ |
| env-113 | SRS014470_C3580284 | + | 17 | 67 | **RNA**→ rpmE (PRK00019)Ribosomal_L31 (pfam01197)→ |

| | | | | | |
|---|---|---|---|---|---|
| env-114 | SRS014684_C3226119 | - | 425 | 375 | RNA→ rpmE (PRK00019)Ribosomal_L31 (pfam01197)→ |
| env-115 | SRS015154_WUGC_scaffold_4241 | + | 17 | 67 | RNA→ rpmE (PRK00019)Ribosomal_L31 (pfam01197)→ |
| env-116 | SRS015650_WUGC_scaffold_11302 | + | 60 | 110 | RNA→ rpmE (PRK00019)Ribosomal_L31 (pfam01197)→ |
| env-117 | SRS016200_WUGC_scaffold_15359 | + | 2090 | 2140 | RNA→ rpmE (PRK00019)Ribosomal_L31 (pfam01197)→ |
| env-118 | SRS016342_Baylor_scaffold_20824 | + | 57 | 107 | RNA→ rpmE (PRK00019)Ribosomal_L31 (pfam01197)→ |
| env-119 | SRS016503_LANL_scaffold_2981 | + | 71 | 121 | RNA→ rpmE (PRK00019)Ribosomal_L31 (pfam01197)→ |
| env-120 | SRS016533_C471857 | + | 71 | 121 | RNA→ rpmE (PRK00019)Ribosomal_L31 (pfam01197)→ |
| env-121 | SRS016575_Baylor_scaffold_14305 | + | 208 | 258 | RNA→ rpmE (PRK00019)Ribosomal_L31 (pfam01197)→ |
| env-122 | SRS017445_Baylor_scaffold_14093 | + | 48 | 98 | RNA→ ←TonB-siderophor (TIGR01783)TonB_dep_Rec (pfam00593) |
| env-123 | SRS018665_WUGC_scaffold_24884 | + | 71 | 121 | RNA→ ←PRK13524 (PRK13524)ligand_gated_channel (cd01347) |
| env-124 | SRS018791_WUGC_scaffold_50431 | + | 17 | 67 | RNA→ rpmE (PRK00019)Ribosomal_L31 (pfam01197)→ |
| env-125 | SRS022621_Baylor_scaffold_64879 | + | 117 | 167 | RNA→ rpmE (PRK00019)Ribosomal_L31 (pfam01197)→ |
| env-126 | SRS022625_Baylor_scaffold_7805 | - | 66 | 16 | RNA→ |
| env-127 | SRS022721_C501186 | - | 318 | 268 | RNA→ rpmE (PRK00019)Ribosomal_L31 (pfam01197)→ |
| env-128 | SRS023617_Baylor_scaffold_71993 | + | 48 | 98 | RNA→ rpmE (PRK00019)Ribosomal_L31 (pfam01197)→ |
| env-129 | SRS023835_Baylor_scaffold_11069 | - | 295 | 245 | RNA→ rpmE (PRK00019)Ribosomal_L31 (pfam01197)→ |
| env-130 | SRS023926_Baylor_scaffold_35367 | + | 75 | 125 | RNA→ rpmE (PRK00019)Ribosomal_L31 (pfam01197)→ |
| env-131 | SRS024318_LANL_scaffold_16452 | + | 48 | 98 | RNA→ rpmE (PRK00019)Ribosomal_L31 (pfam01197)→ |
| env-132 | SRS024381_LANL_scaffold_28820 | + | 71 | 121 | RNA→ rpmE (PRK00019)Ribosomal_L31 (pfam01197)→ |
| env-133 | SRS024557_C267439 | + | 71 | 121 | RNA→ rpmE (PRK00019)Ribosomal_L31 (pfam01197)→ |
| env-134 | SRS042131_WUGC_scaffold_18379 | + | 1363 | 1413 | RNA→ rpmE (PRK00019)Ribosomal_L31 (pfam01197)→ |
| env-135 | SRS048719_WUGC_scaffold_10207 | + | 648 | 698 | RNA→ rpmE (PRK00019)Ribosomal_L31 (pfam01197)→ |
| env-136 | SRS052874_LANL_scaffold_13398 | + | 539 | 589 | RNA→ rpmE (PRK00019)Ribosomal_L31 (pfam01197)→ |
| env-137 | SRS054956_LANL_scaffold_30996 | + | 2601 | 2651 | RNA→ rpmE (PRK00019)Ribosomal_L31 (pfam01197)→ |
| env-138 | SRS064449_LANL_scaffold_6703 | + | 601 | 651 | RNA→ rpmE (PRK00019)Ribosomal_L31 (pfam01197)→ |
| Hpa-2-3 | NZ_JVPX01000007.1 | - | 10911 | 10861 | RNA→ rpmE (PRK00019)Ribosomal_L31 (pfam01197)→ |
| env-139 | 3300001196_1009759 | + | 160 | 210 | RNA→ rpmE (PRK00019)Ribosomal_L31 (pfam01197)→ |
| env-140 | scaffold25950_1_MH0073 | + | 515 | 565 | RNA→ rpmE (PRK00019)Ribosomal_L31 (pfam01197)→ |
| env-141 | SRS011086_WUGC_scaffold_33563 | - | 474 | 424 | RNA→ rpmE (PRK00019)Ribosomal_L31 (pfam01197)→ |
| env-142 | SRS014692_WUGC_scaffold_12598 | + | 345 | 395 | RNA→ rpmE (PRK00019)Ribosomal_L31 (pfam01197)→ |
| env-143 | SRS017088_C1144155 | - | 81 | 31 | RNA→ |
| env-144 | SRS018774_WUGC_scaffold_1636 | - | 68 | 18 | RNA→ |
| env-145 | SRS021986_C925803 | - | 129 | 79 | RNA→ rpmE (PRK00019)rpmE2 (PRK01678)Ribosomal_L31 (pfam01197)→ |
| env-146 | SRS043239_C1968841 | + | 117 | 167 | RNA→ rpmE (PRK00019)Ribosomal_L31 (pfam01197)→ |
| env-147 | SRS044366_C951592 | + | 2864 | 2914 | RNA→ rpmE (PRK00019)Ribosomal_L31 (pfam01197)→ |
| env-148 | SRS053630_LANL_scaffold_5204 | + | 273260 | 273310 | RNA→ rpmE (PRK00019)Ribosomal_L31 (pfam01197)→ |
| env-149 | SRS053917_C3768051 | + | 554 | 604 | RNA→ |
| env-150 | SRS064329_LANL_scaffold_3760 | + | 358 | 408 | RNA→ rpmE (PRK00019)Ribosomal_L31 (pfam01197)→ |
| env-151 | SRS075404_C3577072 | - | 300 | 250 | RNA→ rpmE (PRK00019)Ribosomal_L31 (pfam01197)→ |
| env-152 | Saliva_LANL_C141248 | + | 159 | 209 | RNA→ rpmE (PRK00019)Ribosomal_L31 (pfam01197)→ |
| env-153 | Throat_LANL_C5774541 | + | 179 | 229 | RNA→ rpmE (PRK00019)Ribosomal_L31 (pfam01197)→ |
| env-154 | UnmappedStool_Broad_C252872408 | - | 329 | 279 | RNA→ rpmE (PRK00019)Ribosomal_L31 (pfam01197)→ |
| env-155 | SRS011247_Baylor_scaffold_2642 | - | 68 | 18 | RNA→ |
| env-156 | SRS011529_C4573705 | + | 205 | 255 | RNA→ rpmE (PRK00019)Ribosomal_L31 (pfam01197)→ |
| env-157 | SRS012279_Baylor_scaffold_42064 | - | 295 | 245 | RNA→ rpmE (PRK00019)Ribosomal_L31 (pfam01197)→ |
| env-158 | SRS013234_Baylor_scaffold_25561 | - | 434 | 384 | RNA→ rpmE (PRK00019)Ribosomal_L31 (pfam01197)→ |
| env-159 | SRS013502_Baylor_scaffold_15687 | - | 357 | 307 | RNA→ rpmE (PRK00019)Ribosomal_L31 (pfam01197)→ |
| env-160 | SRS013705_Baylor_scaffold_9676 | - | 304 | 254 | RNA→ rpmE (PRK00019)Ribosomal_L31 (pfam01197)→ |
| env-161 | SRS013711_C1736629 | - | 324 | 274 | RNA→ rpmE (PRK00019)Ribosomal_L31 (pfam01197)→ |
| env-162 | SRS013723_Baylor_scaffold_21329 | - | 159 | 109 | RNA→ rpmE (PRK00019)Ribosomal_L31 (pfam01197)→ |
| env-163 | SRS013818_Baylor_scaffold_39426 | - | 207 | 157 | RNA→ rpmE (PRK00019)Ribosomal_L31 (pfam01197)→ |
| env-164 | SRS013825_Baylor_scaffold_3706 | + | 131 | 181 | RNA→ rpmE (PRK00019)Ribosomal_L31 (pfam01197)→ |

| | | | | | |
|---|---|---|---|---|---|
| env-165 | SRS013836_C4728831 | + | 152 | 202 | **RNA**→ rpmE (PRK00019)Ribosomal_L31 (pfam01197)→ |
| env-166 | SRS013881_WUGC_scaffold_7653 | - | 253 | 203 | **RNA**→ rpmE (PRK00019)Ribosomal_L31 (pfam01197)→ |
| env-167 | SRS013945_WUGC_scaffold_15499 | - | 346 | 296 | **RNA**→ rpmE (PRK00019)Ribosomal_L31 (pfam01197)→ |
| env-168 | SRS013947_WUGC_scaffold_18108 | - | 207 | 157 | **RNA**→ rpmE (PRK00019)Ribosomal_L31 (pfam01197)→ |
| env-169 | SRS013948_WUGC_scaffold_4756 | + | 610 | 660 | **RNA**→ rpmE (PRK00019)Ribosomal_L31 (pfam01197)→ |
| env-170 | SRS013950_C2184733 | + | 2868 | 2918 | **RNA**→ rpmE (PRK00528)rpmE (PRK00019)rpmE2 (PRK01678)Ribosomal_L31 (pfam01197)→ |
| env-171 | SRS014126_C559458 | + | 145 | 195 | **RNA**→ rpmE (PRK00019)Ribosomal_L31 (pfam01197)→ |
| env-172 | SRS014271_WUGC_scaffold_41463 | + | 179 | 229 | **RNA**→ rpmE (PRK00019)Ribosomal_L31 (pfam01197)→ |
| env-173 | SRS014575_C476087 | + | 226 | 276 | **RNA**→ rpmE (PRK00019)Ribosomal_L31 (pfam01197)→ |
| env-174 | SRS014686_C892496 | + | 364 | 414 | **RNA**→ rpmE (PRK00019)Ribosomal_L31 (pfam01197)→ |
| env-175 | SRS014890_C397523 | - | 260 | 210 | **RNA**→ rpmE (PRK00019)Ribosomal_L31 (pfam01197)→ |
| env-176 | SRS015057_C3383085 | + | 266 | 316 | **RNA**→ rpmE (PRK00019)Ribosomal_L31 (pfam01197)→ |
| env-177 | SRS015209_WUGC_scaffold_13430 | - | 873 | 823 | **RNA**→ rpmE (PRK00019)Ribosomal_L31 (pfam01197)→ |
| env-178 | SRS015272_C3511097 | - | 79 | 29 | **RNA**→ |
| env-179 | SRS015395_C2368773 | - | 127 | 77 | **RNA**→ rpmE (PRK00528)rpmE (PRK00019)rpmE2 (PRK01678)Ribosomal_L31 (pfam01197)→ |
| env-180 | SRS015434_WUGC_scaffold_16743 | - | 64 | 14 | **RNA**→ |
| env-181 | SRS015578_WUGC_scaffold_34785 | - | 689 | 639 | **RNA**→ rpmE (PRK00019)Ribosomal_L31 (pfam01197)→ |
| env-182 | SRS015745_WUGC_scaffold_1107 | - | 148 | 98 | **RNA**→ rpmE (PRK00019)Ribosomal_L31 (pfam01197)→ |
| env-183 | SRS015797_WUGC_scaffold_3481 | + | 265 | 315 | **RNA**→ rpmE (PRK00019)Ribosomal_L31 (pfam01197)→ |
| env-184 | SRS015893_C2295370 | - | 293 | 243 | **RNA**→ rpmE (PRK00019)Ribosomal_L31 (pfam01197)→ |
| env-185 | SRS015895_WUGC_scaffold_997 | - | 4030 | 3980 | **RNA**→ rpmE (PRK00019)Ribosomal_L31 (pfam01197)→ |
| env-186 | SRS015899_WUGC_scaffold_31920 | + | 1115 | 1165 | **RNA**→ |
| env-187 | SRS015921_WUGC_scaffold_8625 | - | 2466 | 2416 | **RNA**→ rpmE (PRK00019)Ribosomal_L31 (pfam01197)→ |
| env-188 | SRS015941_WUGC_scaffold_8499 | + | 674 | 724 | **RNA**→ rpmE (PRK00019)Ribosomal_L31 (pfam01197)→ |
| env-189 | SRS015989_WUGC_scaffold_6985 | - | 66 | 16 | **RNA**→ |
| env-190 | SRS016319_WUGC_scaffold_52031 | + | 117 | 167 | **RNA**→ rpmE (PRK00019)Ribosomal_L31 (pfam01197)→ |
| env-191 | SRS016501_LANL_scaffold_19928 | + | 109 | 159 | **RNA**→ rpmE (PRK00019)Ribosomal_L31 (pfam01197)→ |
| env-192 | SRS016600_Baylor_scaffold_790 | - | 153 | 103 | **RNA**→ rpmE (PRK00019)Ribosomal_L31 (pfam01197)→ |
| env-193 | SRS017080_C1218701 | - | 375 | 325 | **RNA**→ rpmE (PRK00019)Ribosomal_L31 (pfam01197)→ |
| env-194 | SRS017120_Baylor_scaffold_43405 | - | 677 | 627 | **RNA**→ rpmE (PRK00019)Ribosomal_L31 (pfam01197)→ |
| env-195 | SRS017209_C2427531 | + | 266 | 316 | **RNA**→ |
| env-196 | SRS017713_Baylor_scaffold_3587 | - | 374 | 324 | **RNA**→ rpmE (PRK00019)Ribosomal_L31 (pfam01197)→ |
| env-197 | SRS018145_Baylor_scaffold_10947 | - | 73 | 23 | **RNA**→ |
| env-198 | SRS018300_Baylor_scaffold_30 | + | 260 | 310 | **RNA**→ |
| env-199 | SRS018359_Baylor_scaffold_4860 | - | 257 | 207 | **RNA**→ rpmE (PRK00019)Ribosomal_L31 (pfam01197)→ |
| env-200 | SRS018439_Baylor_scaffold_16838 | + | 239 | 289 | **RNA**→ |
| env-201 | SRS018573_C2361858 | - | 65 | 15 | **RNA**→ |
| env-202 | SRS018591_WUGC_scaffold_44156 | - | 226 | 176 | **RNA**→ rpmE (PRK00019)Ribosomal_L31 (pfam01197)→ |
| env-203 | SRS018661_WUGC_scaffold_2690 | - | 953 | 903 | **RNA**→ rpmE (PRK00019)Ribosomal_L31 (pfam01197)→ |
| env-204 | SRS018739_C3611004 | - | 374 | 324 | **RNA**→ rpmE (PRK00019)Ribosomal_L31 (pfam01197)→ |
| env-205 | SRS018971_WUGC_scaffold_6404 | + | 266 | 316 | **RNA**→ ←PRK13524 (PRK13524)ligand_gated_channel (cd01347) |
| env-206 | SRS018975_WUGC_scaffold_41000 | - | 368 | 318 | **RNA**→ rpmE (PRK00019)Ribosomal_L31 (pfam01197)→ |
| env-207 | SRS019022_WUGC_scaffold_65126 | + | 326 | 376 | **RNA**→ rpmE (PRK00019)Ribosomal_L31 (pfam01197)→ |
| env-208 | SRS019026_C1969885 | + | 607 | 657 | **RNA**→ rpmE (PRK00019)Ribosomal_L31 (pfam01197)→ |
| env-209 | SRS019027_WUGC_scaffold_18155 | - | 713 | 663 | **RNA**→ rpmE (PRK00019)Ribosomal_L31 (pfam01197)→ |
| env-210 | SRS019045_WUGC_scaffold_19548 | + | 272 | 322 | **RNA**→ rpmE (PRK00019)Ribosomal_L31 (pfam01197)→ |
| env-211 | SRS019120_C1279990 | + | 248 | 298 | **RNA**→ |
| env-212 | SRS019122_WUGC_scaffold_7081 | - | 561 | 511 | **RNA**→ rpmE (PRK00019)Ribosomal_L31 (pfam01197)→ |
| env-213 | SRS019124_WUGC_scaffold_7080 | - | 180 | 130 | **RNA**→ rpmE (PRK00019)Ribosomal_L31 (pfam01197)→ |
| env-214 | SRS019126_WUGC_scaffold_8147 | + | 629 | 679 | **RNA**→ rpmE (PRK00019)Ribosomal_L31 (pfam01197)→ |
| env-215 | SRS019127_C1598465 | - | 374 | 324 | **RNA**→ rpmE (PRK00019)Ribosomal_L31 (pfam01197)→ |
| env-216 | SRS019129_C1955039 | - | 253 | 203 | **RNA**→ rpmE (PRK00019)Ribosomal_L31 (pfam01197)→ |

| | | | | | |
|---|---|---|---|---|---|
| env-217 | SRS019219_C2807201 | + | 89 | 139 | **RNA**→ rpmE (PRK00019)Ribosomal_L31 (pfam01197)→ |
| env-218 | SRS019327_C2615848 | + | 281 | 331 | **RNA**→ |
| env-219 | SRS019389_C2178886 | - | 330 | 280 | **RNA**→ rpmE (PRK00019)Ribosomal_L31 (pfam01197)→ |
| env-220 | SRS019391_WUGC_scaffold_1544 | - | 264 | 214 | **RNA**→ |
| env-221 | SRS019397_C1439804 | - | 920 | 870 | **RNA**→ rpmE (PRK00019)Ribosomal_L31 (pfam01197)→ |
| env-222 | SRS019607_WUGC_scaffold_1072 | + | 1281 | 1331 | **RNA**→ rpmE (PRK00019)Ribosomal_L31 (pfam01197)→ |
| env-223 | SRS019872_C622759 | + | 365 | 415 | **RNA**→ rpmE (PRK00019)Ribosomal_L31 (pfam01197)→ |
| env-224 | SRS019894_WUGC_scaffold_40693 | - | 460 | 410 | **RNA**→ ←PRK13524 (PRK13524)TonB_dep_Rec (pfam00593) |
| env-225 | SRS020334_Baylor_scaffold_37850 | - | 778 | 728 | **RNA**→ rpmE (PRK00019)Ribosomal_L31 (pfam01197)→ |
| env-226 | SRS021477_Baylor_scaffold_70117 | + | 470 | 520 | **RNA**→ rpmE (PRK00019)Ribosomal_L31 (pfam01197)→ |
| env-227 | SRS021496_Baylor_scaffold_549 | - | 394 | 344 | **RNA**→ rpmE (PRK00019)Ribosomal_L31 (pfam01197)→ |
| env-228 | SRS021948_C2754471 | - | 240 | 190 | **RNA**→ rpmE (PRK00019)Ribosomal_L31 (pfam01197)→ |
| env-229 | SRS021954_Baylor_scaffold_5062 | - | 148 | 98 | **RNA**→ rpmE (PRK00019)Ribosomal_L31 (pfam01197)→ |
| env-230 | SRS022077_C2432114 | + | 107 | 157 | **RNA**→ rpmE (PRK00019)Ribosomal_L31 (pfam01197)→ |
| env-231 | SRS022079_Baylor_scaffold_515 | - | 15255 | 15205 | **RNA**→ rpmE (PRK00019)Ribosomal_L31 (pfam01197)→ |
| env-232 | SRS022083_Baylor_scaffold_4111 | + | 1531 | 1581 | **RNA**→ rpmE (PRK00019)Ribosomal_L31 (pfam01197)→ |
| env-233 | SRS022143_WUGC_scaffold_83552 | - | 318 | 268 | **RNA**→ rpmE (PRK00019)Ribosomal_L31 (pfam01197)→ |
| env-234 | SRS022532_LANL_scaffold_10532 | + | 671 | 721 | **RNA**→ ←hypo |
| env-235 | SRS023837_C582069 | + | 172 | 222 | **RNA**→ rpmE (PRK00019)Ribosomal_L31 (pfam01197)→ |
| env-236 | SRS023964_Baylor_scaffold_37730 | - | 207 | 157 | **RNA**→ rpmE (PRK00019)Ribosomal_L31 (pfam01197)→ |
| env-237 | SRS024081_LANL_scaffold_839 | + | 386 | 436 | **RNA**→ rpmE (PRK00019)Ribosomal_L31 (pfam01197)→ |
| env-238 | SRS024375_LANL_scaffold_34090 | - | 295 | 245 | **RNA**→ rpmE (PRK00019)Ribosomal_L31 (pfam01197)→ |
| env-239 | SRS024470_LANL_scaffold_1444 | + | 270 | 320 | **RNA**→ rpmE (PRK00019)Ribosomal_L31 (pfam01197)→ |
| env-240 | SRS043676_WUGC_scaffold_7680 | - | 68 | 18 | **RNA**→ |
| env-241 | SRS043701_C1745554 | - | 295 | 245 | **RNA**→ rpmE (PRK00019)Ribosomal_L31 (pfam01197)→ |
| env-242 | SRS044373_C2721994 | - | 67 | 17 | **RNA**→ |
| env-243 | SRS044486_WUGC_scaffold_2702 | + | 290 | 340 | **RNA**→ rpmE (PRK00019)Ribosomal_L31 (pfam01197)→ |
| env-244 | SRS045127_LANL_scaffold_25629 | - | 213 | 163 | **RNA**→ rpmE (PRK00019)Ribosomal_L31 (pfam01197)→ |
| env-245 | SRS045645_C1878673 | + | 83 | 133 | **RNA**→ rpmE (PRK00019)Ribosomal_L31 (pfam01197)→ |
| env-246 | SRS045715_C4716689 | - | 253 | 203 | **RNA**→ rpmE (PRK00019)Ribosomal_L31 (pfam01197)→ |
| env-247 | SRS045978_C962638 | - | 76 | 26 | **RNA**→ |
| env-248 | SRS047210_WUGC_scaffold_43713 | - | 305 | 255 | **RNA**→ rpmE (PRK00019)Ribosomal_L31 (pfam01197)→ |
| env-249 | SRS047219_WUGC_scaffold_2120 | + | 278 | 328 | **RNA**→ rpmE (PRK00019)Ribosomal_L31 (pfam01197)→ |
| env-250 | SRS047824_WUGC_scaffold_42445 | + | 423 | 473 | **RNA**→ rpmE (PRK00019)Ribosomal_L31 (pfam01197)→ |
| env-251 | SRS048411_C6924320 | - | 360 | 310 | **RNA**→ rpmE (PRK00019)Ribosomal_L31 (pfam01197)→ |
| env-252 | SRS048791_LANL_scaffold_60240 | - | 253 | 203 | **RNA**→ rpmE (PRK00019)Ribosomal_L31 (pfam01197)→ |
| env-253 | SRS049147_LANL_scaffold_50458 | - | 171 | 121 | **RNA**→ rpmE (PRK00019)Ribosomal_L31 (pfam01197)→ |
| env-254 | SRS050029_C660889 | + | 260 | 310 | **RNA**→ |
| env-255 | SRS051244_LANL_scaffold_49385 | + | 361 | 411 | **RNA**→ |
| env-256 | SRS051791_LANL_scaffold_5910 | + | 208 | 258 | **RNA**→ rpmE (PRK00019)Ribosomal_L31 (pfam01197)→ |
| env-257 | SRS052027_C1805428 | - | 348 | 298 | **RNA**→ rpmE (PRK00019)Ribosomal_L31 (pfam01197)→ |
| env-258 | SRS052227_LANL_scaffold_41974 | - | 295 | 245 | **RNA**→ rpmE (PRK00019)Ribosomal_L31 (pfam01197)→ |
| env-259 | SRS052604_C1681687 | + | 239 | 289 | **RNA**→ |
| env-260 | SRS053603_C3252729 | + | 266 | 316 | **RNA**→ rpmE (PRK00019)Ribosomal_L31 (pfam01197)→ |
| env-261 | SRS054430_LANL_scaffold_5925 | - | 66 | 16 | **RNA**→ |
| env-262 | SRS054590_C647943 | - | 247 | 197 | **RNA**→ rpmE (PRK00019)Ribosomal_L31 (pfam01197)→ |
| env-263 | SRS054687_C3379381 | - | 302 | 252 | **RNA**→ rpmE (PRK00019)Ribosomal_L31 (pfam01197)→ |
| env-264 | SRS055401_LANL_scaffold_32442 | - | 252 | 202 | **RNA**→ rpmE (PRK00019)Ribosomal_L31 (pfam01197)→ |
| env-265 | SRS055426_LANL_scaffold_18857 | - | 951 | 901 | **RNA**→ ←PRK13528 (PRK13528)TonB_dep_Rec (pfam00593) |
| env-266 | SRS056323_C2564669 | - | 291 | 241 | **RNA**→ rpmE (PRK00019)Ribosomal_L31 (pfam01197)→ |
| env-267 | SRS056622_LANL_scaffold_38163 | + | 117 | 167 | **RNA**→ rpmE (PRK00019)Ribosomal_L31 (pfam01197)→ |
| env-268 | SRS056892_LANL_scaffold_6433 | + | 339 | 389 | **RNA**→ ←PRK13528 (PRK13528)ligand_gated_channel (cd01347) |

| | | | | | |
|---|---|---|---|---|---|
| env-269 | SRS057539_LANL_scaffold_29320 | + | 406 | 456 | **RNA**→ ←PRK13528 (PRK13528)ligand_gated_channel (cd01347) |
| env-270 | SRS057692_LANL_scaffold_16818 | + | 97 | 147 | **RNA**→ rpmE (PRK00019)Ribosomal_L31 (pfam01197)→ |
| env-271 | SRS057791_C3589113 | + | 145 | 195 | **RNA**→ rpmE (PRK00019)Ribosomal_L31 (pfam01197)→ |
| env-272 | SRS058336_C2100928 | + | 217 | 267 | **RNA**→ rpmE (PRK00019)Ribosomal_L31 (pfam01197)→ |
| env-273 | SRS058723_C6753173 | + | 271 | 321 | **RNA**→ rpmE (PRK00019)Ribosomal_L31 (pfam01197)→ |
| env-274 | SRS062540_LANL_scaffold_77508 | - | 354 | 304 | **RNA**→ rpmE (PRK00019)Ribosomal_L31 (pfam01197)→ |
| env-275 | SRS062544_C2913519 | + | 260 | 310 | **RNA**→ rpmE (PRK00019)Ribosomal_L31 (pfam01197)→ |
| env-276 | SRS063478_C532042 | + | 109 | 159 | **RNA**→ rpmE (PRK00019)Ribosomal_L31 (pfam01197)→ |
| env-277 | SRS064423_LANL_scaffold_7733 | - | 403 | 353 | **RNA**→ rpmE (PRK00019)Ribosomal_L31 (pfam01197)→ |
| env-278 | SRS064774_C2757027 | - | 295 | 245 | **RNA**→ rpmE (PRK00019)Ribosomal_L31 (pfam01197)→ |
| env-279 | SRS065278_LANL_scaffold_48090 | - | 535 | 485 | **RNA**→ rpmE (PRK00019)Ribosomal_L31 (pfam01197)→ |
| env-280 | SRS065335_C936123 | - | 321 | 271 | **RNA**→ rpmE (PRK00019)Ribosomal_L31 (pfam01197)→ |
| env-281 | SRS075406_C811674 | + | 97 | 147 | **RNA**→ rpmE (PRK00019)Ribosomal_L31 (pfam01197)→ |
| env-282 | SRS075410_LANL_scaffold_13122 | + | 361 | 411 | **RNA**→ rpmE (PRK00019)Ribosomal_L31 (pfam01197)→ |
| env-283 | SRS077736_LANL_scaffold_33263 | - | 287 | 237 | **RNA**→ rpmE (PRK00019)Ribosomal_L31 (pfam01197)→ |
| Hpa-2-4 | NZ_JUOR01000021.1 | - | 10901 | 10851 | **RNA**→ rpmE (PRK00019)Ribosomal_L31 (pfam01197)→ |
| Hpa-6-1 | NC_015964.1 | - | 912647 | 912597 | **RNA**→ rpmE (PRK00019)Ribosomal_L31 (pfam01197)→ |
| env-284 | 4491485.3_NODE_13737 | + | 3635 | 3685 | **RNA**→ rpmE (PRK00019)Ribosomal_L31 (pfam01197)→ |
| env-285 | Keratinized_gingiva_LANL_scaffold_2331 | + | 325 | 375 | **RNA**→ rpmE (PRK00019)Ribosomal_L31 (pfam01197)→ |
| env-286 | SRS015540_C1078977 | + | 208 | 258 | **RNA**→ |
| env-287 | SRS016541_Baylor_scaffold_18251 | - | 14291 | 14241 | **RNA**→ rpmE (PRK00019)Ribosomal_L31 (pfam01197)→ |
| env-288 | SRS023841_Baylor_scaffold_26182 | + | 952 | 1002 | **RNA**→ rpmE (PRK00019)Ribosomal_L31 (pfam01197)→ |
| env-289 | SRS056157_C273441 | + | 153 | 203 | **RNA**→ rpmE (PRK00019)Ribosomal_L31 (pfam01197)→ |
| env-290 | NLM007_scaffold6885_2 | - | 257 | 207 | **RNA**→ rpmE (PRK00019)Ribosomal_L31 (pfam01197)→ |
| env-291 | NOM018_scaffold11447_1 | + | 1091 | 1141 | **RNA**→ rpmE (PRK00019)Ribosomal_L31 (pfam01197)→ |
| env-292 | SRS011098_Baylor_scaffold_8386 | - | 2267 | 2217 | **RNA**→ rpmE (PRK00019)Ribosomal_L31 (pfam01197)→ |
| env-293 | SRS011310_Baylor_scaffold_12329 | - | 240 | 190 | **RNA**→ rpmE (PRK00019)Ribosomal_L31 (pfam01197)→ |
| env-294 | SRS011343_Baylor_scaffold_2577 | + | 789 | 839 | **RNA**→ rpmE (PRK00019)Ribosomal_L31 (pfam01197)→ |
| env-295 | SRS013164_Baylor_scaffold_6398 | + | 1941 | 1991 | **RNA**→ hypo→ |
| env-296 | SRS013533_PGA_scaffold_83577 | - | 4677 | 4627 | **RNA**→ rpmE (PRK00019)Ribosomal_L31 (pfam01197)→ |
| env-297 | SRS014477_C1618804 | + | 520 | 570 | **RNA**→ rpmE (PRK00019)rpmE2 (PRK01678)Ribosomal_L31 (pfam01197)→ |
| env-298 | SRS014690_WUGC_scaffold_24846 | + | 2301 | 2351 | **RNA**→ rpmE (PRK00019)Ribosomal_L31 (pfam01197)→ |
| env-299 | SRS014894_WUGC_scaffold_15243 | + | 3458 | 3508 | **RNA**→ rpmE (PRK00019)Ribosomal_L31 (pfam01197)→ |
| env-300 | SRS015040_WUGC_scaffold_3662 | - | 66 | 16 | **RNA**→ |
| env-301 | SRS015044_WUGC_scaffold_680 | - | 16214 | 16164 | **RNA**→ ←PRK13524 (PRK13524)ligand_gated_channel (cd01347) |
| env-302 | SRS015063_WUGC_scaffold_20082 | - | 483 | 433 | **RNA**→ rpmE (PRK00019)Ribosomal_L31 (pfam01197)→ |
| env-303 | SRS015215_WUGC_scaffold_46212 | - | 4914 | 4864 | **RNA**→ rpmE (PRK00019)Ribosomal_L31 (pfam01197)→ |
| env-304 | SRS015374_WUGC_scaffold_2905 | + | 23086 | 23136 | **RNA**→ rpmE (PRK00019)Ribosomal_L31 (pfam01197)→ |
| env-305 | SRS015378_WUGC_scaffold_12166 | + | 554 | 604 | **RNA**→ rpmE (PRK00019)Ribosomal_L31 (pfam01197)→ |
| env-306 | SRS015440_WUGC_scaffold_25622 | + | 512 | 562 | **RNA**→ rpmE (PRK00019)Ribosomal_L31 (pfam01197)→ |
| env-307 | SRS015755_C1590475 | + | 281 | 331 | **RNA**→ |
| env-308 | SRS016043_WUGC_scaffold_17944 | + | 1164 | 1214 | **RNA**→ rpmE (PRK00528)rpmE (PRK00019)rpmE2 (PRK01678)Ribosomal_L31 (pfam01197)→ |
| env-309 | SRS016092_WUGC_scaffold_1540 | - | 14964 | 14914 | **RNA**→ rpmE (PRK00019)Ribosomal_L31 (pfam01197)→ |
| env-310 | SRS016196_C485265 | + | 1027 | 1077 | **RNA**→ |
| env-311 | SRS016331_C4184433 | + | 888 | 938 | **RNA**→ rpmE (PRK00019)Ribosomal_L31 (pfam01197)→ |
| env-312 | SRS016360_Baylor_scaffold_42110 | + | 1164 | 1214 | **RNA**→ |
| env-313 | SRS017227_Baylor_scaffold_17966 | - | 2211 | 2161 | **RNA**→ rpmE (PRK00019)Ribosomal_L31 (pfam01197)→ |
| env-314 | SRS017441_C716525 | - | 122 | 72 | **RNA**→ rpmE (PRK00528)rpmE (PRK00019)rpmE2 (PRK01678)Ribosomal_L31 (pfam01197)→ |
| env-315 | SRS017691_Baylor_scaffold_71029 | + | 1502 | 1552 | **RNA**→ |
| env-316 | SRS018149_Baylor_scaffold_8518 | - | 67 | 17 | **RNA**→ |
| env-317 | SRS018337_Baylor_scaffold_3991 | - | 160 | 110 | **RNA**→ rpmE (PRK00019)Ribosomal_L31 (pfam01197)→ |
| env-318 | SRS018394_Baylor_scaffold_31222 | - | 1643 | 1593 | **RNA**→ rpmE (PRK00019)Ribosomal_L31 (pfam01197)→ |

| | | | | | |
|---|---|---|---|---|---|
| env-319 | SRS019029_WUGC_scaffold_48805 | - | 66 | 16 | RNA→ |
| env-320 | SRS019333_WUGC_scaffold_15291 | + | 2869 | 2919 | RNA→ rpmE (PRK00019)Ribosomal_L31 (pfam01197)→ |
| env-321 | SRS019976_Baylor_scaffold_6230 | - | 243 | 193 | RNA→ rpmE (PRK00019)Ribosomal_L31 (pfam01197)→ |
| env-322 | SRS019980_Baylor_scaffold_31327 | + | 562 | 612 | RNA→ |
| env-323 | SRS020226_Baylor_scaffold_32347 | - | 213 | 163 | RNA→ rpmE (PRK00019)Ribosomal_L31 (pfam01197)→ |
| env-324 | SRS020340_Baylor_scaffold_20184 | + | 496 | 546 | RNA→ rpmE (PRK00019)Ribosomal_L31 (pfam01197)→ |
| env-325 | SRS020858_C769095 | - | 68 | 18 | RNA→ |
| env-326 | SRS020862_Baylor_scaffold_23178 | - | 1984 | 1934 | RNA→ rpmE (PRK00019)Ribosomal_L31 (pfam01197)→ |
| env-327 | SRS021960_Baylor_scaffold_7710 | - | 174 | 124 | RNA→ rpmE (PRK00019)Ribosomal_L31 (pfam01197)→ |
| env-328 | SRS022149_LANL_scaffold_6234 | - | 68 | 18 | RNA→ |
| env-329 | SRS023358_C3482659 | - | 66 | 16 | RNA→ |
| env-330 | SRS023538_Baylor_scaffold_23152 | - | 66 | 16 | RNA→ |
| env-331 | SRS023595_Baylor_scaffold_91416 | - | 66 | 16 | RNA→ |
| env-332 | SRS023987_Baylor_scaffold_1631 | + | 1076 | 1126 | RNA→ rpmE (PRK00019)Ribosomal_L31 (pfam01197)→ |
| env-333 | SRS024017_Baylor_scaffold_15613 | - | 68 | 18 | RNA→ |
| env-334 | SRS024144_LANL_scaffold_20602 | + | 916 | 966 | RNA→ |
| env-335 | SRS024561_C2824542 | - | 66 | 16 | RNA→ |
| env-336 | SRS042457_LANL_scaffold_4939 | + | 713 | 763 | RNA→ |
| env-337 | SRS042984_LANL_scaffold_74580 | + | 421 | 471 | RNA→ |
| env-338 | SRS045313_LANL_scaffold_12892 | - | 1373 | 1323 | RNA→ rpmE (PRK00019)Ribosomal_L31 (pfam01197)→ Glyco_transf_25 (pfam01755)→ PRK10422 (PRK10422)RfaF (COG0859)→ |
| env-339 | SRS047634_LANL_scaffold_115926 | - | 66 | 16 | RNA→ |
| env-340 | SRS049268_C4184946 | + | 288 | 338 | RNA→ |
| env-341 | SRS049318_C6295479 | - | 66 | 16 | RNA→ |
| env-342 | SRS050244_C4383723 | + | 117 | 167 | RNA→ rpmE (PRK00019)Ribosomal_L31 (pfam01197)→ |
| env-343 | SRS050628_LANL_scaffold_4839 | - | 68 | 18 | RNA→ |
| env-344 | SRS052876_LANL_scaffold_39738 | + | 595 | 645 | RNA→ |
| env-345 | SRS053584_LANL_scaffold_42738 | + | 163 | 213 | RNA→ rpmE (PRK00019)Ribosomal_L31 (pfam01197)→ |
| env-346 | SRS054653_C2080848 | - | 159 | 109 | RNA→ rpmE (PRK00019)Ribosomal_L31 (pfam01197)→ |
| env-347 | SRS055450_LANL_scaffold_67168 | + | 1178 | 1228 | RNA→ |
| env-348 | SRS058808_LANL_scaffold_12344 | + | 2015 | 2065 | RNA→ |
| env-349 | SRS063287_LANL_scaffold_6852 | + | 404 | 454 | RNA→ rpmE (PRK00019)Ribosomal_L31 (pfam01197)→ |
| env-350 | SRS064809_LANL_scaffold_5427 | + | 157 | 207 | RNA→ rpmE (PRK00019)Ribosomal_L31 (pfam01197)→ |
| env-351 | SRS065310_C1245580 | - | 68 | 18 | RNA→ |
| Hpa-2-5 | NZ_JVSU01000025.1 | + | 2976 | 3026 | RNA→ rpmE (PRK00019)Ribosomal_L31 (pfam01197)→ |
| Hpa-2-6 | NZ_JWCE01000018.1 | - | 105224 | 105174 | RNA→ rpmE (PRK00019)Ribosomal_L31 (pfam01197)→ |
| Hpa-2-7 | NZ_JVSB01000011.1 | + | 2976 | 3026 | RNA→ rpmE (PRK00019)Ribosomal_L31 (pfam01197)→ |
| Hpa-2-8 | NZ_JVRH01000079.1 | - | 114460 | 114410 | RNA→ rpmE (PRK00019)Ribosomal_L31 (pfam01197)→ |
| Hpa-2-9 | NZ_JUTJ01000002.1 | - | 24649 | 24599 | RNA→ rpmE (PRK00019)Ribosomal_L31 (pfam01197)→ |
| Hpa-2-10 | NZ_JVFE01000001.1 | - | 48872 | 48822 | RNA→ rpmE (PRK00019)Ribosomal_L31 (pfam01197)→ |
| Hpa-5-1 | NZ_AJMW01000057.1 | - | 27592 | 27542 | RNA→ rpmE (PRK00019)Ribosomal_L31 (pfam01197)→ |
| Hpa-4-1 | NZ_AJTC01000031.1 | - | 50368 | 50318 | RNA→ rpmE (PRK00019)Ribosomal_L31 (pfam01197)→ |
| Hpa-3-1 | NZ_GL872339.1 | + | 1999941 | 1999991 | RNA→ rpmE (PRK00019)Ribosomal_L31 (pfam01197)→ |
| env-352 | SRS017076_Baylor_scaffold_26623 | - | 284 | 234 | RNA→ rpmE (PRK00019)Ribosomal_L31 (pfam01197)→ |
| env-353 | SRS015038_C1823634 | + | 113 | 163 | RNA→ rpmE (PRK00019)Ribosomal_L31 (pfam01197)→ |
| env-354 | SRS024637_C2357778 | - | 307 | 257 | RNA→ rpmE (PRK00019)Ribosomal_L31 (pfam01197)→ |
| env-355 | SRS053854_LANL_scaffold_40299 | + | 271 | 321 | RNA→ rpmE (PRK00019)Ribosomal_L31 (pfam01197)→ |
| env-356 | SRS022602_Baylor_scaffold_39274 | - | 1217 | 1167 | RNA→ rpmE (PRK00019)Ribosomal_L31 (pfam01197)→ |
| env-357 | SRS015470_WUGC_scaffold_7219 | + | 442 | 492 | RNA→ rpmE (PRK00019)Ribosomal_L31 (pfam01197)→ |
| env-358 | SRS014578_WUGC_scaffold_4249 | - | 4087 | 4037 | RNA→ rpmE (PRK00019)Ribosomal_L31 (pfam01197)→ |
| env-359 | SRS014474_C733884 | + | 2001 | 2051 | RNA→ rpmE (PRK00019)Ribosomal_L31 (pfam01197)→ |
| env-360 | SRS020220_Baylor_scaffold_39783 | - | 240 | 190 | RNA→ rpmE (PRK00019)Ribosomal_L31 (pfam01197)→ |

| | | | | | |
|---|---|---|---|---|---|
| env-361 | SRS022145_C596044 | + | 247 | 297 | **RNA**→ |
| env-362 | SRS042643_WUGC_scaffold_30660 | + | 182 | 232 | **RNA**→ rpmE (PRK00019)Ribosomal_L31 (pfam01197)→ |
| Hpa-2-11 | NZ_JVYT01000034.1 | + | 2978 | 3028 | **RNA**→ rpmE (PRK00019)Ribosomal_L31 (pfam01197)→ |
| Rpn-1-1 | NZ_BBIX01000005.1 | - | 3001 | 2951 | **RNA**→ rpmE (PRK00019)Ribosomal_L31 (pfam01197)→ |
| Rpn-1-2 | NZ_BBIX01000009.1 | - | 658570 | 658520 | **RNA**→ rpmE (PRK00019)Ribosomal_L31 (pfam01197)→ |
| Ppn-1-1 | NZ_KB904052.1 | + | 167149 | 167199 | **RNA**→ rpmE (PRK00019)Ribosomal_L31 (pfam01197)→ |
| env-363 | SRS014692_C1497409 | + | 379 | 429 | **RNA**→ rpmE (PRK00019)Ribosomal_L31 (pfam01197)→ |
| env-364 | SRS014126_C565578 | - | 84 | 34 | **RNA**→ |
| env-365 | SRS014687_WUGC_scaffold_2169 | - | 3350 | 3300 | **RNA**→ rpmE (PRK00019)Ribosomal_L31 (pfam01197)→ PRK11204 (PRK11204)Glycos_transf_2 (pfam00535)→ |
| | | | | | GT8_A4GalT_like (cd04194)PRK15171 (PRK15171)→ Glyco_transf_25 (pfam01755)→ Glyco_transf_25 (cd06532)→ |
| env-366 | SRS018978_C1739365 | - | 62 | 12 | **RNA**→ |
| env-367 | SRS048719_C812527 | + | 2851 | 2901 | **RNA**→ rpmE (PRK00019)Ribosomal_L31 (pfam01197)→ |
| env-368 | SRS016196_C475885 | + | 466 | 516 | **RNA**→ |
| env-369 | SRS021986_C944337 | - | 84 | 34 | **RNA**→ |
| env-370 | SRS011090_Baylor_scaffold_4448 | - | 1061 | 1011 | **RNA**→ rpmE (PRK00019)Ribosomal_L31 (pfam01197)→ Glyco_transf_25 (pfam01755)POX (pfam07526)→ |
| env-371 | SRS014472_C402330 | - | 548 | 498 | **RNA**→ rpmE (PRK00019)Ribosomal_L31 (pfam01197)→ Glyco_transf_25 (pfam01755)→ |
| env-372 | SRS016039_WUGC_scaffold_8838 | - | 2881 | 2831 | **RNA**→ rpmE (PRK00019)Ribosomal_L31 (pfam01197)→ Glyco_transf_25 (pfam01755)POX (pfam07526)→ |
| | | | | | GT8_A4GalT_like (cd04194)PRK15171 (PRK15171)→ Glyco_transf_25 (pfam01755)→ |
| env-373 | SRS017013_Baylor_scaffold_22515 | + | 291 | 341 | **RNA**→ Glyco_transf_25 (pfam01755)POX (pfam07526)→ |
| env-374 | SRS017810_Baylor_scaffold_7758 | + | 2802 | 2852 | **RNA**→ rpmE (PRK00019)Ribosomal_L31 (pfam01197)→ Glyco_transf_25 (pfam01755)POX (pfam07526)→ |
| env-375 | SRS018329_Baylor_scaffold_7486 | + | 448 | 498 | **RNA**→ rpmE (PRK00019)rpmE2 (PRK01678)Ribosomal_L31 (pfam01197)→ |
| env-376 | SRS018359_C1229806 | + | 1424 | 1474 | **RNA**→ |
| env-377 | SRS019329_C786673 | - | 68 | 18 | **RNA**→ |
| env-378 | SRS019587_C1052142 | + | 652 | 702 | **RNA**→ |
| env-379 | SRS021473_Baylor_scaffold_4924 | + | 2802 | 2852 | **RNA**→ rpmE (PRK00019)Ribosomal_L31 (pfam01197)→ Glyco_transf_25 (pfam01755)POX (pfam07526)→ |
| env-380 | SRS023534_C139914 | - | 324 | 274 | **RNA**→ rpmE (PRK00019)Ribosomal_L31 (pfam01197)→ |
| env-381 | SRS023538_Baylor_scaffold_23260 | + | 216 | 266 | **RNA**→ rpmE (PRK00019)rpmE2 (PRK01678)Ribosomal_L31 (pfam01197)→ |
| env-382 | SRS024377_LANL_scaffold_33 | - | 23058 | 23008 | **RNA**→ rpmE (PRK00019)Ribosomal_L31 (pfam01197)→ Glyco_transf_25 (pfam01755)POX (pfam07526)→ |
| | | | | | GT8_A4GalT_like (cd04194)PRK15171 (PRK15171)→ Glyco_transf_25 (pfam01755)→ |
| env-383 | SRS024381_LANL_scaffold_54075 | - | 559 | 509 | **RNA**→ rpmE (PRK00019)Ribosomal_L31 (pfam01197)→ Glyco_transf_25 (pfam01755)→ |
| env-384 | SRS052668_C1028805 | + | 444 | 494 | **RNA**→ |
| env-385 | SRS015985_WUGC_scaffold_5736 | + | 8804 | 8854 | **RNA**→ rpmE (PRK00019)Ribosomal_L31 (pfam01197)→ RfaB (COG0438)Glyco_trans_1_4 (pfam13692)→ |
| | | | | | GT8_A4GalT_like (cd04194)PRK15171 (PRK15171)→ Glyco_transf_25 (pfam01755)→ |
| | | | | | Glyco_transf_25 (pfam01755)→ Glyco_tranf_GTA_type (cd00761)PRK10073 (PRK10073)→ |
| env-386 | SRS011098_Baylor_scaffold_48474 | - | 745 | 695 | **RNA**→ rpmE (PRK00019)Ribosomal_L31 (pfam01197)→ hypo→ |
| env-387 | SRS011144_C527765 | - | 148 | 98 | **RNA**→ rpmE (PRK00019)Ribosomal_L31 (pfam01197)→ |
| env-388 | SRS016088_C183306 | - | 94 | 44 | **RNA**→ |
| env-389 | SRS017127_Baylor_scaffold_6911 | + | 2934 | 2984 | **RNA**→ rpmE (PRK00019)Ribosomal_L31 (pfam01197)→ Glyco_transf_25 (pfam01755)POX (pfam07526)→ |
| | | | | | GT8_A4GalT_like (cd04194)PRK15171 (PRK15171)→ Glyco_transf_25 (pfam01755)→ |
| env-390 | SRS017139_C3563542 | + | 262 | 312 | **RNA**→ |
| env-391 | SRS017441_C719147 | + | 344 | 394 | **RNA**→ rpmE (PRK00528)rpmE (PRK00019)rpmE2 (PRK01678)Ribosomal_L31 (pfam01197)→ |
| env-392 | SRS022145_C605259 | - | 68 | 18 | **RNA**→ |
| env-393 | SRS051116_LANL_scaffold_1628 | + | 2934 | 2984 | **RNA**→ rpmE (PRK00019)Ribosomal_L31 (pfam01197)→ Glyco_transf_25 (pfam01755)POX (pfam07526)→ |
| | | | | | GT8_A4GalT_like (cd04194)PRK15171 (PRK15171)→ Glyco_transf_25 (pfam01755)→ |
| env-394 | SRS011343_Baylor_scaffold_11773 | - | 68 | 18 | **RNA**→ |
| env-395 | SRS019387_C1774500 | + | 1449 | 1499 | **RNA**→ rpmE (PRK00019)Ribosomal_L31 (pfam01197)→ |
| env-396 | SRS020858_C776329 | + | 1031 | 1081 | **RNA**→ |
| env-397 | SRS020862_Baylor_scaffold_28830 | - | 1159 | 1109 | **RNA**→ rpmE (PRK00019)Ribosomal_L31 (pfam01197)→ Glyco_transf_25 (pfam01755)→ |
| env-398 | SRS022532_LANL_scaffold_11832 | + | 8865 | 8915 | **RNA**→ Glyco_transf_25 (pfam01755)→ |
| env-399 | SRS024281_C986018 | + | 1138 | 1188 | **RNA**→ |
| env-400 | SRS024289_LANL_scaffold_11530 | - | 68 | 18 | **RNA**→ |
| env-401 | SRS013164_C2241878 | + | 310 | 360 | **RNA**→ |
| env-402 | SRS022625_C2147320 | - | 68 | 18 | **RNA**→ |
| env-403 | SRS023987_C1180207 | - | 68 | 18 | **RNA**→ |

| | | | | | |
|---|---|---|---|---|---|
| env-404 | SRS017080_Baylor_scaffold_3076 | - | 1935 | 1885 | **RNA**→ rpmE (PRK00019)Ribosomal_L31 (pfam01197)→ Glyco_transf_25 (pfam01755)POX (pfam07526)→ GT8_A4GalT_like (cd04194)PRK15171 (PRK15171)→ |
| env-405 | SRS024318_LANL_scaffold_15937 | - | 68 | 18 | **RNA**→ |
| env-406 | SRS050628_LANL_scaffold_21056 | + | 835 | 885 | **RNA**→ rpmE (PRK00019)rpmE2 (PRK01678)Ribosomal_L31 (pfam01197)→ |
| env-407 | SRS045262_WUGC_scaffold_11350 | + | 1338 | 1388 | **RNA**→ ←hypo |
| env-408 | SRS045049_WUGC_scaffold_4937 | - | 259 | 209 | **RNA**→ rpmE (PRK00019)Ribosomal_L31 (pfam01197)→ |
| env-409 | SRS013239_Baylor_scaffold_8305 | + | 2802 | 2852 | **RNA**→ |
| env-410 | SRS024017_C890428 | - | 68 | 18 | **RNA**→ |
| env-411 | SRS013818_Baylor_scaffold_30131 | - | 190 | 140 | **RNA**→ rpmE (PRK00019)Ribosomal_L31 (pfam01197)→ |
| env-412 | SRS018300_Baylor_scaffold_30376 | - | 198 | 148 | **RNA**→ rpmE (PRK00019)Ribosomal_L31 (pfam01197)→ |
| env-413 | SRS020856_C2499523 | + | 297 | 347 | **RNA**→ |
| env-414 | SRS024441_LANL_scaffold_23848 | + | 1393 | 1443 | **RNA**→ rpmE (PRK00019)Ribosomal_L31 (pfam01197)→ |
| env-415 | SRS043701_C1809232 | + | 527 | 577 | **RNA**→ rpmE (PRK00019)Ribosomal_L31 (pfam01197)→ |
| env-416 | SRS049389_C4590327 | + | 399 | 449 | **RNA**→ |
| env-417 | SRS017215_Baylor_scaffold_5416 | + | 724 | 774 | **RNA**→ |
| Hha-1-1 | NZ_LCTK01000024.1 | + | 36220 | 36270 | **RNA**→ rpmE (PRK00019)Ribosomal_L31 (pfam01197)→ |
| env-418 | SRS011247_C1322309 | + | 307 | 357 | **RNA**→ |
| env-419 | SRS015574_C3331718 | + | 437 | 487 | **RNA**→ |
| Hha-1-2 | NZ_LCTI01000049.1 | - | 6537 | 6487 | **RNA**→ rpmE (PRK00019)Ribosomal_L31 (pfam01197)→ |
| env-420 | SRS019028_C2927759 | - | 68 | 18 | **RNA**→ |
| env-421 | SRS024087_LANL_scaffold_49246 | - | 237 | 187 | **RNA**→ rpmE (PRK00019)Ribosomal_L31 (pfam01197)→ |
| Hha-1-3 | NZ_LCTH01000053.1 | - | 9141 | 9091 | **RNA**→ rpmE (PRK00019)Ribosomal_L31 (pfam01197)→ CgtA (pfam06306)→ Glyco_trans_1_4 (pfam13692)GT1_WabH_like (cd03811)→ Glyco_transf_25 (pfam01755)→ |
| Hha-1-4 | NZ_LCTJ01000055.1 | - | 6545 | 6495 | **RNA**→ rpmE (PRK00019)Ribosomal_L31 (pfam01197)→ |
| env-422 | SRS023930_Baylor_scaffold_6246 | + | 2806 | 2856 | **RNA**→ ←hypo |
| env-423 | SRS043422_LANL_scaffold_13569 | + | 103 | 153 | **RNA**→ rpmE (PRK00019)Ribosomal_L31 (pfam01197)→ |
| env-424 | SRS017127_C1115545 | + | 1102 | 1152 | **RNA**→ rpmE (PRK00019)Ribosomal_L31 (pfam01197)→ |
| env-425 | SRS019125_WUGC_scaffold_9446 | + | 973 | 1023 | **RNA**→ rpmE (PRK00019)Ribosomal_L31 (pfam01197)→ |
| env-426 | SRS019221_WUGC_scaffold_2235 | - | 213 | 163 | **RNA**→ rpmE (PRK00019)Ribosomal_L31 (pfam01197)→ |
| env-427 | SRS062761_C4146270 | + | 274 | 324 | **RNA**→ |
| Hha-4-1 | NZ_AFQP01000010.1 | + | 312767 | 312817 | **RNA**→ rpmE (PRK00019)Ribosomal_L31 (pfam01197)→ |
| Hha-2-1 | NZ_AJSV01000029.1 | - | 6279 | 6229 | **RNA**→ rpmE (PRK00019)Ribosomal_L31 (pfam01197)→ |
| env-428 | Saliva_LANL_C126650 | + | 119 | 169 | **RNA**→ rpmE (PRK00019)Ribosomal_L31 (pfam01197)→ |
| env-429 | SRS015803_C2530019 | + | 300 | 350 | **RNA**→ |
| Hha-3-1 | NZ_AFQO01000007.1 | - | 7110 | 7060 | **RNA**→ rpmE (PRK00019)Ribosomal_L31 (pfam01197)→ |
| Hsp-2-1 | NZ_JH591072.1 | + | 167214 | 167264 | **RNA**→ rpmE (PRK00019)Ribosomal_L31 (pfam01197)→ |
| env-430 | SRS015436_WUGC_scaffold_10632 | - | 502 | 452 | **RNA**→ rpmE (PRK00019)Ribosomal_L31 (pfam01197)→ |
| Hha-5-1 | NZ_AFQQ01000021.1 | - | 6823 | 6773 | **RNA**→ rpmE (PRK00019)Ribosomal_L31 (pfam01197)→ rpmE (PRK00019)Ribosomal_L31 (pfam01197)→ |
| Hha-6-1 | NZ_AFQR01000017.1 | - | 23162 | 23112 | **RNA**→ rpmE (PRK00019)Ribosomal_L31 (pfam01197)→ |
| Hin-17-1 | NZ_AAZI01000001.1 | + | 341323 | 341373 | **RNA**→ rpmE (PRK00019)Ribosomal_L31 (pfam01197)→ rpmE (PRK00019)Ribosomal_L31 (pfam01197)→ |
| Hin-19-1 | NC_017451.1 | + | 1667269 | 1667319 | **RNA**→ rpmE (PRK00019)Ribosomal_L31 (pfam01197)→ |
| env-431 | SRS014475_C164765 | + | 539 | 589 | **RNA**→ rpmE (PRK00019)Ribosomal_L31 (pfam01197)→ |
| Hin-5-1 | NZ_JXCI01000014.1 | - | 45598 | 45548 | **RNA**→ rpmE (PRK00019)Ribosomal_L31 (pfam01197)→ |
| Hin-7-1 | NZ_ABWV01000001.1 | - | 45931 | 45881 | **RNA**→ rpmE (PRK00019)Ribosomal_L31 (pfam01197)→ rpmE (PRK00019)Ribosomal_L31 (pfam01197)→ |
| Hin-21-1 | NZ_ACSM01000001.1 | - | 45997 | 45947 | **RNA**→ rpmE (PRK00019)Ribosomal_L31 (pfam01197)→ rpmE (PRK00019)Ribosomal_L31 (pfam01197)→ |
| Hin-16-1 | NZ_AAZH01000004.1 | + | 87662 | 87712 | **RNA**→ rpmE (PRK00019)Ribosomal_L31 (pfam01197)→ rpmE (PRK00019)Ribosomal_L31 (pfam01197)→ |
| Hin-20-1 | NC_000907.1 | - | 822480 | 822430 | **RNA**→ rpmE (PRK00019)Ribosomal_L31 (pfam01197)→ |
| Hin-15-1 | NC_009566.1 | + | 1630832 | 1630882 | **RNA**→ rpmE (PRK00019)Ribosomal_L31 (pfam01197)→ |
| Hin-8-1 | NC_007146.2 | - | 887870 | 887820 | **RNA**→ rpmE (PRK00019)Ribosomal_L31 (pfam01197)→ |
| Hin-18-1 | NC_017452.1 | + | 1629437 | 1629487 | **RNA**→ rpmE (PRK00019)Ribosomal_L31 (pfam01197)→ |
| Hin-10-1 | NC_014920.1 | + | 1454654 | 1454704 | **RNA**→ rpmE (PRK00019)Ribosomal_L31 (pfam01197)→ |
| Hae-1-1 | NZ_GL878526.1 | - | 481719 | 481669 | **RNA**→ rpmE (PRK00019)Ribosomal_L31 (pfam01197)→ |
| Hin-2-1 | NC_016809.1 | - | 927122 | 927072 | **RNA**→ rpmE (PRK00019)Ribosomal_L31 (pfam01197)→ |

| | | | | | |
|---|---|---|---|---|---|
| Hin-1-1 | NZ_CP007472.1 | + | 866657 | 866707 | **RNA**→ rpmE (PRK00019)Ribosomal_L31 (pfam01197)→ |
| Hin-9-1 | NZ_CP007805.1 | + | 1761399 | 1761449 | **RNA**→ rpmE (PRK00019)Ribosomal_L31 (pfam01197)→ |
| Hin-1-2 | NZ_CP009610.1 | + | 226803 | 226853 | **RNA**→ rpmE (PRK00019)Ribosomal_L31 (pfam01197)→ |
| Apa-1-1 | NZ_LAEO01000011.1 | + | 134437 | 134487 | **RNA**→ rpmE (PRK00019)Ribosomal_L31 (pfam01197)→ |
| Apa-2-1 | NZ_AOGF01000002.1 | + | 91207 | 91257 | **RNA**→ rpmE (PRK00019)Ribosomal_L31 (pfam01197)→ |
| Apa-3-1 | NZ_AFFP02000002.1 | + | 105702 | 105752 | **RNA**→ rpmE (PRK00019)Ribosomal_L31 (pfam01197)→ |
| Apa-4-1 | NZ_CBMK010000011.1 | - | 233267 | 233217 | **RNA**→ rpmE (PRK00019)Ribosomal_L31 (pfam01197)→ |
| Apa-1-2 | NZ_LAEN01000032.1 | + | 12136 | 12186 | **RNA**→ rpmE (PRK00019)Ribosomal_L31 (pfam01197)→ |
| Nro-1-1 | NZ_CCMQ01000009.1 | - | 101998 | 101948 | **RNA**→ rpmE (PRK00019)Ribosomal_L31 (pfam01197)→ |
| Hma-1-1 | NZ_CCFL01000036.1 | - | 64728 | 64678 | **RNA**→ rpmE (PRK00019)Ribosomal_L31 (pfam01197)→ |
| Hsp-1-1 | NZ_CCNK01000036.1 | - | 64728 | 64678 | **RNA**→ rpmE (PRK00019)Ribosomal_L31 (pfam01197)→ |
| Psp-56-1 | NZ_CDON01000014.1 | - | 178969 | 178919 | **RNA**→ rpmE (PRK00019)Ribosomal_L31 (pfam01197)→ |
| Psp-56-2 | NZ_CDNG01000014.1 | - | 178969 | 178919 | **RNA**→ rpmE (PRK00019)Ribosomal_L31 (pfam01197)→ |
| Psp-56-3 | NZ_CDEX01000018.1 | - | 178969 | 178919 | **RNA**→ rpmE (PRK00019)Ribosomal_L31 (pfam01197)→ |
| Hin-17-2 | NZ_AAZI01000001.1 | + | 243123 | 243173 | **RNA**→ hypo→ Glyco_transf_25 (pfam01755)→ Glyco_transf_25 (pfam01755)→ heptsyl_trn_III (TIGR02201)RfaF (COG0859)→ heptsyl_trn_III (TIGR02201)RfaF (COG0859)→ |
| Hin-19-2 | NC_017451.1 | + | 1569026 | 1569076 | **RNA**→ hypo→ hypo→ hypo→ Glyco_transf_25 (pfam01755)→ heptsyl_trn_III (TIGR02201)RfaF (COG0859)→ |
| env-432 | IMNBL1_c0005061 | + | 1709 | 1758 | **RNA**→ rpmE (PRK00019)Ribosomal_L31 (pfam01197)→ |
| Mmu-1-1 | NZ_JWIZ01000037.1 | - | 22959 | 22904 | **RNA**→ rpmE (PRK00019)Ribosomal_L31 (pfam01197)→ Glyco_transf_25 (cd06532)→ Glyco_transf_4 (pfam13439)Glyco_trans_1_4 (pfam13692)GT1_WabH_like (cd03811)→ PBP2_IlvY (cd08430)PRK11716 (PRK11716)HTH_1 (pfam00126)→ Upp (COG0035)upp (PRK00129)→ |
| Cor-1-1 | NZ_JSUM01000010.1 | + | 3479 | 3528 | **RNA**→ rpmE (PRK00019)Ribosomal_L31 (pfam01197)→ NTP-PPase_MazG_Nterm (cd11528)NTP-PPase_MazG_Cterm (cd11529)mazG (PRK09562)→ |
| Baq-1-1 | NZ_ATYS01000043.1 | - | 32716 | 32666 | **RNA**→ rpmE (PRK00019)Ribosomal_L31 (pfam01197)→ |
| Pfo-1-1 | NZ_CP010423.1 | - | 3853120 | 3853071 | **RNA**→ rpmE (PRK00019)Ribosomal_L31 (pfam01197)→ |
| env-433 | CEPZ01031768.1 | - | 3598 | 3538 | **RNA**→ rpmE (PRK00019)Ribosomal_L31 (pfam01197)→ |
| env-434 | CESI01260480.1 | - | 92 | 32 | **RNA**→ |
| env-435 | SRS018157_Baylor_scaffold_12494 | - | 682 | 619 | **RNA**→ rpmE (PRK00019)Ribosomal_L31 (pfam01197)→ |
| env-436 | SRS047634_LANL_scaffold_94015 | - | 848 | 785 | **RNA**→ ←RfaF (COG0859)heptsyl_trn_I (TIGR02193) |
| env-437 | SRS015440_WUGC_scaffold_33201 | + | 172 | 235 | **RNA**→ rpmE (PRK00019)Ribosomal_L31 (pfam01197)→ |
| env-438 | SRS013949_WUGC_scaffold_20645 | + | 1423 | 1486 | **RNA**→ rpmE (PRK00019)Ribosomal_L31 (pfam01197)→ |
| env-439 | SRS013950_C2020939 | + | 249 | 312 | **RNA**→ |
| env-440 | SRS019129_WUGC_scaffold_8522 | + | 436 | 499 | **RNA**→ rpmE (PRK00019)Ribosomal_L31 (pfam01197)→ |
| env-441 | SRS053917_LANL_scaffold_40839 | - | 230 | 167 | **RNA**→ rpmE (PRK00019)Ribosomal_L31 (pfam01197)→ |
| env-442 | SRS017445_C2959731 | + | 208 | 271 | **RNA**→ |
| env-443 | SRS018394_C3997563 | + | 368 | 431 | **RNA**→ rpmE (PRK00019)Ribosomal_L31 (pfam01197)→ |
| env-444 | SRS045197_C3344225 | + | 229 | 292 | **RNA**→ |
| env-445 | SRS013170_Baylor_scaffold_137087 | + | 944 | 1007 | **RNA**→ rpmE (PRK00019)Ribosomal_L31 (pfam01197)→ |
| env-446 | SRS024087_LANL_scaffold_38158 | + | 3632 | 3695 | **RNA**→ rpmE (PRK00019)Ribosomal_L31 (pfam01197)→ |
| env-447 | SRS055378_C3854289 | + | 246 | 309 | **RNA**→ |
| env-448 | SRS055401_C2303800 | + | 358 | 421 | **RNA**→ rpmE (PRK00019)Ribosomal_L31 (pfam01197)→ |
| env-449 | SRS064449_C4092018 | - | 272 | 209 | **RNA**→ rpmE (PRK00019)Ribosomal_L31 (pfam01197)→ |
| Tvi-1-1 | NZ_JYNJ01000035.1 | - | 80853 | 80804 | **RNA**→ rpmE (PRK00019)Ribosomal_L31 (pfam01197)→ rpmE (PRK00019)Ribosomal_L31 (pfam01197)→ |
| Tac-1-1 | NZ_JYNI01000086.1 | + | 37893 | 37942 | **RNA**→ rpmE (PRK00019)Ribosomal_L31 (pfam01197)→ |
| env-450 | JGI24723J26617_10002644 | + | 13634 | 13692 | **RNA**→ rpmE (PRK00019)Ribosomal_L31 (pfam01197)→ |
| env-451 | JGI24723J26617_10002693 | + | 13459 | 13517 | **RNA**→ rpmE (PRK00019)Ribosomal_L31 (pfam01197)→ |
| env-452 | KGI_S2_ANT06_2345m_c1007050 | + | 3448 | 3506 | **RNA**→ |
| env-453 | CEWQ01046878.1 | + | 431 | 489 | **RNA**→ rpmE (PRK00019)Ribosomal_L31 (pfam01197)→ |
| env-454 | KGI_S1_ANT01_95m_c10073717 | + | 821 | 879 | **RNA**→ rpmE (PRK00019)Ribosomal_L31 (pfam01197)→ |
| env-455 | KGI_S2_ANT04_2345m_c1000317 | + | 1025 | 1083 | **RNA**→ rpmE (PRK00019)Ribosomal_L31 (pfam01197)→ |
| env-456 | KGI_S1_ANT02_95m_c10008266 | + | 756 | 814 | **RNA**→ rpmE (PRK00019)Ribosomal_L31 (pfam01197)→ |
| env-457 | CESO01253667.1 | + | 628 | 686 | **RNA**→ rpmE (PRK00019)Ribosomal_L31 (pfam01197)→ |
| env-458 | JGI24724J26744_10001006 | + | 3321 | 3379 | **RNA**→ rpmE (PRK00019)Ribosomal_L31 (pfam01197)→ |

| | | | | | |
|---|---|---|---|---|---|
| env-459 | KGI_S2_ANT04_2345m_c1002142 | + | 7997 | 8055 | **RNA**→ |
| env-460 | KGI_S1_ANT01_95m_c10018623 | + | 2892 | 2950 | **RNA**→ ←hypo |
| env-461 | KGI_S1_ANT01_95m_c10184467 | - | 71 | 13 | **RNA**→ ←hypo |
| env-462 | JCVI_SCAF_1101668217967 | + | 711 | 759 | **RNA**→ rpmE (PRK00019)Ribosomal_L31 (pfam01197)→ |
| env-463 | CETA01009654.1 | - | 3746 | 3690 | **RNA**→ rpmE (PRK00019)Ribosomal_L31 (pfam01197)→ |
| env-464 | CETX01104058.1 | - | 71 | 15 | **RNA**→ |
| env-465 | CEUK01157528.1 | + | 2126 | 2182 | **RNA**→ rpmE (PRK00019)Ribosomal_L31 (pfam01197)→ |
| Iba-1-1 | NZ_CH672403.1 | + | 337077 | 337133 | **RNA**→ rpmE (PRK00019)Ribosomal_L31 (pfam01197)→ |
| env-466 | CETM01039028.1 | + | 1422 | 1478 | **RNA**→ rpmE (PRK00019)Ribosomal_L31 (pfam01197)→ |
| env-467 | CESO01134195.1 | - | 3172 | 3116 | **RNA**→ rpmE (PRK00019)Ribosomal_L31 (pfam01197)→ |
| env-468 | CETM01039027.1 | + | 30 | 86 | **RNA**→ rpmE (PRK00019)Ribosomal_L31 (pfam01197)→ |
| env-469 | JGI11881J13070_1002314 | + | 722 | 772 | **RNA**→ rpmE (PRK00019)Ribosomal_L31 (pfam01197)→ |
| env-470 | CEQC01048050.1 | - | 211 | 161 | **RNA**→ rpmE (PRK00019)Ribosomal_L31 (pfam01197)→ |
| env-471 | CEQH01152759.1 | + | 421 | 471 | **RNA**→ |
| env-472 | CEQK01044707.1 | - | 2989 | 2939 | **RNA**→ rpmE (PRK00019)Ribosomal_L31 (pfam01197)→ |
| env-473 | CESK01005100.1 | - | 13926 | 13876 | **RNA**→ rpmE (PRK00019)Ribosomal_L31 (pfam01197)→ |
| env-474 | CEVG01126604.1 | - | 447 | 397 | **RNA**→ rpmE (PRK00019)Ribosomal_L31 (pfam01197)→ |
| env-475 | CERU01038670.1 | - | 1641 | 1591 | **RNA**→ rpmE (PRK00019)Ribosomal_L31 (pfam01197)→ |
| env-476 | CETF01102798.1 | + | 365 | 415 | **RNA**→ rpmE (PRK00019)Ribosomal_L31 (pfam01197)→ |
| env-477 | CENV01129508.1 | + | 828 | 878 | **RNA**→ rpmE (PRK00019)Ribosomal_L31 (pfam01197)→ |
| env-478 | CESO01310600.1 | + | 1303 | 1353 | **RNA**→ rpmE (PRK00019)Ribosomal_L31 (pfam01197)→ |
| Isp-1-1 | NZ_BANL01000034.1 | + | 133074 | 133124 | **RNA**→ rpmE (PRK00019)Ribosomal_L31 (pfam01197)→ |
| env-479 | CETA01011209.1 | + | 242 | 292 | **RNA**→ rpmE (PRK00019)Ribosomal_L31 (pfam01197)→ |
| Ilo-2-1 | NC_006512.1 | + | 2636814 | 2636864 | **RNA**→ rpmE (PRK00019)Ribosomal_L31 (pfam01197)→ |
| Ilo-1-1 | NC_021286.1 | + | 2636929 | 2636979 | **RNA**→ rpmE (PRK00019)Ribosomal_L31 (pfam01197)→ |
| env-480 | JGI24724J26744_10119556 | + | 251 | 301 | **RNA**→ ←hypo |
| env-481 | CEGE01001000.1 | - | 1200 | 1150 | **RNA**→ rpmE (PRK00019)Ribosomal_L31 (pfam01197)→ |
| Vru-1-1 | NZ_AJYK01000068.1 | + | 61091 | 61172 | **RNA**→ rpmE (PRK00019)Ribosomal_L31 (pfam01197)→ |
| env-482 | AUXO015586256.1 | - | 5177 | 5128 | **RNA**→ rpmE (PRK00019)Ribosomal_L31 (pfam01197)→ |
| env-483 | CEUX01028586.1 | + | 276 | 324 | **RNA**→ rpmE (PRK00019)Ribosomal_L31 (pfam01197)→ |
| Ama-1-1 | NZ_AUBY01000009.1 | + | 261972 | 262030 | **RNA**→ rpmE (PRK00019)Ribosomal_L31 (pfam01197)→ |
| Ata-2-1 | NZ_AUBZ01000002.1 | + | 259960 | 260018 | **RNA**→ rpmE (PRK00019)Ribosomal_L31 (pfam01197)→ |
| env-484 | topACOD_F95O9CU02IUQBW | + | 144 | 205 | **RNA**→ rpmE (PRK00019)Ribosomal_L31 (pfam01197)→ |
| env-485 | JGI24739J22299_10000025 | + | 17932 | 17991 | **RNA**→ rpmE (PRK00019)Ribosomal_L31 (pfam01197)→ |
| env-486 | JGI24737J22298_10026474 | + | 1377 | 1436 | **RNA**→ rpmE (PRK00019)Ribosomal_L31 (pfam01197)→ |
| Pan-2-1 | NZ_JMJL01000007.1 | - | 24847 | 24788 | **RNA**→ rpmE (PRK00019)Ribosomal_L31 (pfam01197)→ |
| Pan-1-1 | NZ_JMJK01000008.1 | - | 22834 | 22775 | **RNA**→ rpmE (PRK00019)Ribosomal_L31 (pfam01197)→ |
| Pan-3-1 | NZ_JMJJ01000008.1 | - | 24828 | 24769 | **RNA**→ rpmE (PRK00019)Ribosomal_L31 (pfam01197)→ |
| Pan-3-2 | NZ_JFZU01000018.1 | - | 25163 | 25104 | **RNA**→ rpmE (PRK00019)Ribosomal_L31 (pfam01197)→ |
| Pan-1-4 | NZ_CAEI01000100.1 | + | 94980 | 95039 | **RNA**→ rpmE (PRK00019)Ribosomal_L31 (pfam01197)→ |
| Pan-1-5 | NZ_CAEJ01000213.1 | + | 94965 | 95024 | **RNA**→ rpmE (PRK00019)Ribosomal_L31 (pfam01197)→ |
| Pan-4-1 | NZ_AZTE01000093.1 | - | 24860 | 24801 | **RNA**→ rpmE (PRK00019)Ribosomal_L31 (pfam01197)→ |
| Pan-8-1 | NC_013956.2 | - | 4273458 | 4273399 | **RNA**→ rpmE (PRK00019)Ribosomal_L31 (pfam01197)→ |
| Pan-10-1 | NC_017554.1 | + | 198388 | 198447 | **RNA**→ rpmE (PRK00019)Ribosomal_L31 (pfam01197)→ |
| Pan-6-1 | NC_017531.1 | - | 3680829 | 3680770 | **RNA**→ rpmE (PRK00019)Ribosomal_L31 (pfam01197)→ |
| Pan-7-1 | NZ_KI440898.1 | - | 25142 | 25083 | **RNA**→ rpmE (PRK00019)Ribosomal_L31 (pfam01197)→ |
| Pan-9-1 | NC_016816.1 | + | 197910 | 197969 | **RNA**→ rpmE (PRK00019)Ribosomal_L31 (pfam01197)→ |
| Pan-1-2 | NZ_JQZZ01000153.1 | + | 20373 | 20432 | **RNA**→ rpmE (PRK00019)Ribosomal_L31 (pfam01197)→ |
| Psp-49-1 | NC_014837.1 | - | 4221521 | 4221464 | **RNA**→ rpmE (PRK00019)Ribosomal_L31 (pfam01197)→ |
| Psp-45-1 | NZ_LGIS01000015.1 | + | 106087 | 106144 | **RNA**→ rpmE (PRK00019)Ribosomal_L31 (pfam01197)→ |
| Psp-50-1 | NZ_AKIU01000004.1 | + | 105774 | 105831 | **RNA**→ rpmE (PRK00019)Ribosomal_L31 (pfam01197)→ rpmE (PRK00019)Ribosomal_L31 (pfam01197)→ |
| Psp-53-1 | NZ_AKIT01000110.1 | - | 22761 | 22704 | **RNA**→ rpmE (PRK00019)Ribosomal_L31 (pfam01197)→ rpmE (PRK00019)Ribosomal_L31 (pfam01197)→ |

| | | | | | |
|---|---|---|---|---|---|
| env-487 | JGI26055J44792_1011881 | + | 746 | 804 | **RNA**→ rpmE (PRK00019)Ribosomal_L31 (pfam01197)→ |
| env-488 | JGI25162J39368_1004374 | + | 567 | 625 | **RNA**→ rpmE (PRK00019)Ribosomal_L31 (pfam01197)→ |
| env-489 | JGI26052J44787_1009334 | + | 340 | 398 | **RNA**→ rpmE (PRK00019)Ribosomal_L31 (pfam01197)→ |
| env-490 | JGI26053J44789_1014726 | - | 757 | 699 | **RNA**→ rpmE (PRK00019)Ribosomal_L31 (pfam01197)→ |
| env-491 | JGI26057J44791_1022320 | + | 76 | 134 | **RNA**→ rpmE (PRK00019)Ribosomal_L31 (pfam01197)→ |
| env-492 | JGI26054J46599_1001267 | - | 4935 | 4877 | **RNA**→ rpmE (PRK00019)Ribosomal_L31 (pfam01197)→ |
| Psp-41-1 | NZ_JSUT01000007.1 | + | 90918 | 90976 | **RNA**→ rpmE (PRK00019)Ribosomal_L31 (pfam01197)→ |
| Pag-4-1 | NZ_ANKX01000043.1 | - | 23467 | 23409 | **RNA**→ rpmE (PRK00019)Ribosomal_L31 (pfam01197)→ |
| Psp-54-1 | NZ_AEDL01000016.1 | - | 22943 | 22885 | **RNA**→ rpmE (PRK00019)Ribosomal_L31 (pfam01197)→ rpmE (PRK00019)Ribosomal_L31 (pfam01197)→ |
| Pst-4-1 | NZ_AHIE01000032.1 | + | 133115 | 133173 | **RNA**→ rpmE (PRK00019)Ribosomal_L31 (pfam01197)→ rpmE (PRK00019)Ribosomal_L31 (pfam01197)→ |
| Pst-3-1 | NZ_JPKO01000013.1 | + | 92601 | 92659 | **RNA**→ rpmE (PRK00019)Ribosomal_L31 (pfam01197)→ |
| Pst-1-1 | NZ_JRWI01000025.1 | - | 22125 | 22067 | **RNA**→ rpmE (PRK00019)Ribosomal_L31 (pfam01197)→ |
| Pst-1-2 | NZ_JSXF01000016.1 | + | 100240 | 100298 | **RNA**→ rpmE (PRK00019)Ribosomal_L31 (pfam01197)→ |
| Psp-36-1 | NZ_JMRT02000009.1 | + | 90963 | 91021 | **RNA**→ rpmE (PRK00019)Ribosomal_L31 (pfam01197)→ |
| Pva-1-1 | NZ_JPKP01000004.1 | - | 524677 | 524619 | **RNA**→ rpmE (PRK00019)Ribosomal_L31 (pfam01197)→ |
| Pva-2-1 | NC_014562.1 | - | 3321901 | 3321843 | **RNA**→ rpmE (PRK00019)Ribosomal_L31 (pfam01197)→ |
| Psp-37-1 | NZ_JQNO01000001.1 | - | 3712474 | 3712416 | **RNA**→ rpmE (PRK00019)Ribosomal_L31 (pfam01197)→ |
| env-493 | TrFG_GO06Z1102HIXED | + | 45 | 103 | **RNA**→ rpmE (PRK00019)Ribosomal_L31 (pfam01197)→ |
| Pan-5-1 | NZ_JXXL01000002.1 | - | 24189 | 24131 | **RNA**→ rpmE (PRK00019)Ribosomal_L31 (pfam01197)→ |
| Psp-52-1 | NZ_AJFP01000001.1 | - | 750284 | 750226 | **RNA**→ rpmE (PRK00019)Ribosomal_L31 (pfam01197)→ rpmE (PRK00019)Ribosomal_L31 (pfam01197)→ |
| Psp-43-1 | NZ_CP009880.1 | - | 1060052 | 1059996 | **RNA**→ rpmE (PRK00019)Ribosomal_L31 (pfam01197)→ |
| env-494 | JGI26051J44788_1011887 | + | 355 | 411 | **RNA**→ rpmE (PRK00019)Ribosomal_L31 (pfam01197)→ |
| env-495 | JGI26053J44789_1003248 | - | 1683 | 1627 | **RNA**→ rpmE (PRK00019)Ribosomal_L31 (pfam01197)→ |
| env-496 | JGI26052J44787_1008131 | + | 177 | 233 | **RNA**→ rpmE (PRK00019)Ribosomal_L31 (pfam01197)→ |
| env-497 | JGI26055J44792_1005306 | - | 1251 | 1195 | **RNA**→ rpmE (PRK00019)Ribosomal_L31 (pfam01197)→ |
| env-498 | JGI26057J44791_1000727 | + | 3248 | 3304 | **RNA**→ rpmE (PRK00019)Ribosomal_L31 (pfam01197)→ |
| env-499 | JGI26054J46599_1000150 | - | 5814 | 5758 | **RNA**→ rpmE (PRK00019)Ribosomal_L31 (pfam01197)→ |
| Pva-1-2 | NZ_JUQR01000121.1 | + | 95265 | 95324 | **RNA**→ rpmE (PRK00019)Ribosomal_L31 (pfam01197)→ |
| Pse-1-1 | NZ_CCAQ010000025.1 | - | 34969 | 34910 | **RNA**→ rpmE (PRK00019)Ribosomal_L31 (pfam01197)→ rpmE (PRK00019)Ribosomal_L31 (pfam01197)→ |
| Psp-42-1 | NZ_JPQA01000010.1 | + | 91852 | 91911 | **RNA**→ rpmE (PRK00019)Ribosomal_L31 (pfam01197)→ |
| Pro-1-1 | NZ_JTJJ01000012.1 | + | 100019 | 100075 | **RNA**→ rpmE (PRK00019)Ribosomal_L31 (pfam01197)→ |
| Psp-46-1 | NZ_JZRD01000022.1 | - | 22790 | 22734 | **RNA**→ rpmE (PRK00019)Ribosomal_L31 (pfam01197)→ |
| Prw-1-1 | NZ_CP009454.1 | + | 660150 | 660206 | **RNA**→ rpmE (PRK00019)Ribosomal_L31 (pfam01197)→ |
| env-500 | AECF_contig08144 | + | 2575 | 2631 | **RNA**→ rpmE (PRK00019)Ribosomal_L31 (pfam01197)→ |
| Psp-38-1 | NZ_JZRH01000036.1 | - | 22821 | 22765 | **RNA**→ rpmE (PRK00019)Ribosomal_L31 (pfam01197)→ |
| env-501 | APTF_contig25596 | + | 28 | 84 | **RNA**→ rpmE (PRK00019)Ribosomal_L31 (pfam01197)→ |
| env-502 | CLOF_contig32609 | + | 188 | 244 | **RNA**→ rpmE (PRK00019)Ribosomal_L31 (pfam01197)→ |
| Eca-1-1 | NZ_JRUP01000012.1 | - | 22784 | 22728 | **RNA**→ rpmE (PRK00019)Ribosomal_L31 (pfam01197)→ |
| env-503 | ACEF_contig13072 | + | 4745 | 4801 | **RNA**→ rpmE (PRK00019)Ribosomal_L31 (pfam01197)→ |
| env-504 | ACOFG987_contig06308 | - | 345 | 289 | **RNA**→ rpmE (PRK00019)Ribosomal_L31 (pfam01197)→ |
| env-505 | ACOFG988_contig23352 | - | 761 | 705 | **RNA**→ rpmE (PRK00019)Ribosomal_L31 (pfam01197)→ |
| env-506 | CLOF_contig29606 | - | 3568 | 3512 | **RNA**→ rpmE (PRK00019)Ribosomal_L31 (pfam01197)→ |
| Pst-5-1 | NC_022546.1 | + | 2480650 | 2480706 | **RNA**→ rpmE (PRK00019)Ribosomal_L31 (pfam01197)→ |
| Pdi-2-1 | NZ_AVSS01000005.1 | - | 23665 | 23609 | **RNA**→ rpmE (PRK00019)Ribosomal_L31 (pfam01197)→ |
| env-507 | AECF_contig21769 | - | 377 | 320 | **RNA**→ rpmE (PRK00019)Ribosomal_L31 (pfam01197)→ |
| Tmo-1-1 | NZ_JPKR02000005.1 | - | 545319 | 545262 | **RNA**→ rpmE (PRK00019)Ribosomal_L31 (pfam01197)→ |
| Tsp-2-1 | NZ_JFJX01000045.1 | - | 21515 | 21458 | **RNA**→ rpmE (PRK00019)Ribosomal_L31 (pfam01197)→ |
| Tsa-1-1 | NZ_ATMI01000006.1 | + | 58800 | 58857 | **RNA**→ rpmE (PRK00019)Ribosomal_L31 (pfam01197)→ rpmE (PRK00019)Ribosomal_L31 (pfam01197)→ |
| Tpt-1-1 | NZ_ATMJ01000077.1 | + | 69932 | 69989 | **RNA**→ rpmE (PRK00019)Ribosomal_L31 (pfam01197)→ |
| Tpt-1-2 | NZ_JMPR01000011.1 | + | 71180 | 71237 | **RNA**→ rpmE (PRK00019)Ribosomal_L31 (pfam01197)→ |
| env-508 | FGTFG_contig02879 | + | 93249 | 93307 | **RNA**→ rpmE (PRK00019)Ribosomal_L31 (pfam01197)→ |
| Pag-2-1 | NZ_JPKQ01000009.1 | - | 95475 | 95417 | **RNA**→ rpmE (PRK00019)Ribosomal_L31 (pfam01197)→ |

| | | | | | |
|---|---|---|---|---|---|
| Pag-5-1 | NZ_BAEF01000013.1 | - | 23435 | 23377 | **RNA**→ rpmE (PRK00019)Ribosomal_L31 (pfam01197)→ rpmE (PRK00019)Ribosomal_L31 (pfam01197)→ |
| Psp-39-1 | NZ_LGYX01000019.1 | + | 92817 | 92875 | **RNA**→ rpmE (PRK00019)Ribosomal_L31 (pfam01197)→ |
| Pan-1-3 | NZ_JYGW01000002.1 | + | 93322 | 93380 | **RNA**→ rpmE (PRK00019)Ribosomal_L31 (pfam01197)→ |
| Psp-51-1 | NZ_ADWZ01000015.1 | - | 23434 | 23376 | **RNA**→ rpmE (PRK00019)Ribosomal_L31 (pfam01197)→ rpmE (PRK00019)Ribosomal_L31 (pfam01197)→ |
| Pag-2-2 | NZ_JFOK01000007.1 | - | 23564 | 23506 | **RNA**→ rpmE (PRK00019)Ribosomal_L31 (pfam01197)→ |
| Pag-2-6 | NZ_JPOT01000008.1 | + | 93015 | 93073 | **RNA**→ rpmE (PRK00019)Ribosomal_L31 (pfam01197)→ |
| Pag-2-3 | NZ_JNVA01000019.1 | - | 24939 | 24881 | **RNA**→ rpmE (PRK00019)Ribosomal_L31 (pfam01197)→ |
| Pag-2-4 | NZ_JNGC01000002.1 | - | 294027 | 293969 | **RNA**→ rpmE (PRK00019)Ribosomal_L31 (pfam01197)→ |
| Pva-1-3 | NZ_LFQL01000031.1 | + | 91121 | 91179 | **RNA**→ rpmE (PRK00019)Ribosomal_L31 (pfam01197)→ |
| Pag-3-1 | NZ_KK403334.1 | + | 94095 | 94153 | **RNA**→ rpmE (PRK00019)Ribosomal_L31 (pfam01197)→ |
| Pag-6-1 | NZ_KI440944.1 | + | 94622 | 94680 | **RNA**→ rpmE (PRK00019)Ribosomal_L31 (pfam01197)→ |
| Pag-2-5 | NZ_JWLQ01000007.1 | + | 134357 | 134415 | **RNA**→ rpmE (PRK00019)Ribosomal_L31 (pfam01197)→ |
| env-509 | ACOFG987_contig03760 | - | 311 | 253 | **RNA**→ rpmE (PRK00019)Ribosomal_L31 (pfam01197)→ |
| env-510 | ACOFG988_F31SS7V02FQ5RM | + | 210 | 268 | **RNA**→ rpmE (PRK00019)Ribosomal_L31 (pfam01197)→ |
| Psp-48-1 | NZ_ALXE01000044.1 | - | 21482 | 21425 | **RNA**→ rpmE (PRK00019)Ribosomal_L31 (pfam01197)→ |
| env-511 | AUXO014079757.1 | + | 108 | 157 | **RNA**→ ←hypo |
| env-512 | HLSNC_1000009 | - | 72348 | 72298 | **RNA**→ rpmE (PRK00019)Ribosomal_L31 (pfam01197)→ |
| env-513 | SR_TP_S2_1000441 | + | 2919 | 2968 | **RNA**→ rpmE (PRK00019)Ribosomal_L31 (pfam01197)→ |
| env-514 | SR_TTP_S3_1000590 | + | 2911 | 2960 | **RNA**→ rpmE (PRK00019)Ribosomal_L31 (pfam01197)→ |
| env-515 | SS_2KS_010_SOIL_10367424 | + | 142 | 192 | **RNA**→ rpmE (PRK00019)Ribosomal_L31 (pfam01197)→ |
| env-516 | SR_TP_S13sol_1000025 | - | 70388 | 70338 | **RNA**→ rpmE (PRK00019)Ribosomal_L31 (pfam01197)→ |
| Isp-2-1 | NZ_AFPO01000012.1 | + | 277955 | 278005 | **RNA**→ rpmE (PRK00019)Ribosomal_L31 (pfam01197)→ rpmE (PRK00019)Ribosomal_L31 (pfam01197)→ |
| env-517 | SS_3KL_010_SOIL_10380658 | + | 208 | 258 | **RNA**→ ←hypo |
| env-518 | PML_READ_00685738 | + | 151 | 202 | **RNA**→ |
| Asp-8-1 | NC_015554.1 | + | 4361206 | 4361257 | **RNA**→ rpmE (PRK00019)Ribosomal_L31 (pfam01197)→ |
| env-519 | CESF01238860.1 | + | 483 | 534 | **RNA**→ rpmE (PRK00019)Ribosomal_L31 (pfam01197)→ |
| env-520 | TahiMoana_1000136 | - | 193 | 142 | **RNA**→ rpmE (PRK00019)Ribosomal_L31 (pfam01197)→ |
| Vrh-1-1 | NZ_JONG01000015.1 | + | 19747 | 19798 | **RNA**→ rpmE (PRK00019)Ribosomal_L31 (pfam01197)→ |
| env-521 | BMHBC_125854 | - | 1492 | 1439 | **RNA**→ rpmE (PRK00019)Ribosomal_L31 (pfam01197)→ |
| env-522 | BMHBC_2_2794853 | + | 48 | 101 | **RNA**→ |
| env-523 | BMHBC_2_1493885 | + | 48 | 101 | **RNA**→ |
| env-524 | BMHBC_2_2089384 | + | 48 | 101 | **RNA**→ |
| env-525 | BMHBC_2_884118 | - | 73 | 20 | **RNA**→ hypo→ |
| env-526 | BMHBC_2_3380210 | + | 48 | 101 | **RNA**→ |
| Spu-2-1 | NC_009438.1 | + | 601661 | 601714 | **RNA**→ rpmE (PRK00019)Ribosomal_L31 (pfam01197)→ rpmE (PRK00019)Ribosomal_L31 (pfam01197)→ |
| Spu-1-1 | NC_017566.1 | + | 532157 | 532210 | **RNA**→ rpmE (PRK00019)Ribosomal_L31 (pfam01197)→ |
| Ssp-9-1 | NC_008750.1 | - | 4138866 | 4138813 | **RNA**→ rpmE (PRK00019)Ribosomal_L31 (pfam01197)→ rpmE (PRK00019)Ribosomal_L31 (pfam01197)→ |
| Ssp-5-1 | NZ_AFOZ01000042.1 | - | 15569 | 15516 | **RNA**→ rpmE (PRK00019)Ribosomal_L31 (pfam01197)→ rpmE (PRK00019)Ribosomal_L31 (pfam01197)→ rpmE (PRK00019)Ribosomal_L31 (pfam01197)→ |
| Sba-3-1 | NC_009052.1 | - | 4515864 | 4515810 | **RNA**→ rpmE (PRK00019)Ribosomal_L31 (pfam01197)→ |
| Sba-7-1 | NC_011663.1 | + | 571662 | 571716 | **RNA**→ rpmE (PRK00019)Ribosomal_L31 (pfam01197)→ |
| Sba-5-1 | NC_009665.1 | + | 541101 | 541155 | **RNA**→ rpmE (PRK00019)Ribosomal_L31 (pfam01197)→ |
| Sba-2-1 | NC_017579.1 | - | 4615835 | 4615781 | **RNA**→ rpmE (PRK00019)Ribosomal_L31 (pfam01197)→ |
| Sba-1-1 | NC_017571.1 | + | 651813 | 651867 | **RNA**→ rpmE (PRK00019)Ribosomal_L31 (pfam01197)→ |
| Sba-4-1 | NZ_CM001435.1 | - | 4392352 | 4392298 | **RNA**→ rpmE (PRK00019)Ribosomal_L31 (pfam01197)→ rpmE (PRK00019)Ribosomal_L31 (pfam01197)→ |
| Sba-6-1 | NC_009997.1 | + | 570132 | 570186 | **RNA**→ rpmE (PRK00019)Ribosomal_L31 (pfam01197)→ |
| Sba-9-1 | NC_016901.1 | + | 566504 | 566558 | **RNA**→ rpmE (PRK00019)Ribosomal_L31 (pfam01197)→ |
| Sba-8-1 | NZ_AGEX01000011.1 | - | 113918 | 113864 | **RNA**→ rpmE (PRK00019)Ribosomal_L31 (pfam01197)→ |
| env-527 | 3300001239_10036133 | - | 345 | 291 | **RNA**→ rpmE (PRK00019)Ribosomal_L31 (pfam01197)→ |
| env-528 | 3300001605_10743065 | - | 345 | 291 | **RNA**→ rpmE (PRK00019)Ribosomal_L31 (pfam01197)→ |
| Sma-1-1 | NZ_JPEO01000004.1 | + | 103613 | 103665 | **RNA**→ rpmE (PRK00019)Ribosomal_L31 (pfam01197)→ |
| Sco-1-1 | NZ_JAEC01000006.1 | + | 43528 | 43580 | **RNA**→ rpmE (PRK00019)Ribosomal_L31 (pfam01197)→ |

| | | | | | |
|---|---|---|---|---|---|
| Ssp-4-1 | NZ_JWGX01000031.1 | - | 43326 | 43275 | **RNA**→ rpmE (PRK00019)Ribosomal_L31 (pfam01197)→ |
| Sam-2-1 | NC_008700.1 | + | 552817 | 552869 | **RNA**→ ←hypo |
| Son-1-1 | NC_004347.2 | - | 4279346 | 4279296 | **RNA**→ rpmE (PRK00019)Ribosomal_L31 (pfam01197)→ |
| Ssp-10-1 | NZ_JTLE01000094.1 | + | 2661313 | 2661363 | **RNA**→ rpmE (PRK00019)Ribosomal_L31 (pfam01197)→ |
| env-529 | KGI_S1_ANT02_95m_c10000044 | + | 8538 | 8592 | **RNA**→ rpmE (PRK00019)Ribosomal_L31 (pfam01197)→ |
| Swo-1-1 | NC_010506.1 | - | 5359927 | 5359875 | **RNA**→ rpmE (PRK00019)Ribosomal_L31 (pfam01197)→ |
| Sse-1-1 | NC_009831.1 | + | 655635 | 655687 | **RNA**→ rpmE (PRK00019)Ribosomal_L31 (pfam01197)→ |
| Sbe-1-1 | NZ_ABIC01000023.1 | + | 45676 | 45728 | **RNA**→ rpmE (PRK00019)Ribosomal_L31 (pfam01197)→ rpmE (PRK00019)Ribosomal_L31 (pfam01197)→ |
| Svi-1-1 | NC_014012.1 | + | 337588 | 337640 | **RNA**→ rpmE (PRK00019)Ribosomal_L31 (pfam01197)→ |
| env-530 | JCVI_SCAF_1096627387720 | - | 14506 | 14454 | **RNA**→ rpmE (PRK00019)Ribosomal_L31 (pfam01197)→ hypo→ |
| env-531 | GO4KUYH01CXCC6 | + | 142 | 194 | **RNA**→ |
| env-532 | GO4KUYH01C3ZKW | + | 141 | 193 | **RNA**→ rpmE (PRK00019)Ribosomal_L31 (pfam01197)→ |
| env-533 | GooDraft_1002869 | + | 3149 | 3201 | **RNA**→ rpmE (PRK00019)Ribosomal_L31 (pfam01197)→ |
| Ssp-8-1 | NZ_AKZL01000033.1 | - | 34459 | 34407 | **RNA**→ rpmE (PRK00019)Ribosomal_L31 (pfam01197)→ |
| Sxi-1-1 | NZ_JGVI01000084.1 | + | 35320 | 35372 | **RNA**→ rpmE (PRK00019)Ribosomal_L31 (pfam01197)→ |
| Ssp-7-1 | NC_008322.1 | + | 522940 | 522992 | **RNA**→ rpmE (PRK00019)Ribosomal_L31 (pfam01197)→ |
| Sde-1-1 | NZ_AXZL01000063.1 | + | 70809 | 70861 | **RNA**→ rpmE (PRK00019)Ribosomal_L31 (pfam01197)→ |
| env-534 | JCVI_SCAF_1096627387716 | - | 23316 | 23264 | **RNA**→ rpmE (PRK00019)Ribosomal_L31 (pfam01197)→ hypo→ |
| Ssp-2-1 | NC_008577.1 | - | 4408636 | 4408584 | **RNA**→ rpmE (PRK00019)Ribosomal_L31 (pfam01197)→ |
| Ssp-6-1 | NC_008321.1 | - | 4178394 | 4178342 | **RNA**→ rpmE (PRK00019)Ribosomal_L31 (pfam01197)→ |
| Sma-2-1 | NZ_BALM01000013.1 | + | 59899 | 59951 | **RNA**→ rpmE (PRK00019)Ribosomal_L31 (pfam01197)→ |
| Spi-1-1 | NC_011566.1 | + | 499660 | 499712 | **RNA**→ rpmE (PRK00019)Ribosomal_L31 (pfam01197)→ rpmE (PRK00019)Ribosomal_L31 (pfam01197)→ |
| Ssp-3-1 | NZ_JSFF01000009.1 | - | 106570 | 106518 | **RNA**→ rpmE (PRK00019)Ribosomal_L31 (pfam01197)→ |
| Ssp-11-1 | NZ_JPII01000042.1 | + | 12098 | 12150 | **RNA**→ rpmE (PRK00019)Ribosomal_L31 (pfam01197)→ rpmE (PRK00019)Ribosomal_L31 (pfam01197)→ |
| Slo-1-1 | NC_009092.1 | + | 431236 | 431288 | **RNA**→ rpmE (PRK00019)Ribosomal_L31 (pfam01197)→ rpmE (PRK00019)Ribosomal_L31 (pfam01197)→ |
| Swa-1-1 | NZ_KI912497.1 | - | 254535 | 254484 | **RNA**→ rpmE (PRK00019)Ribosomal_L31 (pfam01197)→ |
| Sde-2-1 | NC_007954.1 | - | 3997631 | 3997580 | **RNA**→ rpmE (PRK00019)Ribosomal_L31 (pfam01197)→ |
| env-535 | KGI_S1_ANT01_95m_c10083218 | - | 640 | 588 | **RNA**→ rpmE (PRK00019)Ribosomal_L31 (pfam01197)→ |
| Sha-1-1 | NC_010334.1 | - | 4683887 | 4683836 | **RNA**→ rpmE (PRK00019)Ribosomal_L31 (pfam01197)→ |
| Spe-1-1 | NC_009901.1 | - | 4591286 | 4591235 | **RNA**→ rpmE (PRK00019)Ribosomal_L31 (pfam01197)→ |
| Sfi-1-1 | NZ_JADX01000010.1 | - | 251293 | 251242 | **RNA**→ rpmE (PRK00019)Ribosomal_L31 (pfam01197)→ |
| Sfr-1-1 | NC_008345.1 | - | 4467060 | 4467008 | **RNA**→ rpmE (PRK00019)Ribosomal_L31 (pfam01197)→ |
| CRe-1-1 | NZ_GL379589.1 | - | 502536 | 502482 | **RNA**→ rpmE (PRK00019)Ribosomal_L31 (pfam01197)→ rpmE (PRK00019)Ribosomal_L31 (pfam01197)→ |
| Eba-2-1 | NZ_DF158888.1 | + | 91727 | 91780 | **RNA**→ rpmE (PRK00019)Ribosomal_L31 (pfam01197)→ |
| Sgl-1-1 | NC_007712.1 | - | 3774802 | 3774749 | **RNA**→ rpmE (PRK00019)Ribosomal_L31 (pfam01197)→ |
| CSo-1-1 | NZ_CP006568.1 | + | 939488 | 939543 | **RNA**→ rpmE (PRK00019)Ribosomal_L31 (pfam01197)→ |
| env-536 | XAGC_contig20879 | - | 400 | 347 | **RNA**→ rpmE (PRK00019)Ribosomal_L31 (pfam01197)→ |
| Spr-1-1 | NZ_CP006569.1 | - | 4560293 | 4560240 | **RNA**→ rpmE (PRK00019)Ribosomal_L31 (pfam01197)→ |
| Psp-47-1 | NZ_JRKB01000114.1 | - | 29350 | 29291 | **RNA**→ rpmE (PRK00019)Ribosomal_L31 (pfam01197)→ |
| Psh-1-1 | NZ_JWHQ01000025.1 | - | 1797 | 1738 | **RNA**→ rpmE (PRK00019)Ribosomal_L31 (pfam01197)→ |
| Psh-2-1 | NZ_KB944511.1 | - | 1649396 | 1649337 | **RNA**→ rpmE (PRK00019)Ribosomal_L31 (pfam01197)→ |
| Aal-2-1 | NZ_BARX01000001.1 | - | 16360 | 16301 | **RNA**→ rpmE (PRK00019)Ribosomal_L31 (pfam01197)→ |
| env-537 | GO4KUYH02HSOZH | - | 181 | 129 | **RNA**→ hypo→ |
| Pte-1-1 | NZ_JONU01000008.1 | + | 256804 | 256856 | **RNA**→ rpmE (PRK00019)Ribosomal_L31 (pfam01197)→ rpmE (PRK00019)Ribosomal_L31 (pfam01197)→ hypo |
| env-538 | 3300000052_491248 | + | 102 | 154 | **RNA**→ rpmE (PRK00019)Ribosomal_L31 (pfam01197)→ |
| env-539 | 3300000052_373220 | - | 197 | 145 | **RNA**→ rpmE (PRK00019)Ribosomal_L31 (pfam01197)→ |
| Tli-1-1 | NZ_AZUK01000001.1 | + | 2305027 | 2305077 | **RNA**→ rpmE (PRK00019)Ribosomal_L31 (pfam01197)→ |
| Tau-1-1 | NC_012691.1 | + | 541965 | 542015 | **RNA**→ rpmE (PRK00019)Ribosomal_L31 (pfam01197)→ |
| Ana-1-1 | NZ_AUCC01000060.1 | - | 4940 | 4890 | **RNA**→ rpmE (PRK00019)Ribosomal_L31 (pfam01197)→ |
| Aen-3-1 | NZ_JRLH01000010.1 | + | 51375 | 51425 | **RNA**→ rpmE (PRK00019)Ribosomal_L31 (pfam01197)→ |
| Asa-10-1 | NZ_KB894494.1 | - | 58337 | 58287 | **RNA**→ rpmE (PRK00019)Ribosomal_L31 (pfam01197)→ |
| Iat-1-1 | NZ_JPIN01000002.1 | - | 98575 | 98527 | **RNA**→ rpmE (PRK00019)Ribosomal_L31 (pfam01197)→ |

| | | | | | |
|---|---|---|---|---|---|
| Ise-1-1 | NZ_AUHL01000006.1 | - | 66251 | 66203 | **RNA**→ rpmE (PRK00019)Ribosomal_L31 (pfam01197)→ rpmE (PRK00019)Ribosomal_L31 (pfam01197)→ |
| Isa-1-1 | NZ_JPER01000004.1 | + | 235268 | 235316 | **RNA**→ rpmE (PRK00019)Ribosomal_L31 (pfam01197)→ |
| Ixi-1-1 | NZ_AMRG01000014.1 | - | 30706 | 30654 | **RNA**→ rpmE (PRK00019)Ribosomal_L31 (pfam01197)→ rpmE (PRK00019)Ribosomal_L31 (pfam01197)→ |
| env-540 | AYRE01004792.1 | - | 218 | 167 | **RNA**→ rpmE (PRK00019)Ribosomal_L31 (pfam01197)→ |
| Osm-1-1 | NZ_KB908459.1 | + | 180853 | 180904 | **RNA**→ rpmE (PRK00019)Ribosomal_L31 (pfam01197)→ |
| Osp-1-1 | NC_016745.1 | + | 726676 | 726727 | **RNA**→ rpmE (PRK00019)Ribosomal_L31 (pfam01197)→ |
| env-541 | 2231941556 | + | 4 | 58 | **RNA**→ |
| Ema-2-1 | NZ_JFHN01000029.1 | + | 87873 | 87932 | **RNA**→ rpmE (PRK00019)Ribosomal_L31 (pfam01197)→ |
| Etr-1-1 | NZ_JXNU01000003.1 | + | 3289663 | 3289722 | **RNA**→ rpmE (PRK00019)Ribosomal_L31 (pfam01197)→ |
| Etr-2-1 | NZ_KE136311.1 | - | 127906 | 127847 | **RNA**→ rpmE (PRK00019)Ribosomal_L31 (pfam01197)→ |
| Psp-40-1 | NZ_JFGT01000004.1 | - | 137795 | 137735 | **RNA**→ rpmE (PRK00019)Ribosomal_L31 (pfam01197)→ |
| Ein-1-1 | NZ_JRXF01000015.1 | - | 23356 | 23296 | **RNA**→ rpmE (PRK00019)Ribosomal_L31 (pfam01197)→ |
| Ein-1-2 | NZ_JRXE01000029.1 | - | 23448 | 23388 | **RNA**→ rpmE (PRK00019)Ribosomal_L31 (pfam01197)→ |
| env-542 | JGI26052J44787_1013371 | + | 333 | 393 | **RNA**→ rpmE (PRK00019)Ribosomal_L31 (pfam01197)→ |
| env-543 | JGI26055J44792_1089657 | + | 258 | 318 | **RNA**→ |
| env-544 | JGI26057J44791_1023823 | - | 676 | 616 | **RNA**→ rpmE (PRK00019)Ribosomal_L31 (pfam01197)→ |
| env-545 | JGI26054J46599_1079959 | + | 92 | 152 | **RNA**→ rpmE (PRK00019)Ribosomal_L31 (pfam01197)→ |
| Eto-1-1 | NZ_KB372804.1 | - | 1363173 | 1363113 | **RNA**→ ←hypo |
| Esp-55-1 | NZ_JQNE01000001.1 | + | 4128718 | 4128777 | **RNA**→ rpmE (PRK00019)Ribosomal_L31 (pfam01197)→ |
| Eol-1-1 | NZ_JNVB01000002.1 | + | 112156 | 112215 | **RNA**→ rpmE (PRK00019)Ribosomal_L31 (pfam01197)→ |
| Ebi-2-1 | NC_014306.1 | + | 171661 | 171721 | **RNA**→ rpmE (PRK00019)Ribosomal_L31 (pfam01197)→ |
| Ebi-1-1 | NZ_LHXI01000018.1 | + | 107630 | 107690 | **RNA**→ rpmE (PRK00019)Ribosomal_L31 (pfam01197)→ |
| bsy-1-1 | NZ_JMSO01000006.1 | + | 18997 | 19057 | **RNA**→ rpmE (PRK00019)Ribosomal_L31 (pfam01197)→ |
| Epy-2-1 | NC_012214.1 | + | 161194 | 161254 | **RNA**→ rpmE (PRK00019)Ribosomal_L31 (pfam01197)→ |
| Epy-1-1 | NC_017390.1 | + | 161200 | 161260 | **RNA**→ rpmE (PRK00019)Ribosomal_L31 (pfam01197)→ |
| Esp-74-1 | NC_017445.1 | + | 1434028 | 1434088 | **RNA**→ rpmE (PRK00019)Ribosomal_L31 (pfam01197)→ |
| Eta-9-1 | NC_010694.1 | + | 140209 | 140269 | **RNA**→ rpmE (PRK00019)Ribosomal_L31 (pfam01197)→ |
| Epi-2-1 | NZ_CAHS01000002.1 | + | 113089 | 113149 | **RNA**→ rpmE (PRK00019)Ribosomal_L31 (pfam01197)→ hypo→ |
| Eam-1-1 | NZ_BAYW01000017.1 | - | 21692 | 21632 | **RNA**→ rpmE (PRK00019)Ribosomal_L31 (pfam01197)→ |
| Eam-10-1 | NZ_CAOX01000002.1 | + | 109737 | 109797 | **RNA**→ rpmE (PRK00019)Ribosomal_L31 (pfam01197)→ |
| Eam-8-1 | NC_013961.1 | + | 156303 | 156363 | **RNA**→ rpmE (PRK00019)Ribosomal_L31 (pfam01197)→ |
| Eam-5-1 | NC_013971.1 | + | 156303 | 156363 | **RNA**→ rpmE (PRK00019)Ribosomal_L31 (pfam01197)→ hypo→ hypo→ hypo→ hypo→ |
| Eam-4-1 | NZ_AFHN01000002.1 | + | 109783 | 109843 | **RNA**→ rpmE (PRK00019)Ribosomal_L31 (pfam01197)→ |
| Eam-15-1 | NZ_CAPC01000002.1 | + | 109783 | 109843 | **RNA**→ rpmE (PRK00019)Ribosomal_L31 (pfam01197)→ |
| Eam-11-1 | NZ_CAPD01000002.1 | + | 97809 | 97869 | **RNA**→ rpmE (PRK00019)Ribosomal_L31 (pfam01197)→ |
| Eam-9-1 | NZ_CAOY01000002.1 | + | 109776 | 109836 | **RNA**→ rpmE (PRK00019)Ribosomal_L31 (pfam01197)→ |
| Eam-6-1 | NZ_CAPB01000002.1 | + | 109783 | 109843 | **RNA**→ rpmE (PRK00019)Ribosomal_L31 (pfam01197)→ |
| Eam-7-1 | NZ_CAOZ01000002.1 | + | 109749 | 109809 | **RNA**→ rpmE (PRK00019)Ribosomal_L31 (pfam01197)→ |
| Eam-3-1 | NZ_CAPA01000002.1 | + | 109783 | 109843 | **RNA**→ rpmE (PRK00019)Ribosomal_L31 (pfam01197)→ |
| Eam-13-1 | NZ_CBVT010000002.1 | + | 109584 | 109644 | **RNA**→ rpmE (PRK00019)Ribosomal_L31 (pfam01197)→ |
| Eam-14-1 | NZ_CBVU010000002.1 | + | 109584 | 109644 | **RNA**→ rpmE (PRK00019)Ribosomal_L31 (pfam01197)→ |
| Eam-12-1 | NZ_CBVS010000002.1 | + | 109584 | 109644 | **RNA**→ rpmE (PRK00019)Ribosomal_L31 (pfam01197)→ |
| Ety-1-1 | NZ_JRUQ01000016.1 | + | 133327 | 133387 | **RNA**→ rpmE (PRK00019)Ribosomal_L31 (pfam01197)→ |
| env-546 | XAGC_contig27338 | + | 221 | 281 | **RNA**→ ←hypo |
| Psp-44-1 | NZ_CP009866.1 | - | 3634114 | 3634055 | **RNA**→ rpmE (PRK00019)Ribosomal_L31 (pfam01197)→ |
| env-547 | CETM01121351.1 | - | 624 | 574 | **RNA**→ rpmE (PRK00019)Ribosomal_L31 (pfam01197)→ |
| env-548 | CEVA01063086.1 | - | 14576 | 14526 | **RNA**→ rpmE (PRK00019)Ribosomal_L31 (pfam01197)→ |
| Pru-3-1 | NZ_JXXY01000018.1 | - | 149896 | 149846 | **RNA**→ rpmE (PRK00019)Ribosomal_L31 (pfam01197)→ rpmE (PRK00019)Ribosomal_L31 (pfam01197)→ |
| Pru-3-2 | NZ_JXXZ01000010.1 | - | 626883 | 626833 | **RNA**→ rpmE (PRK00019)Ribosomal_L31 (pfam01197)→ rpmE (PRK00019)Ribosomal_L31 (pfam01197)→ |
| Pru-4-1 | NZ_AOPM01000105.1 | + | 40951 | 41001 | **RNA**→ rpmE (PRK00019)Ribosomal_L31 (pfam01197)→ rpmE (PRK00019)Ribosomal_L31 (pfam01197)→ rpmE (PRK00019)Ribosomal_L31 (pfam01197)→ |
| env-549 | CEPZ01177373.1 | + | 388 | 438 | **RNA**→ rpmE (PRK00019)Ribosomal_L31 (pfam01197)→ |

| | | | | | |
|---|---|---|---|---|---|
| Psp-33-1 | NZ_AHCE02000021.1 | + | 18594 | 18644 | **RNA**→ rpmE (PRK00019)Ribosomal_L31 (pfam01197)→ |
| Ppi-1-1 | NZ_CP009888.1 | - | 346889 | 346839 | **RNA**→ rpmE (PRK00019)Ribosomal_L31 (pfam01197)→ |
| Pci-1-1 | NZ_AHBZ02000178.1 | - | 16984 | 16934 | **RNA**→ rpmE (PRK00019)Ribosomal_L31 (pfam01197)→ |
| Plu-2-1 | NZ_AUSV01000022.1 | - | 2218 | 2166 | **RNA**→ rpmE (PRK00019)Ribosomal_L31 (pfam01197)→ rpmE (PRK00019)Ribosomal_L31 (pfam01197)→ |
| Plu-3-1 | NZ_CAPN01000031.1 | - | 44749 | 44698 | **RNA**→ rpmE (PRK00019)Ribosomal_L31 (pfam01197)→ |
| env-550 | CERC01133101.1 | - | 82 | 31 | **RNA**→ |
| Ppi-3-1 | NZ_KB907371.1 | - | 399235 | 399184 | **RNA**→ rpmE (PRK00019)Ribosomal_L31 (pfam01197)→ rpmE (PRK00019)Ribosomal_L31 (pfam01197)→ rpmE (PRK00019)Ribosomal_L31 (pfam01197)→ |
| Ppi-4-1 | NZ_AHCC02000195.1 | + | 23369 | 23420 | **RNA**→ rpmE (PRK00019)Ribosomal_L31 (pfam01197)→ rpmE (PRK00019)Ribosomal_L31 (pfam01197)→ |
| Ppi-2-1 | NZ_JXXX01000108.1 | - | 44988 | 44937 | **RNA**→ rpmE (PRK00019)Ribosomal_L31 (pfam01197)→ rpmE (PRK00019)Ribosomal_L31 (pfam01197)→ |
| Pfl-1-1 | NZ_JH650748.1 | + | 23482 | 23533 | **RNA**→ rpmE (PRK00019)Ribosomal_L31 (pfam01197)→ |
| Pel-1-1 | NZ_JWIH01000018.1 | + | 24597 | 24648 | **RNA**→ rpmE (PRK00019)Ribosomal_L31 (pfam01197)→ rpmE (PRK00019)Ribosomal_L31 (pfam01197)→ |
| Ppi-2-2 | NZ_JXXW01000011.1 | + | 23330 | 23381 | **RNA**→ rpmE (PRK00019)Ribosomal_L31 (pfam01197)→ rpmE (PRK00019)Ribosomal_L31 (pfam01197)→ |
| Psp-15-1 | NZ_AKXJ01000003.1 | - | 426568 | 426517 | **RNA**→ rpmE (PRK00019)Ribosomal_L31 (pfam01197)→ rpmE (PRK00019)Ribosomal_L31 (pfam01197)→ rpmE (PRK00019)Ribosomal_L31 (pfam01197)→ |
| env-551 | CEPQ01102968.1 | - | 435 | 384 | **RNA**→ rpmE (PRK00019)Ribosomal_L31 (pfam01197)→ |
| env-552 | JGI11829J13087_107135 | - | 139 | 88 | **RNA**→ rpmE (PRK00019)Ribosomal_L31 (pfam01197)→ |
| env-553 | JGI12020J13220_1000475 | - | 5069 | 5018 | **RNA**→ rpmE (PRK00019)Ribosomal_L31 (pfam01197)→ |
| env-554 | CEPV01195072.1 | + | 400 | 451 | **RNA**→ rpmE (PRK00019)Ribosomal_L31 (pfam01197)→ |
| env-555 | CEPZ01204578.1 | + | 666 | 717 | **RNA**→ rpmE (PRK00019)Ribosomal_L31 (pfam01197)→ |
| env-556 | CEQE01276759.1 | + | 560 | 611 | **RNA**→ rpmE (PRK00019)Ribosomal_L31 (pfam01197)→ |
| env-557 | CETV01108507.1 | + | 476 | 527 | **RNA**→ rpmE (PRK00019)Ribosomal_L31 (pfam01197)→ |
| env-558 | CEUK01292042.1 | + | 691 | 742 | **RNA**→ |
| env-559 | CEQG01236191.1 | + | 694 | 745 | **RNA**→ |
| env-560 | CERC01031317.1 | - | 852 | 801 | **RNA**→ rpmE (PRK00019)Ribosomal_L31 (pfam01197)→ |
| env-561 | CERD01031679.1 | + | 5331 | 5382 | **RNA**→ rpmE (PRK00019)Ribosomal_L31 (pfam01197)→ |
| env-562 | CERL01075421.1 | + | 216 | 267 | **RNA**→ rpmE (PRK00019)Ribosomal_L31 (pfam01197)→ |
| env-563 | CETA01065829.1 | - | 448 | 397 | **RNA**→ rpmE (PRK00019)Ribosomal_L31 (pfam01197)→ |
| env-564 | CEUX01031767.1 | + | 2189 | 2240 | **RNA**→ rpmE (PRK00019)Ribosomal_L31 (pfam01197)→ |
| env-565 | CEWB01087664.1 | - | 382 | 331 | **RNA**→ rpmE (PRK00019)Ribosomal_L31 (pfam01197)→ |
| env-566 | JGI11949J13268_1006774 | + | 325 | 376 | **RNA**→ rpmE (PRK00019)Ribosomal_L31 (pfam01197)→ |
| env-567 | CESF01139237.1 | - | 14048 | 13997 | **RNA**→ rpmE (PRK00019)Ribosomal_L31 (pfam01197)→ |
| env-568 | CEUD01330189.1 | + | 1588 | 1639 | **RNA**→ rpmE (PRK00019)Ribosomal_L31 (pfam01197)→ |
| env-569 | CEUO01029621.1 | + | 332 | 383 | **RNA**→ rpmE (PRK00019)Ribosomal_L31 (pfam01197)→ |
| env-570 | CEWI01244144.1 | - | 323 | 272 | **RNA**→ rpmE (PRK00019)Ribosomal_L31 (pfam01197)→ |
| env-571 | _GX4VOVV01DM434 | - | 83 | 32 | **RNA**→ hypo→ |
| Psp-17-1 | NZ_AJTK01000019.1 | - | 4461 | 4410 | **RNA**→ rpmE (PRK00019)Ribosomal_L31 (pfam01197)→ |
| Csp-1-1 | NZ_JONX01000008.1 | + | 64919 | 64970 | **RNA**→ rpmE (PRK00019)Ribosomal_L31 (pfam01197)→ |
| Psp-14-1 | NZ_JQFL01000080.1 | - | 131090 | 131039 | **RNA**→ rpmE (PRK00019)Ribosomal_L31 (pfam01197)→ |
| env-572 | CENU01396304.1 | + | 1029 | 1080 | **RNA**→ rpmE (PRK00019)Ribosomal_L31 (pfam01197)→ |
| Psp-8-1 | NZ_BADW01000079.1 | + | 29656 | 29707 | **RNA**→ rpmE (PRK00019)Ribosomal_L31 (pfam01197)→ rpmE (PRK00019)Ribosomal_L31 (pfam01197)→ |
| env-573 | JGI11838J13071_130222 | - | 91 | 40 | **RNA**→ hypo→ |
| env-574 | JGI11906J13219_100106 | + | 2033 | 2084 | **RNA**→ rpmE (PRK00019)Ribosomal_L31 (pfam01197)→ |
| env-575 | JGI12020J13220_1008982 | - | 426 | 375 | **RNA**→ rpmE (PRK00019)Ribosomal_L31 (pfam01197)→ |
| env-576 | JGI11754J13217_1001331 | - | 2564 | 2513 | **RNA**→ rpmE (PRK00019)Ribosomal_L31 (pfam01197)→ |
| env-577 | JGI12218J13221_1000027 | - | 540 | 489 | **RNA**→ rpmE (PRK00019)Ribosomal_L31 (pfam01197)→ |
| env-578 | JGI12218J13221_1000194 | + | 1789 | 1840 | **RNA**→ rpmE (PRK00019)Ribosomal_L31 (pfam01197)→ |
| env-579 | JGI11931J13222_1012993 | - | 124 | 73 | **RNA**→ rpmE (PRK00019)Ribosomal_L31 (pfam01197)→ |
| env-580 | JGI12156J13080_113928 | + | 216 | 267 | **RNA**→ rpmE (PRK00019)Ribosomal_L31 (pfam01197)→ |
| env-581 | CENZ01163183.1 | + | 102 | 153 | **RNA**→ rpmE (PRK00019)Ribosomal_L31 (pfam01197)→ |
| env-582 | CEQS01011751.1 | - | 21184 | 21133 | **RNA**→ rpmE (PRK00019)Ribosomal_L31 (pfam01197)→ |
| env-583 | CERC01016695.1 | - | 2342 | 2291 | **RNA**→ rpmE (PRK00019)Ribosomal_L31 (pfam01197)→ |
| env-584 | CERD01038299.1 | + | 663 | 714 | **RNA**→ rpmE (PRK00019)Ribosomal_L31 (pfam01197)→ |

| | | | | | |
|---|---|---|---|---|---|
| env-585 | CERE01031172.1 | + | 6395 | 6446 | **RNA**→ rpmE (PRK00019)Ribosomal_L31 (pfam01197)→ |
| env-586 | CESS01027590.1 | - | 156 | 105 | **RNA**→ rpmE (PRK00019)Ribosomal_L31 (pfam01197)→ |
| env-587 | CETM01162887.1 | + | 1686 | 1737 | **RNA**→ rpmE (PRK00019)Ribosomal_L31 (pfam01197)→ |
| env-588 | JGI24025J20009_10001015 | + | 10992 | 11043 | **RNA**→ rpmE (PRK00019)Ribosomal_L31 (pfam01197)→ |
| Pag-1-1 | NZ_APME01000074.1 | + | 65874 | 65925 | **RNA**→ rpmE (PRK00019)Ribosomal_L31 (pfam01197)→ rpmE (PRK00019)Ribosomal_L31 (pfam01197)→ |
| Psp-16-1 | NZ_AZIO01000115.1 | - | 148851 | 148800 | **RNA**→ rpmE (PRK00019)Ribosomal_L31 (pfam01197)→ rpmE (PRK00019)Ribosomal_L31 (pfam01197)→ rpmE (PRK00019)Ribosomal_L31 (pfam01197)→ |
| env-589 | CERE01032072.1 | - | 203 | 152 | **RNA**→ rpmE (PRK00019)Ribosomal_L31 (pfam01197)→ |
| env-590 | CERL01102156.1 | + | 104 | 155 | **RNA**→ rpmE (PRK00019)Ribosomal_L31 (pfam01197)→ |
| Psp-24-1 | NC_014803.1 | + | 329863 | 329914 | **RNA**→ rpmE (PRK00019)Ribosomal_L31 (pfam01197)→ |
| Psp-23-1 | NZ_KK070023.1 | + | 1202960 | 1203011 | **RNA**→ rpmE (PRK00019)Ribosomal_L31 (pfam01197)→ |
| Psp-31-1 | NZ_AUTH01000084.1 | + | 64821 | 64872 | **RNA**→ rpmE (PRK00019)Ribosomal_L31 (pfam01197)→ |
| Psp-2-1 | NZ_JADO01000011.1 | + | 64319 | 64370 | **RNA**→ rpmE (PRK00019)Ribosomal_L31 (pfam01197)→ |
| Psp-3-1 | NZ_JIAM01000011.1 | + | 64044 | 64095 | **RNA**→ rpmE (PRK00019)Ribosomal_L31 (pfam01197)→ |
| Psp-4-1 | NZ_JPMC01000016.1 | + | 210417 | 210468 | **RNA**→ rpmE (PRK00019)Ribosomal_L31 (pfam01197)→ |
| env-591 | CEQG01078170.1 | + | 93 | 144 | **RNA**→ rpmE (PRK00019)Ribosomal_L31 (pfam01197)→ |
| env-592 | CETF01103377.1 | - | 841 | 790 | **RNA**→ rpmE (PRK00019)Ribosomal_L31 (pfam01197)→ |
| Pli-1-1 | NZ_KK070036.1 | - | 168509 | 168458 | **RNA**→ rpmE (PRK00019)Ribosomal_L31 (pfam01197)→ |
| Psp-6-1 | NZ_BADU01000157.1 | + | 29544 | 29595 | **RNA**→ rpmE (PRK00019)Ribosomal_L31 (pfam01197)→ rpmE (PRK00019)Ribosomal_L31 (pfam01197)→ |
| env-593 | LAZR01004165.1 | - | 5394 | 5343 | **RNA**→ rpmE (PRK00019)Ribosomal_L31 (pfam01197)→ |
| Pdi-1-1 | NZ_JWIG01000001.1 | - | 843923 | 843872 | **RNA**→ rpmE (PRK00019)Ribosomal_L31 (pfam01197)→ |
| Pha-1-1 | NZ_ADOP01000005.1 | + | 8477 | 8528 | **RNA**→ rpmE (PRK00019)Ribosomal_L31 (pfam01197)→ rpmE (PRK00019)Ribosomal_L31 (pfam01197)→ |
| Psp-5-1 | NZ_AUTK01000406.1 | + | 27446 | 27497 | **RNA**→ rpmE (PRK00019)Ribosomal_L31 (pfam01197)→ |
| Psp-30-1 | NZ_AUTI01000314.1 | - | 2912 | 2861 | **RNA**→ rpmE (PRK00019)Ribosomal_L31 (pfam01197)→ |
| Pfl-2-1 | NZ_JTDZ01000012.1 | - | 588159 | 588108 | **RNA**→ rpmE (PRK00019)Ribosomal_L31 (pfam01197)→ |
| Psp-26-1 | NZ_AUTN01000149.1 | + | 65662 | 65713 | **RNA**→ rpmE (PRK00019)Ribosomal_L31 (pfam01197)→ |
| Psp-27-1 | NZ_AUTL01000176.1 | - | 12649 | 12598 | **RNA**→ rpmE (PRK00019)Ribosomal_L31 (pfam01197)→ |
| Psp-28-1 | NZ_AUTM01000342.1 | - | 54996 | 54945 | **RNA**→ rpmE (PRK00019)Ribosomal_L31 (pfam01197)→ |
| env-594 | JGI24025J20009_10000510 | + | 30088 | 30139 | **RNA**→ |
| Pun-1-1 | NZ_AHCF02000035.1 | - | 124891 | 124840 | **RNA**→ rpmE (PRK00019)Ribosomal_L31 (pfam01197)→ |
| Aba-1-1 | NZ_AAVS01000013.1 | + | 62431 | 62482 | **RNA**→ rpmE (PRK00019)Ribosomal_L31 (pfam01197)→ rpmE (PRK00019)Ribosomal_L31 (pfam01197)→ |
| env-595 | JCVI_SCAF_1096626478483 | - | 754 | 703 | **RNA**→ rpmE (PRK00019)Ribosomal_L31 (pfam01197)→ |
| env-596 | JGI11906J13219_116351 | + | 271 | 322 | **RNA**→ |
| env-597 | CESU01212683.1 | + | 608 | 659 | **RNA**→ rpmE (PRK00019)Ribosomal_L31 (pfam01197)→ |
| env-598 | A09P201000a_Sequence0000002282 | - | 1478 | 1427 | **RNA**→ rpmE (PRK00019)Ribosomal_L31 (pfam01197)→ |
| Pma-1-1 | NZ_AHCB02000038.1 | + | 62021 | 62072 | **RNA**→ rpmE (PRK00019)Ribosomal_L31 (pfam01197)→ |
| Psp-13-1 | NZ_JWGY01000001.1 | + | 61985 | 62036 | **RNA**→ rpmE (PRK00019)Ribosomal_L31 (pfam01197)→ |
| Psp-9-1 | NZ_BADX01000055.1 | + | 23372 | 23423 | **RNA**→ rpmE (PRK00019)Ribosomal_L31 (pfam01197)→ rpmE (PRK00019)Ribosomal_L31 (pfam01197)→ |
| Pha-2-1 | NZ_AHCA01000009.1 | - | 150446 | 150395 | **RNA**→ rpmE (PRK00019)Ribosomal_L31 (pfam01197)→ |
| Psp-11-1 | NZ_BADT01000142.1 | + | 28576 | 28627 | **RNA**→ rpmE (PRK00019)Ribosomal_L31 (pfam01197)→ rpmE (PRK00019)Ribosomal_L31 (pfam01197)→ |
| Psp-32-1 | NZ_AUTQ01000186.1 | - | 26876 | 26825 | **RNA**→ rpmE (PRK00019)Ribosomal_L31 (pfam01197)→ |
| Psp-19-1 | NZ_JXYD01000037.1 | - | 57348 | 57297 | **RNA**→ rpmE (PRK00019)Ribosomal_L31 (pfam01197)→ |
| Psp-12-1 | NZ_AMYA01000022.1 | - | 2067 | 2016 | **RNA**→ rpmE (PRK00019)Ribosomal_L31 (pfam01197)→ |
| Psp-20-1 | NZ_JJNY01000024.1 | - | 36149 | 36098 | **RNA**→ rpmE (PRK00019)Ribosomal_L31 (pfam01197)→ |
| Psp-10-1 | NZ_BADY01000117.1 | - | 2048 | 1997 | **RNA**→ rpmE (PRK00019)Ribosomal_L31 (pfam01197)→ rpmE (PRK00019)Ribosomal_L31 (pfam01197)→ |
| Pfu-1-1 | NZ_JJNZ01000017.1 | + | 69496 | 69547 | **RNA**→ rpmE (PRK00019)Ribosomal_L31 (pfam01197)→ |
| env-599 | wf3_Contig4170 | + | 986 | 1037 | **RNA**→ |
| env-600 | LV_Brine_h2_0102_1000031 | + | 5692 | 5743 | **RNA**→ rpmE (PRK00019)Ribosomal_L31 (pfam01197)→ |
| Pha-3-1 | NC_007481.1 | - | 2892806 | 2892755 | **RNA**→ rpmE (PRK00019)Ribosomal_L31 (pfam01197)→ rpmE (PRK00019)Ribosomal_L31 (pfam01197)→ |
| Pru-1-1 | NZ_JXYA01000021.1 | + | 73438 | 73489 | **RNA**→ rpmE (PRK00019)Ribosomal_L31 (pfam01197)→ rpmE (PRK00019)Ribosomal_L31 (pfam01197)→ |
| Pru-1-2 | NZ_LFZX01000030.1 | + | 37188 | 37239 | **RNA**→ rpmE (PRK00019)Ribosomal_L31 (pfam01197)→ rpmE (PRK00019)Ribosomal_L31 (pfam01197)→ |
| Pru-2-1 | NZ_AHCD02000031.1 | + | 22677 | 22728 | **RNA**→ rpmE (PRK00019)Ribosomal_L31 (pfam01197)→ rpmE (PRK00019)Ribosomal_L31 (pfam01197)→ |

| | | | | | |
|---|---|---|---|---|---|
| Plu-4-1 | NZ_AUXW01000198.1 | - | 44235 | 44184 | **RNA**→ rpmE (PRK00019)Ribosomal_L31 (pfam01197)→ |
| Plu-1-1 | NZ_JWIC01000001.1 | + | 71814 | 71865 | **RNA**→ rpmE (PRK00019)Ribosomal_L31 (pfam01197)→ |
| Par-1-1 | NZ_AHBY02000081.1 | - | 369 | 318 | **RNA**→ rpmE (PRK00019)Ribosomal_L31 (pfam01197)→ |
| Psp-21-1 | NZ_AUTS01000045.1 | + | 48461 | 48512 | **RNA**→ rpmE (PRK00019)Ribosomal_L31 (pfam01197)→ |
| Psp-22-1 | NZ_AUTR01000020.1 | - | 193290 | 193239 | **RNA**→ rpmE (PRK00019)Ribosomal_L31 (pfam01197)→ |
| Psp-29-1 | NZ_AUTJ01000247.1 | + | 71641 | 71692 | **RNA**→ rpmE (PRK00019)Ribosomal_L31 (pfam01197)→ hypo→ |
| Psp-25-1 | NZ_AUTP01000215.1 | + | 15068 | 15119 | **RNA**→ rpmE (PRK00019)Ribosomal_L31 (pfam01197)→ |
| Psp-7-1 | NZ_BADV01000031.1 | - | 2245 | 2194 | **RNA**→ rpmE (PRK00019)Ribosomal_L31 (pfam01197)→ rpmE (PRK00019)Ribosomal_L31 (pfam01197)→ |
| env-601 | CEUP01039342.1 | - | 391 | 340 | **RNA**→ rpmE (PRK00019)Ribosomal_L31 (pfam01197)→ |
| env-602 | CEVA01501963.1 | - | 276 | 225 | **RNA**→ rpmE (PRK00019)Ribosomal_L31 (pfam01197)→ |
| env-603 | CEPZ01051150.1 | - | 473 | 422 | **RNA**→ rpmE (PRK00019)Ribosomal_L31 (pfam01197)→ |
| env-604 | CEQS01116987.1 | + | 480 | 531 | **RNA**→ |
| env-605 | TCCM__contig09952 | - | 265 | 214 | **RNA**→ rpmE (PRK00019)Ribosomal_L31 (pfam01197)→ |
| Psp-1-1 | NZ_BBIN01000213.1 | - | 19009 | 18959 | **RNA**→ rpmE (PRK00019)Ribosomal_L31 (pfam01197)→ |
| Ptu-1-1 | NZ_AAOH01000003.1 | + | 284768 | 284818 | **RNA**→ rpmE (PRK00019)Ribosomal_L31 (pfam01197)→ rpmE (PRK00019)Ribosomal_L31 (pfam01197)→ |
| env-606 | ABMI01022972.1 | + | 23 | 75 | **RNA**→ |
| Eca-5-1 | NZ_AJYH01000066.1 | - | 21608 | 21558 | **RNA**→ rpmE (PRK00019)Ribosomal_L31 (pfam01197)→ |
| Eca-7-1 | NZ_AJYF01000130.1 | + | 138107 | 138157 | **RNA**→ rpmE (PRK00019)Ribosomal_L31 (pfam01197)→ |
| Eca-4-1 | NZ_AJYG01000029.1 | + | 139048 | 139098 | **RNA**→ rpmE (PRK00019)Ribosomal_L31 (pfam01197)→ |
| Eca-6-1 | NZ_KK211219.1 | + | 164382 | 164432 | **RNA**→ rpmE (PRK00019)Ribosomal_L31 (pfam01197)→ |
| Gho-1-1 | NZ_ADAQ01000008.1 | - | 84673 | 84623 | **RNA**→ rpmE (PRK00019)Ribosomal_L31 (pfam01197)→ rpmE (PRK00019)Ribosomal_L31 (pfam01197)→ |
| Gsp-4-1 | NZ_ANFM02000002.1 | - | 72505 | 72455 | **RNA**→ rpmE (PRK00019)Ribosomal_L31 (pfam01197)→ |
| Gsp-3-1 | NZ_LASY01000133.1 | + | 6375 | 6425 | **RNA**→ rpmE (PRK00019)Ribosomal_L31 (pfam01197)→ |
| Eno-3-1 | NZ_AJWN01000101.1 | + | 97798 | 97847 | **RNA**→ rpmE (PRK00019)Ribosomal_L31 (pfam01197)→ |
| Eno-2-1 | NZ_AJYD01000004.1 | + | 100008 | 100057 | **RNA**→ rpmE (PRK00019)Ribosomal_L31 (pfam01197)→ |
| Eno-1-1 | NZ_AJYE01000099.1 | + | 19319 | 19368 | **RNA**→ rpmE (PRK00019)Ribosomal_L31 (pfam01197)→ |
| env-607 | JGI24724J26744_10298770 | - | 465 | 419 | **RNA**→ rpmE (PRK00019)Ribosomal_L31 (pfam01197)→ |
| env-608 | 3300000513_101996 | - | 1403 | 1355 | **RNA**→ rpmE (PRK00019)Ribosomal_L31 (pfam01197)→ |
| env-609 | 3300001239_10010964 | - | 1403 | 1355 | **RNA**→ rpmE (PRK00019)Ribosomal_L31 (pfam01197)→ |
| env-610 | 3300001605_10069366 | - | 1992 | 1944 | **RNA**→ rpmE (PRK00019)Ribosomal_L31 (pfam01197)→ |
| Tsp-1-1 | NZ_JQDZ01000010.1 | - | 1358 | 1309 | **RNA**→ ←hypo |
| Pha-5-1 | NZ_AWXP01000020.1 | + | 164833 | 164882 | **RNA**→ rpmE (PRK00019)Ribosomal_L31 (pfam01197)→ rpmE (PRK00019)Ribosomal_L31 (pfam01197)→ hypo→ HipA (COG3550)PRK09775 (PRK09775)HTH_ARSR (cd00090)→ |
| Pvu-2-1 | NZ_KN150745.1 | - | 686197 | 686148 | **RNA**→ rpmE (PRK00019)Ribosomal_L31 (pfam01197)→ |
| Pmi-3-1 | NC_022000.1 | + | 3534400 | 3534449 | **RNA**→ rpmE (PRK00019)Ribosomal_L31 (pfam01197)→ rpmE (PRK00019)Ribosomal_L31 (pfam01197)→ |
| env-611 | JGI24707J26582_10420588 | + | 135 | 189 | **RNA**→ rpmE (PRK00019)Ribosomal_L31 (pfam01197)→ |
| Ffu-1-1 | NZ_AUGL01000001.1 | - | 147568 | 147519 | **RNA**→ rpmE (PRK00019)Ribosomal_L31 (pfam01197)→ |
| Fky-1-1 | NZ_AUDJ01000001.1 | + | 302704 | 302753 | **RNA**→ ←hypo |
| Asi-1-1 | NZ_CDBY01000035.1 | + | 3638 | 3691 | **RNA**→ rpmE (PRK00019)Ribosomal_L31 (pfam01197)→ SRPBCC (cd07812)YbjT (COG0702)DUF2867 (pfam11066)NAD_binding_10 (pfam13460)→ |
| Afi-1-1 | NZ_JRGK01000350.1 | - | 15755 | 15703 | **RNA**→ rpmE (PRK00019)Ribosomal_L31 (pfam01197)→ |
| Aja-1-1 | NZ_JWJR01000011.1 | - | 124061 | 124009 | **RNA**→ rpmE (PRK00019)Ribosomal_L31 (pfam01197)→ |
| Adh-1-1 | NZ_CDBP01000004.1 | + | 72490 | 72542 | **RNA**→ rpmE (PRK00019)Ribosomal_L31 (pfam01197)→ |
| Adh-1-2 | NZ_CDBH01000029.1 | + | 72699 | 72751 | **RNA**→ rpmE (PRK00019)Ribosomal_L31 (pfam01197)→ |
| Aen-2-1 | NZ_LDWG01000014.1 | - | 162402 | 162350 | **RNA**→ rpmE (PRK00019)Ribosomal_L31 (pfam01197)→ |
| Adh-1-3 | NZ_JDWD01000154.1 | + | 20498 | 20550 | **RNA**→ rpmE (PRK00019)Ribosomal_L31 (pfam01197)→ |
| Ahy-4-1 | NZ_JEMK01000035.1 | + | 72881 | 72933 | **RNA**→ rpmE (PRK00019)Ribosomal_L31 (pfam01197)→ |
| Ahy-18-1 | NZ_CP007518.1 | + | 4401696 | 4401748 | **RNA**→ rpmE (PRK00019)Ribosomal_L31 (pfam01197)→ |
| Ahy-17-1 | NZ_JH815591.1 | - | 3833332 | 3833280 | **RNA**→ rpmE (PRK00019)Ribosomal_L31 (pfam01197)→ |
| Ahy-2-1 | NZ_ANPN01000010.1 | - | 162590 | 162538 | **RNA**→ rpmE (PRK00019)Ribosomal_L31 (pfam01197)→ |
| Ahy-9-1 | NZ_AOBQ01000006.1 | + | 72707 | 72759 | **RNA**→ rpmE (PRK00019)Ribosomal_L31 (pfam01197)→ |
| Ahy-8-1 | NZ_AOBP01000008.1 | - | 131017 | 130965 | **RNA**→ rpmE (PRK00019)Ribosomal_L31 (pfam01197)→ |
| Ahy-6-1 | NZ_AOBO01000002.1 | - | 186283 | 186231 | **RNA**→ rpmE (PRK00019)Ribosomal_L31 (pfam01197)→ |

| | | | | | |
|---|---|---|---|---|---|
| Ahy-3-1 | NZ_AOBM01000003.1 | - | 130263 | 130211 | **RNA**→ rpmE (PRK00019)Ribosomal_L31 (pfam01197)→ |
| Ahy-5-1 | NZ_AOBN01000020.1 | + | 72684 | 72736 | **RNA**→ rpmE (PRK00019)Ribosomal_L31 (pfam01197)→ |
| Aaq-1-1 | NZ_BAFL01000026.1 | + | 72708 | 72760 | **RNA**→ rpmE (PRK00019)Ribosomal_L31 (pfam01197)→ rpmE (PRK00019)Ribosomal_L31 (pfam01197)→ |
| Aja-1-2 | NZ_CDBV01000003.1 | + | 54015 | 54067 | **RNA**→ rpmE (PRK00019)Ribosomal_L31 (pfam01197)→ |
| Asp-2-1 | NZ_JXIR01000034.1 | - | 113264 | 113212 | **RNA**→ rpmE (PRK00019)Ribosomal_L31 (pfam01197)→ |
| Ala-1-1 | NZ_JRGM01000150.1 | + | 53033 | 53085 | **RNA**→ rpmE (PRK00019)Ribosomal_L31 (pfam01197)→ |
| Aja-2-1 | NZ_JFDL01000010.1 | + | 54457 | 54509 | **RNA**→ rpmE (PRK00019)Ribosomal_L31 (pfam01197)→ |
| Aau-1-1 | NZ_CDDH01000054.1 | - | 10218 | 10166 | **RNA**→ rpmE (PRK00019)Ribosomal_L31 (pfam01197)→ |
| Amo-1-1 | NZ_AQGQ01000086.1 | - | 10566 | 10515 | **RNA**→ rpmE (PRK00019)Ribosomal_L31 (pfam01197)→ |
| Api-1-1 | NZ_CDBL01000057.1 | - | 131319 | 131268 | **RNA**→ rpmE (PRK00019)Ribosomal_L31 (pfam01197)→ |
| Asa-5-1 | NZ_CDDW01000113.1 | - | 28277 | 28226 | **RNA**→ rpmE (PRK00019)Ribosomal_L31 (pfam01197)→ |
| Apo-1-1 | NZ_CDBI01000047.1 | + | 73787 | 73838 | **RNA**→ rpmE (PRK00019)Ribosomal_L31 (pfam01197)→ |
| Asa-5-2 | NZ_JXTA01000018.1 | + | 52404 | 52455 | **RNA**→ rpmE (PRK00019)Ribosomal_L31 (pfam01197)→ |
| Asa-3-1 | NZ_BAWQ01000020.1 | - | 14307 | 14256 | **RNA**→ rpmE (PRK00019)Ribosomal_L31 (pfam01197)→ |
| Asa-4-1 | NZ_ARYZ02000006.1 | - | 52708 | 52657 | **RNA**→ rpmE (PRK00019)Ribosomal_L31 (pfam01197)→ |
| Ahy-16-1 | NZ_AMQA01000007.1 | + | 48674 | 48725 | **RNA**→ rpmE (PRK00019)Ribosomal_L31 (pfam01197)→ |
| Asa-6-1 | NZ_AGVO01000060.1 | + | 52105 | 52156 | **RNA**→ rpmE (PRK00019)Ribosomal_L31 (pfam01197)→ |
| Asa-7-1 | NC_009348.1 | + | 140845 | 140896 | **RNA**→ rpmE (PRK00019)Ribosomal_L31 (pfam01197)→ |
| Asa-2-1 | NZ_AMQG02000058.1 | - | 13927 | 13876 | **RNA**→ rpmE (PRK00019)Ribosomal_L31 (pfam01197)→ |
| Asa-5-3 | NZ_JRYW01000019.1 | + | 52405 | 52456 | **RNA**→ rpmE (PRK00019)Ribosomal_L31 (pfam01197)→ |
| Asa-5-4 | NZ_JRYV01000091.1 | - | 17030 | 16979 | **RNA**→ rpmE (PRK00019)Ribosomal_L31 (pfam01197)→ |
| env-612 | JCVI_SCAF_1096627194609 | - | 4672 | 4621 | **RNA**→ rpmE (PRK00019)Ribosomal_L31 (pfam01197)→ YbjT (COG0702)DUF2867 (pfam11066)NAD_binding_10 (pfam13460)→ |
| Ahy-21-1 | NZ_CDDC01000091.1 | - | 122529 | 122478 | **RNA**→ rpmE (PRK00019)Ribosomal_L31 (pfam01197)→ |
| Ahy-11-1 | NZ_JFJO01000068.1 | - | 10099 | 10048 | **RNA**→ rpmE (PRK00019)Ribosomal_L31 (pfam01197)→ |
| Ahy-20-1 | NC_008570.1 | - | 4646089 | 4646038 | **RNA**→ rpmE (PRK00019)Ribosomal_L31 (pfam01197)→ |
| Ahy-14-1 | NC_021290.1 | - | 4925796 | 4925745 | **RNA**→ rpmE (PRK00019)Ribosomal_L31 (pfam01197)→ |
| Ahy-12-1 | NZ_CP007566.1 | - | 4925161 | 4925110 | **RNA**→ rpmE (PRK00019)Ribosomal_L31 (pfam01197)→ |
| Ahy-19-1 | NZ_CP007576.1 | - | 4925129 | 4925078 | **RNA**→ rpmE (PRK00019)Ribosomal_L31 (pfam01197)→ |
| Ahy-13-1 | NZ_CP006883.1 | + | 142305 | 142356 | **RNA**→ rpmE (PRK00019)Ribosomal_L31 (pfam01197)→ |
| Ahy-1-1 | NZ_CP010947.1 | - | 4784963 | 4784912 | **RNA**→ rpmE (PRK00019)Ribosomal_L31 (pfam01197)→ |
| Ahy-1-2 | NZ_CP011100.1 | + | 204318 | 204369 | **RNA**→ rpmE (PRK00019)Ribosomal_L31 (pfam01197)→ |
| Ahy-15-1 | NZ_CP006870.1 | + | 142342 | 142393 | **RNA**→ rpmE (PRK00019)Ribosomal_L31 (pfam01197)→ |
| Ahy-7-1 | NZ_JEML01000047.1 | + | 52150 | 52201 | **RNA**→ rpmE (PRK00019)Ribosomal_L31 (pfam01197)→ |
| Abi-1-1 | NZ_CDBT01000035.1 | - | 149827 | 149776 | **RNA**→ rpmE (PRK00019)Ribosomal_L31 (pfam01197)→ |
| env-613 | GooDraft_1010177 | + | 247 | 298 | **RNA**→ rpmE (PRK00019)Ribosomal_L31 (pfam01197)→ |
| Aso-1-1 | NZ_CDBW01000004.1 | + | 44945 | 44996 | **RNA**→ rpmE (PRK00019)Ribosomal_L31 (pfam01197)→ |
| Aen-2-2 | NZ_CDDE01000010.1 | + | 51645 | 51696 | **RNA**→ rpmE (PRK00019)Ribosomal_L31 (pfam01197)→ |
| Afl-1-1 | NZ_CDBO01000009.1 | + | 51638 | 51689 | **RNA**→ rpmE (PRK00019)Ribosomal_L31 (pfam01197)→ |
| Ave-1-1 | NZ_CDDU01000053.1 | + | 54632 | 54683 | **RNA**→ rpmE (PRK00019)Ribosomal_L31 (pfam01197)→ |
| Aen-2-3 | NZ_CDCG01000034.1 | - | 93277 | 93226 | **RNA**→ rpmE (PRK00019)Ribosomal_L31 (pfam01197)→ |
| Ave-9-1 | NZ_CDDK01000028.1 | + | 53357 | 53408 | **RNA**→ rpmE (PRK00019)Ribosomal_L31 (pfam01197)→ |
| Ave-1-2 | NZ_CDBU01000031.1 | + | 53055 | 53106 | **RNA**→ rpmE (PRK00019)Ribosomal_L31 (pfam01197)→ |
| Ave-7-1 | NZ_ATFB01000030.1 | - | 126444 | 126393 | **RNA**→ rpmE (PRK00019)Ribosomal_L31 (pfam01197)→ |
| Ave-1-3 | NZ_JRBE01000009.1 | - | 111575 | 111524 | **RNA**→ rpmE (PRK00019)Ribosomal_L31 (pfam01197)→ |
| Ave-6-1 | NC_015424.1 | - | 171080 | 171029 | **RNA**→ rpmE (PRK00019)Ribosomal_L31 (pfam01197)→ |
| Ave-3-1 | NZ_JH815586.1 | + | 52965 | 53016 | **RNA**→ rpmE (PRK00019)Ribosomal_L31 (pfam01197)→ |
| Ave-2-1 | NZ_JH815583.1 | + | 55914 | 55965 | **RNA**→ rpmE (PRK00019)Ribosomal_L31 (pfam01197)→ |
| Ave-4-1 | NZ_JH823256.1 | - | 1129313 | 1129262 | **RNA**→ rpmE (PRK00019)Ribosomal_L31 (pfam01197)→ |
| Ave-5-1 | NZ_JH815589.1 | + | 321999 | 322050 | **RNA**→ rpmE (PRK00019)Ribosomal_L31 (pfam01197)→ |
| Ave-8-1 | NZ_CDBQ01000036.1 | + | 53773 | 53824 | **RNA**→ rpmE (PRK00019)Ribosomal_L31 (pfam01197)→ |
| Asp-3-1 | NZ_JRJY01000041.1 | - | 59091 | 59040 | **RNA**→ rpmE (PRK00019)Ribosomal_L31 (pfam01197)→ |

| | | | | | |
|---|---|---|---|---|---|
| Aal-1-1 | NZ_CDCB01000029.1 | + | 53445 | 53496 | RNA→ rpmE (PRK00019)Ribosomal_L31 (pfam01197)→ |
| Aal-1-2 | NZ_CDBR01000043.1 | + | 32916 | 32967 | RNA→ rpmE (PRK00019)Ribosomal_L31 (pfam01197)→ |
| env-614 | GO4KUYH01ENIZ1 | + | 340 | 391 | RNA→ rpmE (PRK00528)rpmE (PRK00019)rpmE2 (PRK01678)Ribosomal_L31 (pfam01197)→ |
| env-615 | JGI11944J13513_1007729 | + | 807 | 858 | RNA→ rpmE (PRK00019)Ribosomal_L31 (pfam01197)→ |
| Aen-1-1 | NZ_CDDI01000023.1 | - | 131318 | 131267 | RNA→ rpmE (PRK00019)Ribosomal_L31 (pfam01197)→ |
| Ahy-10-1 | NZ_CP006579.1 | - | 4352087 | 4352036 | RNA→ rpmE (PRK00019)Ribosomal_L31 (pfam01197)→ |
| env-616 | JGI26532J50257_10010618 | - | 2542 | 2491 | RNA→ rpmE (PRK00019)Ribosomal_L31 (pfam01197)→ |
| env-617 | JGI26536J50258_1000749 | + | 19423 | 19474 | RNA→ rpmE (PRK00019)Ribosomal_L31 (pfam01197)→ |
| env-618 | JGI26540J51217_10020906 | + | 793 | 844 | RNA→ rpmE (PRK00019)Ribosomal_L31 (pfam01197)→ |
| Asa-8-1 | NZ_CDBN01000071.1 | - | 4965 | 4914 | RNA→ rpmE (PRK00019)Ribosomal_L31 (pfam01197)→ |
| Aca-2-1 | NZ_CACP01000112.1 | - | 130878 | 130827 | RNA→ rpmE (PRK00019)Ribosomal_L31 (pfam01197)→ rpmE (PRK00019)Ribosomal_L31 (pfam01197)→ |
| Asp-1-1 | NZ_JEMQ01000008.1 | + | 76191 | 76242 | RNA→ rpmE (PRK00019)Ribosomal_L31 (pfam01197)→ |
| Aca-1-1 | NZ_KQ087569.1 | - | 134377 | 134326 | RNA→ rpmE (PRK00019)Ribosomal_L31 (pfam01197)→ |
| Asp-4-1 | NZ_JRQJ01000349.1 | - | 10593 | 10542 | RNA→ rpmE (PRK00019)Ribosomal_L31 (pfam01197)→ |
| Ata-1-1 | NZ_BAWK01000091.1 | - | 20016 | 19965 | RNA→ rpmE (PRK00019)Ribosomal_L31 (pfam01197)→ |
| | | | | | SRPBCC (cd07812)YbjT (COG0702)DUF2867 (pfam11066)NAD_binding_10 (pfam13460)→ |
| Ata-1-2 | NZ_CDDD01000058.1 | - | 134875 | 134824 | RNA→ rpmE (PRK00019)Ribosomal_L31 (pfam01197)→ |
| | | | | | SRPBCC (cd07812)YbjT (COG0702)DUF2867 (pfam11066)NAD_binding_10 (pfam13460)→ |
| Ari-1-1 | NZ_CDBJ01000055.1 | + | 6337 | 6388 | RNA→ rpmE (PRK00019)Ribosomal_L31 (pfam01197)→ |
| Ate-1-1 | NZ_CDCA01000034.1 | + | 49892 | 49942 | RNA→ rpmE (PRK00019)Ribosomal_L31 (pfam01197)→ |
| Ame-1-1 | NZ_CDBZ01000101.1 | - | 64689 | 64639 | RNA→ rpmE (PRK00019)Ribosomal_L31 (pfam01197)→ |
| Ame-1-2 | NZ_JRBF01000002.1 | - | 94963 | 94913 | RNA→ rpmE (PRK00019)Ribosomal_L31 (pfam01197)→ |
| Ame-2-1 | NZ_CP007567.1 | - | 4055949 | 4055899 | RNA→ rpmE (PRK00019)Ribosomal_L31 (pfam01197)→ |
| Ame-1-3 | NZ_JRBG01000006.1 | - | 70612 | 70562 | RNA→ rpmE (PRK00019)Ribosomal_L31 (pfam01197)→ |
| Aaq-2-1 | NZ_JRGL01000008.1 | - | 78368 | 78318 | RNA→ rpmE (PRK00019)Ribosomal_L31 (pfam01197)→ |
| env-619 | XAGC_contig09808 | - | 443 | 393 | RNA→ rpmE (PRK00019)Ribosomal_L31 (pfam01197)→ |
| Abe-1-1 | NZ_CDDA01000005.1 | - | 159579 | 159529 | RNA→ rpmE (PRK00019)Ribosomal_L31 (pfam01197)→ |
| Asa-1-1 | NZ_JPWL01000105.1 | - | 218920 | 218870 | RNA→ rpmE (PRK00019)Ribosomal_L31 (pfam01197)→ |
| Aeu-1-1 | NZ_CDDF01000018.1 | - | 151708 | 151658 | RNA→ rpmE (PRK00019)Ribosomal_L31 (pfam01197)→ |
| Asc-1-1 | NZ_CDDB01000021.1 | + | 125280 | 125333 | RNA→ rpmE (PRK00019)Ribosomal_L31 (pfam01197)→ |
| | | | | | SRPBCC (cd07812)YbjT (COG0702)DUF2867 (pfam11066)NAD_binding_10 (pfam13460)→ |
| Adi-2-1 | NZ_CDCE01000027.1 | - | 19201 | 19148 | RNA→ rpmE (PRK00019)Ribosomal_L31 (pfam01197)→ |
| | | | | | SRPBCC_CalC_Aha1-like_GntR-HTH (cd08893)YbjT (COG0702)DUF2867 (pfam11066)NAD_binding_10 (pfam13 |
| Adi-1-1 | NZ_APVG01000009.1 | + | 71044 | 71097 | RNA→ rpmE (PRK00019)Ribosomal_L31 (pfam01197)→ |
| | | | | | SRPBCC_CalC_Aha1-like_GntR-HTH (cd08893)YbjT (COG0702)DUF2867 (pfam11066)NAD_binding_10 (pfam13 |
| env-620 | GSLSAS_contig00578 | - | 2300 | 2248 | RNA→ rpmE (PRK00019)Ribosomal_L31 (pfam01197)→ |
| Sso-15-1 | NZ_AQOD01000036.1 | - | 8278 | 8226 | RNA→ rpmE (PRK00019)Ribosomal_L31 (pfam01197)→ |
| Sso-14-1 | NZ_AQOE01000001.1 | - | 17831 | 17779 | RNA→ rpmE (PRK00019)Ribosomal_L31 (pfam01197)→ |
| env-621 | CEGE01004975.1 | - | 1222 | 1170 | RNA→ rpmE (PRK00019)Ribosomal_L31 (pfam01197)→ |
| env-622 | GSLSAS_contig00020 | + | 11479 | 11531 | RNA→ rpmE (PRK00019)Ribosomal_L31 (pfam01197)→ |
| Sco-3-1 | NZ_AQOF01000004.1 | + | 41661 | 41713 | RNA→ rpmE (PRK00019)Ribosomal_L31 (pfam01197)→ |
| env-623 | GSLSAS_contig02665 | + | 1265 | 1317 | RNA→ rpmE (PRK00019)Ribosomal_L31 (pfam01197)→ |
| Ssp-27-1 | NZ_LAQR01000036.1 | + | 121364 | 121416 | RNA→ rpmE (PRK00019)Ribosomal_L31 (pfam01197)→ |
| Psp-57-1 | NZ_AMZO01000033.1 | - | 59095 | 59042 | RNA→ rpmE (PRK00019)Ribosomal_L31 (pfam01197)→ |
| Pan-11-1 | NZ_JZSW01000011.1 | + | 88087 | 88139 | RNA→ rpmE (PRK00019)Ribosomal_L31 (pfam01197)→ |
| Pan-12-1 | NZ_CH902603.1 | - | 60950 | 60898 | RNA→ rpmE (PRK00019)Ribosomal_L31 (pfam01197)→ |
| Pan-11-2 | NZ_JZSO01000016.1 | + | 86032 | 86084 | RNA→ rpmE (PRK00019)Ribosomal_L31 (pfam01197)→ |
| Pan-11-3 | NZ_JZSN01000009.1 | - | 60994 | 60942 | RNA→ rpmE (PRK00019)Ribosomal_L31 (pfam01197)→ |
| Pan-11-4 | NZ_JZSS01000006.1 | + | 115493 | 115545 | RNA→ rpmE (PRK00019)Ribosomal_L31 (pfam01197)→ |
| Pan-11-5 | NZ_JZSY01000010.1 | + | 90276 | 90328 | RNA→ rpmE (PRK00019)Ribosomal_L31 (pfam01197)→ |
| Pan-11-6 | NZ_JZSZ01000011.1 | + | 90260 | 90312 | RNA→ rpmE (PRK00019)Ribosomal_L31 (pfam01197)→ |
| Pan-11-7 | NZ_JZSU01000008.1 | - | 60993 | 60941 | RNA→ rpmE (PRK00019)Ribosomal_L31 (pfam01197)→ |
| Pan-11-8 | NZ_JZST01000009.1 | + | 115491 | 115543 | RNA→ rpmE (PRK00019)Ribosomal_L31 (pfam01197)→ |

| | | | | | | |
|---|---|---|---|---|---|---|
| Pan-11-9 | NZ_JZSM01000004.1 | - | 135032 | 134980 | **RNA**→ | rpmE (PRK00019)Ribosomal_L31 (pfam01197)→ |
| Pda-2-1 | NZ_JZSI01000010.1 | + | 90276 | 90328 | **RNA**→ | rpmE (PRK00019)Ribosomal_L31 (pfam01197)→ |
| Pan-11-10 | NZ_JZSV01000008.1 | - | 60990 | 60938 | **RNA**→ | rpmE (PRK00019)Ribosomal_L31 (pfam01197)→ |
| Pan-11-11 | NZ_JZSX01000009.1 | + | 90266 | 90318 | **RNA**→ | rpmE (PRK00019)Ribosomal_L31 (pfam01197)→ |
| Pan-11-12 | NZ_JZTA01000010.1 | + | 90591 | 90643 | **RNA**→ | rpmE (PRK00019)Ribosomal_L31 (pfam01197)→ |
| Psp-58-1 | NZ_AAOU01000028.1 | + | 15095 | 15147 | **RNA**→ | rpmE (PRK00019)Ribosomal_L31 (pfam01197)→ rpmE (PRK00019)Ribosomal_L31 (pfam01197)→ |
| Pap-1-1 | NZ_LDOV01000029.1 | - | 41563 | 41511 | **RNA**→ | rpmE (PRK00019)Ribosomal_L31 (pfam01197)→ |
| Psa-1-1 | NZ_JGVO01001562.1 | + | 1691 | 1743 | **RNA**→ | rpmE (PRK00019)Ribosomal_L31 (pfam01197)→ |
| Paq-2-1 | NZ_LDOT01000005.1 | + | 157449 | 157501 | **RNA**→ | rpmE (PRK00019)Ribosomal_L31 (pfam01197)→ |
| Pga-2-1 | NZ_CP005974.1 | + | 545149 | 545201 | **RNA**→ | rpmE (PRK00019)Ribosomal_L31 (pfam01197)→ |
| Pga-1-1 | NZ_JWLZ01000213.1 | - | 58688 | 58636 | **RNA**→ | rpmE (PRK00019)Ribosomal_L31 (pfam01197)→ |
| Ple-1-1 | NZ_JZSL01000002.1 | - | 60040 | 59988 | **RNA**→ | rpmE (PRK00019)Ribosomal_L31 (pfam01197)→ |
| Ple-1-2 | NZ_JZSK01000007.1 | - | 60039 | 59987 | **RNA**→ | rpmE (PRK00019)Ribosomal_L31 (pfam01197)→ |
| Ple-3-1 | NZ_DF093598.1 | - | 93166 | 93114 | **RNA**→ | rpmE (PRK00019)Ribosomal_L31 (pfam01197)→ rpmE (PRK00019)Ribosomal_L31 (pfam01197)→ |
| Ple-2-1 | NZ_DF196819.1 | - | 1468186 | 1468134 | **RNA**→ | rpmE (PRK00019)Ribosomal_L31 (pfam01197)→ |
| Pda-4-1 | NZ_KB405046.1 | + | 159586 | 159638 | **RNA**→ | rpmE (PRK00019)Ribosomal_L31 (pfam01197)→ |
| Pda-3-1 | NZ_ADBS01000001.1 | + | 3048450 | 3048502 | **RNA**→ | rpmE (PRK00019)Ribosomal_L31 (pfam01197)→ rpmE (PRK00019)Ribosomal_L31 (pfam01197)→ |
| Psw-1-1 | NZ_LELC01000008.1 | + | 114365 | 114416 | **RNA**→ | rpmE (PRK00019)Ribosomal_L31 (pfam01197)→ |
| Pph-1-1 | NZ_JZSJ01000013.1 | + | 34239 | 34290 | **RNA**→ | rpmE (PRK00019)Ribosomal_L31 (pfam01197)→ |
| Pki-1-1 | NZ_JZTD01000009.1 | - | 53868 | 53817 | **RNA**→ | rpmE (PRK00019)Ribosomal_L31 (pfam01197)→ |
| Pki-1-2 | NZ_JZTE01000008.1 | - | 53868 | 53817 | **RNA**→ | rpmE (PRK00019)Ribosomal_L31 (pfam01197)→ |
| Pki-1-3 | NZ_JZTC01000007.1 | - | 53870 | 53819 | **RNA**→ | rpmE (PRK00019)Ribosomal_L31 (pfam01197)→ |
| Pki-1-4 | NZ_JZTB01000007.1 | - | 53867 | 53816 | **RNA**→ | rpmE (PRK00019)Ribosomal_L31 (pfam01197)→ |
| Pil-1-1 | NZ_JZSQ01000007.1 | + | 107422 | 107473 | **RNA**→ | rpmE (PRK00019)Ribosomal_L31 (pfam01197)→ |
| Pil-1-2 | NZ_JZSR01000004.1 | + | 109269 | 109320 | **RNA**→ | rpmE (PRK00019)Ribosomal_L31 (pfam01197)→ |
| Pki-1-5 | NZ_JZSP01000008.1 | - | 26274 | 26223 | **RNA**→ | rpmE (PRK00019)Ribosomal_L31 (pfam01197)→ |
| Pha-6-1 | NZ_JWYV01000006.1 | - | 63341 | 63289 | **RNA**→ | rpmE (PRK00019)Ribosomal_L31 (pfam01197)→ |
| Pha-7-1 | NZ_AULG01000012.1 | + | 172893 | 172945 | **RNA**→ | rpmE (PRK00019)Ribosomal_L31 (pfam01197)→ |
| Pga-3-1 | NZ_JMIB01000030.1 | - | 62198 | 62146 | **RNA**→ | rpmE (PRK00019)Ribosomal_L31 (pfam01197)→ |
| Pga-4-1 | NZ_LDOU01000024.1 | - | 60173 | 60119 | **RNA**→ | rpmE (PRK00019)Ribosomal_L31 (pfam01197)→ |
| Ppr-1-1 | NZ_CH724140.1 | - | 69796 | 69744 | **RNA**→ | rpmE (PRK00019)Ribosomal_L31 (pfam01197)→ |
| env-624 | KGI_S1_ANT01_95m_c10724678 | + | 129 | 181 | **RNA**→ | rpmE (PRK00019)rpmE2 (PRK01678)Ribosomal_L31 (pfam01197)→ |
| Ppr-2-1 | NC_006370.1 | + | 266374 | 266426 | **RNA**→ | rpmE (PRK00019)Ribosomal_L31 (pfam01197)→ |
| Vez-1-1 | NZ_BATM01000037.1 | + | 129339 | 129390 | **RNA**→ | rpmE (PRK00019)Ribosomal_L31 (pfam01197)→ |
| Vha-1-1 | NZ_BAUJ01000003.1 | + | 79401 | 79452 | **RNA**→ | rpmE (PRK00019)Ribosomal_L31 (pfam01197)→ |
| env-625 | ASRK01000687.1 | - | 597 | 548 | **RNA**→ | rpmE (PRK00019)Ribosomal_L31 (pfam01197)→ |
| Vpr-1-1 | NZ_BATJ01000004.1 | + | 154354 | 154404 | **RNA**→ | rpmE (PRK00019)Ribosomal_L31 (pfam01197)→ |
| Vbr-2-1 | NZ_AKXW01000019.1 | - | 57344 | 57294 | **RNA**→ | rpmE (PRK00019)Ribosomal_L31 (pfam01197)→ |
| Vbr-3-1 | NZ_AJYM01000057.1 | - | 57302 | 57252 | **RNA**→ | rpmE (PRK00019)Ribosomal_L31 (pfam01197)→ |
| Vbr-4-1 | NZ_AJYL01000130.1 | + | 139283 | 139333 | **RNA**→ | rpmE (PRK00019)Ribosomal_L31 (pfam01197)→ |
| Fse-1-1 | NZ_AUGM01000007.1 | - | 231862 | 231812 | **RNA**→ | rpmE (PRK00019)Ribosomal_L31 (pfam01197)→ |
| Vmy-1-1 | NZ_JXOK01000076.1 | - | 101239 | 101190 | **RNA**→ | rpmE (PRK00019)Ribosomal_L31 (pfam01197)→ |
| Vsp-27-1 | NZ_JZAM01000160.1 | - | 49297 | 49246 | **RNA**→ | rpmE (PRK00019)Ribosomal_L31 (pfam01197)→ |
| Vna-2-1 | NZ_JMCF01000037.1 | - | 19898 | 19847 | **RNA**→ | rpmE (PRK00019)Ribosomal_L31 (pfam01197)→ |
| Vna-2-2 | NZ_JMCI01000042.1 | + | 6951 | 7002 | **RNA**→ | rpmE (PRK00019)Ribosomal_L31 (pfam01197)→ |
| Vna-2-3 | NZ_JMCH01000025.1 | + | 16370 | 16421 | **RNA**→ | rpmE (PRK00019)Ribosomal_L31 (pfam01197)→ |
| Vna-2-4 | NZ_JMCG01000001.1 | + | 496985 | 497036 | **RNA**→ | rpmE (PRK00019)Ribosomal_L31 (pfam01197)→ |
| Vvu-8-1 | NC_014965.1 | + | 326873 | 326924 | **RNA**→ | rpmE (PRK00019)Ribosomal_L31 (pfam01197)→ |
| Vsa-1-1 | NZ_BAOJ01000037.1 | - | 34285 | 34235 | **RNA**→ | rpmE (PRK00019)Ribosomal_L31 (pfam01197)→ |
| Vaz-1-1 | NZ_BAOB01000011.1 | + | 28481 | 28531 | **RNA**→ | rpmE (PRK00019)Ribosomal_L31 (pfam01197)→ |
| Vaz-1-2 | NZ_BATL01000020.1 | - | 35437 | 35387 | **RNA**→ | rpmE (PRK00019)Ribosomal_L31 (pfam01197)→ |
| Vro-3-1 | NZ_AFAJ01000015.1 | + | 3027 | 3077 | **RNA**→ | rpmE (PRK00019)Ribosomal_L31 (pfam01197)→ rpmE (PRK00019)Ribosomal_L31 (pfam01197)→ |

| | | | | | |
|---|---|---|---|---|---|
| Vro-2-1 | NZ_BAOI01000078.1 | - | 79201 | 79151 | **RNA**→ rpmE (PRK00019)Ribosomal_L31 (pfam01197)→ |
| Vhy-1-1 | NZ_BBLF01000022.1 | + | 27647 | 27697 | **RNA**→ rpmE (PRK00019)Ribosomal_L31 (pfam01197)→ |
| Vja-1-1 | NZ_BBKZ01000028.1 | - | 303783 | 303733 | **RNA**→ rpmE (PRK00019)Ribosomal_L31 (pfam01197)→ |
| Vhy-1-2 | NZ_BBLD01000021.1 | - | 69972 | 69922 | **RNA**→ rpmE (PRK00019)Ribosomal_L31 (pfam01197)→ |
| Vja-1-2 | NZ_BBLA01000033.1 | - | 289672 | 289622 | **RNA**→ rpmE (PRK00019)Ribosomal_L31 (pfam01197)→ |
| Vja-3-1 | NZ_BAOG01000025.1 | - | 30346 | 30296 | **RNA**→ rpmE (PRK00019)Ribosomal_L31 (pfam01197)→ |
| Vna-1-1 | NZ_ATFJ01000038.1 | + | 148901 | 148951 | **RNA**→ rpmE (PRK00019)Ribosomal_L31 (pfam01197)→ |
| Vna-1-2 | NZ_ATWU01000061.1 | + | 212711 | 212761 | **RNA**→ rpmE (PRK00019)Ribosomal_L31 (pfam01197)→ |
| Vsp-6-1 | NC_016613.1 | + | 288935 | 288985 | **RNA**→ rpmE (PRK00019)Ribosomal_L31 (pfam01197)→ |
| Vja-4-1 | NZ_BAOA01000021.1 | + | 69740 | 69790 | **RNA**→ rpmE (PRK00019)Ribosomal_L31 (pfam01197)→ |
| Vpa-87-1 | NZ_AWMB01000121.1 | - | 5724 | 5674 | **RNA**→ rpmE (PRK00019)Ribosomal_L31 (pfam01197)→ |
| Vpa-88-1 | NZ_AWMA01000113.1 | + | 9881 | 9931 | **RNA**→ rpmE (PRK00019)Ribosomal_L31 (pfam01197)→ |
| Vpa-156-1 | NZ_AWJK01000001.1 | + | 63708 | 63758 | **RNA**→ rpmE (PRK00019)Ribosomal_L31 (pfam01197)→ |
| Val-2-1 | NZ_LFZC01000032.1 | - | 106376 | 106326 | **RNA**→ rpmE (PRK00019)Ribosomal_L31 (pfam01197)→ |
| Vpa-240-1 | NZ_AVPW01000029.1 | + | 19681 | 19731 | **RNA**→ rpmE (PRK00019)Ribosomal_L31 (pfam01197)→ |
| Vpa-238-1 | NZ_AOOW01000293.1 | + | 7296 | 7346 | **RNA**→ rpmE (PRK00019)Ribosomal_L31 (pfam01197)→ |
| Vpa-94-1 | NZ_AWLU01000147.1 | + | 8348 | 8398 | **RNA**→ rpmE (PRK00019)Ribosomal_L31 (pfam01197)→ |
| Vhy-1-3 | NZ_BBLE01000010.1 | + | 164226 | 164276 | **RNA**→ rpmE (PRK00019)Ribosomal_L31 (pfam01197)→ |
| env-626 | GSLSAS_GLMUJHB02HDDBS | + | 419 | 469 | **RNA**→ |
| Vpa-47-1 | NZ_BBQD01000006.1 | + | 238207 | 238257 | **RNA**→ rpmE (PRK00019)Ribosomal_L31 (pfam01197)→ |
| Vpa-23-1 | NZ_AOOZ01000001.1 | + | 266710 | 266760 | **RNA**→ rpmE (PRK00019)Ribosomal_L31 (pfam01197)→ |
| Vpa-21-1 | NZ_AOOY01000012.1 | + | 32552 | 32602 | **RNA**→ rpmE (PRK00019)Ribosomal_L31 (pfam01197)→ |
| Vpa-237-1 | NZ_AOOV01000001.1 | + | 266710 | 266760 | **RNA**→ rpmE (PRK00019)Ribosomal_L31 (pfam01197)→ |
| Vpa-140-1 | NZ_AWKA01000014.1 | - | 5727 | 5677 | **RNA**→ rpmE (PRK00019)Ribosomal_L31 (pfam01197)→ |
| Vpa-126-1 | NZ_AWKO01000078.1 | - | 5726 | 5676 | **RNA**→ rpmE (PRK00019)Ribosomal_L31 (pfam01197)→ |
| Vpa-70-1 | NZ_AWMS01000089.1 | - | 14428 | 14378 | **RNA**→ rpmE (PRK00019)Ribosomal_L31 (pfam01197)→ |
| Vpa-125-1 | NZ_AWKP01000009.1 | + | 52752 | 52802 | **RNA**→ rpmE (PRK00019)Ribosomal_L31 (pfam01197)→ |
| Vpa-78-1 | NZ_AWMK01000010.1 | + | 51675 | 51725 | **RNA**→ rpmE (PRK00019)Ribosomal_L31 (pfam01197)→ |
| Vpa-83-1 | NZ_AWMF01000124.1 | + | 8335 | 8385 | **RNA**→ rpmE (PRK00019)Ribosomal_L31 (pfam01197)→ |
| Vpa-66-1 | NZ_AWMW01000061.1 | + | 23084 | 23134 | **RNA**→ rpmE (PRK00019)Ribosomal_L31 (pfam01197)→ |
| Vpa-124-1 | NZ_AWKQ01000001.1 | + | 47328 | 47378 | **RNA**→ rpmE (PRK00019)Ribosomal_L31 (pfam01197)→ |
| Vpa-127-1 | NZ_AWKN01000016.1 | + | 40862 | 40912 | **RNA**→ rpmE (PRK00019)Ribosomal_L31 (pfam01197)→ |
| Vpa-24-1 | NZ_AVPV01000004.1 | + | 421744 | 421794 | **RNA**→ rpmE (PRK00019)Ribosomal_L31 (pfam01197)→ |
| Vpa-33-1 | NZ_JFFP01000029.1 | + | 224759 | 224809 | **RNA**→ rpmE (PRK00019)Ribosomal_L31 (pfam01197)→ |
| Vpa-45-1 | NZ_ACKB01000038.1 | - | 1783 | 1733 | **RNA**→ rpmE (PRK00019)Ribosomal_L31 (pfam01197)→ rpmE (PRK00019)Ribosomal_L31 (pfam01197)→ |
| Vpa-71-1 | NZ_AWMR01000251.1 | - | 5731 | 5681 | **RNA**→ rpmE (PRK00019)Ribosomal_L31 (pfam01197)→ |
| Vpa-62-1 | NZ_AWNA01000002.1 | - | 5725 | 5675 | **RNA**→ rpmE (PRK00019)Ribosomal_L31 (pfam01197)→ |
| Vpa-69-1 | NZ_AWMT01000122.1 | - | 5724 | 5674 | **RNA**→ rpmE (PRK00019)Ribosomal_L31 (pfam01197)→ |
| Vpa-65-1 | NZ_AWMX01000034.1 | - | 5716 | 5666 | **RNA**→ rpmE (PRK00019)Ribosomal_L31 (pfam01197)→ |
| Vpa-61-1 | NZ_AWNB01000020.1 | - | 5751 | 5701 | **RNA**→ rpmE (PRK00019)Ribosomal_L31 (pfam01197)→ |
| VPa-1-1 | NZ_AXNJ01000046.1 | + | 227758 | 227808 | **RNA**→ rpmE (PRK00019)Ribosomal_L31 (pfam01197)→ |
| VPa-2-1 | NZ_AXNK01000046.1 | - | 264322 | 264272 | **RNA**→ rpmE (PRK00019)Ribosomal_L31 (pfam01197)→ |
| Vpa-60-1 | NZ_AWNC01000013.1 | + | 26584 | 26634 | **RNA**→ rpmE (PRK00019)Ribosomal_L31 (pfam01197)→ |
| Vpa-48-1 | NZ_AVOM01000003.1 | - | 321091 | 321041 | **RNA**→ rpmE (PRK00019)Ribosomal_L31 (pfam01197)→ |
| Vpa-77-1 | NZ_AWML01000131.1 | + | 8341 | 8391 | **RNA**→ rpmE (PRK00019)Ribosomal_L31 (pfam01197)→ |
| Vpa-63-1 | NZ_AWMZ01000037.1 | - | 5720 | 5670 | **RNA**→ rpmE (PRK00019)Ribosomal_L31 (pfam01197)→ |
| Vpa-64-1 | NZ_AWMY01000005.1 | + | 66806 | 66856 | **RNA**→ rpmE (PRK00019)Ribosomal_L31 (pfam01197)→ |
| Vpa-59-1 | NZ_AWND01000118.1 | - | 5744 | 5694 | **RNA**→ rpmE (PRK00019)Ribosomal_L31 (pfam01197)→ |
| Vpa-27-1 | NZ_JNUE01000003.1 | + | 14971 | 15021 | **RNA**→ rpmE (PRK00019)Ribosomal_L31 (pfam01197)→ |
| Vpa-55-1 | NZ_ACFM01000100.1 | - | 2781 | 2731 | **RNA**→ rpmE (PRK00019)Ribosomal_L31 (pfam01197)→ rpmE (PRK00019)Ribosomal_L31 (pfam01197)→ |
| Vpa-84-1 | NZ_AWME01000007.1 | - | 9036 | 8986 | **RNA**→ rpmE (PRK00019)Ribosomal_L31 (pfam01197)→ |
| Vpa-30-1 | NZ_ACFN01000016.1 | - | 2994 | 2944 | **RNA**→ rpmE (PRK00019)Ribosomal_L31 (pfam01197)→ rpmE (PRK00019)Ribosomal_L31 (pfam01197)→ |

| | | | | | | |
|---|---|---|---|---|---|---|
| Vpa-95-1 | NZ_AWLT01000086.1 | - | 2796 | 2746 | **RNA**→ | rpmE (PRK00019)Ribosomal_L31 (pfam01197)→ |
| Vpa-53-1 | NZ_AOCL01000001.1 | - | 15138 | 15088 | **RNA**→ | rpmE (PRK00019)Ribosomal_L31 (pfam01197)→ |
| Vpa-224-1 | NZ_AXNM01000020.1 | - | 145227 | 145177 | **RNA**→ | rpmE (PRK00019)Ribosomal_L31 (pfam01197)→ |
| Vpa-232-1 | NZ_AVON01000009.1 | + | 26500 | 26550 | **RNA**→ | rpmE (PRK00019)Ribosomal_L31 (pfam01197)→ |
| Vpa-132-1 | NZ_AWKI01000110.1 | - | 15116 | 15066 | **RNA**→ | rpmE (PRK00019)Ribosomal_L31 (pfam01197)→ |
| Vpa-123-1 | NZ_AWKR01000136.1 | - | 4460 | 4410 | **RNA**→ | rpmE (PRK00019)Ribosomal_L31 (pfam01197)→ |
| Vpa-118-1 | NZ_AWKW01000003.1 | + | 77006 | 77056 | **RNA**→ | rpmE (PRK00019)Ribosomal_L31 (pfam01197)→ |
| Vpa-113-1 | NZ_AWLB01000029.1 | + | 25233 | 25283 | **RNA**→ | rpmE (PRK00019)Ribosomal_L31 (pfam01197)→ |
| Vpa-144-1 | NZ_AWJW01000077.1 | - | 5722 | 5672 | **RNA**→ | rpmE (PRK00019)Ribosomal_L31 (pfam01197)→ |
| Vpa-131-1 | NZ_AWKJ01000063.1 | + | 13874 | 13924 | **RNA**→ | rpmE (PRK00019)Ribosomal_L31 (pfam01197)→ |
| Vpa-121-1 | NZ_AWKT01000003.1 | + | 95261 | 95311 | **RNA**→ | rpmE (PRK00019)Ribosomal_L31 (pfam01197)→ |
| Vpa-117-1 | NZ_AWKX01000001.1 | - | 5719 | 5669 | **RNA**→ | rpmE (PRK00019)Ribosomal_L31 (pfam01197)→ |
| Vpa-116-1 | NZ_AWKY01000001.1 | - | 5725 | 5675 | **RNA**→ | rpmE (PRK00019)Ribosomal_L31 (pfam01197)→ |
| Vpa-103-1 | NZ_AWLL01000285.1 | - | 3101 | 3051 | **RNA**→ | rpmE (PRK00019)Ribosomal_L31 (pfam01197)→ |
| Vpa-138-1 | NZ_AWKC01000048.1 | + | 24625 | 24675 | **RNA**→ | rpmE (PRK00019)Ribosomal_L31 (pfam01197)→ |
| Vpa-128-1 | NZ_AWKM01000125.1 | + | 8342 | 8392 | **RNA**→ | rpmE (PRK00019)Ribosomal_L31 (pfam01197)→ |
| Vpa-119-1 | NZ_AWKV01000122.1 | - | 5717 | 5667 | **RNA**→ | rpmE (PRK00019)Ribosomal_L31 (pfam01197)→ |
| Vpa-114-1 | NZ_AWLA01000080.1 | + | 14625 | 14675 | **RNA**→ | rpmE (PRK00019)Ribosomal_L31 (pfam01197)→ |
| Vpa-101-1 | NZ_AWLN01000126.1 | + | 8426 | 8476 | **RNA**→ | rpmE (PRK00019)Ribosomal_L31 (pfam01197)→ |
| Vpa-143-1 | NZ_AWJX01000108.1 | + | 10341 | 10391 | **RNA**→ | rpmE (PRK00019)Ribosomal_L31 (pfam01197)→ |
| Vpa-139-1 | NZ_AWKB01000002.1 | + | 88686 | 88736 | **RNA**→ | rpmE (PRK00019)Ribosomal_L31 (pfam01197)→ |
| Vpa-130-1 | NZ_AWKK01000008.1 | + | 45640 | 45690 | **RNA**→ | rpmE (PRK00019)Ribosomal_L31 (pfam01197)→ |
| Vpa-129-1 | NZ_AWKL01000001.1 | + | 92746 | 92796 | **RNA**→ | rpmE (PRK00019)Ribosomal_L31 (pfam01197)→ |
| Vpa-120-1 | NZ_AWKU01000205.1 | + | 16 | 66 | **RNA**→ | rpmE (PRK00019)Ribosomal_L31 (pfam01197)→ |
| Vpa-115-1 | NZ_AWKZ01000276.1 | + | 16 | 66 | **RNA**→ | rpmE (PRK00019)Ribosomal_L31 (pfam01197)→ |
| Vpa-111-1 | NZ_AWLD01000121.1 | + | 10001 | 10051 | **RNA**→ | rpmE (PRK00019)Ribosomal_L31 (pfam01197)→ |
| Vpa-154-1 | NZ_AWJM01000096.1 | - | 2751 | 2701 | **RNA**→ | rpmE (PRK00019)Ribosomal_L31 (pfam01197)→ |
| Vpa-141-1 | NZ_AWJZ01000006.1 | - | 4398 | 4348 | **RNA**→ | rpmE (PRK00019)Ribosomal_L31 (pfam01197)→ |
| Vpa-155-1 | NZ_AWJL01000004.1 | + | 42147 | 42197 | **RNA**→ | rpmE (PRK00019)Ribosomal_L31 (pfam01197)→ |
| Vpa-151-1 | NZ_AWJP01000158.1 | - | 5710 | 5660 | **RNA**→ | rpmE (PRK00019)Ribosomal_L31 (pfam01197)→ |
| Vpa-137-1 | NZ_AWKD01000009.1 | + | 41289 | 41339 | **RNA**→ | rpmE (PRK00019)Ribosomal_L31 (pfam01197)→ |
| Vpa-153-1 | NZ_AWJN01000089.1 | - | 5718 | 5668 | **RNA**→ | rpmE (PRK00019)Ribosomal_L31 (pfam01197)→ |
| Vpa-98-1 | NZ_AWLQ01000252.1 | + | 22 | 72 | **RNA**→ | rpmE (PRK00019)Ribosomal_L31 (pfam01197)→ |
| Vpa-152-1 | NZ_AWJO01000079.1 | + | 13233 | 13283 | **RNA**→ | rpmE (PRK00019)Ribosomal_L31 (pfam01197)→ |
| Vpa-212-1 | NZ_AWHG01000024.1 | + | 37943 | 37993 | **RNA**→ | rpmE (PRK00019)Ribosomal_L31 (pfam01197)→ |
| Vpa-178-1 | NZ_AWIO01000008.1 | - | 5716 | 5666 | **RNA**→ | rpmE (PRK00019)Ribosomal_L31 (pfam01197)→ |
| Vpa-177-1 | NZ_AWIP01000091.1 | - | 2793 | 2743 | **RNA**→ | rpmE (PRK00019)Ribosomal_L31 (pfam01197)→ |
| Vpa-173-1 | NZ_AWIT01000001.1 | - | 5739 | 5689 | **RNA**→ | rpmE (PRK00019)Ribosomal_L31 (pfam01197)→ |
| Vpa-198-1 | NZ_AWHU01000008.1 | + | 37741 | 37791 | **RNA**→ | rpmE (PRK00019)Ribosomal_L31 (pfam01197)→ |
| Vpa-180-1 | NZ_AWIM01000008.1 | - | 5714 | 5664 | **RNA**→ | rpmE (PRK00019)Ribosomal_L31 (pfam01197)→ |
| Vpa-175-1 | NZ_AWIR01000001.1 | + | 88686 | 88736 | **RNA**→ | rpmE (PRK00019)Ribosomal_L31 (pfam01197)→ |
| Vpa-196-1 | NZ_AWHW01000005.1 | + | 37727 | 37777 | **RNA**→ | rpmE (PRK00019)Ribosomal_L31 (pfam01197)→ |
| Vpa-176-1 | NZ_AWIQ01000002.1 | + | 95269 | 95319 | **RNA**→ | rpmE (PRK00019)Ribosomal_L31 (pfam01197)→ |
| Vpa-171-1 | NZ_AWIV01000001.1 | - | 5726 | 5676 | **RNA**→ | rpmE (PRK00019)Ribosomal_L31 (pfam01197)→ |
| Vpa-164-1 | NZ_AWJC01000010.1 | - | 5713 | 5663 | **RNA**→ | rpmE (PRK00019)Ribosomal_L31 (pfam01197)→ |
| Vpa-167-1 | NZ_AWIZ01000002.1 | - | 5741 | 5691 | **RNA**→ | rpmE (PRK00019)Ribosomal_L31 (pfam01197)→ |
| Vpa-163-1 | NZ_AWJD01000006.1 | - | 5709 | 5659 | **RNA**→ | rpmE (PRK00019)Ribosomal_L31 (pfam01197)→ |
| Vpa-162-1 | NZ_AWJE01000001.1 | + | 50993 | 51043 | **RNA**→ | rpmE (PRK00019)Ribosomal_L31 (pfam01197)→ |
| Vpa-56-1 | NC_004603.1 | + | 266709 | 266759 | **RNA**→ | rpmE (PRK00019)Ribosomal_L31 (pfam01197)→ |
| Vpa-18-1 | NZ_KI912682.1 | - | 314852 | 314802 | **RNA**→ | rpmE (PRK00019)Ribosomal_L31 (pfam01197)→ |
| Vpa-44-1 | NZ_KI929149.1 | - | 315229 | 315179 | **RNA**→ | rpmE (PRK00019)Ribosomal_L31 (pfam01197)→ |
| Vpa-54-1 | NZ_KI929565.1 | + | 25058 | 25108 | **RNA**→ | rpmE (PRK00019)Ribosomal_L31 (pfam01197)→ |

| | | | | | |
|---|---|---|---|---|---|
| Vpa-31-1 | NZ_KI911857.1 | - | 314830 | 314780 | **RNA**→ rpmE (PRK00019)Ribosomal_L31 (pfam01197)→ |
| Vpa-43-1 | NZ_KI929393.1 | - | 315348 | 315298 | **RNA**→ rpmE (PRK00019)Ribosomal_L31 (pfam01197)→ |
| Vpa-36-1 | NZ_KI965925.1 | - | 314934 | 314884 | **RNA**→ rpmE (PRK00019)Ribosomal_L31 (pfam01197)→ |
| Vpa-37-1 | NZ_KI929290.1 | + | 231685 | 231735 | **RNA**→ rpmE (PRK00019)Ribosomal_L31 (pfam01197)→ |
| Vpa-233-1 | NZ_KK020603.1 | + | 185410 | 185460 | **RNA**→ rpmE (PRK00019)Ribosomal_L31 (pfam01197)→ |
| Vpa-222-1 | NZ_KK019419.1 | + | 227930 | 227980 | **RNA**→ rpmE (PRK00019)Ribosomal_L31 (pfam01197)→ |
| Vpa-231-1 | NZ_KK081667.1 | + | 228093 | 228143 | **RNA**→ rpmE (PRK00019)Ribosomal_L31 (pfam01197)→ |
| Vpa-20-1 | NZ_KI965795.1 | - | 315927 | 315877 | **RNA**→ rpmE (PRK00019)Ribosomal_L31 (pfam01197)→ |
| Vpa-29-1 | NZ_KK073793.1 | - | 297930 | 297880 | **RNA**→ rpmE (PRK00019)Ribosomal_L31 (pfam01197)→ |
| Vpa-32-1 | NC_019955.1 | + | 286013 | 286063 | **RNA**→ rpmE (PRK00019)Ribosomal_L31 (pfam01197)→ |
| Vpa-229-1 | NZ_AXNL01000094.1 | + | 4296 | 4346 | **RNA**→ rpmE (PRK00019)Ribosomal_L31 (pfam01197)→ |
| Vpa-46-1 | NZ_JALL01000115.1 | - | 98380 | 98330 | **RNA**→ rpmE (PRK00019)Ribosomal_L31 (pfam01197)→ |
| Vpa-157-1 | NZ_AWJJ01000001.1 | - | 5723 | 5673 | **RNA**→ rpmE (PRK00019)Ribosomal_L31 (pfam01197)→ |
| Vro-1-1 | NZ_BBLC01000048.1 | + | 223563 | 223613 | **RNA**→ rpmE (PRK00019)Ribosomal_L31 (pfam01197)→ |
| Vpa-79-1 | NZ_AWMJ01000046.1 | + | 14638 | 14688 | **RNA**→ rpmE (PRK00019)Ribosomal_L31 (pfam01197)→ |
| Vpa-80-1 | NZ_AWMI01000001.1 | - | 4354 | 4304 | **RNA**→ rpmE (PRK00019)Ribosomal_L31 (pfam01197)→ |
| Vpa-207-1 | NZ_AWHL01000001.1 | - | 14454 | 14404 | **RNA**→ rpmE (PRK00019)Ribosomal_L31 (pfam01197)→ |
| Vpa-219-1 | NZ_BAVJ01000011.1 | + | 245390 | 245440 | **RNA**→ rpmE (PRK00019)Ribosomal_L31 (pfam01197)→ |
| Val-2-2 | NZ_JXIN01000014.1 | - | 106189 | 106139 | **RNA**→ rpmE (PRK00019)Ribosomal_L31 (pfam01197)→ |
| Val-2-3 | NZ_JXIO01000014.1 | - | 106189 | 106139 | **RNA**→ rpmE (PRK00019)Ribosomal_L31 (pfam01197)→ |
| Val-2-4 | NZ_JZWD01000003.1 | - | 319570 | 319520 | **RNA**→ rpmE (PRK00019)Ribosomal_L31 (pfam01197)→ |
| Val-2-5 | NZ_LCSG01000007.1 | + | 32246 | 32296 | **RNA**→ rpmE (PRK00019)Ribosomal_L31 (pfam01197)→ |
| Val-2-6 | NZ_LCUM01000005.1 | + | 16468 | 16518 | **RNA**→ rpmE (PRK00019)Ribosomal_L31 (pfam01197)→ |
| Vdi-1-1 | NZ_CCKK01000023.1 | + | 38258 | 38308 | **RNA**→ rpmE (PRK00019)Ribosomal_L31 (pfam01197)→ |
| Val-4-1 | NZ_ACZB01000006.1 | - | 131835 | 131785 | **RNA**→ rpmE (PRK00019)Ribosomal_L31 (pfam01197)→ rpmE (PRK00019)Ribosomal_L31 (pfam01197)→ |
| Val-5-1 | NZ_AMPD01000006.1 | - | 120699 | 120649 | **RNA**→ rpmE (PRK00019)Ribosomal_L31 (pfam01197)→ |
| Vsp-1-1 | NZ_ANBX01000002.1 | + | 10986 | 11036 | **RNA**→ rpmE (PRK00019)Ribosomal_L31 (pfam01197)→ |
| Vsp-24-1 | NZ_JPIL01000006.1 | + | 222387 | 222437 | **RNA**→ rpmE (PRK00019)Ribosomal_L31 (pfam01197)→ |
| Val-2-7 | NZ_LFWH01000029.1 | + | 40818 | 40868 | **RNA**→ rpmE (PRK00019)Ribosomal_L31 (pfam01197)→ |
| Vpa-199-1 | NZ_AWHT01000012.1 | + | 37150 | 37200 | **RNA**→ rpmE (PRK00019)Ribosomal_L31 (pfam01197)→ |
| Val-2-8 | NZ_JOKY01000004.1 | - | 117745 | 117695 | **RNA**→ rpmE (PRK00019)Ribosomal_L31 (pfam01197)→ |
| Vsp-8-1 | NC_013456.1 | - | 130415 | 130365 | **RNA**→ rpmE (PRK00019)Ribosomal_L31 (pfam01197)→ |
| Val-6-1 | NC_022349.1 | + | 2643213 | 2643263 | **RNA**→ rpmE (PRK00019)Ribosomal_L31 (pfam01197)→ |
| Val-3-1 | NZ_CH902591.1 | + | 483698 | 483748 | **RNA**→ rpmE (PRK00019)Ribosomal_L31 (pfam01197)→ |
| Vha-9-1 | NZ_BAOD01000009.1 | + | 112541 | 112591 | **RNA**→ rpmE (PRK00019)Ribosomal_L31 (pfam01197)→ |
| Vha-9-2 | NZ_AMCT01000001.1 | + | 10582 | 10632 | **RNA**→ rpmE (PRK00019)Ribosomal_L31 (pfam01197)→ |
| Vha-10-1 | NZ_CAUN01000001.1 | + | 221925 | 221975 | **RNA**→ rpmE (PRK00019)Ribosomal_L31 (pfam01197)→ |
| Vow-5-1 | NZ_JPRD01000041.1 | + | 10157 | 10207 | **RNA**→ rpmE (PRK00019)Ribosomal_L31 (pfam01197)→ |
| Vha-7-1 | NZ_AYKI01000011.1 | - | 258359 | 258309 | **RNA**→ rpmE (PRK00019)Ribosomal_L31 (pfam01197)→ |
| Vha-6-1 | NZ_AHHQ01000061.1 | - | 135341 | 135291 | **RNA**→ rpmE (PRK00019)Ribosomal_L31 (pfam01197)→ |
| Vsp-9-1 | NZ_JH976042.1 | - | 105560 | 105510 | **RNA**→ rpmE (PRK00019)Ribosomal_L31 (pfam01197)→ |
| Vha-12-1 | NZ_JH720475.1 | + | 252971 | 253021 | **RNA**→ rpmE (PRK00019)Ribosomal_L31 (pfam01197)→ |
| Vha-4-1 | NZ_AOMR01000019.1 | + | 237184 | 237234 | **RNA**→ rpmE (PRK00019)Ribosomal_L31 (pfam01197)→ |
| Vow-1-1 | NZ_BAOH01000058.1 | + | 10195 | 10245 | **RNA**→ rpmE (PRK00019)Ribosomal_L31 (pfam01197)→ |
| Vha-11-1 | NZ_CAUO01000001.1 | - | 283952 | 283902 | **RNA**→ rpmE (PRK00019)Ribosomal_L31 (pfam01197)→ |
| Vca-3-1 | NC_022269.1 | - | 2006268 | 2006218 | **RNA**→ rpmE (PRK00019)Ribosomal_L31 (pfam01197)→ |
| Vha-5-1 | NC_009783.1 | + | 721255 | 721305 | **RNA**→ rpmE (PRK00019)Ribosomal_L31 (pfam01197)→ |
| Vow-3-1 | NZ_JPRC01000010.1 | + | 104312 | 104362 | **RNA**→ rpmE (PRK00019)Ribosomal_L31 (pfam01197)→ |
| Vow-2-1 | NZ_BBKN01000104.1 | + | 104573 | 104623 | **RNA**→ rpmE (PRK00019)Ribosomal_L31 (pfam01197)→ |
| Vow-2-2 | NZ_BBKO01000049.1 | + | 27652 | 27702 | **RNA**→ rpmE (PRK00019)Ribosomal_L31 (pfam01197)→ |
| Vow-2-3 | NZ_BBPJ01000004.1 | + | 221575 | 221625 | **RNA**→ rpmE (PRK00019)Ribosomal_L31 (pfam01197)→ |
| Vow-2-4 | NZ_BBLB01000056.1 | - | 63133 | 63083 | **RNA**→ rpmE (PRK00019)Ribosomal_L31 (pfam01197)→ |

| | | | | | |
|---|---|---|---|---|---|
| Vow-6-1 | NZ_BAOE01000009.1 | - | 159659 | 159609 | **RNA**→ rpmE (PRK00019)Ribosomal_L31 (pfam01197)→ |
| Vow-4-1 | NZ_BANZ01000007.1 | + | 221354 | 221404 | **RNA**→ rpmE (PRK00019)Ribosomal_L31 (pfam01197)→ |
| Vha-3-1 | NZ_ACZC01000005.1 | + | 75147 | 75197 | **RNA**→ rpmE (PRK00019)Ribosomal_L31 (pfam01197)→ rpmE (PRK00019)Ribosomal_L31 (pfam01197)→ |
| Vsp-10-1 | NZ_JH976131.1 | - | 290388 | 290338 | **RNA**→ rpmE (PRK00019)Ribosomal_L31 (pfam01197)→ |
| env-627 | CEUP01099403.1 | + | 1243 | 1293 | **RNA**→ |
| env-628 | CEUY01075272.1 | + | 1198 | 1248 | **RNA**→ rpmE (PRK00019)Ribosomal_L31 (pfam01197)→ |
| env-629 | CEVA01026273.1 | + | 584 | 634 | **RNA**→ rpmE (PRK00019)Ribosomal_L31 (pfam01197)→ |
| Vca-1-1 | NZ_BBKU01000066.1 | - | 291589 | 291539 | **RNA**→ rpmE (PRK00019)Ribosomal_L31 (pfam01197)→ |
| Vca-1-2 | NZ_BBKX01000044.1 | + | 229368 | 229418 | **RNA**→ rpmE (PRK00019)Ribosomal_L31 (pfam01197)→ |
| Vca-1-3 | NZ_BBKG01000111.1 | - | 228720 | 228670 | **RNA**→ rpmE (PRK00019)Ribosomal_L31 (pfam01197)→ |
| Vca-1-4 | NZ_BBKW01000020.1 | + | 122219 | 122269 | **RNA**→ rpmE (PRK00019)Ribosomal_L31 (pfam01197)→ |
| Vca-1-5 | NZ_BBKV01000034.1 | - | 289838 | 289788 | **RNA**→ rpmE (PRK00019)Ribosomal_L31 (pfam01197)→ |
| Vca-4-1 | NZ_BAOF01000005.1 | - | 213680 | 213630 | **RNA**→ rpmE (PRK00019)Ribosomal_L31 (pfam01197)→ |
| Vca-2-1 | NZ_BANY01000042.1 | - | 156590 | 156540 | **RNA**→ rpmE (PRK00019)Ribosomal_L31 (pfam01197)→ |
| Vca-5-1 | NZ_AGIE01000044.1 | + | 24998 | 25048 | **RNA**→ rpmE (PRK00019)Ribosomal_L31 (pfam01197)→ rpmE (PRK00019)Ribosomal_L31 (pfam01197)→ |
| Vca-1-6 | NZ_JPTJ01000344.1 | - | 60557 | 60507 | **RNA**→ rpmE (PRK00019)Ribosomal_L31 (pfam01197)→ |
| Vca-1-7 | NZ_JPTF01000271.1 | + | 6528 | 6578 | **RNA**→ rpmE (PRK00019)Ribosomal_L31 (pfam01197)→ |
| Vca-1-8 | NZ_JPTG01000526.1 | - | 34186 | 34136 | **RNA**→ rpmE (PRK00019)Ribosomal_L31 (pfam01197)→ |
| Vca-4-2 | NZ_AMDG01000002.1 | + | 21188 | 21238 | **RNA**→ rpmE (PRK00019)Ribosomal_L31 (pfam01197)→ |
| Vha-8-1 | NZ_DS179523.1 | - | 1250 | 1200 | **RNA**→ rpmE (PRK00019)Ribosomal_L31 (pfam01197)→ |
| Vha-2-1 | NZ_DF967934.1 | + | 223234 | 223284 | **RNA**→ rpmE (PRK00019)Ribosomal_L31 (pfam01197)→ |
| Vca-1-9 | NZ_JPTH01000426.1 | + | 2525 | 2575 | **RNA**→ rpmE (PRK00019)Ribosomal_L31 (pfam01197)→ |
| Vca-1-10 | NZ_JSFE01000004.1 | + | 218917 | 218967 | **RNA**→ rpmE (PRK00019)Ribosomal_L31 (pfam01197)→ |
| Vsp-2-1 | NZ_ABGR01000076.1 | + | 1750 | 1800 | **RNA**→ rpmE (PRK00019)Ribosomal_L31 (pfam01197)→ rpmE (PRK00019)Ribosomal_L31 (pfam01197)→ |
| Vsp-44-1 | NZ_AJZF01000019.1 | + | 83395 | 83443 | **RNA**→ rpmE (PRK00019)Ribosomal_L31 (pfam01197)→ |
| Vsp-39-1 | NZ_AJZG01000004.1 | + | 88039 | 88087 | **RNA**→ rpmE (PRK00019)Ribosomal_L31 (pfam01197)→ |
| Vsp-42-1 | NZ_AJZI01000104.1 | + | 130365 | 130413 | **RNA**→ rpmE (PRK00019)Ribosomal_L31 (pfam01197)→ |
| Vsp-37-1 | NZ_AJZJ01000171.1 | - | 55509 | 55461 | **RNA**→ rpmE (PRK00019)Ribosomal_L31 (pfam01197)→ |
| Vsp-36-1 | NZ_AIDS01000064.1 | + | 80876 | 80924 | **RNA**→ rpmE (PRK00019)Ribosomal_L31 (pfam01197)→ |
| Vsp-41-1 | NZ_AJZH01000106.1 | - | 157442 | 157394 | **RNA**→ rpmE (PRK00019)Ribosomal_L31 (pfam01197)→ |
| Vsp-34-1 | NZ_AAMR01000027.1 | - | 57919 | 57871 | **RNA**→ rpmE (PRK00019)Ribosomal_L31 (pfam01197)→ rpmE (PRK00019)Ribosomal_L31 (pfam01197)→ |
| Vsp-33-1 | NZ_AJZK01000123.1 | - | 5581 | 5533 | **RNA**→ rpmE (PRK00019)Ribosomal_L31 (pfam01197)→ |
| Vcy-2-1 | NZ_AIDB01000019.1 | - | 153914 | 153866 | **RNA**→ rpmE (PRK00019)Ribosomal_L31 (pfam01197)→ |
| Vcy-5-1 | NZ_AICZ01000040.1 | + | 82209 | 82257 | **RNA**→ rpmE (PRK00019)Ribosomal_L31 (pfam01197)→ |
| Vcy-15-1 | NZ_AIDQ01000003.1 | - | 64827 | 64779 | **RNA**→ rpmE (PRK00019)Ribosomal_L31 (pfam01197)→ |
| Vsp-45-1 | NZ_AJZE01000045.1 | - | 868 | 820 | **RNA**→ rpmE (PRK00019)Ribosomal_L31 (pfam01197)→ |
| Vcy-13-1 | NZ_AIDN01000109.1 | - | 154821 | 154773 | **RNA**→ rpmE (PRK00019)Ribosomal_L31 (pfam01197)→ |
| Vcy-18-1 | NZ_JH636802.1 | - | 154566 | 154518 | **RNA**→ rpmE (PRK00019)Ribosomal_L31 (pfam01197)→ |
| Vsp-5-1 | NZ_LAUM01000025.1 | - | 7503 | 7455 | **RNA**→ rpmE (PRK00019)Ribosomal_L31 (pfam01197)→ |
| Vcr-1-1 | NZ_CCKB01000054.1 | - | 64767 | 64719 | **RNA**→ rpmE (PRK00019)Ribosomal_L31 (pfam01197)→ |
| Vcr-1-2 | NZ_CCKC01000053.1 | - | 64768 | 64720 | **RNA**→ rpmE (PRK00019)Ribosomal_L31 (pfam01197)→ |
| Vcr-1-3 | NZ_CCJW01000071.1 | - | 64767 | 64719 | **RNA**→ rpmE (PRK00019)Ribosomal_L31 (pfam01197)→ |
| Vcr-1-4 | NZ_CCKE01000137.1 | - | 64767 | 64719 | **RNA**→ rpmE (PRK00019)Ribosomal_L31 (pfam01197)→ |
| Vcr-1-5 | NZ_CCKD01000003.1 | + | 80738 | 80786 | **RNA**→ rpmE (PRK00019)Ribosomal_L31 (pfam01197)→ |
| Vcr-1-6 | NZ_CCJU01000033.1 | - | 64791 | 64743 | **RNA**→ rpmE (PRK00019)Ribosomal_L31 (pfam01197)→ |
| Vcr-2-1 | NZ_AJYY01000105.1 | - | 64875 | 64827 | **RNA**→ rpmE (PRK00019)Ribosomal_L31 (pfam01197)→ |
| Vba-1-1 | NZ_AAZW01000016.1 | + | 72717 | 72765 | **RNA**→ rpmE (PRK00019)Ribosomal_L31 (pfam01197)→ rpmE (PRK00019)Ribosomal_L31 (pfam01197)→ |
| Vsp-21-1 | NZ_CCKL01000023.1 | - | 71200 | 71152 | **RNA**→ rpmE (PRK00019)Ribosomal_L31 (pfam01197)→ |
| Vsp-16-1 | NZ_CCKH01000049.1 | - | 65057 | 65009 | **RNA**→ rpmE (PRK00019)Ribosomal_L31 (pfam01197)→ |
| Vsp-11-1 | NZ_CCJZ01000014.1 | + | 80837 | 80885 | **RNA**→ rpmE (PRK00019)Ribosomal_L31 (pfam01197)→ |
| Vcr-3-1 | NZ_AJYZ01000064.1 | + | 82711 | 82759 | **RNA**→ rpmE (PRK00019)Ribosomal_L31 (pfam01197)→ |
| Vcr-4-1 | NZ_AJZA01000103.1 | + | 8727 | 8775 | **RNA**→ rpmE (PRK00019)Ribosomal_L31 (pfam01197)→ |

| | | | | | |
|---|---|---|---|---|---|
| Vcr-5-1 | NZ_AJZB01000073.1 | - | 64832 | 64784 | **RNA**→ rpmE (PRK00019)Ribosomal_L31 (pfam01197)→ |
| Vcr-6-1 | NZ_AJZC01000134.1 | - | 154849 | 154801 | **RNA**→ rpmE (PRK00019)Ribosomal_L31 (pfam01197)→ |
| Vsp-17-1 | NZ_LK934107.1 | - | 64903 | 64855 | **RNA**→ rpmE (PRK00019)Ribosomal_L31 (pfam01197)→ |
| Vsp-13-1 | NZ_LK934399.1 | - | 64965 | 64917 | **RNA**→ rpmE (PRK00019)Ribosomal_L31 (pfam01197)→ |
| Vsp-12-1 | NZ_LK933700.1 | + | 80899 | 80947 | **RNA**→ rpmE (PRK00019)Ribosomal_L31 (pfam01197)→ |
| Vsp-20-1 | NZ_LK933651.1 | + | 80775 | 80823 | **RNA**→ rpmE (PRK00019)Ribosomal_L31 (pfam01197)→ |
| Vsp-19-1 | NZ_LK933553.1 | + | 81252 | 81300 | **RNA**→ rpmE (PRK00019)Ribosomal_L31 (pfam01197)→ |
| Vsp-18-1 | NZ_LK933855.1 | - | 65082 | 65034 | **RNA**→ rpmE (PRK00019)Ribosomal_L31 (pfam01197)→ |
| Vsp-14-1 | NZ_LK934544.1 | + | 80994 | 81042 | **RNA**→ rpmE (PRK00019)Ribosomal_L31 (pfam01197)→ |
| Vsp-15-1 | NZ_LK934195.1 | - | 52838 | 52790 | **RNA**→ rpmE (PRK00019)Ribosomal_L31 (pfam01197)→ |
| env-630 | CEUP01012127.1 | - | 372 | 324 | **RNA**→ rpmE (PRK00019)Ribosomal_L31 (pfam01197)→ |
| env-631 | CEUY01376693.1 | + | 406 | 454 | **RNA**→ rpmE (PRK00019)Ribosomal_L31 (pfam01197)→ |
| env-632 | CEVA01031311.1 | + | 1334 | 1382 | **RNA**→ rpmE (PRK00019)Ribosomal_L31 (pfam01197)→ |
| Vsp-40-1 | NZ_AFWG01000002.1 | + | 80636 | 80684 | **RNA**→ rpmE (PRK00019)Ribosomal_L31 (pfam01197)→ |
| Vsp-35-1 | NZ_AJZD01000142.1 | - | 7513 | 7465 | **RNA**→ rpmE (PRK00019)Ribosomal_L31 (pfam01197)→ |
| Vka-1-1 | NZ_AJYX01000051.1 | - | 64615 | 64567 | **RNA**→ rpmE (PRK00019)Ribosomal_L31 (pfam01197)→ |
| env-633 | JGI24025J20009_10090133 | - | 649 | 601 | **RNA**→ rpmE (PRK00019)Ribosomal_L31 (pfam01197)→ |
| Vsp-4-1 | NZ_LAUL01000014.1 | - | 65430 | 65382 | **RNA**→ rpmE (PRK00019)Ribosomal_L31 (pfam01197)→ |
| Vcy-11-1 | NZ_AIDM01000034.1 | + | 88163 | 88211 | **RNA**→ rpmE (PRK00019)Ribosomal_L31 (pfam01197)→ |
| Vcy-14-1 | NZ_AIDP01000054.1 | - | 71422 | 71374 | **RNA**→ rpmE (PRK00019)Ribosomal_L31 (pfam01197)→ |
| Vcy-19-1 | NZ_AIDK01000001.1 | - | 102003 | 101955 | **RNA**→ rpmE (PRK00019)Ribosomal_L31 (pfam01197)→ |
| Vcy-4-1 | NZ_AIDD01000066.1 | + | 8925 | 8973 | **RNA**→ rpmE (PRK00019)Ribosomal_L31 (pfam01197)→ |
| Vcy-20-1 | NZ_AIDL01000041.1 | + | 19770 | 19818 | **RNA**→ rpmE (PRK00019)Ribosomal_L31 (pfam01197)→ |
| Vcy-17-1 | NZ_AIDI01000041.1 | + | 52335 | 52383 | **RNA**→ rpmE (PRK00019)Ribosomal_L31 (pfam01197)→ |
| Vcy-8-1 | NZ_AIDG01000061.1 | - | 154606 | 154558 | **RNA**→ rpmE (PRK00019)Ribosomal_L31 (pfam01197)→ |
| Vcy-10-1 | NZ_AIDH01000094.1 | - | 1744 | 1696 | **RNA**→ rpmE (PRK00019)Ribosomal_L31 (pfam01197)→ rpmE (PRK00019)Ribosomal_L31 (pfam01197)→ |
| Vcy-1-1 | NZ_AHTI01000209.1 | + | 27717 | 27765 | **RNA**→ rpmE (PRK00019)Ribosomal_L31 (pfam01197)→ |
| Vcy-3-1 | NZ_AIDC01000227.1 | + | 87986 | 88034 | **RNA**→ rpmE (PRK00019)Ribosomal_L31 (pfam01197)→ |
| Vle-1-1 | NZ_LFQI01000004.1 | + | 80855 | 80903 | **RNA**→ rpmE (PRK00019)Ribosomal_L31 (pfam01197)→ |
| Vcy-7-1 | NZ_JH636741.1 | + | 180960 | 181008 | **RNA**→ rpmE (PRK00019)Ribosomal_L31 (pfam01197)→ |
| Vcy-16-1 | NZ_JH636876.1 | + | 82223 | 82271 | **RNA**→ rpmE (PRK00019)Ribosomal_L31 (pfam01197)→ |
| Vcy-12-1 | NZ_JH636836.1 | + | 88238 | 88286 | **RNA**→ rpmE (PRK00019)Ribosomal_L31 (pfam01197)→ |
| Vcy-9-1 | NZ_JH636656.1 | - | 256575 | 256527 | **RNA**→ rpmE (PRK00019)Ribosomal_L31 (pfam01197)→ |
| Vcy-6-1 | NZ_JH636615.1 | - | 154644 | 154596 | **RNA**→ rpmE (PRK00019)Ribosomal_L31 (pfam01197)→ |
| Vsp-31-1 | NZ_CVNE01000009.1 | + | 88422 | 88470 | **RNA**→ rpmE (PRK00019)Ribosomal_L31 (pfam01197)→ |
| env-634 | AYRE01002890.1 | + | 135 | 183 | **RNA**→ rpmE (PRK00019)Ribosomal_L31 (pfam01197)→ |
| env-635 | AYRF01043169.1 | + | 149 | 197 | **RNA**→ |
| Vsp-38-1 | NZ_AJZL01000194.1 | + | 19737 | 19785 | **RNA**→ rpmE (PRK00019)Ribosomal_L31 (pfam01197)→ |
| Afi-2-1 | NZ_LCYF01000005.1 | + | 152582 | 152635 | **RNA**→ rpmE (PRK00019)Ribosomal_L31 (pfam01197)→ |
| Vfi-1-1 | NC_006840.2 | - | 2555303 | 2555250 | **RNA**→ rpmE (PRK00019)Ribosomal_L31 (pfam01197)→ |
| Vfi-2-1 | NC_011184.1 | - | 2565485 | 2565432 | **RNA**→ rpmE (PRK00019)Ribosomal_L31 (pfam01197)→ |
| Vfi-3-1 | NZ_CM001400.1 | - | 2527750 | 2527697 | **RNA**→ rpmE (PRK00019)Ribosomal_L31 (pfam01197)→ |
| Afi-3-1 | NZ_AJYI01000167.1 | + | 62506 | 62559 | **RNA**→ rpmE (PRK00019)Ribosomal_L31 (pfam01197)→ |
| Vor-1-1 | NZ_AJYV01000135.1 | + | 15084 | 15133 | **RNA**→ rpmE (PRK00019)Ribosomal_L31 (pfam01197)→ Malic_M (smart00919)SfcA (COG0281)→ Malic_M (smart00919)SfcA (COG0281)→ |
| Vor-4-1 | NZ_AJYT01000033.1 | - | 45641 | 45592 | **RNA**→ rpmE (PRK00019)Ribosomal_L31 (pfam01197)→ |
| Vor-5-1 | NZ_AJYU01000056.1 | + | 119483 | 119532 | **RNA**→ rpmE (PRK00019)Ribosomal_L31 (pfam01197)→ Malic_M (smart00919)SfcA (COG0281)→ Malic_M (smart00919)SfcA (COG0281)→ |
| Vge-5-1 | NZ_AJYW01000134.1 | - | 56037 | 55990 | **RNA**→ rpmE (PRK00019)Ribosomal_L31 (pfam01197)→ |
| Vca-7-1 | NZ_AEIU01000066.1 | - | 8794 | 8742 | **RNA**→ rpmE (PRK00019)Ribosomal_L31 (pfam01197)→ rpmE (PRK00019)Ribosomal_L31 (pfam01197)→ |
| Vne-1-1 | NZ_JXXU01000034.1 | + | 24784 | 24834 | **RNA**→ rpmE (PRK00019)Ribosomal_L31 (pfam01197)→ |
| Vco-3-1 | NZ_AEQS01000202.1 | - | 30435 | 30385 | **RNA**→ rpmE (PRK00019)Ribosomal_L31 (pfam01197)→ |
| Vsp-3-1 | NZ_JPQB01000026.1 | - | 151183 | 151133 | **RNA**→ rpmE (PRK00019)Ribosomal_L31 (pfam01197)→ |

| | | | | | |
|---|---|---|---|---|---|
| env-636 | JQGR01000570.1 | - | 2317 | 2267 | **RNA**→ rpmE (PRK00019)Ribosomal_L31 (pfam01197)→ |
| env-637 | ASRM01000514.1 | + | 1915 | 1965 | **RNA**→ rpmE (PRK00019)Ribosomal_L31 (pfam01197)→ |
| env-638 | ASRP01020733.1 | - | 74 | 24 | **RNA**→ hypo→ |
| env-639 | AYRF01005860.1 | - | 150 | 100 | **RNA**→ rpmE (PRK00019)rpmE2 (PRK01678)Ribosomal_L31 (pfam01197)→ |
| Vmi-5-1 | NZ_ADAJ01000008.1 | + | 2127116 | 2127169 | **RNA**→ rpmE (PRK00019)Ribosomal_L31 (pfam01197)→ rpmE (PRK00019)Ribosomal_L31 (pfam01197)→ |
| Vsp-25-1 | NZ_ACZT01000021.1 | + | 188396 | 188449 | **RNA**→ rpmE (PRK00019)Ribosomal_L31 (pfam01197)→ rpmE (PRK00019)Ribosomal_L31 (pfam01197)→ |
| Vch-3-1 | NZ_ACFQ01000008.1 | - | 82042 | 81989 | **RNA**→ rpmE (PRK00019)Ribosomal_L31 (pfam01197)→ rpmE (PRK00019)Ribosomal_L31 (pfam01197)→ |
| Vch-115-1 | NC_017270.1 | + | 2678381 | 2678434 | **RNA**→ rpmE (PRK00019)Ribosomal_L31 (pfam01197)→ RpmE (COG0254)rpmE (PRK00019)→ hypo→ hypo– hypo→ |
| Vmi-1-1 | NZ_AWWL01000147.1 | + | 15291 | 15344 | **RNA**→ rpmE (PRK00019)Ribosomal_L31 (pfam01197)→ |
| Vmi-2-1 | NZ_AOMO01000029.1 | - | 84099 | 84046 | **RNA**→ rpmE (PRK00019)Ribosomal_L31 (pfam01197)→ |
| Vch-7-1 | NZ_AAUT01000067.1 | - | 679 | 626 | **RNA**→ rpmE (PRK00019)Ribosomal_L31 (pfam01197)→ rpmE (PRK00019)Ribosomal_L31 (pfam01197)→ hypo→ hypo→ |
| Vch-196-1 | NZ_JRJX01000170.1 | + | 166028 | 166081 | **RNA**→ rpmE (PRK00019)Ribosomal_L31 (pfam01197)→ |
| Vch-185-1 | NZ_ADAI01000043.1 | - | 82275 | 82222 | **RNA**→ rpmE (PRK00019)Ribosomal_L31 (pfam01197)→ rpmE (PRK00019)Ribosomal_L31 (pfam01197)→ hypo→ hypo→ Malic_M (smart00919)SfcA (COG0281)→ Malic_M (smart00919)SfcA (COG0281)→ |
| Vch-37-1 | NZ_ADAL01000222.1 | - | 11533 | 11480 | **RNA**→ rpmE (PRK00019)Ribosomal_L31 (pfam01197)→ rpmE (PRK00019)Ribosomal_L31 (pfam01197)→ hypo→ hypo→ Malic_M (smart00919)SfcA (COG0281)→ Malic_M (smart00919)SfcA (COG0281)→ |
| Vch-114-1 | NZ_ADAK01000050.1 | + | 83945 | 83998 | **RNA**→ rpmE (PRK00019)Ribosomal_L31 (pfam01197)→ rpmE (PRK00019)Ribosomal_L31 (pfam01197)→ hypo→ hypo→ Malic_M (smart00919)SfcA (COG0281)→ Malic_M (smart00919)SfcA (COG0281)→ |
| Vch-35-1 | NZ_ANNM01000107.1 | + | 15189 | 15242 | **RNA**→ rpmE (PRK00019)Ribosomal_L31 (pfam01197)→ |
| Vch-32-1 | NZ_AMWU01000165.1 | - | 24184 | 24131 | **RNA**→ rpmE (PRK00019)Ribosomal_L31 (pfam01197)→ |
| Vch-33-1 | NZ_AMWV01000181.1 | - | 24184 | 24131 | **RNA**→ rpmE (PRK00019)Ribosomal_L31 (pfam01197)→ |
| Vch-34-1 | NZ_AMWR01000112.1 | - | 24184 | 24131 | **RNA**→ rpmE (PRK00019)Ribosomal_L31 (pfam01197)→ |
| Vch-36-1 | NZ_AMWW01000167.1 | - | 24184 | 24131 | **RNA**→ rpmE (PRK00019)Ribosomal_L31 (pfam01197)→ |
| Vch-30-1 | NZ_AMWS01000172.1 | - | 24184 | 24131 | **RNA**→ rpmE (PRK00019)Ribosomal_L31 (pfam01197)→ |
| Vch-29-1 | NZ_AMWF01000174.1 | - | 24184 | 24131 | **RNA**→ rpmE (PRK00019)Ribosomal_L31 (pfam01197)→ |
| Vch-31-1 | NZ_AMWT01000032.1 | - | 24184 | 24131 | **RNA**→ rpmE (PRK00019)Ribosomal_L31 (pfam01197)→ |
| Vch-184-1 | NZ_AWWF01000005.1 | - | 80007 | 79954 | **RNA**→ rpmE (PRK00019)Ribosomal_L31 (pfam01197)→ |
| Vch-183-1 | NZ_AWWE01000005.1 | - | 80008 | 79955 | **RNA**→ rpmE (PRK00019)Ribosomal_L31 (pfam01197)→ |
| Vch-182-1 | NZ_AWWD01000005.1 | - | 80007 | 79954 | **RNA**→ rpmE (PRK00019)Ribosomal_L31 (pfam01197)→ |
| Vch-54-1 | NZ_AXDR01000012.1 | + | 65426 | 65479 | **RNA**→ rpmE (PRK00019)Ribosomal_L31 (pfam01197)→ RpmE (COG0254)rpmE (PRK00019)→ hypo→ Malic_M (smart00919)SfcA (COG0281)→ |
| Vch-126-1 | NZ_AYNN01000014.1 | + | 166095 | 166148 | **RNA**→ rpmE (PRK00019)Ribosomal_L31 (pfam01197)→ |
| Vch-129-1 | NZ_AYNM01000035.1 | - | 37698 | 37645 | **RNA**→ rpmE (PRK00019)Ribosomal_L31 (pfam01197)→ |
| Vch-130-1 | NZ_AYNL01000117.1 | - | 37688 | 37635 | **RNA**→ rpmE (PRK00019)Ribosomal_L31 (pfam01197)→ |
| Vch-125-1 | NZ_AYOM01000167.1 | + | 140313 | 140366 | **RNA**→ rpmE (PRK00019)Ribosomal_L31 (pfam01197)→ |
| Val-1-1 | NZ_JADA01000005.1 | - | 80513 | 80460 | **RNA**→ rpmE (PRK00019)Ribosomal_L31 (pfam01197)→ |
| Vch-14-1 | NZ_ACIA01000006.1 | - | 83502 | 83449 | **RNA**→ rpmE (PRK00019)Ribosomal_L31 (pfam01197)→ rpmE (PRK00019)Ribosomal_L31 (pfam01197)→ |
| Vch-111-1 | NZ_AFOR01000017.1 | + | 23142 | 23195 | **RNA**→ rpmE (PRK00019)Ribosomal_L31 (pfam01197)→ RpmE (COG0254)rpmE (PRK00019)→ |
| Vch-47-1 | NZ_ALDL01000016.1 | + | 169034 | 169087 | **RNA**→ rpmE (PRK00019)Ribosomal_L31 (pfam01197)→ RpmE (COG0254)rpmE (PRK00019)→ hypo→ Malic_M (smart00919)SfcA (COG0281)→ |
| Vch-108-1 | NZ_ALED01000028.1 | + | 174043 | 174096 | **RNA**→ rpmE (PRK00019)Ribosomal_L31 (pfam01197)→ RpmE (COG0254)rpmE (PRK00019)→ |
| Vch-39-1 | NZ_AKGH01000001.1 | + | 2527089 | 2527142 | **RNA**→ rpmE (PRK00019)Ribosomal_L31 (pfam01197)→ |
| Vch-1-1 | NZ_ACVW01000009.1 | - | 64807 | 64754 | **RNA**→ rpmE (PRK00019)Ribosomal_L31 (pfam01197)→ rpmE (PRK00019)Ribosomal_L31 (pfam01197)→ hypo→ hypo→ Malic_M (smart00919)SfcA (COG0281)→ Malic_M (smart00919)SfcA (COG0281)→ |
| Vch-51-1 | NZ_AGUN01000136.1 | + | 166224 | 166277 | **RNA**→ rpmE (PRK00019)Ribosomal_L31 (pfam01197)→ RpmE (COG0254)rpmE (PRK00019)→ hypo→ Malic_M (smart00919)SfcA (COG0281)→ |
| Vch-65-1 | NZ_ALDQ01000022.1 | + | 171464 | 171517 | **RNA**→ rpmE (PRK00019)Ribosomal_L31 (pfam01197)→ hypo→ Malic_M (smart00919)SfcA (COG0281)→ |
| Vch-63-1 | NZ_ALDP01000017.1 | + | 170302 | 170355 | **RNA**→ rpmE (PRK00019)Ribosomal_L31 (pfam01197)→ RpmE (COG0254)rpmE (PRK00019)→ |
| Vch-16-1 | NZ_ALDA01000016.1 | + | 167150 | 167203 | **RNA**→ rpmE (PRK00019)Ribosomal_L31 (pfam01197)→ RpmE (COG0254)rpmE (PRK00019)→ hypo→ Malic_M (smart00919)SfcA (COG0281)→ |
| Vch-133-1 | NZ_AHFX01000066.1 | + | 166303 | 166356 | **RNA**→ rpmE (PRK00019)Ribosomal_L31 (pfam01197)→ |
| Vch-12-1 | NZ_ACHZ01000015.1 | + | 166071 | 166124 | **RNA**→ rpmE (PRK00019)Ribosomal_L31 (pfam01197)→ rpmE (PRK00019)Ribosomal_L31 (pfam01197)→ hypo→ hypo→ Malic_M (smart00919)SfcA (COG0281)→ Malic_M (smart00919)SfcA (COG0281)→ |

| | | | | | |
|---|---|---|---|---|---|
| Vch-62-1 | NZ_AGUQ01000019.1 | + | 167891 | 167944 | **RNA**→ rpmE (PRK00019)Ribosomal_L31 (pfam01197)→ hypo→ Malic_M (smart00919)SfcA (COG0281)→ |
| Vch-20-1 | NZ_ALDC01000014.1 | + | 167366 | 167419 | **RNA**→ rpmE (PRK00019)Ribosomal_L31 (pfam01197)→ RpmE (COG0254)rpmE (PRK00019)→ hypo→ Malic_M (smart00919)SfcA (COG0281)→ |
| Vch-22-1 | NZ_ALDE01000023.1 | + | 167278 | 167331 | **RNA**→ rpmE (PRK00019)Ribosomal_L31 (pfam01197)→ hypo→ Malic_M (smart00919)SfcA (COG0281)→ |
| Vch-23-1 | NZ_ALDF01000013.1 | + | 167122 | 167175 | **RNA**→ rpmE (PRK00019)Ribosomal_L31 (pfam01197)→ RpmE (COG0254)rpmE (PRK00019)→ hypo→ Malic_M (smart00919)SfcA (COG0281)→ |
| Vch-70-1 | NZ_AFOM01000019.1 | + | 168579 | 168632 | **RNA**→ rpmE (PRK00019)Ribosomal_L31 (pfam01197)→ hypo→ Malic_M (smart00919)SfcA (COG0281)→ |
| Vch-110-1 | NZ_AFOQ01000010.1 | + | 171737 | 171790 | **RNA**→ rpmE (PRK00019)Ribosomal_L31 (pfam01197)→ RpmE (COG0254)rpmE (PRK00019)→ hypo→ hypo→ Malic_M (smart00919)SfcA (COG0281)→ |
| Vch-27-1 | NZ_ALDJ01000015.1 | + | 169778 | 169831 | **RNA**→ rpmE (PRK00019)Ribosomal_L31 (pfam01197)→ RpmE (COG0254)rpmE (PRK00019)→ hypo→ Malic_M (smart00919)SfcA (COG0281)→ |
| Vch-86-1 | NZ_AJRU01000023.1 | + | 172852 | 172905 | **RNA**→ rpmE (PRK00019)Ribosomal_L31 (pfam01197)→ RpmE (COG0254)rpmE (PRK00019)→ hypo→ Malic_M (smart00919)SfcA (COG0281)→ |
| Vch-50-1 | NZ_AGUM01000012.1 | + | 169488 | 169541 | **RNA**→ rpmE (PRK00019)Ribosomal_L31 (pfam01197)→ RpmE (COG0254)rpmE (PRK00019)→ hypo→ Malic_M (smart00919)SfcA (COG0281)→ |
| Vch-106-1 | NZ_ALEC01000008.1 | + | 171021 | 171074 | **RNA**→ rpmE (PRK00019)Ribosomal_L31 (pfam01197)→ rpmE (PRK00019)Ribosomal_L31 (pfam01197)→ hypo→ Malic_M (smart00919)SfcA (COG0281)→ |
| Vch-25-1 | NZ_ALDH01000025.1 | + | 168608 | 168661 | **RNA**→ rpmE (PRK00019)Ribosomal_L31 (pfam01197)→ RpmE (COG0254)rpmE (PRK00019)→ hypo→ Malic_M (smart00919)SfcA (COG0281)→ |
| Vch-103-1 | NZ_AFOO01000020.1 | + | 167567 | 167620 | **RNA**→ rpmE (PRK00019)Ribosomal_L31 (pfam01197)→ RpmE (COG0254)rpmE (PRK00019)→ hypo→ Malic_M (smart00919)SfcA (COG0281)→ |
| Vch-85-1 | NZ_AGUS01000011.1 | + | 167472 | 167525 | **RNA**→ rpmE (PRK00019)Ribosomal_L31 (pfam01197)→ RpmE (COG0254)rpmE (PRK00019)→ hypo→ Malic_M (smart00919)SfcA (COG0281)→ |
| Vch-6-1 | NZ_JNEW01000014.1 | + | 97535 | 97588 | **RNA**→ rpmE (PRK00019)Ribosomal_L31 (pfam01197)→ |
| Vch-17-1 | NZ_AJRL01000032.1 | + | 166987 | 167040 | **RNA**→ rpmE (PRK00019)Ribosomal_L31 (pfam01197)→ RpmE (COG0254)rpmE (PRK00019)→ hypo→ Malic_M (smart00919)SfcA (COG0281)→ |
| Vch-11-1 | NZ_JHXR01000011.1 | - | 80293 | 80240 | **RNA**→ rpmE (PRK00019)Ribosomal_L31 (pfam01197)→ |
| Vch-42-1 | NZ_AGUI01000126.1 | + | 166151 | 166204 | **RNA**→ rpmE (PRK00019)Ribosomal_L31 (pfam01197)→ hypo→ Malic_M (smart00919)SfcA (COG0281)→ |
| Vch-134-1 | NZ_AHFY01000032.1 | - | 1904 | 1851 | **RNA**→ rpmE (PRK00019)Ribosomal_L31 (pfam01197)→ |
| Vch-195-1 | NZ_ATEV02000004.1 | + | 164199 | 164252 | **RNA**→ rpmE (PRK00019)Ribosomal_L31 (pfam01197)→ |
| Vch-138-1 | NZ_AHGC01000053.1 | + | 4310 | 4363 | **RNA**→ rpmE (PRK00019)Ribosomal_L31 (pfam01197)→ |
| Vch-152-1 | NZ_AHGP01000035.1 | - | 1904 | 1851 | **RNA**→ rpmE (PRK00019)Ribosomal_L31 (pfam01197)→ |
| Vch-192-1 | NZ_AMBS01000018.1 | - | 80028 | 79975 | **RNA**→ rpmE (PRK00019)Ribosomal_L31 (pfam01197)→ |
| Vch-197-1 | NZ_ACHV01000001.1 | + | 1368480 | 1368533 | **RNA**→ rpmE (PRK00019)Ribosomal_L31 (pfam01197)→ rpmE (PRK00019)Ribosomal_L31 (pfam01197)→ |
| Vch-194-1 | NZ_AMBT01000021.1 | + | 166169 | 166222 | **RNA**→ rpmE (PRK00019)Ribosomal_L31 (pfam01197)→ |
| Vch-158-1 | NZ_AHGQ01000033.1 | + | 166151 | 166204 | **RNA**→ rpmE (PRK00019)Ribosomal_L31 (pfam01197)→ |
| Vch-157-1 | NZ_AFSV01000006.1 | + | 72274 | 72327 | **RNA**→ rpmE (PRK00019)Ribosomal_L31 (pfam01197)→ |
| Vch-148-1 | NZ_AHGK01000006.1 | + | 166158 | 166211 | **RNA**→ rpmE (PRK00019)Ribosomal_L31 (pfam01197)→ |
| Vch-181-1 | NZ_AIJR01000043.1 | + | 141625 | 141678 | **RNA**→ rpmE (PRK00019)Ribosomal_L31 (pfam01197)→ RpmE (COG0254)rpmE (PRK00019)→ hypo→ Malic_M (smart00919)SfcA (COG0281)→ |
| Vch-137-1 | NZ_AHGB01000061.1 | - | 80003 | 79950 | **RNA**→ rpmE (PRK00019)Ribosomal_L31 (pfam01197)→ |
| Vch-141-1 | NZ_AELI01000040.1 | - | 1908 | 1855 | **RNA**→ rpmE (PRK00019)Ribosomal_L31 (pfam01197)→ |
| Vch-188-1 | NZ_ACHW01000024.1 | + | 173500 | 173553 | **RNA**→ rpmE (PRK00019)Ribosomal_L31 (pfam01197)→ rpmE (PRK00019)Ribosomal_L31 (pfam01197)→ |
| Vch-150-1 | NZ_AHGM01000040.1 | - | 1911 | 1858 | **RNA**→ rpmE (PRK00019)Ribosomal_L31 (pfam01197)→ |
| Vch-149-1 | NZ_AHGL01000070.1 | + | 166303 | 166356 | **RNA**→ rpmE (PRK00019)Ribosomal_L31 (pfam01197)→ |
| Vch-142-1 | NZ_AHGD01000007.1 | - | 1904 | 1851 | **RNA**→ rpmE (PRK00019)Ribosomal_L31 (pfam01197)→ |
| Vch-147-1 | NZ_AHGJ01000008.1 | - | 1906 | 1853 | **RNA**→ rpmE (PRK00019)Ribosomal_L31 (pfam01197)→ |
| Vch-193-1 | NZ_AMBR01000002.1 | + | 166162 | 166215 | **RNA**→ rpmE (PRK00019)Ribosomal_L31 (pfam01197)→ |
| Vch-146-1 | NZ_AHGH01000007.1 | - | 1908 | 1855 | **RNA**→ rpmE (PRK00019)Ribosomal_L31 (pfam01197)→ |
| Vch-151-1 | NZ_AHGN01000009.1 | - | 1903 | 1850 | **RNA**→ rpmE (PRK00019)Ribosomal_L31 (pfam01197)→ |
| Vch-144-1 | NZ_AHGF01000007.1 | - | 1906 | 1853 | **RNA**→ rpmE (PRK00019)Ribosomal_L31 (pfam01197)→ |
| Vch-136-1 | NZ_AHGA01000061.1 | + | 166303 | 166356 | **RNA**→ rpmE (PRK00019)Ribosomal_L31 (pfam01197)→ |
| Vch-145-1 | NZ_AHGG01000060.1 | + | 88775 | 88828 | **RNA**→ rpmE (PRK00019)Ribosomal_L31 (pfam01197)→ |
| Vch-135-1 | NZ_AHFZ01000034.1 | - | 1919 | 1866 | **RNA**→ rpmE (PRK00019)Ribosomal_L31 (pfam01197)→ |
| Vch-143-1 | NZ_AHGE01000007.1 | + | 166319 | 166372 | **RNA**→ rpmE (PRK00019)Ribosomal_L31 (pfam01197)→ |

| | | | | | |
|---|---|---|---|---|---|
| Vch-140-1 | NZ_AELJ01000042.1 | - | 1903 | 1850 | **RNA**→ rpmE (PRK00019)Ribosomal_L31 (pfam01197)→ |
| Vmi-6-1 | NZ_ACYV01000004.1 | - | 168966 | 168913 | **RNA**→ rpmE (PRK00019)Ribosomal_L31 (pfam01197)→ rpmE (PRK00019)Ribosomal_L31 (pfam01197)→ hypo→ Malic_M (smart00919)SfcA (COG0281)→ Malic_M (smart00919)SfcA (COG0281)→ |
| Vmi-3-1 | NZ_ADAF01000001.1 | - | 86644 | 86591 | **RNA**→ rpmE (PRK00019)Ribosomal_L31 (pfam01197)→ rpmE (PRK00019)Ribosomal_L31 (pfam01197)→ hypo→ Malic_M (smart00919)SfcA (COG0281)→ Malic_M (smart00919)SfcA (COG0281)→ |
| Vmi-7-1 | NZ_ACYU01000088.1 | - | 21850 | 21797 | **RNA**→ rpmE (PRK00019)Ribosomal_L31 (pfam01197)→ rpmE (PRK00019)Ribosomal_L31 (pfam01197)→ hypo→ Malic_M (smart00919)SfcA (COG0281)→ Malic_M (smart00919)SfcA (COG0281)→ |
| Vch-128-1 | NC_002505.1 | + | 2847750 | 2847803 | **RNA**→ rpmE (PRK00019)Ribosomal_L31 (pfam01197)→ hypo→ Malic_M (smart00919)SfcA (COG0281)→ |
| Vch-121-1 | NZ_DS178186.1 | + | 88174 | 88227 | **RNA**→ rpmE (PRK00019)Ribosomal_L31 (pfam01197)→ hypo→ Malic_M (smart00919)SfcA (COG0281)→ |
| Vch-10-1 | NZ_DS265229.1 | + | 74064 | 74117 | **RNA**→ rpmE (PRK00019)Ribosomal_L31 (pfam01197)→ rpmE (PRK00019)Ribosomal_L31 (pfam01197)→ |
| Vch-179-1 | NC_012582.1 | + | 2933043 | 2933096 | **RNA**→ rpmE (PRK00019)Ribosomal_L31 (pfam01197)→ hypo→ Malic_M (smart00919)SfcA (COG0281)→ |
| Vch-101-1 | NZ_AGVC02000009.1 | + | 168641 | 168694 | **RNA**→ rpmE (PRK00019)Ribosomal_L31 (pfam01197)→ RpmE (COG0254)rpmE (PRK00019)→ hypo→ Malic_M (smart00919)SfcA (COG0281)→ |
| Vch-8-1 | NZ_AMVL01000016.1 | + | 167875 | 167928 | **RNA**→ rpmE (PRK00019)Ribosomal_L31 (pfam01197)→ RpmE (COG0254)rpmE (PRK00019)→ hypo→ Malic_M (smart00919)SfcA (COG0281)→ |
| Vch-187-1 | NZ_ACHX01000008.1 | + | 168810 | 168863 | **RNA**→ rpmE (PRK00019)Ribosomal_L31 (pfam01197)→ rpmE (PRK00019)Ribosomal_L31 (pfam01197)→ |
| Vch-189-1 | NZ_ACHY01000014.1 | + | 164082 | 164135 | **RNA**→ rpmE (PRK00019)Ribosomal_L31 (pfam01197)→ rpmE (PRK00019)Ribosomal_L31 (pfam01197)→ |
| Vch-186-1 | NZ_GG774560.1 | + | 167617 | 167670 | **RNA**→ rpmE (PRK00019)Ribosomal_L31 (pfam01197)→ hypo→ |
| Vch-117-1 | NZ_GG774581.1 | - | 79982 | 79929 | **RNA**→ hypo→ rpmE (PRK00019)Ribosomal_L31 (pfam01197)→ |
| Vch-119-1 | NZ_DS990139.1 | + | 166078 | 166131 | **RNA**→ rpmE (PRK00019)Ribosomal_L31 (pfam01197)→ rpmE (PRK00019)Ribosomal_L31 (pfam01197)→ hypo→ hypo→ Malic_M (smart00919)SfcA (COG0281)→ Malic_M (smart00919)SfcA (COG0281)→ |
| Vmi-4-1 | NZ_GL989611.1 | - | 83946 | 83893 | **RNA**→ rpmE (PRK00019)Ribosomal_L31 (pfam01197)→ hypo→ Malic_M (smart00919)SfcA (COG0281)→ |
| Vch-56-1 | NZ_GL989441.1 | + | 140393 | 140446 | **RNA**→ rpmE (PRK00019)Ribosomal_L31 (pfam01197)→ RpmE (COG0254)rpmE (PRK00019)→ hypo→ Malic_M (smart00919)SfcA (COG0281)→ |
| Vch-68-1 | NZ_GL988459.1 | + | 96678 | 96731 | **RNA**→ rpmE (PRK00019)Ribosomal_L31 (pfam01197)→ RpmE (COG0254)rpmE (PRK00019)→ hypo→ Malic_M (smart00919)SfcA (COG0281)→ |
| Vch-104-1 | NZ_GL988785.1 | + | 94619 | 94672 | **RNA**→ rpmE (PRK00019)Ribosomal_L31 (pfam01197)→ RpmE (COG0254)rpmE (PRK00019)→ |
| Vch-13-1 | NZ_GL989317.1 | + | 88093 | 88146 | **RNA**→ rpmE (PRK00019)Ribosomal_L31 (pfam01197)→ RpmE (COG0254)rpmE (PRK00019)→ hypo→ Malic_M (smart00919)SfcA (COG0281)→ |
| Vch-112-1 | NZ_GL988901.1 | + | 166145 | 166198 | **RNA**→ rpmE (PRK00019)Ribosomal_L31 (pfam01197)→ RpmE (COG0254)rpmE (PRK00019)→ hypo→ Malic_M (smart00919)SfcA (COG0281)→ |
| Vch-94-1 | NZ_GL988622.1 | + | 166218 | 166271 | **RNA**→ rpmE (PRK00019)Ribosomal_L31 (pfam01197)→ RpmE (COG0254)rpmE (PRK00019)→ hypo→ Malic_M (smart00919)SfcA (COG0281)→ |
| Vch-45-1 | NZ_JH377912.1 | + | 166228 | 166281 | **RNA**→ rpmE (PRK00019)Ribosomal_L31 (pfam01197)→ RpmE (COG0254)rpmE (PRK00019)→ hypo→ Malic_M (smart00919)SfcA (COG0281)→ |
| Vch-52-1 | NZ_JH377642.1 | + | 166224 | 166277 | **RNA**→ rpmE (PRK00019)Ribosomal_L31 (pfam01197)→ RpmE (COG0254)rpmE (PRK00019)→ hypo→ Malic_M (smart00919)SfcA (COG0281)→ |
| Vch-49-1 | NZ_JH377763.1 | + | 166391 | 166444 | **RNA**→ rpmE (PRK00019)Ribosomal_L31 (pfam01197)→ RpmE (COG0254)rpmE (PRK00019)→ hypo→ Malic_M (smart00919)SfcA (COG0281)→ |
| Vch-48-1 | NZ_JH377867.1 | + | 166149 | 166202 | **RNA**→ rpmE (PRK00019)Ribosomal_L31 (pfam01197)→ RpmE (COG0254)rpmE (PRK00019)→ hypo→ Malic_M (smart00919)SfcA (COG0281)→ |
| Vch-53-1 | NZ_JH377415.1 | + | 166220 | 166273 | **RNA**→ rpmE (PRK00019)Ribosomal_L31 (pfam01197)→ RpmE (COG0254)rpmE (PRK00019)→ hypo→ Malic_M (smart00919)SfcA (COG0281)→ |
| Vch-69-1 | NZ_JH377599.1 | + | 96678 | 96731 | **RNA**→ rpmE (PRK00019)Ribosomal_L31 (pfam01197)→ RpmE (COG0254)rpmE (PRK00019)→ hypo→ Malic_M (smart00919)SfcA (COG0281)→ |
| Vch-15-1 | NZ_JH724675.1 | + | 166163 | 166216 | **RNA**→ rpmE (PRK00019)Ribosomal_L31 (pfam01197)→ RpmE (COG0254)rpmE (PRK00019)→ hypo→ Malic_M (smart00919)SfcA (COG0281)→ |
| Vch-79-1 | NZ_JH724533.1 | + | 166212 | 166265 | **RNA**→ rpmE (PRK00019)Ribosomal_L31 (pfam01197)→ RpmE (COG0254)rpmE (PRK00019)→ hypo→ Malic_M (smart00919)SfcA (COG0281)→ |
| Vch-67-1 | NZ_JH724981.1 | + | 166199 | 166252 | **RNA**→ rpmE (PRK00019)Ribosomal_L31 (pfam01197)→ RpmE (COG0254)rpmE (PRK00019)→ hypo→ Malic_M (smart00919)SfcA (COG0281)→ |
| Vch-61-1 | NZ_JH724365.1 | + | 166237 | 166290 | **RNA**→ rpmE (PRK00019)Ribosomal_L31 (pfam01197)→ RpmE (COG0254)rpmE (PRK00019)→ hypo→ Malic_M (smart00919)SfcA (COG0281)→ |
| Vch-81-1 | NZ_JH724637.1 | + | 166235 | 166288 | **RNA**→ rpmE (PRK00019)Ribosomal_L31 (pfam01197)→ RpmE (COG0254)rpmE (PRK00019)→ hypo→ Malic_M (smart00919)SfcA (COG0281)→ |
| Vch-26-1 | NZ_JH724792.1 | + | 166090 | 166143 | **RNA**→ rpmE (PRK00019)Ribosomal_L31 (pfam01197)→ RpmE (COG0254)rpmE (PRK00019)→ hypo→ Malic_M (smart00919)SfcA (COG0281)→ |

| | | | | | |
|---|---|---|---|---|---|
| Vch-57-1 | NZ_JH942113.1 | + | 166129 | 166182 | **RNA**→ rpmE (PRK00019)Ribosomal_L31 (pfam01197)→ RpmE (COG0254)rpmE (PRK00019)→ hypo→ Malic_M (smart00919)SfcA (COG0281)→ |
| Vch-102-1 | NZ_JH941267.1 | + | 166210 | 166263 | **RNA**→ rpmE (PRK00019)Ribosomal_L31 (pfam01197)→ RpmE (COG0254)rpmE (PRK00019)→ hypo→ Malic_M (smart00919)SfcA (COG0281)→ |
| Vch-64-1 | NZ_JH973832.1 | + | 15199 | 15252 | **RNA**→ rpmE (PRK00019)Ribosomal_L31 (pfam01197)→ RpmE (COG0254)rpmE (PRK00019)→ |
| Vch-28-1 | NZ_JH941761.1 | + | 166181 | 166234 | **RNA**→ rpmE (PRK00019)Ribosomal_L31 (pfam01197)→ RpmE (COG0254)rpmE (PRK00019)→ hypo→ Malic_M (smart00919)SfcA (COG0281)→ |
| Vch-21-1 | NZ_JH942554.1 | + | 166156 | 166209 | **RNA**→ rpmE (PRK00019)Ribosomal_L31 (pfam01197)→ RpmE (COG0254)rpmE (PRK00019)→ hypo→ Malic_M (smart00919)SfcA (COG0281)→ |
| Vch-76-1 | NZ_JH973997.1 | + | 64867 | 64920 | **RNA**→ rpmE (PRK00019)Ribosomal_L31 (pfam01197)→ RpmE (COG0254)rpmE (PRK00019)→ hypo→ Malic_M (smart00919)SfcA (COG0281)→ |
| Vch-75-1 | NZ_JH942782.1 | + | 64906 | 64959 | **RNA**→ rpmE (PRK00019)Ribosomal_L31 (pfam01197)→ RpmE (COG0254)rpmE (PRK00019)→ hypo→ Malic_M (smart00919)SfcA (COG0281)→ |
| Vch-73-1 | NZ_JH941360.1 | + | 64867 | 64920 | **RNA**→ rpmE (PRK00019)Ribosomal_L31 (pfam01197)→ RpmE (COG0254)rpmE (PRK00019)→ hypo→ Malic_M (smart00919)SfcA (COG0281)→ |
| Vch-105-1 | NZ_JH941443.1 | + | 164229 | 164282 | **RNA**→ rpmE (PRK00019)Ribosomal_L31 (pfam01197)→ |
| Vch-80-1 | NZ_JH941836.1 | + | 64865 | 64918 | **RNA**→ rpmE (PRK00019)Ribosomal_L31 (pfam01197)→ RpmE (COG0254)rpmE (PRK00019)→ hypo→ Malic_M (smart00919)SfcA (COG0281)→ |
| Vch-59-1 | NZ_JH941105.1 | + | 166233 | 166286 | **RNA**→ rpmE (PRK00019)Ribosomal_L31 (pfam01197)→ RpmE (COG0254)rpmE (PRK00019)→ hypo→ Malic_M (smart00919)SfcA (COG0281)→ |
| Vch-41-1 | NZ_JH973434.1 | + | 64873 | 64926 | **RNA**→ rpmE (PRK00019)Ribosomal_L31 (pfam01197)→ RpmE (COG0254)rpmE (PRK00019)→ hypo→ Malic_M (smart00919)SfcA (COG0281)→ |
| Vch-74-1 | NZ_JH942250.1 | + | 64867 | 64920 | **RNA**→ rpmE (PRK00019)Ribosomal_L31 (pfam01197)→ RpmE (COG0254)rpmE (PRK00019)→ hypo→ Malic_M (smart00919)SfcA (COG0281)→ |
| Vch-78-1 | NZ_JH942415.1 | + | 64859 | 64912 | **RNA**→ rpmE (PRK00019)Ribosomal_L31 (pfam01197)→ RpmE (COG0254)rpmE (PRK00019)→ hypo→ Malic_M (smart00919)SfcA (COG0281)→ |
| Vch-19-1 | NZ_JH942327.1 | + | 97406 | 97459 | **RNA**→ rpmE (PRK00019)Ribosomal_L31 (pfam01197)→ RpmE (COG0254)rpmE (PRK00019)→ hypo→ Malic_M (smart00919)SfcA (COG0281)→ |
| Vch-77-1 | NZ_JH967247.1 | + | 64887 | 64940 | **RNA**→ rpmE (PRK00019)Ribosomal_L31 (pfam01197)→ RpmE (COG0254)rpmE (PRK00019)→ hypo→ Malic_M (smart00919)SfcA (COG0281)→ |
| Vch-93-1 | NZ_JH974133.1 | + | 166143 | 166196 | **RNA**→ rpmE (PRK00019)Ribosomal_L31 (pfam01197)→ hypo→ Malic_M (smart00919)SfcA (COG0281)→ |
| Vch-18-1 | NZ_JH966256.1 | + | 131489 | 131542 | **RNA**→ rpmE (PRK00019)Ribosomal_L31 (pfam01197)→ RpmE (COG0254)rpmE (PRK00019)→ |
| Vch-71-1 | NZ_JH941925.1 | + | 64869 | 64922 | **RNA**→ rpmE (PRK00019)Ribosomal_L31 (pfam01197)→ RpmE (COG0254)rpmE (PRK00019)→ hypo→ Malic_M (smart00919)SfcA (COG0281)→ |
| Vch-24-1 | NZ_JH942693.1 | + | 166218 | 166271 | **RNA**→ rpmE (PRK00019)Ribosomal_L31 (pfam01197)→ RpmE (COG0254)rpmE (PRK00019)→ hypo→ Malic_M (smart00919)SfcA (COG0281)→ |
| Vch-83-1 | NZ_JH973563.1 | + | 64871 | 64924 | **RNA**→ rpmE (PRK00019)Ribosomal_L31 (pfam01197)→ RpmE (COG0254)rpmE (PRK00019)→ hypo→ Malic_M (smart00919)SfcA (COG0281)→ |
| Vch-72-1 | NZ_JH967136.1 | + | 26466 | 26519 | **RNA**→ rpmE (PRK00019)Ribosomal_L31 (pfam01197)→ RpmE (COG0254)rpmE (PRK00019)→ hypo→ Malic_M (smart00919)SfcA (COG0281)→ |
| Vch-82-1 | NZ_JH967405.1 | + | 64871 | 64924 | **RNA**→ rpmE (PRK00019)Ribosomal_L31 (pfam01197)→ RpmE (COG0254)rpmE (PRK00019)→ hypo→ Malic_M (smart00919)SfcA (COG0281)→ |
| Vch-55-1 | NZ_JH973724.1 | + | 166137 | 166190 | **RNA**→ rpmE (PRK00019)Ribosomal_L31 (pfam01197)→ RpmE (COG0254)rpmE (PRK00019)→ hypo→ Malic_M (smart00919)SfcA (COG0281)→ |
| Vch-88-1 | NZ_JH976196.1 | + | 166222 | 166275 | **RNA**→ rpmE (PRK00019)Ribosomal_L31 (pfam01197)→ RpmE (COG0254)rpmE (PRK00019)→ hypo→ Malic_M (smart00919)SfcA (COG0281)→ |
| Vch-60-1 | NZ_JH966759.1 | + | 15217 | 15270 | **RNA**→ rpmE (PRK00019)Ribosomal_L31 (pfam01197)→ RpmE (COG0254)rpmE (PRK00019)→ |
| Vch-107-1 | NZ_JH967832.1 | + | 39583 | 39636 | **RNA**→ rpmE (PRK00019)Ribosomal_L31 (pfam01197)→ RpmE (COG0254)rpmE (PRK00019)→ hypo→ Malic_M (smart00919)SfcA (COG0281)→ |
| Vch-109-1 | NZ_JH967937.1 | + | 39583 | 39636 | **RNA**→ rpmE (PRK00019)Ribosomal_L31 (pfam01197)→ RpmE (COG0254)rpmE (PRK00019)→ hypo→ hypo– Malic_M (smart00919)SfcA (COG0281)→ |
| Vch-87-1 | NZ_JH967583.1 | + | 166149 | 166202 | **RNA**→ rpmE (PRK00019)Ribosomal_L31 (pfam01197)→ hypo→ Malic_M (smart00919)SfcA (COG0281)→ |
| Vch-44-1 | NZ_JH974086.1 | + | 166208 | 166261 | **RNA**→ rpmE (PRK00019)Ribosomal_L31 (pfam01197)→ RpmE (COG0254)rpmE (PRK00019)→ hypo→ Malic_M (smart00919)SfcA (COG0281)→ |
| Vch-84-1 | NZ_JH967537.1 | + | 64883 | 64936 | **RNA**→ rpmE (PRK00019)Ribosomal_L31 (pfam01197)→ hypo→ Malic_M (smart00919)SfcA (COG0281)→ |
| Vch-97-1 | NZ_JH967742.1 | + | 166214 | 166267 | **RNA**→ rpmE (PRK00019)Ribosomal_L31 (pfam01197)→ RpmE (COG0254)rpmE (PRK00019)→ hypo→ Malic_M (smart00919)SfcA (COG0281)→ |

| | | | | | |
|---|---|---|---|---|---|
| Vch-91-1 | NZ_KB454392.1 | + | 166145 | 166198 | **RNA**→ rpmE (PRK00019)Ribosomal_L31 (pfam01197)→ RpmE (COG0254)rpmE (PRK00019)→ hypo→ Malic_M (smart00919)SfcA (COG0281)→ |
| Vch-100-1 | NZ_KB453749.1 | + | 166261 | 166314 | **RNA**→ rpmE (PRK00019)Ribosomal_L31 (pfam01197)→ RpmE (COG0254)rpmE (PRK00019)→ hypo→ Malic_M (smart00919)SfcA (COG0281)→ |
| Vch-89-1 | NZ_KB454200.1 | + | 166220 | 166273 | **RNA**→ rpmE (PRK00019)Ribosomal_L31 (pfam01197)→ RpmE (COG0254)rpmE (PRK00019)→ hypo→ Malic_M (smart00919)SfcA (COG0281)→ |
| Vch-96-1 | NZ_KB453977.1 | + | 96717 | 96770 | **RNA**→ rpmE (PRK00019)Ribosomal_L31 (pfam01197)→ RpmE (COG0254)rpmE (PRK00019)→ hypo→ Malic_M (smart00919)SfcA (COG0281)→ |
| Vch-90-1 | NZ_KB454302.1 | + | 166149 | 166202 | **RNA**→ rpmE (PRK00019)Ribosomal_L31 (pfam01197)→ RpmE (COG0254)rpmE (PRK00019)→ hypo→ Malic_M (smart00919)SfcA (COG0281)→ |
| Vch-95-1 | NZ_KB454023.1 | + | 166216 | 166269 | **RNA**→ rpmE (PRK00019)Ribosomal_L31 (pfam01197)→ RpmE (COG0254)rpmE (PRK00019)→ hypo→ Malic_M (smart00919)SfcA (COG0281)→ |
| Vch-92-1 | NZ_KB454115.1 | + | 166145 | 166198 | **RNA**→ rpmE (PRK00019)Ribosomal_L31 (pfam01197)→ RpmE (COG0254)rpmE (PRK00019)→ hypo→ Malic_M (smart00919)SfcA (COG0281)→ |
| Vch-98-1 | NZ_KB453880.1 | + | 64873 | 64926 | **RNA**→ rpmE (PRK00019)Ribosomal_L31 (pfam01197)→ RpmE (COG0254)rpmE (PRK00019)→ hypo→ Malic_M (smart00919)SfcA (COG0281)→ |
| Vch-153-1 | NZ_KB661509.1 | + | 164143 | 164196 | **RNA**→ rpmE (PRK00019)Ribosomal_L31 (pfam01197)→ RpmE (COG0254)rpmE (PRK00019)→ |
| Vch-171-1 | NZ_KB662560.1 | + | 166228 | 166281 | **RNA**→ rpmE (PRK00019)Ribosomal_L31 (pfam01197)→ RpmE (COG0254)rpmE (PRK00019)→ hypo→ Malic_M (smart00919)SfcA (COG0281)→ |
| Vch-38-1 | NZ_KI530559.1 | + | 166138 | 166191 | **RNA**→ rpmE (PRK00019)Ribosomal_L31 (pfam01197)→ |
| Vch-177-1 | NZ_KK037977.1 | + | 166271 | 166324 | **RNA**→ rpmE (PRK00019)Ribosomal_L31 (pfam01197)→ |
| Vch-163-1 | NZ_KB663382.1 | + | 166153 | 166206 | **RNA**→ rpmE (PRK00019)Ribosomal_L31 (pfam01197)→ hypo→ Malic_M (smart00919)SfcA (COG0281)→ |
| Vch-160-1 | NZ_KB661766.1 | + | 166232 | 166285 | **RNA**→ rpmE (PRK00019)Ribosomal_L31 (pfam01197)→ RpmE (COG0254)rpmE (PRK00019)→ hypo→ Malic_M (smart00919)SfcA (COG0281)→ |
| Vch-165-1 | NZ_KB662943.1 | + | 166233 | 166286 | **RNA**→ rpmE (PRK00019)Ribosomal_L31 (pfam01197)→ RpmE (COG0254)rpmE (PRK00019)→ hypo→ Malic_M (smart00919)SfcA (COG0281)→ |
| Vch-155-1 | NZ_KB662662.1 | + | 166230 | 166283 | **RNA**→ rpmE (PRK00019)Ribosomal_L31 (pfam01197)→ RpmE (COG0254)rpmE (PRK00019)→ hypo→ Malic_M (smart00919)SfcA (COG0281)→ |
| Vch-173-1 | NZ_KB661868.1 | + | 284370 | 284423 | **RNA**→ rpmE (PRK00019)Ribosomal_L31 (pfam01197)→ RpmE (COG0254)rpmE (PRK00019)→ hypo→ Malic_M (smart00919)SfcA (COG0281)→ |
| Vch-162-1 | NZ_KB661631.1 | + | 166230 | 166283 | **RNA**→ rpmE (PRK00019)Ribosomal_L31 (pfam01197)→ RpmE (COG0254)rpmE (PRK00019)→ hypo→ Malic_M (smart00919)SfcA (COG0281)→ |
| Vch-167-1 | NZ_KB662883.1 | + | 166226 | 166279 | **RNA**→ rpmE (PRK00019)Ribosomal_L31 (pfam01197)→ RpmE (COG0254)rpmE (PRK00019)→ hypo→ Malic_M (smart00919)SfcA (COG0281)→ |
| Vch-161-1 | NZ_KB663436.1 | + | 166222 | 166275 | **RNA**→ rpmE (PRK00019)Ribosomal_L31 (pfam01197)→ RpmE (COG0254)rpmE (PRK00019)→ hypo→ Malic_M (smart00919)SfcA (COG0281)→ |
| Vch-164-1 | NZ_KB663305.1 | + | 166229 | 166282 | **RNA**→ rpmE (PRK00019)Ribosomal_L31 (pfam01197)→ RpmE (COG0254)rpmE (PRK00019)→ hypo→ Malic_M (smart00919)SfcA (COG0281)→ |
| Vch-156-1 | NZ_KB663101.1 | + | 166157 | 166210 | **RNA**→ rpmE (PRK00019)Ribosomal_L31 (pfam01197)→ RpmE (COG0254)rpmE (PRK00019)→ hypo→ Malic_M (smart00919)SfcA (COG0281)→ |
| Vch-172-1 | NZ_KB662607.1 | + | 166153 | 166206 | **RNA**→ rpmE (PRK00019)Ribosomal_L31 (pfam01197)→ RpmE (COG0254)rpmE (PRK00019)→ hypo→ Malic_M (smart00919)SfcA (COG0281)→ |
| Vch-132-1 | NZ_KB661281.1 | + | 68216 | 68269 | **RNA**→ rpmE (PRK00019)Ribosomal_L31 (pfam01197)→ |
| Vch-176-1 | NZ_KB662149.1 | + | 166222 | 166275 | **RNA**→ rpmE (PRK00019)Ribosomal_L31 (pfam01197)→ RpmE (COG0254)rpmE (PRK00019)→ hypo→ Malic_M (smart00919)SfcA (COG0281)→ |
| Vch-131-1 | NZ_KB661126.1 | + | 166230 | 166283 | **RNA**→ rpmE (PRK00019)Ribosomal_L31 (pfam01197)→ RpmE (COG0254)rpmE (PRK00019)→ hypo→ Malic_M (smart00919)SfcA (COG0281)→ |
| Vch-178-1 | NZ_KB661943.1 | + | 258269 | 258322 | **RNA**→ rpmE (PRK00019)Ribosomal_L31 (pfam01197)→ hypo→ Malic_M (smart00919)SfcA (COG0281)→ |
| Vch-168-1 | NZ_KB662761.1 | + | 176840 | 176893 | **RNA**→ rpmE (PRK00019)Ribosomal_L31 (pfam01197)→ RpmE (COG0254)rpmE (PRK00019)→ hypo→ Malic_M (smart00919)SfcA (COG0281)→ |
| Vch-174-1 | NZ_KB662328.1 | + | 166153 | 166206 | **RNA**→ rpmE (PRK00019)Ribosomal_L31 (pfam01197)→ RpmE (COG0254)rpmE (PRK00019)→ hypo→ Malic_M (smart00919)SfcA (COG0281)→ |
| Vch-166-1 | NZ_KB663195.1 | + | 166153 | 166206 | **RNA**→ rpmE (PRK00019)Ribosomal_L31 (pfam01197)→ hypo→ Malic_M (smart00919)SfcA (COG0281)→ |
| Vch-169-1 | NZ_KB662391.1 | + | 166157 | 166210 | **RNA**→ rpmE (PRK00019)Ribosomal_L31 (pfam01197)→ RpmE (COG0254)rpmE (PRK00019)→ hypo→ Malic_M (smart00919)SfcA (COG0281)→ |
| Vch-175-1 | NZ_KB661618.1 | + | 166149 | 166202 | **RNA**→ rpmE (PRK00019)Ribosomal_L31 (pfam01197)→ RpmE (COG0254)rpmE (PRK00019)→ hypo→ Malic_M (smart00919)SfcA (COG0281)→ |

| | | | | | |
|---|---|---|---|---|---|
| Vch-159-1 | NZ_KB663522.1 | + | 166151 | 166204 | **RNA**→ rpmE (PRK00019)Ribosomal_L31 (pfam01197)→ RpmE (COG0254)rpmE (PRK00019)→ hypo→ Malic_M (smart00919)SfcA (COG0281)→ |
| Vch-154-1 | NZ_KB662046.1 | + | 97478 | 97531 | **RNA**→ rpmE (PRK00019)Ribosomal_L31 (pfam01197)→ RpmE (COG0254)rpmE (PRK00019)→ hypo→ Malic_M (smart00919)SfcA (COG0281)→ |
| Vch-116-1 | NC_012578.1 | + | 2784875 | 2784928 | **RNA**→ rpmE (PRK00019)Ribosomal_L31 (pfam01197)→ hypo→ Malic_M (smart00919)SfcA (COG0281)→ |
| Vch-179-2 | NC_009457.1 | + | 2401623 | 2401676 | **RNA**→ rpmE (PRK00019)Ribosomal_L31 (pfam01197)→ hypo→ Malic_M (smart00919)SfcA (COG0281)→ |
| Vch-118-1 | NC_012668.1 | - | 746911 | 746858 | **RNA**→ rpmE (PRK00019)Ribosomal_L31 (pfam01197)→ |
| Vch-40-1 | NZ_GL989090.1 | + | 64879 | 64932 | **RNA**→ rpmE (PRK00019)Ribosomal_L31 (pfam01197)→ RpmE (COG0254)rpmE (PRK00019)→ hypo→ Malic_M (smart00919)SfcA (COG0281)→ |
| Vch-139-1 | NC_016445.1 | + | 2317908 | 2317961 | **RNA**→ rpmE (PRK00019)Ribosomal_L31 (pfam01197)→ hypo→ Malic_M (smart00919)SfcA (COG0281)→ |
| Vch-190-1 | NZ_KQ257374.1 | - | 83622 | 83569 | **RNA**→ rpmE (PRK00019)Ribosomal_L31 (pfam01197)→ |
| Vch-123-1 | NZ_LGNW01000019.1 | - | 80238 | 80185 | **RNA**→ rpmE (PRK00019)Ribosomal_L31 (pfam01197)→ |
| Vch-123-2 | NZ_LGNN01000069.1 | + | 166375 | 166428 | **RNA**→ rpmE (PRK00019)Ribosomal_L31 (pfam01197)→ |
| Vch-123-3 | NZ_LGNQ01000044.1 | + | 166376 | 166429 | **RNA**→ rpmE (PRK00019)Ribosomal_L31 (pfam01197)→ |
| Vch-123-4 | NZ_LGOB01000007.1 | + | 166374 | 166427 | **RNA**→ rpmE (PRK00019)Ribosomal_L31 (pfam01197)→ |
| Vch-123-5 | NZ_LGOJ01000066.1 | + | 166366 | 166419 | **RNA**→ rpmE (PRK00019)Ribosomal_L31 (pfam01197)→ |
| Vch-123-6 | NZ_LGOT01000059.1 | + | 166378 | 166431 | **RNA**→ rpmE (PRK00019)Ribosomal_L31 (pfam01197)→ |
| Vch-123-7 | NZ_LGNZ01000002.1 | - | 80248 | 80195 | **RNA**→ rpmE (PRK00019)Ribosomal_L31 (pfam01197)→ |
| Vch-123-8 | NZ_LGON01000072.1 | - | 80254 | 80201 | **RNA**→ rpmE (PRK00019)Ribosomal_L31 (pfam01197)→ |
| Vch-123-9 | NZ_LGOM01000082.1 | + | 166376 | 166429 | **RNA**→ rpmE (PRK00019)Ribosomal_L31 (pfam01197)→ |
| Vch-123-10 | NZ_LGNI01000064.1 | + | 166378 | 166431 | **RNA**→ rpmE (PRK00019)Ribosomal_L31 (pfam01197)→ |
| Vch-123-11 | NZ_LGNL01000021.1 | + | 166350 | 166403 | **RNA**→ rpmE (PRK00019)Ribosomal_L31 (pfam01197)→ |
| Vch-123-12 | NZ_LGNO01000044.1 | + | 166376 | 166429 | **RNA**→ rpmE (PRK00019)Ribosomal_L31 (pfam01197)→ |
| Vch-123-13 | NZ_LGOG01000113.1 | - | 80230 | 80177 | **RNA**→ rpmE (PRK00019)Ribosomal_L31 (pfam01197)→ |
| Vch-123-14 | NZ_LGOH01000029.1 | + | 166376 | 166429 | **RNA**→ rpmE (PRK00019)Ribosomal_L31 (pfam01197)→ |
| Vch-123-15 | NZ_LGOP01000049.1 | - | 80210 | 80157 | **RNA**→ rpmE (PRK00019)Ribosomal_L31 (pfam01197)→ |
| Vch-123-16 | NZ_LGOS01000062.1 | - | 80228 | 80175 | **RNA**→ rpmE (PRK00019)Ribosomal_L31 (pfam01197)→ |
| Vch-123-17 | NZ_LGOE01000049.1 | + | 166376 | 166429 | **RNA**→ rpmE (PRK00019)Ribosomal_L31 (pfam01197)→ |
| Vch-123-18 | NZ_LGOK01000130.1 | + | 166173 | 166226 | **RNA**→ rpmE (PRK00019)Ribosomal_L31 (pfam01197)→ |
| Vch-123-19 | NZ_LGNG01000064.1 | - | 80274 | 80221 | **RNA**→ rpmE (PRK00019)Ribosomal_L31 (pfam01197)→ |
| Vch-123-20 | NZ_LGNK01000118.1 | + | 166342 | 166395 | **RNA**→ rpmE (PRK00019)Ribosomal_L31 (pfam01197)→ |
| Vch-123-21 | NZ_LGNM01000071.1 | + | 166376 | 166429 | **RNA**→ rpmE (PRK00019)Ribosomal_L31 (pfam01197)→ |
| Vch-123-22 | NZ_LGNU01000074.1 | - | 80274 | 80221 | **RNA**→ rpmE (PRK00019)Ribosomal_L31 (pfam01197)→ |
| Vch-123-23 | NZ_LGNS01000082.1 | - | 80264 | 80211 | **RNA**→ rpmE (PRK00019)Ribosomal_L31 (pfam01197)→ |
| Vch-123-24 | NZ_LGNR01000046.1 | - | 37719 | 37666 | **RNA**→ rpmE (PRK00019)Ribosomal_L31 (pfam01197)→ |
| Vch-123-25 | NZ_LGOO01000098.1 | - | 80234 | 80181 | **RNA**→ rpmE (PRK00019)Ribosomal_L31 (pfam01197)→ |
| Vch-123-26 | NZ_LGNV01000021.1 | + | 166350 | 166403 | **RNA**→ rpmE (PRK00019)Ribosomal_L31 (pfam01197)→ |
| Vch-99-1 | NZ_AGVA02000012.1 | + | 168790 | 168843 | **RNA**→ rpmE (PRK00019)Ribosomal_L31 (pfam01197)→ hypo→ Malic_M (smart00919)SfcA (COG0281)→ |
| Vch-180-1 | NZ_ANHS01000115.1 | - | 37694 | 37641 | **RNA**→ rpmE (PRK00019)Ribosomal_L31 (pfam01197)→ |
| Vch-123-27 | NZ_LGNX01000040.1 | - | 80274 | 80221 | **RNA**→ rpmE (PRK00019)Ribosomal_L31 (pfam01197)→ |
| Vch-123-28 | NZ_LGNY01000087.1 | - | 80232 | 80179 | **RNA**→ rpmE (PRK00019)Ribosomal_L31 (pfam01197)→ |
| Vch-123-29 | NZ_LGOL01000001.1 | + | 166376 | 166429 | **RNA**→ rpmE (PRK00019)Ribosomal_L31 (pfam01197)→ |
| Vch-123-30 | NZ_LGOQ01000030.1 | - | 80264 | 80211 | **RNA**→ rpmE (PRK00019)Ribosomal_L31 (pfam01197)→ |
| Vch-123-31 | NZ_LGOR01000066.1 | + | 166378 | 166431 | **RNA**→ rpmE (PRK00019)Ribosomal_L31 (pfam01197)→ |
| Vch-123-32 | NZ_LGNH01000051.1 | - | 80268 | 80215 | **RNA**→ rpmE (PRK00019)Ribosomal_L31 (pfam01197)→ |
| Vch-123-33 | NZ_LGNJ01000075.1 | + | 166368 | 166421 | **RNA**→ rpmE (PRK00019)Ribosomal_L31 (pfam01197)→ |
| Vch-123-34 | NZ_LGNP01000076.1 | + | 166335 | 166388 | **RNA**→ rpmE (PRK00019)Ribosomal_L31 (pfam01197)→ |
| Vch-123-35 | NZ_LGNT01000064.1 | - | 80274 | 80221 | **RNA**→ rpmE (PRK00019)Ribosomal_L31 (pfam01197)→ |
| Vch-123-36 | NZ_LGOA01000065.1 | - | 80266 | 80213 | **RNA**→ rpmE (PRK00019)Ribosomal_L31 (pfam01197)→ |
| Vch-123-37 | NZ_LGOD01000058.1 | + | 166368 | 166421 | **RNA**→ rpmE (PRK00019)Ribosomal_L31 (pfam01197)→ |
| Vch-123-38 | NZ_LGOF01000063.1 | - | 80252 | 80199 | **RNA**→ rpmE (PRK00019)Ribosomal_L31 (pfam01197)→ |
| Vch-123-39 | NZ_LGOI01000018.1 | - | 80230 | 80177 | **RNA**→ rpmE (PRK00019)Ribosomal_L31 (pfam01197)→ |
| Vch-123-40 | NZ_LGOU01000021.1 | - | 80264 | 80211 | **RNA**→ rpmE (PRK00019)Ribosomal_L31 (pfam01197)→ |

| | | | | | |
|---|---|---|---|---|---|
| Vch-122-1 | NZ_DS177959.1 | + | 45142 | 45195 | **RNA**→ rpmE (PRK00019)Ribosomal_L31 (pfam01197)→ hypo→ Malic_M (smart00919)SfcA (COG0281)→ |
| Vch-43-1 | NZ_JH966648.1 | + | 166215 | 166268 | **RNA**→ rpmE (PRK00019)Ribosomal_L31 (pfam01197)→ RpmE (COG0254)rpmE (PRK00019)→ hypo→ Malic_M (smart00919)SfcA (COG0281)→ |
| Vch-120-1 | NZ_AP014524.1 | + | 2708026 | 2708079 | **RNA**→ rpmE (PRK00019)Ribosomal_L31 (pfam01197)→ |
| Vch-2-1 | NZ_CP007634.1 | + | 2318308 | 2318361 | **RNA**→ rpmE (PRK00019)Ribosomal_L31 (pfam01197)→ |
| Vch-124-1 | NZ_CP009042.1 | + | 2932367 | 2932420 | **RNA**→ rpmE (PRK00019)Ribosomal_L31 (pfam01197)→ |
| Vch-2-2 | NZ_CP010811.1 | - | 245075 | 245022 | **RNA**→ rpmE (PRK00019)Ribosomal_L31 (pfam01197)→ |
| Vch-2-3 | NZ_CP010812.1 | - | 232393 | 232340 | **RNA**→ rpmE (PRK00019)Ribosomal_L31 (pfam01197)→ |
| Vch-4-1 | NZ_KQ410628.1 | - | 89751 | 89698 | **RNA**→ rpmE (PRK00019)Ribosomal_L31 (pfam01197)→ |
| Vch-191-1 | NZ_KQ410502.1 | - | 81251 | 81198 | **RNA**→ rpmE (PRK00019)Ribosomal_L31 (pfam01197)→ |
| Vch-9-1 | NZ_KQ410586.1 | + | 511976 | 512029 | **RNA**→ rpmE (PRK00019)Ribosomal_L31 (pfam01197)→ |
| Vch-2-4 | NZ_CP007653.1 | + | 2824210 | 2824263 | **RNA**→ rpmE (PRK00019)Ribosomal_L31 (pfam01197)→ |
| Vch-170-1 | NZ_CM001785.1 | - | 1052063 | 1052010 | **RNA**→ RpmE (COG0254)rpmE (PRK00019)→ rpmE (PRK00019)Ribosomal_L31 (pfam01197)→ hypo→ Malic_M (smart00919)SfcA (COG0281)→ |
| Vch-58-1 | NZ_GL988242.1 | + | 96693 | 96746 | **RNA**→ rpmE (PRK00019)Ribosomal_L31 (pfam01197)→ RpmE (COG0254)rpmE (PRK00019)→ hypo→ Malic_M (smart00919)SfcA (COG0281)→ |
| Vch-113-1 | NC_016944.1 | + | 2894052 | 2894105 | **RNA**→ rpmE (PRK00019)Ribosomal_L31 (pfam01197)→ |
| Vch-5-1 | NZ_AJFN02000034.1 | - | 37684 | 37631 | **RNA**→ rpmE (PRK00019)Ribosomal_L31 (pfam01197)→ |
| Vch-123-41 | NZ_LGOC01000036.1 | + | 166343 | 166396 | **RNA**→ rpmE (PRK00019)Ribosomal_L31 (pfam01197)→ |
| Vch-127-1 | NZ_JDVX01000004.1 | + | 166061 | 166114 | **RNA**→ rpmE (PRK00019)Ribosomal_L31 (pfam01197)→ |
| Vch-46-1 | NZ_AJRO01000013.1 | + | 65177 | 65230 | **RNA**→ rpmE (PRK00019)Ribosomal_L31 (pfam01197)→ RpmE (COG0254)rpmE (PRK00019)→ hypo→ Malic_M (smart00919)SfcA (COG0281)→ |
| Vch-66-1 | NZ_AJSL01000015.1 | + | 15488 | 15541 | **RNA**→ rpmE (PRK00019)Ribosomal_L31 (pfam01197)→ RpmE (COG0254)rpmE (PRK00019)→ |
| Fba-1-1 | NC_014541.1 | - | 3979465 | 3979416 | **RNA**→ rpmE (PRK00019)Ribosomal_L31 (pfam01197)→ |
| Vsi-2-1 | NZ_AEVT01000072.1 | - | 1849 | 1800 | **RNA**→ rpmE (PRK00019)Ribosomal_L31 (pfam01197)→ rpmE (PRK00019)Ribosomal_L31 (pfam01197)→ |
| Val-2-9 | NZ_LFWL01000062.1 | + | 184291 | 184340 | **RNA**→ rpmE (PRK00019)Ribosomal_L31 (pfam01197)→ |
| Vor-7-1 | NZ_ACZV01000005.1 | + | 2348491 | 2348540 | **RNA**→ rpmE (PRK00019)Ribosomal_L31 (pfam01197)→ rpmE (PRK00019)Ribosomal_L31 (pfam01197)→ |
| Vor-8-1 | NZ_AFWH01000025.1 | - | 8828 | 8779 | **RNA**→ rpmE (PRK00019)Ribosomal_L31 (pfam01197)→ |
| Vdi-2-1 | NZ_BBJY01000009.1 | - | 86132 | 86083 | **RNA**→ rpmE (PRK00019)Ribosomal_L31 (pfam01197)→ |
| env-640 | CEGC01006263.1 | - | 190 | 141 | **RNA**→ |
| Vsp-32-1 | NZ_JTLH01000041.1 | + | 156463 | 156512 | **RNA**→ rpmE (PRK00019)Ribosomal_L31 (pfam01197)→ |
| Van-2-1 | NC_015633.1 | - | 544299 | 544250 | **RNA**→ rpmE (PRK00019)Ribosomal_L31 (pfam01197)→ Malic_M (smart00919)SfcA (COG0281)→ Malic_M (smart00919)SfcA (COG0281)→ |
| Lan-1-1 | NC_022223.1 | - | 544215 | 544166 | **RNA**→ rpmE (PRK00019)Ribosomal_L31 (pfam01197)→ Malic_M (smart00919)SfcA (COG0281)→ Malic_M (smart00919)SfcA (COG0281)→ |
| Van-3-1 | NZ_AEZA01000180.1 | + | 175711 | 175760 | **RNA**→ rpmE (PRK00019)Ribosomal_L31 (pfam01197)→ Malic_M (smart00919)SfcA (COG0281)→ Malic_M (smart00919)SfcA (COG0281)→ |
| Van-4-1 | NZ_AEZB01000141.1 | + | 8300 | 8349 | **RNA**→ rpmE (PRK00019)Ribosomal_L31 (pfam01197)→ Malic_M (smart00919)SfcA (COG0281)→ Malic_M (smart00919)SfcA (COG0281)→ |
| Vor-2-1 | NZ_AEZC01000054.1 | - | 18796 | 18747 | **RNA**→ rpmE (PRK00019)Ribosomal_L31 (pfam01197)→ rpmE (PRK00019)Ribosomal_L31 (pfam01197)→ |
| Van-1-1 | NZ_LK021130.1 | + | 2795609 | 2795658 | **RNA**→ rpmE (PRK00019)Ribosomal_L31 (pfam01197)→ |
| Vor-3-1 | NZ_AJYR01000120.1 | - | 13360 | 13311 | **RNA**→ rpmE (PRK00019)Ribosomal_L31 (pfam01197)→ |
| Vor-6-1 | NZ_AJYS01000220.1 | + | 119473 | 119522 | **RNA**→ rpmE (PRK00019)Ribosomal_L31 (pfam01197)→ Malic_M (smart00919)SfcA (COG0281)→ Malic_M (smart00919)SfcA (COG0281)→ |
| Vge-3-1 | NZ_AJYO01000176.1 | + | 32585 | 32634 | **RNA**→ rpmE (PRK00019)Ribosomal_L31 (pfam01197)→ |
| Vge-1-1 | NZ_AJYN01000189.1 | + | 32671 | 32720 | **RNA**→ rpmE (PRK00019)Ribosomal_L31 (pfam01197)→ |
| Vge-4-1 | NZ_AJYQ01000004.1 | - | 155012 | 154963 | **RNA**→ rpmE (PRK00019)Ribosomal_L31 (pfam01197)→ |
| Vge-2-1 | NZ_AJYP01000253.1 | - | 8489 | 8440 | **RNA**→ rpmE (PRK00019)Ribosomal_L31 (pfam01197)→ |
| Vni-9-1 | NZ_CAOC01000112.1 | + | 87830 | 87879 | **RNA**→ rpmE (PRK00019)Ribosomal_L31 (pfam01197)→ |
| Vme-2-1 | NZ_JYJP01000116.1 | - | 87552 | 87503 | **RNA**→ rpmE (PRK00019)Ribosomal_L31 (pfam01197)→ |
| Vsi-1-1 | NZ_JXBJ01000020.1 | + | 64370 | 64420 | **RNA**→ rpmE (PRK00019)Ribosomal_L31 (pfam01197)→ |
| Vpa-8-1 | NZ_DS999347.1 | + | 49201 | 49251 | **RNA**→ rpmE (PRK00019)Ribosomal_L31 (pfam01197)→ rpmE (PRK00019)Ribosomal_L31 (pfam01197)→ |
| Vsi-1-2 | NZ_JRWP01000001.1 | - | 9440 | 9390 | **RNA**→ rpmE (PRK00019)Ribosomal_L31 (pfam01197)→ |
| Vca-6-1 | NZ_JRWR01000026.1 | - | 9377 | 9327 | **RNA**→ rpmE (PRK00019)Ribosomal_L31 (pfam01197)→ |

| | | | | | |
|---|---|---|---|---|---|
| Vsi-1-3 | NZ_JWLV01000015.1 | + | 63119 | 63169 | **RNA**→ rpmE (PRK00019)Ribosomal_L31 (pfam01197)→ |
| Vsi-1-4 | NZ_JWLU01000041.1 | - | 9417 | 9367 | **RNA**→ rpmE (PRK00019)Ribosomal_L31 (pfam01197)→ |
| Vva-1-1 | NZ_JRWM01000039.1 | + | 63092 | 63142 | **RNA**→ rpmE (PRK00019)Ribosomal_L31 (pfam01197)→ |
| Vne-2-1 | NZ_LHPJ01000014.1 | - | 9210 | 9160 | **RNA**→ rpmE (PRK00019)Ribosomal_L31 (pfam01197)→ |
| Vga-1-1 | NZ_JXXV01000019.1 | - | 9082 | 9031 | **RNA**→ rpmE (PRK00019)Ribosomal_L31 (pfam01197)→ |
| Vbr-1-1 | NZ_AEVS01000038.1 | + | 4830 | 4881 | **RNA**→ rpmE (PRK00019)Ribosomal_L31 (pfam01197)→ rpmE (PRK00019)Ribosomal_L31 (pfam01197)→ |
| Vpa-1-1 | NZ_JONH01000021.1 | + | 58049 | 58100 | **RNA**→ rpmE (PRK00019)Ribosomal_L31 (pfam01197)→ |
| Vsp-30-1 | NZ_LBLS01000010.1 | - | 9548 | 9497 | **RNA**→ rpmE (PRK00019)Ribosomal_L31 (pfam01197)→ |
| Vtu-2-2 | NZ_AFWI01000035.1 | - | 9169 | 9118 | **RNA**→ rpmE (PRK00019)Ribosomal_L31 (pfam01197)→ rpmE (PRK00019)Ribosomal_L31 (pfam01197)→ |
| Vtu-2-1 | NZ_CP009354.1 | + | 3076212 | 3076263 | **RNA**→ rpmE (PRK00019)Ribosomal_L31 (pfam01197)→ |
| Vtu-3-1 | NZ_AHHF01000012.1 | - | 9134 | 9083 | **RNA**→ rpmE (PRK00019)Ribosomal_L31 (pfam01197)→ |
| Vtu-1-1 | NZ_JRWQ01000006.1 | - | 11627 | 11576 | **RNA**→ rpmE (PRK00019)Ribosomal_L31 (pfam01197)→ |
| Vhe-1-1 | NZ_LHPI01000017.1 | + | 58937 | 58988 | **RNA**→ rpmE (PRK00019)Ribosomal_L31 (pfam01197)→ |
| env-641 | 2231812703 | - | 323 | 274 | **RNA**→ rpmE (PRK00019)Ribosomal_L31 (pfam01197)→ |
| Vfu-3-1 | NZ_JTLJ01000033.1 | + | 28258 | 28307 | **RNA**→ rpmE (PRK00019)Ribosomal_L31 (pfam01197)→ |
| Vfl-1-1 | NZ_JXXQ01000009.1 | - | 5247 | 5198 | **RNA**→ rpmE (PRK00019)Ribosomal_L31 (pfam01197)→ |
| Vfl-5-1 | NZ_ASXS01000007.1 | - | 93784 | 93735 | **RNA**→ rpmE (PRK00019)Ribosomal_L31 (pfam01197)→ |
| Vfl-3-1 | NZ_JQHW01000024.1 | - | 1142 | 1093 | **RNA**→ rpmE (PRK00019)Ribosomal_L31 (pfam01197)→ |
| Vfl-2-1 | NZ_JQHX01000023.1 | - | 82414 | 82365 | **RNA**→ rpmE (PRK00019)Ribosomal_L31 (pfam01197)→ |
| Vfl-4-1 | NZ_ASXT01000003.1 | + | 178429 | 178478 | **RNA**→ rpmE (PRK00019)Ribosomal_L31 (pfam01197)→ |
| Vfu-2-1 | NC_016602.1 | - | 611055 | 611006 | **RNA**→ rpmE (PRK00019)Ribosomal_L31 (pfam01197)→ |
| Vfu-1-1 | NZ_ACZP01000018.1 | - | 103722 | 103673 | **RNA**→ rpmE (PRK00019)Ribosomal_L31 (pfam01197)→ rpmE (PRK00019)Ribosomal_L31 (pfam01197)→ |
| Vsp-26-1 | NZ_ADBD01000005.1 | + | 179408 | 179459 | **RNA**→ rpmE (PRK00019)Ribosomal_L31 (pfam01197)→ rpmE (PRK00019)Ribosomal_L31 (pfam01197)→ |
| Vre-1-1 | NZ_JTKI01000012.1 | + | 137514 | 137562 | **RNA**→ rpmE (PRK00019)Ribosomal_L31 (pfam01197)→ |
| Vre-1-2 | NZ_JTKH01000005.1 | + | 9661 | 9709 | **RNA**→ rpmE (PRK00019)Ribosomal_L31 (pfam01197)→ |
| Vsc-1-1 | NZ_AFWE01000063.1 | - | 23847 | 23799 | **RNA**→ rpmE (PRK00019)Ribosomal_L31 (pfam01197)→ rpmE (PRK00019)Ribosomal_L31 (pfam01197)→ |
| Vsp-23-1 | NZ_AFWD01000056.1 | + | 110632 | 110680 | **RNA**→ rpmE (PRK00019)Ribosomal_L31 (pfam01197)→ rpmE (PRK00019)Ribosomal_L31 (pfam01197)→ |
| Vic-1-1 | NZ_AFWF01000319.1 | - | 14236 | 14188 | **RNA**→ rpmE (PRK00019)Ribosomal_L31 (pfam01197)→ rpmE (PRK00019)Ribosomal_L31 (pfam01197)→ |
| Awo-1-1 | NZ_LN554846.1 | + | 373049 | 373100 | **RNA**→ rpmE (PRK00019)Ribosomal_L31 (pfam01197)→ |
| Asa-11-1 | NC_011312.1 | + | 2970876 | 2970927 | **RNA**→ rpmE (PRK00019)Ribosomal_L31 (pfam01197)→ |
| Alo-1-1 | NZ_AJYJ01000097.1 | + | 40923 | 40974 | **RNA**→ rpmE (PRK00019)Ribosomal_L31 (pfam01197)→ |
| Alo-2-1 | NZ_ASAH01000010.1 | + | 62670 | 62721 | **RNA**→ rpmE (PRK00019)Ribosomal_L31 (pfam01197)→ |
| env-642 | CEPX01402402.1 | + | 28 | 76 | **RNA**→ rpmE (PRK00019)Ribosomal_L31 (pfam01197)→ |
| Vta-2-1 | NZ_AJZM01000039.1 | + | 89281 | 89329 | **RNA**→ rpmE (PRK00019)Ribosomal_L31 (pfam01197)→ |
| Vsp-22-1 | NZ_CH902613.1 | + | 16168 | 16216 | **RNA**→ rpmE (PRK00019)Ribosomal_L31 (pfam01197)→ |
| env-643 | CEUP01013542.1 | + | 1403 | 1451 | **RNA**→ rpmE (PRK00019)Ribosomal_L31 (pfam01197)→ |
| env-644 | CEUY01108946.1 | - | 745 | 697 | **RNA**→ rpmE (PRK00019)Ribosomal_L31 (pfam01197)→ |
| env-645 | CEVA01131518.1 | - | 743 | 695 | **RNA**→ rpmE (PRK00019)Ribosomal_L31 (pfam01197)→ |
| Vfo-1-1 | NZ_JFFR01000005.1 | + | 71657 | 71705 | **RNA**→ rpmE (PRK00019)Ribosomal_L31 (pfam01197)→ |
| Vta-3-1 | NZ_AJZO01000189.1 | - | 107060 | 107012 | **RNA**→ rpmE (PRK00019)Ribosomal_L31 (pfam01197)→ |
| Vta-1-1 | NZ_AJZN01000372.1 | - | 1937 | 1889 | **RNA**→ rpmE (PRK00019)Ribosomal_L31 (pfam01197)→ |
| Vsp-43-1 | NC_011753.2 | - | 3051148 | 3051100 | **RNA**→ rpmE (PRK00019)Ribosomal_L31 (pfam01197)→ |
| env-646 | CEUP01002391.1 | + | 3518 | 3569 | **RNA**→ rpmE (PRK00019)Ribosomal_L31 (pfam01197)→ |
| Vsh-2-1 | NZ_ABCH01000012.1 | - | 28236 | 28185 | **RNA**→ rpmE (PRK00019)Ribosomal_L31 (pfam01197)→ rpmE (PRK00019)Ribosomal_L31 (pfam01197)→ |
| Vsh-1-1 | NZ_ASHJ01000339.1 | + | 4607 | 4658 | **RNA**→ rpmE (PRK00019)Ribosomal_L31 (pfam01197)→ |
| Vsp-7-1 | NZ_JPJA01000096.1 | - | 67831 | 67780 | **RNA**→ rpmE (PRK00019)Ribosomal_L31 (pfam01197)→ |
| Vma-1-1 | NZ_BBMR01000011.1 | + | 93746 | 93798 | **RNA**→ rpmE (PRK00019)Ribosomal_L31 (pfam01197)→ |
| Vma-1-2 | NZ_BBMT01000020.1 | + | 8867 | 8919 | **RNA**→ rpmE (PRK00019)Ribosomal_L31 (pfam01197)→ |
| Vme-1-1 | NZ_ACZO01000005.1 | - | 90681 | 90631 | **RNA**→ rpmE (PRK00019)Ribosomal_L31 (pfam01197)→ rpmE (PRK00019)Ribosomal_L31 (pfam01197)→ rpmE (PRK00019)Ribosomal_L31 (pfam01197)→ |
| Ymo-1-1 | NZ_CQDS01000022.1 | + | 45976 | 46030 | **RNA**→ rpmE (PRK00019)Ribosomal_L31 (pfam01197)→ |
| Ymo-1-2 | NZ_CPYJ01000009.1 | + | 45908 | 45962 | **RNA**→ rpmE (PRK00019)Ribosomal_L31 (pfam01197)→ |

| | | | | | |
|---|---|---|---|---|---|
| Ymo-1-3 | NZ_CQBM01000006.1 | - | 164691 | 164637 | **RNA**→ rpmE (PRK00019)Ribosomal_L31 (pfam01197)→ |
| Ymo-1-4 | NZ_CTIO01000059.1 | - | 9110 | 9056 | **RNA**→ rpmE (PRK00019)Ribosomal_L31 (pfam01197)→ |
| Ymo-1-5 | NZ_CTKJ01000023.1 | + | 45799 | 45853 | **RNA**→ rpmE (PRK00019)Ribosomal_L31 (pfam01197)→ |
| Ymo-1-6 | NZ_CTRC01000025.1 | + | 46524 | 46578 | **RNA**→ rpmE (PRK00019)Ribosomal_L31 (pfam01197)→ |
| Ymo-2-1 | NZ_AALD02000032.1 | - | 9725 | 9671 | **RNA**→ rpmE (PRK00019)Ribosomal_L31 (pfam01197)→ rpmE (PRK00019)Ribosomal_L31 (pfam01197)→ |
| Ymo-1-7 | NZ_CTEP01000029.1 | + | 23903 | 23957 | **RNA**→ rpmE (PRK00019)Ribosomal_L31 (pfam01197)→ |
| Ybe-1-1 | NZ_CQBU01000019.1 | - | 10230 | 10176 | **RNA**→ rpmE (PRK00019)Ribosomal_L31 (pfam01197)→ |
| Ybe-2-1 | NZ_AALC02000088.1 | + | 2405 | 2459 | **RNA**→ rpmE (PRK00019)Ribosomal_L31 (pfam01197)→ rpmE (PRK00019)Ribosomal_L31 (pfam01197)→ rpmE (PRK00019)Ribosomal_L31 (pfam01197)→ |
| Yma-2-1 | NZ_HE997650.1 | + | 43162 | 43216 | **RNA**→ rpmE (PRK00019)Ribosomal_L31 (pfam01197)→ |
| Yfr-1-1 | NZ_CQEN01000012.1 | - | 112641 | 112587 | **RNA**→ rpmE (PRK00019)Ribosomal_L31 (pfam01197)→ |
| Yfr-1-2 | NZ_CTJA01000014.1 | + | 44765 | 44819 | **RNA**→ rpmE (PRK00019)Ribosomal_L31 (pfam01197)→ |
| Yma-1-1 | NZ_CQBH01000009.1 | + | 43447 | 43501 | **RNA**→ rpmE (PRK00019)Ribosomal_L31 (pfam01197)→ |
| Yfr-1-3 | NZ_CPZP01000012.1 | - | 112412 | 112358 | **RNA**→ rpmE (PRK00019)Ribosomal_L31 (pfam01197)→ |
| Yfr-1-4 | NZ_CTKH01000009.1 | + | 42513 | 42567 | **RNA**→ rpmE (PRK00019)Ribosomal_L31 (pfam01197)→ |
| Yfr-1-5 | NZ_CPYY01000014.1 | - | 33557 | 33503 | **RNA**→ rpmE (PRK00019)Ribosomal_L31 (pfam01197)→ |
| Yfr-1-6 | NZ_CPVN01000012.1 | + | 100741 | 100795 | **RNA**→ rpmE (PRK00019)Ribosomal_L31 (pfam01197)→ |
| Ype-1-1 | NZ_CQAZ01000031.1 | + | 21873 | 21927 | **RNA**→ rpmE (PRK00019)Ribosomal_L31 (pfam01197)→ |
| Ype-1-2 | NZ_CWJL01000021.1 | + | 57627 | 57681 | **RNA**→ rpmE (PRK00019)Ribosomal_L31 (pfam01197)→ |
| Yfr-1-7 | NZ_CQDT01000005.1 | - | 35538 | 35484 | **RNA**→ rpmE (PRK00019)Ribosomal_L31 (pfam01197)→ |
| Yfr-1-8 | NZ_CPWN01000012.1 | - | 35062 | 35008 | **RNA**→ rpmE (PRK00019)Ribosomal_L31 (pfam01197)→ |
| Yfr-1-9 | NZ_CQBR01000013.1 | - | 34560 | 34506 | **RNA**→ rpmE (PRK00019)Ribosomal_L31 (pfam01197)→ |
| Yfr-1-10 | NZ_CQBF01000012.1 | + | 107426 | 107480 | **RNA**→ rpmE (PRK00019)Ribosomal_L31 (pfam01197)→ |
| Yfr-1-11 | NZ_CQEC01000013.1 | + | 114411 | 114465 | **RNA**→ rpmE (PRK00019)Ribosomal_L31 (pfam01197)→ |
| Yfr-1-12 | NZ_CQEP01000014.1 | - | 25751 | 25697 | **RNA**→ rpmE (PRK00019)Ribosomal_L31 (pfam01197)→ |
| Yfr-1-13 | NZ_CQBO01000025.1 | + | 44190 | 44244 | **RNA**→ rpmE (PRK00019)Ribosomal_L31 (pfam01197)→ |
| Yfr-2-2 | NZ_AALE02000018.1 | + | 75774 | 75828 | **RNA**→ rpmE (PRK00019)Ribosomal_L31 (pfam01197)→ rpmE (PRK00019)Ribosomal_L31 (pfam01197)→ |
| Yfr-2-1 | NZ_KN150731.1 | + | 3005078 | 3005132 | **RNA**→ rpmE (PRK00019)Ribosomal_L31 (pfam01197)→ |
| Yfr-1-14 | NZ_CQBG01000011.1 | - | 36689 | 36635 | **RNA**→ rpmE (PRK00019)Ribosomal_L31 (pfam01197)→ |
| Yfr-1-15 | NZ_CQBX01000015.1 | + | 103415 | 103469 | **RNA**→ rpmE (PRK00019)Ribosomal_L31 (pfam01197)→ |
| Yfr-1-16 | NZ_CQDU01000015.1 | + | 94102 | 94156 | **RNA**→ rpmE (PRK00019)Ribosomal_L31 (pfam01197)→ |
| Yfr-1-17 | NZ_CQBW01000011.1 | - | 34252 | 34198 | **RNA**→ rpmE (PRK00019)Ribosomal_L31 (pfam01197)→ |
| Epi-1-1 | NZ_BAYT01000033.1 | + | 32851 | 32903 | **RNA**→ rpmE (PRK00019)Ribosomal_L31 (pfam01197)→ |
| Eta-6-1 | NC_013508.1 | - | 3620131 | 3620079 | **RNA**→ rpmE (PRK00019)Ribosomal_L31 (pfam01197)→ |
| Eta-7-1 | NC_017309.1 | - | 3544275 | 3544223 | **RNA**→ rpmE (PRK00019)Ribosomal_L31 (pfam01197)→ |
| Eta-5-1 | NC_020796.1 | - | 3687618 | 3687566 | **RNA**→ rpmE (PRK00019)Ribosomal_L31 (pfam01197)→ |
| Epi-1-2 | NZ_BAYU01000006.1 | + | 32953 | 33005 | **RNA**→ rpmE (PRK00019)Ribosomal_L31 (pfam01197)→ |
| Ean-3-1 | NZ_CP006664.1 | - | 1629414 | 1629362 | **RNA**→ rpmE (PRK00019)Ribosomal_L31 (pfam01197)→ |
| Esp-56-1 | NZ_CP011364.1 | + | 751576 | 751628 | **RNA**→ rpmE (PRK00019)Ribosomal_L31 (pfam01197)→ |
| Ean-1-1 | NZ_JABY01000008.1 | + | 32857 | 32909 | **RNA**→ rpmE (PRK00019)Ribosomal_L31 (pfam01197)→ |
| Ean-4-1 | NZ_JACP01000017.1 | + | 32830 | 32882 | **RNA**→ rpmE (PRK00019)Ribosomal_L31 (pfam01197)→ |
| Ean-2-1 | NZ_JABS01000011.1 | - | 207016 | 206964 | **RNA**→ rpmE (PRK00019)Ribosomal_L31 (pfam01197)→ |
| Eic-2-1 | NC_012779.2 | + | 3684424 | 3684476 | **RNA**→ rpmE (PRK00019)Ribosomal_L31 (pfam01197)→ |
| Eic-1-1 | NZ_AFJI01000056.1 | - | 19577 | 19525 | **RNA**→ rpmE (PRK00019)Ribosomal_L31 (pfam01197)→ |
| Eta-2-1 | NZ_AFJG01000032.1 | + | 21708 | 21760 | **RNA**→ rpmE (PRK00019)Ribosomal_L31 (pfam01197)→ |
| Eho-5-1 | NZ_BAUC01000010.1 | - | 117607 | 117555 | **RNA**→ rpmE (PRK00019)Ribosomal_L31 (pfam01197)→ |
| Eta-4-1 | NZ_GG739637.1 | - | 235527 | 235475 | **RNA**→ rpmE (PRK00019)Ribosomal_L31 (pfam01197)→ |
| Eta-1-1 | NZ_CP011359.1 | + | 64883 | 64935 | **RNA**→ rpmE (PRK00019)Ribosomal_L31 (pfam01197)→ |
| Eta-3-1 | NZ_AFJJ01000018.1 | + | 109310 | 109362 | **RNA**→ rpmE (PRK00019)Ribosomal_L31 (pfam01197)→ |
| Eta-8-1 | NZ_BANW01000013.1 | - | 156383 | 156331 | **RNA**→ rpmE (PRK00019)Ribosomal_L31 (pfam01197)→ |
| Yal-1-1 | NZ_CQEH01000017.1 | - | 39180 | 39127 | **RNA**→ rpmE (PRK00019)Ribosomal_L31 (pfam01197)→ |
| Yal-1-2 | NZ_CQEJ01000021.1 | - | 38832 | 38779 | **RNA**→ rpmE (PRK00019)Ribosomal_L31 (pfam01197)→ |

| | | | | | |
|---|---|---|---|---|---|
| Yal-1-3 | NZ_CQCP01000020.1 | + | 47144 | 47197 | **RNA**→ rpmE (PRK00019)Ribosomal_L31 (pfam01197)→ |
| Yal-4-1 | NZ_ACCB01000047.1 | - | 1909 | 1856 | **RNA**→ rpmE (PRK00019)Ribosomal_L31 (pfam01197)→ rpmE (PRK00019)Ribosomal_L31 (pfam01197)→ |
| Yen-14-1 | NC_008800.1 | + | 120155 | 120208 | **RNA**→ rpmE (PRK00019)Ribosomal_L31 (pfam01197)→ |
| Yen-1-1 | NZ_CP009846.1 | + | 4065802 | 4065855 | **RNA**→ rpmE (PRK00019)Ribosomal_L31 (pfam01197)→ |
| Yal-1-4 | NZ_CQAX01000010.1 | + | 122176 | 122229 | **RNA**→ rpmE (PRK00019)Ribosomal_L31 (pfam01197)→ |
| Yen-21-1 | NZ_ASHT01000043.1 | - | 36650 | 36597 | **RNA**→ rpmE (PRK00019)Ribosomal_L31 (pfam01197)→ |
| Yen-22-1 | NZ_ASHU01000055.1 | - | 36354 | 36301 | **RNA**→ rpmE (PRK00019)Ribosomal_L31 (pfam01197)→ |
| Yen-19-1 | NZ_ASHV01000059.1 | - | 36237 | 36184 | **RNA**→ rpmE (PRK00019)Ribosomal_L31 (pfam01197)→ |
| Yen-20-1 | NZ_ASHW01000053.1 | - | 36469 | 36416 | **RNA**→ rpmE (PRK00019)Ribosomal_L31 (pfam01197)→ |
| Yen-11-1 | NZ_CAOV01000011.1 | + | 83723 | 83776 | **RNA**→ rpmE (PRK00019)Ribosomal_L31 (pfam01197)→ |
| Yen-17-1 | NZ_AGQO01000036.1 | - | 35756 | 35703 | **RNA**→ rpmE (PRK00019)Ribosomal_L31 (pfam01197)→ |
| Yen-5-1 | NZ_CACU01000005.1 | + | 132412 | 132465 | **RNA**→ rpmE (PRK00019)Ribosomal_L31 (pfam01197)→ |
| Yen-15-1 | NZ_AKKR01000116.1 | - | 27907 | 27854 | **RNA**→ rpmE (PRK00019)Ribosomal_L31 (pfam01197)→ |
| Yen-7-1 | NZ_CACW01000016.1 | - | 5952 | 5899 | **RNA**→ rpmE (PRK00019)Ribosomal_L31 (pfam01197)→ |
| Yen-6-1 | NZ_CACV01000011.1 | - | 200350 | 200297 | **RNA**→ rpmE (PRK00019)Ribosomal_L31 (pfam01197)→ |
| Yen-16-1 | NC_015224.1 | + | 131163 | 131216 | **RNA**→ rpmE (PRK00019)Ribosomal_L31 (pfam01197)→ |
| Yen-18-1 | NC_017564.1 | - | 2886749 | 2886696 | **RNA**→ rpmE (PRK00019)Ribosomal_L31 (pfam01197)→ |
| Yen-1-2 | NZ_CP009367.1 | - | 3690254 | 3690201 | **RNA**→ rpmE (PRK00019)Ribosomal_L31 (pfam01197)→ |
| Yen-1-3 | NZ_CP009838.1 | + | 2005651 | 2005704 | **RNA**→ rpmE (PRK00019)Ribosomal_L31 (pfam01197)→ |
| Yen-9-1 | NZ_HF933424.1 | + | 128752 | 128805 | **RNA**→ rpmE (PRK00019)Ribosomal_L31 (pfam01197)→ |
| Yen-4-1 | NZ_HF933423.1 | + | 127495 | 127548 | **RNA**→ rpmE (PRK00019)Ribosomal_L31 (pfam01197)→ |
| Yen-2-1 | NZ_HF933426.1 | - | 4598604 | 4598551 | **RNA**→ rpmE (PRK00019)Ribosomal_L31 (pfam01197)→ |
| Yen-8-1 | NZ_HF933425.1 | + | 157671 | 157724 | **RNA**→ rpmE (PRK00019)Ribosomal_L31 (pfam01197)→ |
| Yen-10-1 | NZ_HF933206.1 | + | 132388 | 132441 | **RNA**→ rpmE (PRK00019)Ribosomal_L31 (pfam01197)→ |
| Ype-127-1 | NZ_ADPS01000326.1 | + | 13715 | 13768 | **RNA**→ rpmE (PRK00019)Ribosomal_L31 (pfam01197)→ |
| Ype-132-1 | NZ_ABCD01000004.1 | + | 834679 | 834732 | **RNA**→ rpmE (PRK00019)Ribosomal_L31 (pfam01197)→ rpmE (PRK00019)Ribosomal_L31 (pfam01197)→ |
| Ype-76-1 | NZ_AOSA01000185.1 | + | 13890 | 13943 | **RNA**→ rpmE (PRK00019)Ribosomal_L31 (pfam01197)→ |
| Ype-77-1 | NZ_AOMY01000087.1 | - | 7151 | 7098 | **RNA**→ rpmE (PRK00019)Ribosomal_L31 (pfam01197)→ |
| Ype-227-1 | NZ_AAYS01000001.1 | - | 719404 | 719351 | **RNA**→ rpmE (PRK00019)Ribosomal_L31 (pfam01197)→ rpmE (PRK00019)Ribosomal_L31 (pfam01197)→ rpmE (PRK00019)Ribosomal_L31 (pfam01197)→ |
| Ype-221-1 | NZ_AAYT01000010.1 | + | 29809 | 29862 | **RNA**→ rpmE (PRK00019)Ribosomal_L31 (pfam01197)→ rpmE (PRK00019)Ribosomal_L31 (pfam01197)→ |
| Ype-224-1 | NZ_ABAT01000037.1 | - | 8040 | 7987 | **RNA**→ rpmE (PRK00019)Ribosomal_L31 (pfam01197)→ |
| Ype-228-1 | NZ_ACNS01000001.1 | + | 2230658 | 2230711 | **RNA**→ rpmE (PRK00019)Ribosomal_L31 (pfam01197)→ rpmE (PRK00019)Ribosomal_L31 (pfam01197)→ |
| Ype-225-1 | NZ_AAOS02000002.1 | - | 191563 | 191510 | **RNA**→ rpmE (PRK00019)Ribosomal_L31 (pfam01197)→ rpmE (PRK00019)Ribosomal_L31 (pfam01197)→ rpmE (PRK00019)Ribosomal_L31 (pfam01197)→ |
| Ype-226-1 | NZ_ACNR01000002.1 | + | 105009 | 105062 | **RNA**→ rpmE (PRK00019)Ribosomal_L31 (pfam01197)→ rpmE (PRK00019)Ribosomal_L31 (pfam01197)→ |
| Ype-73-1 | NZ_ADSB01001145.1 | - | 769 | 716 | **RNA**→ rpmE (PRK00019)Ribosomal_L31 (pfam01197)→ |
| Ype-114-1 | NZ_JYJW01000062.1 | + | 13935 | 13988 | **RNA**→ rpmE (PRK00019)Ribosomal_L31 (pfam01197)→ |
| Ype-114-2 | NZ_JYJX01000087.1 | + | 13930 | 13983 | **RNA**→ rpmE (PRK00019)Ribosomal_L31 (pfam01197)→ |
| Ype-115-1 | NZ_LBFJ01000039.1 | + | 79507 | 79560 | **RNA**→ rpmE (PRK00019)Ribosomal_L31 (pfam01197)→ |
| Ype-218-1 | NZ_AAYU01000010.1 | - | 36730 | 36677 | **RNA**→ rpmE (PRK00019)Ribosomal_L31 (pfam01197)→ rpmE (PRK00019)Ribosomal_L31 (pfam01197)→ |
| Ype-219-1 | NZ_ADRZ01000574.1 | - | 769 | 716 | **RNA**→ rpmE (PRK00019)Ribosomal_L31 (pfam01197)→ |
| Ype-219-2 | NZ_AAYV01000014.1 | - | 8610 | 8557 | **RNA**→ rpmE (PRK00019)Ribosomal_L31 (pfam01197)→ rpmE (PRK00019)Ribosomal_L31 (pfam01197)→ |
| Ype-105-1 | NZ_AGJZ01000163.1 | - | 7134 | 7081 | **RNA**→ rpmE (PRK00019)Ribosomal_L31 (pfam01197)→ |
| Ype-106-1 | NZ_AGKA01000038.1 | - | 7176 | 7123 | **RNA**→ rpmE (PRK00019)Ribosomal_L31 (pfam01197)→ |
| Ype-109-1 | NZ_AGJV01000211.1 | + | 12389 | 12442 | **RNA**→ rpmE (PRK00019)Ribosomal_L31 (pfam01197)→ |
| Ype-104-1 | NZ_AGJU01000166.1 | - | 7123 | 7070 | **RNA**→ rpmE (PRK00019)Ribosomal_L31 (pfam01197)→ |
| Ype-108-1 | NZ_AGJY01000045.1 | - | 7176 | 7123 | **RNA**→ rpmE (PRK00019)Ribosomal_L31 (pfam01197)→ |
| Ype-107-1 | NZ_AGJT01000136.1 | - | 7132 | 7079 | **RNA**→ rpmE (PRK00019)Ribosomal_L31 (pfam01197)→ |
| Ype-110-1 | NZ_AGJS01000160.1 | - | 7177 | 7124 | **RNA**→ rpmE (PRK00019)Ribosomal_L31 (pfam01197)→ |
| Ype-111-1 | NZ_AGJW01000073.1 | - | 7101 | 7048 | **RNA**→ rpmE (PRK00019)Ribosomal_L31 (pfam01197)→ |
| Ype-112-1 | NZ_AGJX01000102.1 | - | 7175 | 7122 | **RNA**→ rpmE (PRK00019)Ribosomal_L31 (pfam01197)→ |
| Ype-174-1 | NZ_AKRZ01000067.1 | + | 10124 | 10177 | **RNA**→ rpmE (PRK00019)Ribosomal_L31 (pfam01197)→ |

| | | | | | |
|---|---|---|---|---|---|
| Ype-117-1 | NZ_ADOY01000335.1 | - | 769 | 716 | **RNA**→ rpmE (PRK00019)Ribosomal_L31 (pfam01197)→ |
| Ype-170-1 | NZ_AKRW01000074.1 | + | 8810 | 8863 | **RNA**→ rpmE (PRK00019)Ribosomal_L31 (pfam01197)→ |
| Ype-9-1 | NZ_ADOW01000210.1 | - | 335 | 282 | **RNA**→ rpmE (PRK00019)Ribosomal_L31 (pfam01197)→ |
| Ype-175-1 | NZ_AKSA01000062.1 | + | 12442 | 12495 | **RNA**→ rpmE (PRK00019)Ribosomal_L31 (pfam01197)→ |
| Ype-179-1 | NZ_AKSE01000056.1 | + | 8851 | 8904 | **RNA**→ rpmE (PRK00019)Ribosomal_L31 (pfam01197)→ |
| Ype-14-1 | NZ_ADPD01000556.1 | + | 10503 | 10556 | **RNA**→ rpmE (PRK00019)Ribosomal_L31 (pfam01197)→ |
| Ype-189-1 | NZ_AKSO01000061.1 | + | 8798 | 8851 | **RNA**→ rpmE (PRK00019)Ribosomal_L31 (pfam01197)→ |
| Ype-123-1 | NZ_ADPI01000278.1 | + | 13715 | 13768 | **RNA**→ rpmE (PRK00019)Ribosomal_L31 (pfam01197)→ |
| Ype-194-1 | NZ_AKST01000031.1 | + | 8772 | 8825 | **RNA**→ rpmE (PRK00019)Ribosomal_L31 (pfam01197)→ |
| Ype-20-1 | NZ_ADPM01000236.1 | - | 7093 | 7040 | **RNA**→ rpmE (PRK00019)Ribosomal_L31 (pfam01197)→ |
| Ype-201-1 | NZ_AKTA01000022.1 | + | 12445 | 12498 | **RNA**→ rpmE (PRK00019)Ribosomal_L31 (pfam01197)→ |
| Ype-207-1 | NZ_AKTG01000058.1 | + | 8844 | 8897 | **RNA**→ rpmE (PRK00019)Ribosomal_L31 (pfam01197)→ |
| Ype-42-1 | NZ_ADQL01000708.1 | - | 769 | 716 | **RNA**→ rpmE (PRK00019)Ribosomal_L31 (pfam01197)→ |
| Ype-208-1 | NZ_AKTH01000039.1 | + | 12442 | 12495 | **RNA**→ rpmE (PRK00019)Ribosomal_L31 (pfam01197)→ |
| Ype-214-1 | NZ_AKTN01000075.1 | + | 5074 | 5127 | **RNA**→ rpmE (PRK00019)Ribosomal_L31 (pfam01197)→ |
| Ype-166-1 | NZ_AKTO01000032.1 | + | 8856 | 8909 | **RNA**→ rpmE (PRK00019)Ribosomal_L31 (pfam01197)→ |
| Ype-53-1 | NZ_ADRA01001343.1 | + | 9993 | 10046 | **RNA**→ rpmE (PRK00019)Ribosomal_L31 (pfam01197)→ |
| Ype-183-1 | NZ_AKSI01000056.1 | + | 8820 | 8873 | **RNA**→ rpmE (PRK00019)Ribosomal_L31 (pfam01197)→ |
| Ype-159-1 | NZ_AKRP01000066.1 | + | 8787 | 8840 | **RNA**→ rpmE (PRK00019)Ribosomal_L31 (pfam01197)→ |
| Ype-185-1 | NZ_AKSK01000024.1 | + | 8853 | 8906 | **RNA**→ rpmE (PRK00019)Ribosomal_L31 (pfam01197)→ |
| Ype-188-1 | NZ_AKSN01000036.1 | + | 8837 | 8890 | **RNA**→ rpmE (PRK00019)Ribosomal_L31 (pfam01197)→ |
| Ype-15-1 | NZ_ADPE01000834.1 | + | 5067 | 5120 | **RNA**→ rpmE (PRK00019)Ribosomal_L31 (pfam01197)→ |
| Ype-211-1 | NZ_AKTK01000059.1 | + | 8826 | 8879 | **RNA**→ rpmE (PRK00019)Ribosomal_L31 (pfam01197)→ |
| Ype-169-1 | NZ_AKTR01000071.1 | + | 8836 | 8889 | **RNA**→ rpmE (PRK00019)Ribosomal_L31 (pfam01197)→ |
| Ype-21-1 | NZ_ADPO01001311.1 | + | 5067 | 5120 | **RNA**→ rpmE (PRK00019)Ribosomal_L31 (pfam01197)→ |
| Ype-215-1 | NZ_ACNT01000002.1 | + | 99358 | 99411 | **RNA**→ rpmE (PRK00019)Ribosomal_L31 (pfam01197)→ rpmE (PRK00019)Ribosomal_L31 (pfam01197)→ |
| Ype-176-1 | NZ_AKSB01000035.1 | + | 8780 | 8833 | **RNA**→ rpmE (PRK00019)Ribosomal_L31 (pfam01197)→ |
| Ype-172-1 | NZ_AKRX01000066.1 | + | 8773 | 8826 | **RNA**→ rpmE (PRK00019)Ribosomal_L31 (pfam01197)→ |
| Ype-162-1 | NZ_AKRS01000057.1 | + | 8792 | 8845 | **RNA**→ rpmE (PRK00019)Ribosomal_L31 (pfam01197)→ |
| Ype-180-1 | NZ_AKSF01000045.1 | + | 12446 | 12499 | **RNA**→ rpmE (PRK00019)Ribosomal_L31 (pfam01197)→ |
| Ype-164-1 | NZ_AKRU01000036.1 | + | 5005 | 5058 | **RNA**→ rpmE (PRK00019)Ribosomal_L31 (pfam01197)→ |
| Ype-190-1 | NZ_AKSP01000050.1 | + | 9378 | 9431 | **RNA**→ rpmE (PRK00019)Ribosomal_L31 (pfam01197)→ |
| Ype-165-1 | NZ_AKRV01000028.1 | + | 12442 | 12495 | **RNA**→ rpmE (PRK00019)Ribosomal_L31 (pfam01197)→ |
| Ype-191-1 | NZ_AKSQ01000022.1 | + | 9219 | 9272 | **RNA**→ rpmE (PRK00019)Ribosomal_L31 (pfam01197)→ |
| Ype-156-1 | NZ_AKRM01000069.1 | + | 12432 | 12485 | **RNA**→ rpmE (PRK00019)Ribosomal_L31 (pfam01197)→ |
| Ype-195-1 | NZ_AKSU01000058.1 | + | 8770 | 8823 | **RNA**→ rpmE (PRK00019)Ribosomal_L31 (pfam01197)→ |
| Ype-197-1 | NZ_AKSW01000057.1 | + | 12440 | 12493 | **RNA**→ rpmE (PRK00019)Ribosomal_L31 (pfam01197)→ |
| Ype-4-1 | NZ_ADOV01000797.1 | + | 10642 | 10695 | **RNA**→ rpmE (PRK00019)Ribosomal_L31 (pfam01197)→ |
| Ype-202-1 | NZ_AKTB01000030.1 | + | 6451 | 6504 | **RNA**→ rpmE (PRK00019)Ribosomal_L31 (pfam01197)→ |
| Ype-203-1 | NZ_AKTC01000019.1 | + | 8778 | 8831 | **RNA**→ rpmE (PRK00019)Ribosomal_L31 (pfam01197)→ |
| Ype-11-1 | NZ_ADOX01001006.1 | + | 623 | 676 | **RNA**→ rpmE (PRK00019)Ribosomal_L31 (pfam01197)→ |
| Ype-209-1 | NZ_AKTI01000030.1 | + | 12442 | 12495 | **RNA**→ rpmE (PRK00019)Ribosomal_L31 (pfam01197)→ |
| Ype-168-1 | NZ_AKTQ01000048.1 | + | 8785 | 8838 | **RNA**→ rpmE (PRK00019)Ribosomal_L31 (pfam01197)→ |
| Ype-122-1 | NZ_ADPG01000719.1 | - | 798 | 745 | **RNA**→ rpmE (PRK00019)Ribosomal_L31 (pfam01197)→ |
| Ype-186-1 | NZ_AKSL01000056.1 | + | 12446 | 12499 | **RNA**→ rpmE (PRK00019)Ribosomal_L31 (pfam01197)→ |
| Ype-124-1 | NZ_ADPL01000265.1 | + | 62 | 115 | **RNA**→ rpmE (PRK00019)Ribosomal_L31 (pfam01197)→ |
| Ype-196-1 | NZ_AKSV01000044.1 | + | 8842 | 8895 | **RNA**→ rpmE (PRK00019)Ribosomal_L31 (pfam01197)→ |
| Ype-23-1 | NZ_ADPQ01000548.1 | - | 798 | 745 | **RNA**→ rpmE (PRK00019)Ribosomal_L31 (pfam01197)→ |
| Ype-173-1 | NZ_AKRY01000059.1 | + | 8848 | 8901 | **RNA**→ rpmE (PRK00019)Ribosomal_L31 (pfam01197)→ |
| Ype-178-1 | NZ_AKSD01000032.1 | + | 8843 | 8896 | **RNA**→ rpmE (PRK00019)Ribosomal_L31 (pfam01197)→ |
| Ype-128-1 | NZ_ADPU01000583.1 | - | 798 | 745 | **RNA**→ rpmE (PRK00019)Ribosomal_L31 (pfam01197)→ |
| Ype-187-1 | NZ_AKSM01000049.1 | + | 12444 | 12497 | **RNA**→ rpmE (PRK00019)Ribosomal_L31 (pfam01197)→ |

| | | | | | |
|---|---|---|---|---|---|
| Ype-193-1 | NZ_AKSS01000018.1 | + | 9267 | 9320 | **RNA**→ rpmE (PRK00019)Ribosomal_L31 (pfam01197)→ |
| Ype-130-1 | NZ_ADQA01000477.1 | - | 412 | 359 | **RNA**→ rpmE (PRK00019)Ribosomal_L31 (pfam01197)→ |
| Ype-199-1 | NZ_AKSY01000055.1 | + | 12442 | 12495 | **RNA**→ rpmE (PRK00019)Ribosomal_L31 (pfam01197)→ |
| Ype-160-1 | NZ_AKRQ01000033.1 | + | 12432 | 12485 | **RNA**→ rpmE (PRK00019)Ribosomal_L31 (pfam01197)→ |
| Ype-206-1 | NZ_AKTF01000032.1 | + | 12436 | 12489 | **RNA**→ rpmE (PRK00019)Ribosomal_L31 (pfam01197)→ |
| Ype-163-1 | NZ_AKRT01000060.1 | + | 8826 | 8879 | **RNA**→ rpmE (PRK00019)Ribosomal_L31 (pfam01197)→ |
| Ype-213-1 | NZ_AKTM01000034.1 | + | 12438 | 12491 | **RNA**→ rpmE (PRK00019)Ribosomal_L31 (pfam01197)→ |
| Ype-151-1 | NZ_AMQL01000001.1 | + | 119897 | 119950 | **RNA**→ rpmE (PRK00019)Ribosomal_L31 (pfam01197)→ |
| Ype-182-1 | NZ_AKSH01000070.1 | + | 8780 | 8833 | **RNA**→ rpmE (PRK00019)Ribosomal_L31 (pfam01197)→ |
| Ype-158-1 | NZ_AKRO01000081.1 | + | 6527 | 6580 | **RNA**→ rpmE (PRK00019)Ribosomal_L31 (pfam01197)→ |
| Ype-204-1 | NZ_AKTD01000072.1 | + | 8776 | 8829 | **RNA**→ rpmE (PRK00019)Ribosomal_L31 (pfam01197)→ |
| Ype-118-1 | NZ_AKVQ01000089.1 | - | 7230 | 7177 | **RNA**→ rpmE (PRK00019)Ribosomal_L31 (pfam01197)→ |
| Ype-210-1 | NZ_AKTJ01000027.1 | + | 8752 | 8805 | **RNA**→ rpmE (PRK00019)Ribosomal_L31 (pfam01197)→ |
| Ype-120-1 | NZ_ADPA01000150.1 | + | 13715 | 13768 | **RNA**→ rpmE (PRK00019)Ribosomal_L31 (pfam01197)→ |
| Ype-167-1 | NZ_AKTP01000020.1 | + | 8836 | 8889 | **RNA**→ rpmE (PRK00019)Ribosomal_L31 (pfam01197)→ |
| Ype-16-1 | NZ_ADPF01000098.1 | - | 798 | 745 | **RNA**→ rpmE (PRK00019)Ribosomal_L31 (pfam01197)→ |
| Ype-177-1 | NZ_AKSC01000026.1 | + | 8781 | 8834 | **RNA**→ rpmE (PRK00019)Ribosomal_L31 (pfam01197)→ |
| Ype-181-1 | NZ_AKSG01000061.1 | + | 12440 | 12493 | **RNA**→ rpmE (PRK00019)Ribosomal_L31 (pfam01197)→ |
| Ype-184-1 | NZ_AKSJ01000016.1 | + | 12444 | 12497 | **RNA**→ rpmE (PRK00019)Ribosomal_L31 (pfam01197)→ |
| Ype-134-1 | NZ_ADQT01000515.1 | - | 798 | 745 | **RNA**→ rpmE (PRK00019)Ribosomal_L31 (pfam01197)→ |
| Ype-192-1 | NZ_AKSR01000032.1 | + | 6522 | 6575 | **RNA**→ rpmE (PRK00019)Ribosomal_L31 (pfam01197)→ |
| Ype-198-1 | NZ_AKSX01000076.1 | + | 8836 | 8889 | **RNA**→ rpmE (PRK00019)Ribosomal_L31 (pfam01197)→ |
| Ype-40-1 | NZ_ADQI01001024.1 | - | 798 | 745 | **RNA**→ rpmE (PRK00019)Ribosomal_L31 (pfam01197)→ |
| Ype-205-1 | NZ_AKTE01000056.1 | + | 8828 | 8881 | **RNA**→ rpmE (PRK00019)Ribosomal_L31 (pfam01197)→ |
| Ype-161-1 | NZ_AKRR01000069.1 | + | 8837 | 8890 | **RNA**→ rpmE (PRK00019)Ribosomal_L31 (pfam01197)→ |
| Ype-38-1 | NZ_ADQG01000238.1 | - | 769 | 716 | **RNA**→ rpmE (PRK00019)Ribosomal_L31 (pfam01197)→ |
| Ype-212-1 | NZ_AKTL01000068.1 | + | 8860 | 8913 | **RNA**→ rpmE (PRK00019)Ribosomal_L31 (pfam01197)→ |
| Ype-171-1 | NZ_AKTS01000032.1 | + | 8838 | 8891 | **RNA**→ rpmE (PRK00019)Ribosomal_L31 (pfam01197)→ |
| Ype-200-1 | NZ_AKSZ01000056.1 | + | 6521 | 6574 | **RNA**→ rpmE (PRK00019)Ribosomal_L31 (pfam01197)→ |
| Ype-72-1 | NZ_ADSA02000142.1 | - | 7075 | 7022 | **RNA**→ rpmE (PRK00019)Ribosomal_L31 (pfam01197)→ |
| Ype-28-1 | NZ_ADPW01000087.1 | + | 10482 | 10535 | **RNA**→ rpmE (PRK00019)Ribosomal_L31 (pfam01197)→ |
| Ype-131-1 | NZ_ADQB01000681.1 | + | 13715 | 13768 | **RNA**→ rpmE (PRK00019)Ribosomal_L31 (pfam01197)→ |
| Ype-57-1 | NZ_ADRG01000150.1 | - | 769 | 716 | **RNA**→ rpmE (PRK00019)Ribosomal_L31 (pfam01197)→ |
| Ype-63-1 | NZ_ADRQ01000346.1 | - | 543 | 490 | **RNA**→ rpmE (PRK00019)Ribosomal_L31 (pfam01197)→ |
| Ype-67-1 | NZ_ADRU01000229.1 | - | 769 | 716 | **RNA**→ rpmE (PRK00019)Ribosomal_L31 (pfam01197)→ |
| Ype-148-1 | NZ_ADSF01000103.1 | + | 13920 | 13973 | **RNA**→ rpmE (PRK00019)Ribosomal_L31 (pfam01197)→ |
| Ype-80-1 | NZ_ADSK01000857.1 | + | 5067 | 5120 | **RNA**→ rpmE (PRK00019)Ribosomal_L31 (pfam01197)→ |
| Ype-126-1 | NZ_ADPR01000439.1 | - | 798 | 745 | **RNA**→ rpmE (PRK00019)Ribosomal_L31 (pfam01197)→ |
| Ype-27-1 | NZ_ADPV01000112.1 | - | 335 | 282 | **RNA**→ rpmE (PRK00019)Ribosomal_L31 (pfam01197)→ |
| Ype-136-1 | NZ_ADQZ01000375.1 | - | 480 | 427 | **RNA**→ rpmE (PRK00019)Ribosomal_L31 (pfam01197)→ |
| Ype-70-1 | NZ_ADRY01000109.1 | - | 769 | 716 | **RNA**→ rpmE (PRK00019)Ribosomal_L31 (pfam01197)→ |
| Ype-47-1 | NZ_ADQR01000100.1 | - | 769 | 716 | **RNA**→ rpmE (PRK00019)Ribosomal_L31 (pfam01197)→ |
| Ype-135-1 | NZ_ADQX01000781.1 | - | 798 | 745 | **RNA**→ rpmE (PRK00019)Ribosomal_L31 (pfam01197)→ |
| Ype-140-1 | NZ_ADRM01000463.1 | - | 769 | 716 | **RNA**→ rpmE (PRK00019)Ribosomal_L31 (pfam01197)→ |
| Ype-97-1 | NZ_ADTB01000365.1 | + | 5079 | 5132 | **RNA**→ rpmE (PRK00019)Ribosomal_L31 (pfam01197)→ |
| Ype-26-1 | NZ_ADPT01000656.1 | + | 5077 | 5130 | **RNA**→ rpmE (PRK00019)Ribosomal_L31 (pfam01197)→ |
| Ype-35-1 | NZ_ADQD01000442.1 | + | 13715 | 13768 | **RNA**→ rpmE (PRK00019)Ribosomal_L31 (pfam01197)→ |
| Ype-137-1 | NZ_ADRD01000263.1 | + | 13715 | 13768 | **RNA**→ rpmE (PRK00019)Ribosomal_L31 (pfam01197)→ |
| Ype-61-1 | NZ_ADRN01000955.1 | + | 10028 | 10081 | **RNA**→ rpmE (PRK00019)Ribosomal_L31 (pfam01197)→ |
| Ype-91-1 | NZ_ADSX01000354.1 | - | 798 | 745 | **RNA**→ rpmE (PRK00019)Ribosomal_L31 (pfam01197)→ |
| Ype-46-1 | NZ_ADQQ01000130.1 | - | 769 | 716 | **RNA**→ rpmE (PRK00019)Ribosomal_L31 (pfam01197)→ |
| Ype-60-1 | NZ_ADRL01000403.1 | - | 798 | 745 | **RNA**→ rpmE (PRK00019)Ribosomal_L31 (pfam01197)→ |

| | | | | | | | |
|---|---|---|---|---|---|---|---|
| Ype-62-1 | NZ_ADRP01000734.1 | - | 325 | 272 | **RNA**→ | rpmE (PRK00019)Ribosomal_L31 (pfam01197)→ | |
| Ype-103-1 | NZ_ADTI01001185.1 | + | 9163 | 9216 | **RNA**→ | rpmE (PRK00019)Ribosomal_L31 (pfam01197)→ | |
| Ype-68-1 | NZ_ADRV01000666.1 | - | 769 | 716 | **RNA**→ | rpmE (PRK00019)Ribosomal_L31 (pfam01197)→ | |
| Ype-154-1 | NZ_ADSU01000714.1 | + | 13716 | 13769 | **RNA**→ | rpmE (PRK00019)Ribosomal_L31 (pfam01197)→ | |
| Ype-41-1 | NZ_ADQK01000507.1 | - | 769 | 716 | **RNA**→ | rpmE (PRK00019)Ribosomal_L31 (pfam01197)→ | |
| Ype-49-1 | NZ_ADQU01000137.1 | - | 769 | 716 | **RNA**→ | rpmE (PRK00019)Ribosomal_L31 (pfam01197)→ | |
| Ype-139-1 | NZ_ADRJ01000349.1 | - | 769 | 716 | **RNA**→ | rpmE (PRK00019)Ribosomal_L31 (pfam01197)→ | |
| Ype-75-1 | NZ_ADSD01001379.1 | + | 9545 | 9598 | **RNA**→ | rpmE (PRK00019)Ribosomal_L31 (pfam01197)→ | |
| Ype-92-1 | NZ_ADSY01001236.1 | + | 8207 | 8260 | **RNA**→ | rpmE (PRK00019)Ribosomal_L31 (pfam01197)→ | |
| Ype-133-1 | NZ_ADQN01000513.1 | + | 13920 | 13973 | **RNA**→ | rpmE (PRK00019)Ribosomal_L31 (pfam01197)→ | |
| Ype-44-1 | NZ_ADQO01000836.1 | - | 7070 | 7017 | **RNA**→ | rpmE (PRK00019)Ribosomal_L31 (pfam01197)→ | |
| Ype-58-1 | NZ_ADRI01000395.1 | - | 798 | 745 | **RNA**→ | rpmE (PRK00019)Ribosomal_L31 (pfam01197)→ | |
| Ype-65-1 | NZ_ADRS01001043.1 | + | 9838 | 9891 | **RNA**→ | rpmE (PRK00019)Ribosomal_L31 (pfam01197)→ | |
| Ype-87-1 | NZ_ADSS01000896.1 | - | 769 | 716 | **RNA**→ | rpmE (PRK00019)Ribosomal_L31 (pfam01197)→ | |
| Ype-98-1 | NZ_ADTC01000594.1 | - | 769 | 716 | **RNA**→ | rpmE (PRK00019)Ribosomal_L31 (pfam01197)→ | |
| Ype-129-1 | NZ_ADPX01000345.1 | + | 13715 | 13768 | **RNA**→ | rpmE (PRK00019)Ribosomal_L31 (pfam01197)→ | |
| Ype-119-1 | NZ_ADOZ01000454.1 | - | 798 | 745 | **RNA**→ | rpmE (PRK00019)Ribosomal_L31 (pfam01197)→ | |
| Ype-18-1 | NZ_ADPJ01000086.1 | - | 335 | 282 | **RNA**→ | rpmE (PRK00019)Ribosomal_L31 (pfam01197)→ | |
| Ype-51-1 | NZ_ADQW01000954.1 | + | 5067 | 5120 | **RNA**→ | rpmE (PRK00019)Ribosomal_L31 (pfam01197)→ | |
| Ype-54-1 | NZ_ADRB01000030.1 | - | 798 | 745 | **RNA**→ | rpmE (PRK00019)Ribosomal_L31 (pfam01197)→ | |
| Ype-34-1 | NZ_ADQC01000685.1 | + | 5067 | 5120 | **RNA**→ | rpmE (PRK00019)Ribosomal_L31 (pfam01197)→ | |
| Ype-43-1 | NZ_ADQM01000370.1 | - | 781 | 728 | **RNA**→ | rpmE (PRK00019)Ribosomal_L31 (pfam01197)→ | |
| Ype-55-1 | NZ_ADRC01000710.1 | - | 798 | 745 | **RNA**→ | rpmE (PRK00019)Ribosomal_L31 (pfam01197)→ | |
| Ype-138-1 | NZ_ADRH01000811.1 | + | 12403 | 12456 | **RNA**→ | rpmE (PRK00019)Ribosomal_L31 (pfam01197)→ | |
| Ype-48-1 | NZ_ADQS01000710.1 | + | 13920 | 13973 | **RNA**→ | rpmE (PRK00019)Ribosomal_L31 (pfam01197)→ | |
| Ype-64-1 | NZ_ADRR01001210.1 | + | 62 | 115 | **RNA**→ | rpmE (PRK00019)Ribosomal_L31 (pfam01197)→ | |
| Ype-145-1 | NZ_ADRW01000267.1 | + | 12028 | 12081 | **RNA**→ | rpmE (PRK00019)Ribosomal_L31 (pfam01197)→ | |
| Ype-89-1 | NZ_ADSV01001085.1 | - | 769 | 716 | **RNA**→ | rpmE (PRK00019)Ribosomal_L31 (pfam01197)→ | |
| Ype-149-1 | NZ_ADSG01000304.1 | - | 752 | 699 | **RNA**→ | rpmE (PRK00019)Ribosomal_L31 (pfam01197)→ | |
| Ype-81-1 | NZ_ADSL01000341.1 | + | 62 | 115 | **RNA**→ | rpmE (PRK00019)Ribosomal_L31 (pfam01197)→ | |
| Ype-220-1 | NZ_AAYR01000027.1 | - | 8565 | 8512 | **RNA**→ | rpmE (PRK00019)Ribosomal_L31 (pfam01197)→ | rpmE (PRK00019)Ribosomal_L31 (pfam01197)→ |
| Yps-15-1 | NZ_BAUR01000075.1 | + | 71087 | 71140 | **RNA**→ | rpmE (PRK00019)Ribosomal_L31 (pfam01197)→ | |
| Ype-146-1 | NZ_AAUB01000173.1 | + | 6501 | 6554 | **RNA**→ | rpmE (PRK00019)Ribosomal_L31 (pfam01197)→ | rpmE (PRK00019)Ribosomal_L31 (pfam01197)→ |
| Ype-69-1 | NZ_ADRX01000324.1 | - | 769 | 716 | **RNA**→ | rpmE (PRK00019)Ribosomal_L31 (pfam01197)→ | |
| Ype-74-1 | NZ_ADSC01000669.1 | + | 4245 | 4298 | **RNA**→ | rpmE (PRK00019)Ribosomal_L31 (pfam01197)→ | |
| Ype-78-1 | NZ_ADSH01000912.1 | + | 9300 | 9353 | **RNA**→ | rpmE (PRK00019)Ribosomal_L31 (pfam01197)→ | |
| Ype-82-1 | NZ_ADSM01000470.1 | + | 5067 | 5120 | **RNA**→ | rpmE (PRK00019)Ribosomal_L31 (pfam01197)→ | |
| Ype-147-1 | NZ_ADSE01000689.1 | + | 13920 | 13973 | **RNA**→ | rpmE (PRK00019)Ribosomal_L31 (pfam01197)→ | |
| Ype-71-1 | NZ_ATWT01000295.1 | - | 2939 | 2886 | **RNA**→ | rpmE (PRK00019)Ribosomal_L31 (pfam01197)→ | |
| Ype-88-1 | NZ_ADST01000002.1 | + | 13725 | 13778 | **RNA**→ | rpmE (PRK00019)Ribosomal_L31 (pfam01197)→ | |
| Ype-100-1 | NZ_ADTE01000181.1 | + | 9873 | 9926 | **RNA**→ | rpmE (PRK00019)Ribosomal_L31 (pfam01197)→ | |
| Ype-93-1 | NZ_ADSZ01000320.1 | + | 10485 | 10538 | **RNA**→ | rpmE (PRK00019)Ribosomal_L31 (pfam01197)→ | |
| Ype-142-1 | NC_003143.1 | + | 119914 | 119967 | **RNA**→ | rpmE (PRK00019)Ribosomal_L31 (pfam01197)→ | rpmE (PRK00019)Ribosomal_L31 (pfam01197)→ |
| Yps-14-1 | NC_009708.1 | + | 124888 | 124941 | **RNA**→ | rpmE (PRK00019)Ribosomal_L31 (pfam01197)→ | |
| Ype-94-2 | NC_009381.1 | - | 4365187 | 4365134 | **RNA**→ | rpmE (PRK00019)Ribosomal_L31 (pfam01197)→ | |
| Yps-11-2 | NC_010634.1 | + | 110418 | 110471 | **RNA**→ | rpmE (PRK00019)Ribosomal_L31 (pfam01197)→ | |
| Ype-86-1 | NZ_ADSQ01000035.1 | - | 798 | 745 | **RNA**→ | rpmE (PRK00019)Ribosomal_L31 (pfam01197)→ | |
| Ype-101-1 | NZ_ADTG01000548.1 | - | 751 | 698 | **RNA**→ | rpmE (PRK00019)Ribosomal_L31 (pfam01197)→ | |
| Ype-19-1 | NZ_ADPK01000243.1 | + | 5067 | 5120 | **RNA**→ | rpmE (PRK00019)Ribosomal_L31 (pfam01197)→ | |
| Ype-22-1 | NZ_ADPP01000198.1 | + | 13715 | 13768 | **RNA**→ | rpmE (PRK00019)Ribosomal_L31 (pfam01197)→ | |
| Ype-30-1 | NZ_ADPY01001373.1 | + | 13715 | 13768 | **RNA**→ | rpmE (PRK00019)Ribosomal_L31 (pfam01197)→ | |
| Ype-52-1 | NZ_ADQY01000753.1 | - | 798 | 745 | **RNA**→ | rpmE (PRK00019)Ribosomal_L31 (pfam01197)→ | |

| | | | | | |
|---|---|---|---|---|---|
| Ype-83-1 | NZ_ADSN01000864.1 | + | 5067 | 5120 | **RNA**→ rpmE (PRK00019)Ribosomal_L31 (pfam01197)→ |
| Ype-90-1 | NZ_ADSW01000343.1 | + | 62 | 115 | **RNA**→ rpmE (PRK00019)Ribosomal_L31 (pfam01197)→ |
| Ype-102-1 | NZ_ADTH01000245.1 | + | 6953 | 7006 | **RNA**→ rpmE (PRK00019)Ribosomal_L31 (pfam01197)→ |
| Ype-121-1 | NZ_ADPC01000333.1 | - | 769 | 716 | **RNA**→ rpmE (PRK00019)Ribosomal_L31 (pfam01197)→ |
| Ype-125-1 | NZ_ADPN01000484.1 | - | 798 | 745 | **RNA**→ rpmE (PRK00019)Ribosomal_L31 (pfam01197)→ |
| Ype-59-1 | NZ_ADRK01001149.1 | - | 549 | 496 | **RNA**→ rpmE (PRK00019)Ribosomal_L31 (pfam01197)→ |
| Ype-85-1 | NZ_ADSP01001051.1 | + | 5067 | 5120 | **RNA**→ rpmE (PRK00019)Ribosomal_L31 (pfam01197)→ |
| Ype-33-1 | NZ_ADPZ01000150.1 | - | 769 | 716 | **RNA**→ rpmE (PRK00019)Ribosomal_L31 (pfam01197)→ |
| Ype-37-1 | NZ_ADQF01000591.1 | - | 769 | 716 | **RNA**→ rpmE (PRK00019)Ribosomal_L31 (pfam01197)→ |
| Ype-56-1 | NZ_ADRE01000062.1 | + | 12515 | 12568 | **RNA**→ rpmE (PRK00019)Ribosomal_L31 (pfam01197)→ |
| Ype-141-1 | NZ_ADRO01000211.1 | + | 13735 | 13788 | **RNA**→ rpmE (PRK00019)Ribosomal_L31 (pfam01197)→ |
| Ype-66-1 | NZ_ADRT01000420.1 | - | 7070 | 7017 | **RNA**→ rpmE (PRK00019)Ribosomal_L31 (pfam01197)→ |
| Ype-79-1 | NZ_ADSJ01000717.1 | + | 5067 | 5120 | **RNA**→ rpmE (PRK00019)Ribosomal_L31 (pfam01197)→ |
| Ype-84-1 | NZ_ADSO01000781.1 | - | 798 | 745 | **RNA**→ rpmE (PRK00019)Ribosomal_L31 (pfam01197)→ |
| Ype-216-1 | NZ_ADTF01000446.1 | + | 12403 | 12456 | **RNA**→ rpmE (PRK00019)Ribosomal_L31 (pfam01197)→ |
| Yps-10-2 | NC_006155.1 | + | 115862 | 115915 | **RNA**→ rpmE (PRK00019)Ribosomal_L31 (pfam01197)→ |
| Ype-153-1 | NC_004088.1 | + | 319573 | 319626 | **RNA**→ rpmE (PRK00019)Ribosomal_L31 (pfam01197)→ rpmE (PRK00019)Ribosomal_L31 (pfam01197)→ |
| Ype-7-1 | NZ_AYLS01000023.1 | + | 99382 | 99435 | **RNA**→ rpmE (PRK00019)Ribosomal_L31 (pfam01197)→ |
| Ype-10-1 | NZ_AYLR01000006.1 | - | 2608204 | 2608151 | **RNA**→ rpmE (PRK00019)Ribosomal_L31 (pfam01197)→ |
| Yps-12-2 | NC_010465.1 | - | 4530451 | 4530398 | **RNA**→ rpmE (PRK00019)Ribosomal_L31 (pfam01197)→ |
| Ype-31-2 | NC_010159.1 | - | 4044329 | 4044276 | **RNA**→ rpmE (PRK00019)Ribosomal_L31 (pfam01197)→ hypo→ RAYT (COG1943)→ RAYT (COG1943)→ |
| Ype-217-1 | NC_014029.1 | + | 118900 | 118953 | **RNA**→ rpmE (PRK00019)Ribosomal_L31 (pfam01197)→ rpmE (PRK00019)Ribosomal_L31 (pfam01197)→ rpmE (PRK00019)Ribosomal_L31 (pfam01197)→ |
| Ype-29-2 | NC_017168.1 | - | 985179 | 985126 | **RNA**→ rpmE (PRK00019)Ribosomal_L31 (pfam01197)→ |
| Ype-5-1 | NZ_CP006783.1 | + | 3932284 | 3932337 | **RNA**→ rpmE (PRK00019)Ribosomal_L31 (pfam01197)→ |
| Ype-13-1 | NZ_CP006754.1 | - | 1018418 | 1018365 | **RNA**→ rpmE (PRK00019)Ribosomal_L31 (pfam01197)→ |
| Ype-6-1 | NZ_CP006762.1 | - | 1856755 | 1856702 | **RNA**→ rpmE (PRK00019)Ribosomal_L31 (pfam01197)→ |
| Ype-8-1 | NZ_CP006758.1 | - | 626171 | 626118 | **RNA**→ rpmE (PRK00019)Ribosomal_L31 (pfam01197)→ |
| Ype-3-1 | NZ_CP006794.1 | - | 1252367 | 1252314 | **RNA**→ rpmE (PRK00019)Ribosomal_L31 (pfam01197)→ |
| Ype-12-1 | NZ_CP006792.1 | - | 3098142 | 3098089 | **RNA**→ rpmE (PRK00019)Ribosomal_L31 (pfam01197)→ |
| Ype-24-1 | NZ_CP006806.1 | + | 163420 | 163473 | **RNA**→ rpmE (PRK00019)Ribosomal_L31 (pfam01197)→ |
| Ype-17-1 | NZ_CP006751.1 | - | 1158455 | 1158402 | **RNA**→ rpmE (PRK00019)Ribosomal_L31 (pfam01197)→ |
| Ype-25-1 | NZ_CP006748.1 | + | 3470401 | 3470454 | **RNA**→ rpmE (PRK00019)Ribosomal_L31 (pfam01197)→ |
| Ype-152-1 | NZ_ADSR01000499.1 | + | 13715 | 13768 | **RNA**→ rpmE (PRK00019)Ribosomal_L31 (pfam01197)→ |
| Ype-45-1 | NZ_ADQP01000988.1 | + | 4329 | 4382 | **RNA**→ rpmE (PRK00019)Ribosomal_L31 (pfam01197)→ |
| Ype-150-1 | NZ_ADSI01000634.1 | - | 798 | 745 | **RNA**→ rpmE (PRK00019)Ribosomal_L31 (pfam01197)→ |
| Ype-39-1 | NZ_ADQH01001213.1 | + | 5067 | 5120 | **RNA**→ rpmE (PRK00019)Ribosomal_L31 (pfam01197)→ |
| Ype-96-1 | NZ_ADTA01000354.1 | + | 9546 | 9599 | **RNA**→ rpmE (PRK00019)Ribosomal_L31 (pfam01197)→ |
| Ype-50-1 | NZ_ADQV01000561.1 | + | 5067 | 5120 | **RNA**→ rpmE (PRK00019)Ribosomal_L31 (pfam01197)→ |
| Ype-155-1 | NC_008149.1 | - | 4294903 | 4294850 | **RNA**→ rpmE (PRK00019)Ribosomal_L31 (pfam01197)→ |
| Ype-32-2 | NC_008150.1 | + | 302083 | 302136 | **RNA**→ rpmE (PRK00019)Ribosomal_L31 (pfam01197)→ |
| Ype-143-1 | NC_017154.1 | + | 118900 | 118953 | **RNA**→ rpmE (PRK00019)Ribosomal_L31 (pfam01197)→ rpmE (PRK00019)Ribosomal_L31 (pfam01197)→ rpmE (PRK00019)Ribosomal_L31 (pfam01197)→ |
| Ype-144-1 | NC_017160.1 | + | 118961 | 119014 | **RNA**→ rpmE (PRK00019)Ribosomal_L31 (pfam01197)→ rpmE (PRK00019)Ribosomal_L31 (pfam01197)→ rpmE (PRK00019)Ribosomal_L31 (pfam01197)→ |
| Ype-2-1 | NZ_CP009492.1 | + | 2504726 | 2504779 | **RNA**→ rpmE (PRK00019)Ribosomal_L31 (pfam01197)→ |
| Yps-11-1 | NZ_CP009780.1 | + | 4050379 | 4050432 | **RNA**→ rpmE (PRK00019)Ribosomal_L31 (pfam01197)→ |
| Ype-2-2 | NZ_CP009844.1 | - | 4507031 | 4506978 | **RNA**→ rpmE (PRK00019)Ribosomal_L31 (pfam01197)→ |
| Ype-32-1 | NZ_CP009906.1 | + | 3397336 | 3397389 | **RNA**→ rpmE (PRK00019)Ribosomal_L31 (pfam01197)→ |
| Ype-31-1 | NZ_CP009935.1 | - | 536663 | 536610 | **RNA**→ rpmE (PRK00019)Ribosomal_L31 (pfam01197)→ hypo→ RAYT (COG1943)→ |
| Yps-13-1 | NZ_CP010067.1 | + | 4077636 | 4077689 | **RNA**→ rpmE (PRK00019)Ribosomal_L31 (pfam01197)→ |
| Ype-2-3 | NZ_CP010293.1 | + | 484831 | 484884 | **RNA**→ rpmE (PRK00019)Ribosomal_L31 (pfam01197)→ |
| Yps-1-1 | NZ_CP009757.1 | + | 3889992 | 3890045 | **RNA**→ rpmE (PRK00019)Ribosomal_L31 (pfam01197)→ |

| | | | | | | |
|---|---|---|---|---|---|---|
| Ype-2-4 | NZ_CP009704.1 | + | 3323809 | 3323862 | RNA→ | rpmE (PRK00019)Ribosomal_L31 (pfam01197)→ |
| Yps-10-1 | NZ_CP009712.1 | - | 2780604 | 2780551 | RNA→ | rpmE (PRK00019)Ribosomal_L31 (pfam01197)→ |
| Ype-2-5 | NZ_CP009785.1 | - | 886875 | 886822 | RNA→ | rpmE (PRK00019)Ribosomal_L31 (pfam01197)→ |
| Ype-113-1 | NZ_CP010023.1 | + | 4500934 | 4500987 | RNA→ | rpmE (PRK00019)Ribosomal_L31 (pfam01197)→ |
| Ype-2-6 | NZ_CP009723.1 | - | 3814528 | 3814475 | RNA→ | rpmE (PRK00019)Ribosomal_L31 (pfam01197)→ |
| Yps-1-2 | NZ_CP009786.1 | + | 4234015 | 4234068 | RNA→ | rpmE (PRK00019)Ribosomal_L31 (pfam01197)→ |
| Ype-2-7 | NZ_CP009991.1 | - | 1952990 | 1952937 | RNA→ | rpmE (PRK00019)Ribosomal_L31 (pfam01197)→ |
| Yps-12-1 | NZ_CP009792.1 | - | 490330 | 490277 | RNA→ | rpmE (PRK00019)Ribosomal_L31 (pfam01197)→ |
| Ype-29-1 | NZ_CP009840.1 | + | 2556450 | 2556503 | RNA→ | rpmE (PRK00019)Ribosomal_L31 (pfam01197)→ |
| Ype-2-8 | NZ_CP009996.1 | - | 323257 | 323204 | RNA→ | rpmE (PRK00019)Ribosomal_L31 (pfam01197)→ |
| Ype-95-1 | NZ_CP010247.1 | - | 579933 | 579880 | RNA→ | rpmE (PRK00019)Ribosomal_L31 (pfam01197)→ |
| Ype-94-1 | NZ_CP009715.1 | + | 758495 | 758548 | RNA→ | rpmE (PRK00019)Ribosomal_L31 (pfam01197)→ |
| Ype-222-1 | NC_017265.1 | + | 253058 | 253111 | RNA→ | rpmE (PRK00019)Ribosomal_L31 (pfam01197)→ |
| Ype-223-1 | NC_005810.1 | + | 120657 | 120710 | RNA→ | rpmE (PRK00019)Ribosomal_L31 (pfam01197)→ |
| Ype-2-9 | NZ_CP009836.1 | - | 1128920 | 1128867 | RNA→ | rpmE (PRK00019)Ribosomal_L31 (pfam01197)→ |
| Ype-36-1 | NZ_ADQE01000355.1 | + | 62 | 115 | RNA→ | rpmE (PRK00019)Ribosomal_L31 (pfam01197)→ |
| Ype-99-1 | NZ_ADTD01000838.1 | - | 769 | 716 | RNA→ | rpmE (PRK00019)Ribosomal_L31 (pfam01197)→ |
| Yps-9-1 | NZ_CBKR010000111.1 | - | 7080 | 7027 | RNA→ | rpmE (PRK00019)Ribosomal_L31 (pfam01197)→ |
| Yps-8-1 | NZ_CBKS010000081.1 | + | 71103 | 71156 | RNA→ | rpmE (PRK00019)Ribosomal_L31 (pfam01197)→ |
| Ype-116-1 | NZ_AXDG01000093.1 | + | 12398 | 12451 | RNA→ | rpmE (PRK00019)Ribosomal_L31 (pfam01197)→ |
| Ype-157-1 | NZ_AKRN01000061.1 | + | 12430 | 12483 | RNA→ | rpmE (PRK00019)Ribosomal_L31 (pfam01197)→ |
| Yal-2-1 | NZ_CP009781.1 | - | 2568066 | 2568013 | RNA→ | rpmE (PRK00019)Ribosomal_L31 (pfam01197)→ |
| Ynu-1-1 | NZ_CPYD01000015.1 | + | 93996 | 94049 | RNA→ | rpmE (PRK00019)Ribosomal_L31 (pfam01197)→ |
| Yru-1-1 | NZ_CP009539.1 | - | 2020716 | 2020663 | RNA→ | rpmE (PRK00019)Ribosomal_L31 (pfam01197)→ |
| Dze-2-1 | NZ_APMV01000013.1 | + | 179089 | 179142 | RNA→ | rpmE (PRK00019)Ribosomal_L31 (pfam01197)→ |
| Dsp-1-1 | NZ_JSYG01000017.1 | + | 181093 | 181146 | RNA→ | rpmE (PRK00019)Ribosomal_L31 (pfam01197)→ |
| Ssp-23-1 | NZ_AWXH01000002.1 | - | 14589 | 14536 | RNA→ | rpmE (PRK00019)Ribosomal_L31 (pfam01197)→ |
| env-647 | HBC_scaff_10311262 | - | 789 | 736 | RNA→ | hypo→ |
| env-648 | HBC_scaff_10330317 | + | 27 | 80 | RNA→ | metL (PRK09466)Homoserine_dh (pfam00742)→ |
| env-649 | HBC_ctgs_1043907 | - | 789 | 736 | RNA→ | hypo→ |
| env-650 | HBC_ctgs_1006921 | + | 27 | 80 | RNA→ | metL (PRK09466)Homoserine_dh (pfam00742)→ |
| Lqu-1-1 | NZ_JIBP01000021.1 | + | 129696 | 129749 | RNA→ | rpmE (PRK00019)Ribosomal_L31 (pfam01197)→ |
| Lqu-1-2 | NZ_JIBO01000012.1 | - | 15246 | 15193 | RNA→ | rpmE (PRK00019)Ribosomal_L31 (pfam01197)→ |
| Lqu-1-3 | NZ_JIBQ01000013.1 | + | 129634 | 129687 | RNA→ | rpmE (PRK00019)Ribosomal_L31 (pfam01197)→ |
| Dsp-2-1 | NZ_JXBN01000033.1 | + | 48158 | 48211 | RNA→ | rpmE (PRK00019)Ribosomal_L31 (pfam01197)→ |
| Dch-1-1 | NZ_JRWY01000048.1 | + | 61153 | 61206 | RNA→ | rpmE (PRK00019)Ribosomal_L31 (pfam01197)→ |
| Dsp-6-1 | NZ_JXBO01000013.1 | - | 15193 | 15140 | RNA→ | rpmE (PRK00019)Ribosomal_L31 (pfam01197)→ |
| Dso-1-1 | NZ_CP009460.1 | + | 3086245 | 3086298 | RNA→ | rpmE (PRK00019)Ribosomal_L31 (pfam01197)→ |
| Dsp-5-1 | NZ_CM001979.1 | + | 218165 | 218218 | RNA→ | rpmE (PRK00019)Ribosomal_L31 (pfam01197)→ |
| Dsp-4-1 | NZ_CM001984.1 | + | 228039 | 228092 | RNA→ | rpmE (PRK00019)Ribosomal_L31 (pfam01197)→ |
| Dze-7-1 | NZ_APWM01000030.1 | + | 26292 | 26345 | RNA→ | rpmE (PRK00019)Ribosomal_L31 (pfam01197)→ |
| Dze-9-1 | NZ_AJVN01000010.1 | + | 188349 | 188402 | RNA→ | rpmE (PRK00019)Ribosomal_L31 (pfam01197)→ |
| Dda-5-1 | NC_013592.1 | - | 4501180 | 4501127 | RNA→ | rpmE (PRK00019)Ribosomal_L31 (pfam01197)→ |
| Dze-8-1 | NZ_CM001858.1 | - | 4245021 | 4244968 | RNA→ | rpmE (PRK00019)Ribosomal_L31 (pfam01197)→ |
| Dze-5-1 | NZ_CM001977.1 | - | 4240558 | 4240505 | RNA→ | rpmE (PRK00019)Ribosomal_L31 (pfam01197)→ |
| Dze-1-1 | NZ_CM001972.1 | - | 4399809 | 4399756 | RNA→ | rpmE (PRK00019)Ribosomal_L31 (pfam01197)→ |
| Dze-6-1 | NZ_CM001980.1 | - | 4317617 | 4317564 | RNA→ | rpmE (PRK00019)Ribosomal_L31 (pfam01197)→ |
| Dze-4-1 | NZ_CM001985.1 | - | 4353927 | 4353874 | RNA→ | rpmE (PRK00019)Ribosomal_L31 (pfam01197)→ |
| Dze-3-1 | NZ_CP006929.1 | - | 4276325 | 4276272 | RNA→ | rpmE (PRK00019)Ribosomal_L31 (pfam01197)→ |
| Dch-3-1 | NZ_CM001974.1 | + | 156157 | 156210 | RNA→ | rpmE (PRK00019)Ribosomal_L31 (pfam01197)→ |
| Dch-2-1 | NZ_CM001981.1 | + | 138095 | 138148 | RNA→ | rpmE (PRK00019)Ribosomal_L31 (pfam01197)→ |
| Dda-1-1 | NZ_CM001982.1 | + | 212261 | 212314 | RNA→ | rpmE (PRK00019)Ribosomal_L31 (pfam01197)→ |

| | | | | | |
|---|---|---|---|---|---|
| Dda-3-1 | NZ_CM001978.1 | + | 204493 | 204546 | **RNA**→ rpmE (PRK00019)Ribosomal_L31 (pfam01197)→ |
| Dda-4-1 | NC_014500.1 | + | 216804 | 216857 | **RNA**→ rpmE (PRK00019)Ribosomal_L31 (pfam01197)→ |
| Dda-2-1 | NZ_CM001976.1 | + | 246468 | 246521 | **RNA**→ rpmE (PRK00019)Ribosomal_L31 (pfam01197)→ |
| Dch-1-2 | NZ_JSYH01000045.1 | + | 149402 | 149455 | **RNA**→ rpmE (PRK00019)Ribosomal_L31 (pfam01197)→ |
| Dch-4-1 | NZ_CM001904.1 | + | 133855 | 133908 | **RNA**→ rpmE (PRK00019)Ribosomal_L31 (pfam01197)→ |
| Dsp-9-1 | NZ_CM001975.1 | + | 190423 | 190476 | **RNA**→ rpmE (PRK00019)Ribosomal_L31 (pfam01197)→ |
| Xdo-1-1 | NZ_FO704550.1 | + | 206252 | 206305 | **RNA**→ rpmE (PRK00019)Ribosomal_L31 (pfam01197)→ |
| Xpo-1-1 | NZ_FO704551.1 | - | 3528869 | 3528816 | **RNA**→ rpmE (PRK00019)Ribosomal_L31 (pfam01197)→ |
| Xbo-11-1 | NC_013892.1 | - | 4104240 | 4104187 | **RNA**→ rpmE (PRK00019)Ribosomal_L31 (pfam01197)→ rpmE (PRK00019)Ribosomal_L31 (pfam01197)→ |
| Xbo-3-1 | NZ_HG428366.1 | - | 23364 | 23311 | **RNA**→ rpmE (PRK00019)Ribosomal_L31 (pfam01197)→ |
| Xbo-8-1 | NZ_HG427595.1 | + | 21398 | 21451 | **RNA**→ rpmE (PRK00019)Ribosomal_L31 (pfam01197)→ |
| Xbo-2-1 | NZ_HG428558.1 | + | 25827 | 25880 | **RNA**→ rpmE (PRK00019)Ribosomal_L31 (pfam01197)→ |
| Xbo-6-1 | NZ_HG427126.1 | - | 7746 | 7693 | **RNA**→ rpmE (PRK00019)Ribosomal_L31 (pfam01197)→ |
| Xbo-5-1 | NZ_HG426603.1 | - | 23994 | 23941 | **RNA**→ rpmE (PRK00019)Ribosomal_L31 (pfam01197)→ |
| Xbo-10-1 | NZ_HG779933.1 | - | 23971 | 23918 | **RNA**→ rpmE (PRK00019)Ribosomal_L31 (pfam01197)→ |
| Xbo-4-1 | NZ_HG426761.1 | - | 24006 | 23953 | **RNA**→ rpmE (PRK00019)Ribosomal_L31 (pfam01197)→ |
| Xbo-7-1 | NZ_HG428161.1 | - | 23817 | 23764 | **RNA**→ rpmE (PRK00019)Ribosomal_L31 (pfam01197)→ |
| Xbo-9-1 | NZ_HG427430.1 | + | 93568 | 93621 | **RNA**→ rpmE (PRK00019)Ribosomal_L31 (pfam01197)→ |
| Xbo-1-1 | NZ_FO818637.1 | - | 4500575 | 4500522 | **RNA**→ rpmE (PRK00019)Ribosomal_L31 (pfam01197)→ |
| Xsz-1-1 | NZ_CBXF010000099.1 | + | 173156 | 173209 | **RNA**→ rpmE (PRK00019)Ribosomal_L31 (pfam01197)→ |
| Xca-1-1 | NZ_CBXE010000012.1 | - | 24443 | 24390 | **RNA**→ rpmE (PRK00019)Ribosomal_L31 (pfam01197)→ |
| Xkh-1-1 | NZ_LFCV01000027.1 | - | 4729 | 4676 | **RNA**→ ←hypo |
| Xsp-1-1 | NZ_JTHK01000060.1 | - | 23322 | 23269 | **RNA**→ rpmE (PRK00019)Ribosomal_L31 (pfam01197)→ |
| Xne-5-1 | NZ_CCWM01000116.1 | + | 16700 | 16753 | **RNA**→ rpmE (PRK00019)Ribosomal_L31 (pfam01197)→ |
| Xne-5-2 | NZ_CCWM01000334.1 | + | 16700 | 16753 | **RNA**→ rpmE (PRK00019)Ribosomal_L31 (pfam01197)→ |
| Xne-6-1 | NZ_CCWW01000414.1 | - | 7580 | 7527 | **RNA**→ rpmE (PRK00019)Ribosomal_L31 (pfam01197)→ |
| Xne-3-1 | NC_014228.1 | + | 198616 | 198669 | **RNA**→ rpmE (PRK00019)Ribosomal_L31 (pfam01197)→ |
| Xne-4-1 | NZ_HF952048.1 | + | 184374 | 184427 | **RNA**→ rpmE (PRK00019)Ribosomal_L31 (pfam01197)→ |
| Xne-2-1 | NZ_LN681227.1 | + | 198617 | 198670 | **RNA**→ rpmE (PRK00019)Ribosomal_L31 (pfam01197)→ |
| Xne-1-1 | NZ_JRJV01000010.1 | - | 42870 | 42817 | **RNA**→ rpmE (PRK00019)Ribosomal_L31 (pfam01197)→ |
| Plu-5-1 | NZ_JQOC01000024.1 | - | 35411 | 35358 | **RNA**→ rpmE (PRK00019)Ribosomal_L31 (pfam01197)→ |
| Plu-9-1 | NZ_JXSK01000009.1 | - | 70770 | 70717 | **RNA**→ rpmE (PRK00019)Ribosomal_L31 (pfam01197)→ |
| Plu-7-1 | NZ_AZAB02000203.1 | - | 35364 | 35311 | **RNA**→ rpmE (PRK00019)Ribosomal_L31 (pfam01197)→ |
| Plu-8-1 | NZ_JTHJ02000138.1 | + | 22410 | 22463 | **RNA**→ rpmE (PRK00019)Ribosomal_L31 (pfam01197)→ |
| Plu-5-2 | NZ_JXUR02000066.1 | + | 22410 | 22463 | **RNA**→ rpmE (PRK00019)Ribosomal_L31 (pfam01197)→ |
| Plu-6-1 | NZ_JFGV01000042.1 | - | 14184 | 14131 | **RNA**→ rpmE (PRK00019)Ribosomal_L31 (pfam01197)→ |
| Pte-4-1 | NZ_CP011104.1 | - | 1726439 | 1726386 | **RNA**→ rpmE (PRK00019)Ribosomal_L31 (pfam01197)→ |
| Pte-6-1 | NZ_AUXQ01000046.1 | + | 22142 | 22195 | **RNA**→ hypo→ rpmE (PRK00019)Ribosomal_L31 (pfam01197)→ |
| Pte-5-1 | NZ_AXDT01000086.1 | + | 18583 | 18636 | **RNA**→ rpmE (PRK00019)Ribosomal_L31 (pfam01197)→ |
| Pte-3-1 | NZ_JGVH01000076.1 | - | 10895 | 10842 | **RNA**→ rpmE (PRK00019)Ribosomal_L31 (pfam01197)→ |
| Plu-10-1 | NC_005126.1 | - | 5539908 | 5539855 | **RNA**→ rpmE (PRK00019)Ribosomal_L31 (pfam01197)→ |
| Pte-2-1 | NZ_AYSJ01000007.1 | + | 43299 | 43352 | **RNA**→ rpmE (PRK00019)Ribosomal_L31 (pfam01197)→ |
| Ssp-15-1 | NZ_AYKS02000100.1 | - | 30017 | 29965 | **RNA**→ rpmE (PRK00019)Ribosomal_L31 (pfam01197)→ |
| Pca-2-1 | NZ_JRMH01000002.1 | - | 333539 | 333487 | **RNA**→ rpmE (PRK00019)Ribosomal_L31 (pfam01197)→ |
| Pca-1-1 | NZ_JSXC01000044.1 | - | 17660 | 17608 | **RNA**→ rpmE (PRK00019)Ribosomal_L31 (pfam01197)→ |
| Ddi-5-1 | NZ_CM001841.1 | + | 187089 | 187141 | **RNA**→ rpmE (PRK00019)Ribosomal_L31 (pfam01197)→ |
| Ddi-3-1 | NZ_CM001838.1 | + | 162116 | 162168 | **RNA**→ rpmE (PRK00019)Ribosomal_L31 (pfam01197)→ |
| Ddi-4-1 | NZ_CM001840.1 | + | 170048 | 170100 | **RNA**→ rpmE (PRK00019)Ribosomal_L31 (pfam01197)→ |
| Dsp-7-1 | NZ_CM001973.1 | + | 162116 | 162168 | **RNA**→ rpmE (PRK00019)Ribosomal_L31 (pfam01197)→ |
| Ddi-1-1 | NZ_CM002023.1 | + | 192156 | 192208 | **RNA**→ rpmE (PRK00019)Ribosomal_L31 (pfam01197)→ |
| Ddi-2-1 | NZ_KQ046817.1 | - | 4180925 | 4180873 | **RNA**→ rpmE (PRK00019)Ribosomal_L31 (pfam01197)→ |
| Ssp-13-1 | NZ_JQEI01000010.1 | + | 16683 | 16735 | **RNA**→ rpmE (PRK00019)Ribosomal_L31 (pfam01197)→ |

| | | | | | |
|---|---|---|---|---|---|
| Ssp-14-1 | NZ_JQEJ01000064.1 | + | 16684 | 16736 | **RNA**→ rpmE (PRK00019)Ribosomal_L31 (pfam01197)→ |
| Dpa-1-1 | NZ_CM001857.1 | - | 4364511 | 4364459 | **RNA**→ rpmE (PRK00019)Ribosomal_L31 (pfam01197)→ |
| Sfo-1-1 | NZ_CP011254.1 | - | 1590388 | 1590336 | **RNA**→ rpmE (PRK00019)Ribosomal_L31 (pfam01197)→ |
| Sfo-2-1 | NZ_AVAH01000002.1 | - | 32669 | 32617 | **RNA**→ rpmE (PRK00019)Ribosomal_L31 (pfam01197)→ |
| Sfo-5-1 | NZ_ASZB01000023.1 | + | 61525 | 61577 | **RNA**→ rpmE (PRK00019)Ribosomal_L31 (pfam01197)→ |
| Sfo-4-1 | NZ_ASZA01000012.1 | - | 28374 | 28322 | **RNA**→ rpmE (PRK00019)Ribosomal_L31 (pfam01197)→ |
| Sfo-3-1 | NZ_AUZV01000007.1 | + | 146615 | 146667 | **RNA**→ rpmE (PRK00019)Ribosomal_L31 (pfam01197)→ |
| Pca-7-1 | NZ_AODU01000021.1 | - | 23353 | 23301 | **RNA**→ rpmE (PRK00019)Ribosomal_L31 (pfam01197)→ |
| Ppa-2-1 | NZ_JENG01000020.1 | + | 33412 | 33464 | **RNA**→ rpmE (PRK00019)Ribosomal_L31 (pfam01197)→ |
| Pwa-3-1 | NC_013421.1 | + | 198446 | 198498 | **RNA**→ rpmE (PRK00019)Ribosomal_L31 (pfam01197)→ |
| Ppa-1-1 | NZ_KN151108.1 | - | 181617 | 181565 | **RNA**→ rpmE (PRK00019)Ribosomal_L31 (pfam01197)→ |
| Pat-3-1 | NZ_KB933471.1 | + | 739413 | 739465 | **RNA**→ rpmE (PRK00019)Ribosomal_L31 (pfam01197)→ |
| Pat-4-1 | NC_004547.2 | - | 4770742 | 4770690 | **RNA**→ rpmE (PRK00019)Ribosomal_L31 (pfam01197)→ |
| Pat-2-1 | NZ_CP007744.1 | - | 4721893 | 4721841 | **RNA**→ rpmE (PRK00019)Ribosomal_L31 (pfam01197)→ |
| Pat-2-2 | NZ_CP009125.1 | - | 4676226 | 4676174 | **RNA**→ rpmE (PRK00019)Ribosomal_L31 (pfam01197)→ |
| Psp-55-1 | NC_017845.1 | + | 210989 | 211041 | **RNA**→ rpmE (PRK00019)Ribosomal_L31 (pfam01197)→ |
| Pwa-2-1 | NZ_AKVS01000042.1 | + | 29133 | 29185 | **RNA**→ rpmE (PRK00019)Ribosomal_L31 (pfam01197)→ |
| Pwa-1-1 | NZ_JQHP01000018.1 | - | 24420 | 24368 | **RNA**→ rpmE (PRK00019)Ribosomal_L31 (pfam01197)→ |
| Pwa-1-2 | NZ_JQOH01000017.1 | + | 30041 | 30093 | **RNA**→ rpmE (PRK00019)Ribosomal_L31 (pfam01197)→ |
| Pca-8-1 | NZ_JQOF01000004.1 | + | 276707 | 276759 | **RNA**→ rpmE (PRK00019)Ribosomal_L31 (pfam01197)→ |
| Pca-8-2 | NZ_JQOG01000006.1 | - | 24482 | 24430 | **RNA**→ rpmE (PRK00019)Ribosomal_L31 (pfam01197)→ |
| Bgo-1-1 | NZ_CGIG01000001.1 | - | 1386796 | 1386744 | **RNA**→ rpmE (PRK00019)Ribosomal_L31 (pfam01197)→ |
| Bsp-1-1 | NZ_CM001230.1 | - | 4603649 | 4603597 | **RNA**→ rpmE (PRK00019)Ribosomal_L31 (pfam01197)→ rpmE (PRK00019)Ribosomal_L31 (pfam01197)→ rpmE (PRK00019)Ribosomal_L31 (pfam01197)→ |
| Dsp-3-1 | NZ_CM001983.1 | + | 176356 | 176408 | **RNA**→ rpmE (PRK00019)Ribosomal_L31 (pfam01197)→ |
| Pbe-1-1 | NZ_JQHM01000017.1 | + | 44048 | 44100 | **RNA**→ rpmE (PRK00019)Ribosomal_L31 (pfam01197)→ |
| Pbe-1-2 | NZ_JQHL01000014.1 | - | 18376 | 18324 | **RNA**→ rpmE (PRK00019)Ribosomal_L31 (pfam01197)→ |
| Pca-5-11 | NZ_JUJI01000017.1 | - | 23832 | 23780 | **RNA**→ rpmE (PRK00019)Ribosomal_L31 (pfam01197)→ |
| Ppo-1-1 | NZ_JQHN01000020.1 | + | 52227 | 52279 | **RNA**→ rpmE (PRK00019)Ribosomal_L31 (pfam01197)→ |
| env-651 | JGI26054J46599_1244098 | + | 95 | 147 | **RNA**→ ←hypo |
| env-652 | SPBFG_contig14881 | - | 539 | 487 | **RNA**→ rpmE (PRK00019)Ribosomal_L31 (pfam01197)→ |
| Sma-27-1 | NZ_ALOV01000001.1 | + | 94626 | 94678 | **RNA**→ rpmE (PRK00019)Ribosomal_L31 (pfam01197)→ |
| Sma-23-1 | NZ_AVSR01000009.1 | - | 25295 | 25243 | **RNA**→ rpmE (PRK00019)Ribosomal_L31 (pfam01197)→ |
| Sma-20-1 | NZ_JVCS01000035.1 | - | 25362 | 25310 | **RNA**→ rpmE (PRK00019)Ribosomal_L31 (pfam01197)→ |
| Sso-1-1 | NZ_JUSZ01000089.1 | + | 94274 | 94326 | **RNA**→ rpmE (PRK00019)Ribosomal_L31 (pfam01197)→ |
| Sma-9-1 | NZ_AYMO01000018.1 | - | 25741 | 25689 | **RNA**→ rpmE (PRK00019)Ribosomal_L31 (pfam01197)→ |
| Sma-28-1 | NC_020211.1 | - | 5122946 | 5122894 | **RNA**→ rpmE (PRK00019)Ribosomal_L31 (pfam01197)→ rpmE (PRK00019)Ribosomal_L31 (pfam01197)→ |
| Sma-7-1 | NZ_KK214336.1 | + | 24122 | 24174 | **RNA**→ rpmE (PRK00019)Ribosomal_L31 (pfam01197)→ rpmE (PRK00019)Ribosomal_L31 (pfam01197)→ |
| Sma-5-1 | NZ_KI929263.1 | - | 2697435 | 2697383 | **RNA**→ rpmE (PRK00019)Ribosomal_L31 (pfam01197)→ rpmE (PRK00019)Ribosomal_L31 (pfam01197)→ |
| Sma-19-1 | NZ_AP013063.1 | + | 4816609 | 4816661 | **RNA**→ rpmE (PRK00019)Ribosomal_L31 (pfam01197)→ rpmE (PRK00019)Ribosomal_L31 (pfam01197)→ |
| Sma-18-1 | NZ_AYMT01000027.1 | - | 25712 | 25660 | **RNA**→ rpmE (PRK00019)Ribosomal_L31 (pfam01197)→ |
| Sma-3-2 | NZ_JTBR01000001.1 | + | 3163415 | 3163467 | **RNA**→ rpmE (PRK00019)Ribosomal_L31 (pfam01197)→ |
| env-653 | AEQT01000902.1 | - | 25741 | 25689 | **RNA**→ rpmE (PRK00019)Ribosomal_L31 (pfam01197)→ |
| env-654 | APTF_contig00062 | + | 84678 | 84730 | **RNA**→ rpmE (PRK00019)Ribosomal_L31 (pfam01197)→ |
| Sur-1-1 | NZ_JSFB01000001.1 | - | 3972705 | 3972653 | **RNA**→ rpmE (PRK00019)Ribosomal_L31 (pfam01197)→ |
| Ssp-20-1 | NZ_LDEG01000010.1 | - | 25885 | 25833 | **RNA**→ rpmE (PRK00019)Ribosomal_L31 (pfam01197)→ |
| Sma-3-3 | NZ_JVEJ01000246.1 | - | 25311 | 25259 | **RNA**→ rpmE (PRK00019)Ribosomal_L31 (pfam01197)→ |
| Ssp-12-1 | NZ_JVEL01000457.1 | - | 8014 | 7962 | **RNA**→ rpmE (PRK00019)Ribosomal_L31 (pfam01197)→ rpmE (PRK00019)Ribosomal_L31 (pfam01197)→ |
| Sma-21-1 | NZ_JMPQ01000029.1 | + | 94339 | 94391 | **RNA**→ rpmE (PRK00019)Ribosomal_L31 (pfam01197)→ |
| Sne-1-1 | NZ_JPUX01000001.1 | - | 29493 | 29441 | **RNA**→ rpmE (PRK00019)Ribosomal_L31 (pfam01197)→ rpmE (PRK00019)Ribosomal_L31 (pfam01197)→ |
| Ssp-19-1 | NZ_CP003424.1 | - | 4914868 | 4914816 | **RNA**→ rpmE (PRK00019)Ribosomal_L31 (pfam01197)→ rpmE (PRK00019)Ribosomal_L31 (pfam01197)→ |
| Ssp-16-1 | NZ_CP005927.1 | - | 849700 | 849648 | **RNA**→ rpmE (PRK00019)Ribosomal_L31 (pfam01197)→ rpmE (PRK00019)Ribosomal_L31 (pfam01197)→ |

| | | | | | | |
|---|---|---|---|---|---|---|
| Sma-8-1 | NZ_KK214298.1 | - | 25945 | 25893 | **RNA**→ | rpmE (PRK00019)Ribosomal_L31 (pfam01197)→ rpmE (PRK00019)Ribosomal_L31 (pfam01197)→ |
| Sma-6-1 | NZ_KI929261.1 | + | 420108 | 420160 | **RNA**→ | rpmE (PRK00019)Ribosomal_L31 (pfam01197)→ rpmE (PRK00019)Ribosomal_L31 (pfam01197)→ |
| Sma-26-1 | NZ_AORJ01000046.1 | - | 25636 | 25584 | **RNA**→ | rpmE (PRK00019)Ribosomal_L31 (pfam01197)→ |
| Sma-3-4 | NZ_JTBU01000001.1 | + | 2106852 | 2106904 | **RNA**→ | rpmE (PRK00019)Ribosomal_L31 (pfam01197)→ |
| env-655 | F76GVBA01EWCUB | + | 406 | 458 | **RNA**→ | hypo→ |
| env-656 | CLOF_contig22543 | + | 3049 | 3101 | **RNA**→ | |
| Sma-20-2 | NZ_LFJS01000008.1 | - | 181443 | 181391 | **RNA**→ | rpmE (PRK00019)Ribosomal_L31 (pfam01197)→ |
| Sbo-2-1 | NZ_JWAE01000021.1 | - | 25579 | 25527 | **RNA**→ | rpmE (PRK00019)Ribosomal_L31 (pfam01197)→ |
| Sbo-2-2 | NZ_JWBK01000133.1 | + | 17268 | 17320 | **RNA**→ | rpmE (PRK00019)Ribosomal_L31 (pfam01197)→ |
| Sbo-2-3 | NZ_JVWX01000015.1 | - | 25565 | 25513 | **RNA**→ | rpmE (PRK00019)Ribosomal_L31 (pfam01197)→ |
| Sma-22-1 | NZ_HG326223.1 | - | 4286664 | 4286612 | **RNA**→ | rpmE (PRK00019)Ribosomal_L31 (pfam01197)→ rpmE (PRK00019)Ribosomal_L31 (pfam01197)→ |
| Spl-5-1 | NZ_AJTB01000062.1 | + | 34460 | 34512 | **RNA**→ | rpmE (PRK00019)Ribosomal_L31 (pfam01197)→ rpmE (PRK00019)Ribosomal_L31 (pfam01197)→ |
| Ssy-1-1 | NZ_CCES01000004.1 | - | 27914 | 27862 | **RNA**→ | rpmE (PRK00019)Ribosomal_L31 (pfam01197)→ |
| Ssy-2-1 | NZ_GL636109.1 | - | 16479 | 16427 | **RNA**→ | rpmE (PRK00019)Ribosomal_L31 (pfam01197)→ rpmE (PRK00019)Ribosomal_L31 (pfam01197)→ |
| Sgr-1-1 | NZ_JGVP01000045.1 | - | 1623 | 1571 | **RNA**→ | rpmE (PRK00019)Ribosomal_L31 (pfam01197)→ |
| Sod-1-1 | NZ_GG753567.1 | - | 1210885 | 1210833 | **RNA**→ | rpmE (PRK00019)Ribosomal_L31 (pfam01197)→ rpmE (PRK00019)Ribosomal_L31 (pfam01197)→ |
| Spl-1-1 | NZ_ARWD01000001.1 | - | 5361904 | 5361852 | **RNA**→ | rpmE (PRK00019)Ribosomal_L31 (pfam01197)→ |
| Spl-2-1 | NC_021591.1 | - | 5175142 | 5175090 | **RNA**→ | rpmE (PRK00019)Ribosomal_L31 (pfam01197)→ |
| Spl-6-1 | NC_021659.1 | - | 5314607 | 5314555 | **RNA**→ | rpmE (PRK00019)Ribosomal_L31 (pfam01197)→ |
| Spl-3-1 | NZ_AMSV01000076.1 | - | 19552 | 19500 | **RNA**→ | rpmE (PRK00019)Ribosomal_L31 (pfam01197)→ |
| Spr-2-1 | NC_009832.1 | - | 5307340 | 5307288 | **RNA**→ | rpmE (PRK00019)Ribosomal_L31 (pfam01197)→ |
| Ssp-25-1 | NZ_KB661120.1 | - | 5154241 | 5154189 | **RNA**→ | rpmE (PRK00019)Ribosomal_L31 (pfam01197)→ |
| env-657 | JGI24735J21928_10403343 | - | 113 | 61 | **RNA**→ | |
| env-658 | DPOL_contig07438 | - | 12704 | 12652 | **RNA**→ | rpmE (PRK00019)Ribosomal_L31 (pfam01197)→ |
| env-659 | SWWA_contig25429 | - | 3135 | 3083 | **RNA**→ | ←PRK05264 (PRK05264)MetJ (COG3060) |
| Sli-3-1 | NC_021741.1 | - | 5069699 | 5069647 | **RNA**→ | rpmE (PRK00019)Ribosomal_L31 (pfam01197)→ |
| Sli-1-1 | NZ_CP011303.1 | + | 5203476 | 5203528 | **RNA**→ | rpmE (PRK00019)Ribosomal_L31 (pfam01197)→ |
| Sli-2-1 | NZ_DF820433.1 | + | 97289 | 97341 | **RNA**→ | rpmE (PRK00019)Ribosomal_L31 (pfam01197)→ |
| Sma-16-1 | NZ_ATOH01000018.1 | + | 95385 | 95437 | **RNA**→ | rpmE (PRK00019)Ribosomal_L31 (pfam01197)→ |
| Sma-13-1 | NZ_ATOK01000032.1 | + | 95385 | 95437 | **RNA**→ | rpmE (PRK00019)Ribosomal_L31 (pfam01197)→ |
| Sma-12-1 | NZ_ATOL01000045.1 | - | 25725 | 25673 | **RNA**→ | rpmE (PRK00019)Ribosomal_L31 (pfam01197)→ rpmE (PRK00019)Ribosomal_L31 (pfam01197)→ |
| Sma-4-1 | NZ_ATOM01000026.1 | - | 25727 | 25675 | **RNA**→ | rpmE (PRK00019)Ribosomal_L31 (pfam01197)→ rpmE (PRK00019)Ribosomal_L31 (pfam01197)→ |
| Sma-17-1 | NZ_ATOG01000076.1 | - | 25572 | 25520 | **RNA**→ | rpmE (PRK00019)Ribosomal_L31 (pfam01197)→ |
| Sma-15-1 | NZ_KE375711.1 | + | 95619 | 95671 | **RNA**→ | rpmE (PRK00019)Ribosomal_L31 (pfam01197)→ |
| Sma-14-1 | NZ_KE375758.1 | + | 95401 | 95453 | **RNA**→ | rpmE (PRK00019)Ribosomal_L31 (pfam01197)→ rpmE (PRK00019)Ribosomal_L31 (pfam01197)→ |
| Ssp-21-1 | NC_015566.1 | - | 5290302 | 5290250 | **RNA**→ | rpmE (PRK00019)Ribosomal_L31 (pfam01197)→ |
| Ssp-22-1 | NC_017573.1 | - | 5289850 | 5289798 | **RNA**→ | rpmE (PRK00019)Ribosomal_L31 (pfam01197)→ |
| Spl-4-1 | NC_015567.1 | - | 5290173 | 5290121 | **RNA**→ | rpmE (PRK00019)Ribosomal_L31 (pfam01197)→ |
| Sma-24-1 | NC_020064.1 | - | 4697421 | 4697369 | **RNA**→ | rpmE (PRK00019)Ribosomal_L31 (pfam01197)→ rpmE (PRK00019)Ribosomal_L31 (pfam01197)→ |
| Sma-3-1 | NZ_CP011642.1 | + | 2708667 | 2708719 | **RNA**→ | rpmE (PRK00019)Ribosomal_L31 (pfam01197)→ |
| Pca-3-1 | NZ_JPSO01000025.1 | - | 24470 | 24418 | **RNA**→ | rpmE (PRK00019)Ribosomal_L31 (pfam01197)→ |
| Pca-6-1 | NZ_AODT01000022.1 | - | 23586 | 23534 | **RNA**→ | rpmE (PRK00019)Ribosomal_L31 (pfam01197)→ |
| Pca-5-1 | NZ_JQHJ01000008.1 | + | 136473 | 136525 | **RNA**→ | rpmE (PRK00019)Ribosomal_L31 (pfam01197)→ |
| Pca-5-2 | NZ_JUJR01000004.1 | - | 26044 | 25992 | **RNA**→ | rpmE (PRK00019)Ribosomal_L31 (pfam01197)→ |
| Pca-5-3 | NZ_JUJG01000011.1 | - | 26099 | 26047 | **RNA**→ | rpmE (PRK00019)Ribosomal_L31 (pfam01197)→ |
| Pca-5-4 | NZ_JUJS01000003.1 | - | 792139 | 792087 | **RNA**→ | rpmE (PRK00019)Ribosomal_L31 (pfam01197)→ |
| Pca-10-1 | NC_012917.1 | + | 199661 | 199713 | **RNA**→ | rpmE (PRK00019)Ribosomal_L31 (pfam01197)→ |
| Pca-3-2 | NZ_JPSM01000001.1 | - | 24461 | 24409 | **RNA**→ | rpmE (PRK00019)Ribosomal_L31 (pfam01197)→ |
| Pca-3-3 | NZ_JUJJ01000011.1 | - | 24104 | 24052 | **RNA**→ | rpmE (PRK00019)Ribosomal_L31 (pfam01197)→ |
| Pca-5-5 | NZ_JUJT01000020.1 | - | 24009 | 23957 | **RNA**→ | rpmE (PRK00019)Ribosomal_L31 (pfam01197)→ |
| Pca-3-4 | NZ_JUJE01000005.1 | - | 28337 | 28285 | **RNA**→ | rpmE (PRK00019)Ribosomal_L31 (pfam01197)→ |
| Pca-3-5 | NZ_JUJL01000007.1 | - | 24410 | 24358 | **RNA**→ | rpmE (PRK00019)Ribosomal_L31 (pfam01197)→ |

| | | | | | |
|---|---|---|---|---|---|
| Pca-12-1 | NZ_ABVY01000037.1 | - | 14362 | 14310 | **RNA**→ rpmE (PRK00019)Ribosomal_L31 (pfam01197)→ rpmE (PRK00019)Ribosomal_L31 (pfam01197)→ |
| Cmu-2-1 | NZ_CP007044.2 | + | 1090545 | 1090597 | **RNA**→ rpmE (PRK00019)Ribosomal_L31 (pfam01197)→ |
| Pca-5-6 | NZ_JUJO01000019.1 | - | 24319 | 24267 | **RNA**→ rpmE (PRK00019)Ribosomal_L31 (pfam01197)→ |
| Pca-5-7 | NZ_JUJQ01000006.1 | - | 26072 | 26020 | **RNA**→ rpmE (PRK00019)Ribosomal_L31 (pfam01197)→ |
| Pca-5-8 | NZ_JUJP01000007.1 | - | 25212 | 25160 | **RNA**→ rpmE (PRK00019)Ribosomal_L31 (pfam01197)→ |
| Pca-5-9 | NZ_JUJF01000020.1 | - | 24130 | 24078 | **RNA**→ rpmE (PRK00019)Ribosomal_L31 (pfam01197)→ |
| Pca-3-6 | NZ_JUJD01000003.1 | - | 24283 | 24231 | **RNA**→ rpmE (PRK00019)Ribosomal_L31 (pfam01197)→ |
| Pca-3-7 | NZ_JUJB01000021.1 | - | 24944 | 24892 | **RNA**→ rpmE (PRK00019)Ribosomal_L31 (pfam01197)→ |
| Pca-5-10 | NZ_JUJH01000022.1 | - | 26713 | 26661 | **RNA**→ rpmE (PRK00019)Ribosomal_L31 (pfam01197)→ |
| Pca-3-8 | NZ_JUJC01000010.1 | - | 23496 | 23444 | **RNA**→ rpmE (PRK00019)Ribosomal_L31 (pfam01197)→ |
| Pca-9-1 | NZ_ABVX01000016.1 | - | 18047 | 17995 | **RNA**→ rpmE (PRK00019)Ribosomal_L31 (pfam01197)→ rpmE (PRK00019)Ribosomal_L31 (pfam01197)→ |
| Pca-3-9 | NZ_JUJK01000006.1 | - | 1101 | 1049 | **RNA**→ rpmE (PRK00019)Ribosomal_L31 (pfam01197)→ |
| Pca-3-10 | NZ_JUJN01000013.1 | - | 25803 | 25751 | **RNA**→ rpmE (PRK00019)Ribosomal_L31 (pfam01197)→ |
| Pca-3-11 | NZ_JUJM01000006.1 | - | 23692 | 23640 | **RNA**→ rpmE (PRK00019)Ribosomal_L31 (pfam01197)→ |
| Pca-3-12 | NZ_LGRF01000008.1 | + | 135274 | 135326 | **RNA**→ rpmE (PRK00019)Ribosomal_L31 (pfam01197)→ |
| Pca-4-1 | NZ_ALIU01000001.1 | - | 18504 | 18452 | **RNA**→ rpmE (PRK00019)Ribosomal_L31 (pfam01197)→ |
| Pca-11-1 | NC_018525.1 | - | 4586906 | 4586854 | **RNA**→ rpmE (PRK00019)Ribosomal_L31 (pfam01197)→ |
| Pca-3-13 | NZ_JPSN01000031.1 | - | 107877 | 107825 | **RNA**→ rpmE (PRK00019)Ribosomal_L31 (pfam01197)→ |
| Pca-3-14 | NZ_JQOE01000006.1 | + | 275251 | 275303 | **RNA**→ rpmE (PRK00019)Ribosomal_L31 (pfam01197)→ |
| Pca-3-15 | NZ_JQOD01000007.1 | - | 18268 | 18216 | **RNA**→ rpmE (PRK00019)Ribosomal_L31 (pfam01197)→ |
| Ssp-17-1 | NZ_AYMN01000039.1 | + | 151302 | 151354 | **RNA**→ rpmE (PRK00019)Ribosomal_L31 (pfam01197)→ |
| Ssp-18-1 | NZ_AYMQ01000065.1 | + | 114568 | 114620 | **RNA**→ rpmE (PRK00019)Ribosomal_L31 (pfam01197)→ |
| Pas-2-1 | NC_012962.1 | - | 4916832 | 4916778 | **RNA**→ rpmE (PRK00019)Ribosomal_L31 (pfam01197)→ rpmE (PRK00019)Ribosomal_L31 (pfam01197)→ |
| Pas-1-1 | NZ_JONO01000042.1 | + | 23154 | 23208 | **RNA**→ rpmE (PRK00019)Ribosomal_L31 (pfam01197)→ |
| Xgr-1-1 | NZ_LDNM01000030.1 | - | 22309 | 22254 | **RNA**→ rpmE (PRK00019)Ribosomal_L31 (pfam01197)→ |
| Hpa-1-1 | NZ_JWGZ01000009.1 | + | 170462 | 170514 | **RNA**→ rpmE (PRK00019)Ribosomal_L31 (pfam01197)→ |
| Hal-1-1 | NZ_JTET01000010.1 | - | 22434 | 22382 | **RNA**→ rpmE (PRK00019)Ribosomal_L31 (pfam01197)→ |
| Hal-1-2 | NZ_JTEU02000022.1 | + | 113408 | 113460 | **RNA**→ rpmE (PRK00019)Ribosomal_L31 (pfam01197)→ |
| Hal-1-3 | NZ_LAZF01000026.1 | + | 11854 | 11906 | **RNA**→ rpmE (PRK00019)Ribosomal_L31 (pfam01197)→ |
| Hal-1-4 | NZ_LAZG01000008.1 | - | 21781 | 21729 | **RNA**→ rpmE (PRK00019)Ribosomal_L31 (pfam01197)→ |
| Hal-2-1 | NZ_JMPK01000018.1 | + | 58626 | 58678 | **RNA**→ rpmE (PRK00019)Ribosomal_L31 (pfam01197)→ |
| env-660 | DPOL_F64H56M02IXVBN | + | 28 | 80 | **RNA**→ rpmE (PRK00019)Ribosomal_L31 (pfam01197)→ |
| Hpa-1-2 | NZ_JWBZ01000059.1 | - | 434 | 382 | **RNA**→ rpmE (PRK00019)Ribosomal_L31 (pfam01197)→ |
| Hal-4-1 | NZ_JH417535.1 | + | 157332 | 157384 | **RNA**→ rpmE (PRK00019)Ribosomal_L31 (pfam01197)→ |
| Eba-3-1 | NZ_ADCU02000001.1 | + | 4449192 | 4449244 | **RNA**→ rpmE (PRK00019)Ribosomal_L31 (pfam01197)→ |
| Hal-3-1 | NZ_CP009706.1 | + | 299355 | 299407 | **RNA**→ rpmE (PRK00019)Ribosomal_L31 (pfam01197)→ |
| env-661 | SPBFG_contig15870 | - | 5793 | 5739 | **RNA**→ rpmE (PRK00019)Ribosomal_L31 (pfam01197)→ |
| gpr-1-1 | NZ_AMYV01000018.1 | + | 32852 | 32905 | **RNA**→ rpmE (PRK00019)Ribosomal_L31 (pfam01197)→ |
| Ssp-24-1 | NZ_AJHJ01000020.1 | + | 52512 | 52565 | **RNA**→ rpmE (PRK00019)Ribosomal_L31 (pfam01197)→ rpmE (PRK00019)Ribosomal_L31 (pfam01197)→ |
| Rch-1-1 | NZ_JRWU01000009.1 | - | 26174 | 26121 | **RNA**→ rpmE (PRK00019)Ribosomal_L31 (pfam01197)→ rpmE (PRK00019)Ribosomal_L31 (pfam01197)→ |
| Eam-2-1 | NZ_JMPJ01000068.1 | + | 95292 | 95345 | **RNA**→ rpmE (PRK00019)Ribosomal_L31 (pfam01197)→ |
| env-662 | DPOF_contig09155 | - | 2266 | 2213 | **RNA**→ rpmE (PRK00019)Ribosomal_L31 (pfam01197)→ |
| env-663 | DPOLF_contig09092 | - | 6580 | 6527 | **RNA**→ rpmE (PRK00019)Ribosomal_L31 (pfam01197)→ |
| env-664 | DPOL_contig00038 | + | 36878 | 36931 | **RNA**→ rpmE (PRK00019)Ribosomal_L31 (pfam01197)→ |
| env-665 | SPBFG_GA1E3IJ02I46OL | + | 306 | 359 | **RNA**→ |
| env-666 | FGTW_contig30418 | - | 31336 | 31283 | **RNA**→ rpmE (PRK00019)Ribosomal_L31 (pfam01197)→ |
| env-667 | FGTFG_contig02103 | + | 85330 | 85383 | **RNA**→ rpmE (PRK00019)Ribosomal_L31 (pfam01197)→ |
| env-668 | JGI25162J39368_1000075 | - | 25553 | 25500 | **RNA**→ ←hypo |
| env-669 | JGI25163J39215_1000213 | - | 4721 | 4668 | **RNA**→ ←hypo |
| env-670 | JGI25164J39214_1000053 | - | 25553 | 25500 | **RNA**→ ←hypo |
| env-671 | SwRhRL2b_contig_56033 | + | 517 | 570 | **RNA**→ rpmE (PRK00019)Ribosomal_L31 (pfam01197)→ |
| Raq-3-1 | NC_017047.1 | - | 4814745 | 4814692 | **RNA**→ rpmE (PRK00019)Ribosomal_L31 (pfam01197)→ rpmE (PRK00019)Ribosomal_L31 (pfam01197)→ |

| | | | | | |
|---|---|---|---|---|---|
| Rsp-4-1 | NC_015061.1 | - | 4716764 | 4716711 | **RNA**→ rpmE (PRK00019)Ribosomal_L31 (pfam01197)→ rpmE (PRK00019)Ribosomal_L31 (pfam01197)→ |
| Rsp-3-1 | NZ_JQJR01000015.1 | - | 25545 | 25492 | **RNA**→ rpmE (PRK00019)Ribosomal_L31 (pfam01197)→ rpmE (PRK00019)Ribosomal_L31 (pfam01197)→ |
| Raq-1-1 | NZ_JUHL01000031.1 | + | 18582 | 18635 | **RNA**→ rpmE (PRK00019)Ribosomal_L31 (pfam01197)→ |
| Raq-1-2 | NZ_JUHM01000047.1 | - | 5015 | 4962 | **RNA**→ rpmE (PRK00019)Ribosomal_L31 (pfam01197)→ |
| Raq-2-2 | NC_016818.1 | - | 4714466 | 4714413 | **RNA**→ rpmE (PRK00019)Ribosomal_L31 (pfam01197)→ |
| Raq-2-1 | NZ_JMPO01000146.1 | + | 95132 | 95185 | **RNA**→ rpmE (PRK00019)Ribosomal_L31 (pfam01197)→ |
| env-672 | XylAfBL_contig05016 | + | 910 | 963 | **RNA**→ rpmE (PRK00019)Ribosomal_L31 (pfam01197)→ |
| env-673 | XAGC_contig03916 | + | 8510 | 8563 | **RNA**→ ←hypo |
| env-674 | XylAfBA_contig26731 | + | 2773 | 2826 | **RNA**→ rpmE (PRK00019)Ribosomal_L31 (pfam01197)→ |
| Mmo-1-1 | NZ_LGYC01000001.1 | - | 31789 | 31738 | **RNA**→ rpmE (PRK00019)Ribosomal_L31 (pfam01197)→ |
| Mmo-1-2 | NZ_JSWU01000006.1 | + | 171759 | 171810 | **RNA**→ rpmE (PRK00019)Ribosomal_L31 (pfam01197)→ |
| Mmo-3-1 | NZ_AYMP01000005.1 | - | 32378 | 32327 | **RNA**→ rpmE (PRK00019)Ribosomal_L31 (pfam01197)→ |
| Mmo-2-1 | NZ_CCFB01000001.1 | + | 198262 | 198313 | **RNA**→ rpmE (PRK00019)Ribosomal_L31 (pfam01197)→ |
| Mmo-1-3 | NZ_LAGC01000010.1 | - | 143210 | 143159 | **RNA**→ rpmE (PRK00019)Ribosomal_L31 (pfam01197)→ |
| Mmo-1-4 | NZ_JUYZ01000051.1 | - | 52099 | 52048 | **RNA**→ rpmE (PRK00019)Ribosomal_L31 (pfam01197)→ |
| Mmo-6-1 | NC_020418.1 | + | 647912 | 647963 | **RNA**→ rpmE (PRK00019)Ribosomal_L31 (pfam01197)→ |
| Mmo-4-1 | NZ_CBWE010000048.1 | + | 118052 | 118103 | **RNA**→ rpmE (PRK00019)Ribosomal_L31 (pfam01197)→ |
| Msp-5-1 | NZ_AZRH01000004.1 | - | 103144 | 103093 | **RNA**→ rpmE (PRK00019)Ribosomal_L31 (pfam01197)→ |
| Mmo-5-1 | NZ_AMWL02000012.1 | - | 105828 | 105777 | **RNA**→ rpmE (PRK00019)Ribosomal_L31 (pfam01197)→ rpmE (PRK00019)Ribosomal_L31 (pfam01197)→ |
| Mmo-1-5 | NZ_JQGP01000003.1 | - | 77455 | 77404 | **RNA**→ rpmE (PRK00019)Ribosomal_L31 (pfam01197)→ |
| Mmo-1-6 | NZ_JRKW01000009.1 | + | 165680 | 165731 | **RNA**→ rpmE (PRK00019)Ribosomal_L31 (pfam01197)→ |
| Mmo-1-8 | NZ_JTBT01000001.1 | + | 1238015 | 1238066 | **RNA**→ rpmE (PRK00019)Ribosomal_L31 (pfam01197)→ |
| Mmo-1-7 | NZ_JUVX01000049.1 | + | 20761 | 20813 | **RNA**→ rpmE (PRK00019)Ribosomal_L31 (pfam01197)→ |
| Mwi-1-1 | NZ_LDOF01000037.1 | - | 16081 | 16031 | **RNA**→ rpmE (PRK00019)Ribosomal_L31 (pfam01197)→ |
| Psn-1-1 | NZ_CM001773.1 | - | 286368 | 286317 | **RNA**→ rpmE (PRK00019)Ribosomal_L31 (pfam01197)→ |
| Pbu-1-1 | NZ_KB233226.1 | - | 31250 | 31199 | **RNA**→ rpmE (PRK00019)Ribosomal_L31 (pfam01197)→ |
| Pal-5-1 | NZ_AZYZ01000023.1 | + | 170468 | 170519 | **RNA**→ rpmE (PRK00019)Ribosomal_L31 (pfam01197)→ |
| Pal-3-1 | NZ_JALA01000018.1 | - | 30690 | 30639 | **RNA**→ rpmE (PRK00019)Ribosomal_L31 (pfam01197)→ |
| Pre-3-1 | NZ_CM001774.1 | + | 548141 | 548192 | **RNA**→ rpmE (PRK00019)Ribosomal_L31 (pfam01197)→ |
| Pre-1-1 | NZ_CVLT01000011.1 | + | 8877 | 8928 | **RNA**→ rpmE (PRK00019)Ribosomal_L31 (pfam01197)→ |
| Pre-1-2 | NZ_JSEQ01000006.1 | - | 32695 | 32644 | **RNA**→ rpmE (PRK00019)Ribosomal_L31 (pfam01197)→ |
| Pre-2-1 | NZ_GG705271.1 | + | 112866 | 112917 | **RNA**→ rpmE (PRK00019)Ribosomal_L31 (pfam01197)→ |
| Pru-5-1 | NZ_GG703823.1 | - | 35067 | 35016 | **RNA**→ rpmE (PRK00019)Ribosomal_L31 (pfam01197)→ |
| Pal-8-1 | NZ_ABXW01000070.1 | - | 30334 | 30283 | **RNA**→ rpmE (PRK00019)Ribosomal_L31 (pfam01197)→ rpmE (PRK00019)Ribosomal_L31 (pfam01197)→ |
| Pal-9-1 | NZ_KB233235.1 | - | 30888 | 30837 | **RNA**→ rpmE (PRK00019)Ribosomal_L31 (pfam01197)→ |
| Pal-2-1 | NZ_JACS01000017.1 | - | 31209 | 31158 | **RNA**→ rpmE (PRK00019)Ribosomal_L31 (pfam01197)→ |
| Pal-4-1 | NZ_JAKZ01000057.1 | + | 50253 | 50304 | **RNA**→ rpmE (PRK00019)Ribosomal_L31 (pfam01197)→ |
| Pal-7-1 | NZ_JALB01000006.1 | - | 30356 | 30305 | **RNA**→ rpmE (PRK00019)Ribosomal_L31 (pfam01197)→ |
| Pal-6-1 | NZ_JALC01000028.1 | + | 173522 | 173573 | **RNA**→ rpmE (PRK00019)Ribosomal_L31 (pfam01197)→ |
| Pal-1-1 | NZ_JALD01000049.1 | + | 172466 | 172517 | **RNA**→ rpmE (PRK00019)Ribosomal_L31 (pfam01197)→ |
| Eba-4-1 | NZ_AMFN01000021.1 | + | 169629 | 169682 | **RNA**→ rpmE (PRK00019)Ribosomal_L31 (pfam01197)→ |
| Msp-4-1 | NZ_JJMI01000052.1 | + | 104126 | 104179 | **RNA**→ rpmE (PRK00019)Ribosomal_L31 (pfam01197)→ |
| env-675 | HBC_scaff_10000440 | + | 5214 | 5266 | **RNA**→ ←hypo |
| env-676 | HBC_ctgs_1005247 | + | 2966 | 3018 | **RNA**→ ←hypo |
| env-677 | CLOF_contig24321 | - | 704 | 652 | **RNA**→ rpmE (PRK00019)Ribosomal_L31 (pfam01197)→ |
| env-678 | ACEF_contig00049 | - | 523 | 472 | **RNA**→ rpmE (PRK00019)Ribosomal_L31 (pfam01197)→ |
| Cmu-1-1 | NZ_AJKU01000051.1 | - | 3469 | 3418 | **RNA**→ rpmE (PRK00019)Ribosomal_L31 (pfam01197)→ |
| Csa-13-1 | NC_023032.1 | + | 149771 | 149822 | **RNA**→ rpmE (PRK00019)Ribosomal_L31 (pfam01197)→ |
| Csa-10-1 | NZ_CALF01000071.1 | + | 4097945 | 4097996 | **RNA**→ rpmE (PRK00019)Ribosomal_L31 (pfam01197)→ |
| Csa-3-1 | NZ_AWSP01000023.1 | + | 208400 | 208451 | **RNA**→ rpmE (PRK00019)Ribosomal_L31 (pfam01197)→ |
| Csa-11-1 | NZ_CALE01000616.1 | + | 4090594 | 4090645 | **RNA**→ rpmE (PRK00019)Ribosomal_L31 (pfam01197)→ hypo→ |
| Csa-6-1 | NZ_AJLB01000065.1 | - | 2039 | 1988 | **RNA**→ rpmE (PRK00019)Ribosomal_L31 (pfam01197)→ |

| | | | | | |
|---|---|---|---|---|---|
| Csa-4-1 | NZ_AJLA01000020.1 | - | 36481 | 36430 | **RNA**→ rpmE (PRK00019)Ribosomal_L31 (pfam01197)→ |
| Csa-2-1 | NZ_AJKT01000018.1 | + | 66704 | 66755 | **RNA**→ rpmE (PRK00019)Ribosomal_L31 (pfam01197)→ |
| Cdu-3-1 | NZ_AJKY01000079.1 | + | 62855 | 62906 | **RNA**→ rpmE (PRK00019)Ribosomal_L31 (pfam01197)→ |
| Cdu-1-1 | NZ_AJKZ01000025.1 | - | 32358 | 32307 | **RNA**→ rpmE (PRK00019)Ribosomal_L31 (pfam01197)→ |
| Cdu-2-1 | NZ_AJKX01000093.1 | - | 35818 | 35767 | **RNA**→ rpmE (PRK00019)Ribosomal_L31 (pfam01197)→ |
| Csa-8-1 | NZ_AWFW01000041.1 | - | 32602 | 32551 | **RNA**→ rpmE (PRK00019)Ribosomal_L31 (pfam01197)→ |
| Cdu-4-1 | NZ_CAKZ01000164.1 | + | 29188 | 29239 | **RNA**→ rpmE (PRK00019)Ribosomal_L31 (pfam01197)→ |
| Ctu-3-1 | NZ_CALB01000097.1 | + | 3916369 | 3916420 | **RNA**→ rpmE (PRK00019)Ribosomal_L31 (pfam01197)→ |
| Csa-14-1 | NZ_AFMO01000075.1 | - | 17965 | 17914 | **RNA**→ rpmE (PRK00019)Ribosomal_L31 (pfam01197)→ |
| Cdu-5-1 | NZ_CALA01000393.1 | + | 4235981 | 4236032 | **RNA**→ rpmE (PRK00019)Ribosomal_L31 (pfam01197)→ |
| Cco-1-1 | NZ_CAKW01000043.1 | + | 51829 | 51880 | **RNA**→ rpmE (PRK00019)Ribosomal_L31 (pfam01197)→ |
| Ctu-4-1 | NC_013282.2 | + | 183302 | 183353 | **RNA**→ rpmE (PRK00019)Ribosomal_L31 (pfam01197)→ |
| Csa-16-1 | NC_020260.1 | - | 3781421 | 3781370 | **RNA**→ rpmE (PRK00019)Ribosomal_L31 (pfam01197)→ |
| Cun-1-1 | NZ_AJKW01000018.1 | - | 32659 | 32608 | **RNA**→ rpmE (PRK00019)Ribosomal_L31 (pfam01197)→ |
| Cun-1-2 | NZ_CAKX01000221.1 | + | 3809092 | 3809143 | **RNA**→ rpmE (PRK00019)Ribosomal_L31 (pfam01197)→ |
| Csa-7-1 | NZ_BAWU01000020.1 | + | 2500 | 2551 | **RNA**→ rpmE (PRK00019)Ribosomal_L31 (pfam01197)→ |
| Csa-5-1 | NZ_AJLC01000180.1 | + | 62760 | 62811 | **RNA**→ rpmE (PRK00019)Ribosomal_L31 (pfam01197)→ |
| Cma-2-1 | NZ_JXTD01000005.1 | + | 64019 | 64070 | **RNA**→ rpmE (PRK00019)Ribosomal_L31 (pfam01197)→ |
| Cma-3-1 | NZ_AJKV01000030.1 | + | 59963 | 60014 | **RNA**→ rpmE (PRK00019)Ribosomal_L31 (pfam01197)→ |
| Cma-1-1 | NZ_JPZE01000012.1 | + | 76282 | 76333 | **RNA**→ rpmE (PRK00019)Ribosomal_L31 (pfam01197)→ |
| Cma-1-2 | NZ_JPZG01000177.1 | - | 6258 | 6207 | **RNA**→ rpmE (PRK00019)Ribosomal_L31 (pfam01197)→ |
| Cma-1-3 | NZ_JPZH01000130.1 | - | 44040 | 43989 | **RNA**→ rpmE (PRK00019)Ribosomal_L31 (pfam01197)→ |
| Cma-4-1 | NZ_CALD01000014.1 | + | 4023113 | 4023164 | **RNA**→ rpmE (PRK00019)Ribosomal_L31 (pfam01197)→ |
| Csa-9-1 | NZ_CALG01000065.1 | + | 30206 | 30257 | **RNA**→ rpmE (PRK00019)Ribosomal_L31 (pfam01197)→ |
| Cma-1-4 | NZ_JPZD01000152.1 | + | 5905 | 5956 | **RNA**→ rpmE (PRK00019)Ribosomal_L31 (pfam01197)→ |
| Cma-1-5 | NZ_JPZF01000101.1 | - | 6338 | 6287 | **RNA**→ rpmE (PRK00019)Ribosomal_L31 (pfam01197)→ |
| Cma-5-1 | NZ_CALC01000087.1 | - | 3657473 | 3657422 | **RNA**→ rpmE (PRK00019)Ribosomal_L31 (pfam01197)→ |
| Csa-12-1 | NC_009778.1 | - | 3776784 | 3776733 | **RNA**→ rpmE (PRK00019)Ribosomal_L31 (pfam01197)→ |
| Csa-15-1 | NC_017933.1 | - | 4083623 | 4083572 | **RNA**→ rpmE (PRK00019)Ribosomal_L31 (pfam01197)→ |
| Csa-1-1 | NZ_CP011047.1 | - | 827969 | 827918 | **RNA**→ rpmE (PRK00019)Ribosomal_L31 (pfam01197)→ |
| env-679 | FGTW_contig20460 | - | 365 | 314 | **RNA**→ rpmE (PRK00019)Ribosomal_L31 (pfam01197)→ |
| Tod-1-1 | NZ_JNGH01000037.1 | - | 69319 | 69268 | **RNA**→ rpmE (PRK00019)Ribosomal_L31 (pfam01197)→ |
| Tod-1-2 | NZ_JNGI01000158.1 | - | 23681 | 23630 | **RNA**→ rpmE (PRK00019)Ribosomal_L31 (pfam01197)→ |
| env-680 | 2227358547 | + | 86186 | 86237 | **RNA**→ rpmE (PRK00019)Ribosomal_L31 (pfam01197)→ |
| env-681 | TrFG_GO06Z1101E2O72 | + | 74 | 125 | **RNA**→ rpmE (PRK00019)Ribosomal_L31 (pfam01197)→ |
| env-682 | ACOFG987_contig24115 | + | 393 | 444 | **RNA**→ rpmE (PRK00019)rpmE2 (PRK01678)Ribosomal_L31 (pfam01197)→ |
| env-683 | APTF_contig08800 | + | 1648 | 1699 | **RNA**→ rpmE (PRK00019)Ribosomal_L31 (pfam01197)→ |
| env-684 | ACOFG987_contig23961 | - | 394 | 343 | **RNA**→ rpmE (PRK00019)Ribosomal_L31 (pfam01197)→ |
| Eba-5-1 | NC_020063.1 | - | 4635348 | 4635297 | **RNA**→ rpmE (PRK00019)Ribosomal_L31 (pfam01197)→ |
| env-685 | FGTW_GHRKC7402FUCRI | - | 60 | 9 | **RNA**→ |
| Tgu-1-1 | NZ_JMTB01000106.1 | + | 15805 | 15856 | **RNA**→ rpmE (PRK00019)Ribosomal_L31 (pfam01197)→ |
| env-686 | JGI26055J44792_1031982 | - | 374 | 323 | **RNA**→ rpmE (PRK00019)Ribosomal_L31 (pfam01197)→ |
| env-687 | TrFG_contig00049 | - | 5566 | 5515 | **RNA**→ rpmE (PRK00019)Ribosomal_L31 (pfam01197)→ |
| env-688 | ACOFG987_contig40025 | + | 6347 | 6398 | **RNA**→ rpmE (PRK00019)Ribosomal_L31 (pfam01197)→ |
| env-689 | ACOFG988_contig25298 | - | 1504 | 1453 | **RNA**→ rpmE (PRK00019)Ribosomal_L31 (pfam01197)→ |
| env-690 | scaffold610_9_V1.UC-21 | + | 568 | 619 | **RNA**→ rpmE (PRK00019)Ribosomal_L31 (pfam01197)→ |
| env-691 | DOF008_scaffold12770_1 | - | 3518 | 3467 | **RNA**→ rpmE (PRK00019)Ribosomal_L31 (pfam01197)→ |
| env-692 | PaFGMunAill_c011219 | + | 225 | 276 | **RNA**→ rpmE (PRK00019)Ribosomal_L31 (pfam01197)→ |
| env-693 | JGI26057J44791_1008390 | - | 1105 | 1054 | **RNA**→ rpmE (PRK00019)Ribosomal_L31 (pfam01197)→ |
| env-694 | APTF_contig04656 | - | 272 | 221 | **RNA**→ rpmE (PRK00019)Ribosomal_L31 (pfam01197)→ |
| env-695 | ACEF_contig16949 | + | 52822 | 52873 | **RNA**→ rpmE (PRK00019)Ribosomal_L31 (pfam01197)→ |
| env-696 | ACOFG987_contig47692 | + | 281 | 332 | **RNA**→ rpmE (PRK00019)Ribosomal_L31 (pfam01197)→ |

| | | | | | |
|---|---|---|---|---|---|
| env-697 | ACOFG988_contig32966 | + | 7759 | 7810 | **RNA**→ rpmE (PRK00019)Ribosomal_L31 (pfam01197)→ |
| env-698 | CLOF_contig00503 | + | 11843 | 11894 | **RNA**→ rpmE (PRK00019)Ribosomal_L31 (pfam01197)→ |
| env-699 | AECF_contig10395 | + | 1675 | 1726 | **RNA**→ rpmE (PRK00019)Ribosomal_L31 (pfam01197)→ |
| Esp-71-1 | NZ_LEDV01000005.1 | - | 25147 | 25096 | **RNA**→ rpmE (PRK00019)Ribosomal_L31 (pfam01197)→ |
| Eho-7-1 | NZ_ANIA01000010.1 | + | 18731 | 18782 | **RNA**→ rpmE (PRK00019)Ribosomal_L31 (pfam01197)→ |
| Ecl-67-1 | NZ_AMGJ01000018.1 | + | 18729 | 18780 | **RNA**→ rpmE (PRK00019)Ribosomal_L31 (pfam01197)→ |
| Ecl-9-1 | NZ_CP008823.1 | - | 4756273 | 4756222 | **RNA**→ rpmE (PRK00019)Ribosomal_L31 (pfam01197)→ |
| Ecl-15-1 | NZ_KK736253.1 | - | 4808834 | 4808783 | **RNA**→ rpmE (PRK00019)Ribosomal_L31 (pfam01197)→ |
| Ecl-61-1 | NZ_KI535482.1 | - | 1900916 | 1900865 | **RNA**→ rpmE (PRK00019)Ribosomal_L31 (pfam01197)→ |
| Ecl-64-1 | NZ_KI535506.1 | - | 4967519 | 4967468 | **RNA**→ rpmE (PRK00019)Ribosomal_L31 (pfam01197)→ |
| Esp-65-1 | NZ_KI535607.1 | - | 1051094 | 1051043 | **RNA**→ rpmE (PRK00019)Ribosomal_L31 (pfam01197)→ |
| Ecl-55-1 | NZ_KI535567.1 | - | 2547796 | 2547745 | **RNA**→ rpmE (PRK00019)Ribosomal_L31 (pfam01197)→ |
| Ecl-18-1 | NZ_KI973154.1 | + | 546623 | 546674 | **RNA**→ rpmE (PRK00019)Ribosomal_L31 (pfam01197)→ |
| Esp-4-1 | NZ_KI973172.1 | - | 1856201 | 1856150 | **RNA**→ rpmE (PRK00019)Ribosomal_L31 (pfam01197)→ |
| Ecl-21-1 | NZ_KI973143.1 | - | 25734 | 25683 | **RNA**→ rpmE (PRK00019)Ribosomal_L31 (pfam01197)→ |
| Ecl-22-1 | NZ_KI973132.1 | + | 2398583 | 2398634 | **RNA**→ rpmE (PRK00019)Ribosomal_L31 (pfam01197)→ |
| Esp-20-1 | NZ_KI973336.1 | - | 25242 | 25191 | **RNA**→ rpmE (PRK00019)Ribosomal_L31 (pfam01197)→ |
| Esp-31-1 | NZ_KI973278.1 | - | 109546 | 109495 | **RNA**→ rpmE (PRK00019)Ribosomal_L31 (pfam01197)→ |
| Esp-3-1 | NZ_KI973178.1 | - | 690446 | 690395 | **RNA**→ rpmE (PRK00019)Ribosomal_L31 (pfam01197)→ |
| Esp-14-1 | NZ_KI973197.1 | - | 1835108 | 1835057 | **RNA**→ rpmE (PRK00019)Ribosomal_L31 (pfam01197)→ |
| Esp-32-1 | NZ_KI973270.1 | - | 581090 | 581039 | **RNA**→ rpmE (PRK00019)Ribosomal_L31 (pfam01197)→ |
| Ecl-6-1 | NZ_KI973186.1 | - | 3623723 | 3623672 | **RNA**→ rpmE (PRK00019)Ribosomal_L31 (pfam01197)→ |
| Esp-27-1 | NZ_KI973212.1 | - | 103050 | 102999 | **RNA**→ rpmE (PRK00019)Ribosomal_L31 (pfam01197)→ |
| Ecl-23-1 | NZ_KI973128.1 | - | 95766 | 95715 | **RNA**→ rpmE (PRK00019)Ribosomal_L31 (pfam01197)→ |
| Esp-21-1 | NZ_KI973257.1 | - | 383611 | 383560 | **RNA**→ rpmE (PRK00019)Ribosomal_L31 (pfam01197)→ |
| Esp-26-1 | NZ_KI973320.1 | - | 27166 | 27115 | **RNA**→ rpmE (PRK00019)Ribosomal_L31 (pfam01197)→ |
| Esp-24-1 | NZ_KI973217.1 | - | 3681096 | 3681045 | **RNA**→ rpmE (PRK00019)Ribosomal_L31 (pfam01197)→ |
| Eho-6-1 | NZ_GL892086.1 | - | 148706 | 148655 | **RNA**→ rpmE (PRK00019)Ribosomal_L31 (pfam01197)→ |
| Eho-2-1 | NZ_CP010377.1 | - | 4491382 | 4491331 | **RNA**→ rpmE (PRK00019)Ribosomal_L31 (pfam01197)→ |
| Eho-3-1 | NZ_CP010384.1 | - | 4645020 | 4644969 | **RNA**→ rpmE (PRK00019)Ribosomal_L31 (pfam01197)→ |
| Ecl-4-1 | NZ_KK736218.1 | - | 27669 | 27618 | **RNA**→ rpmE (PRK00019)Ribosomal_L31 (pfam01197)→ |
| Ecl-3-1 | NZ_KN912611.1 | - | 181038 | 180987 | **RNA**→ rpmE (PRK00019)Ribosomal_L31 (pfam01197)→ |
| Eho-4-1 | NZ_CP010376.2 | - | 4745095 | 4745044 | **RNA**→ rpmE (PRK00019)Ribosomal_L31 (pfam01197)→ |
| Ecl-1-1 | NZ_CP011584.1 | - | 3888938 | 3888887 | **RNA**→ rpmE (PRK00019)Ribosomal_L31 (pfam01197)→ |
| Ecl-1-2 | NZ_CP011650.1 | - | 3861467 | 3861416 | **RNA**→ rpmE (PRK00019)Ribosomal_L31 (pfam01197)→ |
| Ecl-1-3 | NZ_CP011572.1 | + | 2928705 | 2928756 | **RNA**→ rpmE (PRK00019)Ribosomal_L31 (pfam01197)→ |
| Ecl-1-4 | NZ_CP011581.1 | + | 1042420 | 1042471 | **RNA**→ rpmE (PRK00019)Ribosomal_L31 (pfam01197)→ |
| Esp-12-1 | NZ_KQ089963.1 | - | 1910866 | 1910815 | **RNA**→ rpmE (PRK00019)Ribosomal_L31 (pfam01197)→ |
| Esp-35-1 | NZ_KQ089940.1 | + | 544460 | 544511 | **RNA**→ rpmE (PRK00019)Ribosomal_L31 (pfam01197)→ |
| Esp-34-1 | NZ_KQ089934.1 | - | 36906 | 36855 | **RNA**→ rpmE (PRK00019)Ribosomal_L31 (pfam01197)→ |
| Esp-9-1 | NZ_KQ089956.1 | - | 370660 | 370609 | **RNA**→ rpmE (PRK00019)Ribosomal_L31 (pfam01197)→ |
| Esp-10-1 | NZ_KQ089962.1 | + | 561134 | 561185 | **RNA**→ rpmE (PRK00019)Ribosomal_L31 (pfam01197)→ |
| Esp-18-1 | NZ_KQ089922.1 | + | 443512 | 443563 | **RNA**→ rpmE (PRK00019)Ribosomal_L31 (pfam01197)→ |
| Esp-37-1 | NZ_KQ089949.1 | + | 3599004 | 3599055 | **RNA**→ rpmE (PRK00019)Ribosomal_L31 (pfam01197)→ |
| Esp-16-1 | NZ_KQ089855.1 | - | 35241 | 35190 | **RNA**→ rpmE (PRK00019)Ribosomal_L31 (pfam01197)→ |
| Esp-8-1 | NZ_KQ090002.1 | - | 4088043 | 4087992 | **RNA**→ rpmE (PRK00019)Ribosomal_L31 (pfam01197)→ |
| Esp-17-1 | NZ_KQ089884.1 | - | 31776 | 31725 | **RNA**→ rpmE (PRK00019)Ribosomal_L31 (pfam01197)→ |
| Esp-11-1 | NZ_KQ089967.1 | + | 1947798 | 1947849 | **RNA**→ rpmE (PRK00019)Ribosomal_L31 (pfam01197)→ |
| Esp-7-1 | NZ_KQ089989.1 | + | 4391029 | 4391080 | **RNA**→ rpmE (PRK00019)Ribosomal_L31 (pfam01197)→ |
| Esp-39-1 | NZ_KQ089891.1 | - | 25347 | 25296 | **RNA**→ rpmE (PRK00019)Ribosomal_L31 (pfam01197)→ |
| Eho-4-2 | NZ_CP012167.1 | - | 4548238 | 4548187 | **RNA**→ rpmE (PRK00019)Ribosomal_L31 (pfam01197)→ |
| Eho-3-2 | NZ_CP012165.1 | - | 4790402 | 4790351 | **RNA**→ rpmE (PRK00019)Ribosomal_L31 (pfam01197)→ |

| | | | | | | |
|---|---|---|---|---|---|---|
| Eho-8-1 | NZ_ANID01000005.1 | + | 18734 | 18785 | **RNA**→ | rpmE (PRK00019)Ribosomal_L31 (pfam01197)→ |
| Esp-57-1 | NZ_AXLK01000002.1 | - | 662290 | 662239 | **RNA**→ | rpmE (PRK00019)Ribosomal_L31 (pfam01197)→ |
| Eca-2-1 | NZ_ANIC01000004.1 | - | 513413 | 513362 | **RNA**→ | rpmE (PRK00019)Ribosomal_L31 (pfam01197)→ |
| Ecl-1-9 | NZ_JTBO01000001.1 | + | 4527233 | 4527284 | **RNA**→ | rpmE (PRK00019)Ribosomal_L31 (pfam01197)→ |
| Ecl-1-10 | NZ_JTBN01000004.1 | + | 1394084 | 1394135 | **RNA**→ | rpmE (PRK00019)Ribosomal_L31 (pfam01197)→ |
| env-700 | FGTW_contig23348 | + | 302 | 353 | **RNA**→ | rpmE (PRK00019)Ribosomal_L31 (pfam01197)→ |
| env-701 | TrFG_GO06Z1101EGFU2 | - | 143 | 92 | **RNA**→ | rpmE (PRK00019)Ribosomal_L31 (pfam01197)→ |
| env-702 | JGI26532J50257_10015026 | + | 2407 | 2458 | **RNA**→ | rpmE (PRK00019)Ribosomal_L31 (pfam01197)→ |
| env-703 | JGI26540J51217_10040168 | + | 700 | 751 | **RNA**→ | rpmE (PRK00019)Ribosomal_L31 (pfam01197)→ |
| env-704 | FGTW_contig31546 | + | 3921 | 3972 | **RNA**→ | |
| env-705 | FGTW_contig33102 | + | 679 | 730 | **RNA**→ | rpmE (PRK00019)Ribosomal_L31 (pfam01197)→ |
| Esp-19-1 | NZ_JSZC01000008.1 | + | 81942 | 81993 | **RNA**→ | rpmE (PRK00019)Ribosomal_L31 (pfam01197)→ |
| Ecl-31-1 | NZ_JZZI01000030.1 | + | 11753 | 11804 | **RNA**→ | rpmE (PRK00019)Ribosomal_L31 (pfam01197)→ |
| Ecl-29-1 | NZ_LABD01000016.1 | + | 76834 | 76885 | **RNA**→ | rpmE (PRK00019)Ribosomal_L31 (pfam01197)→ |
| Ecl-38-1 | NZ_JZYC01000011.1 | + | 76772 | 76823 | **RNA**→ | rpmE (PRK00019)Ribosomal_L31 (pfam01197)→ |
| Ecl-32-1 | NZ_JZZF01000011.1 | + | 76674 | 76725 | **RNA**→ | rpmE (PRK00019)Ribosomal_L31 (pfam01197)→ |
| Ecl-30-1 | NZ_JZZX01000013.1 | + | 71383 | 71434 | **RNA**→ | rpmE (PRK00019)Ribosomal_L31 (pfam01197)→ |
| Ecl-34-1 | NZ_JZYX01000023.1 | + | 66329 | 66380 | **RNA**→ | rpmE (PRK00019)Ribosomal_L31 (pfam01197)→ |
| Ecl-35-1 | NZ_JZYR01000012.1 | + | 76673 | 76724 | **RNA**→ | rpmE (PRK00019)Ribosomal_L31 (pfam01197)→ |
| Ecl-37-1 | NZ_JZYJ01000013.1 | - | 34145 | 34094 | **RNA**→ | rpmE (PRK00019)Ribosomal_L31 (pfam01197)→ |
| Ecl-41-1 | NZ_JZKL01000011.1 | + | 77056 | 77107 | **RNA**→ | rpmE (PRK00019)Ribosomal_L31 (pfam01197)→ |
| Ecl-46-1 | NZ_LEET01000001.1 | + | 76857 | 76908 | **RNA**→ | rpmE (PRK00019)Ribosomal_L31 (pfam01197)→ |
| Ecl-51-1 | NZ_LECZ01000019.1 | - | 26537 | 26486 | **RNA**→ | rpmE (PRK00019)Ribosomal_L31 (pfam01197)→ |
| Ecl-48-1 | NZ_LEES01000015.1 | - | 26503 | 26452 | **RNA**→ | rpmE (PRK00019)Ribosomal_L31 (pfam01197)→ |
| Ecl-50-1 | NZ_LEDI01000013.1 | - | 8101 | 8050 | **RNA**→ | rpmE (PRK00019)Ribosomal_L31 (pfam01197)→ |
| Ecl-43-1 | NZ_LEEP01000004.1 | - | 34305 | 34254 | **RNA**→ | rpmE (PRK00019)Ribosomal_L31 (pfam01197)→ |
| Ecl-52-5 | NZ_LFDQ01000011.1 | + | 75190 | 75241 | **RNA**→ | rpmE (PRK00019)Ribosomal_L31 (pfam01197)→ |
| Lsp-1-1 | NZ_LDUO01000006.1 | + | 85683 | 85734 | **RNA**→ | rpmE (PRK00019)Ribosomal_L31 (pfam01197)→ |
| Lad-1-1 | NZ_LDWM01000006.1 | + | 85682 | 85733 | **RNA**→ | rpmE (PRK00019)Ribosomal_L31 (pfam01197)→ |
| Ecl-28-1 | NZ_JWGU01000218.1 | + | 34093 | 34144 | **RNA**→ | rpmE (PRK00019)Ribosomal_L31 (pfam01197)→ |
| Ecl-28-2 | NZ_JWGT01000199.1 | + | 34093 | 34144 | **RNA**→ | rpmE (PRK00019)Ribosomal_L31 (pfam01197)→ |
| Ecl-28-3 | NZ_JWAF01000124.1 | - | 26447 | 26396 | **RNA**→ | rpmE (PRK00019)Ribosomal_L31 (pfam01197)→ |
| Ecl-28-4 | NZ_JVMT01000150.1 | - | 26451 | 26400 | **RNA**→ | rpmE (PRK00019)Ribosomal_L31 (pfam01197)→ |
| Ecl-28-5 | NZ_JVND01000119.1 | + | 71438 | 71489 | **RNA**→ | rpmE (PRK00019)Ribosomal_L31 (pfam01197)→ |
| Esp-40-1 | NZ_JUMS01000116.1 | - | 34207 | 34156 | **RNA**→ | rpmE (PRK00019)Ribosomal_L31 (pfam01197)→ |
| Ecl-28-6 | NZ_JVWV01000106.1 | + | 11834 | 11885 | **RNA**→ | rpmE (PRK00019)Ribosomal_L31 (pfam01197)→ |
| Ecl-1-11 | NZ_JUOF01000010.1 | + | 42106 | 42157 | **RNA**→ | rpmE (PRK00019)Ribosomal_L31 (pfam01197)→ |
| Ecl-57-1 | NZ_ATHX01000018.1 | + | 76731 | 76782 | **RNA**→ | rpmE (PRK00019)Ribosomal_L31 (pfam01197)→ |
| Esp-1-1 | NZ_JFHW01000016.1 | - | 26463 | 26412 | **RNA**→ | rpmE (PRK00019)Ribosomal_L31 (pfam01197)→ |
| Ecl-14-1 | NZ_JDWG01000020.1 | - | 34120 | 34069 | **RNA**→ | rpmE (PRK00019)Ribosomal_L31 (pfam01197)→ |
| Esp-73-1 | NZ_ALNS01000021.1 | - | 26409 | 26358 | **RNA**→ | rpmE (PRK00019)Ribosomal_L31 (pfam01197)→ rpmE (PRK00019)Ribosomal_L31 (pfam01197)→ |
| Ecl-53-1 | NZ_JALR01000031.1 | + | 24192 | 24243 | **RNA**→ | rpmE (PRK00019)Ribosomal_L31 (pfam01197)→ |
| Esp-46-1 | NZ_AZUB01000011.1 | + | 64891 | 64942 | **RNA**→ | rpmE (PRK00019)Ribosomal_L31 (pfam01197)→ |
| Esp-44-1 | NZ_AZUA01000011.1 | + | 71526 | 71577 | **RNA**→ | rpmE (PRK00019)Ribosomal_L31 (pfam01197)→ |
| Eas-2-1 | NZ_JACW01000036.1 | - | 26054 | 26003 | **RNA**→ | rpmE (PRK00019)Ribosomal_L31 (pfam01197)→ |
| Esp-45-1 | NZ_AZXZ01000015.1 | + | 64891 | 64942 | **RNA**→ | rpmE (PRK00019)Ribosomal_L31 (pfam01197)→ |
| Ecl-8-1 | NZ_KK736240.1 | - | 3203895 | 3203844 | **RNA**→ | rpmE (PRK00019)Ribosomal_L31 (pfam01197)→ |
| Ecl-16-1 | NZ_KK736266.1 | - | 37488 | 37437 | **RNA**→ | rpmE (PRK00019)Ribosomal_L31 (pfam01197)→ |
| Esp-64-1 | NZ_KI535616.1 | - | 2057472 | 2057421 | **RNA**→ | rpmE (PRK00019)Ribosomal_L31 (pfam01197)→ |
| Ecl-62-1 | NZ_KI535461.1 | - | 2643968 | 2643917 | **RNA**→ | rpmE (PRK00019)Ribosomal_L31 (pfam01197)→ |
| Ecl-56-1 | NZ_KI535548.1 | - | 1326474 | 1326423 | **RNA**→ | rpmE (PRK00019)Ribosomal_L31 (pfam01197)→ |
| Esp-58-1 | NZ_KI535695.1 | - | 3774199 | 3774148 | **RNA**→ | rpmE (PRK00019)Ribosomal_L31 (pfam01197)→ |

| | | | | | |
|---|---|---|---|---|---|
| Ecl-25-1 | NZ_KI973093.1 | - | 738284 | 738233 | **RNA**→ rpmE (PRK00019)Ribosomal_L31 (pfam01197)→ |
| Ecl-24-1 | NZ_KI973119.1 | - | 3729528 | 3729477 | **RNA**→ rpmE (PRK00019)Ribosomal_L31 (pfam01197)→ |
| Eas-3-1 | NZ_CP007546.1 | + | 4028910 | 4028961 | **RNA**→ rpmE (PRK00019)Ribosomal_L31 (pfam01197)→ |
| Eas-1-1 | NZ_CP011591.1 | + | 3197317 | 3197368 | **RNA**→ rpmE (PRK00019)Ribosomal_L31 (pfam01197)→ |
| Esp-41-1 | NZ_KQ087598.1 | - | 4674371 | 4674320 | **RNA**→ rpmE (PRK00019)Ribosomal_L31 (pfam01197)→ |
| Ecl-36-1 | NZ_CP012162.1 | - | 4845841 | 4845790 | **RNA**→ rpmE (PRK00019)Ribosomal_L31 (pfam01197)→ |
| Ecl-13-1 | NZ_JDWH01000014.1 | - | 34223 | 34172 | **RNA**→ rpmE (PRK00019)Ribosomal_L31 (pfam01197)→ |
| Esp-50-1 | NZ_JRJC01000045.1 | + | 72599 | 72650 | **RNA**→ rpmE (PRK00019)Ribosomal_L31 (pfam01197)→ |
| Esp-69-1 | NZ_JRUR01000040.1 | + | 75694 | 75745 | **RNA**→ rpmE (PRK00019)Ribosomal_L31 (pfam01197)→ |
| Ksp-8-1 | NZ_ALNJ01000088.1 | - | 7833 | 7782 | **RNA**→ rpmE (PRK00019)Ribosomal_L31 (pfam01197)→ rpmE (PRK00019)Ribosomal_L31 (pfam01197)→ |
| Ecl-52-1 | NZ_LDCI01000017.1 | - | 26551 | 26500 | **RNA**→ rpmE (PRK00019)Ribosomal_L31 (pfam01197)→ |
| env-706 | bisonPool14jan08_C4751 | + | 678 | 729 | **RNA**→ |
| env-707 | bisonPool05dec07_BXBC29500_b1 | + | 542 | 593 | **RNA**→ ←hypo |
| env-708 | BISONP_C3973 | - | 751 | 700 | **RNA**→ rpmE (PRK00019)Ribosomal_L31 (pfam01197)→ |
| env-709 | DOM005_scaffold1784_4 | - | 8533 | 8482 | **RNA**→ rpmE (PRK00019)Ribosomal_L31 (pfam01197)→ |
| Eho-1-1 | NZ_JZYQ01000016.1 | - | 26597 | 26546 | **RNA**→ rpmE (PRK00019)Ribosomal_L31 (pfam01197)→ |
| Ecl-33-1 | NZ_JZZB01000010.1 | + | 86850 | 86901 | **RNA**→ rpmE (PRK00019)Ribosomal_L31 (pfam01197)→ |
| Eho-1-2 | NZ_JZZN01000011.1 | - | 26606 | 26555 | **RNA**→ rpmE (PRK00019)Ribosomal_L31 (pfam01197)→ |
| Eho-1-3 | NZ_JZYY01000014.1 | + | 86140 | 86191 | **RNA**→ rpmE (PRK00019)Ribosomal_L31 (pfam01197)→ |
| Eho-1-4 | NZ_JZZQ01000016.1 | + | 86140 | 86191 | **RNA**→ rpmE (PRK00019)Ribosomal_L31 (pfam01197)→ |
| Eho-1-5 | NZ_JZZO01000011.1 | + | 86145 | 86196 | **RNA**→ rpmE (PRK00019)Ribosomal_L31 (pfam01197)→ |
| Eho-1-6 | NZ_JZZT01000013.1 | + | 86145 | 86196 | **RNA**→ rpmE (PRK00019)Ribosomal_L31 (pfam01197)→ |
| Ecl-45-1 | NZ_LEEJ01000002.1 | + | 80227 | 80278 | **RNA**→ rpmE (PRK00019)Ribosomal_L31 (pfam01197)→ |
| Ecl-47-1 | NZ_LEDQ01000007.1 | + | 84661 | 84712 | **RNA**→ rpmE (PRK00019)Ribosomal_L31 (pfam01197)→ |
| Eho-1-7 | NZ_LEDF01000011.1 | - | 26773 | 26722 | **RNA**→ rpmE (PRK00019)Ribosomal_L31 (pfam01197)→ |
| Eho-1-8 | NZ_LEED01000021.1 | - | 26771 | 26720 | **RNA**→ rpmE (PRK00019)Ribosomal_L31 (pfam01197)→ |
| Esp-66-1 | NZ_AXLJ01000001.1 | - | 4555433 | 4555382 | **RNA**→ rpmE (PRK00019)Ribosomal_L31 (pfam01197)→ |
| Ecl-27-1 | NZ_JVNW01000148.1 | + | 11730 | 11781 | **RNA**→ rpmE (PRK00019)Ribosomal_L31 (pfam01197)→ |
| Lam-1-1 | NZ_KK737778.1 | - | 749896 | 749845 | **RNA**→ rpmE (PRK00019)Ribosomal_L31 (pfam01197)→ |
| Ecl-65-1 | NZ_KI535486.1 | - | 1069309 | 1069258 | **RNA**→ rpmE (PRK00019)Ribosomal_L31 (pfam01197)→ |
| Esp-63-1 | NZ_KI535633.1 | - | 97928 | 97877 | **RNA**→ rpmE (PRK00019)Ribosomal_L31 (pfam01197)→ |
| Ecl-63-1 | NZ_KI535513.1 | - | 26618 | 26567 | **RNA**→ rpmE (PRK00019)Ribosomal_L31 (pfam01197)→ |
| Ecl-12-1 | NZ_KI911561.1 | - | 3608360 | 3608309 | **RNA**→ rpmE (PRK00019)Ribosomal_L31 (pfam01197)→ |
| Ecl-26-1 | NZ_KI973090.1 | - | 30534 | 30483 | **RNA**→ rpmE (PRK00019)Ribosomal_L31 (pfam01197)→ |
| Esp-22-1 | NZ_KI973240.1 | + | 4664452 | 4664503 | **RNA**→ rpmE (PRK00019)Ribosomal_L31 (pfam01197)→ |
| Esp-15-1 | NZ_KI973191.1 | - | 26615 | 26564 | **RNA**→ rpmE (PRK00019)Ribosomal_L31 (pfam01197)→ |
| Esp-23-1 | NZ_KI973229.1 | + | 3189653 | 3189704 | **RNA**→ rpmE (PRK00019)Ribosomal_L31 (pfam01197)→ |
| Esp-33-1 | NZ_KI973265.1 | + | 3567038 | 3567089 | **RNA**→ rpmE (PRK00019)Ribosomal_L31 (pfam01197)→ |
| Esp-30-1 | NZ_KI973283.1 | + | 3569631 | 3569682 | **RNA**→ rpmE (PRK00019)Ribosomal_L31 (pfam01197)→ |
| Ecl-1-5 | NZ_CP009854.1 | - | 4534500 | 4534449 | **RNA**→ rpmE (PRK00019)Ribosomal_L31 (pfam01197)→ |
| Esp-25-1 | NZ_KI973322.1 | + | 86166 | 86217 | **RNA**→ rpmE (PRK00019)Ribosomal_L31 (pfam01197)→ |
| Esp-5-1 | NZ_KI973170.1 | + | 4660747 | 4660798 | **RNA**→ rpmE (PRK00019)Ribosomal_L31 (pfam01197)→ |
| Esp-13-1 | NZ_KK214763.1 | + | 100101 | 100152 | **RNA**→ rpmE (PRK00019)Ribosomal_L31 (pfam01197)→ |
| Ecl-10-1 | NZ_CP008897.1 | - | 4541025 | 4540974 | **RNA**→ rpmE (PRK00019)Ribosomal_L31 (pfam01197)→ |
| Ecl-11-1 | NZ_CP008905.1 | - | 4529003 | 4528952 | **RNA**→ rpmE (PRK00019)Ribosomal_L31 (pfam01197)→ |
| Esp-36-1 | NZ_KQ089941.1 | + | 4327417 | 4327468 | **RNA**→ rpmE (PRK00019)Ribosomal_L31 (pfam01197)→ |
| Esp-38-1 | NZ_KQ089925.1 | + | 1012995 | 1013046 | **RNA**→ rpmE (PRK00019)Ribosomal_L31 (pfam01197)→ |
| env-710 | BAAW01000972.1 | - | 3278 | 3227 | **RNA**→ rpmE (PRK00019)Ribosomal_L31 (pfam01197)→ |
| Kox-15-1 | NZ_JH603150.1 | + | 3606885 | 3606936 | **RNA**→ rpmE (PRK00019)Ribosomal_L31 (pfam01197)→ |
| Kva-1-1 | NZ_KK736636.1 | + | 72250 | 72301 | **RNA**→ rpmE (PRK00019)Ribosomal_L31 (pfam01197)→ |
| env-711 | ACOFG988_contig05406 | + | 809 | 860 | **RNA**→ rpmE (PRK00019)Ribosomal_L31 (pfam01197)→ |
| Emo-1-1 | NZ_GL890783.1 | + | 74375 | 74426 | **RNA**→ rpmE (PRK00019)Ribosomal_L31 (pfam01197)→ |

| | | | | | |
|---|---|---|---|---|---|
| env-712 | ACOFG987_contig12924 | + | 6742 | 6793 | **RNA**→ rpmE (PRK00019)Ribosomal_L31 (pfam01197)→ |
| env-713 | TrFG_GO06Z1101DZBFU | - | 245 | 194 | **RNA**→ ←hypo |
| Eae-3-1 | NC_015663.1 | + | 1578215 | 1578266 | **RNA**→ rpmE (PRK00019)Ribosomal_L31 (pfam01197)→ |
| Kae-2-1 | NZ_KK736189.1 | - | 26043 | 25992 | **RNA**→ rpmE (PRK00019)Ribosomal_L31 (pfam01197)→ |
| Kae-8-1 | NZ_KI973151.1 | + | 4566597 | 4566648 | **RNA**→ rpmE (PRK00019)Ribosomal_L31 (pfam01197)→ |
| Kae-10-1 | NZ_KI973115.1 | + | 4912164 | 4912215 | **RNA**→ rpmE (PRK00019)Ribosomal_L31 (pfam01197)→ |
| Eae-2-1 | NZ_KB911089.1 | + | 60962 | 61013 | **RNA**→ rpmE (PRK00019)Ribosomal_L31 (pfam01197)→ |
| Kae-9-1 | NZ_KI973149.1 | - | 26765 | 26714 | **RNA**→ rpmE (PRK00019)Ribosomal_L31 (pfam01197)→ |
| Ecl-39-1 | NZ_JZYA01000020.1 | + | 70697 | 70748 | **RNA**→ rpmE (PRK00019)Ribosomal_L31 (pfam01197)→ |
| Ecl-40-1 | NZ_JZXZ01000015.1 | + | 70697 | 70748 | **RNA**→ rpmE (PRK00019)Ribosomal_L31 (pfam01197)→ |
| Ecl-49-1 | NZ_LEDN01000009.1 | - | 34047 | 33996 | **RNA**→ rpmE (PRK00019)Ribosomal_L31 (pfam01197)→ |
| Eas-1-2 | NZ_LEDT01000010.1 | - | 34185 | 34134 | **RNA**→ rpmE (PRK00019)Ribosomal_L31 (pfam01197)→ |
| env-714 | JCVI_SCAF_1096628076268 | + | 1539 | 1590 | **RNA**→ rpmE (PRK00019)Ribosomal_L31 (pfam01197)→ |
| Ksp-2-1 | NZ_JAND01000018.1 | - | 27347 | 27296 | **RNA**→ rpmE (PRK00019)Ribosomal_L31 (pfam01197)→ |
| Kox-18-1 | NC_016612.1 | + | 1528968 | 1529019 | **RNA**→ rpmE (PRK00019)Ribosomal_L31 (pfam01197)→ |
| Kox-17-1 | NC_018106.1 | + | 102388 | 102439 | **RNA**→ rpmE (PRK00019)Ribosomal_L31 (pfam01197)→ |
| Kox-9-1 | NZ_CP008788.1 | + | 136463 | 136514 | **RNA**→ rpmE (PRK00019)Ribosomal_L31 (pfam01197)→ |
| Kmi-3-1 | NZ_CP004887.1 | - | 549059 | 549008 | **RNA**→ rpmE (PRK00019)Ribosomal_L31 (pfam01197)→ |
| Kox-1-4 | NZ_JTBQ01000003.1 | + | 1597391 | 1597442 | **RNA**→ rpmE (PRK00019)Ribosomal_L31 (pfam01197)→ |
| Ecl-1-6 | NZ_CP009756.1 | - | 4713790 | 4713739 | **RNA**→ rpmE (PRK00019)Ribosomal_L31 (pfam01197)→ |
| env-715 | JGI26055J44792_1064861 | - | 162 | 111 | **RNA**→ |
| env-716 | JGI26054J46599_1155624 | + | 96 | 147 | **RNA**→ rpmE (PRK00019)Ribosomal_L31 (pfam01197)→ |
| Lad-2-1 | NZ_JMPM01000080.1 | - | 5520 | 5469 | **RNA**→ rpmE (PRK00019)Ribosomal_L31 (pfam01197)→ |
| env-717 | BAAW01000278.1 | - | 9404 | 9353 | **RNA**→ rpmE (PRK00019)Ribosomal_L31 (pfam01197)→ |
| env-718 | BAAX01012058.1 | - | 967 | 916 | **RNA**→ rpmE (PRK00019)Ribosomal_L31 (pfam01197)→ |
| env-719 | DLM016_C725662_1 | - | 371 | 320 | **RNA**→ ←hypo |
| Ror-3-1 | NZ_JNPD01000022.1 | + | 72914 | 72965 | **RNA**→ rpmE (PRK00019)Ribosomal_L31 (pfam01197)→ |
| Ror-2-1 | NZ_JNPC01000052.1 | + | 72989 | 73040 | **RNA**→ rpmE (PRK00019)Ribosomal_L31 (pfam01197)→ |
| Ror-4-2 | NC_021066.1 | - | 4009168 | 4009117 | **RNA**→ rpmE (PRK00019)Ribosomal_L31 (pfam01197)→ |
| Kox-14-1 | NZ_JH603146.1 | + | 5383932 | 5383983 | **RNA**→ rpmE (PRK00019)Ribosomal_L31 (pfam01197)→ |
| env-720 | BMHB3a_c17472 | - | 62 | 11 | **RNA**→ |
| env-721 | JGI20225J20221_1000350 | + | 59385 | 59436 | **RNA**→ rpmE (PRK00019)Ribosomal_L31 (pfam01197)→ |
| env-722 | JGI20225J20221_1000514 | - | 18991 | 18940 | **RNA**→ rpmE (PRK00019)Ribosomal_L31 (pfam01197)→ |
| env-723 | JGI2065J20421_1001533 | - | 7784 | 7733 | **RNA**→ rpmE (PRK00019)Ribosomal_L31 (pfam01197)→ |
| env-724 | JGI26051J44788_1097127 | - | 262 | 211 | **RNA**→ rpmE (PRK00019)Ribosomal_L31 (pfam01197)→ |
| env-725 | JGI26052J44787_1053593 | - | 334 | 283 | **RNA**→ rpmE (PRK00019)Ribosomal_L31 (pfam01197)→ |
| env-726 | JGI26055J44792_1036479 | - | 374 | 323 | **RNA**→ rpmE (PRK00019)Ribosomal_L31 (pfam01197)→ |
| env-727 | JGI26054J46599_1011686 | - | 900 | 849 | **RNA**→ rpmE (PRK00019)Ribosomal_L31 (pfam01197)→ |
| Esp-51-1 | NZ_JTLF01000168.1 | + | 39738 | 39789 | **RNA**→ rpmE (PRK00019)Ribosomal_L31 (pfam01197)→ |
| Elu-1-1 | NZ_JTLO01000001.1 | - | 3106148 | 3106097 | **RNA**→ rpmE (PRK00019)Ribosomal_L31 (pfam01197)→ |
| Ecl-52-2 | NZ_JZYG01000016.1 | + | 85427 | 85478 | **RNA**→ rpmE (PRK00019)Ribosomal_L31 (pfam01197)→ |
| Eko-1-1 | NZ_JZYS01000009.1 | + | 92343 | 92394 | **RNA**→ rpmE (PRK00019)Ribosomal_L31 (pfam01197)→ |
| Eko-1-2 | NZ_JZYH01000012.1 | + | 90533 | 90584 | **RNA**→ rpmE (PRK00019)Ribosomal_L31 (pfam01197)→ |
| Eko-1-3 | NZ_JZXR01000011.1 | + | 92366 | 92417 | **RNA**→ rpmE (PRK00019)Ribosomal_L31 (pfam01197)→ |
| Ecl-52-3 | NZ_JZKU01000006.1 | + | 84152 | 84203 | **RNA**→ rpmE (PRK00019)Ribosomal_L31 (pfam01197)→ |
| Ecl-44-1 | NZ_LDCK01000029.1 | + | 87672 | 87723 | **RNA**→ rpmE (PRK00019)Ribosomal_L31 (pfam01197)→ |
| Eko-1-4 | NZ_LDCJ01000023.1 | - | 26538 | 26487 | **RNA**→ rpmE (PRK00019)Ribosomal_L31 (pfam01197)→ |
| Eko-1-5 | NZ_LDCL01000005.1 | + | 90555 | 90606 | **RNA**→ rpmE (PRK00019)Ribosomal_L31 (pfam01197)→ |
| Ecl-54-1 | NZ_LEDR01000013.1 | - | 26431 | 26380 | **RNA**→ rpmE (PRK00019)Ribosomal_L31 (pfam01197)→ |
| Ecl-1-12 | NZ_LEDX01000013.1 | - | 26457 | 26406 | **RNA**→ rpmE (PRK00019)Ribosomal_L31 (pfam01197)→ |
| Eko-1-6 | NZ_LEEC01000023.1 | - | 26443 | 26392 | **RNA**→ rpmE (PRK00019)Ribosomal_L31 (pfam01197)→ |
| Eko-1-7 | NZ_LECY01000021.1 | + | 96372 | 96423 | **RNA**→ rpmE (PRK00019)Ribosomal_L31 (pfam01197)→ |

| | | | | | |
|---|---|---|---|---|---|
| Esp-70-1 | NZ_LEDW01000020.1 | + | 61773 | 61824 | **RNA**→ rpmE (PRK00019)Ribosomal_L31 (pfam01197)→ |
| Elu-1-2 | NZ_LEEO01000026.1 | - | 26445 | 26394 | **RNA**→ rpmE (PRK00019)Ribosomal_L31 (pfam01197)→ |
| Ecl-52-4 | NZ_LEDL01000002.1 | - | 34294 | 34243 | **RNA**→ rpmE (PRK00019)Ribosomal_L31 (pfam01197)→ |
| Eko-1-8 | NZ_LEDC01000018.1 | + | 92385 | 92436 | **RNA**→ rpmE (PRK00019)Ribosomal_L31 (pfam01197)→ |
| Eko-1-9 | NZ_LEEK01000034.1 | + | 75795 | 75846 | **RNA**→ rpmE (PRK00019)Ribosomal_L31 (pfam01197)→ |
| Eko-1-10 | NZ_LEEL01000010.1 | - | 26539 | 26488 | **RNA**→ rpmE (PRK00019)Ribosomal_L31 (pfam01197)→ |
| Elu-1-3 | NZ_LEER01000024.1 | + | 44534 | 44585 | **RNA**→ rpmE (PRK00019)Ribosomal_L31 (pfam01197)→ |
| Eko-1-11 | NZ_LEEQ01000013.1 | + | 94220 | 94271 | **RNA**→ rpmE (PRK00019)Ribosomal_L31 (pfam01197)→ |
| Elu-1-4 | NZ_JVAG01000129.1 | - | 26272 | 26221 | **RNA**→ rpmE (PRK00019)Ribosomal_L31 (pfam01197)→ |
| Ecl-66-1 | NZ_ATCK01000043.1 | - | 26572 | 26521 | **RNA**→ rpmE (PRK00019)Ribosomal_L31 (pfam01197)→ |
| Esp-49-1 | NZ_JALW01000032.1 | - | 34139 | 34088 | **RNA**→ rpmE (PRK00019)Ribosomal_L31 (pfam01197)→ |
| Ecl-68-1 | NC_014121.1 | - | 5180845 | 5180794 | **RNA**→ rpmE (PRK00019)Ribosomal_L31 (pfam01197)→ |
| Ecl-17-1 | NZ_CP006580.1 | + | 191531 | 191582 | **RNA**→ rpmE (PRK00019)Ribosomal_L31 (pfam01197)→ |
| Elu-1-5 | NZ_LGIV01000009.1 | - | 26778 | 26727 | **RNA**→ rpmE (PRK00019)Ribosomal_L31 (pfam01197)→ |
| Ecl-5-1 | NZ_KK736229.1 | - | 126143 | 126092 | **RNA**→ rpmE (PRK00019)Ribosomal_L31 (pfam01197)→ |
| Ecl-20-1 | NZ_KK736277.1 | - | 26447 | 26396 | **RNA**→ rpmE (PRK00019)Ribosomal_L31 (pfam01197)→ |
| Esp-59-1 | NZ_KI535662.1 | - | 3665245 | 3665194 | **RNA**→ rpmE (PRK00019)Ribosomal_L31 (pfam01197)→ |
| Esp-62-1 | NZ_KI535636.1 | - | 2645376 | 2645325 | **RNA**→ rpmE (PRK00019)Ribosomal_L31 (pfam01197)→ |
| Esp-60-1 | NZ_KI535658.1 | - | 31674 | 31623 | **RNA**→ rpmE (PRK00019)Ribosomal_L31 (pfam01197)→ |
| Esp-6-1 | NZ_KI973169.1 | + | 364492 | 364543 | **RNA**→ rpmE (PRK00019)Ribosomal_L31 (pfam01197)→ |
| Ecl-19-1 | NZ_KI973153.1 | + | 4412446 | 4412497 | **RNA**→ rpmE (PRK00019)Ribosomal_L31 (pfam01197)→ |
| Esp-28-1 | NZ_KI973206.1 | - | 1100641 | 1100590 | **RNA**→ rpmE (PRK00019)Ribosomal_L31 (pfam01197)→ |
| Esp-2-1 | NZ_KI973182.1 | - | 3892522 | 3892471 | **RNA**→ rpmE (PRK00019)Ribosomal_L31 (pfam01197)→ |
| Ecl-58-1 | NC_016514.1 | - | 4606548 | 4606497 | **RNA**→ rpmE (PRK00019)Ribosomal_L31 (pfam01197)→ |
| Ecl-69-1 | NC_018405.1 | - | 4625000 | 4624949 | **RNA**→ rpmE (PRK00019)Ribosomal_L31 (pfam01197)→ |
| Ecl-1-7 | NZ_CP009850.1 | - | 4664467 | 4664416 | **RNA**→ rpmE (PRK00019)Ribosomal_L31 (pfam01197)→ |
| Ecl-70-1 | NC_018079.1 | - | 4844043 | 4843992 | **RNA**→ rpmE (PRK00019)Ribosomal_L31 (pfam01197)→ |
| Ecl-1-8 | NZ_CP011798.1 | - | 327444 | 327393 | **RNA**→ rpmE (PRK00019)Ribosomal_L31 (pfam01197)→ |
| env-728 | ACEF_contig00957 | - | 2301 | 2250 | **RNA**→ rpmE (PRK00019)Ribosomal_L31 (pfam01197)→ |
| env-729 | AECF_contig26314 | - | 81 | 30 | **RNA**→ |
| Kae-3-1 | NZ_KK736200.1 | + | 3717888 | 3717939 | **RNA**→ rpmE (PRK00019)Ribosomal_L31 (pfam01197)→ |
| Kae-1-1 | NZ_CP011574.1 | - | 3117235 | 3117184 | **RNA**→ rpmE (PRK00019)Ribosomal_L31 (pfam01197)→ |
| env-730 | XAGC_contig23616 | - | 1010 | 959 | **RNA**→ ←hypo |
| Kmi-2-1 | NZ_AYMI01000011.1 | + | 97857 | 97908 | **RNA**→ rpmE (PRK00019)Ribosomal_L31 (pfam01197)→ |
| env-731 | TrFG_contig00373 | + | 1944 | 1995 | **RNA**→ rpmE (PRK00019)Ribosomal_L31 (pfam01197)→ |
| Eas-4-1 | NC_015968.1 | - | 4661093 | 4661042 | **RNA**→ rpmE (PRK00019)Ribosomal_L31 (pfam01197)→ |
| Lad-1-2 | NZ_JWJV01000033.1 | - | 7978 | 7927 | **RNA**→ rpmE (PRK00019)Ribosomal_L31 (pfam01197)→ |
| env-732 | 4491419.3_NODE_4719 | - | 473 | 422 | **RNA**→ rpmE (PRK00019)Ribosomal_L31 (pfam01197)→ |
| env-733 | 4491419.3_NODE_11764 | - | 474 | 423 | **RNA**→ rpmE (PRK00019)Ribosomal_L31 (pfam01197)→ |
| env-734 | DLM028_scaffold4351_7 | - | 4065 | 4014 | **RNA**→ rpmE (PRK00019)Ribosomal_L31 (pfam01197)→ |
| Kpn-326-1 | NZ_AQOC01000021.1 | - | 29376 | 29325 | **RNA**→ rpmE (PRK00019)Ribosomal_L31 (pfam01197)→ |
| Kqu-2-1 | NZ_JVQN01000176.1 | - | 1038 | 987 | **RNA**→ rpmE (PRK00019)Ribosomal_L31 (pfam01197)→ |
| Kqu-2-2 | NZ_JVJE01000086.1 | + | 33812 | 33863 | **RNA**→ rpmE (PRK00019)Ribosomal_L31 (pfam01197)→ |
| Kqu-2-3 | NZ_JVVC01000018.1 | - | 26482 | 26431 | **RNA**→ rpmE (PRK00019)Ribosomal_L31 (pfam01197)→ |
| Kqu-2-4 | NZ_CBZR010000001.1 | + | 69045 | 69096 | **RNA**→ rpmE (PRK00019)Ribosomal_L31 (pfam01197)→ |
| Kpn-275-1 | NZ_KI535595.1 | + | 3403936 | 3403987 | **RNA**→ rpmE (PRK00019)Ribosomal_L31 (pfam01197)→ |
| Kpn-237-1 | NZ_AOGO01000011.1 | + | 69040 | 69091 | **RNA**→ rpmE (PRK00019)Ribosomal_L31 (pfam01197)→ |
| env-735 | TrFG_contig13861 | - | 281 | 230 | **RNA**→ ←hypo |
| Kva-2-1 | NC_013850.1 | - | 5332563 | 5332512 | **RNA**→ rpmE (PRK00019)Ribosomal_L31 (pfam01197)→ |
| Kpn-232-1 | NC_011283.1 | - | 5516732 | 5516681 | **RNA**→ rpmE (PRK00019)Ribosomal_L31 (pfam01197)→ |
| Kva-1-6 | NZ_CP009274.1 | + | 1816499 | 1816550 | **RNA**→ rpmE (PRK00019)Ribosomal_L31 (pfam01197)→ |
| Rte-1-1 | NZ_LANE01000003.1 | - | 120401 | 120350 | **RNA**→ rpmE (PRK00019)Ribosomal_L31 (pfam01197)→ |

| | | | | | | |
|---|---|---|---|---|---|---|
| Eho-1-9 | NZ_LBLX01000029.1 | + | 85605 | 85656 | **RNA**→ | rpmE (PRK00019)Ribosomal_L31 (pfam01197)→ |
| env-736 | 4491419.3_NODE_12924 | + | 144 | 195 | **RNA**→ | |
| env-737 | 4491488.3_NODE_1236 | + | 328 | 379 | **RNA**→ | rpmE (PRK00019)Ribosomal_L31 (pfam01197)→ |
| Ksp-4-1 | NZ_GG745516.1 | - | 26527 | 26476 | **RNA**→ | rpmE (PRK00019)Ribosomal_L31 (pfam01197)→ rpmE (PRK00019)Ribosomal_L31 (pfam01197)→ |
| Kva-1-2 | NZ_KK737586.1 | + | 69197 | 69248 | **RNA**→ | rpmE (PRK00019)Ribosomal_L31 (pfam01197)→ |
| Kva-1-3 | NZ_KK737692.1 | + | 69506 | 69557 | **RNA**→ | rpmE (PRK00019)Ribosomal_L31 (pfam01197)→ |
| Kva-1-4 | NZ_KK737663.1 | + | 508763 | 508814 | **RNA**→ | rpmE (PRK00019)Ribosomal_L31 (pfam01197)→ |
| Kpn-274-1 | NZ_KI535605.1 | + | 1344252 | 1344303 | **RNA**→ | rpmE (PRK00019)Ribosomal_L31 (pfam01197)→ |
| Kpn-280-1 | NZ_KI535485.1 | + | 75229 | 75280 | **RNA**→ | rpmE (PRK00019)Ribosomal_L31 (pfam01197)→ |
| Kpn-268-1 | NZ_KI535672.1 | + | 4608393 | 4608444 | **RNA**→ | rpmE (PRK00019)Ribosomal_L31 (pfam01197)→ |
| Kva-1-5 | NZ_KK036667.1 | + | 1372107 | 1372158 | **RNA**→ | rpmE (PRK00019)Ribosomal_L31 (pfam01197)→ |
| Ksp-6-1 | NZ_KB976100.1 | + | 5373858 | 5373909 | **RNA**→ | rpmE (PRK00019)Ribosomal_L31 (pfam01197)→ |
| Kva-1-7 | NZ_CP010523.1 | + | 3265862 | 3265913 | **RNA**→ | rpmE (PRK00019)Ribosomal_L31 (pfam01197)→ |
| Kox-8-1 | NZ_JAKX01000043.1 | + | 84191 | 84242 | **RNA**→ | rpmE (PRK00019)Ribosomal_L31 (pfam01197)→ |
| Kox-12-1 | NZ_JH603143.1 | + | 4820997 | 4821048 | **RNA**→ | rpmE (PRK00019)Ribosomal_L31 (pfam01197)→ |
| Kox-2-1 | NZ_KQ235800.1 | + | 1506324 | 1506375 | **RNA**→ | rpmE (PRK00019)Ribosomal_L31 (pfam01197)→ |
| Esp-29-1 | NZ_KI973298.1 | + | 90849 | 90900 | **RNA**→ | rpmE (PRK00019)Ribosomal_L31 (pfam01197)→ |
| Ecl-2-1 | NZ_KI973222.1 | + | 1710483 | 1710534 | **RNA**→ | rpmE (PRK00019)Ribosomal_L31 (pfam01197)→ |
| Ror-1-1 | NZ_CP010557.1 | + | 596820 | 596871 | **RNA**→ | rpmE (PRK00019)Ribosomal_L31 (pfam01197)→ |
| env-738 | TrFG_GO06Z1101DX8CW | + | 410 | 461 | **RNA**→ | |
| Ecl-42-1 | NZ_JZKT01000038.1 | - | 26924 | 26873 | **RNA**→ | rpmE (PRK00019)Ribosomal_L31 (pfam01197)→ |
| Ecl-7-1 | NZ_KK736236.1 | - | 40059 | 40008 | **RNA**→ | rpmE (PRK00019)Ribosomal_L31 (pfam01197)→ |
| env-739 | JCVI_SCAF_1101667396151 | - | 261 | 210 | **RNA**→ | ←PriA (COG1198)PRK05580 (PRK05580) |
| Ror-4-1 | NZ_CVRH01000021.1 | - | 26617 | 26566 | **RNA**→ | rpmE (PRK00019)Ribosomal_L31 (pfam01197)→ |
| Rpl-2-1 | NZ_JMPP01000013.1 | - | 26477 | 26426 | **RNA**→ | rpmE (PRK00019)Ribosomal_L31 (pfam01197)→ |
| Rpl-1-3 | NZ_JTBS01000001.1 | + | 1859515 | 1859566 | **RNA**→ | rpmE (PRK00019)Ribosomal_L31 (pfam01197)→ |
| Rpl-1-1 | NZ_JUZO01000017.1 | - | 26397 | 26346 | **RNA**→ | rpmE (PRK00019)Ribosomal_L31 (pfam01197)→ |
| Rpl-1-2 | NZ_JPRG01000005.1 | - | 29247 | 29196 | **RNA**→ | rpmE (PRK00019)Ribosomal_L31 (pfam01197)→ |
| Eca-3-1 | NZ_GG704867.1 | + | 87486 | 87537 | **RNA**→ | rpmE (PRK00019)Ribosomal_L31 (pfam01197)→ |
| Kqu-1-1 | NZ_KQ089430.1 | + | 80289 | 80340 | **RNA**→ | rpmE (PRK00019)Ribosomal_L31 (pfam01197)→ |
| Esp-67-1 | NC_009436.1 | - | 4390991 | 4390940 | **RNA**→ | rpmE (PRK00019)Ribosomal_L31 (pfam01197)→ |
| env-740 | BAAY01007565.1 | + | 809 | 860 | **RNA**→ | rpmE (PRK00019)Ribosomal_L31 (pfam01197)→ |
| env-741 | NLM027_scaffold15214_2 | - | 8262 | 8211 | **RNA**→ | rpmE (PRK00019)Ribosomal_L31 (pfam01197)→ |
| Kae-5-1 | NZ_KK736212.1 | + | 4447956 | 4448007 | **RNA**→ | rpmE (PRK00019)Ribosomal_L31 (pfam01197)→ |
| Kae-4-1 | NZ_KK736209.1 | + | 59203 | 59254 | **RNA**→ | rpmE (PRK00019)Ribosomal_L31 (pfam01197)→ |
| Kae-11-1 | NZ_KI973112.1 | - | 26144 | 26093 | **RNA**→ | rpmE (PRK00019)Ribosomal_L31 (pfam01197)→ |
| Kae-6-1 | NZ_KI973163.1 | + | 4470166 | 4470217 | **RNA**→ | rpmE (PRK00019)Ribosomal_L31 (pfam01197)→ |
| Kae-7-1 | NZ_KI973160.1 | - | 439123 | 439072 | **RNA**→ | rpmE (PRK00019)Ribosomal_L31 (pfam01197)→ |
| Kae-13-1 | NZ_KI973099.1 | - | 76517 | 76466 | **RNA**→ | rpmE (PRK00019)Ribosomal_L31 (pfam01197)→ |
| Kae-12-1 | NZ_KI973103.1 | + | 5175205 | 5175256 | **RNA**→ | rpmE (PRK00019)Ribosomal_L31 (pfam01197)→ |
| Eae-1-1 | NC_020181.1 | - | 3840695 | 3840644 | **RNA**→ | rpmE (PRK00019)Ribosomal_L31 (pfam01197)→ |
| env-742 | JGI26053J44789_1079225 | + | 118 | 169 | **RNA**→ | rpmE (PRK00019)Ribosomal_L31 (pfam01197)→ |
| env-743 | scaffold31432_4_MH0008 | - | 1152 | 1101 | **RNA**→ | rpmE (PRK00019)Ribosomal_L31 (pfam01197)→ |
| env-744 | scaffold66042_2_MH0028 | + | 325 | 376 | **RNA**→ | rpmE (PRK00019)Ribosomal_L31 (pfam01197)→ |
| env-745 | DLF010_scaffold7891_17 | - | 257 | 206 | **RNA**→ | rpmE (PRK00019)Ribosomal_L31 (pfam01197)→ |
| env-746 | DLM010_scaffold43772_3 | + | 3869 | 3920 | **RNA**→ | rpmE (PRK00019)Ribosomal_L31 (pfam01197)→ |
| env-747 | DLM013_scaffold3780_4 | - | 2536 | 2485 | **RNA**→ | rpmE (PRK00019)Ribosomal_L31 (pfam01197)→ |
| env-748 | DOM025_scaffold154_16 | - | 1239 | 1188 | **RNA**→ | rpmE (PRK00019)Ribosomal_L31 (pfam01197)→ |
| env-749 | NLM006_scaffold9282_3 | + | 1048 | 1099 | **RNA**→ | rpmE (PRK00019)Ribosomal_L31 (pfam01197)→ |
| env-750 | NOM005_scaffold41513_1 | - | 528 | 477 | **RNA**→ | rpmE (PRK00019)Ribosomal_L31 (pfam01197)→ |
| env-751 | NOM020_scaffold17044_1 | - | 530 | 479 | **RNA**→ | rpmE (PRK00019)Ribosomal_L31 (pfam01197)→ |
| env-752 | NOM026_scaffold17616_5 | + | 3875 | 3926 | **RNA**→ | rpmE (PRK00019)Ribosomal_L31 (pfam01197)→ |

| | | | | | | | |
|---|---|---|---|---|---|---|---|
| env-753 | SRS017191_C1645600 | - | 298 | 247 | **RNA**→ | ←hypo | |
| Kpn-336-1 | NZ_AKAI01000019.1 | + | 69325 | 69376 | **RNA**→ | rpmE (PRK00019)Ribosomal_L31 (pfam01197)→ | |
| Kpn-339-1 | NZ_AKAK01000019.1 | + | 69445 | 69496 | **RNA**→ | rpmE (PRK00019)Ribosomal_L31 (pfam01197)→ | |
| Kpn-344-1 | NZ_AJZX01000018.1 | + | 69310 | 69361 | **RNA**→ | rpmE (PRK00019)Ribosomal_L31 (pfam01197)→ | |
| Kpn-342-1 | NZ_AKAN01000030.1 | - | 30770 | 30719 | **RNA**→ | rpmE (PRK00019)Ribosomal_L31 (pfam01197)→ | |
| Kpn-337-1 | NZ_AKAJ01000017.1 | + | 68885 | 68936 | **RNA**→ | rpmE (PRK00019)Ribosomal_L31 (pfam01197)→ | |
| Kpn-341-1 | NZ_AKAM01000021.1 | + | 69493 | 69544 | **RNA**→ | rpmE (PRK00019)Ribosomal_L31 (pfam01197)→ | |
| Kpn-346-1 | NZ_AJZZ01000020.1 | - | 30703 | 30652 | **RNA**→ | rpmE (PRK00019)Ribosomal_L31 (pfam01197)→ | |
| Kpn-130-1 | NZ_JAPZ01000068.1 | - | 30404 | 30353 | **RNA**→ | rpmE (PRK00019)Ribosomal_L31 (pfam01197)→ | |
| Kpn-226-1 | NZ_AFQL01000035.1 | - | 30305 | 30254 | **RNA**→ | rpmE (PRK00019)Ribosomal_L31 (pfam01197)→ | |
| Kpn-343-1 | NZ_AJZW01000019.1 | - | 30594 | 30543 | **RNA**→ | rpmE (PRK00019)Ribosomal_L31 (pfam01197)→ | |
| Kpn-340-1 | NZ_AKAL01000044.1 | - | 30767 | 30716 | **RNA**→ | rpmE (PRK00019)Ribosomal_L31 (pfam01197)→ | |
| Kpn-315-1 | NZ_ANGI02000123.1 | - | 30404 | 30353 | **RNA**→ | rpmE (PRK00019)Ribosomal_L31 (pfam01197)→ | |
| Kpn-331-1 | NZ_AKAD01000019.1 | - | 30622 | 30571 | **RNA**→ | rpmE (PRK00019)Ribosomal_L31 (pfam01197)→ | |
| Kpn-205-4 | NZ_CDJH01000058.1 | - | 30308 | 30257 | **RNA**→ | rpmE (PRK00019)Ribosomal_L31 (pfam01197)→ | |
| Kpn-200-1 | NZ_JNBM01000001.1 | + | 71686 | 71737 | **RNA**→ | rpmE (PRK00019)Ribosomal_L31 (pfam01197)→ | |
| Kpn-300-1 | NZ_ARRU01000117.1 | - | 30307 | 30256 | **RNA**→ | rpmE (PRK00019)Ribosomal_L31 (pfam01197)→ | rpmE (PRK00019)Ribosomal_L31 (pfam01197)→ |
| Kpn-293-1 | NZ_ARSM01000115.1 | - | 30314 | 30263 | **RNA**→ | rpmE (PRK00019)Ribosomal_L31 (pfam01197)→ | rpmE (PRK00019)Ribosomal_L31 (pfam01197)→ |
| Kpn-288-1 | NZ_ARSK01000223.1 | - | 30308 | 30257 | **RNA**→ | rpmE (PRK00019)Ribosomal_L31 (pfam01197)→ | rpmE (PRK00019)Ribosomal_L31 (pfam01197)→ |
| Kpn-204-1 | NZ_JMSW01000001.1 | + | 3639812 | 3639863 | **RNA**→ | rpmE (PRK00019)Ribosomal_L31 (pfam01197)→ | |
| Kpn-203-1 | NZ_JMSX01000001.1 | + | 3139354 | 3139405 | **RNA**→ | rpmE (PRK00019)Ribosomal_L31 (pfam01197)→ | |
| Kpn-3-1 | NZ_JMMY01000001.1 | + | 1290527 | 1290578 | **RNA**→ | rpmE (PRK00019)Ribosomal_L31 (pfam01197)→ | |
| Kpn-10-1 | NZ_AQQH01000032.1 | - | 30385 | 30334 | **RNA**→ | rpmE (PRK00019)Ribosomal_L31 (pfam01197)→ | |
| Kpn-128-1 | NZ_LBLV01000039.1 | + | 64047 | 64098 | **RNA**→ | rpmE (PRK00019)Ribosomal_L31 (pfam01197)→ | |
| Kpn-166-1 | NZ_LBID01000034.1 | - | 30659 | 30608 | **RNA**→ | rpmE (PRK00019)Ribosomal_L31 (pfam01197)→ | |
| Kpn-167-1 | NZ_LBLU01000064.1 | + | 69284 | 69335 | **RNA**→ | rpmE (PRK00019)Ribosomal_L31 (pfam01197)→ | |
| Kpn-165-1 | NZ_LBIE01000001.1 | + | 63787 | 63838 | **RNA**→ | rpmE (PRK00019)Ribosomal_L31 (pfam01197)→ | |
| Kpn-169-1 | NZ_AOPN02000142.1 | + | 33886 | 33937 | **RNA**→ | rpmE (PRK00019)Ribosomal_L31 (pfam01197)→ | |
| Kpn-164-1 | NZ_LBIK01000035.1 | + | 26789 | 26840 | **RNA**→ | rpmE (PRK00019)Ribosomal_L31 (pfam01197)→ | |
| Kpn-262-1 | NZ_AVNZ01000012.1 | + | 63706 | 63757 | **RNA**→ | rpmE (PRK00019)Ribosomal_L31 (pfam01197)→ | rpmE (PRK00019)Ribosomal_L31 (pfam01197)→ |
| Kpn-205-1 | NZ_JVKF01000039.1 | + | 63885 | 63936 | **RNA**→ | rpmE (PRK00019)Ribosomal_L31 (pfam01197)→ | |
| Kpn-241-1 | NZ_AXLF01000003.1 | + | 4686167 | 4686218 | **RNA**→ | rpmE (PRK00019)Ribosomal_L31 (pfam01197)→ | |
| Kpn-240-1 | NZ_AXLG01000003.1 | + | 4750718 | 4750769 | **RNA**→ | rpmE (PRK00019)Ribosomal_L31 (pfam01197)→ | |
| Kpn-242-1 | NZ_AXLE01000001.1 | + | 4741668 | 4741719 | **RNA**→ | rpmE (PRK00019)Ribosomal_L31 (pfam01197)→ | |
| Kpn-209-1 | NZ_AVFC01000077.1 | + | 69005 | 69056 | **RNA**→ | rpmE (PRK00019)Ribosomal_L31 (pfam01197)→ | |
| Kpn-205-2 | NZ_JULI01000116.1 | - | 30352 | 30301 | **RNA**→ | rpmE (PRK00019)Ribosomal_L31 (pfam01197)→ | |
| Kpn-205-3 | NZ_JULH01000061.1 | - | 30362 | 30311 | **RNA**→ | rpmE (PRK00019)Ribosomal_L31 (pfam01197)→ | |
| Kpn-3-2 | NZ_ARSN01000079.1 | - | 30275 | 30224 | **RNA**→ | rpmE (PRK00019)Ribosomal_L31 (pfam01197)→ | rpmE (PRK00019)Ribosomal_L31 (pfam01197)→ |
| Kpn-264-1 | NZ_AQQD01000294.1 | + | 65257 | 65308 | **RNA**→ | rpmE (PRK00019)Ribosomal_L31 (pfam01197)→ | rpmE (PRK00019)Ribosomal_L31 (pfam01197)→ |
| Kpn-233-1 | NZ_APWF01000076.1 | - | 30307 | 30256 | **RNA**→ | rpmE (PRK00019)Ribosomal_L31 (pfam01197)→ | rpmE (PRK00019)Ribosomal_L31 (pfam01197)→ |
| Kpn-294-1 | NZ_ARSH01000011.1 | - | 30308 | 30257 | **RNA**→ | rpmE (PRK00019)Ribosomal_L31 (pfam01197)→ | rpmE (PRK00019)Ribosomal_L31 (pfam01197)→ |
| Kpn-199-2 | NZ_ARVN01000047.1 | - | 30307 | 30256 | **RNA**→ | rpmE (PRK00019)Ribosomal_L31 (pfam01197)→ | rpmE (PRK00019)Ribosomal_L31 (pfam01197)→ |
| Kpn-236-1 | NZ_APWJ01000011.1 | + | 22829 | 22880 | **RNA**→ | rpmE (PRK00019)Ribosomal_L31 (pfam01197)→ | rpmE (PRK00019)Ribosomal_L31 (pfam01197)→ |
| Kpn-319-1 | NZ_APWA01000100.1 | - | 30308 | 30257 | **RNA**→ | rpmE (PRK00019)Ribosomal_L31 (pfam01197)→ | rpmE (PRK00019)Ribosomal_L31 (pfam01197)→ |
| Kpn-295-1 | NZ_APVU01000051.1 | - | 30307 | 30256 | **RNA**→ | rpmE (PRK00019)Ribosomal_L31 (pfam01197)→ | rpmE (PRK00019)Ribosomal_L31 (pfam01197)→ |
| Kpn-308-1 | NZ_ARSE01000152.1 | - | 30308 | 30257 | **RNA**→ | rpmE (PRK00019)Ribosomal_L31 (pfam01197)→ | rpmE (PRK00019)Ribosomal_L31 (pfam01197)→ |
| Kpn-306-1 | NZ_ARPP01000028.1 | - | 30307 | 30256 | **RNA**→ | rpmE (PRK00019)Ribosomal_L31 (pfam01197)→ | rpmE (PRK00019)Ribosomal_L31 (pfam01197)→ |
| Kpn-235-1 | NZ_APWI01000146.1 | - | 30307 | 30256 | **RNA**→ | rpmE (PRK00019)Ribosomal_L31 (pfam01197)→ | rpmE (PRK00019)Ribosomal_L31 (pfam01197)→ |
| Kpn-321-1 | NZ_APVZ01000120.1 | - | 30307 | 30256 | **RNA**→ | rpmE (PRK00019)Ribosomal_L31 (pfam01197)→ | rpmE (PRK00019)Ribosomal_L31 (pfam01197)→ |
| Kpn-231-1 | NZ_AVAN01000034.1 | + | 64640 | 64691 | **RNA**→ | rpmE (PRK00019)Ribosomal_L31 (pfam01197)→ | |
| Kpn-255-1 | NZ_ARPO01000036.1 | - | 30308 | 30257 | **RNA**→ | rpmE (PRK00019)Ribosomal_L31 (pfam01197)→ | rpmE (PRK00019)Ribosomal_L31 (pfam01197)→ |
| Kpn-298-1 | NZ_APVT01000051.1 | - | 30307 | 30256 | **RNA**→ | rpmE (PRK00019)Ribosomal_L31 (pfam01197)→ | rpmE (PRK00019)Ribosomal_L31 (pfam01197)→ |

| | | | | | | | |
|---|---|---|---|---|---|---|---|
| Kpn-200-2 | NZ_ARVM01000045.1 | - | 30308 | 30257 | **RNA**→ | rpmE (PRK00019)Ribosomal_L31 (pfam01197)→ | rpmE (PRK00019)Ribosomal_L31 (pfam01197)→ |
| Kpn-310-1 | NZ_ARSL01000131.1 | + | 68924 | 68975 | **RNA**→ | rpmE (PRK00019)Ribosomal_L31 (pfam01197)→ | rpmE (PRK00019)Ribosomal_L31 (pfam01197)→ |
| Kpn-257-1 | NZ_ARSB01000142.1 | - | 30314 | 30263 | **RNA**→ | rpmE (PRK00019)Ribosomal_L31 (pfam01197)→ | rpmE (PRK00019)Ribosomal_L31 (pfam01197)→ |
| Kpn-314-1 | NZ_ARRY01000131.1 | - | 30308 | 30257 | **RNA**→ | rpmE (PRK00019)Ribosomal_L31 (pfam01197)→ | rpmE (PRK00019)Ribosomal_L31 (pfam01197)→ |
| Kpn-287-1 | NZ_APVP01000044.1 | - | 30308 | 30257 | **RNA**→ | rpmE (PRK00019)Ribosomal_L31 (pfam01197)→ | rpmE (PRK00019)Ribosomal_L31 (pfam01197)→ |
| Kpn-299-1 | NZ_APVR01000150.1 | + | 22831 | 22882 | **RNA**→ | rpmE (PRK00019)Ribosomal_L31 (pfam01197)→ | rpmE (PRK00019)Ribosomal_L31 (pfam01197)→ |
| Kpn-291-1 | NZ_APVO01000145.1 | - | 30307 | 30256 | **RNA**→ | rpmE (PRK00019)Ribosomal_L31 (pfam01197)→ | rpmE (PRK00019)Ribosomal_L31 (pfam01197)→ |
| Kpn-305-1 | NZ_ARRV01000085.1 | - | 30307 | 30256 | **RNA**→ | rpmE (PRK00019)Ribosomal_L31 (pfam01197)→ | rpmE (PRK00019)Ribosomal_L31 (pfam01197)→ |
| Kpn-292-1 | NZ_ARSG01000160.1 | - | 30308 | 30257 | **RNA**→ | rpmE (PRK00019)Ribosomal_L31 (pfam01197)→ | rpmE (PRK00019)Ribosomal_L31 (pfam01197)→ |
| Kpn-296-1 | NZ_AQQA01000109.1 | - | 30307 | 30256 | **RNA**→ | rpmE (PRK00019)Ribosomal_L31 (pfam01197)→ | rpmE (PRK00019)Ribosomal_L31 (pfam01197)→ |
| Kpn-127-1 | NZ_AQQI01000044.1 | - | 30387 | 30336 | **RNA**→ | rpmE (PRK00019)Ribosomal_L31 (pfam01197)→ | |
| Kpn-357-1 | NZ_ATEZ01000131.1 | + | 68945 | 68996 | **RNA**→ | rpmE (PRK00019)Ribosomal_L31 (pfam01197)→ | rpmE (PRK00019)Ribosomal_L31 (pfam01197)→ |
| Kpn-234-1 | NZ_APWG01000108.1 | - | 30318 | 30267 | **RNA**→ | rpmE (PRK00019)Ribosomal_L31 (pfam01197)→ | rpmE (PRK00019)Ribosomal_L31 (pfam01197)→ |
| Kpn-230-1 | NZ_ARSQ01000036.1 | + | 22838 | 22889 | **RNA**→ | rpmE (PRK00019)Ribosomal_L31 (pfam01197)→ | rpmE (PRK00019)Ribosomal_L31 (pfam01197)→ |
| Kpn-229-1 | NZ_ARSP01000052.1 | - | 30307 | 30256 | **RNA**→ | rpmE (PRK00019)Ribosomal_L31 (pfam01197)→ | rpmE (PRK00019)Ribosomal_L31 (pfam01197)→ |
| Kpn-263-1 | NZ_APWE01000184.1 | - | 30309 | 30258 | **RNA**→ | rpmE (PRK00019)Ribosomal_L31 (pfam01197)→ | |
| Kpn-228-1 | NZ_ARSO01000057.1 | - | 30308 | 30257 | **RNA**→ | rpmE (PRK00019)Ribosomal_L31 (pfam01197)→ | rpmE (PRK00019)Ribosomal_L31 (pfam01197)→ |
| Kpn-290-1 | NZ_ARSJ01000101.1 | - | 30307 | 30256 | **RNA**→ | rpmE (PRK00019)Ribosomal_L31 (pfam01197)→ | rpmE (PRK00019)Ribosomal_L31 (pfam01197)→ |
| Kpn-258-1 | NZ_AFQK01000005.1 | - | 30326 | 30275 | **RNA**→ | rpmE (PRK00019)Ribosomal_L31 (pfam01197)→ | |
| Kpn-332-1 | NZ_AKAE01000020.1 | - | 30565 | 30514 | **RNA**→ | rpmE (PRK00019)Ribosomal_L31 (pfam01197)→ | |
| Kpn-355-1 | NZ_CANS01000126.1 | + | 69343 | 69394 | **RNA**→ | rpmE (PRK00019)Ribosomal_L31 (pfam01197)→ | rpmE (PRK00019)Ribosomal_L31 (pfam01197)→ |
| Kpn-328-1 | NZ_AJJI01000017.1 | + | 68833 | 68884 | **RNA**→ | rpmE (PRK00019)Ribosomal_L31 (pfam01197)→ | rpmE (PRK00019)Ribosomal_L31 (pfam01197)→ |
| Kpn-194-1 | NZ_JJNI01000022.1 | + | 60242 | 60293 | **RNA**→ | rpmE (PRK00019)Ribosomal_L31 (pfam01197)→ | |
| Kpn-190-1 | NZ_JJNJ01000018.1 | + | 70166 | 70217 | **RNA**→ | rpmE (PRK00019)Ribosomal_L31 (pfam01197)→ | |
| Kpn-161-1 | NZ_JJNK01000022.1 | + | 63710 | 63761 | **RNA**→ | rpmE (PRK00019)Ribosomal_L31 (pfam01197)→ | |
| Kpn-199-1 | NZ_JMSY01000001.1 | + | 3417357 | 3417408 | **RNA**→ | rpmE (PRK00019)Ribosomal_L31 (pfam01197)→ | |
| Kpn-211-1 | NZ_AYOV01000023.1 | + | 69005 | 69056 | **RNA**→ | rpmE (PRK00019)Ribosomal_L31 (pfam01197)→ | |
| Kpn-212-1 | NZ_AYOW01000027.1 | + | 69005 | 69056 | **RNA**→ | rpmE (PRK00019)Ribosomal_L31 (pfam01197)→ | |
| Kpn-348-1 | NZ_AKAB01000020.1 | + | 69393 | 69444 | **RNA**→ | rpmE (PRK00019)Ribosomal_L31 (pfam01197)→ | |
| Kpn-345-1 | NZ_AJZY01000021.1 | - | 30650 | 30599 | **RNA**→ | rpmE (PRK00019)Ribosomal_L31 (pfam01197)→ | |
| Kpn-354-1 | NZ_CANR01000136.1 | + | 69501 | 69552 | **RNA**→ | rpmE (PRK00019)Ribosomal_L31 (pfam01197)→ | rpmE (PRK00019)Ribosomal_L31 (pfam01197)→ |
| Kpn-350-1 | NZ_AJVY01000169.1 | + | 64800 | 64851 | **RNA**→ | rpmE (PRK00019)Ribosomal_L31 (pfam01197)→ | |
| Kpn-347-1 | NZ_AKAA01000020.1 | - | 30633 | 30582 | **RNA**→ | rpmE (PRK00019)Ribosomal_L31 (pfam01197)→ | |
| Kpn-356-1 | NZ_CAJM02000254.1 | + | 69080 | 69131 | **RNA**→ | rpmE (PRK00019)Ribosomal_L31 (pfam01197)→ | rpmE (PRK00019)Ribosomal_L31 (pfam01197)→ |
| Kpn-353-1 | NZ_ALIS01000023.1 | - | 30146 | 30095 | **RNA**→ | rpmE (PRK00019)Ribosomal_L31 (pfam01197)→ | |
| Kpn-289-1 | NZ_APVV01000006.1 | + | 68928 | 68979 | **RNA**→ | rpmE (PRK00019)Ribosomal_L31 (pfam01197)→ | rpmE (PRK00019)Ribosomal_L31 (pfam01197)→ |
| Kpn-322-1 | NZ_APWC01000036.1 | + | 68928 | 68979 | **RNA**→ | rpmE (PRK00019)Ribosomal_L31 (pfam01197)→ | rpmE (PRK00019)Ribosomal_L31 (pfam01197)→ |
| Kpn-323-1 | NZ_APWD01000119.1 | + | 22834 | 22885 | **RNA**→ | rpmE (PRK00019)Ribosomal_L31 (pfam01197)→ | rpmE (PRK00019)Ribosomal_L31 (pfam01197)→ |
| Kpn-352-1 | NZ_ATAK01000075.1 | + | 68887 | 68938 | **RNA**→ | rpmE (PRK00019)Ribosomal_L31 (pfam01197)→ | rpmE (PRK00019)Ribosomal_L31 (pfam01197)→ |
| Kpn-301-1 | NZ_ARVQ01000045.1 | + | 68928 | 68979 | **RNA**→ | rpmE (PRK00019)Ribosomal_L31 (pfam01197)→ | rpmE (PRK00019)Ribosomal_L31 (pfam01197)→ |
| Kpn-318-1 | NZ_APVY01000110.1 | - | 30307 | 30256 | **RNA**→ | rpmE (PRK00019)Ribosomal_L31 (pfam01197)→ | rpmE (PRK00019)Ribosomal_L31 (pfam01197)→ |
| Kpn-312-1 | NZ_ARRX01000168.1 | - | 30308 | 30257 | **RNA**→ | rpmE (PRK00019)Ribosomal_L31 (pfam01197)→ | rpmE (PRK00019)Ribosomal_L31 (pfam01197)→ |
| Kpn-138-1 | NZ_AYQE01000011.1 | - | 29562 | 29511 | **RNA**→ | rpmE (PRK00019)Ribosomal_L31 (pfam01197)→ | |
| Kpn-317-1 | NZ_APVX01000016.1 | - | 30308 | 30257 | **RNA**→ | rpmE (PRK00019)Ribosomal_L31 (pfam01197)→ | rpmE (PRK00019)Ribosomal_L31 (pfam01197)→ |
| Kpn-316-1 | NZ_APVW01000045.1 | + | 68928 | 68979 | **RNA**→ | rpmE (PRK00019)Ribosomal_L31 (pfam01197)→ | rpmE (PRK00019)Ribosomal_L31 (pfam01197)→ |
| Kpn-297-1 | NZ_APVS01000015.1 | - | 30307 | 30256 | **RNA**→ | rpmE (PRK00019)Ribosomal_L31 (pfam01197)→ | rpmE (PRK00019)Ribosomal_L31 (pfam01197)→ |
| Kpn-311-1 | NZ_ARRW01000042.1 | - | 30307 | 30256 | **RNA**→ | rpmE (PRK00019)Ribosomal_L31 (pfam01197)→ | rpmE (PRK00019)Ribosomal_L31 (pfam01197)→ |
| Kpn-320-1 | NZ_APWB01000051.1 | - | 30308 | 30257 | **RNA**→ | rpmE (PRK00019)Ribosomal_L31 (pfam01197)→ | rpmE (PRK00019)Ribosomal_L31 (pfam01197)→ |
| Kpn-313-1 | NZ_APVQ01000201.1 | - | 30314 | 30263 | **RNA**→ | rpmE (PRK00019)Ribosomal_L31 (pfam01197)→ | rpmE (PRK00019)Ribosomal_L31 (pfam01197)→ |
| Kpn-302-1 | NZ_ARSI01000095.1 | - | 30307 | 30256 | **RNA**→ | rpmE (PRK00019)Ribosomal_L31 (pfam01197)→ | rpmE (PRK00019)Ribosomal_L31 (pfam01197)→ |
| Kpn-309-1 | NZ_ARSC01000038.1 | + | 68928 | 68979 | **RNA**→ | rpmE (PRK00019)Ribosomal_L31 (pfam01197)→ | rpmE (PRK00019)Ribosomal_L31 (pfam01197)→ |
| Kpn-304-1 | NZ_AQOT01000031.1 | + | 68928 | 68979 | **RNA**→ | rpmE (PRK00019)Ribosomal_L31 (pfam01197)→ | rpmE (PRK00019)Ribosomal_L31 (pfam01197)→ |

| | | | | | |
|---|---|---|---|---|---|
| Kpn-203-2 | NZ_ARVO01000169.1 | - | 30310 | 30259 | **RNA**→ rpmE (PRK00019)Ribosomal_L31 (pfam01197)→ rpmE (PRK00019)Ribosomal_L31 (pfam01197)→ |
| Kpn-303-1 | NZ_ARPQ01000202.1 | - | 30308 | 30257 | **RNA**→ rpmE (PRK00019)Ribosomal_L31 (pfam01197)→ rpmE (PRK00019)Ribosomal_L31 (pfam01197)→ |
| Kpn-204-2 | NZ_ARVP01000082.1 | + | 68924 | 68975 | **RNA**→ rpmE (PRK00019)Ribosomal_L31 (pfam01197)→ rpmE (PRK00019)Ribosomal_L31 (pfam01197)→ |
| Kpn-307-1 | NZ_ARPR01000083.1 | + | 22839 | 22890 | **RNA**→ rpmE (PRK00019)Ribosomal_L31 (pfam01197)→ rpmE (PRK00019)Ribosomal_L31 (pfam01197)→ |
| Kpn-219-1 | NZ_AWOM01000016.1 | - | 30326 | 30275 | **RNA**→ rpmE (PRK00019)Ribosomal_L31 (pfam01197)→ |
| Kpn-220-1 | NZ_AWON01000029.1 | + | 35737 | 35788 | **RNA**→ rpmE (PRK00019)Ribosomal_L31 (pfam01197)→ |
| Kpn-210-1 | NZ_AVFD01000075.1 | + | 69293 | 69344 | **RNA**→ rpmE (PRK00019)Ribosomal_L31 (pfam01197)→ |
| Kpn-163-1 | NZ_JSVB01000050.1 | + | 37050 | 37101 | **RNA**→ rpmE (PRK00019)Ribosomal_L31 (pfam01197)→ |
| Kpn-136-1 | NZ_AQRD01000007.1 | - | 3937419 | 3937368 | **RNA**→ rpmE (PRK00019)Ribosomal_L31 (pfam01197)→ |
| Kpn-150-1 | NZ_JJNB01000021.1 | + | 63818 | 63869 | **RNA**→ rpmE (PRK00019)Ribosomal_L31 (pfam01197)→ |
| Kpn-65-1 | NZ_JJNG01000016.1 | + | 68967 | 69018 | **RNA**→ rpmE (PRK00019)Ribosomal_L31 (pfam01197)→ |
| Kpn-59-1 | NZ_JJNH01000017.1 | + | 72322 | 72373 | **RNA**→ rpmE (PRK00019)Ribosomal_L31 (pfam01197)→ |
| Kpn-6-1 | NZ_JDWP01000017.1 | - | 30401 | 30350 | **RNA**→ rpmE (PRK00019)Ribosomal_L31 (pfam01197)→ |
| Kpn-9-1 | NZ_JDWM01000007.1 | - | 29721 | 29670 | **RNA**→ rpmE (PRK00019)Ribosomal_L31 (pfam01197)→ |
| Kpn-7-1 | NZ_JDWO01000009.1 | - | 30401 | 30350 | **RNA**→ rpmE (PRK00019)Ribosomal_L31 (pfam01197)→ |
| Kpn-227-1 | NZ_AFXH01000001.1 | - | 15574 | 15523 | **RNA**→ rpmE (PRK00019)Ribosomal_L31 (pfam01197)→ |
| Kpn-2-1 | NZ_JDWN01000019.1 | + | 64235 | 64286 | **RNA**→ rpmE (PRK00019)Ribosomal_L31 (pfam01197)→ |
| Kpn-134-1 | NZ_APMF01000021.1 | - | 30307 | 30256 | **RNA**→ rpmE (PRK00019)Ribosomal_L31 (pfam01197)→ |
| Kpn-133-1 | NZ_AOSJ01000020.1 | - | 30307 | 30256 | **RNA**→ rpmE (PRK00019)Ribosomal_L31 (pfam01197)→ |
| Kpn-135-1 | NZ_APMG01000019.1 | - | 30307 | 30256 | **RNA**→ rpmE (PRK00019)Ribosomal_L31 (pfam01197)→ |
| Kpn-131-1 | NZ_AQCJ01000019.1 | - | 30310 | 30259 | **RNA**→ rpmE (PRK00019)Ribosomal_L31 (pfam01197)→ |
| Kpn-132-1 | NZ_AQCI01000019.1 | + | 68968 | 69019 | **RNA**→ rpmE (PRK00019)Ribosomal_L31 (pfam01197)→ |
| Kpn-330-1 | NC_016845.1 | + | 90172 | 90223 | **RNA**→ rpmE (PRK00019)Ribosomal_L31 (pfam01197)→ |
| Kpn-351-1 | NC_009648.1 | + | 4632889 | 4632940 | **RNA**→ rpmE (PRK00019)Ribosomal_L31 (pfam01197)→ |
| Kpn-216-1 | NZ_CP007731.1 | + | 92618 | 92669 | **RNA**→ rpmE (PRK00019)Ribosomal_L31 (pfam01197)→ |
| Kpn-239-1 | NZ_CP006659.1 | + | 90175 | 90226 | **RNA**→ rpmE (PRK00019)Ribosomal_L31 (pfam01197)→ |
| Kpn-126-1 | NZ_AUTW02000084.1 | - | 30242 | 30191 | **RNA**→ rpmE (PRK00019)Ribosomal_L31 (pfam01197)→ |
| Kpn-1-1 | NZ_CP010361.1 | - | 5219933 | 5219882 | **RNA**→ rpmE (PRK00019)Ribosomal_L31 (pfam01197)→ |
| Kpn-362-1 | NZ_GG703533.1 | + | 65601 | 65652 | **RNA**→ rpmE (PRK00019)Ribosomal_L31 (pfam01197)→ rpmE (PRK00019)Ribosomal_L31 (pfam01197)→ |
| Kpn-91-1 | NZ_KK214620.1 | + | 69995 | 70046 | **RNA**→ rpmE (PRK00019)Ribosomal_L31 (pfam01197)→ |
| Kpn-96-1 | NZ_KK214479.1 | + | 69063 | 69114 | **RNA**→ rpmE (PRK00019)Ribosomal_L31 (pfam01197)→ |
| Kpn-109-1 | NZ_KK214372.1 | + | 69063 | 69114 | **RNA**→ rpmE (PRK00019)Ribosomal_L31 (pfam01197)→ |
| Kpn-77-1 | NZ_KK736955.1 | + | 581455 | 581506 | **RNA**→ rpmE (PRK00019)Ribosomal_L31 (pfam01197)→ |
| Kpn-123-1 | NZ_KK737461.1 | + | 2876697 | 2876748 | **RNA**→ rpmE (PRK00019)Ribosomal_L31 (pfam01197)→ |
| Kpn-121-1 | NZ_KK737432.1 | + | 73324 | 73375 | **RNA**→ rpmE (PRK00019)Ribosomal_L31 (pfam01197)→ |
| Kpn-155-1 | NZ_KK737589.1 | + | 3182677 | 3182728 | **RNA**→ rpmE (PRK00019)Ribosomal_L31 (pfam01197)→ |
| Kpn-153-1 | NZ_KK737549.1 | - | 634579 | 634528 | **RNA**→ rpmE (PRK00019)Ribosomal_L31 (pfam01197)→ |
| Kpn-154-1 | NZ_KK737562.1 | + | 73376 | 73427 | **RNA**→ rpmE (PRK00019)Ribosomal_L31 (pfam01197)→ |
| Kpn-157-1 | NZ_KK737614.1 | + | 71967 | 72018 | **RNA**→ rpmE (PRK00019)Ribosomal_L31 (pfam01197)→ |
| Kpn-152-1 | NZ_KK737538.1 | + | 63914 | 63965 | **RNA**→ rpmE (PRK00019)Ribosomal_L31 (pfam01197)→ |
| Kpn-196-1 | NZ_KK737766.1 | - | 3724699 | 3724648 | **RNA**→ rpmE (PRK00019)Ribosomal_L31 (pfam01197)→ |
| Kpn-162-1 | NZ_KK737679.1 | + | 68934 | 68985 | **RNA**→ rpmE (PRK00019)Ribosomal_L31 (pfam01197)→ |
| Kpn-156-1 | NZ_KK737605.1 | + | 63679 | 63730 | **RNA**→ rpmE (PRK00019)Ribosomal_L31 (pfam01197)→ |
| Kpn-38-1 | NZ_KK736604.1 | + | 69390 | 69441 | **RNA**→ rpmE (PRK00019)Ribosomal_L31 (pfam01197)→ |
| Kpn-148-1 | NZ_KK737484.1 | + | 583791 | 583842 | **RNA**→ rpmE (PRK00019)Ribosomal_L31 (pfam01197)→ |
| Kpn-278-1 | NZ_KK737503.1 | + | 4119556 | 4119607 | **RNA**→ rpmE (PRK00019)Ribosomal_L31 (pfam01197)→ |
| Kpn-192-1 | NZ_KK737733.1 | + | 3278687 | 3278738 | **RNA**→ rpmE (PRK00019)Ribosomal_L31 (pfam01197)→ |
| Kpn-122-1 | NZ_KK737447.1 | + | 4942116 | 4942167 | **RNA**→ rpmE (PRK00019)Ribosomal_L31 (pfam01197)→ |
| Kpn-158-1 | NZ_KK737629.1 | + | 115507 | 115558 | **RNA**→ rpmE (PRK00019)Ribosomal_L31 (pfam01197)→ |
| Kpn-151-1 | NZ_KK737530.1 | + | 4494417 | 4494468 | **RNA**→ rpmE (PRK00019)Ribosomal_L31 (pfam01197)→ |
| Kpn-188-1 | NZ_KK737707.1 | + | 4500007 | 4500058 | **RNA**→ rpmE (PRK00019)Ribosomal_L31 (pfam01197)→ |
| Kpn-195-1 | NZ_KK737755.1 | + | 1400819 | 1400870 | **RNA**→ rpmE (PRK00019)Ribosomal_L31 (pfam01197)→ |

| | | | | | | |
|---|---|---|---|---|---|---|
| Kpn-160-1 | NZ_KK737646.1 | + | 63830 | 63881 | **RNA**→ | rpmE (PRK00019)Ribosomal_L31 (pfam01197)→ |
| Kpn-149-1 | NZ_KK737520.1 | - | 78478 | 78427 | **RNA**→ | rpmE (PRK00019)Ribosomal_L31 (pfam01197)→ |
| Kpn-187-1 | NZ_KK737696.1 | + | 4798861 | 4798912 | **RNA**→ | rpmE (PRK00019)Ribosomal_L31 (pfam01197)→ |
| Kpn-193-1 | NZ_KK737742.1 | + | 4381090 | 4381141 | **RNA**→ | rpmE (PRK00019)Ribosomal_L31 (pfam01197)→ |
| Kpn-159-1 | NZ_KK737636.1 | + | 3151654 | 3151705 | **RNA**→ | rpmE (PRK00019)Ribosomal_L31 (pfam01197)→ |
| Kpn-120-1 | NZ_KK737419.1 | + | 2864858 | 2864909 | **RNA**→ | rpmE (PRK00019)Ribosomal_L31 (pfam01197)→ |
| Kpn-40-1 | NZ_KK736624.1 | + | 3233701 | 3233752 | **RNA**→ | rpmE (PRK00019)Ribosomal_L31 (pfam01197)→ |
| Kpn-42-1 | NZ_KK736658.1 | + | 577414 | 577465 | **RNA**→ | rpmE (PRK00019)Ribosomal_L31 (pfam01197)→ |
| Kpn-39-1 | NZ_KK736620.1 | + | 63835 | 63886 | **RNA**→ | rpmE (PRK00019)Ribosomal_L31 (pfam01197)→ |
| Kpn-110-1 | NZ_KK737334.1 | + | 69390 | 69441 | **RNA**→ | rpmE (PRK00019)Ribosomal_L31 (pfam01197)→ |
| Kpn-118-1 | NZ_KK737405.1 | + | 2867900 | 2867951 | **RNA**→ | rpmE (PRK00019)Ribosomal_L31 (pfam01197)→ |
| Kpn-50-1 | NZ_KK736666.1 | + | 4827133 | 4827184 | **RNA**→ | rpmE (PRK00019)Ribosomal_L31 (pfam01197)→ |
| Kpn-72-1 | NZ_KK736890.1 | + | 69390 | 69441 | **RNA**→ | rpmE (PRK00019)Ribosomal_L31 (pfam01197)→ |
| Kpn-97-1 | NZ_KK737177.1 | + | 4679594 | 4679645 | **RNA**→ | rpmE (PRK00019)Ribosomal_L31 (pfam01197)→ |
| Kpn-76-1 | NZ_KK736937.1 | + | 3239685 | 3239736 | **RNA**→ | rpmE (PRK00019)Ribosomal_L31 (pfam01197)→ |
| Kpn-41-1 | NZ_KK736641.1 | - | 123068 | 123017 | **RNA**→ | rpmE (PRK00019)Ribosomal_L31 (pfam01197)→ |
| Kpn-55-1 | NZ_KK736735.1 | + | 1426606 | 1426657 | **RNA**→ | rpmE (PRK00019)Ribosomal_L31 (pfam01197)→ |
| Kpn-90-1 | NZ_KK737113.1 | + | 3359104 | 3359155 | **RNA**→ | rpmE (PRK00019)Ribosomal_L31 (pfam01197)→ |
| Kpn-112-1 | NZ_KK737351.1 | + | 69126 | 69177 | **RNA**→ | rpmE (PRK00019)Ribosomal_L31 (pfam01197)→ |
| Kpn-51-1 | NZ_KK736683.1 | + | 72227 | 72278 | **RNA**→ | rpmE (PRK00019)Ribosomal_L31 (pfam01197)→ |
| Kpn-57-1 | NZ_KK736760.1 | + | 3379575 | 3379626 | **RNA**→ | rpmE (PRK00019)Ribosomal_L31 (pfam01197)→ |
| Kpn-67-1 | NZ_KK736836.1 | + | 69126 | 69177 | **RNA**→ | rpmE (PRK00019)Ribosomal_L31 (pfam01197)→ |
| Kpn-73-1 | NZ_KK736899.1 | + | 73376 | 73427 | **RNA**→ | rpmE (PRK00019)Ribosomal_L31 (pfam01197)→ |
| Kpn-104-1 | NZ_KK737273.1 | + | 4895637 | 4895688 | **RNA**→ | rpmE (PRK00019)Ribosomal_L31 (pfam01197)→ |
| Kpn-107-1 | NZ_KK737307.1 | + | 4735918 | 4735969 | **RNA**→ | rpmE (PRK00019)Ribosomal_L31 (pfam01197)→ |
| Kpn-64-1 | NZ_KK736818.1 | + | 69126 | 69177 | **RNA**→ | rpmE (PRK00019)Ribosomal_L31 (pfam01197)→ |
| Kpn-85-1 | NZ_KK737050.1 | + | 4908031 | 4908082 | **RNA**→ | rpmE (PRK00019)Ribosomal_L31 (pfam01197)→ |
| Kpn-94-1 | NZ_KK737157.1 | + | 1431419 | 1431470 | **RNA**→ | rpmE (PRK00019)Ribosomal_L31 (pfam01197)→ |
| Kpn-115-1 | NZ_KK737382.1 | + | 64630 | 64681 | **RNA**→ | rpmE (PRK00019)Ribosomal_L31 (pfam01197)→ |
| Kpn-116-1 | NZ_KK737388.1 | + | 1428472 | 1428523 | **RNA**→ | rpmE (PRK00019)Ribosomal_L31 (pfam01197)→ |
| Kpn-114-1 | NZ_KK737366.1 | + | 1425775 | 1425826 | **RNA**→ | rpmE (PRK00019)Ribosomal_L31 (pfam01197)→ |
| Kpn-102-1 | NZ_KK737235.1 | + | 69390 | 69441 | **RNA**→ | rpmE (PRK00019)Ribosomal_L31 (pfam01197)→ |
| Kpn-92-1 | NZ_KK737128.1 | + | 69033 | 69084 | **RNA**→ | rpmE (PRK00019)Ribosomal_L31 (pfam01197)→ |
| Kpn-103-1 | NZ_KK737268.1 | + | 4887246 | 4887297 | **RNA**→ | rpmE (PRK00019)Ribosomal_L31 (pfam01197)→ |
| Kpn-99-1 | NZ_KK737192.1 | + | 3399123 | 3399174 | **RNA**→ | rpmE (PRK00019)Ribosomal_L31 (pfam01197)→ |
| Kpn-78-1 | NZ_KK736970.1 | + | 576185 | 576236 | **RNA**→ | rpmE (PRK00019)Ribosomal_L31 (pfam01197)→ |
| Kpn-71-1 | NZ_KK736871.1 | + | 582040 | 582091 | **RNA**→ | rpmE (PRK00019)Ribosomal_L31 (pfam01197)→ |
| Kpn-79-1 | NZ_KK736979.1 | + | 5071703 | 5071754 | **RNA**→ | rpmE (PRK00019)Ribosomal_L31 (pfam01197)→ |
| Kpn-68-1 | NZ_KK736847.1 | + | 69390 | 69441 | **RNA**→ | rpmE (PRK00019)Ribosomal_L31 (pfam01197)→ |
| Kpn-87-1 | NZ_KK737072.1 | + | 2869909 | 2869960 | **RNA**→ | rpmE (PRK00019)Ribosomal_L31 (pfam01197)→ |
| Kpn-88-1 | NZ_KK737090.1 | + | 3362502 | 3362553 | **RNA**→ | rpmE (PRK00019)Ribosomal_L31 (pfam01197)→ |
| Kpn-106-1 | NZ_KK737291.1 | + | 583431 | 583482 | **RNA**→ | rpmE (PRK00019)Ribosomal_L31 (pfam01197)→ |
| Kpn-119-1 | NZ_KK737414.1 | + | 4907268 | 4907319 | **RNA**→ | rpmE (PRK00019)Ribosomal_L31 (pfam01197)→ |
| Kpn-111-1 | NZ_KK737340.1 | + | 3355062 | 3355113 | **RNA**→ | rpmE (PRK00019)Ribosomal_L31 (pfam01197)→ |
| Kpn-101-1 | NZ_KK737216.1 | + | 3340842 | 3340893 | **RNA**→ | rpmE (PRK00019)Ribosomal_L31 (pfam01197)→ |
| Kpn-84-1 | NZ_KK737037.1 | + | 2868328 | 2868379 | **RNA**→ | rpmE (PRK00019)Ribosomal_L31 (pfam01197)→ |
| Kpn-95-1 | NZ_KK737167.1 | + | 3253271 | 3253322 | **RNA**→ | rpmE (PRK00019)Ribosomal_L31 (pfam01197)→ |
| Kpn-117-1 | NZ_KK737398.1 | + | 2884275 | 2884326 | **RNA**→ | rpmE (PRK00019)Ribosomal_L31 (pfam01197)→ |
| Kpn-56-1 | NZ_KK736752.1 | + | 73376 | 73427 | **RNA**→ | rpmE (PRK00019)Ribosomal_L31 (pfam01197)→ |
| Kpn-75-1 | NZ_KK736922.1 | + | 3226370 | 3226421 | **RNA**→ | rpmE (PRK00019)Ribosomal_L31 (pfam01197)→ |
| Kpn-66-1 | NZ_KK736824.1 | + | 2870499 | 2870550 | **RNA**→ | rpmE (PRK00019)Ribosomal_L31 (pfam01197)→ |
| Kpn-83-1 | NZ_KK737028.1 | + | 4924426 | 4924477 | **RNA**→ | rpmE (PRK00019)Ribosomal_L31 (pfam01197)→ |

| | | | | | | |
|---|---|---|---|---|---|---|
| Kpn-61-1 | NZ_KK736786.1 | + | 3318945 | 3318996 | **RNA**→ | rpmE (PRK00019)Ribosomal_L31 (pfam01197)→ |
| Kpn-80-1 | NZ_KK736984.1 | + | 4535238 | 4535289 | **RNA**→ | rpmE (PRK00019)Ribosomal_L31 (pfam01197)→ |
| Kpn-89-1 | NZ_KK737103.1 | + | 2869188 | 2869239 | **RNA**→ | rpmE (PRK00019)Ribosomal_L31 (pfam01197)→ |
| Kpn-105-1 | NZ_KK737277.1 | + | 4533715 | 4533766 | **RNA**→ | rpmE (PRK00019)Ribosomal_L31 (pfam01197)→ |
| Kpn-58-1 | NZ_KK736771.1 | + | 578958 | 579009 | **RNA**→ | rpmE (PRK00019)Ribosomal_L31 (pfam01197)→ |
| Kpn-147-1 | NZ_KK737472.1 | + | 1117063 | 1117114 | **RNA**→ | rpmE (PRK00019)Ribosomal_L31 (pfam01197)→ |
| Kpn-63-1 | NZ_KK736804.1 | + | 3369698 | 3369749 | **RNA**→ | rpmE (PRK00019)Ribosomal_L31 (pfam01197)→ |
| Kpn-124-1 | NZ_KK737466.1 | + | 4912013 | 4912064 | **RNA**→ | rpmE (PRK00019)Ribosomal_L31 (pfam01197)→ |
| Kpn-19-1 | NZ_KK736587.1 | + | 3144951 | 3145002 | **RNA**→ | rpmE (PRK00019)Ribosomal_L31 (pfam01197)→ |
| Kpn-98-1 | NZ_KK737183.1 | + | 4638212 | 4638263 | **RNA**→ | rpmE (PRK00019)Ribosomal_L31 (pfam01197)→ |
| Kpn-113-1 | NZ_KK737354.1 | + | 3354410 | 3354461 | **RNA**→ | rpmE (PRK00019)Ribosomal_L31 (pfam01197)→ |
| Kpn-108-1 | NZ_KK737323.1 | + | 4751603 | 4751654 | **RNA**→ | rpmE (PRK00019)Ribosomal_L31 (pfam01197)→ |
| Kpn-69-1 | NZ_KK736861.1 | + | 70162 | 70213 | **RNA**→ | rpmE (PRK00019)Ribosomal_L31 (pfam01197)→ |
| Kpn-81-1 | NZ_KK737002.1 | + | 69390 | 69441 | **RNA**→ | rpmE (PRK00019)Ribosomal_L31 (pfam01197)→ |
| Kpn-189-1 | NZ_KK737711.1 | + | 4562420 | 4562471 | **RNA**→ | rpmE (PRK00019)Ribosomal_L31 (pfam01197)→ |
| Kpn-74-1 | NZ_KK736911.1 | + | 4523291 | 4523342 | **RNA**→ | rpmE (PRK00019)Ribosomal_L31 (pfam01197)→ |
| Kpn-86-1 | NZ_KK737063.1 | + | 69126 | 69177 | **RNA**→ | rpmE (PRK00019)Ribosomal_L31 (pfam01197)→ |
| Kpn-60-1 | NZ_KK736776.1 | + | 4900314 | 4900365 | **RNA**→ | rpmE (PRK00019)Ribosomal_L31 (pfam01197)→ |
| Kpn-54-1 | NZ_KK736708.1 | + | 2866599 | 2866650 | **RNA**→ | rpmE (PRK00019)Ribosomal_L31 (pfam01197)→ |
| Kpn-82-1 | NZ_KK737013.1 | + | 2866351 | 2866402 | **RNA**→ | rpmE (PRK00019)Ribosomal_L31 (pfam01197)→ |
| Kpn-53-1 | NZ_KK736703.1 | + | 63762 | 63813 | **RNA**→ | rpmE (PRK00019)Ribosomal_L31 (pfam01197)→ |
| Kpn-191-1 | NZ_KK737723.1 | + | 73376 | 73427 | **RNA**→ | rpmE (PRK00019)Ribosomal_L31 (pfam01197)→ |
| Kpn-93-1 | NZ_KK737140.1 | + | 2732162 | 2732213 | **RNA**→ | rpmE (PRK00019)Ribosomal_L31 (pfam01197)→ |
| Kpn-62-1 | NZ_KK736796.1 | + | 1430900 | 1430951 | **RNA**→ | rpmE (PRK00019)Ribosomal_L31 (pfam01197)→ |
| Kpn-100-1 | NZ_KK737201.1 | + | 68880 | 68931 | **RNA**→ | rpmE (PRK00019)Ribosomal_L31 (pfam01197)→ |
| Kpn-172-1 | NZ_KL407317.1 | + | 3279007 | 3279058 | **RNA**→ | rpmE (PRK00019)Ribosomal_L31 (pfam01197)→ |
| Kpn-12-1 | NZ_KL407321.1 | + | 2250192 | 2250243 | **RNA**→ | rpmE (PRK00019)Ribosomal_L31 (pfam01197)→ |
| Kpn-24-1 | NZ_KL407345.1 | + | 1374223 | 1374274 | **RNA**→ | rpmE (PRK00019)Ribosomal_L31 (pfam01197)→ |
| Ksp-5-1 | NZ_JH414807.1 | + | 70580 | 70631 | **RNA**→ | rpmE (PRK00019)Ribosomal_L31 (pfam01197)→ |
| Kpn-360-1 | NZ_JH930422.1 | - | 123563 | 123512 | **RNA**→ | rpmE (PRK00019)Ribosomal_L31 (pfam01197)→ |
| Kpn-358-1 | NZ_JH930439.1 | - | 123670 | 123619 | **RNA**→ | rpmE (PRK00019)Ribosomal_L31 (pfam01197)→ |
| Kpn-359-1 | NZ_JH930419.1 | + | 5098754 | 5098805 | **RNA**→ | rpmE (PRK00019)Ribosomal_L31 (pfam01197)→ |
| Kpn-361-1 | NZ_JH930433.1 | + | 97306 | 97357 | **RNA**→ | rpmE (PRK00019)Ribosomal_L31 (pfam01197)→ |
| Kpn-329-1 | NZ_HF536482.1 | + | 5149422 | 5149473 | **RNA**→ | rpmE (PRK00019)Ribosomal_L31 (pfam01197)→ |
| Kpn-245-1 | NZ_KI535426.1 | + | 70377 | 70428 | **RNA**→ | rpmE (PRK00019)Ribosomal_L31 (pfam01197)→ |
| Kpn-243-1 | NZ_KI535444.1 | + | 495195 | 495246 | **RNA**→ | rpmE (PRK00019)Ribosomal_L31 (pfam01197)→ |
| Kpn-285-1 | NZ_KI535498.1 | + | 69087 | 69138 | **RNA**→ | rpmE (PRK00019)Ribosomal_L31 (pfam01197)→ |
| Kpn-283-1 | NZ_KI535509.1 | + | 4696783 | 4696834 | **RNA**→ | rpmE (PRK00019)Ribosomal_L31 (pfam01197)→ |
| Kpn-277-1 | NZ_KI535580.1 | + | 1331131 | 1331182 | **RNA**→ | rpmE (PRK00019)Ribosomal_L31 (pfam01197)→ |
| Kpn-272-1 | NZ_KI535620.1 | + | 73553 | 73604 | **RNA**→ | rpmE (PRK00019)Ribosomal_L31 (pfam01197)→ |
| Kpn-270-1 | NZ_KI535623.1 | - | 32015 | 31964 | **RNA**→ | rpmE (PRK00019)Ribosomal_L31 (pfam01197)→ |
| Kpn-253-1 | NZ_KI535560.1 | + | 69194 | 69245 | **RNA**→ | rpmE (PRK00019)Ribosomal_L31 (pfam01197)→ |
| Kpn-252-1 | NZ_KI535573.1 | - | 30634 | 30583 | **RNA**→ | rpmE (PRK00019)Ribosomal_L31 (pfam01197)→ |
| Kpn-251-1 | NZ_KI535346.1 | + | 730498 | 730549 | **RNA**→ | rpmE (PRK00019)Ribosomal_L31 (pfam01197)→ |
| Kpn-276-1 | NZ_KI535587.1 | + | 1356604 | 1356655 | **RNA**→ | rpmE (PRK00019)Ribosomal_L31 (pfam01197)→ |
| Kpn-282-1 | NZ_KI535517.1 | - | 167340 | 167289 | **RNA**→ | rpmE (PRK00019)Ribosomal_L31 (pfam01197)→ |
| Kpn-247-1 | NZ_KI535416.1 | + | 492678 | 492729 | **RNA**→ | rpmE (PRK00019)Ribosomal_L31 (pfam01197)→ |
| Kpn-250-1 | NZ_KI535359.1 | + | 826395 | 826446 | **RNA**→ | rpmE (PRK00019)Ribosomal_L31 (pfam01197)→ |
| Kpn-266-1 | NZ_KI535679.1 | - | 127694 | 127643 | **RNA**→ | rpmE (PRK00019)Ribosomal_L31 (pfam01197)→ |
| Kpn-286-1 | NZ_KI535492.1 | + | 577955 | 578006 | **RNA**→ | rpmE (PRK00019)Ribosomal_L31 (pfam01197)→ |
| Kpn-267-1 | NZ_KI535678.1 | + | 4357833 | 4357884 | **RNA**→ | rpmE (PRK00019)Ribosomal_L31 (pfam01197)→ |
| Kpn-284-1 | NZ_KI535503.1 | + | 4585522 | 4585573 | **RNA**→ | rpmE (PRK00019)Ribosomal_L31 (pfam01197)→ |

| | | | | | | |
|---|---|---|---|---|---|---|
| Kpn-273-1 | NZ_KI535611.1 | + | 70501 | 70552 | **RNA**→ | rpmE (PRK00019)Ribosomal_L31 (pfam01197)→ |
| Kpn-249-1 | NZ_KI535412.1 | + | 4539853 | 4539904 | **RNA**→ | rpmE (PRK00019)Ribosomal_L31 (pfam01197)→ |
| Kpn-246-1 | NZ_KI535420.1 | + | 4238528 | 4238579 | **RNA**→ | rpmE (PRK00019)Ribosomal_L31 (pfam01197)→ |
| Kpn-269-1 | NZ_KI535663.1 | + | 4648189 | 4648240 | **RNA**→ | rpmE (PRK00019)Ribosomal_L31 (pfam01197)→ |
| Kpn-265-1 | NZ_KI535689.1 | + | 68523 | 68574 | **RNA**→ | rpmE (PRK00019)Ribosomal_L31 (pfam01197)→ |
| Kpn-221-1 | NZ_KI517376.1 | - | 5252285 | 5252234 | **RNA**→ | rpmE (PRK00019)Ribosomal_L31 (pfam01197)→ |
| Kpn-223-1 | NZ_HG518477.1 | - | 4904955 | 4904904 | **RNA**→ | rpmE (PRK00019)Ribosomal_L31 (pfam01197)→ |
| Kpn-22-1 | NZ_KI929271.1 | + | 4503228 | 4503279 | **RNA**→ | rpmE (PRK00019)Ribosomal_L31 (pfam01197)→ |
| Kpn-208-1 | NZ_HG518479.1 | - | 4685746 | 4685695 | **RNA**→ | rpmE (PRK00019)Ribosomal_L31 (pfam01197)→ |
| Kpn-224-1 | NZ_HG518478.1 | - | 4726590 | 4726539 | **RNA**→ | rpmE (PRK00019)Ribosomal_L31 (pfam01197)→ |
| Kpn-175-1 | NZ_KK036648.1 | + | 4561960 | 4562011 | **RNA**→ | rpmE (PRK00019)Ribosomal_L31 (pfam01197)→ |
| Kpn-179-1 | NZ_KK036616.1 | + | 4695172 | 4695223 | **RNA**→ | rpmE (PRK00019)Ribosomal_L31 (pfam01197)→ |
| Kpn-248-1 | NZ_KK036740.1 | + | 1316016 | 1316067 | **RNA**→ | rpmE (PRK00019)Ribosomal_L31 (pfam01197)→ |
| Kpn-182-1 | NZ_KK036586.1 | + | 69272 | 69323 | **RNA**→ | rpmE (PRK00019)Ribosomal_L31 (pfam01197)→ |
| Kpn-45-1 | NZ_KK036932.1 | + | 66222 | 66273 | **RNA**→ | rpmE (PRK00019)Ribosomal_L31 (pfam01197)→ |
| Kpn-184-1 | NZ_KK036571.1 | + | 4342385 | 4342436 | **RNA**→ | rpmE (PRK00019)Ribosomal_L31 (pfam01197)→ |
| Kpn-37-1 | NZ_KK036686.1 | + | 4560269 | 4560320 | **RNA**→ | rpmE (PRK00019)Ribosomal_L31 (pfam01197)→ |
| Kpn-146-1 | NZ_KK036944.1 | + | 2970777 | 2970828 | **RNA**→ | rpmE (PRK00019)Ribosomal_L31 (pfam01197)→ |
| Kpn-271-1 | NZ_KK036969.1 | + | 1997125 | 1997176 | **RNA**→ | rpmE (PRK00019)Ribosomal_L31 (pfam01197)→ |
| Kpn-186-1 | NZ_KK036555.1 | + | 1862408 | 1862459 | **RNA**→ | rpmE (PRK00019)Ribosomal_L31 (pfam01197)→ |
| Kpn-35-1 | NZ_KK036701.1 | + | 4380641 | 4380692 | **RNA**→ | rpmE (PRK00019)Ribosomal_L31 (pfam01197)→ |
| Kpn-27-1 | NZ_KK036732.1 | + | 2646020 | 2646071 | **RNA**→ | rpmE (PRK00019)Ribosomal_L31 (pfam01197)→ |
| Kpn-176-1 | NZ_KK036640.1 | + | 580651 | 580702 | **RNA**→ | rpmE (PRK00019)Ribosomal_L31 (pfam01197)→ |
| Kpn-183-1 | NZ_KK036575.1 | + | 4843690 | 4843741 | **RNA**→ | rpmE (PRK00019)Ribosomal_L31 (pfam01197)→ |
| Kpn-180-1 | NZ_KK036610.1 | + | 1384294 | 1384345 | **RNA**→ | rpmE (PRK00019)Ribosomal_L31 (pfam01197)→ |
| Kpn-36-1 | NZ_KK036697.1 | + | 69043 | 69094 | **RNA**→ | rpmE (PRK00019)Ribosomal_L31 (pfam01197)→ |
| Kpn-44-1 | NZ_KK036866.1 | + | 70380 | 70431 | **RNA**→ | rpmE (PRK00019)Ribosomal_L31 (pfam01197)→ |
| Kpn-26-1 | NZ_KK036882.1 | + | 68932 | 68983 | **RNA**→ | rpmE (PRK00019)Ribosomal_L31 (pfam01197)→ |
| Kpn-21-1 | NZ_KK036888.1 | + | 1217538 | 1217589 | **RNA**→ | rpmE (PRK00019)Ribosomal_L31 (pfam01197)→ |
| Kpn-141-1 | NZ_KK036984.1 | + | 71754 | 71805 | **RNA**→ | rpmE (PRK00019)Ribosomal_L31 (pfam01197)→ |
| Kpn-177-1 | NZ_KK036634.1 | + | 694277 | 694328 | **RNA**→ | rpmE (PRK00019)Ribosomal_L31 (pfam01197)→ |
| Kpn-198-1 | NZ_KK037007.1 | + | 2272584 | 2272635 | **RNA**→ | rpmE (PRK00019)Ribosomal_L31 (pfam01197)→ |
| Kpn-29-1 | NZ_KK036680.1 | + | 326960 | 327011 | **RNA**→ | rpmE (PRK00019)Ribosomal_L31 (pfam01197)→ |
| Kpn-185-1 | NZ_KK036563.1 | + | 4808777 | 4808828 | **RNA**→ | rpmE (PRK00019)Ribosomal_L31 (pfam01197)→ |
| Kpn-23-1 | NZ_KK036758.1 | + | 1229368 | 1229419 | **RNA**→ | rpmE (PRK00019)Ribosomal_L31 (pfam01197)→ |
| Kpn-30-1 | NZ_KK036718.1 | + | 4570752 | 4570803 | **RNA**→ | rpmE (PRK00019)Ribosomal_L31 (pfam01197)→ |
| Kpn-18-1 | NZ_KK036784.1 | + | 1382201 | 1382252 | **RNA**→ | rpmE (PRK00019)Ribosomal_L31 (pfam01197)→ |
| Kpn-13-1 | NZ_KK036863.1 | + | 66760 | 66811 | **RNA**→ | rpmE (PRK00019)Ribosomal_L31 (pfam01197)→ |
| Kpn-32-1 | NZ_KK036710.1 | + | 73184 | 73235 | **RNA**→ | rpmE (PRK00019)Ribosomal_L31 (pfam01197)→ |
| Kpn-181-1 | NZ_KK036593.1 | + | 4760906 | 4760957 | **RNA**→ | rpmE (PRK00019)Ribosomal_L31 (pfam01197)→ |
| Kpn-197-1 | NZ_KK036901.1 | - | 655958 | 655907 | **RNA**→ | rpmE (PRK00019)Ribosomal_L31 (pfam01197)→ |
| Kpn-20-1 | NZ_KK036779.1 | + | 73328 | 73379 | **RNA**→ | rpmE (PRK00019)Ribosomal_L31 (pfam01197)→ |
| Kpn-43-1 | NZ_KK037004.1 | + | 69390 | 69441 | **RNA**→ | rpmE (PRK00019)Ribosomal_L31 (pfam01197)→ |
| Kpn-16-1 | NZ_KK036799.1 | - | 127335 | 127284 | **RNA**→ | rpmE (PRK00019)Ribosomal_L31 (pfam01197)→ |
| Kpn-11-1 | NZ_KK036895.1 | + | 580946 | 580997 | **RNA**→ | rpmE (PRK00019)Ribosomal_L31 (pfam01197)→ |
| Kpn-15-1 | NZ_KK036828.1 | + | 4511150 | 4511201 | **RNA**→ | rpmE (PRK00019)Ribosomal_L31 (pfam01197)→ |
| Kpn-49-1 | NZ_KK036908.1 | + | 5252304 | 5252355 | **RNA**→ | rpmE (PRK00019)Ribosomal_L31 (pfam01197)→ |
| Kpn-47-1 | NZ_KK036921.1 | + | 4589468 | 4589519 | **RNA**→ | rpmE (PRK00019)Ribosomal_L31 (pfam01197)→ |
| Kpn-28-1 | NZ_KK036724.1 | + | 2619848 | 2619899 | **RNA**→ | rpmE (PRK00019)Ribosomal_L31 (pfam01197)→ |
| Kpn-48-1 | NZ_KK036911.1 | + | 3085051 | 3085102 | **RNA**→ | rpmE (PRK00019)Ribosomal_L31 (pfam01197)→ |
| Kpn-25-1 | NZ_KK036744.1 | + | 3484063 | 3484114 | **RNA**→ | rpmE (PRK00019)Ribosomal_L31 (pfam01197)→ |
| Kpn-14-1 | NZ_KK036838.1 | + | 4750317 | 4750368 | **RNA**→ | rpmE (PRK00019)Ribosomal_L31 (pfam01197)→ |

| | | | | | | | |
|---|---|---|---|---|---|---|---|
| Kpn-33-1 | NZ_KK036705.1 | + | 4723059 | 4723110 | **RNA**→ | rpmE (PRK00019)Ribosomal_L31 (pfam01197)→ | |
| Kpn-145-1 | NZ_KK036949.1 | + | 4329695 | 4329746 | **RNA**→ | rpmE (PRK00019)Ribosomal_L31 (pfam01197)→ | |
| Kpn-173-1 | NZ_KK036663.1 | - | 661579 | 661528 | **RNA**→ | rpmE (PRK00019)Ribosomal_L31 (pfam01197)→ | |
| Kpn-144-1 | NZ_KK036953.1 | + | 1102473 | 1102524 | **RNA**→ | rpmE (PRK00019)Ribosomal_L31 (pfam01197)→ | |
| Kpn-174-1 | NZ_KK036655.1 | + | 801825 | 801876 | **RNA**→ | rpmE (PRK00019)Ribosomal_L31 (pfam01197)→ | |
| Kpn-178-1 | NZ_KK037012.1 | + | 1362262 | 1362313 | **RNA**→ | rpmE (PRK00019)Ribosomal_L31 (pfam01197)→ | |
| Kpn-34-1 | NZ_KK037013.1 | - | 608696 | 608645 | **RNA**→ | rpmE (PRK00019)Ribosomal_L31 (pfam01197)→ | |
| Kpn-349-1 | NZ_JH930400.1 | - | 121654 | 121603 | **RNA**→ | rpmE (PRK00019)Ribosomal_L31 (pfam01197)→ | |
| Kpn-139-1 | NZ_KE504626.1 | - | 5255393 | 5255342 | **RNA**→ | rpmE (PRK00019)Ribosomal_L31 (pfam01197)→ | |
| Kpn-140-1 | NZ_KE504629.1 | - | 5515152 | 5515101 | **RNA**→ | rpmE (PRK00019)Ribosomal_L31 (pfam01197)→ | |
| Kpn-279-1 | NC_012731.1 | + | 89967 | 90018 | **RNA**→ | rpmE (PRK00019)Ribosomal_L31 (pfam01197)→ | |
| Kpn-261-1 | NC_017540.1 | + | 5087242 | 5087293 | **RNA**→ | rpmE (PRK00019)Ribosomal_L31 (pfam01197)→ | |
| Kpn-202-1 | NZ_CP011989.1 | - | 5205226 | 5205175 | **RNA**→ | rpmE (PRK00019)Ribosomal_L31 (pfam01197)→ | |
| Kpn-201-1 | NZ_CP011985.1 | - | 5196465 | 5196414 | **RNA**→ | rpmE (PRK00019)Ribosomal_L31 (pfam01197)→ | |
| Kpn-125-1 | NZ_CP011976.1 | - | 5269356 | 5269305 | **RNA**→ | rpmE (PRK00019)Ribosomal_L31 (pfam01197)→ | |
| Kpn-325-1 | NC_018522.1 | - | 5262049 | 5261998 | **RNA**→ | rpmE (PRK00019)Ribosomal_L31 (pfam01197)→ | rpmE (PRK00019)Ribosomal_L31 (pfam01197)→ |
| Kpn-260-1 | NC_022082.1 | - | 5183982 | 5183931 | **RNA**→ | rpmE (PRK00019)Ribosomal_L31 (pfam01197)→ | rpmE (PRK00019)Ribosomal_L31 (pfam01197)→ |
| Kpn-254-1 | NC_022566.1 | + | 4488386 | 4488437 | **RNA**→ | rpmE (PRK00019)Ribosomal_L31 (pfam01197)→ | |
| Kpn-1-2 | NZ_CP008929.1 | + | 1842237 | 1842288 | **RNA**→ | rpmE (PRK00019)Ribosomal_L31 (pfam01197)→ | |
| Kpn-1-3 | NZ_CP009461.1 | - | 4775192 | 4775141 | **RNA**→ | rpmE (PRK00019)Ribosomal_L31 (pfam01197)→ | |
| Kpn-206-1 | NZ_CP009775.1 | + | 90073 | 90124 | **RNA**→ | rpmE (PRK00019)Ribosomal_L31 (pfam01197)→ | |
| Kpn-206-2 | NZ_CP009872.1 | + | 90073 | 90124 | **RNA**→ | rpmE (PRK00019)Ribosomal_L31 (pfam01197)→ | |
| Kpn-206-3 | NZ_CP009771.1 | + | 90073 | 90124 | **RNA**→ | rpmE (PRK00019)Ribosomal_L31 (pfam01197)→ | |
| Kpn-206-4 | NZ_CP009863.1 | + | 85038 | 85089 | **RNA**→ | rpmE (PRK00019)Ribosomal_L31 (pfam01197)→ | |
| Kpn-129-1 | NZ_CP006738.1 | - | 5209835 | 5209784 | **RNA**→ | rpmE (PRK00019)Ribosomal_L31 (pfam01197)→ | |
| Kpn-207-1 | NZ_CP006722.1 | - | 5216292 | 5216241 | **RNA**→ | rpmE (PRK00019)Ribosomal_L31 (pfam01197)→ | |
| Kpn-1-4 | NZ_CP010392.1 | - | 5189077 | 5189026 | **RNA**→ | rpmE (PRK00019)Ribosomal_L31 (pfam01197)→ | |
| Kpn-1-9 | NZ_LK022721.1 | + | 63891 | 63942 | **RNA**→ | rpmE (PRK00019)Ribosomal_L31 (pfam01197)→ | |
| Kpn-222-1 | NZ_CP006798.1 | - | 3463792 | 3463741 | **RNA**→ | rpmE (PRK00019)Ribosomal_L31 (pfam01197)→ | |
| Kpn-218-1 | NZ_CP003999.1 | - | 5187695 | 5187644 | **RNA**→ | rpmE (PRK00019)Ribosomal_L31 (pfam01197)→ | |
| Kpn-17-1 | NZ_KK036791.1 | + | 1461115 | 1461166 | **RNA**→ | rpmE (PRK00019)Ribosomal_L31 (pfam01197)→ | |
| Kpn-4-1 | NZ_CP006923.1 | - | 5138876 | 5138825 | **RNA**→ | rpmE (PRK00019)Ribosomal_L31 (pfam01197)→ | |
| Kpn-5-1 | NZ_CP006918.1 | - | 5168803 | 5168752 | **RNA**→ | rpmE (PRK00019)Ribosomal_L31 (pfam01197)→ | |
| Kpn-214-1 | NZ_CP007727.1 | + | 90073 | 90124 | **RNA**→ | rpmE (PRK00019)Ribosomal_L31 (pfam01197)→ | |
| Kpn-52-1 | NZ_KK736687.1 | + | 69390 | 69441 | **RNA**→ | rpmE (PRK00019)Ribosomal_L31 (pfam01197)→ | |
| Kpn-215-1 | NZ_CP008797.1 | + | 90073 | 90124 | **RNA**→ | rpmE (PRK00019)Ribosomal_L31 (pfam01197)→ | |
| Kpn-217-1 | NZ_CP008831.1 | + | 90073 | 90124 | **RNA**→ | rpmE (PRK00019)Ribosomal_L31 (pfam01197)→ | |
| Kpn-213-1 | NZ_CP008827.1 | + | 90073 | 90124 | **RNA**→ | rpmE (PRK00019)Ribosomal_L31 (pfam01197)→ | |
| Kpn-1-5 | NZ_CP009114.1 | + | 3598750 | 3598801 | **RNA**→ | rpmE (PRK00019)Ribosomal_L31 (pfam01197)→ | |
| Kpn-206-5 | NZ_CP009208.1 | - | 3330554 | 3330503 | **RNA**→ | rpmE (PRK00019)Ribosomal_L31 (pfam01197)→ | |
| Kpn-8-1 | NZ_CP011980.1 | - | 5183712 | 5183661 | **RNA**→ | rpmE (PRK00019)Ribosomal_L31 (pfam01197)→ | |
| Eco-902-1 | NZ_KI522728.1 | + | 10412 | 10463 | **RNA**→ | rpmE (PRK00019)Ribosomal_L31 (pfam01197)→ | rpmE (PRK00019)Ribosomal_L31 (pfam01197)→ |
| Kpn-225-1 | NC_021232.1 | + | 194552 | 194603 | **RNA**→ | rpmE (PRK00019)Ribosomal_L31 (pfam01197)→ | |
| Kpn-1-6 | NZ_CP011578.1 | + | 3572358 | 3572409 | **RNA**→ | rpmE (PRK00019)Ribosomal_L31 (pfam01197)→ | |
| Kpn-1-7 | NZ_CP011647.1 | + | 1269750 | 1269801 | **RNA**→ | rpmE (PRK00019)Ribosomal_L31 (pfam01197)→ | |
| Kpn-1-8 | NZ_CP011624.1 | - | 2090102 | 2090051 | **RNA**→ | rpmE (PRK00019)Ribosomal_L31 (pfam01197)→ | |
| Kpn-324-1 | NZ_AOIZ01000021.1 | - | 8000 | 7949 | **RNA**→ | rpmE (PRK00019)Ribosomal_L31 (pfam01197)→ | |
| Kpn-238-1 | NZ_AOGQ01000046.1 | + | 69011 | 69062 | **RNA**→ | rpmE (PRK00019)Ribosomal_L31 (pfam01197)→ | |
| Kpn-327-1 | NZ_AQCG01001093.1 | + | 1849 | 1900 | **RNA**→ | rpmE (PRK00019)Ribosomal_L31 (pfam01197)→ | |
| Ksp-1-1 | NZ_AKYX01000005.1 | + | 73137 | 73188 | **RNA**→ | rpmE (PRK00019)Ribosomal_L31 (pfam01197)→ | |
| Kpn-137-1 | NZ_AQPG01000012.1 | + | 5338757 | 5338808 | **RNA**→ | rpmE (PRK00019)Ribosomal_L31 (pfam01197)→ | |
| Kpn-170-1 | NZ_JAQL01000017.1 | - | 30330 | 30279 | **RNA**→ | rpmE (PRK00019)Ribosomal_L31 (pfam01197)→ | |

| | | | | | | | |
|---|---|---|---|---|---|---|---|
| Kpn-171-1 | NZ_JAQM01000046.1 | - | 15043 | 14992 | **RNA**→ | rpmE (PRK00019)Ribosomal_L31 (pfam01197)→ | |
| Kpn-259-1 | NZ_ANGH02000122.1 | - | 30407 | 30356 | **RNA**→ | rpmE (PRK00019)Ribosomal_L31 (pfam01197)→ | |
| Kpn-333-1 | NZ_AKAF01000019.1 | + | 69079 | 69130 | **RNA**→ | rpmE (PRK00019)Ribosomal_L31 (pfam01197)→ | |
| Kpn-335-1 | NZ_AKAH01000021.1 | + | 69289 | 69340 | **RNA**→ | rpmE (PRK00019)Ribosomal_L31 (pfam01197)→ | |
| Kpn-338-1 | NZ_AJZV01000021.1 | + | 69354 | 69405 | **RNA**→ | rpmE (PRK00019)Ribosomal_L31 (pfam01197)→ | |
| Kpn-334-1 | NZ_AKAG01000020.1 | - | 30706 | 30655 | **RNA**→ | rpmE (PRK00019)Ribosomal_L31 (pfam01197)→ | |
| env-754 | topACOD_FV90NF401A4K61 | + | 172 | 223 | **RNA**→ | rpmE (PRK00019)Ribosomal_L31 (pfam01197)→ | |
| env-755 | CLOF_F4EQGSL02IC6OG | - | 459 | 408 | **RNA**→ | rpmE (PRK00019)Ribosomal_L31 (pfam01197)→ | |
| env-756 | CLOF_F4EQGSL02FU17A | - | 147 | 96 | **RNA**→ | rpmE (PRK00019)Ribosomal_L31 (pfam01197)→ | |
| Kcr-1-1 | NZ_LGHZ01000022.1 | + | 77178 | 77229 | **RNA**→ | rpmE (PRK00019)Ribosomal_L31 (pfam01197)→ | |
| Kas-1-1 | NZ_JMPL01000010.1 | + | 63578 | 63629 | **RNA**→ | rpmE (PRK00019)Ribosomal_L31 (pfam01197)→ | |
| Fhe-1-1 | NZ_JVIC01000038.1 | + | 35499 | 35550 | **RNA**→ | rpmE (PRK00019)Ribosomal_L31 (pfam01197)→ | |
| Fhe-4-1 | NZ_AWFX01000038.1 | - | 54843 | 54792 | **RNA**→ | rpmE (PRK00019)Ribosomal_L31 (pfam01197)→ | |
| Fhe-2-1 | NZ_AXDL01000043.1 | - | 6798 | 6747 | **RNA**→ | rpmE (PRK00019)Ribosomal_L31 (pfam01197)→ | |
| Fpu-1-1 | NZ_LFEJ01000010.1 | + | 152430 | 152481 | **RNA**→ | rpmE (PRK00019)Ribosomal_L31 (pfam01197)→ | |
| Fpu-3-1 | NZ_AXSY01000052.1 | - | 32389 | 32338 | **RNA**→ | rpmE (PRK00019)Ribosomal_L31 (pfam01197)→ | |
| Fpu-2-1 | NZ_AXSZ01000028.1 | - | 11077 | 11026 | **RNA**→ | rpmE (PRK00019)Ribosomal_L31 (pfam01197)→ | |
| Fpu-4-1 | NZ_JHYZ01000015.1 | - | 32736 | 32685 | **RNA**→ | rpmE (PRK00019)Ribosomal_L31 (pfam01197)→ | |
| Fhe-3-1 | NZ_AXDK01000124.1 | - | 8590 | 8539 | **RNA**→ | rpmE (PRK00019)Ribosomal_L31 (pfam01197)→ | |
| Fpu-5-1 | NZ_AWFY01000004.1 | + | 156607 | 156658 | **RNA**→ | rpmE (PRK00019)Ribosomal_L31 (pfam01197)→ | |
| Cne-1-1 | NZ_CP009459.1 | + | 1199721 | 1199772 | **RNA**→ | rpmE (PRK00019)Ribosomal_L31 (pfam01197)→ | |
| Cne-1-2 | NZ_CP009458.1 | + | 1216774 | 1216825 | **RNA**→ | rpmE (PRK00019)Ribosomal_L31 (pfam01197)→ | |
| Esp-68-1 | NZ_AKXM01000024.1 | + | 88396 | 88447 | **RNA**→ | rpmE (PRK00019)Ribosomal_L31 (pfam01197)→ | rpmE (PRK00019)Ribosomal_L31 (pfam01197)→ |
| Cne-1-3 | NZ_CP009451.1 | - | 2536387 | 2536336 | **RNA**→ | rpmE (PRK00019)Ribosomal_L31 (pfam01197)→ | |
| Cda-1-1 | NZ_KE161030.1 | + | 4763143 | 4763194 | **RNA**→ | rpmE (PRK00019)Ribosomal_L31 (pfam01197)→ | |
| Sco-2-1 | NZ_JMSQ01000013.1 | + | 67335 | 67386 | **RNA**→ | rpmE (PRK00019)Ribosomal_L31 (pfam01197)→ | |
| Bag-2-1 | NZ_JMPI01000074.1 | + | 98246 | 98298 | **RNA**→ | rpmE (PRK00019)Ribosomal_L31 (pfam01197)→ | |
| Ebl-1-1 | NC_017910.1 | - | 3996940 | 3996888 | **RNA**→ | rpmE (PRK00019)Ribosomal_L31 (pfam01197)→ | |
| Sbl-1-1 | NZ_BAHA01000017.1 | + | 91171 | 91223 | **RNA**→ | rpmE (PRK00019)Ribosomal_L31 (pfam01197)→ | |
| Sen-311-1 | NC_010067.1 | - | 3496303 | 3496251 | **RNA**→ | rpmE (PRK00019)Ribosomal_L31 (pfam01197)→ | rpmE (PRK00019)Ribosomal_L31 (pfam01197)→ |
| Sen-3-1 | NZ_CP006693.1 | - | 3463342 | 3463290 | **RNA**→ | rpmE (PRK00019)Ribosomal_L31 (pfam01197)→ | |
| Sen-1013-1 | NZ_ABAN01000010.1 | - | 45116 | 45064 | **RNA**→ | rpmE (PRK00019)Ribosomal_L31 (pfam01197)→ | rpmE (PRK00019)Ribosomal_L31 (pfam01197)→ |
| Sen-89-1 | NZ_JRLP01000010.1 | + | 43143 | 43195 | **RNA**→ | rpmE (PRK00019)Ribosomal_L31 (pfam01197)→ | |
| Sen-729-1 | NZ_JSWQ01000008.1 | - | 49158 | 49106 | **RNA**→ | rpmE (PRK00019)Ribosomal_L31 (pfam01197)→ | |
| Sen-1007-1 | NZ_ABEL01000012.1 | - | 37576 | 37524 | **RNA**→ | rpmE (PRK00019)Ribosomal_L31 (pfam01197)→ | rpmE (PRK00019)Ribosomal_L31 (pfam01197)→ |
| Sen-1030-1 | NZ_ABFF01000013.1 | - | 37462 | 37410 | **RNA**→ | rpmE (PRK00019)Ribosomal_L31 (pfam01197)→ | rpmE (PRK00019)Ribosomal_L31 (pfam01197)→ |
| Sen-1028-1 | NZ_ABAO01000008.1 | + | 112768 | 112820 | **RNA**→ | rpmE (PRK00019)Ribosomal_L31 (pfam01197)→ | rpmE (PRK00019)Ribosomal_L31 (pfam01197)→ |
| Sen-1008-1 | NZ_ABEI01000014.1 | + | 110111 | 110163 | **RNA**→ | rpmE (PRK00019)Ribosomal_L31 (pfam01197)→ | rpmE (PRK00019)Ribosomal_L31 (pfam01197)→ |
| Sen-239-1 | NZ_ARYW01000006.1 | - | 45147 | 45095 | **RNA**→ | rpmE (PRK00019)Ribosomal_L31 (pfam01197)→ | |
| Sen-940-1 | NZ_AOXX01000025.1 | + | 95096 | 95148 | **RNA**→ | rpmE (PRK00019)Ribosomal_L31 (pfam01197)→ | rpmE (PRK00019)Ribosomal_L31 (pfam01197)→ |
| Sen-929-1 | NZ_AHUI01000026.1 | + | 42783 | 42835 | **RNA**→ | rpmE (PRK00019)Ribosomal_L31 (pfam01197)→ | |
| Sen-58-1 | NZ_JRLS01000058.1 | + | 4123 | 4175 | **RNA**→ | rpmE (PRK00019)Ribosomal_L31 (pfam01197)→ | |
| Sen-111-1 | NZ_JXXJ01000012.1 | - | 45085 | 45033 | **RNA**→ | rpmE (PRK00019)Ribosomal_L31 (pfam01197)→ | |
| Sen-329-1 | NZ_CATD01000035.1 | - | 37552 | 37500 | **RNA**→ | rpmE (PRK00019)Ribosomal_L31 (pfam01197)→ | |
| Sen-350-1 | NZ_CATW01000042.1 | - | 37552 | 37500 | **RNA**→ | rpmE (PRK00019)Ribosomal_L31 (pfam01197)→ | |
| Sen-408-1 | NZ_CARI01000040.1 | + | 111839 | 111891 | **RNA**→ | rpmE (PRK00019)Ribosomal_L31 (pfam01197)→ | |
| Sen-413-1 | NZ_CARD01000040.1 | - | 37553 | 37501 | **RNA**→ | rpmE (PRK00019)Ribosomal_L31 (pfam01197)→ | |
| Sen-403-1 | NZ_CARL01000082.1 | + | 25648 | 25700 | **RNA**→ | rpmE (PRK00019)Ribosomal_L31 (pfam01197)→ | |
| Sen-288-1 | NZ_CDLI01000125.1 | - | 39519 | 39467 | **RNA**→ | rpmE (PRK00019)Ribosomal_L31 (pfam01197)→ | |
| Sen-1027-2 | NZ_ABEJ01000012.1 | + | 103003 | 103055 | **RNA**→ | rpmE (PRK00019)Ribosomal_L31 (pfam01197)→ | rpmE (PRK00019)Ribosomal_L31 (pfam01197)→ |
| Sen-1009-1 | NZ_ABAK02000001.1 | + | 3619652 | 3619704 | **RNA**→ | rpmE (PRK00019)Ribosomal_L31 (pfam01197)→ | rpmE (PRK00019)Ribosomal_L31 (pfam01197)→ |
| Sen-1010-1 | NZ_ABEW01000011.1 | + | 105665 | 105717 | **RNA**→ | rpmE (PRK00019)Ribosomal_L31 (pfam01197)→ | rpmE (PRK00019)Ribosomal_L31 (pfam01197)→ |

| | | | | | | | |
|---|---|---|---|---|---|---|---|
| Sen-813-1 | NZ_AESY01000065.1 | - | 35204 | 35152 | **RNA**→ | rpmE (PRK00019)Ribosomal_L31 (pfam01197)→ | rpmE (PRK00019)Ribosomal_L31 (pfam01197)→ |
| Sen-228-1 | NZ_JZWK01000010.1 | - | 45598 | 45546 | **RNA**→ | rpmE (PRK00019)Ribosomal_L31 (pfam01197)→ | |
| Sen-756-1 | NZ_AMNZ01000014.1 | + | 44988 | 45040 | **RNA**→ | rpmE (PRK00019)Ribosomal_L31 (pfam01197)→ | |
| Sen-188-1 | NZ_AMOF01000015.1 | + | 100319 | 100371 | **RNA**→ | rpmE (PRK00019)Ribosomal_L31 (pfam01197)→ | |
| Sen-185-1 | NZ_AMOC01000019.1 | - | 37507 | 37455 | **RNA**→ | rpmE (PRK00019)Ribosomal_L31 (pfam01197)→ | |
| Sen-184-1 | NZ_AMOA01000042.1 | - | 37497 | 37445 | **RNA**→ | rpmE (PRK00019)Ribosomal_L31 (pfam01197)→ | |
| Sen-187-1 | NZ_AMOE01000058.1 | + | 100554 | 100606 | **RNA**→ | rpmE (PRK00019)Ribosomal_L31 (pfam01197)→ | |
| Sen-189-1 | NZ_AMOG01000112.1 | + | 58896 | 58948 | **RNA**→ | rpmE (PRK00019)Ribosomal_L31 (pfam01197)→ | |
| Sen-112-1 | NZ_JZUV01000021.1 | + | 104758 | 104810 | **RNA**→ | rpmE (PRK00019)Ribosomal_L31 (pfam01197)→ | |
| Sen-113-1 | NZ_JZUW01000036.1 | - | 49763 | 49711 | **RNA**→ | rpmE (PRK00019)Ribosomal_L31 (pfam01197)→ | |
| Sen-222-1 | NZ_AMLQ01000020.1 | - | 37602 | 37550 | **RNA**→ | rpmE (PRK00019)Ribosomal_L31 (pfam01197)→ | |
| Sen-220-1 | NZ_AMLS01000127.1 | - | 14386 | 14334 | **RNA**→ | rpmE (PRK00019)Ribosomal_L31 (pfam01197)→ | |
| Sen-186-1 | NZ_AMOD01000033.1 | + | 100556 | 100608 | **RNA**→ | rpmE (PRK00019)Ribosomal_L31 (pfam01197)→ | |
| Sen-192-1 | NZ_AMOJ01000025.1 | - | 37710 | 37658 | **RNA**→ | rpmE (PRK00019)Ribosomal_L31 (pfam01197)→ | |
| Sen-190-1 | NZ_AMOH01000008.1 | - | 37530 | 37478 | **RNA**→ | rpmE (PRK00019)Ribosomal_L31 (pfam01197)→ | |
| Sen-759-1 | NZ_AMOI01000089.1 | - | 37516 | 37464 | **RNA**→ | rpmE (PRK00019)Ribosomal_L31 (pfam01197)→ | |
| Sen-191-1 | NZ_AMOK01000144.1 | + | 100556 | 100608 | **RNA**→ | rpmE (PRK00019)Ribosomal_L31 (pfam01197)→ | |
| Sen-232-1 | NZ_JZUU01000023.1 | + | 129364 | 129416 | **RNA**→ | rpmE (PRK00019)Ribosomal_L31 (pfam01197)→ | |
| Sen-758-1 | NZ_AMOB01000059.1 | + | 100641 | 100693 | **RNA**→ | rpmE (PRK00019)Ribosomal_L31 (pfam01197)→ | |
| Sen-309-1 | NZ_JXTT01000068.1 | + | 95702 | 95754 | **RNA**→ | rpmE (PRK00019)Ribosomal_L31 (pfam01197)→ | |
| Sen-167-1 | NZ_LAPA01000017.1 | - | 45228 | 45176 | **RNA**→ | rpmE (PRK00019)Ribosomal_L31 (pfam01197)→ | |
| Sen-170-1 | NZ_LAPK01000016.1 | - | 37548 | 37496 | **RNA**→ | rpmE (PRK00019)Ribosomal_L31 (pfam01197)→ | |
| Sen-286-1 | NZ_LAPB01000004.1 | - | 45128 | 45076 | **RNA**→ | rpmE (PRK00019)Ribosomal_L31 (pfam01197)→ | |
| Sen-167-2 | NZ_LAPE01000030.1 | + | 100068 | 100120 | **RNA**→ | rpmE (PRK00019)Ribosomal_L31 (pfam01197)→ | |
| Sen-167-3 | NZ_LAOV01000025.1 | - | 45125 | 45073 | **RNA**→ | rpmE (PRK00019)Ribosomal_L31 (pfam01197)→ | |
| Sen-170-2 | NZ_LAPI01000047.1 | + | 100376 | 100428 | **RNA**→ | rpmE (PRK00019)Ribosomal_L31 (pfam01197)→ | |
| Sen-225-1 | NZ_LAOZ01000065.1 | - | 37573 | 37521 | **RNA**→ | rpmE (PRK00019)Ribosomal_L31 (pfam01197)→ | |
| Sen-170-3 | NZ_LAPJ01000049.1 | - | 37548 | 37496 | **RNA**→ | rpmE (PRK00019)Ribosomal_L31 (pfam01197)→ | |
| Sen-225-2 | NZ_LAOT01000033.1 | - | 37573 | 37521 | **RNA**→ | rpmE (PRK00019)Ribosomal_L31 (pfam01197)→ | |
| Sen-225-3 | NZ_LAOW01000010.1 | - | 15419 | 15367 | **RNA**→ | rpmE (PRK00019)Ribosomal_L31 (pfam01197)→ | |
| Sen-170-4 | NZ_LAOS01000078.1 | - | 17555 | 17503 | **RNA**→ | rpmE (PRK00019)Ribosomal_L31 (pfam01197)→ | |
| Sen-225-4 | NZ_LAPL01000035.1 | + | 55787 | 55839 | **RNA**→ | rpmE (PRK00019)Ribosomal_L31 (pfam01197)→ | |
| Sen-98-1 | NZ_JMML02000005.1 | + | 103476 | 103528 | **RNA**→ | rpmE (PRK00019)Ribosomal_L31 (pfam01197)→ | |
| Sen-856-1 | NZ_AOYV01000096.1 | - | 45077 | 45025 | **RNA**→ | rpmE (PRK00019)Ribosomal_L31 (pfam01197)→ | rpmE (PRK00019)Ribosomal_L31 (pfam01197)→ |
| Sen-167-4 | NZ_LHHD01000025.1 | - | 45515 | 45463 | **RNA**→ | rpmE (PRK00019)Ribosomal_L31 (pfam01197)→ | |
| Sen-167-5 | NZ_LHIP01000011.1 | - | 45288 | 45236 | **RNA**→ | rpmE (PRK00019)Ribosomal_L31 (pfam01197)→ | |
| Sen-167-6 | NZ_LHNA01000015.1 | + | 100286 | 100338 | **RNA**→ | rpmE (PRK00019)Ribosomal_L31 (pfam01197)→ | |
| Sen-170-5 | NZ_LHNR01000007.1 | - | 37518 | 37466 | **RNA**→ | rpmE (PRK00019)Ribosomal_L31 (pfam01197)→ | |
| Sen-170-6 | NZ_LHEU01000001.1 | - | 37518 | 37466 | **RNA**→ | rpmE (PRK00019)Ribosomal_L31 (pfam01197)→ | |
| Sen-170-7 | NZ_LHFX01000020.1 | - | 37518 | 37466 | **RNA**→ | rpmE (PRK00019)Ribosomal_L31 (pfam01197)→ | |
| Sen-170-8 | NZ_LHGB01000017.1 | - | 37518 | 37466 | **RNA**→ | rpmE (PRK00019)Ribosomal_L31 (pfam01197)→ | |
| Sen-170-50 | NZ_LHHQ01000072.1 | - | 37518 | 37466 | **RNA**→ | rpmE (PRK00019)Ribosomal_L31 (pfam01197)→ | |
| Sen-170-9 | NZ_LHHT01000023.1 | - | 37518 | 37466 | **RNA**→ | rpmE (PRK00019)Ribosomal_L31 (pfam01197)→ | |
| Sen-167-7 | NZ_LHIZ01000020.1 | + | 100286 | 100338 | **RNA**→ | rpmE (PRK00019)Ribosomal_L31 (pfam01197)→ | |
| Sen-167-8 | NZ_LHJE01000040.1 | + | 100286 | 100338 | **RNA**→ | rpmE (PRK00019)Ribosomal_L31 (pfam01197)→ | |
| Sen-170-10 | NZ_LHKX01000011.1 | + | 43559 | 43611 | **RNA**→ | rpmE (PRK00019)Ribosomal_L31 (pfam01197)→ | |
| Sen-771-1 | NZ_AMMK01000337.1 | - | 34305 | 34253 | **RNA**→ | rpmE (PRK00019)Ribosomal_L31 (pfam01197)→ | |
| Sen-288-2 | NZ_CUZH01000020.1 | - | 39384 | 39332 | **RNA**→ | rpmE (PRK00019)Ribosomal_L31 (pfam01197)→ | |
| Sen-300-1 | NZ_CTNK01000018.1 | + | 42508 | 42560 | **RNA**→ | rpmE (PRK00019)Ribosomal_L31 (pfam01197)→ | |
| Sen-755-1 | NZ_AMMO01000064.1 | + | 1167 | 1219 | **RNA**→ | rpmE (PRK00019)Ribosomal_L31 (pfam01197)→ | |
| Sen-196-1 | NZ_AMON01000006.1 | - | 37717 | 37665 | **RNA**→ | rpmE (PRK00019)Ribosomal_L31 (pfam01197)→ | |
| Sen-288-3 | NZ_CUZP01000016.1 | - | 39536 | 39484 | **RNA**→ | rpmE (PRK00019)Ribosomal_L31 (pfam01197)→ | |

| | | | | | | |
|---|---|---|---|---|---|---|
| Sen-288-4 | NZ_CUZJ01000016.1 | + | 104156 | 104208 | RNA→ | rpmE (PRK00019) Ribosomal_L31 (pfam01197)→ |
| Sen-194-1 | NZ_AMOL01000071.1 | - | 37527 | 37475 | RNA→ | rpmE (PRK00019) Ribosomal_L31 (pfam01197)→ |
| Sen-288-5 | NZ_CUZY01000016.1 | - | 39534 | 39482 | RNA→ | rpmE (PRK00019) Ribosomal_L31 (pfam01197)→ |
| Sen-170-11 | NZ_LHEG01000017.1 | - | 37518 | 37466 | RNA→ | rpmE (PRK00019) Ribosomal_L31 (pfam01197)→ |
| Sen-170-12 | NZ_LHEK01000011.1 | + | 43461 | 43513 | RNA→ | rpmE (PRK00019) Ribosomal_L31 (pfam01197)→ |
| Sen-170-13 | NZ_LHEL01000026.1 | + | 43461 | 43513 | RNA→ | rpmE (PRK00019) Ribosomal_L31 (pfam01197)→ |
| Sen-170-14 | NZ_LHGF01000016.1 | - | 37788 | 37736 | RNA→ | rpmE (PRK00019) Ribosomal_L31 (pfam01197)→ |
| Sen-170-15 | NZ_LHHU01000005.1 | - | 37758 | 37706 | RNA→ | rpmE (PRK00019) Ribosomal_L31 (pfam01197)→ |
| Sen-130-1 | NZ_LHJK01000002.1 | - | 43074 | 43022 | RNA→ | rpmE (PRK00019) Ribosomal_L31 (pfam01197)→ |
| Sen-225-5 | NZ_LHIA01000022.1 | + | 110618 | 110670 | RNA→ | rpmE (PRK00019) Ribosomal_L31 (pfam01197)→ |
| Sen-225-6 | NZ_LHIU01000047.1 | - | 37837 | 37785 | RNA→ | rpmE (PRK00019) Ribosomal_L31 (pfam01197)→ |
| Sen-12-1 | NZ_LHJT01000036.1 | - | 37541 | 37489 | RNA→ | rpmE (PRK00019) Ribosomal_L31 (pfam01197)→ |
| Sen-170-16 | NZ_LHIY01000013.1 | + | 43461 | 43513 | RNA→ | rpmE (PRK00019) Ribosomal_L31 (pfam01197)→ |
| Sen-129-1 | NZ_LHLL01000005.1 | - | 45297 | 45245 | RNA→ | rpmE (PRK00019) Ribosomal_L31 (pfam01197)→ |
| Sen-14-1 | NZ_LHNN01000010.1 | + | 106822 | 106874 | RNA→ | rpmE (PRK00019) Ribosomal_L31 (pfam01197)→ |
| Sen-170-17 | NZ_LHMK01000033.1 | - | 3250 | 3198 | RNA→ | rpmE (PRK00019) Ribosomal_L31 (pfam01197)→ |
| Sen-170-18 | NZ_LHMM01000016.1 | + | 100688 | 100740 | RNA→ | rpmE (PRK00019) Ribosomal_L31 (pfam01197)→ |
| Sen-137-1 | NZ_LHNP01000044.1 | + | 33072 | 33124 | RNA→ | rpmE (PRK00019) Ribosomal_L31 (pfam01197)→ |
| Sen-5-1 | NZ_LHNH01000020.1 | - | 45358 | 45306 | RNA→ | rpmE (PRK00019) Ribosomal_L31 (pfam01197)→ |
| Sen-225-7 | NZ_LHOC01000018.1 | + | 43745 | 43797 | RNA→ | rpmE (PRK00019) Ribosomal_L31 (pfam01197)→ |
| Sen-14-2 | NZ_LHGC01000023.1 | - | 41667 | 41615 | RNA→ | rpmE (PRK00019) Ribosomal_L31 (pfam01197)→ |
| Sen-170-19 | NZ_LHOD01000025.1 | + | 100688 | 100740 | RNA→ | rpmE (PRK00019) Ribosomal_L31 (pfam01197)→ |
| Sen-138-1 | NZ_LHHP01000012.1 | - | 49389 | 49337 | RNA→ | rpmE (PRK00019) Ribosomal_L31 (pfam01197)→ |
| Sen-5-2 | NZ_LHOM01000026.1 | + | 104489 | 104541 | RNA→ | rpmE (PRK00019) Ribosomal_L31 (pfam01197)→ |
| Sen-170-20 | NZ_LHOW01000002.1 | - | 37758 | 37706 | RNA→ | rpmE (PRK00019) Ribosomal_L31 (pfam01197)→ |
| Sen-137-2 | NZ_LHKI01000166.1 | + | 4122 | 4174 | RNA→ | rpmE (PRK00019) Ribosomal_L31 (pfam01197)→ |
| Sen-225-8 | NZ_LHDM01000014.1 | - | 37858 | 37806 | RNA→ | rpmE (PRK00019) Ribosomal_L31 (pfam01197)→ |
| Sen-138-2 | NZ_LHLY01000014.1 | + | 104581 | 104633 | RNA→ | rpmE (PRK00019) Ribosomal_L31 (pfam01197)→ |
| Sen-225-9 | NZ_LHEN01000003.1 | - | 37859 | 37807 | RNA→ | rpmE (PRK00019) Ribosomal_L31 (pfam01197)→ |
| Sen-138-3 | NZ_LHMB01000021.1 | - | 49457 | 49405 | RNA→ | rpmE (PRK00019) Ribosomal_L31 (pfam01197)→ |
| Sen-225-10 | NZ_LHEZ01000010.1 | - | 37859 | 37807 | RNA→ | rpmE (PRK00019) Ribosomal_L31 (pfam01197)→ |
| Sen-225-11 | NZ_LHFO01000006.1 | - | 37858 | 37806 | RNA→ | rpmE (PRK00019) Ribosomal_L31 (pfam01197)→ |
| Sen-130-2 | NZ_LHMR01000007.1 | - | 43074 | 43022 | RNA→ | rpmE (PRK00019) Ribosomal_L31 (pfam01197)→ |
| Sen-225-12 | NZ_LHHH01000031.1 | + | 110443 | 110495 | RNA→ | rpmE (PRK00019) Ribosomal_L31 (pfam01197)→ |
| Sen-225-13 | NZ_LHJC01000006.1 | + | 110627 | 110679 | RNA→ | rpmE (PRK00019) Ribosomal_L31 (pfam01197)→ |
| Sen-225-14 | NZ_LHKJ01000025.1 | + | 110443 | 110495 | RNA→ | rpmE (PRK00019) Ribosomal_L31 (pfam01197)→ |
| Sen-12-2 | NZ_LHET01000013.1 | + | 80459 | 80511 | RNA→ | rpmE (PRK00019) Ribosomal_L31 (pfam01197)→ |
| Sen-15-1 | NZ_LHFH01000010.1 | + | 98924 | 98976 | RNA→ | rpmE (PRK00019) Ribosomal_L31 (pfam01197)→ |
| Sen-225-15 | NZ_LHLB01000016.1 | - | 37859 | 37807 | RNA→ | rpmE (PRK00019) Ribosomal_L31 (pfam01197)→ |
| Sen-225-16 | NZ_LHLC01000015.1 | - | 38115 | 38063 | RNA→ | rpmE (PRK00019) Ribosomal_L31 (pfam01197)→ |
| Sen-130-3 | NZ_LHHJ01000016.1 | - | 43074 | 43022 | RNA→ | rpmE (PRK00019) Ribosomal_L31 (pfam01197)→ |
| Sen-225-17 | NZ_LHLG01000042.1 | - | 37859 | 37807 | RNA→ | rpmE (PRK00019) Ribosomal_L31 (pfam01197)→ |
| Sen-14-3 | NZ_LHIF01000005.1 | - | 41667 | 41615 | RNA→ | rpmE (PRK00019) Ribosomal_L31 (pfam01197)→ |
| Sen-12-3 | NZ_LHKF01000013.1 | + | 141508 | 141560 | RNA→ | rpmE (PRK00019) Ribosomal_L31 (pfam01197)→ |
| Sen-225-18 | NZ_LHLT01000024.1 | + | 110443 | 110495 | RNA→ | rpmE (PRK00019) Ribosomal_L31 (pfam01197)→ |
| Sen-129-2 | NZ_LHKW01000027.1 | + | 134845 | 134897 | RNA→ | rpmE (PRK00019) Ribosomal_L31 (pfam01197)→ |
| Sen-6-1 | NZ_LHMF01000001.1 | - | 45365 | 45313 | RNA→ | rpmE (PRK00019) Ribosomal_L31 (pfam01197)→ |
| Sen-6-2 | NZ_LHNW01000009.1 | + | 104489 | 104541 | RNA→ | rpmE (PRK00019) Ribosomal_L31 (pfam01197)→ |
| Sen-170-21 | NZ_LHOB01000004.1 | - | 37787 | 37735 | RNA→ | rpmE (PRK00019) Ribosomal_L31 (pfam01197)→ |
| Sen-15-2 | NZ_LHFG01000019.1 | + | 98923 | 98975 | RNA→ | rpmE (PRK00019) Ribosomal_L31 (pfam01197)→ |
| Sen-112-2 | NZ_LHGO01000001.1 | - | 49213 | 49161 | RNA→ | rpmE (PRK00019) Ribosomal_L31 (pfam01197)→ |
| Sen-224-1 | NZ_LHOL01000024.1 | + | 118678 | 118730 | RNA→ | rpmE (PRK00019) Ribosomal_L31 (pfam01197)→ |

| | | | | | |
|---|---|---|---|---|---|
| Sen-112-3 | NZ_LHGP01000036.1 | - | 49493 | 49441 | RNA→ rpmE (PRK00019)Ribosomal_L31 (pfam01197)→ |
| Sen-225-19 | NZ_LHOT01000027.1 | - | 38140 | 38088 | RNA→ rpmE (PRK00019)Ribosomal_L31 (pfam01197)→ |
| Sen-5-3 | NZ_LHEA01000014.1 | + | 134515 | 134567 | RNA→ rpmE (PRK00019)Ribosomal_L31 (pfam01197)→ |
| Sen-137-3 | NZ_LHHE01000026.1 | + | 33061 | 33113 | RNA→ rpmE (PRK00019)Ribosomal_L31 (pfam01197)→ |
| Sen-137-4 | NZ_LHKH01000008.1 | + | 4122 | 4174 | RNA→ rpmE (PRK00019)Ribosomal_L31 (pfam01197)→ |
| Sen-138-4 | NZ_LHMP01000018.1 | - | 49457 | 49405 | RNA→ rpmE (PRK00019)Ribosomal_L31 (pfam01197)→ |
| Sen-224-2 | NZ_LHHX01000010.1 | - | 337 | 285 | RNA→ rpmE (PRK00019)Ribosomal_L31 (pfam01197)→ |
| Sen-138-5 | NZ_LHOE01000015.1 | - | 49211 | 49159 | RNA→ rpmE (PRK00019)Ribosomal_L31 (pfam01197)→ |
| Sen-5-4 | NZ_LHKM01000033.1 | + | 43435 | 43487 | RNA→ rpmE (PRK00019)Ribosomal_L31 (pfam01197)→ |
| Sen-15-3 | NZ_LHON01000004.1 | + | 98924 | 98976 | RNA→ rpmE (PRK00019)Ribosomal_L31 (pfam01197)→ |
| Sen-6-3 | NZ_LHMA01000017.1 | + | 104489 | 104541 | RNA→ rpmE (PRK00019)Ribosomal_L31 (pfam01197)→ |
| Sen-137-5 | NZ_LHJB01000007.1 | + | 95635 | 95687 | RNA→ rpmE (PRK00019)Ribosomal_L31 (pfam01197)→ |
| Sen-6-4 | NZ_LHLX01000040.1 | - | 45358 | 45306 | RNA→ rpmE (PRK00019)Ribosomal_L31 (pfam01197)→ |
| Sen-6-5 | NZ_LHLF01000017.1 | + | 104489 | 104541 | RNA→ rpmE (PRK00019)Ribosomal_L31 (pfam01197)→ |
| Sen-14-4 | NZ_LHKN01000021.1 | + | 106828 | 106880 | RNA→ rpmE (PRK00019)Ribosomal_L31 (pfam01197)→ |
| Sen-288-6 | NZ_CUYH01000014.1 | - | 41503 | 41451 | RNA→ rpmE (PRK00019)Ribosomal_L31 (pfam01197)→ |
| Sen-306-1 | NZ_LHFT01000001.1 | - | 45297 | 45245 | RNA→ rpmE (PRK00019)Ribosomal_L31 (pfam01197)→ |
| Sen-306-2 | NZ_LHFW01000015.1 | + | 104298 | 104350 | RNA→ rpmE (PRK00019)Ribosomal_L31 (pfam01197)→ |
| Sen-306-3 | NZ_LHGI01000004.1 | + | 104488 | 104540 | RNA→ rpmE (PRK00019)Ribosomal_L31 (pfam01197)→ |
| Sen-306-4 | NZ_LHHK01000013.1 | - | 45297 | 45245 | RNA→ rpmE (PRK00019)Ribosomal_L31 (pfam01197)→ |
| Sen-306-5 | NZ_LHIW01000004.1 | + | 104298 | 104350 | RNA→ rpmE (PRK00019)Ribosomal_L31 (pfam01197)→ |
| Sen-286-2 | NZ_LHEO01000040.1 | + | 4123 | 4175 | RNA→ rpmE (PRK00019)Ribosomal_L31 (pfam01197)→ |
| Sen-280-1 | NZ_LHOQ01000013.1 | + | 104644 | 104696 | RNA→ rpmE (PRK00019)Ribosomal_L31 (pfam01197)→ |
| Sen-280-2 | NZ_LHDZ01000014.1 | - | 37830 | 37778 | RNA→ rpmE (PRK00019)Ribosomal_L31 (pfam01197)→ |
| Sen-306-6 | NZ_LHED01000008.1 | + | 104298 | 104350 | RNA→ rpmE (PRK00019)Ribosomal_L31 (pfam01197)→ |
| Sen-306-7 | NZ_LHER01000039.1 | + | 104298 | 104350 | RNA→ rpmE (PRK00019)Ribosomal_L31 (pfam01197)→ |
| Sen-306-8 | NZ_LHEX01000022.1 | + | 104409 | 104461 | RNA→ rpmE (PRK00019)Ribosomal_L31 (pfam01197)→ |
| Sen-306-9 | NZ_LHFF01000003.1 | - | 45297 | 45245 | RNA→ rpmE (PRK00019)Ribosomal_L31 (pfam01197)→ |
| Sen-306-10 | NZ_LHFP01000006.1 | + | 43372 | 43424 | RNA→ rpmE (PRK00019)Ribosomal_L31 (pfam01197)→ |
| Sen-306-11 | NZ_LHGJ01000015.1 | + | 104298 | 104350 | RNA→ rpmE (PRK00019)Ribosomal_L31 (pfam01197)→ |
| Sen-306-12 | NZ_LHGL01000067.1 | - | 45407 | 45355 | RNA→ rpmE (PRK00019)Ribosomal_L31 (pfam01197)→ |
| Sen-306-13 | NZ_LHHB01000046.1 | - | 45296 | 45244 | RNA→ rpmE (PRK00019)Ribosomal_L31 (pfam01197)→ |
| Sen-306-14 | NZ_LHID01000028.1 | - | 45297 | 45245 | RNA→ rpmE (PRK00019)Ribosomal_L31 (pfam01197)→ |
| Sen-306-15 | NZ_LHIQ01000056.1 | + | 43376 | 43428 | RNA→ rpmE (PRK00019)Ribosomal_L31 (pfam01197)→ |
| Sen-306-16 | NZ_LHIX01000050.1 | - | 45297 | 45245 | RNA→ rpmE (PRK00019)Ribosomal_L31 (pfam01197)→ |
| Sen-306-17 | NZ_LHJZ01000008.1 | + | 104298 | 104350 | RNA→ rpmE (PRK00019)Ribosomal_L31 (pfam01197)→ |
| Sen-306-18 | NZ_LHKD01000012.1 | - | 45297 | 45245 | RNA→ rpmE (PRK00019)Ribosomal_L31 (pfam01197)→ |
| Sen-306-19 | NZ_LHKL01000021.1 | + | 43403 | 43455 | RNA→ rpmE (PRK00019)Ribosomal_L31 (pfam01197)→ |
| Sen-306-20 | NZ_LHKK01000020.1 | + | 15365 | 15417 | RNA→ rpmE (PRK00019)Ribosomal_L31 (pfam01197)→ |
| Sen-306-21 | NZ_LHNG01000019.1 | + | 104298 | 104350 | RNA→ rpmE (PRK00019)Ribosomal_L31 (pfam01197)→ |
| Sen-306-22 | NZ_LHNL01000012.1 | - | 45297 | 45245 | RNA→ rpmE (PRK00019)Ribosomal_L31 (pfam01197)→ |
| Sen-286-3 | NZ_LHEF01000001.1 | - | 45301 | 45249 | RNA→ rpmE (PRK00019)Ribosomal_L31 (pfam01197)→ |
| Sen-306-23 | NZ_LHEP01000016.1 | + | 104298 | 104350 | RNA→ rpmE (PRK00019)Ribosomal_L31 (pfam01197)→ |
| Sen-225-20 | NZ_LHDL01000036.1 | - | 37859 | 37807 | RNA→ rpmE (PRK00019)Ribosomal_L31 (pfam01197)→ |
| Sen-306-24 | NZ_LHEW01000007.1 | + | 104297 | 104349 | RNA→ rpmE (PRK00019)Ribosomal_L31 (pfam01197)→ |
| Sen-306-25 | NZ_LHEY01000045.1 | - | 45297 | 45245 | RNA→ rpmE (PRK00019)Ribosomal_L31 (pfam01197)→ |
| Sen-225-21 | NZ_LHDS01000006.1 | - | 37859 | 37807 | RNA→ rpmE (PRK00019)Ribosomal_L31 (pfam01197)→ |
| Sen-306-26 | NZ_LHFU01000021.1 | - | 45297 | 45245 | RNA→ rpmE (PRK00019)Ribosomal_L31 (pfam01197)→ |
| Sen-225-22 | NZ_LHEC01000015.1 | - | 37859 | 37807 | RNA→ rpmE (PRK00019)Ribosomal_L31 (pfam01197)→ |
| Sen-306-27 | NZ_LHGH01000007.1 | - | 45297 | 45245 | RNA→ rpmE (PRK00019)Ribosomal_L31 (pfam01197)→ |
| Sen-286-4 | NZ_LHHI01000015.1 | + | 4124 | 4176 | RNA→ rpmE (PRK00019)Ribosomal_L31 (pfam01197)→ |
| Sen-306-28 | NZ_LHIE01000025.1 | - | 45297 | 45245 | RNA→ rpmE (PRK00019)Ribosomal_L31 (pfam01197)→ |

| | | | | | | |
|---|---|---|---|---|---|---|
| Sen-226-1 | NZ_LHIM01000031.1 | - | 45095 | 45043 | RNA→ | rpmE (PRK00019)Ribosomal_L31 (pfam01197)→ |
| Sen-306-29 | NZ_LHIK01000030.1 | - | 45297 | 45245 | RNA→ | rpmE (PRK00019)Ribosomal_L31 (pfam01197)→ |
| Sen-225-23 | NZ_LHJO01000023.1 | + | 110443 | 110495 | RNA→ | rpmE (PRK00019)Ribosomal_L31 (pfam01197)→ |
| Sen-306-30 | NZ_LHJJ01000014.1 | + | 104298 | 104350 | RNA→ | rpmE (PRK00019)Ribosomal_L31 (pfam01197)→ |
| Sen-225-24 | NZ_LHLR01000018.1 | + | 110489 | 110541 | RNA→ | rpmE (PRK00019)Ribosomal_L31 (pfam01197)→ |
| Sen-306-31 | NZ_LHLO01000017.1 | - | 45297 | 45245 | RNA→ | rpmE (PRK00019)Ribosomal_L31 (pfam01197)→ |
| Sen-225-25 | NZ_LHMN01000017.1 | + | 110595 | 110647 | RNA→ | rpmE (PRK00019)Ribosomal_L31 (pfam01197)→ |
| Sen-306-32 | NZ_LHLP01000013.1 | + | 104298 | 104350 | RNA→ | rpmE (PRK00019)Ribosomal_L31 (pfam01197)→ |
| Sen-225-26 | NZ_LHMQ01000007.1 | - | 37859 | 37807 | RNA→ | rpmE (PRK00019)Ribosomal_L31 (pfam01197)→ |
| Sen-233-1 | NZ_LHNX01000010.1 | + | 97858 | 97910 | RNA→ | rpmE (PRK00019)Ribosomal_L31 (pfam01197)→ |
| Sen-306-33 | NZ_LHNI01000006.1 | - | 45297 | 45245 | RNA→ | rpmE (PRK00019)Ribosomal_L31 (pfam01197)→ |
| Sen-230-1 | NZ_LHOI01000018.1 | + | 101470 | 101522 | RNA→ | rpmE (PRK00019)Ribosomal_L31 (pfam01197)→ |
| Sen-226-2 | NZ_LHIN01000021.1 | + | 99690 | 99742 | RNA→ | rpmE (PRK00019)Ribosomal_L31 (pfam01197)→ |
| Sen-306-34 | NZ_LHGY01000018.1 | - | 45297 | 45245 | RNA→ | rpmE (PRK00019)Ribosomal_L31 (pfam01197)→ |
| Sen-306-35 | NZ_LHJU01000006.1 | + | 104298 | 104350 | RNA→ | rpmE (PRK00019)Ribosomal_L31 (pfam01197)→ |
| Sen-227-1 | NZ_LHLZ01000020.1 | + | 104655 | 104707 | RNA→ | rpmE (PRK00019)Ribosomal_L31 (pfam01197)→ |
| Sen-230-2 | NZ_LHOS01000013.1 | - | 37982 | 37930 | RNA→ | rpmE (PRK00019)Ribosomal_L31 (pfam01197)→ |
| Sen-306-36 | NZ_LHOJ01000015.1 | + | 104298 | 104350 | RNA→ | rpmE (PRK00019)Ribosomal_L31 (pfam01197)→ |
| Sen-225-27 | NZ_LHDO01000014.1 | + | 110443 | 110495 | RNA→ | rpmE (PRK00019)Ribosomal_L31 (pfam01197)→ |
| Sen-286-5 | NZ_LHEH01000008.1 | + | 105522 | 105574 | RNA→ | rpmE (PRK00019)Ribosomal_L31 (pfam01197)→ |
| Sen-225-28 | NZ_LHEB01000004.1 | - | 37858 | 37806 | RNA→ | rpmE (PRK00019)Ribosomal_L31 (pfam01197)→ |
| Sen-306-37 | NZ_LHOH01000009.1 | - | 45297 | 45245 | RNA→ | rpmE (PRK00019)Ribosomal_L31 (pfam01197)→ |
| Sen-225-29 | NZ_LHEI01000015.1 | - | 37858 | 37806 | RNA→ | rpmE (PRK00019)Ribosomal_L31 (pfam01197)→ |
| Sen-237-1 | NZ_LHES01000005.1 | - | 45233 | 45181 | RNA→ | rpmE (PRK00019)Ribosomal_L31 (pfam01197)→ |
| Sen-226-3 | NZ_LHGE01000011.1 | - | 45335 | 45283 | RNA→ | rpmE (PRK00019)Ribosomal_L31 (pfam01197)→ |
| Sen-306-38 | NZ_LHJF01000021.1 | + | 104297 | 104349 | RNA→ | rpmE (PRK00019)Ribosomal_L31 (pfam01197)→ |
| Sen-225-30 | NZ_LHGN01000027.1 | - | 37859 | 37807 | RNA→ | rpmE (PRK00019)Ribosomal_L31 (pfam01197)→ |
| Sen-306-39 | NZ_LHKA01000019.1 | + | 104298 | 104350 | RNA→ | rpmE (PRK00019)Ribosomal_L31 (pfam01197)→ |
| Sen-306-40 | NZ_LHLK01000012.1 | - | 45428 | 45376 | RNA→ | rpmE (PRK00019)Ribosomal_L31 (pfam01197)→ |
| Sen-226-4 | NZ_LHIG01000022.1 | + | 99667 | 99719 | RNA→ | rpmE (PRK00019)Ribosomal_L31 (pfam01197)→ |
| Sen-306-41 | NZ_LHMT01000039.1 | + | 104298 | 104350 | RNA→ | rpmE (PRK00019)Ribosomal_L31 (pfam01197)→ |
| Sen-237-2 | NZ_LHIJ01000001.1 | + | 105504 | 105556 | RNA→ | rpmE (PRK00019)Ribosomal_L31 (pfam01197)→ |
| Sen-306-42 | NZ_LHOG01000039.1 | + | 104298 | 104350 | RNA→ | rpmE (PRK00019)Ribosomal_L31 (pfam01197)→ |
| Sen-233-2 | NZ_LHKV01000011.1 | + | 129710 | 129762 | RNA→ | rpmE (PRK00019)Ribosomal_L31 (pfam01197)→ |
| Sen-306-43 | NZ_LHEQ01000030.1 | - | 45300 | 45248 | RNA→ | rpmE (PRK00019)Ribosomal_L31 (pfam01197)→ |
| Sen-225-31 | NZ_LHKZ01000044.1 | + | 110628 | 110680 | RNA→ | rpmE (PRK00019)Ribosomal_L31 (pfam01197)→ |
| Sen-306-44 | NZ_LHFB01000001.1 | - | 45428 | 45376 | RNA→ | rpmE (PRK00019)Ribosomal_L31 (pfam01197)→ |
| Sen-225-32 | NZ_LHLN01000022.1 | + | 110627 | 110679 | RNA→ | rpmE (PRK00019)Ribosomal_L31 (pfam01197)→ |
| Sen-306-45 | NZ_LHFS01000008.1 | + | 104297 | 104349 | RNA→ | rpmE (PRK00019)Ribosomal_L31 (pfam01197)→ |
| Sen-225-33 | NZ_LHMG01000021.1 | - | 37859 | 37807 | RNA→ | rpmE (PRK00019)Ribosomal_L31 (pfam01197)→ |
| Sen-306-46 | NZ_LHFV01000011.1 | + | 43443 | 43495 | RNA→ | rpmE (PRK00019)Ribosomal_L31 (pfam01197)→ |
| Sen-225-34 | NZ_LHOR01000003.1 | + | 110443 | 110495 | RNA→ | rpmE (PRK00019)Ribosomal_L31 (pfam01197)→ |
| Sen-306-47 | NZ_LHGZ01000018.1 | - | 45557 | 45505 | RNA→ | rpmE (PRK00019)Ribosomal_L31 (pfam01197)→ |
| Sen-237-3 | NZ_LHDR01000020.1 | - | 45233 | 45181 | RNA→ | rpmE (PRK00019)Ribosomal_L31 (pfam01197)→ |
| Sen-308-1 | NZ_LHIL01000010.1 | - | 37811 | 37759 | RNA→ | rpmE (PRK00019)Ribosomal_L31 (pfam01197)→ |
| Sen-233-3 | NZ_LHGG01000023.1 | - | 35466 | 35414 | RNA→ | rpmE (PRK00019)Ribosomal_L31 (pfam01197)→ |
| Sen-306-48 | NZ_LHIR01000020.1 | + | 104298 | 104350 | RNA→ | rpmE (PRK00019)Ribosomal_L31 (pfam01197)→ |
| Sen-225-35 | NZ_LHGX01000039.1 | + | 110443 | 110495 | RNA→ | rpmE (PRK00019)Ribosomal_L31 (pfam01197)→ |
| Sen-306-49 | NZ_LHKC01000026.1 | - | 45297 | 45245 | RNA→ | rpmE (PRK00019)Ribosomal_L31 (pfam01197)→ |
| Sen-225-36 | NZ_LHHC01000016.1 | - | 37859 | 37807 | RNA→ | rpmE (PRK00019)Ribosomal_L31 (pfam01197)→ |
| Sen-306-50 | NZ_LHLA01000003.1 | + | 104298 | 104350 | RNA→ | rpmE (PRK00019)Ribosomal_L31 (pfam01197)→ |
| Sen-230-3 | NZ_LHHR01000021.1 | - | 37982 | 37930 | RNA→ | rpmE (PRK00019)Ribosomal_L31 (pfam01197)→ |

| | | | | | | |
|---|---|---|---|---|---|---|
| Sen-308-2 | NZ_LHOK01000024.1 | + | 100381 | 100433 | **RNA**→ | rpmE (PRK00019)Ribosomal_L31 (pfam01197)→ |
| Sen-227-2 | NZ_LHHW01000019.1 | - | 45628 | 45576 | **RNA**→ | rpmE (PRK00019)Ribosomal_L31 (pfam01197)→ |
| Sen-225-37 | NZ_LHJM01000029.1 | - | 37859 | 37807 | **RNA**→ | rpmE (PRK00019)Ribosomal_L31 (pfam01197)→ |
| Sen-225-38 | NZ_LHJP01000060.1 | - | 14715 | 14663 | **RNA**→ | rpmE (PRK00019)Ribosomal_L31 (pfam01197)→ |
| Sen-237-4 | NZ_LHKR01000017.1 | - | 45233 | 45181 | **RNA**→ | rpmE (PRK00019)Ribosomal_L31 (pfam01197)→ |
| Sen-308-3 | NZ_LHGK01000003.1 | - | 37864 | 37812 | **RNA**→ | rpmE (PRK00019)Ribosomal_L31 (pfam01197)→ |
| Sen-233-4 | NZ_LHKY01000034.1 | - | 35226 | 35174 | **RNA**→ | rpmE (PRK00019)Ribosomal_L31 (pfam01197)→ |
| Sen-237-5 | NZ_LHLW01000028.1 | + | 170370 | 170422 | **RNA**→ | rpmE (PRK00019)Ribosomal_L31 (pfam01197)→ |
| Sen-225-39 | NZ_LHNZ01000016.1 | + | 104535 | 104587 | **RNA**→ | rpmE (PRK00019)Ribosomal_L31 (pfam01197)→ |
| Sen-225-40 | NZ_LHOU01000014.1 | - | 37859 | 37807 | **RNA**→ | rpmE (PRK00019)Ribosomal_L31 (pfam01197)→ |
| Sen-237-6 | NZ_LHFZ01000007.1 | - | 45233 | 45181 | **RNA**→ | rpmE (PRK00019)Ribosomal_L31 (pfam01197)→ |
| Sen-228-2 | NZ_LHGD01000004.1 | - | 45717 | 45665 | **RNA**→ | rpmE (PRK00019)Ribosomal_L31 (pfam01197)→ |
| Sen-225-41 | NZ_LHJQ01000014.1 | + | 110332 | 110384 | **RNA**→ | rpmE (PRK00019)Ribosomal_L31 (pfam01197)→ |
| Sen-225-42 | NZ_LHLE01000029.1 | - | 38116 | 38064 | **RNA**→ | rpmE (PRK00019)Ribosomal_L31 (pfam01197)→ |
| Sen-225-43 | NZ_LHNJ01000028.1 | - | 38141 | 38089 | **RNA**→ | rpmE (PRK00019)Ribosomal_L31 (pfam01197)→ |
| Sen-237-7 | NZ_LHDX01000028.1 | - | 45233 | 45181 | **RNA**→ | rpmE (PRK00019)Ribosomal_L31 (pfam01197)→ |
| Sen-237-8 | NZ_LHOV01000014.1 | + | 170530 | 170582 | **RNA**→ | rpmE (PRK00019)Ribosomal_L31 (pfam01197)→ |
| Sen-237-9 | NZ_LHJA01000018.1 | + | 170373 | 170425 | **RNA**→ | rpmE (PRK00019)Ribosomal_L31 (pfam01197)→ |
| Sen-279-1 | NZ_LHEJ01000024.1 | + | 4123 | 4175 | **RNA**→ | rpmE (PRK00019)Ribosomal_L31 (pfam01197)→ |
| Sen-225-44 | NZ_LHJV01000040.1 | + | 110443 | 110495 | **RNA**→ | rpmE (PRK00019)Ribosomal_L31 (pfam01197)→ |
| Sen-237-10 | NZ_LHEM01000027.1 | - | 45233 | 45181 | **RNA**→ | rpmE (PRK00019)Ribosomal_L31 (pfam01197)→ |
| Sen-225-45 | NZ_LHJW01000015.1 | + | 110598 | 110650 | **RNA**→ | rpmE (PRK00019)Ribosomal_L31 (pfam01197)→ |
| Sen-225-46 | NZ_LHMU01000012.1 | - | 37859 | 37807 | **RNA**→ | rpmE (PRK00019)Ribosomal_L31 (pfam01197)→ |
| Sen-270-1 | NZ_LHFR01000039.1 | - | 45189 | 45137 | **RNA**→ | rpmE (PRK00019)Ribosomal_L31 (pfam01197)→ |
| Sen-225-47 | NZ_LHHA01000023.1 | + | 43611 | 43663 | **RNA**→ | rpmE (PRK00019)Ribosomal_L31 (pfam01197)→ |
| Sen-278-1 | NZ_LHFY01000007.1 | + | 104454 | 104506 | **RNA**→ | rpmE (PRK00019)Ribosomal_L31 (pfam01197)→ |
| Sen-237-11 | NZ_LHGR01000017.1 | - | 45233 | 45181 | **RNA**→ | rpmE (PRK00019)Ribosomal_L31 (pfam01197)→ |
| Sen-237-12 | NZ_LHHF01000011.1 | - | 45532 | 45480 | **RNA**→ | rpmE (PRK00019)Ribosomal_L31 (pfam01197)→ |
| Sen-278-2 | NZ_LHIC01000007.1 | + | 104453 | 104505 | **RNA**→ | rpmE (PRK00019)Ribosomal_L31 (pfam01197)→ |
| Sen-278-3 | NZ_LHIH01000022.1 | + | 104454 | 104506 | **RNA**→ | rpmE (PRK00019)Ribosomal_L31 (pfam01197)→ |
| Sen-278-4 | NZ_LHIS01000017.1 | + | 104453 | 104505 | **RNA**→ | rpmE (PRK00019)Ribosomal_L31 (pfam01197)→ |
| Sen-278-5 | NZ_LHKB01000030.1 | - | 45232 | 45180 | **RNA**→ | rpmE (PRK00019)Ribosomal_L31 (pfam01197)→ |
| Sen-279-2 | NZ_LHKE01000013.1 | - | 35503 | 35451 | **RNA**→ | rpmE (PRK00019)Ribosomal_L31 (pfam01197)→ |
| Sen-277-1 | NZ_LHKG01000033.1 | + | 97652 | 97704 | **RNA**→ | rpmE (PRK00019)Ribosomal_L31 (pfam01197)→ |
| Sen-278-6 | NZ_LHLI01000036.1 | + | 43581 | 43633 | **RNA**→ | rpmE (PRK00019)Ribosomal_L31 (pfam01197)→ |
| Sen-237-13 | NZ_LHMH01000002.1 | + | 170373 | 170425 | **RNA**→ | rpmE (PRK00019)Ribosomal_L31 (pfam01197)→ |
| Sen-279-3 | NZ_LHMV01000016.1 | + | 103255 | 103307 | **RNA**→ | rpmE (PRK00019)Ribosomal_L31 (pfam01197)→ |
| Sen-278-7 | NZ_LHDT01000024.1 | + | 104454 | 104506 | **RNA**→ | rpmE (PRK00019)Ribosomal_L31 (pfam01197)→ |
| Sen-278-8 | NZ_LHHN01000005.1 | + | 104454 | 104506 | **RNA**→ | rpmE (PRK00019)Ribosomal_L31 (pfam01197)→ |
| Sen-279-4 | NZ_LHJG01000019.1 | - | 2093 | 2041 | **RNA**→ | rpmE (PRK00019)Ribosomal_L31 (pfam01197)→ |
| Sen-278-9 | NZ_LHJN01000001.1 | - | 45365 | 45313 | **RNA**→ | rpmE (PRK00019)Ribosomal_L31 (pfam01197)→ |
| Sen-278-10 | NZ_LHMJ01000035.1 | + | 8177 | 8229 | **RNA**→ | rpmE (PRK00019)Ribosomal_L31 (pfam01197)→ |
| Sen-279-5 | NZ_LHMX01000008.1 | + | 4123 | 4175 | **RNA**→ | rpmE (PRK00019)Ribosomal_L31 (pfam01197)→ |
| Sen-279-6 | NZ_LHNF01000052.1 | - | 2092 | 2040 | **RNA**→ | rpmE (PRK00019)Ribosomal_L31 (pfam01197)→ |
| Sen-279-7 | NZ_LHNY01000033.1 | - | 2091 | 2039 | **RNA**→ | rpmE (PRK00019)Ribosomal_L31 (pfam01197)→ |
| Sen-278-11 | NZ_LHDY01000004.1 | + | 104454 | 104506 | **RNA**→ | rpmE (PRK00019)Ribosomal_L31 (pfam01197)→ |
| Sen-279-8 | NZ_LHFI01000008.1 | + | 103254 | 103306 | **RNA**→ | rpmE (PRK00019)Ribosomal_L31 (pfam01197)→ |
| Sen-278-12 | NZ_LHGQ01000010.1 | - | 45505 | 45453 | **RNA**→ | rpmE (PRK00019)Ribosomal_L31 (pfam01197)→ |
| Sen-279-9 | NZ_LHHM01000020.1 | - | 35738 | 35686 | **RNA**→ | rpmE (PRK00019)Ribosomal_L31 (pfam01197)→ |
| Sen-278-13 | NZ_LHII01000006.1 | - | 45365 | 45313 | **RNA**→ | rpmE (PRK00019)Ribosomal_L31 (pfam01197)→ |
| Sen-243-1 | NZ_LHIV01000005.1 | - | 45298 | 45246 | **RNA**→ | rpmE (PRK00019)Ribosomal_L31 (pfam01197)→ |
| Sen-308-4 | NZ_LHGV01000009.1 | - | 37811 | 37759 | **RNA**→ | rpmE (PRK00019)Ribosomal_L31 (pfam01197)→ |

| | | | | | |
|---|---|---|---|---|---|
| Sen-277-2 | NZ_LHJD01000020.1 | + | 97390 | 97442 | RNA→ rpmE (PRK00019)Ribosomal_L31 (pfam01197)→ |
| Sen-278-14 | NZ_LHJS01000013.1 | - | 45326 | 45274 | RNA→ rpmE (PRK00019)Ribosomal_L31 (pfam01197)→ |
| Sen-278-15 | NZ_LHKS01000005.1 | + | 19850 | 19902 | RNA→ rpmE (PRK00019)Ribosomal_L31 (pfam01197)→ |
| Sen-278-16 | NZ_LHKT01000094.1 | + | 19849 | 19901 | RNA→ rpmE (PRK00019)Ribosomal_L31 (pfam01197)→ |
| Sen-279-10 | NZ_LHLQ01000028.1 | - | 2093 | 2041 | RNA→ rpmE (PRK00019)Ribosomal_L31 (pfam01197)→ |
| Sen-240-1 | NZ_LHMO01000017.1 | - | 35408 | 35356 | RNA→ rpmE (PRK00019)Ribosomal_L31 (pfam01197)→ |
| Sen-237-14 | NZ_LHNT01000030.1 | + | 34658 | 34710 | RNA→ rpmE (PRK00019)Ribosomal_L31 (pfam01197)→ |
| Sen-237-15 | NZ_LHOO01000015.1 | + | 170400 | 170452 | RNA→ rpmE (PRK00019)Ribosomal_L31 (pfam01197)→ |
| Sen-279-11 | NZ_LHDW01000023.1 | - | 2091 | 2039 | RNA→ rpmE (PRK00019)Ribosomal_L31 (pfam01197)→ |
| Sen-277-3 | NZ_LHGA01000003.1 | + | 97652 | 97704 | RNA→ rpmE (PRK00019)Ribosomal_L31 (pfam01197)→ |
| Sen-278-17 | NZ_LHIT01000015.1 | - | 45365 | 45313 | RNA→ rpmE (PRK00019)Ribosomal_L31 (pfam01197)→ |
| Sen-278-18 | NZ_LHJH01000036.1 | - | 3114 | 3062 | RNA→ rpmE (PRK00019)Ribosomal_L31 (pfam01197)→ |
| Sen-278-19 | NZ_LHLH01000022.1 | - | 45244 | 45192 | RNA→ rpmE (PRK00019)Ribosomal_L31 (pfam01197)→ |
| Sen-277-4 | NZ_LHML01000023.1 | - | 3161 | 3109 | RNA→ rpmE (PRK00019)Ribosomal_L31 (pfam01197)→ |
| Sen-277-5 | NZ_LHMZ01000005.1 | - | 35557 | 35505 | RNA→ rpmE (PRK00019)Ribosomal_L31 (pfam01197)→ |
| Sen-278-20 | NZ_LHOA01000019.1 | + | 104454 | 104506 | RNA→ rpmE (PRK00019)Ribosomal_L31 (pfam01197)→ |
| Sen-279-12 | NZ_LHOP01000008.1 | + | 42069 | 42121 | RNA→ rpmE (PRK00019)Ribosomal_L31 (pfam01197)→ |
| Sen-280-3 | NZ_LHNK01000016.1 | - | 38071 | 38019 | RNA→ rpmE (PRK00019)Ribosomal_L31 (pfam01197)→ |
| Sen-279-13 | NZ_LHGW01000029.1 | - | 35431 | 35379 | RNA→ rpmE (PRK00019)Ribosomal_L31 (pfam01197)→ |
| Sen-278-21 | NZ_LHJR01000005.1 | + | 104169 | 104221 | RNA→ rpmE (PRK00019)Ribosomal_L31 (pfam01197)→ |
| Sen-288-7 | NZ_CHFY01000014.1 | - | 39456 | 39404 | RNA→ rpmE (PRK00019)Ribosomal_L31 (pfam01197)→ |
| Sen-300-2 | NZ_CTGT01000031.1 | + | 13413 | 13465 | RNA→ rpmE (PRK00019)Ribosomal_L31 (pfam01197)→ |
| Sen-300-3 | NZ_CTGI01000018.1 | - | 45153 | 45101 | RNA→ rpmE (PRK00019)Ribosomal_L31 (pfam01197)→ |
| Sen-300-4 | NZ_CTIE01000017.1 | - | 45343 | 45291 | RNA→ rpmE (PRK00019)Ribosomal_L31 (pfam01197)→ |
| Sen-288-8 | NZ_CUYY01000016.1 | - | 39532 | 39480 | RNA→ rpmE (PRK00019)Ribosomal_L31 (pfam01197)→ |
| Sen-288-9 | NZ_CUZT01000016.1 | - | 39535 | 39483 | RNA→ rpmE (PRK00019)Ribosomal_L31 (pfam01197)→ |
| Sen-288-10 | NZ_CUZD01000016.1 | - | 39530 | 39478 | RNA→ rpmE (PRK00019)Ribosomal_L31 (pfam01197)→ |
| Sen-288-11 | NZ_CUYL01000013.1 | - | 41231 | 41179 | RNA→ rpmE (PRK00019)Ribosomal_L31 (pfam01197)→ |
| Sen-288-12 | NZ_CUZG01000016.1 | - | 39532 | 39480 | RNA→ rpmE (PRK00019)Ribosomal_L31 (pfam01197)→ |
| Sen-288-13 | NZ_CUZB01000013.1 | - | 41498 | 41446 | RNA→ rpmE (PRK00019)Ribosomal_L31 (pfam01197)→ |
| Sen-288-14 | NZ_CUZE01000016.1 | - | 39526 | 39474 | RNA→ rpmE (PRK00019)Ribosomal_L31 (pfam01197)→ |
| Sen-288-15 | NZ_CUYP01000016.1 | + | 104146 | 104198 | RNA→ rpmE (PRK00019)Ribosomal_L31 (pfam01197)→ |
| Sen-288-16 | NZ_CUZV01000016.1 | - | 39537 | 39485 | RNA→ rpmE (PRK00019)Ribosomal_L31 (pfam01197)→ |
| Sen-288-17 | NZ_CUZM01000016.1 | + | 103905 | 103957 | RNA→ rpmE (PRK00019)Ribosomal_L31 (pfam01197)→ |
| Sen-288-18 | NZ_CUZK01000016.1 | + | 104178 | 104230 | RNA→ rpmE (PRK00019)Ribosomal_L31 (pfam01197)→ |
| Sen-300-5 | NZ_CTMV01000010.1 | - | 37187 | 37135 | RNA→ rpmE (PRK00019)Ribosomal_L31 (pfam01197)→ |
| Sen-300-6 | NZ_CTGJ01000017.1 | + | 42456 | 42508 | RNA→ rpmE (PRK00019)Ribosomal_L31 (pfam01197)→ |
| Sen-300-7 | NZ_CTHA01000017.1 | - | 45179 | 45127 | RNA→ rpmE (PRK00019)Ribosomal_L31 (pfam01197)→ |
| Sen-300-8 | NZ_CTLT01000019.1 | - | 45112 | 45060 | RNA→ rpmE (PRK00019)Ribosomal_L31 (pfam01197)→ |
| Sen-300-9 | NZ_CTHV01000017.1 | - | 45337 | 45285 | RNA→ rpmE (PRK00019)Ribosomal_L31 (pfam01197)→ |
| Sen-300-10 | NZ_CTQD01000017.1 | - | 45198 | 45146 | RNA→ rpmE (PRK00019)Ribosomal_L31 (pfam01197)→ |
| Sen-288-19 | NZ_CUYM01000016.1 | - | 39531 | 39479 | RNA→ rpmE (PRK00019)Ribosomal_L31 (pfam01197)→ |
| Sen-288-20 | NZ_CUZF01000016.1 | - | 39535 | 39483 | RNA→ rpmE (PRK00019)Ribosomal_L31 (pfam01197)→ |
| Sen-772-1 | NZ_AMME01000310.1 | + | 12820 | 12872 | RNA→ rpmE (PRK00019)Ribosomal_L31 (pfam01197)→ |
| Sen-288-21 | NZ_CUZA01000016.1 | + | 104133 | 104185 | RNA→ rpmE (PRK00019)Ribosomal_L31 (pfam01197)→ |
| Sen-288-22 | NZ_CUZN01000016.1 | - | 39532 | 39480 | RNA→ rpmE (PRK00019)Ribosomal_L31 (pfam01197)→ |
| Sen-288-23 | NZ_CUZO01000017.1 | + | 104197 | 104249 | RNA→ rpmE (PRK00019)Ribosomal_L31 (pfam01197)→ |
| Sen-751-1 | NZ_AMMI01000362.1 | + | 28318 | 28370 | RNA→ rpmE (PRK00019)Ribosomal_L31 (pfam01197)→ |
| Sen-288-1107 | NZ_CUZW01000016.1 | + | 104149 | 104201 | RNA→ rpmE (PRK00019)Ribosomal_L31 (pfam01197)→ |
| Sen-288-24 | NZ_CUYK01000016.1 | + | 104120 | 104172 | RNA→ rpmE (PRK00019)Ribosomal_L31 (pfam01197)→ |
| Sen-288-25 | NZ_CUYV01000016.1 | - | 39527 | 39475 | RNA→ rpmE (PRK00019)Ribosomal_L31 (pfam01197)→ |
| Sen-180-1 | NZ_AMMA01000024.1 | - | 37602 | 37550 | RNA→ rpmE (PRK00019)Ribosomal_L31 (pfam01197)→ |

| | | | | | |
|---|---|---|---|---|---|
| Sen-288-26 | NZ_CUYJ01000015.1 | - | 39535 | 39483 | **RNA**→ rpmE (PRK00019)Ribosomal_L31 (pfam01197)→ |
| Sen-753-1 | NZ_AMMG01000052.1 | + | 100556 | 100608 | **RNA**→ rpmE (PRK00019)Ribosomal_L31 (pfam01197)→ |
| Sen-288-27 | NZ_CUYU01000016.1 | + | 104154 | 104206 | **RNA**→ rpmE (PRK00019)Ribosomal_L31 (pfam01197)→ |
| Sen-750-1 | NZ_AMMH01000191.1 | + | 66756 | 66808 | **RNA**→ rpmE (PRK00019)Ribosomal_L31 (pfam01197)→ |
| Sen-754-1 | NZ_AMMN01000182.1 | - | 32704 | 32652 | **RNA**→ rpmE (PRK00019)Ribosomal_L31 (pfam01197)→ |
| Sen-179-1 | NZ_AMMR01000082.1 | + | 100658 | 100710 | **RNA**→ rpmE (PRK00019)Ribosomal_L31 (pfam01197)→ |
| Sen-300-11 | NZ_CTPQ01000018.1 | - | 46977 | 46925 | **RNA**→ rpmE (PRK00019)Ribosomal_L31 (pfam01197)→ |
| Sen-182-1 | NZ_AMMU01000426.1 | + | 58961 | 59013 | **RNA**→ rpmE (PRK00019)Ribosomal_L31 (pfam01197)→ |
| Sen-300-12 | NZ_CTGR01000022.1 | - | 45111 | 45059 | **RNA**→ rpmE (PRK00019)Ribosomal_L31 (pfam01197)→ |
| Sen-288-28 | NZ_CUZX01000016.1 | + | 104147 | 104199 | **RNA**→ rpmE (PRK00019)Ribosomal_L31 (pfam01197)→ |
| Sen-288-29 | NZ_CUYW01000016.1 | - | 39536 | 39484 | **RNA**→ rpmE (PRK00019)Ribosomal_L31 (pfam01197)→ |
| Sen-288-30 | NZ_CUZS01000016.1 | + | 104156 | 104208 | **RNA**→ rpmE (PRK00019)Ribosomal_L31 (pfam01197)→ |
| Sen-288-31 | NZ_CUYT01000016.1 | - | 39526 | 39474 | **RNA**→ rpmE (PRK00019)Ribosomal_L31 (pfam01197)→ |
| Sen-288-32 | NZ_CUZR01000016.1 | - | 39535 | 39483 | **RNA**→ rpmE (PRK00019)Ribosomal_L31 (pfam01197)→ |
| Sen-288-33 | NZ_CUYZ01000016.1 | - | 39534 | 39482 | **RNA**→ rpmE (PRK00019)Ribosomal_L31 (pfam01197)→ |
| Sen-178-1 | NZ_AMLY01000042.1 | + | 100656 | 100708 | **RNA**→ rpmE (PRK00019)Ribosomal_L31 (pfam01197)→ |
| Sen-183-1 | NZ_AMMD01000057.1 | + | 20766 | 20818 | **RNA**→ rpmE (PRK00019)Ribosomal_L31 (pfam01197)→ |
| Sen-288-34 | NZ_CUYO01000010.1 | + | 133787 | 133839 | **RNA**→ rpmE (PRK00019)Ribosomal_L31 (pfam01197)→ |
| Sen-288-35 | NZ_CUYS01000016.1 | + | 104149 | 104201 | **RNA**→ rpmE (PRK00019)Ribosomal_L31 (pfam01197)→ |
| Sen-288-36 | NZ_CUYX01000016.1 | + | 104141 | 104193 | **RNA**→ rpmE (PRK00019)Ribosomal_L31 (pfam01197)→ |
| Sen-288-37 | NZ_CUZI01000016.1 | - | 39536 | 39484 | **RNA**→ rpmE (PRK00019)Ribosomal_L31 (pfam01197)→ |
| Sen-288-38 | NZ_CUYI01000015.1 | + | 103893 | 103945 | **RNA**→ rpmE (PRK00019)Ribosomal_L31 (pfam01197)→ |
| Sen-195-1 | NZ_AMOM01000044.1 | - | 37502 | 37450 | **RNA**→ rpmE (PRK00019)Ribosomal_L31 (pfam01197)→ |
| Sen-288-39 | NZ_CUZU01000016.1 | - | 39527 | 39475 | **RNA**→ rpmE (PRK00019)Ribosomal_L31 (pfam01197)→ |
| Sen-177-1 | NZ_AMLZ01000013.1 | - | 37630 | 37578 | **RNA**→ rpmE (PRK00019)Ribosomal_L31 (pfam01197)→ |
| Sen-752-1 | NZ_AMMF01000163.1 | - | 21720 | 21668 | **RNA**→ rpmE (PRK00019)Ribosomal_L31 (pfam01197)→ |
| Sen-774-1 | NZ_AMMM01000001.1 | + | 9526 | 9578 | **RNA**→ rpmE (PRK00019)Ribosomal_L31 (pfam01197)→ |
| Sen-176-1 | NZ_AMMS01000110.1 | - | 37699 | 37647 | **RNA**→ rpmE (PRK00019)Ribosomal_L31 (pfam01197)→ |
| Sen-7-1 | NZ_LAPG01000029.1 | + | 104291 | 104343 | **RNA**→ rpmE (PRK00019)Ribosomal_L31 (pfam01197)→ |
| Sen-7-2 | NZ_LAPH01000026.1 | - | 45127 | 45075 | **RNA**→ rpmE (PRK00019)Ribosomal_L31 (pfam01197)→ |
| Sen-7-3 | NZ_LAPQ01000010.1 | + | 104291 | 104343 | **RNA**→ rpmE (PRK00019)Ribosomal_L31 (pfam01197)→ |
| Sen-8-1 | NZ_LHEE01000039.1 | - | 38019 | 37967 | **RNA**→ rpmE (PRK00019)Ribosomal_L31 (pfam01197)→ |
| Sen-288-40 | NZ_CUZQ01000017.1 | - | 39533 | 39481 | **RNA**→ rpmE (PRK00019)Ribosomal_L31 (pfam01197)→ |
| Sen-288-41 | NZ_CUYN01000014.1 | - | 41508 | 41456 | **RNA**→ rpmE (PRK00019)Ribosomal_L31 (pfam01197)→ |
| Sen-288-42 | NZ_CUYR01000016.1 | - | 39531 | 39479 | **RNA**→ rpmE (PRK00019)Ribosomal_L31 (pfam01197)→ |
| Sen-216-1 | NZ_AMLW01000211.1 | - | 1474 | 1422 | **RNA**→ rpmE (PRK00019)Ribosomal_L31 (pfam01197)→ |
| Sen-288-43 | NZ_CUZL01000015.1 | + | 103906 | 103958 | **RNA**→ rpmE (PRK00019)Ribosomal_L31 (pfam01197)→ |
| Sen-288-44 | NZ_CUYQ01000016.1 | - | 39568 | 39516 | **RNA**→ rpmE (PRK00019)Ribosomal_L31 (pfam01197)→ |
| Sen-288-45 | NZ_CUZC01000015.1 | - | 39558 | 39506 | **RNA**→ rpmE (PRK00019)Ribosomal_L31 (pfam01197)→ |
| Sen-866-1 | NZ_AUQP01000004.1 | + | 105451 | 105503 | **RNA**→ rpmE (PRK00019)Ribosomal_L31 (pfam01197)→ |
| Sen-864-1 | NZ_AUQR01000004.1 | - | 45390 | 45338 | **RNA**→ rpmE (PRK00019)Ribosomal_L31 (pfam01197)→ |
| Sen-865-1 | NZ_AUQQ01000005.1 | - | 45390 | 45338 | **RNA**→ rpmE (PRK00019)Ribosomal_L31 (pfam01197)→ |
| Sen-291-5 | NZ_LFDW01000010.1 | - | 588736 | 588684 | **RNA**→ rpmE (PRK00019)Ribosomal_L31 (pfam01197)→ |
| Sen-291-6 | NZ_LFGN01000015.1 | - | 504820 | 504768 | **RNA**→ rpmE (PRK00019)Ribosomal_L31 (pfam01197)→ |
| Sen-137-6 | NZ_LAZB01000004.1 | + | 95741 | 95793 | **RNA**→ rpmE (PRK00019)Ribosomal_L31 (pfam01197)→ |
| Sen-133-1 | NZ_LELI01000015.1 | - | 36659 | 36607 | **RNA**→ rpmE (PRK00019)Ribosomal_L31 (pfam01197)→ |
| Sen-242-1 | NZ_LFIH01000123.1 | - | 559 | 507 | **RNA**→ rpmE (PRK00019)Ribosomal_L31 (pfam01197)→ hypo→ hypo→ |
| Sen-248-1 | NZ_APGQ01000022.1 | - | 16137 | 16085 | **RNA**→ rpmE (PRK00019)Ribosomal_L31 (pfam01197)→ |
| Sen-288-46 | NZ_CVQC01000013.1 | - | 41503 | 41451 | **RNA**→ rpmE (PRK00019)Ribosomal_L31 (pfam01197)→ |
| Sen-249-1 | NZ_APGS01000122.1 | - | 45070 | 45018 | **RNA**→ rpmE (PRK00019)Ribosomal_L31 (pfam01197)→ |
| Sen-288-47 | NZ_CDLH01000019.1 | - | 41652 | 41600 | **RNA**→ rpmE (PRK00019)Ribosomal_L31 (pfam01197)→ |
| Sen-333-1 | NZ_CATY01000043.1 | + | 111327 | 111379 | **RNA**→ rpmE (PRK00019)Ribosomal_L31 (pfam01197)→ |

| | | | | | |
|---|---|---|---|---|---|
| Sen-245-1 | NZ_APGT01000012.1 | - | 45268 | 45216 | **RNA**→ rpmE (PRK00019)Ribosomal_L31 (pfam01197)→ |
| Sen-246-1 | NZ_APGV01000022.1 | + | 105431 | 105483 | **RNA**→ rpmE (PRK00019)Ribosomal_L31 (pfam01197)→ |
| Sen-244-1 | NZ_APGU01000022.1 | + | 105431 | 105483 | **RNA**→ rpmE (PRK00019)Ribosomal_L31 (pfam01197)→ |
| Sen-250-1 | NZ_APGR01000135.1 | - | 6913 | 6861 | **RNA**→ rpmE (PRK00019)Ribosomal_L31 (pfam01197)→ |
| Sen-247-1 | NZ_APHF01000061.1 | + | 105419 | 105471 | **RNA**→ rpmE (PRK00019)Ribosomal_L31 (pfam01197)→ |
| Sen-898-1 | NZ_AYEG01000116.1 | - | 45279 | 45227 | **RNA**→ rpmE (PRK00019)Ribosomal_L31 (pfam01197)→ |
| Sen-930-1 | NZ_AOYA01000083.1 | - | 35287 | 35235 | **RNA**→ rpmE (PRK00019)Ribosomal_L31 (pfam01197)→ rpmE (PRK00019)Ribosomal_L31 (pfam01197)→ |
| Sen-369-1 | NZ_AYDB01000067.1 | + | 111693 | 111745 | **RNA**→ rpmE (PRK00019)Ribosomal_L31 (pfam01197)→ rpmE (PRK00019)Ribosomal_L31 (pfam01197)→ |
| Sen-2-1 | NZ_LFCC01000063.1 | + | 17175 | 17227 | **RNA**→ rpmE (PRK00019)Ribosomal_L31 (pfam01197)→ |
| Sen-261-1 | NZ_APHB01000016.1 | + | 105451 | 105503 | **RNA**→ rpmE (PRK00019)Ribosomal_L31 (pfam01197)→ |
| Sen-288-48 | NZ_CVQF01000016.1 | + | 104155 | 104207 | **RNA**→ rpmE (PRK00019)Ribosomal_L31 (pfam01197)→ |
| Sen-288-49 | NZ_CVQB01000015.1 | + | 104149 | 104201 | **RNA**→ rpmE (PRK00019)Ribosomal_L31 (pfam01197)→ |
| Sen-288-50 | NZ_CVQO01000014.1 | + | 104154 | 104206 | **RNA**→ rpmE (PRK00019)Ribosomal_L31 (pfam01197)→ |
| Sen-251-1 | NZ_APHC01000073.1 | + | 20125 | 20177 | **RNA**→ rpmE (PRK00019)Ribosomal_L31 (pfam01197)→ |
| Sen-288-51 | NZ_CVQN01000010.1 | + | 133799 | 133851 | **RNA**→ rpmE (PRK00019)Ribosomal_L31 (pfam01197)→ |
| Sen-288-52 | NZ_CVPK01000016.1 | + | 104140 | 104192 | **RNA**→ rpmE (PRK00019)Ribosomal_L31 (pfam01197)→ |
| Sen-288-53 | NZ_CVPQ01000016.1 | + | 104149 | 104201 | **RNA**→ rpmE (PRK00019)Ribosomal_L31 (pfam01197)→ |
| Sen-288-54 | NZ_CVQM01000016.1 | - | 39532 | 39480 | **RNA**→ rpmE (PRK00019)Ribosomal_L31 (pfam01197)→ |
| Sen-288-55 | NZ_CVPS01000016.1 | - | 39588 | 39536 | **RNA**→ rpmE (PRK00019)Ribosomal_L31 (pfam01197)→ |
| Sen-269-1 | NZ_APGX01000014.1 | - | 45390 | 45338 | **RNA**→ rpmE (PRK00019)Ribosomal_L31 (pfam01197)→ |
| Sen-262-1 | NZ_APGY01000013.1 | + | 105451 | 105503 | **RNA**→ rpmE (PRK00019)Ribosomal_L31 (pfam01197)→ |
| Sen-259-1 | NZ_APGZ01000013.1 | - | 45274 | 45222 | **RNA**→ rpmE (PRK00019)Ribosomal_L31 (pfam01197)→ |
| Sen-264-1 | NZ_APHD01000133.1 | - | 2882 | 2830 | **RNA**→ rpmE (PRK00019)Ribosomal_L31 (pfam01197)→ |
| Sen-265-1 | NZ_APHE01000099.1 | - | 45389 | 45337 | **RNA**→ rpmE (PRK00019)Ribosomal_L31 (pfam01197)→ |
| Sen-263-1 | NZ_APHG01000048.1 | + | 105398 | 105450 | **RNA**→ rpmE (PRK00019)Ribosomal_L31 (pfam01197)→ |
| Sen-268-1 | NZ_APGW01000014.1 | - | 45390 | 45338 | **RNA**→ rpmE (PRK00019)Ribosomal_L31 (pfam01197)→ |
| Sen-260-1 | NZ_APHA01000015.1 | - | 45273 | 45221 | **RNA**→ rpmE (PRK00019)Ribosomal_L31 (pfam01197)→ |
| Sen-170-22 | NZ_LHMI01000009.1 | + | 100687 | 100739 | **RNA**→ rpmE (PRK00019)Ribosomal_L31 (pfam01197)→ |
| Sen-170-23 | NZ_LHDP01000023.1 | - | 37621 | 37569 | **RNA**→ rpmE (PRK00019)Ribosomal_L31 (pfam01197)→ |
| Sen-170-24 | NZ_LHFK01000021.1 | - | 37518 | 37466 | **RNA**→ rpmE (PRK00019)Ribosomal_L31 (pfam01197)→ |
| Sen-170-25 | NZ_LHFM01000003.1 | + | 100914 | 100966 | **RNA**→ rpmE (PRK00019)Ribosomal_L31 (pfam01197)→ |
| Sen-170-26 | NZ_LHFN01000033.1 | - | 36158 | 36106 | **RNA**→ rpmE (PRK00019)Ribosomal_L31 (pfam01197)→ |
| Sen-170-51 | NZ_LHHL01000008.1 | + | 42794 | 42846 | **RNA**→ rpmE (PRK00019)Ribosomal_L31 (pfam01197)→ |
| Sen-170-27 | NZ_LHHY01000001.1 | - | 37518 | 37466 | **RNA**→ rpmE (PRK00019)Ribosomal_L31 (pfam01197)→ |
| Sen-170-28 | NZ_LHIB01000026.1 | - | 37517 | 37465 | **RNA**→ rpmE (PRK00019)Ribosomal_L31 (pfam01197)→ |
| Sen-170-29 | NZ_LHLM01000008.1 | + | 100688 | 100740 | **RNA**→ rpmE (PRK00019)Ribosomal_L31 (pfam01197)→ |
| Sen-170-30 | NZ_LHMD01000023.1 | + | 100688 | 100740 | **RNA**→ rpmE (PRK00019)Ribosomal_L31 (pfam01197)→ |
| Sen-170-31 | NZ_LHNQ01000030.1 | + | 43462 | 43514 | **RNA**→ rpmE (PRK00019)Ribosomal_L31 (pfam01197)→ |
| Sen-170-32 | NZ_LHNU01000034.1 | + | 100688 | 100740 | **RNA**→ rpmE (PRK00019)Ribosomal_L31 (pfam01197)→ |
| Sen-170-33 | NZ_LHFD01000017.1 | + | 100590 | 100642 | **RNA**→ rpmE (PRK00019)Ribosomal_L31 (pfam01197)→ |
| Sen-170-34 | NZ_LHFE01000007.1 | + | 100590 | 100642 | **RNA**→ rpmE (PRK00019)Ribosomal_L31 (pfam01197)→ |
| Sen-170-52 | NZ_LHME01000039.1 | - | 37788 | 37736 | **RNA**→ rpmE (PRK00019)Ribosomal_L31 (pfam01197)→ |
| Sen-170-35 | NZ_LHGU01000033.1 | + | 42035 | 42087 | **RNA**→ rpmE (PRK00019)Ribosomal_L31 (pfam01197)→ |
| Sen-170-36 | NZ_LHNV01000009.1 | + | 100688 | 100740 | **RNA**→ rpmE (PRK00019)Ribosomal_L31 (pfam01197)→ |
| Sen-170-37 | NZ_LHDQ01000029.1 | + | 42035 | 42087 | **RNA**→ rpmE (PRK00019)Ribosomal_L31 (pfam01197)→ |
| Sen-170-38 | NZ_LHFJ01000018.1 | + | 100688 | 100740 | **RNA**→ rpmE (PRK00019)Ribosomal_L31 (pfam01197)→ |
| Sen-170-39 | NZ_LHFQ01000021.1 | + | 100688 | 100740 | **RNA**→ rpmE (PRK00019)Ribosomal_L31 (pfam01197)→ |
| Sen-170-40 | NZ_LHHO01000035.1 | + | 43729 | 43781 | **RNA**→ rpmE (PRK00019)Ribosomal_L31 (pfam01197)→ |
| Sen-170-41 | NZ_LHJY01000010.1 | + | 100688 | 100740 | **RNA**→ rpmE (PRK00019)Ribosomal_L31 (pfam01197)→ |
| Sen-167-9 | NZ_LHLD01000049.1 | - | 45288 | 45236 | **RNA**→ rpmE (PRK00019)Ribosomal_L31 (pfam01197)→ |
| Sen-170-42 | NZ_LHLV01000033.1 | + | 42046 | 42098 | **RNA**→ rpmE (PRK00019)Ribosomal_L31 (pfam01197)→ |
| Sen-167-10 | NZ_LHND01000022.1 | - | 45288 | 45236 | **RNA**→ rpmE (PRK00019)Ribosomal_L31 (pfam01197)→ |

| | | | | | |
|---|---|---|---|---|---|
| Sen-167-11 | NZ_LHNO01000018.1 | - | 45288 | 45236 | **RNA**→ rpmE (PRK00019)Ribosomal_L31 (pfam01197)→ |
| Sen-170-43 | NZ_LHKU01000027.1 | - | 37758 | 37706 | **RNA**→ rpmE (PRK00019)Ribosomal_L31 (pfam01197)→ |
| Sen-170-44 | NZ_LHHS01000031.1 | - | 37517 | 37465 | **RNA**→ rpmE (PRK00019)Ribosomal_L31 (pfam01197)→ |
| Sen-170-45 | NZ_LHJL01000038.1 | + | 100688 | 100740 | **RNA**→ rpmE (PRK00019)Ribosomal_L31 (pfam01197)→ |
| Sen-288-56 | NZ_CVQG01000009.1 | + | 133788 | 133840 | **RNA**→ rpmE (PRK00019)Ribosomal_L31 (pfam01197)→ |
| Sen-288-57 | NZ_CVQD01000016.1 | - | 41505 | 41453 | **RNA**→ rpmE (PRK00019)Ribosomal_L31 (pfam01197)→ |
| Sen-288-58 | NZ_CVQA01000015.1 | + | 104149 | 104201 | **RNA**→ rpmE (PRK00019)Ribosomal_L31 (pfam01197)→ |
| Sen-288-59 | NZ_CVPU01000016.1 | - | 39566 | 39514 | **RNA**→ rpmE (PRK00019)Ribosomal_L31 (pfam01197)→ |
| Sen-288-60 | NZ_CVPR01000016.1 | + | 104157 | 104209 | **RNA**→ rpmE (PRK00019)Ribosomal_L31 (pfam01197)→ |
| Sen-288-1108 | NZ_CVPY01000015.1 | - | 39536 | 39484 | **RNA**→ rpmE (PRK00019)Ribosomal_L31 (pfam01197)→ |
| Sen-288-61 | NZ_CVPX01000016.1 | + | 104194 | 104246 | **RNA**→ rpmE (PRK00019)Ribosomal_L31 (pfam01197)→ |
| Sen-288-62 | NZ_CVPP01000016.1 | + | 104164 | 104216 | **RNA**→ rpmE (PRK00019)Ribosomal_L31 (pfam01197)→ |
| Sen-288-63 | NZ_CVQK01000016.1 | + | 104154 | 104206 | **RNA**→ rpmE (PRK00019)Ribosomal_L31 (pfam01197)→ |
| Sen-288-64 | NZ_CVPZ01000016.1 | - | 39536 | 39484 | **RNA**→ rpmE (PRK00019)Ribosomal_L31 (pfam01197)→ |
| Sen-288-65 | NZ_CVPM01000015.1 | - | 39556 | 39504 | **RNA**→ rpmE (PRK00019)Ribosomal_L31 (pfam01197)→ |
| Sen-288-66 | NZ_CVPO01000016.1 | + | 104154 | 104206 | **RNA**→ rpmE (PRK00019)Ribosomal_L31 (pfam01197)→ |
| Sen-288-1109 | NZ_CVPW01000016.1 | - | 39512 | 39460 | **RNA**→ rpmE (PRK00019)Ribosomal_L31 (pfam01197)→ |
| Sen-288-67 | NZ_CVPT01000016.1 | + | 104162 | 104214 | **RNA**→ rpmE (PRK00019)Ribosomal_L31 (pfam01197)→ |
| Sen-288-68 | NZ_CVPV01000016.1 | + | 103992 | 104044 | **RNA**→ rpmE (PRK00019)Ribosomal_L31 (pfam01197)→ |
| Sen-288-69 | NZ_CVPL01000014.1 | - | 39534 | 39482 | **RNA**→ rpmE (PRK00019)Ribosomal_L31 (pfam01197)→ |
| Sen-288-70 | NZ_CVPN01000016.1 | - | 39534 | 39482 | **RNA**→ rpmE (PRK00019)Ribosomal_L31 (pfam01197)→ |
| Sen-288-71 | NZ_CVQL01000015.1 | + | 104176 | 104228 | **RNA**→ rpmE (PRK00019)Ribosomal_L31 (pfam01197)→ |
| Sen-288-72 | NZ_CVQE01000015.1 | - | 39535 | 39483 | **RNA**→ rpmE (PRK00019)Ribosomal_L31 (pfam01197)→ |
| Sen-288-73 | NZ_CVQJ01000016.1 | - | 39531 | 39479 | **RNA**→ rpmE (PRK00019)Ribosomal_L31 (pfam01197)→ |
| Sen-167-12 | NZ_LHMY01000012.1 | + | 100286 | 100338 | **RNA**→ rpmE (PRK00019)Ribosomal_L31 (pfam01197)→ |
| Sen-167-13 | NZ_LHNC01000011.1 | - | 45288 | 45236 | **RNA**→ rpmE (PRK00019)Ribosomal_L31 (pfam01197)→ |
| Sen-167-14 | NZ_LHLS01000024.1 | - | 45288 | 45236 | **RNA**→ rpmE (PRK00019)Ribosomal_L31 (pfam01197)→ |
| Sen-167-15 | NZ_LHFC01000001.1 | + | 100283 | 100335 | **RNA**→ rpmE (PRK00019)Ribosomal_L31 (pfam01197)→ |
| Sen-167-16 | NZ_LHGM01000010.1 | + | 55519 | 55571 | **RNA**→ rpmE (PRK00019)Ribosomal_L31 (pfam01197)→ |
| Sen-167-17 | NZ_LHNB01000004.1 | + | 100286 | 100338 | **RNA**→ rpmE (PRK00019)Ribosomal_L31 (pfam01197)→ |
| Sen-300-13 | NZ_CTNF01000019.1 | - | 45179 | 45127 | **RNA**→ rpmE (PRK00019)Ribosomal_L31 (pfam01197)→ |
| Sen-167-18 | NZ_LHFA01000003.1 | + | 100286 | 100338 | **RNA**→ rpmE (PRK00019)Ribosomal_L31 (pfam01197)→ |
| Sen-288-74 | NZ_CEYU01000021.1 | + | 26787 | 26839 | **RNA**→ rpmE (PRK00019)Ribosomal_L31 (pfam01197)→ |
| Sen-288-75 | NZ_CEXT01000016.1 | + | 104150 | 104202 | **RNA**→ rpmE (PRK00019)Ribosomal_L31 (pfam01197)→ |
| Sen-288-76 | NZ_CJBN01000027.1 | + | 19742 | 19794 | **RNA**→ rpmE (PRK00019)Ribosomal_L31 (pfam01197)→ |
| Sen-288-77 | NZ_CJTY01000007.1 | + | 190518 | 190570 | **RNA**→ rpmE (PRK00019)Ribosomal_L31 (pfam01197)→ |
| Sen-288-78 | NZ_CJCM01000016.1 | - | 39535 | 39483 | **RNA**→ rpmE (PRK00019)Ribosomal_L31 (pfam01197)→ |
| Sen-288-79 | NZ_CJRP01000014.1 | + | 109738 | 109790 | **RNA**→ rpmE (PRK00019)Ribosomal_L31 (pfam01197)→ |
| Sen-288-80 | NZ_CJPH01000016.1 | + | 104151 | 104203 | **RNA**→ rpmE (PRK00019)Ribosomal_L31 (pfam01197)→ |
| Sen-288-81 | NZ_CIZR01000016.1 | - | 39534 | 39482 | **RNA**→ rpmE (PRK00019)Ribosomal_L31 (pfam01197)→ |
| Sen-906-1 | NZ_AOYL01000004.1 | + | 98734 | 98786 | **RNA**→ rpmE (PRK00019)Ribosomal_L31 (pfam01197)→ rpmE (PRK00019)Ribosomal_L31 (pfam01197)→ |
| Sen-288-82 | NZ_CJIY01000016.1 | + | 104157 | 104209 | **RNA**→ rpmE (PRK00019)Ribosomal_L31 (pfam01197)→ |
| Sen-288-83 | NZ_CEYF01000015.1 | + | 103788 | 103840 | **RNA**→ rpmE (PRK00019)Ribosomal_L31 (pfam01197)→ |
| Sen-288-84 | NZ_CJLH01000011.1 | - | 39456 | 39404 | **RNA**→ rpmE (PRK00019)Ribosomal_L31 (pfam01197)→ |
| Sen-288-85 | NZ_CIPI01000004.1 | + | 104159 | 104211 | **RNA**→ rpmE (PRK00019)Ribosomal_L31 (pfam01197)→ |
| Sen-288-86 | NZ_CJUT01000014.1 | + | 107995 | 108047 | **RNA**→ rpmE (PRK00019)Ribosomal_L31 (pfam01197)→ |
| Sen-288-87 | NZ_CJOV01000016.1 | - | 39723 | 39671 | **RNA**→ rpmE (PRK00019)Ribosomal_L31 (pfam01197)→ |
| Sen-288-88 | NZ_CEXS01000016.1 | - | 39532 | 39480 | **RNA**→ rpmE (PRK00019)Ribosomal_L31 (pfam01197)→ |
| Sen-288-89 | NZ_CIZY01000011.1 | - | 39456 | 39404 | **RNA**→ rpmE (PRK00019)Ribosomal_L31 (pfam01197)→ |
| Sen-288-90 | NZ_CILO01000016.1 | - | 39601 | 39549 | **RNA**→ rpmE (PRK00019)Ribosomal_L31 (pfam01197)→ |
| Sen-347-1 | NZ_CASQ01000042.1 | + | 111150 | 111202 | **RNA**→ rpmE (PRK00019)Ribosomal_L31 (pfam01197)→ |
| Sen-392-1 | NZ_CART01000035.1 | - | 37552 | 37500 | **RNA**→ rpmE (PRK00019)Ribosomal_L31 (pfam01197)→ |

| | | | | | | | |
|---|---|---|---|---|---|---|---|
| Sen-377-1 | NZ_AYCS01000004.1 | + | 111233 | 111285 | **RNA**→ | rpmE (PRK00019)Ribosomal_L31 (pfam01197)→ | rpmE (PRK00019)Ribosomal_L31 (pfam01197)→ |
| Sen-352-1 | NZ_AYCL01000005.1 | + | 83635 | 83687 | **RNA**→ | rpmE (PRK00019)Ribosomal_L31 (pfam01197)→ | rpmE (PRK00019)Ribosomal_L31 (pfam01197)→ |
| Sen-372-1 | NZ_AOZV01000015.1 | + | 42798 | 42850 | **RNA**→ | rpmE (PRK00019)Ribosomal_L31 (pfam01197)→ | rpmE (PRK00019)Ribosomal_L31 (pfam01197)→ |
| Sen-344-1 | NZ_AYDI01000048.1 | + | 111595 | 111647 | **RNA**→ | rpmE (PRK00019)Ribosomal_L31 (pfam01197)→ | rpmE (PRK00019)Ribosomal_L31 (pfam01197)→ |
| Sen-340-1 | NZ_CASS01000018.1 | + | 111225 | 111277 | **RNA**→ | rpmE (PRK00019)Ribosomal_L31 (pfam01197)→ | |
| Sen-348-1 | NZ_CASP01000044.1 | + | 111150 | 111202 | **RNA**→ | rpmE (PRK00019)Ribosomal_L31 (pfam01197)→ | |
| Sen-349-1 | NZ_CASO01000054.1 | - | 17553 | 17501 | **RNA**→ | rpmE (PRK00019)Ribosomal_L31 (pfam01197)→ | |
| Sen-388-1 | NZ_CARX01000045.1 | - | 37768 | 37716 | **RNA**→ | rpmE (PRK00019)Ribosomal_L31 (pfam01197)→ | |
| Sen-393-1 | NZ_CARS01000036.1 | - | 37552 | 37500 | **RNA**→ | rpmE (PRK00019)Ribosomal_L31 (pfam01197)→ | |
| Sen-379-1 | NZ_AYCQ01000005.1 | + | 111262 | 111314 | **RNA**→ | rpmE (PRK00019)Ribosomal_L31 (pfam01197)→ | rpmE (PRK00019)Ribosomal_L31 (pfam01197)→ |
| Sen-355-1 | NZ_AYCM01000006.1 | + | 111260 | 111312 | **RNA**→ | rpmE (PRK00019)Ribosomal_L31 (pfam01197)→ | rpmE (PRK00019)Ribosomal_L31 (pfam01197)→ |
| Sen-383-1 | NZ_AYCO01000001.1 | - | 37555 | 37503 | **RNA**→ | rpmE (PRK00019)Ribosomal_L31 (pfam01197)→ | rpmE (PRK00019)Ribosomal_L31 (pfam01197)→ |
| Sen-384-1 | NZ_AYCW01000013.1 | + | 111445 | 111497 | **RNA**→ | rpmE (PRK00019)Ribosomal_L31 (pfam01197)→ | rpmE (PRK00019)Ribosomal_L31 (pfam01197)→ |
| Sen-342-1 | NZ_AYDJ01000060.1 | + | 111622 | 111674 | **RNA**→ | rpmE (PRK00019)Ribosomal_L31 (pfam01197)→ | rpmE (PRK00019)Ribosomal_L31 (pfam01197)→ |
| Sen-362-1 | NZ_AYDH01000066.1 | + | 111625 | 111677 | **RNA**→ | rpmE (PRK00019)Ribosomal_L31 (pfam01197)→ | rpmE (PRK00019)Ribosomal_L31 (pfam01197)→ |
| Sen-345-1 | NZ_CATX01000012.1 | + | 112246 | 112298 | **RNA**→ | rpmE (PRK00019)Ribosomal_L31 (pfam01197)→ | |
| Sen-390-1 | NZ_CARV01000038.1 | - | 37552 | 37500 | **RNA**→ | rpmE (PRK00019)Ribosomal_L31 (pfam01197)→ | |
| Sen-391-1 | NZ_CARU01000036.1 | - | 37552 | 37500 | **RNA**→ | rpmE (PRK00019)Ribosomal_L31 (pfam01197)→ | |
| Sen-346-1 | NZ_AYCY01000031.1 | + | 11826 | 11878 | **RNA**→ | rpmE (PRK00019)Ribosomal_L31 (pfam01197)→ | |
| Sen-366-1 | NZ_AYDE01000034.1 | + | 111736 | 111788 | **RNA**→ | rpmE (PRK00019)Ribosomal_L31 (pfam01197)→ | rpmE (PRK00019)Ribosomal_L31 (pfam01197)→ |
| Sen-365-1 | NZ_AYDF01000058.1 | + | 111692 | 111744 | **RNA**→ | rpmE (PRK00019)Ribosomal_L31 (pfam01197)→ | rpmE (PRK00019)Ribosomal_L31 (pfam01197)→ |
| Sen-341-1 | NZ_CASR01000038.1 | + | 111200 | 111252 | **RNA**→ | rpmE (PRK00019)Ribosomal_L31 (pfam01197)→ | |
| Sen-389-1 | NZ_CARW01000047.1 | + | 111473 | 111525 | **RNA**→ | rpmE (PRK00019)Ribosomal_L31 (pfam01197)→ | |
| Sen-363-1 | NZ_AYDG01000035.1 | + | 111679 | 111731 | **RNA**→ | rpmE (PRK00019)Ribosomal_L31 (pfam01197)→ | rpmE (PRK00019)Ribosomal_L31 (pfam01197)→ |
| Sen-368-1 | NZ_AYDC01000067.1 | + | 111674 | 111726 | **RNA**→ | rpmE (PRK00019)Ribosomal_L31 (pfam01197)→ | |
| Sen-367-1 | NZ_AYDD01000027.1 | + | 111684 | 111736 | **RNA**→ | rpmE (PRK00019)Ribosomal_L31 (pfam01197)→ | rpmE (PRK00019)Ribosomal_L31 (pfam01197)→ |
| Sen-376-1 | NZ_CASB01000116.1 | - | 17617 | 17565 | **RNA**→ | rpmE (PRK00019)Ribosomal_L31 (pfam01197)→ | |
| Sen-370-1 | NZ_AYCT01000039.1 | - | 37612 | 37560 | **RNA**→ | rpmE (PRK00019)Ribosomal_L31 (pfam01197)→ | rpmE (PRK00019)Ribosomal_L31 (pfam01197)→ |
| Sen-380-1 | NZ_AYCP01000055.1 | - | 37519 | 37467 | **RNA**→ | rpmE (PRK00019)Ribosomal_L31 (pfam01197)→ | rpmE (PRK00019)Ribosomal_L31 (pfam01197)→ |
| Sen-373-1 | NZ_CASD01000056.1 | + | 111150 | 111202 | **RNA**→ | rpmE (PRK00019)Ribosomal_L31 (pfam01197)→ | |
| Sen-374-1 | NZ_CASC01000035.1 | - | 37552 | 37500 | **RNA**→ | rpmE (PRK00019)Ribosomal_L31 (pfam01197)→ | |
| Sen-387-1 | NZ_CATU01000036.1 | + | 55551 | 55603 | **RNA**→ | rpmE (PRK00019)Ribosomal_L31 (pfam01197)→ | |
| Sen-385-1 | NZ_CAAD010000062.1 | + | 61209 | 61261 | **RNA**→ | rpmE (PRK00019)Ribosomal_L31 (pfam01197)→ | |
| Sen-353-1 | NZ_CASM01000050.1 | + | 111284 | 111336 | **RNA**→ | rpmE (PRK00019)Ribosomal_L31 (pfam01197)→ | |
| Sen-354-1 | NZ_CASL01000044.1 | - | 37552 | 37500 | **RNA**→ | rpmE (PRK00019)Ribosomal_L31 (pfam01197)→ | |
| Sen-359-1 | NZ_CASH01000039.1 | + | 111150 | 111202 | **RNA**→ | rpmE (PRK00019)Ribosomal_L31 (pfam01197)→ | |
| Sen-360-1 | NZ_CASG01000064.1 | + | 55563 | 55615 | **RNA**→ | rpmE (PRK00019)Ribosomal_L31 (pfam01197)→ | |
| Sen-375-1 | NZ_AYCX01000037.1 | + | 111230 | 111282 | **RNA**→ | rpmE (PRK00019)Ribosomal_L31 (pfam01197)→ | rpmE (PRK00019)Ribosomal_L31 (pfam01197)→ |
| Sen-356-1 | NZ_CASK01000036.1 | - | 37552 | 37500 | **RNA**→ | rpmE (PRK00019)Ribosomal_L31 (pfam01197)→ | |
| Sen-378-1 | NZ_AYCR01000008.1 | + | 83965 | 84017 | **RNA**→ | rpmE (PRK00019)Ribosomal_L31 (pfam01197)→ | rpmE (PRK00019)Ribosomal_L31 (pfam01197)→ |
| Sen-364-1 | NZ_CASE01000032.1 | - | 37552 | 37500 | **RNA**→ | rpmE (PRK00019)Ribosomal_L31 (pfam01197)→ | |
| Sen-382-1 | NZ_CARZ01000055.1 | + | 61126 | 61178 | **RNA**→ | rpmE (PRK00019)Ribosomal_L31 (pfam01197)→ | |
| Sen-386-1 | NZ_CARY01000035.1 | - | 37552 | 37500 | **RNA**→ | rpmE (PRK00019)Ribosomal_L31 (pfam01197)→ | |
| Sen-351-1 | NZ_CASN01000036.1 | - | 37552 | 37500 | **RNA**→ | rpmE (PRK00019)Ribosomal_L31 (pfam01197)→ | |
| Sen-358-1 | NZ_CASI01000035.1 | - | 37552 | 37500 | **RNA**→ | rpmE (PRK00019)Ribosomal_L31 (pfam01197)→ | |
| Sen-371-1 | NZ_CASF01000145.1 | - | 17572 | 17520 | **RNA**→ | rpmE (PRK00019)Ribosomal_L31 (pfam01197)→ | |
| Sen-381-1 | NZ_CASA01000057.1 | + | 61126 | 61178 | **RNA**→ | rpmE (PRK00019)Ribosomal_L31 (pfam01197)→ | |
| Sen-357-1 | NZ_CASJ01000039.1 | + | 143491 | 143543 | **RNA**→ | rpmE (PRK00019)Ribosomal_L31 (pfam01197)→ | |
| Sen-288-91 | NZ_CIXI01000014.1 | - | 39456 | 39404 | **RNA**→ | rpmE (PRK00019)Ribosomal_L31 (pfam01197)→ | |
| Sen-288-92 | NZ_CILL01000016.1 | - | 39381 | 39329 | **RNA**→ | rpmE (PRK00019)Ribosomal_L31 (pfam01197)→ | |
| Sen-300-14 | NZ_CQGP01000016.1 | + | 42532 | 42584 | **RNA**→ | rpmE (PRK00019)Ribosomal_L31 (pfam01197)→ | |
| Sen-288-93 | NZ_CIVV01000012.1 | + | 107996 | 108048 | **RNA**→ | rpmE (PRK00019)Ribosomal_L31 (pfam01197)→ | |

| | | | | | |
|---|---|---|---|---|---|
| Sen-288-94 | NZ_CINM01000012.1 | + | 133824 | 133876 | **RNA**→ rpmE (PRK00019)Ribosomal_L31 (pfam01197)→ |
| Sen-288-95 | NZ_CILS01000015.1 | - | 39542 | 39490 | **RNA**→ rpmE (PRK00019)Ribosomal_L31 (pfam01197)→ |
| Sen-288-96 | NZ_CJQE01000015.1 | - | 39601 | 39549 | **RNA**→ rpmE (PRK00019)Ribosomal_L31 (pfam01197)→ |
| Sen-288-97 | NZ_CJVR01000015.1 | - | 41231 | 41179 | **RNA**→ rpmE (PRK00019)Ribosomal_L31 (pfam01197)→ |
| Sen-288-98 | NZ_CJMA01000010.1 | - | 41499 | 41447 | **RNA**→ rpmE (PRK00019)Ribosomal_L31 (pfam01197)→ |
| Sen-288-99 | NZ_CIWL01000015.1 | + | 103992 | 104044 | **RNA**→ rpmE (PRK00019)Ribosomal_L31 (pfam01197)→ |
| Sen-288-100 | NZ_CJNY01000010.1 | + | 104137 | 104189 | **RNA**→ rpmE (PRK00019)Ribosomal_L31 (pfam01197)→ |
| Sen-288-101 | NZ_CJWX01000017.1 | + | 104185 | 104237 | **RNA**→ rpmE (PRK00019)Ribosomal_L31 (pfam01197)→ |
| Sen-288-102 | NZ_CJCL01000016.1 | - | 39536 | 39484 | **RNA**→ rpmE (PRK00019)Ribosomal_L31 (pfam01197)→ |
| Sen-288-103 | NZ_CJJG01000010.1 | + | 133795 | 133847 | **RNA**→ rpmE (PRK00019)Ribosomal_L31 (pfam01197)→ |
| Sen-288-104 | NZ_CIUZ01000017.1 | + | 104176 | 104228 | **RNA**→ rpmE (PRK00019)Ribosomal_L31 (pfam01197)→ |
| Sen-288-105 | NZ_CJIF01000016.1 | - | 39535 | 39483 | **RNA**→ rpmE (PRK00019)Ribosomal_L31 (pfam01197)→ |
| Sen-288-106 | NZ_CJKI01000006.1 | - | 39456 | 39404 | **RNA**→ rpmE (PRK00019)Ribosomal_L31 (pfam01197)→ |
| Sen-288-107 | NZ_CJMB01000015.1 | + | 104141 | 104193 | **RNA**→ rpmE (PRK00019)Ribosomal_L31 (pfam01197)→ |
| Sen-288-108 | NZ_CJAS01000016.1 | + | 104148 | 104200 | **RNA**→ rpmE (PRK00019)Ribosomal_L31 (pfam01197)→ |
| Sen-288-109 | NZ_CIXD01000016.1 | + | 104172 | 104224 | **RNA**→ rpmE (PRK00019)Ribosomal_L31 (pfam01197)→ |
| Sen-288-110 | NZ_CJEU01000016.1 | - | 39537 | 39485 | **RNA**→ rpmE (PRK00019)Ribosomal_L31 (pfam01197)→ |
| Sen-288-111 | NZ_CJUE01000003.1 | - | 216862 | 216810 | **RNA**→ rpmE (PRK00019)Ribosomal_L31 (pfam01197)→ |
| Sen-288-112 | NZ_CJKA01000002.1 | + | 434934 | 434986 | **RNA**→ rpmE (PRK00019)Ribosomal_L31 (pfam01197)→ |
| Sen-288-113 | NZ_CJYE01000015.1 | - | 39456 | 39404 | **RNA**→ rpmE (PRK00019)Ribosomal_L31 (pfam01197)→ |
| Sen-288-114 | NZ_CIUT01000015.1 | + | 107848 | 107900 | **RNA**→ rpmE (PRK00019)Ribosomal_L31 (pfam01197)→ |
| Sen-288-115 | NZ_CIWB01000015.1 | - | 39506 | 39454 | **RNA**→ rpmE (PRK00019)Ribosomal_L31 (pfam01197)→ |
| Sen-288-116 | NZ_CJLA01000015.1 | - | 39535 | 39483 | **RNA**→ rpmE (PRK00019)Ribosomal_L31 (pfam01197)→ |
| Sen-288-117 | NZ_CJTN01000014.1 | - | 39372 | 39320 | **RNA**→ rpmE (PRK00019)Ribosomal_L31 (pfam01197)→ |
| Sen-288-118 | NZ_CIVE01000017.1 | - | 39385 | 39333 | **RNA**→ rpmE (PRK00019)Ribosomal_L31 (pfam01197)→ |
| Sen-288-119 | NZ_CJZC01000014.1 | + | 107832 | 107884 | **RNA**→ rpmE (PRK00019)Ribosomal_L31 (pfam01197)→ |
| Sen-288-120 | NZ_CJSE01000014.1 | + | 104161 | 104213 | **RNA**→ rpmE (PRK00019)Ribosomal_L31 (pfam01197)→ |
| Sen-69-1 | NZ_JMMG01000025.1 | - | 45104 | 45052 | **RNA**→ rpmE (PRK00019)Ribosomal_L31 (pfam01197)→ |
| Sen-288-121 | NZ_CJDT01000016.1 | - | 39535 | 39483 | **RNA**→ rpmE (PRK00019)Ribosomal_L31 (pfam01197)→ |
| Sen-288-122 | NZ_CJVQ01000001.1 | + | 104154 | 104206 | **RNA**→ rpmE (PRK00019)Ribosomal_L31 (pfam01197)→ |
| Sen-423-1 | NZ_ANOT01000041.1 | + | 116944 | 116996 | **RNA**→ rpmE (PRK00019)Ribosomal_L31 (pfam01197)→ rpmE (PRK00019)Ribosomal_L31 (pfam01197)→ |
| Sen-288-123 | NZ_CJWR01000013.1 | - | 41253 | 41201 | **RNA**→ rpmE (PRK00019)Ribosomal_L31 (pfam01197)→ |
| Sen-398-1 | NZ_CARP01000034.1 | - | 37552 | 37500 | **RNA**→ rpmE (PRK00019)Ribosomal_L31 (pfam01197)→ |
| Sen-288-124 | NZ_CJJP01000015.1 | + | 104140 | 104192 | **RNA**→ rpmE (PRK00019)Ribosomal_L31 (pfam01197)→ |
| Sen-288-125 | NZ_CJHA01000015.1 | - | 39531 | 39479 | **RNA**→ rpmE (PRK00019)Ribosomal_L31 (pfam01197)→ |
| Sen-405-1 | NZ_CARK01000035.1 | - | 37552 | 37500 | **RNA**→ rpmE (PRK00019)Ribosomal_L31 (pfam01197)→ |
| Sen-288-126 | NZ_CJHK01000014.1 | - | 39564 | 39512 | **RNA**→ rpmE (PRK00019)Ribosomal_L31 (pfam01197)→ |
| Sen-411-1 | NZ_CARF01000053.1 | + | 40879 | 40931 | **RNA**→ rpmE (PRK00019)Ribosomal_L31 (pfam01197)→ |
| Sen-288-127 | NZ_CIYT01000003.1 | - | 198399 | 198347 | **RNA**→ rpmE (PRK00019)Ribosomal_L31 (pfam01197)→ |
| Sen-288-128 | NZ_CJOD01000016.1 | - | 39537 | 39485 | **RNA**→ rpmE (PRK00019)Ribosomal_L31 (pfam01197)→ |
| Sen-420-1 | NZ_AYDX01000010.1 | + | 111568 | 111620 | **RNA**→ rpmE (PRK00019)Ribosomal_L31 (pfam01197)→ rpmE (PRK00019)Ribosomal_L31 (pfam01197)→ |
| Sen-288-129 | NZ_CIXO01000014.1 | - | 39528 | 39476 | **RNA**→ rpmE (PRK00019)Ribosomal_L31 (pfam01197)→ |
| Sen-394-1 | NZ_AYCV01000014.1 | + | 111281 | 111333 | **RNA**→ rpmE (PRK00019)Ribosomal_L31 (pfam01197)→ rpmE (PRK00019)Ribosomal_L31 (pfam01197)→ |
| Sen-288-130 | NZ_CIVB01000016.1 | - | 39541 | 39489 | **RNA**→ rpmE (PRK00019)Ribosomal_L31 (pfam01197)→ |
| Sen-288-131 | NZ_CJLB01000010.1 | + | 133796 | 133848 | **RNA**→ rpmE (PRK00019)Ribosomal_L31 (pfam01197)→ |
| Sen-433-1 | NZ_AOZP01000119.1 | + | 95342 | 95394 | **RNA**→ rpmE (PRK00019)Ribosomal_L31 (pfam01197)→ rpmE (PRK00019)Ribosomal_L31 (pfam01197)→ |
| Sen-288-132 | NZ_CJPV01000009.1 | - | 39601 | 39549 | **RNA**→ rpmE (PRK00019)Ribosomal_L31 (pfam01197)→ |
| Sen-424-1 | NZ_ANOU01000032.1 | + | 111695 | 111747 | **RNA**→ rpmE (PRK00019)Ribosomal_L31 (pfam01197)→ rpmE (PRK00019)Ribosomal_L31 (pfam01197)→ |
| Sen-288-133 | NZ_CIVX01000015.1 | + | 104162 | 104214 | **RNA**→ rpmE (PRK00019)Ribosomal_L31 (pfam01197)→ |
| Sen-288-134 | NZ_CJQJ01000009.1 | - | 39601 | 39549 | **RNA**→ rpmE (PRK00019)Ribosomal_L31 (pfam01197)→ |
| Sen-399-1 | NZ_CARO01000044.1 | + | 111371 | 111423 | **RNA**→ rpmE (PRK00019)Ribosomal_L31 (pfam01197)→ |
| Sen-288-135 | NZ_CJXW01000016.1 | + | 104185 | 104237 | **RNA**→ rpmE (PRK00019)Ribosomal_L31 (pfam01197)→ |

| | | | | | |
|---|---|---|---|---|---|
| Sen-407-1 | NZ_CARJ01000039.1 | - | 37552 | 37500 | **RNA**→ rpmE (PRK00019)Ribosomal_L31 (pfam01197)→ |
| Sen-288-136 | NZ_CJAD01000016.1 | - | 39569 | 39517 | **RNA**→ rpmE (PRK00019)Ribosomal_L31 (pfam01197)→ |
| Sen-419-1 | NZ_AYDW01000050.1 | + | 48948 | 49000 | **RNA**→ rpmE (PRK00019)Ribosomal_L31 (pfam01197)→ rpmE (PRK00019)Ribosomal_L31 (pfam01197)→ |
| Sen-288-137 | NZ_CIVW01000016.1 | + | 104014 | 104066 | **RNA**→ rpmE (PRK00019)Ribosomal_L31 (pfam01197)→ |
| Sen-417-1 | NZ_AYDN01000011.1 | - | 37792 | 37740 | **RNA**→ rpmE (PRK00019)Ribosomal_L31 (pfam01197)→ rpmE (PRK00019)Ribosomal_L31 (pfam01197)→ |
| Sen-288-138 | NZ_CJNW01000013.1 | + | 107993 | 108045 | **RNA**→ rpmE (PRK00019)Ribosomal_L31 (pfam01197)→ |
| Sen-438-1 | NZ_AOZQ01000041.1 | + | 103718 | 103770 | **RNA**→ rpmE (PRK00019)Ribosomal_L31 (pfam01197)→ rpmE (PRK00019)Ribosomal_L31 (pfam01197)→ |
| Sen-288-1110 | NZ_CIYE01000016.1 | - | 39531 | 39479 | **RNA**→ rpmE (PRK00019)Ribosomal_L31 (pfam01197)→ |
| Sen-288-139 | NZ_CJAU01000016.1 | + | 104154 | 104206 | **RNA**→ rpmE (PRK00019)Ribosomal_L31 (pfam01197)→ |
| Sen-288-140 | NZ_CJGP01000015.1 | + | 104038 | 104090 | **RNA**→ rpmE (PRK00019)Ribosomal_L31 (pfam01197)→ |
| Sen-288-141 | NZ_CJAQ01000017.1 | + | 104151 | 104203 | **RNA**→ rpmE (PRK00019)Ribosomal_L31 (pfam01197)→ |
| Sen-288-142 | NZ_CJFS01000016.1 | - | 39534 | 39482 | **RNA**→ rpmE (PRK00019)Ribosomal_L31 (pfam01197)→ |
| Sen-288-143 | NZ_CJJH01000014.1 | - | 41507 | 41455 | **RNA**→ rpmE (PRK00019)Ribosomal_L31 (pfam01197)→ |
| Sen-288-144 | NZ_CIWD01000015.1 | + | 104162 | 104214 | **RNA**→ rpmE (PRK00019)Ribosomal_L31 (pfam01197)→ |
| Sen-288-145 | NZ_CIVF01000017.1 | - | 39385 | 39333 | **RNA**→ rpmE (PRK00019)Ribosomal_L31 (pfam01197)→ |
| Sen-288-146 | NZ_CJGL01000015.1 | - | 39456 | 39404 | **RNA**→ rpmE (PRK00019)Ribosomal_L31 (pfam01197)→ |
| Sen-288-147 | NZ_CIVT01000016.1 | - | 39387 | 39335 | **RNA**→ rpmE (PRK00019)Ribosomal_L31 (pfam01197)→ |
| Sen-288-148 | NZ_CJIM01000014.1 | + | 104149 | 104201 | **RNA**→ rpmE (PRK00019)Ribosomal_L31 (pfam01197)→ |
| Sen-288-149 | NZ_CJQP01000016.1 | - | 39601 | 39549 | **RNA**→ rpmE (PRK00019)Ribosomal_L31 (pfam01197)→ |
| Sen-288-150 | NZ_CIZD01000016.1 | - | 39533 | 39481 | **RNA**→ rpmE (PRK00019)Ribosomal_L31 (pfam01197)→ |
| Sen-288-151 | NZ_CIXK01000015.1 | + | 103990 | 104042 | **RNA**→ rpmE (PRK00019)Ribosomal_L31 (pfam01197)→ |
| Sen-288-152 | NZ_CJUD01000016.1 | + | 104141 | 104193 | **RNA**→ rpmE (PRK00019)Ribosomal_L31 (pfam01197)→ |
| Sen-288-153 | NZ_CJCS01000016.1 | - | 39573 | 39521 | **RNA**→ rpmE (PRK00019)Ribosomal_L31 (pfam01197)→ |
| Sen-288-154 | NZ_CJOE01000016.1 | + | 104157 | 104209 | **RNA**→ rpmE (PRK00019)Ribosomal_L31 (pfam01197)→ |
| Sen-288-155 | NZ_CIYS01000014.1 | + | 107961 | 108013 | **RNA**→ rpmE (PRK00019)Ribosomal_L31 (pfam01197)→ |
| Sen-288-156 | NZ_CJVA01000015.1 | + | 104154 | 104206 | **RNA**→ rpmE (PRK00019)Ribosomal_L31 (pfam01197)→ |
| Sen-288-157 | NZ_CILV01000012.1 | + | 107993 | 108045 | **RNA**→ rpmE (PRK00019)Ribosomal_L31 (pfam01197)→ |
| Sen-288-158 | NZ_CJBL01000016.1 | + | 104155 | 104207 | **RNA**→ rpmE (PRK00019)Ribosomal_L31 (pfam01197)→ |
| Sen-288-159 | NZ_CJBS01000017.1 | + | 104148 | 104200 | **RNA**→ rpmE (PRK00019)Ribosomal_L31 (pfam01197)→ |
| Sen-288-160 | NZ_CIWK01000014.1 | + | 107980 | 108032 | **RNA**→ rpmE (PRK00019)Ribosomal_L31 (pfam01197)→ |
| Sen-288-161 | NZ_CIZB01000016.1 | - | 39534 | 39482 | **RNA**→ rpmE (PRK00019)Ribosomal_L31 (pfam01197)→ |
| Sen-288-162 | NZ_CJEJ01000015.1 | + | 104154 | 104206 | **RNA**→ rpmE (PRK00019)Ribosomal_L31 (pfam01197)→ |
| Sen-288-163 | NZ_CJLD01000014.1 | - | 39570 | 39518 | **RNA**→ rpmE (PRK00019)Ribosomal_L31 (pfam01197)→ |
| Sen-288-164 | NZ_CIWI01000014.1 | + | 103990 | 104042 | **RNA**→ rpmE (PRK00019)Ribosomal_L31 (pfam01197)→ |
| Sen-288-165 | NZ_CJUM01000016.1 | - | 39371 | 39319 | **RNA**→ rpmE (PRK00019)Ribosomal_L31 (pfam01197)→ |
| Sen-288-166 | NZ_CJAE01000016.1 | + | 104155 | 104207 | **RNA**→ rpmE (PRK00019)Ribosomal_L31 (pfam01197)→ |
| Sen-288-1111 | NZ_CJJU01000015.1 | - | 39572 | 39520 | **RNA**→ rpmE (PRK00019)Ribosomal_L31 (pfam01197)→ |
| Sen-288-167 | NZ_CJUN01000016.1 | - | 39366 | 39314 | **RNA**→ rpmE (PRK00019)Ribosomal_L31 (pfam01197)→ |
| Sen-288-168 | NZ_CJYM01000015.1 | - | 39372 | 39320 | **RNA**→ rpmE (PRK00019)Ribosomal_L31 (pfam01197)→ |
| Sen-288-169 | NZ_CJKT01000014.1 | + | 104147 | 104199 | **RNA**→ rpmE (PRK00019)Ribosomal_L31 (pfam01197)→ |
| Sen-288-170 | NZ_CIXM01000015.1 | + | 103990 | 104042 | **RNA**→ rpmE (PRK00019)Ribosomal_L31 (pfam01197)→ |
| Sen-288-171 | NZ_CILK01000016.1 | - | 39379 | 39327 | **RNA**→ rpmE (PRK00019)Ribosomal_L31 (pfam01197)→ |
| Sen-288-172 | NZ_CJXI01000017.1 | - | 39543 | 39491 | **RNA**→ rpmE (PRK00019)Ribosomal_L31 (pfam01197)→ |
| Sen-288-173 | NZ_CJAO01000016.1 | + | 104150 | 104202 | **RNA**→ rpmE (PRK00019)Ribosomal_L31 (pfam01197)→ |
| Sen-288-174 | NZ_CJTQ01000015.1 | + | 104157 | 104209 | **RNA**→ rpmE (PRK00019)Ribosomal_L31 (pfam01197)→ |
| Sen-288-175 | NZ_CJUA01000016.1 | - | 39370 | 39318 | **RNA**→ rpmE (PRK00019)Ribosomal_L31 (pfam01197)→ |
| Sen-288-176 | NZ_CEZO01000015.1 | + | 104143 | 104195 | **RNA**→ rpmE (PRK00019)Ribosomal_L31 (pfam01197)→ |
| Sen-288-177 | NZ_CJMK01000015.1 | - | 39353 | 39301 | **RNA**→ rpmE (PRK00019)Ribosomal_L31 (pfam01197)→ |
| Sen-288-178 | NZ_CJMR01000016.1 | + | 104017 | 104069 | **RNA**→ rpmE (PRK00019)Ribosomal_L31 (pfam01197)→ |
| Sen-288-179 | NZ_CJTK01000015.1 | + | 104163 | 104215 | **RNA**→ rpmE (PRK00019)Ribosomal_L31 (pfam01197)→ |
| Sen-288-180 | NZ_CJBR01000014.1 | + | 104148 | 104200 | **RNA**→ rpmE (PRK00019)Ribosomal_L31 (pfam01197)→ |
| Sen-288-181 | NZ_CJCP01000015.1 | - | 39456 | 39404 | **RNA**→ rpmE (PRK00019)Ribosomal_L31 (pfam01197)→ |

| | | | | | |
|---|---|---|---|---|---|
| Sen-288-182 | NZ_CJGG01000016.1 | + | 104146 | 104198 | **RNA**→ rpmE (PRK00019)Ribosomal_L31 (pfam01197)→ |
| Sen-288-183 | NZ_CJNK01000014.1 | - | 41504 | 41452 | **RNA**→ rpmE (PRK00019)Ribosomal_L31 (pfam01197)→ |
| Sen-288-184 | NZ_CJIL01000012.1 | + | 135291 | 135343 | **RNA**→ rpmE (PRK00019)Ribosomal_L31 (pfam01197)→ |
| Sen-288-185 | NZ_CJNR01000015.1 | - | 41512 | 41460 | **RNA**→ rpmE (PRK00019)Ribosomal_L31 (pfam01197)→ |
| Sen-288-186 | NZ_CJHU01000015.1 | + | 104200 | 104252 | **RNA**→ rpmE (PRK00019)Ribosomal_L31 (pfam01197)→ |
| Sen-288-187 | NZ_CJAA01000015.1 | - | 39626 | 39574 | **RNA**→ rpmE (PRK00019)Ribosomal_L31 (pfam01197)→ |
| Sen-288-188 | NZ_CJDF01000008.1 | - | 39456 | 39404 | **RNA**→ rpmE (PRK00019)Ribosomal_L31 (pfam01197)→ |
| Sen-288-189 | NZ_CJVD01000013.1 | + | 104003 | 104055 | **RNA**→ rpmE (PRK00019)Ribosomal_L31 (pfam01197)→ |
| Sen-288-190 | NZ_CJRZ01000015.1 | + | 104157 | 104209 | **RNA**→ rpmE (PRK00019)Ribosomal_L31 (pfam01197)→ |
| Sen-288-191 | NZ_CJSU01000015.1 | - | 41231 | 41179 | **RNA**→ rpmE (PRK00019)Ribosomal_L31 (pfam01197)→ |
| Sen-288-192 | NZ_CJJC01000014.1 | - | 41500 | 41448 | **RNA**→ rpmE (PRK00019)Ribosomal_L31 (pfam01197)→ |
| Sen-288-193 | NZ_CJER01000016.1 | - | 39536 | 39484 | **RNA**→ rpmE (PRK00019)Ribosomal_L31 (pfam01197)→ |
| Sen-288-194 | NZ_CJKY01000009.1 | - | 39562 | 39510 | **RNA**→ rpmE (PRK00019)Ribosomal_L31 (pfam01197)→ |
| Sen-288-1112 | NZ_CIXE01000015.1 | - | 39353 | 39301 | **RNA**→ rpmE (PRK00019)Ribosomal_L31 (pfam01197)→ |
| Sen-288-195 | NZ_CJUG01000006.1 | + | 190341 | 190393 | **RNA**→ rpmE (PRK00019)Ribosomal_L31 (pfam01197)→ |
| Sen-288-196 | NZ_CJPN01000008.1 | - | 39601 | 39549 | **RNA**→ rpmE (PRK00019)Ribosomal_L31 (pfam01197)→ |
| Sen-288-197 | NZ_CJJD01000011.1 | + | 136982 | 137034 | **RNA**→ rpmE (PRK00019)Ribosomal_L31 (pfam01197)→ |
| Sen-288-1113 | NZ_CIWW01000016.1 | + | 104212 | 104264 | **RNA**→ rpmE (PRK00019)Ribosomal_L31 (pfam01197)→ |
| Sen-288-198 | NZ_CJHG01000016.1 | - | 39534 | 39482 | **RNA**→ rpmE (PRK00019)Ribosomal_L31 (pfam01197)→ |
| Sen-288-199 | NZ_CIZN01000016.1 | - | 39537 | 39485 | **RNA**→ rpmE (PRK00019)Ribosomal_L31 (pfam01197)→ |
| Sen-288-200 | NZ_CIYX01000013.1 | + | 104153 | 104205 | **RNA**→ rpmE (PRK00019)Ribosomal_L31 (pfam01197)→ |
| Sen-288-201 | NZ_CJUX01000016.1 | + | 104160 | 104212 | **RNA**→ rpmE (PRK00019)Ribosomal_L31 (pfam01197)→ |
| Sen-288-202 | NZ_CIZK01000016.1 | + | 104178 | 104230 | **RNA**→ rpmE (PRK00019)Ribosomal_L31 (pfam01197)→ |
| Sen-288-203 | NZ_CJGJ01000016.1 | + | 104143 | 104195 | **RNA**→ rpmE (PRK00019)Ribosomal_L31 (pfam01197)→ |
| Sen-288-204 | NZ_CJMH01000010.1 | + | 107983 | 108035 | **RNA**→ rpmE (PRK00019)Ribosomal_L31 (pfam01197)→ |
| Sen-288-205 | NZ_CJEM01000015.1 | - | 39537 | 39485 | **RNA**→ rpmE (PRK00019)Ribosomal_L31 (pfam01197)→ |
| Sen-288-206 | NZ_CJNF01000014.1 | - | 41507 | 41455 | **RNA**→ rpmE (PRK00019)Ribosomal_L31 (pfam01197)→ |
| Sen-288-207 | NZ_CJUV01000015.1 | - | 39368 | 39316 | **RNA**→ rpmE (PRK00019)Ribosomal_L31 (pfam01197)→ |
| Sen-288-208 | NZ_CJDU01000016.1 | + | 104148 | 104200 | **RNA**→ rpmE (PRK00019)Ribosomal_L31 (pfam01197)→ |
| Sen-288-209 | NZ_CJYQ01000013.1 | - | 39538 | 39486 | **RNA**→ rpmE (PRK00019)Ribosomal_L31 (pfam01197)→ |
| Sen-288-210 | NZ_CJIP01000019.1 | + | 43218 | 43270 | **RNA**→ rpmE (PRK00019)Ribosomal_L31 (pfam01197)→ |
| Sen-288-211 | NZ_CJHF01000015.1 | + | 103990 | 104042 | **RNA**→ rpmE (PRK00019)Ribosomal_L31 (pfam01197)→ |
| Sen-288-212 | NZ_CJTZ01000015.1 | + | 107417 | 107469 | **RNA**→ rpmE (PRK00019)Ribosomal_L31 (pfam01197)→ |
| Sen-288-213 | NZ_CJQQ01000016.1 | - | 39601 | 39549 | **RNA**→ rpmE (PRK00019)Ribosomal_L31 (pfam01197)→ |
| Sen-288-214 | NZ_CJWC01000014.1 | + | 104157 | 104209 | **RNA**→ rpmE (PRK00019)Ribosomal_L31 (pfam01197)→ |
| Sen-288-215 | NZ_CJEK01000016.1 | + | 104154 | 104206 | **RNA**→ rpmE (PRK00019)Ribosomal_L31 (pfam01197)→ |
| Sen-288-216 | NZ_CILH01000015.1 | - | 39456 | 39404 | **RNA**→ rpmE (PRK00019)Ribosomal_L31 (pfam01197)→ |
| Sen-288-217 | NZ_CJJQ01000011.1 | + | 104148 | 104200 | **RNA**→ rpmE (PRK00019)Ribosomal_L31 (pfam01197)→ |
| Sen-288-218 | NZ_CJBW01000017.1 | - | 39537 | 39485 | **RNA**→ rpmE (PRK00019)Ribosomal_L31 (pfam01197)→ |
| Sen-288-219 | NZ_CJAZ01000015.1 | + | 103986 | 104038 | **RNA**→ rpmE (PRK00019)Ribosomal_L31 (pfam01197)→ |
| Sen-288-220 | NZ_CEYB01000016.1 | - | 39569 | 39517 | **RNA**→ rpmE (PRK00019)Ribosomal_L31 (pfam01197)→ |
| Sen-288-221 | NZ_CIXT01000015.1 | + | 103990 | 104042 | **RNA**→ rpmE (PRK00019)Ribosomal_L31 (pfam01197)→ |
| Sen-288-222 | NZ_CJWE01000015.1 | - | 41258 | 41206 | **RNA**→ rpmE (PRK00019)Ribosomal_L31 (pfam01197)→ |
| Sen-288-223 | NZ_CJAC01000016.1 | - | 39524 | 39472 | **RNA**→ rpmE (PRK00019)Ribosomal_L31 (pfam01197)→ |
| Sen-288-224 | NZ_CIYD01000011.1 | - | 39456 | 39404 | **RNA**→ rpmE (PRK00019)Ribosomal_L31 (pfam01197)→ |
| Sen-288-225 | NZ_CJDQ01000016.1 | - | 39538 | 39486 | **RNA**→ rpmE (PRK00019)Ribosomal_L31 (pfam01197)→ |
| Sen-288-226 | NZ_CJEY01000003.1 | + | 104147 | 104199 | **RNA**→ rpmE (PRK00019)Ribosomal_L31 (pfam01197)→ |
| Sen-288-227 | NZ_CJEV01000016.1 | - | 39535 | 39483 | **RNA**→ rpmE (PRK00019)Ribosomal_L31 (pfam01197)→ |
| Sen-288-228 | NZ_CJFU01000014.1 | + | 104165 | 104217 | **RNA**→ rpmE (PRK00019)Ribosomal_L31 (pfam01197)→ |
| Sen-288-229 | NZ_CJFQ01000015.1 | - | 39536 | 39484 | **RNA**→ rpmE (PRK00019)Ribosomal_L31 (pfam01197)→ |
| Sen-288-230 | NZ_CJPT01000015.1 | - | 39601 | 39549 | **RNA**→ rpmE (PRK00019)Ribosomal_L31 (pfam01197)→ |
| Sen-288-231 | NZ_CJHZ01000015.1 | - | 39534 | 39482 | **RNA**→ rpmE (PRK00019)Ribosomal_L31 (pfam01197)→ |

| | | | | | |
|---|---|---|---|---|---|
| Sen-288-232 | NZ_CILT01000015.1 | - | 41673 | 41621 | **RNA**→ rpmE (PRK00019)Ribosomal_L31 (pfam01197)→ |
| Sen-288-233 | NZ_CJTS01000014.1 | + | 104157 | 104209 | **RNA**→ rpmE (PRK00019)Ribosomal_L31 (pfam01197)→ |
| Sen-288-234 | NZ_CJBY01000016.1 | + | 104160 | 104212 | **RNA**→ rpmE (PRK00019)Ribosomal_L31 (pfam01197)→ |
| Sen-288-235 | NZ_CJKS01000015.1 | - | 39478 | 39426 | **RNA**→ rpmE (PRK00019)Ribosomal_L31 (pfam01197)→ |
| Sen-288-236 | NZ_CJYU01000015.1 | - | 44803 | 44751 | **RNA**→ rpmE (PRK00019)Ribosomal_L31 (pfam01197)→ |
| Sen-288-237 | NZ_CJXE01000011.1 | + | 163323 | 163375 | **RNA**→ rpmE (PRK00019)Ribosomal_L31 (pfam01197)→ |
| Sen-288-238 | NZ_CIWA01000016.1 | - | 39514 | 39462 | **RNA**→ rpmE (PRK00019)Ribosomal_L31 (pfam01197)→ |
| Sen-288-239 | NZ_CJVL01000014.1 | - | 41231 | 41179 | **RNA**→ rpmE (PRK00019)Ribosomal_L31 (pfam01197)→ |
| Sen-288-240 | NZ_CIZP01000016.1 | + | 104152 | 104204 | **RNA**→ rpmE (PRK00019)Ribosomal_L31 (pfam01197)→ |
| Sen-288-241 | NZ_CJQI01000017.1 | + | 106102 | 106154 | **RNA**→ rpmE (PRK00019)Ribosomal_L31 (pfam01197)→ |
| Sen-288-242 | NZ_CJEE01000016.1 | - | 39723 | 39671 | **RNA**→ rpmE (PRK00019)Ribosomal_L31 (pfam01197)→ |
| Sen-288-243 | NZ_CJVY01000015.1 | + | 104146 | 104198 | **RNA**→ rpmE (PRK00019)Ribosomal_L31 (pfam01197)→ |
| Sen-288-244 | NZ_CJQM01000014.1 | + | 107413 | 107465 | **RNA**→ rpmE (PRK00019)Ribosomal_L31 (pfam01197)→ |
| Sen-288-245 | NZ_CEZA01000017.1 | + | 104146 | 104198 | **RNA**→ rpmE (PRK00019)Ribosomal_L31 (pfam01197)→ |
| Sen-288-246 | NZ_CJPZ01000009.1 | - | 39601 | 39549 | **RNA**→ rpmE (PRK00019)Ribosomal_L31 (pfam01197)→ |
| Sen-288-247 | NZ_CIVH01000013.1 | + | 104201 | 104253 | **RNA**→ rpmE (PRK00019)Ribosomal_L31 (pfam01197)→ |
| Sen-288-248 | NZ_CJEX01000016.1 | + | 104156 | 104208 | **RNA**→ rpmE (PRK00019)Ribosomal_L31 (pfam01197)→ |
| Sen-288-249 | NZ_CJIS01000019.1 | + | 43848 | 43900 | **RNA**→ rpmE (PRK00019)Ribosomal_L31 (pfam01197)→ |
| Sen-288-250 | NZ_CJQA01000008.1 | - | 39601 | 39549 | **RNA**→ rpmE (PRK00019)Ribosomal_L31 (pfam01197)→ |
| Sen-288-251 | NZ_CJNG01000014.1 | - | 41511 | 41459 | **RNA**→ rpmE (PRK00019)Ribosomal_L31 (pfam01197)→ |
| Sen-288-252 | NZ_CJGM01000010.1 | - | 39456 | 39404 | **RNA**→ rpmE (PRK00019)Ribosomal_L31 (pfam01197)→ |
| Sen-288-253 | NZ_CJKD01000010.1 | - | 39361 | 39309 | **RNA**→ rpmE (PRK00019)Ribosomal_L31 (pfam01197)→ |
| Sen-288-254 | NZ_CJWL01000015.1 | - | 39531 | 39479 | **RNA**→ rpmE (PRK00019)Ribosomal_L31 (pfam01197)→ |
| Sen-288-255 | NZ_CJUQ01000015.1 | + | 104156 | 104208 | **RNA**→ rpmE (PRK00019)Ribosomal_L31 (pfam01197)→ |
| Sen-288-256 | NZ_CJMW01000013.1 | + | 104152 | 104204 | **RNA**→ rpmE (PRK00019)Ribosomal_L31 (pfam01197)→ |
| Sen-288-257 | NZ_CILN01000016.1 | + | 104151 | 104203 | **RNA**→ rpmE (PRK00019)Ribosomal_L31 (pfam01197)→ |
| Sen-288-258 | NZ_CJBJ01000016.1 | + | 104162 | 104214 | **RNA**→ rpmE (PRK00019)Ribosomal_L31 (pfam01197)→ |
| Sen-288-1114 | NZ_CIWU01000015.1 | + | 104119 | 104171 | **RNA**→ rpmE (PRK00019)Ribosomal_L31 (pfam01197)→ |
| Sen-288-259 | NZ_CJGQ01000015.1 | + | 104149 | 104201 | **RNA**→ rpmE (PRK00019)Ribosomal_L31 (pfam01197)→ |
| Sen-288-260 | NZ_CJMC01000015.1 | - | 41499 | 41447 | **RNA**→ rpmE (PRK00019)Ribosomal_L31 (pfam01197)→ |
| Sen-288-261 | NZ_CIUN01000015.1 | + | 104162 | 104214 | **RNA**→ rpmE (PRK00019)Ribosomal_L31 (pfam01197)→ |
| Sen-288-262 | NZ_CIWR01000015.1 | + | 104111 | 104163 | **RNA**→ rpmE (PRK00019)Ribosomal_L31 (pfam01197)→ |
| Sen-288-263 | NZ_CIUL01000014.1 | - | 41231 | 41179 | **RNA**→ rpmE (PRK00019)Ribosomal_L31 (pfam01197)→ |
| Sen-288-264 | NZ_CJCG01000016.1 | - | 39537 | 39485 | **RNA**→ rpmE (PRK00019)Ribosomal_L31 (pfam01197)→ |
| Sen-288-265 | NZ_CJMO01000013.1 | - | 41484 | 41432 | **RNA**→ rpmE (PRK00019)Ribosomal_L31 (pfam01197)→ |
| Sen-288-1115 | NZ_CIVO01000015.1 | - | 41516 | 41464 | **RNA**→ rpmE (PRK00019)Ribosomal_L31 (pfam01197)→ |
| Sen-288-266 | NZ_CJUU01000015.1 | + | 104153 | 104205 | **RNA**→ rpmE (PRK00019)Ribosomal_L31 (pfam01197)→ |
| Sen-288-267 | NZ_CJOM01000013.1 | + | 104141 | 104193 | **RNA**→ rpmE (PRK00019)Ribosomal_L31 (pfam01197)→ |
| Sen-288-268 | NZ_CJXC01000017.1 | - | 39381 | 39329 | **RNA**→ rpmE (PRK00019)Ribosomal_L31 (pfam01197)→ |
| Sen-288-269 | NZ_CIYC01000016.1 | + | 104218 | 104270 | **RNA**→ rpmE (PRK00019)Ribosomal_L31 (pfam01197)→ |
| Sen-288-270 | NZ_CIUJ01000015.1 | - | 39505 | 39453 | **RNA**→ rpmE (PRK00019)Ribosomal_L31 (pfam01197)→ |
| Sen-288-271 | NZ_CJSA01000015.1 | + | 104163 | 104215 | **RNA**→ rpmE (PRK00019)Ribosomal_L31 (pfam01197)→ |
| Sen-288-272 | NZ_CIWF01000016.1 | - | 39601 | 39549 | **RNA**→ rpmE (PRK00019)Ribosomal_L31 (pfam01197)→ |
| Sen-288-273 | NZ_CJOK01000016.1 | + | 104162 | 104214 | **RNA**→ rpmE (PRK00019)Ribosomal_L31 (pfam01197)→ |
| Sen-288-274 | NZ_CJGV01000016.1 | - | 39536 | 39484 | **RNA**→ rpmE (PRK00019)Ribosomal_L31 (pfam01197)→ |
| Sen-288-275 | NZ_CILE01000015.1 | + | 104154 | 104206 | **RNA**→ rpmE (PRK00019)Ribosomal_L31 (pfam01197)→ |
| Sen-288-276 | NZ_CJYX01000015.1 | + | 104158 | 104210 | **RNA**→ rpmE (PRK00019)Ribosomal_L31 (pfam01197)→ |
| Sen-288-277 | NZ_CJLF01000009.1 | + | 152651 | 152703 | **RNA**→ rpmE (PRK00019)Ribosomal_L31 (pfam01197)→ |
| Sen-848-1 | NZ_AETN01000013.1 | - | 35207 | 35155 | **RNA**→ rpmE (PRK00019)Ribosomal_L31 (pfam01197)→ |
| Sen-288-278 | NZ_CJWF01000015.1 | + | 104158 | 104210 | **RNA**→ rpmE (PRK00019)Ribosomal_L31 (pfam01197)→ |
| Sen-288-279 | NZ_CJNS01000014.1 | - | 39702 | 39650 | **RNA**→ rpmE (PRK00019)Ribosomal_L31 (pfam01197)→ |
| Sen-288-280 | NZ_CJAY01000015.1 | + | 104150 | 104202 | **RNA**→ rpmE (PRK00019)Ribosomal_L31 (pfam01197)→ |

| | | | | | |
|---|---|---|---|---|---|
| Sen-288-281 | NZ_CJIV01000012.1 | - | 41499 | 41447 | **RNA**→ rpmE (PRK00019)Ribosomal_L31 (pfam01197)→ |
| Sen-852-1 | NZ_AESU01000031.1 | - | 42784 | 42732 | **RNA**→ rpmE (PRK00019)Ribosomal_L31 (pfam01197)→ rpmE (PRK00019)Ribosomal_L31 (pfam01197)→ |
| Sen-288-282 | NZ_CJVH01000013.1 | - | 41231 | 41179 | **RNA**→ rpmE (PRK00019)Ribosomal_L31 (pfam01197)→ |
| Sen-288-283 | NZ_CIWH01000015.1 | + | 107416 | 107468 | **RNA**→ rpmE (PRK00019)Ribosomal_L31 (pfam01197)→ |
| Sen-288-284 | NZ_CJFL01000015.1 | - | 39560 | 39508 | **RNA**→ rpmE (PRK00019)Ribosomal_L31 (pfam01197)→ |
| Sen-884-1 | NZ_AHUG01000050.1 | + | 100567 | 100619 | **RNA**→ rpmE (PRK00019)Ribosomal_L31 (pfam01197)→ rpmE (PRK00019)Ribosomal_L31 (pfam01197)→ |
| Sen-288-285 | NZ_CJSY01000015.1 | - | 41516 | 41464 | **RNA**→ rpmE (PRK00019)Ribosomal_L31 (pfam01197)→ |
| Sen-288-286 | NZ_CJXS01000009.1 | - | 39558 | 39506 | **RNA**→ rpmE (PRK00019)Ribosomal_L31 (pfam01197)→ |
| Sen-288-287 | NZ_CJAL01000014.1 | + | 104209 | 104261 | **RNA**→ rpmE (PRK00019)Ribosomal_L31 (pfam01197)→ |
| Sen-288-288 | NZ_CJAI01000016.1 | + | 104143 | 104195 | **RNA**→ rpmE (PRK00019)Ribosomal_L31 (pfam01197)→ |
| Sen-872-1 | NZ_AHUE01000073.1 | - | 45277 | 45225 | **RNA**→ rpmE (PRK00019)Ribosomal_L31 (pfam01197)→ |
| Sen-288-289 | NZ_CJWT01000015.1 | - | 44769 | 44717 | **RNA**→ rpmE (PRK00019)Ribosomal_L31 (pfam01197)→ |
| Sen-288-290 | NZ_CJRX01000010.1 | + | 141198 | 141250 | **RNA**→ rpmE (PRK00019)Ribosomal_L31 (pfam01197)→ |
| Sen-288-291 | NZ_CJVN01000014.1 | - | 41231 | 41179 | **RNA**→ rpmE (PRK00019)Ribosomal_L31 (pfam01197)→ |
| Sen-288-292 | NZ_CJEL01000016.1 | - | 39723 | 39671 | **RNA**→ rpmE (PRK00019)Ribosomal_L31 (pfam01197)→ |
| Sen-862-1 | NZ_APAE01000207.1 | - | 3173 | 3121 | **RNA**→ rpmE (PRK00019)Ribosomal_L31 (pfam01197)→ rpmE (PRK00019)Ribosomal_L31 (pfam01197)→ |
| Sen-288-293 | NZ_CJCV01000014.1 | - | 39539 | 39487 | **RNA**→ rpmE (PRK00019)Ribosomal_L31 (pfam01197)→ |
| Sen-288-294 | NZ_CJKQ01000015.1 | + | 104149 | 104201 | **RNA**→ rpmE (PRK00019)Ribosomal_L31 (pfam01197)→ |
| Sen-288-295 | NZ_CJYN01000014.1 | + | 107792 | 107844 | **RNA**→ rpmE (PRK00019)Ribosomal_L31 (pfam01197)→ |
| Sen-288-296 | NZ_CJUY01000015.1 | + | 104009 | 104061 | **RNA**→ rpmE (PRK00019)Ribosomal_L31 (pfam01197)→ |
| Sen-842-1 | NZ_AHIK01000014.1 | - | 35207 | 35155 | **RNA**→ rpmE (PRK00019)Ribosomal_L31 (pfam01197)→ |
| Sen-288-297 | NZ_CJLE01000002.1 | - | 39456 | 39404 | **RNA**→ rpmE (PRK00019)Ribosomal_L31 (pfam01197)→ |
| Sen-288-298 | NZ_CJNN01000003.1 | - | 68376 | 68324 | **RNA**→ rpmE (PRK00019)Ribosomal_L31 (pfam01197)→ |
| Sen-869-1 | NZ_AHUB01000025.1 | - | 4720 | 4668 | **RNA**→ rpmE (PRK00019)Ribosomal_L31 (pfam01197)→ |
| Sen-288-299 | NZ_CJOL01000016.1 | - | 39377 | 39325 | **RNA**→ rpmE (PRK00019)Ribosomal_L31 (pfam01197)→ |
| Sen-288-300 | NZ_CIZJ01000015.1 | - | 39537 | 39485 | **RNA**→ rpmE (PRK00019)Ribosomal_L31 (pfam01197)→ |
| Sen-288-301 | NZ_CJVB01000016.1 | + | 104163 | 104215 | **RNA**→ rpmE (PRK00019)Ribosomal_L31 (pfam01197)→ |
| Sen-892-1 | NZ_AYDY01000049.1 | + | 100567 | 100619 | **RNA**→ rpmE (PRK00019)Ribosomal_L31 (pfam01197)→ rpmE (PRK00019)Ribosomal_L31 (pfam01197)→ |
| Sen-288-302 | NZ_CJXO01000011.1 | + | 133824 | 133876 | **RNA**→ rpmE (PRK00019)Ribosomal_L31 (pfam01197)→ |
| Sen-288-303 | NZ_CEXQ01000013.1 | + | 104163 | 104215 | **RNA**→ rpmE (PRK00019)Ribosomal_L31 (pfam01197)→ |
| Sen-851-1 | NZ_AESP01000050.1 | + | 97839 | 97891 | **RNA**→ rpmE (PRK00019)Ribosomal_L31 (pfam01197)→ |
| Sen-288-304 | NZ_CIWV01000015.1 | - | 39620 | 39568 | **RNA**→ rpmE (PRK00019)Ribosomal_L31 (pfam01197)→ |
| Sen-288-305 | NZ_CIWT01000016.1 | - | 39601 | 39549 | **RNA**→ rpmE (PRK00019)Ribosomal_L31 (pfam01197)→ |
| Sen-888-1 | NZ_AJMO01000028.1 | - | 45069 | 45017 | **RNA**→ rpmE (PRK00019)Ribosomal_L31 (pfam01197)→ |
| Sen-288-306 | NZ_CJCO01000015.1 | - | 39570 | 39518 | **RNA**→ rpmE (PRK00019)Ribosomal_L31 (pfam01197)→ |
| Sen-288-307 | NZ_CJHW01000016.1 | + | 104154 | 104206 | **RNA**→ rpmE (PRK00019)Ribosomal_L31 (pfam01197)→ |
| Sen-288-308 | NZ_CJVV01000013.1 | - | 41510 | 41458 | **RNA**→ rpmE (PRK00019)Ribosomal_L31 (pfam01197)→ |
| Sen-873-1 | NZ_AHUF01000060.1 | - | 36249 | 36197 | **RNA**→ rpmE (PRK00019)Ribosomal_L31 (pfam01197)→ |
| Sen-288-309 | NZ_CJMG01000014.1 | - | 41503 | 41451 | **RNA**→ rpmE (PRK00019)Ribosomal_L31 (pfam01197)→ |
| Sen-875-1 | NZ_AHUD01000038.1 | + | 100524 | 100576 | **RNA**→ rpmE (PRK00019)Ribosomal_L31 (pfam01197)→ |
| Sen-288-310 | NZ_CJCI01000015.1 | - | 39537 | 39485 | **RNA**→ rpmE (PRK00019)Ribosomal_L31 (pfam01197)→ |
| Sen-288-311 | NZ_CJCJ01000015.1 | - | 39534 | 39482 | **RNA**→ rpmE (PRK00019)Ribosomal_L31 (pfam01197)→ |
| Sen-890-1 | NZ_AYEA01000016.1 | - | 45273 | 45221 | **RNA**→ rpmE (PRK00019)Ribosomal_L31 (pfam01197)→ rpmE (PRK00019)Ribosomal_L31 (pfam01197)→ |
| Sen-288-312 | NZ_CJLL01000015.1 | + | 104153 | 104205 | **RNA**→ rpmE (PRK00019)Ribosomal_L31 (pfam01197)→ |
| Sen-288-313 | NZ_CIZA01000016.1 | + | 104178 | 104230 | **RNA**→ rpmE (PRK00019)Ribosomal_L31 (pfam01197)→ |
| Sen-867-1 | NZ_AYDM01000074.1 | + | 100113 | 100165 | **RNA**→ rpmE (PRK00019)Ribosomal_L31 (pfam01197)→ |
| Sen-288-314 | NZ_CJKP01000013.1 | + | 107998 | 108050 | **RNA**→ rpmE (PRK00019)Ribosomal_L31 (pfam01197)→ |
| Sen-288-315 | NZ_CJXY01000015.1 | + | 104164 | 104216 | **RNA**→ rpmE (PRK00019)Ribosomal_L31 (pfam01197)→ |
| Sen-855-1 | NZ_AOXN01000002.1 | - | 45077 | 45025 | **RNA**→ rpmE (PRK00019)Ribosomal_L31 (pfam01197)→ rpmE (PRK00019)Ribosomal_L31 (pfam01197)→ |
| Sen-288-316 | NZ_CJPU01000015.1 | - | 39601 | 39549 | **RNA**→ rpmE (PRK00019)Ribosomal_L31 (pfam01197)→ |
| Sen-288-317 | NZ_CJHR01000017.1 | - | 39535 | 39483 | **RNA**→ rpmE (PRK00019)Ribosomal_L31 (pfam01197)→ |
| Sen-288-318 | NZ_CJMD01000013.1 | + | 107990 | 108042 | **RNA**→ rpmE (PRK00019)Ribosomal_L31 (pfam01197)→ |

| | | | | | | | |
|---|---|---|---|---|---|---|---|
| Sen-863-1 | NZ_APAD01000053.1 | - | 3017 | 2965 | **RNA**→ | rpmE (PRK00019)Ribosomal_L31 (pfam01197)→ | rpmE (PRK00019)Ribosomal_L31 (pfam01197)→ |
| Sen-288-319 | NZ_CJBX01000015.1 | - | 39589 | 39537 | **RNA**→ | rpmE (PRK00019)Ribosomal_L31 (pfam01197)→ | |
| Sen-288-320 | NZ_CJBK01000015.1 | + | 104149 | 104201 | **RNA**→ | rpmE (PRK00019)Ribosomal_L31 (pfam01197)→ | |
| Sen-288-321 | NZ_CJCK01000016.1 | + | 104153 | 104205 | **RNA**→ | rpmE (PRK00019)Ribosomal_L31 (pfam01197)→ | |
| Sen-845-1 | NZ_AETK01000003.1 | + | 97841 | 97893 | **RNA**→ | rpmE (PRK00019)Ribosomal_L31 (pfam01197)→ | |
| Sen-288-1116 | NZ_CIYI01000014.1 | + | 104286 | 104338 | **RNA**→ | rpmE (PRK00019)Ribosomal_L31 (pfam01197)→ | |
| Sen-844-1 | NZ_AEST01000030.1 | + | 97733 | 97785 | **RNA**→ | rpmE (PRK00019)Ribosomal_L31 (pfam01197)→ | rpmE (PRK00019)Ribosomal_L31 (pfam01197)→ |
| Sen-288-322 | NZ_CJLJ01000016.1 | + | 104043 | 104095 | **RNA**→ | rpmE (PRK00019)Ribosomal_L31 (pfam01197)→ | |
| Sen-288-323 | NZ_CILP01000015.1 | + | 107794 | 107846 | **RNA**→ | rpmE (PRK00019)Ribosomal_L31 (pfam01197)→ | |
| Sen-843-1 | NZ_AETW01000044.1 | - | 35207 | 35155 | **RNA**→ | rpmE (PRK00019)Ribosomal_L31 (pfam01197)→ | |
| Sen-288-324 | NZ_CIYU01000012.1 | - | 39456 | 39404 | **RNA**→ | rpmE (PRK00019)Ribosomal_L31 (pfam01197)→ | |
| Sen-850-1 | NZ_AESO01000043.1 | - | 35285 | 35233 | **RNA**→ | rpmE (PRK00019)Ribosomal_L31 (pfam01197)→ | |
| Sen-288-325 | NZ_CJVC01000015.1 | - | 41231 | 41179 | **RNA**→ | rpmE (PRK00019)Ribosomal_L31 (pfam01197)→ | |
| Sen-887-1 | NZ_AJMN01000011.1 | - | 45390 | 45338 | **RNA**→ | rpmE (PRK00019)Ribosomal_L31 (pfam01197)→ | |
| Sen-288-326 | NZ_CIWO01000015.1 | + | 107983 | 108035 | **RNA**→ | rpmE (PRK00019)Ribosomal_L31 (pfam01197)→ | |
| Sen-883-1 | NZ_AHUC01000010.1 | + | 100542 | 100594 | **RNA**→ | rpmE (PRK00019)Ribosomal_L31 (pfam01197)→ | |
| Sen-288-327 | NZ_CJFW01000015.1 | + | 103990 | 104042 | **RNA**→ | rpmE (PRK00019)Ribosomal_L31 (pfam01197)→ | |
| Sen-891-1 | NZ_AYDZ01000001.1 | - | 45273 | 45221 | **RNA**→ | rpmE (PRK00019)Ribosomal_L31 (pfam01197)→ | |
| Sen-860-1 | NZ_AYCZ01000097.1 | + | 97435 | 97487 | **RNA**→ | rpmE (PRK00019)Ribosomal_L31 (pfam01197)→ | rpmE (PRK00019)Ribosomal_L31 (pfam01197)→ |
| Sen-889-1 | NZ_AYEB01000069.1 | + | 93391 | 93443 | **RNA**→ | rpmE (PRK00019)Ribosomal_L31 (pfam01197)→ | |
| Sen-896-1 | NZ_AYEI01000076.1 | + | 16984 | 17036 | **RNA**→ | rpmE (PRK00019)Ribosomal_L31 (pfam01197)→ | |
| Sen-847-1 | NZ_AETM01000086.1 | + | 2601 | 2653 | **RNA**→ | rpmE (PRK00019)Ribosomal_L31 (pfam01197)→ | |
| Sen-874-1 | NZ_AHTR01000033.1 | - | 45077 | 45025 | **RNA**→ | rpmE (PRK00019)Ribosomal_L31 (pfam01197)→ | |
| Sen-871-1 | NZ_AHTQ01000033.1 | + | 100304 | 100356 | **RNA**→ | rpmE (PRK00019)Ribosomal_L31 (pfam01197)→ | rpmE (PRK00019)Ribosomal_L31 (pfam01197)→ |
| Sen-886-1 | NZ_AOGG01000035.1 | + | 42041 | 42093 | **RNA**→ | rpmE (PRK00019)Ribosomal_L31 (pfam01197)→ | |
| Sen-858-1 | NZ_AOYX01000026.1 | + | 42042 | 42094 | **RNA**→ | rpmE (PRK00019)Ribosomal_L31 (pfam01197)→ | rpmE (PRK00019)Ribosomal_L31 (pfam01197)→ |
| Sen-870-1 | NZ_AHTP01000001.1 | + | 100541 | 100593 | **RNA**→ | rpmE (PRK00019)Ribosomal_L31 (pfam01197)→ | |
| Sen-878-1 | NZ_AHTM01000018.1 | + | 105423 | 105475 | **RNA**→ | rpmE (PRK00019)Ribosomal_L31 (pfam01197)→ | |
| Sen-894-1 | NZ_AOGI01000026.1 | + | 100031 | 100083 | **RNA**→ | rpmE (PRK00019)Ribosomal_L31 (pfam01197)→ | |
| Sen-859-1 | NZ_AOYW01000025.1 | - | 35385 | 35333 | **RNA**→ | rpmE (PRK00019)Ribosomal_L31 (pfam01197)→ | rpmE (PRK00019)Ribosomal_L31 (pfam01197)→ |
| Sen-841-1 | NZ_AETV01000024.1 | - | 35284 | 35232 | **RNA**→ | rpmE (PRK00019)Ribosomal_L31 (pfam01197)→ | |
| Sen-846-1 | NZ_AETL01000105.1 | - | 35284 | 35232 | **RNA**→ | rpmE (PRK00019)Ribosomal_L31 (pfam01197)→ | |
| Sen-861-1 | NZ_AYUP01000215.1 | - | 9770 | 9718 | **RNA**→ | rpmE (PRK00019)Ribosomal_L31 (pfam01197)→ | |
| Sen-849-1 | NZ_AETA01000041.1 | - | 23926 | 23874 | **RNA**→ | rpmE (PRK00019)Ribosomal_L31 (pfam01197)→ | |
| Sen-876-1 | NZ_AHTL01000005.1 | + | 105413 | 105465 | **RNA**→ | rpmE (PRK00019)Ribosomal_L31 (pfam01197)→ | |
| Sen-893-1 | NZ_AOGJ01000036.1 | - | 45120 | 45068 | **RNA**→ | rpmE (PRK00019)Ribosomal_L31 (pfam01197)→ | |
| Sen-266-1 | NZ_CCMR01000001.1 | + | 277430 | 277482 | **RNA**→ | rpmE (PRK00019)Ribosomal_L31 (pfam01197)→ | |
| Sen-880-1 | NZ_AHTN01000018.1 | - | 45387 | 45335 | **RNA**→ | rpmE (PRK00019)Ribosomal_L31 (pfam01197)→ | |
| Sen-877-1 | NZ_AHTO01000001.1 | - | 45075 | 45023 | **RNA**→ | rpmE (PRK00019)Ribosomal_L31 (pfam01197)→ | |
| Sen-882-1 | NZ_AHTS01000001.1 | + | 105415 | 105467 | **RNA**→ | rpmE (PRK00019)Ribosomal_L31 (pfam01197)→ | |
| Sen-881-1 | NZ_AHTK01000004.1 | + | 105417 | 105469 | **RNA**→ | rpmE (PRK00019)Ribosomal_L31 (pfam01197)→ | |
| Sen-885-1 | NZ_AOGH01000031.1 | + | 100387 | 100439 | **RNA**→ | rpmE (PRK00019)Ribosomal_L31 (pfam01197)→ | |
| Sen-288-328 | NZ_CJVG01000014.1 | - | 44930 | 44878 | **RNA**→ | rpmE (PRK00019)Ribosomal_L31 (pfam01197)→ | |
| Sen-288-329 | NZ_CJMP01000015.1 | - | 41498 | 41446 | **RNA**→ | rpmE (PRK00019)Ribosomal_L31 (pfam01197)→ | |
| Sen-288-330 | NZ_CJSO01000015.1 | - | 41513 | 41461 | **RNA**→ | rpmE (PRK00019)Ribosomal_L31 (pfam01197)→ | |
| Sen-288-331 | NZ_CJIO01000014.1 | + | 105490 | 105542 | **RNA**→ | rpmE (PRK00019)Ribosomal_L31 (pfam01197)→ | |
| Sen-853-1 | NZ_AESR01000011.1 | - | 35207 | 35155 | **RNA**→ | rpmE (PRK00019)Ribosomal_L31 (pfam01197)→ | rpmE (PRK00019)Ribosomal_L31 (pfam01197)→ |
| Sen-879-1 | NZ_AHTJ01000074.1 | - | 45390 | 45338 | **RNA**→ | rpmE (PRK00019)Ribosomal_L31 (pfam01197)→ | |
| Sen-857-1 | NZ_AOYT01000022.1 | - | 45150 | 45098 | **RNA**→ | rpmE (PRK00019)Ribosomal_L31 (pfam01197)→ | rpmE (PRK00019)Ribosomal_L31 (pfam01197)→ |
| Sen-288-332 | NZ_CJOB01000016.1 | + | 104145 | 104197 | **RNA**→ | rpmE (PRK00019)Ribosomal_L31 (pfam01197)→ | |
| Sen-288-333 | NZ_CJIE01000017.1 | + | 104161 | 104213 | **RNA**→ | rpmE (PRK00019)Ribosomal_L31 (pfam01197)→ | |
| Sen-288-334 | NZ_CJIH01000014.1 | + | 104152 | 104204 | **RNA**→ | rpmE (PRK00019)Ribosomal_L31 (pfam01197)→ | |

| | | | | | |
|---|---|---|---|---|---|
| Sen-288-335 | NZ_CJMJ01000009.1 | - | 39534 | 39482 | **RNA**→ rpmE (PRK00019)Ribosomal_L31 (pfam01197)→ |
| Sen-288-336 | NZ_CIXA01000011.1 | - | 39556 | 39504 | **RNA**→ rpmE (PRK00019)Ribosomal_L31 (pfam01197)→ |
| Sen-288-337 | NZ_CIXR01000016.1 | - | 39529 | 39477 | **RNA**→ rpmE (PRK00019)Ribosomal_L31 (pfam01197)→ |
| Sen-288-338 | NZ_CJXK01000010.1 | - | 39378 | 39326 | **RNA**→ rpmE (PRK00019)Ribosomal_L31 (pfam01197)→ |
| Sen-288-339 | NZ_CJDM01000016.1 | + | 104156 | 104208 | **RNA**→ rpmE (PRK00019)Ribosomal_L31 (pfam01197)→ |
| Sen-288-340 | NZ_CJMZ01000015.1 | + | 104156 | 104208 | **RNA**→ rpmE (PRK00019)Ribosomal_L31 (pfam01197)→ |
| Sen-288-341 | NZ_CJAG01000017.1 | - | 39532 | 39480 | **RNA**→ rpmE (PRK00019)Ribosomal_L31 (pfam01197)→ |
| Sen-288-342 | NZ_CJIA01000015.1 | + | 104146 | 104198 | **RNA**→ rpmE (PRK00019)Ribosomal_L31 (pfam01197)→ |
| Sen-288-343 | NZ_CJIB01000016.1 | - | 39535 | 39483 | **RNA**→ rpmE (PRK00019)Ribosomal_L31 (pfam01197)→ |
| Sen-288-344 | NZ_CJLV01000015.1 | - | 41505 | 41453 | **RNA**→ rpmE (PRK00019)Ribosomal_L31 (pfam01197)→ |
| Sen-288-345 | NZ_CJFN01000016.1 | - | 39558 | 39506 | **RNA**→ rpmE (PRK00019)Ribosomal_L31 (pfam01197)→ |
| Sen-288-346 | NZ_CJDC01000002.1 | - | 198555 | 198503 | **RNA**→ rpmE (PRK00019)Ribosomal_L31 (pfam01197)→ |
| Sen-288-347 | NZ_CJSB01000016.1 | + | 104174 | 104226 | **RNA**→ rpmE (PRK00019)Ribosomal_L31 (pfam01197)→ |
| Sen-288-348 | NZ_CJGZ01000016.1 | + | 104153 | 104205 | **RNA**→ rpmE (PRK00019)Ribosomal_L31 (pfam01197)→ |
| Sen-288-349 | NZ_CJPL01000015.1 | - | 39601 | 39549 | **RNA**→ rpmE (PRK00019)Ribosomal_L31 (pfam01197)→ |
| Sen-288-1117 | NZ_CIWJ01000014.1 | - | 41502 | 41450 | **RNA**→ rpmE (PRK00019)Ribosomal_L31 (pfam01197)→ |
| Sen-288-350 | NZ_CJZD01000016.1 | + | 104156 | 104208 | **RNA**→ rpmE (PRK00019)Ribosomal_L31 (pfam01197)→ |
| Sen-288-351 | NZ_CJWU01000014.1 | - | 39586 | 39534 | **RNA**→ rpmE (PRK00019)Ribosomal_L31 (pfam01197)→ |
| Sen-288-352 | NZ_CJTF01000015.1 | - | 39367 | 39315 | **RNA**→ rpmE (PRK00019)Ribosomal_L31 (pfam01197)→ |
| Sen-288-353 | NZ_CIVP01000016.1 | + | 104175 | 104227 | **RNA**→ rpmE (PRK00019)Ribosomal_L31 (pfam01197)→ |
| Sen-288-354 | NZ_CJEH01000016.1 | + | 104155 | 104207 | **RNA**→ rpmE (PRK00019)Ribosomal_L31 (pfam01197)→ |
| Sen-288-355 | NZ_CIYM01000016.1 | + | 104162 | 104214 | **RNA**→ rpmE (PRK00019)Ribosomal_L31 (pfam01197)→ |
| Sen-288-356 | NZ_CJDK01000011.1 | + | 104148 | 104200 | **RNA**→ rpmE (PRK00019)Ribosomal_L31 (pfam01197)→ |
| Sen-288-357 | NZ_CIZL01000011.1 | + | 137065 | 137117 | **RNA**→ rpmE (PRK00019)Ribosomal_L31 (pfam01197)→ |
| Sen-288-358 | NZ_CJWI01000014.1 | + | 104153 | 104205 | **RNA**→ rpmE (PRK00019)Ribosomal_L31 (pfam01197)→ |
| Sen-288-359 | NZ_CJDN01000015.1 | - | 39372 | 39320 | **RNA**→ rpmE (PRK00019)Ribosomal_L31 (pfam01197)→ |
| Sen-288-360 | NZ_CJMU01000015.1 | + | 104152 | 104204 | **RNA**→ rpmE (PRK00019)Ribosomal_L31 (pfam01197)→ |
| Sen-288-361 | NZ_CJHB01000016.1 | - | 39535 | 39483 | **RNA**→ rpmE (PRK00019)Ribosomal_L31 (pfam01197)→ |
| Sen-288-362 | NZ_CJTU01000016.1 | + | 104163 | 104215 | **RNA**→ rpmE (PRK00019)Ribosomal_L31 (pfam01197)→ |
| Sen-288-363 | NZ_CJDW01000017.1 | + | 104153 | 104205 | **RNA**→ rpmE (PRK00019)Ribosomal_L31 (pfam01197)→ |
| Sen-288-364 | NZ_CJSW01000016.1 | + | 104153 | 104205 | **RNA**→ rpmE (PRK00019)Ribosomal_L31 (pfam01197)→ |
| Sen-288-365 | NZ_CJQN01000015.1 | + | 107846 | 107898 | **RNA**→ rpmE (PRK00019)Ribosomal_L31 (pfam01197)→ |
| Sen-305-1 | NZ_AUXE01000010.1 | - | 45181 | 45129 | **RNA**→ rpmE (PRK00019)Ribosomal_L31 (pfam01197)→ |
| Sen-288-366 | NZ_CJPR01000016.1 | + | 104151 | 104203 | **RNA**→ rpmE (PRK00019)Ribosomal_L31 (pfam01197)→ |
| Sen-288-367 | NZ_CJRO01000014.1 | - | 39552 | 39500 | **RNA**→ rpmE (PRK00019)Ribosomal_L31 (pfam01197)→ |
| Sen-288-368 | NZ_CILG01000015.1 | - | 39576 | 39524 | **RNA**→ rpmE (PRK00019)Ribosomal_L31 (pfam01197)→ |
| Sen-288-369 | NZ_CJOO01000015.1 | + | 107413 | 107465 | **RNA**→ rpmE (PRK00019)Ribosomal_L31 (pfam01197)→ |
| Sen-986-1 | NZ_AMDZ02000070.1 | + | 104391 | 104443 | **RNA**→ rpmE (PRK00019)Ribosomal_L31 (pfam01197)→ |
| Sen-288-370 | NZ_CJSG01000015.1 | - | 39539 | 39487 | **RNA**→ rpmE (PRK00019)Ribosomal_L31 (pfam01197)→ |
| Sen-984-1 | NZ_AMED02000023.1 | - | 45415 | 45363 | **RNA**→ rpmE (PRK00019)Ribosomal_L31 (pfam01197)→ |
| Sen-288-371 | NZ_CJQS01000015.1 | - | 39556 | 39504 | **RNA**→ rpmE (PRK00019)Ribosomal_L31 (pfam01197)→ |
| Sen-995-1 | NZ_AYVJ01000205.1 | + | 27408 | 27460 | **RNA**→ rpmE (PRK00019)Ribosomal_L31 (pfam01197)→ |
| Sen-288-372 | NZ_CJKN01000015.1 | + | 107411 | 107463 | **RNA**→ rpmE (PRK00019)Ribosomal_L31 (pfam01197)→ |
| Sen-288-373 | NZ_CJDD01000012.1 | + | 109398 | 109450 | **RNA**→ rpmE (PRK00019)Ribosomal_L31 (pfam01197)→ |
| Sen-970-1 | NZ_AOXO01000023.1 | + | 104435 | 104487 | **RNA**→ rpmE (PRK00019)Ribosomal_L31 (pfam01197)→ |
| Sen-288-374 | NZ_CJCD01000016.1 | + | 104152 | 104204 | **RNA**→ rpmE (PRK00019)Ribosomal_L31 (pfam01197)→ |
| Sen-972-1 | NZ_AJTU01000006.1 | + | 145115 | 145167 | **RNA**→ rpmE (PRK00019)Ribosomal_L31 (pfam01197)→ |
| Sen-288-375 | NZ_CJHD01000016.1 | - | 39723 | 39671 | **RNA**→ rpmE (PRK00019)Ribosomal_L31 (pfam01197)→ |
| Sen-987-1 | NZ_AMEA02000019.1 | - | 45415 | 45363 | **RNA**→ rpmE (PRK00019)Ribosomal_L31 (pfam01197)→ |
| Sen-288-376 | NZ_CJIX01000014.1 | - | 41505 | 41453 | **RNA**→ rpmE (PRK00019)Ribosomal_L31 (pfam01197)→ |
| Sen-977-1 | NZ_AMEE02000078.1 | + | 106171 | 106223 | **RNA**→ rpmE (PRK00019)Ribosomal_L31 (pfam01197)→ |
| Sen-288-377 | NZ_CJDA01000014.1 | + | 104149 | 104201 | **RNA**→ rpmE (PRK00019)Ribosomal_L31 (pfam01197)→ |

| | | | | | |
|---|---|---|---|---|---|
| Sen-978-1 | NZ_AMEF02000109.1 | - | 45424 | 45372 | **RNA**→ rpmE (PRK00019)Ribosomal_L31 (pfam01197)→ |
| Sen-989-1 | NZ_AERV01000009.1 | + | 104113 | 104165 | **RNA**→ rpmE (PRK00019)Ribosomal_L31 (pfam01197)→ rpmE (PRK00019)Ribosomal_L31 (pfam01197)→ |
| Sen-288-378 | NZ_CJKU01000014.1 | + | 107993 | 108045 | **RNA**→ rpmE (PRK00019)Ribosomal_L31 (pfam01197)→ |
| Sen-288-379 | NZ_CJCC01000015.1 | + | 104226 | 104278 | **RNA**→ rpmE (PRK00019)Ribosomal_L31 (pfam01197)→ |
| Sen-980-1 | NZ_AMDY01000201.1 | + | 104301 | 104353 | **RNA**→ rpmE (PRK00019)Ribosomal_L31 (pfam01197)→ |
| Sen-288-380 | NZ_CJGS01000015.1 | - | 39567 | 39515 | **RNA**→ rpmE (PRK00019)Ribosomal_L31 (pfam01197)→ |
| Sen-998-1 | NZ_AOXK01000345.1 | - | 38141 | 38089 | **RNA**→ rpmE (PRK00019)Ribosomal_L31 (pfam01197)→ rpmE (PRK00019)Ribosomal_L31 (pfam01197)→ |
| Sen-976-1 | NZ_AMDX02000028.1 | - | 45307 | 45255 | **RNA**→ rpmE (PRK00019)Ribosomal_L31 (pfam01197)→ |
| Sen-982-1 | NZ_AMEC02000007.1 | + | 104391 | 104443 | **RNA**→ rpmE (PRK00019)Ribosomal_L31 (pfam01197)→ |
| Sen-983-1 | NZ_AMEH02000005.1 | + | 2211 | 2263 | **RNA**→ rpmE (PRK00019)Ribosomal_L31 (pfam01197)→ |
| Sen-974-1 | NZ_AOXD01000071.1 | + | 69832 | 69884 | **RNA**→ rpmE (PRK00019)Ribosomal_L31 (pfam01197)→ |
| Sen-1023-1 | NZ_AOXI01000017.1 | + | 17636 | 17688 | **RNA**→ rpmE (PRK00019)Ribosomal_L31 (pfam01197)→ |
| Sen-997-1 | NZ_AOXM01000219.1 | - | 37004 | 36952 | **RNA**→ rpmE (PRK00019)Ribosomal_L31 (pfam01197)→ |
| Sen-981-1 | NZ_AMEB02000038.1 | + | 106171 | 106223 | **RNA**→ rpmE (PRK00019)Ribosomal_L31 (pfam01197)→ |
| Sen-979-1 | NZ_AMEG02000182.1 | - | 45307 | 45255 | **RNA**→ rpmE (PRK00019)Ribosomal_L31 (pfam01197)→ |
| Sen-300-15 | NZ_CQGX01000017.1 | - | 45178 | 45126 | **RNA**→ rpmE (PRK00019)Ribosomal_L31 (pfam01197)→ |
| Sen-300-16 | NZ_CQHG01000017.1 | - | 45269 | 45217 | **RNA**→ rpmE (PRK00019)Ribosomal_L31 (pfam01197)→ |
| Sen-300-17 | NZ_CQHN01000015.1 | - | 47295 | 47243 | **RNA**→ rpmE (PRK00019)Ribosomal_L31 (pfam01197)→ |
| Sen-975-1 | NZ_AUVE01000010.1 | - | 45181 | 45129 | **RNA**→ rpmE (PRK00019)Ribosomal_L31 (pfam01197)→ |
| Sen-304-1 | NZ_AUVD01000058.1 | - | 45077 | 45025 | **RNA**→ rpmE (PRK00019)Ribosomal_L31 (pfam01197)→ |
| Sen-994-1 | NZ_AYUO01000004.1 | + | 57838 | 57890 | **RNA**→ rpmE (PRK00019)Ribosomal_L31 (pfam01197)→ |
| Sen-300-18 | NZ_CQGT01000017.1 | + | 42541 | 42593 | **RNA**→ rpmE (PRK00019)Ribosomal_L31 (pfam01197)→ |
| Sen-300-19 | NZ_CQHH01000015.1 | - | 45254 | 45202 | **RNA**→ rpmE (PRK00019)Ribosomal_L31 (pfam01197)→ |
| Sen-300-20 | NZ_CQGZ01000017.1 | + | 42532 | 42584 | **RNA**→ rpmE (PRK00019)Ribosomal_L31 (pfam01197)→ |
| Sen-300-21 | NZ_CQHR01000015.1 | - | 45343 | 45291 | **RNA**→ rpmE (PRK00019)Ribosomal_L31 (pfam01197)→ |
| Sen-300-22 | NZ_CQGJ01000016.1 | - | 45252 | 45200 | **RNA**→ rpmE (PRK00019)Ribosomal_L31 (pfam01197)→ |
| Sen-300-23 | NZ_CQHD01000016.1 | - | 45196 | 45144 | **RNA**→ rpmE (PRK00019)Ribosomal_L31 (pfam01197)→ |
| Sen-300-24 | NZ_CQGM01000018.1 | - | 45252 | 45200 | **RNA**→ rpmE (PRK00019)Ribosomal_L31 (pfam01197)→ |
| Sen-300-25 | NZ_CQGO01000016.1 | + | 42519 | 42571 | **RNA**→ rpmE (PRK00019)Ribosomal_L31 (pfam01197)→ |
| Sen-300-26 | NZ_CQHW01000017.1 | - | 45178 | 45126 | **RNA**→ rpmE (PRK00019)Ribosomal_L31 (pfam01197)→ |
| Sen-302-1 | NZ_AHVA01000016.1 | + | 104730 | 104782 | **RNA**→ rpmE (PRK00019)Ribosomal_L31 (pfam01197)→ |
| Sen-985-1 | NZ_ATWR01000081.1 | + | 107605 | 107657 | **RNA**→ rpmE (PRK00019)Ribosomal_L31 (pfam01197)→ |
| Sen-300-27 | NZ_CQHC01000015.1 | + | 42549 | 42601 | **RNA**→ rpmE (PRK00019)Ribosomal_L31 (pfam01197)→ |
| Sen-300-28 | NZ_CQGQ01000019.1 | - | 45254 | 45202 | **RNA**→ rpmE (PRK00019)Ribosomal_L31 (pfam01197)→ |
| Sen-300-29 | NZ_CQHA01000015.1 | + | 42549 | 42601 | **RNA**→ rpmE (PRK00019)Ribosomal_L31 (pfam01197)→ |
| Sen-303-1 | NZ_AHUZ01000016.1 | + | 104759 | 104811 | **RNA**→ rpmE (PRK00019)Ribosomal_L31 (pfam01197)→ |
| Sen-288-381 | NZ_CIVR01000009.1 | - | 39382 | 39330 | **RNA**→ rpmE (PRK00019)Ribosomal_L31 (pfam01197)→ |
| Sen-288-382 | NZ_CJNX01000014.1 | - | 41232 | 41180 | **RNA**→ rpmE (PRK00019)Ribosomal_L31 (pfam01197)→ |
| Sen-288-383 | NZ_CJLY01000015.1 | + | 104151 | 104203 | **RNA**→ rpmE (PRK00019)Ribosomal_L31 (pfam01197)→ |
| Sen-288-384 | NZ_CJPW01000010.1 | - | 39601 | 39549 | **RNA**→ rpmE (PRK00019)Ribosomal_L31 (pfam01197)→ |
| Sen-288-385 | NZ_CEZM01000016.1 | - | 39537 | 39485 | **RNA**→ rpmE (PRK00019)Ribosomal_L31 (pfam01197)→ |
| Sen-288-386 | NZ_CJHQ01000017.1 | - | 39537 | 39485 | **RNA**→ rpmE (PRK00019)Ribosomal_L31 (pfam01197)→ |
| Sen-288-387 | NZ_CEXK01000015.1 | - | 39680 | 39628 | **RNA**→ rpmE (PRK00019)Ribosomal_L31 (pfam01197)→ |
| Sen-288-388 | NZ_CIWG01000014.1 | + | 104159 | 104211 | **RNA**→ rpmE (PRK00019)Ribosomal_L31 (pfam01197)→ |
| Sen-288-389 | NZ_CINK01000011.1 | - | 44769 | 44717 | **RNA**→ rpmE (PRK00019)Ribosomal_L31 (pfam01197)→ |
| Sen-288-390 | NZ_CJOT01000015.1 | + | 104156 | 104208 | **RNA**→ rpmE (PRK00019)Ribosomal_L31 (pfam01197)→ |
| Sen-288-391 | NZ_CJIT01000010.1 | - | 39562 | 39510 | **RNA**→ rpmE (PRK00019)Ribosomal_L31 (pfam01197)→ |
| Sen-288-392 | NZ_CJFD01000013.1 | - | 39574 | 39522 | **RNA**→ rpmE (PRK00019)Ribosomal_L31 (pfam01197)→ |
| Sen-288-393 | NZ_CJYH01000004.1 | - | 216848 | 216796 | **RNA**→ rpmE (PRK00019)Ribosomal_L31 (pfam01197)→ |
| Sen-288-394 | NZ_CJCT01000016.1 | + | 104146 | 104198 | **RNA**→ rpmE (PRK00019)Ribosomal_L31 (pfam01197)→ |
| Sen-288-395 | NZ_CJFK01000013.1 | + | 104153 | 104205 | **RNA**→ rpmE (PRK00019)Ribosomal_L31 (pfam01197)→ |
| Sen-288-396 | NZ_CJKF01000007.1 | + | 189409 | 189461 | **RNA**→ rpmE (PRK00019)Ribosomal_L31 (pfam01197)→ |

| | | | | | |
|---|---|---|---|---|---|
| Sen-288-397 | NZ_CILW01000010.1 | - | 39540 | 39488 | RNA→ rpmE (PRK00019) Ribosomal_L31 (pfam01197)→ |
| Sen-288-398 | NZ_CIYJ01000011.1 | + | 137633 | 137685 | RNA→ rpmE (PRK00019) Ribosomal_L31 (pfam01197)→ |
| Sen-288-399 | NZ_CJBA01000015.1 | - | 39566 | 39514 | RNA→ rpmE (PRK00019) Ribosomal_L31 (pfam01197)→ |
| Sen-288-400 | NZ_CIVL01000015.1 | - | 39544 | 39492 | RNA→ rpmE (PRK00019) Ribosomal_L31 (pfam01197)→ |
| Sen-288-401 | NZ_CJDV01000016.1 | - | 39534 | 39482 | RNA→ rpmE (PRK00019) Ribosomal_L31 (pfam01197)→ |
| Sen-288-402 | NZ_CJVE01000015.1 | + | 104132 | 104184 | RNA→ rpmE (PRK00019) Ribosomal_L31 (pfam01197)→ |
| Sen-288-403 | NZ_CJYL01000014.1 | - | 41510 | 41458 | RNA→ rpmE (PRK00019) Ribosomal_L31 (pfam01197)→ |
| Sen-288-404 | NZ_CJIZ01000020.1 | - | 39374 | 39322 | RNA→ rpmE (PRK00019) Ribosomal_L31 (pfam01197)→ |
| Sen-288-405 | NZ_CJFX01000013.1 | - | 39536 | 39484 | RNA→ rpmE (PRK00019) Ribosomal_L31 (pfam01197)→ |
| Sen-288-406 | NZ_CIXL01000015.1 | - | 39556 | 39504 | RNA→ rpmE (PRK00019) Ribosomal_L31 (pfam01197)→ |
| Sen-288-407 | NZ_CEZC01000016.1 | + | 104161 | 104213 | RNA→ rpmE (PRK00019) Ribosomal_L31 (pfam01197)→ |
| Sen-288-408 | NZ_CEXH01000014.1 | + | 103992 | 104044 | RNA→ rpmE (PRK00019) Ribosomal_L31 (pfam01197)→ |
| Sen-288-409 | NZ_CJEG01000016.1 | - | 39652 | 39600 | RNA→ rpmE (PRK00019) Ribosomal_L31 (pfam01197)→ |
| Sen-288-410 | NZ_CJUB01000015.1 | - | 41231 | 41179 | RNA→ rpmE (PRK00019) Ribosomal_L31 (pfam01197)→ |
| Sen-288-411 | NZ_CJUJ01000015.1 | - | 41231 | 41179 | RNA→ rpmE (PRK00019) Ribosomal_L31 (pfam01197)→ |
| Sen-288-412 | NZ_CIWP01000016.1 | + | 104141 | 104193 | RNA→ rpmE (PRK00019) Ribosomal_L31 (pfam01197)→ |
| Sen-288-413 | NZ_CJVJ01000017.1 | - | 39377 | 39325 | RNA→ rpmE (PRK00019) Ribosomal_L31 (pfam01197)→ |
| Sen-288-414 | NZ_CJUW01000012.1 | - | 39355 | 39303 | RNA→ rpmE (PRK00019) Ribosomal_L31 (pfam01197)→ |
| Sen-288-415 | NZ_CJPE01000016.1 | - | 39601 | 39549 | RNA→ rpmE (PRK00019) Ribosomal_L31 (pfam01197)→ |
| Sen-288-416 | NZ_CIWE01000016.1 | + | 103936 | 103988 | RNA→ rpmE (PRK00019) Ribosomal_L31 (pfam01197)→ |
| Sen-288-417 | NZ_CIYW01000008.1 | + | 170860 | 170912 | RNA→ rpmE (PRK00019) Ribosomal_L31 (pfam01197)→ |
| Sen-288-418 | NZ_CJXX01000017.1 | - | 39362 | 39310 | RNA→ rpmE (PRK00019) Ribosomal_L31 (pfam01197)→ |
| Sen-288-419 | NZ_CIUM01000014.1 | - | 39585 | 39533 | RNA→ rpmE (PRK00019) Ribosomal_L31 (pfam01197)→ |
| Sen-288-420 | NZ_CJOC01000016.1 | - | 39536 | 39484 | RNA→ rpmE (PRK00019) Ribosomal_L31 (pfam01197)→ |
| Sen-288-421 | NZ_CJGD01000016.1 | - | 39535 | 39483 | RNA→ rpmE (PRK00019) Ribosomal_L31 (pfam01197)→ |
| Sen-288-422 | NZ_CIVM01000016.1 | - | 39382 | 39330 | RNA→ rpmE (PRK00019) Ribosomal_L31 (pfam01197)→ |
| Sen-288-423 | NZ_CJTR01000016.1 | + | 104150 | 104202 | RNA→ rpmE (PRK00019) Ribosomal_L31 (pfam01197)→ |
| Sen-288-424 | NZ_CIUS01000015.1 | + | 107419 | 107471 | RNA→ rpmE (PRK00019) Ribosomal_L31 (pfam01197)→ |
| Sen-288-425 | NZ_CJND01000015.1 | + | 107419 | 107471 | RNA→ rpmE (PRK00019) Ribosomal_L31 (pfam01197)→ |
| Sen-288-426 | NZ_CJKL01000011.1 | - | 39456 | 39404 | RNA→ rpmE (PRK00019) Ribosomal_L31 (pfam01197)→ |
| Sen-288-427 | NZ_CIWS01000015.1 | + | 104216 | 104268 | RNA→ rpmE (PRK00019) Ribosomal_L31 (pfam01197)→ |
| Sen-288-428 | NZ_CJRT01000016.1 | + | 104154 | 104206 | RNA→ rpmE (PRK00019) Ribosomal_L31 (pfam01197)→ |
| Sen-288-429 | NZ_CJPD01000014.1 | - | 41228 | 41176 | RNA→ rpmE (PRK00019) Ribosomal_L31 (pfam01197)→ |
| Sen-288-430 | NZ_CIXX01000022.1 | + | 43222 | 43274 | RNA→ rpmE (PRK00019) Ribosomal_L31 (pfam01197)→ |
| Sen-288-431 | NZ_CIWX01000016.1 | - | 39527 | 39475 | RNA→ rpmE (PRK00019) Ribosomal_L31 (pfam01197)→ |
| Sen-288-432 | NZ_CJLS01000016.1 | - | 39536 | 39484 | RNA→ rpmE (PRK00019) Ribosomal_L31 (pfam01197)→ |
| Sen-288-433 | NZ_CJBM01000016.1 | + | 104179 | 104231 | RNA→ rpmE (PRK00019) Ribosomal_L31 (pfam01197)→ |
| Sen-288-434 | NZ_CJTA01000015.1 | - | 39577 | 39525 | RNA→ rpmE (PRK00019) Ribosomal_L31 (pfam01197)→ |
| Sen-288-435 | NZ_CJJT01000014.1 | + | 104154 | 104206 | RNA→ rpmE (PRK00019) Ribosomal_L31 (pfam01197)→ |
| Sen-288-436 | NZ_CJPM01000016.1 | - | 39383 | 39331 | RNA→ rpmE (PRK00019) Ribosomal_L31 (pfam01197)→ |
| Sen-288-437 | NZ_CJBF01000011.1 | + | 141125 | 141177 | RNA→ rpmE (PRK00019) Ribosomal_L31 (pfam01197)→ |
| Sen-288-438 | NZ_CJJI01000002.1 | + | 405136 | 405188 | RNA→ rpmE (PRK00019) Ribosomal_L31 (pfam01197)→ |
| Sen-288-439 | NZ_CJIU01000013.1 | - | 41231 | 41179 | RNA→ rpmE (PRK00019) Ribosomal_L31 (pfam01197)→ |
| Sen-288-440 | NZ_CJJM01000014.1 | - | 41506 | 41454 | RNA→ rpmE (PRK00019) Ribosomal_L31 (pfam01197)→ |
| Sen-288-441 | NZ_CJEF01000016.1 | + | 104145 | 104197 | RNA→ rpmE (PRK00019) Ribosomal_L31 (pfam01197)→ |
| Sen-288-442 | NZ_CJAW01000016.1 | - | 39536 | 39484 | RNA→ rpmE (PRK00019) Ribosomal_L31 (pfam01197)→ |
| Sen-288-443 | NZ_CJYK01000014.1 | - | 39370 | 39318 | RNA→ rpmE (PRK00019) Ribosomal_L31 (pfam01197)→ |
| Sen-288-444 | NZ_CJWN01000016.1 | + | 104145 | 104197 | RNA→ rpmE (PRK00019) Ribosomal_L31 (pfam01197)→ |
| Sen-288-445 | NZ_CJSF01000017.1 | - | 39601 | 39549 | RNA→ rpmE (PRK00019) Ribosomal_L31 (pfam01197)→ |
| Sen-288-446 | NZ_CJHO01000016.1 | + | 104160 | 104212 | RNA→ rpmE (PRK00019) Ribosomal_L31 (pfam01197)→ |
| Sen-288-447 | NZ_CJYG01000015.1 | - | 41672 | 41620 | RNA→ rpmE (PRK00019) Ribosomal_L31 (pfam01197)→ |
| Sen-288-448 | NZ_CJDX01000017.1 | - | 39584 | 39532 | RNA→ rpmE (PRK00019) Ribosomal_L31 (pfam01197)→ |

| | | | | | |
|---|---|---|---|---|---|
| Sen-288-449 | NZ_CJLM01000015.1 | + | 103992 | 104044 | **RNA**→ rpmE (PRK00019)Ribosomal_L31 (pfam01197)→ |
| Sen-288-450 | NZ_CJXR01000015.1 | - | 39382 | 39330 | **RNA**→ rpmE (PRK00019)Ribosomal_L31 (pfam01197)→ |
| Sen-288-451 | NZ_CILF01000016.1 | - | 39530 | 39478 | **RNA**→ rpmE (PRK00019)Ribosomal_L31 (pfam01197)→ |
| Sen-288-452 | NZ_CJSC01000016.1 | - | 39549 | 39497 | **RNA**→ rpmE (PRK00019)Ribosomal_L31 (pfam01197)→ |
| Sen-288-453 | NZ_CJJV01000011.1 | - | 39456 | 39404 | **RNA**→ rpmE (PRK00019)Ribosomal_L31 (pfam01197)→ |
| Sen-288-454 | NZ_CIUO01000015.1 | - | 41231 | 41179 | **RNA**→ rpmE (PRK00019)Ribosomal_L31 (pfam01197)→ |
| Sen-288-455 | NZ_CIVD01000014.1 | - | 41227 | 41175 | **RNA**→ rpmE (PRK00019)Ribosomal_L31 (pfam01197)→ |
| Sen-288-456 | NZ_CJVZ01000016.1 | - | 39363 | 39311 | **RNA**→ rpmE (PRK00019)Ribosomal_L31 (pfam01197)→ |
| Sen-288-457 | NZ_CJQH01000016.1 | + | 104156 | 104208 | **RNA**→ rpmE (PRK00019)Ribosomal_L31 (pfam01197)→ |
| Sen-288-458 | NZ_CJGB01000016.1 | + | 104179 | 104231 | **RNA**→ rpmE (PRK00019)Ribosomal_L31 (pfam01197)→ |
| Sen-288-459 | NZ_CJUZ01000013.1 | + | 107861 | 107913 | **RNA**→ rpmE (PRK00019)Ribosomal_L31 (pfam01197)→ |
| Sen-288-460 | NZ_CIZF01000016.1 | - | 39534 | 39482 | **RNA**→ rpmE (PRK00019)Ribosomal_L31 (pfam01197)→ |
| Sen-288-461 | NZ_CJFH01000017.1 | - | 39663 | 39611 | **RNA**→ rpmE (PRK00019)Ribosomal_L31 (pfam01197)→ |
| Sen-288-462 | NZ_CJVU01000016.1 | + | 104011 | 104063 | **RNA**→ rpmE (PRK00019)Ribosomal_L31 (pfam01197)→ |
| Sen-288-463 | NZ_CJIC01000016.1 | + | 104142 | 104194 | **RNA**→ rpmE (PRK00019)Ribosomal_L31 (pfam01197)→ |
| Sen-288-464 | NZ_CJIQ01000013.1 | + | 107705 | 107757 | **RNA**→ rpmE (PRK00019)Ribosomal_L31 (pfam01197)→ |
| Sen-288-465 | NZ_CJFT01000016.1 | + | 104156 | 104208 | **RNA**→ rpmE (PRK00019)Ribosomal_L31 (pfam01197)→ |
| Sen-288-466 | NZ_CJGF01000016.1 | - | 39538 | 39486 | **RNA**→ rpmE (PRK00019)Ribosomal_L31 (pfam01197)→ |
| Sen-288-467 | NZ_CJSP01000016.1 | + | 104164 | 104216 | **RNA**→ rpmE (PRK00019)Ribosomal_L31 (pfam01197)→ |
| Sen-288-468 | NZ_CIWY01000011.1 | - | 39583 | 39531 | **RNA**→ rpmE (PRK00019)Ribosomal_L31 (pfam01197)→ |
| Sen-288-469 | NZ_CIXJ01000016.1 | - | 39527 | 39475 | **RNA**→ rpmE (PRK00019)Ribosomal_L31 (pfam01197)→ |
| Sen-426-1 | NZ_AOZW01000064.1 | + | 84250 | 84302 | **RNA**→ rpmE (PRK00019)Ribosomal_L31 (pfam01197)→ |
| Sen-288-470 | NZ_CJSS01000016.1 | - | 39373 | 39321 | **RNA**→ rpmE (PRK00019)Ribosomal_L31 (pfam01197)→ |
| Sen-288-471 | NZ_CJXZ01000017.1 | - | 39540 | 39488 | **RNA**→ rpmE (PRK00019)Ribosomal_L31 (pfam01197)→ |
| Sen-428-1 | NZ_APAA01000073.1 | + | 98899 | 98951 | **RNA**→ rpmE (PRK00019)Ribosomal_L31 (pfam01197)→ |
| Sen-288-472 | NZ_CJHC01000016.1 | + | 104148 | 104200 | **RNA**→ rpmE (PRK00019)Ribosomal_L31 (pfam01197)→ |
| Sen-18-1 | NZ_AUXR01000014.1 | + | 105391 | 105443 | **RNA**→ rpmE (PRK00019)Ribosomal_L31 (pfam01197)→ |
| Sen-288-473 | NZ_CJAV01000016.1 | - | 39534 | 39482 | **RNA**→ rpmE (PRK00019)Ribosomal_L31 (pfam01197)→ |
| Sen-95-1 | NZ_JMMF01000016.1 | + | 103451 | 103503 | **RNA**→ rpmE (PRK00019)Ribosomal_L31 (pfam01197)→ |
| Sen-288-474 | NZ_CJSN01000016.1 | + | 104157 | 104209 | **RNA**→ rpmE (PRK00019)Ribosomal_L31 (pfam01197)→ |
| Sen-288-475 | NZ_CJJN01000016.1 | + | 104130 | 104182 | **RNA**→ rpmE (PRK00019)Ribosomal_L31 (pfam01197)→ |
| Sen-41-1 | NZ_JMMI01000001.1 | + | 55530 | 55582 | **RNA**→ rpmE (PRK00019)Ribosomal_L31 (pfam01197)→ |
| Sen-288-476 | NZ_CJHV01000015.1 | + | 103986 | 104038 | **RNA**→ rpmE (PRK00019)Ribosomal_L31 (pfam01197)→ |
| Sen-397-1 | NZ_CARQ01000039.1 | - | 37552 | 37500 | **RNA**→ rpmE (PRK00019)Ribosomal_L31 (pfam01197)→ |
| Sen-288-477 | NZ_CJTG01000016.1 | + | 104155 | 104207 | **RNA**→ rpmE (PRK00019)Ribosomal_L31 (pfam01197)→ |
| Sen-288-478 | NZ_CJFG01000016.1 | - | 39536 | 39484 | **RNA**→ rpmE (PRK00019)Ribosomal_L31 (pfam01197)→ |
| Sen-410-1 | NZ_CARG01000040.1 | - | 37550 | 37498 | **RNA**→ rpmE (PRK00019)Ribosomal_L31 (pfam01197)→ |
| Sen-288-479 | NZ_CJGT01000015.1 | - | 39574 | 39522 | **RNA**→ rpmE (PRK00019)Ribosomal_L31 (pfam01197)→ |
| Sen-415-1 | NZ_CARB01000036.1 | - | 37553 | 37501 | **RNA**→ rpmE (PRK00019)Ribosomal_L31 (pfam01197)→ |
| Sen-288-480 | NZ_CJFR01000016.1 | - | 39536 | 39484 | **RNA**→ rpmE (PRK00019)Ribosomal_L31 (pfam01197)→ |
| Sen-422-1 | NZ_AYDL01000045.1 | + | 111493 | 111545 | **RNA**→ rpmE (PRK00019)Ribosomal_L31 (pfam01197)→ rpmE (PRK00019)Ribosomal_L31 (pfam01197)→ |
| Sen-288-481 | NZ_CJSZ01000014.1 | + | 107858 | 107910 | **RNA**→ rpmE (PRK00019)Ribosomal_L31 (pfam01197)→ |
| Sen-288-482 | NZ_CJXM01000010.1 | - | 39370 | 39318 | **RNA**→ rpmE (PRK00019)Ribosomal_L31 (pfam01197)→ |
| Sen-418-1 | NZ_AYDV01000082.1 | + | 111737 | 111789 | **RNA**→ rpmE (PRK00019)Ribosomal_L31 (pfam01197)→ rpmE (PRK00019)Ribosomal_L31 (pfam01197)→ |
| Sen-434-1 | NZ_AOZN01000010.1 | + | 40589 | 40641 | **RNA**→ rpmE (PRK00019)Ribosomal_L31 (pfam01197)→ rpmE (PRK00019)Ribosomal_L31 (pfam01197)→ |
| Sen-288-483 | NZ_CJDH01000017.1 | - | 39536 | 39484 | **RNA**→ rpmE (PRK00019)Ribosomal_L31 (pfam01197)→ |
| Sen-288-484 | NZ_CJNO01000014.1 | - | 41507 | 41455 | **RNA**→ rpmE (PRK00019)Ribosomal_L31 (pfam01197)→ |
| Sen-113-2 | NZ_AOZO01000047.1 | + | 104627 | 104679 | **RNA**→ rpmE (PRK00019)Ribosomal_L31 (pfam01197)→ |
| Sen-288-485 | NZ_CJME01000017.1 | + | 107419 | 107471 | **RNA**→ rpmE (PRK00019)Ribosomal_L31 (pfam01197)→ |
| Sen-435-1 | NZ_AOZT01000071.1 | + | 104272 | 104324 | **RNA**→ rpmE (PRK00019)Ribosomal_L31 (pfam01197)→ rpmE (PRK00019)Ribosomal_L31 (pfam01197)→ |
| Sen-288-486 | NZ_CJNB01000013.1 | - | 41505 | 41453 | **RNA**→ rpmE (PRK00019)Ribosomal_L31 (pfam01197)→ |
| Sen-406-1 | NZ_AOZU01000014.1 | - | 37534 | 37482 | **RNA**→ rpmE (PRK00019)Ribosomal_L31 (pfam01197)→ rpmE (PRK00019)Ribosomal_L31 (pfam01197)→ |

| | | | | | |
|---|---|---|---|---|---|
| Sen-288-487 | NZ_CIVQ01000014.1 | + | 108011 | 108063 | **RNA**→ rpmE (PRK00019)Ribosomal_L31 (pfam01197)→ |
| Sen-404-1 | NZ_AOZY01000035.1 | + | 101428 | 101480 | **RNA**→ rpmE (PRK00019)Ribosomal_L31 (pfam01197)→ rpmE (PRK00019)Ribosomal_L31 (pfam01197)→ |
| Sen-288-488 | NZ_CJJO01000016.1 | + | 104159 | 104211 | **RNA**→ rpmE (PRK00019)Ribosomal_L31 (pfam01197)→ |
| Sen-288-1118 | NZ_CIXY01000014.1 | - | 39556 | 39504 | **RNA**→ rpmE (PRK00019)Ribosomal_L31 (pfam01197)→ |
| Sen-432-1 | NZ_ALPN01000011.1 | - | 45077 | 45025 | **RNA**→ rpmE (PRK00019)Ribosomal_L31 (pfam01197)→ |
| Sen-288-489 | NZ_CJPX01000009.1 | - | 39601 | 39549 | **RNA**→ rpmE (PRK00019)Ribosomal_L31 (pfam01197)→ |
| Sen-425-1 | NZ_ANOS01000058.1 | + | 37363 | 37415 | **RNA**→ rpmE (PRK00019)Ribosomal_L31 (pfam01197)→ rpmE (PRK00019)Ribosomal_L31 (pfam01197)→ |
| Sen-288-490 | NZ_CJZF01000016.1 | - | 39578 | 39526 | **RNA**→ rpmE (PRK00019)Ribosomal_L31 (pfam01197)→ |
| Sen-288-491 | NZ_CIYG01000016.1 | + | 104148 | 104200 | **RNA**→ rpmE (PRK00019)Ribosomal_L31 (pfam01197)→ |
| Sen-395-1 | NZ_CARR01000042.1 | - | 37552 | 37500 | **RNA**→ rpmE (PRK00019)Ribosomal_L31 (pfam01197)→ |
| Sen-288-492 | NZ_CJOW01000016.1 | - | 39706 | 39654 | **RNA**→ rpmE (PRK00019)Ribosomal_L31 (pfam01197)→ |
| Sen-400-1 | NZ_CARN01000056.1 | + | 61476 | 61528 | **RNA**→ rpmE (PRK00019)Ribosomal_L31 (pfam01197)→ |
| Sen-401-1 | NZ_CARM01000014.1 | - | 37552 | 37500 | **RNA**→ rpmE (PRK00019)Ribosomal_L31 (pfam01197)→ |
| Sen-409-1 | NZ_CATS01000038.1 | - | 37552 | 37500 | **RNA**→ rpmE (PRK00019)Ribosomal_L31 (pfam01197)→ |
| Sen-414-1 | NZ_CARC01000038.1 | + | 110915 | 110967 | **RNA**→ rpmE (PRK00019)Ribosomal_L31 (pfam01197)→ |
| Sen-421-1 | NZ_AYDK01000018.1 | + | 47113 | 47165 | **RNA**→ rpmE (PRK00019)Ribosomal_L31 (pfam01197)→ rpmE (PRK00019)Ribosomal_L31 (pfam01197)→ |
| Sen-402-1 | NZ_AYCU01000016.1 | - | 37792 | 37740 | **RNA**→ rpmE (PRK00019)Ribosomal_L31 (pfam01197)→ rpmE (PRK00019)Ribosomal_L31 (pfam01197)→ |
| Sen-437-1 | NZ_AOZR01000102.1 | - | 45537 | 45485 | **RNA**→ rpmE (PRK00019)Ribosomal_L31 (pfam01197)→ rpmE (PRK00019)Ribosomal_L31 (pfam01197)→ |
| Sen-416-1 | NZ_AOZX01000052.1 | + | 111713 | 111765 | **RNA**→ rpmE (PRK00019)Ribosomal_L31 (pfam01197)→ rpmE (PRK00019)Ribosomal_L31 (pfam01197)→ |
| Sen-439-1 | NZ_CQPO01000009.1 | + | 138562 | 138614 | **RNA**→ rpmE (PRK00019)Ribosomal_L31 (pfam01197)→ |
| Sen-439-2 | NZ_CQPH01000012.1 | - | 45285 | 45233 | **RNA**→ rpmE (PRK00019)Ribosomal_L31 (pfam01197)→ |
| Sen-439-3 | NZ_CQPJ01000008.1 | + | 138584 | 138636 | **RNA**→ rpmE (PRK00019)Ribosomal_L31 (pfam01197)→ |
| Sen-16-1 | NZ_ARYP01000011.1 | + | 98887 | 98939 | **RNA**→ rpmE (PRK00019)Ribosomal_L31 (pfam01197)→ |
| Sen-19-1 | NZ_ARYQ01000080.1 | - | 45140 | 45088 | **RNA**→ rpmE (PRK00019)Ribosomal_L31 (pfam01197)→ |
| Sen-49-1 | NZ_JMMM01000017.1 | + | 103448 | 103500 | **RNA**→ rpmE (PRK00019)Ribosomal_L31 (pfam01197)→ |
| Sen-439-4 | NZ_CQPB01000009.1 | + | 106471 | 106523 | **RNA**→ rpmE (PRK00019)Ribosomal_L31 (pfam01197)→ |
| Sen-439-5 | NZ_CQPG01000014.1 | - | 45583 | 45531 | **RNA**→ rpmE (PRK00019)Ribosomal_L31 (pfam01197)→ |
| Sen-21-1 | NZ_JMMK01000030.1 | + | 103456 | 103508 | **RNA**→ rpmE (PRK00019)Ribosomal_L31 (pfam01197)→ |
| Sen-31-1 | NZ_JMMH01000008.1 | - | 45105 | 45053 | **RNA**→ rpmE (PRK00019)Ribosomal_L31 (pfam01197)→ |
| Sen-288-493 | NZ_CJHN01000016.1 | + | 104144 | 104196 | **RNA**→ rpmE (PRK00019)Ribosomal_L31 (pfam01197)→ |
| Sen-288-494 | NZ_CJOZ01000016.1 | + | 104156 | 104208 | **RNA**→ rpmE (PRK00019)Ribosomal_L31 (pfam01197)→ |
| Sen-288-495 | NZ_CJYV01000015.1 | - | 41680 | 41628 | **RNA**→ rpmE (PRK00019)Ribosomal_L31 (pfam01197)→ |
| Sen-288-496 | NZ_CJIW01000014.1 | + | 104151 | 104203 | **RNA**→ rpmE (PRK00019)Ribosomal_L31 (pfam01197)→ |
| Sen-288-497 | NZ_CELU01000016.1 | + | 104133 | 104185 | **RNA**→ rpmE (PRK00019)Ribosomal_L31 (pfam01197)→ |
| Sen-288-498 | NZ_CEYE01000016.1 | - | 39723 | 39671 | **RNA**→ rpmE (PRK00019)Ribosomal_L31 (pfam01197)→ |
| Sen-288-499 | NZ_CEXO01000016.1 | - | 39595 | 39543 | **RNA**→ rpmE (PRK00019)Ribosomal_L31 (pfam01197)→ |
| Sen-288-500 | NZ_CITF01000015.1 | + | 104157 | 104209 | **RNA**→ rpmE (PRK00019)Ribosomal_L31 (pfam01197)→ |
| Sen-288-501 | NZ_CIRA01000016.1 | - | 39572 | 39520 | **RNA**→ rpmE (PRK00019)Ribosomal_L31 (pfam01197)→ |
| Sen-288-502 | NZ_CIUE01000002.1 | + | 404999 | 405051 | **RNA**→ rpmE (PRK00019)Ribosomal_L31 (pfam01197)→ |
| Sen-288-503 | NZ_CIPP01000016.1 | + | 104162 | 104214 | **RNA**→ rpmE (PRK00019)Ribosomal_L31 (pfam01197)→ |
| Sen-288-504 | NZ_CIOU01000014.1 | - | 41231 | 41179 | **RNA**→ rpmE (PRK00019)Ribosomal_L31 (pfam01197)→ |
| Sen-288-505 | NZ_CIPY01000009.1 | - | 39541 | 39489 | **RNA**→ rpmE (PRK00019)Ribosomal_L31 (pfam01197)→ |
| Sen-288-506 | NZ_CIRH01000015.1 | + | 104027 | 104079 | **RNA**→ rpmE (PRK00019)Ribosomal_L31 (pfam01197)→ |
| Sen-288-507 | NZ_CIOJ01000016.1 | - | 39368 | 39316 | **RNA**→ rpmE (PRK00019)Ribosomal_L31 (pfam01197)→ |
| Sen-288-508 | NZ_CIRB01000014.1 | + | 104160 | 104212 | **RNA**→ rpmE (PRK00019)Ribosomal_L31 (pfam01197)→ |
| Sen-288-509 | NZ_CIST01000013.1 | - | 41228 | 41176 | **RNA**→ rpmE (PRK00019)Ribosomal_L31 (pfam01197)→ |
| Sen-288-510 | NZ_CIQL01000015.1 | + | 104145 | 104197 | **RNA**→ rpmE (PRK00019)Ribosomal_L31 (pfam01197)→ |
| Sen-288-511 | NZ_CIQU01000004.1 | + | 104154 | 104206 | **RNA**→ rpmE (PRK00019)Ribosomal_L31 (pfam01197)→ |
| Sen-288-512 | NZ_CITE01000014.1 | + | 103992 | 104044 | **RNA**→ rpmE (PRK00019)Ribosomal_L31 (pfam01197)→ |
| Sen-288-513 | NZ_CITH01000016.1 | - | 39543 | 39491 | **RNA**→ rpmE (PRK00019)Ribosomal_L31 (pfam01197)→ |
| Sen-288-514 | NZ_CJBV01000015.1 | + | 104159 | 104211 | **RNA**→ rpmE (PRK00019)Ribosomal_L31 (pfam01197)→ |
| Sen-288-515 | NZ_CITO01000014.1 | - | 39380 | 39328 | **RNA**→ rpmE (PRK00019)Ribosomal_L31 (pfam01197)→ |

| | | | | | |
|---|---|---|---|---|---|
| Sen-288-516 | NZ_CJLW01000014.1 | + | 104151 | 104203 | RNA→ rpmE (PRK00019)Ribosomal_L31 (pfam01197)→ |
| Sen-288-517 | NZ_CISW01000015.1 | + | 104138 | 104190 | RNA→ rpmE (PRK00019)Ribosomal_L31 (pfam01197)→ |
| Sen-288-518 | NZ_CIYO01000014.1 | + | 107981 | 108033 | RNA→ rpmE (PRK00019)Ribosomal_L31 (pfam01197)→ |
| Sen-288-519 | NZ_CIRU01000011.1 | - | 39576 | 39524 | RNA→ rpmE (PRK00019)Ribosomal_L31 (pfam01197)→ |
| Sen-288-520 | NZ_CJTB01000016.1 | + | 104163 | 104215 | RNA→ rpmE (PRK00019)Ribosomal_L31 (pfam01197)→ |
| Sen-288-521 | NZ_CITL01000016.1 | + | 103986 | 104038 | RNA→ rpmE (PRK00019)Ribosomal_L31 (pfam01197)→ |
| Sen-288-522 | NZ_CIXC01000010.1 | - | 39454 | 39402 | RNA→ rpmE (PRK00019)Ribosomal_L31 (pfam01197)→ |
| Sen-288-523 | NZ_CIPR01000014.1 | + | 109634 | 109686 | RNA→ rpmE (PRK00019)Ribosomal_L31 (pfam01197)→ |
| Sen-288-524 | NZ_CJID01000016.1 | + | 104149 | 104201 | RNA→ rpmE (PRK00019)Ribosomal_L31 (pfam01197)→ |
| Sen-288-525 | NZ_CITK01000013.1 | - | 41230 | 41178 | RNA→ rpmE (PRK00019)Ribosomal_L31 (pfam01197)→ |
| Sen-288-526 | NZ_CINW01000014.1 | + | 104151 | 104203 | RNA→ rpmE (PRK00019)Ribosomal_L31 (pfam01197)→ |
| Sen-288-527 | NZ_CJZB01000016.1 | - | 39372 | 39320 | RNA→ rpmE (PRK00019)Ribosomal_L31 (pfam01197)→ |
| Sen-288-528 | NZ_CIRT01000010.1 | - | 39560 | 39508 | RNA→ rpmE (PRK00019)Ribosomal_L31 (pfam01197)→ |
| Sen-288-529 | NZ_CJCU01000016.1 | - | 39534 | 39482 | RNA→ rpmE (PRK00019)Ribosomal_L31 (pfam01197)→ |
| Sen-288-530 | NZ_CITT01000013.1 | + | 107996 | 108048 | RNA→ rpmE (PRK00019)Ribosomal_L31 (pfam01197)→ |
| Sen-288-531 | NZ_CIVN01000015.1 | - | 39383 | 39331 | RNA→ rpmE (PRK00019)Ribosomal_L31 (pfam01197)→ |
| Sen-288-532 | NZ_CIQA01000015.1 | + | 104162 | 104214 | RNA→ rpmE (PRK00019)Ribosomal_L31 (pfam01197)→ |
| Sen-288-533 | NZ_CJFF01000015.1 | + | 103990 | 104042 | RNA→ rpmE (PRK00019)Ribosomal_L31 (pfam01197)→ |
| Sen-288-534 | NZ_CIMW01000015.1 | - | 41230 | 41178 | RNA→ rpmE (PRK00019)Ribosomal_L31 (pfam01197)→ |
| Sen-288-535 | NZ_CIYV01000016.1 | + | 104156 | 104208 | RNA→ rpmE (PRK00019)Ribosomal_L31 (pfam01197)→ |
| Sen-288-536 | NZ_CISA01000015.1 | - | 39573 | 39521 | RNA→ rpmE (PRK00019)Ribosomal_L31 (pfam01197)→ |
| Sen-288-537 | NZ_CJUL01000015.1 | + | 104157 | 104209 | RNA→ rpmE (PRK00019)Ribosomal_L31 (pfam01197)→ |
| Sen-288-538 | NZ_CITS01000016.1 | - | 39579 | 39527 | RNA→ rpmE (PRK00019)Ribosomal_L31 (pfam01197)→ |
| Sen-288-539 | NZ_CITW01000013.1 | - | 41231 | 41179 | RNA→ rpmE (PRK00019)Ribosomal_L31 (pfam01197)→ |
| Sen-288-540 | NZ_CJMY01000013.1 | + | 107986 | 108038 | RNA→ rpmE (PRK00019)Ribosomal_L31 (pfam01197)→ |
| Sen-288-541 | NZ_CEXI01000016.1 | - | 39535 | 39483 | RNA→ rpmE (PRK00019)Ribosomal_L31 (pfam01197)→ |
| Sen-288-542 | NZ_CJNJ01000013.1 | - | 39536 | 39484 | RNA→ rpmE (PRK00019)Ribosomal_L31 (pfam01197)→ |
| Sen-288-543 | NZ_CIOL01000015.1 | + | 104163 | 104215 | RNA→ rpmE (PRK00019)Ribosomal_L31 (pfam01197)→ |
| Sen-288-544 | NZ_CJJJ01000004.1 | - | 194650 | 194598 | RNA→ rpmE (PRK00019)Ribosomal_L31 (pfam01197)→ |
| Sen-288-545 | NZ_CIRQ01000015.1 | - | 39584 | 39532 | RNA→ rpmE (PRK00019)Ribosomal_L31 (pfam01197)→ |
| Sen-288-546 | NZ_CJYZ01000013.1 | + | 107698 | 107750 | RNA→ rpmE (PRK00019)Ribosomal_L31 (pfam01197)→ |
| Sen-288-547 | NZ_CIPF01000015.1 | - | 39379 | 39327 | RNA→ rpmE (PRK00019)Ribosomal_L31 (pfam01197)→ |
| Sen-288-548 | NZ_CIXQ01000016.1 | + | 104148 | 104200 | RNA→ rpmE (PRK00019)Ribosomal_L31 (pfam01197)→ |
| Sen-288-549 | NZ_CIQG01000006.1 | + | 190326 | 190378 | RNA→ rpmE (PRK00019)Ribosomal_L31 (pfam01197)→ |
| Sen-288-550 | NZ_CJWO01000014.1 | + | 104155 | 104207 | RNA→ rpmE (PRK00019)Ribosomal_L31 (pfam01197)→ |
| Sen-288-551 | NZ_CIUB01000015.1 | + | 104161 | 104213 | RNA→ rpmE (PRK00019)Ribosomal_L31 (pfam01197)→ |
| Sen-288-552 | NZ_CIZQ01000010.1 | - | 41508 | 41456 | RNA→ rpmE (PRK00019)Ribosomal_L31 (pfam01197)→ |
| Sen-288-553 | NZ_CIPU01000003.1 | - | 198388 | 198336 | RNA→ rpmE (PRK00019)Ribosomal_L31 (pfam01197)→ |
| Sen-288-554 | NZ_CIYK01000016.1 | - | 39723 | 39671 | RNA→ rpmE (PRK00019)Ribosomal_L31 (pfam01197)→ |
| Sen-288-555 | NZ_CINV01000016.1 | - | 39601 | 39549 | RNA→ rpmE (PRK00019)Ribosomal_L31 (pfam01197)→ |
| Sen-288-556 | NZ_CJOQ01000014.1 | + | 107408 | 107460 | RNA→ rpmE (PRK00019)Ribosomal_L31 (pfam01197)→ |
| Sen-288-557 | NZ_CISJ01000014.1 | + | 103990 | 104042 | RNA→ rpmE (PRK00019)Ribosomal_L31 (pfam01197)→ |
| Sen-288-558 | NZ_CEYC01000016.1 | + | 104161 | 104213 | RNA→ rpmE (PRK00019)Ribosomal_L31 (pfam01197)→ |
| Sen-288-559 | NZ_CIVG01000015.1 | - | 41690 | 41638 | RNA→ rpmE (PRK00019)Ribosomal_L31 (pfam01197)→ |
| Sen-288-560 | NZ_CIOK01000016.1 | + | 104161 | 104213 | RNA→ rpmE (PRK00019)Ribosomal_L31 (pfam01197)→ |
| Sen-288-561 | NZ_CJHL01000016.1 | + | 104140 | 104192 | RNA→ rpmE (PRK00019)Ribosomal_L31 (pfam01197)→ |
| Sen-288-562 | NZ_CINF01000015.1 | - | 39370 | 39318 | RNA→ rpmE (PRK00019)Ribosomal_L31 (pfam01197)→ |
| Sen-288-563 | NZ_CJXQ01000011.1 | - | 39540 | 39488 | RNA→ rpmE (PRK00019)Ribosomal_L31 (pfam01197)→ |
| Sen-288-564 | NZ_CIRD01000016.1 | - | 39569 | 39517 | RNA→ rpmE (PRK00019)Ribosomal_L31 (pfam01197)→ |
| Sen-288-565 | NZ_CIPQ01000013.1 | + | 107862 | 107914 | RNA→ rpmE (PRK00019)Ribosomal_L31 (pfam01197)→ |
| Sen-288-566 | NZ_CISQ01000015.1 | - | 39591 | 39539 | RNA→ rpmE (PRK00019)Ribosomal_L31 (pfam01197)→ |
| Sen-288-567 | NZ_CIQV01000016.1 | + | 104145 | 104197 | RNA→ rpmE (PRK00019)Ribosomal_L31 (pfam01197)→ |

| | | | | | |
|---|---|---|---|---|---|
| Sen-288-568 | NZ_CIQR01000015.1 | - | 39361 | 39309 | **RNA**→ rpmE (PRK00019)Ribosomal_L31 (pfam01197)→ |
| Sen-288-569 | NZ_CIPT01000015.1 | + | 104157 | 104209 | **RNA**→ rpmE (PRK00019)Ribosomal_L31 (pfam01197)→ |
| Sen-288-570 | NZ_CISD01000015.1 | + | 103990 | 104042 | **RNA**→ rpmE (PRK00019)Ribosomal_L31 (pfam01197)→ |
| Sen-288-571 | NZ_CINU01000015.1 | - | 39376 | 39324 | **RNA**→ rpmE (PRK00019)Ribosomal_L31 (pfam01197)→ |
| Sen-288-572 | NZ_CIPC01000014.1 | + | 104163 | 104215 | **RNA**→ rpmE (PRK00019)Ribosomal_L31 (pfam01197)→ |
| Sen-288-573 | NZ_CIOT01000015.1 | + | 107419 | 107471 | **RNA**→ rpmE (PRK00019)Ribosomal_L31 (pfam01197)→ |
| Sen-288-574 | NZ_CITA01000014.1 | + | 103990 | 104042 | **RNA**→ rpmE (PRK00019)Ribosomal_L31 (pfam01197)→ |
| Sen-288-575 | NZ_CISH01000015.1 | - | 39585 | 39533 | **RNA**→ rpmE (PRK00019)Ribosomal_L31 (pfam01197)→ |
| Sen-288-576 | NZ_CIQB01000016.1 | - | 39591 | 39539 | **RNA**→ rpmE (PRK00019)Ribosomal_L31 (pfam01197)→ |
| Sen-288-577 | NZ_CISF01000015.1 | + | 103990 | 104042 | **RNA**→ rpmE (PRK00019)Ribosomal_L31 (pfam01197)→ |
| Sen-288-578 | NZ_CISS01000014.1 | + | 103990 | 104042 | **RNA**→ rpmE (PRK00019)Ribosomal_L31 (pfam01197)→ |
| Sen-288-579 | NZ_CIPA01000016.1 | - | 39584 | 39532 | **RNA**→ rpmE (PRK00019)Ribosomal_L31 (pfam01197)→ |
| Sen-288-580 | NZ_CIUA01000014.1 | - | 39588 | 39536 | **RNA**→ rpmE (PRK00019)Ribosomal_L31 (pfam01197)→ |
| Sen-288-581 | NZ_CIPV01000004.1 | + | 104146 | 104198 | **RNA**→ rpmE (PRK00019)Ribosomal_L31 (pfam01197)→ |
| Sen-288-582 | NZ_CION01000015.1 | - | 41231 | 41179 | **RNA**→ rpmE (PRK00019)Ribosomal_L31 (pfam01197)→ |
| Sen-288-583 | NZ_CIPW01000013.1 | - | 41509 | 41457 | **RNA**→ rpmE (PRK00019)Ribosomal_L31 (pfam01197)→ |
| Sen-288-584 | NZ_CIOX01000006.1 | - | 39373 | 39321 | **RNA**→ rpmE (PRK00019)Ribosomal_L31 (pfam01197)→ |
| Sen-288-585 | NZ_CINQ01000018.1 | - | 39456 | 39404 | **RNA**→ rpmE (PRK00019)Ribosomal_L31 (pfam01197)→ |
| Sen-288-586 | NZ_CIQK01000016.1 | + | 104142 | 104194 | **RNA**→ rpmE (PRK00019)Ribosomal_L31 (pfam01197)→ |
| Sen-288-587 | NZ_CIQY01000014.1 | + | 107419 | 107471 | **RNA**→ rpmE (PRK00019)Ribosomal_L31 (pfam01197)→ |
| Sen-288-588 | NZ_CIOG01000003.1 | + | 103986 | 104038 | **RNA**→ rpmE (PRK00019)Ribosomal_L31 (pfam01197)→ |
| Sen-288-589 | NZ_CITX01000013.1 | - | 41509 | 41457 | **RNA**→ rpmE (PRK00019)Ribosomal_L31 (pfam01197)→ |
| Sen-288-590 | NZ_CIRZ01000014.1 | + | 103989 | 104041 | **RNA**→ rpmE (PRK00019)Ribosomal_L31 (pfam01197)→ |
| Sen-288-591 | NZ_CIQM01000016.1 | - | 39503 | 39451 | **RNA**→ rpmE (PRK00019)Ribosomal_L31 (pfam01197)→ |
| Sen-288-592 | NZ_CITZ01000012.1 | - | 41231 | 41179 | **RNA**→ rpmE (PRK00019)Ribosomal_L31 (pfam01197)→ |
| Sen-811-1 | NZ_AETB01000027.1 | - | 35284 | 35232 | **RNA**→ rpmE (PRK00019)Ribosomal_L31 (pfam01197)→ |
| Sen-814-1 | NZ_AETE01000046.1 | - | 35284 | 35232 | **RNA**→ rpmE (PRK00019)Ribosomal_L31 (pfam01197)→ |
| Sen-838-1 | NZ_AETS01000077.1 | - | 35207 | 35155 | **RNA**→ rpmE (PRK00019)Ribosomal_L31 (pfam01197)→ |
| Sen-831-1 | NZ_AHHS01000107.1 | + | 97829 | 97881 | **RNA**→ rpmE (PRK00019)Ribosomal_L31 (pfam01197)→ |
| Sen-231-1 | NZ_ARYT01000025.1 | + | 101525 | 101577 | **RNA**→ rpmE (PRK00019)Ribosomal_L31 (pfam01197)→ |
| Sen-803-1 | NZ_AESV01000060.1 | + | 97648 | 97700 | **RNA**→ rpmE (PRK00019)Ribosomal_L31 (pfam01197)→ |
| Sen-821-1 | NZ_AETG01000061.1 | - | 35207 | 35155 | **RNA**→ rpmE (PRK00019)Ribosomal_L31 (pfam01197)→ |
| Sen-801-1 | NZ_ALPQ01000042.1 | + | 8188 | 8240 | **RNA**→ rpmE (PRK00019)Ribosomal_L31 (pfam01197)→ |
| Sen-826-1 | NZ_AESZ01000035.1 | - | 35284 | 35232 | **RNA**→ rpmE (PRK00019)Ribosomal_L31 (pfam01197)→ |
| Sen-807-1 | NZ_AETQ01000019.1 | + | 97839 | 97891 | **RNA**→ rpmE (PRK00019)Ribosomal_L31 (pfam01197)→ |
| Sen-829-1 | NZ_AESQ01000056.1 | + | 97651 | 97703 | **RNA**→ rpmE (PRK00019)Ribosomal_L31 (pfam01197)→ |
| Sen-824-1 | NZ_AETH01000095.1 | + | 97835 | 97887 | **RNA**→ rpmE (PRK00019)Ribosomal_L31 (pfam01197)→ |
| Sen-800-1 | NZ_AOYO01000024.1 | - | 35210 | 35158 | **RNA**→ rpmE (PRK00019)Ribosomal_L31 (pfam01197)→ rpmE (PRK00019)Ribosomal_L31 (pfam01197)→ |
| Sen-819-1 | NZ_AESM01000030.1 | - | 35648 | 35596 | **RNA**→ rpmE (PRK00019)Ribosomal_L31 (pfam01197)→ |
| Sen-808-1 | NZ_AESH01000041.1 | - | 35284 | 35232 | **RNA**→ rpmE (PRK00019)Ribosomal_L31 (pfam01197)→ |
| Sen-815-1 | NZ_AESI01000032.1 | + | 97839 | 97891 | **RNA**→ rpmE (PRK00019)Ribosomal_L31 (pfam01197)→ |
| Sen-234-1 | NZ_AHUX01000027.1 | + | 79706 | 79758 | **RNA**→ rpmE (PRK00019)Ribosomal_L31 (pfam01197)→ |
| Sen-827-1 | NZ_AOYQ01000006.1 | + | 97727 | 97779 | **RNA**→ rpmE (PRK00019)Ribosomal_L31 (pfam01197)→ rpmE (PRK00019)Ribosomal_L31 (pfam01197)→ |
| Sen-804-1 | NZ_AETO01000003.1 | - | 35284 | 35232 | **RNA**→ rpmE (PRK00019)Ribosomal_L31 (pfam01197)→ |
| Sen-288-593 | NZ_CINB01000016.1 | - | 39380 | 39328 | **RNA**→ rpmE (PRK00019)Ribosomal_L31 (pfam01197)→ |
| Sen-812-1 | NZ_AESX01000043.1 | + | 98859 | 98911 | **RNA**→ rpmE (PRK00019)Ribosomal_L31 (pfam01197)→ rpmE (PRK00019)Ribosomal_L31 (pfam01197)→ |
| Sen-288-594 | NZ_CIRJ01000011.1 | - | 39370 | 39318 | **RNA**→ rpmE (PRK00019)Ribosomal_L31 (pfam01197)→ |
| Sen-793-1 | NZ_AUQI01000090.1 | - | 37855 | 37803 | **RNA**→ rpmE (PRK00019)Ribosomal_L31 (pfam01197)→ |
| Sen-288-595 | NZ_CJYW01000015.1 | - | 39580 | 39528 | **RNA**→ rpmE (PRK00019)Ribosomal_L31 (pfam01197)→ |
| Sen-809-1 | NZ_AETC01000022.1 | - | 35310 | 35258 | **RNA**→ rpmE (PRK00019)Ribosomal_L31 (pfam01197)→ |
| Sen-288-596 | NZ_CIWN01000014.1 | + | 104144 | 104196 | **RNA**→ rpmE (PRK00019)Ribosomal_L31 (pfam01197)→ |
| Sen-818-1 | NZ_AESL01000022.1 | + | 97839 | 97891 | **RNA**→ rpmE (PRK00019)Ribosomal_L31 (pfam01197)→ |

| | | | | | |
|---|---|---|---|---|---|
| Sen-288-597 | NZ_CJCA01000015.1 | + | 104180 | 104232 | **RNA**→ rpmE (PRK00019)Ribosomal_L31 (pfam01197)→ |
| Sen-825-1 | NZ_AHHR01000028.1 | - | 35284 | 35232 | **RNA**→ rpmE (PRK00019)Ribosomal_L31 (pfam01197)→ |
| Sen-288-598 | NZ_CJWH01000016.1 | + | 104162 | 104214 | **RNA**→ rpmE (PRK00019)Ribosomal_L31 (pfam01197)→ |
| Sen-832-1 | NZ_AHHT01000024.1 | - | 35284 | 35232 | **RNA**→ rpmE (PRK00019)Ribosomal_L31 (pfam01197)→ |
| Sen-288-599 | NZ_CJST01000016.1 | + | 104158 | 104210 | **RNA**→ rpmE (PRK00019)Ribosomal_L31 (pfam01197)→ |
| Sen-823-1 | NZ_AETF01000059.1 | - | 35284 | 35232 | **RNA**→ rpmE (PRK00019)Ribosomal_L31 (pfam01197)→ |
| Sen-288-600 | NZ_CJGA01000013.1 | + | 104148 | 104200 | **RNA**→ rpmE (PRK00019)Ribosomal_L31 (pfam01197)→ |
| Sen-840-1 | NZ_AETU01000084.1 | - | 35207 | 35155 | **RNA**→ rpmE (PRK00019)Ribosomal_L31 (pfam01197)→ |
| Sen-288-601 | NZ_CJNP01000015.1 | + | 104157 | 104209 | **RNA**→ rpmE (PRK00019)Ribosomal_L31 (pfam01197)→ |
| Sen-839-1 | NZ_AETT01000012.1 | - | 35208 | 35156 | **RNA**→ rpmE (PRK00019)Ribosomal_L31 (pfam01197)→ |
| Sen-288-602 | NZ_CJGY01000016.1 | + | 104142 | 104194 | **RNA**→ rpmE (PRK00019)Ribosomal_L31 (pfam01197)→ |
| Sen-288-603 | NZ_CINP01000009.1 | - | 39456 | 39404 | **RNA**→ rpmE (PRK00019)Ribosomal_L31 (pfam01197)→ |
| Sen-798-1 | NZ_AOYN01000139.1 | - | 45480 | 45428 | **RNA**→ rpmE (PRK00019)Ribosomal_L31 (pfam01197)→ |
| Sen-288-604 | NZ_CIQH01000013.1 | + | 107423 | 107475 | **RNA**→ rpmE (PRK00019)Ribosomal_L31 (pfam01197)→ |
| Sen-799-1 | NZ_AOYS01000026.1 | - | 40814 | 40762 | **RNA**→ rpmE (PRK00019)Ribosomal_L31 (pfam01197)→ |
| Sen-288-605 | NZ_CJIN01000016.1 | + | 104146 | 104198 | **RNA**→ rpmE (PRK00019)Ribosomal_L31 (pfam01197)→ |
| Sen-288-606 | NZ_CILJ01000015.1 | - | 39365 | 39313 | **RNA**→ rpmE (PRK00019)Ribosomal_L31 (pfam01197)→ |
| Sen-794-1 | NZ_APAN01000075.1 | + | 100234 | 100286 | **RNA**→ rpmE (PRK00019)Ribosomal_L31 (pfam01197)→ |
| Sen-288-607 | NZ_CJKX01000014.1 | - | 39456 | 39404 | **RNA**→ rpmE (PRK00019)Ribosomal_L31 (pfam01197)→ |
| Sen-805-1 | NZ_AETP01000016.1 | - | 35207 | 35155 | **RNA**→ rpmE (PRK00019)Ribosomal_L31 (pfam01197)→ |
| Sen-288-608 | NZ_CJTV01000016.1 | + | 104157 | 104209 | **RNA**→ rpmE (PRK00019)Ribosomal_L31 (pfam01197)→ |
| Sen-810-1 | NZ_AETD01000015.1 | + | 97636 | 97688 | **RNA**→ rpmE (PRK00019)Ribosomal_L31 (pfam01197)→ |
| Sen-288-609 | NZ_CJFM01000016.1 | + | 104015 | 104067 | **RNA**→ rpmE (PRK00019)Ribosomal_L31 (pfam01197)→ |
| Sen-816-1 | NZ_AESJ01000001.1 | + | 97835 | 97887 | **RNA**→ rpmE (PRK00019)Ribosomal_L31 (pfam01197)→ |
| Sen-288-610 | NZ_CJDL01000016.1 | + | 104150 | 104202 | **RNA**→ rpmE (PRK00019)Ribosomal_L31 (pfam01197)→ |
| Sen-822-1 | NZ_AETJ01000084.1 | - | 35207 | 35155 | **RNA**→ rpmE (PRK00019)Ribosomal_L31 (pfam01197)→ |
| Sen-288-611 | NZ_CJGW01000016.1 | - | 39535 | 39483 | **RNA**→ rpmE (PRK00019)Ribosomal_L31 (pfam01197)→ |
| Sen-797-1 | NZ_AOYR01000043.1 | - | 37499 | 37447 | **RNA**→ rpmE (PRK00019)Ribosomal_L31 (pfam01197)→ |
| Sen-288-612 | NZ_CJJX01000016.1 | - | 39711 | 39659 | **RNA**→ rpmE (PRK00019)Ribosomal_L31 (pfam01197)→ |
| Sen-288-613 | NZ_CIWC01000015.1 | - | 39456 | 39404 | **RNA**→ rpmE (PRK00019)Ribosomal_L31 (pfam01197)→ |
| Sen-288-614 | NZ_CJPI01000016.1 | + | 104156 | 104208 | **RNA**→ rpmE (PRK00019)Ribosomal_L31 (pfam01197)→ |
| Sen-795-1 | NZ_APAL01000294.1 | - | 2980 | 2928 | **RNA**→ rpmE (PRK00019)Ribosomal_L31 (pfam01197)→ rpmE (PRK00019)Ribosomal_L31 (pfam01197)→ |
| Sen-288-615 | NZ_CJPC01000016.1 | + | 104162 | 104214 | **RNA**→ rpmE (PRK00019)Ribosomal_L31 (pfam01197)→ |
| Sen-288-616 | NZ_CJPQ01000015.1 | + | 104156 | 104208 | **RNA**→ rpmE (PRK00019)Ribosomal_L31 (pfam01197)→ |
| Sen-830-1 | NZ_AYUN01000226.1 | - | 4846 | 4794 | **RNA**→ rpmE (PRK00019)Ribosomal_L31 (pfam01197)→ |
| Sen-288-617 | NZ_CJJA01000014.1 | - | 41503 | 41451 | **RNA**→ rpmE (PRK00019)Ribosomal_L31 (pfam01197)→ |
| Sen-820-1 | NZ_AESN01000032.1 | + | 79699 | 79751 | **RNA**→ rpmE (PRK00019)Ribosomal_L31 (pfam01197)→ |
| Sen-288-618 | NZ_CIVY01000014.1 | - | 41526 | 41474 | **RNA**→ rpmE (PRK00019)Ribosomal_L31 (pfam01197)→ |
| Sen-288-619 | NZ_CIKZ01000013.1 | + | 104165 | 104217 | **RNA**→ rpmE (PRK00019)Ribosomal_L31 (pfam01197)→ |
| Sen-796-1 | NZ_AMRS01000013.1 | - | 37964 | 37912 | **RNA**→ rpmE (PRK00019)Ribosomal_L31 (pfam01197)→ |
| Sen-835-1 | NZ_AHHW01000009.1 | + | 95080 | 95132 | **RNA**→ rpmE (PRK00019)Ribosomal_L31 (pfam01197)→ |
| Sen-288-620 | NZ_CISK01000005.1 | - | 198400 | 198348 | **RNA**→ rpmE (PRK00019)Ribosomal_L31 (pfam01197)→ |
| Sen-288-621 | NZ_CIQC01000013.1 | + | 104161 | 104213 | **RNA**→ rpmE (PRK00019)Ribosomal_L31 (pfam01197)→ |
| Sen-837-1 | NZ_AETR01000029.1 | + | 97844 | 97896 | **RNA**→ rpmE (PRK00019)Ribosomal_L31 (pfam01197)→ |
| Sen-288-622 | NZ_CIPM01000001.1 | - | 39375 | 39323 | **RNA**→ rpmE (PRK00019)Ribosomal_L31 (pfam01197)→ |
| Sen-836-1 | NZ_JMMJ01000005.1 | - | 28770 | 28718 | **RNA**→ rpmE (PRK00019)Ribosomal_L31 (pfam01197)→ |
| Sen-288-623 | NZ_CEWW01000016.1 | - | 39533 | 39481 | **RNA**→ rpmE (PRK00019)Ribosomal_L31 (pfam01197)→ |
| Sen-288-624 | NZ_CILD01000015.1 | + | 104049 | 104101 | **RNA**→ rpmE (PRK00019)Ribosomal_L31 (pfam01197)→ |
| Sen-288-625 | NZ_CJRV01000010.1 | - | 39557 | 39505 | **RNA**→ rpmE (PRK00019)Ribosomal_L31 (pfam01197)→ |
| Sen-828-1 | NZ_AESS01000101.1 | + | 73609 | 73661 | **RNA**→ rpmE (PRK00019)Ribosomal_L31 (pfam01197)→ rpmE (PRK00019)Ribosomal_L31 (pfam01197)→ |
| Sen-806-1 | NZ_AESW01000001.1 | + | 97696 | 97748 | **RNA**→ rpmE (PRK00019)Ribosomal_L31 (pfam01197)→ rpmE (PRK00019)Ribosomal_L31 (pfam01197)→ |
| Sen-288-626 | NZ_CJNA01000014.1 | - | 41509 | 41457 | **RNA**→ rpmE (PRK00019)Ribosomal_L31 (pfam01197)→ |

| | | | | | |
|---|---|---|---|---|---|
| Sen-288-627 | NZ_CING01000015.1 | + | 104008 | 104060 | RNA→ rpmE (PRK00019)Ribosomal_L31 (pfam01197)→ |
| Sen-792-1 | NZ_AOYZ01000072.1 | - | 37522 | 37470 | RNA→ rpmE (PRK00019)Ribosomal_L31 (pfam01197)→ |
| Sen-288-628 | NZ_CIUQ01000016.1 | - | 39541 | 39489 | RNA→ rpmE (PRK00019)Ribosomal_L31 (pfam01197)→ |
| Sen-834-1 | NZ_AHHV01000035.1 | + | 97838 | 97890 | RNA→ rpmE (PRK00019)Ribosomal_L31 (pfam01197)→ |
| Sen-288-629 | NZ_CIUP01000015.1 | + | 108000 | 108052 | RNA→ rpmE (PRK00019)Ribosomal_L31 (pfam01197)→ |
| Sen-791-1 | NZ_AHUU01000022.1 | + | 110824 | 110876 | RNA→ rpmE (PRK00019)Ribosomal_L31 (pfam01197)→ |
| Sen-288-630 | NZ_CJBC01000016.1 | - | 39766 | 39714 | RNA→ rpmE (PRK00019)Ribosomal_L31 (pfam01197)→ |
| Sen-288-631 | NZ_CJRS01000015.1 | - | 39549 | 39497 | RNA→ rpmE (PRK00019)Ribosomal_L31 (pfam01197)→ |
| Sen-817-1 | NZ_AESK01000040.1 | - | 35284 | 35232 | RNA→ rpmE (PRK00019)Ribosomal_L31 (pfam01197)→ |
| Sen-833-1 | NZ_AHHU01000014.1 | + | 97835 | 97887 | RNA→ rpmE (PRK00019)Ribosomal_L31 (pfam01197)→ |
| Sen-288-632 | NZ_CJUF01000016.1 | - | 39702 | 39650 | RNA→ rpmE (PRK00019)Ribosomal_L31 (pfam01197)→ |
| Sen-288-633 | NZ_CITD01000015.1 | - | 39586 | 39534 | RNA→ rpmE (PRK00019)Ribosomal_L31 (pfam01197)→ |
| Sen-288-634 | NZ_CIQI01000003.1 | + | 104147 | 104199 | RNA→ rpmE (PRK00019)Ribosomal_L31 (pfam01197)→ |
| Sen-288-635 | NZ_CJXU01000016.1 | - | 39456 | 39404 | RNA→ rpmE (PRK00019)Ribosomal_L31 (pfam01197)→ |
| Sen-288-636 | NZ_CJCB01000014.1 | - | 39572 | 39520 | RNA→ rpmE (PRK00019)Ribosomal_L31 (pfam01197)→ |
| Sen-288-637 | NZ_CJSI01000014.1 | + | 107419 | 107471 | RNA→ rpmE (PRK00019)Ribosomal_L31 (pfam01197)→ |
| Sen-288-638 | NZ_CJEN01000016.1 | + | 104147 | 104199 | RNA→ rpmE (PRK00019)Ribosomal_L31 (pfam01197)→ |
| Sen-288-639 | NZ_CIZT01000016.1 | + | 104159 | 104211 | RNA→ rpmE (PRK00019)Ribosomal_L31 (pfam01197)→ |
| Sen-288-640 | NZ_CJWB01000014.1 | - | 41231 | 41179 | RNA→ rpmE (PRK00019)Ribosomal_L31 (pfam01197)→ |
| Sen-288-641 | NZ_CJFZ01000016.1 | - | 39537 | 39485 | RNA→ rpmE (PRK00019)Ribosomal_L31 (pfam01197)→ |
| Sen-288-642 | NZ_CJJL01000015.1 | - | 39562 | 39510 | RNA→ rpmE (PRK00019)Ribosomal_L31 (pfam01197)→ |
| Sen-288-643 | NZ_CIXB01000015.1 | - | 39614 | 39562 | RNA→ rpmE (PRK00019)Ribosomal_L31 (pfam01197)→ |
| Sen-288-644 | NZ_CJWY01000016.1 | + | 104160 | 104212 | RNA→ rpmE (PRK00019)Ribosomal_L31 (pfam01197)→ |
| Sen-288-645 | NZ_CJTC01000016.1 | - | 39383 | 39331 | RNA→ rpmE (PRK00019)Ribosomal_L31 (pfam01197)→ |
| Sen-288-646 | NZ_CJWJ01000015.1 | - | 39363 | 39311 | RNA→ rpmE (PRK00019)Ribosomal_L31 (pfam01197)→ |
| Sen-288-647 | NZ_CJNT01000013.1 | + | 107992 | 108044 | RNA→ rpmE (PRK00019)Ribosomal_L31 (pfam01197)→ |
| Sen-288-648 | NZ_CJGX01000014.1 | - | 39575 | 39523 | RNA→ rpmE (PRK00019)Ribosomal_L31 (pfam01197)→ |
| Sen-288-649 | NZ_CILR01000015.1 | + | 109294 | 109346 | RNA→ rpmE (PRK00019)Ribosomal_L31 (pfam01197)→ |
| Sen-288-650 | NZ_CJCH01000015.1 | - | 39573 | 39521 | RNA→ rpmE (PRK00019)Ribosomal_L31 (pfam01197)→ |
| Sen-288-651 | NZ_CIZW01000016.1 | + | 104148 | 104200 | RNA→ rpmE (PRK00019)Ribosomal_L31 (pfam01197)→ |
| Sen-288-652 | NZ_CJRU01000015.1 | + | 104162 | 104214 | RNA→ rpmE (PRK00019)Ribosomal_L31 (pfam01197)→ |
| Sen-288-653 | NZ_CJFE01000016.1 | + | 104137 | 104189 | RNA→ rpmE (PRK00019)Ribosomal_L31 (pfam01197)→ |
| Sen-288-654 | NZ_CJAP01000017.1 | - | 39539 | 39487 | RNA→ rpmE (PRK00019)Ribosomal_L31 (pfam01197)→ |
| Sen-288-655 | NZ_CJKO01000009.1 | + | 133799 | 133851 | RNA→ rpmE (PRK00019)Ribosomal_L31 (pfam01197)→ |
| Sen-288-656 | NZ_CJJS01000013.1 | - | 41505 | 41453 | RNA→ rpmE (PRK00019)Ribosomal_L31 (pfam01197)→ |
| Sen-288-657 | NZ_CJAJ01000016.1 | + | 104153 | 104205 | RNA→ rpmE (PRK00019)Ribosomal_L31 (pfam01197)→ |
| Sen-288-658 | NZ_CJTW01000015.1 | - | 41512 | 41460 | RNA→ rpmE (PRK00019)Ribosomal_L31 (pfam01197)→ |
| Sen-288-659 | NZ_CJNL01000015.1 | + | 104158 | 104210 | RNA→ rpmE (PRK00019)Ribosomal_L31 (pfam01197)→ |
| Sen-288-660 | NZ_CJNU01000016.1 | - | 39708 | 39656 | RNA→ rpmE (PRK00019)Ribosomal_L31 (pfam01197)→ |
| Sen-288-661 | NZ_CJHH01000013.1 | - | 39563 | 39511 | RNA→ rpmE (PRK00019)Ribosomal_L31 (pfam01197)→ |
| Sen-288-662 | NZ_CJDO01000014.1 | - | 39535 | 39483 | RNA→ rpmE (PRK00019)Ribosomal_L31 (pfam01197)→ |
| Sen-288-663 | NZ_CJSQ01000016.1 | - | 39361 | 39309 | RNA→ rpmE (PRK00019)Ribosomal_L31 (pfam01197)→ |
| Sen-288-664 | NZ_CJDE01000014.1 | + | 107811 | 107863 | RNA→ rpmE (PRK00019)Ribosomal_L31 (pfam01197)→ |
| Sen-288-665 | NZ_CIXS01000016.1 | + | 104124 | 104176 | RNA→ rpmE (PRK00019)Ribosomal_L31 (pfam01197)→ |
| Sen-288-666 | NZ_CIXH01000014.1 | + | 104249 | 104301 | RNA→ rpmE (PRK00019)Ribosomal_L31 (pfam01197)→ |
| Sen-288-667 | NZ_CJHX01000015.1 | - | 39530 | 39478 | RNA→ rpmE (PRK00019)Ribosomal_L31 (pfam01197)→ |
| Sen-288-668 | NZ_CJEA01000016.1 | + | 104151 | 104203 | RNA→ rpmE (PRK00019)Ribosomal_L31 (pfam01197)→ |
| Sen-288-669 | NZ_CJFI01000015.1 | - | 39532 | 39480 | RNA→ rpmE (PRK00019)Ribosomal_L31 (pfam01197)→ |
| Sen-288-670 | NZ_CJQL01000016.1 | + | 104172 | 104224 | RNA→ rpmE (PRK00019)Ribosomal_L31 (pfam01197)→ |
| Sen-288-671 | NZ_CJKR01000015.1 | + | 104124 | 104176 | RNA→ rpmE (PRK00019)Ribosomal_L31 (pfam01197)→ |
| Sen-288-672 | NZ_CJHS01000016.1 | + | 104160 | 104212 | RNA→ rpmE (PRK00019)Ribosomal_L31 (pfam01197)→ |
| Sen-288-673 | NZ_CJKC01000013.1 | + | 104143 | 104195 | RNA→ rpmE (PRK00019)Ribosomal_L31 (pfam01197)→ |

| | | | | | |
|---|---|---|---|---|---|
| Sen-288-674 | NZ_CIVA01000015.1 | - | 39386 | 39334 | **RNA**→ rpmE (PRK00019)Ribosomal_L31 (pfam01197)→ |
| Sen-288-675 | NZ_CJPG01000008.1 | + | 159488 | 159540 | **RNA**→ rpmE (PRK00019)Ribosomal_L31 (pfam01197)→ |
| Sen-288-676 | NZ_CJTX01000015.1 | - | 39371 | 39319 | **RNA**→ rpmE (PRK00019)Ribosomal_L31 (pfam01197)→ |
| Sen-288-677 | NZ_CJOA01000016.1 | + | 104153 | 104205 | **RNA**→ rpmE (PRK00019)Ribosomal_L31 (pfam01197)→ |
| Sen-288-678 | NZ_CJEW01000016.1 | - | 39532 | 39480 | **RNA**→ rpmE (PRK00019)Ribosomal_L31 (pfam01197)→ |
| Sen-288-679 | NZ_CJZA01000016.1 | - | 39540 | 39488 | **RNA**→ rpmE (PRK00019)Ribosomal_L31 (pfam01197)→ |
| Sen-288-680 | NZ_CIXN01000015.1 | + | 103986 | 104038 | **RNA**→ rpmE (PRK00019)Ribosomal_L31 (pfam01197)→ |
| Sen-288-681 | NZ_CJWM01000014.1 | + | 104141 | 104193 | **RNA**→ rpmE (PRK00019)Ribosomal_L31 (pfam01197)→ |
| Sen-288-682 | NZ_CJEC01000016.1 | - | 39536 | 39484 | **RNA**→ rpmE (PRK00019)Ribosomal_L31 (pfam01197)→ |
| Sen-288-683 | NZ_CJCN01000015.1 | - | 39538 | 39486 | **RNA**→ rpmE (PRK00019)Ribosomal_L31 (pfam01197)→ |
| Sen-288-684 | NZ_CJOU01000016.1 | + | 104172 | 104224 | **RNA**→ rpmE (PRK00019)Ribosomal_L31 (pfam01197)→ |
| Sen-288-685 | NZ_CJKW01000011.1 | - | 39456 | 39404 | **RNA**→ rpmE (PRK00019)Ribosomal_L31 (pfam01197)→ |
| Sen-288-686 | NZ_CJCR01000016.1 | - | 39597 | 39545 | **RNA**→ rpmE (PRK00019)Ribosomal_L31 (pfam01197)→ |
| Sen-288-687 | NZ_CJVS01000016.1 | - | 39366 | 39314 | **RNA**→ rpmE (PRK00019)Ribosomal_L31 (pfam01197)→ |
| Sen-288-688 | NZ_CJES01000016.1 | + | 104133 | 104185 | **RNA**→ rpmE (PRK00019)Ribosomal_L31 (pfam01197)→ |
| Sen-288-689 | NZ_CJXN01000017.1 | + | 104165 | 104217 | **RNA**→ rpmE (PRK00019)Ribosomal_L31 (pfam01197)→ |
| Sen-288-690 | NZ_CIZG01000016.1 | + | 104156 | 104208 | **RNA**→ rpmE (PRK00019)Ribosomal_L31 (pfam01197)→ |
| Sen-288-691 | NZ_CJCY01000013.1 | - | 39535 | 39483 | **RNA**→ rpmE (PRK00019)Ribosomal_L31 (pfam01197)→ |
| Sen-288-692 | NZ_CJNM01000012.1 | - | 41512 | 41460 | **RNA**→ rpmE (PRK00019)Ribosomal_L31 (pfam01197)→ |
| Sen-288-693 | NZ_CIOP01000015.1 | + | 104153 | 104205 | **RNA**→ rpmE (PRK00019)Ribosomal_L31 (pfam01197)→ |
| Sen-288-694 | NZ_CITB01000013.1 | - | 39583 | 39531 | **RNA**→ rpmE (PRK00019)Ribosomal_L31 (pfam01197)→ |
| Sen-288-695 | NZ_CIRE01000014.1 | - | 41231 | 41179 | **RNA**→ rpmE (PRK00019)Ribosomal_L31 (pfam01197)→ |
| Sen-288-696 | NZ_CIRP01000016.1 | + | 104151 | 104203 | **RNA**→ rpmE (PRK00019)Ribosomal_L31 (pfam01197)→ |
| Sen-288-697 | NZ_CIQQ01000014.1 | + | 104157 | 104209 | **RNA**→ rpmE (PRK00019)Ribosomal_L31 (pfam01197)→ |
| Sen-288-698 | NZ_CIRY01000014.1 | + | 103989 | 104041 | **RNA**→ rpmE (PRK00019)Ribosomal_L31 (pfam01197)→ |
| Sen-288-699 | NZ_CITG01000015.1 | + | 103990 | 104042 | **RNA**→ rpmE (PRK00019)Ribosomal_L31 (pfam01197)→ |
| Sen-288-700 | NZ_CIPO01000007.1 | + | 170843 | 170895 | **RNA**→ rpmE (PRK00019)Ribosomal_L31 (pfam01197)→ |
| Sen-288-701 | NZ_CIPX01000015.1 | + | 104162 | 104214 | **RNA**→ rpmE (PRK00019)Ribosomal_L31 (pfam01197)→ |
| Sen-288-702 | NZ_CITY01000015.1 | + | 104162 | 104214 | **RNA**→ rpmE (PRK00019)Ribosomal_L31 (pfam01197)→ |
| Sen-288-703 | NZ_CIRG01000002.1 | - | 198392 | 198340 | **RNA**→ rpmE (PRK00019)Ribosomal_L31 (pfam01197)→ |
| Sen-288-704 | NZ_CINL01000015.1 | - | 39382 | 39330 | **RNA**→ rpmE (PRK00019)Ribosomal_L31 (pfam01197)→ |
| Sen-288-705 | NZ_CITV01000013.1 | + | 107862 | 107914 | **RNA**→ rpmE (PRK00019)Ribosomal_L31 (pfam01197)→ |
| Sen-288-706 | NZ_CIOV01000014.1 | + | 104206 | 104258 | **RNA**→ rpmE (PRK00019)Ribosomal_L31 (pfam01197)→ |
| Sen-288-707 | NZ_CIOI01000016.1 | + | 104160 | 104212 | **RNA**→ rpmE (PRK00019)Ribosomal_L31 (pfam01197)→ |
| Sen-288-708 | NZ_CISM01000014.1 | + | 103990 | 104042 | **RNA**→ rpmE (PRK00019)Ribosomal_L31 (pfam01197)→ |
| Sen-288-709 | NZ_CIRR01000008.1 | + | 152660 | 152712 | **RNA**→ rpmE (PRK00019)Ribosomal_L31 (pfam01197)→ |
| Sen-288-710 | NZ_CISE01000014.1 | + | 104134 | 104186 | **RNA**→ rpmE (PRK00019)Ribosomal_L31 (pfam01197)→ |
| Sen-288-711 | NZ_CIPB01000016.1 | + | 104163 | 104215 | **RNA**→ rpmE (PRK00019)Ribosomal_L31 (pfam01197)→ |
| Sen-288-712 | NZ_CILB01000015.1 | + | 104166 | 104218 | **RNA**→ rpmE (PRK00019)Ribosomal_L31 (pfam01197)→ |
| Sen-288-713 | NZ_CIQD01000016.1 | + | 104150 | 104202 | **RNA**→ rpmE (PRK00019)Ribosomal_L31 (pfam01197)→ |
| Sen-288-714 | NZ_CITC01000015.1 | - | 39574 | 39522 | **RNA**→ rpmE (PRK00019)Ribosomal_L31 (pfam01197)→ |
| Sen-288-715 | NZ_CISO01000015.1 | + | 103990 | 104042 | **RNA**→ rpmE (PRK00019)Ribosomal_L31 (pfam01197)→ |
| Sen-288-716 | NZ_CIQN01000016.1 | + | 104152 | 104204 | **RNA**→ rpmE (PRK00019)Ribosomal_L31 (pfam01197)→ |
| Sen-288-717 | NZ_CIQX01000015.1 | - | 39367 | 39315 | **RNA**→ rpmE (PRK00019)Ribosomal_L31 (pfam01197)→ |
| Sen-288-718 | NZ_CIRF01000016.1 | - | 39361 | 39309 | **RNA**→ rpmE (PRK00019)Ribosomal_L31 (pfam01197)→ |
| Sen-288-719 | NZ_CIOF01000014.1 | - | 41231 | 41179 | **RNA**→ rpmE (PRK00019)Ribosomal_L31 (pfam01197)→ |
| Sen-288-720 | NZ_CIUG01000016.1 | + | 104161 | 104213 | **RNA**→ rpmE (PRK00019)Ribosomal_L31 (pfam01197)→ |
| Sen-288-721 | NZ_CISX01000014.1 | + | 103990 | 104042 | **RNA**→ rpmE (PRK00019)Ribosomal_L31 (pfam01197)→ |
| Sen-288-722 | NZ_CITJ01000014.1 | - | 39374 | 39322 | **RNA**→ rpmE (PRK00019)Ribosomal_L31 (pfam01197)→ |
| Sen-288-723 | NZ_CIOR01000014.1 | + | 104163 | 104215 | **RNA**→ rpmE (PRK00019)Ribosomal_L31 (pfam01197)→ |
| Sen-288-724 | NZ_CILA01000011.1 | - | 39571 | 39519 | **RNA**→ rpmE (PRK00019)Ribosomal_L31 (pfam01197)→ |
| Sen-288-725 | NZ_CIRW01000015.1 | + | 104133 | 104185 | **RNA**→ rpmE (PRK00019)Ribosomal_L31 (pfam01197)→ |

| | | | | | |
|---|---|---|---|---|---|
| Sen-288-726 | NZ_CIRL01000015.1 | - | 39588 | 39536 | **RNA**→ rpmE (PRK00019)Ribosomal_L31 (pfam01197)→ |
| Sen-288-727 | NZ_CISV01000015.1 | + | 103990 | 104042 | **RNA**→ rpmE (PRK00019)Ribosomal_L31 (pfam01197)→ |
| Sen-288-728 | NZ_CINO01000016.1 | + | 104157 | 104209 | **RNA**→ rpmE (PRK00019)Ribosomal_L31 (pfam01197)→ |
| Sen-288-729 | NZ_CINT01000015.1 | - | 39589 | 39537 | **RNA**→ rpmE (PRK00019)Ribosomal_L31 (pfam01197)→ |
| Sen-288-730 | NZ_CIZU01000016.1 | - | 39536 | 39484 | **RNA**→ rpmE (PRK00019)Ribosomal_L31 (pfam01197)→ |
| Sen-288-731 | NZ_CIUF01000015.1 | + | 104167 | 104219 | **RNA**→ rpmE (PRK00019)Ribosomal_L31 (pfam01197)→ |
| Sen-288-732 | NZ_CJQC01000008.1 | + | 152693 | 152745 | **RNA**→ rpmE (PRK00019)Ribosomal_L31 (pfam01197)→ |
| Sen-288-733 | NZ_CJON01000014.1 | - | 39844 | 39792 | **RNA**→ rpmE (PRK00019)Ribosomal_L31 (pfam01197)→ |
| Sen-288-734 | NZ_CJZE01000016.1 | - | 39361 | 39309 | **RNA**→ rpmE (PRK00019)Ribosomal_L31 (pfam01197)→ |
| Sen-288-735 | NZ_CJUR01000016.1 | - | 39365 | 39313 | **RNA**→ rpmE (PRK00019)Ribosomal_L31 (pfam01197)→ |
| Sen-288-736 | NZ_CJSR01000015.1 | + | 104161 | 104213 | **RNA**→ rpmE (PRK00019)Ribosomal_L31 (pfam01197)→ |
| Sen-288-737 | NZ_CJTT01000014.1 | + | 107817 | 107869 | **RNA**→ rpmE (PRK00019)Ribosomal_L31 (pfam01197)→ |
| Sen-288-738 | NZ_CJUC01000015.1 | + | 104153 | 104205 | **RNA**→ rpmE (PRK00019)Ribosomal_L31 (pfam01197)→ |
| Sen-288-739 | NZ_CIVZ01000016.1 | - | 39549 | 39497 | **RNA**→ rpmE (PRK00019)Ribosomal_L31 (pfam01197)→ |
| Sen-288-740 | NZ_CJMM01000014.1 | + | 107827 | 107879 | **RNA**→ rpmE (PRK00019)Ribosomal_L31 (pfam01197)→ |
| Sen-288-741 | NZ_CIVJ01000014.1 | - | 41231 | 41179 | **RNA**→ rpmE (PRK00019)Ribosomal_L31 (pfam01197)→ |
| Sen-288-742 | NZ_CJCX01000016.1 | + | 104152 | 104204 | **RNA**→ rpmE (PRK00019)Ribosomal_L31 (pfam01197)→ |
| Sen-288-743 | NZ_CJYJ01000015.1 | + | 104154 | 104206 | **RNA**→ rpmE (PRK00019)Ribosomal_L31 (pfam01197)→ |
| Sen-288-744 | NZ_CJJB01000015.1 | + | 107662 | 107714 | **RNA**→ rpmE (PRK00019)Ribosomal_L31 (pfam01197)→ |
| Sen-288-745 | NZ_CJOG01000016.1 | - | 39541 | 39489 | **RNA**→ rpmE (PRK00019)Ribosomal_L31 (pfam01197)→ |
| Sen-288-746 | NZ_CJGN01000016.1 | - | 39537 | 39485 | **RNA**→ rpmE (PRK00019)Ribosomal_L31 (pfam01197)→ |
| Sen-288-747 | NZ_CJGI01000002.1 | + | 405531 | 405583 | **RNA**→ rpmE (PRK00019)Ribosomal_L31 (pfam01197)→ |
| Sen-288-748 | NZ_CJMV01000003.1 | - | 216953 | 216901 | **RNA**→ rpmE (PRK00019)Ribosomal_L31 (pfam01197)→ |
| Sen-288-749 | NZ_CJTP01000015.1 | + | 107862 | 107914 | **RNA**→ rpmE (PRK00019)Ribosomal_L31 (pfam01197)→ |
| Sen-288-750 | NZ_CJOF01000016.1 | + | 104155 | 104207 | **RNA**→ rpmE (PRK00019)Ribosomal_L31 (pfam01197)→ |
| Sen-288-751 | NZ_CJLR01000015.1 | + | 104138 | 104190 | **RNA**→ rpmE (PRK00019)Ribosomal_L31 (pfam01197)→ |
| Sen-288-752 | NZ_CJHJ01000016.1 | - | 39532 | 39480 | **RNA**→ rpmE (PRK00019)Ribosomal_L31 (pfam01197)→ |
| Sen-288-753 | NZ_CJHP01000015.1 | - | 39723 | 39671 | **RNA**→ rpmE (PRK00019)Ribosomal_L31 (pfam01197)→ |
| Sen-288-754 | NZ_CIXZ01000015.1 | + | 104219 | 104271 | **RNA**→ rpmE (PRK00019)Ribosomal_L31 (pfam01197)→ |
| Sen-288-755 | NZ_CJJZ01000016.1 | + | 104142 | 104194 | **RNA**→ rpmE (PRK00019)Ribosomal_L31 (pfam01197)→ |
| Sen-288-756 | NZ_CJCE01000015.1 | + | 103986 | 104038 | **RNA**→ rpmE (PRK00019)Ribosomal_L31 (pfam01197)→ |
| Sen-288-757 | NZ_CIVC01000015.1 | + | 104178 | 104230 | **RNA**→ rpmE (PRK00019)Ribosomal_L31 (pfam01197)→ |
| Sen-288-758 | NZ_CJFO01000016.1 | + | 104149 | 104201 | **RNA**→ rpmE (PRK00019)Ribosomal_L31 (pfam01197)→ |
| Sen-288-759 | NZ_CIZV01000016.1 | + | 104157 | 104209 | **RNA**→ rpmE (PRK00019)Ribosomal_L31 (pfam01197)→ |
| Sen-288-760 | NZ_CJYD01000009.1 | + | 163325 | 163377 | **RNA**→ rpmE (PRK00019)Ribosomal_L31 (pfam01197)→ |
| Sen-288-761 | NZ_CJRR01000014.1 | + | 108666 | 108718 | **RNA**→ rpmE (PRK00019)Ribosomal_L31 (pfam01197)→ |
| Sen-288-762 | NZ_CIUV01000016.1 | + | 104162 | 104214 | **RNA**→ rpmE (PRK00019)Ribosomal_L31 (pfam01197)→ |
| Sen-288-763 | NZ_CJNI01000015.1 | + | 104158 | 104210 | **RNA**→ rpmE (PRK00019)Ribosomal_L31 (pfam01197)→ |
| Sen-288-764 | NZ_CJJK01000003.1 | + | 103208 | 103260 | **RNA**→ rpmE (PRK00019)Ribosomal_L31 (pfam01197)→ |
| Sen-288-765 | NZ_CJXP01000016.1 | + | 104184 | 104236 | **RNA**→ rpmE (PRK00019)Ribosomal_L31 (pfam01197)→ |
| Sen-288-766 | NZ_CJDG01000014.1 | + | 108006 | 108058 | **RNA**→ rpmE (PRK00019)Ribosomal_L31 (pfam01197)→ |
| Sen-288-767 | NZ_CJEQ01000016.1 | + | 104150 | 104202 | **RNA**→ rpmE (PRK00019)Ribosomal_L31 (pfam01197)→ |
| Sen-288-768 | NZ_CJKZ01000014.1 | - | 41231 | 41179 | **RNA**→ rpmE (PRK00019)Ribosomal_L31 (pfam01197)→ |
| Sen-288-769 | NZ_CJYS01000016.1 | + | 104159 | 104211 | **RNA**→ rpmE (PRK00019)Ribosomal_L31 (pfam01197)→ |
| Sen-288-770 | NZ_CJSX01000016.1 | + | 104016 | 104068 | **RNA**→ rpmE (PRK00019)Ribosomal_L31 (pfam01197)→ |
| Sen-288-771 | NZ_CJBZ01000016.1 | + | 104158 | 104210 | **RNA**→ rpmE (PRK00019)Ribosomal_L31 (pfam01197)→ |
| Sen-288-772 | NZ_CJOX01000015.1 | + | 104151 | 104203 | **RNA**→ rpmE (PRK00019)Ribosomal_L31 (pfam01197)→ |
| Sen-288-773 | NZ_CJFA01000015.1 | - | 39536 | 39484 | **RNA**→ rpmE (PRK00019)Ribosomal_L31 (pfam01197)→ |
| Sen-288-774 | NZ_CIYH01000015.1 | + | 104203 | 104255 | **RNA**→ rpmE (PRK00019)Ribosomal_L31 (pfam01197)→ |
| Sen-288-775 | NZ_CJTL01000016.1 | - | 39371 | 39319 | **RNA**→ rpmE (PRK00019)Ribosomal_L31 (pfam01197)→ |
| Sen-288-776 | NZ_CJWA01000015.1 | + | 104148 | 104200 | **RNA**→ rpmE (PRK00019)Ribosomal_L31 (pfam01197)→ |
| Sen-288-777 | NZ_CJUO01000016.1 | + | 104155 | 104207 | **RNA**→ rpmE (PRK00019)Ribosomal_L31 (pfam01197)→ |

| | | | | | |
|---|---|---|---|---|---|
| Sen-288-778 | NZ_CJPB01000015.1 | - | 39706 | 39654 | **RNA**→ rpmE (PRK00019)Ribosomal_L31 (pfam01197)→ |
| Sen-288-779 | NZ_CIWQ01000015.1 | + | 103990 | 104042 | **RNA**→ rpmE (PRK00019)Ribosomal_L31 (pfam01197)→ |
| Sen-288-780 | NZ_CJKH01000010.1 | + | 152666 | 152718 | **RNA**→ rpmE (PRK00019)Ribosomal_L31 (pfam01197)→ |
| Sen-288-781 | NZ_CJFP01000015.1 | - | 39531 | 39479 | **RNA**→ rpmE (PRK00019)Ribosomal_L31 (pfam01197)→ |
| Sen-288-782 | NZ_CEYH01000016.1 | + | 104151 | 104203 | **RNA**→ rpmE (PRK00019)Ribosomal_L31 (pfam01197)→ |
| Sen-288-783 | NZ_CEZU01000016.1 | + | 104138 | 104190 | **RNA**→ rpmE (PRK00019)Ribosomal_L31 (pfam01197)→ |
| Sen-288-784 | NZ_CEXJ01000016.1 | - | 39536 | 39484 | **RNA**→ rpmE (PRK00019)Ribosomal_L31 (pfam01197)→ |
| Sen-288-785 | NZ_CIZC01000015.1 | + | 104148 | 104200 | **RNA**→ rpmE (PRK00019)Ribosomal_L31 (pfam01197)→ |
| Sen-288-786 | NZ_CJYO01000012.1 | + | 107743 | 107795 | **RNA**→ rpmE (PRK00019)Ribosomal_L31 (pfam01197)→ |
| Sen-288-787 | NZ_CJQW01000015.1 | - | 39388 | 39336 | **RNA**→ rpmE (PRK00019)Ribosomal_L31 (pfam01197)→ |
| Sen-288-788 | NZ_CJTM01000016.1 | + | 104157 | 104209 | **RNA**→ rpmE (PRK00019)Ribosomal_L31 (pfam01197)→ |
| Sen-288-789 | NZ_CJPP01000015.1 | - | 39601 | 39549 | **RNA**→ rpmE (PRK00019)Ribosomal_L31 (pfam01197)→ |
| Sen-288-790 | NZ_CJMS01000003.1 | - | 200080 | 200028 | **RNA**→ rpmE (PRK00019)Ribosomal_L31 (pfam01197)→ |
| Sen-288-791 | NZ_CJVO01000016.1 | - | 39365 | 39313 | **RNA**→ rpmE (PRK00019)Ribosomal_L31 (pfam01197)→ |
| Sen-288-792 | NZ_CILI01000008.1 | - | 39456 | 39404 | **RNA**→ rpmE (PRK00019)Ribosomal_L31 (pfam01197)→ |
| Sen-288-793 | NZ_CJLN01000002.1 | + | 405306 | 405358 | **RNA**→ rpmE (PRK00019)Ribosomal_L31 (pfam01197)→ |
| Sen-288-794 | NZ_CIYZ01000016.1 | - | 39535 | 39483 | **RNA**→ rpmE (PRK00019)Ribosomal_L31 (pfam01197)→ |
| Sen-288-795 | NZ_CJRQ01000008.1 | - | 39557 | 39505 | **RNA**→ rpmE (PRK00019)Ribosomal_L31 (pfam01197)→ |
| Sen-288-796 | NZ_CIWZ01000015.1 | + | 104053 | 104105 | **RNA**→ rpmE (PRK00019)Ribosomal_L31 (pfam01197)→ |
| Sen-288-797 | NZ_CJGH01000014.1 | - | 39583 | 39531 | **RNA**→ rpmE (PRK00019)Ribosomal_L31 (pfam01197)→ |
| Sen-288-798 | NZ_CJKG01000006.1 | + | 182313 | 182365 | **RNA**→ rpmE (PRK00019)Ribosomal_L31 (pfam01197)→ |
| Sen-288-799 | NZ_CJKM01000010.1 | - | 39456 | 39404 | **RNA**→ rpmE (PRK00019)Ribosomal_L31 (pfam01197)→ |
| Sen-288-800 | NZ_CIXG01000015.1 | - | 39565 | 39513 | **RNA**→ rpmE (PRK00019)Ribosomal_L31 (pfam01197)→ |
| Sen-288-801 | NZ_CJIR01000019.1 | - | 39361 | 39309 | **RNA**→ rpmE (PRK00019)Ribosomal_L31 (pfam01197)→ |
| Sen-288-802 | NZ_CJDS01000012.1 | + | 107993 | 108045 | **RNA**→ rpmE (PRK00019)Ribosomal_L31 (pfam01197)→ |
| Sen-288-803 | NZ_CJAT01000015.1 | + | 104153 | 104205 | **RNA**→ rpmE (PRK00019)Ribosomal_L31 (pfam01197)→ |
| Sen-288-804 | NZ_CJJF01000002.1 | - | 198048 | 197996 | **RNA**→ rpmE (PRK00019)Ribosomal_L31 (pfam01197)→ |
| Sen-288-805 | NZ_CJTH01000015.1 | - | 41512 | 41460 | **RNA**→ rpmE (PRK00019)Ribosomal_L31 (pfam01197)→ |
| Sen-288-806 | NZ_CJBG01000015.1 | - | 39456 | 39404 | **RNA**→ rpmE (PRK00019)Ribosomal_L31 (pfam01197)→ |
| Sen-288-807 | NZ_CJRY01000016.1 | - | 39506 | 39454 | **RNA**→ rpmE (PRK00019)Ribosomal_L31 (pfam01197)→ |
| Sen-288-1119 | NZ_CILC01000016.1 | + | 104213 | 104265 | **RNA**→ rpmE (PRK00019)Ribosomal_L31 (pfam01197)→ |
| Sen-288-808 | NZ_CJDJ01000015.1 | - | 39456 | 39404 | **RNA**→ rpmE (PRK00019)Ribosomal_L31 (pfam01197)→ |
| Sen-288-809 | NZ_CIZI01000015.1 | - | 39534 | 39482 | **RNA**→ rpmE (PRK00019)Ribosomal_L31 (pfam01197)→ |
| Sen-288-810 | NZ_CJYB01000016.1 | - | 39456 | 39404 | **RNA**→ rpmE (PRK00019)Ribosomal_L31 (pfam01197)→ |
| Sen-288-811 | NZ_CIVI01000014.1 | + | 104162 | 104214 | **RNA**→ rpmE (PRK00019)Ribosomal_L31 (pfam01197)→ |
| Sen-288-812 | NZ_CJAM01000016.1 | - | 39456 | 39404 | **RNA**→ rpmE (PRK00019)Ribosomal_L31 (pfam01197)→ |
| Sen-288-813 | NZ_CJOP01000016.1 | - | 39383 | 39331 | **RNA**→ rpmE (PRK00019)Ribosomal_L31 (pfam01197)→ |
| Sen-288-814 | NZ_CJPJ01000017.1 | - | 39601 | 39549 | **RNA**→ rpmE (PRK00019)Ribosomal_L31 (pfam01197)→ |
| Sen-288-815 | NZ_CJQR01000015.1 | - | 39561 | 39509 | **RNA**→ rpmE (PRK00019)Ribosomal_L31 (pfam01197)→ |
| Sen-288-816 | NZ_CJTO01000016.1 | - | 39371 | 39319 | **RNA**→ rpmE (PRK00019)Ribosomal_L31 (pfam01197)→ |
| Sen-288-817 | NZ_CJMF01000013.1 | - | 41498 | 41446 | **RNA**→ rpmE (PRK00019)Ribosomal_L31 (pfam01197)→ |
| Sen-288-818 | NZ_CJET01000016.1 | + | 104147 | 104199 | **RNA**→ rpmE (PRK00019)Ribosomal_L31 (pfam01197)→ |
| Sen-288-819 | NZ_CEXP01000016.1 | + | 104148 | 104200 | **RNA**→ rpmE (PRK00019)Ribosomal_L31 (pfam01197)→ |
| Sen-288-820 | NZ_CIYA01000015.1 | - | 39556 | 39504 | **RNA**→ rpmE (PRK00019)Ribosomal_L31 (pfam01197)→ |
| Sen-288-821 | NZ_CJCQ01000016.1 | - | 39540 | 39488 | **RNA**→ rpmE (PRK00019)Ribosomal_L31 (pfam01197)→ |
| Sen-288-822 | NZ_CJHI01000016.1 | - | 39537 | 39485 | **RNA**→ rpmE (PRK00019)Ribosomal_L31 (pfam01197)→ |
| Sen-288-823 | NZ_CJXL01000017.1 | - | 39552 | 39500 | **RNA**→ rpmE (PRK00019)Ribosomal_L31 (pfam01197)→ |
| Sen-288-824 | NZ_CIXW01000015.1 | + | 103990 | 104042 | **RNA**→ rpmE (PRK00019)Ribosomal_L31 (pfam01197)→ |
| Sen-288-825 | NZ_CJYR01000014.1 | + | 104157 | 104209 | **RNA**→ rpmE (PRK00019)Ribosomal_L31 (pfam01197)→ |
| Sen-288-826 | NZ_CJDI01000015.1 | - | 39582 | 39530 | **RNA**→ rpmE (PRK00019)Ribosomal_L31 (pfam01197)→ |
| Sen-288-827 | NZ_CJMT01000014.1 | - | 41231 | 41179 | **RNA**→ rpmE (PRK00019)Ribosomal_L31 (pfam01197)→ |
| Sen-288-828 | NZ_CJAK01000014.1 | + | 104156 | 104208 | **RNA**→ rpmE (PRK00019)Ribosomal_L31 (pfam01197)→ |

| | | | | | | | |
|---|---|---|---|---|---|---|---|
| Sen-288-829 | NZ_CIVK01000013.1 | - | 39506 | 39454 | **RNA**→ | rpmE (PRK00019)Ribosomal_L31 (pfam01197)→ | |
| Sen-288-830 | NZ_CJWP01000003.1 | - | 198560 | 198508 | **RNA**→ | rpmE (PRK00019)Ribosomal_L31 (pfam01197)→ | |
| Sen-131-1 | NZ_LIMK01000047.1 | - | 7856 | 7804 | **RNA**→ | rpmE (PRK00019)Ribosomal_L31 (pfam01197)→ | |
| Sen-114-1 | NZ_AZKO02000007.1 | + | 14026 | 14078 | **RNA**→ | rpmE (PRK00019)Ribosomal_L31 (pfam01197)→ | |
| Sen-119-1 | NZ_AZLA01000036.1 | - | 14715 | 14663 | **RNA**→ | rpmE (PRK00019)Ribosomal_L31 (pfam01197)→ | |
| Sen-439-6 | NZ_CQPF01000014.1 | - | 45175 | 45123 | **RNA**→ | rpmE (PRK00019)Ribosomal_L31 (pfam01197)→ | |
| Sen-136-1 | NZ_AYUE01000015.1 | + | 21280 | 21332 | **RNA**→ | rpmE (PRK00019)Ribosomal_L31 (pfam01197)→ | |
| Sen-126-1 | NZ_AZLB01000007.1 | - | 37541 | 37489 | **RNA**→ | rpmE (PRK00019)Ribosomal_L31 (pfam01197)→ | |
| Sen-439-7 | NZ_CQPI01000012.1 | - | 45367 | 45315 | **RNA**→ | rpmE (PRK00019)Ribosomal_L31 (pfam01197)→ | |
| Sen-449-1 | NZ_AYVC01000013.1 | - | 36680 | 36628 | **RNA**→ | rpmE (PRK00019)Ribosomal_L31 (pfam01197)→ | |
| Sen-443-1 | NZ_AYVH01000013.1 | - | 5782 | 5730 | **RNA**→ | rpmE (PRK00019)Ribosomal_L31 (pfam01197)→ | |
| Sen-465-1 | NZ_AOZH01000225.1 | - | 19515 | 19463 | **RNA**→ | rpmE (PRK00019)Ribosomal_L31 (pfam01197)→ | rpmE (PRK00019)Ribosomal_L31 (pfam01197)→ |
| Sen-288-831 | NZ_CIOO01000013.1 | + | 107861 | 107913 | **RNA**→ | rpmE (PRK00019)Ribosomal_L31 (pfam01197)→ | |
| Sen-441-1 | NZ_AOZM01000038.1 | + | 34562 | 34614 | **RNA**→ | rpmE (PRK00019)Ribosomal_L31 (pfam01197)→ | |
| Sen-288-832 | NZ_CIQJ01000015.1 | + | 104150 | 104202 | **RNA**→ | rpmE (PRK00019)Ribosomal_L31 (pfam01197)→ | |
| Sen-288-833 | NZ_CIQW01000016.1 | - | 39536 | 39484 | **RNA**→ | rpmE (PRK00019)Ribosomal_L31 (pfam01197)→ | |
| Sen-471-1 | NZ_APAF01000259.1 | - | 8143 | 8091 | **RNA**→ | rpmE (PRK00019)Ribosomal_L31 (pfam01197)→ | rpmE (PRK00019)Ribosomal_L31 (pfam01197)→ |
| Sen-288-834 | NZ_CIUK01000013.1 | - | 44930 | 44878 | **RNA**→ | rpmE (PRK00019)Ribosomal_L31 (pfam01197)→ | |
| Sen-442-1 | NZ_APAP01000035.1 | - | 45429 | 45377 | **RNA**→ | rpmE (PRK00019)Ribosomal_L31 (pfam01197)→ | |
| Sen-288-835 | NZ_CIMZ01000012.1 | + | 107813 | 107865 | **RNA**→ | rpmE (PRK00019)Ribosomal_L31 (pfam01197)→ | |
| Sen-127-1 | NZ_AZKQ02000008.1 | + | 18889 | 18941 | **RNA**→ | rpmE (PRK00019)Ribosomal_L31 (pfam01197)→ | |
| Sen-439-8 | NZ_CQPK01000013.1 | - | 15585 | 15533 | **RNA**→ | rpmE (PRK00019)Ribosomal_L31 (pfam01197)→ | |
| Sen-288-836 | NZ_CIQF01000016.1 | - | 39576 | 39524 | **RNA**→ | rpmE (PRK00019)Ribosomal_L31 (pfam01197)→ | |
| Sen-439-9 | NZ_CQPE01000017.1 | - | 45348 | 45296 | **RNA**→ | rpmE (PRK00019)Ribosomal_L31 (pfam01197)→ | |
| Sen-288-837 | NZ_CIOS01000015.1 | + | 104160 | 104212 | **RNA**→ | rpmE (PRK00019)Ribosomal_L31 (pfam01197)→ | |
| Sen-288-838 | NZ_CITN01000009.1 | - | 39370 | 39318 | **RNA**→ | rpmE (PRK00019)Ribosomal_L31 (pfam01197)→ | |
| Sen-464-1 | NZ_ATEU01000051.1 | - | 45254 | 45202 | **RNA**→ | rpmE (PRK00019)Ribosomal_L31 (pfam01197)→ | |
| Sen-288-839 | NZ_CINN01000015.1 | + | 104181 | 104233 | **RNA**→ | rpmE (PRK00019)Ribosomal_L31 (pfam01197)→ | |
| Sen-448-1 | NZ_AYVD01000012.1 | + | 95202 | 95254 | **RNA**→ | rpmE (PRK00019)Ribosomal_L31 (pfam01197)→ | |
| Sen-288-840 | NZ_CIQO01000015.1 | - | 39538 | 39486 | **RNA**→ | rpmE (PRK00019)Ribosomal_L31 (pfam01197)→ | |
| Sen-469-1 | NZ_AHUJ01000005.1 | - | 49367 | 49315 | **RNA**→ | rpmE (PRK00019)Ribosomal_L31 (pfam01197)→ | rpmE (PRK00019)Ribosomal_L31 (pfam01197)→ |
| Sen-288-841 | NZ_CINC01000016.1 | + | 104014 | 104066 | **RNA**→ | rpmE (PRK00019)Ribosomal_L31 (pfam01197)→ | |
| Sen-444-1 | NZ_AOZJ01000055.1 | - | 36680 | 36628 | **RNA**→ | rpmE (PRK00019)Ribosomal_L31 (pfam01197)→ | rpmE (PRK00019)Ribosomal_L31 (pfam01197)→ |
| Sen-288-842 | NZ_CIOW01000015.1 | - | 39541 | 39489 | **RNA**→ | rpmE (PRK00019)Ribosomal_L31 (pfam01197)→ | |
| Sen-445-1 | NZ_AYVG01000011.1 | + | 95203 | 95255 | **RNA**→ | rpmE (PRK00019)Ribosomal_L31 (pfam01197)→ | |
| Sen-288-843 | NZ_CISP01000015.1 | - | 39580 | 39528 | **RNA**→ | rpmE (PRK00019)Ribosomal_L31 (pfam01197)→ | |
| Sen-288-844 | NZ_CITM01000016.1 | - | 39541 | 39489 | **RNA**→ | rpmE (PRK00019)Ribosomal_L31 (pfam01197)→ | |
| Sen-461-1 | NZ_AOZI01000030.1 | + | 97354 | 97406 | **RNA**→ | rpmE (PRK00019)Ribosomal_L31 (pfam01197)→ | rpmE (PRK00019)Ribosomal_L31 (pfam01197)→ |
| Sen-288-845 | NZ_CIOH01000015.1 | - | 41231 | 41179 | **RNA**→ | rpmE (PRK00019)Ribosomal_L31 (pfam01197)→ | |
| Sen-125-1 | NZ_AZKU01000012.1 | - | 14715 | 14663 | **RNA**→ | rpmE (PRK00019)Ribosomal_L31 (pfam01197)→ | |
| Sen-124-1 | NZ_AZKT01000038.1 | - | 37597 | 37545 | **RNA**→ | rpmE (PRK00019)Ribosomal_L31 (pfam01197)→ | |
| Sen-288-846 | NZ_CIOE01000015.1 | + | 107384 | 107436 | **RNA**→ | rpmE (PRK00019)Ribosomal_L31 (pfam01197)→ | |
| Sen-121-1 | NZ_AZKV01000024.1 | + | 97957 | 98009 | **RNA**→ | rpmE (PRK00019)Ribosomal_L31 (pfam01197)→ | |
| Sen-288-847 | NZ_CIRI01000016.1 | - | 39564 | 39512 | **RNA**→ | rpmE (PRK00019)Ribosomal_L31 (pfam01197)→ | |
| Sen-288-848 | NZ_CISZ01000015.1 | + | 103990 | 104042 | **RNA**→ | rpmE (PRK00019)Ribosomal_L31 (pfam01197)→ | |
| Sen-118-1 | NZ_AZKZ01000037.1 | + | 104477 | 104529 | **RNA**→ | rpmE (PRK00019)Ribosomal_L31 (pfam01197)→ | |
| Sen-288-849 | NZ_CISU01000014.1 | - | 41249 | 41197 | **RNA**→ | rpmE (PRK00019)Ribosomal_L31 (pfam01197)→ | |
| Sen-123-1 | NZ_AZKS01000081.1 | - | 11432 | 11380 | **RNA**→ | rpmE (PRK00019)Ribosomal_L31 (pfam01197)→ | |
| Sen-288-850 | NZ_CIPL01000016.1 | + | 104154 | 104206 | **RNA**→ | rpmE (PRK00019)Ribosomal_L31 (pfam01197)→ | |
| Sen-117-1 | NZ_AZKY01000039.1 | + | 14027 | 14079 | **RNA**→ | rpmE (PRK00019)Ribosomal_L31 (pfam01197)→ | |
| Sen-288-851 | NZ_CIOM01000017.1 | - | 39569 | 39517 | **RNA**→ | rpmE (PRK00019)Ribosomal_L31 (pfam01197)→ | |
| Sen-115-1 | NZ_AZKW01000008.1 | - | 45095 | 45043 | **RNA**→ | rpmE (PRK00019)Ribosomal_L31 (pfam01197)→ | |

| | | | | | | |
|---|---|---|---|---|---|---|
| Sen-288-852 | NZ_CIPG01000014.1 | - | 44758 | 44706 | **RNA**→ | rpmE (PRK00019)Ribosomal_L31 (pfam01197)→ |
| Sen-288-853 | NZ_CISC01000015.1 | - | 39589 | 39537 | **RNA**→ | rpmE (PRK00019)Ribosomal_L31 (pfam01197)→ |
| Sen-439-10 | NZ_CQPN01000012.1 | - | 45266 | 45214 | **RNA**→ | rpmE (PRK00019)Ribosomal_L31 (pfam01197)→ |
| Sen-288-854 | NZ_CIOB01000018.1 | + | 107995 | 108047 | **RNA**→ | rpmE (PRK00019)Ribosomal_L31 (pfam01197)→ hypo→ |
| Sen-288-855 | NZ_CIND01000016.1 | - | 39601 | 39549 | **RNA**→ | rpmE (PRK00019)Ribosomal_L31 (pfam01197)→ |
| Sen-288-856 | NZ_CJRI01000013.1 | + | 104161 | 104213 | **RNA**→ | rpmE (PRK00019)Ribosomal_L31 (pfam01197)→ |
| Sen-288-857 | NZ_CIPK01000016.1 | + | 104157 | 104209 | **RNA**→ | rpmE (PRK00019)Ribosomal_L31 (pfam01197)→ |
| Sen-453-1 | NZ_AYUY01000011.1 | + | 95202 | 95254 | **RNA**→ | rpmE (PRK00019)Ribosomal_L31 (pfam01197)→ |
| Sen-288-858 | NZ_CIPZ01000010.1 | + | 104165 | 104217 | **RNA**→ | rpmE (PRK00019)Ribosomal_L31 (pfam01197)→ |
| Sen-128-1 | NZ_AZKP01000048.1 | - | 11305 | 11253 | **RNA**→ | rpmE (PRK00019)Ribosomal_L31 (pfam01197)→ |
| Sen-447-1 | NZ_AYVE01000011.1 | + | 95203 | 95255 | **RNA**→ | rpmE (PRK00019)Ribosomal_L31 (pfam01197)→ |
| Sen-288-859 | NZ_CIRC01000015.1 | + | 104144 | 104196 | **RNA**→ | rpmE (PRK00019)Ribosomal_L31 (pfam01197)→ |
| Sen-450-1 | NZ_AYVB01000014.1 | - | 36680 | 36628 | **RNA**→ | rpmE (PRK00019)Ribosomal_L31 (pfam01197)→ |
| Sen-288-860 | NZ_CIOD01000015.1 | - | 41231 | 41179 | **RNA**→ | rpmE (PRK00019)Ribosomal_L31 (pfam01197)→ |
| Sen-288-861 | NZ_CINA01000016.1 | - | 39601 | 39549 | **RNA**→ | rpmE (PRK00019)Ribosomal_L31 (pfam01197)→ |
| Sen-288-862 | NZ_CJRC01000015.1 | - | 41510 | 41458 | **RNA**→ | rpmE (PRK00019)Ribosomal_L31 (pfam01197)→ |
| Sen-288-863 | NZ_CIOY01000014.1 | - | 41231 | 41179 | **RNA**→ | rpmE (PRK00019)Ribosomal_L31 (pfam01197)→ |
| Sen-120-1 | NZ_AZLC01000015.1 | + | 104477 | 104529 | **RNA**→ | rpmE (PRK00019)Ribosomal_L31 (pfam01197)→ |
| Sen-288-864 | NZ_CIOZ01000018.1 | + | 104178 | 104230 | **RNA**→ | rpmE (PRK00019)Ribosomal_L31 (pfam01197)→ |
| Sen-122-1 | NZ_AZKR01000018.1 | + | 18897 | 18949 | **RNA**→ | rpmE (PRK00019)Ribosomal_L31 (pfam01197)→ |
| Sen-288-865 | NZ_CISN01000015.1 | - | 39576 | 39524 | **RNA**→ | rpmE (PRK00019)Ribosomal_L31 (pfam01197)→ |
| Sen-116-1 | NZ_AZKX01000030.1 | + | 104479 | 104531 | **RNA**→ | rpmE (PRK00019)Ribosomal_L31 (pfam01197)→ |
| Sen-288-866 | NZ_CIRV01000014.1 | - | 39588 | 39536 | **RNA**→ | rpmE (PRK00019)Ribosomal_L31 (pfam01197)→ |
| Sen-451-1 | NZ_AYVA01000013.1 | - | 36680 | 36628 | **RNA**→ | rpmE (PRK00019)Ribosomal_L31 (pfam01197)→ |
| Sen-288-867 | NZ_CITQ01000016.1 | - | 39374 | 39322 | **RNA**→ | rpmE (PRK00019)Ribosomal_L31 (pfam01197)→ |
| Sen-456-1 | NZ_AYUV01000014.1 | - | 36603 | 36551 | **RNA**→ | rpmE (PRK00019)Ribosomal_L31 (pfam01197)→ |
| Sen-288-868 | NZ_CIQT01000016.1 | - | 39518 | 39466 | **RNA**→ | rpmE (PRK00019)Ribosomal_L31 (pfam01197)→ |
| Sen-457-1 | NZ_AYUU01000011.1 | - | 36680 | 36628 | **RNA**→ | rpmE (PRK00019)Ribosomal_L31 (pfam01197)→ |
| Sen-288-869 | NZ_CIUD01000012.1 | + | 107998 | 108050 | **RNA**→ | rpmE (PRK00019)Ribosomal_L31 (pfam01197)→ |
| Sen-288-870 | NZ_CIPH01000014.1 | - | 39541 | 39489 | **RNA**→ | rpmE (PRK00019)Ribosomal_L31 (pfam01197)→ |
| Sen-467-1 | NZ_AHUK01000056.1 | - | 49403 | 49351 | **RNA**→ | rpmE (PRK00019)Ribosomal_L31 (pfam01197)→ rpmE (PRK00019)Ribosomal_L31 (pfam01197)→ |
| Sen-288-871 | NZ_CISI01000014.1 | + | 103990 | 104042 | **RNA**→ | rpmE (PRK00019)Ribosomal_L31 (pfam01197)→ |
| Sen-452-1 | NZ_AYUZ01000012.1 | - | 36680 | 36628 | **RNA**→ | rpmE (PRK00019)Ribosomal_L31 (pfam01197)→ |
| Sen-288-872 | NZ_CIKY01000015.1 | + | 107851 | 107903 | **RNA**→ | rpmE (PRK00019)Ribosomal_L31 (pfam01197)→ |
| Sen-459-1 | NZ_AYUS01000043.1 | - | 268 | 216 | **RNA**→ | rpmE (PRK00019)Ribosomal_L31 (pfam01197)→ |
| Sen-288-873 | NZ_CIOA01000015.1 | + | 104165 | 104217 | **RNA**→ | rpmE (PRK00019)Ribosomal_L31 (pfam01197)→ |
| Sen-288-874 | NZ_CJRL01000015.1 | + | 104152 | 104204 | **RNA**→ | rpmE (PRK00019)Ribosomal_L31 (pfam01197)→ |
| Sen-455-1 | NZ_AYUW01000012.1 | + | 95203 | 95255 | **RNA**→ | rpmE (PRK00019)Ribosomal_L31 (pfam01197)→ |
| Sen-288-875 | NZ_CIMY01000014.1 | + | 104030 | 104082 | **RNA**→ | rpmE (PRK00019)Ribosomal_L31 (pfam01197)→ |
| Sen-446-1 | NZ_AYVF01000014.1 | - | 36680 | 36628 | **RNA**→ | rpmE (PRK00019)Ribosomal_L31 (pfam01197)→ |
| Sen-288-876 | NZ_CIQS01000015.1 | + | 104012 | 104064 | **RNA**→ | rpmE (PRK00019)Ribosomal_L31 (pfam01197)→ |
| Sen-458-1 | NZ_AYUT01000012.1 | - | 36680 | 36628 | **RNA**→ | rpmE (PRK00019)Ribosomal_L31 (pfam01197)→ |
| Sen-288-877 | NZ_CIRM01000008.1 | + | 133807 | 133859 | **RNA**→ | rpmE (PRK00019)Ribosomal_L31 (pfam01197)→ |
| Sen-454-1 | NZ_AYUX01000013.1 | + | 70771 | 70823 | **RNA**→ | rpmE (PRK00019)Ribosomal_L31 (pfam01197)→ |
| Sen-288-878 | NZ_CIPE01000015.1 | - | 39501 | 39449 | **RNA**→ | rpmE (PRK00019)Ribosomal_L31 (pfam01197)→ |
| Sen-460-1 | NZ_AYUR01000011.1 | + | 95203 | 95255 | **RNA**→ | rpmE (PRK00019)Ribosomal_L31 (pfam01197)→ |
| Sen-288-879 | NZ_CINR01000014.1 | + | 104181 | 104233 | **RNA**→ | rpmE (PRK00019)Ribosomal_L31 (pfam01197)→ |
| Sen-288-880 | NZ_CINE01000015.1 | - | 41673 | 41621 | **RNA**→ | rpmE (PRK00019)Ribosomal_L31 (pfam01197)→ |
| Sen-288-881 | NZ_CIPJ01000014.1 | - | 39537 | 39485 | **RNA**→ | rpmE (PRK00019)Ribosomal_L31 (pfam01197)→ |
| Sen-288-882 | NZ_CIRK01000012.1 | + | 107987 | 108039 | **RNA**→ | rpmE (PRK00019)Ribosomal_L31 (pfam01197)→ |
| Sen-288-883 | NZ_CJXD01000017.1 | + | 104163 | 104215 | **RNA**→ | rpmE (PRK00019)Ribosomal_L31 (pfam01197)→ |
| Sen-288-884 | NZ_CINX01000016.1 | - | 39596 | 39544 | **RNA**→ | rpmE (PRK00019)Ribosomal_L31 (pfam01197)→ |

| | | | | | | |
|---|---|---|---|---|---|---|
| Sen-288-885 | NZ_CJCW01000016.1 | - | 39532 | 39480 | **RNA**→ | rpmE (PRK00019)Ribosomal_L31 (pfam01197)→ |
| Sen-288-886 | NZ_CIUY01000006.1 | - | 39383 | 39331 | **RNA**→ | rpmE (PRK00019)Ribosomal_L31 (pfam01197)→ |
| Sen-288-1120 | NZ_CIYL01000011.1 | - | 39456 | 39404 | **RNA**→ | rpmE (PRK00019)Ribosomal_L31 (pfam01197)→ |
| Sen-288-887 | NZ_CJPO01000007.1 | + | 152696 | 152748 | **RNA**→ | rpmE (PRK00019)Ribosomal_L31 (pfam01197)→ |
| Sen-288-888 | NZ_CJHT01000016.1 | + | 104150 | 104202 | **RNA**→ | rpmE (PRK00019)Ribosomal_L31 (pfam01197)→ |
| Sen-288-889 | NZ_CJGC01000012.1 | + | 104169 | 104221 | **RNA**→ | rpmE (PRK00019)Ribosomal_L31 (pfam01197)→ |
| Sen-288-1121 | NZ_CIOC01000014.1 | - | 41512 | 41460 | **RNA**→ | rpmE (PRK00019)Ribosomal_L31 (pfam01197)→ |
| Sen-288-890 | NZ_CIUI01000014.1 | + | 103989 | 104041 | **RNA**→ | rpmE (PRK00019)Ribosomal_L31 (pfam01197)→ |
| Sen-288-891 | NZ_CISL01000014.1 | + | 103990 | 104042 | **RNA**→ | rpmE (PRK00019)Ribosomal_L31 (pfam01197)→ |
| Sen-288-892 | NZ_CIQP01000015.1 | - | 41231 | 41179 | **RNA**→ | rpmE (PRK00019)Ribosomal_L31 (pfam01197)→ |
| Sen-288-893 | NZ_CITU01000014.1 | + | 107894 | 107946 | **RNA**→ | rpmE (PRK00019)Ribosomal_L31 (pfam01197)→ |
| Sen-288-894 | NZ_CIUH01000013.1 | + | 104149 | 104201 | **RNA**→ | rpmE (PRK00019)Ribosomal_L31 (pfam01197)→ |
| Sen-288-895 | NZ_CITI01000014.1 | - | 39588 | 39536 | **RNA**→ | rpmE (PRK00019)Ribosomal_L31 (pfam01197)→ |
| Sen-288-896 | NZ_CINZ01000016.1 | - | 39601 | 39549 | **RNA**→ | rpmE (PRK00019)Ribosomal_L31 (pfam01197)→ |
| Sen-288-897 | NZ_CINS01000015.1 | + | 104164 | 104216 | **RNA**→ | rpmE (PRK00019)Ribosomal_L31 (pfam01197)→ |
| Sen-288-898 | NZ_CINH01000015.1 | + | 104148 | 104200 | **RNA**→ | rpmE (PRK00019)Ribosomal_L31 (pfam01197)→ |
| Sen-288-899 | NZ_CIQE01000016.1 | + | 104151 | 104203 | **RNA**→ | rpmE (PRK00019)Ribosomal_L31 (pfam01197)→ |
| Sen-288-900 | NZ_CIRX01000010.1 | + | 133852 | 133904 | **RNA**→ | rpmE (PRK00019)Ribosomal_L31 (pfam01197)→ |
| Sen-288-901 | NZ_CIQZ01000006.1 | - | 39370 | 39318 | **RNA**→ | rpmE (PRK00019)Ribosomal_L31 (pfam01197)→ |
| Sen-288-902 | NZ_CIRO01000014.1 | + | 104138 | 104190 | **RNA**→ | rpmE (PRK00019)Ribosomal_L31 (pfam01197)→ |
| Sen-288-903 | NZ_CISB01000014.1 | - | 39586 | 39534 | **RNA**→ | rpmE (PRK00019)Ribosomal_L31 (pfam01197)→ |
| Sen-288-904 | NZ_CITP01000015.1 | - | 39549 | 39497 | **RNA**→ | rpmE (PRK00019)Ribosomal_L31 (pfam01197)→ |
| Sen-288-905 | NZ_CITR01000003.1 | - | 198388 | 198336 | **RNA**→ | rpmE (PRK00019)Ribosomal_L31 (pfam01197)→ |
| Sen-288-906 | NZ_CINI01000016.1 | - | 39379 | 39327 | **RNA**→ | rpmE (PRK00019)Ribosomal_L31 (pfam01197)→ |
| Sen-288-907 | NZ_CIRS01000017.1 | + | 104157 | 104209 | **RNA**→ | rpmE (PRK00019)Ribosomal_L31 (pfam01197)→ |
| Sen-288-908 | NZ_CIPS01000015.1 | - | 41507 | 41455 | **RNA**→ | rpmE (PRK00019)Ribosomal_L31 (pfam01197)→ |
| Sen-288-909 | NZ_CIPD01000003.1 | + | 104158 | 104210 | **RNA**→ | rpmE (PRK00019)Ribosomal_L31 (pfam01197)→ |
| Sen-288-910 | NZ_CISY01000013.1 | - | 39584 | 39532 | **RNA**→ | rpmE (PRK00019)Ribosomal_L31 (pfam01197)→ |
| Sen-288-911 | NZ_CIUC01000015.1 | + | 104162 | 104214 | **RNA**→ | rpmE (PRK00019)Ribosomal_L31 (pfam01197)→ |
| Sen-288-912 | NZ_CIKX01000016.1 | + | 104173 | 104225 | **RNA**→ | rpmE (PRK00019)Ribosomal_L31 (pfam01197)→ |
| Sen-288-913 | NZ_CISG01000015.1 | + | 103989 | 104041 | **RNA**→ | rpmE (PRK00019)Ribosomal_L31 (pfam01197)→ |
| Sen-288-914 | NZ_CIRN01000016.1 | - | 39588 | 39536 | **RNA**→ | rpmE (PRK00019)Ribosomal_L31 (pfam01197)→ |
| Sen-288-915 | NZ_CINY01000011.1 | + | 107828 | 107880 | **RNA**→ | rpmE (PRK00019)Ribosomal_L31 (pfam01197)→ |
| Sen-288-916 | NZ_CIMX01000015.1 | + | 104154 | 104206 | **RNA**→ | rpmE (PRK00019)Ribosomal_L31 (pfam01197)→ |
| Sen-288-917 | NZ_CISR01000015.1 | - | 39588 | 39536 | **RNA**→ | rpmE (PRK00019)Ribosomal_L31 (pfam01197)→ |
| Sen-288-918 | NZ_CIOQ01000015.1 | + | 104025 | 104077 | **RNA**→ | rpmE (PRK00019)Ribosomal_L31 (pfam01197)→ |
| Sen-288-919 | NZ_CINJ01000015.1 | - | 39388 | 39336 | **RNA**→ | rpmE (PRK00019)Ribosomal_L31 (pfam01197)→ |
| Sen-288-920 | NZ_CIPN01000015.1 | + | 107996 | 108048 | **RNA**→ | rpmE (PRK00019)Ribosomal_L31 (pfam01197)→ |
| Sen-288-921 | NZ_CJNC01000013.1 | - | 41507 | 41455 | **RNA**→ | rpmE (PRK00019)Ribosomal_L31 (pfam01197)→ |
| Sen-288-922 | NZ_CJLQ01000014.1 | + | 104151 | 104203 | **RNA**→ | rpmE (PRK00019)Ribosomal_L31 (pfam01197)→ |
| Sen-288-923 | NZ_CJNV01000007.1 | - | 41508 | 41456 | **RNA**→ | rpmE (PRK00019)Ribosomal_L31 (pfam01197)→ |
| Sen-288-924 | NZ_CJNH01000014.1 | + | 104156 | 104208 | **RNA**→ | rpmE (PRK00019)Ribosomal_L31 (pfam01197)→ |
| Sen-288-925 | NZ_CJBB01000016.1 | - | 39541 | 39489 | **RNA**→ | rpmE (PRK00019)Ribosomal_L31 (pfam01197)→ |
| Sen-288-926 | NZ_CJDY01000016.1 | - | 39710 | 39658 | **RNA**→ | rpmE (PRK00019)Ribosomal_L31 (pfam01197)→ |
| Sen-288-927 | NZ_CJWS01000016.1 | - | 39456 | 39404 | **RNA**→ | rpmE (PRK00019)Ribosomal_L31 (pfam01197)→ |
| Sen-288-928 | NZ_CIYP01000014.1 | + | 104206 | 104258 | **RNA**→ | rpmE (PRK00019)Ribosomal_L31 (pfam01197)→ |
| Sen-288-929 | NZ_CILM01000016.1 | - | 39357 | 39305 | **RNA**→ | rpmE (PRK00019)Ribosomal_L31 (pfam01197)→ |
| Sen-288-930 | NZ_CJNQ01000014.1 | - | 41499 | 41447 | **RNA**→ | rpmE (PRK00019)Ribosomal_L31 (pfam01197)→ |
| Sen-288-1122 | NZ_CIWM01000014.1 | - | 41505 | 41453 | **RNA**→ | rpmE (PRK00019)Ribosomal_L31 (pfam01197)→ |
| Sen-288-931 | NZ_CJBT01000010.1 | + | 141195 | 141247 | **RNA**→ | rpmE (PRK00019)Ribosomal_L31 (pfam01197)→ |
| Sen-288-932 | NZ_CJLT01000014.1 | + | 104151 | 104203 | **RNA**→ | rpmE (PRK00019)Ribosomal_L31 (pfam01197)→ |
| Sen-288-933 | NZ_CJCF01000015.1 | + | 103986 | 104038 | **RNA**→ | rpmE (PRK00019)Ribosomal_L31 (pfam01197)→ |

| | | | | | |
|---|---|---|---|---|---|
| Sen-288-934 | NZ_CJYC01000017.1 | - | 44768 | 44716 | **RNA**→ rpmE (PRK00019)Ribosomal_L31 (pfam01197)→ |
| Sen-288-935 | NZ_CIYY01000016.1 | + | 104137 | 104189 | **RNA**→ rpmE (PRK00019)Ribosomal_L31 (pfam01197)→ |
| Sen-288-936 | NZ_CJWD01000016.1 | + | 104147 | 104199 | **RNA**→ rpmE (PRK00019)Ribosomal_L31 (pfam01197)→ |
| Sen-288-937 | NZ_CJMI01000015.1 | - | 41231 | 41179 | **RNA**→ rpmE (PRK00019)Ribosomal_L31 (pfam01197)→ |
| Sen-288-938 | NZ_CIXF01000016.1 | - | 39588 | 39536 | **RNA**→ rpmE (PRK00019)Ribosomal_L31 (pfam01197)→ |
| Sen-288-939 | NZ_CJBH01000015.1 | + | 103986 | 104038 | **RNA**→ rpmE (PRK00019)Ribosomal_L31 (pfam01197)→ |
| Sen-288-940 | NZ_CJLC01000016.1 | + | 104109 | 104161 | **RNA**→ rpmE (PRK00019)Ribosomal_L31 (pfam01197)→ |
| Sen-288-941 | NZ_CJXA01000016.1 | - | 39378 | 39326 | **RNA**→ rpmE (PRK00019)Ribosomal_L31 (pfam01197)→ |
| Sen-315-1 | NZ_AHUT01000041.1 | + | 42411 | 42463 | **RNA**→ rpmE (PRK00019)Ribosomal_L31 (pfam01197)→ |
| Sen-322-1 | NZ_CATM01000040.1 | + | 111239 | 111291 | **RNA**→ rpmE (PRK00019)Ribosomal_L31 (pfam01197)→ |
| Sen-327-1 | NZ_CATR01000036.1 | - | 37562 | 37510 | **RNA**→ rpmE (PRK00019)Ribosomal_L31 (pfam01197)→ |
| Sen-332-1 | NZ_CATA01000042.1 | + | 42800 | 42852 | **RNA**→ rpmE (PRK00019)Ribosomal_L31 (pfam01197)→ |
| Sen-338-1 | NZ_CASU01000047.1 | + | 42880 | 42932 | **RNA**→ rpmE (PRK00019)Ribosomal_L31 (pfam01197)→ |
| Sen-316-1 | NZ_CATQ01000043.1 | + | 111287 | 111339 | **RNA**→ rpmE (PRK00019)Ribosomal_L31 (pfam01197)→ |
| Sen-323-1 | NZ_CATL01000048.1 | + | 111226 | 111278 | **RNA**→ rpmE (PRK00019)Ribosomal_L31 (pfam01197)→ |
| Sen-328-1 | NZ_CATE01000036.1 | - | 37552 | 37500 | **RNA**→ rpmE (PRK00019)Ribosomal_L31 (pfam01197)→ |
| Sen-334-1 | NZ_CASY01000039.1 | + | 111202 | 111254 | **RNA**→ rpmE (PRK00019)Ribosomal_L31 (pfam01197)→ |
| Sen-335-1 | NZ_CASX01000036.1 | + | 111336 | 111388 | **RNA**→ rpmE (PRK00019)Ribosomal_L31 (pfam01197)→ |
| Sen-339-1 | NZ_CAST01000046.1 | + | 102302 | 102354 | **RNA**→ rpmE (PRK00019)Ribosomal_L31 (pfam01197)→ |
| Sen-9-1 | NZ_JGYE01000109.1 | + | 4194 | 4246 | **RNA**→ rpmE (PRK00019)Ribosomal_L31 (pfam01197)→ |
| Sen-313-1 | NZ_AHUS01000028.1 | - | 45078 | 45026 | **RNA**→ rpmE (PRK00019)Ribosomal_L31 (pfam01197)→ |
| Sen-321-1 | NZ_CATN01000044.1 | + | 111197 | 111249 | **RNA**→ rpmE (PRK00019)Ribosomal_L31 (pfam01197)→ |
| Sen-326-1 | NZ_CATF01000034.1 | - | 37552 | 37500 | **RNA**→ rpmE (PRK00019)Ribosomal_L31 (pfam01197)→ |
| Sen-331-1 | NZ_CATB01000040.1 | - | 37552 | 37500 | **RNA**→ rpmE (PRK00019)Ribosomal_L31 (pfam01197)→ |
| Sen-337-1 | NZ_CASV01000053.1 | + | 55458 | 55510 | **RNA**→ rpmE (PRK00019)Ribosomal_L31 (pfam01197)→ |
| Sen-320-1 | NZ_AYDT01000147.1 | + | 111582 | 111634 | **RNA**→ rpmE (PRK00019)Ribosomal_L31 (pfam01197)→ rpmE (PRK00019)Ribosomal_L31 (pfam01197)→ |
| Sen-317-1 | NZ_CATP01000025.1 | + | 111848 | 111900 | **RNA**→ rpmE (PRK00019)Ribosomal_L31 (pfam01197)→ |
| Sen-319-1 | NZ_CATO01000040.1 | + | 111150 | 111202 | **RNA**→ rpmE (PRK00019)Ribosomal_L31 (pfam01197)→ |
| Sen-324-1 | NZ_CATK01000043.1 | - | 37550 | 37498 | **RNA**→ rpmE (PRK00019)Ribosomal_L31 (pfam01197)→ |
| Sen-325-1 | NZ_CATJ01000034.1 | - | 37552 | 37500 | **RNA**→ rpmE (PRK00019)Ribosomal_L31 (pfam01197)→ |
| Sen-330-1 | NZ_CATC01000039.1 | - | 37552 | 37500 | **RNA**→ rpmE (PRK00019)Ribosomal_L31 (pfam01197)→ |
| Sen-336-1 | NZ_CASW01000025.1 | + | 111111 | 111163 | **RNA**→ rpmE (PRK00019)Ribosomal_L31 (pfam01197)→ |
| Sen-318-1 | NZ_AYDU01000103.1 | - | 37530 | 37478 | **RNA**→ rpmE (PRK00019)Ribosomal_L31 (pfam01197)→ rpmE (PRK00019)Ribosomal_L31 (pfam01197)→ |
| Sen-288-942 | NZ_CJXJ01000016.1 | + | 104162 | 104214 | **RNA**→ rpmE (PRK00019)Ribosomal_L31 (pfam01197)→ |
| Sen-288-943 | NZ_CEZQ01000016.1 | + | 104159 | 104211 | **RNA**→ rpmE (PRK00019)Ribosomal_L31 (pfam01197)→ |
| Sen-288-944 | NZ_CJEO01000016.1 | + | 104148 | 104200 | **RNA**→ rpmE (PRK00019)Ribosomal_L31 (pfam01197)→ |
| Sen-288-945 | NZ_CILQ01000012.1 | - | 39601 | 39549 | **RNA**→ rpmE (PRK00019)Ribosomal_L31 (pfam01197)→ |
| Sen-288-946 | NZ_CJNE01000014.1 | - | 41516 | 41464 | **RNA**→ rpmE (PRK00019)Ribosomal_L31 (pfam01197)→ |
| Sen-288-947 | NZ_CJHM01000016.1 | + | 104139 | 104191 | **RNA**→ rpmE (PRK00019)Ribosomal_L31 (pfam01197)→ |
| Sen-288-948 | NZ_CJLG01000016.1 | + | 103212 | 103264 | **RNA**→ rpmE (PRK00019)Ribosomal_L31 (pfam01197)→ |
| Sen-288-949 | NZ_CJTJ01000014.1 | - | 41231 | 41179 | **RNA**→ rpmE (PRK00019)Ribosomal_L31 (pfam01197)→ |
| Sen-288-950 | NZ_CIZX01000016.1 | + | 104149 | 104201 | **RNA**→ rpmE (PRK00019)Ribosomal_L31 (pfam01197)→ |
| Sen-288-951 | NZ_CJAX01000016.1 | + | 104135 | 104187 | **RNA**→ rpmE (PRK00019)Ribosomal_L31 (pfam01197)→ |
| Sen-288-952 | NZ_CJPF01000015.1 | - | 41228 | 41176 | **RNA**→ rpmE (PRK00019)Ribosomal_L31 (pfam01197)→ |
| Sen-288-953 | NZ_CJTI01000014.1 | - | 41231 | 41179 | **RNA**→ rpmE (PRK00019)Ribosomal_L31 (pfam01197)→ |
| Sen-288-954 | NZ_CJFV01000016.1 | - | 39579 | 39527 | **RNA**→ rpmE (PRK00019)Ribosomal_L31 (pfam01197)→ |
| Sen-288-955 | NZ_CJQT01000015.1 | + | 104170 | 104222 | **RNA**→ rpmE (PRK00019)Ribosomal_L31 (pfam01197)→ |
| Sen-288-956 | NZ_CEYX01000016.1 | - | 39535 | 39483 | **RNA**→ rpmE (PRK00019)Ribosomal_L31 (pfam01197)→ |
| Sen-288-957 | NZ_CIVS01000015.1 | + | 109396 | 109448 | **RNA**→ rpmE (PRK00019)Ribosomal_L31 (pfam01197)→ |
| Sen-288-958 | NZ_CJJE01000015.1 | + | 104136 | 104188 | **RNA**→ rpmE (PRK00019)Ribosomal_L31 (pfam01197)→ |
| Sen-288-959 | NZ_CJPK01000008.1 | - | 39601 | 39549 | **RNA**→ rpmE (PRK00019)Ribosomal_L31 (pfam01197)→ |
| Sen-288-960 | NZ_CJAH01000016.1 | - | 39456 | 39404 | **RNA**→ rpmE (PRK00019)Ribosomal_L31 (pfam01197)→ |

| | | | | | |
|---|---|---|---|---|---|
| Sen-288-961 | NZ_CIYQ01000011.1 | + | 116070 | 116122 | **RNA**→ rpmE (PRK00019)Ribosomal_L31 (pfam01197)→ |
| Sen-288-962 | NZ_CIXU01000014.1 | + | 104140 | 104192 | **RNA**→ rpmE (PRK00019)Ribosomal_L31 (pfam01197)→ |
| Sen-288-963 | NZ_CJPS01000013.1 | - | 39601 | 39549 | **RNA**→ rpmE (PRK00019)Ribosomal_L31 (pfam01197)→ |
| Sen-288-964 | NZ_CJLI01000006.1 | - | 39456 | 39404 | **RNA**→ rpmE (PRK00019)Ribosomal_L31 (pfam01197)→ |
| Sen-288-965 | NZ_CJTE01000016.1 | - | 39371 | 39319 | **RNA**→ rpmE (PRK00019)Ribosomal_L31 (pfam01197)→ |
| Sen-288-966 | NZ_CIVU01000016.1 | + | 104172 | 104224 | **RNA**→ rpmE (PRK00019)Ribosomal_L31 (pfam01197)→ |
| Sen-288-967 | NZ_CJYA01000016.1 | - | 39456 | 39404 | **RNA**→ rpmE (PRK00019)Ribosomal_L31 (pfam01197)→ |
| Sen-288-968 | NZ_CJHY01000015.1 | + | 104142 | 104194 | **RNA**→ rpmE (PRK00019)Ribosomal_L31 (pfam01197)→ |
| Sen-288-969 | NZ_CIUW01000016.1 | + | 104162 | 104214 | **RNA**→ rpmE (PRK00019)Ribosomal_L31 (pfam01197)→ |
| Sen-288-970 | NZ_CIUX01000015.1 | + | 107419 | 107471 | **RNA**→ rpmE (PRK00019)Ribosomal_L31 (pfam01197)→ |
| Sen-288-971 | NZ_CIZH01000015.1 | - | 39531 | 39479 | **RNA**→ rpmE (PRK00019)Ribosomal_L31 (pfam01197)→ |
| Sen-288-972 | NZ_CJJR01000013.1 | - | 41201 | 41149 | **RNA**→ rpmE (PRK00019)Ribosomal_L31 (pfam01197)→ |
| Sen-288-973 | NZ_CJIG01000015.1 | - | 39560 | 39508 | **RNA**→ rpmE (PRK00019)Ribosomal_L31 (pfam01197)→ |
| Sen-288-974 | NZ_CJMX01000014.1 | - | 41231 | 41179 | **RNA**→ rpmE (PRK00019)Ribosomal_L31 (pfam01197)→ |
| Sen-288-975 | NZ_CJQY01000016.1 | - | 39601 | 39549 | **RNA**→ rpmE (PRK00019)Ribosomal_L31 (pfam01197)→ |
| Sen-288-976 | NZ_CJOY01000016.1 | - | 39706 | 39654 | **RNA**→ rpmE (PRK00019)Ribosomal_L31 (pfam01197)→ |
| Sen-288-977 | NZ_CILU01000016.1 | - | 39370 | 39318 | **RNA**→ rpmE (PRK00019)Ribosomal_L31 (pfam01197)→ |
| Sen-288-978 | NZ_CJXV01000017.1 | - | 39374 | 39322 | **RNA**→ rpmE (PRK00019)Ribosomal_L31 (pfam01197)→ |
| Sen-288-979 | NZ_CJAB01000016.1 | - | 39538 | 39486 | **RNA**→ rpmE (PRK00019)Ribosomal_L31 (pfam01197)→ |
| Sen-288-980 | NZ_CJXT01000016.1 | + | 104164 | 104216 | **RNA**→ rpmE (PRK00019)Ribosomal_L31 (pfam01197)→ |
| Sen-288-981 | NZ_CIXP01000022.1 | - | 39353 | 39301 | **RNA**→ rpmE (PRK00019)Ribosomal_L31 (pfam01197)→ |
| Sen-288-982 | NZ_CJTD01000014.1 | + | 107864 | 107916 | **RNA**→ rpmE (PRK00019)Ribosomal_L31 (pfam01197)→ |
| Sen-288-983 | NZ_CJLO01000014.1 | - | 41501 | 41449 | **RNA**→ rpmE (PRK00019)Ribosomal_L31 (pfam01197)→ |
| Sen-288-1123 | NZ_CIYN01000015.1 | - | 39528 | 39476 | **RNA**→ rpmE (PRK00019)Ribosomal_L31 (pfam01197)→ |
| Sen-288-984 | NZ_CJFC01000015.1 | + | 104157 | 104209 | **RNA**→ rpmE (PRK00019)Ribosomal_L31 (pfam01197)→ |
| Sen-288-985 | NZ_CJWG01000017.1 | + | 104159 | 104211 | **RNA**→ rpmE (PRK00019)Ribosomal_L31 (pfam01197)→ |
| Sen-288-986 | NZ_CJAR01000014.1 | + | 104154 | 104206 | **RNA**→ rpmE (PRK00019)Ribosomal_L31 (pfam01197)→ |
| Sen-288-987 | NZ_CJPY01000015.1 | + | 104172 | 104224 | **RNA**→ rpmE (PRK00019)Ribosomal_L31 (pfam01197)→ |
| Sen-288-988 | NZ_CJZG01000016.1 | - | 39601 | 39549 | **RNA**→ rpmE (PRK00019)Ribosomal_L31 (pfam01197)→ |
| Sen-288-989 | NZ_CJQO01000015.1 | + | 104149 | 104201 | **RNA**→ rpmE (PRK00019)Ribosomal_L31 (pfam01197)→ |
| Sen-288-990 | NZ_CJYY01000014.1 | + | 107837 | 107889 | **RNA**→ rpmE (PRK00019)Ribosomal_L31 (pfam01197)→ |
| Sen-175-1 | NZ_AMNI01000018.1 | + | 43440 | 43492 | **RNA**→ rpmE (PRK00019)Ribosomal_L31 (pfam01197)→ |
| Sen-209-1 | NZ_AMNN01000060.1 | + | 16157 | 16209 | **RNA**→ rpmE (PRK00019)Ribosomal_L31 (pfam01197)→ |
| Sen-204-1 | NZ_AMNS01000001.1 | - | 36301 | 36249 | **RNA**→ rpmE (PRK00019)Ribosomal_L31 (pfam01197)→ |
| Sen-211-1 | NZ_AMNJ01000008.1 | + | 43441 | 43493 | **RNA**→ rpmE (PRK00019)Ribosomal_L31 (pfam01197)→ |
| Sen-200-1 | NZ_AMNO01000081.1 | - | 37603 | 37551 | **RNA**→ rpmE (PRK00019)Ribosomal_L31 (pfam01197)→ |
| Sen-205-1 | NZ_AMNT01000032.1 | + | 25569 | 25621 | **RNA**→ rpmE (PRK00019)Ribosomal_L31 (pfam01197)→ |
| Sen-213-1 | NZ_AMMB01000092.1 | + | 131954 | 132006 | **RNA**→ rpmE (PRK00019)Ribosomal_L31 (pfam01197)→ |
| Sen-212-1 | NZ_AMNK01000018.1 | - | 37498 | 37446 | **RNA**→ rpmE (PRK00019)Ribosomal_L31 (pfam01197)→ |
| Sen-306-51 | NZ_LINK01000004.1 | - | 9890 | 9838 | **RNA**→ rpmE (PRK00019)Ribosomal_L31 (pfam01197)→ |
| Sen-198-1 | NZ_AMNP01000004.1 | - | 37494 | 37442 | **RNA**→ rpmE (PRK00019)Ribosomal_L31 (pfam01197)→ |
| Sen-306-52 | NZ_LINL01000061.1 | + | 42496 | 42548 | **RNA**→ rpmE (PRK00019)Ribosomal_L31 (pfam01197)→ |
| Sen-773-1 | NZ_AKYL01000049.1 | + | 100902 | 100954 | **RNA**→ rpmE (PRK00019)Ribosomal_L31 (pfam01197)→ |
| Sen-790-1 | NZ_AUQG01000125.1 | - | 37523 | 37471 | **RNA**→ rpmE (PRK00019)Ribosomal_L31 (pfam01197)→ |
| Sen-788-1 | NZ_AUQJ01000078.1 | - | 37855 | 37803 | **RNA**→ rpmE (PRK00019)Ribosomal_L31 (pfam01197)→ |
| Sen-225-48 | NZ_LINR01000022.1 | + | 61621 | 61673 | **RNA**→ rpmE (PRK00019)Ribosomal_L31 (pfam01197)→ |
| Sen-761-1 | NZ_AMBU01000035.1 | - | 37498 | 37446 | **RNA**→ rpmE (PRK00019)Ribosomal_L31 (pfam01197)→ |
| Sen-170-46 | NZ_LIMW01000021.1 | - | 37518 | 37466 | **RNA**→ rpmE (PRK00019)Ribosomal_L31 (pfam01197)→ |
| Sen-201-1 | NZ_AKYN01000107.1 | - | 37697 | 37645 | **RNA**→ rpmE (PRK00019)Ribosomal_L31 (pfam01197)→ |
| Sen-208-1 | NZ_AMMC01000016.1 | + | 42026 | 42078 | **RNA**→ rpmE (PRK00019)Ribosomal_L31 (pfam01197)→ |
| Sen-306-53 | NZ_LIOG01000018.1 | - | 45387 | 45335 | **RNA**→ rpmE (PRK00019)Ribosomal_L31 (pfam01197)→ |
| Sen-767-1 | NZ_AMMT01000186.1 | - | 37600 | 37548 | **RNA**→ rpmE (PRK00019)Ribosomal_L31 (pfam01197)→ |

| | | | | | |
|---|---|---|---|---|---|
| Sen-306-54 | NZ_LINM01000005.1 | - | 45297 | 45245 | **RNA**→ rpmE (PRK00019)Ribosomal_L31 (pfam01197)→ |
| Sen-225-49 | NZ_LINO01000036.1 | - | 37859 | 37807 | **RNA**→ rpmE (PRK00019)Ribosomal_L31 (pfam01197)→ |
| Sen-769-1 | NZ_AMMV01000445.1 | - | 37768 | 37716 | **RNA**→ rpmE (PRK00019)Ribosomal_L31 (pfam01197)→ |
| Sen-207-1 | NZ_AMNC01000031.1 | + | 43443 | 43495 | **RNA**→ rpmE (PRK00019)Ribosomal_L31 (pfam01197)→ |
| Sen-202-1 | NZ_AMND01000144.1 | - | 23081 | 23029 | **RNA**→ rpmE (PRK00019)Ribosomal_L31 (pfam01197)→ |
| Sen-764-1 | NZ_AMBX01000018.1 | - | 37498 | 37446 | **RNA**→ rpmE (PRK00019)Ribosomal_L31 (pfam01197)→ |
| Sen-221-1 | NZ_AMLR01000008.1 | + | 52031 | 52083 | **RNA**→ rpmE (PRK00019)Ribosomal_L31 (pfam01197)→ |
| Sen-5-5 | NZ_LIMH01000026.1 | - | 45667 | 45615 | **RNA**→ rpmE (PRK00019)Ribosomal_L31 (pfam01197)→ |
| Sen-775-1 | NZ_AMMJ01000088.1 | - | 37496 | 37444 | **RNA**→ rpmE (PRK00019)Ribosomal_L31 (pfam01197)→ |
| Sen-170-47 | NZ_LIMR01000020.1 | + | 100579 | 100631 | **RNA**→ rpmE (PRK00019)Ribosomal_L31 (pfam01197)→ |
| Sen-215-1 | NZ_AMMW01000224.1 | + | 1116 | 1168 | **RNA**→ rpmE (PRK00019)Ribosomal_L31 (pfam01197)→ |
| Sen-306-55 | NZ_LINE01000024.1 | - | 45200 | 45148 | **RNA**→ rpmE (PRK00019)Ribosomal_L31 (pfam01197)→ |
| Sen-210-1 | NZ_AMNE01000034.1 | + | 131957 | 132009 | **RNA**→ rpmE (PRK00019)Ribosomal_L31 (pfam01197)→ |
| Sen-199-1 | NZ_AMNQ01000115.1 | - | 15260 | 15208 | **RNA**→ rpmE (PRK00019)Ribosomal_L31 (pfam01197)→ |
| Sen-197-1 | NZ_AMNR01000090.1 | - | 37498 | 37446 | **RNA**→ rpmE (PRK00019)Ribosomal_L31 (pfam01197)→ |
| Sen-217-1 | NZ_AMLX01000040.1 | + | 100589 | 100641 | **RNA**→ rpmE (PRK00019)Ribosomal_L31 (pfam01197)→ |
| Sen-787-1 | NZ_AUQF01000013.1 | + | 87900 | 87952 | **RNA**→ rpmE (PRK00019)Ribosomal_L31 (pfam01197)→ |
| Sen-779-1 | NZ_ATHK01000082.1 | - | 37777 | 37725 | **RNA**→ rpmE (PRK00019)Ribosomal_L31 (pfam01197)→ |
| Sen-765-1 | NZ_AKYM01000112.1 | + | 100553 | 100605 | **RNA**→ rpmE (PRK00019)Ribosomal_L31 (pfam01197)→ |
| Sen-218-1 | NZ_AMLV01000008.1 | - | 37697 | 37645 | **RNA**→ rpmE (PRK00019)Ribosomal_L31 (pfam01197)→ |
| Sen-279-14 | NZ_LINN01000029.1 | + | 4123 | 4175 | **RNA**→ rpmE (PRK00019)Ribosomal_L31 (pfam01197)→ |
| Sen-768-1 | NZ_AMMQ01000223.1 | - | 36339 | 36287 | **RNA**→ rpmE (PRK00019)Ribosomal_L31 (pfam01197)→ |
| Sen-214-1 | NZ_AMNA01000068.1 | - | 34202 | 34150 | **RNA**→ rpmE (PRK00019)Ribosomal_L31 (pfam01197)→ |
| Sen-306-56 | NZ_LIMZ01000035.1 | - | 45097 | 45045 | **RNA**→ rpmE (PRK00019)Ribosomal_L31 (pfam01197)→ |
| Sen-789-1 | NZ_AUQH01000002.1 | + | 110425 | 110477 | **RNA**→ rpmE (PRK00019)Ribosomal_L31 (pfam01197)→ |
| Sen-762-1 | NZ_AMBV01000030.1 | - | 37701 | 37649 | **RNA**→ rpmE (PRK00019)Ribosomal_L31 (pfam01197)→ |
| Sen-237-16 | NZ_LINT01000029.1 | - | 9659 | 9607 | **RNA**→ rpmE (PRK00019)Ribosomal_L31 (pfam01197)→ |
| Sen-776-1 | NZ_AMLT01000019.1 | + | 97303 | 97355 | **RNA**→ rpmE (PRK00019)Ribosomal_L31 (pfam01197)→ |
| Sen-778-1 | NZ_ASRF01000014.1 | - | 24955 | 24903 | **RNA**→ rpmE (PRK00019)Ribosomal_L31 (pfam01197)→ |
| Sen-219-1 | NZ_AMLU01000019.1 | + | 25396 | 25448 | **RNA**→ rpmE (PRK00019)Ribosomal_L31 (pfam01197)→ |
| Sen-243-2 | NZ_LINY01000015.1 | + | 105471 | 105523 | **RNA**→ rpmE (PRK00019)Ribosomal_L31 (pfam01197)→ |
| Sen-203-1 | NZ_AMNF01000034.1 | - | 37601 | 37549 | **RNA**→ rpmE (PRK00019)Ribosomal_L31 (pfam01197)→ |
| Sen-170-48 | NZ_LIOC01000040.1 | + | 43729 | 43781 | **RNA**→ rpmE (PRK00019)Ribosomal_L31 (pfam01197)→ |
| Sen-777-1 | NZ_AOZC01000026.1 | + | 95441 | 95493 | **RNA**→ rpmE (PRK00019)Ribosomal_L31 (pfam01197)→ |
| Sen-306-57 | NZ_LIOB01000011.1 | + | 104279 | 104331 | **RNA**→ rpmE (PRK00019)Ribosomal_L31 (pfam01197)→ |
| Sen-782-1 | NZ_AOZA01000016.1 | + | 103042 | 103094 | **RNA**→ rpmE (PRK00019)Ribosomal_L31 (pfam01197)→ rpmE (PRK00019)Ribosomal_L31 (pfam01197)→ |
| Sen-770-1 | NZ_AKYK01000046.1 | + | 100992 | 101044 | **RNA**→ rpmE (PRK00019)Ribosomal_L31 (pfam01197)→ |
| Sen-181-1 | NZ_AKYO01000018.1 | - | 37601 | 37549 | **RNA**→ rpmE (PRK00019)Ribosomal_L31 (pfam01197)→ |
| Sen-763-1 | NZ_AMBW01000014.1 | - | 37691 | 37639 | **RNA**→ rpmE (PRK00019)Ribosomal_L31 (pfam01197)→ |
| Sen-225-50 | NZ_LIMQ01000012.1 | - | 37858 | 37806 | **RNA**→ rpmE (PRK00019)Ribosomal_L31 (pfam01197)→ |
| Sen-757-1 | NZ_AMMX01000038.1 | - | 23868 | 23816 | **RNA**→ rpmE (PRK00019)Ribosomal_L31 (pfam01197)→ |
| Sen-781-1 | NZ_AOZD01000012.1 | - | 25990 | 25938 | **RNA**→ rpmE (PRK00019)Ribosomal_L31 (pfam01197)→ rpmE (PRK00019)Ribosomal_L31 (pfam01197)→ |
| Sen-170-49 | NZ_LIOA01000019.1 | - | 37518 | 37466 | **RNA**→ rpmE (PRK00019)Ribosomal_L31 (pfam01197)→ |
| Sen-206-1 | NZ_AMNU01000070.1 | - | 37496 | 37444 | **RNA**→ rpmE (PRK00019)Ribosomal_L31 (pfam01197)→ |
| Sen-786-1 | NZ_AYDR01000087.1 | - | 37855 | 37803 | **RNA**→ rpmE (PRK00019)Ribosomal_L31 (pfam01197)→ |
| Sen-224-3 | NZ_LINA01000049.1 | - | 337 | 285 | **RNA**→ rpmE (PRK00019)Ribosomal_L31 (pfam01197)→ |
| Sen-278-22 | NZ_LING01000005.1 | - | 45365 | 45313 | **RNA**→ rpmE (PRK00019)Ribosomal_L31 (pfam01197)→ |
| Sen-225-51 | NZ_LINU01000054.1 | + | 43650 | 43702 | **RNA**→ rpmE (PRK00019)Ribosomal_L31 (pfam01197)→ |
| Sen-225-52 | NZ_LIOD01000004.1 | + | 110443 | 110495 | **RNA**→ rpmE (PRK00019)Ribosomal_L31 (pfam01197)→ |
| Sen-230-4 | NZ_LIOE01000015.1 | - | 37726 | 37674 | **RNA**→ rpmE (PRK00019)Ribosomal_L31 (pfam01197)→ |
| Sen-233-5 | NZ_LINX01000023.1 | + | 97750 | 97802 | **RNA**→ rpmE (PRK00019)Ribosomal_L31 (pfam01197)→ |
| Sen-288-991 | NZ_CJPA01000016.1 | - | 39706 | 39654 | **RNA**→ rpmE (PRK00019)Ribosomal_L31 (pfam01197)→ |

| | | | | | |
|---|---|---|---|---|---|
| Sen-288-992 | NZ_CJAF01000016.1 | - | 39540 | 39488 | **RNA**→ rpmE (PRK00019)Ribosomal_L31 (pfam01197)→ |
| Sen-288-993 | NZ_CIZS01000014.1 | + | 104154 | 104206 | **RNA**→ rpmE (PRK00019)Ribosomal_L31 (pfam01197)→ |
| Sen-288-1124 | NZ_CIYB01000016.1 | + | 104158 | 104210 | **RNA**→ rpmE (PRK00019)Ribosomal_L31 (pfam01197)→ |
| Sen-288-994 | NZ_CJVP01000015.1 | + | 103906 | 103958 | **RNA**→ rpmE (PRK00019)Ribosomal_L31 (pfam01197)→ |
| Sen-288-995 | NZ_CJSJ01000015.1 | - | 41231 | 41179 | **RNA**→ rpmE (PRK00019)Ribosomal_L31 (pfam01197)→ |
| Sen-288-996 | NZ_CIYR01000014.1 | + | 107420 | 107472 | **RNA**→ rpmE (PRK00019)Ribosomal_L31 (pfam01197)→ |
| Sen-288-997 | NZ_CJXB01000011.1 | - | 39375 | 39323 | **RNA**→ rpmE (PRK00019)Ribosomal_L31 (pfam01197)→ |
| Sen-288-998 | NZ_CJOR01000016.1 | - | 39723 | 39671 | **RNA**→ rpmE (PRK00019)Ribosomal_L31 (pfam01197)→ |
| Sen-288-999 | NZ_CJXG01000016.1 | - | 39456 | 39404 | **RNA**→ rpmE (PRK00019)Ribosomal_L31 (pfam01197)→ |
| Sen-306-58 | NZ_LIMT01000017.1 | - | 45297 | 45245 | **RNA**→ rpmE (PRK00019)Ribosomal_L31 (pfam01197)→ |
| Sen-288-1000 | NZ_CJKE01000013.1 | + | 107689 | 107741 | **RNA**→ rpmE (PRK00019)Ribosomal_L31 (pfam01197)→ |
| Sen-288-1001 | NZ_CJNZ01000016.1 | - | 41231 | 41179 | **RNA**→ rpmE (PRK00019)Ribosomal_L31 (pfam01197)→ |
| Sen-288-1002 | NZ_CJDB01000010.1 | - | 39456 | 39404 | **RNA**→ rpmE (PRK00019)Ribosomal_L31 (pfam01197)→ |
| Sen-288-1003 | NZ_CJOI01000014.1 | - | 39382 | 39330 | **RNA**→ rpmE (PRK00019)Ribosomal_L31 (pfam01197)→ |
| Sen-288-1004 | NZ_CJSV01000015.1 | + | 104172 | 104224 | **RNA**→ rpmE (PRK00019)Ribosomal_L31 (pfam01197)→ |
| Sen-288-1005 | NZ_CJQV01000016.1 | + | 104162 | 104214 | **RNA**→ rpmE (PRK00019)Ribosomal_L31 (pfam01197)→ |
| Sen-288-1006 | NZ_CIXV01000015.1 | + | 104055 | 104107 | **RNA**→ rpmE (PRK00019)Ribosomal_L31 (pfam01197)→ |
| Sen-288-1007 | NZ_CJFB01000014.1 | - | 39456 | 39404 | **RNA**→ rpmE (PRK00019)Ribosomal_L31 (pfam01197)→ |
| Sen-288-1008 | NZ_CIZO01000016.1 | + | 104149 | 104201 | **RNA**→ rpmE (PRK00019)Ribosomal_L31 (pfam01197)→ |
| Sen-288-1009 | NZ_CJQD01000015.1 | + | 107622 | 107674 | **RNA**→ rpmE (PRK00019)Ribosomal_L31 (pfam01197)→ |
| Sen-288-1010 | NZ_CJML01000013.1 | + | 107971 | 108023 | **RNA**→ rpmE (PRK00019)Ribosomal_L31 (pfam01197)→ |
| Sen-288-1011 | NZ_CJQK01000014.1 | + | 104156 | 104208 | **RNA**→ rpmE (PRK00019)Ribosomal_L31 (pfam01197)→ |
| Sen-288-1012 | NZ_CJYI01000016.1 | - | 39587 | 39535 | **RNA**→ rpmE (PRK00019)Ribosomal_L31 (pfam01197)→ |
| Sen-288-1013 | NZ_CJWK01000015.1 | - | 41231 | 41179 | **RNA**→ rpmE (PRK00019)Ribosomal_L31 (pfam01197)→ |
| Sen-288-1014 | NZ_CJAN01000017.1 | - | 39533 | 39481 | **RNA**→ rpmE (PRK00019)Ribosomal_L31 (pfam01197)→ |
| Sen-288-1015 | NZ_CJBI01000011.1 | - | 39535 | 39483 | **RNA**→ rpmE (PRK00019)Ribosomal_L31 (pfam01197)→ |
| Sen-288-1016 | NZ_CJIK01000015.1 | - | 39560 | 39508 | **RNA**→ rpmE (PRK00019)Ribosomal_L31 (pfam01197)→ |
| Sen-288-1017 | NZ_CJWZ01000010.1 | - | 39382 | 39330 | **RNA**→ rpmE (PRK00019)Ribosomal_L31 (pfam01197)→ |
| Sen-288-1018 | NZ_CJUH01000016.1 | + | 104153 | 104205 | **RNA**→ rpmE (PRK00019)Ribosomal_L31 (pfam01197)→ |
| Sen-288-1019 | NZ_CJQU01000014.1 | - | 39534 | 39482 | **RNA**→ rpmE (PRK00019)Ribosomal_L31 (pfam01197)→ |
| Sen-288-1020 | NZ_CJBD01000015.1 | - | 39535 | 39483 | **RNA**→ rpmE (PRK00019)Ribosomal_L31 (pfam01197)→ |
| Sen-288-1021 | NZ_CJLK01000010.1 | - | 39456 | 39404 | **RNA**→ rpmE (PRK00019)Ribosomal_L31 (pfam01197)→ |
| Sen-288-1022 | NZ_CJIJ01000015.1 | + | 104149 | 104201 | **RNA**→ rpmE (PRK00019)Ribosomal_L31 (pfam01197)→ |
| Sen-288-1023 | NZ_CJOS01000016.1 | - | 39601 | 39549 | **RNA**→ rpmE (PRK00019)Ribosomal_L31 (pfam01197)→ |
| Sen-288-1024 | NZ_CJFY01000016.1 | + | 104150 | 104202 | **RNA**→ rpmE (PRK00019)Ribosomal_L31 (pfam01197)→ |
| Sen-288-1025 | NZ_CJKK01000012.1 | + | 107993 | 108045 | **RNA**→ rpmE (PRK00019)Ribosomal_L31 (pfam01197)→ |
| Sen-288-1026 | NZ_CJHE01000016.1 | - | 39535 | 39483 | **RNA**→ rpmE (PRK00019)Ribosomal_L31 (pfam01197)→ |
| Sen-288-1027 | NZ_CJYF01000016.1 | + | 104092 | 104144 | **RNA**→ rpmE (PRK00019)Ribosomal_L31 (pfam01197)→ |
| Sen-288-1028 | NZ_CJKV01000015.1 | - | 41231 | 41179 | **RNA**→ rpmE (PRK00019)Ribosomal_L31 (pfam01197)→ |
| Sen-288-1029 | NZ_CJSK01000014.1 | + | 107994 | 108046 | **RNA**→ rpmE (PRK00019)Ribosomal_L31 (pfam01197)→ |
| Sen-288-1030 | NZ_CJVT01000015.1 | - | 39373 | 39321 | **RNA**→ rpmE (PRK00019)Ribosomal_L31 (pfam01197)→ |
| Sen-288-1031 | NZ_CJBP01000016.1 | + | 104148 | 104200 | **RNA**→ rpmE (PRK00019)Ribosomal_L31 (pfam01197)→ |
| Sen-288-1032 | NZ_CJCZ01000022.1 | + | 43275 | 43327 | **RNA**→ rpmE (PRK00019)Ribosomal_L31 (pfam01197)→ |
| Sen-288-1033 | NZ_CJSD01000015.1 | - | 39361 | 39309 | **RNA**→ rpmE (PRK00019)Ribosomal_L31 (pfam01197)→ |
| Sen-288-1034 | NZ_CJMQ01000013.1 | + | 107855 | 107907 | **RNA**→ rpmE (PRK00019)Ribosomal_L31 (pfam01197)→ |
| Sen-288-1035 | NZ_CJVK01000013.1 | + | 104142 | 104194 | **RNA**→ rpmE (PRK00019)Ribosomal_L31 (pfam01197)→ |
| Sen-288-1036 | NZ_CJVM01000016.1 | + | 104161 | 104213 | **RNA**→ rpmE (PRK00019)Ribosomal_L31 (pfam01197)→ |
| Sen-288-1037 | NZ_CJJW01000015.1 | + | 104139 | 104191 | **RNA**→ rpmE (PRK00019)Ribosomal_L31 (pfam01197)→ |
| Sen-288-1038 | NZ_CJSL01000014.1 | + | 103996 | 104048 | **RNA**→ rpmE (PRK00019)Ribosomal_L31 (pfam01197)→ |
| Sen-288-1039 | NZ_CIYF01000014.1 | + | 104144 | 104196 | **RNA**→ rpmE (PRK00019)Ribosomal_L31 (pfam01197)→ |
| Sen-306-59 | NZ_LIMV01000003.1 | + | 104298 | 104350 | **RNA**→ rpmE (PRK00019)Ribosomal_L31 (pfam01197)→ |
| Sen-306-60 | NZ_LINF01000043.1 | + | 42428 | 42480 | **RNA**→ rpmE (PRK00019)Ribosomal_L31 (pfam01197)→ |

| | | | | | |
|---|---|---|---|---|---|
| Sen-306-61 | NZ_LINJ01000010.1 | + | 55677 | 55729 | **RNA**→ rpmE (PRK00019)Ribosomal_L31 (pfam01197)→ |
| Sen-306-62 | NZ_LINI01000110.1 | + | 22614 | 22666 | **RNA**→ rpmE (PRK00019)Ribosomal_L31 (pfam01197)→ |
| Sen-306-63 | NZ_LIMU01000037.1 | - | 45297 | 45245 | **RNA**→ rpmE (PRK00019)Ribosomal_L31 (pfam01197)→ |
| Sen-293-1 | NZ_JHAH01000023.1 | - | 45098 | 45046 | **RNA**→ rpmE (PRK00019)Ribosomal_L31 (pfam01197)→ |
| Sen-1015-1 | NZ_CAAR01000266.1 | - | 2721 | 2669 | **RNA**→ rpmE (PRK00019)Ribosomal_L31 (pfam01197)→ rpmE (PRK00019)Ribosomal_L31 (pfam01197)→ |
| Sen-296-1 | NZ_JHAE01000047.1 | + | 42429 | 42481 | **RNA**→ rpmE (PRK00019)Ribosomal_L31 (pfam01197)→ |
| Sen-297-1 | NZ_JHAF01000021.1 | - | 45098 | 45046 | **RNA**→ rpmE (PRK00019)Ribosomal_L31 (pfam01197)→ |
| Sen-300-30 | NZ_CQGI01000015.1 | + | 42522 | 42574 | **RNA**→ rpmE (PRK00019)Ribosomal_L31 (pfam01197)→ |
| Sen-300-31 | NZ_CQGG01000016.1 | + | 42512 | 42564 | **RNA**→ rpmE (PRK00019)Ribosomal_L31 (pfam01197)→ |
| Sen-300-32 | NZ_CQHF01000017.1 | - | 45178 | 45126 | **RNA**→ rpmE (PRK00019)Ribosomal_L31 (pfam01197)→ |
| Sen-294-1 | NZ_JHAI01000009.1 | - | 45098 | 45046 | **RNA**→ rpmE (PRK00019)Ribosomal_L31 (pfam01197)→ |
| Sen-300-33 | NZ_CQGW01000017.1 | + | 42551 | 42603 | **RNA**→ rpmE (PRK00019)Ribosomal_L31 (pfam01197)→ |
| Sen-300-34 | NZ_CQHV01000017.1 | - | 45252 | 45200 | **RNA**→ rpmE (PRK00019)Ribosomal_L31 (pfam01197)→ |
| Sen-300-35 | NZ_CQGU01000016.1 | + | 42527 | 42579 | **RNA**→ rpmE (PRK00019)Ribosomal_L31 (pfam01197)→ |
| Sen-951-1 | NZ_AJGK01000058.1 | - | 19387 | 19335 | **RNA**→ rpmE (PRK00019)Ribosomal_L31 (pfam01197)→ |
| Sen-952-1 | NZ_AKZO01000027.1 | - | 19333 | 19281 | **RNA**→ rpmE (PRK00019)Ribosomal_L31 (pfam01197)→ |
| Sen-967-1 | NZ_AUQL01000037.1 | + | 104438 | 104490 | **RNA**→ rpmE (PRK00019)Ribosomal_L31 (pfam01197)→ |
| Sen-959-1 | NZ_AUQT01000002.1 | - | 45277 | 45225 | **RNA**→ rpmE (PRK00019)Ribosomal_L31 (pfam01197)→ |
| Sen-966-1 | NZ_AUQN01000036.1 | + | 104414 | 104466 | **RNA**→ rpmE (PRK00019)Ribosomal_L31 (pfam01197)→ |
| Sen-964-1 | NZ_AUQM01000035.1 | + | 104413 | 104465 | **RNA**→ rpmE (PRK00019)Ribosomal_L31 (pfam01197)→ |
| Sen-961-1 | NZ_AUQV01000004.1 | - | 45184 | 45132 | **RNA**→ rpmE (PRK00019)Ribosomal_L31 (pfam01197)→ |
| Sen-290-1 | NZ_JABZ01000035.1 | - | 39416 | 39364 | **RNA**→ rpmE (PRK00019)Ribosomal_L31 (pfam01197)→ |
| Sen-300-36 | NZ_CQHE01000015.1 | - | 45235 | 45183 | **RNA**→ rpmE (PRK00019)Ribosomal_L31 (pfam01197)→ |
| Sen-288-1040 | NZ_CJRN01000014.1 | - | 41512 | 41460 | **RNA**→ rpmE (PRK00019)Ribosomal_L31 (pfam01197)→ |
| Sen-288-1041 | NZ_CJRJ01000014.1 | - | 39360 | 39308 | **RNA**→ rpmE (PRK00019)Ribosomal_L31 (pfam01197)→ |
| Sen-288-1042 | NZ_CJRD01000011.1 | + | 157688 | 157740 | **RNA**→ rpmE (PRK00019)Ribosomal_L31 (pfam01197)→ |
| Sen-300-37 | NZ_CQHU01000017.1 | + | 42532 | 42584 | **RNA**→ rpmE (PRK00019)Ribosomal_L31 (pfam01197)→ |
| Sen-490-1 | NZ_AKFX01000012.1 | - | 49253 | 49201 | **RNA**→ rpmE (PRK00019)Ribosomal_L31 (pfam01197)→ |
| Sen-963-1 | NZ_AOXC01000032.1 | - | 45190 | 45138 | **RNA**→ rpmE (PRK00019)Ribosomal_L31 (pfam01197)→ |
| Sen-495-1 | NZ_AYVI01000129.1 | - | 34669 | 34617 | **RNA**→ rpmE (PRK00019)Ribosomal_L31 (pfam01197)→ |
| Sen-300-38 | NZ_CQHJ01000016.1 | + | 43122 | 43174 | **RNA**→ rpmE (PRK00019)Ribosomal_L31 (pfam01197)→ |
| Sen-509-1 | NZ_ALEP01000014.1 | + | 103706 | 103758 | **RNA**→ rpmE (PRK00019)Ribosomal_L31 (pfam01197)→ rpmE (PRK00019)Ribosomal_L31 (pfam01197)→ |
| Sen-300-39 | NZ_CQGS01000016.1 | - | 45260 | 45208 | **RNA**→ rpmE (PRK00019)Ribosomal_L31 (pfam01197)→ |
| Sen-513-1 | NZ_ALGM01000023.1 | - | 49346 | 49294 | **RNA**→ rpmE (PRK00019)Ribosomal_L31 (pfam01197)→ rpmE (PRK00019)Ribosomal_L31 (pfam01197)→ |
| Sen-300-40 | NZ_CQHS01000018.1 | - | 45164 | 45112 | **RNA**→ rpmE (PRK00019)Ribosomal_L31 (pfam01197)→ |
| Sen-514-1 | NZ_ALGN01000014.1 | + | 103703 | 103755 | **RNA**→ rpmE (PRK00019)Ribosomal_L31 (pfam01197)→ rpmE (PRK00019)Ribosomal_L31 (pfam01197)→ |
| Sen-300-41 | NZ_CQHQ01000017.1 | + | 42531 | 42583 | **RNA**→ rpmE (PRK00019)Ribosomal_L31 (pfam01197)→ |
| Sen-288-1043 | NZ_CJRH01000015.1 | + | 104161 | 104213 | **RNA**→ rpmE (PRK00019)Ribosomal_L31 (pfam01197)→ |
| Sen-288-1044 | NZ_CJUS01000013.1 | - | 41231 | 41179 | **RNA**→ rpmE (PRK00019)Ribosomal_L31 (pfam01197)→ |
| Sen-484-1 | NZ_AJVD01000011.1 | - | 49252 | 49200 | **RNA**→ rpmE (PRK00019)Ribosomal_L31 (pfam01197)→ |
| Sen-300-42 | NZ_CQGL01000016.1 | + | 42551 | 42603 | **RNA**→ rpmE (PRK00019)Ribosomal_L31 (pfam01197)→ |
| Sen-472-1 | NZ_AKFY01000012.1 | + | 96338 | 96390 | **RNA**→ rpmE (PRK00019)Ribosomal_L31 (pfam01197)→ |
| Sen-300-43 | NZ_CQGY01000016.1 | + | 42551 | 42603 | **RNA**→ rpmE (PRK00019)Ribosomal_L31 (pfam01197)→ |
| Sen-473-1 | NZ_AKFZ01000014.1 | - | 49346 | 49294 | **RNA**→ rpmE (PRK00019)Ribosomal_L31 (pfam01197)→ |
| Sen-957-1 | NZ_AJTE01000099.1 | - | 13364 | 13312 | **RNA**→ rpmE (PRK00019)Ribosomal_L31 (pfam01197)→ |
| Sen-511-1 | NZ_ALGO01000040.1 | + | 103706 | 103758 | **RNA**→ rpmE (PRK00019)Ribosomal_L31 (pfam01197)→ rpmE (PRK00019)Ribosomal_L31 (pfam01197)→ |
| Sen-956-1 | NZ_AJTD01000295.1 | + | 4207 | 4259 | **RNA**→ rpmE (PRK00019)Ribosomal_L31 (pfam01197)→ |
| Sen-288-1045 | NZ_CJRA01000015.1 | + | 108136 | 108188 | **RNA**→ rpmE (PRK00019)Ribosomal_L31 (pfam01197)→ |
| Sen-491-1 | NZ_AKFW01000012.1 | - | 49253 | 49201 | **RNA**→ rpmE (PRK00019)Ribosomal_L31 (pfam01197)→ |
| Sen-954-1 | NZ_AJXA01000168.1 | + | 4121 | 4173 | **RNA**→ rpmE (PRK00019)Ribosomal_L31 (pfam01197)→ |
| Sen-960-1 | NZ_AUQU01000003.1 | - | 45184 | 45132 | **RNA**→ rpmE (PRK00019)Ribosomal_L31 (pfam01197)→ |
| Sen-517-1 | NZ_ALEI01000040.1 | - | 49344 | 49292 | **RNA**→ rpmE (PRK00019)Ribosomal_L31 (pfam01197)→ rpmE (PRK00019)Ribosomal_L31 (pfam01197)→ |

| | | | | | |
|---|---|---|---|---|---|
| Sen-497-1 | NZ_ALEN01000014.1 | - | 49251 | 49199 | **RNA**→ rpmE (PRK00019)Ribosomal_L31 (pfam01197)→ rpmE (PRK00019)Ribosomal_L31 (pfam01197)→ |
| Sen-965-1 | NZ_AUQO01000020.1 | - | 45183 | 45131 | **RNA**→ rpmE (PRK00019)Ribosomal_L31 (pfam01197)→ |
| Sen-508-1 | NZ_ALEO01000005.1 | - | 49252 | 49200 | **RNA**→ rpmE (PRK00019)Ribosomal_L31 (pfam01197)→ rpmE (PRK00019)Ribosomal_L31 (pfam01197)→ |
| Sen-300-44 | NZ_CQHI01000016.1 | + | 42508 | 42560 | **RNA**→ rpmE (PRK00019)Ribosomal_L31 (pfam01197)→ |
| Sen-300-45 | NZ_CQGK01000016.1 | + | 42508 | 42560 | **RNA**→ rpmE (PRK00019)Ribosomal_L31 (pfam01197)→ |
| Sen-300-46 | NZ_CQGN01000018.1 | + | 42526 | 42578 | **RNA**→ rpmE (PRK00019)Ribosomal_L31 (pfam01197)→ |
| Sen-300-47 | NZ_CQGV01000017.1 | + | 42523 | 42575 | **RNA**→ rpmE (PRK00019)Ribosomal_L31 (pfam01197)→ |
| Sen-478-1 | NZ_ALGV01000705.1 | + | 1026 | 1078 | **RNA**→ rpmE (PRK00019)Ribosomal_L31 (pfam01197)→ rpmE (PRK00019)Ribosomal_L31 (pfam01197)→ |
| Sen-1026-1 | NZ_ACBF01000011.1 | + | 90926 | 90978 | **RNA**→ rpmE (PRK00019)Ribosomal_L31 (pfam01197)→ rpmE (PRK00019)Ribosomal_L31 (pfam01197)→ |
| Sen-300-48 | NZ_CQGR01000016.1 | + | 42549 | 42601 | **RNA**→ rpmE (PRK00019)Ribosomal_L31 (pfam01197)→ |
| Sen-300-49 | NZ_CQHB01000017.1 | - | 45260 | 45208 | **RNA**→ rpmE (PRK00019)Ribosomal_L31 (pfam01197)→ |
| Sen-1012-1 | NZ_ABAM02000001.1 | + | 4673479 | 4673531 | **RNA**→ rpmE (PRK00019)Ribosomal_L31 (pfam01197)→ rpmE (PRK00019)Ribosomal_L31 (pfam01197)→ |
| Sen-299-1 | NZ_AHUV01000040.1 | - | 45184 | 45132 | **RNA**→ rpmE (PRK00019)Ribosomal_L31 (pfam01197)→ |
| Sen-479-1 | NZ_ALHB01000061.1 | - | 49392 | 49340 | **RNA**→ rpmE (PRK00019)Ribosomal_L31 (pfam01197)→ rpmE (PRK00019)Ribosomal_L31 (pfam01197)→ |
| Sen-504-1 | NZ_ALHG01000001.1 | - | 49346 | 49294 | **RNA**→ rpmE (PRK00019)Ribosomal_L31 (pfam01197)→ rpmE (PRK00019)Ribosomal_L31 (pfam01197)→ |
| Sen-913-1 | NZ_AFYZ01000020.1 | + | 135012 | 135064 | **RNA**→ rpmE (PRK00019)Ribosomal_L31 (pfam01197)→ |
| Sen-300-50 | NZ_CQGH01000017.1 | - | 45179 | 45127 | **RNA**→ rpmE (PRK00019)Ribosomal_L31 (pfam01197)→ |
| Sen-288-1046 | NZ_CJQZ01000013.1 | - | 39377 | 39325 | **RNA**→ rpmE (PRK00019)Ribosomal_L31 (pfam01197)→ |
| Sen-945-1 | NZ_AOXV01000110.1 | + | 71399 | 71451 | **RNA**→ rpmE (PRK00019)Ribosomal_L31 (pfam01197)→ |
| Sen-526-1 | NZ_ALEK01000010.1 | - | 49346 | 49294 | **RNA**→ rpmE (PRK00019)Ribosomal_L31 (pfam01197)→ rpmE (PRK00019)Ribosomal_L31 (pfam01197)→ |
| Sen-474-1 | NZ_AKGA01000012.1 | - | 49252 | 49200 | **RNA**→ rpmE (PRK00019)Ribosomal_L31 (pfam01197)→ |
| Sen-907-1 | NZ_AROZ01000014.1 | + | 3149 | 3201 | **RNA**→ rpmE (PRK00019)Ribosomal_L31 (pfam01197)→ rpmE (PRK00019)Ribosomal_L31 (pfam01197)→ |
| Sen-548-1 | NZ_ALFS01000030.1 | - | 49346 | 49294 | **RNA**→ rpmE (PRK00019)Ribosomal_L31 (pfam01197)→ rpmE (PRK00019)Ribosomal_L31 (pfam01197)→ |
| Sen-512-1 | NZ_ALEM01000020.1 | - | 49252 | 49200 | **RNA**→ rpmE (PRK00019)Ribosomal_L31 (pfam01197)→ rpmE (PRK00019)Ribosomal_L31 (pfam01197)→ |
| Sen-554-1 | NZ_ALFX01000011.1 | + | 103706 | 103758 | **RNA**→ rpmE (PRK00019)Ribosomal_L31 (pfam01197)→ rpmE (PRK00019)Ribosomal_L31 (pfam01197)→ |
| Sen-492-1 | NZ_ALEX01000033.1 | - | 49252 | 49200 | **RNA**→ rpmE (PRK00019)Ribosomal_L31 (pfam01197)→ rpmE (PRK00019)Ribosomal_L31 (pfam01197)→ |
| Sen-284-1 | NZ_APWL01000003.1 | - | 82310 | 82258 | **RNA**→ rpmE (PRK00019)Ribosomal_L31 (pfam01197)→ |
| Sen-550-1 | NZ_ALGD01000026.1 | + | 103707 | 103759 | **RNA**→ rpmE (PRK00019)Ribosomal_L31 (pfam01197)→ rpmE (PRK00019)Ribosomal_L31 (pfam01197)→ |
| Sen-506-1 | NZ_ALGK01000023.1 | - | 49346 | 49294 | **RNA**→ rpmE (PRK00019)Ribosomal_L31 (pfam01197)→ rpmE (PRK00019)Ribosomal_L31 (pfam01197)→ |
| Sen-914-1 | NZ_AFZA01000020.1 | + | 98771 | 98823 | **RNA**→ rpmE (PRK00019)Ribosomal_L31 (pfam01197)→ |
| Sen-900-1 | NZ_AYEE01000002.1 | - | 45343 | 45291 | **RNA**→ rpmE (PRK00019)Ribosomal_L31 (pfam01197)→ |
| Sen-527-1 | NZ_ALHH01000057.1 | + | 103704 | 103756 | **RNA**→ rpmE (PRK00019)Ribosomal_L31 (pfam01197)→ rpmE (PRK00019)Ribosomal_L31 (pfam01197)→ |
| Sen-515-1 | NZ_ALGP01000021.1 | + | 103706 | 103758 | **RNA**→ rpmE (PRK00019)Ribosomal_L31 (pfam01197)→ rpmE (PRK00019)Ribosomal_L31 (pfam01197)→ |
| Sen-533-1 | NZ_ALHM01000062.1 | - | 49346 | 49294 | **RNA**→ rpmE (PRK00019)Ribosomal_L31 (pfam01197)→ rpmE (PRK00019)Ribosomal_L31 (pfam01197)→ |
| Sen-501-1 | NZ_ALHE01000055.1 | - | 49346 | 49294 | **RNA**→ rpmE (PRK00019)Ribosomal_L31 (pfam01197)→ rpmE (PRK00019)Ribosomal_L31 (pfam01197)→ |
| Sen-937-1 | NZ_AYDP01000046.1 | - | 278 | 226 | **RNA**→ rpmE (PRK00019)Ribosomal_L31 (pfam01197)→ rpmE (PRK00019)Ribosomal_L31 (pfam01197)→ |
| Sen-502-1 | NZ_ALHF01000013.1 | - | 49346 | 49294 | **RNA**→ rpmE (PRK00019)Ribosomal_L31 (pfam01197)→ rpmE (PRK00019)Ribosomal_L31 (pfam01197)→ |
| Sen-543-1 | NZ_ALEL01000033.1 | - | 49252 | 49200 | **RNA**→ rpmE (PRK00019)Ribosomal_L31 (pfam01197)→ rpmE (PRK00019)Ribosomal_L31 (pfam01197)→ |
| Sen-519-1 | NZ_ALEQ01000021.1 | - | 49252 | 49200 | **RNA**→ rpmE (PRK00019)Ribosomal_L31 (pfam01197)→ rpmE (PRK00019)Ribosomal_L31 (pfam01197)→ |
| Sen-934-1 | NZ_AOXF01000013.1 | + | 104433 | 104485 | **RNA**→ rpmE (PRK00019)Ribosomal_L31 (pfam01197)→ |
| Sen-477-1 | NZ_AHUQ01000003.1 | + | 103706 | 103758 | **RNA**→ rpmE (PRK00019)Ribosomal_L31 (pfam01197)→ rpmE (PRK00019)Ribosomal_L31 (pfam01197)→ |
| Sen-544-1 | NZ_ALEV01000015.1 | - | 49252 | 49200 | **RNA**→ rpmE (PRK00019)Ribosomal_L31 (pfam01197)→ rpmE (PRK00019)Ribosomal_L31 (pfam01197)→ |
| Sen-939-1 | NZ_AOXW01000036.1 | + | 104624 | 104676 | **RNA**→ rpmE (PRK00019)Ribosomal_L31 (pfam01197)→ rpmE (PRK00019)Ribosomal_L31 (pfam01197)→ |
| Sen-564-1 | NZ_ALFK01000006.1 | - | 49252 | 49200 | **RNA**→ rpmE (PRK00019)Ribosomal_L31 (pfam01197)→ rpmE (PRK00019)Ribosomal_L31 (pfam01197)→ |
| Sen-288-1047 | NZ_CJRK01000015.1 | - | 41507 | 41455 | **RNA**→ rpmE (PRK00019)Ribosomal_L31 (pfam01197)→ |
| Sen-920-1 | NZ_AOYB01000081.1 | + | 63749 | 63801 | **RNA**→ rpmE (PRK00019)Ribosomal_L31 (pfam01197)→ |
| Sen-559-1 | NZ_ALFO01000008.1 | - | 49346 | 49294 | **RNA**→ rpmE (PRK00019)Ribosomal_L31 (pfam01197)→ rpmE (PRK00019)Ribosomal_L31 (pfam01197)→ |
| Sen-288-1048 | NZ_CJRB01000013.1 | + | 115815 | 115867 | **RNA**→ rpmE (PRK00019)Ribosomal_L31 (pfam01197)→ |
| Sen-552-1 | NZ_ALFP01000008.1 | + | 103716 | 103768 | **RNA**→ rpmE (PRK00019)Ribosomal_L31 (pfam01197)→ rpmE (PRK00019)Ribosomal_L31 (pfam01197)→ |
| Sen-288-1049 | NZ_CJQX01000014.1 | + | 104143 | 104195 | **RNA**→ rpmE (PRK00019)Ribosomal_L31 (pfam01197)→ |
| Sen-915-1 | NZ_AOYF01000047.1 | - | 45153 | 45101 | **RNA**→ rpmE (PRK00019)Ribosomal_L31 (pfam01197)→ rpmE (PRK00019)Ribosomal_L31 (pfam01197)→ |
| Sen-549-1 | NZ_ALFT01000006.1 | - | 49343 | 49291 | **RNA**→ rpmE (PRK00019)Ribosomal_L31 (pfam01197)→ rpmE (PRK00019)Ribosomal_L31 (pfam01197)→ |

| | | | | | |
|---|---|---|---|---|---|
| Sen-288-1050 | NZ_CJRF01000014.1 | - | 41231 | 41179 | **RNA**→ rpmE (PRK00019)Ribosomal_L31 (pfam01197)→ |
| Sen-557-1 | NZ_ALFU01000022.1 | - | 49252 | 49200 | **RNA**→ rpmE (PRK00019)Ribosomal_L31 (pfam01197)→ rpmE (PRK00019)Ribosomal_L31 (pfam01197)→ |
| Sen-919-1 | NZ_AOYG01000081.1 | - | 19765 | 19713 | **RNA**→ rpmE (PRK00019)Ribosomal_L31 (pfam01197)→ rpmE (PRK00019)Ribosomal_L31 (pfam01197)→ |
| Sen-555-1 | NZ_ALFY01000031.1 | - | 49343 | 49291 | **RNA**→ rpmE (PRK00019)Ribosomal_L31 (pfam01197)→ rpmE (PRK00019)Ribosomal_L31 (pfam01197)→ |
| Sen-938-1 | NZ_AOYP01000063.1 | - | 10370 | 10318 | **RNA**→ rpmE (PRK00019)Ribosomal_L31 (pfam01197)→ rpmE (PRK00019)Ribosomal_L31 (pfam01197)→ |
| Sen-560-1 | NZ_ALFZ01000023.1 | - | 49346 | 49294 | **RNA**→ rpmE (PRK00019)Ribosomal_L31 (pfam01197)→ rpmE (PRK00019)Ribosomal_L31 (pfam01197)→ |
| Sen-499-1 | NZ_ALIA01000399.1 | + | 1803 | 1855 | **RNA**→ rpmE (PRK00019)Ribosomal_L31 (pfam01197)→ rpmE (PRK00019)Ribosomal_L31 (pfam01197)→ |
| Sen-556-1 | NZ_ALGE01000038.1 | - | 49191 | 49139 | **RNA**→ rpmE (PRK00019)Ribosomal_L31 (pfam01197)→ rpmE (PRK00019)Ribosomal_L31 (pfam01197)→ |
| Sen-529-1 | NZ_ALHI01000010.1 | + | 103704 | 103756 | **RNA**→ rpmE (PRK00019)Ribosomal_L31 (pfam01197)→ rpmE (PRK00019)Ribosomal_L31 (pfam01197)→ |
| Sen-904-1 | NZ_APAK01000385.1 | - | 9620 | 9568 | **RNA**→ rpmE (PRK00019)Ribosomal_L31 (pfam01197)→ |
| Sen-535-1 | NZ_ALHN01000038.1 | + | 103703 | 103755 | **RNA**→ rpmE (PRK00019)Ribosomal_L31 (pfam01197)→ rpmE (PRK00019)Ribosomal_L31 (pfam01197)→ |
| Sen-924-1 | NZ_AHIA01000022.1 | - | 35206 | 35154 | **RNA**→ rpmE (PRK00019)Ribosomal_L31 (pfam01197)→ rpmE (PRK00019)Ribosomal_L31 (pfam01197)→ |
| Sen-516-1 | NZ_ALIC01000015.1 | - | 49252 | 49200 | **RNA**→ rpmE (PRK00019)Ribosomal_L31 (pfam01197)→ rpmE (PRK00019)Ribosomal_L31 (pfam01197)→ |
| Sen-523-1 | NZ_ALEJ01000020.1 | - | 49346 | 49294 | **RNA**→ rpmE (PRK00019)Ribosomal_L31 (pfam01197)→ rpmE (PRK00019)Ribosomal_L31 (pfam01197)→ |
| Sen-943-1 | NZ_CAGQ01000056.1 | - | 3031 | 2979 | **RNA**→ rpmE (PRK00019)Ribosomal_L31 (pfam01197)→ |
| Sen-288-1051 | NZ_CJRG01000004.1 | + | 104158 | 104210 | **RNA**→ rpmE (PRK00019)Ribosomal_L31 (pfam01197)→ |
| Sen-524-1 | NZ_ALES01000011.1 | - | 49252 | 49200 | **RNA**→ rpmE (PRK00019)Ribosomal_L31 (pfam01197)→ rpmE (PRK00019)Ribosomal_L31 (pfam01197)→ |
| Sen-910-1 | NZ_AFYX01000021.1 | + | 104135 | 104187 | **RNA**→ rpmE (PRK00019)Ribosomal_L31 (pfam01197)→ |
| Sen-525-1 | NZ_ALET01000030.1 | + | 103705 | 103757 | **RNA**→ rpmE (PRK00019)Ribosomal_L31 (pfam01197)→ rpmE (PRK00019)Ribosomal_L31 (pfam01197)→ |
| Sen-912-1 | NZ_AFYY01000022.1 | + | 135205 | 135257 | **RNA**→ rpmE (PRK00019)Ribosomal_L31 (pfam01197)→ |
| Sen-493-1 | NZ_ALFD01000131.1 | + | 8328 | 8380 | **RNA**→ rpmE (PRK00019)Ribosomal_L31 (pfam01197)→ rpmE (PRK00019)Ribosomal_L31 (pfam01197)→ |
| Sen-901-1 | NZ_AYED01000002.1 | - | 45156 | 45104 | **RNA**→ rpmE (PRK00019)Ribosomal_L31 (pfam01197)→ rpmE (PRK00019)Ribosomal_L31 (pfam01197)→ |
| Sen-547-1 | NZ_ALEY01000030.1 | - | 49252 | 49200 | **RNA**→ rpmE (PRK00019)Ribosomal_L31 (pfam01197)→ rpmE (PRK00019)Ribosomal_L31 (pfam01197)→ |
| Sen-553-1 | NZ_ALFM01000022.1 | - | 49252 | 49200 | **RNA**→ rpmE (PRK00019)Ribosomal_L31 (pfam01197)→ rpmE (PRK00019)Ribosomal_L31 (pfam01197)→ |
| Sen-897-1 | NZ_AYEH01000007.1 | + | 55314 | 55366 | **RNA**→ rpmE (PRK00019)Ribosomal_L31 (pfam01197)→ |
| Sen-567-1 | NZ_ALFR01000017.1 | + | 103706 | 103758 | **RNA**→ rpmE (PRK00019)Ribosomal_L31 (pfam01197)→ rpmE (PRK00019)Ribosomal_L31 (pfam01197)→ |
| Sen-510-1 | NZ_ALHZ01000159.1 | - | 6062 | 6010 | **RNA**→ rpmE (PRK00019)Ribosomal_L31 (pfam01197)→ rpmE (PRK00019)Ribosomal_L31 (pfam01197)→ |
| Sen-566-1 | NZ_ALGG01000006.1 | - | 49346 | 49294 | **RNA**→ rpmE (PRK00019)Ribosomal_L31 (pfam01197)→ rpmE (PRK00019)Ribosomal_L31 (pfam01197)→ |
| Sen-921-1 | NZ_AOXH01000205.1 | + | 25845 | 25897 | **RNA**→ rpmE (PRK00019)Ribosomal_L31 (pfam01197)→ rpmE (PRK00019)Ribosomal_L31 (pfam01197)→ |
| Sen-521-1 | NZ_ALGL01000025.1 | - | 49346 | 49294 | **RNA**→ rpmE (PRK00019)Ribosomal_L31 (pfam01197)→ rpmE (PRK00019)Ribosomal_L31 (pfam01197)→ |
| Sen-941-1 | NZ_AOXU01000041.1 | - | 41814 | 41762 | **RNA**→ rpmE (PRK00019)Ribosomal_L31 (pfam01197)→ rpmE (PRK00019)Ribosomal_L31 (pfam01197)→ |
| Sen-518-1 | NZ_ALGQ01000023.1 | - | 49252 | 49200 | **RNA**→ rpmE (PRK00019)Ribosomal_L31 (pfam01197)→ rpmE (PRK00019)Ribosomal_L31 (pfam01197)→ |
| Sen-944-1 | NZ_AOXZ01000060.1 | + | 110031 | 110083 | **RNA**→ rpmE (PRK00019)Ribosomal_L31 (pfam01197)→ rpmE (PRK00019)Ribosomal_L31 (pfam01197)→ |
| Sen-522-1 | NZ_ALHA01000033.1 | - | 49346 | 49294 | **RNA**→ rpmE (PRK00019)Ribosomal_L31 (pfam01197)→ rpmE (PRK00019)Ribosomal_L31 (pfam01197)→ |
| Sen-505-1 | NZ_ALHT01000042.1 | - | 49343 | 49291 | **RNA**→ rpmE (PRK00019)Ribosomal_L31 (pfam01197)→ rpmE (PRK00019)Ribosomal_L31 (pfam01197)→ |
| Sen-918-1 | NZ_AOYE01000040.1 | - | 45074 | 45022 | **RNA**→ rpmE (PRK00019)Ribosomal_L31 (pfam01197)→ rpmE (PRK00019)Ribosomal_L31 (pfam01197)→ |
| Sen-532-1 | NZ_ALHL01000082.1 | - | 49347 | 49295 | **RNA**→ rpmE (PRK00019)Ribosomal_L31 (pfam01197)→ rpmE (PRK00019)Ribosomal_L31 (pfam01197)→ |
| Sen-927-1 | NZ_AOYI01000002.1 | - | 49162 | 49110 | **RNA**→ rpmE (PRK00019)Ribosomal_L31 (pfam01197)→ |
| Sen-494-1 | NZ_ALHX01000060.1 | - | 49252 | 49200 | **RNA**→ rpmE (PRK00019)Ribosomal_L31 (pfam01197)→ rpmE (PRK00019)Ribosomal_L31 (pfam01197)→ |
| Sen-520-1 | NZ_ALER01000036.1 | + | 103705 | 103757 | **RNA**→ rpmE (PRK00019)Ribosomal_L31 (pfam01197)→ rpmE (PRK00019)Ribosomal_L31 (pfam01197)→ |
| Sen-545-1 | NZ_ALEW01000006.1 | - | 49252 | 49200 | **RNA**→ rpmE (PRK00019)Ribosomal_L31 (pfam01197)→ rpmE (PRK00019)Ribosomal_L31 (pfam01197)→ |
| Sen-933-1 | NZ_AQQN01000034.1 | - | 36977 | 36925 | **RNA**→ rpmE (PRK00019)Ribosomal_L31 (pfam01197)→ rpmE (PRK00019)Ribosomal_L31 (pfam01197)→ |
| Sen-558-1 | NZ_ALFL01000006.1 | - | 49346 | 49294 | **RNA**→ rpmE (PRK00019)Ribosomal_L31 (pfam01197)→ rpmE (PRK00019)Ribosomal_L31 (pfam01197)→ |
| Sen-480-1 | NZ_AHUM01000178.1 | + | 53030 | 53082 | **RNA**→ rpmE (PRK00019)Ribosomal_L31 (pfam01197)→ rpmE (PRK00019)Ribosomal_L31 (pfam01197)→ |
| Sen-911-1 | NZ_AFYW01000025.1 | + | 135019 | 135071 | **RNA**→ rpmE (PRK00019)Ribosomal_L31 (pfam01197)→ |
| Sen-551-1 | NZ_ALFQ01000008.1 | + | 103705 | 103757 | **RNA**→ rpmE (PRK00019)Ribosomal_L31 (pfam01197)→ rpmE (PRK00019)Ribosomal_L31 (pfam01197)→ |
| Sen-500-1 | NZ_ALID01000015.1 | - | 49400 | 49348 | **RNA**→ rpmE (PRK00019)Ribosomal_L31 (pfam01197)→ rpmE (PRK00019)Ribosomal_L31 (pfam01197)→ |
| Sen-902-1 | NZ_AYEC01000105.1 | + | 100326 | 100378 | **RNA**→ rpmE (PRK00019)Ribosomal_L31 (pfam01197)→ |
| Sen-561-1 | NZ_ALGA01000017.1 | - | 20301 | 20249 | **RNA**→ rpmE (PRK00019)Ribosomal_L31 (pfam01197)→ rpmE (PRK00019)Ribosomal_L31 (pfam01197)→ |
| Sen-931-1 | NZ_AYDS01000004.1 | - | 45346 | 45294 | **RNA**→ rpmE (PRK00019)Ribosomal_L31 (pfam01197)→ |
| Sen-288-1052 | NZ_CJRE01000016.1 | - | 39355 | 39303 | **RNA**→ rpmE (PRK00019)Ribosomal_L31 (pfam01197)→ |
| Sen-563-1 | NZ_ALGF01000007.1 | - | 49252 | 49200 | **RNA**→ rpmE (PRK00019)Ribosomal_L31 (pfam01197)→ rpmE (PRK00019)Ribosomal_L31 (pfam01197)→ |

| | | | | | | | |
|---|---|---|---|---|---|---|---|
| Sen-476-1 | NZ_AWOI01000031.1 | - | 49718 | 49666 | **RNA**→ | rpmE (PRK00019)Ribosomal_L31 (pfam01197)→ | |
| Sen-922-1 | NZ_AYDQ01000012.1 | + | 111446 | 111498 | **RNA**→ | rpmE (PRK00019)Ribosomal_L31 (pfam01197)→ | rpmE (PRK00019)Ribosomal_L31 (pfam01197)→ |
| Sen-530-1 | NZ_ALHJ01000014.1 | + | 103703 | 103755 | **RNA**→ | rpmE (PRK00019)Ribosomal_L31 (pfam01197)→ | rpmE (PRK00019)Ribosomal_L31 (pfam01197)→ |
| Sen-926-1 | NZ_AYDA01000017.1 | - | 49162 | 49110 | **RNA**→ | rpmE (PRK00019)Ribosomal_L31 (pfam01197)→ | |
| Sen-498-1 | NZ_ALFB01000033.1 | + | 103706 | 103758 | **RNA**→ | rpmE (PRK00019)Ribosomal_L31 (pfam01197)→ | rpmE (PRK00019)Ribosomal_L31 (pfam01197)→ |
| Sen-531-1 | NZ_ALHK01000054.1 | + | 103704 | 103756 | **RNA**→ | rpmE (PRK00019)Ribosomal_L31 (pfam01197)→ | rpmE (PRK00019)Ribosomal_L31 (pfam01197)→ |
| Sen-942-1 | NZ_AYEJ01000008.1 | + | 9768 | 9820 | **RNA**→ | rpmE (PRK00019)Ribosomal_L31 (pfam01197)→ | rpmE (PRK00019)Ribosomal_L31 (pfam01197)→ |
| Sen-507-1 | NZ_ALFC01000017.1 | + | 103666 | 103718 | **RNA**→ | rpmE (PRK00019)Ribosomal_L31 (pfam01197)→ | rpmE (PRK00019)Ribosomal_L31 (pfam01197)→ |
| Sen-537-1 | NZ_ALHO01000024.1 | - | 49346 | 49294 | **RNA**→ | rpmE (PRK00019)Ribosomal_L31 (pfam01197)→ | rpmE (PRK00019)Ribosomal_L31 (pfam01197)→ |
| Sen-923-1 | NZ_AOXG01000019.1 | + | 110333 | 110385 | **RNA**→ | rpmE (PRK00019)Ribosomal_L31 (pfam01197)→ | rpmE (PRK00019)Ribosomal_L31 (pfam01197)→ |
| Sen-503-1 | NZ_ALHU01000058.1 | - | 49345 | 49293 | **RNA**→ | rpmE (PRK00019)Ribosomal_L31 (pfam01197)→ | rpmE (PRK00019)Ribosomal_L31 (pfam01197)→ |
| Sen-538-1 | NZ_ALHP01000022.1 | - | 49345 | 49293 | **RNA**→ | rpmE (PRK00019)Ribosomal_L31 (pfam01197)→ | rpmE (PRK00019)Ribosomal_L31 (pfam01197)→ |
| Sen-948-1 | NZ_AOXQ01000190.1 | - | 18527 | 18475 | **RNA**→ | rpmE (PRK00019)Ribosomal_L31 (pfam01197)→ | rpmE (PRK00019)Ribosomal_L31 (pfam01197)→ |
| Sen-562-1 | NZ_ALFJ01000009.1 | + | 103711 | 103763 | **RNA**→ | rpmE (PRK00019)Ribosomal_L31 (pfam01197)→ | rpmE (PRK00019)Ribosomal_L31 (pfam01197)→ |
| Sen-496-1 | NZ_AHUO01000034.1 | + | 3317 | 3369 | **RNA**→ | rpmE (PRK00019)Ribosomal_L31 (pfam01197)→ | rpmE (PRK00019)Ribosomal_L31 (pfam01197)→ |
| Sen-565-1 | NZ_ALFI01000001.1 | + | 103715 | 103767 | **RNA**→ | rpmE (PRK00019)Ribosomal_L31 (pfam01197)→ | rpmE (PRK00019)Ribosomal_L31 (pfam01197)→ |
| Sen-932-1 | NZ_AOXY01000013.1 | - | 45355 | 45303 | **RNA**→ | rpmE (PRK00019)Ribosomal_L31 (pfam01197)→ | |
| Sen-540-1 | NZ_ALEZ01000020.1 | - | 49346 | 49294 | **RNA**→ | rpmE (PRK00019)Ribosomal_L31 (pfam01197)→ | rpmE (PRK00019)Ribosomal_L31 (pfam01197)→ |
| Sen-917-1 | NZ_AOYC01000001.1 | - | 45077 | 45025 | **RNA**→ | rpmE (PRK00019)Ribosomal_L31 (pfam01197)→ | rpmE (PRK00019)Ribosomal_L31 (pfam01197)→ |
| Sen-541-1 | NZ_ALHR01000009.1 | + | 103734 | 103786 | **RNA**→ | rpmE (PRK00019)Ribosomal_L31 (pfam01197)→ | rpmE (PRK00019)Ribosomal_L31 (pfam01197)→ |
| Sen-916-1 | NZ_AOYD01000029.1 | - | 45153 | 45101 | **RNA**→ | rpmE (PRK00019)Ribosomal_L31 (pfam01197)→ | rpmE (PRK00019)Ribosomal_L31 (pfam01197)→ |
| Sen-288-1053 | NZ_CJRM01000011.1 | + | 140955 | 141007 | **RNA**→ | rpmE (PRK00019)Ribosomal_L31 (pfam01197)→ | |
| Sen-909-1 | NZ_AOYH01000028.1 | + | 73247 | 73299 | **RNA**→ | rpmE (PRK00019)Ribosomal_L31 (pfam01197)→ | rpmE (PRK00019)Ribosomal_L31 (pfam01197)→ |
| Sen-905-1 | NZ_AOYM01000106.1 | - | 36696 | 36644 | **RNA**→ | rpmE (PRK00019)Ribosomal_L31 (pfam01197)→ | rpmE (PRK00019)Ribosomal_L31 (pfam01197)→ |
| Sen-542-1 | NZ_ALHW01000479.1 | + | 6260 | 6312 | **RNA**→ | rpmE (PRK00019)Ribosomal_L31 (pfam01197)→ | rpmE (PRK00019)Ribosomal_L31 (pfam01197)→ |
| Sen-903-1 | NZ_APAM01000189.1 | + | 8581 | 8633 | **RNA**→ | rpmE (PRK00019)Ribosomal_L31 (pfam01197)→ | |
| Sen-536-1 | NZ_ALFA01000040.1 | + | 103705 | 103757 | **RNA**→ | rpmE (PRK00019)Ribosomal_L31 (pfam01197)→ | rpmE (PRK00019)Ribosomal_L31 (pfam01197)→ |
| Sen-528-1 | NZ_ALHS01000040.1 | + | 22613 | 22665 | **RNA**→ | rpmE (PRK00019)Ribosomal_L31 (pfam01197)→ | rpmE (PRK00019)Ribosomal_L31 (pfam01197)→ |
| Sen-946-1 | NZ_APAR01000027.1 | - | 49439 | 49387 | **RNA**→ | rpmE (PRK00019)Ribosomal_L31 (pfam01197)→ | rpmE (PRK00019)Ribosomal_L31 (pfam01197)→ |
| Sen-539-1 | NZ_ALHQ01000071.1 | - | 12300 | 12248 | **RNA**→ | rpmE (PRK00019)Ribosomal_L31 (pfam01197)→ | rpmE (PRK00019)Ribosomal_L31 (pfam01197)→ |
| Sen-288-1054 | NZ_CJVW01000015.1 | + | 107860 | 107912 | **RNA**→ | rpmE (PRK00019)Ribosomal_L31 (pfam01197)→ | |
| Sen-274-1 | NZ_ARYU01000013.1 | + | 97576 | 97628 | **RNA**→ | rpmE (PRK00019)Ribosomal_L31 (pfam01197)→ | |
| Sen-534-1 | NZ_ALHV01000013.1 | + | 19300 | 19352 | **RNA**→ | rpmE (PRK00019)Ribosomal_L31 (pfam01197)→ | rpmE (PRK00019)Ribosomal_L31 (pfam01197)→ |
| Sen-949-1 | NZ_AMSN01000034.1 | - | 2989 | 2937 | **RNA**→ | rpmE (PRK00019)Ribosomal_L31 (pfam01197)→ | |
| Sen-546-1 | NZ_ALIB01000011.1 | - | 49100 | 49048 | **RNA**→ | rpmE (PRK00019)Ribosomal_L31 (pfam01197)→ | rpmE (PRK00019)Ribosomal_L31 (pfam01197)→ |
| Sen-283-1 | NZ_AHUY01000024.1 | - | 37543 | 37491 | **RNA**→ | rpmE (PRK00019)Ribosomal_L31 (pfam01197)→ | |
| Sen-282-1 | NZ_ARYV01000035.1 | + | 42107 | 42159 | **RNA**→ | rpmE (PRK00019)Ribosomal_L31 (pfam01197)→ | |
| Sen-935-1 | NZ_AOYU01000078.1 | + | 104625 | 104677 | **RNA**→ | rpmE (PRK00019)Ribosomal_L31 (pfam01197)→ | rpmE (PRK00019)Ribosomal_L31 (pfam01197)→ |
| Sen-947-1 | NZ_AOXS01000229.1 | + | 15341 | 15393 | **RNA**→ | rpmE (PRK00019)Ribosomal_L31 (pfam01197)→ | |
| Sen-288-1055 | NZ_CIUU01000016.1 | - | 39380 | 39328 | **RNA**→ | rpmE (PRK00019)Ribosomal_L31 (pfam01197)→ | |
| Sen-288-1056 | NZ_CJLP01000012.1 | - | 39527 | 39475 | **RNA**→ | rpmE (PRK00019)Ribosomal_L31 (pfam01197)→ | |
| Sen-288-1057 | NZ_CJWV01000015.1 | - | 39378 | 39326 | **RNA**→ | rpmE (PRK00019)Ribosomal_L31 (pfam01197)→ | |
| Sen-288-1058 | NZ_CJYP01000016.1 | + | 104128 | 104180 | **RNA**→ | rpmE (PRK00019)Ribosomal_L31 (pfam01197)→ | |
| Sen-288-1059 | NZ_CJYT01000016.1 | + | 104158 | 104210 | **RNA**→ | rpmE (PRK00019)Ribosomal_L31 (pfam01197)→ | |
| Sen-288-1060 | NZ_CJGE01000014.1 | - | 41507 | 41455 | **RNA**→ | rpmE (PRK00019)Ribosomal_L31 (pfam01197)→ | |
| Sen-746-1 | NZ_AJGY01000070.1 | + | 20275 | 20327 | **RNA**→ | rpmE (PRK00019)Ribosomal_L31 (pfam01197)→ | |
| Sen-288-1061 | NZ_CJQG01000016.1 | - | 39601 | 39549 | **RNA**→ | rpmE (PRK00019)Ribosomal_L31 (pfam01197)→ | |
| Sen-288-1062 | NZ_CJUK01000016.1 | + | 104154 | 104206 | **RNA**→ | rpmE (PRK00019)Ribosomal_L31 (pfam01197)→ | |
| Sen-573-1 | NZ_ALFN01000042.1 | - | 49346 | 49294 | **RNA**→ | rpmE (PRK00019)Ribosomal_L31 (pfam01197)→ | rpmE (PRK00019)Ribosomal_L31 (pfam01197)→ |
| Sen-288-1063 | NZ_CJXH01000016.1 | - | 39371 | 39319 | **RNA**→ | rpmE (PRK00019)Ribosomal_L31 (pfam01197)→ | |
| Sen-574-1 | NZ_ALGC01000047.1 | - | 49343 | 49291 | **RNA**→ | rpmE (PRK00019)Ribosomal_L31 (pfam01197)→ | rpmE (PRK00019)Ribosomal_L31 (pfam01197)→ |
| Sen-288-1064 | NZ_CJUP01000015.1 | - | 41231 | 41179 | **RNA**→ | rpmE (PRK00019)Ribosomal_L31 (pfam01197)→ | |

| | | | | | |
|---|---|---|---|---|---|
| Sen-568-1 | NZ_ALGH01000012.1 | + | 103706 | 103758 | **RNA**→ rpmE (PRK00019)Ribosomal_L31 (pfam01197)→ rpmE (PRK00019)Ribosomal_L31 (pfam01197)→ |
| Sen-570-1 | NZ_ALGI01000014.1 | + | 103700 | 103752 | **RNA**→ rpmE (PRK00019)Ribosomal_L31 (pfam01197)→ rpmE (PRK00019)Ribosomal_L31 (pfam01197)→ |
| Sen-288-1065 | NZ_CJLU01000013.1 | + | 107988 | 108040 | **RNA**→ rpmE (PRK00019)Ribosomal_L31 (pfam01197)→ |
| Sen-583-1 | NZ_ALGS01000034.1 | + | 103661 | 103713 | **RNA**→ rpmE (PRK00019)Ribosomal_L31 (pfam01197)→ rpmE (PRK00019)Ribosomal_L31 (pfam01197)→ |
| Sen-288-1066 | NZ_CJGK01000016.1 | - | 39535 | 39483 | **RNA**→ rpmE (PRK00019)Ribosomal_L31 (pfam01197)→ |
| Sen-727-1 | NZ_ALGX01000047.1 | - | 49252 | 49200 | **RNA**→ rpmE (PRK00019)Ribosomal_L31 (pfam01197)→ rpmE (PRK00019)Ribosomal_L31 (pfam01197)→ |
| Sen-288-1067 | NZ_CJEP01000016.1 | - | 39536 | 39484 | **RNA**→ rpmE (PRK00019)Ribosomal_L31 (pfam01197)→ |
| Sen-577-1 | NZ_ALHC01000023.1 | - | 49191 | 49139 | **RNA**→ rpmE (PRK00019)Ribosomal_L31 (pfam01197)→ rpmE (PRK00019)Ribosomal_L31 (pfam01197)→ |
| Sen-288-1068 | NZ_CEXR01000016.1 | - | 39537 | 39485 | **RNA**→ rpmE (PRK00019)Ribosomal_L31 (pfam01197)→ |
| Sen-288-1069 | NZ_CJXF01000015.1 | + | 104154 | 104206 | **RNA**→ rpmE (PRK00019)Ribosomal_L31 (pfam01197)→ |
| Sen-723-1 | NZ_AHUN01000085.1 | - | 40121 | 40069 | **RNA**→ rpmE (PRK00019)Ribosomal_L31 (pfam01197)→ rpmE (PRK00019)Ribosomal_L31 (pfam01197)→ |
| Sen-288-1070 | NZ_CJQF01000016.1 | - | 39601 | 39549 | **RNA**→ rpmE (PRK00019)Ribosomal_L31 (pfam01197)→ |
| Sen-733-1 | NZ_AOZG01000039.1 | + | 32849 | 32901 | **RNA**→ rpmE (PRK00019)Ribosomal_L31 (pfam01197)→ rpmE (PRK00019)Ribosomal_L31 (pfam01197)→ |
| Sen-288-1071 | NZ_CIZE01000017.1 | - | 41512 | 41460 | **RNA**→ rpmE (PRK00019)Ribosomal_L31 (pfam01197)→ |
| Sen-739-1 | NZ_APAO01000024.1 | + | 105311 | 105363 | **RNA**→ rpmE (PRK00019)Ribosomal_L31 (pfam01197)→ rpmE (PRK00019)Ribosomal_L31 (pfam01197)→ |
| Sen-288-1072 | NZ_CJMN01000014.1 | + | 107976 | 108028 | **RNA**→ rpmE (PRK00019)Ribosomal_L31 (pfam01197)→ |
| Sen-288-1073 | NZ_CJSH01000016.1 | + | 78112 | 78164 | **RNA**→ rpmE (PRK00019)Ribosomal_L31 (pfam01197)→ |
| Sen-165-1 | NZ_AZRG01000037.1 | - | 14220 | 14168 | **RNA**→ rpmE (PRK00019)Ribosomal_L31 (pfam01197)→ |
| Sen-288-1074 | NZ_CJSM01000016.1 | + | 104155 | 104207 | **RNA**→ rpmE (PRK00019)Ribosomal_L31 (pfam01197)→ |
| Sen-738-1 | NZ_ALPO01000015.1 | - | 21962 | 21910 | **RNA**→ rpmE (PRK00019)Ribosomal_L31 (pfam01197)→ |
| Sen-163-1 | NZ_CAGR02000040.1 | + | 103650 | 103702 | **RNA**→ rpmE (PRK00019)Ribosomal_L31 (pfam01197)→ |
| Sen-288-1075 | NZ_CJKJ01000014.1 | - | 41511 | 41459 | **RNA**→ rpmE (PRK00019)Ribosomal_L31 (pfam01197)→ |
| Sen-288-1076 | NZ_CJLZ01000002.1 | - | 198398 | 198346 | **RNA**→ rpmE (PRK00019)Ribosomal_L31 (pfam01197)→ |
| Sen-166-1 | NZ_JOKK01000017.1 | + | 93924 | 93976 | **RNA**→ rpmE (PRK00019)Ribosomal_L31 (pfam01197)→ |
| Sen-288-1077 | NZ_CJBE01000011.1 | - | 39456 | 39404 | **RNA**→ rpmE (PRK00019)Ribosomal_L31 (pfam01197)→ |
| Sen-745-1 | NZ_AJGZ01000018.1 | - | 37514 | 37462 | **RNA**→ rpmE (PRK00019)Ribosomal_L31 (pfam01197)→ |
| Sen-288-1078 | NZ_CIUR01000015.1 | + | 104168 | 104220 | **RNA**→ rpmE (PRK00019)Ribosomal_L31 (pfam01197)→ |
| Sen-744-1 | NZ_AJHA01000094.1 | - | 3193 | 3141 | **RNA**→ rpmE (PRK00019)Ribosomal_L31 (pfam01197)→ |
| Sen-288-1079 | NZ_CJGU01000016.1 | + | 104144 | 104196 | **RNA**→ rpmE (PRK00019)Ribosomal_L31 (pfam01197)→ |
| Sen-571-1 | NZ_ALGJ01000040.1 | - | 49252 | 49200 | **RNA**→ rpmE (PRK00019)Ribosomal_L31 (pfam01197)→ rpmE (PRK00019)Ribosomal_L31 (pfam01197)→ |
| Sen-288-1080 | NZ_CJQB01000016.1 | - | 39601 | 39549 | **RNA**→ rpmE (PRK00019)Ribosomal_L31 (pfam01197)→ |
| Sen-579-1 | NZ_ALGT01000019.1 | - | 49252 | 49200 | **RNA**→ rpmE (PRK00019)Ribosomal_L31 (pfam01197)→ rpmE (PRK00019)Ribosomal_L31 (pfam01197)→ |
| Sen-288-1081 | NZ_CJII01000015.1 | - | 41504 | 41452 | **RNA**→ rpmE (PRK00019)Ribosomal_L31 (pfam01197)→ |
| Sen-728-1 | NZ_ALGY01000015.1 | + | 103708 | 103760 | **RNA**→ rpmE (PRK00019)Ribosomal_L31 (pfam01197)→ rpmE (PRK00019)Ribosomal_L31 (pfam01197)→ |
| Sen-288-1082 | NZ_CJWQ01000013.1 | + | 107425 | 107477 | **RNA**→ rpmE (PRK00019)Ribosomal_L31 (pfam01197)→ |
| Sen-172-1 | NZ_APIX01000052.1 | + | 43443 | 43495 | **RNA**→ rpmE (PRK00019)Ribosomal_L31 (pfam01197)→ |
| Sen-288-1083 | NZ_CJEZ01000016.1 | + | 104158 | 104210 | **RNA**→ rpmE (PRK00019)Ribosomal_L31 (pfam01197)→ |
| Sen-173-1 | NZ_AMNL01000067.1 | - | 4749 | 4697 | **RNA**→ rpmE (PRK00019)Ribosomal_L31 (pfam01197)→ |
| Sen-288-1084 | NZ_CJDZ01000015.1 | + | 104151 | 104203 | **RNA**→ rpmE (PRK00019)Ribosomal_L31 (pfam01197)→ |
| Sen-742-1 | NZ_AMNM01000007.1 | - | 37497 | 37445 | **RNA**→ rpmE (PRK00019)Ribosomal_L31 (pfam01197)→ |
| Sen-288-1085 | NZ_CIZM01000016.1 | + | 104147 | 104199 | **RNA**→ rpmE (PRK00019)Ribosomal_L31 (pfam01197)→ |
| Sen-736-1 | NZ_ALPP01000010.1 | - | 45323 | 45271 | **RNA**→ rpmE (PRK00019)Ribosomal_L31 (pfam01197)→ |
| Sen-288-1086 | NZ_CJVI01000016.1 | + | 104157 | 104209 | **RNA**→ rpmE (PRK00019)Ribosomal_L31 (pfam01197)→ |
| Sen-749-1 | NZ_AJGW01000005.1 | - | 37498 | 37446 | **RNA**→ rpmE (PRK00019)Ribosomal_L31 (pfam01197)→ |
| Sen-288-1087 | NZ_CJUI01000001.1 | - | 39357 | 39305 | **RNA**→ rpmE (PRK00019)Ribosomal_L31 (pfam01197)→ |
| Sen-743-1 | NZ_AJGX01000011.1 | + | 43277 | 43329 | **RNA**→ rpmE (PRK00019)Ribosomal_L31 (pfam01197)→ |
| Sen-288-1088 | NZ_CJBU01000013.1 | - | 39578 | 39526 | **RNA**→ rpmE (PRK00019)Ribosomal_L31 (pfam01197)→ |
| Sen-288-1089 | NZ_CJGO01000015.1 | - | 39574 | 39522 | **RNA**→ rpmE (PRK00019)Ribosomal_L31 (pfam01197)→ |
| Sen-572-1 | NZ_ALGB01000053.1 | - | 49347 | 49295 | **RNA**→ rpmE (PRK00019)Ribosomal_L31 (pfam01197)→ rpmE (PRK00019)Ribosomal_L31 (pfam01197)→ |
| Sen-288-1090 | NZ_CJVX01000017.1 | + | 104154 | 104206 | **RNA**→ rpmE (PRK00019)Ribosomal_L31 (pfam01197)→ |
| Sen-724-1 | NZ_ALGR01000015.1 | - | 49252 | 49200 | **RNA**→ rpmE (PRK00019)Ribosomal_L31 (pfam01197)→ rpmE (PRK00019)Ribosomal_L31 (pfam01197)→ |
| Sen-288-1091 | NZ_CJKB01000008.1 | - | 39361 | 39309 | **RNA**→ rpmE (PRK00019)Ribosomal_L31 (pfam01197)→ |

| | | | | | |
|---|---|---|---|---|---|
| Sen-582-1 | NZ_ALGW01000052.1 | + | 103706 | 103758 | **RNA**→ rpmE (PRK00019)Ribosomal_L31 (pfam01197)→ rpmE (PRK00019)Ribosomal_L31 (pfam01197)→ |
| Sen-288-1125 | NZ_CJFJ01000015.1 | - | 39564 | 39512 | **RNA**→ rpmE (PRK00019)Ribosomal_L31 (pfam01197)→ |
| Sen-737-1 | NZ_AOZE01000034.1 | - | 45179 | 45127 | **RNA**→ rpmE (PRK00019)Ribosomal_L31 (pfam01197)→ |
| Sen-288-1092 | NZ_CJLX01000013.1 | - | 41508 | 41456 | **RNA**→ rpmE (PRK00019)Ribosomal_L31 (pfam01197)→ |
| Sen-174-1 | NZ_AMNG01000040.1 | + | 43295 | 43347 | **RNA**→ rpmE (PRK00019)Ribosomal_L31 (pfam01197)→ |
| Sen-288-1093 | NZ_CJWW01000016.1 | - | 39366 | 39314 | **RNA**→ rpmE (PRK00019)Ribosomal_L31 (pfam01197)→ |
| Sen-747-1 | NZ_AMNH01000065.1 | + | 43992 | 44044 | **RNA**→ rpmE (PRK00019)Ribosomal_L31 (pfam01197)→ |
| Sen-288-1094 | NZ_CJEB01000016.1 | - | 39534 | 39482 | **RNA**→ rpmE (PRK00019)Ribosomal_L31 (pfam01197)→ |
| Sen-569-1 | NZ_ALFV01000010.1 | - | 49343 | 49291 | **RNA**→ rpmE (PRK00019)Ribosomal_L31 (pfam01197)→ rpmE (PRK00019)Ribosomal_L31 (pfam01197)→ |
| Sen-288-1095 | NZ_CJDP01000015.1 | - | 39373 | 39321 | **RNA**→ rpmE (PRK00019)Ribosomal_L31 (pfam01197)→ |
| Sen-581-1 | NZ_ALGU01000026.1 | + | 103706 | 103758 | **RNA**→ rpmE (PRK00019)Ribosomal_L31 (pfam01197)→ rpmE (PRK00019)Ribosomal_L31 (pfam01197)→ |
| Sen-288-1096 | NZ_CJVF01000015.1 | - | 41231 | 41179 | **RNA**→ rpmE (PRK00019)Ribosomal_L31 (pfam01197)→ |
| Sen-288-1097 | NZ_CJOJ01000016.1 | - | 39387 | 39335 | **RNA**→ rpmE (PRK00019)Ribosomal_L31 (pfam01197)→ |
| Sen-580-1 | NZ_ALGZ01000049.1 | - | 49107 | 49055 | **RNA**→ rpmE (PRK00019)Ribosomal_L31 (pfam01197)→ rpmE (PRK00019)Ribosomal_L31 (pfam01197)→ |
| Sen-288-1098 | NZ_CJOH01000016.1 | + | 104153 | 104205 | **RNA**→ rpmE (PRK00019)Ribosomal_L31 (pfam01197)→ |
| Sen-730-1 | NZ_AHUH01000066.1 | + | 104411 | 104463 | **RNA**→ rpmE (PRK00019)Ribosomal_L31 (pfam01197)→ rpmE (PRK00019)Ribosomal_L31 (pfam01197)→ |
| Sen-288-1099 | NZ_CJDR01000016.1 | + | 104156 | 104208 | **RNA**→ rpmE (PRK00019)Ribosomal_L31 (pfam01197)→ |
| Sen-740-1 | NZ_APAH01000055.1 | - | 2949 | 2897 | **RNA**→ rpmE (PRK00019)Ribosomal_L31 (pfam01197)→ |
| Sen-288-1100 | NZ_CJGR01000016.1 | - | 39535 | 39483 | **RNA**→ rpmE (PRK00019)Ribosomal_L31 (pfam01197)→ |
| Sen-288-1101 | NZ_CJBO01000016.1 | + | 104166 | 104218 | **RNA**→ rpmE (PRK00019)Ribosomal_L31 (pfam01197)→ |
| Sen-171-1 | NZ_AMMZ01000027.1 | - | 37498 | 37446 | **RNA**→ rpmE (PRK00019)Ribosomal_L31 (pfam01197)→ |
| Sen-288-1102 | NZ_CJEI01000015.1 | + | 103986 | 104038 | **RNA**→ rpmE (PRK00019)Ribosomal_L31 (pfam01197)→ |
| Sen-288-1103 | NZ_CJED01000016.1 | + | 104139 | 104191 | **RNA**→ rpmE (PRK00019)Ribosomal_L31 (pfam01197)→ |
| Sen-725-1 | NZ_ALFE01000018.1 | - | 4666 | 4614 | **RNA**→ rpmE (PRK00019)Ribosomal_L31 (pfam01197)→ rpmE (PRK00019)Ribosomal_L31 (pfam01197)→ |
| Sen-726-1 | NZ_AHUL01000075.1 | + | 103648 | 103700 | **RNA**→ rpmE (PRK00019)Ribosomal_L31 (pfam01197)→ rpmE (PRK00019)Ribosomal_L31 (pfam01197)→ |
| Sen-741-1 | NZ_AMMY01000035.1 | + | 8157 | 8209 | **RNA**→ rpmE (PRK00019)Ribosomal_L31 (pfam01197)→ |
| Sen-722-1 | NZ_ALHY01000024.1 | - | 9651 | 9599 | **RNA**→ rpmE (PRK00019)Ribosomal_L31 (pfam01197)→ |
| Sen-169-1 | NZ_ARYS01000012.1 | + | 104604 | 104656 | **RNA**→ rpmE (PRK00019)Ribosomal_L31 (pfam01197)→ |
| Sen-735-1 | NZ_AYUQ01000070.1 | + | 104455 | 104507 | **RNA**→ rpmE (PRK00019)Ribosomal_L31 (pfam01197)→ |
| Sen-1004-1 | NZ_ABFG01000008.1 | - | 54519 | 54467 | **RNA**→ rpmE (PRK00019)Ribosomal_L31 (pfam01197)→ rpmE (PRK00019)Ribosomal_L31 (pfam01197)→ |
| Sen-576-1 | NZ_ALFH01000027.1 | - | 49346 | 49294 | **RNA**→ rpmE (PRK00019)Ribosomal_L31 (pfam01197)→ rpmE (PRK00019)Ribosomal_L31 (pfam01197)→ |
| Sen-679-1 | NZ_AHUR01000081.1 | + | 103706 | 103758 | **RNA**→ rpmE (PRK00019)Ribosomal_L31 (pfam01197)→ rpmE (PRK00019)Ribosomal_L31 (pfam01197)→ |
| Sen-168-1 | NZ_ARYR01000001.1 | + | 100051 | 100103 | **RNA**→ rpmE (PRK00019)Ribosomal_L31 (pfam01197)→ |
| Sen-578-1 | NZ_ALFF01000033.1 | - | 26193 | 26141 | **RNA**→ rpmE (PRK00019)Ribosomal_L31 (pfam01197)→ rpmE (PRK00019)Ribosomal_L31 (pfam01197)→ |
| Sen-575-1 | NZ_ALFG01000008.1 | - | 49346 | 49294 | **RNA**→ rpmE (PRK00019)Ribosomal_L31 (pfam01197)→ rpmE (PRK00019)Ribosomal_L31 (pfam01197)→ |
| Sen-431-1 | NZ_AFCK01001570.1 | + | 1664 | 1716 | **RNA**→ rpmE (PRK00019)Ribosomal_L31 (pfam01197)→ |
| Sen-5-6 | NZ_LIMM01000012.1 | + | 104491 | 104543 | **RNA**→ rpmE (PRK00019)Ribosomal_L31 (pfam01197)→ |
| Sen-1014-1 | NC_003198.1 | - | 3639708 | 3639656 | **RNA**→ rpmE (PRK00019)Ribosomal_L31 (pfam01197)→ |
| Sen-1017-1 | NC_003197.1 | + | 4305705 | 4305757 | **RNA**→ rpmE (PRK00019)Ribosomal_L31 (pfam01197)→ |
| Sen-440-1 | NZ_HF969015.1 | + | 4155499 | 4155551 | **RNA**→ rpmE (PRK00019)Ribosomal_L31 (pfam01197)→ |
| Sen-731-1 | NZ_CM001153.1 | - | 3480716 | 3480664 | **RNA**→ rpmE (PRK00019)Ribosomal_L31 (pfam01197)→ rpmE (PRK00019)Ribosomal_L31 (pfam01197)→ |
| Sen-1000-1 | NZ_CM001062.1 | + | 4224780 | 4224832 | **RNA**→ rpmE (PRK00019)Ribosomal_L31 (pfam01197)→ rpmE (PRK00019)Ribosomal_L31 (pfam01197)→ hypo→ hypo→ |
| Sen-468-1 | NZ_CM001151.1 | + | 4332666 | 4332718 | **RNA**→ rpmE (PRK00019)Ribosomal_L31 (pfam01197)→ rpmE (PRK00019)Ribosomal_L31 (pfam01197)→ rpmE (PRK00019)Ribosomal_L31 (pfam01197)→ |
| Sen-953-1 | NC_016832.1 | - | 3614165 | 3614113 | **RNA**→ rpmE (PRK00019)Ribosomal_L31 (pfam01197)→ |
| Sen-971-1 | NC_016810.1 | + | 4327143 | 4327195 | **RNA**→ rpmE (PRK00019)Ribosomal_L31 (pfam01197)→ |
| Sen-470-1 | NZ_KB731431.1 | + | 30272 | 30324 | **RNA**→ rpmE (PRK00019)Ribosomal_L31 (pfam01197)→ |
| Sen-483-1 | NZ_KE349350.1 | + | 30273 | 30325 | **RNA**→ rpmE (PRK00019)Ribosomal_L31 (pfam01197)→ |
| Sen-489-1 | NZ_KE350075.1 | + | 55984 | 56036 | **RNA**→ rpmE (PRK00019)Ribosomal_L31 (pfam01197)→ |
| Sen-485-1 | NZ_KE349599.1 | + | 30273 | 30325 | **RNA**→ rpmE (PRK00019)Ribosomal_L31 (pfam01197)→ |
| Sen-487-1 | NZ_KE349812.1 | + | 30273 | 30325 | **RNA**→ rpmE (PRK00019)Ribosomal_L31 (pfam01197)→ |
| Sen-482-1 | NZ_KE349293.1 | + | 30273 | 30325 | **RNA**→ rpmE (PRK00019)Ribosomal_L31 (pfam01197)→ |

| | | | | | |
|---|---|---|---|---|---|
| Sen-486-1 | NZ_KE349740.1 | + | 30273 | 30325 | **RNA**→ rpmE (PRK00019)Ribosomal_L31 (pfam01197)→ |
| Sen-475-1 | NZ_KE349031.1 | + | 30273 | 30325 | **RNA**→ rpmE (PRK00019)Ribosomal_L31 (pfam01197)→ |
| Sen-481-1 | NZ_KE349069.1 | + | 30273 | 30325 | **RNA**→ rpmE (PRK00019)Ribosomal_L31 (pfam01197)→ |
| Sen-488-1 | NZ_KE350026.1 | + | 30274 | 30326 | **RNA**→ rpmE (PRK00019)Ribosomal_L31 (pfam01197)→ |
| Sen-463-1 | NC_021818.1 | + | 364880 | 364932 | **RNA**→ rpmE (PRK00019)Ribosomal_L31 (pfam01197)→ |
| Sen-307-1 | NZ_KI911915.1 | + | 3835966 | 3836018 | **RNA**→ rpmE (PRK00019)Ribosomal_L31 (pfam01197)→ |
| Sen-300-330 | NC_022569.1 | + | 4382615 | 4382667 | **RNA**→ rpmE (PRK00019)Ribosomal_L31 (pfam01197)→ |
| Sen-601-1 | NZ_CP007251.1 | + | 4194336 | 4194388 | **RNA**→ rpmE (PRK00019)Ribosomal_L31 (pfam01197)→ |
| Sen-241-1 | NZ_KK082459.1 | - | 37622 | 37570 | **RNA**→ rpmE (PRK00019)Ribosomal_L31 (pfam01197)→ |
| Sen-612-1 | NZ_CP007344.1 | + | 4173178 | 4173230 | **RNA**→ rpmE (PRK00019)Ribosomal_L31 (pfam01197)→ |
| Sen-136-2 | NZ_KI635587.1 | + | 21280 | 21332 | **RNA**→ rpmE (PRK00019)Ribosomal_L31 (pfam01197)→ |
| Sen-621-1 | NZ_CP007353.1 | + | 4173243 | 4173295 | **RNA**→ rpmE (PRK00019)Ribosomal_L31 (pfam01197)→ |
| Sen-699-1 | NZ_CP007301.1 | + | 4173901 | 4173953 | **RNA**→ rpmE (PRK00019)Ribosomal_L31 (pfam01197)→ |
| Sen-626-1 | NZ_CP007335.1 | + | 4173885 | 4173937 | **RNA**→ rpmE (PRK00019)Ribosomal_L31 (pfam01197)→ |
| Sen-748-1 | NC_021810.1 | + | 1275161 | 1275213 | **RNA**→ rpmE (PRK00019)Ribosomal_L31 (pfam01197)→ |
| Sen-271-1 | NZ_CP012038.1 | + | 2994691 | 2994743 | **RNA**→ rpmE (PRK00019)Ribosomal_L31 (pfam01197)→ |
| Sen-1031-1 | NC_011294.1 | + | 4173886 | 4173938 | **RNA**→ rpmE (PRK00019)Ribosomal_L31 (pfam01197)→ |
| Sen-1003-1 | NC_011274.1 | - | 3480716 | 3480664 | **RNA**→ rpmE (PRK00019)Ribosomal_L31 (pfam01197)→ |
| Sen-908-1 | NC_006511.1 | + | 4083750 | 4083802 | **RNA**→ rpmE (PRK00019)Ribosomal_L31 (pfam01197)→ |
| Sen-1027-1 | NC_011094.1 | + | 4182786 | 4182838 | **RNA**→ rpmE (PRK00019)Ribosomal_L31 (pfam01197)→ |
| Sen-266-2 | NC_011080.1 | + | 4297332 | 4297384 | **RNA**→ rpmE (PRK00019)Ribosomal_L31 (pfam01197)→ |
| Sen-1001-1 | NC_011205.1 | + | 4332663 | 4332715 | **RNA**→ rpmE (PRK00019)Ribosomal_L31 (pfam01197)→ |
| Sen-1006-1 | NC_011083.1 | + | 4310089 | 4310141 | **RNA**→ rpmE (PRK00019)Ribosomal_L31 (pfam01197)→ |
| Sen-999-1 | NC_011149.1 | + | 4232095 | 4232147 | **RNA**→ rpmE (PRK00019)Ribosomal_L31 (pfam01197)→ |
| Sen-1-1 | NC_011147.1 | + | 4079434 | 4079486 | **RNA**→ rpmE (PRK00019)Ribosomal_L31 (pfam01197)→ |
| Sen-1032-1 | NC_012125.1 | + | 4278107 | 4278159 | **RNA**→ rpmE (PRK00019)Ribosomal_L31 (pfam01197)→ hypo→ hypo→ |
| Sen-968-1 | NC_016854.1 | + | 4328460 | 4328512 | **RNA**→ rpmE (PRK00019)Ribosomal_L31 (pfam01197)→ |
| Sen-973-1 | NC_016857.1 | + | 4327144 | 4327196 | **RNA**→ rpmE (PRK00019)Ribosomal_L31 (pfam01197)→ |
| Sen-991-1 | NC_016863.1 | + | 4266896 | 4266948 | **RNA**→ rpmE (PRK00019)Ribosomal_L31 (pfam01197)→ |
| Sen-139-1 | NZ_CP011791.1 | + | 4166192 | 4166244 | **RNA**→ rpmE (PRK00019)Ribosomal_L31 (pfam01197)→ |
| Sen-143-1 | NZ_CP007249.2 | + | 4200339 | 4200391 | **RNA**→ rpmE (PRK00019)Ribosomal_L31 (pfam01197)→ |
| Sen-1005-1 | NC_017623.1 | + | 216124 | 216176 | **RNA**→ rpmE (PRK00019)Ribosomal_L31 (pfam01197)→ |
| Sen-988-1 | NC_016860.1 | + | 4410794 | 4410846 | **RNA**→ rpmE (PRK00019)Ribosomal_L31 (pfam01197)→ |
| Sen-969-1 | NC_022544.1 | + | 4264269 | 4264321 | **RNA**→ rpmE (PRK00019)Ribosomal_L31 (pfam01197)→ |
| Sen-990-1 | NC_021151.1 | + | 4300603 | 4300655 | **RNA**→ rpmE (PRK00019)Ribosomal_L31 (pfam01197)→ |
| Sen-1002-1 | NC_016831.1 | - | 3603834 | 3603782 | **RNA**→ rpmE (PRK00019)Ribosomal_L31 (pfam01197)→ |
| Sen-158-1 | NZ_CP007329.2 | + | 4167433 | 4167485 | **RNA**→ rpmE (PRK00019)Ribosomal_L31 (pfam01197)→ |
| Sen-139-2 | NZ_CP011790.1 | + | 4166710 | 4166762 | **RNA**→ rpmE (PRK00019)Ribosomal_L31 (pfam01197)→ |
| Sen-955-1 | NC_021176.1 | - | 3625365 | 3625313 | **RNA**→ rpmE (PRK00019)Ribosomal_L31 (pfam01197)→ |
| Sen-159-1 | NZ_CP007267.2 | + | 4167317 | 4167369 | **RNA**→ rpmE (PRK00019)Ribosomal_L31 (pfam01197)→ |
| Sen-314-1 | NC_021820.1 | - | 93015 | 92963 | **RNA**→ rpmE (PRK00019)Ribosomal_L31 (pfam01197)→ |
| Sen-436-1 | NC_021844.1 | - | 4520505 | 4520453 | **RNA**→ rpmE (PRK00019)Ribosomal_L31 (pfam01197)→ |
| Sen-466-1 | NZ_KE348840.1 | + | 30272 | 30324 | **RNA**→ rpmE (PRK00019)Ribosomal_L31 (pfam01197)→ |
| Sen-732-1 | NC_022221.1 | - | 3597464 | 3597412 | **RNA**→ rpmE (PRK00019)Ribosomal_L31 (pfam01197)→ |
| Sen-928-1 | NC_021984.1 | - | 3651991 | 3651939 | **RNA**→ rpmE (PRK00019)Ribosomal_L31 (pfam01197)→ |
| Sen-895-1 | NC_021902.1 | - | 4667263 | 4667211 | **RNA**→ rpmE (PRK00019)Ribosomal_L31 (pfam01197)→ rpmE (PRK00019)Ribosomal_L31 (pfam01197)→ |
| Sen-300-51 | NZ_CTAM01000028.1 | - | 45107 | 45055 | **RNA**→ rpmE (PRK00019)Ribosomal_L31 (pfam01197)→ |
| Sen-300-52 | NZ_CTAF01000018.1 | - | 45343 | 45291 | **RNA**→ rpmE (PRK00019)Ribosomal_L31 (pfam01197)→ |
| Sen-300-53 | NZ_CTCI01000019.1 | + | 42438 | 42490 | **RNA**→ rpmE (PRK00019)Ribosomal_L31 (pfam01197)→ |
| Sen-300-54 | NZ_CTAS01000021.1 | + | 42473 | 42525 | **RNA**→ rpmE (PRK00019)Ribosomal_L31 (pfam01197)→ |
| Sen-300-55 | NZ_CTBO01000018.1 | + | 42484 | 42536 | **RNA**→ rpmE (PRK00019)Ribosomal_L31 (pfam01197)→ |
| Sen-300-56 | NZ_CTMH01000016.1 | - | 45343 | 45291 | **RNA**→ rpmE (PRK00019)Ribosomal_L31 (pfam01197)→ |

| | | | | | | |
|---|---|---|---|---|---|---|
| Sen-300-57 | NZ_CTOK01000019.1 | - | 45129 | 45077 | **RNA**→ | rpmE (PRK00019)Ribosomal_L31 (pfam01197)→ |
| Sen-300-58 | NZ_CTNR01000018.1 | - | 45179 | 45127 | **RNA**→ | rpmE (PRK00019)Ribosomal_L31 (pfam01197)→ |
| Sen-300-59 | NZ_CTLF01000015.1 | - | 45343 | 45291 | **RNA**→ | rpmE (PRK00019)Ribosomal_L31 (pfam01197)→ |
| Sen-300-60 | NZ_CTMD01000019.1 | - | 45179 | 45127 | **RNA**→ | rpmE (PRK00019)Ribosomal_L31 (pfam01197)→ |
| Sen-300-61 | NZ_CTKB01000021.1 | - | 45179 | 45127 | **RNA**→ | rpmE (PRK00019)Ribosomal_L31 (pfam01197)→ |
| Sen-300-62 | NZ_CTMP01000018.1 | + | 42525 | 42577 | **RNA**→ | rpmE (PRK00019)Ribosomal_L31 (pfam01197)→ |
| Sen-300-63 | NZ_CTGQ01000015.1 | + | 42526 | 42578 | **RNA**→ | rpmE (PRK00019)Ribosomal_L31 (pfam01197)→ |
| Sen-300-64 | NZ_CTHY01000018.1 | - | 45107 | 45055 | **RNA**→ | rpmE (PRK00019)Ribosomal_L31 (pfam01197)→ |
| Sen-300-65 | NZ_CTMO01000028.1 | - | 32531 | 32479 | **RNA**→ | rpmE (PRK00019)Ribosomal_L31 (pfam01197)→ |
| Sen-300-66 | NZ_CTPW01000017.1 | + | 42472 | 42524 | **RNA**→ | rpmE (PRK00019)Ribosomal_L31 (pfam01197)→ |
| Sen-300-67 | NZ_CTNM01000018.1 | - | 45152 | 45100 | **RNA**→ | rpmE (PRK00019)Ribosomal_L31 (pfam01197)→ |
| Sen-300-68 | NZ_CTME01000020.1 | + | 42484 | 42536 | **RNA**→ | rpmE (PRK00019)Ribosomal_L31 (pfam01197)→ |
| Sen-300-69 | NZ_CTQE01000008.1 | + | 42483 | 42535 | **RNA**→ | rpmE (PRK00019)Ribosomal_L31 (pfam01197)→ |
| Sen-300-70 | NZ_CTOJ01000017.1 | - | 45337 | 45285 | **RNA**→ | rpmE (PRK00019)Ribosomal_L31 (pfam01197)→ |
| Sen-950-1 | NC_022525.1 | + | 4159260 | 4159312 | **RNA**→ | rpmE (PRK00019)Ribosomal_L31 (pfam01197)→ |
| Sen-343-1 | NC_022991.1 | + | 4197585 | 4197637 | **RNA**→ | rpmE (PRK00019)Ribosomal_L31 (pfam01197)→ |
| Sen-760-1 | NC_021812.2 | + | 4742642 | 4742694 | **RNA**→ | rpmE (PRK00019)Ribosomal_L31 (pfam01197)→ |
| Sen-193-1 | NZ_CP005995.1 | + | 2820952 | 2821004 | **RNA**→ | rpmE (PRK00019)Ribosomal_L31 (pfam01197)→ |
| Sen-273-1 | NZ_CP009559.1 | - | 1380546 | 1380494 | **RNA**→ | rpmE (PRK00019)Ribosomal_L31 (pfam01197)→ |
| Sen-273-2 | NZ_CP009049.1 | + | 104123 | 104175 | **RNA**→ | rpmE (PRK00019)Ribosomal_L31 (pfam01197)→ |
| Sen-1016-1 | NC_004631.1 | - | 3625368 | 3625316 | **RNA**→ | rpmE (PRK00019)Ribosomal_L31 (pfam01197)→ |
| Sen-958-1 | NC_016856.1 | + | 4319290 | 4319342 | **RNA**→ | rpmE (PRK00019)Ribosomal_L31 (pfam01197)→ |
| Sen-1011-1 | NC_010102.1 | + | 4247096 | 4247148 | **RNA**→ | rpmE (PRK00019)Ribosomal_L31 (pfam01197)→ |
| Sen-780-1 | NZ_CM001274.1 | + | 4283628 | 4283680 | **RNA**→ | rpmE (PRK00019)Ribosomal_L31 (pfam01197)→ |
| Sen-300-71 | NZ_CTJW01000019.1 | + | 42531 | 42583 | **RNA**→ | rpmE (PRK00019)Ribosomal_L31 (pfam01197)→ |
| Sen-300-72 | NZ_CTOX01000017.1 | - | 45201 | 45149 | **RNA**→ | rpmE (PRK00019)Ribosomal_L31 (pfam01197)→ |
| Sen-288-1104 | NZ_CJJY01000016.1 | + | 104152 | 104204 | **RNA**→ | rpmE (PRK00019)Ribosomal_L31 (pfam01197)→ |
| Sen-300-73 | NZ_CTGN01000016.1 | - | 45167 | 45115 | **RNA**→ | rpmE (PRK00019)Ribosomal_L31 (pfam01197)→ |
| Sen-300-74 | NZ_CTLO01000018.1 | + | 42452 | 42504 | **RNA**→ | rpmE (PRK00019)Ribosomal_L31 (pfam01197)→ |
| Sen-300-75 | NZ_CTQN01000018.1 | - | 45337 | 45285 | **RNA**→ | rpmE (PRK00019)Ribosomal_L31 (pfam01197)→ |
| Sen-300-76 | NZ_CTBT01000017.1 | + | 42551 | 42603 | **RNA**→ | rpmE (PRK00019)Ribosomal_L31 (pfam01197)→ |
| Sen-300-77 | NZ_CTBY01000017.1 | + | 42479 | 42531 | **RNA**→ | rpmE (PRK00019)Ribosomal_L31 (pfam01197)→ |
| Sen-300-78 | NZ_CTHE01000019.1 | + | 42480 | 42532 | **RNA**→ | rpmE (PRK00019)Ribosomal_L31 (pfam01197)→ |
| Sen-300-79 | NZ_CTHD01000017.1 | - | 45517 | 45465 | **RNA**→ | rpmE (PRK00019)Ribosomal_L31 (pfam01197)→ |
| Sen-300-80 | NZ_CTBF01000019.1 | - | 45344 | 45292 | **RNA**→ | rpmE (PRK00019)Ribosomal_L31 (pfam01197)→ |
| Sen-300-81 | NZ_CTBQ01000019.1 | + | 42505 | 42557 | **RNA**→ | rpmE (PRK00019)Ribosomal_L31 (pfam01197)→ |
| Sen-300-82 | NZ_CTNQ01000016.1 | + | 42524 | 42576 | **RNA**→ | rpmE (PRK00019)Ribosomal_L31 (pfam01197)→ |
| Sen-300-83 | NZ_CTBE01000020.1 | + | 42438 | 42490 | **RNA**→ | rpmE (PRK00019)Ribosomal_L31 (pfam01197)→ |
| Sen-300-84 | NZ_CTIA01000021.1 | + | 42453 | 42505 | **RNA**→ | rpmE (PRK00019)Ribosomal_L31 (pfam01197)→ |
| Sen-300-85 | NZ_CTNO01000020.1 | + | 42456 | 42508 | **RNA**→ | rpmE (PRK00019)Ribosomal_L31 (pfam01197)→ |
| Sen-300-86 | NZ_CTBR01000011.1 | + | 104187 | 104239 | **RNA**→ | rpmE (PRK00019)Ribosomal_L31 (pfam01197)→ |
| Sen-300-87 | NZ_CTBJ01000020.1 | - | 45107 | 45055 | **RNA**→ | rpmE (PRK00019)Ribosomal_L31 (pfam01197)→ |
| Sen-300-88 | NZ_CTMX01000017.1 | + | 42531 | 42583 | **RNA**→ | rpmE (PRK00019)Ribosomal_L31 (pfam01197)→ |
| Sen-300-89 | NZ_CTBX01000019.1 | - | 45107 | 45055 | **RNA**→ | rpmE (PRK00019)Ribosomal_L31 (pfam01197)→ |
| Sen-300-90 | NZ_CTGF01000018.1 | - | 45179 | 45127 | **RNA**→ | rpmE (PRK00019)Ribosomal_L31 (pfam01197)→ |
| Sen-300-91 | NZ_CTJY01000017.1 | + | 42489 | 42541 | **RNA**→ | rpmE (PRK00019)Ribosomal_L31 (pfam01197)→ |
| Sen-300-92 | NZ_CTBK01000019.1 | - | 45107 | 45055 | **RNA**→ | rpmE (PRK00019)Ribosomal_L31 (pfam01197)→ |
| Sen-300-93 | NZ_CTIL01000018.1 | + | 42496 | 42548 | **RNA**→ | rpmE (PRK00019)Ribosomal_L31 (pfam01197)→ |
| Sen-300-94 | NZ_CTBB01000018.1 | + | 42532 | 42584 | **RNA**→ | rpmE (PRK00019)Ribosomal_L31 (pfam01197)→ |
| Sen-300-95 | NZ_CTNL01000017.1 | - | 46977 | 46925 | **RNA**→ | rpmE (PRK00019)Ribosomal_L31 (pfam01197)→ |
| Sen-300-96 | NZ_CTCJ01000017.1 | - | 45272 | 45220 | **RNA**→ | rpmE (PRK00019)Ribosomal_L31 (pfam01197)→ |
| Sen-300-97 | NZ_CTPA01000018.1 | - | 45141 | 45089 | **RNA**→ | rpmE (PRK00019)Ribosomal_L31 (pfam01197)→ |

| | | | | | |
|---|---|---|---|---|---|
| Sen-300-98 | NZ_CTAZ01000013.1 | + | 104065 | 104117 | **RNA**→ rpmE (PRK00019)Ribosomal_L31 (pfam01197)→ |
| Sen-300-99 | NZ_CTLB01000018.1 | + | 42527 | 42579 | **RNA**→ rpmE (PRK00019)Ribosomal_L31 (pfam01197)→ |
| Sen-300-100 | NZ_CTMY01000021.1 | - | 45179 | 45127 | **RNA**→ rpmE (PRK00019)Ribosomal_L31 (pfam01197)→ |
| Sen-300-101 | NZ_CTLG01000005.1 | - | 45116 | 45064 | **RNA**→ rpmE (PRK00019)Ribosomal_L31 (pfam01197)→ |
| Sen-300-102 | NZ_CTPC01000020.1 | + | 42473 | 42525 | **RNA**→ rpmE (PRK00019)Ribosomal_L31 (pfam01197)→ |
| Sen-300-103 | NZ_CTQU01000018.1 | - | 45179 | 45127 | **RNA**→ rpmE (PRK00019)Ribosomal_L31 (pfam01197)→ |
| Sen-300-104 | NZ_CTMB01000017.1 | + | 42472 | 42524 | **RNA**→ rpmE (PRK00019)Ribosomal_L31 (pfam01197)→ |
| Sen-300-105 | NZ_CTQL01000158.1 | + | 1706 | 1758 | **RNA**→ rpmE (PRK00019)Ribosomal_L31 (pfam01197)→ |
| Sen-300-106 | NZ_CTPE01000019.1 | - | 45110 | 45058 | **RNA**→ rpmE (PRK00019)Ribosomal_L31 (pfam01197)→ |
| Sen-300-107 | NZ_CTQA01000017.1 | + | 42501 | 42553 | **RNA**→ rpmE (PRK00019)Ribosomal_L31 (pfam01197)→ |
| Sen-300-108 | NZ_CTOD01000014.1 | + | 104184 | 104236 | **RNA**→ rpmE (PRK00019)Ribosomal_L31 (pfam01197)→ |
| Sen-300-109 | NZ_CTMU01000020.1 | + | 43122 | 43174 | **RNA**→ rpmE (PRK00019)Ribosomal_L31 (pfam01197)→ |
| Sen-300-110 | NZ_CTOB01000018.1 | - | 45343 | 45291 | **RNA**→ rpmE (PRK00019)Ribosomal_L31 (pfam01197)→ |
| Sen-300-111 | NZ_CTGB01000019.1 | - | 45107 | 45055 | **RNA**→ rpmE (PRK00019)Ribosomal_L31 (pfam01197)→ |
| Sen-300-112 | NZ_CTGS01000018.1 | - | 45214 | 45162 | **RNA**→ rpmE (PRK00019)Ribosomal_L31 (pfam01197)→ |
| Sen-300-113 | NZ_CTAT01000013.1 | - | 45107 | 45055 | **RNA**→ rpmE (PRK00019)Ribosomal_L31 (pfam01197)→ |
| Sen-300-114 | NZ_CTGO01000018.1 | + | 42484 | 42536 | **RNA**→ rpmE (PRK00019)Ribosomal_L31 (pfam01197)→ |
| Sen-300-115 | NZ_CTLS01000021.1 | - | 45179 | 45127 | **RNA**→ rpmE (PRK00019)Ribosomal_L31 (pfam01197)→ |
| Sen-300-116 | NZ_CTNX01000021.1 | + | 42503 | 42555 | **RNA**→ rpmE (PRK00019)Ribosomal_L31 (pfam01197)→ |
| Sen-300-117 | NZ_CTPF01000017.1 | - | 45176 | 45124 | **RNA**→ rpmE (PRK00019)Ribosomal_L31 (pfam01197)→ |
| Sen-300-118 | NZ_CTOC01000018.1 | - | 45107 | 45055 | **RNA**→ rpmE (PRK00019)Ribosomal_L31 (pfam01197)→ |
| Sen-300-119 | NZ_CTNV01000019.1 | - | 45179 | 45127 | **RNA**→ rpmE (PRK00019)Ribosomal_L31 (pfam01197)→ |
| Sen-300-120 | NZ_CTQT01000020.1 | - | 45179 | 45127 | **RNA**→ rpmE (PRK00019)Ribosomal_L31 (pfam01197)→ |
| Sen-300-121 | NZ_CTLJ01000018.1 | + | 42531 | 42583 | **RNA**→ rpmE (PRK00019)Ribosomal_L31 (pfam01197)→ |
| Sen-300-122 | NZ_CTLQ01000018.1 | + | 42485 | 42537 | **RNA**→ rpmE (PRK00019)Ribosomal_L31 (pfam01197)→ |
| Sen-300-123 | NZ_CTCD01000016.1 | - | 45343 | 45291 | **RNA**→ rpmE (PRK00019)Ribosomal_L31 (pfam01197)→ |
| Sen-300-124 | NZ_CTMA01000017.1 | - | 45179 | 45127 | **RNA**→ rpmE (PRK00019)Ribosomal_L31 (pfam01197)→ |
| Sen-300-125 | NZ_CTMI01000019.1 | - | 45107 | 45055 | **RNA**→ rpmE (PRK00019)Ribosomal_L31 (pfam01197)→ |
| Sen-300-126 | NZ_CTAH01000011.1 | - | 46977 | 46925 | **RNA**→ rpmE (PRK00019)Ribosomal_L31 (pfam01197)→ |
| Sen-300-127 | NZ_CTCF01000017.1 | - | 45271 | 45219 | **RNA**→ rpmE (PRK00019)Ribosomal_L31 (pfam01197)→ |
| Sen-300-128 | NZ_CTPG01000019.1 | - | 45179 | 45127 | **RNA**→ rpmE (PRK00019)Ribosomal_L31 (pfam01197)→ |
| Sen-300-129 | NZ_CTAK01000019.1 | - | 45108 | 45056 | **RNA**→ rpmE (PRK00019)Ribosomal_L31 (pfam01197)→ |
| Sen-300-130 | NZ_CTOQ01000019.1 | + | 42472 | 42524 | **RNA**→ rpmE (PRK00019)Ribosomal_L31 (pfam01197)→ |
| Sen-300-131 | NZ_CTCG01000017.1 | - | 45343 | 45291 | **RNA**→ rpmE (PRK00019)Ribosomal_L31 (pfam01197)→ |
| Sen-300-132 | NZ_CTAD01000017.1 | - | 46430 | 46378 | **RNA**→ rpmE (PRK00019)Ribosomal_L31 (pfam01197)→ |
| Sen-300-133 | NZ_CTAP01000017.1 | + | 42490 | 42542 | **RNA**→ rpmE (PRK00019)Ribosomal_L31 (pfam01197)→ |
| Sen-300-134 | NZ_CTAQ01000017.1 | + | 42528 | 42580 | **RNA**→ rpmE (PRK00019)Ribosomal_L31 (pfam01197)→ |
| Sen-962-1 | NC_017046.1 | + | 4325695 | 4325747 | **RNA**→ rpmE (PRK00019)Ribosomal_L31 (pfam01197)→ |
| Sen-620-1 | NZ_CP007352.1 | + | 4173893 | 4173945 | **RNA**→ rpmE (PRK00019)Ribosomal_L31 (pfam01197)→ |
| Sen-606-1 | NZ_CP007327.1 | + | 4173907 | 4173959 | **RNA**→ rpmE (PRK00019)Ribosomal_L31 (pfam01197)→ |
| Sen-160-1 | NZ_CP007245.1 | + | 4196527 | 4196579 | **RNA**→ rpmE (PRK00019)Ribosomal_L31 (pfam01197)→ |
| Sen-706-1 | NZ_CP007306.1 | + | 4173906 | 4173958 | **RNA**→ rpmE (PRK00019)Ribosomal_L31 (pfam01197)→ |
| Sen-610-1 | NZ_CP007433.1 | + | 4173880 | 4173932 | **RNA**→ rpmE (PRK00019)Ribosomal_L31 (pfam01197)→ |
| Sen-648-1 | NZ_CP007380.1 | + | 4173899 | 4173951 | **RNA**→ rpmE (PRK00019)Ribosomal_L31 (pfam01197)→ |
| Sen-663-1 | NZ_CP007382.1 | + | 4173900 | 4173952 | **RNA**→ rpmE (PRK00019)Ribosomal_L31 (pfam01197)→ |
| Sen-696-1 | NZ_CP007295.1 | + | 4173849 | 4173901 | **RNA**→ rpmE (PRK00019)Ribosomal_L31 (pfam01197)→ |
| Sen-301-1 | NZ_AP014565.1 | + | 4500867 | 4500919 | **RNA**→ rpmE (PRK00019)Ribosomal_L31 (pfam01197)→ |
| Sen-597-1 | NZ_CP007359.1 | + | 4158686 | 4158738 | **RNA**→ rpmE (PRK00019)Ribosomal_L31 (pfam01197)→ |
| Sen-595-1 | NZ_CP007358.1 | + | 4155155 | 4155207 | **RNA**→ rpmE (PRK00019)Ribosomal_L31 (pfam01197)→ |
| Sen-139-3 | NZ_CP009085.2 | + | 4167334 | 4167386 | **RNA**→ rpmE (PRK00019)Ribosomal_L31 (pfam01197)→ |
| Sen-139-4 | NZ_CP009084.2 | + | 4167333 | 4167385 | **RNA**→ rpmE (PRK00019)Ribosomal_L31 (pfam01197)→ |
| Sen-255-1 | NZ_CP010280.1 | + | 3658105 | 3658157 | **RNA**→ rpmE (PRK00019)Ribosomal_L31 (pfam01197)→ |

| | | | | | |
|---|---|---|---|---|---|
| Sen-139-5 | NZ_CP007507.1 | + | 4166959 | 4167011 | RNA→ rpmE (PRK00019)Ribosomal_L31 (pfam01197)→ |
| Sen-600-1 | NZ_CP007266.1 | + | 4154819 | 4154871 | RNA→ rpmE (PRK00019)Ribosomal_L31 (pfam01197)→ |
| Sen-10-1 | NZ_CP007532.1 | + | 1359262 | 1359314 | RNA→ rpmE (PRK00019)Ribosomal_L31 (pfam01197)→ |
| Sen-682-1 | NZ_CP007279.1 | + | 4173886 | 4173938 | RNA→ rpmE (PRK00019)Ribosomal_L31 (pfam01197)→ |
| Sen-680-1 | NZ_CP007277.1 | + | 4173900 | 4173952 | RNA→ rpmE (PRK00019)Ribosomal_L31 (pfam01197)→ |
| Sen-704-1 | NZ_CP007304.1 | + | 4173906 | 4173958 | RNA→ rpmE (PRK00019)Ribosomal_L31 (pfam01197)→ |
| Sen-690-1 | NZ_CP007288.1 | + | 4173879 | 4173931 | RNA→ rpmE (PRK00019)Ribosomal_L31 (pfam01197)→ |
| Sen-695-1 | NZ_CP007294.1 | + | 4173888 | 4173940 | RNA→ rpmE (PRK00019)Ribosomal_L31 (pfam01197)→ |
| Sen-705-1 | NZ_CP007305.1 | + | 4173909 | 4173961 | RNA→ rpmE (PRK00019)Ribosomal_L31 (pfam01197)→ |
| Sen-701-1 | NZ_CP007274.1 | - | 4451692 | 4451640 | RNA→ rpmE (PRK00019)Ribosomal_L31 (pfam01197)→ |
| Sen-685-1 | NZ_CP007282.1 | + | 4173887 | 4173939 | RNA→ rpmE (PRK00019)Ribosomal_L31 (pfam01197)→ |
| Sen-623-1 | NZ_CP007362.1 | + | 4173904 | 4173956 | RNA→ rpmE (PRK00019)Ribosomal_L31 (pfam01197)→ |
| Sen-624-1 | NZ_CP007363.1 | + | 4173904 | 4173956 | RNA→ rpmE (PRK00019)Ribosomal_L31 (pfam01197)→ |
| Sen-617-1 | NZ_CP007349.1 | + | 4173904 | 4173956 | RNA→ rpmE (PRK00019)Ribosomal_L31 (pfam01197)→ |
| Sen-703-1 | NZ_CP007303.1 | + | 4173905 | 4173957 | RNA→ rpmE (PRK00019)Ribosomal_L31 (pfam01197)→ |
| Sen-717-1 | NZ_CP007431.1 | + | 4174081 | 4174133 | RNA→ rpmE (PRK00019)Ribosomal_L31 (pfam01197)→ |
| Sen-151-1 | NZ_CP007260.1 | + | 4162468 | 4162520 | RNA→ rpmE (PRK00019)Ribosomal_L31 (pfam01197)→ |
| Sen-720-1 | NZ_CP007372.1 | + | 4173901 | 4173953 | RNA→ rpmE (PRK00019)Ribosomal_L31 (pfam01197)→ |
| Sen-692-1 | NZ_CP007290.1 | + | 4173905 | 4173957 | RNA→ rpmE (PRK00019)Ribosomal_L31 (pfam01197)→ |
| Sen-584-1 | NZ_CP007320.1 | + | 4149921 | 4149973 | RNA→ rpmE (PRK00019)Ribosomal_L31 (pfam01197)→ |
| Sen-149-1 | NZ_CP007262.1 | + | 4166935 | 4166987 | RNA→ rpmE (PRK00019)Ribosomal_L31 (pfam01197)→ |
| Sen-585-1 | NZ_CP007321.1 | + | 4158170 | 4158222 | RNA→ rpmE (PRK00019)Ribosomal_L31 (pfam01197)→ |
| Sen-150-1 | NZ_CP007261.1 | + | 4171043 | 4171095 | RNA→ rpmE (PRK00019)Ribosomal_L31 (pfam01197)→ |
| Sen-709-1 | NZ_CP007309.1 | + | 4173904 | 4173956 | RNA→ rpmE (PRK00019)Ribosomal_L31 (pfam01197)→ |
| Sen-153-1 | NZ_CP007258.1 | + | 4162080 | 4162132 | RNA→ rpmE (PRK00019)Ribosomal_L31 (pfam01197)→ |
| Sen-654-1 | NZ_CP007373.1 | + | 4173904 | 4173956 | RNA→ rpmE (PRK00019)Ribosomal_L31 (pfam01197)→ |
| Sen-684-1 | NZ_CP007281.1 | + | 4173903 | 4173955 | RNA→ rpmE (PRK00019)Ribosomal_L31 (pfam01197)→ |
| Sen-285-1 | NZ_CP007505.1 | + | 909485 | 909537 | RNA→ rpmE (PRK00019)Ribosomal_L31 (pfam01197)→ |
| Sen-235-1 | NZ_CP007530.1 | + | 3875383 | 3875435 | RNA→ rpmE (PRK00019)Ribosomal_L31 (pfam01197)→ |
| Sen-607-1 | NZ_CP007328.1 | + | 4173904 | 4173956 | RNA→ rpmE (PRK00019)Ribosomal_L31 (pfam01197)→ |
| Sen-613-1 | NZ_CP007345.1 | + | 4173898 | 4173950 | RNA→ rpmE (PRK00019)Ribosomal_L31 (pfam01197)→ |
| Sen-132-1 | NZ_CP007533.1 | + | 3681808 | 3681860 | RNA→ rpmE (PRK00019)Ribosomal_L31 (pfam01197)→ |
| Sen-693-1 | NZ_CP007291.1 | + | 4173893 | 4173945 | RNA→ rpmE (PRK00019)Ribosomal_L31 (pfam01197)→ |
| Sen-614-1 | NZ_CP007346.1 | + | 4173899 | 4173951 | RNA→ rpmE (PRK00019)Ribosomal_L31 (pfam01197)→ |
| Sen-700-1 | NZ_CP007334.1 | + | 580417 | 580469 | RNA→ rpmE (PRK00019)Ribosomal_L31 (pfam01197)→ |
| Sen-658-1 | NZ_CP007396.1 | + | 4173900 | 4173952 | RNA→ rpmE (PRK00019)Ribosomal_L31 (pfam01197)→ |
| Sen-659-1 | NZ_CP007397.1 | + | 4173889 | 4173941 | RNA→ rpmE (PRK00019)Ribosomal_L31 (pfam01197)→ |
| Sen-660-1 | NZ_CP007398.1 | + | 4173905 | 4173957 | RNA→ rpmE (PRK00019)Ribosomal_L31 (pfam01197)→ |
| Sen-687-1 | NZ_CP007284.1 | + | 4173899 | 4173951 | RNA→ rpmE (PRK00019)Ribosomal_L31 (pfam01197)→ |
| Sen-702-1 | NZ_CP007302.1 | + | 4173892 | 4173944 | RNA→ rpmE (PRK00019)Ribosomal_L31 (pfam01197)→ |
| Sen-707-1 | NZ_CP007307.1 | + | 4173908 | 4173960 | RNA→ rpmE (PRK00019)Ribosomal_L31 (pfam01197)→ |
| Sen-657-1 | NZ_CP007395.1 | + | 4173913 | 4173965 | RNA→ rpmE (PRK00019)Ribosomal_L31 (pfam01197)→ |
| Sen-638-1 | NZ_CP007405.1 | + | 4173903 | 4173955 | RNA→ rpmE (PRK00019)Ribosomal_L31 (pfam01197)→ |
| Sen-708-1 | NZ_CP007308.1 | + | 4173906 | 4173958 | RNA→ rpmE (PRK00019)Ribosomal_L31 (pfam01197)→ |
| Sen-686-1 | NZ_CP007283.1 | + | 4173887 | 4173939 | RNA→ rpmE (PRK00019)Ribosomal_L31 (pfam01197)→ |
| Sen-619-1 | NZ_CP007351.1 | + | 4173904 | 4173956 | RNA→ rpmE (PRK00019)Ribosomal_L31 (pfam01197)→ |
| Sen-698-1 | NZ_CP007300.1 | + | 4173902 | 4173954 | RNA→ rpmE (PRK00019)Ribosomal_L31 (pfam01197)→ |
| Sen-710-1 | NZ_CP007310.1 | + | 4173912 | 4173964 | RNA→ rpmE (PRK00019)Ribosomal_L31 (pfam01197)→ |
| Sen-718-1 | NZ_CP007317.1 | + | 4173901 | 4173953 | RNA→ rpmE (PRK00019)Ribosomal_L31 (pfam01197)→ |
| Sen-716-1 | NZ_CP007316.1 | + | 4173895 | 4173947 | RNA→ rpmE (PRK00019)Ribosomal_L31 (pfam01197)→ |
| Sen-719-1 | NZ_CP007318.1 | + | 4173895 | 4173947 | RNA→ rpmE (PRK00019)Ribosomal_L31 (pfam01197)→ |
| Sen-713-1 | NZ_CP007313.1 | + | 4173907 | 4173959 | RNA→ rpmE (PRK00019)Ribosomal_L31 (pfam01197)→ |

| | | | | | | |
|---|---|---|---|---|---|---|
| Sen-604-1 | NZ_CP007325.1 | + | 4173909 | 4173961 | **RNA**→ | rpmE (PRK00019)Ribosomal_L31 (pfam01197)→ |
| Sen-652-1 | NZ_CP007370.1 | + | 4173899 | 4173951 | **RNA**→ | rpmE (PRK00019)Ribosomal_L31 (pfam01197)→ |
| Sen-616-1 | NZ_CP007348.1 | + | 4173876 | 4173928 | **RNA**→ | rpmE (PRK00019)Ribosomal_L31 (pfam01197)→ |
| Sen-650-1 | NZ_CP007368.1 | + | 4173906 | 4173958 | **RNA**→ | rpmE (PRK00019)Ribosomal_L31 (pfam01197)→ |
| Sen-634-1 | NZ_CP007401.1 | + | 4173906 | 4173958 | **RNA**→ | rpmE (PRK00019)Ribosomal_L31 (pfam01197)→ |
| Sen-632-1 | NZ_CP007343.1 | + | 4173899 | 4173951 | **RNA**→ | rpmE (PRK00019)Ribosomal_L31 (pfam01197)→ |
| Sen-668-1 | NZ_CP007387.1 | + | 4173904 | 4173956 | **RNA**→ | rpmE (PRK00019)Ribosomal_L31 (pfam01197)→ |
| Sen-694-1 | NZ_CP007293.1 | + | 4173853 | 4173905 | **RNA**→ | rpmE (PRK00019)Ribosomal_L31 (pfam01197)→ |
| Sen-11-1 | NZ_CP007534.1 | + | 1393133 | 1393185 | **RNA**→ | rpmE (PRK00019)Ribosomal_L31 (pfam01197)→ |
| Sen-715-1 | NZ_CP007315.1 | + | 4173902 | 4173954 | **RNA**→ | rpmE (PRK00019)Ribosomal_L31 (pfam01197)→ |
| Sen-625-1 | NZ_CP007364.1 | + | 4173905 | 4173957 | **RNA**→ | rpmE (PRK00019)Ribosomal_L31 (pfam01197)→ |
| Sen-637-1 | NZ_CP007404.1 | + | 4173903 | 4173955 | **RNA**→ | rpmE (PRK00019)Ribosomal_L31 (pfam01197)→ |
| Sen-714-1 | NZ_CP007314.1 | + | 4173860 | 4173912 | **RNA**→ | rpmE (PRK00019)Ribosomal_L31 (pfam01197)→ |
| Sen-618-1 | NZ_CP007350.1 | + | 4173904 | 4173956 | **RNA**→ | rpmE (PRK00019)Ribosomal_L31 (pfam01197)→ |
| Sen-688-1 | NZ_CP007285.1 | + | 4173888 | 4173940 | **RNA**→ | rpmE (PRK00019)Ribosomal_L31 (pfam01197)→ |
| Sen-627-1 | NZ_CP007336.1 | + | 4173855 | 4173907 | **RNA**→ | rpmE (PRK00019)Ribosomal_L31 (pfam01197)→ |
| Sen-691-1 | NZ_CP007289.1 | + | 4173857 | 4173909 | **RNA**→ | rpmE (PRK00019)Ribosomal_L31 (pfam01197)→ |
| Sen-711-1 | NZ_CP007311.1 | + | 4173900 | 4173952 | **RNA**→ | rpmE (PRK00019)Ribosomal_L31 (pfam01197)→ |
| Sen-655-1 | NZ_CP007374.1 | + | 4173906 | 4173958 | **RNA**→ | rpmE (PRK00019)Ribosomal_L31 (pfam01197)→ |
| Sen-142-1 | NZ_CP007528.1 | - | 96083 | 96031 | **RNA**→ | rpmE (PRK00019)Ribosomal_L31 (pfam01197)→ |
| Sen-661-1 | NZ_CP007429.1 | + | 4173888 | 4173940 | **RNA**→ | rpmE (PRK00019)Ribosomal_L31 (pfam01197)→ |
| Sen-689-1 | NZ_CP007286.1 | + | 4173887 | 4173939 | **RNA**→ | rpmE (PRK00019)Ribosomal_L31 (pfam01197)→ |
| Sen-17-1 | NZ_CP007531.1 | - | 2862649 | 2862597 | **RNA**→ | rpmE (PRK00019)Ribosomal_L31 (pfam01197)→ |
| Sen-636-1 | NZ_CP007403.1 | + | 4173904 | 4173956 | **RNA**→ | rpmE (PRK00019)Ribosomal_L31 (pfam01197)→ |
| Sen-640-1 | NZ_CP007407.1 | + | 4173906 | 4173958 | **RNA**→ | rpmE (PRK00019)Ribosomal_L31 (pfam01197)→ |
| Sen-670-1 | NZ_CP007411.1 | + | 4173906 | 4173958 | **RNA**→ | rpmE (PRK00019)Ribosomal_L31 (pfam01197)→ |
| Sen-681-1 | NZ_CP007278.1 | + | 4173904 | 4173956 | **RNA**→ | rpmE (PRK00019)Ribosomal_L31 (pfam01197)→ |
| Sen-641-1 | NZ_CP007408.1 | + | 4173903 | 4173955 | **RNA**→ | rpmE (PRK00019)Ribosomal_L31 (pfam01197)→ |
| Sen-635-1 | NZ_CP007402.1 | + | 4173916 | 4173968 | **RNA**→ | rpmE (PRK00019)Ribosomal_L31 (pfam01197)→ |
| Sen-678-1 | NZ_CP007424.1 | + | 4173893 | 4173945 | **RNA**→ | rpmE (PRK00019)Ribosomal_L31 (pfam01197)→ |
| Sen-712-1 | NZ_CP007312.1 | + | 4173886 | 4173938 | **RNA**→ | rpmE (PRK00019)Ribosomal_L31 (pfam01197)→ |
| Sen-673-1 | NZ_CP007414.1 | + | 4173907 | 4173959 | **RNA**→ | rpmE (PRK00019)Ribosomal_L31 (pfam01197)→ |
| Sen-599-1 | NZ_CP007323.1 | + | 4174039 | 4174091 | **RNA**→ | rpmE (PRK00019)Ribosomal_L31 (pfam01197)→ |
| Sen-697-1 | NZ_CP007297.1 | + | 4173855 | 4173907 | **RNA**→ | rpmE (PRK00019)Ribosomal_L31 (pfam01197)→ |
| Sen-672-1 | NZ_CP007413.1 | + | 4173902 | 4173954 | **RNA**→ | rpmE (PRK00019)Ribosomal_L31 (pfam01197)→ |
| Sen-645-1 | NZ_CP007377.1 | + | 4173902 | 4173954 | **RNA**→ | rpmE (PRK00019)Ribosomal_L31 (pfam01197)→ |
| Sen-667-1 | NZ_CP007386.1 | + | 4173908 | 4173960 | **RNA**→ | rpmE (PRK00019)Ribosomal_L31 (pfam01197)→ |
| Sen-646-1 | NZ_CP007378.1 | + | 4173903 | 4173955 | **RNA**→ | rpmE (PRK00019)Ribosomal_L31 (pfam01197)→ |
| Sen-647-1 | NZ_CP007379.1 | + | 4173894 | 4173946 | **RNA**→ | rpmE (PRK00019)Ribosomal_L31 (pfam01197)→ |
| Sen-683-1 | NZ_CP007280.1 | + | 4173887 | 4173939 | **RNA**→ | rpmE (PRK00019)Ribosomal_L31 (pfam01197)→ |
| Sen-639-1 | NZ_CP007406.1 | + | 4173905 | 4173957 | **RNA**→ | rpmE (PRK00019)Ribosomal_L31 (pfam01197)→ |
| Sen-721-1 | NZ_CP007417.1 | + | 4173901 | 4173953 | **RNA**→ | rpmE (PRK00019)Ribosomal_L31 (pfam01197)→ |
| Sen-664-1 | NZ_CP007383.1 | + | 4173902 | 4173954 | **RNA**→ | rpmE (PRK00019)Ribosomal_L31 (pfam01197)→ |
| Sen-669-1 | NZ_CP007388.1 | + | 4173906 | 4173958 | **RNA**→ | rpmE (PRK00019)Ribosomal_L31 (pfam01197)→ |
| Sen-161-1 | NZ_CP007332.1 | + | 4643968 | 4644020 | **RNA**→ | rpmE (PRK00019)Ribosomal_L31 (pfam01197)→ |
| Sen-671-1 | NZ_CP007412.1 | + | 4173902 | 4173954 | **RNA**→ | rpmE (PRK00019)Ribosomal_L31 (pfam01197)→ |
| Sen-666-1 | NZ_CP007385.1 | + | 4173915 | 4173967 | **RNA**→ | rpmE (PRK00019)Ribosomal_L31 (pfam01197)→ |
| Sen-676-1 | NZ_CP007418.1 | + | 4174075 | 4174127 | **RNA**→ | rpmE (PRK00019)Ribosomal_L31 (pfam01197)→ |
| Sen-642-1 | NZ_CP007375.1 | + | 4173917 | 4173969 | **RNA**→ | rpmE (PRK00019)Ribosomal_L31 (pfam01197)→ |
| Sen-633-1 | NZ_CP007400.1 | + | 4173899 | 4173951 | **RNA**→ | rpmE (PRK00019)Ribosomal_L31 (pfam01197)→ |
| Sen-665-1 | NZ_CP007384.1 | + | 4173908 | 4173960 | **RNA**→ | rpmE (PRK00019)Ribosomal_L31 (pfam01197)→ |
| Sen-631-1 | NZ_CP007341.1 | + | 4173899 | 4173951 | **RNA**→ | rpmE (PRK00019)Ribosomal_L31 (pfam01197)→ |

| | | | | | | |
|---|---|---|---|---|---|---|
| Sen-674-1 | NZ_CP007415.1 | + | 4173996 | 4174048 | RNA→ | rpmE (PRK00019)Ribosomal_L31 (pfam01197)→ |
| Sen-586-1 | NZ_CP007430.1 | + | 4173903 | 4173955 | RNA→ | rpmE (PRK00019)Ribosomal_L31 (pfam01197)→ |
| Sen-677-1 | NZ_CP007419.1 | + | 4173986 | 4174038 | RNA→ | rpmE (PRK00019)Ribosomal_L31 (pfam01197)→ |
| Sen-675-1 | NZ_CP007416.1 | + | 4173982 | 4174034 | RNA→ | rpmE (PRK00019)Ribosomal_L31 (pfam01197)→ |
| Sen-154-1 | NZ_CP007263.1 | + | 4166548 | 4166600 | RNA→ | rpmE (PRK00019)Ribosomal_L31 (pfam01197)→ |
| Sen-603-1 | NZ_CP007324.1 | + | 4173903 | 4173955 | RNA→ | rpmE (PRK00019)Ribosomal_L31 (pfam01197)→ |
| Sen-615-1 | NZ_CP007347.1 | + | 4173902 | 4173954 | RNA→ | rpmE (PRK00019)Ribosomal_L31 (pfam01197)→ |
| Sen-630-1 | NZ_CP007340.1 | + | 4173900 | 4173952 | RNA→ | rpmE (PRK00019)Ribosomal_L31 (pfam01197)→ |
| Sen-656-1 | NZ_CP007464.1 | + | 4178809 | 4178861 | RNA→ | rpmE (PRK00019)Ribosomal_L31 (pfam01197)→ |
| Sen-591-1 | NZ_CP007355.1 | + | 4173897 | 4173949 | RNA→ | rpmE (PRK00019)Ribosomal_L31 (pfam01197)→ |
| Sen-152-1 | NZ_CP007259.1 | + | 4195383 | 4195435 | RNA→ | rpmE (PRK00019)Ribosomal_L31 (pfam01197)→ |
| Sen-164-1 | NZ_CP007465.1 | + | 4174359 | 4174411 | RNA→ | rpmE (PRK00019)Ribosomal_L31 (pfam01197)→ |
| Sen-587-1 | NZ_CP007322.1 | + | 4163250 | 4163302 | RNA→ | rpmE (PRK00019)Ribosomal_L31 (pfam01197)→ |
| Sen-608-1 | NZ_CP007330.1 | + | 4120423 | 4120475 | RNA→ | rpmE (PRK00019)Ribosomal_L31 (pfam01197)→ |
| Sen-588-1 | NZ_CP007298.1 | + | 4173899 | 4173951 | RNA→ | rpmE (PRK00019)Ribosomal_L31 (pfam01197)→ |
| Sen-609-1 | NZ_CP007331.1 | + | 4176562 | 4176614 | RNA→ | rpmE (PRK00019)Ribosomal_L31 (pfam01197)→ |
| Sen-629-1 | NZ_CP007338.1 | + | 4173902 | 4173954 | RNA→ | rpmE (PRK00019)Ribosomal_L31 (pfam01197)→ |
| Sen-596-1 | NZ_CP007432.1 | + | 4173897 | 4173949 | RNA→ | rpmE (PRK00019)Ribosomal_L31 (pfam01197)→ |
| Sen-628-1 | NZ_CP007337.1 | + | 4173902 | 4173954 | RNA→ | rpmE (PRK00019)Ribosomal_L31 (pfam01197)→ |
| Sen-592-1 | NZ_CP007356.1 | + | 4173897 | 4173949 | RNA→ | rpmE (PRK00019)Ribosomal_L31 (pfam01197)→ |
| Sen-605-1 | NZ_CP007326.1 | + | 4173906 | 4173958 | RNA→ | rpmE (PRK00019)Ribosomal_L31 (pfam01197)→ |
| Sen-611-1 | NZ_CP007434.1 | + | 4173889 | 4173941 | RNA→ | rpmE (PRK00019)Ribosomal_L31 (pfam01197)→ |
| Sen-145-1 | NZ_CP007246.1 | + | 4162763 | 4162815 | RNA→ | rpmE (PRK00019)Ribosomal_L31 (pfam01197)→ |
| Sen-146-1 | NZ_CP007247.1 | + | 4193044 | 4193096 | RNA→ | rpmE (PRK00019)Ribosomal_L31 (pfam01197)→ |
| Sen-157-1 | NZ_CP007252.1 | + | 4163237 | 4163289 | RNA→ | rpmE (PRK00019)Ribosomal_L31 (pfam01197)→ |
| Sen-590-1 | NZ_CP007421.1 | + | 4152968 | 4153020 | RNA→ | rpmE (PRK00019)Ribosomal_L31 (pfam01197)→ |
| Sen-144-1 | NZ_CP007248.1 | + | 4165166 | 4165218 | RNA→ | rpmE (PRK00019)Ribosomal_L31 (pfam01197)→ |
| Sen-651-1 | NZ_CP007369.1 | + | 4173851 | 4173903 | RNA→ | rpmE (PRK00019)Ribosomal_L31 (pfam01197)→ |
| Sen-147-1 | NZ_CP007175.1 | + | 4195011 | 4195063 | RNA→ | rpmE (PRK00019)Ribosomal_L31 (pfam01197)→ |
| Sen-598-1 | NZ_CP007360.1 | + | 4173885 | 4173937 | RNA→ | rpmE (PRK00019)Ribosomal_L31 (pfam01197)→ |
| Sen-300-135 | NZ_CTHT01000018.1 | + | 42438 | 42490 | RNA→ | rpmE (PRK00019)Ribosomal_L31 (pfam01197)→ |
| Sen-300-136 | NZ_CTBM01000029.1 | + | 13413 | 13465 | RNA→ | rpmE (PRK00019)Ribosomal_L31 (pfam01197)→ |
| Sen-300-137 | NZ_CTBC01000018.1 | + | 42472 | 42524 | RNA→ | rpmE (PRK00019)Ribosomal_L31 (pfam01197)→ |
| Sen-300-138 | NZ_CTOS01000016.1 | + | 42474 | 42526 | RNA→ | rpmE (PRK00019)Ribosomal_L31 (pfam01197)→ |
| Sen-300-139 | NZ_CTAJ01000017.1 | - | 45157 | 45105 | RNA→ | rpmE (PRK00019)Ribosomal_L31 (pfam01197)→ |
| Sen-300-140 | NZ_CTHL01000017.1 | - | 45272 | 45220 | RNA→ | rpmE (PRK00019)Ribosomal_L31 (pfam01197)→ |
| Sen-300-141 | NZ_CTMZ01000016.1 | + | 42908 | 42960 | RNA→ | rpmE (PRK00019)Ribosomal_L31 (pfam01197)→ |
| Sen-300-142 | NZ_CTHW01000018.1 | + | 42472 | 42524 | RNA→ | rpmE (PRK00019)Ribosomal_L31 (pfam01197)→ |
| Sen-300-143 | NZ_CTHI01000026.1 | + | 13439 | 13491 | RNA→ | rpmE (PRK00019)Ribosomal_L31 (pfam01197)→ |
| Sen-300-144 | NZ_CTPH01000017.1 | + | 42497 | 42549 | RNA→ | rpmE (PRK00019)Ribosomal_L31 (pfam01197)→ |
| Sen-288-1105 | NZ_CJBQ01000016.1 | + | 104151 | 104203 | RNA→ | rpmE (PRK00019)Ribosomal_L31 (pfam01197)→ |
| Sen-300-145 | NZ_CTPR01000021.1 | - | 45179 | 45127 | RNA→ | rpmE (PRK00019)Ribosomal_L31 (pfam01197)→ |
| Sen-300-146 | NZ_CTQM01000016.1 | + | 42910 | 42962 | RNA→ | rpmE (PRK00019)Ribosomal_L31 (pfam01197)→ |
| Sen-300-147 | NZ_CTHG01000017.1 | + | 42531 | 42583 | RNA→ | rpmE (PRK00019)Ribosomal_L31 (pfam01197)→ |
| Sen-300-148 | NZ_CTQF01000017.1 | + | 42517 | 42569 | RNA→ | rpmE (PRK00019)Ribosomal_L31 (pfam01197)→ |
| Sen-300-149 | NZ_CTJJ01000022.1 | + | 42437 | 42489 | RNA→ | rpmE (PRK00019)Ribosomal_L31 (pfam01197)→ |
| Sen-300-150 | NZ_CTOL01000019.1 | + | 42453 | 42505 | RNA→ | rpmE (PRK00019)Ribosomal_L31 (pfam01197)→ |
| Sen-300-151 | NZ_CTPL01000021.1 | - | 45179 | 45127 | RNA→ | rpmE (PRK00019)Ribosomal_L31 (pfam01197)→ |
| Sen-300-152 | NZ_CTPX01000020.1 | - | 45107 | 45055 | RNA→ | rpmE (PRK00019)Ribosomal_L31 (pfam01197)→ |
| Sen-300-153 | NZ_CTNB01000021.1 | - | 35684 | 35632 | RNA→ | rpmE (PRK00019)Ribosomal_L31 (pfam01197)→ |
| Sen-300-154 | NZ_CTHS01000020.1 | + | 42466 | 42518 | RNA→ | rpmE (PRK00019)Ribosomal_L31 (pfam01197)→ |
| Sen-300-155 | NZ_CTQJ01000023.1 | + | 49795 | 49847 | RNA→ | rpmE (PRK00019)Ribosomal_L31 (pfam01197)→ |

| | | | | | |
|---|---|---|---|---|---|
| Sen-300-156 | NZ_CTQY01000017.1 | + | 42507 | 42559 | RNA→ rpmE (PRK00019)Ribosomal_L31 (pfam01197)→ |
| Sen-300-157 | NZ_CTIC01000016.1 | + | 42478 | 42530 | RNA→ rpmE (PRK00019)Ribosomal_L31 (pfam01197)→ |
| Sen-300-158 | NZ_CTMR01000018.1 | + | 42531 | 42583 | RNA→ rpmE (PRK00019)Ribosomal_L31 (pfam01197)→ |
| Sen-300-159 | NZ_CTJT01000018.1 | - | 45270 | 45218 | RNA→ rpmE (PRK00019)Ribosomal_L31 (pfam01197)→ |
| Sen-300-160 | NZ_CTPS01000016.1 | - | 45343 | 45291 | RNA→ rpmE (PRK00019)Ribosomal_L31 (pfam01197)→ |
| Sen-300-161 | NZ_CTMN01000019.1 | - | 45179 | 45127 | RNA→ rpmE (PRK00019)Ribosomal_L31 (pfam01197)→ |
| Sen-300-162 | NZ_CTNP01000017.1 | - | 45343 | 45291 | RNA→ rpmE (PRK00019)Ribosomal_L31 (pfam01197)→ |
| Sen-300-163 | NZ_CTLW01000018.1 | + | 42472 | 42524 | RNA→ rpmE (PRK00019)Ribosomal_L31 (pfam01197)→ |
| Sen-300-164 | NZ_CTLN01000018.1 | + | 42531 | 42583 | RNA→ rpmE (PRK00019)Ribosomal_L31 (pfam01197)→ |
| Sen-300-165 | NZ_CTHP01000017.1 | - | 45194 | 45142 | RNA→ rpmE (PRK00019)Ribosomal_L31 (pfam01197)→ |
| Sen-300-166 | NZ_CTNT01000016.1 | - | 45196 | 45144 | RNA→ rpmE (PRK00019)Ribosomal_L31 (pfam01197)→ |
| Sen-300-167 | NZ_CTBH01000027.1 | + | 13435 | 13487 | RNA→ rpmE (PRK00019)Ribosomal_L31 (pfam01197)→ |
| Sen-300-168 | NZ_CTGL01000018.1 | - | 45178 | 45126 | RNA→ rpmE (PRK00019)Ribosomal_L31 (pfam01197)→ |
| Sen-300-169 | NZ_CTAG01000019.1 | - | 45179 | 45127 | RNA→ rpmE (PRK00019)Ribosomal_L31 (pfam01197)→ |
| Sen-300-170 | NZ_CTQZ01000017.1 | - | 45343 | 45291 | RNA→ rpmE (PRK00019)Ribosomal_L31 (pfam01197)→ |
| Sen-288-1106 | NZ_CTAA01000016.1 | - | 41507 | 41455 | RNA→ rpmE (PRK00019)Ribosomal_L31 (pfam01197)→ |
| Sen-300-171 | NZ_CTAU01000017.1 | - | 45179 | 45127 | RNA→ rpmE (PRK00019)Ribosomal_L31 (pfam01197)→ |
| Sen-300-172 | NZ_CTON01000018.1 | + | 55105 | 55157 | RNA→ rpmE (PRK00019)Ribosomal_L31 (pfam01197)→ |
| Sen-300-173 | NZ_CTAE01000017.1 | + | 42938 | 42990 | RNA→ rpmE (PRK00019)Ribosomal_L31 (pfam01197)→ |
| Sen-300-174 | NZ_CTOM01000021.1 | + | 42452 | 42504 | RNA→ rpmE (PRK00019)Ribosomal_L31 (pfam01197)→ |
| Sen-300-175 | NZ_CTNA01000017.1 | - | 45107 | 45055 | RNA→ rpmE (PRK00019)Ribosomal_L31 (pfam01197)→ |
| Sen-300-176 | NZ_CTBI01000020.1 | - | 45123 | 45071 | RNA→ rpmE (PRK00019)Ribosomal_L31 (pfam01197)→ |
| Sen-300-177 | NZ_CTLD01000019.1 | + | 42548 | 42600 | RNA→ rpmE (PRK00019)Ribosomal_L31 (pfam01197)→ |
| Sen-300-178 | NZ_CTQH01000018.1 | - | 45343 | 45291 | RNA→ rpmE (PRK00019)Ribosomal_L31 (pfam01197)→ |
| Sen-300-179 | NZ_CTIG01000019.1 | - | 45130 | 45078 | RNA→ rpmE (PRK00019)Ribosomal_L31 (pfam01197)→ |
| Sen-300-180 | NZ_CTPO01000019.1 | - | 45343 | 45291 | RNA→ rpmE (PRK00019)Ribosomal_L31 (pfam01197)→ |
| Sen-300-181 | NZ_CTLE01000017.1 | + | 42536 | 42588 | RNA→ rpmE (PRK00019)Ribosomal_L31 (pfam01197)→ |
| Sen-300-182 | NZ_CTLH01000017.1 | - | 45107 | 45055 | RNA→ rpmE (PRK00019)Ribosomal_L31 (pfam01197)→ |
| Sen-300-183 | NZ_CTAN01000021.1 | - | 47096 | 47044 | RNA→ rpmE (PRK00019)Ribosomal_L31 (pfam01197)→ |
| Sen-300-184 | NZ_CTKI01000017.1 | - | 47085 | 47033 | RNA→ rpmE (PRK00019)Ribosomal_L31 (pfam01197)→ |
| Sen-300-185 | NZ_CTBN01000019.1 | + | 42472 | 42524 | RNA→ rpmE (PRK00019)Ribosomal_L31 (pfam01197)→ |
| Sen-300-186 | NZ_CTHN01000005.1 | - | 228766 | 228714 | RNA→ rpmE (PRK00019)Ribosomal_L31 (pfam01197)→ |
| Sen-300-187 | NZ_CTCC01000018.1 | - | 45176 | 45124 | RNA→ rpmE (PRK00019)Ribosomal_L31 (pfam01197)→ |
| Sen-300-188 | NZ_CTPZ01000017.1 | - | 45161 | 45109 | RNA→ rpmE (PRK00019)Ribosomal_L31 (pfam01197)→ |
| Sen-300-189 | NZ_CTBU01000020.1 | + | 42438 | 42490 | RNA→ rpmE (PRK00019)Ribosomal_L31 (pfam01197)→ |
| Sen-300-190 | NZ_CTQK01000018.1 | + | 42476 | 42528 | RNA→ rpmE (PRK00019)Ribosomal_L31 (pfam01197)→ |
| Sen-300-191 | NZ_CTBS01000017.1 | - | 45179 | 45127 | RNA→ rpmE (PRK00019)Ribosomal_L31 (pfam01197)→ |
| Sen-300-192 | NZ_CTFW01000019.1 | - | 45107 | 45055 | RNA→ rpmE (PRK00019)Ribosomal_L31 (pfam01197)→ |
| Sen-300-193 | NZ_CTAR01000018.1 | + | 42531 | 42583 | RNA→ rpmE (PRK00019)Ribosomal_L31 (pfam01197)→ |
| Sen-300-194 | NZ_CTQO01000016.1 | + | 42546 | 42598 | RNA→ rpmE (PRK00019)Ribosomal_L31 (pfam01197)→ |
| Sen-300-195 | NZ_CTAI01000017.1 | - | 45174 | 45122 | RNA→ rpmE (PRK00019)Ribosomal_L31 (pfam01197)→ |
| Sen-300-196 | NZ_CTPY01000020.1 | - | 45179 | 45127 | RNA→ rpmE (PRK00019)Ribosomal_L31 (pfam01197)→ |
| Sen-300-197 | NZ_CTAX01000017.1 | - | 45343 | 45291 | RNA→ rpmE (PRK00019)Ribosomal_L31 (pfam01197)→ |
| Sen-300-198 | NZ_CTAV01000019.1 | + | 42489 | 42541 | RNA→ rpmE (PRK00019)Ribosomal_L31 (pfam01197)→ |
| Sen-300-199 | NZ_CTAY01000018.1 | - | 45343 | 45291 | RNA→ rpmE (PRK00019)Ribosomal_L31 (pfam01197)→ |
| Sen-300-200 | NZ_CTCB01000010.1 | + | 104185 | 104237 | RNA→ rpmE (PRK00019)Ribosomal_L31 (pfam01197)→ |
| Sen-589-1 | NZ_CP007422.1 | + | 4123191 | 4123243 | RNA→ rpmE (PRK00019)Ribosomal_L31 (pfam01197)→ |
| Sen-148-1 | NZ_CP007250.1 | + | 4165607 | 4165659 | RNA→ rpmE (PRK00019)Ribosomal_L31 (pfam01197)→ |
| Sen-594-1 | NZ_CP007420.1 | + | 4122037 | 4122089 | RNA→ rpmE (PRK00019)Ribosomal_L31 (pfam01197)→ |
| Sen-602-1 | NZ_CP007498.1 | + | 4173902 | 4173954 | RNA→ rpmE (PRK00019)Ribosomal_L31 (pfam01197)→ |
| Sen-155-1 | NZ_CP007254.1 | + | 4163479 | 4163531 | RNA→ rpmE (PRK00019)Ribosomal_L31 (pfam01197)→ |
| Sen-644-1 | NZ_CP007463.1 | + | 4113654 | 4113706 | RNA→ rpmE (PRK00019)Ribosomal_L31 (pfam01197)→ |

| | | | | | |
|---|---|---|---|---|---|
| Sen-156-1 | NZ_CP007253.1 | + | 4167210 | 4167262 | **RNA**→ rpmE (PRK00019)Ribosomal_L31 (pfam01197)→ |
| Sen-291-1 | NZ_CP007581.1 | + | 4382613 | 4382665 | **RNA**→ rpmE (PRK00019)Ribosomal_L31 (pfam01197)→ |
| Sen-141-1 | NZ_CP007598.1 | + | 1099087 | 1099139 | **RNA**→ rpmE (PRK00019)Ribosomal_L31 (pfam01197)→ |
| Sen-291-2 | NZ_CP007804.1 | + | 4212582 | 4212634 | **RNA**→ rpmE (PRK00019)Ribosomal_L31 (pfam01197)→ |
| Sen-291-3 | NZ_CP009102.1 | + | 4241573 | 4241625 | **RNA**→ rpmE (PRK00019)Ribosomal_L31 (pfam01197)→ |
| Sen-134-1 | NZ_CP007639.1 | + | 4197939 | 4197991 | **RNA**→ rpmE (PRK00019)Ribosomal_L31 (pfam01197)→ hypo→ hypo→ |
| Sen-139-6 | NZ_CP009083.1 | + | 4166984 | 4167036 | **RNA**→ rpmE (PRK00019)Ribosomal_L31 (pfam01197)→ |
| Sen-139-7 | NZ_CP009087.1 | + | 4166622 | 4166674 | **RNA**→ rpmE (PRK00019)Ribosomal_L31 (pfam01197)→ |
| Sen-139-8 | NZ_CP009091.1 | + | 4159359 | 4159411 | **RNA**→ rpmE (PRK00019)Ribosomal_L31 (pfam01197)→ |
| Sen-139-9 | NZ_CP009093.1 | + | 4190780 | 4190832 | **RNA**→ rpmE (PRK00019)Ribosomal_L31 (pfam01197)→ |
| Sen-139-10 | NZ_CP009088.1 | + | 4166019 | 4166071 | **RNA**→ rpmE (PRK00019)Ribosomal_L31 (pfam01197)→ |
| Sen-139-11 | NZ_CP009089.1 | + | 4199181 | 4199233 | **RNA**→ rpmE (PRK00019)Ribosomal_L31 (pfam01197)→ |
| Sen-139-12 | NZ_CP009092.1 | + | 4199355 | 4199407 | **RNA**→ rpmE (PRK00019)Ribosomal_L31 (pfam01197)→ |
| Sen-139-13 | NZ_CP009086.1 | + | 4166667 | 4166719 | **RNA**→ rpmE (PRK00019)Ribosomal_L31 (pfam01197)→ |
| Sen-139-14 | NZ_CP009090.1 | - | 3533924 | 3533872 | **RNA**→ rpmE (PRK00019)Ribosomal_L31 (pfam01197)→ |
| Sen-162-1 | NZ_CP007269.1 | + | 4163479 | 4163531 | **RNA**→ rpmE (PRK00019)Ribosomal_L31 (pfam01197)→ |
| Sen-139-15 | NZ_CP008928.1 | + | 2471151 | 2471203 | **RNA**→ rpmE (PRK00019)Ribosomal_L31 (pfam01197)→ |
| Sen-1033-1 | NC_006905.1 | + | 4238765 | 4238817 | **RNA**→ rpmE (PRK00019)Ribosomal_L31 (pfam01197)→ rpmE (PRK00019)Ribosomal_L31 (pfam01197)→ hypo→ hypo→ hypo→ hypo→ hypo→ hypo→ |
| Sen-649-1 | NZ_CP007272.1 | + | 2781849 | 2781901 | **RNA**→ hypo→ rpmE (PRK00019)Ribosomal_L31 (pfam01197)→ |
| Sen-649-2 | NZ_CP007272.1 | - | 4860128 | 4860076 | **RNA**→ ←hypo |
| Sen-662-1 | NZ_CP007381.1 | + | 4173906 | 4173958 | **RNA**→ rpmE (PRK00019)Ribosomal_L31 (pfam01197)→ |
| Sen-252-1 | NZ_CP010282.1 | - | 4355853 | 4355801 | **RNA**→ rpmE (PRK00019)Ribosomal_L31 (pfam01197)→ |
| Sen-256-1 | NZ_CP010281.1 | - | 3676986 | 3676934 | **RNA**→ rpmE (PRK00019)Ribosomal_L31 (pfam01197)→ |
| Sen-254-1 | NZ_CP010279.1 | + | 3848446 | 3848498 | **RNA**→ rpmE (PRK00019)Ribosomal_L31 (pfam01197)→ |
| Sen-257-1 | NZ_CP010284.1 | - | 538801 | 538749 | **RNA**→ rpmE (PRK00019)Ribosomal_L31 (pfam01197)→ |
| Sen-253-1 | NZ_CP010283.1 | - | 4242129 | 4242077 | **RNA**→ rpmE (PRK00019)Ribosomal_L31 (pfam01197)→ |
| Sen-275-1 | NZ_LK931482.1 | - | 3404936 | 3404884 | **RNA**→ rpmE (PRK00019)Ribosomal_L31 (pfam01197)→ |
| Sen-223-1 | NZ_LN649235.1 | + | 4201024 | 4201076 | **RNA**→ rpmE (PRK00019)Ribosomal_L31 (pfam01197)→ |
| Sen-258-1 | NZ_CP009561.1 | + | 1415376 | 1415428 | **RNA**→ rpmE (PRK00019)Ribosomal_L31 (pfam01197)→ |
| Sen-429-1 | NZ_CP007483.1 | - | 4514966 | 4514914 | **RNA**→ rpmE (PRK00019)Ribosomal_L31 (pfam01197)→ |
| Sen-993-1 | NC_021814.1 | - | 4121916 | 4121864 | **RNA**→ rpmE (PRK00019)Ribosomal_L31 (pfam01197)→ |
| Sen-13-1 | NZ_CP011259.1 | + | 1265510 | 1265562 | **RNA**→ rpmE (PRK00019)Ribosomal_L31 (pfam01197)→ |
| Sen-593-1 | NZ_CP007357.1 | + | 4173901 | 4173953 | **RNA**→ rpmE (PRK00019)Ribosomal_L31 (pfam01197)→ |
| Sen-298-1 | NZ_CP007523.1 | + | 221910 | 221962 | **RNA**→ rpmE (PRK00019)Ribosomal_L31 (pfam01197)→ |
| Sen-854-1 | NZ_CP007540.1 | - | 4268937 | 4268885 | **RNA**→ rpmE (PRK00019)Ribosomal_L31 (pfam01197)→ |
| Sen-868-1 | NZ_CP007559.1 | - | 4541938 | 4541886 | **RNA**→ rpmE (PRK00019)Ribosomal_L31 (pfam01197)→ |
| Sen-430-1 | NZ_CP007584.1 | + | 4315734 | 4315786 | **RNA**→ rpmE (PRK00019)Ribosomal_L31 (pfam01197)→ |
| Sen-291-4 | NZ_CP011365.1 | + | 4204324 | 4204376 | **RNA**→ rpmE (PRK00019)Ribosomal_L31 (pfam01197)→ |
| Sen-4-1 | NZ_CP011428.1 | + | 4333576 | 4333628 | **RNA**→ rpmE (PRK00019)Ribosomal_L31 (pfam01197)→ |
| Sen-287-1 | NZ_CP011396.1 | - | 4403134 | 4403082 | **RNA**→ rpmE (PRK00019)Ribosomal_L31 (pfam01197)→ |
| Sen-140-1 | NZ_CP011394.1 | - | 4431786 | 4431734 | **RNA**→ rpmE (PRK00019)Ribosomal_L31 (pfam01197)→ |
| Sen-300-201 | NZ_CTCA01000019.1 | + | 42551 | 42603 | **RNA**→ rpmE (PRK00019)Ribosomal_L31 (pfam01197)→ |
| Sen-300-202 | NZ_CTAB01000019.1 | + | 42495 | 42547 | **RNA**→ rpmE (PRK00019)Ribosomal_L31 (pfam01197)→ |
| Sen-300-203 | NZ_CTBA01000020.1 | - | 45179 | 45127 | **RNA**→ rpmE (PRK00019)Ribosomal_L31 (pfam01197)→ |
| Sen-300-204 | NZ_CTAL01000022.1 | - | 45111 | 45059 | **RNA**→ rpmE (PRK00019)Ribosomal_L31 (pfam01197)→ |
| Sen-300-205 | NZ_CTAW01000017.1 | + | 42480 | 42532 | **RNA**→ rpmE (PRK00019)Ribosomal_L31 (pfam01197)→ |
| Sen-300-206 | NZ_CTBZ01000017.1 | + | 42531 | 42583 | **RNA**→ rpmE (PRK00019)Ribosomal_L31 (pfam01197)→ |
| Sen-300-207 | NZ_CTFZ01000021.1 | - | 45343 | 45291 | **RNA**→ rpmE (PRK00019)Ribosomal_L31 (pfam01197)→ |
| Sen-300-208 | NZ_CTBV01000027.1 | + | 13413 | 13465 | **RNA**→ rpmE (PRK00019)Ribosomal_L31 (pfam01197)→ |
| Sen-300-209 | NZ_CTMW01000030.1 | + | 13443 | 13495 | **RNA**→ rpmE (PRK00019)Ribosomal_L31 (pfam01197)→ |
| Sen-300-210 | NZ_CTLI01000017.1 | + | 42508 | 42560 | **RNA**→ rpmE (PRK00019)Ribosomal_L31 (pfam01197)→ |
| Sen-300-211 | NZ_CTGK01000017.1 | - | 45193 | 45141 | **RNA**→ rpmE (PRK00019)Ribosomal_L31 (pfam01197)→ |

| | | | | | |
|---|---|---|---|---|---|
| Sen-300-212 | NZ_CTLK01000017.1 | - | 45343 | 45291 | **RNA**→ rpmE (PRK00019)Ribosomal_L31 (pfam01197)→ |
| Sen-300-213 | NZ_CTLM01000019.1 | + | 42551 | 42603 | **RNA**→ rpmE (PRK00019)Ribosomal_L31 (pfam01197)→ |
| Sen-300-214 | NZ_CTJO01000020.1 | + | 42453 | 42505 | **RNA**→ rpmE (PRK00019)Ribosomal_L31 (pfam01197)→ |
| Sen-300-215 | NZ_CTII01000017.1 | - | 45343 | 45291 | **RNA**→ rpmE (PRK00019)Ribosomal_L31 (pfam01197)→ |
| Sen-300-216 | NZ_CTJK01000018.1 | - | 47016 | 46964 | **RNA**→ rpmE (PRK00019)Ribosomal_L31 (pfam01197)→ |
| Sen-300-217 | NZ_CTMC01000020.1 | - | 45179 | 45127 | **RNA**→ rpmE (PRK00019)Ribosomal_L31 (pfam01197)→ |
| Sen-300-218 | NZ_CTHM01000017.1 | + | 42513 | 42565 | **RNA**→ rpmE (PRK00019)Ribosomal_L31 (pfam01197)→ |
| Sen-300-219 | NZ_CTOO01000018.1 | + | 42496 | 42548 | **RNA**→ rpmE (PRK00019)Ribosomal_L31 (pfam01197)→ |
| Sen-300-220 | NZ_CTJS01000019.1 | - | 45135 | 45083 | **RNA**→ rpmE (PRK00019)Ribosomal_L31 (pfam01197)→ |
| Sen-300-221 | NZ_CTGW01000024.1 | - | 32539 | 32487 | **RNA**→ rpmE (PRK00019)Ribosomal_L31 (pfam01197)→ |
| Sen-300-222 | NZ_CTGX01000021.1 | - | 45124 | 45072 | **RNA**→ rpmE (PRK00019)Ribosomal_L31 (pfam01197)→ |
| Sen-300-223 | NZ_CTOE01000018.1 | + | 42532 | 42584 | **RNA**→ rpmE (PRK00019)Ribosomal_L31 (pfam01197)→ |
| Sen-300-224 | NZ_CTPM01000019.1 | - | 45167 | 45115 | **RNA**→ rpmE (PRK00019)Ribosomal_L31 (pfam01197)→ |
| Sen-300-225 | NZ_CTHX01000019.1 | - | 45107 | 45055 | **RNA**→ rpmE (PRK00019)Ribosomal_L31 (pfam01197)→ |
| Sen-300-226 | NZ_CTNC01000019.1 | - | 45179 | 45127 | **RNA**→ rpmE (PRK00019)Ribosomal_L31 (pfam01197)→ |
| Sen-300-227 | NZ_CTGZ01000022.1 | + | 42532 | 42584 | **RNA**→ rpmE (PRK00019)Ribosomal_L31 (pfam01197)→ |
| Sen-300-228 | NZ_CTMG01000018.1 | - | 46977 | 46925 | **RNA**→ rpmE (PRK00019)Ribosomal_L31 (pfam01197)→ |
| Sen-300-229 | NZ_CTOY01000017.1 | + | 42548 | 42600 | **RNA**→ rpmE (PRK00019)Ribosomal_L31 (pfam01197)→ |
| Sen-300-230 | NZ_CTQP01000019.1 | + | 42536 | 42588 | **RNA**→ rpmE (PRK00019)Ribosomal_L31 (pfam01197)→ |
| Sen-300-231 | NZ_CTNY01000018.1 | - | 45125 | 45073 | **RNA**→ rpmE (PRK00019)Ribosomal_L31 (pfam01197)→ |
| Sen-300-232 | NZ_CTLY01000020.1 | - | 45179 | 45127 | **RNA**→ rpmE (PRK00019)Ribosomal_L31 (pfam01197)→ |
| Sen-300-233 | NZ_CTFY01000017.1 | - | 45179 | 45127 | **RNA**→ rpmE (PRK00019)Ribosomal_L31 (pfam01197)→ |
| Sen-300-234 | NZ_CTKG01000019.1 | - | 45179 | 45127 | **RNA**→ rpmE (PRK00019)Ribosomal_L31 (pfam01197)→ |
| Sen-300-235 | NZ_CTJU01000017.1 | - | 45120 | 45068 | **RNA**→ rpmE (PRK00019)Ribosomal_L31 (pfam01197)→ |
| Sen-300-236 | NZ_CTQQ01000018.1 | + | 42546 | 42598 | **RNA**→ rpmE (PRK00019)Ribosomal_L31 (pfam01197)→ |
| Sen-300-237 | NZ_CTJN01000020.1 | + | 42551 | 42603 | **RNA**→ rpmE (PRK00019)Ribosomal_L31 (pfam01197)→ |
| Sen-300-238 | NZ_CTPT01000017.1 | - | 46977 | 46925 | **RNA**→ rpmE (PRK00019)Ribosomal_L31 (pfam01197)→ |
| Sen-300-239 | NZ_CTHU01000018.1 | + | 42546 | 42598 | **RNA**→ rpmE (PRK00019)Ribosomal_L31 (pfam01197)→ |
| Sen-300-240 | NZ_CTNN01000019.1 | - | 45136 | 45084 | **RNA**→ rpmE (PRK00019)Ribosomal_L31 (pfam01197)→ |
| Sen-300-241 | NZ_CTND01000017.1 | - | 45343 | 45291 | **RNA**→ rpmE (PRK00019)Ribosomal_L31 (pfam01197)→ |
| Sen-300-242 | NZ_CTHJ01000018.1 | - | 45179 | 45127 | **RNA**→ rpmE (PRK00019)Ribosomal_L31 (pfam01197)→ |
| Sen-300-243 | NZ_CTOG01000010.1 | - | 37221 | 37169 | **RNA**→ rpmE (PRK00019)Ribosomal_L31 (pfam01197)→ |
| Sen-300-244 | NZ_CTLX01000011.1 | + | 104185 | 104237 | **RNA**→ rpmE (PRK00019)Ribosomal_L31 (pfam01197)→ |
| Sen-300-245 | NZ_CTQX01000021.1 | - | 45343 | 45291 | **RNA**→ rpmE (PRK00019)Ribosomal_L31 (pfam01197)→ |
| Sen-300-246 | NZ_CTNZ01000020.1 | - | 45114 | 45062 | **RNA**→ rpmE (PRK00019)Ribosomal_L31 (pfam01197)→ |
| Sen-300-247 | NZ_CTJQ01000016.1 | + | 42510 | 42562 | **RNA**→ rpmE (PRK00019)Ribosomal_L31 (pfam01197)→ |
| Sen-300-248 | NZ_CTMF01000017.1 | + | 42505 | 42557 | **RNA**→ rpmE (PRK00019)Ribosomal_L31 (pfam01197)→ |
| Sen-300-249 | NZ_CTHR01000017.1 | + | 42536 | 42588 | **RNA**→ rpmE (PRK00019)Ribosomal_L31 (pfam01197)→ |
| Sen-300-250 | NZ_CTMK01000017.1 | - | 45343 | 45291 | **RNA**→ rpmE (PRK00019)Ribosomal_L31 (pfam01197)→ |
| Sen-300-251 | NZ_CTKA01000017.1 | - | 46977 | 46925 | **RNA**→ rpmE (PRK00019)Ribosomal_L31 (pfam01197)→ |
| Sen-300-252 | NZ_CTOH01000019.1 | + | 42453 | 42505 | **RNA**→ rpmE (PRK00019)Ribosomal_L31 (pfam01197)→ |
| Sen-300-253 | NZ_CTHB01000017.1 | - | 45179 | 45127 | **RNA**→ rpmE (PRK00019)Ribosomal_L31 (pfam01197)→ |
| Sen-300-254 | NZ_CTLA01000020.1 | - | 45256 | 45204 | **RNA**→ rpmE (PRK00019)Ribosomal_L31 (pfam01197)→ |
| Sen-300-255 | NZ_CTLR01000017.1 | + | 42509 | 42561 | **RNA**→ rpmE (PRK00019)Ribosomal_L31 (pfam01197)→ |
| Sen-300-256 | NZ_CTMJ01000019.1 | - | 45107 | 45055 | **RNA**→ rpmE (PRK00019)Ribosomal_L31 (pfam01197)→ |
| Sen-300-257 | NZ_CTLV01000020.1 | - | 45179 | 45127 | **RNA**→ rpmE (PRK00019)Ribosomal_L31 (pfam01197)→ |
| Sen-300-258 | NZ_CTHO01000020.1 | + | 42484 | 42536 | **RNA**→ rpmE (PRK00019)Ribosomal_L31 (pfam01197)→ |
| Sen-300-259 | NZ_CTKD01000016.1 | + | 42478 | 42530 | **RNA**→ rpmE (PRK00019)Ribosomal_L31 (pfam01197)→ |
| Sen-300-260 | NZ_CTMQ01000017.1 | + | 42438 | 42490 | **RNA**→ rpmE (PRK00019)Ribosomal_L31 (pfam01197)→ |
| Sen-427-1 | NZ_CP007211.1 | + | 217000 | 217052 | **RNA**→ rpmE (PRK00019)Ribosomal_L31 (pfam01197)→ |
| Sen-267-1 | NZ_CP007216.1 | + | 222105 | 222157 | **RNA**→ rpmE (PRK00019)Ribosomal_L31 (pfam01197)→ |
| Sen-992-1 | NZ_CP007235.1 | + | 222015 | 222067 | **RNA**→ rpmE (PRK00019)Ribosomal_L31 (pfam01197)→ |

| | | | | | | |
|---|---|---|---|---|---|---|
| Sen-236-1 | NZ_CP007222.1 | + | 219817 | 219869 | **RNA**→ | rpmE (PRK00019)Ribosomal_L31 (pfam01197)→ |
| Sen-273-3 | NZ_CP011967.1 | + | 4083750 | 4083802 | **RNA**→ | rpmE (PRK00019)Ribosomal_L31 (pfam01197)→ |
| Sen-135-1 | NZ_CP012344.1 | - | 4536970 | 4536918 | **RNA**→ | rpmE (PRK00019)Ribosomal_L31 (pfam01197)→ |
| Sen-276-1 | NZ_CP012347.1 | - | 4534664 | 4534612 | **RNA**→ | rpmE (PRK00019)Ribosomal_L31 (pfam01197)→ |
| Sen-272-1 | NZ_CP012346.1 | - | 4305396 | 4305344 | **RNA**→ | rpmE (PRK00019)Ribosomal_L31 (pfam01197)→ |
| Sen-281-1 | NZ_CP012349.1 | - | 4562745 | 4562693 | **RNA**→ | rpmE (PRK00019)Ribosomal_L31 (pfam01197)→ |
| Sen-622-1 | NZ_CP007354.1 | + | 4173831 | 4173883 | **RNA**→ | rpmE (PRK00019)Ribosomal_L31 (pfam01197)→ |
| Sen-643-1 | NZ_CP007376.1 | + | 4173841 | 4173893 | **RNA**→ | rpmE (PRK00019)Ribosomal_L31 (pfam01197)→ |
| Sen-653-1 | NZ_CP007371.1 | + | 4173817 | 4173869 | **RNA**→ | rpmE (PRK00019)Ribosomal_L31 (pfam01197)→ |
| Sen-300-261 | NZ_CTMM01000027.1 | - | 45179 | 45127 | **RNA**→ | rpmE (PRK00019)Ribosomal_L31 (pfam01197)→ |
| Sen-300-262 | NZ_CTPV01000021.1 | + | 42473 | 42525 | **RNA**→ | rpmE (PRK00019)Ribosomal_L31 (pfam01197)→ |
| Sen-300-263 | NZ_CTPB01000017.1 | - | 45179 | 45127 | **RNA**→ | rpmE (PRK00019)Ribosomal_L31 (pfam01197)→ |
| Sen-300-264 | NZ_CTIJ01000012.1 | - | 45337 | 45285 | **RNA**→ | rpmE (PRK00019)Ribosomal_L31 (pfam01197)→ |
| Sen-300-265 | NZ_CTGE01000016.1 | - | 45343 | 45291 | **RNA**→ | rpmE (PRK00019)Ribosomal_L31 (pfam01197)→ |
| Sen-300-266 | NZ_CTGM01000017.1 | - | 45179 | 45127 | **RNA**→ | rpmE (PRK00019)Ribosomal_L31 (pfam01197)→ |
| Sen-300-267 | NZ_CTQB01000021.1 | + | 42438 | 42490 | **RNA**→ | rpmE (PRK00019)Ribosomal_L31 (pfam01197)→ |
| Sen-300-268 | NZ_CTFX01000018.1 | + | 42453 | 42505 | **RNA**→ | rpmE (PRK00019)Ribosomal_L31 (pfam01197)→ |
| Sen-300-269 | NZ_CTGY01000018.1 | + | 42510 | 42562 | **RNA**→ | rpmE (PRK00019)Ribosomal_L31 (pfam01197)→ |
| Sen-300-270 | NZ_CTPK01000017.1 | - | 45179 | 45127 | **RNA**→ | rpmE (PRK00019)Ribosomal_L31 (pfam01197)→ |
| Sen-300-271 | NZ_CTOU01000017.1 | - | 45179 | 45127 | **RNA**→ | rpmE (PRK00019)Ribosomal_L31 (pfam01197)→ |
| Sen-300-272 | NZ_CTLU01000018.1 | + | 42484 | 42536 | **RNA**→ | rpmE (PRK00019)Ribosomal_L31 (pfam01197)→ |
| Sen-300-273 | NZ_CTNW01000020.1 | + | 42453 | 42505 | **RNA**→ | rpmE (PRK00019)Ribosomal_L31 (pfam01197)→ |
| Sen-300-274 | NZ_CTNJ01000021.1 | - | 45107 | 45055 | **RNA**→ | rpmE (PRK00019)Ribosomal_L31 (pfam01197)→ |
| Sen-300-275 | NZ_CTLP01000019.1 | + | 42467 | 42519 | **RNA**→ | rpmE (PRK00019)Ribosomal_L31 (pfam01197)→ |
| Sen-300-276 | NZ_CTHH01000028.1 | - | 32546 | 32494 | **RNA**→ | rpmE (PRK00019)Ribosomal_L31 (pfam01197)→ |
| Sen-300-277 | NZ_CTIB01000016.1 | + | 42438 | 42490 | **RNA**→ | rpmE (PRK00019)Ribosomal_L31 (pfam01197)→ |
| Sen-300-278 | NZ_CTFV01000017.1 | + | 42453 | 42505 | **RNA**→ | rpmE (PRK00019)Ribosomal_L31 (pfam01197)→ |
| Sen-300-279 | NZ_CTPD01000016.1 | - | 50325 | 50273 | **RNA**→ | rpmE (PRK00019)Ribosomal_L31 (pfam01197)→ |
| Sen-300-280 | NZ_CTOF01000017.1 | + | 42479 | 42531 | **RNA**→ | rpmE (PRK00019)Ribosomal_L31 (pfam01197)→ |
| Sen-300-281 | NZ_CTQC01000017.1 | + | 42490 | 42542 | **RNA**→ | rpmE (PRK00019)Ribosomal_L31 (pfam01197)→ |
| Sen-300-282 | NZ_CTLC01000017.1 | - | 45179 | 45127 | **RNA**→ | rpmE (PRK00019)Ribosomal_L31 (pfam01197)→ |
| Sen-300-283 | NZ_CTML01000016.1 | - | 45343 | 45291 | **RNA**→ | rpmE (PRK00019)Ribosomal_L31 (pfam01197)→ |
| Sen-300-284 | NZ_CTPP01000019.1 | - | 45176 | 45124 | **RNA**→ | rpmE (PRK00019)Ribosomal_L31 (pfam01197)→ |
| Sen-300-285 | NZ_CTNH01000018.1 | + | 42532 | 42584 | **RNA**→ | rpmE (PRK00019)Ribosomal_L31 (pfam01197)→ |
| Sen-300-286 | NZ_CTJZ01000017.1 | - | 45247 | 45195 | **RNA**→ | rpmE (PRK00019)Ribosomal_L31 (pfam01197)→ |
| Sen-300-287 | NZ_CTOZ01000017.1 | + | 42472 | 42524 | **RNA**→ | rpmE (PRK00019)Ribosomal_L31 (pfam01197)→ |
| Sen-300-288 | NZ_CTGP01000020.1 | - | 45343 | 45291 | **RNA**→ | rpmE (PRK00019)Ribosomal_L31 (pfam01197)→ |
| Sen-300-289 | NZ_CTJM01000019.1 | - | 45107 | 45055 | **RNA**→ | rpmE (PRK00019)Ribosomal_L31 (pfam01197)→ |
| Sen-300-290 | NZ_CTNI01000017.1 | - | 45150 | 45098 | **RNA**→ | rpmE (PRK00019)Ribosomal_L31 (pfam01197)→ |
| Sen-300-291 | NZ_CTIF01000021.1 | - | 45181 | 45129 | **RNA**→ | rpmE (PRK00019)Ribosomal_L31 (pfam01197)→ |
| Sen-300-292 | NZ_CTQS01000016.1 | + | 42508 | 42560 | **RNA**→ | rpmE (PRK00019)Ribosomal_L31 (pfam01197)→ |
| Sen-300-293 | NZ_CTQR01000017.1 | - | 45179 | 45127 | **RNA**→ | rpmE (PRK00019)Ribosomal_L31 (pfam01197)→ |
| Sen-300-294 | NZ_CTJV01000019.1 | - | 45113 | 45061 | **RNA**→ | rpmE (PRK00019)Ribosomal_L31 (pfam01197)→ |
| Sen-300-295 | NZ_CTNG01000027.1 | + | 22519 | 22571 | **RNA**→ | rpmE (PRK00019)Ribosomal_L31 (pfam01197)→ |
| Sen-300-296 | NZ_CTJP01000007.1 | + | 175854 | 175906 | **RNA**→ | rpmE (PRK00019)Ribosomal_L31 (pfam01197)→ |
| Sen-300-297 | NZ_CTNS01000020.1 | - | 45107 | 45055 | **RNA**→ | rpmE (PRK00019)Ribosomal_L31 (pfam01197)→ |
| Sen-300-298 | NZ_CTPJ01000020.1 | + | 42544 | 42596 | **RNA**→ | rpmE (PRK00019)Ribosomal_L31 (pfam01197)→ |
| Sen-300-299 | NZ_CTJR01000017.1 | - | 45255 | 45203 | **RNA**→ | rpmE (PRK00019)Ribosomal_L31 (pfam01197)→ |
| Sen-300-300 | NZ_CTQI01000017.1 | + | 43122 | 43174 | **RNA**→ | rpmE (PRK00019)Ribosomal_L31 (pfam01197)→ |
| Sen-300-301 | NZ_CTPU01000020.1 | - | 45107 | 45055 | **RNA**→ | rpmE (PRK00019)Ribosomal_L31 (pfam01197)→ |
| Sen-300-302 | NZ_CTKZ01000020.1 | - | 45116 | 45064 | **RNA**→ | rpmE (PRK00019)Ribosomal_L31 (pfam01197)→ |
| Sen-300-303 | NZ_CTOR01000017.1 | - | 45179 | 45127 | **RNA**→ | rpmE (PRK00019)Ribosomal_L31 (pfam01197)→ |

| | | | | | |
|---|---|---|---|---|---|
| Sen-300-304 | NZ_CTID01000016.1 | + | 42774 | 42826 | **RNA**→ rpmE (PRK00019)Ribosomal_L31 (pfam01197)→ |
| Sen-300-305 | NZ_CTHC01000020.1 | - | 45179 | 45127 | **RNA**→ rpmE (PRK00019)Ribosomal_L31 (pfam01197)→ |
| Sen-300-306 | NZ_CTGH01000020.1 | - | 45107 | 45055 | **RNA**→ rpmE (PRK00019)Ribosomal_L31 (pfam01197)→ |
| Sen-300-307 | NZ_CTMS01000017.1 | + | 42506 | 42558 | **RNA**→ rpmE (PRK00019)Ribosomal_L31 (pfam01197)→ |
| Sen-300-308 | NZ_CTHQ01000015.1 | - | 45343 | 45291 | **RNA**→ rpmE (PRK00019)Ribosomal_L31 (pfam01197)→ |
| Sen-300-309 | NZ_CTMT01000018.1 | - | 45107 | 45055 | **RNA**→ rpmE (PRK00019)Ribosomal_L31 (pfam01197)→ |
| Sen-300-310 | NZ_CTOI01000027.1 | - | 32545 | 32493 | **RNA**→ rpmE (PRK00019)Ribosomal_L31 (pfam01197)→ |
| Sen-300-311 | NZ_CTOW01000021.1 | - | 45197 | 45145 | **RNA**→ rpmE (PRK00019)Ribosomal_L31 (pfam01197)→ |
| Sen-300-312 | NZ_CTLZ01000017.1 | + | 42531 | 42583 | **RNA**→ rpmE (PRK00019)Ribosomal_L31 (pfam01197)→ |
| Sen-300-313 | NZ_CTKC01000017.1 | - | 45174 | 45122 | **RNA**→ rpmE (PRK00019)Ribosomal_L31 (pfam01197)→ |
| Sen-300-314 | NZ_CTNU01000017.1 | + | 42517 | 42569 | **RNA**→ rpmE (PRK00019)Ribosomal_L31 (pfam01197)→ |
| Sen-300-315 | NZ_CTIH01000018.1 | - | 45343 | 45291 | **RNA**→ rpmE (PRK00019)Ribosomal_L31 (pfam01197)→ |
| Sen-300-316 | NZ_CTOT01000019.1 | + | 42461 | 42513 | **RNA**→ rpmE (PRK00019)Ribosomal_L31 (pfam01197)→ |
| Sen-300-317 | NZ_CTGD01000017.1 | + | 42532 | 42584 | **RNA**→ rpmE (PRK00019)Ribosomal_L31 (pfam01197)→ |
| Sen-300-318 | NZ_CTOP01000024.1 | + | 13413 | 13465 | **RNA**→ rpmE (PRK00019)Ribosomal_L31 (pfam01197)→ |
| Sen-300-319 | NZ_CTQG01000020.1 | + | 42499 | 42551 | **RNA**→ rpmE (PRK00019)Ribosomal_L31 (pfam01197)→ |
| Sen-300-320 | NZ_CTJX01000017.1 | + | 42438 | 42490 | **RNA**→ rpmE (PRK00019)Ribosomal_L31 (pfam01197)→ |
| Sen-300-321 | NZ_CTPI01000017.1 | - | 45199 | 45147 | **RNA**→ rpmE (PRK00019)Ribosomal_L31 (pfam01197)→ |
| Sen-300-322 | NZ_CTNE01000017.1 | + | 42517 | 42569 | **RNA**→ rpmE (PRK00019)Ribosomal_L31 (pfam01197)→ |
| Sen-300-323 | NZ_CTHK01000017.1 | - | 45343 | 45291 | **RNA**→ rpmE (PRK00019)Ribosomal_L31 (pfam01197)→ |
| Sen-300-324 | NZ_CTHF01000019.1 | + | 42460 | 42512 | **RNA**→ rpmE (PRK00019)Ribosomal_L31 (pfam01197)→ |
| Sen-300-325 | NZ_CTAO01000017.1 | - | 45255 | 45203 | **RNA**→ rpmE (PRK00019)Ribosomal_L31 (pfam01197)→ |
| Sen-300-326 | NZ_CTCH01000021.1 | - | 45107 | 45055 | **RNA**→ rpmE (PRK00019)Ribosomal_L31 (pfam01197)→ |
| Sen-300-327 | NZ_CTBW01000021.1 | - | 45117 | 45065 | **RNA**→ rpmE (PRK00019)Ribosomal_L31 (pfam01197)→ |
| Sen-300-328 | NZ_CTCE01000019.1 | + | 42473 | 42525 | **RNA**→ rpmE (PRK00019)Ribosomal_L31 (pfam01197)→ |
| Sen-300-329 | NZ_CTAC01000018.1 | - | 45179 | 45127 | **RNA**→ rpmE (PRK00019)Ribosomal_L31 (pfam01197)→ |
| Sen-802-1 | NZ_AFCR01001558.1 | + | 1664 | 1716 | **RNA**→ rpmE (PRK00019)Ribosomal_L31 (pfam01197)→ |
| Sen-996-1 | NZ_AFCV01001391.1 | + | 6411 | 6463 | **RNA**→ rpmE (PRK00019)Ribosomal_L31 (pfam01197)→ |
| Sen-734-1 | NZ_AFCM01000234.1 | - | 1925 | 1873 | **RNA**→ rpmE (PRK00019)Ribosomal_L31 (pfam01197)→ |
| Sen-289-1 | NZ_AKIC01000146.1 | + | 3535 | 3587 | **RNA**→ rpmE (PRK00019)Ribosomal_L31 (pfam01197)→ |
| Sen-1025-1 | NZ_CAFD01000090.1 | - | 1029 | 977 | **RNA**→ rpmE (PRK00019)Ribosomal_L31 (pfam01197)→ |
| Sen-439-11 | NZ_CSTH01000013.1 | - | 45242 | 45190 | **RNA**→ rpmE (PRK00019)Ribosomal_L31 (pfam01197)→ |
| Sen-412-1 | NZ_CARE01000039.1 | - | 37552 | 37500 | **RNA**→ rpmE (PRK00019)Ribosomal_L31 (pfam01197)→ |
| Sen-229-1 | NZ_CBKW020000007.1 | + | 103767 | 103819 | **RNA**→ rpmE (PRK00019)Ribosomal_L31 (pfam01197)→ |
| Sen-291-7 | NZ_LFGV01000021.1 | - | 4804 | 4752 | **RNA**→ rpmE (PRK00019)Ribosomal_L31 (pfam01197)→ |
| Sen-396-1 | NZ_CATT01000048.1 | - | 37552 | 37500 | **RNA**→ rpmE (PRK00019)Ribosomal_L31 (pfam01197)→ |
| Sen-361-1 | NZ_CATV01000026.1 | - | 38256 | 38204 | **RNA**→ rpmE (PRK00019)Ribosomal_L31 (pfam01197)→ |
| Sen-238-1 | NZ_ARYX01000002.1 | + | 104268 | 104320 | **RNA**→ rpmE (PRK00019)Ribosomal_L31 (pfam01197)→ |
| Sen-310-1 | NZ_ATFA01000012.1 | - | 37653 | 37601 | **RNA**→ rpmE (PRK00019)Ribosomal_L31 (pfam01197)→ |
| Sen-1018-1 | NZ_ABFH02000001.1 | - | 831201 | 831149 | **RNA**→ rpmE (PRK00019)Ribosomal_L31 (pfam01197)→ rpmE (PRK00019)Ribosomal_L31 (pfam01197)→ |
| Sen-936-1 | NZ_AYDO01000075.1 | - | 37705 | 37653 | **RNA**→ rpmE (PRK00019)Ribosomal_L31 (pfam01197)→ rpmE (PRK00019)Ribosomal_L31 (pfam01197)→ |
| Sen-1024-1 | NZ_AOXE01000061.1 | + | 25782 | 25834 | **RNA**→ rpmE (PRK00019)Ribosomal_L31 (pfam01197)→ rpmE (PRK00019)Ribosomal_L31 (pfam01197)→ |
| Sen-899-1 | NZ_AYEF01000033.1 | - | 45341 | 45289 | **RNA**→ rpmE (PRK00019)Ribosomal_L31 (pfam01197)→ rpmE (PRK00019)Ribosomal_L31 (pfam01197)→ |
| Sen-61-1 | NZ_JRDC01000027.1 | - | 45217 | 45165 | **RNA**→ rpmE (PRK00019)Ribosomal_L31 (pfam01197)→ |
| Sen-50-1 | NZ_JRDM01000013.1 | + | 103447 | 103499 | **RNA**→ rpmE (PRK00019)Ribosomal_L31 (pfam01197)→ |
| Sen-102-1 | NZ_JRBQ01000021.1 | - | 45104 | 45052 | **RNA**→ rpmE (PRK00019)Ribosomal_L31 (pfam01197)→ |
| Sen-77-1 | NZ_JRCK01000003.1 | + | 43610 | 43662 | **RNA**→ rpmE (PRK00019)Ribosomal_L31 (pfam01197)→ |
| Sen-27-1 | NZ_JREG01000026.1 | - | 45105 | 45053 | **RNA**→ rpmE (PRK00019)Ribosomal_L31 (pfam01197)→ |
| Sen-23-1 | NZ_JREK01000016.1 | - | 45182 | 45130 | **RNA**→ rpmE (PRK00019)Ribosomal_L31 (pfam01197)→ |
| Sen-60-1 | NZ_JRDD01000019.1 | + | 4123 | 4175 | **RNA**→ rpmE (PRK00019)Ribosomal_L31 (pfam01197)→ |
| Sen-109-1 | NZ_JRBJ01000021.1 | - | 45105 | 45053 | **RNA**→ rpmE (PRK00019)Ribosomal_L31 (pfam01197)→ |
| Sen-99-1 | NZ_JRBS01000026.1 | - | 45104 | 45052 | **RNA**→ rpmE (PRK00019)Ribosomal_L31 (pfam01197)→ |

| | | | | | |
|---|---|---|---|---|---|
| Sen-45-1 | NZ_JRDQ01000023.1 | + | 98924 | 98976 | **RNA**→ rpmE (PRK00019)Ribosomal_L31 (pfam01197)→ |
| Sen-74-1 | NZ_JRCN01000001.1 | - | 45105 | 45053 | **RNA**→ rpmE (PRK00019)Ribosomal_L31 (pfam01197)→ |
| Sen-75-1 | NZ_JRCM01000022.1 | - | 45105 | 45053 | **RNA**→ rpmE (PRK00019)Ribosomal_L31 (pfam01197)→ |
| Sen-110-1 | NZ_JRBI01000096.1 | - | 3356 | 3304 | **RNA**→ rpmE (PRK00019)Ribosomal_L31 (pfam01197)→ |
| Sen-52-1 | NZ_JRDK01000048.1 | - | 45216 | 45164 | **RNA**→ rpmE (PRK00019)Ribosomal_L31 (pfam01197)→ |
| Sen-100-1 | NZ_JRBR01000031.1 | + | 4124 | 4176 | **RNA**→ rpmE (PRK00019)Ribosomal_L31 (pfam01197)→ |
| Sen-70-1 | NZ_JRCR01000027.1 | - | 45104 | 45052 | **RNA**→ rpmE (PRK00019)Ribosomal_L31 (pfam01197)→ |
| Sen-71-1 | NZ_JRCQ01000018.1 | - | 45105 | 45053 | **RNA**→ rpmE (PRK00019)Ribosomal_L31 (pfam01197)→ |
| Sen-76-1 | NZ_JRCL01000003.1 | + | 104296 | 104348 | **RNA**→ rpmE (PRK00019)Ribosomal_L31 (pfam01197)→ |
| Sen-22-1 | NZ_JREL01000019.1 | - | 45105 | 45053 | **RNA**→ rpmE (PRK00019)Ribosomal_L31 (pfam01197)→ |
| Sen-59-1 | NZ_JRDE01000031.1 | + | 4122 | 4174 | **RNA**→ rpmE (PRK00019)Ribosomal_L31 (pfam01197)→ |
| Sen-44-1 | NZ_JRDR01000023.1 | - | 45217 | 45165 | **RNA**→ rpmE (PRK00019)Ribosomal_L31 (pfam01197)→ |
| Sen-73-1 | NZ_JRCO01000013.1 | - | 45104 | 45052 | **RNA**→ rpmE (PRK00019)Ribosomal_L31 (pfam01197)→ |
| Sen-32-1 | NZ_JREC01000027.1 | + | 103206 | 103258 | **RNA**→ rpmE (PRK00019)Ribosomal_L31 (pfam01197)→ |
| Sen-42-1 | NZ_JRDT01000022.1 | + | 43449 | 43501 | **RNA**→ rpmE (PRK00019)Ribosomal_L31 (pfam01197)→ |
| Sen-47-1 | NZ_JRDO01000005.1 | + | 103439 | 103491 | **RNA**→ rpmE (PRK00019)Ribosomal_L31 (pfam01197)→ |
| Sen-20-1 | NZ_JREM01000014.1 | - | 45104 | 45052 | **RNA**→ rpmE (PRK00019)Ribosomal_L31 (pfam01197)→ |
| Sen-37-1 | NZ_JRDX01000002.1 | - | 45196 | 45144 | **RNA**→ rpmE (PRK00019)Ribosomal_L31 (pfam01197)→ |
| Sen-51-1 | NZ_JRDL01000012.1 | + | 103445 | 103497 | **RNA**→ rpmE (PRK00019)Ribosomal_L31 (pfam01197)→ |
| Sen-65-1 | NZ_JRCY01000019.1 | - | 45105 | 45053 | **RNA**→ rpmE (PRK00019)Ribosomal_L31 (pfam01197)→ |
| Sen-57-1 | NZ_JRDF01000014.1 | - | 45217 | 45165 | **RNA**→ rpmE (PRK00019)Ribosomal_L31 (pfam01197)→ |
| Sen-53-1 | NZ_JRDJ01000006.1 | + | 103457 | 103509 | **RNA**→ rpmE (PRK00019)Ribosomal_L31 (pfam01197)→ |
| Sen-72-1 | NZ_JRCP01000025.1 | - | 45104 | 45052 | **RNA**→ rpmE (PRK00019)Ribosomal_L31 (pfam01197)→ |
| Sen-30-1 | NZ_JRED01000031.1 | + | 43390 | 43442 | **RNA**→ rpmE (PRK00019)Ribosomal_L31 (pfam01197)→ |
| Sen-39-1 | NZ_JRDV01000026.1 | - | 45217 | 45165 | **RNA**→ rpmE (PRK00019)Ribosomal_L31 (pfam01197)→ |
| Sen-24-1 | NZ_JREJ01000027.1 | - | 45104 | 45052 | **RNA**→ rpmE (PRK00019)Ribosomal_L31 (pfam01197)→ |
| Sen-36-1 | NZ_JRDY01000016.1 | - | 45303 | 45251 | **RNA**→ rpmE (PRK00019)Ribosomal_L31 (pfam01197)→ |
| Sen-28-1 | NZ_JREF01000014.1 | - | 45216 | 45164 | **RNA**→ rpmE (PRK00019)Ribosomal_L31 (pfam01197)→ |
| Sen-33-1 | NZ_JREB01000019.1 | + | 4122 | 4174 | **RNA**→ rpmE (PRK00019)Ribosomal_L31 (pfam01197)→ |
| Sen-35-1 | NZ_JRDZ01000195.1 | + | 8320 | 8372 | **RNA**→ rpmE (PRK00019)Ribosomal_L31 (pfam01197)→ |
| Sen-40-1 | NZ_JRDU01000022.1 | - | 45217 | 45165 | **RNA**→ rpmE (PRK00019)Ribosomal_L31 (pfam01197)→ |
| Sen-26-1 | NZ_JREH01000017.1 | + | 8335 | 8387 | **RNA**→ rpmE (PRK00019)Ribosomal_L31 (pfam01197)→ |
| Sen-108-1 | NZ_JRBK01000010.1 | + | 103451 | 103503 | **RNA**→ rpmE (PRK00019)Ribosomal_L31 (pfam01197)→ |
| Sen-56-1 | NZ_JRDG01000018.1 | - | 45217 | 45165 | **RNA**→ rpmE (PRK00019)Ribosomal_L31 (pfam01197)→ |
| Sen-105-1 | NZ_JRBN01000053.1 | - | 45105 | 45053 | **RNA**→ rpmE (PRK00019)Ribosomal_L31 (pfam01197)→ |
| Sen-68-1 | NZ_JRCS01000047.1 | - | 45095 | 45043 | **RNA**→ rpmE (PRK00019)Ribosomal_L31 (pfam01197)→ |
| Sen-79-1 | NZ_JRCI01000083.1 | - | 1205 | 1153 | **RNA**→ rpmE (PRK00019)Ribosomal_L31 (pfam01197)→ |
| Sen-90-1 | NZ_JRBZ01000014.1 | - | 45105 | 45053 | **RNA**→ rpmE (PRK00019)Ribosomal_L31 (pfam01197)→ |
| Sen-94-1 | NZ_JRBV01000025.1 | + | 103450 | 103502 | **RNA**→ rpmE (PRK00019)Ribosomal_L31 (pfam01197)→ |
| Sen-96-1 | NZ_JRBU01000019.1 | - | 45105 | 45053 | **RNA**→ rpmE (PRK00019)Ribosomal_L31 (pfam01197)→ |
| Sen-85-1 | NZ_JRCD01000010.1 | + | 103451 | 103503 | **RNA**→ rpmE (PRK00019)Ribosomal_L31 (pfam01197)→ |
| Sen-925-1 | NZ_AOYK01000093.1 | + | 42029 | 42081 | **RNA**→ rpmE (PRK00019)Ribosomal_L31 (pfam01197)→ rpmE (PRK00019)Ribosomal_L31 (pfam01197)→ |
| Sen-107-1 | NZ_JRBL01000010.1 | - | 45105 | 45053 | **RNA**→ rpmE (PRK00019)Ribosomal_L31 (pfam01197)→ |
| Sen-54-1 | NZ_JRDI01000020.1 | - | 45217 | 45165 | **RNA**→ rpmE (PRK00019)Ribosomal_L31 (pfam01197)→ |
| Sen-55-1 | NZ_JRDH01000008.1 | - | 45105 | 45053 | **RNA**→ rpmE (PRK00019)Ribosomal_L31 (pfam01197)→ |
| Sen-104-1 | NZ_JRBO01000052.1 | + | 15512 | 15564 | **RNA**→ rpmE (PRK00019)Ribosomal_L31 (pfam01197)→ |
| Sen-67-1 | NZ_JRCT01000026.1 | - | 45173 | 45121 | **RNA**→ rpmE (PRK00019)Ribosomal_L31 (pfam01197)→ |
| Sen-78-1 | NZ_JRCJ01000053.1 | - | 45105 | 45053 | **RNA**→ rpmE (PRK00019)Ribosomal_L31 (pfam01197)→ |
| Sen-93-1 | NZ_JRBW01000021.1 | + | 55522 | 55574 | **RNA**→ rpmE (PRK00019)Ribosomal_L31 (pfam01197)→ |
| Sen-34-1 | NZ_JREA01000027.1 | + | 103445 | 103497 | **RNA**→ rpmE (PRK00019)Ribosomal_L31 (pfam01197)→ |
| Sen-43-1 | NZ_JRDS01000015.1 | + | 103420 | 103472 | **RNA**→ rpmE (PRK00019)Ribosomal_L31 (pfam01197)→ |
| Sen-82-1 | NZ_JRCF01000015.1 | + | 103451 | 103503 | **RNA**→ rpmE (PRK00019)Ribosomal_L31 (pfam01197)→ |

| | | | | | |
|---|---|---|---|---|---|
| Sen-84-1 | NZ_JRCE01000022.1 | + | 103451 | 103503 | **RNA**→ rpmE (PRK00019)Ribosomal_L31 (pfam01197)→ |
| Sen-48-1 | NZ_JRDN01000019.1 | - | 45217 | 45165 | **RNA**→ rpmE (PRK00019)Ribosomal_L31 (pfam01197)→ |
| Sen-64-1 | NZ_JRCZ01000043.1 | + | 4122 | 4174 | **RNA**→ rpmE (PRK00019)Ribosomal_L31 (pfam01197)→ |
| Sen-292-1 | NZ_JHAG01000043.1 | + | 42429 | 42481 | **RNA**→ rpmE (PRK00019)Ribosomal_L31 (pfam01197)→ |
| Sen-63-1 | NZ_JRDA01000002.1 | + | 103469 | 103521 | **RNA**→ rpmE (PRK00019)Ribosomal_L31 (pfam01197)→ |
| Sen-66-1 | NZ_JRCW01000065.1 | + | 4122 | 4174 | **RNA**→ rpmE (PRK00019)Ribosomal_L31 (pfam01197)→ |
| Sen-103-1 | NZ_JRBP01000017.1 | + | 4123 | 4175 | **RNA**→ rpmE (PRK00019)Ribosomal_L31 (pfam01197)→ |
| Sen-92-1 | NZ_JRBX01000004.1 | - | 45104 | 45052 | **RNA**→ rpmE (PRK00019)Ribosomal_L31 (pfam01197)→ |
| Sen-81-1 | NZ_JRCG01000038.1 | + | 4122 | 4174 | **RNA**→ rpmE (PRK00019)Ribosomal_L31 (pfam01197)→ |
| Sen-87-1 | NZ_JRCB01000003.1 | + | 103451 | 103503 | **RNA**→ rpmE (PRK00019)Ribosomal_L31 (pfam01197)→ |
| Sen-88-1 | NZ_JRCA01000012.1 | + | 103451 | 103503 | **RNA**→ rpmE (PRK00019)Ribosomal_L31 (pfam01197)→ |
| Sen-462-1 | NZ_APAG01000166.1 | + | 96846 | 96898 | **RNA**→ rpmE (PRK00019)Ribosomal_L31 (pfam01197)→ rpmE (PRK00019)Ribosomal_L31 (pfam01197)→ |
| Sen-766-1 | NZ_AMMP01000013.1 | + | 4125 | 4177 | **RNA**→ rpmE (PRK00019)Ribosomal_L31 (pfam01197)→ |
| Sen-62-1 | NZ_JRDB01000008.1 | + | 103327 | 103379 | **RNA**→ rpmE (PRK00019)Ribosomal_L31 (pfam01197)→ |
| Sen-25-1 | NZ_JREI01000009.1 | - | 45213 | 45161 | **RNA**→ rpmE (PRK00019)Ribosomal_L31 (pfam01197)→ |
| Sen-38-1 | NZ_JRDW01000021.1 | + | 103444 | 103496 | **RNA**→ rpmE (PRK00019)Ribosomal_L31 (pfam01197)→ |
| Sen-106-1 | NZ_JRBM01000066.1 | + | 43379 | 43431 | **RNA**→ rpmE (PRK00019)Ribosomal_L31 (pfam01197)→ |
| Sen-46-1 | NZ_JRDP01000018.1 | - | 9628 | 9576 | **RNA**→ rpmE (PRK00019)Ribosomal_L31 (pfam01197)→ |
| Sen-91-1 | NZ_JRBY01000004.1 | + | 103450 | 103502 | **RNA**→ rpmE (PRK00019)Ribosomal_L31 (pfam01197)→ |
| Sen-29-1 | NZ_JREE01000016.1 | + | 103448 | 103500 | **RNA**→ rpmE (PRK00019)Ribosomal_L31 (pfam01197)→ |
| Sen-97-1 | NZ_JRBT01000001.1 | + | 103451 | 103503 | **RNA**→ rpmE (PRK00019)Ribosomal_L31 (pfam01197)→ |
| Sen-80-1 | NZ_JRCH01000029.1 | + | 103451 | 103503 | **RNA**→ rpmE (PRK00019)Ribosomal_L31 (pfam01197)→ |
| Sen-86-1 | NZ_JRCC01000044.1 | + | 43272 | 43324 | **RNA**→ rpmE (PRK00019)Ribosomal_L31 (pfam01197)→ |
| Sen-101-1 | NZ_JRLO01000034.1 | - | 45105 | 45053 | **RNA**→ rpmE (PRK00019)Ribosomal_L31 (pfam01197)→ |
| Sen-83-1 | NZ_JRLQ01000100.1 | + | 4123 | 4175 | **RNA**→ rpmE (PRK00019)Ribosomal_L31 (pfam01197)→ |
| Sen-2-2 | NZ_JTBM01000001.1 | - | 3487697 | 3487645 | **RNA**→ rpmE (PRK00019)Ribosomal_L31 (pfam01197)→ |
| Sen-295-1 | NZ_JTED01000011.1 | - | 45184 | 45132 | **RNA**→ rpmE (PRK00019)Ribosomal_L31 (pfam01197)→ |
| Sbo-3-1 | NC_021870.1 | + | 4124212 | 4124264 | **RNA**→ rpmE (PRK00019)Ribosomal_L31 (pfam01197)→ |
| Sbo-4-1 | NC_015761.1 | + | 3964037 | 3964089 | **RNA**→ rpmE (PRK00019)Ribosomal_L31 (pfam01197)→ rpmE (PRK00019)Ribosomal_L31 (pfam01197)→ rpmE (PRK00019)Ribosomal_L31 (pfam01197)→ |
| Sbo-1-1 | NZ_CP006692.1 | + | 3836915 | 3836967 | **RNA**→ rpmE (PRK00019)Ribosomal_L31 (pfam01197)→ |
| Sen-1029-1 | NZ_ABEH02000008.1 | + | 207232 | 207284 | **RNA**→ rpmE (PRK00019)Ribosomal_L31 (pfam01197)→ rpmE (PRK00019)Ribosomal_L31 (pfam01197)→ |
| Sen-783-1 | NZ_AHUW01000041.1 | - | 44329 | 44277 | **RNA**→ rpmE (PRK00019)Ribosomal_L31 (pfam01197)→ rpmE (PRK00019)Ribosomal_L31 (pfam01197)→ |
| Sen-785-1 | NZ_AOZB01000105.1 | + | 102730 | 102782 | **RNA**→ rpmE (PRK00019)Ribosomal_L31 (pfam01197)→ rpmE (PRK00019)Ribosomal_L31 (pfam01197)→ |
| Sen-784-1 | NC_020307.1 | - | 2717973 | 2717921 | **RNA**→ rpmE (PRK00019)Ribosomal_L31 (pfam01197)→ rpmE (PRK00019)Ribosomal_L31 (pfam01197)→ |
| Sen-1021-1 | NZ_AOXJ01000015.1 | - | 34309 | 34257 | **RNA**→ rpmE (PRK00019)Ribosomal_L31 (pfam01197)→ |
| Sen-1020-1 | NZ_ANHR01000083.1 | + | 63387 | 63439 | **RNA**→ rpmE (PRK00019)Ribosomal_L31 (pfam01197)→ |
| Sen-1022-1 | NZ_CM001471.1 | + | 4136164 | 4136216 | **RNA**→ rpmE (PRK00019)Ribosomal_L31 (pfam01197)→ |
| Sen-1019-1 | NZ_JWSP01000004.1 | - | 1022116 | 1022064 | **RNA**→ rpmE (PRK00019)Ribosomal_L31 (pfam01197)→ |
| Sen-312-1 | NZ_APAC01000103.1 | + | 85432 | 85484 | **RNA**→ rpmE (PRK00019)Ribosomal_L31 (pfam01197)→ rpmE (PRK00019)Ribosomal_L31 (pfam01197)→ |
| Bag-1-1 | NZ_JPRU01000006.1 | - | 38990 | 38938 | **RNA**→ rpmE (PRK00019)Ribosomal_L31 (pfam01197)→ |
| env-757 | JGI26054J46599_1163256 | + | 160 | 212 | **RNA**→ rpmE (PRK00019)Ribosomal_L31 (pfam01197)→ |
| Stu-1-1 | NZ_AWFZ01000002.1 | + | 111257 | 111309 | **RNA**→ rpmE (PRK00019)Ribosomal_L31 (pfam01197)→ |
| Ctu-2-1 | NZ_AXDM01000005.1 | - | 25734 | 25682 | **RNA**→ rpmE (PRK00019)Ribosomal_L31 (pfam01197)→ |
| Ctu-1-1 | NZ_AVPP01000003.1 | + | 47207 | 47259 | **RNA**→ rpmE (PRK00019)Ribosomal_L31 (pfam01197)→ |
| Cfr-3-1 | NZ_CP007557.1 | + | 1495160 | 1495212 | **RNA**→ rpmE (PRK00019)Ribosomal_L31 (pfam01197)→ |
| Ksp-3-1 | NZ_LGIT01000007.1 | - | 29099 | 29047 | **RNA**→ rpmE (PRK00019)Ribosomal_L31 (pfam01197)→ |
| Eco-1011-1 | NZ_AQCZ01000115.1 | + | 23399 | 23451 | **RNA**→ rpmE (PRK00019)Ribosomal_L31 (pfam01197)→ |
| env-758 | BAAV01033320.1 | + | 305 | 357 | **RNA**→ rpmE (PRK00019)Ribosomal_L31 (pfam01197)→ |
| env-759 | BAAW01007240.1 | - | 565 | 513 | **RNA**→ rpmE (PRK00019)Ribosomal_L31 (pfam01197)→ |
| env-760 | BAAW01011360.1 | + | 754 | 806 | **RNA**→ ←hypo |
| env-761 | BABF01001643.1 | - | 1445 | 1393 | **RNA**→ rpmE (PRK00019)Ribosomal_L31 (pfam01197)→ |

| | | | | | |
|---|---|---|---|---|---|
| env-762 | AEQT01000001.1 | + | 88843 | 88895 | RNA→ rpmE (PRK00019)Ribosomal_L31 (pfam01197)→ |
| Cfr-5-1 | NZ_BBMV01000030.1 | - | 35429 | 35377 | RNA→ rpmE (PRK00019)Ribosomal_L31 (pfam01197)→ |
| Cpa-1-1 | NZ_CDHL01000043.1 | - | 29572 | 29520 | RNA→ rpmE (PRK00019)Ribosomal_L31 (pfam01197)→ |
| Cfr-11-1 | NZ_AOUE01000017.1 | + | 87594 | 87646 | RNA→ rpmE (PRK00019)Ribosomal_L31 (pfam01197)→ |
| Csp-16-1 | NZ_AKTT01000023.1 | + | 87561 | 87613 | RNA→ rpmE (PRK00019)Ribosomal_L31 (pfam01197)→ rpmE (PRK00019)Ribosomal_L31 (pfam01197)→ |
| Cfr-2-2 | NZ_ANAV01000016.1 | + | 78413 | 78465 | RNA→ rpmE (PRK00019)Ribosomal_L31 (pfam01197)→ |
| Cfr-2-1 | NZ_JMTA01000012.1 | + | 78415 | 78467 | RNA→ rpmE (PRK00019)Ribosomal_L31 (pfam01197)→ |
| Cfr-6-1 | NZ_JATT01000019.1 | + | 78434 | 78486 | RNA→ rpmE (PRK00019)Ribosomal_L31 (pfam01197)→ |
| Cfr-7-1 | NZ_KI929269.1 | + | 4901833 | 4901885 | RNA→ rpmE (PRK00019)Ribosomal_L31 (pfam01197)→ |
| Csp-18-1 | NZ_KB976057.1 | + | 90814 | 90866 | RNA→ rpmE (PRK00019)Ribosomal_L31 (pfam01197)→ |
| Cfr-10-1 | NZ_AOMS01000029.1 | - | 35491 | 35439 | RNA→ rpmE (PRK00019)Ribosomal_L31 (pfam01197)→ |
| Cfr-1-1 | NZ_CP011657.1 | - | 2678236 | 2678184 | RNA→ rpmE (PRK00019)Ribosomal_L31 (pfam01197)→ |
| Cfr-1-2 | NZ_CP011612.1 | + | 3379319 | 3379371 | RNA→ rpmE (PRK00019)Ribosomal_L31 (pfam01197)→ |
| Csp-6-1 | NZ_KQ089819.1 | - | 838497 | 838445 | RNA→ rpmE (PRK00019)Ribosomal_L31 (pfam01197)→ |
| Csp-11-1 | NZ_KQ089794.1 | - | 242684 | 242632 | RNA→ rpmE (PRK00019)Ribosomal_L31 (pfam01197)→ |
| Csp-5-1 | NZ_KQ089814.1 | + | 89593 | 89645 | RNA→ rpmE (PRK00019)Ribosomal_L31 (pfam01197)→ |
| Csp-4-1 | NZ_KQ089805.1 | - | 35482 | 35430 | RNA→ rpmE (PRK00019)Ribosomal_L31 (pfam01197)→ |
| Csp-2-1 | NZ_KQ089848.1 | - | 2277940 | 2277888 | RNA→ rpmE (PRK00019)Ribosomal_L31 (pfam01197)→ |
| Csp-9-1 | NZ_KQ089838.1 | - | 444760 | 444708 | RNA→ rpmE (PRK00019)Ribosomal_L31 (pfam01197)→ |
| Csp-20-1 | NZ_AMPE01000010.1 | + | 87560 | 87612 | RNA→ rpmE (PRK00019)Ribosomal_L31 (pfam01197)→ |
| Cfr-1-3 | NZ_JTBJ01000001.1 | + | 4640958 | 4641010 | RNA→ rpmE (PRK00019)Ribosomal_L31 (pfam01197)→ |
| Eco-1781-1 | NZ_ASHB01000053.1 | + | 32271 | 32323 | RNA→ rpmE (PRK00019)Ribosomal_L31 (pfam01197)→ |
| Cam-1-1 | NZ_LAMY01000136.1 | + | 29848 | 29900 | RNA→ rpmE (PRK00019)Ribosomal_L31 (pfam01197)→ |
| Esp-48-1 | NZ_LEDM01000004.1 | - | 45562 | 45510 | RNA→ rpmE (PRK00019)Ribosomal_L31 (pfam01197)→ |
| Cbr-1-1 | NZ_JUKU01000045.1 | + | 60235 | 60287 | RNA→ rpmE (PRK00019)Ribosomal_L31 (pfam01197)→ |
| Cbr-1-2 | NZ_JWAG01000048.1 | + | 60248 | 60300 | RNA→ rpmE (PRK00019)Ribosomal_L31 (pfam01197)→ |
| Cbr-1-3 | NZ_JUYY01000053.1 | + | 60968 | 61020 | RNA→ rpmE (PRK00019)Ribosomal_L31 (pfam01197)→ |
| Cbr-1-4 | NZ_JULD01000069.1 | - | 27815 | 27763 | RNA→ rpmE (PRK00019)Ribosomal_L31 (pfam01197)→ |
| Csp-15-1 | NZ_KI391987.1 | - | 39609 | 39557 | RNA→ rpmE (PRK00019)Ribosomal_L31 (pfam01197)→ |
| Cfr-8-1 | NZ_KI929267.1 | + | 90664 | 90716 | RNA→ rpmE (PRK00019)Ribosomal_L31 (pfam01197)→ |
| Csp-19-1 | NZ_KB976073.1 | + | 200071 | 200123 | RNA→ rpmE (PRK00019)Ribosomal_L31 (pfam01197)→ |
| Csp-17-1 | NZ_KB976068.1 | + | 125011 | 125063 | RNA→ rpmE (PRK00019)Ribosomal_L31 (pfam01197)→ |
| Csp-3-1 | NZ_KQ089873.1 | + | 62426 | 62478 | RNA→ rpmE (PRK00019)Ribosomal_L31 (pfam01197)→ |
| Csp-10-1 | NZ_KQ089833.1 | - | 40759 | 40707 | RNA→ rpmE (PRK00019)Ribosomal_L31 (pfam01197)→ hypo→ KdgM (pfam06178)→ MelB (COG2211)gph (TIGR00792)→ |
| Csp-7-1 | NZ_KQ089824.1 | - | 2325449 | 2325397 | RNA→ rpmE (PRK00019)Ribosomal_L31 (pfam01197)→ |
| Cbr-1-5 | NZ_JRHL01000001.1 | - | 2295239 | 2295187 | RNA→ rpmE (PRK00019)Ribosomal_L31 (pfam01197)→ |
| Cbr-1-6 | NZ_JRHK01000001.1 | + | 107410 | 107462 | RNA→ rpmE (PRK00019)Ribosomal_L31 (pfam01197)→ |
| Sso-1-2 | NZ_CWTS01000028.1 | + | 39051 | 39103 | RNA→ rpmE (PRK00019)Ribosomal_L31 (pfam01197)→ |
| Sfl-70-1 | NZ_AKMW01000069.1 | - | 54097 | 54045 | RNA→ rpmE (PRK00019)Ribosomal_L31 (pfam01197)→ rpmE (PRK00019)Ribosomal_L31 (pfam01197)→ |
| Sfl-71-1 | NZ_AERO01000112.1 | - | 1241 | 1189 | RNA→ rpmE (PRK00019)Ribosomal_L31 (pfam01197)→ |
| Eco-390-1 | NZ_HG977162.1 | + | 91704 | 91756 | RNA→ rpmE (PRK00019)Ribosomal_L31 (pfam01197)→ |
| Sfl-69-1 | NZ_LAJR01000096.1 | - | 1240 | 1188 | RNA→ rpmE (PRK00019)Ribosomal_L31 (pfam01197)→ |
| Ssp-20-2 | NZ_LDEG01000011.1 | - | 34084 | 34032 | RNA→ rpmE (PRK00019)Ribosomal_L31 (pfam01197)→ |
| Cfr-9-1 | NZ_JH414877.1 | - | 39489 | 39437 | RNA→ rpmE (PRK00019)Ribosomal_L31 (pfam01197)→ |
| Cfr-4-1 | NZ_KK736169.1 | + | 94576 | 94628 | RNA→ rpmE (PRK00019)Ribosomal_L31 (pfam01197)→ |
| Cfr-1-4 | NZ_JTBV01000001.1 | - | 898514 | 898462 | RNA→ rpmE (PRK00019)Ribosomal_L31 (pfam01197)→ |
| Eco-1460-1 | NZ_KB732916.1 | + | 3520416 | 3520468 | RNA→ rpmE (PRK00019)Ribosomal_L31 (pfam01197)→ |
| env-763 | DLM009_scaffold16386_3 | + | 718 | 770 | RNA→ rpmE (PRK00019)Ribosomal_L31 (pfam01197)→ |
| env-764 | NLF015_scaffold22851_1 | - | 263 | 211 | RNA→ rpmE (PRK00019)Ribosomal_L31 (pfam01197)→ |
| env-765 | NOM004_scaffold4453_6 | + | 1622 | 1674 | RNA→ rpmE (PRK00019)Ribosomal_L31 (pfam01197)→ |
| Eco-1383-1 | NZ_CBWB010000190.1 | + | 91974 | 92026 | RNA→ rpmE (PRK00019)Ribosomal_L31 (pfam01197)→ |

| | | | | | |
|---|---|---|---|---|---|
| Eco-325-1 | NZ_KK214157.1 | + | 92332 | 92384 | **RNA**→ rpmE (PRK00019)Ribosomal_L31 (pfam01197)→ |
| Eco-316-1 | NZ_KK736352.1 | + | 92567 | 92619 | **RNA**→ rpmE (PRK00019)Ribosomal_L31 (pfam01197)→ |
| Eco-305-1 | NZ_KI929660.1 | + | 3035802 | 3035854 | **RNA**→ rpmE (PRK00019)Ribosomal_L31 (pfam01197)→ |
| Eco-296-1 | NZ_KI929714.1 | + | 4459892 | 4459944 | **RNA**→ rpmE (PRK00019)Ribosomal_L31 (pfam01197)→ |
| Eco-302-1 | NZ_KI929774.1 | - | 205100 | 205048 | **RNA**→ rpmE (PRK00019)Ribosomal_L31 (pfam01197)→ |
| Eco-304-1 | NZ_KI929668.1 | - | 97421 | 97369 | **RNA**→ rpmE (PRK00019)Ribosomal_L31 (pfam01197)→ |
| Eco-310-1 | NZ_KI929804.1 | + | 4463923 | 4463975 | **RNA**→ rpmE (PRK00019)Ribosomal_L31 (pfam01197)→ |
| Eco-298-1 | NZ_KI929698.1 | - | 639989 | 639937 | **RNA**→ rpmE (PRK00019)Ribosomal_L31 (pfam01197)→ |
| Eco-297-1 | NZ_KI929706.1 | - | 636775 | 636723 | **RNA**→ rpmE (PRK00019)Ribosomal_L31 (pfam01197)→ |
| Eco-299-1 | NZ_KI929690.1 | + | 1502179 | 1502231 | **RNA**→ rpmE (PRK00019)Ribosomal_L31 (pfam01197)→ |
| Eco-327-1 | NZ_KI929758.1 | + | 93576 | 93628 | **RNA**→ rpmE (PRK00019)Ribosomal_L31 (pfam01197)→ |
| Eco-303-1 | NZ_KI929771.1 | + | 92568 | 92620 | **RNA**→ rpmE (PRK00019)Ribosomal_L31 (pfam01197)→ |
| Eco-295-1 | NZ_KI929738.1 | + | 206307 | 206359 | **RNA**→ rpmE (PRK00019)Ribosomal_L31 (pfam01197)→ |
| Eco-1015-1 | NZ_AXLI01000001.1 | + | 4460664 | 4460716 | **RNA**→ rpmE (PRK00019)Ribosomal_L31 (pfam01197)→ |
| Eco-258-1 | NZ_AYQW01000048.1 | - | 687 | 635 | **RNA**→ rpmE (PRK00019)Ribosomal_L31 (pfam01197)→ |
| Eco-1647-1 | NZ_AYNI01000026.1 | - | 938 | 886 | **RNA**→ rpmE (PRK00019)Ribosomal_L31 (pfam01197)→ |
| Eco-1646-1 | NZ_AYNH01000020.1 | - | 922 | 870 | **RNA**→ rpmE (PRK00019)Ribosomal_L31 (pfam01197)→ |
| Eco-428-1 | NZ_JDVF01000018.1 | - | 52260 | 52208 | **RNA**→ rpmE (PRK00019)Ribosomal_L31 (pfam01197)→ |
| Cwe-1-1 | NZ_BBMW01000019.1 | - | 35233 | 35181 | **RNA**→ rpmE (PRK00019)Ribosomal_L31 (pfam01197)→ |
| Sso-1-3 | NZ_CWSV01000068.1 | + | 20842 | 20894 | **RNA**→ rpmE (PRK00019)Ribosomal_L31 (pfam01197)→ |
| env-766 | BABF01000073.1 | - | 6870 | 6818 | **RNA**→ rpmE (PRK00019)Ribosomal_L31 (pfam01197)→ |
| env-767 | scaffold134588_2_MH0012 | + | 210 | 262 | **RNA**→ rpmE (PRK00019)Ribosomal_L31 (pfam01197)→ |
| env-768 | 4491407.3_NODE_4519 | - | 2364 | 2312 | **RNA**→ rpmE (PRK00019)Ribosomal_L31 (pfam01197)→ |
| env-769 | longitudinal_136_11 | + | 77179 | 77231 | **RNA**→ rpmE (PRK00019)Ribosomal_L31 (pfam01197)→ |
| env-770 | longitudinal_137_363 | + | 76993 | 77045 | **RNA**→ rpmE (PRK00019)Ribosomal_L31 (pfam01197)→ |
| env-771 | longitudinal_138_145 | + | 77051 | 77103 | **RNA**→ rpmE (PRK00019)Ribosomal_L31 (pfam01197)→ |
| env-772 | longitudinal_382_105 | + | 76965 | 77017 | **RNA**→ rpmE (PRK00019)Ribosomal_L31 (pfam01197)→ |
| env-773 | longitudinal_383_30 | - | 52083 | 52031 | **RNA**→ rpmE (PRK00019)Ribosomal_L31 (pfam01197)→ |
| env-774 | longitudinal_476_25 | - | 52085 | 52033 | **RNA**→ rpmE (PRK00019)Ribosomal_L31 (pfam01197)→ |
| env-775 | longitudinal_477_11 | - | 18180 | 18128 | **RNA**→ rpmE (PRK00019)Ribosomal_L31 (pfam01197)→ |
| env-776 | longitudinal_478_15 | + | 7304 | 7356 | **RNA**→ rpmE (PRK00019)Ribosomal_L31 (pfam01197)→ |
| env-777 | longitudinal_479_7 | - | 52071 | 52019 | **RNA**→ rpmE (PRK00019)Ribosomal_L31 (pfam01197)→ |
| env-778 | longitudinal_480_186 | + | 93226 | 93278 | **RNA**→ rpmE (PRK00019)Ribosomal_L31 (pfam01197)→ |
| env-779 | longitudinal_481_108 | + | 76967 | 77019 | **RNA**→ rpmE (PRK00019)Ribosomal_L31 (pfam01197)→ |
| env-780 | longitudinal_523_386 | + | 775 | 827 | **RNA**→ rpmE (PRK00019)Ribosomal_L31 (pfam01197)→ |
| env-781 | SRS012273_Baylor_scaffold_27707 | - | 310 | 258 | **RNA**→ ←hypo |
| env-782 | SRS013687_C2419310 | + | 106 | 158 | **RNA**→ ←hypo |
| env-783 | SRS064276_LANL_scaffold_67167 | + | 192 | 244 | **RNA**→ rpmE (PRK00019)Ribosomal_L31 (pfam01197)→ |
| Eco-1069-1 | NZ_AIFH01000083.1 | + | 95460 | 95512 | **RNA**→ rpmE (PRK00019)Ribosomal_L31 (pfam01197)→ |
| Eco-932-1 | NZ_ANME01000140.1 | + | 91944 | 91996 | **RNA**→ rpmE (PRK00019)Ribosomal_L31 (pfam01197)→ |
| Eco-1953-1 | NZ_AOEL01000099.1 | + | 92477 | 92529 | **RNA**→ rpmE (PRK00019)Ribosomal_L31 (pfam01197)→ |
| Eco-1949-1 | NZ_AOEK01000130.1 | + | 91968 | 92020 | **RNA**→ rpmE (PRK00019)Ribosomal_L31 (pfam01197)→ |
| Eco-943-1 | NZ_AOEQ01000149.1 | + | 91698 | 91750 | **RNA**→ rpmE (PRK00019)Ribosomal_L31 (pfam01197)→ |
| Eco-935-1 | NZ_AODY01000196.1 | + | 94220 | 94272 | **RNA**→ rpmE (PRK00019)Ribosomal_L31 (pfam01197)→ |
| Eco-1977-1 | NZ_AOEO01000117.1 | + | 44640 | 44692 | **RNA**→ rpmE (PRK00019)Ribosomal_L31 (pfam01197)→ |
| Eco-1075-1 | NZ_AIFN01000076.1 | + | 96431 | 96483 | **RNA**→ rpmE (PRK00019)Ribosomal_L31 (pfam01197)→ |
| Eco-941-1 | NZ_AOER01000130.1 | + | 88384 | 88436 | **RNA**→ rpmE (PRK00019)Ribosomal_L31 (pfam01197)→ |
| Eco-939-1 | NZ_AOEC01000217.1 | + | 88385 | 88437 | **RNA**→ rpmE (PRK00019)Ribosomal_L31 (pfam01197)→ |
| Eco-1972-1 | NZ_AOEM01000150.1 | + | 44656 | 44708 | **RNA**→ rpmE (PRK00019)Ribosomal_L31 (pfam01197)→ |
| Eco-736-1 | NZ_JWZW01000013.1 | - | 52488 | 52436 | **RNA**→ rpmE (PRK00019)Ribosomal_L31 (pfam01197)→ |
| Eco-1802-1 | NZ_ABHT01000011.1 | - | 47240 | 47188 | **RNA**→ rpmE (PRK00019)Ribosomal_L31 (pfam01197)→ rpmE (PRK00019)Ribosomal_L31 (pfam01197)→ |
| Eco-1800-1 | NZ_ABHR01000009.1 | + | 96825 | 96877 | **RNA**→ rpmE (PRK00019)Ribosomal_L31 (pfam01197)→ rpmE (PRK00019)Ribosomal_L31 (pfam01197)→ |

| | | | | | | | |
|---|---|---|---|---|---|---|---|
| Eco-1794-1 | NZ_ABHQ01000006.1 | + | 210561 | 210613 | **RNA**→ | rpmE (PRK00019)Ribosomal_L31 (pfam01197)→ | rpmE (PRK00019)Ribosomal_L31 (pfam01197)→ |
| Eco-1795-1 | NZ_ABHP01000009.1 | - | 47048 | 46996 | **RNA**→ | rpmE (PRK00019)Ribosomal_L31 (pfam01197)→ | rpmE (PRK00019)Ribosomal_L31 (pfam01197)→ |
| Eco-1804-1 | NZ_ABHU01000005.1 | + | 261003 | 261055 | **RNA**→ | rpmE (PRK00019)Ribosomal_L31 (pfam01197)→ | rpmE (PRK00019)Ribosomal_L31 (pfam01197)→ |
| Eco-1803-1 | NZ_ABHW01000012.1 | - | 47087 | 47035 | **RNA**→ | rpmE (PRK00019)Ribosomal_L31 (pfam01197)→ | rpmE (PRK00019)Ribosomal_L31 (pfam01197)→ |
| Eco-1801-1 | NZ_ABHS01000007.1 | + | 209927 | 209979 | **RNA**→ | rpmE (PRK00019)Ribosomal_L31 (pfam01197)→ | rpmE (PRK00019)Ribosomal_L31 (pfam01197)→ |
| Eco-859-1 | NZ_AMTN01000126.1 | + | 91910 | 91962 | **RNA**→ | rpmE (PRK00019)Ribosomal_L31 (pfam01197)→ | |
| Eco-937-1 | NZ_AOEA01000208.1 | + | 94248 | 94300 | **RNA**→ | rpmE (PRK00019)Ribosomal_L31 (pfam01197)→ | |
| Eco-940-1 | NZ_AOED01000229.1 | + | 93715 | 93767 | **RNA**→ | rpmE (PRK00019)Ribosomal_L31 (pfam01197)→ | |
| Eco-569-1 | NZ_ASHA01000104.1 | - | 486 | 434 | **RNA**→ | rpmE (PRK00019)Ribosomal_L31 (pfam01197)→ | |
| Eco-1798-1 | NZ_ABHO01000013.1 | - | 49906 | 49854 | **RNA**→ | rpmE (PRK00019)Ribosomal_L31 (pfam01197)→ | rpmE (PRK00019)Ribosomal_L31 (pfam01197)→ |
| Eco-1792-1 | NZ_ABHM02000001.1 | + | 3295143 | 3295195 | **RNA**→ | rpmE (PRK00019)Ribosomal_L31 (pfam01197)→ | rpmE (PRK00019)Ribosomal_L31 (pfam01197)→ |
| Eco-346-1 | NZ_AIAR01000054.1 | - | 5358 | 5306 | **RNA**→ | rpmE (PRK00019)Ribosomal_L31 (pfam01197)→ | |
| Eco-574-1 | NZ_LAZD01000064.1 | - | 557 | 505 | **RNA**→ | rpmE (PRK00019)Ribosomal_L31 (pfam01197)→ | |
| Eco-574-2 | NZ_LCWU01000211.1 | + | 17316 | 17368 | **RNA**→ | rpmE (PRK00019)Ribosomal_L31 (pfam01197)→ | |
| Eco-574-3 | NZ_LAYW01000197.1 | - | 556 | 504 | **RNA**→ | rpmE (PRK00019)Ribosomal_L31 (pfam01197)→ | |
| Sso-1-4 | NZ_CWTD01000021.1 | - | 555 | 503 | **RNA**→ | rpmE (PRK00019)Ribosomal_L31 (pfam01197)→ | |
| Eco-380-1 | NZ_AIBY01000120.1 | - | 485 | 433 | **RNA**→ | rpmE (PRK00019)Ribosomal_L31 (pfam01197)→ | |
| Eco-351-1 | NZ_AIAW01000183.1 | - | 706 | 654 | **RNA**→ | rpmE (PRK00019)Ribosomal_L31 (pfam01197)→ | |
| Eco-757-1 | NZ_AVRI01000125.1 | + | 46581 | 46633 | **RNA**→ | rpmE (PRK00019)Ribosomal_L31 (pfam01197)→ | |
| Eco-979-1 | NZ_AVQW01000112.1 | + | 91931 | 91983 | **RNA**→ | rpmE (PRK00019)Ribosomal_L31 (pfam01197)→ | |
| Eco-984-1 | NZ_AVRA01000101.1 | + | 91696 | 91748 | **RNA**→ | rpmE (PRK00019)Ribosomal_L31 (pfam01197)→ | |
| Eco-962-1 | NZ_AVQS01000121.1 | + | 94248 | 94300 | **RNA**→ | rpmE (PRK00019)Ribosomal_L31 (pfam01197)→ | |
| Eco-977-1 | NZ_AVQU01000053.1 | + | 95584 | 95636 | **RNA**→ | rpmE (PRK00019)Ribosomal_L31 (pfam01197)→ | |
| Eco-1998-1 | NZ_AVRN01000113.1 | + | 91692 | 91744 | **RNA**→ | rpmE (PRK00019)Ribosomal_L31 (pfam01197)→ | |
| Eco-978-1 | NZ_AVQV01000003.1 | + | 96537 | 96589 | **RNA**→ | rpmE (PRK00019)Ribosomal_L31 (pfam01197)→ | |
| Eco-967-1 | NZ_AVQT01000050.1 | + | 96695 | 96747 | **RNA**→ | rpmE (PRK00019)Ribosomal_L31 (pfam01197)→ | |
| Eco-982-1 | NZ_AVQZ01000113.1 | + | 91945 | 91997 | **RNA**→ | rpmE (PRK00019)Ribosomal_L31 (pfam01197)→ | |
| Eco-772-1 | NZ_AVRQ01000125.1 | + | 94905 | 94957 | **RNA**→ | rpmE (PRK00019)Ribosomal_L31 (pfam01197)→ | |
| Eco-985-1 | NZ_AVRB01000098.1 | + | 91952 | 92004 | **RNA**→ | rpmE (PRK00019)Ribosomal_L31 (pfam01197)→ | |
| Eco-981-1 | NZ_AVQY01000110.1 | + | 91935 | 91987 | **RNA**→ | rpmE (PRK00019)Ribosomal_L31 (pfam01197)→ | |
| Eco-1997-1 | NZ_AVRM01000138.1 | + | 44627 | 44679 | **RNA**→ | rpmE (PRK00019)Ribosomal_L31 (pfam01197)→ | |
| Eco-991-1 | NZ_AVRC01000046.1 | + | 94622 | 94674 | **RNA**→ | rpmE (PRK00019)Ribosomal_L31 (pfam01197)→ | |
| Eco-1002-1 | NZ_AVRF01000049.1 | + | 96837 | 96889 | **RNA**→ | rpmE (PRK00019)Ribosomal_L31 (pfam01197)→ | |
| Eco-995-1 | NZ_AVSH01000114.1 | + | 92149 | 92201 | **RNA**→ | rpmE (PRK00019)Ribosomal_L31 (pfam01197)→ | |
| Eco-965-1 | NZ_AVRU01000122.1 | + | 44646 | 44698 | **RNA**→ | rpmE (PRK00019)Ribosomal_L31 (pfam01197)→ | |
| Eco-760-1 | NZ_AVQR01000061.1 | + | 94786 | 94838 | **RNA**→ | rpmE (PRK00019)Ribosomal_L31 (pfam01197)→ | |
| Eco-1999-1 | NZ_AVRO01000114.1 | + | 29667 | 29719 | **RNA**→ | rpmE (PRK00019)Ribosomal_L31 (pfam01197)→ | |
| Eco-966-1 | NZ_AVRV01000107.1 | + | 92178 | 92230 | **RNA**→ | rpmE (PRK00019)Ribosomal_L31 (pfam01197)→ | |
| Eco-990-1 | NZ_AVSG01000112.1 | + | 91949 | 92001 | **RNA**→ | rpmE (PRK00019)Ribosomal_L31 (pfam01197)→ | |
| Eco-963-1 | NZ_AVRS01000114.1 | + | 91671 | 91723 | **RNA**→ | rpmE (PRK00019)Ribosomal_L31 (pfam01197)→ | |
| Eco-970-1 | NZ_AVRY01000117.1 | + | 94217 | 94269 | **RNA**→ | rpmE (PRK00019)Ribosomal_L31 (pfam01197)→ | |
| Eco-997-1 | NZ_AVSJ01000120.1 | + | 93623 | 93675 | **RNA**→ | rpmE (PRK00019)Ribosomal_L31 (pfam01197)→ | rpmE (PRK00019)Ribosomal_L31 (pfam01197)→ |
| Eco-1003-1 | NZ_AVRG01000051.1 | + | 95891 | 95943 | **RNA**→ | rpmE (PRK00019)Ribosomal_L31 (pfam01197)→ | |
| Eco-992-1 | NZ_AVRD01000047.1 | + | 96895 | 96947 | **RNA**→ | rpmE (PRK00019)Ribosomal_L31 (pfam01197)→ | |
| Eco-964-1 | NZ_AVRT01000114.1 | + | 94254 | 94306 | **RNA**→ | rpmE (PRK00019)Ribosomal_L31 (pfam01197)→ | |
| Eco-969-1 | NZ_AVRX01000111.1 | + | 92141 | 92193 | **RNA**→ | rpmE (PRK00019)Ribosomal_L31 (pfam01197)→ | |
| Eco-972-1 | NZ_AVSA01000108.1 | + | 93637 | 93689 | **RNA**→ | rpmE (PRK00019)Ribosomal_L31 (pfam01197)→ | |
| Eco-968-1 | NZ_AVRW01000116.1 | + | 44654 | 44706 | **RNA**→ | rpmE (PRK00019)Ribosomal_L31 (pfam01197)→ | |
| Eco-1000-1 | NZ_AVSM01000111.1 | + | 93633 | 93685 | **RNA**→ | rpmE (PRK00019)Ribosomal_L31 (pfam01197)→ | |
| Eco-999-1 | NZ_AVSL01000116.1 | + | 92472 | 92524 | **RNA**→ | rpmE (PRK00019)Ribosomal_L31 (pfam01197)→ | |
| Eco-998-1 | NZ_AVSK01000131.1 | + | 91687 | 91739 | **RNA**→ | rpmE (PRK00019)Ribosomal_L31 (pfam01197)→ | |
| Eco-987-1 | NZ_AVSE01000119.1 | + | 44642 | 44694 | **RNA**→ | rpmE (PRK00019)Ribosomal_L31 (pfam01197)→ | |

| | | | | | | |
|---|---|---|---|---|---|---|
| Eco-974-1 | NZ_AVSC01000008.1 | - | 4109 | 4057 | **RNA**→ | rpmE (PRK00019)Ribosomal_L31 (pfam01197)→ |
| Eco-758-1 | NZ_AVRJ01000121.1 | + | 94936 | 94988 | **RNA**→ | rpmE (PRK00019)Ribosomal_L31 (pfam01197)→ |
| Eco-448-1 | NZ_AWPR01000018.1 | + | 91639 | 91691 | **RNA**→ | rpmE (PRK00019)Ribosomal_L31 (pfam01197)→ |
| Eco-759-1 | NZ_AVRK01000132.1 | + | 46570 | 46622 | **RNA**→ | rpmE (PRK00019)Ribosomal_L31 (pfam01197)→ |
| Eco-449-1 | NZ_AWPS01000020.1 | + | 91641 | 91693 | **RNA**→ | rpmE (PRK00019)Ribosomal_L31 (pfam01197)→ |
| Eco-437-1 | NZ_AWQC01000422.1 | + | 91641 | 91693 | **RNA**→ | rpmE (PRK00019)Ribosomal_L31 (pfam01197)→ |
| Eco-1001-1 | NZ_AVRE01000043.1 | + | 94898 | 94950 | **RNA**→ | rpmE (PRK00019)Ribosomal_L31 (pfam01197)→ |
| Eco-447-1 | NZ_AWPQ01000136.1 | + | 91638 | 91690 | **RNA**→ | rpmE (PRK00019)Ribosomal_L31 (pfam01197)→ |
| Eco-438-1 | NZ_AWQD01000028.1 | + | 91639 | 91691 | **RNA**→ | rpmE (PRK00019)Ribosomal_L31 (pfam01197)→ |
| Eco-432-1 | NZ_AWPX01000127.1 | + | 12748 | 12800 | **RNA**→ | rpmE (PRK00019)Ribosomal_L31 (pfam01197)→ |
| Eco-433-1 | NZ_AWPY01000116.1 | + | 12746 | 12798 | **RNA**→ | rpmE (PRK00019)Ribosomal_L31 (pfam01197)→ |
| Eco-421-1 | NZ_AWQJ01000017.1 | + | 91639 | 91691 | **RNA**→ | rpmE (PRK00019)Ribosomal_L31 (pfam01197)→ |
| Eco-430-1 | NZ_AWPV01000015.1 | + | 12744 | 12796 | **RNA**→ | rpmE (PRK00019)Ribosomal_L31 (pfam01197)→ |
| Eco-451-1 | NZ_AWPU01000098.1 | + | 12748 | 12800 | **RNA**→ | rpmE (PRK00019)Ribosomal_L31 (pfam01197)→ |
| Eco-434-1 | NZ_AWPZ01000093.1 | + | 19850 | 19902 | **RNA**→ | rpmE (PRK00019)Ribosomal_L31 (pfam01197)→ |
| Eco-750-1 | NZ_AWQO01000113.1 | + | 12748 | 12800 | **RNA**→ | rpmE (PRK00019)Ribosomal_L31 (pfam01197)→ |
| Eco-440-1 | NZ_AWPN01000045.1 | + | 12746 | 12798 | **RNA**→ | rpmE (PRK00019)Ribosomal_L31 (pfam01197)→ |
| Eco-439-1 | NZ_AWQE01000031.1 | + | 91641 | 91693 | **RNA**→ | rpmE (PRK00019)Ribosomal_L31 (pfam01197)→ |
| Eco-444-1 | NZ_AWQI01000063.1 | + | 12748 | 12800 | **RNA**→ | rpmE (PRK00019)Ribosomal_L31 (pfam01197)→ |
| Eco-436-1 | NZ_AWQB01000157.1 | + | 12747 | 12799 | **RNA**→ | rpmE (PRK00019)Ribosomal_L31 (pfam01197)→ |
| Eco-435-1 | NZ_AWQA01000092.1 | + | 12748 | 12800 | **RNA**→ | rpmE (PRK00019)Ribosomal_L31 (pfam01197)→ |
| Eco-747-1 | NZ_AWQL01000099.1 | + | 12748 | 12800 | **RNA**→ | rpmE (PRK00019)Ribosomal_L31 (pfam01197)→ |
| Eco-749-1 | NZ_AWQN01000168.1 | + | 12746 | 12798 | **RNA**→ | rpmE (PRK00019)Ribosomal_L31 (pfam01197)→ |
| Eco-748-1 | NZ_AWQM01000034.1 | + | 12748 | 12800 | **RNA**→ | rpmE (PRK00019)Ribosomal_L31 (pfam01197)→ |
| Eco-443-1 | NZ_AWQH01000004.1 | + | 2951 | 3003 | **RNA**→ | rpmE (PRK00019)Ribosomal_L31 (pfam01197)→ |
| Eco-278-1 | NZ_AIAK01000020.1 | - | 1021 | 969 | **RNA**→ | rpmE (PRK00019)Ribosomal_L31 (pfam01197)→ |
| Eco-980-1 | NZ_AVQX01000117.1 | + | 92014 | 92066 | **RNA**→ | rpmE (PRK00019)Ribosomal_L31 (pfam01197)→ |
| Eco-973-1 | NZ_AVSB01000121.1 | - | 4116 | 4064 | **RNA**→ | rpmE (PRK00019)Ribosomal_L31 (pfam01197)→ |
| Eco-2045-1 | NZ_AVSO01000133.1 | + | 94217 | 94269 | **RNA**→ | rpmE (PRK00019)Ribosomal_L31 (pfam01197)→ |
| Eco-986-1 | NZ_AVSD01000125.1 | + | 92039 | 92091 | **RNA**→ | rpmE (PRK00019)Ribosomal_L31 (pfam01197)→ |
| Eco-996-1 | NZ_AVSI01000127.1 | + | 94242 | 94294 | **RNA**→ | rpmE (PRK00019)Ribosomal_L31 (pfam01197)→ rpmE (PRK00019)Ribosomal_L31 (pfam01197)→ |
| Eco-1023-1 | NZ_AVRL01000126.1 | + | 91700 | 91752 | **RNA**→ | rpmE (PRK00019)Ribosomal_L31 (pfam01197)→ |
| Eco-2000-1 | NZ_AVRP01000114.1 | + | 94245 | 94297 | **RNA**→ | rpmE (PRK00019)Ribosomal_L31 (pfam01197)→ |
| Eco-971-1 | NZ_AVRZ01000117.1 | + | 94219 | 94271 | **RNA**→ | rpmE (PRK00019)Ribosomal_L31 (pfam01197)→ |
| Eco-989-1 | NZ_AVSF01000119.1 | + | 44639 | 44691 | **RNA**→ | rpmE (PRK00019)Ribosomal_L31 (pfam01197)→ |
| Eco-2036-1 | NZ_AKLW01000069.1 | + | 95016 | 95068 | **RNA**→ | rpmE (PRK00019)Ribosomal_L31 (pfam01197)→ |
| Eco-1968-1 | NZ_AKLQ01000060.1 | + | 23530 | 23582 | **RNA**→ | rpmE (PRK00019)Ribosomal_L31 (pfam01197)→ |
| Eco-2031-1 | NZ_AKLV01000044.1 | + | 95383 | 95435 | **RNA**→ | rpmE (PRK00019)Ribosomal_L31 (pfam01197)→ |
| Eco-1791-1 | NZ_ADMX01000127.1 | + | 91609 | 91661 | **RNA**→ | rpmE (PRK00019)Ribosomal_L31 (pfam01197)→ |
| Eco-1805-1 | NZ_ACXO01000010.1 | + | 30986 | 31038 | **RNA**→ | rpmE (PRK00019)Ribosomal_L31 (pfam01197)→ rpmE (PRK00019)Ribosomal_L31 (pfam01197)→ |
| Eco-620-1 | NZ_ADVC01000558.1 | + | 91610 | 91662 | **RNA**→ | rpmE (PRK00019)Ribosomal_L31 (pfam01197)→ |
| Eco-1790-1 | NZ_AERQ01000011.1 | - | 879247 | 879195 | **RNA**→ | rpmE (PRK00019)Ribosomal_L31 (pfam01197)→ |
| Eco-1788-1 | NZ_AERP01000068.1 | - | 47614 | 47562 | **RNA**→ | rpmE (PRK00019)Ribosomal_L31 (pfam01197)→ |
| Eco-605-1 | NZ_JHKM01000001.1 | + | 91629 | 91681 | **RNA**→ | rpmE (PRK00019)Ribosomal_L31 (pfam01197)→ |
| Eco-579-1 | NZ_JHKZ01000005.1 | - | 451 | 399 | **RNA**→ | rpmE (PRK00019)Ribosomal_L31 (pfam01197)→ |
| Eco-598-1 | NZ_JHKT01000066.1 | + | 91629 | 91681 | **RNA**→ | rpmE (PRK00019)Ribosomal_L31 (pfam01197)→ |
| Eco-622-1 | NZ_JHJT01000041.1 | + | 91630 | 91682 | **RNA**→ | rpmE (PRK00019)Ribosomal_L31 (pfam01197)→ |
| Eco-624-1 | NZ_JHJP01000049.1 | - | 450 | 398 | **RNA**→ | rpmE (PRK00019)Ribosomal_L31 (pfam01197)→ |
| Eco-586-1 | NZ_JHGD01000027.1 | + | 18891 | 18943 | **RNA**→ | rpmE (PRK00019)Ribosomal_L31 (pfam01197)→ |
| Eco-582-1 | NZ_JHHI01000057.1 | + | 91627 | 91679 | **RNA**→ | rpmE (PRK00019)Ribosomal_L31 (pfam01197)→ |
| Eco-609-1 | NZ_JHKI01000039.1 | - | 451 | 399 | **RNA**→ | rpmE (PRK00019)Ribosomal_L31 (pfam01197)→ |
| Eco-1787-1 | NZ_JJOM01000126.1 | + | 10323 | 10375 | **RNA**→ | rpmE (PRK00019)Ribosomal_L31 (pfam01197)→ |

| | | | | | |
|---|---|---|---|---|---|
| Eco-616-1 | NZ_JHKB01000029.1 | - | 450 | 398 | **RNA**→ rpmE (PRK00019)Ribosomal_L31 (pfam01197)→ |
| Eco-603-1 | NZ_JHKO01000068.1 | + | 91629 | 91681 | **RNA**→ rpmE (PRK00019)Ribosomal_L31 (pfam01197)→ |
| Eco-607-1 | NZ_JHKK01000018.1 | + | 91629 | 91681 | **RNA**→ rpmE (PRK00019)Ribosomal_L31 (pfam01197)→ |
| Eco-580-1 | NZ_JHKY01000056.1 | - | 451 | 399 | **RNA**→ rpmE (PRK00019)Ribosomal_L31 (pfam01197)→ |
| Eco-2028-1 | NZ_AKLY01000063.1 | + | 95632 | 95684 | **RNA**→ rpmE (PRK00019)Ribosomal_L31 (pfam01197)→ |
| Eco-623-1 | NZ_JHJQ01000008.1 | + | 91629 | 91681 | **RNA**→ rpmE (PRK00019)Ribosomal_L31 (pfam01197)→ |
| Eco-629-1 | NZ_JHJK01000015.1 | - | 450 | 398 | **RNA**→ rpmE (PRK00019)Ribosomal_L31 (pfam01197)→ |
| Eco-627-1 | NZ_JHJM01000040.1 | - | 450 | 398 | **RNA**→ rpmE (PRK00019)Ribosomal_L31 (pfam01197)→ |
| Eco-617-1 | NZ_JHKA01000046.1 | + | 91629 | 91681 | **RNA**→ rpmE (PRK00019)Ribosomal_L31 (pfam01197)→ |
| Eco-1067-1 | NZ_AIFF01000060.1 | + | 96859 | 96911 | **RNA**→ rpmE (PRK00019)Ribosomal_L31 (pfam01197)→ |
| Eco-611-1 | NZ_JHKG01000053.1 | + | 91627 | 91679 | **RNA**→ rpmE (PRK00019)Ribosomal_L31 (pfam01197)→ |
| Eco-626-1 | NZ_JHJN01000046.1 | - | 450 | 398 | **RNA**→ rpmE (PRK00019)Ribosomal_L31 (pfam01197)→ |
| Eco-1954-1 | NZ_AKLI01000067.1 | + | 93083 | 93135 | **RNA**→ hypo→ rpmE (PRK00019)Ribosomal_L31 (pfam01197)→ |
| Eco-618-1 | NZ_ADUZ01000101.1 | + | 19096 | 19148 | **RNA**→ rpmE (PRK00019)Ribosomal_L31 (pfam01197)→ |
| Eco-2030-1 | NZ_AKMA01000052.1 | + | 92700 | 92752 | **RNA**→ rpmE (PRK00019)Ribosomal_L31 (pfam01197)→ |
| Eco-587-1 | NZ_JHGC01000018.1 | - | 889 | 837 | **RNA**→ rpmE (PRK00019)Ribosomal_L31 (pfam01197)→ |
| Eco-604-1 | NZ_JHKN01000014.1 | - | 450 | 398 | **RNA**→ rpmE (PRK00019)Ribosomal_L31 (pfam01197)→ |
| Eco-628-1 | NZ_JHJL01000028.1 | + | 91627 | 91679 | **RNA**→ rpmE (PRK00019)Ribosomal_L31 (pfam01197)→ |
| Eco-625-1 | NZ_JHJO01000073.1 | + | 91629 | 91681 | **RNA**→ rpmE (PRK00019)Ribosomal_L31 (pfam01197)→ |
| Eco-1786-1 | NZ_JJOL01000613.1 | - | 450 | 398 | **RNA**→ rpmE (PRK00019)Ribosomal_L31 (pfam01197)→ |
| Eco-1797-1 | NZ_ADVA01000255.1 | - | 430 | 378 | **RNA**→ rpmE (PRK00019)Ribosomal_L31 (pfam01197)→ |
| Eco-614-1 | NZ_JHKD01000047.1 | - | 450 | 398 | **RNA**→ rpmE (PRK00019)Ribosomal_L31 (pfam01197)→ |
| Eco-933-1 | NZ_ANMF01000108.1 | + | 92081 | 92133 | **RNA**→ rpmE (PRK00019)Ribosomal_L31 (pfam01197)→ |
| Eco-602-1 | NZ_JHKP01000022.1 | - | 450 | 398 | **RNA**→ rpmE (PRK00019)Ribosomal_L31 (pfam01197)→ |
| Eco-764-1 | NZ_AMTO01000096.1 | + | 92142 | 92194 | **RNA**→ rpmE (PRK00019)Ribosomal_L31 (pfam01197)→ |
| Eco-615-1 | NZ_JHKC01000036.1 | - | 450 | 398 | **RNA**→ rpmE (PRK00019)Ribosomal_L31 (pfam01197)→ |
| Eco-583-1 | NZ_JHGQ01000004.1 | + | 23860 | 23912 | **RNA**→ rpmE (PRK00019)Ribosomal_L31 (pfam01197)→ |
| Eco-584-1 | NZ_JHGF01000028.1 | + | 91945 | 91997 | **RNA**→ rpmE (PRK00019)Ribosomal_L31 (pfam01197)→ |
| Eco-575-1 | NZ_JHNI01000046.1 | - | 450 | 398 | **RNA**→ rpmE (PRK00019)Ribosomal_L31 (pfam01197)→ |
| Eco-1806-1 | NZ_ACXN01000137.1 | + | 91610 | 91662 | **RNA**→ rpmE (PRK00019)Ribosomal_L31 (pfam01197)→ rpmE (PRK00019)Ribosomal_L31 (pfam01197)→ |
| Eco-1072-1 | NZ_AIFK01000050.1 | + | 26780 | 26832 | **RNA**→ rpmE (PRK00019)Ribosomal_L31 (pfam01197)→ |
| Eco-1077-1 | NZ_AIFP01000064.1 | + | 96827 | 96879 | **RNA**→ rpmE (PRK00019)Ribosomal_L31 (pfam01197)→ |
| Eco-639-1 | NZ_JHJA01000020.1 | + | 91780 | 91832 | **RNA**→ rpmE (PRK00019)Ribosomal_L31 (pfam01197)→ |
| Eco-636-1 | NZ_JHJD01000062.1 | + | 28264 | 28316 | **RNA**→ rpmE (PRK00019)Ribosomal_L31 (pfam01197)→ |
| Eco-644-1 | NZ_JHIV01000025.1 | - | 385 | 333 | **RNA**→ rpmE (PRK00019)Ribosomal_L31 (pfam01197)→ |
| Eco-647-1 | NZ_JHIS01000020.1 | - | 1165 | 1113 | **RNA**→ rpmE (PRK00019)Ribosomal_L31 (pfam01197)→ |
| Eco-653-1 | NZ_JHIM01000108.1 | - | 1167 | 1115 | **RNA**→ rpmE (PRK00019)Ribosomal_L31 (pfam01197)→ |
| Eco-660-1 | NZ_JHID01000024.1 | + | 91933 | 91985 | **RNA**→ |
| Eco-665-1 | NZ_JHHY01000031.1 | + | 91851 | 91903 | **RNA**→ rpmE (PRK00019)Ribosomal_L31 (pfam01197)→ |
| Eco-673-1 | NZ_JHHH01000049.1 | + | 91629 | 91681 | **RNA**→ rpmE (PRK00019)Ribosomal_L31 (pfam01197)→ |
| Eco-635-1 | NZ_JHJF01000046.1 | - | 686 | 634 | **RNA**→ rpmE (PRK00019)Ribosomal_L31 (pfam01197)→ |
| Eco-1115-1 | NZ_AKMN01000055.1 | + | 94670 | 94722 | **RNA**→ rpmE (PRK00019)Ribosomal_L31 (pfam01197)→ |
| Eco-633-1 | NZ_JHJH01000029.1 | - | 450 | 398 | **RNA**→ rpmE (PRK00019)Ribosomal_L31 (pfam01197)→ |
| Eco-640-1 | NZ_JHIZ01000031.1 | + | 90872 | 90924 | **RNA**→ rpmE (PRK00019)Ribosomal_L31 (pfam01197)→ |
| Eco-641-1 | NZ_JHIY01000023.1 | + | 90756 | 90808 | **RNA**→ rpmE (PRK00019)Ribosomal_L31 (pfam01197)→ |
| Eco-648-1 | NZ_JHIR01000036.1 | + | 91660 | 91712 | **RNA**→ rpmE (PRK00019)Ribosomal_L31 (pfam01197)→ |
| Eco-645-1 | NZ_JHIU01000032.1 | - | 1552 | 1500 | **RNA**→ rpmE (PRK00019)Ribosomal_L31 (pfam01197)→ |
| Eco-421-2 | NZ_JQFE01000162.1 | - | 5435 | 5383 | **RNA**→ rpmE (PRK00019)Ribosomal_L31 (pfam01197)→ |
| Eco-655-1 | NZ_JHII01000022.1 | - | 1206 | 1154 | **RNA**→ rpmE (PRK00019)Ribosomal_L31 (pfam01197)→ |
| Eco-661-1 | NZ_JHIC01000130.1 | + | 91932 | 91984 | **RNA**→ rpmE (PRK00019)Ribosomal_L31 (pfam01197)→ |
| Eco-666-1 | NZ_JHHX01000127.1 | + | 80205 | 80257 | **RNA**→ rpmE (PRK00019)Ribosomal_L31 (pfam01197)→ |
| Eco-667-1 | NZ_JHHW01000029.1 | + | 92008 | 92060 | **RNA**→ rpmE (PRK00019)Ribosomal_L31 (pfam01197)→ |

| | | | | | |
|---|---|---|---|---|---|
| Eco-652-1 | NZ_JHIN01000027.1 | + | 91794 | 91846 | **RNA**→ rpmE (PRK00019)Ribosomal_L31 (pfam01197)→ |
| Eco-637-1 | NZ_JHJC01000021.1 | - | 1444 | 1392 | **RNA**→ rpmE (PRK00019)Ribosomal_L31 (pfam01197)→ |
| Eco-1111-1 | NZ_AKMO01000049.1 | + | 92352 | 92404 | **RNA**→ rpmE (PRK00019)Ribosomal_L31 (pfam01197)→ |
| Eco-642-1 | NZ_JHIX01000008.1 | + | 38907 | 38959 | **RNA**→ rpmE (PRK00019)Ribosomal_L31 (pfam01197)→ |
| Eco-649-1 | NZ_JHIQ01000016.1 | + | 92060 | 92112 | **RNA**→ rpmE (PRK00019)Ribosomal_L31 (pfam01197)→ |
| Eco-1074-1 | NZ_AIFM01000051.1 | + | 29971 | 30023 | **RNA**→ rpmE (PRK00019)Ribosomal_L31 (pfam01197)→ |
| Eco-646-1 | NZ_JHIT01000029.1 | + | 92053 | 92105 | **RNA**→ rpmE (PRK00019)Ribosomal_L31 (pfam01197)→ |
| Eco-656-1 | NZ_JHIH01000027.1 | + | 91457 | 91509 | **RNA**→ rpmE (PRK00019)Ribosomal_L31 (pfam01197)→ |
| Eco-662-1 | NZ_JHIB01000019.1 | - | 617 | 565 | **RNA**→ rpmE (PRK00019)Ribosomal_L31 (pfam01197)→ |
| Eco-421-3 | NZ_JQFC01000080.1 | - | 5475 | 5423 | **RNA**→ rpmE (PRK00019)Ribosomal_L31 (pfam01197)→ |
| Eco-659-1 | NZ_JHIE01000023.1 | + | 92013 | 92065 | **RNA**→ rpmE (PRK00019)Ribosomal_L31 (pfam01197)→ |
| Eco-669-1 | NZ_JHHK01000045.1 | + | 91704 | 91756 | **RNA**→ rpmE (PRK00019)Ribosomal_L31 (pfam01197)→ |
| Eco-668-1 | NZ_JHHV01000025.1 | - | 1045 | 993 | **RNA**→ rpmE (PRK00019)Ribosomal_L31 (pfam01197)→ |
| Eco-421-4 | NZ_JQFH01000455.1 | - | 9813 | 9761 | **RNA**→ rpmE (PRK00019)Ribosomal_L31 (pfam01197)→ |
| Eco-631-1 | NZ_JHJI01000008.1 | - | 451 | 399 | **RNA**→ rpmE (PRK00019)Ribosomal_L31 (pfam01197)→ |
| Eco-1710-1 | NZ_AJMD01000038.1 | + | 191195 | 191247 | **RNA**→ rpmE (PRK00019)Ribosomal_L31 (pfam01197)→ |
| Eco-634-1 | NZ_JHJG01000006.1 | + | 91629 | 91681 | **RNA**→ rpmE (PRK00019)Ribosomal_L31 (pfam01197)→ |
| Eco-638-1 | NZ_JHJB01000094.1 | + | 20677 | 20729 | **RNA**→ hypo→ rpmE (PRK00019)Ribosomal_L31 (pfam01197)→ |
| Eco-643-1 | NZ_JHIW01000024.1 | - | 648 | 596 | **RNA**→ rpmE (PRK00019)Ribosomal_L31 (pfam01197)→ |
| Eco-650-1 | NZ_JHIP01000024.1 | - | 1458 | 1406 | **RNA**→ rpmE (PRK00019)Ribosomal_L31 (pfam01197)→ |
| Eco-651-1 | NZ_JHIO01000024.1 | + | 91973 | 92025 | **RNA**→ rpmE (PRK00019)Ribosomal_L31 (pfam01197)→ |
| Eco-654-1 | NZ_JHIL01000021.1 | + | 92073 | 92125 | **RNA**→ rpmE (PRK00019)Ribosomal_L31 (pfam01197)→ |
| Eco-657-1 | NZ_JHIG01000029.1 | + | 91857 | 91909 | **RNA**→ rpmE (PRK00019)Ribosomal_L31 (pfam01197)→ |
| Eco-658-1 | NZ_JHIF01000022.1 | - | 702 | 650 | **RNA**→ rpmE (PRK00019)Ribosomal_L31 (pfam01197)→ |
| Eco-663-1 | NZ_JHIA01000018.1 | + | 30623 | 30675 | **RNA**→ rpmE (PRK00019)Ribosomal_L31 (pfam01197)→ |
| Eco-1070-1 | NZ_AIFI01000086.1 | + | 95517 | 95569 | **RNA**→ rpmE (PRK00019)Ribosomal_L31 (pfam01197)→ |
| Eco-1076-1 | NZ_AIFO01000059.1 | + | 96837 | 96889 | **RNA**→ rpmE (PRK00019)Ribosomal_L31 (pfam01197)→ |
| Eco-672-1 | NZ_JJOK01000008.1 | - | 429 | 377 | **RNA**→ rpmE (PRK00019)Ribosomal_L31 (pfam01197)→ |
| Eco-619-1 | NZ_ADUX01000057.1 | + | 91609 | 91661 | **RNA**→ rpmE (PRK00019)Ribosomal_L31 (pfam01197)→ |
| Eco-1976-1 | NZ_AMTQ01000001.1 | + | 95107 | 95159 | **RNA**→ rpmE (PRK00019)Ribosomal_L31 (pfam01197)→ |
| Eco-613-1 | NZ_JHKE01000012.1 | - | 450 | 398 | **RNA**→ rpmE (PRK00019)Ribosomal_L31 (pfam01197)→ |
| Eco-592-1 | NZ_JHLG01000045.1 | - | 450 | 398 | **RNA**→ rpmE (PRK00019)Ribosomal_L31 (pfam01197)→ |
| Eco-593-1 | NZ_JHLF01000079.1 | - | 450 | 398 | **RNA**→ rpmE (PRK00019)Ribosomal_L31 (pfam01197)→ |
| Eco-595-1 | NZ_JHLD01000027.1 | - | 450 | 398 | **RNA**→ rpmE (PRK00019)Ribosomal_L31 (pfam01197)→ |
| Eco-610-1 | NZ_JHKH01000027.1 | + | 91881 | 91933 | **RNA**→ rpmE (PRK00019)Ribosomal_L31 (pfam01197)→ |
| Eco-597-1 | NZ_JHLB01000049.1 | - | 450 | 398 | **RNA**→ rpmE (PRK00019)Ribosomal_L31 (pfam01197)→ |
| Eco-2164-1 | NZ_AEYT01000045.1 | + | 77041 | 77093 | **RNA**→ rpmE (PRK00019)Ribosomal_L31 (pfam01197)→ |
| Eco-1799-1 | NZ_ABHK02000001.1 | + | 2307056 | 2307108 | **RNA**→ rpmE (PRK00019)Ribosomal_L31 (pfam01197)→ rpmE (PRK00019)Ribosomal_L31 (pfam01197)→ |
| Eco-608-1 | NZ_JHKJ01000002.1 | - | 450 | 398 | **RNA**→ rpmE (PRK00019)Ribosomal_L31 (pfam01197)→ |
| Eco-600-1 | NZ_JHKR01000082.1 | - | 450 | 398 | **RNA**→ rpmE (PRK00019)Ribosomal_L31 (pfam01197)→ |
| Eco-594-1 | NZ_JHLE01000058.1 | - | 450 | 398 | **RNA**→ rpmE (PRK00019)Ribosomal_L31 (pfam01197)→ |
| Eco-596-1 | NZ_JHLC01000006.1 | - | 450 | 398 | **RNA**→ rpmE (PRK00019)Ribosomal_L31 (pfam01197)→ |
| Eco-601-1 | NZ_JHKQ01000035.1 | - | 450 | 398 | **RNA**→ rpmE (PRK00019)Ribosomal_L31 (pfam01197)→ |
| Eco-591-1 | NZ_JHLH01000025.1 | - | 450 | 398 | **RNA**→ rpmE (PRK00019)Ribosomal_L31 (pfam01197)→ |
| Eco-1793-1 | NZ_ABHL02000001.1 | + | 2175397 | 2175449 | **RNA**→ rpmE (PRK00019)Ribosomal_L31 (pfam01197)→ rpmE (PRK00019)Ribosomal_L31 (pfam01197)→ |
| Eco-606-1 | NZ_JHKL01000031.1 | - | 450 | 398 | **RNA**→ rpmE (PRK00019)Ribosomal_L31 (pfam01197)→ |
| Eco-599-1 | NZ_JHKS01000060.1 | - | 450 | 398 | **RNA**→ rpmE (PRK00019)Ribosomal_L31 (pfam01197)→ |
| Eco-1789-1 | NZ_AERR01000025.1 | + | 95838 | 95890 | **RNA**→ rpmE (PRK00019)Ribosomal_L31 (pfam01197)→ |
| Eco-589-1 | NZ_JHLK01000122.1 | - | 1188 | 1136 | **RNA**→ rpmE (PRK00019)Ribosomal_L31 (pfam01197)→ |
| Eco-581-1 | NZ_JHKW01000036.1 | + | 91629 | 91681 | **RNA**→ rpmE (PRK00019)Ribosomal_L31 (pfam01197)→ |
| Eco-231-1 | NZ_AYQU01000019.1 | - | 32637 | 32585 | **RNA**→ rpmE (PRK00019)Ribosomal_L31 (pfam01197)→ |
| Eco-577-1 | NZ_JHNF01000058.1 | + | 91629 | 91681 | **RNA**→ rpmE (PRK00019)Ribosomal_L31 (pfam01197)→ |

| | | | | | |
|---|---|---|---|---|---|
| Eco-578-1 | NZ_JHNE01000052.1 | + | 91627 | 91679 | **RNA**→ rpmE (PRK00019)Ribosomal_L31 (pfam01197)→ |
| Eco-570-1 | NZ_JHME01000049.1 | + | 44021 | 44073 | **RNA**→ rpmE (PRK00019)Ribosomal_L31 (pfam01197)→ |
| Eco-576-1 | NZ_JHNG01000027.1 | - | 450 | 398 | **RNA**→ rpmE (PRK00019)Ribosomal_L31 (pfam01197)→ |
| Eco-585-1 | NZ_JHGE01000017.1 | + | 91918 | 91970 | **RNA**→ rpmE (PRK00019)Ribosomal_L31 (pfam01197)→ |
| Eco-590-1 | NZ_JHLI01000054.1 | - | 450 | 398 | **RNA**→ rpmE (PRK00019)Ribosomal_L31 (pfam01197)→ |
| Eco-630-1 | NZ_AVCD01000005.1 | + | 2518806 | 2518858 | **RNA**→ rpmE (PRK00019)Ribosomal_L31 (pfam01197)→ |
| Eco-293-1 | NZ_AYGW02000101.1 | + | 91627 | 91679 | **RNA**→ rpmE (PRK00019)Ribosomal_L31 (pfam01197)→ |
| Eco-445-1 | NZ_AWPO01000180.1 | + | 49800 | 49852 | **RNA**→ rpmE (PRK00019)Ribosomal_L31 (pfam01197)→ |
| Eco-1809-1 | NC_002695.1 | + | 4928446 | 4928498 | **RNA**→ rpmE (PRK00019)Ribosomal_L31 (pfam01197)→ |
| Eco-2156-1 | NC_007946.1 | + | 4403952 | 4404004 | **RNA**→ ←hypo |
| Eco-1496-1 | NZ_ANYQ01000052.1 | + | 77161 | 77213 | **RNA**→ rpmE (PRK00019)Ribosomal_L31 (pfam01197)→ |
| Eco-944-1 | NZ_AOEF01000177.1 | + | 94253 | 94305 | **RNA**→ rpmE (PRK00019)Ribosomal_L31 (pfam01197)→ |
| Eco-930-1 | NZ_AOEV01000127.1 | + | 46610 | 46662 | **RNA**→ rpmE (PRK00019)Ribosomal_L31 (pfam01197)→ |
| Eco-1810-1 | NC_013008.1 | + | 4987619 | 4987671 | **RNA**→ rpmE (PRK00019)Ribosomal_L31 (pfam01197)→ |
| Eco-1381-1 | NC_017628.1 | + | 4516074 | 4516126 | **RNA**→ rpmE (PRK00019)Ribosomal_L31 (pfam01197)→ |
| Eco-2046-1 | NC_017632.1 | + | 4145232 | 4145284 | **RNA**→ rpmE (PRK00019)Ribosomal_L31 (pfam01197)→ |
| Eco-1796-1 | NC_011353.1 | + | 5031559 | 5031611 | **RNA**→ rpmE (PRK00019)Ribosomal_L31 (pfam01197)→ |
| Eco-2162-1 | NC_017906.1 | + | 4855185 | 4855237 | **RNA**→ rpmE (PRK00019)Ribosomal_L31 (pfam01197)→ |
| Eco-950-1 | NC_008563.1 | + | 4435966 | 4436018 | **RNA**→ rpmE (PRK00019)Ribosomal_L31 (pfam01197)→ rpmE (PRK00019)Ribosomal_L31 (pfam01197)→ rpmE (PRK00019)Ribosomal_L31 (pfam01197)→ |
| Eco-670-1 | NZ_CP008805.1 | + | 4983353 | 4983405 | **RNA**→ rpmE (PRK00019)Ribosomal_L31 (pfam01197)→ |
| Eco-1785-1 | NC_002655.2 | + | 4997210 | 4997262 | **RNA**→ rpmE (PRK00019)Ribosomal_L31 (pfam01197)→ |
| Eco-1811-1 | NZ_CM000662.1 | - | 1831197 | 1831145 | **RNA**→ rpmE (PRK00019)Ribosomal_L31 (pfam01197)→ |
| Eco-1679-1 | NZ_GG772990.1 | - | 52128 | 52076 | **RNA**→ rpmE (PRK00019)Ribosomal_L31 (pfam01197)→ |
| Eco-913-1 | NZ_KB001687.1 | + | 91681 | 91733 | **RNA**→ rpmE (PRK00019)Ribosomal_L31 (pfam01197)→ |
| Eco-912-1 | NZ_KB001158.1 | + | 91691 | 91743 | **RNA**→ rpmE (PRK00019)Ribosomal_L31 (pfam01197)→ |
| Eco-916-1 | NZ_KB003763.1 | + | 91671 | 91723 | **RNA**→ rpmE (PRK00019)Ribosomal_L31 (pfam01197)→ |
| Eco-868-1 | NZ_KB007262.1 | + | 91757 | 91809 | **RNA**→ rpmE (PRK00019)Ribosomal_L31 (pfam01197)→ |
| Eco-906-1 | NZ_KB005335.1 | + | 91753 | 91805 | **RNA**→ rpmE (PRK00019)Ribosomal_L31 (pfam01197)→ |
| Eco-872-1 | NZ_KB006068.1 | + | 88410 | 88462 | **RNA**→ rpmE (PRK00019)Ribosomal_L31 (pfam01197)→ |
| Eco-870-1 | NZ_KB006781.1 | + | 91761 | 91813 | **RNA**→ rpmE (PRK00019)Ribosomal_L31 (pfam01197)→ |
| Eco-910-1 | NZ_KB002140.1 | + | 88458 | 88510 | **RNA**→ rpmE (PRK00019)Ribosomal_L31 (pfam01197)→ |
| Eco-922-1 | NZ_KB003073.1 | + | 91734 | 91786 | **RNA**→ rpmE (PRK00019)Ribosomal_L31 (pfam01197)→ |
| Eco-314-1 | NZ_KK736334.1 | + | 79261 | 79313 | **RNA**→ rpmE (PRK00019)Ribosomal_L31 (pfam01197)→ |
| Eco-1172-1 | NZ_GL871984.1 | - | 52350 | 52298 | **RNA**→ rpmE (PRK00019)Ribosomal_L31 (pfam01197)→ |
| Eco-1174-1 | NZ_GL896790.1 | - | 2228748 | 2228696 | **RNA**→ rpmE (PRK00019)Ribosomal_L31 (pfam01197)→ |
| Eco-1171-1 | NZ_GL871899.1 | + | 77435 | 77487 | **RNA**→ rpmE (PRK00019)Ribosomal_L31 (pfam01197)→ |
| Eco-1175-1 | NZ_JH590476.1 | - | 52676 | 52624 | **RNA**→ rpmE (PRK00019)Ribosomal_L31 (pfam01197)→ |
| Eco-1957-1 | NZ_JH706532.1 | + | 91681 | 91733 | **RNA**→ rpmE (PRK00019)Ribosomal_L31 (pfam01197)→ |
| Eco-1160-1 | NZ_JH698273.1 | + | 91691 | 91743 | **RNA**→ rpmE (PRK00019)Ribosomal_L31 (pfam01197)→ |
| Eco-1959-1 | NZ_JH711380.1 | + | 91753 | 91805 | **RNA**→ rpmE (PRK00019)Ribosomal_L31 (pfam01197)→ |
| Eco-1138-1 | NZ_JH696208.1 | + | 91679 | 91731 | **RNA**→ rpmE (PRK00019)Ribosomal_L31 (pfam01197)→ |
| Eco-1969-1 | NZ_JH703729.1 | + | 91757 | 91809 | **RNA**→ rpmE (PRK00019)Ribosomal_L31 (pfam01197)→ |
| Eco-1975-1 | NZ_JH709966.1 | + | 91759 | 91811 | **RNA**→ rpmE (PRK00019)Ribosomal_L31 (pfam01197)→ |
| Eco-1956-1 | NZ_JH707511.1 | + | 91755 | 91807 | **RNA**→ rpmE (PRK00019)Ribosomal_L31 (pfam01197)→ |
| Eco-1970-1 | NZ_JH703250.1 | + | 91689 | 91741 | **RNA**→ rpmE (PRK00019)Ribosomal_L31 (pfam01197)→ |
| Eco-1137-1 | NZ_JH696593.1 | + | 91681 | 91733 | **RNA**→ rpmE (PRK00019)Ribosomal_L31 (pfam01197)→ |
| Eco-904-1 | NZ_JH710989.1 | + | 91681 | 91733 | **RNA**→ rpmE (PRK00019)Ribosomal_L31 (pfam01197)→ |
| Eco-2026-1 | NZ_JH702793.1 | + | 91750 | 91802 | **RNA**→ rpmE (PRK00019)Ribosomal_L31 (pfam01197)→ |
| Eco-1958-1 | NZ_JH706083.1 | + | 91685 | 91737 | **RNA**→ rpmE (PRK00019)Ribosomal_L31 (pfam01197)→ |
| Eco-1154-1 | NZ_JH692587.1 | + | 91689 | 91741 | **RNA**→ rpmE (PRK00019)Ribosomal_L31 (pfam01197)→ |
| Eco-2027-1 | NZ_JH702437.1 | + | 91680 | 91732 | **RNA**→ rpmE (PRK00019)Ribosomal_L31 (pfam01197)→ |

| | | | | | |
|---|---|---|---|---|---|
| Eco-1960-1 | NZ_JH707064.1 | + | 91761 | 91813 | **RNA**→ rpmE (PRK00019)Ribosomal_L31 (pfam01197)→ |
| Eco-1961-1 | NZ_JH705659.1 | + | 91683 | 91735 | **RNA**→ rpmE (PRK00019)Ribosomal_L31 (pfam01197)→ |
| Eco-1962-1 | NZ_JH705097.1 | + | 91689 | 91741 | **RNA**→ rpmE (PRK00019)Ribosomal_L31 (pfam01197)→ |
| Eco-1966-1 | NZ_JH704534.1 | + | 8213 | 8265 | **RNA**→ rpmE (PRK00019)Ribosomal_L31 (pfam01197)→ |
| Eco-1124-1 | NZ_JH692963.1 | + | 91755 | 91807 | **RNA**→ rpmE (PRK00019)Ribosomal_L31 (pfam01197)→ |
| Eco-1978-1 | NZ_JH709459.1 | + | 91686 | 91738 | **RNA**→ rpmE (PRK00019)Ribosomal_L31 (pfam01197)→ |
| Eco-1132-1 | NZ_JH698686.1 | + | 91751 | 91803 | **RNA**→ rpmE (PRK00019)Ribosomal_L31 (pfam01197)→ |
| Eco-1139-1 | NZ_JH694736.1 | + | 91686 | 91738 | **RNA**→ rpmE (PRK00019)Ribosomal_L31 (pfam01197)→ |
| Eco-1135-1 | NZ_JH699668.1 | + | 91685 | 91737 | **RNA**→ rpmE (PRK00019)Ribosomal_L31 (pfam01197)→ |
| Eco-2039-1 | NZ_JH695371.1 | + | 8219 | 8271 | **RNA**→ rpmE (PRK00019)Ribosomal_L31 (pfam01197)→ |
| Eco-1116-1 | NZ_JH693435.1 | + | 91685 | 91737 | **RNA**→ rpmE (PRK00019)Ribosomal_L31 (pfam01197)→ |
| Eco-1141-1 | NZ_JH697067.1 | + | 91685 | 91737 | **RNA**→ rpmE (PRK00019)Ribosomal_L31 (pfam01197)→ |
| Eco-1140-1 | NZ_JH694330.1 | + | 91750 | 91802 | **RNA**→ rpmE (PRK00019)Ribosomal_L31 (pfam01197)→ |
| Eco-1134-1 | NZ_JH699178.1 | + | 88425 | 88477 | **RNA**→ rpmE (PRK00019)Ribosomal_L31 (pfam01197)→ |
| Eco-1963-1 | NZ_JH964280.1 | + | 91688 | 91740 | **RNA**→ rpmE (PRK00019)Ribosomal_L31 (pfam01197)→ |
| Eco-1120-1 | NZ_JH953686.1 | + | 91679 | 91731 | **RNA**→ rpmE (PRK00019)Ribosomal_L31 (pfam01197)→ |
| Eco-1713-1 | NZ_JH957915.1 | + | 91763 | 91815 | **RNA**→ rpmE (PRK00019)Ribosomal_L31 (pfam01197)→ |
| Eco-1153-1 | NZ_JH962205.1 | + | 91766 | 91818 | **RNA**→ rpmE (PRK00019)Ribosomal_L31 (pfam01197)→ |
| Eco-1967-1 | NZ_JH960930.1 | - | 90108 | 90056 | **RNA**→ rpmE (PRK00019)Ribosomal_L31 (pfam01197)→ |
| Eco-1971-1 | NZ_JH959095.1 | + | 91691 | 91743 | **RNA**→ rpmE (PRK00019)Ribosomal_L31 (pfam01197)→ |
| Eco-1657-1 | NZ_JH957193.1 | + | 87134 | 87186 | **RNA**→ rpmE (PRK00019)Ribosomal_L31 (pfam01197)→ |
| Eco-1155-1 | NZ_JH964868.1 | + | 91806 | 91858 | **RNA**→ rpmE (PRK00019)Ribosomal_L31 (pfam01197)→ |
| Eco-1025-1 | NZ_JH956755.1 | + | 86912 | 86964 | **RNA**→ rpmE (PRK00019)Ribosomal_L31 (pfam01197)→ |
| Eco-1112-1 | NZ_JH958301.1 | + | 91756 | 91808 | **RNA**→ rpmE (PRK00019)Ribosomal_L31 (pfam01197)→ |
| Eco-765-1 | NZ_JH969496.1 | + | 91753 | 91805 | **RNA**→ rpmE (PRK00019)Ribosomal_L31 (pfam01197)→ |
| Eco-1156-1 | NZ_JH963939.1 | + | 91765 | 91817 | **RNA**→ rpmE (PRK00019)Ribosomal_L31 (pfam01197)→ |
| Eco-1162-1 | NZ_JH962614.1 | + | 91689 | 91741 | **RNA**→ rpmE (PRK00019)Ribosomal_L31 (pfam01197)→ |
| Eco-1114-1 | NZ_JH963514.1 | + | 88419 | 88471 | **RNA**→ rpmE (PRK00019)Ribosomal_L31 (pfam01197)→ |
| Eco-1121-1 | NZ_JH953257.1 | + | 91761 | 91813 | **RNA**→ rpmE (PRK00019)Ribosomal_L31 (pfam01197)→ |
| Eco-847-1 | NZ_JH971373.1 | + | 91765 | 91817 | **RNA**→ rpmE (PRK00019)Ribosomal_L31 (pfam01197)→ |
| Eco-1965-1 | NZ_JH966096.1 | + | 88442 | 88494 | **RNA**→ rpmE (PRK00019)Ribosomal_L31 (pfam01197)→ |
| Eco-754-1 | NZ_JH943323.1 | + | 91763 | 91815 | **RNA**→ rpmE (PRK00019)Ribosomal_L31 (pfam01197)→ |
| Eco-1163-1 | NZ_JH961354.1 | + | 91823 | 91875 | **RNA**→ rpmE (PRK00019)Ribosomal_L31 (pfam01197)→ |
| Eco-1117-1 | NZ_JH956301.1 | + | 91755 | 91807 | **RNA**→ rpmE (PRK00019)Ribosomal_L31 (pfam01197)→ |
| Eco-838-1 | NZ_JH970961.1 | + | 91677 | 91729 | **RNA**→ rpmE (PRK00019)Ribosomal_L31 (pfam01197)→ |
| Eco-2021-1 | NZ_JH957502.1 | + | 91767 | 91819 | **RNA**→ rpmE (PRK00019)Ribosomal_L31 (pfam01197)→ |
| Eco-1119-1 | NZ_JH955815.1 | + | 91687 | 91739 | **RNA**→ rpmE (PRK00019)Ribosomal_L31 (pfam01197)→ |
| Eco-1129-1 | NZ_JH947101.1 | + | 91763 | 91815 | **RNA**→ rpmE (PRK00019)Ribosomal_L31 (pfam01197)→ |
| Eco-856-1 | NZ_JH970471.1 | + | 91687 | 91739 | **RNA**→ rpmE (PRK00019)Ribosomal_L31 (pfam01197)→ |
| Eco-1955-1 | NZ_JH959984.1 | + | 91758 | 91810 | **RNA**→ rpmE (PRK00019)Ribosomal_L31 (pfam01197)→ |
| Eco-1113-1 | NZ_JH965229.1 | + | 91684 | 91736 | **RNA**→ rpmE (PRK00019)Ribosomal_L31 (pfam01197)→ |
| Eco-1122-1 | NZ_JH952871.1 | + | 91691 | 91743 | **RNA**→ rpmE (PRK00019)Ribosomal_L31 (pfam01197)→ |
| Eco-1123-1 | NZ_JH952449.1 | + | 91753 | 91805 | **RNA**→ rpmE (PRK00019)Ribosomal_L31 (pfam01197)→ |
| Eco-1118-1 | NZ_JH963066.1 | + | 91753 | 91805 | **RNA**→ rpmE (PRK00019)Ribosomal_L31 (pfam01197)→ |
| Eco-1712-1 | NZ_JH946694.1 | + | 91677 | 91729 | **RNA**→ rpmE (PRK00019)Ribosomal_L31 (pfam01197)→ |
| Eco-865-1 | NZ_JH968340.1 | + | 91746 | 91798 | **RNA**→ rpmE (PRK00019)Ribosomal_L31 (pfam01197)→ |
| Eco-1127-1 | NZ_JH944307.1 | + | 91687 | 91739 | **RNA**→ rpmE (PRK00019)Ribosomal_L31 (pfam01197)→ |
| Eco-918-1 | NZ_KB002678.1 | + | 91681 | 91733 | **RNA**→ rpmE (PRK00019)Ribosomal_L31 (pfam01197)→ |
| Eco-866-1 | NZ_KB006465.1 | + | 91685 | 91737 | **RNA**→ rpmE (PRK00019)Ribosomal_L31 (pfam01197)→ |
| Eco-924-1 | NZ_JH999841.1 | + | 93668 | 93720 | **RNA**→ rpmE (PRK00019)Ribosomal_L31 (pfam01197)→ |
| Eco-863-1 | NZ_JH969035.1 | + | 91747 | 91799 | **RNA**→ rpmE (PRK00019)Ribosomal_L31 (pfam01197)→ |
| Eco-1125-1 | NZ_JH946246.1 | + | 88431 | 88483 | **RNA**→ rpmE (PRK00019)Ribosomal_L31 (pfam01197)→ |

| | | | | | | |
|---|---|---|---|---|---|---|
| Eco-908-1 | NZ_KB005700.1 | + | 91689 | 91741 | **RNA**→ | rpmE (PRK00019)Ribosomal_L31 (pfam01197)→ |
| Eco-919-1 | NZ_KB000775.1 | + | 88442 | 88494 | **RNA**→ | rpmE (PRK00019)Ribosomal_L31 (pfam01197)→ |
| Eco-905-1 | NZ_KB004122.1 | + | 91687 | 91739 | **RNA**→ | rpmE (PRK00019)Ribosomal_L31 (pfam01197)→ |
| Eco-929-1 | NZ_KB004465.1 | + | 5425 | 5477 | **RNA**→ | rpmE (PRK00019)Ribosomal_L31 (pfam01197)→ |
| Eco-867-1 | NZ_KB000427.1 | + | 91685 | 91737 | **RNA**→ | rpmE (PRK00019)Ribosomal_L31 (pfam01197)→ |
| Eco-931-1 | NZ_KB004956.1 | - | 85517 | 85465 | **RNA**→ | rpmE (PRK00019)Ribosomal_L31 (pfam01197)→ |
| Eco-1589-1 | NZ_KB732234.1 | + | 199572 | 199624 | **RNA**→ | rpmE (PRK00019)Ribosomal_L31 (pfam01197)→ |
| Eco-1284-1 | NZ_KE700830.1 | + | 3082190 | 3082242 | **RNA**→ | rpmE (PRK00019)Ribosomal_L31 (pfam01197)→ |
| Eco-1363-1 | NZ_KE699552.1 | + | 1425559 | 1425611 | **RNA**→ | rpmE (PRK00019)Ribosomal_L31 (pfam01197)→ |
| Eco-1193-1 | NZ_KE699732.1 | + | 80106 | 80158 | **RNA**→ | rpmE (PRK00019)Ribosomal_L31 (pfam01197)→ |
| Eco-1215-1 | NZ_KE700025.1 | + | 3022754 | 3022806 | **RNA**→ | rpmE (PRK00019)Ribosomal_L31 (pfam01197)→ |
| Eco-1299-1 | NZ_KE700948.1 | + | 80532 | 80584 | **RNA**→ | rpmE (PRK00019)Ribosomal_L31 (pfam01197)→ |
| Eco-2064-1 | NZ_KE701659.1 | + | 2188141 | 2188193 | **RNA**→ | rpmE (PRK00019)Ribosomal_L31 (pfam01197)→ |
| Eco-2090-1 | NZ_KE702030.1 | + | 1589080 | 1589132 | **RNA**→ | rpmE (PRK00019)Ribosomal_L31 (pfam01197)→ |
| Eco-1310-1 | NZ_KE701076.1 | + | 81617 | 81669 | **RNA**→ | rpmE (PRK00019)Ribosomal_L31 (pfam01197)→ |
| Eco-1265-1 | NZ_KE700589.1 | + | 80503 | 80555 | **RNA**→ | rpmE (PRK00019)Ribosomal_L31 (pfam01197)→ |
| Eco-1288-1 | NZ_KE700850.1 | + | 4139451 | 4139503 | **RNA**→ | rpmE (PRK00019)Ribosomal_L31 (pfam01197)→ |
| Eco-1281-1 | NZ_KE700802.1 | + | 82469 | 82521 | **RNA**→ | rpmE (PRK00019)Ribosomal_L31 (pfam01197)→ |
| Eco-2132-1 | NZ_KE702477.1 | + | 721545 | 721597 | **RNA**→ | rpmE (PRK00019)Ribosomal_L31 (pfam01197)→ |
| Eco-1305-1 | NZ_KE701022.1 | + | 429776 | 429828 | **RNA**→ | rpmE (PRK00019)Ribosomal_L31 (pfam01197)→ |
| Eco-1398-1 | NZ_KE701238.1 | + | 81941 | 81993 | **RNA**→ | rpmE (PRK00019)Ribosomal_L31 (pfam01197)→ |
| Eco-1277-1 | NZ_KE700778.1 | + | 2969295 | 2969347 | **RNA**→ | rpmE (PRK00019)Ribosomal_L31 (pfam01197)→ |
| Eco-2050-1 | NZ_KE701458.1 | + | 4128279 | 4128331 | **RNA**→ | rpmE (PRK00019)Ribosomal_L31 (pfam01197)→ |
| Eco-1290-1 | NZ_KE700866.1 | + | 3081361 | 3081413 | **RNA**→ | rpmE (PRK00019)Ribosomal_L31 (pfam01197)→ |
| Eco-1207-1 | NZ_KE699929.1 | + | 81923 | 81975 | **RNA**→ | rpmE (PRK00019)Ribosomal_L31 (pfam01197)→ |
| Eco-1276-1 | NZ_KE700763.1 | + | 40914 | 40966 | **RNA**→ | rpmE (PRK00019)Ribosomal_L31 (pfam01197)→ |
| Eco-2129-1 | NZ_KE702454.1 | + | 3004717 | 3004769 | **RNA**→ | rpmE (PRK00019)Ribosomal_L31 (pfam01197)→ |
| Eco-2089-1 | NZ_KE702022.1 | + | 82164 | 82216 | **RNA**→ | rpmE (PRK00019)Ribosomal_L31 (pfam01197)→ |
| Eco-2145-1 | NZ_KE702629.1 | - | 52361 | 52309 | **RNA**→ | rpmE (PRK00019)Ribosomal_L31 (pfam01197)→ |
| Eco-2147-1 | NZ_KE702650.1 | + | 2279303 | 2279355 | **RNA**→ | rpmE (PRK00019)Ribosomal_L31 (pfam01197)→ |
| Eco-1224-1 | NZ_KE700110.1 | + | 3032185 | 3032237 | **RNA**→ | rpmE (PRK00019)Ribosomal_L31 (pfam01197)→ |
| Eco-1298-1 | NZ_KE700927.1 | + | 81739 | 81791 | **RNA**→ | rpmE (PRK00019)Ribosomal_L31 (pfam01197)→ |
| Eco-1256-1 | NZ_KE700501.1 | + | 1510971 | 1511023 | **RNA**→ | rpmE (PRK00019)Ribosomal_L31 (pfam01197)→ |
| Eco-2074-1 | NZ_KE701779.1 | + | 81844 | 81896 | **RNA**→ | rpmE (PRK00019)Ribosomal_L31 (pfam01197)→ |
| Eco-2138-1 | NZ_KE702517.1 | + | 81885 | 81937 | **RNA**→ | rpmE (PRK00019)Ribosomal_L31 (pfam01197)→ |
| Eco-2109-1 | NZ_KE702295.1 | + | 202832 | 202884 | **RNA**→ | rpmE (PRK00019)Ribosomal_L31 (pfam01197)→ |
| Eco-1191-1 | NZ_KE747793.1 | + | 1499856 | 1499908 | **RNA**→ | rpmE (PRK00019)Ribosomal_L31 (pfam01197)→ |
| Eco-441-1 | NZ_KI303532.1 | + | 91677 | 91729 | **RNA**→ | rpmE (PRK00019)Ribosomal_L31 (pfam01197)→ |
| Eco-422-1 | NZ_KI303976.1 | + | 91676 | 91728 | **RNA**→ | rpmE (PRK00019)Ribosomal_L31 (pfam01197)→ |
| Eco-442-1 | NZ_KI303614.1 | + | 119771 | 119823 | **RNA**→ | rpmE (PRK00019)Ribosomal_L31 (pfam01197)→ |
| Eco-446-1 | NZ_KI302222.1 | + | 91843 | 91895 | **RNA**→ | rpmE (PRK00019)Ribosomal_L31 (pfam01197)→ |
| Eco-431-1 | NZ_KI302783.1 | + | 79232 | 79284 | **RNA**→ | rpmE (PRK00019)Ribosomal_L31 (pfam01197)→ |
| Eco-1566-1 | NZ_KE136719.1 | + | 78966 | 79018 | **RNA**→ | rpmE (PRK00019)Ribosomal_L31 (pfam01197)→ |
| Eco-1235-1 | NZ_KI538716.1 | + | 1125873 | 1125925 | **RNA**→ | rpmE (PRK00019)Ribosomal_L31 (pfam01197)→ |
| Eco-1263-1 | NZ_KI538784.1 | + | 1554820 | 1554872 | **RNA**→ | rpmE (PRK00019)Ribosomal_L31 (pfam01197)→ |
| Eco-1209-1 | NZ_KI538675.1 | + | 3015588 | 3015640 | **RNA**→ | rpmE (PRK00019)Ribosomal_L31 (pfam01197)→ |
| Eco-1302-1 | NZ_KI669364.1 | + | 77079 | 77131 | **RNA**→ | rpmE (PRK00019)Ribosomal_L31 (pfam01197)→ |
| Eco-307-1 | NZ_KI929791.1 | + | 2901756 | 2901808 | **RNA**→ | rpmE (PRK00019)Ribosomal_L31 (pfam01197)→ |
| Eco-306-1 | NZ_KI929797.1 | + | 2884603 | 2884655 | **RNA**→ | rpmE (PRK00019)Ribosomal_L31 (pfam01197)→ |
| Eco-761-1 | NZ_KB453217.1 | + | 91679 | 91731 | **RNA**→ | rpmE (PRK00019)Ribosomal_L31 (pfam01197)→ |
| Eco-1598-1 | NZ_KB732740.1 | + | 81850 | 81902 | **RNA**→ | rpmE (PRK00019)Ribosomal_L31 (pfam01197)→ |
| Eco-1599-1 | NZ_KB733044.1 | + | 202007 | 202059 | **RNA**→ | rpmE (PRK00019)Ribosomal_L31 (pfam01197)→ |

| | | | | | |
|---|---|---|---|---|---|
| Eco-1595-1 | NZ_KB732701.1 | + | 77074 | 77126 | **RNA**→ rpmE (PRK00019)Ribosomal_L31 (pfam01197)→ |
| Eco-1542-1 | NZ_KB733023.1 | + | 4117024 | 4117076 | **RNA**→ rpmE (PRK00019)Ribosomal_L31 (pfam01197)→ |
| Eco-1627-1 | NZ_KB733107.1 | + | 91628 | 91680 | **RNA**→ rpmE (PRK00019)Ribosomal_L31 (pfam01197)→ |
| Eco-1552-1 | NZ_KB733235.1 | + | 3048568 | 3048620 | **RNA**→ rpmE (PRK00019)Ribosomal_L31 (pfam01197)→ |
| Eco-1603-1 | NZ_KB732751.1 | - | 1572177 | 1572125 | **RNA**→ rpmE (PRK00019)Ribosomal_L31 (pfam01197)→ |
| Eco-1606-1 | NZ_KB733066.1 | + | 3009872 | 3009924 | **RNA**→ rpmE (PRK00019)Ribosomal_L31 (pfam01197)→ |
| Eco-1433-1 | NZ_KB733111.1 | + | 201057 | 201109 | **RNA**→ rpmE (PRK00019)Ribosomal_L31 (pfam01197)→ |
| Eco-1578-1 | NZ_KB732224.1 | + | 200445 | 200497 | **RNA**→ rpmE (PRK00019)Ribosomal_L31 (pfam01197)→ |
| Eco-1439-1 | NZ_KB733071.1 | + | 1502061 | 1502113 | **RNA**→ rpmE (PRK00019)Ribosomal_L31 (pfam01197)→ |
| Eco-1550-1 | NZ_KB733358.1 | + | 391531 | 391583 | **RNA**→ rpmE (PRK00019)Ribosomal_L31 (pfam01197)→ |
| Eco-1569-1 | NZ_KE136581.1 | + | 2924005 | 2924057 | **RNA**→ rpmE (PRK00019)Ribosomal_L31 (pfam01197)→ |
| Eco-1611-1 | NZ_KE136591.1 | + | 1576213 | 1576265 | **RNA**→ rpmE (PRK00019)Ribosomal_L31 (pfam01197)→ |
| Eco-1563-1 | NZ_KE137004.1 | + | 4441094 | 4441146 | **RNA**→ rpmE (PRK00019)Ribosomal_L31 (pfam01197)→ |
| Eco-1274-1 | NZ_KE698996.1 | + | 82952 | 83004 | **RNA**→ rpmE (PRK00019)Ribosomal_L31 (pfam01197)→ |
| Eco-1341-1 | NZ_KE698838.1 | + | 202726 | 202778 | **RNA**→ rpmE (PRK00019)Ribosomal_L31 (pfam01197)→ |
| Eco-1359-1 | NZ_KE699460.1 | + | 1555547 | 1555599 | **RNA**→ rpmE (PRK00019)Ribosomal_L31 (pfam01197)→ |
| Eco-1348-1 | NZ_KE699341.1 | + | 38316 | 38368 | **RNA**→ rpmE (PRK00019)Ribosomal_L31 (pfam01197)→ |
| Eco-1357-1 | NZ_KE699437.1 | + | 1585903 | 1585955 | **RNA**→ rpmE (PRK00019)Ribosomal_L31 (pfam01197)→ |
| Eco-1188-1 | NZ_KE698778.1 | + | 746991 | 747043 | **RNA**→ rpmE (PRK00019)Ribosomal_L31 (pfam01197)→ |
| Eco-1327-1 | NZ_KE699160.1 | + | 205160 | 205212 | **RNA**→ rpmE (PRK00019)Ribosomal_L31 (pfam01197)→ |
| Eco-1325-1 | NZ_KE699137.1 | + | 196255 | 196307 | **RNA**→ rpmE (PRK00019)Ribosomal_L31 (pfam01197)→ |
| Eco-1340-1 | NZ_KE699296.1 | + | 79965 | 80017 | **RNA**→ rpmE (PRK00019)Ribosomal_L31 (pfam01197)→ |
| Eco-1322-1 | NZ_KE698817.1 | + | 223232 | 223284 | **RNA**→ rpmE (PRK00019)Ribosomal_L31 (pfam01197)→ |
| Eco-1323-1 | NZ_KE699123.1 | + | 204897 | 204949 | **RNA**→ rpmE (PRK00019)Ribosomal_L31 (pfam01197)→ |
| Eco-1335-1 | NZ_KE699242.1 | + | 741196 | 741248 | **RNA**→ rpmE (PRK00019)Ribosomal_L31 (pfam01197)→ |
| Eco-1214-1 | NZ_KE700005.1 | - | 52165 | 52113 | **RNA**→ rpmE (PRK00019)Ribosomal_L31 (pfam01197)→ |
| Eco-1951-1 | NZ_JH708009.1 | + | 91689 | 91741 | **RNA**→ rpmE (PRK00019)Ribosomal_L31 (pfam01197)→ |
| Eco-1159-1 | NZ_JH710487.1 | + | 91681 | 91733 | **RNA**→ rpmE (PRK00019)Ribosomal_L31 (pfam01197)→ |
| Eco-1947-1 | NZ_JH709052.1 | + | 91683 | 91735 | **RNA**→ rpmE (PRK00019)Ribosomal_L31 (pfam01197)→ |
| Eco-1136-1 | NZ_JH697709.1 | + | 91677 | 91729 | **RNA**→ rpmE (PRK00019)Ribosomal_L31 (pfam01197)→ |
| Eco-1142-1 | NZ_JH693924.1 | + | 91683 | 91735 | **RNA**→ rpmE (PRK00019)Ribosomal_L31 (pfam01197)→ |
| Eco-1157-1 | NZ_JH701983.1 | + | 91683 | 91735 | **RNA**→ rpmE (PRK00019)Ribosomal_L31 (pfam01197)→ |
| Eco-1950-1 | NZ_JH708588.1 | + | 91759 | 91811 | **RNA**→ rpmE (PRK00019)Ribosomal_L31 (pfam01197)→ |
| Eco-2029-1 | NZ_JH700687.1 | + | 8195 | 8247 | **RNA**→ rpmE (PRK00019)Ribosomal_L31 (pfam01197)→ |
| Eco-1130-1 | NZ_JH945202.1 | + | 91757 | 91809 | **RNA**→ rpmE (PRK00019)Ribosomal_L31 (pfam01197)→ |
| Eco-1711-1 | NZ_JH960537.1 | + | 91749 | 91801 | **RNA**→ rpmE (PRK00019)Ribosomal_L31 (pfam01197)→ |
| Eco-1128-1 | NZ_JH944742.1 | + | 88417 | 88469 | **RNA**→ rpmE (PRK00019)Ribosomal_L31 (pfam01197)→ |
| Eco-917-1 | NZ_KB009442.1 | + | 91687 | 91739 | **RNA**→ rpmE (PRK00019)Ribosomal_L31 (pfam01197)→ |
| Eco-871-1 | NZ_KB007779.1 | + | 88442 | 88494 | **RNA**→ rpmE (PRK00019)Ribosomal_L31 (pfam01197)→ |
| Eco-926-1 | NZ_KB008854.1 | + | 88429 | 88481 | **RNA**→ rpmE (PRK00019)Ribosomal_L31 (pfam01197)→ |
| Eco-1161-1 | NZ_JH961766.1 | + | 91689 | 91741 | **RNA**→ rpmE (PRK00019)Ribosomal_L31 (pfam01197)→ |
| Eco-1164-1 | NZ_JH943934.1 | + | 91767 | 91819 | **RNA**→ rpmE (PRK00019)Ribosomal_L31 (pfam01197)→ |
| Eco-862-1 | NZ_JH970100.1 | + | 91671 | 91723 | **RNA**→ rpmE (PRK00019)Ribosomal_L31 (pfam01197)→ |
| Eco-1974-1 | NZ_JH959572.1 | + | 91693 | 91745 | **RNA**→ rpmE (PRK00019)Ribosomal_L31 (pfam01197)→ |
| Eco-839-1 | NZ_AOET01000116.1 | + | 91981 | 92033 | **RNA**→ rpmE (PRK00019)Ribosomal_L31 (pfam01197)→ |
| Eco-1498-1 | NZ_ANYR01000076.1 | + | 77184 | 77236 | **RNA**→ rpmE (PRK00019)Ribosomal_L31 (pfam01197)→ |
| Eco-953-1 | NZ_AOEH01000082.1 | - | 4113 | 4061 | **RNA**→ rpmE (PRK00019)Ribosomal_L31 (pfam01197)→ |
| Eco-1493-1 | NZ_ANYO01000058.1 | + | 29022 | 29074 | **RNA**→ rpmE (PRK00019)Ribosomal_L31 (pfam01197)→ |
| Eco-938-1 | NZ_AOEB01000174.1 | + | 23013 | 23065 | **RNA**→ rpmE (PRK00019)Ribosomal_L31 (pfam01197)→ |
| Eco-1952-1 | NZ_AOEJ01000126.1 | + | 91844 | 91896 | **RNA**→ rpmE (PRK00019)Ribosomal_L31 (pfam01197)→ |
| Eco-1948-1 | NZ_AOEI01000113.1 | + | 91787 | 91839 | **RNA**→ rpmE (PRK00019)Ribosomal_L31 (pfam01197)→ |
| Eco-909-1 | NZ_AOEU01000182.1 | + | 55101 | 55153 | **RNA**→ rpmE (PRK00019)Ribosomal_L31 (pfam01197)→ |

| | | | | | |
|---|---|---|---|---|---|
| Eco-936-1 | NZ_AODZ01000257.1 | + | 44648 | 44700 | **RNA**→ rpmE (PRK00019)Ribosomal_L31 (pfam01197)→ |
| Eco-1964-1 | NZ_AOES01000010.1 | - | 4113 | 4061 | **RNA**→ rpmE (PRK00019)Ribosomal_L31 (pfam01197)→ |
| Eco-671-1 | NZ_CP010304.1 | + | 4948099 | 4948151 | **RNA**→ rpmE (PRK00019)Ribosomal_L31 (pfam01197)→ |
| Eco-290-1 | NZ_CP005930.1 | + | 2266617 | 2266669 | **RNA**→ rpmE (PRK00019)Ribosomal_L31 (pfam01197)→ |
| Eco-1158-1 | NZ_JH701513.1 | + | 91691 | 91743 | **RNA**→ rpmE (PRK00019)Ribosomal_L31 (pfam01197)→ |
| Eco-1979-1 | NC_022370.1 | + | 4435942 | 4435994 | **RNA**→ rpmE (PRK00019)Ribosomal_L31 (pfam01197)→ |
| Eco-621-1 | NZ_CP008957.1 | + | 5016072 | 5016124 | **RNA**→ rpmE (PRK00019)Ribosomal_L31 (pfam01197)→ |
| Eco-899-1 | NZ_KI534620.1 | + | 8157 | 8209 | **RNA**→ rpmE (PRK00019)Ribosomal_L31 (pfam01197)→ |
| Esp-52-1 | NZ_KQ235739.1 | + | 77226 | 77278 | **RNA**→ rpmE (PRK00019)Ribosomal_L31 (pfam01197)→ |
| Eco-393-1 | NZ_KK583188.1 | - | 330264 | 330212 | **RNA**→ rpmE (PRK00019)Ribosomal_L31 (pfam01197)→ |
| Eco-1184-1 | NZ_HE572566.1 | - | 2016950 | 2016898 | **RNA**→ rpmE (PRK00019)Ribosomal_L31 (pfam01197)→ |
| Eco-632-1 | NZ_CAOT01000021.1 | + | 91956 | 92008 | **RNA**→ rpmE (PRK00019)Ribosomal_L31 (pfam01197)→ |
| Eco-834-1 | NZ_AVRR01000119.1 | + | 44656 | 44708 | **RNA**→ rpmE (PRK00019)Ribosomal_L31 (pfam01197)→ |
| Eco-429-1 | NZ_AWPM01000182.1 | + | 12739 | 12791 | **RNA**→ rpmE (PRK00019)Ribosomal_L31 (pfam01197)→ |
| Eco-450-1 | NZ_AWPT01000222.1 | + | 13300 | 13352 | **RNA**→ rpmE (PRK00019)Ribosomal_L31 (pfam01197)→ |
| Eco-612-1 | NZ_JHKF01000045.1 | + | 91627 | 91679 | **RNA**→ rpmE (PRK00019)Ribosomal_L31 (pfam01197)→ |
| Eco-664-1 | NZ_JHHZ01000024.1 | - | 911 | 859 | **RNA**→ rpmE (PRK00019)Ribosomal_L31 (pfam01197)→ |
| Eco-571-1 | NZ_JHMD01000021.1 | + | 91540 | 91592 | **RNA**→ rpmE (PRK00019)Ribosomal_L31 (pfam01197)→ |
| Eco-1165-1 | NZ_AMTP01000117.1 | + | 92274 | 92326 | **RNA**→ rpmE (PRK00019)Ribosomal_L31 (pfam01197)→ |
| Eco-1066-1 | NZ_AIFE01000058.1 | + | 96821 | 96873 | **RNA**→ rpmE (PRK00019)Ribosomal_L31 (pfam01197)→ |
| Eco-1068-1 | NZ_AIFG01000065.1 | + | 94325 | 94377 | **RNA**→ rpmE (PRK00019)Ribosomal_L31 (pfam01197)→ |
| Eco-1073-1 | NZ_AIFL01000073.1 | + | 95249 | 95301 | **RNA**→ rpmE (PRK00019)Ribosomal_L31 (pfam01197)→ |
| Eco-925-1 | NZ_ANMC01000254.1 | + | 4802 | 4854 | **RNA**→ rpmE (PRK00019)Ribosomal_L31 (pfam01197)→ |
| Eco-942-1 | NZ_AOEE01000236.1 | + | 7965 | 8017 | **RNA**→ rpmE (PRK00019)Ribosomal_L31 (pfam01197)→ |
| env-784 | NLF014_scaffold17740_1 | + | 423 | 475 | **RNA**→ rpmE (PRK00019)Ribosomal_L31 (pfam01197)→ |
| env-785 | 2222084013.827_7 | - | 639 | 587 | **RNA**→ ←hypo |
| Esp-79-1 | NZ_AEME01000001.1 | + | 879790 | 879842 | **RNA**→ rpmE (PRK00019)Ribosomal_L31 (pfam01197)→ rpmE (PRK00019)Ribosomal_L31 (pfam01197)→ |
| Esp-77-1 | NZ_AEJW01000018.1 | + | 15607 | 15659 | **RNA**→ rpmE (PRK00019)Ribosomal_L31 (pfam01197)→ |
| Esp-53-1 | NZ_AEJZ01000171.1 | - | 17689 | 17637 | **RNA**→ rpmE (PRK00019)Ribosomal_L31 (pfam01197)→ |
| Esp-78-1 | NZ_AEJV01000033.1 | + | 22157 | 22209 | **RNA**→ rpmE (PRK00019)Ribosomal_L31 (pfam01197)→ |
| Eco-15-1 | NZ_JHDF01000032.1 | - | 51954 | 51902 | **RNA**→ rpmE (PRK00019)Ribosomal_L31 (pfam01197)→ |
| Eco-1425-1 | NZ_KB732249.1 | + | 3306906 | 3306958 | **RNA**→ rpmE (PRK00019)Ribosomal_L31 (pfam01197)→ |
| Eco-1410-1 | NZ_KE701379.1 | - | 4263807 | 4263755 | **RNA**→ rpmE (PRK00019)Ribosomal_L31 (pfam01197)→ |
| Eco-1475-1 | NZ_KE137293.1 | - | 53399 | 53347 | **RNA**→ rpmE (PRK00019)Ribosomal_L31 (pfam01197)→ |
| Eco-1571-1 | NZ_KB976077.1 | - | 52047 | 51995 | **RNA**→ rpmE (PRK00019)Ribosomal_L31 (pfam01197)→ |
| Eco-1639-1 | NZ_KE137149.1 | - | 39308 | 39256 | **RNA**→ rpmE (PRK00019)Ribosomal_L31 (pfam01197)→ |
| Eco-1429-1 | NZ_KE136626.1 | - | 1021432 | 1021380 | **RNA**→ rpmE (PRK00019)Ribosomal_L31 (pfam01197)→ |
| Eco-1570-1 | NZ_KE136726.1 | - | 1008527 | 1008475 | **RNA**→ rpmE (PRK00019)Ribosomal_L31 (pfam01197)→ |
| Eco-1592-1 | NZ_KE138680.1 | + | 1352301 | 1352353 | **RNA**→ rpmE (PRK00019)Ribosomal_L31 (pfam01197)→ |
| Ema-1-1 | NZ_KN714332.1 | - | 33004 | 32952 | **RNA**→ rpmE (PRK00019)Ribosomal_L31 (pfam01197)→ |
| Cro-2-1 | NZ_BBNA01000079.1 | + | 8219 | 8271 | **RNA**→ rpmE (PRK00019)Ribosomal_L31 (pfam01197)→ |
| Cro-1-1 | NZ_JXUN01000174.1 | + | 2944 | 2996 | **RNA**→ rpmE (PRK00019)Ribosomal_L31 (pfam01197)→ |
| Cro-3-1 | NC_013716.1 | - | 4016831 | 4016779 | **RNA**→ rpmE (PRK00019)Ribosomal_L31 (pfam01197)→ |
| Eal-2-1 | NZ_BBMY01000009.1 | - | 42980 | 42928 | **RNA**→ rpmE (PRK00019)Ribosomal_L31 (pfam01197)→ |
| Eal-4-1 | NZ_AEJU01000166.1 | + | 12329 | 12381 | **RNA**→ rpmE (PRK00019)Ribosomal_L31 (pfam01197)→ |
| Eal-5-1 | NZ_AEJY01000264.1 | - | 1037 | 985 | **RNA**→ rpmE (PRK00019)Ribosomal_L31 (pfam01197)→ |
| Eal-1-1 | NZ_CP007025.1 | - | 2095964 | 2095912 | **RNA**→ rpmE (PRK00019)Ribosomal_L31 (pfam01197)→ |
| Eal-3-1 | NZ_CH991859.1 | + | 1832703 | 1832755 | **RNA**→ rpmE (PRK00019)Ribosomal_L31 (pfam01197)→ |
| Sso-1-5 | NZ_CWUD01000027.1 | + | 39075 | 39127 | **RNA**→ rpmE (PRK00019)Ribosomal_L31 (pfam01197)→ |
| Eco-835-1 | NZ_AFAE02000014.1 | - | 44473 | 44421 | **RNA**→ rpmE (PRK00019)Ribosomal_L31 (pfam01197)→ |
| Ban-1-1 | NZ_ABOT01000022.1 | - | 97 | 45 | **RNA**→ hypo→ |
| Eco-1442-1 | NZ_KE137230.1 | + | 202992 | 203044 | **RNA**→ rpmE (PRK00019)Ribosomal_L31 (pfam01197)→ |

| | | | | | |
|---|---|---|---|---|---|
| Eco-1586-1 | NZ_KB732639.1 | + | 82570 | 82622 | RNA→ rpmE (PRK00019)Ribosomal_L31 (pfam01197)→ |
| Cyo-1-1 | NZ_GG730300.1 | + | 190031 | 190083 | RNA→ rpmE (PRK00019)Ribosomal_L31 (pfam01197)→ |
| Eco-763-1 | NZ_AEZI02000021.1 | + | 1102773 | 1102825 | RNA→ rpmE (PRK00019)Ribosomal_L31 (pfam01197)→ |
| Efe-1-1 | NZ_JZWP01000017.1 | - | 45156 | 45104 | RNA→ rpmE (PRK00019)Ribosomal_L31 (pfam01197)→ |
| Efe-1-2 | NZ_JZWQ01000024.1 | + | 86602 | 86654 | RNA→ rpmE (PRK00019)Ribosomal_L31 (pfam01197)→ |
| Efe-1-3 | NZ_JZWN01000028.1 | + | 83564 | 83616 | RNA→ rpmE (PRK00019)Ribosomal_L31 (pfam01197)→ |
| Esp-54-1 | NZ_AEJX01000050.1 | + | 12223 | 12275 | RNA→ rpmE (PRK00019)Ribosomal_L31 (pfam01197)→ |
| Eco-84-1 | NZ_JNPK01000044.1 | + | 90831 | 90883 | RNA→ rpmE (PRK00019)Ribosomal_L31 (pfam01197)→ |
| Eco-1655-1 | NZ_GL872050.1 | + | 91674 | 91726 | RNA→ rpmE (PRK00019)Ribosomal_L31 (pfam01197)→ |
| Efe-3-1 | NZ_GL870859.1 | - | 45478 | 45426 | RNA→ rpmE (PRK00019)Ribosomal_L31 (pfam01197)→ |
| Eco-1989-1 | NZ_AEXD01000024.1 | + | 95154 | 95206 | RNA→ rpmE (PRK00019)Ribosomal_L31 (pfam01197)→ |
| Efe-2-1 | NC_011740.1 | - | 3932957 | 3932905 | RNA→ rpmE (PRK00019)Ribosomal_L31 (pfam01197)→ |
| Efe-4-1 | NZ_CM001142.1 | - | 3881886 | 3881834 | RNA→ rpmE (PRK00019)Ribosomal_L31 (pfam01197)→ |
| Sdy-5-1 | NZ_LAIG01000046.1 | - | 686 | 634 | RNA→ rpmE (PRK00019)Ribosomal_L31 (pfam01197)→ |
| Sdy-6-1 | NZ_AERM01000110.1 | + | 14260 | 14312 | RNA→ rpmE (PRK00019)Ribosomal_L31 (pfam01197)→ |
| env-786 | scaffold88732_1_MH0014 | + | 177 | 229 | RNA→ rpmE (PRK00019)Ribosomal_L31 (pfam01197)→ |
| env-787 | scaffold94036_1_MH0025 | - | 440 | 388 | RNA→ rpmE (PRK00019)Ribosomal_L31 (pfam01197)→ |
| env-788 | C1998588_1_MH0079 | - | 203 | 151 | RNA→ ←hypo |
| env-789 | scaffold1530_9_O2.UC-18 | + | 19529 | 19581 | RNA→ ←hypo |
| env-790 | scaffold8769_24_O2.UC-23 | + | 453 | 505 | RNA→ rpmE (PRK00019)Ribosomal_L31 (pfam01197)→ |
| env-791 | scaffold39006_2_V1.UC-17 | + | 567 | 619 | RNA→ ←hypo |
| env-792 | 4491404.3_NODE_67640 | - | 319 | 267 | RNA→ rpmE (PRK00019)Ribosomal_L31 (pfam01197)→ |
| env-793 | 4491406.3_NODE_5404 | + | 85961 | 86013 | RNA→ rpmE (PRK00019)Ribosomal_L31 (pfam01197)→ |
| env-794 | 4491409.3_NODE_11159 | - | 722 | 670 | RNA→ rpmE (PRK00019)Ribosomal_L31 (pfam01197)→ |
| env-795 | 4491410.3_NODE_23735 | + | 163 | 215 | RNA→ ←hypo |
| env-796 | 4491411.3_NODE_14623 | + | 66007 | 66059 | RNA→ rpmE (PRK00019)Ribosomal_L31 (pfam01197)→ |
| env-797 | 4491420.3_NODE_260 | + | 83392 | 83444 | RNA→ rpmE (PRK00019)Ribosomal_L31 (pfam01197)→ |
| env-798 | 4491479.3_NODE_3717 | + | 11526 | 11578 | RNA→ rpmE (PRK00019)Ribosomal_L31 (pfam01197)→ |
| env-799 | 4491482.3_NODE_114102 | + | 1175 | 1227 | RNA→ rpmE (PRK00019)Ribosomal_L31 (pfam01197)→ |
| env-800 | 4491484.3_NODE_12754 | + | 80216 | 80268 | RNA→ rpmE (PRK00019)Ribosomal_L31 (pfam01197)→ |
| env-801 | MC6.MG3.AS1.C42510 | + | 670 | 722 | RNA→ rpmE (PRK00019)Ribosomal_L31 (pfam01197)→ |
| env-802 | SRS018313_C7874553 | - | 263 | 211 | RNA→ rpmE (PRK00019)Ribosomal_L31 (pfam01197)→ |
| env-803 | SRS057478_LANL_scaffold_18323 | + | 764 | 816 | RNA→ ←hypo |
| env-804 | DLF002_scaffold11095_3 | - | 441 | 389 | RNA→ rpmE (PRK00019)Ribosomal_L31 (pfam01197)→ |
| env-805 | DLF006_scaffold8915_21 | - | 999 | 947 | RNA→ rpmE (PRK00019)Ribosomal_L31 (pfam01197)→ |
| env-806 | DLF008_scaffold4013_1 | - | 452 | 400 | RNA→ rpmE (PRK00019)Ribosomal_L31 (pfam01197)→ |
| env-807 | DLF009_scaffold6274_1 | + | 32314 | 32366 | RNA→ rpmE (PRK00019)Ribosomal_L31 (pfam01197)→ |
| env-808 | DLF012_C220366_1 | + | 276 | 328 | RNA→ rpmE (PRK00019)Ribosomal_L31 (pfam01197)→ |
| env-809 | DLF013_scaffold50084_2 | + | 132 | 184 | RNA→ rpmE (PRK00019)Ribosomal_L31 (pfam01197)→ |
| env-810 | DLM005_scaffold10654_5 | + | 248 | 300 | RNA→ rpmE (PRK00019)Ribosomal_L31 (pfam01197)→ |
| env-811 | DLM011_C451971_1 | + | 271 | 323 | RNA→ rpmE (PRK00019)Ribosomal_L31 (pfam01197)→ |
| env-812 | DLM012_scaffold26432_1 | + | 784 | 836 | RNA→ rpmE (PRK00019)Ribosomal_L31 (pfam01197)→ |
| env-813 | DLM022_scaffold90450_1 | - | 6267 | 6215 | RNA→ rpmE (PRK00019)Ribosomal_L31 (pfam01197)→ |
| env-814 | DLM023_scaffold8878_5 | - | 933 | 881 | RNA→ rpmE (PRK00019)Ribosomal_L31 (pfam01197)→ |
| env-815 | DLM024_C119635_1 | - | 578 | 526 | RNA→ rpmE (PRK00019)Ribosomal_L31 (pfam01197)→ |
| env-816 | DLM028_scaffold4352_1 | - | 39427 | 39375 | RNA→ rpmE (PRK00019)Ribosomal_L31 (pfam01197)→ |
| env-817 | DOF004_scaffold50245_1 | - | 473 | 421 | RNA→ rpmE (PRK00019)Ribosomal_L31 (pfam01197)→ |
| env-818 | DOF006_C278658_1 | + | 276 | 328 | RNA→ rpmE (PRK00019)Ribosomal_L31 (pfam01197)→ |
| env-819 | DOF007_C334852_1 | - | 319 | 267 | RNA→ rpmE (PRK00019)Ribosomal_L31 (pfam01197)→ |
| env-820 | DOF010_scaffold4818_8 | + | 1645 | 1697 | RNA→ rpmE (PRK00019)Ribosomal_L31 (pfam01197)→ |
| env-821 | DOF012_scaffold101_4 | - | 364 | 312 | RNA→ rpmE (PRK00019)Ribosomal_L31 (pfam01197)→ |
| env-822 | DOM003_scaffold11449_1 | - | 39427 | 39375 | RNA→ rpmE (PRK00019)Ribosomal_L31 (pfam01197)→ |

| | | | | | |
|---|---|---|---|---|---|
| env-823 | DOM008_scaffold1034_13 | + | 1472 | 1524 | **RNA**→ rpmE (PRK00019)Ribosomal_L31 (pfam01197)→ |
| env-824 | DOM016_scaffold5749_2 | + | 1289 | 1341 | **RNA**→ rpmE (PRK00019)Ribosomal_L31 (pfam01197)→ |
| env-825 | NLF008_scaffold4821_2 | - | 516 | 464 | **RNA**→ rpmE (PRK00019)Ribosomal_L31 (pfam01197)→ |
| env-826 | NLM002_C331626_1 | + | 159 | 211 | **RNA**→ rpmE (PRK00019)Ribosomal_L31 (pfam01197)→ |
| env-827 | NLM003_scaffold29307_1 | - | 621 | 569 | **RNA**→ rpmE (PRK00019)Ribosomal_L31 (pfam01197)→ |
| env-828 | NLM006_scaffold26360_2 | + | 30174 | 30226 | **RNA**→ rpmE (PRK00019)Ribosomal_L31 (pfam01197)→ |
| env-829 | NLM015_scaffold10348_11 | + | 1248 | 1300 | **RNA**→ rpmE (PRK00019)Ribosomal_L31 (pfam01197)→ |
| env-830 | NLM016_scaffold4185_1 | + | 346 | 398 | **RNA**→ rpmE (PRK00019)Ribosomal_L31 (pfam01197)→ |
| env-831 | NLM017_scaffold15483_4 | + | 491 | 543 | **RNA**→ rpmE (PRK00019)Ribosomal_L31 (pfam01197)→ |
| env-832 | NLM022_C583280_1 | + | 276 | 328 | **RNA**→ rpmE (PRK00019)Ribosomal_L31 (pfam01197)→ |
| env-833 | NLM024_C270022_1 | + | 816 | 868 | **RNA**→ rpmE (PRK00019)Ribosomal_L31 (pfam01197)→ |
| env-834 | NLM028_scaffold24732_2 | + | 337 | 389 | **RNA**→ rpmE (PRK00019)Ribosomal_L31 (pfam01197)→ |
| env-835 | NOF002_scaffold5884_5 | + | 505 | 557 | **RNA**→ rpmE (PRK00019)Ribosomal_L31 (pfam01197)→ |
| env-836 | NOF006_scaffold6742_1 | - | 452 | 400 | **RNA**→ rpmE (PRK00019)Ribosomal_L31 (pfam01197)→ |
| env-837 | NOF010_scaffold32693_2 | + | 187 | 239 | **RNA**→ rpmE (PRK00019)Ribosomal_L31 (pfam01197)→ |
| env-838 | NOM008_scaffold26361_2 | + | 421 | 473 | **RNA**→ rpmE (PRK00019)Ribosomal_L31 (pfam01197)→ |
| env-839 | NOM018_scaffold38037_2 | + | 115 | 167 | **RNA**→ rpmE (PRK00019)Ribosomal_L31 (pfam01197)→ |
| env-840 | NOM020_scaffold104_1 | - | 1043 | 991 | **RNA**→ rpmE (PRK00019)Ribosomal_L31 (pfam01197)→ |
| env-841 | NOM025_scaffold24626_1 | - | 474 | 422 | **RNA**→ rpmE (PRK00019)Ribosomal_L31 (pfam01197)→ |
| env-842 | NOM028_scaffold47489_1 | + | 23473 | 23525 | **RNA**→ rpmE (PRK00019)Ribosomal_L31 (pfam01197)→ |
| env-843 | NOM029_scaffold36166_1 | - | 466 | 414 | **RNA**→ rpmE (PRK00019)Ribosomal_L31 (pfam01197)→ |
| env-844 | longitudinal_69_558 | - | 44412 | 44360 | **RNA**→ rpmE (PRK00019)Ribosomal_L31 (pfam01197)→ |
| env-845 | longitudinal_70_173 | - | 546960 | 546908 | **RNA**→ rpmE (PRK00019)Ribosomal_L31 (pfam01197)→ |
| env-846 | longitudinal_71_356 | + | 21299 | 21351 | **RNA**→ rpmE (PRK00019)Ribosomal_L31 (pfam01197)→ |
| env-847 | longitudinal_72_263 | - | 44442 | 44390 | **RNA**→ rpmE (PRK00019)Ribosomal_L31 (pfam01197)→ |
| env-848 | longitudinal_115_82 | + | 20819 | 20871 | **RNA**→ rpmE (PRK00019)Ribosomal_L31 (pfam01197)→ |
| env-849 | longitudinal_116_26 | + | 20918 | 20970 | **RNA**→ rpmE (PRK00019)Ribosomal_L31 (pfam01197)→ |
| env-850 | longitudinal_117_29 | - | 44410 | 44358 | **RNA**→ rpmE (PRK00019)Ribosomal_L31 (pfam01197)→ |
| env-851 | longitudinal_303_178 | - | 44417 | 44365 | **RNA**→ rpmE (PRK00019)Ribosomal_L31 (pfam01197)→ |
| env-852 | longitudinal_304_63 | - | 44415 | 44363 | **RNA**→ rpmE (PRK00019)Ribosomal_L31 (pfam01197)→ |
| env-853 | longitudinal_348_423 | + | 1372 | 1424 | **RNA**→ rpmE (PRK00019)Ribosomal_L31 (pfam01197)→ |
| env-854 | longitudinal_349_45 | + | 32353 | 32405 | **RNA**→ rpmE (PRK00019)Ribosomal_L31 (pfam01197)→ |
| env-855 | PaFGMunAill_c006456 | - | 489 | 437 | **RNA**→ rpmE (PRK00019)Ribosomal_L31 (pfam01197)→ |
| env-856 | 3300001567_10000498 | - | 39514 | 39462 | **RNA**→ rpmE (PRK00019)Ribosomal_L31 (pfam01197)→ |
| env-857 | SRS017191_C1719102 | - | 442 | 390 | **RNA**→ rpmE (PRK00019)Ribosomal_L31 (pfam01197)→ |
| env-858 | SRS017521_Baylor_scaffold_53627 | + | 39694 | 39746 | **RNA**→ rpmE (PRK00019)Ribosomal_L31 (pfam01197)→ |
| env-859 | SRS019601_C1965939 | + | 486 | 538 | **RNA**→ ←hypo |
| env-860 | FGTW_contig17927 | + | 168 | 220 | **RNA**→ rpmE (PRK00019)Ribosomal_L31 (pfam01197)→ |
| Eco-1931-1 | NZ_AQBH01000074.1 | + | 34085 | 34137 | **RNA**→ rpmE (PRK00019)Ribosomal_L31 (pfam01197)→ |
| Eco-1010-1 | NZ_APXM01000157.1 | + | 11559 | 11611 | **RNA**→ rpmE (PRK00019)Ribosomal_L31 (pfam01197)→ |
| Eco-1935-1 | NZ_AQBL01000072.1 | + | 32496 | 32548 | **RNA**→ rpmE (PRK00019)Ribosomal_L31 (pfam01197)→ |
| Eco-1012-1 | NZ_APXN01000126.1 | + | 30077 | 30129 | **RNA**→ rpmE (PRK00019)Ribosomal_L31 (pfam01197)→ |
| Eco-1937-1 | NZ_AQBM01000080.1 | + | 34161 | 34213 | **RNA**→ rpmE (PRK00019)Ribosomal_L31 (pfam01197)→ |
| Eco-1007-1 | NZ_AQFJ01000025.1 | + | 86615 | 86667 | **RNA**→ rpmE (PRK00019)Ribosomal_L31 (pfam01197)→ |
| Eco-1941-1 | NZ_AQBQ01000078.1 | + | 34132 | 34184 | **RNA**→ rpmE (PRK00019)Ribosomal_L31 (pfam01197)→ |
| Eco-1006-1 | NZ_APZM01000121.1 | + | 29881 | 29933 | **RNA**→ rpmE (PRK00019)Ribosomal_L31 (pfam01197)→ |
| Eco-1942-1 | NZ_AQBR01000109.1 | + | 34119 | 34171 | **RNA**→ rpmE (PRK00019)Ribosomal_L31 (pfam01197)→ |
| Eco-1086-1 | NZ_AIGA01000058.1 | + | 91006 | 91058 | **RNA**→ rpmE (PRK00019)Ribosomal_L31 (pfam01197)→ |
| Eco-2167-1 | NZ_AQBU01000262.1 | + | 18685 | 18737 | **RNA**→ rpmE (PRK00019)Ribosomal_L31 (pfam01197)→ |
| Eco-1091-1 | NZ_AIGF01000049.1 | + | 93764 | 93816 | **RNA**→ rpmE (PRK00019)Ribosomal_L31 (pfam01197)→ |
| Eco-2174-1 | NZ_AQBZ01000220.1 | + | 18684 | 18736 | **RNA**→ rpmE (PRK00019)Ribosomal_L31 (pfam01197)→ |
| Eco-831-1 | NZ_AQDC01000103.1 | + | 87700 | 87752 | **RNA**→ rpmE (PRK00019)Ribosomal_L31 (pfam01197)→ |

| | | | | | |
|---|---|---|---|---|---|
| Eco-1060-1 | NZ_AIEY01000064.1 | + | 90155 | 90207 | **RNA**→ rpmE (PRK00019)Ribosomal_L31 (pfam01197)→ |
| Eco-1084-1 | NZ_AIFW01000046.1 | + | 91475 | 91527 | **RNA**→ rpmE (PRK00019)Ribosomal_L31 (pfam01197)→ rpmE (PRK00019)Ribosomal_L31 (pfam01197)→ |
| Eco-1049-1 | NZ_AIHL01000077.1 | + | 101568 | 101620 | **RNA**→ rpmE (PRK00019)Ribosomal_L31 (pfam01197)→ |
| Eco-1943-1 | NZ_AQBS01000074.1 | + | 34100 | 34152 | **RNA**→ rpmE (PRK00019)Ribosomal_L31 (pfam01197)→ |
| Eco-1926-1 | NZ_AQBD01000151.1 | + | 87569 | 87621 | **RNA**→ rpmE (PRK00019)Ribosomal_L31 (pfam01197)→ |
| Eco-1932-1 | NZ_AQBI01000070.1 | + | 34119 | 34171 | **RNA**→ rpmE (PRK00019)Ribosomal_L31 (pfam01197)→ |
| Eco-1005-1 | NZ_AQDA01000087.1 | + | 18694 | 18746 | **RNA**→ rpmE (PRK00019)Ribosomal_L31 (pfam01197)→ |
| Eco-1938-1 | NZ_AQBN01000075.1 | + | 34114 | 34166 | **RNA**→ rpmE (PRK00019)Ribosomal_L31 (pfam01197)→ |
| Eco-1087-1 | NZ_AIGB01000037.1 | + | 93728 | 93780 | **RNA**→ rpmE (PRK00019)Ribosomal_L31 (pfam01197)→ |
| Eco-2168-1 | NZ_AQBV01000151.1 | + | 18698 | 18750 | **RNA**→ rpmE (PRK00019)Ribosomal_L31 (pfam01197)→ |
| Eco-1063-1 | NZ_AIFB01000068.1 | + | 89616 | 89668 | **RNA**→ rpmE (PRK00019)Ribosomal_L31 (pfam01197)→ |
| Eco-2178-1 | NZ_AQCD01000147.1 | + | 18690 | 18742 | **RNA**→ rpmE (PRK00019)Ribosomal_L31 (pfam01197)→ |
| Eco-1064-1 | NZ_AIFC01000041.1 | + | 89813 | 89865 | **RNA**→ rpmE (PRK00019)Ribosomal_L31 (pfam01197)→ |
| Eco-2177-1 | NZ_AQCC01000175.1 | + | 18742 | 18794 | **RNA**→ rpmE (PRK00019)Ribosomal_L31 (pfam01197)→ |
| Eco-1071-1 | NZ_AIFJ01000088.1 | + | 96866 | 96918 | **RNA**→ rpmE (PRK00019)Ribosomal_L31 (pfam01197)→ |
| Eco-1050-1 | NZ_AIHM01000068.1 | + | 30233 | 30285 | **RNA**→ rpmE (PRK00019)Ribosomal_L31 (pfam01197)→ |
| Eco-1927-1 | NZ_AQBE01000065.1 | + | 87685 | 87737 | **RNA**→ rpmE (PRK00019)Ribosomal_L31 (pfam01197)→ |
| Eco-833-1 | NZ_APXK01000246.1 | + | 48768 | 48820 | **RNA**→ rpmE (PRK00019)Ribosomal_L31 (pfam01197)→ |
| Eco-1933-1 | NZ_AQBJ01000075.1 | + | 34205 | 34257 | **RNA**→ rpmE (PRK00019)Ribosomal_L31 (pfam01197)→ |
| Eco-1089-1 | NZ_AIGD01000026.1 | + | 90023 | 90075 | **RNA**→ rpmE (PRK00019)Ribosomal_L31 (pfam01197)→ |
| Eco-1939-1 | NZ_AQBO01000076.1 | + | 32750 | 32802 | **RNA**→ rpmE (PRK00019)Ribosomal_L31 (pfam01197)→ |
| Eco-1057-1 | NZ_AIEV01000058.1 | + | 88320 | 88372 | **RNA**→ rpmE (PRK00019)Ribosomal_L31 (pfam01197)→ |
| Eco-1058-1 | NZ_AIEW01000062.1 | + | 89743 | 89795 | **RNA**→ rpmE (PRK00019)Ribosomal_L31 (pfam01197)→ |
| Eco-2171-1 | NZ_AQBW01000092.1 | + | 18599 | 18651 | **RNA**→ rpmE (PRK00019)Ribosomal_L31 (pfam01197)→ rpmE (PRK00019)Ribosomal_L31 (pfam01197)→ |
| Eco-1065-1 | NZ_AIFD01000051.1 | + | 91758 | 91810 | **RNA**→ rpmE (PRK00019)Ribosomal_L31 (pfam01197)→ |
| Eco-418-1 | NZ_JAQA01000142.1 | - | 430 | 378 | **RNA**→ rpmE (PRK00019)Ribosomal_L31 (pfam01197)→ |
| Sso-7-1 | NZ_AMKL01000066.1 | + | 88372 | 88424 | **RNA**→ rpmE (PRK00019)Ribosomal_L31 (pfam01197)→ |
| Sso-3-1 | NZ_AMKH01000064.1 | + | 1250 | 1302 | **RNA**→ rpmE (PRK00019)Ribosomal_L31 (pfam01197)→ |
| Sdy-7-1 | NZ_AMJS01000102.1 | + | 139660 | 139712 | **RNA**→ rpmE (PRK00019)Ribosomal_L31 (pfam01197)→ |
| Sso-4-1 | NZ_AMKI01000053.1 | + | 675911 | 675963 | **RNA**→ rpmE (PRK00019)Ribosomal_L31 (pfam01197)→ |
| Sbo-17-1 | NZ_AMJX01000076.1 | + | 186189 | 186241 | **RNA**→ rpmE (PRK00019)Ribosomal_L31 (pfam01197)→ |
| Sso-5-1 | NZ_AMKJ01000065.1 | + | 11296 | 11348 | **RNA**→ rpmE (PRK00019)Ribosomal_L31 (pfam01197)→ |
| Sbo-6-1 | NZ_AMKA01000088.1 | + | 82733 | 82785 | **RNA**→ rpmE (PRK00019)Ribosomal_L31 (pfam01197)→ |
| Sbo-11-1 | NZ_AMKG01000054.1 | + | 186704 | 186756 | **RNA**→ rpmE (PRK00019)Ribosomal_L31 (pfam01197)→ |
| Sbo-5-1 | NZ_AMJZ01000079.1 | + | 82295 | 82347 | **RNA**→ rpmE (PRK00019)Ribosomal_L31 (pfam01197)→ |
| Sso-6-1 | NZ_AMKK01000080.1 | + | 88121 | 88173 | **RNA**→ rpmE (PRK00019)Ribosomal_L31 (pfam01197)→ |
| Sbo-10-1 | NZ_AMKE01000113.1 | + | 199814 | 199866 | **RNA**→ rpmE (PRK00019)Ribosomal_L31 (pfam01197)→ |
| Sbo-9-1 | NZ_AMKD01000099.1 | + | 44335 | 44387 | **RNA**→ rpmE (PRK00019)Ribosomal_L31 (pfam01197)→ |
| Sfl-73-1 | NZ_AMKF01000096.1 | + | 14330 | 14382 | **RNA**→ rpmE (PRK00019)Ribosomal_L31 (pfam01197)→ |
| Sso-11-1 | NZ_AMJR01000162.1 | + | 48321 | 48373 | **RNA**→ rpmE (PRK00019)Ribosomal_L31 (pfam01197)→ |
| Sbo-7-1 | NZ_AMKB01000096.1 | + | 114428 | 114480 | **RNA**→ rpmE (PRK00019)Ribosomal_L31 (pfam01197)→ |
| Sfl-77-1 | NZ_AMJT01000150.1 | + | 49168 | 49220 | **RNA**→ rpmE (PRK00019)Ribosomal_L31 (pfam01197)→ |
| Sfl-78-1 | NZ_AMJV01000159.1 | + | 49169 | 49221 | **RNA**→ rpmE (PRK00019)Ribosomal_L31 (pfam01197)→ |
| Sfl-80-1 | NZ_AMJY01000153.1 | + | 131442 | 131494 | **RNA**→ rpmE (PRK00019)Ribosomal_L31 (pfam01197)→ |
| Sfl-75-1 | NZ_AMJP01000130.1 | + | 49169 | 49221 | **RNA**→ rpmE (PRK00019)Ribosomal_L31 (pfam01197)→ |
| Sbo-8-1 | NZ_AMKC01000199.1 | + | 34062 | 34114 | **RNA**→ rpmE (PRK00019)Ribosomal_L31 (pfam01197)→ |
| Sfl-79-1 | NZ_AMJW01000030.1 | + | 35330 | 35382 | **RNA**→ rpmE (PRK00019)Ribosomal_L31 (pfam01197)→ |
| Sbo-16-1 | NZ_AMJU01000250.1 | + | 48467 | 48519 | **RNA**→ rpmE (PRK00019)Ribosomal_L31 (pfam01197)→ |
| Sdy-1-1 | NZ_AMJQ01000036.1 | - | 5806 | 5754 | **RNA**→ rpmE (PRK00019)Ribosomal_L31 (pfam01197)→ |
| Sfl-76-1 | NZ_ANAN01000865.1 | + | 33995 | 34047 | **RNA**→ rpmE (PRK00019)Ribosomal_L31 (pfam01197)→ |
| Eco-994-1 | NZ_AAJT02000014.1 | + | 76738 | 76790 | **RNA**→ rpmE (PRK00019)Ribosomal_L31 (pfam01197)→ rpmE (PRK00019)Ribosomal_L31 (pfam01197)→ |
| Eco-1152-1 | NZ_AAJU02000017.1 | + | 31540 | 31592 | **RNA**→ rpmE (PRK00019)Ribosomal_L31 (pfam01197)→ rpmE (PRK00019)Ribosomal_L31 (pfam01197)→ |

| | | | | | |
|---|---|---|---|---|---|
| Eco-1107-1 | NZ_AAJV02000012.1 | - | 49537 | 49485 | **RNA**→ ←hypo |
| Eco-1103-1 | NZ_AAJW02000005.1 | + | 149007 | 149059 | **RNA**→ rpmE (PRK00019)Ribosomal_L31 (pfam01197)→ rpmE (PRK00019)Ribosomal_L31 (pfam01197)→ |
| Eco-767-1 | NZ_AAMK02000013.1 | - | 39845 | 39793 | **RNA**→ rpmE (PRK00019)Ribosomal_L31 (pfam01197)→ rpmE (PRK00019)Ribosomal_L31 (pfam01197)→ |
| Eco-837-1 | NZ_AFAC02000012.1 | - | 1611362 | 1611310 | **RNA**→ rpmE (PRK00019)Ribosomal_L31 (pfam01197)→ |
| Eco-849-1 | NZ_AAKB02000001.1 | - | 1814208 | 1814156 | **RNA**→ rpmE (PRK00019)Ribosomal_L31 (pfam01197)→ rpmE (PRK00019)Ribosomal_L31 (pfam01197)→ |
| Eco-975-1 | NZ_AAJX02000008.1 | - | 1237 | 1185 | **RNA**→ rpmE (PRK00019)Ribosomal_L31 (pfam01197)→ rpmE (PRK00019)Ribosomal_L31 (pfam01197)→ |
| Eco-1768-1 | NZ_AKAX01000157.1 | - | 651 | 599 | **RNA**→ rpmE (PRK00019)Ribosomal_L31 (pfam01197)→ |
| Eco-468-1 | NZ_JHLL01000039.1 | - | 605 | 553 | **RNA**→ rpmE (PRK00019)Ribosomal_L31 (pfam01197)→ |
| Eco-40-1 | NZ_JOSK01000099.1 | + | 84179 | 84231 | **RNA**→ rpmE (PRK00019)Ribosomal_L31 (pfam01197)→ |
| Eco-956-1 | NZ_AZBZ01000002.1 | - | 450 | 398 | **RNA**→ rpmE (PRK00019)Ribosomal_L31 (pfam01197)→ |
| Eco-2172-1 | NZ_AQBX01000136.1 | + | 18739 | 18791 | **RNA**→ rpmE (PRK00019)Ribosomal_L31 (pfam01197)→ |
| Eco-2179-1 | NZ_AQCE01000128.1 | + | 18671 | 18723 | **RNA**→ rpmE (PRK00019)Ribosomal_L31 (pfam01197)→ |
| Eco-1082-1 | NZ_AIFU01000024.1 | + | 31669 | 31721 | **RNA**→ rpmE (PRK00019)Ribosomal_L31 (pfam01197)→ |
| Eco-1924-1 | NZ_AQBB01000153.1 | + | 87690 | 87742 | **RNA**→ rpmE (PRK00019)Ribosomal_L31 (pfam01197)→ |
| Eco-1037-1 | NZ_AIGZ01000027.1 | + | 125333 | 125385 | **RNA**→ rpmE (PRK00019)Ribosomal_L31 (pfam01197)→ |
| Eco-873-1 | NZ_AFAI02000031.1 | + | 737717 | 737769 | **RNA**→ rpmE (PRK00019)Ribosomal_L31 (pfam01197)→ |
| Eco-1973-1 | NZ_AOEN01000114.1 | + | 91791 | 91843 | **RNA**→ rpmE (PRK00019)Ribosomal_L31 (pfam01197)→ |
| Eco-2169-1 | NZ_APZI01000449.1 | + | 13475 | 13527 | **RNA**→ rpmE (PRK00019)Ribosomal_L31 (pfam01197)→ |
| Eco-2170-1 | NZ_APZZ01000144.1 | + | 18534 | 18586 | **RNA**→ rpmE (PRK00019)Ribosomal_L31 (pfam01197)→ |
| Eco-1014-1 | NZ_AQCY01000139.1 | + | 23759 | 23811 | **RNA**→ rpmE (PRK00019)Ribosomal_L31 (pfam01197)→ |
| Eco-1928-1 | NZ_AQBF01000062.1 | + | 87675 | 87727 | **RNA**→ rpmE (PRK00019)Ribosomal_L31 (pfam01197)→ |
| Eco-1934-1 | NZ_AQBK01000072.1 | + | 34111 | 34163 | **RNA**→ rpmE (PRK00019)Ribosomal_L31 (pfam01197)→ |
| Eco-1940-1 | NZ_AQBP01000082.1 | + | 34095 | 34147 | **RNA**→ rpmE (PRK00019)Ribosomal_L31 (pfam01197)→ |
| Eco-1008-1 | NZ_APXL01000532.1 | + | 7809 | 7861 | **RNA**→ rpmE (PRK00019)Ribosomal_L31 (pfam01197)→ |
| Eco-365-1 | NZ_JXBV01000104.1 | + | 79967 | 80019 | **RNA**→ rpmE (PRK00019)Ribosomal_L31 (pfam01197)→ |
| Sfl-74-1 | NZ_AMWM01000231.1 | + | 43473 | 43525 | **RNA**→ rpmE (PRK00019)Ribosomal_L31 (pfam01197)→ |
| Eco-282-1 | NZ_AEZP02000004.1 | - | 469851 | 469799 | **RNA**→ rpmE (PRK00019)Ribosomal_L31 (pfam01197)→ |
| Eco-2166-1 | NZ_AQBT01000172.1 | + | 18701 | 18753 | **RNA**→ rpmE (PRK00019)Ribosomal_L31 (pfam01197)→ |
| Eco-1085-1 | NZ_AIFZ01000036.1 | + | 91384 | 91436 | **RNA**→ rpmE (PRK00019)Ribosomal_L31 (pfam01197)→ |
| Eco-2173-1 | NZ_AQBY01000146.1 | + | 18635 | 18687 | **RNA**→ rpmE (PRK00019)Ribosomal_L31 (pfam01197)→ |
| Eco-1090-1 | NZ_AIGE01000037.1 | + | 90172 | 90224 | **RNA**→ rpmE (PRK00019)Ribosomal_L31 (pfam01197)→ |
| Eco-1925-1 | NZ_AQBC01000182.1 | + | 87680 | 87732 | **RNA**→ rpmE (PRK00019)Ribosomal_L31 (pfam01197)→ |
| Eco-2175-1 | NZ_AQCA01000232.1 | + | 18733 | 18785 | **RNA**→ rpmE (PRK00019)Ribosomal_L31 (pfam01197)→ |
| Sso-13-1 | NZ_AGRD01000092.1 | + | 94585 | 94637 | **RNA**→ rpmE (PRK00019)Ribosomal_L31 (pfam01197)→ |
| Eco-1185-1 | NZ_APNX01000009.1 | - | 52622 | 52570 | **RNA**→ rpmE (PRK00019)Ribosomal_L31 (pfam01197)→ |
| Eco-1182-1 | NZ_APNU01000001.1 | - | 52129 | 52077 | **RNA**→ rpmE (PRK00019)Ribosomal_L31 (pfam01197)→ |
| Eco-1186-1 | NZ_APNV01000024.1 | + | 64755 | 64807 | **RNA**→ rpmE (PRK00019)Ribosomal_L31 (pfam01197)→ |
| Eco-2013-1 | NZ_AOQY01000020.1 | - | 57102 | 57050 | **RNA**→ rpmE (PRK00019)Ribosomal_L31 (pfam01197)→ |
| Eco-2006-1 | NZ_AOQQ01000030.1 | + | 32479 | 32531 | **RNA**→ rpmE (PRK00019)Ribosomal_L31 (pfam01197)→ |
| Eco-2005-1 | NZ_AORB01000013.1 | + | 32411 | 32463 | **RNA**→ rpmE (PRK00019)Ribosomal_L31 (pfam01197)→ |
| Eco-2010-1 | NZ_AORA01000044.1 | - | 52396 | 52344 | **RNA**→ rpmE (PRK00019)Ribosomal_L31 (pfam01197)→ |
| Eco-2015-1 | NZ_AOQC01000042.1 | - | 52225 | 52173 | **RNA**→ rpmE (PRK00019)Ribosomal_L31 (pfam01197)→ |
| Eco-2018-1 | NZ_AOQM01000007.1 | - | 52226 | 52174 | **RNA**→ rpmE (PRK00019)Ribosomal_L31 (pfam01197)→ |
| Eco-2019-1 | NZ_AOQN01000054.1 | - | 52224 | 52172 | **RNA**→ rpmE (PRK00019)Ribosomal_L31 (pfam01197)→ |
| Eco-2011-1 | NZ_AOQW01000017.1 | - | 28856 | 28804 | **RNA**→ rpmE (PRK00019)Ribosomal_L31 (pfam01197)→ |
| Eco-2012-1 | NZ_AOQX01000001.1 | - | 28848 | 28796 | **RNA**→ rpmE (PRK00019)Ribosomal_L31 (pfam01197)→ |
| Eco-2017-1 | NZ_AOQE01000030.1 | - | 21619 | 21567 | **RNA**→ rpmE (PRK00019)Ribosomal_L31 (pfam01197)→ |
| Eco-2020-1 | NZ_AOQO01000021.1 | + | 8155 | 8207 | **RNA**→ rpmE (PRK00019)Ribosomal_L31 (pfam01197)→ |
| Eco-2005-2 | NZ_AOQP01000122.1 | + | 48352 | 48404 | **RNA**→ rpmE (PRK00019)Ribosomal_L31 (pfam01197)→ |
| Eco-2009-1 | NZ_AOQU01000249.1 | - | 443 | 391 | **RNA**→ rpmE (PRK00019)Ribosomal_L31 (pfam01197)→ |
| Eco-2014-1 | NZ_AOQZ01000031.1 | - | 28877 | 28825 | **RNA**→ rpmE (PRK00019)Ribosomal_L31 (pfam01197)→ |
| Eco-2007-1 | NZ_AOQR01000094.1 | + | 18381 | 18433 | **RNA**→ rpmE (PRK00019)Ribosomal_L31 (pfam01197)→ |

| | | | | | |
|---|---|---|---|---|---|
| Eco-2008-1 | NZ_AOQS01000188.1 | + | 18322 | 18374 | **RNA**→ rpmE (PRK00019)Ribosomal_L31 (pfam01197)→ |
| Eco-1770-1 | NZ_AJVT01000257.1 | + | 35439 | 35491 | **RNA**→ rpmE (PRK00019)Ribosomal_L31 (pfam01197)→ |
| Eco-1776-1 | NZ_AKAW01000246.1 | - | 430 | 378 | **RNA**→ rpmE (PRK00019)Ribosomal_L31 (pfam01197)→ |
| Eco-461-1 | NZ_JDRV01000378.1 | - | 506 | 454 | **RNA**→ rpmE (PRK00019)Ribosomal_L31 (pfam01197)→ |
| Eco-920-1 | NZ_AEZQ02000013.1 | + | 1490930 | 1490982 | **RNA**→ rpmE (PRK00019)Ribosomal_L31 (pfam01197)→ rpmE (PRK00019)Ribosomal_L31 (pfam01197)→ |
| Eco-923-1 | NZ_AEZW02000001.1 | + | 2765917 | 2765969 | **RNA**→ rpmE (PRK00019)Ribosomal_L31 (pfam01197)→ |
| Eco-845-1 | NZ_AEZK02000026.1 | + | 2868888 | 2868940 | **RNA**→ rpmE (PRK00019)Ribosomal_L31 (pfam01197)→ |
| Eco-928-1 | NZ_AEZL02000030.1 | - | 10772 | 10720 | **RNA**→ rpmE (PRK00019)Ribosomal_L31 (pfam01197)→ |
| Eco-1702-1 | NZ_ADWR01000068.1 | + | 21376 | 21428 | **RNA**→ rpmE (PRK00019)Ribosomal_L31 (pfam01197)→ |
| Eco-762-1 | NZ_AEZO02000020.1 | + | 311836 | 311888 | **RNA**→ rpmE (PRK00019)Ribosomal_L31 (pfam01197)→ |
| Eco-843-1 | NZ_AEZU02000105.1 | - | 167468 | 167416 | **RNA**→ rpmE (PRK00019)Ribosomal_L31 (pfam01197)→ rpmE (PRK00019)Ribosomal_L31 (pfam01197)→ |
| Eco-844-1 | NZ_AFAA02000008.1 | - | 253538 | 253486 | **RNA**→ rpmE (PRK00019)Ribosomal_L31 (pfam01197)→ |
| Eco-934-1 | NZ_AEZR02000014.1 | - | 89280 | 89228 | **RNA**→ rpmE (PRK00019)Ribosomal_L31 (pfam01197)→ |
| Eco-1684-1 | NZ_ADWT01000012.1 | + | 89502 | 89554 | **RNA**→ rpmE (PRK00019)Ribosomal_L31 (pfam01197)→ rpmE (PRK00019)Ribosomal_L31 (pfam01197)→ |
| Eco-927-1 | NZ_AEZJ02000011.1 | - | 47842 | 47790 | **RNA**→ rpmE (PRK00019)Ribosomal_L31 (pfam01197)→ |
| Eco-2016-1 | NZ_AOQD01000036.1 | - | 57103 | 57051 | **RNA**→ rpmE (PRK00019)Ribosomal_L31 (pfam01197)→ |
| Eco-728-1 | NZ_AOUQ01000152.1 | + | 41502 | 41554 | **RNA**→ rpmE (PRK00019)Ribosomal_L31 (pfam01197)→ |
| Eco-381-1 | NZ_AIBZ01000106.1 | - | 52248 | 52196 | **RNA**→ rpmE (PRK00019)Ribosomal_L31 (pfam01197)→ |
| Eco-291-2 | NZ_ASHD01000050.1 | + | 80292 | 80344 | **RNA**→ rpmE (PRK00019)Ribosomal_L31 (pfam01197)→ |
| Eco-287-1 | NZ_ATOA01000041.1 | + | 64688 | 64740 | **RNA**→ rpmE (PRK00019)Ribosomal_L31 (pfam01197)→ |
| Eco-281-1 | NZ_AEZY02000004.1 | + | 82291 | 82343 | **RNA**→ rpmE (PRK00019)Ribosomal_L31 (pfam01197)→ |
| Eco-392-1 | NZ_JRYM01000024.1 | + | 86369 | 86421 | **RNA**→ rpmE (PRK00019)Ribosomal_L31 (pfam01197)→ |
| Eco-1831-1 | NZ_AICG01000014.1 | + | 88929 | 88981 | **RNA**→ rpmE (PRK00019)Ribosomal_L31 (pfam01197)→ |
| Eco-344-1 | NZ_AIAP01000056.1 | - | 52126 | 52074 | **RNA**→ rpmE (PRK00019)Ribosomal_L31 (pfam01197)→ |
| Eco-341-1 | NZ_AIAL01000036.1 | - | 39155 | 39103 | **RNA**→ rpmE (PRK00019)Ribosomal_L31 (pfam01197)→ |
| Eco-361-1 | NZ_AIBF01000024.1 | + | 80060 | 80112 | **RNA**→ rpmE (PRK00019)Ribosomal_L31 (pfam01197)→ |
| Sso-1-6 | NZ_CXEQ01000019.1 | - | 645 | 593 | **RNA**→ rpmE (PRK00019)Ribosomal_L31 (pfam01197)→ |
| Sso-1-7 | NZ_CXOD01000022.1 | + | 39082 | 39134 | **RNA**→ rpmE (PRK00019)Ribosomal_L31 (pfam01197)→ |
| Eco-369-1 | NZ_AIBN01000020.1 | - | 505 | 453 | **RNA**→ rpmE (PRK00019)Ribosomal_L31 (pfam01197)→ |
| Eco-357-1 | NZ_AIBC01000257.1 | - | 613 | 561 | **RNA**→ rpmE (PRK00019)Ribosomal_L31 (pfam01197)→ |
| Eco-371-1 | NZ_AIBP01000011.1 | - | 18225 | 18173 | **RNA**→ rpmE (PRK00019)Ribosomal_L31 (pfam01197)→ |
| Eco-377-1 | NZ_AIBU01000027.1 | - | 690 | 638 | **RNA**→ rpmE (PRK00019)Ribosomal_L31 (pfam01197)→ |
| Eco-383-1 | NZ_AICB01000019.1 | + | 11304 | 11356 | **RNA**→ rpmE (PRK00019)Ribosomal_L31 (pfam01197)→ |
| Eco-386-1 | NZ_AICE01000057.1 | + | 80383 | 80435 | **RNA**→ rpmE (PRK00019)Ribosomal_L31 (pfam01197)→ |
| Eco-333-1 | NZ_AIAE01000024.1 | + | 91910 | 91962 | **RNA**→ rpmE (PRK00019)Ribosomal_L31 (pfam01197)→ |
| Eco-354-1 | NZ_AIAZ01000228.1 | - | 505 | 453 | **RNA**→ rpmE (PRK00019)Ribosomal_L31 (pfam01197)→ |
| Eco-342-1 | NZ_AIAN01000137.1 | + | 23487 | 23539 | **RNA**→ rpmE (PRK00019)Ribosomal_L31 (pfam01197)→ |
| Eco-382-1 | NZ_AICA01000012.1 | - | 487 | 435 | **RNA**→ rpmE (PRK00019)Ribosomal_L31 (pfam01197)→ |
| Eco-347-1 | NZ_AIAS01000067.1 | + | 39822 | 39874 | **RNA**→ rpmE (PRK00019)Ribosomal_L31 (pfam01197)→ |
| Eco-370-1 | NZ_AIBO01000227.1 | - | 39194 | 39142 | **RNA**→ rpmE (PRK00019)Ribosomal_L31 (pfam01197)→ |
| Eco-376-1 | NZ_AIBV01000160.1 | - | 486 | 434 | **RNA**→ rpmE (PRK00019)Ribosomal_L31 (pfam01197)→ |
| Eco-543-1 | NZ_ASZR01000033.1 | + | 85134 | 85186 | **RNA**→ rpmE (PRK00019)Ribosomal_L31 (pfam01197)→ |
| Eco-355-1 | NZ_AIBA01000470.1 | + | 46666 | 46718 | **RNA**→ rpmE (PRK00019)Ribosomal_L31 (pfam01197)→ |
| Eco-340-1 | NZ_AIAM01000047.1 | - | 52250 | 52198 | **RNA**→ rpmE (PRK00019)Ribosomal_L31 (pfam01197)→ |
| Eco-544-1 | NZ_ASZS01000023.1 | - | 18236 | 18184 | **RNA**→ rpmE (PRK00019)Ribosomal_L31 (pfam01197)→ |
| Eco-332-1 | NZ_AIAD01000260.1 | - | 547 | 495 | **RNA**→ rpmE (PRK00019)Ribosomal_L31 (pfam01197)→ |
| Eco-363-1 | NZ_AIBI01000108.1 | + | 84228 | 84280 | **RNA**→ rpmE (PRK00019)Ribosomal_L31 (pfam01197)→ |
| Eco-373-1 | NZ_AIBS01000001.1 | + | 76094 | 76146 | **RNA**→ rpmE (PRK00019)Ribosomal_L31 (pfam01197)→ |
| Eco-334-1 | NZ_AIAF01000014.1 | - | 486 | 434 | **RNA**→ rpmE (PRK00019)Ribosomal_L31 (pfam01197)→ |
| Eco-385-1 | NZ_AICD01000096.1 | - | 52255 | 52203 | **RNA**→ rpmE (PRK00019)Ribosomal_L31 (pfam01197)→ |
| Eco-360-1 | NZ_AIBG01000067.1 | - | 52126 | 52074 | **RNA**→ rpmE (PRK00019)Ribosomal_L31 (pfam01197)→ |
| Eco-289-1 | NZ_CWHW01000022.1 | - | 39312 | 39260 | **RNA**→ rpmE (PRK00019)Ribosomal_L31 (pfam01197)→ |

| | | | | | |
|---|---|---|---|---|---|
| Eco-45-1 | NZ_JHRA01000033.1 | - | 526 | 474 | **RNA**→ rpmE (PRK00019)Ribosomal_L31 (pfam01197)→ |
| Eco-195-1 | NZ_JHRC01000049.1 | + | 8435 | 8487 | **RNA**→ rpmE (PRK00019)Ribosomal_L31 (pfam01197)→ |
| Eco-44-1 | NZ_JHQZ01000062.1 | - | 5988 | 5936 | **RNA**→ rpmE (PRK00019)Ribosomal_L31 (pfam01197)→ |
| Eco-196-1 | NZ_JHRG01000048.1 | + | 11856 | 11908 | **RNA**→ rpmE (PRK00019)Ribosomal_L31 (pfam01197)→ |
| Eco-211-1 | NZ_JHRM01000023.1 | - | 39670 | 39618 | **RNA**→ rpmE (PRK00019)Ribosomal_L31 (pfam01197)→ |
| Eco-212-1 | NZ_JHRQ01000039.1 | + | 23521 | 23573 | **RNA**→ rpmE (PRK00019)Ribosomal_L31 (pfam01197)→ |
| Eco-43-1 | NZ_JHRT01000008.1 | - | 39527 | 39475 | **RNA**→ rpmE (PRK00019)Ribosomal_L31 (pfam01197)→ |
| Sso-1-8 | NZ_CXLF01000021.1 | + | 39082 | 39134 | **RNA**→ rpmE (PRK00019)Ribosomal_L31 (pfam01197)→ |
| Sso-1-9 | NZ_CXDE01000021.1 | + | 39007 | 39059 | **RNA**→ rpmE (PRK00019)Ribosomal_L31 (pfam01197)→ |
| Sso-1-10 | NZ_CXMH01000026.1 | + | 39089 | 39141 | **RNA**→ rpmE (PRK00019)Ribosomal_L31 (pfam01197)→ |
| Sso-1-11 | NZ_CXLA01000024.1 | + | 39089 | 39141 | **RNA**→ rpmE (PRK00019)Ribosomal_L31 (pfam01197)→ |
| Sso-1-12 | NZ_CXDX01000024.1 | + | 39030 | 39082 | **RNA**→ rpmE (PRK00019)Ribosomal_L31 (pfam01197)→ |
| Sso-1-13 | NZ_CXKH01000025.1 | + | 39089 | 39141 | **RNA**→ rpmE (PRK00019)Ribosomal_L31 (pfam01197)→ |
| Sso-1-14 | NZ_CWYN01000026.1 | - | 660 | 608 | **RNA**→ rpmE (PRK00019)Ribosomal_L31 (pfam01197)→ |
| Sso-1-15 | NZ_CXBZ01000018.1 | - | 636 | 584 | **RNA**→ rpmE (PRK00019)Ribosomal_L31 (pfam01197)→ |
| Sso-1-16 | NZ_CXDU01000021.1 | + | 47902 | 47954 | **RNA**→ rpmE (PRK00019)Ribosomal_L31 (pfam01197)→ |
| Sso-1-17 | NZ_CXGT01000025.1 | - | 1211 | 1159 | **RNA**→ rpmE (PRK00019)Ribosomal_L31 (pfam01197)→ |
| Sso-1-18 | NZ_CXAN01000023.1 | + | 39089 | 39141 | **RNA**→ rpmE (PRK00019)Ribosomal_L31 (pfam01197)→ |
| Sso-1-19 | NZ_CXJC01000027.1 | + | 39089 | 39141 | **RNA**→ rpmE (PRK00019)Ribosomal_L31 (pfam01197)→ |
| Sso-1-20 | NZ_CXAI01000025.1 | + | 39045 | 39097 | **RNA**→ rpmE (PRK00019)Ribosomal_L31 (pfam01197)→ |
| Sso-1-21 | NZ_CXGW01000022.1 | - | 3806 | 3754 | **RNA**→ rpmE (PRK00019)Ribosomal_L31 (pfam01197)→ |
| Sso-1-22 | NZ_CXIW01000023.1 | + | 39104 | 39156 | **RNA**→ rpmE (PRK00019)Ribosomal_L31 (pfam01197)→ |
| Sso-1-23 | NZ_CWYW01000026.1 | + | 39257 | 39309 | **RNA**→ rpmE (PRK00019)Ribosomal_L31 (pfam01197)→ |
| Sso-1-24 | NZ_CXCZ01000017.1 | - | 630 | 578 | **RNA**→ rpmE (PRK00019)Ribosomal_L31 (pfam01197)→ |
| Sso-1-25 | NZ_CXLY01000025.1 | - | 1734 | 1682 | **RNA**→ rpmE (PRK00019)Ribosomal_L31 (pfam01197)→ |
| Sso-1-26 | NZ_CXBY01000020.1 | - | 1174 | 1122 | **RNA**→ rpmE (PRK00019)Ribosomal_L31 (pfam01197)→ |
| Sso-1-27 | NZ_CXMR01000026.1 | - | 1627 | 1575 | **RNA**→ rpmE (PRK00019)Ribosomal_L31 (pfam01197)→ |
| Sso-1-28 | NZ_CXID01000025.1 | - | 793 | 741 | **RNA**→ rpmE (PRK00019)Ribosomal_L31 (pfam01197)→ |
| Sso-1-29 | NZ_CXJG01000027.1 | + | 39082 | 39134 | **RNA**→ rpmE (PRK00019)Ribosomal_L31 (pfam01197)→ |
| Sso-1-30 | NZ_CXJN01000027.1 | + | 39082 | 39134 | **RNA**→ rpmE (PRK00019)Ribosomal_L31 (pfam01197)→ |
| Sso-1-31 | NZ_CXFQ01000025.1 | - | 664 | 612 | **RNA**→ rpmE (PRK00019)Ribosomal_L31 (pfam01197)→ |
| Sso-1-32 | NZ_CXOC01000023.1 | + | 39089 | 39141 | **RNA**→ rpmE (PRK00019)Ribosomal_L31 (pfam01197)→ |
| Sso-1-33 | NZ_CWZL01000026.1 | + | 39089 | 39141 | **RNA**→ rpmE (PRK00019)Ribosomal_L31 (pfam01197)→ |
| Sso-1-34 | NZ_CWWR01000026.1 | + | 39082 | 39134 | **RNA**→ rpmE (PRK00019)Ribosomal_L31 (pfam01197)→ |
| Sso-1-35 | NZ_CXDO01000020.1 | + | 39023 | 39075 | **RNA**→ rpmE (PRK00019)Ribosomal_L31 (pfam01197)→ |
| Sso-1-36 | NZ_CXLN01000027.1 | + | 39089 | 39141 | **RNA**→ rpmE (PRK00019)Ribosomal_L31 (pfam01197)→ |
| Sso-1-37 | NZ_CXMW01000027.1 | - | 683 | 631 | **RNA**→ rpmE (PRK00019)Ribosomal_L31 (pfam01197)→ |
| Sso-1-38 | NZ_CXEG01000022.1 | + | 39030 | 39082 | **RNA**→ rpmE (PRK00019)Ribosomal_L31 (pfam01197)→ |
| Sso-1-39 | NZ_CXBG01000024.1 | + | 39023 | 39075 | **RNA**→ rpmE (PRK00019)Ribosomal_L31 (pfam01197)→ |
| Sso-1-40 | NZ_CXKW01000025.1 | - | 1627 | 1575 | **RNA**→ rpmE (PRK00019)Ribosomal_L31 (pfam01197)→ |
| Sfl-25-1 | NZ_LAIB01000061.1 | - | 31124 | 31072 | **RNA**→ rpmE (PRK00019)Ribosomal_L31 (pfam01197)→ |
| Sfl-25-2 | NZ_LAIF01000103.1 | - | 31124 | 31072 | **RNA**→ rpmE (PRK00019)Ribosomal_L31 (pfam01197)→ |
| Sfl-25-3 | NZ_LAHW01000133.1 | - | 31124 | 31072 | **RNA**→ rpmE (PRK00019)Ribosomal_L31 (pfam01197)→ |
| Eco-460-1 | NZ_LBLW01000064.1 | + | 82857 | 82909 | **RNA**→ rpmE (PRK00019)Ribosomal_L31 (pfam01197)→ |
| Sfl-25-9 | NZ_LAHX01000020.1 | - | 31124 | 31072 | **RNA**→ rpmE (PRK00019)Ribosomal_L31 (pfam01197)→ |
| Sso-1-41 | NZ_CXEY01000021.1 | - | 1726 | 1674 | **RNA**→ rpmE (PRK00019)Ribosomal_L31 (pfam01197)→ |
| Sso-1-42 | NZ_CXCW01000015.1 | - | 378 | 326 | **RNA**→ rpmE (PRK00019)Ribosomal_L31 (pfam01197)→ |
| Sfl-25-4 | NZ_LAIA01000057.1 | + | 24623 | 24675 | **RNA**→ rpmE (PRK00019)Ribosomal_L31 (pfam01197)→ |
| Sso-1-43 | NZ_CXDA01000016.1 | - | 428 | 376 | **RNA**→ rpmE (PRK00019)Ribosomal_L31 (pfam01197)→ hypo→ |
| Sso-1-44 | NZ_CWZJ01000023.1 | - | 1174 | 1122 | **RNA**→ rpmE (PRK00019)Ribosomal_L31 (pfam01197)→ |
| Sso-1-45 | NZ_CWSX01000023.1 | + | 38756 | 38808 | **RNA**→ rpmE (PRK00019)Ribosomal_L31 (pfam01197)→ |
| Sso-1-46 | NZ_CWUJ01000026.1 | - | 660 | 608 | **RNA**→ rpmE (PRK00019)Ribosomal_L31 (pfam01197)→ |

| | | | | | | |
|---|---|---|---|---|---|---|
| Sfl-25-5 | NZ_LAHY01000049.1 | + | 24753 | 24805 | **RNA**→ | rpmE (PRK00019)Ribosomal_L31 (pfam01197)→ |
| Sso-1-47 | NZ_CWTE01000026.1 | - | 529 | 477 | **RNA**→ | rpmE (PRK00019)Ribosomal_L31 (pfam01197)→ |
| Sso-1-48 | NZ_CWVE01000028.1 | - | 800 | 748 | **RNA**→ | rpmE (PRK00019)Ribosomal_L31 (pfam01197)→ |
| Sso-1-49 | NZ_CWUI01000024.1 | + | 38914 | 38966 | **RNA**→ | rpmE (PRK00019)Ribosomal_L31 (pfam01197)→ |
| Sso-1-50 | NZ_CWUS01000027.1 | - | 1730 | 1678 | **RNA**→ | rpmE (PRK00019)Ribosomal_L31 (pfam01197)→ |
| Sso-1-51 | NZ_CXNJ01000026.1 | + | 39089 | 39141 | **RNA**→ | rpmE (PRK00019)Ribosomal_L31 (pfam01197)→ |
| Sso-1-52 | NZ_CWUR01000020.1 | - | 495 | 443 | **RNA**→ | rpmE (PRK00019)Ribosomal_L31 (pfam01197)→ |
| Sso-1-53 | NZ_CWZX01000025.1 | - | 1539 | 1487 | **RNA**→ | rpmE (PRK00019)Ribosomal_L31 (pfam01197)→ |
| Sso-1-54 | NZ_CWVJ01000024.1 | + | 39082 | 39134 | **RNA**→ | rpmE (PRK00019)Ribosomal_L31 (pfam01197)→ |
| Sso-1-55 | NZ_CXFL01000025.1 | - | 481 | 429 | **RNA**→ | rpmE (PRK00019)Ribosomal_L31 (pfam01197)→ |
| Sso-1-56 | NZ_CWSU01000026.1 | - | 529 | 477 | **RNA**→ | rpmE (PRK00019)Ribosomal_L31 (pfam01197)→ |
| Sso-1-57 | NZ_CXIK01000024.1 | + | 39082 | 39134 | **RNA**→ | rpmE (PRK00019)Ribosomal_L31 (pfam01197)→ |
| Sso-1-58 | NZ_CWVL01000028.1 | + | 39089 | 39141 | **RNA**→ | rpmE (PRK00019)Ribosomal_L31 (pfam01197)→ |
| Sso-1-59 | NZ_CWYU01000024.1 | + | 39039 | 39091 | **RNA**→ | rpmE (PRK00019)Ribosomal_L31 (pfam01197)→ |
| Sso-1-60 | NZ_CWVI01000027.1 | + | 39099 | 39151 | **RNA**→ | rpmE (PRK00019)Ribosomal_L31 (pfam01197)→ |
| Sso-1-61 | NZ_CXHA01000022.1 | + | 39104 | 39156 | **RNA**→ | rpmE (PRK00019)Ribosomal_L31 (pfam01197)→ |
| Sso-1-62 | NZ_CWTL01000023.1 | - | 1776 | 1724 | **RNA**→ | rpmE (PRK00019)Ribosomal_L31 (pfam01197)→ |
| Sso-1-63 | NZ_CXER01000023.1 | + | 39030 | 39082 | **RNA**→ | rpmE (PRK00019)Ribosomal_L31 (pfam01197)→ |
| Sso-1-64 | NZ_CWTV01000027.1 | + | 39076 | 39128 | **RNA**→ | rpmE (PRK00019)Ribosomal_L31 (pfam01197)→ |
| Sso-1-65 | NZ_CXNI01000023.1 | - | 1174 | 1122 | **RNA**→ | rpmE (PRK00019)Ribosomal_L31 (pfam01197)→ |
| Sso-1-66 | NZ_CWWF01000027.1 | + | 39093 | 39145 | **RNA**→ | rpmE (PRK00019)Ribosomal_L31 (pfam01197)→ |
| Sso-1-67 | NZ_CXJV01000026.1 | - | 695 | 643 | **RNA**→ | rpmE (PRK00019)Ribosomal_L31 (pfam01197)→ |
| Sso-1-68 | NZ_CWTG01000022.1 | - | 1137 | 1085 | **RNA**→ | rpmE (PRK00019)Ribosomal_L31 (pfam01197)→ |
| Sso-1-69 | NZ_CWZQ01000026.1 | + | 39082 | 39134 | **RNA**→ | rpmE (PRK00019)Ribosomal_L31 (pfam01197)→ |
| Sso-1-70 | NZ_CWXI01000025.1 | - | 1137 | 1085 | **RNA**→ | rpmE (PRK00019)Ribosomal_L31 (pfam01197)→ |
| Sso-1-71 | NZ_CWWE01000028.1 | - | 662 | 610 | **RNA**→ | rpmE (PRK00019)Ribosomal_L31 (pfam01197)→ |
| Sso-1-72 | NZ_CXIA01000027.1 | + | 39089 | 39141 | **RNA**→ | rpmE (PRK00019)Ribosomal_L31 (pfam01197)→ |
| Sso-1-73 | NZ_CXHM01000025.1 | - | 782 | 730 | **RNA**→ | rpmE (PRK00019)Ribosomal_L31 (pfam01197)→ |
| Sso-1-74 | NZ_CWVP01000027.1 | - | 1234 | 1182 | **RNA**→ | rpmE (PRK00019)Ribosomal_L31 (pfam01197)→ |
| Sso-1-75 | NZ_CXGI01000026.1 | - | 668 | 616 | **RNA**→ | rpmE (PRK00019)Ribosomal_L31 (pfam01197)→ |
| Sso-1-76 | NZ_CWUZ01000025.1 | - | 1627 | 1575 | **RNA**→ | rpmE (PRK00019)Ribosomal_L31 (pfam01197)→ |
| Sso-1-77 | NZ_CXAE01000028.1 | - | 529 | 477 | **RNA**→ | rpmE (PRK00019)Ribosomal_L31 (pfam01197)→ |
| Sso-1-78 | NZ_CXMN01000028.1 | + | 39082 | 39134 | **RNA**→ | rpmE (PRK00019)Ribosomal_L31 (pfam01197)→ |
| Sso-1-79 | NZ_CXKR01000024.1 | + | 39089 | 39141 | **RNA**→ | rpmE (PRK00019)Ribosomal_L31 (pfam01197)→ |
| Sso-1-80 | NZ_CXIV01000028.1 | - | 1627 | 1575 | **RNA**→ | rpmE (PRK00019)Ribosomal_L31 (pfam01197)→ |
| Sso-1-81 | NZ_CXLI01000025.1 | - | 1539 | 1487 | **RNA**→ | rpmE (PRK00019)Ribosomal_L31 (pfam01197)→ |
| Sso-1-82 | NZ_CWVQ01000028.1 | - | 1224 | 1172 | **RNA**→ | rpmE (PRK00019)Ribosomal_L31 (pfam01197)→ |
| Sso-1-83 | NZ_CXCX01000002.1 | + | 80210 | 80262 | **RNA**→ | rpmE (PRK00019)Ribosomal_L31 (pfam01197)→ |
| Sso-1-84 | NZ_CWTU01000026.1 | + | 39106 | 39158 | **RNA**→ | rpmE (PRK00019)Ribosomal_L31 (pfam01197)→ |
| Sso-1-85 | NZ_CXGM01000023.1 | - | 867 | 815 | **RNA**→ | rpmE (PRK00019)Ribosomal_L31 (pfam01197)→ |
| Sso-1-86 | NZ_CXHR01000027.1 | + | 39089 | 39141 | **RNA**→ | rpmE (PRK00019)Ribosomal_L31 (pfam01197)→ |
| Sso-1-87 | NZ_CWTX01000029.1 | - | 660 | 608 | **RNA**→ | rpmE (PRK00019)Ribosomal_L31 (pfam01197)→ |
| Sso-1-88 | NZ_CXIN01000026.1 | - | 786 | 734 | **RNA**→ | rpmE (PRK00019)Ribosomal_L31 (pfam01197)→ |
| Sso-1-89 | NZ_CXDH01000024.1 | - | 632 | 580 | **RNA**→ | rpmE (PRK00019)Ribosomal_L31 (pfam01197)→ |
| Sso-1-90 | NZ_CWTW01000025.1 | - | 556 | 504 | **RNA**→ | rpmE (PRK00019)Ribosomal_L31 (pfam01197)→ |
| Sso-1-91 | NZ_CXAH01000027.1 | + | 39082 | 39134 | **RNA**→ | rpmE (PRK00019)Ribosomal_L31 (pfam01197)→ |
| Sso-1-92 | NZ_CWXW01000022.1 | + | 39104 | 39156 | **RNA**→ | rpmE (PRK00019)Ribosomal_L31 (pfam01197)→ |
| Sso-1-93 | NZ_CXMO01000027.1 | + | 39082 | 39134 | **RNA**→ | rpmE (PRK00019)Ribosomal_L31 (pfam01197)→ |
| Sso-1-94 | NZ_CWUA01000024.1 | - | 1539 | 1487 | **RNA**→ | rpmE (PRK00019)Ribosomal_L31 (pfam01197)→ |
| Sso-1-95 | NZ_CXAY01000023.1 | - | 1137 | 1085 | **RNA**→ | rpmE (PRK00019)Ribosomal_L31 (pfam01197)→ |
| Sso-1-96 | NZ_CWYH01000026.1 | - | 664 | 612 | **RNA**→ | rpmE (PRK00019)Ribosomal_L31 (pfam01197)→ |
| Sso-1-97 | NZ_CXNC01000028.1 | - | 800 | 748 | **RNA**→ | rpmE (PRK00019)Ribosomal_L31 (pfam01197)→ |

| | | | | | |
|---|---|---|---|---|---|
| Sso-1-98 | NZ_CWTT01000028.1 | + | 38880 | 38932 | **RNA**→ rpmE (PRK00019)Ribosomal_L31 (pfam01197)→ |
| Sso-1-99 | NZ_CXBA01000025.1 | - | 1137 | 1085 | **RNA**→ rpmE (PRK00019)Ribosomal_L31 (pfam01197)→ |
| Sso-1-100 | NZ_CXHG01000028.1 | - | 695 | 643 | **RNA**→ rpmE (PRK00019)Ribosomal_L31 (pfam01197)→ |
| Sso-1-101 | NZ_CXIH01000027.1 | - | 768 | 716 | **RNA**→ rpmE (PRK00019)Ribosomal_L31 (pfam01197)→ |
| Sso-1-102 | NZ_CWST01000068.1 | - | 660 | 608 | **RNA**→ rpmE (PRK00019)Ribosomal_L31 (pfam01197)→ |
| Sso-1-103 | NZ_CXGQ01000025.1 | - | 1174 | 1122 | **RNA**→ rpmE (PRK00019)Ribosomal_L31 (pfam01197)→ |
| Sso-1-104 | NZ_CWYP01000024.1 | - | 1696 | 1644 | **RNA**→ rpmE (PRK00019)Ribosomal_L31 (pfam01197)→ |
| Sso-1-105 | NZ_CWVT01000028.1 | - | 660 | 608 | **RNA**→ rpmE (PRK00019)Ribosomal_L31 (pfam01197)→ |
| Sso-1-106 | NZ_CXNR01000022.1 | - | 630 | 578 | **RNA**→ rpmE (PRK00019)Ribosomal_L31 (pfam01197)→ |
| Sso-1-107 | NZ_CXHY01000027.1 | - | 1627 | 1575 | **RNA**→ rpmE (PRK00019)Ribosomal_L31 (pfam01197)→ |
| Sso-1-108 | NZ_CXHB01000028.1 | + | 39089 | 39141 | **RNA**→ rpmE (PRK00019)Ribosomal_L31 (pfam01197)→ |
| Sso-1-109 | NZ_CWUO01000021.1 | - | 660 | 608 | **RNA**→ rpmE (PRK00019)Ribosomal_L31 (pfam01197)→ |
| Sso-1-110 | NZ_CXCC01000002.1 | + | 80327 | 80379 | **RNA**→ rpmE (PRK00019)Ribosomal_L31 (pfam01197)→ |
| Sso-1-111 | NZ_CWXK01000292.1 | + | 787 | 839 | **RNA**→ rpmE (PRK00019)Ribosomal_L31 (pfam01197)→ |
| Sso-1-112 | NZ_CWTH01000026.1 | + | 39059 | 39111 | **RNA**→ rpmE (PRK00019)Ribosomal_L31 (pfam01197)→ |
| Sso-1-113 | NZ_CXHL01000028.1 | + | 39045 | 39097 | **RNA**→ rpmE (PRK00019)Ribosomal_L31 (pfam01197)→ |
| Sso-1-114 | NZ_CXAQ01000027.1 | - | 6653 | 6601 | **RNA**→ rpmE (PRK00019)Ribosomal_L31 (pfam01197)→ |
| Sso-1-115 | NZ_CWTZ01000023.1 | + | 39040 | 39092 | **RNA**→ rpmE (PRK00019)Ribosomal_L31 (pfam01197)→ |
| Sso-1-116 | NZ_CWZU01000029.1 | - | 664 | 612 | **RNA**→ rpmE (PRK00019)Ribosomal_L31 (pfam01197)→ |
| Sso-1-117 | NZ_CWZH01000027.1 | + | 39089 | 39141 | **RNA**→ rpmE (PRK00019)Ribosomal_L31 (pfam01197)→ |
| Sso-1-118 | NZ_CWUU01000028.1 | + | 39082 | 39134 | **RNA**→ rpmE (PRK00019)Ribosomal_L31 (pfam01197)→ |
| Sso-1-119 | NZ_CWXP01000027.1 | + | 39089 | 39141 | **RNA**→ rpmE (PRK00019)Ribosomal_L31 (pfam01197)→ |
| Sso-1-120 | NZ_CXFX01000021.1 | - | 1174 | 1122 | **RNA**→ rpmE (PRK00019)Ribosomal_L31 (pfam01197)→ |
| Sso-1-121 | NZ_CWWB01000027.1 | + | 39093 | 39145 | **RNA**→ rpmE (PRK00019)Ribosomal_L31 (pfam01197)→ |
| Sso-1-122 | NZ_CXBJ01000069.1 | + | 20736 | 20788 | **RNA**→ rpmE (PRK00019)Ribosomal_L31 (pfam01197)→ |
| Sso-1-123 | NZ_CXFR01000024.1 | - | 481 | 429 | **RNA**→ rpmE (PRK00019)Ribosomal_L31 (pfam01197)→ |
| Sso-1-124 | NZ_CWVM01000027.1 | + | 39093 | 39145 | **RNA**→ rpmE (PRK00019)Ribosomal_L31 (pfam01197)→ |
| Sso-1-125 | NZ_CXDC01000015.1 | + | 47850 | 47902 | **RNA**→ rpmE (PRK00019)Ribosomal_L31 (pfam01197)→ |
| Sso-1-126 | NZ_CXLJ01000026.1 | + | 39089 | 39141 | **RNA**→ rpmE (PRK00019)Ribosomal_L31 (pfam01197)→ |
| Sso-1-127 | NZ_CWUE01000025.1 | - | 1535 | 1483 | **RNA**→ rpmE (PRK00019)Ribosomal_L31 (pfam01197)→ |
| Sso-1-128 | NZ_CXIU01000022.1 | - | 1137 | 1085 | **RNA**→ rpmE (PRK00019)Ribosomal_L31 (pfam01197)→ |
| Sso-1-129 | NZ_CXFT01000022.1 | - | 481 | 429 | **RNA**→ rpmE (PRK00019)Ribosomal_L31 (pfam01197)→ |
| Sso-1-130 | NZ_CWTO01000027.1 | - | 660 | 608 | **RNA**→ rpmE (PRK00019)Ribosomal_L31 (pfam01197)→ |
| Sso-1-131 | NZ_CXKJ01000023.1 | - | 1627 | 1575 | **RNA**→ rpmE (PRK00019)Ribosomal_L31 (pfam01197)→ |
| Sso-1-132 | NZ_CWXM01000026.1 | + | 39089 | 39141 | **RNA**→ rpmE (PRK00019)Ribosomal_L31 (pfam01197)→ |
| Sso-1-133 | NZ_CWWC01000028.1 | + | 39082 | 39134 | **RNA**→ rpmE (PRK00019)Ribosomal_L31 (pfam01197)→ |
| Sso-1-134 | NZ_CWXV01000020.1 | - | 854 | 802 | **RNA**→ rpmE (PRK00019)Ribosomal_L31 (pfam01197)→ |
| Sso-1-135 | NZ_CXGY01000028.1 | - | 529 | 477 | **RNA**→ rpmE (PRK00019)Ribosomal_L31 (pfam01197)→ |
| Sso-1-136 | NZ_CWVU01000028.1 | + | 39089 | 39141 | **RNA**→ rpmE (PRK00019)Ribosomal_L31 (pfam01197)→ |
| Sso-1-137 | NZ_CXMZ01000018.1 | - | 6792 | 6740 | **RNA**→ rpmE (PRK00019)Ribosomal_L31 (pfam01197)→ |
| Sso-1-138 | NZ_CXKS01000028.1 | - | 800 | 748 | **RNA**→ rpmE (PRK00019)Ribosomal_L31 (pfam01197)→ |
| Sso-1-139 | NZ_CWUX01000023.1 | - | 2669 | 2617 | **RNA**→ rpmE (PRK00019)Ribosomal_L31 (pfam01197)→ |
| Sso-1-140 | NZ_CWXS01000023.1 | + | 39104 | 39156 | **RNA**→ rpmE (PRK00019)Ribosomal_L31 (pfam01197)→ |
| Sso-1-141 | NZ_CXLD01000025.1 | + | 39089 | 39141 | **RNA**→ rpmE (PRK00019)Ribosomal_L31 (pfam01197)→ |
| Sso-1-142 | NZ_CWVF01000027.1 | + | 39082 | 39134 | **RNA**→ rpmE (PRK00019)Ribosomal_L31 (pfam01197)→ |
| Sso-1-143 | NZ_CXJB01000028.1 | - | 790 | 738 | **RNA**→ rpmE (PRK00019)Ribosomal_L31 (pfam01197)→ |
| Sso-1-144 | NZ_CXBF01000020.1 | + | 39023 | 39075 | **RNA**→ rpmE (PRK00019)Ribosomal_L31 (pfam01197)→ |
| Sso-1-145 | NZ_CWTF01000026.1 | - | 660 | 608 | **RNA**→ rpmE (PRK00019)Ribosomal_L31 (pfam01197)→ |
| Sso-1-146 | NZ_CXMS01000026.1 | - | 795 | 743 | **RNA**→ rpmE (PRK00019)Ribosomal_L31 (pfam01197)→ |
| Sso-1-147 | NZ_CWWT01000020.1 | - | 843 | 791 | **RNA**→ rpmE (PRK00019)Ribosomal_L31 (pfam01197)→ |
| Sso-1-148 | NZ_CWVK01000026.1 | + | 39089 | 39141 | **RNA**→ rpmE (PRK00019)Ribosomal_L31 (pfam01197)→ |
| Sso-1-149 | NZ_CWUN01000018.1 | - | 495 | 443 | **RNA**→ rpmE (PRK00019)Ribosomal_L31 (pfam01197)→ |

| | | | | | |
|---|---|---|---|---|---|
| Sso-1-150 | NZ_CXLM01000026.1 | + | 39089 | 39141 | **RNA**→ rpmE (PRK00019)Ribosomal_L31 (pfam01197)→ |
| Sso-1-151 | NZ_CWVO01000028.1 | + | 39093 | 39145 | **RNA**→ rpmE (PRK00019)Ribosomal_L31 (pfam01197)→ |
| Sso-1-152 | NZ_CXEW01000014.1 | + | 39108 | 39160 | **RNA**→ rpmE (PRK00019)Ribosomal_L31 (pfam01197)→ |
| Sso-1-153 | NZ_CWVA01000027.1 | - | 684 | 632 | **RNA**→ rpmE (PRK00019)Ribosomal_L31 (pfam01197)→ |
| Sso-1-154 | NZ_CWUB01000022.1 | - | 555 | 503 | **RNA**→ rpmE (PRK00019)Ribosomal_L31 (pfam01197)→ |
| Sso-1-155 | NZ_CXIL01000022.1 | + | 39082 | 39134 | **RNA**→ rpmE (PRK00019)Ribosomal_L31 (pfam01197)→ |
| Sso-1-156 | NZ_CWUF01000024.1 | + | 39068 | 39120 | **RNA**→ rpmE (PRK00019)Ribosomal_L31 (pfam01197)→ |
| Sso-1-157 | NZ_CXJW01000027.1 | + | 39089 | 39141 | **RNA**→ rpmE (PRK00019)Ribosomal_L31 (pfam01197)→ |
| Sso-1-158 | NZ_CWVZ01000025.1 | - | 1223 | 1171 | **RNA**→ rpmE (PRK00019)Ribosomal_L31 (pfam01197)→ |
| Sso-1-159 | NZ_CXEJ01000023.1 | - | 644 | 592 | **RNA**→ rpmE (PRK00019)Ribosomal_L31 (pfam01197)→ |
| Sso-1-160 | NZ_CXMU01000027.1 | + | 39094 | 39146 | **RNA**→ rpmE (PRK00019)Ribosomal_L31 (pfam01197)→ |
| Sso-1-161 | NZ_CWVH01000025.1 | + | 39093 | 39145 | **RNA**→ rpmE (PRK00019)Ribosomal_L31 (pfam01197)→ |
| Sso-1-162 | NZ_CXLC01000025.1 | + | 39094 | 39146 | **RNA**→ rpmE (PRK00019)Ribosomal_L31 (pfam01197)→ |
| Sso-1-163 | NZ_CXJE01000028.1 | + | 39089 | 39141 | **RNA**→ rpmE (PRK00019)Ribosomal_L31 (pfam01197)→ |
| Sso-1-164 | NZ_CWTR01000029.1 | - | 495 | 443 | **RNA**→ rpmE (PRK00019)Ribosomal_L31 (pfam01197)→ |
| Sso-1-165 | NZ_CXDR01000004.1 | - | 630 | 578 | **RNA**→ rpmE (PRK00019)Ribosomal_L31 (pfam01197)→ |
| Sso-1-166 | NZ_CXNM01000026.1 | - | 1539 | 1487 | **RNA**→ rpmE (PRK00019)Ribosomal_L31 (pfam01197)→ |
| Sso-1-167 | NZ_CXMX01000027.1 | + | 39082 | 39134 | **RNA**→ rpmE (PRK00019)Ribosomal_L31 (pfam01197)→ |
| Sso-1-168 | NZ_CWZD01000024.1 | + | 39089 | 39141 | **RNA**→ rpmE (PRK00019)Ribosomal_L31 (pfam01197)→ |
| Sso-1-169 | NZ_CXEF01000015.1 | + | 39030 | 39082 | **RNA**→ rpmE (PRK00019)Ribosomal_L31 (pfam01197)→ |
| Sso-1-170 | NZ_CXBM01000061.1 | - | 2767 | 2715 | **RNA**→ rpmE (PRK00019)Ribosomal_L31 (pfam01197)→ |
| Sso-1-171 | NZ_CXLL01000024.1 | + | 39082 | 39134 | **RNA**→ rpmE (PRK00019)Ribosomal_L31 (pfam01197)→ |
| Sso-1-172 | NZ_CXMF01000026.1 | + | 39089 | 39141 | **RNA**→ rpmE (PRK00019)Ribosomal_L31 (pfam01197)→ |
| Sso-1-173 | NZ_CXFP01000025.1 | - | 644 | 592 | **RNA**→ rpmE (PRK00019)Ribosomal_L31 (pfam01197)→ |
| Sso-1-174 | NZ_CXKA01000026.1 | - | 662 | 610 | **RNA**→ rpmE (PRK00019)Ribosomal_L31 (pfam01197)→ |
| Sso-1-175 | NZ_CXEZ01000023.1 | + | 39123 | 39175 | **RNA**→ rpmE (PRK00019)Ribosomal_L31 (pfam01197)→ |
| Sso-1-176 | NZ_CXEM01000023.1 | + | 39031 | 39083 | **RNA**→ rpmE (PRK00019)Ribosomal_L31 (pfam01197)→ |
| Sso-1-177 | NZ_CXLS01000028.1 | + | 39082 | 39134 | **RNA**→ rpmE (PRK00019)Ribosomal_L31 (pfam01197)→ |
| Sso-1-178 | NZ_CXDT01000023.1 | - | 630 | 578 | **RNA**→ rpmE (PRK00019)Ribosomal_L31 (pfam01197)→ |
| Sso-1-179 | NZ_CXEE01000022.1 | - | 650 | 598 | **RNA**→ rpmE (PRK00019)Ribosomal_L31 (pfam01197)→ |
| Sso-1-180 | NZ_CWZR01000026.1 | + | 39041 | 39093 | **RNA**→ rpmE (PRK00019)Ribosomal_L31 (pfam01197)→ |
| Sso-1-181 | NZ_CXKY01000024.1 | - | 1137 | 1085 | **RNA**→ rpmE (PRK00019)Ribosomal_L31 (pfam01197)→ |
| Sso-1-182 | NZ_CXFI01000026.1 | - | 1174 | 1122 | **RNA**→ rpmE (PRK00019)Ribosomal_L31 (pfam01197)→ |
| Sso-1-183 | NZ_CXKO01000028.1 | - | 776 | 724 | **RNA**→ rpmE (PRK00019)Ribosomal_L31 (pfam01197)→ |
| Sso-1-184 | NZ_CXMV01000027.1 | + | 39082 | 39134 | **RNA**→ rpmE (PRK00019)Ribosomal_L31 (pfam01197)→ |
| Sso-1-185 | NZ_CXLO01000028.1 | - | 695 | 643 | **RNA**→ rpmE (PRK00019)Ribosomal_L31 (pfam01197)→ |
| Sso-1-186 | NZ_CXJM01000025.1 | + | 39082 | 39134 | **RNA**→ rpmE (PRK00019)Ribosomal_L31 (pfam01197)→ |
| Sso-1-187 | NZ_CXNG01000028.1 | - | 529 | 477 | **RNA**→ rpmE (PRK00019)Ribosomal_L31 (pfam01197)→ |
| Sso-1-188 | NZ_CXGZ01000014.1 | - | 627 | 575 | **RNA**→ rpmE (PRK00019)Ribosomal_L31 (pfam01197)→ |
| Sso-1-189 | NZ_CWWZ01000021.1 | - | 846 | 794 | **RNA**→ rpmE (PRK00019)Ribosomal_L31 (pfam01197)→ |
| Sso-1-190 | NZ_CXNB01000027.1 | + | 39089 | 39141 | **RNA**→ rpmE (PRK00019)Ribosomal_L31 (pfam01197)→ |
| Sso-1-191 | NZ_CXGO01000022.1 | - | 850 | 798 | **RNA**→ rpmE (PRK00019)Ribosomal_L31 (pfam01197)→ |
| Sso-1-192 | NZ_CWXT01000023.1 | + | 39089 | 39141 | **RNA**→ rpmE (PRK00019)Ribosomal_L31 (pfam01197)→ |
| Sso-1-193 | NZ_CXLZ01000024.1 | - | 1703 | 1651 | **RNA**→ rpmE (PRK00019)Ribosomal_L31 (pfam01197)→ |
| Sso-1-194 | NZ_CWZV01000027.1 | + | 39089 | 39141 | **RNA**→ rpmE (PRK00019)Ribosomal_L31 (pfam01197)→ |
| Sso-1-195 | NZ_CXDY01000020.1 | - | 644 | 592 | **RNA**→ rpmE (PRK00019)Ribosomal_L31 (pfam01197)→ |
| Sso-1-196 | NZ_CXCQ01000030.1 | - | 405 | 353 | **RNA**→ rpmE (PRK00019)Ribosomal_L31 (pfam01197)→ |
| Sso-1-197 | NZ_CXAM01000038.1 | - | 630 | 578 | **RNA**→ rpmE (PRK00019)Ribosomal_L31 (pfam01197)→ |
| Sso-1-198 | NZ_CXCU01000026.1 | - | 636 | 584 | **RNA**→ rpmE (PRK00019)Ribosomal_L31 (pfam01197)→ |
| Eco-353-1 | NZ_AIAY01000004.1 | + | 76542 | 76594 | **RNA**→ rpmE (PRK00019)Ribosomal_L31 (pfam01197)→ |
| Eco-343-1 | NZ_AIAO01000005.1 | - | 39155 | 39103 | **RNA**→ rpmE (PRK00019)Ribosomal_L31 (pfam01197)→ |
| Eco-348-1 | NZ_AIAT01000054.1 | + | 18409 | 18461 | **RNA**→ rpmE (PRK00019)Ribosomal_L31 (pfam01197)→ |

| | | | | | | |
|---|---|---|---|---|---|---|
| Eco-345-1 | NZ_AIAQ01000283.1 | + | 29275 | 29327 | **RNA**→ | rpmE (PRK00019)Ribosomal_L31 (pfam01197)→ |
| Eco-384-1 | NZ_AICC01000004.1 | + | 46666 | 46718 | **RNA**→ | rpmE (PRK00019)Ribosomal_L31 (pfam01197)→ |
| Eco-356-1 | NZ_AIBB01000072.1 | - | 39155 | 39103 | **RNA**→ | rpmE (PRK00019)Ribosomal_L31 (pfam01197)→ |
| Eco-349-1 | NZ_AIAU01000029.1 | - | 486 | 434 | **RNA**→ | rpmE (PRK00019)Ribosomal_L31 (pfam01197)→ |
| Eco-378-1 | NZ_AIBX01000062.1 | + | 47741 | 47793 | **RNA**→ | rpmE (PRK00019)Ribosomal_L31 (pfam01197)→ |
| Eco-339-1 | NZ_AIAJ01000152.1 | - | 640 | 588 | **RNA**→ | rpmE (PRK00019)Ribosomal_L31 (pfam01197)→ |
| Eco-336-1 | NZ_AIAH01000234.1 | - | 547 | 495 | **RNA**→ | rpmE (PRK00019)Ribosomal_L31 (pfam01197)→ |
| Eco-337-1 | NZ_AIAI01000001.1 | - | 41795 | 41743 | **RNA**→ | rpmE (PRK00019)Ribosomal_L31 (pfam01197)→ |
| Eco-358-1 | NZ_AIBD01000070.1 | + | 46666 | 46718 | **RNA**→ | rpmE (PRK00019)Ribosomal_L31 (pfam01197)→ |
| Eco-379-1 | NZ_AIBW01000118.1 | - | 640 | 588 | **RNA**→ | rpmE (PRK00019)Ribosomal_L31 (pfam01197)→ |
| Eco-50-1 | NZ_ATOE01000063.1 | + | 39109 | 39161 | **RNA**→ | rpmE (PRK00019)Ribosomal_L31 (pfam01197)→ |
| Eco-366-1 | NZ_AIBK01000054.1 | + | 28476 | 28528 | **RNA**→ | rpmE (PRK00019)Ribosomal_L31 (pfam01197)→ |
| Eco-338-1 | NZ_AICF01000195.1 | - | 486 | 434 | **RNA**→ | rpmE (PRK00019)Ribosomal_L31 (pfam01197)→ |
| Eco-372-1 | NZ_AIBR01000067.1 | + | 47726 | 47778 | **RNA**→ | rpmE (PRK00019)Ribosomal_L31 (pfam01197)→ |
| Eco-359-1 | NZ_AIBE01000033.1 | - | 52126 | 52074 | **RNA**→ | rpmE (PRK00019)Ribosomal_L31 (pfam01197)→ |
| Eco-350-1 | NZ_AIAV01000087.1 | + | 60721 | 60773 | **RNA**→ | rpmE (PRK00019)Ribosomal_L31 (pfam01197)→ |
| Eco-375-1 | NZ_AIBQ01000159.1 | + | 11322 | 11374 | **RNA**→ | rpmE (PRK00019)Ribosomal_L31 (pfam01197)→ |
| Sso-1-199 | NZ_CXAJ01000024.1 | + | 39089 | 39141 | **RNA**→ | rpmE (PRK00019)Ribosomal_L31 (pfam01197)→ |
| Sso-1-200 | NZ_CWXR01000025.1 | - | 848 | 796 | **RNA**→ | rpmE (PRK00019)Ribosomal_L31 (pfam01197)→ |
| Sso-1-201 | NZ_CXML01000027.1 | + | 39089 | 39141 | **RNA**→ | rpmE (PRK00019)Ribosomal_L31 (pfam01197)→ |
| Sso-1-202 | NZ_CWXU01000023.1 | - | 855 | 803 | **RNA**→ | rpmE (PRK00019)Ribosomal_L31 (pfam01197)→ |
| Sso-1-203 | NZ_CXBD01000022.1 | - | 1137 | 1085 | **RNA**→ | rpmE (PRK00019)Ribosomal_L31 (pfam01197)→ |
| Sfl-25-6 | NZ_LAHZ01000037.1 | + | 24424 | 24476 | **RNA**→ | rpmE (PRK00019)Ribosomal_L31 (pfam01197)→ |
| Sso-1-204 | NZ_CXJT01000025.1 | + | 39089 | 39141 | **RNA**→ | rpmE (PRK00019)Ribosomal_L31 (pfam01197)→ |
| Sso-1-205 | NZ_CWXZ01000023.1 | - | 865 | 813 | **RNA**→ | rpmE (PRK00019)Ribosomal_L31 (pfam01197)→ |
| Sso-1-206 | NZ_CXIX01000027.1 | - | 786 | 734 | **RNA**→ | rpmE (PRK00019)Ribosomal_L31 (pfam01197)→ |
| Sso-1-207 | NZ_CWXD01000025.1 | + | 39104 | 39156 | **RNA**→ | rpmE (PRK00019)Ribosomal_L31 (pfam01197)→ |
| Sfl-25-10 | NZ_LAID01000111.1 | - | 31124 | 31072 | **RNA**→ | rpmE (PRK00019)Ribosomal_L31 (pfam01197)→ |
| Sso-1-208 | NZ_CXFH01000026.1 | - | 1174 | 1122 | **RNA**→ | rpmE (PRK00019)Ribosomal_L31 (pfam01197)→ |
| Sfl-51-1 | NZ_LAIC01000109.1 | + | 24423 | 24475 | **RNA**→ | rpmE (PRK00019)Ribosomal_L31 (pfam01197)→ |
| Sso-1-209 | NZ_CXHU01000023.1 | - | 1137 | 1085 | **RNA**→ | rpmE (PRK00019)Ribosomal_L31 (pfam01197)→ |
| Sso-1-210 | NZ_CXEH01000026.1 | - | 1137 | 1085 | **RNA**→ | rpmE (PRK00019)Ribosomal_L31 (pfam01197)→ |
| Sfl-25-7 | NZ_LAIE01000028.1 | + | 30022 | 30074 | **RNA**→ | rpmE (PRK00019)Ribosomal_L31 (pfam01197)→ |
| Sso-1-211 | NZ_CXKP01000022.1 | - | 1137 | 1085 | **RNA**→ | rpmE (PRK00019)Ribosomal_L31 (pfam01197)→ |
| Sso-1-212 | NZ_CWWJ01000027.1 | + | 39093 | 39145 | **RNA**→ | rpmE (PRK00019)Ribosomal_L31 (pfam01197)→ |
| Sso-1-213 | NZ_CXBI01000018.1 | + | 39023 | 39075 | **RNA**→ | rpmE (PRK00019)Ribosomal_L31 (pfam01197)→ |
| Sso-1-214 | NZ_CXIY01000027.1 | - | 793 | 741 | **RNA**→ | rpmE (PRK00019)Ribosomal_L31 (pfam01197)→ |
| Sso-1-215 | NZ_CXHS01000025.1 | + | 39045 | 39097 | **RNA**→ | rpmE (PRK00019)Ribosomal_L31 (pfam01197)→ |
| Sso-1-216 | NZ_CWUK01000025.1 | - | 555 | 503 | **RNA**→ | rpmE (PRK00019)Ribosomal_L31 (pfam01197)→ |
| Sso-1-217 | NZ_CWYX01000027.1 | + | 39115 | 39167 | **RNA**→ | rpmE (PRK00019)Ribosomal_L31 (pfam01197)→ |
| Sso-1-218 | NZ_CWWW01000023.1 | + | 39099 | 39151 | **RNA**→ | rpmE (PRK00019)Ribosomal_L31 (pfam01197)→ |
| Sso-1-219 | NZ_CXJH01000024.1 | + | 39082 | 39134 | **RNA**→ | rpmE (PRK00019)Ribosomal_L31 (pfam01197)→ |
| Sso-1-220 | NZ_CWUL01000025.1 | + | 38789 | 38841 | **RNA**→ | rpmE (PRK00019)Ribosomal_L31 (pfam01197)→ |
| Sso-1-221 | NZ_CXIM01000022.1 | + | 39082 | 39134 | **RNA**→ | rpmE (PRK00019)Ribosomal_L31 (pfam01197)→ |
| Sfl-25-8 | NZ_LAHV01000005.1 | + | 24423 | 24475 | **RNA**→ | rpmE (PRK00019)Ribosomal_L31 (pfam01197)→ |
| Sso-1-222 | NZ_CXGR01000027.1 | - | 695 | 643 | **RNA**→ | rpmE (PRK00019)Ribosomal_L31 (pfam01197)→ |
| Sso-1-223 | NZ_CXLK01000027.1 | + | 39082 | 39134 | **RNA**→ | rpmE (PRK00019)Ribosomal_L31 (pfam01197)→ |
| Sso-1-224 | NZ_CXCD01000021.1 | + | 39007 | 39059 | **RNA**→ | rpmE (PRK00019)Ribosomal_L31 (pfam01197)→ |
| Sso-1-225 | NZ_CWZP01000023.1 | - | 664 | 612 | **RNA**→ | rpmE (PRK00019)Ribosomal_L31 (pfam01197)→ |
| Sso-1-226 | NZ_CWZK01000029.1 | + | 39099 | 39151 | **RNA**→ | rpmE (PRK00019)Ribosomal_L31 (pfam01197)→ |
| Sso-1-227 | NZ_CWYC01000025.1 | + | 39084 | 39136 | **RNA**→ | rpmE (PRK00019)Ribosomal_L31 (pfam01197)→ |
| Sso-1-228 | NZ_CXCS01000004.1 | + | 76716 | 76768 | **RNA**→ | rpmE (PRK00019)Ribosomal_L31 (pfam01197)→ |

| | | | | | |
|---|---|---|---|---|---|
| Sso-1-229 | NZ_CXFO01000025.1 | - | 1174 | 1122 | **RNA**→ rpmE (PRK00019)Ribosomal_L31 (pfam01197)→ |
| Sso-1-230 | NZ_CXJF01000027.1 | - | 1539 | 1487 | **RNA**→ rpmE (PRK00019)Ribosomal_L31 (pfam01197)→ |
| Sso-1-231 | NZ_CXFF01000024.1 | + | 39045 | 39097 | **RNA**→ rpmE (PRK00019)Ribosomal_L31 (pfam01197)→ |
| Sso-1-232 | NZ_CWYB01000025.1 | + | 39089 | 39141 | **RNA**→ rpmE (PRK00019)Ribosomal_L31 (pfam01197)→ |
| Sso-1-233 | NZ_CWZY01000026.1 | + | 39082 | 39134 | **RNA**→ rpmE (PRK00019)Ribosomal_L31 (pfam01197)→ |
| Sso-1-234 | NZ_CXMG01000026.1 | - | 792 | 740 | **RNA**→ rpmE (PRK00019)Ribosomal_L31 (pfam01197)→ |
| Sso-1-235 | NZ_CXBS01000019.1 | - | 630 | 578 | **RNA**→ rpmE (PRK00019)Ribosomal_L31 (pfam01197)→ |
| Sso-1-236 | NZ_CXBX01000019.1 | + | 39027 | 39079 | **RNA**→ rpmE (PRK00019)Ribosomal_L31 (pfam01197)→ |
| Sso-1-237 | NZ_CXKF01000028.1 | + | 39089 | 39141 | **RNA**→ rpmE (PRK00019)Ribosomal_L31 (pfam01197)→ |
| Sso-1-238 | NZ_CWWP01000024.1 | + | 39099 | 39151 | **RNA**→ rpmE (PRK00019)Ribosomal_L31 (pfam01197)→ |
| Sso-1-239 | NZ_CXBW01000012.1 | + | 47420 | 47472 | **RNA**→ rpmE (PRK00019)Ribosomal_L31 (pfam01197)→ |
| Sso-1-240 | NZ_CXEI01000023.1 | - | 1137 | 1085 | **RNA**→ rpmE (PRK00019)Ribosomal_L31 (pfam01197)→ |
| Sso-1-241 | NZ_CXJZ01000027.1 | - | 695 | 643 | **RNA**→ rpmE (PRK00019)Ribosomal_L31 (pfam01197)→ |
| Sso-1-242 | NZ_CXKU01000020.1 | - | 1174 | 1122 | **RNA**→ rpmE (PRK00019)Ribosomal_L31 (pfam01197)→ |
| Sso-1-243 | NZ_CXLB01000016.1 | + | 39082 | 39134 | **RNA**→ rpmE (PRK00019)Ribosomal_L31 (pfam01197)→ |
| Sso-1-244 | NZ_CXJP01000026.1 | + | 39089 | 39141 | **RNA**→ rpmE (PRK00019)Ribosomal_L31 (pfam01197)→ |
| Sso-1-245 | NZ_CXGK01000028.1 | - | 556 | 504 | **RNA**→ rpmE (PRK00019)Ribosomal_L31 (pfam01197)→ |
| Sso-1-246 | NZ_CXEO01000018.1 | - | 644 | 592 | **RNA**→ rpmE (PRK00019)Ribosomal_L31 (pfam01197)→ |
| Sso-1-247 | NZ_CWYG01000025.1 | - | 1680 | 1628 | **RNA**→ rpmE (PRK00019)Ribosomal_L31 (pfam01197)→ |
| Sso-1-248 | NZ_CXLG01000028.1 | + | 39082 | 39134 | **RNA**→ rpmE (PRK00019)Ribosomal_L31 (pfam01197)→ |
| Sso-1-249 | NZ_CWWU01000025.1 | + | 39104 | 39156 | **RNA**→ rpmE (PRK00019)Ribosomal_L31 (pfam01197)→ |
| Sso-1-250 | NZ_CWZO01000026.1 | - | 662 | 610 | **RNA**→ rpmE (PRK00019)Ribosomal_L31 (pfam01197)→ |
| Sso-1-251 | NZ_CXGC01000028.1 | - | 1627 | 1575 | **RNA**→ rpmE (PRK00019)Ribosomal_L31 (pfam01197)→ |
| Sso-1-252 | NZ_CWUC01000026.1 | - | 660 | 608 | **RNA**→ rpmE (PRK00019)Ribosomal_L31 (pfam01197)→ |
| Sso-1-253 | NZ_CXEB01000017.1 | + | 47821 | 47873 | **RNA**→ rpmE (PRK00019)Ribosomal_L31 (pfam01197)→ |
| Sso-1-254 | NZ_CXAG01000028.1 | - | 664 | 612 | **RNA**→ rpmE (PRK00019)Ribosomal_L31 (pfam01197)→ |
| Sso-1-255 | NZ_CWWG01000023.1 | + | 39082 | 39134 | **RNA**→ rpmE (PRK00019)Ribosomal_L31 (pfam01197)→ |
| Sso-1-256 | NZ_CXES01000023.1 | + | 39030 | 39082 | **RNA**→ rpmE (PRK00019)Ribosomal_L31 (pfam01197)→ |
| Sso-1-257 | NZ_CWYR01000024.1 | + | 39089 | 39141 | **RNA**→ rpmE (PRK00019)Ribosomal_L31 (pfam01197)→ |
| Sso-1-258 | NZ_CWVN01000027.1 | - | 799 | 747 | **RNA**→ rpmE (PRK00019)Ribosomal_L31 (pfam01197)→ |
| Sso-1-259 | NZ_CWWS01000022.1 | - | 833 | 781 | **RNA**→ rpmE (PRK00019)Ribosomal_L31 (pfam01197)→ |
| Sso-1-260 | NZ_CXFD01000026.1 | - | 652 | 600 | **RNA**→ rpmE (PRK00019)Ribosomal_L31 (pfam01197)→ |
| Sso-1-261 | NZ_CWYD01000027.1 | + | 39100 | 39152 | **RNA**→ rpmE (PRK00019)Ribosomal_L31 (pfam01197)→ |
| Sso-1-262 | NZ_CWUW01000028.1 | - | 1768 | 1716 | **RNA**→ rpmE (PRK00019)Ribosomal_L31 (pfam01197)→ |
| Sso-1-263 | NZ_CXCO01000033.1 | - | 632 | 580 | **RNA**→ rpmE (PRK00019)Ribosomal_L31 (pfam01197)→ |
| Sso-1-264 | NZ_CWZB01000027.1 | - | 672 | 620 | **RNA**→ rpmE (PRK00019)Ribosomal_L31 (pfam01197)→ |
| Sso-1-265 | NZ_CWVG01000028.1 | - | 1732 | 1680 | **RNA**→ rpmE (PRK00019)Ribosomal_L31 (pfam01197)→ |
| Sso-1-266 | NZ_CXJQ01000027.1 | + | 39082 | 39134 | **RNA**→ rpmE (PRK00019)Ribosomal_L31 (pfam01197)→ |
| Sso-1-267 | NZ_CXCR01000016.1 | - | 465 | 413 | **RNA**→ rpmE (PRK00019)Ribosomal_L31 (pfam01197)→ |
| Sso-1-268 | NZ_CWTP01000028.1 | + | 39087 | 39139 | **RNA**→ rpmE (PRK00019)Ribosomal_L31 (pfam01197)→ |
| Sso-1-269 | NZ_CXAR01000002.1 | - | 1174 | 1122 | **RNA**→ rpmE (PRK00019)Ribosomal_L31 (pfam01197)→ |
| Sso-1-270 | NZ_CXIG01000026.1 | - | 1539 | 1487 | **RNA**→ rpmE (PRK00019)Ribosomal_L31 (pfam01197)→ |
| Sso-1-271 | NZ_CWTA01000027.1 | + | 39144 | 39196 | **RNA**→ rpmE (PRK00019)Ribosomal_L31 (pfam01197)→ |
| Sso-1-272 | NZ_CWXC01000024.1 | + | 39104 | 39156 | **RNA**→ rpmE (PRK00019)Ribosomal_L31 (pfam01197)→ |
| Sso-1-273 | NZ_CXBH01000062.1 | + | 20736 | 20788 | **RNA**→ rpmE (PRK00019)Ribosomal_L31 (pfam01197)→ |
| Sso-1-274 | NZ_CWTK01000015.1 | - | 660 | 608 | **RNA**→ rpmE (PRK00019)Ribosomal_L31 (pfam01197)→ |
| Sso-1-275 | NZ_CXBC01000024.1 | - | 1174 | 1122 | **RNA**→ rpmE (PRK00019)Ribosomal_L31 (pfam01197)→ |
| Sso-1-276 | NZ_CXCI01000022.1 | - | 630 | 578 | **RNA**→ rpmE (PRK00019)Ribosomal_L31 (pfam01197)→ |
| Sso-1-277 | NZ_CWWD01000024.1 | - | 1202 | 1150 | **RNA**→ rpmE (PRK00019)Ribosomal_L31 (pfam01197)→ |
| Sso-1-278 | NZ_CXDJ01000008.1 | - | 630 | 578 | **RNA**→ rpmE (PRK00019)Ribosomal_L31 (pfam01197)→ |
| Sso-1-279 | NZ_CXFS01000026.1 | + | 39031 | 39083 | **RNA**→ rpmE (PRK00019)Ribosomal_L31 (pfam01197)→ |
| Sso-1-280 | NZ_CWTI01000027.1 | + | 39109 | 39161 | **RNA**→ rpmE (PRK00019)Ribosomal_L31 (pfam01197)→ |

| | | | | | |
|---|---|---|---|---|---|
| Sso-1-281 | NZ_CXDV01000023.1 | + | 47867 | 47919 | **RNA**→ rpmE (PRK00019)Ribosomal_L31 (pfam01197)→ |
| Sso-1-282 | NZ_CXAW01000024.1 | - | 1174 | 1122 | **RNA**→ rpmE (PRK00019)Ribosomal_L31 (pfam01197)→ |
| Sso-1-283 | NZ_CWTC01000022.1 | + | 39082 | 39134 | **RNA**→ rpmE (PRK00019)Ribosomal_L31 (pfam01197)→ |
| Sso-1-284 | NZ_CXLP01000026.1 | - | 794 | 742 | **RNA**→ rpmE (PRK00019)Ribosomal_L31 (pfam01197)→ |
| Sso-1-285 | NZ_CXIQ01000021.1 | - | 2679 | 2627 | **RNA**→ rpmE (PRK00019)Ribosomal_L31 (pfam01197)→ |
| Sso-1-286 | NZ_CWTY01000023.1 | + | 39069 | 39121 | **RNA**→ rpmE (PRK00019)Ribosomal_L31 (pfam01197)→ |
| Sso-1-287 | NZ_CXEX01000023.1 | + | 39045 | 39097 | **RNA**→ rpmE (PRK00019)Ribosomal_L31 (pfam01197)→ |
| Sso-1-288 | NZ_CXAV01000016.1 | + | 39007 | 39059 | **RNA**→ rpmE (PRK00019)Ribosomal_L31 (pfam01197)→ |
| Sso-1-289 | NZ_CWUT01000025.1 | + | 39093 | 39145 | **RNA**→ rpmE (PRK00019)Ribosomal_L31 (pfam01197)→ |
| Sso-1-290 | NZ_CXLQ01000026.1 | - | 1279 | 1227 | **RNA**→ rpmE (PRK00019)Ribosomal_L31 (pfam01197)→ |
| Sso-1-291 | NZ_CXND01000027.1 | - | 915 | 863 | **RNA**→ rpmE (PRK00019)Ribosomal_L31 (pfam01197)→ |
| Sso-1-292 | NZ_CWWH01000021.1 | + | 39093 | 39145 | **RNA**→ rpmE (PRK00019)Ribosomal_L31 (pfam01197)→ |
| Sso-1-293 | NZ_CWWX01000020.1 | - | 835 | 783 | **RNA**→ rpmE (PRK00019)Ribosomal_L31 (pfam01197)→ |
| Sso-1-294 | NZ_CXFK01000023.1 | + | 39045 | 39097 | **RNA**→ rpmE (PRK00019)Ribosomal_L31 (pfam01197)→ |
| Sso-1-295 | NZ_CWVD01000031.1 | + | 39089 | 39141 | **RNA**→ rpmE (PRK00019)Ribosomal_L31 (pfam01197)→ |
| Sso-1-296 | NZ_CWTN01000022.1 | - | 660 | 608 | **RNA**→ rpmE (PRK00019)Ribosomal_L31 (pfam01197)→ |
| Sso-1-297 | NZ_CXHI01000028.1 | - | 792 | 740 | **RNA**→ rpmE (PRK00019)Ribosomal_L31 (pfam01197)→ |
| Sso-1-298 | NZ_CWTQ01000027.1 | + | 38832 | 38884 | **RNA**→ rpmE (PRK00019)Ribosomal_L31 (pfam01197)→ |
| Sso-1-299 | NZ_CWVV01000028.1 | + | 39099 | 39151 | **RNA**→ rpmE (PRK00019)Ribosomal_L31 (pfam01197)→ |
| Sso-1-300 | NZ_CWZI01000025.1 | - | 1692 | 1640 | **RNA**→ rpmE (PRK00019)Ribosomal_L31 (pfam01197)→ |
| Sso-1-301 | NZ_CWTB01000024.1 | - | 660 | 608 | **RNA**→ rpmE (PRK00019)Ribosomal_L31 (pfam01197)→ |
| Sso-1-302 | NZ_CXAB01000024.1 | - | 1174 | 1122 | **RNA**→ rpmE (PRK00019)Ribosomal_L31 (pfam01197)→ |
| Sso-1-303 | NZ_CWUM01000015.1 | + | 39156 | 39208 | **RNA**→ rpmE (PRK00019)Ribosomal_L31 (pfam01197)→ |
| Sso-1-304 | NZ_CWVX01000027.1 | + | 39093 | 39145 | **RNA**→ rpmE (PRK00019)Ribosomal_L31 (pfam01197)→ |
| Sso-1-305 | NZ_CXKV01000028.1 | - | 682 | 630 | **RNA**→ rpmE (PRK00019)Ribosomal_L31 (pfam01197)→ |
| Sso-1-306 | NZ_CWSW01000026.1 | - | 660 | 608 | **RNA**→ rpmE (PRK00019)Ribosomal_L31 (pfam01197)→ |
| Sso-1-307 | NZ_CWWM01000029.1 | + | 39082 | 39134 | **RNA**→ rpmE (PRK00019)Ribosomal_L31 (pfam01197)→ |
| Sso-1-308 | NZ_CXDI01000014.1 | - | 480 | 428 | **RNA**→ rpmE (PRK00019)Ribosomal_L31 (pfam01197)→ |
| Sso-1-309 | NZ_CWVC01000026.1 | + | 39093 | 39145 | **RNA**→ rpmE (PRK00019)Ribosomal_L31 (pfam01197)→ |
| Sso-1-310 | NZ_CWTJ01000023.1 | - | 1539 | 1487 | **RNA**→ rpmE (PRK00019)Ribosomal_L31 (pfam01197)→ |
| Sso-1-311 | NZ_CXBL01000018.1 | + | 39135 | 39187 | **RNA**→ rpmE (PRK00019)Ribosomal_L31 (pfam01197)→ |
| Sso-1-312 | NZ_CWVY01000029.1 | + | 39082 | 39134 | **RNA**→ rpmE (PRK00019)Ribosomal_L31 (pfam01197)→ |
| Sso-1-313 | NZ_CXEL01000014.1 | - | 644 | 592 | **RNA**→ rpmE (PRK00019)Ribosomal_L31 (pfam01197)→ |
| Sso-1-314 | NZ_CXHH01000027.1 | - | 695 | 643 | **RNA**→ rpmE (PRK00019)Ribosomal_L31 (pfam01197)→ |
| Sso-1-315 | NZ_CWVW01000029.1 | - | 1397 | 1345 | **RNA**→ rpmE (PRK00019)Ribosomal_L31 (pfam01197)→ |
| Sso-1-316 | NZ_CWZW01000026.1 | - | 559 | 507 | **RNA**→ rpmE (PRK00019)Ribosomal_L31 (pfam01197)→ |
| Sso-1-317 | NZ_CXHJ01000027.1 | - | 695 | 643 | **RNA**→ rpmE (PRK00019)Ribosomal_L31 (pfam01197)→ |
| Sso-1-318 | NZ_CWUH01000025.1 | + | 39104 | 39156 | **RNA**→ rpmE (PRK00019)Ribosomal_L31 (pfam01197)→ |
| Sso-1-319 | NZ_CXNS01000027.1 | - | 1294 | 1242 | **RNA**→ rpmE (PRK00019)Ribosomal_L31 (pfam01197)→ |
| Sso-1-320 | NZ_CXGV01000027.1 | + | 39094 | 39146 | **RNA**→ rpmE (PRK00019)Ribosomal_L31 (pfam01197)→ |
| Sso-1-321 | NZ_CXHO01000023.1 | + | 39089 | 39141 | **RNA**→ rpmE (PRK00019)Ribosomal_L31 (pfam01197)→ |
| Sso-1-322 | NZ_CWTM01000025.1 | + | 39158 | 39210 | **RNA**→ rpmE (PRK00019)Ribosomal_L31 (pfam01197)→ |
| Sso-1-323 | NZ_CXMK01000027.1 | + | 39089 | 39141 | **RNA**→ rpmE (PRK00019)Ribosomal_L31 (pfam01197)→ |
| Sso-1-324 | NZ_CXBU01000023.1 | + | 39030 | 39082 | **RNA**→ rpmE (PRK00019)Ribosomal_L31 (pfam01197)→ |
| Sso-1-325 | NZ_CWVR01000030.1 | + | 39093 | 39145 | **RNA**→ rpmE (PRK00019)Ribosomal_L31 (pfam01197)→ |
| Sso-1-326 | NZ_CXIF01000024.1 | - | 776 | 724 | **RNA**→ rpmE (PRK00019)Ribosomal_L31 (pfam01197)→ |
| Sso-1-327 | NZ_CXDZ01000014.1 | - | 1137 | 1085 | **RNA**→ rpmE (PRK00019)Ribosomal_L31 (pfam01197)→ |
| Sso-1-328 | NZ_CWWO01000025.1 | + | 39089 | 39141 | **RNA**→ rpmE (PRK00019)Ribosomal_L31 (pfam01197)→ |
| Sso-1-329 | NZ_CWYT01000026.1 | - | 664 | 612 | **RNA**→ rpmE (PRK00019)Ribosomal_L31 (pfam01197)→ |
| Sso-1-330 | NZ_CXHQ01000026.1 | + | 39082 | 39134 | **RNA**→ rpmE (PRK00019)Ribosomal_L31 (pfam01197)→ |
| Sso-1-331 | NZ_CWYY01000025.1 | - | 1539 | 1487 | **RNA**→ rpmE (PRK00019)Ribosomal_L31 (pfam01197)→ |
| Sso-1-332 | NZ_CXKM01000025.1 | - | 771 | 719 | **RNA**→ rpmE (PRK00019)Ribosomal_L31 (pfam01197)→ |

| | | | | | |
|---|---|---|---|---|---|
| Sso-1-333 | NZ_CWZF01000025.1 | - | 662 | 610 | **RNA**→ rpmE (PRK00019)Ribosomal_L31 (pfam01197)→ |
| Sso-1-334 | NZ_CWWA01000027.1 | - | 800 | 748 | **RNA**→ rpmE (PRK00019)Ribosomal_L31 (pfam01197)→ |
| Sso-1-335 | NZ_CXHE01000025.1 | - | 2674 | 2622 | **RNA**→ rpmE (PRK00019)Ribosomal_L31 (pfam01197)→ |
| Sso-1-336 | NZ_CXBN01000024.1 | + | 39023 | 39075 | **RNA**→ rpmE (PRK00019)Ribosomal_L31 (pfam01197)→ |
| Sso-1-337 | NZ_CXEN01000022.1 | - | 821 | 769 | **RNA**→ rpmE (PRK00019)Ribosomal_L31 (pfam01197)→ |
| Sso-1-338 | NZ_CXAT01000016.1 | + | 39023 | 39075 | **RNA**→ rpmE (PRK00019)Ribosomal_L31 (pfam01197)→ |
| Sso-1-339 | NZ_CXJA01000025.1 | + | 39089 | 39141 | **RNA**→ rpmE (PRK00019)Ribosomal_L31 (pfam01197)→ |
| Sso-1-340 | NZ_CWUP01000024.1 | - | 555 | 503 | **RNA**→ rpmE (PRK00019)Ribosomal_L31 (pfam01197)→ |
| Sso-1-341 | NZ_CXBO01000021.1 | - | 1174 | 1122 | **RNA**→ rpmE (PRK00019)Ribosomal_L31 (pfam01197)→ |
| Sso-1-342 | NZ_CXCY01000005.1 | + | 80319 | 80371 | **RNA**→ rpmE (PRK00019)Ribosomal_L31 (pfam01197)→ |
| Sso-1-343 | NZ_CWUG01000016.1 | + | 38772 | 38824 | **RNA**→ rpmE (PRK00019)Ribosomal_L31 (pfam01197)→ |
| Sso-1-344 | NZ_CXHW01000028.1 | + | 39089 | 39141 | **RNA**→ rpmE (PRK00019)Ribosomal_L31 (pfam01197)→ |
| Sso-1-345 | NZ_CXIR01000026.1 | + | 39082 | 39134 | **RNA**→ rpmE (PRK00019)Ribosomal_L31 (pfam01197)→ |
| Sso-1-346 | NZ_CWVB01000026.1 | - | 794 | 742 | **RNA**→ rpmE (PRK00019)Ribosomal_L31 (pfam01197)→ |
| Sso-1-347 | NZ_CXLR01000027.1 | + | 39089 | 39141 | **RNA**→ rpmE (PRK00019)Ribosomal_L31 (pfam01197)→ |
| Sso-1-348 | NZ_CXCL01000027.1 | - | 632 | 580 | **RNA**→ rpmE (PRK00019)Ribosomal_L31 (pfam01197)→ |
| Sso-1-349 | NZ_CWSY01000022.1 | - | 555 | 503 | **RNA**→ rpmE (PRK00019)Ribosomal_L31 (pfam01197)→ |
| Sso-1-350 | NZ_CXDN01000021.1 | - | 1174 | 1122 | **RNA**→ rpmE (PRK00019)Ribosomal_L31 (pfam01197)→ |
| Sso-1-351 | NZ_CWXB01000016.1 | + | 39104 | 39156 | **RNA**→ rpmE (PRK00019)Ribosomal_L31 (pfam01197)→ |
| Sso-1-352 | NZ_CWXA01000021.1 | + | 39104 | 39156 | **RNA**→ rpmE (PRK00019)Ribosomal_L31 (pfam01197)→ |
| Sso-1-353 | NZ_CWSZ01000022.1 | - | 660 | 608 | **RNA**→ rpmE (PRK00019)Ribosomal_L31 (pfam01197)→ |
| Sso-1-354 | NZ_CXAL01000025.1 | - | 676 | 624 | **RNA**→ rpmE (PRK00019)Ribosomal_L31 (pfam01197)→ |
| Sso-1-355 | NZ_CXLT01000025.1 | + | 39089 | 39141 | **RNA**→ rpmE (PRK00019)Ribosomal_L31 (pfam01197)→ |
| Sso-1-356 | NZ_CWWK01000028.1 | - | 800 | 748 | **RNA**→ rpmE (PRK00019)Ribosomal_L31 (pfam01197)→ |
| Sso-1-357 | NZ_CXDS01000022.1 | - | 5350 | 5298 | **RNA**→ rpmE (PRK00019)Ribosomal_L31 (pfam01197)→ |
| Sso-1-358 | NZ_CWUV01000023.1 | + | 39082 | 39134 | **RNA**→ rpmE (PRK00019)Ribosomal_L31 (pfam01197)→ |
| Sso-1-359 | NZ_CXGN01000021.1 | - | 628 | 576 | **RNA**→ rpmE (PRK00019)Ribosomal_L31 (pfam01197)→ |
| Sso-1-360 | NZ_CWYK01000022.1 | + | 39082 | 39134 | **RNA**→ rpmE (PRK00019)Ribosomal_L31 (pfam01197)→ |
| Sso-1-361 | NZ_CWUQ01000021.1 | - | 660 | 608 | **RNA**→ rpmE (PRK00019)Ribosomal_L31 (pfam01197)→ |
| Sso-1-362 | NZ_CXKG01000049.1 | + | 22507 | 22559 | **RNA**→ rpmE (PRK00019)Ribosomal_L31 (pfam01197)→ |
| Sso-1-363 | NZ_CXFC01000024.1 | - | 1701 | 1649 | **RNA**→ rpmE (PRK00019)Ribosomal_L31 (pfam01197)→ |
| Sso-1-364 | NZ_CWWL01000026.1 | - | 1227 | 1175 | **RNA**→ rpmE (PRK00019)Ribosomal_L31 (pfam01197)→ |
| Sso-1-365 | NZ_CXGH01000025.1 | + | 39045 | 39097 | **RNA**→ rpmE (PRK00019)Ribosomal_L31 (pfam01197)→ |
| Sso-1-366 | NZ_CXEU01000023.1 | - | 768 | 716 | **RNA**→ rpmE (PRK00019)Ribosomal_L31 (pfam01197)→ |
| Sso-1-367 | NZ_CWVS01000024.1 | - | 3757 | 3705 | **RNA**→ rpmE (PRK00019)Ribosomal_L31 (pfam01197)→ |
| Sso-1-368 | NZ_CXHD01000027.1 | + | 39082 | 39134 | **RNA**→ rpmE (PRK00019)Ribosomal_L31 (pfam01197)→ |
| Sso-1-369 | NZ_CXJR01000027.1 | + | 39089 | 39141 | **RNA**→ rpmE (PRK00019)Ribosomal_L31 (pfam01197)→ |
| Sso-1-370 | NZ_CWWI01000027.1 | - | 1221 | 1169 | **RNA**→ rpmE (PRK00019)Ribosomal_L31 (pfam01197)→ |
| Sso-1-371 | NZ_CXIS01000027.1 | - | 669 | 617 | **RNA**→ rpmE (PRK00019)Ribosomal_L31 (pfam01197)→ |
| Sso-1-372 | NZ_CXFW01000022.1 | - | 1174 | 1122 | **RNA**→ rpmE (PRK00019)Ribosomal_L31 (pfam01197)→ |
| Sso-1-373 | NZ_CXIC01000002.1 | + | 80379 | 80431 | **RNA**→ rpmE (PRK00019)Ribosomal_L31 (pfam01197)→ |
| Sso-1-374 | NZ_CWWQ01000023.1 | + | 39082 | 39134 | **RNA**→ rpmE (PRK00019)Ribosomal_L31 (pfam01197)→ |
| Sso-1-375 | NZ_CXMJ01000027.1 | - | 798 | 746 | **RNA**→ rpmE (PRK00019)Ribosomal_L31 (pfam01197)→ |
| Sso-1-376 | NZ_CXDP01000021.1 | - | 433 | 381 | **RNA**→ rpmE (PRK00019)Ribosomal_L31 (pfam01197)→ hypo→ |
| Sso-1-377 | NZ_CXLE01000027.1 | + | 39089 | 39141 | **RNA**→ rpmE (PRK00019)Ribosomal_L31 (pfam01197)→ |
| Sso-1-378 | NZ_CWZG01000025.1 | + | 39089 | 39141 | **RNA**→ rpmE (PRK00019)Ribosomal_L31 (pfam01197)→ |
| Sso-1-379 | NZ_CXHC01000027.1 | - | 695 | 643 | **RNA**→ rpmE (PRK00019)Ribosomal_L31 (pfam01197)→ |
| Sso-1-380 | NZ_CWYM01000024.1 | + | 39089 | 39141 | **RNA**→ rpmE (PRK00019)Ribosomal_L31 (pfam01197)→ |
| Sso-1-381 | NZ_CXDB01000009.1 | + | 33320 | 33372 | **RNA**→ rpmE (PRK00019)Ribosomal_L31 (pfam01197)→ |
| Sso-1-382 | NZ_CXMM01000026.1 | + | 39089 | 39141 | **RNA**→ rpmE (PRK00019)Ribosomal_L31 (pfam01197)→ |
| Sso-1-383 | NZ_CWYI01000029.1 | - | 664 | 612 | **RNA**→ rpmE (PRK00019)Ribosomal_L31 (pfam01197)→ |
| Sso-1-384 | NZ_CWXN01000026.1 | + | 39106 | 39158 | **RNA**→ rpmE (PRK00019)Ribosomal_L31 (pfam01197)→ |

| | | | | | |
|---|---|---|---|---|---|
| Sso-1-385 | NZ_CXNN01000025.1 | - | 1539 | 1487 | **RNA**→ rpmE (PRK00019)Ribosomal_L31 (pfam01197)→ |
| Sso-1-386 | NZ_CXBT01000023.1 | + | 39007 | 39059 | **RNA**→ rpmE (PRK00019)Ribosomal_L31 (pfam01197)→ |
| Sso-1-387 | NZ_CXFA01000020.1 | + | 47728 | 47780 | **RNA**→ rpmE (PRK00019)Ribosomal_L31 (pfam01197)→ |
| Sso-1-388 | NZ_CXGF01000027.1 | - | 556 | 504 | **RNA**→ rpmE (PRK00019)Ribosomal_L31 (pfam01197)→ |
| Sso-1-389 | NZ_CXGD01000020.1 | + | 39045 | 39097 | **RNA**→ rpmE (PRK00019)Ribosomal_L31 (pfam01197)→ |
| Sso-1-390 | NZ_CXDG01000045.1 | - | 488 | 436 | **RNA**→ rpmE (PRK00019)Ribosomal_L31 (pfam01197)→ |
| Sso-1-391 | NZ_CXGE01000027.1 | + | 39045 | 39097 | **RNA**→ rpmE (PRK00019)Ribosomal_L31 (pfam01197)→ |
| Sso-1-392 | NZ_CWXQ01000020.1 | - | 846 | 794 | **RNA**→ rpmE (PRK00019)Ribosomal_L31 (pfam01197)→ |
| Sso-1-393 | NZ_CXFB01000026.1 | - | 695 | 643 | **RNA**→ rpmE (PRK00019)Ribosomal_L31 (pfam01197)→ |
| Sso-1-394 | NZ_CXDK01000016.1 | + | 47867 | 47919 | **RNA**→ rpmE (PRK00019)Ribosomal_L31 (pfam01197)→ |
| Sso-1-395 | NZ_CXKI01000002.1 | + | 67788 | 67840 | **RNA**→ rpmE (PRK00019)Ribosomal_L31 (pfam01197)→ |
| Sso-1-396 | NZ_CXCT01000016.1 | - | 479 | 427 | **RNA**→ rpmE (PRK00019)Ribosomal_L31 (pfam01197)→ |
| Sso-1-397 | NZ_CXCN01000005.1 | + | 70940 | 70992 | **RNA**→ rpmE (PRK00019)Ribosomal_L31 (pfam01197)→ |
| Sso-1-398 | NZ_CXCA01000025.1 | - | 632 | 580 | **RNA**→ rpmE (PRK00019)Ribosomal_L31 (pfam01197)→ |
| Sso-1-399 | NZ_CXCE01000021.1 | - | 632 | 580 | **RNA**→ rpmE (PRK00019)Ribosomal_L31 (pfam01197)→ |
| Sso-1-400 | NZ_CXNQ01000027.1 | + | 39089 | 39141 | **RNA**→ rpmE (PRK00019)Ribosomal_L31 (pfam01197)→ |
| Sso-1-401 | NZ_CXCH01000001.1 | - | 632 | 580 | **RNA**→ rpmE (PRK00019)Ribosomal_L31 (pfam01197)→ |
| Sso-1-402 | NZ_CXAZ01000023.1 | + | 39023 | 39075 | **RNA**→ rpmE (PRK00019)Ribosomal_L31 (pfam01197)→ |
| Sso-1-403 | NZ_CXET01000019.1 | - | 646 | 594 | **RNA**→ rpmE (PRK00019)Ribosomal_L31 (pfam01197)→ |
| Sso-1-404 | NZ_CWYZ01000027.1 | - | 660 | 608 | **RNA**→ rpmE (PRK00019)Ribosomal_L31 (pfam01197)→ |
| Sso-1-405 | NZ_CXIT01000027.1 | - | 786 | 734 | **RNA**→ rpmE (PRK00019)Ribosomal_L31 (pfam01197)→ |
| Sso-1-406 | NZ_CWXG01000022.1 | + | 39104 | 39156 | **RNA**→ rpmE (PRK00019)Ribosomal_L31 (pfam01197)→ |
| Sso-1-407 | NZ_CXBP01000067.1 | - | 401 | 349 | **RNA**→ rpmE (PRK00019)Ribosomal_L31 (pfam01197)→ |
| Sso-1-408 | NZ_CXHK01000020.1 | - | 3534 | 3482 | **RNA**→ rpmE (PRK00019)Ribosomal_L31 (pfam01197)→ |
| Sso-1-409 | NZ_CXFE01000028.1 | - | 695 | 643 | **RNA**→ rpmE (PRK00019)Ribosomal_L31 (pfam01197)→ |
| Sso-1-410 | NZ_CWXY01000026.1 | - | 862 | 810 | **RNA**→ rpmE (PRK00019)Ribosomal_L31 (pfam01197)→ |
| Sso-1-411 | NZ_CXAU01000013.1 | + | 43096 | 43148 | **RNA**→ rpmE (PRK00019)Ribosomal_L31 (pfam01197)→ |
| Sso-1-412 | NZ_CXBV01000021.1 | + | 39007 | 39059 | **RNA**→ rpmE (PRK00019)Ribosomal_L31 (pfam01197)→ |
| Sso-1-413 | NZ_CXNH01000020.1 | - | 2678 | 2626 | **RNA**→ rpmE (PRK00019)Ribosomal_L31 (pfam01197)→ |
| Sso-1-414 | NZ_CXDM01000029.1 | + | 37772 | 37824 | **RNA**→ rpmE (PRK00019)Ribosomal_L31 (pfam01197)→ |
| Sso-1-415 | NZ_CXIB01000026.1 | + | 39089 | 39141 | **RNA**→ rpmE (PRK00019)Ribosomal_L31 (pfam01197)→ |
| Sso-1-416 | NZ_CXGA01000026.1 | + | 39045 | 39097 | **RNA**→ rpmE (PRK00019)Ribosomal_L31 (pfam01197)→ |
| Sso-1-417 | NZ_CXJU01000026.1 | - | 779 | 727 | **RNA**→ rpmE (PRK00019)Ribosomal_L31 (pfam01197)→ |
| Sso-1-418 | NZ_CXMA01000023.1 | + | 39089 | 39141 | **RNA**→ rpmE (PRK00019)Ribosomal_L31 (pfam01197)→ |
| Sso-1-419 | NZ_CXHF01000026.1 | - | 695 | 643 | **RNA**→ rpmE (PRK00019)Ribosomal_L31 (pfam01197)→ |
| Sso-1-420 | NZ_CWZT01000023.1 | + | 39094 | 39146 | **RNA**→ rpmE (PRK00019)Ribosomal_L31 (pfam01197)→ |
| Sso-1-421 | NZ_CXGP01000022.1 | - | 2682 | 2630 | **RNA**→ rpmE (PRK00019)Ribosomal_L31 (pfam01197)→ |
| Sso-1-422 | NZ_CXAK01000025.1 | - | 662 | 610 | **RNA**→ rpmE (PRK00019)Ribosomal_L31 (pfam01197)→ |
| Sso-1-423 | NZ_CWXL01000027.1 | - | 664 | 612 | **RNA**→ rpmE (PRK00019)Ribosomal_L31 (pfam01197)→ |
| Sso-1-424 | NZ_CXBB01000103.1 | - | 1174 | 1122 | **RNA**→ rpmE (PRK00019)Ribosomal_L31 (pfam01197)→ |
| Sso-1-425 | NZ_CXMC01000027.1 | + | 39089 | 39141 | **RNA**→ rpmE (PRK00019)Ribosomal_L31 (pfam01197)→ |
| Sso-1-426 | NZ_CXKD01000028.1 | - | 812 | 760 | **RNA**→ rpmE (PRK00019)Ribosomal_L31 (pfam01197)→ |
| Sso-1-427 | NZ_CWYL01000026.1 | - | 666 | 614 | **RNA**→ rpmE (PRK00019)Ribosomal_L31 (pfam01197)→ |
| Sso-1-428 | NZ_CXGG01000026.1 | - | 496 | 444 | **RNA**→ rpmE (PRK00019)Ribosomal_L31 (pfam01197)→ |
| Sso-1-429 | NZ_CWZM01000026.1 | - | 668 | 616 | **RNA**→ rpmE (PRK00019)Ribosomal_L31 (pfam01197)→ |
| Sso-1-430 | NZ_CXIO01000028.1 | + | 39045 | 39097 | **RNA**→ rpmE (PRK00019)Ribosomal_L31 (pfam01197)→ |
| Sso-1-431 | NZ_CXJJ01000026.1 | + | 39082 | 39134 | **RNA**→ rpmE (PRK00019)Ribosomal_L31 (pfam01197)→ |
| Sso-1-432 | NZ_CXHT01000025.1 | - | 779 | 727 | **RNA**→ rpmE (PRK00019)Ribosomal_L31 (pfam01197)→ |
| Sso-1-433 | NZ_CXIJ01000028.1 | - | 793 | 741 | **RNA**→ rpmE (PRK00019)Ribosomal_L31 (pfam01197)→ |
| Sso-1-434 | NZ_CXAD01000025.1 | - | 1137 | 1085 | **RNA**→ rpmE (PRK00019)Ribosomal_L31 (pfam01197)→ |
| Sso-1-435 | NZ_CXOH01000026.1 | + | 39089 | 39141 | **RNA**→ rpmE (PRK00019)Ribosomal_L31 (pfam01197)→ |
| Sso-1-436 | NZ_CXFJ01000027.1 | - | 637 | 585 | **RNA**→ rpmE (PRK00019)Ribosomal_L31 (pfam01197)→ |

| | | | | | |
|---|---|---|---|---|---|
| Sso-1-437 | NZ_CXJL01000025.1 | + | 39089 | 39141 | **RNA**→ rpmE (PRK00019)Ribosomal_L31 (pfam01197)→ |
| Sso-1-438 | NZ_CXFY01000027.1 | + | 39045 | 39097 | **RNA**→ rpmE (PRK00019)Ribosomal_L31 (pfam01197)→ |
| Sso-1-439 | NZ_CXNL01000026.1 | - | 799 | 747 | **RNA**→ rpmE (PRK00019)Ribosomal_L31 (pfam01197)→ |
| Sso-1-440 | NZ_CWZS01000023.1 | + | 39089 | 39141 | **RNA**→ rpmE (PRK00019)Ribosomal_L31 (pfam01197)→ |
| Sso-1-441 | NZ_CXAC01000026.1 | - | 661 | 609 | **RNA**→ rpmE (PRK00019)Ribosomal_L31 (pfam01197)→ |
| Sso-1-442 | NZ_CXKN01000026.1 | - | 769 | 717 | **RNA**→ rpmE (PRK00019)Ribosomal_L31 (pfam01197)→ |
| Sso-1-443 | NZ_CXCP01000015.1 | + | 39185 | 39237 | **RNA**→ rpmE (PRK00019)Ribosomal_L31 (pfam01197)→ |
| Sso-1-444 | NZ_CXJS01000017.1 | - | 1137 | 1085 | **RNA**→ rpmE (PRK00019)Ribosomal_L31 (pfam01197)→ |
| Sso-1-445 | NZ_CXKC01000028.1 | + | 39082 | 39134 | **RNA**→ rpmE (PRK00019)Ribosomal_L31 (pfam01197)→ |
| Sso-1-446 | NZ_CXBK01000055.1 | + | 20736 | 20788 | **RNA**→ rpmE (PRK00019)Ribosomal_L31 (pfam01197)→ |
| Sso-1-447 | NZ_CXME01000026.1 | + | 39082 | 39134 | **RNA**→ rpmE (PRK00019)Ribosomal_L31 (pfam01197)→ |
| Sso-1-448 | NZ_CXLX01000028.1 | + | 39089 | 39141 | **RNA**→ rpmE (PRK00019)Ribosomal_L31 (pfam01197)→ |
| Sso-1-449 | NZ_CXLU01000028.1 | - | 695 | 643 | **RNA**→ rpmE (PRK00019)Ribosomal_L31 (pfam01197)→ |
| Sso-1-450 | NZ_CXHV01000025.1 | + | 39089 | 39141 | **RNA**→ rpmE (PRK00019)Ribosomal_L31 (pfam01197)→ |
| Sso-1-451 | NZ_CXHX01000026.1 | - | 1627 | 1575 | **RNA**→ rpmE (PRK00019)Ribosomal_L31 (pfam01197)→ |
| Sso-1-452 | NZ_CXNE01000028.1 | + | 39082 | 39134 | **RNA**→ rpmE (PRK00019)Ribosomal_L31 (pfam01197)→ |
| Sso-1-453 | NZ_CXAO01000026.1 | - | 529 | 477 | **RNA**→ rpmE (PRK00019)Ribosomal_L31 (pfam01197)→ |
| Sso-1-454 | NZ_CXNO01000025.1 | - | 1174 | 1122 | **RNA**→ rpmE (PRK00019)Ribosomal_L31 (pfam01197)→ |
| Sso-1-455 | NZ_CXGX01000025.1 | - | 1174 | 1122 | **RNA**→ rpmE (PRK00019)Ribosomal_L31 (pfam01197)→ |
| Sso-1-456 | NZ_CXEC01000022.1 | - | 644 | 592 | **RNA**→ rpmE (PRK00019)Ribosomal_L31 (pfam01197)→ |
| Sso-1-457 | NZ_CWZE01000026.1 | - | 664 | 612 | **RNA**→ rpmE (PRK00019)Ribosomal_L31 (pfam01197)→ |
| Sso-1-458 | NZ_CXOB01000028.1 | - | 804 | 752 | **RNA**→ rpmE (PRK00019)Ribosomal_L31 (pfam01197)→ |
| Sso-1-459 | NZ_CWXF01000027.1 | + | 39104 | 39156 | **RNA**→ rpmE (PRK00019)Ribosomal_L31 (pfam01197)→ |
| Sso-1-460 | NZ_CXCV01000008.1 | - | 27546 | 27494 | **RNA**→ rpmE (PRK00019)Ribosomal_L31 (pfam01197)→ |
| Sso-1-461 | NZ_CXEV01000018.1 | - | 2655 | 2603 | **RNA**→ rpmE (PRK00019)Ribosomal_L31 (pfam01197)→ |
| Sso-1-462 | NZ_CXGB01000025.1 | - | 695 | 643 | **RNA**→ rpmE (PRK00019)Ribosomal_L31 (pfam01197)→ |
| Sso-1-463 | NZ_CXMT01000026.1 | - | 1627 | 1575 | **RNA**→ rpmE (PRK00019)Ribosomal_L31 (pfam01197)→ |
| Sso-1-464 | NZ_CXLH01000027.1 | - | 800 | 748 | **RNA**→ rpmE (PRK00019)Ribosomal_L31 (pfam01197)→ |
| Sso-1-465 | NZ_CXDW01000020.1 | - | 2418 | 2366 | **RNA**→ rpmE (PRK00019)Ribosomal_L31 (pfam01197)→ |
| Sso-1-466 | NZ_CXAP01000002.1 | + | 80220 | 80272 | **RNA**→ rpmE (PRK00019)Ribosomal_L31 (pfam01197)→ |
| Sso-1-467 | NZ_CXNK01000026.1 | - | 812 | 760 | **RNA**→ rpmE (PRK00019)Ribosomal_L31 (pfam01197)→ |
| Sso-1-468 | NZ_CWYQ01000027.1 | + | 39041 | 39093 | **RNA**→ rpmE (PRK00019)Ribosomal_L31 (pfam01197)→ |
| Sso-1-469 | NZ_CXMY01000022.1 | + | 39089 | 39141 | **RNA**→ rpmE (PRK00019)Ribosomal_L31 (pfam01197)→ |
| Sso-1-470 | NZ_CWZC01000027.1 | - | 666 | 614 | **RNA**→ rpmE (PRK00019)Ribosomal_L31 (pfam01197)→ |
| Sso-1-471 | NZ_CXIZ01000027.1 | + | 39045 | 39097 | **RNA**→ rpmE (PRK00019)Ribosomal_L31 (pfam01197)→ |
| Sso-1-472 | NZ_CXMP01000024.1 | - | 812 | 760 | **RNA**→ rpmE (PRK00019)Ribosomal_L31 (pfam01197)→ |
| Sso-1-473 | NZ_CWYA01000024.1 | - | 853 | 801 | **RNA**→ rpmE (PRK00019)Ribosomal_L31 (pfam01197)→ |
| Sso-1-474 | NZ_CWYJ01000023.1 | - | 1174 | 1122 | **RNA**→ rpmE (PRK00019)Ribosomal_L31 (pfam01197)→ |
| Sso-1-475 | NZ_CXED01000015.1 | + | 39085 | 39137 | **RNA**→ rpmE (PRK00019)Ribosomal_L31 (pfam01197)→ |
| Sso-1-476 | NZ_CWXJ01000027.1 | - | 736 | 684 | **RNA**→ rpmE (PRK00019)Ribosomal_L31 (pfam01197)→ |
| Sso-1-477 | NZ_CXII01000026.1 | + | 39089 | 39141 | **RNA**→ rpmE (PRK00019)Ribosomal_L31 (pfam01197)→ |
| Sso-1-478 | NZ_CXHN01000027.1 | + | 39082 | 39134 | **RNA**→ rpmE (PRK00019)Ribosomal_L31 (pfam01197)→ |
| Sso-1-479 | NZ_CXKX01000025.1 | + | 39089 | 39141 | **RNA**→ rpmE (PRK00019)Ribosomal_L31 (pfam01197)→ |
| Sso-1-480 | NZ_CXKT01000026.1 | - | 1174 | 1122 | **RNA**→ rpmE (PRK00019)Ribosomal_L31 (pfam01197)→ |
| Sso-1-481 | NZ_CXJO01000026.1 | + | 39082 | 39134 | **RNA**→ rpmE (PRK00019)Ribosomal_L31 (pfam01197)→ |
| Sso-1-482 | NZ_CXJX01000030.1 | + | 39082 | 39134 | **RNA**→ rpmE (PRK00019)Ribosomal_L31 (pfam01197)→ |
| Sso-1-483 | NZ_CXLV01000031.1 | + | 39082 | 39134 | **RNA**→ rpmE (PRK00019)Ribosomal_L31 (pfam01197)→ |
| Sso-1-484 | NZ_CWYE01000019.1 | - | 4838 | 4786 | **RNA**→ rpmE (PRK00019)Ribosomal_L31 (pfam01197)→ |
| Sso-1-485 | NZ_CWXO01000026.1 | + | 39089 | 39141 | **RNA**→ rpmE (PRK00019)Ribosomal_L31 (pfam01197)→ |
| Sso-1-486 | NZ_CXFV01000025.1 | + | 39045 | 39097 | **RNA**→ rpmE (PRK00019)Ribosomal_L31 (pfam01197)→ |
| Sso-1-487 | NZ_CXIE01000027.1 | - | 1627 | 1575 | **RNA**→ rpmE (PRK00019)Ribosomal_L31 (pfam01197)→ |
| Sso-1-488 | NZ_CXKL01000026.1 | - | 662 | 610 | **RNA**→ rpmE (PRK00019)Ribosomal_L31 (pfam01197)→ |

| | | | | | |
|---|---|---|---|---|---|
| Sso-1-489 | NZ_CWXE01000031.1 | - | 855 | 803 | **RNA**→ rpmE (PRK00019)Ribosomal_L31 (pfam01197)→ |
| Sso-1-490 | NZ_CXAF01000026.1 | + | 39082 | 39134 | **RNA**→ rpmE (PRK00019)Ribosomal_L31 (pfam01197)→ |
| Sso-1-491 | NZ_CWZN01000027.1 | - | 664 | 612 | **RNA**→ rpmE (PRK00019)Ribosomal_L31 (pfam01197)→ |
| Sso-1-492 | NZ_CXEK01000014.1 | - | 644 | 592 | **RNA**→ rpmE (PRK00019)Ribosomal_L31 (pfam01197)→ |
| Sso-1-493 | NZ_CXFN01000026.1 | - | 481 | 429 | **RNA**→ rpmE (PRK00019)Ribosomal_L31 (pfam01197)→ |
| Sso-1-494 | NZ_CXFZ01000023.1 | + | 39045 | 39097 | **RNA**→ rpmE (PRK00019)Ribosomal_L31 (pfam01197)→ |
| Sso-1-495 | NZ_CXCG01000013.1 | - | 334 | 282 | **RNA**→ rpmE (PRK00019)Ribosomal_L31 (pfam01197)→ |
| Sso-1-496 | NZ_CXNF01000028.1 | + | 39089 | 39141 | **RNA**→ rpmE (PRK00019)Ribosomal_L31 (pfam01197)→ |
| Sso-1-497 | NZ_CXKQ01000026.1 | + | 39045 | 39097 | **RNA**→ rpmE (PRK00019)Ribosomal_L31 (pfam01197)→ |
| Sso-1-498 | NZ_CWYV01000027.1 | + | 39041 | 39093 | **RNA**→ rpmE (PRK00019)Ribosomal_L31 (pfam01197)→ |
| Sso-1-499 | NZ_CXCM01000057.1 | + | 20736 | 20788 | **RNA**→ rpmE (PRK00019)Ribosomal_L31 (pfam01197)→ |
| Sso-1-500 | NZ_CXAX01000027.1 | + | 39023 | 39075 | **RNA**→ rpmE (PRK00019)Ribosomal_L31 (pfam01197)→ |
| Sso-1-501 | NZ_CWXX01000025.1 | + | 39089 | 39141 | **RNA**→ rpmE (PRK00019)Ribosomal_L31 (pfam01197)→ |
| Sso-1-502 | NZ_CXMB01000026.1 | - | 773 | 721 | **RNA**→ rpmE (PRK00019)Ribosomal_L31 (pfam01197)→ |
| Sso-1-503 | NZ_CXJD01000027.1 | - | 1539 | 1487 | **RNA**→ rpmE (PRK00019)Ribosomal_L31 (pfam01197)→ |
| Sso-1-504 | NZ_CWWY01000017.1 | - | 799 | 747 | **RNA**→ rpmE (PRK00019)Ribosomal_L31 (pfam01197)→ |
| Sso-1-505 | NZ_CXNA01000027.1 | + | 39082 | 39134 | **RNA**→ rpmE (PRK00019)Ribosomal_L31 (pfam01197)→ |
| Sso-1-506 | NZ_CXGL01000026.1 | - | 855 | 803 | **RNA**→ rpmE (PRK00019)Ribosomal_L31 (pfam01197)→ |
| Sso-1-507 | NZ_CXJI01000026.1 | - | 1627 | 1575 | **RNA**→ rpmE (PRK00019)Ribosomal_L31 (pfam01197)→ |
| Sso-1-508 | NZ_CWYS01000026.1 | - | 679 | 627 | **RNA**→ rpmE (PRK00019)Ribosomal_L31 (pfam01197)→ |
| Sso-1-509 | NZ_CWWN01000026.1 | - | 751 | 699 | **RNA**→ rpmE (PRK00019)Ribosomal_L31 (pfam01197)→ |
| Sso-1-510 | NZ_CWYO01000025.1 | + | 39093 | 39145 | **RNA**→ rpmE (PRK00019)Ribosomal_L31 (pfam01197)→ |
| Sso-1-511 | NZ_CXMI01000027.1 | + | 39089 | 39141 | **RNA**→ rpmE (PRK00019)Ribosomal_L31 (pfam01197)→ |
| Sso-1-512 | NZ_CXBR01000018.1 | + | 39007 | 39059 | **RNA**→ rpmE (PRK00019)Ribosomal_L31 (pfam01197)→ |
| Sso-1-513 | NZ_CXOF01000024.1 | - | 1627 | 1575 | **RNA**→ rpmE (PRK00019)Ribosomal_L31 (pfam01197)→ |
| Sso-1-514 | NZ_CXFU01000025.1 | + | 39045 | 39097 | **RNA**→ rpmE (PRK00019)Ribosomal_L31 (pfam01197)→ |
| Sso-1-515 | NZ_CXFG01000031.1 | - | 481 | 429 | **RNA**→ rpmE (PRK00019)Ribosomal_L31 (pfam01197)→ |
| Sso-1-516 | NZ_CWYF01000025.1 | + | 39089 | 39141 | **RNA**→ rpmE (PRK00019)Ribosomal_L31 (pfam01197)→ |
| Sso-1-517 | NZ_CXBQ01000065.1 | + | 20750 | 20802 | **RNA**→ rpmE (PRK00019)Ribosomal_L31 (pfam01197)→ |
| Sso-1-518 | NZ_CXMD01000027.1 | - | 794 | 742 | **RNA**→ rpmE (PRK00019)Ribosomal_L31 (pfam01197)→ |
| Sso-1-519 | NZ_CXHZ01000022.1 | - | 2667 | 2615 | **RNA**→ rpmE (PRK00019)Ribosomal_L31 (pfam01197)→ |
| Sso-1-520 | NZ_CXGS01000026.1 | - | 867 | 815 | **RNA**→ rpmE (PRK00019)Ribosomal_L31 (pfam01197)→ |
| Sso-1-521 | NZ_CXKE01000028.1 | - | 779 | 727 | **RNA**→ rpmE (PRK00019)Ribosomal_L31 (pfam01197)→ |
| Sso-1-522 | NZ_CXHP01000025.1 | + | 39082 | 39134 | **RNA**→ rpmE (PRK00019)Ribosomal_L31 (pfam01197)→ |
| Sso-1-523 | NZ_CXLW01000026.1 | + | 39082 | 39134 | **RNA**→ rpmE (PRK00019)Ribosomal_L31 (pfam01197)→ |
| Sso-1-524 | NZ_CXEA01000023.1 | - | 646 | 594 | **RNA**→ rpmE (PRK00019)Ribosomal_L31 (pfam01197)→ |
| Sso-1-525 | NZ_CWZA01000021.1 | - | 1539 | 1487 | **RNA**→ rpmE (PRK00019)Ribosomal_L31 (pfam01197)→ |
| Sso-1-526 | NZ_CXKZ01000025.1 | + | 39089 | 39141 | **RNA**→ rpmE (PRK00019)Ribosomal_L31 (pfam01197)→ |
| Sso-1-527 | NZ_CXGJ01000025.1 | + | 39045 | 39097 | **RNA**→ rpmE (PRK00019)Ribosomal_L31 (pfam01197)→ |
| Sso-1-528 | NZ_CXOG01000062.1 | + | 23400 | 23452 | **RNA**→ rpmE (PRK00019)Ribosomal_L31 (pfam01197)→ |
| Sso-1-529 | NZ_CXDF01000021.1 | + | 39593 | 39645 | **RNA**→ rpmE (PRK00019)Ribosomal_L31 (pfam01197)→ |
| Sso-1-530 | NZ_CXDL01000052.1 | - | 634 | 582 | **RNA**→ rpmE (PRK00019)Ribosomal_L31 (pfam01197)→ |
| Sso-1-531 | NZ_CXMQ01000026.1 | + | 39089 | 39141 | **RNA**→ rpmE (PRK00019)Ribosomal_L31 (pfam01197)→ |
| Sso-1-532 | NZ_CWXH01000034.1 | - | 855 | 803 | **RNA**→ rpmE (PRK00019)Ribosomal_L31 (pfam01197)→ |
| Sso-1-533 | NZ_CXAA01000028.1 | + | 39082 | 39134 | **RNA**→ rpmE (PRK00019)Ribosomal_L31 (pfam01197)→ |
| Sso-1-534 | NZ_CWWV01000019.1 | + | 38339 | 38391 | **RNA**→ rpmE (PRK00019)Ribosomal_L31 (pfam01197)→ |
| Sso-1-535 | NZ_CXJK01000023.1 | + | 39082 | 39134 | **RNA**→ rpmE (PRK00019)Ribosomal_L31 (pfam01197)→ |
| Sso-1-536 | NZ_CXGU01000032.1 | - | 621 | 569 | **RNA**→ rpmE (PRK00019)Ribosomal_L31 (pfam01197)→ |
| Sso-1-537 | NZ_CXIP01000030.1 | + | 39045 | 39097 | **RNA**→ rpmE (PRK00019)Ribosomal_L31 (pfam01197)→ |
| Sso-1-538 | NZ_CXKB01000026.1 | + | 39089 | 39141 | **RNA**→ rpmE (PRK00019)Ribosomal_L31 (pfam01197)→ |
| Sso-1-539 | NZ_CXAS01000019.1 | - | 2634 | 2582 | **RNA**→ rpmE (PRK00019)Ribosomal_L31 (pfam01197)→ |
| Eco-2024-1 | NZ_AVRH01000067.1 | + | 46925 | 46977 | **RNA**→ rpmE (PRK00019)Ribosomal_L31 (pfam01197)→ |

| | | | | | |
|---|---|---|---|---|---|
| Sdy-2-1 | NZ_AURX01000023.1 | + | 37576 | 37628 | **RNA**→ rpmE (PRK00019)Ribosomal_L31 (pfam01197)→ |
| Eco-2044-1 | NZ_AVSN01000121.1 | + | 44651 | 44703 | **RNA**→ rpmE (PRK00019)Ribosomal_L31 (pfam01197)→ |
| Sso-2-1 | NZ_AURV01000019.1 | - | 39143 | 39091 | **RNA**→ rpmE (PRK00019)Ribosomal_L31 (pfam01197)→ |
| Eco-1983-1 | NZ_ATKA01000044.1 | - | 507 | 455 | **RNA**→ rpmE (PRK00019)Ribosomal_L31 (pfam01197)→ |
| Cmi-1-1 | NZ_AEJF01000307.1 | - | 141 | 89 | **RNA**→ rpmE (PRK00019)Ribosomal_L31 (pfam01197)→ |
| Eco-1830-1 | NZ_AGTI01000011.1 | - | 722 | 670 | **RNA**→ rpmE (PRK00019)Ribosomal_L31 (pfam01197)→ |
| Eco-1778-1 | NZ_AGTJ01000006.1 | - | 39164 | 39112 | **RNA**→ rpmE (PRK00019)Ribosomal_L31 (pfam01197)→ |
| Eco-1021-1 | NZ_AXLH01000003.1 | + | 4432498 | 4432550 | **RNA**→ rpmE (PRK00019)Ribosomal_L31 (pfam01197)→ |
| Eco-728-2 | NZ_AOUP01000502.1 | + | 305 | 357 | **RNA**→ rpmE (PRK00019)Ribosomal_L31 (pfam01197)→ |
| Eco-414-1 | NZ_ASXQ01000020.1 | - | 44445 | 44393 | **RNA**→ rpmE (PRK00019)Ribosomal_L31 (pfam01197)→ |
| Eco-1919-1 | NZ_AQAW01000068.1 | + | 87681 | 87733 | **RNA**→ rpmE (PRK00019)Ribosomal_L31 (pfam01197)→ |
| Eco-362-1 | NZ_AIBH01000033.1 | + | 107851 | 107903 | **RNA**→ rpmE (PRK00019)Ribosomal_L31 (pfam01197)→ |
| Eco-1879-1 | NZ_APYX01000072.1 | + | 118012 | 118064 | **RNA**→ rpmE (PRK00019)Ribosomal_L31 (pfam01197)→ |
| Eco-2176-1 | NZ_AQCB01000200.1 | + | 18734 | 18786 | **RNA**→ rpmE (PRK00019)Ribosomal_L31 (pfam01197)→ |
| Eco-706-1 | NZ_JHLN01000031.1 | - | 655 | 603 | **RNA**→ rpmE (PRK00019)Ribosomal_L31 (pfam01197)→ |
| Eco-914-1 | NZ_AWFJ01000105.1 | - | 1379 | 1327 | **RNA**→ rpmE (PRK00019)Ribosomal_L31 (pfam01197)→ |
| Sbo-2-4 | NZ_JWBP01000001.1 | - | 474 | 422 | **RNA**→ rpmE (PRK00019)Ribosomal_L31 (pfam01197)→ |
| Sbo-2-5 | NZ_JVYE01000206.1 | + | 32322 | 32374 | **RNA**→ rpmE (PRK00019)Ribosomal_L31 (pfam01197)→ |
| Sbo-2-6 | NZ_JVNX01000122.1 | + | 32322 | 32374 | **RNA**→ rpmE (PRK00019)Ribosomal_L31 (pfam01197)→ |
| Sbo-2-7 | NZ_JVJU01000018.1 | + | 32335 | 32387 | **RNA**→ rpmE (PRK00019)Ribosomal_L31 (pfam01197)→ |
| Sbo-2-8 | NZ_JVYF01000453.1 | - | 1032 | 980 | **RNA**→ rpmE (PRK00019)Ribosomal_L31 (pfam01197)→ |
| Sso-1-540 | NZ_JVNM01000130.1 | + | 32400 | 32452 | **RNA**→ rpmE (PRK00019)Ribosomal_L31 (pfam01197)→ |
| Sbo-2-9 | NZ_JVNL01000131.1 | + | 32406 | 32458 | **RNA**→ rpmE (PRK00019)Ribosomal_L31 (pfam01197)→ |
| Eco-1723-1 | NZ_AMVT01000005.1 | + | 85592 | 85644 | **RNA**→ rpmE (PRK00019)Ribosomal_L31 (pfam01197)→ |
| Eco-1424-1 | NZ_ANXO01000041.1 | + | 11336 | 11388 | **RNA**→ rpmE (PRK00019)Ribosomal_L31 (pfam01197)→ |
| Eco-1477-1 | NZ_ANYI01000065.1 | + | 32593 | 32645 | **RNA**→ rpmE (PRK00019)Ribosomal_L31 (pfam01197)→ |
| Eco-1491-1 | NZ_ANYN01000098.1 | + | 32501 | 32553 | **RNA**→ rpmE (PRK00019)Ribosomal_L31 (pfam01197)→ |
| Eco-1440-1 | NZ_ANXU01000077.1 | + | 32410 | 32462 | **RNA**→ rpmE (PRK00019)Ribosomal_L31 (pfam01197)→ |
| Eco-1728-1 | NZ_AMVY01000010.1 | + | 85592 | 85644 | **RNA**→ rpmE (PRK00019)Ribosomal_L31 (pfam01197)→ |
| Eco-776-1 | NZ_APWZ01000454.1 | + | 33662 | 33714 | **RNA**→ rpmE (PRK00019)Ribosomal_L31 (pfam01197)→ |
| Eco-1878-1 | NZ_APYW01000114.1 | + | 117712 | 117764 | **RNA**→ rpmE (PRK00019)Ribosomal_L31 (pfam01197)→ |
| Eco-1907-1 | NZ_AQAM01000194.1 | + | 46822 | 46874 | **RNA**→ rpmE (PRK00019)Ribosomal_L31 (pfam01197)→ |
| Eco-915-1 | NZ_AVDU01000093.1 | - | 2645 | 2593 | **RNA**→ rpmE (PRK00019)Ribosomal_L31 (pfam01197)→ |
| Eco-409-1 | NZ_AXOC01000008.1 | + | 49059 | 49111 | **RNA**→ rpmE (PRK00019)Ribosomal_L31 (pfam01197)→ |
| Eco-1982-1 | NZ_ATJZ01000076.1 | - | 954 | 902 | **RNA**→ rpmE (PRK00019)Ribosomal_L31 (pfam01197)→ |
| Eco-1144-1 | NZ_AHHN01000007.1 | + | 85918 | 85970 | **RNA**→ rpmE (PRK00019)Ribosomal_L31 (pfam01197)→ |
| Eco-410-1 | NZ_AXOD01000008.1 | + | 49059 | 49111 | **RNA**→ rpmE (PRK00019)Ribosomal_L31 (pfam01197)→ |
| Eco-1918-1 | NZ_AQAV01000071.1 | + | 87581 | 87633 | **RNA**→ rpmE (PRK00019)Ribosomal_L31 (pfam01197)→ |
| Eco-1870-1 | NZ_APYR01000172.1 | + | 87716 | 87768 | **RNA**→ rpmE (PRK00019)Ribosomal_L31 (pfam01197)→ |
| Eco-1923-1 | NZ_AQBA01000071.1 | + | 87684 | 87736 | **RNA**→ rpmE (PRK00019)Ribosomal_L31 (pfam01197)→ |
| Eco-1945-1 | NZ_AHHO01000057.1 | + | 58857 | 58909 | **RNA**→ rpmE (PRK00019)Ribosomal_L31 (pfam01197)→ |
| Eco-424-1 | NZ_AYLY01000017.1 | - | 44955 | 44903 | **RNA**→ rpmE (PRK00019)Ribosomal_L31 (pfam01197)→ |
| Eco-229-1 | NZ_AYQQ01000021.1 | + | 32271 | 32323 | **RNA**→ rpmE (PRK00019)Ribosomal_L31 (pfam01197)→ |
| Eco-526-1 | NZ_JHFC01000048.1 | + | 83133 | 83185 | **RNA**→ rpmE (PRK00019)Ribosomal_L31 (pfam01197)→ |
| Eco-555-1 | NZ_JHFW01000009.1 | + | 113689 | 113741 | **RNA**→ rpmE (PRK00019)Ribosomal_L31 (pfam01197)→ |
| Eco-523-1 | NZ_JHFM01000019.1 | - | 39110 | 39058 | **RNA**→ rpmE (PRK00019)Ribosomal_L31 (pfam01197)→ |
| Eco-480-1 | NZ_JHKU01000042.1 | - | 451 | 399 | **RNA**→ rpmE (PRK00019)Ribosomal_L31 (pfam01197)→ |
| Eco-540-1 | NZ_JHIJ01000018.1 | + | 83133 | 83185 | **RNA**→ rpmE (PRK00019)Ribosomal_L31 (pfam01197)→ |
| Eco-499-1 | NZ_JHHU01000047.1 | - | 451 | 399 | **RNA**→ rpmE (PRK00019)Ribosomal_L31 (pfam01197)→ |
| Eco-500-1 | NZ_JHHT01000020.1 | - | 655 | 603 | **RNA**→ rpmE (PRK00019)Ribosomal_L31 (pfam01197)→ |
| Eco-504-1 | NZ_JHHP01000019.1 | - | 451 | 399 | **RNA**→ rpmE (PRK00019)Ribosomal_L31 (pfam01197)→ |
| Eco-505-1 | NZ_JHHO01000094.1 | + | 88567 | 88619 | **RNA**→ rpmE (PRK00019)Ribosomal_L31 (pfam01197)→ |

| | | | | | |
|---|---|---|---|---|---|
| Eco-486-1 | NZ_JHFE01000034.1 | - | 451 | 399 | RNA→ rpmE (PRK00019)Ribosomal_L31 (pfam01197)→ |
| Eco-488-1 | NZ_JHMX01000046.1 | + | 93655 | 93707 | RNA→ rpmE (PRK00019)Ribosomal_L31 (pfam01197)→ |
| Eco-483-1 | NZ_JHFZ01000080.1 | + | 88516 | 88568 | RNA→ rpmE (PRK00019)Ribosomal_L31 (pfam01197)→ |
| Eco-546-1 | NZ_JASO01000010.1 | + | 2235040 | 2235092 | RNA→ rpmE (PRK00019)Ribosomal_L31 (pfam01197)→ |
| Eco-531-1 | NZ_JHMG01000006.1 | - | 39184 | 39132 | RNA→ rpmE (PRK00019)Ribosomal_L31 (pfam01197)→ |
| Eco-519-1 | NZ_JHGN01000005.1 | - | 39111 | 39059 | RNA→ rpmE (PRK00019)Ribosomal_L31 (pfam01197)→ |
| Eco-485-1 | NZ_JHFF01000047.1 | - | 451 | 399 | RNA→ rpmE (PRK00019)Ribosomal_L31 (pfam01197)→ |
| Eco-527-1 | NZ_JHMV01000020.1 | - | 39111 | 39059 | RNA→ rpmE (PRK00019)Ribosomal_L31 (pfam01197)→ |
| Eco-520-1 | NZ_JHGL01000022.1 | + | 83133 | 83185 | RNA→ rpmE (PRK00019)Ribosomal_L31 (pfam01197)→ |
| Eco-524-1 | NZ_JHFL01000061.1 | + | 83052 | 83104 | RNA→ rpmE (PRK00019)Ribosomal_L31 (pfam01197)→ |
| Eco-501-1 | NZ_JHHS01000028.1 | + | 7404 | 7456 | RNA→ rpmE (PRK00019)Ribosomal_L31 (pfam01197)→ |
| Eco-507-1 | NZ_JHHM01000024.1 | - | 723 | 671 | RNA→ rpmE (PRK00019)Ribosomal_L31 (pfam01197)→ |
| Eco-492-1 | NZ_JHMR01000017.1 | + | 88567 | 88619 | RNA→ rpmE (PRK00019)Ribosomal_L31 (pfam01197)→ |
| Eco-511-1 | NZ_JASP01000001.1 | + | 2904157 | 2904209 | RNA→ rpmE (PRK00019)Ribosomal_L31 (pfam01197)→ |
| Eco-1777-1 | NZ_AGTH01000174.1 | + | 20821 | 20873 | RNA→ rpmE (PRK00019)Ribosomal_L31 (pfam01197)→ |
| Eco-530-1 | NZ_JHMJ01000018.1 | - | 39359 | 39307 | RNA→ rpmE (PRK00019)Ribosomal_L31 (pfam01197)→ |
| Eco-522-1 | NZ_JHGG01000033.1 | + | 83133 | 83185 | RNA→ rpmE (PRK00019)Ribosomal_L31 (pfam01197)→ |
| Eco-525-1 | NZ_JHFK01000039.1 | + | 83560 | 83612 | RNA→ rpmE (PRK00019)Ribosomal_L31 (pfam01197)→ |
| Eco-512-1 | NZ_JHKX01000025.1 | - | 912 | 860 | RNA→ rpmE (PRK00019)Ribosomal_L31 (pfam01197)→ |
| Eco-489-1 | NZ_JHMW01000016.1 | - | 451 | 399 | RNA→ rpmE (PRK00019)Ribosomal_L31 (pfam01197)→ |
| Eco-538-1 | NZ_JHJR01000002.1 | - | 39112 | 39060 | RNA→ rpmE (PRK00019)Ribosomal_L31 (pfam01197)→ |
| Eco-482-1 | NZ_JHGS01000024.1 | + | 88528 | 88580 | RNA→ rpmE (PRK00019)Ribosomal_L31 (pfam01197)→ |
| Eco-513-1 | NZ_JHGP01000041.1 | + | 79309 | 79361 | RNA→ rpmE (PRK00019)Ribosomal_L31 (pfam01197)→ |
| Eco-521-1 | NZ_JHGH01000021.1 | - | 39111 | 39059 | RNA→ rpmE (PRK00019)Ribosomal_L31 (pfam01197)→ |
| Eco-481-1 | NZ_JHGU01000024.1 | - | 451 | 399 | RNA→ rpmE (PRK00019)Ribosomal_L31 (pfam01197)→ |
| Eco-502-1 | NZ_JHHR01000021.1 | - | 783 | 731 | RNA→ rpmE (PRK00019)Ribosomal_L31 (pfam01197)→ |
| Eco-506-1 | NZ_JHHN01000035.1 | - | 451 | 399 | RNA→ rpmE (PRK00019)Ribosomal_L31 (pfam01197)→ |
| Eco-97-1 | NZ_JNPZ01000021.1 | + | 85592 | 85644 | RNA→ rpmE (PRK00019)Ribosomal_L31 (pfam01197)→ |
| Eco-493-1 | NZ_JHMQ01000031.1 | - | 451 | 399 | RNA→ rpmE (PRK00019)Ribosomal_L31 (pfam01197)→ |
| Eco-537-1 | NZ_JASV01000004.1 | + | 2866260 | 2866312 | RNA→ rpmE (PRK00019)Ribosomal_L31 (pfam01197)→ |
| Eco-102-1 | NZ_JNQA01000020.1 | + | 33535 | 33587 | RNA→ rpmE (PRK00019)Ribosomal_L31 (pfam01197)→ |
| Eco-548-2 | NZ_AGTL01000019.1 | + | 67580 | 67632 | RNA→ rpmE (PRK00019)Ribosomal_L31 (pfam01197)→ |
| Eco-494-1 | NZ_JHMA01000070.1 | + | 88568 | 88620 | RNA→ rpmE (PRK00019)Ribosomal_L31 (pfam01197)→ |
| Eco-86-1 | NZ_JNPJ01000150.1 | + | 28740 | 28792 | RNA→ rpmE (PRK00019)Ribosomal_L31 (pfam01197)→ |
| Eco-484-1 | NZ_JHFG01000005.1 | + | 88567 | 88619 | RNA→ rpmE (PRK00019)Ribosomal_L31 (pfam01197)→ |
| Eco-91-1 | NZ_JNPM01000070.1 | + | 87598 | 87650 | RNA→ rpmE (PRK00019)Ribosomal_L31 (pfam01197)→ |
| Eco-554-1 | NZ_JHKV01000042.1 | + | 113307 | 113359 | RNA→ rpmE (PRK00019)Ribosomal_L31 (pfam01197)→ |
| Eco-99-1 | NZ_JNPO01000013.1 | - | 27928 | 27876 | RNA→ rpmE (PRK00019)Ribosomal_L31 (pfam01197)→ |
| Eco-515-1 | NZ_JHNX01000006.1 | + | 83133 | 83185 | RNA→ rpmE (PRK00019)Ribosomal_L31 (pfam01197)→ |
| Eco-106-1 | NZ_JNQK01000038.1 | - | 3888 | 3836 | RNA→ rpmE (PRK00019)Ribosomal_L31 (pfam01197)→ |
| Eco-516-1 | NZ_JHNR01000001.1 | + | 83133 | 83185 | RNA→ rpmE (PRK00019)Ribosomal_L31 (pfam01197)→ |
| Eco-118-1 | NZ_JNQQ01000016.1 | + | 88828 | 88880 | RNA→ rpmE (PRK00019)Ribosomal_L31 (pfam01197)→ |
| Eco-539-1 | NZ_JHIK01000019.1 | + | 83133 | 83185 | RNA→ rpmE (PRK00019)Ribosomal_L31 (pfam01197)→ |
| Eco-111-1 | NZ_JNQN01000020.1 | + | 62519 | 62571 | RNA→ rpmE (PRK00019)Ribosomal_L31 (pfam01197)→ |
| Eco-541-1 | NZ_JHHC01000012.1 | + | 87650 | 87702 | RNA→ rpmE (PRK00019)Ribosomal_L31 (pfam01197)→ |
| Eco-149-1 | NZ_JNRO01000012.1 | - | 41639 | 41587 | RNA→ rpmE (PRK00019)Ribosomal_L31 (pfam01197)→ |
| Eco-547-1 | NZ_JHHD01000017.1 | + | 85958 | 86010 | RNA→ rpmE (PRK00019)Ribosomal_L31 (pfam01197)→ |
| Eco-503-1 | NZ_JHHQ01000058.1 | - | 451 | 399 | RNA→ rpmE (PRK00019)Ribosomal_L31 (pfam01197)→ |
| Eco-146-1 | NZ_JNRM01000006.1 | - | 129387 | 129335 | RNA→ rpmE (PRK00019)Ribosomal_L31 (pfam01197)→ |
| Eco-508-1 | NZ_JHHL01000028.1 | + | 88625 | 88677 | RNA→ rpmE (PRK00019)Ribosomal_L31 (pfam01197)→ |
| Eco-152-1 | NZ_JNRR01000018.1 | + | 86382 | 86434 | RNA→ rpmE (PRK00019)Ribosomal_L31 (pfam01197)→ |
| Eco-528-1 | NZ_JHMO01000008.1 | + | 83133 | 83185 | RNA→ rpmE (PRK00019)Ribosomal_L31 (pfam01197)→ |

| | | | | | |
|---|---|---|---|---|---|
| Eco-155-1 | NZ_JNRS01000020.1 | + | 88027 | 88079 | RNA→ rpmE (PRK00019)Ribosomal_L31 (pfam01197)→ |
| Eco-159-1 | NZ_JNRW01000009.1 | + | 89304 | 89356 | RNA→ rpmE (PRK00019)Ribosomal_L31 (pfam01197)→ |
| Eco-160-1 | NZ_JNRX01000009.1 | - | 39867 | 39815 | RNA→ rpmE (PRK00019)Ribosomal_L31 (pfam01197)→ |
| Eco-154-1 | NZ_JONC01000118.1 | + | 28684 | 28736 | RNA→ rpmE (PRK00019)Ribosomal_L31 (pfam01197)→ |
| Eco-145-1 | NZ_JOSL01000055.1 | + | 87855 | 87907 | RNA→ rpmE (PRK00019)Ribosomal_L31 (pfam01197)→ |
| Eco-161-1 | NZ_JNMT01000023.1 | - | 39691 | 39639 | RNA→ rpmE (PRK00019)Ribosomal_L31 (pfam01197)→ |
| Eco-89-1 | NZ_JNPH01000081.1 | + | 88998 | 89050 | RNA→ rpmE (PRK00019)Ribosomal_L31 (pfam01197)→ |
| Eco-82-1 | NZ_JNPE01000121.1 | + | 86220 | 86272 | RNA→ rpmE (PRK00019)Ribosomal_L31 (pfam01197)→ |
| Eco-78-1 | NZ_JNOZ01000062.1 | + | 87039 | 87091 | RNA→ rpmE (PRK00019)Ribosomal_L31 (pfam01197)→ |
| Eco-90-1 | NZ_JNPL01000135.1 | + | 88947 | 88999 | RNA→ rpmE (PRK00019)Ribosomal_L31 (pfam01197)→ |
| Eco-105-1 | NZ_JNQJ01000010.1 | - | 39999 | 39947 | RNA→ rpmE (PRK00019)Ribosomal_L31 (pfam01197)→ |
| Eco-2161-1 | NZ_AFVX01000011.1 | - | 25826 | 25774 | RNA→ rpmE (PRK00019)Ribosomal_L31 (pfam01197)→ |
| Eco-119-1 | NZ_JNQR01000040.1 | + | 32316 | 32368 | RNA→ rpmE (PRK00019)Ribosomal_L31 (pfam01197)→ |
| Eco-113-1 | NZ_JNQO01000012.1 | + | 89995 | 90047 | RNA→ rpmE (PRK00019)Ribosomal_L31 (pfam01197)→ |
| Eco-147-1 | NZ_JNRN01000027.1 | + | 57743 | 57795 | RNA→ rpmE (PRK00019)Ribosomal_L31 (pfam01197)→ |
| Eco-2035-1 | NZ_AELC01000028.1 | + | 23717 | 23769 | RNA→ rpmE (PRK00019)Ribosomal_L31 (pfam01197)→ |
| Eco-156-1 | NZ_JNRT01000008.1 | - | 79980 | 79928 | RNA→ rpmE (PRK00019)Ribosomal_L31 (pfam01197)→ |
| Eco-123-1 | NZ_JNQV01000020.1 | + | 88561 | 88613 | RNA→ rpmE (PRK00019)Ribosomal_L31 (pfam01197)→ |
| Esp-80-1 | NZ_AEKA01000063.1 | + | 12224 | 12276 | RNA→ rpmE (PRK00019)Ribosomal_L31 (pfam01197)→ |
| Eco-115-1 | NZ_JOMW01000186.1 | + | 65886 | 65938 | RNA→ rpmE (PRK00019)Ribosomal_L31 (pfam01197)→ |
| Eco-114-1 | NZ_JOSP01000088.1 | + | 88483 | 88535 | RNA→ rpmE (PRK00019)Ribosomal_L31 (pfam01197)→ |
| Eco-81-1 | NZ_JNPB01000106.1 | + | 38018 | 38070 | RNA→ rpmE (PRK00019)Ribosomal_L31 (pfam01197)→ |
| Eco-2158-1 | NZ_AGIA01000015.1 | - | 39247 | 39195 | RNA→ rpmE (PRK00019)Ribosomal_L31 (pfam01197)→ rpmE (PRK00019)Ribosomal_L31 (pfam01197)→ |
| Eco-83-1 | NZ_JNPF01000067.1 | + | 87496 | 87548 | RNA→ rpmE (PRK00019)Ribosomal_L31 (pfam01197)→ |
| Eco-96-1 | NZ_JNPN01000049.1 | + | 29332 | 29384 | RNA→ rpmE (PRK00019)Ribosomal_L31 (pfam01197)→ |
| Eco-98-1 | NZ_JNPV01000011.1 | - | 41495 | 41443 | RNA→ rpmE (PRK00019)Ribosomal_L31 (pfam01197)→ |
| Eco-93-1 | NZ_JNQF01000020.1 | - | 1220 | 1168 | RNA→ rpmE (PRK00019)Ribosomal_L31 (pfam01197)→ |
| Eco-88-1 | NZ_JNQE01000113.1 | + | 83333 | 83385 | RNA→ rpmE (PRK00019)Ribosomal_L31 (pfam01197)→ |
| Eco-104-1 | NZ_JNQI01000008.1 | - | 41038 | 40986 | RNA→ rpmE (PRK00019)Ribosomal_L31 (pfam01197)→ |
| Eco-100-1 | NZ_JNQH01000013.1 | - | 26588 | 26536 | RNA→ rpmE (PRK00019)Ribosomal_L31 (pfam01197)→ |
| Eco-141-1 | NZ_JNRK01000010.1 | - | 40492 | 40440 | RNA→ rpmE (PRK00019)Ribosomal_L31 (pfam01197)→ |
| Eco-150-1 | NZ_JNRP01000018.1 | - | 41639 | 41587 | RNA→ rpmE (PRK00019)Ribosomal_L31 (pfam01197)→ |
| Eco-2040-1 | NZ_AELE01000001.1 | - | 107913 | 107861 | RNA→ rpmE (PRK00019)Ribosomal_L31 (pfam01197)→ |
| Eco-157-1 | NZ_JNRU01000011.1 | + | 79676 | 79728 | RNA→ rpmE (PRK00019)Ribosomal_L31 (pfam01197)→ |
| Eco-137-1 | NZ_JONA01000068.1 | + | 86290 | 86342 | RNA→ rpmE (PRK00019)Ribosomal_L31 (pfam01197)→ |
| Eco-110-1 | NZ_JOSN01000103.1 | + | 61758 | 61810 | RNA→ rpmE (PRK00019)Ribosomal_L31 (pfam01197)→ |
| Eco-131-1 | NZ_JOSS01000074.1 | + | 85558 | 85610 | RNA→ rpmE (PRK00019)Ribosomal_L31 (pfam01197)→ |
| Eco-79-1 | NZ_JNNG01000165.1 | + | 26964 | 27016 | RNA→ rpmE (PRK00019)Ribosomal_L31 (pfam01197)→ |
| Eco-80-1 | NZ_JNPA01000051.1 | + | 87539 | 87591 | RNA→ rpmE (PRK00019)Ribosomal_L31 (pfam01197)→ |
| Eco-92-1 | NZ_JNPI01000134.1 | + | 88008 | 88060 | RNA→ rpmE (PRK00019)Ribosomal_L31 (pfam01197)→ |
| Eco-85-1 | NZ_JNPG01000074.1 | + | 49989 | 50041 | RNA→ rpmE (PRK00019)Ribosomal_L31 (pfam01197)→ |
| Eco-103-1 | NZ_JNPW01000018.1 | - | 1405 | 1353 | RNA→ rpmE (PRK00019)Ribosomal_L31 (pfam01197)→ |
| Eco-1993-1 | NZ_AFDZ01000026.1 | + | 94461 | 94513 | RNA→ rpmE (PRK00019)Ribosomal_L31 (pfam01197)→ |
| Eco-1990-1 | NZ_AFDQ01000030.1 | + | 120795 | 120847 | RNA→ rpmE (PRK00019)Ribosomal_L31 (pfam01197)→ |
| Eco-1995-1 | NZ_AFEA01000021.1 | + | 89529 | 89581 | RNA→ rpmE (PRK00019)Ribosomal_L31 (pfam01197)→ |
| Eco-2037-1 | NZ_AELD01000027.1 | - | 2133 | 2081 | RNA→ rpmE (PRK00019)Ribosomal_L31 (pfam01197)→ |
| Eco-1826-1 | NZ_AEUB01000054.1 | + | 92443 | 92495 | RNA→ rpmE (PRK00019)Ribosomal_L31 (pfam01197)→ |
| Eco-2042-1 | NZ_AFEB01000062.1 | + | 82453 | 82505 | RNA→ rpmE (PRK00019)Ribosomal_L31 (pfam01197)→ |
| Eco-1817-1 | NZ_AKBA01000257.1 | + | 47707 | 47759 | RNA→ rpmE (PRK00019)Ribosomal_L31 (pfam01197)→ |
| Eco-2025-1 | NZ_AFAG02000007.1 | + | 81627 | 81679 | RNA→ rpmE (PRK00019)Ribosomal_L31 (pfam01197)→ |
| Eco-2023-1 | NZ_AKLT01000016.1 | - | 3775 | 3723 | RNA→ rpmE (PRK00019)Ribosomal_L31 (pfam01197)→ |
| Eco-707-1 | NZ_JHFB01000041.1 | - | 655 | 603 | RNA→ rpmE (PRK00019)Ribosomal_L31 (pfam01197)→ |

| | | | | | |
|---|---|---|---|---|---|
| Eco-947-1 | NZ_AFET01000003.1 | + | 367780 | 367832 | **RNA**→ rpmE (PRK00019)Ribosomal_L31 (pfam01197)→ |
| Eco-1031-1 | NZ_AIGT01000034.1 | - | 907 | 855 | **RNA**→ rpmE (PRK00019)Ribosomal_L31 (pfam01197)→ |
| Eco-732-1 | NZ_JHGO01000021.1 | + | 115572 | 115624 | **RNA**→ rpmE (PRK00019)Ribosomal_L31 (pfam01197)→ |
| Eco-1032-1 | NZ_AIGU01000065.1 | - | 4128 | 4076 | **RNA**→ rpmE (PRK00019)Ribosomal_L31 (pfam01197)→ |
| Eco-712-1 | NZ_JHGY01000036.1 | + | 46647 | 46699 | **RNA**→ rpmE (PRK00019)Ribosomal_L31 (pfam01197)→ |
| Eco-1036-1 | NZ_AIGY01000052.1 | + | 9545 | 9597 | **RNA**→ rpmE (PRK00019)Ribosomal_L31 (pfam01197)→ |
| Eco-708-1 | NZ_JHMS01000030.1 | + | 47743 | 47795 | **RNA**→ rpmE (PRK00019)Ribosomal_L31 (pfam01197)→ |
| Eco-1041-1 | NZ_AIHD01000074.1 | + | 106498 | 106550 | **RNA**→ rpmE (PRK00019)Ribosomal_L31 (pfam01197)→ |
| Eco-698-1 | NZ_JHMP01000084.1 | - | 655 | 603 | **RNA**→ rpmE (PRK00019)Ribosomal_L31 (pfam01197)→ |
| Eco-1047-1 | NZ_AIHJ01000045.1 | + | 51483 | 51535 | **RNA**→ rpmE (PRK00019)Ribosomal_L31 (pfam01197)→ |
| Eco-1052-1 | NZ_AIHO01000050.1 | - | 3640 | 3588 | **RNA**→ rpmE (PRK00019)Ribosomal_L31 (pfam01197)→ |
| Eco-700-1 | NZ_JHMI01000040.1 | + | 47743 | 47795 | **RNA**→ rpmE (PRK00019)Ribosomal_L31 (pfam01197)→ |
| Eco-1782-1 | NZ_AGTG01000310.1 | + | 91692 | 91744 | **RNA**→ rpmE (PRK00019)Ribosomal_L31 (pfam01197)→ |
| Eco-949-1 | NZ_AJPQ01000119.1 | + | 11893 | 11945 | **RNA**→ rpmE (PRK00019)Ribosomal_L31 (pfam01197)→ |
| Eco-691-1 | NZ_JHGI01000054.1 | + | 47743 | 47795 | **RNA**→ rpmE (PRK00019)Ribosomal_L31 (pfam01197)→ |
| Eco-1061-1 | NZ_AIEZ01000034.1 | + | 89055 | 89107 | **RNA**→ rpmE (PRK00019)Ribosomal_L31 (pfam01197)→ |
| Eco-724-1 | NZ_JFBE01000009.1 | + | 23451 | 23503 | **RNA**→ rpmE (PRK00019)Ribosomal_L31 (pfam01197)→ |
| Eco-1033-1 | NZ_AIGV01000053.1 | + | 90417 | 90469 | **RNA**→ rpmE (PRK00019)Ribosomal_L31 (pfam01197)→ |
| Eco-715-1 | NZ_JASS01000001.1 | - | 2254243 | 2254191 | **RNA**→ rpmE (PRK00019)Ribosomal_L31 (pfam01197)→ |
| Eco-1038-1 | NZ_AIHA01000041.1 | + | 93933 | 93985 | **RNA**→ rpmE (PRK00019)Ribosomal_L31 (pfam01197)→ |
| Eco-1043-1 | NZ_AIHF01000032.1 | + | 18538 | 18590 | **RNA**→ rpmE (PRK00019)Ribosomal_L31 (pfam01197)→ |
| Eco-727-1 | NZ_JHJX01000130.1 | - | 5325 | 5273 | **RNA**→ rpmE (PRK00019)Ribosomal_L31 (pfam01197)→ |
| Eco-1048-1 | NZ_AIHK01000029.1 | - | 52984 | 52932 | **RNA**→ rpmE (PRK00019)Ribosomal_L31 (pfam01197)→ |
| Eco-573-1 | NZ_JHJY01000046.1 | - | 39120 | 39068 | **RNA**→ rpmE (PRK00019)Ribosomal_L31 (pfam01197)→ |
| Eco-1053-1 | NZ_AIHP01000031.1 | + | 126889 | 126941 | **RNA**→ rpmE (PRK00019)Ribosomal_L31 (pfam01197)→ |
| Eco-716-1 | NZ_JHNP01000054.1 | + | 47846 | 47898 | **RNA**→ rpmE (PRK00019)Ribosomal_L31 (pfam01197)→ |
| Eco-907-1 | NZ_AEZT02000041.1 | - | 3273 | 3221 | **RNA**→ rpmE (PRK00019)Ribosomal_L31 (pfam01197)→ |
| Eco-720-1 | NZ_JHHA01000004.1 | - | 605 | 553 | **RNA**→ rpmE (PRK00019)Ribosomal_L31 (pfam01197)→ |
| Eco-869-1 | NZ_AEZZ02000031.1 | + | 33807 | 33859 | **RNA**→ ←hypo |
| Eco-733-1 | NZ_JHGW01000029.1 | + | 84488 | 84540 | **RNA**→ rpmE (PRK00019)Ribosomal_L31 (pfam01197)→ |
| Eco-719-1 | NZ_JHHG01000014.1 | - | 512 | 460 | **RNA**→ rpmE (PRK00019)Ribosomal_L31 (pfam01197)→ |
| Eco-242-1 | NZ_JORD01000185.1 | + | 18741 | 18793 | **RNA**→ rpmE (PRK00019)Ribosomal_L31 (pfam01197)→ |
| Eco-697-1 | NZ_JHND01000051.1 | + | 47743 | 47795 | **RNA**→ rpmE (PRK00019)Ribosomal_L31 (pfam01197)→ |
| Eco-253-1 | NZ_JORX01000060.1 | + | 89329 | 89381 | **RNA**→ rpmE (PRK00019)Ribosomal_L31 (pfam01197)→ |
| Eco-701-1 | NZ_JHMF01000044.1 | - | 655 | 603 | **RNA**→ rpmE (PRK00019)Ribosomal_L31 (pfam01197)→ |
| Eco-559-1 | NZ_JHFS01000024.1 | + | 113298 | 113350 | **RNA**→ rpmE (PRK00019)Ribosomal_L31 (pfam01197)→ |
| Eco-263-1 | NZ_JORY01000190.1 | + | 32400 | 32452 | **RNA**→ rpmE (PRK00019)Ribosomal_L31 (pfam01197)→ |
| Eco-1813-1 | NZ_JAST01000013.1 | + | 1682622 | 1682674 | **RNA**→ rpmE (PRK00019)Ribosomal_L31 (pfam01197)→ |
| Eco-250-1 | NZ_JOSC01000067.1 | + | 77673 | 77725 | **RNA**→ rpmE (PRK00019)Ribosomal_L31 (pfam01197)→ |
| Eco-737-1 | NZ_AQGP01000014.1 | + | 46642 | 46694 | **RNA**→ rpmE (PRK00019)Ribosomal_L31 (pfam01197)→ |
| Eco-251-1 | NZ_JOSF01000157.1 | + | 117812 | 117864 | **RNA**→ rpmE (PRK00019)Ribosomal_L31 (pfam01197)→ |
| Eco-738-1 | NZ_AQGO01000053.1 | - | 450 | 398 | **RNA**→ rpmE (PRK00019)Ribosomal_L31 (pfam01197)→ |
| Eco-244-1 | NZ_JORF01000038.1 | + | 101362 | 101414 | **RNA**→ rpmE (PRK00019)Ribosomal_L31 (pfam01197)→ |
| Eco-739-1 | NZ_AQGN01000068.1 | - | 469 | 417 | **RNA**→ rpmE (PRK00019)Ribosomal_L31 (pfam01197)→ |
| Eco-558-1 | NZ_JHFT01000023.1 | + | 113389 | 113441 | **RNA**→ rpmE (PRK00019)Ribosomal_L31 (pfam01197)→ |
| Eco-1832-1 | NZ_ADUQ01000074.1 | + | 324503 | 324555 | **RNA**→ rpmE (PRK00019)Ribosomal_L31 (pfam01197)→ rpmE (PRK00019)Ribosomal_L31 (pfam01197)→ |
| Eco-269-1 | NZ_JORZ01000117.1 | + | 64110 | 64162 | **RNA**→ rpmE (PRK00019)Ribosomal_L31 (pfam01197)→ |
| Eco-1833-1 | NZ_ADUR01000080.1 | + | 18632 | 18684 | **RNA**→ rpmE (PRK00019)Ribosomal_L31 (pfam01197)→ |
| Eco-271-1 | NZ_JOSB01000055.1 | + | 80836 | 80888 | **RNA**→ rpmE (PRK00019)Ribosomal_L31 (pfam01197)→ |
| Eco-1821-1 | NZ_AJQW01000016.1 | - | 39344 | 39292 | **RNA**→ rpmE (PRK00019)Ribosomal_L31 (pfam01197)→ |
| Eco-265-1 | NZ_JOSD01000063.1 | + | 87627 | 87679 | **RNA**→ rpmE (PRK00019)Ribosomal_L31 (pfam01197)→ |
| Eco-239-1 | NZ_JORC01000094.1 | + | 89119 | 89171 | **RNA**→ rpmE (PRK00019)Ribosomal_L31 (pfam01197)→ |

| | | | | | | |
|---|---|---|---|---|---|---|
| Eco-1818-1 | NZ_AKBB01000286.1 | + | 47707 | 47759 | RNA→ | rpmE (PRK00019)Ribosomal_L31 (pfam01197)→ |
| Eco-391-1 | NZ_CCCR010000424.1 | - | 319 | 267 | RNA→ | rpmE (PRK00019)Ribosomal_L31 (pfam01197)→ |
| Eco-1994-1 | NZ_AFEX01000042.1 | + | 24076 | 24128 | RNA→ | rpmE (PRK00019)Ribosomal_L31 (pfam01197)→ |
| Eco-709-1 | NZ_JHNY01000076.1 | + | 42083 | 42135 | RNA→ | rpmE (PRK00019)Ribosomal_L31 (pfam01197)→ |
| Eco-1030-1 | NZ_AJWV01000005.1 | - | 430 | 378 | RNA→ | rpmE (PRK00019)Ribosomal_L31 (pfam01197)→ |
| Eco-556-1 | NZ_JHFV01000020.1 | - | 42199 | 42147 | RNA→ | rpmE (PRK00019)Ribosomal_L31 (pfam01197)→ |
| Eco-710-1 | NZ_JHHJ01000123.1 | - | 470 | 418 | RNA→ | rpmE (PRK00019)Ribosomal_L31 (pfam01197)→ |
| Eco-713-1 | NZ_JHGJ01000012.1 | - | 1410 | 1358 | RNA→ | rpmE (PRK00019)Ribosomal_L31 (pfam01197)→ |
| Eco-1034-1 | NZ_AIGW01000044.1 | + | 124125 | 124177 | RNA→ | rpmE (PRK00019)Ribosomal_L31 (pfam01197)→ |
| Eco-730-1 | NZ_JHHF01000016.1 | - | 470 | 418 | RNA→ | rpmE (PRK00019)Ribosomal_L31 (pfam01197)→ |
| Eco-1039-1 | NZ_AIHB01000071.1 | + | 255720 | 255772 | RNA→ | rpmE (PRK00019)Ribosomal_L31 (pfam01197)→ |
| Eco-696-1 | NZ_JHFD01000005.1 | + | 47743 | 47795 | RNA→ | rpmE (PRK00019)Ribosomal_L31 (pfam01197)→ |
| Eco-1044-1 | NZ_AIHG01000032.1 | + | 124962 | 125014 | RNA→ | rpmE (PRK00019)Ribosomal_L31 (pfam01197)→ |
| Eco-588-1 | NZ_JHMK01000020.1 | - | 470 | 418 | RNA→ | rpmE (PRK00019)Ribosomal_L31 (pfam01197)→ |
| Eco-1054-1 | NZ_AIHQ01000029.1 | + | 28373 | 28425 | RNA→ | rpmE (PRK00019)Ribosomal_L31 (pfam01197)→ |
| Eco-561-1 | NZ_JHFQ01000010.1 | - | 41718 | 41666 | RNA→ | rpmE (PRK00019)Ribosomal_L31 (pfam01197)→ |
| Eco-731-1 | NZ_JASQ01000001.1 | + | 2087491 | 2087543 | RNA→ | rpmE (PRK00019)Ribosomal_L31 (pfam01197)→ |
| Eco-289-2 | NZ_AMSK01000022.1 | - | 39312 | 39260 | RNA→ | rpmE (PRK00019)Ribosomal_L31 (pfam01197)→ |
| Eco-1822-1 | NZ_AGTK01000047.1 | - | 658 | 606 | RNA→ | rpmE (PRK00019)Ribosomal_L31 (pfam01197)→ |
| Eco-562-1 | NZ_JHFP01000015.1 | + | 113491 | 113543 | RNA→ | rpmE (PRK00019)Ribosomal_L31 (pfam01197)→ |
| Eco-988-1 | NZ_AFAH02000005.1 | + | 348627 | 348679 | RNA→ | rpmE (PRK00019)Ribosomal_L31 (pfam01197)→ |
| Eco-243-1 | NZ_JORE01000086.1 | + | 18702 | 18754 | RNA→ | rpmE (PRK00019)Ribosomal_L31 (pfam01197)→ |
| Eco-1984-1 | NZ_AJFG01000039.1 | + | 32298 | 32350 | RNA→ | rpmE (PRK00019)Ribosomal_L31 (pfam01197)→ |
| Eco-264-1 | NZ_JORV01000162.1 | + | 86327 | 86379 | RNA→ | rpmE (PRK00019)Ribosomal_L31 (pfam01197)→ |
| Eco-273-1 | NZ_JOSA01000051.1 | + | 84041 | 84093 | RNA→ | rpmE (PRK00019)Ribosomal_L31 (pfam01197)→ |
| Eco-721-1 | NZ_JHJW01000066.1 | + | 34177 | 34229 | RNA→ | rpmE (PRK00019)Ribosomal_L31 (pfam01197)→ |
| Eco-256-1 | NZ_JOSG01000075.1 | + | 49467 | 49519 | RNA→ | rpmE (PRK00019)Ribosomal_L31 (pfam01197)→ |
| Eco-722-1 | NZ_JHJU01000002.1 | - | 39350 | 39298 | RNA→ | rpmE (PRK00019)Ribosomal_L31 (pfam01197)→ |
| Eco-270-1 | NZ_JOSQ01000141.1 | + | 63861 | 63913 | RNA→ | rpmE (PRK00019)Ribosomal_L31 (pfam01197)→ |
| Eco-714-1 | NZ_JHFI01000013.1 | - | 1185 | 1133 | RNA→ | rpmE (PRK00019)Ribosomal_L31 (pfam01197)→ |
| Eco-858-1 | NZ_AJWT01000023.1 | + | 86064 | 86116 | RNA→ | rpmE (PRK00019)Ribosomal_L31 (pfam01197)→ |
| Eco-690-1 | NZ_JHGK01000050.1 | + | 47735 | 47787 | RNA→ | rpmE (PRK00019)Ribosomal_L31 (pfam01197)→ |
| Eco-564-1 | NZ_JHFN01000056.1 | - | 42067 | 42015 | RNA→ | rpmE (PRK00019)Ribosomal_L31 (pfam01197)→ |
| Eco-1062-1 | NZ_AFJB01000037.1 | + | 88476 | 88528 | RNA→ | rpmE (PRK00019)Ribosomal_L31 (pfam01197)→ |
| Eco-1035-1 | NZ_AIGX01000036.1 | - | 2477 | 2425 | RNA→ | rpmE (PRK00019)Ribosomal_L31 (pfam01197)→ |
| Eco-717-1 | NZ_JASN01000012.1 | - | 1273280 | 1273228 | RNA→ | rpmE (PRK00019)Ribosomal_L31 (pfam01197)→ |
| Eco-1784-1 | NZ_AHZD02000014.1 | + | 121770 | 121822 | RNA→ | rpmE (PRK00019)Ribosomal_L31 (pfam01197)→ |
| Eco-1040-1 | NZ_AIHC01000084.1 | + | 46886 | 46938 | RNA→ | rpmE (PRK00019)Ribosomal_L31 (pfam01197)→ |
| Eco-836-1 | NZ_AEZS02000010.1 | + | 2298721 | 2298773 | RNA→ | rpmE (PRK00019)Ribosomal_L31 (pfam01197)→ |
| Eco-1045-1 | NZ_AIHH01000059.1 | + | 123112 | 123164 | RNA→ | ←hypo |
| Eco-846-1 | NZ_AEZX02000008.1 | - | 166424 | 166372 | RNA→ | rpmE (PRK00019)Ribosomal_L31 (pfam01197)→ |
| Eco-1046-1 | NZ_AIHI01000030.1 | + | 123842 | 123894 | RNA→ | rpmE (PRK00019)Ribosomal_L31 (pfam01197)→ |
| Eco-172-1 | NZ_JNMD01000017.1 | + | 34663 | 34715 | RNA→ | rpmE (PRK00019)Ribosomal_L31 (pfam01197)→ |
| Eco-1051-1 | NZ_AIHN01000068.1 | + | 65831 | 65883 | RNA→ | rpmE (PRK00019)Ribosomal_L31 (pfam01197)→ |
| Eco-1055-1 | NZ_AIHR01000030.1 | + | 127087 | 127139 | RNA→ | rpmE (PRK00019)Ribosomal_L31 (pfam01197)→ |
| Eco-181-1 | NZ_JNMI01000027.1 | + | 34751 | 34803 | RNA→ | rpmE (PRK00019)Ribosomal_L31 (pfam01197)→ |
| Eco-1056-1 | NZ_AIHS01000050.1 | + | 28401 | 28453 | RNA→ | rpmE (PRK00019)Ribosomal_L31 (pfam01197)→ |
| Eco-187-1 | NZ_JNMS01000012.1 | + | 85669 | 85721 | RNA→ | rpmE (PRK00019)Ribosomal_L31 (pfam01197)→ |
| Eco-560-1 | NZ_JHFR01000019.1 | + | 113461 | 113513 | RNA→ | rpmE (PRK00019)Ribosomal_L31 (pfam01197)→ |
| Eco-186-1 | NZ_JNMW01000014.1 | - | 40226 | 40174 | RNA→ | rpmE (PRK00019)Ribosomal_L31 (pfam01197)→ |
| Eco-6-1 | NZ_JHDM01000084.1 | + | 28050 | 28102 | RNA→ | rpmE (PRK00019)Ribosomal_L31 (pfam01197)→ |
| Eco-165-1 | NZ_JNSA01000020.1 | + | 34427 | 34479 | RNA→ | rpmE (PRK00019)Ribosomal_L31 (pfam01197)→ |

| | | | | | |
|---|---|---|---|---|---|
| Eco-3-1 | NZ_JHDP01000068.1 | + | 89417 | 89469 | RNA→ rpmE (PRK00019)Ribosomal_L31 (pfam01197)→ |
| Eco-178-1 | NZ_JOND01000226.1 | + | 84724 | 84776 | RNA→ rpmE (PRK00019)Ribosomal_L31 (pfam01197)→ |
| Eco-4-1 | NZ_JHDO01000123.1 | + | 86211 | 86263 | RNA→ rpmE (PRK00019)Ribosomal_L31 (pfam01197)→ |
| Eco-567-1 | NZ_JHMH01000011.1 | + | 46647 | 46699 | RNA→ rpmE (PRK00019)Ribosomal_L31 (pfam01197)→ |
| Eco-26-1 | NZ_JJLM01000079.1 | + | 84269 | 84321 | RNA→ rpmE (PRK00019)Ribosomal_L31 (pfam01197)→ |
| Eco-227-1 | NZ_JOST01000207.1 | + | 86014 | 86066 | RNA→ rpmE (PRK00019)Ribosomal_L31 (pfam01197)→ |
| Eco-557-1 | NZ_JHFU01000017.1 | + | 113389 | 113441 | RNA→ rpmE (PRK00019)Ribosomal_L31 (pfam01197)→ |
| Eco-208-1 | NZ_JOQH01000012.1 | + | 90856 | 90908 | RNA→ rpmE (PRK00019)Ribosomal_L31 (pfam01197)→ |
| Eco-16-1 | NZ_JJLQ01000099.1 | + | 22129 | 22181 | RNA→ rpmE (PRK00019)Ribosomal_L31 (pfam01197)→ |
| Eco-188-1 | NZ_JOQL01000012.1 | + | 86189 | 86241 | RNA→ rpmE (PRK00019)Ribosomal_L31 (pfam01197)→ |
| Eco-70-1 | NZ_JMGU01000105.1 | + | 89454 | 89506 | RNA→ rpmE (PRK00019)Ribosomal_L31 (pfam01197)→ |
| Eco-217-1 | NZ_JOQQ01000012.1 | + | 89454 | 89506 | RNA→ rpmE (PRK00019)Ribosomal_L31 (pfam01197)→ |
| Eco-66-1 | NZ_JMGQ01000119.1 | + | 100759 | 100811 | RNA→ rpmE (PRK00019)Ribosomal_L31 (pfam01197)→ |
| Eco-237-1 | NZ_JORA01000046.1 | + | 102409 | 102461 | RNA→ rpmE (PRK00019)Ribosomal_L31 (pfam01197)→ |
| Eco-75-1 | NZ_JNNE01000139.1 | + | 56087 | 56139 | RNA→ rpmE (PRK00019)Ribosomal_L31 (pfam01197)→ |
| Eco-38-1 | NZ_JNPP01000004.1 | - | 40299 | 40247 | RNA→ rpmE (PRK00019)Ribosomal_L31 (pfam01197)→ |
| Eco-175-1 | NZ_JNME01000058.1 | + | 23766 | 23818 | RNA→ rpmE (PRK00019)Ribosomal_L31 (pfam01197)→ |
| Eco-167-1 | NZ_JNMA01000015.1 | + | 93888 | 93940 | RNA→ rpmE (PRK00019)Ribosomal_L31 (pfam01197)→ |
| Eco-35-1 | NZ_JNPT01000021.1 | - | 857 | 805 | RNA→ rpmE (PRK00019)Ribosomal_L31 (pfam01197)→ |
| Eco-189-1 | NZ_JNMJ01000009.1 | + | 86006 | 86058 | RNA→ rpmE (PRK00019)Ribosomal_L31 (pfam01197)→ |
| Eco-32-1 | NZ_JOSR01000127.1 | + | 33305 | 33357 | RNA→ rpmE (PRK00019)Ribosomal_L31 (pfam01197)→ |
| Eco-169-1 | NZ_JNMO01000007.1 | + | 57000 | 57052 | RNA→ rpmE (PRK00019)Ribosomal_L31 (pfam01197)→ |
| Eco-9-1 | NZ_JHDJ01000187.1 | + | 22118 | 22170 | RNA→ rpmE (PRK00019)Ribosomal_L31 (pfam01197)→ |
| Eco-194-1 | NZ_JNMX01000008.1 | + | 70796 | 70848 | RNA→ rpmE (PRK00019)Ribosomal_L31 (pfam01197)→ |
| Eco-2-1 | NZ_JHDQ01000081.1 | + | 87608 | 87660 | RNA→ rpmE (PRK00019)Ribosomal_L31 (pfam01197)→ |
| Eco-203-1 | NZ_JNMY01000024.1 | + | 32464 | 32516 | RNA→ rpmE (PRK00019)Ribosomal_L31 (pfam01197)→ |
| Eco-18-1 | NZ_JJLI01000182.1 | + | 22109 | 22161 | RNA→ rpmE (PRK00019)Ribosomal_L31 (pfam01197)→ |
| Eco-166-1 | NZ_JNSB01000008.1 | + | 70954 | 71006 | RNA→ rpmE (PRK00019)Ribosomal_L31 (pfam01197)→ |
| Eco-31-1 | NZ_JJLU01000129.1 | + | 67734 | 67786 | RNA→ rpmE (PRK00019)Ribosomal_L31 (pfam01197)→ |
| Eco-23-1 | NZ_JJLT01000167.1 | + | 11783 | 11835 | RNA→ rpmE (PRK00019)Ribosomal_L31 (pfam01197)→ |
| Eco-190-1 | NZ_JOQC01000020.1 | - | 721 | 669 | RNA→ rpmE (PRK00019)Ribosomal_L31 (pfam01197)→ |
| Eco-20-1 | NZ_JMGS01000101.1 | + | 86002 | 86054 | RNA→ rpmE (PRK00019)Ribosomal_L31 (pfam01197)→ |
| Eco-228-1 | NZ_JOQM01000023.1 | - | 39867 | 39815 | RNA→ rpmE (PRK00019)Ribosomal_L31 (pfam01197)→ |
| Eco-72-1 | NZ_JNNC01000114.1 | + | 14514 | 14566 | RNA→ rpmE (PRK00019)Ribosomal_L31 (pfam01197)→ |
| Eco-222-1 | NZ_JOQU01000013.1 | + | 88168 | 88220 | RNA→ rpmE (PRK00019)Ribosomal_L31 (pfam01197)→ |
| Eco-76-1 | NZ_JNNF01000065.1 | + | 87059 | 87111 | RNA→ rpmE (PRK00019)Ribosomal_L31 (pfam01197)→ |
| Eco-60-1 | NZ_JNPQ01000048.1 | + | 19619 | 19671 | RNA→ rpmE (PRK00019)Ribosomal_L31 (pfam01197)→ |
| Eco-225-1 | NZ_JOQW01000020.1 | + | 88702 | 88754 | RNA→ rpmE (PRK00019)Ribosomal_L31 (pfam01197)→ |
| Eco-36-1 | NZ_JNPU01000018.1 | - | 2206 | 2154 | RNA→ rpmE (PRK00019)Ribosomal_L31 (pfam01197)→ |
| Eco-14-1 | NZ_JHDG01000181.1 | + | 40836 | 40888 | RNA→ rpmE (PRK00019)Ribosomal_L31 (pfam01197)→ |
| Eco-851-1 | NZ_AJWQ01000257.1 | - | 598 | 546 | RNA→ rpmE (PRK00019)Ribosomal_L31 (pfam01197)→ |
| Eco-10-1 | NZ_JHDI01000087.1 | + | 22108 | 22160 | RNA→ rpmE (PRK00019)Ribosomal_L31 (pfam01197)→ |
| Eco-854-1 | NZ_AJWS01000002.1 | + | 64656 | 64708 | RNA→ rpmE (PRK00019)Ribosomal_L31 (pfam01197)→ |
| Eco-7-1 | NZ_JHDL01000086.1 | + | 28654 | 28706 | RNA→ rpmE (PRK00019)Ribosomal_L31 (pfam01197)→ |
| Eco-25-1 | NZ_JJLP01000055.1 | + | 84896 | 84948 | RNA→ rpmE (PRK00019)Ribosomal_L31 (pfam01197)→ |
| Eco-850-1 | NZ_AJWR01000026.1 | + | 46611 | 46663 | RNA→ rpmE (PRK00019)Ribosomal_L31 (pfam01197)→ |
| Eco-41-1 | NZ_JJLK01000087.1 | + | 85941 | 85993 | RNA→ rpmE (PRK00019)Ribosomal_L31 (pfam01197)→ |
| Eco-24-1 | NZ_JJLS01000289.1 | + | 11674 | 11726 | RNA→ rpmE (PRK00019)Ribosomal_L31 (pfam01197)→ |
| Eco-176-1 | NZ_JNMF01000033.1 | + | 22119 | 22171 | RNA→ rpmE (PRK00019)Ribosomal_L31 (pfam01197)→ |
| Eco-168-1 | NZ_JNMB01000027.1 | - | 39867 | 39815 | RNA→ rpmE (PRK00019)Ribosomal_L31 (pfam01197)→ |
| Eco-71-1 | NZ_JMGV01000071.1 | + | 89398 | 89450 | RNA→ rpmE (PRK00019)Ribosomal_L31 (pfam01197)→ |
| Eco-183-1 | NZ_JNMG01000025.1 | - | 39785 | 39733 | RNA→ rpmE (PRK00019)Ribosomal_L31 (pfam01197)→ |

| | | | | | | |
|---|---|---|---|---|---|---|
| Eco-74-1 | NZ_JMGR01000079.1 | + | 88902 | 88954 | **RNA**→ | rpmE (PRK00019)Ribosomal_L31 (pfam01197)→ |
| Eco-193-1 | NZ_JNMK01000014.1 | - | 41267 | 41215 | **RNA**→ | rpmE (PRK00019)Ribosomal_L31 (pfam01197)→ |
| Eco-69-1 | NZ_JNNB01000156.1 | + | 86845 | 86897 | **RNA**→ | rpmE (PRK00019)Ribosomal_L31 (pfam01197)→ |
| Eco-170-1 | NZ_JNML01000017.1 | + | 87682 | 87734 | **RNA**→ | rpmE (PRK00019)Ribosomal_L31 (pfam01197)→ |
| Eco-73-1 | NZ_JNND01000109.1 | + | 14502 | 14554 | **RNA**→ | rpmE (PRK00019)Ribosomal_L31 (pfam01197)→ |
| Eco-173-1 | NZ_JNMP01000063.1 | + | 24046 | 24098 | **RNA**→ | rpmE (PRK00019)Ribosomal_L31 (pfam01197)→ |
| Eco-62-1 | NZ_JNPR01000049.1 | + | 19639 | 19691 | **RNA**→ | rpmE (PRK00019)Ribosomal_L31 (pfam01197)→ |
| Eco-184-1 | NZ_JNMQ01000015.1 | + | 86809 | 86861 | **RNA**→ | rpmE (PRK00019)Ribosomal_L31 (pfam01197)→ |
| Eco-182-1 | NZ_JNMU01000024.1 | + | 34648 | 34700 | **RNA**→ | rpmE (PRK00019)Ribosomal_L31 (pfam01197)→ |
| Eco-210-1 | NZ_JNMZ01000011.1 | + | 86362 | 86414 | **RNA**→ | rpmE (PRK00019)Ribosomal_L31 (pfam01197)→ |
| Eco-162-1 | NZ_JNRY01000050.1 | - | 2050 | 1998 | **RNA**→ | rpmE (PRK00019)Ribosomal_L31 (pfam01197)→ |
| Eco-5-1 | NZ_JHDN01000076.1 | + | 28050 | 28102 | **RNA**→ | rpmE (PRK00019)Ribosomal_L31 (pfam01197)→ |
| Eco-233-1 | NZ_JOMS01000137.1 | + | 91874 | 91926 | **RNA**→ | rpmE (PRK00019)Ribosomal_L31 (pfam01197)→ |
| Eco-206-1 | NZ_JOQJ01000013.1 | - | 27977 | 27925 | **RNA**→ | rpmE (PRK00019)Ribosomal_L31 (pfam01197)→ |
| Eco-11-1 | NZ_JHDH01000162.1 | + | 32464 | 32516 | **RNA**→ | rpmE (PRK00019)Ribosomal_L31 (pfam01197)→ |
| Eco-8-1 | NZ_JHDK01000152.1 | + | 93103 | 93155 | **RNA**→ | rpmE (PRK00019)Ribosomal_L31 (pfam01197)→ |
| Eco-221-1 | NZ_JOQT01000016.1 | + | 86416 | 86468 | **RNA**→ | rpmE (PRK00019)Ribosomal_L31 (pfam01197)→ |
| Eco-65-1 | NZ_JJLJ01000063.1 | + | 26558 | 26610 | **RNA**→ | rpmE (PRK00019)Ribosomal_L31 (pfam01197)→ |
| Eco-238-1 | NZ_JORB01000073.1 | + | 90314 | 90366 | **RNA**→ | rpmE (PRK00019)Ribosomal_L31 (pfam01197)→ |
| Eco-27-1 | NZ_JJLH01000058.1 | + | 85415 | 85467 | **RNA**→ | rpmE (PRK00019)Ribosomal_L31 (pfam01197)→ |
| Eco-842-1 | NZ_ADUM01000088.1 | + | 46158 | 46210 | **RNA**→ | rpmE (PRK00019)Ribosomal_L31 (pfam01197)→ |
| Eco-840-1 | NZ_AFAF02000002.1 | - | 554932 | 554880 | **RNA**→ | rpmE (PRK00019)Ribosomal_L31 (pfam01197)→ |
| Eco-64-1 | NZ_JJLN01000275.1 | + | 26565 | 26617 | **RNA**→ | rpmE (PRK00019)Ribosomal_L31 (pfam01197)→ |
| Eco-164-1 | NZ_JNLZ01000007.1 | - | 39917 | 39865 | **RNA**→ | rpmE (PRK00019)Ribosomal_L31 (pfam01197)→ |
| Eco-39-1 | NZ_JJLR01000144.1 | + | 78890 | 78942 | **RNA**→ | rpmE (PRK00019)Ribosomal_L31 (pfam01197)→ |
| Eco-171-1 | NZ_JNMC01000016.1 | - | 40179 | 40127 | **RNA**→ | rpmE (PRK00019)Ribosomal_L31 (pfam01197)→ |
| Eco-17-1 | NZ_JMGW01000087.1 | + | 22117 | 22169 | **RNA**→ | rpmE (PRK00019)Ribosomal_L31 (pfam01197)→ |
| Eco-185-1 | NZ_JNMH01000014.1 | + | 86284 | 86336 | **RNA**→ | rpmE (PRK00019)Ribosomal_L31 (pfam01197)→ |
| Eco-61-1 | NZ_JMGT01000069.1 | + | 65843 | 65895 | **RNA**→ | rpmE (PRK00019)Ribosomal_L31 (pfam01197)→ |
| Eco-177-1 | NZ_JNMM01000020.1 | - | 792 | 740 | **RNA**→ | rpmE (PRK00019)Ribosomal_L31 (pfam01197)→ |
| Eco-179-1 | NZ_JNMR01000020.1 | + | 88701 | 88753 | **RNA**→ | rpmE (PRK00019)Ribosomal_L31 (pfam01197)→ |
| Eco-37-1 | NZ_JNPY01000004.1 | - | 40945 | 40893 | **RNA**→ | rpmE (PRK00019)Ribosomal_L31 (pfam01197)→ |
| Eco-63-1 | NZ_JNPS01000013.1 | + | 89385 | 89437 | **RNA**→ | rpmE (PRK00019)Ribosomal_L31 (pfam01197)→ |
| Eco-180-1 | NZ_JNMV01000024.1 | + | 32368 | 32420 | **RNA**→ | rpmE (PRK00019)Ribosomal_L31 (pfam01197)→ |
| Eco-1759-1 | NZ_AFWP01000188.1 | + | 84134 | 84186 | **RNA**→ | rpmE (PRK00019)Ribosomal_L31 (pfam01197)→ |
| Eco-1707-1 | NZ_AEFA01000012.1 | - | 52072 | 52020 | **RNA**→ | rpmE (PRK00019)Ribosomal_L31 (pfam01197)→ rpmE (PRK00019)Ribosomal_L31 (pfam01197)→ |
| Eco-1775-1 | NZ_AKAV01000113.1 | + | 88546 | 88598 | **RNA**→ | rpmE (PRK00019)Ribosomal_L31 (pfam01197)→ |
| Eco-685-1 | NZ_JHOE01000044.1 | - | 821 | 769 | **RNA**→ | rpmE (PRK00019)Ribosomal_L31 (pfam01197)→ |
| Eco-679-1 | NZ_JHOC01000057.1 | + | 22349 | 22401 | **RNA**→ | rpmE (PRK00019)Ribosomal_L31 (pfam01197)→ |
| Eco-676-1 | NZ_JHJJ01000015.1 | - | 3338 | 3286 | **RNA**→ | rpmE (PRK00019)Ribosomal_L31 (pfam01197)→ |
| Eco-1100-1 | NZ_AIGO01000057.1 | - | 3488 | 3436 | **RNA**→ | rpmE (PRK00019)Ribosomal_L31 (pfam01197)→ |
| Eco-1704-1 | NZ_ADWQ01000019.1 | + | 85972 | 86024 | **RNA**→ | rpmE (PRK00019)Ribosomal_L31 (pfam01197)→ |
| Eco-681-1 | NZ_JHJV01000007.1 | + | 82800 | 82852 | **RNA**→ | rpmE (PRK00019)Ribosomal_L31 (pfam01197)→ |
| Eco-1105-1 | NZ_ADUO01000104.1 | - | 534 | 482 | **RNA**→ | rpmE (PRK00019)Ribosomal_L31 (pfam01197)→ |
| Eco-1717-1 | NZ_AJVQ01000020.1 | + | 3811 | 3863 | **RNA**→ | rpmE (PRK00019)Ribosomal_L31 (pfam01197)→ |
| Eco-1774-1 | NZ_AJVV01000390.1 | - | 417 | 365 | **RNA**→ | rpmE (PRK00019)Ribosomal_L31 (pfam01197)→ |
| Eco-1390-1 | NZ_AJWO01000023.1 | + | 65109 | 65161 | **RNA**→ | rpmE (PRK00019)Ribosomal_L31 (pfam01197)→ |
| Eco-1391-1 | NZ_AJWP01000137.1 | - | 430 | 378 | **RNA**→ | rpmE (PRK00019)Ribosomal_L31 (pfam01197)→ |
| Eco-1096-1 | NZ_AIGK01000064.1 | - | 794 | 742 | **RNA**→ | rpmE (PRK00019)Ribosomal_L31 (pfam01197)→ |
| Eco-1097-1 | NZ_AIGL01000055.1 | - | 4074 | 4022 | **RNA**→ | rpmE (PRK00019)Ribosomal_L31 (pfam01197)→ |
| Eco-470-1 | NZ_JHJZ01000022.1 | - | 1543 | 1491 | **RNA**→ | rpmE (PRK00019)Ribosomal_L31 (pfam01197)→ |
| Eco-476-1 | NZ_JHJS01000024.1 | + | 88763 | 88815 | **RNA**→ | rpmE (PRK00019)Ribosomal_L31 (pfam01197)→ |

| | | | | | |
|---|---|---|---|---|---|
| Eco-467-1 | NZ_JHNC01000043.1 | - | 1014 | 962 | **RNA**→ rpmE (PRK00019)Ribosomal_L31 (pfam01197)→ |
| Eco-473-1 | NZ_JHGR01000007.1 | - | 451 | 399 | **RNA**→ rpmE (PRK00019)Ribosomal_L31 (pfam01197)→ |
| Eco-477-1 | NZ_JFGU01000002.1 | - | 450 | 398 | **RNA**→ rpmE (PRK00019)Ribosomal_L31 (pfam01197)→ |
| Eco-1807-1 | NZ_AETX01000108.1 | + | 33794 | 33846 | **RNA**→ rpmE (PRK00019)Ribosomal_L31 (pfam01197)→ |
| Eco-678-1 | NZ_JHNW01000023.1 | + | 24132 | 24184 | **RNA**→ rpmE (PRK00019)Ribosomal_L31 (pfam01197)→ |
| Eco-1683-1 | NZ_ADWU01000058.1 | - | 1253 | 1201 | **RNA**→ rpmE (PRK00019)Ribosomal_L31 (pfam01197)→ rpmE (PRK00019)Ribosomal_L31 (pfam01197)→ |
| Eco-1716-1 | NZ_AJVR01000251.1 | + | 46629 | 46681 | **RNA**→ rpmE (PRK00019)Ribosomal_L31 (pfam01197)→ |
| Eco-1146-1 | NZ_AKNI01000042.1 | + | 23288 | 23340 | **RNA**→ rpmE (PRK00019)Ribosomal_L31 (pfam01197)→ |
| Eco-1386-1 | NZ_AEZV02000042.1 | - | 584 | 532 | **RNA**→ rpmE (PRK00019)Ribosomal_L31 (pfam01197)→ |
| Eco-680-1 | NZ_JHMZ01000045.1 | - | 1193 | 1141 | **RNA**→ rpmE (PRK00019)Ribosomal_L31 (pfam01197)→ |
| Eco-465-1 | NZ_JHFY01000094.1 | - | 1174 | 1122 | **RNA**→ rpmE (PRK00019)Ribosomal_L31 (pfam01197)→ |
| Eco-1708-1 | NZ_AJMB01000085.1 | - | 3692 | 3640 | **RNA**→ rpmE (PRK00019)Ribosomal_L31 (pfam01197)→ |
| Eco-689-1 | NZ_JHGV01000084.1 | + | 47743 | 47795 | **RNA**→ rpmE (PRK00019)Ribosomal_L31 (pfam01197)→ |
| Eco-1709-1 | NZ_AJLU01000016.1 | + | 86588 | 86640 | **RNA**→ rpmE (PRK00019)Ribosomal_L31 (pfam01197)→ |
| Eco-1718-1 | NZ_AGSG01000243.1 | - | 3943 | 3891 | **RNA**→ rpmE (PRK00019)Ribosomal_L31 (pfam01197)→ |
| Eco-1685-1 | NZ_ADWS01000023.1 | + | 66459 | 66511 | **RNA**→ rpmE (PRK00019)Ribosomal_L31 (pfam01197)→ rpmE (PRK00019)Ribosomal_L31 (pfam01197)→ |
| Eco-675-1 | NZ_JHMM01000008.1 | - | 42766 | 42714 | **RNA**→ rpmE (PRK00019)Ribosomal_L31 (pfam01197)→ |
| Eco-1678-1 | NZ_ADWV01000042.1 | + | 23555 | 23607 | **RNA**→ rpmE (PRK00019)Ribosomal_L31 (pfam01197)→ rpmE (PRK00019)Ribosomal_L31 (pfam01197)→ |
| Eco-1168-1 | NZ_AFDX01000043.1 | + | 38823 | 38875 | **RNA**→ rpmE (PRK00019)Ribosomal_L31 (pfam01197)→ |
| Eco-684-1 | NZ_JHNT01000082.1 | - | 502 | 450 | **RNA**→ rpmE (PRK00019)Ribosomal_L31 (pfam01197)→ |
| Eco-1147-1 | NZ_AKNH01000066.1 | + | 82725 | 82777 | **RNA**→ rpmE (PRK00019)Ribosomal_L31 (pfam01197)→ |
| Eco-686-1 | NZ_JHNO01000045.1 | - | 512 | 460 | **RNA**→ rpmE (PRK00019)Ribosomal_L31 (pfam01197)→ |
| Eco-1099-1 | NZ_AIGN01000060.1 | + | 126535 | 126587 | **RNA**→ rpmE (PRK00019)Ribosomal_L31 (pfam01197)→ |
| Eco-158-1 | NZ_JNRV01000013.1 | + | 57668 | 57720 | **RNA**→ rpmE (PRK00019)Ribosomal_L31 (pfam01197)→ |
| Eco-464-1 | NZ_JHOD01000035.1 | + | 86530 | 86582 | **RNA**→ rpmE (PRK00019)Ribosomal_L31 (pfam01197)→ |
| Eco-479-1 | NZ_JHNS01000069.1 | + | 88567 | 88619 | **RNA**→ rpmE (PRK00019)Ribosomal_L31 (pfam01197)→ |
| Eco-163-1 | NZ_JNRZ01000023.1 | - | 25355 | 25303 | **RNA**→ rpmE (PRK00019)Ribosomal_L31 (pfam01197)→ |
| Eco-395-1 | NZ_JDFV01000023.1 | + | 87347 | 87399 | **RNA**→ rpmE (PRK00019)Ribosomal_L31 (pfam01197)→ |
| Eco-475-1 | NZ_JHLU01000058.1 | - | 451 | 399 | **RNA**→ rpmE (PRK00019)Ribosomal_L31 (pfam01197)→ |
| Eco-223-1 | NZ_JONE01000058.1 | + | 88201 | 88253 | **RNA**→ rpmE (PRK00019)Ribosomal_L31 (pfam01197)→ |
| Eco-245-1 | NZ_JORG01000123.1 | + | 101921 | 101973 | **RNA**→ rpmE (PRK00019)Ribosomal_L31 (pfam01197)→ |
| Eco-247-1 | NZ_JORH01000200.1 | + | 86694 | 86746 | **RNA**→ rpmE (PRK00019)Ribosomal_L31 (pfam01197)→ |
| Eco-240-1 | NZ_JORK01000124.1 | + | 87639 | 87691 | **RNA**→ rpmE (PRK00019)Ribosomal_L31 (pfam01197)→ |
| Eco-252-1 | NZ_JORP01000053.1 | + | 90001 | 90053 | **RNA**→ rpmE (PRK00019)Ribosomal_L31 (pfam01197)→ |
| Eco-267-1 | NZ_JOSE01000067.1 | + | 87144 | 87196 | **RNA**→ rpmE (PRK00019)Ribosomal_L31 (pfam01197)→ |
| Eco-272-1 | NZ_JOSI01000129.1 | + | 81087 | 81139 | **RNA**→ rpmE (PRK00019)Ribosomal_L31 (pfam01197)→ |
| Eco-204-1 | NZ_JOQF01000011.1 | + | 90131 | 90183 | **RNA**→ rpmE (PRK00019)Ribosomal_L31 (pfam01197)→ |
| Eco-219-1 | NZ_JOQP01000016.1 | + | 89457 | 89509 | **RNA**→ rpmE (PRK00019)Ribosomal_L31 (pfam01197)→ |
| Eco-235-1 | NZ_JOQZ01000040.1 | + | 101316 | 101368 | **RNA**→ rpmE (PRK00019)Ribosomal_L31 (pfam01197)→ |
| Eco-1131-1 | NZ_AMFM01000015.1 | + | 87827 | 87879 | **RNA**→ rpmE (PRK00019)Ribosomal_L31 (pfam01197)→ |
| Eco-1148-1 | NZ_ADUN01000112.1 | - | 1762 | 1710 | **RNA**→ rpmE (PRK00019)Ribosomal_L31 (pfam01197)→ |
| Eco-261-1 | NZ_JORU01000067.1 | + | 89179 | 89231 | **RNA**→ rpmE (PRK00019)Ribosomal_L31 (pfam01197)→ |
| Eco-218-1 | NZ_JOQO01000026.1 | - | 26793 | 26741 | **RNA**→ rpmE (PRK00019)Ribosomal_L31 (pfam01197)→ |
| Eco-216-1 | NZ_JOQS01000013.1 | - | 41073 | 41021 | **RNA**→ rpmE (PRK00019)Ribosomal_L31 (pfam01197)→ |
| Eco-1094-1 | NZ_AIGI01000048.1 | + | 48588 | 48640 | **RNA**→ rpmE (PRK00019)Ribosomal_L31 (pfam01197)→ |
| Eco-241-1 | NZ_JORT01000089.1 | + | 19614 | 19666 | **RNA**→ rpmE (PRK00019)Ribosomal_L31 (pfam01197)→ |
| Eco-220-1 | NZ_JOQN01000016.1 | + | 89415 | 89467 | **RNA**→ rpmE (PRK00019)Ribosomal_L31 (pfam01197)→ |
| Eco-276-1 | NZ_JOMV01000046.1 | + | 83575 | 83627 | **RNA**→ rpmE (PRK00019)Ribosomal_L31 (pfam01197)→ |
| Eco-236-1 | NZ_JONF01000114.1 | + | 103257 | 103309 | **RNA**→ rpmE (PRK00019)Ribosomal_L31 (pfam01197)→ |
| Eco-266-1 | NZ_JORW01000076.1 | + | 87173 | 87225 | **RNA**→ rpmE (PRK00019)Ribosomal_L31 (pfam01197)→ |
| Eco-207-1 | NZ_JOQK01000011.1 | - | 35712 | 35660 | **RNA**→ rpmE (PRK00019)Ribosomal_L31 (pfam01197)→ |
| Eco-1083-1 | NZ_AIFV01000036.1 | + | 89358 | 89410 | **RNA**→ rpmE (PRK00019)Ribosomal_L31 (pfam01197)→ |

| | | | | | | |
|---|---|---|---|---|---|---|
| Eco-1088-1 | NZ_AIGC01000024.1 | + | 89715 | 89767 | **RNA**→ | rpmE (PRK00019)Ribosomal_L31 (pfam01197)→ |
| Eco-417-1 | NZ_CBWC010000015.1 | + | 4177 | 4229 | **RNA**→ | rpmE (PRK00019)Ribosomal_L31 (pfam01197)→ |
| Eco-255-1 | NZ_JORS01000088.1 | + | 49497 | 49549 | **RNA**→ | rpmE (PRK00019)Ribosomal_L31 (pfam01197)→ |
| Eco-192-1 | NZ_JOQE01000031.1 | - | 1379 | 1327 | **RNA**→ | rpmE (PRK00019)Ribosomal_L31 (pfam01197)→ |
| Eco-209-1 | NZ_JOQI01000010.1 | - | 40093 | 40041 | **RNA**→ | rpmE (PRK00019)Ribosomal_L31 (pfam01197)→ |
| Eco-215-1 | NZ_JDWT01000016.1 | + | 52138 | 52190 | **RNA**→ | rpmE (PRK00019)Ribosomal_L31 (pfam01197)→ |
| Eco-215-2 | NZ_JDWT01000047.1 | - | 39821 | 39769 | **RNA**→ | rpmE (PRK00019)Ribosomal_L31 (pfam01197)→ |
| Eco-246-1 | NZ_JORM01000046.1 | + | 101824 | 101876 | **RNA**→ | rpmE (PRK00019)Ribosomal_L31 (pfam01197)→ |
| Eco-394-1 | NZ_JDFU02000010.1 | + | 2149849 | 2149901 | **RNA**→ | rpmE (PRK00019)Ribosomal_L31 (pfam01197)→ |
| Eco-268-1 | NZ_JDWS01000004.1 | - | 53977 | 53925 | **RNA**→ | rpmE (PRK00019)Ribosomal_L31 (pfam01197)→ |
| Eco-1133-1 | NZ_AERU01000031.1 | - | 830 | 778 | **RNA**→ | rpmE (PRK00019)Ribosomal_L31 (pfam01197)→ |
| Eco-1081-1 | NZ_AIFT01000041.1 | + | 95082 | 95134 | **RNA**→ | rpmE (PRK00019)Ribosomal_L31 (pfam01197)→ rpmE (PRK00019)Ribosomal_L31 (pfam01197)→ |
| Eco-248-1 | NZ_JORJ01000139.1 | + | 86559 | 86611 | **RNA**→ | rpmE (PRK00019)Ribosomal_L31 (pfam01197)→ |
| Eco-262-1 | NZ_JORO01000129.1 | + | 89396 | 89448 | **RNA**→ | rpmE (PRK00019)Ribosomal_L31 (pfam01197)→ |
| Eco-191-1 | NZ_JOQD01000045.1 | + | 30376 | 30428 | **RNA**→ | rpmE (PRK00019)Ribosomal_L31 (pfam01197)→ |
| Eco-214-1 | NZ_JDWU01000026.1 | - | 39906 | 39854 | **RNA**→ | rpmE (PRK00019)Ribosomal_L31 (pfam01197)→ |
| Eco-677-1 | NZ_APWX01000006.1 | - | 577 | 525 | **RNA**→ | rpmE (PRK00019)Ribosomal_L31 (pfam01197)→ |
| Eco-274-1 | NZ_JOMT01000130.1 | + | 83847 | 83899 | **RNA**→ | rpmE (PRK00019)Ribosomal_L31 (pfam01197)→ |
| Eco-254-1 | NZ_JORQ01000056.1 | + | 89943 | 89995 | **RNA**→ | rpmE (PRK00019)Ribosomal_L31 (pfam01197)→ |
| Eco-232-1 | NZ_JOQR01000016.1 | - | 19180 | 19128 | **RNA**→ | rpmE (PRK00019)Ribosomal_L31 (pfam01197)→ |
| Eco-1079-1 | NZ_AIFR01000051.1 | + | 95532 | 95584 | **RNA**→ | rpmE (PRK00019)Ribosomal_L31 (pfam01197)→ |
| Eco-275-1 | NZ_JOMU01000122.1 | + | 80876 | 80928 | **RNA**→ | rpmE (PRK00019)Ribosomal_L31 (pfam01197)→ |
| Eco-249-1 | NZ_JORI01000142.1 | + | 119549 | 119601 | **RNA**→ | rpmE (PRK00019)Ribosomal_L31 (pfam01197)→ |
| Eco-260-1 | NZ_JORN01000052.1 | + | 89277 | 89329 | **RNA**→ | rpmE (PRK00019)Ribosomal_L31 (pfam01197)→ |
| Eco-205-1 | NZ_JOQG01000029.1 | + | 32464 | 32516 | **RNA**→ | rpmE (PRK00019)Ribosomal_L31 (pfam01197)→ |
| Eco-687-1 | NZ_JHGZ01000099.1 | - | 578 | 526 | **RNA**→ | rpmE (PRK00019)Ribosomal_L31 (pfam01197)→ |
| Eco-2033-1 | NZ_AELA01000036.1 | + | 35234 | 35286 | **RNA**→ | rpmE (PRK00019)Ribosomal_L31 (pfam01197)→ |
| Eco-718-1 | NZ_JHLA01000019.1 | - | 579 | 527 | **RNA**→ | rpmE (PRK00019)Ribosomal_L31 (pfam01197)→ |
| Eco-726-1 | NZ_JHNZ01000062.1 | + | 26430 | 26482 | **RNA**→ | rpmE (PRK00019)Ribosomal_L31 (pfam01197)→ |
| Eco-711-1 | NZ_JHOA01000076.1 | + | 46647 | 46699 | **RNA**→ | rpmE (PRK00019)Ribosomal_L31 (pfam01197)→ |
| Eco-683-1 | NZ_JHGM01000054.1 | - | 606 | 554 | **RNA**→ | rpmE (PRK00019)Ribosomal_L31 (pfam01197)→ |
| Eco-1823-1 | NZ_AEUA01000081.1 | + | 92414 | 92466 | **RNA**→ | rpmE (PRK00019)Ribosomal_L31 (pfam01197)→ |
| Eco-2154-1 | NZ_AGTD01000001.1 | + | 4670150 | 4670202 | **RNA**→ | rpmE (PRK00019)Ribosomal_L31 (pfam01197)→ |
| Eco-2159-1 | NZ_AERT01000012.1 | - | 53530 | 53478 | **RNA**→ | rpmE (PRK00019)Ribosomal_L31 (pfam01197)→ |
| Eco-1992-1 | NZ_AFDW01000036.1 | + | 90212 | 90264 | **RNA**→ | rpmE (PRK00019)Ribosomal_L31 (pfam01197)→ |
| Eco-1980-1 | NZ_ADUS01000024.1 | - | 43866 | 43814 | **RNA**→ | rpmE (PRK00019)Ribosomal_L31 (pfam01197)→ |
| Eco-729-1 | NZ_JHNJ01000021.1 | - | 736 | 684 | **RNA**→ | rpmE (PRK00019)Ribosomal_L31 (pfam01197)→ |
| Eco-695-1 | NZ_JHFH01000073.1 | + | 47743 | 47795 | **RNA**→ | rpmE (PRK00019)Ribosomal_L31 (pfam01197)→ |
| Eco-705-1 | NZ_JHLS01000064.1 | - | 1472 | 1420 | **RNA**→ | rpmE (PRK00019)Ribosomal_L31 (pfam01197)→ |
| Eco-699-1 | NZ_JHMN01000069.1 | - | 583 | 531 | **RNA**→ | rpmE (PRK00019)Ribosomal_L31 (pfam01197)→ |
| Eco-688-1 | NZ_JHGX01000054.1 | - | 576 | 524 | **RNA**→ | rpmE (PRK00019)Ribosomal_L31 (pfam01197)→ |
| Eco-702-1 | NZ_JHLY01000043.1 | - | 1068 | 1016 | **RNA**→ | rpmE (PRK00019)Ribosomal_L31 (pfam01197)→ |
| Eco-1808-1 | NZ_AEUC01000085.1 | + | 33893 | 33945 | **RNA**→ | rpmE (PRK00019)Ribosomal_L31 (pfam01197)→ |
| Eco-101-1 | NZ_JNPX01000067.1 | - | 682 | 630 | **RNA**→ | rpmE (PRK00019)Ribosomal_L31 (pfam01197)→ |
| Eco-725-1 | NZ_JHNM01000060.1 | + | 23451 | 23503 | **RNA**→ | rpmE (PRK00019)Ribosomal_L31 (pfam01197)→ |
| Eco-87-1 | NZ_JNQD01000204.1 | + | 80611 | 80663 | **RNA**→ | rpmE (PRK00019)Ribosomal_L31 (pfam01197)→ |
| Eco-694-1 | NZ_JHFJ01000058.1 | - | 533 | 481 | **RNA**→ | rpmE (PRK00019)Ribosomal_L31 (pfam01197)→ |
| Eco-117-1 | NZ_JNQP01000014.1 | - | 27805 | 27753 | **RNA**→ | rpmE (PRK00019)Ribosomal_L31 (pfam01197)→ |
| Eco-2160-1 | NZ_AFYG01000012.1 | - | 39376 | 39324 | **RNA**→ | rpmE (PRK00019)Ribosomal_L31 (pfam01197)→ |
| Eco-107-1 | NZ_JNQL01000039.1 | - | 855 | 803 | **RNA**→ | rpmE (PRK00019)Ribosomal_L31 (pfam01197)→ |
| Eco-692-1 | NZ_JHGA01000065.1 | + | 47743 | 47795 | **RNA**→ | rpmE (PRK00019)Ribosomal_L31 (pfam01197)→ |
| Eco-144-1 | NZ_JNRL01000006.1 | + | 89192 | 89244 | **RNA**→ | rpmE (PRK00019)Ribosomal_L31 (pfam01197)→ |

| | | | | | |
|---|---|---|---|---|---|
| Eco-693-1 | NZ_JHFX01000110.1 | + | 47743 | 47795 | **RNA**→ rpmE (PRK00019)Ribosomal_L31 (pfam01197)→ |
| Eco-151-1 | NZ_JNRQ01000020.1 | + | 39986 | 40038 | **RNA**→ rpmE (PRK00019)Ribosomal_L31 (pfam01197)→ |
| Eco-703-1 | NZ_JHLX01000056.1 | + | 47743 | 47795 | **RNA**→ rpmE (PRK00019)Ribosomal_L31 (pfam01197)→ |
| Eco-77-1 | NZ_JOSJ01000091.1 | + | 87648 | 87700 | **RNA**→ rpmE (PRK00019)Ribosomal_L31 (pfam01197)→ |
| Eco-109-1 | NZ_JOSM01000256.1 | + | 61749 | 61801 | **RNA**→ rpmE (PRK00019)Ribosomal_L31 (pfam01197)→ |
| Eco-94-1 | NZ_JOSO01000082.1 | + | 87720 | 87772 | **RNA**→ rpmE (PRK00019)Ribosomal_L31 (pfam01197)→ |
| Eco-19-1 | NZ_JJMD01000081.1 | + | 88208 | 88260 | **RNA**→ rpmE (PRK00019)Ribosomal_L31 (pfam01197)→ |
| Eco-22-1 | NZ_JJLV01000287.1 | + | 11761 | 11813 | **RNA**→ rpmE (PRK00019)Ribosomal_L31 (pfam01197)→ |
| Eco-129-1 | NZ_JNRA01000017.1 | - | 46298 | 46246 | **RNA**→ rpmE (PRK00019)Ribosomal_L31 (pfam01197)→ |
| Eco-138-1 | NZ_JNRE01000016.1 | + | 86382 | 86434 | **RNA**→ rpmE (PRK00019)Ribosomal_L31 (pfam01197)→ |
| Eco-139-1 | NZ_JNRF01000030.1 | + | 21849 | 21901 | **RNA**→ rpmE (PRK00019)Ribosomal_L31 (pfam01197)→ |
| Eco-59-1 | NZ_JJLZ01000070.1 | + | 85773 | 85825 | **RNA**→ rpmE (PRK00019)Ribosomal_L31 (pfam01197)→ |
| Eco-13-1 | NZ_JJLY01000090.1 | + | 89272 | 89324 | **RNA**→ rpmE (PRK00019)Ribosomal_L31 (pfam01197)→ |
| Eco-136-1 | NZ_JNRD01000020.1 | + | 34619 | 34671 | **RNA**→ rpmE (PRK00019)Ribosomal_L31 (pfam01197)→ |
| Eco-143-1 | NZ_JNRI01000010.1 | + | 89453 | 89505 | **RNA**→ rpmE (PRK00019)Ribosomal_L31 (pfam01197)→ |
| Eco-116-1 | NZ_JOMX01000149.1 | + | 65251 | 65303 | **RNA**→ rpmE (PRK00019)Ribosomal_L31 (pfam01197)→ |
| Eco-12-1 | NZ_JJMA01000084.1 | + | 86866 | 86918 | **RNA**→ rpmE (PRK00019)Ribosomal_L31 (pfam01197)→ |
| Eco-135-1 | NZ_JNRC01000019.1 | + | 34683 | 34735 | **RNA**→ rpmE (PRK00019)Ribosomal_L31 (pfam01197)→ |
| Eco-142-1 | NZ_JNRH01000010.1 | - | 34510 | 34458 | **RNA**→ rpmE (PRK00019)Ribosomal_L31 (pfam01197)→ |
| Eco-133-1 | NZ_JOMZ01000112.1 | + | 82463 | 82515 | **RNA**→ rpmE (PRK00019)Ribosomal_L31 (pfam01197)→ |
| Eco-125-1 | NZ_JNRJ01000025.1 | - | 1790 | 1738 | **RNA**→ rpmE (PRK00019)Ribosomal_L31 (pfam01197)→ |
| Eco-112-1 | NZ_JORR01000098.1 | + | 83115 | 83167 | **RNA**→ rpmE (PRK00019)Ribosomal_L31 (pfam01197)→ |
| Eco-48-1 | NZ_JDWR01000004.1 | + | 79506 | 79558 | **RNA**→ rpmE (PRK00019)Ribosomal_L31 (pfam01197)→ |
| Eco-57-1 | NZ_JJME01000073.1 | + | 85793 | 85845 | **RNA**→ rpmE (PRK00019)Ribosomal_L31 (pfam01197)→ |
| Eco-121-1 | NZ_JNQT01000021.1 | - | 40206 | 40154 | **RNA**→ rpmE (PRK00019)Ribosomal_L31 (pfam01197)→ |
| Eco-127-1 | NZ_JNQY01000020.1 | + | 85463 | 85515 | **RNA**→ rpmE (PRK00019)Ribosomal_L31 (pfam01197)→ |
| Eco-67-1 | NZ_JJMB01000084.1 | + | 90394 | 90446 | **RNA**→ rpmE (PRK00019)Ribosomal_L31 (pfam01197)→ |
| Eco-58-1 | NZ_JJMC01000070.1 | + | 88903 | 88955 | **RNA**→ rpmE (PRK00019)Ribosomal_L31 (pfam01197)→ |
| Eco-128-1 | NZ_JNQZ01000017.1 | - | 39492 | 39440 | **RNA**→ rpmE (PRK00019)Ribosomal_L31 (pfam01197)→ |
| Eco-132-1 | NZ_JOSU01000161.1 | + | 79731 | 79783 | **RNA**→ rpmE (PRK00019)Ribosomal_L31 (pfam01197)→ |
| Eco-47-1 | NZ_JDWQ01000542.1 | + | 14061 | 14113 | **RNA**→ rpmE (PRK00019)Ribosomal_L31 (pfam01197)→ |
| Eco-788-1 | NZ_AFDS01000072.1 | + | 396474 | 396526 | **RNA**→ rpmE (PRK00019)Ribosomal_L31 (pfam01197)→ rpmE (PRK00019)Ribosomal_L31 (pfam01197)→ |
| Eco-21-1 | NZ_JJLX01000084.1 | + | 87592 | 87644 | **RNA**→ rpmE (PRK00019)Ribosomal_L31 (pfam01197)→ |
| Eco-126-1 | NZ_JNQX01000020.1 | + | 84700 | 84752 | **RNA**→ rpmE (PRK00019)Ribosomal_L31 (pfam01197)→ |
| Eco-134-1 | NZ_JNRB01000021.1 | - | 40829 | 40777 | **RNA**→ rpmE (PRK00019)Ribosomal_L31 (pfam01197)→ |
| Eco-140-1 | NZ_JNRG01000015.1 | + | 56974 | 57026 | **RNA**→ rpmE (PRK00019)Ribosomal_L31 (pfam01197)→ |
| Eco-787-1 | NZ_ADUL01000081.1 | - | 7831 | 7779 | **RNA**→ rpmE (PRK00019)Ribosomal_L31 (pfam01197)→ rpmE (PRK00019)Ribosomal_L31 (pfam01197)→ |
| Eco-108-1 | NZ_JNQM01000040.1 | + | 22863 | 22915 | **RNA**→ rpmE (PRK00019)Ribosomal_L31 (pfam01197)→ |
| Eco-124-1 | NZ_JNQW01000036.1 | - | 713 | 661 | **RNA**→ rpmE (PRK00019)Ribosomal_L31 (pfam01197)→ |
| Eco-130-1 | NZ_JOMY01000091.1 | + | 84423 | 84475 | **RNA**→ rpmE (PRK00019)Ribosomal_L31 (pfam01197)→ |
| Eco-1761-1 | NZ_AFSO01000036.1 | + | 85014 | 85066 | **RNA**→ rpmE (PRK00019)Ribosomal_L31 (pfam01197)→ |
| Eco-1782-2 | NZ_AETY01000099.1 | + | 33905 | 33957 | **RNA**→ rpmE (PRK00019)Ribosomal_L31 (pfam01197)→ |
| Eco-1725-1 | NZ_AMVV01000014.1 | + | 85592 | 85644 | **RNA**→ rpmE (PRK00019)Ribosomal_L31 (pfam01197)→ |
| Eco-201-1 | NZ_AYQP01000023.1 | + | 21352 | 21404 | **RNA**→ rpmE (PRK00019)Ribosomal_L31 (pfam01197)→ |
| Eco-1946-1 | NZ_AHHP01000059.1 | - | 430 | 378 | **RNA**→ rpmE (PRK00019)Ribosomal_L31 (pfam01197)→ |
| Eco-412-1 | NZ_AXOA01000007.1 | + | 49060 | 49112 | **RNA**→ rpmE (PRK00019)Ribosomal_L31 (pfam01197)→ |
| Eco-51-1 | NZ_AYQI01000015.1 | - | 487 | 435 | **RNA**→ rpmE (PRK00019)Ribosomal_L31 (pfam01197)→ |
| Eco-56-1 | NZ_AYQX01000041.1 | - | 449 | 397 | **RNA**→ rpmE (PRK00019)Ribosomal_L31 (pfam01197)→ |
| Eco-1783-1 | NZ_AETZ01000105.1 | + | 33906 | 33958 | **RNA**→ rpmE (PRK00019)Ribosomal_L31 (pfam01197)→ |
| Eco-1724-1 | NZ_AMVU01000007.1 | + | 85592 | 85644 | **RNA**→ rpmE (PRK00019)Ribosomal_L31 (pfam01197)→ |
| Eco-1730-1 | NZ_AMVZ01000013.1 | + | 85592 | 85644 | **RNA**→ rpmE (PRK00019)Ribosomal_L31 (pfam01197)→ |
| Eco-487-1 | NZ_JHMY01000014.1 | - | 451 | 399 | **RNA**→ rpmE (PRK00019)Ribosomal_L31 (pfam01197)→ |

| | | | | | |
|---|---|---|---|---|---|
| Eco-474-1 | NZ_JHGB01000047.1 | - | 451 | 399 | **RNA**→ rpmE (PRK00019)Ribosomal_L31 (pfam01197)→ |
| Eco-1758-1 | NZ_AFWO01000116.1 | - | 450 | 398 | **RNA**→ rpmE (PRK00019)Ribosomal_L31 (pfam01197)→ |
| Eco-1729-1 | NZ_AMWA01000009.1 | + | 85592 | 85644 | **RNA**→ rpmE (PRK00019)Ribosomal_L31 (pfam01197)→ |
| Eco-471-1 | NZ_JHLJ01000007.1 | - | 470 | 418 | **RNA**→ rpmE (PRK00019)Ribosomal_L31 (pfam01197)→ |
| Eco-565-1 | NZ_JHNA01000056.1 | - | 42099 | 42047 | **RNA**→ rpmE (PRK00019)Ribosomal_L31 (pfam01197)→ |
| Eco-497-1 | NZ_JHLO01000076.1 | + | 88545 | 88597 | **RNA**→ rpmE (PRK00019)Ribosomal_L31 (pfam01197)→ |
| Eco-466-1 | NZ_JHHE01000034.1 | - | 605 | 553 | **RNA**→ rpmE (PRK00019)Ribosomal_L31 (pfam01197)→ |
| Eco-568-1 | NZ_JHLP01000031.1 | + | 47300 | 47352 | **RNA**→ rpmE (PRK00019)Ribosomal_L31 (pfam01197)→ |
| Eco-498-1 | NZ_JHLM01000051.1 | + | 88567 | 88619 | **RNA**→ rpmE (PRK00019)Ribosomal_L31 (pfam01197)→ |
| Eco-1722-1 | NZ_AMVS01000017.1 | + | 85592 | 85644 | **RNA**→ rpmE (PRK00019)Ribosomal_L31 (pfam01197)→ |
| Eco-1727-1 | NZ_AMVX01000007.1 | + | 85592 | 85644 | **RNA**→ rpmE (PRK00019)Ribosomal_L31 (pfam01197)→ |
| Eco-533-1 | NZ_JHMB01000069.1 | - | 39184 | 39132 | **RNA**→ rpmE (PRK00019)Ribosomal_L31 (pfam01197)→ |
| Eco-469-1 | NZ_JHNB01000020.1 | - | 1265 | 1213 | **RNA**→ rpmE (PRK00019)Ribosomal_L31 (pfam01197)→ |
| Eco-532-1 | NZ_JHMC01000004.1 | - | 39111 | 39059 | **RNA**→ rpmE (PRK00019)Ribosomal_L31 (pfam01197)→ |
| Eco-478-1 | NZ_JHNU01000036.1 | + | 88567 | 88619 | **RNA**→ rpmE (PRK00019)Ribosomal_L31 (pfam01197)→ |
| Eco-534-1 | NZ_JHLZ01000024.1 | + | 83133 | 83185 | **RNA**→ rpmE (PRK00019)Ribosomal_L31 (pfam01197)→ |
| Eco-495-1 | NZ_JHLW01000035.1 | + | 88567 | 88619 | **RNA**→ rpmE (PRK00019)Ribosomal_L31 (pfam01197)→ |
| Eco-490-1 | NZ_JHMU01000039.1 | + | 88567 | 88619 | **RNA**→ rpmE (PRK00019)Ribosomal_L31 (pfam01197)→ |
| Eco-1384-1 | NZ_AICK01000040.1 | - | 39344 | 39292 | **RNA**→ rpmE (PRK00019)Ribosomal_L31 (pfam01197)→ |
| Eco-1432-1 | NZ_ANXR01000072.1 | + | 32531 | 32583 | **RNA**→ rpmE (PRK00019)Ribosomal_L31 (pfam01197)→ |
| Eco-536-1 | NZ_JHLR01000009.1 | - | 39111 | 39059 | **RNA**→ rpmE (PRK00019)Ribosomal_L31 (pfam01197)→ |
| Eco-510-1 | NZ_JHNK01000139.1 | + | 28051 | 28103 | **RNA**→ rpmE (PRK00019)Ribosomal_L31 (pfam01197)→ |
| Eco-529-1 | NZ_JHML01000050.1 | + | 83085 | 83137 | **RNA**→ rpmE (PRK00019)Ribosomal_L31 (pfam01197)→ |
| Eco-517-1 | NZ_JHNH01000023.1 | - | 39183 | 39131 | **RNA**→ rpmE (PRK00019)Ribosomal_L31 (pfam01197)→ |
| Eco-518-1 | NZ_JHGT01000031.1 | - | 39111 | 39059 | **RNA**→ rpmE (PRK00019)Ribosomal_L31 (pfam01197)→ |
| Eco-496-1 | NZ_JHLQ01000067.1 | - | 450 | 398 | **RNA**→ rpmE (PRK00019)Ribosomal_L31 (pfam01197)→ |
| Eco-548-1 | NZ_JHEY01000018.1 | + | 113832 | 113884 | **RNA**→ rpmE (PRK00019)Ribosomal_L31 (pfam01197)→ |
| Eco-535-1 | NZ_JHLT01000018.1 | - | 39183 | 39131 | **RNA**→ rpmE (PRK00019)Ribosomal_L31 (pfam01197)→ |
| Eco-553-1 | NZ_JHNN01000013.1 | + | 113307 | 113359 | **RNA**→ rpmE (PRK00019)Ribosomal_L31 (pfam01197)→ |
| Eco-491-1 | NZ_JHMT01000080.1 | - | 451 | 399 | **RNA**→ rpmE (PRK00019)Ribosomal_L31 (pfam01197)→ |
| Eco-404-1 | NZ_AYRC01000021.1 | + | 87436 | 87488 | **RNA**→ rpmE (PRK00019)Ribosomal_L31 (pfam01197)→ |
| Eco-425-1 | NZ_AYOO01000016.1 | - | 44972 | 44920 | **RNA**→ rpmE (PRK00019)Ribosomal_L31 (pfam01197)→ |
| Eco-413-1 | NZ_AXOB01000006.1 | + | 49060 | 49112 | **RNA**→ rpmE (PRK00019)Ribosomal_L31 (pfam01197)→ |
| Eco-197-1 | NZ_AYQM01000023.1 | - | 26396 | 26344 | **RNA**→ rpmE (PRK00019)Ribosomal_L31 (pfam01197)→ |
| Eco-277-1 | NZ_AYQR01000023.1 | + | 20893 | 20945 | **RNA**→ rpmE (PRK00019)Ribosomal_L31 (pfam01197)→ |
| Eco-53-1 | NZ_AYQJ01000020.1 | - | 711 | 659 | **RNA**→ rpmE (PRK00019)Ribosomal_L31 (pfam01197)→ |
| Eco-54-1 | NZ_AYQK01000016.1 | + | 82842 | 82894 | **RNA**→ rpmE (PRK00019)Ribosomal_L31 (pfam01197)→ |
| Eco-199-1 | NZ_AYQL01000015.1 | - | 491 | 439 | **RNA**→ rpmE (PRK00019)Ribosomal_L31 (pfam01197)→ |
| Eco-257-1 | NZ_AYQV01000040.1 | + | 41491 | 41543 | **RNA**→ rpmE (PRK00019)Ribosomal_L31 (pfam01197)→ |
| Eco-1645-1 | NZ_AYNG01000007.1 | - | 46298 | 46246 | **RNA**→ rpmE (PRK00019)Ribosomal_L31 (pfam01197)→ |
| Eco-1644-1 | NZ_AYNF01000011.1 | - | 52703 | 52651 | **RNA**→ rpmE (PRK00019)Ribosomal_L31 (pfam01197)→ |
| Eco-200-1 | NZ_AYQO01000044.1 | + | 32235 | 32287 | **RNA**→ rpmE (PRK00019)Ribosomal_L31 (pfam01197)→ |
| Eco-148-1 | NZ_AYQZ01000048.1 | - | 449 | 397 | **RNA**→ rpmE (PRK00019)Ribosomal_L31 (pfam01197)→ |
| Eco-213-1 | NZ_AYQT01000039.1 | - | 417 | 365 | **RNA**→ rpmE (PRK00019)Ribosomal_L31 (pfam01197)→ |
| Eco-411-1 | NZ_AXOE01000008.1 | + | 49059 | 49111 | **RNA**→ rpmE (PRK00019)Ribosomal_L31 (pfam01197)→ |
| Eco-283-1 | NZ_AYQY01000014.1 | + | 43050 | 43102 | **RNA**→ rpmE (PRK00019)Ribosomal_L31 (pfam01197)→ |
| Eco-294-1 | NZ_AYMX02000444.1 | + | 47725 | 47777 | **RNA**→ rpmE (PRK00019)Ribosomal_L31 (pfam01197)→ |
| Eco-958-1 | NZ_AYRW01000368.1 | - | 39239 | 39187 | **RNA**→ rpmE (PRK00019)Ribosomal_L31 (pfam01197)→ |
| Eco-259-1 | NZ_AYQN01000026.1 | + | 32272 | 32324 | **RNA**→ rpmE (PRK00019)Ribosomal_L31 (pfam01197)→ |
| Eco-1643-1 | NZ_AYNJ01000061.1 | + | 92332 | 92384 | **RNA**→ rpmE (PRK00019)Ribosomal_L31 (pfam01197)→ |
| Eco-401-1 | NZ_AYJY01000016.1 | + | 113253 | 113305 | **RNA**→ rpmE (PRK00019)Ribosomal_L31 (pfam01197)→ |
| Sfl-85-1 | NZ_AFGW01000061.1 | - | 44611 | 44559 | **RNA**→ rpmE (PRK00019)Ribosomal_L31 (pfam01197)→ |

| | | | | | | |
|---|---|---|---|---|---|---|
| Sbo-12-1 | NZ_AKNB01000294.1 | + | 56283 | 56335 | **RNA**→ | rpmE (PRK00019)Ribosomal_L31 (pfam01197)→ |
| Sfl-61-1 | NZ_AFHD01000041.1 | - | 49743 | 49691 | **RNA**→ | rpmE (PRK00019)Ribosomal_L31 (pfam01197)→ |
| Sfl-65-1 | NZ_AFHC01000055.1 | - | 49661 | 49609 | **RNA**→ | rpmE (PRK00019)Ribosomal_L31 (pfam01197)→ |
| Sbo-14-1 | NZ_AERN01000044.1 | - | 696 | 644 | **RNA**→ | rpmE (PRK00019)Ribosomal_L31 (pfam01197)→ |
| Sso-8-1 | NZ_AKND01000057.1 | + | 87100 | 87152 | **RNA**→ | rpmE (PRK00019)Ribosomal_L31 (pfam01197)→ rpmE (PRK00019)Ribosomal_L31 (pfam01197)→ |
| Sdy-4-1 | NZ_AKNG01000073.1 | + | 16736 | 16788 | **RNA**→ | rpmE (PRK00019)Ribosomal_L31 (pfam01197)→ |
| Sso-9-1 | NZ_AKNE01000079.1 | + | 32321 | 32373 | **RNA**→ | rpmE (PRK00019)Ribosomal_L31 (pfam01197)→ |
| Sfl-25-11 | NZ_CELV01000020.1 | - | 31220 | 31168 | **RNA**→ | rpmE (PRK00019)Ribosomal_L31 (pfam01197)→ |
| Sbo-13-1 | NZ_AFGE01000068.1 | + | 90513 | 90565 | **RNA**→ | rpmE (PRK00019)Ribosomal_L31 (pfam01197)→ |
| Eco-403-1 | NZ_AYKB01000016.1 | + | 113255 | 113307 | **RNA**→ | rpmE (PRK00019)Ribosomal_L31 (pfam01197)→ |
| Eco-405-1 | NZ_AYKC01000030.1 | - | 39439 | 39387 | **RNA**→ | rpmE (PRK00019)Ribosomal_L31 (pfam01197)→ |
| Eco-400-1 | NZ_AYJZ01000015.1 | + | 113245 | 113297 | **RNA**→ | rpmE (PRK00019)Ribosomal_L31 (pfam01197)→ |
| Eco-407-1 | NZ_AYKE01000042.1 | - | 919 | 867 | **RNA**→ | rpmE (PRK00019)Ribosomal_L31 (pfam01197)→ |
| Eco-198-1 | NZ_AYQS01000013.1 | - | 28061 | 28009 | **RNA**→ | rpmE (PRK00019)Ribosomal_L31 (pfam01197)→ |
| Eco-55-1 | NZ_AYJX01000022.1 | + | 32282 | 32334 | **RNA**→ | rpmE (PRK00019)Ribosomal_L31 (pfam01197)→ |
| Eco-406-1 | NZ_AYKD01000019.1 | - | 489 | 437 | **RNA**→ | rpmE (PRK00019)Ribosomal_L31 (pfam01197)→ |
| Eco-957-1 | NZ_AZCA01000201.1 | + | 46647 | 46699 | **RNA**→ | rpmE (PRK00019)Ribosomal_L31 (pfam01197)→ |
| Eco-279-1 | NZ_AYRB01000533.1 | + | 1654 | 1706 | **RNA**→ | rpmE (PRK00019)Ribosomal_L31 (pfam01197)→ |
| Eco-292-1 | NZ_AWXM02000304.1 | + | 113311 | 113363 | **RNA**→ | rpmE (PRK00019)Ribosomal_L31 (pfam01197)→ |
| Eco-1382-1 | NZ_CBWH010000046.1 | + | 79652 | 79704 | **RNA**→ | rpmE (PRK00019)Ribosomal_L31 (pfam01197)→ |
| Eco-416-1 | NZ_CBWG010000182.1 | + | 65101 | 65153 | **RNA**→ | rpmE (PRK00019)Ribosomal_L31 (pfam01197)→ |
| Sfl-32-1 | NZ_AZOI01000027.1 | - | 31127 | 31075 | **RNA**→ | rpmE (PRK00019)Ribosomal_L31 (pfam01197)→ |
| Sfl-30-1 | NZ_AZOS01000026.1 | + | 24428 | 24480 | **RNA**→ | rpmE (PRK00019)Ribosomal_L31 (pfam01197)→ |
| Sfl-22-1 | NZ_AZPX01000049.1 | + | 24428 | 24480 | **RNA**→ | rpmE (PRK00019)Ribosomal_L31 (pfam01197)→ |
| Sfl-67-1 | NZ_AZPG01000200.1 | - | 27825 | 27773 | **RNA**→ | rpmE (PRK00019)Ribosomal_L31 (pfam01197)→ |
| Sfl-59-1 | NZ_AZPW01000280.1 | - | 31364 | 31312 | **RNA**→ | rpmE (PRK00019)Ribosomal_L31 (pfam01197)→ |
| Sfl-34-1 | NZ_AZOO01000003.1 | - | 31133 | 31081 | **RNA**→ | rpmE (PRK00019)Ribosomal_L31 (pfam01197)→ |
| Sfl-48-1 | NZ_AZOP01000079.1 | - | 31135 | 31083 | **RNA**→ | rpmE (PRK00019)Ribosomal_L31 (pfam01197)→ |
| Sfl-37-1 | NZ_AZOU01000070.1 | + | 24432 | 24484 | **RNA**→ | rpmE (PRK00019)Ribosomal_L31 (pfam01197)→ |
| Sfl-49-1 | NZ_AZOY01000093.1 | - | 31123 | 31071 | **RNA**→ | rpmE (PRK00019)Ribosomal_L31 (pfam01197)→ |
| Sfl-27-1 | NZ_AZPE01000102.1 | - | 31292 | 31240 | **RNA**→ | rpmE (PRK00019)Ribosomal_L31 (pfam01197)→ |
| Sfl-60-1 | NZ_AZPY01000287.1 | - | 31486 | 31434 | **RNA**→ | rpmE (PRK00019)Ribosomal_L31 (pfam01197)→ |
| Sfl-84-1 | NZ_AZQD01000281.1 | - | 31338 | 31286 | **RNA**→ | rpmE (PRK00019)Ribosomal_L31 (pfam01197)→ |
| Sfl-41-1 | NZ_AZOQ01000079.1 | + | 24434 | 24486 | **RNA**→ | rpmE (PRK00019)Ribosomal_L31 (pfam01197)→ |
| Sfl-21-1 | NZ_AZOV01000067.1 | + | 24434 | 24486 | **RNA**→ | rpmE (PRK00019)Ribosomal_L31 (pfam01197)→ |
| Sfl-13-1 | NZ_AZPP01000004.1 | - | 31121 | 31069 | **RNA**→ | rpmE (PRK00019)Ribosomal_L31 (pfam01197)→ |
| Sfl-36-1 | NZ_AZOJ01000046.1 | + | 24432 | 24484 | **RNA**→ | rpmE (PRK00019)Ribosomal_L31 (pfam01197)→ |
| Sfl-4-1 | NZ_AZPK01000077.1 | + | 24436 | 24488 | **RNA**→ | rpmE (PRK00019)Ribosomal_L31 (pfam01197)→ |
| Sfl-43-1 | NZ_AZQE01000091.1 | + | 24436 | 24488 | **RNA**→ | rpmE (PRK00019)Ribosomal_L31 (pfam01197)→ |
| Sfl-24-1 | NZ_AZPZ01000091.1 | - | 31117 | 31065 | **RNA**→ | rpmE (PRK00019)Ribosomal_L31 (pfam01197)→ |
| Sfl-2-1 | NZ_AZPH01000073.1 | + | 24422 | 24474 | **RNA**→ | rpmE (PRK00019)Ribosomal_L31 (pfam01197)→ |
| Sfl-83-1 | NZ_AZQC01000174.1 | - | 31492 | 31440 | **RNA**→ | rpmE (PRK00019)Ribosomal_L31 (pfam01197)→ |
| Sfl-15-1 | NZ_AZOZ01000072.1 | - | 31133 | 31081 | **RNA**→ | rpmE (PRK00019)Ribosomal_L31 (pfam01197)→ |
| Sfl-23-1 | NZ_AZON01000035.1 | - | 31133 | 31081 | **RNA**→ | rpmE (PRK00019)Ribosomal_L31 (pfam01197)→ |
| Sfl-7-1 | NZ_AZPL01000292.1 | + | 24472 | 24524 | **RNA**→ | rpmE (PRK00019)Ribosomal_L31 (pfam01197)→ |
| Sfl-28-1 | NZ_AZPF01000209.1 | + | 24439 | 24491 | **RNA**→ | rpmE (PRK00019)Ribosomal_L31 (pfam01197)→ |
| Sfl-16-1 | NZ_AZPS01000215.1 | - | 31471 | 31419 | **RNA**→ | rpmE (PRK00019)Ribosomal_L31 (pfam01197)→ |
| Sfl-40-1 | NZ_AZOX01000009.1 | - | 31131 | 31079 | **RNA**→ | rpmE (PRK00019)Ribosomal_L31 (pfam01197)→ |
| Sfl-10-1 | NZ_AZPC01000017.1 | - | 31131 | 31079 | **RNA**→ | rpmE (PRK00019)Ribosomal_L31 (pfam01197)→ |
| Sfl-8-1 | NZ_AZPM01000105.1 | + | 24432 | 24484 | **RNA**→ | rpmE (PRK00019)Ribosomal_L31 (pfam01197)→ |
| Sfl-11-1 | NZ_AZPN01000265.1 | - | 31305 | 31253 | **RNA**→ | rpmE (PRK00019)Ribosomal_L31 (pfam01197)→ |
| Sfl-6-1 | NZ_AZQH01000008.1 | + | 24424 | 24476 | **RNA**→ | rpmE (PRK00019)Ribosomal_L31 (pfam01197)→ |

| | | | | | |
|---|---|---|---|---|---|
| Sfl-54-1 | NZ_AZPO01000232.1 | + | 24480 | 24532 | **RNA**→ rpmE (PRK00019)Ribosomal_L31 (pfam01197)→ |
| Sfl-18-1 | NZ_AZPT01000020.1 | + | 24494 | 24546 | **RNA**→ rpmE (PRK00019)Ribosomal_L31 (pfam01197)→ |
| Sfl-9-1 | NZ_AZQI01000099.1 | + | 24519 | 24571 | **RNA**→ rpmE (PRK00019)Ribosomal_L31 (pfam01197)→ |
| Sfl-42-1 | NZ_AZOM01000009.1 | - | 31135 | 31083 | **RNA**→ rpmE (PRK00019)Ribosomal_L31 (pfam01197)→ |
| Sfl-50-1 | NZ_AZPB01000027.1 | - | 31121 | 31069 | **RNA**→ rpmE (PRK00019)Ribosomal_L31 (pfam01197)→ |
| Sfl-14-1 | NZ_AZPQ01000223.1 | - | 31360 | 31308 | **RNA**→ rpmE (PRK00019)Ribosomal_L31 (pfam01197)→ |
| Sfl-5-1 | NZ_AZQG01000007.1 | + | 24424 | 24476 | **RNA**→ rpmE (PRK00019)Ribosomal_L31 (pfam01197)→ |
| Sfl-58-1 | NZ_AZPV01000133.1 | + | 24550 | 24602 | **RNA**→ rpmE (PRK00019)Ribosomal_L31 (pfam01197)→ |
| Sfl-81-1 | NZ_AZQA01000184.1 | - | 31349 | 31297 | **RNA**→ rpmE (PRK00019)Ribosomal_L31 (pfam01197)→ |
| Sfl-35-1 | NZ_AZOG01000027.1 | - | 31129 | 31077 | **RNA**→ rpmE (PRK00019)Ribosomal_L31 (pfam01197)→ |
| Sfl-47-1 | NZ_AZOR01000045.1 | + | 24434 | 24486 | **RNA**→ rpmE (PRK00019)Ribosomal_L31 (pfam01197)→ |
| Sfl-29-1 | NZ_AZPA01000041.1 | - | 31191 | 31139 | **RNA**→ rpmE (PRK00019)Ribosomal_L31 (pfam01197)→ |
| Sfl-33-1 | NZ_AZOW01000075.1 | + | 24434 | 24486 | **RNA**→ rpmE (PRK00019)Ribosomal_L31 (pfam01197)→ |
| Sfl-31-1 | NZ_AZQF01000166.1 | - | 31137 | 31085 | **RNA**→ rpmE (PRK00019)Ribosomal_L31 (pfam01197)→ |
| Sfl-12-1 | NZ_AZQJ01000145.1 | + | 24571 | 24623 | **RNA**→ rpmE (PRK00019)Ribosomal_L31 (pfam01197)→ |
| Eco-397-1 | NZ_AONF01000012.1 | - | 41676 | 41624 | **RNA**→ rpmE (PRK00019)Ribosomal_L31 (pfam01197)→ |
| Sfl-56-1 | NZ_AZPU01000275.1 | + | 24653 | 24705 | **RNA**→ rpmE (PRK00019)Ribosomal_L31 (pfam01197)→ |
| Sfl-52-1 | NZ_AZPJ01000292.1 | - | 31282 | 31230 | **RNA**→ rpmE (PRK00019)Ribosomal_L31 (pfam01197)→ |
| Sfl-38-1 | NZ_AZOH01000090.1 | + | 24430 | 24482 | **RNA**→ rpmE (PRK00019)Ribosomal_L31 (pfam01197)→ |
| Sfl-39-1 | NZ_AZOK01000053.1 | + | 24432 | 24484 | **RNA**→ rpmE (PRK00019)Ribosomal_L31 (pfam01197)→ |
| Sfl-3-1 | NZ_AZPI01000293.1 | + | 24420 | 24472 | **RNA**→ rpmE (PRK00019)Ribosomal_L31 (pfam01197)→ |
| Sfl-46-1 | NZ_AZOT01000006.1 | + | 24434 | 24486 | **RNA**→ rpmE (PRK00019)Ribosomal_L31 (pfam01197)→ |
| Sfl-44-1 | NZ_AZOL01000016.1 | - | 31123 | 31071 | **RNA**→ rpmE (PRK00019)Ribosomal_L31 (pfam01197)→ |
| Sfl-19-1 | NZ_AZQL01000102.1 | + | 24428 | 24480 | **RNA**→ rpmE (PRK00019)Ribosomal_L31 (pfam01197)→ |
| Sfl-45-1 | NZ_AZQB01000275.1 | - | 31288 | 31236 | **RNA**→ rpmE (PRK00019)Ribosomal_L31 (pfam01197)→ |
| Sfl-26-1 | NZ_AZPD01000034.1 | - | 31130 | 31078 | **RNA**→ rpmE (PRK00019)Ribosomal_L31 (pfam01197)→ |
| Sfl-20-1 | NZ_AZQM01000042.1 | + | 24424 | 24476 | **RNA**→ rpmE (PRK00019)Ribosomal_L31 (pfam01197)→ |
| Eco-459-1 | NZ_JEMI01000046.1 | - | 52620 | 52568 | **RNA**→ rpmE (PRK00019)Ribosomal_L31 (pfam01197)→ |
| Sfl-17-1 | NZ_AZQK01000022.1 | - | 31129 | 31077 | **RNA**→ rpmE (PRK00019)Ribosomal_L31 (pfam01197)→ |
| Sfl-55-1 | NZ_AZPR01000289.1 | + | 24486 | 24538 | **RNA**→ rpmE (PRK00019)Ribosomal_L31 (pfam01197)→ |
| Sso-1-541 | NZ_CXFM01000027.1 | + | 39045 | 39097 | **RNA**→ rpmE (PRK00019)Ribosomal_L31 (pfam01197)→ |
| Sso-1-542 | NZ_CXKK01000025.1 | - | 790 | 738 | **RNA**→ rpmE (PRK00019)Ribosomal_L31 (pfam01197)→ |
| Sso-1-543 | NZ_CXBE01000023.1 | + | 39030 | 39082 | **RNA**→ rpmE (PRK00019)Ribosomal_L31 (pfam01197)→ |
| Sso-1-544 | NZ_CXJY01000028.1 | + | 39089 | 39141 | **RNA**→ rpmE (PRK00019)Ribosomal_L31 (pfam01197)→ |
| Sdy-3-1 | NZ_AAMJ02000013.1 | - | 49902 | 49850 | **RNA**→ rpmE (PRK00019)Ribosomal_L31 (pfam01197)→ rpmE (PRK00019)Ribosomal_L31 (pfam01197)→ |
| Eco-752-2 | NC_000913.3 | + | 4126954 | 4127006 | **RNA**→ rpmE (PRK00019)Ribosomal_L31 (pfam01197)→ |
| Eco-2153-1 | NC_011751.1 | + | 4613769 | 4613821 | **RNA**→ rpmE (PRK00019)Ribosomal_L31 (pfam01197)→ |
| Eco-1829-1 | NC_017634.1 | + | 4166613 | 4166665 | **RNA**→ rpmE (PRK00019)Ribosomal_L31 (pfam01197)→ |
| Eco-1380-1 | NC_011750.1 | - | 3185961 | 3185909 | **RNA**→ rpmE (PRK00019)Ribosomal_L31 (pfam01197)→ |
| Eco-1742-1 | NC_018658.1 | - | 5023166 | 5023114 | **RNA**→ rpmE (PRK00019)Ribosomal_L31 (pfam01197)→ |
| Sdy-8-1 | NC_007606.1 | - | 3539800 | 3539748 | **RNA**→ rpmE (PRK00019)Ribosomal_L31 (pfam01197)→ |
| Sfl-63-1 | NC_004337.2 | + | 4142031 | 4142083 | **RNA**→ rpmE (PRK00019)Ribosomal_L31 (pfam01197)→ |
| Sfl-66-1 | NC_008258.1 | + | 4113206 | 4113258 | **RNA**→ rpmE (PRK00019)Ribosomal_L31 (pfam01197)→ rpmE (PRK00019)Ribosomal_L31 (pfam01197)→ rpmE (PRK00019)Ribosomal_L31 (pfam01197)→ |
| Sbo-18-1 | NC_007613.1 | + | 3980530 | 3980582 | **RNA**→ rpmE (PRK00019)Ribosomal_L31 (pfam01197)→ |
| Eco-1986-1 | NC_013654.1 | + | 4115387 | 4115439 | **RNA**→ rpmE (PRK00019)Ribosomal_L31 (pfam01197)→ |
| Eco-1985-1 | NC_011415.1 | + | 4388453 | 4388505 | **RNA**→ rpmE (PRK00019)Ribosomal_L31 (pfam01197)→ |
| Eco-1020-1 | NC_012759.1 | + | 4014646 | 4014698 | **RNA**→ rpmE (PRK00019)Ribosomal_L31 (pfam01197)→ |
| Sfl-53-1 | NC_017328.1 | + | 4181705 | 4181757 | **RNA**→ rpmE (PRK00019)Ribosomal_L31 (pfam01197)→ |
| Eco-1988-1 | NC_010498.1 | + | 4465819 | 4465871 | **RNA**→ rpmE (PRK00019)Ribosomal_L31 (pfam01197)→ |
| Eco-1019-1 | NC_012947.1 | - | 4307595 | 4307543 | **RNA**→ rpmE (PRK00019)Ribosomal_L31 (pfam01197)→ |
| Eco-1445-1 | NZ_ANXX01000089.1 | + | 65724 | 65776 | **RNA**→ rpmE (PRK00019)Ribosomal_L31 (pfam01197)→ |

| | | | | | | | |
|---|---|---|---|---|---|---|---|
| Eco-1448-1 | NZ_ANXY01000069.1 | + | 80032 | 80084 | **RNA**→ | rpmE (PRK00019)Ribosomal_L31 (pfam01197)→ | |
| Eco-1466-1 | NZ_ANYE01000075.1 | + | 93209 | 93261 | **RNA**→ | rpmE (PRK00019)Ribosomal_L31 (pfam01197)→ | |
| Eco-1455-1 | NZ_ANYB01000068.1 | + | 87346 | 87398 | **RNA**→ | rpmE (PRK00019)Ribosomal_L31 (pfam01197)→ | |
| Eco-1444-1 | NZ_ANXW01000101.1 | + | 98443 | 98495 | **RNA**→ | rpmE (PRK00019)Ribosomal_L31 (pfam01197)→ | |
| Eco-1642-1 | NZ_ANZA01000057.1 | + | 32586 | 32638 | **RNA**→ | rpmE (PRK00019)Ribosomal_L31 (pfam01197)→ | |
| Eco-1637-1 | NZ_ANYX01000066.1 | + | 32374 | 32426 | **RNA**→ | rpmE (PRK00019)Ribosomal_L31 (pfam01197)→ | |
| Eco-1468-1 | NZ_ANYF01000041.1 | + | 32366 | 32418 | **RNA**→ | rpmE (PRK00019)Ribosomal_L31 (pfam01197)→ | |
| Eco-1463-1 | NZ_ANYD01000058.1 | + | 32368 | 32420 | **RNA**→ | rpmE (PRK00019)Ribosomal_L31 (pfam01197)→ | |
| Eco-1494-1 | NZ_ANYP01000077.1 | + | 65180 | 65232 | **RNA**→ | rpmE (PRK00019)Ribosomal_L31 (pfam01197)→ | |
| Eco-1480-1 | NZ_ANYJ01000072.1 | + | 85888 | 85940 | **RNA**→ | rpmE (PRK00019)Ribosomal_L31 (pfam01197)→ | |
| Eco-1631-1 | NZ_ANYV01000057.1 | + | 77047 | 77099 | **RNA**→ | rpmE (PRK00019)Ribosomal_L31 (pfam01197)→ | |
| Eco-1556-1 | NZ_ANYS01000075.1 | + | 27826 | 27878 | **RNA**→ | rpmE (PRK00019)Ribosomal_L31 (pfam01197)→ | |
| Eco-1019-2 | NC_012971.2 | + | 4034877 | 4034929 | **RNA**→ | rpmE (PRK00019)Ribosomal_L31 (pfam01197)→ | |
| Eco-948-1 | NC_017631.1 | + | 4515506 | 4515558 | **RNA**→ | ←hypo | |
| Eco-1019-3 | NC_012892.2 | + | 4034871 | 4034923 | **RNA**→ | rpmE (PRK00019)Ribosomal_L31 (pfam01197)→ | |
| Eco-2157-1 | NC_017635.1 | + | 4385130 | 4385182 | **RNA**→ | rpmE (PRK00019)Ribosomal_L31 (pfam01197)→ | |
| Sso-12-1 | NC_007384.1 | + | 4340994 | 4341046 | **RNA**→ | rpmE (PRK00019)Ribosomal_L31 (pfam01197)→ | |
| Eco-755-1 | NC_017626.1 | + | 4599214 | 4599266 | **RNA**→ | rpmE (PRK00019)Ribosomal_L31 (pfam01197)→ | |
| Eco-1392-1 | NC_016902.1 | - | 4641114 | 4641062 | **RNA**→ | rpmE (PRK00019)Ribosomal_L31 (pfam01197)→ | |
| Eco-2180-1 | NC_017652.1 | + | 4468480 | 4468532 | **RNA**→ | ←hypo | |
| Eco-2181-1 | NC_017651.1 | + | 4468480 | 4468532 | **RNA**→ | ←hypo | |
| Sso-10-1 | NC_016822.1 | + | 4496466 | 4496518 | **RNA**→ | rpmE (PRK00019)Ribosomal_L31 (pfam01197)→ | |
| Eco-1944-1 | NC_017663.1 | + | 4432216 | 4432268 | **RNA**→ | rpmE (PRK00019)Ribosomal_L31 (pfam01197)→ | |
| Eco-2157-2 | NC_017664.1 | + | 4381611 | 4381663 | **RNA**→ | rpmE (PRK00019)Ribosomal_L31 (pfam01197)→ | |
| Eco-1706-1 | NC_017644.1 | + | 4189500 | 4189552 | **RNA**→ | rpmE (PRK00019)Ribosomal_L31 (pfam01197)→ | |
| Eco-1825-1 | NC_017656.1 | + | 4724417 | 4724469 | **RNA**→ | rpmE (PRK00019)Ribosomal_L31 (pfam01197)→ | |
| Eco-1740-1 | NC_018650.1 | - | 5003301 | 5003249 | **RNA**→ | rpmE (PRK00019)Ribosomal_L31 (pfam01197)→ | |
| Eco-1392-2 | NC_017660.1 | - | 593611 | 593559 | **RNA**→ | rpmE (PRK00019)Ribosomal_L31 (pfam01197)→ | |
| Eco-1741-1 | NC_018661.1 | - | 5062731 | 5062679 | **RNA**→ | rpmE (PRK00019)Ribosomal_L31 (pfam01197)→ | |
| Eco-951-1 | NC_020163.1 | + | 85695 | 85747 | **RNA**→ | rpmE (PRK00019)Ribosomal_L31 (pfam01197)→ | |
| Eco-1101-1 | NC_017625.1 | - | 4387516 | 4387464 | **RNA**→ | rpmE (PRK00019)Ribosomal_L31 (pfam01197)→ | |
| Eco-1652-1 | NC_022364.1 | + | 4304568 | 4304620 | **RNA**→ | rpmE (PRK00019)Ribosomal_L31 (pfam01197)→ | |
| Eco-1109-1 | NC_009801.1 | + | 4454628 | 4454680 | **RNA**→ | rpmE (PRK00019)Ribosomal_L31 (pfam01197)→ | |
| Eco-1387-1 | NC_022648.1 | + | 4479116 | 4479168 | **RNA**→ | rpmE (PRK00019)Ribosomal_L31 (pfam01197)→ | |
| Eco-723-1 | NZ_AUZS01000108.1 | + | 34252 | 34304 | **RNA**→ | rpmE (PRK00019)Ribosomal_L31 (pfam01197)→ | |
| Sfl-68-1 | NZ_CM001474.1 | + | 4119177 | 4119229 | **RNA**→ | rpmE (PRK00019)Ribosomal_L31 (pfam01197)→ | rpmE (PRK00019)Ribosomal_L31 (pfam01197)→ |
| Eco-2155-1 | NC_017641.1 | + | 4645410 | 4645462 | **RNA**→ | rpmE (PRK00019)Ribosomal_L31 (pfam01197)→ | |
| Eco-2184-1 | NZ_CM000960.1 | + | 4124944 | 4124996 | **RNA**→ | rpmE (PRK00019)Ribosomal_L31 (pfam01197)→ | |
| Eco-983-1 | NZ_GG749342.1 | + | 87874 | 87926 | **RNA**→ | rpmE (PRK00019)Ribosomal_L31 (pfam01197)→ | rpmE (PRK00019)Ribosomal_L31 (pfam01197)→ |
| Eco-976-1 | NZ_GG749188.1 | + | 127059 | 127111 | **RNA**→ | rpmE (PRK00019)Ribosomal_L31 (pfam01197)→ | rpmE (PRK00019)Ribosomal_L31 (pfam01197)→ |
| Eco-1699-1 | NZ_GG774371.1 | + | 80684 | 80736 | **RNA**→ | rpmE (PRK00019)Ribosomal_L31 (pfam01197)→ | |
| Ssp-26-1 | NZ_GG657392.1 | + | 88633 | 88685 | **RNA**→ | rpmE (PRK00019)Ribosomal_L31 (pfam01197)→ | rpmE (PRK00019)Ribosomal_L31 (pfam01197)→ |
| Eco-1166-1 | NZ_GG774918.1 | + | 89061 | 89113 | **RNA**→ | rpmE (PRK00019)Ribosomal_L31 (pfam01197)→ | rpmE (PRK00019)Ribosomal_L31 (pfam01197)→ |
| Eco-1691-1 | NZ_GG774248.1 | - | 52196 | 52144 | **RNA**→ | rpmE (PRK00019)Ribosomal_L31 (pfam01197)→ | |
| Eco-1698-1 | NZ_GG774461.1 | + | 64498 | 64550 | **RNA**→ | rpmE (PRK00019)Ribosomal_L31 (pfam01197)→ | |
| Eco-1694-1 | NZ_GG771479.1 | + | 22528 | 22580 | **RNA**→ | rpmE (PRK00019)Ribosomal_L31 (pfam01197)→ | |
| Eco-960-1 | NZ_GG749150.1 | + | 85259 | 85311 | **RNA**→ | rpmE (PRK00019)Ribosomal_L31 (pfam01197)→ | rpmE (PRK00019)Ribosomal_L31 (pfam01197)→ |
| Eco-1682-1 | NZ_GG772852.1 | - | 462 | 410 | **RNA**→ | rpmE (PRK00019)Ribosomal_L31 (pfam01197)→ | |
| Eco-1703-1 | NZ_GG771679.1 | + | 85822 | 85874 | **RNA**→ | rpmE (PRK00019)Ribosomal_L31 (pfam01197)→ | |
| Eco-1697-1 | NZ_GG772206.1 | + | 32504 | 32556 | **RNA**→ | rpmE (PRK00019)Ribosomal_L31 (pfam01197)→ | |
| Eco-1700-1 | NZ_GG772413.1 | + | 25779 | 25831 | **RNA**→ | rpmE (PRK00019)Ribosomal_L31 (pfam01197)→ | |
| Eco-864-1 | NZ_GG669103.1 | - | 52064 | 52012 | **RNA**→ | rpmE (PRK00019)Ribosomal_L31 (pfam01197)→ | |

| | | | | | | |
|---|---|---|---|---|---|---|
| Eco-1167-1 | NZ_GG749232.1 | + | 86937 | 86989 | **RNA**→ | rpmE (PRK00019)Ribosomal_L31 (pfam01197)→ rpmE (PRK00019)Ribosomal_L31 (pfam01197)→ |
| Eco-1689-1 | NZ_GG773609.1 | + | 32438 | 32490 | **RNA**→ | rpmE (PRK00019)Ribosomal_L31 (pfam01197)→ |
| Eco-1695-1 | NZ_GG773824.1 | + | 80487 | 80539 | **RNA**→ | rpmE (PRK00019)Ribosomal_L31 (pfam01197)→ |
| Eco-1680-1 | NZ_GG771822.1 | + | 86729 | 86781 | **RNA**→ | rpmE (PRK00019)Ribosomal_L31 (pfam01197)→ |
| Eco-1696-1 | NZ_GG772729.1 | - | 13634 | 13582 | **RNA**→ | rpmE (PRK00019)Ribosomal_L31 (pfam01197)→ |
| Eco-1687-1 | NZ_GG773153.1 | + | 81002 | 81054 | **RNA**→ | rpmE (PRK00019)Ribosomal_L31 (pfam01197)→ |
| Eco-1701-1 | NZ_GG773309.1 | + | 104840 | 104892 | **RNA**→ | rpmE (PRK00019)Ribosomal_L31 (pfam01197)→ |
| Eco-1690-1 | NZ_GG771886.1 | - | 460 | 408 | **RNA**→ | rpmE (PRK00019)Ribosomal_L31 (pfam01197)→ |
| Eco-1686-1 | NZ_GG772092.1 | + | 44158 | 44210 | **RNA**→ | rpmE (PRK00019)Ribosomal_L31 (pfam01197)→ |
| Eco-1692-1 | NZ_GG772571.1 | - | 108651 | 108599 | **RNA**→ | rpmE (PRK00019)Ribosomal_L31 (pfam01197)→ |
| Eco-420-1 | NZ_KK097726.1 | + | 80911 | 80963 | **RNA**→ | rpmE (PRK00019)Ribosomal_L31 (pfam01197)→ |
| Eco-326-1 | NZ_KK214059.1 | + | 80697 | 80749 | **RNA**→ | rpmE (PRK00019)Ribosomal_L31 (pfam01197)→ |
| Eco-1027-1 | NZ_KI543994.1 | + | 87495 | 87547 | **RNA**→ | rpmE (PRK00019)Ribosomal_L31 (pfam01197)→ |
| Eco-1026-1 | NZ_KI543649.1 | + | 4300 | 4352 | **RNA**→ | rpmE (PRK00019)Ribosomal_L31 (pfam01197)→ |
| Eco-1028-1 | NZ_KI544375.1 | + | 34333 | 34385 | **RNA**→ | rpmE (PRK00019)Ribosomal_L31 (pfam01197)→ |
| Eco-1746-1 | NZ_KB021558.1 | + | 84299 | 84351 | **RNA**→ | rpmE (PRK00019)Ribosomal_L31 (pfam01197)→ |
| Eco-921-1 | NZ_KB003425.1 | + | 91671 | 91723 | **RNA**→ | rpmE (PRK00019)Ribosomal_L31 (pfam01197)→ |
| Eco-1747-1 | NZ_KB021571.1 | + | 180804 | 180856 | **RNA**→ | rpmE (PRK00019)Ribosomal_L31 (pfam01197)→ |
| Eco-1752-1 | NZ_KB021472.1 | + | 181474 | 181526 | **RNA**→ | rpmE (PRK00019)Ribosomal_L31 (pfam01197)→ |
| Eco-1649-1 | NZ_KB223477.1 | + | 2193206 | 2193258 | **RNA**→ | rpmE (PRK00019)Ribosomal_L31 (pfam01197)→ |
| Eco-1756-1 | NZ_KB021464.1 | + | 85592 | 85644 | **RNA**→ | rpmE (PRK00019)Ribosomal_L31 (pfam01197)→ |
| Eco-1750-1 | NZ_KB021485.1 | + | 184834 | 184886 | **RNA**→ | rpmE (PRK00019)Ribosomal_L31 (pfam01197)→ |
| Eco-308-1 | NZ_KK736286.1 | + | 196918 | 196970 | **RNA**→ | rpmE (PRK00019)Ribosomal_L31 (pfam01197)→ |
| Eco-458-1 | NZ_KK736506.1 | + | 3269686 | 3269738 | **RNA**→ | rpmE (PRK00019)Ribosomal_L31 (pfam01197)→ |
| Eco-315-1 | NZ_KK736338.1 | + | 4058460 | 4058512 | **RNA**→ | rpmE (PRK00019)Ribosomal_L31 (pfam01197)→ |
| Eco-389-1 | NZ_KK736486.1 | + | 209675 | 209727 | **RNA**→ | rpmE (PRK00019)Ribosomal_L31 (pfam01197)→ |
| Eco-322-1 | NZ_KK736423.1 | + | 93266 | 93318 | **RNA**→ | rpmE (PRK00019)Ribosomal_L31 (pfam01197)→ |
| Eco-321-1 | NZ_KK736416.1 | + | 82463 | 82515 | **RNA**→ | rpmE (PRK00019)Ribosomal_L31 (pfam01197)→ |
| Eco-741-1 | NZ_KK736534.1 | + | 86741 | 86793 | **RNA**→ | rpmE (PRK00019)Ribosomal_L31 (pfam01197)→ |
| Eco-323-1 | NZ_KK736437.1 | + | 2266431 | 2266483 | **RNA**→ | rpmE (PRK00019)Ribosomal_L31 (pfam01197)→ |
| Eco-313-1 | NZ_KK736322.1 | + | 80906 | 80958 | **RNA**→ | rpmE (PRK00019)Ribosomal_L31 (pfam01197)→ |
| Eco-740-1 | NZ_KK736517.1 | + | 486722 | 486774 | **RNA**→ | rpmE (PRK00019)Ribosomal_L31 (pfam01197)→ |
| Eco-319-1 | NZ_KK736385.1 | + | 87501 | 87553 | **RNA**→ | rpmE (PRK00019)Ribosomal_L31 (pfam01197)→ |
| Eco-324-1 | NZ_KK736448.1 | + | 115918 | 115970 | **RNA**→ | rpmE (PRK00019)Ribosomal_L31 (pfam01197)→ |
| Eco-318-1 | NZ_KK736372.1 | + | 2909588 | 2909640 | **RNA**→ | rpmE (PRK00019)Ribosomal_L31 (pfam01197)→ |
| Eco-744-1 | NZ_KK736563.1 | + | 1241877 | 1241929 | **RNA**→ | rpmE (PRK00019)Ribosomal_L31 (pfam01197)→ |
| Eco-457-1 | NZ_KK736497.1 | + | 90449 | 90501 | **RNA**→ | rpmE (PRK00019)Ribosomal_L31 (pfam01197)→ |
| Eco-387-1 | NZ_KK736462.1 | + | 115174 | 115226 | **RNA**→ | rpmE (PRK00019)Ribosomal_L31 (pfam01197)→ |
| Eco-311-1 | NZ_KK736301.1 | + | 195414 | 195466 | **RNA**→ | rpmE (PRK00019)Ribosomal_L31 (pfam01197)→ |
| Eco-742-1 | NZ_KK736542.1 | + | 4150583 | 4150635 | **RNA**→ | rpmE (PRK00019)Ribosomal_L31 (pfam01197)→ |
| Eco-743-1 | NZ_KK736552.1 | + | 132495 | 132547 | **RNA**→ | rpmE (PRK00019)Ribosomal_L31 (pfam01197)→ |
| Eco-309-1 | NZ_KK736297.1 | + | 87560 | 87612 | **RNA**→ | rpmE (PRK00019)Ribosomal_L31 (pfam01197)→ |
| Eco-745-1 | NZ_KK736578.1 | + | 88541 | 88593 | **RNA**→ | rpmE (PRK00019)Ribosomal_L31 (pfam01197)→ |
| Eco-388-1 | NZ_KK736473.1 | + | 2466433 | 2466485 | **RNA**→ | rpmE (PRK00019)Ribosomal_L31 (pfam01197)→ |
| Eco-320-1 | NZ_KK736404.1 | + | 81196 | 81248 | **RNA**→ | rpmE (PRK00019)Ribosomal_L31 (pfam01197)→ |
| Eco-312-1 | NZ_KK736313.1 | + | 194713 | 194765 | **RNA**→ | rpmE (PRK00019)Ribosomal_L31 (pfam01197)→ |
| Eco-317-1 | NZ_KK736365.1 | + | 88105 | 88157 | **RNA**→ | rpmE (PRK00019)Ribosomal_L31 (pfam01197)→ |
| Eco-1170-1 | NZ_GL871851.1 | + | 43112 | 43164 | **RNA**→ | rpmE (PRK00019)Ribosomal_L31 (pfam01197)→ |
| Eco-911-1 | NZ_GL878028.1 | + | 724541 | 724593 | **RNA**→ | rpmE (PRK00019)Ribosomal_L31 (pfam01197)→ |
| Eco-1180-1 | NZ_GL883798.1 | + | 86450 | 86502 | **RNA**→ | rpmE (PRK00019)Ribosomal_L31 (pfam01197)→ rpmE (PRK00019)Ribosomal_L31 (pfam01197)→ |
| Eco-1176-1 | NZ_GL872255.1 | - | 40223 | 40171 | **RNA**→ | rpmE (PRK00019)Ribosomal_L31 (pfam01197)→ |
| Eco-1178-1 | NZ_GL884468.1 | + | 87820 | 87872 | **RNA**→ | rpmE (PRK00019)Ribosomal_L31 (pfam01197)→ rpmE (PRK00019)Ribosomal_L31 (pfam01197)→ |

| | | | | | |
|---|---|---|---|---|---|
| Eco-1110-1 | NZ_GL871816.1 | + | 87804 | 87856 | **RNA**→ rpmE (PRK00019)Ribosomal_L31 (pfam01197)→ |
| Eco-1653-1 | NZ_GL883924.1 | + | 84931 | 84983 | **RNA**→ rpmE (PRK00019)Ribosomal_L31 (pfam01197)→ rpmE (PRK00019)Ribosomal_L31 (pfam01197)→ |
| Eco-1104-1 | NZ_GL871751.1 | - | 651 | 599 | **RNA**→ rpmE (PRK00019)Ribosomal_L31 (pfam01197)→ |
| Eco-2032-1 | NZ_GL872210.1 | + | 93460 | 93512 | **RNA**→ rpmE (PRK00019)Ribosomal_L31 (pfam01197)→ |
| Eco-2004-1 | NZ_GL884294.1 | + | 87814 | 87866 | **RNA**→ rpmE (PRK00019)Ribosomal_L31 (pfam01197)→ rpmE (PRK00019)Ribosomal_L31 (pfam01197)→ |
| Eco-1764-1 | NZ_GL989523.1 | + | 84226 | 84278 | **RNA**→ rpmE (PRK00019)Ribosomal_L31 (pfam01197)→ |
| Eco-2002-1 | NZ_GL884232.1 | + | 83146 | 83198 | **RNA**→ rpmE (PRK00019)Ribosomal_L31 (pfam01197)→ rpmE (PRK00019)Ribosomal_L31 (pfam01197)→ |
| Eco-1106-1 | NZ_GL871776.1 | + | 87176 | 87228 | **RNA**→ rpmE (PRK00019)Ribosomal_L31 (pfam01197)→ |
| Eco-1654-1 | NZ_GL884138.1 | + | 94025 | 94077 | **RNA**→ rpmE (PRK00019)Ribosomal_L31 (pfam01197)→ rpmE (PRK00019)Ribosomal_L31 (pfam01197)→ |
| Eco-2003-1 | NZ_GL884196.1 | + | 81379 | 81431 | **RNA**→ rpmE (PRK00019)Ribosomal_L31 (pfam01197)→ rpmE (PRK00019)Ribosomal_L31 (pfam01197)→ |
| Eco-2001-1 | NZ_JH590773.1 | - | 42161 | 42109 | **RNA**→ rpmE (PRK00019)Ribosomal_L31 (pfam01197)→ |
| Eco-1177-1 | NZ_JH590617.1 | + | 88767 | 88819 | **RNA**→ rpmE (PRK00019)Ribosomal_L31 (pfam01197)→ |
| Eco-1102-1 | NZ_JH590524.1 | + | 92133 | 92185 | **RNA**→ rpmE (PRK00019)Ribosomal_L31 (pfam01197)→ |
| Eco-961-1 | NZ_JH590664.1 | - | 9874 | 9822 | **RNA**→ rpmE (PRK00019)Ribosomal_L31 (pfam01197)→ |
| Eco-1656-1 | NZ_JH659572.1 | + | 695309 | 695361 | **RNA**→ rpmE (PRK00019)Ribosomal_L31 (pfam01197)→ |
| Eco-2038-1 | NZ_JH695848.1 | + | 91681 | 91733 | **RNA**→ rpmE (PRK00019)Ribosomal_L31 (pfam01197)→ |
| Esp-76-1 | NZ_JH660542.1 | - | 40026 | 39974 | **RNA**→ rpmE (PRK00019)Ribosomal_L31 (pfam01197)→ |
| Eco-993-1 | NZ_JH768572.1 | + | 1165956 | 1166008 | **RNA**→ rpmE (PRK00019)Ribosomal_L31 (pfam01197)→ |
| Eco-1179-1 | NZ_JH768585.1 | + | 1121171 | 1121223 | **RNA**→ rpmE (PRK00019)Ribosomal_L31 (pfam01197)→ |
| Esp-75-1 | NZ_JH815282.1 | + | 114130 | 114182 | **RNA**→ rpmE (PRK00019)Ribosomal_L31 (pfam01197)→ |
| Eco-2022-1 | NZ_JH954133.1 | + | 32834 | 32886 | **RNA**→ rpmE (PRK00019)Ribosomal_L31 (pfam01197)→ |
| Eco-841-1 | NZ_JH953850.1 | + | 85845 | 85897 | **RNA**→ rpmE (PRK00019)Ribosomal_L31 (pfam01197)→ |
| Eco-952-1 | NZ_JH954331.1 | + | 76681 | 76733 | **RNA**→ rpmE (PRK00019)Ribosomal_L31 (pfam01197)→ |
| Eco-1143-1 | NZ_JH955422.1 | + | 91755 | 91807 | **RNA**→ rpmE (PRK00019)Ribosomal_L31 (pfam01197)→ |
| Eco-1705-1 | NZ_JH965420.1 | + | 47331 | 47383 | **RNA**→ rpmE (PRK00019)Ribosomal_L31 (pfam01197)→ |
| Eco-860-1 | NZ_JH970602.1 | + | 81850 | 81902 | **RNA**→ rpmE (PRK00019)Ribosomal_L31 (pfam01197)→ |
| Eco-852-1 | NZ_JH954938.1 | + | 91698 | 91750 | **RNA**→ rpmE (PRK00019)Ribosomal_L31 (pfam01197)→ |
| Eco-756-1 | NZ_JH964479.1 | + | 22701 | 22753 | **RNA**→ rpmE (PRK00019)Ribosomal_L31 (pfam01197)→ |
| Eco-766-1 | NZ_JH968717.1 | + | 91759 | 91811 | **RNA**→ rpmE (PRK00019)Ribosomal_L31 (pfam01197)→ |
| Eco-861-1 | NZ_JH969598.1 | + | 81857 | 81909 | **RNA**→ rpmE (PRK00019)Ribosomal_L31 (pfam01197)→ |
| Eco-2041-1 | NZ_JH954585.1 | + | 47568 | 47620 | **RNA**→ rpmE (PRK00019)Ribosomal_L31 (pfam01197)→ |
| Eco-753-1 | NZ_JH943432.1 | + | 46671 | 46723 | **RNA**→ rpmE (PRK00019)Ribosomal_L31 (pfam01197)→ |
| Eco-855-1 | NZ_JH958335.1 | - | 512 | 460 | **RNA**→ rpmE (PRK00019)Ribosomal_L31 (pfam01197)→ |
| Eco-1755-1 | NZ_KB021584.1 | + | 87738 | 87790 | **RNA**→ rpmE (PRK00019)Ribosomal_L31 (pfam01197)→ |
| Eco-2163-1 | NZ_HE962388.1 | + | 4198777 | 4198829 | **RNA**→ rpmE (PRK00019)Ribosomal_L31 (pfam01197)→ |
| Eco-1726-1 | NZ_KB021620.1 | + | 85592 | 85644 | **RNA**→ rpmE (PRK00019)Ribosomal_L31 (pfam01197)→ |
| Eco-1754-1 | NZ_KB021513.1 | + | 85592 | 85644 | **RNA**→ rpmE (PRK00019)Ribosomal_L31 (pfam01197)→ |
| Eco-1757-1 | NZ_KB021454.1 | + | 1171507 | 1171559 | **RNA**→ rpmE (PRK00019)Ribosomal_L31 (pfam01197)→ |
| Eco-1749-1 | NZ_KB021497.1 | + | 1170984 | 1171036 | **RNA**→ rpmE (PRK00019)Ribosomal_L31 (pfam01197)→ |
| Eco-1748-1 | NZ_KB021525.1 | + | 427364 | 427416 | **RNA**→ rpmE (PRK00019)Ribosomal_L31 (pfam01197)→ |
| Eco-1751-1 | NZ_KB021544.1 | + | 85116 | 85168 | **RNA**→ rpmE (PRK00019)Ribosomal_L31 (pfam01197)→ |
| Eco-1567-1 | NZ_KB732351.1 | + | 97932 | 97984 | **RNA**→ rpmE (PRK00019)Ribosomal_L31 (pfam01197)→ |
| Eco-1505-1 | NZ_KB732415.1 | + | 113044 | 113096 | **RNA**→ rpmE (PRK00019)Ribosomal_L31 (pfam01197)→ |
| Eco-1522-1 | NZ_KB732473.1 | + | 743128 | 743180 | **RNA**→ rpmE (PRK00019)Ribosomal_L31 (pfam01197)→ |
| Eco-1414-1 | NZ_KB732238.1 | + | 1437519 | 1437571 | **RNA**→ rpmE (PRK00019)Ribosomal_L31 (pfam01197)→ |
| Eco-1436-1 | NZ_KB731785.1 | + | 32367 | 32419 | **RNA**→ rpmE (PRK00019)Ribosomal_L31 (pfam01197)→ |
| Eco-1441-1 | NZ_KB732177.1 | + | 84318 | 84370 | **RNA**→ rpmE (PRK00019)Ribosomal_L31 (pfam01197)→ |
| Eco-1476-1 | NZ_KB732283.1 | + | 81703 | 81755 | **RNA**→ rpmE (PRK00019)Ribosomal_L31 (pfam01197)→ |
| Eco-1564-1 | NZ_KB732326.1 | + | 87030 | 87082 | **RNA**→ rpmE (PRK00019)Ribosomal_L31 (pfam01197)→ |
| Eco-1577-1 | NZ_KB732362.1 | + | 80466 | 80518 | **RNA**→ rpmE (PRK00019)Ribosomal_L31 (pfam01197)→ |
| Eco-1495-1 | NZ_KB732396.1 | + | 195853 | 195905 | **RNA**→ rpmE (PRK00019)Ribosomal_L31 (pfam01197)→ |
| Eco-1510-1 | NZ_KB732448.1 | + | 782725 | 782777 | **RNA**→ rpmE (PRK00019)Ribosomal_L31 (pfam01197)→ |

| | | | | | |
|---|---|---|---|---|---|
| Eco-1527-1 | NZ_KB732498.1 | + | 193073 | 193125 | **RNA**→ rpmE (PRK00019)Ribosomal_L31 (pfam01197)→ |
| Eco-1554-1 | NZ_KB732596.1 | + | 199313 | 199365 | **RNA**→ rpmE (PRK00019)Ribosomal_L31 (pfam01197)→ |
| Eco-1561-1 | NZ_KB733385.1 | + | 2566546 | 2566598 | **RNA**→ rpmE (PRK00019)Ribosomal_L31 (pfam01197)→ |
| Eco-1479-1 | NZ_KB732953.1 | + | 2179276 | 2179328 | **RNA**→ rpmE (PRK00019)Ribosomal_L31 (pfam01197)→ |
| Eco-1483-1 | NZ_KB731709.1 | + | 32524 | 32576 | **RNA**→ rpmE (PRK00019)Ribosomal_L31 (pfam01197)→ |
| Eco-2106-1 | NZ_KE702260.1 | + | 90983 | 91035 | **RNA**→ rpmE (PRK00019)Ribosomal_L31 (pfam01197)→ |
| Eco-2131-1 | NZ_KE702466.1 | - | 52307 | 52255 | **RNA**→ rpmE (PRK00019)Ribosomal_L31 (pfam01197)→ |
| Eco-2104-1 | NZ_KE702230.1 | + | 115511 | 115563 | **RNA**→ rpmE (PRK00019)Ribosomal_L31 (pfam01197)→ |
| Eco-2113-1 | NZ_KE702352.1 | + | 89164 | 89216 | **RNA**→ rpmE (PRK00019)Ribosomal_L31 (pfam01197)→ |
| Eco-1270-1 | NZ_KE700660.1 | + | 403149 | 403201 | **RNA**→ rpmE (PRK00019)Ribosomal_L31 (pfam01197)→ |
| Eco-1336-1 | NZ_KE699249.1 | + | 188168 | 188220 | **RNA**→ rpmE (PRK00019)Ribosomal_L31 (pfam01197)→ |
| Eco-2121-1 | NZ_KE702395.1 | + | 1075165 | 1075217 | **RNA**→ rpmE (PRK00019)Ribosomal_L31 (pfam01197)→ |
| Eco-1199-1 | NZ_KE699807.1 | + | 1128823 | 1128875 | **RNA**→ rpmE (PRK00019)Ribosomal_L31 (pfam01197)→ |
| Eco-1297-1 | NZ_KE699028.1 | + | 79986 | 80038 | **RNA**→ rpmE (PRK00019)Ribosomal_L31 (pfam01197)→ |
| Eco-1345-1 | NZ_KE699321.1 | + | 4351742 | 4351794 | **RNA**→ rpmE (PRK00019)Ribosomal_L31 (pfam01197)→ |
| Eco-1243-1 | NZ_KE700363.1 | + | 85160 | 85212 | **RNA**→ rpmE (PRK00019)Ribosomal_L31 (pfam01197)→ |
| Eco-1369-1 | NZ_KE699618.1 | + | 84335 | 84387 | **RNA**→ rpmE (PRK00019)Ribosomal_L31 (pfam01197)→ |
| Eco-1280-1 | NZ_KE700796.1 | + | 4000295 | 4000347 | **RNA**→ rpmE (PRK00019)Ribosomal_L31 (pfam01197)→ |
| Eco-1337-1 | NZ_KE699258.1 | + | 743182 | 743234 | **RNA**→ rpmE (PRK00019)Ribosomal_L31 (pfam01197)→ |
| Eco-1220-1 | NZ_KE700087.1 | + | 3167873 | 3167925 | **RNA**→ rpmE (PRK00019)Ribosomal_L31 (pfam01197)→ |
| Eco-1234-1 | NZ_KE700242.1 | + | 2622626 | 2622678 | **RNA**→ rpmE (PRK00019)Ribosomal_L31 (pfam01197)→ |
| Eco-1233-1 | NZ_KE700237.1 | + | 1196136 | 1196188 | **RNA**→ rpmE (PRK00019)Ribosomal_L31 (pfam01197)→ |
| Eco-1253-1 | NZ_KE700478.1 | + | 4545085 | 4545137 | **RNA**→ rpmE (PRK00019)Ribosomal_L31 (pfam01197)→ |
| Eco-1344-1 | NZ_KE699313.1 | + | 97329 | 97381 | **RNA**→ rpmE (PRK00019)Ribosomal_L31 (pfam01197)→ |
| Eco-1249-1 | NZ_KE700419.1 | + | 742045 | 742097 | **RNA**→ rpmE (PRK00019)Ribosomal_L31 (pfam01197)→ |
| Eco-1296-1 | NZ_KE700922.1 | - | 41764 | 41712 | **RNA**→ rpmE (PRK00019)Ribosomal_L31 (pfam01197)→ |
| Eco-1197-1 | NZ_KE699787.1 | + | 349152 | 349204 | **RNA**→ rpmE (PRK00019)Ribosomal_L31 (pfam01197)→ |
| Eco-1221-1 | NZ_KE700096.1 | + | 88104 | 88156 | **RNA**→ rpmE (PRK00019)Ribosomal_L31 (pfam01197)→ |
| Eco-1373-1 | NZ_KE699643.1 | + | 3534137 | 3534189 | **RNA**→ rpmE (PRK00019)Ribosomal_L31 (pfam01197)→ |
| Eco-1222-1 | NZ_KE700100.1 | + | 4500805 | 4500857 | **RNA**→ rpmE (PRK00019)Ribosomal_L31 (pfam01197)→ |
| Eco-1291-1 | NZ_KE700872.1 | + | 80483 | 80535 | **RNA**→ rpmE (PRK00019)Ribosomal_L31 (pfam01197)→ |
| Eco-1248-1 | NZ_KE700414.1 | + | 199577 | 199629 | **RNA**→ rpmE (PRK00019)Ribosomal_L31 (pfam01197)→ |
| Eco-2094-1 | NZ_KE702091.1 | + | 85614 | 85666 | **RNA**→ rpmE (PRK00019)Ribosomal_L31 (pfam01197)→ |
| Eco-1229-1 | NZ_KE700205.1 | + | 1104705 | 1104757 | **RNA**→ rpmE (PRK00019)Ribosomal_L31 (pfam01197)→ |
| Eco-1321-1 | NZ_KE699105.1 | + | 2180086 | 2180138 | **RNA**→ rpmE (PRK00019)Ribosomal_L31 (pfam01197)→ |
| Eco-1208-1 | NZ_KE699939.1 | + | 88102 | 88154 | **RNA**→ rpmE (PRK00019)Ribosomal_L31 (pfam01197)→ |
| Eco-1202-1 | NZ_KE699849.1 | + | 1590970 | 1591022 | **RNA**→ rpmE (PRK00019)Ribosomal_L31 (pfam01197)→ |
| Eco-1301-1 | NZ_KE700974.1 | + | 245573 | 245625 | **RNA**→ rpmE (PRK00019)Ribosomal_L31 (pfam01197)→ |
| Eco-1269-1 | NZ_KE700653.1 | + | 325873 | 325925 | **RNA**→ rpmE (PRK00019)Ribosomal_L31 (pfam01197)→ |
| Eco-2061-1 | NZ_KE701627.1 | + | 121771 | 121823 | **RNA**→ rpmE (PRK00019)Ribosomal_L31 (pfam01197)→ |
| Eco-1356-1 | NZ_KE699409.1 | + | 204658 | 204710 | **RNA**→ rpmE (PRK00019)Ribosomal_L31 (pfam01197)→ |
| Eco-1210-1 | NZ_KE699949.1 | + | 742245 | 742297 | **RNA**→ rpmE (PRK00019)Ribosomal_L31 (pfam01197)→ |
| Eco-1308-1 | NZ_KE701068.1 | + | 83463 | 83515 | **RNA**→ rpmE (PRK00019)Ribosomal_L31 (pfam01197)→ |
| Eco-1266-1 | NZ_KE700607.1 | + | 185536 | 185588 | **RNA**→ rpmE (PRK00019)Ribosomal_L31 (pfam01197)→ |
| Eco-1405-1 | NZ_KE701348.1 | + | 85496 | 85548 | **RNA**→ rpmE (PRK00019)Ribosomal_L31 (pfam01197)→ |
| Eco-1260-1 | NZ_KE700567.1 | + | 85985 | 86037 | **RNA**→ rpmE (PRK00019)Ribosomal_L31 (pfam01197)→ |
| Eco-2047-1 | NZ_KE701430.1 | + | 85390 | 85442 | **RNA**→ rpmE (PRK00019)Ribosomal_L31 (pfam01197)→ |
| Eco-1227-1 | NZ_KE700176.1 | + | 2293232 | 2293284 | **RNA**→ rpmE (PRK00019)Ribosomal_L31 (pfam01197)→ |
| Eco-1190-1 | NZ_KE699700.1 | + | 467270 | 467322 | **RNA**→ rpmE (PRK00019)Ribosomal_L31 (pfam01197)→ |
| Eco-1371-1 | NZ_KE699635.1 | + | 2173713 | 2173765 | **RNA**→ rpmE (PRK00019)Ribosomal_L31 (pfam01197)→ |
| Eco-1254-1 | NZ_KE700490.1 | + | 2177479 | 2177531 | **RNA**→ rpmE (PRK00019)Ribosomal_L31 (pfam01197)→ |
| Eco-1295-1 | NZ_KE700916.1 | + | 86774 | 86826 | **RNA**→ rpmE (PRK00019)Ribosomal_L31 (pfam01197)→ |

| | | | | | | |
|---|---|---|---|---|---|---|
| Eco-1300-1 | NZ_KE700953.1 | + | 84308 | 84360 | **RNA**→ | rpmE (PRK00019)Ribosomal_L31 (pfam01197)→ |
| Eco-1194-1 | NZ_KE699743.1 | + | 181707 | 181759 | **RNA**→ | rpmE (PRK00019)Ribosomal_L31 (pfam01197)→ |
| Eco-2055-1 | NZ_KE701518.1 | + | 92772 | 92824 | **RNA**→ | rpmE (PRK00019)Ribosomal_L31 (pfam01197)→ |
| Eco-1201-1 | NZ_KE699839.1 | + | 87380 | 87432 | **RNA**→ | rpmE (PRK00019)Ribosomal_L31 (pfam01197)→ |
| Eco-1370-1 | NZ_KE699628.1 | + | 319058 | 319110 | **RNA**→ | rpmE (PRK00019)Ribosomal_L31 (pfam01197)→ |
| Eco-2069-1 | NZ_KE701708.1 | + | 227760 | 227812 | **RNA**→ | rpmE (PRK00019)Ribosomal_L31 (pfam01197)→ |
| Eco-1397-1 | NZ_KE701232.1 | + | 79950 | 80002 | **RNA**→ | rpmE (PRK00019)Ribosomal_L31 (pfam01197)→ |
| Eco-1230-1 | NZ_KE700212.1 | + | 80100 | 80152 | **RNA**→ | rpmE (PRK00019)Ribosomal_L31 (pfam01197)→ |
| Eco-1231-1 | NZ_KE700228.1 | + | 85258 | 85310 | **RNA**→ | rpmE (PRK00019)Ribosomal_L31 (pfam01197)→ |
| Eco-1394-1 | NZ_KE701414.1 | + | 192950 | 193002 | **RNA**→ | rpmE (PRK00019)Ribosomal_L31 (pfam01197)→ |
| Eco-1232-1 | NZ_KE700230.1 | - | 42749 | 42697 | **RNA**→ | rpmE (PRK00019)Ribosomal_L31 (pfam01197)→ |
| Eco-2057-1 | NZ_KE701560.1 | - | 44526 | 44474 | **RNA**→ | rpmE (PRK00019)Ribosomal_L31 (pfam01197)→ |
| Eco-1237-1 | NZ_KE700264.1 | + | 2639992 | 2640044 | **RNA**→ | rpmE (PRK00019)Ribosomal_L31 (pfam01197)→ |
| Eco-1268-1 | NZ_KE700644.1 | + | 80782 | 80834 | **RNA**→ | rpmE (PRK00019)Ribosomal_L31 (pfam01197)→ |
| Eco-2068-1 | NZ_KE701687.1 | + | 85680 | 85732 | **RNA**→ | rpmE (PRK00019)Ribosomal_L31 (pfam01197)→ |
| Eco-1198-1 | NZ_KE699798.1 | + | 1589000 | 1589052 | **RNA**→ | rpmE (PRK00019)Ribosomal_L31 (pfam01197)→ |
| Eco-2081-1 | NZ_KE701872.1 | + | 85305 | 85357 | **RNA**→ | rpmE (PRK00019)Ribosomal_L31 (pfam01197)→ |
| Eco-1278-1 | NZ_KE700785.1 | + | 204693 | 204745 | **RNA**→ | rpmE (PRK00019)Ribosomal_L31 (pfam01197)→ |
| Eco-1292-1 | NZ_KE700883.1 | + | 117631 | 117683 | **RNA**→ | rpmE (PRK00019)Ribosomal_L31 (pfam01197)→ |
| Eco-1283-1 | NZ_KE700822.1 | + | 1471785 | 1471837 | **RNA**→ | rpmE (PRK00019)Ribosomal_L31 (pfam01197)→ |
| Eco-1396-1 | NZ_KE701181.1 | + | 281387 | 281439 | **RNA**→ | rpmE (PRK00019)Ribosomal_L31 (pfam01197)→ |
| Eco-1242-1 | NZ_KE700346.1 | + | 87380 | 87432 | **RNA**→ | rpmE (PRK00019)Ribosomal_L31 (pfam01197)→ |
| Eco-1211-1 | NZ_KE699962.1 | + | 179632 | 179684 | **RNA**→ | rpmE (PRK00019)Ribosomal_L31 (pfam01197)→ |
| Eco-1216-1 | NZ_KE700044.1 | + | 91616 | 91668 | **RNA**→ | rpmE (PRK00019)Ribosomal_L31 (pfam01197)→ |
| Eco-1403-1 | NZ_KE701325.1 | + | 81714 | 81766 | **RNA**→ | rpmE (PRK00019)Ribosomal_L31 (pfam01197)→ |
| Eco-1271-1 | NZ_KE700720.1 | - | 747529 | 747477 | **RNA**→ | rpmE (PRK00019)Ribosomal_L31 (pfam01197)→ |
| Eco-2052-1 | NZ_KE701490.1 | + | 743787 | 743839 | **RNA**→ | rpmE (PRK00019)Ribosomal_L31 (pfam01197)→ |
| Eco-1273-1 | NZ_KE700745.1 | + | 99117 | 99169 | **RNA**→ | rpmE (PRK00019)Ribosomal_L31 (pfam01197)→ |
| Eco-1294-1 | NZ_KE700903.1 | + | 748033 | 748085 | **RNA**→ | rpmE (PRK00019)Ribosomal_L31 (pfam01197)→ |
| Eco-1313-1 | NZ_KE701161.1 | - | 495077 | 495025 | **RNA**→ | rpmE (PRK00019)Ribosomal_L31 (pfam01197)→ |
| Eco-1204-1 | NZ_KE699870.1 | + | 21397 | 21449 | **RNA**→ | rpmE (PRK00019)Ribosomal_L31 (pfam01197)→ |
| Eco-2082-1 | NZ_KE701899.1 | + | 1059289 | 1059341 | **RNA**→ | rpmE (PRK00019)Ribosomal_L31 (pfam01197)→ |
| Eco-1226-1 | NZ_KE700140.1 | + | 194311 | 194363 | **RNA**→ | rpmE (PRK00019)Ribosomal_L31 (pfam01197)→ |
| Eco-2071-1 | NZ_KE701737.1 | + | 84955 | 85007 | **RNA**→ | rpmE (PRK00019)Ribosomal_L31 (pfam01197)→ |
| Eco-1206-1 | NZ_KE699898.1 | + | 84775 | 84827 | **RNA**→ | rpmE (PRK00019)Ribosomal_L31 (pfam01197)→ |
| Eco-1212-1 | NZ_KE699972.1 | + | 118706 | 118758 | **RNA**→ | rpmE (PRK00019)Ribosomal_L31 (pfam01197)→ |
| Eco-1406-1 | NZ_KE701351.1 | + | 1610660 | 1610712 | **RNA**→ | rpmE (PRK00019)Ribosomal_L31 (pfam01197)→ |
| Eco-1309-1 | NZ_KE701070.1 | + | 739347 | 739399 | **RNA**→ | rpmE (PRK00019)Ribosomal_L31 (pfam01197)→ |
| Eco-2093-1 | NZ_KE702074.1 | + | 774010 | 774062 | **RNA**→ | rpmE (PRK00019)Ribosomal_L31 (pfam01197)→ |
| Eco-1213-1 | NZ_KE699994.1 | + | 85300 | 85352 | **RNA**→ | rpmE (PRK00019)Ribosomal_L31 (pfam01197)→ |
| Eco-1311-1 | NZ_KE701117.1 | + | 27360 | 27412 | **RNA**→ | rpmE (PRK00019)Ribosomal_L31 (pfam01197)→ |
| Eco-1312-1 | NZ_KE701119.1 | + | 91804 | 91856 | **RNA**→ | rpmE (PRK00019)Ribosomal_L31 (pfam01197)→ |
| Eco-1393-1 | NZ_KE701223.1 | + | 202372 | 202424 | **RNA**→ | rpmE (PRK00019)Ribosomal_L31 (pfam01197)→ |
| Eco-2059-1 | NZ_KE701583.1 | + | 199908 | 199960 | **RNA**→ | rpmE (PRK00019)Ribosomal_L31 (pfam01197)→ |
| Eco-1267-1 | NZ_KE700619.1 | + | 3121410 | 3121462 | **RNA**→ | rpmE (PRK00019)Ribosomal_L31 (pfam01197)→ |
| Eco-1404-1 | NZ_KE701335.1 | + | 318980 | 319032 | **RNA**→ | rpmE (PRK00019)Ribosomal_L31 (pfam01197)→ |
| Eco-2150-1 | NZ_KE702713.1 | + | 199335 | 199387 | **RNA**→ | rpmE (PRK00019)Ribosomal_L31 (pfam01197)→ |
| Eco-2125-1 | NZ_KE702411.1 | + | 1536533 | 1536585 | **RNA**→ | rpmE (PRK00019)Ribosomal_L31 (pfam01197)→ |
| Eco-2048-1 | NZ_KE701438.1 | + | 1586317 | 1586369 | **RNA**→ | rpmE (PRK00019)Ribosomal_L31 (pfam01197)→ |
| Eco-1408-1 | NZ_KE701207.1 | + | 121266 | 121318 | **RNA**→ | rpmE (PRK00019)Ribosomal_L31 (pfam01197)→ |
| Eco-1261-1 | NZ_KE700575.1 | + | 83633 | 83685 | **RNA**→ | rpmE (PRK00019)Ribosomal_L31 (pfam01197)→ |
| Eco-2065-1 | NZ_KE701670.1 | + | 2925945 | 2925997 | **RNA**→ | rpmE (PRK00019)Ribosomal_L31 (pfam01197)→ |

| | | | | | |
|---|---|---|---|---|---|
| Eco-1293-1 | NZ_KE700892.1 | + | 1709668 | 1709720 | RNA→ rpmE (PRK00019)Ribosomal_L31 (pfam01197)→ |
| Eco-2067-1 | NZ_KE701675.1 | + | 192386 | 192438 | RNA→ rpmE (PRK00019)Ribosomal_L31 (pfam01197)→ |
| Eco-2139-1 | NZ_KE702536.1 | + | 29590 | 29642 | RNA→ rpmE (PRK00019)Ribosomal_L31 (pfam01197)→ |
| Eco-2053-1 | NZ_KE701494.1 | + | 3675398 | 3675450 | RNA→ rpmE (PRK00019)Ribosomal_L31 (pfam01197)→ |
| Eco-1400-1 | NZ_KE701267.1 | + | 213951 | 214003 | RNA→ rpmE (PRK00019)Ribosomal_L31 (pfam01197)→ |
| Eco-1258-1 | NZ_KE700536.1 | + | 1137068 | 1137120 | RNA→ rpmE (PRK00019)Ribosomal_L31 (pfam01197)→ |
| Eco-2084-1 | NZ_KE701945.1 | + | 84267 | 84319 | RNA→ rpmE (PRK00019)Ribosomal_L31 (pfam01197)→ |
| Eco-2099-1 | NZ_KE702161.1 | + | 85444 | 85496 | RNA→ rpmE (PRK00019)Ribosomal_L31 (pfam01197)→ |
| Eco-2085-1 | NZ_KE701948.1 | + | 78493 | 78545 | RNA→ rpmE (PRK00019)Ribosomal_L31 (pfam01197)→ |
| Eco-1395-1 | NZ_KE701420.1 | + | 88540 | 88592 | RNA→ rpmE (PRK00019)Ribosomal_L31 (pfam01197)→ |
| Eco-2143-1 | NZ_KE702605.1 | + | 25421 | 25473 | RNA→ rpmE (PRK00019)Ribosomal_L31 (pfam01197)→ |
| Eco-2112-1 | NZ_KE702337.1 | + | 88193 | 88245 | RNA→ rpmE (PRK00019)Ribosomal_L31 (pfam01197)→ |
| Eco-1240-1 | NZ_KE700291.1 | + | 117994 | 118046 | RNA→ rpmE (PRK00019)Ribosomal_L31 (pfam01197)→ |
| Eco-2063-1 | NZ_KE701654.1 | + | 185838 | 185890 | RNA→ rpmE (PRK00019)Ribosomal_L31 (pfam01197)→ |
| Eco-1246-1 | NZ_KE700403.1 | + | 85286 | 85338 | RNA→ rpmE (PRK00019)Ribosomal_L31 (pfam01197)→ |
| Eco-2100-1 | NZ_KE702175.1 | + | 80475 | 80527 | RNA→ rpmE (PRK00019)Ribosomal_L31 (pfam01197)→ |
| Eco-1289-1 | NZ_KE700861.1 | + | 85906 | 85958 | RNA→ rpmE (PRK00019)Ribosomal_L31 (pfam01197)→ |
| Eco-2056-1 | NZ_KE701523.1 | + | 2947159 | 2947211 | RNA→ rpmE (PRK00019)Ribosomal_L31 (pfam01197)→ |
| Eco-1306-1 | NZ_KE701027.1 | + | 85359 | 85411 | RNA→ rpmE (PRK00019)Ribosomal_L31 (pfam01197)→ |
| Eco-1402-1 | NZ_KE701300.1 | + | 85206 | 85258 | RNA→ rpmE (PRK00019)Ribosomal_L31 (pfam01197)→ |
| Eco-2108-1 | NZ_KE702265.1 | + | 93283 | 93335 | RNA→ rpmE (PRK00019)Ribosomal_L31 (pfam01197)→ |
| Eco-2151-1 | NZ_KE702724.1 | + | 83451 | 83503 | RNA→ rpmE (PRK00019)Ribosomal_L31 (pfam01197)→ |
| Eco-2095-1 | NZ_KE702095.1 | - | 52165 | 52113 | RNA→ rpmE (PRK00019)Ribosomal_L31 (pfam01197)→ |
| Eco-1314-1 | NZ_KE701170.1 | + | 196467 | 196519 | RNA→ rpmE (PRK00019)Ribosomal_L31 (pfam01197)→ |
| Eco-2092-1 | NZ_KE702067.1 | + | 92478 | 92530 | RNA→ rpmE (PRK00019)Ribosomal_L31 (pfam01197)→ |
| Eco-2133-1 | NZ_KE702487.1 | + | 95092 | 95144 | RNA→ rpmE (PRK00019)Ribosomal_L31 (pfam01197)→ |
| Eco-1401-1 | NZ_KE701274.1 | + | 85390 | 85442 | RNA→ rpmE (PRK00019)Ribosomal_L31 (pfam01197)→ |
| Eco-2149-1 | NZ_KE702705.1 | + | 78536 | 78588 | RNA→ rpmE (PRK00019)Ribosomal_L31 (pfam01197)→ |
| Eco-2078-1 | NZ_KE701848.1 | + | 119686 | 119738 | RNA→ rpmE (PRK00019)Ribosomal_L31 (pfam01197)→ |
| Eco-2058-1 | NZ_KE701578.1 | + | 196466 | 196518 | RNA→ rpmE (PRK00019)Ribosomal_L31 (pfam01197)→ |
| Eco-2135-1 | NZ_KE702499.1 | + | 204063 | 204115 | RNA→ rpmE (PRK00019)Ribosomal_L31 (pfam01197)→ |
| Eco-2117-1 | NZ_KE702369.1 | + | 199988 | 200040 | RNA→ rpmE (PRK00019)Ribosomal_L31 (pfam01197)→ |
| Eco-2137-1 | NZ_KE702509.1 | + | 81443 | 81495 | RNA→ rpmE (PRK00019)Ribosomal_L31 (pfam01197)→ |
| Eco-1304-1 | NZ_KE701008.1 | + | 741430 | 741482 | RNA→ rpmE (PRK00019)Ribosomal_L31 (pfam01197)→ |
| Eco-2049-1 | NZ_KE701450.1 | + | 329620 | 329672 | RNA→ rpmE (PRK00019)Ribosomal_L31 (pfam01197)→ |
| Eco-1303-1 | NZ_KE700987.1 | + | 4378097 | 4378149 | RNA→ rpmE (PRK00019)Ribosomal_L31 (pfam01197)→ |
| Eco-2070-1 | NZ_KE701712.1 | + | 766321 | 766373 | RNA→ rpmE (PRK00019)Ribosomal_L31 (pfam01197)→ |
| Eco-1219-1 | NZ_KE700086.1 | + | 83699 | 83751 | RNA→ rpmE (PRK00019)Ribosomal_L31 (pfam01197)→ |
| Eco-2080-1 | NZ_KE701868.1 | + | 90346 | 90398 | RNA→ rpmE (PRK00019)Ribosomal_L31 (pfam01197)→ |
| Eco-1409-1 | NZ_KE701377.1 | + | 200139 | 200191 | RNA→ rpmE (PRK00019)Ribosomal_L31 (pfam01197)→ |
| Eco-1287-1 | NZ_KE700845.1 | + | 299546 | 299598 | RNA→ rpmE (PRK00019)Ribosomal_L31 (pfam01197)→ |
| Eco-2062-1 | NZ_KE701633.1 | + | 192993 | 193045 | RNA→ rpmE (PRK00019)Ribosomal_L31 (pfam01197)→ |
| Eco-2083-1 | NZ_KE701910.1 | + | 317539 | 317591 | RNA→ rpmE (PRK00019)Ribosomal_L31 (pfam01197)→ |
| Eco-2103-1 | NZ_KE702202.1 | + | 81678 | 81730 | RNA→ rpmE (PRK00019)Ribosomal_L31 (pfam01197)→ |
| Eco-1195-1 | NZ_KE699763.1 | + | 91882 | 91934 | RNA→ rpmE (PRK00019)Ribosomal_L31 (pfam01197)→ |
| Eco-2118-1 | NZ_KE702382.1 | + | 82835 | 82887 | RNA→ rpmE (PRK00019)Ribosomal_L31 (pfam01197)→ |
| Eco-1251-1 | NZ_KE700461.1 | - | 43511 | 43459 | RNA→ rpmE (PRK00019)Ribosomal_L31 (pfam01197)→ |
| Eco-1412-1 | NZ_KE701406.1 | + | 1581102 | 1581154 | RNA→ rpmE (PRK00019)Ribosomal_L31 (pfam01197)→ |
| Eco-2142-1 | NZ_KE702584.1 | + | 2386278 | 2386330 | RNA→ rpmE (PRK00019)Ribosomal_L31 (pfam01197)→ |
| Eco-1411-1 | NZ_KE701395.1 | + | 3203329 | 3203381 | RNA→ rpmE (PRK00019)Ribosomal_L31 (pfam01197)→ |
| Eco-1244-1 | NZ_KE700366.1 | + | 83391 | 83443 | RNA→ rpmE (PRK00019)Ribosomal_L31 (pfam01197)→ |
| Eco-2126-1 | NZ_KE702423.1 | + | 86433 | 86485 | RNA→ rpmE (PRK00019)Ribosomal_L31 (pfam01197)→ |

| | | | | | |
|---|---|---|---|---|---|
| Eco-2054-1 | NZ_KE701500.1 | + | 32308 | 32360 | **RNA**→ rpmE (PRK00019)Ribosomal_L31 (pfam01197)→ |
| Eco-1282-1 | NZ_KE700813.1 | + | 197775 | 197827 | **RNA**→ rpmE (PRK00019)Ribosomal_L31 (pfam01197)→ |
| Eco-2128-1 | NZ_KE702432.1 | + | 79824 | 79876 | **RNA**→ rpmE (PRK00019)Ribosomal_L31 (pfam01197)→ |
| Eco-2096-1 | NZ_KE702100.1 | + | 85155 | 85207 | **RNA**→ rpmE (PRK00019)Ribosomal_L31 (pfam01197)→ |
| Eco-1259-1 | NZ_KE700541.1 | - | 376622 | 376570 | **RNA**→ rpmE (PRK00019)Ribosomal_L31 (pfam01197)→ |
| Eco-2140-1 | NZ_KE702556.1 | + | 204424 | 204476 | **RNA**→ rpmE (PRK00019)Ribosomal_L31 (pfam01197)→ |
| Eco-2088-1 | NZ_KE701999.1 | + | 305573 | 305625 | **RNA**→ rpmE (PRK00019)Ribosomal_L31 (pfam01197)→ |
| Eco-2073-1 | NZ_KE701764.1 | + | 85198 | 85250 | **RNA**→ rpmE (PRK00019)Ribosomal_L31 (pfam01197)→ |
| Eco-1399-1 | NZ_KE701264.1 | + | 90963 | 91015 | **RNA**→ rpmE (PRK00019)Ribosomal_L31 (pfam01197)→ |
| Eco-2115-1 | NZ_KE702359.1 | + | 89025 | 89077 | **RNA**→ rpmE (PRK00019)Ribosomal_L31 (pfam01197)→ |
| Eco-2060-1 | NZ_KE701597.1 | + | 798535 | 798587 | **RNA**→ rpmE (PRK00019)Ribosomal_L31 (pfam01197)→ |
| Eco-2152-1 | NZ_KE702739.1 | + | 85432 | 85484 | **RNA**→ rpmE (PRK00019)Ribosomal_L31 (pfam01197)→ |
| Eco-1407-1 | NZ_KE701357.1 | + | 3796106 | 3796158 | **RNA**→ rpmE (PRK00019)Ribosomal_L31 (pfam01197)→ |
| Eco-2051-1 | NZ_KE701474.1 | + | 183061 | 183113 | **RNA**→ rpmE (PRK00019)Ribosomal_L31 (pfam01197)→ |
| Eco-2087-1 | NZ_KE701982.1 | + | 191708 | 191760 | **RNA**→ rpmE (PRK00019)Ribosomal_L31 (pfam01197)→ |
| Eco-2097-1 | NZ_KE702123.1 | + | 198791 | 198843 | **RNA**→ rpmE (PRK00019)Ribosomal_L31 (pfam01197)→ |
| Eco-2072-1 | NZ_KE701745.1 | + | 90258 | 90310 | **RNA**→ rpmE (PRK00019)Ribosomal_L31 (pfam01197)→ |
| Eco-2111-1 | NZ_KE702333.1 | + | 184802 | 184854 | **RNA**→ rpmE (PRK00019)Ribosomal_L31 (pfam01197)→ |
| Eco-2098-1 | NZ_KE702128.1 | + | 81858 | 81910 | **RNA**→ rpmE (PRK00019)Ribosomal_L31 (pfam01197)→ |
| Eco-2110-1 | NZ_KE702301.1 | + | 87657 | 87709 | **RNA**→ rpmE (PRK00019)Ribosomal_L31 (pfam01197)→ |
| Eco-2076-1 | NZ_KE701799.1 | + | 85001 | 85053 | **RNA**→ rpmE (PRK00019)Ribosomal_L31 (pfam01197)→ |
| Eco-2077-1 | NZ_KE701803.1 | + | 85072 | 85124 | **RNA**→ rpmE (PRK00019)Ribosomal_L31 (pfam01197)→ |
| Eco-2091-1 | NZ_KE702043.1 | + | 203223 | 203275 | **RNA**→ rpmE (PRK00019)Ribosomal_L31 (pfam01197)→ |
| Eco-2134-1 | NZ_KE702490.1 | + | 4289533 | 4289585 | **RNA**→ rpmE (PRK00019)Ribosomal_L31 (pfam01197)→ |
| Eco-2086-1 | NZ_KE701969.1 | + | 712304 | 712356 | **RNA**→ rpmE (PRK00019)Ribosomal_L31 (pfam01197)→ |
| Eco-2120-1 | NZ_KE702386.1 | + | 3870113 | 3870165 | **RNA**→ rpmE (PRK00019)Ribosomal_L31 (pfam01197)→ |
| Eco-2075-1 | NZ_KE701784.1 | + | 94679 | 94731 | **RNA**→ rpmE (PRK00019)Ribosomal_L31 (pfam01197)→ |
| Eco-2079-1 | NZ_KE701857.1 | + | 763071 | 763123 | **RNA**→ rpmE (PRK00019)Ribosomal_L31 (pfam01197)→ |
| Eco-2124-1 | NZ_KE702407.1 | + | 2192167 | 2192219 | **RNA**→ rpmE (PRK00019)Ribosomal_L31 (pfam01197)→ |
| Eco-2127-1 | NZ_KE702425.1 | + | 3699233 | 3699285 | **RNA**→ rpmE (PRK00019)Ribosomal_L31 (pfam01197)→ |
| Eco-2146-1 | NZ_KE702642.1 | + | 185296 | 185348 | **RNA**→ rpmE (PRK00019)Ribosomal_L31 (pfam01197)→ |
| Eco-2148-1 | NZ_KE702686.1 | + | 818241 | 818293 | **RNA**→ rpmE (PRK00019)Ribosomal_L31 (pfam01197)→ |
| Eco-2144-1 | NZ_KE702621.1 | + | 80610 | 80662 | **RNA**→ rpmE (PRK00019)Ribosomal_L31 (pfam01197)→ |
| Eco-2141-1 | NZ_KE702572.1 | + | 83823 | 83875 | **RNA**→ rpmE (PRK00019)Ribosomal_L31 (pfam01197)→ |
| Eco-1275-1 | NZ_KE747799.1 | + | 206535 | 206587 | **RNA**→ rpmE (PRK00019)Ribosomal_L31 (pfam01197)→ |
| Eco-2130-1 | NZ_KE705271.1 | + | 3310513 | 3310565 | **RNA**→ rpmE (PRK00019)Ribosomal_L31 (pfam01197)→ |
| Eco-1374-1 | NZ_KE705265.1 | + | 175598 | 175650 | **RNA**→ rpmE (PRK00019)Ribosomal_L31 (pfam01197)→ |
| Eco-771-1 | NZ_KI521938.1 | + | 26072 | 26124 | **RNA**→ rpmE (PRK00019)Ribosomal_L31 (pfam01197)→ |
| Eco-874-1 | NZ_KI522050.1 | - | 2343 | 2291 | **RNA**→ rpmE (PRK00019)Ribosomal_L31 (pfam01197)→ |
| Eco-1375-1 | NZ_KE705252.1 | + | 1104995 | 1105047 | **RNA**→ rpmE (PRK00019)Ribosomal_L31 (pfam01197)→ |
| Eco-883-1 | NZ_KI522363.1 | + | 79390 | 79442 | **RNA**→ rpmE (PRK00019)Ribosomal_L31 (pfam01197)→ |
| Eco-901-1 | NZ_KI522509.1 | + | 32268 | 32320 | **RNA**→ rpmE (PRK00019)Ribosomal_L31 (pfam01197)→ |
| Eco-2107-1 | NZ_KI530646.1 | + | 1159405 | 1159457 | **RNA**→ rpmE (PRK00019)Ribosomal_L31 (pfam01197)→ |
| Eco-769-1 | NZ_KI521734.1 | - | 41852 | 41800 | **RNA**→ rpmE (PRK00019)Ribosomal_L31 (pfam01197)→ |
| Eco-894-1 | NZ_KI533778.1 | + | 64774 | 64826 | **RNA**→ rpmE (PRK00019)Ribosomal_L31 (pfam01197)→ |
| Eco-1342-1 | NZ_KI530595.1 | - | 145459 | 145407 | **RNA**→ rpmE (PRK00019)Ribosomal_L31 (pfam01197)→ |
| Eco-881-1 | NZ_KI522310.1 | + | 60402 | 60454 | **RNA**→ rpmE (PRK00019)Ribosomal_L31 (pfam01197)→ |
| Eco-768-1 | NZ_KI521711.1 | - | 2401 | 2349 | **RNA**→ rpmE (PRK00019)Ribosomal_L31 (pfam01197)→ |
| Eco-903-1 | NZ_KI535003.1 | - | 52232 | 52180 | **RNA**→ rpmE (PRK00019)Ribosomal_L31 (pfam01197)→ |
| Eco-1378-1 | NZ_KI530619.1 | + | 85133 | 85185 | **RNA**→ rpmE (PRK00019)Ribosomal_L31 (pfam01197)→ |
| Eco-889-1 | NZ_KI532955.1 | + | 79292 | 79344 | **RNA**→ rpmE (PRK00019)Ribosomal_L31 (pfam01197)→ |
| Eco-880-1 | NZ_KI531695.1 | + | 148651 | 148703 | **RNA**→ rpmE (PRK00019)Ribosomal_L31 (pfam01197)→ |

| | | | | | |
|---|---|---|---|---|---|
| Eco-895-1 | NZ_KI534085.1 | + | 64817 | 64869 | **RNA**→ rpmE (PRK00019)Ribosomal_L31 (pfam01197)→ |
| Eco-2119-1 | NZ_KI530691.1 | + | 742265 | 742317 | **RNA**→ rpmE (PRK00019)Ribosomal_L31 (pfam01197)→ |
| Eco-2122-1 | NZ_KI530639.1 | + | 84693 | 84745 | **RNA**→ rpmE (PRK00019)Ribosomal_L31 (pfam01197)→ |
| Eco-1016-1 | NZ_KI535409.1 | + | 2483096 | 2483148 | **RNA**→ rpmE (PRK00019)Ribosomal_L31 (pfam01197)→ |
| Eco-2136-1 | NZ_KI530662.1 | + | 1099626 | 1099678 | **RNA**→ rpmE (PRK00019)Ribosomal_L31 (pfam01197)→ |
| Eco-875-1 | NZ_KI530989.1 | + | 32307 | 32359 | **RNA**→ rpmE (PRK00019)Ribosomal_L31 (pfam01197)→ |
| Eco-2116-1 | NZ_KI530626.1 | + | 87523 | 87575 | **RNA**→ rpmE (PRK00019)Ribosomal_L31 (pfam01197)→ |
| Eco-882-1 | NZ_KI531752.1 | + | 64690 | 64742 | **RNA**→ rpmE (PRK00019)Ribosomal_L31 (pfam01197)→ |
| Eco-898-1 | NZ_KI534543.1 | + | 28730 | 28782 | **RNA**→ rpmE (PRK00019)Ribosomal_L31 (pfam01197)→ |
| Eco-946-1 | NZ_KI535243.1 | + | 80068 | 80120 | **RNA**→ rpmE (PRK00019)Ribosomal_L31 (pfam01197)→ |
| Eco-879-1 | NZ_KI531489.1 | - | 44412 | 44360 | **RNA**→ rpmE (PRK00019)Ribosomal_L31 (pfam01197)→ |
| Eco-1423-1 | NZ_KE137211.1 | + | 1128617 | 1128669 | **RNA**→ rpmE (PRK00019)Ribosomal_L31 (pfam01197)→ |
| Eco-1613-1 | NZ_KE137107.1 | + | 1403747 | 1403799 | **RNA**→ rpmE (PRK00019)Ribosomal_L31 (pfam01197)→ |
| Eco-1449-1 | NZ_KE137273.1 | + | 87029 | 87081 | **RNA**→ rpmE (PRK00019)Ribosomal_L31 (pfam01197)→ |
| Eco-1489-1 | NZ_KE137316.1 | + | 3244486 | 3244538 | **RNA**→ rpmE (PRK00019)Ribosomal_L31 (pfam01197)→ |
| Eco-1641-1 | NZ_KE137153.1 | - | 88837 | 88785 | **RNA**→ rpmE (PRK00019)Ribosomal_L31 (pfam01197)→ |
| Eco-900-1 | NZ_KI534784.1 | + | 79620 | 79672 | **RNA**→ rpmE (PRK00019)Ribosomal_L31 (pfam01197)→ |
| Eco-1017-1 | NZ_KI535398.1 | + | 309573 | 309625 | **RNA**→ rpmE (PRK00019)Ribosomal_L31 (pfam01197)→ |
| Eco-1555-1 | NZ_KE136642.1 | + | 179829 | 179881 | **RNA**→ rpmE (PRK00019)Ribosomal_L31 (pfam01197)→ |
| Eco-1541-1 | NZ_KE136951.1 | + | 2950291 | 2950343 | **RNA**→ rpmE (PRK00019)Ribosomal_L31 (pfam01197)→ |
| Eco-1544-1 | NZ_KE136956.1 | + | 180294 | 180346 | **RNA**→ rpmE (PRK00019)Ribosomal_L31 (pfam01197)→ |
| Eco-1504-1 | NZ_KE136876.1 | + | 2069968 | 2070020 | **RNA**→ rpmE (PRK00019)Ribosomal_L31 (pfam01197)→ |
| Eco-1514-1 | NZ_KE136897.1 | + | 314032 | 314084 | **RNA**→ rpmE (PRK00019)Ribosomal_L31 (pfam01197)→ |
| Eco-876-1 | NZ_KI531098.1 | - | 44414 | 44362 | **RNA**→ rpmE (PRK00019)Ribosomal_L31 (pfam01197)→ |
| Eco-1579-1 | NZ_KE136820.1 | + | 1390963 | 1391015 | **RNA**→ rpmE (PRK00019)Ribosomal_L31 (pfam01197)→ |
| Eco-1413-1 | NZ_KE136565.1 | + | 183210 | 183262 | **RNA**→ rpmE (PRK00019)Ribosomal_L31 (pfam01197)→ |
| Eco-1562-1 | NZ_KE136702.1 | + | 123684 | 123736 | **RNA**→ rpmE (PRK00019)Ribosomal_L31 (pfam01197)→ |
| Eco-1521-1 | NZ_KE136944.1 | + | 743534 | 743586 | **RNA**→ rpmE (PRK00019)Ribosomal_L31 (pfam01197)→ |
| Eco-1575-1 | NZ_KE136772.1 | + | 84349 | 84401 | **RNA**→ rpmE (PRK00019)Ribosomal_L31 (pfam01197)→ |
| Eco-1503-1 | NZ_KE136872.1 | - | 345466 | 345414 | **RNA**→ rpmE (PRK00019)Ribosomal_L31 (pfam01197)→ |
| Eco-1548-1 | NZ_KE136980.1 | + | 750943 | 750995 | **RNA**→ rpmE (PRK00019)Ribosomal_L31 (pfam01197)→ |
| Eco-1417-1 | NZ_KE137162.1 | + | 713495 | 713547 | **RNA**→ rpmE (PRK00019)Ribosomal_L31 (pfam01197)→ |
| Eco-1610-1 | NZ_KE137068.1 | + | 1539326 | 1539378 | **RNA**→ rpmE (PRK00019)Ribosomal_L31 (pfam01197)→ |
| Eco-1612-1 | NZ_KE137074.1 | + | 222135 | 222187 | **RNA**→ rpmE (PRK00019)Ribosomal_L31 (pfam01197)→ |
| Eco-1418-1 | NZ_KE137186.1 | + | 26461 | 26513 | **RNA**→ rpmE (PRK00019)Ribosomal_L31 (pfam01197)→ |
| Eco-1443-1 | NZ_KE137233.1 | + | 216568 | 216620 | **RNA**→ rpmE (PRK00019)Ribosomal_L31 (pfam01197)→ |
| Eco-2114-1 | NZ_KI530681.1 | + | 1627520 | 1627572 | **RNA**→ rpmE (PRK00019)Ribosomal_L31 (pfam01197)→ |
| Eco-1018-1 | NZ_KI535383.1 | + | 3646346 | 3646398 | **RNA**→ rpmE (PRK00019)Ribosomal_L31 (pfam01197)→ |
| Eco-1520-1 | NZ_KE136683.1 | + | 92806 | 92858 | **RNA**→ rpmE (PRK00019)Ribosomal_L31 (pfam01197)→ |
| Eco-1574-1 | NZ_KE136752.1 | + | 116804 | 116856 | **RNA**→ rpmE (PRK00019)Ribosomal_L31 (pfam01197)→ |
| Eco-1518-1 | NZ_KE136939.1 | + | 827982 | 828034 | **RNA**→ rpmE (PRK00019)Ribosomal_L31 (pfam01197)→ |
| Eco-1457-1 | NZ_KE136655.1 | + | 1741425 | 1741477 | **RNA**→ rpmE (PRK00019)Ribosomal_L31 (pfam01197)→ |
| Eco-1572-1 | NZ_KE136745.1 | - | 62476 | 62424 | **RNA**→ rpmE (PRK00019)Ribosomal_L31 (pfam01197)→ |
| Eco-1517-1 | NZ_KE136913.1 | + | 2263750 | 2263802 | **RNA**→ rpmE (PRK00019)Ribosomal_L31 (pfam01197)→ |
| Eco-1609-1 | NZ_KE137053.1 | + | 1171162 | 1171214 | **RNA**→ rpmE (PRK00019)Ribosomal_L31 (pfam01197)→ |
| Eco-1328-1 | NZ_KI538668.1 | + | 125792 | 125844 | **RNA**→ rpmE (PRK00019)Ribosomal_L31 (pfam01197)→ |
| Eco-1616-1 | NZ_KE137133.1 | + | 84349 | 84401 | **RNA**→ rpmE (PRK00019)Ribosomal_L31 (pfam01197)→ |
| Eco-1239-1 | NZ_KI538738.1 | + | 93530 | 93582 | **RNA**→ rpmE (PRK00019)Ribosomal_L31 (pfam01197)→ |
| Eco-2066-1 | NZ_KI538796.1 | + | 717176 | 717228 | **RNA**→ rpmE (PRK00019)Ribosomal_L31 (pfam01197)→ |
| Eco-1022-1 | NZ_KI535541.1 | + | 768557 | 768609 | **RNA**→ rpmE (PRK00019)Ribosomal_L31 (pfam01197)→ |
| Eco-1368-1 | NZ_KI538698.1 | + | 356734 | 356786 | **RNA**→ rpmE (PRK00019)Ribosomal_L31 (pfam01197)→ |
| Eco-1615-1 | NZ_KE137109.1 | + | 1090822 | 1090874 | **RNA**→ rpmE (PRK00019)Ribosomal_L31 (pfam01197)→ |

| | | | | | |
|---|---|---|---|---|---|
| Eco-1223-1 | NZ_KI538713.1 | + | 84208 | 84260 | **RNA**→ rpmE (PRK00019)Ribosomal_L31 (pfam01197)→ |
| Eco-1196-1 | NZ_KI538815.1 | + | 3031145 | 3031197 | **RNA**→ rpmE (PRK00019)Ribosomal_L31 (pfam01197)→ |
| Eco-896-1 | NZ_KI534157.1 | + | 79484 | 79536 | **RNA**→ rpmE (PRK00019)Ribosomal_L31 (pfam01197)→ |
| Eco-888-1 | NZ_KI532739.1 | - | 42075 | 42023 | **RNA**→ rpmE (PRK00019)Ribosomal_L31 (pfam01197)→ |
| Eco-892-1 | NZ_KI533438.1 | - | 479 | 427 | **RNA**→ rpmE (PRK00019)Ribosomal_L31 (pfam01197)→ |
| Eco-897-1 | NZ_KI534353.1 | - | 44556 | 44504 | **RNA**→ rpmE (PRK00019)Ribosomal_L31 (pfam01197)→ |
| Eco-945-1 | NZ_KI535081.1 | - | 52219 | 52167 | **RNA**→ rpmE (PRK00019)Ribosomal_L31 (pfam01197)→ |
| Eco-887-1 | NZ_KI532548.1 | + | 64822 | 64874 | **RNA**→ rpmE (PRK00019)Ribosomal_L31 (pfam01197)→ |
| Eco-1365-1 | NZ_KI669338.1 | + | 85482 | 85534 | **RNA**→ rpmE (PRK00019)Ribosomal_L31 (pfam01197)→ |
| Eco-752-1 | NZ_KI912161.1 | + | 85941 | 85993 | **RNA**→ rpmE (PRK00019)Ribosomal_L31 (pfam01197)→ |
| Eco-1315-1 | NZ_KI669335.1 | + | 81184 | 81236 | **RNA**→ rpmE (PRK00019)Ribosomal_L31 (pfam01197)→ |
| Eco-2123-1 | NZ_KI669372.1 | + | 3497209 | 3497261 | **RNA**→ rpmE (PRK00019)Ribosomal_L31 (pfam01197)→ |
| Eco-1262-1 | NZ_KI669356.1 | + | 1449482 | 1449534 | **RNA**→ rpmE (PRK00019)Ribosomal_L31 (pfam01197)→ |
| Eco-300-1 | NZ_KI929682.1 | + | 4436190 | 4436242 | **RNA**→ rpmE (PRK00019)Ribosomal_L31 (pfam01197)→ |
| Eco-329-1 | NZ_KI929787.1 | + | 4431189 | 4431241 | **RNA**→ rpmE (PRK00019)Ribosomal_L31 (pfam01197)→ |
| Eco-396-1 | NZ_KI894099.1 | + | 18404 | 18456 | **RNA**→ rpmE (PRK00019)Ribosomal_L31 (pfam01197)→ |
| Eco-301-1 | NZ_KI929678.1 | + | 708660 | 708712 | **RNA**→ rpmE (PRK00019)Ribosomal_L31 (pfam01197)→ |
| Eco-330-1 | NZ_KI929782.1 | + | 4994418 | 4994470 | **RNA**→ rpmE (PRK00019)Ribosomal_L31 (pfam01197)→ |
| Eco-2034-1 | NZ_AELB01000053.1 | + | 19626 | 19678 | **RNA**→ rpmE (PRK00019)Ribosomal_L31 (pfam01197)→ |
| Eco-1467-1 | NZ_KB732280.1 | + | 715829 | 715881 | **RNA**→ rpmE (PRK00019)Ribosomal_L31 (pfam01197)→ |
| Eco-1532-1 | NZ_KB732523.1 | + | 91133 | 91185 | **RNA**→ rpmE (PRK00019)Ribosomal_L31 (pfam01197)→ |
| Eco-1543-1 | NZ_KB733374.1 | + | 739997 | 740049 | **RNA**→ rpmE (PRK00019)Ribosomal_L31 (pfam01197)→ |
| Eco-1597-1 | NZ_KB732717.1 | + | 126864 | 126916 | **RNA**→ rpmE (PRK00019)Ribosomal_L31 (pfam01197)→ |
| Eco-1608-1 | NZ_KB732770.1 | + | 390084 | 390136 | **RNA**→ rpmE (PRK00019)Ribosomal_L31 (pfam01197)→ |
| Eco-1519-1 | NZ_KB732206.1 | + | 87364 | 87416 | **RNA**→ rpmE (PRK00019)Ribosomal_L31 (pfam01197)→ |
| Eco-1435-1 | NZ_KB732269.1 | + | 769148 | 769200 | **RNA**→ rpmE (PRK00019)Ribosomal_L31 (pfam01197)→ |
| Eco-1499-1 | NZ_KB732407.1 | + | 185493 | 185545 | **RNA**→ rpmE (PRK00019)Ribosomal_L31 (pfam01197)→ |
| Eco-1506-1 | NZ_KB732426.1 | + | 386694 | 386746 | **RNA**→ rpmE (PRK00019)Ribosomal_L31 (pfam01197)→ |
| Eco-1512-1 | NZ_KB732454.1 | + | 2967821 | 2967873 | **RNA**→ rpmE (PRK00019)Ribosomal_L31 (pfam01197)→ |
| Eco-1525-1 | NZ_KB732469.1 | + | 217122 | 217174 | **RNA**→ rpmE (PRK00019)Ribosomal_L31 (pfam01197)→ |
| Eco-1529-1 | NZ_KB732511.1 | + | 2112749 | 2112801 | **RNA**→ rpmE (PRK00019)Ribosomal_L31 (pfam01197)→ |
| Eco-1534-1 | NZ_KB732530.1 | + | 4480523 | 4480575 | **RNA**→ rpmE (PRK00019)Ribosomal_L31 (pfam01197)→ |
| Eco-1547-1 | NZ_KB732561.1 | + | 81703 | 81755 | **RNA**→ rpmE (PRK00019)Ribosomal_L31 (pfam01197)→ |
| Eco-1591-1 | NZ_KB732652.1 | + | 89385 | 89437 | **RNA**→ rpmE (PRK00019)Ribosomal_L31 (pfam01197)→ |
| Eco-1531-1 | NZ_KB732305.1 | + | 92502 | 92554 | **RNA**→ rpmE (PRK00019)Ribosomal_L31 (pfam01197)→ |
| Eco-1583-1 | NZ_KB732388.1 | + | 93133 | 93185 | **RNA**→ rpmE (PRK00019)Ribosomal_L31 (pfam01197)→ |
| Eco-1507-1 | NZ_KB732429.1 | + | 37417 | 37469 | **RNA**→ rpmE (PRK00019)Ribosomal_L31 (pfam01197)→ |
| Eco-1526-1 | NZ_KB732489.1 | + | 247369 | 247421 | **RNA**→ rpmE (PRK00019)Ribosomal_L31 (pfam01197)→ |
| Eco-1535-1 | NZ_KB732535.1 | + | 3871288 | 3871340 | **RNA**→ rpmE (PRK00019)Ribosomal_L31 (pfam01197)→ |
| Eco-1601-1 | NZ_KB732723.1 | + | 80818 | 80870 | **RNA**→ rpmE (PRK00019)Ribosomal_L31 (pfam01197)→ |
| Eco-1565-1 | NZ_KB732337.1 | + | 1098559 | 1098611 | **RNA**→ rpmE (PRK00019)Ribosomal_L31 (pfam01197)→ |
| Eco-1538-1 | NZ_KB732546.1 | + | 210180 | 210232 | **RNA**→ rpmE (PRK00019)Ribosomal_L31 (pfam01197)→ |
| Eco-1614-1 | NZ_KB733402.1 | + | 81830 | 81882 | **RNA**→ rpmE (PRK00019)Ribosomal_L31 (pfam01197)→ |
| Eco-1426-1 | NZ_KB732931.1 | + | 4693303 | 4693355 | **RNA**→ rpmE (PRK00019)Ribosomal_L31 (pfam01197)→ |
| Eco-1472-1 | NZ_KB733424.1 | + | 88823 | 88875 | **RNA**→ rpmE (PRK00019)Ribosomal_L31 (pfam01197)→ |
| Eco-1462-1 | NZ_KB733119.1 | + | 748025 | 748077 | **RNA**→ rpmE (PRK00019)Ribosomal_L31 (pfam01197)→ |
| Eco-1536-1 | NZ_KB732539.1 | + | 203898 | 203950 | **RNA**→ rpmE (PRK00019)Ribosomal_L31 (pfam01197)→ |
| Eco-1629-1 | NZ_KB732876.1 | + | 79585 | 79637 | **RNA**→ rpmE (PRK00019)Ribosomal_L31 (pfam01197)→ |
| Eco-1633-1 | NZ_KB732995.1 | + | 192553 | 192605 | **RNA**→ rpmE (PRK00019)Ribosomal_L31 (pfam01197)→ |
| Eco-1587-1 | NZ_KB733034.1 | + | 217425 | 217477 | **RNA**→ rpmE (PRK00019)Ribosomal_L31 (pfam01197)→ |
| Eco-1438-1 | NZ_KB731947.1 | + | 64751 | 64803 | **RNA**→ rpmE (PRK00019)Ribosomal_L31 (pfam01197)→ |
| Eco-1551-1 | NZ_KB732574.1 | + | 3169698 | 3169750 | **RNA**→ rpmE (PRK00019)Ribosomal_L31 (pfam01197)→ |

| | | | | | | |
|---|---|---|---|---|---|---|
| Eco-1593-1 | NZ_KB732678.1 | + | 80428 | 80480 | RNA→ | rpmE (PRK00019)Ribosomal_L31 (pfam01197)→ |
| Eco-1607-1 | NZ_KB732766.1 | + | 1078458 | 1078510 | RNA→ | rpmE (PRK00019)Ribosomal_L31 (pfam01197)→ |
| Eco-1619-1 | NZ_KB732824.1 | + | 86495 | 86547 | RNA→ | rpmE (PRK00019)Ribosomal_L31 (pfam01197)→ |
| Eco-1630-1 | NZ_KB732885.1 | + | 312298 | 312350 | RNA→ | rpmE (PRK00019)Ribosomal_L31 (pfam01197)→ |
| Eco-1478-1 | NZ_KB732934.1 | + | 1549529 | 1549581 | RNA→ | rpmE (PRK00019)Ribosomal_L31 (pfam01197)→ |
| Eco-1485-1 | NZ_KB732981.1 | + | 79252 | 79304 | RNA→ | rpmE (PRK00019)Ribosomal_L31 (pfam01197)→ |
| Eco-1620-1 | NZ_KB733059.1 | + | 89746 | 89798 | RNA→ | rpmE (PRK00019)Ribosomal_L31 (pfam01197)→ |
| Eco-1515-1 | NZ_KB733178.1 | + | 185235 | 185287 | RNA→ | rpmE (PRK00019)Ribosomal_L31 (pfam01197)→ |
| Eco-1588-1 | NZ_KB732644.1 | + | 117671 | 117723 | RNA→ | rpmE (PRK00019)Ribosomal_L31 (pfam01197)→ |
| Eco-1623-1 | NZ_KB732835.1 | + | 90079 | 90131 | RNA→ | rpmE (PRK00019)Ribosomal_L31 (pfam01197)→ |
| Eco-1636-1 | NZ_KB732900.1 | + | 81758 | 81810 | RNA→ | rpmE (PRK00019)Ribosomal_L31 (pfam01197)→ |
| Eco-1621-1 | NZ_KB733069.1 | + | 1528753 | 1528805 | RNA→ | rpmE (PRK00019)Ribosomal_L31 (pfam01197)→ |
| Eco-1635-1 | NZ_KB733085.1 | + | 90282 | 90334 | RNA→ | rpmE (PRK00019)Ribosomal_L31 (pfam01197)→ |
| Eco-1474-1 | NZ_KB733136.1 | + | 89188 | 89240 | RNA→ | rpmE (PRK00019)Ribosomal_L31 (pfam01197)→ |
| Eco-1511-1 | NZ_KB733148.1 | + | 201599 | 201651 | RNA→ | rpmE (PRK00019)Ribosomal_L31 (pfam01197)→ |
| Eco-1557-1 | NZ_KB732580.1 | + | 2122614 | 2122666 | RNA→ | rpmE (PRK00019)Ribosomal_L31 (pfam01197)→ |
| Eco-1492-1 | NZ_KB733163.1 | + | 215587 | 215639 | RNA→ | rpmE (PRK00019)Ribosomal_L31 (pfam01197)→ |
| Eco-1624-1 | NZ_KB732854.1 | + | 179054 | 179106 | RNA→ | rpmE (PRK00019)Ribosomal_L31 (pfam01197)→ |
| Eco-1523-1 | NZ_KB733190.1 | + | 84884 | 84936 | RNA→ | rpmE (PRK00019)Ribosomal_L31 (pfam01197)→ |
| Eco-1600-1 | NZ_KB732993.1 | + | 1144242 | 1144294 | RNA→ | rpmE (PRK00019)Ribosomal_L31 (pfam01197)→ |
| Eco-1584-1 | NZ_KB733366.1 | + | 85959 | 86011 | RNA→ | rpmE (PRK00019)Ribosomal_L31 (pfam01197)→ |
| Eco-1431-1 | NZ_KB732889.1 | + | 87725 | 87777 | RNA→ | rpmE (PRK00019)Ribosomal_L31 (pfam01197)→ |
| Eco-1537-1 | NZ_KB733209.1 | + | 1116317 | 1116369 | RNA→ | rpmE (PRK00019)Ribosomal_L31 (pfam01197)→ |
| Eco-1430-1 | NZ_KB733437.1 | + | 194575 | 194627 | RNA→ | rpmE (PRK00019)Ribosomal_L31 (pfam01197)→ |
| Eco-1509-1 | NZ_KB733164.1 | + | 4861972 | 4862024 | RNA→ | rpmE (PRK00019)Ribosomal_L31 (pfam01197)→ |
| Eco-1530-1 | NZ_KB733361.1 | + | 2206480 | 2206532 | RNA→ | rpmE (PRK00019)Ribosomal_L31 (pfam01197)→ |
| Eco-1560-1 | NZ_KB732614.1 | + | 95899 | 95951 | RNA→ | rpmE (PRK00019)Ribosomal_L31 (pfam01197)→ |
| Eco-1622-1 | NZ_KB732987.1 | + | 728728 | 728780 | RNA→ | rpmE (PRK00019)Ribosomal_L31 (pfam01197)→ |
| Eco-1434-1 | NZ_KB732923.1 | + | 4068090 | 4068142 | RNA→ | rpmE (PRK00019)Ribosomal_L31 (pfam01197)→ |
| Eco-1590-1 | NZ_KB733431.1 | + | 1094753 | 1094805 | RNA→ | rpmE (PRK00019)Ribosomal_L31 (pfam01197)→ |
| Eco-1416-1 | NZ_KB733078.1 | + | 2993520 | 2993572 | RNA→ | rpmE (PRK00019)Ribosomal_L31 (pfam01197)→ |
| Eco-1546-1 | NZ_KB733355.1 | + | 80038 | 80090 | RNA→ | rpmE (PRK00019)Ribosomal_L31 (pfam01197)→ |
| Eco-1453-1 | NZ_KB733095.1 | + | 123923 | 123975 | RNA→ | rpmE (PRK00019)Ribosomal_L31 (pfam01197)→ |
| Eco-1513-1 | NZ_KB733153.1 | + | 196949 | 197001 | RNA→ | rpmE (PRK00019)Ribosomal_L31 (pfam01197)→ |
| Eco-1490-1 | NZ_KB733443.1 | + | 729974 | 730026 | RNA→ | rpmE (PRK00019)Ribosomal_L31 (pfam01197)→ |
| Eco-1501-1 | NZ_KB733173.1 | + | 3320254 | 3320306 | RNA→ | rpmE (PRK00019)Ribosomal_L31 (pfam01197)→ |
| Eco-1618-1 | NZ_KB732816.1 | + | 95428 | 95480 | RNA→ | rpmE (PRK00019)Ribosomal_L31 (pfam01197)→ |
| Eco-1516-1 | NZ_KB733185.1 | + | 182340 | 182392 | RNA→ | rpmE (PRK00019)Ribosomal_L31 (pfam01197)→ |
| Eco-1553-1 | NZ_KB733021.1 | + | 88442 | 88494 | RNA→ | rpmE (PRK00019)Ribosomal_L31 (pfam01197)→ |
| Eco-1528-1 | NZ_KB733194.1 | + | 81745 | 81797 | RNA→ | rpmE (PRK00019)Ribosomal_L31 (pfam01197)→ |
| Eco-1533-1 | NZ_KB733226.1 | + | 97399 | 97451 | RNA→ | rpmE (PRK00019)Ribosomal_L31 (pfam01197)→ |
| Eco-1540-1 | NZ_KB733215.1 | + | 81328 | 81380 | RNA→ | rpmE (PRK00019)Ribosomal_L31 (pfam01197)→ |
| Eco-1461-1 | NZ_KB732947.1 | + | 1431844 | 1431896 | RNA→ | rpmE (PRK00019)Ribosomal_L31 (pfam01197)→ |
| Eco-1488-1 | NZ_KB732965.1 | + | 89272 | 89324 | RNA→ | rpmE (PRK00019)Ribosomal_L31 (pfam01197)→ |
| Eco-1497-1 | NZ_KB733007.1 | + | 87248 | 87300 | RNA→ | rpmE (PRK00019)Ribosomal_L31 (pfam01197)→ |
| Eco-1452-1 | NZ_KB733086.1 | + | 1415738 | 1415790 | RNA→ | rpmE (PRK00019)Ribosomal_L31 (pfam01197)→ |
| Eco-1459-1 | NZ_KB733104.1 | + | 118416 | 118468 | RNA→ | rpmE (PRK00019)Ribosomal_L31 (pfam01197)→ |
| Eco-1502-1 | NZ_KB733172.1 | + | 700985 | 701037 | RNA→ | rpmE (PRK00019)Ribosomal_L31 (pfam01197)→ |
| Eco-1558-1 | NZ_KB732598.1 | + | 181845 | 181897 | RNA→ | rpmE (PRK00019)Ribosomal_L31 (pfam01197)→ |
| Eco-1524-1 | NZ_KB733201.1 | + | 697684 | 697736 | RNA→ | rpmE (PRK00019)Ribosomal_L31 (pfam01197)→ |
| Eco-1568-1 | NZ_KB733016.1 | + | 701071 | 701123 | RNA→ | rpmE (PRK00019)Ribosomal_L31 (pfam01197)→ |
| Eco-1585-1 | NZ_KB733032.1 | + | 1465991 | 1466043 | RNA→ | rpmE (PRK00019)Ribosomal_L31 (pfam01197)→ |

| | | | | | | |
|---|---|---|---|---|---|---|
| Eco-1594-1 | NZ_KB733040.1 | + | 738674 | 738726 | **RNA**→ | rpmE (PRK00019)Ribosomal_L31 (pfam01197)→ |
| Eco-1604-1 | NZ_KB733048.1 | + | 85198 | 85250 | **RNA**→ | rpmE (PRK00019)Ribosomal_L31 (pfam01197)→ |
| Eco-1539-1 | NZ_KB733228.1 | + | 206290 | 206342 | **RNA**→ | rpmE (PRK00019)Ribosomal_L31 (pfam01197)→ |
| Eco-1465-1 | NZ_KB733135.1 | + | 2714412 | 2714464 | **RNA**→ | rpmE (PRK00019)Ribosomal_L31 (pfam01197)→ |
| Eco-1481-1 | NZ_KB733141.1 | + | 1453826 | 1453878 | **RNA**→ | rpmE (PRK00019)Ribosomal_L31 (pfam01197)→ |
| Eco-1628-1 | NZ_KB731625.1 | + | 65760 | 65812 | **RNA**→ | rpmE (PRK00019)Ribosomal_L31 (pfam01197)→ |
| Eco-1559-1 | NZ_KB732610.1 | + | 3217660 | 3217712 | **RNA**→ | rpmE (PRK00019)Ribosomal_L31 (pfam01197)→ |
| Eco-1486-1 | NZ_KB733000.1 | + | 4628049 | 4628101 | **RNA**→ | rpmE (PRK00019)Ribosomal_L31 (pfam01197)→ |
| Eco-1596-1 | NZ_KB732702.1 | + | 2093565 | 2093617 | **RNA**→ | rpmE (PRK00019)Ribosomal_L31 (pfam01197)→ |
| Eco-1458-1 | NZ_KB733090.1 | + | 345664 | 345716 | **RNA**→ | rpmE (PRK00019)Ribosomal_L31 (pfam01197)→ |
| Eco-1582-1 | NZ_KB733426.1 | + | 3687548 | 3687600 | **RNA**→ | rpmE (PRK00019)Ribosomal_L31 (pfam01197)→ |
| Eco-1464-1 | NZ_KB733128.1 | + | 185480 | 185532 | **RNA**→ | rpmE (PRK00019)Ribosomal_L31 (pfam01197)→ |
| Eco-1419-1 | NZ_KB733317.1 | + | 21535 | 21587 | **RNA**→ | rpmE (PRK00019)Ribosomal_L31 (pfam01197)→ |
| Eco-1581-1 | NZ_KB733011.1 | + | 86410 | 86462 | **RNA**→ | rpmE (PRK00019)Ribosomal_L31 (pfam01197)→ |
| Eco-1996-1 | NZ_KB822529.1 | - | 52192 | 52140 | **RNA**→ | rpmE (PRK00019)Ribosomal_L31 (pfam01197)→ |
| Eco-1602-1 | NZ_KE137015.1 | + | 1385265 | 1385317 | **RNA**→ | rpmE (PRK00019)Ribosomal_L31 (pfam01197)→ |
| Eco-1580-1 | NZ_KE136843.1 | + | 716940 | 716992 | **RNA**→ | rpmE (PRK00019)Ribosomal_L31 (pfam01197)→ |
| Eco-1471-1 | NZ_KE137290.1 | + | 178813 | 178865 | **RNA**→ | rpmE (PRK00019)Ribosomal_L31 (pfam01197)→ |
| Eco-1573-1 | NZ_KE136621.1 | - | 63621 | 63569 | **RNA**→ | rpmE (PRK00019)Ribosomal_L31 (pfam01197)→ |
| Eco-1415-1 | NZ_KE137155.1 | + | 114384 | 114436 | **RNA**→ | rpmE (PRK00019)Ribosomal_L31 (pfam01197)→ |
| Eco-1605-1 | NZ_KE137028.1 | + | 4278970 | 4279022 | **RNA**→ | rpmE (PRK00019)Ribosomal_L31 (pfam01197)→ |
| Eco-1437-1 | NZ_KE137222.1 | + | 1659902 | 1659954 | **RNA**→ | rpmE (PRK00019)Ribosomal_L31 (pfam01197)→ |
| Eco-1451-1 | NZ_KE137276.1 | - | 548492 | 548440 | **RNA**→ | rpmE (PRK00019)Ribosomal_L31 (pfam01197)→ |
| Eco-1626-1 | NZ_KB733418.1 | + | 88751 | 88803 | **RNA**→ | rpmE (PRK00019)Ribosomal_L31 (pfam01197)→ |
| Eco-1545-1 | NZ_KE136972.1 | + | 1439611 | 1439663 | **RNA**→ | rpmE (PRK00019)Ribosomal_L31 (pfam01197)→ |
| Eco-1446-1 | NZ_KE136600.1 | + | 2913667 | 2913719 | **RNA**→ | rpmE (PRK00019)Ribosomal_L31 (pfam01197)→ |
| Eco-1549-1 | NZ_KE136988.1 | + | 80699 | 80751 | **RNA**→ | rpmE (PRK00019)Ribosomal_L31 (pfam01197)→ |
| Eco-1508-1 | NZ_KE136675.1 | + | 2773214 | 2773266 | **RNA**→ | rpmE (PRK00019)Ribosomal_L31 (pfam01197)→ |
| Eco-1632-1 | NZ_KE137143.1 | + | 1102318 | 1102370 | **RNA**→ | rpmE (PRK00019)Ribosomal_L31 (pfam01197)→ |
| Eco-1422-1 | NZ_KE137205.1 | + | 191756 | 191808 | **RNA**→ | rpmE (PRK00019)Ribosomal_L31 (pfam01197)→ |
| Eco-1447-1 | NZ_KE137259.1 | + | 537843 | 537895 | **RNA**→ | rpmE (PRK00019)Ribosomal_L31 (pfam01197)→ |
| Eco-455-1 | NZ_KE375261.1 | - | 41954 | 41902 | **RNA**→ | rpmE (PRK00019)Ribosomal_L31 (pfam01197)→ |
| Eco-454-1 | NZ_KE375303.1 | + | 64690 | 64742 | **RNA**→ | rpmE (PRK00019)Ribosomal_L31 (pfam01197)→ |
| Eco-456-1 | NZ_KE375225.1 | - | 44622 | 44570 | **RNA**→ | rpmE (PRK00019)Ribosomal_L31 (pfam01197)→ |
| Eco-453-1 | NZ_KE375333.1 | - | 42079 | 42027 | **RNA**→ | rpmE (PRK00019)Ribosomal_L31 (pfam01197)→ |
| Eco-284-1 | NZ_KE375526.1 | - | 44499 | 44447 | **RNA**→ | rpmE (PRK00019)Ribosomal_L31 (pfam01197)→ |
| Eco-49-1 | NZ_KE375613.1 | + | 24839 | 24891 | **RNA**→ | rpmE (PRK00019)Ribosomal_L31 (pfam01197)→ |
| Eco-1576-1 | NZ_KE136794.1 | + | 89196 | 89248 | **RNA**→ | rpmE (PRK00019)Ribosomal_L31 (pfam01197)→ |
| Eco-1487-1 | NZ_KE137305.1 | + | 2743351 | 2743403 | **RNA**→ | rpmE (PRK00019)Ribosomal_L31 (pfam01197)→ |
| Eco-1500-1 | NZ_KE136847.1 | + | 123541 | 123593 | **RNA**→ | rpmE (PRK00019)Ribosomal_L31 (pfam01197)→ |
| Eco-452-1 | NZ_KE375397.1 | + | 24841 | 24893 | **RNA**→ | rpmE (PRK00019)Ribosomal_L31 (pfam01197)→ |
| Eco-1285-1 | NZ_KE698801.1 | + | 2222813 | 2222865 | **RNA**→ | rpmE (PRK00019)Ribosomal_L31 (pfam01197)→ |
| Eco-285-1 | NZ_KE375504.1 | - | 44622 | 44570 | **RNA**→ | rpmE (PRK00019)Ribosomal_L31 (pfam01197)→ |
| Eco-1108-1 | NZ_KE353061.1 | - | 469 | 417 | **RNA**→ | rpmE (PRK00019)Ribosomal_L31 (pfam01197)→ |
| Eco-1333-1 | NZ_KE699213.1 | + | 2261174 | 2261226 | **RNA**→ | rpmE (PRK00019)Ribosomal_L31 (pfam01197)→ |
| Eco-1350-1 | NZ_KE699361.1 | + | 85199 | 85251 | **RNA**→ | rpmE (PRK00019)Ribosomal_L31 (pfam01197)→ |
| Eco-1372-1 | NZ_KE698882.1 | + | 3962636 | 3962688 | **RNA**→ | rpmE (PRK00019)Ribosomal_L31 (pfam01197)→ |
| Eco-1346-1 | NZ_KE699328.1 | + | 765355 | 765407 | **RNA**→ | rpmE (PRK00019)Ribosomal_L31 (pfam01197)→ |
| Eco-1247-1 | NZ_KE698925.1 | + | 741829 | 741881 | **RNA**→ | rpmE (PRK00019)Ribosomal_L31 (pfam01197)→ |
| Eco-1332-1 | NZ_KE698833.1 | + | 1212504 | 1212556 | **RNA**→ | rpmE (PRK00019)Ribosomal_L31 (pfam01197)→ |
| Eco-1339-1 | NZ_KE699290.1 | + | 203429 | 203481 | **RNA**→ | rpmE (PRK00019)Ribosomal_L31 (pfam01197)→ |
| Eco-1351-1 | NZ_KE699371.1 | + | 1139517 | 1139569 | **RNA**→ | rpmE (PRK00019)Ribosomal_L31 (pfam01197)→ |

| | | | | | |
|---|---|---|---|---|---|
| Eco-1354-1 | NZ_KE699399.1 | - | 46140 | 46088 | **RNA**→ rpmE (PRK00019)Ribosomal_L31 (pfam01197)→ |
| Eco-1355-1 | NZ_KE698860.1 | + | 85312 | 85364 | **RNA**→ rpmE (PRK00019)Ribosomal_L31 (pfam01197)→ |
| Eco-1352-1 | NZ_KE699379.1 | + | 3129427 | 3129479 | **RNA**→ rpmE (PRK00019)Ribosomal_L31 (pfam01197)→ |
| Eco-1349-1 | NZ_KE698854.1 | + | 87029 | 87081 | **RNA**→ rpmE (PRK00019)Ribosomal_L31 (pfam01197)→ |
| Eco-1347-1 | NZ_KE699336.1 | + | 2182479 | 2182531 | **RNA**→ rpmE (PRK00019)Ribosomal_L31 (pfam01197)→ |
| Eco-1364-1 | NZ_KE699566.1 | - | 582415 | 582363 | **RNA**→ rpmE (PRK00019)Ribosomal_L31 (pfam01197)→ |
| Eco-1330-1 | NZ_KE699189.1 | + | 197919 | 197971 | **RNA**→ rpmE (PRK00019)Ribosomal_L31 (pfam01197)→ |
| Eco-1361-1 | NZ_KE699534.1 | + | 78550 | 78602 | **RNA**→ rpmE (PRK00019)Ribosomal_L31 (pfam01197)→ |
| Eco-1316-1 | NZ_KE699037.1 | + | 2979691 | 2979743 | **RNA**→ rpmE (PRK00019)Ribosomal_L31 (pfam01197)→ |
| Eco-1367-1 | NZ_KE699590.1 | + | 2993670 | 2993722 | **RNA**→ rpmE (PRK00019)Ribosomal_L31 (pfam01197)→ |
| Eco-1358-1 | NZ_KE699456.1 | + | 85185 | 85237 | **RNA**→ rpmE (PRK00019)Ribosomal_L31 (pfam01197)→ |
| Eco-1317-1 | NZ_KE699041.1 | + | 92815 | 92867 | **RNA**→ rpmE (PRK00019)Ribosomal_L31 (pfam01197)→ |
| Eco-1376-1 | NZ_KE699686.1 | + | 204988 | 205040 | **RNA**→ rpmE (PRK00019)Ribosomal_L31 (pfam01197)→ |
| Eco-1331-1 | NZ_KE699194.1 | + | 85115 | 85167 | **RNA**→ rpmE (PRK00019)Ribosomal_L31 (pfam01197)→ |
| Eco-1326-1 | NZ_KE699142.1 | + | 90612 | 90664 | **RNA**→ rpmE (PRK00019)Ribosomal_L31 (pfam01197)→ |
| Eco-1366-1 | NZ_KE699581.1 | + | 1099205 | 1099257 | **RNA**→ rpmE (PRK00019)Ribosomal_L31 (pfam01197)→ |
| Eco-1324-1 | NZ_KE699129.1 | + | 204686 | 204738 | **RNA**→ rpmE (PRK00019)Ribosomal_L31 (pfam01197)→ |
| Eco-1353-1 | NZ_KE699395.1 | + | 745950 | 746002 | **RNA**→ rpmE (PRK00019)Ribosomal_L31 (pfam01197)→ |
| Eco-1255-1 | NZ_KE698942.1 | + | 83897 | 83949 | **RNA**→ rpmE (PRK00019)Ribosomal_L31 (pfam01197)→ |
| Eco-1360-1 | NZ_KE699480.1 | + | 298169 | 298221 | **RNA**→ rpmE (PRK00019)Ribosomal_L31 (pfam01197)→ |
| Eco-1377-1 | NZ_KE699698.1 | + | 203590 | 203642 | **RNA**→ rpmE (PRK00019)Ribosomal_L31 (pfam01197)→ |
| Eco-1205-1 | NZ_KE699895.1 | + | 84426 | 84478 | **RNA**→ rpmE (PRK00019)Ribosomal_L31 (pfam01197)→ |
| Eco-1318-1 | NZ_KE699079.1 | + | 24904 | 24956 | **RNA**→ rpmE (PRK00019)Ribosomal_L31 (pfam01197)→ |
| Eco-1320-1 | NZ_KE699099.1 | + | 1146447 | 1146499 | **RNA**→ rpmE (PRK00019)Ribosomal_L31 (pfam01197)→ |
| Eco-1307-1 | NZ_KE699031.1 | + | 181978 | 182030 | **RNA**→ rpmE (PRK00019)Ribosomal_L31 (pfam01197)→ |
| Eco-1189-1 | NZ_KE698890.1 | + | 88102 | 88154 | **RNA**→ rpmE (PRK00019)Ribosomal_L31 (pfam01197)→ |
| Eco-1319-1 | NZ_KE699094.1 | + | 85356 | 85408 | **RNA**→ rpmE (PRK00019)Ribosomal_L31 (pfam01197)→ |
| Eco-1264-1 | NZ_KE698975.1 | + | 86763 | 86815 | **RNA**→ rpmE (PRK00019)Ribosomal_L31 (pfam01197)→ |
| Eco-1217-1 | NZ_KE698897.1 | + | 125488 | 125540 | **RNA**→ rpmE (PRK00019)Ribosomal_L31 (pfam01197)→ |
| Eco-1250-1 | NZ_KE700457.1 | + | 935564 | 935616 | **RNA**→ rpmE (PRK00019)Ribosomal_L31 (pfam01197)→ |
| Eco-1334-1 | NZ_KE699236.1 | + | 329829 | 329881 | **RNA**→ rpmE (PRK00019)Ribosomal_L31 (pfam01197)→ |
| Eco-1245-1 | NZ_KE700389.1 | + | 1530219 | 1530271 | **RNA**→ rpmE (PRK00019)Ribosomal_L31 (pfam01197)→ |
| Eco-1329-1 | NZ_KE699173.1 | + | 769817 | 769869 | **RNA**→ rpmE (PRK00019)Ribosomal_L31 (pfam01197)→ |
| Eco-1228-1 | NZ_KE700182.1 | + | 196885 | 196937 | **RNA**→ rpmE (PRK00019)Ribosomal_L31 (pfam01197)→ |
| Eco-1225-1 | NZ_KE700123.1 | + | 78535 | 78587 | **RNA**→ rpmE (PRK00019)Ribosomal_L31 (pfam01197)→ |
| Eco-1286-1 | NZ_KE699009.1 | + | 201650 | 201702 | **RNA**→ rpmE (PRK00019)Ribosomal_L31 (pfam01197)→ |
| Eco-1238-1 | NZ_KE700271.1 | + | 121843 | 121895 | **RNA**→ rpmE (PRK00019)Ribosomal_L31 (pfam01197)→ |
| Eco-1343-1 | NZ_KE699306.1 | + | 1570515 | 1570567 | **RNA**→ rpmE (PRK00019)Ribosomal_L31 (pfam01197)→ |
| Eco-1257-1 | NZ_KE700520.1 | + | 318247 | 318299 | **RNA**→ rpmE (PRK00019)Ribosomal_L31 (pfam01197)→ |
| Eco-1203-1 | NZ_KE699873.1 | + | 35069 | 35121 | **RNA**→ rpmE (PRK00019)Ribosomal_L31 (pfam01197)→ |
| Eco-1338-1 | NZ_KE699267.1 | + | 3281857 | 3281909 | **RNA**→ rpmE (PRK00019)Ribosomal_L31 (pfam01197)→ |
| Eco-1279-1 | NZ_KE700788.1 | + | 1208073 | 1208125 | **RNA**→ rpmE (PRK00019)Ribosomal_L31 (pfam01197)→ |
| Eco-1192-1 | NZ_KE699721.1 | + | 84995 | 85047 | **RNA**→ rpmE (PRK00019)Ribosomal_L31 (pfam01197)→ |
| Eco-1252-1 | NZ_KE700471.1 | + | 728585 | 728637 | **RNA**→ rpmE (PRK00019)Ribosomal_L31 (pfam01197)→ |
| Eco-1236-1 | NZ_KE700249.1 | - | 53043 | 52991 | **RNA**→ rpmE (PRK00019)Ribosomal_L31 (pfam01197)→ |
| Eco-1200-1 | NZ_KE699826.1 | + | 194261 | 194313 | **RNA**→ rpmE (PRK00019)Ribosomal_L31 (pfam01197)→ |
| Eco-1218-1 | NZ_KE700049.1 | + | 122728 | 122780 | **RNA**→ rpmE (PRK00019)Ribosomal_L31 (pfam01197)→ |
| Eco-1241-1 | NZ_KE700299.1 | + | 53857 | 53909 | **RNA**→ rpmE (PRK00019)Ribosomal_L31 (pfam01197)→ |
| Eco-2105-1 | NZ_KE702238.1 | + | 772837 | 772889 | **RNA**→ rpmE (PRK00019)Ribosomal_L31 (pfam01197)→ |
| Eco-2102-1 | NZ_KE702186.1 | + | 83352 | 83404 | **RNA**→ rpmE (PRK00019)Ribosomal_L31 (pfam01197)→ |
| Eco-2101-1 | NZ_KE702179.1 | + | 486494 | 486546 | **RNA**→ rpmE (PRK00019)Ribosomal_L31 (pfam01197)→ |
| Eco-848-1 | NC_008253.1 | + | 4326358 | 4326410 | **RNA**→ rpmE (PRK00019)Ribosomal_L31 (pfam01197)→ |

| | | | | | | |
|---|---|---|---|---|---|---|
| Sfl-25-12 | NZ_LAJS01000047.1 | + | 24423 | 24475 | **RNA**→ | rpmE (PRK00019)Ribosomal_L31 (pfam01197)→ |
| Sfl-25-13 | NZ_LAJU01000056.1 | + | 24788 | 24840 | **RNA**→ | rpmE (PRK00019)Ribosomal_L31 (pfam01197)→ |
| Sso-1-545 | NZ_LAJT01000097.1 | + | 39222 | 39274 | **RNA**→ | rpmE (PRK00019)Ribosomal_L31 (pfam01197)→ |
| Sfl-64-1 | NZ_LAJW01000048.1 | + | 34707 | 34759 | **RNA**→ | rpmE (PRK00019)Ribosomal_L31 (pfam01197)→ |
| Sfl-64-2 | NZ_LAJP01000066.1 | + | 34638 | 34690 | **RNA**→ | rpmE (PRK00019)Ribosomal_L31 (pfam01197)→ |
| Sfl-51-2 | NZ_LAJV01000117.1 | - | 31124 | 31072 | **RNA**→ | rpmE (PRK00019)Ribosomal_L31 (pfam01197)→ |
| Eco-1991-1 | NZ_AFDR01000029.1 | + | 123981 | 124033 | **RNA**→ | rpmE (PRK00019)Ribosomal_L31 (pfam01197)→ |
| Eco-1126-1 | NZ_JH945811.1 | + | 47754 | 47806 | **RNA**→ | rpmE (PRK00019)Ribosomal_L31 (pfam01197)→ |
| Eco-1470-1 | NZ_KB733130.1 | + | 2157129 | 2157181 | **RNA**→ | rpmE (PRK00019)Ribosomal_L31 (pfam01197)→ |
| Eco-286-1 | NZ_KE375480.1 | - | 44497 | 44445 | **RNA**→ | rpmE (PRK00019)Ribosomal_L31 (pfam01197)→ |
| Eco-1450-1 | NZ_ANXZ01000039.1 | + | 77024 | 77076 | **RNA**→ | rpmE (PRK00019)Ribosomal_L31 (pfam01197)→ |
| Eco-1625-1 | NZ_ANYT01000117.1 | + | 64751 | 64803 | **RNA**→ | rpmE (PRK00019)Ribosomal_L31 (pfam01197)→ |
| Eco-1827-1 | NZ_CAPL01000021.1 | - | 39907 | 39855 | **RNA**→ | rpmE (PRK00019)Ribosomal_L31 (pfam01197)→ |
| Eco-1721-1 | NZ_AMVR01000013.1 | + | 85592 | 85644 | **RNA**→ | rpmE (PRK00019)Ribosomal_L31 (pfam01197)→ |
| Eco-1421-1 | NZ_ANXN01000074.1 | + | 78602 | 78654 | **RNA**→ | rpmE (PRK00019)Ribosomal_L31 (pfam01197)→ |
| Eco-1469-1 | NZ_ANYG01000054.1 | + | 32597 | 32649 | **RNA**→ | rpmE (PRK00019)Ribosomal_L31 (pfam01197)→ |
| Eco-1428-1 | NZ_ANXQ01000047.1 | + | 79108 | 79160 | **RNA**→ | rpmE (PRK00019)Ribosomal_L31 (pfam01197)→ |
| Eco-1765-1 | NZ_CAPK01000190.1 | - | 39191 | 39139 | **RNA**→ | rpmE (PRK00019)Ribosomal_L31 (pfam01197)→ |
| Eco-1420-1 | NZ_ANXM01000069.1 | + | 64751 | 64803 | **RNA**→ | rpmE (PRK00019)Ribosomal_L31 (pfam01197)→ |
| Eco-1456-1 | NZ_ANYC01000070.1 | + | 41451 | 41503 | **RNA**→ | rpmE (PRK00019)Ribosomal_L31 (pfam01197)→ |
| Eco-1484-1 | NZ_ANYM01000060.1 | + | 77056 | 77108 | **RNA**→ | rpmE (PRK00019)Ribosomal_L31 (pfam01197)→ |
| Eco-1640-1 | NZ_ANYZ01000053.1 | + | 65240 | 65292 | **RNA**→ | rpmE (PRK00019)Ribosomal_L31 (pfam01197)→ |
| Eco-1634-1 | NZ_ANYW01000072.1 | + | 18519 | 18571 | **RNA**→ | rpmE (PRK00019)Ribosomal_L31 (pfam01197)→ |
| Eco-1473-1 | NZ_ANYH01000078.1 | + | 66273 | 66325 | **RNA**→ | rpmE (PRK00019)Ribosomal_L31 (pfam01197)→ |
| Eco-1482-1 | NZ_ANYK01000069.1 | + | 85888 | 85940 | **RNA**→ | rpmE (PRK00019)Ribosomal_L31 (pfam01197)→ |
| Eco-463-4 | NZ_CAPM01000059.1 | - | 52489 | 52437 | **RNA**→ | rpmE (PRK00019)Ribosomal_L31 (pfam01197)→ |
| Eco-857-1 | NZ_AOEP01000134.1 | + | 94220 | 94272 | **RNA**→ | rpmE (PRK00019)Ribosomal_L31 (pfam01197)→ |
| Eco-1454-1 | NZ_ANYA01000061.1 | + | 32592 | 32644 | **RNA**→ | rpmE (PRK00019)Ribosomal_L31 (pfam01197)→ |
| Eco-1427-1 | NZ_ANXP01000124.1 | + | 34000 | 34052 | **RNA**→ | rpmE (PRK00019)Ribosomal_L31 (pfam01197)→ |
| Eco-1638-1 | NZ_ANYY01000082.1 | + | 65676 | 65728 | **RNA**→ | rpmE (PRK00019)Ribosomal_L31 (pfam01197)→ |
| Eco-751-1 | NZ_HG738867.1 | - | 2653144 | 2653092 | **RNA**→ | rpmE (PRK00019)Ribosomal_L31 (pfam01197)→ |
| Eco-402-1 | NZ_CP009578.1 | + | 4817517 | 4817569 | **RNA**→ | rpmE (PRK00019)Ribosomal_L31 (pfam01197)→ |
| Eco-399-1 | NZ_CP009644.1 | + | 4098610 | 4098662 | **RNA**→ | rpmE (PRK00019)Ribosomal_L31 (pfam01197)→ |
| Sfl-62-1 | NC_004741.1 | - | 3631186 | 3631134 | **RNA**→ | rpmE (PRK00019)Ribosomal_L31 (pfam01197)→ |
| Eco-1029-1 | NC_004431.1 | + | 4646648 | 4646700 | **RNA**→ | ←hypo |
| Eco-2185-1 | NC_007779.1 | - | 3509727 | 3509675 | **RNA**→ | rpmE (PRK00019)Ribosomal_L31 (pfam01197)→ |
| Eco-1715-1 | NC_013353.1 | - | 4722457 | 4722405 | **RNA**→ | rpmE (PRK00019)Ribosomal_L31 (pfam01197)→ |
| Eco-1766-1 | NC_013364.1 | + | 4809048 | 4809100 | **RNA**→ | rpmE (PRK00019)Ribosomal_L31 (pfam01197)→ |
| Eco-959-1 | NC_012967.1 | + | 4106437 | 4106489 | **RNA**→ | rpmE (PRK00019)Ribosomal_L31 (pfam01197)→ |
| Eco-954-1 | NC_010468.1 | - | 4508346 | 4508294 | **RNA**→ | rpmE (PRK00019)Ribosomal_L31 (pfam01197)→ |
| Sbo-15-1 | NC_010658.1 | + | 4123060 | 4123112 | **RNA**→ | rpmE (PRK00019)Ribosomal_L31 (pfam01197)→ |
| Eco-1379-1 | NC_011741.1 | + | 4208059 | 4208111 | **RNA**→ | rpmE (PRK00019)Ribosomal_L31 (pfam01197)→ |
| Eco-1812-1 | NC_013361.1 | - | 4648170 | 4648118 | **RNA**→ | rpmE (PRK00019)Ribosomal_L31 (pfam01197)→ |
| Eco-2182-1 | NC_010473.1 | + | 4224674 | 4224726 | **RNA**→ | rpmE (PRK00019)Ribosomal_L31 (pfam01197)→ |
| Eco-1145-1 | NC_011745.1 | + | 4578369 | 4578421 | **RNA**→ | rpmE (PRK00019)Ribosomal_L31 (pfam01197)→ |
| Eco-853-1 | NC_011748.1 | + | 4508708 | 4508760 | **RNA**→ | rpmE (PRK00019)Ribosomal_L31 (pfam01197)→ |
| Eco-1187-1 | NC_009800.1 | + | 4154963 | 4155015 | **RNA**→ | rpmE (PRK00019)Ribosomal_L31 (pfam01197)→ |
| Eco-1780-1 | NC_011601.1 | + | 4442040 | 4442092 | **RNA**→ | rpmE (PRK00019)Ribosomal_L31 (pfam01197)→ |
| Eco-1735-1 | NZ_JH378015.1 | + | 84387 | 84439 | **RNA**→ | rpmE (PRK00019)Ribosomal_L31 (pfam01197)→ |
| Eco-1824-1 | NC_013941.1 | + | 4816088 | 4816140 | **RNA**→ | rpmE (PRK00019)Ribosomal_L31 (pfam01197)→ |
| Eco-1743-1 | NZ_JH378081.1 | + | 329552 | 329604 | **RNA**→ | rpmE (PRK00019)Ribosomal_L31 (pfam01197)→ |
| Eco-1719-1 | NZ_JH378073.1 | + | 1107751 | 1107803 | **RNA**→ | rpmE (PRK00019)Ribosomal_L31 (pfam01197)→ |

| | | | | | |
|---|---|---|---|---|---|
| Eco-1739-1 | NZ_JH377982.1 | + | 84391 | 84443 | **RNA**→ rpmE (PRK00019)Ribosomal_L31 (pfam01197)→ |
| Eco-472-2 | NZ_JH378089.1 | + | 1162708 | 1162760 | **RNA**→ rpmE (PRK00019)Ribosomal_L31 (pfam01197)→ |
| Eco-1720-1 | NZ_JH378068.1 | + | 84396 | 84448 | **RNA**→ rpmE (PRK00019)Ribosomal_L31 (pfam01197)→ |
| Eco-1736-1 | NZ_JH378006.1 | + | 84397 | 84449 | **RNA**→ rpmE (PRK00019)Ribosomal_L31 (pfam01197)→ |
| Eco-1737-1 | NZ_JH377999.1 | + | 88840 | 88892 | **RNA**→ rpmE (PRK00019)Ribosomal_L31 (pfam01197)→ |
| Eco-1828-1 | NC_017646.1 | + | 4694260 | 4694312 | **RNA**→ rpmE (PRK00019)Ribosomal_L31 (pfam01197)→ |
| Eco-1733-1 | NZ_JH378032.1 | + | 183333 | 183385 | **RNA**→ rpmE (PRK00019)Ribosomal_L31 (pfam01197)→ |
| Eco-1738-1 | NZ_JH377991.1 | + | 1168004 | 1168056 | **RNA**→ rpmE (PRK00019)Ribosomal_L31 (pfam01197)→ |
| Eco-1731-1 | NZ_JH378054.1 | + | 775608 | 775660 | **RNA**→ rpmE (PRK00019)Ribosomal_L31 (pfam01197)→ |
| Eco-1732-1 | NZ_JH378041.1 | + | 181741 | 181793 | **RNA**→ rpmE (PRK00019)Ribosomal_L31 (pfam01197)→ |
| Eco-1734-1 | NZ_JH378023.1 | + | 1163099 | 1163151 | **RNA**→ rpmE (PRK00019)Ribosomal_L31 (pfam01197)→ |
| Eco-1101-2 | NC_017638.1 | + | 4106059 | 4106111 | **RNA**→ rpmE (PRK00019)Ribosomal_L31 (pfam01197)→ |
| Eco-2183-1 | NC_020518.1 | + | 3556167 | 3556219 | **RNA**→ rpmE (PRK00019)Ribosomal_L31 (pfam01197)→ |
| Eco-1651-1 | NC_011993.1 | + | 4191555 | 4191607 | **RNA**→ rpmE (PRK00019)Ribosomal_L31 (pfam01197)→ |
| Eco-1753-1 | NZ_KB021597.1 | + | 86049 | 86101 | **RNA**→ rpmE (PRK00019)Ribosomal_L31 (pfam01197)→ |
| Eco-551-1 | NZ_CP006027.1 | + | 4927607 | 4927659 | **RNA**→ rpmE (PRK00019)Ribosomal_L31 (pfam01197)→ |
| Eco-572-1 | NZ_CP007592.1 | + | 4700635 | 4700687 | **RNA**→ rpmE (PRK00019)Ribosomal_L31 (pfam01197)→ |
| Eco-52-1 | NZ_CP009166.1 | + | 4408327 | 4408379 | **RNA**→ rpmE (PRK00019)Ribosomal_L31 (pfam01197)→ |
| Eco-398-1 | NZ_CP010344.1 | + | 4283291 | 4283343 | **RNA**→ rpmE (PRK00019)Ribosomal_L31 (pfam01197)→ |
| Eco-552-1 | NZ_CP006262.1 | + | 4745473 | 4745525 | **RNA**→ rpmE (PRK00019)Ribosomal_L31 (pfam01197)→ |
| Eco-549-1 | NZ_CP007136.1 | + | 4927604 | 4927656 | **RNA**→ rpmE (PRK00019)Ribosomal_L31 (pfam01197)→ |
| Eco-550-1 | NZ_CP007133.1 | + | 4745477 | 4745529 | **RNA**→ rpmE (PRK00019)Ribosomal_L31 (pfam01197)→ |
| Eco-463-1 | NZ_CP007799.1 | + | 4549787 | 4549839 | **RNA**→ hypo→ hypo→ hypo→ rpmE (PRK00019)Ribosomal_L31 (pfam01197)→ rpmE (PRK00019)Ribosomal_L31 (pfam01197)→ rpmE (PRK00019)Ribosomal_L31 (pfam01197)→ |
| Eco-463-2 | NZ_CP007799.1 | + | 4552090 | 4552142 | **RNA**→ hypo→ hypo→ hypo→ rpmE (PRK00019)Ribosomal_L31 (pfam01197)→ rpmE (PRK00019)Ribosomal_L31 (pfam01197)→ rpmE (PRK00019)Ribosomal_L31 (pfam01197)→ |
| Eco-463-3 | NZ_CP007799.1 | + | 4554393 | 4554445 | **RNA**→ hypo→ hypo→ hypo→ rpmE (PRK00019)Ribosomal_L31 (pfam01197)→ rpmE (PRK00019)Ribosomal_L31 (pfam01197)→ rpmE (PRK00019)Ribosomal_L31 (pfam01197)→ |
| Eco-423-1 | NZ_CP008801.1 | + | 4203810 | 4203862 | **RNA**→ rpmE (PRK00019)Ribosomal_L31 (pfam01197)→ |
| Sfl-57-1 | NZ_CP004056.1 | + | 4130700 | 4130752 | **RNA**→ rpmE (PRK00019)Ribosomal_L31 (pfam01197)→ |
| Sfl-82-1 | NZ_CP004057.1 | + | 4155779 | 4155831 | **RNA**→ rpmE (PRK00019)Ribosomal_L31 (pfam01197)→ |
| Eco-291-1 | NZ_CP009072.1 | - | 587468 | 587416 | **RNA**→ rpmE (PRK00019)Ribosomal_L31 (pfam01197)→ |
| Eco-328-1 | NZ_CP009273.1 | + | 4116882 | 4116934 | **RNA**→ rpmE (PRK00019)Ribosomal_L31 (pfam01197)→ |
| Eco-893-1 | NZ_KI533530.1 | + | 25625 | 25677 | **RNA**→ rpmE (PRK00019)Ribosomal_L31 (pfam01197)→ |
| Eco-1693-1 | NZ_GG774009.1 | + | 32525 | 32577 | **RNA**→ rpmE (PRK00019)Ribosomal_L31 (pfam01197)→ |
| Eco-878-1 | NZ_KI531365.1 | + | 79602 | 79654 | **RNA**→ rpmE (PRK00019)Ribosomal_L31 (pfam01197)→ |
| Eco-885-1 | NZ_KI532294.1 | + | 32305 | 32357 | **RNA**→ rpmE (PRK00019)Ribosomal_L31 (pfam01197)→ |
| Eco-890-1 | NZ_KI533169.1 | + | 22217 | 22269 | **RNA**→ rpmE (PRK00019)Ribosomal_L31 (pfam01197)→ |
| Eco-1688-1 | NZ_GG773540.1 | - | 52256 | 52204 | **RNA**→ rpmE (PRK00019)Ribosomal_L31 (pfam01197)→ |
| Eco-770-1 | NZ_KI530777.1 | + | 119139 | 119191 | **RNA**→ rpmE (PRK00019)Ribosomal_L31 (pfam01197)→ |
| Eco-886-1 | NZ_KI532500.1 | - | 52076 | 52024 | **RNA**→ rpmE (PRK00019)Ribosomal_L31 (pfam01197)→ |
| Eco-891-1 | NZ_KI533323.1 | + | 82900 | 82952 | **RNA**→ rpmE (PRK00019)Ribosomal_L31 (pfam01197)→ |
| Eco-1149-1 | NC_017633.1 | + | 4549461 | 4549513 | **RNA**→ rpmE (PRK00019)Ribosomal_L31 (pfam01197)→ |
| Eco-877-1 | NZ_KI531308.1 | + | 126494 | 126546 | **RNA**→ rpmE (PRK00019)Ribosomal_L31 (pfam01197)→ |
| Eco-884-1 | NZ_KI532131.1 | - | 549 | 497 | **RNA**→ rpmE (PRK00019)Ribosomal_L31 (pfam01197)→ |
| Eco-1681-1 | NZ_GG773415.1 | - | 39384 | 39332 | **RNA**→ rpmE (PRK00019)Ribosomal_L31 (pfam01197)→ |
| Eco-1-4 | NZ_LK931544.1 | - | 47804 | 47752 | **RNA**→ rpmE (PRK00019)Ribosomal_L31 (pfam01197)→ |
| Sfl-1-2 | NZ_LM651928.1 | + | 4058590 | 4058642 | **RNA**→ rpmE (PRK00019)Ribosomal_L31 (pfam01197)→ |
| Eco-1-1 | NZ_LK985426.1 | + | 27751 | 27803 | **RNA**→ rpmE (PRK00019)Ribosomal_L31 (pfam01197)→ |
| Eco-746-1 | NZ_CP011134.1 | + | 4469729 | 4469781 | **RNA**→ rpmE (PRK00019)Ribosomal_L31 (pfam01197)→ |
| Eco-682-1 | NZ_HG941718.1 | + | 4499005 | 4499057 | **RNA**→ rpmE (PRK00019)Ribosomal_L31 (pfam01197)→ |
| Eco-419-1 | NZ_LN832404.1 | + | 4123427 | 4123479 | **RNA**→ rpmE (PRK00019)Ribosomal_L31 (pfam01197)→ |
| Eco-472-1 | NZ_CP011331.1 | - | 4330723 | 4330671 | **RNA**→ rpmE (PRK00019)Ribosomal_L31 (pfam01197)→ |

| | | | | | |
|---|---|---|---|---|---|
| Sso-1-547 | NZ_CP011422.1 | + | 966341 | 966393 | RNA→ rpmE (PRK00019)Ribosomal_L31 (pfam01197)→ |
| Eco-1-2 | NZ_LK999974.1 | - | 48085 | 48033 | RNA→ rpmE (PRK00019)Ribosomal_L31 (pfam01197)→ |
| Eco-1-3 | NZ_LK999940.1 | - | 49482 | 49430 | RNA→ rpmE (PRK00019)Ribosomal_L31 (pfam01197)→ |
| Eco-1169-1 | NZ_CP011343.1 | + | 365674 | 365726 | RNA→ rpmE (PRK00019)Ribosomal_L31 (pfam01197)→ |
| Eco-1169-2 | NZ_CP011342.1 | - | 2842199 | 2842147 | RNA→ rpmE (PRK00019)Ribosomal_L31 (pfam01197)→ |
| Sfl-72-1 | NZ_CP007037.1 | + | 4126943 | 4126995 | RNA→ rpmE (PRK00019)Ribosomal_L31 (pfam01197)→ |
| Sbo-2-10 | NZ_CP011511.1 | + | 4030216 | 4030268 | RNA→ rpmE (PRK00019)Ribosomal_L31 (pfam01197)→ |
| Eco-735-1 | NZ_CP006636.1 | + | 4090689 | 4090741 | RNA→ rpmE (PRK00019)Ribosomal_L31 (pfam01197)→ |
| Eco-734-1 | NZ_CP006632.1 | + | 4399932 | 4399984 | RNA→ rpmE (PRK00019)Ribosomal_L31 (pfam01197)→ |
| Eco-288-1 | NZ_CP007442.1 | + | 4329255 | 4329307 | RNA→ rpmE (PRK00019)Ribosomal_L31 (pfam01197)→ |
| Eco-1760-1 | NZ_GL988050.1 | + | 4993697 | 4993749 | RNA→ rpmE (PRK00019)Ribosomal_L31 (pfam01197)→ |
| Eco-1648-1 | NZ_JH650819.1 | + | 699275 | 699327 | RNA→ rpmE (PRK00019)Ribosomal_L31 (pfam01197)→ |
| Eco-1763-1 | NZ_JH648168.1 | + | 1166678 | 1166730 | RNA→ rpmE (PRK00019)Ribosomal_L31 (pfam01197)→ |
| Eco-1650-1 | NZ_KB000001.1 | + | 2277793 | 2277845 | RNA→ rpmE (PRK00019)Ribosomal_L31 (pfam01197)→ |
| Eco-1762-1 | NZ_JH648165.1 | + | 4457835 | 4457887 | RNA→ rpmE (PRK00019)Ribosomal_L31 (pfam01197)→ |
| Sfl-1-1 | NZ_KL503838.1 | - | 31116 | 31064 | RNA→ rpmE (PRK00019)Ribosomal_L31 (pfam01197)→ |
| Eco-1981-1 | NZ_AOGN01000115.1 | + | 13965 | 14017 | RNA→ rpmE (PRK00019)Ribosomal_L31 (pfam01197)→ |
| Eco-1714-1 | NZ_AOGM01000086.1 | + | 28059 | 28111 | RNA→ rpmE (PRK00019)Ribosomal_L31 (pfam01197)→ |
| Eco-1744-1 | NZ_AHAV01000106.1 | + | 32410 | 32462 | RNA→ rpmE (PRK00019)Ribosomal_L31 (pfam01197)→ |
| Eco-1834-1 | NZ_AHAX01000061.1 | + | 86405 | 86457 | RNA→ rpmE (PRK00019)Ribosomal_L31 (pfam01197)→ |
| Eco-1745-1 | NZ_AHAU01000082.1 | + | 84181 | 84233 | RNA→ rpmE (PRK00019)Ribosomal_L31 (pfam01197)→ |
| Eco-1389-1 | NZ_AQFA01000051.1 | + | 88710 | 88762 | RNA→ rpmE (PRK00019)Ribosomal_L31 (pfam01197)→ |
| Eco-1388-1 | NZ_AQFB01000026.1 | + | 46854 | 46906 | RNA→ rpmE (PRK00019)Ribosomal_L31 (pfam01197)→ |
| Eco-774-1 | NZ_AQGI01000087.1 | + | 88674 | 88726 | RNA→ rpmE (PRK00019)Ribosomal_L31 (pfam01197)→ |
| Eco-1093-1 | NZ_AIGH01000071.1 | + | 97778 | 97830 | RNA→ rpmE (PRK00019)Ribosomal_L31 (pfam01197)→ |
| Eco-1183-1 | NZ_APNY01000014.1 | + | 87903 | 87955 | RNA→ rpmE (PRK00019)Ribosomal_L31 (pfam01197)→ |
| Sfl-25-14 | NZ_LAJQ01000116.1 | - | 31124 | 31072 | RNA→ rpmE (PRK00019)Ribosomal_L31 (pfam01197)→ |
| Eco-1181-1 | NZ_APNW01000010.1 | + | 91909 | 91961 | RNA→ rpmE (PRK00019)Ribosomal_L31 (pfam01197)→ |
| Eco-2010-2 | NZ_AOQV01000032.1 | - | 52231 | 52179 | RNA→ rpmE (PRK00019)Ribosomal_L31 (pfam01197)→ |
| Eco-1042-1 | NZ_AIHE01000043.1 | + | 47119 | 47171 | RNA→ rpmE (PRK00019)Ribosomal_L31 (pfam01197)→ |
| Eco-784-1 | NZ_AFAB02000228.1 | + | 211717 | 211769 | RNA→ rpmE (PRK00019)Ribosomal_L31 (pfam01197)→ |
| Eco-368-1 | NZ_AIBM01000066.1 | - | 39338 | 39286 | RNA→ rpmE (PRK00019)Ribosomal_L31 (pfam01197)→ |
| Eco-1675-1 | NZ_AQEV01000020.1 | + | 89153 | 89205 | RNA→ rpmE (PRK00019)Ribosomal_L31 (pfam01197)→ |
| Eco-790-1 | NZ_AQFF01000029.1 | + | 24026 | 24078 | RNA→ rpmE (PRK00019)Ribosomal_L31 (pfam01197)→ |
| Eco-2043-1 | NZ_AQEJ01000024.1 | + | 119619 | 119671 | RNA→ rpmE (PRK00019)Ribosomal_L31 (pfam01197)→ |
| Eco-1004-1 | NZ_AQFK01000021.1 | + | 91846 | 91898 | RNA→ rpmE (PRK00019)Ribosomal_L31 (pfam01197)→ |
| Eco-1660-1 | NZ_AQEY01000025.1 | + | 21685 | 21737 | RNA→ rpmE (PRK00019)Ribosomal_L31 (pfam01197)→ |
| Eco-1009-1 | NZ_AQFI01000025.1 | + | 90459 | 90511 | RNA→ rpmE (PRK00019)Ribosomal_L31 (pfam01197)→ |
| Eco-1915-1 | NZ_AQEM01000049.1 | + | 91353 | 91405 | RNA→ rpmE (PRK00019)Ribosomal_L31 (pfam01197)→ |
| Eco-1674-1 | NZ_AQEW01000025.1 | + | 121217 | 121269 | RNA→ rpmE (PRK00019)Ribosomal_L31 (pfam01197)→ |
| Eco-1677-1 | NZ_AQEU01000023.1 | + | 86825 | 86877 | RNA→ rpmE (PRK00019)Ribosomal_L31 (pfam01197)→ |
| Eco-789-1 | NZ_AQFG01000030.1 | + | 117781 | 117833 | RNA→ rpmE (PRK00019)Ribosomal_L31 (pfam01197)→ |
| Eco-1930-1 | NZ_AQEL01000014.1 | + | 89670 | 89722 | RNA→ rpmE (PRK00019)Ribosomal_L31 (pfam01197)→ |
| Eco-1900-1 | NZ_AQEN01000035.1 | + | 90256 | 90308 | RNA→ rpmE (PRK00019)Ribosomal_L31 (pfam01197)→ |
| Eco-1150-1 | NZ_AQFD01000047.1 | + | 91814 | 91866 | RNA→ rpmE (PRK00019)Ribosomal_L31 (pfam01197)→ |
| Eco-1872-1 | NZ_AQEQ01000029.1 | + | 121013 | 121065 | RNA→ rpmE (PRK00019)Ribosomal_L31 (pfam01197)→ |
| Eco-1661-1 | NZ_AQEX01000030.1 | + | 90110 | 90162 | RNA→ rpmE (PRK00019)Ribosomal_L31 (pfam01197)→ |
| Eco-2165-1 | NZ_AQEK01000042.1 | + | 27965 | 28017 | RNA→ rpmE (PRK00019)Ribosomal_L31 (pfam01197)→ |
| Eco-1837-1 | NZ_APXR01000184.1 | + | 23817 | 23869 | RNA→ rpmE (PRK00019)Ribosomal_L31 (pfam01197)→ |
| Eco-786-1 | NZ_AQFH01000031.1 | + | 29182 | 29234 | RNA→ rpmE (PRK00019)Ribosomal_L31 (pfam01197)→ |
| Eco-822-1 | NZ_APXI01000107.1 | + | 87655 | 87707 | RNA→ rpmE (PRK00019)Ribosomal_L31 (pfam01197)→ |
| Eco-1151-1 | NZ_AQFC01000057.1 | + | 92055 | 92107 | RNA→ rpmE (PRK00019)Ribosomal_L31 (pfam01197)→ |

| | | | | | |
|---|---|---|---|---|---|
| Eco-1013-1 | NZ_APXO01000114.1 | + | 30089 | 30141 | **RNA**→ ←hypo |
| Eco-1850-1 | NZ_AQES01000017.1 | + | 91820 | 91872 | **RNA**→ rpmE (PRK00019)Ribosomal_L31 (pfam01197)→ |
| Eco-367-1 | NZ_AIBL01000107.1 | - | 690 | 638 | **RNA**→ rpmE (PRK00019)Ribosomal_L31 (pfam01197)→ |
| Eco-335-1 | NZ_AIAG01000120.1 | + | 841 | 893 | **RNA**→ rpmE (PRK00019)Ribosomal_L31 (pfam01197)→ |
| Eco-352-1 | NZ_AIAX01000027.1 | + | 85782 | 85834 | **RNA**→ rpmE (PRK00019)Ribosomal_L31 (pfam01197)→ |
| Eco-364-1 | NZ_AIBJ01000226.1 | + | 53110 | 53162 | **RNA**→ rpmE (PRK00019)Ribosomal_L31 (pfam01197)→ |
| Eco-331-1 | NZ_AIAC01000054.1 | + | 88135 | 88187 | **RNA**→ rpmE (PRK00019)Ribosomal_L31 (pfam01197)→ |
| Eco-374-1 | NZ_AIBT01000104.1 | - | 486 | 434 | **RNA**→ rpmE (PRK00019)Ribosomal_L31 (pfam01197)→ |
| Eco-427-1 | NZ_AYOG01000027.1 | + | 72663 | 72715 | **RNA**→ rpmE (PRK00019)Ribosomal_L31 (pfam01197)→ |
| Sdy-10-1 | NZ_AXUU01000203.1 | - | 464 | 412 | **RNA**→ rpmE (PRK00019)Ribosomal_L31 (pfam01197)→ |
| Sdy-9-1 | NZ_AXUT01000390.1 | + | 8180 | 8232 | **RNA**→ rpmE (PRK00019)Ribosomal_L31 (pfam01197)→ |
| Eco-426-1 | NZ_AYOP01000027.1 | + | 72663 | 72715 | **RNA**→ rpmE (PRK00019)Ribosomal_L31 (pfam01197)→ |
| Eco-955-1 | NZ_AYOF01000070.1 | - | 556 | 504 | **RNA**→ rpmE (PRK00019)Ribosomal_L31 (pfam01197)→ |
| Eco-415-1 | NZ_CBWA010000372.1 | + | 4180 | 4232 | **RNA**→ rpmE (PRK00019)Ribosomal_L31 (pfam01197)→ |
| Eco-674-1 | NZ_JHJE01000048.1 | - | 42020 | 41968 | **RNA**→ rpmE (PRK00019)Ribosomal_L31 (pfam01197)→ |
| Eco-563-1 | NZ_JHFO01000028.1 | + | 113553 | 113605 | **RNA**→ rpmE (PRK00019)Ribosomal_L31 (pfam01197)→ |
| Eco-34-1 | NZ_JJLL01000159.1 | + | 83742 | 83794 | **RNA**→ rpmE (PRK00019)Ribosomal_L31 (pfam01197)→ |
| Eco-801-1 | NZ_AQEA01000073.1 | + | 18586 | 18638 | **RNA**→ rpmE (PRK00019)Ribosomal_L31 (pfam01197)→ |
| Eco-802-1 | NZ_AQDZ01000090.1 | + | 18323 | 18375 | **RNA**→ rpmE (PRK00019)Ribosomal_L31 (pfam01197)→ |
| Eco-823-1 | NZ_AQDJ01000065.1 | + | 87862 | 87914 | **RNA**→ rpmE (PRK00019)Ribosomal_L31 (pfam01197)→ |
| Eco-812-1 | NZ_AQDQ01000090.1 | + | 32876 | 32928 | **RNA**→ rpmE (PRK00019)Ribosomal_L31 (pfam01197)→ |
| Eco-807-1 | NZ_AQDU01000131.1 | + | 66030 | 66082 | **RNA**→ rpmE (PRK00019)Ribosomal_L31 (pfam01197)→ |
| Eco-1890-1 | NZ_AQEO01000037.1 | + | 87406 | 87458 | **RNA**→ rpmE (PRK00019)Ribosomal_L31 (pfam01197)→ |
| Eco-808-1 | NZ_AQDT01000070.1 | + | 18684 | 18736 | **RNA**→ rpmE (PRK00019)Ribosomal_L31 (pfam01197)→ |
| Eco-1885-1 | NZ_AQAC01000120.1 | + | 87618 | 87670 | **RNA**→ rpmE (PRK00019)Ribosomal_L31 (pfam01197)→ |
| Eco-799-1 | NZ_AQGH01000063.1 | + | 119118 | 119170 | **RNA**→ rpmE (PRK00019)Ribosomal_L31 (pfam01197)→ |
| Eco-1884-1 | NZ_AQAB01000058.1 | + | 87798 | 87850 | **RNA**→ rpmE (PRK00019)Ribosomal_L31 (pfam01197)→ |
| Eco-791-1 | NZ_AQEE01000140.1 | + | 11616 | 11668 | **RNA**→ rpmE (PRK00019)Ribosomal_L31 (pfam01197)→ |
| Eco-1865-1 | NZ_APYM01000235.1 | + | 33620 | 33672 | **RNA**→ rpmE (PRK00019)Ribosomal_L31 (pfam01197)→ |
| Eco-811-1 | NZ_AQGG01000114.1 | + | 46224 | 46276 | **RNA**→ rpmE (PRK00019)Ribosomal_L31 (pfam01197)→ |
| Eco-1866-1 | NZ_APYN01000077.1 | + | 41465 | 41517 | **RNA**→ rpmE (PRK00019)Ribosomal_L31 (pfam01197)→ |
| Eco-778-1 | NZ_APZJ01000082.1 | + | 31396 | 31448 | **RNA**→ rpmE (PRK00019)Ribosomal_L31 (pfam01197)→ |
| Eco-1871-1 | NZ_APYS01000184.1 | + | 87770 | 87822 | **RNA**→ rpmE (PRK00019)Ribosomal_L31 (pfam01197)→ |
| Eco-797-1 | NZ_AQEB01000177.1 | + | 82811 | 82863 | **RNA**→ rpmE (PRK00019)Ribosomal_L31 (pfam01197)→ |
| Eco-1892-1 | NZ_APZA01000064.1 | + | 84354 | 84406 | **RNA**→ rpmE (PRK00019)Ribosomal_L31 (pfam01197)→ |
| Eco-818-1 | NZ_AQDL01000080.1 | + | 18684 | 18736 | **RNA**→ rpmE (PRK00019)Ribosomal_L31 (pfam01197)→ |
| Eco-1898-1 | NZ_APZG01000363.1 | + | 80717 | 80769 | **RNA**→ rpmE (PRK00019)Ribosomal_L31 (pfam01197)→ |
| Eco-803-1 | NZ_AQDY01000177.1 | + | 74715 | 74767 | **RNA**→ rpmE (PRK00019)Ribosomal_L31 (pfam01197)→ |
| Eco-33-1 | NZ_JJLO01000155.1 | + | 60700 | 60752 | **RNA**→ rpmE (PRK00019)Ribosomal_L31 (pfam01197)→ |
| Eco-30-1 | NZ_JJLW01000160.1 | + | 84245 | 84297 | **RNA**→ rpmE (PRK00019)Ribosomal_L31 (pfam01197)→ |
| Eco-704-1 | NZ_JHLV01000044.1 | - | 835 | 783 | **RNA**→ rpmE (PRK00019)Ribosomal_L31 (pfam01197)→ |
| Eco-542-1 | NZ_JHHB01000007.1 | - | 605 | 553 | **RNA**→ rpmE (PRK00019)Ribosomal_L31 (pfam01197)→ |
| Eco-509-1 | NZ_JHNQ01000014.1 | + | 28051 | 28103 | **RNA**→ rpmE (PRK00019)Ribosomal_L31 (pfam01197)→ |
| Eco-122-1 | NZ_JNQU01000023.1 | - | 46318 | 46266 | **RNA**→ rpmE (PRK00019)Ribosomal_L31 (pfam01197)→ |
| Eco-174-1 | NZ_JNMN01000019.1 | + | 85263 | 85315 | **RNA**→ rpmE (PRK00019)Ribosomal_L31 (pfam01197)→ |
| Eco-29-1 | NZ_JNQC01000014.1 | + | 85569 | 85621 | **RNA**→ rpmE (PRK00019)Ribosomal_L31 (pfam01197)→ |
| Eco-120-1 | NZ_JNQS01000019.1 | - | 29219 | 29167 | **RNA**→ rpmE (PRK00019)Ribosomal_L31 (pfam01197)→ |
| Eco-68-1 | NZ_JNNA01000147.1 | + | 11560 | 11612 | **RNA**→ rpmE (PRK00019)Ribosomal_L31 (pfam01197)→ |
| Eco-545-1 | NZ_JASU01000001.1 | - | 1459090 | 1459038 | **RNA**→ rpmE (PRK00019)Ribosomal_L31 (pfam01197)→ |
| Eco-28-1 | NZ_JNQB01000014.1 | - | 43513 | 43461 | **RNA**→ rpmE (PRK00019)Ribosomal_L31 (pfam01197)→ |
| Eco-566-1 | NZ_JASR01000001.1 | - | 2462015 | 2461963 | **RNA**→ rpmE (PRK00019)Ribosomal_L31 (pfam01197)→ |
| Eco-226-1 | NZ_JOQV01000010.1 | + | 124927 | 124979 | **RNA**→ rpmE (PRK00019)Ribosomal_L31 (pfam01197)→ |

| | | | | | |
|---|---|---|---|---|---|
| Eco-224-1 | NZ_JOQY01000010.1 | - | 43045 | 42993 | **RNA**→ rpmE (PRK00019)Ribosomal_L31 (pfam01197)→ |
| Eco-234-1 | NZ_JORL01000069.1 | + | 93206 | 93258 | **RNA**→ rpmE (PRK00019)Ribosomal_L31 (pfam01197)→ |
| Eco-153-1 | NZ_JONB01000076.1 | + | 87678 | 87730 | **RNA**→ rpmE (PRK00019)Ribosomal_L31 (pfam01197)→ |
| Eco-42-1 | NZ_JOSH01000256.1 | + | 86293 | 86345 | **RNA**→ rpmE (PRK00019)Ribosomal_L31 (pfam01197)→ |
| Eco-1902-1 | NZ_AQAH01000120.1 | + | 46756 | 46808 | **RNA**→ rpmE (PRK00019)Ribosomal_L31 (pfam01197)→ |
| Eco-809-1 | NZ_AQDS01000152.1 | + | 89137 | 89189 | **RNA**→ rpmE (PRK00019)Ribosomal_L31 (pfam01197)→ |
| Eco-805-1 | NZ_AQDW01000073.1 | + | 18711 | 18763 | **RNA**→ rpmE (PRK00019)Ribosomal_L31 (pfam01197)→ |
| Eco-1903-1 | NZ_AQAI01000051.1 | + | 46814 | 46866 | **RNA**→ rpmE (PRK00019)Ribosomal_L31 (pfam01197)→ |
| Eco-796-1 | NZ_APXE01000196.1 | + | 29919 | 29971 | **RNA**→ rpmE (PRK00019)Ribosomal_L31 (pfam01197)→ |
| Eco-1905-1 | NZ_AQAK01000105.1 | + | 46790 | 46842 | **RNA**→ rpmE (PRK00019)Ribosomal_L31 (pfam01197)→ |
| Eco-1911-1 | NZ_AQAQ01000057.1 | + | 46776 | 46828 | **RNA**→ rpmE (PRK00019)Ribosomal_L31 (pfam01197)→ |
| Eco-781-1 | NZ_APXC01000301.1 | + | 18755 | 18807 | **RNA**→ rpmE (PRK00019)Ribosomal_L31 (pfam01197)→ |
| Eco-824-1 | NZ_APXJ01000122.1 | + | 18695 | 18747 | **RNA**→ rpmE (PRK00019)Ribosomal_L31 (pfam01197)→ rpmE (PRK00019)Ribosomal_L31 (pfam01197)→ |
| Eco-1891-1 | NZ_APYZ01000056.1 | + | 84492 | 84544 | **RNA**→ rpmE (PRK00019)Ribosomal_L31 (pfam01197)→ |
| Eco-792-1 | NZ_AQFE01000042.1 | + | 88874 | 88926 | **RNA**→ rpmE (PRK00019)Ribosomal_L31 (pfam01197)→ rpmE (PRK00019)Ribosomal_L31 (pfam01197)→ |
| Eco-1814-1 | NZ_AMXO01000238.1 | - | 638 | 586 | **RNA**→ rpmE (PRK00019)Ribosomal_L31 (pfam01197)→ |
| Eco-1894-1 | NZ_APZC01000326.1 | + | 84351 | 84403 | **RNA**→ rpmE (PRK00019)Ribosomal_L31 (pfam01197)→ |
| Eco-798-1 | NZ_APZK01000191.1 | + | 23771 | 23823 | **RNA**→ rpmE (PRK00019)Ribosomal_L31 (pfam01197)→ |
| Eco-1779-1 | NZ_AHAW01000062.1 | + | 86201 | 86253 | **RNA**→ rpmE (PRK00019)Ribosomal_L31 (pfam01197)→ |
| Eco-1877-1 | NZ_APYV01000111.1 | + | 117684 | 117736 | **RNA**→ rpmE (PRK00019)Ribosomal_L31 (pfam01197)→ |
| Eco-775-1 | NZ_APWY01000309.1 | + | 14050 | 14102 | **RNA**→ rpmE (PRK00019)Ribosomal_L31 (pfam01197)→ |
| Eco-785-1 | NZ_AFAD02000007.1 | + | 29056 | 29108 | **RNA**→ rpmE (PRK00019)Ribosomal_L31 (pfam01197)→ |
| Eco-1867-1 | NZ_APYO01000081.1 | + | 87500 | 87552 | **RNA**→ rpmE (PRK00019)Ribosomal_L31 (pfam01197)→ |
| Eco-1662-1 | NZ_AQCW01000153.1 | + | 7794 | 7846 | **RNA**→ rpmE (PRK00019)Ribosomal_L31 (pfam01197)→ |
| Eco-828-1 | NZ_AQDF01000073.1 | + | 87908 | 87960 | **RNA**→ rpmE (PRK00019)Ribosomal_L31 (pfam01197)→ |
| Eco-1658-1 | NZ_AQEZ01000029.1 | + | 87870 | 87922 | **RNA**→ rpmE (PRK00019)Ribosomal_L31 (pfam01197)→ |
| Eco-1873-1 | NZ_APYT01000068.1 | + | 52672 | 52724 | **RNA**→ rpmE (PRK00019)Ribosomal_L31 (pfam01197)→ |
| Eco-1874-1 | NZ_APZT01000083.1 | + | 30333 | 30385 | **RNA**→ rpmE (PRK00019)Ribosomal_L31 (pfam01197)→ |
| Eco-1849-1 | NZ_APYB01000069.1 | + | 12751 | 12803 | **RNA**→ rpmE (PRK00019)Ribosomal_L31 (pfam01197)→ |
| Eco-782-1 | NZ_AQEG01000132.1 | + | 46739 | 46791 | **RNA**→ rpmE (PRK00019)Ribosomal_L31 (pfam01197)→ |
| Eco-1881-1 | NZ_APZV01000166.1 | + | 33953 | 34005 | **RNA**→ rpmE (PRK00019)Ribosomal_L31 (pfam01197)→ |
| Eco-1841-1 | NZ_APXV01000269.1 | + | 23763 | 23815 | **RNA**→ rpmE (PRK00019)Ribosomal_L31 (pfam01197)→ |
| Eco-1899-1 | NZ_APZH01000117.1 | + | 84317 | 84369 | **RNA**→ rpmE (PRK00019)Ribosomal_L31 (pfam01197)→ |
| Eco-773-1 | NZ_AQEI01000288.1 | + | 2100 | 2152 | **RNA**→ rpmE (PRK00019)Ribosomal_L31 (pfam01197)→ |
| Eco-1092-1 | NZ_AIGG01000081.1 | + | 93774 | 93826 | **RNA**→ rpmE (PRK00019)Ribosomal_L31 (pfam01197)→ |
| Eco-1904-1 | NZ_AQAJ01000054.1 | + | 46831 | 46883 | **RNA**→ rpmE (PRK00019)Ribosomal_L31 (pfam01197)→ |
| Eco-825-1 | NZ_AQDI01000113.1 | + | 87539 | 87591 | **RNA**→ rpmE (PRK00019)Ribosomal_L31 (pfam01197)→ |
| Eco-1666-1 | NZ_AQCS01000124.1 | + | 7818 | 7870 | **RNA**→ rpmE (PRK00019)Ribosomal_L31 (pfam01197)→ |
| Eco-1906-1 | NZ_AQAL01000124.1 | + | 46820 | 46872 | **RNA**→ rpmE (PRK00019)Ribosomal_L31 (pfam01197)→ |
| Eco-1671-1 | NZ_AQCN01000186.1 | + | 47879 | 47931 | **RNA**→ rpmE (PRK00019)Ribosomal_L31 (pfam01197)→ |
| Eco-815-1 | NZ_AQDN01000103.1 | + | 119323 | 119375 | **RNA**→ rpmE (PRK00019)Ribosomal_L31 (pfam01197)→ |
| Eco-1912-1 | NZ_AQAR01000135.1 | + | 46818 | 46870 | **RNA**→ rpmE (PRK00019)Ribosomal_L31 (pfam01197)→ |
| Eco-1851-1 | NZ_APYC01000065.1 | + | 78035 | 78087 | **RNA**→ rpmE (PRK00019)Ribosomal_L31 (pfam01197)→ |
| Eco-820-1 | NZ_AQDK01000143.1 | + | 7833 | 7885 | **RNA**→ rpmE (PRK00019)Ribosomal_L31 (pfam01197)→ |
| Eco-1920-1 | NZ_AQAX01000145.1 | + | 87587 | 87639 | **RNA**→ rpmE (PRK00019)Ribosomal_L31 (pfam01197)→ |
| Eco-1859-1 | NZ_APZR01000088.1 | + | 84547 | 84599 | **RNA**→ rpmE (PRK00019)Ribosomal_L31 (pfam01197)→ |
| Eco-1875-1 | NZ_APZU01000076.1 | + | 30357 | 30409 | **RNA**→ rpmE (PRK00019)Ribosomal_L31 (pfam01197)→ |
| Eco-814-1 | NZ_AQDO01000114.1 | + | 87792 | 87844 | **RNA**→ rpmE (PRK00019)Ribosomal_L31 (pfam01197)→ |
| Eco-1820-1 | NZ_AKBC01000335.1 | - | 584 | 532 | **RNA**→ rpmE (PRK00019)Ribosomal_L31 (pfam01197)→ |
| Eco-1883-1 | NZ_AQEP01000022.1 | + | 90682 | 90734 | **RNA**→ rpmE (PRK00019)Ribosomal_L31 (pfam01197)→ |
| Eco-810-1 | NZ_AQDR01000106.1 | + | 122355 | 122407 | **RNA**→ rpmE (PRK00019)Ribosomal_L31 (pfam01197)→ |
| Eco-1886-1 | NZ_AQAD01000068.1 | + | 87873 | 87925 | **RNA**→ rpmE (PRK00019)Ribosomal_L31 (pfam01197)→ |

| | | | | | |
|---|---|---|---|---|---|
| Eco-1098-1 | NZ_AIGM01000052.1 | - | 51199 | 51147 | **RNA**→ rpmE (PRK00019)Ribosomal_L31 (pfam01197)→ |
| Eco-1868-1 | NZ_APYP01000074.1 | + | 87717 | 87769 | **RNA**→ rpmE (PRK00019)Ribosomal_L31 (pfam01197)→ |
| Eco-804-1 | NZ_AQDX01000116.1 | + | 82883 | 82935 | **RNA**→ rpmE (PRK00019)Ribosomal_L31 (pfam01197)→ |
| Eco-1664-1 | NZ_AQCU01000277.1 | + | 7814 | 7866 | **RNA**→ rpmE (PRK00019)Ribosomal_L31 (pfam01197)→ |
| Eco-1869-1 | NZ_APYQ01000075.1 | + | 23398 | 23450 | **RNA**→ rpmE (PRK00019)Ribosomal_L31 (pfam01197)→ |
| Eco-1672-1 | NZ_AQCM01000086.1 | + | 85975 | 86027 | **RNA**→ rpmE (PRK00019)Ribosomal_L31 (pfam01197)→ |
| Eco-783-1 | NZ_AQEF01000111.1 | + | 46744 | 46796 | **RNA**→ ←hypo |
| Eco-1887-1 | NZ_APZW01000065.1 | + | 87850 | 87902 | **RNA**→ rpmE (PRK00019)Ribosomal_L31 (pfam01197)→ |
| Eco-1676-1 | NZ_APXP01000199.1 | + | 89949 | 90001 | **RNA**→ rpmE (PRK00019)Ribosomal_L31 (pfam01197)→ |
| Eco-1896-1 | NZ_APZE01000057.1 | + | 84083 | 84135 | **RNA**→ rpmE (PRK00019)Ribosomal_L31 (pfam01197)→ rpmE (PRK00019)Ribosomal_L31 (pfam01197)→ |
| Eco-793-1 | NZ_AQED01000166.1 | + | 32326 | 32378 | **RNA**→ rpmE (PRK00019)Ribosomal_L31 (pfam01197)→ rpmE (PRK00019)Ribosomal_L31 (pfam01197)→ |
| Eco-1836-1 | NZ_APXQ01000157.1 | + | 23781 | 23833 | **RNA**→ rpmE (PRK00019)Ribosomal_L31 (pfam01197)→ |
| Eco-1889-1 | NZ_AQAF01000060.1 | + | 87791 | 87843 | **RNA**→ rpmE (PRK00019)Ribosomal_L31 (pfam01197)→ |
| Eco-800-1 | NZ_APXF01000193.1 | + | 29965 | 30017 | **RNA**→ rpmE (PRK00019)Ribosomal_L31 (pfam01197)→ |
| Eco-1854-1 | NZ_APYF01000071.1 | + | 78099 | 78151 | **RNA**→ rpmE (PRK00019)Ribosomal_L31 (pfam01197)→ |
| Eco-1908-1 | NZ_AQAN01000124.1 | + | 46749 | 46801 | **RNA**→ rpmE (PRK00019)Ribosomal_L31 (pfam01197)→ |
| Eco-795-1 | NZ_APXD01000154.1 | + | 28153 | 28205 | **RNA**→ rpmE (PRK00019)Ribosomal_L31 (pfam01197)→ |
| Eco-1844-1 | NZ_APXW01000179.1 | + | 23741 | 23793 | **RNA**→ rpmE (PRK00019)Ribosomal_L31 (pfam01197)→ |
| Eco-1913-1 | NZ_AQAS01000085.1 | + | 46815 | 46867 | **RNA**→ rpmE (PRK00019)Ribosomal_L31 (pfam01197)→ |
| Eco-1847-1 | NZ_APXZ01000347.1 | + | 23890 | 23942 | **RNA**→ rpmE (PRK00019)Ribosomal_L31 (pfam01197)→ |
| Eco-1921-1 | NZ_AQAY01000065.1 | + | 87679 | 87731 | **RNA**→ rpmE (PRK00019)Ribosomal_L31 (pfam01197)→ |
| Eco-777-1 | NZ_APXA01000319.1 | + | 44177 | 44229 | **RNA**→ rpmE (PRK00019)Ribosomal_L31 (pfam01197)→ rpmE (PRK00019)Ribosomal_L31 (pfam01197)→ |
| Eco-1659-1 | NZ_APZN01000151.1 | + | 20924 | 20976 | **RNA**→ rpmE (PRK00019)Ribosomal_L31 (pfam01197)→ |
| Eco-1922-1 | NZ_AQAZ01000159.1 | + | 75652 | 75704 | **RNA**→ rpmE (PRK00019)Ribosomal_L31 (pfam01197)→ |
| Eco-1838-1 | NZ_APXS01000116.1 | + | 23729 | 23781 | **RNA**→ rpmE (PRK00019)Ribosomal_L31 (pfam01197)→ |
| Eco-1893-1 | NZ_APZB01000101.1 | + | 84367 | 84419 | **RNA**→ rpmE (PRK00019)Ribosomal_L31 (pfam01197)→ |
| Eco-1667-1 | NZ_AQCR01000138.1 | + | 7883 | 7935 | **RNA**→ rpmE (PRK00019)Ribosomal_L31 (pfam01197)→ |
| Eco-1773-1 | NZ_AJVU01000157.1 | - | 430 | 378 | **RNA**→ rpmE (PRK00019)Ribosomal_L31 (pfam01197)→ |
| Eco-462-1 | NZ_ASRI01000044.1 | - | 39367 | 39315 | **RNA**→ rpmE (PRK00019)Ribosomal_L31 (pfam01197)→ |
| Eco-408-1 | NZ_JPKI01000004.1 | - | 820306 | 820254 | **RNA**→ rpmE (PRK00019)Ribosomal_L31 (pfam01197)→ |
| Eco-1876-1 | NZ_APYU01000063.1 | + | 117681 | 117733 | **RNA**→ rpmE (PRK00019)Ribosomal_L31 (pfam01197)→ |
| Eco-819-1 | NZ_APXG01000161.1 | + | 18777 | 18829 | **RNA**→ rpmE (PRK00019)Ribosomal_L31 (pfam01197)→ |
| Eco-1852-1 | NZ_APYD01000069.1 | + | 88151 | 88203 | **RNA**→ rpmE (PRK00019)Ribosomal_L31 (pfam01197)→ |
| Eco-1895-1 | NZ_APZD01000070.1 | + | 80732 | 80784 | **RNA**→ rpmE (PRK00019)Ribosomal_L31 (pfam01197)→ |
| Eco-1860-1 | NZ_APZS01000115.1 | + | 84337 | 84389 | **RNA**→ rpmE (PRK00019)Ribosomal_L31 (pfam01197)→ |
| Eco-817-1 | NZ_APZL01000142.1 | + | 46914 | 46966 | **RNA**→ rpmE (PRK00019)Ribosomal_L31 (pfam01197)→ rpmE (PRK00019)Ribosomal_L31 (pfam01197)→ |
| Eco-1882-1 | NZ_AQAA01000396.1 | + | 47121 | 47173 | **RNA**→ rpmE (PRK00019)Ribosomal_L31 (pfam01197)→ |
| Eco-827-1 | NZ_AQDG01000067.1 | + | 87789 | 87841 | **RNA**→ rpmE (PRK00019)Ribosomal_L31 (pfam01197)→ |
| Eco-1772-1 | NZ_AMXQ01000200.1 | + | 58855 | 58907 | **RNA**→ rpmE (PRK00019)Ribosomal_L31 (pfam01197)→ |
| Eco-1888-1 | NZ_AQAE01000110.1 | + | 75775 | 75827 | **RNA**→ rpmE (PRK00019)Ribosomal_L31 (pfam01197)→ rpmE (PRK00019)Ribosomal_L31 (pfam01197)→ |
| Eco-826-1 | NZ_AQDH01000122.1 | + | 87807 | 87859 | **RNA**→ rpmE (PRK00019)Ribosomal_L31 (pfam01197)→ |
| Eco-1385-1 | NZ_ALIN02000018.1 | - | 54267 | 54215 | **RNA**→ rpmE (PRK00019)Ribosomal_L31 (pfam01197)→ |
| Eco-813-1 | NZ_AQDP01000060.1 | + | 87872 | 87924 | **RNA**→ rpmE (PRK00019)Ribosomal_L31 (pfam01197)→ |
| Eco-1914-1 | NZ_APZX01000452.1 | + | 87677 | 87729 | **RNA**→ rpmE (PRK00019)Ribosomal_L31 (pfam01197)→ |
| Eco-1673-1 | NZ_AQCL01000237.1 | + | 86159 | 86211 | **RNA**→ rpmE (PRK00019)Ribosomal_L31 (pfam01197)→ |
| Eco-816-1 | NZ_AQDM01000074.1 | + | 87795 | 87847 | **RNA**→ rpmE (PRK00019)Ribosomal_L31 (pfam01197)→ |
| Eco-1897-1 | NZ_APZF01000116.1 | + | 84315 | 84367 | **RNA**→ rpmE (PRK00019)Ribosomal_L31 (pfam01197)→ |
| Eco-1855-1 | NZ_APYG01000106.1 | + | 78045 | 78097 | **RNA**→ rpmE (PRK00019)Ribosomal_L31 (pfam01197)→ |
| Eco-1901-1 | NZ_AQAG01000128.1 | + | 46798 | 46850 | **RNA**→ ←hypo |
| Eco-829-1 | NZ_AQDE01000110.1 | + | 34384 | 34436 | **RNA**→ rpmE (PRK00019)Ribosomal_L31 (pfam01197)→ |
| Eco-1842-1 | NZ_APZO01000122.1 | + | 23706 | 23758 | **RNA**→ rpmE (PRK00019)Ribosomal_L31 (pfam01197)→ |
| Eco-806-1 | NZ_AQDV01000047.1 | + | 87714 | 87766 | **RNA**→ rpmE (PRK00019)Ribosomal_L31 (pfam01197)→ |

| | | | | | |
|---|---|---|---|---|---|
| Eco-1858-1 | NZ_APZQ01000074.1 | + | 84729 | 84781 | **RNA**→ rpmE (PRK00019)Ribosomal_L31 (pfam01197)→ |
| Eco-1910-1 | NZ_AQAP01000479.1 | + | 46858 | 46910 | **RNA**→ rpmE (PRK00019)Ribosomal_L31 (pfam01197)→ rpmE (PRK00019)Ribosomal_L31 (pfam01197)→ |
| Eco-1845-1 | NZ_APXX01000117.1 | + | 23667 | 23719 | **RNA**→ rpmE (PRK00019)Ribosomal_L31 (pfam01197)→ |
| Eco-1909-1 | NZ_AQAO01000051.1 | + | 46817 | 46869 | **RNA**→ rpmE (PRK00019)Ribosomal_L31 (pfam01197)→ |
| Eco-780-1 | NZ_AQEH01000121.1 | + | 88400 | 88452 | **RNA**→ rpmE (PRK00019)Ribosomal_L31 (pfam01197)→ |
| Eco-1909-2 | NZ_AQAO01000113.1 | - | 589 | 537 | **RNA**→ rpmE (PRK00019)Ribosomal_L31 (pfam01197)→ |
| Eco-1916-1 | NZ_AQAT01000072.1 | + | 87698 | 87750 | **RNA**→ rpmE (PRK00019)Ribosomal_L31 (pfam01197)→ |
| Eco-794-1 | NZ_AQEC01000081.1 | + | 18602 | 18654 | **RNA**→ rpmE (PRK00019)Ribosomal_L31 (pfam01197)→ |
| Eco-1846-1 | NZ_APXY01000411.1 | + | 23795 | 23847 | **RNA**→ rpmE (PRK00019)Ribosomal_L31 (pfam01197)→ |
| Eco-1917-1 | NZ_AQAU01000164.1 | + | 87687 | 87739 | **RNA**→ rpmE (PRK00019)Ribosomal_L31 (pfam01197)→ |
| Eco-1839-1 | NZ_APXT01000186.1 | + | 18324 | 18376 | **RNA**→ rpmE (PRK00019)Ribosomal_L31 (pfam01197)→ |
| Eco-779-1 | NZ_APXB01000350.1 | + | 58819 | 58871 | **RNA**→ rpmE (PRK00019)Ribosomal_L31 (pfam01197)→ |
| Eco-1663-1 | NZ_AQCV01000086.1 | + | 7832 | 7884 | **RNA**→ rpmE (PRK00019)Ribosomal_L31 (pfam01197)→ |
| Eco-1880-1 | NZ_APYY01000581.1 | + | 14000 | 14052 | **RNA**→ rpmE (PRK00019)Ribosomal_L31 (pfam01197)→ rpmE (PRK00019)Ribosomal_L31 (pfam01197)→ |
| Eco-1668-1 | NZ_AQCQ01000084.1 | + | 7879 | 7931 | **RNA**→ rpmE (PRK00019)Ribosomal_L31 (pfam01197)→ |
| Eco-821-1 | NZ_APXH01000145.1 | + | 23860 | 23912 | **RNA**→ rpmE (PRK00019)Ribosomal_L31 (pfam01197)→ |
| Eco-1864-1 | NZ_APYL01000074.1 | + | 87767 | 87819 | **RNA**→ rpmE (PRK00019)Ribosomal_L31 (pfam01197)→ |
| Eco-1853-1 | NZ_APYE01000072.1 | + | 78009 | 78061 | **RNA**→ rpmE (PRK00019)Ribosomal_L31 (pfam01197)→ rpmE (PRK00019)Ribosomal_L31 (pfam01197)→ |
| Eco-1815-1 | NZ_AKAZ01000268.1 | - | 584 | 532 | **RNA**→ rpmE (PRK00019)Ribosomal_L31 (pfam01197)→ |
| Eco-1095-1 | NZ_AIGJ01000066.1 | + | 92486 | 92538 | **RNA**→ rpmE (PRK00019)Ribosomal_L31 (pfam01197)→ |
| Eco-1835-1 | NZ_AQET01000058.1 | + | 61158 | 61210 | **RNA**→ rpmE (PRK00019)Ribosomal_L31 (pfam01197)→ |
| Eco-1059-1 | NZ_AIEX01000073.1 | + | 23889 | 23941 | **RNA**→ rpmE (PRK00019)Ribosomal_L31 (pfam01197)→ |
| Eco-1856-1 | NZ_APYH01000109.1 | + | 78061 | 78113 | **RNA**→ rpmE (PRK00019)Ribosomal_L31 (pfam01197)→ |
| Eco-1857-1 | NZ_APYI01000070.1 | + | 34181 | 34233 | **RNA**→ rpmE (PRK00019)Ribosomal_L31 (pfam01197)→ |
| Eco-1840-1 | NZ_APXU01000193.1 | + | 23762 | 23814 | **RNA**→ rpmE (PRK00019)Ribosomal_L31 (pfam01197)→ |
| Eco-1848-1 | NZ_APYA01000065.1 | + | 87764 | 87816 | **RNA**→ rpmE (PRK00019)Ribosomal_L31 (pfam01197)→ |
| Eco-1665-1 | NZ_AQCT01000137.1 | + | 7887 | 7939 | **RNA**→ rpmE (PRK00019)Ribosomal_L31 (pfam01197)→ |
| Eco-1669-1 | NZ_AQCP01000088.1 | + | 7856 | 7908 | **RNA**→ rpmE (PRK00019)Ribosomal_L31 (pfam01197)→ |
| Eco-1670-1 | NZ_AQCO01000090.1 | + | 7829 | 7881 | **RNA**→ rpmE (PRK00019)Ribosomal_L31 (pfam01197)→ |
| Eco-1078-1 | NZ_AIFQ01000032.1 | + | 93309 | 93361 | **RNA**→ rpmE (PRK00019)Ribosomal_L31 (pfam01197)→ |
| Eco-1843-1 | NZ_APZP01000199.1 | + | 23764 | 23816 | **RNA**→ rpmE (PRK00019)Ribosomal_L31 (pfam01197)→ rpmE (PRK00019)Ribosomal_L31 (pfam01197)→ |
| Eco-1863-1 | NZ_APYK01000175.1 | + | 87879 | 87931 | **RNA**→ rpmE (PRK00019)Ribosomal_L31 (pfam01197)→ |
| Eco-832-1 | NZ_AQDB01000114.1 | + | 87811 | 87863 | **RNA**→ rpmE (PRK00019)Ribosomal_L31 (pfam01197)→ |
| Eco-1024-1 | NZ_AQCX01000178.1 | + | 86186 | 86238 | **RNA**→ rpmE (PRK00019)Ribosomal_L31 (pfam01197)→ |
| Eco-1936-1 | NZ_APZY01000103.1 | + | 34233 | 34285 | **RNA**→ rpmE (PRK00019)Ribosomal_L31 (pfam01197)→ |
| Eco-830-1 | NZ_AQDD01000104.1 | + | 34455 | 34507 | **RNA**→ rpmE (PRK00019)Ribosomal_L31 (pfam01197)→ |
| Eco-1929-1 | NZ_AQBG01000072.1 | + | 87685 | 87737 | **RNA**→ rpmE (PRK00019)Ribosomal_L31 (pfam01197)→ |
| Eco-1769-1 | NZ_AJVS01000201.1 | + | 47721 | 47773 | **RNA**→ rpmE (PRK00019)Ribosomal_L31 (pfam01197)→ |
| Eco-1816-1 | NZ_AJVX01000290.1 | - | 585 | 533 | **RNA**→ rpmE (PRK00019)Ribosomal_L31 (pfam01197)→ |
| Eco-1771-1 | NZ_AKAY01000190.1 | + | 47720 | 47772 | **RNA**→ rpmE (PRK00019)Ribosomal_L31 (pfam01197)→ |
| Eco-1819-1 | NZ_AJVW01000268.1 | + | 47709 | 47761 | **RNA**→ rpmE (PRK00019)Ribosomal_L31 (pfam01197)→ |
| Eco-1080-1 | NZ_AIFS01000049.1 | + | 28249 | 28301 | **RNA**→ rpmE (PRK00019)Ribosomal_L31 (pfam01197)→ |
| Eco-1767-1 | NZ_AMXP01000360.1 | - | 492 | 440 | **RNA**→ rpmE (PRK00019)Ribosomal_L31 (pfam01197)→ |
| Eco-1862-1 | NZ_AQER01000021.1 | + | 90838 | 90890 | **RNA**→ ←hypo |
| Eco-1861-1 | NZ_APYJ01000184.1 | + | 87723 | 87775 | **RNA**→ ←hypo |
| env-861 | TrFG_contig04772 | + | 590 | 642 | **RNA**→ rpmE (PRK00019)Ribosomal_L31 (pfam01197)→ |
| Cko-1-2 | NZ_JTAV01000001.1 | - | 2039494 | 2039442 | **RNA**→ rpmE (PRK00019)Ribosomal_L31 (pfam01197)→ |
| Cko-1-1 | NZ_JUYC01000147.1 | - | 1421 | 1369 | **RNA**→ rpmE (PRK00019)Ribosomal_L31 (pfam01197)→ |
| Cko-2-1 | NC_009792.1 | - | 2839161 | 2839109 | **RNA**→ ←hypo |
| Csp-14-1 | NZ_DF830265.1 | + | 5041211 | 5041263 | **RNA**→ rpmE (PRK00019)Ribosomal_L31 (pfam01197)→ |
| Csp-8-1 | NZ_KQ089823.1 | + | 657952 | 658004 | **RNA**→ rpmE (PRK00019)Ribosomal_L31 (pfam01197)→ |
| env-862 | HBC_scaff_10316882 | - | 972 | 920 | **RNA**→ rpmE (PRK00019)Ribosomal_L31 (pfam01197)→ |

| | | | | | |
|---|---|---|---|---|---|
| env-863 | HBC_ctgs_1033478 | - | 972 | 920 | **RNA**→ rpmE (PRK00019)Ribosomal_L31 (pfam01197)→ |
| env-864 | JGI25152J39213_1000244 | - | 11208 | 11156 | **RNA**→ rpmE (PRK00019)Ribosomal_L31 (pfam01197)→ PRK14515 (PRK14515)→ |
| env-865 | JGI26052J44787_1048675 | + | 102 | 154 | **RNA**→ rpmE (PRK00019)Ribosomal_L31 (pfam01197)→ |
| env-866 | JGI26055J44792_1064428 | + | 109 | 161 | **RNA**→ rpmE (PRK00019)Ribosomal_L31 (pfam01197)→ |
| Ecl-59-1 | NZ_AXOM01000028.1 | + | 73061 | 73113 | **RNA**→ ←hypo |
| Cam-2-1 | NZ_CP011132.1 | + | 4667907 | 4667959 | **RNA**→ rpmE (PRK00019)Ribosomal_L31 (pfam01197)→ |
| env-867 | GooDraft_1000552 | - | 4465 | 4413 | **RNA**→ rpmE (PRK00019)Ribosomal_L31 (pfam01197)→ |
| Cse-1-1 | NZ_BBNB01000017.1 | + | 87854 | 87906 | **RNA**→ rpmE (PRK00019)Ribosomal_L31 (pfam01197)→ |
| env-868 | RicEn_C1233 | - | 705 | 653 | **RNA**→ ←PRK05264 (PRK05264)MetJ (COG3060) |
| Esp-43-1 | NZ_JXAF01000019.1 | - | 42097 | 42045 | **RNA**→ rpmE (PRK00019)Ribosomal_L31 (pfam01197)→ |
| Kor-1-1 | NZ_JZLI01000016.1 | - | 25625 | 25573 | **RNA**→ rpmE (PRK00019)Ribosomal_L31 (pfam01197)→ |
| Esp-72-1 | NC_021500.1 | + | 3129679 | 3129731 | **RNA**→ rpmE (PRK00019)Ribosomal_L31 (pfam01197)→ |
| Ksa-1-1 | NZ_CP007215.1 | - | 2992737 | 2992685 | **RNA**→ rpmE (PRK00019)Ribosomal_L31 (pfam01197)→ |
| Mma-2-1 | NZ_HE578950.1 | + | 83372 | 83424 | **RNA**→ rpmE (PRK00019)Ribosomal_L31 (pfam01197)→ |
| Pur-1-1 | NZ_CP011602.1 | - | 4453563 | 4453511 | **RNA**→ rpmE (PRK00019)Ribosomal_L31 (pfam01197)→ |
| Cfa-1-1 | NZ_BBMX01000021.1 | + | 105611 | 105663 | **RNA**→ rpmE (PRK00019)Ribosomal_L31 (pfam01197)→ |
| env-869 | GO4KUYH01D3UFU | + | 163 | 215 | **RNA**→ ←hypo |
| env-870 | JGI26054J46599_1125534 | + | 31 | 83 | **RNA**→ ←hypo |
| Kcf-1-1 | NZ_BADH01000654.1 | + | 6305 | 6357 | **RNA**→ ←hypo |
| env-871 | AECF_contig04828 | - | 82 | 30 | **RNA**→ |
| Esp-42-1 | NZ_JXAG01000025.1 | - | 24558 | 24506 | **RNA**→ rpmE (PRK00019)Ribosomal_L31 (pfam01197)→ |
| Era-1-1 | NZ_JH725435.1 | - | 117259 | 117207 | **RNA**→ rpmE (PRK00019)Ribosomal_L31 (pfam01197)→ hypo→ |
| Kra-1-1 | NZ_JDYJ01000019.1 | + | 81305 | 81357 | **RNA**→ rpmE (PRK00019)Ribosomal_L31 (pfam01197)→ |
| Kra-2-1 | NZ_JSFC01000001.1 | - | 4090078 | 4090026 | **RNA**→ rpmE (PRK00019)Ribosomal_L31 (pfam01197)→ |
| Csp-12-1 | NZ_JAQB01000052.1 | + | 46953 | 47005 | **RNA**→ rpmE (PRK00019)Ribosomal_L31 (pfam01197)→ |
| Cam-1-2 | NZ_JMQQ01000019.1 | + | 108204 | 108256 | **RNA**→ rpmE (PRK00019)Ribosomal_L31 (pfam01197)→ |
| Cam-1-3 | NZ_CDQV01000565.1 | - | 351 | 299 | **RNA**→ rpmE (PRK00019)Ribosomal_L31 (pfam01197)→ |
| Csp-13-1 | NZ_KK736183.1 | + | 1511889 | 1511941 | **RNA**→ rpmE (PRK00019)Ribosomal_L31 (pfam01197)→ |
| Ecl-60-1 | NC_014618.1 | - | 4692425 | 4692373 | **RNA**→ rpmE (PRK00019)Ribosomal_L31 (pfam01197)→ |
| env-872 | ACOFG988_contig03604 | - | 434 | 382 | **RNA**→ rpmE (PRK00019)Ribosomal_L31 (pfam01197)→ |
| env-873 | CLOF_contig01917 | - | 175 | 123 | **RNA**→ rpmE (PRK00019)Ribosomal_L31 (pfam01197)→ |
| env-874 | DPOLF_F46JATG01CXPZU | + | 132 | 184 | **RNA**→ hypo→ |
| env-875 | TrFG_GO06Z1101D3LWN | - | 296 | 244 | **RNA**→ ←hypo |
| Pvu-1-1 | NZ_BBMZ01000025.1 | + | 73082 | 73133 | **RNA**→ rpmE (PRK00019)Ribosomal_L31 (pfam01197)→ |
| Gpe-1-1 | NZ_LFWC01000015.1 | + | 70062 | 70109 | **RNA**→ ←PriA (COG1198)PRK05580 (PRK05580) |
| Gxi-1-1 | NZ_AMRI01000003.1 | + | 50253 | 50300 | **RNA**→ ←PriA (COG1198)PRK05580 (PRK05580) |
| env-876 | CEPH01076795.1 | - | 934 | 882 | **RNA**→ rpmE (PRK00019)Ribosomal_L31 (pfam01197)→ |
| env-877 | CEUT01138684.1 | + | 360 | 412 | **RNA**→ rpmE (PRK00019)Ribosomal_L31 (pfam01197)→ |
| env-878 | CEUX01102889.1 | + | 2080 | 2132 | **RNA**→ rpmE (PRK00019)Ribosomal_L31 (pfam01197)→ |
| env-879 | CEVB01010953.1 | + | 2587 | 2639 | **RNA**→ rpmE (PRK00019)Ribosomal_L31 (pfam01197)→ |
| env-880 | 3300000053_316318 | - | 326 | 276 | **RNA**→ rpmE (PRK00019)Ribosomal_L31 (pfam01197)→ |
| Pme-1-1 | NZ_LAVS01000089.1 | - | 2227 | 2177 | **RNA**→ rpmE (PRK00019)Ribosomal_L31 (pfam01197)→ rpmE (PRK00019)Ribosomal_L31 (pfam01197)→ |
| Rsp-2-1 | NZ_LAMX01000027.1 | + | 22231 | 22281 | **RNA**→ rpmE (PRK00019)Ribosomal_L31 (pfam01197)→ rpmE (PRK00019)Ribosomal_L31 (pfam01197)→ |
| env-881 | JCVI_SCAF_1096627203505 | + | 760 | 810 | **RNA**→ rpmE (PRK00019)Ribosomal_L31 (pfam01197)→ |
| Rsp-1-1 | NZ_AFHI01000019.1 | + | 59978 | 60028 | **RNA**→ rpmE (PRK00019)Ribosomal_L31 (pfam01197)→ rpmE (PRK00019)Ribosomal_L31 (pfam01197)→ rpmE (PRK00019)Ribosomal_L31 (pfam01197)→ hypo→ |
| Aag-1-1 | NZ_AKKU01000026.1 | + | 135962 | 136016 | **RNA**→ rpmE (PRK00019)Ribosomal_L31 (pfam01197)→ rpmE (PRK00019)Ribosomal_L31 (pfam01197)→ rpmE (PRK00019)Ribosomal_L31 (pfam01197)→ |
| Aae-1-1 | NZ_ALAB01000027.1 | - | 258817 | 258762 | **RNA**→ rpmE (PRK00019)Ribosomal_L31 (pfam01197)→ rpmE (PRK00019)Ribosomal_L31 (pfam01197)→ rpmE (PRK00019)Ribosomal_L31 (pfam01197)→ |
| env-882 | 3300000053_006648 | + | 1670 | 1725 | **RNA**→ rpmE (PRK00019)Ribosomal_L31 (pfam01197)→ |
| env-883 | 3300000052_128519 | - | 369 | 314 | **RNA**→ rpmE (PRK00019)Ribosomal_L31 (pfam01197)→ |

| | | | | | |
|---|---|---|---|---|---|
| Aje-1-1 | NZ_AHTH01000040.1 | + | 47811 | 47866 | **RNA**→ rpmE (PRK00019)Ribosomal_L31 (pfam01197)→ rpmE (PRK00019)Ribosomal_L31 (pfam01197)→ rpmE (PRK00019)Ribosomal_L31 (pfam01197)→ |
| Aik-1-1 | NZ_LAHO01000004.1 | + | 191768 | 191818 | **RNA**→ rpmE (PRK00019)Ribosomal_L31 (pfam01197)→ rpmE (PRK00019)Ribosomal_L31 (pfam01197)→ |
| env-884 | CEWF01130100.1 | - | 390 | 340 | **RNA**→ rpmE (PRK00019)Ribosomal_L31 (pfam01197)→ |
| env-885 | CESO01326287.1 | + | 25250 | 25300 | **RNA**→ rpmE (PRK00019)Ribosomal_L31 (pfam01197)→ |
| Asp-9-1 | NZ_LAHP01000005.1 | + | 145176 | 145226 | **RNA**→ rpmE (PRK00019)Ribosomal_L31 (pfam01197)→ rpmE (PRK00019)Ribosomal_L31 (pfam01197)→ |
| Rpe-1-1 | NZ_KB907708.1 | - | 105042 | 104992 | **RNA**→ rpmE (PRK00019)Ribosomal_L31 (pfam01197)→ |
| Rba-1-1 | NZ_AUDG01000008.1 | + | 95570 | 95622 | **RNA**→ rpmE (PRK00019)Ribosomal_L31 (pfam01197)→ |
| Rna-1-1 | NZ_BAFK01000001.1 | + | 221629 | 221679 | **RNA**→ rpmE (PRK00019)Ribosomal_L31 (pfam01197)→ rpmE (PRK00019)Ribosomal_L31 (pfam01197)→ rpmE (PRK00019)Ribosomal_L31 (pfam01197)→ |
| env-886 | P_2C_Liq_3_UnCty_100070 | + | 17818 | 17869 | **RNA**→ rpmE (PRK00019)Ribosomal_L31 (pfam01197)→ |
| env-887 | P_2C_Liq_1_UnCty_1000092 | + | 12358 | 12409 | **RNA**→ rpmE (PRK00019)Ribosomal_L31 (pfam01197)→ |
| env-888 | P_2C_Liq_2_UnCty_1000391 | - | 4101 | 4050 | **RNA**→ rpmE (PRK00019)Ribosomal_L31 (pfam01197)→ |
| Asa-9-1 | NZ_AUBH01000006.1 | + | 241719 | 241770 | **RNA**→ rpmE (PRK00019)Ribosomal_L31 (pfam01197)→ |
| Gli-1-1 | NZ_BAEN01000076.1 | - | 267505 | 267455 | **RNA**→ rpmE (PRK00019)Ribosomal_L31 (pfam01197)→ rpmE (PRK00019)Ribosomal_L31 (pfam01197)→ hypo→ hypo→ |
| Gpu-1-1 | NZ_BAET01000031.1 | + | 85003 | 85056 | **RNA**→ rpmE (PRK00019)Ribosomal_L31 (pfam01197)→ rpmE (PRK00019)Ribosomal_L31 (pfam01197)→ |
| env-889 | P_2C_Liq_1_UnCty_1151257 | + | 164 | 215 | **RNA**→ rpmE (PRK00019)Ribosomal_L31 (pfam01197)→ |
| Gsp-2-1 | NZ_DS989813.1 | - | 29050 | 28999 | **RNA**→ rpmE (PRK00019)Ribosomal_L31 (pfam01197)→ |
| env-890 | JGI20152J14361_10002432 | - | 5019 | 4967 | **RNA**→ rpmE (PRK00019)Ribosomal_L31 (pfam01197)→ |
| env-891 | JGI20151J14362_10056971 | + | 509 | 561 | **RNA**→ rpmE (PRK00019)Ribosomal_L31 (pfam01197)→ |
| env-892 | JGI20153J14295_10036242 | - | 781 | 729 | **RNA**→ rpmE (PRK00019)Ribosomal_L31 (pfam01197)→ |
| env-893 | JGI20153J14318_10036242 | - | 781 | 729 | **RNA**→ rpmE (PRK00019)Ribosomal_L31 (pfam01197)→ |
| Gni-1-1 | NC_016041.1 | + | 3221564 | 3221615 | **RNA**→ rpmE (PRK00019)Ribosomal_L31 (pfam01197)→ |
| Gpa-1-1 | NZ_KE386814.1 | - | 95601 | 95549 | **RNA**→ rpmE (PRK00019)Ribosomal_L31 (pfam01197)→ |
| env-894 | _GX4VOVV01EZ7Q7 | - | 122 | 71 | **RNA**→ hypo→ |
| Gpo-1-1 | NZ_BAER01000045.1 | + | 100823 | 100873 | **RNA**→ rpmE (PRK00019)Ribosomal_L31 (pfam01197)→ rpmE (PRK00019)Ribosomal_L31 (pfam01197)→ rpmE (PRK00019)Ribosomal_L31 (pfam01197)→ rpmE (PRK00019)Ribosomal_L31 (pfam01197)→ |
| env-895 | JGI20157J14317_10084041 | - | 798 | 748 | **RNA**→ rpmE (PRK00019)Ribosomal_L31 (pfam01197)→ |
| Gar-1-1 | NZ_BAEO01000065.1 | - | 149160 | 149110 | **RNA**→ rpmE (PRK00019)Ribosomal_L31 (pfam01197)→ rpmE (PRK00019)Ribosomal_L31 (pfam01197)→ |
| env-896 | JGI20153J14295_10040431 | - | 589 | 539 | **RNA**→ rpmE (PRK00019)Ribosomal_L31 (pfam01197)→ |
| env-897 | JGI20153J14318_10040431 | - | 589 | 539 | **RNA**→ rpmE (PRK00019)Ribosomal_L31 (pfam01197)→ |
| Gps-1-1 | NC_020514.1 | + | 218373 | 218423 | **RNA**→ ←hypo |
| env-898 | CEWJ01088929.1 | - | 6335 | 6285 | **RNA**→ rpmE (PRK00019)Ribosomal_L31 (pfam01197)→ |
| env-899 | CERD01001178.1 | + | 488 | 538 | **RNA**→ rpmE (PRK00019)Ribosomal_L31 (pfam01197)→ |
| env-900 | JGI12097J13213_1000038 | - | 8918 | 8868 | **RNA**→ rpmE (PRK00019)Ribosomal_L31 (pfam01197)→ |
| env-901 | CEUB01117245.1 | + | 201 | 251 | **RNA**→ rpmE (PRK00019)Ribosomal_L31 (pfam01197)→ |
| env-902 | CENJ01188181.1 | + | 311 | 361 | **RNA**→ rpmE (PRK00019)Ribosomal_L31 (pfam01197)→ |
| env-903 | CEQI01175305.1 | - | 182 | 132 | **RNA**→ rpmE (PRK00019)Ribosomal_L31 (pfam01197)→ |
| env-904 | CEQJ01016256.1 | - | 1813 | 1763 | **RNA**→ rpmE (PRK00019)Ribosomal_L31 (pfam01197)→ |
| env-905 | CEQQ01064592.1 | + | 268 | 318 | **RNA**→ rpmE (PRK00019)Ribosomal_L31 (pfam01197)→ |
| env-906 | CEQU01072249.1 | + | 4389 | 4439 | **RNA**→ rpmE (PRK00019)Ribosomal_L31 (pfam01197)→ |
| env-907 | CESF01168137.1 | + | 311 | 361 | **RNA**→ rpmE (PRK00019)Ribosomal_L31 (pfam01197)→ |
| env-908 | CESJ01134068.1 | + | 721 | 771 | **RNA**→ rpmE (PRK00019)Ribosomal_L31 (pfam01197)→ |
| env-909 | CESL01022392.1 | + | 8352 | 8402 | **RNA**→ rpmE (PRK00019)Ribosomal_L31 (pfam01197)→ |
| env-910 | CESM01050812.1 | + | 387 | 437 | **RNA**→ rpmE (PRK00019)Ribosomal_L31 (pfam01197)→ |
| env-911 | CESR01320609.1 | - | 725 | 675 | **RNA**→ rpmE (PRK00019)Ribosomal_L31 (pfam01197)→ |
| env-912 | CETH01013924.1 | + | 3079 | 3129 | **RNA**→ rpmE (PRK00019)Ribosomal_L31 (pfam01197)→ |
| env-913 | CETJ01046849.1 | + | 3068 | 3118 | **RNA**→ rpmE (PRK00019)Ribosomal_L31 (pfam01197)→ |
| env-914 | CETO01002955.1 | + | 1403 | 1453 | **RNA**→ rpmE (PRK00019)Ribosomal_L31 (pfam01197)→ |
| env-915 | CETQ01029354.1 | - | 3342 | 3292 | **RNA**→ rpmE (PRK00019)Ribosomal_L31 (pfam01197)→ |
| env-916 | CETS01286873.1 | - | 271 | 221 | **RNA**→ rpmE (PRK00019)Ribosomal_L31 (pfam01197)→ |
| env-917 | CETT01166069.1 | + | 280 | 330 | **RNA**→ rpmE (PRK00019)Ribosomal_L31 (pfam01197)→ |

| | | | | | |
|---|---|---|---|---|---|
| env-918 | CETZ01011127.1 | - | 189 | 139 | **RNA**→ rpmE (PRK00019)Ribosomal_L31 (pfam01197)→ |
| env-919 | CEUC01278456.1 | - | 212 | 162 | **RNA**→ rpmE (PRK00019)Ribosomal_L31 (pfam01197)→ |
| env-920 | CEUL01103588.1 | - | 129 | 79 | **RNA**→ rpmE (PRK00528)rpmE (PRK00019)Ribosomal_L31 (pfam01197)→ |
| env-921 | CEUM01053021.1 | - | 9973 | 9923 | **RNA**→ rpmE (PRK00019)Ribosomal_L31 (pfam01197)→ |
| env-922 | CEUQ01106239.1 | + | 1010 | 1060 | **RNA**→ rpmE (PRK00019)Ribosomal_L31 (pfam01197)→ |
| env-923 | CEUU01007342.1 | + | 10738 | 10788 | **RNA**→ rpmE (PRK00019)Ribosomal_L31 (pfam01197)→ |
| env-924 | CEUW01253280.1 | + | 3470 | 3520 | **RNA**→ rpmE (PRK00019)Ribosomal_L31 (pfam01197)→ |
| env-925 | CEUX01014635.1 | + | 306 | 356 | **RNA**→ rpmE (PRK00019)Ribosomal_L31 (pfam01197)→ |
| env-926 | CEUY01025164.1 | - | 498 | 448 | **RNA**→ rpmE (PRK00019)Ribosomal_L31 (pfam01197)→ |
| env-927 | CEVF01153062.1 | - | 1235 | 1185 | **RNA**→ rpmE (PRK00019)Ribosomal_L31 (pfam01197)→ |
| env-928 | CEWF01048426.1 | + | 795 | 845 | **RNA**→ rpmE (PRK00019)Ribosomal_L31 (pfam01197)→ |
| env-929 | CEWI01078971.1 | + | 376 | 426 | **RNA**→ rpmE (PRK00019)Ribosomal_L31 (pfam01197)→ |
| env-930 | CEQS01114527.1 | + | 267 | 317 | **RNA**→ rpmE (PRK00019)Ribosomal_L31 (pfam01197)→ |
| env-931 | CERB01320821.1 | - | 434 | 384 | **RNA**→ rpmE (PRK00019)Ribosomal_L31 (pfam01197)→ |
| env-932 | CESW01183859.1 | - | 333 | 283 | **RNA**→ rpmE (PRK00019)Ribosomal_L31 (pfam01197)→ |
| env-933 | CENT01094743.1 | - | 7856 | 7806 | **RNA**→ rpmE (PRK00019)Ribosomal_L31 (pfam01197)→ |
| env-934 | CETX01054649.1 | + | 40 | 90 | **RNA**→ hypo→ rpmE (PRK00019)Ribosomal_L31 (pfam01197)→ |
| Pat-1-1 | NC_008228.1 | - | 5045228 | 5045178 | **RNA**→ rpmE (PRK00019)Ribosomal_L31 (pfam01197)→ |
| Gsp-1-1 | NC_015497.1 | - | 4815194 | 4815144 | **RNA**→ rpmE (PRK00019)Ribosomal_L31 (pfam01197)→ rpmE (PRK00019)Ribosomal_L31 (pfam01197)→ |
| Gme-1-1 | NZ_BAEP01000006.1 | + | 5181 | 5231 | **RNA**→ ←hypo |
| Gch-1-1 | NZ_BAEM01000019.1 | - | 23453 | 23403 | **RNA**→ rpmE (PRK00019)Ribosomal_L31 (pfam01197)→ rpmE (PRK00019)Ribosomal_L31 (pfam01197)→ |
| Psp-18-1 | NZ_JRKG01000003.1 | - | 204430 | 204380 | **RNA**→ rpmE (PRK00019)Ribosomal_L31 (pfam01197)→ rpmE (PRK00019)Ribosomal_L31 (pfam01197)→ |
| Gag-1-1 | NZ_BAEK01000035.1 | - | 25147 | 25097 | **RNA**→ rpmE (PRK00019)Ribosomal_L31 (pfam01197)→ rpmE (PRK00019)Ribosomal_L31 (pfam01197)→ |
| env-935 | CENT01097505.1 | - | 7154 | 7104 | **RNA**→ rpmE (PRK00019)Ribosomal_L31 (pfam01197)→ |
| Asp-7-1 | NZ_JQHG01000007.1 | + | 134541 | 134591 | **RNA**→ rpmE (PRK00019)Ribosomal_L31 (pfam01197)→ |
| Asp-6-1 | NZ_JQFW01000050.1 | + | 134541 | 134591 | **RNA**→ rpmE (PRK00019)Ribosomal_L31 (pfam01197)→ |
| env-936 | CEPZ01032191.1 | - | 667 | 617 | **RNA**→ rpmE (PRK00019)Ribosomal_L31 (pfam01197)→ |
| env-937 | CEPV01037448.1 | + | 598 | 648 | **RNA**→ rpmE (PRK00019)Ribosomal_L31 (pfam01197)→ |
| env-938 | CEQH01105122.1 | + | 558 | 608 | **RNA**→ |
| env-939 | CERD01127562.1 | - | 11742 | 11692 | **RNA**→ rpmE (PRK00019)Ribosomal_L31 (pfam01197)→ |
| env-940 | CERQ01120834.1 | + | 1165 | 1215 | **RNA**→ rpmE (PRK00019)Ribosomal_L31 (pfam01197)→ |
| env-941 | CESC01037646.1 | - | 1488 | 1438 | **RNA**→ rpmE (PRK00019)Ribosomal_L31 (pfam01197)→ |
| env-942 | CESQ01222567.1 | + | 1419 | 1469 | **RNA**→ rpmE (PRK00019)Ribosomal_L31 (pfam01197)→ |
| env-943 | CETH01076277.1 | - | 632 | 582 | **RNA**→ rpmE (PRK00019)Ribosomal_L31 (pfam01197)→ |
| env-944 | CETO01078733.1 | + | 15481 | 15531 | **RNA**→ rpmE (PRK00019)Ribosomal_L31 (pfam01197)→ |
| env-945 | CETZ01121052.1 | + | 1200 | 1250 | **RNA**→ rpmE (PRK00019)Ribosomal_L31 (pfam01197)→ |
| env-946 | CEUK01183503.1 | + | 5639 | 5689 | **RNA**→ rpmE (PRK00019)Ribosomal_L31 (pfam01197)→ |
| env-947 | CEUL01119818.1 | + | 637 | 687 | **RNA**→ rpmE (PRK00019)Ribosomal_L31 (pfam01197)→ |
| env-948 | CEUN01029593.1 | + | 2483 | 2533 | **RNA**→ rpmE (PRK00019)Ribosomal_L31 (pfam01197)→ |
| env-949 | CEUP01063689.1 | + | 379 | 429 | **RNA**→ rpmE (PRK00019)Ribosomal_L31 (pfam01197)→ |
| env-950 | CEUQ01293793.1 | - | 193 | 143 | **RNA**→ rpmE (PRK00019)Ribosomal_L31 (pfam01197)→ |
| env-951 | CEUS01268477.1 | + | 379 | 429 | **RNA**→ rpmE (PRK00019)Ribosomal_L31 (pfam01197)→ |
| env-952 | CEUY01052883.1 | + | 4169 | 4219 | **RNA**→ rpmE (PRK00019)Ribosomal_L31 (pfam01197)→ |
| env-953 | CEVA01307699.1 | - | 183 | 133 | **RNA**→ rpmE (PRK00019)Ribosomal_L31 (pfam01197)→ |
| Sch-1-1 | NZ_KB899393.1 | - | 40924 | 40874 | **RNA**→ rpmE (PRK00019)Ribosomal_L31 (pfam01197)→ |
| env-954 | CETX01177667.1 | + | 221 | 272 | **RNA**→ rpmE (PRK00019)Ribosomal_L31 (pfam01197)→ |
| env-955 | JGI12097J13213_1001517 | - | 1180 | 1129 | **RNA**→ rpmE (PRK00019)Ribosomal_L31 (pfam01197)→ |
| Asp-5-1 | NZ_JFBK01000001.1 | - | 4465586 | 4465534 | **RNA**→ rpmE (PRK00019)Ribosomal_L31 (pfam01197)→ |
| env-956 | CESM01022288.1 | + | 175 | 227 | **RNA**→ rpmE (PRK00019)Ribosomal_L31 (pfam01197)→ |
| Ama-14-1 | NZ_JWLW01000010.1 | + | 95215 | 95267 | **RNA**→ rpmE (PRK00019)Ribosomal_L31 (pfam01197)→ hypo→ |
| env-957 | JGI12120J13076_102920 | - | 400 | 348 | **RNA**→ rpmE (PRK00019)Ribosomal_L31 (pfam01197)→ |
| env-958 | JGI12020J13220_1000023 | + | 611 | 663 | **RNA**→ rpmE (PRK00019)Ribosomal_L31 (pfam01197)→ |

| | | | | | |
|---|---|---|---|---|---|
| env-959 | JGI11800J13271_1000309 | + | 1785 | 1837 | **RNA**→ rpmE (PRK00019)Ribosomal_L31 (pfam01197)→ |
| env-960 | CENO01130258.1 | + | 217 | 269 | **RNA**→ rpmE (PRK00019)Ribosomal_L31 (pfam01197)→ |
| env-961 | CENX01098222.1 | + | 441 | 493 | **RNA**→ rpmE (PRK00019)Ribosomal_L31 (pfam01197)→ |
| env-962 | CEOC01158531.1 | + | 359 | 411 | **RNA**→ rpmE (PRK00019)Ribosomal_L31 (pfam01197)→ |
| env-963 | CEPA01172521.1 | - | 2899 | 2847 | **RNA**→ rpmE (PRK00019)Ribosomal_L31 (pfam01197)→ |
| env-964 | CEPH01005174.1 | - | 133 | 81 | **RNA**→ |
| env-965 | CEPK01169822.1 | - | 3662 | 3610 | **RNA**→ rpmE (PRK00019)Ribosomal_L31 (pfam01197)→ |
| env-966 | CEPU01151402.1 | + | 251 | 303 | **RNA**→ rpmE (PRK00019)Ribosomal_L31 (pfam01197)→ |
| env-967 | CEPV01200560.1 | + | 250 | 302 | **RNA**→ rpmE (PRK00019)Ribosomal_L31 (pfam01197)→ |
| env-968 | CEPW01224637.1 | - | 461 | 409 | **RNA**→ rpmE (PRK00019)Ribosomal_L31 (pfam01197)→ |
| env-969 | CEPY01054066.1 | + | 243 | 295 | **RNA**→ rpmE (PRK00019)Ribosomal_L31 (pfam01197)→ |
| env-970 | CERS01171998.1 | + | 621 | 673 | **RNA**→ rpmE (PRK00019)Ribosomal_L31 (pfam01197)→ |
| env-971 | CETB01056501.1 | - | 13247 | 13195 | **RNA**→ rpmE (PRK00019)Ribosomal_L31 (pfam01197)→ |
| env-972 | CETI01039859.1 | - | 2772 | 2720 | **RNA**→ rpmE (PRK00019)Ribosomal_L31 (pfam01197)→ |
| env-973 | CETR01033810.1 | - | 294 | 242 | **RNA**→ rpmE (PRK00019)Ribosomal_L31 (pfam01197)→ |
| env-974 | CEVK01195631.1 | - | 373 | 321 | **RNA**→ rpmE (PRK00019)Ribosomal_L31 (pfam01197)→ |
| env-975 | CEVM01241445.1 | + | 721 | 773 | **RNA**→ ←hypo |
| env-976 | CEVQ01117480.1 | + | 28172 | 28224 | **RNA**→ rpmE (PRK00019)Ribosomal_L31 (pfam01197)→ |
| env-977 | CEVZ01044906.1 | - | 3231 | 3179 | **RNA**→ rpmE (PRK00019)Ribosomal_L31 (pfam01197)→ |
| env-978 | CEWI01013184.1 | - | 2889 | 2837 | **RNA**→ rpmE (PRK00019)Ribosomal_L31 (pfam01197)→ |
| env-979 | CEWJ01032880.1 | + | 1770 | 1822 | **RNA**→ rpmE (PRK00019)Ribosomal_L31 (pfam01197)→ |
| env-980 | Mariner_1003560 | + | 602 | 654 | **RNA**→ rpmE (PRK00019)Ribosomal_L31 (pfam01197)→ |
| env-981 | TahiMoana_1000279 | + | 110727 | 110779 | **RNA**→ rpmE (PRK00019)Ribosomal_L31 (pfam01197)→ |
| env-982 | CEQG01016892.1 | - | 90 | 38 | **RNA**→ |
| env-983 | CEQH01004879.1 | - | 232 | 180 | **RNA**→ rpmE (PRK00019)Ribosomal_L31 (pfam01197)→ |
| env-984 | CEQM01102687.1 | - | 3030 | 2978 | **RNA**→ rpmE (PRK00019)Ribosomal_L31 (pfam01197)→ |
| env-985 | CESC01010137.1 | - | 90 | 38 | **RNA**→ |
| env-986 | CERG01114847.1 | + | 251 | 303 | **RNA**→ rpmE (PRK00019)Ribosomal_L31 (pfam01197)→ |
| env-987 | CERR01251468.1 | + | 128 | 180 | **RNA**→ rpmE (PRK00019)Ribosomal_L31 (pfam01197)→ |
| env-988 | CERV01054361.1 | - | 2998 | 2946 | **RNA**→ rpmE (PRK00019)Ribosomal_L31 (pfam01197)→ |
| env-989 | CESI01008217.1 | + | 5871 | 5923 | **RNA**→ rpmE (PRK00019)Ribosomal_L31 (pfam01197)→ |
| env-990 | CETE01023379.1 | + | 39511 | 39563 | **RNA**→ rpmE (PRK00019)Ribosomal_L31 (pfam01197)→ |
| env-991 | CETQ01016815.1 | + | 345 | 397 | **RNA**→ rpmE (PRK00019)Ribosomal_L31 (pfam01197)→ |
| env-992 | JGI11881J13070_1018350 | - | 364 | 312 | **RNA**→ rpmE (PRK00019)Ribosomal_L31 (pfam01197)→ |
| env-993 | JGI12025J13092_100507 | + | 516 | 568 | **RNA**→ rpmE (PRK00019)Ribosomal_L31 (pfam01197)→ |
| env-994 | JGI12107J13081_100823 | + | 1838 | 1890 | **RNA**→ rpmE (PRK00019)Ribosomal_L31 (pfam01197)→ |
| env-995 | JGI11755J13075_102346 | - | 67 | 15 | **RNA**→ |
| env-996 | JGI11765J13083_105572 | + | 198 | 250 | **RNA**→ rpmE (PRK00019)Ribosomal_L31 (pfam01197)→ |
| env-997 | JGI11805J13103_105660 | + | 220 | 272 | **RNA**→ rpmE (PRK00019)Ribosomal_L31 (pfam01197)→ |
| env-998 | JGI11754J13217_1010636 | + | 234 | 286 | **RNA**→ rpmE (PRK00019)Ribosomal_L31 (pfam01197)→ |
| env-999 | JGI12207J13218_1017989 | + | 147 | 199 | **RNA**→ rpmE (PRK00019)Ribosomal_L31 (pfam01197)→ |
| env-1000 | JGI12218J13221_1001289 | - | 3331 | 3279 | **RNA**→ rpmE (PRK00019)Ribosomal_L31 (pfam01197)→ |
| env-1001 | JGI11978J13257_1000948 | - | 2287 | 2235 | **RNA**→ rpmE (PRK00019)Ribosomal_L31 (pfam01197)→ |
| env-1002 | JGI12148J13107_102525 | + | 1320 | 1372 | **RNA**→ rpmE (PRK00019)Ribosomal_L31 (pfam01197)→ |
| env-1003 | JGI11949J13268_1023255 | + | 99 | 151 | **RNA**→ rpmE (PRK00019)Ribosomal_L31 (pfam01197)→ |
| env-1004 | JGI11931J13222_1000347 | + | 5042 | 5094 | **RNA**→ rpmE (PRK00019)Ribosomal_L31 (pfam01197)→ |
| env-1005 | JGI12156J13080_107206 | + | 180 | 232 | **RNA**→ rpmE (PRK00019)Ribosomal_L31 (pfam01197)→ |
| env-1006 | CEOW01017245.1 | - | 1797 | 1745 | **RNA**→ rpmE (PRK00019)Ribosomal_L31 (pfam01197)→ |
| env-1007 | CEPC01258783.1 | - | 681 | 629 | **RNA**→ rpmE (PRK00019)Ribosomal_L31 (pfam01197)→ |
| env-1008 | CEPD01089067.1 | + | 250 | 302 | **RNA**→ rpmE (PRK00019)Ribosomal_L31 (pfam01197)→ |
| env-1009 | CEPM01022322.1 | - | 136 | 84 | **RNA**→ |
| env-1010 | CEQL01158694.1 | + | 267 | 319 | **RNA**→ rpmE (PRK00019)Ribosomal_L31 (pfam01197)→ |

| | | | | | |
|---|---|---|---|---|---|
| env-1011 | CEQP01301609.1 | - | 558 | 506 | **RNA**→ rpmE (PRK00019)Ribosomal_L31 (pfam01197)→ |
| env-1012 | CEQX01011574.1 | + | 291 | 343 | **RNA**→ rpmE (PRK00019)Ribosomal_L31 (pfam01197)→ |
| env-1013 | CERL01235058.1 | - | 936 | 884 | **RNA**→ rpmE (PRK00019)Ribosomal_L31 (pfam01197)→ |
| env-1014 | CESG01011657.1 | + | 212 | 264 | **RNA**→ rpmE (PRK00019)Ribosomal_L31 (pfam01197)→ |
| env-1015 | CESR01104947.1 | + | 324 | 376 | **RNA**→ rpmE (PRK00019)Ribosomal_L31 (pfam01197)→ |
| env-1016 | CESX01007994.1 | - | 514 | 462 | **RNA**→ rpmE (PRK00019)Ribosomal_L31 (pfam01197)→ |
| env-1017 | CESZ01006041.1 | - | 1040 | 988 | **RNA**→ rpmE (PRK00019)Ribosomal_L31 (pfam01197)→ |
| env-1018 | CETH01095267.1 | - | 295 | 243 | **RNA**→ rpmE (PRK00019)Ribosomal_L31 (pfam01197)→ |
| env-1019 | CETM01005225.1 | - | 232 | 180 | **RNA**→ rpmE (PRK00019)Ribosomal_L31 (pfam01197)→ |
| env-1020 | CETV01018356.1 | - | 1037 | 985 | **RNA**→ rpmE (PRK00019)Ribosomal_L31 (pfam01197)→ |
| env-1021 | CEUC01310578.1 | + | 502 | 554 | **RNA**→ rpmE (PRK00019)Ribosomal_L31 (pfam01197)→ |
| env-1022 | CEUK01103776.1 | - | 854 | 802 | **RNA**→ rpmE (PRK00019)Ribosomal_L31 (pfam01197)→ |
| env-1023 | CEUS01001549.1 | + | 1749 | 1801 | **RNA**→ rpmE (PRK00019)Ribosomal_L31 (pfam01197)→ |
| env-1024 | PRT_FXEMIOB02I0Y6Q | + | 336 | 388 | **RNA**→ |
| Ama-2-1 | NZ_JWLY01000014.1 | + | 124090 | 124142 | **RNA**→ rpmE (PRK00019)Ribosomal_L31 (pfam01197)→ hypo→ |
| Ama-2-2 | NZ_JWLX01000009.1 | + | 137955 | 138007 | **RNA**→ rpmE (PRK00019)Ribosomal_L31 (pfam01197)→ hypo→ |
| Ama-2-3 | NZ_JXRW01000067.1 | + | 4135 | 4187 | **RNA**→ rpmE (PRK00019)Ribosomal_L31 (pfam01197)→ hypo→ |
| Ama-6-1 | NC_018679.1 | + | 4142089 | 4142141 | **RNA**→ rpmE (PRK00019)Ribosomal_L31 (pfam01197)→ hypo→ |
| Ama-3-1 | NC_018632.1 | + | 4038733 | 4038785 | **RNA**→ rpmE (PRK00019)Ribosomal_L31 (pfam01197)→ hypo→ |
| Ama-9-1 | NC_018678.1 | + | 4082894 | 4082946 | **RNA**→ rpmE (PRK00019)Ribosomal_L31 (pfam01197)→ hypo→ hypo→ hypo→ |
| env-1025 | JGI12097J13213_1002839 | + | 279 | 331 | **RNA**→ rpmE (PRK00019)Ribosomal_L31 (pfam01197)→ |
| env-1026 | CESQ01018836.1 | - | 3232 | 3180 | **RNA**→ rpmE (PRK00019)Ribosomal_L31 (pfam01197)→ |
| env-1027 | CENU01132442.1 | - | 183 | 131 | **RNA**→ |
| env-1028 | CESS01183283.1 | - | 394 | 342 | **RNA**→ rpmE (PRK00019)Ribosomal_L31 (pfam01197)→ |
| env-1029 | CENY01193254.1 | - | 408 | 356 | **RNA**→ rpmE (PRK00019)Ribosomal_L31 (pfam01197)→ |
| env-1030 | CEVI01152444.1 | - | 189 | 138 | **RNA**→ rpmE (PRK00019)Ribosomal_L31 (pfam01197)→ |
| env-1031 | CENG01015084.1 | - | 330 | 278 | **RNA**→ rpmE (PRK00019)Ribosomal_L31 (pfam01197)→ |
| Ama-7-1 | NC_018692.1 | + | 3943384 | 3943436 | **RNA**→ rpmE (PRK00019)Ribosomal_L31 (pfam01197)→ |
| env-1032 | CENH01017879.1 | + | 4473 | 4525 | **RNA**→ rpmE (PRK00019)Ribosomal_L31 (pfam01197)→ |
| env-1033 | CENT01009082.1 | + | 1961 | 2013 | **RNA**→ rpmE (PRK00019)Ribosomal_L31 (pfam01197)→ |
| env-1034 | CEOF01054473.1 | + | 162 | 214 | **RNA**→ rpmE (PRK00019)Ribosomal_L31 (pfam01197)→ |
| Ama-4-1 | NC_019393.1 | + | 4144827 | 4144879 | **RNA**→ rpmE (PRK00019)Ribosomal_L31 (pfam01197)→ |
| Ama-8-1 | NC_011138.3 | + | 4003832 | 4003884 | **RNA**→ rpmE (PRK00019)Ribosomal_L31 (pfam01197)→ |
| Ama-12-1 | NC_021714.1 | + | 3960995 | 3961047 | **RNA**→ rpmE (PRK00019)Ribosomal_L31 (pfam01197)→ |
| Ama-11-1 | NC_021712.1 | + | 3975520 | 3975572 | **RNA**→ rpmE (PRK00019)Ribosomal_L31 (pfam01197)→ |
| Ama-5-1 | NC_023045.1 | + | 3921718 | 3921770 | **RNA**→ rpmE (PRK00019)Ribosomal_L31 (pfam01197)→ |
| Ama-13-1 | NC_021713.1 | + | 4126897 | 4126949 | **RNA**→ rpmE (PRK00019)Ribosomal_L31 (pfam01197)→ |
| Ama-10-1 | NC_021717.1 | + | 4004559 | 4004611 | **RNA**→ rpmE (PRK00019)Ribosomal_L31 (pfam01197)→ |
| env-1035 | CEQI01061962.1 | - | 175 | 123 | **RNA**→ rpmE (PRK00019)Ribosomal_L31 (pfam01197)→ |
| env-1036 | CEPQ01008655.1 | - | 175 | 123 | **RNA**→ rpmE (PRK00019)Ribosomal_L31 (pfam01197)→ |
| env-1037 | CENH01008023.1 | - | 1725 | 1674 | **RNA**→ rpmE (PRK00019)Ribosomal_L31 (pfam01197)→ |
| env-1038 | CEPO01000325.1 | - | 741 | 690 | **RNA**→ rpmE (PRK00019)Ribosomal_L31 (pfam01197)→ |
| env-1039 | CEPX01405457.1 | - | 564 | 513 | **RNA**→ rpmE (PRK00019)Ribosomal_L31 (pfam01197)→ |
| env-1040 | CEPY01060952.1 | - | 1146 | 1095 | **RNA**→ rpmE (PRK00019)Ribosomal_L31 (pfam01197)→ |
| env-1041 | CEQF01170735.1 | - | 268 | 217 | **RNA**→ rpmE (PRK00019)Ribosomal_L31 (pfam01197)→ |
| env-1042 | CEQG01249219.1 | - | 343 | 292 | **RNA**→ rpmE (PRK00019)Ribosomal_L31 (pfam01197)→ |
| env-1043 | CEQH01275712.1 | - | 508 | 457 | **RNA**→ rpmE (PRK00019)Ribosomal_L31 (pfam01197)→ |
| env-1044 | CEQI01021998.1 | - | 423 | 372 | **RNA**→ rpmE (PRK00019)Ribosomal_L31 (pfam01197)→ |
| env-1045 | CEQQ01141891.1 | + | 1236 | 1287 | **RNA**→ rpmE (PRK00019)Ribosomal_L31 (pfam01197)→ |
| env-1046 | CEQS01013898.1 | + | 914 | 965 | **RNA**→ rpmE (PRK00019)Ribosomal_L31 (pfam01197)→ |
| env-1047 | CEQU01246787.1 | - | 692 | 641 | **RNA**→ rpmE (PRK00019)Ribosomal_L31 (pfam01197)→ |
| env-1048 | CEQW01065128.1 | + | 15681 | 15732 | **RNA**→ rpmE (PRK00019)Ribosomal_L31 (pfam01197)→ |

| | | | | | |
|---|---|---|---|---|---|
| env-1049 | CEQX01342402.1 | + | 853 | 904 | **RNA**→ rpmE (PRK00019)Ribosomal_L31 (pfam01197)→ |
| env-1050 | CERB01035857.1 | + | 914 | 965 | **RNA**→ rpmE (PRK00019)Ribosomal_L31 (pfam01197)→ |
| env-1051 | CERC01009557.1 | + | 9311 | 9362 | **RNA**→ rpmE (PRK00019)Ribosomal_L31 (pfam01197)→ |
| env-1052 | CERD01017474.1 | - | 551 | 500 | **RNA**→ rpmE (PRK00019)Ribosomal_L31 (pfam01197)→ |
| env-1053 | CERE01022366.1 | - | 692 | 641 | **RNA**→ rpmE (PRK00019)Ribosomal_L31 (pfam01197)→ |
| env-1054 | CERG01021368.1 | + | 2982 | 3033 | **RNA**→ rpmE (PRK00019)Ribosomal_L31 (pfam01197)→ |
| env-1055 | CERL01022088.1 | - | 6990 | 6939 | **RNA**→ rpmE (PRK00019)Ribosomal_L31 (pfam01197)→ |
| env-1056 | CERQ01243698.1 | - | 1660 | 1609 | **RNA**→ rpmE (PRK00019)Ribosomal_L31 (pfam01197)→ |
| env-1057 | CERR01097688.1 | - | 14701 | 14650 | **RNA**→ rpmE (PRK00019)Ribosomal_L31 (pfam01197)→ |
| env-1058 | CERS01310048.1 | + | 853 | 904 | **RNA**→ rpmE (PRK00019)Ribosomal_L31 (pfam01197)→ |
| env-1059 | CERV01201358.1 | - | 463 | 412 | **RNA**→ rpmE (PRK00019)Ribosomal_L31 (pfam01197)→ |
| env-1060 | CERX01037102.1 | + | 914 | 965 | **RNA**→ rpmE (PRK00019)Ribosomal_L31 (pfam01197)→ |
| env-1061 | CESC01002520.1 | + | 28189 | 28240 | **RNA**→ rpmE (PRK00019)Ribosomal_L31 (pfam01197)→ |
| env-1062 | CESL01235759.1 | + | 660 | 711 | **RNA**→ rpmE (PRK00019)Ribosomal_L31 (pfam01197)→ |
| env-1063 | CESM01253004.1 | + | 346 | 397 | **RNA**→ rpmE (PRK00019)rpmE2 (PRK01678)Ribosomal_L31 (pfam01197)→ |
| env-1064 | CESO01220634.1 | + | 374 | 425 | **RNA**→ rpmE (PRK00019)Ribosomal_L31 (pfam01197)→ |
| env-1065 | CESQ01033314.1 | - | 39300 | 39249 | **RNA**→ rpmE (PRK00019)Ribosomal_L31 (pfam01197)→ |
| env-1066 | CESR01169867.1 | + | 15683 | 15734 | **RNA**→ rpmE (PRK00019)Ribosomal_L31 (pfam01197)→ |
| env-1067 | CESY01007781.1 | - | 786 | 735 | **RNA**→ rpmE (PRK00019)Ribosomal_L31 (pfam01197)→ |
| env-1068 | CETG01034723.1 | - | 509 | 458 | **RNA**→ rpmE (PRK00019)Ribosomal_L31 (pfam01197)→ |
| env-1069 | CETH01358830.1 | + | 328 | 379 | **RNA**→ rpmE (PRK00019)Ribosomal_L31 (pfam01197)→ |
| env-1070 | CETJ01151418.1 | + | 432 | 483 | **RNA**→ rpmE (PRK00019)Ribosomal_L31 (pfam01197)→ |
| env-1071 | CETM01434228.1 | - | 509 | 458 | **RNA**→ rpmE (PRK00019)Ribosomal_L31 (pfam01197)→ |
| env-1072 | CETN01152556.1 | + | 995 | 1046 | **RNA**→ rpmE (PRK00019)Ribosomal_L31 (pfam01197)→ |
| env-1073 | CETO01022433.1 | + | 15683 | 15734 | **RNA**→ rpmE (PRK00019)Ribosomal_L31 (pfam01197)→ |
| env-1074 | CETR01195172.1 | - | 320 | 269 | **RNA**→ rpmE (PRK00019)Ribosomal_L31 (pfam01197)→ |
| env-1075 | CETS01064668.1 | + | 666 | 717 | **RNA**→ rpmE (PRK00019)Ribosomal_L31 (pfam01197)→ |
| env-1076 | CETV01229244.1 | + | 266 | 317 | **RNA**→ rpmE (PRK00019)Ribosomal_L31 (pfam01197)→ |
| env-1077 | CETY01197886.1 | + | 15683 | 15734 | **RNA**→ rpmE (PRK00019)Ribosomal_L31 (pfam01197)→ |
| env-1078 | CEUC01145376.1 | - | 2776 | 2725 | **RNA**→ rpmE (PRK00019)Ribosomal_L31 (pfam01197)→ |
| env-1079 | CEUD01121151.1 | + | 15683 | 15734 | **RNA**→ rpmE (PRK00019)Ribosomal_L31 (pfam01197)→ |
| env-1080 | CEUM01133227.1 | - | 1141 | 1090 | **RNA**→ rpmE (PRK00019)Ribosomal_L31 (pfam01197)→ |
| env-1081 | CEUN01034438.1 | - | 509 | 458 | **RNA**→ rpmE (PRK00019)Ribosomal_L31 (pfam01197)→ |
| env-1082 | CEUP01008540.1 | + | 914 | 965 | **RNA**→ rpmE (PRK00019)Ribosomal_L31 (pfam01197)→ |
| env-1083 | CEUS01321896.1 | - | 272 | 221 | **RNA**→ rpmE (PRK00019)Ribosomal_L31 (pfam01197)→ |
| env-1084 | CEUY01032768.1 | - | 2465 | 2414 | **RNA**→ rpmE (PRK00019)Ribosomal_L31 (pfam01197)→ |
| env-1085 | CEVA01041336.1 | + | 14542 | 14593 | **RNA**→ rpmE (PRK00019)Ribosomal_L31 (pfam01197)→ |
| env-1086 | CEVG01310261.1 | + | 713 | 764 | **RNA**→ rpmE (PRK00019)Ribosomal_L31 (pfam01197)→ |
| env-1087 | CEVI01003491.1 | - | 39966 | 39915 | **RNA**→ rpmE (PRK00019)Ribosomal_L31 (pfam01197)→ |
| env-1088 | CEVK01179875.1 | - | 373 | 322 | **RNA**→ rpmE (PRK00019)Ribosomal_L31 (pfam01197)→ |
| env-1089 | CEVZ01370195.1 | - | 341 | 290 | **RNA**→ rpmE (PRK00019)Ribosomal_L31 (pfam01197)→ |
| env-1090 | CEWB01191093.1 | + | 853 | 904 | **RNA**→ rpmE (PRK00019)Ribosomal_L31 (pfam01197)→ |
| env-1091 | CEWI01259732.1 | + | 1163 | 1214 | **RNA**→ rpmE (PRK00019)Ribosomal_L31 (pfam01197)→ |
| env-1092 | CEWJ01143993.1 | + | 15683 | 15734 | **RNA**→ rpmE (PRK00019)Ribosomal_L31 (pfam01197)→ |
| Aau-2-1 | NZ_CP008849.1 | + | 3814040 | 3814091 | **RNA**→ rpmE (PRK00019)Ribosomal_L31 (pfam01197)→ |
| Aau-2-2 | NZ_CP010912.1 | + | 3941841 | 3941892 | **RNA**→ rpmE (PRK00019)Ribosomal_L31 (pfam01197)→ |
| Pha-4-1 | NZ_ATUO01000067.1 | + | 19196 | 19259 | **RNA**→ rpmE (PRK00019)Ribosomal_L31 (pfam01197)→ |
| env-1093 | BS_KBA_SWE02_21m_10551754 | + | 152 | 216 | **RNA**→ rpmE (PRK00019)Ribosomal_L31 (pfam01197)→ |
| env-1094 | KGI_S1_ANT01_95m_c10012892 | - | 141 | 77 | **RNA**→ rpmE (PRK00019)Ribosomal_L31 (pfam01197)→ |
| env-1095 | KGI_S2_ANT05_2345m_c1007829 | + | 2549 | 2613 | **RNA**→ rpmE (PRK00019)Ribosomal_L31 (pfam01197)→ |
| env-1096 | KGI_S1_ANT01_95m_c10008728 | - | 288 | 224 | **RNA**→ rpmE (PRK00019)Ribosomal_L31 (pfam01197)→ |
| env-1097 | KGI_S2_ANT04_2345m_c1009121 | - | 1653 | 1589 | **RNA**→ rpmE (PRK00019)Ribosomal_L31 (pfam01197)→ |

| | | | | | |
|---|---|---|---|---|---|
| env-1098 | KGI_S2_ANT05_2345m_c1003874 | + | 4157 | 4221 | **RNA**→ rpmE (PRK00019)Ribosomal_L31 (pfam01197)→ |
| Pin-1-1 | NC_008709.1 | - | 3538323 | 3538259 | **RNA**→ rpmE (PRK00019)Ribosomal_L31 (pfam01197)→ |
| env-1099 | SA_S2_NOR13_50m_c1137819 | + | 116 | 180 | **RNA**→ rpmE (PRK00019)Ribosomal_L31 (pfam01197)→ |
| env-1100 | SA_S1_NOR05_45m_c10035075 | + | 932 | 996 | **RNA**→ rpmE (PRK00019)Ribosomal_L31 (pfam01197)→ |
| env-1101 | SA_S1_NOR08_45m_c10037975 | - | 506 | 442 | **RNA**→ rpmE (PRK00019)Ribosomal_L31 (pfam01197)→ |
| env-1102 | SA_S2_NOR15_50m_c10183128 | + | 221 | 285 | **RNA**→ rpmE (PRK00019)Ribosomal_L31 (pfam01197)→ |
| env-1103 | SA_S1_NOR05_45m_c10141124 | - | 371 | 307 | **RNA**→ rpmE (PRK00019)Ribosomal_L31 (pfam01197)→ |
| env-1104 | ASRP01007864.1 | - | 203 | 139 | **RNA**→ rpmE (PRK00019)Ribosomal_L31 (pfam01197)→ |
| Par-2-1 | NZ_AXWP01000041.1 | - | 63330 | 63266 | **RNA**→ rpmE (PRK00019)Ribosomal_L31 (pfam01197)→ |
| Paq-1-1 | NZ_AUAM01000062.1 | - | 1573 | 1509 | **RNA**→ rpmE (PRK00019)Ribosomal_L31 (pfam01197)→ |
| Pos-1-1 | NZ_KB907022.1 | + | 11035 | 11099 | **RNA**→ rpmE (PRK00019)Ribosomal_L31 (pfam01197)→ rpmE (PRK00019)Ribosomal_L31 (pfam01197)→ rpmE (PRK00019)Ribosomal_L31 (pfam01197)→ hypo→ |
| Psp-35-1 | NZ_CBRF010000116.1 | - | 14588 | 14523 | **RNA**→ rpmE (PRK00019)Ribosomal_L31 (pfam01197)→ |
| env-1105 | KGI_S1_ANT01_95m_c10000182 | - | 2970 | 2905 | **RNA**→ rpmE (PRK00019)Ribosomal_L31 (pfam01197)→ |
| env-1106 | KGI_S2_ANT05_2345m_c1036643 | - | 656 | 591 | **RNA**→ rpmE (PRK00019)Ribosomal_L31 (pfam01197)→ |
| env-1107 | KGI_S1_ANT03_95m_c1000085 | + | 397 | 462 | **RNA**→ rpmE (PRK00019)Ribosomal_L31 (pfam01197)→ |
| env-1108 | KGI_S1_ANT02_95m_c10004749 | - | 2245 | 2180 | **RNA**→ rpmE (PRK00019)Ribosomal_L31 (pfam01197)→ |
| env-1109 | KGI_S2_ANT04_2345m_c1001133 | + | 149 | 214 | **RNA**→ rpmE (PRK00019)Ribosomal_L31 (pfam01197)→ |
| env-1110 | SA_S1_NOR05_45m_c10067223 | - | 515 | 451 | **RNA**→ rpmE (PRK00019)Ribosomal_L31 (pfam01197)→ |
| env-1111 | SA_S1_NOR08_45m_c10078152 | - | 527 | 463 | **RNA**→ rpmE (PRK00019)Ribosomal_L31 (pfam01197)→ |
| env-1112 | SA_S2_NOR15_50m_c10042781 | + | 1628 | 1692 | **RNA**→ rpmE (PRK00019)Ribosomal_L31 (pfam01197)→ |
| Psp-34-1 | NC_020802.1 | - | 2578482 | 2578419 | **RNA**→ rpmE (PRK00019)Ribosomal_L31 (pfam01197)→ |
| Sam-1-1 | NZ_KB899636.1 | + | 535889 | 535941 | **RNA**→ rpmE (PRK00019)Ribosomal_L31 (pfam01197)→ |
| env-1113 | CETY01166192.1 | + | 4690 | 4742 | **RNA**→ rpmE (PRK00019)Ribosomal_L31 (pfam01197)→ |
| Pfl-3-1 | NZ_JAEE01000006.1 | - | 27868 | 27808 | **RNA**→ rpmE (PRK00019)Ribosomal_L31 (pfam01197)→ rpmE (PRK00019)Ribosomal_L31 (pfam01197)→ |
| Ykr-2-1 | NZ_ACCA01000030.1 | + | 28292 | 28346 | **RNA**→ rpmE (PRK00019)Ribosomal_L31 (pfam01197)→ rpmE (PRK00019)Ribosomal_L31 (pfam01197)→ |
| Pge-1-1 | NZ_CP009450.1 | + | 982043 | 982094 | **RNA**→ rpmE (PRK00019)Ribosomal_L31 (pfam01197)→ |
| Kpn-281-1 | NZ_KI535451.1 | + | 4373213 | 4373264 | **RNA**→ rpmE (PRK00019)Ribosomal_L31 (pfam01197)→ |
| Vja-2-1 | NZ_BAOC01000004.1 | + | 79462 | 79512 | **RNA**→ rpmE (PRK00019)Ribosomal_L31 (pfam01197)→ |
| Kpn-256-1 | NZ_ARRZ01000059.1 | + | 70097 | 70148 | **RNA**→ rpmE (PRK00019)Ribosomal_L31 (pfam01197)→ rpmE (PRK00019)Ribosomal_L31 (pfam01197)→ |
| Kpn-70-1 | NZ_JJNF01000020.1 | + | 68880 | 68931 | **RNA**→ rpmE (PRK00019)Ribosomal_L31 (pfam01197)→ |
| Ksp-7-1 | NZ_GL882802.1 | - | 7317 | 7266 | **RNA**→ rpmE (PRK00019)Ribosomal_L31 (pfam01197)→ rpmE (PRK00019)Ribosomal_L31 (pfam01197)→ |
| Kpn-244-1 | NZ_KI535433.1 | + | 4551638 | 4551689 | **RNA**→ rpmE (PRK00019)Ribosomal_L31 (pfam01197)→ |
| Kpn-31-1 | NZ_KK036716.1 | + | 4570872 | 4570923 | **RNA**→ rpmE (PRK00019)Ribosomal_L31 (pfam01197)→ |
| Kpn-143-1 | NZ_KK036961.1 | + | 1586585 | 1586636 | **RNA**→ rpmE (PRK00019)Ribosomal_L31 (pfam01197)→ |
| Kpn-142-1 | NZ_KK036965.1 | + | 4373208 | 4373259 | **RNA**→ rpmE (PRK00019)Ribosomal_L31 (pfam01197)→ |
| Kpn-206-6 | NZ_CP009876.1 | + | 90246 | 90297 | **RNA**→ rpmE (PRK00019)Ribosomal_L31 (pfam01197)→ |
| Hin-6-1 | NZ_ABWW01000003.1 | - | 44774 | 44724 | **RNA**→ rpmE (PRK00019)Ribosomal_L31 (pfam01197)→ rpmE (PRK00019)Ribosomal_L31 (pfam01197)→ |
| Hin-11-1 | NC_014922.1 | + | 379270 | 379320 | **RNA**→ rpmE (PRK00019)Ribosomal_L31 (pfam01197)→ |
| Eba-1-1 | NZ_KN150796.1 | + | 4226043 | 4226094 | **RNA**→ rpmE (PRK00019)Ribosomal_L31 (pfam01197)→ |
| Ysi-1-1 | NZ_CP007230.1 | + | 1742261 | 1742314 | **RNA**→ rpmE (PRK00019)Ribosomal_L31 (pfam01197)→ |
| Hin-1-3 | NZ_CP007470.1 | + | 969362 | 969412 | **RNA**→ rpmE (PRK00019)Ribosomal_L31 (pfam01197)→ |
| Kox-4-2 | NZ_AKCF01000001.1 | - | 987481 | 987430 | **RNA**→ rpmE (PRK00019)Ribosomal_L31 (pfam01197)→ |
| Kox-6-1 | NZ_ARVT01000001.1 | + | 5597804 | 5597855 | **RNA**→ rpmE (PRK00019)Ribosomal_L31 (pfam01197)→ |
| Kox-10-1 | NZ_JAKY01000077.1 | + | 76452 | 76503 | **RNA**→ rpmE (PRK00019)Ribosomal_L31 (pfam01197)→ |
| Kox-7-1 | NZ_KK097713.1 | + | 287765 | 287816 | **RNA**→ rpmE (PRK00019)Ribosomal_L31 (pfam01197)→ |
| Kox-13-1 | NZ_JH603137.1 | + | 5368788 | 5368839 | **RNA**→ rpmE (PRK00019)Ribosomal_L31 (pfam01197)→ |
| Kox-19-1 | NZ_KI535631.1 | + | 608704 | 608755 | **RNA**→ rpmE (PRK00019)Ribosomal_L31 (pfam01197)→ |
| Kox-20-1 | NZ_KI535598.1 | + | 3002323 | 3002374 | **RNA**→ rpmE (PRK00019)Ribosomal_L31 (pfam01197)→ |
| Kox-1-1 | NZ_CP011618.1 | - | 3481846 | 3481795 | **RNA**→ rpmE (PRK00019)Ribosomal_L31 (pfam01197)→ |
| Kox-1-2 | NZ_CP011597.1 | - | 3460832 | 3460781 | **RNA**→ rpmE (PRK00019)Ribosomal_L31 (pfam01197)→ |
| Kox-5-1 | NZ_KQ235791.1 | + | 607562 | 607613 | **RNA**→ rpmE (PRK00019)Ribosomal_L31 (pfam01197)→ |

| | | | | | |
|---|---|---|---|---|---|
| Kox-4-1 | NZ_KQ235784.1 | - | 5111984 | 5111933 | **RNA**→ rpmE (PRK00019)Ribosomal_L31 (pfam01197)→ |
| Kox-3-1 | NZ_KQ235774.1 | - | 2026518 | 2026467 | **RNA**→ rpmE (PRK00019)Ribosomal_L31 (pfam01197)→ |
| Kox-16-1 | NZ_AIEM01000124.1 | + | 33854 | 33905 | **RNA**→ rpmE (PRK00019)Ribosomal_L31 (pfam01197)→ |
| Hin-13-1 | NZ_ACSL01000001.1 | - | 4352 | 4302 | **RNA**→ rpmE (PRK00019)Ribosomal_L31 (pfam01197)→ rpmE (PRK00019)Ribosomal_L31 (pfam01197)→ |
| Hin-1-4 | NZ_CP007471.1 | + | 209664 | 209714 | **RNA**→ rpmE (PRK00019)Ribosomal_L31 (pfam01197)→ |
| Sma-10-1 | NZ_KE560346.1 | + | 418205 | 418257 | **RNA**→ rpmE (PRK00019)Ribosomal_L31 (pfam01197)→ rpmE (PRK00019)Ribosomal_L31 (pfam01197)→ |
| Sma-11-1 | NZ_KE560341.1 | + | 214674 | 214726 | **RNA**→ rpmE (PRK00019)Ribosomal_L31 (pfam01197)→ |
| Sma-25-1 | NZ_JH670250.1 | + | 3952174 | 3952226 | **RNA**→ rpmE (PRK00019)Ribosomal_L31 (pfam01197)→ rpmE (PRK00019)Ribosomal_L31 (pfam01197)→ rpmE (PRK00019)Ribosomal_L31 (pfam01197)→ |
| Yru-2-2 | NZ_ACCC01000012.1 | - | 37748 | 37695 | **RNA**→ rpmE (PRK00019)Ribosomal_L31 (pfam01197)→ rpmE (PRK00019)Ribosomal_L31 (pfam01197)→ |
| Yru-2-1 | NZ_KN150747.1 | - | 419490 | 419437 | **RNA**→ rpmE (PRK00019)Ribosomal_L31 (pfam01197)→ |
| Yru-1-2 | NZ_CP011078.1 | - | 2139842 | 2139789 | **RNA**→ rpmE (PRK00019)Ribosomal_L31 (pfam01197)→ |
| Yru-1-3 | NZ_JRWX01000001.1 | + | 89041 | 89094 | **RNA**→ rpmE (PRK00019)Ribosomal_L31 (pfam01197)→ |
| Vni-11-1 | NZ_CAOB01000008.1 | - | 6262 | 6213 | **RNA**→ rpmE (PRK00019)Ribosomal_L31 (pfam01197)→ |
| Vni-8-1 | NZ_CAOA01000077.1 | + | 57549 | 57598 | **RNA**→ rpmE (PRK00019)Ribosomal_L31 (pfam01197)→ |
| Vni-10-1 | NZ_CAOD01000077.1 | + | 4423 | 4472 | **RNA**→ rpmE (PRK00019)Ribosomal_L31 (pfam01197)→ |
| Vni-6-1 | NZ_CANW01000076.1 | + | 57549 | 57598 | **RNA**→ rpmE (PRK00019)Ribosomal_L31 (pfam01197)→ |
| Vni-4-1 | NZ_CANU01000076.1 | - | 6171 | 6122 | **RNA**→ rpmE (PRK00019)Ribosomal_L31 (pfam01197)→ |
| Vni-5-1 | NZ_CANV01000028.1 | - | 6171 | 6122 | **RNA**→ rpmE (PRK00019)Ribosomal_L31 (pfam01197)→ |
| Vni-2-1 | NZ_CANT01000080.1 | + | 44452 | 44501 | **RNA**→ rpmE (PRK00019)Ribosomal_L31 (pfam01197)→ |
| Vni-3-1 | NZ_AFWJ01000267.1 | - | 88872 | 88823 | **RNA**→ rpmE (PRK00019)Ribosomal_L31 (pfam01197)→ rpmE (PRK00019)Ribosomal_L31 (pfam01197)→ |
| Vni-13-1 | NZ_LK391965.1 | + | 1712210 | 1712259 | **RNA**→ rpmE (PRK00019)Ribosomal_L31 (pfam01197)→ |
| Vni-1-1 | NC_022528.1 | - | 3799901 | 3799852 | **RNA**→ rpmE (PRK00019)Ribosomal_L31 (pfam01197)→ |
| Vni-14-1 | NZ_LK391967.1 | - | 747727 | 747678 | **RNA**→ rpmE (PRK00019)Ribosomal_L31 (pfam01197)→ |
| Vni-12-1 | NZ_LK391963.1 | - | 4084011 | 4083962 | **RNA**→ rpmE (PRK00019)Ribosomal_L31 (pfam01197)→ |
| Vni-7-1 | NZ_LK054223.1 | + | 2036267 | 2036316 | **RNA**→ rpmE (PRK00019)Ribosomal_L31 (pfam01197)→ |
| Yin-2-1 | NZ_AALF02000041.1 | - | 35133 | 35079 | **RNA**→ rpmE (PRK00019)Ribosomal_L31 (pfam01197)→ |
| Yin-1-1 | NZ_CP009801.1 | + | 2155185 | 2155239 | **RNA**→ rpmE (PRK00019)Ribosomal_L31 (pfam01197)→ |
| Hin-18-2 | NC_017452.1 | + | 1536435 | 1536485 | **RNA**→ hypo→ hypo→ hypo→ hypo→ hypo→ COG3306 (COG3306)→ COG3306 (COG3306)→ heptsyl_trn_III (TIGR02201)RfaF (COG0859)→ |
| Hin-14-1 | NZ_AAZG01000004.1 | + | 103226 | 103276 | **RNA**→ rpmE (PRK00019)Ribosomal_L31 (pfam01197)→ rpmE (PRK00019)Ribosomal_L31 (pfam01197)→ |
| Hin-4-1 | NZ_AAZF01000001.1 | - | 45759 | 45709 | **RNA**→ rpmE (PRK00019)Ribosomal_L31 (pfam01197)→ rpmE (PRK00019)Ribosomal_L31 (pfam01197)→ |
| Hin-3-1 | NZ_CP008740.1 | - | 919236 | 919186 | **RNA**→ rpmE (PRK00019)Ribosomal_L31 (pfam01197)→ |
| Vvu-10-1 | NZ_CM001799.1 | + | 1209887 | 1209938 | **RNA**→ rpmE (PRK00019)Ribosomal_L31 (pfam01197)→ |
| Vvu-3-1 | NZ_JDSE01000046.1 | + | 2549 | 2600 | **RNA**→ rpmE (PRK00019)Ribosomal_L31 (pfam01197)→ |
| Ywa-1-1 | NZ_CBLI010000383.1 | - | 346 | 293 | **RNA**→ rpmE (PRK00019)Ribosomal_L31 (pfam01197)→ |
| Ywa-1-2 | NZ_CBLG010000169.1 | + | 14348 | 14401 | **RNA**→ rpmE (PRK00019)Ribosomal_L31 (pfam01197)→ |
| Dso-4-1 | NZ_CM001839.1 | + | 191962 | 192015 | **RNA**→ rpmE (PRK00019)Ribosomal_L31 (pfam01197)→ |
| Dso-3-1 | NZ_CM001859.1 | + | 192153 | 192206 | **RNA**→ rpmE (PRK00019)Ribosomal_L31 (pfam01197)→ |
| Dso-5-1 | NZ_CM001842.1 | + | 191961 | 192014 | **RNA**→ rpmE (PRK00019)Ribosomal_L31 (pfam01197)→ |
| Dso-2-1 | NZ_CM001860.1 | + | 192284 | 192337 | **RNA**→ rpmE (PRK00019)Ribosomal_L31 (pfam01197)→ |
| Dso-6-1 | NZ_KI866531.1 | - | 2542514 | 2542461 | **RNA**→ rpmE (PRK00019)Ribosomal_L31 (pfam01197)→ |
| Dsp-8-1 | NZ_AMWE01000004.1 | - | 1998925 | 1998872 | **RNA**→ rpmE (PRK00019)Ribosomal_L31 (pfam01197)→ |
| Yps-1-3 | NZ_CP008943.1 | - | 2566211 | 2566158 | **RNA**→ rpmE (PRK00019)Ribosomal_L31 (pfam01197)→ |
| Yps-1-4 | NZ_CP009759.1 | - | 2277500 | 2277447 | **RNA**→ rpmE (PRK00019)Ribosomal_L31 (pfam01197)→ |
| Yps-5-1 | NZ_CAQW01000045.1 | + | 24851 | 24904 | **RNA**→ rpmE (PRK00019)Ribosomal_L31 (pfam01197)→ |
| Yps-2-1 | NZ_CAQT01000042.1 | - | 7087 | 7034 | **RNA**→ rpmE (PRK00019)Ribosomal_L31 (pfam01197)→ |
| Yps-7-1 | NZ_CAQY01000083.1 | + | 13728 | 13781 | **RNA**→ rpmE (PRK00019)Ribosomal_L31 (pfam01197)→ |
| Yps-3-1 | NZ_CAQU01000109.1 | - | 7077 | 7024 | **RNA**→ rpmE (PRK00019)Ribosomal_L31 (pfam01197)→ |
| Yps-4-1 | NZ_CAQV01000017.1 | + | 71002 | 71055 | **RNA**→ rpmE (PRK00019)Ribosomal_L31 (pfam01197)→ |
| Yps-6-1 | NZ_CAQX01000032.1 | + | 13731 | 13784 | **RNA**→ rpmE (PRK00019)Ribosomal_L31 (pfam01197)→ |
| Sso-1-546 | NZ_CXJA01000293.1 | + | 448 | 499 | **RNA**→ rpmE (PRK00019)Ribosomal_L31 (pfam01197)→ |
| Yal-3-1 | NZ_CP011975.1 | - | 2144109 | 2144055 | **RNA**→ rpmE (PRK00019)Ribosomal_L31 (pfam01197)→ |

| | | | | | |
|---|---|---|---|---|---|
| Sal-1-1 | NZ_LN810019.1 | + | 59430 | 59481 | RNA→ rpmE (PRK00019)Ribosomal_L31 (pfam01197)→ |
| Ssp-1-1 | NZ_JADP01000004.1 | + | 57303 | 57354 | RNA→ rpmE (PRK00019)Ribosomal_L31 (pfam01197)→ |
| Sha-2-1 | NZ_BALL01000002.1 | + | 56991 | 57042 | RNA→ rpmE (PRK00019)Ribosomal_L31 (pfam01197)→ |
| Sal-2-1 | NZ_BALO01000003.1 | + | 42263 | 42314 | RNA→ rpmE (PRK00019)Ribosomal_L31 (pfam01197)→ |
| Kpn-168-1 | NZ_AZAP01000001.1 | + | 69358 | 69409 | RNA→ rpmE (PRK00019)Ribosomal_L31 (pfam01197)→ |
| Kpn-46-1 | NZ_KK036936.1 | - | 673181 | 673130 | RNA→ rpmE (PRK00019)Ribosomal_L31 (pfam01197)→ |
| Kpn-206-7 | NZ_CP011313.1 | - | 5158789 | 5158738 | RNA→ rpmE (PRK00019)Ribosomal_L31 (pfam01197)→ |
| Vvu-9-1 | NZ_AMQV01000002.1 | - | 49207 | 49156 | RNA→ rpmE (PRK00019)Ribosomal_L31 (pfam01197)→ |
| Vvu-2-1 | NZ_AMQR01000003.1 | + | 15395 | 15446 | RNA→ rpmE (PRK00019)Ribosomal_L31 (pfam01197)→ |
| Vvu-7-1 | NZ_AFSW01000084.1 | + | 15849 | 15900 | RNA→ rpmE (PRK00019)Ribosomal_L31 (pfam01197)→ |
| Vvu-6-1 | NZ_AFSX01000029.1 | - | 40054 | 40003 | RNA→ rpmE (PRK00019)Ribosomal_L31 (pfam01197)→ |
| Yro-2-1 | NZ_ACCD01000008.1 | + | 93813 | 93867 | RNA→ rpmE (PRK00019)Ribosomal_L31 (pfam01197)→ rpmE (PRK00019)Ribosomal_L31 (pfam01197)→ rpmE (PRK00019)Ribosomal_L31 (pfam01197)→ |
| Yro-1-1 | NZ_CP009787.1 | + | 3005532 | 3005586 | RNA→ rpmE (PRK00019)Ribosomal_L31 (pfam01197)→ |
| Yen-13-1 | NZ_CACY01000042.1 | - | 36129 | 36075 | RNA→ rpmE (PRK00019)Ribosomal_L31 (pfam01197)→ |
| Yen-12-1 | NZ_CACZ01000062.1 | + | 13330 | 13384 | RNA→ rpmE (PRK00019)Ribosomal_L31 (pfam01197)→ |
| Yen-3-1 | NZ_HF571988.1 | + | 128432 | 128486 | RNA→ rpmE (PRK00019)Ribosomal_L31 (pfam01197)→ |
| Yen-1-4 | NZ_CP011118.1 | - | 4578762 | 4578708 | RNA→ rpmE (PRK00019)Ribosomal_L31 (pfam01197)→ |
| Pmi-2-1 | NZ_GG668582.1 | - | 34951 | 34902 | RNA→ rpmE (PRK00019)Ribosomal_L31 (pfam01197)→ |
| Pmi-5-1 | NC_010554.1 | + | 3535542 | 3535591 | RNA→ rpmE (PRK00019)Ribosomal_L31 (pfam01197)→ |
| Pmi-8-1 | NZ_JH815508.1 | - | 34281 | 34232 | RNA→ rpmE (PRK00019)Ribosomal_L31 (pfam01197)→ |
| Pmi-7-1 | NZ_JH815533.1 | - | 71312 | 71263 | RNA→ rpmE (PRK00019)Ribosomal_L31 (pfam01197)→ |
| Pmi-4-1 | NZ_KB206032.1 | - | 37018 | 36969 | RNA→ rpmE (PRK00019)Ribosomal_L31 (pfam01197)→ |
| Pmi-1-1 | NZ_KN150749.1 | - | 3010571 | 3010522 | RNA→ rpmE (PRK00019)Ribosomal_L31 (pfam01197)→ |
| Pmi-6-1 | NZ_AORN01000002.1 | + | 155202 | 155251 | RNA→ rpmE (PRK00019)Ribosomal_L31 (pfam01197)→ |
| Pmi-1-2 | NZ_JTBA01000001.1 | + | 4050178 | 4050227 | RNA→ rpmE (PRK00019)Ribosomal_L31 (pfam01197)→ |
| Pmi-1-3 | NZ_JTBP01000001.1 | + | 2513335 | 2513384 | RNA→ rpmE (PRK00019)Ribosomal_L31 (pfam01197)→ |
| Pmi-1-4 | NZ_JTBW01000001.1 | - | 3605112 | 3605063 | RNA→ rpmE (PRK00019)Ribosomal_L31 (pfam01197)→ |
| Pmi-1-5 | NZ_JTBB01000001.1 | + | 3920452 | 3920501 | RNA→ rpmE (PRK00019)Ribosomal_L31 (pfam01197)→ |
| Pmi-1-6 | NZ_JTAW01000004.1 | + | 2330106 | 2330155 | RNA→ rpmE (PRK00019)Ribosomal_L31 (pfam01197)→ |
| Vpa-183-1 | NZ_AWIJ01000088.1 | - | 5725 | 5675 | RNA→ rpmE (PRK00019)Ribosomal_L31 (pfam01197)→ |
| Vpa-182-1 | NZ_AWIK01000006.1 | - | 14426 | 14376 | RNA→ rpmE (PRK00019)Ribosomal_L31 (pfam01197)→ |
| Vpa-217-1 | NZ_BAVG01000031.1 | + | 252975 | 253025 | RNA→ rpmE (PRK00019)Ribosomal_L31 (pfam01197)→ |
| Vpa-7-1 | NZ_AOPF01000052.1 | + | 256268 | 256318 | RNA→ rpmE (PRK00019)Ribosomal_L31 (pfam01197)→ |
| Vpa-19-1 | NZ_AOOX01000058.1 | + | 213943 | 213993 | RNA→ rpmE (PRK00019)Ribosomal_L31 (pfam01197)→ |
| Vpa-25-1 | NZ_AOOU01000022.1 | + | 256268 | 256318 | RNA→ rpmE (PRK00019)Ribosomal_L31 (pfam01197)→ |
| Vpa-38-1 | NZ_AOPH01000018.1 | + | 256269 | 256319 | RNA→ rpmE (PRK00019)Ribosomal_L31 (pfam01197)→ |
| Vpa-11-1 | NZ_AOPA01000042.1 | + | 256268 | 256318 | RNA→ rpmE (PRK00019)Ribosomal_L31 (pfam01197)→ |
| Vpa-41-1 | NZ_AOPK01000019.1 | + | 256269 | 256319 | RNA→ rpmE (PRK00019)Ribosomal_L31 (pfam01197)→ |
| Vpa-42-1 | NZ_AOPL01000021.1 | + | 256268 | 256318 | RNA→ rpmE (PRK00019)Ribosomal_L31 (pfam01197)→ |
| Vpa-14-1 | NZ_AOPD01000389.1 | - | 14501 | 14451 | RNA→ rpmE (PRK00019)Ribosomal_L31 (pfam01197)→ |
| Vpa-39-1 | NZ_AOPI01000388.1 | + | 10709 | 10759 | RNA→ rpmE (PRK00019)Ribosomal_L31 (pfam01197)→ |
| Vpa-12-1 | NZ_AOPB01000323.1 | + | 40203 | 40253 | RNA→ rpmE (PRK00019)Ribosomal_L31 (pfam01197)→ |
| Vpa-40-1 | NZ_AOPJ01000367.1 | - | 86970 | 86920 | RNA→ rpmE (PRK00019)Ribosomal_L31 (pfam01197)→ |
| Vpa-16-1 | NZ_AONA01000310.1 | - | 14503 | 14453 | RNA→ rpmE (PRK00019)Ribosomal_L31 (pfam01197)→ |
| Vpa-22-1 | NZ_AOPG01000006.1 | + | 43335 | 43385 | RNA→ rpmE (PRK00019)Ribosomal_L31 (pfam01197)→ |
| Vpa-15-1 | NZ_AOPE01000363.1 | - | 33905 | 33855 | RNA→ rpmE (PRK00019)Ribosomal_L31 (pfam01197)→ |
| Vpa-13-1 | NZ_AOPC01000423.1 | - | 12383 | 12333 | RNA→ rpmE (PRK00019)Ribosomal_L31 (pfam01197)→ |
| Vsp-28-1 | NZ_JYOQ01000177.1 | + | 238961 | 239011 | RNA→ rpmE (PRK00019)Ribosomal_L31 (pfam01197)→ |
| Vsp-29-1 | NZ_JYOR01000005.1 | + | 238961 | 239011 | RNA→ rpmE (PRK00019)Ribosomal_L31 (pfam01197)→ |
| Vpa-234-1 | NZ_AVOI01000008.1 | - | 16042 | 15992 | RNA→ rpmE (PRK00019)Ribosomal_L31 (pfam01197)→ |
| Vpa-174-1 | NZ_AWIS01000006.1 | + | 25917 | 25967 | RNA→ rpmE (PRK00019)Ribosomal_L31 (pfam01197)→ |

| | | | | | | | |
|---|---|---|---|---|---|---|---|
| Vpa-108-1 | NZ_AWLG01000122.1 | - | 5118 | 5068 | **RNA**→ | rpmE (PRK00019)Ribosomal_L31 (pfam01197)→ | |
| Vpa-68-1 | NZ_AWMU01000003.1 | - | 5712 | 5662 | **RNA**→ | rpmE (PRK00019)Ribosomal_L31 (pfam01197)→ | |
| Vpa-209-1 | NZ_AWHJ01000001.1 | - | 14442 | 14392 | **RNA**→ | rpmE (PRK00019)Ribosomal_L31 (pfam01197)→ | |
| Vpa-203-1 | NZ_AWHP01000127.1 | + | 9049 | 9099 | **RNA**→ | rpmE (PRK00019)Ribosomal_L31 (pfam01197)→ | |
| Vpa-170-1 | NZ_AWIW01000029.1 | + | 31633 | 31683 | **RNA**→ | rpmE (PRK00019)Ribosomal_L31 (pfam01197)→ | |
| Vpa-106-1 | NZ_AWLI01000091.1 | + | 13347 | 13397 | **RNA**→ | rpmE (PRK00019)Ribosomal_L31 (pfam01197)→ | |
| Vpa-146-1 | NZ_AWJU01000061.1 | + | 14639 | 14689 | **RNA**→ | rpmE (PRK00019)Ribosomal_L31 (pfam01197)→ | |
| Vpa-205-1 | NZ_AWHN01000015.1 | - | 5723 | 5673 | **RNA**→ | rpmE (PRK00019)Ribosomal_L31 (pfam01197)→ | |
| Vpa-49-1 | NZ_AVPX01000004.1 | - | 2971905 | 2971855 | **RNA**→ | rpmE (PRK00019)Ribosomal_L31 (pfam01197)→ | |
| Vpa-3-1 | NZ_AVOH01000008.1 | + | 32867 | 32917 | **RNA**→ | rpmE (PRK00019)Ribosomal_L31 (pfam01197)→ | |
| Vpa-76-1 | NZ_AWMM01000015.1 | - | 5718 | 5668 | **RNA**→ | rpmE (PRK00019)Ribosomal_L31 (pfam01197)→ | |
| Vpa-57-1 | NZ_AWNF01000119.1 | - | 5725 | 5675 | **RNA**→ | rpmE (PRK00019)Ribosomal_L31 (pfam01197)→ | |
| Vpa-75-1 | NZ_AWMN01000075.1 | + | 9602 | 9652 | **RNA**→ | rpmE (PRK00019)Ribosomal_L31 (pfam01197)→ | |
| Vpa-10-1 | NZ_AVOL01000005.1 | - | 2951504 | 2951454 | **RNA**→ | rpmE (PRK00019)Ribosomal_L31 (pfam01197)→ | |
| Vpa-72-1 | NZ_AWMQ01000039.1 | + | 27138 | 27188 | **RNA**→ | rpmE (PRK00019)Ribosomal_L31 (pfam01197)→ | |
| Vpa-67-1 | NZ_AWMV01000005.1 | + | 45007 | 45057 | **RNA**→ | rpmE (PRK00019)Ribosomal_L31 (pfam01197)→ | |
| Vpa-58-1 | NZ_AWNE01000131.1 | - | 5731 | 5681 | **RNA**→ | rpmE (PRK00019)Ribosomal_L31 (pfam01197)→ | |
| Vpa-74-1 | NZ_AWMO01000129.1 | + | 8347 | 8397 | **RNA**→ | rpmE (PRK00019)Ribosomal_L31 (pfam01197)→ | |
| Vpa-73-1 | NZ_AWMP01000001.1 | + | 105903 | 105953 | **RNA**→ | rpmE (PRK00019)Ribosomal_L31 (pfam01197)→ | |
| Vpa-85-1 | NZ_AWMD01000134.1 | - | 3102 | 3052 | **RNA**→ | rpmE (PRK00019)Ribosomal_L31 (pfam01197)→ | |
| Vpa-52-1 | NZ_JNUD01000062.1 | + | 94002 | 94052 | **RNA**→ | rpmE (PRK00019)Ribosomal_L31 (pfam01197)→ | |
| Vpa-28-1 | NZ_ACFO01000016.1 | + | 250127 | 250177 | **RNA**→ | rpmE (PRK00019)Ribosomal_L31 (pfam01197)→ | rpmE (PRK00019)Ribosomal_L31 (pfam01197)→ |
| Vpa-86-1 | NZ_AWMC01000105.1 | - | 5744 | 5694 | **RNA**→ | rpmE (PRK00019)Ribosomal_L31 (pfam01197)→ | |
| Vpa-81-1 | NZ_AWMH01000023.1 | - | 41439 | 41389 | **RNA**→ | rpmE (PRK00019)Ribosomal_L31 (pfam01197)→ | |
| Vpa-89-1 | NZ_AWLZ01000119.1 | + | 9880 | 9930 | **RNA**→ | rpmE (PRK00019)Ribosomal_L31 (pfam01197)→ | |
| Vpa-235-1 | NZ_AVOJ01000007.1 | - | 62093 | 62043 | **RNA**→ | rpmE (PRK00019)Ribosomal_L31 (pfam01197)→ | |
| Vpa-90-1 | NZ_AWLY01000125.1 | - | 2796 | 2746 | **RNA**→ | rpmE (PRK00019)Ribosomal_L31 (pfam01197)→ | |
| Vpa-93-1 | NZ_AWLV01000109.1 | - | 5719 | 5669 | **RNA**→ | rpmE (PRK00019)Ribosomal_L31 (pfam01197)→ | |
| Vpa-96-1 | NZ_AWLS01000176.1 | - | 2590 | 2540 | **RNA**→ | rpmE (PRK00019)Ribosomal_L31 (pfam01197)→ | |
| Vpa-92-1 | NZ_AWLW01000157.1 | + | 8341 | 8391 | **RNA**→ | rpmE (PRK00019)Ribosomal_L31 (pfam01197)→ | |
| Vpa-91-1 | NZ_AWLX01000155.1 | + | 8339 | 8389 | **RNA**→ | rpmE (PRK00019)Ribosomal_L31 (pfam01197)→ | |
| Vpa-82-1 | NZ_AWMG01000090.1 | - | 5722 | 5672 | **RNA**→ | rpmE (PRK00019)Ribosomal_L31 (pfam01197)→ | |
| Vpa-227-1 | NZ_AXNP01000089.1 | + | 9009 | 9059 | **RNA**→ | rpmE (PRK00019)Ribosomal_L31 (pfam01197)→ | |
| Vpa-228-1 | NZ_AXNR01000069.1 | - | 232409 | 232359 | **RNA**→ | rpmE (PRK00019)Ribosomal_L31 (pfam01197)→ | |
| Vpa-223-1 | NZ_AXNQ01000037.1 | - | 114511 | 114461 | **RNA**→ | rpmE (PRK00019)Ribosomal_L31 (pfam01197)→ | |
| Vpa-236-1 | NZ_AVOK01000001.1 | - | 2881819 | 2881769 | **RNA**→ | rpmE (PRK00019)Ribosomal_L31 (pfam01197)→ | |
| Vpa-220-1 | NZ_BAVK01000001.1 | - | 282471 | 282421 | **RNA**→ | rpmE (PRK00019)Ribosomal_L31 (pfam01197)→ | |
| Vpa-226-1 | NZ_AXNN01000032.1 | - | 232504 | 232454 | **RNA**→ | rpmE (PRK00019)Ribosomal_L31 (pfam01197)→ | |
| Vpa-214-1 | NZ_AMRZ01000013.1 | + | 20471 | 20521 | **RNA**→ | rpmE (PRK00019)Ribosomal_L31 (pfam01197)→ | |
| Vpa-149-1 | NZ_AWJR01000059.1 | + | 15133 | 15183 | **RNA**→ | rpmE (PRK00019)Ribosomal_L31 (pfam01197)→ | |
| Vpa-145-1 | NZ_AWJV01000051.1 | + | 19667 | 19717 | **RNA**→ | rpmE (PRK00019)Ribosomal_L31 (pfam01197)→ | |
| Vpa-136-1 | NZ_AWKE01000259.1 | - | 3103 | 3053 | **RNA**→ | rpmE (PRK00019)Ribosomal_L31 (pfam01197)→ | |
| Vpa-109-1 | NZ_AWLF01000108.1 | + | 338 | 388 | **RNA**→ | rpmE (PRK00019)Ribosomal_L31 (pfam01197)→ | |
| Vpa-105-1 | NZ_AWLJ01000343.1 | + | 1646 | 1696 | **RNA**→ | rpmE (PRK00019)Ribosomal_L31 (pfam01197)→ | |
| Vpa-104-1 | NZ_AWLK01000105.1 | - | 2793 | 2743 | **RNA**→ | rpmE (PRK00019)Ribosomal_L31 (pfam01197)→ | |
| Vpa-99-1 | NZ_AWLP01000087.1 | + | 9324 | 9374 | **RNA**→ | rpmE (PRK00019)Ribosomal_L31 (pfam01197)→ | |
| Vpa-148-1 | NZ_AWJS01000117.1 | - | 5712 | 5662 | **RNA**→ | rpmE (PRK00019)Ribosomal_L31 (pfam01197)→ | |
| Vpa-135-1 | NZ_AWKF01000118.1 | + | 11247 | 11297 | **RNA**→ | rpmE (PRK00019)Ribosomal_L31 (pfam01197)→ | |
| Vpa-122-1 | NZ_AWKS01000010.1 | + | 46837 | 46887 | **RNA**→ | rpmE (PRK00019)Ribosomal_L31 (pfam01197)→ | |
| Vpa-112-1 | NZ_AWLC01000077.1 | + | 16312 | 16362 | **RNA**→ | rpmE (PRK00019)Ribosomal_L31 (pfam01197)→ | |
| Vpa-142-1 | NZ_AWJY01000001.1 | - | 5717 | 5667 | **RNA**→ | rpmE (PRK00019)Ribosomal_L31 (pfam01197)→ | |
| Vpa-133-1 | NZ_AWKH01000066.1 | + | 16328 | 16378 | **RNA**→ | rpmE (PRK00019)Ribosomal_L31 (pfam01197)→ | |

| | | | | | |
|---|---|---|---|---|---|
| Vpa-110-1 | NZ_AWLE01000231.1 | + | 1597 | 1647 | **RNA**→ rpmE (PRK00019)Ribosomal_L31 (pfam01197)→ |
| Vpa-100-1 | NZ_AWLO01000088.1 | - | 5717 | 5667 | **RNA**→ rpmE (PRK00019)Ribosomal_L31 (pfam01197)→ |
| Vpa-147-1 | NZ_AWJT01000151.1 | + | 6962 | 7012 | **RNA**→ rpmE (PRK00019)Ribosomal_L31 (pfam01197)→ |
| Vpa-134-1 | NZ_AWKG01000077.1 | - | 5725 | 5675 | **RNA**→ rpmE (PRK00019)Ribosomal_L31 (pfam01197)→ |
| Vpa-102-1 | NZ_AWLM01000036.1 | + | 11274 | 11324 | **RNA**→ rpmE (PRK00019)Ribosomal_L31 (pfam01197)→ |
| Vpa-150-1 | NZ_AWJQ01000056.1 | - | 5722 | 5672 | **RNA**→ rpmE (PRK00019)Ribosomal_L31 (pfam01197)→ |
| Vpa-97-1 | NZ_AWLR01000092.1 | + | 12388 | 12438 | **RNA**→ rpmE (PRK00019)Ribosomal_L31 (pfam01197)→ |
| Vpa-218-1 | NZ_BAVI01000019.1 | - | 18992 | 18942 | **RNA**→ rpmE (PRK00019)Ribosomal_L31 (pfam01197)→ |
| Vpa-215-1 | NZ_BAVH01000008.1 | + | 253593 | 253643 | **RNA**→ rpmE (PRK00019)Ribosomal_L31 (pfam01197)→ |
| Vpa-206-1 | NZ_AWHM01000082.1 | - | 5734 | 5684 | **RNA**→ rpmE (PRK00019)Ribosomal_L31 (pfam01197)→ |
| Vpa-201-1 | NZ_AWHR01000011.1 | - | 5725 | 5675 | **RNA**→ rpmE (PRK00019)Ribosomal_L31 (pfam01197)→ |
| Vpa-193-1 | NZ_AWHZ01000070.1 | - | 5709 | 5659 | **RNA**→ rpmE (PRK00019)Ribosomal_L31 (pfam01197)→ |
| Vpa-192-1 | NZ_AWIA01000080.1 | + | 13363 | 13413 | **RNA**→ rpmE (PRK00019)Ribosomal_L31 (pfam01197)→ |
| Vpa-188-1 | NZ_AWIE01000003.1 | + | 51026 | 51076 | **RNA**→ rpmE (PRK00019)Ribosomal_L31 (pfam01197)→ |
| Vpa-179-1 | NZ_AWIN01000218.1 | - | 5712 | 5662 | **RNA**→ rpmE (PRK00019)Ribosomal_L31 (pfam01197)→ |
| Vpa-169-1 | NZ_AWIX01000099.1 | + | 11334 | 11384 | **RNA**→ rpmE (PRK00019)Ribosomal_L31 (pfam01197)→ |
| Vpa-187-1 | NZ_AWIF01000009.1 | - | 5723 | 5673 | **RNA**→ rpmE (PRK00019)Ribosomal_L31 (pfam01197)→ |
| Vpa-172-1 | NZ_AWIU01000005.1 | - | 14448 | 14398 | **RNA**→ rpmE (PRK00019)Ribosomal_L31 (pfam01197)→ |
| Vpa-210-1 | NZ_AWHI01000010.1 | + | 37406 | 37456 | **RNA**→ rpmE (PRK00019)Ribosomal_L31 (pfam01197)→ |
| Vpa-208-1 | NZ_AWHK01000122.1 | + | 13363 | 13413 | **RNA**→ rpmE (PRK00019)Ribosomal_L31 (pfam01197)→ |
| Vpa-202-1 | NZ_AWHQ01000003.1 | - | 5715 | 5665 | **RNA**→ rpmE (PRK00019)Ribosomal_L31 (pfam01197)→ |
| Vpa-194-1 | NZ_AWHY01000088.1 | - | 5707 | 5657 | **RNA**→ rpmE (PRK00019)Ribosomal_L31 (pfam01197)→ |
| Vpa-189-1 | NZ_AWID01000085.1 | - | 5723 | 5673 | **RNA**→ rpmE (PRK00019)Ribosomal_L31 (pfam01197)→ |
| Vpa-184-1 | NZ_AWII01000060.1 | + | 13360 | 13410 | **RNA**→ rpmE (PRK00019)Ribosomal_L31 (pfam01197)→ |
| Vpa-211-1 | NZ_AWHH01000001.1 | + | 114662 | 114712 | **RNA**→ rpmE (PRK00019)Ribosomal_L31 (pfam01197)→ |
| Vpa-204-1 | NZ_AWHO01000125.1 | + | 8341 | 8391 | **RNA**→ rpmE (PRK00019)Ribosomal_L31 (pfam01197)→ |
| Vpa-195-1 | NZ_AWHX01000081.1 | - | 5710 | 5660 | **RNA**→ rpmE (PRK00019)Ribosomal_L31 (pfam01197)→ |
| Vpa-191-1 | NZ_AWIB01000016.1 | - | 5711 | 5661 | **RNA**→ rpmE (PRK00019)Ribosomal_L31 (pfam01197)→ |
| Vpa-190-1 | NZ_AWIC01000085.1 | - | 5710 | 5660 | **RNA**→ rpmE (PRK00019)Ribosomal_L31 (pfam01197)→ |
| Vpa-186-1 | NZ_AWIG01000001.1 | - | 5744 | 5694 | **RNA**→ rpmE (PRK00019)Ribosomal_L31 (pfam01197)→ |
| Vpa-185-1 | NZ_AWIH01000001.1 | - | 5711 | 5661 | **RNA**→ rpmE (PRK00019)Ribosomal_L31 (pfam01197)→ |
| Vpa-181-1 | NZ_AWIL01000049.1 | + | 14828 | 14878 | **RNA**→ rpmE (PRK00019)Ribosomal_L31 (pfam01197)→ |
| Vpa-159-1 | NZ_AWJH01000003.1 | - | 5713 | 5663 | **RNA**→ rpmE (PRK00019)Ribosomal_L31 (pfam01197)→ |
| Vpa-158-1 | NZ_AWJI01000003.1 | - | 5708 | 5658 | **RNA**→ rpmE (PRK00019)Ribosomal_L31 (pfam01197)→ |
| Vpa-168-1 | NZ_AWIY01000019.1 | + | 31982 | 32032 | **RNA**→ rpmE (PRK00019)Ribosomal_L31 (pfam01197)→ |
| Vpa-160-1 | NZ_AWJG01000249.1 | - | 396 | 346 | **RNA**→ rpmE (PRK00019)Ribosomal_L31 (pfam01197)→ |
| Vpa-200-1 | NZ_AWHS01000092.1 | - | 5726 | 5676 | **RNA**→ rpmE (PRK00019)Ribosomal_L31 (pfam01197)→ |
| Vpa-165-1 | NZ_AWJB01000007.1 | - | 396 | 346 | **RNA**→ rpmE (PRK00019)Ribosomal_L31 (pfam01197)→ |
| Vpa-161-1 | NZ_AWJF01000049.1 | - | 5721 | 5671 | **RNA**→ rpmE (PRK00019)Ribosomal_L31 (pfam01197)→ |
| Vpa-197-1 | NZ_AWHV01000125.1 | - | 396 | 346 | **RNA**→ rpmE (PRK00019)Ribosomal_L31 (pfam01197)→ |
| Vpa-166-1 | NZ_AWJA01000056.1 | - | 14431 | 14381 | **RNA**→ rpmE (PRK00019)Ribosomal_L31 (pfam01197)→ |
| Vpa-4-1 | NZ_KI549257.1 | - | 292241 | 292191 | **RNA**→ rpmE (PRK00019)Ribosomal_L31 (pfam01197)→ |
| Vpa-9-1 | NZ_KI912894.1 | + | 256465 | 256515 | **RNA**→ rpmE (PRK00019)Ribosomal_L31 (pfam01197)→ |
| Vpa-6-1 | NZ_KI601405.1 | - | 292231 | 292181 | **RNA**→ rpmE (PRK00019)Ribosomal_L31 (pfam01197)→ |
| Vpa-213-1 | NZ_KI929493.1 | - | 292480 | 292430 | **RNA**→ rpmE (PRK00019)Ribosomal_L31 (pfam01197)→ |
| Vpa-34-1 | NZ_KI783133.1 | - | 291972 | 291922 | **RNA**→ rpmE (PRK00019)Ribosomal_L31 (pfam01197)→ |
| Vpa-17-1 | NZ_KI912778.1 | + | 236877 | 236927 | **RNA**→ rpmE (PRK00019)Ribosomal_L31 (pfam01197)→ |
| Vpa-26-1 | NZ_KI783213.1 | - | 196217 | 196167 | **RNA**→ rpmE (PRK00019)Ribosomal_L31 (pfam01197)→ |
| Vpa-35-1 | NZ_KK042409.1 | + | 247870 | 247920 | **RNA**→ rpmE (PRK00019)Ribosomal_L31 (pfam01197)→ |
| Vpa-5-1 | NZ_AFBW01000018.1 | + | 256269 | 256319 | **RNA**→ rpmE (PRK00019)Ribosomal_L31 (pfam01197)→ |
| Vpa-241-1 | NZ_KK081893.1 | + | 21893 | 21943 | **RNA**→ rpmE (PRK00019)Ribosomal_L31 (pfam01197)→ |
| Vpa-239-1 | NZ_KK081837.1 | - | 103889 | 103839 | **RNA**→ rpmE (PRK00019)Ribosomal_L31 (pfam01197)→ |

| | | | | | |
|---|---|---|---|---|---|
| Vpa-243-1 | NZ_KK082063.1 | - | 295164 | 295114 | **RNA**→ rpmE (PRK00019)Ribosomal_L31 (pfam01197)→ |
| Vpa-242-1 | NZ_KK081954.1 | + | 246629 | 246679 | **RNA**→ rpmE (PRK00019)Ribosomal_L31 (pfam01197)→ |
| Vpa-244-1 | NZ_KB932197.1 | - | 292184 | 292134 | **RNA**→ hypo→ |
| Vpa-50-1 | NC_021848.1 | - | 2637243 | 2637193 | **RNA**→ rpmE (PRK00019)Ribosomal_L31 (pfam01197)→ |
| Vpa-51-1 | NC_021847.1 | + | 655200 | 655250 | **RNA**→ rpmE (PRK00019)Ribosomal_L31 (pfam01197)→ |
| Vpa-221-1 | NZ_CP007004.1 | + | 305279 | 305329 | **RNA**→ rpmE (PRK00019)Ribosomal_L31 (pfam01197)→ |
| Vpa-2-1 | NZ_CP009982.1 | - | 2981797 | 2981747 | **RNA**→ rpmE (PRK00019)Ribosomal_L31 (pfam01197)→ |
| Vpa-107-1 | NZ_AWLH01000247.1 | + | 1797 | 1847 | **RNA**→ rpmE (PRK00019)Ribosomal_L31 (pfam01197)→ |
| Vpa-230-1 | NZ_AXNS01000054.1 | - | 232477 | 232427 | **RNA**→ rpmE (PRK00019)Ribosomal_L31 (pfam01197)→ |
| Vpa-225-1 | NZ_AXNO01000047.1 | + | 185816 | 185866 | **RNA**→ rpmE (PRK00019)Ribosomal_L31 (pfam01197)→ |
| Vpa-216-1 | NZ_BAVF01000010.1 | + | 243480 | 243530 | **RNA**→ rpmE (PRK00019)Ribosomal_L31 (pfam01197)→ |
| Ehe-1-1 | NZ_BAFF01000008.1 | + | 130616 | 130667 | **RNA**→ rpmE (PRK00019)Ribosomal_L31 (pfam01197)→ rpmE (PRK00019)Ribosomal_L31 (pfam01197)→ |
| Eco-230-1 | NZ_JXMX01000003.1 | - | 847619 | 847567 | **RNA**→ rpmE (PRK00019)Ribosomal_L31 (pfam01197)→ |
| Eco-95-1 | NZ_JNQG01000012.1 | + | 113897 | 113949 | **RNA**→ rpmE (PRK00019)Ribosomal_L31 (pfam01197)→ |
| Eco-202-1 | NZ_AYQH01000026.1 | - | 32643 | 32591 | **RNA**→ rpmE (PRK00019)Ribosomal_L31 (pfam01197)→ |
| Eco-514-1 | NZ_JHNV01000011.1 | - | 49699 | 49647 | **RNA**→ rpmE (PRK00019)Ribosomal_L31 (pfam01197)→ |
| Eco-46-1 | NZ_AYQG01000014.1 | - | 32634 | 32582 | **RNA**→ rpmE (PRK00019)Ribosomal_L31 (pfam01197)→ |
| Eco-230-2 | NZ_AYRA01000032.1 | - | 32648 | 32596 | **RNA**→ rpmE (PRK00019)Ribosomal_L31 (pfam01197)→ |
| Eco-280-1 | NZ_AYQF01000011.1 | - | 32645 | 32593 | **RNA**→ rpmE (PRK00019)Ribosomal_L31 (pfam01197)→ |
| Eco-1173-1 | NZ_GL884547.1 | + | 79856 | 79908 | **RNA**→ rpmE (PRK00019)Ribosomal_L31 (pfam01197)→ rpmE (PRK00019)Ribosomal_L31 (pfam01197)→ |
| Eco-1617-1 | NZ_KB732779.1 | + | 1174695 | 1174747 | **RNA**→ rpmE (PRK00019)Ribosomal_L31 (pfam01197)→ |
| Eco-1362-1 | NZ_KE699546.1 | + | 733437 | 733489 | **RNA**→ rpmE (PRK00019)Ribosomal_L31 (pfam01197)→ |
| Eco-1272-1 | NZ_KE700731.1 | + | 4435409 | 4435461 | **RNA**→ rpmE (PRK00019)Ribosomal_L31 (pfam01197)→ |
| Eco-1987-1 | NZ_AOGL01000074.1 | + | 21073 | 21125 | **RNA**→ rpmE (PRK00019)Ribosomal_L31 (pfam01197)→ |
| Vvu-5-1 | NZ_LBNN01000028.1 | + | 15413 | 15464 | **RNA**→ rpmE (PRK00019)Ribosomal_L31 (pfam01197)→ |
| Vvu-11-1 | NC_005139.1 | - | 3092393 | 3092342 | **RNA**→ rpmE (PRK00019)Ribosomal_L31 (pfam01197)→ |
| Vvu-4-1 | NC_004459.3 | + | 1340353 | 1340404 | **RNA**→ rpmE (PRK00019)Ribosomal_L31 (pfam01197)→ |
| Vvu-1-1 | NZ_CP009261.1 | - | 3027015 | 3026964 | **RNA**→ rpmE (PRK00019)Ribosomal_L31 (pfam01197)→ |
| Cbr-1-7 | NZ_JUYY01000023.1 | + | 59204 | 59254 | **RNA**→ rpmE (PRK00019)Ribosomal_L31 (pfam01197)→ |
| Esp-47-1 | NZ_AVOP01000007.1 | + | 72571 | 72622 | **RNA**→ rpmE (PRK00019)Ribosomal_L31 (pfam01197)→ |
| Esp-61-1 | NZ_KI535657.1 | - | 1293528 | 1293477 | **RNA**→ rpmE (PRK00019)Ribosomal_L31 (pfam01197)→ |
| Hin-12-1 | NC_022356.1 | - | 434942 | 434892 | **RNA**→ rpmE (PRK00019)Ribosomal_L31 (pfam01197)→ |
| Kox-11-1 | NZ_JH603153.1 | + | 4689088 | 4689139 | **RNA**→ rpmE (PRK00019)Ribosomal_L31 (pfam01197)→ |
| Kmi-1-1 | NZ_KK036986.1 | + | 403430 | 403481 | **RNA**→ rpmE (PRK00019)Ribosomal_L31 (pfam01197)→ |
| Pst-7-1 | NC_017731.1 | + | 2402863 | 2402914 | **RNA**→ rpmE (PRK00019)Ribosomal_L31 (pfam01197)→ |
| Pst-6-1 | NZ_DS607672.1 | - | 59943 | 59892 | **RNA**→ rpmE (PRK00019)Ribosomal_L31 (pfam01197)→ |
| Pst-2-1 | NZ_CP008920.1 | - | 1023294 | 1023243 | **RNA**→ rpmE (PRK00019)Ribosomal_L31 (pfam01197)→ |
| Vco-2-1 | NZ_ACZN01000020.1 | + | 156744 | 156794 | **RNA**→ rpmE (PRK00019)Ribosomal_L31 (pfam01197)→ rpmE (PRK00019)Ribosomal_L31 (pfam01197)→ |
| Vco-1-1 | NZ_CP009617.1 | + | 3234588 | 3234638 | **RNA**→ rpmE (PRK00019)Ribosomal_L31 (pfam01197)→ |
| Vco-1-2 | NZ_CP009264.1 | + | 3063229 | 3063279 | **RNA**→ rpmE (PRK00019)Ribosomal_L31 (pfam01197)→ |
| Yfr-3-1 | NZ_CP009364.1 | - | 3370416 | 3370362 | **RNA**→ rpmE (PRK00019)Ribosomal_L31 (pfam01197)→ |
| Ykr-1-1 | NZ_CP009997.1 | + | 2908986 | 2909040 | **RNA**→ rpmE (PRK00019)Ribosomal_L31 (pfam01197)→ |
| Ykr-1-2 | NZ_CP008955.1 | + | 3824913 | 3824967 | **RNA**→ rpmE (PRK00019)Ribosomal_L31 (pfam01197)→ |
| Kmi-1-2 | NZ_KK036954.1 | + | 23422 | 23473 | **RNA**→ rpmE (PRK00019)Ribosomal_L31 (pfam01197)→ |
| Kox-1-3 | NZ_CP011636.1 | - | 259495 | 259444 | **RNA**→ rpmE (PRK00019)Ribosomal_L31 (pfam01197)→ |

#### 12.3 Conserved domains

Conserved domains found in protein-coding genes listed in Section 12.2 are shown below, with the first sentence in their description from the Conserved Domain Database (if any). Conserved domains associated with more than one L31-Gammaproteobacteria RNA are assigned a color, while others are shown in gray. The number in parentheses after the colored domain name is the number of occurrences in Section 12.2.

cd00090 (1) Arsenical Resistance Operon Repressor and similar prokaryotic, metal regulated homodimeric repressors.
cd00761 (18) Glycosyltransferase family A (GT-A) includes diverse families of glycosyl transferases with a common GT-A type structural fold.
cd01347 (5) TonB dependent/Ligand-Gated channels are created by a monomeric 22 strand (22,24) anti-parallel beta-barrel.
cd03811 (12) This family is most closely related to the GT1 family of glycosyltransferases.
cd04194 (11) A4GalT_like proteins catalyze the addition of galactose or glucose residues to the lipooligosaccharide (LOS) or lipopolysaccharide (LPS) of the bacterial cell surface.
cd05399 (1) Nucleotidyltransferase (NT) domain of RelA- and SpoT-like ppGpp synthetases and hydrolases. [D/E]
cd06532 (18) Glycosyltransferase family 25 [lipooligosaccharide (LOS) biosynthesis protein]
cd07812 (4) START/RHO_alpha_C/PITP/Bet_v1/CoxG/CalC (SRPBCC) ligand-binding domain superfamily.
cd08430 (12) The C-terminal substrate binding of LysR-type transcriptional regulator IlvY, which activates the expression of ilvC gene that encoding acetohydroxy acid isomeroreductase for the biosynthesis of branched amino acids; contains the type 2 periplasmic binding fold.
cd08893 (2) Putative hydrophobic ligand-binding SRPBCC domain of an uncharacterized subgroup of CalC- and Aha1-like proteins; some contain an N-terminal GntR family winged HTH DNA-binding domain.
cd11528 (1) Nucleoside Triphosphate Pyrophosphohydrolase (EC 3.
cd11529 (1) Nucleoside Triphosphate Pyrophosphohydrolase (EC 3.
COG0035 (1) Uracil phosphoribosyltransferase [Nucleotide transport and metabolism]
COG0254 (101) Ribosomal protein L31 [Translation, ribosomal structure and biogenesis]
COG0281 (144) Malic enzyme [Energy production and conversion]
COG0438 (11) Glycosyltransferase involved in cell wall bisynthesis [Cell wall/membrane/envelope biogenesis]
COG0702 (7) Uncharacterized conserved protein YbjT, contains NAD(P)-binding and DUF2867 domains [General function prediction only]
COG0859 (133) ADP-heptose:LPS heptosyltransferase [Cell wall/membrane/envelope biogenesis]
COG1198 (3) Primosomal protein N' (replication factor Y) - superfamily II helicase [Replication, recombination and repair]
COG1368 (1) Phosphoglycerol transferase MdoB or a related enzyme of AlkP superfamily [Cell wall/membrane/envelope biogenesis]
COG1943 (3) REP element-mobilizing transposase RayT [Mobilome: prophages, transposons]
COG2194 (2) Phosphoethanolamine transferase for periplasmic glucans (OPG), alkaline phosphatase superfamily [Cell wall/membrane/envelope biogenesis]
COG2211 (1) Na+/melibiose symporter or related transporter [Carbohydrate transport and metabolism]
COG3060 (2) Transcriptional regulator of met regulon [Transcription, Amino acid transport and metabolism]
COG3306 (4) Glycosyltransferase involved in LPS biosynthesis, GR25 family [Cell wall/membrane/envelope biogenesis]
COG3550 (1) Serine/threonine protein kinase HipA, toxin component of the HipAB toxin-antitoxin module [Signal transduction mechanisms]
pfam00126 (12) Bacterial regulatory helix-turn-helix protein, lysR family.
pfam00535 (2) Glycosyl transferase family 2.
pfam00593 (3) TonB dependent receptor.
pfam00742 (2) Homoserine dehydrogenase.
pfam01197 (10003) Ribosomal protein L31.
pfam01755 (73) Glycosyltransferase family 25 (LPS biosynthesis protein).
pfam06178 (1) Oligogalacturonate-specific porin protein (KdgM).
pfam06306 (3) Beta-1,4-N-acetylgalactosaminyltransferase (CgtA).
pfam07526 (9) Associated with HOX.
pfam11066 (7) Protein of unknown function (DUF2867).
pfam13439 (9) Glycosyltransferase Family 4.
pfam13460 (7) NAD(P)H-binding.
pfam13641 (14) Glycosyltransferase like family 2.
pfam13692 (23) Glycosyl transferases group 1.
PRK00019 (10104) 50S ribosomal protein L31; Reviewed
PRK00129 (1) uracil phosphoribosyltransferase; Reviewed
PRK00528 (8) 50S ribosomal protein L31; Reviewed
PRK01678 (16) 50S ribosomal protein L31 type B; Reviewed
PRK05264 (2) transcriptional repressor protein MetJ; Provisional
PRK05580 (3) primosome assembly protein PriA; Validated
PRK09466 (2) bifunctional aspartate kinase II/homoserine dehydrogenase II; Provisional
PRK09562 (1) nucleoside triphosphate pyrophosphohydrolase; Reviewed
PRK09775 (1) putative DNA-binding transcriptional regulator; Provisional
PRK10073 (2) putative glycosyl transferase; Provisional
PRK10422 (2) lipopolysaccharide core biosynthesis protein; Provisional
PRK11204 (2) N-glycosyltransferase; Provisional
PRK11560 (2) phosphoethanolamine transferase; Provisional
PRK11716 (12) DNA-binding transcriptional regulator IlvY; Provisional
PRK13524 (4) outer membrane receptor FepA; Provisional
PRK13528 (3) outer membrane receptor FepA; Provisional
PRK14515 (1) aspartate ammonia-lyase; Provisional
PRK15171 (11) lipopolysaccharide 1,3-galactosyltransferase; Provisional
smart00919 (144) Malic enzyme, NAD binding domain.
smart00954 (1) Region found in RelA / SpoT proteins.
TIGR00792 (1) sugar (Glycoside-Pentoside-Hexuronide) transporter. [Transport and binding proteins, Carbohydrates, organic alcohols, and acids]
TIGR01783 (1) TonB-dependent siderophore receptor. [Transport and binding proteins, Cations and iron carrying compounds, Transport and binding proteins, Porins]
TIGR02193 (53) lipopolysaccharide heptosyltransferase I. [Cell envelope, Biosynthesis and degradation of surface polysaccharides and lipopolysaccharides]
TIGR02195 (9) lipopolysaccharide heptosyltransferase II. [Cell envelope, Biosynthesis and degradation of surface polysaccharides and lipopolysaccharides]
TIGR02201 (63) lipopolysaccharide heptosyltransferase III, putative. [Cell envelope, Biosynthesis and degradation of surface polysaccharides and lipopolysaccharides]
TIGR03937 (2) poly-beta-1,6 N-acetyl-D-glucosamine synthase.

#### 12.4 Multiple-sequence alignment

Each L31-Gammaproteobacteria RNA is denoted by an abbreviation (like "Eco-1-1") that refers to a taxonomy given in Section 12.1. The alignment may include sequences containing the RNA motif, as well as flanking sequence. The L31-Gammaproteobacteria RNA itself is denoted by the line underneath marked 5′ and 3′ on either end. Nucleotides in flanking sequences (i.e., not part of the motif) are written in gray letters. Stems of predicted rho-independent transcription terminators, if any, are shaded yellow. (Note: terminator predictions have not been analyzed manually, and many are likely to be false positives. Terminator predictions are those of the RNIE software.) Nucleotides predicted to function (as DNA) as transcription-factor binding sites, if any, are shaded in green. (But note: these predictions are manually annotated, so they might be under-predicted.) Annotated start codons, if any, are shaded green. (Note: start codons are frequently misannotated, especially in environmental samples.) Nucleotides proposed to basepair as part of the consensus structure are shaded in color when they comprise Watson-Crick or G-U pairs. Otherwise they are shaded gray. Conserved stems are also indicated at the bottom of the alignment by angle brackets, where matching $<$ and $>$ denote base-paired columns. Below these angle brackets, the symbol "2" denotes base pairs exhibiting covariation according to the statistically well-founded R-scape method. "1" denotes base pairs exhibiting covariation according to R2R's simplistic method. "0" denotes base pairs that are not observed to mutate and "?" denotes base pairs that have a significant frequency of non-canonical nucleotides for Watson-Crick or G-U pairs ($>$ 5%). Below these base pair annotation is the consensus sequence: "R" = "A" or "G", "Y" = "C" or "U", red nucleotides: nucleotide identity conserved more than 97% of the time, black nucleotides: 90%, gray nucleotides: 75%, red circle ($\odot$): nucleotide is present 97% of the time, black circle ($\odot$): 90%, gray circle ($\odot$): 75%, white circle ($\odot$): 50%. All percentages of sequences just described (e.g. 97% conserved) assume that sequences have been weighted by the GSC algorithm implemented by the Infernal software package.

Duplicate sequences: the following putative homologs are not shown in the alignment because their sequences are identical to a homolog already shown: Aac-10-1, Aac-12-1, Aac-13-1, Aac-15-1, Aac-16-1, Aac-19-1, Aac-2-1, Aac-20-1, Aac-3-1, Aac-4-1, Aac-5-1, Aac-5-2, Aac-6-2, Aac-7-1, Aac-8-1, Aac-9-1, Aal-1-2, Aap-1-1, Aap-2-1, Aap-3-1, Aap-4-1, Aaq-1-1, Aau-2-1, Aau-2-2, Abe-1-1, Aca-1-1, Aca-2-1, Adh-1-1, Adh-1-2, Adh-1-3, Adi-1-1, Aen-1-1, Aen-2-1, Aen-2-2, Aen-2-3, Afi-3-1, Afl-1-1, Ahy-1-1, Ahy-1-2, Ahy-10-1, Ahy-11-1, Ahy-12-1, Ahy-13-1, Ahy-14-1, Ahy-15-1, Ahy-16-1, Ahy-17-1, Ahy-18-1, Ahy-19-1, Ahy-2-1, Ahy-20-1, Ahy-21-1, Ahy-3-1, Ahy-4-1, Ahy-5-1, Ahy-6-1, Ahy-7-1, Ahy-8-1, Ahy-9-1, Aja-2-1, Aje-1-1, Ala-1-1, Alo-1-1, Alo-2-1, Ama-10-1, Ama-11-1, Ama-12-1, Ama-13-1, Ama-2-1, Ama-2-2, Ama-2-3, Ama-3-1, Ama-4-1, Ama-5-1, Ama-6-1, Ama-8-1, Ama-9-1, Ame-1-2, Ame-1-3, Ame-2-1, Apa-2-1, Apa-3-1, Apa-4-1, Apl-1-2, Apl-1-3, Apl-1-4, Apl-1-5, Apl-1-6, Apl-10-1, Apl-11-1, Apl-12-1, Apl-13-2, Apl-14-1, Apl-15-1, Apl-2-1, Apl-3-1, Apl-4-1, Apl-4-2, Apl-5-1, Apl-7-1, Apl-9-1, Apo-1-1, Asa-1-1, Asa-2-1, Asa-3-1, Asa-4-1, Asa-5-1, Asa-5-2, Asa-5-3, Asa-5-4, Asa-6-1, Asa-7-1, Asa-8-1, Ase-1-1, Ase-2-1, Aso-1-1, Asp-1-1, Asp-2-1, Asp-3-1, Asp-4-1, Asp-6-1, Asu-2-1, Ata-1-2, Ave-1-1, Ave-1-2, Ave-1-3, Ave-2-1, Ave-3-1, Ave-4-1, Ave-5-1, Ave-6-1, Ave-7-1, Ave-9-1, Cam-1-2, Cam-1-3, Cbr-1-1, Cbr-1-2, Cbr-1-3, Cbr-1-4, Cbr-1-5, Cbr-1-6, Cco-1-1, Cdu-1-1, Cdu-2-1, Cdu-3-1, Cdu-4-1, Cdu-5-1, Cfr-1-1, Cfr-1-2, Cfr-1-3, Cfr-1-4, Cfr-10-1, Cfr-11-1, Cfr-2-1, Cfr-2-2, Cfr-4-1, Cfr-5-1, Cfr-6-1, Cfr-7-1, Cfr-8-1, Cfr-9-1, Cko-1-1, Cko-1-2, Cma-1-1, Cma-1-2, Cma-1-3, Cma-1-4, Cma-1-5, Cma-2-1, Cma-3-1, Cma-4-1, Cma-5-1, Cmi-1-1, Cne-1-2, Cpa-1-1, Cro-1-1, Cro-3-1, Csa-1-1, Csa-11-1, Csa-12-1, Csa-14-1, Csa-15-1, Csa-16-1, Csa-2-1, Csa-3-1, Csa-4-1, Csa-5-1, Csa-6-1, Csa-8-1, Csa-9-1, Csp-1-1, Csp-10-1, Csp-11-1, Csp-13-1, Csp-15-1, Csp-16-1, Csp-17-1, Csp-18-1, Csp-19-1, Csp-2-1, Csp-20-1, Csp-3-1, Csp-4-1, Csp-5-1, Csp-6-1, Csp-7-1, Csp-8-1, Csp-9-1, Ctu-1-1, Ctu-2-1, Ctu-3-1, Ctu-4-1, Cun-1-2, Dch-1-1, Dch-4-1, Dda-3-1, Dda-4-1, Dda-5-1, Ddi-1-1, Ddi-2-1, Ddi-3-1, Ddi-4-1, Dso-1-1, Dso-2-1, Dso-3-1, Dso-5-1, Dso-6-1, Dsp-4-1, Dsp-5-1, Dsp-6-1, Dsp-7-1, Dsp-8-1, Dze-1-1, Dze-3-1, Dze-4-1, Dze-5-1, Dze-6-1, Dze-8-1, Dze-9-1, Eae-1-1, Eae-2-1, Eae-3-1, Eal-1-1, Eal-3-1, Eal-4-1, Eal-5-1, Eam-10-1, Eam-11-1, Eam-12-1, Eam-13-1, Eam-14-1, Eam-15-1, Eam-3-1, Eam-4-1, Eam-5-1, Eam-6-1, Eam-7-1, Eam-8-1, Eam-9-1, Ean-1-1, Ean-2-1, Ean-3-1, Ean-4-1, Eas-1-1, Eas-1-2, Eas-2-1, Eas-3-1, Eas-4-1, Eba-3-1, Eba-5-1, Eca-1-1, Eca-2-1, Eca-4-1, Eca-7-1, Ecl-1-1, Ecl-1-10, Ecl-1-11, Ecl-1-12, Ecl-1-2, Ecl-1-3, Ecl-1-4, Ecl-1-5, Ecl-1-7, Ecl-1-8, Ecl-1-9, Ecl-10-1, Ecl-11-1, Ecl-12-1, Ecl-13-1, Ecl-14-1, Ecl-15-1, Ecl-16-1, Ecl-17-1, Ecl-18-1, Ecl-19-1, Ecl-2-1, Ecl-20-1, Ecl-21-1, Ecl-22-1, Ecl-23-1, Ecl-24-1, Ecl-25-1, Ecl-26-1, Ecl-27-1, Ecl-28-1, Ecl-28-2, Ecl-28-3, Ecl-28-4, Ecl-28-5, Ecl-28-6, Ecl-29-1, Ecl-3-1, Ecl-30-1, Ecl-31-1, Ecl-32-1, Ecl-33-1, Ecl-34-1, Ecl-35-1, Ecl-36-1, Ecl-37-1, Ecl-38-1, Ecl-4-1, Ecl-40-1, Ecl-41-1, Ecl-43-1, Ecl-44-1, Ecl-45-1, Ecl-46-1, Ecl-47-1, Ecl-48-1, Ecl-49-1, Ecl-5-1, Ecl-50-1, Ecl-51-1, Ecl-52-2, Ecl-52-3, Ecl-52-4, Ecl-52-5, Ecl-53-1, Ecl-54-1, Ecl-55-1, Ecl-56-1, Ecl-57-1, Ecl-58-1, Ecl-59-1, Ecl-6-1, Ecl-61-1, Ecl-62-1, Ecl-63-1, Ecl-64-1, Ecl-65-1, Ecl-66-1, Ecl-67-1, Ecl-68-1, Ecl-69-1, Ecl-7-1, Ecl-70-1, Ecl-8-1, Ecl-9-1, Eco-1-1, Eco-1-2, Eco-1-3, Eco-1-4, Eco-10-1, Eco-100-1, Eco-1000-1, Eco-1001-1, Eco-1002-1, Eco-1003-1, Eco-1004-1, Eco-1005-1, Eco-1006-1, Eco-1007-1, Eco-1008-1, Eco-1009-1, Eco-101-1, Eco-1010-1, Eco-1012-1, Eco-1013-1, Eco-1014-1, Eco-1015-1, Eco-1016-1, Eco-1017-1, Eco-1018-1, Eco-1019-1, Eco-1019-2, Eco-1019-3, Eco-102-1, Eco-1020-1, Eco-1021-1, Eco-1022-1, Eco-1023-1, Eco-1024-1, Eco-1025-1, Eco-1026-1, Eco-1027-1, Eco-1028-1, Eco-1029-1, Eco-103-1, Eco-1030-1, Eco-1031-1, Eco-1032-1, Eco-1033-1, Eco-1034-1, Eco-1035-1, Eco-1036-1, Eco-1037-1, Eco-1038-1, Eco-1039-1, Eco-104-1, Eco-1040-1, Eco-1041-1, Eco-1042-1, Eco-1043-1, Eco-1044-1, Eco-1045-1, Eco-1046-1, Eco-1047-1, Eco-1048-1, Eco-1049-1, Eco-105-1, Eco-1050-1,

Eco-1051-1, Eco-1052-1, Eco-1053-1, Eco-1054-1, Eco-1055-1, Eco-1056-1, Eco-1057-1, Eco-1058-1, Eco-1059-1, Eco-106-1, Eco-1060-1, Eco-1061-1, Eco-1062-1, Eco-
1063-1, Eco-1064-1, Eco-1065-1, Eco-1066-1, Eco-1067-1, Eco-1068-1, Eco-1069-1, Eco-107-1, Eco-1070-1, Eco-1071-1, Eco-1072-1, Eco-1073-1, Eco-1074-1, Eco-1075-1,
Eco-1076-1, Eco-1077-1, Eco-1078-1, Eco-1079-1, Eco-108-1, Eco-1080-1, Eco-1081-1, Eco-1082-1, Eco-1083-1, Eco-1084-1, Eco-1085-1, Eco-1086-1, Eco-1087-1, Eco-
1088-1, Eco-1089-1, Eco-109-1, Eco-1090-1, Eco-1091-1, Eco-1092-1, Eco-1093-1, Eco-1094-1, Eco-1095-1, Eco-1096-1, Eco-1097-1, Eco-1098-1, Eco-1099-1, Eco-11-1,
Eco-110-1, Eco-1100-1, Eco-1101-1, Eco-1101-2, Eco-1102-1, Eco-1103-1, Eco-1104-1, Eco-1105-1, Eco-1106-1, Eco-1107-1, Eco-1108-1, Eco-1109-1, Eco-111-1, Eco-
1110-1, Eco-1111-1, Eco-1112-1, Eco-1113-1, Eco-1114-1, Eco-1115-1, Eco-1116-1, Eco-1117-1, Eco-1118-1, Eco-1119-1, Eco-112-1, Eco-1120-1, Eco-1121-1, Eco-1122-1,
Eco-1123-1, Eco-1124-1, Eco-1125-1, Eco-1126-1, Eco-1127-1, Eco-1128-1, Eco-1129-1, Eco-113-1, Eco-1130-1, Eco-1131-1, Eco-1132-1, Eco-1133-1, Eco-1134-1, Eco-
1135-1, Eco-1136-1, Eco-1137-1, Eco-1138-1, Eco-1139-1, Eco-114-1, Eco-1140-1, Eco-1141-1, Eco-1142-1, Eco-1143-1, Eco-1144-1, Eco-1145-1, Eco-1146-1, Eco-1147-1,
Eco-1148-1, Eco-1149-1, Eco-115-1, Eco-1150-1, Eco-1151-1, Eco-1152-1, Eco-1153-1, Eco-1154-1, Eco-1155-1, Eco-1156-1, Eco-1157-1, Eco-1158-1, Eco-1159-1, Eco-
116-1, Eco-1160-1, Eco-1161-1, Eco-1162-1, Eco-1163-1, Eco-1164-1, Eco-1165-1, Eco-1166-1, Eco-1167-1, Eco-1168-1, Eco-1169-1, Eco-1169-2, Eco-117-1, Eco-1170-1,
Eco-1171-1, Eco-1172-1, Eco-1173-1, Eco-1174-1, Eco-1175-1, Eco-1176-1, Eco-1177-1, Eco-1178-1, Eco-1179-1, Eco-118-1, Eco-1180-1, Eco-1181-1, Eco-1182-1, Eco-
1183-1, Eco-1184-1, Eco-1185-1, Eco-1186-1, Eco-1187-1, Eco-1188-1, Eco-1189-1, Eco-119-1, Eco-1190-1, Eco-1191-1, Eco-1192-1, Eco-1193-1, Eco-1194-1, Eco-1195-1,
Eco-1196-1, Eco-1197-1, Eco-1198-1, Eco-1199-1, Eco-12-1, Eco-120-1, Eco-1200-1, Eco-1201-1, Eco-1202-1, Eco-1203-1, Eco-1204-1, Eco-1205-1, Eco-1206-1, Eco-
1207-1, Eco-1208-1, Eco-1209-1, Eco-121-1, Eco-1210-1, Eco-1211-1, Eco-1212-1, Eco-1213-1, Eco-1214-1, Eco-1215-1, Eco-1216-1, Eco-1217-1, Eco-1218-1, Eco-1219-1,
Eco-122-1, Eco-1220-1, Eco-1221-1, Eco-1222-1, Eco-1223-1, Eco-1224-1, Eco-1225-1, Eco-1226-1, Eco-1227-1, Eco-1228-1, Eco-1229-1, Eco-123-1, Eco-1230-1, Eco-
1231-1, Eco-1232-1, Eco-1233-1, Eco-1234-1, Eco-1235-1, Eco-1236-1, Eco-1237-1, Eco-1238-1, Eco-1239-1, Eco-124-1, Eco-1240-1, Eco-1241-1, Eco-1242-1, Eco-1243-1,
Eco-1244-1, Eco-1245-1, Eco-1246-1, Eco-1247-1, Eco-1248-1, Eco-1249-1, Eco-125-1, Eco-1250-1, Eco-1251-1, Eco-1252-1, Eco-1253-1, Eco-1254-1, Eco-1255-1, Eco-
1256-1, Eco-1257-1, Eco-1258-1, Eco-1259-1, Eco-126-1, Eco-1260-1, Eco-1261-1, Eco-1262-1, Eco-1263-1, Eco-1264-1, Eco-1265-1, Eco-1266-1, Eco-1267-1, Eco-1268-1,
Eco-1269-1, Eco-127-1, Eco-1270-1, Eco-1271-1, Eco-1272-1, Eco-1273-1, Eco-1274-1, Eco-1275-1, Eco-1276-1, Eco-1277-1, Eco-1278-1, Eco-1279-1, Eco-128-1, Eco-
1280-1, Eco-1281-1, Eco-1282-1, Eco-1283-1, Eco-1284-1, Eco-1285-1, Eco-1286-1, Eco-1287-1, Eco-1288-1, Eco-1289-1, Eco-129-1, Eco-1290-1, Eco-1291-1, Eco-1292-1,
Eco-1293-1, Eco-1294-1, Eco-1295-1, Eco-1296-1, Eco-1297-1, Eco-1298-1, Eco-1299-1, Eco-13-1, Eco-130-1, Eco-1300-1, Eco-1301-1, Eco-1302-1, Eco-1303-1, Eco-
1304-1, Eco-1305-1, Eco-1306-1, Eco-1307-1, Eco-1308-1, Eco-1309-1, Eco-131-1, Eco-1310-1, Eco-1311-1, Eco-1312-1, Eco-1313-1, Eco-1314-1, Eco-1315-1, Eco-1316-1,
Eco-1317-1, Eco-1318-1, Eco-1319-1, Eco-132-1, Eco-1320-1, Eco-1321-1, Eco-1322-1, Eco-1323-1, Eco-1324-1, Eco-1325-1, Eco-1326-1, Eco-1327-1, Eco-1328-1, Eco-
1329-1, Eco-133-1, Eco-1330-1, Eco-1331-1, Eco-1332-1, Eco-1333-1, Eco-1334-1, Eco-1335-1, Eco-1336-1, Eco-1337-1, Eco-1338-1, Eco-1339-1, Eco-134-1, Eco-1340-1,
Eco-1341-1, Eco-1342-1, Eco-1343-1, Eco-1344-1, Eco-1345-1, Eco-1346-1, Eco-1347-1, Eco-1348-1, Eco-1349-1, Eco-135-1, Eco-1350-1, Eco-1351-1, Eco-1352-1, Eco-
1353-1, Eco-1354-1, Eco-1355-1, Eco-1356-1, Eco-1357-1, Eco-1358-1, Eco-1359-1, Eco-136-1, Eco-1360-1, Eco-1361-1, Eco-1362-1, Eco-1363-1, Eco-1364-1, Eco-1365-1,
Eco-1366-1, Eco-1367-1, Eco-1368-1, Eco-1369-1, Eco-137-1, Eco-1370-1, Eco-1371-1, Eco-1372-1, Eco-1373-1, Eco-1374-1, Eco-1375-1, Eco-1376-1, Eco-1377-1, Eco-
1378-1, Eco-1379-1, Eco-138-1, Eco-1380-1, Eco-1381-1, Eco-1382-1, Eco-1383-1, Eco-1384-1, Eco-1385-1, Eco-1386-1, Eco-1387-1, Eco-1388-1, Eco-1389-1, Eco-139-1,
Eco-1390-1, Eco-1391-1, Eco-1392-1, Eco-1392-2, Eco-1393-1, Eco-1394-1, Eco-1395-1, Eco-1396-1, Eco-1397-1, Eco-1398-1, Eco-1399-1, Eco-14-1, Eco-140-1, Eco-
1400-1, Eco-1401-1, Eco-1402-1, Eco-1403-1, Eco-1404-1, Eco-1405-1, Eco-1406-1, Eco-1407-1, Eco-1408-1, Eco-1409-1, Eco-141-1, Eco-1410-1, Eco-1411-1, Eco-1412-1,
Eco-1413-1, Eco-1414-1, Eco-1415-1, Eco-1416-1, Eco-1417-1, Eco-1418-1, Eco-1419-1, Eco-142-1, Eco-1420-1, Eco-1421-1, Eco-1422-1, Eco-1423-1, Eco-1424-1, Eco-
1425-1, Eco-1426-1, Eco-1427-1, Eco-1428-1, Eco-1429-1, Eco-143-1, Eco-1430-1, Eco-1431-1, Eco-1432-1, Eco-1433-1, Eco-1434-1, Eco-1435-1, Eco-1436-1, Eco-1437-1,
Eco-1438-1, Eco-1439-1, Eco-144-1, Eco-1440-1, Eco-1441-1, Eco-1443-1, Eco-1444-1, Eco-1445-1, Eco-1446-1, Eco-1447-1, Eco-1448-1, Eco-1449-1, Eco-145-1, Eco-
1450-1, Eco-1451-1, Eco-1452-1, Eco-1453-1, Eco-1454-1, Eco-1455-1, Eco-1456-1, Eco-1457-1, Eco-1458-1, Eco-1459-1, Eco-146-1, Eco-1461-1, Eco-1462-1, Eco-1463-1,
Eco-1464-1, Eco-1465-1, Eco-1466-1, Eco-1467-1, Eco-1468-1, Eco-1469-1, Eco-147-1, Eco-1470-1, Eco-1471-1, Eco-1472-1, Eco-1473-1, Eco-1474-1, Eco-1475-1, Eco-
1476-1, Eco-1477-1, Eco-1478-1, Eco-1479-1, Eco-148-1, Eco-1480-1, Eco-1481-1, Eco-1482-1, Eco-1483-1, Eco-1484-1, Eco-1485-1, Eco-1486-1, Eco-1487-1, Eco-1488-1,
Eco-1489-1, Eco-149-1, Eco-1490-1, Eco-1491-1, Eco-1492-1, Eco-1493-1, Eco-1494-1, Eco-1495-1, Eco-1496-1, Eco-1497-1, Eco-1498-1, Eco-1499-1, Eco-15-1, Eco-
150-1, Eco-1500-1, Eco-1501-1, Eco-1502-1, Eco-1503-1, Eco-1504-1, Eco-1505-1, Eco-1506-1, Eco-1507-1, Eco-1508-1, Eco-1509-1, Eco-151-1, Eco-1510-1, Eco-1511-1,
Eco-1512-1, Eco-1513-1, Eco-1514-1, Eco-1515-1, Eco-1516-1, Eco-1517-1, Eco-1518-1, Eco-1519-1, Eco-152-1, Eco-1520-1, Eco-1521-1, Eco-1522-1, Eco-1523-1, Eco-
1524-1, Eco-1525-1, Eco-1526-1, Eco-1527-1, Eco-1528-1, Eco-1529-1, Eco-153-1, Eco-1530-1, Eco-1531-1, Eco-1532-1, Eco-1533-1, Eco-1534-1, Eco-1535-1, Eco-1536-1,
Eco-1537-1, Eco-1538-1, Eco-1539-1, Eco-154-1, Eco-1540-1, Eco-1541-1, Eco-1542-1, Eco-1543-1, Eco-1544-1, Eco-1545-1, Eco-1546-1, Eco-1547-1, Eco-1548-1, Eco-
1549-1, Eco-155-1, Eco-1550-1, Eco-1551-1, Eco-1552-1, Eco-1553-1, Eco-1554-1, Eco-1555-1, Eco-1556-1, Eco-1557-1, Eco-1558-1, Eco-1559-1, Eco-156-1, Eco-1560-1,
Eco-1561-1, Eco-1562-1, Eco-1563-1, Eco-1564-1, Eco-1565-1, Eco-1566-1, Eco-1567-1, Eco-1568-1, Eco-1569-1, Eco-157-1, Eco-1570-1, Eco-1571-1, Eco-1572-1, Eco-
1573-1, Eco-1574-1, Eco-1575-1, Eco-1576-1, Eco-1577-1, Eco-1578-1, Eco-1579-1, Eco-158-1, Eco-1580-1, Eco-1581-1, Eco-1582-1, Eco-1583-1, Eco-1584-1, Eco-1585-1,

Eco-1586-1, Eco-1587-1, Eco-1588-1, Eco-1589-1, Eco-159-1, Eco-1590-1, Eco-1591-1, Eco-1592-1, Eco-1593-1, Eco-1594-1, Eco-1595-1, Eco-1596-1, Eco-1597-1, Eco-1598-1, Eco-1599-1, Eco-16-1, Eco-160-1, Eco-1600-1, Eco-1601-1, Eco-1602-1, Eco-1603-1, Eco-1604-1, Eco-1605-1, Eco-1606-1, Eco-1607-1, Eco-1608-1, Eco-1609-1, Eco-161-1, Eco-1610-1, Eco-1611-1, Eco-1612-1, Eco-1613-1, Eco-1614-1, Eco-1615-1, Eco-1616-1, Eco-1617-1, Eco-1618-1, Eco-1619-1, Eco-162-1, Eco-1620-1, Eco-1621-1, Eco-1622-1, Eco-1623-1, Eco-1624-1, Eco-1625-1, Eco-1626-1, Eco-1627-1, Eco-1628-1, Eco-1629-1, Eco-163-1, Eco-1630-1, Eco-1631-1, Eco-1632-1, Eco-1633-1, Eco-1634-1, Eco-1635-1, Eco-1636-1, Eco-1637-1, Eco-1638-1, Eco-1639-1, Eco-164-1, Eco-1640-1, Eco-1641-1, Eco-1642-1, Eco-1643-1, Eco-1644-1, Eco-1645-1, Eco-1646-1, Eco-1647-1, Eco-1648-1, Eco-1649-1, Eco-165-1, Eco-1650-1, Eco-1651-1, Eco-1652-1, Eco-1653-1, Eco-1654-1, Eco-1655-1, Eco-1656-1, Eco-1657-1, Eco-1658-1, Eco-1659-1, Eco-166-1, Eco-1660-1, Eco-1661-1, Eco-1662-1, Eco-1663-1, Eco-1664-1, Eco-1665-1, Eco-1666-1, Eco-1667-1, Eco-1668-1, Eco-1669-1, Eco-167-1, Eco-1670-1, Eco-1671-1, Eco-1672-1, Eco-1673-1, Eco-1674-1, Eco-1675-1, Eco-1676-1, Eco-1677-1, Eco-1678-1, Eco-1679-1, Eco-168-1, Eco-1680-1, Eco-1681-1, Eco-1682-1, Eco-1683-1, Eco-1684-1, Eco-1685-1, Eco-1686-1, Eco-1687-1, Eco-1688-1, Eco-1689-1, Eco-169-1, Eco-1690-1, Eco-1691-1, Eco-1692-1, Eco-1693-1, Eco-1694-1, Eco-1695-1, Eco-1696-1, Eco-1697-1, Eco-1698-1, Eco-1699-1, Eco-17-1, Eco-170-1, Eco-1700-1, Eco-1701-1, Eco-1702-1, Eco-1703-1, Eco-1704-1, Eco-1705-1, Eco-1706-1, Eco-1707-1, Eco-1708-1, Eco-1709-1, Eco-171-1, Eco-1710-1, Eco-1711-1, Eco-1712-1, Eco-1713-1, Eco-1714-1, Eco-1715-1, Eco-1716-1, Eco-1717-1, Eco-1718-1, Eco-1719-1, Eco-172-1, Eco-1720-1, Eco-1721-1, Eco-1722-1, Eco-1723-1, Eco-1724-1, Eco-1725-1, Eco-1726-1, Eco-1727-1, Eco-1728-1, Eco-1729-1, Eco-173-1, Eco-1730-1, Eco-1731-1, Eco-1732-1, Eco-1733-1, Eco-1734-1, Eco-1735-1, Eco-1736-1, Eco-1737-1, Eco-1738-1, Eco-1739-1, Eco-174-1, Eco-1740-1, Eco-1741-1, Eco-1742-1, Eco-1743-1, Eco-1744-1, Eco-1745-1, Eco-1746-1, Eco-1747-1, Eco-1748-1, Eco-1749-1, Eco-175-1, Eco-1750-1, Eco-1751-1, Eco-1752-1, Eco-1753-1, Eco-1754-1, Eco-1755-1, Eco-1756-1, Eco-1757-1, Eco-1758-1, Eco-1759-1, Eco-176-1, Eco-1760-1, Eco-1761-1, Eco-1762-1, Eco-1763-1, Eco-1764-1, Eco-1765-1, Eco-1766-1, Eco-1767-1, Eco-1768-1, Eco-1769-1, Eco-177-1, Eco-1770-1, Eco-1771-1, Eco-1772-1, Eco-1773-1, Eco-1774-1, Eco-1775-1, Eco-1776-1, Eco-1777-1, Eco-1778-1, Eco-1779-1, Eco-178-1, Eco-1780-1, Eco-1782-1, Eco-1782-2, Eco-1783-1, Eco-1784-1, Eco-1785-1, Eco-1786-1, Eco-1787-1, Eco-1788-1, Eco-1789-1, Eco-179-1, Eco-1790-1, Eco-1791-1, Eco-1792-1, Eco-1793-1, Eco-1794-1, Eco-1795-1, Eco-1796-1, Eco-1797-1, Eco-1798-1, Eco-1799-1, Eco-18-1, Eco-180-1, Eco-1800-1, Eco-1801-1, Eco-1802-1, Eco-1803-1, Eco-1804-1, Eco-1805-1, Eco-1806-1, Eco-1807-1, Eco-1808-1, Eco-1809-1, Eco-181-1, Eco-1810-1, Eco-1811-1, Eco-1812-1, Eco-1813-1, Eco-1814-1, Eco-1815-1, Eco-1816-1, Eco-1817-1, Eco-1818-1, Eco-1819-1, Eco-182-1, Eco-1820-1, Eco-1821-1, Eco-1822-1, Eco-1823-1, Eco-1824-1, Eco-1825-1, Eco-1826-1, Eco-1827-1, Eco-1828-1, Eco-1829-1, Eco-183-1, Eco-1830-1, Eco-1831-1, Eco-1832-1, Eco-1833-1, Eco-1834-1, Eco-1835-1, Eco-1836-1, Eco-1837-1, Eco-1838-1, Eco-1839-1, Eco-184-1, Eco-1840-1, Eco-1841-1, Eco-1842-1, Eco-1843-1, Eco-1844-1, Eco-1845-1, Eco-1846-1, Eco-1847-1, Eco-1848-1, Eco-1849-1, Eco-185-1, Eco-1850-1, Eco-1851-1, Eco-1852-1, Eco-1853-1, Eco-1854-1, Eco-1855-1, Eco-1856-1, Eco-1857-1, Eco-1858-1, Eco-1859-1, Eco-186-1, Eco-1860-1, Eco-1861-1, Eco-1863-1, Eco-1864-1, Eco-1865-1, Eco-1866-1, Eco-1867-1, Eco-1868-1, Eco-1869-1, Eco-187-1, Eco-1870-1, Eco-1871-1, Eco-1872-1, Eco-1873-1, Eco-1874-1, Eco-1875-1, Eco-1876-1, Eco-1877-1, Eco-1878-1, Eco-1879-1, Eco-188-1, Eco-1880-1, Eco-1881-1, Eco-1882-1, Eco-1883-1, Eco-1884-1, Eco-1885-1, Eco-1886-1, Eco-1887-1, Eco-1888-1, Eco-1889-1, Eco-189-1, Eco-1890-1, Eco-1891-1, Eco-1892-1, Eco-1893-1, Eco-1894-1, Eco-1895-1, Eco-1896-1, Eco-1897-1, Eco-1898-1, Eco-1899-1, Eco-19-1, Eco-190-1, Eco-1900-1, Eco-1901-1, Eco-1902-1, Eco-1903-1, Eco-1904-1, Eco-1905-1, Eco-1906-1, Eco-1907-1, Eco-1908-1, Eco-1909-1, Eco-1909-2, Eco-191-1, Eco-1910-1, Eco-1911-1, Eco-1912-1, Eco-1913-1, Eco-1914-1, Eco-1915-1, Eco-1916-1, Eco-1917-1, Eco-1918-1, Eco-1919-1, Eco-192-1, Eco-1920-1, Eco-1921-1, Eco-1922-1, Eco-1923-1, Eco-1924-1, Eco-1925-1, Eco-1926-1, Eco-1927-1, Eco-1928-1, Eco-1929-1, Eco-193-1, Eco-1930-1, Eco-1931-1, Eco-1932-1, Eco-1933-1, Eco-1934-1, Eco-1935-1, Eco-1936-1, Eco-1937-1, Eco-1938-1, Eco-1939-1, Eco-194-1, Eco-1940-1, Eco-1941-1, Eco-1942-1, Eco-1943-1, Eco-1944-1, Eco-1945-1, Eco-1946-1, Eco-1947-1, Eco-1948-1, Eco-1949-1, Eco-195-1, Eco-1950-1, Eco-1951-1, Eco-1952-1, Eco-1953-1, Eco-1954-1, Eco-1955-1, Eco-1956-1, Eco-1957-1, Eco-1958-1, Eco-1959-1, Eco-196-1, Eco-1960-1, Eco-1961-1, Eco-1962-1, Eco-1963-1, Eco-1964-1, Eco-1965-1, Eco-1966-1, Eco-1967-1, Eco-1968-1, Eco-1969-1, Eco-197-1, Eco-1970-1, Eco-1971-1, Eco-1972-1, Eco-1973-1, Eco-1974-1, Eco-1975-1, Eco-1976-1, Eco-1977-1, Eco-1978-1, Eco-1979-1, Eco-198-1, Eco-1980-1, Eco-1981-1, Eco-1982-1, Eco-1983-1, Eco-1984-1, Eco-1985-1, Eco-1986-1, Eco-1987-1, Eco-1988-1, Eco-1989-1, Eco-199-1, Eco-1990-1, Eco-1991-1, Eco-1992-1, Eco-1993-1, Eco-1994-1, Eco-1995-1, Eco-1996-1, Eco-1997-1, Eco-1998-1, Eco-1999-1, Eco-2-1, Eco-20-1, Eco-200-1, Eco-2000-1, Eco-2001-1, Eco-2002-1, Eco-2003-1, Eco-2004-1, Eco-2005-1, Eco-2005-2, Eco-2006-1, Eco-2007-1, Eco-2008-1, Eco-2009-1, Eco-201-1, Eco-2010-1, Eco-2010-2, Eco-2011-1, Eco-2012-1, Eco-2013-1, Eco-2014-1, Eco-2015-1, Eco-2016-1, Eco-2017-1, Eco-2018-1, Eco-2019-1, Eco-202-1, Eco-2020-1, Eco-2021-1, Eco-2022-1, Eco-2023-1, Eco-2024-1, Eco-2025-1, Eco-2026-1, Eco-2027-1, Eco-2028-1, Eco-2029-1, Eco-203-1, Eco-2030-1, Eco-2031-1, Eco-2032-1, Eco-2033-1, Eco-2034-1, Eco-2035-1, Eco-2036-1, Eco-2037-1, Eco-2038-1, Eco-2039-1, Eco-204-1, Eco-2040-1, Eco-2041-1, Eco-2042-1, Eco-2043-1, Eco-2044-1, Eco-2045-1, Eco-2046-1, Eco-2047-1, Eco-2048-1, Eco-2049-1, Eco-205-1, Eco-2050-1, Eco-2051-1, Eco-2052-1, Eco-2053-1, Eco-2054-1, Eco-2055-1, Eco-2056-1, Eco-2057-1, Eco-2058-1, Eco-2059-1, Eco-206-1, Eco-2060-1, Eco-2061-1, Eco-2062-1, Eco-2063-1, Eco-2064-1, Eco-2065-1, Eco-2066-1, Eco-2067-1, Eco-2068-1, Eco-2069-1, Eco-207-1, Eco-2070-1, Eco-2071-1, Eco-2072-1, Eco-2073-1, Eco-2074-1, Eco-2075-1, Eco-2076-1, Eco-2077-1, Eco-2078-1, Eco-2079-1, Eco-208-1, Eco-2080-1, Eco-2081-1, Eco-2082-1, Eco-2083-1, Eco-2084-1, Eco-2085-1, Eco-2086-1, Eco-2087-1, Eco-2088-1, Eco-2089-1, Eco-209-1, Eco-2090-1, Eco-2091-1, Eco-2092-1, Eco-2093-1, Eco-2094-1, Eco-2095-1, Eco-2096-1, Eco-2097-1, Eco-2098-1, Eco-2099-1, Eco-21-1, Eco-210-1, Eco-2100-1, Eco-2101-1, Eco-2102-1, Eco-2103-1, Eco-2104-1, Eco-2105-1, Eco-2106-1, Eco-2107-1, Eco-2108-1, Eco-2109-1, Eco-211-1, Eco-2110-1, Eco-2111-1, Eco-2112-1, Eco-2113-1, Eco-2114-1, Eco-2115-1, Eco-2116-1, Eco-2117-1, Eco-

2118-1, Eco-2119-1, Eco-212-1, Eco-2120-1, Eco-2121-1, Eco-2122-1, Eco-2123-1, Eco-2124-1, Eco-2125-1, Eco-2126-1, Eco-2127-1, Eco-2128-1, Eco-2129-1, Eco-213-1, Eco-2130-1, Eco-2131-1, Eco-2132-1, Eco-2133-1, Eco-2134-1, Eco-2135-1, Eco-2136-1, Eco-2137-1, Eco-2138-1, Eco-2139-1, Eco-214-1, Eco-2140-1, Eco-2141-1, Eco-2142-1, Eco-2143-1, Eco-2144-1, Eco-2145-1, Eco-2146-1, Eco-2147-1, Eco-2148-1, Eco-2149-1, Eco-215-1, Eco-215-2, Eco-2150-1, Eco-2151-1, Eco-2152-1, Eco-2153-1, Eco-2154-1, Eco-2155-1, Eco-2156-1, Eco-2157-1, Eco-2157-2, Eco-2158-1, Eco-2159-1, Eco-216-1, Eco-2160-1, Eco-2161-1, Eco-2162-1, Eco-2163-1, Eco-2164-1, Eco-2165-1, Eco-2166-1, Eco-2167-1, Eco-2168-1, Eco-2169-1, Eco-217-1, Eco-2170-1, Eco-2171-1, Eco-2172-1, Eco-2173-1, Eco-2174-1, Eco-2175-1, Eco-2176-1, Eco-2177-1, Eco-2178-1, Eco-2179-1, Eco-218-1, Eco-2180-1, Eco-2181-1, Eco-2182-1, Eco-2183-1, Eco-2184-1, Eco-2185-1, Eco-219-1, Eco-22-1, Eco-220-1, Eco-221-1, Eco-222-1, Eco-223-1, Eco-224-1, Eco-225-1, Eco-226-1, Eco-227-1, Eco-228-1, Eco-229-1, Eco-23-1, Eco-230-2, Eco-231-1, Eco-232-1, Eco-233-1, Eco-234-1, Eco-235-1, Eco-236-1, Eco-237-1, Eco-238-1, Eco-239-1, Eco-24-1, Eco-240-1, Eco-241-1, Eco-242-1, Eco-243-1, Eco-244-1, Eco-245-1, Eco-246-1, Eco-247-1, Eco-248-1, Eco-249-1, Eco-25-1, Eco-250-1, Eco-251-1, Eco-252-1, Eco-253-1, Eco-254-1, Eco-255-1, Eco-256-1, Eco-257-1, Eco-258-1, Eco-259-1, Eco-26-1, Eco-260-1, Eco-261-1, Eco-262-1, Eco-263-1, Eco-264-1, Eco-265-1, Eco-266-1, Eco-267-1, Eco-268-1, Eco-269-1, Eco-27-1, Eco-270-1, Eco-271-1, Eco-272-1, Eco-273-1, Eco-274-1, Eco-275-1, Eco-276-1, Eco-277-1, Eco-278-1, Eco-279-1, Eco-28-1, Eco-280-1, Eco-281-1, Eco-282-1, Eco-283-1, Eco-284-1, Eco-285-1, Eco-286-1, Eco-287-1, Eco-288-1, Eco-289-1, Eco-289-2, Eco-29-1, Eco-290-1, Eco-291-1, Eco-291-2, Eco-292-1, Eco-293-1, Eco-294-1, Eco-295-1, Eco-296-1, Eco-297-1, Eco-298-1, Eco-299-1, Eco-3-1, Eco-30-1, Eco-300-1, Eco-301-1, Eco-302-1, Eco-303-1, Eco-304-1, Eco-305-1, Eco-306-1, Eco-307-1, Eco-308-1, Eco-309-1, Eco-31-1, Eco-310-1, Eco-311-1, Eco-312-1, Eco-313-1, Eco-314-1, Eco-315-1, Eco-316-1, Eco-317-1, Eco-318-1, Eco-319-1, Eco-32-1, Eco-320-1, Eco-321-1, Eco-322-1, Eco-323-1, Eco-324-1, Eco-325-1, Eco-326-1, Eco-327-1, Eco-328-1, Eco-329-1, Eco-33-1, Eco-330-1, Eco-331-1, Eco-332-1, Eco-333-1, Eco-334-1, Eco-335-1, Eco-336-1, Eco-337-1, Eco-338-1, Eco-339-1, Eco-34-1, Eco-340-1, Eco-341-1, Eco-342-1, Eco-343-1, Eco-344-1, Eco-345-1, Eco-346-1, Eco-347-1, Eco-348-1, Eco-349-1, Eco-35-1, Eco-350-1, Eco-351-1, Eco-352-1, Eco-353-1, Eco-354-1, Eco-355-1, Eco-356-1, Eco-357-1, Eco-358-1, Eco-359-1, Eco-36-1, Eco-360-1, Eco-361-1, Eco-362-1, Eco-363-1, Eco-364-1, Eco-365-1, Eco-366-1, Eco-367-1, Eco-368-1, Eco-369-1, Eco-37-1, Eco-370-1, Eco-371-1, Eco-372-1, Eco-373-1, Eco-374-1, Eco-375-1, Eco-376-1, Eco-377-1, Eco-378-1, Eco-379-1, Eco-38-1, Eco-380-1, Eco-381-1, Eco-382-1, Eco-383-1, Eco-384-1, Eco-385-1, Eco-386-1, Eco-387-1, Eco-388-1, Eco-389-1, Eco-39-1, Eco-390-1, Eco-391-1, Eco-392-1, Eco-393-1, Eco-394-1, Eco-395-1, Eco-396-1, Eco-397-1, Eco-398-1, Eco-399-1, Eco-4-1, Eco-40-1, Eco-400-1, Eco-401-1, Eco-402-1, Eco-403-1, Eco-404-1, Eco-405-1, Eco-406-1, Eco-407-1, Eco-408-1, Eco-409-1, Eco-41-1, Eco-410-1, Eco-411-1, Eco-412-1, Eco-413-1, Eco-414-1, Eco-415-1, Eco-416-1, Eco-417-1, Eco-418-1, Eco-419-1, Eco-42-1, Eco-420-1, Eco-421-1, Eco-421-2, Eco-421-3, Eco-421-4, Eco-422-1, Eco-423-1, Eco-424-1, Eco-425-1, Eco-426-1, Eco-427-1, Eco-428-1, Eco-429-1, Eco-43-1, Eco-430-1, Eco-431-1, Eco-432-1, Eco-433-1, Eco-434-1, Eco-435-1, Eco-436-1, Eco-437-1, Eco-438-1, Eco-439-1, Eco-44-1, Eco-440-1, Eco-441-1, Eco-442-1, Eco-443-1, Eco-444-1, Eco-445-1, Eco-446-1, Eco-447-1, Eco-448-1, Eco-449-1, Eco-45-1, Eco-450-1, Eco-451-1, Eco-452-1, Eco-453-1, Eco-454-1, Eco-455-1, Eco-456-1, Eco-457-1, Eco-458-1, Eco-459-1, Eco-46-1, Eco-460-1, Eco-461-1, Eco-462-1, Eco-463-1, Eco-463-2, Eco-463-3, Eco-463-4, Eco-464-1, Eco-465-1, Eco-466-1, Eco-467-1, Eco-468-1, Eco-469-1, Eco-47-1, Eco-470-1, Eco-471-1, Eco-472-1, Eco-472-2, Eco-473-1, Eco-474-1, Eco-475-1, Eco-476-1, Eco-477-1, Eco-478-1, Eco-479-1, Eco-48-1, Eco-480-1, Eco-481-1, Eco-482-1, Eco-483-1, Eco-484-1, Eco-485-1, Eco-486-1, Eco-487-1, Eco-488-1, Eco-489-1, Eco-49-1, Eco-490-1, Eco-491-1, Eco-492-1, Eco-493-1, Eco-494-1, Eco-495-1, Eco-496-1, Eco-497-1, Eco-498-1, Eco-499-1, Eco-5-1, Eco-50-1, Eco-500-1, Eco-501-1, Eco-502-1, Eco-503-1, Eco-504-1, Eco-505-1, Eco-506-1, Eco-507-1, Eco-508-1, Eco-509-1, Eco-51-1, Eco-510-1, Eco-511-1, Eco-512-1, Eco-513-1, Eco-514-1, Eco-515-1, Eco-516-1, Eco-517-1, Eco-518-1, Eco-519-1, Eco-52-1, Eco-520-1, Eco-521-1, Eco-522-1, Eco-523-1, Eco-524-1, Eco-525-1, Eco-526-1, Eco-527-1, Eco-528-1, Eco-529-1, Eco-53-1, Eco-530-1, Eco-531-1, Eco-532-1, Eco-533-1, Eco-534-1, Eco-535-1, Eco-536-1, Eco-537-1, Eco-538-1, Eco-539-1, Eco-54-1, Eco-540-1, Eco-541-1, Eco-542-1, Eco-543-1, Eco-544-1, Eco-545-1, Eco-546-1, Eco-547-1, Eco-548-1, Eco-548-2, Eco-549-1, Eco-55-1, Eco-550-1, Eco-551-1, Eco-552-1, Eco-553-1, Eco-554-1, Eco-555-1, Eco-556-1, Eco-557-1, Eco-558-1, Eco-559-1, Eco-56-1, Eco-560-1, Eco-561-1, Eco-562-1, Eco-563-1, Eco-564-1, Eco-565-1, Eco-566-1, Eco-567-1, Eco-568-1, Eco-569-1, Eco-57-1, Eco-570-1, Eco-571-1, Eco-572-1, Eco-573-1, Eco-574-1, Eco-574-2, Eco-574-3, Eco-575-1, Eco-576-1, Eco-577-1, Eco-578-1, Eco-579-1, Eco-58-1, Eco-580-1, Eco-581-1, Eco-582-1, Eco-583-1, Eco-584-1, Eco-585-1, Eco-586-1, Eco-587-1, Eco-588-1, Eco-589-1, Eco-59-1, Eco-590-1, Eco-591-1, Eco-592-1, Eco-593-1, Eco-594-1, Eco-595-1, Eco-596-1, Eco-597-1, Eco-598-1, Eco-599-1, Eco-6-1, Eco-60-1, Eco-600-1, Eco-601-1, Eco-602-1, Eco-603-1, Eco-604-1, Eco-605-1, Eco-606-1, Eco-607-1, Eco-608-1, Eco-609-1, Eco-61-1, Eco-610-1, Eco-611-1, Eco-612-1, Eco-613-1, Eco-614-1, Eco-615-1, Eco-616-1, Eco-617-1, Eco-618-1, Eco-619-1, Eco-62-1, Eco-620-1, Eco-621-1, Eco-622-1, Eco-623-1, Eco-624-1, Eco-625-1, Eco-626-1, Eco-627-1, Eco-628-1, Eco-629-1, Eco-63-1, Eco-630-1, Eco-631-1, Eco-632-1, Eco-633-1, Eco-634-1, Eco-635-1, Eco-636-1, Eco-637-1, Eco-638-1, Eco-639-1, Eco-64-1, Eco-640-1, Eco-641-1, Eco-642-1, Eco-643-1, Eco-644-1, Eco-645-1, Eco-646-1, Eco-647-1, Eco-648-1, Eco-649-1, Eco-65-1, Eco-650-1, Eco-651-1, Eco-652-1, Eco-653-1, Eco-654-1, Eco-655-1, Eco-656-1, Eco-657-1, Eco-658-1, Eco-659-1, Eco-66-1, Eco-660-1, Eco-661-1, Eco-662-1, Eco-663-1, Eco-664-1, Eco-665-1, Eco-666-1, Eco-667-1, Eco-668-1, Eco-669-1, Eco-67-1, Eco-670-1, Eco-671-1, Eco-672-1, Eco-673-1, Eco-674-1, Eco-675-1, Eco-676-1, Eco-677-1, Eco-678-1, Eco-679-1, Eco-68-1, Eco-680-1, Eco-681-1, Eco-682-1, Eco-683-1, Eco-684-1, Eco-685-1, Eco-686-1, Eco-687-1, Eco-688-1, Eco-689-1, Eco-69-1, Eco-690-1, Eco-691-1, Eco-692-1, Eco-693-1, Eco-694-1, Eco-695-1, Eco-696-1, Eco-697-1, Eco-698-1, Eco-699-1, Eco-7-1, Eco-70-1, Eco-700-1, Eco-701-1, Eco-702-1, Eco-703-1, Eco-704-1, Eco-705-1, Eco-706-1, Eco-707-1, Eco-708-1, Eco-709-1, Eco-71-1, Eco-710-1, Eco-711-1, Eco-712-1, Eco-713-1, Eco-714-1, Eco-715-1, Eco-716-1, Eco-717-1, Eco-718-1, Eco-719-1, Eco-72-1, Eco-720-1, Eco-721-1, Eco-722-1, Eco-723-1, Eco-724-1, Eco-725-1,

Eco-726-1, Eco-727-1, Eco-728-1, Eco-728-2, Eco-729-1, Eco-73-1, Eco-730-1, Eco-731-1, Eco-732-1, Eco-733-1, Eco-734-1, Eco-735-1, Eco-736-1, Eco-737-1, Eco-738-1, Eco-739-1, Eco-74-1, Eco-740-1, Eco-741-1, Eco-742-1, Eco-743-1, Eco-744-1, Eco-745-1, Eco-746-1, Eco-747-1, Eco-748-1, Eco-749-1, Eco-75-1, Eco-750-1, Eco-751-1, Eco-752-1, Eco-752-2, Eco-753-1, Eco-754-1, Eco-755-1, Eco-756-1, Eco-757-1, Eco-758-1, Eco-759-1, Eco-76-1, Eco-760-1, Eco-761-1, Eco-762-1, Eco-764-1, Eco-765-1, Eco-766-1, Eco-767-1, Eco-768-1, Eco-769-1, Eco-77-1, Eco-770-1, Eco-771-1, Eco-772-1, Eco-773-1, Eco-774-1, Eco-775-1, Eco-776-1, Eco-777-1, Eco-778-1, Eco-779-1, Eco-78-1, Eco-780-1, Eco-781-1, Eco-782-1, Eco-783-1, Eco-784-1, Eco-785-1, Eco-786-1, Eco-787-1, Eco-788-1, Eco-789-1, Eco-79-1, Eco-790-1, Eco-791-1, Eco-792-1, Eco-793-1, Eco-794-1, Eco-795-1, Eco-796-1, Eco-797-1, Eco-798-1, Eco-799-1, Eco-8-1, Eco-80-1, Eco-800-1, Eco-801-1, Eco-802-1, Eco-803-1, Eco-804-1, Eco-805-1, Eco-806-1, Eco-807-1, Eco-808-1, Eco-809-1, Eco-81-1, Eco-810-1, Eco-811-1, Eco-812-1, Eco-813-1, Eco-814-1, Eco-815-1, Eco-816-1, Eco-817-1, Eco-818-1, Eco-819-1, Eco-82-1, Eco-820-1, Eco-821-1, Eco-822-1, Eco-823-1, Eco-824-1, Eco-825-1, Eco-826-1, Eco-827-1, Eco-828-1, Eco-829-1, Eco-83-1, Eco-830-1, Eco-831-1, Eco-832-1, Eco-833-1, Eco-834-1, Eco-836-1, Eco-837-1, Eco-838-1, Eco-839-1, Eco-84-1, Eco-840-1, Eco-841-1, Eco-842-1, Eco-843-1, Eco-844-1, Eco-845-1, Eco-846-1, Eco-847-1, Eco-848-1, Eco-849-1, Eco-85-1, Eco-850-1, Eco-851-1, Eco-852-1, Eco-853-1, Eco-854-1, Eco-855-1, Eco-856-1, Eco-857-1, Eco-858-1, Eco-859-1, Eco-86-1, Eco-860-1, Eco-861-1, Eco-862-1, Eco-863-1, Eco-864-1, Eco-865-1, Eco-866-1, Eco-867-1, Eco-868-1, Eco-869-1, Eco-87-1, Eco-870-1, Eco-871-1, Eco-872-1, Eco-873-1, Eco-874-1, Eco-875-1, Eco-876-1, Eco-877-1, Eco-878-1, Eco-879-1, Eco-88-1, Eco-880-1, Eco-881-1, Eco-882-1, Eco-883-1, Eco-884-1, Eco-885-1, Eco-886-1, Eco-887-1, Eco-888-1, Eco-889-1, Eco-89-1, Eco-890-1, Eco-891-1, Eco-892-1, Eco-893-1, Eco-894-1, Eco-895-1, Eco-896-1, Eco-897-1, Eco-898-1, Eco-899-1, Eco-9-1, Eco-90-1, Eco-900-1, Eco-901-1, Eco-902-1, Eco-903-1, Eco-904-1, Eco-905-1, Eco-906-1, Eco-907-1, Eco-908-1, Eco-909-1, Eco-91-1, Eco-910-1, Eco-911-1, Eco-912-1, Eco-913-1, Eco-914-1, Eco-915-1, Eco-916-1, Eco-917-1, Eco-918-1, Eco-919-1, Eco-92-1, Eco-920-1, Eco-921-1, Eco-922-1, Eco-923-1, Eco-924-1, Eco-925-1, Eco-926-1, Eco-927-1, Eco-928-1, Eco-929-1, Eco-93-1, Eco-930-1, Eco-931-1, Eco-932-1, Eco-933-1, Eco-934-1, Eco-935-1, Eco-936-1, Eco-937-1, Eco-938-1, Eco-939-1, Eco-94-1, Eco-940-1, Eco-941-1, Eco-942-1, Eco-943-1, Eco-944-1, Eco-945-1, Eco-946-1, Eco-947-1, Eco-948-1, Eco-949-1, Eco-95-1, Eco-950-1, Eco-951-1, Eco-952-1, Eco-953-1, Eco-954-1, Eco-955-1, Eco-956-1, Eco-957-1, Eco-958-1, Eco-959-1, Eco-96-1, Eco-960-1, Eco-961-1, Eco-962-1, Eco-963-1, Eco-964-1, Eco-965-1, Eco-966-1, Eco-967-1, Eco-968-1, Eco-969-1, Eco-97-1, Eco-970-1, Eco-971-1, Eco-972-1, Eco-973-1, Eco-974-1, Eco-975-1, Eco-976-1, Eco-977-1, Eco-978-1, Eco-979-1, Eco-98-1, Eco-980-1, Eco-981-1, Eco-982-1, Eco-983-1, Eco-984-1, Eco-985-1, Eco-986-1, Eco-987-1, Eco-988-1, Eco-989-1, Eco-99-1, Eco-990-1, Eco-991-1, Eco-992-1, Eco-993-1, Eco-994-1, Eco-995-1, Eco-996-1, Eco-997-1, Eco-998-1, Eco-999-1, Efe-1-1, Efe-1-2, Efe-1-3, Efe-2-1, Efe-3-1, Efe-4-1, Eho-1-1, Eho-1-2, Eho-1-3, Eho-1-4, Eho-1-5, Eho-1-6, Eho-1-7, Eho-1-8, Eho-2-1, Eho-3-1, Eho-3-2, Eho-4-1, Eho-4-2, Eho-6-1, Eho-7-1, Eho-8-1, Eic-1-1, Ein-1-2, Eko-1-1, Eko-1-10, Eko-1-11, Eko-1-2, Eko-1-3, Eko-1-4, Eko-1-5, Eko-1-6, Eko-1-7, Eko-1-8, Eko-1-9, Elu-1-1, Elu-1-2, Elu-1-3, Elu-1-4, Elu-1-5, Ema-1-1, Emo-1-1, Eno-1-1, Eno-2-1, Epy-1-1, Esp-1-1, Esp-10-1, Esp-11-1, Esp-12-1, Esp-13-1, Esp-14-1, Esp-15-1, Esp-16-1, Esp-17-1, Esp-18-1, Esp-19-1, Esp-2-1, Esp-20-1, Esp-21-1, Esp-22-1, Esp-23-1, Esp-24-1, Esp-25-1, Esp-26-1, Esp-27-1, Esp-28-1, Esp-3-1, Esp-30-1, Esp-31-1, Esp-32-1, Esp-33-1, Esp-34-1, Esp-35-1, Esp-36-1, Esp-37-1, Esp-38-1, Esp-39-1, Esp-4-1, Esp-40-1, Esp-41-1, Esp-44-1, Esp-45-1, Esp-46-1, Esp-48-1, Esp-49-1, Esp-5-1, Esp-50-1, Esp-51-1, Esp-52-1, Esp-53-1, Esp-54-1, Esp-56-1, Esp-57-1, Esp-58-1, Esp-59-1, Esp-6-1, Esp-60-1, Esp-61-1, Esp-62-1, Esp-63-1, Esp-64-1, Esp-65-1, Esp-66-1, Esp-68-1, Esp-69-1, Esp-7-1, Esp-70-1, Esp-71-1, Esp-72-1, Esp-73-1, Esp-75-1, Esp-76-1, Esp-77-1, Esp-78-1, Esp-79-1, Esp-8-1, Esp-80-1, Esp-9-1, Eta-1-1, Eta-3-1, Eta-5-1, Eta-6-1, Eta-7-1, Eta-8-1, Etr-2-1, Fhe-4-1, Fpu-2-1, Fpu-3-1, Fpu-4-1, Gag-1-1, Gch-1-1, Gme-1-1, Gsp-1-1, Gsp-2-1, Hae-1-1, Hal-1-2, Hal-1-3, Hal-1-4, Hal-2-1, Hal-4-1, Hha-2-1, Hha-3-1, Hha-4-1, Hha-5-1, Hin-1-1, Hin-1-2, Hin-1-4, Hin-10-1, Hin-11-1, Hin-15-1, Hin-16-1, Hin-18-1, Hin-19-1, Hin-19-2, Hin-2-1, Hin-20-1, Hin-21-1, Hin-3-1, Hin-4-1, Hin-5-1, Hin-7-1, Hin-8-1, Hin-9-1, Hpa-1-2, Hpa-2-1, Hpa-2-10, Hpa-2-3, Hpa-2-4, Hpa-2-5, Hpa-2-6, Hpa-2-7, Hpa-2-8, Hpa-2-9, Hpa-3-1, Hpa-4-1, Hpa-5-1, Hpa-6-1, Hpa-7-1, Hsp-1-1, Hsp-2-1, Hsp-4-1, Iba-1-1, Ilo-1-1, Ilo-2-1, Isp-1-1, Kae-1-1, Kae-10-1, Kae-11-1, Kae-12-1, Kae-13-1, Kae-2-1, Kae-3-1, Kae-4-1, Kae-5-1, Kae-6-1, Kae-7-1, Kae-8-1, Kae-9-1, Kmi-1-1, Kmi-2-1, Kmi-3-1, Kox-1-1, Kox-1-2, Kox-1-3, Kox-1-4, Kox-10-1, Kox-12-1, Kox-13-1, Kox-14-1, Kox-15-1, Kox-16-1, Kox-17-1, Kox-18-1, Kox-19-1, Kox-2-1, Kox-20-1, Kox-3-1, Kox-4-1, Kox-5-1, Kox-6-1, Kox-7-1, Kox-9-1, Kpn-1-1, Kpn-1-2, Kpn-1-3, Kpn-1-4, Kpn-1-5, Kpn-1-6, Kpn-1-7, Kpn-1-8, Kpn-1-9, Kpn-10-1, Kpn-100-1, Kpn-101-1, Kpn-102-1, Kpn-103-1, Kpn-104-1, Kpn-105-1, Kpn-106-1, Kpn-107-1, Kpn-108-1, Kpn-109-1, Kpn-11-1, Kpn-110-1, Kpn-111-1, Kpn-112-1, Kpn-113-1, Kpn-114-1, Kpn-115-1, Kpn-116-1, Kpn-117-1, Kpn-118-1, Kpn-119-1, Kpn-12-1, Kpn-120-1, Kpn-121-1, Kpn-122-1, Kpn-123-1, Kpn-124-1, Kpn-125-1, Kpn-126-1, Kpn-127-1, Kpn-128-1, Kpn-129-1, Kpn-13-1, Kpn-130-1, Kpn-131-1, Kpn-132-1, Kpn-133-1, Kpn-134-1, Kpn-135-1, Kpn-136-1, Kpn-137-1, Kpn-138-1, Kpn-139-1, Kpn-14-1, Kpn-140-1, Kpn-141-1, Kpn-142-1, Kpn-143-1, Kpn-144-1, Kpn-145-1, Kpn-146-1, Kpn-147-1, Kpn-148-1, Kpn-149-1, Kpn-15-1, Kpn-150-1, Kpn-151-1, Kpn-152-1, Kpn-153-1, Kpn-154-1, Kpn-155-1, Kpn-156-1, Kpn-157-1, Kpn-158-1, Kpn-159-1, Kpn-16-1, Kpn-160-1, Kpn-161-1, Kpn-162-1, Kpn-163-1, Kpn-164-1, Kpn-165-1, Kpn-166-1, Kpn-167-1, Kpn-169-1, Kpn-17-1, Kpn-170-1, Kpn-171-1, Kpn-172-1, Kpn-173-1, Kpn-174-1, Kpn-175-1, Kpn-176-1, Kpn-177-1, Kpn-178-1, Kpn-179-1, Kpn-18-1, Kpn-180-1, Kpn-181-1, Kpn-182-1, Kpn-183-1, Kpn-184-1, Kpn-185-1, Kpn-186-1, Kpn-187-1, Kpn-188-1, Kpn-189-1, Kpn-19-1, Kpn-190-1, Kpn-191-1, Kpn-192-1, Kpn-193-1, Kpn-194-1, Kpn-195-1, Kpn-196-1, Kpn-197-1, Kpn-198-1, Kpn-199-1, Kpn-199-2, Kpn-2-1, Kpn-20-1, Kpn-200-1, Kpn-200-2, Kpn-201-1, Kpn-202-1, Kpn-203-1, Kpn-203-2, Kpn-204-1, Kpn-204-2, Kpn-205-1, Kpn-205-2, Kpn-205-3, Kpn-205-4, Kpn-206-1, Kpn-206-2, Kpn-206-3, Kpn-206-4, Kpn-206-5, Kpn-206-6, Kpn-206-7, Kpn-207-1, Kpn-208-1, Kpn-209-1, Kpn-21-1, Kpn-210-1, Kpn-211-1, Kpn-212-1, Kpn-213-1, Kpn-214-1, Kpn-215-1, Kpn-216-1, Kpn-217-1, Kpn-218-1, Kpn-219-1, Kpn-22-1, Kpn-220-1, Kpn-221-1,

Kpn-222-1, Kpn-223-1, Kpn-224-1, Kpn-225-1, Kpn-226-1, Kpn-227-1, Kpn-228-1, Kpn-229-1, Kpn-23-1, Kpn-230-1, Kpn-231-1, Kpn-232-1, Kpn-233-1, Kpn-234-1, Kpn-235-1, Kpn-236-1, Kpn-237-1, Kpn-238-1, Kpn-239-1, Kpn-24-1, Kpn-240-1, Kpn-241-1, Kpn-242-1, Kpn-243-1, Kpn-244-1, Kpn-245-1, Kpn-246-1, Kpn-247-1, Kpn-248-1, Kpn-249-1, Kpn-25-1, Kpn-250-1, Kpn-251-1, Kpn-252-1, Kpn-253-1, Kpn-254-1, Kpn-255-1, Kpn-257-1, Kpn-258-1, Kpn-259-1, Kpn-26-1, Kpn-260-1, Kpn-261-1, Kpn-262-1, Kpn-263-1, Kpn-264-1, Kpn-265-1, Kpn-266-1, Kpn-267-1, Kpn-268-1, Kpn-269-1, Kpn-27-1, Kpn-270-1, Kpn-271-1, Kpn-272-1, Kpn-273-1, Kpn-274-1, Kpn-275-1, Kpn-276-1, Kpn-277-1, Kpn-278-1, Kpn-279-1, Kpn-28-1, Kpn-280-1, Kpn-282-1, Kpn-283-1, Kpn-284-1, Kpn-285-1, Kpn-286-1, Kpn-287-1, Kpn-288-1, Kpn-289-1, Kpn-29-1, Kpn-290-1, Kpn-291-1, Kpn-292-1, Kpn-293-1, Kpn-294-1, Kpn-295-1, Kpn-296-1, Kpn-297-1, Kpn-298-1, Kpn-299-1, Kpn-3-1, Kpn-3-2, Kpn-30-1, Kpn-300-1, Kpn-301-1, Kpn-302-1, Kpn-303-1, Kpn-304-1, Kpn-305-1, Kpn-306-1, Kpn-307-1, Kpn-308-1, Kpn-309-1, Kpn-31-1, Kpn-310-1, Kpn-311-1, Kpn-312-1, Kpn-313-1, Kpn-314-1, Kpn-315-1, Kpn-316-1, Kpn-317-1, Kpn-318-1, Kpn-319-1, Kpn-32-1, Kpn-320-1, Kpn-321-1, Kpn-322-1, Kpn-323-1, Kpn-324-1, Kpn-325-1, Kpn-326-1, Kpn-327-1, Kpn-328-1, Kpn-329-1, Kpn-33-1, Kpn-330-1, Kpn-331-1, Kpn-332-1, Kpn-333-1, Kpn-334-1, Kpn-335-1, Kpn-336-1, Kpn-337-1, Kpn-338-1, Kpn-339-1, Kpn-34-1, Kpn-340-1, Kpn-341-1, Kpn-342-1, Kpn-343-1, Kpn-344-1, Kpn-345-1, Kpn-346-1, Kpn-347-1, Kpn-348-1, Kpn-349-1, Kpn-35-1, Kpn-350-1, Kpn-351-1, Kpn-352-1, Kpn-353-1, Kpn-354-1, Kpn-355-1, Kpn-356-1, Kpn-357-1, Kpn-358-1, Kpn-359-1, Kpn-36-1, Kpn-360-1, Kpn-361-1, Kpn-362-1, Kpn-37-1, Kpn-38-1, Kpn-39-1, Kpn-4-1, Kpn-40-1, Kpn-41-1, Kpn-42-1, Kpn-43-1, Kpn-44-1, Kpn-45-1, Kpn-46-1, Kpn-47-1, Kpn-48-1, Kpn-49-1, Kpn-5-1, Kpn-50-1, Kpn-51-1, Kpn-52-1, Kpn-53-1, Kpn-54-1, Kpn-55-1, Kpn-56-1, Kpn-57-1, Kpn-58-1, Kpn-59-1, Kpn-6-1, Kpn-60-1, Kpn-61-1, Kpn-62-1, Kpn-63-1, Kpn-64-1, Kpn-65-1, Kpn-66-1, Kpn-67-1, Kpn-68-1, Kpn-69-1, Kpn-7-1, Kpn-70-1, Kpn-71-1, Kpn-72-1, Kpn-73-1, Kpn-74-1, Kpn-75-1, Kpn-76-1, Kpn-77-1, Kpn-78-1, Kpn-79-1, Kpn-8-1, Kpn-80-1, Kpn-81-1, Kpn-82-1, Kpn-83-1, Kpn-84-1, Kpn-85-1, Kpn-86-1, Kpn-87-1, Kpn-88-1, Kpn-89-1, Kpn-9-1, Kpn-90-1, Kpn-91-1, Kpn-92-1, Kpn-93-1, Kpn-94-1, Kpn-95-1, Kpn-96-1, Kpn-97-1, Kpn-98-1, Kpn-99-1, Kqu-2-1, Kqu-2-2, Kqu-2-3, Kqu-2-4, Kra-1-1, Ksp-1-1, Ksp-2-1, Ksp-4-1, Ksp-5-1, Ksp-6-1, Ksp-7-1, Kva-1-2, Kva-1-3, Kva-1-4, Kva-1-5, Kva-1-6, Kva-1-7, Kva-2-1, Lad-1-1, Lad-2-1, Lam-1-1, Lan-1-1, Lgr-1-2, Lqu-1-1, Lqu-1-2, Lsp-1-1, Maq-1-1, Mhy-1-1, Mmo-1-2, Mmo-1-3, Mmo-1-4, Mmo-1-5, Mmo-1-6, Mmo-1-8, Mmo-4-1, Mmo-5-1, Mmo-6-1, Msp-2-1, Msp-3-1, Msp-5-1, Pag-1-1, Pag-2-1, Pag-2-2, Pag-2-3, Pag-2-4, Pag-2-6, Pag-3-1, Pag-4-1, Pag-6-1, Pal-1-1, Pal-2-1, Pal-3-1, Pal-4-1, Pal-6-1, Pal-7-1, Pal-9-1, Pan-1-1, Pan-1-2, Pan-1-3, Pan-1-4, Pan-1-5, Pan-10-1, Pan-11-10, Pan-11-11, Pan-11-12, Pan-11-3, Pan-11-4, Pan-11-5, Pan-11-6, Pan-11-7, Pan-11-8, Pan-11-9, Pan-2-1, Pan-3-1, Pan-3-2, Pan-4-1, Pan-6-1, Pan-7-1, Pan-8-1, Pan-9-1, Pat-2-1, Pat-2-2, Pat-4-1, Pbe-1-2, Pbe-2-1, Pca-11-1, Pca-3-10, Pca-3-11, Pca-3-12, Pca-3-13, Pca-3-14, Pca-3-15, Pca-3-4, Pca-3-5, Pca-3-6, Pca-3-7, Pca-3-8, Pca-3-9, Pca-4-1, Pca-5-1, Pca-5-10, Pca-5-3, Pca-5-5, Pca-5-7, Pca-5-9, Pca-8-2, Pca-9-1, Pda-2-1, Pda-3-1, Pdi-1-1, Pdi-2-1, Pfl-2-1, Pfu-1-1, Pha-1-1, Pha-3-1, Pha-7-1, Pil-1-2, Pki-1-2, Pki-1-3, Pki-1-4, Ple-1-2, Ple-2-1, Ple-3-1, Pli-1-1, Plu-5-2, Plu-8-1, Pma-1-1, Pmi-1-1, Pmi-1-2, Pmi-1-3, Pmi-1-4, Pmi-1-5, Pmi-1-6, Pmi-4-1, Pmi-5-1, Pmi-6-1, Pmi-7-1, Pmi-8-1, Pmu-1-1, Pmu-10-1, Pmu-11-1, Pmu-13-1, Pmu-13-10, Pmu-13-11, Pmu-13-12, Pmu-13-2, Pmu-13-3, Pmu-13-4, Pmu-13-5, Pmu-13-6, Pmu-13-7, Pmu-13-8, Pmu-13-9, Pmu-15-1, Pmu-16-1, Pmu-17-1, Pmu-18-1, Pmu-19-1, Pmu-20-1, Pmu-21-1, Pmu-4-1, Pmu-5-1, Pmu-7-1, Pmu-9-1, Ppa-1-1, Ppi-2-2, Ppi-4-1, Ppn-1-1, Ppo-1-1, Pre-1-2, Pru-3-1, Pru-3-2, Pru-4-1, Prw-1-1, Psh-1-1, Psh-2-1, Psp-10-1, Psp-12-1, Psp-13-1, Psp-14-1, Psp-15-1, Psp-16-1, Psp-18-1, Psp-2-1, Psp-20-1, Psp-21-1, Psp-22-1, Psp-23-1, Psp-24-1, Psp-25-1, Psp-27-1, Psp-28-1, Psp-29-1, Psp-3-1, Psp-30-1, Psp-31-1, Psp-33-1, Psp-37-1, Psp-4-1, Psp-41-1, Psp-42-1, Psp-5-1, Psp-50-1, Psp-51-1, Psp-52-1, Psp-54-1, Psp-56-2, Psp-56-3, Psp-7-1, Psp-8-1, Psp-9-1, Pst-1-2, Pst-2-1, Pst-5-1, Pst-6-1, Pte-3-1, Pte-5-1, Pun-1-1, Pva-1-3, Pva-2-1, Pwa-1-1, Pwa-1-2, Pwa-3-1, Raq-1-2, Raq-2-1, Raq-2-2, Raq-3-1, Ror-2-1, Ror-3-1, Ror-4-1, Ror-4-2, Rpl-1-2, Rpl-1-3, Rpl-2-1, Rpn-1-2, Rsp-3-1, Rsp-4-1, Sal-2-1, Sba-1-1, Sba-2-1, Sba-4-1, Sba-5-1, Sba-6-1, Sba-7-1, Sba-8-1, Sba-9-1, Sbl-1-1, Sbo-1-1, Sbo-10-1, Sbo-11-1, Sbo-12-1, Sbo-13-1, Sbo-14-1, Sbo-15-1, Sbo-16-1, Sbo-17-1, Sbo-18-1, Sbo-2-1, Sbo-2-10, Sbo-2-2, Sbo-2-3, Sbo-2-4, Sbo-2-5, Sbo-2-6, Sbo-2-7, Sbo-2-8, Sbo-2-9, Sbo-4-1, Sbo-5-1, Sbo-6-1, Sbo-7-1, Sbo-8-1, Sbo-9-1, Sco-3-1, Sdy-1-1, Sdy-10-1, Sdy-2-1, Sdy-3-1, Sdy-4-1, Sdy-6-1, Sdy-7-1, Sdy-8-1, Sdy-9-1, Sen-1-1, Sen-10-1, Sen-100-1, Sen-1000-1, Sen-1001-1, Sen-1002-1, Sen-1003-1, Sen-1004-1, Sen-1005-1, Sen-1006-1, Sen-1007-1, Sen-1008-1, Sen-1009-1, Sen-101-1, Sen-1010-1, Sen-1011-1, Sen-1012-1, Sen-1014-1, Sen-1015-1, Sen-1016-1, Sen-1017-1, Sen-1018-1, Sen-1019-1, Sen-102-1, Sen-1020-1, Sen-1022-1, Sen-1023-1, Sen-1024-1, Sen-1025-1, Sen-1026-1, Sen-1027-1, Sen-1027-2, Sen-1028-1, Sen-103-1, Sen-1030-1, Sen-1031-1, Sen-1032-1, Sen-1033-1, Sen-104-1, Sen-105-1, Sen-106-1, Sen-107-1, Sen-108-1, Sen-109-1, Sen-11-1, Sen-110-1, Sen-111-1, Sen-112-1, Sen-112-2, Sen-112-3, Sen-113-1, Sen-113-2, Sen-114-1, Sen-115-1, Sen-116-1, Sen-117-1, Sen-118-1, Sen-119-1, Sen-12-1, Sen-12-2, Sen-12-3, Sen-120-1, Sen-121-1, Sen-122-1, Sen-123-1, Sen-124-1, Sen-125-1, Sen-126-1, Sen-127-1, Sen-128-1, Sen-129-1, Sen-129-2, Sen-13-1, Sen-130-1, Sen-130-2, Sen-130-3, Sen-131-1, Sen-132-1, Sen-133-1, Sen-134-1, Sen-135-1, Sen-136-1, Sen-136-2, Sen-137-1, Sen-137-2, Sen-137-3, Sen-137-4, Sen-137-5, Sen-137-6, Sen-138-1, Sen-138-2, Sen-138-3, Sen-138-4, Sen-138-5, Sen-139-1, Sen-139-10, Sen-139-11, Sen-139-12, Sen-139-13, Sen-139-14, Sen-139-15, Sen-139-2, Sen-139-3, Sen-139-4, Sen-139-5, Sen-139-6, Sen-139-7, Sen-139-8, Sen-139-9, Sen-14-1, Sen-14-2, Sen-14-3, Sen-14-4, Sen-140-1, Sen-141-1, Sen-142-1, Sen-143-1, Sen-144-1, Sen-145-1, Sen-146-1, Sen-147-1, Sen-148-1, Sen-149-1, Sen-15-1, Sen-15-2, Sen-15-3, Sen-150-1, Sen-151-1, Sen-152-1, Sen-153-1, Sen-154-1, Sen-155-1, Sen-156-1, Sen-157-1, Sen-158-1, Sen-159-1, Sen-16-1, Sen-160-1, Sen-161-1, Sen-162-1, Sen-163-1, Sen-164-1, Sen-165-1, Sen-166-1, Sen-167-1, Sen-167-10, Sen-167-11, Sen-167-12, Sen-167-13, Sen-167-14, Sen-167-15, Sen-167-16, Sen-167-17, Sen-167-18, Sen-167-2, Sen-167-3, Sen-167-4, Sen-167-5, Sen-167-6, Sen-167-7, Sen-167-8, Sen-167-9, Sen-168-1, Sen-169-1, Sen-17-1, Sen-170-1, Sen-170-10, Sen-170-11, Sen-170-12, Sen-170-13, Sen-170-14, Sen-170-15, Sen-170-16, Sen-170-17, Sen-170-18, Sen-170-19, Sen-170-2, Sen-170-20, Sen-170-21, Sen-

170-22, Sen-170-23, Sen-170-24, Sen-170-25, Sen-170-26, Sen-170-27, Sen-170-28, Sen-170-29, Sen-170-3, Sen-170-30, Sen-170-31, Sen-170-32, Sen-170-33, Sen-170-34, Sen-170-35, Sen-170-36, Sen-170-37, Sen-170-38, Sen-170-39, Sen-170-4, Sen-170-40, Sen-170-41, Sen-170-42, Sen-170-43, Sen-170-44, Sen-170-45, Sen-170-46, Sen-170-47, Sen-170-48, Sen-170-49, Sen-170-5, Sen-170-50, Sen-170-51, Sen-170-52, Sen-170-6, Sen-170-7, Sen-170-8, Sen-170-9, Sen-171-1, Sen-172-1, Sen-173-1, Sen-174-1, Sen-175-1, Sen-176-1, Sen-177-1, Sen-178-1, Sen-179-1, Sen-18-1, Sen-180-1, Sen-181-1, Sen-182-1, Sen-183-1, Sen-184-1, Sen-185-1, Sen-186-1, Sen-187-1, Sen-188-1, Sen-189-1, Sen-19-1, Sen-190-1, Sen-191-1, Sen-192-1, Sen-193-1, Sen-194-1, Sen-195-1, Sen-196-1, Sen-197-1, Sen-198-1, Sen-199-1, Sen-2-1, Sen-2-2, Sen-20-1, Sen-200-1, Sen-201-1, Sen-202-1, Sen-203-1, Sen-204-1, Sen-205-1, Sen-206-1, Sen-207-1, Sen-208-1, Sen-209-1, Sen-21-1, Sen-210-1, Sen-211-1, Sen-212-1, Sen-213-1, Sen-214-1, Sen-215-1, Sen-216-1, Sen-217-1, Sen-218-1, Sen-219-1, Sen-22-1, Sen-220-1, Sen-221-1, Sen-222-1, Sen-223-1, Sen-224-1, Sen-224-2, Sen-224-3, Sen-225-1, Sen-225-10, Sen-225-11, Sen-225-12, Sen-225-13, Sen-225-14, Sen-225-15, Sen-225-16, Sen-225-17, Sen-225-18, Sen-225-19, Sen-225-2, Sen-225-20, Sen-225-21, Sen-225-22, Sen-225-23, Sen-225-24, Sen-225-25, Sen-225-26, Sen-225-27, Sen-225-28, Sen-225-29, Sen-225-3, Sen-225-30, Sen-225-31, Sen-225-32, Sen-225-33, Sen-225-34, Sen-225-35, Sen-225-36, Sen-225-37, Sen-225-38, Sen-225-39, Sen-225-4, Sen-225-40, Sen-225-41, Sen-225-42, Sen-225-43, Sen-225-44, Sen-225-45, Sen-225-46, Sen-225-47, Sen-225-48, Sen-225-49, Sen-225-5, Sen-225-50, Sen-225-51, Sen-225-52, Sen-225-6, Sen-225-7, Sen-225-8, Sen-225-9, Sen-226-1, Sen-226-2, Sen-226-3, Sen-226-4, Sen-227-1, Sen-227-2, Sen-228-1, Sen-228-2, Sen-229-1, Sen-23-1, Sen-230-1, Sen-230-2, Sen-230-3, Sen-230-4, Sen-231-1, Sen-232-1, Sen-233-1, Sen-233-2, Sen-233-3, Sen-233-4, Sen-233-5, Sen-234-1, Sen-235-1, Sen-236-1, Sen-237-1, Sen-237-10, Sen-237-11, Sen-237-12, Sen-237-13, Sen-237-14, Sen-237-15, Sen-237-16, Sen-237-2, Sen-237-3, Sen-237-4, Sen-237-5, Sen-237-6, Sen-237-7, Sen-237-8, Sen-237-9, Sen-238-1, Sen-239-1, Sen-24-1, Sen-240-1, Sen-241-1, Sen-242-1, Sen-243-1, Sen-243-2, Sen-244-1, Sen-245-1, Sen-246-1, Sen-247-1, Sen-248-1, Sen-249-1, Sen-25-1, Sen-250-1, Sen-251-1, Sen-252-1, Sen-253-1, Sen-254-1, Sen-255-1, Sen-256-1, Sen-257-1, Sen-258-1, Sen-259-1, Sen-26-1, Sen-260-1, Sen-261-1, Sen-262-1, Sen-263-1, Sen-264-1, Sen-265-1, Sen-266-1, Sen-266-2, Sen-267-1, Sen-268-1, Sen-269-1, Sen-27-1, Sen-270-1, Sen-271-1, Sen-272-1, Sen-273-1, Sen-273-2, Sen-273-3, Sen-274-1, Sen-275-1, Sen-276-1, Sen-277-1, Sen-277-2, Sen-277-3, Sen-277-4, Sen-277-5, Sen-278-1, Sen-278-10, Sen-278-11, Sen-278-12, Sen-278-13, Sen-278-14, Sen-278-15, Sen-278-16, Sen-278-17, Sen-278-18, Sen-278-19, Sen-278-2, Sen-278-20, Sen-278-21, Sen-278-22, Sen-278-3, Sen-278-4, Sen-278-5, Sen-278-6, Sen-278-7, Sen-278-8, Sen-278-9, Sen-279-1, Sen-279-10, Sen-279-11, Sen-279-12, Sen-279-13, Sen-279-14, Sen-279-2, Sen-279-3, Sen-279-4, Sen-279-5, Sen-279-6, Sen-279-7, Sen-279-8, Sen-279-9, Sen-28-1, Sen-280-1, Sen-280-2, Sen-280-3, Sen-281-1, Sen-282-1, Sen-283-1, Sen-284-1, Sen-285-1, Sen-286-1, Sen-286-2, Sen-286-3, Sen-286-4, Sen-286-5, Sen-287-1, Sen-288-1, Sen-288-10, Sen-288-100, Sen-288-1000, Sen-288-1001, Sen-288-1002, Sen-288-1003, Sen-288-1004, Sen-288-1005, Sen-288-1006, Sen-288-1007, Sen-288-1008, Sen-288-1009, Sen-288-101, Sen-288-1010, Sen-288-1011, Sen-288-1012, Sen-288-1013, Sen-288-1014, Sen-288-1015, Sen-288-1016, Sen-288-1017, Sen-288-1018, Sen-288-1019, Sen-288-102, Sen-288-1020, Sen-288-1021, Sen-288-1022, Sen-288-1023, Sen-288-1024, Sen-288-1025, Sen-288-1026, Sen-288-1027, Sen-288-1028, Sen-288-1029, Sen-288-103, Sen-288-1030, Sen-288-1031, Sen-288-1032, Sen-288-1033, Sen-288-1034, Sen-288-1035, Sen-288-1036, Sen-288-1037, Sen-288-1038, Sen-288-1039, Sen-288-104, Sen-288-1040, Sen-288-1041, Sen-288-1042, Sen-288-1043, Sen-288-1044, Sen-288-1045, Sen-288-1046, Sen-288-1047, Sen-288-1048, Sen-288-1049, Sen-288-105, Sen-288-1050, Sen-288-1051, Sen-288-1052, Sen-288-1053, Sen-288-1054, Sen-288-1055, Sen-288-1056, Sen-288-1057, Sen-288-1058, Sen-288-1059, Sen-288-106, Sen-288-1060, Sen-288-1061, Sen-288-1062, Sen-288-1063, Sen-288-1064, Sen-288-1065, Sen-288-1066, Sen-288-1067, Sen-288-1068, Sen-288-1069, Sen-288-107, Sen-288-1070, Sen-288-1071, Sen-288-1072, Sen-288-1073, Sen-288-1074, Sen-288-1075, Sen-288-1076, Sen-288-1077, Sen-288-1078, Sen-288-1079, Sen-288-108, Sen-288-1080, Sen-288-1081, Sen-288-1082, Sen-288-1083, Sen-288-1084, Sen-288-1085, Sen-288-1086, Sen-288-1087, Sen-288-1088, Sen-288-1089, Sen-288-109, Sen-288-1090, Sen-288-1091, Sen-288-1092, Sen-288-1093, Sen-288-1094, Sen-288-1095, Sen-288-1096, Sen-288-1097, Sen-288-1098, Sen-288-1099, Sen-288-11, Sen-288-110, Sen-288-1100, Sen-288-1101, Sen-288-1102, Sen-288-1103, Sen-288-1104, Sen-288-1105, Sen-288-1106, Sen-288-1107, Sen-288-1108, Sen-288-1109, Sen-288-111, Sen-288-1110, Sen-288-1111, Sen-288-1112, Sen-288-1113, Sen-288-1114, Sen-288-1115, Sen-288-1116, Sen-288-1117, Sen-288-1118, Sen-288-1119, Sen-288-112, Sen-288-1120, Sen-288-1121, Sen-288-1122, Sen-288-1123, Sen-288-1124, Sen-288-1125, Sen-288-113, Sen-288-114, Sen-288-115, Sen-288-116, Sen-288-117, Sen-288-118, Sen-288-119, Sen-288-12, Sen-288-120, Sen-288-121, Sen-288-122, Sen-288-123, Sen-288-124, Sen-288-125, Sen-288-126, Sen-288-127, Sen-288-128, Sen-288-129, Sen-288-13, Sen-288-130, Sen-288-131, Sen-288-132, Sen-288-133, Sen-288-134, Sen-288-135, Sen-288-136, Sen-288-137, Sen-288-138, Sen-288-139, Sen-288-14, Sen-288-140, Sen-288-141, Sen-288-142, Sen-288-143, Sen-288-144, Sen-288-145, Sen-288-146, Sen-288-147, Sen-288-148, Sen-288-149, Sen-288-15, Sen-288-150, Sen-288-151, Sen-288-152, Sen-288-153, Sen-288-154, Sen-288-155, Sen-288-156, Sen-288-157, Sen-288-158, Sen-288-159, Sen-288-16, Sen-288-160, Sen-288-161, Sen-288-162, Sen-288-163, Sen-288-164, Sen-288-165, Sen-288-166, Sen-288-167, Sen-288-168, Sen-288-169, Sen-288-17, Sen-288-170, Sen-288-171, Sen-288-172, Sen-288-173, Sen-288-174, Sen-288-175, Sen-288-176, Sen-288-177, Sen-288-178, Sen-288-179, Sen-288-18, Sen-288-180, Sen-288-181, Sen-288-182, Sen-288-183, Sen-288-184, Sen-288-185, Sen-288-186, Sen-288-187, Sen-288-188, Sen-288-189, Sen-288-19, Sen-288-190, Sen-288-191, Sen-288-192, Sen-288-193, Sen-288-194, Sen-288-195, Sen-288-196, Sen-288-197, Sen-288-198, Sen-288-199, Sen-288-2, Sen-288-20, Sen-288-200, Sen-288-201, Sen-288-202, Sen-288-203, Sen-288-204, Sen-288-205, Sen-288-206, Sen-288-207, Sen-288-208, Sen-288-209, Sen-288-21, Sen-288-210, Sen-288-211, Sen-288-212, Sen-288-213, Sen-288-214, Sen-288-215, Sen-288-216, Sen-288-217, Sen-288-218, Sen-288-219, Sen-288-22, Sen-288-220, Sen-288-221, Sen-288-222, Sen-288-223, Sen-288-224, Sen-288-225, Sen-288-226, Sen-288-227, Sen-288-228, Sen-288-229, Sen-288-23, Sen-288-230, Sen-288-231, Sen-288-232, Sen-288-233, Sen-288-234, Sen-288-235, Sen-288-236, Sen-288-237, Sen-288-238, Sen-288-239,

Sen-288-24, Sen-288-240, Sen-288-241, Sen-288-242, Sen-288-243, Sen-288-244, Sen-288-245, Sen-288-246, Sen-288-247, Sen-288-248, Sen-288-249, Sen-288-25, Sen-288-250, Sen-288-251, Sen-288-252, Sen-288-253, Sen-288-254, Sen-288-255, Sen-288-256, Sen-288-257, Sen-288-258, Sen-288-259, Sen-288-26, Sen-288-260, Sen-288-261, Sen-288-262, Sen-288-263, Sen-288-264, Sen-288-265, Sen-288-266, Sen-288-267, Sen-288-268, Sen-288-269, Sen-288-27, Sen-288-270, Sen-288-271, Sen-288-272, Sen-288-273, Sen-288-274, Sen-288-275, Sen-288-276, Sen-288-277, Sen-288-278, Sen-288-279, Sen-288-28, Sen-288-280, Sen-288-281, Sen-288-282, Sen-288-283, Sen-288-284, Sen-288-285, Sen-288-286, Sen-288-287, Sen-288-288, Sen-288-289, Sen-288-29, Sen-288-290, Sen-288-291, Sen-288-292, Sen-288-293, Sen-288-294, Sen-288-295, Sen-288-296, Sen-288-297, Sen-288-298, Sen-288-299, Sen-288-3, Sen-288-30, Sen-288-300, Sen-288-301, Sen-288-302, Sen-288-303, Sen-288-304, Sen-288-305, Sen-288-306, Sen-288-307, Sen-288-308, Sen-288-309, Sen-288-31, Sen-288-310, Sen-288-311, Sen-288-312, Sen-288-313, Sen-288-314, Sen-288-315, Sen-288-316, Sen-288-317, Sen-288-318, Sen-288-319, Sen-288-32, Sen-288-320, Sen-288-321, Sen-288-322, Sen-288-323, Sen-288-324, Sen-288-325, Sen-288-326, Sen-288-327, Sen-288-328, Sen-288-329, Sen-288-33, Sen-288-330, Sen-288-331, Sen-288-332, Sen-288-333, Sen-288-334, Sen-288-335, Sen-288-336, Sen-288-337, Sen-288-338, Sen-288-339, Sen-288-34, Sen-288-340, Sen-288-341, Sen-288-342, Sen-288-343, Sen-288-344, Sen-288-345, Sen-288-346, Sen-288-347, Sen-288-348, Sen-288-349, Sen-288-35, Sen-288-350, Sen-288-351, Sen-288-352, Sen-288-353, Sen-288-354, Sen-288-355, Sen-288-356, Sen-288-357, Sen-288-358, Sen-288-359, Sen-288-36, Sen-288-360, Sen-288-361, Sen-288-362, Sen-288-363, Sen-288-364, Sen-288-365, Sen-288-366, Sen-288-367, Sen-288-368, Sen-288-369, Sen-288-37, Sen-288-370, Sen-288-371, Sen-288-372, Sen-288-373, Sen-288-374, Sen-288-375, Sen-288-376, Sen-288-377, Sen-288-378, Sen-288-379, Sen-288-38, Sen-288-380, Sen-288-381, Sen-288-382, Sen-288-383, Sen-288-384, Sen-288-385, Sen-288-386, Sen-288-387, Sen-288-388, Sen-288-389, Sen-288-39, Sen-288-390, Sen-288-391, Sen-288-392, Sen-288-393, Sen-288-394, Sen-288-395, Sen-288-396, Sen-288-397, Sen-288-398, Sen-288-399, Sen-288-4, Sen-288-40, Sen-288-400, Sen-288-401, Sen-288-402, Sen-288-403, Sen-288-404, Sen-288-405, Sen-288-406, Sen-288-407, Sen-288-408, Sen-288-409, Sen-288-41, Sen-288-410, Sen-288-411, Sen-288-412, Sen-288-413, Sen-288-414, Sen-288-415, Sen-288-416, Sen-288-417, Sen-288-418, Sen-288-419, Sen-288-42, Sen-288-420, Sen-288-421, Sen-288-422, Sen-288-423, Sen-288-424, Sen-288-425, Sen-288-426, Sen-288-427, Sen-288-428, Sen-288-429, Sen-288-43, Sen-288-430, Sen-288-431, Sen-288-432, Sen-288-433, Sen-288-434, Sen-288-435, Sen-288-436, Sen-288-437, Sen-288-438, Sen-288-439, Sen-288-44, Sen-288-440, Sen-288-441, Sen-288-442, Sen-288-443, Sen-288-444, Sen-288-445, Sen-288-446, Sen-288-447, Sen-288-448, Sen-288-449, Sen-288-45, Sen-288-450, Sen-288-451, Sen-288-452, Sen-288-453, Sen-288-454, Sen-288-455, Sen-288-456, Sen-288-457, Sen-288-458, Sen-288-459, Sen-288-46, Sen-288-460, Sen-288-461, Sen-288-462, Sen-288-463, Sen-288-464, Sen-288-465, Sen-288-466, Sen-288-467, Sen-288-468, Sen-288-469, Sen-288-47, Sen-288-470, Sen-288-471, Sen-288-472, Sen-288-473, Sen-288-474, Sen-288-475, Sen-288-476, Sen-288-477, Sen-288-478, Sen-288-479, Sen-288-48, Sen-288-480, Sen-288-481, Sen-288-482, Sen-288-483, Sen-288-484, Sen-288-485, Sen-288-486, Sen-288-487, Sen-288-488, Sen-288-489, Sen-288-49, Sen-288-490, Sen-288-491, Sen-288-492, Sen-288-493, Sen-288-494, Sen-288-495, Sen-288-496, Sen-288-497, Sen-288-498, Sen-288-499, Sen-288-5, Sen-288-50, Sen-288-500, Sen-288-501, Sen-288-502, Sen-288-503, Sen-288-504, Sen-288-505, Sen-288-506, Sen-288-507, Sen-288-508, Sen-288-509, Sen-288-51, Sen-288-510, Sen-288-511, Sen-288-512, Sen-288-513, Sen-288-514, Sen-288-515, Sen-288-516, Sen-288-517, Sen-288-518, Sen-288-519, Sen-288-52, Sen-288-520, Sen-288-521, Sen-288-522, Sen-288-523, Sen-288-524, Sen-288-525, Sen-288-526, Sen-288-527, Sen-288-528, Sen-288-529, Sen-288-53, Sen-288-530, Sen-288-531, Sen-288-532, Sen-288-533, Sen-288-534, Sen-288-535, Sen-288-536, Sen-288-537, Sen-288-538, Sen-288-539, Sen-288-54, Sen-288-540, Sen-288-541, Sen-288-542, Sen-288-543, Sen-288-544, Sen-288-545, Sen-288-546, Sen-288-547, Sen-288-548, Sen-288-549, Sen-288-55, Sen-288-550, Sen-288-551, Sen-288-552, Sen-288-553, Sen-288-554, Sen-288-555, Sen-288-556, Sen-288-557, Sen-288-558, Sen-288-559, Sen-288-56, Sen-288-560, Sen-288-561, Sen-288-562, Sen-288-563, Sen-288-564, Sen-288-565, Sen-288-566, Sen-288-567, Sen-288-568, Sen-288-569, Sen-288-57, Sen-288-570, Sen-288-571, Sen-288-572, Sen-288-573, Sen-288-574, Sen-288-575, Sen-288-576, Sen-288-577, Sen-288-578, Sen-288-579, Sen-288-58, Sen-288-580, Sen-288-581, Sen-288-582, Sen-288-583, Sen-288-584, Sen-288-585, Sen-288-586, Sen-288-587, Sen-288-588, Sen-288-589, Sen-288-59, Sen-288-590, Sen-288-591, Sen-288-592, Sen-288-593, Sen-288-594, Sen-288-595, Sen-288-596, Sen-288-597, Sen-288-598, Sen-288-599, Sen-288-6, Sen-288-60, Sen-288-600, Sen-288-601, Sen-288-602, Sen-288-603, Sen-288-604, Sen-288-605, Sen-288-606, Sen-288-607, Sen-288-608, Sen-288-609, Sen-288-61, Sen-288-610, Sen-288-611, Sen-288-612, Sen-288-613, Sen-288-614, Sen-288-615, Sen-288-616, Sen-288-617, Sen-288-618, Sen-288-619, Sen-288-62, Sen-288-620, Sen-288-621, Sen-288-622, Sen-288-623, Sen-288-624, Sen-288-625, Sen-288-626, Sen-288-627, Sen-288-628, Sen-288-629, Sen-288-63, Sen-288-630, Sen-288-631, Sen-288-632, Sen-288-633, Sen-288-634, Sen-288-635, Sen-288-636, Sen-288-637, Sen-288-638, Sen-288-639, Sen-288-64, Sen-288-640, Sen-288-641, Sen-288-642, Sen-288-643, Sen-288-644, Sen-288-645, Sen-288-646, Sen-288-647, Sen-288-648, Sen-288-649, Sen-288-65, Sen-288-650, Sen-288-651, Sen-288-652, Sen-288-653, Sen-288-654, Sen-288-655, Sen-288-656, Sen-288-657, Sen-288-658, Sen-288-659, Sen-288-66, Sen-288-660, Sen-288-661, Sen-288-662, Sen-288-663, Sen-288-664, Sen-288-665, Sen-288-666, Sen-288-667, Sen-288-668, Sen-288-669, Sen-288-67, Sen-288-670, Sen-288-671, Sen-288-672, Sen-288-673, Sen-288-674, Sen-288-675, Sen-288-676, Sen-288-677, Sen-288-678, Sen-288-679, Sen-288-68, Sen-288-680, Sen-288-681, Sen-288-682, Sen-288-683, Sen-288-684, Sen-288-685, Sen-288-686, Sen-288-687, Sen-288-688, Sen-288-689, Sen-288-69, Sen-288-690, Sen-288-691, Sen-288-692, Sen-288-693, Sen-288-694, Sen-288-695, Sen-288-696, Sen-288-697, Sen-288-698, Sen-288-699, Sen-288-7, Sen-288-70, Sen-288-700, Sen-288-701, Sen-288-702, Sen-288-703, Sen-288-704, Sen-288-705, Sen-288-706, Sen-288-707, Sen-288-708, Sen-288-709, Sen-288-71, Sen-288-710, Sen-288-711, Sen-288-712, Sen-288-713, Sen-288-714, Sen-288-715, Sen-288-716, Sen-288-717, Sen-288-718, Sen-288-719, Sen-288-72, Sen-288-720, Sen-288-721, Sen-288-722, Sen-288-723, Sen-288-724, Sen-288-725, Sen-288-726, Sen-288-727, Sen-288-728, Sen-288-729, Sen-288-73, Sen-288-730, Sen-288-731, Sen-288-732, Sen-288-733, Sen-288-734,

Sen-288-735, Sen-288-736, Sen-288-737, Sen-288-738, Sen-288-739, Sen-288-74, Sen-288-740, Sen-288-741, Sen-288-742, Sen-288-743, Sen-288-744, Sen-288-745, Sen-288-746, Sen-288-747, Sen-288-748, Sen-288-749, Sen-288-75, Sen-288-750, Sen-288-751, Sen-288-752, Sen-288-753, Sen-288-754, Sen-288-755, Sen-288-756, Sen-288-757, Sen-288-758, Sen-288-759, Sen-288-76, Sen-288-760, Sen-288-761, Sen-288-762, Sen-288-763, Sen-288-764, Sen-288-765, Sen-288-766, Sen-288-767, Sen-288-768, Sen-288-769, Sen-288-77, Sen-288-770, Sen-288-771, Sen-288-772, Sen-288-773, Sen-288-774, Sen-288-775, Sen-288-776, Sen-288-777, Sen-288-778, Sen-288-779, Sen-288-78, Sen-288-780, Sen-288-781, Sen-288-782, Sen-288-783, Sen-288-784, Sen-288-785, Sen-288-786, Sen-288-787, Sen-288-788, Sen-288-789, Sen-288-79, Sen-288-790, Sen-288-791, Sen-288-792, Sen-288-793, Sen-288-794, Sen-288-795, Sen-288-796, Sen-288-797, Sen-288-798, Sen-288-799, Sen-288-8, Sen-288-80, Sen-288-800, Sen-288-801, Sen-288-802, Sen-288-803, Sen-288-804, Sen-288-805, Sen-288-806, Sen-288-807, Sen-288-808, Sen-288-809, Sen-288-81, Sen-288-810, Sen-288-811, Sen-288-812, Sen-288-813, Sen-288-814, Sen-288-815, Sen-288-816, Sen-288-817, Sen-288-818, Sen-288-819, Sen-288-82, Sen-288-820, Sen-288-821, Sen-288-822, Sen-288-823, Sen-288-824, Sen-288-825, Sen-288-826, Sen-288-827, Sen-288-828, Sen-288-829, Sen-288-83, Sen-288-830, Sen-288-831, Sen-288-832, Sen-288-833, Sen-288-834, Sen-288-835, Sen-288-836, Sen-288-837, Sen-288-838, Sen-288-839, Sen-288-84, Sen-288-840, Sen-288-841, Sen-288-842, Sen-288-843, Sen-288-844, Sen-288-845, Sen-288-846, Sen-288-847, Sen-288-848, Sen-288-849, Sen-288-85, Sen-288-850, Sen-288-851, Sen-288-852, Sen-288-853, Sen-288-854, Sen-288-855, Sen-288-856, Sen-288-857, Sen-288-858, Sen-288-859, Sen-288-86, Sen-288-860, Sen-288-861, Sen-288-862, Sen-288-863, Sen-288-864, Sen-288-865, Sen-288-866, Sen-288-867, Sen-288-868, Sen-288-869, Sen-288-87, Sen-288-870, Sen-288-871, Sen-288-872, Sen-288-873, Sen-288-874, Sen-288-875, Sen-288-876, Sen-288-877, Sen-288-878, Sen-288-879, Sen-288-88, Sen-288-880, Sen-288-881, Sen-288-882, Sen-288-883, Sen-288-884, Sen-288-885, Sen-288-886, Sen-288-887, Sen-288-888, Sen-288-889, Sen-288-89, Sen-288-890, Sen-288-891, Sen-288-892, Sen-288-893, Sen-288-894, Sen-288-895, Sen-288-896, Sen-288-897, Sen-288-898, Sen-288-899, Sen-288-9, Sen-288-90, Sen-288-900, Sen-288-901, Sen-288-902, Sen-288-903, Sen-288-904, Sen-288-905, Sen-288-906, Sen-288-907, Sen-288-908, Sen-288-909, Sen-288-91, Sen-288-910, Sen-288-911, Sen-288-912, Sen-288-913, Sen-288-914, Sen-288-915, Sen-288-916, Sen-288-917, Sen-288-918, Sen-288-919, Sen-288-92, Sen-288-920, Sen-288-921, Sen-288-922, Sen-288-923, Sen-288-924, Sen-288-925, Sen-288-926, Sen-288-927, Sen-288-928, Sen-288-929, Sen-288-93, Sen-288-930, Sen-288-931, Sen-288-932, Sen-288-933, Sen-288-934, Sen-288-935, Sen-288-936, Sen-288-937, Sen-288-938, Sen-288-939, Sen-288-94, Sen-288-940, Sen-288-941, Sen-288-942, Sen-288-943, Sen-288-944, Sen-288-945, Sen-288-946, Sen-288-947, Sen-288-948, Sen-288-949, Sen-288-95, Sen-288-950, Sen-288-951, Sen-288-952, Sen-288-953, Sen-288-954, Sen-288-955, Sen-288-956, Sen-288-957, Sen-288-958, Sen-288-959, Sen-288-96, Sen-288-960, Sen-288-961, Sen-288-962, Sen-288-963, Sen-288-964, Sen-288-965, Sen-288-966, Sen-288-967, Sen-288-968, Sen-288-969, Sen-288-97, Sen-288-970, Sen-288-971, Sen-288-972, Sen-288-973, Sen-288-974, Sen-288-975, Sen-288-976, Sen-288-977, Sen-288-978, Sen-288-979, Sen-288-98, Sen-288-980, Sen-288-981, Sen-288-982, Sen-288-983, Sen-288-984, Sen-288-985, Sen-288-986, Sen-288-987, Sen-288-988, Sen-288-989, Sen-288-99, Sen-288-990, Sen-288-991, Sen-288-992, Sen-288-993, Sen-288-994, Sen-288-995, Sen-288-996, Sen-288-997, Sen-288-998, Sen-288-999, Sen-289-1, Sen-29-1, Sen-290-1, Sen-291-1, Sen-291-2, Sen-291-3, Sen-291-4, Sen-291-5, Sen-291-6, Sen-291-7, Sen-292-1, Sen-293-1, Sen-294-1, Sen-295-1, Sen-296-1, Sen-297-1, Sen-298-1, Sen-299-1, Sen-3-1, Sen-30-1, Sen-300-1, Sen-300-10, Sen-300-100, Sen-300-101, Sen-300-102, Sen-300-103, Sen-300-104, Sen-300-105, Sen-300-106, Sen-300-107, Sen-300-108, Sen-300-109, Sen-300-11, Sen-300-110, Sen-300-111, Sen-300-112, Sen-300-113, Sen-300-114, Sen-300-115, Sen-300-116, Sen-300-117, Sen-300-118, Sen-300-119, Sen-300-12, Sen-300-120, Sen-300-121, Sen-300-122, Sen-300-123, Sen-300-124, Sen-300-125, Sen-300-126, Sen-300-127, Sen-300-128, Sen-300-129, Sen-300-13, Sen-300-130, Sen-300-131, Sen-300-132, Sen-300-133, Sen-300-134, Sen-300-135, Sen-300-136, Sen-300-137, Sen-300-138, Sen-300-139, Sen-300-14, Sen-300-140, Sen-300-141, Sen-300-142, Sen-300-143, Sen-300-144, Sen-300-145, Sen-300-146, Sen-300-147, Sen-300-148, Sen-300-149, Sen-300-15, Sen-300-150, Sen-300-151, Sen-300-152, Sen-300-153, Sen-300-154, Sen-300-155, Sen-300-156, Sen-300-157, Sen-300-158, Sen-300-159, Sen-300-16, Sen-300-160, Sen-300-161, Sen-300-162, Sen-300-163, Sen-300-164, Sen-300-165, Sen-300-166, Sen-300-167, Sen-300-168, Sen-300-169, Sen-300-17, Sen-300-170, Sen-300-171, Sen-300-172, Sen-300-173, Sen-300-174, Sen-300-175, Sen-300-176, Sen-300-177, Sen-300-178, Sen-300-179, Sen-300-18, Sen-300-180, Sen-300-181, Sen-300-182, Sen-300-183, Sen-300-184, Sen-300-185, Sen-300-186, Sen-300-187, Sen-300-188, Sen-300-189, Sen-300-19, Sen-300-190, Sen-300-191, Sen-300-192, Sen-300-193, Sen-300-194, Sen-300-195, Sen-300-196, Sen-300-197, Sen-300-198, Sen-300-199, Sen-300-2, Sen-300-20, Sen-300-200, Sen-300-201, Sen-300-202, Sen-300-203, Sen-300-204, Sen-300-205, Sen-300-206, Sen-300-207, Sen-300-208, Sen-300-209, Sen-300-21, Sen-300-210, Sen-300-211, Sen-300-212, Sen-300-213, Sen-300-214, Sen-300-215, Sen-300-216, Sen-300-217, Sen-300-218, Sen-300-219, Sen-300-22, Sen-300-220, Sen-300-221, Sen-300-222, Sen-300-223, Sen-300-224, Sen-300-225, Sen-300-226, Sen-300-227, Sen-300-228, Sen-300-229, Sen-300-23, Sen-300-230, Sen-300-231, Sen-300-232, Sen-300-233, Sen-300-234, Sen-300-235, Sen-300-236, Sen-300-237, Sen-300-238, Sen-300-239, Sen-300-24, Sen-300-240, Sen-300-241, Sen-300-242, Sen-300-243, Sen-300-244, Sen-300-245, Sen-300-246, Sen-300-247, Sen-300-248, Sen-300-249, Sen-300-25, Sen-300-250, Sen-300-251, Sen-300-252, Sen-300-253, Sen-300-254, Sen-300-255, Sen-300-256, Sen-300-257, Sen-300-258, Sen-300-259, Sen-300-26, Sen-300-260, Sen-300-261, Sen-300-262, Sen-300-263, Sen-300-264, Sen-300-265, Sen-300-266, Sen-300-267, Sen-300-268, Sen-300-269, Sen-300-27, Sen-300-270, Sen-300-271, Sen-300-272, Sen-300-273, Sen-300-274, Sen-300-275, Sen-300-276, Sen-300-277, Sen-300-278, Sen-300-279, Sen-300-28, Sen-300-280, Sen-300-281, Sen-300-282, Sen-300-283, Sen-300-284, Sen-300-285, Sen-300-286, Sen-300-287, Sen-300-288, Sen-300-289, Sen-300-29, Sen-300-290, Sen-300-291, Sen-300-292, Sen-300-293, Sen-300-294, Sen-300-295, Sen-300-296, Sen-300-297, Sen-300-298, Sen-300-299, Sen-300-3, Sen-300-30, Sen-300-300, Sen-300-301, Sen-300-302, Sen-300-303, Sen-300-304, Sen-300-305, Sen-300-306, Sen-300-307, Sen-300-308, Sen-300-309, Sen-300-31, Sen-300-310, Sen-300-311, Sen-300-312, Sen-300-313, Sen-300-314, Sen-300-

315, Sen-300-316, Sen-300-317, Sen-300-318, Sen-300-319, Sen-300-32, Sen-300-320, Sen-300-321, Sen-300-322, Sen-300-323, Sen-300-324, Sen-300-325, Sen-300-326, Sen-300-327, Sen-300-328, Sen-300-329, Sen-300-33, Sen-300-330, Sen-300-34, Sen-300-35, Sen-300-36, Sen-300-37, Sen-300-38, Sen-300-39, Sen-300-4, Sen-300-40, Sen-300-41, Sen-300-42, Sen-300-43, Sen-300-44, Sen-300-45, Sen-300-46, Sen-300-47, Sen-300-48, Sen-300-49, Sen-300-5, Sen-300-50, Sen-300-51, Sen-300-52, Sen-300-53, Sen-300-54, Sen-300-55, Sen-300-56, Sen-300-57, Sen-300-58, Sen-300-59, Sen-300-6, Sen-300-60, Sen-300-61, Sen-300-62, Sen-300-63, Sen-300-64, Sen-300-65, Sen-300-66, Sen-300-67, Sen-300-68, Sen-300-69, Sen-300-7, Sen-300-70, Sen-300-71, Sen-300-72, Sen-300-73, Sen-300-74, Sen-300-75, Sen-300-76, Sen-300-77, Sen-300-78, Sen-300-79, Sen-300-8, Sen-300-80, Sen-300-81, Sen-300-82, Sen-300-83, Sen-300-84, Sen-300-85, Sen-300-86, Sen-300-87, Sen-300-88, Sen-300-89, Sen-300-9, Sen-300-90, Sen-300-91, Sen-300-92, Sen-300-93, Sen-300-94, Sen-300-95, Sen-300-96, Sen-300-97, Sen-300-98, Sen-300-99, Sen-301-1, Sen-302-1, Sen-303-1, Sen-304-1, Sen-305-1, Sen-306-1, Sen-306-10, Sen-306-11, Sen-306-12, Sen-306-13, Sen-306-14, Sen-306-15, Sen-306-16, Sen-306-17, Sen-306-18, Sen-306-19, Sen-306-2, Sen-306-20, Sen-306-21, Sen-306-22, Sen-306-23, Sen-306-24, Sen-306-25, Sen-306-26, Sen-306-27, Sen-306-28, Sen-306-29, Sen-306-3, Sen-306-30, Sen-306-31, Sen-306-32, Sen-306-33, Sen-306-34, Sen-306-35, Sen-306-36, Sen-306-37, Sen-306-38, Sen-306-39, Sen-306-4, Sen-306-40, Sen-306-41, Sen-306-42, Sen-306-43, Sen-306-44, Sen-306-45, Sen-306-46, Sen-306-47, Sen-306-48, Sen-306-49, Sen-306-5, Sen-306-50, Sen-306-51, Sen-306-52, Sen-306-53, Sen-306-54, Sen-306-55, Sen-306-56, Sen-306-57, Sen-306-58, Sen-306-59, Sen-306-6, Sen-306-60, Sen-306-61, Sen-306-62, Sen-306-63, Sen-306-7, Sen-306-8, Sen-306-9, Sen-307-1, Sen-308-1, Sen-308-2, Sen-308-3, Sen-308-4, Sen-309-1, Sen-31-1, Sen-310-1, Sen-313-1, Sen-314-1, Sen-315-1, Sen-316-1, Sen-317-1, Sen-318-1, Sen-319-1, Sen-32-1, Sen-320-1, Sen-321-1, Sen-322-1, Sen-323-1, Sen-324-1, Sen-325-1, Sen-326-1, Sen-327-1, Sen-328-1, Sen-329-1, Sen-33-1, Sen-330-1, Sen-331-1, Sen-332-1, Sen-333-1, Sen-334-1, Sen-335-1, Sen-336-1, Sen-337-1, Sen-338-1, Sen-339-1, Sen-34-1, Sen-340-1, Sen-341-1, Sen-342-1, Sen-343-1, Sen-344-1, Sen-345-1, Sen-346-1, Sen-347-1, Sen-348-1, Sen-349-1, Sen-35-1, Sen-350-1, Sen-351-1, Sen-352-1, Sen-353-1, Sen-354-1, Sen-355-1, Sen-356-1, Sen-357-1, Sen-358-1, Sen-359-1, Sen-36-1, Sen-360-1, Sen-361-1, Sen-362-1, Sen-363-1, Sen-364-1, Sen-365-1, Sen-366-1, Sen-367-1, Sen-368-1, Sen-369-1, Sen-37-1, Sen-370-1, Sen-371-1, Sen-372-1, Sen-373-1, Sen-374-1, Sen-375-1, Sen-376-1, Sen-377-1, Sen-378-1, Sen-379-1, Sen-38-1, Sen-380-1, Sen-381-1, Sen-382-1, Sen-383-1, Sen-384-1, Sen-385-1, Sen-386-1, Sen-387-1, Sen-388-1, Sen-389-1, Sen-39-1, Sen-390-1, Sen-391-1, Sen-392-1, Sen-393-1, Sen-394-1, Sen-395-1, Sen-396-1, Sen-397-1, Sen-398-1, Sen-399-1, Sen-4-1, Sen-40-1, Sen-400-1, Sen-401-1, Sen-402-1, Sen-403-1, Sen-404-1, Sen-405-1, Sen-406-1, Sen-407-1, Sen-408-1, Sen-409-1, Sen-41-1, Sen-410-1, Sen-411-1, Sen-412-1, Sen-413-1, Sen-414-1, Sen-415-1, Sen-416-1, Sen-417-1, Sen-418-1, Sen-419-1, Sen-42-1, Sen-420-1, Sen-421-1, Sen-422-1, Sen-423-1, Sen-424-1, Sen-425-1, Sen-426-1, Sen-427-1, Sen-428-1, Sen-429-1, Sen-43-1, Sen-430-1, Sen-431-1, Sen-432-1, Sen-433-1, Sen-434-1, Sen-435-1, Sen-436-1, Sen-437-1, Sen-438-1, Sen-439-1, Sen-439-10, Sen-439-11, Sen-439-2, Sen-439-3, Sen-439-4, Sen-439-5, Sen-439-6, Sen-439-7, Sen-439-8, Sen-439-9, Sen-44-1, Sen-440-1, Sen-441-1, Sen-442-1, Sen-443-1, Sen-444-1, Sen-445-1, Sen-446-1, Sen-447-1, Sen-448-1, Sen-449-1, Sen-45-1, Sen-450-1, Sen-451-1, Sen-452-1, Sen-453-1, Sen-454-1, Sen-455-1, Sen-456-1, Sen-457-1, Sen-458-1, Sen-459-1, Sen-46-1, Sen-460-1, Sen-461-1, Sen-462-1, Sen-463-1, Sen-464-1, Sen-465-1, Sen-466-1, Sen-467-1, Sen-468-1, Sen-469-1, Sen-47-1, Sen-470-1, Sen-471-1, Sen-472-1, Sen-473-1, Sen-474-1, Sen-475-1, Sen-476-1, Sen-477-1, Sen-478-1, Sen-479-1, Sen-48-1, Sen-480-1, Sen-481-1, Sen-482-1, Sen-483-1, Sen-484-1, Sen-485-1, Sen-486-1, Sen-487-1, Sen-488-1, Sen-489-1, Sen-49-1, Sen-490-1, Sen-491-1, Sen-492-1, Sen-493-1, Sen-494-1, Sen-495-1, Sen-496-1, Sen-497-1, Sen-498-1, Sen-499-1, Sen-5-1, Sen-5-2, Sen-5-3, Sen-5-4, Sen-5-5, Sen-5-6, Sen-50-1, Sen-500-1, Sen-501-1, Sen-502-1, Sen-503-1, Sen-504-1, Sen-505-1, Sen-506-1, Sen-507-1, Sen-508-1, Sen-509-1, Sen-51-1, Sen-510-1, Sen-511-1, Sen-512-1, Sen-513-1, Sen-514-1, Sen-515-1, Sen-516-1, Sen-517-1, Sen-518-1, Sen-519-1, Sen-52-1, Sen-520-1, Sen-521-1, Sen-522-1, Sen-523-1, Sen-524-1, Sen-525-1, Sen-526-1, Sen-527-1, Sen-528-1, Sen-529-1, Sen-53-1, Sen-530-1, Sen-531-1, Sen-532-1, Sen-533-1, Sen-534-1, Sen-535-1, Sen-536-1, Sen-537-1, Sen-538-1, Sen-539-1, Sen-54-1, Sen-540-1, Sen-541-1, Sen-542-1, Sen-543-1, Sen-544-1, Sen-545-1, Sen-546-1, Sen-547-1, Sen-548-1, Sen-549-1, Sen-55-1, Sen-550-1, Sen-551-1, Sen-552-1, Sen-553-1, Sen-554-1, Sen-555-1, Sen-556-1, Sen-557-1, Sen-558-1, Sen-559-1, Sen-56-1, Sen-560-1, Sen-561-1, Sen-562-1, Sen-563-1, Sen-564-1, Sen-565-1, Sen-566-1, Sen-567-1, Sen-568-1, Sen-569-1, Sen-57-1, Sen-570-1, Sen-571-1, Sen-572-1, Sen-573-1, Sen-574-1, Sen-575-1, Sen-576-1, Sen-577-1, Sen-578-1, Sen-579-1, Sen-58-1, Sen-580-1, Sen-581-1, Sen-582-1, Sen-583-1, Sen-584-1, Sen-585-1, Sen-586-1, Sen-587-1, Sen-588-1, Sen-589-1, Sen-59-1, Sen-590-1, Sen-591-1, Sen-592-1, Sen-593-1, Sen-594-1, Sen-595-1, Sen-596-1, Sen-597-1, Sen-598-1, Sen-599-1, Sen-6-1, Sen-6-2, Sen-6-3, Sen-6-4, Sen-6-5, Sen-60-1, Sen-600-1, Sen-601-1, Sen-602-1, Sen-603-1, Sen-604-1, Sen-605-1, Sen-606-1, Sen-607-1, Sen-608-1, Sen-609-1, Sen-61-1, Sen-610-1, Sen-611-1, Sen-612-1, Sen-613-1, Sen-614-1, Sen-615-1, Sen-616-1, Sen-617-1, Sen-618-1, Sen-619-1, Sen-62-1, Sen-620-1, Sen-621-1, Sen-622-1, Sen-623-1, Sen-624-1, Sen-625-1, Sen-626-1, Sen-627-1, Sen-628-1, Sen-629-1, Sen-63-1, Sen-630-1, Sen-631-1, Sen-632-1, Sen-633-1, Sen-634-1, Sen-635-1, Sen-636-1, Sen-637-1, Sen-638-1, Sen-639-1, Sen-64-1, Sen-640-1, Sen-641-1, Sen-642-1, Sen-643-1, Sen-644-1, Sen-645-1, Sen-646-1, Sen-647-1, Sen-648-1, Sen-649-1, Sen-649-2, Sen-65-1, Sen-650-1, Sen-651-1, Sen-652-1, Sen-653-1, Sen-654-1, Sen-655-1, Sen-656-1, Sen-657-1, Sen-658-1, Sen-659-1, Sen-66-1, Sen-660-1, Sen-661-1, Sen-662-1, Sen-663-1, Sen-664-1, Sen-665-1, Sen-666-1, Sen-667-1, Sen-668-1, Sen-669-1, Sen-67-1, Sen-670-1, Sen-671-1, Sen-672-1, Sen-673-1, Sen-674-1, Sen-675-1, Sen-676-1, Sen-677-1, Sen-678-1, Sen-679-1, Sen-68-1, Sen-680-1, Sen-681-1, Sen-682-1, Sen-683-1, Sen-684-1, Sen-685-1, Sen-686-1, Sen-687-1, Sen-688-1, Sen-689-1, Sen-69-1, Sen-690-1, Sen-691-1, Sen-692-1, Sen-693-1, Sen-694-1, Sen-695-1, Sen-696-1, Sen-697-1, Sen-698-1, Sen-699-1, Sen-7-1, Sen-7-2, Sen-7-3, Sen-70-1, Sen-700-1, Sen-701-1, Sen-702-1, Sen-703-1, Sen-704-1, Sen-705-1, Sen-706-1, Sen-707-1, Sen-708-1, Sen-709-1, Sen-71-1, Sen-710-1, Sen-711-1, Sen-712-1, Sen-713-1, Sen-714-1, Sen-715-1, Sen-716-1, Sen-717-1, Sen-718-1, Sen-719-1, Sen-72-1, Sen-720-1, Sen-721-1, Sen-722-1, Sen-723-1, Sen-724-1, Sen-725-1,

Sen-726-1, Sen-727-1, Sen-728-1, Sen-729-1, Sen-73-1, Sen-730-1, Sen-731-1, Sen-732-1, Sen-733-1, Sen-734-1, Sen-735-1, Sen-736-1, Sen-737-1, Sen-738-1, Sen-739-1, Sen-74-1, Sen-740-1, Sen-741-1, Sen-742-1, Sen-743-1, Sen-744-1, Sen-745-1, Sen-746-1, Sen-747-1, Sen-748-1, Sen-749-1, Sen-75-1, Sen-750-1, Sen-751-1, Sen-752-1, Sen-753-1, Sen-754-1, Sen-755-1, Sen-756-1, Sen-757-1, Sen-758-1, Sen-759-1, Sen-76-1, Sen-760-1, Sen-761-1, Sen-762-1, Sen-763-1, Sen-764-1, Sen-765-1, Sen-766-1, Sen-767-1, Sen-768-1, Sen-769-1, Sen-77-1, Sen-770-1, Sen-771-1, Sen-772-1, Sen-773-1, Sen-774-1, Sen-775-1, Sen-776-1, Sen-777-1, Sen-778-1, Sen-779-1, Sen-78-1, Sen-780-1, Sen-781-1, Sen-782-1, Sen-783-1, Sen-784-1, Sen-785-1, Sen-786-1, Sen-787-1, Sen-788-1, Sen-789-1, Sen-79-1, Sen-790-1, Sen-791-1, Sen-792-1, Sen-793-1, Sen-794-1, Sen-795-1, Sen-796-1, Sen-797-1, Sen-798-1, Sen-799-1, Sen-8-1, Sen-80-1, Sen-800-1, Sen-801-1, Sen-802-1, Sen-803-1, Sen-804-1, Sen-805-1, Sen-806-1, Sen-807-1, Sen-808-1, Sen-809-1, Sen-81-1, Sen-810-1, Sen-811-1, Sen-812-1, Sen-813-1, Sen-814-1, Sen-815-1, Sen-816-1, Sen-817-1, Sen-818-1, Sen-819-1, Sen-82-1, Sen-820-1, Sen-821-1, Sen-822-1, Sen-823-1, Sen-824-1, Sen-825-1, Sen-826-1, Sen-827-1, Sen-828-1, Sen-829-1, Sen-83-1, Sen-830-1, Sen-831-1, Sen-832-1, Sen-833-1, Sen-834-1, Sen-835-1, Sen-836-1, Sen-837-1, Sen-838-1, Sen-839-1, Sen-84-1, Sen-840-1, Sen-841-1, Sen-842-1, Sen-843-1, Sen-844-1, Sen-845-1, Sen-846-1, Sen-847-1, Sen-848-1, Sen-849-1, Sen-85-1, Sen-850-1, Sen-851-1, Sen-852-1, Sen-853-1, Sen-854-1, Sen-855-1, Sen-856-1, Sen-857-1, Sen-858-1, Sen-859-1, Sen-86-1, Sen-860-1, Sen-861-1, Sen-862-1, Sen-863-1, Sen-864-1, Sen-865-1, Sen-866-1, Sen-867-1, Sen-868-1, Sen-869-1, Sen-87-1, Sen-870-1, Sen-871-1, Sen-872-1, Sen-873-1, Sen-874-1, Sen-875-1, Sen-876-1, Sen-877-1, Sen-878-1, Sen-879-1, Sen-88-1, Sen-880-1, Sen-881-1, Sen-882-1, Sen-883-1, Sen-884-1, Sen-885-1, Sen-886-1, Sen-887-1, Sen-888-1, Sen-889-1, Sen-890-1, Sen-891-1, Sen-892-1, Sen-893-1, Sen-894-1, Sen-895-1, Sen-896-1, Sen-897-1, Sen-898-1, Sen-899-1, Sen-9-1, Sen-90-1, Sen-900-1, Sen-901-1, Sen-902-1, Sen-903-1, Sen-904-1, Sen-905-1, Sen-906-1, Sen-907-1, Sen-908-1, Sen-909-1, Sen-91-1, Sen-910-1, Sen-911-1, Sen-912-1, Sen-913-1, Sen-914-1, Sen-915-1, Sen-916-1, Sen-917-1, Sen-918-1, Sen-919-1, Sen-92-1, Sen-920-1, Sen-921-1, Sen-922-1, Sen-923-1, Sen-924-1, Sen-925-1, Sen-926-1, Sen-927-1, Sen-928-1, Sen-929-1, Sen-93-1, Sen-930-1, Sen-931-1, Sen-932-1, Sen-933-1, Sen-934-1, Sen-935-1, Sen-936-1, Sen-937-1, Sen-938-1, Sen-939-1, Sen-94-1, Sen-940-1, Sen-941-1, Sen-942-1, Sen-943-1, Sen-944-1, Sen-945-1, Sen-946-1, Sen-947-1, Sen-948-1, Sen-949-1, Sen-95-1, Sen-950-1, Sen-951-1, Sen-952-1, Sen-953-1, Sen-954-1, Sen-955-1, Sen-956-1, Sen-957-1, Sen-958-1, Sen-959-1, Sen-96-1, Sen-960-1, Sen-961-1, Sen-962-1, Sen-963-1, Sen-964-1, Sen-965-1, Sen-966-1, Sen-967-1, Sen-968-1, Sen-969-1, Sen-97-1, Sen-970-1, Sen-971-1, Sen-972-1, Sen-973-1, Sen-974-1, Sen-975-1, Sen-976-1, Sen-977-1, Sen-978-1, Sen-979-1, Sen-98-1, Sen-980-1, Sen-981-1, Sen-982-1, Sen-983-1, Sen-984-1, Sen-985-1, Sen-986-1, Sen-987-1, Sen-988-1, Sen-989-1, Sen-99-1, Sen-990-1, Sen-991-1, Sen-992-1, Sen-993-1, Sen-994-1, Sen-995-1, Sen-996-1, Sen-997-1, Sen-998-1, Sen-999-1, Sfl-1-1, Sfl-1-2, Sfl-10-1, Sfl-11-1, Sfl-12-1, Sfl-13-1, Sfl-14-1, Sfl-15-1, Sfl-16-1, Sfl-17-1, Sfl-18-1, Sfl-19-1, Sfl-2-1, Sfl-20-1, Sfl-21-1, Sfl-22-1, Sfl-23-1, Sfl-24-1, Sfl-25-1, Sfl-25-10, Sfl-25-11, Sfl-25-12, Sfl-25-13, Sfl-25-14, Sfl-25-2, Sfl-25-3, Sfl-25-4, Sfl-25-5, Sfl-25-6, Sfl-25-7, Sfl-25-8, Sfl-25-9, Sfl-26-1, Sfl-27-1, Sfl-28-1, Sfl-29-1, Sfl-3-1, Sfl-30-1, Sfl-31-1, Sfl-32-1, Sfl-33-1, Sfl-34-1, Sfl-35-1, Sfl-36-1, Sfl-37-1, Sfl-38-1, Sfl-39-1, Sfl-4-1, Sfl-40-1, Sfl-41-1, Sfl-42-1, Sfl-43-1, Sfl-44-1, Sfl-45-1, Sfl-46-1, Sfl-47-1, Sfl-48-1, Sfl-49-1, Sfl-5-1, Sfl-50-1, Sfl-51-1, Sfl-51-2, Sfl-52-1, Sfl-53-1, Sfl-54-1, Sfl-55-1, Sfl-56-1, Sfl-57-1, Sfl-58-1, Sfl-59-1, Sfl-6-1, Sfl-60-1, Sfl-61-1, Sfl-62-1, Sfl-63-1, Sfl-64-1, Sfl-64-2, Sfl-65-1, Sfl-66-1, Sfl-67-1, Sfl-68-1, Sfl-69-1, Sfl-7-1, Sfl-71-1, Sfl-72-1, Sfl-73-1, Sfl-74-1, Sfl-75-1, Sfl-76-1, Sfl-77-1, Sfl-78-1, Sfl-79-1, Sfl-8-1, Sfl-80-1, Sfl-81-1, Sfl-82-1, Sfl-83-1, Sfl-84-1, Sfl-85-1, Sfl-9-1, Sfo-2-1, Sfo-3-1, Sfo-4-1, Sfo-5-1, Sha-2-1, Sli-1-1, Sli-2-1, Sli-3-1, Sma-11-1, Sma-12-1, Sma-13-1, Sma-14-1, Sma-15-1, Sma-17-1, Sma-18-1, Sma-19-1, Sma-20-1, Sma-20-2, Sma-21-1, Sma-22-1, Sma-23-1, Sma-25-1, Sma-26-1, Sma-27-1, Sma-28-1, Sma-3-2, Sma-3-3, Sma-3-4, Sma-4-1, Sma-5-1, Sma-6-1, Sma-7-1, Sma-8-1, Sma-9-1, Sne-1-1, Spl-2-1, Spl-3-1, Spl-4-1, Spl-6-1, Spu-1-1, Sso-1-1, Sso-1-10, Sso-1-100, Sso-1-101, Sso-1-102, Sso-1-103, Sso-1-104, Sso-1-105, Sso-1-106, Sso-1-107, Sso-1-108, Sso-1-109, Sso-1-11, Sso-1-110, Sso-1-111, Sso-1-112, Sso-1-113, Sso-1-114, Sso-1-115, Sso-1-116, Sso-1-117, Sso-1-118, Sso-1-119, Sso-1-12, Sso-1-120, Sso-1-121, Sso-1-122, Sso-1-123, Sso-1-124, Sso-1-125, Sso-1-126, Sso-1-127, Sso-1-128, Sso-1-129, Sso-1-13, Sso-1-130, Sso-1-131, Sso-1-132, Sso-1-133, Sso-1-134, Sso-1-135, Sso-1-136, Sso-1-137, Sso-1-138, Sso-1-139, Sso-1-14, Sso-1-140, Sso-1-141, Sso-1-142, Sso-1-143, Sso-1-144, Sso-1-145, Sso-1-146, Sso-1-147, Sso-1-148, Sso-1-149, Sso-1-15, Sso-1-150, Sso-1-151, Sso-1-152, Sso-1-153, Sso-1-154, Sso-1-155, Sso-1-156, Sso-1-157, Sso-1-158, Sso-1-159, Sso-1-16, Sso-1-160, Sso-1-161, Sso-1-162, Sso-1-163, Sso-1-164, Sso-1-165, Sso-1-166, Sso-1-167, Sso-1-168, Sso-1-169, Sso-1-17, Sso-1-170, Sso-1-171, Sso-1-172, Sso-1-173, Sso-1-174, Sso-1-175, Sso-1-176, Sso-1-177, Sso-1-178, Sso-1-179, Sso-1-18, Sso-1-180, Sso-1-181, Sso-1-182, Sso-1-183, Sso-1-184, Sso-1-185, Sso-1-186, Sso-1-187, Sso-1-188, Sso-1-189, Sso-1-19, Sso-1-190, Sso-1-191, Sso-1-192, Sso-1-193, Sso-1-194, Sso-1-195, Sso-1-196, Sso-1-197, Sso-1-198, Sso-1-199, Sso-1-20, Sso-1-200, Sso-1-201, Sso-1-202, Sso-1-203, Sso-1-204, Sso-1-205, Sso-1-206, Sso-1-207, Sso-1-208, Sso-1-209, Sso-1-21, Sso-1-210, Sso-1-211, Sso-1-212, Sso-1-213, Sso-1-214, Sso-1-215, Sso-1-216, Sso-1-217, Sso-1-218, Sso-1-219, Sso-1-22, Sso-1-220, Sso-1-221, Sso-1-222, Sso-1-223, Sso-1-224, Sso-1-225, Sso-1-226, Sso-1-227, Sso-1-228, Sso-1-229, Sso-1-23, Sso-1-230, Sso-1-231, Sso-1-232, Sso-1-233, Sso-1-234, Sso-1-235, Sso-1-236, Sso-1-237, Sso-1-238, Sso-1-239, Sso-1-24, Sso-1-240, Sso-1-241, Sso-1-242, Sso-1-243, Sso-1-244, Sso-1-245, Sso-1-246, Sso-1-247, Sso-1-248, Sso-1-249, Sso-1-25, Sso-1-250, Sso-1-251, Sso-1-252, Sso-1-253, Sso-1-254, Sso-1-255, Sso-1-256, Sso-1-257, Sso-1-258, Sso-1-259, Sso-1-26, Sso-1-260, Sso-1-261, Sso-1-262, Sso-1-263, Sso-1-264, Sso-1-265, Sso-1-266, Sso-1-267, Sso-1-268, Sso-1-269, Sso-1-27, Sso-1-270, Sso-1-271, Sso-1-272, Sso-1-273, Sso-1-274, Sso-1-275, Sso-1-276, Sso-1-277, Sso-1-278, Sso-1-279, Sso-1-28, Sso-1-280, Sso-1-281, Sso-1-282, Sso-1-283, Sso-1-284, Sso-1-285, Sso-1-286, Sso-1-287, Sso-1-288, Sso-1-289, Sso-1-29, Sso-1-290, Sso-1-291, Sso-1-292, Sso-1-293, Sso-1-294, Sso-1-295, Sso-1-296, Sso-1-297, Sso-1-298, Sso-1-299, Sso-1-30, Sso-1-300, Sso-1-301, Sso-1-302, Sso-1-303, Sso-1-304, Sso-1-305, Sso-1-306, Sso-1-307, Sso-1-308, Sso-1-309, Sso-1-31, Sso-1-310, Sso-1-311, Sso-1-312, Sso-1-313, Sso-1-314, Sso-1-315, Sso-1-316, Sso-1-317, Sso-1-318, Sso-1-319, Sso-1-32, Sso-1-320, Sso-1-321, Sso-1-322, Sso-1-323, Sso-1-324, Sso-1-325, Sso-1-326, Sso-

1-327, Sso-1-328, Sso-1-329, Sso-1-33, Sso-1-330, Sso-1-331, Sso-1-332, Sso-1-333, Sso-1-334, Sso-1-335, Sso-1-336, Sso-1-337, Sso-1-338, Sso-1-339, Sso-1-34, Sso-1-340, Sso-1-341, Sso-1-342, Sso-1-343, Sso-1-344, Sso-1-345, Sso-1-346, Sso-1-347, Sso-1-348, Sso-1-349, Sso-1-35, Sso-1-350, Sso-1-351, Sso-1-352, Sso-1-353, Sso-1-354, Sso-1-355, Sso-1-356, Sso-1-357, Sso-1-358, Sso-1-359, Sso-1-36, Sso-1-360, Sso-1-361, Sso-1-362, Sso-1-363, Sso-1-364, Sso-1-365, Sso-1-366, Sso-1-367, Sso-1-368, Sso-1-369, Sso-1-37, Sso-1-370, Sso-1-371, Sso-1-372, Sso-1-373, Sso-1-374, Sso-1-375, Sso-1-376, Sso-1-377, Sso-1-378, Sso-1-379, Sso-1-38, Sso-1-380, Sso-1-381, Sso-1-382, Sso-1-383, Sso-1-384, Sso-1-385, Sso-1-386, Sso-1-387, Sso-1-388, Sso-1-389, Sso-1-39, Sso-1-390, Sso-1-391, Sso-1-392, Sso-1-393, Sso-1-394, Sso-1-395, Sso-1-396, Sso-1-397, Sso-1-398, Sso-1-399, Sso-1-4, Sso-1-40, Sso-1-400, Sso-1-401, Sso-1-402, Sso-1-403, Sso-1-404, Sso-1-405, Sso-1-406, Sso-1-407, Sso-1-408, Sso-1-409, Sso-1-41, Sso-1-410, Sso-1-411, Sso-1-412, Sso-1-413, Sso-1-414, Sso-1-415, Sso-1-416, Sso-1-417, Sso-1-418, Sso-1-419, Sso-1-42, Sso-1-420, Sso-1-421, Sso-1-422, Sso-1-423, Sso-1-424, Sso-1-425, Sso-1-426, Sso-1-427, Sso-1-428, Sso-1-429, Sso-1-43, Sso-1-430, Sso-1-431, Sso-1-432, Sso-1-433, Sso-1-434, Sso-1-435, Sso-1-436, Sso-1-437, Sso-1-438, Sso-1-439, Sso-1-44, Sso-1-440, Sso-1-441, Sso-1-442, Sso-1-443, Sso-1-444, Sso-1-445, Sso-1-446, Sso-1-447, Sso-1-448, Sso-1-449, Sso-1-45, Sso-1-450, Sso-1-451, Sso-1-452, Sso-1-453, Sso-1-454, Sso-1-455, Sso-1-456, Sso-1-457, Sso-1-458, Sso-1-459, Sso-1-46, Sso-1-460, Sso-1-461, Sso-1-462, Sso-1-463, Sso-1-464, Sso-1-465, Sso-1-466, Sso-1-467, Sso-1-468, Sso-1-469, Sso-1-47, Sso-1-470, Sso-1-471, Sso-1-472, Sso-1-473, Sso-1-474, Sso-1-475, Sso-1-476, Sso-1-477, Sso-1-478, Sso-1-479, Sso-1-48, Sso-1-480, Sso-1-481, Sso-1-482, Sso-1-483, Sso-1-484, Sso-1-485, Sso-1-486, Sso-1-487, Sso-1-488, Sso-1-489, Sso-1-49, Sso-1-490, Sso-1-491, Sso-1-492, Sso-1-493, Sso-1-494, Sso-1-495, Sso-1-496, Sso-1-497, Sso-1-498, Sso-1-499, Sso-1-50, Sso-1-500, Sso-1-501, Sso-1-502, Sso-1-503, Sso-1-504, Sso-1-505, Sso-1-506, Sso-1-507, Sso-1-508, Sso-1-509, Sso-1-51, Sso-1-510, Sso-1-511, Sso-1-512, Sso-1-513, Sso-1-514, Sso-1-515, Sso-1-516, Sso-1-517, Sso-1-518, Sso-1-519, Sso-1-52, Sso-1-520, Sso-1-521, Sso-1-522, Sso-1-523, Sso-1-524, Sso-1-525, Sso-1-526, Sso-1-527, Sso-1-528, Sso-1-529, Sso-1-53, Sso-1-530, Sso-1-531, Sso-1-532, Sso-1-533, Sso-1-534, Sso-1-535, Sso-1-536, Sso-1-537, Sso-1-538, Sso-1-539, Sso-1-54, Sso-1-540, Sso-1-541, Sso-1-542, Sso-1-543, Sso-1-544, Sso-1-545, Sso-1-547, Sso-1-55, Sso-1-56, Sso-1-57, Sso-1-58, Sso-1-59, Sso-1-6, Sso-1-60, Sso-1-61, Sso-1-62, Sso-1-63, Sso-1-64, Sso-1-65, Sso-1-66, Sso-1-67, Sso-1-68, Sso-1-69, Sso-1-7, Sso-1-70, Sso-1-71, Sso-1-72, Sso-1-73, Sso-1-74, Sso-1-75, Sso-1-76, Sso-1-77, Sso-1-78, Sso-1-79, Sso-1-8, Sso-1-80, Sso-1-81, Sso-1-82, Sso-1-83, Sso-1-84, Sso-1-85, Sso-1-86, Sso-1-87, Sso-1-88, Sso-1-89, Sso-1-9, Sso-1-90, Sso-1-91, Sso-1-92, Sso-1-93, Sso-1-94, Sso-1-95, Sso-1-96, Sso-1-97, Sso-1-98, Sso-1-99, Sso-10-1, Sso-11-1, Sso-12-1, Sso-13-1, Sso-14-1, Sso-15-1, Sso-2-1, Sso-3-1, Sso-4-1, Sso-5-1, Sso-6-1, Sso-7-1, Sso-8-1, Sso-9-1, Ssp-1-1, Ssp-12-1, Ssp-14-1, Ssp-16-1, Ssp-18-1, Ssp-19-1, Ssp-2-1, Ssp-20-1, Ssp-22-1, Ssp-25-1, Ssp-26-1, Ssp-8-1, Ssp-9-1, Stu-1-1, Sur-1-1, Sxi-1-1, Tmo-1-1, Tod-1-1, Tod-1-2, Tpt-1-2, VPa-1-1, VPa-2-1, Val-1-1, Val-2-2, Val-2-3, Val-2-4, Val-2-5, Val-2-6, Val-2-7, Val-2-8, Val-3-1, Val-4-1, Val-5-1, Val-6-1, Van-1-1, Van-2-1, Van-3-1, Van-4-1, Vaz-1-2, Vbr-3-1, Vca-1-1, Vca-1-10, Vca-1-2, Vca-1-3, Vca-1-4, Vca-1-5, Vca-1-6, Vca-1-7, Vca-1-8, Vca-1-9, Vca-2-1, Vca-4-1, Vca-4-2, Vca-5-1, Vca-6-1, Vch-1-1, Vch-10-1, Vch-100-1, Vch-101-1, Vch-102-1, Vch-103-1, Vch-104-1, Vch-105-1, Vch-106-1, Vch-107-1, Vch-108-1, Vch-109-1, Vch-11-1, Vch-110-1, Vch-111-1, Vch-112-1, Vch-113-1, Vch-114-1, Vch-115-1, Vch-116-1, Vch-117-1, Vch-118-1, Vch-119-1, Vch-12-1, Vch-120-1, Vch-121-1, Vch-122-1, Vch-123-1, Vch-123-10, Vch-123-11, Vch-123-12, Vch-123-13, Vch-123-14, Vch-123-15, Vch-123-16, Vch-123-17, Vch-123-18, Vch-123-19, Vch-123-2, Vch-123-20, Vch-123-21, Vch-123-22, Vch-123-23, Vch-123-24, Vch-123-25, Vch-123-26, Vch-123-27, Vch-123-28, Vch-123-29, Vch-123-3, Vch-123-30, Vch-123-31, Vch-123-32, Vch-123-33, Vch-123-34, Vch-123-35, Vch-123-36, Vch-123-37, Vch-123-38, Vch-123-39, Vch-123-4, Vch-123-40, Vch-123-41, Vch-123-5, Vch-123-6, Vch-123-7, Vch-123-8, Vch-123-9, Vch-124-1, Vch-125-1, Vch-126-1, Vch-127-1, Vch-128-1, Vch-129-1, Vch-13-1, Vch-130-1, Vch-131-1, Vch-132-1, Vch-133-1, Vch-134-1, Vch-135-1, Vch-136-1, Vch-137-1, Vch-138-1, Vch-139-1, Vch-14-1, Vch-140-1, Vch-141-1, Vch-142-1, Vch-143-1, Vch-144-1, Vch-145-1, Vch-146-1, Vch-147-1, Vch-148-1, Vch-149-1, Vch-15-1, Vch-150-1, Vch-151-1, Vch-152-1, Vch-153-1, Vch-154-1, Vch-155-1, Vch-156-1, Vch-157-1, Vch-158-1, Vch-159-1, Vch-16-1, Vch-160-1, Vch-161-1, Vch-162-1, Vch-163-1, Vch-164-1, Vch-165-1, Vch-166-1, Vch-167-1, Vch-168-1, Vch-169-1, Vch-17-1, Vch-170-1, Vch-171-1, Vch-172-1, Vch-173-1, Vch-174-1, Vch-175-1, Vch-176-1, Vch-177-1, Vch-178-1, Vch-179-1, Vch-179-2, Vch-18-1, Vch-180-1, Vch-181-1, Vch-182-1, Vch-183-1, Vch-184-1, Vch-185-1, Vch-186-1, Vch-187-1, Vch-188-1, Vch-189-1, Vch-19-1, Vch-190-1, Vch-191-1, Vch-192-1, Vch-193-1, Vch-194-1, Vch-195-1, Vch-196-1, Vch-197-1, Vch-2-1, Vch-2-2, Vch-2-3, Vch-2-4, Vch-20-1, Vch-21-1, Vch-22-1, Vch-23-1, Vch-24-1, Vch-25-1, Vch-26-1, Vch-27-1, Vch-28-1, Vch-29-1, Vch-30-1, Vch-31-1, Vch-32-1, Vch-33-1, Vch-34-1, Vch-35-1, Vch-36-1, Vch-37-1, Vch-38-1, Vch-39-1, Vch-4-1, Vch-40-1, Vch-41-1, Vch-42-1, Vch-43-1, Vch-44-1, Vch-45-1, Vch-46-1, Vch-47-1, Vch-48-1, Vch-49-1, Vch-5-1, Vch-50-1, Vch-51-1, Vch-52-1, Vch-53-1, Vch-54-1, Vch-55-1, Vch-56-1, Vch-57-1, Vch-58-1, Vch-59-1, Vch-6-1, Vch-60-1, Vch-61-1, Vch-62-1, Vch-63-1, Vch-64-1, Vch-65-1, Vch-66-1, Vch-67-1, Vch-68-1, Vch-69-1, Vch-70-1, Vch-71-1, Vch-72-1, Vch-73-1, Vch-74-1, Vch-75-1, Vch-76-1, Vch-77-1, Vch-78-1, Vch-79-1, Vch-8-1, Vch-80-1, Vch-81-1, Vch-82-1, Vch-83-1, Vch-84-1, Vch-85-1, Vch-86-1, Vch-87-1, Vch-88-1, Vch-89-1, Vch-9-1, Vch-90-1, Vch-91-1, Vch-92-1, Vch-93-1, Vch-94-1, Vch-95-1, Vch-96-1, Vch-97-1, Vch-98-1, Vch-99-1, Vco-1-1, Vco-1-2, Vcr-1-2, Vcr-1-3, Vcr-1-4, Vcr-1-5, Vcr-1-6, Vcr-2-1, Vcr-3-1, Vcr-4-1, Vcr-5-1, Vcr-6-1, Vcy-1-1, Vcy-10-1, Vcy-11-1, Vcy-12-1, Vcy-14-1, Vcy-16-1, Vcy-17-1, Vcy-18-1, Vcy-19-1, Vcy-20-1, Vcy-3-1, Vcy-4-1, Vcy-5-1, Vcy-6-1, Vcy-7-1, Vcy-8-1, Vcy-9-1, Vdi-1-1, Vfi-1-1, Vfi-3-1, Vfl-1-1, Vfl-2-1, Vfl-3-1, Vfl-5-1, Vfo-1-1, Vfu-1-1, Vfu-3-1, Vge-2-1, Vge-4-1, Vha-10-1, Vha-11-1, Vha-12-1, Vha-2-1, Vha-3-1, Vha-4-1, Vha-5-1, Vha-6-1, Vha-7-1, Vha-8-1, Vha-9-2, Vja-1-2, Vja-3-1, Vle-1-1, Vmi-1-1, Vmi-3-1, Vmi-4-1, Vmi-6-1, Vmi-7-1, Vna-1-2, Vna-2-2, Vna-2-4, Vni-1-1, Vni-10-1, Vni-12-1, Vni-13-1, Vni-14-1, Vni-2-1, Vni-3-1, Vni-4-1, Vni-5-1, Vni-6-1, Vni-7-1, Vni-8-1, Vor-2-1, Vor-3-1, Vor-4-1, Vor-5-1, Vor-6-1, Vor-8-1, Vow-1-1, Vow-2-1, Vow-2-2, Vow-2-3, Vow-2-4, Vow-4-1, Vow-6-1, Vpa-10-1, Vpa-100-1, Vpa-101-1, Vpa-102-1, Vpa-103-1, Vpa-104-1, Vpa-105-1, Vpa-106-1, Vpa-107-1, Vpa-108-1, Vpa-

109-1, Vpa-11-1, Vpa-110-1, Vpa-111-1, Vpa-112-1, Vpa-113-1, Vpa-114-1, Vpa-115-1, Vpa-116-1, Vpa-117-1, Vpa-118-1, Vpa-119-1, Vpa-12-1, Vpa-120-1, Vpa-121-1, Vpa-122-1, Vpa-123-1, Vpa-124-1, Vpa-125-1, Vpa-126-1, Vpa-127-1, Vpa-128-1, Vpa-129-1, Vpa-13-1, Vpa-130-1, Vpa-131-1, Vpa-132-1, Vpa-133-1, Vpa-134-1, Vpa-135-1, Vpa-136-1, Vpa-137-1, Vpa-138-1, Vpa-139-1, Vpa-14-1, Vpa-140-1, Vpa-141-1, Vpa-142-1, Vpa-143-1, Vpa-144-1, Vpa-145-1, Vpa-146-1, Vpa-147-1, Vpa-148-1, Vpa-149-1, Vpa-15-1, Vpa-150-1, Vpa-151-1, Vpa-152-1, Vpa-153-1, Vpa-154-1, Vpa-155-1, Vpa-156-1, Vpa-158-1, Vpa-159-1, Vpa-16-1, Vpa-160-1, Vpa-161-1, Vpa-162-1, Vpa-163-1, Vpa-164-1, Vpa-165-1, Vpa-166-1, Vpa-167-1, Vpa-168-1, Vpa-169-1, Vpa-17-1, Vpa-170-1, Vpa-171-1, Vpa-172-1, Vpa-173-1, Vpa-174-1, Vpa-175-1, Vpa-176-1, Vpa-177-1, Vpa-178-1, Vpa-179-1, Vpa-18-1, Vpa-180-1, Vpa-181-1, Vpa-182-1, Vpa-184-1, Vpa-185-1, Vpa-186-1, Vpa-187-1, Vpa-188-1, Vpa-189-1, Vpa-19-1, Vpa-190-1, Vpa-191-1, Vpa-192-1, Vpa-193-1, Vpa-194-1, Vpa-195-1, Vpa-196-1, Vpa-197-1, Vpa-198-1, Vpa-199-1, Vpa-2-1, Vpa-20-1, Vpa-200-1, Vpa-201-1, Vpa-202-1, Vpa-203-1, Vpa-204-1, Vpa-205-1, Vpa-206-1, Vpa-208-1, Vpa-209-1, Vpa-21-1, Vpa-210-1, Vpa-211-1, Vpa-212-1, Vpa-213-1, Vpa-214-1, Vpa-215-1, Vpa-216-1, Vpa-217-1, Vpa-218-1, Vpa-22-1, Vpa-220-1, Vpa-221-1, Vpa-222-1, Vpa-223-1, Vpa-224-1, Vpa-225-1, Vpa-226-1, Vpa-227-1, Vpa-228-1, Vpa-229-1, Vpa-23-1, Vpa-230-1, Vpa-231-1, Vpa-232-1, Vpa-233-1, Vpa-234-1, Vpa-235-1, Vpa-236-1, Vpa-237-1, Vpa-239-1, Vpa-24-1, Vpa-241-1, Vpa-242-1, Vpa-243-1, Vpa-244-1, Vpa-25-1, Vpa-26-1, Vpa-27-1, Vpa-28-1, Vpa-29-1, Vpa-3-1, Vpa-30-1, Vpa-31-1, Vpa-32-1, Vpa-33-1, Vpa-34-1, Vpa-35-1, Vpa-36-1, Vpa-37-1, Vpa-38-1, Vpa-39-1, Vpa-4-1, Vpa-40-1, Vpa-41-1, Vpa-42-1, Vpa-43-1, Vpa-44-1, Vpa-45-1, Vpa-46-1, Vpa-47-1, Vpa-48-1, Vpa-49-1, Vpa-5-1, Vpa-50-1, Vpa-51-1, Vpa-52-1, Vpa-53-1, Vpa-54-1, Vpa-55-1, Vpa-56-1, Vpa-57-1, Vpa-58-1, Vpa-59-1, Vpa-6-1, Vpa-60-1, Vpa-61-1, Vpa-62-1, Vpa-63-1, Vpa-64-1, Vpa-65-1, Vpa-66-1, Vpa-67-1, Vpa-68-1, Vpa-69-1, Vpa-7-1, Vpa-70-1, Vpa-71-1, Vpa-72-1, Vpa-73-1, Vpa-74-1, Vpa-75-1, Vpa-76-1, Vpa-77-1, Vpa-78-1, Vpa-8-1, Vpa-80-1, Vpa-81-1, Vpa-82-1, Vpa-83-1, Vpa-84-1, Vpa-85-1, Vpa-86-1, Vpa-88-1, Vpa-89-1, Vpa-9-1, Vpa-90-1, Vpa-91-1, Vpa-92-1, Vpa-93-1, Vpa-95-1, Vpa-96-1, Vpa-97-1, Vpa-98-1, Vpa-99-1, Vre-1-2, Vsh-1-1, Vsi-1-2, Vsi-1-4, Vsp-1-1, Vsp-11-1, Vsp-12-1, Vsp-13-1, Vsp-14-1, Vsp-15-1, Vsp-16-1, Vsp-17-1, Vsp-18-1, Vsp-19-1, Vsp-20-1, Vsp-22-1, Vsp-23-1, Vsp-24-1, Vsp-28-1, Vsp-29-1, Vsp-33-1, Vsp-34-1, Vsp-36-1, Vsp-37-1, Vsp-38-1, Vsp-39-1, Vsp-4-1, Vsp-40-1, Vsp-41-1, Vsp-42-1, Vsp-43-1, Vsp-6-1, Vsp-7-1, Vsp-8-1, Vsp-9-1, Vta-1-1, Vtu-2-1, Vtu-3-1, Vva-1-1, Vvu-1-1, Vvu-11-1, Vvu-2-1, Vvu-3-1, Vvu-4-1, Vvu-6-1, Vvu-7-1, Xbo-1-1, Xbo-10-1, Xbo-2-1, Xbo-3-1, Xbo-4-1, Xbo-5-1, Xbo-7-1, Xbo-9-1, Xne-1-1, Xne-2-1, Xne-3-1, Xne-4-1, Xne-5-2, Xne-6-1, Yal-1-2, Yal-1-3, Yal-4-1, Ybe-2-1, Yen-1-1, Yen-1-2, Yen-1-3, Yen-1-4, Yen-10-1, Yen-11-1, Yen-12-1, Yen-15-1, Yen-16-1, Yen-17-1, Yen-18-1, Yen-19-1, Yen-2-1, Yen-20-1, Yen-22-1, Yen-3-1, Yen-4-1, Yen-5-1, Yen-6-1, Yen-7-1, Yen-8-1, Yen-9-1, Yfr-1-10, Yfr-1-11, Yfr-1-12, Yfr-1-13, Yfr-1-16, Yfr-1-17, Yfr-1-2, Yfr-1-3, Yfr-1-4, Yfr-1-6, Yfr-1-9, Yfr-2-1, Yin-1-1, Ykr-1-1, Ykr-1-2, Ymo-1-2, Ymo-1-3, Ymo-1-4, Ymo-1-5, Ymo-1-6, Ymo-1-7, Ymo-2-1, Ype-1-2, Ype-10-1, Ype-100-1, Ype-101-1, Ype-102-1, Ype-103-1, Ype-104-1, Ype-105-1, Ype-106-1, Ype-107-1, Ype-108-1, Ype-109-1, Ype-11-1, Ype-110-1, Ype-111-1, Ype-112-1, Ype-113-1, Ype-114-1, Ype-114-2, Ype-115-1, Ype-116-1, Ype-117-1, Ype-118-1, Ype-119-1, Ype-12-1, Ype-120-1, Ype-121-1, Ype-122-1, Ype-123-1, Ype-124-1, Ype-125-1, Ype-126-1, Ype-128-1, Ype-129-1, Ype-13-1, Ype-130-1, Ype-131-1, Ype-132-1, Ype-133-1, Ype-134-1, Ype-135-1, Ype-136-1, Ype-137-1, Ype-138-1, Ype-139-1, Ype-14-1, Ype-140-1, Ype-141-1, Ype-142-1, Ype-143-1, Ype-144-1, Ype-145-1, Ype-146-1, Ype-147-1, Ype-148-1, Ype-149-1, Ype-15-1, Ype-150-1, Ype-151-1, Ype-152-1, Ype-153-1, Ype-154-1, Ype-155-1, Ype-156-1, Ype-157-1, Ype-158-1, Ype-159-1, Ype-16-1, Ype-160-1, Ype-161-1, Ype-162-1, Ype-163-1, Ype-164-1, Ype-165-1, Ype-166-1, Ype-167-1, Ype-168-1, Ype-169-1, Ype-17-1, Ype-170-1, Ype-171-1, Ype-172-1, Ype-173-1, Ype-174-1, Ype-175-1, Ype-176-1, Ype-177-1, Ype-178-1, Ype-179-1, Ype-18-1, Ype-180-1, Ype-181-1, Ype-182-1, Ype-183-1, Ype-184-1, Ype-185-1, Ype-186-1, Ype-187-1, Ype-188-1, Ype-189-1, Ype-19-1, Ype-190-1, Ype-191-1, Ype-192-1, Ype-193-1, Ype-194-1, Ype-195-1, Ype-196-1, Ype-197-1, Ype-198-1, Ype-199-1, Ype-2-1, Ype-2-2, Ype-2-3, Ype-2-4, Ype-2-5, Ype-2-6, Ype-2-7, Ype-2-8, Ype-2-9, Ype-20-1, Ype-200-1, Ype-201-1, Ype-202-1, Ype-203-1, Ype-204-1, Ype-205-1, Ype-206-1, Ype-207-1, Ype-208-1, Ype-209-1, Ype-21-1, Ype-210-1, Ype-211-1, Ype-212-1, Ype-213-1, Ype-214-1, Ype-215-1, Ype-216-1, Ype-217-1, Ype-218-1, Ype-219-1, Ype-219-2, Ype-22-1, Ype-220-1, Ype-221-1, Ype-222-1, Ype-223-1, Ype-224-1, Ype-225-1, Ype-226-1, Ype-227-1, Ype-228-1, Ype-23-1, Ype-24-1, Ype-25-1, Ype-26-1, Ype-27-1, Ype-28-1, Ype-29-1, Ype-29-2, Ype-3-1, Ype-30-1, Ype-31-1, Ype-31-2, Ype-32-1, Ype-32-2, Ype-33-1, Ype-34-1, Ype-35-1, Ype-36-1, Ype-37-1, Ype-38-1, Ype-39-1, Ype-4-1, Ype-40-1, Ype-41-1, Ype-42-1, Ype-43-1, Ype-44-1, Ype-45-1, Ype-46-1, Ype-47-1, Ype-48-1, Ype-49-1, Ype-5-1, Ype-50-1, Ype-51-1, Ype-52-1, Ype-53-1, Ype-54-1, Ype-55-1, Ype-56-1, Ype-57-1, Ype-58-1, Ype-59-1, Ype-6-1, Ype-60-1, Ype-61-1, Ype-62-1, Ype-63-1, Ype-64-1, Ype-65-1, Ype-66-1, Ype-67-1, Ype-68-1, Ype-69-1, Ype-7-1, Ype-70-1, Ype-71-1, Ype-72-1, Ype-73-1, Ype-74-1, Ype-75-1, Ype-76-1, Ype-77-1, Ype-78-1, Ype-79-1, Ype-8-1, Ype-80-1, Ype-81-1, Ype-82-1, Ype-83-1, Ype-84-1, Ype-85-1, Ype-86-1, Ype-87-1, Ype-88-1, Ype-89-1, Ype-9-1, Ype-90-1, Ype-91-1, Ype-92-1, Ype-93-1, Ype-94-1, Ype-94-2, Ype-95-1, Ype-96-1, Ype-97-1, Ype-98-1, Ype-99-1, Yps-1-1, Yps-1-2, Yps-1-4, Yps-10-1, Yps-10-2, Yps-11-1, Yps-11-2, Yps-12-1, Yps-12-2, Yps-13-1, Yps-14-1, Yps-15-1, Yps-2-1, Yps-3-1, Yps-4-1, Yps-5-1, Yps-6-1, Yps-7-1, Yps-8-1, Yps-9-1, Yro-1-1, Yru-1-2, Yru-1-3, Yru-2-1, Ywa-1-2, env-100, env-1000, env-1001, env-1002, env-1003, env-1004, env-1005, env-1006, env-1007, env-1008, env-1009, env-1010, env-1011, env-1012, env-1013, env-1014, env-1015, env-1016, env-1017, env-1018, env-1019, env-1020, env-1021, env-1022, env-1023, env-1024, env-1026, env-1028, env-1033, env-1034, env-1038, env-1039, env-104, env-1040, env-1041, env-1042, env-1043, env-1044, env-1045, env-1046, env-1047, env-1048, env-1049, env-1050, env-1051, env-1052, env-1053, env-1054, env-1055, env-1056, env-1057, env-1058, env-1059, env-1060, env-1061, env-1062, env-1063, env-1064, env-1065, env-1066, env-1067, env-1068, env-1069, env-107, env-1070, env-1071, env-1072, env-1073, env-1074, env-1075, env-1076, env-1077, env-1078, env-1079, env-108, env-1080, env-1081, env-1082, env-1083, env-1084, env-1085, env-1086, env-1087, env-1088, env-1089, env-1090, env-1091, env-1092, env-1095, env-1098, env-110, env-1100, env-1101, env-1102, env-1106, env-1107, env-1108, env-111, env-1111, env-1112, env-112, env-113, env-114, env-115, env-116, env-117, env-118, env-119, env-120, env-121, env-122, env-123, env-124, env-125, env-126, env-127, env-128, env-129, env-130, env-131, env-132, env-133, env-134, env-135, env-136, env-137, env-138, env-141, env-142, env-143, env-144, env-145, env-146, env-147, env-148, env-149, env-150, env-151, env-152, env-153, env-154, env-155, env-156, env-157, env-158, env-159, env-160, env-161, env-162, env-163, env-164, env-165, env-166, env-167, env-168, env-169, env-17, env-170, env-171, env-172, env-173, env-174, env-175, env-176, env-177, env-178, env-179, env-18, env-180, env-181, env-182, env-183, env-184, env-185, env-186, env-187, env-188, env-189, env-190, env-191, env-192, env-193, env-194, env-195, env-196, env-197, env-198, env-199, env-2, env-200, env-201, env-202, env-203, env-204, env-205, env-206, env-207, env-208, env-209, env-210, env-211, env-212, env-213, env-214, env-215, env-216, env-217, env-218, env-219, env-22, env-220, env-221, env-222, env-223, env-224, env-225, env-226, env-227, env-228, env-229, env-230, env-231, env-232, env-233, env-234, env-235, env-236, env-237, env-238, env-239, env-240, env-241, env-242, env-243, env-244, env-245, env-246, env-247, env-248, env-249, env-25, env-250, env-251, env-252, env-253, env-254, env-255, env-256, env-257, env-258, env-259, env-26, env-260, env-261, env-262, env-263, env-264, env-265, env-266, env-267, env-268, env-269, env-27, env-270, env-271, env-272, env-273, env-274, env-275, env-276, env-277, env-278, env-279, env-28, env-280, env-281, env-282, env-283, env-285, env-286, env-287, env-288, env-289, env-29, env-290, env-291, env-292, env-293, env-294, env-295, env-296, env-297, env-298, env-299, env-3, env-30, env-300, env-301, env-302, env-303, env-304, env-305, env-306, env-307, env-308, env-309, env-31, env-310, env-311, env-312, env-313, env-314, env-315, env-316, env-317, env-318, env-319, env-32, env-320, env-321, env-322, env-323, env-324, env-325, env-326, env-327, env-328, env-329, env-33, env-330, env-331, env-332, env-333, env-334, env-335, env-336, env-337, env-338, env-339, env-34, env-340, env-341, env-342, env-343, env-344, env-345, env-346, env-347, env-348, env-349, env-35, env-350, env-351, env-353, env-354, env-355, env-36, env-364, env-365, env-366, env-367, env-37, env-370, env-371, env-372, env-373, env-374, env-375, env-376, env-377, env-378, env-379, env-38, env-380, env-381, env-382, env-383, env-384, env-386, env-387, env-388, env-389, env-39, env-390, env-391, env-392, env-393, env-395, env-396, env-397, env-398, env-4, env-400, env-402, env-403, env-405, env-406, env-41, env-412, env-413, env-414, env-415, env-416, env-42, env-421, env-424, env-425, env-426, env-427, env-429, env-43, env-436, env-439, env-44, env-440, env-442, env-444, env-446, env-447, env-448, env-449, env-451, env-455, env-456, env-464, env-465, env-468, env-470, env-471, env-472, env-473, env-474, env-476, env-478, env-48, env-491, env-492, env-495, env-496, env-497, env-498, env-499, env-5, env-502, env-505, env-506, env-510, env-514, env-523, env-524, env-525, env-526, env-528, env-53, env-531, env-532, env-533, env-54, env-543, env-544, env-545, env-548, env-55, env-553, env-554, env-555, env-556, env-557, env-558, env-560, env-561, env-562, env-563, env-564, env-565, env-567, env-568, env-569, env-570, env-571, env-574, env-575, env-576, env-577, env-578, env-579, env-580, env-581, env-582, env-583, env-584, env-585, env-586, env-587, env-588, env-590, env-592, env-596, env-597, env-598, env-6, env-600, env-602, env-604, env-605, env-609, env-610, env-615, env-617, env-618, env-62, env-622, env-628, env-629, env-631, env-632, env-635, env-638, env-639, env-64, env-644, env-645, env-648, env-649, env-65, env-650, env-652, env-654, env-659, env-66, env-663, env-664, env-665, env-666, env-667, env-669, env-67, env-670, env-671, env-673, env-676, env-68, env-684, env-689, env-69, env-691, env-692, env-693, env-694, env-695, env-696, env-697, env-698, env-699, env-7, env-70, env-703, env-704, env-705, env-707, env-708, env-709, env-716, env-718, env-719, env-72, env-721, env-722, env-723, env-727, env-733, env-734, env-737, env-744, env-745, env-746, env-747, env-748, env-749, env-75, env-750, env-751, env-752, env-753, env-759, env-760, env-761, env-762, env-764, env-765, env-767, env-768, env-769, env-770, env-771, env-772, env-773, env-774, env-775, env-776, env-777, env-778, env-779, env-78, env-780, env-781, env-782, env-783, env-787, env-788, env-789, env-79, env-790, env-791, env-792, env-793, env-794, env-795, env-796, env-797, env-798, env-799, env-80, env-800, env-801, env-802, env-803, env-804, env-805, env-806, env-807, env-808, env-809, env-81, env-810, env-811, env-812, env-813, env-814, env-815, env-816, env-817, env-818, env-819, env-82, env-820, env-821, env-822, env-823, env-824, env-825, env-826, env-827, env-828, env-829, env-83, env-830, env-831, env-832, env-833, env-834, env-835, env-836, env-837, env-838, env-839, env-84, env-840, env-841, env-842, env-843, env-844, env-845, env-846, env-847, env-848, env-849, env-85, env-850, env-851, env-852, env-853, env-854, env-855, env-856, env-857, env-858, env-859, env-860, env-863, env-864, env-865, env-866, env-873, env-877, env-878, env-879, env-887, env-888, env-891, env-892, env-893, env-897, env-901, env-903, env-904, env-905, env-906, env-907, env-908, env-909, env-910, env-911, env-912, env-913, env-914, env-915, env-916, env-917, env-918, env-919, env-92, env-920, env-921, env-922, env-923, env-924, env-925, env-926, env-927, env-928, env-929, env-931, env-932, env-938, env-939, env-940, env-941, env-942, env-943, env-944, env-945, env-946, env-947, env-948, env-949, env-950, env-951, env-952, env-953, env-958, env-959, env-960, env-961, env-962, env-963, env-964, env-965, env-966, env-967, env-968, env-969, env-970, env-971, env-972, env-973, env-974, env-975, env-976, env-977, env-978, env-979, env-98, env-980, env-983, env-984, env-985, env-987, env-988, env-989, env-99, env-990, env-991, env-993, env-994, env-995, env-996, env-997, env-998, env-999

alignment positions $1 \cdots 79$

```
CBa-1-1   UGCAUUCUGUAUUAUCCGUCGCCUUUGAUGUCAUAUCACAUAUCAGACGACUUCAAAGUGAUGAAACUUAACAUUUAAU
Apl-1-1   AUUGAUCUUCCUUGCCAAAUACCAUAAAAUACGCACUCUUUUUUGGCGUCGAGUAACUGCUCGACAUAUUAUUUAUUUC

          ...............................................................................
          ...............................................................................
          ●●●●●●●●●●●●●●●●●●●●●●●●●●●●●●●●●●●●●●●●●●●●●●●●●●●●●●●●●●●●●●●●●●●●●●●●●●●●RY●
```

```
Apl-6-1     UAUUGAUCUUUGUUGCCAAAUGCCGUAGAAUACGCACUCUUUUUUGGCGUCGAGUAACUGCUCGGCAUAUUAUUUAUUC
Apl-13-1    UAUUGAUCUUUGUUGCCAAAUGCCGUAGAAUACGCAAUCUUUUUUGGCGUCGAGUAACUGCUCGGCAUAUUAUUUAUUC
Apl-8-1     UAUUGAUCUUCCUUGCCAAAUACCAUAAAAUACACGCUCUUUUUUGGCGUCGAGUAAAUGCUCGGCAUAUUAUUUAUUC
Aur-1-1     GUAUUGAUCUUAAUGACUAAAUACCAUAGAAUACGCACUCUUUUUGGCGUCGAGCAGAUGCUCGGCAUAUUAUUUAUUC
Aca-3-1     GUAUUGAUCUUAAUGGCUAAAUGCCAUAGAAUACGCACUCUUUUUGGCGUCGAGUAAAUACUCGGCAUAUUAUUUAUUC
Aeq-1-1     AGUAUUGAUCUUAAUUGCCAAAUACCAUAGAAUAUUGGCUCUUUUGGCGUCGAGCAAAUGCUCGGCAUAUUAUUUAUUC
Asu-3-1     GUAUUGAUCUUAAUGACUAAAUACCAUAGAAUACGCACUCUUUUUGGCGUCGAGUAAAUACUCGGCAUAUUAUUUAUUC
Mgr-1-1     GGUUGAUCUCGGCUACAAAUUUAUCUAGAAUAUGCACUCUUUUUAGGCGUUGAGCAAAUGCUCAAUAUUUUUAUUAAUC
Mha-1-1     UCUCGGCUACAAAUUUAUCUAGAAUAUGCACUCUUUUUAUAGGCGUUGAGCAGCAAAUGCUCAAUAUUUUUAUUAAUC
Ami-1-1     CUAUUGAUCUCUGAAAUCAUUUAGCCUAAAAUACGCACUCCUUUAGGCGUUAAACAGAUGUUUAACAUUUUUAUUAAUU
env-1       GCGGUUGAUCUUCGCUCAACUUUCACUUAGAAUACGCACUCUUUUAGGCGUUAGGCAUAAGCUUAACAUAUUAUCAAUC
env-8       AAAUUCUGCAUUAAAUUAGCGGUUGAUCUUCGCUCAACUUUCACUUAGAAUACGCACUCUUUUAGGCGUUAGGNCAAUC
env-9       UUGGUUGAUCUCCACAGGAAAUUCACAUAAAAUACGAACUCUUUUUGGUGCUAAAUCUAGAUUUAGCGUAUUAUUAAUC
env-10      UUGGUUGAUCUCCACAGGAAAUUCACAUAAAAUACGGACUCUUUUUGGUGCUAAAUUUAGAUUUAGCGUAUUAUUAAUC
Hsp-3-1     UUGGUUGAUCUCCACAGGAAAUUCACAUAAAAUACGGGCUCUUUUUGGUGCUAAAUUUAGAUUUAGCGUAUUAUUAAUC
Ami-2-1     GCUUGAUCUCUUUACUUAUCUCGUCUAAAAUACGCACUCUUUUUAGGCGUUAAACAUCUGUUUAACAUUUUUAUUAAUU
Lgr-1-1     UGCGUGGUUCAUCUCUAGCCUGUAUAAUCUGCGGCCUUUGAUUCUCAAUGAGGGUCAAACCACUCACCCAUUAUUUAUC
Asu-1-1     ACAGUCUUGAAUAAUGUAUUUUUUUCGGUAUAAUUUGCCGUCUAAUUUCCGUGCUGCACCGUGCAGCGUAAUAUUAACU
env-11      UGCGUCCGGGAUGCAAUCUGAUAGACUUCUGCGUCCAUUUGAGCAGUGCAUGAUUGUGCAUUGCUUUAGCGAUUGAUUC
env-12      GCUAUGGAUCUGAUAUUAUUCGCCGUCCGAAAUCAGCAUGGCGCAAGCGCAGAGUUUGCACCGGUUGGACNCACGAUUC
env-13      GCUAUGGAUCUGAUAUUAUUCGCCGUCCGAAAUCAGCAUGGCGCAAGCGCAGAGUUUGCACCGGUUGGACCCACGAUUC
env-14      UGAUGCCACAAUUGCAAGUAAGAUUCGCGGUCUGUUUCCGGCAGGGCAUGAUUGUGCCCCGUCUUGUCAAUUGAUUCAA
env-15      .......................................CGGCAGGGCAUGAUUGUGCCCGGUCUUCUCAAUUGAUUCAA
env-16      UGCGACACUAAUUGCAAGUAAGAUUCGCGGUCUGUUUCCGGCUGGGCAUGAUUGUGCCCCGUCUUCUCAAUUGAUUCAA
env-19      UGCGACACUAAUUGCAAGUAAGAUUCGCGGUCUGUUUCCGGCAGGGCAUGAUUGUGCCCCGUCUUCUCAAUUGAUUCAA
env-20      UGCGACACUAAUUGCAAGUAAGAUUCGCGAUCUGUUUCCGGCUGGGCAUGAUUGUGCCCCGUCUUCUCAAUUGAUUCAA
env-21      UGCAGCAAUAAUUGCAAGUAAGAUUCGCGGUCUGUUUCCGGCAGGGCAUGAUUGUGCCCCGUCUUCUCAAUUGAUUCAA
Msu-1-1     UGUGGCAAUAAUUGCAAGUAAGAUUCGCGGUCUGUUUCCGGCAGGGCAUGAUUGUGCCCCGUCUUCUCAAUUGAUUCAA
env-23      UGUGGUCGGGUUUGUCAGUAAGAUUCGCGGUCUGUUUCUGGCCGGGCUUGAUUGUGCACGGUCUUUUCAAUUGAUUCAA
env-24      UGUGGUCGGGUUUGUCAGUAAGAUUCGCGGUCUGUUUCUGGCCGGGCUUGAUUGUGCACGGUCUUUUCAAUUGAUUCAA
env-40      UGUGGUCGGGUUUGUCAGUAAGAUUCGCGGUCUGUUUCUGGCCGGGCUUGAUUGUGCACGGUCUUUUCAAUUGAUUCAA
Msp-1-1     UGUGGUCGGGUUUGUCAGUAAGAUUCGCGGUCUGUUUCUGGCCGGGCUUGAUUGUGCACGGUCUUUUCAAUUGAUUCAA
Vli-1-1     UGUGUUGAUCGAUGGAGACGGAUUCACUAUUAUACUGCGCCUCUAACGAUGUAUAUUUUUACAUCAUUUUUUAGUAACA
Mma-1-1     AUGUAGUUUAUUUAUGCAUCUGCUAUUGAUAUUAGGGGGUAAGAUCCUAUAUUAUUCGCAGCCUAAAAAUUCUAUAACU
env-45      CUUGAACUACAUGAAUUUUUUCUGUAAAAUUUGCCCCCUUACACACAAGCGUGCCAUUUUAAAUGGCGUAAUAUUAAUC
env-46      CUUGAACUACAACUAUUUUUUCUGUACAAUUCCGAGCCUUACACACAAGCGUGCCAUUUAAAAUGGCGUAAUAUUAAUC
env-47      GUUGAAUGUCAUUUUCUUUUUCUGUAAAAUUCGCAUCCUUACACACAAGCGUGCCAUUUAUAAUGGCGUAAUAUUAAUC
env-49      UUGAAUGUCAUUUUCUUUUUCUGUAAAAUUCCCAGCCUUACACACAAGCGUGCCAUUUUAUAAUGGCGUAAUAUUAAUC
env-50      GUUGAAUGUCAUUUUCUUUUUCUGUAAAAUUCACAACCUUACACACAAGCGUGCCAUUUAUAAUGGCGUAAUAUUAAUC
Asp-10-1    GUUGAAUGUCAUUUUCUUUUUCUGUAAAAUUCGCAAACUUACACACAAGCGUGCCAUUUAUAAUGGCGUAAUAUUAAUC
env-51      GUUGAAUGUCAUUUUCUUUUUCUGUAAAAUCCGCAACCUUACACACAAGCGUGCCAUUUAUGAUGGCGUAAUAUUAAUC
env-52      GUUGAAUGUCAUUUUCUUUUUCUGUAAAAUUCGCAACCUUACACACAAGCGUGCCAUUUAUAAUGGCGUAAUAUUAAUC
env-56      GUUGAAUGUCAUUUUCUUUUUCUGUAAAAUUUGCACCCUUACACACAAGCGUGCCAUUUAUAAUGGCGUAAUAUUAAUC
env-57      GUUGAAUGUCAUUUUCUUUUUCUGUAAAAUUUGCACCCUUACACACAAGCGUGCCAUUUAUAAUGGCGUAAUAUUAAUC
env-58      GUUGAAUGUCAUUUUCUUUUUCUGUAAAAUUCGCAACCUUACACACAAGCGUGCCAUUUAUAAUGGCGUAAUAUUAAUC
env-59      UUGAACUACGCUAAUUUUUUCUGUAAAAUUCGCAACCUUACACACAAGCGUGCCAUUUUGUAAUGGCGUAAUAUUAAUC
env-60      UUGAACUACGCUAAUUUUUUCUGUAAAAUUCGCAACCUUACACACAAGCGUGCCAUUUUGUAAUGGCGUAAUAUUAAUC
env-61      UGAACUGCACUAAUUUUUUCUGUAGAAUUUGCAACCUUACACACAAGGGUGUCAUUUUGUAAUGGCAUAAAUAUUAAUC
env-63      UGAACUACACUAAUUUUUUCUGUAGAAUUUGCAACCUUACACACAAGGGUGUCAUUUUGUAAUGGCAUAAAUAUUAAUC
env-71      UGAACUACACUAAUUUUUUCUGUAAAAUUUGCAACCCUACACACAAGGGUGUCAUUUUGUAAUGGCAUAAAUAUUAAUC
env-73      ............................................................................AAUC
Aac-6-1     UUGAAUUGCGCUUUUUUUUUCUGUAAAAUUCGCAGCCUUACACACAAGCGUGCCAUUUUGAAAUGGCGUAAUAUUAAUC
Aac-18-1    UUGAAUUGCGCUUUUUUUUUCUGUAAAAUUCGCAACCUUACACACAAGCGUGCCAUUUUGAAAUGGCGUAAUAUUAAUC
env-74      UUGAACUACGCUAAUUUUUUCUGUAAAAUUCGCAACCUUACACACAAGCGUGCCAUUUUGUAAUGGCGUAAUAUUAAUC
Aac-11-1    UUGAAUUGCACUUUUUUUUUCUGUAAAAUUCGCAGCCUUACACACAAACGUGCCAUUUUAGAAUGGCGUAAUAUUAAUC
env-76      UUGAACUACGCUAAUUUUUUCUGUAAAAUUUGCAACCUUACACACAAGCGUGCCAUUUUGUAAUGGCGUAAUAUUAAUC
Aac-17-1    UUGAAUUGCACUUUUUUUUUCUGUAAAAUUCGCAGCCUUACACACAAGCGUGCCAUUUUAGAAUGGCGUAAUAUUAAUC
Aac-1-1     CUUGAAUUGCACUUUUUUUUCUGUAAAAUUCGCAGCCUUACACACAAACGUGCCAUUUUAGAAUGGCGUAAUAUUAAUC
Aac-14-1    UUGAAUUGCGCUUUUUUUUUCUGUAAAAUUCGCAGCCUUACACACAAACGUGCCAUUUUAGAAUGGCGUAAUAUUAAUC
env-77      .................................GCAUCCUUACACACAAGCGUGCCAUUUAUAAUGGCGUAAUAUUAAUC
env-86      GUUGAAUGUCAUUUUCUUUUUCUGUAAAAUUCGCAUCCUUACACACAAGCGUGCCAUUUAUAAUGGCGUAAUAUUAAUC
env-87      GUUGAAUGUCAUUUUCUUUUUCUGUAAAAUUCGCACCCUUACACACAAGCGUGCCAUUUAUAAUGGCGUAAUAUUAAUC
env-88      CUUGAACUACAUGGAUUUUUUCUGUAAAAUUCGCAUCCUUACACACAAGCGUGCCAUUUAUAAUGGCGUAAUAUUAAUC
env-89      UUGAACUACGCUAAUUUUUUCUGUAAAAUUCGCAACCUUACACACAAGCGUGCCAUUUUGUAAUGGCGUAAUAUUAAUC
env-90      GUUGAAUGUCAUUUUCUUUUUCUGUACAAUUCGCAACCUUACACACAAGCGUGCCAUUUUAAAUGGCGUAAUAUUAAUC
env-91      GUUGAAUGUCAUUUUCUUUUUCUGUAAAAUUCGCAUCCUUACACACAAGCGUGCCAUUUUAAAUGGCGUAAUAUUAAUC
env-93      CUUGAACUAUGCUAAUUUUUUCUGUAGAAUUUGCACCCUUACACACAAGCGUGCCAUUUAUAAUGGCGUAAUAUUAAUC
env-94      GUUGAAUGUCAUUUUCUUUUUCUGUAAAAUUUGCAUCCUUACACACAAGCGUGCCAUUUAUAAUGGCGUAAUAUUAAUC
env-95      CUUGAACUACAUGGAUUUUUUCUGUAAAAUUCGCAUCCUUACACACAAGCGUGCCAUUUAUGAUGGCGUAAUAUUAAUC
Pda-1-1     AUCGGACUUGCAACUUUAUCGCUUUUUUUGUACAAUUCGCCAUCUUACAUACAAACUGCCUUUUGGCGUAAUAUUAACU
Pmu-14-1    UUUGGCUUGAAUAUUCGAUCAUUUUUUUGUAGAAUUUGCCACCUUACACAUAACUUGCCAUUUUGGCGUAAUAUUAACU
Pmu-2-1     CUUCUACUUGCAACUUUAUCUGUUUUUUUGUACAAUCCUGCCUCUUACAUACAAACUGCCAUACGGCGUAAUAUUAACA
Pmu-8-1     AAAAUUGUUGAAAAUUACUUUAGUAUUUUGUAGAAUCCAACAGCUUACAUACAAACUGCCAUAUGGCGUAAUAUUAACU
Pmu-12-1    GGCUUGAAUAUUCUGCUUUUUUUUUGUAGAAUUUGCCACCUUACAUACAAACUGCUACUUAUGUGGCGUAAUAUUAAUU
Pmu-3-1     UACACUUGCAAGACUGUCCCUUUUUUUGUACAAUCUGCACCCUUACAUACAAACUGCCAUUUAUGGCGUAAUAUUAAUU
Pmu-6-1     UCUGCUUGCAACUUUAUCCCUUUUUUUGUACAAUCUGCACCCUUACAUACAAACUGCCAUUUAUGGCGUAAUAUUAAUU
Mma-3-1     AUUUUUUAAGGUAUAAUUUGCGCUCUUACUAACGAGCCUUCAGAAAGUUGUUGAUACUUUCGAUGGCGUAAUAUUAAUC
Msu-2-1     UUUUUUACGGUAUAAUUUGCUCUCUUUUACACAGAGUCUUCAGAAAAUCAUAUUGAUUUUCGAUGACAUAAUAUUAAUC
env-96      AAGCACACCUCUUGAUCAAUUUGAUUUUUUCAGUAUAAUUCGCAGUCUAUUUGGUGCCGGAUUGUCGGCGAUUAUUAAC
env-97      AAGCACACCUCUUGAUCAAUUUGAUUUUUUCAGUAUAAUUCGCAGUCUAUUUGGUGCCGGAUUGUCGGCGAUUAUUAAC
Hpa-2-2     AAGCACACCUCUUGAUCAAUUUGAUUUUUUCAGUAUAAUUCGCAGUCUAUUUGGUGCCGGAUUGUCGGCGAUUAUUAAC
env-101     UGUANNNNNNNNNGAUCAAUUUGAUUUUUUCAGUAUAAUUCGCAGUCUAUUUGGUGCCGGAUUGUCGGCGAUUAUUAAC
env-102     AAGCACGCCUCUUGAUCAAUUUGAUUUUUUCAGUAUAAUUCGCAGUCUAUUUGGUGCCGGAUUGUCGGCGAUUAUUAAC
env-103     AAGCACACCACUUGAUCAAUUUGAUUUUUUCAGUAUAAUUCGCAGUCUAUUUGGUGCCGGAUUGUCGGCGGUUAUUAAC

            ................................................................................
            ................................................................................
            ●●●●●●●●●●●●●●●●●●●●●●●●●●●●●●●●●●●●●●●●●●●●●●●●●●●●●●●●●●●●●●●●●●●●●●●●●●●●●RY●
```

```
env-105   CCAUUGUAGAACUGAAUGGGCAAANNNNNNNNNNNNNNNNNNNNNNNNNNNNUUGGUGCCGGAUUGUCGGCGAUUAUUAAC
env-106   AAGCACACCACUUGAUCAAUUUGAUUUUUUCAGUAUAAUUCGCAGUCUAUUUGGUGCCGGAUUGUCGGUGAUUAUUAAC
env-109   .................................................................UCGGCGAUUAUUAAC
env-139   AGCACACCACUUGAUCAAUUUGAUUUUUUCAGUAUAAUUCGCAGUCUAAUUUGGUGCCGGAUUGUCGGCGAUUAUUAAC
env-140   AAGCACACCUCUUGAUCAAUUUGAUUUUUUCAGUAUAAUUCGCAGUCUAUUUGGUGCCGGAUUGUCGGCGAUUAUUAAC
env-284   AAGCACACCACUUGAUCAAUUUGAUUUUUUCAGUAUAAUUCGCAGUCUAUUUGGUGCCGGAUUGUCGGCGAUUAUUAAC
env-352   AAGCAUACCUCUUGAUCAAUUUGAUUUUUUCAGUAUAAUUCGCAGUCUAUUUGGUGCCGGAUUGUCGGCGAUUAUUAAC
env-356   AAGCACACCACUUGAUCAAUCUGAUUUUUUCAGUAUAAUUCGCAGUCUAUUUGGUGCCGGAUUGUCGGCGAUUAUUAAC
env-357   NNNNNNNNNACUUGAUCAAUUUGAUUUUUUCAGUAUAAUUCGCAGUCUAUUUGGUGCCGGAUUGUCGGCGAUUAUUAAC
env-358   UUCCUAAAGUGCGGUCNNNNNNNNNNNNNNNNNNUAUAAUUCGCAGUCUAUUUGGUGCCGGAUUGUCGGCGAUUAUUAAC
env-359   AAGCACACCUCUUGAUCAAUUUGAUUUUUUUAGUAUAAUUCGCAGUCUAUUUGGUGCCGGAUUAUCGGCGAUUAUUAAC
env-360   CUUUCCUAAAGUGCGGUCAAANNNUUUUUUCAGUAUAAUUCGCAAUCUAUUUGGUGCCGGAUUGUCGGCGAUUAUUAAC
env-361   AAGCGAACCUCUUGAUCAAUUUGAUUUUUUCAGUAUAAUUCGCAGUCUAUUUGGUGCCGGAUUGUCGGCGAUUAUUAAC
env-362   GACAGAGGAUUGGGCUAACAUAACUUUCCUAAAGUGCGGUNNNNNNNNNNNNNNNNNNNNNNNNNNNNNNAUUAUUAAC
Hpa-2-11  AAGCACACCUCUUGAUCAAUUUGAUUUUUUCAGUAUAAUUCGCAGUCUAUUUGGUGCCGGAUUGUCGGCAAUUAUUAAC
Rpn-1-1   AUCUCCCCCCUCUUGAUCAAUAUCAUUUUUUCAGUAUAAUUUGCGGUCUAUUUGGUGCCGAAUAACGGCGAUUAUUAAC
env-363   CUAAAAUGGGGCUUGUUUAUCUAUGUUUUUUCAGUAUAAUUCGCAGUCUAUUUGGUGCCAAAUAUGGCGUAUUAUUCAC
env-368   CUAAAAUGGGGCUUGUUUAUCUAUGUUUUUUCAGUAUAAUUCGCAGUCUAUUUGGUGCCAGAUAUGGCGUAUUAUUCAC
env-369   CUAAAAUGGGGCUUGUUUAUCUAUGUUUUUUCAGUAUAAUUCACAGCCUAUUUGGUGCCAGAUAUGGCGUAUUAUUCAC
env-385   CUAAAAUGGGGCUUGUUUAUCUAUGUUUUUUCAGUAUAAUUCGCAGUCUAUUUGGUGCCAAAUAUGGCGUAUUAUUCAU
env-394   CUAAAAUGGGGCUUGUUUAUCUAUGUUUUUUCAGUAUAAUUCGCAGUCUAUUUGGUGCCAAAUAUGGCGUGUUAUUCAC
env-399   CUAAAAUGGGGCUUGUUUAUCUAUGUUUUUUCAGUAUAAUUCGCAGUCUAUUUGGUGCCAAAUAUGGCGUAUUAUUCAC
env-401   CUAAAAUGGGGCUUGUUUAUCUAUGUUUUUUCAGUAUAAUUCGCAGCCUAUUUGGUGCCAAAUAUGGCGUAUUAUUCAC
env-404   CUAAAAUGGGACUUGUUUAUCUAUGUUUUUUCAGUAUAAUUCGCAGUCUAUUUGGUGCCAAAUAUGGCGUAUUAUUCAU
env-407   CUAAAAUGGGGCUUGUUUAUCUAUGUUUUUUCAGUAUAAUUCACAUCCUAUUUGGUGCCAGAUAUGGCGUAUUAUUCAC
env-408   CUAAAAUGGGGCUUGUUUAUCUAUGUUUUUUCAGUAUAAUUCACAGCCUAUUUGGUGCCAGAUAUGGCGUAUUAUUCAU
env-409   CGAAAAUAGGGCUUGUUUAUCUGUAUUUUUUCAGUAUAAUCCGCGAUCUAUUUGGUGCCAAUUAUGGCGUAUUAUUAAC
env-410   CGAAAAUAGGGCUUGUUUAUCUGUAUUUUUUCAGUAUAAUCCGCGAUCUAUUUGGUGCCAAAUAUGGCGUAUUAUUCAC
env-411   AAAUCCGGACGCUUGAUCAAUUCUCAUUUUUUAGUAUAAUUUGCGCCCUUUGGUGUCGCUUGAGCGACGUAUUAUUAAC
env-417   CUAAAAUUGGGCUUGUUUAUCUAUGUUUUUUCAGUAUAAUUCACAGCCUAUUUGGUGCCAGAUAUGGCGUAUUAUUAAC
Hha-1-1   CGAAAAUAGGGCUUGUUUAUCUGUAUUUUUUCAGUAUAAUCCGCGAUCUAUUUGGUGCCAAUUAUGGCGUAUUAUUAAC
env-418   CGAAAAUAGGGCUUGUUUAUCUGUAUUUUUUCAGUAUAAUCCGCGAUCUAUUUGGUGCCAAUUAUGGCGUAUUAUUCAC
env-419   AAAACGGAGGGCUUGUUUAUCUUUGUUUUUUCGGUAUAAUUCGCGGCCUAUUUGGUGCCGGUUACGGCGUAUUAUUCAC
Hha-1-2   CGAAAAUAAGGCUUGUUUAUCUGUAUUUUUUCAGUAUAAUCCACGAUCUAUUUGGUGCCAAUUAUGGCGUAUUAUUAAC
env-420   AAAACGGAGGGCUUGUUUAUCUUUGUUUUUUCGGUAUAAUUCGCGGCCUAUUUGGUGCCGAUUACGGCGUAUUAUUCAC
Hha-1-3   CGAAAAUAAGGCUUGUUUAUCUGUAUUUUUUCAGUAUAAUCCGCGAUCUAUUUGGUGCCAAUUAUGGCGUAUUAUUAAC
Hha-1-4   CGAAAAUAAGGCUUGUUUAUCUGUAUUUUUUCAGUAUAAUCCGCGAUCUAUUUGGUGCCAAUUAUGGCGUAUUAUUAAC
env-422   AUAAGGCUUGUUUAUCUGUAUUUUUUCAGUAUAAUCCGCGAUCUAAUCUAUUUGGUGCCAAUUAUGGCGUAUUAUUAAC
env-423   CGAAAAUAGGGCUUGUUUAUCUGUAUUUUUUCAGUAUAAUCCGCGAUCUAUUUGGUGCCAAUUAUGGCGUAUUAUUAAC
env-428   CGAAAAUAAGGCUUGUUUAUCUGUAUUUUUUCAGUAUAAUCCGCGAUCUAUUUGGUGCCAAUUAUGGCGUAUUAUUAAC
env-430   CGAAAAUAAGGCUUGUUUAUCUGUAUUUUUUCAGUAUAAUCCGCAAUCUAUUUGGUGCCAAUUAUGGCGUAUUAUUAAC
Hha-6-1   CGAAAAUAGGGCUUGUUUAUCUGUAUUUUUUCAGUAUAAUCCGCGAUCUAUUUGGUGCCAAUUAUGGCGUAUUAUUAAC
Hin-17-1  CGAAAAUAGAGCUUGUUUAUCUGCAUUUUUUCAGUAUAAUUCGCAGCCUAUUUGGUGCCAAUUAUGGCGUAUUAUUCAC
env-431   CGAAAAUAGAGCUUGUUUAUCUGCAUUUUUUCAGUAUAAUUCGCAACCUAUUUGGUGCCAAUUAUGGCGUAUUAUUCAC
Apa-1-1   UUCCUUUUUUUUCGGUAUAAUUCGCGGCUUGAAUAAACGAUCCUGUUUAUGGUGAUUCACUAUAAACGUAAUAUUAACC
Apa-1-2   UUCCUUUUUUUUCGGUAUAAUUCGCGACUUGAAUAAACGAUCCUGUUUAUGGUGAUUCACUAUAAACGUAAUAUUAACC
Nro-1-1   CACCCCCUUGAAUAUUCCUUUUUUUCCGGUAUAAUCUGCCUCCUGACAAAAGCUGCCGUUGUACGGCGUAAUAUUAACC
Hma-1-1   CCACCCCUUGAAUAUUCCUUUUUUUCGGGUAUAAUCUGCCUCCUUAUUAGCGUUGCCGUUAUACGGCGUAAUAUUAACC
Psp-56-1  CCGCCCCUUGAAUAUUCCUUUUUUUCGGGUAUAAUCUGCCUCCUUAUUAGCAUUGCCGUUAUACGGCGUAAUAUUAACC
Hin-17-2  CCUACACUUGCAAGACUGUCCUUUUUUGUAUAAUCUGCACUCUUACAUACAAACUACCAUUUAUGGCAUAAUAUUAAUU
env-432   CAUUUCCAACUAAAGGGCGUAUAAUUCGCCACCUUAAUACUCAUGUAGUAUUGAAUCAUCAUUGAUUUUUUAAUAAAUU
Mmu-1-1   AAAGAUCUUGAAUUUCCCCUGAAUUUAACCUAGAAUAUGCAACCUUUAUGGCGCUAGGAUCACCUAGCAUUUUAUCAAU
Cor-1-1   CUAUUGCUUAACCGGUUUUUAUUCUGUAUAAUCUUCCGUCUUUAAAAGACCUGCGCAGCUGCGCAUAAUUAUUAACAAU
Baq-1-1   AAUAUCAACUCUGUAUAAUGCGCCUCCUUUGGUUCAUUUAGGUGAAUCCAAAACACGACUCAUCAUCAUUAUUUAUAUU
Pfo-1-1   UGCGUUGAUUAUAUGGCAUCUGUAUAAUACGCCACCUUUGGUUCAAAACAUGAAUCAAACCACUCAACCAUUUUUUACU
env-433   UAUUUUUUAGCACUUUUUACUUGUUGAGUGUUGCAGUUAAGGACUUGAUCCAAUACAAUGCGCGCCCAUUAAAUUGAAU
env-434   UAAAAUUGAUUAAUUAAUUACCACAUAGGCUUGCCUUGCCUGUCGCGAUACCUUAUUAUUCGCCACCAUUAUUUUUAAC
env-435   AAUAAUCUGUUUUUUACGGUAUAAUUUGCCGUCUAUUUUCAACGUGUCGCCAAUUAAAAACGGCGGUGUAAUAUUAACU
env-437   CUGCAUAAAAAUCCGAAUUUAAGAUAAAUUGGCUUAUUANNNNNNNNNNNNNNNNNNNNNNNNNNNNNNNNNNNNNACU
env-438   AAUAAUCUGUUUUUUACGGUAUAAUUUACCGUCUAUUUUCAACGUGUCGCCAAUUAAAAACGGCGGUGUAAUAUUAACU
env-441   AAUAAUCUGUUUUUUACGGUAUAAUUUGCCGUCUAUUUUGAUCGUGCCGCCAAUUGAAAACGGCGGUGUAAUAUUAACU
env-443   AUAAUCUGUUUUUUACGGUAUAAUUUACCGUCUAUUUUCAACGUGUCGCCAAUUAAAAAACGGCGGUGUAAUAUUAACU
env-445   GAAUAAUCUGUUUUUUACGGUAUAAUUUGCCGUCUAUUUUCAACGUGUCGCCAAUUAAAACGGCGGUGUAAUAUUAACU
Tvi-1-1   AUUUUUUCCCGGCGGAUCGCCAUCUUAGUGUUGAACUUCUGCCCCCUAUGCUAUAUUAUUCGCCACCAUUAUUUUUAAC
Tac-1-1   AUUUUUUCCUGCCUGAUUGCGAAGUUAAUGUUGAACUUCAUGGCUUGAUGCUUUAUUAUUCGGCCCAUUAUUUUUAAC
env-450   CUGAUUGCUUAAUUUUUUAGCAGUUUUUUAUUGCUAUGGCAGUAAUGAUGCUAUAACAUUCGCCUCCAUUAUUUUUAAC
env-452   CUCGUUGCUUAAUUUUUUAGCAGUUUUUUAUUGCUAUGGAGGAAAUGAUCCUAUAACAUUCGCCUCCAUUAUUUUUAAC
env-453   AUUUAUUGCUUAAAUUUCUAGCAAUUUUUAUUGCUAUGGUGGUAAUGAUCCUAUAACAUUCGCCUCCAUUAUUUUUAAC
env-454   CAAAAAUGCUCAAAUUUUUAGCAAUUUUUAUUGCUAUGGGGGUGAUGAUCCUAUAACAUUCGCCUCCAUUAUUUUUAAC
env-457   GUAAAUUGCUUAUUUUUUUAGCGGUUUUUAUUGCUAUGCGGGUGGGGAUCUUAUAAUAUUCGCCUCCAUUAUUUUUAAC
env-458   ACAAAUUGCUUAUUUUUUUAGCGGUUUUUAUUGCUAUGCGAAGGGGGAUCUUAUAAUAUUCGCCUCCAUUAUUUUUAAC
env-459   ACCUGAUUGUUCAAUUUAUAGCUUUUUUUAUUGCUAUAGAGAGCAUCAUGCUAUAACAUUCGCCUCCAUUAUUUUUAAC
env-460   CUUCAUUGCUUAUUUUUUUAGCAAUUUUUAUUGCUAUGGGGUAAAUGAUCCUAUAACAUUCGCCUCCAUUAUUUUUAAC
env-461   CUUUAUAGCUUAGUUUUGUAGCAAUUUUUAUUGCUAUGGGAGAGAUGAUGCUAUAACAUUCGCCUCCAUUAUUUUUAAC
env-462   AUUUAAUGAUUUUUAAGGGGUGUUUUAUUGUAAAUAACGCAGCAAACCUGUAUGAUUCGCCAGUUUUUUUAAUAUUAAU
env-463   AAAACAAGCGUAAUAAGCCUCUUGAAUGAUCAGUGAAGAUCGCAUAAAAUGCCGCGCUCUUAAACAUUGUUGUUUUUUU
env-466   AAAACAAGCGUAAUAAGCCUCUUGAAUGAUCAGCGAAGAUCGCAUAAAAUGCCGCGCUCUUAAACAUUGUUGUUUUUUU
env-467   ....................................................CCGCGCUCUUAAACAUUGUUGUUUUUUU
env-469   UACAAGGCAAUAGAUUCUGCUUGAUCACAAGCCGUUGAUCGCUUAAAAUGCGCGGCUCUUAAAUCAUUGAUGUUUUUUU
env-475   UACAAGGCAGCAGAUUCUGCUUGAUCACAGGCUCAGGAUCACAUAAAAUGCGCGGCUCUUAAUUCAUUGAUGUUUUUUU
env-477   UACAAGGCAGCAGAUUCUGCUUGAUCACAGGCGCAGGAUCACAUAAAAUGCGCGGCUCUUAAUUCAUUGAUGUUUUUUU
env-479   UACAAGGCAGCAGAUUCUACUUGAUCACAGGCUCAGGAUCACAUAAAAUGCGCGGCUCUUAAUUCAUUGAUGUUUUUUU
env-480   UACAAGGCAGCAGAUUCUGCUUGAUCACANGCUCAGGAUCACAUAAAAUGCGCGGCUCUUAAUUCAUUGAUGUUUUUUU
env-481   UACAAGGCAAUAGAUUAUGCUUGAUCACAAGCCGUGGAUCGCAUAAAAUGCGCGGCUCUUAAAUCAUUGAUGUUUUUUU

          ................................................................................
          ................................................................................
          ●●●●●●●●●●●●●●●●●●●●●●●●●●●●●●●●●●●●●●●●●●●●●●●●●●●●●●●●●●●●●●●●●●●●●●●●●●●●●RY●
```

```
Vru-1-1     AUGUGUUGAUCGAUUGCGGCGGAUUCAUUACUAUACUGCGCCUCGAACGAUGCAGAUUUCUGUAUCAUUAUUAAAAACU
env-482     AAAAAAUAGAUCUUGCACAAAUUGAUAAGUGGUUAUAUUAUGCGUGAGUUGCACGAUGCGGCGGUUUGUUAGUUUAAUU
env-483     CGGCGCAAGCGCGAUCCUGUGCUUGAUCAUUGCACAGGGUUCAGGUAGAAUCCGCGCCCUAAUUAUUCCAUUAUUAACC
Ama-1-1     UCAGGGAUCCUUGCUUGGGUGAGCGGCUUACUAUAGAAUUCGCGACCUAAAAUUUGUUAAGGCUCUGAUAGAGUUUAAU
Ata-2-1     UCGGGGAUCCUUGCUUGGUUGAGCGGCUUACUAUAGAAUUCGCGACCUAAAAAUUUGUUAAGGCUCUGAUAGAGUUUAAU
env-484     CCGCCGUCGAUACGGUAUUUGCGUAUCACGAUACCUGAAAAUCACUGAGAUUUUAGGUUUAUCAUUUAACUAUUAAUUC
env-485     CAGCAUUAAACAUUAGCCUGUAUAAUGUGCCGCCUUCGGUAAGAAAGACCUUGCCGGAGUUCACCAUUAAUGUUCAAUC
env-486     CAGCAUUAAACAUUAGCCUGUAUAAUAUGCCGCCUUCGGUAAGAAAGACCUUGCCGGAGUUCACCAUUAAUGUUCAAUC
Psp-49-1    CAAAAGCGCUCAUUUUCUGUAUACUACGCCGCCUUUGAUGAGAAUCGCUCGUCAAAGAGUACACAUUAAUUGUUCAACC
Psp-45-1    UGCUAAAAGUGGUUAAUUUCUGUAUACUAUGCCGCCUUCGAUGAGAUCGCUUGUCGGAGCAUACACUUAAUGUUCAAUC
Psp-53-1    UGCUAAAACCGGUUAAUUUCUGUAUACUAUGCCGCCUUCGAUGAGAUCGCUUGUCGGAGCAUACACUUAAUGUUCAACC
env-487     AGCAAUCGAAACUUACCUGUAUAAUAUGCCGCCUUCGGUGAGAAAUAUNUGGCCGGAGUACACCCUAUAAUGUUCAACC
env-488     AGCAAUCGAAACUUACCUGUAUAAUAUGCCGCCUUCGGUGAGAAAUAUUUGGCCGGAGUACACCCUAUAAUGUUCAACC
env-489     AGCAAUCGAAACUUACCUGUAUAAUAUGCCGCCUUCGGUGAGAAAUAUCUGGCCGGAGUACACCCUAUAAUGUUCAACC
env-490     ...........................................AAUAUCUGGCCGGAGUACACCCUAUAAUGUUCAACC
Pst-4-1     CAGCAGCAAACAUUAGCCUGUAUAAUAUGCCGCCUUCGGUAGGAAAGACCUUACCGGAGUACACCAUUAAUGUUCAAUC
Pst-3-1     CAGCAGCAAACAUUAGCCUGUAUAAUAUGCCGCCUUCGGUAGGAAAGACCUUACCGGAGUACACCAUUAAUGUUCAAUC
Pst-1-1     CACCAGCAAACAUUAGCCUGUAUAAUAUGCCGCCUUCGGUAGGAAAGACCUUACCGGAGUACACCAUUAAUGUUCAAUC
Psp-36-1    CAGCAAUCAGAAGUUAGCUGUAUAAUGUUGCGCCUUCGGUGAGAAAGAUCUUGCCGGAGUCCACCCUUUUUGUUCAACC
Pva-1-1     AGCAAUCGAAACUUACCUGUAUAAUAUGCCGCCUUCGGUGAGAAAUAUCUGGCCGGAGUACACCCUAUAAUGUUCAACC
env-493     ....................................GUGAGAAACGACCUUGCCGGAGUACACCCUAUAAUGUUCAACC
Pan-5-1     AGCAAUGAUAACUUCUCUGUAUAAUAUGCCGCCUUCGGUGAGAAAGACCUUGCCGGAGUACACCCUAUAAUGUUCAACC
Psp-43-1    UUGCCUCGGGCGGUUAAAUUCUGUAUACUAUGCCGCCUCUGGUGAGAUAGCUCGUCAGAGAUACAUAUAAUGUUCAACC
env-494     UUGCCUUGGGCGGUUAAAUUCUGUAUACUAUGCCGCCUUUGGUGAGAUGGCUCGCCAGAGAUACAUAUAAUGUUCAACC
Pva-1-2     UAAAAGCGGUUAUUUUCUGUAUAGUAUGCCGCCUUUGGUGAGAAAGGUCUCGCCAGAGCCUAAACAUUAAUGUUCAACC
Pse-1-1     UAAAAGCGGUUAUUUUCUGUAUAGUAUGCCGCCUUUGGUGAGAAAGGUCUCGCCAGAGCUUAAACAUUAAUGUUCAACC
Pro-1-1     UGCCUGGUGCGGUCAAAUUCUGUAUACUAUGCCGCCUUUGAUGAGACGACUCGUCAAAGAUACACUUAAUUGUUCAACC
Psp-46-1    UGCCUGGUGCGGUCAAUUUCUGUAUACUAUGCCGCCUUUGAUGAGAUGACUUGUCAAAGAUACACUUAAUUGUUCAACC
env-500     UGCCUGUAGCGGUCAAUUUCUGUAUACUAUGCCGCCUUUGAUGAGAUGACUCGUCAAAGCAUACACAUAUUGUUCAACC
Psp-38-1    UGCCUGGUGCGGUCAAUUUCUGUAUACUAUGCCGCCUUUGAUGAGAUGGCUUGUCAAAGAUACACUUAAUUGUUCAACC
env-501     .....................................................GUCAAAGAUACACUUAAUUAUUAACC
env-503     UUGCCUGGUGCGGUCAAUUUCUGUAUACUAUGCCGCCUUUGAUGAGAUGACUUGUCAAAGAUACACUUAAUUAUUAACC
env-504     ..................UUUCUGUAUACUAUGCCGCCUUUGAUGAGAUGACUCGUCAAAGAUACACCAUUUAUUAACC
env-507     UCAUGCCGGUGGGGUUCUGUAUAAUACAGCGCCUCUGUUGCCGCGAGUGGCUGCAGAGAUACGAAAAAUUCUUUUAACC
Tsp-2-1     UCCGGCGAACUUCUGUAUAAUAUAGCGCCUCUGUUGCCUGAAAGGUGGCAGGGAACUAAAAAAACAUUCUUUUUUAACC
Tsa-1-1     GGGGAUAGUCUGUAUAAUAUAGCGCCUCUGGUGCUGUAGCCGGCGAACAGAGAGCUAAAUAAAACAUUCUUUUUUAACC
Tpt-1-1     CCGGCGAACUUCUGUAUAAUAUAGCGCCUCUGUUGCCUGAAAGGUGGCAGGGAACUAAAAAAAACAUUCUUUUUUAACC
env-508     AGCAAUCGAAACUUACCUGUAUAAUAUACCGCCUUCGGUGAGAAAUAUCUGGCCGGAGUACACCCUAUAAUGUUCAAUC
Pag-5-1     AGCAAUCGAAACUUACCUGUAUAAUGUGCCGCCUUCGGUGAGAAAUAUCUGGCCGGAGUACACCCUAUAAUGUUCAAUC
Psp-39-1    AGCAAUCGAAACUUACCUGUAUAAUAUGCCGCCUUCGGUGAGAAAUAUCUGGCCGGAGUACACCCUAUAAUGUUCAAUC
Pag-2-5     AGCAAUCGAAACUUACCUGUAUAAUAUGCCGCCUUCGGUGAGAAAUAUCUGGCCGAAGUACACCCUAUAAUGUUCAAUC
env-509     GUCUAUUUUCUGUAUAAUAUGACGCCUUCGUAAGACAGCGUAAUUAUUACGCGGCGAAGACAUACACUUAUUAUUAAUC
Psp-48-1    CCAGGCGGGCUGGGCUGUAUAAUAUGCCGCCUUUGGUGAGGUGUUCCUCGUCAAAGAAAACAUAACUUAUUGUUCAAUC
env-511     UGUUGAAUAACAUUUUAAGAAGUGCUAGUAUGCACCGUCACAGUUUGCAAAGACUGUUGCAGAAGAUUUUGUUAAUCCU
env-512     GUAAGUGAACGGCAAAUGAGCGUGUGUUCUGCCAAUCUACAGAACAAAUAGCAAGCUCAGGCGCCUAGAUUAUGAAACA
env-513     GCUCUACUGGCCUUAUCUCAGAAUGCUUGAUCAAUAACCAACACUGGGGUAAAAUCCGCGCUCCGAAUUUAUAGUUAAU
env-515     CACUAGAAUUUGUUGACGUGUUGCUUGAUCAUUAGUCGAUAGCAAGGUAAAAUCCGCCGCUCUUUUAACCAAGCUAGUU
env-516     UGAUCGCGCUUGAUCCCACCGGGUGACUGGGGUAGAAUUCGCGACCUUAAUUUCGUAUUAAGCGCAGAUUUUUUAAACA
Isp-2-1     GAUGUUGCUUGAUCCUUAGCCAGCUCUGCGCUAGAAUCCGCGUCCUUAAAUUAGAUUUAAGUACGAAUUUUAUUAACAA
env-517     AAAUGACUCUGAAAAAAGCACGGCUUGAUCCUUGGCCUGCUUCAGGGUAAAAUCCGCCGCUCUUUUAACCAAGCUUGUU
env-518     CUUUAGUUGCUUCCUUUAUUAUGCGCCGCUCUUUUGCGGCUGACUGCAAUUUCUGCGGUUGGUCAUUGAAUUUAAAGUU
Asp-8-1     CUUUAGUUGCUUCCUUUAUUAUGCGCCGCUCUUUUGCGGCUGACUGCAAUUACUGCGGUUGGUCAUUGAAUUUAAAGUU
env-519     CUUACGAUGCUUCCUUUAUUAUGCGCCGCUCUUUUGCGGCUGACUGCAAUUUCUGCGGUCGGUCAUUGAAUUUAAAGUU
env-520     GAUCCCCGAAAGACGAUACAUUAAUAUGCGCGCCUCUUUGCCUCAAGGGCAAAAUGAGAAAGAAUUUUUGUAAUUAACA
Vrh-1-1     UCAUGGAUUCUUUAAUAUAGCGCGCCUUGAUGAUGCGGAUACCCGUAUCACAACCGAUGUAAGACGAUAUGAUUAACUG
env-521     CCUGUUUUUUCUGCGAUCAUACUUGCGGCUGUAGGGAUCGAUCCUAUAUUAUUCGCACCCUCAAAACAUCCGUUCCAAU
env-522     .................................GGGAUCGAUCCUAUAUUAUUCGCACCCUCAAAACAUCCGUUCCAAU
Spu-2-1     GUGAAAUGCUCAGAUAAUCUACUUGUAUCCGCAGUGACGAUACUGUACCAUUCUGCGCCCUAACAUAUCGUUAUUAACU
Ssp-5-1     GUUAAACGCUCAGAUAAUCUACUUGUAUCCGCCGUGGUGAUACUGUACUAUUCUGCGCCCUAACAUAUCGUUAUUAACU
Sba-3-1     CAAACGCUCAGAUAAUCUACUUGUAUCCCUUGUGGGUAUGCUGUACCAUUCUGCGCCCUAAACAUAUUGUUAUUAACAU
env-527     CAACCGCUCAGAUAAUCUACUUGUAUCCCUUGUAGGUAUGCUGUACCAUUCUGCGCCCUAAACAUAUUGUUAUUAACAU
Sma-1-1     AGUUUGACGCGCUGCUUUAAUGCUUGUAUUGCCCCCAAUGAUCCGGUAAUAUCCUGCGCCCUAAACAACUGUAUUAACC
Sco-1-1     GUUACUUAAUUAAUAGUGUGGGUGCUUGUAUCUAUUGUUGCUAUCCUGUAGUAUCUCGCGCCUGAAUAUUCUAUUAAUU
Ssp-4-1     UCGAGCAGAUGUAGUCACCUACUUGUCAGCGCUGGGCAGAUACUGUACUAUUCUGCGCCCUGAACUUUUGAUAUUAACC
Sam-2-1     GCUUUUUAAGCGACGAUCUGCUUGUCAGUAUGGGUGAGAUCCUGUACCAUUCUGCGCCCUAACAUACUUGUUAUUAACU
Son-1-1     AGAAAAUGCGCAGAUAAUCUACUUGUAUCCACUUAGGCGAUACUGUAAUAUCCUGCGCCCUAACAUAUCGUUAUUAACU
Ssp-10-1    AGAAAAUGCGCAGAUAAUCUACUUGUAUCCACUUGGGUGAUACUGUAAUAUCCUGCGCCCUAACAUAUCGUUAUUAACU
env-529     CGUUGAUUCCUUUAGCUGUUUUUUAUCUUGUAUCUAUUUUUACUUCAUGUACAAUUUCGCGCCUAAUAUUUUAUUAACU
Swo-1-1     UGUAAUAUUAAGCGUUAUAUUUCUCUUGUAACCCUUGAAUAGUUACUGUACUAUCUUGCGCCUAAAUAUUUUAUUAACA
Sse-1-1     CGUUGAAUCUUUGCCCUGCUUUUCUCUUGUAUCUCUUUUCCAUAUCCUGUAGUAUUUCGCGCCUAAUAUUUUAUUAACU
Sbe-1-1     CUAGGAACUAAAGAUCUGUUCCUCUCUUGCAUCUUUUUUUAAAUCCUGUAUCAUUUUGCGCCUGAAUAUUAUAUUAACC
Svi-1-1     GUCAGUGCGAAGUUAGGUUUUUACUCUUGCAUAUUUUUUUAAAUCCUGUAGUAUUUUGCGCCUGAAUAUUCUAUUAAUC
env-530     UAAAAUGCUCAGAUAAUCUACUUGUAUCCGCUUAGGUGAUACUGUAAUAUCCUGCGCCCUAACAUAUCCGUUAUUAACU
Ssp-7-1     UGAAAAUGCUCAGAUAAUCUACUUGUAUCCGCGUAGGUGAUACUGUACUAUUCUGCGCCCUAACAUAUCGUUAUUAACU
Sde-1-1     UGAAAAUGCUCAGAUAAUCUACUUGUGUCUGCGUGGGCGAUACUGUAGUAUUCUGCGCCCUAACAUAUCGUUAUUAACU
env-534     UGAAAAUGCUCAGAUAAUCUACUUGUGUCUGCGUGGGUGAUACUGUAGUAUUCUGCGCCCUAACAUAUCGUUAUUAACU
Ssp-6-1     UGAAAAUGCUCAGAUAAUCUACUUGUAUCCGCGUAGGUGGUACUGUAUUAUUCUGCGCCCUAACAUAUCGUUAUUAACU
Sma-2-1     GCAUUAGUUAAUGGUCAAUAAUUAGCUUGUAUCGCUGUGGUAGAUACUGUACUAUUGCGCGUCAAAAUUUCUAUUAACC
Spi-1-1     CUUGGAGUUUUAUUUACAGGUCGCUUGAAUCUCAGGGGCAGUUACUGUAUCAUCCUGCGCCCUAAACUAUCUAUUAACU
Ssp-3-1     GGGGGUGAAUAGUUGCCGGGGAUGCUUGUAUCUAUUGCUGCUAUCCUGUACUAUCUCGCGCCUAAAUAUUUUAUUAACU
Ssp-11-1    GCAGCAGAAUAGUUACAGGCUAUGCUUGUAUCUAUUGGUGGUAUCCUGUACUAUCUCGCGCCUAAAUAUUUUAUUAACU
Slo-1-1     GCAGGAGAAUAGUUGCAGGCUAUGCUUGUAACUAUUGAUGGUAUCCUGUACUAUCUCGCGCCUAAAUAUUCUAUUAACU
Swa-1-1     CAAAAAGCACGGGUUAAUUUCGAUGCUUGUAUCUUGUUGGCAGUUGCUGUAAUAUCCGCCGCCUAAUUAUUUAUUAACU
Sde-2-1     GCUAUCGCUGAUAAAUUCUACUUGUUUGCCAUGGGGGCUUACUGUAUUAUUGCGCGCCUAAAACACUCAGUUAUUAACU

            ...............................................................................
            ...............................................................................
            ●●●●●●●●●●●●●●●●●●●●●●●●●●●●●●●●●●●●●●●●●●●●●●●●●●●●●●●●●●●●●●●●●●●●●●●●●●●RY●
```

| | |
|---|---|
| env-535 | AUUUAUGGUGAUUCAUGACGGGUAGCUGGAUCUAUUGCUUAAAUGCUGUAUAAUCCUGCGCCCUAAACUAUUAUUAACU |
| Sha-1-1 | GAUUAAUCGUACAUUGUGGCGCUUGAAUCUACAAAUGAGAUCCUGUAGUAUUCGGCGCCUAAAUAAUUUUAUUUUAACU |
| Spe-1-1 | UUUAUUGAUCACGUAUUGCAUUGCUUGAAUCUAUCAUCUAUAUGCUGUAGUAUUCGCCGCCUAAAUAAUUUUAUUAACU |
| Sfi-1-1 | GAUUAAUCGUCCACUUUGUUGCUUGAAACUGCUAUAGAGUUGCUGUACUAUUCCGCGCCUAAAUAAUUUUAUUUUAACC |
| Sfr-1-1 | AGUAUCGAUUAGAAAUUCUACUUGUUUGUGUAUUAGAGACACUGUAUUAUCUCGCGCCUAAAACAUAUUGUUAUUAACU |
| CRe-1-1 | AACUUCAAAGGCGAAGGGCAUAUAUGGCCAGAUUGCGUCGAAGGUUAACAUCUUGUAUGAUCCGCGACUAUUGAUAACU |
| Eba-2-1 | UCGAUUGCAUGGUGAAAACAGAUUCUGUAUGAUUCGCCACCUUUGAUAUAAUUCCAUAUCAUUGGAUACAUUAUCCACC |
| Sgl-1-1 | UUCUGUUUGCAAAGAUUGGCGGCAUUCUGUAUGAUUCGCCGCCUUUGAUGUUAUCCGGCAUUAAAGCGAUAAACUCCAU |
| CSo-1-1 | UUGUGUUUGCAAAGAUUGGCGGCAUUCUGUAUGAUUCGCCGCCUUUGAUGUCAUCCGGCAUUAAAGCGAUAAACUCCAU |
| env-536 | UGUGUUUGCAAAUAGCCCCGGCAUUCUGUAUGAUUCGCCGCCUUUGCUGCCAUCCGGCAUCAGAGAAACAAUACACCCU |
| Spr-1-1 | UUGUGUUUGCAAAGAUUGGCGGCAUUCUGUAUGAUUCGCCGCCUUUGAUGUCAUCCGGCAUUAAAGCGAUAAACGCCAU |
| Psp-47-1 | GUGAAAAAGACGGCAGAUGGCUUGAAAGGGGUAGGCGAAUGCCGUAUCAUUCUGCGCCUUAUAACUGAGUUAUUAACCC |
| Aal-2-1 | GUUGUUGGGUGCUUGCUAUGCUAGGUGGCUUCUUAUAUGAUCCGCCACCUAAAAUUUAUAGGCUCUGAUAGAGUUGCUU |
| env-537 | ...........................GUUGAACCGGCAGGGCAGAUCUAUUAAGCUACGCGCCCUAAUUUAACUGAAAU |
| Pte-1-1 | UUUUUUCUGCGAAAUCGGAAUAAAAAGUUGAACCGGCAAGGCAGAUCUAUUAAGCUACGCGCCCUAAUUUAACUGAAAU |
| env-538 | GUUUUUUUCUGAAGGCGGUUUUUUUUGUUGAAUAAGGCCUGCUGAUCUAUUAAGCUACGCGCCCUAAUUUAACUGAAAU |
| env-539 | CUUUUUUUCCGAAGCCGGUUUUUUUUGUUGAAUAAGGCCUGCUGAUCUAUUAAGCUACGCGCCCUAAUUUAACUGAAAU |
| Tli-1-1 | UGUUUUUUCUGCGUCUUUACUUGCGGGUGUGAUGCCUAUCCUAUAUCAUUCUGACCCUUGAAAAUAUCCGUUUAAUGUU |
| Tau-1-1 | UGUUUUUUCUGCGCAUAUGCUUGCGGGGGUGGGGUCUAUCCUAUAUCAUUCUGACCCUUGAAAAUAUCCGUUUAACGUU |
| Ana-1-1 | UAUAAGCGGAUUUUCUGUAUGAUCCGCCACCUUUAUUAGUAAUUUUGAUGAUUACUAAGCACUAUUCUUAAUCGAUUUG |
| Aen-3-1 | GCGCUAUAAACAGAUUUUCUGUAUGAUCCGCCACCUUUGUUAGUAAUUUUGAUUACUAACACUAUUCUUAAUCGAUUUG |
| Asa-10-1 | CACGCGCAAAAUGAUCGAUUAAAUAUUGAUUAUUUGGUCGGCUUCAUUUAGAAUUCGCGCCUUUAUUUUAAGUUAUUAA |
| Iat-1-1 | CGAUCCAAUGCUUGAUCAUUACUUUGCCUUCGGGUAGAAUUCGCGCCCUAUUUAAUGACGAUAAGUCGUAAUUUUAACC |
| Ise-1-1 | GAUCCUGCCGUUGAUCCUUAUGCUUGAUCAAUGACCAGUGAUGGGGUAAAAUCCGCGCCCUAUUAAUUUGACAUUAACC |
| Isa-1-1 | CGCUUCAGGGUACUUGAUCAGCGACACCCGUUAGGGUAGAAUUCGCGCCCUUGUUAAACGAAUCGAUUAACUAUUAACC |
| Ixi-1-1 | GCUCUUAAAUGCCGCUGCUUUUUUAAGCUGAUACUGUCAUGCUUAGUCAUUUAGCCCGGUGUUUGUUAUUGAUACUUUU |
| env-540 | GGUGAUUUUUCCCCCGUUUGACCUUGCCUGUAAAAAGGCAAUCCUAUAUUAUUCGCACCCUUCACAAAACGUGUAUCAC |
| Osm-1-1 | UGAUUUUUUCAUCCGCAUUUUCCUUGCCUGAAAAAAAGGCGAUCCUAUAUUAUUCGCACCCUUCACAAAACGUGUAACUC |
| Osp-1-1 | UGGUUUUUUCAUCCCCAUUGUCCUUGCCUGAAAAAAAGGCGAUCCUAUAUUAUUCGCACCCUUCACAAAACGUGUAUCAC |
| env-541 | ..............................................................................AC |
| Ema-2-1 | GGCGCGUGGUUAAUUUCUGUAUAGUUCGCCGCCGUUGAUACGUUAAGUGGUAUCAGCAUACAGCUGUUAUUUUUAAUCU |
| Etr-1-1 | AGUGGUCGGUUAAUUUCUGUAUAAUUCGCCGCCGUUGAUACCUUUUUAUAGUAUCACGUACACCUGAAUUUUUUAAUUU |
| Psp-40-1 | UGCAGCGAGCGGUUACUUUCUGUAUAGUUCGCCGCCGUGAUACGGCAUUUUGGUAUCAUUACACCUGAAAUAUUAAUCU |
| Ein-1-1 | AGAGGGUGGUGAUUUUCUGUAUAGUUCGCCGCCUUUGAUACCGCGUUUCAGUAUCAAUGACACCUAAUUUUUUUAAUCU |
| env-542 | AGAGGAUGGUGAUUUUCUGUAUAGUUCGCCGCCUUUGAUACCGCGUUUCAGUAUCAAUGACACCUAAUUUUUUUAAUCU |
| Eto-1-1 | AGGCGCUGGCGACUUUCUGUAUAGUUCGCCGCCUUUGAUACCGUGUUUUAGUAUCAAUGACACCUAAUUUUUUUAAUCU |
| Esp-55-1 | UUGCAGCGAACGGUUACUUUCUGUAUAGUUCGCCGCCUUGAUACUGCAUUCAGGUAUCAAUUCAUCUGAAUUUUAAUCA |
| Eol-1-1 | UUGCAGCGAACGGUUACUUUCUGUAUAGUUCGCCGCCUUGAUGCUGCAUUCAGGUAUCAAUUCAUCUGAAUUUUAAUCA |
| Ebi-2-1 | AGCAGACGUUACCUUUCUGUAUAGUUUGCCGCCGUUGAUACGGCAUUAUUGGUAUCAAAACACCUGUUAUUAUUAAUCC |
| Ebi-1-1 | AGCAGACGUUACCUUUCUGUAUAGUUUGCCGCCGUUGAUACGGCAUUAUUGGUAUCAAAACACCUGUUAUUAUUAAUCC |
| bsy-1-1 | CGGUGUGUGGUUAAUUUCUGUAUAGUUCGCCGCCGUUGGUACGGCAUUUUGGUAUCAACUUCACUUGAAACAUUAAUCU |
| Epy-2-1 | GCAGCGGGUGGUUAAUUUCUGUAUAGUCCGCCGCCGUUGGUACGGCAUUUUAGUAUCAACUUCACCUAAAUAUUAAUCU |
| Esp-74-1 | GCAGCGGGUGGUUAAUUUCUGUAUAGUCCGCCGCCGUUGGUACGGCAUUUUGGUAUCAACUUCACCUAAAUAUUAAUCU |
| Eta-9-1 | GCAGCGGGUGGUUAUUUUCUGUAUAGUCCGCCGCCGUUGGUACGGCAUUUUGGUAUCAACUUCACCUAAUUAUUAAUUU |
| Epi-2-1 | GCAACGGGUGGUUAAUUUCUGUAUAAUUCGCCGCCGUUGGUACGACAUUUUGGUAUCAACUUCACUUAACUAUUAAUUU |
| Eam-1-1 | GCAGCGGAUGGUUAUUUUCUGUAUAGUCCGCCGCCGUUGGUACGGCAUUUUGGUAUCUACUUCACCUAAAUAUUAAUCU |
| Ety-1-1 | UGCAGCGGGCGGUUACUUUCUGUAUAGUUCGCCGCCGUUGAUGCGGCAUUUUGGUAUCAACACACCUGAAUUUUAAUCU |
| env-546 | UGCAGCGAGCGGUUACUUUCUGUAUAGUUCGCCGCCGUUGAUGCGGCAUUUUGGUAUCAACACACCUGAAUUUUAAUCU |
| Psp-44-1 | CAUCAGCCGGUUAUUUUCUGUAUAGUUCGCCGCCUUUGGUGCAAAACAGCGCAUCAAAAAUCACCAGAAUUUUUAAUCC |
| env-547 | UAAUCAGGAGUCAAAUAAAAUACGCGGCCUUAAGGGUGCUGAGCGGGUUAUAUCCAUGCUUUGGCACAUUGAAACUAAC |
| env-549 | UUAUCUGAAGAUGGUUAAAGAUCUUGCUAUUGGCAUGUUAGUUGGCUAGAAUUCGCGCCCUAUAUAUUUUUGUUUAACA |
| Ppi-1-1 | UUAAAUCACGGUGGUUAAAGAUCUUGCUAUUACGCAGUUUGUUGGCUAGAAUUCGCGCCCUAUAUAUUUUUGUUUAACA |
| Pci-1-1 | AUUUAGCUAAAUAGUGUAUUUAGUGCUUGUAGAUUGUGCUGUAGUUACAUAAAAUACGCGUCCAAUUAUUUGUAUAACU |
| Plu-2-1 | AUCUAGUUAGAUACUAAUUUAUUGCUUGUAGUUUGAGCAUGAAAGUCCUAAAAUACGCGUCCAAUUUAUUUGUAUAACG |
| Plu-3-1 | AGGGAAUAAAAGAUCAUUUUCUGCUUGUCGAUUGUACAGAUCAUGCAUUAGAAUACGCGACCAAUUAUUUUGUUUAACC |
| env-550 | UUUUAGCGAAAGAUGCAUUUAAAGCUUGUCGCAGAGCGUAUUGGCCUAUAAAAUACGCGGCCUAUUAAUUGUAUUAACG |
| Ppi-3-1 | GGUGAUAAAAAACUCUGAAAUAUUGCUUGUAGAUUGUUCAGUUAGGCACUAAAAUACGCGUCCAUAUUUUUGUAUAACU |
| Ppi-2-1 | GGUGACAAAAAACUCUGAAAUAUUGCUUGUAGAUUGUUCAGAUAGGCACUAAAAUACGCGUCCACAUUUUUGUAUAACU |
| Pfl-1-1 | GGUGAUAAAAAACUCUGAAAUAUCGCUUGUAGAUUGUUCAGUUAAGCACUAAAAUACGCGUCCACAUUUUUGUAUAACU |
| Pel-1-1 | GGCGAUAAAAAACUCUGAAAUAUUGCUUGUAGAUUGUUCAGUUAAGCACUAAAAUACGCGUCCACAUUUUUGUAUAACU |
| env-551 | UUUUAGCGAAAGAUGUAUUUAAAGCUUGUCGUAAGGCGUCUUGAGCUAUAAAAUACGCGGCCUAUUAAUUGUAUUAACU |
| env-552 | UUUUAGCGAAAGAUGUAUUUAAAGCUUGUCGUAAGGCUUCUUGAGCUAUAAAAUACGCGGCCUAUUAAUUGUAUUAACU |
| env-559 | UUUUAGCGAAAUAUGUAUUUAAAGCUUGUCGCAAGGUCACUUGAGCUAUAAAAUACGCGGCCUAUUAAUUGUAUUAACU |
| env-566 | ..........................UUGUCGCAAGGUCACUUGAGCUAUAAAAUACGCGGCCUAUUAAUUGUAUUAACU |
| Psp-17-1 | UUUAGCAAAAAGGGUAAUUAAAGCUUGUGUGCGUGCGGGUUGUUGGAUAAAAUUCGCGGCCAAUAAAUUUGUUUUAACG |
| env-572 | UUUAGCAAAAAGAAUAAUUAAAGCUUGUGUGCGUGCUGGUUGUUGGAUAAAAUUCGCGACCAAUAAAUUUGUUUUAACG |
| env-573 | UUUGGCAAAAAGCGUAAUUAAAGCUUGUGCGGGGCAAGCUAGUUGGAUAAAAUCCGCGGCCUAUAAAUUUGUUUUAACG |
| env-589 | UUUAGCAAAAAGGGUAAUUAAAGCUUGUGUGCGUGCGGGUUGUUGGAUAAAAUUCGCGACCAAUAAAUUUGUUUUAACG |
| env-591 | UUUUAGCUAAAGAUGAAUUUAAAGCUUGUCGAGAGGCUGCUUGAGCUAUAAAAUACGCGACCUAUUAAUUGUAUUAACG |
| Psp-6-1 | UUUAGCAAAAAGGGUAAUUAAAGCUUGUGUGUGUGUGGGUUGUUGGAUAAAAUUCGCGACCAAUAAAUUUGUUUUAACG |
| env-593 | UUUAGCGAAAAGAUAAAUAAAUACUUGUGCAGGGGAAGGUUGUUGGAUAAAAUCCGCCACCAACAAAUUUGUUUUAACG |
| Psp-26-1 | UUUGGCUAAAACAUAAAUAAAUACUUGUGUAGGGGAACGUUGUUGGAUAAAAUCCGCCACCAACAAAUUUGUUUUAACG |
| env-594 | UUUAGUAAAAAGAAUAAUUAAAGCUUGUGUGCGUGCGGGUUGUUGGAUAAAAUUCGCGACCAAUAAAUUUGUUUUAACG |
| Aba-1-1 | UUUAGCAAAACAGUUAAUAAUAGCUUGUGUGUUUGAUGGUAGUUGGAUAAAAUCCGCGACCAACAAAUUUGUUUUAAUG |
| env-595 | UUUAGCAAAACAGUUAAUAAUAGCUUGUGUGUUUGAUGGUAGUUGGAUAAAAUCCGCGACCAACAAAUUUGUUUUAACG |
| Pha-2-1 | UUAGCAAAAACGCUAUUUAAAGCUUGUAGCAAGCGGGGUUGUUGCAUAGAAUACGCGUCCUAAAUAAUUUGUUUUAACG |
| Psp-11-1 | UUUAGCAAAAACAUAAAUAAAUACUUGUGUAGGGCUUGGUUGUUGGAUAAAAUCCGCGACCAACAAAUUUGUUUUAACG |
| Psp-32-1 | UUUUAGCAAAAAAUAUAAUUAAAACUUGUGUGCGGGGUUGUUGUUGGAUAAAAUCCGGCUCCAACAAUUUGUUUUAACG |
| Psp-19-1 | UUAGCAAAAAAUUAUAAUUAAACCUUGUGUGCUCGGUAGUUGUUGGAUAAAAUCCGCCACCAACAAAUUUGUUUUAACG |
| env-599 | UUUAGCAAAAAUACUAAUUAAUGCUUGUGUGCUUGAUGGUUGUUGGAUAAAAUCCGCCACCAAUAAAUUUGUUUUAACG |
| Pru-1-1 | AUAGCAGUUAUUUUGCAUUUAUUGCUUGUGGAUCGCCCGGAUUGACCCUAAAAUACGCGUCCACAAUUUUUGUAUAACU |
| Pru-1-2 | AUAACAGUUAAAUUGUAUUUAUUGCUUGUGGAUCGUCCCGAUUGACCCUAAAAUACGCGUCCACAAUUUUUGUAUAACU |
| Pru-2-1 | AUAACAGUUAAAUUGCAUUUAUUGCUUGUGGAUCAUUCCGAUUGACCCUAAAAUACGCGUCCACAAUUUUUGUAUAACU |
| Plu-4-1 | AUCUAAUGGAUUACUAAUUUAUUGCUUGUAGUUUGAGCAUGAAAGACCUAAAAUACGCGUCCAAUUUAUUUGUAUAACG |
| | ................................................................................ |
| | ................................................................................ |
| | ●●●●●●●●●●●●●●●●●●●●●●●●●●●●●●●●●●●●●●●●●●●●●●●●●●●●●●●●●●●●●●●●●●●●●●●●●●●●●RY● |

```
Plu-1-1    AUCUAAUGGAAUACUAAUUUAUUGCUUGUAGUUUGAGCAUGAAAGACCUAAAAUACGCGUCCAAUUUAUUUGUAUAACG
Par-1-1    UUUGGCAAAAAUAUAAAUAAAUACUUGUGUGGGGGAGGUUGUUGGAUAAAAUCCGCCACCAACAAAUUUGUUUUAACG
env-601    AUUUAGUUUUUGUGCAAUUAUUGCUUGUAGAUCUCAGGCGACUCCAAUAAAAUACGCGUCCACUAAUUUGAAUAUAACU
env-603    AUUUACUUUUUGUGCAAUUAUUGCUUGUAGAUUUCGGACGACUCCAAUAAAAUACGCGUCCACUAAUUUGAAUAUAACU
Psp-1-1    UAAUGCUUGAAGAUCUUGGGUGAUUUGUCUAAAAUACGCGCCCUAACUUAUCAAUCAAUGAUUGAUGUAUAAAUUAACA
Ptu-1-1    UAACUUGACUUAAGAUAUAAAUCUUGUGAAGUGCGUCGGAUUUGUCUAAUAUACGCGCCCUAACUACAUUUGUUUAACA
env-606    ............................................................CGUCCACAAUUUUGUAUAACU
Eca-5-1    UGAAAUGAAUUGAACAAAAAAGUUGCGAUCCUAGGUCUCCUUGAUUAGAAUUUCGCGCCUUACAAUUUGUUUGUUAACG
Eca-6-1    AAUAAAUGAAUUGAACAAAAAAGUUGCUAUCACAGGUCUCCUUGAUUAGAAUUUCGCGCCUUACAAAUUUUUGUCAACG
Gho-1-1    GAAAAUGAAUUGAACAAAAAAGUUGCGAUCGACGGUCACCUUGAUUAGAAUUUCGCGCCUUACAAACUGUUUGUUAACG
Gsp-4-1    UGGAAAUGAAUUGAACAAAAAAGUUGCGAUCGCAGGUCACCUUGAUUAGAAUCCGCGCCUUACAAACAGUUUGUUAACG
Gsp-3-1    UGGAAAUGAAUUGAACAAAAAAGUUGCGAUCGCAGGUCACCUUGAUUAGAAUCCGCGCCUUACAAACAGUUUGUUAACG
Eno-3-1    UCAAAUGAAUUGAACAAAAAAGUUGCUAUCAAAGGUCUCCUUGAUUAGAAUUUCGCGCCUUACAAAAUGUUUGUUAACG
env-607    .........................................ACCGAUUAUAAUUCGCGCCCUAAAUUUAUUUAAACU
env-608    AACCCGCAGCGCGAUACGCGAAGGUGCUUGAUCAAAAUGCCGCUGUGCAGUACAAUUCGCGCCCUUAUUAACAUUAAUC
Tsp-1-1    GAUAAAUUUGACUGUUUUUAACGGUUUUAGUUGUACAUUGGCCAAUGUUCCCGUAUUAUUCGCCACCUUAAAUUGAAAC
Pha-5-1    UUGCGCGUUUAGUGGAUAUUCUGUAUGAUCCGCCCCCUUUGGUAGAUAAAUUACCAAACACUUAAUUUCAAUCAACACG
Pvu-2-1    UUGCGCGUUUAGUAGAUAUUCUGUAUGAUCCGCCCCCUUUGGUAGAUAAAUUACCAGACACUUAAUUUCAAUCAACACG
Pmi-3-1    UUGCGCGUUUGGCGGAUAUUCUGUAUGAUCCGCCCCCUUUGGUAGAUAAAAUACCAAACAAUUUAAUUCAAUCAACACG
env-611    GGUGAUUUAUCCUCCAUUUAUCCUUGCCUGUAAAAAAGCGAUCCUAUAUUAUUCGCACCCUUCACUAAAUGUGAAUCAU
Ffu-1-1    UUACUAUGCAGAGCAUUUCUCUCCUUGCGGGGGUUGAUUGCAUCGUGUACUAUUACGCGCCUUUAAUUUAUCACUAACG
Fky-1-1    GAUACUAUGGAGGCGGAUUGAUGCUUGCGGCGGUUGGUUACAUCAUGUACCAUUACGCGCCUUAAAAUUGUCAAUAACG
Asi-1-1    CUAUUUUUUCUGCGAAGACACUUGCCUGAGCUCAGGCGUUUCCUAUAUUAUUCGCUGCCUUUGAAAUUGUGUUCAAUAU
Afi-1-1    CUAUUUUUUCUGCGAAGCCACUUGCCUGAUCUCAGGCAUUUCCUAUAUUAUUCGCAGCCUUUGAAACUGUGUUCAACAA
Aja-1-1    CUAUUUUUUCUGCGAAGCCACUUGCCUGAUCUCAGGCAUUUCCUAUAUUAUUCGCAGCCUUUGAAACUGUGUUCAACAA
Aja-1-2    CUAUUUUUUCUGCGAAGCCACUUGCCUGAUCUCAGGCAUUUCCUAUAUUAUUCGCAGCCUUUGAAACUGUGUUCAACAA
Aau-1-1    CUAUUUUUUCUGCGAAGCCACUUGCCUGAACUCAGGCAUUUCCUAUAUUAUUCGCAGCCUUUGAAACUGUGUUCAACAA
Amo-1-1    CUAUUUUUUCUGCGAAGCCACUUGCCUGCUAUCAGGCAUUUCCUAUAUUAUUCGCAGCCUUUGAAACUGUGUUCAACAA
Api-1-1    CUAUUUUUUCUGCGAAGCCACUUGCCUGAUCUCAGGCAUUUCCUAUAUUAUUCGCAGCCUUUGAAACUGUGUUCAACAA
env-612    CUAUUUUUUCUGCGAAGCCACUUGCCUGAACUCAGGCAUUUCCUAUAUUAUUCGCAGCCUUUGAAACUGUGUUCAACAA
Abi-1-1    CUAUUUUUUCUGCGAAGCCACUUGCCUGCCAUCAGGCAUUUCCUAUAUUAUUCGCAGCCUUUGAAACUGUGUUCAACAA
env-613    CUAUUUUUUCUGCGAAGCCACUUGCCUGAUCUCAGGCAUUUCCUAUAUUAUUCGCAGCCUUUGAAACUGUGUUCAACAA
Ave-8-1    CUAUUUUUUCUGCGAAGCCACUUGCCUGAUAUCAGGCAUUUCCUAUAUUAUUCGCAGCCUUUGAAACUGUGUUCAACAA
Aal-1-1    CUAUUUUUUCUGCGAAGCCACUUGCCUGAUCUCAGGCAUUUCCUAUAUUAUUCGCAGCCUUUGAAACUGUGUUCAACAA
env-614    CUAUUUUUUCUGCGAAGCCACUUGCCUGAACUCAGGCAUUUCCUAUAUUAUUCGCAGCCUUUGAAACUGUGUUCAACAA
env-616    CUAUUUUUUCUGCGAAGCCACUUGCCUGAUCUCAGGCGUUUCCUAUAUUAUUCGCAGCCUUUGAAACUGUGUUCAACAA
Ata-1-1    CUAUUUUUUCUGCGAAGCCACUUGCCUGAUCUCAGGCAUUUCCUAUAUUAUUCGCAGCCUUUGAAACUGUGUUCAACAA
Ari-1-1    CUAUUUUUUCUGCGAAGCCACUUGCCUGCUAUCAGGCAUUUCCUAUAUUAUUCGCAGCCUUUGAAACUGUGUUCAACAA
Ate-1-1    CUAUUUUUUCUGCGAAGCCACUUGCCUGAACUCAGGCAUUUCCUAUAUUAUUCGCAGCCUUUGAAACUGUGUUCAACAA
Ame-1-1    CUAUUUUUUCUGCGAAGCCACUUGCCUGAACUCAGGCAUUUCCUAUAUUAUUCGCAGCCUUUGAAACUGUGUUCAACAA
Aaq-2-1    CUAUUUUUUCUGCGAAGCUACUUGCCUGAUCUCAGGCAUUUCCUAUAUUAUUCGCAGCCUUUGAAACUGUGUUCAACAA
env-619    .........CUGCGAAGCCACUUGCCUGAUCUCAGGCAUUUCCUAUAUUAUUCGCAGCCUUUGAAACUGUGUUCAACAA
Aeu-1-1    CUAUUUUUUCUGCGAAGCCACUUGCCUGAUCUCAGGCAUUUCCUAUAUUAUUCGCAGCCUUUGAAACUGUGUUCAACAA
Asc-1-1    CUAUUUUUUCCGCGGGGCCACUUGCCUGAUCUCAGGCGUUUCCUAUAUUAUUCGCUGCCUUUGAAACUGUGUUCAAUAU
Adi-2-1    CUAUUUUUUCCGCGGGGCCACUUGCCUGAGCUCAGGCGUUUCCUAUAUUAUUCGCUGCCUUUGAAACUGUGUUCAAUAA
env-620    GCGUGGUGGUUGCACAAAAAAGUUGCGAAACCGGAGGGGCUUGAUUAGAAUUGCGCGCCUUAGAAAAUAUUUGUUAACG
env-621    GCGUGGUGAUUGCACAAAAAAGUUGCGAAACCGAGGGGGCUUGAUUAGAAUUGCGCGCCUUAGAAAAUAUUUGUUAACG
env-623    GCGCUAUGAUUGCACAAAAAAGUUGCGAAAAAGAGGGCGCUUGAUUAGAAUUGCGCGCCUUAGAAAAUGUUUGUUAACG
Ssp-27-1   ACACGCGGAUUGCACAAAAAAGUUGCGAAAAAGAGGGCGCUUGAUUAGAAUUGCGCGCCUUAGAAAAUGUUUGUUAACG
Psp-57-1   GAUUGAUUACUAUACGCCGCCUGAAUUUAGCGUAUUGAUAUCAGCUUGCUGAUGGUAAUACGUUAUCCUGUUAUUAAUG
Pan-11-1   CUGUAUUGAUCUCCGCGGGAUGAUUGAUUACUAUACGCCGCCUGAUUUGUCGCAUAAUGUGGCAAGCUAUUUUUUAACG
Pan-12-1   CUGUAUUGAUCUCCGCAGGAUGAUUGAUUACUAUACCCCGCCUGAUUUGUCACAUAAUGUGGCAAGCUAUUUUUUAACG
Pan-11-2   CUGUAUUGAUCUCCGCGGGAUGAUUGAUUACUAUACGCCGCCUGAUUUGUCACAUAAUGUGGCAAGCUAUUUUUUAACG
Psp-58-1   CUGUAUUGAUCUCCGCGGUAUGAUUGAUUACUAUACGCCGCCUGAUUUGUCACAUAAUGUGGCAAGCUAUUUUUUAACG
Pap-1-1    UGGACGAGACGAUUGAUUACUAUACGCCGCCUAAUUUGAUGGAUUGGUCACGGCCUGUCCAUUGACUAUUUUUUUAAUG
Psa-1-1    AGGGACGAUUGAUUACUAUACGCCGCCUGAAUUCGGCGUAUUGGCUUAUUUGCUGCCGUUACGCUUACUGUUAUUAACG
Paq-2-1    CUGGUGGACAUGAUUGAUUACUAUACGCCGCCUGAAUUUAGCAUAUCGGCAGUCGCCAAUAUGCUUACUUUUAUUAACG
Pga-2-1    UCUGGGCUGCGCGAUUGAUUACUAUACGCCGCCUGAAUUUGGCGUAUUGGCAAUGUCGCCGUUACGCUUGUUAUUAACG
Pga-1-1    UCUGGGCUGCGCGAUUGAUUACUAUACGCCGCCUGAAUUUGGCGUAUUGGCAAUGUUGCCGUUACGCUUGUUAUUAACG
Ple-1-1    CUCUGUUGAUCUCCGUGAGAUGAUUGAUUAUUAUACGCCGCCUGAUUUGUCGCAUAAUGUGGCAAGAUAUUUUUUAACG
Pda-4-1    CGUAUUAGUUGAUCUGUGACGUGUGAUUGAUUACUAUACGCCGCCUAAUUUGUCACAUCAGUGGCAACUAUUUUUAACG
Psw-1-1    UGAUUAGUUGAUCUGUGCGGUGUGAUUGAUUACUAUACGCCGCCUGAAUUAUCGCCUUGCGCGGUAACUAUUUUUUAAC
Pph-1-1    GAGUUGAUCUAUUGAGGAUGAUUGAUUACUAUACGCCGCCUGAUUUGUUGUAAAUUUACGACAGACUAUUUUAUUAACC
Pki-1-1    GAGUUGAUCUAUGGGGAAUGAUUGAUUACUAUACGCCGCCUGAUUUGUUGUAAAUCUACGACAGACUAUUUUAUUAACC
Pil-1-1    GAGUUGAUCUCUUGAUAAUGAUUGAUUACUAUACGCCGCCUGAUUUGUCGUAAAUUUACGGCAGACUAUUUUAUUAAUC
Pki-1-5    GAGUUGAUCUAUGGGGAAUGAUUGAUUACUAUACGCCGCCUGAUUUGUCGUAAAUCUACGACAGACUAUUUUAUUAACC
Pha-6-1    GGUUGAUCUGUGUCGGCUGAUUGAUUAUCAUACGCCGCCUAUUUGUCCAGCGCCGUUGGCCUGGCAGUAUUUAUUAACC
Pga-3-1    GGUUGAUCUGUUUUGACUGAUUGAUUAUCAUACGCCGCCUAUUUGUCCAGUGCCGUCGGCCUGGCAGUAUUUAUUAACC
Pga-4-1    UUUGGAGGGCGUGAUUGAUUACUAUACGCGGCCUGAAUUUGGCGUAUUGGCCAUACCUAUACGCUGUUUGUUAUUAACG
Ppr-1-1    GUGUCUGUUGAUCUGAGCGUAAUGAUUGAUUAUCAUACGCGCCCUAAUUUGUCGUUUAUGCGGCAUACUAUUUUUAACG
env-624    GUGUCGGUUGAUCUGAGCGUAAUGAUUGAUUACUAUACGCGGCCUAAUUUGUCGUUUAUGCGGCAUACUAUUUUUAAUG
Ppr-2-1    GUGUCUGUUGAUCUCCGUGGAAUGAUUGAUUAUCAUACGCGCCCUAAUUUGUCGUUUAUGCGGCAUACUAUUUUUAACG
Vez-1-1    CAGUUGAUCCCUAAAGACUAAUUCAUUACUAUACUGCGCCUUGAAGAUGCUGUAUUACGCGGUGUCACUGUUAGAAACU
Vha-1-1    CAGUUGAUCCUUAAAGACUAAUUCAUUACUAUACUGCGCCUUGAAGAUGCUGUAUUACGCGGUGUCACUGUUAGAAACU
env-625    AAUGAGUUGAUUCCAAUGUCUUGUUUCAUUAAUAUACUGCGCCUUGACGGUAUAUCAAUUUAUGCCAAUUUGUGUAAAU
Vpr-1-1    CGAGUGAUUCAUUACUAUAUCGCGCCUUAGGAUAUGGCUUGUCAUACGACAGCGGUAUCCGAACUGUGUUUUUUUAACU
Vbr-2-1    AAGUUGAUCCCAACAGACUAUUUCAUUAGUAUACUGCGCCUUGAAGAUGUCGUAUUAAACGAUAUCACUGUUAGAAACU
Vbr-4-1    AAGUUGAUCCCAACAGACUAUUUCAUUACUAUACUGCGCCUUGAAGAUGUCGUAUUAAACGAUAUCACUGUUAGAAACU
Fse-1-1    GCAGAAAAUCGCCAGCGAUGAUUGCUCUGAACCGGGCUUUACUAUAGAAUCCUGCGCCCUGAACCUAUUUGUAAACCAU
Vmy-1-1    AUGCGAUUCAUUAGUAUAUCGCGCCUUGAGAUAUGGCUUCGGCAAUCAGCCAAAGCGGUAUCGACUUUUGAUGUUAAUU
Vsp-27-1   AGCCGAUUCAUUACUAUAUCGCGCCUUAAGAUAUGGCUUCGCAUCAGCGAGCGCGGUAUCUAAGAGUUCAUUUUUAACU
Vna-2-1    AGCCGAUUCAUUACUAUAUCGCGCCUUAAGAUAUGGCUUCGCAUUAGCGAGCGCGGUAUCUAAGAGUUCAUUUUUAACU
Vna-2-3    AGUCGAUUCAUUACUAUAUCGCGCCUUAAGAUAUGGCUUCGCAUUAGCGAUCGCGGUAUCUAAGAGUUCAUUUUUAACU
Vvu-8-1    GAGACGAUUCAUUACUAUAUCGCGCCUUAAGAUAUGGCUUUAAUUCUGUUUCAGCGGUAUCUGAGUUUUAUUUUUAACU

           ...............................................................................
           ...............................................................................
           ............................................................................RY.
```

```
Vsa-1-1      CAGUGAUUCAUUAUCAUACUGCGCCUUAAGAUAUGGCUUCAGCUAUAAGCAGAUGCGGUAUCAUAUUUUUAUAUUAAUU
Vaz-1-1      CAGUGAUUCAUUAUUAUACUGCGCCUUGAGAUAUGGCUUCAGCUAUAAGCAGAUGCGGUAUCACAUUUUUAUAUUAAUU
Vro-3-1      GGUGAUUCAUUACUAUACUGCGCCUUGAGAUAUGGCUUCGGCAAUUGGUCGAUGCGGUAUCGCAUUUUUGAUAUUAAUU
Vro-2-1      GGUGAUUCAUUACUAUACUGCGCCUUGAGAUAUGGCUUCGGCAAUUGGUCGAUGCGGUAUCGCAUUUUUGAUAUUAAUU
Vhy-1-1      AAGUGAUUCAUUACUAUACUGCGCCUUGAGAUAUGGCUUUGGCAAUUGGCCGAUGCGGUACUCAUUUCCAUUAUUAAUU
Vja-1-1      AAGUGAUUCAUUACUAUACUGCACCUUGAGAUAUGGCUUCGGCAAUUGGCCGAUGCGGUACUCAUUUCUAUUAUUAAUU
Vhy-1-2      AAGUGAUUCAUUACUAUACUGCGCCUUGAGAUAUGGCUUCGGCAAUUGGCCGAUGCGGUACUCAUUUCCAUUAUUAAUU
Vna-1-1      UGCGAUUCAUUAGUAUAUCGCGCCUUGAGAUAUGGCUUCAGCAUUAUGCUGAUGCGGUAUCGCACUUUUGAUAUUAAUU
Vja-4-1      AAGUGAUUCAUUACUAUACUGCGCCUUGAGAUAUGGCUUCGACAAUUGGCCGAUGCGGUACUCAUUUCUAUUAUUAAUU
Vpa-87-1     AGGGGAUUCAUUACUAUAUUGCGCCUUGAGAUAUGGCUUCAGCAAUUGGUUGGUGCGGUAUCCCAUUUUAUUAUUAAUU
Val-2-1      AUGUGAUUCUUUACUAUAUCGCGCCUUGAGAUAUGGCUUCGGCAAUUAGUUGAUGCAGUAUCCCAUUUUAUUAUUAAUU
Vpa-240-1    AGGGGAUUCAUUACUAUAUCGCGCCUUGAGAUAUGGCUUCAGCAACUGGUUGGUGCGGUAUCCCAUUUUAUUAUUAAUU
Vpa-238-1    AUGUGAUUCUUUACUAUAUCGCGCCUUGAGAUAUGGCUUCAGCAAUUGGUUGGUGCGGUAUCCCAUUUUAUUAUUAAUU
Vpa-94-1     AGGGGAUUCAUUACUAUAUCUCGCCUUGAGAUAUGGCUUCAGCAAUUGGUUGGUGCGGUAUCCCAUUUUAUUAUUAAUU
Vhy-1-3      AAGUGAUUCAUUACUAUACUGCGCCUUGAGAUAUGGCUUCGGCAAUUGGCCGAUGCGGUACUCAUUUCCAUUAUUAAUU
env-626      .................................................................UUUUAUUAUUAAUU
Vpa-157-1    AGGGGAUUCAUUACUAUAUCGCGCCUUGAGAUAUGGCUUCAGCAACAGGUUGGUGCGGUAUCCCAUUUUAUUAUUAAUU
Vro-1-1      GGUGAUUCAUUACUAUACUGCGCCUUGAGAUAUGGCUUCGGCAAUUGGUCGAUGCGGUAUCGCAUUUUUGAUAUUAAUU
Vpa-79-1     AGGGGAUUCAUUACUAUAUCGCGCCUUGAGAUAUGGCUUCAGCAAUUGGUUGGUGCGGUAUCCCAUUUUAUUGUUAAUU
Vpa-207-1    AGGGGAUUCAUUACUAUAUCGCGCCUUGAGAUAUGGCUUCAGCAAUUAGUUGGUGCGGUAUCCCAUUUUAUUAUUAAUU
Vpa-219-1    AUGUGAUUCUUUACUAUAUCGCGCCUUGAGAUAUGGCUUCGGCAAUUAGUUGAUGCGGUAUCCCAUUUUAUUAUUAAUU
Vha-9-1      AGUGAUUCAUUACUAUACUGCGCCUUGAGAUAUGGCUUCGGCGAUUGGCCGAUGCGGUAUCGCACUUUUAUUAUUAAUU
Vow-5-1      AGUGAUUCAUUACUAUACUGCGCCUUGAGAUAUGGCUUCGGCGAUUGGUCGAUGCGGUAUCGCACUUUUAUUAUUAAUU
Vca-3-1      UAGUGAUUCAUUACUAUACUGCGCCUUAGGUAUGGCUUCGGCAUUUGGUCGAUGCGGUAUCGCACUUUUAUUUUUCAUU
Vow-3-1      UAGCGAUUCAUUAGUAUACUGCGCCUUGAGAUAUGGCUUCGGCAAAUGGUCGAUGCGGUAUCGCAUUUUAUUAUUAAUU
Vsp-10-1     UAGCGAUUCAUUAGUAUACUGCGCCUUGAGAUAUGGCUUCGGCAAAUAGUCGAUGCGGUAUCGCAUUUUAUUAUUAAUU
env-627      UAGUGAUUCAUUACUAUACUGCGCCUUAGAUAUGGCUUCGGCAUUUGGUCGAUGCGGUAUCGCACUUUUAUUUUUCAUU
Vsp-2-1      UAGUGAUUCAUUAAUAUACUGCGCCUUAGAUAUGACUUCGGCACUUAGCCGAUGCGGUAUCGCACUUUUAUUAUUCAUU
Vsp-44-1     CAUUACUAUACUGCGCCUUAAAGGUAUGGCUUUAUCGCAGUUUAGUGAUGCAGCGGUGUCUUUAUUUAUUUUUUAACU
Vcy-2-1      UUCAUUAUUAUACUGCGCCUUAAAGGUAUGGCUUUAUCUCUAUUUAGUGAUACAGCGGUGUCUUUAUUUAUUUUUUAACU
Vcy-15-1     CAUUACUAUACUGCGCCUUAAAGGUAUGGCUUUAUCUCUAUUUAGUGAUAUAGCGGUGUCUUUAUUUAUUUUUUAACU
Vsp-45-1     CAUUAAUAUACUGCGCCUUAAAGGUAUGGCUUUAUCUCUAUUUAGUGAUGCAGCGGUGUCUUUAUUUAUUUUUUAACU
Vcy-13-1     UCAUUAUUAUACUGCGCCUUAAAGGUAUGGCUUUAUCUCUAUUUAGUGAUACAGCGGUGUCUUUAUUUAUUUUUUAACU
Vsp-5-1      AUUACUAUACUGCGCCUUAAAGGUAUGGCUUCGUCACUUUUUUGUACGGAUGCGGUGUCUUUAUUUCUUUUUUUAAACU
Vcr-1-1      CAUUACUAUACUGCGCCUUAAAGGUAUGGCUUCAUCACUCUUUUGUGUCGAUGCGGUGUCUUUAUUUCUUUUUUUAAACU
Vba-1-1      UCAUUACUAUACUGCGCCUUAAAGGUAUGGCUUCGUCGCGCUUUAGUGAUGAGCGGUGUCUUUAUUUCUUUUUUUAAACU
Vsp-21-1     AUUACUAUACUGCGCCUUAAAGGUAUGGCUUCAUCACUCUUUUGUGUCGAUGCGGUGUCUUUAUUUCUUUUUUUAAACU
env-630      UUCAUUACUAUACUGCGCCUUAAAGGUAUGGCUUCAUCGCGUUUUAGUGAUAAGCGGUGUCUUUAUUUCUUUUUUAAACU
Vsp-35-1     CAUUACUAUACUGCGCCUUAAAGGUAUGGCUUCGUCACUCUUUUGUGCGAAUGCGGUGUCUUUAUUUCUUUUUUUAAACU
Vka-1-1      CAUUACUAUACUGCGCCUUAAAGGUAUGGCUUCGUCACUUUUUUGUACGGAUGCGGUGUCUUUAUUUCUUUUUUUAAACU
env-633      CAUUACUAUACUGCGCCUUAAAGGUAUGGCUUUAUCUCUAUUUAGUGAUACAGCGGUGUCUUUAUUUAUUUUUUAACU
Vsp-31-1     AUUACUAUACUGCGCCUUAAAGGUAUGGCUUCAUCACUCUUUUGUGUCGAUGCGGUGUCUUUAUUUCUUUUUUUAAACU
env-634      CAUUACUAUACUGCGCCUUAAAGGUAUGGCUUUAUCGCAGUUUAGUGAUGCAGCGGUGUCUUUAUUUAUUUUUUAACU
Afi-2-1      GAUUGAUCGCUCCAGAUUCAUUACUAUACGCGACCUUAAGUAAGUGUAAUCCAUUUACGCUUACUAUUUUUGUUAAACU
Vfi-2-1      GAUUGAUCGCUCCAGAUUCAUUACUAUACGCGACCUUAAGUAAGUGUAAUCCAUUUAUGCUUACUAUUUUUGUUAAACU
Vor-1-1      AUUAAUAUAUCGCGCCUUGAUGAUAUGGCUUCUUGCUUGAUGUUAAGCAGUCAGCGGUAUCAACAUUUUUUUAUUAACU
Vge-5-1      UUCAUUACUAUACUGCGCCUUAACGUCAUGGCUUAUUUUACCUUUGAGUAAUUUAGCGGUGGCAAUAAUUUUUUAAACU
Vca-7-1      UCCGCAGCUUAGUAUUCAUUAAAAUAGCGCGCCUUGAUGAUAUGGCUCACAAGCGGUAUACAUCAAUUGACAAUUAACU
Vne-1-1      UCUGUAUUCAUUACUAUAUCGCGCCUUAGUGAUAUGGCUUGCUUUUGAGCAGCGGUAUCCAAAGGAAAUUUUAUUAACU
Vco-3-1      CCGUAUUCAUUACUAUAUCGCGCCUUAGUGAUAUGGCUUGUUUUUGAACAGCGGUAUCCCAAGUGAAAUUUUAUUAACU
Vsp-3-1      CCGUAUUCAUUACUAUAUCGCGCCUUAGUGAUAUGGCUUGUUUUUGAACAGCGAUAUCCCAAGUGAAAUUUUAUUAACU
env-636      GGUUGAUCCGAACUAAGCUAUUCAUUAAUAUACUGCGCCUUAAUGGUGUAGCAUUGCUAUGCUUUUUACUUUUGAACAC
env-637      AGUUGAUCCUAGCCUUGGGAUUCAUUACUAUACUGCGCCUUAAUGGUGUAGCUACGCUAUACUUUUUACUUACGAACAC
Vmi-5-1      GAUUCAUUAAUAUAUCGCGCCUUACGGCAUCCUAGAGGGUUGACCUCUCUGGAUAUGUAAACUCGUUAUUUUUAACACU
Vsp-25-1     GAUUCAUUAAUAUAUCGCGCCUUACGGCAUCCUAGAGGGUUGACCUCUCUGGAUAUGCAAAAUCGUUAUUUUUAACACU
Vch-3-1      GAUUCAUUAAUAUAUUGCGCCUUACGGCAUCCUAGAGGGUUGACCUCUCUGGAUAUGCAAACUCGUUAUUUUUAACACU
Vmi-2-1      AGAUUCAUUAAUAUAUCGCGCCUUACGGCAUCCUAGAGGGUUGACCCUCUGGAUAUGCAAACUCGUUAUUUUUAACACU
Vch-7-1      GAUUCAUUAAUAUAUCGCGCCUUACGGCAUCCUAGAGGGUUGACCUCUCUGGAUAUGCAAACUCGUUAUUUUUAACACU
Fba-1-1      GCGUAGAAAUUGUACAGAAUGCUUGUGGGCAUUGCGGCACUGCUGUAUCAUCUCGCGCCCUGAAAGUAUCACCAACCUA
Vsi-2-1      ACAUGAUUCAUUACUAUAUCGCGCCUUAGAGAUAUGGCUCGCCAAACGGCAGCGGUAUCCUAGCAAUUUUUUAUUCACU
Val-2-9      CAUUAAUAUAUCGCGCCUUGAUGAUAUGGCUUAGUGCUCACUAUUGAACAUGAAGCGGUAUCAACAUUUUUUAUUAACU
Vor-7-1      GCUCACGUAUUCAUUACUAUAUCGCGCCUUAGGAUAUGGCUCGCCUUAGUGUAGCGGUAUCUAAGUUAUUUUUUUCACU
Vdi-2-1      CAGGAUUCAUUAAUAUAUCGCGCCUUGAUGAGACGCUUGUAAGAACGCUCUUCAAGUGUCACCAAUUAUUUAUUAAUUU
env-640      CAGGAUUCAUUAAUAUAUCGCGCCUUGACGAGACGCUUGUAAGAACGCUCUUCAAGUGUUACCAAUUAUUUAUUAAUUU
Vsp-32-1     AUUAAUAUAUCGCGCCUUGAUGAUAUGGCUUCUUGCUUGAUGUUAAGCAGUCAGCGGUAUCAACAUUUUUUUAUUAACU
Vge-3-1      AGUGAUAGGGCUUUCUGAUGGGGUAUUGGUCAUCACGACCCGCCAAGAAGAGAGUGUUAUCCAUACUGUUUUAUUAACU
Vge-1-1      AGUGAUAGGGCUUUCUGAUAGGGUAUUGGUCAUCACGACCCGCCAAGAAGAGAGUGUUAUCCAUACUGUUUUAUUAACU
Vni-9-1      UAGGUUGAUCCUACCCUCAUGUUUCAUUACUAUACUGCGCCUCGAUGAUGUGGUCAUUCUGCAUCGAAUUAUCUAAACU
Vme-2-1      ACUAUACUGCGCCUUAGCGUUAUGGCUCACUUCAUCUUUGGAUGUCGUAGCGGUACGCAAUAAGUAAAUUUUAUUAACU
Vsi-1-1      GAGAGAUGGAUUCAUUACUAUAGCGCGCCUUAGAGAUAUGGCUCGCCAAUGGCAGCGGUAUCCAAGCUAUUUUUUAACU
Vsi-1-3      GGGAGAUGGAUUCAUUACUAUAGCGCGCCUUAGAGAUAUGGCUCGCCAAUGGCAGCGGUAUCCAAGCUAUUUUUUAACU
Vne-2-1      GCGUAUUCAUUACUAUAUCGCGCCUUAGUGAUAUGGCGGCGUUACGCGACGCGGUAUCAAAGAUAAUUUUUUAUUCACU
Vga-1-1      AGCUAUUCAUUAAUAUAUCGCGCCUUAGAGAUAUGGCUCUCGCUGUAGUAGCGGUAUCUAGAAAGGUAUUUUAUUAACU
Vbr-1-1      GACGUAUUCAUUACUAUAUCGCGCCUUAGAGAUAUGGCUUGCUCUUUUGCAGCGGUAUCCUAAAUACUUUUUAUUAACU
Vpa-1-1      UGGCGCGAUUCAUUACUAUAUCGCGCCUUAGAGAUAUGGCUGUCAUUUGGCGGCGGUAUCCAAGCAACUUUUAUUAACU
Vsp-30-1     AGCUAUUCAUUACUAUAUCGCGCCUUAGUGAUAUGGCUGGCUCUAGAGCAGCGGUAUCCAAAAAGGUUUUUUAUUAACU
Vtu-2-2      AGGUAUUCAUUACUAUAUCGCGCCUUAGUGAUAUGGCUAGCUUUACAGCAGCGGUAUCCGAAAAGGUUUUUUAUUAACU
Vtu-1-1      AGCUAUUCAUUACUAUAUCGCGCCUUAGUGAUAUGGCUAGCUUAAUAGCAGCGGUAUCCGAAAAGGUUUUUUAUUAACU
Vhe-1-1      UGGAUUCAUUACUAUAUCGCGCCUUAGAGAUAUGGCUUGCUUUUUGAGCAGCGGUAUCCAAGCUAAUUUUUUAUUAACU
env-641      AGCGGUGGAUUCAUUAAUAUAUCGCGCCUUGAAGAUGCUUGGUCAACAAGGCUGAGUAUCGAACAGUAUCAUUAACAUU
Vfl-4-1      AGCGGGGGAUUCAUUAAUAUAUCGCGCCUUGAAGAUGCUUGGUCAACAAGGCUGAGUAUCGAACAGUAUCAUUAACAUU
Vfu-2-1      CGCGGGGGAUUCAUUAAUAUAUCGCGCCUUGAAGAUGCUUGGUCACCAAGGCUGAGUAUCGAACAGUAUCAUUAACAUU
Vsp-26-1     GAUUCAUUAAUAUAUCGCGCCUUACGGCAUCCUAGAGGGUUGACCUCUCUGGAUAUGCAAACUCGUUAUUUUUAACACU
Vre-1-1      AUUCAUUACUAUACUGCGCCUUAGCACUAUGGCUCGAUGAUUCAUUCAUCAAGCGGUAGCGCGAAAUUUUUUAUUAACU

             ...............................................................................
             ...............................................................................
             ●●●●●●●●●●●●●●●●●●●●●●●●●●●●●●●●●●●●●●●●●●●●●●●●●●●●●●●●●●●●●●●●●●●●●●●●●●●●RY●
```

```
Vsc-1-1      UUCAUUACUAUACUGCGCCUUAGCACUAUGGCUCGGUGAUUUAUUCAUUAAGCGGUAGAGCGAAAUUUUUUUAUUAACC
Vic-1-1      AUUCAUUACUAUACUGCGCCUUAGCACUAUGGCUGAUGAUUCGUUCAUUCGGCGGUAGAGCGAAAUUUUUUUAUUAACU
Awo-1-1      GAUUGAUCGCUCCGGAUUCAUUAUUAUACGCGACCUUAAGUAAGUGUAAUCUGUUUACGCUUAUUAUUUUUGUUAAACU
Asa-11-1     GAUUGAUUGGCCGCGAUUCAUUAUUAUACGCGACCUUAAGUAGGUGUAAUUUAUUUAUGCUUAUUAUUUUUGUUACACG
env-642      ......................................................CGGUGUCUUUAAUCUAUUUUUUAACA
Vta-2-1      UCAUUACUAUACUGCGCCUUAAAGGUAUGGCUUUAUCGCGGUUUAGUGAUACAGCGGUGUCUUUAUUUAUUUUUUAACU
env-643      UCAUUACUAUACGGCGCCUUAAAGGUAUGGCUUCAUCACGUUUUCGUGAGAUGCGGUGUCUUUAAUCUAUUUUUUAACA
Vta-3-1      UCAUUACUAUACUGCGCCUUAAAGGUAUGGCUUUAUCGCGGUUUAGUGAUACAGCGGUGUCUUUAUUUAUUUUUUAACU
env-646      UGGAUUCAUUACUAUAUCGCGCCUUAGAGAUAUGGCUCGCCUGAUGGUAGCGGUAUCCCUAAAUACUAUUUUUUAAACU
Vsh-2-1      UACUAUACUGCGCCUUGAUGGUAUGGCUUGUUCUACAUUUAUUGCAGUAACAGCGAUAUCAUAAGUCUUUUAUUAAACU
Vma-1-1      UACUAUACUGCGCCUUGAUGGCAUGGCUUCUUGCUUUCAUAUUGAAAUGCAAAGCGAUGGUAUCAUCUAUUUAUUAACU
Vma-1-2      UACUAUACUGCGCCUUGAUGGCAUGGCUUCUUGCUUUCUUAUUAAAGUGUGAAGCGAUGGUAUCGACUAUUUAUUAACU
Vme-1-1      UGGCGUGGAUUCAUUAAUAUAUCGCGCCUUAAAGAUGCUUGGCAAAUCGGUUUAGCAUCUGCAUAUUAUCAUUAACAUU
Ymo-1-1      UUCGCCGCCUUUGGUAUAAUUCGAUAUCAAAUCCUGUUAACAACCGUUGGUUACUUUAUCAGCGAUUGUUAGUUAUCAA
Ybe-1-1      UUCGCCGCCUUUGAUAUAAUUCGAUAUCAAAUCCUGUUAACAACCGUUGGUUACUUUAUCAGCGAUUGUUAGUUAUCAA
Yma-2-1      UUCGCCGCCUUUGAUAUAAUUCGAUAUCAAAUCCUGUUAACAAGCGUUGGUUACUUUAUCAGCGAUUGUUAGUUAUCAA
Yfr-1-1      UUCGCCGCCUUUGAUAUAAUUCGAUAUCAAAUCCUGUUAACGAGCGUUGGUUACUUUAUCAGCGAUUGUUAGUUAUCAA
Yma-1-1      UUCGCCGCCUUUGAUAUAAUUCGAUAUCAAAUCCUGUUAACAAGCGUUGGUUACUUUAUCAGCGAUUGUUAGUUAUCAA
Yfr-1-5      UUCGAUAUCAAACUCUGUUAACAGCCGUUGCUUACUUUACCAGCGAUAGUUAGUUAUCAAUAAAAUUAUUAUUUAUCAA
Ype-1-1      UUCGCCGCCUUUGGUAUAAUUUGAUAUCAAAUCCUAUGAACAACCGUUGCUUGUUUUAUCAGCGAUUGUUAUUUCUUAA
Yfr-1-7      UCGAUAUCAAACUCUGUUAACAGCCGUUGCUUAUUUUACCAGUGAUAGUUAGUUAUCAAUAAAAUUAAUUAUUUAUCAA
Yfr-1-8      UCGAUAUCAAACUCUGUUAACAGCCGUUGCUUAUUUUACCAGCGAUAGUUAGUUAUCAAUAAAAUUAAUUAUUUAUCAA
Yfr-2-2      UUCGAUAUCAAACUCUGUUAACAGCCGUUGCUUACUUUACCAGCGAUAGUUAGUUAUCAAUAAAAUUAUUAUUUAUCAA
Yfr-1-14     UUCGAUAUCAAAUCCUGUUAACAACCGUUGCUCAGUUUAUCAGCGGCUGUUCGUUAAAAAUCAAAUUGUUAUUUAUCAA
Yfr-1-15     UUCGCCGCCUUUGAUAGAAUUCGAUAUCAAAUCCUGUUAACAAGCGUUGUUUAUUUUAUCGACGAUUGUUAUUUAUCAA
Epi-1-1      GUGAUUGCGUUGGUCGGCUAUUUUCUGUAUGAUUCGCCGCCUUUGAUAUAAUUUUGUAUCAAACCCGAUUUUAUCAACC
Epi-1-2      GUGAUUGCGUUAGGCGGCUAUUUUCUGUAUGAUUCGCCGCCUUUGAUAUAAUUUUGUAUCAAACCCGAUUUUAUCAACC
Eic-2-1      GUGAUUGCGCUGGACGGCUAUUUUCUGUAUGAUUCGCCGCCUUUGAUAUAAUUUUGUAUCAAACCCGAUUUUAUCAACC
Eta-2-1      GUGAUUGCGUUAGGCGGCUAUUUUCUGUAUGAUUCGCCGCCUUUGAUAUCGUUUUGUAUCAAACCCAAUUUUAUCAACC
Eho-5-1      GUGAUUGCGUUAGUCGGCUAUUUUCUGUAUGAUUCGCCGCCUUUGAUAUCGUUUUGUAUCAAACCCGAUUUUAUCAACC
Eta-4-1      GUGAUUGCGUUAGGCGGCUAUUUUCUGUAUGAUUCGCCGCCUUUGAUAUCGUUUUGUAUCAAACCCAAUUUUAUCAACC
Yal-1-1      UUCGCCGCCUUUGGUAUAAUUUGAUAUCAAAUCCUGUUAACAGCCGUUGUUUGUUUUAUCAGCGCUUGUUAUUUUCUAA
Yen-14-1     UUCGCCGCCUUUGAUAUAAUUUGAUAUCAAAUUCUGUUAACAAUCGUUGUUUGUUUUAUCAGCGCUUGUUAUUUUCUAA
Yal-1-4      UUCGCCGCCUUUGGUAUAAUUUGAUAUCAAAUCCAGUUAACAACCGUUGUUUGUUUUAUCAGCGCUUGUUAUUUUCUAA
Yen-21-1     UUCGCCGCCUUUGAUAUAAUUUGAUAUCAAAUUCUGUUAACAACCGUUGUUUGUUUUAUCAGCGCUUGUUAUUUUCUAA
Ype-127-1    ....................UUUGAUAUCAAAUCCUGUUAACCAUCGUUGCUUAUUUUAUCAACGAUGGUUAUUUAUCAA
Yal-2-1      UUCGCCGCCUUUGGUAUAAUUUGAUAUCAAAUCCUGUUAACAGCCGUUGUUUGUUUUAUCAACGCUUGUUAUUUUCUAA
Ynu-1-1      AUUCGAUUGCGAGGGCUGCUCAAAUUCUGUAUGAUUCGCCGCCUUUGAUAUAAUCCGAUAUCAAAUCCUAUUAUCAACU
Yru-1-1      AUUCGAUUGCGGGGGCUGCUUAAAUUCUGUAUGAUUCGCCGCCUUUGAUAUAAUCCGAUAUCAAAUCUUAUUAUCAACU
Dze-2-1      UCCAAUUGCAUCGGGUGGUUAAUUUCUGUAUGAUCCGCCGCCUUUAUACCUCGUAGUAUAAAGACUAUUUCACUCAACU
Dsp-1-1      UCCAAUUGCAUCGGUUGGUUAAUUUCUGUAUGAUCCGCCGCCUUUAUACCUUGCGGUGUAAAGAACAUUUCAUUCAACU
Ssp-23-1     AUGUGAUUGCAUCAGCUGACUAAUUUCUGUAUGAUUCGCCGCCUUUAUACCUCGUGGUAUAAUGAUAUUUUACUCAACU
env-647      .......................................................AGAUAAAACGAAUUCAACUUCAACC
Lqu-1-3      UGCAUCGGCUGGNUAAUUUCUGUAUGAUUCGCCGCCUUUAUACCGCGUGUAUAAAGAUAAAACGAAUUCAACUUCAACC
Dsp-2-1      UCCAAUUGCAUCGGGUGGUUAAUUUCUGUAUGAUCCGCCGCCUUUAUACCUUGUGGUGUAAAGAACAUUUCACUCAACU
Dze-7-1      UCCAAUUGCAUCGGGUGGUUAAUUUCUGUAUGAUCCGCCGCCUUUAUACCUCGUAGUAUAAAGACUAUUUCACUCAACU
Dch-3-1      CCAAUUUGCAUCAGCUGGUUAAUUUCUGUAUGAUCCGCCGCCUUUAUACCUUGUGGUAUGAAGAACAUUUCACUCAACU
Dch-2-1      UCCAAUUGCAUCAGGUGGUUAAUUUCUGUAUGAUCCGCCGCCUUUGUACCUUGUGGUAUGAAGAACAUUUCACUCAACU
Dda-1-1      UCCAAUUGCAUCAGGUGGUUAAUUUCUGUAUGAUCCGCCGCCUUUAUACCUUGUGGUGUAAAGAACAUUUCACUCAACU
Dda-2-1      UCCAAUUGCAUCAGGUGGUUAAUUUCUGUAUGAUUCGCCGCCUUUAUACCUUGUGGUGUAAAGAACAUUUCACUCAACU
Dch-1-2      UCCAAUUGCAUCAGGUGGUUAAUUUCUGUAUGAUCCGCCGCCUUUAUACCUUGUGGUAUGAAGAACAUUUCACUCAACU
Dsp-9-1      UCUGAUUGCAUCAUAGGGUUAAUUUCUGUAUGAUCCGCCGCCUUUAUACCUUGUAGUAUAAAGAACAUUUCACUCAACU
Xdo-1-1      CAUUGCGCAAUAAACAGAUAUUCUGUAUGAUCCGCCGCCUUUGGUAUAGAAAUUACCGAUACUUAUUUUUAUCAACACG
Xpo-1-1      GCGCAAUAAACCGAUAUUCUGUAUGAUCCGCCGCCUUUGGUAUAGAAAGAAAUUCCCGAGAAUUAUUUUCAUCAACACG
Xbo-11-1     UCAUUGCGCAAUAAACCGAUAUUCUGUAUGAUCCGCCGCCUUUGGUACAGAAAUUACCGAGCUUAUUUCAAUCAACACG
Xbo-8-1      UUAUUGCGCAAUAAACCGAUAUUCUGUAUGAUCCGCCGCCUUUGGUACAGAAAUUACCGAACUUAUUUCAAUCAACACG
Xbo-6-1      UCAUUGCGCAAUAAACCGAUAUUCUGUAUGAUCCGCCGCCUUUGGUACAGAAAUUACCGAACUUAUUUCAAUCAACACG
Xsz-1-1      CAUUGCACAAUAACCAGAUAUUCUGUAUGAUCCGCCGCCUUUGGUAUAGAAAUUACCGAUACUUAUUUUAAUCAACACG
Xca-1-1      UCAUUGCGCAAUAAAACGAUAUUCUGUAUGAUCCGCCGCCUUUGGUAUAAAAAUUACCGAACUUAUUUUAAUCAACACG
Xkh-1-1      AUUUUCUGUAUGAUCCGCCGCCUUUGGUAUAGAAAUUACCGAUACUGAUUUUUAUCAACACGACAUUUUUAUCAACAUG
Xsp-1-1      UCAUUGCGCAAUAAAACGAUAUUCUGUAUGAUUCGCCGCCUUUGGUAUGAAAAUUACCGAACUUAUUUUAAUCAACACG
Xne-5-1      CAUUGCGCAAUAAACCGAUAUUCUGUAAGAUCCGCCGCCUUUGGUAUAGAAAUUACCGAUACUUAUUUUUAUCAACACG
Plu-5-1      UCAUUGCGAGAUAGGCAGAUCUUCUGUAUGAUCCGCCGCCUUUGGUAUAAGAAGAUACCAAACUCAUUUUAUCAACACG
Plu-9-1      UUCAUUGCGGGAUAGGCAGAUCUUCUGUAUGAUCCGCCGCCUUUGGUAUGGAAUAUACCAAACUCAUUUUAUCAACACG
Plu-7-1      UUCAUUGCGAGAUAGGCAGAUCUUCUGUAUGAUCCGCCGCCUUUGGUAUGGAAGAUACCAAACUCAUUUUAUCAACACG
Plu-6-1      UUCAUUGCGAGAUAGGCAGAUCUUCUGUAUGAUCCGCCGCCUUUGGUAUAGAAGAUACCAAACUCAUUUUAUCAACACG
Pte-4-1      UUCAUUGCGGGAUAGGCAGAUAUUCUGUAUGAUCCGCCGCCUUUGGUACAAAAGAUACCAAACUUAUUUUAUCAACACG
Pte-6-1      UUCAUUGCGGGAUGGGCAGAUAUUCUGUAUGAUCCGCCGCCUUUGGUACAAAAGAUACCAAACUUAUUUUAUCAACACG
Plu-10-1     UUCAUUGCGAGAUAGGUAGAUAUUCUGUAUGAUCCGCCGCCUUUGGUACGGAAGAUACCAAACUCAUUUUAUCAACACG
Pte-2-1      UUCAUUGCGGGAUAGGUAGAUAUUCUGUAUGAUCCGCCGCCUUUGGUACAAAAGAUACCAAACUUAUUUUAUCAACACG
Ssp-15-1     GAUCCGAUUGCGCAGUAUAUUUUAUUCUGUAUAAUUUGCCGCCUUUGAUAUAAUUCCAUAUCAAAUAUGAUGAUUAACU
Pca-2-1      AUUCUGUAUGAUCCGCCGCCACUAUGCUGUUCGCGCAAACGCUUGUGCCGUUCGCAUAAUGAACUCACUUACUCAACUU
Pca-1-1      CUAUGCUGUUCGCACAAGUGAUGGUUCGUUCGCGAAAAUCAUUGCGCCGUUCGCAUAAUGAACUUACUUAACUCAACUU
Ddi-5-1      UCCAAUUGCACUGGGUGGUUAAUUUCUGUAUGAUUCGCCGCCUUUAUACCUUGUGGUGUAAAGAACAUUUCACUCAACU
Ssp-13-1     GCAGAUCCGUUUGCGCGGUGCAUUUAAUCCUGUAUAAUUUGCCGCCUUUGAUAUAAUUCCGUAUCAAAAUUUAUCAAUC
Dpa-1-1      UAUGAUUGCAUCAUCUGGUUAAUUUCUGUAUGAUCCGCCGCCUUUGUACCGUGUGGUAUAAAGCUCGUUUAACUCAACU
Sfo-1-1      GAUCCGAUUGCGCGCUACAUUUUAUUCUGUAUAAUUUGCCGCCUUUGGUAUAAUCCCAUAUCAAAACCUAUUAUCAACC
Pca-7-1      GCGAUUGCAUCGGGGUAUCAGAUUCUGUAUGAUCCGCCGCCAUUGUGCCGUUCGCAUAAUGAACUUACUUACUCAACUU
Ppa-2-1      AUUCUGUAUGAUCCGCCGCCACUGUGUUGUUCGCGCAAAUGACUGUGCCGUUCGCAUAAUGAACUUACUUACUCAACUU
Pat-3-1      UGUGUCGUUAGAAGAAAUGAUCGUGUUGUUCGCGCAAAUGAUGAUUUCGUUCGCAUAAUGAACUUAACUUACUCAACUU
Psp-55-1     UUCUGUAUGAUCCGCCGCCACUGUGUUGUUCGCGCAAAAUGACUGUGCCGUUCGCAUAAUGAACUUACUUACUCAACUU
Pwa-2-1      UUGUGUCGUUAGAAGAAAUGAUCGUGUUGCUCGCGCAAAUGAUGAUUUCGUUCGCAUAAUGAACUUACUUACUCAACUU
Pca-8-1      UCUGUAUGAUCCGCCGCCACUAUGCUGUUCGCGCGAAUUAUUUGUGCCGUCCGCAUAAUGCACUUAACUUACUCAACUU
Bgo-1-1      GGAGAUGAGUUUCUGUAUGAUUCGCCGCCAUUAUACUGCGUUGUAUAAUGGUUGAUUGACUCAACUUCUUUACUCACUU

             ................................................................................
             ................................................................................
             •••••••••••••••••••••••••••••••••••••••••••••••••••••••••••••••••••••••••••••RY•
```

| | |
|---|---|
| Bsp-1-1 | GCUGGUUGGAUUCUGUAUGAUUCGCCGCCAUUAUACUGCGUGGUAUAAUGGUUGAUUGACUCAACUUCUUACUCAACUU |
| Dsp-3-1 | ACACCUUGUGUUGCAGAAAGGCAUGUACCAGUGCGCAUGUUUUGCUGUCAGGGCGUGAACAUUCACUAUAUAGACAACU |
| Pbe-1-1 | UGUGUCGUUAGAAGAAAUGAUCGUGCCGUUCGCGCCAAUGGUUAUUUCGUUCGCAUAAUGAACUUAACUUACUCAACUU |
| Pca-5-11 | UCUGUAUGAUUCGCCGCCAUUAUGCUGUUCGCGAAAAUCAUUGCGCCGUUUGCAUAAUGAACUUACUCAUAUACAACUU |
| env-651 | UUCCGAUUGCGCGCCCCGGCUAAUGCUGUAUAAUUUGCCGCCUUUGGUAUAAUUCGAUACCAAAAUUCUAUUAUCAACC |
| env-653 | UUCCGAUUGCGCGCCCCGACUAAUGCUGUAUAAUUUGCCGCCUUUGGUAUAAUUCGAUACCAAAAUUCUAUUAUCAACC |
| env-655 | UUCCGAUUGCGCGGUACUUUUAAUUCUGUAUAAUUUGCCGCCUUUGGUAUAAUUCGAUACCAAAUUUUUUUAUCAACCC |
| env-656 | UUCCGAUUGCGCGCCUCGGCUAAUGCUGUAUAAUUUGCCGCCUUUGGUAUAAUUCGAUACCAAAAUUCUAUUAUCAACC |
| Spl-5-1 | AUCCAAUUGCGCGCUAGGUUGAAUUUCUGUAUAGUUUGCCGCCUUUGGUAUAAUCCGAUACCAAAAUAUAUUAUCAACC |
| Ssy-1-1 | GUUCCGAUUGCGUAGUACUUUUAAUUCUGUAUAAUUUAGCGCCUUUGGUAUAAUUCGAUACCAAAUUCUAUUAUCAACC |
| Ssy-2-1 | GUUCCGAUUGCGUAGUACUUUUAAUUCUGUAUAAUUUAGCGCCUUUGGUAUAAUUCGAUACCAAAUUCUACUAUCAACC |
| Sgr-1-1 | GUUCCGAUUGCGCGGCACUUUUAAUUCUGUAUAAUUUGCCGCCUUUGGUAUAAUUCGAUACCAAAUUUUAUUAUCAACC |
| Sod-1-1 | AUCCGAUUGCGCGGGUGAUAUUAUUCUGUAUAAUUUGCCGCCUUUGGUAUAAUUCGAUACCAAAAAAUUAUCAUCAAUC |
| Spl-1-1 | UUCCAAUUGCGCGCUACAUUGAAUUUCUGUAUAGUUUGCCGCCUUUGGUAUAAUCCGAUACCAAAAUAUAUUAUCAACC |
| Spr-2-1 | GGUUCCGAUUGCGCGGUACUUUUAAUUCUGUAUAAUUUGCCGCCUUUGGUAUAAUUCGAUACCAAAUUCUUUAUCAACC |
| env-657 | GUUCCGAUUGCGCGGCACUUUUAAUUCUGUAUAAUUUGCCGCCUUUGGUAUAAUUCGAUACCAAAUUCUAUUAUCAACC |
| env-658 | GUUCCGAUUGCGUGGUACUUUUAAUUCUGUAUAAUUUGCCGCCUUUGGUAUAAUUCGAUACCAAAUUUUAUUAUCAACC |
| Sma-16-1 | UUCCGAUUGCGCGCCUCGACUAAUGCUGUAUAAUUUGCCGCCUUUGGUAUAAUUCGAUACCAAAAUUCUAUUAUCAACC |
| Ssp-21-1 | AUCCAAUUGCGCGCCACAUUGAAUUUCUGUAUAAUUUGCCGCCUUUGGUAUAAUCCGAUACCAAAAUAUAUUAUCAACC |
| Sma-24-1 | AUCCGAUUGCGCGGUGUAUUUAAUUCUGUAUAAUUUGCCGCCUUUGGUAUAAUUCGAUACCAAUAAAUUAUCAUCAACC |
| Sma-3-1 | UUCCGAUUGCGCGCCUCGACUAAUACUGUAUAAUUUGCCGCCUUUGGUAUAAUUCGAUACCAAAAUUCUAUUAUCAACC |
| Pca-3-1 | CGAUUGCAUCAGGGUAGCGGAUUCUGUAUGAUCCGCCGCCAUUGUGCCAUUUGCAUAAUGAAACUUACUUAUACAACUU |
| Pca-6-1 | CGAUUGCAUCGGGGUAGCGGAUUCUGUAUGAUCCGCCGCCAUUAUGCCAUUUGCAUAAUGAAACUUACUUAUACAACUU |
| Pca-5-2 | CGAUUGCAUCGGGGUAGCGGAUUCUGUAUGAUCCGCCGCCAUUAUGCCAUUUGCAUAAUGAAACUUACUUAUACAACUU |
| Pca-5-4 | CGAUUGCAUCGUGGUAGCGGAUUCUGUAUGAUCCGCCGCCAUUAUGCCAUUUGCAUAAUGAAACUUACUUAUACAACUU |
| Pca-10-1 | CUGUGCUGUUCGCGCAAAUGAUAGGUCGUUCGCGAAAAUCAUUGCGCCGUUCGCAUAAUGAAACUUACUUAUACAACUU |
| Pca-3-2 | CGAUUGCAUCAGGGUAGCGGAUUCUGUAUGAUCCGCCGCCAUUGUGCCGUUUGCACAAUGAAACUUACUUAUACAACUU |
| Pca-3-3 | CGAUUGCAUCAGGGUAGCGGAUUCUGUAUGAUCCGCCGCCAUUGUGCCGUUCGCAUAAUGAAACUUACUUAUACAACUU |
| Pca-12-1 | CGAUUGCAUCAGGGUGGCGGAUUCUGUAUGAUCCGCCGCCAUUAUGCCAUUUGCAUAAUGAAACUUACUUAUACAACUU |
| Cmu-2-1 | GUUCCGUUUGCGUGGCCCAUUUAAUUCUGUAUAAUUUACCGCCUUUGAUAUAAUUCCAUAUCGAAGUUGAUUAUCAACC |
| Pca-5-6 | CGAUUGCAUCAGGGUGACGGAUUCUGUAUGAUCCGCCGCCAUUAUGCCAUUUGCAUAAUGAAACUUACUUAUACAACUU |
| Pca-5-8 | CGAUUGCAUCAGGGUAGCGGAUUCUGUAUGAUCCGCCGCCAUUGUGCCGUUUGCAUAAUGAAACUUACUUAUACAACUU |
| Ssp-17-1 | GAUCCGAUUGCGUGCUACAUUUUAUUCUGUAUAAUUUGCCGCCUUUGGUAUAAUUCCAUAUCAAAACCUAUUAUCAACC |
| Pas-2-1 | UUCAUUGCGAGAGAGGCAGAUAUUCUGUAUGAUCCGCCGCCUUUGGUACGGAAGAUACCAAACUUAUUUUAUCAACGUG |
| Pas-1-1 | UUCAUUGCGAGAUAGAUGGAUAUUCUGUAUGAUCCGCCGCCUUUGGUACGGAAGAUGCCAAACUUAUUUUAUCAACGCG |
| Xgr-1-1 | AUUGCGCAAUAAACAGAUAUUCUGUAUGAUCCGCCGCCUUUGGUAUAGAAAUUACCGAUACAUAUUUUUUAUCAACACG |
| Hpa-1-1 | AUUCGAUUGCGGUGUAUGUUUGAUCCCUGUAUAAUUCGCCGCCUUUGAUAGGCUUUUUAUCAAACCAAAUUUAUCAACC |
| Hal-1-1 | UUCGAUUGCGGUGGAUGUUUUAUCUCUGUAUAAUUCGCCGCCUUUGAUAGUCUUUUUUAUCAAACCAAAUUUAUCAACC |
| env-660 | ......................................................UUUUUAUCAAACCAAAUUUAUCAACC |
| Hal-3-1 | UUCGAUUGCGGUGGAUGUUUUAUCUCUGUAUAAUUCGCCGCCUUUGAUAGGCUUUUUUAUCAAACCAAAUUUAUCAACC |
| env-661 | UGAUUGCGCGCUCAGAUUAUAUUCUGUAUGAUUCGCCGCCUUUGAUGCAUUCCGUAUCAGAGACCUAAUCAUUAACACC |
| gpr-1-1 | UGAUUGCGCGGUCACUUUAUAUUCUGUAUGAUUCGCCGCCUUUGAUGCAUUCCGCAUCAGAGACCUAAUCAUUUACACC |
| Ssp-24-1 | UGAUUGCGCGGUCAGAUUUUAUUCUGUAUGAUUCGCCGCCUUUGAUGCAUUCCGUAUCAGAGACCUAAUCAUUAACACC |
| Rch-1-1 | UGAUUGCGCGGUCAGAUUAUAUUCUGUAUGAUUCGCCGCCUUUGAUGCAUUCCGUAUCAGAGAACUAAUCAUUAACACU |
| Eam-2-1 | ACAUCAUUGCGCGGCAUAAAUAGAUUCUGUAUGAUUCGCCGCCUUUGGUGCAUUUCGCAUCAAAACCUAAUCAUCCACU |
| env-662 | AACAUCAUUGCGCGGUUCUCAGAGAUUCUGUAUGAUUCGCCGCCUUUAGUGCAUUUCGUACUAAAGCCUAUCAUCCACU |
| env-668 | ACAUGAUUGCGCGGUUGUCAGAGAUUCUGUAUGAUUCGCCGCCUUUGGUGCAUUUCGUACUCAAAGCCUAUCAUCCACU |
| Raq-1-1 | ACAUCAUUGCGCGGUUGUCAGAGAUUCUGUAUGAUUCGCCGCCUUUGGUGCAUUUCGUACUCAAAGCCUAUCAUCCACU |
| env-672 | ACAUCAUUGCGCGGUUCUCAGGGAUUCUGUAUGAUUCGCCGCCUUUGGUGCAUUUCGCACUCAAAGCCUAUCAUCCACU |
| env-674 | ACAUCAUUGCGCGGUUCUCAGAGAUUCUGUAUGAUUCGCCGCCUUUGGUGCAUUUCGUACUCAAAGCCUAUCAUCCACU |
| Mmo-1-1 | AUUGCAUGGUUCCGGGAUUUUCUGUAUGAUCCGCCGCCUUUGGUAAUUAAAAUACCAGCGAUAACUUAUCAUCAACACG |
| Mmo-3-1 | AUUGCAUGGUUCAGGGAUUUUCUGUAUGAUCCGCCGCCUUUGGUAAUUAAAAUACCACCGAUAACUUAUCAUCAACACG |
| Mmo-2-1 | AUUGCAUGGUUCAGGGAAUUUCUGUAUGAUCCGCCGCCUUUGGUAAUUAAAAUACCACCGAUAACUUAUCAUCAACACG |
| Mmo-1-7 | AUUGCAUGGUUCAGGGAAUUUCUGUAUGAUCCGCCGCCUUUGGUAAUUAAAAUACCACCGAUAACUUAUCAUCAACACG |
| Mwi-1-1 | UAAAUAGUUUUUCUGUAUGAUCCGCCGCCUUUGGUAUCGAUUUUUACCAAUAAAAAUAUCCUAAUUUAUAAUUAACACG |
| Psn-1-1 | UUGCGUAAUAAAUAGAUUUUCUGUAUGAUCCGCCGCCUUUGGUAGCAUUUUUACCGAUACGAUAUUUUUAAUCAACACG |
| Pbu-1-1 | GUAUGUGAUUCGGGUAGCUAAGUGAUGAUAACAUCUAUGCAGCUUCAAUUAUGAAGAGUAAUAAUUUCUAAUCAACACG |
| Pal-5-1 | AAACAGAUUUUCUGUAUGAUCCGCCGCCUUUGGUAUCAAUUAAUUUUGUUUUGCCAAUAAACUAAAUUUAAUCAACACG |
| Pre-3-1 | UGCGUGAUAAACAGAAUUUCUGUAUGAUCCGCCGCCUUUGGUAUUUAUUUUUACCAAUAUAUUAAAUUCAAUCAACACG |
| Pre-1-1 | GCGUGAUAAACAGAAUUUCUGUAUGAUCCGCCGCCUUUGGUAGUUAUUUUUGCCAAUAAUAUUAAAUUCAAUCAACACG |
| Pre-2-1 | UGCGUGAUAAACAGAAUUUCUGUAUGAUCCGCCGCCUUUGGUAUCUAUUUUUACCAAUAUAUUAAAUUCAAUCAACACG |
| Pru-5-1 | UUGGUAUCAUUAAUUUUGCGAAAAUAUUUGUUUUUGUGAAAACCUAAUCCUUGCCAAUAUACUAAAUUUAAUCAACACG |
| Pal-8-1 | AAACAGAUUUUCUGUAUGAUCCGCCGCCUUUGGUAUCAAUUAAUUUUGUUUUGCCAAUAAACUAAAUUUAAUCAACACG |
| Eba-4-1 | AUUCGCUAUCAGGGUUCUGUAUGAUCCGCCCCCUUUGAUGCGUUAUUUUCGCGCAGUCAAAGAACUCACUUAUAUUAAC |
| Msp-4-1 | UUUGCAUACCCUGGUUAAUUUCUGUAUGAUUCGCCGCCUUUGAUGUGCAAUUUCGCACUCAAAAACAUUACUAUCAACU |
| env-675 | CGCUCAUAAAUCCCUGUAUGAUUCGCCGCCUUCGAUGCGCUUUUCUGCCAUCAAGAAACCUGAAUUAUUCAUAUUCACA |
| env-677 | GAUUGCAUCGGCUGUUCAAAUUCUGUAUGAUUCGCCGCCUUUGGUGCGCACAUCCGCCAUCAAAGAUUCACUUUUAACA |
| env-678 | UGCAUAGCGCAUAAAGAUUCUGUAUGAUCCUCGCCUUUGAUGCGAAGAAAUCGUAUCAAUGACCUAAACUUUAUCAACA |
| Cmu-1-1 | GUUUGCAUGGGGCGGUUAUUUUCUGUAUGAUUCGCCGCCUUUGAUGCCCUUUAAAAGUAUCAAACCGACUUUAUCAACA |
| Csa-13-1 | GUUUGCAUGAGGCGGUUAUUUUCUGUAUGAUUCACCGCCUUUGAUGCCCUUUAAAAGUAUCAAACCGACUUUAUCAACA |
| Csa-10-1 | GUUUGCAUGGGGCGGUUAUUUUCUGUAUGAUUCGCCGCCUUUGAUGCCCUUUAAAAGUAUCAAACCGACUUUAUCAACA |
| Cun-1-1 | GUUUGCAUGGGGCGGUUAUUUUCUGUAUGAUUCUCCGCCUUUGAUGCCCUUUAAAAGUAUCAAACCGACUUUAUCAACA |
| Csa-7-1 | GUUUGCAUGAGGCGGUUAUUUUCUGUAUGAUUCGCCGCCUUUGAUGCCCUUUAAAAGUAUCAAACCGACUUUAUCAACA |
| env-679 | UGCAACCGCUGGCUAAAUUCUGUAUGAUUCGCCGCCUUUAAUGCGAAAUAAUCGCGUUAAACUACCAUUAUCACUAACA |
| env-680 | GCAUACGCUUCUCAGAUUCUGUAUGAUCCGCCGCCUUUGGUGCAAAUAAUUGCAUCAAGAGACCAAUACAUUAUCAACA |
| env-681 | .......UGGUUAAAUUCUGUAUGAUUCGCCGCCUUUGAUGCGCAUUAAUGCGGUAUCAAAUGAACUAUACUAUUAACU |
| env-682 | CAACCCCUGGUUAAAUUCUGUAUGAUUCGCCGCCUUUGAUGCGCAUUAAUGCGGUAUCAAAGUAACUAUACUAUUAACU |
| env-683 | ACCCUGCUCAGAUUCUGUAUGAUCCGCCGCCUUUGAUGCAGAAAAAAUGCGUCGAAAGCCGAGAAGUUUACUAUCAACA |
| env-685 | GCAUCUGCUGGUUAAAUUCUGUAUGAUUCGCCGCCUUUGGUGCUGAUCGGCUUUGCGCCAAAAUACAAUACAUUUAACA |
| Tgu-1-1 | UGCAUCCGAUGGUUAAAUUCUGUAUGAUUCGCCGCCUUUAAUGCGAAAUAGUCGCGUUAAACUACCUAUAUUAUUAACA |
| env-686 | UGUUGACCGCUUCAACAAGCGUUUCAGCAUUGAUUCGCCGCCUUUGGUGUACUUUACGCCAAAGAACUUACAUUCAACA |
| env-687 | UCAGUUUGCAUAUGCUGGUCAAAUUCUGUAUGAUUCGCCGCCUUUGGUGUAUUUUACGCCAAAGAACUUACAUUCAACA |
| env-688 | UCAGUUUGCAUACGCUGGUCAAAUUCUGUAUGAUUCGCCGCCUUUGGUGUAUUUUACGCCAAAGAACUUACAUUCAACU |
| env-690 | UCAGUUUGCAUACGUUGGUCAAAUUCUGUAUGAUUCGCCGCCUUUGGUGUAUUUUACGCCAAAGAACUUACAUUCAACA |
| | ................................................................................ |
| | ................................................................................ |
| | ●●●●●●●●●●●●●●●●●●●●●●●●●●●●●●●●●●●●●●●●●●●●●●●●●●●●●●●●●●●●●●●●●●●●●●●●●●●●●RY● |

```
env-700      CAUCAAACUCCCAAAACCCGGGAAAUUCGCUAAGCCGGUUUACGCCAGCGGCGCGAAGGUCGUCGGGAAAUUAUUAAUA
env-701      UUUGCAUACGCAGGUCAAAUUCUGUAUGAUUCGCCGCCUUUGAUGCGCUUUAUGCAGCGUCAAAACUAUACUAUUAACA
env-702      UCAGUUUGCAUACGUUGGUCAAAUUCUGUAUGAUUCGCCGCCUUUGGUGUACUUUACGCCAAAGAACUUACAAUCAACA
Ksp-8-1      UUUGAUGCGCUUUGUGCGCCAUCAAAAUCUAUUACCCGGGAAUUUCGCUUCGCGCAAUCGCACCGGGAAAUUAUUAACA
Ecl-52-1     UCAGUUUGCAUACGUUGGUCAAAUUCUGUAUGAUUCGCCGCCUUUGGUGUACUUUAUGCCAAAGAACUUACAUUCAACA
env-706      UCAGUUUGCAUACGUUGGUUAAAUUCUGUAUGAUUCGCCGCCUUUGGUGUACUUUACGCCAAAGAACUUACAUUCAACA
env-710      UUUGAUGCGCUUUAUGCGGUAUCAAAAUCUAUUACCCGGCAAUUUCGCUUUGCGCAAUCGCACCGGGAAAUUAUUAACA
Kva-1-1      UAUACCGCAUUGGAGGAUUGACCAUUACCGUACGCUUCGCGCUGUCAGCAGCGUGGAAGACGACGGGAAAUUAUUAAUA
env-711      UCAGUUUGCAUACGCUGGUCAAAUUCUGUAUGAUUCGCCGCCUUUGGUGUAUAUUACGCCAAAGAAAUUACAUUCAACA
env-712      CAUCAAACUCCCAAAACCCGGGAAAUUCGUUAAGCCGGUAUGCGCCAGCUGUGCGAAGGUCGUCGGGAAAUUAUUAAUA
env-713      AUCAAACAUCCCAAAACCCGGGAAAUUCGCUAAGCCGGUUUACGCCAGCGGCGCGAAGGUCGUCGGGAAAUUAUUAAUA
Ecl-39-1     UCAGUUUGCAUACACUGGUUAAAUUCUGUAUGAUUCGCCGCCUUUGGUGUAGAUUACGCCAGAGAAAUUACAUUCAACA
env-714      GAUUCGCCGCCUUUGAUGCGCUUUGUGCGACAUCAAAAUCUAUUACCCGGUAGUGUAAUCGCUCCGGGAAUUAUUAACA
Ecl-1-6      UCAGUUUGCAUACGUUGGUCAAAUUCUGUAUGAUUCGCCGCCUUUGGUGUACUUUACGUCAAAGAACUUACAUUCAACA
env-715      UCAGUUUGCAUACCCUGGUUAAAUUCUGUAUGAUUCGCCGCCUUUGGUGCACUUUGCAUCAAAGAACUUACAUUCAACA
env-717      UUUGGUGCGCUUCGUGCAGCAUCAGAAUCUAUUACCCGGGGAUUACGCUUAGCGGAAUUGCGCCGGGAAAUUAUUAACA
env-720      UUGCAUACGGUGGUCAAAUUCUGUAUGAUUCGCCGCCUUUGAUGCGCUUUAUGCAGCGUCAAAACCUAUACUAUUAACA
env-724      ..............................................................AGAANUUACAUUCAACA
env-725      ..................UCAAAUUCUGUAUGAUUCGCCGCCUUUGGUGUACUUUACGCCAAAGAACUUACAUUCAACA
env-726      UCAGUUUGCAUACGUUGGUCAAAUUCUGUAUGAUUCGCCGCCUUUGGUGUACUUUACGCCAAAGAACUUACAUUCAACA
env-728      AUCAAACUCCCAAAACCCGGGAAAUUCGCUAAGCCGGUUUACGCUCAGCUGUGCGAAGGUCGUCGGGAAAUUAUUAAUA
env-729      AUCAAACAUCCCAAAACCCGGGAAAUUCGCUAAGCCGGUUUACGCCAGCUGCGCGAAGGUCGUCGGGAAAUUAUUAAUA
env-730      UUUGAUGCGCUUUGUGCGGCAUCAAAAUCUAUUACCCGGCAAUUUCGCUUUGCGCAAUCGCACCGGGAAAUUAUUAACA
env-731      UUGCAUACACAGGUCAAAUUCUGUAUGAUUCGCCGCCUUUGAUGCGCUUUAUGCAGCGUCAAAACCUAUACUAUUAACA
Lad-1-2      UCAGUUUGCAUACCCUGGUUAAAUUCUGUAUGAUUCGCCGCCUUUGGUGCACUUUGCAUCAAAGAACUUACAUUCAACA
env-732      UUAUAUCGCAUUGAAGAUUGACCAUUACCGUGUGCUUCACGCUGUCAGCAGCGUAGAGGGCGACGGGAAAUUACUAAUA
env-735      UUAUACCGCAUUGGAGAUUGACCAUUACCGUACGCUUCGCGCUGUCAGCAGUGUGGAAGACGACGGGAAAUUAUUAAUA
Rte-1-1      UGUGCUUCGUGCAGUGCCAAAGCCUAUACUAUUACCCAUGAGUUCCGCACGGCGGAAAGUCUACGGGAAAUUAUUAACA
Eho-1-9      UCAGUUUGCAUACGUUGGUUAAAUUCUGUAUGAUUCGCCGCCUUUGGUGUACUUUAUGCCAAAGAACUUACAUUCAACA
env-736      UUAUACCGCAUUGGAGAUUGACCAUUACCGUACGCUUCGCGCUGUCAGCAGCGUGGAAGACGACGGGAAAUUAUUAAUA
Kox-8-1      GAUUCGCCGCCUUUGAUGUGCUUCGUGUAGCAUCAAAAUCUAUUACCCGGUAGUGUAAUCGCUCCGGGAAUUAUUAACA
Esp-29-1     UCAGUUUGCAUACGUUGGUUAAAUUCUGUAUGAUUCUCCGCCUUUGGUGUACUUUACGCCAAAGAACUUACAUUCAACA
Ror-1-1      UGUGCUUCGUGCAGCAUCAAAGCGUAUACUAUUACCCAUGAGUUCCGCACGGCGGAAAGUCUACGGGAAAUUAUUAACA
env-738      UGCAUACGCAGGUCAAAUUCUGUAUGAUGCGCCGCCUUUGAUGCGCUUUAUGCAGCGUCAAAAACCUNUACUAUUAACA
Ecl-42-1     UCAGUUUGCAUACACUGGUUAAAUUCUGUAUGAUUCGCCGCCUUUGGUGUAUAUUACGCCAGAGAAAUUACAUUCAACA
env-739      UUUGGUGCGCUUCGUGCAGCAUCAGAAUCUAUUACCCGGGGAUUGCGUUUAGCGGAAUUGCGCCGGGAAAUUAUUAACA
Rpl-1-1      UUUGGUGCGCUUCGUGCAGCAUCAGAAUCUAUUACCCGGGGAUUACGUUUAGCGGAAUUGCGCCGGGAAAUUAUUAACA
Eca-3-1      UCAGUUUGCAUACGCUGGUUAAAUUCUGUAUGAUUCGCCGCCUUUGGUGUACUUUACGCCAAAGAACUUACAUUCAACA
Kqu-1-1      UUAUAUCGCAUUGAAGAUUGACCAUUACCGCGUGCUUCACGCUAUCAACAGCGUAGAGGGCGACGGGAAAUUACUAAUA
Esp-67-1     CAGUUUGCAUCCACCGAAUGGAUUCUGUAUGAUUCGCCGCCUUUGGUGUAACAUUACGCCAAAGAAUUUACUAUCAACA
env-740      UUUGGUGCGCUUCGUGCAGCAUCAGAAUCUAUUACCCGGGGAUUACGCUUAGCGGAAUUGCGCCGGGAAAUUAUAAACA
env-741      AUCAAACAUCCCAAAACCCGGGAAAUUCGCUAAGCCGGUUUACGCUAGCUGCGCGAAGGUCGUCGGGAAAUUAUUAAUA
env-742      UCAGUUUGCAUCGCCUGGUUAAAUUCUGUAUGAUUCGCCGCCUUUGGUGUACUUUACGCCAAAGAACUUACAUUCAACA
env-743      UUAUAUCGCAUUGAAGAUUGACCAUUACCGCGUGCUUCACGCUGUCAGCAGCGUAGAAGGCGACGGGAAAUUACUAAUA
env-754      CAGAUUGCAUCUGCUGGUUAAAUUCUGUAUGAUUCGCCGCCUUUGAUGUAUAUUUGCAUCAUCAACUUUAUCAUUAACA
env-755      ..............................................................................A
env-756      UCAGUUUGCAUAGCCUGGUUAAUUUCUGUAUGAUUCGCCGCCUUUGGUGUAAUUUACGCCAGAGAACUUACAAUCAACA
Kcr-1-1      GUUUGCAUCUGCUGGUCAAAUUCUGUAUGAUUCGCCGCCUUUGAUGCAUUUUGGCAUCAGGAAAUAUACUACAUUAACA
Kas-1-1      AGUUUGCAUCUGUUGGUCAAAUUCUGUAUGAUUCGCGGCCUUUGAUGCAUCUUUGUAUCAUCGACCUAUACUAUUAACA
Fhe-1-1      AGUUUGCAUACGGCGGUUAUUUUCUGUAUGAUUCGCCGCCUUUGAUGUGCAGUUUCGCCAUCAAACCUUUCUAUCAACA
Fhe-2-1      AGUUUGCAUACGGCGGUUAUUUUUUGUAUGAUUUGCCGCCUUUGAUGUGCAGUUUUGCCAUCAAACCUUUCUAUCAACA
Fpu-1-1      GAUUUGCAUUGGGUGGUUAUUUUCUGUAUGAUUCGCCGCCUUUGAUGUGCAGUUCCGCCAUCAAACCUUACUAUCAACA
Fhe-3-1      AGUUUGCAUACGGCGGUUAUUUUUUGUAUGAUUUGCCGCCUUUGAUGUGCAGUUUCGCCAUCAAACCUUUCUAUCAACA
Fpu-5-1      GGUUUGCAUUGGGCGGUUAUUUUCUGUAUGAUUCGCCGCCUUUGAUGUGCAGUUCCGCCAUCAAACCUUACUAUCAACA
Cne-1-1      UGCACCGGGUGAGCAAUUUCUGUAUGAUUCGCCGCCUUUGAUGCGCAGUUUUGGUAUCAAAAAACCUAAUUUAUUAACA
Cne-1-3      GCACCGGCUGAGUAAGUUCUGUAUGAUUCGCCGCCUUUGAUGCGCAGUUUUGGUAUCGAAAAAACCUAAUUUAUUAACU
Cda-1-1      UGCACCGGCAGGUUAAUCUCUGUAUGAUUCGCCGCCUUUGAUGCGCAGUUUUGGUAUCGAAAAAACCUAAUUAUUAACA
Sco-2-1      AUCCGGUGGCUAUUUUCUGUAUGAUUCGCCGCCUUUGAUGCGAUUUAUUCGCGUUGAAGCUCACCAAAAUACAUUAACA
Bag-2-1      GCAUCCACUCGCGGGGUUCUGUAUGAUUCGCCGCCUUUGAUGCGCAGUUUUUGGUGUCAAAAAACCUAAUUUAUCAACU
Ebl-1-1      UGCACAGGGCGGAAACUCUCUGUAUAAUUCGCCGCCUUUGAUGCGCGAUUUUCGCCUUCAAUAUACAUCAUUAUCAACC
Sen-311-1    UUUGCAUACAGAGGCCAAAUUCUGUAUGAUUCGCCGCCUUUGGUGCGUGUUAUGCGCAUCAAACCCUUUACUAUUAACU
Sen-1013-1   UUUGCAUACGGUGGUCAAAUUCUGUAUGAUUCGCCGCCUUUGGUGUGUGUUGUGAGCAUCAAACCCUUUACUAUUAACU
Sen-89-1     UUUGCAUACGGUGGUCAAAUUCUGUAUGAUUCGCCGCCUUUGGUGCGUGUUGUGAGCAUCAAACCCUUUACUAUUAACU
Sbo-3-1      UUUGCAUACGGUGGUCAAAUUCUGUAUGAUUCGCCGCCUUUGGUGCGUAUUGCGAGCAUCAAACCCUUUACUAUUAACU
Sen-1029-1   UUUGCAUACGGUGGUCUAAUUCUGUAUGAUUCGCCGCCUUUGGUGCGUGUUGUGAGCAUCAAACCCUUUACUAUUAACU
Sen-1021-1   UUUGCAUACGGUGGUCAAAUUCUGUAUGAUUCGCCGCCUUUGGUGCGUGUUGUGAGCAUCAAACCCUUUACUAUUAACU
Sen-312-1    UUUGCAUACGGUGGUCAAAUUCUGUAUGAUUCGCCGCCUUUGGUGCGUGUUAUGAGCAUCAAACCCUUUACUAUUAACU
Bag-1-1      GCAUCCAACGGUGGGGUUCUGUAUGAUUCGCCGCCUUUGAUGCGCAGUUUUUGGUAUCAAAAAACCUAAUUUAUCAACU
env-757      CAGGGCGGUUAUUUUCUGUAUGAUUCGCCGCCUUCGAUGCGAUUAAUUCGUGUUGAAGCUAACCAAAAUACAUUCAACA
Cfr-3-1      GAUUGCGUCAAACAAUGACCAUUACCCGAGAAUUUCAAGCUGUUGAUGCAGUGUGAAAGGUGACGGGGAAUAAUUAACA
Ksp-3-1      UUUGCAUCCGCUGGUUAAAUUCUGUAUGAUUCGCCCCCUUUGAUGCGCUAAUCGCGUGUCAAAAAAAUUACCAUUAACA
Eco-1011-1   AGUUUGCAUACGCUGGUUAAUUUCUGUAUGAUUUCGCGCCUUCGUACGAAAUGAUCGUAUUGAAGCUAUACUUUUAACA
env-758      AGUUUGCAUGAGCAGGUUAAUUUCUGUAUGAUUCGCCGCCUUUGAUACAAAUUUGUAUCACUGACCUAUACUAUUAACA
Eco-1781-1   AGUUUGCAUACGCUGGUUAAUUUCUGUAUAAUUUCGCGCCUUCGUACGAAAUGAUCGUAUUGAAGCUAUACUUUUAACA
Cam-1-1      GAUUGCGUCAAACAAUGACCAUUACCCGAGAGUUUCAGGCUGUUGAUGCAGCGUGAAAGGUGACGGGGAAUAAUUAACA
Sso-1-2      NNNNNNNNNNNNNNNNNNNNNNNNNNAUGAUUUCGCGCCUUCGUACGAAAUGAUCGUAUUGAAGCUAUACUUUUAACA
Sfl-70-1     AGUUUGCAUACGUUGGUUAAUUUCUGUAUGAUUUCGCGCCUUCGUACGAAAUGAUCGUAUUGAAGCUAUACUUUUAACA
Ssp-20-2     CAUAGUGAACUAUGUGACCAGAGUGAGUGAACGCAGGCAACGCAUCAGCAGUGUGAAAGGUGACGGGGAAUAAUUAACA
Eco-1460-1   AGUUUGCAUACGCUGGUUAAUUUCUGUAUGAUUUCGCGCCUUCUUACGAAAUGAUCGUAUUGAAGCUAUACUUUUAACA
env-763      AGUUUGCAUCCGCUGGUUAAUUUCUGUAUGAUUUCGCGCCUUCGUACGAAAUGAUCGUAUUGAAGCUAUACUUUUAACA
Cwe-1-1      GAUUGCGUCAAACAAUGACCAUUACCCGAGAAUUUCAAGCUGUUGAUGCAGUGUGAAAGGUGACGGGGAAUAAUUAACA
Sso-1-3      NNNNNNNNNNNNNNNNNNNNNNNNNNNNNNNNNNNNNNCCUUCGUACGAAAUGAUCGUAUUGAAGCUAUACUUUUAACA
env-766      AGUUUGCAUAUGCUGGUUAAUUUCUGUAUGAUUUCGCGCCUUCGUACGAAAUGAUCGUAUUGAAGCUAUACUUUUAACA
env-784      CAGUUUGCAUCCGCUGGUUAAUUUCUGUAUGAUUUCGCGCCUUGUACGAAAUGAUCGUAUUGAAGCUAUACUUUUAACA

             ...............................................................................
             ...............................................................................
             ••••••••••••••••••••••••••••••••••••••••••••••••••••••••••••••••••••••••••••RY•
```

```
env-785       GUUUGCAUACGCUGGUUAAUUUCUGUAUGAUUCGGCGCCUUUGAUACGAAAUGAUCGUAUUGAAACUAUACUUUAAACA
Cro-2-1       GAUUGUGUCGAAAAUUGACCAUUACCCGAGAAUUUCGCGCUGCUGGUGUAGAGUGGAAGGCGACGGGAAAUAAUUAACA
Eal-2-1       AGUUUGCAUACGCUGGUUAAUUUCUGUAUGAUUCGGCGCCUUUGAUGCGAAAUGAUCGUGUCAAACUAUACUUUAAACA
Sso-1-5       NNNNNNNNNNNNNNNNNNNNNNNNNNNNNNNNNNNNNNNGCCUUCGUACGAAAUGAUCGUAUUGAAGCUAUACUUUUAACA
Eco-835-1     AGUUUGCAUACGCUGGUUAAUUUCUGUAUGAUUUCACGCCUUCGUACGAAAUGAUCGUAUUGAAGCUAUACUUUUAACA
Ban-1-1       GUUUUGCAUACGCUGGUUAAUUUCUGUAUGAUUUCGCGCCUUCGUACGAAAUGAUCGUAUUGAAGCUAUACUUUUAACA
Eco-1442-1    AGUUUGCAUACGCUGGUUAAUUUCUGUAUGAUUUCGCGCCUUCGUACGAAAUGAUCGUAUUGAAGCUCUACUUUUAACA
Cyo-1-1       GAUUGCGUCAAAUAAUGACCAUUACCCGAGAAUUUCAAGCUGUUGAUGCAGUGUGAAAGGUGACGGGGAAUAAUUAACA
Eco-763-1     AGUUUGCAUACGCUGGUUAAUUUCUGUAUGAUUCGGCGCCUUUGAUACGAAAUGAUCGUGUCAAACUAUACUUUAAACA
Sdy-5-1       AGUUUGCAUACGCUGGUUAAUUUCUGUAUGAUUUCGCGCCUUUGUACGAAAUGAUCGUAUUGAAGCUAUACUUUUAACA
env-786       AGUUUGCAUACGCUGGUUAAUUUCUGUAUGAUUUCGCGCCUUCGUACGAAAUGAUCGUAUUGAAGCUAUACUUUUAACA
Eco-1862-1    AGUUUGCAUACGCUGGUUAAUUUCUGUAUGAUUUCGCGCCUUCGUACGAAAUGAUCGUAUUGAAGCUAUACUUUUAACA
env-861       AGUUUGCAUACGCUGGUCAAAUUCUGUAUGAUUCGCCGCCUUUGAUACACAUUUGUAUCAUCGACCUAUACUAUCAACA
Cko-2-1       AGUUUGCAUACGCUGGUCAAAUUCUGUAUGAUUCGCCGCCUUUGAUACACAUUUGUAUCAUCGACCUAUAUUAUCAACA
Csp-14-1      AGUUUGCAUGAGCAGGUUAAUUUCUGUAUGAUUCGCCGCCUUUGAUACAAAUUUGUAUCACUGACCUAUACUAUUAACA
env-862       UGCAUACGCUGGUUAAUUUCUGUAUGAUUCGCCGCCUUUGAUGCUCAUUGCGGCGUCAAAGAUUCACUUACAUAUCAAC
Cam-2-1       GAUGCAAAUUGAUUGUGUCAAAACUUGACCGUUACCCGAGAAUUUCACGCUGCUGGAAGGCGACGGGUAAAAAUUAACA
env-867       GUUUGCAUGAGCGGGCUAAUUUCUGUAUGAUUCGCCGCCUUUGAUGCAAAUUGCUUGUGUCAAAACUAUACUAUCAACA
Cse-1-1       GAUCGCGUCAAAAAUUGACCAUUACCCGAGAAUUUCGCGCUGCUGGUGUAGAGUGGAAGGCGACGGGAAAUAAUUAACA
env-868       GCAUGUGCUGGUCAAUUUCUGUAUGAUUCGCCGCCUUUAGUGCGGAUUUAGCGCGCCAAAGAUACACUUAUUAUUUAAC
Esp-43-1      UGCAUACGUUGGUUAAUUUCUGUAUGAUUCGCCGCCUUUGGUGCGUAUUCACGUGCCAGAGAUCCACUAAUACAUUAAC
Kor-1-1       GCAUGUGCUGGCUAAUUUCUGUAUGAUUCGCCGCCUUUGAUGCGUUUUGAACGUGCCAAAGAUACACUUACAUAUUAAC
Ksa-1-1       GCAUGUGCUGGCUAAUUUCUGUAUGAUUCGUCGCCUUUGAUGCGUUUUGAACGUGCCAAAGAUACACUUACAUAUUAAC
Mma-2-1       CUUAUUGCCCGAGAAUUUCCCGCUUCCUGAAUGUGGCUAACGCAUCAGCGGUGUGAGAGACAACGGGGAAAUUAUUAAC
Pur-1-1       UGCAUACGUUGGUUAAUUUCUGUAUGAUUCGCCGCCUUUGGUACGUAUUCACGCGCCAGAGAUCCACUAAUACAUUAAC
Cfa-1-1       GUUUGCAUGAGCGGGCUAAUUUCUGUAUGAUUCGCCGCCUUUGAUGCAAAUUACUUGUGUCAAAACUAUACUAUCAACA
env-869       GCAUACGCUGGCUAAUUUCUGUAUGAUUCGCCGCCUUUGGUGCGUAUUCACGCAGCAGAGAUCCACUAAUACAUUAACU
env-870       ..................................................CGGCGUCAGAGAUUCACUUACAUAUCAAC
Kcf-1-1       UGCAUACGCUGGUUAAUUUCUGUAUGAUUCGCCGCCUUUGAUGCUCAUUGCGGCGUCAGAGAUUCACUUACAUAUCAAC
env-871       GUUUGCAUGAGCAGGCUAAUUUCUGUAUGAUUCGCCGCCUUUGAUGCAAAUUGAUUGUGUCAAAACUAUACUAUCAACA
Esp-42-1      UGCAUGUGCUGGUUAAUUUCUGUAUGAUUCGCCGCCUUUGAUGCGUUUUAGCGCGCCAAAGAUACACUUACAUAUUAAC
Era-1-1       GCAUGUGCUGGUUAAUUUCUGUAUGAUUCGCCGCCUUUAGUGCGGAUUUAGCGCGCCAAAGAUACACUUAUUAUUUAAC
Kra-2-1       CAUGUGCUGGUUAAUUUCUGUAUGAUUCGCCGCCUUUAGUGCGGAUUUAGCGCGCCAAAGAUACACUUAAUAUAUUAAC
Csp-12-1      AGUUUGCAUGAGUGGGCUAAUUUCUGUAUGAUUCGCCGCCUUUGAUGCAAAUUGAUUGUGUCAAACUAUACUAUCAACA
Ecl-60-1      UUGCAUACGGUGGUUAAAUUCUGUAUGAUUCGCCGCCUUUGAUGCGCUAUCUGCGCGGCAAAAAAUCCUAAUAUCAACU
env-872       CAGUUUGCAUAGACUGGCUAAAUUCUGUAUGAUUCGCCGCCUUUGAUACAGAUUCGUAUCAGGGACUAAUCAAUCAACA
env-874       UCAUCGGUUAAUUUCUGUAUGAUUCGCCGCCUUUGAUGCGCAGUUUUGGCAUGCAAAGCAAAAAAACCGUUUAUUAACA
env-875       GAUUGCAUGCGCUGGUUAAAUUCUGUAUGAUCCGCCGCCUUUGAUACAGAUUUGUAUCACUGACACUAUACUAUCAACU
Pvu-1-1       UUUGCAUAUGGCGGUUAAUUUCUGUAUGAUUCGCCGCCUUUGGUGUUGCAUUCUGCCGCCAGAGCUUAAAUUAAUUAUC
Gpe-1-1       AUAACCGGAGGGGGGGAUUGAUUAUUGAACAACUGAUCCGCUAUGAUGCGCGCCUUCGAAUUUUUGAACGUUAACACAA
Gxi-1-1       AUCACGCGGGGCGGGGAUUGAUUAUUGAGCAAGGGAUCCGGUAUUAUGCGCGCCUUCGAAUUCCCGAACGUUAACACAA
env-876       AAAUAUCAGCGUAAAGACAGAAGCUGUUGCAACUUAAGCGGCUAUCAAUUAAGCUACGCGCCCUAAAUUAACUGUAAUU
env-880       CCAUUAAAAAUAAAUUAGCCAAUUUGUUGAAACUUUAAGCCUUAGCAAUUAAACUACGCGCCCUAAUUUAACUGAAAUU
Pme-1-1       CCAUUAAAAAUAAAUCAGCCAUUUUGUUGAAACUUUAAGCCCUAGCCAUUAAACUACGCGCCCUAAUUUAACUGAAAUU
Rsp-2-1       CCAUUAAAAAUAAAUCAGCCAAUUUGUUGAAACUUUAAGCCUUAGCCAUUAAACUACGCGCCCUAAUUUAACUGAAAUU
env-881       CCAUUAAAAAUAAAUUAGUCAAUUUGUUGAAACUUUGAGCCUUACCAAUUAAACUACGCGCCCUAAUUUAACUGAAAUU
Rsp-1-1       CCAUUAAAAAUAAAUUGGUCAAUUUGUUGAAACUUUGAGCCUUACCAAUUAAACUACGCGCCCUAAUUUAACUGAAAUU
Aag-1-1       AAAUGUCGGCUUUUUCAGUUUUACGGUUGCAACUUUAAUGGCUAUCUAUUAAGCUACGCGCCCUUAAUUAACUGUAAUU
Aae-1-1       AAUUCUGUGCUUUUCCGGUUUUCUGGUUGCAACUUCAGCGGCUAUCUAUUAAGCUACGCGCCCUAAAUUAACUGUAAUU
env-882       AANUCUGUGCUUUUCNGGUUUUCUGGUUGCAACUUCAGCGGCUAUCUAUUAAGCUACGCGCCCUAAAUUAACUGUAAUU
env-883       AAAUCUGUGCUUUUCUGGUUUUCUGGUUGCAACUUCAGCGGCUAUCUAUUAAGCUACGCGCCCUAAAUUAACUGUAAUU
Aik-1-1       AAGUUGUAAAUAAGCCUGCCUGGUAGUUGCAACUUUGCUGGCUAUCUAUUAAGCUACGCGCCCUUAAUUAACUGUAAUU
env-884       AAAUGCUAAAUAUCUGUGGAUGCCAGUUGCAACUUUGCUGGCUAUCUAUUAAGCUACGCGCCCUUAAUUAACUGUAAUU
env-885       AAGUGCUAAAUAAGUCAGGUUGGCAGUUGCAACUUGGCUGGCUAUCUAUUAAGCUACGCGCCCUUAAUUAACUGUAAUU
Asp-9-1       AAGCGGCAAAUAGGUGGGUUUGCCAGUUGAAACUUUGAUGGCUAUCUAUUAAGCUACGCGCCCUUAAUAAACUGUAAUU
Rpe-1-1       AAGUUGUAAAUAAGCCGGUAUGACAGUUGCAACUUUGCUGGCUAUCUAUUAAGCUACGCGCCCUUAAUUAACUGUAAUU
Rba-1-1       AAUUUAUCCUGCAACGGCUUCAAGUAUUGCAACUUAAGCGGCUAUCAAUUAAGCUACGCGCCCUAAUUUAACUGUAAUU
Rna-1-1       AAUUUGCCGUAACAGCGUCUGGCCUGUUGCAACUUGACCGGCUAUCUAUUAAGCUACGCGCCCUUAAUUAACUGUAAUU
env-886       UGCACAAUCAGUUGUUUUACUUUAAUAUGCGCCUCCCUUAACAGCAAGUCGAAUGUUCGAUUUGUAUAAAAAUUUAAUU
Asa-9-1       UACAUUAAUAUGCGCGCCUCUUUGCGGCAAGCCCUGGAAAUGGAAACCUUUUCAUGACCAGGCGGCCAACAUUAUAAUC
Gli-1-1       UCAAAACAGCUUGAUCAAUACAGACCUUUACAUUAAUAUGCGCGCCUCUUUGCGGCAAGCCGCUAUUUAAAAUUAAUUU
Gpu-1-1       GCUCCACAGGUUUAAUUUGUAUAGUAUUCGUGCUCAAAUUUUCAAGUAUAGAAUAAUCUGUGCUUACAUUUAAUUAACA
env-889       UGAUCAUUUAAGGAAGUUACAUUAUUAUGCGCGUCCAUUUUGCGAGCUACUGCACGUGCGGUAGGUCAUGUUAAUUAUU
env-890       UGAUCAAAGGGCUAAGUUACAUUAUUAUGCGCGUCCAUUUUGCGACUCAUUGCGUGUGCGGUGAGUCAUGUUAAUUAUU
Gni-1-1       CUUGAUUAAGGGCCUCAACUCCAUUAUUAUGCGCGUCCAAAUUUCUGUAGUUGAAAACUCAGCUGCUUUUUAAUUAAUU
Gpa-1-1       CUUGAUUAAGGAGUGUAAGUCCAUUAUCAUGCCGGUCCAAAUUUCUGUAGUUGAUUAUUCAGCUGCUUUUUAAUUAAUU
env-894       AAGGCCCUGCCAUCCUUUAUUAUGCGCCGCUCUUUUGCGAUAGGCUGUUCUUUGAACGACUUAUCAUUUGAAUUAAGUA
Gpo-1-1       UGAUCAGGGCAAGGUGUUACAUUAAUAUGCCCGGCUCUUUGCGGCAAGCCGCAUUGAUCGCAUAAUUUGAAUUUAAUUU
env-895       AACCCUUGAUCGCCUAGGCCCGUUACAUUAAUAUGCGCGCCACUUUGCGGCAAGCCACAAAGCUUUUUUAAUUUAAUUU
Gar-1-1       AAGCCUUGAUCGCCCAAGCCCGUUACAUUAAUAUGCGCGCCACUUUGCGGCAAGCCACAAAGCUUUUUUAAUUUAAUUU
env-896       AAACCCUUGAUCGCCCCGGCCCGUUACAUUAAUAUGCGCGCCACUUUGCGGCAAGCCACAAAGCUUUUUAAUUUAAUUU
Gps-1-1       ACCCUUGAUCGCCCAAGCCCGUUACAUUAAUAUGCGCGCCACUUUGCGGCAAGCCACAAAGUCUUUUUUAAUUUAAUUU
env-898       CUUGAUUGAUUAUUCGGCAUCCAUUAAUAUGCGCCGCUCCAAAAAGCAGGCGGUAUGUCGUCUGUAGUGAAAAUUAGUC
env-899       CUUGAUUGAUUAUUCGGCAUCCAUUAAUAUGCGCCGCUCUAAAAAGCAGGCGGUAUGUUGCCUGUAGUGAAAAUUAGUC
env-900       CUUGAUUGAUUAUUCGGCAUCCAUUAAUAUGCGCCGCUCUAAAAAGCGGGCGGUAUGUCGCCUGUAGUGAAAAUUAGUC
env-902       CUUGAUUGAUUAUUCGGCAUCCAUUAAUAUGCGCCGCUCUAAAAAGCAGGCGGUAUGUUGCCUGUAGUGAAAAUUAGUC
env-930       CUUGAUUGAUUAUUCGGCAUCCAUUAAUAUGCGCCGCUCCAAAAAGCAGGCGGUAUGUCGUCUGUAGUGAAAAUUAGUC
env-933       CUUGAUUGAUUAUUCGGCAUCCAUUAAUAUGCGCCGCUCGAAAAAGCAGGCGGUAUGUCGCUUGUAGUGAAAAUUAGUC
env-934       .........................................AAAAGCGGGCGGUAUGUUGCCUGUAGUGAAAAUUAGUC
Pat-1-1       UGAUCAGGGCAAGGCGUUACAUUAAUAUGCCCGGCUCUUUGCGGCAAGCCGCAUUGAUCGCAUAAAUCGAAUUUAAUUU
env-935       CUGGCGCGGCUUCCUUUAUUAUCCGCGCUCUUUUGCGAUGGGCUGUUCCAAGAACUGCUCAUCAUUUGAAUAUUAAGUU
Asp-7-1       UGGCGCAGCUUCCUUUAUUAUCCGCCGCUCUUUUGCGAUGGGUUGUUCUAAGAACCGCUCAUCAUUUGAAUAUUAAGUU
env-936       CCUUGAUUGAUUAAUCGACAUCCAGUAAUAUGCGCCGCUCUUAAAAACCGAGCGAUUAUCGCUUGUACUGAAUUUAGCA
env-937       CCUUGAUUGAUUAAUCGACAUCCAGUAAUAUGCGCCGCUCUUAAAAACCGAGCGAUUAUCGCUUGUACUGAAUUUAGCA

              ...............................................................................
              ...............................................................................
              ............................................................................RY.
```

| | |
|---|---|
| Sch-1-1 | CUUGAUCAGCGAGGCUACUUACAUUAAUAUGCGCCGCUCUUUACGGCAAGGUCGUUUUGACUUUGUAAAUAUUUAAGUA |
| env-954 | UGAUCAGCAGGACAUCAUACAUUAAUAUGCGCGCCUCUUUACGGCAAGGCCAGUUUGGUUUUGCCACUAUUAUUGAAUA |
| env-955 | UUGAUCAAACGGGCGCCAUACAUUAAUAUGCGCCGCUCUUUACGGCAAGGCCAGUUGUGUCUUGCAUUUGAAUAUAGUU |
| Asp-5-1 | AAGCCACUCCCAUCCUUUAUUAUGCGCCGCUCUUUUGCGAUAGGCUGUUCUUUGAACGACUUAUCAUUUGAAUUAAGUA |
| env-956 | ACCCUCUGCCAUCCUUUAUUAUGCGCCGCUCUUUUUGCGAUAGGCUGUUCUUUGAACGACUUAUCAUUUGAAUUAAGUA |
| Ama-14-1 | AAGGCUUUGCCAUCCUUUAUUAUGCGCCGCUCUUUUGCGAUAGGCUGUUCUUUGAACGACUUAUCAUUUGAAUUAAGUA |
| env-957 | AAGGUCCUGCCAUCCUUUAUUAUGCGCCGCUCUUUUGCGAUAGGCUGUUCUUUGAACGACUUAUCAUUUGAAUUAAGUA |
| env-981 | AAGGCCCUGCCAUCCUUUAUUAUGCGCCGCUGUUUUGCGAUAGGCUGUUCUUUGAACGACUUAUCAUUUGAAUUAAGUA |
| env-982 | AGCCCCUGCCAUCCUUUAUUAUGCGCCGCUCUUUUUGCGAUAGGCUGUUCUUUGAACGACUUAUCAUUUGAAUUAAGUA |
| env-986 | AAGGCUCUGCCAUCCUUUAUUAUGCGCCGCUCUUUUGCGAUAGGCUGUUCUUUGAACGACUUAUCAUUUGAAUUAAGUA |
| env-992 | AAGGCCCUGCCAUCCUUUAUUAUGCGCCGCUCUUUUGCGAUAGGCUGUUCUUUGAACGACUUAUCAUUUGAAUUAAGUA |
| env-1025 | AAGGCCCUGCCAUCCUUUAUUAUGCGCCGCUCUUUUGCGAUAGGCUGUUCUUUGAACAACUUAUCAUUUGAAUUAAGUA |
| env-1027 | AAAGCCCUGCCAUCCUUUAUUAUGCGCCGCUCUUUUGCGAUAGGCUGUUCUUUGAACGACUUAUCAUUUGAAUUAAGUA |
| env-1029 | AGCCCUUCCCAUCCUUUAUUAUGCGCCGCUCUUUUUGCGAUAGGCUGUUCUUUGAACGACUUAUCAUUUGAAUUAAGUA |
| env-1030 | GCGCCUCCAUCUAUUAUUAUGCGCCGCUCUUUUGCGAUGGGCUGUUUAUGUAAACGACUCAUCAUUUGAAUUUUAAGUU |
| env-1031 | CGAGCACGCAUACAUUAUUAUGCGCCGCUCUUUUGCGAUAGGUUGUUAUUUAAACCACUUAUCAUUUGAAUAUUAAGUA |
| Ama-7-1 | AGCACCUGCCAUCCUUUAUUAUGCGCCGCUCUUUUUGCGAUAGGCUGUUCUUUGAACGACUUAUCAUUUGAAUUAAGUA |
| env-1032 | ACCCUCUGCCAUCCUUUAUUAUGCGCCGCUCUUUUUGCGAUAGGCUGUUCUUUGAACGACUUAUCAUUUGAAUUAAGUA |
| env-1035 | AGCCCCUGCCAUCCUUUAUUAUGCGCCGCUCUUUUUGCGAUAGGCUGUUCUUUGAACGACUUAUCAUUUGAAUUAAGUA |
| env-1036 | AGGCACUUCCAUCCUAUAUCAUGCGCCGCUCUUUUUGCGAUAGGUUGUUCUUUAAACGACUUAUCAUUUGAACUAAGUA |
| env-1037 | GCAGCCCUCUUCCUUUAAUAUGCGCCGCUCUUUUUACGGCUGCCUGCACUUAUGCGGUCGGUCAUUUGAAUAUUAAGUU |
| Pha-4-1 | UGUUAUAUUGAUUAUAUUUAUUGAUUGUAUUUAUUGAUUGGCUUGUUUAUAAUUUGCGCCUUAAUUUUUAAUGAUAAAU |
| env-1093 | CGGUAAAAUACACUGUUUUUAUGUUGUAUUUUUAUUGUUCAUUCCUAUAUAAUUCGCACCUUGAAAUUUAAUGAUAAAU |
| env-1094 | AGUGCUAAGAAUUCAUUUUAUAGUUGUAUUUUUAUUAAACAUUGCUAUACAAUUUGCAACCUGAAAUUUAACGAUAAAU |
| env-1096 | AGUGCUAAGAAUUCACUUUAUAGUUGUAUUUUUAUUAAACAUUGCUAUACAAUUUGCAACCUGAAAUUUAACGAUAAAU |
| env-1097 | ACUGCUAAGAAUACACUUUUUAGUUGUAUUUUUAUUAAACAUUACUAUACAAUUUGCAACCUGAAAUUUAACGAUAAAU |
| Pin-1-1 | AGUGCUGAGAAUACACUUUAUAGUUGUAUUUUUAUUAAACAUUGCUAUACAAUUUGCAACCUGAAAUUUAAUGAUAAAU |
| env-1099 | AAUAAUAAUAAUGGUGUUUAUUGUUGUAUUUUUAUUAAACAUUCCUAUACAAUUCGCCACCUAAAUUUUAAUGAUAAAU |
| env-1103 | AGUAAUAAAAAUAGCGAUUAUUGUUGUCUUUUUAUUAAUCAUUCCUAUACAAUUCGUCACCUAAAUUUUAAUGAUAAAU |
| env-1104 | AACUGAAAAAAACAGUUUUUAUAUUGUCUUUUUAUUGAACGUUCCUAUAUAAUUUGCAACCUGAAAUUUUAUGAUAAAU |
| Par-2-1 | UUGUUAUUUUUGGUUAUUUUAAGUUGUACUUUUAUUGCCCAUUACUAUACAAUCCGCAACCUGAAAUUUAAUGAAAAAU |
| Paq-1-1 | AGUGGAAAUAUCCUAUAUUUUGGUUGUAUUUUUAUUGCACAUCCUUAUAUAAUUCGCCACCUAAAAAUUUACGAUUAAU |
| Pos-1-1 | AGUCGAAAUAUCCCAUUUUUUGGUUGUAUUUUUAUUGCACAUCCUUAUAUAAUUCGCCACCUAAAAAUUUACGAUAAAU |
| Psp-35-1 | UGUUAAAUUUUAGUUAUUUUAAGUUGUACUUUUAUUAUUCGUUACUAUACAAUCCGCAACCUGUAAUACAAUGAUAAAU |
| env-1105 | CGAUUGAUUUUAGUUAUUUUAAGUUGUACUUUUAUUGUUCGUUGCUAUACAAUCCGCAACUUGAAAUUUAAUGAUAAAU |
| env-1109 | CGAUUGAUUUUAGUUAUUUUAAGUUGUACUUUUAUUGGUCGUUGCUAUACAAUCCGCAACUUGAAAUUUAAUGAUAAAU |
| env-1110 | ACUAAUAAAAAUACAGUUUAUAGUUGUAUUUUUAUUAAACAUUCCUAUACAAUUCGCCACCUGAAAUUCAAUGAUAAAU |
| Psp-34-1 | UUUAGGUAAAAAAACAGAUAUAAGUUACUUUUUGGACUUAACUUCUAUAUAAUUCGCAACCUGAAAUUUUAUGAUAAAU |
| Sam-1-1 | CAUUAUUUUUAUUACACCUAUUGCACAUUCUCUCUCAAUAAUAUAUCAUGCGACCCGUUUUUUGACUUUUACUUUAACU |
| env-1113 | UUUUAUCAUGAUUCACAAAGGGUUUUAGUUGAAUUUUUAGGGUUUGAAGGUUAGAAUAGCGGCCUAUUUUUACACAUAU |
| Pfl-3-1 | CAGUACGGAAUUGUUCGUUAUACUGUGUUGCCCUGAAUCAUAUUAAGUUAAUUACAGGGUAACACUAUUUUAUUUAAUC |
| Ykr-2-1 | CUAACAACCGUUGUUUAUUUUAUCGGCGAUUGUUCGUUAGUAAUCAACUAUUCGUUAACGAUAAAUUGUUAUUUAUCAA |
| Pge-1-1 | UCAGUUUGCAUUAGCUGCCCAAAUUCUGUAUGAUUCGCCGCCUUUGGUGCAUCCGGCAUCAGAGAUCUAUUAUUCAACA |
| Kpn-281-1 | UUAUAUCGCAUUGAAGAUUGACCAUUACCGCGUGCUUCACGCUGUCAACAGCGUAGAGGGCGACGGGAAAUUACUAAUA |
| Vja-2-1 | AAGUGAUUCAUUACUAUACUGCGCCUUGAGAUAUGGCUUCGGCAAUUGGCCGAUGCGGUACUCAUUUCUAUUAUUAAUU |
| Kpn-256-1 | UUAUAUCGCAUUGAAGAUUGACCAUUACCGCGUGCUUCAUGCUGUCAGCAGCGUAGAAGGCGACGGGAAAUUACUAAUA |
| Hin-6-1 | CGAAAAUAGAGCUUGUUUAUCUGCAUUUUUUCAGUAUAAUUCGCAACCUAUUUGGUGCCAAUUCUGGCGUAUUAUUCAC |
| Eba-1-1 | UCAGUUUGCAUCGCCUGGUUAAAUUCUGUAUGAUUCGCCGCCUUUGGUGUACUUUACGCCAAAGAACUUACAUUCAACA |
| Ysi-1-1 | UUCGCCGCCUUUGAUAUAAUUUGAUAUCAAAUCCUGUUAACAAGCGUUGCUUAUUUUAUCAGCGAUUGUUAUUUGUCAA |
| Hin-1-3 | CGAAAAUAGAGCUUGUUUAUCUGUAUUUUUUCAGUAUAAUUCGCAACCUAUUUGGUGCCAAUUAUGGCGUAUUAUUCAC |
| Kox-4-2 | GAUUCGCCGCCUUUGAUGUGCUUCGUGCAGCAUCAAAAUCUAUUACCCGGUAGUGUAAUCGCUCCGGGAAUUAUUAACA |
| Hin-13-1 | CGAAAAUAGAGCUUGUUUAUCUGCAUUUUUUCAGUAUAAUUCGCAGCCUAUUUGGUGCCAAUUAUGGCGUAUUAUUCAC |
| Sma-10-1 | UUCCGAUUGCGCGUCCCGACUAAUGCUGUAUAAUUUGCCGCCUUUGGUAUAAUUCGAUACCAAAAUUCUAUUAUCAACC |
| Yru-2-2 | AUUCGAUUGCGAGGGCUGCUUAAAUUCUGUAUGAUUCGCCGCCUUUGAUAUAAUCCGAUAUCAAAUCUUAUUAUCAACU |
| Vni-11-1 | UAGGUUGAUCCUACCCUCAUGUUUCAUUACUAUACUGCGCCUUGAUGAUGUGGUCAUUCUGCAUCGAAUUAUCUAAACU |
| Yin-2-1 | UUCGAUAUCAAAUCCUGUUAACAACCGUUGCUUAGUUUAUCAGCGGCUGUUCGUUAACAAUCAAAUUGUUAUUUAUCAA |
| Hin-18-2 | UCCAUAUUUUUGAAUAUCUCUAAUAAAUAUUUAGCUGCCCCCUUACAUACAAACUGCCAUUUAUGGUGUAAUAUUAAUU |
| Hin-14-1 | CGAAAAUAGAGCUUGUUUAUCUGCAUUUUUUCAGUAUAAUUCGCAACCUAUUUGGUGCCAAUUAUGGCGUAUUAUUCAC |
| Vvu-10-1 | GAGACGAUUCAUUACUAUAUCGCGCCUUAAGAUAUGGCUUUAAUUCUGUUUUAGCGGUAUCUGAGUUUUAUUUUUAACU |
| Ywa-1-1 | UUCGCCGCCUUUGAUAUAAUUUGAUAUCAAAUCCUGUUAACAAACGUUGCUUGUUUUAUCAACGAUUGUUAUUUAUCAA |
| Dso-4-1 | UUCAAUUGCAUCGGGUGGUUAAUUUCUGUAUGAUCCGCCGCCUUUAUACCUUGUGGUGUAAAGAACAUUUCACUCAACU |
| Yps-1-3 | UUCGCCGCCUUUGAUAUAAUUUGAUAUCAAAUCCUGUUAACUAUCGUUGCUUAUUUUAUCAACGAUAGUUAUUUAUCAA |
| Sso-1-546 | AAUCGCUUGCAGCUAAAACUGACAGGCAGUAAGAUGCAGCGCCAUUUUUGCAAUUCUGCAACUGGCUUAAUUUUUAAUC |
| Yal-3-1 | UUCGCCGCCUUUGGUAUAAUUCGAUAUCAAAUCCUGUUAACAAGCGUUGGUUACUUUAUCAGCGAUUGUUAGUUAUCAA |
| Sal-1-1 | UCGAGCAGAUGUAGUCGCCUACUUGUCAGCGCUGGGUAGAUACUGUACUAUUCUGCGCCCUGAACUUUUGAUAUUAACC |
| Kpn-168-1 | UUAUAUGGCAUUGAAGAUUGACCAUUACCGCGUGCUUCACGCUGUCAGCAGCGUAGAAGGCGACGGGAAAUUACUAAUA |
| Vvu-9-1 | GAGACGAUUCAUUACUAUAUCGCGCCUUAAGAUAUGGCUUUAAUUCUAUUUUAGCGGUAUCUGAGUUUUAUUUUUAACU |
| Yro-2-1 | UUCGAUAUCAAAUCCUGUUAACAGCCGUUGCUUACUUUAUCAGCGAUAGUUAGUUAUCAAUAAAAUUAUUAUUUUUCAA |
| Yen-13-1 | CUAACAACCGUUGUUUAUUUUAUCAGCGAUUGUUCGUUAGCAAUCAACUAUUCGUUAACGAUAAAUUCUUAUUUAUCAA |
| Pmi-2-1 | UUGCGCGUUUGGCGGAUAUUCUGUAUGAUCCGCCCCCUUUGGUAGAUAAAAUACCAAACAAUUUAAUUCAAUCAACACG |
| Vpa-183-1 | AGGGGAUUCAUUACUAUAUCGCGCCUUGAGAUAUGGCUUCAGCAAUUGGUUGGUGCGGUAUCCCAUUUUAUUAUUAAUU |
| Ehe-1-1 | AAAUCGACCUAUUUCCCCGGCGCGCCGUACUGCAGCGAAUAUACCCGCAGCACGGAAUGAGACGGGGAAUUUAUCAACU |
| Eco-230-1 | AGUUUGCAUACGCUGGUUAAUUUCUGUAUGAUUUCGCGCCUUCAUACGAAAUGAUCGUAUUGAAGCUAUACUUUUAACA |
| Vvu-5-1 | GAGACGAUUCAUUACUAUAUCGCGCCUUAAGAUAUGGCUUUAAUUCUGUUUCAGCGGUAUCUGAGUUUUAUUUUUAACU |
| Cbr-1-7 | GUUAUCUCCAUUUAAAACGGCUAUAAUGCGCGGCCUUUGUUUGUUAAGAAAACAGGCAAAGUAUCACAUUACAUUAAUC |
| Esp-47-1 | UCAGUUUGCAUACGUUGGUCAAAUUCUGUAUGAUUCGCCGCCUUUGGUGUAUAUUACGCCAGAGAAAUUACAUUCAACA |
| Hin-12-1 | CGAAAAUAGAGCUUGUUUAUCUGUAUUUUUUCAGUAUAAUUCGCAACCUAUUUGGUGCCAAUUAUGGCGUAUUAUUCAC |
| Kox-11-1 | UUUGAUGCGCUUUGUGCGGCAUCAAAAUCUAUUACCCGGGAAUUUCGCUUCGCGCAAUCGCACCGGGAAAUUAUUAACA |
| Pst-7-1 | UGCGUGAUACAGAGAUUUUCUGUAUGAUCCGCCGCCUUUGGUAUCUAAUUUUGCCAAUAAAACAUUUUUAAUCAACACG |
| Vco-2-1 | GCCGUAUUCAUUACUAUAUCGCGCCUUAGUGAUAUGGCUUGUUUUUGAACAGCGGUAUCCAAAGGAAAUUUUAUUAACU |
| Yfr-3-1 | UUUAUCAGCGAUUGUUCGUUAGCAAUCAACUAUUCGUUAACGAUAAAUGAUUAUUUAACGACAAAUGGUUAUUUAUCAA |
| Kmi-1-2 | UUUGAUGCGCUUUGUGCGACAUCAAAAUCUAUUACCCGGGAAUUUCGCUUCGCGCAAUCGCACCGGGAAAUUAUUAACA |
| | ................................................................................ |
| | ................................................................................ |
| | ●●●●●●●●●●●●●●●●●●●●●●●●●●●●●●●●●●●●●●●●●●●●●●●●●●●●●●●●●●●●●●●●●●●●●●●●●●●●●RY● |

alignment positions 80 · · · 200

```
CBa-1-1    UCGUGUGGCU...........................UAUAGUUAAAAUCGUCG..........................AUAGCGACACAAUCUGACGAA....UAGGUGUCCC...AUG
Apl-1-1    ACAUAUGGUACUUAUC.AGCGGAUAUAC.........AAAU......................GUAUAACAA..GGUAA.GUGGCGAUAUGGCUUCAUUA.....GAGGUUUUCCA..AUG
Apl-6-1    ACAUAUGGUACUUAUC.AGCGGAUAUAC.........GAAA......................GUAUAACAA..GAUAA.GUGGCGAUAUGGCUUCAUUA.....GAGGUUUUCCA..AUG
Apl-13-1   ACAUAUGGUACUUAUC.AGCGGAUAUAC.........GAAA......................GUAUAACAA..GAUAA.GUGGCGAUAUGGCUUCAUUA.....GAGGUUUUCCA..AUG
Apl-8-1    ACAUAUGGUACUUAUC.AGCGGAUAUAC.........GAAA......................GUAUAACAA..GAUAA.GUGGCGAUAUGGCUUCAUUA.....GAGGUUUUCCA..AUG
Aur-1-1    ACAUAUGGUACUUAUC.AGCGGAUAUAC.........GUAU......................GUAUAACAA..GAUAA.GUGGCGAUAUGGCUUCAUUA.....GAGGUUUUCCA..AUG
Aca-3-1    ACAUAUGGUACUUAUC.AGCGGAUAUAC.........GAAU......................GUAUAACAA..GAUAA.GUGGCGAUAUGGCUUCAUUA.....GAGGUUUUCCA..AUG
Aeq-1-1    ACAUAUGGUACUUAUC.AGCGGAUAUAC.........GAAU......................GUAUAACAA..GAUAA.GUGGCGAUAUGGCUUCAUUA.....GAGGUUUUCCA..AUG
Asu-3-1    ACAUAUGGUACUUAUC.AGCGGAUAUAC.........GAAU......................GUAUAACAA..GAUAA.GUGGCGAUAUGGCUUCAUUA.....GAGGUUUUCCA..AUG
Mgr-1-1    ACAUAUGGUACCUAUU..AGCGGUAG...........AGAGU.......................CUAACAA..GAUAG.GUGGCGAUAUGGCUUCUUAU.....GAGGUUCCCCA..AUG
Mha-1-1    ACAUAUGGUACCUAUC..AGCGGUAG...........AGAGU.......................CUAACAA..GAUAG.GUGGCGAUAUGGCUUCUUAU.....GAGGUUCCCCA..AUG
Ami-1-1    ACAUAUGGUACUUAUC..AGCG.GUU...........CGUGAUGA....................AACAA....GAUAA.GCAGCGAUAUGGCUUCUUAU.....GAGGUUCCCCA..AUG
env-1      ACAUAUGGUACUUAUC..AGCGGUUCAAU........UGAAAUAAUA...............GUUGAAACAA..GAUAA.GCAGCGAUAUGGCUUCUUAU.....GAGGUUCUCCA..AUG
env-8      ACAUAUGGUACUUAUC..AGCGGUUCAAU........UGAAAUAAUA...............GUUGAAACAA..GAUAA.GCAGCGAUAUGGCUUCUUAU.....GAGGUUCUCCA..AUG
env-9      ACAUAUGGUACUUAUC..AGCGGUUC...........UUUU........................GAAACAA..GAUAA.GCAGCGAUAUGGCUUCUUAC.....GAGGUUCUCC...AUG
env-10     ACAUAUGGUACUUAUC..AGCGGUUC...........UUUU........................GAAACAA..GAUAA.GCAGCGAUAUGGCUUCUUAC.....GAGGUUCUCC...AUG
Hsp-3-1    ACAUAUGGUACUUAUC..AGCGGUUC...........UUUU........................GAAACAA..GAUAA.GCAGCGAUAUGGCUUCUUAC.....GAGGUUCUCC...AUG
Ami-2-1    ACAUAUGGUACUUAUC..AGCGGUUC...........AUCAU.......................GAAACAA..GAUAA.GCAGCGAUAUGGCUUCUUAU.....GAGGUUCUCCA..AUG
Lgr-1-1    UCGUAUGGUGUACGG........CA............AAUCGG.......................UG......ACCGU.AUAGCGAUACGACUUUAGA......GAGGUUUCAUC..AUG
Asu-1-1    UCGUAUGGUGUAAGGU.......GGCU..........UGCG.......................GGCC....G.ACGUU.AUGGCGAUACGGCUUCUGACA....GAGGUUUUCCA..AUG
env-11     AAAUGCGGUGCCCGG.C......AG............CGAUUCAAUAA..................CU....G.AUCGG.GUGGCGGCAUUGCCUGAAAAU....GAGGUCUGCC...AUG
env-12     AAAUGCGGUGCCCGG.C....................AGAA...............................G.AUCGG.GUAGCGGCAUUGCCCUGGA......GAGGUUAACC...AUG
env-13     AAAUGCGGUGCCCGG.C....................AGAA...............................G.AUCGG.GUAGCGGCAUUGCCCUGGA......GAGGUUAACC...AUG
env-14     ACGUGCGGUGCCUGG.C..ACCUGGAGC.........GAUUUAA...................GCUCCCGU.G.AUCAG.GUAGCGGCACGACCCGACA......GAGGUUCGCC...AUG
env-15     ACAUGCGGUGCCUGG.C..GCCUGGUA..........AUUCAGA....................UGCCCCGCG.AUCAG.GUAGCGGCAUGACCUAAC.......GAGGUUCCCC...AUG
env-16     ACGUGCGGUGCCUGG.C..GCCUGGGUA.........AUCAAGA...................UGCCCCGC.G.AUCAG.GUAGCGGCAUGACCUAAC.......GAGGUUCACC...AUG
env-19     ACGUGCGGUGCCUGG.C..GCCUGGGUA.........AUCAAGA...................UGCCCCGC.G.AUCAG.GUAGCGGCAUGACCUAAC.......GAGGUUCACC...AUG
env-20     ACGUGCGGUGCCUGG.C..GCCUGGGUA.........AUCAAGA...................UGCCCCGC.G.AUCAG.GUAGCGGCAUGACCUAAC.......GAGGUUCACC...AUG
env-21     ACAUGCGGUGCCUGG.C..GCCUGGGUA.........AUCAAGA...................UACCCCGC.G.AUCAG.GUAGCGGCAUGACCUAAC.......GAGGUUCGCC...AUG
Msu-1-1    ACAUGCGGUGCCUGG.C..GCCAGGGUA.........GUUCAGA...................UGCCCCGU.G.AUCAG.GUAGCGGCAUGACCUAAC.......GAGGUUCGCC...AUG
env-23     ACAUGCGGUGCCUGG.C..ACCUGGGCA.........GCUAAGA...................UGCCCCGU.G.AUCAG.GUAGCGGCAUGACCUAAC.......GAGGUUCGCC...AUG
env-24     ACAUGCGGUGCCUGG.C..ACCUGGGCA.........GCAAAGA...................UGCCCCGU.G.AUCAG.GUAGCGGCAUGACCUAAC.......GAGGUUCGCC...AUG
env-40     ACAUGCGGUGCCUGG.C..ACCUGGGCA.........GCCAAAA...................UGCCCCGU.G.AUCAG.GUAGCGGCAUGACCUAAC.......GAGGUUCGCC...AUG
Msp-1-1    ACAUGCGGUGCCUGG.C..ACCUGGGCA.........GCCAAAA...................UGCCCCGU.G.AUCAG.GUAGCGGCAUGACCUAAA.......GAGGUUCGCC...AUG
Vli-1-1    ACGUAUGGUGUCUGACU......GCUUUGAUA.CUC.GCAA..............GAGAUAUUUGAGUAAGAA.UUCAG.AUAGCGAUACGGACCAAUAA....AAGGUUUGUCCU.AUG
Mma-1-1    UUAUGUGGUGUCUGG.C....................AGU................................GUUUCAG.AUGGCGACAUAACCUAAAUU.....GAGGUUUCU....AUG
env-45     UCAUGCGGUGUCAGG........GAUUGC........UUUC.....................GCAAUU....G.GCUUG.AUAGCGGCAUGGCUUCAAAUUU...GAGGUUUUCU...AUG
env-46     UCAUGCGGUGUCAGG........GAUUGC........UUUC.....................GCAAUU....G.GCUUG.AUAGCGGCAUGGCUUCAAAUUU...GAGGUUUUCU...AUG
env-47     UCAUGCGGUGUCAGG........GAUUGC........UUUC.....................GCAAUU....G.GCUUG.AUAGCGGCAUGGCUUCAAAUUU...GAGGUUUUCU...AUG
env-49     UCAUGCGGUGUCAGG........GAUUGC........UUUU.....................GCGAUU....G.GCUUG.AUAGCGGCAUGGCUUCAACUUU...GAGGUUUUCU...AUG
env-50     UCAUGCGGUGUCAGG........GAUUGC........GUUU.....................GCGAUU....G.GCUUG.AUAGCGGCAUGGCUUCAACUUU...GAGGUUUUCU...AUG
Asp-10-1   UCAUGCGGUGUCAGG........GAUUGC........GUUU.....................GCGAUU....G.GCUUG.AUAGCGGCAUGGCUUCAACUUU...GAGGUUUUCU...AUG
env-51     UCAUGCGGUGUCAGG........GAUUGC........GUUU.....................GCGAUU....G.GCUUG.AUAGCGGCAUGGCUUCAACUUU...GAGGUUUUCU...AUG
env-52     UCAUGCGGUGUCAGG........GAUUGC........GUUU.....................GCGAUU....G.GCUUG.AUAGCGGCAUGGCUUCAACUUU...GAGGUUUUCU...AUG
env-56     UCAUGCGGUGUCAGG........GAUUGC........GUUU.....................GCGAUU....G.GCUUG.AUAGCGGCAUGGCUUCAACUUU...GAGGUUUUCU...AUG
env-57     UCAUGCGGUGUCAGG........GAUUGC........GUUU.....................GCGAUU....G.GCUUG.AUAGCGGCAUGGCUUCAAAUUU...GAGGUUUUCU...AUG
env-58     UCAUGCGGUGUCAGG........GAUUGC........GUUU.....................GCGAUU....G.GCUUG.AUAGCGGCAUGGCUUCAAAUUU...GAGGUUUUCU...AUG
env-59     UCAUGCGGUGUCAGG........GAUUGC........UUCU.....................GCAAUU....G.GCUUG.AUAGCGGCAUGGCUUCAAAUUU...GAGGUUUUCU...AUG
env-60     UCAUGCGGUGUCAGG........GAUUGC........UUCU.....................GCAAUU....G.GCUUG.AUAGCGGCAUGGCUUCCAAUUU...GAGGUUUUCU...AUG
env-61     UCAUGCGGUGUCAAG........GAUUGC........UUCU.....................GCGAUU....G.GCUUG.AUAGCGGUAUGGCUUCAAAUUU...GAGGUUUUCU...AUG
env-63     UCAUGCGGUGUCAAG........GAUUGC........UUCU.....................GCGAUU....G.GCUUG.AUAGCGGCAUGGCUUCAAAUUU...GAGGUUUUCU...AUG
env-71     UCAUGCGGUGUCAAG........GAUUGC........UUCU.....................GCGAUU....G.GCUUG.AUAGCGGCAUGGCUUCAAAUUU...GAGGUUUUCU...AUG
env-73     UCAUGCGGUGUCAGG........GAUUGC........UUCG.....................GCGAUU....G.GCUUG.AUAGCGGCAUGGCUUCAAAUUU...GAGGUUUUCU...AUG
Aac-6-1    UCAUGCGGUGUCAGG........GAUUGC........UUCU.....................GCGAUU....G.GCUUG.AUAGCGGCAUGGCUUCAAAUUU...GAGGUUUUCU...AUG
Aac-18-1   UCAUGCGGUGUCAGG........GAUUGC........UUCU.....................GCGAUU....G.GCUUG.AUAGCGGCAUGGCUUCAAAUUU...GAGGUUUUCU...AUG
env-74     UCAUGCGGUGUCAGG........GAUUGC........UUCU.....................GCGAUU....G.GCUUG.AUAGCGGCAUGGCUUCAAAUUU...GAGGUUUUCU...AUG
Aac-11-1   UCAUGCGGUGUCAGG........GAUUGC........UUCU.....................GCGAUU....G.GCUUG.AUAGCGGCAUGGCUUCAAAUUU...GAGGUUUUCU...AUG
env-76     UCAUGCGGUGUCAGG........GAUUGC........UUCU.....................GCGAUU....G.GCUUG.AUAGCGGCAUGGCUUCAAAUUU...GAGGUUUUCU...AUG
Aac-17-1   UCAUGCGGUGUCAGG........GAUUGC........UUCU.....................GCGAUU....G.GCUUG.AUAGCGGCAUGGCUUCAAAUUU...GAGGUUUUCU...AUG
Aac-1-1    UCAUGCGGUGUCAGG........GAUUGC........UUCU.....................GCGAUU....G.GCUUG.AUAGCGGCAUGGCUUCAAAUUU...GAGGUUUUCU...AUG
Aac-14-1   UCGUGCGGUGUCAGG........GAUUGC........UUCU.....................GCGAUU....G.GCUUG.AUAGCGGCAUGGCUUCAAAUUU...GAGGUUUUCU...AUG
env-77     UCAUGCGGUGUCAGG........GAUUGC........UUUU.....................GCGAUU....G.GCUUG.AUAGCGGCAUGGCUUCAAAUUU...GAGGUUUUCU...AUG
env-86     UCAUGCGGUGUCAGG........GAUUGC........UUUU.....................GCGAUU....G.GCUUG.AUAGCGGCAUGGCUUCAAAUUU...GAGGUUUUCU...AUG
env-87     UCAUGCGGUGUCAGG........GAUUGC........UUUU.....................GCGAUU....G.GCUUG.AUAGCGGCAUGGCUUCAAAUUU...GAGGUUUUCU...AUG
env-88     UCAUGCGGUGUCAGG........GAUUGC........UUUU.....................GCGAUU....G.GCUUG.AUAGCGGCAUGGCUUCAAAUUU...GAGGUUUUCU...AUG
env-89     UCAUGCGGUGUCAGG........GAUUGC........UUUU.....................GCGAUU....G.GCUUG.AUAGCGGCAUGGCUUCAAAUUU...GAGGUUUUCU...AUG
env-90     UCAUGCGGUGUCAGG........GAUUGC........UUUU.....................GCGAUU....G.GCUUG.AUAGCGGCAUGGCUUCAAAUUU...GAGGUUUUCU...AUG
env-91     UCAUGCGGUGUCAGG........GAUUGC........UUUU.....................GCGAUU....G.GCUUG.AUAGCGGCAUGGCUUCAAAUUU...GAGGUUUUCU...AUG
env-93     UCAUGCGGUGUCAGG........GAUUGC........UUUU.....................GCGAUU....G.GCUUG.AUAGCGGCAUGGCUUCAAAUUU...GAGGUUUUCU...AUG
env-94     UCAUGCGGUGUCAGG........GAUUGC........UUUU.....................GCGAUU....G.GCUUG.AUAGCGGCAUGGCUUCAAAUUU...GAGGUUUUCU...AUG
env-95     UCAUGCGGUGUCAGG........GAUUGC........UUUU.....................GCGAUU....G.GCUUG.AUAGCGGCAUGGCUUCAAAUUU...GAGGUUUUCU...AUG
Pda-1-1    ACACGCGGUGUCGG.U.....................UUUGA................................GCUUG.AUAGCGGCGUGGCUUCAAUUU....GAGGUUUUCU...AUG
Pmu-14-1   ACACGCGGUGUC.AGUC....................UUU................................GAACUCG.AUAGCGGCGUGGCUUCAAAUC....GAGGUUUUCC...AUG
Pmu-2-1    ACAUGCGGUGUCAG.........UCU...........AAUA........................AGA......ACUUG.AUAGCGGCAUGGCUUCAAAUU....GAGGUUUUCU...AUG
Pmu-8-1    ACAUGCGGUGUCAG.U.....................CUAAGU...............................ACUUG.AUAGCGGCAUGGCUUCAAAUU....GAGGUUUUCU...AUG
Pmu-12-1   ACAUGCGGUGUC.AGU.....................CUGUGU...............................ACUCG.AUAGCGGCAUGGCUUCAAAUU....GAGGUUUUCU...AUG
Pmu-3-1    ACACGCGGUGUCAG.........UC............CUU..........................GA......ACUUG.AUAGCGGCGUGGCUUCAAAUC....GAGGUUUUCU...AUG
Pmu-6-1    ACACGCGGUGUCAG.........UC............CUU..........................GA......ACUUG.AUAGCGGCGUGGCUUCAAAUC....GAGGUUUUCU...AUG
Mma-3-1    ACUCGUGGUGUUUGGA.......AACUC.........UUUA......................GAGUUG...G.UCCUU.AUAGCGGCGAGGCUUCAAUUU....GAGGUUUUCU...AUG
Msu-2-1    ACUCGUGGUGUUUGGA.......AGCUC.........UCA.......................GAGCUG...G.UCCUU.AUAGCGGCGAGGCUUCAUUUU....GAGGUUUUCU...AUG

           .<<<<<.<<..<<<<<<......<<<<<<<<<<<<<<.................>>>>>>>>>>>>>>....>.>>>>>....>>.>>>>>..............................
           .22222.0...222222......1.1111.11.0010.................0100.11.1111.1....2.22222.....0.22222..............................
           •CGURUGGUG∘YYRR∘Y--------------------••••∘------------------------------G-∘YYRR-A•AGCGAYRCGGCCU•••••∘∘----GAGGU••YCC∘--AUG
            5′                                                                                                      3′
```

```
env-96      UCACGCGGUGUAAGUG.....................GAAAG................................CAUUU.AUAGCGGCGUGGCUUCAAUUU....GAGGUUUUNA...ACG
env-97      UCACGCGGUGUAAGUG.....................GAGAG................................CAUUU.AUAGCGGCGUGGCUUCAAUUU....GAGGUUUUCU...AUG
Hpa-2-2     UCACGCGGUGUAAGUG.....................GAAAG................................CAUUU.AUUGCGGCGUGGCUUCAAUUU....GAGGUUUUCU...AUG
env-101     UCACGCGGUGUAAGUG.....................GAAAG................................CAUUU.AUAGCGGCGUGGCUUCAAUUU....GAGGUUUUCU...AUG
env-102     UCACGCGGUGUAAGUG.....................GAAAG................................CAUUU.AUAGCGGCGUGGCUUCAAUUU....GAGGUUUUCU...AUG
env-103     UCACGCGGUGUAAGUG.....................GAAAG................................CAUUU.AUAGCGGCGUGGCUUCAAUUU....GAGGUUUUCU...AUG
env-105     UCACGCGGUGUAAGUG.....................GAAAG................................CAUUU.AUAGCGGCGUGGCUUCAAUUU....GAGGUUUUCU...AUG
env-106     UCACGCGGUGUAAGUG.....................GAAAG................................CAUUU.AUAGCGGCGUGGCUUCAAUUU....GAGGUUUUCU...AUG
env-109     UCACGCGGUGUAAGUG.....................GAAAG................................CAUUU.AUAGCGGCGUGGCUUCAAUUU....GAGGUUUUCU...AUG
env-139     UCACGCGGUGUAAGUG.....................GAAAG................................CAUUU.AUAGCGGCGUGGCUUCAAUUU....GAGGUUUUCU...AUG
env-140     UCACGCGGUGUAAGUG.....................GAAAG................................CAUUU.AUAGCGGCGUGGCUUCAAUUU....GAGGUUUUCU...AUG
env-284     UCACGCGGUGUAAGUG.....................GAAAG................................CAUUU.AUAGCGGCGUGGCUUCAAUUU....GAGGUUUUCU...AUG
env-352     UCACGCGGUGUAAGUG.....................GAAAG................................CAUUU.AUAGCGGCGUGGCUUCAAUUU....GAGGUUUUCU...AUG
env-356     UCACGCGGUGUAAGUG.....................GAAAG................................CAUUU.AUAGCGGCGUGGCUUCAAUUU....GAGGUUUUCU...AUG
env-357     UCACGCGGUGUAAGUG.....................GAAAG................................CAUUU.AUAGCGGCGUGGCUUCAAUUU....GAGGUUUUCU...AUG
env-358     UCACGCGGUGUAAGUG.....................GAAAG................................CAUUU.AUAGCGGCGUGGCUUCAAUUU....GAGGUUUUCU...AUG
env-359     UCACGCGGUGUAAGUG.....................GAAAG................................CAUUU.AUAGCGGCGUGGCUUCAAUUU....GAGGUUUUCU...AUG
env-360     UCACGCGGUGUAAGUG.....................GAAAG................................CAUUU.AUAGCGGCGUGGCUUCAAUUU....GAGGUUUUCU...AUG
env-361     UCACGCGGUGUAAGUG.....................GAAAG................................CAUUU.AUAGCGGCGUGGCUUCAAUUU....GAGGUUUUCU...AUG
env-362     UCACGCGGUGUAAGUG.....................GAAAG................................CAUUU.AUAGCGGCGUGGCUUCAAUUU....GAGGUUUUCU...AUG
Hpa-2-11    UCACGCGGUGUAAGUG.....................GAAAG................................CAUUU.AUAGCGGCGUGGCUUCAAUUU....GAGGUUUUCU...AUG
Rpn-1-1     UCACGCGGUGUGAGAA.....................GAGAG................................UUCUU.AUAGCGGCGUGGCUUCAUUUU....GAGGUUUUCU...AUG
env-363     UCACGCGGUGUAGGAA.....................GAAAG................................UUCUU.AUAGCGGCGUGGCUUCAUUUU....GAGGUUUUCU...AUG
env-368     UCACGCGGUGUAGGGA.....................GAAAG................................UUCUU.AUAGCGGCGUGGCUUCGUUUU....GAGGUUUUCU...AUG
env-369     UCACGCGGUGUAGGGA.....................GAAAG................................UUCUU.AUAGCGGCGUGGCUUCAUUUU....GAGGUUUUCU...AUG
env-385     UCACGCGGUGUAGGGA.....................GAAAG................................UUCUU.AUAGCGGCGUGGCUUCAUUUU....GAGGUUUUCU...AUG
env-394     UCACGCGGUGUAGGGA.....................GAAAG................................UUCUU.AUAGCGGCGUGGCUUCAUUUU....GAGGUUUUCU...AUG
env-399     UCACGCGGUGUAGGGA.....................GAAAG................................UUCUU.AUAGCGGCGUGGCUUCAUUUU....GAGGUUUUCU...AUG
env-401     UCACGCGGUGUAGGGA.....................GAAAG................................UUCUU.AUAGCGGCGUGGCUUCAUUUU....GAGGUUUUCU...AUG
env-404     UCACGCGGUGUAGGGA.....................GAAAG................................UUCUU.AUAGCGGCGUGGCUUCAUUUU....GAGGUUUUCU...AUG
env-407     UCACGCGGUGUAGGGA.....................GAAAG................................UUCUU.AUAGCGGCGUGGCUUCAUUUU....GAGGUUUUCU...AUG
env-408     UCACGCGGUGUAGGGA.....................GAAAG................................UUCUU.AUAGCGGCGUGGCUUCAUUUU....GAGGUUUUCU...AUG
env-409     UCACGCGGUGUAGGGA.....................GAAAG................................UUCUU.AUAGCGGCGUGGCUUCAUUUU....GAGGUUUUCU...AUG
env-410     UCACGCGGUGUAGGGA.....................GAAAG................................UUCUU.AUAGCGGCGUGGCUUCAUUUU....GAGGUUUUCU...AUG
env-411     UCACGCGGUGUAAGGA.....................GAAAG................................UUCUU.AUAGCGGCGUGGCUUCAUUUU....GAGGUUUUCU...AUG
env-417     UCACGCGGUGUAAGGA.....................GAAAG................................UUCUU.AUAGCGGCGUGGCUUCAUUUU....GAGGUUUUCU...AUG
Hha-1-1     UCACGCGGUGUGAGGA.....................GAAAG................................UUCUU.AUAGCGGCGUGGCUUCAUUUU....GAGGUUUUCU...AUG
env-418     UCACGCGGUGUAGGGA.....................GAAAG................................UUCCU.AUAGCGGCGUGGCUUCAUUUU....GAGGUUUUCU...AUG
env-419     UCACGCGGUGUAGGGA.....................GAAAG................................UUCCU.AUAGCGGCGUGGCUUCAUUUU....GAGGUUUUCU...AUG
Hha-1-2     UCACGCGGUGUGGGGA.....................GAAAG................................UUCCU.AUAGCGGCGUGGCUUCAUUUU....GAGGUUUUCU...AUG
env-420     UCACGCGGUGUGGGGA.....................GAAAG................................UUCCU.AUAGCGGCGUGGCUUCAUUUU....GAGGUUUUCU...AUG
Hha-1-3     UCACGCGGUGUGGGGA.....................GAAAG................................UUCUC.AUAGCGGCGUGGCUUCAUUUU....GAGGUUUUCU...AUG
Hha-1-4     UCACGCGGUGUGGGGA.....................GAAAG................................UUCUU.AUGGCGGCGUGGCUUCAUUUU....GAGGUUUUCU...AUG
env-422     UCACGCGGUGUGGGGA.....................GAAAG................................UUCUU.AUAGCGGCGUGGCUUCAUUUU....GAGGUUUUCU...AUG
env-423     UCACGCGGUGUGGGGA.....................GAAAG................................UUCUU.AUAGCGGCGUGGCUUCAUUUU....GAGGUUUUCU...AUG
env-428     UCACGCGGUGUGGGGA.....................GAAAG................................UUCUU.AUAGCGGCGUGGCUUCAUUUU....GAGGUUUUCU...AUG
env-430     UCACGCGGUGUGGGGA.....................GAAAG................................UUCUU.AUAGCGGCGUGGCUUCAUUUU....GAGGUUUUCU...AUG
Hha-6-1     UCACGCGGUGUGGGAA.....................GAAAG................................UUCUU.AUAGCGGCGUGGCUUCAUUUU....GAGGUUUUCU...AUG
Hin-17-1    UCACGCGGUGUAGGGA.....................GAAAG................................UUCCU.AUAGCGGCGUGGCUUCAUUUU....GAGGUUUUUCU..AUG
env-431     UCACGCGGUGUGGGGA.....................GAAAG................................UUCCU.AUAGCGGCGUGGCUUCAUUUU....GAGGUUUUUCU..AUG
Apa-1-1     ACACUCGGUG.CUAAA.....................UUCAGU...............................UUUAG.AGAGCGGAGUGGCUUCAAAUU....GAGGUUUUCU...AUG
Apa-1-2     ACACUCGGUG.CUAAA.....................UUCAGU...............................UUUAG.AGAGCGGAGUGGCUUCAAAUU....GAGGUUUUCU...AUG
Nro-1-1     ACACUCGGUG.CCGAA.....................UUCAGU...............................UUCGG.AAAGCGGAGUGGCUUCAGAUU....GAGGUUUUCU...AUG
Hma-1-1     ACACUCGGUG.CCGAA.....................UUCAGU...............................UUCGG.AAAGCGGAGUGGCUUCAGAUU....GAGGUUUUCU...AUG
Psp-56-1    ACACUCGGUG.CCGAA.....................UUCAGU...............................UUCGG.AAAGCGGAGUGGCUUCAGAUU....GAGGUUUUCU...AUG
Hin-17-2    ACGCGAGGCGUCAG.......................UCCUUGAAC..............................UUG.AUAGCGGCGUGGCUUUAAAUU....GAGGUUUUCU...AUG
env-432     ACGUGUGGUAUCAGG.U....................GAAAA..............................A.UCCUG.AUGGCGACGCGGCCCAAGA......GAGGUUUUCC...AUG
Mmu-1-1     ACAUAUGGUGUCAGA.C......A.............AAAUAGA.......................U....G.CUCUG.AUGGCGAUAUGGCUUCUAAAU....GAGGUUUUCUAUAAUG
Cor-1-1     ACAUAUGGUGUCGGA......................CUGGUU................................UCCG.AUAGCGAUAUGGCUUCAAUCC....GAGGUUUUCC...AUG
Baq-1-1     UCGCGUGGUGUUCGG.C....................AAUAGU.............................G.ACCGG.AUAGCGACGCGGCUUAAGA......GAGGCCUACC...AUG
Pfo-1-1     UCGCGUGGUGUUCGG.C....................AAUAGU.............................G.ACCGU.AUAGCGACGCGACAUAGA.......GAGGCUAAU....AUG
env-433     UCGUAUGGUGCUUAG.C......AAUU..........AAUAGG.....................AAUU....G.ACUAA.GUGGCGAUACGGCCUAAUACA....GAGGUUUCCC...AUG
env-434     UCGUAUGGUGCUUUG.C......AGC...........AAUAAAUA....................GCU....G.ACUAA.GUGGCGAUGCGGCCUUAAACA....GAGGUUUCC....AUG
env-435     UCGUAUGGUGUUAAG.C......GGUUG.........CAAAGU....................CAAUC....GGGCUUA.AUAGCGAUACGGCUUCUUAAU....GAGGUUCUCCC..AUG
env-437     UCGUAUGGUGUUAAG.C......GGUUG.........CAAAGU....................CAAUC....GGGCUUA.AUAGCGAUACGGCUUCUUAAU....GAGGUUCUCCC..AUG
env-438     UCGUAUGGUGUUAAG.C......GGUUG.........CAAAGU....................CAAUC....GGGCUUA.AUAGCGAUACGGCUUCUUAAU....GAGGUUCUCCC..AUG
env-441     UCGUAUGGUGUUAAG.C......GGUUG.........CAAAGU....................CAAUC....GGGCUUA.AUAGCGAUACGGCUUCUUAAU....GAGGUUCUCCC..AUG
env-443     UCGUAUGGUGUUAAG.C......GGUUG.........CAAAGU....................CAAUC....GGGCUUA.AUAGCGAUACGGCUUCUUAAU....GAGGUUCUCCC..AUG
env-445     UCGUAUGGUGUUAAG.C......GGUUG.........CAAAGU....................CAAUC....GGGCUUA.AUAGCGAUACGGCUUCUUAAU....GAGGUUCUCCC..AUG
Tvi-1-1     UCGUAUGGUACCUGG.C....................AAAA...............................G.AUCAG.GUGGCGAUACGGCCUUAACA.....GAGGUUCCC....AUG
Tac-1-1     UCGUAUGGUACCUGG.C....................AAAA...............................G.AUCAG.GUGGCGAUACGGCCUUAACA.....GAGGUUCCC....AUG
env-450     UCGUAUGGUGCUUGG.C......AG............CAAUAAUGA....................CU....G.AUCAA.GUGGCGAUGCGGCCUUAACA.....GAGGUUCCCC...AUG
env-452     UCGUAUGGUGCUUGG.C......AG............CAAUAAUGA....................CU....G.AUCAA.GUGGCGAUACGGCCUUAACA.....GAGGUUUCCC...AUG
env-453     UCGUAUGGUGCUUGG.C......AG............CAAUAAUGA....................CU....G.AUCAA.GUGGCGAUACGGCCUUAACA.....GAGGUUUCCC...AUG
env-454     UCGUAUGGUGCUUGG.C......AG............CAAUAAUGA....................CU....G.AUCAA.GUGGCGAUACGGCCUUAACA.....GAGGUUUCCC...AUG
env-457     UCGUAUGGUGCUUGG.C......AG............CAAUAAUGA....................CU....G.AUCAA.GUGGCGAUACGGCCUUAACA.....GAGGUUUCCC...AUG
env-458     UCGUAUGGUGCUUGG.C......AG............CAAUAAUGA....................CU....G.AUCAA.GUGGCGAUACGGCCUUAACA.....GAGGUUUCCC...AUG
env-459     UCGUAUGGUGCUUGA.C......AG............CAAUAAUGA....................CU....G.AUCAA.GUGGCGAUACGGCCUUAACA.....GAGGUUUCCC...AUG
env-460     UCGUAUGGUGCUUGG.C......AG............CAAUAAUGA....................CU....G.AUCAA.GUGGCGAUGCGGCCUUAACA.....GAGGUUUCCC...AUG
env-461     UCGUAUGGUGCUUGG.C......AG............CAAUAAUGA....................CU....G.AUCAA.GUGGCGAUGCGGCCUUAACA.....GAGGUUUCCC...AUG
env-462     GCGUGUGGUGUCUGUU.....................GAAACA...............................CAUAG.AUAGCGACACGGCCGACA.......GAGGUAACCCCA.AUG
env-463     ACACAUGGUGUCCUG.C......GC............AGACGAG......................GC....G.ACUGG.AUAGCGAUGUGGCCUAACUC.....GAGGUAUUCCC..AUG
env-466     ACACAUGGUGUCCUG.C......GC............AGACGAG......................GC....G.ACUGG.AUAGCGAUGUGGCCUAACUC.....GAGGUAUUCCC..AUG

            .<<<<<.<<..<<<<<<......<<<<<<<<<<<<<<.................>>>>>>>>>>>>>>....>.>>>>>....>>.>>>>>..............................
            .22222.0...222222......1.1111.11.0010.................0100.11.1111.1....2.22222.....0.22222..............................
            •CGURUGGUG∘YYRR∘Y--------------------•••∘∘------------------------------G-∘YYRR-A•AGCGAYRCGGCU••••∘∘----GAGGU••YCC∘--AUG
             5′                                                                                                  3′
```

```
env-467   ACACAUGGUGUCCUG.C......GC............AAAAGAG......................GC....G.ACUGG.AUAGCGAUGUGGCCCAACUU.....GAGGUAUUCCC..AUG
env-469   GCGCAUGGUGUUCGGCU....................UUG................................G.AUCGA.AUGGCGAUGCGGCCCAUAACU....GAGGUAACCCC..AUG
env-475   GCGCAUGGUGUUCGGCU....................UUG................................G.AUCGA.AUGGCGAUGCGGCCCAUAACU....GAGGUAACCCC..AUG
env-477   GCGCAUGGUGUUCGGCU....................UUG................................G.AUCGA.AUGGCGAUGCGGCCCAUAACU....GAGGUAACCCC..AUG
env-479   GCGCAUGGUGUUCGGCU....................UUG................................G.AUCGA.AUGGCGAUGCGGCCCAUAACU....GAGGUAACCCC..AUG
env-480   GCGCAUGGUGUUCGGCU....................UUG................................G.AUCGA.AUGGCGAUGCGGCCCAUAACU....GAGGUAACCCC..AUG
env-481   GCGCAUGGUGUUCGGCU....................UUG................................G.AUCGA.AUGGCGAUGCGGCCCAUAACU....GAGGUAACCCC..AUG
Vru-1-1   ACGUAUGGUGUCUGGCC.....ACUUUUU.AU.CUUCUUGUA............GAAGUAUAAAGAGGAA..GAACUAG.AUAGCGAUACGGACCAAUAU.....AAGGUUUGUCCU.AUG
env-482   ACGUGUGGUGUCCGACU....................UCG................................G.GUUUG.AUAGCGACACGGCCUAAAUU.....GAGGUUUCCUAA.AUG
env-483   ACGUAUGGUGUGCGGA.....................GAGAG................................UCCGU.AUAGCGAUGCGGCCCAAUU......GAGGUAACCCUU.AUG
Ama-1-1   UCGUGUGGUGUUUGGU.......GCUG..........GUAA.......................CGGUG.....AUCAA.AUGGCGACACGGCCCAUGAUU....GAGGUAAUCCC..AUG
Ata-2-1   UCGUGUGGUGUUUGGU.......GCUG..........GUUA.......................CAGUG.....AUCAA.AUGGCGACACGGCCCAUGAUU....GAGGUAAUCCC..AUG
env-484   UCGUGUGGUGCCGGUGC......AGG...........UGAAUU......................CCUG...G.CACGGGAGAGCGACGCGGCCCUUUACU....GAGGUUUUCC...AUG
env-485   GCGUGUGGUG.CUGCGC......AGG...........UUCACGU.....................CCUG...G.CGCGG.AGAGCGACACGGCCUUAUAU.....GAGGUUUCCC...AUG
env-486   GCGUGUGGUG.CUGCGC......AGG...........UUCACGU.....................CCUG...G.CGCGG.AGAGCGACACGGCCUUAUAU.....GAGGUUUCCC...AUG
Psp-49-1  GCGUGUGGUG.CUGUGC......AGG...........UUUGG.......................CCUG...G.CACGG.AGAGCGACACGGCCUUUUAU.....GAGGUUCUCCC..AUG
Psp-45-1  GCGUGUGGUG.CUGUGC......AGGC..........UGA........................GCCUG...G.CACGG.AGAGCGACACGGCCUUUUAU.....GAGGUUCUCCC..AUG
Psp-53-1  GCGUGUGGUG.CUGUGC......AGGC..........UAA........................GCCU....GGCACGG.AGAGCGACACGGCCUUUUAU.....GAGGUUCUCCC..AUG
env-487   GCGUGUGGUG.CUGUGC......AGGU..........CAAA.......................GCCUG...G.CACGG.AGAGCGACACGGCCUUCUAU.....GAGGUUUCCC...AUG
env-488   GCGUGUGGUG.CUGUGC......AGGU..........CAAA.......................GCCUG...G.CACGG.AGAGCGACACGGCCUUCUAU.....GAGGUUUCCC...AUG
env-489   GCGUGUGGUG.CUGUGC......AGGU..........CAAA.......................GCCUG...G.CACGG.AGAGCGACACGGCCUUCUAU.....GAGGUUUCCC...AUG
env-490   GCGUGUGGUG.CUGUGC......AGGU..........CAAA.......................GCCUG...G.CACGG.AGAGCGACACGGCCUUCUAU.....GAGGUUUCCC...AUG
Pst-4-1   GCGUGUGGUG.CUGCGC......AGG...........UUUCGU......................CCUA...G.CGCGG.AGAGCGACACGGCCUUCUAU.....GAGGUUUCCC...AUG
Pst-3-1   GCGUGUGGUG.CUGCGC......AGG...........UUUCGU......................CCUG...G.CGCGG.AGAGCGACACGGCCUUCUAU.....GAGGUUUCCC...AUG
Pst-1-1   GCGUGUGGUG.CUGCGC......AGG...........UUUCGU......................CCUG...G.CGCGG.AGAGCGACACGGCCUUCUAU.....GAGGUUUCCC...AUG
Psp-36-1  GCGUGUGGUG.CUGUGC......GGG...........UUACGA......................CCUG...G.CACGG.AGAGCGACACGGCCUUAUCU.....GAGGUUUCCC...AUG
Pva-1-1   GCGUGUGGUG.CUGUGC......AGGU..........UAAC.......................GCCUG...G.CACGG.AGAGCGACACGGCCUUCUAU.....GAGGUUUCCC...AUG
env-493   GCGUGUGGUG.CUGUGC......AGGU..........UUAC.......................GCCUG...G.CACGG.AGAGCGACACGGCCUUCUAU.....GAGGUUUCCC...AUG
Pan-5-1   GCGUGUGGUG.CUGUGC......AGGU..........UUAC.......................GCCUG...G.CACGG.AGAGCGACACGGCCUUCUAU.....GAGGUUUCCC...AUG
Psp-43-1  GCGUGUGGUG.CUGUGC......AGG...........UUCG........................CCUG...G.CAUGG.AGAGCGACACGGCCUUUUAU.....GAGGUUUUCCC..AUG
env-494   GCGUGUGGUG.CUGUGC......AGG...........UUCG........................CCUG...G.CAUGG.AGAGCGACACGGCCUUUUAU.....GAGGUUUUCCC..AUG
Pva-1-2   GCGUGUGGUG.CUGUGC......GGGU..........UUUUA......................GCCUG...G.CACGG.AGAGCGACACGGCCUUUUAU.....GAGGUUCUCCC..AUG
Pse-1-1   GCGUGUGGUG.CUGUGC......AGGU..........UUUUA......................GCCUG...G.CACGG.AGAGCGACACGGCCUUUUAU.....GAGGUUCUCCC..AUG
Pro-1-1   GCGUGUGGUG.CUGUGC......AGG...........UUCG........................CCUG...G.CACGG.AGAGCGACACGGCCUUUUAU.....GAGGUUCUCCC..AUG
Psp-46-1  GCGUGUGGUG.CUGUGC......AGG...........UUCG........................CCUG...G.CACGG.AGAGCGACACGGCCUUUUAU.....GAGGUUCUCCC..AUG
env-500   GCGUGUGGUG.CUGUGC......AGG...........UUCG........................CCUG...G.CACGG.AGAGCGACACGGCCUUUUAU.....GAGGUUCUCCC..AUG
Psp-38-1  GCGUGUGGUG.CUGUGC......AGG...........UUCG........................CCUG...G.CACGG.AGAGCGACACGGCCUUUUAU.....GAGGUUCUCCC..AUG
env-501   GCGUGUGGUG.CUGUGC......AGG...........UUCG........................CCUG...G.CACGG.AGAGCGACACGGCCUUUUAU.....GAGGUUCUCCC..AUG
env-503   GCGUGUGGUG.CUGUGC......AGG...........UUCG........................CCUG...G.CACGG.AGAGCGACACGGCCUUUUAU.....GAGGUUCUCCC..AUG
env-504   GCGUGUGGUG.CUGUGC......AGG...........UUCG........................CCUG...G.CACGG.AGAGCGACACGGCCUUUUAU.....GAGGUUCUCCC..AUG
env-507   GCGUGUGGUG.CUGUGC......AGG...........UUCG........................CCU....GGCACGG.AGAGCGACACGGCCUUUUACA....GAGGUUCUCCC..AUG
Tsp-2-1   GCGUGUGGUG.CUGUGC......AGG...........UUCG........................CCU....GGCACGG.AGAGCGACACGGCCUUCUACU....GAGGUUCUCCC..AUG
Tsa-1-1   GCGUGUGGUG.CUGUGC......AGG...........UUCG........................CCU....GGCACGG.AGAGCGACACGGCCUUCUACU....GAGGUUCUCCC..AUG
Tpt-1-1   GCGUGUGGUG.CUGUGC......AGG...........UUCG........................CCU....GGCACGG.AGAGCGACACGGCCUUCUACU....GAGGUUCUCCC..AUG
env-508   GCGUGUGGUG.CUGUGC......AGGU..........UAAA.......................GCCUG...G.CACGG.AGAGCGACACGGCCUUCAAU.....GAGGUUUCCC...AUG
Pag-5-1   GCGUGUGGUG.CUGUGC......AGGU..........UAAA.......................GCCUG...G.CACGG.AGAGCGACACGGCCUUCAAU.....GAGGUUUCCC...AUG
Psp-39-1  GCGUGUGGUG.CUGUGC......AGGU..........UAAA.......................GCCUG...G.CACGG.AGAGCGACACGGCCUUCAAU.....GAGGUUUCCC...AUG
Pag-2-5   GCGUGUGGUG.CUGUGC......AGGU..........UAAA.......................GCCUG...G.CACGG.AGAGCGACACGGCCUUCAAU.....GAGGUUUCCC...AUG
env-509   GCGUGUGGUG.CUGUGC......AAG...........GUAAAC......................UUG....G.CACGG.AGAGCGACACGGCCUUCUACU....GAGGUUAUCCC..AUG
Psp-48-1  GCGUGUGGUG.CUGUGC......AGGG..........UUAA.......................CCUG....G.CACGG.AGAGCGACACGGCCUUUUCU.....GAGGUUUCCC...AUG
env-511   UCGUGUGGUGUCCGG.C....................UCCGG..............................G.CCUGG.AUAGCGACACGGCCUUUGA......GAGGUUUUCC...AUG
env-512   ACGUAUGGUGUUCGCG.....................GAGAG................................CGCGG.AUGGCGAUACGGCCUGAAACU....GAGGUUACCC...AUG
env-513   GCGCAUGGUGUUCGUG.....................GAAAG................................CACGG.AUGGCGAUGCGGCCUGAUAU.....GAGGUAAUCCC..AUG
env-515   GCGCAUGGUGUUCGUG.....................GAAAG................................CACGG.AUGGCGAUGCGGCCCAUAUAU....GAGGUAACCCCC.AUG
env-516   GCGCAUGGUGUUCGUG.....................GAGAG................................CACGG.AUGGCGAUGCGGCCUUCAACU....GAGGUAACCCCUAAUG
Isp-2-1   GCGCGUGGUGUUCGUG.....................GAAAG................................CACGG.AUGGCGAUGCGGCCUUCAACU....GAGGUAACCCCC.AUG
env-517   ACGUAUGGUGUUCGUG.....................GAAAG................................CACGG.AUGGCGAUGCGGCCUACUAUU....GAGGUAACCCCUAAUG
env-518   GCGUAUGGUGUUUGG.C....................UUCGG..............................G..CCAA.AUAGCGAUACGGCCUCAUAUUG...AAGGUAAUCCAAAAUG
Asp-8-1   GCGUAUGGUGUUUGG.C....................UUCGG..............................G..CCAA.AUAGCGAUACGGCCUCAUAUUG...AAGGUAAUCCAAAAUG
env-519   GCGUAUGGUGUUUGGCU....................UCG................................G.GCCAG.AUAGCGAUACGGCCUCAUAUUG...AAGGUAAUCCAAAAUG
env-520   UCGUAUGGUGGCCAG.C....................UUCGG..............................G..UUGG.UUAGCGAUACGGCCUUAAUAUU...GAGGUGCCCC...AUG
Vrh-1-1   ACAUGUGGUGUCCGG.C....................AAC................................GAUCCGG.AUAGCGACAUGGCCCGAUUUUU...GAGGUUCUCCC..AUG
env-521   UCGUGUGGUGUCUGG.C....................GUGAG..............................G..CUGG.AUGGCGACACGGCCUAAUAUAUUA.GAGGUUGUCC...AUG
env-522   UCGUGUGGUGUCUGG.C....................GUGAG..............................G..CUGG.AUGGCGACACGGCCUAAUAUAUUA.GAGGUUGUCC...AUG
Spu-2-1   GCAUGUGGUG.UCCGAC....................UAAGG..............................G.UUGGA.AGAGCGACAUGGCCUUACAUUUU..GAGGUAACCCCA.AUG
Ssp-5-1   GCAUGUGGUG.UCCGAC....................UAAGG..............................G.UUGGA.AGAGCGACAUGGCCUUACAUUUU..GAGGUAACCCCA.AUG
Sba-3-1   GCAUGUGGUG.UCCGAC....................UACGG..............................G.UUGGA.AGAGCGACAUGGCCUUACACUUUU.GAGGUAACCCCA.AUG
env-527   GCAUGUGGUG.UCCGAC....................UACGG..............................G.UUGGA.AGAGCGACAUGGCCUUACACUUUU.GAGGUAACCCCA.AUG
Sma-1-1   GCAUGUGGUG.CUCAAC....................UUCGG..............................G.UUGGG.AGAGCGACAUGGCCUUAAACUU...GAGGUAAUCCCA.AUG
Sco-1-1   ACAUGUGGUG.CCUGAC....................UUCGG..............................G.UUGGG.AGUGCGACGUGGCCUUAAACUU...GAGGUAAUCCAA.AUG
Ssp-4-1   GCAUGUGGUG.UCCGAC....................UUCGG..............................G.UUGGA.AGAGCGACAUGGCCUUAACUU....GAGGUAACCCCA.AUG
Sam-2-1   GCAUGUGGUG.UCCGAC....................UUCGG..............................G.UUGGA.AGAGCGACAUGGCCUUUACUUU...GAGGUAAUCCCG.AUG
Son-1-1   GCAUGUGGUG.UCCGAC....................UUCGG..............................G.UUGGA.AGAGCGACAUGGCCUUACUU.....GAGGUAACCCAA.AUG
Ssp-10-1  GCAUGUGGUG.UCCGAC....................UUCGG..............................G.UUGGA.AGAGCGACAUGGCCUUACUU.....GAGGUAACCCAA.AUG
env-529   UCAUGUGGUG.UCCGAC....................UUCGG..............................G.UUGGG.AGAGCGACAUGGCCUUCAACUAUA.AAGGUAAUCCCA.AUG
Swo-1-1   UCAUGUGGUG.UCCGAC....................UUCGG..............................G.UUGGA.AGAGCGACAUGGCCUUCAACUA...GAGGUAAUCCCA.AUG
Sse-1-1   UCAUGUGGUG.UCCGAC....................UUCGG..............................G.UUGGG.AGAGCGACAUGGCCUUCAACUA...GAGGUAAUCCCA.AUG
Sbe-1-1   GCAUGUGGUG.UCCGAC....................UUCGG..............................G.UUGGG.AGAGCGACAUGGCCUUCAACUU...GAGGUAAUCCAA.AUG
Svi-1-1   GCAUGUGGUG.UCCGAC....................UUCGG..............................G.UUGGG.AGAGCGACAUGGCCUUCAACUU...GAGGUAAUCCAA.AUG
env-530   [illegible]CAUGUGGUG.UCCGAC....................UUCGG..............................G.UUGGA.AGAGCGACAUGGCCUUACUUUU...GAGGUAACCCCA.AUG
Ssp-7-1   GCAUGUGGUG.UCCGAC....................UUCGG..............................G.UUGGA.AGAGCGACAUGGCCUUACAUUU...GAGGUAAUCCCA.AUG
Sde-1-1   GCAUGUGGUG.UCCGAC....................UUCGG..............................G.UUGGA.AGAGCGACAUGGCCUUACAUUU...GAGGUAAUCCCA.AUG
env-534   [illegible]CAUGUGGUG.UCCGAC....................UUCGG..............................G.UUGGA.AGAGCGACAUGGCCUUACAUUU...GAGGUAAUCCCA.AUG

          .<<<<<.<<..<<<<<<......<<<<<<<<<<<<<<.................>>>>>>>>>>>>>>....>.>>>>>....>>.>>>>>..............................
          .22222.0...222222......1.1111.11.0010.................0100.11.1111.1....2.22222.....0.22222..............................
          •CGURUGGUG∘YYRR∘Y--------------------••••∘-------------------------------G-∘YYRR-A•AGCGAYRCGGCCU•••••∘----GAGGU••YCC∘--AUG
          5′                                                                                                             3′
```

```
Ssp-6-1    GCAUGUGGUG.UCCGAC....................UUCGG..............................G.UUGGA.AGAGCGACAUGGCCUUACAUUU...GAGGUAAUCCCA.AUG
Sma-2-1    GCAUGUGGUG.UCCGAC....................UUCGG..............................G.UUGGA.AGAGCGACAUGGCCUUAAACUU...GAGGUAAUUCCA.AUG
Spi-1-1    GCAUGUGGUG.UCCGAC....................UUCGG..............................G.UUGGA.AGAGCGACAUGGCCUUAAACUU...GAGGUAAUUCCA.AUG
Ssp-3-1    GCAUGUGGUG.UCCGAC....................UUCGG..............................G.UUGGA.AGUGCGACAUGGCCUUAAACUU...GAGGUAAUCCCA.AUG
Ssp-11-1   GCAUGUGGUG.UCCGAC....................UUCGG..............................G.UUGGA.AGUGCGACAUGGCCUUAAACUU...GAGGUAAUCCCA.AUG
Slo-1-1    GCAUGUGGUG.UCCGAC....................UUCGG..............................G.UUGGA.AGUGCGACAUGGCCUUAAACUU...GAGGUAAUCCCA.AUG
Swa-1-1    GCAUGUGGUG.CCUGAC....................UUCGG..............................G.UUGGG.AGAGCGACAUGGCCUUCAACU....GAGGUAAUCCAA.AUG
Sde-2-1    UCAUGUGGUG.UCCGAC....................UUCGG..............................G.UUGGA.AGAGCGACAUGGCCUUCAAUU....AAGGUAUCCCAAAAUG
env-535    GCAUGUGGUG.UCCGAC....................UUCGG..............................G.UUGGA.AGAGCGACAUGGCCUUAACCUU...GAGGUAAUUCCA.AUG
Sha-1-1    GCAUGUGGUG.UCCGAC....................UUCGG..............................G.UUGGA.AGAGCGACAUGGCCUUUUACU....GAGGUAAUUCCA.AUG
Spe-1-1    GCAUGUGGUG.UCCGAC....................UUCGG..............................G.UUGGA.AGAGCGACAUGGCCUUUUACU....GAGGUAAUUCCA.AUG
Sfi-1-1    GCAUGUGGUG.UCCGAC....................UUCGG..............................G.UUGGA.AGAGCGACAUGGCCUUUUACU....GAGGUAAUUCCA.AUG
Sfr-1-1    GCAUGUGGUG.CUCAAC....................UUCGG..............................G.UUGGG.AGAGCGACAUGGCCUUCAAUUU...AAGGUAGCCCAAAAUG
CRe-1-1    ACGUGUGGUGUCUGG.C....................GGGAUAAAG..........................G..CCCG.AUAGCGACACGGUCUUAAAGA....GAGCUUUGCU...AUG
Eba-2-1    UCGCGUGGUGUGCGG.C....................GAAAAAU............................G.ACCGU.AUAGCGACGCGGCCUUAACUA....GAGGUUUCCC...AUG
Sgl-1-1    UCGCGUGGUGUCUGG.C......UA............AAG..........................UA....G.ACUGG.AUAGCGACGCGGCCUUAUGAA....GAGGUUUUCC...AUG
CSo-1-1    UCGCGUGGUGUCUGGUU....................AUAAAGUA...........................G.ACCGG.AUAGCGACGCGGCCUUACGAA....GAGGUUUUCC...AUG
env-536    UCGCGUGGUGUCCGG.C....................GUAAGCA............................G.ACCGG.AUAGCGACGCGGCCUUACGAU....GAGGUUUUCC...AUG
Spr-1-1    UCGCGUGGUGUCUGG.C....................UAAAGUA............................G.ACCGG.AUAGCGACGCGGCCUUACGAA....GAGGUUUUC....AUG
Psp-47-1   UCGUGUGGUGUUUGG.C......UGGC..........AGUA.......................GCUGA...G.ACCAA.AUAGCGACACGGCCUUUACUU....GAGGUUUCCCC..AUG
Aal-2-1    UCGUAUGGUGUUUGGU.......GUCA..........AUUG.......................UGGC....G.ACCAG.AUGGCGAUACGGCCCAAGUAUU...GAGGUAAUCC...AUG
env-537    GCGUAUGGUGUCUGA.C......AG............UAAG.........................CU....G.AUCAG.AUAGCGAUACGGCCCUAAC......GAGGUAAGCC...AUG
Pte-1-1    GCGUAUGGUGUCUGA.C......AG............UAAG.........................CU....G.AUCAG.AUAGCGAUACGGCCCUAAC......GAGGUAAGCC...AUG
env-538    ACGUAUGGUGUUUGA.C......AG............CAAGA........................CU....G.AUCAG.AUUGCGAUACGGCCCAAC.......GAGGUAAGCC...AUG
env-539    ACGUAUGGUGUUUGA.C......AG............CAAGA........................CU....G.AUCAG.AUUGCGAUACGGCCCAAC.......GAGGUAAGCC...AUG
Tli-1-1    UCGUGUGGUGUCUGG.C....................UUCG...............................G.GUCAG.AUGGCGACACGGCCUGAAUCA....GAGGUUUCCC...AUG
Tau-1-1    UCGUGUGGUGUCUGGCU....................UCG................................G.GUCAG.AUGGCGACACGGCCUGAAUCA....GAGGUUCUCC...AUG
Ana-1-1    ACGUAUGGUGUCUAG.C....................GUUAGG.............................G..UUAG.AUAGCGAUGCGGCCUAACUA.....GAGGUUUUCU...AUG
Aen-3-1    ACGUAUGGUGUCUAG.C....................GUUAGG.............................G..UUAG.AUAGCGAUGCGGCCUAAGUA.....GAGGUUUUCU...AUG
Asa-10-1   UCGUAUGGUGUUCGG.C....................UUAG...............................G.AUCGA.AUAGCGAUACGGCCUGAAAAA....GAGGUUUUCC...AUG
Iat-1-1    ACGUAUGGUGUCCGG.C....................AACA...............................GA.UCGG.AUGGCGAUGCGGCCUAACU......GAGGUAACCCCU.AUG
Ise-1-1    ACGUAUGGUGUCCGG.C....................ACAG...............................G.AUCGG.AUGGCGAUGCGGCCCAAUU......GAGGUAACCCUU.AUG
Isa-1-1    ACGUGUGGUAUUCGG.C....................AUAA...............................G.AUCGG.AUGGCGAUGCGGCCCAAUU......GAGGUAACCCUU.AUG
Ixi-1-1    ACGUAUGGUGUCUGG.C......GU............UCA..........................GC....G.ACCAG.AUAGCGAUGCGGCCUAACUU.....GAGGUUACCCCU.AUG
env-540    UCGUGUGGUGUCUGGCU....................CUG................................G.AUCAG.AUAGCGACACGGCCUACAGAUA...GAGGUUGUCC...AUG
Osm-1-1    UCGUGUGGUGUCUGG.C....................UCAG...............................G.AUCAG.AUAGCGACACGGCCUACUGACA...GAGGUUGUCC...AUG
Osp-1-1    UCGUGUGGUGUCUGG.C....................UCAG...............................G.AUCAG.AUAGCGACACGGCCUACUGACA...GAGGUUGUCC...AUG
env-541    UCGUGUGGUGUCUGGAC....................UACAG..............................G.AUCAG.AUAGCGACACGGCCUACUGACUA..GAGGUUGUCC...AUG
Ema-2-1    UCGUGUGGUA.CCUGUG......C.AGGA........GUG......................ACCUGG....C.ACGGG.AGAGCGACACGGCCUUCACU.....GAGGUUUCCC...AUG
Etr-1-1    UCGUGUGGUG.CCUGUG......C.AGGU........UUU......................UCCUGG....C.AUGGG.AGAGCGACACGGCCUUAACU.....GAGGUUUCCC...AUG
Psp-40-1   UCGUGUGGUG.CCGGUG......C.AGGG........UCAA.....................CCCUGG....C.ACGGG.AGAGCGACACGGCCUUAACU.....GAGGUUUCCC...AUG
Ein-1-1    UCGUGUGGUG.CCGGUG......C.AGGA........AAUU.....................UCCUGG....C.ACGGG.AGAGCGACGCGGCCUUAACU.....GAGGUUUUCCC..AUG
env-542    UCGUGUGGUG.CCGGUG......C.AGGA........AUUU.....................UCCUGG....C.ACGGG.AGAGCGACGCGGCCUUAACU.....GAGGUUUUCCC..AUG
Eto-1-1    UCGUGUGGUG.CCGGUG......C.AGGA........AUUU.....................UCCUGG....C.ACGGG.AGAGCGACGCGGCCUUAACU.....GAGGUUUUCCC..AUG
Esp-55-1   UCGUGUGGUG.CCGGUG......C.AGGC........GUU......................UCCUGG....C.ACGGG.AGAGCGACACGGCCUUAACU.....GAGGUUUCCC...AUG
Eol-1-1    UCGUGUGGUG.CCGGUG......C.AGGA........AUU......................UCCUGG....C.ACGGG.AGAGCGACACGGCCUUAACU.....GAGGUUUCCC...AUG
Ebi-2-1    GCGUGUGGUG.CCGGUG......C.AGGA........AUAA.....................UCCUGG....C.ACGGG.AGAGCGACACGGCCUUAACU.....GAGGUUUCCC...AUG
Ebi-1-1    GCGUGUGGUG.CCGGUG......C.AGGA........AUGA.....................UCCUGG....C.ACGGG.AGAGCGACACGGCCUUAACU.....GAGGUUUCCC...AUG
bsy-1-1    UCGUGUGGUG.CCGGUG......C.AGGA........GUUA.....................UCCUGG....C.GCGGG.AGAGCGACACGGCCUUAACU.....GAGGUUUUCCC..AUG
Epy-2-1    UCGUGUGGUG.CCGGUG......C.AGGA........UUAU.....................UCCUGG....C.GCGGG.AGAGCGACACGGCCUUAACU.....GAGGUUUUCCC..AUG
Esp-74-1   UCGUGUGGUG.CCGGUG......C.AGGA........UUAU.....................UCCUGG....C.GCGGG.AGAGCGACACGGCCUUAACU.....GAGGUUUUCCC..AUG
Eta-9-1    UCGUGUGGUG.CCGGUG......C.AGGA........GAAU.....................UCCUGG....C.GCGGG.AGAGCGACACGGCCUUAACU.....GAGGUUUUCCC..AUG
Epi-2-1    UCGUGUGGUG.CCGGUG......C.AGGA........GAAU.....................UCCUGG....C.GCGGG.AGAGCGACACGGCCUUAACU.....GAGGUUUUCCC..AUG
Eam-1-1    UCGUGUGGUG.CCGGUG......C.AGGA........GUAU.....................UCCUGG....C.GCGGG.AGAGCGACACGGCCUUAACU.....GAGGUUUUCCC..AUG
Ety-1-1    UCGUGUGGUG.CUUGUG......C.AGGG........CUAA.....................UCCUGG....C.ACGGG.AGAGCGACACGGCCUUAACU.....GAGGUUUUCCC..AUG
env-546    UCGUGUGGUG.CUUGUG......C.AGGG........CUAA.....................UCCUGG....C.ACGGG.AGAGCGACACGGCCUUAACU.....GAGGUUUUCCC..AUG
Psp-44-1   GCGUGUGGUG.CCGGCG......C.AGGU........UUU......................ACCUGG....C.GCGGG.AGAGCGACACGGCCUUUACU.....GAGGUUAUCCC..AUG
env-547    GCGUAUGGUG.CCAAAC....................UUCGG..............................G.UUUGG.AGAGCGAUACGGCCUAUUUA.....GAGGUAACCCCC.AUG
env-549    ACGUAUGGUG.CCAGAC....................UUCGG..............................G.UUUGG.AGAGCGAUGCGGCCUAAUUA.....GAGGUUUACU...AUG
Ppi-1-1    ACGUAUGGUG.CCAGAC....................UUCGG..............................G.UUUGG.AGAGCGAUACGGCCUAAUUA.....GAGGUUCACU...AUG
Pci-1-1    ACGUAUGGUG.CCAAAC....................UUCGG..............................G.UUUGG.AGAGCGAUACGGCCUAACCA.....GAGGUUGCU....AUG
Plu-2-1    ACGUAUGGUG.CCAAAC....................UUCGG..............................G.UUUGG.AGAGCGAUACGGCCUUAAAUUA...GAGGUUUCCU...AUG
Plu-3-1    ACGUAUGGUG.CCAGAC....................UUCGG..............................G.UUUGG.AGAGCGAUACGGCCUUACUUA....GAGGUUCCU....AUG
env-550    ACGUAUGGUG.CCAGAC....................UACGG..............................G.UUUGG.AGAGCGAUGCGGCCUUAACUA....GAGGUUCCCU...AUG
Ppi-3-1    ACGUAUGGUG.CCAGAC....................UUCGG..............................G.UUUGG.AGAGCGAUACGGCCUUAACUA....GAGGUUGCU....AUG
Ppi-2-1    ACGUAUGGUG.CCAGAC....................UUCGG..............................G.UUUGG.AGAGCGAUACGGCCUUAACUA....GAGGUUGCU....AUG
Pfl-1-1    ACGUAUGGUG.CCAGAC....................UUCGG..............................G.UUUGG.AGAGCGAUACGGCCUUAACUA....GAGGUUGCU....AUG
Pel-1-1    ACGUAUGGUG.CCAGAC....................UUCGG..............................G.UUUGG.AGAGCGAUACGGCCUUAACUA....GAGGUUGCU....AUG
env-551    GCGUAUGGUG.CCAGAC....................UUCGG..............................G.UUUGG.AGAGCGAUACGGCCUUAACUA....GAGGUUCCCU...AUG
env-552    GCGUAUGGUG.CCAGAC....................UUCGG..............................G.UUUGG.AGAGCGAUACGGCCUUAACUA....GAGGUUCCCU...AUG
env-559    GCGUAUGGUG.CCAGAC....................UUCGG..............................G.UUUGG.AGAGCGAUACGGCCUUAACUA....GAGGUUCCCU...AUG
env-566    GCGUAUGGUG.CCAGAC....................UUCGG..............................G.UUUGG.AGAGCGAUACGGCCUUAACUA....GAGGUUCCCU...AUG
Psp-17-1   ACGUAUGGUG.CCAGAC....................UUCGG..............................G.UUUGG.AGAGCGAUGCGGCCUUAACUA....GAGGUUCCU....AUG
env-572    ACGUAUGGUG.CCAGAC....................UUCGG..............................G.UUUGG.AGAGCGAUGCGGCCUUAACUA....GAGGUUCCU....AUG
env-573    ACGUAUGGUG.CCAGAC....................UUCGG..............................G.UUUGG.AGAGCGAUGCGGCCUUAACUA....GAGGUUCCU....AUG
env-589    ACGUAUGGUG.CCAGAC....................UUCGG..............................G.UUUGG.AGAGCGAUGCGGCCUUAACUA....GAGGUUCCU....AUG
env-591    ACGUAUGGUG.CCAGAC....................UUCGG..............................G.UUUGG.AGAGCGAUGCGGCCUUAACUA....GAGGUUCCU....AUG
Psp-6-1    ACGUAUGGUG.CCAGAC....................UUCGG..............................G.UUUGG.AGAGCGAUGCGGCCUUAACUA....GAGGUUCCU....AUG
env-593    ACGUAUGGUG.CCAGAC....................UUCGG..............................G.UUUGG.AGAGCGAUGCGGCCUUAACUA....GAGGUUCCU....AUG
Psp-26-1   ACGUAUGGUG.CCAGAC....................UUCGG..............................G.UUUGG.AGAGCGAUGCGGCCUUAACUA....GAGGUUCCU....AUG
env-594    ACGUAUGGUG.CCAGAC....................UUCGG..............................G.UUUGG.AGAGCGAUGCGGCCUUAACUA....GAGGUUCCU....AUG
Aba-1-1    ACGUAUGGUG.CCAGAC....................UUCGG..............................G.UUUGG.AGAGCGAUGCGGCCUUAACUA....GAGGUUUCU....AUG
env-595    ACGUAUGGUG.CCAGAC....................UUCGG..............................G.UUUGG.AGAGCGAUGCGGCCUUAACUA....GAGGUUUCU....AUG

           .<<<<<.<<...<<<<<<......<<<<<<<<<<<<<<................>>>>>>>>>>>>>>....>.>>>>>....>>.>>>>>.............................
           .22222.0...222222......1.1111.11.0010.................0100.11.1111.1....2.22222....0.22222.............................
           ●CGURUGGUG○YYRR○Y------------------●●●○○-----------------------------G-○YYRR-A●AGCGAYRCGGCCU●●●●○○----GAGGU●●YCC○--AUG
           5′                                                                                                          3′
```

```
Pha-2-1   ACGUAUGGUA.CCAGAC...................UUCGG.............................G.UUUGG.AGAGCGAUGCGGCCUUAACUA....GAGGUUUCU....AUG
Psp-11-1  ACGUAUGGUA.CCAGAC...................UACGG.............................G.UUUGG.AGAGCGAUGCGGCCUUAACUA....GAGGUUUCU....AUG
Psp-32-1  ACGUAUGGUA.CCAAAC...................UACGG.............................G.UUUGG.AGAGCGAUGCGGCCUUAACUA....GAGGUUUCU....AUG
Psp-19-1  ACGUAUGGUA.CCAAAC...................UACGG.............................G.UUUGG.AGAGCGAUGCGGCCUUAACUA....GAGGUUUCU....AUG
env-599   ACGUAUGGUG.CCAGAC...................UCCGG.............................G.UUUGG.AGAGCGAUGCGGCCUUAACUA....GAGGUUUCU....AUG
Pru-1-1   ACGUAUGGUG.CCAGAC...................UUCGG.............................G.UUUGG.AGAGCGAUACGGCCUUAACUA....GAGGUGUCCU...AUG
Pru-1-2   ACGUAUGGUG.CCAGAC...................UUCGG.............................G.UUUGG.AGAGCGAUACGGCCUUAACUA....GAGGUGUCCU...AUG
Pru-2-1   ACGUAUGGUG.CCAGAC...................UUCGG.............................G.UUUGG.AGAGCGAUACGGCCUUAACUA....GAGGUGUCCU...AUG
Plu-4-1   ACGUAUGGUG.CCAAAC...................UUCGG.............................G.UUUGG.AGAGCGAUACGGCCUUAACUA....GAGGUUUCCU...AUG
Plu-1-1   ACGUAUGGUG.CCAAAC...................UUCGG.............................G.UUUGG.AGAGCGAUACGGCCUUAACUA....GAGGUUUCCU...AUG
Par-1-1   ACGUAUGGUG.CCAAAC...................UCCGG.............................G.UUUGG.AGAGCGAUGCGGCCUUAACUA....GAGGUUCCU....AUG
env-601   ACGUAUGGUG.CCAAAC...................UUCGG.............................G.UUUGG.AGAGCGAUACGGCCUAACCUA....GAGGUUUACU...AUG
env-603   ACGUAUGGUG.CCAAAC...................UUCGG.............................G.UUUGG.AGAGCGAUACGGCCUAACCUU....GAGGUUUACU...AUG
Psp-1-1   ACGUAUGGUA.CCAGAC...................UUCGG.............................G.UUUGG.AGAGCGAUGCGGCCUUAACU.....GAGGUUUUCCAAAAUG
Ptu-1-1   ACGUAUGGUG.CCAGAC...................UUCGG.............................G.UUUGG.AGAGCGAUACGGCCUUAACU.....GAGGUUUGCU...AUG
env-606   ACGUAUGGUAGCCAGGC...................UUCGG.............................GUUUAGG.AGAGCGAUACGGCCUAAUUA.....GAGGUUGCU....AUG
Eca-5-1   ACGUGUGGUG.CCCGA....................UAGGC...............................UCGGG.AGAGCGACACGGCCUUAUAUUA...GAGGUUAUCCCU.AUG
Eca-6-1   ACGUGUGGUG.CCCGA....................UAGGC...............................UCGGG.AGAGCGACACGGCCUUAUAUUA...GAGGUUAUCCCU.AUG
Gho-1-1   ACGUGUGGUG.CCCGA....................UAGGC...............................UCGGG.AGAGCGACACGGCCUUAGAUUA...GAGGUUAUCCCU.AUG
Gsp-4-1   ACGUGUGGUG.CCCGA....................UAGGC...............................UCGGG.AGAGCGACACGGCCUUAGAUAA...GAGGUAACCCCU.AUG
Gsp-3-1   ACGUGUGGUG.CCCGA....................UAGGC...............................UCGGG.AGAGCGACACGGCCUUAGAUAA...GAGGUAAUCCCU.AUG
Eno-3-1   ACGUGUGGUG.CCCGA....................UAGGC...............................UCGGG.AGAGCGACACGGCCUAUAUUA....GAGGUUAUCCCU.AUG
env-607   UCGUAUGGUAUUCGG.C...................AAU...............................G.ACCGG.AUGGCGAUACGGCCUAAU.......GAGGUAAUCCC..AUG
env-608   ACGCAUGGUAUUCGG.C...................AACA..............................G.AUCGG.AUGGCGAUGCGGCCUGACA......GAGGUAAUCCC..AUG
Tsp-1-1   UCGCAUGGUAUCUGG.C...................AAAA..............................G.AUCAG.AUGGCGAUGCGGCCUAACAC.....AAGGUAAUCCC..AUG
Pha-5-1   ACGUGUGGUGUUCGGCU...................GUG...............................G.GCUGA.AUGGCGACACGGCCUUAGUA.....GAGGUUUUCC...AUG
Pvu-2-1   ACGUGUGGUGUUCGGCU...................GUG...............................G.GCUGA.AUGGCGACACGGCCUUAGUA.....GAGGUUUUCC...AUG
Pmi-3-1   ACGUGUGGUGUUCGGCU...................GAG...............................G.GCUGA.AUAGCGACACGGCCUUUGCA.....GAGGUUUUCC...AUG
env-611   ACGUGUGGUGUCUGG.C...................AUAG..............................G.AUCAG.AUAGCGACACGGCCUACAAAACACAGAGGUUUUCCC..AUG
Ffu-1-1   ACGUAUGGUGUCCGG.C...................UUUG..............................G.AUCGG.AUAGCGAUACGGCCUUAACU.....GAGGUUUUCCCC.AUG
Fky-1-1   ACGUAUGGUGUCCGG.C...................UUUG..............................G.AUCGG.AUAGCGAUACGGCCUUAACU.....GAGGUUUUCCCA.AUG
Asi-1-1   UCGUGUGGUGUCCGACC...................UUUG..............................G.CUCGG.AUAGCGACACGGCCCAAAUCAUU..GAGGUUGUCC...AUG
Afi-1-1   UCGUGUGGUGUCCGA.C...................AAGA..............................G.AUCGG.AUGGCGACACGGCCUAUUAUAUU..GAGGUUGUCC...AUG
Aja-1-1   UCGUGUGGUGUCCGA.C...................AGAA..............................G.AUCGG.AUGGCGACACGGCCUAUUACUUU..GAGGUUGUCC...AUG
Aja-1-2   UCGUGUGGUGUCCGA.C...................AAGA..............................G.AUCGG.AUGGCGACACGGCCUAUUACUUU..GAGGUUGUCC...AUG
Aau-1-1   UCGUGUGGUGUCCGA.C...................AAGA..............................G.AUCGG.AUGGCGACACGGCCUAUUACUUU..GAGGUUGUCC...AUG
Amo-1-1   UCGUGUGGUGUCCGA.C...................AGAA..............................G.AUCGG.AUGGCGACACGGCCUGAUAAUU...GAGGUUGUCC...AUG
Api-1-1   UCGUGUGGUGUCCGA.C...................AGAA..............................G.AUCGG.AUAGCGACACGGCCUGAUAAUU...GAGGUUGUCC...AUG
env-612   UCGUGUGGUGUCCGA.C...................AGAA..............................G.AUCGG.AUAGCGACACGGCCUGAUAAUU...GAGGUUGUCC...AUG
Abi-1-1   UCGUGUGGUGUCCGA.C...................AGAA..............................G.AUCGG.AUAGCGACACGGCCUGAUAAUU...GAGGUUGUCC...AUG
env-613   UCGUGUGGUGUCCGA.C...................AAGA..............................G.AUCGG.AUGGCGACACGGCCUAUUAAUU...GAGGUUGUCC...AUG
Ave-8-1   UCGUGUGGUGUCCGA.C...................AAGA..............................G.AUCGG.AUGGCGACACGGCCUAUUAAUU...GAGGUUGUCC...AUG
Aal-1-1   UCGUGUGGUGUCCGA.C...................GAGA..............................G.AUCGG.AUGGCGACACGGCCUAUUAAUU...GAGGUUGUCC...AUG
env-614   UCGUGUGGUGUCCGA.C...................AAGA..............................G.AUCGG.AUGGCGACACGGCCUGAUAAUC...GAGGUUGUCC...AUG
env-616   UCGUGUGGUGUCCGA.C...................AAGA..............................G.AUCGG.AUGGCGACACGGCCUGAUAAUU...GAGGUUGUCC...AUG
Ata-1-1   UCGUGUGGUGUCCGA.C...................AAGA..............................G.AUCGG.AUGGCGACACGGCCUGAUAAUU...GAGGUUGUCC...AUG
Ari-1-1   UCGUGUGGUGUCCGA.C...................AAGA..............................G.AUCGG.AUGGCGACACGGCCUGAUAAUU...GAGGUUGUCC...AUG
Ate-1-1   UCGUGUGGUGUCCGA.C...................AACA..............................G.AUCGG.AUGGCGACACGGCCUAUAAUC....GAGGUUGUCC...AUG
Ame-1-1   UCGUGUGGUGUCCGA.C...................AAGA..............................G.AUCGG.AUGGCGACACGGCCUAUAAUC....GAGGUUGUCC...AUG
Aaq-2-1   UCGUGUGGUGUCCGA.C...................AAGA..............................G.AUCGG.AUGGCGACACGGCCUAUUAUC....GAGGUUGUCC...AUG
env-619   UCGUGUGGUGUCCGA.C...................AAGA..............................G.AUCGG.AUGGCGACACGGCCUAUUAUC....GAGGUUGUCC...AUG
Aeu-1-1   UCGUGUGGUGUCCGA.C...................AAGA..............................G.AUCGG.AUGGCGACACGGCCUAUUAUC....GAGGUUGUCC...AUG
Asc-1-1   UCGUGUGGUGUCCGACC...................CAAG..............................G.CUCGG.AUGGCGACACGGCCUAUAUCUUU..GAGGUUGUCC...AUG
Adi-2-1   UCGUGUGGUGUCCGACC...................CAAG..............................G.CUCGG.AUGGCGACACGGCUUAUAUCUUA..GAGGUUGUCC...AUG
env-620   ACGUGUGGUG.CCCGGC...................UUAG..............................GAUCGGG.AGAGCGACACGGCCUUUAAAAU...GAGGUUAUCCCU.AUG
env-621   ACGUGUGGUG.CCCGGC...................UUAG..............................GAUCGGG.AGAGCGACACGGCCUUUAAAAU...GAGGUUAUCCCU.AUG
env-623   ACGUGUGGUG.CCCGGC...................UUAG..............................GAUCGGG.AGAGCGACACGGCCUUCAAGAU...GAGGUUAUCCCU.AUG
Ssp-27-1  ACGUGUGGUG.CCCGGC...................UUAG..............................GAUCGGG.AGAGCGACACGGCCUUUAAGAU...GAGGUUAUCCCU.AUG
Psp-57-1  ACGUGUGGUG.CCCGGC...................UUUG..............................GAUCGGG.AGAGCGACACGGCCUUACAUAUU..GAGGUUAUCCC..AUG
Pan-11-1  ACGUGUGGUG.CCCGGC...................UUUG..............................GAUCGGG.AGAGCGACACGGCCUUUAUUUU...GAGGUUAUCCC..AUG
Pan-12-1  ACGUGUGGUG.CCCGGC...................UUUG..............................GAUCGGG.AGAGCGACACGGCCUUUAUUUU...GAGGUUAUCCC..AUG
Pan-11-2  ACGUGUGGUG.CCCGGC...................UUUG..............................GAUCGGG.AGAGCGACACGGCCUUUAUUUU...GAGGUUAUCCC..AUG
Psp-58-1  ACGUGUGGUG.CCCGGC...................UUUG..............................GAUCGGG.AGAGCGACACGGCCUUUAUUUU...GAGGUUAUCCC..AUG
Pap-1-1   ACGUGUGGUG.CCCGGC...................UUAG..............................GAUCGGG.AGAGCGACACGGCCUUACUUUU...GAGGUUAUCCC..AUG
Psa-1-1   ACGUGUGGUG.CCCGGC...................UUUG..............................GAUCGGG.AGAGCGACACGGCCUUACUAUU...GAGGUUAUCCC..AUG
Paq-2-1   ACGUGUGGUG.CCCGGC...................UUUG..............................GAUCGGG.AGAGCGACACGGCCUUACUUUU...GAGGUUAUCCC..AUG
Pga-2-1   ACGUGUGGUG.CCCGGC...................UUUG..............................GAUCGGG.AGAGCGACACGGCCUUACUUUU...GAGGUUAUCCC..AUG
Pga-1-1   ACGUGUGGUG.CCCGGC...................UUUG..............................GAUCGGG.AGAGCGACACGGCCUUACUUUU...GAGGUUAUCCC..AUG
Ple-1-1   ACGUGUGGUG.CCCGGC...................UUUG..............................GAUCGGG.AGAGCGACACGGCCUUAAUUUU...GAGGUUAUCCC..AUG
Pda-4-1   ACGUGUGGUG.CCCGGC...................UUUG..............................GAUCGGG.AGAGCGACACGGCCUUACUACU...GAGGUUAUCCC..AUG
Psw-1-1   GCGUGUGGUG.CCCGGC...................UUUG..............................GAUCGGG.AGAGCGACACGGCCUUUAUUU....GAGGUUAUCCCA.AUG
Pph-1-1   ACGUGUGGUG.CCCGGC...................UUUG..............................GAUCGGG.AGAGCGACACGGCCUUAACUU....GAGGUUUCCCC..AUG
Pki-1-1   ACGUGUGGUG.CCCGGC...................UUUG..............................GAUCGGG.AGAGCGACACGGCCUUAACUU....GAGGUUUCCCC..AUG
Pil-1-1   ACGUGUGGUG.CCCGGC...................UUUG..............................GAUCGGG.AGAGCGACACGGCCUUAACUU....GAGGUUUCCCC..AUG
Pki-1-5   ACGUGUGGUG.CCCGGC...................UUUG..............................GAUCGGG.AGAGCGACACGGCCUUAACUU....GAGGUUUCCCC..AUG
Pha-6-1   ACGUGUGGUG.CCCGGC...................UUUG..............................GAUCGGG.AGAGCGACACGGCCUUAGCUUA...GAGGUUUCCC...AUG
Pga-3-1   ACGUGUGGUG.CCCGGC...................UUUG..............................GAUCGGG.AGAGCGACACGGCCUUAGCUAU...GAGGUUUCCCC..AUG
Pga-4-1   ACGUGUGGUG.CCCGGC...................UUAG..............................GAUCGGG.AGAGCGACACGGCCUUAAAUGUUU.GAGGUUAUCCC..AUG
Ppr-1-1   ACGUGUGGUG.CCCGGC...................UUUG..............................GAUCGGG.AGAGCGACGCGGCCUUAUACUU...GAGGUUUUCCC..AUG
env-624   ACGUGUGGUG.CCCGGC...................UUUG..............................GAUCGGG.AGAGCGACGCGGCCUUAUACUU...GAGGUUUUCCC..AUG
Ppr-2-1   ACGUGUGGUG.CCCGGC...................UUUG..............................GAUCGGG.AGAGCGACGCGGCCUUAUACUU...GAGGUUUUCCC..AUG
Vez-1-1   ACGUGUGGUGUCCAA.C...................CUUA..............................GAUUUGG.AUAGCGACGCGGCCUAUAAAU....GAGGUUUUCCC..AUG
Vha-1-1   ACGUGUGGUGUCCAA.C...................CUUA..............................GAUUUGG.AUAGCGACGCGGCCUAUAAAU....GAGGUUUUCCC..AUG

          .<<<<<.<<..<<<<<<.....<<<<<<<<<<<<<<.................>>>>>>>>>>>>>>....>.>>>>>....>>.>>>>>..............................
          .22222.0...222222.....1.1111.11.0010.................0100.11.1111.1....2.22222.....0.22222..............................
          ●CGURUGGUG○YYRR○Y-------------------●●●●○-----------------------------G-○YYRR-A●AGCGAYRCGGCCU●●●●○○----GAGGU●●YCC○--AUG
           5′                                                                                              3′
```

```
env-625    ACGUGUGGUGUUCGG.C....................UUA................................GAAUCGA.AUGGCGACACGGCCUAUAAU.....GAGGUUUUCCC..AUG
Vpr-1-1    ACGUGUGGUGUCCGG.C....................UCC................................GAUUCGG.AUAGCGACACGGCCCAAUAUU....GAGGUUCUCCC..AUG
Vbr-2-1    ACGUGUGGUGUUCAG.C....................AUA................................GAACUGG.AUAGCGACACGGCCUAUAAAU....GAGGUUUUCCC..AUG
Vbr-4-1    ACGUGUGGUGUUCAG.C....................AUA................................GAACUGG.AUAGCGACACGGCCUAUAAAU....GAGGUUUUCCC..AUG
Fse-1-1    ACGUGUGGUGUCCGG.C....................UUU................................GAAUCGG.AUAGCGACACGGCCUUUGAUU....GAGGUUUUUCC..AUG
Vmy-1-1    GCGUGUGGUGUGCGG.C....................AUA................................GACUCGU.AUAGCGACACGGCCUACUUU.....GAGGUUAUCCU..AUG
Vsp-27-1   ACGUGUGGUGUGCGG.C....................AAAG...............................G.CUCGU.AUAGCGACACGGCCUACUAUUU...GAGGUUAUCCC..AUG
Vna-2-1    ACGUGUGGUGUGCGG.C....................AAAG...............................G.CUCGU.AUAGCGACACGGCCUACUAUUU...GAGGUUAUCCC..AUG
Vna-2-3    ACGUGUGGUGUGCGG.C....................AAAG...............................G.CUCGU.AUAGCGACACGGCCUACUAUUU...GAGGUUAUCCC..AUG
Vvu-8-1    GCGUGUGGUGUGCGG.C....................AAAG...............................G.CUCGU.AUAGCGACACGGCCUACUAUUU...GAGGUUAUCCC..AUG
Vsa-1-1    ACGUGUGGUGUACGGCU....................UUG................................A.UUCGU.AUAGCGACACGGCCUACUAUU....GAGGUUAUCCC..AUG
Vaz-1-1    ACGUGUGGUGUGCGGCU....................UUG................................A.UUCGU.AUAGCGACACGGCCUACUAUU....GAGGUUAUCCC..AUG
Vro-3-1    ACGUGUGGUGUACGG.C....................UUA................................GAUUCGU.AUAGCGACACGGCCUACUAUU....GAGGUUAUCCC..AUG
Vro-2-1    ACGUGUGGUGUGCGG.C....................AUA................................GAUUCGU.AUAGCGACACGGCCUACUAUU....GAGGUUAUCCC..AUG
Vhy-1-1    ACGUGUGGUGUGCGG.C....................UUA................................GACUCGU.AUAGCGACACGGCCUACUAUU....GAGGUUAUCCC..AUG
Vja-1-1    ACGUGUGGUGUGCGG.C....................UUA................................GACUCGU.AUAGCGACACGGCCUACUAUU....GAGGUUAUCCC..AUG
Vhy-1-2    ACGUGUGGUGUGCGG.C....................UUA................................GACUCGU.AUAGCGACACGGCCUACUAUU....GAGGUUAUCCC..AUG
Vna-1-1    ACGUGUGGUGUGCGG.C....................UUA................................GACUCGU.AUAGCGACACGGCCUACUAUU....GAGGUUAUCCC..AUG
Vja-4-1    ACGUGUGGUGUGCGG.C....................UUA................................GACUCGU.AUAGCGACACGGCCUACUAUU....GAGGUUAUCCC..AUG
Vpa-87-1   GCGUGUGGUGUGCGG.C....................UUA................................GAUUCGU.AUAGCGACACGGCCUACUAUU....GAGGUUAUCCC..AUG
Val-2-1    GCGUGUGGUGUGCGG.C....................UUA................................GAUUCGU.AUAGCGACACGGCCUACUAUU....GAGGUUAUCCC..AUG
Vpa-240-1  GCGUGUGGUGUGCGG.C....................UUA................................GAUUCGU.AUAGCGACACGGCCUACUAUU....GAGGUUAUCCC..AUG
Vpa-238-1  GCGUGUGGUGUGCGG.C....................UUA................................GAUUCGU.AUAGCGACACGGCCUACUAUU....GAGGUUAUCCC..AUG
Vpa-94-1   GCGUGUGGUGUGCGG.C....................UUA................................GAUUCGU.AUAGCGACACGGCCUACUAUU....GAGGUUAUCCC..AUG
Vhy-1-3    ACGUGUGGUGUGCGG.C....................UUA................................GAUUCGU.AUAGCGACACGGCCUACUAUU....GAGGUUAUCCC..AUG
env-626    GCGUGUGGUGUGCGG.C....................UUA................................GAUUCGU.AUAGCGACACGGCCUACUAUU....GAGGUUAUCCC..AUG
Vpa-157-1  GCGUGUGGUGUGCGG.C....................UUA................................GAUUCGU.AUAGCGACACGGCCUACUAUU....GAGGUUAUCCC..AUG
Vro-1-1    ACGUGUGGUGUGCGG.C....................UUA................................GAUUCGU.AUAGCGACACGGCCUACUAUU....GAGGUUAUCCC..AUG
Vpa-79-1   GCGUGUGGUGUGCGG.C....................UUA................................GAUUCGU.AUAGCGACACGGCCUACUAUU....GAGGUUAUCCC..AUG
Vpa-207-1  GCGUGUGGUGUGCGG.C....................UUA................................GAUUCGU.AUAGCGACACGGCCUACUAUU....GAGGUUAUCCC..AUG
Vpa-219-1  GCGUGUGGUGUGCGG.C....................UUA................................GAUUCGU.AUAGCGACACGGCCUACUAUU....GAGGUUAUCCC..AUG
Vha-9-1    ACGUGUGGUGUGCGG.C....................UUA................................GACUCGU.AUAGCGACACGGCCUACAAUU....GAGGUUAUCCC..AUG
Vow-5-1    ACGUGUGGUGUGCGG.C....................UUA................................GACUCGU.AUAGCGACACGGCCUACAAUU....GAGGUUAUCCC..AUG
Vca-3-1    ACGUGUGGUGUGCGG.C....................UUA................................GAUUCGU.AUAGCGACACGGCCUACAAUU....GAGGUUAUCCC..AUG
Vow-3-1    ACGUGUGGUGUGCGG.C....................UUA................................GAUUCGU.AUAGCGACACGGCCUACAAUU....GAGGUUAUCCC..AUG
Vsp-10-1   ACGUGUGGUGUGCGG.C....................UUA................................GAUUCGU.AUAGCGACACGGCCUACAAUU....GAGGUUAUCCC..AUG
env-627    ACGUGUGGUGUGCGG.C....................UUA................................GAUUCGU.AUAGCGACACGGCCUACAAUU....GAGGUUAUCCC..AUG
Vsp-2-1    ACGUGUGGUGUGCGG.C....................AUA................................GAUUCGU.AUAGCGACACGGCCUACAAUU....GAGGUUAUCCC..AUG
Vsp-44-1   ACGUGUGGUGUGCGG.C....................UUA................................GAAUCGU.AUAGCGACACGGCCUACUU......GAGGUUAUCCC..AUG
Vcy-2-1    ACGUGUGGUGUGCGG.C....................UUA................................GAAUCGU.AUAGCGACACGGCCUACUU......GAGGUUAUCCC..AUG
Vcy-15-1   ACGUGUGGUGUGCGG.C....................UUA................................GAAUCGU.AUAGCGACACGGCCUACUU......GAGGUUAUCCC..AUG
Vsp-45-1   ACGUGUGGUGUGCGG.C....................UUA................................GAAUCGU.AUAGCGACACGGCCUACUU......GAGGUUAUCCC..AUG
Vcy-13-1   ACGUGUGGUGUGCGG.C....................UUA................................GAAUCGU.AUAGCGACACGGCCUACUU......GAGGUUAUCCC..AUG
Vsp-5-1    ACGUGUGGUGUGCGG.C....................UUA................................GAAUCGU.AUAGCGACACGGCCUACUU......GAGGUUAUCCC..AUG
Vcr-1-1    ACGUGUGGUGUGCGG.C....................UUA................................GAAUCGU.AUAGCGACACGGCCUACUU......GAGGUUAUCCC..AUG
Vba-1-1    ACGUGUGGUGUGCGG.C....................UUA................................GAAUCGU.AUAGCGACACGGCCUACUU......GAGGUUAUCCC..AUG
Vsp-21-1   ACGUGUGGUGUGCGG.C....................UUA................................GAAUCGU.AUAGCGACACGGCCUACUU......GAGGUUAUCCC..AUG
env-630    ACGUGUGGUGUGCGG.C....................UUA................................GAAUCGU.AUAGCGACACGGCCUACUU......GAGGUUAUCCC..AUG
Vsp-35-1   ACGUGUGGUGUGCGG.C....................UUA................................GAAUCGU.AUAGCGACACGGCCUACUU......GAGGUUAUCCC..AUG
Vka-1-1    ACGUGUGGUGUGCGG.C....................UUA................................GAAUCGU.AUAGCGACACGGCCUACUU......GAGGUUAUCCC..AUG
env-633    ACGUGUGGUGUGCGG.C....................UUA................................GAAUCGU.AUAGCGACACGGCCUACUU......GAGGUUAUCCC..AUG
Vsp-31-1   ACGUGUGGUGUGCGG.C....................UUA................................GAAUCGC.AUAGCGACACGGCCUACUU......GAGGUUAUCCC..AUG
env-634    ACGUGUGGUGUACGG.C....................UUA................................GAAUCGU.AUAGCGACACGGCCUACUU......GAGGUUAUCCC..AUG
Afi-2-1    ACGUGUGGUGUCCGG.C....................AUUA...............................GAAUCGG.AUAGCGACACGGCCUAAACUAUU..AAGGUUUUCCC..AUG
Vfi-2-1    ACGUGUGGUGUCCGG.C....................AUUA...............................GAAUCGG.AUAGCGACACGGCCUAAACUAUU..AAGGUUUUCCC..AUG
Vor-1-1    ACGUGUGGUGUCCGGCU....................UCG................................A.UUCGG.AUAGCGACGCGGCCCAAUUU.....GAGGUUAUCCC..AUG
Vge-5-1    ACGUGUGGUGUCCGG.C....................UUA................................GAUUCGG.AUGGCGACACGGCCUAAC.......GAGGUUAUCCC..AUG
Vca-7-1    ACGUGUGGUGUCCGG.C....................AAA................................GAAUCGG.AUAGCGACACGGCCUAUAUUUUA..GAGGUUCUCCC..AUG
Vne-1-1    ACGUGUGGUGUCCGG.C....................AUA................................GAAUCGG.AUAGCGACACGGCCUAUUUUA....GAGGUUCUCCC..AUG
Vco-3-1    ACGUGUGGUGUCCGG.C....................UUA................................GAAUCGG.AUAGCGACACGGCCUAUUUUA....GAGGUUUUCCC..AUG
Vsp-3-1    ACGUGUGGUGUCCGG.C....................UUA................................GAAUCGG.AUAGCGACACGGCCUAUUUUA....GAGGUUUUCCC..AUG
env-636    ACGUGUGGUGUCCGG.C....................UUA................................GAAUCGG.AUAGCGACACGGCCUAUAUAU....GAGGUUUUCCC..AUG
env-637    ACGUGUGGUGUCCGG.C....................UUA................................GAAUCGG.AUGGCGACACGGCCUAUAUAU....GAGGUUCUCCC..AUG
Vmi-5-1    ACGUGUGGUGUCCGA.C....................GACAAUC............................GAUUCGG.AUGGCGACACGGCCUAAAUU.....GAGGUUAUCCC..AUG
Vsp-25-1   ACGUGUGGUGUCCGA.C....................GACAAUC............................GAUUCGG.AUGGCGACACGGCCUAAAUU.....GAGGUUAUCCC..AUG
Vch-3-1    ACGUGUGGUGUCCGA.C....................GACAAUC............................GAUUCGG.AUGGCGACACGGCCUAAAUU.....GAGGUUAUCCC..AUG
Vmi-2-1    ACGUGUGGUGUCCGA.C....................GACAAUC............................GAUUCGG.AUGGCGACACGGCCUAAAUU.....GAGGUUAUCCC..AUG
Vch-7-1    ACGUGUGGUGUCCGA.C....................GACAAUC............................GAUUCGG.AUGGCGACACGGCCUAAAUU.....GAGGUUAUCCC..AUG
Fba-1-1    ACGUGUGGUGUCCGG.C....................AUA................................GAAUCGG.AUAGCGACACGGCCUAAUUU.....GAGGUUUCCC...AUG
Vsi-2-1    ACGUGUGGUGUCCGG.C....................UUA................................GAUUCGG.AUAGCGACACGGCCUAUAUA.....GAGGUUAUCCC..AUG
Val-2-9    ACGUGUGGUGUCCGG.C....................UCA................................GAUUCGG.AUAGCGACACGGCCCAAUUU.....GAGGUUAUCCC..AUG
Vor-7-1    ACGUGUGGUGUCCGG.C....................UUA................................GAUUCGG.AUAGCGACACGGCCUAAUUU.....GAGGUUAUCCC..AUG
Vdi-2-1    ACGUGUGGUGUCCGG.C....................UCA................................GAUUCGG.AUAGCGACACGGCCUAAUUU.....GAGGUUAUCCC..AUG
env-640    ACGUGUGGUGUCCGG.C....................UCA................................GAUUCGG.AUAGCGACACGGCCUAAUUU.....GAGGUUAUCCC..AUG
Vsp-32-1   ACGUGUGGUGUCCGG.C....................AUA................................GAUUCGG.AUAGCGACGCGGCCCAAUUU.....GAGGUUAUCCC..AUG
Vge-3-1    ACGUGUGGUGUCCGG.C....................AUA................................GAAUCGG.AUAGCGACACGGCCUAAAUU.....GAGGUUAUCCC..AUG
Vge-1-1    ACGUGUGGUGUCCGG.C....................AUA................................GAAUCGG.AUAGCGACACGGCCUAAAUU.....GAGGUUAUCCC..AUG
Vni-9-1    ACGUGUGGUGUCCGG.C....................AUA................................GAAUCGG.AUAGCGACACGGCCCAAUUU.....GAGGUUAUCCC..AUG
Vme-2-1    ACGUGUGGUGUCCGG.C....................AUA................................GAAUCGG.AUGGCGACACGGCCCAAUUU.....GAGGUUAUCCC..AUG
Vsi-1-1    ACGUGUGGUGUCCGG.C....................UAA................................GAUUCGG.AUAGCGACACGGCCUAUAUUA....GAGGUUCUCCC..AUG
Vsi-1-3    ACGUGUGGUGUCCGG.C....................UAA................................GAUUCGG.AUAGCGACACGGCCUAUAUUA....GAGGUUCUCCC..AUG
Vne-2-1    ACGUGUGGUGUCCGG.C....................CAA................................GAUUCGG.AUAGCGACACGGCCUAUAUUA....GAGGUUCUCCC..AUG
Vga-1-1    ACGUGUGGUGUCCGG.C....................UUA................................GAAUCGG.AUAGCGACACGGCCUAUACAUA...GAGGUUCUCCC..AUG

           .<<<<<.<<..<<<<<<......<<<<<<<<<<<<<<.................>>>>>>>>>>>>>>....>.>>>>>....>>.>>>>>..............................
           .22222.0...222222......1.1111.11.0010.................0100.11.1111.1....2.22222.....0.22222..............................
           ●CGURUGGUG○YYRR○Y--------------------●●●●○------------------------------G-●YYRR-A●AGCGAYRCGGCCU●●●●●○----GAGGU●●YCC○--AUG
            5′                                                                                                  3′
```

```
Vbr-1-1     ACGUGUGGUGUCCGG.C...................UUA...............................GAAUCGG.AUAGCGACACGGCCUAUAUUCA...GAGGUUCUCCC..AUG
Vpa-1-1     ACGUGUGGUGUCCGG.C...................UUA...............................GAAUCGG.AUAGCGACACGGCCUAUAUUCA...GAGGUUCUCCC..AUG
Vsp-30-1    ACGUGUGGUGUCCGG.C...................UUA...............................GAAUCGG.AUAGCGACACGGCCUAUAUUUA...GAGGUUCUCCC..AUG
Vtu-2-2     ACGUGUGGUGUCCGG.C...................UUA...............................GAAUCGG.AUAGCGACACGGCCUAUAUUUA...GAGGUUCUCCC..AUG
Vtu-1-1     ACGUGUGGUGUCCGG.C...................AUA...............................GAAUCGG.AUAGCGACACGGCCUAUAUUUA...GAGGUUCUCCC..AUG
Vhe-1-1     ACGUGUGGUGUCCGG.C...................UAA...............................GAUUCGG.AUAGCGACACGGCCUAUAUUUA...GAGGUUCUCCC..AUG
env-641     ACGUGUGGUGUCCGA.C...................GUU...............................GAUUCGG.AUAGCGACACGGCCUAAAUU.....GAGGUUAUCCC..AUG
Vfl-4-1     ACGUGUGGUGUCCGA.C...................GUU...............................GAUUCGG.AUAGCGACACGGCCUAAAUU.....GAGGUUAUCCC..AUG
Vfu-2-1     ACGUGUGGUGUCCGA.C...................AUU...............................GAUUCGG.AUAGCGACACGGCCUAAAUU.....GAGGUUAUCCC..AUG
Vsp-26-1    ACGUGUGGUGUCCGA.C...................GAAUC.............................GAUUCGG.AUGGCGACACGGCCUAAAUU.....GAGGUUAUCCC..AUG
Vre-1-1     ACGUGUGGUGUCCGG.C...................UUA...............................GAUUCGG.AUAGCGACACGGCCUAUUU......GAGGUUCUCCC..AUG
Vsc-1-1     ACGUGUGGUGUCCGG.C...................UUA...............................GAUUCGG.AUAGCGACACGGCCUAUUU......GAGGUUCUCCC..AUG
Vic-1-1     ACGUGUGGUGUCCGG.C...................UUA...............................GAUUCGG.AUAGCGACACGGCCUAUUU......GAGGUUCUCCC..AUG
Awo-1-1     ACGUGUGGUGUUCGG.C...................UUA...............................GAUUCGG.AUAGCGACACGGCCUAAAUUAU...AAGGUUUUCCC..AUG
Asa-11-1    ACGUGUGGUGUCCGG.C...................UUA...............................GAUUCGG.AUGGCGACACGGCCUAAAUUAU...AAGGUUUUCCC..AUG
env-642     ACGUGUGGUGUCCGG.C...................UUA...............................GAAUCGG.AUGGCGACACGGCCUAAUU......GAGGUUUUCCC..AUG
Vta-2-1     ACGUGUGGUGUCCGG.C...................UUA...............................GAAUCGG.AUGGCGACACGGCCUAAUU......GAGGUUUUCCC..AUG
env-643     ACGUGUGGUGUCCGG.C...................UUA...............................GAAUCGG.AUGGCGACACGGCCUAAUU......GAGGUUUUCCC..AUG
Vta-3-1     ACGUGUGGUGUCCGG.C...................UUA...............................GAAUCGG.AUGGCGACGCGGCCUAAUU......GAGGUUUUCCC..AUG
env-646     ACGUGUGGUGUCCGG.C...................AUA...............................GAAUCGG.AUGGCGACACGGCCCAUAUAUU...GAGGUUAUCCC..AUG
Vsh-2-1     ACGUGUGGUGUCCGG.C...................UUA...............................GAAUCGG.AUGGCGACACGGCCUAAACUUU...GAGGUUUCCCC..AUG
Vma-1-1     ACGUGUGGUGUCCGG.C...................UUA...............................GAAUCGG.AUGGCGACACGGCCUAAAUUAUU..GAGGUUUCCC...AUG
Vma-1-2     ACGUGUGGUGUCCGG.C...................UUA...............................GAAUCGG.AUGGCGACACGGCCUAAAUUAUU..GAGGUUUCCC...AUG
Vme-1-1     ACGUGUGGUGUCCGG.C...................AUU...............................GAUCCGG.AUAGCGACGCGGCCUACAUUU....GAGGUUAUCCC..AUG
Ymo-1-1     UCGUGUGGUGUCUGG.C...................GGAACAGG..........................G..CUGG.AUAGCGACACGGCCUAAAUAUA...GAGGUUUCC....AUG
Ybe-1-1     UCGUGUGGUGUCUGG.C...................GGAACAGG..........................G..CUGG.AUAGCGACACGGCCUAAAUAUA...GAGGUUUCC....AUG
Yma-2-1     UCGUGUGGUGUCUGG.C...................GAAUCAGG..........................G..CUGG.AUAGCGACACGGCCUAAAUAUA...GAGGUUUUCC...AUG
Yfr-1-1     UCGUGUGGUGUCUGG.C...................GAAACAGG..........................G..CUGG.AUAGCGACACGGCCUAAAUAUA...GAGGUUUUCC...AUG
Yma-1-1     UCGUGUGGUGUCUGG.C...................GAAACAGG..........................G..CUGG.AUAGCGACACGGCCUAAAUAUA...GAGGUUUUCC...AUG
Yfr-1-5     UCGUGUGGUGUCUGG.C...................GAAACAGG..........................G..CUGG.AUAGCGACACGGCCUAAAUAUA...GAGGUUUUCC...AUG
Ype-1-1     UCGUGUGGUGUCUGG.C...................GGAACAGG..........................G..CUGG.AUAGCGACACGGCCUAAAUAUA...GAGGUUUUCC...AUG
Yfr-1-7     UCGUGUGGUGUCUGG.C...................GGAACAGG..........................G..CUGG.AUAGCGACACGGCCUAAAUAUA...GAGGUUUUCC...AUG
Yfr-1-8     UCGUGUGGUGUCUGG.C...................GGAACAGG..........................G..CUGG.AUAGCGACACGGCCUAAAUAUA...GAGGUUUUCC...AUG
Yfr-2-2     UCGUGUGGUGUCUGG.C...................GGAACAGG..........................G..CUGG.AUAGCGACACGGCCUAAAUAUA...GAGGUUUUCC...AUG
Yfr-1-14    UCGUGUGGUGUCUGG.C...................GGAACAGG..........................G..CUGG.AUAGCGACACGGCCUAAAUAUA...GAGGUUUUCC...AUG
Yfr-1-15    UCGUGUGGUGUCUGG.C...................GGAACAGG..........................G..CUGG.AUAGCGACACGGCCUAAAUAUA...GAGGUUUUCC...AUG
Epi-1-1     ACGCGUGGUGUCUGG.C...................GGAAAAGG..........................G..CCGG.AUAGCGACGCGGCCUUAACA.....GAGGUUUUCCCC.AUG
Epi-1-2     ACGCGUGGUGUCUGG.C...................GGAAAAGG..........................G..CCGG.AUAGCGACGCGGCCUUAACA.....GAGGUUUUCCCC.AUG
Eic-2-1     ACGCGUGGUGUCUGG.C...................GGAAAAGG..........................G..CCGG.AUAGCGACGCGGCCUUAACA.....GAGGUUUUCCCC.AUG
Eta-2-1     ACGCGUGGUGUCUGG.C...................GGAAGAGG..........................G..CCGG.AUAGCGACGCGGCCUUAAUA.....GAGGUUUUCCC..AUG
Eho-5-1     ACGCGUGGUGUCUGG.C...................GGAAGAGG..........................G..CCGG.AUAGCGACGCGGCCUUAACA.....GAGGUUUUCCC..AUG
Eta-4-1     ACGCGUGGUGUCUGG.C...................GGAAGAGG..........................G..CCGG.AUAGCGACGCGGCCUUAACA.....GAGGUUUUCCC..AUG
Yal-1-1     UCGUGUGGUGUCUGG.C...................GGAACAGG..........................G..CUGG.AUAGCGACACGGCCUAAUAUA....GAGGUUUCC....AUG
Yen-14-1    UCGUGUGGUGUCUGG.C...................GGAACAGG..........................G..CUGG.AUAGCGACACGGCCUAAUAUA....GAGGUUUCC....AUG
Yal-1-4     UCGUGUGGUGUCUGG.C...................GGAACAGG..........................G..CUGG.AUAGCGACACGGCCUAAUAUA....GAGGUUUCC....AUG
Yen-21-1    UCGUGUGGUGUCUGG.C...................GGAACAGG..........................G..CUGG.AUAGCGACACGGCCUAAUAUA....GAGGUUUCC....AUG
Ype-127-1   UCGUGUGGUGUCUGG.C...................GGAACAGG..........................G..CUGG.AUAGCGACACGGCCUAAUAUA....GAGGUUUCC....AUG
Yal-2-1     UCGUGUGGUGUCUGG.C...................GGAACAGG..........................G..CUGG.AUAGCGACACGGCCUAAUAUA....GAGGUUUCC....AUG
Ynu-1-1     ACGUGUGGUGUCUGG.C...................GGAACAGG..........................G..CUGG.AUAGCGACACGGCCUAAAUUA....GAGGUUUCCC...AUG
Yru-1-1     ACGUGUGGUGUCUGG.C...................GGAACAGG..........................G..CUGG.AUAGCGACACGGCCUUAAUUA....GAGGUUUCCC...AUG
Dze-2-1     UCGUGUGGUGUCUGG.C...................GGAACAGG..........................G..CCGG.AUAGCGACGCGGCCUUAAACA....GAGGUUUCCC...AUG
Dsp-1-1     UCGUGUGGUGUCUGG.C...................GGAAUAGG..........................G..CCGG.AUAGCGACGCGGCCUUAAACA....GAGGUUUCCC...AUG
Ssp-23-1    UCGUGUGGUGUCUGG.C...................GGAAUAGG..........................G..CUGG.AUAGCGACACGGCCUUAAACA....GAGGUUUCCC...AUG
env-647     UCGUGUGGUGUCUGG.C...................GGAAUAGG..........................G..CUGG.AUAGCGACACGGCCUUAAUCA....GAGGUUUCCC...AUG
Lqu-1-3     UCGUGUGGUGUCUGG.C...................GGAAUAGG..........................G..CUGG.AUAGCGACACGGCCUUAAUCA....GAGGUUUCCC...AUG
Dsp-2-1     UCGUGUGGUGUCUGG.C...................GGAACAGG..........................G..CUGG.AUAGCGACACGGCCUUAAACA....GAGGUUUCCC...AUG
Dze-7-1     UCGUGUGGUGUCUGG.C...................GGAACAGG..........................G..CCGG.AUAGCGACACGGCCUUAAACA....GAGGUUUCCC...AUG
Dch-3-1     UCGUGUGGUGUCUGG.C...................GGAACAGG..........................G..CCGG.AUAGCGACACGGCCUUAAACA....GAGGUUUCCC...AUG
Dch-2-1     UCGUGUGGUGUCUGG.C...................GGAACAGG..........................G..CCGG.AUAGCGACACGGCCUUAAACA....GAGGUUUCCC...AUG
Dda-1-1     UCGUGUGGUGUCUGG.C...................GGAACAGG..........................G..CCGG.AUAGCGACACGGCCUUAAACA....GAGGUUUCCC...AUG
Dda-2-1     UCGUGUGGUGUCUGG.C...................GGAACAGG..........................G..CCGG.AUAGCGACACGGCCUUAAACA....GAGGUUUCCC...AUG
Dch-1-2     UCGUGUGGUGUCUGG.C...................GGAACAGG..........................G..CCGG.AUAGCGACACGGCCUUAAACA....GAGGUUUCCC...AUG
Dsp-9-1     UCGUGUGGUGUCUGG.C...................GGAACAAG..........................G..CUGG.AUAGCGACGCGGCCUUAAACA....GAGGUUUCCC...AUG
Xdo-1-1     ACGUGUGGUGUCUGG.C...................GGUCACAAG.........................G..CUGG.AUAGCGACACGGCCUGAACA.....GAGGUUUCCC...AUG
Xpo-1-1     ACGUGUGGUGUCUGG.C...................GAUAACAAG.........................G..CUGG.AUAGCGACACGGCCUGAACA.....GAGGUUUUCC...AUG
Xbo-11-1    ACGUGUGGUGUCUGG.C...................GGCAACAAG.........................G..CUGG.AUAGCGACACGGCCUUAACA.....GAGGUUUCCU...AUG
Xbo-8-1     ACGUGUGGUGUCUGG.C...................GGCAACAAG.........................G..CUGG.AUAGCGACACGGCCUUAACA.....GAGGUUUCCU...AUG
Xbo-6-1     ACGUGUGGUGUCUGG.C...................GGCAACAAG.........................G..CUGG.AUAGCGACACGGCCUUAACA.....GAGGUUUCCU...AUG
Xsz-1-1     ACGUGUGGUAUCUGG.C...................GGCAACAGG.........................G..CUGG.AUGGCGACACGGCCUUAACA.....GAGGUUUUCCC..AUG
Xca-1-1     ACGUGUGGUGUCUGG.C...................GGCGAUAGG.........................G..CUGG.AUAGCGACGCGACCUUAACA.....GAGGUUUUCCCC.AUG
Xkh-1-1     ACGUGUGGUGUCUGG.C...................GGCAACAGG.........................G..CUGG.AUAGCGACACGGCCUGAACA.....GAGGUUUCCC...AUG
Xsp-1-1     ACGUGUGGUGUCUGG.C...................GGCUAUAGG.........................G..CUGG.AUAGCGACACGGCCUUAACA.....GAGGUUUUCCCC.AUG
Xne-5-1     ACGUGUGGUGUCUGG.C...................GGCAAUAGG.........................G..CUGG.AUAGCGACACGGCCUUAACA.....GAGGUUUCCC...AUG
Plu-5-1     ACGUGUGGUGUCUGG.C...................GGCAACAGG.........................G..CUGG.AUAGCGACGCGGCCUUAACA.....GAGGUUAUCCC..AUG
Plu-9-1     ACGUGUGGUGUCUGG.C...................GGCAACAAG.........................G..CUGG.AUAGCGACACGGCCUUAACA.....GAGGUUAUCCC..AUG
Plu-7-1     ACGUGUGGUGUCUGG.C...................GGCAACAGG.........................G..CUGG.AUAGCGACACGGCCUUAACA.....GAGGUUAUCCC..AUG
Plu-6-1     ACGUGUGGUGUCUGG.C...................GGCAACAGG.........................G..CUGG.AUAGCGACACGGCCUUAACA.....GAGGUUAUCCC..AUG
Pte-4-1     ACGUGUGGUGUCUGG.C...................GGCAACAGG.........................G..CUGG.AUAGCGACACGGCCUUAACA.....GAGGUUAUCCC..AUG
Pte-6-1     ACGUGUGGUGUCUGG.C...................GGCAACAGG.........................G..CUGG.AUAGCGACACGGCCUUAACA.....GAGGUUAUCCC..AUG
Plu-10-1    ACGUGUGGUGUCUGG.C...................GGCAACAGG.........................G..CUGG.AUAGCGACACGGCCUUAACA.....GAGGUUAUCCC..AUG
Pte-2-1     ACGUGUGGUGUCUGG.C...................GGCAACAGG.........................G..CUGG.AUAGCGACACGGCCUUAACA.....GAGGUUAUCCC..AUG
Ssp-15-1    ACGUGUGGUGUCUGG.C...................GGAACACG..........................G..CUGG.AUAGCGACACGGCCUUAACU.....GAGGUUCCCCC..AUG
Pca-2-1     UCGUGUGGUGUCUGG.C...................GGAACAAG..........................G..CUGG.AUAGCGACACGGCCUUAACA.....GAGGUUUCCC...AUG

            .<<<<<.<<..<<<<<<......<<<<<<<<<<<<<<..................>>>>>>>>>>>>>>....>.>>>>>....>>.>>>>>..............................
            .22222.0...222222......1.1111.11.0010..................0100.11.1111.1....2.22222.....0.22222..............................
            •CGURUGGUG∘YYRR∘Y--------------------••••∘----------------------------G-∘YYRR-A•AGCGAYRCGGCU••••∘∘----GAGGU••YCC∘--AUG
             5′                                                                                               3′
```

```
Pca-1-1   UCGUGUGGUGUCUGG.C....................GGAACAAG...........................G..CUGG.AUAGCGACACGGCCUUAACA.....GAGGUUUCCC...AUG
Ddi-5-1   UCGUGUGGUGUCUGG.C....................GGAACAAG...........................G..CUGG.AUAGCGACACGGCCUUAACA.....GAGGUUUCCC...AUG
Ssp-13-1  ACGUGUGGUGUCUGG.C....................GAAACAGG...........................G..CUGG.AUAGCGACACGGCCUUAACA.....GAGGUUUCCC...AUG
Dpa-1-1   UCGUGUGGUGUCUGG.C....................GGAAAAGG...........................G..CUGG.AUAGCGACACGGCCUUAACA.....GAGGUUUCCC...AUG
Sfo-1-1   ACGUGUGGUGUCUGG.C....................GGAACAGG...........................G..CUGG.AUAGCGACACGGCCUUAACU.....GAGGUUCUCCC..AUG
Pca-7-1   UCGUGUGGUGUCUGG.C....................GGAACAGG...........................G..CUGG.AUAGCGACACGGCCUUAACA.....GAGGUUUCCC...AUG
Ppa-2-1   UCGUGUGGUGUCUGG.C....................GGAACAGG...........................G..CUGG.AUAGCGACACGGCCUUAACA.....GAGGUUUCCC...AUG
Pat-3-1   UCGUGUGGUGUCUGG.C....................GGAACAGG...........................G..CUGG.AUAGCGACACGGCCUUAACA.....GAGGUUUCCC...AUG
Psp-55-1  UCGUGUGGUGUCUGG.C....................GGAACAGG...........................G..CUGG.AUAGCGACACGGCCUUAACA.....GAGGUUUCCC...AUG
Pwa-2-1   UCGUGUGGUGUCUGG.C....................GGAACAGG...........................G..CUGG.AUAGCGACACGGCCUUAACA.....GAGGUUUCCC...AUG
Pca-8-1   UCGUGUGGUGUCUGG.C....................GGAACAGG...........................G..CUGG.AUAGCGACACGGCCUUAACA.....GAGGUUUCCC...AUG
Bgo-1-1   UCGUGUGGUGUCUGG.C....................GGAACAGG...........................G..CUGG.AUAGCGACACGGCCUUAACA.....GAGGUUUCCC...AUG
Bsp-1-1   UCGUGUGGUGUCUGG.C....................GGAACAGG...........................G..CUGG.AUAGCGACACGGCCUUAACA.....GAGGUUUCCC...AUG
Dsp-3-1   UCGUGUGGUGUCUGG.C....................GGAACAGG...........................G..CUGG.AUAGCGACACGGCCUUAACA.....GAGGUUUCCC...AUG
Pbe-1-1   UCGUGUGGUGUCUGG.C....................GGAACAGG...........................G..CUGG.AUAGCGACACGGCCUUAACA.....GAGGUUUCCC...AUG
Pca-5-11  UCGUGUGGUGUCUGG.C....................GGAACAGG...........................G..CUGG.AUAGCGACACGGCCUUAACA.....GAGGUUUCCCC..AUG
env-651   ACGUGUGGUGUCUGG.C....................GAAACAGG...........................G..CUGG.AUAGCGACACGGCCUUAACU.....GAGGUUUCCC...AUG
env-653   ACGUGUGGUGUCUGG.C....................GAAACAGG...........................G..CUGG.AUAGCGACACGGCCUUAACU.....GAGGUUUCCC...AUG
env-655   ACGUGUGGUGUCUGG.C....................GAAACAGG...........................G..CUGG.AUAGCGACACGGCCUUAACU.....GAGGUUUCCC...AUG
env-656   ACGUGUGGUGUCUGG.C....................GAAACAGG...........................G..CUGG.AUAGCGACACGGCCUUAACU.....GAGGUUUCCC...AUG
Spl-5-1   ACGUGUGGUGUCUGG.C....................GAAACAGG...........................G..CUGG.AUAGCGACACGGCCUUAACU.....GAGGUUUCCC...AUG
Ssy-1-1   UCGUGUGGUGUCUGG.C....................GAAACAGG...........................G..CUGG.AUAGCGACACGGCCUUAACU.....GAGGUUUCCC...AUG
Ssy-2-1   ACGUGUGGUGUCUGG.C....................GAAACAGG...........................G..CUGG.AUAGCGACACGGCCUUAACU.....GAGGUUUCCC...AUG
Sgr-1-1   ACGUGUGGUGUCUGG.C....................GAAACAGG...........................G..CUGG.AUAGCGACACGGCCUUAACU.....GAGGUUUCCC...AUG
Sod-1-1   ACGUGUGGUGUCUGG.C....................GAAACAGG...........................G..CUGG.AUAGCGACACGGCCUUAACU.....GAGGUUUCCCC..AUG
Spl-1-1   ACGUGUGGUGUCUGG.C....................GAAACAGG...........................G..CUGG.AUAGCGACACGGCCUUAACU.....GAGGUUUCCC...AUG
Spr-2-1   ACGUGUGGUGUCUGG.C....................GAAACAGG...........................G..CUGG.AUAGCGACACGGCCUUAACU.....GAGGUUUCCC...AUG
env-657   ACGUGUGGUGUCUGG.C....................GAAACAGG...........................G..CUGG.AUAGCGACACGGCCUUAACU.....GAGGUUUCCC...AUG
env-658   ACGUGUGGUGUCUGG.C....................GAAACAGG...........................G..CUGG.AUAGCGACACGGCCUUAACU.....GAGGUUUCCC...AUG
Sma-16-1  ACGUGUGGUGUCUGG.C....................GAAACAGG...........................G..CUGG.AUAGCGACACGGCCUUAACU.....GAGGUUUCCC...AUG
Ssp-21-1  ACGUGUGGUGUCUGG.C....................GAAACAGG...........................G..CUGG.AUAGCGACACGGCCUUAACU.....GAGGUUUCCC...AUG
Sma-24-1  ACGUGUGGUGUCUGG.C....................GAAACAGG...........................G..CUGG.AUAGCGACACGGCCUUAACU.....GAGGUUUCCCC..AUG
Sma-3-1   ACGUGUGGUGUCUGG.C....................GAAACAGG...........................G..CUGG.AUAGCGACACGGCCUUAACU.....GAGGUUUCCC...AUG
Pca-3-1   UCGUGUGGUGUCUGG.C....................GGAACAGG...........................G..CUGG.AUAGCGACACGGCCUUAACU.....GAGGUUUCCCC..AUG
Pca-6-1   UCGUGUGGUGUCUGG.C....................GGAACAGG...........................G..CUGG.AUAGCGACACGGCCUUAACU.....GAGGUUUCCCC..AUG
Pca-5-2   UCGUGUGGUGUCUGG.C....................GGAACAGG...........................G..CUGG.AUAGCGACACGGCCUUAACU.....GAGGUUUCCC...AUG
Pca-5-4   UCGUGUGGUGUCUGG.C....................GGAACAGG...........................G..CUGG.AUAGCGACACGGCCUUAACU.....GAGGUUUCCC...AUG
Pca-10-1  UCGUGUGGUGUCUGG.C....................GGAACAGG...........................G..CUGG.AUAGCGACACGGCCUUAACU.....GAGGUUUCCCC..AUG
Pca-3-2   UCGUGUGGUGUCUGG.C....................GGAACAGG...........................G..CUGG.AUAGCGACACGGCCUUAACU.....GAGGUUUCCCC..AUG
Pca-3-3   UCGUGUGGUGUCUGG.C....................GGAACAGG...........................G..CUGG.AUAGCGACACGGCCUUAACU.....GAGGUUUCCCC..AUG
Pca-12-1  UCGUGUGGUGUCUGG.C....................GGAACAGG...........................G..CUGG.AUAGCGACACGGCCUUAACU.....GAGGUUUCCCC..AUG
Cmu-2-1   ACGUGUGGUGUCUGG.C....................GGAACAGG...........................G..CUGG.AUAGCGACACGGCCUUAACU.....GAGGUUUCCC...AUG
Pca-5-6   UCGUGUGGUGUCUGG.C....................GGAACAGG...........................G..CUGG.AUAGCGACACGGCCUUAACU.....GAGGUUUCCCC..AUG
Pca-5-8   UCGUGUGGUGUCUGG.C....................GGAACAGG...........................G..CUGG.AUAGCGACACGGCCUUAACU.....GAGGUUUCCCC..AUG
Ssp-17-1  ACGUGUGGUGUCUGG.C....................GGAACAGG...........................G..CUGG.AUAGCGACACGGCCUUAACU.....GAGGUUUUCCC..AUG
Pas-2-1   ACGUGUGGUGUCUGG.C....................GGCAACAGG..........................G..CUGG.AUAGCGACACGGCCUUAAACA....GAGGUUAUCCC..AUG
Pas-1-1   ACGUGUGGUGUUUGG.C....................GGCAACAGG..........................G..CUGG.AUAGCGACACGGCCUUAAAUA....GAGGUUAUCCC..AUG
Xgr-1-1   ACGUGUGGUGUCUGG.C....................AGCAAAACAA.........................G.GCUGG.AUAGCGACACGGCCUGAACA.....GAGGUUUCCC...AUG
Hpa-1-1   ACGCGUGGUGUCUGG.C....................GGAAUAGG...........................G..CUGG.AUAGCGACGCGGCCUUAACU.....GAGGUUAUCCC..AUG
Hal-1-1   ACGCGUGGUGUCUGG.C....................GGAAUAGG...........................G..CUGG.AUAGCGACGCGGCCUUAACU.....GAGGUUAUCCC..AUG
env-660   ACGCGUGGUGUCUGG.C....................GGAAUAGG...........................G..CUGG.AUAGCGACGCGGCCUUAACU.....GAGGUUAUCCC..AUG
Hal-3-1   ACGCGUGGUGUCUGG.C....................GGAAUAGG...........................G..CUGG.AUAGCGACGCGGCCUUAACU.....GAGGUUAUCCC..AUG
env-661   ACGCGUGGUGUCCGA.C....................GGAAAACAC..........................G.GUUGG.AUAGCGACGCGGCCUAAACA.....GAGGUUUUCCC..AUG
gpr-1-1   ACGCGUGGUGUCCGA.C....................GGCAAAACG..........................G..UUGG.AUAGCGACGCGGCCUAAACA.....GAGGUUUUCCC..AUG
Ssp-24-1  ACGCGUGGUGUCCGG.C....................GGUAACACG..........................G..UUGG.AUAGCGACGCGGCCUAAACA.....GAGGUUUUCCC..AUG
Rch-1-1   ACGUGUGGUGUCCGA.C....................GGAAACACG..........................G..UUGG.AUAGCGACACGGCCUAAACA.....GAGGUUUUCUC..AU.
Eam-2-1   ACGCGUGGUGUCCGA.C....................GAAACAGG...........................G..UUGG.AUAGCGACGCGGCCUUAAGAA....GAGGUUUUCCC..AUG
env-662   ACGCGUGGUGUCUGA.C....................GGAACAGG...........................G..UUGG.AUAGCGACGCGGCCUUAAGAA....GAGGUUUUCC...AUG
env-668   ACGCGUGGUGUCUGA.C....................AGAACAGG...........................G..UUGG.AUAGCGACGCGGCCUUAAGAA....GAGGUUUUCC...AUG
Raq-1-1   ACGCGUGGUGUCUGA.C....................AGAACAGG...........................G..UUGG.AUAGCGACGCGGCCUUAAGAA....GAGGUUUUCC...AUG
env-672   ACGCGUGGUGUCUGA.C....................AGAACAGG...........................G..UUGG.AUAGCGACGCGGCCUUAAGAA....GAGGUUUUCC...AUG
env-674   ACGCGUGGUGUCUGA.C....................AGAACAGG...........................G..UUGG.AUAGCGACGCGGCCUUAAGAA....GAGGUUUUCC...AUG
Mmo-1-1   ACGUGUGGUGUCCGG.C....................AAAAAG.............................GC.CCGG.AUAGCGACACGGCCUUAACC.....GAGGUUUUCCCC.AUG
Mmo-3-1   ACGUGUGGUGUCCGG.C....................AAAAAG.............................G.CCCGG.AUAGCGACACGGCCUUAACC.....GAGGUUUUCCC..AUG
Mmo-2-1   ACGUGUGGUGUCCGG.C....................AAAAAG.............................G.CCCGG.AUAGCGACACGGCCUUAACC.....GAGGUUUUCCC..AUG
Mmo-1-7   ACGUGUGGUGUCCGG.C....................AAAAAG.............................G.CCCGG.AUAGCGACACGGCCAAAACCU....GAGGUUUUCCC..AUG
Mwi-1-1   ACGUGUGGUGUCCGG.C....................GUUAGG.............................G..CGGG.AUGGCGACACGGCCUUAAUA.....GAGGUUUUCCC..AUG
Psn-1-1   ACGUGUGGUG.UCCAGC....................GUUAG..............................GCCUGGA.AUGGCGACACGGCCUUAAUC.....GAGGUUUUCCC..AUG
Pbu-1-1   ACGUGUGGUG.UCCGGC....................GUCAGG.............................G.CUGGA.AUGGCGACACGGCCUUAAUA.....GAGGUUUUCCC..AUG
Pal-5-1   ACGUGUGGUG.UCCGGC....................GUUAGG.............................G.CUGGA.AUGGCGACACGGCCUUAUUA.....GAGGUUCUUCC..AUG
Pre-3-1   ACGUGUGGUG.UCCGGC....................GUUAGG.............................G.CUGGA.AUGGCGACACGGCCUUAAUA.....GAGGUUCUCCC..AUG
Pre-1-1   ACGUGUGGUG.UCCGGC....................GUUAGG.............................G.CUGGA.AUGGCGACACGGCCUUAAUA.....GAGGUUCUCCC..AUG
Pre-2-1   ACGUGUGGUG.UCCGGC....................GUUAGG.............................G.CUGGA.AUGGCGACACGGCCUUAAUA.....GAGGUUCUCCC..AUG
Pru-5-1   ACGUGUGGUG.UCCGGC....................GUUAGG.............................G.CUGGA.AUGGCGACACGGCCUUAUUA.....GAGGUUCUCCC..AUG
Pal-8-1   ACGUGUGGUG.UCCGGC....................GUUAGG.............................G.CUGGA.AUGGCGACACGGCCUUAUUA.....GAGGUUCUCCC..AUG
Eba-4-1   UCGCGUGGUG.UCUGGC....................GUUAGG.............................G.CUGGA.AGAGCGACGCGGCCUUCAGAUU...GAGGUUUCCC...AUG
Msp-4-1   UCGCGUGGUG.UCUGGC....................GUUAGG.............................G.CUGGA.AGAGCGACGCGGCCUUAGACCU...GAGGUUUUCC...AUG
env-675   UCGCGUGGUG.UCUGGC....................GUAGGG.............................G.CCGGA.AGAGCGACGCGGCCUUAUACA....GAGGUUUUCCC..AUG
env-677   UCACGCGGUG.UCUGGC....................GUUAGG.............................G.CUGGA.AGAGCGGCGUGGCCUUACACA....GAGGUUUUCC...AUG
env-678   UCGCGUGGUG.UCUGGC....................GUUAGG.............................G.CCGGA.AGAGCGACGCGGCCUUAACU.....GAGGUUUCCC...AUG
Cmu-1-1   UCGCGUGGUG.UCUGGC....................GUAAGG.............................G.CCGGA.AGAGCGACGCGGCCUUAACU.....GAGGUUUCCC...AUG
Csa-13-1  UCGCGUGGUG.UCUGGC....................GUAAGG.............................G.CUGGA.AGAGCGACGCGGCCUUAACU.....GAGGUUUCCC...AUG
Csa-10-1  UCGCGUGGUG.UCUGGC....................GUAAGG.............................G.CUGGA.AGAGCGACGCGGCCUUAACU.....GAGGUUUCCC...AUG

          .<<<<<.<<..<<<<<<......<<<<<<<<<<<<<<.................>>>>>>>>>>>>>>....>.>>>>>....>>.>>>>>..............................
          .22222.0...222222......1.1111.11.0010.................0100.11.1111.1....2.22222....0.22222..............................
          •CGURUGGUG∘YYRR∘Y--------------------●●●∘∘------------------------------G-∘YYRR-A•AGCGAYRCGGCCU●●●●∘∘----GAGGU••YCC∘--AUG
           5′                                                                                                 3′
```

```
Cun-1-1      UCGCGUGGUG.UCUGGC....................GUAAGG.............................G.CUGGA.AGAGCGACGCGGCCUUAACU.....GAGGUUUCCC...AUG
Csa-7-1      UCGCGUGGUG.UCUGGC....................GUAAGG.............................G.CUGGA.AGAGCGACGCGGCCUUAACU.....GAGGUUUCCC...AUG
env-679      UCGCGUGGUG.UCCGGC....................GUUGGG.............................G.CUGGA.AGAGCGACGCGGCCUUAACU.....GAGGUUUCCC...AUG
env-680      UCGCGUGGUG.UCUGGC....................GUUAGG.............................G.CUGGA.AGAGCGACGCGGCCUUUACU.....GAGGUUUCCCC..AUG
env-681      UCGCGUGGUG.UCUGGC....................GUUAGG.............................G.CUGGA.AGAGCGACGCGGCCUUCACU.....GAGGUUUCCC...AUG
env-682      UCGCGUGGUG.UCUGGC....................GUUAGG.............................G.CUGGA.AGAGCGACGCGGCCUUCACU.....GAGGUUUCCC...AUG
env-683      UCGCGUGGUG.UCUGGC....................GUUAGG.............................G.CUGGA.AGAGCGACGCGGCCUUAACU.....GAGGUUUCCC...AUG
env-685      UCGCGUGGUG.UCUGGC....................GUUAGG.............................G.CUGGA.AGAGCGACGCGGCCUUAACU.....GAGGUUUCCC...AUG
Tgu-1-1      UCGCGUGGUG.UCCGGC....................GUUAGG.............................G.CUGGA.AGAGCGACGCGGCCUUAACU.....GAGGUUUCCC...AUG
env-686      UCGCGUGGUG.UCUGGC....................GUUAGG.............................G.CUGGA.AGAGCGACGCGGCCUUUACU.....GAGGUUCUCCC..AUG
env-687      UCGCGUGGUG.UCUGGC....................GUUAGG.............................G.CUGGA.AGAGCGACGCGGCCUUUACU.....GAGGUUCUCCC..AUG
env-688      UCGCGUGGUG.UCUGGC....................GUUAGG.............................G.CUGGA.AGAGCGACGCGGCCUUUACU.....GAGGUUCUCCC..AUG
env-690      UCGCGUGGUG.UCUGGC....................GUUAGG.............................G.CUGGA.AGAGCGACGCGGCCUUAACU.....GAGGUUCUCCC..AUG
env-700      UCGCGUGGUG.UCUGGC....................GUUAGG.............................G.CUGGA.AGAGCGACGCGGCCUUAACU.....GAGGUUCUCCC..AUG
env-701      UCGCGUGGUG.UCUGGC....................GUUAGG.............................G.CUGGA.AGAGCGACGCGGCCUUAACU.....GAGGUUCUCCC..AUG
env-702      UCGCGUGGUG.UCUGGC....................GUUAGG.............................G.CUGGA.AGAGCGACGCGGCCUUAACU.....GAGGUUCUCCC..AUG
Ksp-8-1      UCGCGUGGUG.UCUGGC....................GUUAGG.............................G.CUGGA.AGAGCGACGCGGCCUUAACU.....GAGGUUCUCCC..AUG
Ecl-52-1     UCGCGUGGUG.UCUGGC....................GUUAGG.............................G.CUGGA.AGAGCGACGCGGCCUUAACU.....GAGGUUCUCCC..AUG
env-706      UCGCGUGGUG.UCUGGC....................GUUAGG.............................G.CUGGA.AGAGCGACGCGGCCUUAACU.....GAGGUUCUCCC..AUG
env-710      UCGCGUGGUG.UCUGGC....................GUUAGG.............................G.CUGGA.AGAGCGACGCGGCCUUAACU.....GAGGUUCUCCC..AUG
Kva-1-1      UCGCGUGGUG.UCUGGC....................GUUAGG.............................G.CUGGA.AGAGCGACGCGGCCUUAACU.....GAGGUUCUCCC..AUG
env-711      UCGCGUGGUG.UCUGGC....................GUUAGG.............................G.CUGGA.AGAGCGACGCGGCCUUAACU.....GAGGUUCUCCC..AUG
env-712      UCGCGUGGUG.UCUGGC....................GUUAGG.............................G.CUGGA.AGAGCGACGCGGCCUUAACU.....GAGGUUCUCCC..AUG
env-713      UCGCGUGGUG.UCUGGC....................GUUAGG.............................G.CUGGA.AGAGCGACGCGGCCUUAACU.....GAGGUUCUCCC..AUG
Ecl-39-1     UCGCGUGGUG.UCUGGC....................GUUAGG.............................G.CUGGA.AGAGCGACGCGGCCUUAACU.....GAGGUUCUCCC..AUG
env-714      UCGCGUGGUG.UCUGGC....................GUUAGG.............................G.CUGGA.AGAGCGACGCGGCCUUAACU.....GAGGUUCUCCC..AUG
Ecl-1-6      UCGCGUGGUG.UCUGGC....................GUUAGG.............................G.CUGGA.AGAGCGACGCGGCCUUAACU.....GAGGUUCUCCC..AUG
env-715      UCGCGUGGUG.UCUGGC....................GUUAGG.............................G.CUGGA.AGAGCGACGCGGCCUUAACU.....GAGGUUCUCCCC.AUG
env-717      UCGCGUGGUG.UCUGGC....................GUUAGG.............................G.CUGGA.AGAGCGACGCGGCCUUAACU.....GAGGUUCUCCC..AUG
env-720      UCGCGUGGUG.UCUGGC....................GUUAGG.............................G.CUGGA.AGAGCGACGCGGCCUUAACU.....GAGGUUCUCCC..AUG
env-724      UCGCGUGGUG.UCUGGC....................GUUAGG.............................G.CUGGA.AGAGCGACGCGGCCUUAACU.....GAGGUUCUCCC..AUG
env-725      UCGCGUGGUG.UCUGGC....................GUUAGG.............................G.CUGGA.AGAGCGACGCGGCCUUAACU.....GAGGUUCUCCC..AUG
env-726      UCGCGUGGUG.UCUGGC....................GUUAGG.............................G.CUGGA.AGAGCGACGCGGCCUUAACU.....GAGGUUCUCCC..AUG
env-728      UCGCGUGGUG.UCUGGC....................GUUAGG.............................G.CUGGA.AGAGCGACGCGGCCUUAACU.....GAGGUUCUCCC..AUG
env-729      UCGCGUGGUG.UCUGGC....................GUUAGG.............................G.CUGGA.AGAGCGACGCGGCCUUAACU.....GAGGUUCUCCC..AUG
env-730      UCGCGUGGUG.UCUGGC....................GUUAGG.............................G.CUGGA.AGAGCGACGCGGCCUUAACU.....GAGGUUCUCCC..AUG
env-731      UCGCGUGGUG.UCUGGC....................GUUAGG.............................G.CUGGA.AGAGCGACGCGGCCUUAACU.....GAGGUUCUCCC..AUG
Lad-1-2      UCGCGUGGUG.UCUGGC....................GUUAGG.............................G.CUGGA.AGAGCGACGCGGCCUUAACU.....GAGGUUCUCCC..AUG
env-732      UCGCGUGGUG.UCUGGC....................GUUAGG.............................G.CUGGA.AGAGCGACGCGGCCUUAACU.....GAGGUUCUCCC..AUG
env-735      UCGCGUGGUG.UCUGGC....................GUUAGG.............................G.CUGGA.AGAGCGACGCGGCCUUAACU.....GAGGUUCUCCC..AUG
Rte-1-1      UCGCGUGGUG.UCUGGC....................GUUAGG.............................G.CUGGA.AGAGCGACGCGGCCUUAACU.....GAGGUUCUCCC..AUG
Eho-1-9      UCGCGUGGUG.UCUGGC....................GUUAGG.............................G.CUGGA.AGAGCGACGCGGCCUUAACU.....GAGGUUCUCCC..AUG
env-736      UCGCGUGGUG.UCUGGC....................GUUAGG.............................G.CUGGA.AGAGCGACGCGGCCUUAACU.....GAGGUUCUCCC..AUG
Kox-8-1      UCGCGUGGUG.UCUGGC....................GUUAGG.............................G.CUGGA.AGAGCGACGCGGCCUUAACU.....GAGGUUCUCCC..AUG
Esp-29-1     UCGCGUGGUG.UCUGGC....................GUUAGG.............................G.CUGGA.AGAGCGACGCGGCCUUAACU.....GAGGUUCUCCC..AUG
Ror-1-1      UCGCGUGGUG.UCUGGC....................GUUAGG.............................G.CUGGA.AGAGCGACGCGGCCUUAACU.....GAGGUUCUCCC..AUG
env-738      UCGCGUGGUG.UCUGGC....................GUUAGG.............................G.CUGGA.AGAGCGACGCGGCCUUAACU.....GAGGUUCUCCC..AUG
Ecl-42-1     UCGCGUGGUG.UCUGGC....................GUUAGG.............................G.CUGGA.AGAGCGACGCGGCCUUAACU.....GAGGUUCUCCC..AUG
env-739      UCGCGUGGUG.UCUGGC....................GUUAGG.............................G.CUGGA.AGAGCGACGCGGCCUUAACU.....GAGGUUCUCCC..AUG
Rpl-1-1      UCGCGUGGUG.UCUGGC....................GUUAGG.............................G.CUGGA.AGAGCGACGCGGCCUUAACU.....GAGGUUCUCCC..AUG
Eca-3-1      UCGCGUGGUG.UCUGGC....................GUUAGG.............................G.CUGGA.AGAGCGACGCGGCCUUAACU.....GAGGUUCUCCC..AUG
Kqu-1-1      UCGCGUGGUG.UCUGGC....................GUUAGG.............................G.CUGGA.AGAGCGACGCGGCCUUAACU.....GAGGUUCUCCC..AUG
Esp-67-1     UCGCGUGGUG.UCUGGC....................GUUAGG.............................G.CUGGA.AGAGCGACGCGGCCUUAACU.....GAGGUUCUCCCC.AUG
env-740      UCGCGUGGUG.UCUGGC....................GUUAGG.............................G.CUGGA.AGAGCGACGCGGCCUUAACU.....GAGGUUCUCCC..AUG
env-741      UCGCGUGGUG.UCUGGC....................GUUAGG.............................G.CUGGA.AGAGCGACGCGGCCUUAACU.....GAGGUUCUCCC..AUG
env-742      UCGCGUGGUG.UCUGGC....................GUUAGG.............................G.CUGGA.AGAGCGACGCGGCCUUAACU.....GAGGUUCUCCC..AUG
env-743      UCGCGUGGUG.UCUGGC....................GUUAGG.............................G.CUGGA.AGAGCGACGCGGCCUUAACU.....GAGGUUCUCCC..AUG
env-754      UCGCGUGGUG.UCUGGC....................GUUAGG.............................G.CUGGA.AGAGCGACGCGGCCUUAACU.....GAGGUUUUCCC..AUG
env-755      UCGCGUGGUG.UCUGGC....................GUUAGG.............................G.CUGGA.AGAGCGACGCGGCCUUAACU.....GAGGUUUUCCC..AUG
env-756      UCGCGUGGUG.UCUGGC....................GUUAGG.............................G.CUGGA.AGAGCGACGCGGCCUUAACU.....GAGGUUUUCCC..AUG
Kcr-1-1      UCGCGUGGUG.UCUGGC....................GUUAGG.............................G.CUGGA.AGAGCGACGCGGCCUUAACC.....GAGGUUUUCCCC.AUG
Kas-1-1      UCGCGUGGUG.UCUGGC....................GUUAGG.............................G.CUGGA.AGAGCGACGCGGCCUUAACC.....GAGGUUUUCCCC.AUG
Fhe-1-1      UCGCGUGGUG.UCUGGC....................GUCAGG.............................G.CCGGA.AGAGCGACGCGGCCUUAACC.....GAGGUUUUCCC..AUG
Fhe-2-1      UCGCGUGGUG.UCUGGC....................GUCAGG.............................G.CCGGA.AGAGCGACGCGGCCUUAACC.....GAGGUUUUCCC..AUG
Fpu-1-1      UCGCGUGGUG.UCUGGC....................GUCAGG.............................G.CCGGA.AGAGCGACGCGGCCUUAACC.....GAGGUUUUCCC..AUG
Fhe-3-1      UCGCGUGGUG.UCUGGC....................GUCAGG.............................G.CCGGA.AGAGCGACGCGGCCUUAACC.....GAGGUUUUCCC..AUG
Fpu-5-1      UCGCGUGGUG.UCUGGC....................GUCAGG.............................G.CCGGA.AGAGCGACGCGGCCUUAACC.....GAGGUUUUCCC..AUG
Cne-1-1      UCGCGUGGUG.UCUGGC....................GUUAGG.............................G.CUGGA.AGAGCGACGCGGCCUUUACC.....GAGGUUUACC...AUG
Cne-1-3      UCGCGUGGUG.UCUGGC....................GUUAGG.............................G.CUGGA.AGAGCGACGCGGCCUUUACC.....GAGGUUUACC...AUG
Cda-1-1      UCGCGUGGUG.UCUGGC....................GUUAGG.............................G.CUGGA.AGAGCGACGCGGCCUUUACC.....GAGGUUUACC...AUG
Sco-2-1      UCGCGUGGUG.UCUGGC....................GUCAGG.............................G.CUGGA.AGAGCGACGCGGCCUUAACA.....GAGGUUUCCC...AUG
Bag-2-1      UCGCGCGGUG.UCUGGC....................GUUAGG.............................G.CUGGA.AGAGCGGCGCGGCCUUAUACA....GAGGUUUUCC...AUG
Ebl-1-1      ACGCGUGGUG.UCUGGC....................GUUAGG.............................G.CCGGA.AGAGCGACGCGGCCUUAGACA....GAGGUUUCCC...AUG
Sen-311-1    UCGCGUGGUG.UCUGGC....................GUUAGG.............................G.CUGGA.AGAGCGACGCGGCCUUACACU....GAGGUUCCCC...AUG
Sen-1013-1   UCGCGUGGUG.UCUGGC....................GUUAGG.............................G.CUGGA.AGAGCGACGCGGCCUUACACU....GAGGUUCCCC...AUG
Sen-89-1     UCGCGUGGUG.UCUGGC....................GUUAGG.............................G.CUGGA.AGAGCGACGCGGCCUUACACU....GAGGUUCCCC...AUG
Sbo-3-1      UCGCGUGGUG.UCUGGC....................GUUAGG.............................G.CUGGA.AGAGCGACGCGGCCUUACACU....GAGGUUCCCC...AUG
Sen-1029-1   UCGCGUGGUG.UCUGGC....................GUUAGG.............................G.CUGGA.AGAGCGACGCGGCCUUACACU....GAGGUUCCCC...AUG
Sen-1021-1   UCGCGUGGUG.UCUGGC....................GUUAGG.............................G.CUGGA.AGAGCGACGCGGCCUUACACU....GAGGUUCCCCC..AUG
Sen-312-1    UCGCGUGGUG.UCUGGC....................GUUAGG.............................G.CUGGA.AGAGCGACGCGGCCUUACACU....GAGGUUCCCC...AUG
Bag-1-1      UCGCGUGGUG.UCUGGC....................GUUAGG.............................G.CUGGA.AGAGCGACGCGGCCUUAUACA....GAGGUUUUCC...AUG
env-757      UCGCGUGGUG.UCUGGC....................GUUAGG.............................G.CUGGA.AGAGCGACGCGGCCUUAAACU....GAGGUUUUCC...AUG
Cfr-3-1      UCGCGUGGUG.UCUGGC....................GUUAGG.............................G.CUGGA.AGAGCGACGCGGCCUUAAACU....GAGGUUUUCCC..AUG

             .<<<<<.<<..<<<<<<......<<<<<<<<<<<<<<.................>>>>>>>>>>>>>>....>.>>>>>....>>.>>>>>..............................
             .22222.0...222222......1.1111.11.0010.................0100.11.1111.1....2.22222.....0.22222..............................
             •CGURUGGUG∘YYRR∘Y--------------------••••∘∘-----------------------------G-∘YYRR-A•AGCGAYRCGGCCU••••∘∘----GAGGU••YCC∘--AUG
              5′                                                                                                   3′
```

```
Ksp-3-1     UCGCGUGGUG.UCUGGC....................GUUAGG.............................G.CUGGA.AGAGCGACGCGGCCUUAAACU....GAGGUUUUCCCG.AUG
Eco-1011-1  UCGCGUGGGG.UCUGGC....................GUUAGG.............................G.CUGGA.AGAGCGACGCGGCCUUAAACC....GAGGUUUUCCC..AUG
env-758     UCGCGUGGUG.UCUGGC....................GUUAGG.............................G.CUGGA.AGAGCGACGCGGCCUUAAACC....GAGGUUUUCCC..AUG
Eco-1781-1  UCGCGUGGUG.UCUGGC....................GUUAGG.............................G.CUGGA.AGAGCGACGCGGCCUUAAACC....GAGGUUUUCCC..AUG
Cam-1-1     UCGCGUGGUG.UCUGGC....................GUUAGG.............................G.CUGGA.AGAGCGACGCGGCCUUAAACC....GAGGUUUUCCC..AUG
Sso-1-2     UCGCGUGGUG.UCUGGC....................GUUAGG.............................G.CUGGA.AGAGCGACGCGGCCUUAAACC....GAGGUUUUCCC..AUG
Sfl-70-1    UCGCGUGGUG.UCUGGC....................GUUAGG.............................G.CUGGA.AGAGCGACGCGGCCUUAAACC....GAGGUUUUCCC..AUG
Ssp-20-2    UCGCGUGGUG.UCUGGC....................GUUAGG.............................G.CUGGA.AGAGCGACGCGGCCUUAAACC....GAGGUUUUCCC..AUG
Eco-1460-1  UCGCGUGGUG.UCUGGC....................GUUAGG.............................G.CUGGA.AGAGCGACGCGGCCUUAAACC....GAGGUUUUCCC..AUG
env-763     UCGCGUGGUG.UCUGGC....................GUUAGG.............................G.CUGGA.AGAGCGACGCGGCCUUAAACC....GAGGUUUUCCC..AUG
Cwe-1-1     UCGCGUGGUG.UCUGGC....................GUUAGG.............................G.CUGGA.AGAGCGACGCGGCCUUAAACC....GAGGUUUUCCC..AUG
Sso-1-3     UCGCGUGGUG.UCUGGC....................GUUAGG.............................G.CUGGA.AGAGCGACGCGGCCUUAAACC....GAGGUUUUCCC..AUG
env-766     UCGCGUGGUG.UCUGGC....................GUUAGG.............................G.CUGGA.AGAGCGACGCGGCCUUAAACC....GAGGUUUUCCC..AUG
env-784     UCGCGUGGUG.UCUGGC....................GUUAGG.............................G.CUGGA.AGAGCGACGCGGCCUUAAACC....GAGGUUUUCCC..AUG
env-785     UCGCGUGGUG.UCUGGC....................GUUAGG.............................G.CUGGA.AGAGCGACGCGGCCUUAAACC....GAGGUUUUCCC..AUG
Cro-2-1     UCGCGUGGUG.UCUGGC....................GUUAGG.............................G.CUGGA.AGAGCGACGCGGCCUUAAACC....GAGGUUUUCCC..AUG
Eal-2-1     UCGCGUGGUG.UCUGGC....................GUUAGG.............................G.CUGGA.AGAGCGACGCGGCCUUAAACC....GAGGUUUUCCC..AUG
Sso-1-5     UCGCGUGGUG.UCUGGC....................GUUAGG.............................G.CUGGA.AGAGCGACGCGGCCUUAAACC....GAGGUUUUCCC..AUG
Eco-835-1   UCGCGUGGUG.UCUGGC....................GUUAGG.............................G.CUGGA.AGAGCGACGCGGCCUUAAACC....GAGGUUUUCCC..AUG
Ban-1-1     UCGCGUGGUG.UCUGGC....................GUUAGG.............................G.CUGGA.AGAGCGACGCGGCCUUAAACC....GAGGUUUUCCC..AUG
Eco-1442-1  UCGCGUGGUG.UCUGGC....................GUUAGG.............................G.CUGGA.AGAGCGACGCGGCCUUAAACC....GAGGUUUUCCC..AUG
Cyo-1-1     UCGCGUGGUG.UCUGGC....................GUUAGG.............................G.CUGGA.AGAGCGACGCGGCCUUAAACC....GAGGUUUUCCC..AUG
Eco-763-1   UCGCGUGGUG.UCUGGC....................GUUAGG.............................G.CUGGA.AGAGCGACGCGGCCUUAAACC....GAGGUUUUCCC..AUG
Sdy-5-1     UCGCGUGGUG.UCUGGC....................GUUAGG.............................G.CUGGA.AGAGCGACGCGGCCUUAAACC....GAGGUUUUCCC..AUG
env-786     UCGCGUGGUG.UCUGGC....................GUUAGG.............................G.CUGGA.AGAGCGACGCGGCCUUAAACC....GAGGUUUUCCC..AUG
Eco-1862-1  UCGCGUGGUG.UCUGGC....................GUUAGG.............................G.CUGGA.AGAGCGACGCGGCCUUAAAUC....GAGGUUUUCCC..AUG
env-861     UCGCGUGGUG.UCUGGC....................GUUAGG.............................G.CUGGA.AGAGCGACGCGGCCUUAAACC....GAGGUUUCCCC..AUG
Cko-2-1     UCGCGUGGUG.UCUGGC....................GUUAGG.............................G.CUGGA.AGAGCGACGCGGCCUUAAACC....GAGGUUUCCCC..AUG
Csp-14-1    UCGCGUGGUG.UCUGGC....................GUUAGG.............................G.CUGGA.AGAGCGACGCGGCCUUAAACC....GAGGUUUCCCC..AUG
env-862     UCGCGUGGUG.UCUGGC....................GUUAGG.............................G.CUGGA.AGAGCGACGCGGCCUUAAACU....GAGGUUUCCCC..AUG
Cam-2-1     UCGCGUGGUG.UCUGGC....................GUUAGG.............................G.CUGGA.AGAGCGACGCGGCCUUAAACU....GAGGUUUCCCC..AUG
env-867     UCGCGUGGUG.UCUGGC....................GUUAGG.............................G.CUGGA.AGAGCGACGCGGCCUUAAACU....GAGGUUUCCCC..AUG
Cse-1-1     UCGCGUGGUG.UCUGGC....................GUUAGG.............................G.CUGGA.AGAGCGACGCGGCCUUAAACU....GAGGUUUCCCC..AUG
env-868     UCGCGUGGUG.UCUGGC....................GUUAGG.............................G.CUGGA.AGAGCGACGCGGCCUUAAACU....GAGGUUUCCC...AUG
Esp-43-1    UCGCGUGGUG.UCUGGC....................GUUAGG.............................G.CUGGA.AGAGCGACGCGGCCUUAAACU....GAGGUUUCCC...AUG
Kor-1-1     UCGCGUGGUG.UCUGGC....................GUUAGG.............................G.CUGGA.AGAGCGACGCGGCCUUAAACU....GAGGUUUCCC...AUG
Ksa-1-1     UCGCGUGGUG.UCUGGC....................GUUAGG.............................G.CUGGA.AGAGCGACGCGGCCUUAAACU....GAGGUUUCCC...AUG
Mma-2-1     UCGCGUGGUG.UCUGGC....................GUUAGG.............................G.CUGGA.AGAGCGACGCGGCCUUAAACU....GAGGUUUCCC...AUG
Pur-1-1     UCGCGUGGUG.UCUGGC....................GUUAGG.............................G.CUGGA.AGAGCGACGCGGCCUUAAACU....GAGGUUUCCC...AUG
Cfa-1-1     UCGCGUGGUG.UCUGGC....................GUUAGG.............................G.CUGGA.AGAGCGACGCGGCCUUAAACU....GAGGUUUCCCC..AUG
env-869     UCGCGUGGUG.UCUGGC....................GUUAGG.............................G.CUGGA.AGAGCGACGCGGCCUUAAACU....GAGGUUUCCC...AUG
env-870     UCGCGUGGUG.UCUGGC....................GUUAGG.............................G.CUGGA.AGAGCGACGCGGCCUUAAACU....GAGGUUUCCCC..AUG
Kcf-1-1     UCGCGUGGUG.UCUGGC....................GUUAGG.............................G.CUGGA.AGAGCGACGCGGCCUUAAACU....GAGGUUUCCCC..AUG
env-871     UCGCGUGGUG.UCUGGC....................GUUAGG.............................G.CUGGA.AGAGCGACGCGGCCUUAAACU....GAGGUUUCCC...AUG
Esp-42-1    UCGCGUGGUG.UCUGGC....................GUUAGG.............................G.CUGGA.AGAGCGACGCGGCCUUAAACU....GAGGUUUCCC...AUG
Era-1-1     UCGCGUGGUG.UCUGGC....................GUUAGG.............................G.CUGGA.AGAGCGACGCGGCCUUAAACU....GAGGUUUCCC...AUG
Kra-2-1     UCGCGUGGUG.UCUGGC....................GUUAGG.............................G.CUGGA.AGAGCGACGCGGCCUUAAACU....GAGGUUUCCC...AUG
Csp-12-1    UCGCGUGGUG.UCUGGC....................GUUAGG.............................G.CUGGA.AGAGCGACGCGGCCUUAAACU....GAGGUUUCCC...AUG
Ecl-60-1    UCGCGUGGUG.UCUGGC....................GUUAGG.............................G.CCGGA.AGAGCGACGCGGCCUUAAACU....GAGGUUUCCC...AUG
env-872     UCGCGUGGUG.UCUGGC....................GUUAGG.............................G.CCGGA.AGAGCGACGCGGCCUUAAACU....GAGGUUUCCCC..AUG
env-874     UCGCGUGGUG.UCUGGC....................GUUAGG.............................G.CUGGA.AGAGCGACGCGGCCUUAUACC....GAGGUUUUCC...AUG
env-875     UCGCGCGGUG.UCUGGC....................GUUAGG.............................G.CUGGA.AGAGCGGCGCGGCCUUAAUCA....GAGGUUUCCC...AUG
Pvu-1-1     UCGCAUGGUG.UCCGGC....................GUUAGG.............................G.CUGGA.AGAGCGAUGCGGCCUUAACU.....GAGGUUUCUCCC.AUG
Gpe-1-1     UCGUGUGGUGUCUGG.C....................ACUG...............................G.ACCAG.AUGGCGACACGGCCCACU.......GAGGGAAUCCCC.AUG
Gxi-1-1     UCGUGUGGUGUCUGG.C....................ACUG...............................G.ACCAG.AUGGCGACACGGCCCAAU.......GAGGUAAUCCC..AUG
env-876     ACGUAUGGUGUCUGG.C......AAA...........UUG.........................UUU....G.ACCAG.AUAGCGAUACGGCCUAAC.......GAGGUAACCC...AUG
env-880     GCGUAUGGUGUCUGA.C....................AACAGUU............................G.AUCAG.AUAGCGAUACGGCCUAAC.......GAGGUAACCC...AUG
Pme-1-1     GCGUAUGGUGUCUGA.C....................AACAGUU............................G.AUCAG.AUAGCGAUACGGCCUAAC.......GAGGUAACCC...AUG
Rsp-2-1     GCGUAUGGUGUCUGA.C....................AACAGUU............................G.AUCAG.AUAGCGAUACGGCCUAAC.......GAGGUAACCC...AUG
env-881     GCGUAUGGUGUCUGA.C....................AAAAGUU............................G.AUCAG.AUAGCGAUACGGCCUAAC.......GAGGUAACCC...AUG
Rsp-1-1     GCGUAUGGUGUCUGA.C....................AAAAGUU............................G.AUCAG.AUAGCGAUACGGCCUAAC.......GAGGUAACCC...AUG
Aag-1-1     ACGUAUGGUGUCUGG.C......G.............AUAAAAAUG.....................C....G.ACCAG.AUAGCGAUGCGGCCUAAC.......GAGGUAACCC...AUG
Aae-1-1     ACGUAUGGUGUCUGG.C......G.............AUAAAAAAUG....................C....G.ACCAG.AUGGCGAUACGGCCUAAC.......GAGGUAACCC...AUG
env-882     ACGUAUGGUGUCUGG.C......G.............AUAAAAAAUG....................C....G.ACCAG.AUGGCGAUACGGCCUAAC.......GAGGUAACCC...AUG
env-883     ACGUAUGGUGUCUGG.C......G.............AUAAAAAAUG....................C....G.ACCAG.AUGGCGAUACGGCCUAAC.......GAGGUAACCC...AUG
Aik-1-1     ACGUAUGGUGUCCGG.C....................ACAAGAU............................G.ACCGG.AUAGCGAUACGGCCUAAC.......GAGGUAACCC...AUG
env-884     ACGUAUGGUGUCUGG.C....................ACAUGAU............................G.ACCAG.AUAGCGAUACGGCCUAAC.......GAGGUAAUCC...AUG
env-885     ACGUAUGGUGUCUGG.C....................ACACGAU............................G.ACCAG.AUAGCGAUACGGCCUAAC.......GAGGUAACCC...AUG
Asp-9-1     ACGUAUGGUGUCUGG.C....................ACAUGAU............................G.ACCAG.AUAGCGAUACGGCCUAAC.......GAGGUAACCC...AUG
Rpe-1-1     ACGUAUGGUGUCUGG.C....................ACAUGAU............................G.ACCAG.AUAGCGAUACGGCCUAAC.......GAGGUAACCC...AUG
Rba-1-1     ACGUAUGGUGUCUGG.C....................AACUAAUUU..........................G.ACCAG.AUAGCGAUACGGCCUAAC.......GAGGUAAGCC...AUG
Rna-1-1     ACGUAUGGUGUCUGG.C....................AAUAAUU............................G.ACCAG.AUAGCGAUACGGCCUAAC.......GAGGUAAGCC...AUG
env-886     GCGUAUGGUGUGCAACU....................CCG................................G.GUUGU.AUAGCGAUACGGCCUUAGCUUA...GAGGUAAACC...AUG
Asa-9-1     ACGUAUGGUGGCCAGCU....................UCG................................G.GCUGG.UUAGCGAUGCGGCCUUGAUGAU...GAGGUGACCC...AUG
Gli-1-1     GCGUAUGGUGUGCAA.C....................UUCGG..............................G..UUGU.AUAGCGAUACGGCCCAGAAAU....GAGGUAAUCCCU.AUG
Gpu-1-1     UCGCUUGGUGUGCAA.C....................UUCGG..............................G..UUGU.AUAGCGAGGCGGCCUUAAAUAAGA.AAGGUAUCCC...AUG
env-889     GCGUAUGGUGUGCAA.C....................UUCGG..............................G..UUGU.AUAGCGAUGCGGCCUACAUUUA...AAGGUAUCCC...AUG
env-890     GCGUAUGGUGUACAG.C....................UUCGG..............................G..UUGU.AUAGCGAUACGGCCUAAAUUUAU..AAGGUAACCCCU.AUG
Gni-1-1     UCGUUUGGUGUGCAA.C....................UUCGG..............................G..UUGU.AUAGCGAGACGGCCUAAAAAAU...AAGGUAUCCCA..AUG
Gpa-1-1     UCGUUUGGUGUACAA.C....................UUCGG..............................G..UUGU.AUAGCGAGACGGCCUAAAUAAAU..AAGGUAUCCCA..AUG
env-894     ACGUAUGGUGUGCGA.C....................UUCGG..............................G..UUGU.AUAGCGAUACGGCCUAUUAUGU...GAGGUAACCC...AUG
Gpo-1-1     GCGUAUGGUGUUCAACU....................UCG................................G.GUUGG.AUAGCGAUACGGCCUUACUUU....AAGGGUAAUCC..AUG
env-895     UCGUAUGGUGUUCAACU....................UCG................................G.GUUGG.AUAGCGAUGCGGCCUGACGAA....AAGGUAACCC...AUG

            .<<<<<.<<..<<<<<<......<<<<<<<<<<<<<<.................>>>>>>>>>>>>>>....>.>>>>>....>>.>>>>>..............................
            .22222.0...222222......1.1111.11.0010.................0100.11.1111.1....2.22222.....0.22222..............................
            ●CGURUGGUG○YYRR○Y--------------------●●●●○------------------------------G-○YYRR-A●AGCGAYRCGGCCU●●●●●○----GAGGU●●YCC○--AUG
             5′                                                                                                        3′
```

```
Gar-1-1     UCGUAUGGUGUUCAACU....................UCG................................G.GUUGA.AUAGCGAUACGGCCUGACGAA....GAGGUAACCC...AUG
env-896     UCGUAUGGUGUUCAACU....................UCG................................G.GUUGA.AUAGCGAUGCGGCCUGACGAA....GAGGUAACCC...AUG
Gps-1-1     UCGUAUGGUGUUCAACU....................UCG................................G.GUUGA.AUAGCGAUGCGGCCUGACGAA....GAGGUAACCC...AUG
env-898     ACGUAUGGUGGCCAACU....................CCG................................G.GUUGG.UUAGCGAUACGGCCUUAUUUU....GAGGUAAUCCC..AUG
env-899     ACGUAUGGUGGCCAACU....................UCG................................G.GUUGG.UUAGCGAUACGGCCUUAUUUU....GAGGUAAUCCC..AUG
env-900     ACGUAUGGUGGCCAACU....................UCG................................G.GUUGG.UUAGCGAUGCGGCCUUAUUUU....GAGGUAAUCCC..AUG
env-902     ACGUAUGGUGGCCAACU....................UCG................................G.GUUGG.UUAGCGAUGCGGCCUUAUUUU....GAGGUAAUCCC..AUG
env-930     ACGUAUGGUGGCCAACU....................UCG................................G.GUUGG.UUAGCGAUGCGGCCUUAUUUU....GAGGUAAUCCC..AUG
env-933     ACGUAUGGUGGCCAACU....................UCG................................G.GUUGG.UUAGCGAUGCGGCCUUAUUUU....GAGGUAAUCCC..AUG
env-934     ACGUAUGGUGGCCAACU....................UCG................................G.GUUGG.UUAGCGAUGCGGCCUUAUUUU....GAGGUAAUCCC..AUG
Pat-1-1     UCGUAUGGUGUUCAACU....................UCG................................G.GUUGG.AUAGCGAUACGGCCUUACUUA....GGGGUAAUCC...AUG
env-935     ACGUAUGGUGUUUGGCU....................UCG................................G.GCGAA.AUAGCGAUACGGCCUUAUUGU....GAGGUAAUCCC..AUG
Asp-7-1     ACGUAUGGUGUUUGGCU....................UCG................................G.GCUGA.AUAGCGAUACGGCCUUAUUGU....GAGGUAAUCCC..AUG
env-936     ACGUAUGGUGUUCAGCU....................UCG................................G.GCUGG.AUAGCGAUACGGCCUUUUUGU....GAGGUAACCCA..AUG
env-937     ACGUAUGGUGUUCAGCU....................UCG................................G.GCUGG.AUAGCGAUACGGCCUUUUUUU....GAGGUAACCCA..AUG
Sch-1-1     GCGUAUGGUGUUCAGCU....................UCG................................G.GCUGG.AUAGCGAUACGGCCUUUUUUU....GAGGUAACCC...AUG
env-954     GCGUAUGGUGUUCAGCU....................UCG................................G.GCUGG.AUGGCGAUACGGCCUUUUUUAU...GAGGUAACCC...AUG
env-955     GCGUAUGGUGUUCAGCU....................UCG................................G.GCUGG.AUAGCGAUACGGCCUUAAGAUU...GAGGUAACCC...AUG
Asp-5-1     ACGUAUGGUGUGCGA.C....................UUCGG..............................G..UUGU.AUAGCGAUACGGCCUUAUUAUGU..GAGGUAACCCA..AUG
env-956     ACGUAUGGUGUGCGA.C....................UUCGG..............................G..UUGU.AUAGCGAUACGGCCUUAUUAUGU..GAGGUAACCCA..AUG
Ama-14-1    ACGUAUGGUGUGCGA.C....................UUCGG..............................G..UUGU.AUAGCGAUACGGCCUUAUUAUGU..GAGGUAACCCA..AUG
env-957     ACGUAUGGUGUGCGA.C....................UUCGG..............................G..UUGU.AUAGCGAUACGGCCUUAUUAUGU..GAGGUAACCCA..AUG
env-981     ACGUAUGGUGUGCGA.C....................UUCGG..............................G..UUGU.AUAGCGAUACGGCCUUAUUAUGU..GAGGUAACCCA..AUG
env-982     ACGUAUGGUGUGCGA.C....................UUCGG..............................G..UUGU.AUAGCGAUACGGCCUUAUUAUGU..GAGGUAACCCA..AUG
env-986     ACGUAUGGUGUGCGA.C....................UUCGG..............................G..UUGU.AUAGCGAUACGGCCUUAUUAUGU..GAGGUAACCCA..AUG
env-992     ACGUAUGGUGUGCGA.C....................UUCGG..............................G..UUGU.AUAGCGAUACGGCCUUAUUAUGU..GAGGUAACCCA..AUG
env-1025    ACGUAUGGUGUGCGA.C....................UUCGG..............................G..UUGU.AUAGCGAUACGGCCUUAUUAUGU..GAGGUAACCCA..AUG
env-1027    ACGUAUGGUGUGCGA.C....................UUCGG..............................G..UUGU.AUAGCGAUACGGCCUUAUUAUGU..GAGGUAACCCA..AUG
env-1029    ACGUAUGGUGUGCGA.C....................UUCGG..............................G..UUGU.AUAGCGAUACGGCCUUAUUAUGU..GAGGUAACCCA..AUG
env-1030    ACGUAUGGUGUGCGGCU....................UCG................................G.GCUGU.AUAGCGAUACGGCCUUAUUAAU...GAGGUAAUCCC..AUG
env-1031    ACGUAUGGUGUGCAACU....................UCG................................G.GUUGU.AUAGCGAUACGGCCUUAUAAUGU..GAGGUAAUCCC..AUG
Ama-7-1     ACGUAUGGUGUGCGACU....................UCG................................G.GUUGU.AUAGCGAUACGGCCUUAUUAUGU..GAGGUAAUCCC..AUG
env-1032    ACGUAUGGUGUGCGACU....................UCG................................G.GUUGU.AUAGCGAUACGGCCUUAUUAUGU..GAGGUAAUCCC..AUG
env-1035    ACGUAUGGUGUGCGACU....................UCG................................G.GUUGU.AUAGCGAUACGGCCUUAUUAUGU..GAGGUAAUCCC..AUG
env-1036    GCGUAUGGUGUGCGACU....................UCG................................G.GUCGU.AUAGCGAUACGGCCUUAUUAUUU..GAGGUAACCCA..AUG
env-1037    ACGUAUGGUGUGCAACU....................UCG................................G.GUUGU.AUAGCGAUACGGCCUUUUACGU...GAGGUAAUCCC..AUG
Pha-4-1     ACGUGUGGUACUCUGUU......GAUUGA........AAUAA....................UUCAUCA...A.GCUGA.GCAGCGACACGGACCAAUA......GAGGUUUUCUAACAUG
env-1093    ACGUGUGGUACUUUGUU......GAGUGA........AAUGAU...................UCUCUCA...A.ACAAA.GCAGCGAUGCGGCCCAAUA......GAGGUUUUCCAA.AUG
env-1094    ACGUAUGGUACUUUGUU......GGAUGA........AAUGAU...................UCAUUC....AUGCAGA.GCAGCGAUGCGGCCCAAUA......GAGGUUUUCCCA.AUG
env-1096    ACGUAUGGUACUUUGUU......GAAUGA........AAUGAU...................UCAUUCA...A.GCAGA.GCAGCGAUGCGGCCCAAUA......GAGGUUUUCCCA.AUG
env-1097    ACGUAUGGUACUUUGUU......GGAUGA........AAUGAU...................UCAUUCA...A.GCAGA.GCAGCGAUGCGGCCCAAUA......GAGGUUUUCCCA.AUG
Pin-1-1     ACGUAUGGUACUUUGUU......GAAUGA........AAUGAU...................UCAUUCA...A.GCAGA.GCAGCGAUACGGCCCAAUA......GAGGUUUUCCAA.AUG
env-1099    ACGUGUGGUACUUUGUU......GGAUGA........AAUGAU...................UCAUUCA...A.ACAGA.GCAGCGAUGCGGCCCAAUA......GAGGUUUUCCUU.AUG
env-1103    ACGUGUGGUACUUUGUU......GGAUGA........AAUGAU...................UCAUUCA...A.ACAGA.GCAGCGAUGCGGCCCAAUA......GAGGUUUUCCUU.AUG
env-1104    ACGUGUGGUACUUUGUU......GAGUAA........AAUAAU...................UUAUUCA...A.ACAGA.GCAGCGACACGGACCAAUA......GAGGUUUUCCCA.AUG
Par-2-1     ACGUGUGGUACUUUGUU......GAGUGA........AAUAAU...................UCAUUCA...A.GCUGA.GCAGCGAUACGGCCUAAAA......GAGGUUUUCCCU.AUG
Paq-1-1     ACGUGUGGUACUUUGUU......GAGUGA........AAUAAU...................UCAUUCA...A.ACAGA.GCAGCGACACGGCCCAAUA......GAGGUUUUCCCU.AUG
Pos-1-1     ACGUGUGGUACUUUGUU......GAGUGA........AAUAAU...................UCAUUCA...A.ACUGA.GCAGCGACACGGCCCAAUA......GAGGUUUUCCCU.AUG
Psp-35-1    ACGUGUGGUACUUUGUU......GAAUGA........AAUAAU...................UCAUUC....AAGCUGA.GCAGCGACACGGCCUAAAUA.....GAGGUUUUCCCA.AUG
env-1105    ACGUGUGGUACUUUGUU......GGAUGA........AAUAAU...................UCAUUCA...A.GCUGA.GCAGCGACACGGCCUAAAUA.....GAGGUUUUCCUA.AUG
env-1109    ACGUGUGGUACUUUGUU......GGAUGA........AAUAAU...................UCAUUCA...A.GCUGA.GCAGCGACACGGCCUAAAUA.....GAGGUUUUCCUA.AUG
env-1110    ACGUGUGGUACUUUGUU......GGAUGA........AAUAAU...................UCAUUC....AUGCUGA.GCAGCGACACGGCCCAAUA......GAGGUUUCCCAA.AUG
Psp-34-1    ACGUGUGGUAUUUUGUU......GAGAGA........AAUGCU...................UCUUUCA...A.GCAAG.AUGGCGACACGGCCUAAC.......GAGGUUUUCCCU.AUG
Sam-1-1     AUGUGUGGUGUCAGGC.......GC............GAAA.........................GUG.....GUCUG.AUGGCGACACAGCCUGAAU......GAGGUUUUUCC..AUG
env-1113    ACGUAUGGUGUUAAGC.......U.............UUAG...........................A...G.CUUAA.AUGGCGAUACGGCCGAAAUA.....AAGGUAACCCCU.AUG
Pfl-3-1     ACGUGUGGUG.CCGUGC......GGAU..........CCUGU......................AUCUG...G.UGCGG.AGAGCGACGCGGACUUCUAUC....GAGGUUUUCC...AUG
Ykr-2-1     UCGUGUGGUGUCUGG.C....................GGAGUAGG...........................G..CUGG.AUAGCGACACGGCCUAAAUAUA...GAGGUUUUCC...AUG
Pge-1-1     UCGCGUGGUG.UCUGGC....................GUCAGG.............................G.CUGGA.AGAGCGACGCGGCCUUAAUU.....GAGGUUCUCCC..AUG
Kpn-281-1   UCGCGUGGUG.UCUGGC....................GUUAGG.............................G.CUGGA.AGAGCGACGCGGCCUUAACU.....GAGGUUCUCCC..AUG
Vja-2-1     ACGUGUGGUGUGCGG.C....................UUA................................GACUCGU.AUAGCGACACGGCCUACUAUU....GAGGUUAUCCC..AUG
Kpn-256-1   UCGCGUGGUG.UCUGGC....................GUUAGG.............................G.CUGGA.AGAGCGACGCGGCCUUAACU.....GAGGUUCUCCC..AUG
Hin-6-1     UCACGCGGUGUGGGGA.....................GAAAG................................UUCCU.AUAGCGGCGUGGCUUCAUUUU....GAGGUUUUUCU..AUG
Eba-1-1     UCGCGUGGUG.UCUGGC....................GUUAGG.............................G.CUGGA.AGAGCGACGCGGCCUUUACU.....GAGGUUCUCCC..AUG
Ysi-1-1     UCGUGUGGUGUCUAG.C....................GGAACAGG...........................G..CUGG.AUAGCGACACGGCCUAAUAUA....GAGGUUUCC....AUG
Hin-1-3     UCACGCGGUGUGGGAA.....................GAAAG................................UUCUU.AUAGCGGCGUGGCUUCAUUUU....GAGGUUUUUCU..AUG
Kox-4-2     UCGCGUGGUG.UCUGGC....................GUUAGG.............................G.CUGGA.AGAGCGACGCGGCCUUAACU.....GAGGUUCUCCC..AUG
Hin-13-1    UCACGCGGUGUGGGGA.....................GAAAG................................UUCCU.AUAGCGGCGUGGCUUCAUUUU....GAGGUUUUUCU..AUG
Sma-10-1    ACGUGUGGUGUCUGG.C....................GAAACAGG...........................G..CUGG.AUAGCGACACGGCCUUAACU.....GAGGUUUCCC...AUG
Yru-2-2     ACGUGUGGUGUCUGG.C....................GGAACAGG...........................G..CUGG.AUAGCGACACGGCCUUAAUUA....GAGGUUUCCC...AUG
Vni-11-1    ACGUGUGGUGUCCGG.C....................AUA................................GAAUCGG.AUAGCGACACGGCCCAAUUU.....GAGGUUAUCCC..AUG
Yin-2-1     UCGUGUGGUGUCUGG.C....................GGAACAGG...........................G..CUGG.AUAGCGACACGGCCUAAAUAUA...GAGGUUUUCC...AUG
Hin-18-2    ACACGCGGUGUCAG.........UC............CUU...........................GA.....ACUUG.AUAGCGGCGUGGCUUCAAAUC....GAGGUUUUCU...AUG
Hin-14-1    UCACGCGGUGUAGGGA.....................GAAAG................................UUCCU.AUAGCGGCGUGGCUUCAUUUU....GAGGUUUUUCU..AUG
Vvu-10-1    GCGUGUGGUGUGCGG.C....................AAA................................GACUCGU.AUAGCGACACGGCCUACUAUUU...GAGGUUAUCCC..AUG
Ywa-1-1     UCGUGUGGUGUCUGG.C....................GGAACAGG...........................G..CUGG.AUAGCGACACGGCCUAAUAUA....GAGGUUUCC....AUG
Dso-4-1     UCGUGUGGUGUCUGG.C....................GGAACAGG...........................G..CCGG.AUAGCGACACGGCCUUAAACA....GAGGUUUCCC...AUG
Yps-1-3     UCGUGUGGUGUCUGG.C....................GGAACAGG...........................G..CUGG.AUAGCGACACGGCCUAAUAUA....GAGGUUUCC....AUG
Sso-1-546   UCGCGUGGUG.UCCAGC....................GUUAGG.............................G.CUGGA.AGGGCGACGCGGCCUAUUUU.....AAGGUUUUCCC..AUG
Yal-3-1     UCGUGUGGUGUCUGG.C....................GGAACAGG...........................G..CUGG.AUAGCGACACGGCCUAAAUAUA...GAGGUUUCC....AUG
Sal-1-1     GCAUGUGGUG.UCCGAC....................UUCGG..............................G.UUGGA.AGAGCGACAUGGCCUUAACUU....GAGGUAACCCCA.AUG
Kpn-168-1   UCGCGUGGUG.UCUGGC....................GUUAGG.............................G.CUGGA.AGAGCGACGCGGCCUUAACU.....GAGGUUCUCCC..AUG
Vvu-9-1     GCGUGUGGUGUGCGG.C....................AAA................................GACUCGU.AUAGCGACACGGCCUACUAUUU...GAGGUUAUCCC..AUG
Yro-2-1     UCGUGUGGUGUCUGG.C....................GGAACAGG...........................G..CUGG.AUAGCGACACGGCCUAAAUAUA...GAGGUUUUUCC..AUG

            .<<<<<.<<..<<<<<<......<<<<<<<<<<<<<<................>>>>>>>>>>>>>>....>.>>>>>....>>.>>>>>................................
            .22222.0...222222......1.1111.11.0010................0100.11.1111.1....2.22222....0.22222................................
            •CGURUGGUG∘YYRR∘Y------------------●●●∘∘----------------------------G-∘YYRR-A•AGCGAYRCGGCCU●●●●∘∘----GAGGU••YCC∘--AUG
            5′                                                                                                    3′
```

```
Yen-13-1   UCGUGUGGUGUCUGG.C....................GGAGUAGG...........................G..CUGG.AUAGCGACACGGCCUAAAUAUA...GAGGUUUUCC...AUG
Pmi-2-1    ACGUGUGGUGUUCGGCU....................GAG................................G.GCUGA.AUAGCGACACGGCCUUAGCA.....GAGGUUUUCC...AUG
Vpa-183-1  GCGUGUGGUGUGCGG.C....................UUA................................GAUUCGU.AUAGCGACACGGCCUACUAUU....GAGGUUAUCCC..AUG
Ehe-1-1    UCGCGUGGUG.UCUGGC....................GUUAGG.............................G.CCGGA.AGAGCGACGCGGCCUUAACU.....GAGGUUUCCC...AUG
Eco-230-1  UCGCGUGGUG.UCUGGC....................GUUAGG.............................G.CUGGA.AGAGCGACGCGGCCUUAAACC....GAGGUUUUCCC..AUG
Vvu-5-1    GCGUGUGGUGUGCGG.C....................AAA................................GACUCGU.AUAGCGACACGGCCUACUAUUU...GAGGUUAUCCC..AUG
Cbr-1-7    UCGCGUGGUG.UCUGGC....................GAAAG..............................G.CUGGA.AGGGCGACGCGGCCUAACUU.....AAGGUUUCCC...AUG
Esp-47-1   UCGCGUGGUG.UCUGGC....................GUUAGG.............................G.CUGGA.AGAGCGACGCGGCCUUAACU.....GAGGUUCUCCC..AUG
Hin-12-1   UCACGCGGUGUAGGGA.....................GAAAG................................UUCCU.AUAGCGGCGUGGCUUCAUUUU....GAGGUUUUCU...AUG
Kox-11-1   UCGCGUGGUG.UCUGGC....................GUUAGG.............................G.CUGGA.AGAGCGACGCGGCCUUAACU.....GAGGUUCUCCC..AUG
Pst-7-1    ACGUGUGGUG.UCCGGC....................GUUAGG.............................G.CUGGA.AUGGCGACACGGCCUUAAUA.....GAGGUUUUCCC..AUG
Vco-2-1    ACGUGUGGUGUCCGG.C....................AUA................................GAAUCGG.AUAGCGACACGGCCUAUUUUA....GAGGUUCUCCC..AUG
Yfr-3-1    UCGUGUGGUGUCUGG.C....................GGAGUAGG...........................G..CUGG.AUAGCGACGCGGCCUAAAUAUA...GAGGUUUUUCC..AUG
Kmi-1-2    UCGCGUGGUG.UCUGGC....................GUUAGG.............................G.CUGGA.AGAGCGACGCGGCCUUAACU.....GAGGUUCUCCC..AUG
           .<<<<<.<<..<<<<<<......<<<<<<<<<<<<<<.................>>>>>>>>>>>>>>....>.>>>>>....>>.>>>>>..............................
           .22222.0...222222......1.1111.11.0010.................0100.11.1111.1....2.22222.....0.22222..............................
           ●CGURUGGUG○YYRR○Y--------------------●●●●○------------------------------G-●YYRR-A●AGCGAYRCGGCU●●●●●○-----GAGGU●●YCC○--AUG
            5′                                                                                                          3′
```

### 13 L31-Corynebacteriaceae

#### 13.1 Taxa

The taxonomy of each organism containing a putative L31-Corynebacteriaceae RNA is listed, with abbreviations identifying each hit (e.g., "Eco-1-1" and "Eco-1-2" might hypothetically represent two distinct RNAs in *E. coli*). The abbreviations will be used to identify each individual L31-Corynebacteriaceae RNA in Sections 13.2 and 13.4:.

| abbrev. of hits | taxonomy of species |
|---|---|
| Cac-1-1 | Bacteria Actinobacteria Actinobacteridae Actinomycetales Corynebacterineae Corynebacteriaceae *Corynebacterium accolens* ATCC 49725 |
| Cac-2-1 | Bacteria Actinobacteria Actinobacteridae Actinomycetales Corynebacterineae Corynebacteriaceae *Corynebacterium accolens* ATCC 49726 |
| Cam-1-1 | Bacteria Actinobacteria Actinobacteridae Actinomycetales Corynebacterineae Corynebacteriaceae *Corynebacterium ammoniagenes* DSM 20306 |
| Cam-2-1 | Bacteria Actinobacteria Actinobacteridae Actinomycetales Corynebacterineae Corynebacteriaceae *Corynebacterium amycolatum* SK46 |
| Cau-1-1 | Bacteria Actinobacteria Actinobacteridae Actinomycetales Corynebacterineae Corynebacteriaceae *Corynebacterium aurimucosum* ATCC 700975 |
| Cca-1-1 to Cca-1-2 | Bacteria Actinobacteria Actinobacteridae Actinomycetales Corynebacterineae Corynebacteriaceae *Corynebacterium callunae* DSM 20147 |
| Cca-2-1 | Bacteria Actinobacteria Actinobacteridae Actinomycetales Corynebacterineae Corynebacteriaceae *Corynebacterium capitovis* DSM 44611 |
| Cca-3-1 | Bacteria Actinobacteria Actinobacteridae Actinomycetales Corynebacterineae Corynebacteriaceae *Corynebacterium casei* UCMA 3821 |
| Cca-4-1 | Bacteria Actinobacteria Actinobacteridae Actinomycetales Corynebacterineae Corynebacteriaceae *Corynebacterium caspium* DSM 44850 |
| Cci-1-1 | Bacteria Actinobacteria Actinobacteridae Actinomycetales Corynebacterineae Corynebacteriaceae *Corynebacterium ciconiae* DSM 44920 |
| Ccr-1-1 | Bacteria Actinobacteria Actinobacteridae Actinomycetales Corynebacterineae Corynebacteriaceae *Corynebacterium crenatum* MT |
| Cdi-1-1 | Bacteria Actinobacteria Actinobacteridae Actinomycetales Corynebacterineae Corynebacteriaceae *Corynebacterium diphtheriae* 241 |
| Cdi-2-1 | Bacteria Actinobacteria Actinobacteridae Actinomycetales Corynebacterineae Corynebacteriaceae *Corynebacterium diphtheriae* 31A |
| Cdi-3-1 | Bacteria Actinobacteria Actinobacteridae Actinomycetales Corynebacterineae Corynebacteriaceae *Corynebacterium diphtheriae* BH8 |
| Cdi-15-1 | Bacteria Actinobacteria Actinobacteridae Actinomycetales Corynebacterineae Corynebacteriaceae *Corynebacterium diphtheriae bv. intermedius str.* NCTC 5011 |
| Cdi-16-1 | Bacteria Actinobacteria Actinobacteridae Actinomycetales Corynebacterineae Corynebacteriaceae *Corynebacterium diphtheriae bv. mitis str.* NC03529 |
| Cdi-4-1 | Bacteria Actinobacteria Actinobacteridae Actinomycetales Corynebacterineae Corynebacteriaceae *Corynebacterium diphtheriae* C7 (beta) |
| Cdi-5-1 | Bacteria Actinobacteria Actinobacteridae Actinomycetales Corynebacterineae Corynebacteriaceae *Corynebacterium diphtheriae* CDCE 8392 |
| Cdi-6-1 | Bacteria Actinobacteria Actinobacteridae Actinomycetales Corynebacterineae Corynebacteriaceae *Corynebacterium diphtheriae* DSM 43988 |
| Cdi-7-1 | Bacteria Actinobacteria Actinobacteridae Actinomycetales Corynebacterineae Corynebacteriaceae *Corynebacterium diphtheriae* HC01 |
| Cdi-8-1 | Bacteria Actinobacteria Actinobacteridae Actinomycetales Corynebacterineae Corynebacteriaceae *Corynebacterium diphtheriae* HC02 |
| Cdi-9-1 | Bacteria Actinobacteria Actinobacteridae Actinomycetales Corynebacterineae Corynebacteriaceae *Corynebacterium diphtheriae* HC03 |
| Cdi-10-1 | Bacteria Actinobacteria Actinobacteridae Actinomycetales Corynebacterineae Corynebacteriaceae *Corynebacterium diphtheriae* HC04 |
| Cdi-11-1 | Bacteria Actinobacteria Actinobacteridae Actinomycetales Corynebacterineae Corynebacteriaceae *Corynebacterium diphtheriae* INCA 402 |
| Cdi-12-1 | Bacteria Actinobacteria Actinobacteridae Actinomycetales Corynebacterineae Corynebacteriaceae *Corynebacterium diphtheriae* NCTC 13129 |
| Cdi-13-1 | Bacteria Actinobacteria Actinobacteridae Actinomycetales Corynebacterineae Corynebacteriaceae *Corynebacterium diphtheriae* PW8 |
| Cdi-17-1 | Bacteria Actinobacteria Actinobacteridae Actinomycetales Corynebacterineae Corynebacteriaceae *Corynebacterium diphtheriae str.* Aberdeen |
| Cdi-14-1 | Bacteria Actinobacteria Actinobacteridae Actinomycetales Corynebacterineae Corynebacteriaceae *Corynebacterium diphtheriae* VA01 |
| Cdo-1-1 to Cdo-1-2 | Bacteria Actinobacteria Actinobacteridae Actinomycetales Corynebacterineae Corynebacteriaceae *Corynebacterium doosanense* CAU 212 = DSM 45436 |
| Cef-1-1 | Bacteria Actinobacteria Actinobacteridae Actinomycetales Corynebacterineae Corynebacteriaceae *Corynebacterium efficiens* YS-314 |
| Cge-1-1 | Bacteria Actinobacteria Actinobacteridae Actinomycetales Corynebacterineae Corynebacteriaceae *Corynebacterium genitalium* ATCC 33030 |
| Cgl-1-1 | Bacteria Actinobacteria Actinobacteridae Actinomycetales Corynebacterineae Corynebacteriaceae *Corynebacterium glucuronolyticum* ATCC 51866 |
| Cgl-2-1 | Bacteria Actinobacteria Actinobacteridae Actinomycetales Corynebacterineae Corynebacteriaceae *Corynebacterium glucuronolyticum* ATCC 51867 |

| | |
|---|---|
| Cgl-3-1 to Cgl-3-2 | Bacteria Actinobacteria Actinobacteridae Actinomycetales Corynebacterineae Corynebacteriaceae *Corynebacterium glutamicum* ATCC 13032 |
| Cgl-4-1 | Bacteria Actinobacteria Actinobacteridae Actinomycetales Corynebacterineae Corynebacteriaceae *Corynebacterium glutamicum* K051 |
| Cgl-5-1 | Bacteria Actinobacteria Actinobacteridae Actinomycetales Corynebacterineae Corynebacteriaceae *Corynebacterium glutamicum* MB001 |
| Cgl-6-1 | Bacteria Actinobacteria Actinobacteridae Actinomycetales Corynebacterineae Corynebacteriaceae *Corynebacterium glutamicum* R |
| Cgl-7-1 | Bacteria Actinobacteria Actinobacteridae Actinomycetales Corynebacterineae Corynebacteriaceae *Corynebacterium glutamicum* S9114 |
| Cgl-8-1 | Bacteria Actinobacteria Actinobacteridae Actinomycetales Corynebacterineae Corynebacteriaceae *Corynebacterium glutamicum* SCgG1 |
| Cgl-9-1 | Bacteria Actinobacteria Actinobacteridae Actinomycetales Corynebacterineae Corynebacteriaceae *Corynebacterium glutamicum* SCgG2 |
| Cgl-10-1 | Bacteria Actinobacteria Actinobacteridae Actinomycetales Corynebacterineae Corynebacteriaceae *Corynebacterium glutamicum* Z188 |
| Cha-1-1 to Cha-1-2 | Bacteria Actinobacteria Actinobacteridae Actinomycetales Corynebacterineae Corynebacteriaceae *Corynebacterium halotolerans* YIM 70093 = DSM 44683 |
| Cje-1-1 | Bacteria Actinobacteria Actinobacteridae Actinomycetales Corynebacterineae Corynebacteriaceae *Corynebacterium jeikeium* ATCC 43734 |
| Cje-2-1 | Bacteria Actinobacteria Actinobacteridae Actinomycetales Corynebacterineae Corynebacteriaceae *Corynebacterium jeikeium* K411 |
| Cli-1-1 | Bacteria Actinobacteria Actinobacteridae Actinomycetales Corynebacterineae Corynebacteriaceae *Corynebacterium lipophiloflavum* DSM 44291 |
| Clu-1-1 | Bacteria Actinobacteria Actinobacteridae Actinomycetales Corynebacterineae Corynebacteriaceae *Corynebacterium lubricantis* DSM 45231 |
| Cma-1-1 | Bacteria Actinobacteria Actinobacteridae Actinomycetales Corynebacterineae Corynebacteriaceae *Corynebacterium maris* DSM 45190 |
| Cma-2-1 | Bacteria Actinobacteria Actinobacteridae Actinomycetales Corynebacterineae Corynebacteriaceae *Corynebacterium massiliense* DSM 45435 |
| Cma-3-1 | Bacteria Actinobacteria Actinobacteridae Actinomycetales Corynebacterineae Corynebacteriaceae *Corynebacterium mastitidis* DSM 44356 |
| Cma-4-1 | Bacteria Actinobacteria Actinobacteridae Actinomycetales Corynebacterineae Corynebacteriaceae *Corynebacterium matruchotii* ATCC 14266 |
| Cma-5-1 | Bacteria Actinobacteria Actinobacteridae Actinomycetales Corynebacterineae Corynebacteriaceae *Corynebacterium matruchotii* ATCC 33806 |
| Cnu-1-1 | Bacteria Actinobacteria Actinobacteridae Actinomycetales Corynebacterineae Corynebacteriaceae *Corynebacterium nuruki* S6-4 |
| Cpi-1-1 | Bacteria Actinobacteria Actinobacteridae Actinomycetales Corynebacterineae Corynebacteriaceae *Corynebacterium pilosum* DSM 20521 |
| Cpr-1-1 | Bacteria Actinobacteria Actinobacteridae Actinomycetales Corynebacterineae Corynebacteriaceae *Corynebacterium propinquum* DSM 44285 |
| Cps-1-1 | Bacteria Actinobacteria Actinobacteridae Actinomycetales Corynebacterineae Corynebacteriaceae *Corynebacterium pseudodiphtheriticum* 090104 |
| Cps-2-1 | Bacteria Actinobacteria Actinobacteridae Actinomycetales Corynebacterineae Corynebacteriaceae *Corynebacterium pseudogenitalium* ATCC 33035 |
| Cps-3-1 | Bacteria Actinobacteria Actinobacteridae Actinomycetales Corynebacterineae Corynebacteriaceae *Corynebacterium pseudotuberculosis* 1/06-A |
| Cps-4-1 | Bacteria Actinobacteria Actinobacteridae Actinomycetales Corynebacterineae Corynebacteriaceae *Corynebacterium pseudotuberculosis* 1002 |
| Cps-5-1 | Bacteria Actinobacteria Actinobacteridae Actinomycetales Corynebacterineae Corynebacteriaceae *Corynebacterium pseudotuberculosis* 267 |
| Cps-6-1 | Bacteria Actinobacteria Actinobacteridae Actinomycetales Corynebacterineae Corynebacteriaceae *Corynebacterium pseudotuberculosis* 3/99-5 |
| Cps-7-1 | Bacteria Actinobacteria Actinobacteridae Actinomycetales Corynebacterineae Corynebacteriaceae *Corynebacterium pseudotuberculosis* 316 |
| Cps-8-1 | Bacteria Actinobacteria Actinobacteridae Actinomycetales Corynebacterineae Corynebacteriaceae *Corynebacterium pseudotuberculosis* 42/02-A |
| Cps-9-1 | Bacteria Actinobacteria Actinobacteridae Actinomycetales Corynebacterineae Corynebacteriaceae *Corynebacterium pseudotuberculosis* C231 |
| Cps-10-1 | Bacteria Actinobacteria Actinobacteridae Actinomycetales Corynebacterineae Corynebacteriaceae *Corynebacterium pseudotuberculosis* CIP 52.97 |
| Cps-11-1 | Bacteria Actinobacteria Actinobacteridae Actinomycetales Corynebacterineae Corynebacteriaceae *Corynebacterium pseudotuberculosis* Cp162 |
| Cps-12-1 | Bacteria Actinobacteria Actinobacteridae Actinomycetales Corynebacterineae Corynebacteriaceae *Corynebacterium pseudotuberculosis* FRC41 |
| Cps-13-1 | Bacteria Actinobacteria Actinobacteridae Actinomycetales Corynebacterineae Corynebacteriaceae *Corynebacterium pseudotuberculosis* I19 |
| Cps-14-1 | Bacteria Actinobacteria Actinobacteridae Actinomycetales Corynebacterineae Corynebacteriaceae *Corynebacterium pseudotuberculosis* P54B96 |
| Cps-15-1 | Bacteria Actinobacteria Actinobacteridae Actinomycetales Corynebacterineae Corynebacteriaceae *Corynebacterium pseudotuberculosis* PAT10 |
| Cpy-1-1 | Bacteria Actinobacteria Actinobacteridae Actinomycetales Corynebacterineae Corynebacteriaceae *Corynebacterium pyruviciproducens* ATCC BAA-1742 |
| Cre-1-1 | Bacteria Actinobacteria Actinobacteridae Actinomycetales Corynebacterineae Corynebacteriaceae *Corynebacterium resistens* DSM 45100 |
| Csp-1-1 | Bacteria Actinobacteria Actinobacteridae Actinomycetales Corynebacterineae Corynebacteriaceae *Corynebacterium sp.* HFH0082 |
| Csp-2-1 | Bacteria Actinobacteria Actinobacteridae Actinomycetales Corynebacterineae Corynebacteriaceae *Corynebacterium sp.* KPL1814 |
| Csp-3-1 | Bacteria Actinobacteria Actinobacteridae Actinomycetales Corynebacterineae Corynebacteriaceae *Corynebacterium sp.* KPL1817 |
| Csp-4-1 | Bacteria Actinobacteria Actinobacteridae Actinomycetales Corynebacterineae Corynebacteriaceae *Corynebacterium sp.* KPL1818 |
| Csp-5-1 | Bacteria Actinobacteria Actinobacteridae Actinomycetales Corynebacterineae Corynebacteriaceae *Corynebacterium sp.* KPL1821 |
| Csp-6-1 | Bacteria Actinobacteria Actinobacteridae Actinomycetales Corynebacterineae Corynebacteriaceae *Corynebacterium sp.* KPL1824 |
| Csp-7-1 | Bacteria Actinobacteria Actinobacteridae Actinomycetales Corynebacterineae Corynebacteriaceae *Corynebacterium sp.* KPL1855 |
| Csp-8-1 | Bacteria Actinobacteria Actinobacteridae Actinomycetales Corynebacterineae Corynebacteriaceae *Corynebacterium sp.* KPL1856 |
| Csp-9-1 | Bacteria Actinobacteria Actinobacteridae Actinomycetales Corynebacterineae Corynebacteriaceae *Corynebacterium sp.* KPL1857 |
| Csp-10-1 | Bacteria Actinobacteria Actinobacteridae Actinomycetales Corynebacterineae Corynebacteriaceae *Corynebacterium sp.* KPL1859 |
| Csp-11-1 | Bacteria Actinobacteria Actinobacteridae Actinomycetales Corynebacterineae Corynebacteriaceae *Corynebacterium sp.* KPL1860 |
| Csp-12-1 | Bacteria Actinobacteria Actinobacteridae Actinomycetales Corynebacterineae Corynebacteriaceae *Corynebacterium sp.* KPL1986 |
| Csp-13-1 | Bacteria Actinobacteria Actinobacteridae Actinomycetales Corynebacterineae Corynebacteriaceae *Corynebacterium sp.* KPL1989 |
| Csp-14-1 | Bacteria Actinobacteria Actinobacteridae Actinomycetales Corynebacterineae Corynebacteriaceae *Corynebacterium sp.* KPL1995 |
| Csp-15-1 | Bacteria Actinobacteria Actinobacteridae Actinomycetales Corynebacterineae Corynebacteriaceae *Corynebacterium sp.* KPL1996 |
| Csp-16-1 | Bacteria Actinobacteria Actinobacteridae Actinomycetales Corynebacterineae Corynebacteriaceae *Corynebacterium sp.* KPL1998 |
| Csp-17-1 | Bacteria Actinobacteria Actinobacteridae Actinomycetales Corynebacterineae Corynebacteriaceae *Corynebacterium sp.* KPL2004 |
| Cst-1-1 | Bacteria Actinobacteria Actinobacteridae Actinomycetales Corynebacterineae Corynebacteriaceae *Corynebacterium striatum* ATCC 6940 |
| Cte-1-1 | Bacteria Actinobacteria Actinobacteridae Actinomycetales Corynebacterineae Corynebacteriaceae *Corynebacterium terpenotabidum* Y-11 |
| Ctu-1-1 | Bacteria Actinobacteria Actinobacteridae Actinomycetales Corynebacterineae Corynebacteriaceae *Corynebacterium tuberculostearicum* SK141 |
| Cul-1-1 | Bacteria Actinobacteria Actinobacteridae Actinomycetales Corynebacterineae Corynebacteriaceae *Corynebacterium ulcerans* 0102 |
| Cul-2-1 | Bacteria Actinobacteria Actinobacteridae Actinomycetales Corynebacterineae Corynebacteriaceae *Corynebacterium ulcerans* 809 |
| Cul-3-1 | Bacteria Actinobacteria Actinobacteridae Actinomycetales Corynebacterineae Corynebacteriaceae *Corynebacterium ulcerans* BR-AD22 |
| Cul-4-1 | Bacteria Actinobacteria Actinobacteridae Actinomycetales Corynebacterineae Corynebacteriaceae *Corynebacterium ulcerans* NCTC 12077 |
| Cul-5-1 | Bacteria Actinobacteria Actinobacteridae Actinomycetales Corynebacterineae Corynebacteriaceae *Corynebacterium ulceribovis* DSM 45146 |
| Cur-1-1 | Bacteria Actinobacteria Actinobacteridae Actinomycetales Corynebacterineae Corynebacteriaceae *Corynebacterium urealyticum* DSM 7109 |
| Cur-2-1 | Bacteria Actinobacteria Actinobacteridae Actinomycetales Corynebacterineae Corynebacteriaceae *Corynebacterium urealyticum* DSM 7111 |

| | |
|---|---|
| Cva-1-1 | Bacteria Actinobacteria Actinobacteridae Actinomycetales Corynebacterineae Corynebacteriaceae *Corynebacterium variabile* DSM 44702 |
| Tot-1-1 | Bacteria Actinobacteria Actinobacteridae Actinomycetales Corynebacterineae Corynebacteriaceae *Turicella otitidis* ATCC 51513 |
| [fl-1-1 | Bacteria Actinobacteria Corynebacteriales Corynebacteriaceae Corynebacterium [Brevibacterium] flavum |
| Cat-1-1 | Bacteria Actinobacteria Corynebacteriales Corynebacteriaceae *Corynebacterium atypicum* |
| Cau-2-1 to Cau-2-4 | Bacteria Actinobacteria Corynebacteriales Corynebacteriaceae *Corynebacterium aurimucosum* |
| Cau-3-1 | Bacteria Actinobacteria Corynebacteriales Corynebacteriaceae *Corynebacterium auriscanis* |
| Cca-1-1 to Cca-1-2 | Bacteria Actinobacteria Corynebacteriales Corynebacteriaceae *Corynebacterium callunae* DSM 20147 |
| Cca-5-1 | Bacteria Actinobacteria Corynebacteriales Corynebacteriaceae *Corynebacterium camporealensis* |
| Cca-6-1 | Bacteria Actinobacteria Corynebacteriales Corynebacteriaceae *Corynebacterium casei* LMG S-19264 |
| Ccr-2-1 | Bacteria Actinobacteria Corynebacteriales Corynebacteriaceae *Corynebacterium crenatum* |
| Cdi-18-1 to Cdi-18-2 | Bacteria Actinobacteria Corynebacteriales Corynebacteriaceae *Corynebacterium diphtheriae* |
| Cdi-19-1 | Bacteria Actinobacteria Corynebacteriales Corynebacteriaceae *Corynebacterium diphtheriae bv. gravis str.* ISS 4060 |
| Cdi-20-1 | Bacteria Actinobacteria Corynebacteriales Corynebacteriaceae *Corynebacterium diphtheriae bv. gravis str.* ISS 4746 |
| Cdi-21-1 | Bacteria Actinobacteria Corynebacteriales Corynebacteriaceae *Corynebacterium diphtheriae bv. gravis str.* ISS 4749 |
| Cdi-22-1 | Bacteria Actinobacteria Corynebacteriales Corynebacteriaceae *Corynebacterium diphtheriae bv. mitis str.* ISS 3319 |
| Cdo-1-1 to Cdo-1-2 | Bacteria Actinobacteria Corynebacteriales Corynebacteriaceae *Corynebacterium doosanense* CAU 212 = DSM 45436 |
| Cep-1-1 | Bacteria Actinobacteria Corynebacteriales Corynebacteriaceae *Corynebacterium epidermidicanis* |
| Cfa-1-1 | Bacteria Actinobacteria Corynebacteriales Corynebacteriaceae *Corynebacterium falsenii* |
| Cfa-2-1 | Bacteria Actinobacteria Corynebacteriales Corynebacteriaceae *Corynebacterium falsenii* DSM 44353 |
| Cfr-1-1 | Bacteria Actinobacteria Corynebacteriales Corynebacteriaceae *Corynebacterium freneyi* DNF00450 |
| Cgl-11-1 to Cgl-11-3 | Bacteria Actinobacteria Corynebacteriales Corynebacteriaceae *Corynebacterium glutamicum* |
| Cgl-12-1 | Bacteria Actinobacteria Corynebacteriales Corynebacteriaceae *Corynebacterium glutamicum* ATCC 14067 |
| Cgl-13-1 | Bacteria Actinobacteria Corynebacteriales Corynebacteriaceae *Corynebacterium glyciniphilum* AJ 3170 |
| Cha-2-1 | Bacteria Actinobacteria Corynebacteriales Corynebacteriaceae *Corynebacterium halotolerans* |
| Cha-1-1 to Cha-1-2 | Bacteria Actinobacteria Corynebacteriales Corynebacteriaceae *Corynebacterium halotolerans* YIM 70093 = DSM 44683 |
| Chu-1-1 | Bacteria Actinobacteria Corynebacteriales Corynebacteriaceae *Corynebacterium humireducens* NBRC 106098 = DSM 45392 |
| Cih-1-1 | Bacteria Actinobacteria Corynebacteriales Corynebacteriaceae *Corynebacterium ihumii* |
| Cim-1-1 | Bacteria Actinobacteria Corynebacteriales Corynebacteriaceae *Corynebacterium imitans* |
| Cje-3-1 | Bacteria Actinobacteria Corynebacteriales Corynebacteriaceae *Corynebacterium jeddahense* |
| Cje-4-1 to Cje-4-18 | Bacteria Actinobacteria Corynebacteriales Corynebacteriaceae *Corynebacterium jeikeium* |
| Cku-1-1 | Bacteria Actinobacteria Corynebacteriales Corynebacteriaceae *Corynebacterium kutscheri* |
| Cla-1-1 | Bacteria Actinobacteria Corynebacteriales Corynebacteriaceae *Corynebacterium lactis* RW2-5 |
| Cma-6-1 | Bacteria Actinobacteria Corynebacteriales Corynebacteriaceae *Corynebacterium marinum* DSM 44953 |
| Cmi-1-1 to Cmi-1-2 | Bacteria Actinobacteria Corynebacteriales Corynebacteriaceae *Corynebacterium minutissimum* |
| Cmu-1-1 | Bacteria Actinobacteria Corynebacteriales Corynebacteriaceae *Corynebacterium mustelae* |
| Cpi-2-1 | Bacteria Actinobacteria Corynebacteriales Corynebacteriaceae *Corynebacterium pilosum* |
| Cpr-2-1 to Cpr-2-3 | Bacteria Actinobacteria Corynebacteriales Corynebacteriaceae *Corynebacterium propinquum* |
| Cps-16-1 | Bacteria Actinobacteria Corynebacteriales Corynebacteriaceae *Corynebacterium pseudodiphtheriticum* DSM 44287 |
| Cps-17-1 to Cps-17-9 | Bacteria Actinobacteria Corynebacteriales Corynebacteriaceae *Corynebacterium pseudotuberculosis* |
| Cps-18-1 | Bacteria Actinobacteria Corynebacteriales Corynebacteriaceae *Corynebacterium pseudotuberculosis* 258 |
| Cps-19-1 | Bacteria Actinobacteria Corynebacteriales Corynebacteriaceae *Corynebacterium pseudotuberculosis* 31 |
| Cre-2-1 | Bacteria Actinobacteria Corynebacteriales Corynebacteriaceae *Corynebacterium renale* |
| Cri-1-1 | Bacteria Actinobacteria Corynebacteriales Corynebacteriaceae *Corynebacterium riegelii* |
| Csi-1-1 | Bacteria Actinobacteria Corynebacteriales Corynebacteriaceae *Corynebacterium singulare* |
| Csp-18-1 | Bacteria Actinobacteria Corynebacteriales Corynebacteriaceae *Corynebacterium sp.* ATCC 6931 |
| Csp-19-1 | Bacteria Actinobacteria Corynebacteriales Corynebacteriaceae *Corynebacterium sputi* DSM 45148 |
| Cst-2-1 to Cst-2-8 | Bacteria Actinobacteria Corynebacteriales Corynebacteriaceae *Corynebacterium striatum* |
| Cte-2-1 | Bacteria Actinobacteria Corynebacteriales Corynebacteriaceae *Corynebacterium testudinoris* |
| Cti-1-1 | Bacteria Actinobacteria Corynebacteriales Corynebacteriaceae *Corynebacterium timonense* 5401744 |
| Ctu-2-1 | Bacteria Actinobacteria Corynebacteriales Corynebacteriaceae *Corynebacterium tuscaniense* DNF00037 |
| Cul-6-1 to Cul-6-4 | Bacteria Actinobacteria Corynebacteriales Corynebacteriaceae *Corynebacterium ulcerans* |
| Cul-7-1 | Bacteria Actinobacteria Corynebacteriales Corynebacteriaceae *Corynebacterium ulcerans* FRC11 |
| Cul-8-1 | Bacteria Actinobacteria Corynebacteriales Corynebacteriaceae *Corynebacterium ulcerans* FRC58 |
| Cur-3-1 | Bacteria Actinobacteria Corynebacteriales Corynebacteriaceae *Corynebacterium urealyticum* |
| Cur-4-1 | Bacteria Actinobacteria Corynebacteriales Corynebacteriaceae *Corynebacterium ureicelerivorans* |
| Cut-1-1 | Bacteria Actinobacteria Corynebacteriales Corynebacteriaceae *Corynebacterium uterequi* |
| Cvi-1-1 | Bacteria Actinobacteria Corynebacteriales Corynebacteriaceae *Corynebacterium vitaeruminis* |
| Cvi-2-1 | Bacteria Actinobacteria Corynebacteriales Corynebacteriaceae *Corynebacterium vitaeruminis* DSM 20294 |
| Cvi-3-1 | Bacteria Actinobacteria Corynebacteriales Corynebacteriaceae *Corynebacterium vitaeruminis* Ga6A13 |
| Cxe-1-1 | Bacteria Actinobacteria Corynebacteriales Corynebacteriaceae *Corynebacterium xerosis* |
| Tot-2-1 | Bacteria Actinobacteria Corynebacteriales Corynebacteriaceae *Turicella otitidis* |
| env-1 to env-159 | environmental samples |

#### 13.2 Gene contexts

Each L31-Corynebacteriaceae RNA (indicated by “RNA→”) is listed. For each hit, the downstream genes predicted to reside in a regulated operon are listed. If the nearest downstream gene is encoding in the opposite strand (and therefore presumed to not be a part of a regulated operon), then that gene is still depicted. Some environmental sequences and some RefSeq entries lack gene annotations, and so no genes are available for such sequences. The direction of each gene is indicated with an arrow (→), and each predicted conserved domain in the gene is named. Conserved domains associated with more than one L31-Corynebacteriaceae RNA are assigned a color; other domains are gray. Information about these conserved domains is given in Section 13.3. The accession of the sequence containing each L31-Corynebacteriaceae RNA is given in the column named “Seq. accession”. Accessions beginning with “NC_”, “NS_”, “NW_” or “NZ_” are contained in RefSeq. Other accession refer to environmental samples. Nucleotide coordinates are given for the 5′ and 3′ boundaries of each L31-Corynebacteriaceae RNA. If the 5′ coordinate is greater than the 3′ coordinate, the RNA is present on the reverse-complement strand of the containing genomic DNA sequence. Each hit is denoted by an abbreviation (like “Eco-1-1”) that refers to a taxonomy given in Section 13.1.

| abbrev. | Seq. accession | | 5′ at | 3′ at | genes |
|---|---|---|---|---|---|
| Cut-1-1 | NZ_CP011546.1 | + | 713235 | 713316 | RNA→ rpmE2 (PRK01678)Ribosomal_L31 (pfam01197)→ rpmF (PRK01110)Ribosomal_L32p (pfam01783)→ |
| Cfa-1-1 | NZ_JVKJ01000005.1 | - | 667570 | 667497 | RNA→ rpmE2 (PRK01678)Ribosomal_L31 (pfam01197)→ rpmF (PRK01110)Ribosomal_L32p (pfam01783)→ |
| Cau-3-1 | NZ_JRVJ01000003.1 | + | 120589 | 120653 | RNA→ rpmE2 (PRK01678)Ribosomal_L31 (pfam01197)→ hypo→ hypo→ |
| Tot-2-1 | NZ_LBNF01000055.1 | - | 8790 | 8700 | RNA→ rpmE2 (PRK01678)Ribosomal_L31 (pfam01197)→ rpmF (PRK01110)Ribosomal_L32p (pfam01783)→ |
| Tot-1-1 | NZ_JH815193.1 | - | 531899 | 531809 | RNA→ rpmE2 (PRK01678)Ribosomal_L31 (pfam01197)→ rpmF (PRK01110)Ribosomal_L32p (pfam01783)→ |
| Cnu-1-1 | NZ_AFIZ01000013.1 | - | 97797 | 97736 | RNA→ rpmE2 (PRK01678)Ribosomal_L31 (pfam01197)→ rpmE2 (PRK01678)Ribosomal_L31 (pfam01197)→ rpmF (PRK01110)Ribosomal_L32p (pfam01783)→ rpmF (PRK01110)Ribosomal_L32p (pfam01783)→ |
| Cgl-13-1 | NZ_CP006842.1 | + | 1057041 | 1057122 | RNA→ rpmE2 (PRK01678)Ribosomal_L31 (pfam01197)→ rpmF (PRK01110)Ribosomal_L32p (pfam01783)→ |
| Cte-1-1 | NC_021663.1 | - | 1923752 | 1923653 | RNA→ rpmE2 (PRK01678)Ribosomal_L31 (pfam01197)→ rpmF (PRK01110)Ribosomal_L32p (pfam01783)→ |
| Cva-1-1 | NC_015859.1 | - | 2360714 | 2360640 | RNA→ rpmE2 (PRK01678)Ribosomal_L31 (pfam01197)→ rpmF (PRK01110)Ribosomal_L32p (pfam01783)→ |
| Cps-16-1 | NZ_JIAH01000006.1 | + | 56886 | 57003 | RNA→ rpmE2 (PRK01678)Ribosomal_L31 (pfam01197)→ rpmF (PRK01110)Ribosomal_L32p (pfam01783)→ |
| Csp-14-1 | NZ_KI515715.1 | - | 2247633 | 2247516 | RNA→ rpmE2 (PRK01678)Ribosomal_L31 (pfam01197)→ rpmF (PRK01110)Ribosomal_L32p (pfam01783)→ |
| Csp-13-1 | NZ_KI515718.1 | + | 60683 | 60800 | RNA→ rpmE2 (PRK01678)Ribosomal_L31 (pfam01197)→ rpmF (PRK01110)Ribosomal_L32p (pfam01783)→ |
| env-1 | SRS053437_C431471 | - | 1608 | 1491 | RNA→ rpmE2 (PRK01678)Ribosomal_L31 (pfam01197)→ rpmF (PRK01110)Ribosomal_L32p (pfam01783)→ |
| env-2 | SRS013876_WUGC_scaffold_993 | - | 41899 | 41782 | RNA→ rpmE2 (PRK01678)Ribosomal_L31 (pfam01197)→ rpmF (PRK01110)Ribosomal_L32p (pfam01783)→ |
| Cps-1-1 | NZ_AVFF01000012.1 | - | 4417 | 4300 | RNA→ rpmE2 (PRK01678)Ribosomal_L31 (pfam01197)→ rpmF (PRK01110)Ribosomal_L32p (pfam01783)→ |
| env-3 | SRS017044_C179127 | + | 297 | 403 | RNA→ rpmE2 (PRK01678)Ribosomal_L31 (pfam01197)→ rpmF (PRK01110)Ribosomal_L32p (pfam01783)→ |
| Cpr-2-1 | NZ_JVVY01000068.1 | + | 69983 | 70089 | RNA→ rpmE2 (PRK01678)Ribosomal_L31 (pfam01197)→ rpmF (PRK01110)Ribosomal_L32p (pfam01783)→ |
| Cpr-1-1 | NZ_AQXC01000010.1 | + | 64394 | 64500 | RNA→ rpmE2 (PRK01678)Ribosomal_L31 (pfam01197)→ rpmF (PRK01110)Ribosomal_L32p (pfam01783)→ |
| env-4 | SRS047254_WUGC_scaffold_9884 | + | 208 | 314 | RNA→ |
| env-5 | SRS015752_C119743 | + | 276 | 382 | RNA→ |
| Cpr-2-2 | NZ_JVSI01000039.1 | - | 39215 | 39109 | RNA→ rpmE2 (PRK01678)Ribosomal_L31 (pfam01197)→ rpmF (PRK01110)Ribosomal_L32p (pfam01783)→ |
| Cpr-2-3 | NZ_JVSN01000024.1 | - | 56709 | 56603 | RNA→ rpmE2 (PRK01678)Ribosomal_L31 (pfam01197)→ rpmF (PRK01110)Ribosomal_L32p (pfam01783)→ |
| Cfa-2-1 | NZ_CP007156.1 | - | 2014629 | 2014563 | RNA→ rpmE2 (PRK01678)Ribosomal_L31 (pfam01197)→ rpmF (PRK01110)Ribosomal_L32p (pfam01783)→ |
| Cre-1-1 | NC_015673.1 | - | 1990937 | 1990874 | RNA→ rpmE2 (PRK01678)Ribosomal_L31 (pfam01197)→ rpmF (PRK01110)Ribosomal_L32p (pfam01783)→ |
| Cje-2-1 | NC_007164.1 | - | 1807476 | 1807415 | RNA→ rpmE2 (PRK01678)Ribosomal_L31 (pfam01197)→ rpmF (PRK01110)Ribosomal_L32p (pfam01783)→ |
| Cje-4-1 | NZ_JFCM01000017.1 | - | 3528 | 3467 | RNA→ rpmE2 (PRK01678)Ribosomal_L31 (pfam01197)→ rpmF (PRK01110)Ribosomal_L32p (pfam01783)→ |
| Cje-4-2 | NZ_JFCO01000025.1 | + | 54212 | 54273 | RNA→ rpmE2 (PRK01678)Ribosomal_L31 (pfam01197)→ rpmF (PRK01110)Ribosomal_L32p (pfam01783)→ |
| Cje-4-3 | NZ_JFCR01000012.1 | - | 86567 | 86506 | RNA→ rpmE2 (PRK01678)Ribosomal_L31 (pfam01197)→ rpmF (PRK01110)Ribosomal_L32p (pfam01783)→ |
| Cje-4-4 | NZ_JUNC01000013.1 | - | 142683 | 142622 | RNA→ rpmE2 (PRK01678)Ribosomal_L31 (pfam01197)→ rpmF (PRK01110)Ribosomal_L32p (pfam01783)→ |
| Cje-1-1 | NZ_GG700814.1 | - | 359951 | 359890 | RNA→ rpmE2 (PRK01678)Ribosomal_L31 (pfam01197)→ rpmF (PRK01110)Ribosomal_L32p (pfam01783)→ |
| Cje-4-5 | NZ_JFCH01000061.1 | + | 41014 | 41075 | RNA→ rpmE2 (PRK01678)Ribosomal_L31 (pfam01197)→ rpmF (PRK01110)Ribosomal_L32p (pfam01783)→ |
| Cje-4-6 | NZ_JFCQ01000001.1 | - | 319495 | 319435 | RNA→ rpmE2 (PRK01678)Ribosomal_L31 (pfam01197)→ rpmF (PRK01110)Ribosomal_L32p (pfam01783)→ |
| Cje-4-7 | NZ_JFCL01000010.1 | - | 81768 | 81708 | RNA→ rpmE2 (PRK01678)Ribosomal_L31 (pfam01197)→ rpmF (PRK01110)Ribosomal_L32p (pfam01783)→ |
| Cje-4-8 | NZ_JFCF01000035.1 | + | 36866 | 36926 | RNA→ rpmE2 (PRK01678)Ribosomal_L31 (pfam01197)→ rpmF (PRK01110)Ribosomal_L32p (pfam01783)→ |
| Cje-4-9 | NZ_JFCG01000013.1 | - | 140206 | 140146 | RNA→ rpmE2 (PRK01678)Ribosomal_L31 (pfam01197)→ rpmF (PRK01110)Ribosomal_L32p (pfam01783)→ |

| | | | | | |
|---|---|---|---|---|---|
| Cje-4-10 | NZ_JFCI01000035.1 | + | 37385 | 37445 | **RNA**→ rpmE2 (PRK01678)Ribosomal_L31 (pfam01197)→ rpmF (PRK01110)Ribosomal_L32p (pfam01783)→ |
| Cje-4-11 | NZ_JFCJ01000077.1 | - | 132850 | 132790 | **RNA**→ rpmE2 (PRK01678)Ribosomal_L31 (pfam01197)→ rpmF (PRK01110)Ribosomal_L32p (pfam01783)→ |
| Cje-4-12 | NZ_JFCN01000025.1 | + | 2814 | 2874 | **RNA**→ rpmE2 (PRK01678)Ribosomal_L31 (pfam01197)→ rpmF (PRK01110)Ribosomal_L32p (pfam01783)→ |
| Cje-4-13 | NZ_JFCK01000168.1 | - | 59670 | 59610 | **RNA**→ rpmE2 (PRK01678)Ribosomal_L31 (pfam01197)→ rpmF (PRK01110)Ribosomal_L32p (pfam01783)→ |
| Cje-4-14 | NZ_JFCP01000015.1 | - | 88000 | 87940 | **RNA**→ rpmE2 (PRK01678)Ribosomal_L31 (pfam01197)→ rpmF (PRK01110)Ribosomal_L32p (pfam01783)→ |
| Cur-2-1 | NC_020230.1 | + | 634362 | 634421 | **RNA**→ rpmE2 (PRK01678)Ribosomal_L31 (pfam01197)→ rpmF (PRK01110)Ribosomal_L32p (pfam01783)→ |
| Cur-1-1 | NC_010545.1 | + | 654798 | 654857 | **RNA**→ rpmE2 (PRK01678)Ribosomal_L31 (pfam01197)→ rpmF (PRK01110)Ribosomal_L32p (pfam01783)→ |
| Cdi-18-1 | NZ_JRUZ01000001.1 | - | 346898 | 346839 | **RNA**→ rpmE2 (PRK01678)Ribosomal_L31 (pfam01197)→ rpmF (PRK01110)Ribosomal_L32p (pfam01783)→ |
| Cdi-18-2 | NZ_JZUJ01000002.1 | + | 4530 | 4589 | **RNA**→ rpmE2 (PRK01678)Ribosomal_L31 (pfam01197)→ rpmF (PRK01110)Ribosomal_L32p (pfam01783)→ |
| Cdi-19-1 | NZ_JAQN01000016.1 | + | 59508 | 59567 | **RNA**→ hypo→ rpmE2 (PRK01678)Ribosomal_L31 (pfam01197)→ rpmF (PRK01110)Ribosomal_L32p (pfam01783)→ |
| Cdi-22-1 | NZ_JAQO01000007.1 | + | 59841 | 59900 | **RNA**→ rpmE2 (PRK01678)Ribosomal_L31 (pfam01197)→ rpmF (PRK01110)Ribosomal_L32p (pfam01783)→ |
| Cdi-5-1 | NC_016785.1 | + | 776418 | 776477 | **RNA**→ rpmE2 (PRK01678)Ribosomal_L31 (pfam01197)→ rpmF (PRK01110)Ribosomal_L32p (pfam01783)→ |
| Cdi-16-1 | NZ_AJGI01000001.1 | - | 267143 | 267084 | **RNA**→ PriB (COG2965)→ rpmE2 (PRK01678)Ribosomal_L31 (pfam01197)→ rpmF (PRK01110)Ribosomal_L32p (pfam01783)→ |
| Cdi-1-1 | NC_016782.1 | + | 774515 | 774574 | **RNA**→ hypo→ rpmE2 (PRK01678)Ribosomal_L31 (pfam01197)→ rpmF (PRK01110)Ribosomal_L32p (pfam01783)→ |
| Cdi-9-1 | NC_016787.1 | + | 789776 | 789835 | **RNA**→ rpmE2 (PRK01678)Ribosomal_L31 (pfam01197)→ rpmF (PRK01110)Ribosomal_L32p (pfam01783)→ |
| Cdi-2-1 | NC_016799.1 | + | 854037 | 854096 | **RNA**→ PriB (COG2965)→ rpmE2 (PRK01678)Ribosomal_L31 (pfam01197)→ rpmF (PRK01110)Ribosomal_L32p (pfam01783)→ |
| Cdi-11-1 | NC_016783.1 | + | 770903 | 770962 | **RNA**→ hypo→ rpmE2 (PRK01678)Ribosomal_L31 (pfam01197)→ rpmF (PRK01110)Ribosomal_L32p (pfam01783)→ |
| Cdi-7-1 | NC_016786.1 | + | 774486 | 774545 | **RNA**→ hypo→ rpmE2 (PRK01678)Ribosomal_L31 (pfam01197)→ rpmF (PRK01110)Ribosomal_L32p (pfam01783)→ |
| Cdi-13-1 | NC_016789.1 | + | 835204 | 835263 | **RNA**→ rpmE2 (PRK01678)Ribosomal_L31 (pfam01197)→ rpmF (PRK01110)Ribosomal_L32p (pfam01783)→ |
| Cdi-4-1 | NC_016801.1 | + | 781925 | 781984 | **RNA**→ rpmE2 (PRK01678)Ribosomal_L31 (pfam01197)→ rpmF (PRK01110)Ribosomal_L32p (pfam01783)→ |
| Cdi-12-1 | NC_002935.2 | + | 826926 | 826985 | **RNA**→ rpmE2 (PRK01678)Ribosomal_L31 (pfam01197)→ rpmF (PRK01110)Ribosomal_L32p (pfam01783)→ |
| Cdi-3-1 | NC_016800.1 | + | 806019 | 806078 | **RNA**→ hypo→ rpmE2 (PRK01678)Ribosomal_L31 (pfam01197)→ rpmF (PRK01110)Ribosomal_L32p (pfam01783)→ |
| Cdi-8-1 | NC_016802.1 | + | 795035 | 795094 | **RNA**→ rpmE2 (PRK01678)Ribosomal_L31 (pfam01197)→ rpmF (PRK01110)Ribosomal_L32p (pfam01783)→ |
| Cdi-6-1 | NZ_AUZN01000046.1 | + | 17970 | 18029 | **RNA**→ hypo→ rpmE2 (PRK01678)Ribosomal_L31 (pfam01197)→ rpmF (PRK01110)Ribosomal_L32p (pfam01783)→ |
| Cdi-15-1 | NZ_AJVH01000012.1 | + | 61533 | 61592 | **RNA**→ rpmE2 (PRK01678)Ribosomal_L31 (pfam01197)→ rpmF (PRK01110)Ribosomal_L32p (pfam01783)→ |
| Cdi-20-1 | NZ_JAQP01000015.1 | + | 32179 | 32238 | **RNA**→ hypo→ rpmE2 (PRK01678)Ribosomal_L31 (pfam01197)→ rpmF (PRK01110)Ribosomal_L32p (pfam01783)→ |
| Cdi-21-1 | NZ_JAQQ01000010.1 | + | 32192 | 32251 | **RNA**→ hypo→ rpmE2 (PRK01678)Ribosomal_L31 (pfam01197)→ rpmF (PRK01110)Ribosomal_L32p (pfam01783)→ |
| Cdi-10-1 | NC_016788.1 | + | 799786 | 799845 | **RNA**→ hypo→ rpmE2 (PRK01678)Ribosomal_L31 (pfam01197)→ rpmF (PRK01110)Ribosomal_L32p (pfam01783)→ |
| Cdi-14-1 | NC_016790.1 | + | 767184 | 767243 | **RNA**→ hypo→ rpmE2 (PRK01678)Ribosomal_L31 (pfam01197)→ rpmF (PRK01110)Ribosomal_L32p (pfam01783)→ |
| Cdi-17-1 | NZ_AUZO01000012.1 | + | 32203 | 32262 | **RNA**→ hypo→ rpmE2 (PRK01678)Ribosomal_L31 (pfam01197)→ rpmF (PRK01110)Ribosomal_L32p (pfam01783)→ |
| Cca-4-1 | NZ_KB903376.1 | + | 440662 | 440743 | **RNA**→ rpmE2 (PRK01678)Ribosomal_L31 (pfam01197)→ rpmF (PRK01110)Ribosomal_L32p (pfam01783)→ |
| Cgl-2-1 | NZ_GG667130.1 | + | 67973 | 68043 | **RNA**→ rpmE2 (PRK01678)Ribosomal_L31 (pfam01197)→ rpmF (PRK01110)Ribosomal_L32p (pfam01783)→ |
| Cgl-1-1 | NZ_GG667038.1 | + | 72480 | 72550 | **RNA**→ rpmE2 (PRK01678)Ribosomal_L31 (pfam01197)→ rpmF (PRK01110)Ribosomal_L32p (pfam01783)→ |
| Cpy-1-1 | NZ_KE150446.1 | + | 69361 | 69419 | **RNA**→ rpmE2 (PRK01678)Ribosomal_L31 (pfam01197)→ rpmF (PRK01110)Ribosomal_L32p (pfam01783)→ |
| Cre-2-1 | NZ_LDYE01000007.1 | + | 45841 | 45903 | **RNA**→ rpmE2 (PRK01678)Ribosomal_L31 (pfam01197)→ rpmF (PRK01110)Ribosomal_L32p (pfam01783)→ |
| Cdo-1-2 | NZ_CP006764.1 | + | 832115 | 832176 | **RNA**→ rpmE2 (PRK01678)Ribosomal_L31 (pfam01197)→ rpmF (PRK01110)Ribosomal_L32p (pfam01783)→ |
| Cdo-1-1 | NZ_AQUX01000004.1 | - | 131828 | 131767 | **RNA**→ rpmE2 (PRK01678)Ribosomal_L31 (pfam01197)→ rpmF (PRK01110)Ribosomal_L32p (pfam01783)→ |
| Cca-5-1 | NZ_CP011311.1 | + | 791325 | 791385 | **RNA**→ rpmE2 (PRK01678)Ribosomal_L31 (pfam01197)→ rpmF (PRK01110)Ribosomal_L32p (pfam01783)→ |
| Cma-3-1 | NZ_AQXB01000007.1 | - | 161751 | 161703 | **RNA**→ rpmE2 (PRK01678)Ribosomal_L31 (pfam01197)→ rpmF (PRK01110)Ribosomal_L32p (pfam01783)→ |
| Cmi-1-1 | NZ_JSEF01000008.1 | + | 14324 | 14454 | **RNA**→ rpmE2 (PRK01678)Ribosomal_L31 (pfam01197)→ rpmF (PRK01110)Ribosomal_L32p (pfam01783)→ |
| Cau-1-1 | NC_012590.1 | + | 901957 | 902077 | **RNA**→ rpmE2 (PRK01678)Ribosomal_L31 (pfam01197)→ rpmF (PRK01110)Ribosomal_L32p (pfam01783)→ |
| Cmi-1-2 | NZ_LAYQ01000001.1 | - | 68637 | 68555 | **RNA**→ rpmE2 (PRK01678)Ribosomal_L31 (pfam01197)→ rpmF (PRK01110)Ribosomal_L32p (pfam01783)→ |

| | | | | | |
|---|---|---|---|---|---|
| Cau-2-1 | NZ_JUMN01000003.1 | + | 19608 | 19690 | **RNA**→ rpmE2 (PRK01678)Ribosomal_L31 (pfam01197)→ rpmF (PRK01110)Ribosomal_L32p (pfam01783)→ |
| Csi-1-1 | NZ_CP010827.1 | + | 955732 | 955835 | **RNA**→ rpmE2 (PRK01678)Ribosomal_L31 (pfam01197)→ rpmF (PRK01110)Ribosomal_L32p (pfam01783)→ |
| Cci-1-1 | NZ_AQUW01000002.1 | - | 253907 | 253785 | **RNA**→ rpmE2 (PRK01678)Ribosomal_L31 (pfam01197)→ rpmF (PRK01110)Ribosomal_L32p (pfam01783)→ |
| Cat-1-1 | NZ_CP008944.1 | + | 701875 | 702000 | **RNA**→ rpmE2 (PRK01678)Ribosomal_L31 (pfam01197)→ rpmF (PRK01110)Ribosomal_L32p (pfam01783)→ |
| Cje-4-15 | NZ_JUSD01000006.1 | + | 58418 | 58480 | **RNA**→ rpmE2 (PRK01678)Ribosomal_L31 (pfam01197)→ rpmF (PRK01110)Ribosomal_L32p (pfam01783)→ |
| Csp-18-1 | NZ_CP008913.1 | - | 697524 | 697462 | **RNA**→ rpmE2 (PRK01678)Ribosomal_L31 (pfam01197)→ rpmF (PRK01110)Ribosomal_L32p (pfam01783)→ |
| Cje-4-16 | NZ_JVOW01000015.1 | - | 32940 | 32878 | **RNA**→ rpmE2 (PRK01678)Ribosomal_L31 (pfam01197)→ rpmF (PRK01110)Ribosomal_L32p (pfam01783)→ |
| Cje-4-17 | NZ_JVIP01000011.1 | + | 50025 | 50087 | **RNA**→ rpmE2 (PRK01678)Ribosomal_L31 (pfam01197)→ rpmF (PRK01110)Ribosomal_L32p (pfam01783)→ |
| Csp-1-1 | NZ_KE150404.1 | - | 2427073 | 2427011 | **RNA**→ rpmE2 (PRK01678)Ribosomal_L31 (pfam01197)→ rpmF (PRK01110)Ribosomal_L32p (pfam01783)→ |
| Cam-2-1 | NZ_ABZU01000013.1 | - | 40816 | 40754 | **RNA**→ rpmE2 (PRK01678)Ribosomal_L31 (pfam01197)→ rpmE2 (PRK01678)Ribosomal_L31 (pfam01197)→ rpmF (PRK01110)Ribosomal_L32p (pfam01783)→ rpmF (PRK01110)Ribosomal_L32p (pfam01783)→ |
| Cur-3-1 | NZ_JWEP01000048.1 | - | 32027 | 31965 | **RNA**→ rpmE2 (PRK01678)Ribosomal_L31 (pfam01197)→ rpmF (PRK01110)Ribosomal_L32p (pfam01783)→ |
| Cje-4-18 | NZ_JVPT01000044.1 | - | 37181 | 37119 | **RNA**→ rpmE2 (PRK01678)Ribosomal_L31 (pfam01197)→ rpmF (PRK01110)Ribosomal_L32p (pfam01783)→ |
| Cvi-1-1 | NZ_JVVM01000020.1 | - | 305775 | 305713 | **RNA**→ rpmE2 (PRK01678)Ribosomal_L31 (pfam01197)→ rpmF (PRK01110)Ribosomal_L32p (pfam01783)→ |
| Cla-1-1 | NZ_CP006841.1 | - | 2125937 | 2125813 | **RNA**→ rpmE2 (PRK01678)Ribosomal_L31 (pfam01197)→ rpmF (PRK01110)Ribosomal_L32p (pfam01783)→ |
| Csp-19-1 | NZ_ATYV01000011.1 | + | 77096 | 77159 | **RNA**→ rpmE2 (PRK01678)Ribosomal_L31 (pfam01197)→ rpmF (PRK01110)Ribosomal_L32p (pfam01783)→ DUF3887 (pfam13026)→ |
| env-6 | SRS016292_C136299 | + | 434 | 503 | **RNA**→ rpmE2 (PRK01678)Ribosomal_L31 (pfam01197)→ |
| env-7 | SRS047225_C235885 | - | 2784 | 2715 | **RNA**→ rpmE2 (PRK01678)Ribosomal_L31 (pfam01197)→ rpmF (PRK01110)Ribosomal_L32p (pfam01783)→ |
| Csp-4-1 | NZ_KI515760.1 | + | 78324 | 78393 | **RNA**→ rpmE2 (PRK01678)Ribosomal_L31 (pfam01197)→ rpmF (PRK01110)Ribosomal_L32p (pfam01783)→ |
| env-8 | SRS017451_C217497 | + | 873 | 939 | **RNA**→ rpmE2 (PRK01678)Ribosomal_L31 (pfam01197)→ |
| env-9 | SRS015937_WUGC_scaffold_1576 | - | 2210 | 2144 | **RNA**→ rpmE2 (PRK01678)Ribosomal_L31 (pfam01197)→ rpmF (PRK01110)Ribosomal_L32p (pfam01783)→ |
| env-10 | SRS019986_C257788 | + | 75 | 141 | **RNA**→ rpmE2 (PRK01678)Ribosomal_L31 (pfam01197)→ rpmF (PRK01110)Ribosomal_L32p (pfam01783)→ |
| env-11 | SRS058213_LANL_scaffold_5004 | - | 2005 | 1939 | **RNA**→ rpmE2 (PRK01678)Ribosomal_L31 (pfam01197)→ rpmF (PRK01110)Ribosomal_L32p (pfam01783)→ |
| Cac-1-1 | NZ_GG666998.1 | + | 69464 | 69530 | **RNA**→ rpmE2 (PRK01678)Ribosomal_L31 (pfam01197)→ rpmF (PRK01110)Ribosomal_L32p (pfam01783)→ |
| Csp-15-1 | NZ_KI515712.1 | + | 921794 | 921860 | **RNA**→ rpmE2 (PRK01678)Ribosomal_L31 (pfam01197)→ rpmF (PRK01110)Ribosomal_L32p (pfam01783)→ |
| Csp-17-1 | NZ_KI515704.1 | + | 80492 | 80558 | **RNA**→ rpmE2 (PRK01678)Ribosomal_L31 (pfam01197)→ rpmF (PRK01110)Ribosomal_L32p (pfam01783)→ |
| Csp-16-1 | NZ_KI515710.1 | - | 418615 | 418549 | **RNA**→ rpmE2 (PRK01678)Ribosomal_L31 (pfam01197)→ rpmF (PRK01110)Ribosomal_L32p (pfam01783)→ |
| Csp-12-1 | NZ_KI515719.1 | + | 80492 | 80558 | **RNA**→ rpmE2 (PRK01678)Ribosomal_L31 (pfam01197)→ rpmF (PRK01110)Ribosomal_L32p (pfam01783)→ |
| env-12 | SRS013876_WUGC_scaffold_1100 | - | 378 | 312 | **RNA**→ rpmE2 (PRK01678)Ribosomal_L31 (pfam01197)→ |
| Cac-2-1 | NZ_GL397138.1 | + | 875571 | 875637 | **RNA**→ rpmE2 (PRK01678)Ribosomal_L31 (pfam01197)→ rpmF (PRK01110)Ribosomal_L32p (pfam01783)→ |
| env-13 | SRS014682_WUGC_scaffold_5134 | - | 9672 | 9606 | **RNA**→ rpmE2 (PRK01678)Ribosomal_L31 (pfam01197)→ rpmF (PRK01110)Ribosomal_L32p (pfam01783)→ |
| env-14 | SRS014901_C267752 | + | 75 | 141 | **RNA**→ rpmE2 (PRK01678)Ribosomal_L31 (pfam01197)→ rpmF (PRK01110)Ribosomal_L32p (pfam01783)→ |
| env-15 | SRS017820_C151105 | - | 125 | 59 | **RNA**→ hypo→ |
| env-16 | SRS022006_Baylor_scaffold_3374 | - | 2342 | 2276 | **RNA**→ rpmE2 (PRK01678)Ribosomal_L31 (pfam01197)→ rpmF (PRK01110)Ribosomal_L32p (pfam01783)→ |
| env-17 | SRS047254_WUGC_scaffold_8049 | - | 515 | 449 | **RNA**→ rpmE2 (PRK01678)Ribosomal_L31 (pfam01197)→ rpmF (PRK01110)Ribosomal_L32p (pfam01783)→ |
| Csp-6-1 | NZ_KI515749.1 | + | 937901 | 937967 | **RNA**→ rpmE2 (PRK01678)Ribosomal_L31 (pfam01197)→ rpmF (PRK01110)Ribosomal_L32p (pfam01783)→ |
| env-18 | SRS044474_LANL_scaffold_86 | + | 548 | 614 | **RNA**→ rpmE2 (PRK01678)Ribosomal_L31 (pfam01197)→ rpmF (PRK01110)Ribosomal_L32p (pfam01783)→ |
| Csp-7-1 | NZ_KI515742.1 | - | 405839 | 405773 | **RNA**→ rpmE2 (PRK01678)Ribosomal_L31 (pfam01197)→ rpmF (PRK01110)Ribosomal_L32p (pfam01783)→ |
| Csp-10-1 | NZ_KI515730.1 | + | 76326 | 76392 | **RNA**→ rpmE2 (PRK01678)Ribosomal_L31 (pfam01197)→ rpmF (PRK01110)Ribosomal_L32p (pfam01783)→ |
| Csp-2-1 | NZ_KI515775.1 | + | 77719 | 77785 | **RNA**→ rpmE2 (PRK01678)Ribosomal_L31 (pfam01197)→ rpmF (PRK01110)Ribosomal_L32p (pfam01783)→ |
| Cau-2-2 | NZ_JVXO01000017.1 | - | 651 | 592 | **RNA**→ rpmE2 (PRK01678)Ribosomal_L31 (pfam01197)→ rpmF (PRK01110)Ribosomal_L32p (pfam01783)→ |
| env-19 | L_ear_crease_WUGI_scaffold_15403 | + | 353 | 412 | **RNA**→ rpmE2 (PRK01678)Ribosomal_L31 (pfam01197)→ |
| env-20 | SRS019015_C240949 | + | 347 | 406 | **RNA**→ rpmE2 (PRK01678)Ribosomal_L31 (pfam01197)→ |
| env-21 | SRS019063_C605280 | - | 264 | 205 | **RNA**→ rpmE2 (PRK01678)Ribosomal_L31 (pfam01197)→ |
| env-22 | SRS019867_C148862 | - | 188 | 129 | **RNA**→ rpmE2 (PRK01678)Ribosomal_L31 (pfam01197)→ |
| env-23 | SRS058213_LANL_scaffold_8330 | - | 1796 | 1737 | **RNA**→ rpmE2 (PRK01678)Ribosomal_L31 (pfam01197)→ rpmF (PRK01110)Ribosomal_L32p (pfam01783)→ |
| Csp-11-1 | NZ_KI515722.1 | - | 400130 | 400071 | **RNA**→ rpmE2 (PRK01678)Ribosomal_L31 (pfam01197)→ rpmF (PRK01110)Ribosomal_L32p (pfam01783)→ |
| Csp-9-1 | NZ_KI515731.1 | - | 2393670 | 2393611 | **RNA**→ rpmE2 (PRK01678)Ribosomal_L31 (pfam01197)→ rpmF (PRK01110)Ribosomal_L32p (pfam01783)→ |
| Csp-5-1 | NZ_KI515752.1 | + | 62595 | 62654 | **RNA**→ rpmE2 (PRK01678)Ribosomal_L31 (pfam01197)→ rpmF (PRK01110)Ribosomal_L32p (pfam01783)→ |
| Csp-8-1 | NZ_KI515735.1 | + | 1155727 | 1155786 | **RNA**→ rpmE2 (PRK01678)Ribosomal_L31 (pfam01197)→ rpmF (PRK01110)Ribosomal_L32p (pfam01783)→ |
| Csp-3-1 | NZ_KI515767.1 | + | 58512 | 58571 | **RNA**→ rpmE2 (PRK01678)Ribosomal_L31 (pfam01197)→ rpmF (PRK01110)Ribosomal_L32p (pfam01783)→ |
| env-24 | SRS054061_C156877 | + | 428 | 487 | **RNA**→ |

| | | | | | |
|---|---|---|---|---|---|
| env-25 | SRS056906_LANL_scaffold_9004 | - | 11347 | 11278 | **RNA**→ rpmE2 (PRK01678)Ribosomal_L31 (pfam01197)→ rpmF (PRK01110)Ribosomal_L32p (pfam01783)→ |
| env-26 | SRS014682_C298104 | + | 265 | 334 | **RNA**→ |
| Cau-2-3 | NZ_JUZU01000034.1 | + | 26228 | 26285 | **RNA**→ rpmE2 (PRK01678)Ribosomal_L31 (pfam01197)→ rpmF (PRK01110)Ribosomal_L32p (pfam01783)→ |
| Cau-2-4 | NZ_JUOH01000050.1 | - | 648 | 591 | **RNA**→ rpmE2 (PRK01678)Ribosomal_L31 (pfam01197)→ rpmF (PRK01110)Ribosomal_L32p (pfam01783)→ |
| Ctu-1-1 | NZ_ACVP01000016.1 | + | 27341 | 27398 | **RNA**→ rpmE2 (PRK01678)Ribosomal_L31 (pfam01197)→ rpmE2 (PRK01678)Ribosomal_L31 (pfam01197)→ rpmF (PRK01110)Ribosomal_L32p (pfam01783)→ rpmF (PRK01110)Ribosomal_L32p (pfam01783)→ |
| Cps-2-1 | NZ_GL542875.1 | - | 416624 | 416567 | **RNA**→ rpmE2 (PRK01678)Ribosomal_L31 (pfam01197)→ rpmF (PRK01110)Ribosomal_L32p (pfam01783)→ |
| Cst-2-1 | NZ_LAYR01000011.1 | + | 31347 | 31452 | **RNA**→ rpmE2 (PRK01678)Ribosomal_L31 (pfam01197)→ rpmF (PRK01110)Ribosomal_L32p (pfam01783)→ |
| Cst-2-2 | NZ_JVTN01000092.1 | - | 89483 | 89378 | **RNA**→ rpmE2 (PRK01678)Ribosomal_L31 (pfam01197)→ rpmF (PRK01110)Ribosomal_L32p (pfam01783)→ |
| Cst-2-3 | NZ_JVTL01000033.1 | - | 36207 | 36102 | **RNA**→ rpmE2 (PRK01678)Ribosomal_L31 (pfam01197)→ rpmF (PRK01110)Ribosomal_L32p (pfam01783)→ |
| Cst-2-4 | NZ_JVBA01000054.1 | + | 47752 | 47857 | **RNA**→ rpmE2 (PRK01678)Ribosomal_L31 (pfam01197)→ rpmF (PRK01110)Ribosomal_L32p (pfam01783)→ |
| Cst-2-5 | NZ_JULW01000062.1 | + | 47806 | 47911 | **RNA**→ rpmE2 (PRK01678)Ribosomal_L31 (pfam01197)→ rpmF (PRK01110)Ribosomal_L32p (pfam01783)→ |
| Cst-2-6 | NZ_JULV01000025.1 | + | 47803 | 47908 | **RNA**→ rpmE2 (PRK01678)Ribosomal_L31 (pfam01197)→ rpmF (PRK01110)Ribosomal_L32p (pfam01783)→ |
| Cst-2-7 | NZ_JVCW01000084.1 | + | 75365 | 75470 | **RNA**→ rpmE2 (PRK01678)Ribosomal_L31 (pfam01197)→ rpmF (PRK01110)Ribosomal_L32p (pfam01783)→ |
| Cst-1-1 | NZ_GG667522.1 | - | 145531 | 145426 | **RNA**→ rpmE2 (PRK01678)Ribosomal_L31 (pfam01197)→ rpmF (PRK01110)Ribosomal_L32p (pfam01783)→ |
| Cst-2-8 | NZ_JUSN01000092.1 | - | 950 | 845 | **RNA**→ rpmE2 (PRK01678)Ribosomal_L31 (pfam01197)→ rpmF (PRK01110)Ribosomal_L32p (pfam01783)→ |
| env-27 | AglaG_GBVDVLI01BUUUY | - | 106 | 52 | **RNA**→ hypo→ |
| env-28 | AglaG_contig04927 | - | 168 | 114 | **RNA**→ rpmE2 (PRK01678)Ribosomal_L31 (pfam01197)→ |
| env-29 | AglaG_GDN60OX02HVGVM | + | 158 | 212 | **RNA**→ |
| env-30 | AglaG_GDN60OX02HYEDF | - | 352 | 298 | **RNA**→ rpmE2 (PRK01678)Ribosomal_L31 (pfam01197)→ |
| Cur-4-1 | NZ_CP009215.1 | + | 707148 | 707228 | **RNA**→ rpmE2 (PRK01678)Ribosomal_L31 (pfam01197)→ rpmF (PRK01110)Ribosomal_L32p (pfam01783)→ |
| Cih-1-1 | NZ_HG001322.1 | + | 57297 | 57372 | **RNA**→ rpmE2 (PRK01678)Ribosomal_L31 (pfam01197)→ rpmF (PRK01110)Ribosomal_L32p (pfam01783)→ |
| Cha-2-1 | NZ_JVMD01000015.1 | + | 131313 | 131426 | **RNA**→ rpmE2 (PRK01678)Ribosomal_L31 (pfam01197)→ rpmF (PRK01110)Ribosomal_L32p (pfam01783)→ |
| Cfr-1-1 | NZ_JRNE01000004.1 | - | 33488 | 33375 | **RNA**→ rpmE2 (PRK01678)Ribosomal_L31 (pfam01197)→ rpmF (PRK01110)Ribosomal_L32p (pfam01783)→ |
| env-31 | JGI24707J26582_10105430 | + | 592 | 705 | **RNA**→ rpmE2 (PRK01678)Ribosomal_L31 (pfam01197)→ |
| Cje-3-1 | NZ_CBYN010000125.1 | - | 2960 | 2903 | **RNA**→ rpmE2 (PRK01678)Ribosomal_L31 (pfam01197)→ rpmF (PRK01110)Ribosomal_L32p (pfam01783)→ |
| Cul-6-1 | NZ_CP011095.1 | + | 738588 | 738694 | **RNA**→ rpmE2 (PRK01678)Ribosomal_L31 (pfam01197)→ rpmF (PRK01110)Ribosomal_L32p (pfam01783)→ |
| Ctu-2-1 | NZ_JRNG01000018.1 | - | 53232 | 53172 | **RNA**→ rpmE2 (PRK01678)Ribosomal_L31 (pfam01197)→ rpmF (PRK01110)Ribosomal_L32p (pfam01783)→ |
| Cim-1-1 | NZ_CP009211.1 | + | 754015 | 754070 | **RNA**→ rpmE2 (PRK01678)Ribosomal_L31 (pfam01197)→ rpmF (PRK01110)Ribosomal_L32p (pfam01783)→ |
| Cri-1-1 | NZ_CP012342.1 | - | 1228345 | 1228290 | **RNA**→ rpmE2 (PRK01678)Ribosomal_L31 (pfam01197)→ rpmF (PRK01110)Ribosomal_L32p (pfam01783)→ |
| Cge-1-1 | NZ_CM000961.1 | - | 2235924 | 2235848 | **RNA**→ rpmE2 (PRK01678)Ribosomal_L31 (pfam01197)→ rpmF (PRK01110)Ribosomal_L32p (pfam01783)→ |
| Cul-5-1 | NZ_AQUY01000002.1 | + | 105764 | 105849 | **RNA**→ rpmE2 (PRK01678)Ribosomal_L31 (pfam01197)→ rpmF (PRK01110)Ribosomal_L32p (pfam01783)→ |
| Cca-2-1 | NZ_AQUV01000003.1 | - | 337464 | 337414 | **RNA**→ rpmE2 (PRK01678)Ribosomal_L31 (pfam01197)→ rpmF (PRK01110)Ribosomal_L32p (pfam01783)→ REC (smart00448)Trans_reg_C (smart00862)OmpR (COG0745)→ Trans_reg_C (smart00862)OmpR (COG0745)→ DegQ (COG0265)PDZ_2 (pfam13180)Trypsin_2 (pfam13365)→ DegQ (COG0265)PDZ_2 (pfam13180)Trypsin_2 (pfam13365)→ MoCF_biosynth (smart00852)MogA_MoaB (cd00886)→ |
| Cti-1-1 | NZ_HE998586.1 | - | 222294 | 222239 | **RNA**→ rpmE2 (PRK01678)Ribosomal_L31 (pfam01197)→ rpmF (PRK01110)Ribosomal_L32p (pfam01783)→ |
| Cli-1-1 | NZ_GG667196.1 | - | 108598 | 108535 | **RNA**→ rpmE2 (PRK01678)Ribosomal_L31 (pfam01197)→ rpmF (PRK01110)Ribosomal_L32p (pfam01783)→ |
| Cps-17-1 | NZ_CP012022.1 | + | 703166 | 703274 | **RNA**→ rpmE2 (PRK01678)Ribosomal_L31 (pfam01197)→ rpmF (PRK01110)Ribosomal_L32p (pfam01783)→ |
| Cps-12-1 | NC_014329.1 | + | 696776 | 696884 | **RNA**→ rpmE2 (PRK01678)Ribosomal_L31 (pfam01197)→ rpmF (PRK01110)Ribosomal_L32p (pfam01783)→ |
| Cps-13-1 | NC_017303.1 | + | 696588 | 696696 | **RNA**→ rpmE2 (PRK01678)Ribosomal_L31 (pfam01197)→ rpmF (PRK01110)Ribosomal_L32p (pfam01783)→ |
| Cps-15-1 | NC_017305.1 | + | 696644 | 696752 | **RNA**→ rpmE2 (PRK01678)Ribosomal_L31 (pfam01197)→ rpmE2 (PRK01678)Ribosomal_L31 (pfam01197)→ rpmF (PRK01110)Ribosomal_L32p (pfam01783)→ |
| Cps-8-1 | NC_017306.1 | + | 696771 | 696879 | **RNA**→ rpmE2 (PRK01678)Ribosomal_L31 (pfam01197)→ rpmF (PRK01110)Ribosomal_L32p (pfam01783)→ |
| Cps-6-1 | NC_016781.1 | + | 696768 | 696876 | **RNA**→ rpmE2 (PRK01678)Ribosomal_L31 (pfam01197)→ rpmF (PRK01110)Ribosomal_L32p (pfam01783)→ |
| Cps-14-1 | NC_017031.1 | + | 696696 | 696804 | **RNA**→ rpmE2 (PRK01678)Ribosomal_L31 (pfam01197)→ rpmE2 (PRK01678)Ribosomal_L31 (pfam01197)→ rpmF (PRK01110)Ribosomal_L32p (pfam01783)→ |
| Cps-5-1 | NC_017462.1 | + | 696591 | 696699 | **RNA**→ rpmE2 (PRK01678)Ribosomal_L31 (pfam01197)→ rpmF (PRK01110)Ribosomal_L32p (pfam01783)→ |
| Cps-17-2 | NZ_CP009927.1 | + | 696736 | 696844 | **RNA**→ rpmE2 (PRK01678)Ribosomal_L31 (pfam01197)→ rpmF (PRK01110)Ribosomal_L32p (pfam01783)→ |
| Cps-9-1 | NC_017301.1 | + | 696442 | 696550 | **RNA**→ rpmE2 (PRK01678)Ribosomal_L31 (pfam01197)→ rpmF (PRK01110)Ribosomal_L32p (pfam01783)→ |
| Cps-4-1 | NC_017300.1 | + | 696689 | 696797 | **RNA**→ rpmE2 (PRK01678)Ribosomal_L31 (pfam01197)→ rpmF (PRK01110)Ribosomal_L32p (pfam01783)→ |
| Cps-17-3 | NZ_CP008924.1 | + | 1234105 | 1234213 | **RNA**→ rpmE2 (PRK01678)Ribosomal_L31 (pfam01197)→ rpmF (PRK01110)Ribosomal_L32p (pfam01783)→ |

| | | | | | |
|---|---|---|---|---|---|
| Cps-17-4 | NZ_CP008922.1 | + | 1250687 | 1250795 | RNA→ rpmE2 (PRK01678)Ribosomal_L31 (pfam01197)→ rpmF (PRK01110)Ribosomal_L32p (pfam01783)→ |
| Cps-17-5 | NZ_CP008923.1 | - | 345645 | 345537 | RNA→ rpmE2 (PRK01678)Ribosomal_L31 (pfam01197)→ rpmF (PRK01110)Ribosomal_L32p (pfam01783)→ |
| Cps-17-6 | NZ_CP010889.1 | + | 696683 | 696791 | RNA→ rpmE2 (PRK01678)Ribosomal_L31 (pfam01197)→ rpmF (PRK01110)Ribosomal_L32p (pfam01783)→ |
| Cps-17-7 | NZ_CP011474.1 | + | 696758 | 696866 | RNA→ rpmE2 (PRK01678)Ribosomal_L31 (pfam01197)→ rpmF (PRK01110)Ribosomal_L32p (pfam01783)→ |
| Cps-17-8 | NZ_CP010795.1 | - | 2156721 | 2156613 | RNA→ rpmE2 (PRK01678)Ribosomal_L31 (pfam01197)→ rpmF (PRK01110)Ribosomal_L32p (pfam01783)→ |
| Cku-1-1 | NZ_CP011312.1 | - | 1807067 | 1807007 | RNA→ rpmE2 (PRK01678)Ribosomal_L31 (pfam01197)→ rpmF (PRK01110)Ribosomal_L32p (pfam01783)→ |
| Cam-1-1 | NZ_GG771288.1 | - | 137801 | 137737 | RNA→ rpmE2 (PRK01678)Ribosomal_L31 (pfam01197)→ rpmF (PRK01110)Ribosomal_L32p (pfam01783)→ |
| Cvi-2-1 | NZ_CP004353.1 | + | 974041 | 974128 | RNA→ rpmE2 (PRK01678)Ribosomal_L31 (pfam01197)→ rpmF (PRK01110)Ribosomal_L32p (pfam01783)→ |
| Cvi-3-1 | NZ_JNKV01000010.1 | + | 35324 | 35411 | RNA→ rpmE2 (PRK01678)Ribosomal_L31 (pfam01197)→ rpmF (PRK01110)Ribosomal_L32p (pfam01783)→ |
| env-32 | JGI24707J26582_10216171 | - | 482 | 418 | RNA→ rpmE2 (PRK01678)Ribosomal_L31 (pfam01197)→ rpmF (PRK01110)Ribosomal_L32p (pfam01783)→ |
| Cmu-1-1 | NZ_CP011542.1 | + | 1034276 | 1034329 | RNA→ rpmE2 (PRK01678)Ribosomal_L31 (pfam01197)→ rpmF (PRK01110)Ribosomal_L32p (pfam01783)→ |
| Cca-3-1 | NZ_CAFW01000081.1 | + | 66998 | 67062 | RNA→ rpmE2 (PRK01678)Ribosomal_L31 (pfam01197)→ rpmE2 (PRK01678)Ribosomal_L31 (pfam01197)→ rpmF (PRK01110)Ribosomal_L32p (pfam01783)→ rpmF (PRK01110)Ribosomal_L32p (pfam01783)→ |
| Cca-6-1 | NZ_CP004350.1 | - | 2180252 | 2180188 | RNA→ rpmE2 (PRK01678)Ribosomal_L31 (pfam01197)→ rpmF (PRK01110)Ribosomal_L32p (pfam01783)→ |
| env-33 | 2205737659 | + | 155 | 219 | RNA→ rpmE2 (PRK01678)Ribosomal_L31 (pfam01197)→ rpmF (PRK01110)Ribosomal_L32p (pfam01783)→ |
| env-34 | 2158166148 | + | 155 | 219 | RNA→ rpmE2 (PRK01678)Ribosomal_L31 (pfam01197)→ rpmF (PRK01110)Ribosomal_L32p (pfam01783)→ |
| env-35 | Buccal_mucosa_LANL_scaffold_147023 | + | 1044 | 1107 | RNA→ rpmE2 (PRK01678)Ribosomal_L31 (pfam01197)→ rpmF (PRK01110)Ribosomal_L32p (pfam01783)→ |
| env-36 | SRS015064_C1740324 | - | 3475 | 3412 | RNA→ rpmE2 (PRK01678)Ribosomal_L31 (pfam01197)→ rpmF (PRK01110)Ribosomal_L32p (pfam01783)→ |
| env-37 | SRS017076_C2591228 | - | 210 | 147 | RNA→ rpmE2 (PRK01678)Ribosomal_L31 (pfam01197)→ |
| env-38 | SRS022602_Baylor_scaffold_15996 | + | 4561 | 4624 | RNA→ rpmE2 (PRK01678)Ribosomal_L31 (pfam01197)→ rpmF (PRK01110)Ribosomal_L32p (pfam01783)→ |
| env-39 | SRS023841_C1528757 | - | 216 | 153 | RNA→ rpmE2 (PRK01678)Ribosomal_L31 (pfam01197)→ |
| env-40 | SRS044366_C952506 | + | 2004 | 2067 | RNA→ rpmE2 (PRK01678)Ribosomal_L31 (pfam01197)→ rpmF (PRK01110)Ribosomal_L32p (pfam01783)→ |
| env-41 | SRS053630_LANL_scaffold_21375 | - | 55473 | 55410 | RNA→ rpmE2 (PRK01678)Ribosomal_L31 (pfam01197)→ rpmF (PRK01110)Ribosomal_L32p (pfam01783)→ |
| env-42 | Subgingival_plaque_LANL_C11219698 | - | 389 | 326 | RNA→ rpmE2 (PRK01678)Ribosomal_L31 (pfam01197)→ |
| env-43 | SRS011098_Baylor_scaffold_21411 | - | 9309 | 9246 | RNA→ rpmE2 (PRK01678)Ribosomal_L31 (pfam01197)→ rpmF (PRK01110)Ribosomal_L32p (pfam01783)→ |
| env-44 | SRS011152_Baylor_scaffold_27991 | + | 1045 | 1108 | RNA→ rpmE2 (PRK01678)Ribosomal_L31 (pfam01197)→ rpmF (PRK01110)Ribosomal_L32p (pfam01783)→ |
| env-45 | SRS011343_C2262393 | + | 14035 | 14098 | RNA→ rpmE2 (PRK01678)Ribosomal_L31 (pfam01197)→ rpmF (PRK01110)Ribosomal_L32p (pfam01783)→ |
| env-46 | SRS012285_Baylor_scaffold_4751 | + | 7940 | 8003 | RNA→ rpmE2 (PRK01678)Ribosomal_L31 (pfam01197)→ rpmF (PRK01110)Ribosomal_L32p (pfam01783)→ |
| env-47 | SRS013170_Baylor_scaffold_53613 | - | 476 | 413 | RNA→ rpmE2 (PRK01678)Ribosomal_L31 (pfam01197)→ rpmF (PRK01110)Ribosomal_L32p (pfam01783)→ |
| env-48 | SRS013533_C5500247 | + | 357 | 420 | RNA→ rpmE2 (PRK01678)Ribosomal_L31 (pfam01197)→ rpmF (PRK01110)→ |
| env-49 | SRS013723_Baylor_scaffold_39673 | + | 2404 | 2467 | RNA→ rpmE2 (PRK01678)Ribosomal_L31 (pfam01197)→ rpmF (PRK01110)Ribosomal_L32p (pfam01783)→ |
| env-50 | SRS013836_C4755765 | + | 181 | 244 | RNA→ rpmE2 (PRK01678)Ribosomal_L31 (pfam01197)→ |
| env-51 | SRS013949_WUGC_scaffold_27456 | - | 199 | 136 | RNA→ rpmE2 (PRK01678)Ribosomal_L31 (pfam01197)→ |
| env-52 | SRS013950_C2187510 | + | 1205 | 1268 | RNA→ rpmE2 (PRK01678)Ribosomal_L31 (pfam01197)→ rpmF (PRK01110)Ribosomal_L32p (pfam01783)→ |
| env-53 | SRS014107_C1342401 | - | 235 | 172 | RNA→ rpmE2 (PRK01678)Ribosomal_L31 (pfam01197)→ |
| env-54 | SRS014473_C1089635 | + | 295 | 358 | RNA→ rpmE2 (PRK01678)Ribosomal_L31 (pfam01197)→ |
| env-55 | SRS014476_WUGC_scaffold_41168 | - | 1375 | 1312 | RNA→ rpmE2 (PRK01678)Ribosomal_L31 (pfam01197)→ rpmF (PRK01110)Ribosomal_L32p (pfam01783)→ |
| env-56 | SRS014477_C1645642 | - | 1867 | 1804 | RNA→ rpmE2 (PRK01678)Ribosomal_L31 (pfam01197)→ rpmF (PRK01110)Ribosomal_L32p (pfam01783)→ |
| env-57 | SRS014578_WUGC_scaffold_44820 | + | 551 | 614 | RNA→ rpmE2 (PRK01678)Ribosomal_L31 (pfam01197)→ |
| env-58 | SRS014690_C1892883 | - | 401 | 338 | RNA→ rpmE2 (PRK01678)Ribosomal_L31 (pfam01197)→ |
| env-59 | SRS014691_WUGC_scaffold_10572 | + | 275 | 338 | RNA→ rpmE2 (PRK01678)Ribosomal_L31 (pfam01197)→ |
| env-60 | SRS014894_WUGC_scaffold_28738 | + | 22786 | 22849 | RNA→ rpmE2 (PRK01678)Ribosomal_L31 (pfam01197)→ rpmF (PRK01110)Ribosomal_L32p (pfam01783)→ |
| env-61 | SRS015044_C2942561 | + | 357 | 420 | RNA→ rpmE2 (PRK01678)Ribosomal_L31 (pfam01197)→ |
| env-62 | SRS015060_C1560553 | + | 198 | 261 | RNA→ rpmE2 (PRK01678)Ribosomal_L31 (pfam01197)→ rpmF (PRK01110)Ribosomal_L32p (pfam01783)→ |
| env-63 | SRS015063_WUGC_scaffold_11905 | + | 3939 | 4002 | RNA→ rpmE2 (PRK01678)Ribosomal_L31 (pfam01197)→ rpmF (PRK01110)→ |
| env-64 | SRS015158_WUGC_scaffold_58214 | + | 860 | 923 | RNA→ rpmE2 (PRK01678)Ribosomal_L31 (pfam01197)→ rpmF (PRK01110)Ribosomal_L32p (pfam01783)→ |
| env-65 | SRS015215_WUGC_scaffold_18972 | + | 1864 | 1927 | RNA→ rpmE2 (PRK01678)Ribosomal_L31 (pfam01197)→ rpmF (PRK01110)Ribosomal_L32p (pfam01783)→ |
| env-66 | SRS015278_WUGC_scaffold_11862 | - | 58024 | 57961 | RNA→ rpmE2 (PRK01678)Ribosomal_L31 (pfam01197)→ rpmF (PRK01110)Ribosomal_L32p (pfam01783)→ |
| env-67 | SRS015378_WUGC_scaffold_7793 | + | 1275 | 1338 | RNA→ rpmE2 (PRK01678)Ribosomal_L31 (pfam01197)→ rpmF (PRK01110)Ribosomal_L32p (pfam01783)→ |
| env-68 | SRS015440_WUGC_scaffold_49340 | - | 2220 | 2157 | RNA→ rpmE2 (PRK01678)Ribosomal_L31 (pfam01197)→ rpmF (PRK01110)Ribosomal_L32p (pfam01783)→ |
| env-69 | SRS015470_WUGC_scaffold_45392 | - | 1480 | 1417 | RNA→ rpmE2 (PRK01678)Ribosomal_L31 (pfam01197)→ rpmF (PRK01110)Ribosomal_L32p (pfam01783)→ |
| env-70 | SRS015574_WUGC_scaffold_2816 | + | 74953 | 75016 | RNA→ rpmE2 (PRK01678)Ribosomal_L31 (pfam01197)→ rpmF (PRK01110)Ribosomal_L32p (pfam01783)→ |

| | | | | | | |
|---|---|---|---|---|---|---|
| env-71 | SRS015755_WUGC_scaffold_44984 | + | 1196 | 1259 | RNA→ | rpmE2 (PRK01678)Ribosomal_L31 (pfam01197)→ rpmF (PRK01110)Ribosomal_L32p (pfam01783)→ |
| env-72 | SRS015989_WUGC_scaffold_32775 | - | 5102 | 5039 | RNA→ | rpmE2 (PRK01678)Ribosomal_L31 (pfam01197)→ rpmF (PRK01110)Ribosomal_L32p (pfam01783)→ |
| env-73 | SRS016043_WUGC_scaffold_12513 | - | 56767 | 56704 | RNA→ | rpmE2 (PRK01678)Ribosomal_L31 (pfam01197)→ rpmF (PRK01110)Ribosomal_L32p (pfam01783)→ |
| env-74 | SRS016200_WUGC_scaffold_38664 | - | 3995 | 3932 | RNA→ | rpmE2 (PRK01678)Ribosomal_L31 (pfam01197)→ rpmF (PRK01110)Ribosomal_L32p (pfam01783)→ |
| env-75 | SRS016360_Baylor_scaffold_23915 | - | 10968 | 10905 | RNA→ | rpmE2 (PRK01678)Ribosomal_L31 (pfam01197)→ rpmF (PRK01110)Ribosomal_L32p (pfam01783)→ |
| env-76 | SRS016575_Baylor_scaffold_1786 | - | 7196 | 7133 | RNA→ | rpmE2 (PRK01678)Ribosomal_L31 (pfam01197)→ rpmF (PRK01110)Ribosomal_L32p (pfam01783)→ |
| env-77 | SRS016746_Baylor_scaffold_72724 | + | 7808 | 7871 | RNA→ | rpmE2 (PRK01678)Ribosomal_L31 (pfam01197)→ rpmF (PRK01110)Ribosomal_L32p (pfam01783)→ |
| env-78 | SRS017025_Baylor_scaffold_19305 | - | 682 | 619 | RNA→ | rpmE2 (PRK01678)Ribosomal_L31 (pfam01197)→ rpmF (PRK01110)Ribosomal_L32p (pfam01783)→ |
| env-79 | SRS017227_C5283816 | + | 851 | 914 | RNA→ | ←hypo |
| env-80 | SRS017445_Baylor_scaffold_45380 | - | 6544 | 6481 | RNA→ | rpmE2 (PRK01678)Ribosomal_L31 (pfam01197)→ rpmF (PRK01110)Ribosomal_L32p (pfam01783)→ |
| env-81 | SRS017511_Baylor_scaffold_42338 | + | 1176 | 1239 | RNA→ | rpmE2 (PRK01678)Ribosomal_L31 (pfam01197)→ rpmF (PRK01110)Ribosomal_L32p (pfam01783)→ |
| env-82 | SRS017687_C884952 | + | 126 | 189 | RNA→ | rpmE2 (PRK01678)Ribosomal_L31 (pfam01197)→ rpmF (PRK01110)Ribosomal_L32p (pfam01783)→ |
| env-83 | SRS018337_C2391064 | + | 863 | 926 | RNA→ | rpmE2 (PRK01678)Ribosomal_L31 (pfam01197)→ rpmF (PRK01110)Ribosomal_L32p (pfam01783)→ |
| env-84 | SRS018573_WUGC_scaffold_24099 | - | 238 | 175 | RNA→ | rpmE2 (PRK01678)Ribosomal_L31 (pfam01197)→ |
| env-85 | SRS018665_WUGC_scaffold_21586 | - | 2301 | 2238 | RNA→ | rpmE2 (PRK01678)Ribosomal_L31 (pfam01197)→ rpmF (PRK01110)Ribosomal_L32p (pfam01783)→ |
| env-86 | SRS019025_WUGC_scaffold_5953 | - | 310 | 247 | RNA→ | rpmE2 (PRK01678)Ribosomal_L31 (pfam01197)→ |
| env-87 | SRS019028_WUGC_scaffold_36029 | - | 523 | 460 | RNA→ | rpmE2 (PRK01678)Ribosomal_L31 (pfam01197)→ rpmF (PRK01110)Ribosomal_L32p (pfam01783)→ |
| env-88 | SRS019029_C3309553 | + | 2423 | 2486 | RNA→ | rpmE2 (PRK01678)Ribosomal_L31 (pfam01197)→ rpmF (PRK01110)Ribosomal_L32p (pfam01783)→ |
| env-89 | SRS019128_WUGC_scaffold_42508 | + | 1205 | 1268 | RNA→ | rpmE2 (PRK01678)Ribosomal_L31 (pfam01197)→ rpmF (PRK01110)Ribosomal_L32p (pfam01783)→ |
| env-90 | SRS019129_WUGC_scaffold_20241 | - | 3656 | 3593 | RNA→ | rpmE2 (PRK01678)Ribosomal_L31 (pfam01197)→ rpmF (PRK01110)Ribosomal_L32p (pfam01783)→ |
| env-91 | SRS019225_WUGC_scaffold_23108 | + | 4861 | 4924 | RNA→ | rpmE2 (PRK01678)Ribosomal_L31 (pfam01197)→ rpmF (PRK01110)Ribosomal_L32p (pfam01783)→ |
| env-92 | SRS019333_C1208867 | + | 219 | 282 | RNA→ | |
| env-93 | SRS019387_WUGC_scaffold_16046 | + | 3939 | 4002 | RNA→ | rpmE2 (PRK01678)Ribosomal_L31 (pfam01197)→ rpmF (PRK01110)Ribosomal_L32p (pfam01783)→ |
| env-94 | SRS019591_WUGC_scaffold_40585 | + | 1176 | 1239 | RNA→ | rpmE2 (PRK01678)Ribosomal_L31 (pfam01197)→ rpmF (PRK01110)Ribosomal_L32p (pfam01783)→ |
| env-95 | SRS019906_WUGC_scaffold_28644 | - | 3887 | 3824 | RNA→ | rpmE2 (PRK01678)Ribosomal_L31 (pfam01197)→ rpmF (PRK01110)Ribosomal_L32p (pfam01783)→ |
| env-96 | SRS019980_Baylor_scaffold_30280 | - | 2562 | 2499 | RNA→ | rpmE2 (PRK01678)Ribosomal_L31 (pfam01197)→ rpmF (PRK01110)Ribosomal_L32p (pfam01783)→ |
| env-97 | SRS020220_C3511826 | + | 271 | 334 | RNA→ | ←hypo |
| env-98 | SRS020226_Baylor_scaffold_55927 | - | 3611 | 3548 | RNA→ | rpmE2 (PRK01678)Ribosomal_L31 (pfam01197)→ rpmF (PRK01110)Ribosomal_L32p (pfam01783)→ |
| env-99 | SRS020334_C2557741 | + | 249 | 312 | RNA→ | |
| env-100 | SRS020862_Baylor_scaffold_27689 | - | 8855 | 8792 | RNA→ | rpmE2 (PRK01678)Ribosomal_L31 (pfam01197)→ rpmF (PRK01110)Ribosomal_L32p (pfam01783)→ |
| env-101 | SRS021477_Baylor_scaffold_22850 | - | 15915 | 15852 | RNA→ | rpmE2 (PRK01678)Ribosomal_L31 (pfam01197)→ rpmF (PRK01110)Ribosomal_L32p (pfam01783)→ |
| env-102 | SRS021960_Baylor_scaffold_7131 | - | 2351 | 2288 | RNA→ | rpmE2 (PRK01678)Ribosomal_L31 (pfam01197)→ rpmF (PRK01110)Ribosomal_L32p (pfam01783)→ |
| env-103 | SRS022083_Baylor_scaffold_11512 | - | 55855 | 55792 | RNA→ | rpmE2 (PRK01678)Ribosomal_L31 (pfam01197)→ rpmF (PRK01110)Ribosomal_L32p (pfam01783)→ |
| env-104 | SRS022149_LANL_scaffold_20816 | + | 1063 | 1126 | RNA→ | rpmE2 (PRK01678)Ribosomal_L31 (pfam01197)→ rpmF (PRK01110)Ribosomal_L32p (pfam01783)→ |
| env-105 | SRS022536_LANL_scaffold_114388 | - | 1321 | 1258 | RNA→ | rpmE2 (PRK01678)Ribosomal_L31 (pfam01197)→ rpmF (PRK01110)Ribosomal_L32p (pfam01783)→ |
| env-106 | SRS022725_LANL_scaffold_19220 | + | 1891 | 1954 | RNA→ | rpmE2 (PRK01678)Ribosomal_L31 (pfam01197)→ |
| env-107 | SRS023358_C3452087 | - | 361 | 298 | RNA→ | rpmE2 (PRK01678)Ribosomal_L31 (pfam01197)→ |
| env-108 | SRS023538_C1310182 | - | 1371 | 1308 | RNA→ | rpmE2 (PRK01678)Ribosomal_L31 (pfam01197)→ rpmF (PRK01110)Ribosomal_L32p (pfam01783)→ |
| env-109 | SRS023595_Baylor_scaffold_84979 | + | 919 | 982 | RNA→ | rpmE2 (PRK01678)Ribosomal_L31 (pfam01197)→ rpmF (PRK01110)→ |
| env-110 | SRS023938_Baylor_scaffold_15844 | - | 12477 | 12414 | RNA→ | rpmE2 (PRK01678)Ribosomal_L31 (pfam01197)→ rpmF (PRK01110)Ribosomal_L32p (pfam01783)→ |
| env-111 | SRS023964_Baylor_scaffold_34977 | + | 1176 | 1239 | RNA→ | rpmE2 (PRK01678)Ribosomal_L31 (pfam01197)→ rpmF (PRK01110)Ribosomal_L32p (pfam01783)→ |
| env-112 | SRS024021_Baylor_scaffold_16881 | + | 11059 | 11122 | RNA→ | rpmE2 (PRK01678)Ribosomal_L31 (pfam01197)→ rpmF (PRK01110)Ribosomal_L32p (pfam01783)→ |
| env-113 | SRS024087_LANL_scaffold_41039 | - | 238 | 175 | RNA→ | rpmE2 (PRK01678)Ribosomal_L31 (pfam01197)→ |
| env-114 | SRS024289_LANL_scaffold_12931 | - | 91217 | 91154 | RNA→ | rpmE2 (PRK01678)Ribosomal_L31 (pfam01197)→ rpmF (PRK01110)Ribosomal_L32p (pfam01783)→ |
| env-115 | SRS024381_C2286130 | + | 294 | 357 | RNA→ | |
| env-116 | SRS024447_LANL_scaffold_37014 | - | 3896 | 3833 | RNA→ | rpmE2 (PRK01678)Ribosomal_L31 (pfam01197)→ rpmF (PRK01110)Ribosomal_L32p (pfam01783)→ |
| env-117 | SRS024561_LANL_scaffold_31049 | + | 1456 | 1519 | RNA→ | rpmE2 (PRK01678)Ribosomal_L31 (pfam01197)→ rpmF (PRK01110)Ribosomal_L32p (pfam01783)→ |
| env-118 | SRS024649_LANL_scaffold_19762 | - | 2266 | 2203 | RNA→ | rpmE2 (PRK01678)Ribosomal_L31 (pfam01197)→ rpmF (PRK01110)Ribosomal_L32p (pfam01783)→ |
| env-119 | SRS042984_LANL_scaffold_1303 | - | 1490 | 1427 | RNA→ | rpmE2 (PRK01678)Ribosomal_L31 (pfam01197)→ rpmF (PRK01110)Ribosomal_L32p (pfam01783)→ |
| env-120 | SRS043018_WUGC_scaffold_35426 | + | 991 | 1054 | RNA→ | rpmE2 (PRK01678)Ribosomal_L31 (pfam01197)→ rpmF (PRK01110)Ribosomal_L32p (pfam01783)→ |
| env-121 | SRS043772_WUGC_scaffold_25647 | + | 1553 | 1616 | RNA→ | rpmE2 (PRK01678)Ribosomal_L31 (pfam01197)→ rpmF (PRK01110)Ribosomal_L32p (pfam01783)→ |
| env-122 | SRS047100_WUGC_scaffold_20208 | + | 1176 | 1239 | RNA→ | rpmE2 (PRK01678)Ribosomal_L31 (pfam01197)→ rpmF (PRK01110)Ribosomal_L32p (pfam01783)→ |

| | | | | | |
|---|---|---|---|---|---|
| env-123 | SRS049268_LANL_scaffold_56284 | + | 1050 | 1113 | **RNA**→ rpmE2 (PRK01678)Ribosomal_L31 (pfam01197)→ rpmF (PRK01110)Ribosomal_L32p (pfam01783)→ |
| env-124 | SRS049318_LANL_scaffold_74226 | + | 1275 | 1338 | **RNA**→ ←hypo |
| env-125 | SRS051244_LANL_scaffold_15573 | - | 9216 | 9153 | **RNA**→ rpmE2 (PRK01678)Ribosomal_L31 (pfam01197)→ rpmF (PRK01110)Ribosomal_L32p (pfam01783)→ |
| env-126 | SRS051378_LANL_scaffold_28522 | - | 3634 | 3571 | **RNA**→ rpmE2 (PRK01678)Ribosomal_L31 (pfam01197)→ rpmF (PRK01110)Ribosomal_L32p (pfam01783)→ |
| env-127 | SRS051930_LANL_scaffold_9996 | - | 7943 | 7880 | **RNA**→ rpmE2 (PRK01678)Ribosomal_L31 (pfam01197)→ rpmF (PRK01110)Ribosomal_L32p (pfam01783)→ |
| env-128 | SRS052604_LANL_scaffold_15789 | - | 3462 | 3399 | **RNA**→ rpmE2 (PRK01678)Ribosomal_L31 (pfam01197)→ rpmF (PRK01110)Ribosomal_L32p (pfam01783)→ |
| env-129 | SRS052876_LANL_scaffold_11131 | - | 4560 | 4497 | **RNA**→ rpmE2 (PRK01678)Ribosomal_L31 (pfam01197)→ rpmF (PRK01110)Ribosomal_L32p (pfam01783)→ |
| env-130 | SRS054430_LANL_scaffold_8962 | - | 182 | 119 | **RNA**→ ←hypo |
| env-131 | SRS054653_LANL_scaffold_17280 | - | 2173 | 2110 | **RNA**→ rpmE2 (PRK01678)Ribosomal_L31 (pfam01197)→ rpmF (PRK01110)Ribosomal_L32p (pfam01783)→ |
| env-132 | SRS055378_LANL_scaffold_71521 | + | 1515 | 1578 | **RNA**→ rpmE2 (PRK01678)Ribosomal_L31 (pfam01197)→ rpmF (PRK01110)Ribosomal_L32p (pfam01783)→ |
| env-133 | SRS055401_LANL_scaffold_40435 | + | 268 | 331 | **RNA**→ rpmE2 (PRK01678)Ribosomal_L31 (pfam01197)→ rpmF (PRK01110)Ribosomal_L32p (pfam01783)→ |
| env-134 | SRS055450_LANL_scaffold_48342 | - | 746 | 683 | **RNA**→ rpmE2 (PRK01678)Ribosomal_L31 (pfam01197)→ rpmF (PRK01110)Ribosomal_L32p (pfam01783)→ |
| env-135 | SRS058053_LANL_scaffold_20722 | - | 3271 | 3208 | **RNA**→ rpmE2 (PRK01678)Ribosomal_L31 (pfam01197)→ rpmF (PRK01110)Ribosomal_L32p (pfam01783)→ |
| env-136 | SRS058808_LANL_scaffold_76023 | - | 908 | 845 | **RNA**→ rpmE2 (PRK01678)Ribosomal_L31 (pfam01197)→ rpmF (PRK01110)Ribosomal_L32p (pfam01783)→ |
| env-137 | SRS063215_C2240857 | + | 181 | 244 | **RNA**→ rpmE2 (PRK01678)Ribosomal_L31 (pfam01197)→ rpmF (PRK01110)Ribosomal_L32p (pfam01783)→ |
| env-138 | SRS063932_C3715652 | - | 110 | 47 | **RNA**→ hypo→ |
| env-139 | SRS063999_C3497605 | + | 134 | 197 | **RNA**→ rpmE2 (PRK01678)Ribosomal_L31 (pfam01197)→ |
| env-140 | SRS065099_LANL_scaffold_2670 | + | 5358 | 5421 | **RNA**→ rpmE2 (PRK01678)Ribosomal_L31 (pfam01197)→ rpmF (PRK01110)Ribosomal_L32p (pfam01783)→ |
| env-141 | SRS065310_LANL_scaffold_16984 | - | 1076 | 1013 | **RNA**→ rpmE2 (PRK01678)Ribosomal_L31 (pfam01197)→ rpmF (PRK01110)Ribosomal_L32p (pfam01783)→ |
| Cma-4-1 | NZ_ACSH02000008.1 | - | 471190 | 471127 | **RNA**→ rpmE2 (PRK01678)Ribosomal_L31 (pfam01197)→ rpmE2 (PRK01678)Ribosomal_L31 (pfam01197)→<br>rpmF (PRK01110)Ribosomal_L32p (pfam01783)→ rpmF (PRK01110)Ribosomal_L32p (pfam01783)→ |
| Cma-5-1 | NZ_EQ973332.1 | - | 470329 | 470266 | **RNA**→ rpmE2 (PRK01678)Ribosomal_L31 (pfam01197)→ rpmF (PRK01110)Ribosomal_L32p (pfam01783)→<br>hypo→ REC (smart00448)Trans_reg_C (smart00862)OmpR (COG0745)→<br>HAMP (smart00304)HATPase_c (smart00387)HisKA (smart00388)BaeS (COG0642)→ |
| env-142 | SRS017139_Baylor_scaffold_46230 | + | 1104 | 1167 | **RNA**→ rpmE2 (PRK01678)Ribosomal_L31 (pfam01197)→ rpmF (PRK01110)Ribosomal_L32p (pfam01783)→ |
| env-143 | SRS024318_LANL_scaffold_64348 | - | 95 | 32 | **RNA**→ hypo→ |
| env-144 | SRS015803_WUGC_scaffold_47328 | - | 8195 | 8132 | **RNA**→ rpmE2 (PRK01678)Ribosomal_L31 (pfam01197)→ rpmF (PRK01110)Ribosomal_L32p (pfam01783)→ |
| env-145 | SRS018394_Baylor_scaffold_43567 | + | 1126 | 1189 | **RNA**→ rpmE2 (PRK01678)Ribosomal_L31 (pfam01197)→ |
| env-146 | SRS019077_C2525265 | - | 265 | 202 | **RNA**→ rpmE2 (PRK01678)Ribosomal_L31 (pfam01197)→ |
| env-147 | SRS022719_LANL_scaffold_10847 | - | 240 | 177 | **RNA**→ rpmE2 (PRK01678)Ribosomal_L31 (pfam01197)→ |
| env-148 | SRS045049_C1493001 | + | 93 | 156 | **RNA**→ rpmE2 (PRK01678)Ribosomal_L31 (pfam01197)→ |
| env-149 | SRS045197_C3343939 | + | 181 | 244 | **RNA**→ rpmE (PRK00019)rpmE2 (PRK01678)Ribosomal_L31 (pfam01197)→ |
| env-150 | SRS047634_LANL_scaffold_108263 | + | 2234 | 2297 | **RNA**→ |
| env-151 | SRS051941_LANL_scaffold_38428 | + | 2568 | 2631 | **RNA**→ rpmE2 (PRK01678)Ribosomal_L31 (pfam01197)→ rpmF (PRK01110)Ribosomal_L32p (pfam01783)→ |
| env-152 | SRS053584_LANL_scaffold_32609 | + | 521 | 584 | **RNA**→ |
| env-153 | SRS064449_LANL_scaffold_23248 | + | 2184 | 2247 | **RNA**→ rpmE2 (PRK01678)Ribosomal_L31 (pfam01197)→ rpmF (PRK01110)Ribosomal_L32p (pfam01783)→ |
| env-154 | SRS017304_Baylor_scaffold_4558 | - | 5996 | 5933 | **RNA**→ rpmE2 (PRK01678)Ribosomal_L31 (pfam01197)→ rpmF (PRK01110)Ribosomal_L32p (pfam01783)→ |
| env-155 | SRS024144_C1817014 | - | 4955 | 4892 | **RNA**→ rpmE2 (PRK01678)Ribosomal_L31 (pfam01197)→ rpmF (PRK01110)Ribosomal_L32p (pfam01783)→ |
| env-156 | SRS053917_LANL_scaffold_4723 | - | 52992 | 52939 | **RNA**→ rpmE2 (PRK01678)Ribosomal_L31 (pfam01197)→ rpmF (PRK01110)Ribosomal_L32p (pfam01783)→ |
| Cma-1-1 | NC_021915.1 | + | 861557 | 861618 | **RNA**→ rpmE2 (PRK01678)Ribosomal_L31 (pfam01197)→ rpmF (PRK01110)Ribosomal_L32p (pfam01783)→ |
| Cxe-1-1 | NZ_LAYS01000015.1 | - | 47956 | 47840 | **RNA**→ rpmE2 (PRK01678)Ribosomal_L31 (pfam01197)→ rpmF (PRK01110)Ribosomal_L32p (pfam01783)→ |
| Chu-1-1 | NZ_CP005286.1 | + | 766188 | 766271 | **RNA**→ rpmE2 (PRK01678)Ribosomal_L31 (pfam01197)→ rpmF (PRK01110)Ribosomal_L32p (pfam01783)→ |
| env-157 | JGI24707J26582_10124387 | - | 594 | 528 | **RNA**→ rpmE2 (PRK01678)Ribosomal_L31 (pfam01197)→ rpmF (PRK01110)Ribosomal_L32p (pfam01783)→ |
| env-158 | JGI24708J26588_10046528 | - | 1384 | 1318 | **RNA**→ rpmE2 (PRK01678)Ribosomal_L31 (pfam01197)→ rpmF (PRK01110)Ribosomal_L32p (pfam01783)→<br>REC (smart00448)Trans_reg_C (smart00862)OmpR (COG0745)→ |
| Cte-2-1 | NZ_CP011545.1 | + | 759300 | 759365 | **RNA**→ rpmE2 (PRK01678)Ribosomal_L31 (pfam01197)→ rpmF (PRK01110)Ribosomal_L32p (pfam01783)→ |
| Cma-2-1 | NZ_ATVG01000004.1 | - | 122985 | 122912 | **RNA**→ rpmE2 (PRK01678)Ribosomal_L31 (pfam01197)→ rpmE2 (PRK01678)Ribosomal_L31 (pfam01197)→<br>rpmF (PRK01110)Ribosomal_L32p (pfam01783)→ |
| Cma-6-1 | NZ_CP007790.1 | + | 691873 | 691940 | **RNA**→ rpmE2 (PRK01678)Ribosomal_L31 (pfam01197)→ rpmF (PRK01110)Ribosomal_L32p (pfam01783)→<br>REC (smart00448)Trans_reg_C (smart00862)OmpR (COG0745)→<br>HAMP (smart00304)HATPase_c (smart00387)HisKA (smart00388)BaeS (COG0642)→<br>DegQ (COG0265)PDZ_2 (pfam13180)Trypsin_2 (pfam13365)→<br>DegQ (COG0265)PDZ_2 (pfam13180)Trypsin_2 (pfam13365)→<br>MoCF_biosynth (smart00852)MogA_MoaB (cd00886)→ hypo→ |

| | | | | | |
|---|---|---|---|---|---|
| env-159 | JGI24499J29688_1051025 | + | 20 | 74 | **RNA**→ rpmE2 (PRK01678)Ribosomal_L31 (pfam01197)→ |
| Cef-1-1 | NC_004369.1 | + | 1007841 | 1007895 | **RNA**→ rpmE2 (PRK01678)Ribosomal_L31 (pfam01197)→ rpmF (PRK01110)Ribosomal_L32p (pfam01783)→ |
| Cul-3-1 | NC_015683.1 | + | 772809 | 772915 | **RNA**→ rpmE2 (PRK01678)Ribosomal_L31 (pfam01197)→ rpmF (PRK01110)Ribosomal_L32p (pfam01783)→ |
| Cul-1-1 | NC_018101.1 | + | 844033 | 844139 | **RNA**→ rpmE2 (PRK01678)Ribosomal_L31 (pfam01197)→ rpmF (PRK01110)Ribosomal_L32p (pfam01783)→ |
| Cul-2-1 | NC_017317.1 | + | 770052 | 770158 | **RNA**→ rpmE2 (PRK01678)Ribosomal_L31 (pfam01197)→ rpmF (PRK01110)Ribosomal_L32p (pfam01783)→ |
| Cul-6-2 | NZ_CP009583.1 | + | 759297 | 759403 | **RNA**→ rpmE2 (PRK01678)Ribosomal_L31 (pfam01197)→ rpmF (PRK01110)Ribosomal_L32p (pfam01783)→ |
| Cul-6-3 | NZ_CP009500.1 | + | 756232 | 756338 | **RNA**→ rpmE2 (PRK01678)Ribosomal_L31 (pfam01197)→ rpmF (PRK01110)Ribosomal_L32p (pfam01783)→ |
| Cul-8-1 | NZ_CP011913.1 | + | 800557 | 800663 | **RNA**→ rpmE2 (PRK01678)Ribosomal_L31 (pfam01197)→ rpmF (PRK01110)Ribosomal_L32p (pfam01783)→ |
| Cul-6-4 | NZ_CP009716.1 | + | 764195 | 764301 | **RNA**→ rpmE2 (PRK01678)Ribosomal_L31 (pfam01197)→ rpmF (PRK01110)Ribosomal_L32p (pfam01783)→ |
| Cul-7-1 | NZ_CP009622.1 | + | 751173 | 751279 | **RNA**→ rpmE2 (PRK01678)Ribosomal_L31 (pfam01197)→ rpmF (PRK01110)Ribosomal_L32p (pfam01783)→ |
| Cul-4-1 | NZ_AYUJ01000003.1 | + | 75253 | 75359 | **RNA**→ rpmE2 (PRK01678)Ribosomal_L31 (pfam01197)→ rpmF (PRK01110)Ribosomal_L32p (pfam01783)→ |
| Cca-1-1 | NC_020506.1 | + | 891721 | 891777 | **RNA**→ rpmE2 (PRK01678)Ribosomal_L31 (pfam01197)→ rpmF (PRK01110)Ribosomal_L32p (pfam01783)→ |
| Cca-1-2 | NZ_ATVF01000011.1 | + | 57211 | 57267 | **RNA**→ rpmE2 (PRK01678)Ribosomal_L31 (pfam01197)→ rpmF (PRK01110)Ribosomal_L32p (pfam01783)→ |
| Cha-1-1 | NC_020302.1 | + | 926392 | 926485 | **RNA**→ rpmE2 (PRK01678)Ribosomal_L31 (pfam01197)→ rpmF (PRK01110)Ribosomal_L32p (pfam01783)→ |
| Cha-1-2 | NZ_JIAJ01000009.1 | - | 52009 | 51916 | **RNA**→ rpmE2 (PRK01678)Ribosomal_L31 (pfam01197)→ rpmF (PRK01110)Ribosomal_L32p (pfam01783)→ |
| Cpi-2-1 | NZ_LDYD01000009.1 | + | 112150 | 112215 | **RNA**→ rpmE2 (PRK01678)Ribosomal_L31 (pfam01197)→ rpmF (PRK01110)Ribosomal_L32p (pfam01783)→ |
| Cpi-1-1 | NZ_KB892441.1 | + | 29289 | 29354 | **RNA**→ rpmE2 (PRK01678)Ribosomal_L31 (pfam01197)→ rpmF (PRK01110)Ribosomal_L32p (pfam01783)→ |
| Clu-1-1 | NZ_KB902226.1 | - | 119258 | 119193 | **RNA**→ rpmE2 (PRK01678)Ribosomal_L31 (pfam01197)→ rpmF (PRK01110)Ribosomal_L32p (pfam01783)→ |
| Cep-1-1 | NZ_CP011541.1 | + | 815491 | 815557 | **RNA**→ rpmE2 (PRK01678)Ribosomal_L31 (pfam01197)→ rpmF (PRK01110)Ribosomal_L32p (pfam01783)→ |
| Cps-10-1 | NC_017307.1 | + | 688204 | 688312 | **RNA**→ rpmE2 (PRK01678)Ribosomal_L31 (pfam01197)→ rpmF (PRK01110)Ribosomal_L32p (pfam01783)→ |
| Cps-3-1 | NC_017308.1 | + | 675056 | 675164 | **RNA**→ rpmE2 (PRK01678)Ribosomal_L31 (pfam01197)→ rpmF (PRK01110)Ribosomal_L32p (pfam01783)→ |
| Cps-7-1 | NC_016932.1 | + | 685279 | 685387 | **RNA**→ rpmE2 (PRK01678)Ribosomal_L31 (pfam01197)→ rpmF (PRK01110)Ribosomal_L32p (pfam01783)→ |
| Cps-17-9 | NZ_CP012136.1 | + | 720478 | 720586 | **RNA**→ rpmE2 (PRK01678)Ribosomal_L31 (pfam01197)→ rpmF (PRK01110)Ribosomal_L32p (pfam01783)→ |
| Cps-11-1 | NC_018019.1 | + | 680119 | 680227 | **RNA**→ rpmE2 (PRK01678)Ribosomal_L31 (pfam01197)→ rpmF (PRK01110)Ribosomal_L32p (pfam01783)→ |
| Cps-18-1 | NC_017945.2 | + | 720561 | 720669 | **RNA**→ rpmE2 (PRK01678)Ribosomal_L31 (pfam01197)→ rpmF (PRK01110)Ribosomal_L32p (pfam01783)→ |
| Cps-19-1 | NC_017730.2 | + | 745407 | 745515 | **RNA**→ rpmE2 (PRK01678)Ribosomal_L31 (pfam01197)→ rpmF (PRK01110)Ribosomal_L32p (pfam01783)→ |
| Cgl-3-1 | NC_003450.3 | + | 927407 | 927462 | **RNA**→ rpmE2 (PRK01678)Ribosomal_L31 (pfam01197)→ rpmF (PRK01110)Ribosomal_L32p (pfam01783)→ |
| Cgl-7-1 | NZ_AFYA01000024.1 | + | 65956 | 66011 | **RNA**→ rpmE2 (PRK01678)Ribosomal_L31 (pfam01197)→ rpmF (PRK01110)Ribosomal_L32p (pfam01783)→ |
| Cgl-6-1 | NC_009342.1 | + | 1091957 | 1092012 | **RNA**→ rpmE2 (PRK01678)Ribosomal_L31 (pfam01197)→ rpmF (PRK01110)Ribosomal_L32p (pfam01783)→ |
| Cgl-4-1 | NC_020519.1 | + | 927406 | 927461 | **RNA**→ rpmE2 (PRK01678)Ribosomal_L31 (pfam01197)→ rpmF (PRK01110)Ribosomal_L32p (pfam01783)→ |
| Cgl-9-1 | NC_021352.1 | + | 1128888 | 1128943 | **RNA**→ rpmE2 (PRK01678)Ribosomal_L31 (pfam01197)→ rpmF (PRK01110)Ribosomal_L32p (pfam01783)→ |
| Cgl-8-1 | NC_021351.1 | + | 1128889 | 1128944 | **RNA**→ rpmE2 (PRK01678)Ribosomal_L31 (pfam01197)→ rpmF (PRK01110)Ribosomal_L32p (pfam01783)→ |
| Cgl-5-1 | NC_022040.1 | + | 930302 | 930357 | **RNA**→ rpmE2 (PRK01678)Ribosomal_L31 (pfam01197)→ rpmF (PRK01110)Ribosomal_L32p (pfam01783)→ |
| Cgl-11-1 | NZ_CP010451.1 | + | 991151 | 991206 | **RNA**→ rpmE2 (PRK01678)Ribosomal_L31 (pfam01197)→ rpmF (PRK01110)Ribosomal_L32p (pfam01783)→ |
| Cgl-11-2 | NZ_CP007722.1 | + | 1016675 | 1016730 | **RNA**→ rpmE2 (PRK01678)Ribosomal_L31 (pfam01197)→ rpmF (PRK01110)Ribosomal_L32p (pfam01783)→ |
| Cgl-11-3 | NZ_CP007724.1 | + | 985858 | 985913 | **RNA**→ rpmE2 (PRK01678)Ribosomal_L31 (pfam01197)→ rpmF (PRK01110)Ribosomal_L32p (pfam01783)→ |
| Cgl-3-2 | NC_006958.1 | + | 928877 | 928932 | **RNA**→ rpmE2 (PRK01678)Ribosomal_L31 (pfam01197)→ rpmF (PRK01110)Ribosomal_L32p (pfam01783)→ |
| [fl-1-1 | NZ_CP011309.1 | + | 983067 | 983122 | **RNA**→ rpmE2 (PRK01678)Ribosomal_L31 (pfam01197)→ rpmF (PRK01110)Ribosomal_L32p (pfam01783)→ |
| Cgl-10-1 | NZ_AKXP01000016.1 | + | 65984 | 66039 | **RNA**→ rpmE2 (PRK01678)Ribosomal_L31 (pfam01197)→ rpmF (PRK01110)Ribosomal_L32p (pfam01783)→ |
| Cgl-12-1 | NZ_AGQQ02000002.1 | + | 1168400 | 1168455 | **RNA**→ rpmE2 (PRK01678)Ribosomal_L31 (pfam01197)→ rpmF (PRK01110)Ribosomal_L32p (pfam01783)→ |
| Ccr-2-1 | NZ_JPDH01000040.1 | - | 9017 | 8962 | **RNA**→ rpmE2 (PRK01678)Ribosomal_L31 (pfam01197)→ rpmF (PRK01110)Ribosomal_L32p (pfam01783)→ |
| Ccr-1-1 | NZ_AQPS01000045.1 | + | 65984 | 66039 | **RNA**→ rpmE2 (PRK01678)Ribosomal_L31 (pfam01197)→ rpmF (PRK01110)Ribosomal_L32p (pfam01783)→ |

#### 13.3 Conserved domains

Conserved domains found in protein-coding genes listed in Section 13.2 are shown below, with the first sentence in their description from the Conserved Domain Database (if any). Conserved domains associated with more than one L31-Corynebacteriaceae RNA are assigned a color, while others are shown in gray. The number in parentheses after the colored domain name is the number of occurrences in Section 13.2.

cd00886 (2) MogA_MoaB family.
COG0265 (4) Periplasmic serine protease, S1-C subfamily, contain C-terminal PDZ domain [Posttranslational modification, protein turnover, chaperones]
COG0642 (2) Signal transduction histidine kinase [Signal transduction mechanisms]
COG0745 (5) DNA-binding response regulator, OmpR family, contains REC and winged-helix (wHTH) domain [Signal transduction mechanisms, Transcription]
COG2965 (2) Primosomal replication protein N [Replication, recombination and repair]
pfam01197 (353) Ribosomal protein L31.
pfam01783 (313) Ribosomal L32p protein family.
pfam13026 (1) Protein of unknown function (DUF3887).
pfam13180 (4) PDZ domain.
pfam13365 (4) Trypsin-like peptidase domain.
PRK00019 (1) 50S ribosomal protein L31; Reviewed
PRK01110 (316) 50S ribosomal protein L32; Validated
PRK01678 (353) 50S ribosomal protein L31 type B; Reviewed
smart00304 (2) HAMP (Histidine kinases, Adenylyl cyclases, Methyl binding proteins, Phosphatases) domain.
smart00387 (2) Histidine kinase-like ATPases.
smart00388 (2) His Kinase A (phosphoacceptor) domain.
smart00448 (4) cheY-homologous receiver domain.
smart00852 (2) Probable molybdopterin binding domain.
smart00862 (5) Transcriptional regulatory protein, C terminal.

#### 13.4 Multiple-sequence alignment

Each L31-Corynebacteriaceae RNA is denoted by an abbreviation (like "Eco-1-1") that refers to a taxonomy given in Section 13.1. The alignment may include sequences containing the RNA motif, as well as flanking sequence. The L31-Corynebacteriaceae RNA itself is denoted by the line underneath marked 5′ and 3′ on either end. Nucleotides in flanking sequences (i.e., not part of the motif) are written in gray letters. Stems of predicted rho-independent transcription terminators, if any, are shaded yellow. (Note: terminator predictions have not been analyzed manually, and many are likely to be false positives. Terminator predictions are those of the RNIE software.) Nucleotides predicted to function (as DNA) as transcription-factor binding sites, if any, are shaded in green. (But note: these predictions are manually annotated, so they might be under-predicted.) Annotated start codons, if any, are shaded green. (Note: start codons are frequently misannotated, especially in environmental samples.) Nucleotides proposed to basepair as part of the consensus structure are shaded in color when they comprise Watson-Crick or G-U pairs. Otherwise they are shaded gray. Conserved stems are also indicated at the bottom of the alignment by angle brackets, where matching $<$ and $>$ denote base-paired columns. Below these angle brackets, the symbol "2" denotes base pairs exhibiting covariation according to the statistically well-founded R-scape method. "1" denotes base pairs exhibiting covariation according to R2R's simplistic method. "0" denotes base pairs that are not observed to mutate and "?" denotes base pairs that have a significant frequency of non-canonical nucleotides for Watson-Crick or G-U pairs ($> 5\%$). Below these base pair annotation is the consensus sequence: "R" = "A" or "G", "Y" = "C" or "U", red nucleotides: nucleotide identity conserved more than 97% of the time, black nucleotides: 90%, gray nucleotides: 75%, red circle (•): nucleotide is present 97% of the time, black circle (•): 90%, gray circle (•): 75%, white circle (◦): 50%. All percentages of sequences just described (e.g. 97% conserved) assume that sequences have been weighted by the GSC algorithm implemented by the Infernal software package.

Duplicate sequences: the following putative homologs are not shown in the alignment because their sequences are identical to a homolog already shown: Cac-1-1, Cac-2-1, Cau-2-4, Cca-1-2, Ccr-1-1, Ccr-2-1, Cdi-1-1, Cdi-10-1, Cdi-11-1, Cdi-12-1, Cdi-13-1, Cdi-14-1, Cdi-16-1, Cdi-17-1, Cdi-18-2, Cdi-19-1, Cdi-2-1, Cdi-21-1, Cdi-22-1, Cdi-3-1, Cdi-4-1, Cdi-5-1, Cdi-6-1, Cdi-7-1, Cdi-8-1, Cdi-9-1, Cdo-1-1, Cgl-10-1, Cgl-11-1, Cgl-11-2, Cgl-11-3, Cgl-12-1, Cgl-3-2, Cgl-4-1, Cgl-5-1, Cgl-6-1, Cgl-7-1, Cgl-8-1, Cgl-9-1, Cha-1-2, Cje-4-1, Cje-4-10, Cje-4-12, Cje-4-14, Cje-4-17, Cje-4-18, Cje-4-2, Cje-4-7, Cje-4-9, Cma-4-1, Cma-5-1, Cpi-1-1, Cpr-1-1, Cpr-2-1, Cpr-2-3, Cps-1-1, Cps-11-1, Cps-13-1, Cps-14-1, Cps-15-1, Cps-17-2, Cps-17-3, Cps-17-4, Cps-17-5, Cps-17-6, Cps-17-7, Cps-17-8, Cps-17-9, Cps-18-1, Cps-19-1, Cps-2-1, Cps-3-1, Cps-4-1, Cps-5-1, Cps-6-1, Cps-7-1, Cps-8-1, Cps-9-1, Csp-1-1, Csp-10-1, Csp-11-1, Csp-12-1, Csp-13-1, Csp-15-1, Csp-16-1, Csp-17-1, Csp-18-1, Csp-2-1, Csp-3-1, Csp-4-1, Csp-5-1, Csp-9-1, Cst-1-1, Cst-2-2, Cst-2-3, Cst-2-4, Cst-2-5, Cst-2-6, Cst-2-7, Cul-1-1, Cul-4-1, Cul-6-3, Cul-8-1, Cvi-3-1, Tot-1-1, [fl-1-1, env-10, env-100, env-101, env-102, env-103, env-104, env-105, env-106, env-107, env-108, env-109, env-11, env-110, env-111, env-112, env-113, env-114, env-115, env-116, env-117, env-118, env-119, env-120, env-121, env-122, env-123, env-124, env-125, env-126, env-127, env-128, env-129, env-130, env-131, env-132, env-133, env-134, env-135, env-136, env-137, env-138, env-139, env-140, env-141, env-143, env-145, env-146, env-147, env-148, env-149, env-15, env-150, env-151, env-152, env-153, env-155, env-158, env-16, env-20, env-21, env-22, env-23, env-26, env-28, env-34, env-36, env-37, env-38, env-39, env-40, env-41, env-42, env-43, env-44, env-45, env-46, env-47, env-48, env-49, env-50, env-51, env-52, env-53, env-54, env-55, env-56, env-57, env-58, env-59, env-60, env-61, env-62, env-63, env-64, env-65, env-66, env-67, env-68, env-69, env-7, env-70, env-71, env-72, env-73, env-74, env-75, env-76, env-77, env-78, env-79, env-80, env-81, env-82, env-83, env-84, env-85, env-86, env-87, env-88, env-89, env-90, env-91, env-92, env-93, env-94, env-95, env-96, env-97, env-98, env-99

alignment positions 1 · · · 180

```
Cut-1-1   GGCUUCUGUAUGCUUACCCGAGUUGUCUGCCGCCGCCGCAUAGCGGUCGGGUGGCAGGCCCGAAUAAGAGUAGCAACCCAACUCGAGCGCGAUCGCCCG..............GCC.C..GCCGGC......................CUCUU...........................
Cfa-1-1   CGUAUUGCCGUAGCCAGCUAGUACUAUCACUACGUUGUGCAUUGUCCUGCGCAAGGACUGUGCUUAGCAAUCGCAACCCAACUCGGGCACGACCU................UAGGGGUGCGG..........................CACUCGUA........................
Cau-3-1   AAUUGGGGGCAGGAAGAACUAAGAUUCAUCGAUUGUACGAAUCCUCGGUAACACCCGGGGAGAGUUGUAAUUCUAACCCGACUCGGGCACGACCA..............CGCUGUCA...............................GAAA............................
Tot-2-1   GGGAGCCGCCGGGAUGGCGGCGGCGGCCGGGCCCCUGACCCUAGGGAAUCGGCCACCUUAACGACCUGUCCAUCGACCCGACUCGGGCACGAUCCCGCC..............CGGGGC.GCGCACAGU...................UAACU...........................
Cnu-1-1   GUCCAGGGGAGGAGGGGUUACGAUAUGCGGUUGUGUCUGCCCGCGCGAGGCGCUGUGGCGGAUCCACUACUCCAAACCCGACCCAGGUACGACCCGGCACC...............................................CGCAU...........................
Cgl-13-1  GGAUAGACUGGCCGAUCGUGUUGCGACCUGUCGUCUGUGUUGGUUACGGACGACCCGGUAGCGCCCGCACUCCAAACCCGACCCAGGCACGACUCGC.C..............GUCCCGGUCGCCG......................ACUA............................
Cte-1-1   CCGCGCGCGGGGUCGCGGGUUAUGGUGGUUCGCUGUGUUACUGCCCCGCGAAGACGUGGCAGGCCCACACUCCAAACCCGACCCAGGCACGACCUACGG.........CCUCACCGGUG.CCCGGUCC....................UCAUC...........................
Cva-1-1   CGGAUCGGGGUGGUGCGGGUUAUGGUGGAUCGCUGUGUCACUGCCGCGUUCGGUCGCGGUGUGCCCACACUCCAAACCCGACCCAGGCACGACCUACGG...........UACGGACGG.............................AUCA............................
Cps-16-1  GAAAAACCAAGUGUAGUGGAUAGUUUUUCAUAAUUCGCUUAUGGCUGUAAGAUGUUCAGAGUUGUCCGUAAUCGAACCCAACCUGGGUGCGAUCUUCGG.......UGUUUUGCACUGC.GUGGUGUGAC.UGACCU.CUGU......CAU.......................GCAGUG
Csp-14-1  AAAAACCAAGUGUAGUGGAUAGUUUUUUUAUAAUUCGCUUAUGGCUGUAAGAUGUUCAGAGUUGUCCGUAAUCGAACCCAACCUGGGUGCGAUCUUCGG.......UGUUUUGCACUGC.GUGGUGUGACUGACCUC...........UGUCAUGUAGU....................G
env-1     ....................................GCUUAUGGCUGUAAGAUGUUCAGAGUUGUCCGUAAUCGAACCCAACCUGGGUGCGAUCUUCGG.......UGUUUUGCACUGC.GUGGUGUGACUGACCUC...........UGUCAUGUAGU....................G
env-2     GAAAAACCAAGUGUAGUGGAUAGUUUUUUAUAAUUCGCUUAUGGCUGUAAGAUGUUCAGAGUUGUCCGUAAUCGAACCCAACCUGGGUGCGAUCUUCGG.......UGUUUUGCACUGCGUGGUGUGACUGAUCUCUGU.........CAUGUA.......................GUG
env-3     GAAAAACCAAGUGUAGCGGGCGGUUUUCUAAAACUAGCCUUGAUCUGUAGGAUAACUAGAGUUGUCCAUUGUCGAACCCAACCUGGGUGCGAUCUUCGA.............AAGAUUUACU..GCGCAA.AACACG...........AGAGUA.........................C
env-4     CCAAGUGUAGCGGGCGGUUUUNNNNNNNNNNNNNNNNNNNNNNNNNNUAGGAUAACUAGAGUUGUCCAUUGUCGAACCCAACCUGGGUGCGAUCUUCGA.............AAGAUUUACU..GCGCAA.AACACG...........AGAGUA.........................C
env-5     GAAAAACCAAGUGUAGCGGGCGGUUUUCUAAAAUUAGCUUUGAUCUGUAGGAUAACUAGAGUUGUCCAUUGUCGAACCCAACCUGGGUGCGAUCUUCGA.............AAGAUUUACU..GCGCAA.GACACG...........AGAGUA.........................C
Cpr-2-2   GAAAAACCAAGUGUAGCGGGCGGUUUUCUAAAAUUAGCCUUGGUCUGUAGGAUAACUAGAGUUGUCCAUUGUCGAACCCAACCUGGGUGCGAUCUUCGA.............AAGAUUUACU..GCGCAA.GACACG...........AGAGUA.........................C
Cfa-2-1   UUCGGAAGGUGGGGCACUUUCCGCUAAGGUAUUCAGACUGUACGUAUUUGGCCCACCGGGUCAGUGCGAAUUCUUGCCCGACUCAGGCACGACCACC...............UGCG................................CAA.............................
Cre-1-1   AAUUGGGAGGCUAAGGAAGUAUGAUUCUUCGAUUGUACGAAUCCUCGGCCACCGUCGAGGAGUGCUGUAAUUCUAACCCGACUCGGGCACGACCG.....................................................CAUUUCACAUUCGAUUUA..............
Cje-2-1   GGCGCGCCCAAAAACCACUCGCUUAGCGAGCUGGCCUAAGGGGCGUCAAUAGAAGAAGGCGCGCUGCUUUUUAUGACCCGACUCAGGCACGACCCGCGC.................................................ACGUAAC.........................
Cje-4-3   UCGCGCCCAAAAUCCACCCGCUUAGCGGAGUGGCCUAAAGGGCGUCGGUAAAAAGAAGGCGCGCUGCUUUUUAUGACCCGACUCAGGCACGACCCGCGC.................................................AGGUAAC.........................
Cje-4-4   UCGCGCCCAAAACCCACCCGCUUAGCGGAGUGGCCUAAAGGGCGUCGGUAAAAAGAAGGCGCGCUGCUUUUUAUGACCCGACUCAGGCACGACCCGCGC.................................................AGGUAAC.........................
Cje-1-1   UCACGCCCAAAAUCCACCUGCUUAGCGGGCUGGCCUAAAGGGCGUCAAUAAAAAGAAGGCGCGCUGCUUUUUAUGACCCGACUCAGGCACGACCCGCGC.................................................AAGUAAC.........................
Cje-4-5   UCACGCCCAAAAUCCACCUGCUUAGCGGGCUGGCCUAAAGGGCGUCAGUAAAAAGAAGGCGCGCUGCUUUUUAUGACCCGACUCAGGCACGACCCGCGC.................................................AAGUAAC.........................
Cje-4-6   UUGUCGCGCCCUCGUGGGGCUCGGUAGACGAGUGGCCUAAGGGGCGUCAAUGAAGAAGGCGCGCUGCUUUUUAUGACCCGACUCAGGCACGACCCGCGC.................................................AAGGGAU.........................
Cje-4-8   UUGUCGCGCCCUCGUGGGGCUCGGUUGACGAGUGGCCUAAGGGGCGUCAAUAAAGAAGGCGCGCUGCUUUUUAUGACCCGACUCAGGCACGACCCGCGC.................................................AAGAGAU.........................
Cje-4-11  UUGUCGCGCCCUUGUGGGGCUCGGUAGACGAGUGGCCUAAGGGGCGUCAAUAAAGAAGGCGCGCUGCUUUUUAUGACCCGACUCAGGCACGACCCGCGC.................................................AAGAGAU.........................
Cje-4-13  UUGUCGCGCCCUUGUGGGGCUCGGUAGACGAGUGGCCUAAGGGGCGUCAAUAAAGAAGGCGCGCUGCUUUUUAUGACCCGACUCAGGCACGACCCGCGC.................................................AAGAGAU.........................
Cur-2-1   CGGUCGACAUGCGAUAGACUUUUGCGGUCGUACGUGCUGCCAUCGUGGCAUCUCGAACGUGAAGACAAUGUUUCGCCCCGACUCGGGCACGACCCGCGC.................................................UGUCAU..........................
Cur-1-1   CGGUCGACAUGCGAUAGACUUUUGCGGUCGUACGUGCUGCCAUCGUGGCAUCUCGAACGUGAAGACAAUGUUUCGCCCCGACUCGGGCACGACCCGCGC.................................................UCAUGC..........................
Cdi-18-1  UUUUCUCAAAGGAGUAGUUGGGUGUACACUUUGGCAGGUUGCUUACACUUGUAUAGUGUCGGCGAGCAUCGCGAAACCCAACUCGAGCACGAUCUUCAC.................................................CGUGAU..........................
Cdi-15-1  UUUUCUCAAAGGAGUAGUCGGGUGUACACUUUGUCAGGUUGCUUACACUUGUAUAGUGUCGGCGAGCAUCGCGAAACCCAACUCGAGCACGAUCUUCAC.................................................CGUGAU..........................
Cdi-20-1  UUUUCUCAAAGGAGUAGUUGGGUGUACACUUUGGCAGGUUGCUUACACUUGUAUAGUGUCGGCGAGCAUCGCGAAACCCAACUCGAGCACGAUCUUCAU.................................................CGUGAU..........................
Cca-4-1   UUCCGUGCUAGCUGCUGAUGGUGUACCCUCGUUAAAGUUCACUUUCUAUUGAAGUGAGAAGUCCAUUUUUUAGAAACCCAACUCAGGCACGAUCUGCU...UUCCUUAAUAUAUUUAGA.............................AUUU............................
Cgl-2-1   CCUCCGGCGUGUACUAUGCCAAAGGUUGCUCGAUGUGCUCCUCAGUGGGCGCUGAGAACACUGUAAGAAAUAGGUACCCAACUCAGGCACGACCCGUU..............UCGCUC..............................AUGGC...........................
Cgl-1-1   UCUCUGGCGUGUACUAUGCCAAAGGUUGCUCGAUGUGCUCCCCUGCGGGCGCUGAGAACACUGUAAGAAAUAAGUACCCAACUCAGGCACGACCCGUU..............UCGCUC..............................AUGGC...........................
Cpy-1-1   CCCAGGGGUUGCUCGAUGUGCUGCUGAGAAACAUAUUCCUUUUCUGGUUGCACGAAGCACAGGUAAGAAAUAAGUACCCAACUCAGGCACGACCUACGC.................................................CGGGA...........................
Cre-2-1   GUGGGGAGGUGGUCUAUAGCCGUAUUGCUUUUAGGGGCUGGGUCCUGUACUAUCGAUCGGGUUAUUGUCCAAGAAACCCAACUCAGGCACGAUUCACGC..............GGU................................GUA.............................
Cdo-1-2   GCCGAGGACAUGUAGGCUGUCUGAGGUUCAUCGUUUGUACCGCAGGGCACGUGCGAUGCUGAAUGUUUACACCGACCCCAACUCAGGCACGAUCGCCG..................................................CACAUACCGC......................
Cca-5-1   ACCAGACCUGCAUCUGUAUGAUUUAUCGAGUUGUUGACUCGCCUUACGCGCGGUUCAGCUUAAAUGUAAGUGUAAUCCCAACUCGGGCACGAUCUUCG..................................................ACAGUAUUCGAG....................
Cma-3-1   GCGGCGGUGAUCCUGCGCAGGCGGGGAUUUUCGCAACGCAAGCCCCAUAUAGUAGGGUGCUAAAGGAUUUGACCGAGCCAACCUAGGCACGAUCCC....................................................CCUG............................
Cmi-1-1   UUUAGGUUUUCGGGCCGGUGGCCUGUAGGAUAUUCCGAGUUGUCGGCGCGAGCCGCACAGUUUGUAAGAAUUGAGACCCAACUCGGGCACGAUCCUU.............ACGCGCACAGU.UAAUCUUUGCCAGUGCAGCACUACGCAGCGAUAC............GCAUAGUG.UGC
Cau-1-1   UUCGAUUUUAAUGCCGCUGGCCUGUAGGAUAUUUCGAGUUGUCGGCGCGAGCCGCACAAUUUUGUAAGUAUAGAGACCCAACUCGGGCACGAUCCUUUC..............ACGCU.CCC.AGUUGU..CAUCGGGCGCCGUC...AUAGC..................GACGGCGCG
Cmi-1-2   UUUCGGUUUUAAUGCUGGUGGCCUGUAGGAUAUGUCAAGUUGUCGGCACGAGCCGCACAGUUUGUAAGUAUAGAGACCCAACUCGGGCACGAUCCUUUC..............ACACUUCCUUCC.......................ACGCC...........................
Cau-2-1   UUUCGGUUUUAAUGCUGGUGGCCUGUAGGAUAUGUCAAGUUGUCGGCACGAGCCGCACAGUUUGUAAGUAUAGAGACCCAACUCGGGCACGAUCCUUUC..............ACACUUCCUUCC.......................ACGCC...........................
Csi-1-1   UUUAGGUUUUGGUACCUCGGCCCUGUAGGAUAUUCCGAGUUGUCGGCGAGAGCCGCACAGUUUGUAAGCAUUGAGACCCAACUCGGGCACGAUCCUU.............ACGCACGCUGUGGCGUUGCGACGUGUA...........GUGA...........................U
Cci-1-1   CGCUCUCUUCUAUAGGUGCUCACCCCGCUUGCUGUGGGGAUGGGUGCUGAGGCGCAGAACUGCACACCUAAAGAAACCCAACUCAGGCACGACGCGCGC.......CAUAAGGGUGGU...AACCGUUUUUCG.AGCUG.........AUCGGAAGGUGCCACCUG...........CGG
Cat-1-1   CUGGCGUUUCCCCCGAGUAGGCCGCCGAUUUCGGCUCGGCGAUCUUACCGUGUACAGUGUCUAAAGUUCGUCGAAGCCCAACUCAGGCACGAUCCUGCC....CGGCGUGGUGCGUGU...GCG...CACGCGCC.............AGGGCGUGGUCCUAAGAAUA............
Cje-4-15  UAAUAUUCCAUCGAGUGUCACAUGCCGGUUUGUAGAAAUAUUGACCUGGUGCGUGUCAUUGUAAGUCGUGUUCGUACCCAACUCAGGCACGAUUCUCCC.................................................GCUGUUAGCU......................
Cje-4-16  UAAUAUUCCAUCGAGUGUCACAUGCCGGUUUAUAGAAAUAUUGACCCGGUACGUGUCAUUGUAAGUCGUGUUCGUACCCAACUCAGGCACGAUUCUCUC.................................................ACUCUUAGCU......................
Cam-2-1   UAAUAUUCCAUCGAGUGUCACAUGCCGGUUUGUAGAAGUAUUGAGCUGGUGCGUGUCAUUGUAAGUCGUGUUCGUACCCAACUCAGGCACGAUUCUCUC.................................................ACUCUUAGCU......................
Cur-3-1   UAAUAUUCCAUCGAGUGUCACAUGCCGGUUUAUAGAAAUAUUGACCCGGUGCGUGUCAUUGUAAGUCGUGUUCGUACCCAACUCAGGCACGAUUCUCUC.................................................ACUCUUAGCU......................
Cvi-1-1   UAACAUUCCAUCGAGUGUCACAUGCCGGUUUAUAGAGAUAUUGACCUGGUGCGUGUCAUUGUAAGUCGUAUUCGUACCCAACUCAGGCACGAUUCAUCC..............GC.................................AGUCA...........................
Cla-1-1   UCAAUUUCAGGUACUUAGCCUGUAAUAUUCCCACGAGUGUCACCUCUAAGUAUUUAGGUGCCAUAUUUUGUUCGUACCCAACUCAGGCACGAUUCGCCC........GUAUUCCUA...UG.....UUUUAUAGGA.GAAGUAAG...UAGU...................CUCGCUCAG
Csp-19-1  UUAGGAUUCGGCAUAUCACGUGUACAGUAUUGAAGGUUCAUCGUCGGUCCGCUGGCGGUGCAAACUCAUAAACGAACCCAACUCAGGCACGGUCCCGUU..............GAG................................CAUG............................
env-6     AUGGCGGUGGAAUGUAGGAUAAUUCAGGUUAUCGGUCGGCGCGCUUAGUGGCGCCGGGCGAGCUGUAACUGUACAUCCCAACUCGGGCACGAUCCGUAC.................................................CGUCAUAGUUGUUCAGU...............
env-8     AUGACGGUGGAAUGUAGGAUAAUUCAGGUUAUCGGUCGGCGCGCUUAGUGCCGCCGGGCGAGCUGUAACUGUACAUCCCAACUCGGGCACGAUCCGUAC.................................................CGUCAUUAUUCAGU..................
env-9     ......GUGGAAUGUAGGAUAAUUCAGGUUAUCGGUCGGCGCGCUUAGUGGCGCCGGGCGAGCUGUAACUGUACAUCCCAACUCGGGCACGAUCCGUAC.................................................CGUCAUUAUUCAGU..................
env-12    AUGGCGGUGGAAUGUAGGAUAAUUCAGGUUAUCGGUCGGCGCGCUUAGUGGCGCCGGGCGAGCUGUAACUGUUCAUCCCAACUCGGGCACGAUCCGUAC.................................................CGUCAUUAUUCAGU..................
env-13    AUGGCCGUGGAAUGUAGGAUAAUUCAGGUUAUCGGUCGGCGCGCUGUGUGGCGCCGGGCGAGCUGUAACUGUACAUCCCAACUCGGGCACGAUCCGUAC.................................................CGUCAUUAUUCAGU..................
env-14    ......GUGGAAUGUAGGAUAAUUCAGGUUAUCGGUCGGCGCGCUUAGUGCCGCCGGGCGAGCUGUAACUGUACAUCCCAACUCGGGCACGAUCCGUAC.................................................CGUCAUUAUUCAGU..................
env-17    .........................................GGCUUAGUGGCGCCGGGCGAGCUGUAACUGUACAUCCCAACUCGGGCACGAUCCGUAC.................................................CGUCAUUAUUCAGU..................
Csp-6-1   AUGGCGGUGGAAUGUAGGAUAAUUCAGGUUAUCGGUCGGCGCGCUUUGUGGCGCCGGGCGAGCUGUAACUGUACAUCCCAACUCGGGCACGAUCCGUAC.................................................CGUCAUUAUUCAGU..................
env-18    AUGACGGUGGAAUGUAGGAUAAUUCAGGUUAUCGGUCGGCGCGCUUAGUGCCGCCGGGCGAGCUGUAACUGUACAUCCCAACUCGGGCACGAUCCGUAC.................................................CGUCAUUAUUCAGU..................
Csp-7-1   AAGGCAGUGGAAUGUAGGAUAAUUCAGGUUAUCGGUCGGCGCAUUGUGUGGCGUCGGGCGAGCUGUAACUGUACAUCCCAACUCGGGCACGAUCCGUAC.................................................CGGCAUAAUUCAGU..................
Cau-2-2   GUAGUUGACUGUAGGAUAAUUCAGGUUGUUGACCCGUGCCGCACGGCACUCGGUUAGCAGAGUGAAAUUUGUACGUCCCAACUCGGGCACGAUCCGCUC.................................................CAUAAGU.........................
env-19    GUAGUUGACUGUAGGAUAAUUCAGGUUGUUGACCCGUGCCGCACGGCACGCGGUUAGCAGAGUGAAAUUUGUACGUCCCAACUCGGGCACGAUCCGCUC.................................................CAUAAGU.........................
Csp-8-1   GUAGUUGACUGUAGGAUAAUUCGGGUUGUUGACCCGUGCCGCACGGCACGCGGUUAGCAGAGUGAAAUUUGUACGUCCCAACUCGGGCACGAUCCGCUC.................................................CAUAAGU.........................
env-24    GUAGUUGACUGUAGGAUAAUUCAGGUUGUUGACCCAUACCGCUCGGUACGGGGUCGACAGAGCAAGAACUAUACGUCCCAACUCGGGCACGAUCCAUGC.................................................CGAAUGA.........................
env-25    AUGGCCGUGGAAUGUAGGAUAAUUCAGGUUAUCGGUCGGCGCGCAUUGUGGCGCCGGGCGAGCUGUAACUGUACAUCCCAACUCGGGCACGAUCCGUAC.............CGGC................................AUAGCA..........................
Cau-2-3   UUAGUUGACUGUAGGAUAAUUCAGGUUGUUGACCCGUGCCACGCGGCGCGCGGUUAACAGAGUGAAAUUUGUACGUCCCAACUCGGGCACGAUCCUCAC.................................................CCAGC...........................
Ctu-1-1   UUAGUUGACUGUAGGAUAAUUCAGGUUGUUGACCCGUGCCACGCGGCGCGCGGUUAACAGAGUGAAAUUUGUACGUCCCAACUCGGGCACGAUCCUCAC.................................................UAAGC...........................
Cst-2-1   UGUAAUAUGUUCCGAGUUGUCGACGUCUGCGCCCAGCAGGGGCGGAGGAGUUGACGCCGUUUAUUAAUCUGCACAUCCCAACUCGGGCACGAUCUGCGC....CGCCGAUUACUGUAA..............................GUAAG...........................
Cst-2-8   UGUAGUAUGUUCCGAGUUGUCGACGUCUGCGCCCAGCAGGGGCGGAGGAGUUGACGCCGUUUAUUAAUCUGCACAUCCCAACUCGGGCACGAUCUGCGC....CGCCGAUUACUGUAA..............................GUAAG...........................
env-27    GUUUUCCAUUUGCUGAGUGCGCCGUGUAAGGUAUUGGAGGUUCAUCGCCCGUCGUGGGUGGUGCAAACUUAACGAGCCCAACUCAGGCACGAUCCAU...................................................CACU............................
env-29    GUUUUCCAUUUGCUGAAUGCGCCGUGUAAGGUAUUGGAGGUUCAUCGCCCGUCGUGGGUGGUGCAAACUUAACGAGCCCAACUCAGGCACGAUCCAU...................................................CACU............................
env-30    GUUUUCCAUUUGCUGAGUGCGCCGUGUAAGGUGUUGGAGGUUCAUCGCCCGUCGUGGGUGGUGCAAACUUAACGAGCCCAACUUAGGCACGAUCC.....................................................AUCACUGC........................
Cur-4-1   AUUAGGUCAACGCGCAAGUGAACUGUAAGAUACUGCAGGUUAUUCGGCGACCGCCGGAUGAAACGACCAUUUCGCACCCAACUCGGGCACGAUCUCCG..............CGCUUUGCUUCGCG......................UCG.............................
Cih-1-1   AUUCGGAAAAGGCGAACUAAGCAAGUAUAAUUCUGCGGGUUAUCCGGCGACCGCCGGAUGCGUAGACCAUUUCGCACCCAACUCGGGCACGAUCUUCGC..........GCUCGCUUCUCG...........................UCG.............................
Cha-2-1   UUCCUUAUGGGCGGCCGCGGGUGUAGGGUGCUAGAGGUUCAUCGCCGAUUCAGCGGCGAUGUUGCACGCAUACGAACCCAACUCGGGCACGAUCCGGGC...........UCCGGCGCGAUCCACGUUCCGCGGCG............AAUCAUA.........................
Cfr-1-1   UCCUUAUCGGCGCCCCUGCGUGUAGGGUGCUAGGGGUUCAUCGCCGACAGCAGCGGCGAUGGUGCACGCAUACGAACCCAACUCGGGCACGAUCCGGGC...........CCCGGCGUGACUCACGCUCCACGGCG............AAUGAUA.........................

          ................................................................................<<<<<<<.....<<<<<<<<<............<<<<<<<<<<<<<<<<<<<<<<<<<<<<<<<<<<<....................>>>>>>>>>>>>
          ................................................................................00.2210.....2122222..............111...........1............11.11.00....................00.11.11....
          ●●●●●●●●●●●●●●●●●●●●●●●●●●●●●●●●●●●●●●●●●●●●●●●●●●●●●●●●●●●●●●●●●●●●●●●●●●RACCCAACUCRGGCACGAUCY●○○○--------------○○○--------------------------------●●●●○---------------------------
                                                                                  5′
```

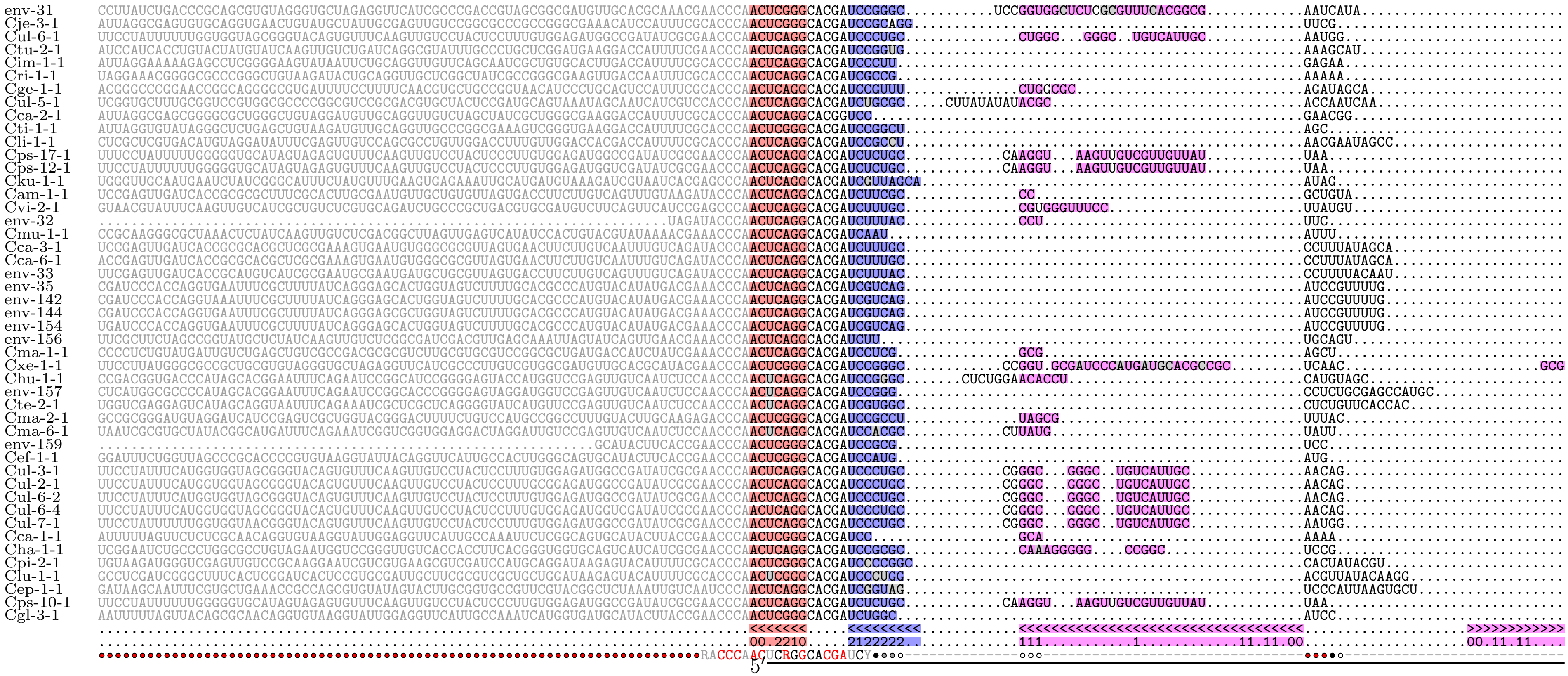

alignment positions 181 · · · 282

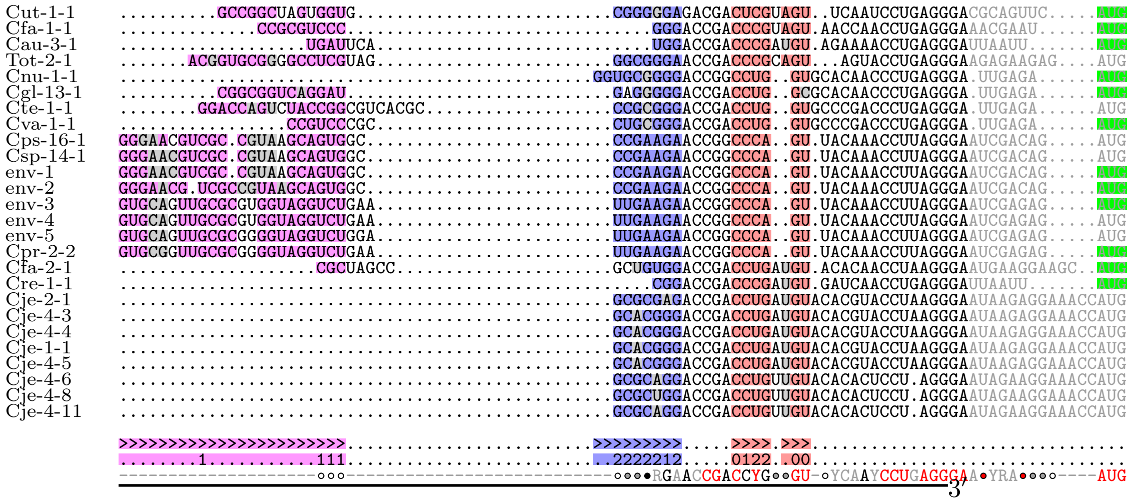

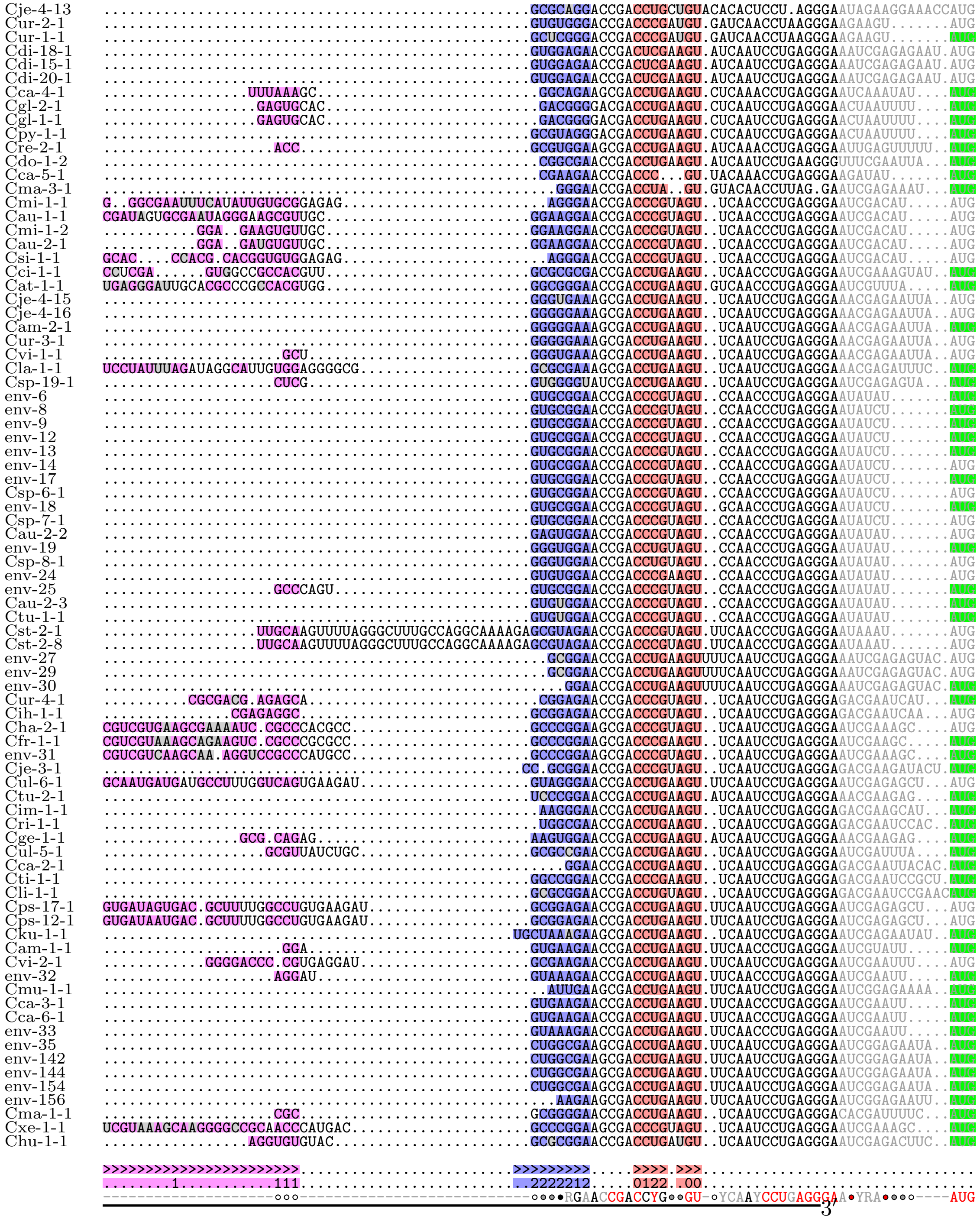
Cje-4-13
Cur-2-1
Cur-1-1
Cdi-18-1
Cdi-15-1
Cdi-20-1
Cca-4-1
Cgl-2-1
Cgl-1-1
Cpy-1-1
Cre-2-1
Cdo-1-2
Cca-5-1
Cma-3-1
Cmi-1-1
Cau-1-1
Cmi-1-2
Cau-2-1
Csi-1-1
Cci-1-1
Cat-1-1
Cje-4-15
Cje-4-16
Cam-2-1
Cur-3-1
Cvi-1-1
Cla-1-1
Csp-19-1
env-6
env-8
env-9
env-12
env-13
env-14
env-17
Csp-6-1
env-18
Csp-7-1
Cau-2-2
env-19
Csp-8-1
env-24
env-25
Cau-2-3
Ctu-1-1
Cst-2-1
Cst-2-8
env-27
env-29
env-30
Cur-4-1
Cih-1-1
Cha-2-1
Cfr-1-1
env-31
Cje-3-1
Cul-6-1
Ctu-2-1
Cim-1-1
Cri-1-1
Cge-1-1
Cul-5-1
Cca-2-1
Cti-1-1
Cli-1-1
Cps-17-1
Cps-12-1
Cku-1-1
Cam-1-1
Cvi-2-1
env-32
Cmu-1-1
Cca-3-1
Cca-6-1
env-33
env-35
env-142
env-144
env-154
env-156
Cma-1-1
Cxe-1-1
Chu-1-1
3'

```
env-157    ...................................................CCUGGAACCGACCUGAUGU..UCAAUCCUGAGGGAAUCGAGACUUC..AUG
Cte-2-1    ..................................................GCCGCGAACCGACCUGAUGU..UCCAUCCUGAGGGAAUCGAGAAUUU..AUG
Cma-2-1    ..................CGCUGUGAGC......................AGGUGGAACCGACCCGAAGU.UUCAACCUGAGGGAAUCGAUA......AUG
Cma-6-1    ...................CAUGC..........................GCGCGGAACCGACCUGAUGU..UCUAUCCUGAGGGAAUCGAGAGUAA..AUG
env-159    ...................................................UGUGGAACCGACCCGUAGU.UUCAAUCCUGAGGGAAUCGAGAGUAA..AUG
Cef-1-1    ...................................................UGUGGAACCGACCCGUAGU.UUCAAUCCUGAGGGAAUCGAGAGUAA..AUG
Cul-3-1    ..GCGAUGAUGAUGCCUCUGGCCAGUGAAGAC..................GUAGGGAACCGACCUGAAGU.UUCAAUCCUGAGGGAAUCGAGAGCU...AUG
Cul-2-1    ..GCGAUGAUGAUGCCUCUGGCCAGUGAAGAU..................GUAGGGAACCGACCUGAAGU.UUCAAUCCUGAGGGAAUCGAGAGCU...AUG
Cul-6-2    ..GCGAUGAUGAUGCCUCUGGCCAGUGAAGAU..................GUAGGGAACCGACCUGAAGU.UUCAAUCCUGAGGGAAUCGAGAGCU...AUG
Cul-6-4    ..GCGAUGAUGAUGCCUCUGGCCAGUGAAGAU..................GUAGGGAACCGACCUGAAGU.UUCAAUCCUGAGGGAAUCGAGAGCU...AUG
Cul-7-1    ..GCAAUGAUGAUGCCUUUGGCCAGUGAAGAU..................GUAGGGAACCGACCUGAAGU.UUCAAUCCUGAGGGAAUCGAGAGCU...AUG
Cca-1-1    ....................UGC..............................UGGAACCGACCUGAAGU.UUCUAUCCUGAGGGAAUCGAGAGUAA..AUG
Cha-1-1    .....GCCGGUGCUCCCCCUCUGAACGG......................GUGCGGAACCGACCUGAAGU..GCAAUCCUGAGGGAAUCGAGACUUA..AUG
Cpi-2-1    .................................................GCCGGAGAACCGACCCGUAGU.CUCAAACCUGAGGGAAACGAAUCAAG..AUG
Clu-1-1    ..................................................CCUUGGAACCGACCCGAUGU..UCAAUCCUGAGGGAAACGAAUAUUU..AUG
Cep-1-1    ..................................................ACGCAGAAGCGACCUGAAGU..UCAAUCCUGAGGGAAUAAACAU.....AUG
Cps-10-1   GUGAUAAUGAC.GCUUUUGGCCUGUGAAGAU...................GCGGAGAACCGACCUGAAGU.UUCAAUCCUGAGGGAAUCGAGAGCU...AUG
Cgl-3-1    ...................................................GCUGGAACCGACCUGAAGU.UUCAAUCCUGAGGGAAUCGAGAGUAA..AUG
           >>>>>>>>>>>>>>>>>>>>>>>.........................>>>>>>>>>.....>>>>.>>>................................
           ........1...........111...........................2222212.....0122..00................................
           --------------------○○○---------------------------○●●●RGAACCGACCYG●●GU-○YCAAYCCUGAGGGAA●YRA●●●○----AUG
                                                                                                    3′
```

### 14 S4-Bacteroidia

#### 14.1 Taxa

The taxonomy of each organism containing a putative S4-Bacteroidia RNA is listed, with abbreviations identifying each hit (e.g., "Eco-1-1" and "Eco-1-2" might hypothetically represent two distinct RNAs in *E. coli*). The abbreviations will be used to identify each individual S4-Bacteroidia RNA in Sections 14.2 and 14.4:.

| **abbrev. of hits** | **taxonomy of species** |
|---|---|
| Bba-1-1 | Bacteria *Bacteroidetes bacterium oral taxon* 272 str. F0290 |
| Bac-1-1 | Bacteria Bacteroidetes Bacteroidia Bacteroidales Bacteroidaceae *Bacteroides acidifaciens* |
| Bba-2-1 | Bacteria Bacteroidetes Bacteroidia Bacteroidales Bacteroidaceae *Bacteroides barnesiae* DSM 18169 |
| Bco-1-1 | Bacteria Bacteroidetes Bacteroidia Bacteroidales Bacteroidaceae *Bacteroides coprosuis* DSM 18011 |
| Bfa-1-1 | Bacteria Bacteroidetes Bacteroidia Bacteroidales Bacteroidaceae *Bacteroides faecichinchillae* JCM 17102 |
| Bfi-1-1 | Bacteria Bacteroidetes Bacteroidia Bacteroidales Bacteroidaceae *Bacteroides finegoldii* DSM 17565 |
| Bfl-1-1 | Bacteria Bacteroidetes Bacteroidia Bacteroidales Bacteroidaceae *Bacteroides fluxus* YIT 12057 |
| Bga-1-1 | Bacteria Bacteroidetes Bacteroidia Bacteroidales Bacteroidaceae *Bacteroides gallinarum* DSM 18171 |
| Bhe-1-1 | Bacteria Bacteroidetes Bacteroidia Bacteroidales Bacteroidaceae *Bacteroides helcogenes* P 36-108 |
| Bma-1-1 | Bacteria Bacteroidetes Bacteroidia Bacteroidales Bacteroidaceae *Bacteroides massiliensis dn*LKV3 |
| Bne-1-1 | Bacteria Bacteroidetes Bacteroidia Bacteroidales Bacteroidaceae *Bacteroides neonati* |
| Bno-1-1 | Bacteria Bacteroidetes Bacteroidia Bacteroidales Bacteroidaceae *Bacteroides nordii* CL02T12C05 |
| Bno-2-1 | Bacteria Bacteroidetes Bacteroidia Bacteroidales Bacteroidaceae *Bacteroides nordii* WAL 11050 = JCM 12987 |
| Bol-1-1 | Bacteria Bacteroidetes Bacteroidia Bacteroidales Bacteroidaceae *Bacteroides oleiciplenus* YIT 12058 |
| Bpa-1-1 | Bacteria Bacteroidetes Bacteroidia Bacteroidales Bacteroidaceae *Bacteroides paurosaccharolyticus* JCM 15092 |
| Bpr-1-1 | Bacteria Bacteroidetes Bacteroidia Bacteroidales Bacteroidaceae *Bacteroides propionicifaciens* DSM 19291 |
| Bpy-1-1 | Bacteria Bacteroidetes Bacteroidia Bacteroidales Bacteroidaceae *Bacteroides pyogenes* F0041 |
| Bpy-2-1 | Bacteria Bacteroidetes Bacteroidia Bacteroidales Bacteroidaceae *Bacteroides pyogenes* JCM 10003 |
| Bre-1-1 | Bacteria Bacteroidetes Bacteroidia Bacteroidales Bacteroidaceae *Bacteroides reticulotermitis* JCM 10512 |
| Bsa-1-1 | Bacteria Bacteroidetes Bacteroidia Bacteroidales Bacteroidaceae *Bacteroides salanitronis* DSM 18170 |
| Bst-1-1 | Bacteria Bacteroidetes Bacteroidia Bacteroidales Bacteroidaceae *Bacteroides stercorirosoris* JCM 17103 |
| Bti-1-1 | Bacteria Bacteroidetes Bacteroidia Bacteroidales Bacteroidaceae *Bacteroides timonensis* |
| Bvi-1-1 | Bacteria Bacteroidetes Bacteroidia Bacteroidales Barnesiellaceae *Barnesiella viscericola* DSM 18177 |
| Dca-1-1 | Bacteria Bacteroidetes Bacteroidia Bacteroidales Dysgonamonadaceae *Dysgonomonas capnocytophagoides* DSM 22835 |
| Dsp-1-1 | Bacteria Bacteroidetes Bacteroidia Bacteroidales Dysgonamonadaceae *Dysgonomonas sp.* BGC7 |
| Dsp-2-1 | Bacteria Bacteroidetes Bacteroidia Bacteroidales Dysgonamonadaceae *Dysgonomonas sp.* HGC4 |
| Dga-1-1 | Bacteria Bacteroidetes Bacteroidia Bacteroidales Porphyromonadaceae *Dysgonomonas gadei* ATCC BAA-286 |
| Ppr-1-1 | Bacteria Bacteroidetes Bacteroidia Bacteroidales Porphyromonadaceae *Paludibacter propionicigenes* WB4 |
| Pca-1-1 | Bacteria Bacteroidetes Bacteroidia Bacteroidales Porphyromonadaceae *Porphyromonas catoniae* ATCC 51270 |
| Pgi-1-1 | Bacteria Bacteroidetes Bacteroidia Bacteroidales Porphyromonadaceae *Porphyromonas gingivalis* F0566 |
| Pgi-2-1 | Bacteria Bacteroidetes Bacteroidia Bacteroidales Porphyromonadaceae *Porphyromonas gingivalis* SJD2 |
| Pgi-3-1 | Bacteria Bacteroidetes Bacteroidia Bacteroidales Porphyromonadaceae *Porphyromonas gingivalis* TDC60 |
| Pgu-1-1 to Pgu-1-2 | Bacteria Bacteroidetes Bacteroidia Bacteroidales Porphyromonadaceae *Porphyromonas gulae* |

| | |
|---|---|
| Pgu-2-1 | Bacteria Bacteroidetes Bacteroidia Bacteroidales Porphyromonadaceae *Porphyromonas gulae* DSM 15663 |
| Psp-1-1 | Bacteria Bacteroidetes Bacteroidia Bacteroidales Porphyromonadaceae *Porphyromonas sp.* COT-239 OH1446 |
| Psp-2-1 | Bacteria Bacteroidetes Bacteroidia Bacteroidales Porphyromonadaceae *Porphyromonas sp.* COT-290 OH860 |
| Psp-3-1 | Bacteria Bacteroidetes Bacteroidia Bacteroidales Porphyromonadaceae *Porphyromonas sp. oral taxon* 278 str. W7784 |
| Pac-1-1 | Bacteria Bacteroidetes Bacteroidia Bacteroidales Porphyromonadaceae *Proteiniphilum acetatigenes* DSM 18083 |
| Pxy-1-1 | Bacteria Bacteroidetes Bacteroidia Bacteroidales Prevotellaceae *Paraprevotella xylaniphila* YIT 11841 |
| Psp-4-1 | Bacteria Bacteroidetes Bacteroidia Bacteroidales Prevotellaceae *Prevotella sp.* 10(H) |
| env-1 to env-517 | environmental samples |

#### 14.2 Gene contexts

Each S4-Bacteroidia RNA (indicated by "RNA→") is listed. For each hit, the downstream genes predicted to reside in a regulated operon are listed. If the nearest downstream gene is encoding in the opposite strand (and therefore presumed to not be a part of a regulated operon), then that gene is still depicted. Some environmental sequences and some RefSeq entries lack gene annotations, and so no genes are available for such sequences. The direction of each gene is indicated with an arrow (→), and each predicted conserved domain in the gene is named. Conserved domains associated with more than one S4-Bacteroidia RNA are assigned a color; other domains are gray. Information about these conserved domains is given in Section 14.3. The accession of the sequence containing each S4-Bacteroidia RNA is given in the column named "Seq. accession". Accessions beginning with "NC_", "NS_", "NW_" or "NZ_" are contained in RefSeq. Other accession refer to environmental samples. Nucleotide coordinates are given for the 5′ and 3′ boundaries of each S4-Bacteroidia RNA. If the 5′ coordinate is greater than the 3′ coordinate, the RNA is present on the reverse-complement strand of the containing genomic DNA sequence. Each hit is denoted by an abbreviation (like "Eco-1-1") that refers to a taxonomy given in Section 14.1.

| abbrev. | Seq. accession | | 5′ at | 3′ at | genes |
|---|---|---|---|---|---|
| env-1 | SRS023468_C1206091 | + | 582 | 641 | **RNA**→ RpsD (COG0522)rpsD (PRK05327)→ |
| env-2 | DLM014_scaffold9702_6 | - | 4201 | 4154 | **RNA**→ RpsD (COG0522)rpsD (PRK05327)→ RpoA (COG0202)PRK05182 (PRK05182)→ rplQ (PRK05591)Ribosomal_L17 (pfam01196)→ |
| env-3 | Buccal_mucosa_LANL_scaffold_41001 | - | 950 | 892 | **RNA**→ RpsD (COG0522)rpsD (PRK05327)→ PRK05182 (PRK05182)RNA_pol_L (pfam01193)→ |
| env-4 | SRS017127_Baylor_scaffold_6196 | + | 1625 | 1683 | **RNA**→ RpsD (COG0522)rpsD (PRK05327)→ |
| env-5 | SRS018149_Baylor_scaffold_12321 | - | 2101 | 2043 | **RNA**→ RpsD (COG0522)rpsD (PRK05327)→ RNAP_alpha_NTD (cd06928)PRK05182 (PRK05182)→ RplQ (COG0203)rplQ (PRK05591)→ |
| env-6 | SRS023964_C2539807 | - | 564 | 506 | **RNA**→ RpsD (COG0522)rpsD (PRK05327)→ |
| env-7 | SRS011310_Baylor_scaffold_16376 | - | 5774 | 5716 | **RNA**→ RpsD (COG0522)rpsD (PRK05327)→ RNAP_alpha_NTD (cd06928)PRK05182 (PRK05182)RNA_pol_A_CTD (pfam03118)→ RplQ (COG0203)rplQ (PRK05591)→ |
| env-8 | SRS065310_C1361795 | + | 968 | 1026 | **RNA**→ RpsD (COG0522)rpsD (PRK05327)→ PRK05182 (PRK05182)RNA_pol_L (pfam01193)→ |
| env-9 | SRS055378_LANL_scaffold_29718 | + | 311 | 362 | **RNA**→ ←hypo |
| env-10 | SRS075410_LANL_scaffold_50382 | + | 626 | 687 | **RNA**→ RpsD (COG0522)rpsD (PRK05327)→ RNAP_alpha_NTD (cd06928)PRK05182 (PRK05182)→ |
| env-11 | SRS016043_WUGC_scaffold_3810 | + | 4122 | 4183 | **RNA**→ RpsD (COG0522)rpsD (PRK05327)→ PRK05182 (PRK05182)RNA_pol_L (pfam01193)→ |
| env-12 | SRS017227_Baylor_scaffold_151231 | + | 598 | 656 | **RNA**→ ←hypo |
| Pca-1-1 | NZ_JDFF01000008.1 | + | 98648 | 98709 | **RNA**→ RpsD (COG0522)rpsD (PRK05327)→ RNAP_alpha_NTD (cd06928)PRK05182 (PRK05182)RNA_pol_A_CTD (pfam03118)→ RplQ (COG0203)rplQ (PRK05591)→ |
| env-13 | SRS023841_C1551522 | - | 144 | 93 | **RNA**→ ←hypo |
| env-14 | SRS011255_Baylor_scaffold_64659 | + | 946 | 1007 | **RNA**→ RpsD (COG0522)rpsD (PRK05327)→ RNAP_alpha_NTD (cd06928)PRK05182 (PRK05182)RNA_pol_A_CTD (pfam03118)→ RplQ (COG0203)rplQ (PRK05591)→ |
| env-15 | SRS022602_Baylor_scaffold_70156 | + | 694 | 755 | **RNA**→ RpsD (COG0522)rpsD (PRK05327)→ RNAP_alpha_NTD (cd06928)PRK05182 (PRK05182)RNA_pol_A_CTD (pfam03118)→ RplQ (COG0203)rplQ (PRK05591)→ |
| env-16 | SRS011343_Baylor_scaffold_39133 | + | 5921 | 5982 | **RNA**→ RpsD (COG0522)rpsD (PRK05327)→ RNAP_alpha_NTD (cd06928)PRK05182 (PRK05182)RNA_pol_A_CTD (pfam03118)→ RplQ (COG0203)rplQ (PRK05591)→ |

| | | | | | |
|---|---|---|---|---|---|
| env-17 | SRS011126_Baylor_scaffold_62966 | + | 893 | 954 | **RNA**→ ←hypo |
| env-18 | SRS020226_Baylor_scaffold_54603 | + | 798 | 857 | **RNA**→ ←hypo |
| env-19 | SRS015574_WUGC_scaffold_12528 | - | 913 | 852 | **RNA**→ RpsD (COG0522)rpsD (PRK05327)→ PRK05182 (PRK05182)RNA_pol_L (pfam01193)→ |
| env-20 | SRS015989_C2177697 | + | 1102 | 1163 | **RNA**→ RpsD (COG0522)rpsD (PRK05327)→ RNAP_alpha_NTD (cd06928)PRK05182 (PRK05182)RNA_pol_A_CTD (pfam03118)→ RplQ (COG0203)rplQ (PRK05591)→ |
| env-21 | SRS017511_Baylor_scaffold_65931 | + | 476 | 535 | **RNA**→ ←hypo |
| env-22 | SRS015899_C1784559 | - | 159 | 98 | **RNA**→ ←hypo |
| env-23 | SRS063999_LANL_scaffold_63196 | - | 141 | 90 | **RNA**→ ←hypo |
| env-24 | SRS023964_C2553979 | - | 311 | 250 | **RNA**→ rpsD (PRK05327)Ribosomal_S4 (pfam00163)→ |
| env-25 | SRS049389_WUGC_scaffold_5829 | - | 2425 | 2364 | **RNA**→ RpsD (COG0522)rpsD (PRK05327)→ RNAP_alpha_NTD (cd06928)PRK05182 (PRK05182)RNA_pol_A_CTD (pfam03118)→ RplQ (COG0203)rplQ (PRK05591)→ |
| env-26 | SRS049268_LANL_scaffold_58337 | + | 435 | 493 | **RNA**→ |
| env-27 | SRS050029_WUGC_scaffold_1469 | + | 6482 | 6543 | **RNA**→ ←hypo |
| env-28 | SRS050628_C1841465 | - | 367 | 306 | **RNA**→ ←hypo |
| Psp-3-1 | NZ_KI259258.1 | + | 268982 | 269043 | **RNA**→ RpsD (COG0522)rpsD (PRK05327)→ RNAP_alpha_NTD (cd06928)PRK05182 (PRK05182)RNA_pol_A_CTD (pfam03118)→ RplQ (COG0203)rplQ (PRK05591)→ |
| env-29 | SRS017808_C3810014 | + | 216 | 277 | **RNA**→ ←hypo |
| env-30 | SRS017810_C1297543 | - | 2305 | 2244 | **RNA**→ RpsD (COG0522)rpsD (PRK05327)→ RNAP_alpha_NTD (cd06928)PRK05182 (PRK05182)RNA_pol_A_CTD (pfam03118)→ RplQ (COG0203)rplQ (PRK05591)→ |
| env-31 | SRS022719_C2873881 | + | 69 | 130 | **RNA**→ rpsD (PRK05327)Ribosomal_S4 (pfam00163)→ |
| env-32 | SRS018157_Baylor_scaffold_50143 | + | 336 | 394 | **RNA**→ |
| env-33 | SRS022077_Baylor_scaffold_39462 | + | 1738 | 1799 | **RNA**→ ←hypo |
| env-34 | SRS019980_Baylor_scaffold_49109 | + | 185 | 246 | **RNA**→ ←hypo |
| env-35 | SRS019122_C5826086 | + | 224 | 285 | **RNA**→ ←hypo |
| env-36 | SRS011098_Baylor_scaffold_35155 | + | 393 | 454 | **RNA**→ |
| env-37 | SRS064329_LANL_scaffold_22656 | - | 2368 | 2307 | **RNA**→ RpsD (COG0522)rpsD (PRK05327)→ RNAP_alpha_NTD (cd06928)PRK05182 (PRK05182)RNA_pol_A_CTD (pfam03118)→ RplQ (COG0203)rplQ (PRK05591)→ |
| env-38 | SRS011243_Baylor_scaffold_40900 | + | 7069 | 7130 | **RNA**→ rpsD (PRK05327)Ribosomal_S4 (pfam00163)→ |
| env-39 | SRS023938_Baylor_scaffold_61057 | + | 322 | 380 | **RNA**→ |
| env-40 | SRS013711_C1768261 | - | 1026 | 965 | **RNA**→ RpsD (COG0522)rpsD (PRK05327)→ PRK05182 (PRK05182)RNA_pol_L (pfam01193)→ |
| env-41 | SRS047100_C2068538 | + | 529 | 590 | **RNA**→ |
| env-42 | SRS016575_Baylor_scaffold_31788 | + | 726 | 784 | **RNA**→ |
| env-43 | SRS017227_Baylor_scaffold_36455 | - | 153 | 95 | **RNA**→ hypo→ |
| env-44 | SRS054569_LANL_scaffold_25042 | + | 6465 | 6526 | **RNA**→ RpsD (COG0522)rpsD (PRK05327)→ RNAP_alpha_NTD (cd06928)PRK05182 (PRK05182)RNA_pol_A_CTD (pfam03118)→ rplQ (PRK05591)Ribosomal_L17 (pfam01196)→ |
| env-45 | SRS015985_WUGC_scaffold_33109 | + | 1425 | 1486 | **RNA**→ RpsD (COG0522)rpsD (PRK05327)→ RNAP_alpha_NTD (cd06928)PRK05182 (PRK05182)RNA_pol_A_CTD (pfam03118)→ RplQ (COG0203)rplQ (PRK05591)→ |
| env-46 | SRS016569_Baylor_scaffold_10142 | + | 7054 | 7115 | **RNA**→ RpsD (COG0522)rpsD (PRK05327)→ RNAP_alpha_NTD (cd06928)PRK05182 (PRK05182)RNA_pol_A_CTD (pfam03118)→ RplQ (COG0203)rplQ (PRK05591)→ |
| env-47 | SRS062878_LANL_scaffold_84002 | + | 2264 | 2325 | **RNA**→ RpsD (COG0522)rpsD (PRK05327)→ RNAP_alpha_NTD (cd06928)PRK05182 (PRK05182)RNA_pol_A_CTD (pfam03118)→ rplQ (PRK05591)Ribosomal_L17 (pfam01196)→ |
| env-48 | SRS043663_LANL_scaffold_33609 | + | 3838 | 3899 | **RNA**→ RpsD (COG0522)rpsD (PRK05327)→ RNAP_alpha_NTD (cd06928)PRK05182 (PRK05182)RNA_pol_A_CTD (pfam03118)→ RplQ (COG0203)rplQ (PRK05591)→ |

| | | | | | |
|---|---|---|---|---|---|
| env-49 | SRS017814_Baylor_scaffold_28373 | + | 1124 | 1185 | **RNA**→ RpsD (COG0522)rpsD (PRK05327)→ RNAP_alpha_NTD (cd06928)PRK05182 (PRK05182)RNA_pol_A_CTD (pfam03118)→ RplQ (COG0203)rplQ (PRK05591)→ |
| env-50 | SRS022725_LANL_scaffold_46531 | + | 399 | 460 | **RNA**→ ←hypo |
| env-51 | SRS016200_WUGC_scaffold_48533 | + | 1527 | 1588 | **RNA**→ RpsD (COG0522)rpsD (PRK05327)→ RNAP_alpha_NTD (cd06928)PRK05182 (PRK05182)RNA_pol_A_CTD (pfam03118)→ RplQ (COG0203)rplQ (PRK05591)→ |
| env-52 | Palatine_Tonsils_LANL_C6173648 | + | 416 | 469 | **RNA**→ |
| env-53 | SRS063932_LANL_scaffold_17762 | - | 154 | 95 | **RNA**→ |
| env-54 | SRS048411_C6928200 | - | 302 | 241 | **RNA**→ rpsD (PRK05327)Ribosomal_S4 (pfam00163)→ |
| env-55 | SRS013164_Baylor_scaffold_10065 | + | 5429 | 5490 | **RNA**→ RpsD (COG0522)rpsD (PRK05327)→ PRK05182 (PRK05182)RNA_pol_L (pfam01193)→ |
| env-56 | SRS015941_C3521747 | - | 131 | 80 | **RNA**→ |
| env-57 | SRS017139_Baylor_scaffold_54946 | - | 310 | 249 | **RNA**→ rpsD (PRK05327)Ribosomal_S4 (pfam00163)→ |
| env-58 | SRS016600_C812171 | + | 234 | 295 | **RNA**→ RpsD (COG0522)rpsD (PRK05327)→ RpoA (COG0202)PRK05182 (PRK05182)→ |
| env-59 | SRS016746_Baylor_scaffold_51447 | - | 792 | 731 | **RNA**→ RpsD (COG0522)rpsD (PRK05327)→ |
| env-60 | SRS055378_LANL_scaffold_10432 | - | 143 | 92 | **RNA**→ |
| env-61 | SRS017439_Baylor_scaffold_45491 | + | 416 | 477 | **RNA**→ ←hypo |
| env-62 | SRS049147_LANL_scaffold_12273 | - | 2261 | 2200 | **RNA**→ RpsD (COG0522)rpsD (PRK05327)→ RNAP_alpha_NTD (cd06928)PRK05182 (PRK05182)RNA_pol_A_CTD (pfam03118)→ RplQ (COG0203)rplQ (PRK05591)→ |
| env-63 | SRS013881_WUGC_scaffold_12223 | - | 5093 | 5032 | **RNA**→ RpsD (COG0522)rpsD (PRK05327)→ RNAP_alpha_NTD (cd06928)PRK05182 (PRK05182)RNA_pol_A_CTD (pfam03118)→ RplQ (COG0203)rplQ (PRK05591)→ |
| env-64 | SRS012279_Baylor_scaffold_38188 | + | 6346 | 6407 | **RNA**→ ←hypo |
| env-65 | SRS044662_C1620928 | + | 41 | 102 | **RNA**→ rpsD (PRK05327)Ribosomal_S4 (pfam00163)→ |
| env-66 | SRS011310_Baylor_scaffold_13983 | + | 2829 | 2890 | **RNA**→ RpsD (COG0522)rpsD (PRK05327)→ RNAP_alpha_NTD (cd06928)PRK05182 (PRK05182)RNA_pol_A_CTD (pfam03118)→ RplQ (COG0203)rplQ (PRK05591)→ |
| env-67 | SRS024355_LANL_scaffold_34750 | - | 442 | 381 | **RNA**→ S4 (smart00363)RpsD (COG0522)→ |
| env-68 | SRS024381_C2388111 | - | 313 | 252 | **RNA**→ rpsD (PRK05327)Ribosomal_S4 (pfam00163)→ |
| env-69 | SRS042131_WUGC_scaffold_5469 | - | 6250 | 6189 | **RNA**→ RpsD (COG0522)rpsD (PRK05327)→ RNAP_alpha_NTD (cd06928)PRK05182 (PRK05182)RNA_pol_A_CTD (pfam03118)→ RplQ (COG0203)rplQ (PRK05591)→ |
| env-70 | SRS011306_Baylor_scaffold_188 | - | 2407 | 2346 | **RNA**→ RpsD (COG0522)rpsD (PRK05327)→ RNAP_alpha_NTD (cd06928)PRK05182 (PRK05182)RNA_pol_A_CTD (pfam03118)→ RplQ (COG0203)rplQ (PRK05591)→ |
| env-71 | SRS011126_Baylor_scaffold_20302 | - | 154 | 95 | **RNA**→ |
| env-72 | SRS011140_Baylor_scaffold_48076 | - | 341 | 280 | **RNA**→ rpsD (PRK05327)Ribosomal_S4 (pfam00163)→ |
| env-73 | SRS013947_WUGC_scaffold_13455 | - | 2590 | 2529 | **RNA**→ RpsD (COG0522)rpsD (PRK05327)→ RNAP_alpha_NTD (cd06928)PRK05182 (PRK05182)RNA_pol_A_CTD (pfam03118)→ RplQ (COG0203)rplQ (PRK05591)→ |
| env-74 | SRS075404_C3684462 | + | 624 | 685 | **RNA**→ RpsD (COG0522)rpsD (PRK05327)→ RNAP_alpha_NTD (cd06928)PRK05182 (PRK05182)RNA_pol_A_CTD (pfam03118)→ RplQ (COG0203)rplQ (PRK05591)→ |
| env-75 | SRS013946_WUGC_scaffold_140 | - | 21413 | 21352 | **RNA**→ RpsD (COG0522)rpsD (PRK05327)→ RNAP_alpha_NTD (cd06928)PRK05182 (PRK05182)RNA_pol_A_CTD (pfam03118)→ RplQ (COG0203)rplQ (PRK05591)→ |
| env-76 | SRS014124_WUGC_scaffold_61000 | + | 4884 | 4945 | **RNA**→ RpsD (COG0522)rpsD (PRK05327)→ RNAP_alpha_NTD (cd06928)PRK05182 (PRK05182)RNA_pol_A_CTD (pfam03118)→ RplQ (COG0203)rplQ (PRK05591)→ |
| env-77 | SRS022602_Baylor_scaffold_69535 | + | 758 | 816 | **RNA**→ |
| env-78 | SRS043422_C2652059 | - | 166 | 104 | **RNA**→ |
| env-79 | SRS014573_WUGC_scaffold_53341 | - | 2102 | 2041 | **RNA**→ RpsD (COG0522)rpsD (PRK05327)→ RNAP_alpha_NTD (cd06928)PRK05182 (PRK05182)RNA_pol_A_CTD (pfam03118)→ rplQ (PRK05591)Ribosomal_L17 (pfam01196)→ |

| | | | | | |
|---|---|---|---|---|---|
| env-80 | SRS013836_C4764451 | - | 172 | 111 | **RNA**→ |
| env-81 | SRS013879_WUGC_scaffold_28433 | - | 386 | 325 | **RNA**→ rpsD (PRK05327)Ribosomal_S4 (pfam00163)→ |
| env-82 | SRS022621_Baylor_scaffold_21872 | - | 2177 | 2116 | **RNA**→ RpsD (COG0522)rpsD (PRK05327)→ RNAP_alpha_NTD (cd06928)PRK05182 (PRK05182)RNA_pol_A_CTD (pfam03118)→ RplQ (COG0203)rplQ (PRK05591)→ |
| env-83 | SRS018439_Baylor_scaffold_23532 | + | 9031 | 9092 | **RNA**→ RpsD (COG0522)rpsD (PRK05327)→ RNAP_alpha_NTD (cd06928)PRK05182 (PRK05182)RNA_pol_A_CTD (pfam03118)→ |
| env-84 | SRS017533_Baylor_scaffold_41553 | + | 217 | 278 | **RNA**→ RpsD (COG0522)rpsD (PRK05327)→ RNAP_alpha_NTD (cd06928)PRK05182 (PRK05182)RNA_pol_A_CTD (pfam03118)→ RplQ (COG0203)rplQ (PRK05591)→ |
| env-85 | SRS019126_WUGC_scaffold_4619 | - | 2437 | 2376 | **RNA**→ RpsD (COG0522)rpsD (PRK05327)→ RNAP_alpha_NTD (cd06928)PRK05182 (PRK05182)RNA_pol_A_CTD (pfam03118)→ RplQ (COG0203)rplQ (PRK05591)→ |
| env-86 | SRS023926_Baylor_scaffold_26039 | + | 6277 | 6338 | **RNA**→ RpsD (COG0522)rpsD (PRK05327)→ RNAP_alpha_NTD (cd06928)PRK05182 (PRK05182)RNA_pol_A_CTD (pfam03118)→ RplQ (COG0203)rplQ (PRK05591)→ |
| env-87 | SRS043772_WUGC_scaffold_22755 | + | 281 | 342 | **RNA**→ RpsD (COG0522)rpsD (PRK05327)→ RNAP_alpha_NTD (cd06928)PRK05182 (PRK05182)RNA_pol_A_CTD (pfam03118)→ |
| env-88 | SRS024375_LANL_scaffold_10736 | - | 3452 | 3391 | **RNA**→ RpsD (COG0522)rpsD (PRK05327)→ RNAP_alpha_NTD (cd06928)PRK05182 (PRK05182)RNA_pol_A_CTD (pfam03118)→ RplQ (COG0203)rplQ (PRK05591)→ |
| env-89 | SRS024441_LANL_scaffold_44794 | + | 3740 | 3801 | **RNA**→ RpsD (COG0522)rpsD (PRK05327)→ RNAP_alpha_NTD (cd06928)PRK05182 (PRK05182)RNA_pol_A_CTD (pfam03118)→ RplQ (COG0203)rplQ (PRK05591)→ |
| env-90 | SRS049318_LANL_scaffold_50729 | - | 142 | 91 | **RNA**→ |
| env-91 | SRS052876_LANL_scaffold_40470 | + | 1404 | 1465 | **RNA**→ RpsD (COG0522)rpsD (PRK05327)→ RNAP_alpha_NTD (cd06928)PRK05182 (PRK05182)RNA_pol_A_CTD (pfam03118)→ RplQ (COG0203)rplQ (PRK05591)→ |
| env-92 | SRS063272_C748000 | - | 2302 | 2241 | **RNA**→ RpsD (COG0522)rpsD (PRK05327)→ RNAP_alpha_NTD (cd06928)PRK05182 (PRK05182)RNA_pol_A_CTD (pfam03118)→ RplQ (COG0203)rplQ (PRK05591)→ |
| Psp-2-1 | NZ_JRAR01000001.1 | + | 21162 | 21223 | **RNA**→ RpsD (COG0522)rpsD (PRK05327)→ RpoA (COG0202)PRK05182 (PRK05182)→ RplQ (COG0203)rplQ (PRK05591)→ |
| Psp-1-1 | NZ_JRAO01000015.1 | - | 141435 | 141374 | **RNA**→ RpsD (COG0522)rpsD (PRK05327)→ RpoA (COG0202)PRK05182 (PRK05182)→ RplQ (COG0203)rplQ (PRK05591)→ |
| env-93 | SRS065099_LANL_scaffold_75481 | + | 444 | 494 | **RNA**→ |
| env-94 | SRS024087_LANL_scaffold_10975 | - | 146 | 89 | **RNA**→ ←hypo |
| env-95 | SRS051941_C4050112 | + | 680 | 740 | **RNA**→ |
| env-96 | SRS013252_C3518045 | - | 147 | 89 | **RNA**→ ←hypo |
| env-97 | SRS020340_C2851743 | + | 298 | 348 | **RNA**→ |
| env-98 | SRS022602_Baylor_scaffold_25508 | + | 442 | 502 | **RNA**→ RpsD (COG0522)rpsD (PRK05327)→ RpoA (COG0202)PRK05182 (PRK05182)→ RplQ (COG0203)rplQ (PRK05591)→ |
| env-99 | SRS013533_C5285183 | - | 174 | 114 | **RNA**→ hypo→ |
| env-100 | SRS064449_LANL_scaffold_59809 | + | 901 | 951 | **RNA**→ |
| env-101 | SRS018394_Baylor_scaffold_53532 | + | 485 | 543 | **RNA**→ |
| env-102 | SRS018443_C3747335 | - | 147 | 89 | **RNA**→ |
| env-103 | SRS055378_LANL_scaffold_13183 | - | 302 | 242 | **RNA**→ rpsD (PRK05327)Ribosomal_S4 (pfam00163)→ |
| env-104 | SRS011343_Baylor_scaffold_40465 | + | 3459 | 3519 | **RNA**→ RpsD (COG0522)rpsD (PRK05327)→ RpoA (COG0202)PRK05182 (PRK05182)→ RplQ (COG0203)rplQ (PRK05591)→ |
| env-105 | SRS022536_LANL_scaffold_94018 | + | 614 | 674 | **RNA**→ rpsD (PRK05327)Ribosomal_S4 (pfam00163)→ |
| env-106 | SRS018394_Baylor_scaffold_31406 | - | 141 | 81 | **RNA**→ |
| env-107 | SRS063932_LANL_scaffold_4135 | + | 370 | 430 | **RNA**→ rpsD (PRK05327)Ribosomal_S4 (pfam00163)→ |
| env-108 | SRS021484_C2188525 | + | 543 | 588 | **RNA**→ RpsD (COG0522)rpsD (PRK05327)→ RpoA (COG0202)PRK05182 (PRK05182)→ |
| env-109 | 4491423.3_NODE_109420 | - | 5213 | 5168 | **RNA**→ RpsD (COG0522)rpsD (PRK05327)→ RNAP_alpha_NTD (cd06928)PRK05182 (PRK05182)→ |
| env-110 | 07_22_CONTROL_DNA_scaffold544_5 | - | 2208 | 2162 | **RNA**→ RpsD (COG0522)rpsD (PRK05327)→ RpoA (COG0202)PRK05182 (PRK05182)→ rplQ (PRK05591)Ribosomal_L17 (pfam01196)→ |

| | | | | | |
|---|---|---|---|---|---|
| env-111 | DLM014_scaffold16166_1 | - | 2097 | 2051 | **RNA**→ RpsD (COG0522)rpsD (PRK05327)→ RpoA (COG0202)PRK05182 (PRK05182)→ RplQ (COG0203)rplQ (PRK05591)→ |
| env-112 | SRS011134_C4924861 | - | 107 | 61 | **RNA**→ |
| env-113 | 08_07_CONTROL_DNA_scaffold19619_1 | + | 164 | 212 | **RNA**→ RpsD (COG0522)rpsD (PRK05327)→ |
| env-114 | 4491417.3_NODE_65482 | + | 43644 | 43690 | **RNA**→ RpsD (COG0522)rpsD (PRK05327)→ RNAP_alpha_NTD (cd06928)PRK05182 (PRK05182)→ PRK05182 (PRK05182)RNA_pol_A_CTD (pfam03118)→ |
| env-115 | UnmappedStool_Broad_scaffold_480295 | + | 268 | 314 | **RNA**→ |
| env-116 | scaffold5632_5_MH0082 | + | 533 | 579 | **RNA**→ RpsD (COG0522)rpsD (PRK05327)→ RNAP_alpha_NTD (cd06928)PRK05182 (PRK05182)RNA_pol_A_CTD (pfam03118)→ |
| env-117 | SRS011134_C4984436 | - | 407 | 361 | **RNA**→ RpsD (COG0522)rpsD (PRK05327)→ |
| env-118 | SRS013476_C3097759 | - | 391 | 347 | **RNA**→ RpsD (COG0522)rpsD (PRK05327)→ |
| env-119 | 08_07_GOS_DNA_scaffold4617_3 | - | 1220 | 1172 | **RNA**→ RpsD (COG0522)rpsD (PRK05327)→ RNAP_alpha_NTD (cd06928)PRK05182 (PRK05182)→ |
| env-120 | 2227505183 | - | 3361 | 3319 | **RNA**→ RpsD (COG0522)rpsD (PRK05327)→ RNAP_alpha_NTD (cd06928)PRK05182 (PRK05182)RNA_pol_A_CTD (pfam03118)→ RplQ (COG0203)rplQ (PRK05591)→ |
| env-121 | AUXO013899695.1 | - | 2595 | 2553 | **RNA**→ RpsD (COG0522)rpsD (PRK05327)→ RpoA (COG0202)PRK05182 (PRK05182)→ RplQ (COG0203)rplQ (PRK05591)→ |
| env-122 | SRMUA_GNVGR9Q01B8SVR | - | 224 | 182 | **RNA**→ hypo→ |
| env-123 | UnmappedStool_Broad_C252248201 | + | 129 | 180 | **RNA**→ rpsD (PRK05327)Ribosomal_S4 (pfam00163)→ |
| env-124 | scaffold47652_4_V1.CD-9 | - | 200 | 152 | **RNA**→ rpsD (PRK05327)Ribosomal_S4 (pfam00163)→ |
| env-125 | scaffold45765_1_V1.CD-14 | - | 234 | 186 | **RNA**→ rpsD (PRK05327)Ribosomal_S4 (pfam00163)→ |
| env-126 | TB_AS07_7DRAFT_10003506 | - | 9850 | 9788 | **RNA**→ AAA (smart00382)COG4185 (COG4185)→ hypo→ |
| env-127 | AUXO012589404.1 | - | 241 | 195 | **RNA**→ rpsD (PRK05327)Ribosomal_S4 (pfam00163)→ |
| Pgu-2-1 | NZ_KB899151.1 | + | 86853 | 86906 | **RNA**→ RpsD (COG0522)rpsD (PRK05327)→ RpoA (COG0202)PRK05182 (PRK05182)→ RplQ (COG0203)rplQ (PRK05591)→ |
| env-128 | SRS015650_WUGC_scaffold_28866 | - | 104 | 54 | **RNA**→ ←hypo |
| Pgi-3-1 | NC_015571.1 | - | 203700 | 203647 | **RNA**→ RpsD (COG0522)rpsD (PRK05327)→ RpoA (COG0202)PRK05182 (PRK05182)→ RplQ (COG0203)rplQ (PRK05591)→ |
| Pgu-1-1 | NZ_KN294118.1 | - | 191559 | 191506 | **RNA**→ RpsD (COG0522)rpsD (PRK05327)→ RpoA (COG0202)PRK05182 (PRK05182)→ RplQ (COG0203)rplQ (PRK05591)→ |
| env-129 | Supragingival_plaque_LANL_C182748641 | + | 464 | 517 | **RNA**→ rpsD (PRK05327)Ribosomal_S4 (pfam00163)→ |
| Pgi-2-1 | NZ_KI629949.1 | - | 30096 | 30043 | **RNA**→ RpsD (COG0522)rpsD (PRK05327)→ RpoA (COG0202)PRK05182 (PRK05182)→ RplQ (COG0203)rplQ (PRK05591)→ hypo→ DUF1661 (pfam07877)→ hypo→ |
| Pgu-1-2 | NZ_KN300353.1 | - | 26757 | 26704 | **RNA**→ RpsD (COG0522)rpsD (PRK05327)→ RpoA (COG0202)PRK05182 (PRK05182)→ RplQ (COG0203)rplQ (PRK05591)→ |
| Pgi-1-1 | NZ_KI260120.1 | + | 16050 | 16103 | **RNA**→ RpsD (COG0522)rpsD (PRK05327)→ RpoA (COG0202)PRK05182 (PRK05182)→ RplQ (COG0203)rplQ (PRK05591)→ hypo→ |
| env-130 | IMNBL1_c0170688 | - | 319 | 279 | **RNA**→ rpsD (PRK05327)Ribosomal_S4 (pfam00163)→ |
| env-131 | scaffold5422_1_MH0052 | - | 2264 | 2218 | **RNA**→ RpsD (COG0522)rpsD (PRK05327)→ RpoA (COG0202)PRK05182 (PRK05182)→ rplQ (PRK05591)Ribosomal_L17 (pfam01196)→ |
| env-132 | 4491404.3_NODE_83097 | + | 488 | 534 | **RNA**→ rpsD (PRK05327)Ribosomal_S4 (pfam00163)→ S4 (smart00363)RpsD (COG0522)→ |
| env-133 | scaffold41480_1_MH0050 | + | 576 | 622 | **RNA**→ RpsD (COG0522)rpsD (PRK05327)→ RNAP_alpha_NTD (cd06928)PRK05182 (PRK05182)→ |
| env-134 | scaffold60668_1_V1.CD-4 | - | 2185 | 2139 | **RNA**→ RpsD (COG0522)rpsD (PRK05327)→ RpoA (COG0202)PRK05182 (PRK05182)→ rplQ (PRK05591)Ribosomal_L17 (pfam01196)→ |
| env-135 | scaffold2465_4_MH0009 | + | 398 | 444 | **RNA**→ RpsD (COG0522)rpsD (PRK05327)→ RpoA (COG0202)PRK05182 (PRK05182)→ rplQ (PRK05591)Ribosomal_L17 (pfam01196)→ |
| env-136 | scaffold8235_1_O2.UC-1 | + | 66 | 112 | **RNA**→ RpsD (COG0522)rpsD (PRK05327)→ RNAP_alpha_NTD (cd06928)PRK05182 (PRK05182)RNA_pol_A_CTD (pfam03118)→ rplQ (PRK05591)Ribosomal_L17 (pfam01196)→ |
| env-137 | RUMENNODE_2120037_111501 | - | 19738 | 19691 | **RNA**→ rpsD (PRK05327)Ribosomal_S4 (pfam00163)→ |
| env-138 | 2226980356 | + | 42770 | 42818 | **RNA**→ RpsD (COG0522)rpsD (PRK05327)→ RpoA (COG0202)PRK05182 (PRK05182)→ RplQ (COG0203)rplQ (PRK05591)→ |
| env-139 | 2226950443 | - | 4600 | 4552 | **RNA**→ RpsD (COG0522)rpsD (PRK05327)→ RpoA (COG0202)PRK05182 (PRK05182)→ RplQ (COG0203)rplQ (PRK05591)→ |
| env-140 | AUXO015494899.1 | - | 713 | 671 | **RNA**→ hypo→ RpsD (COG0522)rpsD (PRK05327)→ |
| env-141 | AUXO012655293.1 | - | 833 | 792 | **RNA**→ RpsD (COG0522)rpsD (PRK05327)→ PRK05182 (PRK05182)RNA_pol_L (pfam01193)→ |

| | | | | | |
|---|---|---|---|---|---|
| env-142 | RUMENNODE_3549585_6500 | + | 14797 | 14838 | RNA→ RpsD (COG0522)rpsD (PRK05327)→ RpoA (COG0202)PRK05182 (PRK05182)→ rplQ (PRK05591)Ribosomal_L17 (pfam01196)→ |
| env-143 | IMNBL1_c0000682 | - | 10946 | 10899 | RNA→ RpsD (COG0522)rpsD (PRK05327)→ RpoA (COG0202)PRK05182 (PRK05182)→ RplQ (COG0203)rplQ (PRK05591)→ |
| env-144 | BABD01035948.1 | + | 534 | 582 | RNA→ rpsD (PRK05327)Ribosomal_S4 (pfam00163)→ |
| env-145 | IMNBL1_c0200996 | + | 120 | 168 | RNA→ rpsD (PRK05327)Ribosomal_S4 (pfam00163)→ |
| env-146 | 4491423.3_NODE_100943 | + | 2001 | 2049 | RNA→ PRK05309 (PRK05309)RpsK (COG0100)→ RpsD (COG0522)rpsD (PRK05327)→ PRK05182 (PRK05182)RNA_pol_L (pfam01193)→ |
| env-147 | scaffold61064_2_MH0006 | + | 191 | 239 | RNA→ RpsD (COG0522)rpsD (PRK05327)→ |
| env-148 | SRS049900_LANL_scaffold_12941 | + | 867 | 915 | RNA→ rpsD (PRK05327)Ribosomal_S4 (pfam00163)→ S4 (smart00363)RpsD (COG0522)→ RpoA (COG0202)PRK05182 (PRK05182)→ RplQ (COG0203)rplQ (PRK05591)→ |
| env-149 | scaffold72032_1_V1.CD-14 | + | 421 | 469 | RNA→ rpsD (PRK05327)Ribosomal_S4 (pfam00163)→ |
| env-150 | ADJT01001361.1 | + | 384 | 432 | RNA→ RpsD (COG0522)rpsD (PRK05327)→ RpoA (COG0202)PRK05182 (PRK05182)→ RplQ (COG0203)rplQ (PRK05591)→ |
| env-151 | scaffold31110_2_MH0019 | + | 232 | 280 | RNA→ hypo→ rpsD (PRK05327)Ribosomal_S4 (pfam00163)→ |
| env-152 | scaffold5909_2_O2.UC-14 | + | 57 | 105 | RNA→ RpsD (COG0522)rpsD (PRK05327)→ |
| env-153 | BABB01007995.1 | + | 205 | 253 | RNA→ RpsD (COG0522)rpsD (PRK05327)→ |
| env-154 | contig19424 | - | 1081 | 1033 | RNA→ RpsD (COG0522)rpsD (PRK05327)→ PRK05182 (PRK05182)RNA_pol_L (pfam01193)→ PRK05182 (PRK05182)RNA_pol_L (pfam01193)→ |
| env-155 | scaffold56457_1_MH0055 | + | 255 | 303 | RNA→ RpsD (COG0522)rpsD (PRK05327)→ |
| env-156 | NLM015_scaffold12871_4 | + | 134 | 182 | RNA→ RpsD (COG0522)rpsD (PRK05327)→ PRK05182 (PRK05182)RNA_pol_L (pfam01193)→ |
| env-157 | scaffold93277_1_MH0053 | + | 467 | 505 | RNA→ hypo→ |
| env-158 | scaffold17281_3_MH0030 | + | 410 | 458 | RNA→ RpsD (COG0522)rpsD (PRK05327)→ RpoA (COG0202)PRK05182 (PRK05182)→ RplQ (COG0203)rplQ (PRK05591)→ |
| env-159 | scaffold738_1_V1.CD-6 | - | 143 | 95 | RNA→ rpsD (PRK05327)Ribosomal_S4 (pfam00163)→ |
| env-160 | scaffold26767_2_MH0040 | + | 57 | 105 | RNA→ RpsD (COG0522)rpsD (PRK05327)→ |
| env-161 | SRS019968_Baylor_scaffold_6283 | + | 260 | 308 | RNA→ RpsD (COG0522)rpsD (PRK05327)→ |
| env-162 | SRS045645_Baylor_scaffold_14899 | + | 400 | 448 | RNA→ RpsD (COG0522)rpsD (PRK05327)→ |
| env-163 | scaffold4241_1_MH0044 | - | 389 | 341 | RNA→ RpsD (COG0522)rpsD (PRK05327)→ |
| env-164 | SRS051882_Baylor_scaffold_18491 | - | 625 | 577 | RNA→ RpsD (COG0522)rpsD (PRK05327)→ |
| env-165 | scaffold33930_1_O2.UC-18 | - | 952 | 904 | RNA→ RpsD (COG0522)rpsD (PRK05327)→ PRK05182 (PRK05182)RNA_pol_L (pfam01193)→ |
| env-166 | BAAZ01001952.1 | + | 529 | 577 | RNA→ RpsD (COG0522)rpsD (PRK05327)→ RNAP_alpha_NTD (cd06928)PRK05182 (PRK05182)→ |
| env-167 | SRS024435_LANL_scaffold_22712 | + | 73 | 121 | RNA→ hypo→ RpsD (COG0522)rpsD (PRK05327)→ |
| env-168 | SRS064557_LANL_scaffold_25854 | - | 129 | 81 | RNA→ |
| env-169 | SRS015782_WUGC_scaffold_11293 | + | 369 | 417 | RNA→ RpsD (COG0522)rpsD (PRK05327)→ |
| env-170 | 4448814.3_Contig42594 | - | 675 | 627 | RNA→ S4 (smart00363)RpsD (COG0522)→ |
| env-171 | DOM019_scaffold22337_4 | - | 435 | 387 | RNA→ S4 (smart00363)RpsD (COG0522)→ |
| env-172 | BABD01010849.1 | - | 1356 | 1308 | RNA→ RpsD (COG0522)rpsD (PRK05327)→ RNAP_alpha_NTD (cd06928)PRK05182 (PRK05182)→ |
| env-173 | 4491416.3_NODE_237 | - | 10253 | 10205 | RNA→ RpsD (COG0522)rpsD (PRK05327)→ RpoA (COG0202)PRK05182 (PRK05182)→ RplQ (COG0203)rplQ (PRK05591)→ |
| env-174 | SRS020869_Baylor_scaffold_2934 | - | 190 | 142 | RNA→ rpsD (PRK05327)Ribosomal_S4 (pfam00163)→ |
| env-175 | 4491421.3_NODE_99843 | + | 4017 | 4065 | RNA→ RpsD (COG0522)rpsD (PRK05327)→ RpoA (COG0202)PRK05182 (PRK05182)→ RplQ (COG0203)rplQ (PRK05591)→ rplQ (PRK05591)Ribosomal_L17 (pfam01196)→ |
| env-176 | JGI994J12354_10027568 | + | 2149 | 2191 | RNA→ RpsD (COG0522)rpsD (PRK05327)→ RNAP_alpha_NTD (cd06928)PRK05182 (PRK05182)→ |
| env-177 | JGI24707J26582_10000222 | - | 2326 | 2281 | RNA→ RpsD (COG0522)rpsD (PRK05327)→ RpoA (COG0202)PRK05182 (PRK05182)→ RplQ (COG0203)rplQ (PRK05591)→ |
| env-178 | BAAZ01024853.1 | - | 919 | 881 | RNA→ RpsD (COG0522)rpsD (PRK05327)→ PRK05182 (PRK05182)RNA_pol_L (pfam01193)→ |
| env-179 | AUXO015025041.1 | - | 398 | 345 | RNA→ rpsD (PRK05327)Ribosomal_S4 (pfam00163)→ |
| env-180 | AUXO016702402.1 | + | 524 | 595 | RNA→ rpsD (PRK05327)Ribosomal_S4 (pfam00163)→ |
| env-181 | NLM006_scaffold52536_3 | + | 16553 | 16592 | RNA→ RpsD (COG0522)rpsD (PRK05327)→ RpoA (COG0202)PRK05182 (PRK05182)→ RplQ (COG0203)rplQ (PRK05591)→ |
| env-182 | DLM007_scaffold5141_17 | + | 197 | 236 | RNA→ RpsD (COG0522)rpsD (PRK05327)→ |
| env-183 | scaffold27115_3_MH0036 | + | 146 | 185 | RNA→ RpsD (COG0522)rpsD (PRK05327)→ RpoA (COG0202)PRK05182 (PRK05182)→ rplQ (PRK05591)Ribosomal_L17 (pfam01196)→ |
| env-184 | scaffold335_4_MH0020 | - | 659 | 620 | RNA→ RpsD (COG0522)rpsD (PRK05327)→ |

| | | | | | |
|---|---|---|---|---|---|
| env-185 | scaffold9913_1_MH0050 | - | 682 | 643 | **RNA**→ RpsD (COG0522)rpsD (PRK05327)→ |
| Pxy-1-1 | NZ_GL883867.1 | - | 21352 | 21313 | **RNA**→ RpsD (COG0522)rpsD (PRK05327)→ RpoA (COG0202)PRK05182 (PRK05182)→ |
| | | | | | RplQ (COG0203)rplQ (PRK05591)→ rplQ (PRK05591)Ribosomal_L17 (pfam01196)→ hypo→ |
| env-186 | SRS048164_WUGC_scaffold_34643 | + | 433 | 472 | **RNA**→ RpsD (COG0522)rpsD (PRK05327)→ |
| env-187 | scaffold10151_5_MH0073 | - | 692 | 653 | **RNA**→ RpsD (COG0522)rpsD (PRK05327)→ |
| env-188 | IMNBL1_c0003450 | + | 7491 | 7530 | **RNA**→ RpsD (COG0522)rpsD (PRK05327)→ |
| | | | | | RNAP_alpha_NTD (cd06928)PRK05182 (PRK05182)RNA_pol_A_CTD (pfam03118)→ |
| | | | | | RplQ (COG0203)rplQ (PRK05591)→ |
| env-189 | SRS011529_WUGC_scaffold_6343 | - | 1690 | 1641 | **RNA**→ RpsD (COG0522)rpsD (PRK05327)→ |
| | | | | | RNAP_alpha_NTD (cd06928)PRK05182 (PRK05182)RNA_pol_A_CTD (pfam03118)→ |
| env-190 | scaffold39757_1_MH0036 | - | 40 | 1 | **RNA**→ |
| env-191 | scaffold56046_2_MH0046 | - | 982 | 933 | **RNA**→ RpsD (COG0522)rpsD (PRK05327)→ PRK05182 (PRK05182)RNA_pol_L (pfam01193)→ |
| env-192 | AUXO013662250.1 | - | 353 | 304 | **RNA**→ rpsD (PRK05327)Ribosomal_S4 (pfam00163)→ |
| env-193 | AUXO011716848.1 | + | 267 | 316 | **RNA**→ S4 (smart00363)RpsD (COG0522)→ |
| env-194 | AUXO012267364.1 | + | 526 | 565 | **RNA**→ |
| env-195 | AUXO014724871.1 | - | 701 | 652 | **RNA**→ RpsD (COG0522)rpsD (PRK05327)→ |
| env-196 | AUXO015059357.1 | + | 29 | 78 | **RNA**→ RpsD (COG0522)rpsD (PRK05327)→ |
| env-197 | AUXO018761249.1 | - | 194 | 146 | **RNA**→ rpsD (PRK05327)Ribosomal_S4 (pfam00163)→ |
| env-198 | 2227339052 | - | 167 | 127 | **RNA**→ rpsD (PRK05327)Ribosomal_S4 (pfam00163)→ |
| env-199 | JGI994J12354_10199883 | - | 251 | 211 | **RNA**→ rpsD (PRK05327)Ribosomal_S4 (pfam00163)→ |
| env-200 | 2227053765 | + | 147 | 187 | **RNA**→ |
| env-201 | JGI994J12354_10460523 | + | 203 | 243 | **RNA**→ rpsD (PRK05327)Ribosomal_S4 (pfam00163)→ |
| env-202 | 2227095262 | - | 330 | 290 | **RNA**→ rpsD (PRK05327)Ribosomal_S4 (pfam00163)→ |
| env-203 | IMNBL1_c0184244 | - | 414 | 374 | **RNA**→ S4 (smart00363)RpsD (COG0522)→ |
| env-204 | JGI994J12354_10281577 | + | 401 | 441 | **RNA**→ rpsD (PRK05327)Ribosomal_S4 (pfam00163)→ |
| env-205 | IMNBL1_c0088446 | - | 33 | 3 | **RNA**→ |
| env-206 | IMNBL1_c0076852 | + | 129 | 169 | **RNA**→ RpsD (COG0522)rpsD (PRK05327)→ PRK05182 (PRK05182)RNA_pol_L (pfam01193)→ |
| env-207 | JGI994J12354_10160833 | - | 704 | 664 | **RNA**→ RpsD (COG0522)rpsD (PRK05327)→ |
| env-208 | 2227437209 | + | 431 | 471 | **RNA**→ RpsD (COG0522)rpsD (PRK05327)→ |
| env-209 | 2227588074 | + | 10 | 50 | **RNA**→ RpsD (COG0522)rpsD (PRK05327)→ |
| env-210 | JGI996J12353_1011164 | - | 683 | 643 | **RNA**→ hypo→ RpsD (COG0522)rpsD (PRK05327)→ |
| env-211 | JGI994J12354_10602131 | - | 192 | 152 | **RNA**→ rpsD (PRK05327)Ribosomal_S4 (pfam00163)→ |
| env-212 | 2227213828 | - | 194 | 154 | **RNA**→ rpsD (PRK05327)Ribosomal_S4 (pfam00163)→ |
| env-213 | JGI994J12354_10069051 | + | 854 | 894 | **RNA**→ RpsD (COG0522)rpsD (PRK05327)→ RpoA (COG0202)PRK05182 (PRK05182)→ |
| env-214 | 2227339122 | + | 4240 | 4280 | **RNA**→ RpsD (COG0522)rpsD (PRK05327)→ |
| | | | | | RNAP_alpha_NTD (cd06928)PRK05182 (PRK05182)RNA_pol_A_CTD (pfam03118)→ |
| | | | | | RplQ (COG0203)rplQ (PRK05591)→ |
| env-215 | IMNBL1_c0070425 | + | 630 | 670 | **RNA**→ RpsD (COG0522)rpsD (PRK05327)→ |
| env-216 | JGI994J12354_10305054 | + | 142 | 182 | **RNA**→ rpsD (PRK05327)Ribosomal_S4 (pfam00163)→ |
| env-217 | JGI994J12354_10567243 | + | 174 | 214 | **RNA**→ rpsD (PRK05327)Ribosomal_S4 (pfam00163)→ |
| env-218 | IMNBL1_c0000677 | - | 14388 | 14348 | **RNA**→ RpsD (COG0522)rpsD (PRK05327)→ |
| | | | | | RNAP_alpha_NTD (cd06928)PRK05182 (PRK05182)RNA_pol_A_CTD (pfam03118)→ |
| | | | | | RplQ (COG0203)rplQ (PRK05591)→ |
| env-219 | AUXO014137175.1 | - | 93 | 54 | **RNA**→ ←hypo |
| env-220 | RUMENNODE_1342001_1 | + | 2330 | 2377 | **RNA**→ rpsD (PRK05327)Ribosomal_S4 (pfam00163)→ |
| env-221 | HCF14C_3_8265_1 | - | 4204 | 4157 | **RNA**→ RpsD (COG0522)rpsD (PRK05327)→ RpoA (COG0202)PRK05182 (PRK05182)→ |
| | | | | | RplQ (COG0203)rplQ (PRK05591)→ |
| env-222 | SRS023604_C783111 | - | 1719 | 1672 | **RNA**→ RpsD (COG0522)rpsD (PRK05327)→ RpoA (COG0202)PRK05182 (PRK05182)→ |
| env-223 | AUXO012249817.1 | - | 397 | 349 | **RNA**→ rpsD (PRK05327)Ribosomal_S4 (pfam00163)→ |
| env-224 | 4491407.3_NODE_8402 | - | 3326 | 3285 | **RNA**→ RpsD (COG0522)rpsD (PRK05327)→ |
| | | | | | RNAP_alpha_NTD (cd06928)PRK05182 (PRK05182)RNA_pol_A_CTD (pfam03118)→ |
| | | | | | RplQ (COG0203)rplQ (PRK05591)→ |
| env-225 | JGI20225J20221_1000009 | + | 207606 | 207653 | **RNA**→ RpsD (COG0522)rpsD (PRK05327)→ RpoA (COG0202)PRK05182 (PRK05182)→ |
| | | | | | RplQ (COG0203)rplQ (PRK05591)→ |

| | | | | | |
|---|---|---|---|---|---|
| env-226 | BMHB3a_c40323 | + | 123 | 170 | **RNA**→ rpsD (PRK05327)Ribosomal_S4 (pfam00163)→ |
| env-227 | JGI26523J50269_1000101 | + | 49816 | 49858 | **RNA**→ RpsD (COG0522)rpsD (PRK05327)→ RNAP_alpha_NTD (cd06928)PRK05182 (PRK05182)RNA_pol_A_CTD (pfam03118)→ RplQ (COG0203)rplQ (PRK05591)→ |
| env-228 | JGI24499J29688_1097403 | + | 252 | 294 | **RNA**→ RpsD (COG0522)rpsD (PRK05327)→ |
| env-229 | JGI24707J26582_10042111 | - | 641 | 599 | **RNA**→ RpsD (COG0522)rpsD (PRK05327)→ |
| env-230 | 2211024960 | + | 90 | 132 | **RNA**→ |
| env-231 | 3300000408_102220 | - | 1904 | 1862 | **RNA**→ RpsD (COG0522)rpsD (PRK05327)→ RNAP_alpha_NTD (cd06928)PRK05182 (PRK05182)RNA_pol_A_CTD (pfam03118)→ rplQ (PRK05591)Ribosomal_L17 (pfam01196)→ |
| env-232 | PBDCA2_contig35478 | - | 23100 | 23058 | **RNA**→ RpsD (COG0522)rpsD (PRK05327)→ RNAP_alpha_NTD (cd06928)PRK05182 (PRK05182)RNA_pol_A_CTD (pfam03118)→ PLN02967 (PLN02967)RplQ (COG0203)rplQ (PRK05591)→ |
| env-233 | 3300000032_0074524 | + | 1154 | 1196 | **RNA**→ RpsD (COG0522)rpsD (PRK05327)→ |
| env-234 | ASA126_GJFDWOI01ADAYG | + | 127 | 159 | **RNA**→ |
| env-235 | ASA129_GJG7ZZE02HS6CS | - | 131 | 89 | **RNA**→ ←hypo |
| env-236 | JGI24707J26582_10313961 | - | 186 | 144 | **RNA**→ rpsD (PRK05327)Ribosomal_S4 (pfam00163)→ |
| env-237 | JGI25321J50212_10130561 | - | 85 | 43 | **RNA**→ |
| env-238 | 3300000568_10484048 | - | 237 | 195 | **RNA**→ rpsD (PRK05327)Ribosomal_S4 (pfam00163)→ |
| env-239 | JGI26523J50269_1018190 | - | 360 | 318 | **RNA**→ rpsD (PRK05327)Ribosomal_S4 (pfam00163)→ |
| Pac-1-1 | NZ_KB905713.1 | - | 5768 | 5726 | **RNA**→ RpsD (COG0522)rpsD (PRK05327)→ RNAP_alpha_NTD (cd06928)PRK05182 (PRK05182)RNA_pol_A_CTD (pfam03118)→ hypo→ RplQ (COG0203)rplQ (PRK05591)→ |
| env-240 | 3300001580_10221456 | - | 676 | 634 | **RNA**→ RpsD (COG0522)rpsD (PRK05327)→ |
| env-241 | 3300001592_10979244 | - | 128 | 86 | **RNA**→ ←hypo |
| env-242 | 4444165.3.06214 | + | 723 | 765 | **RNA**→ RpsD (COG0522)rpsD (PRK05327)→ RpoA (COG0202)PRK05182 (PRK05182)→ PRK05182 (PRK05182)RNA_pol_L (pfam01193)→ |
| env-243 | PRSSGFe2_Sequence0000000352 | - | 11315 | 11273 | **RNA**→ RpsD (COG0522)rpsD (PRK05327)→ RNAP_alpha_NTD (cd06928)PRK05182 (PRK05182)RNA_pol_A_CTD (pfam03118)→ rplQ (PRK05591)Ribosomal_L17 (pfam01196)→ |
| env-244 | BMHBC_1_1_newblercontig05403 | - | 204 | 162 | **RNA**→ hypo→ |
| env-245 | JFJP01044180.1 | + | 7301 | 7343 | **RNA**→ RpsD (COG0522)rpsD (PRK05327)→ RpoA (COG0202)PRK05182 (PRK05182)→ RplQ (COG0203)rplQ (PRK05591)→ |
| Ppr-1-1 | NC_014734.1 | + | 2990427 | 2990469 | **RNA**→ RpsD (COG0522)rpsD (PRK05327)→ RpoA (COG0202)PRK05182 (PRK05182)→ RplQ (COG0203)rplQ (PRK05591)→ |
| env-246 | SRS016095_WUGC_scaffold_14591 | + | 280 | 322 | **RNA**→ RpsD (COG0522)rpsD (PRK05327)→ |
| env-247 | SRS014235_WUGC_scaffold_63059 | + | 873 | 915 | **RNA**→ RpsD (COG0522)rpsD (PRK05327)→ |
| env-248 | JGI24728J21555_1099436 | - | 413 | 371 | **RNA**→ hypo→ RpsD (COG0522)rpsD (PRK05327)→ |
| env-249 | JGI24731J21663_1020569 | + | 1048 | 1090 | **RNA**→ RpsD (COG0522)rpsD (PRK05327)→ |
| env-250 | AUXO012304620.1 | + | 104 | 149 | **RNA**→ rpsD (PRK05327)Ribosomal_S4 (pfam00163)→ |
| env-251 | scaffold9573_4_O2.UC-24 | - | 2883 | 2838 | **RNA**→ RpsD (COG0522)rpsD (PRK05327)→ RNAP_alpha_NTD (cd06928)PRK05182 (PRK05182)RNA_pol_A_CTD (pfam03118)→ rplQ (PRK05591)Ribosomal_L17 (pfam01196)→ |
| env-252 | scaffold28280_3_MH0054 | + | 981 | 1024 | **RNA**→ RpsD (COG0522)rpsD (PRK05327)→ |
| env-253 | 4491423.3_NODE_100453 | + | 1380 | 1423 | **RNA**→ RpsD (COG0522)rpsD (PRK05327)→ |
| env-254 | 2014748740 | + | 450 | 493 | **RNA**→ rpsD (PRK05327)Ribosomal_S4 (pfam00163)→ |
| env-255 | PRSSGFe2_Sequence0000029464 | - | 952 | 909 | **RNA**→ RpsD (COG0522)rpsD (PRK05327)→ PRK05182 (PRK05182)RNA_pol_L (pfam01193)→ |
| env-256 | JGI20225J20221_1000111 | - | 93620 | 93577 | **RNA**→ RpsD (COG0522)rpsD (PRK05327)→ RpoA (COG0202)PRK05182 (PRK05182)→ RplQ (COG0203)rplQ (PRK05591)→ |
| env-257 | BMHBC_193261 | - | 2611 | 2568 | **RNA**→ RpsD (COG0522)rpsD (PRK05327)→ RpoA (COG0202)PRK05182 (PRK05182)→ rplQ (PRK05591)Ribosomal_L17 (pfam01196)→ |
| env-258 | 2210576792 | + | 262 | 295 | **RNA**→ |
| env-259 | JGI24500J29687_10051329 | + | 16282 | 16325 | **RNA**→ RpsD (COG0522)rpsD (PRK05327)→ |

| | | | | | |
|---|---|---|---|---|---|
| Dga-1-1 | NZ_GL891985.1 | - | 188070 | 188027 | **RNA**→ RpsD (COG0522)rpsD (PRK05327)→ RNAP_alpha_NTD (cd06928)PRK05182 (PRK05182)RNA_pol_A_CTD (pfam03118)→ RplQ (COG0203)rplQ (PRK05591)→ |
| env-260 | 2227035932 | + | 1370 | 1413 | **RNA**→ RpsD (COG0522)rpsD (PRK05327)→ RNAP_alpha_NTD (cd06928)PRK05182 (PRK05182)RNA_pol_A_CTD (pfam03118)→ RplQ (COG0203)rplQ (PRK05591)→ |
| Psp-4-1 | NZ_JOMI01000002.1 | + | 192990 | 193033 | **RNA**→ RpsD (COG0522)rpsD (PRK05327)→ RNAP_alpha_NTD (cd06928)PRK05182 (PRK05182)RNA_pol_A_CTD (pfam03118)→ RplQ (COG0203)rplQ (PRK05591)→ |
| env-261 | SRS019267_WUGC_scaffold_20085 | + | 866 | 910 | **RNA**→ RpsD (COG0522)rpsD (PRK05327)→ RpoA (COG0202)PRK05182 (PRK05182)→ rplQ (PRK05591)Ribosomal_L17 (pfam01196)→ |
| env-262 | DOM012_scaffold34729_1 | - | 2272 | 2228 | **RNA**→ RpsD (COG0522)rpsD (PRK05327)→ RpoA (COG0202)PRK05182 (PRK05182)→ rplQ (PRK05591)Ribosomal_L17 (pfam01196)→ |
| env-263 | Subgingival_plaque_LANL_C11140467 | - | 365 | 321 | **RNA**→ rpsD (PRK05327)Ribosomal_S4 (pfam00163)→ |
| env-264 | SRS015650_WUGC_scaffold_24783 | - | 465 | 421 | **RNA**→ RpsD (COG0522)rpsD (PRK05327)→ |
| env-265 | AUXO013584032.1 | + | 2699 | 2747 | **RNA**→ RpsD (COG0522)rpsD (PRK05327)→ RNAP_alpha_NTD (cd06928)PRK05182 (PRK05182)RNA_pol_A_CTD (pfam03118)→ rplQ (PRK05591)Ribosomal_L17 (pfam01196)→ |
| env-266 | AUXO015681238.1 | - | 390 | 342 | **RNA**→ RpsD (COG0522)rpsD (PRK05327)→ |
| env-267 | AUXO016113898.1 | - | 81 | 43 | **RNA**→ |
| env-268 | AUXO017288013.1 | + | 160 | 208 | **RNA**→ RpsD (COG0522)rpsD (PRK05327)→ PRK05182 (PRK05182)RNA_pol_L (pfam01193)→ |
| env-269 | HCF14C_3_8874_1 | + | 1171 | 1219 | **RNA**→ RpsD (COG0522)rpsD (PRK05327)→ |
| env-270 | AUXO013556098.1 | - | 428 | 380 | **RNA**→ RpsD (COG0522)rpsD (PRK05327)→ |
| env-271 | AUXO016146758.1 | - | 533 | 485 | **RNA**→ RpsD (COG0522)rpsD (PRK05327)→ |
| env-272 | BMHB3a_c152975 | + | 10118 | 10161 | **RNA**→ RpsD (COG0522)rpsD (PRK05327)→ RpoA (COG0202)PRK05182 (PRK05182)→ RplQ (COG0203)rplQ (PRK05591)→ |
| env-273 | MA40A_contig27223 | - | 222 | 179 | **RNA**→ rpsD (PRK05327)Ribosomal_S4 (pfam00163)→ |
| env-274 | HCE12Call500_c0170529 | - | 10250 | 10193 | **RNA**→ RpsD (COG0522)rpsD (PRK05327)→ RNAP_alpha_NTD (cd06928)PRK05182 (PRK05182)RNA_pol_A_CTD (pfam03118)→ rplQ (PRK05591)PRK13108 (PRK13108)Ribosomal_L17 (pfam01196)→ |
| env-275 | HCF12C_392227 | + | 332 | 389 | **RNA**→ RpsD (COG0522)rpsD (PRK05327)→ hypo→ |
| env-276 | AUXO017050186.1 | - | 255 | 205 | **RNA**→ rpsD (PRK05327)Ribosomal_S4 (pfam00163)→ |
| env-277 | HCE12Call500_c0119666 | - | 1945 | 1895 | **RNA**→ RpsD (COG0522)rpsD (PRK05327)→ RpoA (COG0202)PRK05182 (PRK05182)→ rplQ (PRK05591)Ribosomal_L17 (pfam01196)→ |
| env-278 | DLM004_scaffold13460_1 | + | 661 | 692 | **RNA**→ |
| env-279 | DOF009_scaffold34853_1 | + | 760 | 801 | **RNA**→ RpsD (COG0522)rpsD (PRK05327)→ |
| env-280 | SRS011239_Baylor_scaffold_2110 | + | 507 | 538 | **RNA**→ |
| env-281 | SRS016095_WUGC_scaffold_12148 | - | 67 | 36 | **RNA**→ |
| env-282 | SRS064276_LANL_scaffold_7093 | + | 4193 | 4234 | **RNA**→ RNAP_alpha_NTD (cd06928)RpsD (COG0522)PRK05182 (PRK05182)rpsD (PRK05327)→ |
| env-283 | NLM015_scaffold155_4_1 | + | 381 | 422 | **RNA**→ RpsD (COG0522)rpsD (PRK05327)→ |
| env-284 | SRS013687_Baylor_scaffold_7394 | + | 426 | 457 | **RNA**→ |
| env-285 | SRS049164_WUGC_scaffold_32178 | - | 432 | 394 | **RNA**→ S4 (smart00363)RpsD (COG0522)→ |
| env-286 | BAAU01005463.1 | - | 1319 | 1278 | **RNA**→ RpsD (COG0522)rpsD (PRK05327)→ RNAP_alpha_NTD (cd06928)PRK05182 (PRK05182)→ |
| env-287 | DLF004_scaffold26868_1 | + | 661 | 692 | **RNA**→ |
| env-288 | 4491423.3_NODE_37638 | - | 96 | 55 | **RNA**→ |
| env-289 | NLM017_scaffold62_7 | - | 557 | 516 | **RNA**→ RpsD (COG0522)rpsD (PRK05327)→ |
| env-290 | SRS014235_WUGC_scaffold_35424 | + | 572 | 603 | **RNA**→ |
| env-291 | SRS022137_WUGC_scaffold_15988 | - | 582 | 541 | **RNA**→ RpsD (COG0522)rpsD (PRK05327)→ |
| env-292 | BABD01000634.1 | - | 4603 | 4562 | **RNA**→ RpsD (COG0522)rpsD (PRK05327)→ RpoA (COG0202)PRK05182 (PRK05182)→ RplQ (COG0203)rplQ (PRK05591)→ |
| env-293 | NOM029_scaffold258_6 | - | 188 | 147 | **RNA**→ rpsD (PRK05327)Ribosomal_S4 (pfam00163)→ |
| env-294 | SRS019582_WUGC_scaffold_26735 | - | 412 | 371 | **RNA**→ S4 (smart00363)RpsD (COG0522)→ |
| env-295 | AJWZ01010988.1 | - | 301 | 260 | **RNA**→ rpsD (PRK05327)Ribosomal_S4 (pfam00163)→ |
| env-296 | pre-rum103-03185 | + | 251 | 292 | **RNA**→ RpsD (COG0522)rpsD (PRK05327)→ PRK05182 (PRK05182)RNA_pol_L (pfam01193)→ |

| | | | | | |
|---|---|---|---|---|---|
| env-297 | longitudinal_403_1247 | - | 4435 | 4394 | **RNA**→ RpsD (COG0522)rpsD (PRK05327)→ RpoA (COG0202)PRK05182 (PRK05182)→ RplQ (COG0203)rplQ (PRK05591)→ |
| env-298 | NOF010_scaffold29793_5 | - | 3201 | 3160 | **RNA**→ RpsD (COG0522)rpsD (PRK05327)→ RpoA (COG0202)PRK05182 (PRK05182)→ RplQ (COG0203)rplQ (PRK05591)→ |
| env-299 | 4491406.3_NODE_31370 | - | 670 | 629 | **RNA**→ RpsD (COG0522)rpsD (PRK05327)→ |
| Dsp-2-1 | NZ_CVRU01000083.1 | + | 63519 | 63561 | **RNA**→ RpsD (COG0522)rpsD (PRK05327)→ RNAP_alpha_NTD (cd06928)PRK05182 (PRK05182)RNA_pol_A_CTD (pfam03118)→ RplQ (COG0203)rplQ (PRK05591)→ |
| env-300 | IMNBL1_c0075309 | - | 465 | 423 | **RNA**→ RpsD (COG0522)rpsD (PRK05327)→ |
| env-301 | IMNBL1_c0368275 | - | 321 | 279 | **RNA**→ rpsD (PRK05327)Ribosomal_S4 (pfam00163)→ |
| env-302 | JGI994J12354_10022051 | + | 3246 | 3288 | **RNA**→ RpsD (COG0522)rpsD (PRK05327)→ |
| env-303 | 2227309966 | + | 229 | 271 | **RNA**→ RpsD (COG0522)rpsD (PRK05327)→ PRK05182 (PRK05182)RNA_pol_L (pfam01193)→ |
| env-304 | IMNBL1_c0230034 | + | 279 | 321 | **RNA**→ rpsD (PRK05327)Ribosomal_S4 (pfam00163)→ |
| env-305 | IMNBL1_c0105304 | - | 213 | 171 | **RNA**→ rpsD (PRK05327)Ribosomal_S4 (pfam00163)→ |
| Dca-1-1 | NZ_AUFL01000019.1 | + | 53200 | 53242 | **RNA**→ RpsD (COG0522)rpsD (PRK05327)→ RNAP_alpha_NTD (cd06928)PRK05182 (PRK05182)RNA_pol_A_CTD (pfam03118)→ RplQ (COG0203)rplQ (PRK05591)→ |
| env-306 | 2210995755 | - | 175 | 133 | **RNA**→ rpsD (PRK05327)Ribosomal_S4 (pfam00163)→ |
| env-307 | JGI24728J21555_1006805 | - | 154 | 112 | **RNA**→ |
| Bpy-2-1 | NZ_BAIU01000011.1 | - | 29809 | 29767 | **RNA**→ RpsD (COG0522)rpsD (PRK05327)→ RpoA (COG0202)PRK05182 (PRK05182)→ RplQ (COG0203)rplQ (PRK05591)→ |
| env-308 | pre-rum103-00894 | - | 9641 | 9599 | **RNA**→ RpsD (COG0522)rpsD (PRK05327)→ RpoA (COG0202)PRK05182 (PRK05182)→ RplQ (COG0203)rplQ (PRK05591)→ |
| Bpy-1-1 | NZ_KE993121.1 | + | 42744 | 42786 | **RNA**→ RpsD (COG0522)rpsD (PRK05327)→ RpoA (COG0202)PRK05182 (PRK05182)→ RplQ (COG0203)rplQ (PRK05591)→ rplQ (PRK05591)Ribosomal_L17 (pfam01196)→ |
| env-309 | IMNBL1_c0157636 | + | 159 | 202 | **RNA**→ RpsD (COG0522)rpsD (PRK05327)→ |
| env-310 | 4491406.3_NODE_15769 | + | 177 | 209 | **RNA**→ |
| env-311 | DLM010_scaffold16098_1 | + | 260 | 302 | **RNA**→ RpsD (COG0522)rpsD (PRK05327)→ |
| env-312 | 4491419.3_NODE_2960 | + | 462 | 504 | **RNA**→ RpsD (COG0522)rpsD (PRK05327)→ RpoA (COG0202)PRK05182 (PRK05182)→ RplQ (COG0203)rplQ (PRK05591)→ |
| env-313 | DOM010_scaffold7295_1 | - | 660 | 618 | **RNA**→ RpsD (COG0522)rpsD (PRK05327)→ |
| env-314 | contig107354 | + | 383 | 425 | **RNA**→ rpsD (PRK05327)Ribosomal_S4 (pfam00163)→ |
| env-315 | NOF012_scaffold53092_1 | - | 107 | 65 | **RNA**→ |
| env-316 | 4491485.3_NODE_4702 | - | 97 | 55 | **RNA**→ |
| env-317 | 4491410.3_NODE_989 | - | 129 | 87 | **RNA**→ |
| env-318 | 4444165.3.13720 | + | 60 | 102 | **RNA**→ rpsD (PRK05327)Ribosomal_S4 (pfam00163)→ S4 (smart00363)S4 (cd00165)→ |
| Bst-1-1 | NZ_BAKL01000061.1 | - | 5724 | 5682 | **RNA**→ RpsD (COG0522)rpsD (PRK05327)→ RpoA (COG0202)PRK05182 (PRK05182)→ RplQ (COG0203)rplQ (PRK05591)→ |
| env-319 | 4491423.3_NODE_71282 | - | 97 | 55 | **RNA**→ hypo→ |
| Bti-1-1 | NZ_CBVI010000016.1 | - | 5424 | 5382 | **RNA**→ RpsD (COG0522)rpsD (PRK05327)→ RpoA (COG0202)PRK05182 (PRK05182)→ RplQ (COG0203)rplQ (PRK05591)→ |
| env-320 | 4491408.3_NODE_13301 | + | 9879 | 9921 | **RNA**→ RpsD (COG0522)rpsD (PRK05327)→ PRK05182 (PRK05182)RNA_pol_A_bac (pfam01000)→ RpoA (COG0202)PRK05182 (PRK05182)→ RplQ (COG0203)rplQ (PRK05591)→ |
| env-321 | 4491420.3_NODE_10528 | + | 459 | 491 | **RNA**→ |
| env-322 | 4491490.3_NODE_12841 | - | 663 | 621 | **RNA**→ RpsD (COG0522)rpsD (PRK05327)→ |
| env-323 | NLM007_scaffold12180_2 | - | 617 | 575 | **RNA**→ RpsD (COG0522)rpsD (PRK05327)→ |
| Bac-1-1 | NZ_CDPR01000609.1 | - | 2777 | 2735 | **RNA**→ RpsD (COG0522)rpsD (PRK05327)→ RpoA (COG0202)PRK05182 (PRK05182)→ RplQ (COG0203)rplQ (PRK05591)→ |
| env-324 | BABA01006018.1 | - | 202 | 160 | **RNA**→ rpsD (PRK05327)Ribosomal_S4 (pfam00163)→ |
| env-325 | NLF013_scaffold515_5 | - | 614 | 572 | **RNA**→ RpsD (COG0522)rpsD (PRK05327)→ |
| Bol-1-1 | NZ_JH992946.1 | + | 318016 | 318058 | **RNA**→ RpsD (COG0522)rpsD (PRK05327)→ RpoA (COG0202)PRK05182 (PRK05182)→ RplQ (COG0203)rplQ (PRK05591)→ |
| env-326 | 08_07_GOS_DNA_scaffold96420_2 | - | 86 | 48 | **RNA**→ |
| env-327 | BMHBC_1_1_newblercontig337321 | + | 91 | 124 | **RNA**→ |
| env-328 | DLM003_C759402_1 | - | 450 | 412 | **RNA**→ S4 (smart00363)RpsD (COG0522)→ |

| | | | | | |
|---|---|---|---|---|---|
| env-329 | SRS022137_WUGC_scaffold_2662 | - | 466 | 423 | **RNA**→ rpsD (PRK05327)Ribosomal_S4 (pfam00163)→ |
| env-330 | NOF008_scaffold26002_3 | + | 1309 | 1342 | **RNA**→ |
| env-331 | BABA01023720.1 | + | 559 | 602 | **RNA**→ hypo→ |
| env-332 | 4491402.3_NODE_2368 | - | 33517 | 33474 | **RNA**→ RpsD (COG0522)rpsD (PRK05327)→ RpoA (COG0202)PRK05182 (PRK05182)→ RplQ (COG0203)rplQ (PRK05591)→ |
| env-333 | longitudinal_29_3 | + | 29258 | 29301 | **RNA**→ RpsD (COG0522)rpsD (PRK05327)→ RpoA (COG0202)PRK05182 (PRK05182)→ RplQ (COG0203)rplQ (PRK05591)→ |
| env-334 | 4491406.3_NODE_26822 | + | 457 | 490 | **RNA**→ |
| Bfa-1-1 | NZ_BAKK01000025.1 | - | 22207 | 22164 | **RNA**→ RpsD (COG0522)rpsD (PRK05327)→ hypo→ hypo→ RplQ (COG0203)rplQ (PRK05591)→ |
| env-335 | NOM012_scaffold23_1 | - | 506 | 463 | **RNA**→ RpsD (COG0522)rpsD (PRK05327)→ |
| env-336 | GO4KUYH01EWM5T | - | 370 | 327 | **RNA**→ rpsD (PRK05327)Ribosomal_S4 (pfam00163)→ |
| Bre-1-1 | NZ_BAIV01000005.1 | + | 187594 | 187637 | **RNA**→ RpsD (COG0522)rpsD (PRK05327)→ RpoA (COG0202)PRK05182 (PRK05182)→ RplQ (COG0203)rplQ (PRK05591)→ |
| env-337 | 4491408.3_NODE_8793 | - | 1690 | 1647 | **RNA**→ RpoA (COG0202)RpsD (COG0522)PRK05182 (PRK05182)rpsD (PRK05327)→ |
| Bno-1-1 | NZ_JH724317.1 | - | 84405 | 84362 | **RNA**→ RpsD (COG0522)rpsD (PRK05327)→ RpoA (COG0202)PRK05182 (PRK05182)→ RplQ (COG0203)rplQ (PRK05591)→ |
| env-338 | 2205287234 | - | 44 | 11 | **RNA**→ hypo→ |
| Bno-2-1 | NZ_BAJA01000009.1 | + | 40118 | 40161 | **RNA**→ RpsD (COG0522)rpsD (PRK05327)→ RpoA (COG0202)PRK05182 (PRK05182)→ RplQ (COG0203)rplQ (PRK05591)→ |
| env-339 | NOF007_scaffold44_16 | - | 1160 | 1117 | **RNA**→ RpsD (COG0522)rpsD (PRK05327)→ RNAP_alpha_NTD (cd06928)PRK05182 (PRK05182)→ |
| env-340 | 4491479.3_NODE_1257 | - | 110 | 67 | **RNA**→ |
| env-341 | DLM020_scaffold13017_1 | + | 262 | 305 | **RNA**→ ←hypo |
| env-342 | DLF001_scaffold15389_1 | + | 209 | 252 | **RNA**→ RpsD (COG0522)rpsD (PRK05327)→ PRK05182 (PRK05182)RNA_pol_L (pfam01193)→ |
| env-343 | BAAY01022580.1 | - | 570 | 527 | **RNA**→ RpsD (COG0522)rpsD (PRK05327)→ |
| env-344 | NLM003_scaffold34637_1 | + | 211 | 254 | **RNA**→ RpsD (COG0522)rpsD (PRK05327)→ |
| env-345 | DLM001_scaffold23897_1 | + | 340 | 383 | **RNA**→ ←hypo |
| Bne-1-1 | NZ_HG726019.1 | + | 504025 | 504068 | **RNA**→ RpsD (COG0522)rpsD (PRK05327)→ RpoA (COG0202)PRK05182 (PRK05182)→ RplQ (COG0203)rplQ (PRK05591)→ |
| env-346 | 4491421.3_NODE_13085 | + | 481 | 519 | **RNA**→ hypo→ |
| env-347 | 4448820.3_Contig_2741 | - | 878 | 840 | **RNA**→ rpsD (PRK05327)→ S4 (smart00363)RpsD (COG0522)→ |
| env-348 | DOF009_scaffold33995_5 | + | 427 | 465 | **RNA**→ RpsD (COG0522)rpsD (PRK05327)→ |
| env-349 | 4491404.3_NODE_40545 | + | 40773 | 40812 | **RNA**→ RpsD (COG0522)rpsD (PRK05327)→ RpoA (COG0202)PRK05182 (PRK05182)→ RplQ (COG0203)rplQ (PRK05591)→ |
| env-350 | NOM026_scaffold21137_8 | + | 135 | 174 | **RNA**→ RpsD (COG0522)rpsD (PRK05327)→ |
| env-351 | CAM_READ_0099620383 | + | 156 | 195 | **RNA**→ rpsD (PRK05327)Ribosomal_S4 (pfam00163)→ |
| env-352 | NOM020_scaffold32164_1 | - | 530 | 491 | **RNA**→ RpsD (COG0522)rpsD (PRK05327)→ |
| env-353 | CAM_READ_0099614309 | - | 253 | 214 | **RNA**→ rpsD (PRK05327)Ribosomal_S4 (pfam00163)→ |
| env-354 | CAM_READ_0100028819 | + | 179 | 218 | **RNA**→ rpsD (PRK05327)Ribosomal_S4 (pfam00163)→ |
| Bba-2-1 | NZ_KB894657.1 | + | 44135 | 44174 | **RNA**→ RpsD (COG0522)rpsD (PRK05327)→ RpsD (COG0522)rpsD (PRK05327)→ RpoA (COG0202)PRK05182 (PRK05182)→ RplQ (COG0203)rplQ (PRK05591)rpsP (PRK14521)→ rplQ (PRK05591)rpsP (PRK14521)Ribosomal_L17 (pfam01196)→ |
| env-355 | CAM_READ_0100328893 | - | 402 | 362 | **RNA**→ rpsD (PRK05327)Ribosomal_S4 (pfam00163)→ |
| env-356 | 4448808.3_Contig2785 | - | 1614 | 1574 | **RNA**→ S4 (smart00363)RpsD (COG0522)→ |
| env-357 | 4448807.3_Contig36355 | + | 56 | 95 | **RNA**→ rpsD (PRK05327)Ribosomal_S4 (pfam00163)→ S4 (smart00363)RpsD (COG0522)→ |
| env-358 | CAM_READ_0100031775 | - | 293 | 253 | **RNA**→ rpsD (PRK05327)Ribosomal_S4 (pfam00163)→ |
| env-359 | 4448817.3_Contig4536 | - | 2122 | 2083 | **RNA**→ rpsD (PRK05327)Ribosomal_S4 (pfam00163)→ S4 (smart00363)RpsD (COG0522)→ RpoA (COG0202)PRK05182 (PRK05182)→ RNAP_alpha_NTD (cd06928)rpoA (TIGR02027)→ |
| env-360 | DLF003_scaffold27229_5 | - | 2090 | 2051 | **RNA**→ RpsD (COG0522)rpsD (PRK05327)→ RpoA (COG0202)PRK05182 (PRK05182)→ RplQ (COG0203)rplQ (PRK05591)→ |
| env-361 | NLM032_scaffold69263_2 | - | 652 | 613 | **RNA**→ RpsD (COG0522)rpsD (PRK05327)→ |
| env-362 | CAM_READ_0099610915 | - | 85 | 46 | **RNA**→ |
| env-363 | NLM015_scaffold45_1 | + | 4595 | 4634 | **RNA**→ RpsD (COG0522)rpsD (PRK05327)→ RpoA (COG0202)PRK05182 (PRK05182)→ RplQ (COG0203)rplQ (PRK05591)→ |
| env-364 | CAM_READ_0099599333 | + | 212 | 241 | **RNA**→ |
| env-365 | CAM_READ_0099868351 | - | 223 | 184 | **RNA**→ rpsD (PRK05327)Ribosomal_S4 (pfam00163)→ |

| | | | | | |
|---|---|---|---|---|---|
| env-366 | 4448815.3_Contig20136 | + | 237 | 276 | RNA→ REC (smart00448)REC (cd00156)→ |
| env-367 | 4448815.3_Contig4225 | + | 1077 | 1117 | RNA→ rpsD (PRK05327)Ribosomal_S4 (pfam00163)→ S4 (smart00363)RpsD (COG0522)→ |
| env-368 | CAM_READ_0100032315 | - | 355 | 315 | RNA→ rpsD (PRK05327)Ribosomal_S4 (pfam00163)→ |
| env-369 | CAM_READ_0099824453 | - | 464 | 423 | RNA→ RpsD (COG0522)rpsD (PRK05327)→ |
| env-370 | 4448811.3_Contig24646 | + | 402 | 441 | RNA→ rpsD (PRK05327)Ribosomal_S4 (pfam00163)→ |
| env-371 | 4448816.3_Contig535 | + | 7083 | 7122 | RNA→ PRK05309 (PRK05309)RpsK (COG0100)→ |
| env-372 | 4491477.3_NODE_13450 | + | 289 | 328 | RNA→ |
| env-373 | BABD01000252.1 | + | 1870 | 1909 | RNA→ RpsD (COG0522)rpsD (PRK05327)→ RpoA (COG0202)PRK05182 (PRK05182)→ RplQ (COG0203)rplQ (PRK05591)→ |
| env-374 | BAAY01002401.1 | - | 744 | 705 | RNA→ RpsD (COG0522)rpsD (PRK05327)→ |
| env-375 | 4448812.3_Contig31152 | + | 433 | 472 | RNA→ rpsD (PRK05327)Ribosomal_S4 (pfam00163)→ |
| env-376 | DOM014_scaffold10613_2 | + | 79 | 118 | RNA→ RpsD (COG0522)rpsD (PRK05327)→ |
| env-377 | longitudinal_9_1858 | - | 1614 | 1575 | RNA→ RpsD (COG0522)rpsD (PRK05327)→ |
| Bma-1-1 | NZ_KE159493.1 | + | 945671 | 945710 | RNA→ RpsD (COG0522)rpsD (PRK05327)→ RpoA (COG0202)PRK05182 (PRK05182)→ RplQ (COG0203)rplQ (PRK05591)→ |
| env-378 | SRS011586_WUGC_scaffold_3940 | + | 681 | 720 | RNA→ |
| env-379 | contig03653 | - | 603 | 564 | RNA→ rpsD (PRK05327)Ribosomal_S4 (pfam00163)→ |
| env-380 | AMCI01000278.1 | - | 2219 | 2180 | RNA→ RpsD (COG0522)rpsD (PRK05327)→ RpoA (COG0202)PRK05182 (PRK05182)→ RplQ (COG0203)rplQ (PRK05591)→ hypo→ |
| env-381 | NLF011_scaffold2976_3 | - | 407 | 368 | RNA→ RpsD (COG0522)rpsD (PRK05327)→ |
| env-382 | SRS018656_C1624822 | - | 39 | 10 | RNA→ |
| env-383 | 4444165.3.59663 | - | 347 | 308 | RNA→ rpsD (PRK05327)Ribosomal_S4 (pfam00163)→ |
| Bsa-1-1 | NC_015164.1 | - | 2309295 | 2309256 | RNA→ RpsD (COG0522)rpsD (PRK05327)→ RpoA (COG0202)PRK05182 (PRK05182)→ RplQ (COG0203)rplQ (PRK05591)PRK10905 (PRK10905)→ |
| env-384 | 4491421.3_NODE_98220 | - | 2451 | 2411 | RNA→ RpsD (COG0522)rpsD (PRK05327)→ RpoA (COG0202)PRK05182 (PRK05182)→ RplQ (COG0203)rplQ (PRK05591)→ |
| env-385 | 4491405.3_NODE_15725 | - | 729 | 690 | RNA→ RpsD (COG0522)rpsD (PRK05327)→ |
| env-386 | 4491405.3_NODE_3691 | - | 706 | 667 | RNA→ RpsD (COG0522)rpsD (PRK05327)→ |
| Bfi-1-1 | NZ_GG688317.1 | + | 227372 | 227415 | RNA→ RpsD (COG0522)rpsD (PRK05327)→ RpoA (COG0202)PRK05182 (PRK05182)→ RplQ (COG0203)rplQ (PRK05591)→ |
| env-387 | DOF008_scaffold9719_2 | + | 318 | 361 | RNA→ rpsD (PRK05327)Ribosomal_S4 (pfam00163)→ |
| env-388 | longitudinal_208_91 | + | 6255 | 6298 | RNA→ hypo→ RpsD (COG0522)rpsD (PRK05327)→ RpoA (COG0202)PRK05182 (PRK05182)→ RplQ (COG0203)rplQ (PRK05591)→ |
| env-389 | SRS022713_LANL_scaffold_2857 | - | 541 | 498 | RNA→ RpsD (COG0522)rpsD (PRK05327)→ |
| env-390 | DLM011_scaffold111_2 | + | 788 | 821 | RNA→ |
| env-391 | SRS023346_LANL_scaffold_1997 | + | 289 | 322 | RNA→ |
| env-392 | scaffold5201_4_MH0074 | - | 616 | 573 | RNA→ RpsD (COG0522)rpsD (PRK05327)→ |
| env-393 | 4491407.3_NODE_11023 | - | 669 | 626 | RNA→ RpsD (COG0522)rpsD (PRK05327)→ |
| env-394 | UnmappedStool_Broad_scaffold_100816 | + | 284 | 318 | RNA→ |
| env-395 | SRS015264_WUGC_scaffold_68015 | - | 1367 | 1323 | RNA→ RpsD (COG0522)rpsD (PRK05327)→ RNAP_alpha_NTD (cd06928)PRK05182 (PRK05182)→ |
| env-396 | SRS011586_C3383160 | + | 59 | 103 | RNA→ RpsD (COG0522)rpsD (PRK05327)→ |
| env-397 | DLM001_scaffold28826_29 | + | 13800 | 13834 | RNA→ |
| env-398 | SRS013476_C2866296 | - | 270 | 226 | RNA→ rpsD (PRK05327)Ribosomal_S4 (pfam00163)→ |
| Bhe-1-1 | NC_014933.1 | + | 1368692 | 1368731 | RNA→ hypo→ RpsD (COG0522)rpsD (PRK05327)→ RpoA (COG0202)PRK05182 (PRK05182)→ RplQ (COG0203)rplQ (PRK05591)→ |
| env-399 | 4491411.3_NODE_7528 | + | 1171 | 1210 | RNA→ RpsD (COG0522)rpsD (PRK05327)→ RpoA (COG0202)PRK05182 (PRK05182)→ RplQ (COG0203)rplQ (PRK05591)→ |
| Bfl-1-1 | NZ_GL882630.1 | + | 110961 | 111000 | RNA→ RpsD (COG0522)rpsD (PRK05327)→ RNAP_alpha_NTD (cd06928)PRK05182 (PRK05182)RNA_pol_A_CTD (pfam03118)→ RplQ (COG0203)rplQ (PRK05591)→ hypo→ |
| env-400 | IMNBL1_c0000119 | + | 20286 | 20326 | RNA→ RpsD (COG0522)rpsD (PRK05327)→ RpoA (COG0202)PRK05182 (PRK05182)→ RplQ (COG0203)rplQ (PRK05591)→ |
| env-401 | 2226960311 | - | 510 | 470 | RNA→ RpsD (COG0522)rpsD (PRK05327)→ |
| env-402 | JGI26534J51046_1128446 | - | 112 | 72 | RNA→ ←hypo |
| env-403 | JGI994J12354_10651307 | - | 257 | 217 | RNA→ rpsD (PRK05327)Ribosomal_S4 (pfam00163)→ |

| | | | | | |
|---|---|---|---|---|---|
| env-404 | 2227529292 | + | 543 | 583 | RNA→ ←hypo |
| env-405 | JGI994J12354_10765493 | + | 91 | 133 | RNA→ rpsD (PRK05327)Ribosomal_S4 (pfam00163)→ |
| env-406 | IMNBL1_c0050277 | - | 364 | 322 | RNA→ rpsD (PRK05327)Ribosomal_S4 (pfam00163)→ |
| Bpa-1-1 | NZ_BAJR01000002.1 | + | 224743 | 224783 | RNA→ RpsD (COG0522)rpsD (PRK05327)→ RpoA (COG0202)PRK05182 (PRK05182)→ RplQ (COG0203)rplQ (PRK05591)→ |
| env-407 | BABD01002563.1 | + | 2434 | 2474 | RNA→ rpsD (PRK05327)Ribosomal_S4 (pfam00163)→ |
| env-408 | 4491404.3_NODE_23173 | - | 1622 | 1582 | RNA→ RpsD (COG0522)rpsD (PRK05327)→ RpoA (COG0202)PRK05182 (PRK05182)→ |
| Bba-1-1 | NZ_AUTU01000007.1 | + | 71664 | 71704 | RNA→ RpsD (COG0522)rpsD (PRK05327)→ RpoA (COG0202)PRK05182 (PRK05182)→ RplQ (COG0203)rplQ (PRK05591)→ |
| env-409 | AUXO016092034.1 | - | 828 | 788 | RNA→ RpsD (COG0522)rpsD (PRK05327)→ PRK05182 (PRK05182)RNA_pol_L (pfam01193)→ |
| env-410 | DLM007_scaffold43740_1 | + | 868 | 908 | RNA→ RpsD (COG0522)rpsD (PRK05327)→ PRK05182 (PRK05182)RNA_pol_L (pfam01193)→ |
| env-411 | BMHB3a_c153300 | + | 16662 | 16702 | RNA→ RpsD (COG0522)rpsD (PRK05327)→ RpoA (COG0202)PRK05182 (PRK05182)→ RplQ (COG0203)rplQ (PRK05591)→ |
| env-412 | JGI24728J21555_1016111 | - | 1883 | 1843 | RNA→ RpsD (COG0522)rpsD (PRK05327)→ RpoA (COG0202)PRK05182 (PRK05182)→ rplQ (PRK05591)Ribosomal_L17 (pfam01196)→ |
| env-413 | BMHBC_922 | + | 4020 | 4060 | RNA→ RpsD (COG0522)rpsD (PRK05327)→ RpoA (COG0202)PRK05182 (PRK05182)→ RplQ (COG0203)rplQ (PRK05591)→ |
| env-414 | HCF12C_147729 | + | 276 | 306 | RNA→ |
| Bpr-1-1 | NZ_AQWS01000016.1 | + | 67128 | 67168 | RNA→ RpsD (COG0522)rpsD (PRK05327)→ RpoA (COG0202)PRK05182 (PRK05182)→ RplQ (COG0203)rplQ (PRK05591)→ |
| Bco-1-1 | NZ_CM001167.1 | - | 250156 | 250116 | RNA→ RpsD (COG0522)rpsD (PRK05327)→ RpsD (COG0522)rpsD (PRK05327)→ RpoA (COG0202)PRK05182 (PRK05182)→ RpoA (COG0202)PRK05182 (PRK05182)→ RplQ (COG0203)rplQ (PRK05591)→ RplQ (COG0203)rplQ (PRK05591)→ |
| env-415 | JGI26523J50269_1004409 | + | 2776 | 2819 | RNA→ RpsD (COG0522)rpsD (PRK05327)→ RNAP_alpha_NTD (cd06928)PRK05182 (PRK05182)→ |
| env-416 | 3300000408_102680 | + | 393 | 436 | RNA→ RpsD (COG0522)rpsD (PRK05327)→ RNAP_alpha_NTD (cd06928)PRK05182 (PRK05182)RNA_pol_A_CTD (pfam03118)→ |
| env-417 | scaffold27906_1_MH0037 | - | 715 | 672 | RNA→ rpsD (PRK05327)Ribosomal_S4 (pfam00163)→ RpsD (COG0522)rpsD (PRK05327)→ |
| Bvi-1-1 | NZ_CP007034.1 | + | 1999562 | 1999605 | RNA→ RpsD (COG0522)rpsD (PRK05327)→ RNAP_alpha_NTD (cd06928)PRK05182 (PRK05182)RNA_pol_A_CTD (pfam03118)→ RplQ (COG0203)rplQ (PRK05591)→ |
| env-418 | scaffold5855_4_MH0006 | - | 1933 | 1890 | RNA→ RpsD (COG0522)rpsD (PRK05327)→ RNAP_alpha_NTD (cd06928)PRK05182 (PRK05182)RNA_pol_A_CTD (pfam03118)→ rplQ (PRK05591)Ribosomal_L17 (pfam01196)→ |
| env-419 | 4491415.3_NODE_16450 | + | 1929 | 1972 | RNA→ RpsD (COG0522)rpsD (PRK05327)→ RpoA (COG0202)PRK05182 (PRK05182)→ |
| env-420 | scaffold13158_6_MH0012 | + | 400 | 443 | RNA→ RpsD (COG0522)rpsD (PRK05327)→ RNAP_alpha_NTD (cd06928)PRK05182 (PRK05182)RNA_pol_A_CTD (pfam03118)→ rplQ (PRK05591)Ribosomal_L17 (pfam01196)→ rplQ (PRK05591)Ribosomal_L17 (pfam01196)→ |
| env-421 | BMHB3a_c149458 | - | 1857 | 1814 | RNA→ RpsD (COG0522)rpsD (PRK05327)→ RNAP_alpha_NTD (cd06928)PRK05182 (PRK05182)RNA_pol_A_CTD (pfam03118)→ rplQ (PRK05591)Ribosomal_L17 (pfam01196)→ |
| env-422 | RUMENNODE_488125_148000 | - | 776 | 733 | RNA→ RpsD (COG0522)rpsD (PRK05327)→ PRK05182 (PRK05182)→ |
| env-423 | scaffold8032_1_MH0020 | - | 100 | 57 | RNA→ |
| env-424 | UnmappedStool_Broad_C251130659 | - | 263 | 220 | RNA→ rpsD (PRK05327)Ribosomal_S4 (pfam00163)→ |
| env-425 | HCE12Call500_c0131960 | + | 95 | 141 | RNA→ RpsD (COG0522)rpsD (PRK05327)→ RNAP_alpha_NTD (cd06928)PRK05182 (PRK05182)RNA_pol_A_CTD (pfam03118)→ rplQ (PRK05591)Ribosomal_L17 (pfam01196)→ |
| env-426 | 2013403773 | + | 632 | 678 | RNA→ ←hypo |
| env-427 | JGI20163J15578_10124444 | - | 233 | 192 | RNA→ rpsD (PRK05327)Ribosomal_S4 (pfam00163)→ |
| env-428 | JGI20163J15578_10757792 | - | 551 | 510 | RNA→ RpsD (COG0522)rpsD (PRK05327)→ |
| env-429 | Incfw_1000509 | - | 63510 | 63469 | RNA→ RpsD (COG0522)rpsD (PRK05327)→ RNAP_alpha_NTD (cd06928)PRK05182 (PRK05182)RNA_pol_A_CTD (pfam03118)→ RplQ (COG0203)rplQ (PRK05591)→ |
| env-430 | IMNBL1_c0128642 | - | 512 | 471 | RNA→ RpsD (COG0522)rpsD (PRK05327)→ |
| env-431 | 2227561212 | - | 532 | 491 | RNA→ RpsD (COG0522)rpsD (PRK05327)→ |
| env-432 | 2227599765 | - | 41 | 10 | RNA→ |

| | | | | | |
|---|---|---|---|---|---|
| Dsp-1-1 | NZ_CVRP01000013.1 | - | 125218 | 125177 | **RNA**→ RpsD (COG0522)rpsD (PRK05327)→ RNAP_alpha_NTD (cd06928)PRK05182 (PRK05182)RNA_pol_A_CTD (pfam03118)→ RplQ (COG0203)rplQ (PRK05591)→ |
| env-433 | 2210517547 | - | 191 | 150 | **RNA**→ rpsD (PRK05327)Ribosomal_S4 (pfam00163)→ |
| env-434 | JGI26523J50269_1102284 | - | 319 | 278 | **RNA**→ rpsD (PRK05327)Ribosomal_S4 (pfam00163)→ |
| Bga-1-1 | NZ_KB894134.1 | - | 31988 | 31948 | **RNA**→ RpsD (COG0522)rpsD (PRK05327)→ RpoA (COG0202)PRK05182 (PRK05182)→ RplQ (COG0203)rplQ (PRK05591)→ rplQ (PRK05591)Ribosomal_L17 (pfam01196)→ |
| env-435 | NLF006_scaffold28489_1 | + | 1163 | 1203 | **RNA**→ |
| env-436 | BABD01000996.1 | + | 324 | 364 | **RNA**→ rpsD (PRK05327)Ribosomal_S4 (pfam00163)→ S4 (smart00363)RpsD (COG0522)→ RpoA (COG0202)PRK05182 (PRK05182)→ RplQ (COG0203)rplQ (PRK05591)→ |
| env-437 | CAM_READ_0099683363 | + | 274 | 314 | **RNA**→ rpsD (PRK05327)Ribosomal_S4 (pfam00163)→ |
| env-438 | CAM_READ_0100181161 | + | 263 | 303 | **RNA**→ |
| env-439 | CAM_READ_0100124625 | + | 293 | 334 | **RNA**→ |
| env-440 | CAM_READ_0099376081 | + | 46 | 86 | **RNA**→ rpsD (PRK05327)Ribosomal_S4 (pfam00163)→ S4 (smart00363)S4 (pfam01479)→ |
| env-441 | CAM_READ_0099881367 | + | 377 | 407 | **RNA**→ |
| env-442 | 4448820.3_Contig_2096 | + | 70 | 110 | **RNA**→ rpsD (PRK05327)Ribosomal_S4 (pfam00163)→ S4 (smart00363)RpsD (COG0522)→ |
| env-443 | 4491406.3_NODE_1651 | - | 106 | 66 | **RNA**→ |
| env-444 | SRS057478_LANL_scaffold_3450 | + | 67 | 107 | **RNA**→ ←hypo |
| env-445 | BABB01003051.1 | + | 162 | 202 | **RNA**→ RpsD (COG0522)rpsD (PRK05327)→ RNAP_alpha_NTD (cd06928)PRK05182 (PRK05182)→ |
| env-446 | BAAY01002159.1 | + | 2011 | 2051 | **RNA**→ RpsD (COG0522)rpsD (PRK05327)→ hypo→ |
| env-447 | 4448808.3_Contig27637 | - | 313 | 273 | **RNA**→ |
| env-448 | DOM001_scaffold38696_1 | - | 521 | 481 | **RNA**→ RpsD (COG0522)rpsD (PRK05327)→ |
| env-449 | NOM016_scaffold2065_1 | + | 772 | 812 | **RNA**→ RpsD (COG0522)rpsD (PRK05327)→ |
| env-450 | SRS023176_Baylor_scaffold_1186 | - | 45 | 15 | **RNA**→ |
| env-451 | 4448815.3_Contig37632 | - | 426 | 386 | **RNA**→ S4 (smart00363)RpsD (COG0522)→ |
| env-452 | 4491401.3_NODE_53198 | + | 747 | 787 | **RNA**→ RpsD (COG0522)rpsD (PRK05327)→ |
| env-453 | 4491419.3_NODE_6796 | - | 720 | 680 | **RNA**→ RpsD (COG0522)rpsD (PRK05327)→ |
| env-454 | DLF004_scaffold28173_1 | + | 663 | 703 | **RNA**→ RpsD (COG0522)rpsD (PRK05327)→ |
| env-455 | NOM012_scaffold42779_1 | - | 646 | 606 | **RNA**→ RpsD (COG0522)rpsD (PRK05327)→ |
| env-456 | NOM020_scaffold27835_3 | - | 612 | 572 | **RNA**→ RpsD (COG0522)rpsD (PRK05327)→ |
| env-457 | 4448814.3_Contig417 | - | 3940 | 3900 | **RNA**→ RpsD (COG0522)rpsD (PRK05327)→ RpoA (COG0202)PRK05182 (PRK05182)→ RplQ (COG0203)rplQ (PRK05591)→ hypo→ |
| env-458 | NOM015_scaffold155_1 | + | 161 | 201 | **RNA**→ RpsD (COG0522)rpsD (PRK05327)→ |
| env-459 | SRS015960_WUGC_scaffold_3709 | - | 45 | 15 | **RNA**→ |
| env-460 | SRS049900_LANL_scaffold_2992 | - | 369 | 329 | **RNA**→ rpsD (PRK05327)Ribosomal_S4 (pfam00163)→ |
| env-461 | 4448813.3_Contig19274 | - | 59 | 29 | **RNA**→ |
| env-462 | SRS018665_C3523114 | + | 23 | 70 | **RNA**→ rpsD (PRK05327)Ribosomal_S4 (pfam00163)→ |
| env-463 | SRS019128_C4417707 | - | 737 | 690 | **RNA**→ RpsD (COG0522)rpsD (PRK05327)→ |
| env-464 | SRS022536_LANL_scaffold_112290 | + | 4797 | 4844 | **RNA**→ S4 (smart00363)RpsD (COG0522)→ PRK05182 (PRK05182)RNA_pol_L (pfam01193)→ |
| env-465 | SRS016575_Baylor_scaffold_36465 | - | 658 | 611 | **RNA**→ RpsD (COG0522)rpsD (PRK05327)→ |
| env-466 | SRS050628_C1820033 | + | 47 | 94 | **RNA**→ rpsD (PRK05327)Ribosomal_S4 (pfam00163)→ |
| env-467 | SRS052876_LANL_scaffold_11771 | + | 2902 | 2949 | **RNA**→ RpsD (COG0522)rpsD (PRK05327)→ RNAP_alpha_NTD (cd06928)PRK05182 (PRK05182)→ |
| env-468 | SRS063603_LANL_scaffold_98718 | - | 50 | 13 | **RNA**→ |
| env-469 | SRS017227_Baylor_scaffold_160228 | + | 643 | 690 | **RNA**→ rpsD (PRK05327)Ribosomal_S4 (pfam00163)→ |
| env-470 | SRS018394_Baylor_scaffold_73299 | + | 6083 | 6130 | **RNA**→ rpsD (PRK05327)Ribosomal_S4 (pfam00163)→ |
| env-471 | SRS021496_Baylor_scaffold_84955 | - | 737 | 690 | **RNA**→ RpsD (COG0522)rpsD (PRK05327)→ |
| env-472 | SRS023595_Baylor_scaffold_87344 | - | 329 | 282 | **RNA**→ rpsD (PRK05327)Ribosomal_S4 (pfam00163)→ |
| env-473 | SRS024289_LANL_scaffold_49264 | + | 21972 | 22019 | **RNA**→ RpsD (COG0522)rpsD (PRK05327)→ RNAP_alpha_NTD (cd06928)PRK05182 (PRK05182)RNA_pol_A_CTD (pfam03118)→ RplQ (COG0203)rplQ (PRK05591)→ |
| env-474 | SRS052604_C1840067 | + | 1153 | 1200 | **RNA**→ RpsD (COG0522)rpsD (PRK05327)→ RNAP_alpha_NTD (cd06928)PRK05182 (PRK05182)RNA_pol_A_CTD (pfam03118)→ RplQ (COG0203)rplQ (PRK05591)→ |
| env-475 | SRS058808_C2801786 | + | 507 | 554 | **RNA**→ |

| | | | | | |
|---|---|---|---|---|---|
| env-476 | SRS065099_C2629417 | - | 284 | 237 | **RNA**→ rpsD (PRK05327)Ribosomal_S4 (pfam00163)→ |
| env-477 | Buccal_mucosa_LANL_scaffold_111229 | - | 710 | 663 | **RNA**→ RpsD (COG0522)rpsD (PRK05327)→ |
| env-478 | SRS013705_Baylor_scaffold_103305 | - | 435 | 388 | **RNA**→ RpsD (COG0522)rpsD (PRK05327)→ |
| env-479 | SRS018443_C3874981 | + | 304 | 351 | **RNA**→ RpsD (COG0522)rpsD (PRK05327)→ RpoA (COG0202)PRK05182 (PRK05182)→ |
| env-480 | SRS017304_Baylor_scaffold_31533 | + | 6469 | 6516 | **RNA**→ RpsD (COG0522)rpsD (PRK05327)→ RNAP_alpha_NTD (cd06928)PRK05182 (PRK05182)RNA_pol_A_CTD (pfam03118)→ RplQ (COG0203)rplQ (PRK05591)→ |
| env-481 | SRS013252_Baylor_scaffold_26940 | + | 960 | 1007 | **RNA**→ RpsD (COG0522)rpsD (PRK05327)→ |
| env-482 | SRS019028_WUGC_scaffold_45751 | - | 423 | 376 | **RNA**→ RpsD (COG0522)rpsD (PRK05327)→ |
| env-483 | SRS012279_Baylor_scaffold_66400 | - | 3211 | 3164 | **RNA**→ RpsD (COG0522)rpsD (PRK05327)→ RNAP_alpha_NTD (cd06928)PRK05182 (PRK05182)RNA_pol_A_CTD (pfam03118)→ RplQ (COG0203)rplQ (PRK05591)→ |
| env-484 | SRS015893_C2389359 | - | 740 | 693 | **RNA**→ RpsD (COG0522)rpsD (PRK05327)→ |
| env-485 | SRS049268_C4370284 | + | 2539 | 2576 | **RNA**→ |
| env-486 | SRS051930_C3081289 | - | 76 | 39 | **RNA**→ |
| env-487 | SRS055378_LANL_scaffold_88414 | - | 2987 | 2940 | **RNA**→ RpsD (COG0522)rpsD (PRK05327)→ RNAP_alpha_NTD (cd06928)PRK05182 (PRK05182)RNA_pol_A_CTD (pfam03118)→ RplQ (COG0203)rplQ (PRK05591)→ |
| env-488 | SRS016360_C2594021 | - | 74 | 37 | **RNA**→ ←hypo |
| env-489 | SRS017511_Baylor_scaffold_44051 | - | 4149 | 4102 | **RNA**→ RpsD (COG0522)rpsD (PRK05327)→ RNAP_alpha_NTD (cd06928)PRK05182 (PRK05182)RNA_pol_A_CTD (pfam03118)→ RplQ (COG0203)rplQ (PRK05591)→ |
| env-490 | SRS018739_WUGC_scaffold_35509 | + | 66 | 113 | **RNA**→ RpsD (COG0522)rpsD (PRK05327)→ |
| env-491 | SRS022143_WUGC_scaffold_58148 | + | 62 | 109 | **RNA**→ RpsD (COG0522)rpsD (PRK05327)→ RNAP_alpha_NTD (cd06928)PRK05182 (PRK05182)RNA_pol_A_CTD (pfam03118)→ rplQ (PRK05591)Ribosomal_L17 (pfam01196)→ |
| env-492 | SRS024087_LANL_scaffold_25676 | + | 1061 | 1108 | **RNA**→ RpsD (COG0522)rpsD (PRK05327)→ RNAP_alpha_NTD (cd06928)PRK05182 (PRK05182)RNA_pol_A_CTD (pfam03118)→ RplQ (COG0203)rplQ (PRK05591)→ |
| env-493 | SRS047113_C5300540 | - | 121 | 74 | **RNA**→ |
| env-494 | SRS055450_LANL_scaffold_68076 | + | 1281 | 1328 | **RNA**→ RpsD (COG0522)rpsD (PRK05327)→ RNAP_alpha_NTD (cd06928)PRK05182 (PRK05182)RNA_pol_A_CTD (pfam03118)→ RplQ (COG0203)rplQ (PRK05591)→ |
| env-495 | SRS013723_Baylor_scaffold_78094 | + | 134 | 181 | **RNA**→ RpsD (COG0522)rpsD (PRK05327)→ |
| env-496 | SRS015574_WUGC_scaffold_47042 | + | 110 | 157 | **RNA**→ RpsD (COG0522)rpsD (PRK05327)→ PRK05182 (PRK05182)RNA_pol_L (pfam01193)→ |
| env-497 | SRS013533_C5397762 | + | 347 | 394 | **RNA**→ |
| env-498 | SRS013170_C3989364 | + | 202 | 245 | **RNA**→ |
| env-499 | SRS015215_C2525062 | - | 60 | 23 | **RNA**→ |
| env-500 | SRS015762_C3609878 | + | 796 | 843 | **RNA**→ |
| env-501 | SRS015434_C5621575 | + | 192 | 237 | **RNA**→ |
| env-502 | SRS013170_Baylor_scaffold_120567 | + | 426 | 471 | **RNA**→ |
| env-503 | SRS023595_C7159178 | + | 773 | 808 | **RNA**→ |
| env-504 | scaffold11811_2_MH0082 | + | 201 | 253 | **RNA**→ RpsD (COG0522)rpsD (PRK05327)→ |
| env-505 | 08_07_XOS_DNA_scaffold14885_1 | - | 241 | 196 | **RNA**→ rpsD (PRK05327)Ribosomal_S4 (pfam00163)→ |
| env-506 | SRS022071_C2972092 | + | 67 | 112 | **RNA**→ RpsD (COG0522)rpsD (PRK05327)→ |
| env-507 | 08_19_CONTROL_DNA_scaffold3842_6 | - | 647 | 602 | **RNA**→ RpsD (COG0522)rpsD (PRK05327)→ |
| env-508 | 07_22_CONTROL_DNA_scaffold5573_1 | + | 364 | 409 | **RNA**→ RpsD (COG0522)rpsD (PRK05327)→ |
| env-509 | 07_22_CONTROL_DNA_scaffold54317_1 | + | 232 | 277 | **RNA**→ RpsD (COG0522)rpsD (PRK05327)→ RNAP_alpha_NTD (cd06928)PRK05182 (PRK05182)RNA_pol_A_CTD (pfam03118)→ rplQ (PRK05591)Ribosomal_L17 (pfam01196)→ |
| env-510 | 07_22_INULIN_DNA_scaffold23708_1 | + | 190 | 235 | **RNA**→ RpsD (COG0522)rpsD (PRK05327)→ |
| env-511 | scaffold11208_4_MH0083 | + | 4936 | 4981 | **RNA**→ RpsD (COG0522)rpsD (PRK05327)→ RpoA (COG0202)PRK05182 (PRK05182)→ RplQ (COG0203)rplQ (PRK05591)→ |
| env-512 | 07_22_INULIN_DNA_scaffold131_3 | - | 1214 | 1169 | **RNA**→ RpsD (COG0522)rpsD (PRK05327)→ RNAP_alpha_NTD (cd06928)PRK05182 (PRK05182)→ |
| env-513 | scaffold80135_1_MH0077 | - | 1023 | 978 | **RNA**→ RpsD (COG0522)rpsD (PRK05327)→ PRK05182 (PRK05182)RNA_pol_L (pfam01193)→ |

| | | | | | |
|---|---|---|---|---|---|
| env-514 | O8_19_POLYDEXT_DNA_scaffold5982_10 | + | 115 | 160 | **RNA**→ RpsD (COG0522)rpsD (PRK05327)→ |
| env-515 | mgutOb2_U_BM_aaa76b05_b1 | - | 766 | 721 | **RNA**→ RpsD (COG0522)rpsD (PRK05327)→ |
| env-516 | SRS022137_WUGC_scaffold_28771 | + | 14373 | 14420 | **RNA**→ RpsD (COG0522)rpsD (PRK05327)→ RNAP_alpha_NTD (cd06928)PRK05182 (PRK05182)RNA_pol_A_CTD (pfam03118)→ RplQ (COG0203)rplQ (PRK05591)→ |
| env-517 | UnmappedStool_Broad_C251091185 | - | 208 | 165 | **RNA**→ rpsD (PRK05327)Ribosomal_S4 (pfam00163)→ |

#### 14.3 Conserved domains

Conserved domains found in protein-coding genes listed in Section 14.2 are shown below, with the first sentence in their description from the Conserved Domain Database (if any). Conserved domains associated with more than one S4-Bacteroidia RNA are assigned a color, while others are shown in gray. The number in parentheses after the colored domain name is the number of occurrences in Section 14.2.

cd00156 (1) Signal receiver domain; originally thought to be unique to bacteria (CheY, OmpR, NtrC, and PhoB), now recently identified in eukaroytes ETR1 Arabidopsis thaliana; this domain receives the signal from the sensor partner in a two-component systems; contains a phosphoacceptor site that is phosphorylated by histidine kinase homologs; usually found N-terminal to a DNA binding effector domain; forms homodimers
cd00165 (1) S4/Hsp/ tRNA synthetase RNA-binding domain; The domain surface is populated by conserved, charged residues that define a likely RNA-binding site; Found in stress proteins, ribosomal proteins and tRNA synthetases; This may imply a hitherto unrecognized functional similarity between these three protein classes.
cd06928 (96) N-terminal domain of the Alpha subunit of Bacterial RNA polymerase.
COG0100 (2) Ribosomal protein S11 [Translation, ribosomal structure and biogenesis]
COG0202 (94) DNA-directed RNA polymerase, alpha subunit/40 kD subunit [Transcription]
COG0203 (132) Ribosomal protein L17 [Translation, ribosomal structure and biogenesis]
COG0522 (341) Ribosomal protein S4 or related protein [Translation, ribosomal structure and biogenesis]
COG4185 (1) Predicted ABC-type ATPase [General function prediction only]
pfam00163 (100) Ribosomal protein S4/S9 N-terminal domain.
pfam01000 (1) RNA polymerase Rpb3/RpoA insert domain.
pfam01193 (26) RNA polymerase Rpb3/Rpb11 dimerisation domain.
pfam01196 (33) Ribosomal protein L17.
pfam01479 (1) S4 domain.
pfam03118 (79) Bacterial RNA polymerase, alpha chain C terminal domain.
pfam07877 (1) Protein of unknown function (DUF1661).
PLN02967 (1) kinase
PRK05182 (218) DNA-directed RNA polymerase subunit alpha; Provisional
PRK05309 (2) 30S ribosomal protein S11; Validated
PRK05327 (423) 30S ribosomal protein S4; Validated
PRK05591 (165) 50S ribosomal protein L17; Validated
PRK10905 (1) cell division protein DamX; Validated
PRK13108 (1) prolipoprotein diacylglyceryl transferase; Reviewed
PRK14521 (2) 30S ribosomal protein S16; Provisional
smart00363 (21) S4 RNA-binding domain.
smart00382 (1) ATPases associated with a variety of cellular activities.
smart00448 (1) cheY-homologous receiver domain.
TIGR02027 (1) DNA-directed RNA polymerase, alpha subunit, bacterial and chloroplast-type. [Transcription, DNA-dependent RNA polymerase]

#### 14.4 Multiple-sequence alignment

Each S4-Bacteroidia RNA is denoted by an abbreviation (like "Eco-1-1") that refers to a taxonomy given in Section 14.1. The alignment may include sequences containing the RNA motif, as well as flanking sequence. The S4-Bacteroidia RNA itself is denoted by the line underneath marked 5′ and 3′ on either end. Nucleotides in flanking sequences (i.e., not part of the motif) are written in gray letters. Stems of predicted rho-independent transcription terminators, if any, are shaded yellow. (Note: terminator predictions have not been analyzed manually, and many are likely to be false positives. Terminator predictions are those of the RNIE software.) Nucleotides predicted to function (as DNA) as transcription-factor binding sites, if any, are shaded in green. (But note: these predictions are manually annotated, so they might be under-predicted.) Annotated start codons, if any, are shaded green. (Note: start codons are frequently misannotated, especially in environmental samples.) Nucleotides proposed to basepair as part of the consensus structure are shaded in color when they comprise Watson-Crick or G-U pairs. Otherwise they are shaded gray. Conserved stems are also indicated at the bottom of the alignment by angle brackets, where matching < and > denote base-paired columns. Below these angle brackets, the symbol "2" denotes base pairs exhibiting covariation according to the statistically well-founded R-scape method. "1" denotes base pairs exhibiting covariation according to R2R's simplistic method. "0" denotes base pairs that are not observed to mutate and "?" denotes base pairs that have a significant frequency of non-canonical nucleotides for Watson-Crick or G-U pairs (> 5%). Below these base pair annotation is the consensus sequence: "R" = "A" or "G", "Y" = "C" or "U", red nucleotides: nucleotide identity conserved more than 97% of the time, black nucleotides: 90%, gray nucleotides: 75%, red circle (•): nucleotide is present 97% of the time, black circle (•): 90%, gray circle (•): 75%, white circle (◦): 50%. All percentages of sequences just described (e.g. 97% conserved) assume that sequences have been weighted by the GSC algorithm implemented by the Infernal software package.

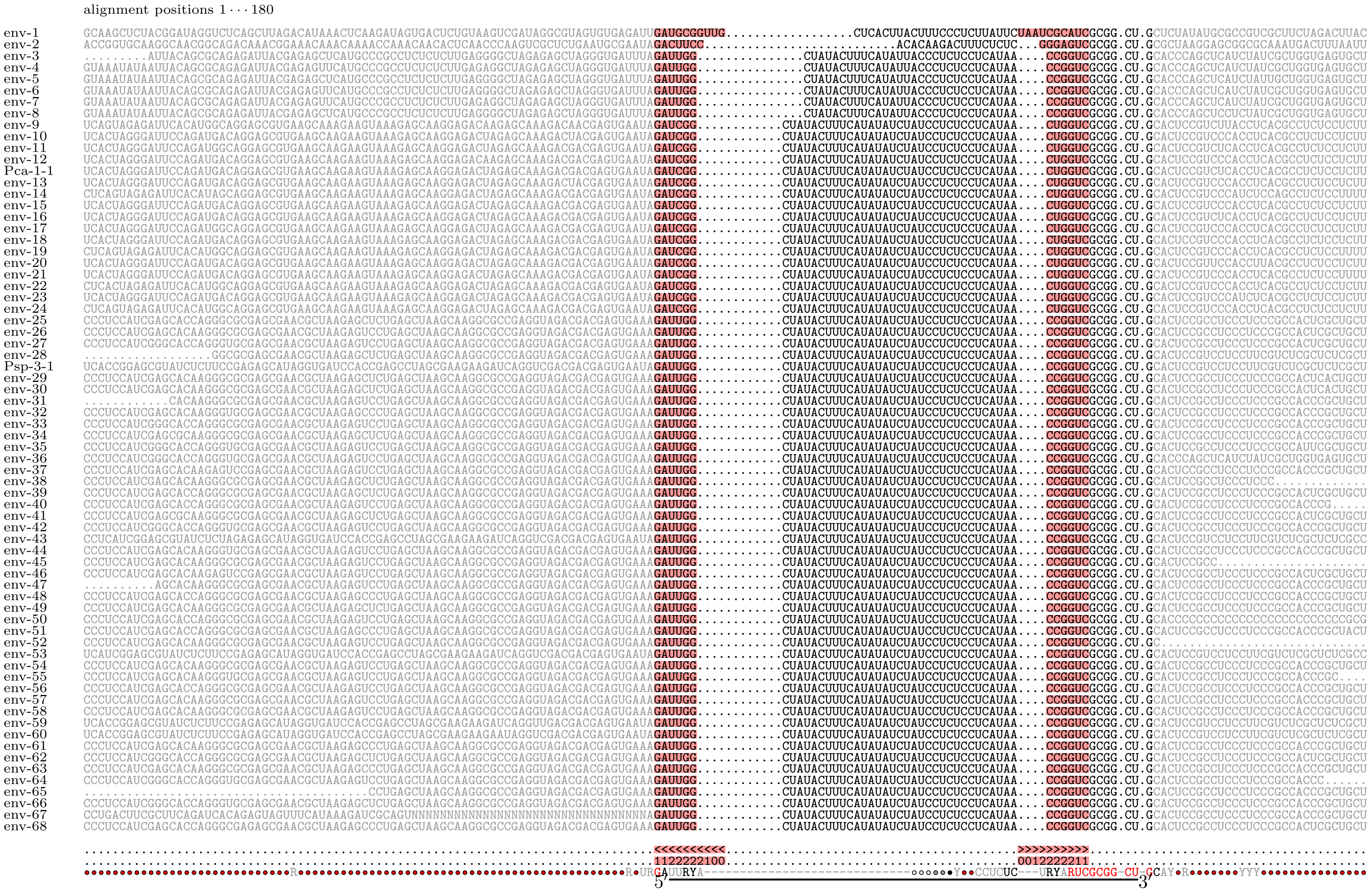

```
env-69    CCCUCCAUCGAGCACCAGGGCGCGAGCGAACGCUAAGAGUCCUGAGCUAAGCAAGGCGCCGAGGUAGACGACGAGUGAAAGAUUGG............CUAUACUUUCAUAUAUCUAUCCUCUCCUCAUAA....CCGGUCGCGG.CU.GCACUCCGCCUUCCUCCCGCCACUCGCUGCU
env-70    CCCUCCAUCGAGCACAAGGGCGCGAGCGAACGCUAAGAGCUCUGAGCUAAGCAAGGCGCCGAGGUAGACGACGAGUGAAAGAUUGG............CUAUACUUUCAUAUAUCUAUCCUCUCCUCAUAA....CCGGUCGCGG.CU.GCACUCCGCCUCCCUCCCGCCACCCGCUGCU
env-71    UCAUCGGAGCGUAUCUCUUCCGAGAGCAUAGGUGAUCCACCGAGCCUAGCGAAGAAGAUCAGGUCGACGACGAGUGAAUAGAUUGG............CUAUACUUUCAUAUAUCUAUCCUCUCCUCAUAA....CCGGUCGCGG.CU.GCACUCCGUCCUCCUUCGUCUCGCUCUCGCC
env-72    CCCUCCAUCGAGCACCAGGGCGCGAGCGAACGCUAAGAGCUCUGAGCUAAGCAAGGCGCCGAGGUAGACGACGAGUGAAAGAUUGG............CUAUACUUUCAUAUAUCUAUCCUCUCCUCAUAA....CCGGUCGCGG.CU.GCACUCCGCCUCCCUCCCGCCACCCGCUGCU
env-73    CCCUCCAUCGAGCACAAGGGCGCGAGCGAACGCUAAGAGUCCUGAGCUAAGCAAGGCGCCGAGGUAGACGACGAGUGAAAGAUUGG............CUAUACUUUCAUAUAUCUAUCCUCUCCUCAUAA....CCGGUCGCGG.CU.GCACUCCGCCUCCCUCCCGCCACUCGCUGCU
env-74    CCCUCCAUCGAGCACAAGGGCGCGAGCGAACGCUAAGAGUCCUGAGCUAAGCAAGGCGCCGAGGUAGACGACGAGUGAAAGAUUGG............CUAUACUUUCAUAUAUCUAUCCUCUCCUCAUAA....CCGGUCGCGG.CU.GCACUCCGCCUCCCUCCCGCCACCCGCUGCU
env-75    CCUCCAUCGAGUGUCGUGAGACAAGAGCGAAUGCUAAGAGCCUGAGCUAAGCAAGACGCCGAGGUAGACGACGAGUGAAAGAUUGG............CUAUACUUUCAUAUAUCUAUCCUCUCCUCAUAA....CCGGUCGCGG.CU.GCACUCCGCCUGCCUCCCGUCCCGCCCACUC
env-76    CCCUCCAUCGAGCACCAGGGCGCGAGCGAACGCUAAGAGUCCUGAGCUAAGCAAGGCGCCGAGGUAGACGACGAGUGAAAGAUUGG............CUAUACUUUCAUAUAUCUAUCCUCUCCUCAUAA....CCGGUCGCGG.CU.GCACUCCGCCUCCCUCCCGCCACUCGCUGCU
env-77    CCCUCCAUCGAGCGCAAGGGCGCGAGCGAACGCUAAGAGUCCUGAGCUAAGCAAGGCGCCGAGGUAGACGACGAGUGAAAGAUUGG............CUAUACUUUCAUAUAUCUAUCCUCUCCUCAUAA....CCGGUCGCGG.CU.GCACUCCGCCUCCCUCCCGCCACCCGCUGCU
env-78    CCUCCAUCGAGUGCCGCAAGACAAGAGCGAAUGCUAAGAGCCUGAGCUAAGCAAGACGCCGAGGUAGACGACGAGUGAAAGAUUGG............CUAUACUUUCAUAUAUCUAUCCUCUCCUCAUAA....CCGGUCGCGG.CU.GCACUCCGCCUGCCUCCCGUCCCGCCCGCUC
env-79    CCCUCCAUCGAGCCCCAGGGCGCGAGCGAACGCUAAGAGUCCUGAGCUAAGCAAGGCGCCGAGGUAGACGACGAGUGAAAGAUUGG............CUAUACUUUCAUAUAUCUAUCCUCUCCUCAUAA....CCGGUCGCGG.CU.GCACUCCGCCUCCCUCCCGCCACUCGCUGCU
env-80    UCAUCGGAGCGUAUCUCUCUAGAGAGCAUAGGUGAUCCACCGAGCCUAGCGAAGAAGAUCAGGUCGACGACGAGUGAAUAGAUUGG............CUAUACUUUCAUAUAUCUAUCCUCUCCUCAUAA....CCGGUCGCGG.CU.GCACUCCGUCCUCCUUCGUCUCGCUCUCGCC
env-81    CCCUCCAUCGGGCACCAGGGUGCGAGCGAACGCUAAGAGUCCUGAGCUAAGCAAGGCGCCGAGGUAGACGACGAGUGAAAGAUUGG............CUAUACUUUCAUAUAUCUAUCCUCUCCUCAUAA....CCGGUCGCGG.CU.GCACUCCGCCUCCCUCCCGCCACCCGCUGCU
env-82    CCCUCCAUCGAGCACCAGGGCGCGAGCGAACGCUAAGAGUCCUGAGCUAAGCAAGGCGCCGAGGUAGACGACGAGUGAAAGAUUGG............CUAUACUUUCAUAUAUCUAUCCUCUCCUCAUAA....CCGGUCGCGG.CU.GCACUCCGCCUCCCUCCCGCCACUCGCUGCU
env-83    CCCUCCAUCGAGCACCAGGGCGCGAGCGAACGCUAAGAGUCCUGAGCUAAGCAAGGCGCCGAGGUAGACGACGAGUGAAAGAUUGG............CUAUACUUUCAUAUAUCUAUCCUCUCCUCAUAA....CCGGUCGCGG.CU.GCACUCCGCCUCCCUCCCGCCACUCGCUGCU
env-84    CCCUCCAUCGAGCACAAGGGCGCGAGCGAACGCUAAGAGCCCUGAGCUAAGCAAGGCGCCGAGGUAGACGACGAGUGAAAGAUUGG............CUAUACUUUCAUAUAUCUAUCCUCUCCUCAUAA....CCGGUCGCGG.CU.GCACUCCGCCUCCCUCCCGCCACUCGCUGCU
env-85    CCCUCCAUCGAGCACAAGGGUGCAAGCGAACGCUAAGAGUCCUGAGCUAAGCAAGGCGCCGAGGUAGACGACGAGUGAAAGAUUGG............CUAUACUUUCAUAUAUCUAUCCUCUCCUCAUAA....CCGGUCGCGG.CU.GCACUCCGCCUCCCUCCCGCCACCCGCUGCU
env-86    CCCUCCAUCGGGCACCAGGGUGCGAGCGAACGCUAAGAGUCCUGAGCUAAGCAAGGCGCCGAGGUAGACGACGAGUGAAAGAUUGG............CUAUACUUUCAUAUAUCUAUCCUCUCCUCAUAA....CCGGUCGCGG.CU.GCACUCCGCCUCCCUCCCGCCACCCGCUGCU
env-87    UCACCGGAGCGUAUCUCUUCCGAGAGUAUAGGUGAUCCACCGAGCCUAGCGAAGAAGAUCAGGUCGACGACGAGUGAAUAGAUUGG............CUAUACUUUCAUAUAUCUAUCCUCUCCUCAUAA....CCGGUCGCGG.CU.GCACUCCGUCCUCCUUCGUCUCGCUCUCGCU
env-88    CCCUCCAUCGAGCACAAGGGCGCGAGCGAAUGCUAAGAGUCCUGAGCUAAGCAAGGCGCCGAGGUAGACGACGAGUGAAAGAUUGG............CUAUACUUUCAUAUAUCUAUCCUCUCCUCAUAA....CCGGUCGCGG.CU.GCACUCCGCCUCCCUCCCGCCACCCGCUGCU
env-89    CCCUCCAUCGAGCACCAGGGCGCGAGCGAACGCUAAGAGUCCUGAGCAAAGCAAGGCGCCGAGGUAGACGACGAGUGAAAGAUUGG............CUAUACUUUCAUAUAUCUAUCCUCUCCUCAUAA....CCGGUCGCGG.CU.GCACUCCGCCUCCCUCCCGCCACCCGCUGCU
env-90    UCAUCGGAGCGUAUCUCUCUAGAGAGCAUAGGUGAUCCGCUGAGCCUAGCGAAGAAGAUCAGGUCGACGACGAGUGAAUAGAUUGG............CUAUACUUUCAUAUAUCUAUCCUCUCCUCAUAA....CCGGUCGCGG.CU.GCACUCCGUCCUCCUUCGUCUCGCUCUCGCC
env-91    CCCUCCAUCGAGCACAAGGGCGCUAGCGAACACUAAGAGUCCUGAGCUAAGCAAGGCGCCGAGGUAGACGACGAGUGAAAGAUUGG............CUAUACUUUCAUAUAUCUAUCCUCUCCUCAUAA....CCGGUCGCGG.CU.GCACUCCGCCUCCCUCCCGCCACCUGCUGCU
env-92    CCUCCAUCGAGUCCCGCGAGACAAGAGCGAAUGCUAAGAGCCUGAGCUAAGCAAGACGCCGAGGUAGACGACGAGUGAAAGAUUGG............CUAUACUUUCAUAUAUCUAUCCUCUCCUCAUAA....CCGGUCGCGG.CU.GCACUCCGCCUGCCUCCCGUCCCGCCCACUC
Psp-2-1   UAAUUCAACGAUGCCGAUGAGAUAGAUUGAGCUCAAGGUCAAAAGACGGAGGAUACAUCAGGCUGGAGAUUUGGUGAAUAGAUUGG............CUAUACUUUCAUAUUUCUAUCCUCUCCUCAUAA....CCGGUCGCGG.CU.GCACCGCUCUGGCUCUCUCCGCACAAAUACU
Psp-1-1   CGAGAGACUCGACACAUGAUUUAUUGAUGCGGUAGGGCGGGGAUGACCCCUCGCAGCUGGAGUCUGAGACUGGGCGAAUAGAUUGG............CUAUACUUUCAUAUUUCUAUCCUCUCCUCAUAA....CCGGUCGCGG.CU.GCGCCACUCAGACACUCUCGAGGCUUCUGAC
env-93    UAGGAGCUAAUGGUCGCUAGAGCAUGACAGAGAAGUAGCAGAAUAGGUCAGGAUCCCCUAGGAGGGACGAUGGGUGAAUAGAUUGG.............CUAUACUUUCAUUAUCUAUCCUCUCCUCAUAA....CCGGUCGCGG.CU.GCACUCCGCCUCCCUCCCGCCACCCGCUGCU
env-94    AAGGAGCUGAUGGUCGCUAGAGCAUGACAGAGAAGUAGCAGAAUAGGUCAGGAUCCCCAAGGAGGGACGAUGGGUGAAUAGAUUGG.............CUAUACUUUCAUUAUCUAUCCUCUCCUCAUAA....CCGGUCGCGG.CU.GCACUCCGCCCUCCUACCCUCGCCUCCUGCC
env-95    UAGGAGCUAAUGGUCGCUAGAGCAUGACAGAGAAGUAGCAGAAUAGGUCAGGAUCCCCUAGGAGGGACGAUGGGUAAAUAGAUUGG.............CUAUACUUUCAUUAUCUAUCCUCUCCUCAUAA....CCGGUCGCGG.CU.GCACUCCGCCCUCCUACCCUCACCUCCUGCC
env-96    AAGGAGCUGAUGGGUGCUAGAGCAUGACAGAGAAGUAGCAGAAUAGGUCAGGAUCCUUCAGGAGGGACGAUGGGUGAAUAGAUUGG.............CUAUACUUUCAUUAUCUAUCCUCUCCUCAUAA....CCGGUCGCGG.CU.GCACUCCGCCCUCCUACCCUCGCCUCCUGCC
env-97    AAGGAGCUGAUGGUCGCUAGAGCAUGACAGAGAAGUAGCAGAAUAGGUCAGGAUCCCCAAGGAGGGACGAUGGGUGAAUAGAUUGG.............CUAUACUUUCAUUAUCUAUCCUCUCCUCAUAA....CCGGUCGCGG.CU.GCACUCCGCCUCCCUCCCGCCACCCGCUGCU
env-98    AAGGAGCUGAUGGUCGCUAGAGCAUGACAGAGAAGUAGCAGAAUAGGUCAGGAUCCCCAAGGAGGGACGAUGGGUGAAUAGAUUGG.............CUAUACUUUCAUUAUCUAUCCUCUCCUCAUAA....CCGGUCGCGG.CU.GCACUCCGCCCUCCUACCCUCGCCUCCUGCC
env-99    AAGGAGCUGAUGGUCGCUAGAGCAUGACAGAGAAGUAGCAGAAUAGGUCAGGAUCCUUCAGGAGGGACGAUGGGUGAAUAGAUUGG.............CUAUACUUUCAUUAUCUAUCCUCUCCUCAUAA....CCGGUCGCGG.CU.GCACUCCGCCCUCCUACCCUCGCCUCCUGCC
env-100   CUAGGAGCUAAUGACGCUAGAGCAUGACAGAGACGUAGCAGAAUAGGUCAGGAUCCCCUAGGAGGGACGAUGGGUGAAUAGAUUGG.............CUAUACUUUCAUUAUCUAUCCUCUCCUCAUAA....CCGGUCGCGG.CU.GCACUCCGCCCUCCUCCCCUCACCUCCUGCC
env-101   AAGGAGCUGAUGGUCUCUAGAGCAUGACAGAGAAGUAGCAGAAUAGGUCAGGAUCCUUCAGGAGGGACGAUGGGUGAAUAGAUUGG.............CUAUACUUUCAUUAUCUAUCCUCUCCUCAUAA....CCGGUCGCGG.CU.GCACUCCGCCCUCCUAUCCUCGCCUCCUACC
env-102   AAGGAACUGAUGGUCGCUAGAGCAUGACAGAGAAGUAGCAGAAUAGGUCAGGAUCCCCUAGGAGGGACGAUGGGUGAAUAGAUUGG.............CUAUACUUUCAUUAUCUAUCCUCUCCUCAUAA....CCGGUCGCGG.CU.GCACUCCGCCCUCCUACCCUCGCCUCCUGAC
env-103   UAGGAGCUAAUGGACGCUAGAGCAUGACAGAGAAGUAGCAGAAUAGGUCAGGAUCCCCUAGGAGGGACGAUGGGUGAAUAGAUUGG.............CUAUACUUUCAUUAUCUAUCCUCUCCUCAUAA....CCGGUCGCGG.CU.GCACUCCGCCCUCCUCCCCUCACCUCCUGUC
env-104   AAAGGAGCUGAUGGUCACUAGAGCAUGACAGAGAAGUAGCAGAUAGGUCAGGAUCCUUCAGGAGGGACGAUGGGUGAAUAGAUUGG.............CUAUACUUUCAUUAUCUAUCCUCUCCUCAUAA....CCGGUCGCGG.CU.GCACUCCGCCCUCCUACCCUCGCCUCCUGCC
env-105   UAGGAGCUAAUGGUCGCUAGAGCAUGACAGAGAAGUAGCAGAAUAGGUCAGGAUCCCCUAGGAGGGACGAUGGGUGAAUAGAUUGG.............CUAUACUUUCAUUAUCUAUCCUCUCCUCAUAA....CCGGUCGCGG.CU.GCACUCCGCCCUCCUCCCCUCCCCUCCUGCC
env-106   CCCGCCAUAGAGCCCUGCUCUCGGACGUGUCGGGGCCUAGCCCUAGGGAUGAUGCUCGGAUGAGGUACGAUGAGUGAAUAGAUUGG.............CUAUACUUUCAAUAUCUAUCCUCUCCUCAUAA....CCGGUCGCGG.CU.GCACUCCGUCCUCCAUCCCAGCUAUCCACCU
env-107   CCCGCCAUAGAGCCCUGCUCUCGGACGUGUCGGGGCCUAGCCCUAGGGAUGAUGCUCGGAUGAGGUACGAUGAGUGAAUAGAUUGG.............CUAUACUUUCAAUAUCUAUCCUCUCCUCAUAA....CCGGUCGCGG.CU.GCACUCCGUCCUCCAUCCCAGCUAUCCACCU
env-108   GCGCCGUCGCGUAUAAUACAGGAAGACAAACAAUACAGCAUACAAGACUUCCGUCCCGCUGAGCGCCCUGAAUGCGAAUAGACCCUC...........................CCAUAACUACCUCUC...GAGGGUCGCGG.CU.GCGCUAAGGGGAGAGCCUCAAGGACACAACA
env-109   GCGCCGUCGCGUAUAAUACAGGAAGACAAACAAAACAGCAUACAAGACUUCCGUCCCGCUGAGCGCCCUGAAUGCGAAUAGACCCUC...........................CCAUAACUACCUCUC...GAGGGUCGCGG.CU.GCGCUAAGGGGAGAGCCUCAAGGACACAACA
env-110   CGAAGAGAAGAAGAGUAUAAUCCGAAGAGGCAUCCAAAAAGAACACGACUUCAGCCGACGAACCGUCCUGAAUGCGAAUAGACCCUC..........................CCAUCACCUUUCUCUC...GAGGGUCGCGG.CU.GCGCUAAGGGAUAGGCGCCGGCACGAAAAAA
env-111   AGAAGAGUCUAAUCGGAAUUUUUUCAAACAAACAGAAUAUUCUGAACAAAGUCCGCCUCCGGUUGUCCUGAAUGCGAAUUGACCGUC..........................ACUUGCACUUUCUCUC...GACGGUCGCGG.CU.GCGCUAAGGGAUAUGCUCAGCGGAUGAGCAA
env-112   CCCGCACAACGGAUGUCGUCCUCCGAAAAGAAGAAGAGUAUAACAUCAAAGCCCCUCGCGCCCGUCUCUGAAUGCGAAUAGACUGCC..........................ACAGCAUCUUUCUCUC...GGCAGUCGCGG.CU.GCGCUAAGGAGCGGGCAGAAAGGCCCACUUU
env-113   GCCUAACUGAUUUGACGACAAGCUGCGCCGACAUCCGUUUUCAUAUCUUUUAACGUGGUGAACGGGCAUGGAAUGUGAAAGACUGCU........................UUCAAAACAUCACCUCUC...AGCAGUCGCGG.CU.GCACUAACGUGUCAAGCACCCCACCCUCGAG
env-114   AACUAUAAGCAUUCACAUAUUGUUUUAUUCUCGCAGCAUCCUAUACGUUAUUGACACCGGCAUCGCGGCUAUGUGAUUUUGAUCAUA..........................UCUUUUUUCACCUCUC...UAUGAUCGCGG.CU.GCACUACACCGCCUAUAAGCCUCAUAACUCU
env-115   UGUAUAAUCCGUCAAUUCCUGUAAAUUUCACUAAUUUAUAAAAUUACGGCUUGACUGCGGAUAAGCAUUGAAUGUGAAUAGACCACA..........................AUUUUUAUCACCUCUC...UGUGGUCGCGG.CU.GCACUAAGGUGUCAAUCCUCAAUCAAACCUA
env-116   AAUAAGAUCUUUAUUUAAAGUUACUUUCACUUAUCUAUCAACUGAGGCUUUUGACUGUGAAGCGGCAUUGAAUGUGAAUAGACCACA..........................AUUUUUAUCACCUCUC...UGUGGUCGCGG.CU.GCACUAAGAUGAACGAACCAUAGUCAAUCCC
env-117   CUUUCGAGCUUUUAGUAAAUAAACAAAUUCUUAUUACCGAGAUUUUCUCCCCCGCGGAGUCAUAAGAGCAAACGUGAAUAGACCAUA..........................AUUUUUAUCACCUCUC...UAUGGUCGCGG.CU.GCACUAAAAGCCUUAUGAAACAAUCUCAACA
env-118   UAAUCUCAGAGAGAAAUUUUUAGAGCAGAAUACUUUUAUCAACACAAACGACUGUGGCAGACAGCGCCGCCGAGUGAAUAGACUAUA............................AUUUAUCACCUCUC...UAUAGUCGCGG.CU.GCACAAAGCGGGCAUAGACUGCCUCGGGUCA
env-119   CGGAUGCCGUCCUCCCAAACGCAGAAGAGUCUAAUAACAAGUAGCGUCCUUUUUUGUCCGGACUGUUCUGAAUGCGAAAAGAUUGUC........................ACCUAUACCUGCCCUCUC...GACAAUCGCGG.CU.GCGCUAAG......................
env-120   CAUUACCUCAUAAUGGUUGUCGUCCUCCAAAGAAAAGAAGAGUUUAAUAAUUCAUAUAACGGAUGCUAAGUUGUGAUAUAGAUUAU.............................ACCUAAAUCCUCUC....AUAAUCGCGG.CU.ACACGACUUUAAUUCAUUAAUAAAAAAGCAA
env-121   GUCCACCUAAGAGAAGAAGAGUAUAAUCAAUCACAAUUCCGACGUCUCCAGUCCAUCCCCUCAGUUCGGUGUCCGAUUUUGGAUGUA..............................UCUUUUCCUUUC...UACAUCCGCUG.CU.GUAAGGACAACGGCUGGUGAGGAGCGACGCU
env-122   ........................AAUCAAUCACAAUUCCGACGUCUCUUGCUCAUCGCGCCAGUUCUGUGUCCGAUUUUGGAUGUA..............................UCUUUUCCUUUC...UACAUCCGCUG.CU.GUAAGGACCGCAGACUGACAUGAUCUGCCGA
env-123   CCCGGGGAAACUUUCCCGGCGGCAAAGCGGCAAGGACACGGCAGGGACAGACGCAUAACAGAGACCGGACCGGCGCGAAAGAUUGCC........................UUUCAAGCCUUUUCUCUC...GGCAAUCGCGG.CU.GCGCGCCCCCGGCUCCAGAGCGCCCACUUUA
env-124   GCAACCACACAGCGUUGCGGACAAAGCGACAAGGACACGGCGGAGACAGACGCAUAAAAGAGACCGGCCAGGUGCGAAAAGAUUGCC........................UUUUAAGCCUUUUCUCUC...GGCAAUCGCGG.CU.GCGCGCCCCCGGCUCUGAAAGCGCCCACUUU
env-125   ACAACCACACAGCGUUGCGGACAAAGCGACAAGGACACGGCGGAGACAGACGCAUAAAAGAGACCGGCCAGGUGCGAAAAGAUUGCC........................UUUUAAGCCUUUUCUCUC...GGCAAUCGCGG.CU.GCGCGCCCCCGGCUCUGAAAGCGCCCACUUU
env-126   AAUGGUGCUGGGAAAACGACCUUUGCAAGAGAAUUUUUAACAAGGGAAGCGGAUUGUCCAGAAUUUAUAAACGCUGAUUUGAUUGCG............CAAGGAUUAUCUCCCUUUGACCCUGAUAAAGC...UGCAAUCGCGG.CU.GCAAAAUUG.....................
env-127   ...........................AAAACAACCAUCGCUCAGGAAACAACCUGACGGACGCCCCGCUACAGUGAAUAGAUUGCA..........................CACAUACCACCCUCUC...UGCAAUCGCGG.CU.GCACUGUACGAGGCCGAAACACCAACCCAAC
Pgu-2-1   AAAGAGAAGAGUUUAAGAGAGAAUAUCUUCUAACAUUUCGGAUGUUUUAUUUUGUGGAAUCCGAAUUCUUUCGGUGUAUGGAUUGC...................CUAUAAACUAUCAAUCUCUCACAUA....GCAAUCGCGG.CU.GUACCGGAAGUAUUCUGUUCAUAGAAAAGCA
env-128   AAAGAGAAGAGUUUAAGAGAGAAGAUCUUCUAACAUUUCGGACGCUCUAUUUUAUGGGAUCCGAAUUCUUUCGGUGUAUGGAUUGC...................CUAUAAACUAUCAAUCUCUCACAUA....GCAAUCGCGG.CU.GUACCGGAAGCAUUCUGUUCAUAGAAAAGCA
Pgi-3-1   AAAGAGAAGAGUUUAAGAGAGAAGAUCUUCUAACAUUUCGGACGCUUUAUUUUAUGGGAUCCGAAUUCUUUCGGUGUAUGGAUUGC...................CUAUAAACUAUCAAUCUCUCACAUA....GCAAUCGCGG.CU.GUACCGGAAGCAUUCUGUUCAUAGAAAAGCA
Pgu-1-1   AAAGAGAAGAGUUUAAGAGAGAAUAUCUUCUAACAUUUCGGAUGUUUUAUUUUGUGGAAUCCGAAUUCUUUCGGUGUAUGGAUUGC...................CUAUAAACUAUCAAUCUCUCACAUA....GCAAUCGCGG.CU.GUACCGGAAGCAUUCUGUUCAUAGAAAAGCA
env-129   AAAGAGAAGAGUUUAAGAGAGAAGAUCUUCUAACAUUUCGGACGCUCUAUUUUAUGGGAUCCGAAUUCUUUCGGUGUAUGGAUUGC...................CUAUAAACUAUCAAUCUCUCACAUA....GCAAUCGCGG.CU.GUACCGGAAGCAUUCUGUUCAUAGAAAAGCA
Pgi-2-1   AAAGAGAAGAGUUUAAGAGAGAAGAUCUUCUAACAUUUCGGACGCUCUAUUUUAUGGGAUCCGAAUUCUUUCGGUGUAUGGAUUGC...................CUAUAAACUAUCAAUCUCUCACAUA....GCAAUCGCGG.CU.GUACCGGAAGCAUUCUGUUCAUAGAAAAGCA
Pgu-1-2   AAAGAGAAGAGUCUAAGAGAGAAUAUCUUCUAACAUUUCGGAUGUUUUAUUUUGUGGAAUCCGAAUUCUUUCGGUGUAUGGAUUGC...................CUAUAAACUAUCAAUCUCUCACAUA....GCAAUCGCGG.CU.GUACCGGAAGCAUUCUGUUCAUAGAAAAGCA
Pgi-1-1   AAAGAGAAGAGUUUAAGAGAGAAGAUCUUCUAACAUUUCGGACGCUCUAUUUUAUGGGAUCUGAAUUCUUUCGGUGUAUGGAUUGC...................CUAUAAACUAUCAAUCUCUCACACA....GCAAUCGCGG.CU.GUACCGGAAGCAUUCUGUUCAUAGAAAAGCA
env-130   ACACAACGGAUGUCGUCCUCCUAAAAGAAGAAGAGUUUAAUUAUUCAUUUAUCUUUAAAAGGAGCGCGGAUAAGUGAAUAGAUUACC................................CAACCCUCUC...GGUAAUCGCGG.CU.GCACAAAUCCAUGUUCCCAAAGAACAAUAAA
env-131   ACAAUGGAUGCCGUCCUCCAAAAAGAAGAAGAGUAUAAUCUCUGAAUAAACAAAACUAAGGAUCCAGCCGAUGUGAAAUAGAUUGCA..........................CCAACAAUCACCUCUC...UGCAAUCGCGG.CU.GCACUAAGCGGAUUCGAUAAAAACUUUAAUU
env-132   NNNNNNNNNNNNNNNNNNNNNNNNNNNNNNNNNNNNNNNNNNNNNNNNNNNNNNNNNNNNNNNNNNNNNNNNNNNNNNNAAUAGAUUGCA..........................CCAACAACCACCUCUC...UGCAAUCGCGG.CU.GCACUAAGCGGAUUCGAUAAAAACUUUAAUU
env-133   ACAAUGGAUGCCGUCCUCCAAAAAGAAGAAGAGUAUAAUCUCUGAAUAAAUAAAACUAAGGAUCCAGCCGAUGUGAAAUAGAUUGCA..........................CCAACAACCACCUCUC...UGCAAUCGCGG.CU.GCACUAAGCGGAUUCGAUAAAAACUUUAAUU
env-134   ACAAUGGAUGCCGUCCUCCAAAAAGAAGAAGAGUAUAAUCUCUGAAUAAACAAAACUAAGGAUCCUGCCGAUGUGAAAUAGAUUGCA..........................CCAACAACCACCUCUC...UGCAAUCGCGG.CU.GCACUAAGCGGAUUCGAUAAAAACUUUAAUU
env-135   ACAAUGGAUGCCGUCCUCCAAAAAGAAGAAGAGUAUAAUCUCUGAAUAAACAAAACUAAGGAUCCAGCCGAUGUGAAAUAGAUUGCA..........................CCAACAACCACCUCUC...UGCAAUCGCGG.CU.GCACUAAGCGGAUUCGAUAAAAACUUUAAUU
env-136   ...............CACCAAAAAGAAGAAGAGUAUAAUCUCUGAAUAAACAAAACUAAGGAUCCAGCCGAUGUGAAAUAGAUUGCA..........................CCAACAACCACCUCUC...UGCAAUCGCGG.CU.GCACUAAGCGGAUUCGAUAAAAACUUUAAUU
env-137   CAACAUUUCUAAAAGAUUUAUUUUUUAUUACAAUCACACAAUUUGCCAGAGGUCGAGCCACAGUGAAUUUAUUCAUUAAGGAUUGCA.........................CAUUCACACUACCUCUC...UGUAAUCGCGG.CU.GCACUGUGCCGGCUCACUGCACAACACUUUA
env-138   AUAAUGGUUGUCGUCCUCCAAAGAAAAGAAGAGUUUAAUAUUAUUUCUUUAAUUUAAAGAAUUCGGUAUAAUAGUGAAUAGAUUAUA........................AAAGACCCUUAUCCUCUC...UAUAAUCGCGG.CU.GCACUAUUCCUUAUUCGUAGAUAAAACAAAA
env-139   CAUAAUGGUUGUCGUCCUCCAAAGAAAAGAAGAGUUUAAUAUUAUUUCUUUAAUUAAAGAAUUCGGUAUAAUAGUGAAUAGAUUAUA........................AAAGACCCUUAUCCUCUC...UAUAAUCGCGG.CU.GCACUAUUCCUUAUUCGUAG..........
env-140   CCGUUCAAUGGCUGUCGUCCUCCCAAGAGAAGAAGAGUUUAAUUAUUAUUUCCCUGCCGUCCCCUCAUACAUUAUUUUAGGACUGCA..............................UCUUUUCCUUUC...UGCAGUCGCGG.CU.GCAAUGUCAGGAGACGGCAAAACAAAAACAA
env-141   CACAACGGAUGUCGUCCUCCAAAGAGAAGAAGAGUUUAAUUAUUAUUUUUUGCCGACAAGCCCGCUCAAAAUAUUUUUGGACUGCA...............................UCUUUCCUUUC...UGCAGUCGCGG.CU.GCUAACCAGCGGGUGCGUCACUUUAAACAAC
env-142   CACAACGGAUGUCGUCCUCCAAAGAGAAGAAGAGUUUAAUUAUUAUUUUUUGCCGACAAGCCCGCUCAAAAUAUAUUUUGGACUGCA...............................UCUUUCCUUUC...UGCAGUCGCGG.CU.GCUAACCAGCGGGUGCGUCACUUUAAACAAC
env-143   GCCUCAUAAUGGAUGUCGCCCUCCAAAGAAAAGAAGAGUUUAAAAUAAUUCCUAAAAAAGAAUUGGAUUUAUAGUGAAUAGAUUAU.........................AAAUAGACCUUGUCCUCUC....AUAAUCGCGG.CU.GCACUAUCCUCCUAUUCAAAGAAAAAAUAAA

          ...................................................................................<<<<<<<<<<...................................>>>>>>>>>>.............................
          ...................................................................................1122222100...................................0012222211.............................
          ••••••••••••••••••••••••••••R••••••••••••••••••••••••••••••••••••••••••••••••••R•URGAUURYA---------------------------○○○○•Y••CCUCUC---URYARUCGCGG-CU-GCAY•R•••••••YYY••••••••••••••••
                                                                                    5′                                                        3′
```

```
env-144   GGAUUGUCGUCCCUCCGAUCAAAGAGAAUAGUCAUGUCUGACUAUCGUUUCUAGAACGUUGAACUCCGGAAUCGUGAAUCGAUUACA..........................CCGCGACCUUAUCCACUC...UGUAGACGCGG.CU.GCUGGGUUCCAGAUUCCGAGCAGAGCGUUAA
env-145   CACAAUGGUUGCCGUCCUCCUAAAAAAAGAAGAGUAUAAUUCAUUUAGUUUCUUGAAUAUUGAAUCGGGAUUUGUGAUAAGAUUGCA..........................UAUUUACCUUACCCUCUC...UGUAAUCGCGG.CU.GCAUGAGUUCAGAUUCAAACAGAACAAUUAA
env-146   NNNNNNNNNNNNNNNNNNNNNNNNNNNNNNNNNNNNNNNNNNNUAUAGUUUCUUAAACAUUGAAUCCGGAAUUGUGAAUAGAUUGCA..........................UAUUGACCUUAUCCUCUC...UGUAAUCGCGG.CU.GCACGGUUCCAGAUUCAAACAGAACAUUUAA
env-147   AUAACGGGUGUCGUCCUCCGAAAAAGAGAAGAGUAUAAUUAAUUAUAGUUUCUUAAACAUUGAAUCCGGAAUUGUGAAUAGAUUGCA..........................UAUUGACCUUAUCCUCUC...UGUAAUCGCGG.CU.GCACGGUUCCAGAUUCAAACAGAACAUUUAA
env-148   AUAACGGGUGUCGUCCUCCGAAAAAGAGAAGAGUAUAAUUAAUUAUAGUUUCUUAAACAUUGAAUCCGGAAUUGUGAAUAGAUUGCA..........................UAUUGACCUUAUCCUCUC...UGUAAUCGCGG.CU.GCACGGUUCCAGAUUCAAACAGAACAUUUAA
env-149   AUAACGGAUGUCGUCCUCCGAAAAAGAGAAGAGUAUAAUUAACUAUAGUUUCUUGAACAUUGAAUCCGGAAUUGUGAAUAGAUUGCA..........................UAUUGACCUUAUCCUCUC...UGUAAUCGCGG.CU.GCACGGUUCCAGAUUCAAACAGAACAUUUAA
env-150   AUAACGGGUGUCGUCCUCCGAAAAAGAGAAGAGUAUAAUUAAUUAUAGUUUCUUAAACAUUGAAUCCGGAAUUGUGAAUAGAUUGCA..........................UAUUGACCUUAUCCUCUC...UGUAAUCGCGG.CU.GCACGGUUCCAGAUUCAAACAGAACAUUUAA
env-151   AUAACGGAUGUCGUCCUCCGAAAAAGAGAAGAGUAUAAUUAAUUAUAGUUUCUUAAACAUUGAAUCCGGAAUUGUGAAUAGAUUGCA..........................UAUUGACCUUAUCCUCUC...UGUAAUCGCGG.CU.GCACGGUUCCAGAUUCAAACAGAACAUUUAA
env-152   ........................AGAGAAGAGUAUAAUUAAUUAUAGUUUCUUAAACAUUGAAUCCGGAAUUGUGAAUAGAUUGCA..........................UAUUGACCUUAUCCUCUC...UGUAAUCGCGG.CU.GCACGGUUCCAGAUUCAGAACAGAACAUUAA
env-153   ..............................................................................UAGAUUGCA..........................UAUUGACCUUAUCCUCUC...UGUAAUCGCGG.CU.GCACGGUUCCAGAUUCAAACAGAACAUUUAA
env-154   AUAACGGAUGUCGUCCUCCGAAAAAGAGAAGAGUAUAAUUAAUUAUAGUUUCUUAAACAUUGAAUCCGGAAUUGUGAAUAGAUUGCA..........................UAUUGACCUUAUCCUCUC...UGUAAUCGCGG.CU.GCACGGUUCCAGAUUCA..............
env-155   AUAACGGAUGUCGUCCUCCGUAAAAGAGAAGAGUAUAAUUAAUUAUAGUUUCUUAAACAUUGAAUCCGGAAUUGUGAAUAGAUUGCA..........................UAUUGACCUUAUCCUCUC...UGUAAUCGCGG.CU.GCACGGUUCCAGAUUCAAACAGAACAUUUAA
env-156   AUAACGGAUGUCGUCCUCCGAAAAAGAGAAGAGUAUAAUUAAUUAUAGUUUCUUAAACAUUGAAUCCGGAAUUGUGAAUAGAUUGCA..........................UAUUGACCUUAUCCUCUC...UGUAAUCGCGG.CU.GCACGGUUCCAGAUUCAAACAAACAUUUAAA
env-157   ACAAUGGUUGUCGUCCUCCGAAAAAGAGAAGAGUAUAAUUAAUUAUAGUUUCUUAAACAUUGAAUCCGGAAUUGUGAAUAGAUUGCA..........................UAUUGACCUUAUCCUCUC...UGUAAUCGCGG.CU.GCACGGUUCCAGAUUCAAACAGAAC......
env-158   CACAAUGGUUGUCGUCCUCCUAAAAAGAGAAGAGUUUAAUAACUUUAGUUUCUUGAAUUUUGAAUCCGGAAUUGUGAAUAGAUUGCA..........................AAUAGACCUUAUCCUCUC...UGUAAUCGCGG.CU.GCACGGUUCCAGAUUCUAUAGAACAUUUAAA
env-159   ACAAUGGUUGUCGUCCUCCGAAAAAGAGAAGAGUAUAAUUAACUAUAGUUUCUUGAACAUUGAAUCCGGAAUUGAGAAUAGAUUAC.........................UCAAAGACCUUAUCCUCUCU....GUAGUCGCGG.CU.GCACGGUUCCAGAUUCAGAACAGAACAUUAA
env-160   ........................AGAGAAGAGUAUAAUUAACUAUAGUUUCUUGAACAUUGAAUCCGGAAUUGUGAAUAGAUUACA..........................CAAAGACCUUAUCCUCUC...UGUAGUCCCGG.CU.GCACGGUUCCAGAUUCAGAACAGAACAUUAA
env-161   GUAUUCGUAAAGGGACCGGGUAACGGACGUGAGUCUGCUANNNNNNNNNNNNNNGAACAUUGAAUCCGGAAUUGUGAAUAGAUUACA..........................CAAAGACCUUAUCCUCUC...UGUAGUCGCGG.CU.GCACGGUUCCAGAUUCAGAACAGAACAUUAA
env-162   ANNNNNNNNNNNNNNNNNNNNAAAGAGAAGAGUAUAAUUAACUAUAGUUUCUUGAACAUUGAAUCCGGAAUUGUGAAUAGAUUACA..........................CAAAGACCUUAUCCUCUC...UGUAGUCGCGG.CU.GCACGGUUCCAGAUUCAGAACAGAACAUUAA
env-163   ACAAUGGUUGUCGUCCUCCGAAAAAGAGAAGAGUAUAAUUAACUAUAGUUUCUUGAACAUUGAAUCCGGAAUUGUGAAUAGAUUACA..........................CAAAGACCUUAUCCUCUC...UGUAGUCGCGG.CU.GCACAGUUCCAGAUUCAGAACAGAACAUUAA
env-164   AUCCACGGUGCAGGUAUUGAAGUAACGGAAAUCGUAGACGUUNNNNNNNNNNNNNNNNNUUGAAUCCGGAAUUGUGAAUAGAUUACA..........................CAAAGACCUUAUCCUCUC...UGUAGUCGCGG.CU.GCACGGUUCCAGAUUCAGAACAGAACAUUAA
env-165   .UAAUGGUUGUCGUCCUCCGAAAAAGAGAAGAGUAUAAUUAACUAUAGUUUCUUGAACAUUGAAUCCGGAAUUGUGAAUAGAUUACA..........................CAAAGACCUUAUCCUCUC...UGUAGUCGCGG.CU.GCACGGUUCCAGAUUCAGAACAGAACAUUAA
env-166   ACAAUGGUUGUCGUCCUCCGAAAAAGAGAAGAGUAUAAUUAACUAUAGUUUCUUGAACAUUGAAUCCGGAAUUGUGAAUAGAUUACA..........................CAAAGACCUUAUCCUCUC...UGUAGUCGCGG.CU.GCACGGUUCCAGAUUCAGAACAGAACAUUAA
env-167   ........UGUCGUCCUCCGAAAAAGAGAAGAGUAUAAUUAACUAUAGUUUCUUGAACAUUGAAUCCGGAAUUGUGAAUAGAUUACA..........................CAAAGACCUUAUCCUCUC...UGUAGUCGCGG.CU.GCACGGUUCCAGAUUCAAACAGAACAUUUAA
env-168   NNNNNNNNNNNNNNNNNCCGAAAAAGAGAAGAGUAUAAUUAACUAUAGUUUCUUGAACAUUGAAUCCGGAAUUGUGAAUAGAUUACA..........................CAAAGACCUUAUCCUCUC...UGUAGUCGCGG.CU.GCACGGUUCCAGAUUCAGAACAGAACAUUAA
env-169   UAAAGGGACCGGGUAACGGACGUGAGUCUGCUANNNNNNNNNCUAUAGUUUCUUGAACAUUGAAUCCGGAAUUGUGAAUAGAUUACA..........................CAAAGACCUUAUCCUCUC...UGUAGUCGCGG.CU.GCACGGUUCCAGAUUCAGAACAGAACAUUAA
env-170   CACAAUGGUUGUCGUCCUCCGAAAAGAGAAGAGUAUAAUUAACUAUAGUUUCUUGAACAUUGAAUCCGGAAUUGUGAAUAGAUUACA..........................CAAAGACCUUAUCCUCUC...UGUAGUCGCGG.CU.GCACGGUUCCAGAUUCAGAACAGAACAUUAA
env-171   ACAAUGGUUGUCGUCCUCCGAAAAAGAGAAGAGUAUAAUUAACUAUAGUUUCUUGAACAUUGAAUCCGGAAUUGUGAAUAGAUUACA..........................CAAAGACCUUAUCCUCUC...UGUAAUCGCGG.CU.GCACGGUUCCAGAUUCAGAACAGAACAUUAA
env-172   ACAAUGGUUGUCGUCCUCCGAAAAAGAGAAGAGUAUAAUUAACUAUAGUUUCUUGAACAUUGAAUCCGGAAUUGUGAAUAGAUUGCA..........................UAAAGACCUUAUCCUCUC...UGUAAUCGCGG.CU.GCACGGUUCCAGAUUCAGACAGAAAAAUUAA
env-173   ACAAUGGUUGUCGUCCUCCGAAAAAGAGAAGAGUAUAAUUAACUAUAGUUUCUUGAACAUUGAAUCCGGAAUUGUGAAUAGAUUGCA..........................UAAAGACCUUAUCCUCUC...UGUAAUCGCGG.CU.GCACGGUUCCAGAUUCAGACAGAAAAAUUAA
env-174   ACAAUGGUUGUCGUCCUCCGAAAAAGAGAAGAGUAUAAUUAACUAUAGUUUCUUGAACAUUGAAUCCGGAAUUGUGAAUAGAUUGCA..........................CAAAGACCUUAUCCUCUC...UGUAAUCGCGG.CU.GCACGGUUCCAGAUUCAGACAGAAAAAUUAA
env-175   ACAAUGGUUGUCGUCCUCCUAAAAAGAGAAGAGUAUAAUUAACUUUAGUUUCUUGAAUAGGGAAUCCGGAAUUGUGAAUAGAUUGCA..........................UUUAGACCUUAUCCUCUC...UGUAAUCGCGG.CU.GCACGGUUCCAGGUUCUUUUAGAACAUUAAA
env-176   AACGGAUGUCGCCCUCCAAAACACAGAAGAGUUUAAUCAAAACAACCGAUUUUUAAAUUGGAAUCUGAAAUUGUGAUUUUGAUUACA................................CAACGACCUUUC...UGUGAUCGCGG.CU.GCACAAUUUGGGCUUUCAUUUAAUUAUUUUA
env-177   AUGUAGACCUCCAAAAAGAAGGAGAGUAUAAUAUUAAGAUUUAUAUAAUCUAAAAAGCGAUGAAACUUGAUUUUGUUUUGGAUUGCA.............................UCUAUUAAUCUUCUC...UGUAAUCGCGG.CU.GUAACAAAUUAAGUUCAUGUAUUAAAAAAUC
env-178   ....................................................UUUAUAGUAACUUGAUUUGUUAAUAAUGGAUGCC...................................ACUCCUUUCC....GUAAUCGCGG.CU.GCAACAAAUAAAGUUCUUUAGUAACAAUUAA
env-179   .......................................AAAAAACGAGUUACAGCUAAAGAGUGAUUAUAUUUUUUAUUGAUUGCA.....................AUACUUUUUAUUAUCAACCUCUC...UGCAAUCGCGG.CU.GCUCACUCCUGUAAUUCAAAACCACUGUUUU
env-180   UUUUGAUACCCGCGAUUUACUCCAACAUCUCUUUCGCAAGUUUUAUUUCUAAAUUUUGCGAACCCGAAUCAGUGAAUAAUGAUUGCA...ACUUUUUUACCAUUCGCAAUAAAAAAGAUACCAUACCUCUC...UGCAAUCGCGG.CU.GCACUGUAGUGGUUCACUACUACAACAACUU
env-181   ACGGUUGUCGUCCUCCUAAGAGAAGAAGAGUUUAAUAAGAGAUAACCCAUAUAACAGUAGGGAUUCCAACGAUUUUCAAGGGCUUCA...................................UUUCCUUUC...UGUAGUCGCGG.CU.GCAAUGACUUGGGAUUCCUGCAAGAACCUUU
env-182   CGGUUGUCGUCCUCCUAAGAGAAGAAGAGUUUAAUAAGAGAUAACCCAUAAUAACAGUAGGGAUUCCAAUGAUUUUCAAGGGCUUCA...................................UUUCCUUUC...UGUAGUCGCGG.CU.GCAAUGACUUGGGGUUCCUGCAAGAACCUUU
env-183   CGGUUGUCGUCCUCCUAAGAGAAGAAGAGUUUAAUAAGAGAUAACCCAUAAUAACAGUAGGGAUUCCAAUGAUUUUCAAGGGCUUCA...................................UUUCCUUUC...UGUAGUCGCGG.CU.GCAAUGACUUGGGAUUCCUACAAGAACCUUU
env-184   ..............................UUAAUAAGAGAUAACCCAUAAUAACAGUAGGGAUUCCAACGAUUUUCAAGGGCUUCA...................................UUUCCUUUC...UGUAGUCGCGG.CU.GCAAUGACUUGGGAUUCCUGCAAGAACCUUU
env-185   ............UCCUAAGAGAAGAAGAGUUUAAUAAGAGAUAACCCAUAAUAACAGUGGGGAUUCCAACGAUUUUCAAGGGCUUCA...................................UUUCCUUUC...UGUAGUCGCGG.CU.GCAAUGACUUGGGAUUCCUGCAAGAACCUUU
Pxy-1-1   CGGUUGUCGUCCUCCUAAGAGAAGAAGAGUUUAAUAAGAGAUAACCCAUAAUAACAGUAGGGAUUCCAAUGAUUUUCAAGGGCUUCA...................................UUUCCUUUC...UGUAGUCGCGG.CU.GCAAUGACUUGGGAUUCCUGCAAGAACCUUU
env-186   CGGUUGUCGUCCUCCUAAGAGAAGAAGAGUUUAAUAAGAGAUAACUCAUAAUAACAGUAGGGAUUCCAACGAUUUUCAAGGGCUUCA...................................UUUCCUUUC...UGUAGUCGCGG.CU.GCAAUGACUUGGGAUUCCUGCAAGAACCUUU
env-187   CGGUUGUCGUCCUCCUAAGAGAAGAAGAGUAUAAUAAGAGAUAACCCAUACUAACAGUAGGGAUUCCAACGAUUUUCAAGGGCUUCA...................................UUUCCUUUC...UGUAGUCGCGG.CU.GCAAUGACUUGGGAUUCCUGCAAGAACCUUU
env-188   CAACGGAUGUCGUCCUCCUAAAAGAAGAAGAGUUUAAUUAUUCAUUUAUUUCUUUACAAAGGAGCGCGGAUAAGUGAAUAGAUUACA...................................AACCCUCUC...UGUAAUCGCGG.CU.GCACAAGUCCAUGUUCCCAAAGAACAAUAAA
env-189   AGAGUUUAAUCACUUCAUUUCACCUUUUUAAACUUUUCGUUUUUUACAUUACAUUUUUCGCGAAUCUCUGUCAGUGAUAAGAUUGCA.........................CUUUAUUUAACAACCUCUC...UGCAAUCGCGG.CU.GCACUGAAGCGAUUCAACAACACAACUUUAA
env-190   AGAAGAGUUUAAUGAAUCAAAUUUAGUUUAAGCUCUUAAAUUAUUUCAUUUAUUUUUUCGCGAAUCUCUGUCAGUGAAUAGAUUGCA.........................UUUUAUUUAACAACCUCUC...UGCAAUCGCGG.CU.G..............................
env-191   GAAGAGUUUAAUGAAUCAAAUUUUAGUUUAAGCUCUUAAAUUAUUUCAUUUAUUUUUUCGCGAAUCUCUGUCAGUGAAUAGAUUGCA.........................UUUUAUUUAACAACCUCUC...UGCAAUCGCGG.CU.GCACUGAAGCGAUUCAUAAACAAACUUUAAU
env-192   CCCCACACUCGGCAGGAGCCUGCAAAGAAAAACAAUUCGUUUUUUAAAAAUUUUUCGCGAAUCACGACCAGUGAAUUUUUGAUUGCA.........................UCUUUUUUAUCAACCUCUC...UGCAAUCGCGG.CU.GCACACUGCGGUGAUUCAUUAUCACACAACA
env-193   UAAUCGGCAGUUUACACCCGAUCACGCUUUUCUGAUUUAUUUUUCACAACAUUUUUUUCGCGAAUCUCUGUCAGUGAUUGGAUUGCA.........................UCUUUCUUAACAACCUCUC...UGCAAUCGCGG.CU.GCACUGAAGCGAUUCUGCAACCCCAACAUCA
env-194   UAUAAUUUCACUACACACUACUUCGAUUAGAGUUUUUAUCAUUAUUAAUCAUUUUUUUCGCGAAUCUCUGUCAGUGAUUGGAUUGCA.........................CCUUUUUUAACAACCUCUC...UGCAAUCGCGG.CU.GCACUGAAGCGAUUC................
env-195   ....................................................UUUUUUCGCGAAUCUCUGUCAGUGAUUGGAUUGCA.........................CCUUUUUUAACAACCUCUC...UGCAAUCGCGG.CU.GCACUGAAGCGAUUCAUCCAGAACUUAACUU
env-196   ....................................................UUUUUUCGCGAAUCUCUGUCAGUGAUUGGAUUGCA.........................CCUUUUUUAACAACCUCUC...UGCAAUCGCGG.CU.GCACUGAAGCGAUUCAUUACAUGAACUUAAC
env-197   AAGAGAAGGAGAGUGUAAUCACAAUCUUAAUCUUUUCAAAUCAUUCAUUAAUUAAUUAGUCGAAUCGAUUAUAGUGAAUGGAUUGCAA...........................CUUUCUUCAACCUCUC..UUGCAAUCGCGG.CU.GCACUAUAGUGAUUCUCGCAUCACAACUUUA
env-198   ACUUCCUCAUAACGGAUGUCGUCCACCAAAAAGAAGAAGAGUAUAAUAAUUAAUAAUGAUAAAUCCCGAAGGUGUAAAUAGAUUGC.................................ACUAAUCCUCUC....GCAAUCGCGG.CU.GCACGGUUCAGGUUUAUAGAAAAAAUUUAA
env-199   AUUAAAAUCAAUGCACAAUGCACAAUUCAAAAUGCACAAUGUAUAAUUAUUAAUAAAUAUGAAUCCCGAAAGUGUAAAUAGAUUGC.................................ACCAAUCCUCUC....GCAAUCGCGG.CU.GCACGGUUCAGGUUUUGUAAAAAAAACUUUU
env-200   UUAAAAUCAAUGCACAAUGCACAAUUCAAAAUGCAAAAUGUAUAAUUAAUAAUAAAAGUGAAUCUUGAAAGUGUAAAAUAGAUUGC.................................AUCAAUCCUCUC....GCAAUCGCGG.CU.GCACGGUUCUGGGUUUAUAAUAAAUUUAAAA
env-201   CACUUCCGCAUAACGGAUGUCGUCCGCCUAAAAGAAGAAGAGUAUAAUAUUAAUAAAUAUGAAUCCAGAAAGUGUAAAUAGAUUGC.................................AUCAAUCCUCUC....GCAAUCGCGG.CU.GCACGGUUCAGGGUUUGUUAUAAAAUUUAAA
env-202   ....................GUCCUCCUAAAAGAAGAAGAGUAUAAUUAUUAAUAACUACGAAUCCCGGAAGUGUAAAUAGAUUGC.................................AUCAAUCCUCUC....GCAAUCGCGG.CU.GCACGGUUCAGGGUUUGUGAUAAAAUUUAAA
env-203   CUUCCGCACAACGGAUGUCGUCCACCGAAAAGAAGGAGAGUAUAAUUAAUUAUAAAAGUGAAUCUUGAAAGUGUAAAAUAGAUUGC.................................AUCAAUCCUCUC....GCAAUCGCGG.CU.GCACGGUUCAGGGUUUAUAAUAAAUUUAAAA
env-204   CUUCCGCAUAACGGAUGUCGUCCACCGAAAAGAAGGAGAGUAUAAUUAAUUAUAAAAGUGAAUCUUGAAAGUGUAAAAUAGAUUGC.................................AUCAAUCCUCUC....GCAAUCGCGG.CU.GCACGGUUCGAGGUUUAUAAUAAAUUUAAAA
env-205   GCUUCCGCACAAUGGAUGCCGUCCACCGAAAAGAAGAAGAGUAUAAUUAAUUAUAAAAGUGAAUUCUGAGAGUGUAAAUAGAUUGC.................................AUCAAUCCUCUC....GCAAUCGCGG.CU.GCA............................
env-206   ACUUCCGCAUAAUGGAUGUCGUCCGCCUAAAAGAAGAAGAGUAUAAUUAAUAAUAAAUAUGAAUCCCGAAAGUGUAAAUAGAUUGC.................................AUCAAUCCUCUC....GCAAUCGCGG.CU.GCACGGUUCAGGUUUCGAAAUAAAAUUUUAA
env-207   CUUCCGCACAACGGAUGUCGUCCACCGAAAAGAAGAAGAGUAUAAUUAAUAAUAAAAAUGAAUCUUGAGAGUGUAAAAUAGAUUGC.................................AUCAAUCCUCUC....GCAAUCGCGG.CU.GCACGGUUCAGGGUUUAUAAGAAAUUUAAAA
env-208   UGAAAAUUGAAAAUUGAAAAUUAAAAAUCAAUGCACAAUGUAUAAUUAAUAAUAAAUAUGGAUUCUGAAAGUGUAAAAUAGAUUGC.................................AUCAAUCCUCUC....GCAAUCGCGG.CU.GCACGACUCGGGAUACAUAAAAAACUUAAAA
env-209   .......................................................................UGUAAAAUAGAUUGC.................................AUCAAUCCUCUC....GCAAUCGCGG.CU.GCACGGUUCGAGGUUUACAAGAAAUUUAAAA
env-210   CGCUUCCGCACAACGGAUGCCGUCCACCGAAAAGAAGAAGAGUAUAAUAAUUAUAAAAGUGAAUCCUGAAAGUGUAAAUAGAUUGC.................................AUCAAUCCUCUC....GCAAUCGCGG.CU.GCACGGUUUGGGGUUUAUAAAAAUUUAAAAC
env-211   GCUUCCGCAUAAUGGAUGUCGUCCGCCUAAAAGAAGAAGAGUAUAAUUAUUAAUAAAUAAGAAUCCCGAAAGUGUAAAUAGAUUGC.................................AUCAAUCCUCUC....GCAAUCGCGG.CU.GCACGGUUCAGGUUUCGAAAUAAAAUUUUAA
env-212   ACUUCCACAUAACGGAUGUCGUCCUCCUAAAAGAAGAAGAGUAUAAUAAUUAAUAAAUAUGAAUCCGGAAAGUGUAAAUAGAUUGC.................................AUCAAUCCUCUC....GCAAUCGCGG.CU.GCACGGUUCAGGUUUCGUAAUAAAAUUUAAA
env-213   ACUACCGCACAACGGAUGUCGUCCUCCUAAAAGAAGAAGAGUAUAAUUAUUAAUAACUAUGAAUCCCGGAAGUGUAAAUAGAUUGC.................................AACAAUCCUCUC....GCAAUCGCGG.CU.GCACGGUUCAGGGUUUGUAAUAAAAUUUAAA
env-214   ACUACCGCAUAACGGAUGUCGUCCUCCUAAAAGAAGAAGAGUAUAAUUAUUAAUAACUAUGAAUCCCGGAAGUGUAAAUAGAUUGC.................................AACAAUCCUCUC....GCAAUCGCGG.CU.GCACGGUUCAGGGUUUAUAAUAAAAUUUAAA
env-215   ACUUCCGCAUAACGGAUGUCGUCCGCCUAAAAGAAGAAGAGUAUAAUUAAUAAUAAAUAUGAAUCCCGAAAGUGUAAAUAGAUUGC.................................CUCAAUCCUCUC....GCAAUCGCGG.CU.GCACGGUUCAGGUUUUGUAAUAAAUUUAAAA
env-216   ACUUCCGCACAACGGAUGUCGUCCGCCUAAAAGAAGAAGAGUAUAAUUAAUAAUAAAUAUGAAUCCCGAAAGUGUAAAUAGAUUGC.................................CUCAAUCCUCUC....GCAAUCGCGG.CU.GCACGGUUCAGGUUUUGUAAUAAAUUUAAAA
env-217   ACAUCAAGGAACGUUGACAAUUAAAAAUCAAUUCACAAUGUAUAAUUAAUCAAUAAAUAUGAAUCCGGAAAGUGUAAAUAGAUUGC.................................AUAAAUCCUCUC....GCAAUCGCGG.CA.GCACGGUUCAGGGUUUGUAAUAAAAAUUUAA
env-218   GCUUCCACAUAAUGGAUGUCGCCCGCCGAAAAGAAGAAGAGUAUAAUUAAUAAUAAAUGUGAAUCCUGGAAGUGUAAAUAGAUUGU.................................AUUAAACCUCUC....GCAAUCGCGG.CU.GCACGGUUCAUGAUUUAUCGUAAACUUAAAA
env-219   CCCAAGAGAAGAAGGGUUUAAGCUUCUUGUGCAUCAGAAUUCGUUAUUAAUUUUUUUCAACGAAUCUCUCAUAGUGAAUAGAUUGCA...........................CCAAUCAACAACCUCUC...UGCAAUCGCGG.CU.GCACUAUCAGUGAUUCCAGACAUCACUUCAA
env-220   UUUAGGCGCUUCCCUUGUCAACAAUUCUGUAUUAACAGUUUUUAGUAACAACAUUUUUCGCGAAUCUCUAUCAGUGAAUGGAUUGCA...........................UUUUUUAACGACCUCUC...UGCAAUCGCGG.CU.GCACUGAAGCGAUUCCAAUUCACAUUAAUCU
env-221   GCGUUUCCUAAACCUCAGUGUGUGUAAUUAAUCACUCUUUAAAAUUUUAUUUCGUUUUCGCGAAUCUCUAUCAGUGAAUGGAUUGCA...........................UUUUUUAACGACCUCUC...UGCAAUCGCGG.CU.GCACUGAAGCGAUUCCAAUUCACACCGUUAA
env-222   UCCUCCCAAGAGAAGAAGAGUUUAAUGGUUUAAACAAUCCUUCAAACAAACAUUUUUUCGCGAAUCUCUGUCAGUGAAUAGAUUGCA...........................CUAUAUAACAACCUCUC...UGCAAUCGCGG.CU.GCACUGAAGCGAUUCAUCAAACAAACUUUAA
env-223   ................................UCAAUUCUCCUUUUUCAUUCAUUUUUUUCGAAUCUUCGUCAGUGAAUAGAUUGCA..........................ACACUUUAACAACCUCUC...UGCAGUCGCGG.CU.GCACUGGCUUGAUUCAGUUUUUCAACUUUAA
env-224   CAAUGGAUGUCGUCCUCCUAAAAGACGUAGAGUUUAAGAUUUACCUUUAUUACUAAGCAUGAAACUUGAUUUUGUUAUUGGAUUGUA.................................CAAUUCUUCUC...UGCAAUCGCGG.CU.GCAACAAAUUGAGUUCAUCAAUAAACAAUUA
env-225   AUAACGGAUGUCGUCCUCCUAAAAAGAGAAGAGUUUAAUUAACUAUUGUUUCUUGUGCAUUGAAUUGAGAAACGUGAAUAGAUUACA...........................AAAGACCUUAUCCUCUC...UGUAAUCGCGG.CU.GCACAGUUCUUUAUUCAUACAGAACAUUUAA

          ................................................................................&lt;&lt;&lt;&lt;&lt;&lt;&lt;&lt;&lt;&lt;.........................................&gt;&gt;&gt;&gt;&gt;&gt;&gt;&gt;&gt;&gt;.......................................
          ................................................................................1122222100.........................................0012222211.......................................
          ●●●●●●●●●●●●●●●●●●●●●●●●●●●●●R●●●●●●●●●●●●●●●●●●●●●●●●●●●●●●●●●●●●●●●●●●●●●●●R●URGAUURYA-----------------------------○○○○●Y●●CCUCUC---URYARUCGCGG-CU-GCAY●R●●●●●●●YYY●●●●●●●●●●●●●●●●
                                                                                5′                                                                 3′
```

```
env-226 AUAACGGGUGUCGUCCUCCUAAAAAGAGAAGAGUUUAAUUAACUAUUGUUCUUCGUGCAUUGAAUUAAGAAUAGUGAAUAGAUUACA...........................AAAGACCUUAUCCUCUC...UGUAAUCGCGG.CU.GCACAGUUCUUUAUUCAUACAGAACAUUUAA
env-227 ACAAUCGAAUUCUAACUGAAUAUAAAGAAAGAGUAAAUAAUCAUAAGUUUCAAUCAUACAGAAACUUGAGCUGUGAUUUUGGUUACA................................CAACAACUUCUC...UGUAAUCGCGG.CA.GCACAGUUUAGGCUUCAAAAAGUUUAAUUUG
env-228 GAUGUCGUCCUCCCAAAGCAAGAAAAGUUUAAUUUGAAUCAACCGUAAGUAUCAUUUCCUGUUAACUUGAGCUGUGAAUUGAUUACA................................CUUCAACUUCUC...UGUAAUCGCGG.CU.GCACAGUUUAGGCUUCAGAAUAAAAGUUUUA
env-229 AUGCCGUCCACCAAAAGCCAGAAAAGUUUAAUCAACAGAUAAUUUAAGUUUCAUUUACCUUGAGACUUGAGCUGUGAAAUGAUUACA................................AUACAGCUUCUC...UGUAAUCGCGG.CU.GCACAGUUUAGGCUUCAGAAAAAAAAGUUUA
env-230 UCCGGUAGUUAUUCGUCUGAAAAUCAAAAAAGAGUAAAUAAUCAUAAGUUUCAAUCAUACAGAAACUUGAGCUGUGAUUUGAUUACA................................AUACAACUUCUC...UGUAAUCGCGG.CU.GCACAGUUUAGGCUUCAGUAGAAAAAGUUUA
env-231 ACAGGAAGGCGGGGAAUAGUCAAACGUUCAUGAAUAAUUAAUUUAAGUUUCAUUUCACUUGAAACUUGAGCUGUGAUAUUGAUUACA................................AUACAACUUCUC...UGUAAUCGCGG.CU.GCACAGUUUAGGCUUCAGUAAAAGUUUAAAA
env-232 UAUGUCGUUGCGAGAAAUAGUCUAAUGAUAUUGAAUAAAUAAUUUAAGUUUCAUUAUCUUGAAACUUGAGCUGUGAUUUUGAUUACA................................ACAAAACUUCUC...UGUAAUCGCGG.CU.GCACAGUUUAGGCUUCAGUAAAAAGUUUAAU
env-233 AACGGAUGCCGUCCCCCGAAACAGAGAAAAGUUUAAUUAAUAAUUUAAGUUUCAUUUAUCUUGAAACUUGAGCUGUGAUUGAUUACA................................ACACAACUUCUC...UGUAAUCGCGG.CU.GCACAGUUUAGGCUUCAGUAAAGUUUAAUUG
env-234 GAUGUCGUCCCCCGAAACAGAGAAAAGUUUAAUCAACAAAUAAUUUAUGUUUCAUUUAUCCUGAAACUUGGGCUGUGAUUGAUUACA................................ACACAACUUCUC...UGUAAUCGCGG.CU.GCACAGUUUAGGC..................
env-235 GAUGUCGUCCCCCGAAACAGAGAAAAGUUUAAUCAACAAAUAAUUUAUGUUUCAUUUAUCCUGAAACUUGGGCUGUGAUUGAUUACA................................ACACAACUUCUC...UGUAAUCGCGG.CU.GCACAGUUUAGGCUUCAGUAAAGUUUAAAAG
env-236 CGGCUGUCGUCCUCCGAAACAAAGAAAAGUUUAAUCAAUCAAUUCAGUUUCAUUUACCUUGAAACUUGAGCUGUGAAUUUGAUUACA................................ACACAACUUCUC...UGUAAUCGCGG.CU.GCACAGUUUAGGCUUCAGUAAAGUUUAAAUU
env-237 CCCACCGAAAGUGAGAAAAGUUUAAUGUGAAUCAAAUAAUUAUUUAAGUUUCAUUUACCUUGAAACUUGAGCUGUGAUUUGAUUACA................................ACACAACUUCUC...UGUAAUCGCGG.CU.GCACAGUUUAGGCUUCAGUAAUAGUUUAAAA
env-238 GUUGCCGUCCCCCGAAACAGCGAAAAGUUUAAUCAUCAAAAAUAAUAAGUUUCAUUGAUCUUGAAACUUGAGCUGUGAUUGAUUACA................................CCACAACUUCUC...UGUAAUCGCGG.CU.GCACAGUUUAGGCUUCAGUAAAGUUUAAUUG
env-239 GUCCACCCAAAGCAAGAAAAGUGUAAGUUUAAUCGAGAAAUAAUUUAAGUUUCAUUUAUCUUGAGACUUGAGCUGUGAUUGAUUACA................................CCACAACUUCUC...UGUAAUCGCGG.CU.GCACAGUUUAGGCUUCAGUUUUUAAAGUUUA
Pac-1-1 AUUAUGCCAUGGCGAGAAAUAGUGCAACGUGUUGAAAAAUAAUUUAAGUUUCAUUUAUCUUGGAACUUGAGCUGUGAUUUGAUUACA................................CCACAACUUCUC...UGUAAUCGCGG.CU.GCACAGUUUAGGCUUCAGUUUUUAAAAGUUU
env-240 GGUGCCGUCCCCCGAAACAGCGAAAAGUUUAAUCAUCAAAAAUAAUAAGUUUCAUUGAUCUUGAAACUUGAGCUGUGAUUGAUUACA................................CCACAACUUCUC...UGUAAUCGCGG.CU.GCACAGUUUAGGCUUCAGUAAAGUUUAAUUG
env-241 CGGAUGUCGCCCCCCGAAACAAAGAAAAGUAUAAUCAAUCAAUUCAGUUUCAUUUACAACGAAACUUGAGCUGUGAAAUUGAUUACA................................CCACAACUUCUC...UGUAAUCGCGG.CU.GCACAGUUUAGGCUUCAGUAAAGUUUAAAAU
env-242 UCAUAAUGGUUGUCGUCCUCCUAAAAGACGUCGUGUAUAAGAUUUACCUUUAUUAUAGAUGAAACUUGAUUUUGUUAUAGGAUUGUA................................UAAUUUCUUCUC...UACAAUCGCGG.CU.GCAACAAAUUAAGUUCAUGAUUAAAACAAUU
env-243 ACCACACAAUGGUUGUCGUCCUCCCAAAAGAAGAAGAGUUUAAUUCAAAAUCCGAAGUUUAGAGUCAUAAUUUGUGAAAAGGUUGCA................................CCAAAUCCUCUC...UGCAAUCGCGG.CU.GCACUGAUUGGGAUUCUAAGUUGACACUAAU
env-244 GUUGUCGUCCACCAAAACGUCGUAGAGUUUAAUUCGUAAUCGAGUUUCCUCGGCAGUCGAAUAGGCGAAGCUAAGAAAUAGGUUGCA................................UUUAAACCUCUC...UGCAAUCGCGG.CU.GUAAUAAGUUUCGGCCGAUUUGAGCACGUGG
env-245 UUGUCGUCCUCCAAAACGUCGUAGAGUUUAAUUCUAAUUAGAGUUUCCUCGGCAACGAGAAUGGGUGAAAUUGAAGAUUAGAUUGCA................................UUUAAACCUCUC...UGCAAUCGCGG.CU.GUGCCAAGUUUCAACCAAUCACUAAACGUGG
Ppr-1-1 GUUGUCGUCCUCCUAAACGUCGUAGAGUGUAAUUCUUAAAAGGGUUUCCUCGGUAGUCGAAUAGGUGAACUAAAGAAAAAGAUUGCA................................UUUAAACCUCUC...UGCAAUCGCGG.CU.GUAAAAAGUUUCGACCCAUUUGUGUACGUGG
env-246 CAACGGAUGUCGUCCCCCGAAAAGAAGAAGAGUAUAAUACAUAUUCUUAUUUAGAACAAUAGAAUCUUGAUCAGUGAAUAGAUUGCA................................CUUAAUCCUCUC...UGCAAUCGCGG.CU.GCACUCCUCGGGUUCGUAGAAAAUUAUAAUU
env-247 .........GUCGUCCUCCGAAAAGAAGAAGAGUCUAACUUAUUCUUUUUAUUAGAAAACUGAAUCUCGAAUAGUGAAUAGAUUGCA................................CUUAAUCCUCUC...UGCAAUCGCGG.CU.GCACUCUUCCGGGUUCGCAACUUAUAAUUUA
env-248 .....................................................................AAUUGUGAUAAGAUUGCA................................CAUUAUCCUCUC...UGCAGUCGCGG.CU.GCACGAUUCCAGAUUCAUACAGAACAUUUAA
env-249 CACAACGGAUGUCGUCCUCCUAAAAAGAGAAGAGUUUAAUUAACAAUAAUUUGCUGCACAUGAAUCCGGAAUUGUGAUAAGAUUGCA................................CAUUAUCCUCUC...UGCAGUCGCGG.CU.GCACGAUUCCAGAUUCAUACAGAACAUUUAA
env-250 GCCGAUGCUCUCAAGCCCUACCUUAACGACGCCUCUGGCUUUUAUCAUUUUAUUACUUCUGAUCUUUCCUGUUGUGAAUAGAUUGCA.............................UUUCUACCACCUCUC...UGCAAUCGCGG.CU.GCACUAAGCUAAGAUCGAAUUCUAAUUCAAU
env-251 AUAAUGGAUGUCGUCCUCCUAAAAGACGGAGAGUAUAAAAUUUACAAGUAAUAAUCAAGAAUCUGUCUGACGUGAAAUAAGAUUGCA.............................UUCAAACCACCUCUC...UGCAAUCGCGG.CU.GCACUAAGAAGAUUCGUAGGACACUUUAAUU
env-252 CCCAUAACGGUUGUCGUCCUCCUAAAAGACGUAGAGUAUAAUUCACUCUUAAACCUUGUCGAGUCACAGGCGGUGAAUAAGAUUGCA...............................AUCAAAUCCUCUC...UGCAAUCGCGG.CU.GCACCUAUUGCGAUUCACAACAAUAUAAAAU
env-253 NNNNNNNNNNNNNNNNNNNNNCUAAAAGACGUAGAGUAUAAUUCACUCUUAAACCUUGUCGAGUCACAGGCGGUGAAUAAGAUUGCA...............................AUCAAAUCCUCUC...UGCAAUCGCGG.CU.GCACCUAUUGCGAUUCACAACAAUAUAAAAU
env-254 GAUGUCGUCCUCCUAAAGCAAGAAGAGUUUAAUCGACAAUUAUUUAAGUUUCAUUACAUCGAAACUUGAGCUGUGAUUUUGAUUACA...............................AUAUUCACUUCUC...UGUAAUCGCGG.CU.GCACAGUUUAGGCUUCAGUAAAAAAGUUUAU
env-255 CACAACGGAUGUCGCCCAGCUAAACGCAGAAGAGUAUAAUAAUCACAAACUUGUUCAUUACCCGGCUAAAAGUGGAUAAAGAUUACA...............................AUUCUAACCUCUC...UGUAAUCGCGG.CU.GCACGGUAAGUCGAGUGUGAUAUAAGCAAUA
env-256 CAUAACGGAUGUCGCCCAGCUAAACGCAGAAGAGUAUAAUAAUCACCAACUUGUUCAUUACCCGGCUAAAAGUGGAUAAAGAUUACA...............................AUUCUAACCUCUC...UGUAAUCGCGG.CU.GCACGGUAGGUCGAGUGUGAUAUAAGCAAUA
env-257 CACAACGGAUGUCGCCCAGCUAAACGCAGAAGAGUAUAAUAAUCACCAACUUGUUCAUUACCCGGCUAAAAGUGGAUAAAGAUUACA...............................AUUCUAACCUCUC...UGUAAUCGCGG.CU.GCACGGUAAGUCGAGUGUGAUAUAAGCAAUA
env-258 AUAACCGUAGACCUAAAUGAAUUCAAAAAAGUGUAAAUAAUCAUAAGUUUCAAUCAUACAGAAACUUGAGCUGUGAUUUUGAUUACA...............................ACACUAACUUCUC...UGUAAUCGCGG.CU.GACACAGUUUAGGCUUCAGU...........
env-259 UAACCGUAGACCUAAAUGAAUUCAAAAAAGUAGUAAAUAAUCAUAAGUUUCAAUCAUACAGAAACUUGAGCUGUGAUUUUGAUUACA...............................ACACUAACUUCUC...UGUAAUCGCGG.CU.GACACAGUUUAGGCUUCAGUAGAAAAAAGUU
Dga-1-1 UUCCUCACAACGGAUGCCGUCCUCCAAAACACAGAAGAGUUUAAUCAAAAUUUUAAAUUUGGAACUUAAAUUGUGAUUUUGAUUGCA...............................AGUUUACCCUCUC...UGUAAUCGCGG.CU.GCACAAUUUCAGCUUUCAUCAAUUAUUAAUU
env-260 CUUCCUCACAACGGAUGUCGUCCUCCAAAACACAGAAGAGUUUAAUCAAAUUUGAAAAUUGGAACUUGAAUUGUGAAUUUGAUUGCA...............................AGUAAACCCUCUC...UGUGAUCGCGG.CU.GCACAAUUUUAGCUUUCGUCAAUUAUUAAUU
Psp-4-1 CUUCCUCACAAUGGAUGUCGUCCUCCUAAACACAGAAGAGUUUAAUCAAAAUUUAAAAUUGGAACUUAAAUUGUGAUUUUGAUUGCA...............................AGUCAACCCUCUC...UGUAAUCGCGG.CU.GCACAAUUUCAGCUUUCAUCAAUUAUUAAUU
env-261 ACACAAUGGAUGUCGUCCUCCUAAGAGAAGAAGAGUAUAAUCUCAGAAAUUAGUAUUUACCAGAAUCUUUUGAUGUGAUUGAUUGCA..............................UUAUACCACCUCUC...UGCAAUCGCGG.CU.GCACUAGAAAGAUUCGUAUUCAACUUUAAUU
env-262 CGCUUCCACACAAUGGUUGUCGUCCUCCUAAAAGAAGAAGAGUAUAAUCAUUUAAAUUAAAAAGAGUCUUAAACGUGAAAGAUUGCA..............................UUCAACCACCUCUC...UGCAAUCGCGG.CU.GCACUGAUAAGAUUCGUUAUCAACUUUAAUA
env-263 CUGCCGCACAAUGGCUGCCGUCCGCCCAAAAGAAGAAAAGUAUAAUUAACAGAUCCGGCGAAUCCCCGAAGAAGUGAUUAGAUUGCA..............................CUAUUCACCCUCUC...UGCAAUCGCGG.CU.GCACGAUUCCGGGAUUCCGAAGAACGAUUAA
env-264 .......................CCCAAAAGAAGAAAAGUAUAAUUAACAGAUCAAGUGAAUUCUCUGAAGACGUGAUUAGAUUGCA..............................CUAUUCACCCUCUC...UGCAAUCGCGG.CU.GCACGAUUCCGGGAUUCCAAAGAACGAUUAA
env-265 GUUUUAGCCCCUUGUUCGGAACCAAAUUUUAUCAAUUUUAUUUACCAAACAAUUUUUUCGCGAGUCUCUUACAGUGAUUGGAUUGCA..........................CUUUAUCAUCGACCUCUC...UGCAAUCGCGG.CU.GCACUGUAGCGAUUCUGCUUUCAAAACUUUA
env-266 UAGUCUACACUUCGACACUAGGUUUAAUUACAUUCAUUUAAACUCACUUUUUAUAACAAGCGAAUUAUUCACAGUGAAUAGAUUGCA..........................UCUCUUACACGACCUCUC...UGCAAUCGCGG.CU.GCACUGUGAUGGUUCUGUAUCACAUCUUUAA
env-267 UGAUCGUCACUUUUCGACAUUUGGUUUAUAACUUCUUUUUUAUAUUUUUUAUUUUCAAGCGAAUUACGCACAGUGAAUUAGAUUGCA..........................UCUUUUACACGACCUCUC...UGCAAUCGCGG.CU.GCACUGUGAUGAUUCUGUAUCACAUCUUUAA
env-268 AGGCGCUUCACGCAUCGUAACCACUGUUUUUAACAGAUUUUUUAUUAACAAACUUUUUCGCGAAUCUCUAUCAGUGAAUGGAUUGCA..........................UUUUUUUGACGACCUCUC...UGCAAUCGCGG.CU.GCACUGAAGCGAUUCCAAUUCACAAAAGCUA
env-269 AAGAGUUUAAUUGCACAACUUCUUGUUUUUAAUCAUCACGAUGAUUUUUUGAUUUUUUCGCGAAUCUCUGUCAGUGAAUGGAUUGCA..........................AUCAUUUAUCGACCUCUC...UGCAAUCGCGG.CU.GCACUGAAGCGAUUCCAAUUCACAACUUGUU
env-270 CCAAGAGAAGAAGAGUCUAAUUCACUUGGAAGAUUUUAUUCGUUUUUUUACUUACACAACGAAUCUCGUAUAGUGAAUAGAUUGCA..........................CUUAAUUAACAACCUCUC...UGCAAUCGCGG.CU.GCACUAUAGUGAUUCUCAACAUCACAAAACU
env-271 AGAAGAGUAUAAUCUAACGCGUGAUUUCUAUUCAACAAUUCGUUUUAAUUUAUUUUUCAACGAGUCUCUUAUAGUGAAUAGAUUGCA..........................UUCAUUUAACAACCUCUC...UGCAAUCGCGG.CU.GCACUAUAGUGAUUCACAGCAUCACACAACU
env-272 UUGCCUCAUAAUGGAUGUCGUCCUCCUAAAAGACGUAGAGUUUAAUUUUUCUAAAUUUGAAAACUUGUUUUUGUUUUUUGAUUACA...............................UUAUCUUCUUCUU...UGUGAUCGCGG.CU.GCAACAAAAUAAGUUUUGAGUAAACAAUUUA
env-273 AUUGCCUCAUAAUGGGUGUCGUCCUCCUAAAAGACGUAGAGUUUAAUUUUUCUAAAUUUGAAAACUUGUUUUUGUUUUUUGAUUACA...............................UUAUCUUCUUCUU...UGUGAUCGCGG.CU.GCAACAAAAUAAGUUUUGAGUAAACAAUUUA
env-274 CGUAGUUUUGAGUGUGAUAACUCCUAACUCCUAAUUCCUAACUCCUAACUUUUAAAAAAAAGAAUCAUUCACAGUGAUUGGAUUGCA.................AUAUCUAUCAUUUAUGCACGACCUCUC...UGCAAUCGCGG.CU.GCACUGUGACGGUUCCUUACCAAAAAAAGAU
env-275 GGAAACGAGCUGAAGAAAAAACUCCACUCUCAACUCUUAACUCUCAAAACUAAUUAAAAAAGAAUCAUUCACAGUGAUUGGAUUGCA.................AUAUCUAUCAUUUAUGCACGACCUCUC...UGCAAUCGCGG.CU.GCACUGUGACGGUUCCUUACCAAAAAAAGAU
env-276 AAGAGAAGAAGAGUUUAAUGUCACUUCUCACUACAUGAUUUUUUAUUAAUUAUUUUUUCGCGAAUCUCUGUCAGUGAUUGGAUUGCA........................CUUUUUCUUAACAACCUCUC...UGCAAUCGCGG.CU.GCACUGAUGCGAUUCAAAACUUAACUUUUUA
env-277 CUCUUGUUCACAAGAAACAAUCACAUUUCUUGAUUCAUUACAAUUAACAAACUUUUUUCGCGAAUCUCUAUCAGUGAUUGGAUUGCA........................CAAUUUUUUAACGACCUCUC...UGCAAUCGCGG.CU.GCACUGUAGCGGUUCCAAUUCACAAUGUUAA
env-278 ACAUAACGGUUGUCGUCCUCCGAAAAGACGUAGAGUUUAAAAUUUACCUUUAAUAGAAGUGAGACUUGAUUUUGUUAUUGGAUUGUA.................................AACUUUCUCUC...UGCAAUCGCGG.CU.GCAACAAACCCAAGUUCAUGAAUAAACAAUU
env-279 ACAUAAUGGUUGUCGUCCUCCGAAAAGACGUAGAGUUUAAAAUUUACCUUUAAUAGAAGUGAGACUUGAUUUUGUUAUUGGAUUGUA.................................AACUUUCUCUC...UGCAAUCGCGG.CU.GCAACAAACCCAAGUUCAUGAAUAAACAAUU
env-280 UGAGAAAGGUAAAAGCAUAUGUGAANNNNNNNNNNNNNNNAAUUUACCUUUAAUAGAAGUGAGACUUGAUUUUGUUAUUGGAUUGUA.................................AACUUUCUCUC...UGCAAUCGCGG.CU.GCAACAAACCCAAGUUCAUGAAUAAACAAUU
env-281 NNNNNNNNNNNNNNNUCCUCCGAAAAGACGUAGAGUUUAAAAUUUACCUUUAAUAGAAGUGAGACUUGAUUUUGUUAUUGGAUUGUA.................................AACUUUCUCUC...UGCAAUCGCGG.CU.GCAACAAACCCAAGUUCAUGAAUAAACAAUU
env-282 NNNNNNNNNNNNNNNNNNNNNNNNNNNNNNNNNNNNNNNNAAUUUACCUUUAAUAGAAGUGAGACUUGAUUUUGUUAUUGGAUUGUA.................................AACUUUCUCUC...UGCAAUCGCGG.CU.GCAACAAACCCAAGUUCAUGAAUAAACAAUU
env-283 CGCACAACGGUUGUCGUCCUCCGAAAAGACGUAGAGUUAAAAUUUACCUUUAAUAGAAGUGAGACUUGAUUUUGUUAUUGGAUUGUA.................................AACUUUCUCUC...UGCAAUCGCGG.CU.GCAACAAACCCAAGUUCAUGAAUAAACAAUU
env-284 AANNNNNNNNNNNNNNNNNNNNNNNNNNNNNNNNNNNNNNAAUUUACCUUUAAUAGAAGUGAGACUUGAUUUUGUUAUUGGAUUGUA.................................AACUUUCUCUC...UGCAAUCGCGG.CU.GCAACAAACCCAAGUUCAUGAAUAAACAAUU
env-285 NNNNNNNNNNNNNNNUCCUCCGAAAAGACGUAGAGUUUAAAAUUUACCUUUAAUAGAAGUGAGACUUGAUUUUGUUAUUGGAUUGUA.................................AACUUUCUCUC...UGCAAUCGCGG.CU.GCAACAAACCC....................
env-286 .................CUCCGAAAAGACGUAGAGUUUAAAAUUUACCUUUAAUAGAAGUGAGACUUGAUUUUGUUAUUGGAUUGUA.................................AACUUUCUCUC...UGCAAUCGCGG.CU.GCAACAAACCCAAGUUCAUGAAUAAACAAUU
env-287 ACAUAAUGGUUGUCGUCCUCCGAAAAGACGUAGAGUUUAAAAUUUACCUUUAAUAGAAGUGAGACUUGAUUUUGUUAUUGGAUUGUA.................................AACUUUCUCUC...UGCAAUCGCGG.CU.GCAACAAACCCAAGUUCAUGAAUAAACAAUU
env-288 GCACAAUGGUUGUCGUCCUCCGAAAAGACGUAGAGUUUAAAAUUUACCUUUAAUAGAAGUGAGACUUGAUUUUGUUAUUGGAUUGUA.................................AACUUUCUCUC...UGCAAUCGCGG.CU.GCAACAAACCC....................
env-289 ACAUAACGGUUGUCGUCCUCCGAAAAGACGUAGAGUUUAAAAUUUACCUUUAAUAGAAGUGAGACUUGAUUUUGUUAUUGGAUUGUA.................................AACUUUCUCUC...UGCAAUCGCGG.CU.GCAACAAACCCAAGUUCAUGAAUAAACAAUU
env-290 NNNNNNNNNNNNNNNNNNNNNNNNNNNNNNNNNNNNNNNNAAUUUACCUUUAAUAGAAGUGAGACUUGAUUUUGUUAUUGGAUUGUA.................................AACUUUCUCUC...UGCAAUCGCGG.CU.GCAACAAACCCAAGUUCAUGAAUAAACAAUU
env-291 NNNNNNNNNNNNNNNUCCUCCGAAAAGACGUAGAGUUUAAAAUUUACCUUUAAUAGAAGUGAGACUUGAUUUUGUUAUUGGAUUGUA.................................AACUUUCUCUC...UGCAAUCGCGG.CU.GCAACAAACCCAAGUUCAUGAAUAAACAAUU
env-292 GCACAAUGGUUGUCGUCCUCCGAAAAGACGUAGAGUUUAAAAUUUACCUUUGAUAGAAGUGAGACUUGAUUUUGUAAUUGGAUUGUA.................................AACUUUCUCUC...UGCAAUCGCGG.CU.GCAACAAACCCAAGUUCAUGAAUAAACAAUU
env-293 GCACAACGGUUGUCGUCCUCCGAAAAGACGUAGAGUUUAAGAUUUACCUUUAAUAGAAGUGAGACUUGAUUUUGUUAUUGGAUUGUA.................................AACUUUCUCUC...UGCAAUCGCGG.CU.GCAACAAACCCAAGUUCAUGAAUAAACAAUU
env-294 GGCUNNNNNNNNNNNUCCUCCGAAAAGACGUAGAGUUUAAAAUUUACCUUUAAUAGAAGUGAGACUUGAUUUUGUUAUUGGAUUGUA.................................AACUUUCUCUC...UGCAAUCGCGG.CU.GCAACAAACCCAAGUUCAUGAAUAAACAAUU
env-295 GCACAACGGUUGUCGUCCUCCGAAAAGACGUAGAGUUUAAAAUUUACCUUUAAUAGAAGUGAGACUUGAUUUUGUUAUUGGAUUGUA.................................AACUUUCUCUC...UGCAAUCGCGG.CU.GCAACAAAUUAAGUUCAUGAAUAAACAAUUA
env-296 GCAUAAUGGUUGUCGUCCUCCGAAAAGACGUAGAGUUUAAGAUUUACCUUUAAUAAGUAUGACACUUGAUUUUGUUUUUGGAUUGUA.................................AACUUUCUCUC...UACAAUCGCGG.CU.GCAACAAAUUGAGUUCAUGAAUAAACAAUUA
env-297 ACAUAAUGGUUGUCGUCCUCCGAAAAGACGUAGAGUUUAAGAUUUACCUUUAAUAAAAAUGAUACUUGAUUUUGUUUUUGGAUUGUA.................................AAAUUUCUCUC...UACAAUCGCGG.CU.GCAACAAAUUGAGUUCAUGAAUAAACAAUUA
env-298 GCAUAAUGGAUGUCGUCCUCCGAAAAGACGUAGAGUUUAAGAUUUACCUUUAAUACAAAUGAUACUUGAUUUUGUUUUUGGAUUACA.................................UUUUUUCUCUC...UGUAAUCGCGG.CU.GCAACAAAUUAAGUUCAUGAAUAAACAAUUA
env-299 ACAUAACGGUUGUCGUCCUCCGAAAAGACGUAGAGUUUAAUUUACCUUUAACUUACACAAUGAUCUUGAUUUUGUUUUUGGAUUGCA.................................UAAUUUCUCUC...UGUAAUCGCGG.CU.GCAACAAAUUGAGUUCAUGAAUAAACAAUUA
Dsp-2-1 ACUUCCUCACAAUGGUUGUCGUCCUCCAAAACACAGAAGAGUUUAAUCCCAAUUUCUAAGUAGAAACAUGAGUUGUGAUUGAUUGCA................................AGUAACCCUCUC...UGUAAUCGCGG.CU.GCACAAUUCUCGCUUCUUUUUAAUGUUUUAA
env-300 AUAACGGAUGUCGUCCUCCUAAACAAAGAAGAGUUUAAUUUUAACGUAAGUAAGAGAGAUUUGAAACUUAAAUUGUGAAUGAUUGCA................................ACAACCCCUCUC...UGUAAUCGCGG.CU.GCACAAUUUCAGAUUCAUUUAAUUUAAUGUA
env-301 .............................................UUUAAUGUAGAGAGUAGGUUGCGGAUUUAUGAUUAGAUUACA................................UAUCAUCCUCUC...UGUAAUCGCGG.CU.GCAUAAACGAUGACUCUACUCUGAAAUAGUA
env-302 UCCCNCACAACGGAUGNCGUCCUCCGAAAAAGAGAAGAGUAUAAUUUUAAUGUAGAGAGUAGGUUGCGGAUUUAUGAUUAGAUUACA................................UAUCAUCCUCUC...UGUGAUCGCGG.CU.GCAUAAACGGAUAACUCUACUCUGAAAUAGU
env-303 UCCCGCACAACGGAUGCCGUCCCCCGAAAAAGAGAAGAGUAUAAUUUUAAUGUAGAGAGUAGGUUGCGGAUUUAUGAUUAGAUUACA................................UAUCAUCCUCUC...UGUGAUCGCGG.CU.GCAUAAACGGAUAACUCUACUCUGAAAUAGU

        ................................................................................<<<<<<<<<<.........................................>>>>>>>>>>.......................................
        ................................................................................1122222100.........................................0012222211.......................................
        •••••••••••••••••••••••••••••R••••••••••••••••••••••••••••••••••••••••••••••R•URGAUURYA-----------------------------○○○○○•Y••CCUCUC---URYARUCGCGG-CU-GCAY•R•••••••YYY•••••••••••••••
                                                                                5′                                                                               3′
```

```
env-304  UUCCGCACAACGGAUGUCGUCCUCCGAAAAAGAGAAGAGUAUAAUUUUAAUGUAGAGAGUAGGUUGCGGAUUUAUGAUUAGAUUACA................................UAUCAUCCUCUC...UGUGAUCGCGG.CU.GCAUAAACGGANAACUCUACUCUNAAAUAGU
env-305  GCACAACGGAUGUCGUCCACCAAAAAGAAGAAGAGUUUAAUUAAUUUAUUUACUGUAUAGGGCAGGAAAUGCGUGGAAUAGAUUACA................................UAAAACCCUCUC...UGUAAUCGCGG.CU.GCACAAGUUUCUUUUCCUUCAAAAUUAACAA
Dca-1-1  CUUCCUCAUAAUGGAUGUCGUCCUCCAAAACACAGAAGAGUUUAAUCCCAAUUUUAAAGUUUGAAACUUGAGUUGUGAUUGAUUGCA................................AAACACCCUCUC...UGUAAUCGCGG.CU.GCACAAUUCUCGCUUCUUUUUAAUGUAUUAA
env-306  .............................................................GACUUAAUUUUGUUUUAUGGAUUGCA................................CUCUUUCUUCUC...UGUAAUCGCGG.CU.GCAACAAAUUAAGUUGAUGAUUACUAUAAUU
env-307  CUCCAUUGCCACAUAAUGGAUGUCGUCCUCCUAAAAGACGUAGAGUUUAAUUAAUAAAAUGAGACUUAAUUUUGUUUUAUGGAUUGCA................................CUCUUUCUUCUC...UGUAAUCGCGG.CU.GCAACAAAUUAAGUUGAUGAUUACUAUAAUU
Bpy-2-1  CGCAUAAUGGUUGUCGUCCUCCUAAAAGACGUAGAGUUUAAUUUACCUUUAACUUAAAAAUGAUCUUGAUUUUGUUUUAUGGAUUGCA................................UUAUUUCUUCUC...UGUAAUCGCGG.CU.GCAACAAAUUGAGUUCAUGAAUAAACAAUUA
env-308  CCCAUAAUGGUUGUCGUCCUCCUAAAAGACGUAGAGUUUAAUUUACCUUUAACUUAAAAAUGAUCUUGAUUUUGUUUUAUGGAUUGCA................................UUAUUUCUUCUC...UGUAAUCGCGG.CU.GCAACAAAUUAAGUUCAUGAAUAAACAAUUA
Bpy-1-1  CGCAUAAUGGUUGUCGUCCUCCUAAAAGACGUAGAGUUUAAUUUACCUUUAACUUAAAAAUGAUCUUGAUUUUGUUUUAUGGAUUGCA................................UUAUUUCUUCUC...UGUAAUCGCGG.CU.GCAACAAAUUGAGUUCAUGAAUAAACAAUUA
env-309  UGCCGCAUAAUGGCUGUCGCCCACCGAAAAGACGUAGAGUAUAAGAAUUACUUUUAAUAAUUGAAACUUGACUUGUUAUGGAUUACA...............................CUAAAUUCUUCUC...UGUAAUCGCGG.CU.GCAACAAAUCGGGGUUCAUAAGUAACAAUUA
env-310  ACAUAAUGGUUGUCGUCCUCCGAAAAGACGUAGAGUUUAAUUUACCUUUAACUUACACAAUGAUCUUGAUUUUGUUUUUGGAUUGCA................................UUAAUUUCUCUC...UGUAAUCGCGG.CU.GCAACAAAUUAAGUUCAUGAAUAAACAAUAA
env-311  GCAUAACGGUUGUCGUCCUCCGAAAAGACGUAGAGUUUAAUUUACCUUUAACUUACACAAUGAUCUUGAUUUUGUUUUUGGAUUGCA................................UUAAUUUCUCUC...UGUAAUCGCGG.CU.GCAACAAAUUAAGUUCAUGAAUAAACAAUUA
env-312  ACAUAAUGGUUGUCGUCCUCCGAAAAGACGUAGAGUUUAAUUUACCUUUAACUUACACAAUGAUCUUGAUUUUGUUUUUGGAUUGCA................................UUAAUUUCUCUC...UGUAAUCGCGG.CU.GCAACAAAUUAAGUUCAUGAAUAAACAAUUA
env-313  CCACACAAUGGUUGUCGUCCUCCGAAAAGACGUAGAGUUUAUUACCUUUAACUUACACAAUGAUCUUGAUUUUGUUUUUGGAUUGCA................................UUAAUUUCUCUC...UGUAAUCGCGG.CU.GCAACAAAUUAAGUUCAUGAAUAAACAAUUA
env-314  ACAUAACGGUUGUCGUCCUCCGAAAAGACGUAGAGUUUAAUUUACCUUUAACUUACACAAUGAUCUUGAUUUUGUUUUUGGAUUGCA................................UUAAUUUCUCUC...UGUAAUCGCGG.CU.GCAACAAAUUAAGUUCAUGAAUAAACAAUUA
env-315  ACAUAAUGGUUGUCGUCCUCCGAAAAGACGUAGAGUUUAAGAUUUACCUUUAAUAAAAAUGAUCUUGAUUUUGUUAUUUGGAUUACA................................UAAAUUUCUCUC...UGUAAUCGCGG.CU.GCAACAAAUUGAGUUCAUGAAUAAAACAAUU
env-316  ACAUAAUGGUUGUCGUCCUCCGAAAAGACGUAGAGUUUAAUUUACCUUUAACUUACACAAUGAUCUUGAUUUUGUUUUUGGAUUACA................................UUAAUUUCUCUC...UGUAAUCGCGG.CU.GCAACAAAUUGAGUUCAUGAAUAAACAAUUA
env-317  ACAUAAUGGUUGUCGUCCUCCGAAAAGACGUAGAGUUUAAGAUUUACCUUUAAUAAAAAUGAUCUUGAUUUUGUUAUUUGGAUUACA................................UUAAUUUCUCUC...UGUAAUCGCGG.CU.GCAACAAAUUGAGUUCAUGAAUAAAACAAUU
env-318  .....................CCUAAAAGACGUAGAGUUUAAUUUACCUUUAAUUUAGAAUGAUCUUGAUUUUGUUUUUGGAUUACA................................UUAAUUUCUCUC...UGUAAUCGCGG.CU.GCAACAAAUUGAGUUCAUCAGUAAACAAUUA
Bst-1-1  CACAUAAUGGUUGUCGUCCUCCGAAAAGACGUAGAGUUUAAGAUUUACCUUUAAUAAAAAUGAUCUUGAUUUUGUUAUUGGAUUACA................................UUAAUUUCUCUC...UGUAAUCGCGG.CU.GCAACAAAUUGAGUUCAUGAAUAAACAAUUA
env-319  ....................................................................AUUUUGUUUUUGGAUUACA................................UUAAUUUCUCUC...UGUAAUCGCGG.CU.GCAACAAAUUGAGUUCAUGAAUAAACAAUUA
Bti-1-1  CACAUAAUGGUUGUCGUCCUCCGAAAAGACGUAGAGUUUAAGAUUUACCUUUAAUAAAAAUGAUCUUGAUUUUGUUUUUGGAUUACA................................UUAAUUUCUCUC...UGUAAUCGCGG.CU.GCAACAAAUUGAGUUCAUGAAUAAAACAAUU
env-320  CACAUAAUGGUUGUCGUCCUCCGAAAAGACGUAGAGUUUAAGAUUUACCUUUAAUAAAAAUGAUCUUGAUUUUGUUUUUGGAUUACA................................UUAAUUUCUCUC...UGUAAUCGCGG.CU.GCAACAAAUUAAGUUCAUGAAUAAAACAAUU
env-321  ACAUAAUGGUUGUCGUCCUCCGAAAAGACGUAGAGUUUAAUUUACCUUUAACUUACACAAUGAUCUUGAUUUUGUUUUUGGAUUACA................................UUAAUUUCUCUC...UGUAAUCGCGG.CU.GCAACAAAUUGAGUUCAUGAAUAAACAAUUA
env-322  ......GGUUGUCGUCCUCCGAAAAGACGUAGAGUUUAAGAUUUACCUUUAAUAAAAAUGAUACUUGAAUUUUGUUUUUGGAUUACA................................UUAAUUUCUCUC...UGUAAUCGCGG.CU.GCAACAAAUUAAGUUCAUGAAUAAAACAAUU
env-323  AUAACAUGGUUGUCGUCCUCCGAAAAGACGUAGAGUUUAAUUUACCUUUAACUUACACAAUGAUCUUGAUUUUGUUUUUGGAUUACA................................UUAAUUUCUCUC...UGUAAUCGCGG.CU.GCAACAAAUUGAGUUCAUGAAUAAACAAUUA
Bac-1-1  CCACAUAAUGGUUGUCGUCCUCCAAAAAGACGUAGAGUUUAAUUUACCUUUAACUUACAAUGAUCUUGAUUUUGUUUUUGGAUUACA................................UUAAUUUCUCUC...UGUAAUCGCGG.CU.GCAACAAAUUGAGUUCAUGAAUAAACAAUUA
env-324  ACAUAAUGGUUGUCGUCCUCCGAAAAGACGUAGAGUUUAAGAUUUACCUUUAAUAAAAAUGAUCUUGAUUUUGUUAUUUGGAUUACA................................UUAAUUUCUCUC...UGUAAUCGCGG.CU.GCAACAAAUUGAGUUCAUGAAUAAAACAAUU
env-325  ACACAAUGGUUGUCGUCCUCCGAAAAGACGUAGAGUUUAAUUUACCUUUAACUUACACAAUGAUCUUGAUUUUGUUUUUGGAUUACA................................UUAAUUUCUCUC...UGUAAUCGCGG.CU.GCAACAAAUUGAGUUCAUGAAUAAACAAUUA
Bol-1-1  CACAUAAUGGUUGUCGUCCGCCGAAAAGACGUAGAGUUUAAGAUUUACCUUUAAUAAAAAUGAUCUUGAUUUUGUUAUUGGAUUACA................................UUAAUUUCUCUC...UGUAAUCGCGG.CU.GCAACAAAUUGAGUUCAUGAAUAAACAAUUA
env-326  ....................................................................AUUUUGUUUUUGGAUUACA................................UUAAUUUCUCUC...UGUAAUCGCGG.CU.GCAACAAAUUAAGUUCAUGAAUAAAACAAUU
env-327  ACAUAAUGGUUGUCGUCCUCCUAAAAGAAGGAGAGUUUAAUAUCACCUAAUUUGAAAUAUGACACUUGAUUUUGUUUUUGGAUUGCA...............................UUUAUUUCCUUUU...UGUAAUCGCGG.CU.GCAACAAAUUAAGUUCAUUAAUAAACAAUUA
env-328  AACGGUUGUCGUCCUCCUAAAAGAAGAAGAGUAUAAUAAAGAUAACCUAUUUAGAUGUAACUUGAUUUUGUUAAAUAAUGGAUUACA....................................UUCCUUUC...UGUGAUCGCGG.CU.GCAACAAAUUGAGUUCAUUAGUAACAAUUAA
env-329  NNNNNNNNNNNNNNNNNNNNNNNNNNNNNNNNNNNNNNNNNNNNNNNNNNNNNNNNNNNNNNAUCUUGAUUUUGUUUUUGGAUUGCA...............................AUAAUUUCUUCUC...UGCAAUCGCGG.CU.GCAACAAAUUGAGUUCAUGAAUAAACAAUUA
env-330  CCACAUAAUGGUUGUCGUCCUCCGAAAAGACGUAGAGUUUAAUUUACCUUUAACUUAAAAUGAUCUUGAUUUUGUUUUUGGAUUGCA...............................AUAAUUUCUUCUC...UGCAAUCGCGG.CU.GCAACAAAUUGAGUUCAUGAAUAAACAAUUA
env-331  CCACACAAUGGUUGUCGUCCUCCGAAAAGACGUAGAGUUUAAUUUACCUUUAACUUAAAAUGAUCUUGAUUUUGUUUUUGGAUUGCA...............................AUAAUUUCUUCUC...UGCAAUCGCGG.CU.GCAACAAAUUGAGUUCAUGAAUAAACAAUUA
env-332  CCACACAAUGGUUGUCGUCCUCCGAAAAGACGUAGAGUUUAAUUUACCUUUAACUUAAAAUGAUCUUGAUUUUGUUUUUGGAUUGCA...............................AUAAUUUCUUCUC...UGCAAUCGCGG.CU.GCAACAAAUUGAGUUCAUGAAUAAACAAUUA
env-333  CCACACAAUGGUUGUCGUCCUCCGAAAAGACGUAGAGUUUAAUUUACCUUUAACUUAAAAUGAUCUUGAUUUUGUUUUUGGAUUGCA...............................AUAAUUUCUUCUC...UGCAAUCGCGG.CU.GCAACAAAUUAAGUUCAUGAAUAAACAAUUA
env-334  CCACACAAUGGUUGUCGUCCUCCGAAAAGACGUAGAGUUUAAUUUACCUUUAACUUAAAAUGAUCUUGAUUUUGUUUUUGGAUUGCA...............................AUAAUUUCUUCUC...UGCAAUCGCGG.CU.GCAACAAAUUAAGUUCAUGAAUAAACAAUUA
Bfa-1-1  CCGCACAAUGGUUGUCGUCCUCCGAAAAGACGUAGAGUUUAAUUUACCUUUAACUUAAAAUGAUCUUGAUUUUGUUUUUGGAUUGCA...............................AUAAUUUCUUCUC...UGCAAUCGCGG.CU.GCAACAAAUUGAGUUCAUGAAUAAACAAUUA
env-335  CCACAUAAUGGUUGUCGUCCUCCGAAAAGACGUAGAGUUUAAUUUACCUUUAACUUAAAAUGAUCUUGAUUUUGUUUUUGGAUUGCA...............................AUAAUUUCUUCUC...UGCAAUCGCGG.CU.GCAACAAAUUGAGUUCAUGAAUAAACAAUUA
env-336  .................................................................UUGACUUUGUUUUUGGAUUGCA...............................UUAAUUUCUUCUC...UGCAAUCGCGG.CU.GCAACAAAUUAAGUUCAUGAAUAAACAAUUA
Bre-1-1  CCACAUAAUGGUUGUCGUCCUCCAAAAAGACGUAGAGUUUAAUUUACCUUUAACUUUAAAUGAUCUUGACUUUGUUUUUGGAUUGCA...............................UUAAUUUCUUCUC...UGCAAUCGCGG.CU.GCAACAAAUUAAGUUCAUGAAUAAACAAUUA
env-337  ACAUAAUGGUUGUCGCCCUCCGAAAAGACGUAGAGUUUAAAAUUUACCUUUAAUGAAAAUGAUACUUGAUUUUGUUUUUGGAUUGCA...............................UCAAAUUCUUCUC...UGUAAUCGCGG.CU.GCAACAAAUUUAAGUUCAUGAAUAAACAAUU
Bno-1-1  ACAUAAUGGUUGUCGUCCUCCAAAAAGACGUAGAGUUUAAGAUUUACCUUUAAUUAAAAUGACACUUGAUUUUGUUUUUGGAUUGCA...............................UCAAUUUCUUCUC...UGUAAUCGCGG.CU.GCAACAAAUUAAGUUCAUGAAUAAACAAUUA
env-338  ACAUAAUGGUUGUCGUCCUCCAAAAAGACGUAGAGUUUAAGAUUUACCUUUAAUUAAAAUGACACUUGAUUUUGUUUUUGGAUUGCA...............................UCAAUUUCUUCUC...UGUAAUCGCGG.CU.GCAACAAAUUA....................
Bno-2-1  ACAUAAUGGUUGUCGUCCUCCAAAAAGACGUAGAGUUUAAGAUUUACCUUUAAUUAAAAUGACACUUGAUUUUGUUUUUGGAUUGCA...............................UCAAUUUCUUCUC...UGUAAUCGCGG.CU.GCAACAAAUUAAGUUCAUGAAUAAACAAUUA
env-339  ACAUAAUGGUUGUCGUCCUCCGAAAAGACGUAGAGUUUAAGAUUUACCUUUAAUAAAAAUGAUACUUGAUUUUGUUUUUGGAUUGCA...............................UUAAUUUCUUCUC...UGUAAUCGCGG.CU.GCAACAAAUUGAGUUCAUGAAUAAACAAUUA
env-340  GCAUAACGGCUGUCGUCCUCCGAAAAGACGUAGAGUUUAAGAUUUACCUUUAAUAAAAAUGAUACUUGAUUUUGUUUUUGGAUUGCA...............................UUAAUUUCUUCUC...UGUAAUCGCGG.CU.GCAACAAAUUGAGUUCAUGAAUAAACAAUUA
env-341  GCAUAACGGUUGUCGUCCUCCGAAAAGACGUAGAGUUUAAGAUUUACCUUUAAUAAAAAUGAUACUUGAUUUUGUUUUUGGAUUGCA...............................UUAAUUUCUUCUC...UGUAAUCGCGG.CU.GCAACAAAUUGAGUUCAUGAAUAAACAAUUA
env-342  GCAUAACGGCUGUCGUCCUCCGAAAAGACGUAGAGUUUAAGAUUUACCUUUAAUAAAAAUGAUACUUGAUUUUGUUUUUGGAUUGCA...............................UUAAUUUCUUCUC...UGUAAUCGCGG.CU.GCAACAAAUUGAGUUCAUGAAUAAACAAUUA
env-343  GCAUAACGGUUGUCGUCCUCCGAAAAGACGUAGAGUUUAAGAUUUACCUUUAAUAAAAAUGAUACUUGAUUUUGUUUUUGGAUUGCA...............................UUAAUUUCUUCUC...UGUAAUCGCGG.CU.GCAACAAAUUGAGUUCAUGAAUAAACAAUUA
env-344  GCAUAACGGUUGUCGUCCUCCGAAAAGACGUAGAGUUUAAGAUUUACCUUUAAUAAAAAUGAUACUUGAUUUUGUUUUUGGAUUGCA...............................UUAAUUUCUUCUC...UGUAAUCGCGG.CU.GCAACAAAUUGAGUUCAUGAAUAAACAUUAA
env-345  GCAUAACGGUUGUCGUCCUCCGAAAAGACGUAGAGUUUAAGAUUUACCUUUAAUAAAAAUGAUACUUGAUUUUGUUUUUGGAUUGCA...............................UUAAUUUCUUCUC...UGUAAUCGCGG.CU.GCAACAAAUUAAGUUCAUGAAUAAACAAUUA
Bne-1-1  CACACAACGGAUGUCGUCCUCCCAAAAGACGUAGAGUGUAAGAUUUACCUUUAAUUAAAUGAAACUUGAUUUUGUUUUUGGAUUGCA...............................AAAAUUUCUUCUC...UGUAAUCGCGG.CU.GCAACAAUAUUGAGUUCAUGAAUAAACAAUU
env-346  AACGGUUGUCGUCCUCCGAAAAGAAGAAGAGUAUAAUAAAGAUAACCUAUUCAGAUGUAACUUGAUUUUGUUAAAUAAUGGAUUGCA....................................UUCCUUUC...UGUAAUCGCGG.CU.GCAACAAAUUGAGUUCAUUAAUAACAAUUAA
env-347  ACAACGGUUGUCGUCCUCCGAAAGAAGAAGAGUAUAAUAAAGAUAACCUAUUCAGAUGUAACUUGAUUUGUUAAAUAAUGGAUUGCA....................................UUCCUUUC...UGUAAUCGCGG.CU.GCAACAAAUUGAGUUCAUUAAUAACAAUUAA
env-348  AACGGUUGUCGUCCUCCGAAAAGAAGAAGAGUAUAAUAAAGAUAACCUAUUCAGAUGUAACUUGAUUUUGUUAAAUAAUGGAUUGCA....................................UUCCUUUC...UGUAAUCGCGG.CU.GCAACAAAUUGAGUUCAUGAAUAACAAUUAA
env-349  GUUGUCGUCCUCCAAAGAGAAGAAGAGUAUAAUUCAGAUAACCUAAUAAUAAGUGUGAUACUUGAUUUUGUUUUAUAAUGGAUUGCA...................................AUUCCUUUC...UGCAAUCGCGG.CU.GCAACAAAUUGAGUACCUUAGUAACAAUUAA
env-350  CAACGGUUGUCGUCCUCCUAAGAGAAGAAGAGUAUAAUAAAGAUAACCUAUUUAGAUGUAACUUGAUUUUGUUAAUAAUGGAUUGCA...................................AUUCCUUUC...UGUAAUCGCGG.CU.GCAACAAAUUAAGUUCAUUAAUAACAAUUAA
env-351  CAAUGGUUGUCGUCCUCCGAAAAGAAGAAGAGUAUAAUAAAGAUAACCUUAUUUGGAUGUAACUUGAUUUGGUUAAUUUGGAUUGCA...................................AUUCCUUUC...UGUAAUCGCGG.CU.GCAACAAAUUGAGUUCAUUAGUAACAAUUAA
env-352  AACGGUUGUCGUCCUCCUAAGAGAAGAAGAGUAUAAUAAAGAUAACCUAUUUAGAUGUAACUUGAUUUUGUUAAAUAAUGGAUUGCA...................................AUUCCUUUC...UGUAAUCGCGG.CU.GCAACAAAUUGAGUUCAUGAAUAACAAUUAA
env-353  CAAUGGUUGUCGUCCUCCGAAAAGAAGAAGAGUAUAAUAAAGAUAACCUUAUUUGGAUGUAACUUGAUUUGGUUAAUUUGGAUUGCA...................................AUUCCUUUC...UGUAAUCGCGG.CU.GCAACAAAUUGAGUUCAUUAGUAACAAUUAA
env-354  AACGGUUGUCGUCCUCCUAAGAGAAGAAGAGUAUAAUAAAGAUAACCUAUUUAGAUGUAACUUGAUUUUGUUAAAUAAUGGAUUGCA...................................AUUCCUUUC...UGUAAUCGCGG.CU.GCAACAAAUUAAGUUCAUUAAUAACAAUUAA
Bba-2-1  ACGGUUGUCGUCCUCCGAAAAGAAGAAGAGUAUAAUACAGAUAACCUAUUUCGAUGUCACUUGAAUUUGUUAAAUAAUUGGAUUGCA...................................AUUCCUUUC...UGUAAUCGCGG.CU.GCAACAAAUUGAGUACAUUAUAACAAUUAAA
env-355  ...............................................CCUUAUUUGGAUGUAACUUGAUUUGGUAAUUUGGAU.UGCA..................................AUUCCUUUC..UGUAAAUCGCGG.CU.GCAACAAAUUGAGUUCAUUAGUAACAAUUAA
env-356  ACGGUUGUCGUCCUCCUAAAGAGAAGAAGAGUAUAAUAAAGAUAACCUAUUUAGAUGUAACUUGAUUUUGUUAAAUAAUGGAUUGCA...................................AUUCCUUUC...UGUAAUCGCGGACU.GCAACAAAUUAAGUUCAUUAAUUAACAAUUA
env-357  .......................AGAAGAGUAUAAUAAAGAUAACCUAUUUAGAUGUAACUUGAUUUUGUUAAAUAAAUGGAUUGCA...................................AUUCCUUUC...UGUAAUCGCGG.CU.GCAACAAAUUAAGUUCAUUAAUAACAAUUAA
env-358  CAACGGUUGUCGUCCUCCUAAAAGAAGAAGAGUAUAAUAAAGAUAACCUAUUUAGAUGUAACUUGAUUUUGUUAAUAAUGGAU.UGCA..................................AUUCCUUUC..UGUAAAUCGCGG.CU.GCAACAAAUUGAGUUCAUGAAUAACAAUUAA
env-359  AACGGUUGUCGUCCUCCUAAGAGAAGAAGAGUAUAAUAAAGAUAACCUAUUUAGAUGUAACUUGAUUUUGUUAAAUAAUGGAUUGCA...................................AUUCCUUUC...UGUAAUCGCGG.CU.GCAACAAAUUAAGUUCAUUAAUAACAAUUAA
env-360  AACGGUUGUCGUCCUCCUAAAAGAAGAAGAGUAUAAUAAAGAUAACCUAUUUAGAUGUAACUUGAUUUUGUUAAAUAAUGGAUUGCA...................................AUUCCUUUC...UGUAAUCGCGG.CU.GCAACAAAUUGAGUUCAUGAAUAACAAUUAA
env-361  AACGGUUGUCGUCCUCCGAAAAGAAGAAGAGUAUAAUAAAGAUAACCUAUUUAGAUGUAACUUGAAUUUGUUAAUAAAUGGAUUGCA...................................AUUCCUUUC...UGUAAUCGCGG.CU.GCAACAAAUUGAGUUCAUUAAUAACAAUUAA
env-362  CAAUGGUUGUCGUCCUCCGAAAAGAAGAAGAGUAUAAUAAAGAUAACCUUAUUUGGAUGUAACUUGAUUUGGUUAAUUUGGAUUGCA...................................AUUCCUUUC...UGUAAUCGCGG.CU.GCAACAAAUUGAGUUCAUUAGUAACAAUUAA
env-363  AACGGUUGUCGUCCUCCUAAAAGAAGAAGAGUAUAAUAAAGAUAACCUAUUUAGAUGUAACUUGAUUUUGUUAAAUAAUGGAUUGCA...................................AUUCCUUUC...UGUAAUCGCGG.CU.GCAACAAAUUAAGUUCAUUAGUAACAAUUAA
env-364  CAAUGGUUGUCGUCCUCCGAAAAGAAGAAGAGUAUAAUAAAGAUAACCUUAUUUGGAUGUAACUUGAUUUGGUUAAUUUGGAUUGCA...................................AUUCCUUUC...UGUAAUCGCGG.CU.GCAACAAAUUGAGUUCAUU............
env-365  CAAUGGUUGUCGUCCUACCGAAAAGAAGAAGAGUAUAAUAAAGAUAACCUUAUUUGGAUGUAACUUGAUUUGGUAAUUUGGAUUGCA...................................AUUCCUUUC...UGUAAUCGCGG.CU.GCAACAAAUUGAGUUCAUUAGUAACAAUUAA
env-366  AACGGUUGUCGUCCUCCUAAAAGAAGAAGAGUAUAAUAAAGAUAACCUAUUUAGAUGUAACUUGAUUUUGUUAAAUAAUGGAUUGCA...................................AUUCCUUUC...UGUAAUCGCGG.CU.GCAA...........................
env-367  AACGGUUGUCGUCCUCCUAAGAGAAGAAGAGUAUAAUAAAGAUAACCUAUUUAGAUGUAACUUGAUUUUGUUAAAUAAUGGAU.UGCA..................................AUUCCUUUC..UGUAAAUCGCGG.CU.GCAAACUAAAUUAAAGUUCAUUAAUAACAAU
env-368  ...............................AAGAGUAUAUAAAGAUACCUAUUUGGAUGUAACUUGAUUUGGUAAUUUGGAUUG.CA..................................AUUCCUUUC..UGUUAAUCGCGG.CU.GCAACAAAUUGAGUUCAUUAGUAACAAUUAA
env-369  .........................................................AUGUAACUUGAUUUGGUAAUUUGGAUUGCA...................................AUUCCUUUC...UGUAAUCGCGGACUAGCAACAAAUUGAGUUCAUUAGUAACAAUUAA
env-370  AACGGUUGUCGUCCUCCUAAGAGAAGAAGAGUAUAAUAAAGAUAACCUAUUUAGAUGUAACUUGAUUUUGUUAAAUAAUGGAUUGCA...................................AUUCCUUUC...UGUAAUCGCGG.CU.GCAACAAAUUAAGUUCAUUAAUAACAAUUAA
env-371  AACGGUUGUCGUCCUCCKAAAAGAAGAAGAGUAUAAUAAAGAUAACCUAUUUAGAUGUAACUUGAUUUUGUUAAAUAAUGGAUUGCA...................................AUUCCUUUC...UGUAAUCGCGG.CU.GCAACAAAUUGAGUUCAUKAAUAACAAUUAA
env-372  AACGGUUGUCGUCCUCCUAAGAGAAGAAGAGUAUAAUAAAGAUAACCUAUUUAGAUGUAACUUGAUUUUGUUAAAUAAUGGAUUGCA...................................AUUCCUUUC...UGUAAUCGCGG.CU.GCAACAAAUUAAGUUCAUUAAUAACAAUUAA
env-373  AACGGUUGUCGUCCUCCUAAGAGAAGAAGAGUAUAAUAAAGAUAACCUAUUUAGAUGUAACUUGAUUUUGUUAAAUAAUGGAUUGCA...................................AUUCCUUUC...UGUAAUCGCGG.CU.GCAACAAAUU.....................

         ................................................................................<<<<<<<<<<.......................................>>>>>>>>>>..........................................
         ................................................................................1122222100.......................................0012222211..........................................
         ●●●●●●●●●●●●●●●●●●●●●●●●●●●●●●R●●●●●●●●●●●●●●●●●●●●●●●●●●●●●●●●●●●●●●●●●●●●●●●●●R●URGAUURYA------------------------------○○○○○●Y●●CCUCUC---URYARUCGCGG-CU-GCAY●R●●●●●●●YYY●●●●●●●●●●●●●●●
                                                                                    5′                                                                     3′
```

```
env-374  CAACGGUUGUCGUCCUCCGAAAAGAAGAAGAGUUUAAUAGAUAACCUAUUUUUAUAUGUAACUUGAUUUUGUUAAUAAUGGAUUGCA.................................CUUCCUUUC...UGUAAUCGCGG.CU.GCAACAAAUUAAGUUCAUUAGUAACAAUUAA
env-375  CAACGGUUGUCGUCCUCCGAAAAGAAGAAGAGUUUAAUAGAUAACCUAUUUUUAUAUGUAACUUGAUUUUGUUAAUAAUGGAUUGCA.................................CUUCCUUUC...UGUAAUCGCGG.CU.GCAACAAAUUAAGUUCAUUAGUAACAAUUAA
env-376  ..UGGUUGUCGUCCUCCGAAGAGAAGAAGAGUAUAAAAGAUAACCCUAUUUUUAUAUGUAACUUGAUUUUGUUAAUAAUGGAUUGCA.................................CUUCCUUUC...UGUAAUCGCGG.CU.GCAACAAAUUAAGUUCAUUAGUAACAAUUAA
env-377  CGCACAACGGUUGUCGUCCUCCGAAAAGANNNNNNNNNNGAUAACCUAUUUUUAUAUGUAACUUGAUUUUGUUAAUAAUGGAUUGCA.................................CUUCCUUUC...UGUAAUCGCGG.CU.GCAACAAAUUAAGUUCAUUAGUAACAAUUAA
Bma-1-1  CAACGGUUGUCGUCCUCCAAAGAGAAGAAGAGUUUAAUAGAUAACCUAUUUUUAUAUGUAACUUGAUUUUGUUAAUAAUGGAUUGCA.................................CUUCCUUUC...UGUAAUCGCGG.CU.GCAACAAAUUAAGUUCAUUAGUAACAAUUAA
env-378  UGCUAUCAGAANNNNNNNNNNNNNNNNAGAAGAGUUUAAUAGAUAACCUAUUUUUAUAUGUAACUUGAUUUUGUUAAUAAUGGAUUGCA.................................CUUCCUUUC...UGUAAUCGCGG.CU.GCAACAAAUUAAGUUCAUUAGUAACAAUUAA
env-379  ACGGUUGUCGUCCUCCGAAAAAGAAGAAGAGUUUAAUAGAUAACCUAUUUUUAUAUGUAACUUGAUUUUGUUAAUAAUGGAUUGCA.................................CUUCCUUUC...UGUAAUCGCGG.CU.GCAACAAAUUAAGUUCAUUAGUAACAAUUAA
env-380  ACAACGGAUGUCGUCCUCCGAAGAGAAGAAGAGUUUAAUAGAUAACCUAUUUUUAUGUAACUUGAUUUUGUUAAUAAUGGAUUGCA.................................CUUCCUUUC...UGUAAUCGCGG.CU.GCAACAAAUUAAGUUCAUUAGUAACAAUUAA
env-381  ACAACGGUUGUCGUCCUCCGAAAAGAAGAAGAGUUUAAUAGAUAACCUAUUUUUAUAUGUAACUUGAUUUUGUUAAUAAGGAUUGCA.................................CUUCCUUUC...UGUAAUCGCGG.CU.GAACAAAUUAAGUUCAUUAGUAACAAUUAAA
env-382  CAACGGUUGUCGUCCUCCAAAGAGAAGAAGAGUUUAAUAGAUAACCUAUUUUUAUAUGUAACUUGAUUUUGUUAAUAAUGGAUUGCA.................................CUUCCUUUC...UGUAAUCGCGG.CU.GCAACAAAUU.....................
env-383  ACACAAUGGUUGUCGUCCUCCAAAAGAGAAGAAGAGUAUAAUACAGAUAACCUAAUUUAUGUAUUCUGAAUUUGUUAAUGGAUUACA.................................UUUCCUUUC...UGUAAUCGCGG.CU.GCAACAAAUUGGGUUUACAAGAACUUUAAA
Bsa-1-1  GUUGUCGUCCUCCGAAAAGAAGAAGAGUAUAAUUUUAGAUAACUUAAUUUAAAGUGUAACUUGAUUUUGUUAGAUACAUGGAUUGCA.................................UUUCCUUUC...UGUAAUCGCGG.CU.GCAACAAAUUGAGUUCAUGAAUAACAAUUAA
env-384  CGGUUGUCGUCCUCCCAAAAGAAGAAGAGUAUAAUACAGAUAACCUAAUAGAAUGAUGUAACUUGAAUUGGUUAAUAAUGG.AUUGCA................................UUUCCUUUC..UGUAAUUCGCGG.CU.GCUACAAUUUAAGUUCAUUAGUAAACCAAUU
env-385  GUUGUCGUCCUCCGAAAAGAAGAAGAGUAUAAUUUUGAUAACCUAAUUUAAAGUGUAACUUGAUUUUGUUAGAUACAUGGAUUGCA.................................UUUCCUUUC...UGUAAUCGCGG.CU.GCAACAAAUUGAGUUCAUGAAUAACAAUUAA
env-386  GUUGUCGUCCUCCGAAGAGAAGAAGAGUAUAAUAAAGGUAACCUAUUUUAAUUAGAUGUAACUUGAUUUUGUUAAUAAUGGAUUGCA.................................UUUCCUUUC...UGUAAUCGCGG.CU.GCAACAAAUUGAGUUCAUGAGUAACAAUUAA
Bfi-1-1  ACAUAAUGGUUGUCGUCCUCCGAAAAGACGUAGAGUUUAAUUUACCUUUAACUUACACAAUGAUCUUGAUUUUGUUUUUGGAUUACA.............................UUUAAUUUCUCUC...UGUAAACGCGG.CU.GCAACAAAUUAAGUUCAUGAAUAAACAAUUA
env-387  ACAUAAUGGUUGUCGUCCUCCGAAAAGACGUAGAGUUUAAUUUACCUUUAGCUUACACAAUGAUCUUGAUUUUGCUUUUGGAUUACA.............................UUUAAUUUCUCUC...UGUAAUCGCGG.CU.GCAACAAAUUAAGUUCAUGAAUAAACAAUUA
env-388  CCACAUAAUGGUUGUCGUCCUCCAAAAAGACGUAGAGUUUAAUUUACCUUUAACUUACAAUGAUCUUGAUUUUGUUUUUGGAUUACA.............................UUUAAUUUCUCUC...UGUAAUCGCGG.CU.GCAACAAAUUAAGUUCAUGAAUAAACAAUUA
env-389  ACAUAAUGAUUGUCGUCCUCCGAAAAGACGUAGAGUUUAAUUUACCUUUAACUUACACAAUGAUCUUGAUUUUGUUUUUGGAUUACA.............................UUUAAUUUCUCUC...UGUAAUCGCGG.CU.GCAACAAAUUAAGUUCAUGAAUAAACAAUUA
env-390  ACACAAUGGUUGUCGUCCUCCGAAAAGACGUAGAGUUUAAUUUACCUUUAACUUACACAAUGAUCUUGAUUUUGUUUUUGGAUUACA.............................UUUAAUUUCUCUC...UGUAAUCGCGG.CU.GCAACAAAUUAAGUUCAUGA...........
env-391  AAAGAAUACUCCUUAUGCAGCNNNNNNNNNNGAGUUUAAUUUACCUUUAACUUACACAAUGAUCUUGAUUUUGUUUUUGGAUUACA.............................UUUAAUUUCUCUC...UGUAAUCGCGG.CU.GCAACAAAUU.....................
env-392  .................CUCCGAAAAGACGUAGAGUUUAAUUUACCUUUAACUUACACAAUGAUCUUGAUUUUGUUUUUGGAUUACA.............................UUUAAUUUCUCUC...UGUAAUCGCGG.CU.GCAACAAAUUGAGUUCAUGAAUAAAACAAUU
env-393  ACAUAAUGGUUGUCGUCCUCCGAAAAGACGUAGAGUUUAAUUUACCUUUAACUUACACAAUGAUCUUGAUUUUGUUUUUGGAUUACA.............................UUUAAUUUCUCUC...UGUAAUCGCGG.CU.GCAACAAAUUAAGUUCAUGAAUAAACAAUUA
env-394  CCACAUAAUGGAUGUCGUCCCCCUAAAAGAAGAAGAGUUUAAUUUUUCUUUUGAAACUGGUGAAUCUAGAAUUGUGAAUAGAUUGCA............................AUUUCUCUCCUCUC...UGCAAUCGCGG.CU.GCA............................
env-395  CCACAUAAUGGAUGUCGUCCCCCUAAAAGAAGAAGAGUUUAAUUUUUCUUUUGAAACUGGUGAAUCUAGAAUUGUGAAUAGAUUGCA............................AUUUCUCUCCUCUC...UGCAAUCGCGG.CU.GCACUAUUCCGGGUUCCAUAAUUUAUAAAUU
env-396  ......................UAAAAGAAGAAGAGUUUAAUCUCUUUUUUAAUUGAAUUUGAAUCUCGAAUGGUGAAUAGAUUGCA............................CUUUUUCUCCUCUC...UGCAAUCGCGG.CU.GCACCAAUCCGGGUUCGUUAAACUAUAAAUU
env-397  CGCAUAAUGGAUGUCGUCCUCCUAAAAGAAGAAGAGUUUAAUCUCUUUUUUAAUUGAAUUUGAAUCUCGAAUGGUGAAUAGAUUGCA............................CUUUUUCUCCUCUC...UGCAAUCGCGG.CU.GCACCAAUCCGGGUU................
env-398  CGCAUAAUGGAUGUCGUCCUCCUAAAAGAAGAAGAGUUUAAUCUCUUUUUUAAUUGAAUUUGAAUCUCGAAUGGUGAAUAGAUUGCA............................CUUUUUCUCCUCUC...UGCAAUCGCGG.CU.GCACCAAUCCGGGUUCGUUAAACUAUAAAUU
Bhe-1-1  CAUAAUGGUUGUCGUCCUCCGAAAAGACGUAGAGUUUAAAAAAUUACCUUUAAUAAAAGUGAUACUUGAUUUUGUUAUUGGAUUGUA.................................CUUCUUCUC...UGCAAUCGCGG.CU.GCAACAAAUUGAGUUCAUGAAUAAACAAUUA
env-399  UAAACAAUUAAAUAUUUAAAAGAAAUGGCUAGAUAUACUGGACNNNNNNNNNNUAAAAAUGACACUUGAUUUUGUUAUUGGAUUGUA.................................AUUCUUCUC...UGCAAUCGCGG.CU.GCAACAAAUUGAGUUCAUGAAUAAACAAUUA
Bfl-1-1  CCACAUAAUGGUUGUCGUCCUCCGAAAAGACGUAGAGUUUAAUUUACCUUUAAUAAAAAAUGACACUUGAUUUUGUUAUUGGAUUGUA.................................AUUCUUCUC...UGCAAUCGCGG.CU.GCAACAAAUUGAGUUCAUGAAUAAACAAUUA
env-400  CACAUAAUGGAUGUCGUCCUCCUAAAAGAAGAAGAGUUUAAUUAAUUUAUUUUCCUGUAUAGGAUGAGAAUACGUGAAUAGAUUGCA................................AUACCCUCUC...UGUAAUCGCGG.CU.GCACAAGUUCCUAUCCACAGAACUAUUAACA
env-401  ..................................................................AGAAUACGUGAAUAGAUUGCA................................AUACCCUCUC...UGUAAUCGCGG.CU.GCACAAGUUCCUAUCCACAGAACUAUUAACA
env-402  CACACAACGGAUGCCGUCCGCCUAAACACAGAAGAGUUUAAUUACUUAUUUUUAAUUUGGUUGAAACUGAGAUUGUGAUUGAUUGCA................................AUAUCCUCUC...UGCAAUCGCGG.CU.GCACAAUUUGAGCUUCACAAAAAUGUUUUAA
env-403  ........GAUGUCGUCCUCCUAAAAGAAGAAGAGUUUAAUUAAUUUACUGUUAAGUAUUUGAGAUAAGAAUAUGUGAAAGAUUGCA................................AGAUCCUCUC...UGUAAUCGCGG.CU.GCACAAGUUCCUGUCUUCAGAACUUUUUAAU
env-404  AUGGAUGUCGUCCUCCUAAAAGAAGAAGAGUUUAAUUAUUAAGUUUUCUUGGUUCUCUUUCGGGGCUUGGAAGGUGAAUAGAUUGCA................................AGAUCCUCUC...UGUAAUCGCGG.CU.GCACAAUCCAAAGUUUCUAAUGGAACAAAAU
env-405  GCACAACGGAUGUCGUCCUCCUAAAAGAAGAAGAGUUUAAUUAAUUUAUUUUCCUGUAUCCGGAUAAGAAUACGUGAAUAGAUUACA..............................UUUUGAUCUCUC...UGUAAUCGCGG.CU.GCACAAGUUCCUGUCCAACAGAACAUUUAAU
env-406  GCACAACGGAUGUCGUCCUCCUAAAAGAAGAAGAGUUUAAUUAAUUUAUUUUCCUGUAUCCGGAUAAGAAUACGUGAAUAGAUUACA..............................UUUCGAUCUCUC...UGUAAUCGCGG.CU.GCACAAGUUCCUGUCCAACAGAACAUUUAAU
Bpa-1-1  ACAAUGGUUGUCGUCCUCCAAAGAGAAGAAGAGUUUAAUAGAUAACCAAAAUUUAAGUAACUUGAUUUUGUUAAUAAAUGGAUUGCA................................CUUUCCUUUC...UGUAAUCGCGG.CU.GCAACAAAUUAAGUUCAUUUAUAACAAUUAA
env-407  AAUGGUUGUCGUCCUCCGAAGAGAAGAAGAGUAUAAAAGAUAACCCUAUUUUUAUAUGUAACUUGAUUUUGUUAAUAAUGGAUUGCA................................CUUUCCUUUC...UGUAAUCGCGG.CU.GCAACAAAUUAAGUUCAUUAGUAACAAUUAA
env-408  AAUGGUUGUCGUCCUCCGAAGAGAAGAAGAGUAUAAAAGAUAACCCUAUUUUUAUAUGUAACUUGAUUUUGUUAAUAAUGGAUUGCA................................CUUUCCUUUC...UGUAAUCGCGG.CU.GCAACAAAUUAAGUUCAUUAGUAACAAUUAA
Bba-1-1  GCAUAAUGGCUGUCGUCCUCCCAAGAGAAGAAGAGUAUAAGAUAAGCUUAAUGUAAAUGUUACUUGAUUCGGUUGAUUUGGAUUACA................................UCUUCCUUUC...UGUAAUCGCGG.CU.GUAACGGAUUAAGUGCAUGAUUAACAAUUUA
env-409  ACAAUGGUUGUCGUCCUCCUAAGAGAAGAAGAGUAUAAGAGAAUAAUCUUUAUUUAUAUGUGACUUAAUCUUGUUGAUUGGAUUACA................................UCUUCCUUUC...UGUAAUCGCGG.CU.GCAACAAAUUAAGUACAUAUGUUUAACAAUU
env-410  ....UAAUGGUUGUCGUCCUCCUAAAAGAAGGAGAGUUUAAGUUUCCUUAUUAUAAGAUGAAACUUGGUUUAGUUAAUGGAUUACA................................AUUUCCUUUC...UGUAAUCGCGG.CU.GCAACAAAUUGAGUUCAUUAGUAACAAUUUA
env-411  ACAUAAUGGUUGUCGUCCUCCAAAAAGAAGGAGAGUCUAAAAUUUCUUUAUUUUAAAUAUGAUACUUGAUUUUGUUAUUGGAUUACA................................UUUUCCUUUC...UGUAAUCGCGG.CU.GCAACAGAUUAAGUUCAUUAACAAACAAUUA
env-412  ACACAAUGGUUGUCGUCCUCCAAAAAGAAGGAGAGUCUAAGAUUUCUUUAUUUUAAAUAUGAUACUUGAUUUUGUUAUUGGAUUACA................................UAUUCCUUUC...UGUAAUCGCGG.CU.GCAACAGAUUAAGUUCAUUAGCAAACAAUUA
env-413  CACAUAAUGGUUGUCGUCCUCCAAAAAGAAGGAGAGUCUAAAUUUCUUUAUUUUAAAUAUGAUACUUGAUUUUGUUAUUGGAUUACA................................UAUUCCUUUC...UGUAAUCGCGG.CU.GCAACAGAUUAAGUUCAUUAACAAACAAUUA
env-414  CCCCGCUUCCCCACAACGGAUGUCGUCCUCCUAAAAGACGUAGAGUUUAAACCAAAUAAGCGUGCCAUGUCCUUGUUUAGGAUUGCA................................UUUUCUUCUC...UGUAAUCGCGG.CU.GUAACAAGGCAUUGCGCGACAAACAACAAAA
Bpr-1-1  UCCACACAAUGGUUGUCGUCCACCAAAAAGACGUCGAGUUUAAUCACUUAUUCUUAUUUAUGAAGCUUGAAUUUGUUUUGGAUUACA................................CCUUCUUCUC...UGUAGUCGCGG.CU.GCAACAAACUUAAGUUCAUAUUUUUACAAUU
Bco-1-1  GCCACACAAUGGUUGUCGUCCUCCAAAAAGACGUAGAGUUUAAUCACUUGAUUCUUACAAUGAGACUUGAUUUUGUUUUGGAUUACA................................UCUUCUUCUC...UGUAAUCGCGG.CU.GCAACAAUUUUAAGUUCAUGUAUAAACAAUU
env-415  CACAACGGAUGUCGUCCUCCUAAAAAGAGAAGAGUUUAAUUAACAAUAGUUUCCUGCACAUGGAUCCGGAAUUGUGAUAAGAUUGCA.............................AAUUAUCCCUCUC...UGCAAUCGCGG.CU.GCACGAUUCCAGAUUCAUACAGAACAUUUAA
env-416  CACAACGGAUGUCGUCCUCCUAAAAAGAGAAGAGUUUAAUUAACAAUAGUUUCCUGCACAUGGAUCCGGAAUUGUGAUAAGAUUGCA.............................CAUUAUCCCUCUC...UGCAGUCGCGG.CU.GCACGAUUCCAGAUUCAUACAGAACAUUUAA
env-417  ACAAUGGUUGUCGUCCUCCUAAGAGAAGAAGAGUUUAAUUUUUCUUUAUUUGUAAACAGAGAAUCCCGAACAGUGAAUAAGAUUGCA.............................UAAUAAUCCUCUC...UGCAAUCGCGG.CU.GCACUCUUCUCGGUUCGCAAUAUUUAAUAAU
Bvi-1-1  CACAAUGGUUGUCGUCCUCCUAAAAGAAGAAGAGUUUAAUUUUCUUUAUUUGUAAACAGAGAAUCCCGAACAGUGAAUAAGAUUGCA.............................UAUUAAUCCUCUC...UGCAAUCGCGG.CU.GCACUCUUCUCGGUUCGCAAUAUUAAUUAUU
env-418  CACAAUGGUUGUCGUCCUCCUAAAAGAAGAAGAGUUUAAUUUUUCUUUAUUUGUAAACAGAGAAUCCCGAACAGUGAAUAGAUUGCA.............................UAUUAAUCCUCUC...UGCAAUCGCGG.CU.GCACUUUUCUCGGUUCGCAAUAUUAUAAUUA
env-419  CACAAUGGUUGUCGUCCUCCCAAAAGAAGAAGAGUUUAAUUUUUCUUUAUUUGUAAACAGAGAAUCCCGAACAGUGAAUAGAUUGCA.............................UAUUAAUCCUCUC...UGCAAUCGCGG.CU.GCACUUUUCUCGGUUCGCAAUAUUAUAAUUA
env-420  CACAAUGGUUGUCGUCCUCCUAAAAGAAGAAGAGUUUAAUUUUUCUUUAUUUGUAAACAGAGAAUCCCGAACAGUGAAUAGAUUGCA.............................UAUUAAUCCUCUC...UGCAAUCGCGG.CU.GCACUUUUCUCGGUUCGCAAUAUUAUAAUUA
env-421  AUGGAUGUCGUCCUCCGAAAAGACGUAGAGUUUAAUAUUUACCUUUAUACAAAAGACAGGCUUUAAUGUCGUUAAAAUAAGAUUGCA.............................UUUAUAUUCUCUC...UGCAAUCGCGG.CU.GCAACGAUUAAGGCUGUCUAGAAAACAAAUU
env-422  AUAACGGUUGUCGACCACCUAAAAAACGUAGAGUAUAAAAAAUACAGUAUUAUAACAACUUUCGAAUCUCGCAUUGUGAUAAUUGCA.............................UUUUAAACCUCUC...UGCAAUUGCGG.CU.GCACGGUUCAAGGUUUGGAAAUAAACAGUAA
env-423  GCAUAAUGGAUGUCGUCCUCCUAAAAGAAGAAGAGUUUAAUCUCUUUUUUAAUUGAAUUUGAAUCUCGAAUGGUGAAUAGAUUGCA.............................UUUUUCUCCUCUC...UGCAAUCGCGG.CU.GCACCAAUCCGGGUUCGUUAAAUUAUAAAUU
env-424  GUCCUCCAAAGAGAAGAAGAGUUUAAUUAUUAUCUCUUAUUUAUCUGAAUUAGAACAAACGGGAUCUUUUAACGUGAAUAGAUUGCA.............................UUUUUAACCUCUC...UGCAAUCGCGG.CU.GCACUAAGGAAGAUUCAAAAAAACUUUAAUU
env-425  CAAGCGCGAGUGGCGUCUCAUCUCACAUUAAACUCAAUCUUUUUUAUUCAACUUCCUGUGUGAAUCUCUGCCAGUGAUUGGAUUGCA..........................AUUUUCACGACCUCUC...UGCAAUCGCGG.CU.GCACUGUA.......................
env-426  GUUUAAUCAACCAGUUCUUCUUGUAAGAUUAAAGUAAUCCUUUUUUAAUUAUUUUUUUCGCGAAUCUCUGUCAGUGAAUAGAUUGCA..........................UUUUUAACAACCUCUC...UGCAAUCGCGG.CU.GCACUGCAGCGAUUCUGCAACACCAAAUUUU
env-427  GNCUCGUUUACUUGUUUACUAAAAAACUCGUCAACUAAAUAAAAAAAGUUUCAUUCACACAGAAACUGAACUGUGAUUUUGAUUACA...............................CACAACUUCUC...UGUAAUCGCGG.CU.GCACAGUAAGGAUUCGGUAAAAAGUUUUAAU
env-428  ....................................................................AACUGUGAUUUUGAUUACA...............................CACAACUUCUC...UGUAAUCGCGG.CU.GCACAGUAAGGAUUCGGUAAAAAGUUUUAAU
env-429  ACUUCCUCACAACGGAUGUCGUCCUCCAAAACACAGAAGAGUUUAAUCAAAUUUAAAAAUUGGAUCUUAAAUUGUGAUUUGAUUACA...............................AAAACCCUCUC...UGUAGUCGCGG.CU.GCACAAUUUUAGCUUCAUUUCAUUAUUAAUU
env-430  UUCCGCAUAAUGGAUGUCGCCCUCCUAAGAGAAGAAGAGUUUAAUUUAUUUAUUUACUUAGGGUAGAAAUACGUGGAAUAGAUUACA...............................ACAACCCUCUC...UGUAAUCGCGG.CU.GCACAAGUUUCUUUUCCUGCAAAAUUAUUAA
env-431  .......................................................GUAUAGGGUGGAAAUACGUGGAAUAGAUUACA...............................AUAACCCUCUC...UGUAAUCGCGG.CU.GCACAAGUUUCUUUUCCUUCAAAAAUUAACA
env-432  CACAUAAUGGAUGUCGUCCUCCUAAGAGAAGAAGAGUUUAAUUAAUUUAUUUACUGUAUAGGGUGGAAAUACGUGGAAUAGAUUACA...............................AUAACCCUCUC...UGUAAUCGCGG.CU.GCACAAGUUC.....................
Dsp-1-1  CUUCCUCACAAUGGUUGUCGUCCUCCAAAACACAGAAGAGUUUAAUCAAAUUUAAAAAUUGGAACUUGAAUUGUGAAUUUGAUUGCA...............................ACGACCCUCUC...UGUAAUCGCGG.CU.GCACAAUUUUAGCUUUCAUCAAUUAUUAAUU
env-433  CUUCCACACAACGGAUGUCGUCCUCCUAAACACAGAAGAGUUUAAUCAAUUUUUGUAUUGAAAUCUUGAAUUGUGAUUAGAUUGCA...............................CGUACCCUCUC...UGUAAUCGCGG.CU.GCACAAUUUUAGCUUUCA.............
env-434  UCCGGUAGUUAUUCGUCUGAACAUCAAAAAAAAGUAAAUAAUCAUAAGUUUCAAUCAUACAGAAACUUGAGCUGUGAUUUGAUUACA...............................AUAUACUUCUC...UGUAAUCGCGG.CU.GCACAGUUUAGGCUUCAGUAGAAAAAGUUUA
Bga-1-1  ACAUAAUGGUUGUCGUCCUCCGAAAAGACGUAGAGUUUAAGAUUUACCUUUAAUAAAAAAUGAAACUUGAUAUUGUUAUUGGAUUGUA................................AAUUCUUCUC...UGCAAUCGCGG.CU.GCAACAAAUUAAGUUCAUGAAUAACAAUUAA
env-435  ..................CCGAAAAGACGUAGAGUUUAAGAUUUACCUUUAAUAAAAAAUGAAACUUGAUAUUGUUAUUGGAUUGUA................................AAUUCUUCUC...UACAAUCGCGG.CU.GCAACAAAUUAAGUUCAUGAAUAACAAUUAA
env-436  ..........................................................AUGAAACUUGAUAUUGUUAUUGGAUUGUA................................AUUUCUUUUC...UACAAUCGCGG.CU.GCAACAAAUUAAGUUCAUGAAUAACAAUUAA
env-437  GCAUAAUGGAUGUCGUCCUCCGAAAAGACGUCGUGUAUAAGAUUUACCUUUAUUAUAUGUGAAACUUGAUUUUGUUAUUGGAUUGUA................................AUUUCUUCUC...UACAGUCGCGG.CU.GCAACAAAUUAAGUUCAUAAUUAAAACAUUU
env-438  GCAUAAUGGAUGUCGUCCUCCGAAAAGACGUCGUGUAUAAGAUUUACCUUUAUUAUAUGUGAAACUUGAUUUUGUUAUUGGAUUGUA................................AUUUCUUCUC...UACAGUCGCGG.CU.GCAACAAAUUAAGUUCAUAAUUAAAACAAUU
env-439  GCAUAAUGGAUGUCGUCCUCCGAAAAGACGUCGUGUAUAAGAUUUACCUUUAUUAUAUGUGAAACUUGAUUUUGUUAUUGG.AUUGUA...............................AUUUCUUCUC..UACAGUUCGCGG.CU.GCAACAAAUUAAGUUCAUAAUUAAAACAAUU
env-440  ...................................UAUAAGAUUUACCUUUAUUAUAUGUGAAACUUGAUUUUGUUAUUGGAUUGUA................................AUUUCUUCUC...UACAGUCGCGG.CU.GCAACAAAUUAAGUUCAUAAUUAAAACAAUU
env-441  GCAUAAUGGAUGUCGUCCUCCGAAAAGACGUCGUGUAUAAGAUUUACCUUUAUUAUAUGUGAAACUUGAUUUUGUUAUUGGAUUGUA................................AUUUCUUCUC...UACAGUCGCGG.CU.GCAACAAAUUAAGUUCAUAAUUAAAACAAUU
env-442  ..........GUCGUCCUCCGAAAAGACGUCGUGUAUAAGAUUUACCUUUAUUAUAUGUGAAACUUGAUUUUGUUAUUGGAUUGUA................................AUUUCUUCUC...UACAGUCGCGG.CU.GCAACAAAUUAAGUUCAUAAUUAAAAACAAU
env-443  ACAUAAUGGUUGUCGUCCUCCGAAAAGACGUAGAGUUUAAGAUUUACCUUUAAUAAAAAAUGAAACUUGAUAUUGUUAUUGGAUUGUA................................AUUUCUUCUC...UACAAUCGCGG.CU.GCAACAAAUUAAGUUCAUGAAUAACAAUUAA
env-444  .............UCCUCCGAAAAGACGUAGAGUUUAAGAUUUACCUUUAAUAAAAAAUGAAACUUGAUAUUGUUAUUGGAUUGUA................................AUUUCUUCUC...UACAAUCGCGG.CU.GCAACAAAUUAAGUUCAUGAAGAACAAUUAA

         ................................................................................<<<<<<<<<<........................................>>>>>>>>>>.......................................
         ................................................................................1122222100........................................0012222211.......................................
         ●●●●●●●●●●●●●●●●●●●●●●●●●●●●●R●●●●●●●●●●●●●●●●●●●●●●●●●●●●●●●●●●●●●●●●●●●●●●R●URGAUURYA--------------------------○○○○○●Y●●CCUCUC---URYARUCGCGG-CU-GCAY●R●●●●●●●YYY●●●●●●●●●●●●●●●
                                                                                5′                                                          3′
```

```
env-445  CAUAAUGGUUGUCGUCCUCCGAAAAGACGUAGAGUUUAAGAUUUACCUUUAAUAAAAAAUGAAACUUGAUAUUGUUAUUGGAUUGUA..................................AUUUCUUCUC...UACAAUCGCGG.CU.GCAACAAAUUAAGUUCAUGAAU.........
env-446  ..............................................................AACUUGAUAUUGUUAUUGGAUUGUA..................................AUUUCUUCUC...UACAAUCGCGG.CU.GCAACAAAUUAAGUUCAUGAAUAACAAUUAA
env-447  ACAUAAUGGUUGUCGUCCUCCGAAAAGACGUAGAGUUUAAGAUUUACCUUUAAUAAAAAUGAAACUUGAUAUUGUUAUUGGAUUGUA..................................AUUUCUUCUC...UACAAUCGCGG.CU.GCAACAAAUUAAGUUCAUGAAUAACAAUUAA
env-448  ACAAUGGUUGUCGUCCUUCCGAAAAGACGUAGAGUUUAAGAUUUACCUUUAAUAAAAAAUGAAACUUGAUAUUGUUAUUGGAUUGUA..................................AUUUCUUCUC...UACAAUCGCGG.CU.GCAACAAAUUAAGUUCAUGAAUAACAAUUAA
env-449  CACAAUGGUUGUCGUCCUCCGAAAAGACGUAGAGUUUAAUUUACCUUUAACUUACACAAUGAAACUUGAUAUUGUUAUUGGAUUGUA..................................AUUUCUUCUC...UACAAUCGCGG.CU.GCAACAAAUUAAGUUCAUGAAUAACAAUUAA
env-450  NNNNNNNNNNNNNNNNNNNNNNNNNNNNNNNNGUUUAAGAUUUACCUUUAAUAAAAAAUGAAACUUGAUAUUGUUAUUGGAUUGUA..................................AUUUCUUCUC...UACAAUCGCGG.CU.GCAACAAAUUAAGUU................
env-451  ACAUAAUGGUUGUCGUCCUCCGAAAAGACGUAGAGUUUAAGAUUUACCUUUAAUAAAAAUGAAACUUGAUAUUGUUAUUGGAUUGUA..................................AUUUCUUCUC...UACAAUCGCGG.CU.GCAACAAAUUAAGUUCAUGAAUAACAAAUUA
env-452  NNNNNNNNNNNNNNNNNNNNNNNNNNNNNNNNNNNNNNNNNNNNNNNNNNNNNNNNNNNNNNAACUUGAUAUUGUUAUUGGAUUGUA..................................AUUUCUUCUC...UACAAUCGCGG.CU.GCAACAAAUUAAGUUCAUGAAUAACAAUUAA
env-453  NNNNAUGGUUGUCGUCCUCCGAAAAGACGUAGAGUUUAAGAUUUACCUUUAAUAAAAAAUGAAACUUGAUAUUGUUAUUGGAUUGUA..................................AUUUCUUCUC...UACAAUCGCGG.CU.GCAACAAAUUAAGUUCAUGAAUAACAAUUAA
env-454  AUAAUGGUUGUCGUCCUCCGAAAAGACGUAGAGUUUAAUUUACCUUUAACUUACACAAAUGAAACUUGAUAUUGUUAUUGGAUUGUA..................................AUUUCUUCUC...UACAAUCGCGG.CU.GCAACAAAUUAAGUUCAUGAAUAACAAUUAA
env-455  CACAACGGUUGUCGUCCUCCGAAAAGACGUAGAGUUUAAGAUUUACCUUUAAUAAAAAAUGAAACUUGAUAUUGUUAUUGGAUUGUA..................................AUUUCUUCUC...UACAAUCGCGG.CU.GCAACAAAUUAAGUUCAUGAAUAACAAUUAA
env-456  CAUAAUGGUUGUCGUCCUCCGAAAAGACGUAGAGUUUAAGAUUUACCUUUAAUAAAAAAUGAAACUUGAUAUUGUUAUUGGAUUGUA..................................AUUUCUUCUC...UACAAUCGCGG.CU.GCAACAAAUUAAGUUCAUGAAUAACAAUUAA
env-457  ........................................................AAAUGAAACUUGAUAUUGUUAUUGGAUUGUA..................................AUUUCUUCUC...UACAAUCGCGG.CU.GCAACAAAUUAAGUUCAUGAAUAACAAUUAA
env-458  CAUAACGGUUGUCGUCCUCCGAAAAGACGUAGAGUUUAAUUUACCUUUAACUUACACAAUGAAACUUGAUAUUGUUAUUGGAUUGUA..................................AUUUCUUCUC...UACAAUCGCGG.CU.GCAACAAAUUAAGUUCAUGAAUAACAAUUAA
env-459  CUUGGCUUGAGAAAGGUAAAGNNNNNNNNNNNGUUUAAGAUUUACCUUUAAUAAAAAAUGAAACUUGAUAUUGUUAUUGGAUUGUA..................................AUUUCUUCUC...UACAAUCGCGG.CU.GCAACAAAUUAAGUU................
env-460  NNNNNNNNNNNNNUCCUCCGAAAAGACGUAGAGUUUAAGAUUUACCUUUAAUAAAAAAUGAAACUUGAUAUUGUUAUUGGAUUGUA..................................AUUUCUUCUC...UACAAUCGCGG.CU.GCAACAAAUUAAGUUCAUGAAUAACAAUUAA
env-461  CAUAAUGGUUGUCGUCCUCCGAAAAGACGUAGAGUUUAAGAUUUACCUUUAAUAAAAAUAGAAACUUGAUAUUGUUAUUGGAUUGUA..................................AUUUCUUCUC...UACAAUCGCGG.CU.GCAACAAAUUAAGUUCAUGAAUAACAAUU..
env-462  ..........................................................CUUUCGGAAAGGGGGCGUGAUAGAUUGCA...........................CUAUUCACAACCCUCUC...UGCAAUCGCGG.CU.GAACAGUUCCCCCUCCGAAAUCGCCUCACC
env-463  AGGAAAGUAUAACAGACAGAGAGAUCAGAAUGGCAGCACAUGGACGUGUGAGAAUCUGCUUUCGGAAAGGGGGCGUGAUAGAUUGCA...........................CUAUUCACAACCCUCUC...UGCAAUCGCGG.CU.GAACAGUUCCCCUCCGGAAUCGCCUCACCAC
env-464  AAAGUAUAACAGACAGAGAGAUCAGAAUGGCAGCACACGGACGUGUGAGACUUUGCUUUCGGGAAAGGGGGGCGUGAUAGAUUGCA...........................CUAUUCACAACCCUCUC...UGCAAUCGCGG.CU.GAACAGUUCCCCCUCCGGAAUCGCCUCACCA
env-465  AGGAAAGUAUAACAGACAGAGAGAUCAGAAUGGCAGCACACGGACGUGUGAGAAUCUACUUUCGGAAAGGGGGCGUGAUAGAUUGCA...........................CUAUUCACAACCCUCUC...UGCAAUCGCGG.CU.GAACAGUUCCCCCUCCGGAAUCGCCUCACCA
env-466  ..................................AGCACAUGGACGUGUGAGAAUCUGCUUUCGGAAAGGGGACGUGAUAGAUUGCA...........................CUAUUCACAACCCUCUC...UGCAAUCGCGG.CU.GAACAGUUCCCCCUCCGGAAUCGCCUCACCA
env-467  AGGAAAGUAUAACAGACAGAGAGAUCAGAAUGGCAGCACAUGGAUGUGUGAGAAUCUACUUUCGGAAAGGGGACGUGAUAGAUUGCA...........................CUAUUCACAACCCUCUC...UGCAAUCGCGG.CU.GAACAGUUCCCCCUCCGGAAUCGCCUCACCA
env-468  AGGAAAGUAUAACAGACAGAGAGAUCAGAAUGGCAGCACAUGGAUGUGUGAGAAUCUGCUUUCGGAAAGGGGGCGUGAUAGAUUGCA...........................CUAUUCACAACCCUCUC...UGCAAUCGCGG.CU.GAACAGUUCCCC...................
env-469  AGGAAAGUAUAACAGACAGAGAGAUCAGAAUGGCAGCACAUGGACGUGUGAGAAUCUGCUUUCGGAAAGGGGGCGUGAUAGAUUGCA...........................CUAUUCACAACCCUCUC...UGCAAUCGCGG.CU.GAACAGUUCCCUCUCCGAAAUCGCCUCACC
env-470  GAAAGUAUAACAGACAGAGAGAUCAGAAUGGCAGCACACGGGCGUGUGAGACUUUGCUUUCGGAAAAGGGGGGCGUGAUAGAUUGCA...........................CUAUUCACAACCCUCUC...UGCAAUCGCGG.CU.GAACAGUUCCCCCUCCGGAAUCGCCUCACCA
env-471  AGGAAAGUAUAACAGACAGAGAGAUCAGAAUGGCAGCACACGGACGUGUGAGAAUCUACUUUCGGAAAGGAGGCGUGAUAGAUUGCA...........................CUAUUCACAACCCUCUC...UGCAAUCGCGG.CU.GAACAGUUCCCCCUCCGGAAUCGCCUCACC
env-472  AGGAAAGUAUAACAGACAGAGAGAUCAGAAUGGCAGCAUACGGACGUGUGAGAAUCUGCUUUCGGAAAGGGGACAUGAUAGAUUGCA...........................CUAUUCACAACCCUCUC...UGCAAUCGCGG.CU.GAACAGUUCCCCCUCCGGAAUCGCCUCACCA
env-473  AGGAAAGUAUAACAGACAGAGAGAUCAGAAUGGCAGCACAUGGACGUGUGAGAAUCUGCUUUCGGAAAGGGGGCAUGAUAGAUUGCA...........................CUAUUCACAACCCUCUC...UGCAAUCGCGG.CU.GAACAGUUCCCCCUCCAAAAUCGCCUCACCA
env-474  AGGAAAGUAUAACAGACAGAGAGAUCAGAAUGGCAGCACACGGACGUGUGAGAAUCUACUUUCGGAAAGGAGGCGUGAUAGAUUGCA...........................CUAUUCACAACCCUCUC...UGCAAUCGCGG.CU.GAACAGUUCCCCCUCCGAAAUCGCCUCACC
env-475  GGAAAGUAUAACAGAUAGAGAGAUCAGAAUGGCAGCACACGGGGCGUGUGAGAAUCUACUUUCGGAAAGGGGGCGUGAUAGAUUGCA...........................CUAUUCACAACCCUCUC...UGCAAUCGCGG.CU.GAACAGUUCCCCCUCCGGAAUCGCCUCACCA
env-476  .................................................GACUUUGCUUUCGGAAAAGGGGGGCGUGAUAGAUUGCA...........................CUAUUCACAACCCUCUC...UGCAAUCGCGG.CU.GAACAGCUCCCCCUCCGAAAUCGCCUCACCA
env-477  .............................................GUGUGAGAAUCUGCUUUCGGAAAGGGGGCGUGAUAGAUUGCA...........................CUAUUCACAACCCUCUC...UGCAAUCGCGG.CU.GAACAGUUCCCCCUCCGGAAUCGCCUCACCA
env-478  AGGAAAGUAUAACAGACAGAGAGAUCAGAAUGGCAGCACAUGGACGUGUGAGAAUCUGCUUUCGGAAAGGGGGCGUGAUAGAUUGCA...........................CUAUUCACAACCCUCUC...UGCAAUCGCGG.CU.GAACAGUUCCCCCUCCGAAAUCGCCUCACCA
env-479  AGGAAAGUAUAACAGACAGAGAGAUCAGAAUGGCAGCACACGGACGUGUGAGACUUUGCUUUCGGAAAGGGGGCGUGAUAGAUUGCA...........................CUAUUCACAACCCUCUC...UGCAAUCGCGG.CU.GAACAGUUUCCCCUCCGGAAUCGCCUCACCA
env-480  AGGAAAGUAUAACAGACAGAGAGAUCAGAAUGGCAGCACACGGACGUGUGAGAAUCUACUUUCGGAAAGGGGACGUGAUAGAUUGCA...........................CUAUUCACAACCCUCUC...UGCAAUCGCGG.CU.GAACAGUUCCCCCUCCGGAAUCGCCUCACCA
env-481  AGGAAAGUAUAACAGACAGAGAGAUCAGAAUGGCAGCACACGGACGUGUGAGAAUCUACUUUCGGAAAGGAGGCGUGAUAGAUUGCA...........................CUAUUCACAACCCUCUC...UGCAAUCGCGG.CU.GAACAGUUCCCCCUCCGGAAUCGCCUCACCA
env-482  GAAAGUAUAACAGACAGAGAGAUCAGAAUGGUAGCACACGGACGUGUGAGACUUUGCUUUCGGAAAGGGGGGCGUGAUAGAUUGCA...........................CUAUUCACAACCCUCUC...UGCAAUCGCGG.CU.GAACAGUUCCCCCUCCGGAAUCGCCUCACCA
env-483  AGGAAAGUAUAACAGACAGAGAGAUCAGAAUGGCAGCACAUGGAUGUGUGAGAAUCUGCUUUCGGAAAGGGGGCGUGAUAGAUUGCA...........................CUAUUCACAACCCUCUC...UGCAAUCGCGG.CU.GAACAGCUCCCCCUCCGGAAUCGCCUCACCA
env-484  AGGAAAGUAUAACAGACAGAGAGAUCAGAAUGGCAGCACACGGACGUGUGAGAAUCUGCUUUCGGAAAGGGGGCGUGAUAGAUUGCA...........................CUAUUCACAACCCUCUC...UGCAAUCGCGG.CU.GAACAGUUCCCCCCUCCGGAAUCGCCUCAC
env-485  AAAGUAUAACAGACAGAGAGAUCAGAAUGGCAGCACACGGACGUGUGAGACUUUGCUUUCGGAAAAGGGGGGCGUGAUAGAUUGCA...........................CUAUUCACAACCCUCUC...UGCAAUCGCGG.CU.GAACAGUUCCCCCUCCGGAAUCGCCUCACCA
env-486  AGGAAAGUAUAACAGACAGAGAGAUCAGAAUGGCAGCACAUGGACGUGUGAGAAUCUGCUUUCGGAAAGGGGGCGUGAUAGAUUGCA...........................CUAUUCACAACCCUCUC...UGCAAUCGCGG.CU.GAACAGUUCCCCCCUCCGGAAUCGCCUCACC
env-487  AGGAAAGUAUAACAGACAGAGAGAUCAGAAUGGCAGCACACGGACGUGUGAGAAUCUGCUUUCGGAAAGGGGGCGUGAUAGAUUGCA...........................CUAUUCACAACCCUCUC...UGCAAUCGCGG.CU.GAACAGUUCCCCCUCCGGAAUCGCCUCACCA
env-488  GGAAAGUAUAACAGAUAGAGAGAUCAGAAUGGCAGCACACGGGGCGUGUGAGAAUCUACUUUCGGAAAGGGGGCGUGAUAGAUUGCA...........................CUAUUCACAACCCUCUC...UGCAAUCGCGG.CU.GAACAGUUCCCCCUCCGGAAUCGCCUCACCA
env-489  AAGGAAAGUAUAACAGACAGAGAGAUCAGAAUGGCAGCACAUGGACGUGUGAGAAUCUGCUUUCGAAAGGGGACGUGAUAGAUUGCA...........................CUAUUCACAACCCUCUC...UGCAAUCGCGG.CU.GAACAGCUCCCCCUCCGGAAUCGCCUCACCA
env-490  ...............ACAGAGAGAUCAGAAUGGCAGCACACGGACGUGUGAGAAUCUGCUUUCGGAAAGGGGGCGUGAUAGAUUGCA...........................CUAUUCACAACCCUCUC...UGCAAUCGCGG.CU.GAACAGUUCCCCCCUCCGAAAUCGCCUCACC
env-491  ...................AGAGAUCAGAAUGGCAGCACAUGGACGUAUGAGAAUCUGCUUUCGGAAAGGGGGCGUGAUAGAUUGCA...........................CUAUUCACAACCCUCUC...UGCAAUCGCGG.CU.GAACAGUUCCCCCUCCGGAAUAGCCUCACCA
env-492  AGGAAAGUAUAACAGACAGAGAGAUCAGAAUGGCAGCACAUGGACGUGUGAGAAUCUGCUUUCGGAAAGGGGACGUGAUAGAUUGCA...........................CUAUUCACAACCCUCUC...UGCAAUCGCGG.CU.GAACAGCUCCCCCUCCGGAAUCGCCUCACCA
env-493  AGGAAAGUAUAACAGACAGAGAGAUCAGAAUGGCAGCACAUGGCCGUGUGAGACUUUGCUUUCGGAAAGGGGGCGUGAUAGAUUGCA...........................CUAUUCACAACCCUCUC...UGCAAUCGCGG.CU.GAACAGUUCCCUCUCCGAAAUCGCCUCACC
env-494  AAAGUAUAACAGACAGAGAGAUCAGAAUGGCAGCACACGGACGUGUGAGACUUUGCUUUCGGAAAAGGGGGGCGUGAUAGAUUGCA...........................CUAUUCACAACCCUCUC...UGCAAUCGCGG.CU.GAACAGUUCCCCCUCCGGAAUCGCCUCACCA
env-495  AGGAAAGUAUAACAGACAGAGAGAUCAGAAUGGCAGCACAUGGACGUGUGAGAAUCUGCUUUCGGAAAGGGGACGUGAUAGAUUGCA...........................CUAUUCACAACCCUCUC...UGCAAUCGCGG.CU.GAACAGCUCCCCCUCCGAAAUCGCCUCACCA
env-496  GAAAGUAUAACAGACAGAGAGAUCAGAAUGGCAGCACACGGACGUGUGAGACUUUGCUUUCGGAAAAGGGGGGCGUGAUAGAUUGCA...........................CUAUUCACAACCCUCUC...UGCAAUCGCGG.CU.GAACAGUUCCCCCCUCCGGAAUCGCCUCACC
env-497  AGGAAAGUAUAACAGACAGAGAGAUCAGAAUGGAAGCACACGGACGUGUGAGACUUCGCUUUCGGAAAGGGGGCGUGAUAGAUUGCA...........................CUAUUCACAACCCUCUC...UGCAAUCGCGG.CU.GAACAGUUCCCCCCUCCGGAAUCGCCUCACC
env-498  GAAAGUAUAACAGACAGAGAGAUCAGAAUGGCAGCACACGGACGUGUGAGACUUUGCUUUCGGAAAAGGGGGGCGUGAUAGAUUGCA...........................CUAUUCACAACCCUCUC...UGCAAUCGCGG.CU.GAACAGUUCCCCCUCCGGAAUCGCCUCACCA
env-499  AGGAAAGUAUAACAGACAGAGAGAUCAGAAUGGCAGCACAUGGACGUGUGAGAAUCUGCUUUCGGAAAGGGGGCGUGAUAGAUUGCA...........................CUAUUCACAACCCUCUC...UGCAAUCGCGG.CU.GAACAGUUCCCCCCUCCGGAAUC........
env-500  ...................................................CUUUGCUUUCGGAAAAGGGGGGCGUGAUAGAUUGCA...........................CUAUUCACAACCCUCUC...UGCAAUCGCGG.CU.GAACAGUUCCCCCCUCCGAAAUCGCCUCACC
env-501  CUAAAAGAAGGAAAGUAUAACACACAAAAAGAGUGGCAGCACACGGCGUGUGAGUGUAUGCUUUCGGGUAGGGCGUAAUAGAUUGCA.............................CUAUUCAACCCUCUC...UGCAAUCGCGG.CU.GAACCGAUCCUCCCCGAAGCCCCUAUCGCCC
env-502  CUAAAAGAAGGAAAGUAUAACACACAAAAAGAGUGGCAGCAUACGGCGUGUGAGUGUAUGCUUUCGGGAAGGGCGUAAUAGAUUGCA.............................CUAUUCAACCCUCUC...UGCAAUCGCGG.CU.GAACCGAUCCUCCCCGACGCCCCCAUCGCCC
env-503  UGUCGUCCUCCUAAAAGAAGGAAAGUAUAACAUAGGAGAACCGCAUACGGAGGUGUGCAUAAUUUGAGAAGGCGUAAUAAGAUUGCA.............................CUAUUCAACCCUCUC...UGCAAUCGCGG.CU.GA.............................
env-504  UCUUACACUGAAAUAUAACGCGUAUCCCCGGCAAUUAUCCUUGACCUGCGACCGGACGGAAACUUACCCGCGGUGAAAUAGACCAUA......................ACUUUUUCGAUACACACCUCUC...UAUGGUCGCGG.CU.GCAC...........................
env-505  AUAACUCAAUUCUCUAAAAACAACCAAUCCGGGAACAACAAAGGCUCACCUUCGAGCCGGAGGAUCCAAAACCAUGAAUAGACCAUU.............................AUCGAAUCACCUCUC...AAUGGUCGCGG.CU.GCAUGACAAAGAACCUCUUCAACGCCCCCGG
env-506  .............CAAGCCAGUGCCGCAGGCUUCAGUUCCGGAAAGCGGGGAUACCUCAGGCCACGAAAAGGGCGAAUAGACCAUA.............................AAUAUUUCACCUCUC...UAUGGUCGCGG.CU.GCGCCCGGACCGACGCCUCUCCCCGGACCCC
env-507  GUCGCGUUUAAUAUCACUAUUUGGAUUUAAUUCAUUUUUUAAGAAAGUAACAUUGUUCAAUAACGCAUUGAAUGUGAAUAGACCACA.............................AUUUUAUCACCUCUC...UGUGGUCGCGG.CU.GCACUAAAAUGAGGGCUUGAACAUGAAGCCU
env-508  GUCGCGUUUAAUAUCACUAUUUGGAUUUAAUUCAUUUUUUAAGAAAGUAACAUUGUUCAAUAACGCAUUGAAUGUGAAUAGACCACA.............................AUUUUAUCACCUCUC...UGUGGUCGCGG.CU.GCACUAAAAUGAGGGUUUGAACAUGAAGCCU
env-509  CCUGCAACCUGUUGCUGCCGGACAUCGCUCGGAGAAAUAUCGACACAACUCAAUGUCAUAGCCUGUGAGCCUCUGUGAAUGACCAUA.............................UUUUCAUCACCUCUC...UAUGGUCGCGG.CU.GCACACGCUCCCUUAGGCGAAAUGACUGAAG
env-510  AAUCAUCUCAACUACAUUUCAUUUUAUUCAACAUCCGGUUUCAUAACUCAGAGAGCAGCGAGAUUCCCGAACGUGAAUUUGACCAUA.............................UUUCUAUCACCUCUC...UAUGGUCGCGG.CU.GCACUGACUGACUCCUCCAAGCUCCCUGACU
env-511  GCAAAGACAUAACUAAAUUUCAUUUACAAUAGGGGCGAGCUUCCGACGAGGCUCUUACGGGCGAAGGCAAACGUGAAAUAGACCAUA.............................UUUCUAUCACCUCUC...UAUGGUCGCGG.CU.GCACUAAACCUUCCCUGCGCAUCGCAAGUGA
env-512  CUGACGAAAAAGUUAAUUAACCCCAACAAGCAUCGAGGGUUCGGGAAGAGGUUACUUGAAAAAACUGAAGAAGUGAAAUAGACCAUA.............................UUUUUAUCACCUCUC...UAUGGUCGCGG.CU.GCACAGACUUGGUUUUAGUAAAGUGAACGAU
env-513  GUGCUUUUGUGACGUCAUUCAUAAACACAACCAACGCCUCGCACCAAGCCUACGGCCACUCCACACGGCAACGUGAAAUAGACCAUA.............................UUUUUAUCACCUCUC...UAUGGUCGCGG.CU.GCACUAAACCGGUGACAGAGUCCACAGACGC
env-514  CAGCUCUUCAUAAUAAAAGCAACUCAAGCAAUCACUUUAUUUCAGGAACGCCUCUAUAGUCUAUCCGGCAGAGUGAAAUUGACCAUA.............................UUUAAUUCACCUCUC...UAUGGUCGCGG.CU.GCACCACCGGAUAAACUGAAGGCAAUCCAAC
env-515  CGCCGGGCACCCAAAGCCCGGAAAGUGCGAACACAGCGACUGCCGGCGGCAGACCCUCGAAGCAUUCCUUAAGGCGAAUAGACCAUA.............................UUCAAUCAGCCUCUC...UAUGGUCGCGG.CU.GCGCCAAAGGCAGGCACCGGCAGCUAACCAA
env-516  ACAGUAUUUCCAAUCAAUUAUAUUGUUUUUUCAUAGCGUAAAUCGUAGUCCAGCUGUCAUUGUGACGGUAAUGUGAUUUUGACCAUA...........................UUUUUUCUUCACCUCUC...UAUGGUCGCGG.CU.GCACUAAGCCUCAUGAGGUCGGUAAGGAUAC
env-517  UUGGCACUUACGCCCCGUAAAGUUAAUUCAAAAUCAAAAGCCAUCAGGCGCCGAAAGCCCGAGCACGGCAAUGUGAAAUAGAUCAUA...............................UUUAUCACCUCUC...UAUGAUCGCGG.CU.GCACUAAACCGACUCAGCACGGCACCGGCAA
         ................................................................................&lt;&lt;&lt;&lt;&lt;&lt;&lt;&lt;&lt;&lt;.........................................&gt;&gt;&gt;&gt;&gt;&gt;&gt;&gt;&gt;&gt;.......................................
         ................................................................................1122222100.........................................0012222211.......................................
         •••••••••••••••••••••••••••••R•••••••••••••••••••••••••••••••••••••••••••••••R•URGAUURYA--------------------------○○○◉◉•Y••CCUCUC---URYARUCGCGG-CU-GCAY•R•••••••YYY•••••••••••••••
                                                                                 5′                                                            3′
```

alignment positions 181 · · · 245

```
env-1    UACUUGAGAUAUGAUCCCAGCCUUGACCACCACAUAGCAGAGAGUGGCGCAU..........AUG

         .................................................................
         .................................................................
         •••••••••○○○○○○----------------------------------------------AUG
```

```
env-2     AAAAAAGUAAACUUUAGAA...........................................AUG
env-3     CUCAACGUAAGACAGCGCAUUAUAAUAAGAAGAAA...........................AUG
env-4     CUCAACGUAAGACAGCGCAUUAUAAUAAGAAGAAA...........................AUG
env-5     CUCAACGUAAGACAGCGCAUUAUAAUAAGAAGAAA...........................AUG
env-6     CUCAACGUAAGACAGCGCAUUAUAAUAAGAAGAAA...........................AUG
env-7     CUCAACGUAAGAUAGCGCAUUAUAAUAAGAAGAAA...........................AUG
env-8     CUCAACGUAAGACAGCGCAUUAUAAUAAGAAGAAA...........................AUG
env-9     UACCACCCAGCAUCACCGACCUCUCUGAGGCUCGCUCGUACUGCCAAACAAUAGAAGAC......
env-10    UACCACCCAGCAUCACCGACCUCUCUGAGGCUCGCUCGUACUGCCAAACAAUAGAAGACA..AUG
env-11    UACCACCCAGCAUCACCGACCUCUCUGAGGCUCGCUCGUACUGCCAAACAAUAGAAGACA..AUG
env-12    UACCACCCAGCAUCACCGACCUCUCUGAGGCUCGCUCGUACUGCCAAACAAUAGAAGACA..AUG
Pca-1-1   UACCACCCAGCAUCACCGACCUCUCUGAGGCUCGCUCGUACUGCCAAACAAAAGAAGACA..AUG
env-13    UACCACCCAGCAUCACCGACCUCUCUGAGGCUCGCUCGUACUGCCAAACAAUAGAAGACA..AU.
env-14    UACCACCUAGCAUCACCGACCUCUCUGAGGCUCGCUCGUACUGCCAAACAAUAGAAGACA..AUG
env-15    UACCACCGAGCAUCACCGACCUCUCUGAGGCUCGCUCGUACUGCCAAACAAUAGAAGAC......
env-16    UACCACCCAGCAUCACCGACCUCUCUGAGGCUCGCUCGUACUGCCAAACAAUAGAAGACA..AUG
env-17    UACCACCCAGCAUCACCGACCUCUCUGAGGCUCGCUCGUACUGCCAAACAAUAGAAAACA..AUG
env-18    UACCACCCAGCAUCACCGACCUCUCUGAGGCUCGCUCGUACUGCCAAACAAUAGAAGACA..AUG
env-19    UACCACCUAGCAUCACCGACCUCUCUGAGGCUCGCUCGUACUGCCAAACAAUAGAAGACA..AUG
env-20    UACCGCCCAGCAUCACCGACCUCUCUGAGGCUCGCUCGUACUGCCAAACAAUAGAAGACA..AUG
env-21    UACCACCCAGCAUCACCGACCUCUCUGAGGCUCGCUCGUACUGCCAAACAAUAGAAGACA..AUG
env-22    UACCACCCAGCAUCACCGACCUCUCUGAGGCUCGCUCGUACUGCCAAACA...............
env-23    UACCACCUAUCAUCACCGACCUCUCUGAGGCUCGCUCGUACUGCCAAACAAUAGAAGAC......
env-24    UACCACCCAGCAUCACCGACCUCUCUGAGGCUCGCUCGUACUGCCAAACAAUAGAAGACA..AUG
env-25    CUCUGCCUCCAGCGUUGCCUCUCCCCGAGAGGUCGCUCGUACUGCUUCAACAACAGAAGACAAUG
env-26    CUUUGCCUCCAGCGUUGCCUCUCCCAGAGAGGUCGCUCGUACUGCUUCAACAAUAGAAGACAAUG
env-27    CUCUGCCUCCAGCGUUGCCUCUCCCCGAGAGGUCGCUCGUACUGCUUCAACAAUAGAAGACAAUG
env-28    CGCCUAAGGGUCCCACACCUGACGCUCGUGAGGUGUCUCGUACUGCCAAACAAUAGAAGACAAUG
Psp-3-1   CGCCUAAGGAUCCCACACCUGACACAUGUGAGGUGUCUCGUACUGCCAAACAAUAGAAGACAAUG
env-29    CUCUGCCUCCAGCGUUGCCUCUCCCCGAGAGGUCACUCGUACUGCUUCAACAAUAGAAGACAAUG
env-30    CUCUGCCUCCAGCGUUGCCUCUCCCCGAGAGGUCGCUCGUACUGCUUCAACAAUAGAAGACAAUG
env-31    CUCUGCCUCCAGCGUUGCCUCUCCCCGAGAGGUCACUCGUACUGCUUCAACAAUAGAAGACAAUG
env-32    CUCUGCCUCCAGCGUUGCCUCUCCCCGAGAGGUCGCUCGUACUGCUUCAACAAUAGAAGACAAUG
env-33    CUUUGCCUCCAGCGUUGCCUCUCCCCGAGAGGUCGCUCGUACUGCUUCAACAAUAGAAGACAAUG
env-34    CUCUGCCUCCAGCGUUGCCUCUCCCAGAGAGGUCGCUCGUACUGCUUCAACAAUAGAAGACAAUG
env-35    CUCUGCCUCCAGCGUUGCCUCUCCCCGAGAGGUCGCUCGUACUGCUUCAACAAUAGAAGACAAUG
env-36    CUCAACGUAAGACAGCGCAUUAUAAUAAGAAGAA...............................
env-37    CUCUGCCUCCAGCGUUGCCUCUCCCCGAGAGGUCGCUCGUACUGCUUCAACAAUAGAAGACAAUG
env-38    .................................................................
env-39    CUCUGCCUCCAGCGUUGCCUCUCCCCGAGAGGUCGCUCGUACUGCUUCAACAAUAGAAGACAAUG
env-40    .................................................................
env-41    CUCUGCCUCCAGCGUUGCCUCUCCCCGAGAGGUCGCUCGUACUGCUUCAACAAUAGAAGACAAUG
env-42    CUUUGCCUCCAGCGUUGCCUCUCCCCGAGAGGUCGCUCGUACUGCUUCAACAAUAGAAGACAAUG
env-43    CUCUUAGGAUUCCACACCUGACGCUCGUGAGGUGUCUCGUACUGCUAAACAAUAGAAGACA.AUG
env-44    CUCUGCCUCCAGCGUUGCCUCUCCCCGAGAGGUCACUCGUACUGCUUCAACAAUAGAAGACAAUG
env-45    .................................................................
env-46    CUCUGCCUCCAGCGUUGCCUCUCCCCGAGAGGUCGCUCGUACUGCUUCAACAAUAGAAGACAAUG
env-47    CUCUGCCUCCAGCGUUGCCUCUCCCCGAGAGGUCGCUCGUACUGCUUCAACAAUAGAAGACAAUG
env-48    CUCUGCCUCCAGCGUUGCCUCUCCCCGAG....................................
env-49    CUCUGCCUCCAGCGUUGCCUCUCCCAGAGAGGUCGCUCGUACUGCUUCAACAAUAGAAGACAAUG
env-50    GGAGCUCGCUCGUACUGCUUCAACAAUAGAAGACA...........................AUG
env-51    CUCUGCCUCCAGCGUUGCCUCUUCCCGAGAGGUCGCUCGUACUGCUUCAACAAUAGAAGACAAUG
env-52    .................................................................
env-53    CUCUUAGGAUCCCGCACCUGACGCUCGUGAGGUGUCUCGUACUGCCAAACAAUAGAAGACA.AUG
env-54    CUCUGCCUCCAGCGUUGCCUCUCUCCGAGAGGUCGCUCGUACUGCUUCAACAAUAGAAGACAAUG
env-55    .................................................................
env-56    CUCUGCCUCCAGCGUUGCCUCUCUCCGAGAGGGCGCUCGUACUGCUUCA................
env-57    CUUUGCCUCCAGCGUUGCCUCUCCCAGAGAGGUCGCUCGUACUGCUUCAACAAUAGAAGACAAUG
env-58    CUUUGCCUCCAGCGUUGCCUCUCCCCGAGAGGUCGCUCGUACUGCUUCAACAAUAGAAGACAAUG
env-59    CGCCUAAGGAUCCCACACCUGACACAUGUGAGGUGUCUCGUACUGCCAAACAAUAGAAGACAAUG
env-60    CGCCUAAUGGUCCCACACCUGACGCUUGUGAGGUGUCUCGUACUGCCAAACAAUAGAAGAC....
env-61    CUCUGCCUCCAGCGUUGCCUCUCCCCAAGAGGUCGCUCGUACUGCUUCAACAAUAGAAGACAAUG
env-62    CUCUGCCUCCAGCGUUGCCUCUCCCCGAGAGGUCGCUCGUACUGCUUCAACAAUAGAAGACAAUG
env-63    CUCUGCCUCCAGCGUUGCCUCUCCCCGAGAGGUCGCUCGUACUGCUUCAACAAUAGAAGACAAUG
env-64    .................................................................
env-65    CUCUGCCUCCAGCGUUGCCUCUUCCCGAGAGGUCGCUCGUACUGCUUCAACAAUAGAAGACAAUG
env-66    CUUUGCCUCCAGCGUUGCCUCUCCCCG......................................
env-67    CUCUGCCUCCAGCGUUGCCUCUCCCAGAGAGGUCGCUCGUACUGCUUCAACAAUAGAAGACAAUG
env-68    CUCUGCCUCCAGCGUUGCCUCUCUCCGAGAGGUCGCUCGUACUGCUUCAACAAUAGAAGACAAUG
env-69    CUCUGCCUCCAGCGUUGCCUCUCCCCGAGAGGUCGCUCGUACUGCUUCAACAAUAGAAGACAAUG
env-70    CUCUGCCUCCAGCGUUGCCUCUCCCCGAGAGGUCGCUCGUACUGCUUCAACAAUAGAAGACAAUG
env-71    CUCCUAGGAUCCCGCACCUGACGCUCGUGAGGUGUCUCGUACUGCCAAACAAUAGAAGACA.AUG
env-72    CUCUGCCUCCAGCGUUGCCUCUCCCCGAGAGGUCGCUCGUACUGCUUCAACAAUAGAAGACAAUG
env-73    CUUUGCCUCCAGCGUUGCCUCUCCCCAAGAGGUCGCUCGUACUGCUUCAACAAUAGAAGACAAUG
env-74    CUUUGCCUCCAGCGUUGCCUCUCCCCAAGAGGUC...............................
env-75    ACUGCCUCCAGCAUUGCCCUCCUCGAGAGGUCGCUCGUACUGCUUCAACAAUAGAAGACA..AUG
env-76    CUCUGCCUCCAGCGUUGCCUCUCCCCGAGAGGUCGCUCGUACUGCUUCAACAAUAGAAGACAAUG
env-77    CUCUGCCUCCAGCGUUGCCUCUCCCCGAGAGGUCGCUCGUACUGCUUCAACAAUAGAAGACAAUG
env-78    ACUGCCUCCAGCAUUGCCCUCCUCGAGAGGUCGCUCGUACUGCUUCAACAAUAGAAGACA..AUG
env-79    CUCUGCCUCCAGCGUUGCCUCUCCCCGAGAGGUCGCUCGUACUGCUUCAACAAUAGAAGACAAUG
env-80    CUCCUAGGAUCCCACACCUGACGCUUGUGAGGUGUCUCGUACUGCCAAACAAUAGAAGACA.AUG
env-81    CUUUGCCUCCAGCGUUGCCUCUCCCCAAGAGGUCGCUCGUACUGCUUCAACAAUAGAAGACAAUG
env-82    CUUUGCCUCCAGCGUUGCCUCUCCCCGAGAGGUCGCUCGUACUGCUUCAACAAUAGAAGACAAUG
env-83    CUCUGCCUCCAGCGUUGCCUCUCCCUGAGAGGUCGCUCGUACUGCUUCAACAAUAGAAGACAAUG

          .................................................................
          .................................................................
          ●●●●●●●●●●●●○○○-----------------------------------------------AUG
```

```
env-84    CUCUGCCUCCAGCGUUGCCUCUCCCCGAGAGGUCGCUCGUACUGCUUCAACAAUAGAAGACAAUG
env-85    CUCUGCCUCCAGCGUUGCCUCUCCCCGAGAGGUCGCUCGUACUGCUUCAACAAUAGAAGACAAUG
env-86    CUCUGCCUCCAGCGUUGCCUCUCCCCGAGAGGUCGCUCGUACUGCUUCAACAAUAGAAGACAAUG
env-87    CGCCUAAGGGUCCCACACCUGACACAUGUGAGGUGUCUCGUACUGCCAAACAAUAGAAGACAAUG
env-88    CUCUGCCUCCAGCGUUGCCUCUCCCCGAGAGGUCGCUCGUACUGCUUCAACAAUAGAAGACAAUG
env-89    CUCUGCCUCCAGCGUUGCCUCUCCCCGAGAGGUCGCUCGUACUGCUUCAACAAUAGAAGACAAUG
env-90    CUCUUAGGAUCCCACACCUGACGCUCGUGAGGUGUCUCGUACUGCCAAACAAUAGAAGAC.....
env-91    CUUUGUCUCCAGCGUUGCCUCUCCCCGAGAGGUCGCUCGUACUGCUUCAACAAUAGAAGACAAUG
env-92    ACUGCCUCCAGCAUUGCCCUCCUCGAGAGGUCGCUCGUACUGCUUCAACAAUAGAAGACA..AUG
Psp-2-1   GAGCUCGUCCUCUCUCUCGAGGCGCUAAACAAGAAAGAAUAAA...................AUG
Psp-1-1   CCCACAGAGCCCCAUCGCCCAAAUUUAAGAUAACA...........................AUG
env-93    CUUUGCCUCC.......................................................
env-94    ACGCUCUCAUCCCCCGACUCGUCGGGAGGCUCGUACUGCCCAACAAUAGAAGACA.......AUG
env-95    ACACUCUCAUCCCCCGACUCGUCGGGUGGCUCGUACUGCCCAACAAUAGAAGACA.......AUG
env-96    ACGCUCCCAUCCCCCGACUCGUCGGGAGGCUCGUACUGCCCAACAAUAGAAGACA.......AUG
env-97    .................................................................
env-98    ACGCUCCCAUCCCCCGACUCGUCGGGAGGCUCGUACUGCCCAACAAUAGAAGACA.......AUG
env-99    ACGCUCCCAUCCCCCGACUCGUCGGGAGGCUCGUACUGCCCAACAAUAGAAGACA.......AUG
env-100   ACACUCCCAUCCCCCGA................................................
env-101   ACGCUCCCAUCCCCCGACUCGUCGGGAGGCUCGUACUGCCCAACAAUAGAAGACA.......AUG
env-102   ACGCUCCCAUCCCCCGACUCUUCGGGAGGCUCGUACUGCCCAACAAUAGAAGACA.......AUG
env-103   ACACUCCCAUCCCCCGACUCGUCGGGUGGCUCGUACUGCCCAACAAUAGAAGACA.......AUG
env-104   ACGCUCCCAUCCCCCGACUCGUCGGGAGGCUCGUACUGCCCAACAAUAGAAGACA.......AUG
env-105   ACACUCCCAUCCCCCGACUCGUCGGGAGGCUCGUACUGCCCAACAAUAGAAGACA.......AUG
env-106   CCCGCGCUACUCCCGCGGCGGAGCUGCUACUCAAUCAAUAGAAGACA...............AUG
env-107   CCCGCGCUACUCCCGCGGUGGAGCUGCUACUCAAUCAAUAGAAGACA...............AUG
env-108   CAGAAGACAAAA..................................................AUG
env-109   CAGAAGACAAAA..................................................AUG
env-110   GAAGACAAAA....................................................AUG
env-111   AAAAGAUUAAACAA................................................AUG
env-112   AAUUAAACAUUAGAAAAAA...........................................AUG
env-113   AG............................................................AUG
env-114   CCGACGCUGCAUUACACAAAUUCAAUUGACAAACUAACAAU........................
env-115   AAAUUCAAGACAAAAAA.............................................AUG
env-116   AAAAUUUAAGAUUUUACAAAAA........................................AUG
env-117   AAACUGAUUUAAACUACA............................................AUG
env-118   AGAUUAAAAACAGAAUAAA...........................................AUG
env-119   .................................................................
env-120   U.............................................................AUG
env-121   GAGAUGUCACAAAAAACAAGGACAAUAAU.................................AUG
env-122   GAGAGGUCACAA..AUAAAAACAAGGACAAUAAU............................AUG
env-123   AUUACCAACCCGGACUUCCGCCAGGCU......................................
env-124   AAUUACCAAUACGCCGG.............................................AUG
env-125   AAUUACCAACACGCCGG.............................................AUG
env-126   ..............................................................AUG
env-127   AUCACUUUAAUUUUAAUUUAGCAAACA...................................AUG
Pgu-2-1   UCUGACACAAAACAAAUAAUA.........................................AUG
env-128   UCUGACACAAAACAAAAAUA..........................................AUG
Pgi-3-1   UCUGACACAAAACAAACAAUA.........................................AUG
Pgu-1-1   UCUGACACAAAACAAAUAAUA.........................................AUG
env-129   UCUGACACAAAACAAACAAUA.........................................AUG
Pgi-2-1   UCUGACUCAAAACAAACAAUA.........................................AUG
Pgu-1-2   UCUGACACAAAACAAAUAAUA.........................................AUG
Pgi-1-1   UCUGACACAAAACAAACAAUA.........................................AUG
env-130   CACAAAAAAAA...................................................AUG
env-131   UUUAAAAGAAUAAAAA..............................................AUG
env-132   UUUAAAAGAAUAAAAA..............................................AUG
env-133   UUUAAAAGAAUAAAAA..............................................AUG
env-134   UUUAAAAGAAUAAAAA..............................................AUG
env-135   UUUAAAAGAAUAAAAA..............................................AUG
env-136   UUUAAAAGAAUAAAAA..............................................AUG
env-137   AUUUUUUUAACAACA...............................................AUG
env-138   AAA...........................................................AUG
env-139   .................................................................
env-140   AAGAUU........................................................AUG
env-141   AAAAAGAAAU....................................................AUG
env-142   AAAAAGAAAU....................................................AUG
env-143   AAA...........................................................AUG
env-144   GACAUUUGUAGG..................................................AUG
env-145   ACACUUAAAAUAAA................................................AUG
env-146   AAAAAUAAAAAA..................................................AUG
env-147   AAAAAUAAAAAA..................................................AUG
env-148   AAAAAUAAAAAAAG...................................................
env-149   AAAAAUAAAAAA..................................................AUG
env-150   AAAAUAAAA.....................................................AUG
env-151   AAAAAUAAAAAA..................................................AUG
env-152   AACAAUAAAAAA..................................................AUG
env-153   AAAAAUAAAAAA..................................................AUG
env-154   .................................................................
env-155   AAAAAUAAAAAA..................................................AUG
env-156   AAAAUAAAAAA...................................................AUG
env-157   .................................................................
env-158   AUAAUAAUAAA...................................................AUG
env-159   AAAAAUAAAAAA..................................................AUG

          .................................................................
          .................................................................
          ●●●●●●●●●●●●○○○-----------------------------------------------AUG
```

| | | |
|---|---|---|
| env-160 | AACAAUAAAAAA... | AUG |
| env-161 | AACAAUAAAAAA... | AUG |
| env-162 | AACAAUAAAAAA... | AUG |
| env-163 | AACAAUAAAAAA... | AUG |
| env-164 | AACAAUAAAAAA... | AUG |
| env-165 | AACAAUAAAAAA... | AUG |
| env-166 | AACAAUAAAAAA... | AUG |
| env-167 | AAAAAUAAAAAA... | AUG |
| env-168 | AACAAUAAAAAA... | AUG |
| env-169 | AACAAUAAAAAA... | AUG |
| env-170 | AACAAUAAAAAUAGG... | ... |
| env-171 | AACAAUAAAAAA... | AUG |
| env-172 | AACAAUAAAAAA... | AUG |
| env-173 | AACAAUAAAAA... | AUG |
| env-174 | AACAAUAAAAAA... | AUG |
| env-175 | AAUAAUAGAAAA... | AUG |
| env-176 | AACUAAAGAAAA... | AUG |
| env-177 | AAUCAAAUAGAAAAAA... | AUG |
| env-178 | AUAGAAAAAAGAA... | AUG |
| env-179 | AAACAACCGAUUUUAAAAAGAACAUUUUUACAAGAUU... | AUG |
| env-180 | UUCUUAACUUUACUAACAACAGAC... | AUG |
| env-181 | AAAAAUUAGAACUUAAA... | AUG |
| env-182 | AAAAAUUAGAAAUUAAA... | AUG |
| env-183 | AAAAAUUAGAAAUUAAA... | AUG |
| env-184 | AAAAAUUAGAACUUAAA... | AUG |
| env-185 | AAAAAUUAGAACUUAAA... | AUG |
| Pxy-1-1 | AAAAAUUAGAAAUUAAA... | AUG |
| env-186 | AAAAAUUAGAACUUAAA... | AUG |
| env-187 | AAAAAUUAGAACUUAAA... | AUG |
| env-188 | CACAAAAAAAA... | AUG |
| env-189 | UAAUUUAGAUUUAAA... | AUG |
| env-190 | ... | ... |
| env-191 | UAUUUAAAAA... | AUG |
| env-192 | CUUUAAUUACUUUCUUUACACACACACUUAGAUU... | AUG |
| env-193 | AUUUAUUUUAUUUAUUUUAAAAACAAGAAA... | AUG |
| env-194 | ... | ... |
| env-195 | UUUUAAAUUUUAAUCAAAACA... | AUG |
| env-196 | UUUUUAAAUUAUUUAUUAAAA... | AUG |
| env-197 | AUUAUUAGAAACA... | AUG |
| env-198 | AAAAAACAGACAAA... | AUG |
| env-199 | AAACACAAACAAA... | AUG |
| env-200 | CACAAACAAAA... | AUG |
| env-201 | AAACACAUACAAA... | AUG |
| env-202 | AACAAACAAAA... | AUG |
| env-203 | CACAAACAAAA... | AUG |
| env-204 | CACAAACAAAA... | AUG |
| env-205 | ... | ... |
| env-206 | AAACACAAACAAA... | AUG |
| env-207 | AACAAACAAA... | AUG |
| env-208 | ACAAA... | AUG |
| env-209 | ACACAAACAAAA... | AUG |
| env-210 | ACAAACAAAA... | AUG |
| env-211 | AAACACAAACAAA... | AUG |
| env-212 | AACACGAACAAA... | AUG |
| env-213 | ACAAACAAAA... | AUG |
| env-214 | AACAAACAAAA... | AUG |
| env-215 | ACACAAACAAA... | AUG |
| env-216 | CACAAACAAA... | AUG |
| env-217 | AAAACAAACAAAA... | AUG |
| env-218 | ACACAAACAAAA... | AUG |
| env-219 | CUUUAAUUUCAAUUAGAACA... | AUG |
| env-220 | AUUUUUAAGAAAAAGACU... | AUG |
| env-221 | UAAUUAAAAAAAGACU... | AUG |
| env-222 | UAAUAAAAAAA... | AUG |
| env-223 | UAAUUUAGAAAA... | AUG |
| env-224 | AAUAUUUAAAAGAA... | AUG |
| env-225 | AAUAAUAAAAAA... | AUG |
| env-226 | AUAUAAUAAAAAA... | AUG |
| env-227 | UAAAA... | AUG |
| env-228 | AUUGUAGACA... | AUG |
| env-229 | AUUGUAAA... | AUG |
| env-230 | AUUUGUAAAA... | AUG |
| env-231 | UUGUAACA... | AUG |
| env-232 | UUGUAAAA... | AUG |
| env-233 | UAAAG... | AUG |
| env-234 | ... | ... |
| env-235 | UUUAAUUGUAAAA... | AUG |
| env-236 | UGUAACA... | AUG |
| env-237 | UUGUAACA... | AUG |
| env-238 | UAAAA... | AUG |
| env-239 | AUUUGUAAA... | AUG |
| Pac-1-1 | AAUUUGUAAA... | AUG |
| env-240 | UAAAA... | AUG |
| env-241 | UCUAACA... | AUG |
| | ... | ... |
| | ... | ... |
|  | ●●●●●●●●●●○○○-------------------------------- | AUG |

```
env-242  AAAAUUUAAAAGAA................................................AUG
env-243  UUUAAAAACAA...................................................AUG
env-244  AACGCAUAAUAAAUUAAAAUAAAUAAAUAAAA..............................AUG
env-245  AACACAUAAACAAAUUUAAAUAUAAAAAAAA...............................AUG
Ppr-1-1  AACACAUAAAUAAUUUUAAAUUUAAAUAAUAAAAA...........................AUG
env-246  AUUAAAAAUUUUUAGAA.............................................AUG
env-247  UAAAAAAAAAUUAGAA..............................................AUG
env-248  AACAAUAGAAAA..................................................AUG
env-249  AACAAUAGAAAA..................................................AUG
env-250  CUUUAAAACGAAUUACCAAA..........................................AUG
env-251  UUAAAAGUAUUUUAGAA.............................................AUG
env-252  UUAAAAAUCUACUAAA..............................................AUG
env-253  UUAAAAAUCUACUAAA..............................................AUG
env-254  UUUAAAAA......................................................AUG
env-255  AUAAUCACAAAA..AUUUUGAAAUA.....................................AUG
env-256  AUAAUCACAAAAACUUUGAAAUA.......................................AUG
env-257  AUAAUCACAAAAACUUUGAAAUA.......................................AUG
env-258  .................................................................
env-259  UAAUUUGUAAAA..................................................AUG
Dga-1-1  AGAAAAGAA.....................................................AUG
env-260  AGAAAAGAAA....................................................AUG
Psp-4-1  AGAAAAGAA.....................................................AUG
env-261  AUUAUUAAAAGUAUUAGAA...........................................AUG
env-262  UUAUUAAAAGAAUUAGAA............................................AUG
env-263  AAAUAAAGAAUAAA................................................AUG
env-264  AAAUAAAGAAUAAA................................................AUG
env-265  AUUAUUUAAAAA..................................................AUG
env-266  UAAUUUUAGAAAA.................................................AUG
env-267  UAAUUUUAGAAA.....................................................
env-268  AUUAUUUUUAAGAAAAGACU..........................................AUG
env-269  AUUUAAUUUUUUAAGAAAAGAAA.......................................AUG
env-270  UUAAUAAUUAGAAAAA..............................................AUG
env-271  UUAAUUAUUAGAAAAA..............................................AUG
env-272  AUAUAAAAAGAAA.................................................AUG
env-273  AUAUAAAAAGAAA.................................................AUG
env-274  UUUUAAUUCAUUAAAUUUUUAGAUUAAA..................................AUG
env-275  UUUUAAUUCAUUAAAUUUUUAGAUUAAA..................................AUG
env-276  AAUUAUUUCAAUUCAAA.............................................AUG
env-277  CUAUCAUUUAAAAAAAGACU..........................................AUG
env-278  AAAUAUUUAAAAGA................................................A..
env-279  AAAUAUUUAAAAGAA...............................................AUG
env-280  AAAUA............................................................
env-281  AAAUA............................................................
env-282  AAAUAUUUAAAAGAA...............................................AUG
env-283  AAAUAUUUAAAAGAA..................................................
env-284  AAAUA............................................................
env-285  .................................................................
env-286  AAAUAUUUAAAAGAA...............................................AUG
env-287  AAAUAUUUAAAAGAA..................................................
env-288  .................................................................
env-289  AAAUAUUUAAAAGAA...............................................AUG
env-290  AAAUA............................................................
env-291  AAAUAUUUAAAAGAA...............................................AUG
env-292  AAAUAUUUAAAAGAA...............................................AUG
env-293  AAAUAUUUAAAAGAA...............................................AUG
env-294  AAAUAUUUAAAAGAA...............................................AUG
env-295  AAUAUUUAAAAGAA................................................AUG
env-296  AAUAUUUAAAAGAA................................................AUG
env-297  AAUAUUUAAAAGAA................................................AUG
env-298  AAUAUUUAAAAAGAA...............................................AUG
env-299  AAUAUUUAAAAGAA................................................AUG
Dsp-2-1  UUUAAAAAA.....A...............................................AUG
env-300  AAUUAAUUAAAAGAAA..............................................AUG
env-301  AAUUUUUAAACA..................................................AUG
env-302  AAAUUUUAAAAACA................................................AUG
env-303  AAAUUUUAAAAACA................................................AUG
env-304  AAAUUUUAAAAACA................................................AUG
env-305  AUAAAAAA......................................................AUG
Dca-1-1  UUUAAAAAAA....................................................AUG
env-306  UAAUAAAAAUAAUA................................................AUG
env-307  UAAUAAAAAUAAUA................................................AUG
Bpy-2-1  AAUAUUUAAAAGAA................................................AUG
env-308  AAUAUUUAAAAGAA................................................AUG
Bpy-1-1  AAUAUUUAAAGAA.................................................AUG
env-309  AAUAAUAGAAAAAA................................................AUG
env-310  AUAUUUAAAA.......................................................
env-311  AAUAUUUAAAAGAA................................................AUG
env-312  AAUAUUUAAAAGAA................................................AUG
env-313  AAUAUUUAAAAGAA................................................AUG
env-314  AAUAUUUAAAAGAA................................................AUG
env-315  AAAAUUUAAAAGAA................................................AUG
env-316  AAUAUUUAAAAGAA................................................AUG
env-317  AAAAUUUAAAAAAA................................................AUG
env-318  AAUAUUUAAAAAGAA...............................................AUG

         .................................................................
         .................................................................
         ●●●●●●●●●●●○○○------------------------------------------------AUG
```

| | | |
|---|---|---|
| Bst-1-1 | AAAUUUAAAAGAA... | AUG |
| env-319 | AAUAUUUAAAAGAA... | AUG |
| Bti-1-1 | AAAAUUUAAAAGAA... | AUG |
| env-320 | AAAAUUUAAAAGAA... | AUG |
| env-321 | AAUAUUUAAAAGA... | ... |
| env-322 | AAAAUUUAAAAGAA... | AUG |
| env-323 | AAUAUUUAAAAGAA... | AUG |
| Bac-1-1 | AAUAUUUAAAAAGAA... | AUG |
| env-324 | AAAAUUUAAAAGAA... | AUG |
| env-325 | AAUAUUUAAAAGAA... | AUG |
| Bol-1-1 | AAAUUUAAAAGAA... | AUG |
| env-326 | AAAAUUUAAAAGAA... | AUG |
| env-327 | AAUAUUUAAAAAAA... | ... |
| env-328 | AUUUAAAGAAGAAA... | AUG |
| env-329 | AAUAUUUAAAAGAA... | AUG |
| env-330 | AA... | ... |
| env-331 | AAUAUUUUAAAGAA... | AUG |
| env-332 | AAUAUUUAAAAGAA... | AUG |
| env-333 | AAUAUUUAAAAGAA... | AUG |
| env-334 | AAUAUUUAAAAGA... | ... |
| Bfa-1-1 | AAUAUUUAAAAGAA... | AUG |
| env-335 | AAUAUUUAAAAGAA... | AUG |
| env-336 | AAUAUUUAAAAGAAA... | AUG |
| Bre-1-1 | AAUAUUUAAAAGAAA... | AUG |
| env-337 | AAAUAUUUAAAAGAA... | AUG |
| Bno-1-1 | AAUAUUUAAAAAAAAA... | AUG |
| env-338 | ... | ... |
| Bno-2-1 | AAUAUUUAAAAAGAAA... | AUG |
| env-339 | AAUAUUUAAAAGAA... | AUG |
| env-340 | AAUAUUUAAAAAGAA... | AUG |
| env-341 | AAUAUUUAAAAGAA... | AUG |
| env-342 | AAUAUUUAAAAGAA... | AUG |
| env-343 | AAUAUUUAAAAAGAA... | AUG |
| env-344 | AUAUUUAAAAAGAA... | AUG |
| env-345 | AAUAUUUAAAAAGAA... | AUG |
| Bne-1-1 | AAAUAUUUAAAAAGAA... | AUG |
| env-346 | AUUAGAAAAGAA... | AUG |
| env-347 | AUUAGAAAAGAA... | AUG |
| env-348 | AUUAAGAAAGAAA... | AUG |
| env-349 | AUUAGAAUAAGAA... | AUG |
| env-350 | AUUUAAAAGUAAGAAA... | AUG |
| env-351 | AUUAAAGAA... | AUG |
| env-352 | AUUAAGAAAGAAA... | AUG |
| env-353 | AUUUAAAGAAA... | AUG |
| env-354 | AUUUAAAAAGUAAGAAA... | AUG |
| Bba-2-1 | UCAGAAAAUAAA... | AUG |
| env-355 | AUUUAAAAGAAA... | AUG |
| env-356 | AAUUUAAAAGUAAGAAA... | AUG |
| env-357 | AUUUAAAAGUAAGAAA... | AUG |
| env-358 | AUUAAGAAAGAAA... | AUG |
| env-359 | AUUUAAAAAGAA... | AUG |
| env-360 | AUUAAGAAAGAAA... | AUG |
| env-361 | AUUAGAAAAGAAA... | AUG |
| env-362 | AUUUAAAAGAAA... | AUG |
| env-363 | AUUAAGAAAGAAA... | AUG |
| env-364 | ... | ... |
| env-365 | UUUAAAAGAAA... | AUG |
| env-366 | ... | ... |
| env-367 | UAAAUUUAAAAGUAAGAAA... | AUG |
| env-368 | AUUUAAAGAAA... | AUG |
| env-369 | AUUUAAAAGAAA... | AUG |
| env-370 | UUUAAAAGUAAGAAA... | AUG |
| env-371 | AUUAAGAAAAGAAA... | AUG |
| env-372 | AUUUAAAAUAAGAAA... | AUG |
| env-373 | ... | ... |
| env-374 | AUUAGAAAAAGAA... | AUG |
| env-375 | AUUAGAAAAGAA... | AUG |
| env-376 | AUUAGAAAAAGAA... | AUG |
| env-377 | AUUAGAAAAAGAA... | AUG |
| Bma-1-1 | AUUAGAAAAAGAA... | AUG |
| env-378 | AUUAGAAAAAGAA... | AUG |
| env-379 | AUUAGAAAAGAA... | AUG |
| env-380 | AUUAGAAAAAAGAA... | AUG |
| env-381 | UUAGAAAAAGAA... | AUG |
| env-382 | ... | ... |
| env-383 | AUUAAGAUUAAAA... | AUG |
| Bsa-1-1 | UUUUAAAUUAAAGAAA... | AUG |
| env-384 | AAAUCAAAGAAGUAAAA... | AUG |
| env-385 | UUUAAAUUAAAGAAA... | AUG |
| env-386 | ACAUUAAAAAGAA... | AUG |
| Bfi-1-1 | AAUAUUUAAAAAGAA... | AUG |
| env-387 | AAUAUUUAAAAGAA... | AUG |
| env-388 | AAUAUUUAAAAAGAA... | AUG |
| env-389 | AAUAUUUAAAAGAA... | AUG |
| | ... | ... |
| | ... | ... |
|  | ●●●●●●●●●●●○○○------ | AUG |

| | |
|---|---|
| env-390 | ... |
| env-391 | ... |
| env-392 | AAAAUUUAAAAGAA...AUG |
| env-393 | AAUAUUUAAAAGAA...AUG |
| env-394 | ... |
| env-395 | UAAAAAAAUUUAGAA...AUG |
| env-396 | UUAAAAGAUUUAGAA...AUG |
| env-397 | ... |
| env-398 | UUAAAAGAUUUAGAA...AUG |
| Bhe-1-1 | AAUAUUUAAAAGAA...AUG |
| env-399 | AAUAUUUAAAAGAA...AUG |
| Bfl-1-1 | AAUAUUUAAAAGAA...AUG |
| env-400 | AAUAAAAAA...AUG |
| env-401 | AAUAAAAAA...AUG |
| env-402 | UUUAAAAAACAAA...AUG |
| env-403 | AAAAAAUAAAA...AUG |
| env-404 | AAAAAAAGUAAAAA...AUG |
| env-405 | AAGAAAAAAA...AUG |
| env-406 | AAGAAAAAAA...AUG |
| Bpa-1-1 | AAUUAAUAGAA...AUG |
| env-407 | AUUAGAAAAAGAA...AUG |
| env-408 | AUUAGAAAAAGAAA...AUG |
| Bba-1-1 | AUUACUAAAGAAA...AUG |
| env-409 | UAAUAAAAAGUAAA...AUG |
| env-410 | AUUUUAAAAAUAAA...AUG |
| env-411 | AAUAUUAAAAAAAA...AUG |
| env-412 | AAUAUUAAAAAAA...AUG |
| env-413 | AAUAUUAAAAAAA...AUG |
| env-414 | AA... |
| Bpr-1-1 | AAAUAUUAAAAUACA...AUG |
| Bco-1-1 | AAAUAUUUAUAAAUAAA...AUG |
| env-415 | AACAAUAGAAAA...AUG |
| env-416 | AACAAUAGAAAA...AUG |
| env-417 | UAAAAAAUUUUAGAA...AUG |
| Bvi-1-1 | AAAAAUUUUUAGAA...AUG |
| env-418 | UUAAAAAUUUUAGAA...AUG |
| env-419 | UUAAAAAUUUUUAGAA...AUG |
| env-420 | UUAAAAAUUUUUAGAA...AUG |
| env-421 | UAAAAAUUAUAGAAA...AUG |
| env-422 | UUAAA...AUG |
| env-423 | UUAAAAGAUUUAGAA...AUG |
| env-424 | AUUAAAAGAAUUUAAA...AUG |
| env-425 | ... |
| env-426 | AAAUUUCAGACA...AUG |
| env-427 | UGUAAAA...AUG |
| env-428 | UGUAAA...AUG |
| env-429 | AGAAAGAAA...AUG |
| env-430 | CAAAUAAAAAA...AUG |
| env-431 | AAUAAAAAA...AUG |
| env-432 | ... |
| Dsp-1-1 | AGAAAAGAA...AUG |
| env-433 | ... |
| env-434 | AUUUGUAAAA...AUG |
| Bga-1-1 | AUAUUUAAAAAGAA...AUG |
| env-435 | AUAUUUAAAAAGAA...AUG |
| env-436 | AUAUUUAAAAGAA...AUG |
| env-437 | AAAUUUAAAAGAA...AUG |
| env-438 | UAAAUUUAAAAGAA...AUG |
| env-439 | UAAUUUAAAAGA...AUG |
| env-440 | UAAAUUUAAAAGAA...AUG |
| env-441 | UAAAUUUAAAAGAAUAGGCUAGAUAUACU... |
| env-442 | UUAAAUUUAAAAGAA...AUG |
| env-443 | AUAUUUAAAAAGAA...AUG |
| env-444 | AUAUUUAAAAGAA...AUG |
| env-445 | ... |
| env-446 | AUAUUUAAAAAGAA...AUG |
| env-447 | AUAUUUAAAAGAA...AUG |
| env-448 | AUAUUUAAAAGAA...AUG |
| env-449 | AUAUUUAAAAAGAA...AUG |
| env-450 | ... |
| env-451 | AAUAUUUAAAAGAA...AUG |
| env-452 | AUAUUUAAAAAGAA...AUG |
| env-453 | AUAUUUAAAAAGAA...AUG |
| env-454 | AUAUUUAAAAGAA...AUG |
| env-455 | AUAUUUAAAAGAA...AUG |
| env-456 | AUAUUUAAAAAGAA...AUG |
| env-457 | AUAUUUAAAGAA...AUG |
| env-458 | AUAUUUAAAAAGAA...AUG |
| env-459 | ... |
| env-460 | AUAUUUAAAAGAA...AUG |
| env-461 | ... |
| env-462 | ACAGCGCUGCUAAUUCAAUAAACAAAGAAGAAA...AUG |
| env-463 | GGCGCUGCUAAUUCAAUAAACAAAGAAGAAA...AUG |
| env-464 | CGGCGCUGCUAAUUCAAUAAACAAAGAAGAAA...AUG |
| | ... |
| | ... |
| | ...AUG |

```
env-465   CGGCGCUGCUAAUUCAAUAAACAAAGAAGAAA............................AUG
env-466   CGGCGCUGCUAAUUCAAUAAACAAAGAAGAAA............................AUG
env-467   CGGCGCUGCUAAUUCAAUAAACAAAGAAGAAA............................AUG
env-468   ...............................................................
env-469   ACGGCGCUGCUAAUUCAAUAAACAAAGAAGAAA...........................AUG
env-470   CGGCGCUGCUAAUUCAAUAAACAAAGAAGAAA............................AUG
env-471   ACGGCGCUGCUAAUUCAAUAAACAAAGAAGAAA...........................AUG
env-472   CGGCGCUGCUAAUUCAAUAAACAAAGAAGAAA............................AUG
env-473   CGGCGCUGCUAAUUCAAUAAACAAAGAAGAAA............................AUG
env-474   ACGGCGCUGCUAAUUCAAUAAACAAAGAAGAAA...........................AUG
env-475   CGGCGCUGCUAAUUCAAUAAACAAAGAAGAAA............................AUG
env-476   CGGCGCUGCUAAUUCAAUAAACAAAGAAGAAA............................AUG
env-477   CGGCGCUGCUAAUUCAAUAAACAAAGAAGAAA............................AUG
env-478   CGGCGCUGCUAAUUCAAUAAACAAAGAAGAAA............................AUG
env-479   CGGCGCUGCUAAUUCAAUAAACAAAGAAGAAA............................AUG
env-480   CGGCGCUGCUAAUUCAAUAAACAAAGAAGAAA............................AUG
env-481   CGGCGCUGCUAAUUCAAUAAACAAAGAAGAAA............................AUG
env-482   CGGCGCUGCUAAUUCAAUAAACAAAGAAGAAA............................AUG
env-483   CGGCGCUGCUAAUUCAAUAAACAAAGAAGAAA............................AUG
env-484   CACGGCGCUGCUAAUUCAAUAAACAAAGAAGAAA..........................AUG
env-485   C..............................................................
env-486   ACGGCGCU.......................................................
env-487   CGGCGCUGCUAAUUCAAUAAACAAAGAAGAAA............................AUG
env-488   CGGCGC.........................................................
env-489   CGGCGCUGCUAAUUCAAUAAACAAAGAAGAAA............................AUG
env-490   ACAGCGCUGCUAAUUCAAUAAACAAAGAAGAAA...........................AUG
env-491   GGGCGCUGCUAAUUCAAUAAACAAAGAAGAAA............................AUG
env-492   CGGCGCUGCUAAUUCAAUAAACAAAGAAGAAA............................AUG
env-493   ACGGCGCUGCUAAUUCAAUAAACAAAGAAGAAA...........................AUG
env-494   CGGCGCUGCUAAUUCAAUAAACAAAGAAGAAA............................AUG
env-495   CGGCGCUGCUAAUUCAAUAAACAAAGAAGAAA............................AUG
env-496   ACGGCGCUGCUAAUUCAAUAAACAAAGAAGAAA...........................AUG
env-497   ACGGCGCUGCUA..AUUCAAUAAACAAAGAAGAAA.........................AUG
env-498   CGGCGCUGCUAAUUCAAUAAACAAAGAAGAAA............................AUG
env-499   ...............................................................
env-500   ACGGCGCUGCUAAUUCAAUAAACAAAGAAGAAA...........................AUG
env-501   CGCUGUUAUUUCAAUAAACAAACGAAGAAA..............................AUG
env-502   CGCUGUUAUUUCAAUAAACAAACGAAGAAA..............................AUG
env-503   ...............................................................
env-504   ...............................................................
env-505   UAACAAAAAAAGAAAAUAAA........................................AUG
env-506   UGAACCGCUUCGCAAACGUAACUGAUUUUAACUACA........................AUG
env-507   UUCACUAAAUUCUGAAACUUAUU.....................................AUG
env-508   UUCACUAAAUUCUGAAACUUAUU.....................................AUG
env-509   UUUUUUUCACAUCCACCCAACAAACCCAAAACUGAAAAAACUAAA...............AUG
env-510   AUCCGGCACAUAUCAUUAACCCGAAAACCAGAC...........................AUG
env-511   CCCCCGCCACUCUUUUUAACCUCAACAUCGAUUAACAACA....................AUG
env-512   UUCCGUGUUUAAAAGGAAG.........................................AUG
env-513   CAGGCGAAGCACUUCAAACCUUCAUUCAACCCGAAACAAAACA.................AUG
env-514   CAAAAACAAAAAGCAAAA..........................................AUG
env-515   GCCCCGACGCAAAAACUGAUUUUAAUACA...............................AUG
env-516   GGAUACGCAACAUAUAAUCACAAUAGACAAAAAUAAAA......................AUG
env-517   CAACCACAACAAUAAUCAGAAAAUAAA.................................AUG
          ...............................................................
          ...............................................................
          ●●●●●●●●●●●●○○○---------------------------------------------AUG
```

### 15 S4-Flavobacteria

#### 15.1 Taxa

The taxonomy of each organism containing a putative S4-Flavobacteria RNA is listed, with abbreviations identifying each hit (e.g., "Eco-1-1" and "Eco-1-2" might hypothetically represent two distinct RNAs in *E. coli*). The abbreviations will be used to identify each individual S4-Flavobacteria RNA in Sections 15.2 and 15.4:.

| abbrev. of hits | taxonomy of species |
|---|---|
| Fba-1-1 | Bacteria Bacteroidetes *Flavobacteria bacterium* BBFL7 |
| Fba-2-1 | Bacteria Bacteroidetes Flavobacteria Flavobacteriales *Flavobacteria bacterium* BAL38 |
| Fjo-1-1 | Bacteria Bacteroidetes Flavobacteria Flavobacteriales Flavobacteriaceae *Flavobacterium johnsoniae* UW101 |
| Gfo-1-1 | Bacteria Bacteroidetes Flavobacteria Flavobacteriales Flavobacteriaceae *Gramella forsetii* KT0803 |
| Kal-1-1 | Bacteria Bacteroidetes Flavobacteria Flavobacteriales Flavobacteriaceae *Kordia algicida* OT-1 |
| Pir-1-1 | Bacteria Bacteroidetes Flavobacteria Flavobacteriales Flavobacteriaceae *Polaribacter irgensii* 23-P |

| | |
|---|---|
| Rbi-1-1 | Bacteria Bacteroidetes Flavobacteria Flavobacteriales Flavobacteriaceae *Robiginitalea biformata* HTCC2501 |
| Fba-3-1 | Bacteria Bacteroidetes Flavobacteria *Flavobacteriales bacterium* HTCC2170 |
| Ale-1-1 | Bacteria Bacteroidetes Flavobacteriia Flavobacteriales Flavobacteriaceae *Algibacter lectus* |
| Aat-1-1 | Bacteria Bacteroidetes Flavobacteriia Flavobacteriales Flavobacteriaceae *Aquimarina atlantica* |
| Ace-1-1 | Bacteria Bacteroidetes Flavobacteriia Flavobacteriales Flavobacteriaceae *Arenibacter certesii* DSM 19833 |
| Ala-1-1 | Bacteria Bacteroidetes Flavobacteriia Flavobacteriales Flavobacteriaceae *Arenibacter latericius* DSM 15913 |
| Cca-1-1 | Bacteria Bacteroidetes Flavobacteriia Flavobacteriales Flavobacteriaceae *Capnocytophaga canimorsus* |
| Cca-2-1 | Bacteria Bacteroidetes Flavobacteriia Flavobacteriales Flavobacteriaceae *Capnocytophaga canis* |
| Ccy-1-1 | Bacteria Bacteroidetes Flavobacteriia Flavobacteriales Flavobacteriaceae *Capnocytophaga cynodegmi* |
| Csp-1-1 | Bacteria Bacteroidetes Flavobacteriia Flavobacteriales Flavobacteriaceae *Capnocytophaga sp. oral taxon* 335 str. F0486 |
| Csp-2-1 | Bacteria Bacteroidetes Flavobacteriia Flavobacteriales Flavobacteriaceae *Capnocytophaga sp. oral taxon* 863 str. F0517 |
| Cal-1-1 | Bacteria Bacteroidetes Flavobacteriia Flavobacteriales Flavobacteriaceae *Cellulophaga algicola* DSM 14237 |
| Cba-1-1 | Bacteria Bacteroidetes Flavobacteriia Flavobacteriales Flavobacteriaceae *Cellulophaga baltica* NN016038 |
| Cly-1-1 | Bacteria Bacteroidetes Flavobacteriia Flavobacteriales Flavobacteriaceae *Cellulophaga lytica* DSM 7489 |
| Csp-3-1 | Bacteria Bacteroidetes Flavobacteriia Flavobacteriales Flavobacteriaceae *Cellulophaga sp.* Hel_I_12 |
| Ddo-1-1 | Bacteria Bacteroidetes Flavobacteriia Flavobacteriales Flavobacteriaceae *Dokdonia donghaensis* DSW-1 |
| Dsp-1-1 | Bacteria Bacteroidetes Flavobacteriia Flavobacteriales Flavobacteriaceae *Dokdonia sp.* MED134 |
| Ead-1-1 | Bacteria Bacteroidetes Flavobacteriia Flavobacteriales Flavobacteriaceae *Eudoraea adriatica* DSM 19308 |
| Fic-1-1 | Bacteria Bacteroidetes Flavobacteriia Flavobacteriales Flavobacteriaceae *Flaviramulus ichthyoenteri* Th78 |
| Fba-4-1 | Bacteria Bacteroidetes Flavobacteriia Flavobacteriales *Flavobacteriaceae bacterium* P7-3-5 |
| Fch-1-1 | Bacteria Bacteroidetes Flavobacteriia Flavobacteriales Flavobacteriaceae *Flavobacterium chungangense* LMG 26729 |
| Fda-1-1 | Bacteria Bacteroidetes Flavobacteriia Flavobacteriales Flavobacteriaceae *Flavobacterium daejeonense* DSM 17708 |
| Ffr-1-1 | Bacteria Bacteroidetes Flavobacteriia Flavobacteriales Flavobacteriaceae *Flavobacterium frigidarium* DSM 17623 |
| Ffr-2-1 | Bacteria Bacteroidetes Flavobacteriia Flavobacteriales Flavobacteriaceae *Flavobacterium frigoris* PS1 |
| Fgi-1-1 | Bacteria Bacteroidetes Flavobacteriia Flavobacteriales Flavobacteriaceae *Flavobacterium gilvum* |
| Fhy-1-1 | Bacteria Bacteroidetes Flavobacteriia Flavobacteriales Flavobacteriaceae *Flavobacterium hydatis* |
| Fin-1-1 | Bacteria Bacteroidetes Flavobacteriia Flavobacteriales Flavobacteriaceae *Flavobacterium indicum* GPTSA100-9 |
| Fps-1-1 | Bacteria Bacteroidetes Flavobacteriia Flavobacteriales Flavobacteriaceae *Flavobacterium psychrophilum* |
| Fre-1-1 | Bacteria Bacteroidetes Flavobacteriia Flavobacteriales Flavobacteriaceae *Flavobacterium reichenbachii* |
| Fsa-1-1 | Bacteria Bacteroidetes Flavobacteriia Flavobacteriales Flavobacteriaceae *Flavobacterium sasangense* DSM 21067 |
| Fsp-1-1 | Bacteria Bacteroidetes Flavobacteriia Flavobacteriales Flavobacteriaceae *Flavobacterium sp.* ABG |
| Fsp-2-1 | Bacteria Bacteroidetes Flavobacteriia Flavobacteriales Flavobacteriaceae *Flavobacterium sp.* ACAM 123 |
| Fsp-3-1 | Bacteria Bacteroidetes Flavobacteriia Flavobacteriales Flavobacteriaceae *Flavobacterium sp.* Fl |
| Fsp-4-1 | Bacteria Bacteroidetes Flavobacteriia Flavobacteriales Flavobacteriaceae *Flavobacterium sp.* JRM |
| Fsp-5-1 | Bacteria Bacteroidetes Flavobacteriia Flavobacteriales Flavobacteriaceae *Flavobacterium sp.* KJJ |
| Fsp-6-1 | Bacteria Bacteroidetes Flavobacteriia Flavobacteriales Flavobacteriaceae *Flavobacterium sp.* MEB061 |
| Fsu-1-1 | Bacteria Bacteroidetes Flavobacteriia Flavobacteriales Flavobacteriaceae *Flavobacterium succinicans* LMG 10402 |
| Fte-1-1 | Bacteria Bacteroidetes Flavobacteriia Flavobacteriales Flavobacteriaceae *Flavobacterium tegetincola* DSM 22377 |
| Fag-1-1 | Bacteria Bacteroidetes Flavobacteriia Flavobacteriales Flavobacteriaceae *Formosa agariphila* KMM 3901 |
| Gsa-1-1 | Bacteria Bacteroidetes Flavobacteriia Flavobacteriales Flavobacteriaceae *Gaetbulibacter saemankumensis* DSM 17032 |
| Gme-1-1 | Bacteria Bacteroidetes Flavobacteriia Flavobacteriales Flavobacteriaceae *Gelidibacter mesophilus* DSM 14095 |
| Gli-1-1 | Bacteria Bacteroidetes Flavobacteriia Flavobacteriales Flavobacteriaceae *Gillisia limnaea* DSM 15749 |
| Gsp-1-1 | Bacteria Bacteroidetes Flavobacteriia Flavobacteriales Flavobacteriaceae *Gillisia sp.* CAL575 |
| Gsp-2-1 | Bacteria Bacteroidetes Flavobacteriia Flavobacteriales Flavobacteriaceae *Gillisia sp.* CBA3202 |
| Gsp-3-1 | Bacteria Bacteroidetes Flavobacteriia Flavobacteriales Flavobacteriaceae *Gillisia sp.* Hel_I_29 |
| Gsp-4-1 | Bacteria Bacteroidetes Flavobacteriia Flavobacteriales Flavobacteriaceae *Gillisia sp.* JM1 |
| Gec-1-1 | Bacteria Bacteroidetes Flavobacteriia Flavobacteriales Flavobacteriaceae *Gramella echinicola* DSM 19838 |
| Gpo-1-1 | Bacteria Bacteroidetes Flavobacteriia Flavobacteriales Flavobacteriaceae *Gramella portivictoriae* DSM 23547 |
| Jpa-1-1 | Bacteria Bacteroidetes Flavobacteriia Flavobacteriales Flavobacteriaceae *Jejuia pallidilutea* |
| Jma-1-1 | Bacteria Bacteroidetes Flavobacteriia Flavobacteriales Flavobacteriaceae *Joostella marina* DSM 19592 |
| Kje-1-1 | Bacteria Bacteroidetes Flavobacteriia Flavobacteriales Flavobacteriaceae *Kordia jejudonensis* |
| Lsp-1-1 | Bacteria Bacteroidetes Flavobacteriia Flavobacteriales Flavobacteriaceae *Lacinutrix sp.* Hel_I_90 |
| Lsp-2-1 | Bacteria Bacteroidetes Flavobacteriia Flavobacteriales Flavobacteriaceae *Leeuwenhoekiella sp.* Hel_I_48 |
| Man-1-1 | Bacteria Bacteroidetes Flavobacteriia Flavobacteriales Flavobacteriaceae *Maribacter antarcticus* DSM 21422 |
| Mfo-1-1 | Bacteria Bacteroidetes Flavobacteriia Flavobacteriales Flavobacteriaceae *Maribacter forsetii* DSM 18668 |
| Msp-1-1 | Bacteria Bacteroidetes Flavobacteriia Flavobacteriales Flavobacteriaceae *Maribacter sp.* Hel_I_7 |
| Mth-1-1 | Bacteria Bacteroidetes Flavobacteriia Flavobacteriales Flavobacteriaceae *Maribacter thermophilus* |
| Mze-1-1 | Bacteria Bacteroidetes Flavobacteriia Flavobacteriales Flavobacteriaceae *Mesoflavibacter zeaxanthinifaciens* DSM 18436 |
| Mze-2-1 | Bacteria Bacteroidetes Flavobacteriia Flavobacteriales Flavobacteriaceae *Mesoflavibacter zeaxanthinifaciens* S86 |
| Mlu-1-1 | Bacteria Bacteroidetes Flavobacteriia Flavobacteriales Flavobacteriaceae *Muricauda lutaonensis* |
| Mru-1-1 | Bacteria Bacteroidetes Flavobacteriia Flavobacteriales Flavobacteriaceae *Muricauda ruestringensis* DSM 13258 |
| Msp-2-1 | Bacteria Bacteroidetes Flavobacteriia Flavobacteriales Flavobacteriaceae *Muricauda sp.* MAR_2010_75 |
| Nul-1-1 to Nul-1-4 | Bacteria Bacteroidetes Flavobacteriia Flavobacteriales Flavobacteriaceae *Nonlabens ulvanivorans* |
| Oma-1-1 | Bacteria Bacteroidetes Flavobacteriia Flavobacteriales Flavobacteriaceae *Olleya marilimosa* CAM030 |
| Osp-1-1 | Bacteria Bacteroidetes Flavobacteriia Flavobacteriales Flavobacteriaceae *Olleya sp.* VCSA23 |

| | |
|---|---|
| Osp-2-1 | Bacteria Bacteroidetes Flavobacteriia Flavobacteriales Flavobacteriaceae *Olleya sp.* VCSM12 |
| Psp-1-1 | Bacteria Bacteroidetes Flavobacteriia Flavobacteriales Flavobacteriaceae *Polaribacter sp.* Hel1_85 |
| Psp-2-1 | Bacteria Bacteroidetes Flavobacteriia Flavobacteriales Flavobacteriaceae *Polaribacter sp.* Hel_I_88 |
| Psp-3-1 | Bacteria Bacteroidetes Flavobacteriia Flavobacteriales Flavobacteriaceae *Polaribacter sp.* MED152 |
| Pgo-1-1 | Bacteria Bacteroidetes Flavobacteriia Flavobacteriales Flavobacteriaceae *Psychroflexus gondwanensis* ACAM 44 |
| Pto-1-1 | Bacteria Bacteroidetes Flavobacteriia Flavobacteriales Flavobacteriaceae *Psychroflexus torquis* ATCC 700755 |
| Ptr-1-1 | Bacteria Bacteroidetes Flavobacteriia Flavobacteriales Flavobacteriaceae *Psychroflexus tropicus* DSM 15496 |
| Pja-1-1 | Bacteria Bacteroidetes Flavobacteriia Flavobacteriales Flavobacteriaceae *Psychroserpens jangbogonensis* |
| Pme-1-1 | Bacteria Bacteroidetes Flavobacteriia Flavobacteriales Flavobacteriaceae *Psychroserpens mesophilus* |
| Ssp-1-1 | Bacteria Bacteroidetes Flavobacteriia Flavobacteriales Flavobacteriaceae *Salegentibacter sp.* Hel_I_6 |
| Ssp-2-1 | Bacteria Bacteroidetes Flavobacteriia Flavobacteriales Flavobacteriaceae *Sediminibacter sp.* Hel_I_10 |
| Sze-1-1 | Bacteria Bacteroidetes Flavobacteriia Flavobacteriales Flavobacteriaceae *Siansivirga zeaxanthinifaciens* CC-SAMT-1 |
| Tna-1-1 | Bacteria Bacteroidetes Flavobacteriia Flavobacteriales Flavobacteriaceae *Tamlana nanhaiensis* |
| Tov-1-1 | Bacteria Bacteroidetes Flavobacteriia Flavobacteriales Flavobacteriaceae *Tenacibaculum ovolyticum* DSM 18103 |
| Zga-1-1 | Bacteria Bacteroidetes Flavobacteriia Flavobacteriales Flavobacteriaceae *Zobellia galactanivorans* |
| Zul-1-1 | Bacteria Bacteroidetes Flavobacteriia Flavobacteriales Flavobacteriaceae *Zobellia uliginosa* |
| env-1 to env-841 | environmental samples |

#### 15.2 Gene contexts

Each S4-Flavobacteria RNA (indicated by "RNA→") is listed. For each hit, the downstream genes predicted to reside in a regulated operon are listed. If the nearest downstream gene is encoding in the opposite strand (and therefore presumed to not be a part of a regulated operon), then that gene is still depicted. Some environmental sequences and some RefSeq entries lack gene annotations, and so no genes are available for such sequences. The direction of each gene is indicated with an arrow (→), and each predicted conserved domain in the gene is named. Conserved domains associated with more than one S4-Flavobacteria RNA are assigned a color; other domains are gray. Information about these conserved domains is given in Section 15.3. The accession of the sequence containing each S4-Flavobacteria RNA is given in the column named "Seq. accession". Accessions beginning with "NC_", "NS_", "NW_" or "NZ_" are contained in RefSeq. Other accession refer to environmental samples. Nucleotide coordinates are given for the 5′ and 3′ boundaries of each S4-Flavobacteria RNA. If the 5′ coordinate is greater than the 3′ coordinate, the RNA is present on the reverse-complement strand of the containing genomic DNA sequence. Each hit is denoted by an abbreviation (like "Eco-1-1") that refers to a taxonomy given in Section 15.1.

| abbrev. | Seq. accession | | 5′ at | 3′ at | genes |
|---|---|---|---|---|---|
| env-1 | JCVI_SCAF_1096627086414 | + | 372 | 420 | **RNA**→ RpsD (COG0522)rpsD (PRK05327)→ RNAP_alpha_NTD (cd06928)PRK05182 (PRK05182)→ |
| env-2 | JCVI_SCAF_1096626661165 | + | 1143 | 1191 | **RNA**→ RpsD (COG0522)rpsD (PRK05327)→ |
| env-3 | TrichMG_c101151 | - | 3047 | 3010 | **RNA**→ RpsD (COG0522)rpsD (PRK05327)→ RpoA (COG0202)PRK05182 (PRK05182)→ RplQ (COG0203)rplQ (PRK05591)→ CPSase_sm_chain (smart01097)CPSase_sm_chain (pfam00988)→ |
| env-4 | KGI_S1_ANT02_95m_c10350754 | + | 179 | 224 | **RNA**→ rpsD (PRK05327)Ribosomal_S4 (pfam00163)→ |
| env-5 | CESE01138582.1 | + | 526 | 571 | **RNA**→ RpsD (COG0522)rpsD (PRK05327)→ |
| env-6 | CESE01260814.1 | + | 526 | 571 | **RNA**→ RpsD (COG0522)rpsD (PRK05327)→ |
| env-7 | JGI20154J14316_10002684 | + | 5028 | 5073 | **RNA**→ RpsD (COG0522)rpsD (PRK05327)→ RpoA (COG0202)PRK05182 (PRK05182)→ RplQ (COG0203)rplQ (PRK05591)→ |
| env-8 | CENR01029873.1 | - | 14623 | 14578 | **RNA**→ RpsD (COG0522)rpsD (PRK05327)→ RpoA (COG0202)PRK05182 (PRK05182)→ |
| env-9 | PML_READ_00292105 | - | 291 | 246 | **RNA**→ rpsD (PRK05327)Ribosomal_S4 (pfam00163)→ |
| env-10 | CETR01015277.1 | - | 2393 | 2348 | **RNA**→ RpsD (COG0522)rpsD (PRK05327)→ RpoA (COG0202)PRK05182 (PRK05182)→ RplQ (COG0203)rplQ (PRK05591)→ |
| env-11 | JGI24723J26617_10003273 | + | 8066 | 8111 | **RNA**→ RpsD (COG0522)rpsD (PRK05327)→ RpoA (COG0202)PRK05182 (PRK05182)→ RplQ (COG0203)rplQ (PRK05591)→ |
| env-12 | DelMOSum2010_c10014645 | - | 2436 | 2391 | **RNA**→ RpsD (COG0522)rpsD (PRK05327)→ RpoA (COG0202)PRK05182 (PRK05182)→ RplQ (COG0203)rplQ (PRK05591)→ |
| env-13 | 2236876010_0077995 | + | 182 | 227 | **RNA**→ rpsD (PRK05327)Ribosomal_S4 (pfam00163)→ |
| env-14 | 2236876010_0613686 | + | 157 | 202 | **RNA**→ rpsD (PRK05327)Ribosomal_S4 (pfam00163)→ rpsD (PRK05327)→ |
| env-15 | DelMOSum2011_c10012933 | + | 1809 | 1854 | **RNA**→ RpsD (COG0522)rpsD (PRK05327)→ RpoA (COG0202)PRK05182 (PRK05182)→ RplQ (COG0203)rplQ (PRK05591)→ |
| env-16 | 2236876010_0135624 | + | 280 | 325 | **RNA**→ |

| | | | | | |
|---|---|---|---|---|---|
| env-17 | 2236876011_290389 | - | 127 | 82 | RNA→ rpsD (PRK05327)Ribosomal_S4 (pfam00163)→ |
| env-18 | JGI20152J14361_10041961 | - | 729 | 684 | RNA→ RpsD (COG0522)rpsD (PRK05327)→ |
| env-19 | JGI20152J14361_10060010 | + | 800 | 845 | RNA→ |
| env-20 | JGI20152J14361_10066480 | + | 773 | 818 | RNA→ |
| env-21 | 2236876009_001520 | - | 435 | 389 | RNA→ S4 (smart00363)RpsD (COG0522)→ |
| env-22 | 2236876010_0329218 | - | 266 | 218 | RNA→ rpsD (PRK05327)Ribosomal_S4 (pfam00163)→ |
| env-23 | P_1C_Liq_1_UnCty_1010446 | + | 1314 | 1348 | RNA→ |
| env-24 | P_2C_Liq_1_UnCty_1056530 | + | 207 | 251 | RNA→ RpsD (COG0522)rpsD (PRK05327)→ |
| Nul-1-1 | NZ_BBMJ01000004.1 | - | 87420 | 87376 | RNA→ RpsD (COG0522)rpsD (PRK05327)→ hypo→ RNAP_alpha_NTD (cd06928)PRK05182 (PRK05182)RNA_pol_A_CTD (pfam03118)→ hypo→ hypo→ CarA (COG0505)PRK12564 (PRK12564)→ hypo→ hypo→ |
| Nul-1-2 | NZ_BBNT01000002.1 | - | 87529 | 87485 | RNA→ RpsD (COG0522)rpsD (PRK05327)→ RNAP_alpha_NTD (cd06928)PRK05182 (PRK05182)RNA_pol_A_CTD (pfam03118)→ RplQ (COG0203)rplQ (PRK05591)→ CarA (COG0505)PRK12564 (PRK12564)→ eno (PRK00077)enolase (cd03313)→ |
| Nul-1-3 | NZ_BBMK01000003.1 | + | 230947 | 230991 | RNA→ RpsD (COG0522)rpsD (PRK05327)→ RNAP_alpha_NTD (cd06928)PRK05182 (PRK05182)RNA_pol_A_CTD (pfam03118)→ RplQ (COG0203)rplQ (PRK05591)→ CarA (COG0505)PRK12564 (PRK12564)→ hypo→ hypo→ |
| Fba-1-1 | NZ_CH672374.1 | - | 1074107 | 1074063 | RNA→ RpsD (COG0522)rpsD (PRK05327)→ RNAP_alpha_NTD (cd06928)PRK05182 (PRK05182)RNA_pol_A_CTD (pfam03118)→ RplQ (COG0203)rplQ (PRK05591)→ CarA (COG0505)PRK12564 (PRK12564)→ eno (PRK00077)enolase (cd03313)→ |
| Nul-1-4 | NZ_BBMM01000006.1 | + | 108642 | 108686 | RNA→ RpsD (COG0522)rpsD (PRK05327)→ RNAP_alpha_NTD (cd06928)PRK05182 (PRK05182)RNA_pol_A_CTD (pfam03118)→ RplQ (COG0203)rplQ (PRK05591)→ hypo→ hypo→ eno (PRK00077)enolase (cd03313)→ |
| env-25 | P_2C_Liq_1_UnCty_1000014 | - | 67042 | 66997 | RNA→ RpsD (COG0522)rpsD (PRK05327)→ RNAP_alpha_NTD (cd06928)PRK05182 (PRK05182)RNA_pol_A_CTD (pfam03118)→ RplQ (COG0203)rplQ (PRK05591)→ CarA (COG0505)PRK12564 (PRK12564)→ eno (PRK00077)enolase (cd03313)→ |
| env-26 | 2210087396 | + | 206 | 248 | RNA→ rpsD (PRK05327)Ribosomal_S4 (pfam00163)→ |
| env-27 | 2210711694 | + | 456 | 488 | RNA→ |
| env-28 | CEVW01139398.1 | + | 983 | 1023 | RNA→ RpsD (COG0522)rpsD (PRK05327)→ PRK05182 (PRK05182)RNA_pol_L (pfam01193)→ |
| env-29 | JCVI_SCAF_1096628258595 | - | 518 | 478 | RNA→ RpsD (COG0522)rpsD (PRK05327)→ |
| env-30 | CERR01133960.1 | - | 144 | 104 | RNA→ rps4 (CHL00113)rpsD (PRK05327)Ribosomal_S4 (pfam00163)→ |
| env-31 | JCVI_READ_1101740017077 | + | 208 | 256 | RNA→ RpsD (COG0522)rpsD (PRK05327)→ |
| Ddo-1-1 | NZ_JSAQ01000001.1 | + | 2692094 | 2692135 | RNA→ RpsD (COG0522)rpsD (PRK05327)→ RNAP_alpha_NTD (cd06928)PRK05182 (PRK05182)RNA_pol_A_CTD (pfam03118)→ RplQ (COG0203)rplQ (PRK05591)→ |
| Dsp-1-1 | NZ_CP009301.1 | + | 2792470 | 2792511 | RNA→ RpsD (COG0522)rpsD (PRK05327)→ RNAP_alpha_NTD (cd06928)PRK05182 (PRK05182)RNA_pol_A_CTD (pfam03118)→ RplQ (COG0203)rplQ (PRK05591)→ |
| env-32 | CEWG01194218.1 | + | 1094 | 1136 | RNA→ |
| env-33 | JCVI_SCAF_1096626222411 | + | 324 | 368 | RNA→ RpsD (COG0522)rpsD (PRK05327)→ |
| env-34 | LPjun09P1210m_c1017679 | - | 357 | 314 | RNA→ rpsD (PRK05327)Ribosomal_S4 (pfam00163)→ |
| env-35 | CESI01021004.1 | - | 629 | 586 | RNA→ RpsD (COG0522)rpsD (PRK05327)→ |
| env-36 | CERI01207796.1 | + | 53 | 96 | RNA→ RpsD (COG0522)rpsD (PRK05327)→ |
| env-37 | CERM01149072.1 | - | 47 | 14 | RNA→ |
| env-38 | DelMOSum2010_c10216602 | + | 506 | 549 | RNA→ rpsD (PRK05327)→ |
| env-39 | CESU01140860.1 | - | 526 | 483 | RNA→ RpsD (COG0522)rpsD (PRK05327)→ |
| env-40 | LPaug08P2610m_c1036481 | - | 442 | 399 | RNA→ S4 (smart00363)RpsD (COG0522)→ |
| env-41 | JGI20160J14292_10002802 | + | 11050 | 11093 | RNA→ RpsD (COG0522)rpsD (PRK05327)→ RpoA (COG0202)PRK05182 (PRK05182)→ RplQ (COG0203)rplQ (PRK05591)→ CPSase_sm_chain (smart01097)CPSase_sm_chain (pfam00988)→ |

| | | | | | |
|---|---|---|---|---|---|
| env-42 | CENJ01167914.1 | + | 816 | 859 | **RNA**→ RpsD (COG0522)rpsD (PRK05327)→ RpoA (COG0202)PRK05182 (PRK05182)→ RplQ (COG0203)rplQ (PRK05591)→ CarA (COG0505)PRK12564 (PRK12564)→ eno (PRK00077)enolase (cd03313)→ GH25_muramidase (cd00599)Glyco_25 (smart00641)→ monomer_idh (TIGR00178)IDH (pfam03971)→ |
| env-43 | CEWK01129157.1 | - | 675 | 632 | **RNA**→ RpsD (COG0522)rpsD (PRK05327)→ |
| env-44 | CENG01098372.1 | + | 851 | 894 | **RNA**→ RpsD (COG0522)rpsD (PRK05327)→ |
| env-45 | JCVI_SCAF_1101668543230 | - | 496 | 453 | **RNA**→ S4 (smart00363)RpsD (COG0522)→ |
| env-46 | DelMOSum2010_c10000017 | - | 35841 | 35798 | **RNA**→ RpsD (COG0522)rpsD (PRK05327)→ RpoA (COG0202)PRK05182 (PRK05182)→ RplQ (COG0203)rplQ (PRK05591)→ CarA (COG0505)PRK12564 (PRK12564)→ eno (PRK00077)enolase (cd03313)→ GH25_muramidase (cd00599)Glyco_25 (smart00641)→ monomer_idh (TIGR00178)IDH (pfam03971)→ |
| env-47 | DelMOSum2010_c10109581 | - | 317 | 274 | **RNA**→ rpsD (PRK05327)Ribosomal_S4 (pfam00163)→ |
| env-48 | JCVI_SCAF_1096627396816 | - | 557 | 514 | **RNA**→ RpsD (COG0522)rpsD (PRK05327)→ |
| env-49 | 2236876001_514958 | - | 290 | 247 | **RNA**→ rpsD (PRK05327)Ribosomal_S4 (pfam00163)→ |
| env-50 | SI34jun09_10m_1018348 | + | 1056 | 1099 | **RNA**→ rpsD (PRK05327)→ |
| env-51 | CESJ01081808.1 | - | 167 | 124 | **RNA**→ rpsD (PRK05327)Ribosomal_S4 (pfam00163)→ |
| env-52 | CESP01137938.1 | - | 184 | 141 | **RNA**→ rpsD (PRK05327)Ribosomal_S4 (pfam00163)→ |
| env-53 | CESQ01357094.1 | + | 349 | 392 | **RNA**→ RpsD (COG0522)rpsD (PRK05327)→ |
| env-54 | CESU01004307.1 | + | 415 | 458 | **RNA**→ RpsD (COG0522)rpsD (PRK05327)→ |
| env-55 | JCVI_SCAF_1096627372950 | + | 1399 | 1442 | **RNA**→ RpsD (COG0522)rpsD (PRK05327)→ PRK05182 (PRK05182)RNA_pol_L (pfam01193)→ |
| env-56 | SR_TTP_S3_1005704 | - | 528 | 487 | **RNA**→ RpsD (COG0522)rpsD (PRK05327)→ |
| env-57 | CENQ01142578.1 | + | 1112 | 1153 | **RNA**→ RpsD (COG0522)rpsD (PRK05327)→ PRK05182 (PRK05182)RNA_pol_L (pfam01193)→ |
| env-58 | JCVI_SCAF_1096627751190 | + | 390 | 431 | **RNA**→ RpsD (COG0522)rpsD (PRK05327)→ RpoA (COG0202)PRK05182 (PRK05182)→ |
| env-59 | SI34jun09_10m_1026568 | + | 377 | 426 | **RNA**→ RpsD (COG0522)rpsD (PRK05327)→ |
| env-60 | 2236876010_0420853 | - | 113 | 64 | **RNA**→ ←hypo |
| env-61 | CERM01058458.1 | - | 47 | 8 | **RNA**→ |
| env-62 | CESF01120564.1 | + | 2547 | 2596 | **RNA**→ |
| env-63 | DelMOWin2010_c10006726 | - | 4916 | 4867 | **RNA**→ RpsD (COG0522)rpsD (PRK05327)→ RNAP_alpha_NTD (cd06928)PRK05182 (PRK05182)RNA_pol_A_CTD (pfam03118)→ RplQ (COG0203)rplQ (PRK05591)→ |
| env-64 | 2236876003_095850 | + | 200 | 249 | **RNA**→ rpsD (PRK05327)Ribosomal_S4 (pfam00163)→ |
| env-65 | JGI20151J14362_10198578 | - | 460 | 411 | **RNA**→ RpsD (COG0522)rpsD (PRK05327)→ |
| env-66 | DelMOSum2011_c10006805 | + | 2258 | 2307 | **RNA**→ RpsD (COG0522)rpsD (PRK05327)→ RpoA (COG0202)PRK05182 (PRK05182)→ RplQ (COG0203)rplQ (PRK05591)→ |
| env-67 | CERM01027048.1 | - | 54 | 15 | **RNA**→ |
| env-68 | SA_S1_NOR05_45m_c10094569 | - | 509 | 460 | **RNA**→ RpsD (COG0522)rpsD (PRK05327)→ |
| env-69 | CESJ01125754.1 | - | 683 | 634 | **RNA**→ RpsD (COG0522)rpsD (PRK05327)→ |
| env-70 | JCVI_SCAF_1096627289576 | + | 1243 | 1292 | **RNA**→ S4 (smart00363)RpsD (COG0522)→ |
| env-71 | SI54feb11_120m_c1075046 | + | 205 | 254 | **RNA**→ |
| env-72 | DelMOSum2010_c10001162 | - | 9097 | 9048 | **RNA**→ RpsD (COG0522)rpsD (PRK05327)→ RNAP_alpha_NTD (cd06928)PRK05182 (PRK05182)RNA_pol_A_CTD (pfam03118)→ RplQ (COG0203)rplQ (PRK05591)→ |
| env-73 | LPaug09P1610m_c1010800 | - | 1174 | 1132 | **RNA**→ RpsD (COG0522)rpsD (PRK05327)→ RNAP_alpha_NTD (cd06928)PRK05182 (PRK05182)→ |
| env-74 | JGI20157J14317_10009143 | + | 851 | 893 | **RNA**→ RpsD (COG0522)rpsD (PRK05327)→ RNAP_alpha_NTD (cd06928)PRK05182 (PRK05182)RNA_pol_A_CTD (pfam03118)→ rplQ (PRK05591)Ribosomal_L17 (pfam01196)→ CarA (COG0505)PRK12564 (PRK12564)→ eno (PRK00077)enolase (cd03313)→ |
| env-75 | DelMOSum2010_c10023112 | + | 1855 | 1897 | **RNA**→ RpsD (COG0522)rpsD (PRK05327)→ RNAP_alpha_NTD (cd06928)PRK05182 (PRK05182)RNA_pol_A_CTD (pfam03118)→ |
| env-76 | JGI20154J14316_10037599 | + | 554 | 596 | **RNA**→ RpsD (COG0522)rpsD (PRK05327)→ RNAP_alpha_NTD (cd06928)PRK05182 (PRK05182)RNA_pol_A_CTD (pfam03118)→ |
| env-77 | LPaug09P1610m_c1025405 | + | 502 | 545 | **RNA**→ rpsD (PRK05327)Ribosomal_S4 (pfam00163)→ |

| | | | | | |
|---|---|---|---|---|---|
| env-78 | CESF01024170.1 | + | 3852 | 3895 | RNA→ RpsD (COG0522)rpsD (PRK05327)→ RNAP_alpha_NTD (cd06928)PRK05182 (PRK05182)RNA_pol_A_CTD (pfam03118)→ RplQ (COG0203)rplQ (PRK05591)→ CarA (COG0505)PRK12564 (PRK12564)→ eno (PRK00077)enolase (cd03313)→ gltA (PRK05614)EcCS_like (cd06114)→ |
| env-79 | CESI01111408.1 | - | 294 | 251 | RNA→ rpsD (PRK05327)Ribosomal_S4 (pfam00163)→ |
| env-80 | CEVX01268119.1 | - | 403 | 360 | RNA→ RpsD (COG0522)rpsD (PRK05327)→ |
| env-81 | CEOP01127922.1 | - | 573 | 530 | RNA→ RpsD (COG0522)rpsD (PRK05327)→ |
| env-82 | CENT01082708.1 | + | 1142 | 1185 | RNA→ RpsD (COG0522)rpsD (PRK05327)→ RNAP_alpha_NTD (cd06928)PRK05182 (PRK05182)→ |
| env-83 | CERI01005517.1 | + | 12209 | 12252 | RNA→ RpsD (COG0522)rpsD (PRK05327)→ RNAP_alpha_NTD (cd06928)PRK05182 (PRK05182)RNA_pol_A_CTD (pfam03118)→ RplQ (COG0203)rplQ (PRK05591)→ GATase1_CPSase (cd01744)PRK12564 (PRK12564)CPSase_sm_chain (pfam00988)→ |
| env-84 | NCBI_BBAY_READ_1105731223526 | - | 451 | 408 | RNA→ RpsD (COG0522)rpsD (PRK05327)→ |
| env-85 | JCVI_SCAF_1096626910973 | + | 1202 | 1246 | RNA→ RpsD (COG0522)rpsD (PRK05327)→ RNAP_alpha_NTD (cd06928)PRK05182 (PRK05182)RNA_pol_A_CTD (pfam03118)→ |
| env-86 | CENJ01088625.1 | - | 632 | 588 | RNA→ RpsD (COG0522)rpsD (PRK05327)→ |
| env-87 | JGI20151J14362_10002063 | + | 4119 | 4163 | RNA→ RpsD (COG0522)rpsD (PRK05327)→ RNAP_alpha_NTD (cd06928)PRK05182 (PRK05182)RNA_pol_A_CTD (pfam03118)→ RplQ (COG0203)rplQ (PRK05591)rpsP (PRK14521)→ CarA (COG0505)PRK12564 (PRK12564)→ eno (PRK00077)enolase (cd03313)→ gltA (PRK05614)EcCS_like (cd06114)→ |
| env-88 | JCVI_SCAF_1096627345251 | + | 852 | 896 | RNA→ RpsD (COG0522)rpsD (PRK05327)→ |
| env-89 | CENU01250245.1 | - | 488 | 444 | RNA→ RpsD (COG0522)rpsD (PRK05327)→ |
| env-90 | JCVI_SCAF_1101668117291 | + | 939 | 982 | RNA→ RpsD (COG0522)rpsD (PRK05327)→ |
| env-91 | NCBI_READ_1112493636597 | - | 239 | 197 | RNA→ |
| env-92 | NCBI_READ_1112637788282 | + | 45 | 87 | RNA→ RpsD (COG0522)rpsD (PRK05327)→ |
| env-93 | DelMOSum2011_c10065322 | + | 1218 | 1261 | RNA→ rpsD (PRK05327)Ribosomal_S4 (pfam00163)→ |
| env-94 | DelMOSum2010_c10001866 | + | 12778 | 12821 | RNA→ RpsD (COG0522)rpsD (PRK05327)→ RNAP_alpha_NTD (cd06928)PRK05182 (PRK05182)RNA_pol_A_CTD (pfam03118)→ RplQ (COG0203)rplQ (PRK05591)→ CarA (COG0505)PRK12564 (PRK12564)→ eno (PRK00077)Enolase_N (pfam03952)→ |
| env-95 | DelMOSum2010_c10000646 | - | 20267 | 20225 | RNA→ RpsD (COG0522)rpsD (PRK05327)→ RNAP_alpha_NTD (cd06928)PRK05182 (PRK05182)RNA_pol_A_CTD (pfam03118)→ RplQ (COG0203)rplQ (PRK05591)→ CarA (COG0505)PRK12564 (PRK12564)→ eno (PRK00077)enolase (cd03313)→ gltA (PRK05614)EcCS_like (cd06114)→ |
| env-96 | CERI01138193.1 | + | 441 | 483 | RNA→ |
| env-97 | JGI20157J14317_10000863 | - | 19631 | 19589 | RNA→ RpsD (COG0522)rpsD (PRK05327)→ RNAP_alpha_NTD (cd06928)PRK05182 (PRK05182)RNA_pol_A_CTD (pfam03118)→ RplQ (COG0203)rplQ (PRK05591)→ CPSase_sm_chain (smart01097)CPSase_sm_chain (pfam00988)→ |
| env-98 | CETY01154757.1 | - | 449 | 405 | RNA→ RpsD (COG0522)rpsD (PRK05327)→ |
| env-99 | JCVI_SCAF_1096627797848 | + | 643 | 687 | RNA→ rpsD (PRK05327)Ribosomal_S4 (pfam00163)→ S4 (smart00363)RpsD (COG0522)→ PRK05182 (PRK05182)RNA_pol_L (pfam01193)→ |
| env-100 | CERI01106383.1 | - | 1156 | 1112 | RNA→ RpsD (COG0522)rpsD (PRK05327)→ RNAP_alpha_NTD (cd06928)PRK05182 (PRK05182)→ |
| env-101 | JGI20156J14371_10258465 | - | 393 | 349 | RNA→ RpsD (COG0522)rpsD (PRK05327)→ |
| env-102 | JGI20156J14371_10000849 | - | 14189 | 14147 | RNA→ RpsD (COG0522)rpsD (PRK05327)→ RNAP_alpha_NTD (cd06928)PRK05182 (PRK05182)RNA_pol_A_CTD (pfam03118)→ RplQ (COG0203)rplQ (PRK05591)→ |
| env-103 | JCVI_SCAF_1101668015840 | - | 1336 | 1289 | RNA→ RpsD (COG0522)rpsD (PRK05327)→ RNAP_alpha_NTD (cd06928)PRK05182 (PRK05182)→ |
| env-104 | CESW01058310.1 | - | 375 | 328 | RNA→ RpsD (COG0522)rpsD (PRK05327)→ |
| env-105 | CEQI01167214.1 | + | 466 | 513 | RNA→ rpsD (PRK05327)Ribosomal_S4 (pfam00163)→ |
| env-106 | KGI_S2_ANT04_2345m_c1444089 | + | 66 | 115 | RNA→ rpsD (PRK05327)Ribosomal_S4 (pfam00163)→ |
| env-107 | KGI_S1_ANT01_95m_c10042760 | - | 553 | 504 | RNA→ RpsD (COG0522)rpsD (PRK05327)→ |
| env-108 | JGI24025J20009_10098185 | - | 82 | 33 | RNA→ |
| env-109 | SA_S1_NOR05_45m_c10522404 | - | 164 | 115 | RNA→ rpsD (PRK05327)Ribosomal_S4 (pfam00163)→ |
| env-110 | SA_S2_NOR15_50m_c10034879 | + | 1070 | 1119 | RNA→ RpsD (COG0522)rpsD (PRK05327)→ |

| | | | | | |
|---|---|---|---|---|---|
| env-111 | KGI_S1_ANT01_95m_c10014341 | - | 2364 | 2315 | RNA→ RpsD (COG0522)rpsD (PRK05327)→ RpoA (COG0202)PRK05182 (PRK05182)→ RplQ (COG0203)rplQ (PRK05591)→ |
| env-112 | SA_S1_NOR08_45m_c10025816 | - | 1459 | 1410 | RNA→ RpsD (COG0522)rpsD (PRK05327)→ |
| env-113 | JGI25923J51411_1187453 | - | 75 | 35 | RNA→ |
| env-114 | LWFCAn_GLQU6AQ01D6DBV | - | 91 | 51 | RNA→ |
| env-115 | CEUH01136830.1 | + | 421 | 461 | RNA→ RpsD (COG0522)rpsD (PRK05327)→ PRK05182 (PRK05182)RNA_pol_L (pfam01193)→ |
| env-116 | JGI24768J34885_10100695 | - | 432 | 392 | RNA→ S4 (smart00363)RpsD (COG0522)→ |
| env-117 | JGI25924J51412_1020305 | - | 684 | 644 | RNA→ RpsD (COG0522)rpsD (PRK05327)→ |
| env-118 | CEPL01069899.1 | - | 397 | 357 | RNA→ RpsD (COG0522)rpsD (PRK05327)→ |
| env-119 | CEUU01249842.1 | + | 1479 | 1519 | RNA→ S4 (smart00363)RpsD (COG0522)→ |
| env-120 | JGI12421J11937_10021494 | + | 1953 | 1993 | RNA→ S4 (smart00363)RpsD (COG0522)→ |
| env-121 | JGI24768J34885_10000210 | + | 12918 | 12958 | RNA→ RpsD (COG0522)rpsD (PRK05327)→ RNAP_alpha_NTD (cd06928)PRK05182 (PRK05182)RNA_pol_A_CTD (pfam03118)→ RplQ (COG0203)rplQ (PRK05591)→ |
| env-122 | JGI25914J50564_10010364 | - | 2244 | 2204 | RNA→ RpsD (COG0522)rpsD (PRK05327)→ RNAP_alpha_NTD (cd06928)PRK05182 (PRK05182)RNA_pol_A_CTD (pfam03118)→ RplQ (COG0203)rplQ (PRK05591)→ |
| env-123 | JGI12421J11937_10125481 | - | 368 | 328 | RNA→ rpsD (PRK05327)Ribosomal_S4 (pfam00163)→ |
| env-124 | JGI20156J14371_10002312 | + | 12798 | 12838 | RNA→ RpsD (COG0522)rpsD (PRK05327)→ RpoA (COG0202)PRK05182 (PRK05182)→ RplQ (COG0203)rplQ (PRK05591)→ |
| env-125 | M3P_c10141944 | + | 153 | 193 | RNA→ rpsD (PRK05327)Ribosomal_S4 (pfam00163)→ |
| env-126 | JGI25914J50564_10045055 | - | 818 | 778 | RNA→ RpsD (COG0522)rpsD (PRK05327)→ PRK05182 (PRK05182)RNA_pol_L (pfam01193)→ |
| env-127 | DelMOSum2010_c10006624 | - | 704 | 664 | RNA→ RpsD (COG0522)rpsD (PRK05327)→ |
| env-128 | JGI25920J50251_10092017 | - | 50 | 20 | RNA→ |
| env-129 | DelMOSum2010_c10002074 | - | 11710 | 11668 | RNA→ RpsD (COG0522)rpsD (PRK05327)→ RpoA (COG0202)PRK05182 (PRK05182)→ PLN02967 (PLN02967)RplQ (COG0203)rplQ (PRK05591)→ |
| env-130 | CESA01021238.1 | - | 2261 | 2219 | RNA→ RpsD (COG0522)rpsD (PRK05327)→ RpoA (COG0202)PRK05182 (PRK05182)→ RplQ (COG0203)rplQ (PRK05591)→ |
| env-131 | NCBI_BBAY_READ_1106105275911 | - | 208 | 155 | RNA→ |
| env-132 | CENG01041113.1 | + | 3621 | 3663 | RNA→ |
| env-133 | CESF01027428.1 | - | 816 | 774 | RNA→ RpsD (COG0522)rpsD (PRK05327)→ PRK05182 (PRK05182)RNA_pol_L (pfam01193)→ |
| env-134 | CEWG01153018.1 | + | 405 | 447 | RNA→ RpsD (COG0522)rpsD (PRK05327)→ |
| env-135 | CETE01270809.1 | - | 1047 | 1005 | RNA→ RpsD (COG0522)rpsD (PRK05327)→ rpoA (CHL00013)PRK05182 (PRK05182)→ |
| env-136 | CEUH01072115.1 | - | 421 | 379 | RNA→ S4 (smart00363)RpsD (COG0522)→ |
| env-137 | CESI01305241.1 | - | 2124 | 2082 | RNA→ RpsD (COG0522)rpsD (PRK05327)→ RpoA (COG0202)PRK05182 (PRK05182)→ RplQ (COG0203)rplQ (PRK05591)→ |
| env-138 | JCVI_SCAF_1101668716688 | + | 592 | 634 | RNA→ RpsD (COG0522)rpsD (PRK05327)→ |
| env-139 | 2236876001_320422 | + | 288 | 342 | RNA→ |
| env-140 | 2236876001_014240 | - | 85 | 44 | RNA→ |
| env-141 | 2236876001_374803 | + | 19 | 60 | RNA→ rpsD (PRK05327)Ribosomal_S4 (pfam00163)→ |
| env-142 | 2236876001_181177 | - | 254 | 213 | RNA→ rpsD (PRK05327)Ribosomal_S4 (pfam00163)→ |
| env-143 | JGI20156J14371_10000248 | + | 11432 | 11480 | RNA→ RpsD (COG0522)rpsD (PRK05327)→ RpoA (COG0202)PRK05182 (PRK05182)→ RplQ (COG0203)rplQ (PRK05591)→ |
| env-144 | JGI20154J14316_10020138 | - | 582 | 534 | RNA→ RpsD (COG0522)rpsD (PRK05327)→ |
| env-145 | JGI20157J14317_10238135 | - | 207 | 166 | RNA→ rpsD (PRK05327)Ribosomal_S4 (pfam00163)→ |
| env-146 | ElkS_mat_MD6A_1006090 | + | 1597 | 1645 | RNA→ RpsD (COG0522)rpsD (PRK05327)→ |
| env-147 | DelMOSum2010_c10001012 | + | 14798 | 14841 | RNA→ RpsD (COG0522)rpsD (PRK05327)→ RNAP_alpha_NTD (cd06928)PRK05182 (PRK05182)RNA_pol_A_CTD (pfam03118)→ RplQ (COG0203)rplQ (PRK05591)→ GATase1_CPSase (cd01744)PRK12564 (PRK12564)CPSase_sm_chain (pfam00988)→ eno (PRK00077)enolase (cd03313)→ |
| env-148 | JGI20159J14440_10208187 | - | 331 | 288 | RNA→ rpsD (PRK05327)Ribosomal_S4 (pfam00163)→ |

| | | | | | |
|---|---|---|---|---|---|
| env-149 | JGI20160J14292_10003529 | - | 10322 | 10279 | **RNA**→ RpsD (COG0522)rpsD (PRK05327)→ |
| | | | | | RNAP_alpha_NTD (cd06928)PRK05182 (PRK05182)RNA_pol_A_CTD (pfam03118)→ |
| | | | | | RplQ (COG0203)rplQ (PRK05591)→ |
| | | | | | GATase1_CPSase (cd01744)PRK12564 (PRK12564)CPSase_sm_chain (pfam00988)→ |
| | | | | | eno (PRK00077)enolase (cd03313)→ |
| env-150 | JGI20156J14371_10005217 | - | 7554 | 7511 | **RNA**→ RpsD (COG0522)rpsD (PRK05327)→ |
| | | | | | RNAP_alpha_NTD (cd06928)PRK05182 (PRK05182)RNA_pol_A_CTD (pfam03118)→ |
| | | | | | RplQ (COG0203)rplQ (PRK05591)→ |
| | | | | | GATase1_CPSase (cd01744)PRK12564 (PRK12564)CPSase_sm_chain (pfam00988)→ |
| | | | | | eno (PRK00077)enolase (cd03313)→ |
| env-151 | JGI20160J14292_10000862 | - | 24569 | 24526 | **RNA**→ RpsD (COG0522)rpsD (PRK05327)→ RpoA (COG0202)PRK05182 (PRK05182)→ |
| | | | | | RplQ (COG0203)rplQ (PRK05591)→ CarA (COG0505)PRK12564 (PRK12564)→ |
| | | | | | eno (PRK00077)enolase (cd03313)→ gltA (PRK05614)EcCS_like (cd06114)→ |
| env-152 | JGI20152J14361_10001272 | + | 11027 | 11070 | **RNA**→ RpsD (COG0522)rpsD (PRK05327)→ PRK05182 (PRK05182)RNA_pol_L (pfam01193)→ |
| env-153 | 2236876001_447696 | - | 404 | 361 | **RNA**→ S4 (smart00363)RpsD (COG0522)→ |
| env-154 | NCBI_READ_1112533167397 | - | 766 | 723 | **RNA**→ RpsD (COG0522)rpsD (PRK05327)→ |
| env-155 | SI34jun09_10m_1127658 | + | 229 | 272 | **RNA**→ ←hypo |
| env-156 | JGI20160J14292_10182352 | + | 404 | 447 | **RNA**→ rpsD (PRK05327)Ribosomal_S4 (pfam00163)→ |
| env-157 | CESJ01259508.1 | - | 513 | 470 | **RNA**→ RpsD (COG0522)rpsD (PRK05327)→ |
| env-158 | 2236876001_073076 | + | 29 | 72 | **RNA**→ rpsD (PRK05327)Ribosomal_S4 (pfam00163)→ |
| env-159 | JGI20154J14316_10148802 | + | 420 | 463 | **RNA**→ rpsD (PRK05327)Ribosomal_S4 (pfam00163)→ |
| env-160 | SA_S1_NOR08_45m_c10566989 | + | 97 | 141 | **RNA**→ rpsD (PRK05327)Ribosomal_S4 (pfam00163)→ |
| Lsp-2-1 | NZ_JHZY01000004.1 | - | 470876 | 470832 | **RNA**→ RpsD (COG0522)rpsD (PRK05327)→ RpoA (COG0202)PRK05182 (PRK05182)→ |
| | | | | | RplQ (COG0203)rplQ (PRK05591)→ |
| env-161 | CEVO01222084.1 | + | 846 | 880 | **RNA**→ |
| env-162 | CESL01055523.1 | + | 471 | 515 | **RNA**→ RpsD (COG0522)rpsD (PRK05327)→ RpoA (COG0202)PRK05182 (PRK05182)→ |
| | | | | | RplQ (COG0203)rplQ (PRK05591)→ |
| env-163 | CEVJ01029531.1 | + | 846 | 890 | **RNA**→ rpsD (PRK05327)Ribosomal_S4 (pfam00163)→ |
| env-164 | CETR01135856.1 | + | 845 | 889 | **RNA**→ RpsD (COG0522)rpsD (PRK05327)→ RpoA (COG0202)PRK05182 (PRK05182)→ |
| | | | | | RplQ (COG0203)rplQ (PRK05591)→ |
| env-165 | CEOS01000692.1 | - | 8747 | 8703 | **RNA**→ RpsD (COG0522)rpsD (PRK05327)→ RpoA (COG0202)PRK05182 (PRK05182)→ |
| | | | | | RplQ (COG0203)rplQ (PRK05591)→ hypo→ CarA (COG0505)PRK12564 (PRK12564)→ |
| env-166 | 2236876011_134002 | + | 294 | 326 | **RNA**→ |
| env-167 | JGI20154J14316_10001276 | - | 5032 | 4990 | **RNA**→ RpsD (COG0522)rpsD (PRK05327)→ |
| | | | | | RNAP_alpha_NTD (cd06928)PRK05182 (PRK05182)RNA_pol_A_CTD (pfam03118)→ |
| env-168 | 2231695257 | - | 173 | 131 | **RNA**→ rpsD (PRK05327)Ribosomal_S4 (pfam00163)→ |
| env-169 | JGI25910J50241_10205082 | - | 41 | 9 | **RNA**→ |
| Fba-2-1 | NZ_AAXX01000001.1 | - | 580267 | 580223 | **RNA**→ RpsD (COG0522)rpsD (PRK05327)→ RpsD (COG0522)rpsD (PRK05327)→ |
| | | | | | RpoA (COG0202)PRK05182 (PRK05182)→ RpoA (COG0202)PRK05182 (PRK05182)→ |
| | | | | | RplQ (COG0203)rplQ (PRK05591)→ RplQ (COG0203)rplQ (PRK05591)→ |
| env-170 | M3P_c10252825 | + | 117 | 160 | **RNA**→ rpsD (PRK05327)Ribosomal_S4 (pfam00163)→ |
| Fsa-1-1 | NZ_JMLU01000009.1 | - | 18897 | 18854 | **RNA**→ RpsD (COG0522)rpsD (PRK05327)→ RpoA (COG0202)PRK05182 (PRK05182)→ |
| | | | | | RplQ (COG0203)rplQ (PRK05591)→ |
| env-171 | JGI25921J50272_10187943 | + | 186 | 229 | **RNA**→ rpsD (PRK05327)Ribosomal_S4 (pfam00163)→ |
| env-172 | 3300000052_174905 | + | 114 | 157 | **RNA**→ |
| env-173 | 3300001592_10384027 | + | 364 | 407 | **RNA**→ rpsD (PRK05327)Ribosomal_S4 (pfam00163)→ |
| env-174 | TB_FS06_10DRAFT_1000051 | - | 30943 | 30900 | **RNA**→ RpsD (COG0522)rpsD (PRK05327)→ |
| | | | | | RNAP_alpha_NTD (cd06928)PRK05182 (PRK05182)RNA_pol_A_CTD (pfam03118)→ |
| | | | | | RplQ (COG0203)rplQ (PRK05591)→ |
| env-175 | 3300001592_10325872 | + | 220 | 263 | **RNA**→ rpsD (PRK05327)Ribosomal_S4 (pfam00163)→ |
| env-176 | CETR01093469.1 | + | 1000 | 1048 | **RNA**→ RpsD (COG0522)rpsD (PRK05327)→ RpoA (COG0202)PRK05182 (PRK05182)→ |
| | | | | | RplQ (COG0203)rplQ (PRK05591)→ |
| env-177 | CEVJ01168016.1 | - | 208 | 160 | **RNA**→ rpsD (PRK05327)Ribosomal_S4 (pfam00163)→ |
| env-178 | CETA01172228.1 | - | 195 | 151 | **RNA**→ rpsD (PRK05327)Ribosomal_S4 (pfam00163)→ |
| env-179 | CEVI01047151.1 | + | 2582 | 2626 | **RNA**→ RpsD (COG0522)rpsD (PRK05327)→ RpoA (COG0202)PRK05182 (PRK05182)→ |
| | | | | | RplQ (COG0203)rplQ (PRK05591)→ |

| | | | | | |
|---|---|---|---|---|---|
| env-180 | JGI24815J26687_1079477 | + | 331 | 375 | RNA→ rpsD (PRK05327)Ribosomal_S4 (pfam00163)→ |
| env-181 | CEUC01056669.1 | + | 895 | 939 | RNA→ rpsD (PRK05327)Ribosomal_S4 (pfam00163)→ |
| env-182 | CESR01033026.1 | + | 887 | 931 | RNA→ RpsD (COG0522)rpsD (PRK05327)→ |
| env-183 | CETO01024088.1 | - | 1009 | 965 | RNA→ RpsD (COG0522)rpsD (PRK05327)→ rpoA (CHL00013)PRK05182 (PRK05182)→ |
| env-184 | CEUS01122815.1 | - | 657 | 613 | RNA→ RpsD (COG0522)rpsD (PRK05327)→ |
| env-185 | CEWR01047726.1 | - | 680 | 636 | RNA→ RpsD (COG0522)rpsD (PRK05327)→ |
| env-186 | CETS01039292.1 | - | 1863 | 1819 | RNA→ RpsD (COG0522)rpsD (PRK05327)→ RpoA (COG0202)PRK05182 (PRK05182)→ RplQ (COG0203)rplQ (PRK05591)→ |
| env-187 | CEVM01006958.1 | - | 238 | 194 | RNA→ rpsD (PRK05327)Ribosomal_S4 (pfam00163)→ |
| env-188 | SI54feb11_120m_c1042447 | - | 137 | 93 | RNA→ rpsD (PRK05327)Ribosomal_S4 (pfam00163)→ |
| env-189 | NCBI_BBAY_READ_1106073052947 | - | 860 | 816 | RNA→ RpsD (COG0522)rpsD (PRK05327)→ |
| env-190 | CENT01023057.1 | - | 131 | 87 | RNA→ rpsD (PRK05327)Ribosomal_S4 (pfam00163)→ |
| env-191 | CEVG01299689.1 | + | 820 | 854 | RNA→ |
| env-192 | JCVI_SCAF_1096627857065 | - | 1611 | 1567 | RNA→ RpsD (COG0522)rpsD (PRK05327)→ |
| env-193 | CERI01112825.1 | - | 699 | 655 | RNA→ RpsD (COG0522)rpsD (PRK05327)→ |
| env-194 | CEWG01256266.1 | - | 432 | 388 | RNA→ RpsD (COG0522)rpsD (PRK05327)→ |
| env-195 | CENG01111780.1 | + | 9651 | 9695 | RNA→ RpsD (COG0522)rpsD (PRK05327)→ |
| env-196 | CENG01073265.1 | + | 732 | 776 | RNA→ RpsD (COG0522)rpsD (PRK05327)→ RNAP_alpha_NTD (cd06928)PRK05182 (PRK05182)→ |
| env-197 | CETY01269978.1 | + | 203 | 247 | RNA→ RpsD (COG0522)rpsD (PRK05327)→ |
| env-198 | CEWG01042362.1 | + | 503 | 547 | RNA→ rpsD (PRK05327)Ribosomal_S4 (pfam00163)→ |
| env-199 | JCVI_SCAF_1096627052264 | - | 632 | 588 | RNA→ RpsD (COG0522)rpsD (PRK05327)→ |
| env-200 | CETK01070832.1 | + | 5009 | 5053 | RNA→ RpsD (COG0522)rpsD (PRK05327)→ RNAP_alpha_NTD (cd06928)PRK05182 (PRK05182)RNA_pol_A_CTD (pfam03118)→ RplQ (COG0203)rplQ (PRK05591)→ CarA (COG0505)PRK12564 (PRK12564)→ eno (PRK00077)enolase (cd03313)→ gltA (PRK05614)EcCS_like (cd06114)→ |
| env-201 | JCVI_SCAF_1101667257495 | - | 457 | 413 | RNA→ RpsD (COG0522)rpsD (PRK05327)→ |
| env-202 | JGI20157J14317_10000316 | - | 23037 | 22993 | RNA→ RpsD (COG0522)rpsD (PRK05327)→ RNAP_alpha_NTD (cd06928)PRK05182 (PRK05182)RNA_pol_A_CTD (pfam03118)→ RplQ (COG0203)rplQ (PRK05591)→ CarA (COG0505)PRK12564 (PRK12564)→ eno (PRK00077)enolase (cd03313)→ gltA (PRK05614)EcCS_like (cd06114)→ |
| env-203 | JGI20156J14371_10020545 | - | 289 | 239 | RNA→ rpsD (PRK05327)Ribosomal_S4 (pfam00163)→ |
| env-204 | JCVI_SCAF_1096626856226 | + | 1215 | 1265 | RNA→ RpsD (COG0522)rpsD (PRK05327)→ RpoA (COG0202)PRK05182 (PRK05182)→ |
| env-205 | JGI20159J14440_10001069 | - | 2266 | 2216 | RNA→ RpsD (COG0522)rpsD (PRK05327)→ RNAP_alpha_NTD (cd06928)PRK05182 (PRK05182)RNA_pol_A_CTD (pfam03118)→ RplQ (COG0203)rplQ (PRK05591)→ |
| env-206 | DelMOSum2010_c10144771 | + | 169 | 218 | RNA→ RpsD (COG0522)rpsD (PRK05327)→ |
| env-207 | SI48aug10_10m_c1005592 | - | 1044 | 995 | RNA→ RpsD (COG0522)rpsD (PRK05327)→ RpoA (COG0202)PRK05182 (PRK05182)→ |
| env-208 | NCBI_BBAY_READ_1106073039522 | - | 543 | 494 | RNA→ RpsD (COG0522)rpsD (PRK05327)→ |
| env-209 | NCBI_READ_1112493604101 | + | 450 | 499 | RNA→ RpsD (COG0522)rpsD (PRK05327)→ |
| env-210 | DelMOSum2010_c10000401 | - | 31221 | 31172 | RNA→ RpsD (COG0522)rpsD (PRK05327)→ RNAP_alpha_NTD (cd06928)PRK05182 (PRK05182)RNA_pol_A_CTD (pfam03118)→ RplQ (COG0203)rplQ (PRK05591)rne (PRK10811)→ hypo→ |
| env-211 | JGI20152J14361_10035130 | - | 700 | 651 | RNA→ RpsD (COG0522)rpsD (PRK05327)→ |
| env-212 | CEWO01191481.1 | + | 536 | 585 | RNA→ RpsD (COG0522)rpsD (PRK05327)→ |
| env-213 | JGI20154J14316_10392850 | - | 59 | 20 | RNA→ |
| env-214 | JGI20160J14292_10012841 | - | 839 | 790 | RNA→ RpsD (COG0522)rpsD (PRK05327)→ PRK05182 (PRK05182)RNA_pol_L (pfam01193)→ |
| Jpa-1-1 | NZ_BBNS01000014.1 | + | 75674 | 75722 | RNA→ RpsD (COG0522)rpsD (PRK05327)→ RpoA (COG0202)PRK05182 (PRK05182)→ |
| env-215 | ElkS_mat_MD6A_1048345 | - | 175 | 127 | RNA→ rpsD (PRK05327)Ribosomal_S4 (pfam00163)→ |
| env-216 | ElkS_mat_MD6A_1011575 | - | 1844 | 1796 | RNA→ RpsD (COG0522)rpsD (PRK05327)→ |
| env-217 | ElkS_mat_MD6A_1054401 | - | 382 | 334 | RNA→ RpsD (COG0522)rpsD (PRK05327)→ |
| env-218 | JGI20160J14292_10190844 | + | 186 | 227 | RNA→ S4 (smart00363)RpsD (COG0522)→ |
| env-219 | JGI12547J11936_1003731 | + | 2052 | 2092 | RNA→ RpsD (COG0522)rpsD (PRK05327)→ RpoA (COG0202)PRK05182 (PRK05182)→ RplQ (COG0203)rplQ (PRK05591)→ |
| env-220 | JGI12421J11937_10050354 | + | 1249 | 1289 | RNA→ ←hypo |

| | | | | | |
|---|---|---|---|---|---|
| env-221 | Keratinized_gingiva_LANL_C5716438 | - | 329 | 287 | **RNA**→ rpsD (PRK05327)Ribosomal_S4 (pfam00163)→ |
| env-222 | SRS017691_Baylor_scaffold_68 | + | 1241 | 1283 | **RNA**→ RpsD (COG0522)rpsD (PRK05327)→ RNAP_alpha_NTD (cd06928)PRK05182 (PRK05182)→ |
| env-223 | SRS018443_Baylor_scaffold_6145 | + | 2897 | 2939 | **RNA**→ rpsD (PRK05327)Ribosomal_S4 (pfam00163)→ |
| env-224 | Palatine_Tonsils_LANL_C6151646 | - | 152 | 110 | **RNA**→ rpsD (PRK05327)Ribosomal_S4 (pfam00163)→ |
| env-225 | SRS012285_Baylor_scaffold_4163 | + | 558 | 600 | **RNA**→ RpsD (COG0522)rpsD (PRK05327)→ |
| | | | | | RNAP_alpha_NTD (cd06928)PRK05182 (PRK05182)RNA_pol_A_CTD (pfam03118)→ |
| | | | | | RplQ (COG0203)rplQ (PRK05591)→ |
| env-226 | SRS016331_WUGC_scaffold_136 | - | 738 | 696 | **RNA**→ RpsD (COG0522)rpsD (PRK05327)→ |
| env-227 | SRS011343_Baylor_scaffold_55324 | - | 267 | 225 | **RNA**→ rpsD (PRK05327)Ribosomal_S4 (pfam00163)→ |
| env-228 | SRS011255_Baylor_scaffold_65092 | + | 718 | 760 | **RNA**→ RpsD (COG0522)rpsD (PRK05327)→ |
| env-229 | SRS022602_Baylor_scaffold_81026 | - | 257 | 216 | **RNA**→ rpsD (PRK05327)Ribosomal_S4 (pfam00163)→ |
| env-230 | SRS047634_LANL_scaffold_130560 | + | 829 | 870 | **RNA**→ RpsD (COG0522)rpsD (PRK05327)→ |
| env-231 | SRS052876_LANL_scaffold_39269 | + | 1190 | 1231 | **RNA**→ |
| env-232 | SRS013836_Baylor_scaffold_28602 | - | 487 | 446 | **RNA**→ RpsD (COG0522)rpsD (PRK05327)→ |
| Csp-1-1 | NZ_AKFR01000015.1 | - | 81600 | 81559 | **RNA**→ RpsD (COG0522)rpsD (PRK05327)→ |
| env-233 | SRS014888_WUGC_scaffold_19943 | + | 291 | 332 | **RNA**→ RpsD (COG0522)rpsD (PRK05327)→ |
| env-234 | SRS013723_Baylor_scaffold_19316 | + | 2562 | 2603 | **RNA**→ |
| env-235 | SRS017511_Baylor_scaffold_85167 | + | 438 | 479 | **RNA**→ |
| env-236 | SRS017209_C2451925 | - | 303 | 262 | **RNA**→ rpsD (PRK05327)Ribosomal_S4 (pfam00163)→ |
| env-237 | Throat_LANL_scaffold_30529 | + | 70 | 111 | **RNA**→ rpsD (PRK05327)Ribosomal_S4 (pfam00163)→ |
| env-238 | SRS014573_C3620607 | - | 364 | 323 | **RNA**→ RpsD (COG0522)rpsD (PRK05327)→ |
| env-239 | SRS015989_WUGC_scaffold_3203 | + | 436 | 477 | **RNA**→ rpsD (PRK05327)Ribosomal_S4 (pfam00163)→ |
| env-240 | SRS051244_LANL_scaffold_57202 | + | 436 | 477 | **RNA**→ |
| env-241 | SRS024381_LANL_scaffold_48261 | + | 437 | 478 | **RNA**→ RpsD (COG0522)rpsD (PRK05327)→ |
| env-242 | SRS053630_LANL_scaffold_15954 | + | 1148 | 1189 | **RNA**→ RpsD (COG0522)rpsD (PRK05327)→ |
| | | | | | RNAP_alpha_NTD (cd06928)PRK05182 (PRK05182)RNA_pol_A_CTD (pfam03118)→ |
| | | | | | RplQ (COG0203)rplQ (PRK05591)→ |
| env-243 | Buccal_mucosa_LANL_scaffold_33054 | + | 15 | 56 | **RNA**→ rpsD (PRK05327)Ribosomal_S4 (pfam00163)→ RpsD (COG0522)rpsD (PRK05327)→ |
| | | | | | RNAP_alpha_NTD (cd06928)PRK05182 (PRK05182)→ |
| env-244 | SRS022083_Baylor_scaffold_14787 | + | 71 | 112 | **RNA**→ RpsD (COG0522)rpsD (PRK05327)→ |
| env-245 | SRS022536_LANL_scaffold_45737 | + | 394 | 435 | **RNA**→ |
| env-246 | SRS049318_LANL_scaffold_22062 | - | 640 | 599 | **RNA**→ rpsD (PRK05327)Ribosomal_S4 (pfam00163)→ S4 (smart00363)RpsD (COG0522)→ rpsD (PRK05327)– |
| env-247 | SRS011343_Baylor_scaffold_68646 | + | 588 | 629 | **RNA**→ |
| env-248 | TDF_OR_ARG05_123m_c1010977 | - | 701 | 661 | **RNA**→ RpsD (COG0522)rpsD (PRK05327)→ |
| env-249 | SA_S1_NOR08_45m_c10064607 | - | 790 | 750 | **RNA**→ RpsD (COG0522)rpsD (PRK05327)→ RNAP_alpha_NTD (cd06928)PRK05182 (PRK05182)→ |
| env-250 | TDF_MC_ARG01_113m_c1000650 | + | 230 | 270 | **RNA**→ RpsD (COG0522)rpsD (PRK05327)→ |
| | | | | | RNAP_alpha_NTD (cd06928)PRK05182 (PRK05182)RNA_pol_A_CTD (pfam03118)→ |
| | | | | | RplQ (COG0203)rplQ (PRK05591)→ |
| env-251 | SA_S2_NOR13_50m_c1156445 | - | 121 | 81 | **RNA**→ rpsD (PRK05327)Ribosomal_S4 (pfam00163)→ |
| env-252 | TDF_OR_ARG04_113m_c1025301 | + | 530 | 570 | **RNA**→ rpsD (PRK05327)Ribosomal_S4 (pfam00163)→ |
| env-253 | JGI24723J26617_10208337 | - | 127 | 87 | **RNA**→ |
| env-254 | TDF_OR_ARG04_113m_c1103690 | - | 163 | 123 | **RNA**→ rpsD (PRK05327)Ribosomal_S4 (pfam00163)→ |
| env-255 | BS_KBA_SWE12_21m_c10302711 | - | 190 | 150 | **RNA**→ rpsD (PRK05327)Ribosomal_S4 (pfam00163)→ |
| env-256 | JGI24723J26617_10789467 | + | 66 | 106 | **RNA**→ rpsD (PRK05327)Ribosomal_S4 (pfam00163)→ |
| env-257 | KGI_S2_ANT05_2345m_c1010433 | - | 779 | 739 | **RNA**→ RpsD (COG0522)rpsD (PRK05327)→ PRK05182 (PRK05182)→ |
| env-258 | KGI_S1_ANT01_95m_c10010832 | + | 492 | 532 | **RNA**→ rpsD (PRK05327)Ribosomal_S4 (pfam00163)→ |
| env-259 | SA_S2_NOR13_50m_c1000962 | + | 5215 | 5255 | **RNA**→ RpsD (COG0522)rpsD (PRK05327)→ rpoA (CHL00013)PRK05182 (PRK05182)→ |
| env-260 | KGI_S1_ANT01_95m_c10110441 | + | 179 | 219 | **RNA**→ RpsD (COG0522)rpsD (PRK05327)→ |
| env-261 | KGI_S1_ANT01_95m_c10085982 | + | 480 | 520 | **RNA**→ RpsD (COG0522)rpsD (PRK05327)→ |
| env-262 | TB_FS06_10DRAFT_1000211 | + | 18911 | 18956 | **RNA**→ RpsD (COG0522)rpsD (PRK05327)→ RpoA (COG0202)PRK05182 (PRK05182)→ |
| | | | | | RplQ (COG0203)rplQ (PRK05591)→ |
| env-263 | DelMOSum2010_c10125148 | - | 972 | 931 | **RNA**→ RpsD (COG0522)rpsD (PRK05327)→ PRK05182 (PRK05182)RNA_pol_L (pfam01193)→ |

| | | | | | |
|---|---|---|---|---|---|
| env-264 | JGI20157J14317_10000116 | + | 22436 | 22477 | **RNA**→ RpsD (COG0522)rpsD (PRK05327)→ RNAP_alpha_NTD (cd06928)PRK05182 (PRK05182)RNA_pol_A_CTD (pfam03118)→ RplQ (COG0203)rplQ (PRK05591)→ CarA (COG0505)PRK12564 (PRK12564)→ eno (PRK00077)enolase (cd03313)→ |
| env-265 | JGI20159J14440_10000121 | + | 30004 | 30045 | **RNA**→ RpsD (COG0522)rpsD (PRK05327)→ RNAP_alpha_NTD (cd06928)PRK05182 (PRK05182)RNA_pol_A_CTD (pfam03118)→ RplQ (COG0203)rplQ (PRK05591)→ CarA (COG0505)PRK12564 (PRK12564)→ eno (PRK00077)enolase (cd03313)→ |
| env-266 | CESE01008383.1 | + | 79060 | 79101 | **RNA**→ RpsD (COG0522)rpsD (PRK05327)→ RNAP_alpha_NTD (cd06928)PRK05182 (PRK05182)RNA_pol_A_CTD (pfam03118)→ RplQ (COG0203)rplQ (PRK05591)→ CarA (COG0505)PRK12564 (PRK12564)→ eno (PRK00077)enolase (cd03313)→ |
| env-267 | JGI20152J14361_10000300 | + | 3945 | 3986 | **RNA**→ RpsD (COG0522)rpsD (PRK05327)→ RNAP_alpha_NTD (cd06928)PRK05182 (PRK05182)RNA_pol_A_CTD (pfam03118)→ RplQ (COG0203)rplQ (PRK05591)→ CarA (COG0505)PRK12564 (PRK12564)→ eno (PRK00077)enolase (cd03313)→ |
| env-268 | DelMOSum2011_c10467853 | + | 40 | 81 | **RNA**→ rpsD (PRK05327)Ribosomal_S4 (pfam00163)→ |
| env-269 | JGI20156J14371_10000212 | - | 49678 | 49637 | **RNA**→ RpsD (COG0522)rpsD (PRK05327)→ RNAP_alpha_NTD (cd06928)PRK05182 (PRK05182)RNA_pol_A_CTD (pfam03118)→ RplQ (COG0203)rplQ (PRK05591)→ CarA (COG0505)PRK12564 (PRK12564)→ eno (PRK00077)enolase (cd03313)→ |
| env-270 | CETE01228191.1 | - | 644 | 603 | **RNA**→ RpsD (COG0522)rpsD (PRK05327)→ |
| env-271 | CEVG01151379.1 | + | 315 | 356 | **RNA**→ RpsD (COG0522)rpsD (PRK05327)→ |
| env-272 | CENO01018210.1 | - | 657 | 616 | **RNA**→ RpsD (COG0522)rpsD (PRK05327)→ |
| env-273 | JCVI_SCAF_1096627080427 | + | 1062 | 1103 | **RNA**→ RpsD (COG0522)rpsD (PRK05327)→ rpoA (CHL00013)PRK05182 (PRK05182)→ |
| env-274 | CEUH01222072.1 | + | 246 | 287 | **RNA**→ RpsD (COG0522)rpsD (PRK05327)→ |
| env-275 | SA_S1_NOR08_45m_c10087543 | - | 795 | 753 | **RNA**→ RpsD (COG0522)rpsD (PRK05327)→ hypo→ |
| env-276 | JGI20156J14371_10002845 | + | 7598 | 7640 | **RNA**→ RpsD (COG0522)rpsD (PRK05327)→ RNAP_alpha_NTD (cd06928)PRK05182 (PRK05182)RNA_pol_A_CTD (pfam03118)→ RplQ (COG0203)rplQ (PRK05591)→ |
| env-277 | TB_GS09_5_10006339 | + | 7277 | 7320 | **RNA**→ RpsD (COG0522)rpsD (PRK05327)→ RNAP_alpha_NTD (cd06928)PRK05182 (PRK05182)RNA_pol_A_CTD (pfam03118)→ RplQ (COG0203)rplQ (PRK05591)→ |
| env-278 | TB_FS06_10DRAFT_1000970 | - | 7745 | 7703 | **RNA**→ RpsD (COG0522)rpsD (PRK05327)→ RNAP_alpha_NTD (cd06928)PRK05182 (PRK05182)RNA_pol_A_CTD (pfam03118)→ RplQ (COG0203)rplQ (PRK05591)→ hypo→ AmyAc_arch_bac_AmyA (cd11313)treS_nterm (TIGR02456)→ TIGR00275 (TIGR00275)HI0933_like (pfam03486)→ TIGR00275 (TIGR00275)HI0933_like (pfam03486)→ |
| env-279 | wf2_AHAA19916_y1 | - | 435 | 392 | **RNA**→ S4 (smart00363)RpsD (COG0522)→ |
| env-280 | JGI20152J14361_10000009 | - | 112324 | 112282 | **RNA**→ RpsD (COG0522)rpsD (PRK05327)→ |
| env-281 | GSLSAAI_GFP0ZZ302IYKTC | - | 96 | 54 | **RNA**→ ←hypo |
| env-282 | SL_9KL_010_SED_10016857 | + | 55 | 99 | **RNA**→ RpsD (COG0522)rpsD (PRK05327)→ RpoA (COG0202)PRK05182 (PRK05182)→ RplQ (COG0203)rplQ (PRK05591)→ |
| env-283 | P_1C_Liq_1_UnCty_1074733 | - | 40 | 6 | **RNA**→ |
| env-284 | JGI11876J14442_10000071 | - | 34341 | 34297 | **RNA**→ RpsD (COG0522)rpsD (PRK05327)→ RpoA (COG0202)PRK05182 (PRK05182)→ rplQ (PRK05591)Ribosomal_L17 (pfam01196)→ CarA (COG0505)PRK12564 (PRK12564)→ |
| Pgo-1-1 | NZ_APLF01000014.1 | - | 34253 | 34209 | **RNA**→ RpsD (COG0522)rpsD (PRK05327)→ RpoA (COG0202)PRK05182 (PRK05182)→ RplQ (COG0203)rplQ (PRK05591)→ rplQ (PRK05591)Ribosomal_L17 (pfam01196)→ CarA (COG0505)PRK12564 (PRK12564)→ |
| env-285 | P_2C_Liq_1_UnCty_1067107 | + | 546 | 578 | **RNA**→ |
| env-286 | ElkS_mat_CD2A_1181468 | - | 226 | 182 | **RNA**→ rpsD (PRK05327)Ribosomal_S4 (pfam00163)→ |
| env-287 | P_1C_Liq_1_UnCty_1015490 | + | 448 | 492 | **RNA**→ RpsD (COG0522)rpsD (PRK05327)→ |
| env-288 | JGI1221J11331_1000639 | + | 5910 | 5954 | **RNA**→ RpsD (COG0522)rpsD (PRK05327)→ RpoA (COG0202)PRK05182 (PRK05182)→ RplQ (COG0203)rplQ (PRK05591)→ CPSase_sm_chain (smart01097)CPSase_sm_chain (pfam00988)→ PRK12564 (PRK12564)GATase (pfam00117)→ |
| Pto-1-1 | NC_018721.1 | - | 1068584 | 1068540 | **RNA**→ RpsD (COG0522)rpsD (PRK05327)→ RpoA (COG0202)PRK05182 (PRK05182)→ RplQ (COG0203)rplQ (PRK05591)→ CarA (COG0505)PRK12564 (PRK12564)→ |

| | | | | | |
|---|---|---|---|---|---|
| Ptr-1-1 | NZ_KB905346.1 | + | 104039 | 104083 | RNA→ RpsD (COG0522)rpsD (PRK05327)→ RpoA (COG0202)PRK05182 (PRK05182)→ RplQ (COG0203)rplQ (PRK05591)→ CarA (COG0505)PRK12564 (PRK12564)→ |
| env-289 | SA_S1_NOR05_45m_c10375136 | + | 157 | 201 | RNA→ rpsD (PRK05327)Ribosomal_S4 (pfam00163)→ |
| env-290 | SA_S2_NOR15_50m_c10192611 | + | 393 | 437 | RNA→ rpsD (PRK05327)Ribosomal_S4 (pfam00163)→ |
| env-291 | SA_S1_NOR05_45m_c10080223 | + | 427 | 471 | RNA→ rpsD (PRK05327)Ribosomal_S4 (pfam00163)→ |
| env-292 | SL_9KL_010_SED_10000028 | - | 31849 | 31807 | RNA→ RpsD (COG0522)rpsD (PRK05327)→ RNAP_alpha_NTD (cd06928)PRK05182 (PRK05182)RNA_pol_A_CTD (pfam03118)→ RplQ (COG0203)rplQ (PRK05591)→ CarA (COG0505)PRK12564 (PRK12564)→ |
| env-293 | SL_7KL_010_BRINE_10000083 | + | 29384 | 29426 | RNA→ RpsD (COG0522)rpsD (PRK05327)→ RNAP_alpha_NTD (cd06928)PRK05182 (PRK05182)RNA_pol_A_CTD (pfam03118)→ RplQ (COG0203)rplQ (PRK05591)→ GATase1_CPSase (cd01744)PRK12564 (PRK12564)CPSase_sm_chain (pfam00988)→ |
| Rbi-1-1 | NC_013222.1 | + | 1475311 | 1475353 | RNA→ RpsD (COG0522)rpsD (PRK05327)→ RNAP_alpha_NTD (cd06928)PRK05182 (PRK05182)RNA_pol_A_CTD (pfam03118)→ RplQ (COG0203)rplQ (PRK05591)→ |
| Gfo-1-1 | NC_008571.1 | - | 2932356 | 2932313 | RNA→ RpsD (COG0522)rpsD (PRK05327)→ RpoA (COG0202)PRK05182 (PRK05182)→ RplQ (COG0203)rplQ (PRK05591)→ |
| Gpo-1-1 | NZ_KE383971.1 | + | 95062 | 95105 | RNA→ RpsD (COG0522)rpsD (PRK05327)→ RpoA (COG0202)PRK05182 (PRK05182)→ RplQ (COG0203)rplQ (PRK05591)→ |
| Gec-1-1 | NZ_AUHG01000011.1 | + | 90775 | 90818 | RNA→ RpsD (COG0522)rpsD (PRK05327)→ RpoA (COG0202)PRK05182 (PRK05182)→ RplQ (COG0203)rplQ (PRK05591)→ hypo→ CarA (COG0505)PRK12564 (PRK12564)→ |
| env-294 | CENZ01218708.1 | - | 288 | 245 | RNA→ rpsD (PRK05327)Ribosomal_S4 (pfam00163)→ |
| env-295 | CESL01104349.1 | + | 2289 | 2332 | RNA→ RpsD (COG0522)rpsD (PRK05327)→ RpoA (COG0202)PRK05182 (PRK05182)→ |
| Fjo-1-1 | NC_009441.1 | - | 396087 | 396045 | RNA→ RpsD (COG0522)rpsD (PRK05327)→ RNAP_alpha_NTD (cd06928)PRK05182 (PRK05182)RNA_pol_A_CTD (pfam03118)→ RplQ (COG0203)rplQ (PRK05591)→ |
| env-296 | KGI_S1_ANT02_95m_c10161500 | - | 420 | 371 | RNA→ S4 (smart00363)RpsD (COG0522)→ |
| env-297 | JGI24023J19991_10010536 | + | 3459 | 3508 | RNA→ RpsD (COG0522)rpsD (PRK05327)→ RpoA (COG0202)PRK05182 (PRK05182)→ |
| env-298 | 2236876001_453531 | - | 209 | 161 | RNA→ rpsD (PRK05327)Ribosomal_S4 (pfam00163)→ |
| env-299 | DelMOSum2010_c10108153 | - | 712 | 664 | RNA→ RpsD (COG0522)rpsD (PRK05327)→ |
| env-300 | JGI20154J14316_10182624 | - | 93 | 45 | RNA→ |
| env-301 | JGI20154J14316_10239595 | - | 85 | 37 | RNA→ |
| env-302 | JGI20160J14292_10255730 | + | 444 | 482 | RNA→ |
| env-303 | CESI01041543.1 | + | 405 | 453 | RNA→ rpsD (PRK05327)Ribosomal_S4 (pfam00163)→ |
| env-304 | DelMOSum2010_c10015854 | - | 4138 | 4090 | RNA→ RpsD (COG0522)rpsD (PRK05327)→ RNAP_alpha_NTD (cd06928)PRK05182 (PRK05182)RNA_pol_A_CTD (pfam03118)→ RplQ (COG0203)rplQ (PRK05591)→ CarA (COG0505)PRK12564 (PRK12564)→ |
| env-305 | JGI20154J14316_10192329 | - | 65 | 24 | RNA→ |
| env-306 | DelMOSum2010_c10003210 | + | 452 | 500 | RNA→ RpsD (COG0522)rpsD (PRK05327)→ RNAP_alpha_NTD (cd06928)PRK05182 (PRK05182)RNA_pol_A_CTD (pfam03118)→ RplQ (COG0203)rplQ (PRK05591)→ |
| env-307 | DelMOSum2010_c10379381 | + | 340 | 378 | RNA→ |
| env-308 | DelMOSum2010_c10492344 | - | 94 | 46 | RNA→ |
| env-309 | JGI20154J14316_10188950 | - | 67 | 24 | RNA→ |
| env-310 | JGI20160J14292_10241200 | + | 448 | 486 | RNA→ |
| env-311 | JGI20153J14295_10182765 | + | 414 | 461 | RNA→ |
| env-312 | CESU01090380.1 | + | 400 | 448 | RNA→ RpsD (COG0522)rpsD (PRK05327)→ RNAP_alpha_NTD (cd06928)PRK05182 (PRK05182)RNA_pol_A_CTD (pfam03118)→ RplQ (COG0203)rplQ (PRK05591)→ CarA (COG0505)PRK12564 (PRK12564)→ |
| env-313 | JGI20154J14316_10183818 | - | 76 | 28 | RNA→ |
| env-314 | JGI20154J14316_10249246 | + | 339 | 386 | RNA→ |
| env-315 | DelMOSum2011_c10107659 | + | 808 | 856 | RNA→ |
| env-316 | KGI_S1_ANT02_95m_c10000070 | - | 9736 | 9688 | RNA→ RpsD (COG0522)rpsD (PRK05327)→ RpoA (COG0202)PRK05182 (PRK05182)→ RplQ (COG0203)rplQ (PRK05591)→ CarA (COG0505)PRK12564 (PRK12564)→ |
| env-317 | DelMOSum2011_c10000248 | - | 37631 | 37583 | RNA→ RpsD (COG0522)rpsD (PRK05327)→ RpoA (COG0202)PRK05182 (PRK05182)→ RplQ (COG0203)rplQ (PRK05591)→ hypo→ CarA (COG0505)PRK12564 (PRK12564)→ |

| | | | | | |
|---|---|---|---|---|---|
| env-318 | JGI20157J14317_10074550 | - | 1039 | 991 | **RNA**→ RpsD (COG0522)rpsD (PRK05327)→ PRK05182 (PRK05182)RNA_pol_L (pfam01193)→ |
| env-319 | JGI20157J14317_10082923 | + | 461 | 509 | **RNA**→ RpsD (COG0522)rpsD (PRK05327)→ |
| Tna-1-1 | NZ_JTDV01000011.1 | - | 67639 | 67591 | **RNA**→ RpsD (COG0522)rpsD (PRK05327)→ RpoA (COG0202)PRK05182 (PRK05182)→ |
| | | | | | RplQ (COG0203)rplQ (PRK05591)→ hypo→ CarA (COG0505)PRK12564 (PRK12564)→ |
| env-320 | JGI12097J13213_1000640 | - | 3883 | 3835 | **RNA**→ RpsD (COG0522)rpsD (PRK05327)→ RpoA (COG0202)PRK05182 (PRK05182)→ |
| | | | | | RplQ (COG0203)rplQ (PRK05591)→ CarA (COG0505)PRK12564 (PRK12564)→ |
| env-321 | NCBI_READ_1112290411083 | - | 940 | 901 | **RNA**→ rpsD (PRK05327)→ |
| env-322 | SRS024087_LANL_scaffold_26415 | + | 693 | 733 | **RNA**→ RpsD (COG0522)rpsD (PRK05327)→ |
| | | | | | RNAP_alpha_NTD (cd06928)PRK05182 (PRK05182)RNA_pol_A_CTD (pfam03118)→ |
| | | | | | RplQ (COG0203)rplQ (PRK05591)→ |
| env-323 | SRS022602_Baylor_scaffold_55726 | - | 1590 | 1550 | **RNA**→ RpsD (COG0522)rpsD (PRK05327)→ |
| | | | | | RNAP_alpha_NTD (cd06928)PRK05182 (PRK05182)RNA_pol_A_CTD (pfam03118)→ |
| env-324 | SRS011343_C2229337 | - | 665 | 625 | **RNA**→ RpsD (COG0522)rpsD (PRK05327)→ |
| env-325 | SRS064449_LANL_scaffold_29232 | - | 83 | 43 | **RNA**→ |
| env-326 | KGI_S1_ANT01_95m_c10069223 | + | 445 | 488 | **RNA**→ RpsD (COG0522)rpsD (PRK05327)→ |
| env-327 | KGI_S1_ANT01_95m_c10437516 | + | 51 | 94 | **RNA**→ rpsD (PRK05327)Ribosomal_S4 (pfam00163)→ |
| env-328 | KGI_S1_ANT01_95m_c10191473 | - | 99 | 56 | **RNA**→ |
| env-329 | KGI_S1_ANT02_95m_c10154153 | - | 555 | 512 | **RNA**→ RpsD (COG0522)rpsD (PRK05327)→ |
| env-330 | KGI_S1_ANT02_95m_c10018669 | - | 104 | 61 | **RNA**→ |
| env-331 | JGI1221J11331_1000195 | - | 5494 | 5451 | **RNA**→ RpsD (COG0522)rpsD (PRK05327)→ RpoA (COG0202)PRK05182 (PRK05182)→ |
| Gli-1-1 | NZ_JH594606.1 | - | 1313497 | 1313454 | **RNA**→ RpsD (COG0522)rpsD (PRK05327)→ RpsD (COG0522)rpsD (PRK05327)→ |
| | | | | | RNAP_alpha_NTD (cd06928)PRK05182 (PRK05182)RNA_pol_A_CTD (pfam03118)→ |
| | | | | | RpoA (COG0202)PRK05182 (PRK05182)→ RplQ (COG0203)rplQ (PRK05591)Mtc1 (pfam10310)→ |
| | | | | | RplQ (COG0203)rplQ (PRK05591)Mtc1 (pfam10310)→ |
| env-332 | ElkS_mat_MD6A_1190446 | + | 69 | 109 | **RNA**→ rpsD (PRK05327)Ribosomal_S4 (pfam00163)→ |
| env-333 | JGI1221J11331_1000080 | + | 24062 | 24105 | **RNA**→ RpsD (COG0522)rpsD (PRK05327)→ RpoA (COG0202)PRK05182 (PRK05182)→ |
| Aat-1-1 | NZ_AQRA01000007.1 | + | 133451 | 133494 | **RNA**→ RpsD (COG0522)rpsD (PRK05327)→ RpoA (COG0202)PRK05182 (PRK05182)→ |
| | | | | | RplQ (COG0203)rplQ (PRK05591)→ |
| env-334 | JGI11876J14442_10876046 | - | 214 | 174 | **RNA**→ rpsD (PRK05327)Ribosomal_S4 (pfam00163)→ |
| Gsp-3-1 | NZ_JQLP01000005.1 | + | 613543 | 613586 | **RNA**→ RpsD (COG0522)rpsD (PRK05327)→ RpoA (COG0202)PRK05182 (PRK05182)→ |
| | | | | | RplQ (COG0203)rplQ (PRK05591)→ |
| env-335 | KGI_S1_ANT01_95m_c10674661 | + | 62 | 105 | **RNA**→ rpsD (PRK05327)Ribosomal_S4 (pfam00163)→ |
| env-336 | KGI_S1_ANT01_95m_c10076701 | - | 615 | 572 | **RNA**→ RpsD (COG0522)rpsD (PRK05327)→ |
| env-337 | KGI_S1_ANT01_95m_c10067832 | + | 463 | 506 | **RNA**→ RpsD (COG0522)rpsD (PRK05327)→ |
| env-338 | KGI_S2_ANT04_2345m_c1337795 | - | 82 | 39 | **RNA**→ |
| env-339 | KGI_S1_ANT02_95m_c10321982 | + | 98 | 141 | **RNA**→ rpsD (PRK05327)Ribosomal_S4 (pfam00163)→ |
| env-340 | KGI_S1_ANT02_95m_c10332312 | + | 96 | 139 | **RNA**→ rpsD (PRK05327)Ribosomal_S4 (pfam00163)→ |
| Gsp-4-1 | NZ_KE384219.1 | - | 887176 | 887133 | **RNA**→ RpsD (COG0522)rpsD (PRK05327)→ RpoA (COG0202)PRK05182 (PRK05182)→ |
| | | | | | RplQ (COG0203)rplQ (PRK05591)→ |
| Gsp-2-1 | NZ_AJLT01000027.1 | - | 3532 | 3489 | **RNA**→ RpsD (COG0522)rpsD (PRK05327)→ RpsD (COG0522)rpsD (PRK05327)→ |
| | | | | | RpoA (COG0202)PRK05182 (PRK05182)→ RpoA (COG0202)PRK05182 (PRK05182)→ |
| | | | | | RplQ (COG0203)rplQ (PRK05591)→ RplQ (COG0203)rplQ (PRK05591)→ |
| env-341 | ElkS_mat_MD6A_1172629 | + | 144 | 187 | **RNA**→ rpsD (PRK05327)Ribosomal_S4 (pfam00163)→ |
| env-342 | CEVI01208716.1 | - | 665 | 622 | **RNA**→ RpsD (COG0522)rpsD (PRK05327)→ |
| Ssp-1-1 | NZ_JQNQ01000001.1 | + | 2652711 | 2652754 | **RNA**→ RpsD (COG0522)rpsD (PRK05327)→ RpoA (COG0202)PRK05182 (PRK05182)→ |
| | | | | | RplQ (COG0203)rplQ (PRK05591)→ |
| Kal-1-1 | NZ_DS544873.1 | - | 2713104 | 2713064 | **RNA**→ RpsD (COG0522)rpsD (PRK05327)→ RpoA (COG0202)PRK05182 (PRK05182)→ |
| | | | | | RplQ (COG0203)rplQ (PRK05591)→ |
| Gsp-1-1 | NZ_APHJ01000040.1 | - | 65659 | 65616 | **RNA**→ RpsD (COG0522)rpsD (PRK05327)→ RpoA (COG0202)PRK05182 (PRK05182)→ |
| | | | | | RplQ (COG0203)rplQ (PRK05591)→ |
| env-343 | P_1C_Liq_3_UnCty_1004385 | - | 1057 | 1016 | **RNA**→ RpsD (COG0522)rpsD (PRK05327)→ PRK05182 (PRK05182)RNA_pol_L (pfam01193)→ |
| env-344 | P_1C_Liq_1_UnCty_1000030 | - | 6951 | 6910 | **RNA**→ RpsD (COG0522)rpsD (PRK05327)→ |
| | | | | | RNAP_alpha_NTD (cd06928)PRK05182 (PRK05182)RNA_pol_A_CTD (pfam03118)→ |
| | | | | | RplQ (COG0203)rplQ (PRK05591)→ CarA (COG0505)PRK12564 (PRK12564)→ |
| env-345 | JGI25913J50563_1051608 | + | 106 | 147 | **RNA**→ RpsD (COG0522)rpsD (PRK05327)→ |

| | | | | | |
|---|---|---|---|---|---|
| env-346 | ABOG01001487.1 | - | 91 | 50 | **RNA**→ hypo→ |
| env-347 | 2204886463 | - | 72 | 31 | **RNA**→ |
| env-348 | GSLSAS_GLMUJHB02GRDZW | - | 120 | 79 | **RNA**→ rpsD (PRK05327)Ribosomal_S4 (pfam00163)→ |
| env-349 | JGI24770J26754_10034354 | + | 1291 | 1332 | **RNA**→ RpsD (COG0522)rpsD (PRK05327)→ RNAP_alpha_NTD (cd06928)PRK05182 (PRK05182)→ |
| Fin-1-1 | NC_017025.1 | + | 672442 | 672483 | **RNA**→ RpsD (COG0522)rpsD (PRK05327)→ RNAP_alpha_NTD (cd06928)PRK05182 (PRK05182)RNA_pol_A_CTD (pfam03118)→ RplQ (COG0203)rplQ (PRK05591)→ |
| env-350 | JGI25920J50251_10035976 | - | 1266 | 1225 | **RNA**→ RpsD (COG0522)rpsD (PRK05327)→ RNAP_alpha_NTD (cd06928)PRK05182 (PRK05182)→ |
| env-351 | P_2C_Liq_2_UnCty_1074319 | + | 58 | 98 | **RNA**→ RpsD (COG0522)rpsD (PRK05327)→ |
| env-352 | DelMOSum2010_c10001779 | - | 4923 | 4883 | **RNA**→ RpsD (COG0522)rpsD (PRK05327)→ RpoA (COG0202)PRK05182 (PRK05182)→ RplQ (COG0203)rplQ (PRK05591)→ CarA (COG0505)PRK12564 (PRK12564)→ |
| env-353 | DelMOWin2010_c10201283 | - | 365 | 325 | **RNA**→ RpsD (COG0522)rpsD (PRK05327)→ |
| env-354 | ABOE01005844.1 | - | 55 | 25 | **RNA**→ ←hypo |
| env-355 | P_2C_Liq_1_UnCty_1005695 | - | 2283 | 2243 | **RNA**→ RpsD (COG0522)rpsD (PRK05327)→ RpoA (COG0202)PRK05182 (PRK05182)→ RplQ (COG0203)rplQ (PRK05591)→ |
| env-356 | CENG01018837.1 | - | 3006 | 2965 | **RNA**→ RpsD (COG0522)rpsD (PRK05327)→ RNAP_alpha_NTD (cd06928)PRK05182 (PRK05182)RNA_pol_A_CTD (pfam03118)→ RplQ (COG0203)rplQ (PRK05591)→ hypo→ |
| env-357 | CENP01317983.1 | + | 486 | 527 | **RNA**→ rpsD (PRK05327)Ribosomal_S4 (pfam00163)→ |
| env-358 | CENW01033667.1 | + | 574 | 605 | **RNA**→ hypo→ |
| env-359 | CEOP01025191.1 | - | 597 | 556 | **RNA**→ RpsD (COG0522)rpsD (PRK05327)→ |
| env-360 | CEPL01236006.1 | - | 586 | 545 | **RNA**→ RpsD (COG0522)rpsD (PRK05327)→ |
| env-361 | CERK01003498.1 | - | 2219 | 2178 | **RNA**→ RpsD (COG0522)rpsD (PRK05327)→ RNAP_alpha_NTD (cd06928)PRK05182 (PRK05182)RNA_pol_A_CTD (pfam03118)→ RplQ (COG0203)rplQ (PRK05591)rpsP (PRK14521)→ |
| env-362 | CESZ01017440.1 | + | 239 | 280 | **RNA**→ RpsD (COG0522)rpsD (PRK05327)→ |
| env-363 | CETQ01116547.1 | + | 277 | 318 | **RNA**→ RpsD (COG0522)rpsD (PRK05327)→ |
| env-364 | CEUG01164120.1 | + | 218 | 259 | **RNA**→ RpsD (COG0522)rpsD (PRK05327)→ |
| env-365 | CEVL01101790.1 | + | 577 | 618 | **RNA**→ rpsD (PRK05327)Ribosomal_S4 (pfam00163)→ |
| env-366 | CEVU01164851.1 | - | 2198 | 2157 | **RNA**→ RpsD (COG0522)rpsD (PRK05327)→ RpoA (COG0202)PRK05182 (PRK05182)→ RplQ (COG0203)rplQ (PRK05591)→ |
| env-367 | CEWG01182900.1 | + | 735 | 776 | **RNA**→ |
| env-368 | CEWK01130899.1 | - | 556 | 515 | **RNA**→ RpsD (COG0522)rpsD (PRK05327)→ |
| env-369 | JCVI_SCAF_1101668693344 | - | 606 | 566 | **RNA**→ RpsD (COG0522)rpsD (PRK05327)→ |
| env-370 | JCVI_SCAF_1096626974148 | - | 920 | 879 | **RNA**→ RpsD (COG0522)rpsD (PRK05327)→ PRK05182 (PRK05182)RNA_pol_L (pfam01193)→ |
| env-371 | CENQ01077134.1 | - | 192 | 151 | **RNA**→ rpsD (PRK05327)Ribosomal_S4 (pfam00163)→ |
| env-372 | CEOM01200204.1 | + | 808 | 849 | **RNA**→ RpsD (COG0522)rpsD (PRK05327)→ RpoA (COG0202)PRK05182 (PRK05182)→ RplQ (COG0203)rplQ (PRK05591)→ |
| env-373 | CEQX01059206.1 | - | 38 | 7 | **RNA**→ |
| env-374 | CESP01186676.1 | + | 437 | 478 | **RNA**→ |
| env-375 | CEUA01031995.1 | + | 888 | 929 | **RNA**→ RpsD (COG0522)rpsD (PRK05327)→ RNAP_alpha_NTD (cd06928)PRK05182 (PRK05182)RNA_pol_A_CTD (pfam03118)→ RplQ (COG0203)rplQ (PRK05591)→ |
| env-376 | CEVG01008164.1 | + | 288 | 329 | **RNA**→ rpsD (PRK05327)Ribosomal_S4 (pfam00163)→ |
| env-377 | CEVR01251554.1 | + | 296 | 337 | **RNA**→ rpsD (PRK05327)Ribosomal_S4 (pfam00163)→ |
| env-378 | CEVX01164589.1 | + | 416 | 457 | **RNA**→ RpsD (COG0522)rpsD (PRK05327)→ |
| env-379 | CEWO01084334.1 | + | 5393 | 5434 | **RNA**→ RpsD (COG0522)rpsD (PRK05327)→ RNAP_alpha_NTD (cd06928)PRK05182 (PRK05182)RNA_pol_A_CTD (pfam03118)→ RplQ (COG0203)rplQ (PRK05591)→ |
| env-380 | CEOV01038829.1 | - | 255 | 212 | **RNA**→ rpsD (PRK05327)Ribosomal_S4 (pfam00163)→ |
| env-381 | CEQQ01099035.1 | + | 406 | 449 | **RNA**→ RpsD (COG0522)rpsD (PRK05327)→ RpoA (COG0202)PRK05182 (PRK05182)→ |
| env-382 | CEQO01051374.1 | + | 920 | 963 | **RNA**→ RpsD (COG0522)rpsD (PRK05327)→ |
| env-383 | CESQ01024469.1 | + | 341 | 384 | **RNA**→ RpsD (COG0522)rpsD (PRK05327)→ RNAP_alpha_NTD (cd06928)PRK05182 (PRK05182)→ |
| env-384 | CEUM01272025.1 | + | 217 | 260 | **RNA**→ RpsD (COG0522)rpsD (PRK05327)→ |

| | | | | | |
|---|---|---|---|---|---|
| env-385 | JCVI_READ_1103770020066 | - | 619 | 576 | **RNA**→ RpsD (COG0522)rpsD (PRK05327)→ |
| env-386 | CESD01077857.1 | - | 283 | 240 | **RNA**→ rpsD (PRK05327)Ribosomal_S4 (pfam00163)→ |
| env-387 | CETB01090522.1 | - | 501 | 458 | **RNA**→ RpsD (COG0522)rpsD (PRK05327)→ |
| env-388 | JCVI_READ_1103769862037 | - | 370 | 327 | **RNA**→ rpsD (PRK05327)Ribosomal_S4 (pfam00163)→ |
| env-389 | JCVI_SCAF_1096627876296 | + | 547 | 588 | **RNA**→ RpsD (COG0522)rpsD (PRK05327)→ RpsD (COG0522)rpsD (PRK05327)→ |
| env-390 | JCVI_SCAF_1101668435598 | + | 664 | 705 | **RNA**→ rpsD (PRK05327)Ribosomal_S4 (pfam00163)→ |
| env-391 | JCVI_SCAF_1096627108452 | - | 156 | 115 | **RNA**→ |
| env-392 | CENJ01155291.1 | - | 444 | 403 | **RNA**→ RpsD (COG0522)rpsD (PRK05327)→ |
| env-393 | CENT01170625.1 | - | 233 | 192 | **RNA**→ rpsD (PRK05327)Ribosomal_S4 (pfam00163)→ |
| env-394 | CENY01156917.1 | + | 415 | 456 | **RNA**→ RpsD (COG0522)rpsD (PRK05327)→ PRK05182 (PRK05182)→ |
| env-395 | CEPC01245844.1 | + | 902 | 933 | **RNA**→ hypo→ |
| env-396 | CEQO01025468.1 | + | 291 | 332 | **RNA**→ RpsD (COG0522)rpsD (PRK05327)→ |
| env-397 | CESN01038593.1 | - | 2422 | 2381 | **RNA**→ RpsD (COG0522)rpsD (PRK05327)→ RpoA (COG0202)PRK05182 (PRK05182)→ RplQ (COG0203)rplQ (PRK05591)→ |
| env-398 | JCVI_SCAF_1096627965867 | + | 486 | 527 | **RNA**→ S4 (smart00363)RpsD (COG0522)→ PRK05182 (PRK05182)RNA_pol_L (pfam01193)→ |
| env-399 | CETY01194632.1 | + | 6186 | 6227 | **RNA**→ RpsD (COG0522)rpsD (PRK05327)→ PRK05182 (PRK05182)RNA_pol_L (pfam01193)→ |
| env-400 | CEVG01161451.1 | + | 1799 | 1840 | **RNA**→ RpsD (COG0522)rpsD (PRK05327)→ PRK05182 (PRK05182)RNA_pol_L (pfam01193)→ |
| env-401 | CEVP01363226.1 | + | 1728 | 1769 | **RNA**→ rpsD (PRK05327)Ribosomal_S4 (pfam00163)→ |
| env-402 | CEVX01015396.1 | + | 5254 | 5295 | **RNA**→ RpsD (COG0522)rpsD (PRK05327)→ |
| env-403 | JCVI_SCAF_1096627738902 | - | 327 | 286 | **RNA**→ rpsD (PRK05327)Ribosomal_S4 (pfam00163)→ |
| env-404 | JCVI_SCAF_1096627987680 | + | 87 | 128 | **RNA**→ RpsD (COG0522)rpsD (PRK05327)→ RpoA (COG0202)PRK05182 (PRK05182)→ |
| env-405 | JCVI_SCAF_1101668751912 | - | 811 | 770 | **RNA**→ RpsD (COG0522)rpsD (PRK05327)→ |
| env-406 | CENP01173649.1 | + | 17909 | 17950 | **RNA**→ RpsD (COG0522)rpsD (PRK05327)→ RNAP_alpha_NTD (cd06928)PRK05182 (PRK05182)RNA_pol_A_CTD (pfam03118)→ RplQ (COG0203)rplQ (PRK05591)→ hypo→ CarA (COG0505)PRK12564 (PRK12564)→ eno (PRK00077)enolase (cd03313)→ gltA (PRK05614)Citrate_synt (pfam00285)→ DdaH (COG1834)→ Amidinotransf (pfam02274)→ Arg_tRNA_synt_N (smart01016)Arg_tRNA_synt_N (pfam03485)→ |
| env-407 | CENX01050295.1 | - | 973 | 932 | **RNA**→ RpsD (COG0522)rpsD (PRK05327)→ PRK05182 (PRK05182)RNA_pol_L (pfam01193)→ |
| env-408 | CEPW01055046.1 | + | 441 | 482 | **RNA**→ RpsD (COG0522)rpsD (PRK05327)→ RpoA (COG0202)PRK05182 (PRK05182)→ RplQ (COG0203)rplQ (PRK05591)→ |
| env-409 | CESF01181460.1 | - | 96 | 55 | **RNA**→ |
| env-410 | CETY01166093.1 | + | 795 | 836 | **RNA**→ |
| env-411 | CEUN01220636.1 | + | 306 | 347 | **RNA**→ rpsD (PRK05327)Ribosomal_S4 (pfam00163)→ |
| env-412 | CEVP01121720.1 | - | 660 | 619 | **RNA**→ RpsD (COG0522)rpsD (PRK05327)→ |
| env-413 | CEVW01163295.1 | - | 43 | 12 | **RNA**→ |
| env-414 | CEWH01074398.1 | + | 631 | 672 | **RNA**→ RpsD (COG0522)rpsD (PRK05327)→ |
| env-415 | 85126595 | - | 708 | 665 | **RNA**→ RpsD (COG0522)rpsD (PRK05327)→ |
| env-416 | CEQM01038432.1 | - | 514 | 471 | **RNA**→ RpsD (COG0522)rpsD (PRK05327)→ |
| env-417 | CEQW01046056.1 | - | 288 | 245 | **RNA**→ rpsD (PRK05327)Ribosomal_S4 (pfam00163)→ |
| env-418 | CESF01121287.1 | + | 799 | 842 | **RNA**→ |
| env-419 | CEUF01015480.1 | + | 459 | 502 | **RNA**→ rpsD (PRK05327)Ribosomal_S4 (pfam00163)→ |
| env-420 | CAM_READ_0234809867 | - | 478 | 435 | **RNA**→ S4 (smart00363)RpsD (COG0522)→ |
| env-421 | CENF01049817.1 | + | 70 | 113 | **RNA**→ RpsD (COG0522)rpsD (PRK05327)→ |
| env-422 | CESW01020303.1 | + | 802 | 845 | **RNA**→ RpsD (COG0522)rpsD (PRK05327)→ |
| env-423 | CEUH01049850.1 | + | 363 | 406 | **RNA**→ rpsD (PRK05327)Ribosomal_S4 (pfam00163)→ |
| env-424 | JCVI_SCAF_1096627517448 | - | 1401 | 1360 | **RNA**→ RpsD (COG0522)rpsD (PRK05327)→ |
| env-425 | JCVI_SCAF_1101668741877 | - | 1146 | 1105 | **RNA**→ RpsD (COG0522)rpsD (PRK05327)→ RNAP_alpha_NTD (cd06928)PRK05182 (PRK05182)→ |
| env-426 | JGI20153J14295_10198722 | + | 211 | 255 | **RNA**→ rpsD (PRK05327)Ribosomal_S4 (pfam00163)→ |
| env-427 | SA_S1_NOR05_45m_c10264163 | + | 183 | 227 | **RNA**→ rpsD (PRK05327)Ribosomal_S4 (pfam00163)→ |
| env-428 | SI60aug11_100m_c1216415 | - | 37 | 5 | **RNA**→ |
| env-429 | SI34jun09_10m_1002524 | - | 5240 | 5198 | **RNA**→ RpsD (COG0522)rpsD (PRK05327)→ |
| env-430 | JGI1221J11331_1019349 | - | 1294 | 1245 | **RNA**→ RpsD (COG0522)rpsD (PRK05327)→ RNAP_alpha_NTD (cd06928)PRK05182 (PRK05182)→ |
| env-431 | P_2C_Liq_1_UnCty_1022676 | + | 198 | 240 | **RNA**→ RpsD (COG0522)rpsD (PRK05327)→ RpoA (COG0202)PRK05182 (PRK05182)→ |

| | | | | | |
|---|---|---|---|---|---|
| env-432 | SA_S1_NOR08_45m_c10303734 | - | 256 | 213 | RNA→ rpsD (PRK05327)Ribosomal_S4 (pfam00163)→ |
| env-433 | JGI24025J20009_10186048 | - | 381 | 339 | RNA→ RpsD (COG0522)rpsD (PRK05327)→ |
| env-434 | TDF_OR_ARG04_113m_c1035950 | - | 334 | 292 | RNA→ rpsD (PRK05327)Ribosomal_S4 (pfam00163)→ |
| env-435 | TDF_MC_ARG02_113m_c1087976 | + | 171 | 213 | RNA→ rpsD (PRK05327)Ribosomal_S4 (pfam00163)→ |
| env-436 | KGI_S1_ANT02_95m_c10052129 | + | 703 | 745 | RNA→ RpsD (COG0522)rpsD (PRK05327)→ |
| env-437 | P_2C_Liq_1_UnCty_1009183 | + | 649 | 691 | RNA→ RpsD (COG0522)rpsD (PRK05327)→ |
| env-438 | CESF01214207.1 | + | 538 | 580 | RNA→ ←hypo |
| env-439 | PML_READ_00354242 | + | 42 | 84 | RNA→ rpsD (PRK05327)Ribosomal_S4 (pfam00163)→ |
| env-440 | KGI_S2_ANT04_2345m_c1245894 | - | 300 | 258 | RNA→ rpsD (PRK05327)Ribosomal_S4 (pfam00163)→ |
| env-441 | KGI_S1_ANT02_95m_c10002151 | - | 4971 | 4929 | RNA→ RpsD (COG0522)rpsD (PRK05327)→ RNAP_alpha_NTD (cd06928)PRK05182 (PRK05182)RNA_pol_A_CTD (pfam03118)→ RplQ (COG0203)rplQ (PRK05591)→ |
| env-442 | JGI24724J26744_10001772 | + | 14502 | 14544 | RNA→ rpsD (PRK05327)Ribosomal_S4 (pfam00163)→ |
| env-443 | KGI_S1_ANT01_95m_c10016085 | - | 758 | 716 | RNA→ RpsD (COG0522)rpsD (PRK05327)→ hypo→ |
| env-444 | KGI_S1_ANT03_95m_c1000649 | - | 2219 | 2177 | RNA→ RpsD (COG0522)rpsD (PRK05327)→ RNAP_alpha_NTD (cd06928)PRK05182 (PRK05182)RNA_pol_A_CTD (pfam03118)→ RplQ (COG0203)rplQ (PRK05591)→ |
| env-445 | SI48aug10_10m_c1075429 | + | 155 | 197 | RNA→ |
| env-446 | JGI20152J14361_10073408 | - | 103 | 61 | RNA→ ←hypo |
| env-447 | JGI24023J19991_10048106 | - | 1743 | 1701 | RNA→ RpsD (COG0522)rpsD (PRK05327)→ RNAP_alpha_NTD (cd06928)PRK05182 (PRK05182)RNA_pol_A_CTD (pfam03118)→ |
| env-448 | SA_S1_NOR05_45m_c10501087 | + | 134 | 176 | RNA→ rpsD (PRK05327)Ribosomal_S4 (pfam00163)→ |
| env-449 | KGI_S2_ANT05_2345m_c1026215 | - | 758 | 716 | RNA→ RpsD (COG0522)rpsD (PRK05327)→ hypo→ |
| env-450 | JGI24025J20009_10005367 | - | 6255 | 6213 | RNA→ RpsD (COG0522)rpsD (PRK05327)→ RNAP_alpha_NTD (cd06928)PRK05182 (PRK05182)RNA_pol_A_CTD (pfam03118)→ RplQ (COG0203)rplQ (PRK05591)→ CarA (COG0505)PRK12564 (PRK12564)→ |
| Fba-3-1 | NC_014472.1 | - | 807022 | 806980 | RNA→ RpsD (COG0522)rpsD (PRK05327)→ RNAP_alpha_NTD (cd06928)PRK05182 (PRK05182)RNA_pol_A_CTD (pfam03118)→ RplQ (COG0203)rplQ (PRK05591)→ CarA (COG0505)PRK12564 (PRK12564)→ eno (PRK00077)enolase (cd03313)→ eno (PRK00077)enolase (cd03313)→ |
| env-451 | SA_S1_NOR08_45m_c10423185 | + | 103 | 145 | RNA→ rpsD (PRK05327)Ribosomal_S4 (pfam00163)→ |
| env-452 | SA_S1_NOR08_45m_c10214572 | + | 57 | 99 | RNA→ RpsD (COG0522)rpsD (PRK05327)→ |
| env-453 | JGI20156J14371_10031151 | + | 222 | 264 | RNA→ RpsD (COG0522)rpsD (PRK05327)→ |
| env-454 | NCBI_READ_1112777383475 | - | 732 | 690 | RNA→ RpsD (COG0522)rpsD (PRK05327)→ |
| env-455 | NCBI_READ_1112788432221 | - | 789 | 747 | RNA→ RpsD (COG0522)rpsD (PRK05327)→ |
| env-456 | NCBI_READ_1112494047845 | + | 599 | 641 | RNA→ rpsD (PRK05327)Ribosomal_S4 (pfam00163)→ |
| env-457 | 2236876010_0236984 | - | 372 | 324 | RNA→ RpsD (COG0522)rpsD (PRK05327)→ |
| env-458 | JGI20160J14292_10208151 | + | 144 | 192 | RNA→ RpsD (COG0522)rpsD (PRK05327)→ |
| env-459 | JGI20155J14468_10404061 | - | 156 | 108 | RNA→ rpsD (PRK05327)Ribosomal_S4 (pfam00163)→ |
| env-460 | SA_S1_NOR08_45m_c10274690 | + | 87 | 129 | RNA→ RpsD (COG0522)rpsD (PRK05327)→ |
| env-461 | 2236876010_0455091 | - | 371 | 329 | RNA→ RpsD (COG0522)rpsD (PRK05327)→ |
| env-462 | 2236876010_0562821 | - | 158 | 116 | RNA→ rpsD (PRK05327)Ribosomal_S4 (pfam00163)→ |
| env-463 | DelMOWin2010_c10188673 | - | 373 | 331 | RNA→ RpsD (COG0522)rpsD (PRK05327)→ |
| env-464 | SA_S1_NOR02_45m_c1174201 | - | 144 | 102 | RNA→ rpsD (PRK05327)Ribosomal_S4 (pfam00163)→ |
| env-465 | JCVI_SCAF_1096626910919 | - | 950 | 908 | RNA→ RpsD (COG0522)rpsD (PRK05327)→ |
| env-466 | DelMOSum2010_c10659421 | - | 71 | 29 | RNA→ |
| env-467 | 2236876010_0311705 | + | 440 | 472 | RNA→ |
| env-468 | 2236876010_0138022 | + | 324 | 366 | RNA→ |
| env-469 | 2236876010_0157521 | + | 63 | 105 | RNA→ rpsD (PRK05327)Ribosomal_S4 (pfam00163)→ |
| env-470 | SA_S1_NOR05_45m_c10203477 | + | 305 | 347 | RNA→ ←hypo |
| env-471 | JGI20157J14317_10005668 | - | 2515 | 2473 | RNA→ RpsD (COG0522)rpsD (PRK05327)→ |
| env-472 | CEWR01000864.1 | + | 572 | 614 | RNA→ rpsD (PRK05327)Ribosomal_S4 (pfam00163)→ |
| env-473 | 2236876010_0363029 | + | 290 | 332 | RNA→ rpsD (PRK05327)Ribosomal_S4 (pfam00163)→ |
| env-474 | 2236876010_0242303 | + | 265 | 307 | RNA→ |

| | | | | | |
|---|---|---|---|---|---|
| env-475 | 2236876010_0101955 | - | 172 | 130 | RNA→ rpsD (PRK05327)Ribosomal_S4 (pfam00163)→ |
| env-476 | DelMOSum2011_c10001719 | - | 610 | 568 | RNA→ RpsD (COG0522)rpsD (PRK05327)→ |
| env-477 | SA_S1_NOR05_45m_c10530560 | - | 224 | 182 | RNA→ rpsD (PRK05327)Ribosomal_S4 (pfam00163)→ |
| env-478 | DelMOSum2011_c10529643 | - | 209 | 167 | RNA→ rpsD (PRK05327)Ribosomal_S4 (pfam00163)→ |
| env-479 | JGI20156J14371_10646125 | - | 87 | 45 | RNA→ |
| Psp-1-1 | NZ_JPDS01000002.1 | + | 1347599 | 1347641 | RNA→ RpsD (COG0522)rpsD (PRK05327)→ |
| | | | | | RNAP_alpha_NTD (cd06928)PRK05182 (PRK05182)RNA_pol_A_CTD (pfam03118)→ |
| | | | | | RplQ (COG0203)rplQ (PRK05591)→ hypo→ |
| env-480 | 2236876010_0517953 | - | 211 | 169 | RNA→ hypo→ |
| env-481 | 2236876010_0238370 | - | 424 | 382 | RNA→ rpsD (PRK05327)Ribosomal_S4 (pfam00163)→ S4 (smart00363)RpsD (COG0522)→ |
| env-482 | 2236876010_0136465 | - | 50 | 18 | RNA→ hypo→ |
| env-483 | SI60aug11_200m_c1106132 | - | 340 | 298 | RNA→ rpsD (PRK05327)Ribosomal_S4 (pfam00163)→ |
| env-484 | CERM01023755.1 | + | 1651 | 1693 | RNA→ RpsD (COG0522)rpsD (PRK05327)→ hypo→ |
| Tov-1-1 | NZ_AUMF01000009.1 | - | 79066 | 79024 | RNA→ RpsD (COG0522)rpsD (PRK05327)→ RpoA (COG0202)PRK05182 (PRK05182)→ |
| | | | | | RplQ (COG0203)rplQ (PRK05591)→ CarA (COG0505)PRK12564 (PRK12564)→ |
| env-485 | 2236876010_0403020 | + | 10 | 52 | RNA→ rpsD (PRK05327)Ribosomal_S4 (pfam00163)→ hypo→ |
| env-486 | CEPZ01013735.1 | - | 141998 | 141956 | RNA→ RpsD (COG0522)rpsD (PRK05327)→ |
| | | | | | RNAP_alpha_NTD (cd06928)PRK05182 (PRK05182)RNA_pol_A_CTD (pfam03118)→ |
| | | | | | RplQ (COG0203)rplQ (PRK05591)→ hypo→ CarA (COG0505)PRK12564 (PRK12564)→ |
| env-487 | KGI_S2_ANT04_2345m_c1133551 | - | 346 | 304 | RNA→ rpsD (PRK05327)Ribosomal_S4 (pfam00163)→ |
| env-488 | SA_S1_NOR05_45m_c10020030 | + | 1304 | 1346 | RNA→ RpsD (COG0522)rpsD (PRK05327)→ |
| env-489 | JGI20152J14361_10000017 | + | 7046 | 7088 | RNA→ RpsD (COG0522)rpsD (PRK05327)→ RpoA (COG0202)PRK05182 (PRK05182)→ |
| | | | | | RplQ (COG0203)rplQ (PRK05591)→ CarA (COG0505)PRK12564 (PRK12564)→ |
| env-490 | DelMOSum2011_c10004810 | - | 4239 | 4197 | RNA→ RpsD (COG0522)rpsD (PRK05327)→ RpoA (COG0202)PRK05182 (PRK05182)→ |
| | | | | | RplQ (COG0203)rplQ (PRK05591)→ CarA (COG0505)PRK12564 (PRK12564)→ |
| env-491 | JGI20154J14316_10079454 | - | 235 | 193 | RNA→ rpsD (PRK05327)Ribosomal_S4 (pfam00163)→ |
| Kje-1-1 | NZ_LBMG01000016.1 | - | 14572 | 14530 | RNA→ RpsD (COG0522)rpsD (PRK05327)→ RpoA (COG0202)PRK05182 (PRK05182)→ |
| | | | | | RplQ (COG0203)rplQ (PRK05591)→ |
| env-492 | KGI_S1_ANT01_95m_c10057790 | - | 1051 | 1009 | RNA→ RpsD (COG0522)rpsD (PRK05327)→ PRK05182 (PRK05182)RNA_pol_L (pfam01193)→ |
| env-493 | JGI20160J14292_10195876 | - | 317 | 275 | RNA→ rpsD (PRK05327)Ribosomal_S4 (pfam00163)→ |
| Psp-2-1 | NZ_JHZZ01000001.1 | - | 237303 | 237261 | RNA→ RpsD (COG0522)rpsD (PRK05327)→ |
| | | | | | RNAP_alpha_NTD (cd06928)PRK05182 (PRK05182)RNA_pol_A_CTD (pfam03118)→ |
| env-494 | JGI20154J14316_10123778 | + | 416 | 458 | RNA→ rpsD (PRK05327)Ribosomal_S4 (pfam00163)→ |
| env-495 | JGI20157J14317_10556627 | + | 133 | 175 | RNA→ rpsD (PRK05327)Ribosomal_S4 (pfam00163)→ |
| env-496 | 2236876010_0174516 | - | 142 | 100 | RNA→ |
| env-497 | BS_KBA_SWE12_21m_c10001102 | + | 2457 | 2499 | RNA→ RpsD (COG0522)rpsD (PRK05327)→ RpoA (COG0202)PRK05182 (PRK05182)→ |
| | | | | | RplQ (COG0203)rplQ (PRK05591)→ hypo→ CarA (COG0505)PRK12564 (PRK12564)→ |
| env-498 | CESE01006475.1 | - | 1107 | 1065 | RNA→ RpsD (COG0522)rpsD (PRK05327)→ RpoA (COG0202)PRK05182 (PRK05182)→ |
| env-499 | wf3_Contig994 | - | 742 | 700 | RNA→ RpsD (COG0522)rpsD (PRK05327)→ |
| env-500 | SA_S2_NOR13_50m_c1005577 | - | 652 | 610 | RNA→ RpsD (COG0522)rpsD (PRK05327)→ |
| env-501 | P_2C_Liq_1_UnCty_1273058 | - | 86 | 45 | RNA→ |
| env-502 | P_2C_Liq_3_UnCty_100504 | + | 604 | 645 | RNA→ RpsD (COG0522)rpsD (PRK05327)→ |
| | | | | | RNAP_alpha_NTD (cd06928)PRK05182 (PRK05182)RNA_pol_A_CTD (pfam03118)→ |
| | | | | | RplQ (COG0203)rplQ (PRK05591)→ CarA (COG0505)PRK12564 (PRK12564)→ |
| env-503 | P_2C_Liq_3_UnCty_100665 | - | 8850 | 8809 | RNA→ RpsD (COG0522)rpsD (PRK05327)→ |
| env-504 | KGI_S1_ANT01_95m_c10123548 | + | 293 | 341 | RNA→ RpsD (COG0522)rpsD (PRK05327)→ |
| env-505 | JGI11876J14442_10581827 | + | 196 | 237 | RNA→ rpsD (PRK05327)Ribosomal_S4 (pfam00163)→ |
| env-506 | JGI24023J19991_10009789 | - | 99 | 58 | RNA→ |
| env-507 | JGI24023J19991_10077987 | + | 1088 | 1129 | RNA→ |
| env-508 | JGI24023J19991_10260636 | - | 75 | 34 | RNA→ |
| env-509 | JGI24723J26617_10023895 | - | 2577 | 2536 | RNA→ RpsD (COG0522)rpsD (PRK05327)→ RNAP_alpha_NTD (cd06928)PRK05182 (PRK05182)→ |
| | | | | | PRK05182 (PRK05182)RNA_pol_A_CTD (pfam03118)→ RplQ (COG0203)rplQ (PRK05591)→ |
| | | | | | PRK12564 (PRK12564)CPSase_sm_chain (pfam00988)→ |
| env-510 | JGI24723J26617_10679119 | - | 223 | 182 | RNA→ rpsD (PRK05327)Ribosomal_S4 (pfam00163)→ |
| env-511 | SRS022602_Baylor_scaffold_12322 | + | 70 | 111 | RNA→ RpsD (COG0522)rpsD (PRK05327)→ |

| | | | | | |
|---|---|---|---|---|---|
| env-512 | ASA120_GJFD58A02GLQWA | + | 249 | 290 | **RNA**→ ←hypo |
| Mze-2-1 | NZ_AFOE01000041.1 | + | 7910 | 7951 | **RNA**→ RpsD (COG0522)rpsD (PRK05327)→ RpsD (COG0522)rpsD (PRK05327)→<br>RNAP_alpha_NTD (cd06928)PRK05182 (PRK05182)RNA_pol_A_CTD (pfam03118)→<br>RNAP_alpha_NTD (cd06928)PRK05182 (PRK05182)RNA_pol_A_CTD (pfam03118)→<br>RplQ (COG0203)rplQ (PRK05591)→ RplQ (COG0203)rplQ (PRK05591)→<br>CarA (COG0505)PRK12564 (PRK12564)→ CarA (COG0505)PRK12564 (PRK12564)→ |
| Csp-2-1 | NZ_KE992086.1 | + | 67634 | 67675 | **RNA**→ RpsD (COG0522)rpsD (PRK05327)→<br>RNAP_alpha_NTD (cd06928)PRK05182 (PRK05182)RNA_pol_A_CTD (pfam03118)→<br>RplQ (COG0203)rplQ (PRK05591)→ |
| Man-1-1 | NZ_KK211214.1 | + | 408244 | 408285 | **RNA**→ RpsD (COG0522)rpsD (PRK05327)→<br>RNAP_alpha_NTD (cd06928)PRK05182 (PRK05182)RNA_pol_A_CTD (pfam03118)→<br>RplQ (COG0203)rplQ (PRK05591)→ rplQ (PRK05591)Ribosomal_L17 (pfam01196)→<br>CarA (COG0505)PRK12564 (PRK12564)→ |
| Msp-1-1 | NZ_JHZW01000002.1 | - | 322870 | 322829 | **RNA**→ RpsD (COG0522)rpsD (PRK05327)→<br>RNAP_alpha_NTD (cd06928)PRK05182 (PRK05182)RNA_pol_A_CTD (pfam03118)→<br>RplQ (COG0203)rplQ (PRK05591)→ CarA (COG0505)PRK12564 (PRK12564)→ |
| env-513 | CEVF01017747.1 | + | 20379 | 20420 | **RNA**→ RpsD (COG0522)rpsD (PRK05327)→<br>RNAP_alpha_NTD (cd06928)PRK05182 (PRK05182)RNA_pol_A_CTD (pfam03118)→ |
| env-514 | LAZR01000718.1 | + | 21183 | 21224 | **RNA**→ RpsD (COG0522)rpsD (PRK05327)→<br>RNAP_alpha_NTD (cd06928)PRK05182 (PRK05182)RNA_pol_A_CTD (pfam03118)→<br>RplQ (COG0203)rplQ (PRK05591)→ |
| env-515 | JGI11876J14442_10577013 | - | 154 | 113 | **RNA**→ rpsD (PRK05327)Ribosomal_S4 (pfam00163)→ |
| env-516 | JGI24023J19991_10007103 | - | 261 | 220 | **RNA**→ rpsD (PRK05327)Ribosomal_S4 (pfam00163)→ |
| env-517 | JGI24023J19991_10074980 | - | 367 | 326 | **RNA**→ RpsD (COG0522)rpsD (PRK05327)→ |
| env-518 | JGI24023J19991_10226678 | - | 76 | 35 | **RNA**→ |
| env-519 | JGI24723J26617_10030345 | + | 1847 | 1888 | **RNA**→ RpsD (COG0522)rpsD (PRK05327)→ |
| env-520 | JGI24723J26617_10764797 | + | 51 | 92 | **RNA**→ rpsD (PRK05327)Ribosomal_S4 (pfam00163)→ |
| env-521 | SRS055378_LANL_scaffold_51092 | - | 215 | 174 | **RNA**→ rpsD (PRK05327)Ribosomal_S4 (pfam00163)→ |
| Csp-3-1 | NZ_JUHB01000001.1 | - | 1541536 | 1541495 | **RNA**→ RpsD (COG0522)rpsD (PRK05327)→<br>RNAP_alpha_NTD (cd06928)PRK05182 (PRK05182)RNA_pol_A_CTD (pfam03118)→<br>RplQ (COG0203)rplQ (PRK05591)→ CarA (COG0505)PRK12564 (PRK12564)→ |
| Cba-1-1 | NZ_CP009887.1 | - | 225786 | 225745 | **RNA**→ RpsD (COG0522)rpsD (PRK05327)→<br>RNAP_alpha_NTD (cd06928)PRK05182 (PRK05182)RNA_pol_A_CTD (pfam03118)→<br>RplQ (COG0203)rplQ (PRK05591)→ CarA (COG0505)PRK12564 (PRK12564)→ |
| Mlu-1-1 | NZ_CP011071.1 | - | 769099 | 769058 | **RNA**→ RpsD (COG0522)rpsD (PRK05327)→<br>RNAP_alpha_NTD (cd06928)PRK05182 (PRK05182)RNA_pol_A_CTD (pfam03118)→ |
| Msp-2-1 | NZ_JQNJ01000001.1 | - | 660265 | 660224 | **RNA**→ RpsD (COG0522)rpsD (PRK05327)→<br>RNAP_alpha_NTD (cd06928)PRK05182 (PRK05182)RNA_pol_A_CTD (pfam03118)→<br>RplQ (COG0203)rplQ (PRK05591)→ CarA (COG0505)PRK12564 (PRK12564)→ |
| env-522 | KGI_S1_ANT01_95m_c10061602 | - | 1117 | 1069 | **RNA**→ RpsD (COG0522)rpsD (PRK05327)→ RpoA (COG0202)PRK05182 (PRK05182)→ |
| env-523 | KGI_S1_ANT01_95m_c10514893 | + | 216 | 264 | **RNA**→ |
| env-524 | SA_S2_NOR13_50m_c1195435 | + | 57 | 105 | **RNA**→ rpsD (PRK05327)Ribosomal_S4 (pfam00163)→ |
| env-525 | SA_S1_NOR08_45m_c10507688 | + | 279 | 317 | **RNA**→ |
| env-526 | SA_S1_NOR08_45m_c10126322 | + | 241 | 289 | **RNA**→ RpsD (COG0522)rpsD (PRK05327)→ |
| env-527 | KGI_S2_ANT04_2345m_c1034044 | + | 979 | 1027 | **RNA**→ rpsD (PRK05327)Ribosomal_S4 (pfam00163)→ |
| env-528 | SA_S1_NOR02_45m_c1006351 | - | 213 | 165 | **RNA**→ rpsD (PRK05327)Ribosomal_S4 (pfam00163)→ |
| env-529 | ElkS_mat_MD6A_1003011 | - | 1378 | 1330 | **RNA**→ RpsD (COG0522)rpsD (PRK05327)→ RNAP_alpha_NTD (cd06928)PRK05182 (PRK05182)→ |
| env-530 | CESG01186264.1 | + | 76 | 124 | **RNA**→ RpsD (COG0522)rpsD (PRK05327)→ |
| env-531 | JGI24023J19991_10302847 | + | 95 | 143 | **RNA**→ rpsD (PRK05327)Ribosomal_S4 (pfam00163)→ |
| Ale-1-1 | NZ_BBNQ01000011.1 | - | 90098 | 90050 | **RNA**→ RpsD (COG0522)rpsD (PRK05327)→ RpoA (COG0202)PRK05182 (PRK05182)→<br>RplQ (COG0203)rplQ (PRK05591)→ |
| Fag-1-1 | NZ_HG315671.1 | + | 689063 | 689111 | **RNA**→ RpsD (COG0522)rpsD (PRK05327)→ RpoA (COG0202)PRK05182 (PRK05182)→<br>RplQ (COG0203)rplQ (PRK05591)→ |
| Osp-1-1 | NZ_JADS01000010.1 | + | 109557 | 109605 | **RNA**→ RpsD (COG0522)rpsD (PRK05327)→ RpoA (COG0202)PRK05182 (PRK05182)→<br>RplQ (COG0203)rplQ (PRK05591)→ CarA (COG0505)PRK12564 (PRK12564)→ |
| env-532 | CENJ01244751.1 | - | 131 | 90 | **RNA**→ |

| | | | | | |
|---|---|---|---|---|---|
| env-533 | KGI_S1_ANT01_95m_c10140035 | + | 608 | 646 | **RNA**→ |
| env-534 | SA_S1_NOR08_45m_c10026887 | - | 62 | 24 | **RNA**→ |
| env-535 | SA_S1_NOR08_45m_c10548832 | - | 224 | 176 | **RNA**→ rpsD (PRK05327)Ribosomal_S4 (pfam00163)→ |
| env-536 | SA_S2_NOR15_50m_c10053365 | + | 1565 | 1613 | **RNA**→ ←hypo |
| env-537 | JGI24024J18818_10001114 | + | 7715 | 7763 | **RNA**→ RpsD (COG0522)rpsD (PRK05327)→ RpoA (COG0202)PRK05182 (PRK05182)→ |
| | | | | | RplQ (COG0203)rplQ (PRK05591)→ PRK12564 (PRK12564)GATase (pfam00117)CPSase_sm_chain (pfam00988)→ |
| env-538 | CEUD01129050.1 | + | 159 | 207 | **RNA**→ rpsD (PRK05327)Ribosomal_S4 (pfam00163)→ |
| env-539 | JGI24724J26744_10024136 | - | 2300 | 2252 | **RNA**→ RpsD (COG0522)rpsD (PRK05327)→ RpoA (COG0202)PRK05182 (PRK05182)→ |
| | | | | | RplQ (COG0203)rplQ (PRK05591)→ |
| Fic-1-1 | NZ_KN525716.1 | + | 2656682 | 2656730 | **RNA**→ RpsD (COG0522)rpsD (PRK05327)→ |
| | | | | | RNAP_alpha_NTD (cd06928)PRK05182 (PRK05182)RNA_pol_A_CTD (pfam03118)→ |
| | | | | | RplQ (COG0203)rplQ (PRK05591)→ CarA (COG0505)PRK12564 (PRK12564)→ |
| Mze-1-1 | NZ_AULQ01000008.1 | - | 44044 | 43996 | **RNA**→ RpsD (COG0522)rpsD (PRK05327)→ |
| | | | | | RNAP_alpha_NTD (cd06928)PRK05182 (PRK05182)RNA_pol_A_CTD (pfam03118)→ |
| | | | | | RplQ (COG0203)rplQ (PRK05591)→ CarA (COG0505)PRK12564 (PRK12564)→ |
| env-540 | KGI_S1_ANT01_95m_c10270403 | + | 123 | 171 | **RNA**→ rpsD (PRK05327)Ribosomal_S4 (pfam00163)→ |
| env-541 | DelMOSum2010_c10000497 | - | 33517 | 33469 | **RNA**→ RpsD (COG0522)rpsD (PRK05327)→ |
| | | | | | RNAP_alpha_NTD (cd06928)PRK05182 (PRK05182)RNA_pol_A_CTD (pfam03118)→ |
| | | | | | RplQ (COG0203)rplQ (PRK05591)→ CarA (COG0505)PRK12564 (PRK12564)→ |
| env-542 | KGI_S1_ANT02_95m_c10388256 | - | 163 | 115 | **RNA**→ rpsD (PRK05327)Ribosomal_S4 (pfam00163)→ |
| Oma-1-1 | NZ_JADR01000010.1 | + | 109053 | 109101 | **RNA**→ RpsD (COG0522)rpsD (PRK05327)→ RpoA (COG0202)PRK05182 (PRK05182)→ |
| | | | | | RplQ (COG0203)rplQ (PRK05591)→ hypo→ hypo→ CarA (COG0505)PRK12564 (PRK12564)→ |
| env-543 | wf3_Contig160 | - | 193 | 152 | **RNA**→ |
| env-544 | KGI_S1_ANT01_95m_c10151354 | - | 449 | 408 | **RNA**→ RpsD (COG0522)rpsD (PRK05327)→ |
| env-545 | SA_S1_NOR05_45m_c10248728 | - | 299 | 258 | **RNA**→ rpsD (PRK05327)Ribosomal_S4 (pfam00163)→ |
| env-546 | SA_S1_NOR08_45m_c10038794 | + | 22 | 63 | **RNA**→ RpsD (COG0522)rpsD (PRK05327)→ |
| | | | | | RNAP_alpha_NTD (cd06928)PRK05182 (PRK05182)RNA_pol_A_CTD (pfam03118)→ |
| | | | | | RplQ (COG0203)rplQ (PRK05591)→ |
| env-547 | SA_S1_NOR08_45m_c10391914 | + | 152 | 193 | **RNA**→ rpsD (PRK05327)Ribosomal_S4 (pfam00163)→ |
| env-548 | JGI24724J26744_10233624 | - | 582 | 534 | **RNA**→ RpsD (COG0522)rpsD (PRK05327)→ |
| env-549 | ElkS_mat_CD2A_1000229 | - | 2419 | 2378 | **RNA**→ RpsD (COG0522)rpsD (PRK05327)→ |
| | | | | | RNAP_alpha_NTD (cd06928)PRK05182 (PRK05182)RNA_pol_A_CTD (pfam03118)→ |
| | | | | | RplQ (COG0203)rplQ (PRK05591)→ PRK12564 (PRK12564)CPSase_sm_chain (pfam00988)→ |
| env-550 | KGI_S1_ANT03_95m_c1144743 | - | 169 | 121 | **RNA**→ rpsD (PRK05327)Ribosomal_S4 (pfam00163)→ |
| env-551 | CEOE01026498.1 | + | 404 | 445 | **RNA**→ RpsD (COG0522)rpsD (PRK05327)→ |
| env-552 | CEQG01271447.1 | - | 1058 | 1017 | **RNA**→ RpsD (COG0522)rpsD (PRK05327)→ rpoA (CHL00013)PRK05182 (PRK05182)→ |
| env-553 | CEVF01017472.1 | - | 1828 | 1787 | **RNA**→ RpsD (COG0522)rpsD (PRK05327)→ |
| | | | | | RNAP_alpha_NTD (cd06928)PRK05182 (PRK05182)RNA_pol_A_CTD (pfam03118)→ |
| env-554 | LAZR01000011.1 | + | 276612 | 276653 | **RNA**→ RpsD (COG0522)rpsD (PRK05327)→ |
| | | | | | RNAP_alpha_NTD (cd06928)PRK05182 (PRK05182)RNA_pol_A_CTD (pfam03118)→ |
| | | | | | RplQ (COG0203)rplQ (PRK05591)→ CarA (COG0505)PRK12564 (PRK12564)→ |
| env-555 | ElkS_mat_CD6A_1038608 | + | 159 | 200 | **RNA**→ rpsD (PRK05327)Ribosomal_S4 (pfam00163)→ |
| env-556 | JGI24023J19991_10001527 | + | 11806 | 11847 | **RNA**→ RpsD (COG0522)rpsD (PRK05327)→ |
| | | | | | RNAP_alpha_NTD (cd06928)PRK05182 (PRK05182)RNA_pol_A_CTD (pfam03118)→ |
| | | | | | RplQ (COG0203)rplQ (PRK05591)→ |
| env-557 | JGI24023J19991_10073144 | - | 242 | 201 | **RNA**→ rpsD (PRK05327)Ribosomal_S4 (pfam00163)→ |
| env-558 | JGI24023J19991_10182654 | - | 76 | 35 | **RNA**→ |
| env-559 | JGI24723J26617_10000010 | - | 143173 | 143132 | **RNA**→ RpsD (COG0522)rpsD (PRK05327)→ |
| | | | | | RNAP_alpha_NTD (cd06928)PRK05182 (PRK05182)RNA_pol_A_CTD (pfam03118)→ |
| | | | | | RplQ (COG0203)rplQ (PRK05591)→ |
| env-560 | JGI24723J26617_10310440 | + | 372 | 403 | **RNA**→ |
| env-561 | SRS013705_Baylor_scaffold_46745 | - | 1715 | 1674 | **RNA**→ RpsD (COG0522)rpsD (PRK05327)→ |
| | | | | | RNAP_alpha_NTD (cd06928)PRK05182 (PRK05182)RNA_pol_A_CTD (pfam03118)→ |
| env-562 | SRS014691_C2644950 | - | 228 | 187 | **RNA**→ rpsD (PRK05327)Ribosomal_S4 (pfam00163)→ |

| | | | | | |
|---|---|---|---|---|---|
| Mth-1-1 | NZ_LDAS01000001.1 | - | 2178073 | 2178032 | **RNA**→ RpsD (COG0522)rpsD (PRK05327)→ RNAP_alpha_NTD (cd06928)PRK05182 (PRK05182)RNA_pol_A_CTD (pfam03118)→ RplQ (COG0203)rplQ (PRK05591)→ CarA (COG0505)PRK12564 (PRK12564)→ |
| Ead-1-1 | NZ_KB907546.1 | + | 154121 | 154162 | **RNA**→ RpsD (COG0522)rpsD (PRK05327)→ RNAP_alpha_NTD (cd06928)PRK05182 (PRK05182)RNA_pol_A_CTD (pfam03118)→ RplQ (COG0203)rplQ (PRK05591)→ rplQ (PRK05591)Ribosomal_L17 (pfam01196)→ CarA (COG0505)PRK12564 (PRK12564)→ eno (PRK00077)enolase (cd03313)→ |
| Cal-1-1 | NC_014934.1 | - | 701743 | 701702 | **RNA**→ RpsD (COG0522)rpsD (PRK05327)→ RNAP_alpha_NTD (cd06928)PRK05182 (PRK05182)RNA_pol_A_CTD (pfam03118)→ RplQ (COG0203)rplQ (PRK05591)→ |
| Zga-1-1 | NC_015844.1 | + | 2988024 | 2988065 | **RNA**→ RpsD (COG0522)rpsD (PRK05327)→ RNAP_alpha_NTD (cd06928)PRK05182 (PRK05182)RNA_pol_A_CTD (pfam03118)→ RplQ (COG0203)rplQ (PRK05591)→ CarA (COG0505)PRK12564 (PRK12564)→ |
| Mfo-1-1 | NZ_JQLH01000001.1 | + | 1025162 | 1025203 | **RNA**→ RpsD (COG0522)rpsD (PRK05327)→ RNAP_alpha_NTD (cd06928)PRK05182 (PRK05182)RNA_pol_A_CTD (pfam03118)→ RplQ (COG0203)rplQ (PRK05591)→ CarA (COG0505)PRK12564 (PRK12564)→ |
| env-563 | CEWJ01225499.1 | - | 661 | 620 | **RNA**→ RpsD (COG0522)rpsD (PRK05327)→ |
| env-564 | JGI11876J14442_10085363 | - | 384 | 343 | **RNA**→ RpsD (COG0522)rpsD (PRK05327)→ |
| env-565 | JGI11876J14442_11014628 | - | 113 | 72 | **RNA**→ |
| env-566 | JGI24023J19991_10047227 | - | 1363 | 1322 | **RNA**→ RpsD (COG0522)rpsD (PRK05327)→ RNAP_alpha_NTD (cd06928)PRK05182 (PRK05182)→ |
| env-567 | JGI24023J19991_10138758 | - | 690 | 649 | **RNA**→ RpsD (COG0522)rpsD (PRK05327)→ |
| env-568 | JGI24723J26617_10014588 | + | 1099 | 1140 | **RNA**→ RpsD (COG0522)rpsD (PRK05327)→ RpoA (COG0202)PRK05182 (PRK05182)→ RplQ (COG0203)rplQ (PRK05591)→ |
| env-569 | JGI24723J26617_10602636 | - | 242 | 201 | **RNA**→ rpsD (PRK05327)Ribosomal_S4 (pfam00163)→ |
| env-570 | Subgingival_plaque_LANL_scaffold_16751 | - | 129 | 88 | **RNA**→ rpsD (PRK05327)Ribosomal_S4 (pfam00163)→ |
| env-571 | ATED_F4WRAXD01DYY4F | + | 189 | 230 | **RNA**→ rpsD (PRK05327)Ribosomal_S4 (pfam00163)→ |
| Mru-1-1 | NC_015945.1 | - | 713998 | 713957 | **RNA**→ RpsD (COG0522)rpsD (PRK05327)→ RNAP_alpha_NTD (cd06928)PRK05182 (PRK05182)RNA_pol_A_CTD (pfam03118)→ RplQ (COG0203)rplQ (PRK05591)→ CarA (COG0505)PRK12564 (PRK12564)→ |
| Cly-1-1 | NC_015167.1 | + | 2435971 | 2436012 | **RNA**→ RpsD (COG0522)rpsD (PRK05327)→ RNAP_alpha_NTD (cd06928)PRK05182 (PRK05182)RNA_pol_A_CTD (pfam03118)→ RplQ (COG0203)rplQ (PRK05591)→ |
| Zul-1-1 | NZ_JQMD01000002.1 | + | 4824073 | 4824114 | **RNA**→ RpsD (COG0522)rpsD (PRK05327)→ RNAP_alpha_NTD (cd06928)PRK05182 (PRK05182)RNA_pol_A_CTD (pfam03118)→ RplQ (COG0203)rplQ (PRK05591)→ CarA (COG0505)PRK12564 (PRK12564)→ |
| env-572 | KGI_S1_ANT01_95m_c10057626 | - | 442 | 394 | **RNA**→ RpsD (COG0522)rpsD (PRK05327)→ |
| env-573 | KGI_S1_ANT01_95m_c10621365 | + | 226 | 264 | **RNA**→ |
| env-574 | SA_S2_NOR13_50m_c1011088 | - | 1383 | 1335 | **RNA**→ RpsD (COG0522)rpsD (PRK05327)→ RNAP_alpha_NTD (cd06928)PRK05182 (PRK05182)→ |
| env-575 | SA_S1_NOR05_45m_c10208693 | + | 159 | 207 | **RNA**→ rpsD (PRK05327)Ribosomal_S4 (pfam00163)→ |
| env-576 | SA_S1_NOR05_45m_c10289234 | + | 223 | 271 | **RNA**→ rpsD (PRK05327)Ribosomal_S4 (pfam00163)→ |
| env-577 | KGI_S2_ANT04_2345m_c1009836 | + | 891 | 939 | **RNA**→ RpsD (COG0522)rpsD (PRK05327)→ RNAP_alpha_NTD (cd06928)PRK05182 (PRK05182)RNA_pol_A_CTD (pfam03118)→ RplQ (COG0203)rplQ (PRK05591)→ |
| env-578 | SA_S2_NOR15_50m_c10285015 | + | 159 | 207 | **RNA**→ rpsD (PRK05327)Ribosomal_S4 (pfam00163)→ |
| env-579 | KGI_S1_ANT03_95m_c1055981 | - | 201 | 153 | **RNA**→ rpsD (PRK05327)Ribosomal_S4 (pfam00163)→ |
| env-580 | JGI24024J18818_10780925 | + | 148 | 196 | **RNA**→ |
| env-581 | CEUX01091020.1 | + | 141 | 189 | **RNA**→ RpsD (COG0522)rpsD (PRK05327)→ RpoA (COG0202)PRK05182 (PRK05182)→ |
| env-582 | KVRMV2_101047941 | + | 60 | 108 | **RNA**→ rpsD (PRK05327)Ribosomal_S4 (pfam00163)→ |
| Gsa-1-1 | NZ_KE384430.1 | + | 1061256 | 1061304 | **RNA**→ RpsD (COG0522)rpsD (PRK05327)→ RpoA (COG0202)PRK05182 (PRK05182)→ RplQ (COG0203)rplQ (PRK05591)→ CarA (COG0505)PRK12564 (PRK12564)→ |
| Fba-4-1 | NZ_ALIH01000001.1 | + | 391313 | 391361 | **RNA**→ RpsD (COG0522)rpsD (PRK05327)→ RpoA (COG0202)PRK05182 (PRK05182)→ RplQ (COG0203)rplQ (PRK05591)→ hypo→ hypo→ CarA (COG0505)PRK12564 (PRK12564)→ |
| Ssp-2-1 | NZ_JHZX01000001.1 | - | 1257672 | 1257624 | **RNA**→ RpsD (COG0522)rpsD (PRK05327)→ RpoA (COG0202)PRK05182 (PRK05182)→ RplQ (COG0203)rplQ (PRK05591)→ |
| env-583 | KGI_S1_ANT01_95m_c10261611 | - | 344 | 296 | **RNA**→ rpsD (PRK05327)Ribosomal_S4 (pfam00163)→ |

| | | | | | |
|---|---|---|---|---|---|
| env-584 | SA_S1_NOR05_45m_c10000531 | + | 10203 | 10251 | **RNA**→ ←hypo |
| env-585 | SA_S1_NOR08_45m_c10128342 | - | 449 | 401 | **RNA**→ RpsD (COG0522)rpsD (PRK05327)→ |
| env-586 | KGI_S2_ANT04_2345m_c1022193 | + | 1360 | 1408 | **RNA**→ rpsD (PRK05327)Ribosomal_S4 (pfam00163)→ |
| env-587 | KGI_S1_ANT02_95m_c10022183 | + | 483 | 531 | **RNA**→ RpsD (COG0522)rpsD (PRK05327)→ RpoA (COG0202)PRK05182 (PRK05182)→ RplQ (COG0203)rplQ (PRK05591)→ |
| env-588 | CEQS01129325.1 | - | 418 | 370 | **RNA**→ S4 (smart00363)RpsD (COG0522)→ |
| env-589 | LGVF01581283.1 | - | 913 | 865 | **RNA**→ RpsD (COG0522)rpsD (PRK05327)→ PRK05182 (PRK05182)RNA_pol_L (pfam01193)→ |
| Lsp-1-1 | NZ_JYNQ01000001.1 | - | 1525387 | 1525339 | **RNA**→ RpsD (COG0522)rpsD (PRK05327)→ RpoA (COG0202)PRK05182 (PRK05182)→ RplQ (COG0203)rplQ (PRK05591)→ |
| Sze-1-1 | NZ_CP007202.1 | - | 225594 | 225546 | **RNA**→ RpsD (COG0522)rpsD (PRK05327)→ RpoA (COG0202)PRK05182 (PRK05182)→ RplQ (COG0203)rplQ (PRK05591)→ CarA (COG0505)PRK12564 (PRK12564)→ |
| Osp-2-1 | NZ_JADT01000008.1 | + | 109566 | 109614 | **RNA**→ RpsD (COG0522)rpsD (PRK05327)→ RpoA (COG0202)PRK05182 (PRK05182)→ RplQ (COG0203)rplQ (PRK05591)→ CarA (COG0505)PRK12564 (PRK12564)→ |
| env-590 | SA_S1_NOR08_45m_c10544044 | + | 126 | 174 | **RNA**→ rpsD (PRK05327)Ribosomal_S4 (pfam00163)→ |
| env-591 | KGI_S1_ANT02_95m_c10037840 | - | 206 | 165 | **RNA**→ rpsD (PRK05327)Ribosomal_S4 (pfam00163)→ |
| env-592 | CESF01075296.1 | + | 14667 | 14715 | **RNA**→ RpsD (COG0522)rpsD (PRK05327)→ RpoA (COG0202)PRK05182 (PRK05182)→ RplQ (COG0203)rplQ (PRK05591)→ hypo→ hypo→ CarA (COG0505)PRK12564 (PRK12564)→ |
| env-593 | ACQI01007085.1 | - | 679 | 639 | **RNA**→ RpsD (COG0522)rpsD (PRK05327)→ |
| env-594 | TDF_OR_ARG05_123m_c1016145 | - | 527 | 486 | **RNA**→ RpsD (COG0522)rpsD (PRK05327)→ |
| env-595 | TDF_OR_ARG04_113m_c1149045 | - | 173 | 132 | **RNA**→ rpsD (PRK05327)Ribosomal_S4 (pfam00163)→ |
| env-596 | SA_S1_NOR05_45m_c10029529 | + | 805 | 846 | **RNA**→ RpsD (COG0522)rpsD (PRK05327)→ hypo→ |
| env-597 | SA_S1_NOR08_45m_c10216906 | - | 145 | 104 | **RNA**→ rpsD (PRK05327)Ribosomal_S4 (pfam00163)→ |
| env-598 | KGI_S2_ANT04_2345m_c1307712 | - | 223 | 182 | **RNA**→ rpsD (PRK05327)Ribosomal_S4 (pfam00163)→ |
| env-599 | KGI_S1_ANT02_95m_c10521979 | + | 127 | 168 | **RNA**→ rpsD (PRK05327)Ribosomal_S4 (pfam00163)→ |
| env-600 | BS_KBA_SWE12_21m_c10766585 | - | 160 | 112 | **RNA**→ rpsD (PRK05327)Ribosomal_S4 (pfam00163)→ |
| env-601 | CENV01048908.1 | + | 2405 | 2446 | **RNA**→ RpsD (COG0522)rpsD (PRK05327)→ PRK05182 (PRK05182)RNA_pol_L (pfam01193)→ |
| env-602 | CEOE01058112.1 | - | 502 | 461 | **RNA**→ RpsD (COG0522)rpsD (PRK05327)→ |
| env-603 | CERQ01210477.1 | - | 335 | 294 | **RNA**→ rpsD (PRK05327)Ribosomal_S4 (pfam00163)→ |
| env-604 | JGI12210J13797_10891149 | + | 219 | 250 | **RNA**→ |
| Psp-3-1 | NC_020830.1 | + | 2776178 | 2776219 | **RNA**→ RpsD (COG0522)rpsD (PRK05327)→ RpoA (COG0202)PRK05182 (PRK05182)→ RplQ (COG0203)rplQ (PRK05591)→ |
| env-605 | SA_S1_NOR05_45m_c10021376 | - | 1404 | 1356 | **RNA**→ RpsD (COG0522)rpsD (PRK05327)→ RNAP_alpha_NTD (cd06928)PRK05182 (PRK05182)→ |
| env-606 | JCVI_SCAF_1101668093316 | - | 1811 | 1770 | **RNA**→ RpsD (COG0522)rpsD (PRK05327)→ PRK05182 (PRK05182)RNA_pol_L (pfam01193)→ RpoA (COG0202)PRK05182 (PRK05182)→ |
| env-607 | JGI20152J14361_10000058 | + | 48115 | 48163 | **RNA**→ RpsD (COG0522)rpsD (PRK05327)→ RpoA (COG0202)PRK05182 (PRK05182)→ RplQ (COG0203)rplQ (PRK05591)→ |
| env-608 | _GVICOHK01D9LRN | + | 86 | 126 | **RNA**→ |
| env-609 | LWAeNNiAF_GBUVFP102I049T | - | 173 | 131 | **RNA**→ rpsD (PRK05327)Ribosomal_S4 (pfam00163)→ |
| env-610 | JGI25909J50240_1174521 | + | 96 | 138 | **RNA**→ rpsD (PRK05327)Ribosomal_S4 (pfam00163)→ |
| env-611 | M3P_c10616965 | + | 93 | 134 | **RNA**→ |
| env-612 | 3300000052_007306 | + | 713 | 755 | **RNA**→ RpsD (COG0522)rpsD (PRK05327)→ RNAP_alpha_NTD (cd06928)PRK05182 (PRK05182)RNA_pol_A_CTD (pfam03118)→ RplQ (COG0203)rplQ (PRK05591)→ |
| Fgi-1-1 | NZ_JNCP01000052.1 | + | 79268 | 79310 | **RNA**→ RpsD (COG0522)rpsD (PRK05327)→ RNAP_alpha_NTD (cd06928)PRK05182 (PRK05182)RNA_pol_A_CTD (pfam03118)→ RplQ (COG0203)rplQ (PRK05591)→ |
| env-613 | JGI12272J11983_1656627 | - | 104 | 62 | **RNA**→ ←hypo |
| env-614 | KGI_S2_ANT04_2345m_c1000254 | - | 19083 | 19041 | **RNA**→ RpsD (COG0522)rpsD (PRK05327)→ RNAP_alpha_NTD (cd06928)PRK05182 (PRK05182)RNA_pol_A_CTD (pfam03118)→ RplQ (COG0203)rplQ (PRK05591)→ |
| env-615 | JGI25908J49247_10003866 | + | 1771 | 1813 | **RNA**→ rpsD (PRK05327)Ribosomal_S4 (pfam00163)→ S4 (smart00363)RpsD (COG0522)→ RNAP_alpha_NTD (cd06928)PRK05182 (PRK05182)RNA_pol_A_CTD (pfam03118)→ RplQ (COG0203)rplQ (PRK05591)→ |
| env-616 | JGI12421J11937_10089621 | - | 766 | 724 | **RNA**→ RpsD (COG0522)rpsD (PRK05327)→ hypo→ |

| | | | | | |
|---|---|---|---|---|---|
| env-617 | JGI25912J50252_10007817 | - | 2296 | 2254 | RNA→ RpsD (COG0522)rpsD (PRK05327)→ RNAP_alpha_NTD (cd06928)PRK05182 (PRK05182)RNA_pol_A_CTD (pfam03118)→ |
| Fps-1-1 | NZ_JRWC01000005.1 | - | 680844 | 680802 | RNA→ RpsD (COG0522)rpsD (PRK05327)→ hypo→ RplQ (COG0203)rplQ (PRK05591)→ |
| env-618 | ARMCC_Contig_363473 | + | 473 | 515 | RNA→ |
| env-619 | JGI20153J14295_10238628 | + | 268 | 310 | RNA→ ←hypo |
| env-620 | JGI25907J50239_1282084 | + | 32 | 74 | RNA→ rpsD (PRK05327)Ribosomal_S4 (pfam00163)→ |
| Ffr-1-1 | NZ_AUDO01000001.1 | - | 396813 | 396771 | RNA→ RpsD (COG0522)rpsD (PRK05327)→ RNAP_alpha_NTD (cd06928)PRK05182 (PRK05182)RNA_pol_A_CTD (pfam03118)→ RplQ (COG0203)rplQ (PRK05591)→ |
| env-621 | KGI_S1_ANT01_95m_c10061422 | + | 544 | 586 | RNA→ RpsD (COG0522)rpsD (PRK05327)→ |
| env-622 | JGI24770J26754_10196236 | + | 572 | 614 | RNA→ |
| env-623 | JGI25909J50240_1035901 | - | 911 | 869 | RNA→ rpsD (PRK05327)Ribosomal_S4 (pfam00163)→ S4 (smart00363)RpsD (COG0522)→ PRK05182 (PRK05182)RNA_pol_L (pfam01193)→ |
| env-624 | JGI25911J50253_10049044 | + | 118 | 160 | RNA→ rpsD (PRK05327)Ribosomal_S4 (pfam00163)→ RpsD (COG0522)rpsD (PRK05327)→ RNAP_alpha_NTD (cd06928)PRK05182 (PRK05182)→ |
| env-625 | 3300000052_115516 | - | 282 | 240 | RNA→ rpsD (PRK05327)Ribosomal_S4 (pfam00163)→ |
| Ffr-2-1 | NZ_AHKF01000008.1 | - | 69789 | 69747 | RNA→ RpsD (COG0522)rpsD (PRK05327)→ RpsD (COG0522)rpsD (PRK05327)→ RNAP_alpha_NTD (cd06928)PRK05182 (PRK05182)RNA_pol_A_CTD (pfam03118)→ RNAP_alpha_NTD (cd06928)PRK05182 (PRK05182)RNA_pol_A_CTD (pfam03118)→ |
| env-626 | KGI_S1_ANT01_95m_c10048862 | + | 770 | 812 | RNA→ RpsD (COG0522)rpsD (PRK05327)→ |
| env-627 | JGI24770J26754_10143512 | - | 394 | 352 | RNA→ RpsD (COG0522)rpsD (PRK05327)→ |
| env-628 | MRS1b_contig_1810326 | - | 411 | 369 | RNA→ S4 (smart00363)RpsD (COG0522)→ |
| Fsp-2-1 | NZ_AJXL01000028.1 | - | 8777 | 8735 | RNA→ RpsD (COG0522)rpsD (PRK05327)→ RNAP_alpha_NTD (cd06928)PRK05182 (PRK05182)RNA_pol_A_CTD (pfam03118)→ RplQ (COG0203)rplQ (PRK05591)→ |
| env-629 | 3300001605_10110064 | + | 1045 | 1087 | RNA→ RpsD (COG0522)rpsD (PRK05327)→ PRK05182 (PRK05182)RNA_pol_L (pfam01193)→ |
| env-630 | JGI25910J50241_10439814 | - | 102 | 60 | RNA→ |
| env-631 | MRS2a_Contig_10143 | + | 194 | 236 | RNA→ RpsD (COG0522)rpsD (PRK05327)→ rpoA (CHL00013)PRK05182 (PRK05182)→ |
| Fch-1-1 | NZ_JASY01000008.1 | - | 165622 | 165580 | RNA→ RpsD (COG0522)rpsD (PRK05327)→ RNAP_alpha_NTD (cd06928)PRK05182 (PRK05182)RNA_pol_A_CTD (pfam03118)→ RplQ (COG0203)rplQ (PRK05591)→ |
| env-632 | M3P_c10026069 | + | 470 | 512 | RNA→ RpsD (COG0522)rpsD (PRK05327)→ |
| env-633 | Y607_1006252 | + | 157633 | 157675 | RNA→ RpsD (COG0522)rpsD (PRK05327)→ RNAP_alpha_NTD (cd06928)PRK05182 (PRK05182)RNA_pol_A_CTD (pfam03118)→ rplQ (PRK05591)Ribosomal_L17 (pfam01196)→ |
| env-634 | AglaG_GDN60OX02GVF13 | + | 112 | 154 | RNA→ rpsD (PRK05327)Ribosomal_S4 (pfam00163)→ |
| Fsp-6-1 | NZ_JXQR01000040.1 | + | 224224 | 224266 | RNA→ RpsD (COG0522)rpsD (PRK05327)→ RNAP_alpha_NTD (cd06928)PRK05182 (PRK05182)RNA_pol_A_CTD (pfam03118)→ RplQ (COG0203)rplQ (PRK05591)→ |
| env-635 | SA_S1_NOR08_45m_c10825333 | - | 194 | 152 | RNA→ rpsD (PRK05327)Ribosomal_S4 (pfam00163)→ |
| env-636 | PBR_1001305 | - | 1487 | 1445 | RNA→ RpsD (COG0522)rpsD (PRK05327)→ RNAP_alpha_NTD (cd06928)PRK05182 (PRK05182)→ |
| env-637 | JGI25908J49247_10195454 | - | 436 | 394 | RNA→ S4 (smart00363)RpsD (COG0522)→ |
| env-638 | JGI25914J50564_10084240 | - | 445 | 403 | RNA→ RpsD (COG0522)rpsD (PRK05327)→ |
| Fte-1-1 | NZ_KE384384.1 | - | 340469 | 340427 | RNA→ RpsD (COG0522)rpsD (PRK05327)→ RNAP_alpha_NTD (cd06928)PRK05182 (PRK05182)RNA_pol_A_CTD (pfam03118)→ RplQ (COG0203)rplQ (PRK05591)→ |
| env-639 | 3300000053_265566 | - | 115 | 73 | RNA→ |
| env-640 | 3300000558_10106130 | + | 1517 | 1559 | RNA→ rpsD (PRK05327)Ribosomal_S4 (pfam00163)→ |
| env-641 | JGI25912J50252_10474158 | + | 162 | 204 | RNA→ |
| env-642 | LM15JUN2010E_c5792 | - | 283 | 241 | RNA→ rpsD (PRK05327)Ribosomal_S4 (pfam00163)→ |
| env-643 | BS_KBA_SWE12_21m_c10390926 | - | 240 | 198 | RNA→ rpsD (PRK05327)Ribosomal_S4 (pfam00163)→ |
| env-644 | KGI_S1_ANT01_95m_c10748070 | - | 50 | 18 | RNA→ |
| env-645 | BS_KBA_SWE21_205m_10011318 | + | 306 | 348 | RNA→ RpsD (COG0522)rpsD (PRK05327)→ RNAP_alpha_NTD (cd06928)PRK05182 (PRK05182)RNA_pol_A_CTD (pfam03118)→ RplQ (COG0203)rplQ (PRK05591)→ |

| | | | | | |
|---|---|---|---|---|---|
| env-646 | BS_KBA_SWE02_21m_10026740 | + | 218 | 260 | RNA→ RpsD (COG0522)rpsD (PRK05327)→ RNAP_alpha_NTD (cd06928)PRK05182 (PRK05182)RNA_pol_A_CTD (pfam03118)→ |
| env-647 | BS_KBA_SWE12_21m_c10002755 | - | 5914 | 5872 | RNA→ RpsD (COG0522)rpsD (PRK05327)→ RNAP_alpha_NTD (cd06928)PRK05182 (PRK05182)RNA_pol_A_CTD (pfam03118)→ RplQ (COG0203)rplQ (PRK05591)→ hypo→ CarA (COG0505)PRK12564 (PRK12564)→ |
| env-648 | TB_FS06_10DRAFT_1022891 | - | 1909 | 1861 | RNA→ RpsD (COG0522)rpsD (PRK05327)→ RpoA (COG0202)PRK05182 (PRK05182)→ RplQ (COG0203)rplQ (PRK05591)→ |
| env-649 | TB_FS08_3DRAFT_1007294 | + | 1041 | 1089 | RNA→ RpsD (COG0522)rpsD (PRK05327)→ RpoA (COG0202)PRK05182 (PRK05182)→ RplQ (COG0203)rplQ (PRK05591)→ |
| env-650 | TB_FS06_10DRAFT_1003825 | - | 2918 | 2870 | RNA→ RpsD (COG0522)rpsD (PRK05327)→ RpoA (COG0202)PRK05182 (PRK05182)→ RplQ (COG0203)rplQ (PRK05591)→ |
| env-651 | TB_PC08_3_1003356 | + | 8254 | 8302 | RNA→ rpsD (PRK05327)Ribosomal_S4 (pfam00163)→ |
| env-652 | BS_KBA_SWE12_21m_c10074507 | - | 439 | 397 | RNA→ RpsD (COG0522)rpsD (PRK05327)→ |
| Fda-1-1 | NZ_AUDK01000020.1 | - | 17618 | 17576 | RNA→ RpsD (COG0522)rpsD (PRK05327)→ RNAP_alpha_NTD (cd06928)PRK05182 (PRK05182)RNA_pol_A_CTD (pfam03118)→ RplQ (COG0203)rplQ (PRK05591)→ |
| env-653 | TB_FS08_3DRAFT_1037856 | - | 82 | 40 | RNA→ |
| env-654 | JGI24770J26754_10248674 | - | 226 | 184 | RNA→ rpsD (PRK05327)Ribosomal_S4 (pfam00163)→ |
| env-655 | JGI24770J26754_10101330 | - | 620 | 578 | RNA→ RpsD (COG0522)rpsD (PRK05327)→ |
| env-656 | JGI25908J49247_10434406 | - | 89 | 47 | RNA→ |
| env-657 | M3P_c10614138 | + | 93 | 134 | RNA→ |
| Fsp-1-1 | NZ_LEKS01000003.1 | - | 349739 | 349697 | RNA→ RpsD (COG0522)rpsD (PRK05327)→ RNAP_alpha_NTD (cd06928)PRK05182 (PRK05182)RNA_pol_A_CTD (pfam03118)→ RplQ (COG0203)rplQ (PRK05591)→ |
| Fhy-1-1 | NZ_JPRM01000006.1 | - | 152624 | 152582 | RNA→ RpsD (COG0522)rpsD (PRK05327)→ RNAP_alpha_NTD (cd06928)PRK05182 (PRK05182)RNA_pol_A_CTD (pfam03118)→ RplQ (COG0203)rplQ (PRK05591)→ |
| env-658 | TB_PC08_66_10110603 | + | 272 | 314 | RNA→ RpsD (COG0522)rpsD (PRK05327)→ |
| env-659 | TB_GS09_5_10159295 | + | 73 | 115 | RNA→ RpsD (COG0522)rpsD (PRK05327)→ |
| env-660 | PBR_1151212 | - | 8605 | 8563 | RNA→ RpsD (COG0522)rpsD (PRK05327)→ RNAP_alpha_NTD (cd06928)PRK05182 (PRK05182)RNA_pol_A_CTD (pfam03118)→ rplQ (PRK05591)Ribosomal_L17 (pfam01196)→ |
| Fsu-1-1 | NZ_JATV01000008.1 | + | 149522 | 149564 | RNA→ RpsD (COG0522)rpsD (PRK05327)→ RNAP_alpha_NTD (cd06928)PRK05182 (PRK05182)RNA_pol_A_CTD (pfam03118)→ RplQ (COG0203)rplQ (PRK05591)→ |
| Fre-1-1 | NZ_JPRL01000001.1 | - | 1417992 | 1417950 | RNA→ RpsD (COG0522)rpsD (PRK05327)→ RNAP_alpha_NTD (cd06928)PRK05182 (PRK05182)RNA_pol_A_CTD (pfam03118)→ RplQ (COG0203)rplQ (PRK05591)→ |
| env-661 | TB_FS08_3DRAFT_1132580 | - | 307 | 265 | RNA→ rpsD (PRK05327)Ribosomal_S4 (pfam00163)→ |
| env-662 | JGI24770J26754_10259849 | + | 431 | 473 | RNA→ rpsD (PRK05327)Ribosomal_S4 (pfam00163)→ |
| env-663 | JGI24770J26754_10222341 | - | 478 | 436 | RNA→ RpsD (COG0522)rpsD (PRK05327)→ |
| Fsp-4-1 | NZ_JSYO01000007.1 | - | 99004 | 98962 | RNA→ RpsD (COG0522)rpsD (PRK05327)→ RNAP_alpha_NTD (cd06928)PRK05182 (PRK05182)RNA_pol_A_CTD (pfam03118)→ RplQ (COG0203)rplQ (PRK05591)→ |
| env-664 | LM20APR2010E_c11148 | + | 62 | 104 | RNA→ RpsD (COG0522)rpsD (PRK05327)→ |
| Fsp-5-1 | NZ_JARP01000003.1 | + | 95196 | 95238 | RNA→ RpsD (COG0522)rpsD (PRK05327)→ RNAP_alpha_NTD (cd06928)PRK05182 (PRK05182)RNA_pol_A_CTD (pfam03118)→ RplQ (COG0203)rplQ (PRK05591)→ |
| Fsp-3-1 | NZ_JQJY01000005.1 | - | 476639 | 476597 | RNA→ RpsD (COG0522)rpsD (PRK05327)→ RNAP_alpha_NTD (cd06928)PRK05182 (PRK05182)RNA_pol_A_CTD (pfam03118)→ RplQ (COG0203)rplQ (PRK05591)→ |
| env-665 | TB_PC08_66_10000572 | + | 14643 | 14685 | RNA→ RpsD (COG0522)rpsD (PRK05327)→ RNAP_alpha_NTD (cd06928)PRK05182 (PRK05182)RNA_pol_A_CTD (pfam03118)→ RplQ (COG0203)rplQ (PRK05591)→ hypo→ CarA (COG0505)PRK12564 (PRK12564)→ |

| | | | | | |
|---|---|---|---|---|---|
| env-666 | JGI12421J11937_10000073 | + | 8864 | 8906 | **RNA**→ RpsD (COG0522)rpsD (PRK05327)→ RNAP_alpha_NTD (cd06928)PRK05182 (PRK05182)RNA_pol_A_CTD (pfam03118)→ RplQ (COG0203)rplQ (PRK05591)→ |
| env-667 | TB_GS09_5_10001765 | + | 16237 | 16284 | **RNA**→ RpsD (COG0522)rpsD (PRK05327)→ RpoA (COG0202)PRK05182 (PRK05182)→ RplQ (COG0203)rplQ (PRK05591)→ |
| env-668 | 2236876009_117224 | - | 159 | 111 | **RNA**→ rpsD (PRK05327)Ribosomal_S4 (pfam00163)→ |
| env-669 | JGI20151J14362_10046846 | - | 1758 | 1710 | **RNA**→ RpsD (COG0522)rpsD (PRK05327)→ RpoA (COG0202)PRK05182 (PRK05182)→ |
| env-670 | JGI20160J14292_10098055 | - | 386 | 338 | **RNA**→ RpsD (COG0522)rpsD (PRK05327)→ |
| env-671 | KGI_S1_ANT01_95m_c10216591 | + | 168 | 209 | **RNA**→ rpsD (PRK05327)Ribosomal_S4 (pfam00163)→ |
| Ace-1-1 | NZ_AUCB01000005.1 | - | 135671 | 135630 | **RNA**→ RpsD (COG0522)rpsD (PRK05327)→ RNAP_alpha_NTD (cd06928)PRK05182 (PRK05182)RNA_pol_A_CTD (pfam03118)→ RplQ (COG0203)rplQ (PRK05591)→ |
| env-672 | CENR01072972.1 | + | 678 | 719 | **RNA**→ RpsD (COG0522)rpsD (PRK05327)→ RNAP_alpha_NTD (cd06928)PRK05182 (PRK05182)RNA_pol_A_CTD (pfam03118)→ RplQ (COG0203)rplQ (PRK05591)→ PRK12564 (PRK12564)GATase (pfam00117)CPSase_sm_chain (pfam00988)→ |
| Ala-1-1 | NZ_AUKX01000021.1 | + | 84226 | 84267 | **RNA**→ RpsD (COG0522)rpsD (PRK05327)→ RNAP_alpha_NTD (cd06928)PRK05182 (PRK05182)RNA_pol_A_CTD (pfam03118)→ RplQ (COG0203)rplQ (PRK05591)→ |
| env-673 | JGI24025J20009_10444090 | + | 195 | 232 | **RNA**→ |
| env-674 | JGI24724J26744_10000005 | + | 153007 | 153048 | **RNA**→ RpsD (COG0522)rpsD (PRK05327)→ RNAP_alpha_NTD (cd06928)PRK05182 (PRK05182)RNA_pol_A_CTD (pfam03118)→ RplQ (COG0203)rplQ (PRK05591)→ CarA (COG0505)PRK12564 (PRK12564)→ |
| env-675 | NCBI_READ_1112308291129 | + | 600 | 641 | **RNA**→ |
| env-676 | SA_S1_NOR05_45m_c10047360 | - | 956 | 915 | **RNA**→ RpsD (COG0522)rpsD (PRK05327)→ PRK05182 (PRK05182)RNA_pol_L (pfam01193)→ |
| env-677 | SRS011255_Baylor_scaffold_31377 | - | 4549 | 4508 | **RNA**→ RpsD (COG0522)rpsD (PRK05327)→ RNAP_alpha_NTD (cd06928)PRK05182 (PRK05182)RNA_pol_A_CTD (pfam03118)→ rplQ (PRK05591)Ribosomal_L17 (pfam01196)→ |
| env-678 | SRS014578_C2475820 | + | 446 | 487 | **RNA**→ |
| env-679 | SRS023841_Baylor_scaffold_6234 | - | 80 | 39 | **RNA**→ |
| env-680 | SRS011126_Baylor_scaffold_35350 | + | 407 | 448 | **RNA**→ |
| env-681 | SRS023841_Baylor_scaffold_1278 | + | 2872 | 2913 | **RNA**→ RpsD (COG0522)rpsD (PRK05327)→ RNAP_alpha_NTD (cd06928)PRK05182 (PRK05182)RNA_pol_A_CTD (pfam03118)→ |
| env-682 | KGI_S2_ANT04_2345m_c1182776 | + | 183 | 224 | **RNA**→ rpsD (PRK05327)Ribosomal_S4 (pfam00163)→ |
| env-683 | SRS015064_WUGC_scaffold_6319 | - | 214 | 173 | **RNA**→ rpsD (PRK05327)Ribosomal_S4 (pfam00163)→ |
| env-684 | SRS015899_WUGC_scaffold_15247 | + | 600 | 641 | **RNA**→ rpsD (PRK05327)Ribosomal_S4 (pfam00163)→ |
| env-685 | SRS043755_WUGC_scaffold_16606 | - | 81 | 40 | **RNA**→ |
| env-686 | SRS053917_LANL_scaffold_21451 | - | 613 | 572 | **RNA**→ RpsD (COG0522)rpsD (PRK05327)→ |
| env-687 | NCBI_READ_1112493645660 | - | 765 | 724 | **RNA**→ RpsD (COG0522)rpsD (PRK05327)→ |
| env-688 | JGI11881J13070_1024505 | - | 310 | 258 | **RNA**→ rpsD (PRK05327)Ribosomal_S4 (pfam00163)→ |
| env-689 | NCBI_READ_1112493894880 | - | 314 | 273 | **RNA**→ rpsD (PRK05327)Ribosomal_S4 (pfam00163)→ |
| env-690 | 2236876002_134331 | + | 288 | 329 | **RNA**→ |
| env-691 | KGI_S1_ANT01_95m_c10003318 | + | 5674 | 5715 | **RNA**→ RpsD (COG0522)rpsD (PRK05327)→ RNAP_alpha_NTD (cd06928)PRK05182 (PRK05182)RNA_pol_A_CTD (pfam03118)→ RplQ (COG0203)rplQ (PRK05591)→ |
| env-692 | TDF_OR_ARG04_113m_c1009262 | + | 1093 | 1134 | **RNA**→ rpsD (PRK05327)Ribosomal_S4 (pfam00163)→ |
| env-693 | BS_KBA_SWE12_21m_c10441963 | - | 48 | 17 | **RNA**→ |
| env-694 | SA_S1_NOR05_45m_c10029776 | + | 910 | 951 | **RNA**→ RpsD (COG0522)rpsD (PRK05327)→ |
| env-695 | SI60aug11_100m_c1001696 | + | 199 | 240 | **RNA**→ RpsD (COG0522)rpsD (PRK05327)→ RNAP_alpha_NTD (cd06928)PRK05182 (PRK05182)RNA_pol_A_CTD (pfam03118)→ RplQ (COG0203)rplQ (PRK05591)→ |
| env-696 | BS_KBA_SWE02_21m_10007670 | - | 3008 | 2967 | **RNA**→ RpsD (COG0522)rpsD (PRK05327)→ RNAP_alpha_NTD (cd06928)PRK05182 (PRK05182)RNA_pol_A_CTD (pfam03118)→ RplQ (COG0203)rplQ (PRK05591)→ |
| env-697 | CESL01223066.1 | - | 581 | 540 | **RNA**→ RpsD (COG0522)rpsD (PRK05327)→ |

| | | | | | |
|---|---|---|---|---|---|
| env-698 | TFO_HY__Contig_1415 | + | 14727 | 14768 | **RNA**→ RpsD (COG0522)rpsD (PRK05327)→ RNAP_alpha_NTD (cd06928)PRK05182 (PRK05182)RNA_pol_A_CTD (pfam03118)→ RplQ (COG0203)rplQ (PRK05591)→ |
| env-699 | 3300001605_10731583 | + | 414 | 455 | **RNA**→ |
| env-700 | KVWG_1_1_1_newblercontig19647 | + | 110 | 151 | **RNA**→ rpsD (PRK05327)Ribosomal_S4 (pfam00163)→ |
| env-701 | JCVI_SCAF_1101668559196 | - | 977 | 935 | **RNA**→ RpsD (COG0522)rpsD (PRK05327)→ PRK05182 (PRK05182)RNA_pol_L (pfam01193)→ |
| env-702 | NCBI_READ_1112494091561 | + | 60 | 101 | **RNA**→ RpsD (COG0522)rpsD (PRK05327)→ |
| env-703 | NCBI_READ_1112532275014 | - | 690 | 649 | **RNA**→ RpsD (COG0522)rpsD (PRK05327)→ |
| env-704 | NCBI_READ_1112777438177 | - | 394 | 353 | **RNA**→ RpsD (COG0522)rpsD (PRK05327)→ |
| env-705 | DelMOSum2011_c10076159 | + | 881 | 922 | **RNA**→ rpsD (PRK05327)Ribosomal_S4 (pfam00163)→ |
| env-706 | TDF_OR_ARG04_113m_c1015903 | - | 887 | 846 | **RNA**→ RpsD (COG0522)rpsD (PRK05327)→ PRK05182 (PRK05182)RNA_pol_L (pfam01193)→ |
| env-707 | BS_KBA_SWE12_21m_c10362886 | + | 6 | 47 | **RNA**→ rpsD (PRK05327)Ribosomal_S4 (pfam00163)→ |
| env-708 | SA_S1_NOR05_45m_c10001294 | - | 776 | 735 | **RNA**→ RpsD (COG0522)rpsD (PRK05327)→ PRK05182 (PRK05182)→ |
| env-709 | KGI_S1_ANT03_95m_c1044459 | - | 439 | 398 | **RNA**→ RpsD (COG0522)rpsD (PRK05327)→ |
| env-710 | JGI20154J14316_10006580 | + | 6557 | 6598 | **RNA**→ RpsD (COG0522)rpsD (PRK05327)→ RNAP_alpha_NTD (cd06928)PRK05182 (PRK05182)RNA_pol_A_CTD (pfam03118)→ |
| env-711 | CESI01170757.1 | + | 255 | 296 | **RNA**→ rpsD (PRK05327)Ribosomal_S4 (pfam00163)→ |
| env-712 | 3300001592_11040489 | + | 52 | 93 | **RNA**→ rpsD (PRK05327)Ribosomal_S4 (pfam00163)→ |
| env-713 | JGI25321J39296_1043397 | + | 52 | 93 | **RNA**→ RpsD (COG0522)rpsD (PRK05327)→ PRK05182 (PRK05182)RNA_pol_L (pfam01193)→ |
| Pir-1-1 | NZ_CH724148.1 | - | 107482 | 107441 | **RNA**→ RpsD (COG0522)rpsD (PRK05327)→ RpoA (COG0202)PRK05182 (PRK05182)→ RplQ (COG0203)rplQ (PRK05591)→ |
| env-714 | NCBI_READ_1112494062227 | - | 199 | 158 | **RNA**→ |
| env-715 | NCBI_READ_1112493702487 | - | 839 | 798 | **RNA**→ RpsD (COG0522)rpsD (PRK05327)→ |
| env-716 | NCBI_READ_1112290461315 | + | 55 | 96 | **RNA**→ RpsD (COG0522)rpsD (PRK05327)→ |
| env-717 | DelMOSum2010_c10003497 | + | 7046 | 7087 | **RNA**→ RpsD (COG0522)rpsD (PRK05327)→ |
| env-718 | SA_S2_NOR13_50m_c1014240 | - | 48 | 17 | **RNA**→ |
| env-719 | BS_KBA_SWE12_21m_c10004498 | - | 3174 | 3133 | **RNA**→ RpsD (COG0522)rpsD (PRK05327)→ RNAP_alpha_NTD (cd06928)PRK05182 (PRK05182)RNA_pol_A_CTD (pfam03118)→ RplQ (COG0203)rplQ (PRK05591)→ |
| env-720 | BS_KBA_SWE12_21m_c10128822 | - | 291 | 250 | **RNA**→ rpsD (PRK05327)Ribosomal_S4 (pfam00163)→ |
| env-721 | SA_S1_NOR08_45m_c10718266 | + | 33 | 74 | **RNA**→ rpsD (PRK05327)Ribosomal_S4 (pfam00163)→ |
| env-722 | JGI20154J14316_10014338 | - | 4356 | 4315 | **RNA**→ RpsD (COG0522)rpsD (PRK05327)→ |
| env-723 | BS_KBA_SWE02_21m_10461828 | + | 81 | 122 | **RNA**→ rpsD (PRK05327)Ribosomal_S4 (pfam00163)→ |
| env-724 | CESU01046469.1 | + | 91 | 132 | **RNA**→ RpsD (COG0522)rpsD (PRK05327)→ PRK05182 (PRK05182)RNA_pol_L (pfam01193)→ |
| env-725 | 3300001580_10549671 | + | 137 | 178 | **RNA**→ rpsD (PRK05327)Ribosomal_S4 (pfam00163)→ |
| env-726 | JGI24724J26744_10022788 | - | 166 | 125 | **RNA**→ rpsD (PRK05327)Ribosomal_S4 (pfam00163)→ |
| Jma-1-1 | NZ_JH651379.1 | - | 2626446 | 2626405 | **RNA**→ RpsD (COG0522)rpsD (PRK05327)→ RpsD (COG0522)rpsD (PRK05327)→ RNAP_alpha_NTD (cd06928)PRK05182 (PRK05182)RNA_pol_A_CTD (pfam03118)→ RpoA (COG0202)PRK05182 (PRK05182)→ RplQ (COG0203)rplQ (PRK05591)→ RplQ (COG0203)rplQ (PRK05591)→ CarA (COG0505)PRK12564 (PRK12564)→ CarA (COG0505)PRK12564 (PRK12564)→ eno (PRK00077)enolase (cd03313)→ eno (PRK00077)enolase (cd03313)→ |
| env-727 | NCBI_READ_1112493915188 | - | 912 | 874 | **RNA**→ hypo→ |
| env-728 | NCBI_READ_1112290401224 | + | 25 | 66 | **RNA**→ RpsD (COG0522)rpsD (PRK05327)→ |
| env-729 | NCBI_READ_1112493792322 | + | 649 | 690 | **RNA**→ |
| env-730 | AMQJ01014928.1 | + | 561 | 602 | **RNA**→ rpsD (PRK05327)Ribosomal_S4 (pfam00163)→ |
| env-731 | SA_S2_NOR13_50m_c1000098 | + | 13594 | 13635 | **RNA**→ RpsD (COG0522)rpsD (PRK05327)→ |
| env-732 | BS_KBA_SWE12_21m_c10395815 | - | 39 | 8 | **RNA**→ |
| env-733 | BS_KBA_SWE12_21m_c10510804 | + | 111 | 152 | **RNA**→ rpsD (PRK05327)Ribosomal_S4 (pfam00163)→ |
| env-734 | SA_S2_NOR15_50m_c10003768 | - | 582 | 541 | **RNA**→ RpsD (COG0522)rpsD (PRK05327)→ |
| env-735 | SI34jun09_10m_1017310 | + | 995 | 1036 | **RNA**→ rpsD (PRK05327)Ribosomal_S4 (pfam00163)→ |
| env-736 | CEQS01147054.1 | - | 337 | 296 | **RNA**→ rpsD (PRK05327)Ribosomal_S4 (pfam00163)→ |

| | | | | | |
|---|---|---|---|---|---|
| env-737 | 3300000558_10037104 | - | 1932 | 1891 | **RNA**→ RpsD (COG0522)rpsD (PRK05327)→<br>RNAP_alpha_NTD (cd06928)PRK05182 (PRK05182)RNA_pol_A_CTD (pfam03118)→<br>rplQ (PRK05591)Ribosomal_L17 (pfam01196)→ |
| env-738 | JGI24025J20009_10008494 | + | 2996 | 3037 | **RNA**→ RpsD (COG0522)rpsD (PRK05327)→<br>RNAP_alpha_NTD (cd06928)PRK05182 (PRK05182)RNA_pol_A_CTD (pfam03118)→<br>RpoA (COG0202)PRK05182 (PRK05182)→ RplQ (COG0203)rplQ (PRK05591)→<br>CPSase_sm_chain (smart01097)CPSase_sm_chain (pfam00988)→ |
| env-739 | JGI25321J50212_10028228 | + | 778 | 819 | **RNA**→ RpsD (COG0522)rpsD (PRK05327)→ RNAP_alpha_NTD (cd06928)PRK05182 (PRK05182)→ |
| env-740 | NCBI_READ_1112493671175 | + | 717 | 758 | **RNA**→ |
| env-741 | NCBI_READ_1112493447629 | - | 601 | 560 | **RNA**→ RpsD (COG0522)rpsD (PRK05327)→ |
| env-742 | SA_S1_NOR08_45m_c10473502 | + | 160 | 200 | **RNA**→ rpsD (PRK05327)Ribosomal_S4 (pfam00163)→ |
| env-743 | SA_S2_NOR15_50m_c10652927 | - | 82 | 42 | **RNA**→ |
| env-744 | SA_S1_NOR05_45m_c10011813 | + | 2157 | 2197 | **RNA**→ RpsD (COG0522)rpsD (PRK05327)→ |
| env-745 | TB_PC08_66_10003287 | - | 3261 | 3221 | **RNA**→ RpsD (COG0522)rpsD (PRK05327)→<br>RNAP_alpha_NTD (cd06928)PRK05182 (PRK05182)RNA_pol_A_CTD (pfam03118)→<br>RplQ (COG0203)rplQ (PRK05591)→ |
| env-746 | LGVC01026408.1 | - | 246 | 206 | **RNA**→ rpsD (PRK05327)Ribosomal_S4 (pfam00163)→ |
| env-747 | LGVF01085840.1 | + | 3439 | 3479 | **RNA**→ RpsD (COG0522)rpsD (PRK05327)→ RNAP_alpha_NTD (cd06928)PRK05182 (PRK05182)→ |
| env-748 | JGI24023J19991_10058338 | + | 371 | 411 | **RNA**→ RpsD (COG0522)rpsD (PRK05327)→ RNAP_alpha_NTD (cd06928)PRK05182 (PRK05182)→ |
| Cca-2-1 | NZ_CDOI01000154.1 | + | 49748 | 49788 | **RNA**→ RpsD (COG0522)rpsD (PRK05327)→<br>RNAP_alpha_NTD (cd06928)PRK05182 (PRK05182)RNA_pol_A_CTD (pfam03118)→<br>RplQ (COG0203)rplQ (PRK05591)→ |
| env-749 | TDF_OR_ARG04_113m_c1311978 | + | 76 | 116 | **RNA**→ rpsD (PRK05327)Ribosomal_S4 (pfam00163)→ |
| env-750 | SA_S1_NOR08_45m_c10021922 | - | 775 | 735 | **RNA**→ RpsD (COG0522)rpsD (PRK05327)→ PRK05182 (PRK05182)→ |
| env-751 | SA_S2_NOR15_50m_c10674918 | + | 250 | 284 | **RNA**→ |
| env-752 | JGI24024J18818_10005433 | - | 2993 | 2953 | **RNA**→ RpsD (COG0522)rpsD (PRK05327)→<br>RNAP_alpha_NTD (cd06928)PRK05182 (PRK05182)RNA_pol_A_CTD (pfam03118)→ |
| env-753 | LGVD01055874.1 | + | 4328 | 4368 | **RNA**→ RpsD (COG0522)rpsD (PRK05327)→<br>RNAP_alpha_NTD (cd06928)PRK05182 (PRK05182)RNA_pol_A_CTD (pfam03118)→ |
| env-754 | JGI26437J51864_10085571 | + | 322 | 362 | **RNA**→ RpsD (COG0522)rpsD (PRK05327)→ |
| env-755 | BS_KBA_SWE12_21m_c10012235 | - | 428 | 388 | **RNA**→ S4 (smart00363)RpsD (COG0522)→ |
| env-756 | BS_KBA_SWE12_21m_c10357750 | + | 138 | 178 | **RNA**→ rpsD (PRK05327)Ribosomal_S4 (pfam00163)→ |
| env-757 | SA_S1_NOR08_45m_c10216310 | + | 259 | 299 | **RNA**→ rpsD (PRK05327)Ribosomal_S4 (pfam00163)→ |
| env-758 | TDF_MC_ARG02_113m_c1073765 | + | 140 | 180 | **RNA**→ rpsD (PRK05327)Ribosomal_S4 (pfam00163)→ |
| env-759 | KGI_S1_ANT01_95m_c10034647 | + | 846 | 886 | **RNA**→ RpsD (COG0522)rpsD (PRK05327)→ RpoA (COG0202)PRK05182 (PRK05182)→ |
| env-760 | JGI24024J18818_10127678 | - | 133 | 93 | **RNA**→ rpsD (PRK05327)Ribosomal_S4 (pfam00163)→ |
| env-761 | LGVD01081681.1 | - | 710 | 670 | **RNA**→ rpsD (PRK05327)Ribosomal_S4 (pfam00163)→ RpsD (COG0522)rpsD (PRK05327)→ |
| env-762 | JGI24023J19991_10002979 | - | 2498 | 2458 | **RNA**→ RpsD (COG0522)rpsD (PRK05327)→<br>RNAP_alpha_NTD (cd06928)PRK05182 (PRK05182)RNA_pol_A_CTD (pfam03118)→<br>RplQ (COG0203)rplQ (PRK05591)→ |
| Ccy-1-1 | NZ_CDOF01000060.1 | - | 6612 | 6572 | **RNA**→ RpsD (COG0522)rpsD (PRK05327)→<br>RNAP_alpha_NTD (cd06928)PRK05182 (PRK05182)RNA_pol_A_CTD (pfam03118)→<br>RplQ (COG0203)rplQ (PRK05591)→ |
| Cca-1-1 | NZ_CDOE01000064.1 | + | 15288 | 15328 | **RNA**→ RpsD (COG0522)rpsD (PRK05327)→<br>RNAP_alpha_NTD (cd06928)PRK05182 (PRK05182)RNA_pol_A_CTD (pfam03118)→<br>RplQ (COG0203)rplQ (PRK05591)→ CarA (COG0505)PRK12564 (PRK12564)→ Methyltransf_23 (pfam13489)→ |
| env-763 | TDF_MC_ARG01_113m_c1021061 | + | 112 | 152 | **RNA**→ S4 (smart00363)RpsD (COG0522)→ |
| env-764 | SA_S2_NOR15_50m_c10115366 | - | 840 | 800 | **RNA**→ RpsD (COG0522)rpsD (PRK05327)→ PRK05182 (PRK05182)RNA_pol_L (pfam01193)→ |
| env-765 | SA_S2_NOR13_50m_c1004409 | + | 1238 | 1278 | **RNA**→ RpsD (COG0522)rpsD (PRK05327)→ |
| env-766 | JGI24024J18818_10558133 | - | 216 | 176 | **RNA**→ rpsD (PRK05327)Ribosomal_S4 (pfam00163)→ |
| env-767 | LGVF01258235.1 | + | 471 | 511 | **RNA**→ RpsD (COG0522)rpsD (PRK05327)→ |
| env-768 | TDF_MC_ARG01_113m_c1001894 | - | 1149 | 1109 | **RNA**→ RpsD (COG0522)rpsD (PRK05327)→ RNAP_alpha_NTD (cd06928)PRK05182 (PRK05182)→ |
| Pja-1-1 | NZ_JSWG01000012.1 | - | 78120 | 78072 | **RNA**→ RpsD (COG0522)rpsD (PRK05327)→ RpoA (COG0202)PRK05182 (PRK05182)→<br>RplQ (COG0203)rplQ (PRK05591)→ CarA (COG0505)PRK12564 (PRK12564)→ |

| | | | | | |
|---|---|---|---|---|---|
| env-769 | BS_KBA_SWE12_21m_c10006531 | + | 2159 | 2207 | RNA→ RpsD (COG0522)rpsD (PRK05327)→ RpoA (COG0202)PRK05182 (PRK05182)→<br>RplQ (COG0203)rplQ (PRK05591)→ |
| env-770 | BS_KBA_SWE02_21m_10206790 | + | 72 | 120 | RNA→ RpsD (COG0522)rpsD (PRK05327)→ |
| Gme-1-1 | NZ_AUHD01000007.1 | + | 277770 | 277818 | RNA→ RpsD (COG0522)rpsD (PRK05327)→ RpoA (COG0202)PRK05182 (PRK05182)→ |
| env-771 | BS_KBA_SWE12_21m_c10002655 | - | 2863 | 2815 | RNA→ RpsD (COG0522)rpsD (PRK05327)→ rpoA (CHL00013)PRK05182 (PRK05182)→<br>RpoA (COG0202)PRK05182 (PRK05182)→ RplQ (COG0203)rplQ (PRK05591)→ |
| Pme-1-1 | NZ_JXJP01000002.1 | + | 289435 | 289483 | RNA→ RpsD (COG0522)rpsD (PRK05327)→ RpoA (COG0202)PRK05182 (PRK05182)→<br>RplQ (COG0203)rplQ (PRK05591)→ CarA (COG0505)PRK12564 (PRK12564)→ |
| env-772 | TDF_OR_ARG04_113m_c1048254 | + | 325 | 373 | RNA→ rpsD (PRK05327)Ribosomal_S4 (pfam00163)→ |
| env-773 | SA_S1_NOR08_45m_c10228226 | + | 189 | 237 | RNA→ rpsD (PRK05327)Ribosomal_S4 (pfam00163)→ |
| env-774 | 3300001592_10000538 | - | 31145 | 31097 | RNA→ RpsD (COG0522)rpsD (PRK05327)→ PRK05182 (PRK05182)RNA_pol_L (pfam01193)→ |
| env-775 | BS_KBA_SWE21_205m_10078219 | + | 596 | 644 | RNA→ rpsD (PRK05327)Ribosomal_S4 (pfam00163)→ |
| env-776 | KGI_S1_ANT02_95m_c10197724 | + | 147 | 195 | RNA→ rpsD (PRK05327)Ribosomal_S4 (pfam00163)→ |
| env-777 | JGI24723J26617_10003269 | + | 213 | 261 | RNA→ RpsD (COG0522)rpsD (PRK05327)→ RpoA (COG0202)PRK05182 (PRK05182)→<br>RplQ (COG0203)rplQ (PRK05591)→ CarA (COG0505)PRK12564 (PRK12564)→ |
| env-778 | JGI24724J26744_10230191 | - | 344 | 296 | RNA→ rpsD (PRK05327)Ribosomal_S4 (pfam00163)→ |
| env-779 | KGI_S2_ANT04_2345m_c1317179 | - | 230 | 182 | RNA→ rpsD (PRK05327)Ribosomal_S4 (pfam00163)→ |
| env-780 | JCVI_SCAF_1096628386002 | - | 1571 | 1525 | RNA→ RpsD (COG0522)rpsD (PRK05327)→ |
| env-781 | LPaug08P2610m_c1098718 | + | 109 | 157 | RNA→ rpsD (PRK05327)Ribosomal_S4 (pfam00163)→ |
| env-782 | DelMOSum2010_c10000647 | - | 14055 | 14007 | RNA→ RpsD (COG0522)rpsD (PRK05327)→<br>RNAP_alpha_NTD (cd06928)PRK05182 (PRK05182)RNA_pol_A_CTD (pfam03118)→<br>RplQ (COG0203)rplQ (PRK05591)→ CarA (COG0505)PRK12564 (PRK12564)→<br>eno (PRK00077)enolase (cd03313)→ gltA (PRK05614)EcCS_like (cd06114)→ |
| env-783 | KGI_S1_ANT01_95m_c10031960 | + | 223 | 266 | RNA→ RpsD (COG0522)rpsD (PRK05327)→<br>RNAP_alpha_NTD (cd06928)PRK05182 (PRK05182)RNA_pol_A_CTD (pfam03118)→<br>RplQ (COG0203)rplQ (PRK05591)→ |
| env-784 | 2236876010_0182615 | - | 474 | 428 | RNA→ rpsD (PRK05327)Ribosomal_S4 (pfam00163)→ S4 (smart00363)RpsD (COG0522)→ |
| env-785 | JCVI_SCAF_1096626851900 | + | 1510 | 1551 | RNA→ RpsD (COG0522)rpsD (PRK05327)→ |
| env-786 | JGI20156J14371_10002853 | - | 815 | 774 | RNA→ RpsD (COG0522)rpsD (PRK05327)→ PRK05182 (PRK05182)RNA_pol_L (pfam01193)→ |
| env-787 | JCVI_SCAF_1096627961966 | + | 1328 | 1370 | RNA→ RpsD (COG0522)rpsD (PRK05327)→ |
| env-788 | JCVI_READ_1103242618048 | - | 562 | 520 | RNA→ RpsD (COG0522)rpsD (PRK05327)→ |
| env-789 | CEWO01110425.1 | + | 727 | 768 | RNA→ RpsD (COG0522)rpsD (PRK05327)→ |
| env-790 | CESJ01017871.1 | + | 773 | 804 | RNA→ |
| env-791 | CESU01012824.1 | + | 1083 | 1124 | RNA→ RpsD (COG0522)rpsD (PRK05327)→ |
| env-792 | CESE01007018.1 | - | 3023 | 2982 | RNA→ RpsD (COG0522)rpsD (PRK05327)→ RpoA (COG0202)PRK05182 (PRK05182)→<br>RplQ (COG0203)rplQ (PRK05591)→ CPSase_sm_chain (smart01097)CPSase_sm_chain (pfam00988)→ |
| env-793 | CESF01014444.1 | + | 686 | 717 | RNA→ |
| env-794 | LPjun09P1210m_c1061424 | - | 156 | 115 | RNA→ rpsD (PRK05327)Ribosomal_S4 (pfam00163)→ |
| env-795 | SI34jun09_10m_1065651 | + | 164 | 205 | RNA→ rpsD (PRK05327)Ribosomal_S4 (pfam00163)→ |
| env-796 | DelMOSum2010_c10031049 | + | 744 | 785 | RNA→ RpsD (COG0522)rpsD (PRK05327)→<br>RNAP_alpha_NTD (cd06928)PRK05182 (PRK05182)RNA_pol_A_CTD (pfam03118)→<br>RplQ (COG0203)rplQ (PRK05591)→ |
| env-797 | JGI20152J14361_10004992 | + | 6162 | 6203 | RNA→ RpsD (COG0522)rpsD (PRK05327)→ RpoA (COG0202)PRK05182 (PRK05182)→ |
| env-798 | JCVI_SCAF_1096626908974 | - | 951 | 910 | RNA→ RpsD (COG0522)rpsD (PRK05327)→ PRK05182 (PRK05182)RNA_pol_L (pfam01193)→ |
| env-799 | 2236876010_0124125 | + | 466 | 497 | RNA→ |
| env-800 | JCVI_SCAF_1096627300879 | - | 885 | 844 | RNA→ RpsD (COG0522)rpsD (PRK05327)→ PRK05182 (PRK05182)RNA_pol_L (pfam01193)→ |
| env-801 | DelMOWin2010_c10056093 | - | 56 | 21 | RNA→ |
| env-802 | DelMOSum2010_c10171752 | + | 62 | 103 | RNA→ RpsD (COG0522)rpsD (PRK05327)→ |
| env-803 | DelMOSum2011_c10049863 | - | 1275 | 1234 | RNA→ RpsD (COG0522)rpsD (PRK05327)→ RNAP_alpha_NTD (cd06928)PRK05182 (PRK05182)→ |
| env-804 | SI47jul10_100m_c1000543 | - | 1464 | 1423 | RNA→ RpsD (COG0522)rpsD (PRK05327)→ RNAP_alpha_NTD (cd06928)PRK05182 (PRK05182)→ |
| env-805 | SI36aug09_120m_1267228 | + | 135 | 176 | RNA→ rpsD (PRK05327)Ribosomal_S4 (pfam00163)→ |
| env-806 | JGI20156J14371_10141859 | + | 313 | 354 | RNA→ RpsD (COG0522)rpsD (PRK05327)→ |
| env-807 | CENT01045813.1 | - | 316 | 275 | RNA→ rpsD (PRK05327)Ribosomal_S4 (pfam00163)→ |
| env-808 | CESI01020841.1 | + | 455 | 496 | RNA→ RpsD (COG0522)rpsD (PRK05327)→ |

| | | | | | |
|---|---|---|---|---|---|
| env-809 | CERW01035365.1 | + | 573 | 614 | **RNA**→ RpsD (COG0522)rpsD (PRK05327)→ |
| env-810 | CESE01023512.1 | - | 2167 | 2126 | **RNA**→ RpsD (COG0522)rpsD (PRK05327)→<br>RNAP_alpha_NTD (cd06928)PRK05182 (PRK05182)RNA_pol_A_CTD (pfam03118)→<br>RplQ (COG0203)rplQ (PRK05591)→ |
| env-811 | CESI01115170.1 | + | 394 | 435 | **RNA**→ rpsD (PRK05327)Ribosomal_S4 (pfam00163)→ |
| env-812 | CESM01447978.1 | - | 644 | 603 | **RNA**→ RpsD (COG0522)rpsD (PRK05327)→ |
| env-813 | CESI01285347.1 | - | 647 | 606 | **RNA**→ RpsD (COG0522)rpsD (PRK05327)→ |
| env-814 | CESJ01035100.1 | + | 13520 | 13561 | **RNA**→ RpsD (COG0522)rpsD (PRK05327)→ RpoA (COG0202)PRK05182 (PRK05182)→<br>RplQ (COG0203)rplQ (PRK05591)→ CarA (COG0505)PRK12564 (PRK12564)→<br>eno (PRK00077)Enolase_N (pfam03952)→ |
| env-815 | CESI01140995.1 | - | 648 | 607 | **RNA**→ RpsD (COG0522)rpsD (PRK05327)→ |
| env-816 | CESL01085606.1 | + | 7356 | 7397 | **RNA**→ RpsD (COG0522)rpsD (PRK05327)→<br>RNAP_alpha_NTD (cd06928)PRK05182 (PRK05182)RNA_pol_A_CTD (pfam03118)→<br>RplQ (COG0203)rplQ (PRK05591)→ |
| env-817 | CETY01336660.1 | - | 667 | 626 | **RNA**→ RpsD (COG0522)rpsD (PRK05327)→ |
| env-818 | CERW01115847.1 | - | 343 | 302 | **RNA**→ rpsD (PRK05327)Ribosomal_S4 (pfam00163)→ |
| env-819 | JCVI_SCAF_1096627085079 | + | 1100 | 1141 | **RNA**→ RpsD (COG0522)rpsD (PRK05327)→ |
| env-820 | 2236876007_0120103 | - | 371 | 330 | **RNA**→ hypo→ |
| env-821 | DelMOWin2010_c10054074 | + | 28 | 69 | **RNA**→ RpsD (COG0522)rpsD (PRK05327)→<br>RNAP_alpha_NTD (cd06928)PRK05182 (PRK05182)RNA_pol_A_CTD (pfam03118)→ |
| env-822 | 2236876010_0562604 | - | 92 | 51 | **RNA**→ ←hypo |
| env-823 | DelMOSum2010_c10187590 | - | 554 | 513 | **RNA**→ RpsD (COG0522)rpsD (PRK05327)→ |
| env-824 | CEOP01438579.1 | + | 571 | 612 | **RNA**→ RpsD (COG0522)rpsD (PRK05327)→ |
| env-825 | CESI01092574.1 | - | 661 | 620 | **RNA**→ RpsD (COG0522)rpsD (PRK05327)→ |
| env-826 | JCVI_SCAF_1096628391443 | + | 527 | 568 | **RNA**→ rpsD (PRK05327)Ribosomal_S4 (pfam00163)→ PRK05182 (PRK05182)RNA_pol_A_CTD (pfam03118)→<br>RplQ (COG0203)rplQ (PRK05591)→ |
| env-827 | CEOK01218656.1 | + | 103 | 144 | **RNA**→ RpsD (COG0522)rpsD (PRK05327)→ |
| env-828 | CETE01044921.1 | - | 657 | 616 | **RNA**→ RpsD (COG0522)rpsD (PRK05327)→ |
| env-829 | CEUF01269544.1 | + | 331 | 372 | **RNA**→ RpsD (COG0522)rpsD (PRK05327)→ PRK05182 (PRK05182)RNA_pol_L (pfam01193)→ |
| env-830 | CENO01020607.1 | - | 670 | 629 | **RNA**→ RpsD (COG0522)rpsD (PRK05327)→ |
| env-831 | CEVH01290424.1 | - | 399 | 358 | **RNA**→ S4 (smart00363)RpsD (COG0522)→ |
| env-832 | CEWG01002875.1 | + | 131 | 172 | **RNA**→ RpsD (COG0522)rpsD (PRK05327)→ RNAP_alpha_NTD (cd06928)PRK05182 (PRK05182)→ |
| env-833 | CEPJ01070513.1 | + | 64 | 105 | **RNA**→ RpsD (COG0522)rpsD (PRK05327)→ |
| env-834 | 2236876001_043944 | - | 301 | 260 | **RNA**→ rpsD (PRK05327)Ribosomal_S4 (pfam00163)→ |
| env-835 | SA_S2_NOR13_50m_c1052491 | - | 136 | 87 | **RNA**→ rpsD (PRK05327)Ribosomal_S4 (pfam00163)→ |
| env-836 | LPaug09P1610m_c1023406 | + | 657 | 699 | **RNA**→ |
| env-837 | CENJ01019829.1 | - | 355 | 313 | **RNA**→ rpsD (PRK05327)Ribosomal_S4 (pfam00163)→ |
| env-838 | CESW01149347.1 | - | 2171 | 2129 | **RNA**→ RpsD (COG0522)rpsD (PRK05327)→ RpoA (COG0202)PRK05182 (PRK05182)→<br>RplQ (COG0203)rplQ (PRK05591)→ |
| env-839 | JCVI_SCAF_1096627389932 | + | 1334 | 1376 | **RNA**→ RpsD (COG0522)rpsD (PRK05327)→ RpoA (COG0202)PRK05182 (PRK05182)→<br>RplQ (COG0203)rplQ (PRK05591)→ CarA (COG0505)PRK12564 (PRK12564)→<br>eno (PRK00077)enolase (cd03313)→ gltA (PRK05614)EcCS_like (cd06114)→<br>DdaH (COG1834)arcA (TIGR01078)→ Amidinotransf (pfam02274)→ |
| env-840 | P_1C_Liq_3_UnCty_1071622 | + | 198 | 240 | **RNA**→ |
| env-841 | P_2C_Liq_1_UnCty_1012564 | + | 496 | 538 | **RNA**→ RpsD (COG0522)rpsD (PRK05327)→<br>RNAP_alpha_NTD (cd06928)PRK05182 (PRK05182)RNA_pol_A_CTD (pfam03118)→ |

#### 15.3 Conserved domains

Conserved domains found in protein-coding genes listed in Section 15.2 are shown below, with the first sentence in their description from the Conserved Domain Database (if any). Conserved domains associated with more than one S4-Flavobacteria RNA are assigned a color, while others are shown in gray. The number in parentheses after the colored domain name is the number of occurrences in Section 15.2.

cd00599 (2) Endo-N-acetylmuramidases (muramidases) are lysozymes (also referred to as peptidoglycan hydrolases) that degrade bacterial cell walls by catalyzing the hydrolysis of 1,4-beta-linkages between N-acetylmuramic acid and N-acetyl-D-glucosamine residues.
cd01744 (5) Small chain of the glutamine-dependent form of carbamoyl phosphate synthase, CPSase II.
cd03313 (29) Enolase: Enolases are homodimeric enzymes that catalyse the reversible dehydration of 2-phospho-D-glycerate to phosphoenolpyruvate as part of the glycolytic and gluconeogenesis pathways.
cd06114 (8) Escherichia coli (Ec) citrate synthase (CS) GltA_like.
cd06928 (183) N-terminal domain of the Alpha subunit of Bacterial RNA polymerase.
cd11313 (1) Alpha amylase catalytic domain found in archaeal and bacterial Alpha-amylases (also called 1,4-alpha-D-glucan-4-glucanohydrolase).
CHL00013 (7) RNA polymerase alpha subunit
CHL00113 (1) ribosomal protein S4; Reviewed
COG0202 (115) DNA-directed RNA polymerase, alpha subunit/40 kD subunit [Transcription]
COG0203 (224) Ribosomal protein L17 [Translation, ribosomal structure and biogenesis]
COG0505 (80) Carbamoylphosphate synthase small subunit [Amino acid transport and metabolism, Nucleotide transport and metabolism]
COG0522 (578) Ribosomal protein S4 or related protein [Translation, ribosomal structure and biogenesis]
COG1834 (2) N-Dimethylarginine dimethylaminohydrolase [Amino acid transport and metabolism]
pfam00117 (3) Glutamine amidotransferase class-I.
pfam00163 (226) Ribosomal protein S4/S9 N-terminal domain.
pfam00285 (1) Citrate synthase, C-terminal domain.
pfam00988 (15) Carbamoyl-phosphate synthase small chain, CPSase domain.
pfam01193 (34) RNA polymerase Rpb3/Rpb11 dimerisation domain.
pfam01196 (9) Ribosomal protein L17.
pfam02274 (2) Amidinotransferase.
pfam03118 (157) Bacterial RNA polymerase, alpha chain C terminal domain.
pfam03485 (1) Arginyl tRNA synthetase N terminal domain.
pfam03486 (2) HI0933-like protein.
pfam03952 (2) Enolase, N-terminal domain.
pfam03971 (2) Monomeric isocitrate dehydrogenase.
pfam10310 (2) Maintenance of telomere capping protein 1.
pfam13489 (1) Methyltransferase domain.
PLN02967 (1) kinase
PRK00077 (31) enolase; Provisional
PRK05182 (345) DNA-directed RNA polymerase subunit alpha; Provisional
PRK05327 (783) 30S ribosomal protein S4; Validated
PRK05591 (233) 50S ribosomal protein L17; Validated
PRK05614 (9) type II citrate synthase; Reviewed
PRK10811 (1) ribonuclease E; Reviewed
PRK12564 (90) carbamoyl phosphate synthase small subunit; Reviewed
PRK14521 (2) 30S ribosomal protein S16; Provisional
smart00363 (26) S4 RNA-binding domain.
smart00641 (2) Glycosyl hydrolases family 25.
smart01016 (1) Arginyl tRNA synthetase N terminal dom.
smart01097 (6) Carbamoyl-phosphate synthase small chain, CPSase domain.
TIGR00178 (2) isocitrate dehydrogenase, NADP-dependent, monomeric type. [Energy metabolism, TCA cycle]
TIGR00275 (2) flavoprotein, HI0933 family. [Unknown function, Enzymes of unknown specificity]
TIGR01078 (1) arginine deiminase. [Energy metabolism, Amino acids and amines]
TIGR02456 (1) trehalose synthase. [Energy metabolism, Biosynthesis and degradation of polysaccharides]

#### 15.4 Multiple-sequence alignment

Each S4-Flavobacteria RNA is denoted by an abbreviation (like “Eco-1-1”) that refers to a taxonomy given in Section 15.1. The alignment may include sequences containing the RNA motif, as well as flanking sequence. The S4-Flavobacteria RNA itself is denoted by the line underneath marked 5′ and 3′ on either end. Nucleotides in flanking sequences (i.e., not part of the motif) are written in gray letters. Stems of predicted rho-independent transcription terminators, if any, are shaded yellow. (Note: terminator predictions have not been analyzed manually, and many are likely to be false positives. Terminator predictions are those of the RNIE software.) Nucleotides predicted to function (as DNA) as transcription-factor binding sites, if any, are shaded in green. (But note: these predictions are manually annotated, so they might be under-predicted.) Annotated start codons, if any, are shaded green. (Note: start codons are frequently misannotated, especially in environmental samples.) Nucleotides proposed to basepair as part of the consensus structure are shaded in color when they comprise Watson-Crick or G-U pairs. Otherwise they are shaded gray. Conserved stems are also indicated at the bottom of the alignment by angle brackets, where matching < and > denote base-paired columns. Below these angle brackets, the symbol “2” denotes base pairs exhibiting covariation according to the statistically well-founded R-scape method. “1” denotes base pairs exhibiting covariation according to R2R’s simplistic method. “0” denotes base pairs that are not observed to mutate and “?” denotes base pairs that have a significant frequency of non-canonical nucleotides for Watson-Crick or G-U pairs (> 5%). Below these base pair annotation is the consensus sequence: “R” = “A” or “G”, “Y” = “C” or “U”, red nucleotides: nucleotide identity conserved more than 97% of the time, black nucleotides: 90%, gray nucleotides: 75%, red circle (•): nucleotide is present 97% of the time, black circle (•): 90%, gray circle (•): 75%, white circle (○): 50%. All percentages of sequences just described (e.g. 97% conserved) assume that sequences have been weighted by the GSC algorithm implemented by the Infernal software package.

alignment positions 1 · · · 165

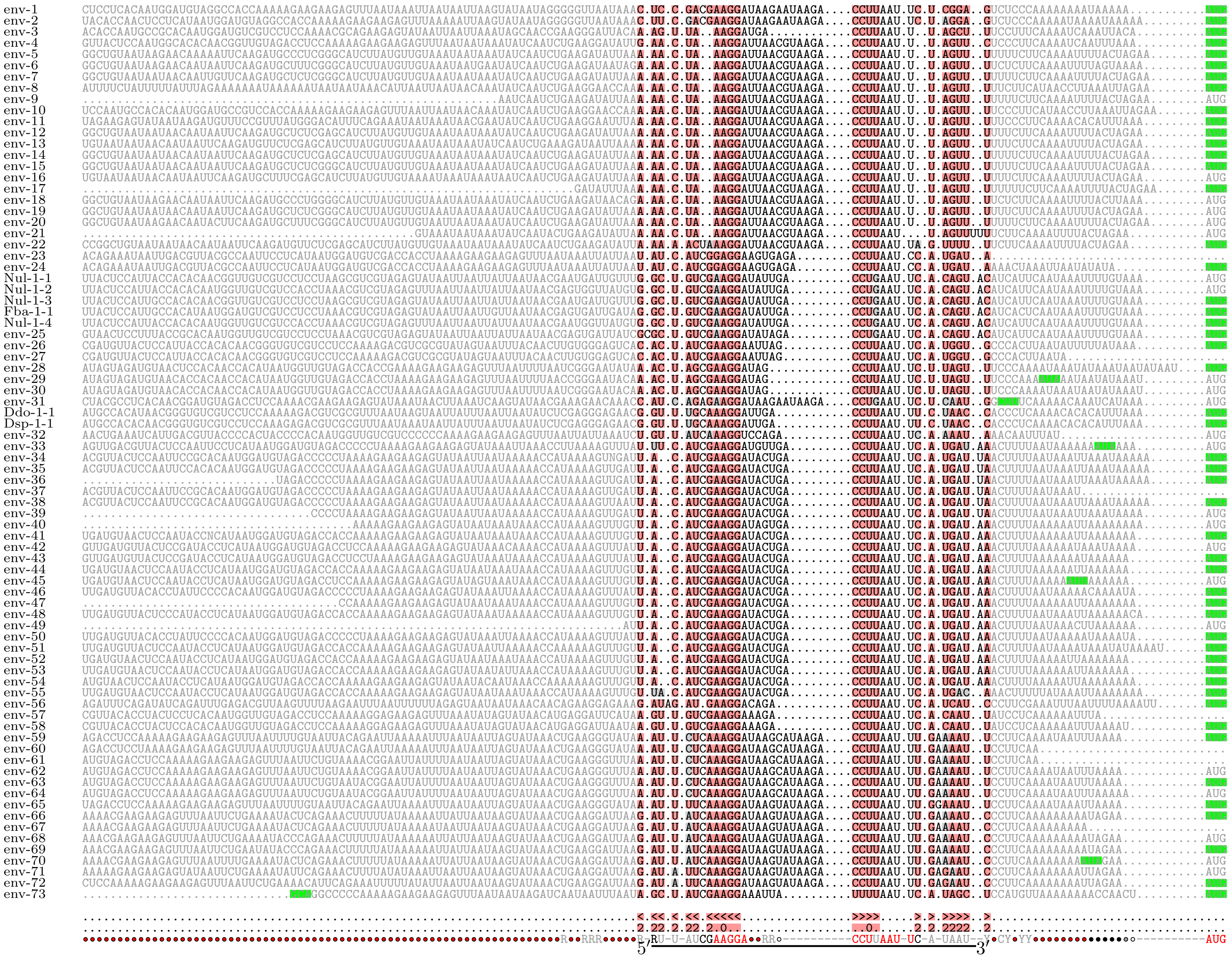

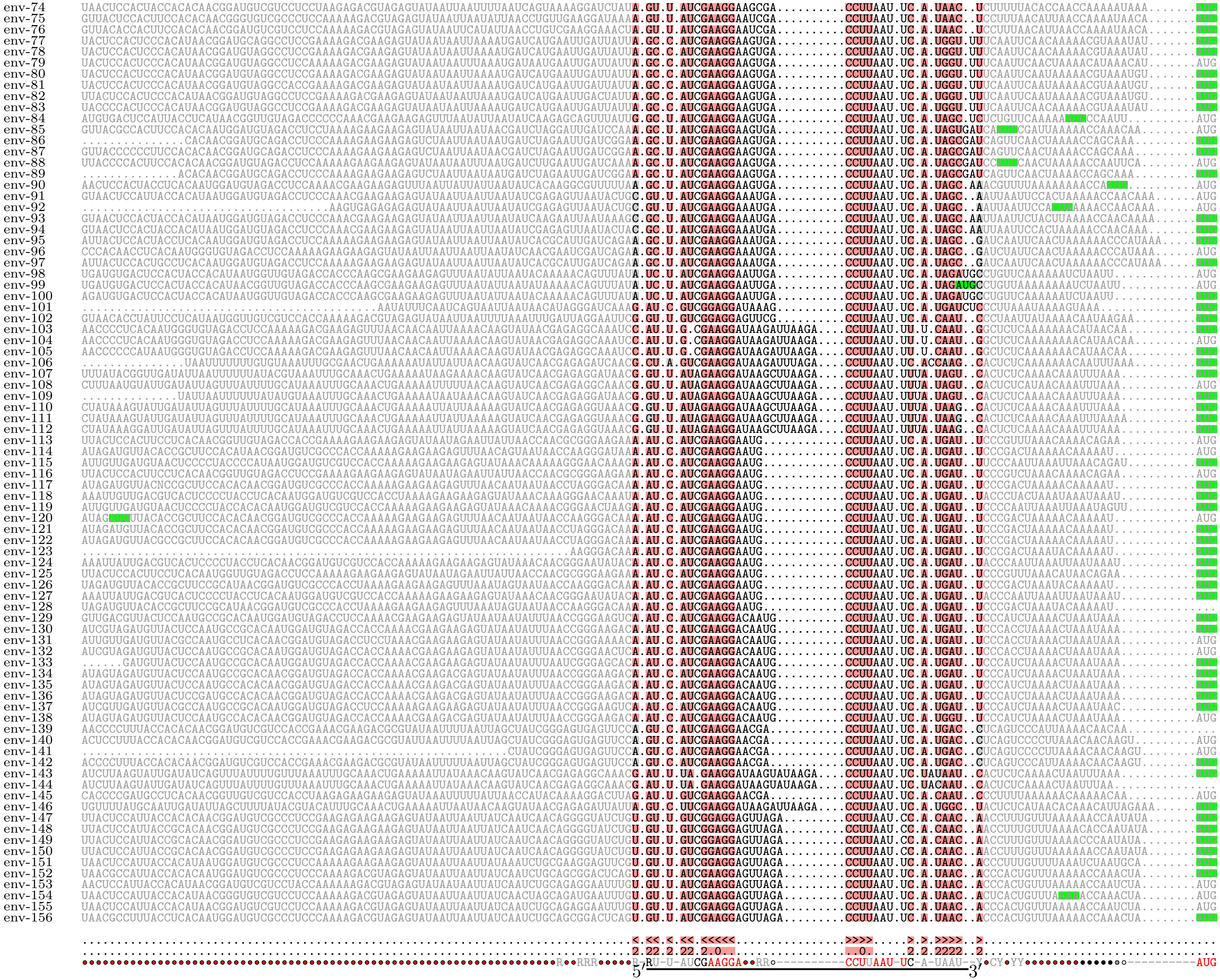
env-74
env-75
env-76
env-77
env-78
env-79
env-80
env-81
env-82
env-83
env-84
env-85
env-86
env-87
env-88
env-89
env-90
env-91
env-92
env-93
env-94
env-95
env-96
env-97
env-98
env-99
env-100
env-101
env-102
env-103
env-104
env-105
env-106
env-107
env-108
env-109
env-110
env-111
env-112
env-113
env-114
env-115
env-116
env-117
env-118
env-119
env-120
env-121
env-122
env-123
env-124
env-125
env-126
env-127
env-128
env-129
env-130
env-131
env-132
env-133
env-134
env-135
env-136
env-137
env-138
env-139
env-140
env-141
env-142
env-143
env-144
env-145
env-146
env-147
env-148
env-149
env-150
env-151
env-152
env-153
env-154
env-155
env-156
5′
3′

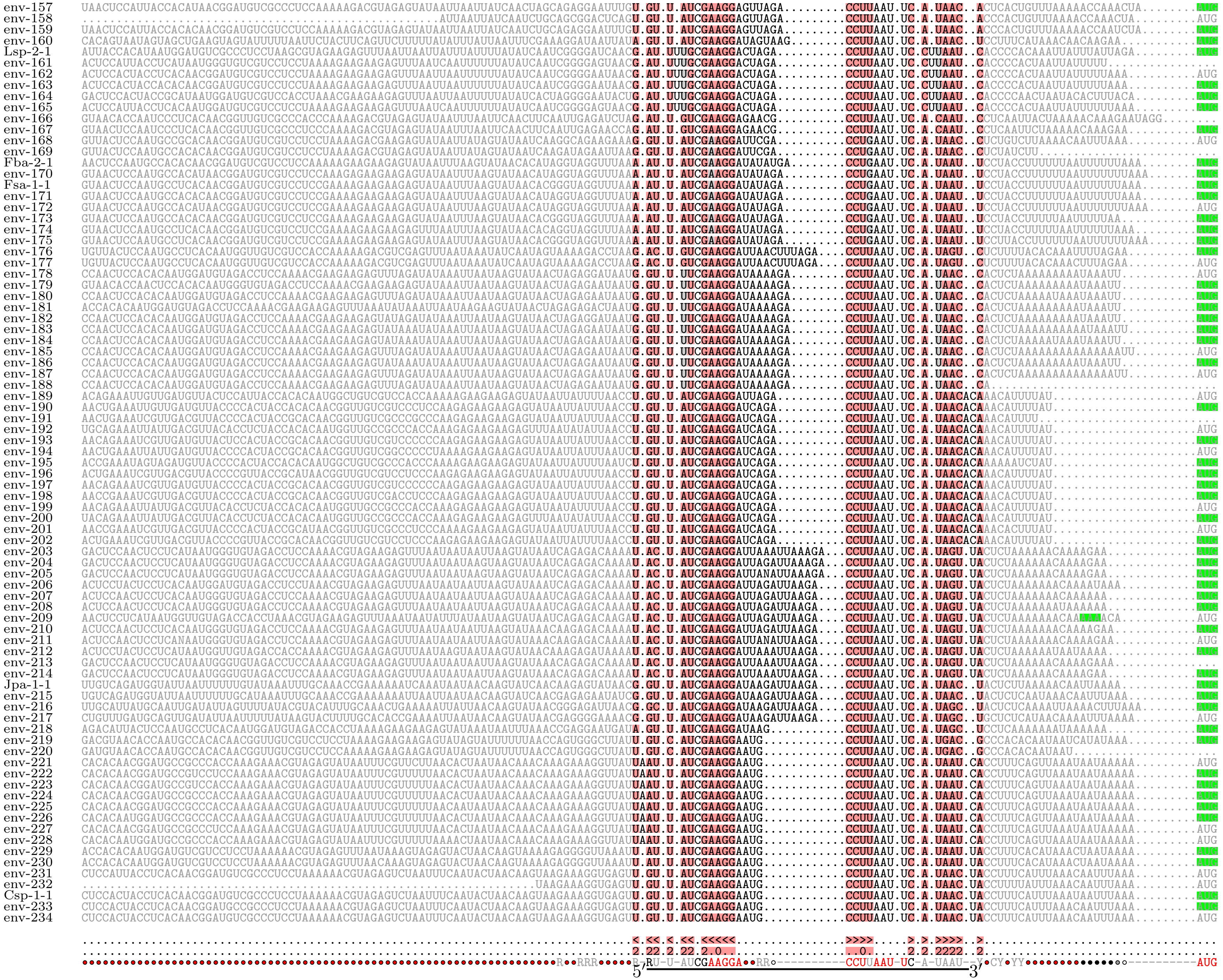

env-157
env-158
env-159
env-160
Lsp-2-1
env-161
env-162
env-163
env-164
env-165
env-166
env-167
env-168
env-169
Fba-2-1
env-170
Fsa-1-1
env-171
env-172
env-173
env-174
env-175
env-176
env-177
env-178
env-179
env-180
env-181
env-182
env-183
env-184
env-185
env-186
env-187
env-188
env-189
env-190
env-191
env-192
env-193
env-194
env-195
env-196
env-197
env-198
env-199
env-200
env-201
env-202
env-203
env-204
env-205
env-206
env-207
env-208
env-209
env-210
env-211
env-212
env-213
env-214
Jpa-1-1
env-215
env-216
env-217
env-218
env-219
env-220
env-221
env-222
env-223
env-224
env-225
env-226
env-227
env-228
env-229
env-230
env-231
env-232
Csp-1-1
env-233
env-234
5′
3′
AUG

```
env-235  CUCCAUUACCUCACAACGGAUGUCGCCCUCCUAAAAAACGUAGAGUCUAAUUUCAAUACUAACAAGUAAGAAAGGUGAGUU.GU.U.AUCGAAGGAAUG............CCUUAAU.UC.A.UAAC.UACCUUUUAUUUAAACAAUUUAAA.........AUG
env-236  .............CACAACGGAUGCCGCCCUCCUAAAAAACGUAGAGUAUAAUUUUUAAUAACAUUUAAGAAAGGUGAGUU.GU.U.AUCGAAGGAAUG............CCUUAAU.UC.A.UAAC.UACCUUUCAUUUAAACAAUUUAAA.........AUG
env-237  ...........CACAACGGAUGCCGCCCUCCUAAAAAACGUAGAGUCUAAUUUCAAUACUAACAAGUAAGAAAGGUGAGUU.GU.U.AUCGAAGGAAUG............CCUUAAU.UC.A.UAAC.UACCUUUCAUUUAAACAAUUUAAA.........AUG
env-238  CUCCACUGCCUCACAACGGAUGCCGCCCUCCUAAAAAACGUAGAGUCUAAUUUCAAUACUAACAAGUAAGAAAGGUGAGUU.GU.U.AUCGAAGGAAUG............CCUUAAU.UC.A.UAAC.UACCUUUCAUUUAAACAAUUUAAA.........AUG
env-239  UACUCCUCUUCCUCACAACGGAUGCCGCCCUCCUAAAAAACGUAGAGUAUAAUUUUUAAUAAUAUUUAAGAAAGGUGAGUU.GU.U.AUCGAAGGAAUG............CCUUAAU.UC.A.UAAC.UGCCUUUCAUUUUAAACAAUUCAAA........AUG
env-240  UACUCCCCUUCCUCACAACGGAUGCCGCCCUCCUAAAAAACGUAGAGUAUGAUUUAAUAUAACAAGUAAGAAAGGUGAGUU.GU.U.AUCGAAGGAAUG............CCUUAAU.UC.A.UAAC.UGCCUUUCAUUUUAAACAAUUCAAA........AUG
env-241  ACUCCUCUUCCUCACAACGGAUGCCGCCCUCCUAAAAAACGUAGAGUAUAAUUUUUAAUAACAUUUAAGAAAGGUGAGUU.GU.U.AUCGAAGGAAUG............CCUUAAU.UC.A.UAAC.UGCCUUUCAUUUUAAACAAUUCAAA........AUG
env-242  .............CACAACGGAUGCCGCCCUCCUAAAAAACGUAGAGUAUAAUUUUUAAUAACAUUUAAGAAAGGUGAGUU.GU.U.AUCGAAGGAAUG............CCUUAAU.UC.A.UAAC.UGCCUUUCAUUUUAAACAAUUCAAA........AUG
env-243  ..............................................................UAAGAAAGGUGAGUU.GU.U.AUCGAAGGAAUG............CCUUAAU.UC.A.UAAC.UGCCUUUCAUUUUAAACAAUUCAAA........AUG
env-244  ..........UCAUAACGGAUGUCGCCCUCCUAAAAAACGUAGAGUCUAAUUUCAAUACUAACAAGUAAGAAAGGUGAGUU.GU.U.AUCGAAGGAAUG............CCUUAAU.UC.A.UAAC.UGCCUUUCAUUUUAAACAAUUCAAA........AUG
env-245  UACUCCCCUUCCUCACAACGGAUGCCGCCCUCCUAAAAAACGUAGAGUAUAAUUUAAUAUAACAAGUAAGAAAGGUGAGUU.GU.U.AUCGAAGGAAUG............CCUUAAU.UC.A.UAAC.UGCCUUUCAUUUUAAACAAUUCAAA........AUG
env-246  UCCUCUACCUCACAACGGAUGCCGUCCUCCUAAAAAACGUCGCGUUUAAUUUCUGAAAUAUUAACAAGAGGAAAGGCUGUU.GU.U.AUCGAAGGAAUG............CCUUAAU.UC.A.UAAC.UGUCUUUCCUACUAAACAUUAAAAA........AUG
env-247  ACUCCUCUUCCUCACAACGGAUGCCGCCCCCCUAAAAAACGUAGAGUAUAAAUUUAAUAUAACAAGUAAGAAAGGUGAGUU.GU.U.AUCGAAGGAAUG............CCUUAAU.UC.A.UAAC.UGCCUUUCAUUUUAAACAAUUCAAA........AUG
env-248  .............................................UUAAUAAAUUUUUUUAAUAACUAAGGAUAAAG.AU.C.AGCGAAGGACUG............CCUUAAU.UC.A.UGAU..CCCUUACAUAAAUAUAAAAG............AUG
env-249  GUAACACCGAUCGCACACAACGGAUGUAGACCUCCGAAAAGAAGAAGAGUUUAAUAUAUUUUUUAACAAGUAAGGAUAAAG.AU.C.AGCGAAGGACUG............CCUUAAU.UC.A.UGAU..CCCUUACAUAAAAACUAAAG............AUG
env-250  GUAACACCGAUCGCACACAACGGGUGUAGACCUCCGAAAAGAAGAAGAGUUUAAUAUAUUUUUUAAUAACUAAGGAUAAAG.AU.C.AGCGAAGGACUG............CCUUAAU.UC.A.UGAU..CCCUUACAUAAAAACUAAAG............AUG
env-251  GUAACACCGAUCGCACACAACGGAUGUAGACCUCCGAAAAGAAGAAGAGUUUAAUAUAUUUUUUAAUAACUAAGGAUAAAG.AU.C.AGCGAAGGACUG............CCUUAAU.UC.A.UGAU..CCCUUACAUAAAAACUAAAG............AUG
env-252  UAACACCUAUCGCACACAACGGAUGUAGACCUCCGAAAAGAAGAAGAGUUUAAUAAAUUUUUUUAAUAACUAAGGAUAAAG.AU.C.AGCGAAGGACUG............CCUUAAU.UC.A.UGAU..CCCUUACAUAAAUAUAAAAG............AUG
env-253  UAACCCCAAUCGCACAUAACGGAUGUAGACCUCCGAAAAGAAGAAGAGUUUAAUAAAUUUUUUUAAUAACUAAGGAUAAAG.AU.C.AGCGAAGGACUG............CCUUAAU.UC.A.UGAU..CCCUUACAUAAAUAUAAAAG............AUG
env-254  UAACUCCGAUUGCUCACAAUGGAUGUAGACCUCCAAAAAGAAGAAGAGUUUAAUAAUAUUUUUUAAUAACUAAGGAUAAAG.AU.C.AGCGAAGGACUG............CCUUAAU.UC.A.UGAU..CCCUUACAUAAAUACAAAAG............AUG
env-255  .....................................................UUUUUAAUAACUAAGGAUUAAG.AU.U.AUCGAAGGACUG............CCUUAAU.UC.A.UGAU..CCCUUACAUAAAUAUAAAAAG...........AUG
env-256  ...............CAUAACGGAUGUAGACCUCCGAAAAGAAGAAGAGUUUAAUAAAUUUUUUAAUAACUAAGGAUAAAG.AU.C.AUCGAAGGACUG............CCUUAAU.UC.A.UGAU..CCCUUACAUAAAUAAUAAAG............AUG
env-257  UAACUCCAAUUGCACACAACGGUUGUAGACCUCCGAAAAGAAGAAGAGUUUAAUAAAUUUUUUUAAUAACUAAGGAUCAAG.AU.C.AUCGAAGGACUG............CCUUAAU.UC.A.UGAU..CCCUUACAAAAAAUAUAAAG............AUG
env-258  UAACUCCAAUUGCACAUAACGGUUGUAGACCUCCGAAAAGAAGAAGAGUUUAAUAAAUUUUUUUAAUAACUAAGGAUCAAG.AU.C.AUCGAAGGACUG............CCUUAAU.UC.A.UGAU..CCCUUACAAAAAAUAUAAAG............AUG
env-259  ..................................AAAGAAGAAGAGUAUAAUUAAAUUUUUAAUAACUGAGGAUUAAA.AU.U.AUCGAAGGACUG............CCUUAAU.UC.A.UAGU..UCCUCAAAUAAAUAUAAAAG............AUG
env-260  UAACACCAAUUGCACAUAACGGAUGUAGACCUCCGAAAAGAAGAAGAGUUUAAUAAAUUUUAUUAAUAACUGGGGAUAAAA.AU.U.AUCGAAGGACUG............CCUUAAU.UC.A.UAGU..UCCUCAAAUAAAUAUAAAAG............AUG
env-261  GUAACACCAAUCGCUCAUAACGGAUGUAGACCUCCGAAAAGAAGAAGAGUAUAAUUAAAUUUUUAAUAACUGAGGAUUAAA.AU.U.AUCGAAGGACUG............CCUUAAU.UC.A.UAGU..UCCUCAAAUAAAUAUAUAAG............AUG
env-262  CCACUUCCUCAUAAUGGAUGUCGUCCACCUAAACGAAGAAGAGUUUAAUUUUUAUAUUUAUUAACCCAGUUAGUACUUUAG.AU.U.AUCGAAGGAUUACUAGA.......CCUGAAU.UC.A.UAAU..CGUACUUAACUUAAUUAUUUUAAUUU......AUG
env-263  ............................................UAAUUAAUUAUUAAUCUAGAAAGGAAUUCG.AU.U.GUCGAAGGAUAUG...........CCUUAAU.UC.A.UAAU..CCCUUUCAAAAAAAAGAAAA............AUG
env-264  GUUACGCCAUUGCCACACAAUGGUUGUAGACCUCCAAAGAGAAGAAGAGUAUAAUUAAUUAUUAAUCAAGAAAGGAAUUCG.AU.U.GUCGAAGGAUAUG...........CCUUAAU.UC.A.UAAU..CCCUUUCAAAAAAAAGAAAA............AUG
env-265  GUUACGCCAUUGCCACACAAUGGUUGUAGACCUCCAAAGAGAAGAAGAGUAUAAUUAAUUAUUAAUCNAGAAAGGAAUUCG.AU.U.GUCGAAGGAUAUG...........CCUUAAU.UC.A.UAAU..CCCUUUCAAAAAAAAGAAAA............AUG
env-266  GUGACACCAUUGCCACAUAAUGGUUGUCGCCCUCCAAAAAGAAGAAGAGUAUAAUUAAUAAUUAAUCUAGAAAGGAAUUCG.AU.U.GUCGAAGGAUAUG...........CCUUAAU.UC.A.UAAU..CCCUUUCAAAAAAAACAAAA............AUG
env-267  GUUACGCCANUGCCACACAAUGGUUGUAGACCUCCAAAGAGAAGAAGAGUAUAAUUAAUUAUUAAUCUAGAAAGGAAUUCG.AU.U.GUCGAAGGAUAUG...........CCUUAAU.UC.A.UAAU..CCCUUUCAAAAAAAAGAAAA............AUG
env-268  ......................................AAGAAGAGUAUAAUUAAUUAUUAAUCUAGAAAGGAAUUCG.AU.U.GUCGAAGGAUAUG...........CCUUAAU.UC.A.UAAU..CCCUUUCAAAAAAAAGAAAA............AUG
env-269  GUUACGCCAUUGCCACACAAUGGUUGUAGACCUCCAAAGAGAAGAAGAGUAUAAUUAAUUAUUAAUCUAGAAAGGAAUUCG.AU.U.GUCGAAGGAUAUG...........CCUUAAU.UC.A.UAAU..CCCUUUCAAAAAAAAGAAAA............AUG
env-270  ...CCUACUCCUCAUAAUGGAUGUAGACCGCCAAAGAGAAGAAGAGUUUAAUAAUAAAUAAGUAUAACCAAAGAGAUAAUA.AU.U.AUCGAAGGAUUGA...........CCUUAAU.UC.A.UAAU..UACUCUUUAAAAAAAUAAAAAUA.........AUG
env-271  ACUCCUACUCCUCAUAAUGGAUGUAGACCACCAAAGAGAAGAAGAGUUUAAUAAUAAAUAAGUAUAACCAAAGAGAUAACA.AU.U.AUCGAAGGAUUGA...........CCUUAAU.UC.A.UAAU..UGCUCUUUAAAAAAUAAAAAAA..........AUG
env-272  GUUACUCCUACCCCUCACAAUGGAUGUAGACCACCUAAAAGAAGAAGAGUUUAAUAAAUAAGUAUAAUAAUAGAGGUUUUA.AU.U.AUCGAAGGAUUGA...........CCUUAAU.UC.A.UAAU..UACUCUAUAAAAAUUUAAAAA...........AUG
env-273  ACACCUACUCCUCAUAAUGGAUGUAGACCGCCAAAGAGAAGAAGAGUUUAAUAAUAAAUAAGUAUAACCAAAGAGAUAAUA.AU.U.AUCGAAGGAUUGA...........CCUUAAU.UC.A.UAAU..UACUCUUUAAAAAAUUAAAAUA..........AUG
env-274  ACACCUACUCCUCAUAAUGGAUGUAGACCGCCAAAGAGAAGAAGAGUUUAAUAAUAAAUAAGUAUAACCAAAGAGAUAAUA.AU.U.AUCGAAGGAUUGA...........CCUUAAU.UC.A.UAAU..UACUCUUUAAAAAAAUAAAAAUA.........AUG
env-275  ACUCCAAUGCCACAUAACGGAUGUCGUCCUCCGAAGAGAAGAAGAGUUUAAUAAGUAUAGUAAAUAACAUAAGAGAUAAAG.AG.U.AUCGGAGGAGGAAG..........CCUUAAU.UC.A.UAUU..CUCUCUUUAAUAAAAGUAAAAA..........AUG
env-276  UUACUCCAAUGCCACAUAACGGAUGUCGUCCUCCGAAAAGAAGAAGAGUUUAAUUAAAUUAAUAACAAAAGGGAAUUAACG.AA.U.AUCGAAGGAGGAAG..........CCUUAAU.UC.A.UAUU..CUCCCUUUAAAUAACAAACAAG..........AUG
env-277  AGUAUAAAUAUUCAUGAUGUAUUUUGUUUUAAAAACACAUCAUUUUUAUAAUUAAAUUAAGUAUAACAUUGGUAGAUAAAAAAU.U.AUCGGAGGAUUUGA..........CCUGAAU.UC.A.UAAU..UACUACCUAAAAAAAUUAAAA...........AUG
env-278  UCCUCAUAAUGGAUGUCGUCCUCCAAAAAGACGUAGAGUUUAAUUUAUAAUAAUAAUUUAAGUAUAAAUAGGUAAAUAUAA.AU.U.AUCGAAGGAUUUGA..........CCUGAAU.UC.A.UAAU..UUUUACCUUAAAAAAUUUUAAAA.........AUG
env-279  AACUCCAGCUACACACAAUGGAUGUAGACCUCCAAAAAGAAGAAGAGUAUAAUUUAAUUUUUAACUAAGUUGGAUUUUAG.AU.U.AUCGAAGGAGUUUCG.........CCUUAAU.UC.A.UAAU..CCCAGCUUUAAUAACUAAGAA...........AUG
env-280  GUAACUCCAAUACCACACAAUGGAUGUAGACCUCCAAAAAGAAGAAGAGUAUAAUUAAAUUUUAACUAAGUUGGAUUUUAG.AU.U.AUCGAAGGAGUUCG..........CCUUAAU.UC.A.UAAU..CCCAACUUUAAUAACUAAGAA...........AUG
env-281  AAUUAUCGAUGUUACUCCUAUUCCUCACAAUGGAUGCAGACCUCCAAAGCGGAGAAGAGUAUAAUACAAUUAAUCAUUACU.AU.U.AUCGAAGGAUAAGA..........CCUUAAU.UC.A.UAAU..AGCACACUUAAACAAUCAACA...........AUG
env-282  ...........................CACAAUGGAUGUAGACCACCAAAACGUAGAAGAGUCUAAUUUAUAACUUUUUACU.AU.U.AUCGAAGGAUAAGA..........CCUUAAU.UC.A.UAAUAGAAAUUAACAAAAACAAAACA...........AUG
env-283  AUCGAUGUUACUCCUAUUCCUCACAACGGAUGUAGACCUCCAAAACGAAGAAGAGUCUAAUUAUAAUAAUCAACUAUUACU.AU.U.AUCGAAGGAUAAGA..........CCUUAAU.UC.A.UAAUAGAGAUAU...........................
env-284  UUAUCGACGUAACACCUAUACCACAUAAUGGAUGUAGACCUCCAAAACGAAGAAGAGUCUAAUUAUAAAUCAACGCUUACU.AU.U.AUCGAAGGAUAAGA..........CCUUAAU.UC.A.UAAUAGAAAAACAAUAAAUACAUAACA..........AUG
Pgo-1-1  UUGAUGUAACUCCAAUUCCACAUAAUGGUUGUAGACCACCAAAACGAAGAAGAGUCUAAUUUUUAUAAAUCAAUUGAUACU.AU.U.AUCGAAGGAUAAGA..........CCUUAAU.UC.A.UAAUAGAGUAAAUUAAAUACACGACA...........AUG
env-285  AAUUAUUGAUGUUACUCCUAUCCCUCACAAUGGAUGCAGACCUCCAAAGAGGAGAAGAGUAUAAUUUAAUUAAUCAUUACU.AU.U.AUCGAAGGAUAAGA..........CCUUAAU.UC.A.UAAU..AGCACACUUAAACAAUCAACA...........AUG
env-286  ...........................AUGGAUGUAGACCUCCAAAACGAAGAAGAGUCUAAUUAUAAAUCAACGCUUACU.AU.U.AUCGAAGGAUAAGA..........CCUUAAU.UC.A.UAAUAGAAAAACAAUAAAUACAUAACA..........AUG
env-287  UUAUCGAUGUAACACCUAUACCACAUAAUGGAUGUAGACCUCCAAAAAGAAGAAGAGUCUAAUUAUAAAUCAAUGCUUACU.AU.U.AUCGAAGGAUAAGA..........CCUUAAU.UC.A.UAAUAGAAAAACAAUAAAUACAUAACA..........AUG
env-288  AUUGAUGUAACUCCAAUACCACAUAAUGGUUGUAGACCACCAAAACGAAGAAGAGUCUAAUUAUAUAAAUCAAUUGAUACU.AU.U.AUCGAAGGAUAAGA..........CCUUAAU.UC.A.UAAUAGAGUAAAUUAAAUACACGACA...........AUG
Pto-1-1  AUUGAUGUAACUCCAAUACCACAUAAUGGUUGUAGACCACCAAAACGAAGAAGAGUCUAAUUAUAUAAAUCAAUUGAUACU.AU.U.AUCGAAGGAUAAGA..........CCUUAAU.UC.A.UAAUAGAGUAAAUUAAAUACAAGACA...........AUG
Ptr-1-1  AUCGAUGUAACUCCUAUCCCACACAAUGGAUGUAGACCUCCUAAAAGAAGAAGAGUCUAAUUAUAUAAAUCAUAGCUUACU.AU.U.AUCGAAGGAUAAGA..........CCUUAAU.UC.A.UAAUAGACAAACGAUAAAUACAUAACA..........AUG
env-289  UACACCAAUGCCACAUAAUGGAUGUAGACCUCCAAAAAGAAGAAGAGUAUAAUUAAAUUAAUUCUAAUUAAUAGGAAUAAG.AU.U.AUCGAAGGAGAUUAAG........CCUUAAU.UC.A.UAAU..CCCUAUUUAAAUAAACAAAGAA..........AUG
env-290  GUUACACCAAUGCCACAUAAUGGAUGUCGCCCUCCAAAAAGAAGAAGAGUAUAAUUAAUAAUUUUAACUAAUAGGAAUUCG.AU.U.AUCGAAGGAGAUUAAG........CCUUAAU.UC.A.UAAU..CCCUAUUUAUAAUAACAUAGAA..........AUG
env-291  GUUACACCAAUGCCGCAUAAUGGAUGUCGCCCUCCAAAAAGAAGAAGAGUAUAAUUAAUAAUUUUAACUAAUAGGAAUUCG.AU.U.AUCGAAGGAGAUUAAG........CCUUAAU.UC.A.UAAU..CCCUAUUUAUAAUAACAUAGAA..........AUG
env-292  AUUAUUGAUGUAACUCCAUUACCACAUAACGGAUGUCGUCCUCCUAAAAGACGUAGAGUUUAAUUAAUUAAAAUUAUAUAG.GU.U.AUCGAAGGAUUGA...........CCUUAAU.UC.A.UAAC.CCAAUUAAAUACAAAUAGA..............AUG
env-293  AGAGAUUAUUGAUGUAACUCCACUACCACACAAUGGAUGUCGACCUCCUAAAAGAAGAAGAGUUUAACUUUGAACGUAUAG.GU.U.AUCGAAGGAUUGA...........CCUUAAU.UC.A.UAAC.CCAAUUAAAUUAUAAUAAA..............AUG
Rbi-1-1  GUGCCGCCCGCCCAAGCGACGCAGAGUCUGAUACGAGCAAGAAACACCAAUAAUUAAUUUUCACUGGGGGUGUUGUUACG.GU.U.AUCGAAGGAUUAAG..........CCUUAAU.UC.A.UAAC..CACAUCCUCAAAAUUUCUGAAA..........AUG
Gfo-1-1  UUACUCCACUUCCACACAACGGUUGUCGUCCACCAAAGAGACGUAGAGUUUAAUAAAUUUUAAUAAUAAGAGAGGAUUUAG.AU.U.AUCGAAGGACUUAUG.........CCUUAAU.UC.A.UAAU..CCCUUUCUAAAAUAACUAUAGUA.........AUG
Gpo-1-1  UCACUCCACUUCCACAUAAUGGAUGUCGUCCUCCUAAGAGACGUAGAGUUUAAUAAAAUUUAAUCAUAGAGAAGGAUUAAG.AU.U.AUCGAAGGACUUAUG.........CCUUAAU.UC.A.UAAU..CCCUUUUCAAAACCACAUUAGUA.........AUG
Gec-1-1  UAACACCACUUCCACAUAACGGUUGUCGUCCACCAAAGAGACGUAGAGUUUAAUAAUUUUAAUUAAUAGGAGAGGAAUUAG.AU.U.AUCGAAGGACUUAUG.........CCUUAAU.UC.A.UAAU..CCCUUUCUAAAACAACUAUAGUA.........AUG
env-294  UCACUCCACUUCCACAUAAUGGAUGUCGUCCUCCUAAGAGACGUAGAGUUUAAAAAAAUUUAAUCAUAGAGAAGGAUUAAG.AU.U.AUCGAAGGACUUAUG.........CCUUAAU.UC.A.UAAU..CCCUUUUCAAAACAACUUAAGUA.........AUG
env-295  UCACUCCACUUCCACACAAUGGAUGUCGUCCUCCUAAGAGACGUAGAGUUUAAUAAAAUUUAAUCAUAGAGAAGGAUUAAG.AU.U.AUCGAAGGACUUAUG.........CCUUAAU.UC.A.UAAU..CCCUUUUCAAAACAACUUAAGUA.........AUG
Fjo-1-1  GCCUCACAACGGAUGUCGUCCUCCAAAAAGACGUAGAGUUUAAUUUUUUUAUUUUUAUAGUAUAAUCAAGGUAGAAUAUAG.AU.U.AUCGGAGGAUUAGA..........CCUGAAU.UC.A.UAAU..CUCUACCUUAAAAUUUUUUUAAA.........AUG
env-296  ACAUUUAUGUGCUGAUAUUAGUUUAUUUUGUAUAAAUUUGCAAACUGAAAAAUAUUAACAAGUAUCAACAAGAGAUCAACG.AU.U.UUUGAAGGAUAAGAUUAAGA....CCUUAAU.UC.AUUUAU..CACUCUUAAAAAACAAUUUAAA..........AUG
env-297  AUCCAAUUAUGGUUUGAUAUUAGUUUAUUUUGCAUAAAUUUGCAAACUGAAAAAUAACAAGUAUCAAUCGAGAGAUCAACG.AU.U.UUUGAAGGAUAAGAUUAAGA....CCUUAAU.UC.AUUUAU..CACUCUCUAAACAAAUUUAAA...........AUG
env-298  UUACUCCAACUCCACACAAUGGAUGUAGACCUCCCAAAAGAAGAAGAGUUUAAUUUUAAUAGUAUAAACGAGAGAUUAAUG.AU.U.GUCGAAGGAUAAGUUUAAGA....CCUUAAU.UC.A.CAAU..CACUUUCAAAAACAAAACAAA...........AUG
env-299  AACUCCUACGCCUCAUAACGGAUGUAGACCACCAAAAAGAAGAAGAGUUUAAUUUUUUAAUAGUAUAAAUAGAGAUCAACG.AC.U.GUCGAAGGAUAAGCUUAAGA....CCUUAAU.UC.A.CAGU..CACUCUUAAAAACAAAACGAA...........AUG
env-300  UAACUCCUACGCCUCACAACGGAUGUAGACCACCAAAAAGAAGAAGAGUUUAAUUUUAAUAGUAUAAACGAGAGAACAACG.AC.U.GUCGAAGGAUAAGCUUAAGA....CCUUAAU.UC.A.CAAU..CACUCUCAAAAACAAAACGAA...........AUG
env-301  CUCCAACGCCUCACAAUGGAUGUAGACCUCCAAAACGAAGAAGAGUUUAAUAAUUUUAAUAGUAUAAACGAGAGAUCAACG.AC.U.GUCGAAGGAUAAGCUUAAGA....CCUUAAU.UC.A.CAAU..CACUCUCAAAAACAAAACAAA...........AUG
env-302  UAACUCCUACACCUCAUAACGGGUGUAGACCACCAAAAAGAAGAAGAGUUUAAUUUUAAUAGUAUAAACGAGAGAUCAACG.AC.U.GUCGAAGGAUAAGCUUAAGA....CCUUAAU.UC.A.CAAU..CA..............................
env-303  ACUCCUACACCUCACAACGGAUGUAGGCCACCAAAAAGAAGAAGAGUUUAAUUAUUUAUUAGUAUAAACGAGAGAUCAACG.AC.U.GUCGAAGGAUAAGCUUAAGA....CCUUAAU.UC.A.CAAU..CACUCUCAAAAACAAAACUAA...........AUG
env-304  UAACUCCUACUCCCCACAAUGGAUGUAGACCACCAAAAAGAAGAAGAGUUUAAUUUUAAUAGUAUAAACGAGAGAUCAACG.AC.U.GUCGAAGGAUAAGCUUAAGA....CCUUAAU.UC.A.CAAU..CACUCUCAAAAACAAAACGAA...........AUG
env-305  ACUCCUACACCUCACAACGGUUGUAGACCUCCAAAACGAAGAAGAGUAUAAAUUUAUAAUAGUAUAAACGAGAGAUCAACG.AC.U.GUCGAAGGAUAAGCUUAAGA....CCUUAAU.UC.A.CAAU..CACUCUCAAAAACAAAACGAA...........AUG
env-306  UAACUCCUACGCCUCACAACGGAUGUAGACCACCAAAAAGAAGAAGAGUUUAAUUUUAAUAGUAUAAACGAGAGAACAACG.AC.U.GUCGAAGGAUAAGCUUAAGA....CCUUAAU.UC.A.CAAU..CACUCUUAAAAACAAAACGAA...........AUG
env-307  ...........................................................................CAACG.AC.U.GUCGAAGGAUAAGCUUAAGA....CCUUAAU.UC.A.CAAU..CACUCUCAAAAACAAAACGAA...........AUG
env-308  ...................................................................CGAGAGAUCAACG.AC.U.GUCGAAGGAUAAGCUUAAGA....CCUUAAU.UC.A.CAAU..CACUCUCAAAAACAAAACAAA...........AUG
env-309  UCCUACACCUCACAAUGGAUGUAGACCUCCAAAACGAAGAAGAGUUUAAUUUUAUUUAAUAGUAUAAACGAGAGAUCAACG.AC.U.GUCGAAGGAUAAGCUUAAGA....CCUUAAU.UC.A.CAAU..CACUCUCAAAAACAAAACGAA...........AUG

         ................................................................................<.<<.<.<<.<<<<<................>>>>.....>.>.>>>>..>..................................
         ................................................................................2.22.2.22.2.0..................0.....2.2.2222..2..................................
         ●●●●●●●●●●●●●●●●●●●●●●●●●●●●●●●●●●●●●●●●●●●●●●●●●●●●●●●●●●●●●●●●●●●R●●RRR●●●●R-RU-U-AUCGAAGGA●●RR○----------CCUUAAU-UC-A-UAAU--Y●CY●YY●●●●●●●●●●●●○○----------AUG
                                                                                 5′                                                3′
```

```
env-310  UCCUACUCCUCACAACGGAUGUAGACCUCCAAAGCGAAGAAGAGUUUAAUUUAUUUUAAUAGUAUAAACGAGAGAUCAACG.AC.U.GUCGAAGGAUAAGCUUAAGA....CCUUAAU.UC.A.CAAU..CACUCUCAAAAACAAAACG................
env-311  CUCCUACUCCUCACAACGGAUGUAGACCUCCAAAGCGAAGAAGAGUUUAAUUUAUUUUAAUAGUAUAAACGAGAGAUCAACG.AC.U.GUCGAAGGAUAAGCUUAAGA....CCUUAAU.UC.A.CAAU..CACUCUCAAAAACAAAACGAA..........AUG
env-312  AACUCCUACUCCUCACAACGGAUGUAGACCACCAAAAAGAAGAAGAGUUUAAUUUUUAAUAGUAUAAACGAGAGAUCAACG.AC.U.GUCGAAGGAUAAGCUUAAGA....CCUUAAU.UC.A.CAAU..CACUCUCAAAAACAAAACGAA..........AUG
env-313  ACUCCUACACCUCACAACGGAUGUAGACCACCAAAAAGAAGAAGAGUUUAAUUAUUUAUUAGUAUAAACGAGAGAUCAACG.AC.U.GUCGAAGGAUAAGCUUAAGA....CCUUAAU.UC.A.CAAU..CACUCUCAAAAACAAAACGAA..........AUG
env-314  UAACUCCUACUCCUCACAAUGGAUGUAGACCACCAAAAAGAAGAAGAGUUUAAUUUUAAUAGUAUAAACGAGAGAUCAACG.AC.U.GUCGAAGGAUAAGCUUAAGA....CCUUAAU.UC.A.CAAU..CACUCUCAAAAACAAAACGAA..........AUG
env-315  CUCCUACACCUCACAACGGUUGUAGACCUCCAAAGAGAAGAAGAGUUUAAUUUAUUUUAAUAGUAUAAACGAGAGAUCAACG.AC.U.GUCGAAGGAUAAGCUUAAGA....CCUUAAU.UC.A.CAAU..CACUCUCAAAAACAAAACGAA..........AUG
env-316  GUCUGUUAUACAGAUGGUAUUAAUUUUUUUGUAUAAAUUUGCAAACUGAAAAUAUUAACAAGUAUCAACAAAGAGAUCAACG.AU.U.GUCGAAGGAUAAGAUUAAGA....CCUUAAU.UC.A.CAAU..CACUCUUAAAUCAAUUAAAGAA........AUG
env-317  CUCCAACGCCUCACAAUGGAUGUAGACCUCCAAAACGAAGAAGAGUUUAAUAAUUUUAAUAGUAUAAACGAGAGAUCAACG.AU.U.GUCGAAGGAUAAGCUUAAGA....CCUUAAU.UC.A.CAAU..CACUCUCAAAAACAAAACAAA..........AUG
env-318  CUCCUACACCUCACAAUGGAUGUAGACCUCCAAAGCGAAGAAGAGUUUAAUUUAUUUUAAUAGUAUAAACGAGAGAUCAACG.AU.U.GUCGAAGGAUAAGCUUAAGA....CCUUAAU.UC.A.CAAU..CACUCUCAAAAACAAAACAAA..........AUG
env-319  CCCCUACGCCUCACAACGGGUGUAGACCUCCAAAACGAAGAAGAGUAUAAUUUAUUUUAAUAGUAUAAACGAGAGAUCAACG.GU.U.GUCGAAGGAUAAGCUUAAGA....CCUUAAU.UC.A.CAAG..CACUCUCAAAAACAAAACAAA..........AUG
Tna-1-1  UUUAUAGCUGAUAUUUGUUUUUUUAUGUACAUUUGCAAUCCGAAAAAAAUAAUAUAAACAAGUAUCAACGAGAGAUUAACG.UU.U.GUCGGAGGAUAAGAUUAAGA....CCUUAAU.UC.A.CAAA..CACUCUCAUAAUUUAAUUUUAAA........AUG
env-320  .................AACUUUUUAUGUGUAAAUUUGCAAACUGAAAAAAUAAACAAGUAUCAAUUAGAGAGAUCAACG.UU.U.GUCGGAGGAUAAGAUUAAGA....CCUUAAU.UC.A.CAAA..CACUCUCAAAAACAAUUAAGAA........AUG
env-321  CAGAAAUAUGAUGUACACCAUCCUCAUAAUGGUGUCGUCUCAAGAGAGAAGAGUAUAAUAAAUAUUUAACCAUAGGAUUAG.A..U.AUCGAAGGAUUA...........GCUUAAU.UC.A.UAAU..CCAUGAUAAAUAAACACGAA..........AUG
env-322  AACUCCACUUCCACAUAAUGGAUGUCGCCCUCCUAAAAGACGUAAAGUAUAACAACUAAUCUAACUCAAAGAGAGAGGCUU.AU.U.AUCGAAGGACAG...........CCUUAAU.UC.A.UAAU..CUCCUCUCUCCAUUUAAAUUAAACGAAAAAA.AUG
env-323  ...........................UCCUAAAAGACGUAAAGUAUAACAACUAAUCUAACUCAAAGAGAGAGGCUU.AU.U.AUCGAAGGACAG...........CCUUAAU.UC.A.UAAU..CUCCUCUCUCCAUUUAAAUUAAACGAAGAAA.AUG
env-324  AACUCCACUACCUCAUAAUGGAUGUCGCCCUCCUAAAAGACGUAAAGUAUAACAACUAAUCUAACUCAAAGAGAGAGGCUU.AU.U.AUCGAAGGACAG...........CCUUAAU.UC.A.UAAU..CUCCUCUCUCCAUUUAAAUUAAACGAAGAAA.AUG
env-325  AACUCCACUUCCACAUAAUGGAUGUCGCCCUCCUAAAAGACGUAAAGUAUAACAACUAAUCUAACUCAAAGAGAGAGGCUU.AU.U.AUCGAAGGACAG...........CCUUAAU.UC.A.UAAU..CUCCUCUCUCCAUUUAAAUUAAACGAAGAAA.AUG
env-326  CCUUUACCACACAAUGGAUGUAGACCUCCUAAAAGACGUAGAGUUUAAUUAAUUAUUAAGUAUAAACGAGAAUGCAAUAAG.AU.U.AUCGAAGGACAGUGA........CCUUAAU.UC.A.UAAU..CGCAUUCUUAAAAAAAACAAUUUUCA......AUG
env-327  ............................AAAAGACGUAGAGUUUAAUUAAUUAUAAUAGUAUAAACGAGAAGGCAACAAG.AU.U.AUCGAAGGACAGUGA........CCUUAAU.UC.A.UAAU..CGCUUUUUCAAAAAAACAUUUUCA.......AUG
env-328  CCUUUGCCACACAAUGGGUGUAGACCUCCAAAAAGACGUAGAGUUUAAUUAAUUAUUAAGUAUAAACGGGAAGGCAAGAAG.AU.U.AUCGAAGGACAGUGA........CCUUAAU.UC.A.UAAU..CGCAUUCUCACAAAAAACAAUUUUCA......AUG
env-329  ..................................................................AAGCAAAAG.AU.U.AUCGAAGGACAGUGA........CCUUAAU.UC.A.UAAU..CGCUUUCUUAUAAAACACAAUUUAA.......AUG
env-330  CCAAUGCCNCACAAUGGAUGUAGNGACCUCCAAAAAGACGUAGAGUUUAAUUAAUUAUUAAGUAUAACAAGGAAAGCAAAAG.AU.U.AUCGAAGGACAGUGA........CCUUAAU.UC.A.UAAU..CGCUUUCUUAUAAAAACACAUUUAA.......AUG
env-331  CCAAUGCCUCACAAUGGAUGCAGACCUCCAAAAAGACGUAGAGUUUAAUUAAUUAUUAAGUAUAACAAGGAAAGCAAAAG.AU.U.AUCGAAGGACAGUGA........CCUUAAU.UC.A.UAAU..CGCUUUCUUAUAAAAACACAAUUUAA.......AUG
Gli-1-1  CCAUUACCACACAAUGGAUGUAGACCUCCAAAAAGACGUAGAGUUUAAUUAACAAUAAGUAUAAACAGGGAAAGCACCCAG.AU.U.AUCGAAGGACAGUGA........CCUUAAU.UC.A.UAAU..CGCUUUCUUAAACACAAAAUUUAAA......AUG
env-332  ...........AACGGAUGCCGUCCCCCGAAACGAAGAAGAGUUUAAUUAAUAUUUAUUGUUUAAUUGAAGGGUUUCCCG.AU.U.AUCGAAGGACAG...........CCUUAAU.UC.A.UAAU..CACCUCAAAAUUAAAAAAAAAC.........AUG
env-333  UCCAUUACCACACAACGGAUGUCGUCCUCCAAAGAGACGUAGAGUUUAAUAAUUAGUAUAACAAACAGGAAAGUAGUUAAG.AU.U.AUCGAAGGACAGAGA........CCUUAAU.UC.A.UAAU..CGCUUUCUAAUAAAAAUUUAAGUUA........AUG
Aat-1-1  GUAUAAAUUUUAUCCCGAAUUUUAACUUCGGGAUCUUUUUGUAUAUAAAUAACAAUAAGUAAAUCUAAAUUGGAGAAUUAG.AU.U.AUCGAAGGACAGAGA........CCUUAAU.UC.A.UAAU..CACCCUAAGUAACUAAAUUAUUAAUA.....AUG
env-334  ......CCGCAUAACGGAUGCCGUCCCCCGAAACGAAGAAGAGUUUAAUUAAUAUUUAUUGUUUAAUUGAAGGGUUUCCCG.AU.U.AUCGAAGGACAG...........CCUUAAU.UC.A.UAAU..CACCUCAAAAUUAAAAAAAAAC.........AUG
Gsp-3-1  CUCCAUUACCGCACAACGGAUGUAGACCUCCAAAAAGACGUAGAGUUUAAUUAUUAAGUAUAAACAAAGAAAGCACUAUAG.AU.U.AUCGAAGGACAGUGA........CCUUAAU.UC.A.UAAU..CGCUUUUUUUUAAAAACAAUUUUCA.....AUG
env-335  ..................AUGGACCUCCAAAAAGACGUAGAGUUUAAUUAAUUAUUAAGUAUAACAAGGAAAGCAAAGAG.AU.U.AUCGAAGGACAGUGA........CCUUAAU.UC.A.UAAU..CGCUUUCUUAAAAAACACAAUUUAA.......AUG
env-336  CCAUUGCCACACAAUGGAUGUAGACCUCCAAAAAGACGUAGAGUUUAAUUAAUUAUUAAGUAUAAACGGGAAGGCAAUAAG.AU.U.AUCGAAGGACAGUGA........CCUUAAU.UC.A.UAAU..CGCAUUCUCACAAAAAACAAUUUUCA......AUG
env-337  CCAUUGCCACACAAUGGGUGUAGACCUCCUAAAAGACGUAGAGUUUAAUUAAUUAUUAAGUAUAAACGAGAAGGCAAUAAG.AU.U.AUCGAAGGACAGUGA........CCUUAAU.UC.A.UAAU..CGCAUUCUUAAAAAAAACAAUUUUCA......AUG
env-338  CCUUUGCCACACAAUGGGUGUAGACCUCCAAAAAGACGUAGAGUUUAAUUAAUUAUUAAGUAUAAACGGGAAGGCAAUAAG.AU.U.AUCGAAGGACAGUGA........CCUUAAU.UC.A.UAAU..CGCAUUCUCACAAAAAACAAUUUUCA......AUG
env-339  CCAAUGCCUCACAAUGGAUGCAGACCUCCAAAAAGACGUAGAGUUUAAUUAAUUAUUAAGUAUAACAAGGAAAGCAAAAG.AU.U.AUCGAAGGACAGUGA........CCUUAAU.UC.A.UAAU..CGCUUUCUUAUAAAAACACAAUUUAA.......AUG
env-340  CCAAUGCCUCACAAUGGAUGCAGACCUCCAAAAAGACGUAGAGUUUAAUUAAUUAUUAAGUAUAACAAGGAAAGCAAAGAG.AU.U.AUCGAAGGACAGUGA........CCUUAAU.UC.A.UAAU..CGCUUUCUUAUAAAAACACAAUUUAA.......AUG
Gsp-4-1  CCUUUGCCACACAAUGGAUGUAGACCUCCAAAAAGACGUAGAGUUUAAUUAAUUAUUAAGUAUAAACGAGAAGGCAAUGAG.AU.U.AUCGAAGGACAGUGA........CCUUAAU.UC.A.UAAU..CGCUUUCUUAAAAAAACAAUUUUCA......AUG
Gsp-2-1  UCCAUUACCUCACAAUGGAUGUAGACCUCCUAAAAGACGUAGAGUUUAAUAAUUAUUAAGUAUAAACGAGAUUGCAAUAAG.AU.U.AUCGAAGGACAGUGA........CCUUAAU.UC.A.UAAU..CGCAUUCUCACAAAAAACAAUUUUCA......AUG
env-341  CCAUUACCACACAACGGAUGUCGUCCUCCUAAGAGACGUAGAGUUUAAUAAUAAGUAUAAAUAAAUAAGAGAGGAGUUAAG.AU.U.AUCGAAGGACAGAGA........CCUUAAU.UC.A.UAAU..CCCUUUCUUCUAAAUUCAAAGUUA.......AUG
env-342  CCACUACCACACAACGGAUGUCGUCCUCCUAAGAGACGUAGAGUUUAAUAAUAAGUAUAAAUAAAUAAGAGAGGAGUUAAG.AU.U.AUCGAAGGACAGAGA........CCUUAAU.UC.A.UAAU..CCCUUUCUUCUAAAUUCAAAGUUA.......AUG
Ssp-1-1  UCCAUUACCACACAAUGGAUGUCGUCCUCCUAAGAGACGUAGAGUUUAAUAAUAAGUAUAAAUAAAUAAGAGAGGAGUUAAG.AU.U.AUCGAAGGACAGAGA........CCUUAAU.UC.A.UAAU..CCCUUUCUACUAAAUUAAAAGUUA.......AUG
Kal-1-1  ACUCCAUUACCACACAAUGGAUGUAGACCUCCAAAAAGAAGAAGAGUAUAAUAAAUUAAUUUUCACAUUGAGAGACUAACG.AU.U.AUCGAAGGACAG...........CCUUAAU.UC.A.UAAU..CACUCUCAUACAACUAUUAGAA.........AUG
Gsp-1-1  CCAUUGCCACACAAUGGAUGUAGACCUCCAAAAAGACGUAGAGUUUAAUUAAUUAUUAAGUAUAAACGGGAACGCAAUAAG.AU.U.AUCGAAGGACAGUGA........CCUUAAU.UC.A.UAAU..CGCAUUCUCACAAAAAACAAUUUUCA......AUG
env-343  UUAAUGAAAUAAUUGAUGUAACGCCUAUACCGCAUAAUGGUUGCAGACCACCAAAAAGAAGAAGAGUAUAAUAAUAUAUAG.AU.U.AUCGAAGGAAAGA..........CCUUAAU.UC.G.UAAU..CAAACAACUAAAAUUUAAAU...........AUG
env-344  UUGAAGUAACUGAAAUAAUCGACGUAACACCAAUACCUCAUAAUGGAUGUAGGCCACCUAAACGUAGAAGAGUAUAAAAAG.AU.U.AUCGAAGGAAAGA..........CCUUAAU.UC.A.UAAU..CAAUAAUUUAAAAAUCAUAAU.........AUG
env-345  CUCCAAUGCCGCAUAAUGGAUGUCGUCCUCCAAAGAGAAGAAGAGUAUAACAAUUAGUAUAAAACAAAGGUAGGAAUUUAG.AU.U.AUCGAAGGAAAGA..........CCUUAAU.UC.A.UAAU..CCUAACUUAAAUUAAAAUUUGUAAA......AUG
env-346  ..............................................UAUGCCACCUAAACGUAGAAGAGUAUAAAAAG.AU.U.AUCGAAGGAAAGA..........CCUUAAU.UC.A.UAAU..CAAAAAUUUUAAAAAAUCAUAAU.........AUG
env-347  CUCCAAUGCCUCAUAAUGGAUGUCGUCCUCCAAAAAGAAGAAGAGUAUAAUUUAGUAUAAAACAAAGGUAGGAAUUUAG.AU.U.AUCGAAGGAAAGA..........CCUUAAU.UC.A.UAAU..CCCUACCUUU........................
env-348  CAGAAAUAUCGAUGUAACUCCUAUACCUCAUAAUGGUUGCCGACCACCAAAAAGAAGAAGAGUUUAAUAACUGAAUAUAG.AU.U.AUCGAAGGAAAGA..........CCUUAAU.UC.A.UAAU..CAAACAAUAAAAAAUAAUU...........AUG
env-349  CUCCGAUGCCACACAAUGGAUGUCGUCCUCCAAAAAGAAGAAGAGUAUAACAAUUAGUAUAAAACAAAGGUAGGAAUUUAG.AU.U.AUCGAAGGAAAGA..........CCUUAAU.UC.A.UAAU..CCUACCUUAAAUUAAAAAUUUGUAAA......AUG
Fin-1-1  UCCAAUGCCUCACAACGGAUGUCGUCCUCCAAAGAGAAGAAGAGUUUAAUAAUUAUUAAGUAUAAACAAAGGUAGGAAUUAG.AU.U.AUCGAAGGAAAGA..........CCUUAAU.UC.A.UAAU..CCUACCUUAUUUUAAAAUUUUAGAA......AUG
env-350  CUCCAAUGCCGCAUAAUGGAUGUCGUCCUCCAAAGAGAAGAAGAGUAUAACAAUUAGUAUAAAACAAAGGUAGGAAUUUAG.AU.U.AUCGAAGGAAAGA..........CCUUAAU.UC.A.UAAU..CCUAUCUUAAAUUAAAAUUUGUAAA......AUG
env-351  .......................AAGAAGAAGAGUUUAAUUAAUUAACGCCCAAGGGCAUUUCACAGAUACUGUAGUUACG.AU.U.AUCGAAGGAUAG...........CCUUAAU.UC.A.UGAU..CACAGUAUCAAAUUUUUUAGAAA.........AUG
env-352  CACAACGGUUGUAGACCUCCAAAAAGAAGAAGAGUUUAAUUAAUUAACGCCCAAGGGCAUUUCACAGAUACUGUAGAUACG.AU.U.AUCGAAGGAUAG...........CCUUAAU.UC.A.UGAU..CACAGUAUCAAAUUUUUUAGAAA.........AUG
env-353  CAUAACGGUUGUAGACCUCCAAAAAGAAGAAGAGUUUAAUUAAUUAACGCCCAAGGGCAUUUCACAGAUACUGUAGAUACG.AU.U.AUCGAAGGAUAG...........CCUUAAU.UC.A.UGAU..CACAGUAUCAAAUUUUUUAGAAA.........AUG
env-354  .............CCUACCAAAACGAAGAAGAGUUUAAUUAAUUAACGCCCAAGGGCAUUUCACAGAUACUGUAGUUACG.AU.U.AUCGAAGGAUAG...........CCUUAAU.UC.A.UGAU..CACAGUAUACUAAAUUUUUUAAGAAA.........
env-355  CANAACGGUUGUAGACCUCCAAAAAGAAGAAGAGUUUAAUUAAUUAACGCCCAAGGGCAUUUCACAGAUACUGUAGNUACG.AU.U.AUCGAAGGAUAG...........CCUUAAU.UC.A.UGAU..CACAGUAUCAAAUUUUUUAGAAA.........AUG
env-356  GAUGUUACUCCAACACCGCACAUAAUGGAUGUAGGCCACCUAAAAGAAGAAGAGUUUAAAAGAAAGUAUAAUUGAGAAAUUG.AU.U.AUCGAAGGAUUGA..........CCUUAAU.UC.A.UAAU..CACUCAAAUAAAAAUUAAAUAAUU.........AUG
env-357  AUCACUCCUACUCCUCACAAUGGAUGUAGACCGCCAAAAAGAAGAAGAGUUUAAUAAUAAGUAUAACAAAAAGAGAUUAUG.AU.U.AUCGAAGGAUUGA..........CCUUAAU.UC.A.UAAU..CACUCUAAAAAAAAAAUUUAAAA.........AUG
env-358  AUGUUACUCCUACUCCACAUAAUGGAUGUAGACCACCAAAAAGAAGAAGAGUAUAAUAUAAGUAUAACUAUAGAGGUUUG.AU.U.AUCGAAGGAUUGA..........CCUUAAU.UC.A.UAAU..CAC...........................
env-359  UGUUACUCCUACUCCGCAUAAUGGAUGUAGGCCACCUAAAAGAAGAAGAGUUUAAUAAUAAGUAUAACAAGAGAGGUAUUG.AU.U.AUCGAAGGAUUGA..........CCUUAAU.UC.A.UAAU..CGCUCUUAAAAAAAUUUAAAAA..........AUG
env-360  AUGUUACUCCUACUCCACAUAAUGGAUGUAGACCACCAAAAAGAAGAAGAGUAUAAUAUAAGUAUAACUAUAGAGGUUUG.AU.U.AUCGAAGGAUUGA..........CCUUAAU.UC.A.UAAU..CACUCUAAUAAAAAUUAAAAA..........AUG
env-361  CACCAACACCUCAUAAUGGUUGUAGACCACCUAAGAGAAGAAGAGUAUAAUUAUUAAAUAGUAUAACAAGGAGAAUAUUG.AU.U.AUCGAAGGAUUGA..........CCUUAAU.UC.A.UAAU..CACUCUUAAAAAAAUAAAAAAU.........AUG
env-362  GAUGUUACUCCGACACCACACAACGGAUGUAGACCACCUAAAAGAAGAAGAGUUUAAAAGAAAGUAUAAUUGAGAAAUUG.AU.U.AUCGAAGGAUUGA..........CCUUAAU.UC.A.UAAU..CACUCAAAAAAAUAAAUAAUU.........AUG
env-363  UGUUACUCCUACUCCGCAUAAUGGAUGUAGGCCACCUAAAAGAAGAAGAGUUUAAUAAUAAGUAUAACAAGAGAGGUAUUG.AU.U.AUCGAAGGAUUGA..........CCUUAAU.UC.A.UAAU..CACUCUUAAAAAAAUUUAAAAA..........AUG
env-364  GAUGUUACUCCGACACCACACAAUGGAUGUAGGCCGCCUAAAAGAAGAAGAGUUUAGAAAAAAGUAUAAUUGAGAAAUUG.AU.U.AUCGAAGGAUUGA..........CCUUAAU.UC.A.UAAU..CACUCAAAAAAAAUAAAUAAUU.........AUG
env-365  AUGUUACUCCUACUCCGCACAACGGAUGUAGACCACCAAAAAGGAGAAGAGUUUAAUAAUAAGUAUAACAAUAGAGAUUUG.AU.U.AUCGAAGGAUUGA..........CCUUAAU.UC.A.UAAU..CACUCUAAUAAAAAAUUAAAAA..........AUG
env-366  AUUACUCCUACUCCUCAUAAUGGAUGUAGACCACCAAAAAGAAGAAGAGUUUAAUAUAAGUAUAACAAAAAGAGAUUAUG.AU.U.AUCGAAGGAUUGA..........CCUUAAU.UC.A.UAAU..CACUCUCUAAAAAAAUUUAAAAA.........AUG
env-367  UGUUACUCCUACUCCUCAUAAUGGAUGUAGACCACCAAAAAGAAGAAGAGUAUAAUAUAAGUAUAACCAGAGGAGAUUUG.AU.U.AUCGAAGGAUUGA..........CCUUAAU.UC.A.UAAU..CACUCUCAAAAAAUUUAAAAA..........AUG
env-368  GAUGUUACCCCAACACCACACAAUGGAUGUAGACCACCUAAAAGAAGAAGAGUUUAAAAGAAAGUAUAAUUGAGAAAUUG.AU.U.AUCGAAGGAUUGA..........CCUUAAU.UC.A.UAAU..CACUCAAAAAAAAAAUAAAUAAUU.........AUG
env-369  AUCACUCCUACUCCUCACAAUGGAUGUAGACCACCAAAAAGAAGAAGAGUUUAAUAUAAGUAUAACAAAAAGAGAUUAUG.AU.U.AUCGAAGGAUUGA..........CCUUAAU.UC.A.UAAU..CACUCUUAAAAAAAAAAUUUAAAA.........AAUG
env-370  AUGUUACUCCUACUCCACACAAUGGAUGUAGACCACCAAAAAGAAGAAGAGUUUAAUAUAAGUAUAACUAUAGAGAUUUG.AU.U.AUCGAAGGAUUGA..........CCUUAAU.UC.A.UAAU..CACUCUAAUAAAAAAUUAAAAA..........AUG
env-371  AUGUUACUCCUACUCCACAUAAUGGAUGUAGACCACCAAAAAGAAGAAGAGUUUAAUAUAAGUAUAACUAUAGAGAUUUG.AU.U.AUCGAAGGAUUGA..........CCUUAAU.UC.A.UAAU..CACUCUAAUAAAAAUUUAAAAA..........AUG
env-372  AUGUUACUCCGACACCACAACGGAUGUAGACCACCAAAAAGAAGAAGAGUAUAAUAUAAGUAUAACAAUAGAGGUUUG.AU.U.AUCGAAGGAUUGA..........CCUUAAU.UC.A.UAAU..CACUCUAAUAAAAAUUAAAAA..........AUG
env-373  UGUUACUCCUACUCCGCAUAAUGGAUGUAGGCCACCUAAAAGAAGAAGAGUUUAAUAAUAAGUAUAACAAGAGAGGUAUUG.AU.U.AUCGAAGGAUUGA..........CCUUAAU.UC.A.UAAU..CACUCUU..........................
env-374  GAUGUUACCCCAACACCACACAAUGGAUGUAGACCACCUAAAAGAAGAAGAGUUUAAAAGAAAGUAUAAUUGAGAAAUUG.AU.U.AUCGAAGGAUUGA..........CCUUAAU.UC.A.UAAU..CACUCAAAAAAUAAAUAAAUAAUU........AUG
env-375  UGUUACUCCUACUCCGCAUAAUGGAUGUAGGCCACCUAAAAGAAGAAGAGUUUAAUAAUAAGUAUAACAAGAGAGGUAUUG.AU.U.AUCGAAGGAUUGA..........CCUUAAU.UC.A.UAAU..CACUCUUAAAAAAAUUUAAAAA..........AUG
env-376  AUGUUACUCCAACUCCACAUAACGGCUGUAGACCACCAAAAAGAAGAAGAGUUUAAUAUAAGUAUAACAAUAGAGAUUUG.AU.U.AUCGAAGGAUUGA..........CCUUAAU.UC.A.UAAU..CACUCUUAUAAAAAAUUAAAAA..........AUG
env-377  GAUGUUACCCCGACACCUCACAAUGGAUGUAGGCCACCUAAAAGAAGAAGAGUUUAGAAGAAAGUAUAAUUGAGAUAUUG.AU.U.AUCGAAGGAUUGA..........CCUUAAU.UC.A.UAAU..CACUCAAAAAAAAAAUAAAUAAUU.........AUG
env-378  AUGUUACCCCUACUCCCACAUAACGGAUGUAGGCCACCAAAAAGAAGAAGAGUUUAAUAUAAGUAUAACAAUAGAGGUUUG.AU.U.AUCGAAGGAUUGA..........CCUUAAU.UC.A.UAAU..CACUCUAAUAAAAAAUUAAAAA..........AUG
env-379  CACCAACACCUCAUAAUGGUUGUAGACCACCUAAGAGAAGAAGAGUAUAAUUAUUAAAUAGUAUAACAAGGAGAAUAUUG.AU.U.AUCGAAGGAUUGA..........CCUUAAU.UC.A.UAAU..CACUCUUAAAAAAAUAAAAAAU..........AUG
env-380  UGUUACUCCUACUCCUCAUAACGGAUGUAGACCCCCUAAAAGAAGAAGAGUUUAAUAAUAAGUAUAAUAAUAGAGAUUUG.AU.U.AUCGAAGGAUUGA..........CCUUAAU.UC.A.UAAUUACUCUAUAAAAAAUUUAAAAA...........AUG
env-381  UGUUACACCUACUCCUCACAACGGAUGUAGACCACCUAAAAGAAGAAGAGUUUAAUAAUAAGUAUAACAAUAGAGGUUUG.AU.U.AUCGAAGGAUUGA..........CCUUAAU.UC.A.UAAUUACUCUAUAAAAAAUUUAAAAA...........AUG
env-382  UGUUACCCCUACUCCUCAUAACGGAUGUAGACCACCUAAAAGAAGAAGAGUUUAAUAAUAAGUAUAACAAUAGAGGUUUG.AU.U.AUCGAAGGAUUGA..........CCUUAAU.UC.A.UAAUUACUCUAUAAAAAAUUUAAAAA...........AUG

         .................................................................................<.<<.<.<<.<<<<<...............>>>>.....>.>.>>>>..>...................................
         .................................................................................2.22.2.22.2.0.................0.....2.2.2222..2...................................
         ●●●●●●●●●●●●●●●●●●●●●●●●●●●●●●●●●●●●●●●●●●●●●●●●●●●●●●●●●●●●●●●●●●●R●●RRR●●●●●R-RU-U-AUCGAAGGA●●RR○---------CCUUAAU-UC-A-UAAU--Y●CY●YY●●●●●●●●●●●●●○○---------AUG
                                                                                  5′                                                     3′
```

```
env-383  UGUUACACCUACUCCUCAUAAUGGAUGUAGACCACCUAAAAGAAGAAGAGUUUAAUAAUAAGUAUAACAAUAGAGGUUUUG.AU.U.AUCGAAGGAUUGA...........CCUUAAU.UC.A.UAAUUACUCUAUAAAAAUUUAAAAA..............AUG
env-384  UGUUACUCCUACUCCCCAUAACGGUUGUAGACCCCCUAAAAGAAGAAGAGUUUAAUAAUAAGUAUAAUAAUAGAGAUUUUG.AU.U.AUCGAAGGAUUGA...........CCUUAAU.UC.A.UAAUUACUCUAUAAAAAUUUAAAAA..............AUG
env-385  UGUUACCCCUACUCCUCACAACGGAUGUAGACCGCCUAAAAGAAGAAGAGUUUAAUAAUAAGUAUAAUAAUAGAGAUUUUG.AU.U.AUCGAAGGAUUGA...........CCUUAAU.UC.A.UAAUUACUCUAUAAAAAUUUAAAAA..............AUG
env-386  UGUUACACCUACUCCUCAUAACGGAUGUAGACCACCUAAAAGAAGAAGAGUUUAAUAAUAAGUAUAACAAUAGGGGUUUUG.AU.U.AUCGAAGGAUUGA...........CCUUAAU.UC.A.UAAUUACUCUAUAAAAAUUUAAAAA..............AUG
env-387  ....ACACCUACUCCUCAUAACGGAUGUAGACCACCUAAAAGAAGAAGAGUUUAAUAAUAAGUAUAACAAUAGGGGCUUAG.AU.U.AUCGAAGGAUUGA...........CCUUAAU.UC.A.UAAUUACUCUAUAAAAAUUUAAAAA..............AUG
env-388  UGUUACUCCUACUCCUCAUAACGGAUGUAGACCACCUAAAAGAAGAAGAGUUUAAUAAUAAGUAUAAUAAUAGAGAUUUUG.AU.U.AUCGAAGGAUUGA...........CCUUAAU.UC.A.UAAUUACUCUAUAAAAAUUUAAAAA..............AUG
env-389  AUCACUCCUACUCCUCAUAAUGGAUGUAGACCACCGAAAAGAAGAAGAGUUUAAUAUAAGUAUAACAAAAAGAGAUUAUUG.AU.U.AUCGAAGGAUUGA...........CCUUAAU.UC.A.UAAU..CACUCUUAAAAAAAAAUUUAAAA..........AUG
env-390  AUCACUCCUACUCCUCAUAAUGGAUGUAGACCACCAAAAAGAAGAAGAGUUUAAUAUAAGUAUAACAAAAAGAGAUUAUUG.AU.U.AUCGAAGGAUUGA...........CCUUAAU.UC.A.UAAU..CACUCUUAAAAAAAAAAUUAAAA..........AUG
env-391  GAUGUUACUCCCACACCACACAAUGGAUGUAGACCACCUAAAAGAAGAAGAGUUUAAAAGAAAGUAUAAUUGAGAAAUAUG.AU.U.AUCGAAGGAUUGA...........CCUUAAU.UC.A.UAAU..CACUCAAAAAAAAAUUAAAUAAUU.........AUG
env-392  UAUCACUCCUACUCCUCAUAAUGGAUGUAGACCACCAAAAAGAAGAAGAGUUUAAUAUAAGUAUAACAAAAGAGAUUAUUG.AU.U.AUCGAAGGAUUGA...........CCUUAAU.UC.A.UAAU..CACUCUUAAAAAAAAAUUUAAAA..........AUG
env-393  AUCACUCCUACUCCUCAUAAUGGAUGUAGACCACCAAAAAGAAGAAGAGUUUAAUAUAAGUAUAACAAAAAGAGAUUAUUG.AU.U.AUCGAAGGAUUGA...........CCUUAAU.UC.A.UAAU..CACUCUUAAAAAAAAAAUUUAAAA.........AUG
env-394  AUGUUACUCCUACUCCACAUAAUGGAUGUAGACCACCAAAAAGAAGAAGAGUUUAAUAUAAGUAUAACUAUAGAGAUUUUG.AU.U.AUCGAAGGAUUGA...........CCUUAAU.UC.A.UAAU..CACUCUAAUAAAAAAUUAAAAA...........AUG
env-395  AUGUUACUCCUACUCCACAUAAUGGAUGUAGACCACCAAAAAGAAGAAGAGUAUAAUAUAAGUAUAACUAUAGAGGUUUUG.AU.U.AUCGAAGGAUUGA...........CCUUAAU.UC.A.UAAU..CACUC...............................
env-396  AUUACUCCUACUCCUCAUAAUGGAUGUAGACCACCUAAAAGAAGAAGAGUUUAAUAUAAGUAUAACAAAAAGAGAUUAUUG.AU.U.AUCGAAGGAUUGA...........CCUUAAU.UC.A.UAAU..CACUCUUAAAAAAAAAUUUAAAA..........AUG
env-397  GAUGUUACCCCAACACCACACAAUGGAUGUAGACCACCUAAAAGAAGAAGAGUUUAAAAAAAAGUAUAAUUGAGAAAUUUG.AU.U.AUCGAAGGAUUGA...........CCUUAAU.UC.A.UAAU..CACUCAAAAAAUAAAUAAAUAAUU.........AUG
env-398  UGUUACUCCAACUCCUCACAAUGGAUGUAGACCACCUAAAAGAAGAAGAGUUUAAUUAUAAGUAUAACCAAAGAGGUAUUG.AU.U.AUCGAAGGAUUGA...........CCUUAAU.UC.A.UAAU..CACUCUUAAAAAUUUUAAAAA............AUG
env-399  GACGUUACUCCGACACCACACAAUGGAUGUAGGCCACCUAAAAGAAGAAGAGUUUAAAACAAAGUAUAAUUGAGAAAUAUG.AU.U.AUCGAAGGAUUGA...........CCUUAAU.UC.A.UAAU..CACUCAAAUAAAAAUUAAAUAAUU.........AUG
env-400  GAUGUUACCCCGACACCACACAAUGGAUGUAGGCCACCUAAAAGAAGAAGAGUUUAGAAGAAAGUAUAAUUGAGAUAUUUG.AU.U.AUCGAAGGAUUGA...........CCUUAAU.UC.A.UAAU..CACUCAAAAAAAAAUAAAUAAUU..........AUG
env-401  GAUGUUACUCCGACACCACACAACGGAUGUAGACCACCUAAAAGAAGAAGAGUUUAAAAUAAAGUAUAAUUGAGAAAUAUG.AU.U.AUCGAAGGAUUGA...........CCUUAAU.UC.A.UAAU..CACUCAAAAAAAAAUAAAUAAUU..........AUG
env-402  CACCAACACCUCAUAAUGGUUGUAGACCACCUAAGAGAAGAAGAGUAUAAUUAUUAAUUAAGUAUAACAAGGAGAAUAUUG.AU.U.AUCGAAGGAUUGA...........CCUUAAU.UC.A.UAAU..CACUCUUAAAAAAAAAUAUAAAAU.........AUG
env-403  GAUGUUACUCCGACACCACACAACGGAUGUAGACCACCUAAAAGAAGAAGAGUUUAAAAGAAAGUAUAAUUGAGAAAUAUG.AU.U.AUCGAAGGAUUGA...........CCUUAAU.UC.A.UAAU..CACUCAAAAAAAAAAUUAAAUAAUU........AUG
env-404  AUGUUACUCCUACUCCACAUAAUGGCUGUAGACCACCAAAAAGAAGAAGAGUUUAAUAUAAGUAUAACAAUAGAGAUUUUG.AU.U.AUCGAAGGAUUGA...........CCUUAAU.UC.A.UAAU..CACUCUAAUAAAAAAUUAAAAA...........AUG
env-405  AUGUUACUCCUACUCCACACAAUGGAUGUAGACCACCAAAAAGAAGAAGAGUAUAAUAUAAGUAUAACUAUAGAGAUUUUG.AU.U.AUCGAAGGAUUGA...........CCUUAAU.UC.A.UAAU..CACUCUAAUAAAAAAUUAAAAA...........AUG
env-406  GAUGUUACUCCAACACCACACAACGGAUGUAGACCACCUAAAAGAAGAAGAGUUUAAAAGAAAGUAUAAUUGAGAAAUUUG.AU.U.AUCGAAGGAUUGA...........CCUUAAU.UC.A.UAAU..CACUCAAAUAAAAAUUAAAUAAUU.........AUG
env-407  GAUGUUACACCCACACCACAUAACGGGUGUAGACCACCUAAAAGAAGAAGAGUUUAAAAUAAAGUAUAAUUGAGAAAUAUG.AU.U.AUCGAAGGAUUGA...........CCUUAAU.UC.A.UAAU..CGCUCAAAUAAAAUUAAAUAAUU..........AUG
env-408  AUCACUCCUACUCCUCAUAACGGAUGUAGACCGCCAAAAAGAAGAAGAGUUUAAUAUAAGUAUAACAAAAAGAGAUUAUUG.AU.U.AUCGAAGGAUUGA...........CCUUAAU.UC.A.UAAU..CACUCUUAAAAAAAAAAUUAAAA..........AUG
env-409  AUCACUCCUACUCCUCACAAUGGAUGUAGACCGCCAAAAAGAAGAAGAGUUUAAUAUAAGUAUAACAAAAAGAGAUUAUUG.AU.U.AUCGAAGGAUUGA...........CCUUAAU.UC.A.UAAU..CACUCUUAAAAAAAAAUUUAAAA..........AUG
env-410  AUCACUCCUACUCCUCAUAAUGGAUGUAGACCACCAAAAAGAAGAAGAGUUUAAUAUAAGUAUAACAAAAAGAGAUUAUUG.AU.U.AUCGAAGGAUUGA...........CCUUAAU.UC.A.UAAU..CACUCUAAAAAAAAAAUUUAAAA..........AUG
env-411  AUGUUACUCCAACUCCACAUAAUGGAUGUAGACCACCAAAAAGAAGAAGAGUUUAAUAUAAGUAUAACUAUAGAGAUUUUG.AU.U.AUCGAAGGAUUGA...........CCUUAAU.UC.A.UAAU..CACUCUAAUAAAAAAUUAAAAA...........AUG
env-412  AUUACUCCUACUCCUCAUAAUGGAUGUAGACCACCAAAAAGAAGAAGAGUUUAAUAUAAGUAUAACAAAAAGAGAUUAUUG.AU.U.AUCGAAGGAUUGA...........CCUUAAU.UC.A.UAAU..CACUCUUAAAAAAAUUUAAAAA...........AUG
env-413  CACCAACACCUCAUAAUGGUUGUAGACCACCUAAGAGAAGAAGAGUAUAAUUAUUAAUUAAGUAUAACAAGGAGAAUAUUG.AU.U.AUCGAAGGAUUGA...........CCUUAAU.UC.A.UAAU..CACUCUUAAAAA........................
env-414  UGUUACUCCAACUCCUCACAAUGGAUGUAGACCACCUAAAAGAAGAAGAGUUUAAUUAUAAGUAUAACCAAAGAGGCAUUG.AU.U.AUCGAAGGAUUGA...........CCUUAAU.UC.A.UAAU..CACUCUUAAAAAUUUUAAAAA............AUG
env-415  UGUUACUCCUACUCCUCACAAUGGAUGUAGACCCCCUAAAAGAAGAAGAGUUUAAUAAUAAGUAUAAUAAUAGAGAUUUUG.AU.U.AUCGAAGGAUUGA...........CCUUAAU.UC.A.UAAUUACUCUAUAAAAAUUUAAAAA..............AUG
env-416  UGUUACACCUACUCCUCAUAACGGAUGUAGACCACCUAAAAGAAGAAGAGUUUAAUAAUAAGUAUAACAAUAGAGGUUUUG.AU.U.AUCGAAGGAUUGA...........CCUUAAU.UC.A.UAAUUACUCUAUAAAAAUUUAAAAA..............AUG
env-417  UGUUACUCCUACUCCUCAUAACGGAUGUAGACCUCCUAAAAGAAGAAGAGUUUAAUAAUAAGUAUAAUAAUAGAGAUUUUG.AU.U.AUCGAAGGAUUGA...........CCUUAAU.UC.A.UAAUUACUCUAUAAAAAUUUAAAAA..............AUG
env-418  CCCCUACUCCCCACAAUGGGUGUAGACCACCUAAAAGAAGAAGAGUAUAAUUAAUUUAAGUAUAACAAAAAGAGACUAUUG.AU.U.AUCGAAGGAUUGA...........CCUUAAU.UC.A.UAAUUACUCUUUAUAAAAUUUAAAAA............AUG
env-419  UGUUACUCCUACUCCCCAUAACGGAUGUAGACCCCCUAAAAGAAGAAGAGUUUAAUAAUAAGUAUAAUAAUAGAGAUUUUG.AU.U.AUCGAAGGAUUGA...........CCUUAAU.UC.A.UAAUUACUCUAUAAAAAUUUAAAAA..............AUG
env-420  .....................................................AAUAAUAAGUAUAAUAAUAGAGAUUUUG.AU.U.AUCGAAGGAUUGA...........CCUUAAU.UC.A.UAAUUACUCUAAUAAAAAUUUAAAAA.............AUG
env-421  ...........CUCCUCAUAACGGAUGUAGACCACCUAAAAGAAGAAGAGUUUAAUAAUAAGUAUAACAAUAGAGGUUUUG.AU.U.AUCGAAGGAUUGA...........CCUUAAU.UC.A.UAAUUACUCUAUAAAAAUUUAAAAA..............AUG
env-422  UGUUACACCUACACCCCAUAACGGAUGUAGACCCCCUAAAAGAAGAAGAGUUUAAUAAUAAGUAUAAUGAUAGAGAUUUUG.AU.U.AUCGAAGGAUUGA...........CCUUAAU.UC.A.UAAUUACUCUAUAAAAAUUUAAAAA..............AUG
env-423  UGUUACUCCUACUCCUCAUAAUGGAUGUAGACCCCCUAAAAGAAGAAGAGUUUAAUAAUAAGUAUAAUAAUAGAGAUUUUG.AU.U.AUCGAAGGAUUGA...........CCUUAAU.UC.A.UAAUUACUCUAUAAAAAUUUAAAAA..............AUG
env-424  AUGUAACCCCUACUCCACAUAAUGGAUGUAGACCACCAAAAAGAAGAAGAGUAUAAUAUAAGUAUAACUAUAGAGGUUUUG.AU.U.AUCGAAGGAUUGA...........CCUUAAU.UC.A.UAAU..CACUCUAAUAAAAAAUUAAAAA...........AUG
env-425  AUGUUACUCCUACUCCACAUAACGGCUGUAGACCACCAAAAAGAAGAAGAGUUUAAUAUAAGUAUAACAAUAGAGAUUUUG.AU.U.AUCGAAGGAUUGA...........CCUUAAU.UC.A.UAAU..CACUCUUAUAAAAAUUAAAAA............AUG
env-426  ACUCCACUUCCACAUAAUGGUUGUAGACCUCCAAAAAGGAGAAGAGUAUAAUUAAAUAAUUUUUAACCGAUUGGAAUUUG.AU.U.AUCGAAGGAUUUAG..........CCUUAAU.UC.A.UAAUCCCCCUAUCAAUUUAACAAAUAAAG..........AUG
env-427  ........................................................UAAUUUUUAACCGAUUAGGAAUUCG.AU.U.AUCGAAGGAUUUAG..........CCUUAAU.UC.A.UAAUCCCUAUCAAUUAACAACUAAAG.............AUG
env-428  UGUUACUCCUACUCCUCACAAUGGUUGUCGUCCACCAAAAAGAAGACGAGUAUAAUUAAAUUUUUAAUAUAUAGGAAACAG.AU.U.AUCGAAGGAUUAAG..........CCUUAAU.UC.A.UAGU..CCCUA...............................
env-429  UGUUACUCCUACUCCUCACAAUGGUUGUCGUCCACCAAAAAGAAGACGAGUAUAAUUAAAUUUUUAAUAUAUAGGAAACAG.AU.U.AUCGAAGGAUUAAG..........CCUUAAU.UC.A.UAGU..CCCUAUUGUAACAAAAACAGAA...........AUG
env-430  CUCCAAUUCCUCACAAUGGAUGUCGUCCUCCUAAGAGACGUAGAGUAUAAUUUUAAAAGUAUAAACAAGAGGGAGACUACG.AU.U.AUCGAAGGAUUAAUAGUAAGA...CCUUAAU.UC.A.UAAU..CACCUUCUAAAAAAAUCAAAUAA..........AUG
env-431  UGUUACAGAAAUUAUCGAUGUAACUCCUUUACCACACAAUGGGUGUAGACCACCUAAACGUAGAAGAGUUUAAUUAAUACG.AU.U.AUCGAAGGAAUAGA..........CCUUAAU.UC.A.UAAU..CACAAAUCAAAUAGA..................AUG
env-432  UACGCCAAUGCCACACAAUGGAUGUCGUCCUCCUAAGAGAAGAAGAGUAUAAUUAAAUUAAUUCUAAACAAUAGGAAUUAG.AU.U.AUCGAAGGAAUAAAG.........CCUUAAU.UC.A.UAAU..CCCUAUUAACAAACAAAGAA............AUG
env-433  CUAUGCCGCAUAAUGGCUGUAGACCUCCAAAAAGAAGAAGAGUUUAAUUAAUAUAAAUAUUUCAUAGGAAAUGUAGUUACG.AU.U.AUCGAAGGAUUAAG..........CCUUAAU.UC.A.UGAU..CACAUUUCCAAAUUAAAAGAAG...........AUG
env-434  CUAUGCCGCAUAAUGGCUGUAGACCCCCAAAAAGAAGAAGAGUUUAAUCAAUAUAAAUAUUUCAUAGGAAAUGUAGUUACG.AU.U.AUCGAAGGAUUAAG..........CCUUAAU.UC.A.UGAU..CACAUUUCCAAAUUAAAAGAAG...........AUG
env-435  CUAUGCCACAUAAUGGCUGUAGACCUCCAAAAAGAAGAAGAGUUUAAUCAAUAUAAAUAUUUCAUAGGAAGUGUAGUUACG.AU.U.AUCGAAGGAUUAAG..........CCUUAAU.UC.A.UGAU..CACAUUUCCAAAUUAAAAGAAG...........AUG
env-436  CAAUGCCACAUAAUGGUUGUAGACCUCCAAAAAGAAGAAGAGUUUAAUCAAUAUAAAUAUUUCAUAGGAAGUGUAGUUACG.AU.U.AUCGAAGGAUUAAG..........CCUUAAU.UC.A.UAAU..CACAUUUCCAAAUUUAAAGAAG...........AUG
env-437  UACACCAAUUCCACAUAACGGUUGUCGUCCACCAAAAAGAAGAAGAGUAUAAUUAAAUAUUUUAACCAAUUAGGAAACAUG.AU.U.AUCGAAGGAUUAAG..........CCUUAAU.UC.A.UAAU..CCCUAAUUAUAACAAAAACAAA...........AUG
env-438  GUUACUCCUACUCCUCACAAUGGUUGUCGUCCACCAAAAAGAAGACGAGUAUAAUUAAAUUUUUAAUAUAUAGGAAACAG.AU.U.AUCGAAGGAUUAAG..........CCUUAAU.UC.A.UAAU..CCCUAUUGUAACAAAAACAGAA...........AUG
env-439  .......................................AAAGAAGACGAGUAUAAUUAAAUUUUAAUAUAUAGGAAACAG.AU.U.AUCGAAGGAUUAAG..........CCUUAAU.UC.A.UAAU..CCCUAUUGUAACAAAACAGAA............AUG
env-440  ............................................UAAUCAAUAUAAAUAUUUCAUAGGAAGUGUAGUUACG.AU.U.AUCGAAGGAUUAAG..........CCUUAAU.UC.A.UAAU..CACAUUUCCAAAUUAAAAGAAG...........AUG
env-441  CAAUGCCACAUAAUGGUUGUAGACCUCCAAAAAGAAGAAGAGUUUAAUAAAUAUAAAUAUUUCAUAGGAAGUGUAGUUACG.AU.U.AUCGAAGGAUUAAG..........CCUUAAU.UC.A.UAAU..CACAUUUCCAAAUUUAAAGAAG...........AUG
env-442  UACACCUAUGCCACACAACGGGUGUAGACCUCCAAAAAGAAGAAGAGUAUAAUUAAUUAAAUUUUAACCGUUAGGAAUUAG.AU.U.AUCGAAGGAUUAAG..........CCUUAAU.UC.A.UAAU..CCCUAUCAUUAACAAAAUAAG............AUG
env-443  CUAUGCCACAUAAUGGCUGUAGACCUCCAAAAAGAAGAAGAGUUUAAUCAAUAUAAAUAUUUCAUAGGAAGUGUAGUUACG.AU.U.AUCGAAGGAUUAAG..........CCUUAAU.UC.A.UAAU..CACAUUUCCAAAUUUAAAGAAG...........AUG
env-444  ACACCUAUGCCACACAACGGUUGUAGACCUCCAAAAAGAAGAAGAGUAUAAUUAAUUUAAAUUUUAACCGAUAGGAAUUUG.AU.U.AUCGAAGGAUUAAG..........CCUUAAU.UC.A.UAAU..CCCUAUCAUUAACAAAAAGAA............AUG
env-445  GUUACUCCUACUCCUCAUAAUGGUUGUCGUCCACCAAAAAGAAGACGAGUAUAAUUAAAUUUUUAAUAUAUAGGAAACAG.AU.U.AUCGAAGGAUUAAG..........CCUUAAU.UC.A.UAAU..CCCUAUUGUAACAAAAACAGAA...........AUG
env-446  UGUUACUCCUACUCCUCACAAUGGUUGUCGUCCACCAAAAAGAAGACGAGUAUAAUUAAAUUUUUAAUAUAUAGGAAACAG.AU.U.AUCGAAGGAUUAAG..........CCUUAAU.UC.A.UAAU..CCCUAUUGUAACAAAAACAGAA...........AUG
env-447  .............................................................UCAUAGGGGGUGUAGUUACG.AU.U.AUCGAAGGAUUAAG..........CCUUAAU.UC.A.UAAU..CACAUUUCCAAAUUUAAAGAAG...........AUG
env-448  CACCUAUGCCACAUAACGGUUGUAGACCUCCAAAAAGAAGAAGAGUAUAAUUAAUUAAAUUUUAACCGAUAGGAUUGUUUG.AU.U.AUCGAAGGAUUAAG..........CCUUAAU.UC.A.UAAU..CCCUAUCAUUAACAAAAAGAA............AUG
env-449  CNAUGCCACAUAAUGGCUGUAGACCUCCAAAAAGAAGAAGAGUUUAAUCAAUAUAAAUAUUUCAUAGGAAGUGUAGUUACG.AU.U.AUCGAAGGAUUAAG..........CCUUAAU.UC.A.UAAU..CACAUUUCCAAAUUUAAAGAAG...........AUG
env-450  UACACCUAUGCCACACAACGGGUGUAGACCUCCAAAAAGAAGAAGAGUAUAAUUAAUUAAAUUUUAACCGUUAGGAAUUAG.AU.U.AUCGAAGGAUUAAG..........CCUUAAU.UC.A.UAAU..CCCUAUCAUUAACAAAACAAG............AUG
Fba-3-1  CAAUGCCACAUAACGGGUGUAGACCUCCAAAAAGAAGAAGAGUUUAAUUAAUAUUAAUAUUUCAAAGGAGGUGUAGUUACG.AU.U.AUCGAAGGAUUAAG..........CCUUAAU.UC.A.UAAU..CACAUUUCCAAAUUAAAAGAAG...........AUG
env-451  CAAUGCCACAUAAUGGCUGUAGACCUCCAAAAAGAAGAAGAGUUUAAUCAAUAUAAAUAUUUCAUAGGAAGUGUAGUUACG.AU.U.AUCGAAGGAUUAAG..........CCUUAAU.UC.A.UAAU..CACAUUUCCAAAUUUAAAGAAG...........AUG
env-452  ........................CUCCAAAAAGAAGAAGAGUUUAAUCAAUAUAAAUAUUUCAUAGGAAGUGUAGUUACG.AU.U.AUCGAAGGAUUAAG..........CCUUAAU.UC.A.UNAU..CACAUUUCCAAAUUAAAAGAAG...........AUG
env-453  UGUUACUCCNACUCCNCANAAUGGUUGUCGUCCACCAAAAAGAAGACGAGUAUAAUUAAAUUUUUAAUAUAUAGGAANCAG.AU.U.AUCGAAGGANUAAG..........CCUUAAU.UC.A.UAAU..CCCUAUUGUAACAAAAACAGAA...........AUG
env-454  UUACACCAGUACCUCACAAUGGUGUCGUCCACUAAGAGAAGAAGAGUAUAAUUAAAUAUUUUAACUAAGUAGGAUUUUUAG.AU.U.AUCGAAGGAAUAAG..........CCUUAAU.UC.A.UAAU..CCCUACUUUAAUAACAACGAA............AUG
env-455  UACACCAGUUCCUCACAAUGGUUGUCGUCCACUAAGAGAAGAAGAGUAUAAUUAAAUAUUUUAACUAAGUAGGAUUUUUAG.AU.U.AUCGAAGGAAUAAG..........CCUUAAU.UC.A.UAAU..CCCUACUUUAAUAACAACGAA............AUG
env-456  ACACCAGUUCCUCACAAUGGUUGUCGUCCACCUAAGAGAAGAAGAGUAUAAUUAAAUAUUUUAACUAAGUAGGAUUUUUAG.AU.U.AUCGAAGGAAUAAG..........CCUUAAU.UC.A.UAAU..CCCUACUUUAAUAACAACGAA............AUG
env-457  UAAUAGAUGUUACUCCAUUACCACACAAUGGUUGUCGACCUCCUAAAAGAAGAAGAGUAUAAUUAAAUAACAAUCGCUGAG.GU.U.AUCGAAGGAUUAAAUUAAGA....CCUUAAU.UC.A.UAAU..CGUUGAAGCAAAAACUACUAUAAA.........AUG
env-458  UAAUAGAUGUUACUCCAUUACCACACAAUGGUUGUCGACCUCCUAAAAGAAGAAGAGUAUAAUUAAUUAACAAUCGCUGAG.AU.U.AUCGAAGGAUUAAAUUAAGA....CCUUAAU.UC.A.UAAU..CGUUGAAGCAAAAACUACUAUAAA.........AUG
env-459  UAAUAGAUGUUACUCCAUUGCCACACAAUGGUUGUCGACCUCCUAAAAGAAGAAGAGUAUAAUUAAAUAACAAUCGCUGAG.AU.U.AUCGAAGGAUUAAAUUAAGA....CCUUAAU.UC.A.UAAU..CGUUGAAGCAAAAACUACUAUAAA.........AUG
env-460  UUACACCAAUGCCACAUAAUGGAUGUCGUCCUCCUAAGAGAAGAAGAGUAUAAUUAAAUUAAUUCUAACAAUAGGAACUAG.AU.U.AUCGAAGGACUAAG..........CCUUAAU.UC.A.UAAU..CCCUAUUAAUAAACAAAGAA............AUG
env-461  AUGUUACUCCAACACCUCACAAUGGUUGUCGUCCACCAAAAAGAAGACGAGUAUAAUAAAUUUUUAAUUAAUAGGAAUCAG.AU.U.AUCGAAGGAGUAAG..........CCUUAAU.UC.A.UAAU..CCCUAUUACAAAACAAAGAA............AUG
env-462  ...........ACCUCAACAAUUGGUUNUCGUCCACCAAAAAGAAGACGAGUAUAAUAAAUUUUAAUUAAUAGGAAUCAG.AU.U.AUCGAAGGAGUAAG..........CCUUAAU.UC.A.UAAU..CCCUAUUACAAAACAAAAGAA...........AUG
env-463  AUGUUACUCCAACGCCUCAUAAUGGUUGUCGUCCACCAAAAAGAAGACGAGUAUAAUUUAAUUUUAAUUGAUAGGAAUCAG.AU.U.AUCGAAGGAGUAAG..........CCUUAAU.UC.A.UAAU..CCCUAUUGUAACAUAAAUAGAA...........AUG
env-464  UACUCCAAUGCCACACAAUGGUUGUAGACCACCAAAGAGAAGAAGAGUAUAAUUAAUUAAUUUUAACCGACAGGAGUUAG.AU.U.AUCGAAGGAGUAAG..........CCUUAAU.UC.A.UAAU..CCCUGUCAAUAACAUAAAAAA............AUG

         ................................................................................<.<<.<.<<.<<<<<................>>>>.....>.>.>>>>..>...................................
         ................................................................................2.22.2.22.2.0..................0......2.2.2222..2...................................
         •••••••••••••••••••••••••••••••••••••••••••••••••••••••••••••••••••••R••RRR••••R-RU-U-AUCGAAGGA••RR○---------CCUUAAU-UC-A-UAAU--Y•CY•YY•••••••••••••◦○---------AUG
                                                                                 5′                                               3′
```

```
env-465  GAUGUUACUCCAACACCUCACAAUGGUUGUCGUCCACCAAAAAGAAGACGAGUAUAAUAAAUUUUAAUUGAUAGGAAUCAG.AU.U.AUCGAAGGAGUAAG.........CCUUAAU.UC.A.UAAU..CCCUAUUACAAAACAAAAGAA..........AUG
env-466  UGUUACUCCAACUCCUCAUAAUGGUUGUCGUCCACCAAAAAGAAGACGAGUAUAAUAAAUUUUAAUAUAUAGGAAUCAG.AU.U.AUCGAAGGAGUAAG.........CCUUAAU.UC.A.UAAU..CCCUAUUGUAACAAAACAGAA..........AUG
env-467  AUGUUACUCCAACACCUCACAAUGGUUGUCGUCCACCAAAAAAGAAGACGAGUAUAAUAAAUUUUAAUUAAUAGGAAUCAG.AU.U.AUCGAAGGAGUAAG.........CCUUAAU.UC.A.UAAU..CCCUA.............................
env-468  ACGUUACUCCAACUCCUCAUAAUGGUUGUCGUCCACCAAAAAGAAGACGAGUAUAAUUAAAUUUUAAUAUAUAGGAACCAG.AU.U.AUCGAAGGAGUAAG.........CCUUAAU.UC.A.UAAU..CCCUAUUGUAACAAAACAGAA..........AUG
env-469  ..................UCACAAUGGUUGUCGUCCACCAAAAAGAAGACGAGUAUAAUAAAUUUUAAUUGAUAGGAAUCAG.AU.U.AUCGAAGGAGUAAG.........CCUUAAU.UC.A.UAAU..CCCUAUUACAAAACAAAAGAA..........AUG
env-470  UACUCCAAUGCCACACAAUGGUUGUAGACCACCAAAGAGAAGAAGAGUAUAAUUAAAUUUUAACCGAUAGGAUGAAG.AU.U.AUCGAAGGAGUAAG.........CCUUAAU.UC.A.UAAU..CCCUAUCAAUUAACAAAAGAA..........AUG
env-471  GAUGUUACUCCAGCACCUCACAAUGGUUGUCGUCCACCAAAAAGAAGACGAGUAUAAUAAAUUUUAAUUGAUAGGAAUCAG.AU.U.AUCGAAGGAGUAAG.........CCUUAAU.UC.A.UAAU..CCCUAUUACAAAACAAAAGAA..........AUG
env-472  AGAAGAAGAGUUUAAGCUGAAUUCAUUUCAGUUUCUAUUUGAGUAUUAGUAGAAUUUAUAUUUUAACUUAUAGGAAUCAG.AU.U.AUCGAAGGAGUAAG.........CCUUAAU.UC.A.UAAU..CCCUAUUUAUAACAAAAAGAA..........AUG
env-473  GAUGUUACUCCAACACCUCACAAUGGUUGUCGUCCACCAAAAAGAAGACGAGUAUAAUAAAUUUUAAUUAAUAGGAAUCAG.AU.U.AUCGAAGGAGUAAG.........CCUUAAU.UC.A.UAAU..CCCUAUU...........................
env-474  ACGUUACUCCAACUCCUCAUAAUGGUUGUCGUCCACCAAAAAGAAGACGAGUAUAAUAAAUUUUAAUAUAUAGGAACCAG.AU.U.AUCGAAGGAGUAAG.........CCUUAAU.UC.A.UAAU..CCCUAUUGUAACAAAACAGAA..........AUG
env-475  GAUGUUACUCCAACUCCUCAUAAUGGUUGUCGUCCACCAAAAGAAGACGAGUAUAAUUAAUUUUAAUAUAUAGGAAUCAG.AU.U.AUCGAAGGAGUAAG.........CCUUAAU.UC.A.UAAU..CCCUAUUGUAACAAAACAGAA..........AUG
env-476  GAUGUUACUCCAACACCUCACAAUGGUUGUCGUCCACCAAAAAGAAGACGAGUAUAAUAAAUUUUAAUUGAUAGGAAUCAG.AU.U.AUCGAAGGAGUAAG.........CCUUAAU.UC.A.UAAU..CCCUAUUGCAAAACAAAAGAA..........AUG
env-477  .............................................GAGUAUAAUUAAUUAAUUUUUAACCGACAGGAGUUAG.AU.U.AUCGAAGGAGUAAG.........CCUUAAU.UC.A.UAAU..CCCUGUCAAUAACAUAAAAAA..........AUG
env-478  UGUUACUCCAACUCCUCAUAAUGGUUGUCGUCCACCAAAAAGAAGACGAGUAUAAUUAAAUUUUAAUAUAUAGGAAUCAG.AU.U.AUCGAAGGAGUAAG.........CCUUAAU.UC.A.UAAU..CCCUAUUGUAACAAAACAGAA..........AUG
env-479  UGUAACUCCAAUUCCUCAUAAUGGUUGUCGUCCACCAAAAAGAAGACGAGUAUAAUAAAUAUUUUAAUAGAUAGGAGUUAG.AU.U.AUCGAAGGAGUAAG.........CCUUAAU.UC.A.UAAU..CCCUAUCAUAACUAAAUUAGAA..........AUG
Psp-1-1  UUACACCUAUUCCUCAUAAUGGAUGUCGUCCACCAAAGAGAAGAAGAGUAUAAUUAAAUAUUUUAACUUGAUAGGAAUCAG.AU.U.AUCGAAGGAGUAAG.........CCUUAAU.UC.A.UAAU..CCCUAUCUAUAACAAAAAAGAA..........AUG
env-480  GUUACUCCAACACCUCACAAUGGUUGUCGUACCACCAAAAAGAAGACGAGUAUAAUAAAUUUUAAUUAAUAGGAAUCAG.AU.U.AUCGAAGGAGUAAG.........CCUUAAU.UC.A.UAAU..CCCUAUUACAAAACAAAAGAA..........AUG
env-481  ..........................................................................AAUCAG.AU.U.AUCGAAGGAGUAAG.........CCUUAAU.UC.A.UAAU..CCCUAUUAACAAAACAAAAGAA..........AUG
env-482  GUUACUCCAACCACCUCACAAUGGUUGUCGUCCACCAAAAAGAAGACGAGUAUAAUAAAUUUUAAUUGAUAGGAACCAG.AU.U.AUCGAAGGAGUAAG.........CCUUAAU.UC.A.UAAU..CCCUAUCUGAGUCGGAGA.................
env-483  ................................UAAAAAGAAUAUCAAAAUUAAUUAUAUUUUAACUUGAUAGGAACUAAG.AU.U.AUCGAAGGAGUAAG.........CCUUAAU.UC.A.UAAU..CCCUAUCUAUAACAAAAAAGAA..........AUG
env-484  GAUGUUACUCCAACACCUCACAAUGGUUGUCGUCCACCAAAAAGAAGACGAGUAUAAUAAAUUUUAAUUGAUAGGAAUCAG.AU.U.AUCGAAGGAGUAAG.........CCUUAAU.UC.A.UAAU..CCCUAUUAUAAAACAAAAGAA..........AUG
Tov-1-1  AGUUGAAUUAACUUCAACUUAUAUUUUAUUUAGCGUACUUUUGUGCGCAAUUAAAAAUAUUUUAACUAAGGUAGGAACAAG.AU.U.AUCGAAGGAGUAAG.........CCUUAAU.UC.A.UAAU..CCCUACUAAAACAAUAAGAA..........AUG
env-485  .................................................................UAGGAAUCAG.AU.U.AUCGAAGGAGUAAG.........CCUUAAU.UC.A.UAAU..CCCUAUUACAAAACAAAAGAA..........AUG
env-486  UACGCCAAUGCCACACAAUGGAUGUCGCCCUCCUAAAAGAAGAAGAGUUUAAUUUAUAUACAUAUUUCAUUUGGGAAUCG.AU.U.AUCGAAGGAGGAAG.........CCUUAAU.UC.A.UAAU..CCCCGAAGCAAAAAGAAAAUCAA..........AUG
env-487  UGCCACACAAUGGUUGUAGACCUCCAAAGAGAAGAAGAGUUUAAUAAAUCACUAUAUAAUUUCACAGAAGUGUACNANACG.AU.U.AUCGAAGGAGGAAG.........CCUUAAU.UC.A.UAAU..CACACUUCAAACUAAAAGAAA..........AUG
env-488  UGCCGCACAACGGUUGUAGACCACCAAAAAGAAGAAGAGUUUAAUAAAUCACUAUAUAAUUUCACAGAGGUGUACUAAACG.AU.U.AUCGAAGGAGGAAG.........CCUUAAU.UC.A.UAAU..CACACUUCAAACUAAAAGAAA..........AUG
env-489  UUACACCAAUGCCGCACAACGGUUGUCGUCCACCAAAGCGAAGAAGAGUAUAAUUAAAUAUUUUAACUAAGUAGGAUCCG.AU.U.AUCGAAGGAGAUAG.........CCUUAAU.UC.A.UAAU..CCCUACUUUAAUAACAACGAA..........AUG
env-490  UUACACCAAUGCCGCACAACGGUUGUCGUCCACCAAAGCGAAGAAGAGUAUAAGUAAAUAUUUUAACUAAGUAGGAUUCG.AU.U.AUCGAAGGAGAUAG.........CCUUAAU.UC.A.UAAU..CCCUACUUUAAUAACAACGAA..........AUG
env-491  AUUUUUCUGAUUAAAUGCUGAAUUUAUUUCAGCAACAAAAAAGAAUAUCAAUUAUUAUAUUUUAACUAAGUAGGAAUUCG.AU.U.AUCGAAGGAGAUAG.........CCUUAAU.UC.A.UAAU..CCCUGCUUUAAUAACAACGAA..........AUG
Kje-1-1  UACUCCAUUACCACACAAUGGAUGUAGACCUCCAAAAAGAAGAAGAGUAUAAUUAAUUUAAUUUUCAUUGAGAGAUUAACG.AU.U.AUCGAAGGAGAAAG.........CCUUAAU.UC.A.UAAU..CACUCUCAUAAUAACUAUUAGAA..........AUG
env-492  CCUAUGCCUCAUAAUGGGUGUAGACCUCCUAAAAGAAGAAGAGUAUAAUUUAAUAAAAUUUUUAACAGAUAGGAUUGUUUG.AU.U.AUCGAAGGAGAAAG.........CCUUAAU.UC.A.UAAU..CCCUAUCAAUAAACAAUAAGAA..........AUG
env-493  ACUAGUUUCAGUUGACGUCGAAUUAAUUUCGUCACCUCUUAUCAGAAAAAUAAUUAUAUUUUAACGAUAGGAUCUAAG.AU.U.AUCGAAGGAGAAAG.........CCUUAAU.UC.A.UAAU..CCCUAUCUAUAACACAAAAGAA..........AUG
Psp-2-1  UUUAGAUUUUUUUUUCUAAAUCUACUUUGUUAUAUACACAAUUAGUAUUUAUUUAUAUUUUAAUCAAUUAGGAUUUAAG.AU.U.AUCGAAGGAGAAAG.........CCUUAAU.UC.A.UAAU..CCCUAAUUUUACAACAACGAA..........AUG
env-494  AUGUUACACCUAUUCCUCACAAUGGUUGUCGUCCACCAAAAAGAAGACGAGUAUAAUAAAUUUUAAUUGAUAGGAAUCAG.AU.U.AUCGAAGGAGAAAG.........CCUUAAU.UC.A.UAAU..CCCUAUUUCUAACAAAAAAGAA..........AUG
env-495  ACUAGUUUCAGUUGAUGUUGAAUUAAUUUCGACACCUCUUAUCAGAAGAAUAAUUAUAUUUUAACAAGAUAGGAUCAAAG.AU.U.AUCGAAGGAGAAAG.........CCUUAAU.UC.A.UAAU..CCCUAUCUAUAACACAAAAGAA..........AUG
env-496  UUACACCUAUGCCUCACAAUGGAUGUCGUCCACCAAAGAGAAGAAGAGUAUAAUUAAAUAUUUUAACUUGAUAGGAAUAAG.AU.U.AUCGAAGGAGAAAG.........CCUUAAU.UC.A.UAAU..CCCUAUCUAUAAUACAAAAAGAA..........AUG
env-497  CACCUAUGCCACACAAUGGUUGUAGACCACCAAAAAGAAGAAGAGUAUAAUUAAAUAAUUUUAACCGAUAGGAUUGUUAG.AU.U.AUCGAAGGAGAAAG.........CCUUAAU.UC.A.UAAU..CCCUAUCAUUAAUCAAUUAAAA..........AUG
env-498  UGUAACUCCAACACCUCACAAUGGUUGUCGUCCACCAAAAAGAAGACGAGUAUAAAUUAAUUUUUAAUUGAUAGGAAUAAG.AU.U.AUCGAAGGAGAAAG.........CCUUAAU.UC.A.UAAU..CCCUAUUAUAAUAAAAAAAGAA..........AUG
env-499  GAUUUAGAAUUUUUUCUAAAUCUACUUUGCUAUGUAUAAAAGUUAGUAAUUAUUUAUAUUUUAAUCAAUUAGGAUUUAAG.AU.U.AUCGAAGGAGAAAG.........CCUUAAU.UC.A.UAAU..CCCUAAUUUAAUAACAACGAA..........AUG
env-500  CCUAUGCCUCAUAAUGGGUGUAGACCUCCAAAAAGAAGAAGAGUAUAAUUUAAUUAAUAUUUUUAACAGAUAGGAUUGUUUG.AU.U.AUCGAAGGAGAAAG.........CCUUAAU.UC.A.UAAU..CCCUAUCAAUAAACAAUAAGAA..........AUG
env-501  UUUGAUACAUACAAAUUAUAUGCUGAAUCAUAUUCAGCAUCCUUAUUAACAAAUAUUAUAUUUUAAUCUGAUAGGAAUACG.AU.U.AGCGAAGGAUAAG..........CCUUAAU.UC.A.UAAU..CCCUAUCUAUAACAAAAACAAA..........AUG
env-502  UUUGAUACAUUCAAAUUAUAUGCUGAAUUAUGUUCAGCAUCCUUAUUAAUAAAUAUUAUAUUUUAAUUUGAUAGGAGUACG.AU.U.AGCGAAGGAUAAG..........CCUUAAU.UC.A.UAAU..CCCUAUCUAUAACAAAAACAAA..........AUG
env-503  UUGAUUGUCUAUCAAAUUAUAUGCUGACUUUGUUUCAGCAUCUUAAUAAUAAAAAUUAUAUUUUAACUUGAUAGGAUUCAG.AU.U.AGCGAAGGAUAAG..........CCUUAAU.UC.A.UAAU..CCCUAUCUAUAACACAAAAGAA..........AUG
env-504  AGAAGAGUUUAAGUCGAACUUGUUUCAGCUUCUUUUUAUGAAAAGUAAAUUAAUUAUUAUAAGUAUAACCGAGAGAUCAACG.AU.U.UUCGAAGGAUAAGCAUAAGA....CCUUAAU.UC.A.UUAU..CACUCUCAAAAUAAAUUAAAAAA..........AUG
env-505  UGCCGCAUAAUGGCUGUAGACCACCUAAAAGAAGAAGAGUUUAAUUAAUCACUUAUAUAUUUCACGGAAGGUGUAGUUACG.AU.U.AUCGAAGGAUAAG..........CCUUAAU.UC.A.UAAU..CUCACUUUCAAAUUAAAAGAAA..........AUG
env-506  GUUACGCCAUUACCACAUAACGGAUGCCGCCCUCCCAAAAGAAGAAGAGUAUAAUAAAUAUUUUAAUCAAUGUGGGAAUUAG.AU.U.AUCGAAGGAUAAG..........CCUUAAU.UC.A.UAAU..CCCUGCUUUAAUAACUAAGAA..........AUG
env-507  UUAGUUUAUAUAAAAUAAAUAAUUACUGAAACUCCUGAAACGUAAGGACAGUAUUUUAAUUAAUUCAUGAAAGGAUAUUAG.AU.U.AUCGAAGGAUAAG..........CCUUAAU.UC.A.UAAU..CCCUUUCUAAAAAACAAAAA..........AUG
env-508  AUAAAAUUGAUAAAAAUAUAUAAAUACUGAAAUCCUGAAAUGUAAGAACAGUAUUAUAAUUAAUUCAUGAAAGGAUAUUAG.AU.U.AUCGAAGGAUAAG..........CCUUAAU.UC.A.UAAU..CCCUUUCUAAAAAACAAAAA..........AUG
env-509  .....CAUAACGGAUGUAGACCACCUAAAAGAAGAAGAGUUUAAUUAAUUACUAAUAUAUUUCAUCGAAAGUGCAGUUACG.AU.U.AUCGAAGGAUAAG..........CCUUAAU.UC.A.UAAU..CUCACUUUCAAAUUUAUAGAAA..........AUG
env-510  ........................................GUUUAAUUAAUUACUAAUAUAUUUCACCGAAAGUGUAGUUACG.AU.U.AUCGAAGGAUAAG..........CCUUAAU.UC.A.UAAU..CUCACUUUCAAAUCAAAAGAAA..........AUG
env-511  ................ACAAUGGUUGUCGCCUCCAAAACGUCGUAAAGUAUAACACUUAUCAACCCCAGAGAGAGGCACAG.AU.U.AUCGAAGGAUAAG..........CCUUAAU.UC.A.UAAU..CCUCUCUCCUUUUAAUCAAAAUAGUAUAAA..AUG
env-512  GGUGUAGACCACCCAAAAGAAGAAGAGUUUAAUUAAGUUACAUAUUUAUAAAAGGUUAAUCACAGGAAGUGCAAUUACG.AU.U.AUCGAAGGAUAAG..........CCUUAAU.UC.A.UAAU..CACAUUUCCAAAUUAAAAGAAG..........AUG
Mze-2-1  CCUCCAAAAAGAAGAGUUUAAUUAUUUUUAUAGUCUGCCCAUAAAGGCAGGCUUUUUAUACAUCGAAUGGUGCAUAAUG.AU.U.AUCGAAGGAUAAG..........CCUUAAU.UC.A.UAAU..CGUACAUUCAAAUAAAUUAGAAA..........AUG
Csp-2-1  CGUAACUCCACUACCACACAAUGGUGUCGCCCUCCUAAAAGACGUAAAGUAUAACACUUAACCCCAAGAGAGAGGCUACG.AU.U.AUCGAAGGAUAAG..........CCUUAAU.UC.A.UAAU..CCUCUCUCCUUUUAAUUAAAAAAAUUCAAA..AUG
Man-1-1  CUAUGCCACAUAAUGGAUGUAGGCCUCCAAAGAGAAGAAGAGUUUAAUUAAUUAAUAUAUUUCACAGAAGGUGCAGUUACG.AU.U.AUCGAAGGAUAAG..........CCUUAAU.UC.A.UAAU..CUCACUUUCAUAACUUAAGAAA..........AUG
Msp-1-1  UGCCUCAUAACGGUUGUAGACCUCCUAAGAGAAGAAGAGUUUAAUUAAUUACUAAUAUAUUUCACCGAAAGUGUAGUUACG.AU.U.AUCGAAGGAUAAG..........CCUUAAU.UC.A.UAAU..CUCACUUUCAAACUUAAAGAAA..........AUG
env-513  CCUCCAAAAAGAAGAAGAGUUUAAUUUAUUUAUUGAUAACCUGUCCGCCGCAGGCGGAUAUCUCAGGAAGGUGCAGUGAAG.AU.U.AUCGAAGGAUAAG..........CCUUAAU.UC.A.UAAU..CGCACUUUCAAAACUAAGAAG..........AUG
env-514  CAGCCGCACAACGGUUGUAGACCACCUAAAAGAAGAAGAGUUUAAUCAAUUUUCAUAUUUUUCACGAAAGUGCAUUGACG.AU.U.AUCGAAGGAUAAG..........CCUUAAU.UC.A.UAAU..CACACUUCAAACCUAAAGAAG..........AUG
env-515  GGAUGCAGACCUCCAAAAAGAAGAAGAGUAUAAUUAAUCAAAUACCCGAUUGGGUGUAAUCAAUAGAGAGUGUACUAGAUG.AU.U.AUCGAAGGAUAAG..........CCUUAAU.UC.A.UAAU..CGCACUUUCAUAAAUAUAAAAG..........AUG
env-516  UUCCUUUAACAUAUAUAGGAAAUAAUAUUAACAGAUGCUGAAACGAGUUCAGUAUUAUAUUUUAAUCAAGUAGGAAAAUAG.AU.U.AUCGAAGGAUAAG..........CCUUAAU.UC.A.UAAU..CCCUGCUUUAAUAACUAAGAA..........AUG
env-517  CAUUGCCACACAAUGGCUGUCGUCCACCAAAGAGAAGAAGAGUUUAAUAAUAAUUAUAUUUUUAAUCAAGUAGGAAAAUAG.AU.U.AUCGAAGGAUAAG..........CCUUAAU.UC.A.UAAU..CCCUACUUUAAUAACAAAGAA..........AUG
env-518  CAUUACCACAUAAUGGUUGUCGUCCACCAAAAAGAAGAAGAGUUUAAUAAUAAUUAUAUUUUAAUCAAGUAGGAAAAUAG.AU.U.AUCGAAGGAUAAG..........CCUUAAU.UC.A.UAAU..CCCUGCUUUAAUAACAAAGAA..........AUG
env-519  CUAAAAGACGAAGAGUAUAAGUUUUCUAAUCGUAUAGGAAACAAACUGUAAUAAUUAUAUUAAUUUGAAAGGAGUAUUG.AU.U.AUCGAAGGAUAAG..........CCUUAAU.UC.A.UAAU..CCCUUUCACUAAUCAACAGAA..........AUG
env-520  ................................CCUAAGAGAAGAAGAGUAUAAUUAAAUUAAUUUUUAACAGGUAGGAGUUUG.AU.U.AUCGAAGGAUAAG..........CCUUAAU.UC.A.UAAU..CCCUAUCAAAUAAUAAUAAAAA..........AUG
env-521  UGUAACUCCACUUCCACACAAUGGUUGUCGCCCUCCAAAACGUCGUAAAGUAUAACACUUAACCCCAGAGAGAAGCACAG.AU.U.AUCGAAGGAUAAG..........CCUUAAU.UC.A.UAAU..CCUCUCUCCUUUUAAUCAAAAUAGUAUAAA..AUG
Csp-3-1  GUAGACCACCAAAAGAAGAAGAGUUUAAUUAUAUAUACAAUUAUCCGAAAACGGAUGUUUCACAGAAAGUAUUGAAUACG.AU.U.AUCGAAGGAUAAG..........CCUUAAU.UC.A.UAAU..CAUACUUUCAAAUUUAAGAAA..........AUG
Cba-1-1  ACCACCAAAAAGAAGAAGAGUUUAAUUAUAAUUACUAAUAGUCCUUUUAAAAUAGGAUAUUUCACAGAAAGUAUUGAUACG.AU.U.AUCGAAGGAUAAG..........CCUUAAU.UC.A.UAAU..CAUACUUUCAAACAACAAGAAG..........AUG
Mlu-1-1  CCUCCAAAAAGAAGAAGAGUUUAAUUUAUUUAUUGAUAACCUGUCCGCCGCAGGCGGAUAUCUAAGGAAGGUGCAGUGAAG.AU.U.AUCGAAGGAUAAG..........CCUUAAU.UC.A.UAAU..CGCACUUUCAAAACUAAGAAG..........AUG
Msp-2-1  GUAGACCGCCCAAAAGAAGAAGAGUUUAAUAAUCAGUUUUAAUAAUGGCCCAAAAGGGUCUUCACCCGAAGGUGCAGUUAG.AU.U.AUCGAAGGAUAAG..........CCUUAAU.UC.A.UAAU..CGCACUUUCAAACUAAUAGAAG..........AUG
env-522  UACACCUUUACCACAUAAUGGAUGUAGACCUCCAAAACGUAGAAGAGUUUAAUAUUAACAAGUAUCAACAGAGAGAUCAACG.AU.U.AUCGAAGGAUAAGAUUAAGA....CCUUAAU.UC.A.UAAU..CACUCUUAAAACAAAUUUAAA..........AUG
env-523  GAAGAAGAGUUUAAUCUGAAUUUAUUUCAGAUACUUUGAAAAUAGAAAUUAUUAAUAACAAGUAUCAACAAGAGAUCAAUG.AU.U.AUCGAAGGAUAAGAUUAAGA....CCUUAAU.UC.A.UAAU..CACUCUUAAAACAAUUUAAA..........AUG
env-524  ..........................................................................CG.AU.U.AUCGAAGGAUAAGAUUAAGA....CCUUAAU.UC.A.UAAU..CACUCUCAAAACAAAUUUAAAA..........AUG
env-525  CGUAGAAGAGUUUAAACUGAACUAGUUUCGGUUACUUUUAAAUAAAAAUUAAUUAAUAACAAGUAUCACAGAGAGAUCAACG.AU.U.AUCGAAGGAUAAGAUUAAGA....CCUUAAU.UC.A.UAAU..CACUCUCAUAA...........................
env-526  UAGAAGAGUUUAAGUUCUAUACUGAAUNUAUUUCAGUNUUUUAUAAUAAUUAAUUUAACAAGUAUCAACGAGAGAUCAACG.AU.U.AUCGAAGGAUAAGAUUAAGA....CCUUAAU.UC.A.UAAU..CACUCUCAAAACAAAUUUAAAA..........AUG
env-527  AAAACGUAGAAGAGUUUAAUCUGAAUUUAAGAUUCAGACUAAUUAAUAAAAAUUAUUAACAAGUAUCAACAGAGAGAUCAACG.AU.U.AUCGAAGGAUAAGAUUAAGA....CCUUAAU.UC.A.UAAU..CACUCUCAUAACUCAAUUUAAGAA.........AUG
env-528  UAGAAGAGUUUAAGUUCUAUACUGAAUAUAUUUCAGUAUUUUAUAAUAAUUAAUUUAACAAGUAUCAACGAGAGAUCAACG.AU.U.AUCGAAGGAUAAGAUUAAGA....CCUUAAU.UC.A.UAAU..CACUCUCAAAACAAAUUUAAAA..........AUG
env-529  UCUGUUUAUCAGGUGGUGUUAAUUUUUUGCAUAAAUUUGCAAACUGAAAAAUAAUUAACAAGUAUCAACGAGAGAUCAACG.AU.U.AUCGAAGGAUAAGAUUAAGA....CCUUAAU.UC.A.UAAU..CACUCUCACAAAACAAUUUAAA..........AUG
env-530  .....GUCGAAGAGUCUUUUAAGAUUUUAAGUAAACUUUUAAAUAAAGAUUAUAUAAACAAGUAUCGACGAGAGAUCAACG.AU.U.AUCGAAGGAUAAGAUUAAGA....CCUUAAU.UC.A.UAAU..CACUCUCAAAACAAAUUUAAAA..........AUG
env-531  AGAGUUUAAAGCGAACACUGAANUUGCUUCAGUGUCUUUUAAAUAAAAAAUUAUUAACAAGUAUCAACCGAGAGAUCAACG.AU.U.AUCGAAGGAUAAGAUUAAGA....CCUUAAU.UC.A.UAAU..CACUCUUAAAACAAAUUUAAA..........AUG
Ale-1-1  ACCUCGUAUGUAGGUGGUAUUAAUUUUUUGUGUAAAUUUGCAAACUGAAAUUAAUUAACAAGUAUCAAUACGGGAGAUCAACG.AU.U.AUCGAAGGAUAAGAUUAAGA....CCUUAAU.UC.A.UAAU..CACUCUUAAAAACAAUUUAUU..........AUG
Fag-1-1  UACAGUUGGUAUUAAUUUUUUUAUAUGUAAAUUUGCACACUUUUAAAAAAUAAUUUAACAAGUAUAACAAGAGAUUAACG.AU.U.AUCGAAGGAUAAGAUUAAGA....CCUUAAU.UC.A.UAAU..CACUCUUUAAAAUCAAUUUAUU..........AUG
Osp-1-1  UCAUCUGUAUUCAGAUGAUAUUAAUUUUUUGUGUAUAUUUGCAAACUGAAUUAAAAAUCAAGUAUCAACAAGAGAUUAACG.AU.U.AUCGAAGGAUAAGAUUAAGA....CCUUAAU.UC.A.UAAU..CACUCUUAAAACAAAUUAAAA..........AUG
env-532  AUGCCUCACAACGGGUGUAGACCUCCAAAAAGAAGAAGAGUUUAAUUAAUUACAAAUAUUUCACCGAAAGUGUAAUUACG.AU.U.AUCGAAGGAUAAG..........CCUUAAU.UC.A.UAAU..CUCACUUUCAAAUUUUUAGAAA..........AUG

         ....................................................................................<.<<.<.<<.<<<<<...............>>>>.....>.>.>>>>..>..................................
         ....................................................................................2.22.2.22.2.0.................0.....2.2.2222..2..................................
         ••••••••••••••••••••••••••••••••••••••••••••••••••••••••••••••R••RRR•••••R-RU-U-AUCGAAGGA••RR○---------CCUUAAU-UC-A-UAAU--Y•CY•YY••••••••••••○○---------AUG
                                                                                  5′                                                3′
```

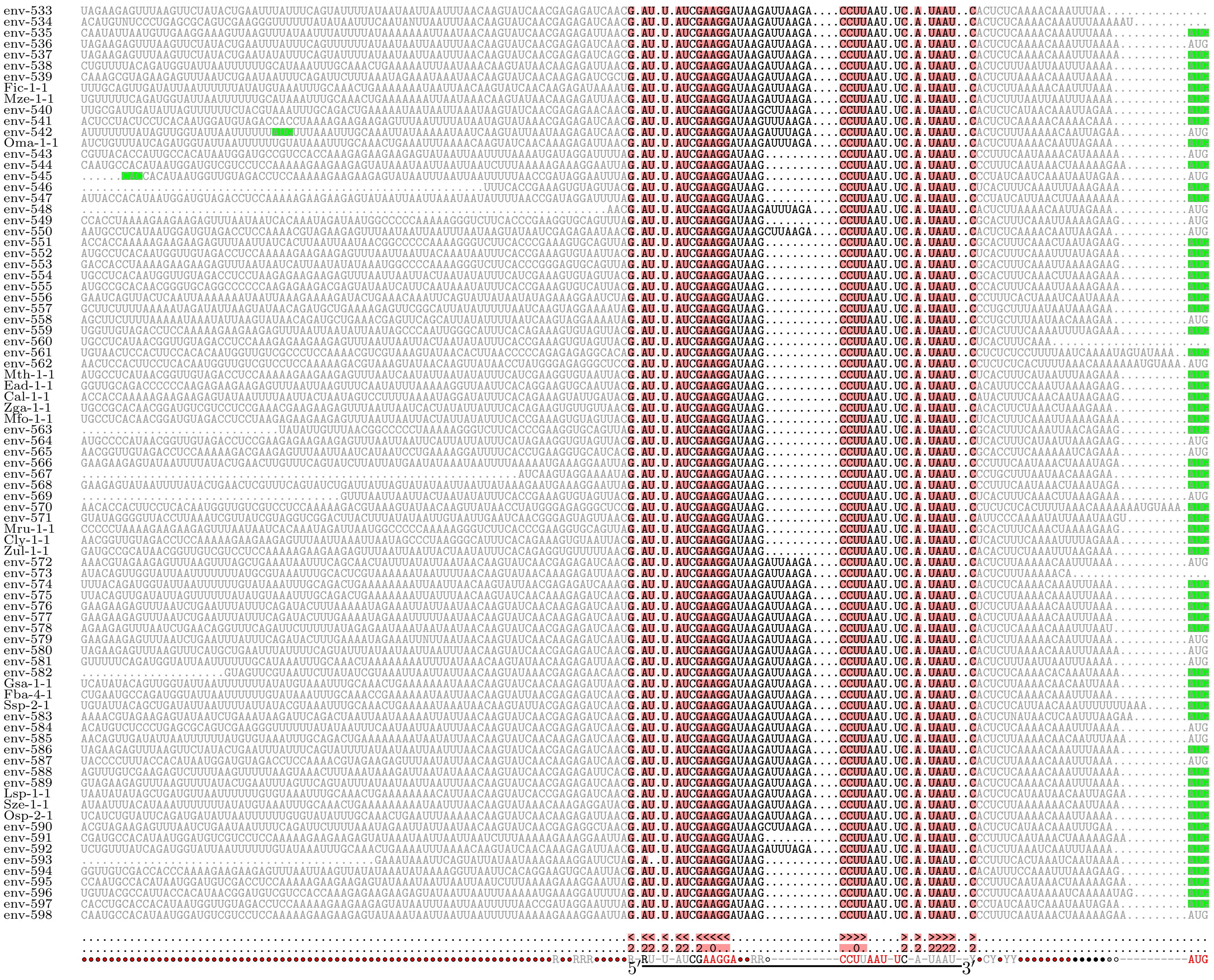
env-533
env-534
env-535
env-536
env-537
env-538
env-539
Fic-1-1
Mze-1-1
env-540
env-541
env-542
Oma-1-1
env-543
env-544
env-545
env-546
env-547
env-548
env-549
env-550
env-551
env-552
env-553
env-554
env-555
env-556
env-557
env-558
env-559
env-560
env-561
env-562
Mth-1-1
Ead-1-1
Cal-1-1
Zga-1-1
Mfo-1-1
env-563
env-564
env-565
env-566
env-567
env-568
env-569
env-570
env-571
Mru-1-1
Cly-1-1
Zul-1-1
env-572
env-573
env-574
env-575
env-576
env-577
env-578
env-579
env-580
env-581
env-582
Gsa-1-1
Fba-4-1
Ssp-2-1
env-583
env-584
env-585
env-586
env-587
env-588
env-589
Lsp-1-1
Sze-1-1
Osp-2-1
env-590
env-591
env-592
env-593
env-594
env-595
env-596
env-597
env-598
5′
3′
AUG

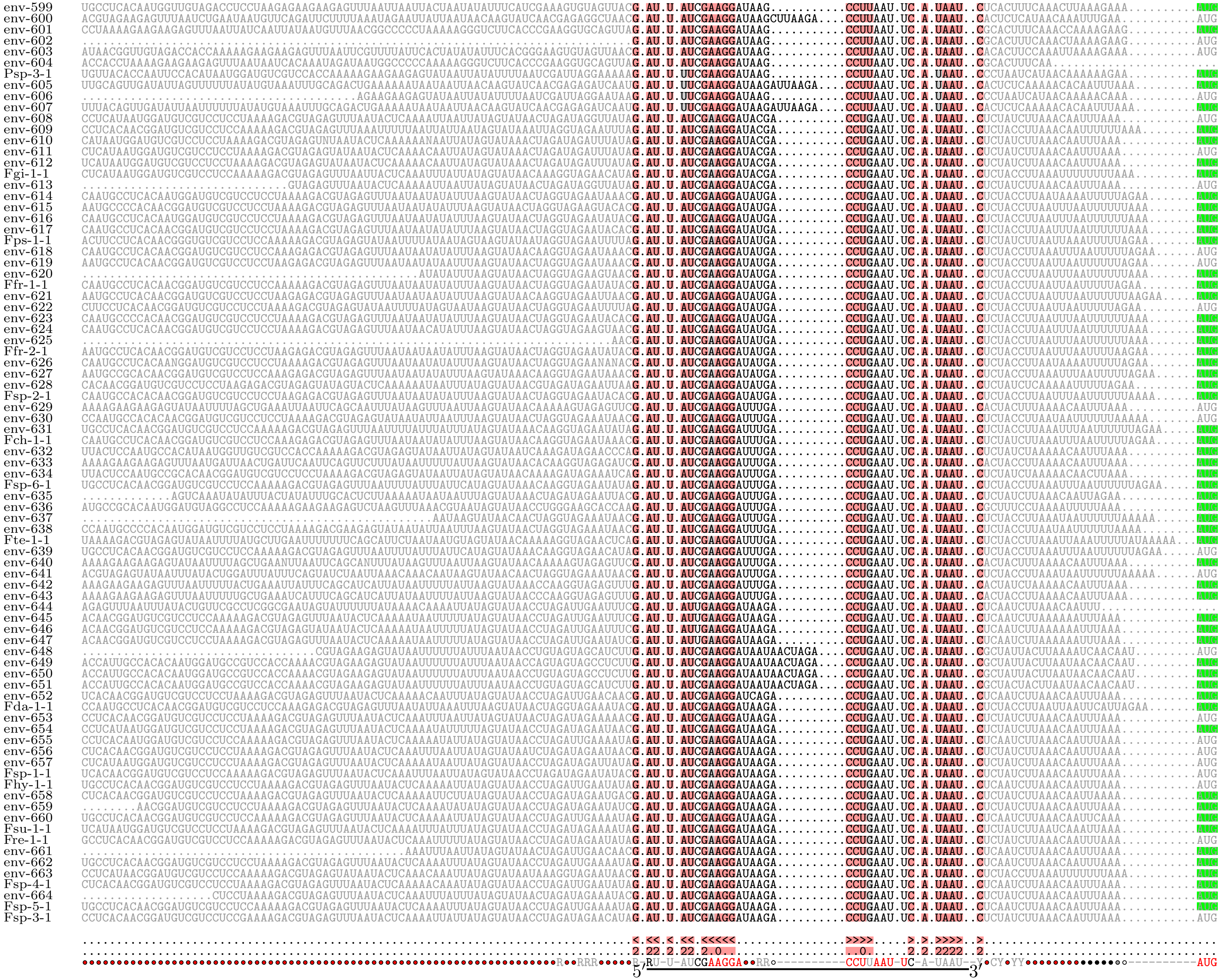
env-599
env-600
env-601
env-602
env-603
env-604
Psp-3-1
env-605
env-606
env-607
env-608
env-609
env-610
env-611
env-612
Fgi-1-1
env-613
env-614
env-615
env-616
env-617
Fps-1-1
env-618
env-619
env-620
Ffr-1-1
env-621
env-622
env-623
env-624
env-625
Ffr-2-1
env-626
env-627
env-628
Fsp-2-1
env-629
env-630
env-631
Fch-1-1
env-632
env-633
env-634
Fsp-6-1
env-635
env-636
env-637
env-638
Fte-1-1
env-639
env-640
env-641
env-642
env-643
env-644
env-645
env-646
env-647
env-648
env-649
env-650
env-651
env-652
Fda-1-1
env-653
env-654
env-655
env-656
env-657
Fsp-1-1
Fhy-1-1
env-658
env-659
env-660
Fsu-1-1
Fre-1-1
env-661
env-662
env-663
Fsp-4-1
env-664
Fsp-5-1
Fsp-3-1
5′
3′
AUG

```
env-665  CCUCACAACGGAUGUCGUCCUCCUAAAAGACGUAGAGUUUAAUACUCAAAAAUUAUUAUAGUAUAACUAGAUUGAAAAUAG.AU.U.AUCGAAGGAUAAGA.........CCUGAAU.UC.A.UAAU..CUCAAUCUUAAACAAUUAAAA..........AUG
env-666  CCUCACAACGGAUGUCGUCCUCCAAAAAGACGUAGAGUAUAAUACUCAAAAUUUAAUUAUAGUAUAACUAGAUUGUAUUACG.AU.U.AUCGAAGGAUAAGA.........CCUGAAU.UC.A.UAAU..CUCAAUCUUAAACAAUUUAAA..........AUG
env-667  CUCCACUUCCACACAACGGGUGUCGUCCACCUAAACGCAGAAGAGUAUAAUUUUUUUUUAUUAACCACGUAAGUACACUUG.AU.U.AUCGAAGGAUAAUACUAGA....CCUGAAU.UC.A.UAAU..CGUACUUACCCAAUAUUUUAAUUU.......AUG
env-668  AAAGAAGAAGAGUGUAAUCUGAUUUUUUUUCAGAUACUCAAAUUAUAAAUUUUAAUAACAAGUAUAAACGAGAGAUUGACG.AU.U.GUCGAAGGAUAAGAUUAAGA....CCUGAAU.UC.A.UAAU..CACUCUCACAACAAAUUUAAA..........AUG
env-669  AAAGAAGAAAAGAGAGUGUAAUCUGAUUUUUUCAGAUGCUCAAAUUAUAAAUUUUAAUAACAAGUAUAAACGAGAGAUUGACG.AU.U.GUCGAAGGAUAAGAUUAAGA....CCUGAAU.UC.A.UAAU..CACUCUCACAACAAAUUUAAA..........AUG
env-670  GAAAAGAAGAAGAGUGUAAUCUGAUUUUUUCAGAUACUCAAAUUAUAAAUUUUAAUAACAAGUAUAAACGAGAGAUUGACG.AU.U.GUCGAAGGAUAAGAUUAAGA....CCUGAAU.UC.A.UAAU..CACUCUCACAACAAAUUUAAA..........AUG
env-671  UGCCACACAAUGGUUGUAGACCUCCUAAAAGAAGAAGAGUUUAAUUAAUUACUAAUAUAUUUCACCGAAAGUGUAGUUACG.AU.U.AUCGAAGGAUAAG..........CCUUAAU.UC.A.UGAU..CUCACUUUCAAAUUAAAAGAAA.........AUG
Ace-1-1  UAGACCACCAAAAAGAAGAAGAGUAUAAUUUAUAUUAAUUAAUAAUCCUAAUAAAAGGGUUUUCAAAGAGGUGUAGUUACG.AU.U.AUCGAAGGAUAAG..........CCUUAAU.UC.A.UGAU..CACACUUCUAAAAUAAAAGAAA.........AUG
env-672  UGUAGACCACCAAAAAGAAGAAGAGUAUAAUUAAUAGUAAUCAAUUGCCCCAAUGGGGUGUUUCAAGGAGAUGUAGUUACG.AU.U.AUCGAAGGAUAAG..........CCUUAAU.UC.A.UGAU..CACGUUUCCAAACUAAAAGAAA.........AUG
Ala-1-1  CCACCAAAAAGAAGAGUUUAAUUAUAUAGUUAUAAUAAUCCUAAUACAAGGAUUUUCAAGGAGUGUAGUUACG.AU.U.AUCGAAGGAUAAG..........CCUUAAU.UC.A.UGAU..CACACUUCUAAAAUAAAAGAAA.........AUG
env-673  UUGUAGACCACCAAAAAGAAGAAGAGUAUAAUUAAUAGUAAUCAAUUGCCCCAUGGGGUAUUUCAAGGAGAUGUAGUUACG.AU.U.AUCGAAGGAUAAG..........CCUUAAU.UC.A.UGAU..CACGUUUCCAAACUAAAAGAAA.........AUG
env-674  UUGUAGACCACCAAAAAGAAGAAGAGUAUAAUUUUUAUUAAUCAAUUGCCCCAAAGGGGUAUUUCAAGGAAGUGUAGUUACG.AU.U.AUCGAAGGAUAAG..........CCUUAAU.UC.A.UGAU..CACGUUUCCAAACUAAAAGAAA.........AUG
env-675  CAAUGGAUGUAGACCUCCAAAAAGAAGAAGAGUUUAAUUAUUUAUAACGCCCAAGGGCAUUUCACAGAUACUGUAGUUACG.AU.U.AUCGAAGGAUUAG..........CCUUAAU.UC.A.UGAU..CACAGUAUCAAAUUUUUAGAAA.........AUG
env-676  ACUCCAAUGCGCACAAUGGAUGUAGACCUCCAAAAAGAAGAAGAGUAUAAUUAAAUAUUUAACCGAUAGGAUUGUAACG.AU.U.AUCGAAGGACUAG..........CCUUAAU.UC.A.UAAU..CCCAUCCAAAUAACUAAAAUAG.........AUG
env-677  GACGUAACCCCACUUCCACACAAUGGUUGUCGCCCUCCAAAAAGACGUAAAGUAUAACACCUAACCCUGAGAGAGGCUCCG.AU.U.AUCGAAGGACAAG..........CCUUAAU.UC.A.UAAU..CCUCUCUCCACUAAUUUAAAAUUAUAAA....AUG
env-678  GACGUAACCCCACUUCCACACAAUGGUUGUCGCCCUCCUAAAAGACGUAAAGUAUAACACUUAACCCUGAGAGAGGCUCCG.AU.U.AUCGAAGGACAAG..........CCUUAAU.UC.A.UAAU..CCUCUCUCCACUAAUUUAAAAUUAUAAA....AUG
env-679  GACGUAACCCCACUUCCACACAAUGGUUGUCGCCCUCCUAAAAGACGUAAAGUAUAACACUUAACCCCGAGAGAGGCUCCG.AU.U.AUCGAAGGACAAG..........CCUUAAU.UC.A.UAAU..CCUCUCUCCACUAAUUUAAAAUUAUAAA....AUG
env-680  GACGUAACCCCACUUCCACACAAUGGUUGUCGCCCUCCAAAAAGACGUAAAGUAUAACACUUAACCCCGAGAGAGGCUCCG.AU.U.AUCGAAGGACAAG..........CCUUAAU.UC.A.UAAU..CCUCUCUCCACUAAUUUAAAAUUAUAAA....AUG
env-681  UACCCCUCUUCCUCACAACGGAUGCCGCCCUCCUAAAAGACGUAAAGUAUAACAAAUGUAACCCUUCAGAGAGAGGCCACG.AU.U.AUCGAAGGACAAG..........CCUUAAU.UC.A.UAAU..CCUCUCUCACUUUAAUUCAAAAUUGUAAA...AUG
env-682  CCACAUAACGGUUGUAGACCUCCGAAAAGAAGAAGAGUUUAAUACAUUCACUAAUUAUAUUUCAUAGAAGUGUAGUUUAAG.AU.U.AUCGAAGGACAAG..........CCUUAAU.UC.A.UAAU..CACACUUCAAAUUAAAAGAAA..........AUG
env-683  UACUCCUCUUCCUCACAACGGAUGCCGCCCUCCUAAAAGACGUAAAGUAUAACAAAUGUAACCCUUCAGAGAGAGGCCACG.AU.U.AUCGAAGGACAAG..........CCUUAAU.UC.A.UAAU..CCUCUCUCACUUUAAUUUAAAAUUGUAAA...AUG
env-684  UACUCCCCUUCCUCACAACGGAUGCCGCCCUCCUAAAAGACGUAAAGUAUAACAAAUGUAACCCUUCAGAGAGAGGCCACG.AU.U.AUCGAAGGACAAG..........CCUUAAU.UC.A.UAAU..CCUCUCUCACUUUAAUUCAAAAUUGUAAA...AUG
env-685  GCCAUCCGUAGUAUUCACAAUGCAGGANNNUCCUAAAAGACGUAAAGUAUAACAAAUGUAACCCUUCAGAGAGAGGCCACG.AU.U.AUCGAAGGACAAG..........CCUUAAU.UC.A.UAAU..CCUCUCUCACUUUAAUUUAAAAUUGUAAA...AUG
env-686  UACUCCUCUUCCUCACAACGGAUGCCGCCCUCCUAAAAGACGUAAAGUAUAACAAAUGUAACCCUUCAGAGAGAGGCCACG.AU.U.AUCGAAGGACAAG..........CCUUAAU.UC.A.UAAU..CCUCUCUCACUUUAAUUCAAAAUUGUAAA...AUG
env-687  GAUGUACACCAAUUCUCAUAAUGGUGUCGUCUCCAAAGAGAAGAAGAGUAUAAAUAAAUAUUUUAACCAAUUAGGAAUUAG.AU.U.AUCGAAGGAUUAG..........CGUUAAU.UC.A.UAAU..CCUAAUUAAUGAAACACGAA..........AUG
env-688  CACCAAUUCCACACAACGGUUGUCGUCCACCAAAAAGACGUAGAGUUUAAUUAAUAGUAAAUAUCAACAAAGAGUAUUACG.AU.U.AUCGAAGGAUUAUAUUUUCAAAGACCUUAAU.UC.A.UAAU..CACUCUUUAAAAUUUAAAUAA..........AUG
env-689  UUACCCCAAUUCCUCAUAACGGUUGUCGUCCACCAAAGAAGAAGAAGAGUAUAAUUAAAUAUUUUAACCAAUUGGGAAUUAG.AU.U.AUCGAAGGAUUAG..........CCUUAAU.UC.A.UAAU..CCCUGAUUAUAACAACAAAGAA.........AUG
env-690  UGUUACCCCAAUACCACACAAUGGAUGUCGUCCACCAAAAAGAAGAAGAGUAUAAUUAAAUUUUAACCUUGAUAGGAAUUAG.AU.U.AUCGAAGGAUUAG..........CCUUAAU.UC.A.UAAU..CCCUAUCUACAAUAUAAAAGA..........AUG
env-691  UUACACCAUUGCCACAUAAUGGAUGUAGACCACCCAAGAGAAGAAGAGUAUAAUUAAUUUUAAAAAGAAGGAUUUUUUAG.AU.U.AUCGAAGGAUUAG..........CCUUAAU.UC.A.UAAU..CCCUUUCCCUAAACAAAUAAAA.........AUG
env-692  GUUACACCAUUGCCACAUAAUGGAUGUAGACCACCCAAGAGAGAAGAAGAGUAUAAUUAAUUUUAAAAAGAAGGAUUUUUUAG.AU.U.AUCGAAGGAUUAG..........CCUUAAU.UC.A.UAAU..CCCUUUCCCUAAACAAAAAAAA.........AUG
env-693  CACCUGCACCACAUAAUGGAUGUAGACCUCCAAAAAGAAGAAGAGUAUAAUUAAUUUUAAUUUUAACCGAUGGGAAUUAG.AU.U.AUCGAAGGAUUAG..........CCUUAAU.UC.A.UAAU..CCCUAUCAAUCAAAUAA..............
env-694  CAUUGCCACACAAUGGUUGCAGACCACCGAAAAGAAGAAGAGUAUAAUUAAAUUAAAUUUUUAACCGAUAGGAUUUUGUAAG.AU.U.AUCGAAGGAUUAG..........CCUUAAU.UC.A.UAAU..CCCUAUCAUCUAAUAUUAAAA..........AUG
env-695  UUACACCAAUGCCUCACAAUGGAUGUCGUCCACCGAAAAGAAGAAGAGUAUAAUUAAAUAUUUUAACCUUGAUAGGAAUUAG.AU.U.AUCGAAGGAUUAG..........CCUUAAU.UC.A.UAAU..CCCUAUCUAUAACAAAAAAGAA.........AUG
env-696  CAUUACCGCACAAUGGUUGCAGACCACCGAAAAGAAGAAGAGUAUAAUUAAAUUAAAUUUUUAACCGAUAGGAUUUUGUAAG.AU.U.AUCGAAGGAUUAG..........CCUUAAU.UC.A.UAAU..CCCUAUCAUCUAAUAUUAAAA..........AUG
env-697  CUCCAUUACCACACAAUGGAUGUCGUCCACCAAAAAGAAGAAGAGUUUAAUAACAUUUAUGUUUAAAAGAAGGUUUUUACG.AU.U.AUCGAAGGAUUAG..........CCUUAAU.UC.A.UAAU..CACCUUCAAAAUAUUACAAAAC.........AUG
env-698  GUAGACCUCCAAAAAGAAGAAGAGUUUAAUCAAUUUUAAUAAUGGCCCGAUAAGGGUCUUCACCCAGAAGGCGCAGUUUAG.AU.U.AUCGAAGGAUUAG..........CCUUAAU.UC.A.UAAU..CGCGCAUUCCAAUUUUAAGAAG.........AUG
env-699  AUUGCCACACAAUGGUUGCAGACCACCGAAACGAAGAAGAGUAUAAUUAAAUUAAAUUUUUAACCGAUAGGAUUUUUCUAAG.AU.U.AUCGAAGGAUUAG..........CCUUAAU.UC.A.UAAU..CCCUAUCAUCUAAUAUAAAAA..........AUG
env-700  UUACGCCAAUUCCUCAUAAUGGAUGUCGUCCACCUAAAAGAAGAAGAGUAUAAUUAAAUAUUUUAACCUUGAUAGGAAAUAG.AU.U.AUCGAAGGAUUAG..........CCUUAAU.UC.A.UAAU..CCCUAUCUAUAACAACAAAGAA.........AUG
env-701  UUACACCGAUACCUCACAAUGGAUGUCGUCCACCUAAAAGAAGAAGAGUAUAAUUAAAUAUUUUAACCUUGAUAGGAAUUAG.AU.U.AUCGAAGGAUUAG..........CCUUAAUAUC.A.UAAU..CCCUAUCUAUAACAAAAAAGAA.........AUG
env-702  .....................GGGGGACGUUGGAGCGUAAGAAGAGAGUAUAAUUAAAUAUUUUAACCUUGAUAGGAAUUAG.AU.U.AUCGAAGGAUUAG..........CCUUAAU.UC.A.UAAU..CCCUAUCUAUAAUAAAAAAGAA.........AUG
env-703  UUACACCAAUUCCUCAUAAUGGAUGUCGUCCACCAUAAAAGAAGAAGAGUAUAAUUAAAUAUUUUAACCUUGAUAGGAAUUAG.AU.U.AUCGAAGGAUUAG..........CCUUAAU.UC.A.UAAU..CCCUAUCUAUAAUAAAAAAGAA.........AUG
env-704  UUACACCAAUUCCUCAUAAUGGUUGUCGUCCACCAAAAAGAAGAAGAGUAUAAUUAAAUAUUUUAACCAAUUAGGAAUUAG.AU.U.AUCGAAGGAUUAG..........CCUUAAU.UC.A.UAAU..CCUAAUUAAACAAACACGAA..........AUG
env-705  UUACACCAAUUCCUCAUAACGGUUGUCGUCCUCCAAAGAGAAGAAGAGUAUAAUUAAAUAUUUUAACCAAUUAGGAAUUAG.AU.U.AUCGAAGGAUUAG..........CCUUAAU.UC.A.UAAU..CCUAAUUAAACAAACACGAA..........AUG
env-706  .......................................................................GAAAGGAUUUUAG.AU.U.AUCGAAGGAUUAG..........CCUUAAU.UC.A.UAAU..CCCUUUCUGUAAACAAUAUAAA.........AUG
env-707  .......................................................................................AUUUAG.AU.U.AUCGAAGGAUUAG..........CCUUAAU.UC.A.UAAU..CCCUAUCAAUCAAAUAAUAGAA.........AUG
env-708  UACACCUGCACCACACAAUGGAUGUAGACCUCCUAAGAGAAGAAGAGUAUAAUUAAUUAUUUUAAAACAGAUAGGAAUUUAG.AU.U.AUCGAAGGAUUAG..........CCUUAAU.UC.A.UAAU..CCCUAUCAUACCAAUAUAAAAA.........AUG
env-709  GUUACACCAUUGCCACAUAAUGGAUGUAGACCACCCAAGAGAGAAGAAGAGUAUAAUUAAUUUUAAAAAGAAGGAUUUUUUAG.AU.U.AUCGAAGGAUUAG..........CCUUAAU.UC.A.UAAU..CCCUUUCCCUAAACAAAUAAAA.........AUG
env-710  UUACCCCAAUACCUCACAAUGGAUGUCGUCCACCUAAAAGAAGAAGAGUAUAAUUAAAUAUUUUAACCUUGAUAGGAAUUAG.AU.U.AUCGAAGGAUUAG..........CCUUAAU.UC.A.UAAU..CCCUAUCUACAAUAUAAAAGAA.........AUG
env-711  UUACCCCAAUACCUCACAAUGGAUGUCGUCCACCUAAAAGAAGAAGAGUAUAAUUAAAUAUUUUAACCUUGAUAGGAAUUAG.AU.U.AUCGAAGGAUUAG..........CCUUAAU.UC.A.UAAU..CCCUAUCUACAAUAUAAAAGAA.........AUG
env-712  ..............................AAAAGACGAAGAGUAUAAUUAAUUUAAAUUUUAACCGAUAGGAUUUUGUAAG.AU.U.AUCGAAGGAUUAG..........CCUUAAU.UC.A.UAAU..CCCUAUCAUCUAAUAUAAAAA..........AUG
env-713  ..............................AACGUAGAAGAGUAUAAUUAAAUAAUUUUAACCGAUAGGAUUUUGUAAG.AU.U.AUCGAAGGAUUAG..........CCUUAAU.UC.A.UAAU..CCCUAUCAUCUAAUAUAAAUA..........AUG
Pir-1-1  UUACCCCAAUUCCUCACAACGGAUGUCGCCCACCAAAAAGAAGAAGAGUAUAAUUAAAUAUUUUAACCUAUUGGGAAUUAG.AU.U.AUCGAAGGAUUAG..........CCUUAAU.UC.A.UAAU..CCUAAUUAUAACAACAAAGAA..........AUG
env-714  UUACACCAAUUCCUCAUAAUGGAUGUCGUCCACCUAAAAGAAGAAGAGUAUAAUUAAAUAUUUUAACCUUGAUAGGAAUUAG.AU.U.AUCGAAGGAUUAG..........CCUUAAU.UC.A.UAAU..CCCUAUCUAUAACAAAAAAGAA.........AUG
env-715  UGUUACCCCAAUCCUCAUAACGGUGUCGUCCACCAAAGAGAAGAAGAGUAUAAUUAAAUAUUUUAACCAAUUGGGAAUUAG.AU.U.AUCGAAGGAUUAG..........CCUUAAU.UC.A.UAAU..CCCUAAUUAUAACAACAAAGAA.........AUG
env-716  ..........................CUUACGGUCUGAGAAGAGAGAGUAUAAUUAAAUAUUUUAACCAAUUAGGAAUUAG.AU.U.AUCGAAGGAUUAG..........CCUUAAU.UC.A.UAAU..CCUAAUUAAUGAAACACGAA..........AUG
env-717  UUACCCCAAUACCACACAAUGGAUGUCGUCCACCAAAAAGAAGAAGAGUAUAAUUAAAUAUUUUAACCUUGAUAGGAAUUAG.AU.U.AUCGAAGGAUUAG..........CCUUAAU.UC.A.UAAU..CCCUAUCUACAAUAUAAAAGAA.........AUG
env-718  GUUACACCAUUGCCACAUAAUGGAUGUAGACCACCCAAGAGAGAAGAAGAGUAUAAUUAAUUUUAAAAAGAAGGAUUUUUUAG.AU.U.AUCGAAGGAUUAG..........CCUUAAU.UC.A.UAAU..CCCUUUCCCUAAACAAA...............
env-719  CAUUGCCACACAAUGGUUGCAGACCUCCGAAAAGAAGAAGAGUAUAAUUAAAUUAAAUUUUUAACCGAUAGGAUUUUGUAAG.AU.U.AUCGAAGGAUUAG..........CCUUAAU.UC.A.UAAU..CCCUAUCAUCUAAUAUUAAAA..........AUG
env-720  UUACCACAUAAUGGCUGUAGACCUCCAAAAAGAAGAAGAGUAUAAUUAAAUUAAAUUAAAUUUUUAACCGAUAGGAGUUGAG.AU.U.AUCGAAGGAUUAG..........CCUUAAU.UC.A.UAAU..CCCUAUCAUUUAACUAAUAGAA.........AUG
env-721  ..................................................AAUUAAUUUAAUUUUAAUAGAUAGGAUUUUAG.AU.U.AUCGAAGGAUUAG..........CCUUAAU.UC.A.UAAU..CCCUAUCAAACAACUAAAAAAA.........AUG
env-722  UUACCCCAAUACCACACAAUGGAUGUCGUCCACCAAAAAGAAGAAGAGUAUAAUUAAAUAUUUUAACCUUGAUAGGAAUUAG.AU.U.AUCGAAGGAUUAG..........CCUUAAU.UC.A.UAAU..CCCUAUCUACAAUACAAAAGAA.........AUG
env-723  CUCCAUUACCACAUAAUGGAUGUAGACCACCAAAAAGAAGAAGAGUAUAAUUAAAUUAAAUUUUUAACCGAUAGGAGUUGAG.AU.U.AUCGAAGGAUUAG..........CCUUAAU.UC.A.UAAU..CCCUAUCAUUUAACUAAUAGAA.........AUG
env-724  UUACACCAAUUCCUCAUAAUGGAUGUCGUCCACCGAAAAGAAGAAGAGUAUAAUUAAAUAUUUUAACCAUUUAGGAAUUAG.AU.U.AUCGAAGGAUUAG..........CCUUAAU.UC.A.UAAU..CCCUAUCUGUAACAACUAAGAA.........AUG
env-725  CAUUGCCACACAAUGGUUGCAGACCACCGAAAAGAAGAAGAGUAUAAUUAAAUUAAAUUUUUAACCGAUAGGAUUUUGUAAG.AU.U.AUCGAAGGAUUAG..........CCUUAAU.UC.A.UAAU..CCCUAUCAUCUAAUAUUAGAA..........AUG
env-726  ACUCCUGCACCUCAUAACGGAUGUAGACCUCCGAAAAGAAGAAGAGUAUAAUUAAAUAUUUUAAACAGAUAGGAUUUAG.AU.U.AUCGAAGGAUUAG..........CCUUAAU.UC.A.UAAU..CCCUAUCAUAUAACAAUAAAAA.........AUG
Jma-1-1  CUCCACUACCACACAAUGGAUGUCGUCCACCAAAAAGAAGAAGAGUUUAAUAACAUUUAUGUUUAAAAGAAGGUUUUUACG.AU.U.AUCGAAGGAUUAG..........CCUUAAU.UC.A.UAAU..CACCUUCAAAAUAUUACAAAAC.........AUG
env-727  ...............................................AGAGAGUAUAUUAAUAUUUUACCAUUAGGAUUAG.AU.U.AUCGAAG.AUUAG...........CUUAAU.UC.A.UA.U..CCUAUUAAUACACGAA..............AUG
env-728  .........................................................AUUUAACUGGCUACUGAUGGAUUAG.AU.U.AUCGAAGGAUUAG..........CCUUAAU.UC.A.UAAU..CCCUAUCUAUAAUAAAAAAGAA.........AUG
env-729  UUACACCAAUUCCUCAUAAUGGUUGUCGUCCUCCAAAGAGAAGAAGAGUAUAAUAAAUAUUUUAACCAAUUGGGAAUUAG.AU.U.AUCGAAGGAUUAG..........CCUUAAU.UC.A.UAAU..CCUAAUUAAAUAAACACGAA..........AUG
env-730  CAUUGCCACACAAUGGUUGCAGACCACCGAAAAGAAGAAGAGUAUAAUUAAAUUAAAUUUUUAACCGAUAGGAUUUUGUAAG.AU.U.AUCGAAGGAUUAG..........CCUUAAU.UC.A.UAAU..CCCUAUCAUAUAAUAUUAGAA..........AUG
env-731  ACACCUGCACCACACAAUGGAUGUAGACCUCCUAAGAGAAGAAGAGUAUAAUUAAUUAUUUUAACAGAUAGGAAUUUAG.AU.U.AUCGAAGGAUUAG..........CCUUAAU.UC.A.UAAU..CCCUAUCAUACCAAUAUAAAAA.........AUG
env-732  CAUUACCGCACAAUGGUUGCAGACCACCGAAAAGAAGAAGAGUAUAAUUAAAUUAAAUUUUUAACCGAUAGGAUUUUGUAAG.AU.U.AUCGAAGGAUUAG..........CCUUAAU.UC.A.UAAU..CCCUAUCA........................
env-733  CUCCAUUACCACAUAAUGGNUGUAGACCNCCAAAAAGAAGAAGAGUAUAAUUAAAUUAAAUUUUUAACCGAUAGGAGUUGAG.AU.U.AUCGAAGGAUUAG..........CCUUAAU.UC.A.UAAU..CCCUAUCAUUUAACUAAUAGAA.........AUG
env-734  CUGCACCACACAAUGGAUGUAGACCUCCUAAGAGAAGAAGAGUAUAAUUAAUUAUUUUAACAGAUAGGAAUUUAG.AU.U.AUCGAAGGAUUAG..........CCUUAAU.UC.A.UAAU..CCCUAUCAUACCAAUAUAAAAA.........AUG
env-735  UGUUACCCCAAUACCACACAAUGGAUGUCGUCCACCAAAAAGAAGAAGAGUAUAAUUAAAUAUUUUAACCUUGAUAGGAAUUAG.AU.U.AUCGAAGGAUUAG..........CCUUAAU.UC.A.UAAU..CCCUAUCUACAAUAUAAAAGAA.........AUG
env-736  CUCCAUUACCACACAAUGGUUGUAGACCACCAAAAAGAAGAAGAGUAUAAUUAAAUUAAAUUUUUAACCGAUAGGAUUUUAG.AU.U.AUCGAAGGAUUAG..........CCUUAAU.UC.A.UAAU..CCCUAUCAAUUAAUAUAAAAA..........AUG
env-737  AUUGCCNCACAAUGGUUGCAGACCACCGAAACGUAGAAGAGUAUAAUUAAAUUAAAUUUUUAACCGAUAGGAUUUUUGUAAG.AU.U.AUCGAAGGAUUAG..........CCUUAAU.UC.A.UAAU..CCCUAUCAUCUAAUAUAAAAA..........AUG
env-738  ACUCCUGCACCUCAUAACGGAUGUAGACCUCCGAAAAGAAGAAGAGUAUAAUUAAUUAUUUUAAUUUUAACAGAUAGGAUUUAG.AU.U.AUCGAAGGAUUAG..........CCUUAAU.UC.A.UAAU..CCCUAUCAUAUAACAAUAAAAA.........AUG
env-739  AUUACCACACAAUGGUUGCAGACCACCGAAACGUAGAAGAGUAUAAUUAAAUUAAAUUUUUAACCGAUAGGAUUUUUGUAAG.AU.U.AUCGAAGGAUUAG..........CCUUAAU.UC.A.UAAU..CCCUAUCAUCUAAUAUAAAUA..........AUG
env-740  UUACACCAAUUCCUCACAACGGUUGUCGUCCUCCAAAGAGAAGAAGAGUAUAAUUAAAUAUUUUAACCAAUUAGGAAUUAG.AU.U.AUCGAAGGAUUAG..........CCUUAAU.UC.A.UAAU..CCCUAAUUAAUAAUACACGAA.........AUG
env-741  UUACACCAAUUCCUCAUAACGGUUGUCGUCCUCCAAAGAGAAGAAGAGUAUAAUUAAAUAUUUUAACCAAUUAGGAAUUAG.AU.U.AUCGAAGGAUUAG..........CCUUAAU.UC.A.UAAU..CCUAAUUAAUGAAACACGAA..........AUG
env-742  GUUGUAGACCACCUAAAAGAAGAAGAGUUUAAUAAAUCGAAUACUAUUCGAAUAAUAUAUUUUAAUUAACUAAGGAUAAAG.AU.U.AUCGAAGGAAUG...........CCUUAAU.UC.A.UAAU..CCCUUAAAUUUUUAUAAAG............AUG
env-743  CGUAACACCAAUUGCUCAUAACGGUUGCAGACCUCCAAAAAGAAGAAGAGUUUAAUUAAUUUUAAUAACUAAGGAUAAAG.AU.U.AUCGAAGGAAUG...........CCUUAAU.UC.A.UAAU..CCCUUAAACAAUUAUAUAAAG..........AUG

         .................................................................................<.<<.<.<<.<<<<<................>>>>.....>.>.>>>>..>................................
         .................................................................................2.22.2.22.2.0..................0......2.2.2222..2................................
         ••••••••••••••••••••••••••••••••••••••••••••••••••••••••••••••R••RRR•••••R RU-U-AUCGAAGGA••RR○---------CCUUAAU-UC-A-UAAU--Y•CY•YY•••••••••••••○○---------AUG
                                                                                  5′                                                   3′
```

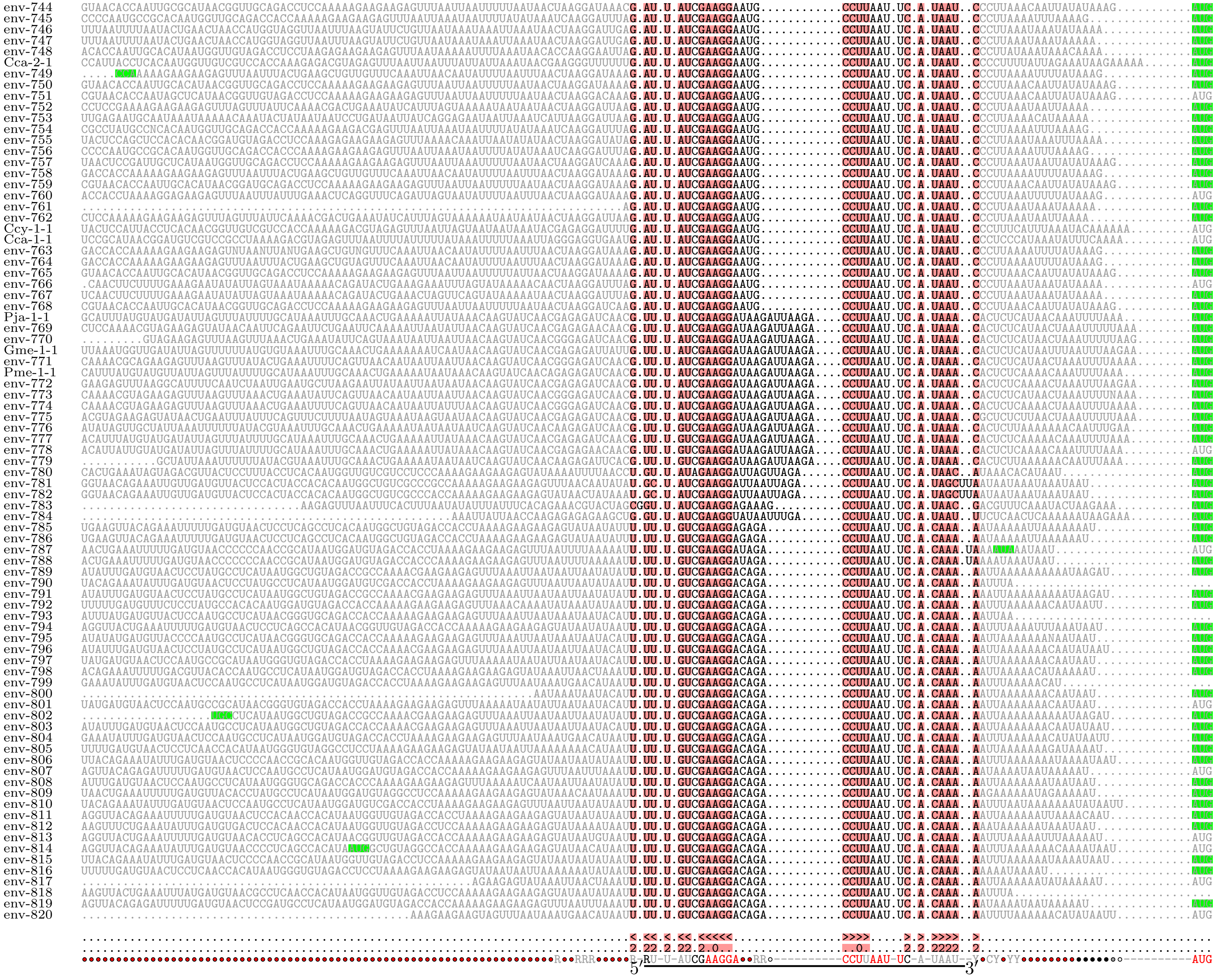
env-744
env-745
env-746
env-747
env-748
Cca-2-1
env-749
env-750
env-751
env-752
env-753
env-754
env-755
env-756
env-757
env-758
env-759
env-760
env-761
env-762
Ccy-1-1
Cca-1-1
env-763
env-764
env-765
env-766
env-767
env-768
Pja-1-1
env-769
env-770
Gme-1-1
env-771
Pme-1-1
env-772
env-773
env-774
env-775
env-776
env-777
env-778
env-779
env-780
env-781
env-782
env-783
env-784
env-785
env-786
env-787
env-788
env-789
env-790
env-791
env-792
env-793
env-794
env-795
env-796
env-797
env-798
env-799
env-800
env-801
env-802
env-803
env-804
env-805
env-806
env-807
env-808
env-809
env-810
env-811
env-812
env-813
env-814
env-815
env-816
env-817
env-818
env-819
env-820
5′
3′
AUG

```
env-821  .................................................AGUAUAAAAAUAAUAUUAAUAAUACAUU.UU.U.GUCGAAGGACAGA..........CCUUAAU.UC.A.CAAA..AAUUAAAAAAACAAUAAU...............AUG
env-822  UAUAUGAUGUUACCCCAAUGCCUCAUAACGGGUGCAGACCACCAAAAGAAGAAGAGUUUAAAUUAACAAAUAAUACAUUU.UU.U.GUCGAAGGACAGA..........CCUUAAU.UC.A.CAAA..AAUUAAAAAAACAAUAAU...............AUG
env-823  AUAUUUGAUGUAACUCCUAUGCCUCACAAUGGGUGUAGACCACCAAAACGAAGAAGAGUUUAAAUUAAUAAUUAAUACAUU.UU.U.GUCGAAGGACAGA..........CCUUAAU.UC.A.CAAA..AAUUAAAAAAAAAAUAUAAU............AUG
env-824  UAUGAUGUAACUCCAAUGCCGCAUAACGGGUGUAGACCACCUAAAAGAAGAAGAGUAUAAAAAUAAUAUUAAUAAUACAUU.UU.U.GUCGAAGGACAGA..........CCUUAAU.UC.A.CAAA..AAUUAAAAAAACAAUAAU...............AUG
env-825  UAUUUGAUGUAACUCCUAUGCCUCAUAAUGGCUGUAGACCGCCAAAACGAAGAAGAGUUUAAAUUCAAUAAUUAAUAUAUU.UU.U.GUCGAAGGACAGA..........CCUUAAU.UC.A.CAAA..AAUUAAAAAAAAAAUAAGAU............AUG
env-826  UCGUUGAUGUUACACCACUUCCUCACAACGGAUGUAGACCACCGAAAAGAAGAAGAGUAUAAAUAAUAAGGAGGGAAUUUG.UU.U.GACGAAGGAUAGA..........CCUUAAU.UC.U.CAUU..CCCCUUCACUAAAUAAUAACU...........AUG
env-827  UUGUUGAUGUUACACCACUUCCUCACAACGGAUGUAGACCACCGAAAAGAAGAAGAGUAUAAAUAAUAAGGAGGGAAUUUG.UU.U.GACGAAGGAUAGA..........CCUUAAU.UC.U.CAUU..CCCCUUCACUAAAUAAUAACU...........AUG
env-828  UUGUUGAUGUUACACCACUUCCUCACAACGGAUGUAGACCACCGAAAAGAAGAAGAGUAUAAAUAAUAAGGAGGGAAUUUG.UU.U.GACGAAGGAUAGA..........CCUUAAU.UC.U.CAUU..CCCCUUCACUAAAUAUAAACU...........AUG
env-829  UUGUUGAUGUUACACCACUUCCUCACAACGGAUGUAGACCACCGAAAAGAAGAAGAGUAUAAAUAACAAGGAGGGAAUUUG.UU.U.GACGAAGGAUAGA..........CCUUAAU.UC.U.CAUU..CCCCUUCACUAAAUAUAAACU...........AUG
env-830  GUUGAUGUUACCCCACUUCCUCACAACGGAUGUAGACCACCGAAAAGAAGAAGAGUGUAAAAAUUAAAAGAAGGAAUUAG.UU.U.GACGAAGGAUAGA..........CCUUAAU.UC.U.CAUU..CCCCUUCAACUAAAAUAAAACU..........AUG
env-831  UCGUUGAUGUUACACCACUUCCUCACAACGGAUGUAGACCACCGAAAAGAAGAAGAGUAUAAAUAAUAAGGAGGGAAUUUG.UU.U.GACGAAGGAUAGA..........CCUUAAU.UC.U.CAUU..CCCCUUCACUAAAUAUAAACU...........AUG
env-832  CGUUGAUGUUACACCACUUCCUCACAACGGAUGUAGACCACCGAAAAGAAGAAGAGUAUAAAUAAUCAAGGAGGGAAUUUG.UU.U.GACGAAGGAUAGA..........CCUUAAU.UC.U.CAUU..CCCCUUCACUAAAUAUAAACU...........AUG
env-833  .................CUUCCUCACAACGGAUGUAGACCACCGAAAAGAAGAAGAGUAUAAAUAACAAGGAGGGAAUUUG.UU.U.GACGAAGGAUAGA..........CCUUAAU.UC.U.CAUU..CCCCUUCACUAAAUAUAAACU...........AUG
env-834  .............CACUUCCUCACAACGGAUGUAGACCACCGAAAAGAAGAAGAGUAUAAUAAUCAAUAGAAGGGAAUUUG.UU.U.GACGAAGGAUAGA..........CCUUAAU.UC.U.CAUA..CCCCUUCCACUAAUUUAGAACU..........AUG
env-835  .............ACAUAAUGGAUGUAGACCUCCAAAAAGAAGAAGAGUAUAAUUAAUUAAGUAUAAACUGAAGGAAAUAG.AA.A.ACCAAAGGAUAAGCAUAAGA...CCUUAAU.UUGG.UAUU..CCCUUCAAAAUAAAAUUUUAAG..........AUG
env-836  UCACCGAAAUAGUUGACGUGACGCCUAUGCCGCACAAUGGAUGUCGUCCACCAAAAAGACGUAGAGUUUAAUACUAACCUC.GC.U.GGCGAAGGAUUGA..........CCUUAAU.UC.C.CAGC.ACUUAAUUUUUAUAAU.................AUG
env-837  UCACCGAAAUAGUUGACGUGACGCCUAUACCGCACAACGGAUGUCGUCCACCAAAAAGACGUAGAGUUUAAUACUAACCCC.GC.U.GGCGAAGGAUUGA..........CCUUAAU.UC.C.CAGC.ACUUAAUUUUUAUAAU.................AUG
env-838  UCACCGAAAUAGUUGACGUGACGCCUAUGCCGCACAACGGAUGUCGUCCACCAAAAAGACGUAGAGUUUAAUACUAACCCC.GC.U.GGCGAAGGAUUGA..........CCUUAAU.UC.C.CAGC.ACUUAAUUUUUAUAAU.................AUG
env-839  UCACCGAAAUAGUUGACGUGACGCCUAUGCCGCAUAACGGAUGUCGUCCACCAAAAAGACGUAGAGUUUAAUACUAACCCC.GC.U.GGCGAAGGAUUGA..........CCUUAAU.UC.C.CAGC.ACUUAAUUUUUAUAAU.................AUG
env-840  UGAUGUUACUCCUUUGCCGCACAAUGGGUGUCGCCCACCGAAAAGAAGAAGAGUAUAAUUAUUUCAAACCCGAGGAGUCAC.GC.U.GGCGAAGGAGGAUG.........CCUUAAU.UC.C.CAGU..GCCUCAUAAUAUUUAAAUAGA...........AUG
env-841  GUAACUCCACUUCCACACAAUGGAUGUCGCCCUCCAAAAAGAAGAAGAGUAUAAUUUAAAUUUAACAUAACGGGGAGUUAC.AU.U.GGCGAAGGAGUAUG.........CCUUAAU.UC.U.CAGU..GCCCCUAUAAAAAACAAGUA............AUG
         ..........................................................................<.<<.<.<<.<<<<<...................>>>>.....>.>.>>>>..>.................................
         ..........................................................................2.22.2.22.2.0.....................0.......2.2.2222..2.................................
         •••••••••••••••••••••••••••••••••••••••••••••••••••••••••••••••••••••R••RRR••••R-RU-U-AUCGAAGGA••RR○----------CCUUAAU-UC-A-UAAU--Y•CY•YY••••••••••••○----------AUG
                                                                              5′                                              3′
```

### 16 S4-Clostridia

#### 16.1 Taxa

The taxonomy of each organism containing a putative S4-Clostridia RNA is listed, with abbreviations identifying each hit (e.g., "Eco-1-1" and "Eco-1-2" might hypothetically represent two distinct RNAs in *E. coli*). The abbreviations will be used to identify each individual S4-Clostridia RNA in Sections 16.2 and 16.4:.

| **abbrev. of hits** | **taxonomy of species** |
|---|---|
| Bha-1-1 | Bacteria Firmicutes Clostridia Clostridiales *Blautia hansenii* DSM 20583 |
| Bhy-1-1 | Bacteria Firmicutes Clostridia Clostridiales *Blautia hydrogenotrophica* DSM 10507 (Ruminococcus |
| Chy-1-1 | Bacteria Firmicutes Clostridia Clostridiales Clostridiaceae *Clostridium hylemonae* DSM 15053 |
| Cne-1-1 | Bacteria Firmicutes Clostridia Clostridiales Clostridiaceae *Clostridium nexile* DSM 1787 |
| Csc-1-1 | Bacteria Firmicutes Clostridia Clostridiales Clostridiaceae *Clostridium scindens* ATCC 35704 |
| Cba-1-1 | Bacteria Firmicutes Clostridia *Clostridiales bacterium* VE202-06 |
| Cba-2-1 | Bacteria Firmicutes Clostridia *Clostridiales bacterium* VE202-14 |
| Cba-3-1 | Bacteria Firmicutes Clostridia *Clostridiales bacterium* VE202-26 |
| [ce-1-1 | Bacteria Firmicutes Clostridia Clostridiales Eubacteriaceae Eubacterium [Eubacterium] cellulosolvens LD2006 |
| Ece-1-1 | Bacteria Firmicutes Clostridia Clostridiales Eubacteriaceae *Eubacterium cellulosolvens* 6 |
| Esp-1-1 | Bacteria Firmicutes Clostridia Clostridiales Eubacteriaceae *Eubacterium sp.* ER2 |
| [sc-1-1 | Bacteria Firmicutes Clostridia Clostridiales Lachnospiraceae [Clostridium] scindens VE202-05 |
| Bpr-1-1 | Bacteria Firmicutes Clostridia Clostridiales Lachnospiraceae *Blautia producta* |
| Bpr-2-1 | Bacteria Firmicutes Clostridia Clostridiales Lachnospiraceae *Blautia producta* ATCC 27340 = DSM 2950 |
| Bsc-1-1 | Bacteria Firmicutes Clostridia Clostridiales Lachnospiraceae *Blautia schinkii* DSM 10518 |
| Bsp-1-1 | Bacteria Firmicutes Clostridia Clostridiales Lachnospiraceae *Blautia sp.* KLE 1732 |
| Bwe-1-1 | Bacteria Firmicutes Clostridia Clostridiales Lachnospiraceae *Blautia wexlerae* AGR2146 |
| Bwe-2-1 | Bacteria Firmicutes Clostridia Clostridiales Lachnospiraceae *Blautia wexlerae* DSM 19850 |
| Rgn-1-1 | Bacteria Firmicutes Clostridia Clostridiales Lachnospiraceae Blautia *Ruminococcus gnavus* AGR2154 |
| Rgn-2-1 | Bacteria Firmicutes Clostridia Clostridiales Lachnospiraceae Blautia *Ruminococcus gnavus* CC55_001C |
| Bfo-1-1 | Bacteria Firmicutes Clostridia Clostridiales Lachnospiraceae *Bryantella formatexigens* DSM 14469 |
| Bsp-2-1 | Bacteria Firmicutes Clostridia Clostridiales Lachnospiraceae *Butyrivibrio sp.* AD3002 |
| Bsp-3-1 | Bacteria Firmicutes Clostridia Clostridiales Lachnospiraceae *Butyrivibrio sp.* AE3004 |
| Bsp-4-1 | Bacteria Firmicutes Clostridia Clostridiales Lachnospiraceae *Butyrivibrio sp.* AE3006 |
| Bsp-5-1 | Bacteria Firmicutes Clostridia Clostridiales Lachnospiraceae *Butyrivibrio sp.* MB2005 |
| Bsp-6-1 | Bacteria Firmicutes Clostridia Clostridiales Lachnospiraceae *Butyrivibrio sp.* NC2002 |
| Bsp-7-1 | Bacteria Firmicutes Clostridia Clostridiales Lachnospiraceae *Butyrivibrio sp.* VCD2006 |
| Bsp-8-1 | Bacteria Firmicutes Clostridia Clostridiales Lachnospiraceae *Butyrivibrio sp.* WCD2001 |
| Bsp-9-1 | Bacteria Firmicutes Clostridia Clostridiales Lachnospiraceae *Butyrivibrio sp.* WCD3002 |
| Csp-1-1 | Bacteria Firmicutes Clostridia Clostridiales Lachnospiraceae *Coprococcus sp.* HPP0048 |

| | |
|---|---|
| Csp-2-1 | Bacteria Firmicutes Clostridia Clostridiales Lachnospiraceae *Coprococcus sp.* HPP0074 |
| Dfo-1-1 | Bacteria Firmicutes Clostridia Clostridiales Lachnospiraceae *Dorea formicigenerans* 4_6_53AFAA |
| Dfo-2-1 | Bacteria Firmicutes Clostridia Clostridiales Lachnospiraceae *Dorea formicigenerans* ATCC 27755 |
| Dlo-1-1 | Bacteria Firmicutes Clostridia Clostridiales Lachnospiraceae *Dorea longicatena* AGR2136 |
| Dlo-2-1 | Bacteria Firmicutes Clostridia Clostridiales Lachnospiraceae *Dorea longicatena* DSM 13814 |
| Dsp-1-1 | Bacteria Firmicutes Clostridia Clostridiales Lachnospiraceae *Dorea sp.* 5-2 |
| Dsp-2-1 | Bacteria Firmicutes Clostridia Clostridiales Lachnospiraceae *Dorea sp.* AGR2135 |
| Dsp-3-1 | Bacteria Firmicutes Clostridia Clostridiales Lachnospiraceae *Dorea sp.* D27 |
| Lba-1-1 | Bacteria Firmicutes Clostridia Clostridiales *Lachnospiraceae bacterium* 1_1_57FAA |
| Lba-2-1 | Bacteria Firmicutes Clostridia Clostridiales *Lachnospiraceae bacterium* 2_1_46FAA |
| Lba-3-1 | Bacteria Firmicutes Clostridia Clostridiales *Lachnospiraceae bacterium* 2_1_58FAA |
| Lba-4-1 | Bacteria Firmicutes Clostridia Clostridiales *Lachnospiraceae bacterium* 3-2 |
| Lba-5-1 | Bacteria Firmicutes Clostridia Clostridiales *Lachnospiraceae bacterium* 3_1_46FAA |
| Lba-6-1 | Bacteria Firmicutes Clostridia Clostridiales *Lachnospiraceae bacterium* 4_1_37FAA |
| Lba-7-1 | Bacteria Firmicutes Clostridia Clostridiales *Lachnospiraceae bacterium* 5_1_57FAA |
| Lba-8-1 | Bacteria Firmicutes Clostridia Clostridiales *Lachnospiraceae bacterium* 6_1_63FAA |
| Lba-9-1 | Bacteria Firmicutes Clostridia Clostridiales *Lachnospiraceae bacterium* 8_1_57FAA |
| Lba-10-1 | Bacteria Firmicutes Clostridia Clostridiales *Lachnospiraceae bacterium* 9_1_43BFAA |
| Lba-11-1 | Bacteria Firmicutes Clostridia Clostridiales *Lachnospiraceae bacterium* FE2018 |
| Rpe-1-1 | Bacteria Firmicutes Clostridia Clostridiales Lachnospiraceae *Robinsoniella peoriensis* |
| Rsp-1-1 | Bacteria Firmicutes Clostridia Clostridiales Lachnospiraceae *Robinsoniella sp.* KNHs210 |
| Dma-1-1 | Bacteria Firmicutes Clostridia Clostridiales Ruminococcaceae *Drancourtella massiliensis* |
| Rga-1-1 | Bacteria Firmicutes Clostridia Clostridiales Ruminococcaceae *Ruminococcus gauvreauii* DSM 19829 |
| Rgn-3-1 | Bacteria Firmicutes Clostridia Clostridiales Ruminococcaceae *Ruminococcus gnavus* ATCC 29149 |
| Rob-1-1 | Bacteria Firmicutes Clostridia Clostridiales Ruminococcaceae *Ruminococcus obeum* ATCC 29174 |
| Rsp-2-1 | Bacteria Firmicutes Clostridia Clostridiales Ruminococcaceae *Ruminococcus sp.* 5_1_39BFAA |
| Rsp-3-1 | Bacteria Firmicutes Clostridia Clostridiales Ruminococcaceae *Ruminococcus sp.* JC304 |
| Rto-1-1 | Bacteria Firmicutes Clostridia Clostridiales Ruminococcaceae *Ruminococcus torques* ATCC 27756 |
| env-1 to env-438 | environmental samples |

#### 16.2 Gene contexts

Each S4-Clostridia RNA (indicated by "RNA→") is listed. For each hit, the downstream genes predicted to reside in a regulated operon are listed. If the nearest downstream gene is encoding in the opposite strand (and therefore presumed to not be a part of a regulated operon), then that gene is still depicted. Some environmental sequences and some RefSeq entries lack gene annotations, and so no genes are available for such sequences. The direction of each gene is indicated with an arrow (→), and each predicted conserved domain in the gene is named. Conserved domains associated with more than one S4-Clostridia RNA are assigned a color; other domains are gray. Information about these conserved domains is given in Section 16.3. The accession of the sequence containing each S4-Clostridia RNA is given in the column named "Seq. accession". Accessions beginning with "NC_", "NS_", "NW_" or "NZ_" are contained in RefSeq. Other accession refer to environmental samples. Nucleotide coordinates are given for the 5′ and 3′ boundaries of each S4-Clostridia RNA. If the 5′ coordinate is greater than the 3′ coordinate, the RNA is present on the reverse-complement strand of the containing genomic DNA sequence. Each hit is denoted by an abbreviation (like "Eco-1-1") that refers to a taxonomy given in Section 16.1.

| abbrev. | Seq. accession | | 5′ at | 3′ at | genes |
|---|---|---|---|---|---|
| env-1 | AUXO015936627.1 | + | 257 | 309 | **RNA**→ RpsM (COG0099)rpsM (PRK05179)→ |
| env-2 | SRS016095_C2391063 | + | 242 | 301 | **RNA**→ rps13 (CHL00137)rpsM (PRK05179)uS13_bact (TIGR03631)→ |
| env-3 | SRS052697_LANL_scaffold_5093 | + | 388 | 447 | **RNA**→ rps13 (CHL00137)RpsM (COG0099)rpsM (PRK05179)uS13_bact (TIGR03631)→ |
| env-4 | scaffold46699_2_MH0044 | + | 217 | 276 | **RNA**→ RpsM (COG0099)rpsM (PRK05179)→ |
| env-5 | scaffold83773_1_MH0059 | - | 378 | 319 | **RNA**→ RpsM (COG0099)rpsM (PRK05179)→ |
| env-6 | NLF007_scaffold4057_7 | + | 767 | 826 | **RNA**→ rps13 (CHL00137)rpsM (PRK05179)uS13_bact (TIGR03631)→ |
| env-7 | scaffold201303_1_MH0006 | - | 337 | 278 | **RNA**→ RpsM (COG0099)rpsM (PRK05179)→ |
| env-8 | scaffold3229_1_MH0038 | - | 339 | 280 | **RNA**→ RpsM (COG0099)rpsM (PRK05179)→ |
| env-9 | UnmappedStool_Broad_scaffold_1083211 | + | 240 | 299 | **RNA**→ RpsM (COG0099)rpsM (PRK05179)→ |
| env-10 | NOM007_scaffold46220_1 | + | 19654 | 19713 | **RNA**→ RpsM (COG0099)rpsM (PRK05179)→ PRK05309 (PRK05309)RpsK (COG0100)→ RpsD (COG0522)rpsD (PRK05327)→ RpoA (COG0202)PRK05182 (PRK05182)→ |

| | | | | | |
|---|---|---|---|---|---|
| env-11 | SRS014923_C3228657 | + | 242 | 301 | **RNA**→ RpsM (COG0099)rpsM (PRK05179)→ |
| env-12 | NLF009_scaffold14126_1 | - | 1235 | 1176 | **RNA**→ RpsM (COG0099)rpsM (PRK05179)→ PRK05309 (PRK05309)RpsK (COG0100)→ RpsD (COG0522)rpsD (PRK05327)→ |
| env-13 | NOF014_scaffold21218_11 | + | 6928 | 6987 | **RNA**→ RpsM (COG0099)rpsM (PRK05179)→ PRK05309 (PRK05309)RpsK (COG0100)→ RpsD (COG0522)rpsD (PRK05327)→ RpoA (COG0202)PRK05182 (PRK05182)→ |
| env-14 | SRS014459_C2557888 | + | 432 | 491 | **RNA**→ rps13 (CHL00137)rpsM (PRK05179)uS13_bact (TIGR03631)→ |
| env-15 | SRS017191_C1739427 | + | 3572 | 3631 | **RNA**→ rps13 (CHL00137)rpsM (PRK05179)uS13_bact (TIGR03631)→ |
| env-16 | SRS015065_WUGC_scaffold_29501 | - | 380 | 321 | **RNA**→ RpsM (COG0099)rpsM (PRK05179)→ |
| env-17 | 08_07_GOS_DNA_scaffold2232_4 | - | 501 | 447 | **RNA**→ RpsM (COG0099)rpsM (PRK05179)→ |
| env-18 | O8_19_POLYDEXT_DNA_scaffold1532_15 | + | 208 | 262 | **RNA**→ RpsM (COG0099)rpsM (PRK05179)→ |
| env-19 | JGI994J12354_10009703 | - | 3666 | 3612 | **RNA**→ RpsM (COG0099)rpsM (PRK05179)→ |
| Bsp-6-1 | NZ_KL370881.1 | + | 27319 | 27373 | **RNA**→ RpsM (COG0099)rpsM (PRK05179)→ PRK05309 (PRK05309)RpsK (COG0100)→ |
| env-20 | JGI20225J20221_1018968 | - | 1486 | 1434 | **RNA**→ RpsM (COG0099)rpsM (PRK05179)→ PRK05309 (PRK05309)RpsK (COG0100)→ RpsD (COG0522)rpsD (PRK05327)→ |
| env-21 | AUXO017144234.1 | + | 5176 | 5235 | **RNA**→ RpsM (COG0099)rpsM (PRK05179)→ PRK05309 (PRK05309)RpsK (COG0100)→ |
| env-22 | AUXO010388311.1 | + | 129 | 199 | **RNA**→ ←hypo |
| env-23 | AUXO013310740.1 | - | 8305 | 8254 | **RNA**→ RpsM (COG0099)rpsM (PRK05179)→ PRK05309 (PRK05309)RpsK (COG0100)→ RpsD (COG0522)rpsD (PRK05327)→ RpoA (COG0202)PRK05182 (PRK05182)→ |
| env-24 | 2227071526 | - | 270 | 181 | **RNA**→ rps13 (CHL00137)rpsM (PRK05179)→ |
| env-25 | 2227624636 | + | 10763 | 10852 | **RNA**→ RpsM (COG0099)rpsM (PRK05179)→ PRK05309 (PRK05309)RpsK (COG0100)→ |
| env-26 | JGI996J12353_1001490 | - | 756 | 667 | **RNA**→ RpsM (COG0099)rpsM (PRK05179)→ PRK05309 (PRK05309)RpsK (COG0100)→ |
| env-27 | JGI994J12354_10000212 | + | 33654 | 33743 | **RNA**→ RpsM (COG0099)rpsM (PRK05179)→ PRK05309 (PRK05309)RpsK (COG0100)→ RpsD (COG0522)rpsD (PRK05327)→ RpoA (COG0202)PRK05182 (PRK05182)→ rplQ (PRK05591)Ribosomal_L17 (pfam01196)→ |
| env-28 | IMNBL1_c0003830 | + | 98 | 187 | **RNA**→ RpsM (COG0099)rpsM (PRK05179)→ PRK05309 (PRK05309)RpsK (COG0100)→ RpsD (COG0522)rpsD (PRK05327)→ RpoA (COG0202)PRK05182 (PRK05182)→ rplQ (PRK05591)Ribosomal_L17 (pfam01196)→ |
| env-29 | 2226950468 | - | 265 | 181 | **RNA**→ rps13 (CHL00137)rpsM (PRK05179)→ |
| env-30 | 2227073852 | - | 2087 | 2000 | **RNA**→ RpsM (COG0099)rpsM (PRK05179)→ PRK05309 (PRK05309)RpsK (COG0100)→ RpsD (COG0522)rpsD (PRK05327)→ RNAP_alpha_NTD (cd06928)PRK05182 (PRK05182)→ |
| env-31 | 2227078579 | + | 1162 | 1249 | **RNA**→ RpsM (COG0099)rpsM (PRK05179)→ PRK05309 (PRK05309)RpsK (COG0100)→ |
| env-32 | IMNBL1_c0000921 | - | 22643 | 22556 | **RNA**→ RpsM (COG0099)rpsM (PRK05179)→ PRK05309 (PRK05309)RpsK (COG0100)→ RpsD (COG0522)rpsD (PRK05327)→ |
| env-33 | 2227013722 | + | 78 | 165 | **RNA**→ RpsM (COG0099)rpsM (PRK05179)→ PRK05309 (PRK05309)RpsK (COG0100)→ RpsD (COG0522)rpsD (PRK05327)→ |
| env-34 | UnmappedStool_Broad_scaffold_1059721 | + | 666 | 716 | **RNA**→ |
| env-35 | SRS014459_WUGC_scaffold_78018 | - | 406 | 356 | **RNA**→ RpsM (COG0099)rpsM (PRK05179)→ |
| env-36 | SRS015065_C2231612 | - | 187 | 137 | **RNA**→ hypo→ |
| env-37 | AUXO016905584.1 | + | 229 | 278 | **RNA**→ rps13 (CHL00137)rpsM (PRK05179)→ |
| env-38 | scaffold82255_1_V1.UC-14 | + | 430 | 480 | **RNA**→ |
| env-39 | SRS014459_WUGC_scaffold_74344 | + | 465 | 518 | **RNA**→ |
| env-40 | SRS015065_C2236339 | - | 162 | 109 | **RNA**→ hypo→ |
| env-41 | SRS056259_LANL_scaffold_44254 | - | 151 | 97 | **RNA**→ ←hypo |
| env-42 | HCF12C_401724 | + | 232 | 297 | **RNA**→ hypo→ RpsM (COG0099)rpsM (PRK05179)→ |
| env-43 | longitudinal_220_106 | + | 5954 | 6044 | **RNA**→ RpsM (COG0099)rpsM (PRK05179)→ PRK05309 (PRK05309)RpsK (COG0100)→ |
| env-44 | longitudinal_221_153 | + | 3442 | 3532 | **RNA**→ RpsM (COG0099)rpsM (PRK05179)→ PRK05309 (PRK05309)RpsK (COG0100)→ |
| env-45 | AJWZ01010485.1 | + | 81 | 143 | **RNA**→ hypo→ |
| env-46 | AUXO010018345.1 | + | 2698 | 2750 | **RNA**→ RpsM (COG0099)rpsM (PRK05179)→ PRK05309 (PRK05309)RpsK (COG0100)→ RpsD (COG0522)rpsD (PRK05327)→ |
| env-47 | AUXO016373287.1 | - | 737 | 678 | **RNA**→ RpsM (COG0099)rpsM (PRK05179)→ |
| env-48 | AUXO016002689.1 | - | 1204 | 1155 | **RNA**→ RpsM (COG0099)rpsM (PRK05179)→ PRK05309 (PRK05309)RpsK (COG0100)→ rpsD (PRK05327)Ribosomal_S4 (pfam00163)→ |
| env-49 | JGI2065J20421_1001636 | + | 4338 | 4413 | **RNA**→ RpsM (COG0099)rpsM (PRK05179)→ PRK05309 (PRK05309)RpsK (COG0100)→ RpsD (COG0522)rpsD (PRK05327)→ RpoA (COG0202)PRK05182 (PRK05182)→ rplQ (PRK05591)Ribosomal_L17 (pfam01196)→ |

| | | | | | |
|---|---|---|---|---|---|
| env-50 | UnmappedStool_Broad_C250603932 | + | 179 | 253 | **RNA**→ |
| Dma-1-1 | NZ_CVPG01000005.1 | + | 471925 | 471999 | **RNA**→ RpsM (COG0099)rpsM (PRK05179)→ PRK05309 (PRK05309)RpsK (COG0100)→ RpsD (COG0522)rpsD (PRK05327)→ RpoA (COG0202)PRK05182 (PRK05182)→ |
| Cba-2-1 | NZ_BAIB02000003.1 | + | 231885 | 231959 | **RNA**→ RpsM (COG0099)rpsM (PRK05179)→ PRK05309 (PRK05309)RpsK (COG0100)→ RpsD (COG0522)rpsD (PRK05327)→ RpoA (COG0202)PRK05182 (PRK05182)→ |
| env-51 | 2227358603 | - | 1575 | 1505 | **RNA**→ RpsM (COG0099)rpsM (PRK05179)→ |
| Lba-2-1 | NZ_KE150017.1 | - | 1725164 | 1725115 | **RNA**→ RpsM (COG0099)rpsM (PRK05179)→ PRK05309 (PRK05309)RpsK (COG0100)→ RpsD (COG0522)rpsD (PRK05327)→ RpoA (COG0202)PRK05182 (PRK05182)→ |
| [ce-1-1 | NZ_JHXY01000007.1 | - | 130151 | 130097 | **RNA**→ RpsM (COG0099)rpsM (PRK05179)→ PRK05309 (PRK05309)RpsK (COG0100)→ RpsD (COG0522)rpsD (PRK05327)→ RpoA (COG0202)PRK05182 (PRK05182)→ |
| Ece-1-1 | NZ_CM001487.1 | + | 2588850 | 2588903 | **RNA**→ RpsM (COG0099)rpsM (PRK05179)→ RpsM (COG0099)rpsM (PRK05179)→ PRK05309 (PRK05309)RpsK (COG0100)→ PRK05309 (PRK05309)RpsK (COG0100)→ RpsD (COG0522)rpsD (PRK05327)→ RpsD (COG0522)rpsD (PRK05327)→ RpoA (COG0202)PRK05182 (PRK05182)→ RpoA (COG0202)PRK05182 (PRK05182)→ |
| env-52 | longitudinal_411_28 | + | 45651 | 45704 | **RNA**→ RpsM (COG0099)rpsM (PRK05179)→ PRK05309 (PRK05309)RpsK (COG0100)→ RpsD (COG0522)rpsD (PRK05327)→ RpoA (COG0202)PRK05182 (PRK05182)→ rplQ (PRK05591)Ribosomal_L17 (pfam01196)→ |
| env-53 | JGI20163J15578_10027836 | - | 917 | 824 | **RNA**→ hypo→ RpsM (COG0099)rpsM (PRK05179)→ PRK05309 (PRK05309)RpsK (COG0100)→ |
| env-54 | scaffold18531_1_V1.CD-15 | + | 542 | 602 | **RNA**→ |
| env-55 | 4491403.3_NODE_1641 | + | 19531 | 19591 | **RNA**→ RpsM (COG0099)rpsM (PRK05179)→ PRK05309 (PRK05309)RpsK (COG0100)→ |
| env-56 | 4491482.3_NODE_7819 | - | 589 | 529 | **RNA**→ RpsM (COG0099)rpsM (PRK05179)→ |
| env-57 | longitudinal_52_17 | - | 410211 | 410151 | **RNA**→ RpsM (COG0099)rpsM (PRK05179)→ PRK05309 (PRK05309)RpsK (COG0100)→ |
| env-58 | longitudinal_231_54 | - | 190275 | 190215 | **RNA**→ RpsM (COG0099)rpsM (PRK05179)→ PRK05309 (PRK05309)RpsK (COG0100)→ |
| env-59 | DOF012_scaffold14355_2 | + | 135 | 195 | **RNA**→ RpsM (COG0099)rpsM (PRK05179)→ PRK05309 (PRK05309)RpsK (COG0100)→ |
| env-60 | longitudinal_230_114 | - | 42745 | 42685 | **RNA**→ RpsM (COG0099)rpsM (PRK05179)→ PRK05309 (PRK05309)RpsK (COG0100)→ |
| env-61 | longitudinal_232_15 | - | 36662 | 36602 | **RNA**→ RpsM (COG0099)rpsM (PRK05179)→ PRK05309 (PRK05309)RpsK (COG0100)→ |
| env-62 | scaffold82035_1_MH0014 | - | 119 | 59 | **RNA**→ |
| env-63 | scaffold818_2_V1.CD-12 | + | 285 | 345 | **RNA**→ RpsM (COG0099)rpsM (PRK05179)→ |
| env-64 | SRS075398_C2026444 | + | 301 | 361 | **RNA**→ ←hypo |
| env-65 | SRS012902_C526759 | + | 741 | 801 | **RNA**→ |
| env-66 | SRS023346_C1056305 | - | 107 | 47 | **RNA**→ hypo→ |
| env-67 | BABA01009209.1 | + | 367 | 427 | **RNA**→ RpsM (COG0099)rpsM (PRK05179)→ PRK05309 (PRK05309)RpsK (COG0100)→ |
| env-68 | scaffold33086_2_V1.CD-1 | - | 516 | 456 | **RNA**→ RpsM (COG0099)rpsM (PRK05179)→ |
| env-69 | DLM009_scaffold11270_3 | + | 4628 | 4688 | **RNA**→ RpsM (COG0099)rpsM (PRK05179)→ PRK05309 (PRK05309)RpsK (COG0100)→ |
| env-70 | NLM017_scaffold20089_2 | + | 813 | 873 | **RNA**→ RpsM (COG0099)rpsM (PRK05179)→ PRK05309 (PRK05309)RpsK (COG0100)→ |
| env-71 | NLM027_scaffold10047_1 | - | 48785 | 48725 | **RNA**→ RpsM (COG0099)rpsM (PRK05179)→ PRK05309 (PRK05309)RpsK (COG0100)→ |
| env-72 | NLM028_scaffold839_7 | - | 1356 | 1296 | **RNA**→ RpsM (COG0099)rpsM (PRK05179)→ PRK05309 (PRK05309)RpsK (COG0100)→ |
| env-73 | NOF005_scaffold31692_1 | - | 1199 | 1139 | **RNA**→ RpsM (COG0099)rpsM (PRK05179)→ PRK05309 (PRK05309)RpsK (COG0100)→ |
| env-74 | NOM002_scaffold8624_11 | - | 5573 | 5513 | **RNA**→ RpsM (COG0099)rpsM (PRK05179)→ PRK05309 (PRK05309)RpsK (COG0100)→ |
| Rgn-3-1 | NZ_AAYG02000020.1 | - | 118252 | 118192 | **RNA**→ RpsM (COG0099)rpsM (PRK05179)→ RpsM (COG0099)rpsM (PRK05179)→ RpsM (COG0099)rpsM (PRK05179)→ PRK05309 (PRK05309)RpsK (COG0100)→ PRK05309 (PRK05309)RpsK (COG0100)→ |
| env-75 | SRS012273_Baylor_scaffold_747 | + | 3356 | 3416 | **RNA**→ RpsM (COG0099)rpsM (PRK05179)→ |
| env-76 | DLM024_scaffold1675_1 | + | 19438 | 19498 | **RNA**→ RpsM (COG0099)rpsM (PRK05179)→ PRK05309 (PRK05309)RpsK (COG0100)→ |
| env-77 | DOM026_scaffold2679_6 | + | 580 | 640 | **RNA**→ RpsM (COG0099)rpsM (PRK05179)→ |
| env-78 | SRS023176_C906831 | - | 179 | 119 | **RNA**→ rps13 (CHL00137)rpsM (PRK05179)→ |
| env-79 | longitudinal_525_50 | + | 44720 | 44780 | **RNA**→ RpsM (COG0099)rpsM (PRK05179)→ PRK05309 (PRK05309)RpsK (COG0100)→ |
| Lba-3-1 | NZ_GL945339.1 | - | 5397 | 5337 | **RNA**→ RpsM (COG0099)rpsM (PRK05179)→ PRK05309 (PRK05309)RpsK (COG0100)→ |
| Rgn-2-1 | NZ_KI669415.1 | - | 296050 | 295990 | **RNA**→ RpsM (COG0099)rpsM (PRK05179)→ PRK05309 (PRK05309)RpsK (COG0100)→ |
| env-80 | NLM007_scaffold2834_4 | + | 3478 | 3538 | **RNA**→ RpsM (COG0099)rpsM (PRK05179)→ |
| env-81 | SRS015578_WUGC_scaffold_47109 | - | 107 | 47 | **RNA**→ |
| Rgn-1-1 | NZ_JAGQ01000004.1 | + | 1161487 | 1161547 | **RNA**→ RpsM (COG0099)rpsM (PRK05179)→ PRK05309 (PRK05309)RpsK (COG0100)→ |
| env-82 | SRS057717_LANL_scaffold_8560 | + | 38 | 98 | **RNA**→ RpsM (COG0099)rpsM (PRK05179)→ |
| env-83 | 4491488.3_NODE_4718 | - | 3548 | 3488 | **RNA**→ RpsM (COG0099)rpsM (PRK05179)→ PRK05309 (PRK05309)RpsK (COG0100)→ |

| | | | | | |
|---|---|---|---|---|---|
| env-84 | NOM026_C930720_1 | + | 365 | 425 | **RNA**→ RpsM (COG0099)rpsM (PRK05179)→ |
| env-85 | NOM027_scaffold1888_6 | + | 683 | 743 | **RNA**→ RpsM (COG0099)rpsM (PRK05179)→ PRK05309 (PRK05309)RpsK (COG0100)→ |
| env-86 | NOM028_scaffold2883_9 | + | 12792 | 12852 | **RNA**→ RpsM (COG0099)rpsM (PRK05179)→ PRK05309 (PRK05309)RpsK (COG0100)→ |
| env-87 | SRS017521_Baylor_scaffold_33616 | - | 298 | 238 | **RNA**→ rps13 (CHL00137)rpsM (PRK05179)→ |
| env-88 | longitudinal_460_95 | + | 247645 | 247697 | **RNA**→ RpsM (COG0099)rpsM (PRK05179)→ PRK05309 (PRK05309)RpsK (COG0100)→ |
| env-89 | 4448814.3_Contig36431 | + | 203 | 262 | **RNA**→ RpsM (COG0099)rpsM (PRK05179)→ |
| env-90 | JGI20163J15578_10056589 | + | 1720 | 1809 | **RNA**→ RpsM (COG0099)rpsM (PRK05179)→ |
| env-91 | JGI20163J15578_10006853 | + | 4205 | 4343 | **RNA**→ RpsM (COG0099)rpsM (PRK05179)→ |
| env-92 | HCF14C_2_1_1_newblercontig58175 | + | 339 | 418 | **RNA**→ |
| env-93 | AUXO011545607.1 | - | 224 | 130 | **RNA**→ ←hypo |
| env-94 | AUXO017992350.1 | + | 172 | 224 | **RNA**→ RpsM (COG0099)rpsM (PRK05179)→ hypo→ |
| Lba-11-1 | NZ_JNKK01000010.1 | - | 51200 | 51148 | **RNA**→ RpsM (COG0099)rpsM (PRK05179)→ PRK05309 (PRK05309)RpsK (COG0100)→ RpsD (COG0522)rpsD (PRK05327)→ RpoA (COG0202)PRK05182 (PRK05182)→ |
| env-95 | AUXO011286893.1 | + | 221 | 272 | **RNA**→ RpsM (COG0099)rpsM (PRK05179)→ |
| env-96 | AUXO012939603.1 | + | 907 | 958 | **RNA**→ |
| env-97 | longitudinal_461_56 | + | 109667 | 109723 | **RNA**→ RpsM (COG0099)rpsM (PRK05179)→ PRK05309 (PRK05309)RpsK (COG0100)→ RpsD (COG0522)rpsD (PRK05327)→ RpoA (COG0202)PRK05182 (PRK05182)→ |
| env-98 | longitudinal_462_123 | + | 109674 | 109730 | **RNA**→ RpsM (COG0099)rpsM (PRK05179)→ PRK05309 (PRK05309)RpsK (COG0100)→ RpsD (COG0522)rpsD (PRK05327)→ RpoA (COG0202)PRK05182 (PRK05182)→ |
| env-99 | longitudinal_463_7 | + | 109694 | 109750 | **RNA**→ RpsM (COG0099)rpsM (PRK05179)→ PRK05309 (PRK05309)RpsK (COG0100)→ RpsD (COG0522)rpsD (PRK05327)→ RpoA (COG0202)PRK05182 (PRK05182)→ |
| env-100 | longitudinal_522_3104 | + | 4325 | 4381 | **RNA**→ RpsM (COG0099)rpsM (PRK05179)→ PRK05309 (PRK05309)RpsK (COG0100)→ RpsD (COG0522)rpsD (PRK05327)→ |
| Dsp-3-1 | NZ_KQ236746.1 | + | 368740 | 368792 | **RNA**→ RpsM (COG0099)rpsM (PRK05179)→ PRK05309 (PRK05309)RpsK (COG0100)→ RpsD (COG0522)rpsD (PRK05327)→ RpoA (COG0202)PRK05182 (PRK05182)→ |
| Dsp-1-1 | NZ_KE159794.1 | - | 157768 | 157723 | **RNA**→ RpsM (COG0099)rpsM (PRK05179)→ PRK05309 (PRK05309)RpsK (COG0100)→ RpsD (COG0522)rpsD (PRK05327)→ RpoA (COG0202)PRK05182 (PRK05182)→ |
| env-101 | SRS011134_C4860384 | + | 270 | 323 | **RNA**→ |
| env-102 | SRS016095_C2345639 | - | 169 | 116 | **RNA**→ rps13 (CHL00137)rpsM (PRK05179)→ |
| Dsp-2-1 | NZ_ATVU01000008.1 | + | 63049 | 63102 | **RNA**→ RpsM (COG0099)rpsM (PRK05179)→ PRK05309 (PRK05309)RpsK (COG0100)→ RpsD (COG0522)rpsD (PRK05327)→ RpoA (COG0202)PRK05182 (PRK05182)→ |
| env-103 | SRS018313_C7830548 | - | 121 | 68 | **RNA**→ |
| env-104 | SRS011271_C2920016 | + | 238 | 291 | **RNA**→ |
| env-105 | SRS015133_C3278084 | + | 230 | 283 | **RNA**→ |
| env-106 | longitudinal_66_246 | + | 13912 | 13965 | **RNA**→ RpsM (COG0099)rpsM (PRK05179)→ PRK05309 (PRK05309)RpsK (COG0100)→ RpsD (COG0522)rpsD (PRK05327)→ RpoA (COG0202)PRK05182 (PRK05182)→ |
| env-107 | longitudinal_67_37 | + | 13912 | 13965 | **RNA**→ RpsM (COG0099)rpsM (PRK05179)→ PRK05309 (PRK05309)RpsK (COG0100)→ RpsD (COG0522)rpsD (PRK05327)→ RpoA (COG0202)PRK05182 (PRK05182)→ |
| env-108 | longitudinal_111_61 | + | 12242 | 12295 | **RNA**→ RpsM (COG0099)rpsM (PRK05179)→ PRK05309 (PRK05309)RpsK (COG0100)→ RpsD (COG0522)rpsD (PRK05327)→ RpoA (COG0202)PRK05182 (PRK05182)→ |
| env-109 | longitudinal_112_225 | + | 13912 | 13965 | **RNA**→ RpsM (COG0099)rpsM (PRK05179)→ PRK05309 (PRK05309)RpsK (COG0100)→ RpsD (COG0522)rpsD (PRK05327)→ RpoA (COG0202)PRK05182 (PRK05182)→ |
| env-110 | longitudinal_187_175 | + | 41148 | 41201 | **RNA**→ RpsM (COG0099)rpsM (PRK05179)→ PRK05309 (PRK05309)RpsK (COG0100)→ RpsD (COG0522)rpsD (PRK05327)→ RpoA (COG0202)PRK05182 (PRK05182)→ |
| env-111 | longitudinal_295_1 | - | 30869 | 30816 | **RNA**→ RpsM (COG0099)rpsM (PRK05179)→ PRK05309 (PRK05309)RpsK (COG0100)→ RpsD (COG0522)rpsD (PRK05327)→ RpoA (COG0202)PRK05182 (PRK05182)→ |
| env-112 | longitudinal_296_203 | - | 16114 | 16061 | **RNA**→ RpsM (COG0099)rpsM (PRK05179)→ PRK05309 (PRK05309)RpsK (COG0100)→ RpsD (COG0522)rpsD (PRK05327)→ RpoA (COG0202)PRK05182 (PRK05182)→ |
| env-113 | longitudinal_297_58 | - | 30869 | 30816 | **RNA**→ RpsM (COG0099)rpsM (PRK05179)→ PRK05309 (PRK05309)RpsK (COG0100)→ RpsD (COG0522)rpsD (PRK05327)→ RpoA (COG0202)PRK05182 (PRK05182)→ |
| env-114 | longitudinal_298_248 | + | 4540 | 4593 | **RNA**→ RpsM (COG0099)rpsM (PRK05179)→ PRK05309 (PRK05309)RpsK (COG0100)→ RpsD (COG0522)rpsD (PRK05327)→ |
| env-115 | longitudinal_421_71 | + | 56051 | 56104 | **RNA**→ RpsM (COG0099)rpsM (PRK05179)→ PRK05309 (PRK05309)RpsK (COG0100)→ RpsD (COG0522)rpsD (PRK05327)→ RpoA (COG0202)PRK05182 (PRK05182)→ |

| | | | | | |
|---|---|---|---|---|---|
| Dfo-2-1 | NZ_AAXA02000012.1 | - | 126306 | 126253 | **RNA**→ RpsM (COG0099)rpsM (PRK05179)→ RpsM (COG0099)rpsM (PRK05179)→ RpsM (COG0099)rpsM (PRK05179)→ PRK05309 (PRK05309)RpsK (COG0100)→ PRK05309 (PRK05309)RpsK (COG0100)→ RpsD (COG0522)rpsD (PRK05327)→ RpsD (COG0522)rpsD (PRK05327)→ RpoA (COG0202)PRK05182 (PRK05182)→ RpoA (COG0202)PRK05182 (PRK05182)→ |
| Dfo-1-1 | NZ_JH126494.1 | - | 91730 | 91677 | **RNA**→ RpsM (COG0099)rpsM (PRK05179)→ PRK05309 (PRK05309)RpsK (COG0100)→ RpsD (COG0522)rpsD (PRK05327)→ RpoA (COG0202)PRK05182 (PRK05182)→ |
| env-116 | SRS015217_C2148859 | - | 111 | 58 | **RNA**→ hypo→ |
| env-117 | SRS023914_C1983791 | - | 382 | 329 | **RNA**→ RpsM (COG0099)rpsM (PRK05179)→ |
| env-118 | DLM022_scaffold5029_5 | - | 1050 | 997 | **RNA**→ RpsM (COG0099)rpsM (PRK05179)→ PRK05309 (PRK05309)RpsK (COG0100)→ hypo→ |
| env-119 | NOM001_scaffold35657_1 | + | 984 | 1035 | **RNA**→ RpsM (COG0099)rpsM (PRK05179)→ |
| env-120 | SRS015217_C2114294 | - | 125 | 74 | **RNA**→ hypo→ |
| env-121 | SRS017521_Baylor_scaffold_12819 | - | 125 | 74 | **RNA**→ hypo→ |
| env-122 | SRS064276_LANL_scaffold_58360 | + | 453 | 504 | **RNA**→ |
| env-123 | 4491413.3_NODE_12466 | - | 145 | 94 | **RNA**→ rps13 (CHL00137)rpsM (PRK05179)→ |
| env-124 | 4491421.3_NODE_115793 | + | 17730 | 17781 | **RNA**→ RpsM (COG0099)rpsM (PRK05179)→ PRK05309 (PRK05309)RpsK (COG0100)→ RpsD (COG0522)rpsD (PRK05327)→ |
| env-125 | longitudinal_113_28 | - | 33326 | 33275 | **RNA**→ RpsM (COG0099)rpsM (PRK05179)→ PRK05309 (PRK05309)RpsK (COG0100)→ RpsD (COG0522)rpsD (PRK05327)→ RpoA (COG0202)PRK05182 (PRK05182)→ rplQ (PRK05591)Ribosomal_L17 (pfam01196)→ |
| env-126 | longitudinal_114_32 | - | 20351 | 20300 | **RNA**→ RpsM (COG0099)rpsM (PRK05179)→ PRK05309 (PRK05309)RpsK (COG0100)→ RpsD (COG0522)rpsD (PRK05327)→ RpoA (COG0202)PRK05182 (PRK05182)→ rplQ (PRK05591)Ribosomal_L17 (pfam01196)→ |
| Dlo-2-1 | NZ_DS264399.1 | + | 39450 | 39501 | **RNA**→ RpsM (COG0099)rpsM (PRK05179)→ PRK05309 (PRK05309)RpsK (COG0100)→ RpsD (COG0522)rpsD (PRK05327)→ RpoA (COG0202)PRK05182 (PRK05182)→ RpoA (COG0202)PRK05182 (PRK05182)→ rplQ (PRK05591)Ribosomal_L17 (pfam01196)→ |
| env-127 | AUXO014831574.1 | - | 23382 | 23331 | **RNA**→ RpsM (COG0099)rpsM (PRK05179)→ PRK05309 (PRK05309)RpsK (COG0100)→ RpsD (COG0522)rpsD (PRK05327)→ RpoA (COG0202)PRK05182 (PRK05182)→ rplQ (PRK05591)Ribosomal_L17 (pfam01196)→ |
| env-128 | SRS015782_C3128808 | + | 194 | 245 | **RNA**→ |
| env-129 | scaffold84298_1_MH0050 | - | 369 | 318 | **RNA**→ RpsM (COG0099)rpsM (PRK05179)→ |
| env-130 | longitudinal_299_23 | - | 26288 | 26235 | **RNA**→ RpsM (COG0099)rpsM (PRK05179)→ PRK05309 (PRK05309)RpsK (COG0100)→ RpsD (COG0522)rpsD (PRK05327)→ RpoA (COG0202)PRK05182 (PRK05182)→ rplQ (PRK05591)Ribosomal_L17 (pfam01196)→ |
| env-131 | longitudinal_300_84 | - | 26610 | 26557 | **RNA**→ RpsM (COG0099)rpsM (PRK05179)→ PRK05309 (PRK05309)RpsK (COG0100)→ RpsD (COG0522)rpsD (PRK05327)→ RpoA (COG0202)PRK05182 (PRK05182)→ rplQ (PRK05591)Ribosomal_L17 (pfam01196)→ |
| env-132 | longitudinal_301_116 | - | 26432 | 26379 | **RNA**→ RpsM (COG0099)rpsM (PRK05179)→ PRK05309 (PRK05309)RpsK (COG0100)→ RpsD (COG0522)rpsD (PRK05327)→ RpoA (COG0202)PRK05182 (PRK05182)→ rplQ (PRK05591)Ribosomal_L17 (pfam01196)→ |
| env-133 | longitudinal_302_50 | + | 53065 | 53118 | **RNA**→ RpsM (COG0099)rpsM (PRK05179)→ PRK05309 (PRK05309)RpsK (COG0100)→ RpsD (COG0522)rpsD (PRK05327)→ RpoA (COG0202)PRK05182 (PRK05182)→ rplQ (PRK05591)Ribosomal_L17 (pfam01196)→ |
| env-134 | longitudinal_346_79 | + | 20368 | 20421 | **RNA**→ RpsM (COG0099)rpsM (PRK05179)→ PRK05309 (PRK05309)RpsK (COG0100)→ RpsD (COG0522)rpsD (PRK05327)→ RpoA (COG0202)PRK05182 (PRK05182)→ rplQ (PRK05591)Ribosomal_L17 (pfam01196)→ |
| env-135 | longitudinal_347_101 | - | 54042 | 53989 | **RNA**→ RpsM (COG0099)rpsM (PRK05179)→ PRK05309 (PRK05309)RpsK (COG0100)→ RpsD (COG0522)rpsD (PRK05327)→ RpoA (COG0202)PRK05182 (PRK05182)→ rplQ (PRK05591)Ribosomal_L17 (pfam01196)→ |
| env-136 | SRS016095_WUGC_scaffold_24083 | - | 346 | 293 | **RNA**→ RpsM (COG0099)rpsM (PRK05179)→ |
| env-137 | NLM027_scaffold13626_3 | - | 4716 | 4663 | **RNA**→ RpsM (COG0099)rpsM (PRK05179)→ PRK05309 (PRK05309)RpsK (COG0100)→ RpsD (COG0522)rpsD (PRK05327)→ RpoA (COG0202)PRK05182 (PRK05182)→ rplQ (PRK05591)Ribosomal_L17 (pfam01196)→ |
| env-138 | NOM029_scaffold5480_12 | - | 185 | 132 | **RNA**→ rps13 (CHL00137)rpsM (PRK05179)→ |

| | | | | | |
|---|---|---|---|---|---|
| env-139 | longitudinal_464_28 | + | 52346 | 52399 | RNA→ RpsM (COG0099)rpsM (PRK05179)→ PRK05309 (PRK05309)RpsK (COG0100)→<br>RpsD (COG0522)rpsD (PRK05327)→ RpoA (COG0202)PRK05182 (PRK05182)→<br>rplQ (PRK05591)Ribosomal_L17 (pfam01196)→ |
| Dlo-1-1 | NZ_KE384171.1 | - | 39745 | 39692 | RNA→ RpsM (COG0099)rpsM (PRK05179)→ PRK05309 (PRK05309)RpsK (COG0100)→<br>RpsD (COG0522)rpsD (PRK05327)→ RpoA (COG0202)PRK05182 (PRK05182)→<br>rplQ (PRK05591)Ribosomal_L17 (pfam01196)→ |
| env-140 | DLM009_scaffold5025_5 | + | 13073 | 13126 | RNA→ RpsM (COG0099)rpsM (PRK05179)→ PRK05309 (PRK05309)RpsK (COG0100)→<br>RpsD (COG0522)rpsD (PRK05327)→ RpoA (COG0202)PRK05182 (PRK05182)→<br>rplQ (PRK05591)Ribosomal_L17 (pfam01196)→ |
| env-141 | longitudinal_134_7 | + | 68515 | 68568 | RNA→ RpsM (COG0099)rpsM (PRK05179)→ PRK05309 (PRK05309)RpsK (COG0100)→<br>RpsD (COG0522)rpsD (PRK05327)→ RpoA (COG0202)PRK05182 (PRK05182)→<br>rplQ (PRK05591)Ribosomal_L17 (pfam01196)→ hypo→<br>GltD (COG0493)NuoF (COG1894)NADH_4Fe-4S (pfam10589)NAD_binding_8 (pfam13450)→<br>Fe_hyd_SSU (smart00902)PreA (COG1146)fer2 (cd00207)Fe_hyd_SSU (pfam02256)Fe_hyd_lg_C (pfam02906)NADH-C |
| env-142 | longitudinal_135_8 | - | 31400 | 31347 | RNA→ RpsM (COG0099)rpsM (PRK05179)→ PRK05309 (PRK05309)RpsK (COG0100)→<br>RpsD (COG0522)rpsD (PRK05327)→ RpoA (COG0202)PRK05182 (PRK05182)→<br>rplQ (PRK05591)Ribosomal_L17 (pfam01196)→ hypo→<br>GltD (COG0493)NuoF (COG1894)NADH_4Fe-4S (pfam10589)NAD_binding_8 (pfam13450)→<br>Fe_hyd_SSU (smart00902)PreA (COG1146)fer2 (cd00207)Fe_hyd_SSU (pfam02256)Fe_hyd_lg_C (pfam02906)NADH-C |
| env-143 | longitudinal_188_28 | + | 68515 | 68568 | RNA→ RpsM (COG0099)rpsM (PRK05179)→ PRK05309 (PRK05309)RpsK (COG0100)→<br>RpsD (COG0522)rpsD (PRK05327)→ RpoA (COG0202)PRK05182 (PRK05182)→<br>rplQ (PRK05591)Ribosomal_L17 (pfam01196)→ hypo→<br>GltD (COG0493)NuoF (COG1894)NADH_4Fe-4S (pfam10589)NAD_binding_8 (pfam13450)→<br>Fe_hyd_SSU (smart00902)PreA (COG1146)fer2 (cd00207)Fe_hyd_SSU (pfam02256)Fe_hyd_lg_C (pfam02906)NADH-C |
| env-144 | longitudinal_189_5 | - | 31403 | 31350 | RNA→ RpsM (COG0099)rpsM (PRK05179)→ PRK05309 (PRK05309)RpsK (COG0100)→<br>RpsD (COG0522)rpsD (PRK05327)→ RpoA (COG0202)PRK05182 (PRK05182)→<br>rplQ (PRK05591)Ribosomal_L17 (pfam01196)→ hypo→<br>GltD (COG0493)NuoF (COG1894)NADH_4Fe-4S (pfam10589)NAD_binding_8 (pfam13450)→<br>Fe_hyd_SSU (smart00902)PreA (COG1146)fer2 (cd00207)Fe_hyd_SSU (pfam02256)Fe_hyd_lg_C (pfam02906)NADH-C |
| env-145 | longitudinal_190_130 | + | 68514 | 68567 | RNA→ RpsM (COG0099)rpsM (PRK05179)→ PRK05309 (PRK05309)RpsK (COG0100)→<br>RpsD (COG0522)rpsD (PRK05327)→ RpoA (COG0202)PRK05182 (PRK05182)→<br>rplQ (PRK05591)Ribosomal_L17 (pfam01196)→ hypo→<br>GltD (COG0493)NuoF (COG1894)NADH_4Fe-4S (pfam10589)NAD_binding_8 (pfam13450)→<br>Fe_hyd_SSU (smart00902)PreA (COG1146)fer2 (cd00207)Fe_hyd_SSU (pfam02256)Fe_hyd_lg_C (pfam02906)NADH-C |
| env-146 | longitudinal_465_52 | + | 52346 | 52399 | RNA→ RpsM (COG0099)rpsM (PRK05179)→ PRK05309 (PRK05309)RpsK (COG0100)→<br>RpsD (COG0522)rpsD (PRK05327)→ RpoA (COG0202)PRK05182 (PRK05182)→<br>rplQ (PRK05591)Ribosomal_L17 (pfam01196)→ |
| env-147 | longitudinal_466_3 | - | 31944 | 31891 | RNA→ RpsM (COG0099)rpsM (PRK05179)→ PRK05309 (PRK05309)RpsK (COG0100)→<br>RpsD (COG0522)rpsD (PRK05327)→ RpoA (COG0202)PRK05182 (PRK05182)→<br>rplQ (PRK05591)Ribosomal_L17 (pfam01196)→ |
| env-148 | longitudinal_467_77 | + | 52344 | 52397 | RNA→ RpsM (COG0099)rpsM (PRK05179)→ PRK05309 (PRK05309)RpsK (COG0100)→<br>RpsD (COG0522)rpsD (PRK05327)→ RpoA (COG0202)PRK05182 (PRK05182)→<br>rplQ (PRK05591)Ribosomal_L17 (pfam01196)→ |
| env-149 | longitudinal_414_23 | + | 90211 | 90261 | RNA→ RpsM (COG0099)rpsM (PRK05179)→ PRK05309 (PRK05309)RpsK (COG0100)→<br>RpsD (COG0522)rpsD (PRK05327)→ RpoA (COG0202)PRK05182 (PRK05182)→ |
| env-150 | SRS017191_Baylor_scaffold_17384 | - | 388 | 337 | RNA→ RpsM (COG0099)rpsM (PRK05179)→ |
| env-151 | SRS015217_WUGC_scaffold_28345 | - | 149 | 98 | RNA→ |
| Chy-1-1 | NZ_GG657761.1 | + | 327237 | 327288 | RNA→ RpsM (COG0099)rpsM (PRK05179)→ RpsM (COG0099)rpsM (PRK05179)→<br>PRK05309 (PRK05309)RpsK (COG0100)→ RpsD (COG0522)rpsD (PRK05327)→<br>RpsD (COG0522)rpsD (PRK05327)→ RpoA (COG0202)PRK05182 (PRK05182)→ |
| env-152 | SRS043411_C1566362 | - | 324 | 270 | RNA→ hypo→ |
| env-153 | SRS016495_LANL_scaffold_8908 | - | 253 | 200 | RNA→ rps13 (CHL00137)rpsM (PRK05179)→ |
| env-154 | SRS016517_C20253773 | - | 120 | 67 | RNA→ hypo→ |
| Esp-1-1 | NZ_JPJE01000006.1 | + | 154152 | 154252 | RNA→ RpsM (COG0099)rpsM (PRK05179)→ |
| env-155 | SRS023971_Baylor_scaffold_879 | - | 742 | 602 | RNA→ RpsM (COG0099)rpsM (PRK05179)→ rps11 (CHL00041)PRK05309 (PRK05309)→ |

| | | | | | |
|---|---|---|---|---|---|
| env-156 | longitudinal_21_17 | - | 9462 | 9322 | **RNA**→ RpsM (COG0099)rpsM (PRK05179)→ PRK05309 (PRK05309)RpsK (COG0100)→ RpsD (COG0522)rpsD (PRK05327)→ |
| env-157 | longitudinal_22_92 | - | 2112 | 1972 | **RNA**→ RpsM (COG0099)rpsM (PRK05179)→ PRK05309 (PRK05309)RpsK (COG0100)→ RpsD (COG0522)rpsD (PRK05327)→ |
| env-158 | longitudinal_23_385 | - | 106549 | 106409 | **RNA**→ RpsM (COG0099)rpsM (PRK05179)→ PRK05309 (PRK05309)RpsK (COG0100)→ RpsD (COG0522)rpsD (PRK05327)→ |
| env-159 | DLM016_scaffold59795_1 | - | 189 | 49 | **RNA**→ |
| env-160 | SRS048870_WUGC_scaffold_43350 | - | 502 | 362 | **RNA**→ RpsM (COG0099)rpsM (PRK05179)→ |
| env-161 | NLM032_scaffold39271_3 | + | 153 | 293 | **RNA**→ RpsM (COG0099)rpsM (PRK05179)→ PRK05309 (PRK05309)RpsK (COG0100)→ RpsD (COG0522)rpsD (PRK05327)→ |
| env-162 | SRS016095_WUGC_scaffold_16877 | - | 640 | 500 | **RNA**→ RpsM (COG0099)rpsM (PRK05179)→ |
| env-163 | SRS049959_C3775241 | - | 206 | 66 | **RNA**→ hypo→ |
| env-164 | scaffold9684_3_O2.UC-11 | + | 3170 | 3310 | **RNA**→ |
| env-165 | longitudinal_441_49 | + | 144537 | 144677 | **RNA**→ RpsM (COG0099)rpsM (PRK05179)→ PRK05309 (PRK05309)RpsK (COG0100)→ RpsD (COG0522)rpsD (PRK05327)→ |
| env-166 | SRS018351_C2776780 | - | 763 | 623 | **RNA**→ RpsM (COG0099)rpsM (PRK05179)→ PRK05309 (PRK05309)RpsK (COG0100)→ |
| env-167 | SRS011586_WUGC_scaffold_26070 | - | 188 | 48 | **RNA**→ hypo→ |
| env-168 | NOF014_scaffold14549_2 | + | 5949 | 6089 | **RNA**→ RpsM (COG0099)rpsM (PRK05179)→ PRK05309 (PRK05309)RpsK (COG0100)→ RpsD (COG0522)rpsD (PRK05327)→ |
| env-169 | DOF008_scaffold13044_2 | - | 703 | 563 | **RNA**→ RpsM (COG0099)rpsM (PRK05179)→ rps11 (CHL00041)PRK05309 (PRK05309)→ |
| env-170 | NLM028_scaffold5808_6 | + | 527 | 667 | **RNA**→ RpsM (COG0099)rpsM (PRK05179)→ PRK05309 (PRK05309)RpsK (COG0100)→ RpsD (COG0522)rpsD (PRK05327)→ |
| env-171 | DOF009_scaffold30020_7 | + | 1575 | 1715 | **RNA**→ RpsM (COG0099)rpsM (PRK05179)→ PRK05309 (PRK05309)RpsK (COG0100)→ hypo→ |
| env-172 | DLM010_scaffold23309_2 | + | 360 | 500 | **RNA**→ RpsM (COG0099)rpsM (PRK05179)→ |
| env-173 | DOM003_scaffold5836_26 | + | 3132 | 3272 | **RNA**→ RpsM (COG0099)rpsM (PRK05179)→ PRK05309 (PRK05309)RpsK (COG0100)→ |
| env-174 | NLM026_scaffold16599_9 | + | 5961 | 6101 | **RNA**→ RpsM (COG0099)rpsM (PRK05179)→ PRK05309 (PRK05309)RpsK (COG0100)→ RpsD (COG0522)rpsD (PRK05327)→ |
| env-175 | DLM027_scaffold4257_3 | - | 2404 | 2264 | **RNA**→ RpsM (COG0099)rpsM (PRK05179)→ PRK05309 (PRK05309)RpsK (COG0100)→ RpsD (COG0522)rpsD (PRK05327)→ |
| env-176 | SRS065504_LANL_scaffold_20275 | - | 380 | 240 | **RNA**→ RpsM (COG0099)rpsM (PRK05179)→ |
| env-177 | NOF010_C660913_1 | + | 68 | 208 | **RNA**→ RpsM (COG0099)rpsM (PRK05179)→ PRK05309 (PRK05309)RpsK (COG0100)→ |
| env-178 | DLM022_scaffold4596_9 | - | 438 | 298 | **RNA**→ RpsM (COG0099)rpsM (PRK05179)→ |
| Lba-4-1 | NZ_KE159795.1 | - | 1371013 | 1370958 | **RNA**→ RpsM (COG0099)rpsM (PRK05179)→ |
| env-179 | DLF012_scaffold283_24 | + | 205 | 261 | **RNA**→ RpsM (COG0099)rpsM (PRK05179)→ PRK05309 (PRK05309)RpsK (COG0100)→ S4 (smart00363)RpsD (COG0522)→ |
| env-180 | longitudinal_337_87 | + | 40213 | 40269 | **RNA**→ RpsM (COG0099)rpsM (PRK05179)→ PRK05309 (PRK05309)RpsK (COG0100)→ RpsD (COG0522)rpsD (PRK05327)→ RpoA (COG0202)PRK05182 (PRK05182)→ |
| Csc-1-1 | NZ_DS499705.1 | - | 60598 | 60542 | **RNA**→ RpsM (COG0099)rpsM (PRK05179)→ PRK05309 (PRK05309)RpsK (COG0100)→ RpsD (COG0522)rpsD (PRK05327)→ RpsD (COG0522)rpsD (PRK05327)→ RpoA (COG0202)PRK05182 (PRK05182)→ |
| Cba-3-1 | NZ_BAII02000008.1 | + | 191914 | 191970 | **RNA**→ RpsM (COG0099)rpsM (PRK05179)→ PRK05309 (PRK05309)RpsK (COG0100)→ RpsD (COG0522)rpsD (PRK05327)→ RpoA (COG0202)PRK05182 (PRK05182)→ |
| [sc-1-1 | NZ_BAHS01000033.1 | + | 21391 | 21446 | **RNA**→ RpsM (COG0099)rpsM (PRK05179)→ PRK05309 (PRK05309)RpsK (COG0100)→ hypo→ RpsD (COG0522)rpsD (PRK05327)→ RpoA (COG0202)PRK05182 (PRK05182)→ |
| Lba-7-1 | NZ_GL945245.1 | - | 61088 | 61032 | **RNA**→ RpsM (COG0099)rpsM (PRK05179)→ PRK05309 (PRK05309)RpsK (COG0100)→ RpsD (COG0522)rpsD (PRK05327)→ RpoA (COG0202)PRK05182 (PRK05182)→ |
| env-181 | DLF009_C396903_1 | + | 387 | 443 | **RNA**→ rps13 (CHL00137)rpsM (PRK05179)→ |
| env-182 | AUXO014627591.1 | + | 809 | 856 | **RNA**→ |
| env-183 | AUXO010674837.1 | - | 397 | 351 | **RNA**→ RpsM (COG0099)rpsM (PRK05179)→ |
| env-184 | 4448819.3_Contig14521 | + | 180 | 277 | **RNA**→ rps13 (CHL00137)rpsM (PRK05179)→ |
| env-185 | scaffold93_3_V1.CD-12 | + | 256 | 353 | **RNA**→ RpsM (COG0099)rpsM (PRK05179)→ |
| env-186 | SRS017521_Baylor_scaffold_37560 | + | 56 | 153 | **RNA**→ RpsM (COG0099)rpsM (PRK05179)→ |
| env-187 | SRS024132_Baylor_scaffold_32332 | + | 1683 | 1780 | **RNA**→ |
| env-188 | SRS042284_C1690458 | - | 139 | 42 | **RNA**→ hypo→ |
| env-189 | BABA01032099.1 | + | 74 | 171 | **RNA**→ RpsM (COG0099)rpsM (PRK05179)→ |

| | | | | | |
|---|---|---|---|---|---|
| env-190 | NOF005_scaffold49548_1 | + | 65 | 162 | RNA→ RpsM (COG0099)rpsM (PRK05179)→ |
| env-191 | DLF006_scaffold33559_1 | + | 534 | 631 | RNA→ ←hypo |
| env-192 | DLM012_scaffold29270_1 | + | 539 | 636 | RNA→ RpsM (COG0099)rpsM (PRK05179)→ |
| env-193 | DLM022_scaffold333_4 | + | 12625 | 12722 | RNA→ RpsM (COG0099)rpsM (PRK05179)→ |
| env-194 | scaffold43093_1_V1.CD-15 | - | 398 | 301 | RNA→ RpsM (COG0099)rpsM (PRK05179)→ |
| env-195 | SRS057478_LANL_scaffold_2364 | + | 6630 | 6727 | RNA→ RpsM (COG0099)rpsM (PRK05179)→ |
| env-196 | SRS014287_WUGC_scaffold_32769 | - | 803 | 706 | RNA→ RpsM (COG0099)rpsM (PRK05179)→ |
| env-197 | SRS014459_WUGC_scaffold_353 | - | 704 | 607 | RNA→ RpsM (COG0099)rpsM (PRK05179)→ |
| env-198 | SRS016495_LANL_scaffold_14198 | + | 1644 | 1741 | RNA→ RpsM (COG0099)rpsM (PRK05179)→ |
| env-199 | SRS064276_LANL_scaffold_67145 | - | 653 | 556 | RNA→ RpsM (COG0099)rpsM (PRK05179)→ |
| Lba-5-1 | NZ_GL890521.1 | - | 320951 | 320854 | RNA→ RpsM (COG0099)rpsM (PRK05179)→ |
| Lba-1-1 | NZ_GL945299.1 | - | 118724 | 118627 | RNA→ RpsM (COG0099)rpsM (PRK05179)→ |
| Lba-9-1 | NZ_GL622455.1 | + | 76223 | 76320 | RNA→ RpsM (COG0099)rpsM (PRK05179)→ |
| Rto-1-1 | NZ_DS264365.1 | - | 22031 | 21934 | RNA→ RpsM (COG0099)rpsM (PRK05179)→ |
| env-200 | UnmappedStool_Broad_scaffold_762794 | + | 200 | 297 | RNA→ rps13 (CHL00137)rpsM (PRK05179)→ |
| env-201 | NLM031_scaffold9790_1 | + | 8440 | 8537 | RNA→ ←hypo |
| env-202 | SRS012273_C3528184 | - | 135 | 38 | RNA→ hypo→ |
| env-203 | SRS014923_C3148149 | - | 135 | 38 | RNA→ hypo→ |
| env-204 | SRS015217_WUGC_scaffold_6745 | + | 283 | 380 | RNA→ |
| env-205 | SRS019397_WUGC_scaffold_29890 | - | 135 | 38 | RNA→ hypo→ |
| env-206 | SRS019601_WUGC_scaffold_3920 | + | 4642 | 4739 | RNA→ hypo→ |
| env-207 | SRS020869_C3206849 | + | 287 | 384 | RNA→ |
| env-208 | SRS042628_C2611262 | + | 367 | 464 | RNA→ hypo→ |
| env-209 | SRS050422_C2650805 | + | 284 | 381 | RNA→ |
| env-210 | SRS077730_LANL_scaffold_23057 | - | 135 | 38 | RNA→ hypo→ |
| env-211 | SRS014613_C1466654 | - | 148 | 51 | RNA→ hypo→ |
| env-212 | SRS015065_WUGC_scaffold_7196 | + | 3193 | 3290 | RNA→ |
| env-213 | SRS015133_WUGC_scaffold_39311 | - | 153 | 56 | RNA→ |
| env-214 | SRS016989_C1459026 | - | 153 | 56 | RNA→ hypo→ |
| env-215 | SRS064645_C414785 | + | 200 | 297 | RNA→ |
| env-216 | SRS058723_C6722821 | + | 142 | 239 | RNA→ RpsM (COG0099)rpsM (PRK05179)→ |
| env-217 | AUXO012083149.1 | - | 437 | 384 | RNA→ hypo→ RpsM (COG0099)rpsM (PRK05179)→ |
| Rpe-1-1 | NZ_JTGN01000002.1 | + | 415740 | 415795 | RNA→ RpsM (COG0099)rpsM (PRK05179)→ PRK05309 (PRK05309)RpsK (COG0100)→ RpsD (COG0522)rpsD (PRK05327)→ RpoA (COG0202)PRK05182 (PRK05182)→ |
| Rsp-1-1 | NZ_JMLZ01000002.1 | + | 4582826 | 4582881 | RNA→ RpsM (COG0099)rpsM (PRK05179)→ PRK05309 (PRK05309)RpsK (COG0100)→ RpsD (COG0522)rpsD (PRK05327)→ RpoA (COG0202)PRK05182 (PRK05182)→ |
| env-218 | HCF12C_72825 | - | 388 | 337 | RNA→ RpsM (COG0099)uS13_bact (TIGR03631)→ |
| env-219 | AUXO015776767.1 | + | 78 | 133 | RNA→ RpsM (COG0099)rpsM (PRK05179)→ |
| env-220 | AUXO011164680.1 | - | 268 | 221 | RNA→ rps13 (CHL00137)rpsM (PRK05179)→ |
| env-221 | SRS050925_LANL_scaffold_36095 | - | 324 | 265 | RNA→ rps13 (CHL00137)rpsM (PRK05179)→ |
| env-222 | NOM009_scaffold13140_13 | - | 2089 | 2030 | RNA→ RpsM (COG0099)rpsM (PRK05179)→ PRK05309 (PRK05309)RpsK (COG0100)→ RpsD (COG0522)rpsD (PRK05327)→ RNAP_alpha_NTD (cd06928)PRK05182 (PRK05182)→ |
| env-223 | SRS023526_C3111217 | + | 472 | 531 | RNA→ |
| env-224 | SRS023583_C1297360 | - | 201 | 142 | RNA→ |
| env-225 | SRS055982_C1612735 | + | 1309 | 1368 | RNA→ |
| env-226 | SRS024075_LANL_scaffold_24329 | - | 214 | 155 | RNA→ |
| env-227 | SRS015578_WUGC_scaffold_35682 | - | 383 | 324 | RNA→ RpsM (COG0099)rpsM (PRK05179)→ |
| env-228 | SRS016095_WUGC_scaffold_27178 | - | 399 | 340 | RNA→ RpsM (COG0099)rpsM (PRK05179)→ |
| env-229 | scaffold6694_1_MH0045 | - | 199 | 140 | RNA→ hypo→ |
| env-230 | scaffold51326_2_MH0086 | + | 2401 | 2460 | RNA→ rps13 (CHL00137)rpsM (PRK05179)→ |
| env-231 | 4491423.3_NODE_156416 | + | 1556 | 1615 | RNA→ rps13 (CHL00137)rpsM (PRK05179)→ |
| env-232 | SRS014459_WUGC_scaffold_16825 | + | 226 | 285 | RNA→ |
| env-233 | SRS015065_C2405519 | + | 836 | 895 | RNA→ |

| | | | | | |
|---|---|---|---|---|---|
| env-234 | scaffold82058_3_MH0002 | + | 1327 | 1386 | **RNA**→ |
| env-235 | scaffold49358_1_MH0011 | + | 406 | 465 | **RNA**→ RpsM (COG0099)rpsM (PRK05179)→ |
| env-236 | scaffold6610_2_V1.UC-6 | - | 583 | 524 | **RNA**→ RpsM (COG0099)rpsM (PRK05179)→ |
| env-237 | UnmappedStool_Broad_scaffold_708525 | + | 266 | 325 | **RNA**→ RpsM (COG0099)rpsM (PRK05179)→ |
| env-238 | SRS017521_C2601561 | - | 183 | 124 | **RNA**→ |
| env-239 | SRS020869_Baylor_scaffold_12378 | - | 384 | 325 | **RNA**→ RpsM (COG0099)rpsM (PRK05179)→ |
| env-240 | NOM013_scaffold46563_2 | - | 763 | 704 | **RNA**→ RpsM (COG0099)rpsM (PRK05179)→ PRK05309 (PRK05309)RpsK (COG0100)→ |
| env-241 | AUXO011855295.1 | + | 1265 | 1312 | **RNA**→ RpsM (COG0099)rpsM (PRK05179)→ hypo→ |
| env-242 | SRS019601_C1835248 | - | 186 | 135 | **RNA**→ rpsM (PRK05179)→ |
| env-243 | DOF006_scaffold6616_2 | - | 320 | 269 | **RNA**→ rps13 (CHL00137)rpsM (PRK05179)→ |
| env-244 | SRS075398_C2013874 | - | 190 | 139 | **RNA**→ rpsM (PRK05179)→ |
| env-245 | UnmappedStool_Broad_scaffold_729622 | - | 165 | 114 | **RNA**→ hypo→ |
| env-246 | DLM024_scaffold288_2 | - | 17086 | 17035 | **RNA**→ RpsM (COG0099)rpsM (PRK05179)→ |
| env-247 | DOM008_scaffold9546_10 | - | 10248 | 10197 | **RNA**→ RpsM (COG0099)rpsM (PRK05179)→ |
| env-248 | NLM023_scaffold268_5 | + | 2617 | 2668 | **RNA**→ RpsM (COG0099)rpsM (PRK05179)→ |
| env-249 | NLM027_scaffold19174_2 | + | 3081 | 3132 | **RNA**→ RpsM (COG0099)rpsM (PRK05179)→ |
| env-250 | NOM002_scaffold15570_5 | + | 2668 | 2719 | **RNA**→ RpsM (COG0099)rpsM (PRK05179)→ |
| env-251 | SRS057717_LANL_scaffold_2277 | + | 5351 | 5402 | **RNA**→ RpsM (COG0099)rpsM (PRK05179)→ |
| Cne-1-1 | NZ_DS995339.4 | + | 445348 | 445399 | **RNA**→ RpsM (COG0099)rpsM (PRK05179)→ RpsM (COG0099)rpsM (PRK05179)→ |
| env-252 | scaffold35150_1_MH0047 | + | 861 | 913 | **RNA**→ RpsM (COG0099)rpsM (PRK05179)→ |
| env-253 | 4491482.3_NODE_6411 | + | 1979 | 2031 | **RNA**→ RpsM (COG0099)rpsM (PRK05179)→ |
| env-254 | NLM027_scaffold9_3 | + | 1877 | 1943 | **RNA**→ RpsM (COG0099)rpsM (PRK05179)→ |
| Lba-6-1 | NZ_GL872351.1 | - | 15002 | 14936 | **RNA**→ RpsM (COG0099)rpsM (PRK05179)→ PRK05309 (PRK05309)RpsK (COG0100)→ RpsD (COG0522)rpsD (PRK05327)→ |
| Csp-1-1 | NZ_KE150443.1 | + | 296207 | 296273 | **RNA**→ RpsM (COG0099)rpsM (PRK05179)→ PRK05309 (PRK05309)RpsK (COG0100)→ RpsD (COG0522)rpsD (PRK05327)→ |
| env-255 | CAM_READ_0099546121 | + | 82 | 148 | **RNA**→ RpsM (COG0099)rpsM (PRK05179)→ |
| Lba-10-1 | NZ_GL890574.1 | - | 15367 | 15301 | **RNA**→ RpsM (COG0099)rpsM (PRK05179)→ PRK05309 (PRK05309)RpsK (COG0100)→ RpsD (COG0522)rpsD (PRK05327)→ |
| Csp-2-1 | NZ_KE150436.1 | + | 369549 | 369615 | **RNA**→ RpsM (COG0099)rpsM (PRK05179)→ PRK05309 (PRK05309)RpsK (COG0100)→ RpsD (COG0522)rpsD (PRK05327)→ |
| env-256 | longitudinal_520_20 | - | 15080 | 15014 | **RNA**→ RpsM (COG0099)rpsM (PRK05179)→ PRK05309 (PRK05309)RpsK (COG0100)→ RpsD (COG0522)rpsD (PRK05327)→ |
| env-257 | longitudinal_521_5 | - | 15036 | 14970 | **RNA**→ RpsM (COG0099)rpsM (PRK05179)→ PRK05309 (PRK05309)RpsK (COG0100)→ RpsD (COG0522)rpsD (PRK05327)→ |
| env-258 | SRS023176_C915644 | + | 1011 | 1077 | **RNA**→ rps13 (CHL00137)rpsM (PRK05179)→ |
| env-259 | DOF006_scaffold3143_2 | - | 554 | 488 | **RNA**→ RpsM (COG0099)rpsM (PRK05179)→ |
| env-260 | AUXO015662166.1 | - | 135 | 84 | **RNA**→ ←hypo |
| env-261 | AUXO010716198.1 | - | 157 | 108 | **RNA**→ hypo→ |
| env-262 | UnmappedStool_Broad_scaffold_1072535 | - | 139 | 50 | **RNA**→ hypo→ |
| env-263 | longitudinal_415_5 | - | 192317 | 192267 | **RNA**→ RpsM (COG0099)rpsM (PRK05179)→ PRK05309 (PRK05309)RpsK (COG0100)→ RpsD (COG0522)rpsD (PRK05327)→ RpoA (COG0202)PRK05182 (PRK05182)→ rplQ (PRK05591)Ribosomal_L17 (pfam01196)→ |
| Bfo-1-1 | NZ_ACCL02000023.1 | - | 56498 | 56409 | **RNA**→ hypo→ hypo→ RpsM (COG0099)rpsM (PRK05179)→ RpsM (COG0099)rpsM (PRK05179)→ PRK05309 (PRK05309)RpsK (COG0100)→ PRK05309 (PRK05309)RpsK (COG0100)→ RpsD (COG0522)rpsD (PRK05327)→ RpsD (COG0522)rpsD (PRK05327)→ |
| Bpr-1-1 | NZ_JPJF01000101.1 | - | 10439 | 10368 | **RNA**→ RpsM (COG0099)rpsM (PRK05179)→ PRK05309 (PRK05309)RpsK (COG0100)→ RpsD (COG0522)rpsD (PRK05327)→ RpoA (COG0202)PRK05182 (PRK05182)→ rplQ (PRK05591)Ribosomal_L17 (pfam01196)→ |
| Cba-1-1 | NZ_BAHT02000178.1 | - | 3508 | 3437 | **RNA**→ RpsM (COG0099)rpsM (PRK05179)→ PRK05309 (PRK05309)RpsK (COG0100)→ RpsD (COG0522)rpsD (PRK05327)→ RpoA (COG0202)PRK05182 (PRK05182)→ rplQ (PRK05591)Ribosomal_L17 (pfam01196)→ |
| env-264 | AUXO012679497.1 | + | 121 | 180 | **RNA**→ |
| env-265 | AUXO010229542.1 | - | 194 | 137 | **RNA**→ |
| env-266 | AUXO015380717.1 | - | 461 | 400 | **RNA**→ RpsM (COG0099)rpsM (PRK05179)→ |

| | | | | | |
|---|---|---|---|---|---|
| env-267 | scaffold5991_3_MH0071 | + | 105 | 161 | **RNA**→ RpsM (COG0099)rpsM (PRK05179)→ |
| env-268 | scaffold2528_1_V1.UC-13 | - | 739 | 683 | **RNA**→ RpsM (COG0099)rpsM (PRK05179)→ PRK05309 (PRK05309)RpsK (COG0100)→ |
| env-269 | ADJT01013136.1 | + | 1342 | 1398 | **RNA**→ RpsM (COG0099)rpsM (PRK05179)→ |
| env-270 | SRS054590_LANL_scaffold_5907 | - | 732 | 676 | **RNA**→ RpsM (COG0099)rpsM (PRK05179)→ PRK05309 (PRK05309)RpsK (COG0100)→ |
| Rob-1-1 | NZ_DS264342.1 | + | 2400 | 2456 | **RNA**→ RpsM (COG0099)rpsM (PRK05179)→ RpsM (COG0099)rpsM (PRK05179)→ PRK05309 (PRK05309)RpsK (COG0100)→ RpsD (COG0522)rpsD (PRK05327)→ RpoA (COG0202)PRK05182 (PRK05182)→ RpoA (COG0202)PRK05182 (PRK05182)→ rplQ (PRK05591)Ribosomal_L17 (pfam01196)→ |
| env-271 | NOM005_scaffold4156_9 | - | 399 | 343 | **RNA**→ RpsM (COG0099)rpsM (PRK05179)→ |
| env-272 | BAAX01036074.1 | + | 131 | 187 | **RNA**→ RpsM (COG0099)rpsM (PRK05179)→ PRK05309 (PRK05309)RpsK (COG0100)→ hypo→ |
| env-273 | longitudinal_278_81 | - | 19214 | 19157 | **RNA**→ RpsM (COG0099)rpsM (PRK05179)→ |
| env-274 | longitudinal_409_317 | - | 113247 | 113190 | **RNA**→ RpsM (COG0099)rpsM (PRK05179)→ |
| env-275 | NLM027_scaffold38083_1 | - | 974 | 918 | **RNA**→ RpsM (COG0099)rpsM (PRK05179)→ PRK05309 (PRK05309)RpsK (COG0100)→ |
| env-276 | AJWZ01005537.1 | + | 83 | 139 | **RNA**→ RpsM (COG0099)rpsM (PRK05179)→ |
| env-277 | scaffold14428_2_V1.CD-12 | - | 284 | 228 | **RNA**→ rps13 (CHL00137)rpsM (PRK05179)→ |
| env-278 | scaffold2111_3_O2.UC-18 | + | 4475 | 4531 | **RNA**→ RpsM (COG0099)rpsM (PRK05179)→ PRK05309 (PRK05309)RpsK (COG0100)→ S4 (smart00363)RpsD (COG0522)→ |
| env-279 | 4448820.3_Contig_54992 | - | 235 | 179 | **RNA**→ rps13 (CHL00137)rpsM (PRK05179)→ |
| env-280 | SRS018351_C2808969 | + | 3733 | 3789 | **RNA**→ |
| env-281 | longitudinal_309_72 | + | 12520 | 12576 | **RNA**→ hypo→ RpsM (COG0099)rpsM (PRK05179)→ PRK05309 (PRK05309)RpsK (COG0100)→ RpsD (COG0522)rpsD (PRK05327)→ RpoA (COG0202)PRK05182 (PRK05182)→ rplQ (PRK05591)Ribosomal_L17 (pfam01196)→ |
| env-282 | longitudinal_205_113 | + | 60575 | 60631 | **RNA**→ hypo→ RpsM (COG0099)rpsM (PRK05179)→ PRK05309 (PRK05309)RpsK (COG0100)→ RpsD (COG0522)rpsD (PRK05327)→ RpoA (COG0202)PRK05182 (PRK05182)→ rplQ (PRK05591)Ribosomal_L17 (pfam01196)→ |
| Bsp-1-1 | NZ_KE993318.1 | + | 57041 | 57097 | **RNA**→ RpsM (COG0099)rpsM (PRK05179)→ PRK05309 (PRK05309)RpsK (COG0100)→ RpsD (COG0522)rpsD (PRK05327)→ RpoA (COG0202)PRK05182 (PRK05182)→ rplQ (PRK05591)Ribosomal_L17 (pfam01196)→ rplQ (PRK05591)Ribosomal_L17 (pfam01196)→ hypo→ |
| env-283 | NOM009_scaffold24969_6 | + | 1081 | 1137 | **RNA**→ RpsM (COG0099)rpsM (PRK05179)→ |
| env-284 | DOF008_scaffold7612_1 | + | 695 | 751 | **RNA**→ RpsM (COG0099)rpsM (PRK05179)→ PRK05309 (PRK05309)RpsK (COG0100)→ RpsD (COG0522)rpsD (PRK05327)→ |
| env-285 | NLM027_scaffold2056_4 | + | 1652 | 1708 | **RNA**→ RpsM (COG0099)rpsM (PRK05179)→ PRK05309 (PRK05309)RpsK (COG0100)→ RpsD (COG0522)rpsD (PRK05327)→ RpoA (COG0202)PRK05182 (PRK05182)→ rplQ (PRK05591)Ribosomal_L17 (pfam01196)→ |
| env-286 | NOM005_scaffold35543_1 | - | 16967 | 16911 | **RNA**→ RpsM (COG0099)rpsM (PRK05179)→ PRK05309 (PRK05309)RpsK (COG0100)→ RpsD (COG0522)rpsD (PRK05327)→ RpoA (COG0202)PRK05182 (PRK05182)→ rplQ (PRK05591)Ribosomal_L17 (pfam01196)→ |
| env-287 | SRS043411_C1591614 | + | 354 | 410 | **RNA**→ RpsM (COG0099)rpsM (PRK05179)→ |
| env-288 | ADJT01013276.1 | - | 15776 | 15720 | **RNA**→ RpsM (COG0099)rpsM (PRK05179)→ PRK05309 (PRK05309)RpsK (COG0100)→ RpsD (COG0522)rpsD (PRK05327)→ RpoA (COG0202)PRK05182 (PRK05182)→ rplQ (PRK05591)Ribosomal_L17 (pfam01196)→ |
| env-289 | DLM012_scaffold20960_1 | + | 3763 | 3819 | **RNA**→ RpsM (COG0099)rpsM (PRK05179)→ |
| env-290 | NOM002_C380510_1 | - | 222 | 166 | **RNA**→ hypo→ |
| env-291 | NOM027_scaffold11759_1 | + | 1905 | 1961 | **RNA**→ RpsM (COG0099)rpsM (PRK05179)→ |
| env-292 | NLM007_scaffold12767_10 | + | 9772 | 9828 | **RNA**→ RpsM (COG0099)rpsM (PRK05179)→ |
| env-293 | SRS018351_C2796305 | + | 1883 | 1939 | **RNA**→ |
| env-294 | SRS050752_C2652917 | - | 205 | 149 | **RNA**→ hypo→ |
| env-295 | SRS017521_Baylor_scaffold_5298 | - | 324 | 268 | **RNA**→ rps13 (CHL00137)rpsM (PRK05179)→ |
| env-296 | DLF012_scaffold19007_1 | - | 35604 | 35548 | **RNA**→ RpsM (COG0099)rpsM (PRK05179)→ |
| env-297 | NLM028_scaffold1532_6 | - | 9451 | 9395 | **RNA**→ RpsM (COG0099)rpsM (PRK05179)→ |
| env-298 | DOM018_scaffold5452_9 | - | 1253 | 1197 | **RNA**→ RpsM (COG0099)rpsM (PRK05179)→ |
| env-299 | NLM002_scaffold3727_2 | - | 562 | 506 | **RNA**→ RpsM (COG0099)rpsM (PRK05179)→ |
| env-300 | NLM032_scaffold28_2 | - | 5783 | 5727 | **RNA**→ RpsM (COG0099)rpsM (PRK05179)→ |
| env-301 | NOM017_scaffold25928_6 | - | 5948 | 5892 | **RNA**→ RpsM (COG0099)rpsM (PRK05179)→ |
| env-302 | ADJS01016621.1 | - | 17385 | 17329 | **RNA**→ RpsM (COG0099)rpsM (PRK05179)→ |

| | | | | | |
|---|---|---|---|---|---|
| env-303 | SRS014313_C2608362 | + | 174 | 230 | **RNA**→ |
| env-304 | SRS018817_C1977109 | + | 122 | 178 | **RNA**→ rps13 (CHL00137)rpsM (PRK05179)→ |
| env-305 | SRS055982_C1571339 | - | 222 | 166 | **RNA**→ |
| Bwe-2-1 | NZ_AXVN01000046.1 | - | 11240 | 11184 | **RNA**→ RpsM (COG0099)rpsM (PRK05179)→ |
| env-306 | SRS018656_C1701680 | + | 38 | 94 | **RNA**→ RpsM (COG0099)rpsM (PRK05179)→ |
| env-307 | scaffold21840_1_V1.UC-7 | + | 171 | 227 | **RNA**→ RpsM (COG0099)rpsM (PRK05179)→ |
| env-308 | SRS018313_C7804088 | + | 115 | 171 | **RNA**→ |
| env-309 | DLM023_scaffold15800_4 | + | 58 | 114 | **RNA**→ RpsM (COG0099)rpsM (PRK05179)→ |
| env-310 | NLF008_scaffold4177_4 | - | 264 | 208 | **RNA**→ rps13 (CHL00137)rpsM (PRK05179)→ |
| env-311 | NLM031_scaffold8043_12 | - | 1012 | 956 | **RNA**→ RpsM (COG0099)rpsM (PRK05179)→ |
| env-312 | NOM009_scaffold9194_1 | - | 1168 | 1112 | **RNA**→ RpsM (COG0099)rpsM (PRK05179)→ |
| env-313 | NOM005_scaffold12219_1 | - | 576 | 520 | **RNA**→ RpsM (COG0099)rpsM (PRK05179)→ |
| env-314 | NOM026_scaffold20027_3 | - | 755 | 699 | **RNA**→ RpsM (COG0099)rpsM (PRK05179)→ |
| env-315 | ADJT01012175.1 | - | 11015 | 10959 | **RNA**→ RpsM (COG0099)rpsM (PRK05179)→ |
| env-316 | SRS050422_LANL_scaffold_5427 | + | 241 | 297 | **RNA**→ RpsM (COG0099)rpsM (PRK05179)→ |
| env-317 | SRS015782_C3373209 | + | 501 | 557 | **RNA**→ |
| env-318 | SRS049712_C15436136 | + | 274 | 330 | **RNA**→ |
| Bwe-1-1 | NZ_AUJF01000003.1 | + | 23323 | 23379 | **RNA**→ RpsM (COG0099)rpsM (PRK05179)→ |
| env-319 | NLM017_scaffold23522_6 | + | 25 | 81 | **RNA**→ RpsM (COG0099)rpsM (PRK05179)→ |
| env-320 | DOF008_scaffold4797_1 | - | 4686 | 4630 | **RNA**→ RpsM (COG0099)rpsM (PRK05179)→ |
| env-321 | scaffold37948_1_O2.UC-19 | - | 326 | 270 | **RNA**→ rps13 (CHL00137)rpsM (PRK05179)→ |
| env-322 | SRS058770_C1599846 | - | 411 | 355 | **RNA**→ RpsM (COG0099)rpsM (PRK05179)→ |
| env-323 | DOF010_scaffold104_3 | - | 22883 | 22827 | **RNA**→ RpsM (COG0099)rpsM (PRK05179)→ |
| Rsp-2-1 | NZ_ACII02000003.1 | - | 3053178 | 3053122 | **RNA**→ RpsM (COG0099)rpsM (PRK05179)→ |
| env-324 | SRS016495_C1820935 | - | 194 | 138 | **RNA**→ hypo→ |
| env-325 | NOM028_scaffold3211_4 | - | 15906 | 15850 | **RNA**→ RpsM (COG0099)rpsM (PRK05179)→ |
| env-326 | scaffold437_1_V1.CD-11 | - | 717 | 661 | **RNA**→ RpsM (COG0099)rpsM (PRK05179)→ |
| env-327 | CAM_READ_0099454135 | + | 66 | 121 | **RNA**→ ←hypo |
| env-328 | SRS014923_WUGC_scaffold_34760 | + | 156 | 211 | **RNA**→ rps13 (CHL00137)rpsM (PRK05179)→ |
| env-329 | longitudinal_439_8 | + | 13623 | 13679 | **RNA**→ RpsM (COG0099)rpsM (PRK05179)→ PRK05309 (PRK05309)RpsK (COG0100)→ RpsD (COG0522)rpsD (PRK05327)→ RpoA (COG0202)PRK05182 (PRK05182)→ |
| env-330 | longitudinal_440_169 | - | 13322 | 13266 | **RNA**→ RpsM (COG0099)rpsM (PRK05179)→ PRK05309 (PRK05309)RpsK (COG0100)→ RpsD (COG0522)rpsD (PRK05327)→ RpoA (COG0202)PRK05182 (PRK05182)→ |
| env-331 | NLM008_scaffold23270_1 | - | 432 | 376 | **RNA**→ RpsM (COG0099)rpsM (PRK05179)→ |
| env-332 | NLM027_scaffold39514_1 | - | 666 | 610 | **RNA**→ RpsM (COG0099)rpsM (PRK05179)→ hypo→ |
| env-333 | longitudinal_20_244 | - | 8699 | 8643 | **RNA**→ RpsM (COG0099)rpsM (PRK05179)→ PRK05309 (PRK05309)RpsK (COG0100)→ RpsD (COG0522)rpsD (PRK05327)→ RpoA (COG0202)PRK05182 (PRK05182)→ |
| env-334 | longitudinal_53_558 | + | 42312 | 42368 | **RNA**→ RpsM (COG0099)rpsM (PRK05179)→ PRK05309 (PRK05309)RpsK (COG0100)→ RpsD (COG0522)rpsD (PRK05327)→ RpoA (COG0202)PRK05182 (PRK05182)→ |
| env-335 | ADJT01008964.1 | + | 1183 | 1239 | **RNA**→ RpsM (COG0099)rpsM (PRK05179)→ |
| env-336 | SRS015217_WUGC_scaffold_18288 | + | 246 | 302 | **RNA**→ RpsM (COG0099)rpsM (PRK05179)→ |
| env-337 | 4491417.3_NODE_73162 | - | 21524 | 21468 | **RNA**→ RpsM (COG0099)rpsM (PRK05179)→ PRK05309 (PRK05309)RpsK (COG0100)→ RpsD (COG0522)rpsD (PRK05327)→ PRK05182 (PRK05182)RNA_pol_L (pfam01193)→ RpoA (COG0202)PRK05182 (PRK05182)→ |
| env-338 | DOM026_scaffold12352_13 | - | 3568 | 3512 | **RNA**→ RpsM (COG0099)rpsM (PRK05179)→ PRK05309 (PRK05309)RpsK (COG0100)→ RpsD (COG0522)rpsD (PRK05327)→ RpoA (COG0202)PRK05182 (PRK05182)→ |
| env-339 | longitudinal_122_170 | + | 39841 | 39897 | **RNA**→ RpsM (COG0099)rpsM (PRK05179)→ PRK05309 (PRK05309)RpsK (COG0100)→ RpsD (COG0522)rpsD (PRK05327)→ RpoA (COG0202)PRK05182 (PRK05182)→ |
| env-340 | SRS014235_C3388785 | + | 157 | 213 | **RNA**→ RpsM (COG0099)rpsM (PRK05179)→ |
| env-341 | SRS014923_WUGC_scaffold_42123 | + | 33 | 89 | **RNA**→ RpsM (COG0099)rpsM (PRK05179)→ |
| env-342 | scaffold34343_3_MH0002 | + | 128 | 184 | **RNA**→ RpsM (COG0099)rpsM (PRK05179)→ |
| Bsc-1-1 | NZ_JNKJ01000057.1 | + | 1161 | 1217 | **RNA**→ RpsM (COG0099)rpsM (PRK05179)→ PRK05309 (PRK05309)RpsK (COG0100)→ RpsD (COG0522)rpsD (PRK05327)→ RpoA (COG0202)PRK05182 (PRK05182)→ |

| | | | | | |
|---|---|---|---|---|---|
| Rsp-3-1 | NZ_HE978651.1 | - | 2529298 | 2529242 | **RNA**→ RpsM (COG0099)rpsM (PRK05179)→ PRK05309 (PRK05309)RpsK (COG0100)→<br>RpsD (COG0522)rpsD (PRK05327)→ RpoA (COG0202)PRK05182 (PRK05182)→ |
| env-343 | AUXO012074612.1 | + | 242 | 300 | **RNA**→ RpsM (COG0099)rpsM (PRK05179)→ PRK05309 (PRK05309)→ |
| env-344 | DLM009_scaffold871_6 | - | 3189 | 3114 | **RNA**→ RpsM (COG0099)rpsM (PRK05179)→ PRK05309 (PRK05309)RpsK (COG0100)→<br>RpsD (COG0522)rpsD (PRK05327)→ RpoA (COG0202)PRK05182 (PRK05182)→<br>rplQ (PRK05591)Ribosomal_L17 (pfam01196)→ |
| env-345 | NLM027_scaffold5861_18 | - | 15285 | 15210 | **RNA**→ RpsM (COG0099)rpsM (PRK05179)→ PRK05309 (PRK05309)RpsK (COG0100)→<br>RpsD (COG0522)rpsD (PRK05327)→ RpoA (COG0202)PRK05182 (PRK05182)→<br>rplQ (PRK05591)Ribosomal_L17 (pfam01196)→ PRK13337 (PRK13337)LCB5 (COG1597)→<br>MntP (COG1971)spore_YtaF (TIGR02840)→ |
| env-346 | CAM_READ_0099493925 | + | 211 | 286 | **RNA**→ rps13 (CHL00137)rpsM (PRK05179)→ |
| env-347 | 4444164.3.49013 | + | 211 | 286 | **RNA**→ rps13 (CHL00137)rpsM (PRK05179)→ |
| env-348 | scaffold12778_1_V1.CD-12 | + | 467 | 542 | **RNA**→ |
| env-349 | SRS018313_C7798604 | + | 179 | 254 | **RNA**→ |
| env-350 | DLF012_scaffold123_4 | + | 22132 | 22207 | **RNA**→ RpsM (COG0099)rpsM (PRK05179)→ PRK05309 (PRK05309)RpsK (COG0100)→<br>RpsD (COG0522)rpsD (PRK05327)→ RpoA (COG0202)PRK05182 (PRK05182)→<br>rplQ (PRK05591)Ribosomal_L17 (pfam01196)→ PRK13337 (PRK13337)LCB5 (COG1597)→<br>MntP (COG1971)spore_YtaF (TIGR02840)→ |
| Lba-8-1 | NZ_GL890549.1 | - | 179387 | 179312 | **RNA**→ RpsM (COG0099)rpsM (PRK05179)→ PRK05309 (PRK05309)RpsK (COG0100)→<br>RpsD (COG0522)rpsD (PRK05327)→ RpoA (COG0202)PRK05182 (PRK05182)→<br>rplQ (PRK05591)Ribosomal_L17 (pfam01196)→ PRK13337 (PRK13337)LCB5 (COG1597)→<br>MntP (COG1971)spore_YtaF (TIGR02840)→ |
| Bha-1-1 | NZ_GG698590.1 | - | 537224 | 537149 | **RNA**→ RpsM (COG0099)rpsM (PRK05179)→ PRK05309 (PRK05309)RpsK (COG0100)→<br>RpsD (COG0522)rpsD (PRK05327)→ RpoA (COG0202)PRK05182 (PRK05182)→<br>rplQ (PRK05591)Ribosomal_L17 (pfam01196)→ PRK13337 (PRK13337)LCB5 (COG1597)→<br>MntP (COG1971)spore_YtaF (TIGR02840)→ |
| env-351 | CAM_READ_0100338075 | + | 116 | 191 | **RNA**→ rps13 (CHL00137)rpsM (PRK05179)→ |
| env-352 | 4448819.3_Contig30791 | + | 5 | 75 | **RNA**→ rpsM (PRK05179)→ |
| env-353 | longitudinal_51_247 | - | 34515 | 34458 | **RNA**→ hypo→ RpsM (COG0099)rpsM (PRK05179)→ |
| Rga-1-1 | NZ_AUDP01000014.1 | - | 27539 | 27482 | **RNA**→ RpsM (COG0099)rpsM (PRK05179)→ |
| env-354 | AUXO015947882.1 | - | 158 | 103 | **RNA**→ rps13 (CHL00137)rpsM (PRK05179)→ |
| env-355 | JGI994J12354_10010207 | + | 5480 | 5533 | **RNA**→ RpsM (COG0099)rpsM (PRK05179)→ |
| env-356 | IMNBL1_c0108913 | - | 287 | 234 | **RNA**→ rps13 (CHL00137)rpsM (PRK05179)→ |
| env-357 | JGI994J12354_10000026 | + | 7090 | 7145 | **RNA**→ RpsM (COG0099)rpsM (PRK05179)→ |
| env-358 | 2227591299 | - | 3740 | 3685 | **RNA**→ RpsM (COG0099)rpsM (PRK05179)→ |
| env-359 | IMNBL1_c0000019 | + | 10768 | 10823 | **RNA**→ RpsM (COG0099)rpsM (PRK05179)→ |
| env-360 | JGI994J12354_10094008 | + | 12 | 69 | **RNA**→ ←hypo |
| env-361 | IMNBL1_c0036661 | + | 988 | 1043 | **RNA**→ RpsM (COG0099)rpsM (PRK05179)→ PRK05309 (PRK05309)RpsK (COG0100)→ |
| env-362 | JGI994J12354_10002700 | - | 15440 | 15385 | **RNA**→ ←hypo |
| env-363 | IMNBL1_c0023149 | + | 2251 | 2307 | **RNA**→ |
| env-364 | 2227528282 | + | 222 | 278 | **RNA**→ RpsM (COG0099)rpsM (PRK05179)→ |
| env-365 | JGI994J12354_10567539 | - | 328 | 272 | **RNA**→ hypo→ rps13 (CHL00137)rpsM (PRK05179)→ |
| env-366 | 2227243856 | + | 905 | 961 | **RNA**→ RpsM (COG0099)rpsM (PRK05179)→ |
| env-367 | scaffold568_3_MH0038 | - | 330 | 279 | **RNA**→ RpsM (COG0099)rpsM (PRK05179)→ |
| env-368 | scaffold119982_1_MH0056 | - | 332 | 281 | **RNA**→ RpsM (COG0099)rpsM (PRK05179)→ |
| env-369 | scaffold11215_5_MH0063 | + | 300 | 351 | **RNA**→ RpsM (COG0099)rpsM (PRK05179)→ |
| env-370 | scaffold3544_6_MH0074 | + | 523 | 574 | **RNA**→ RpsM (COG0099)rpsM (PRK05179)→ |
| env-371 | scaffold17454_3_O2.UC-22 | + | 3387 | 3438 | **RNA**→ rps13 (CHL00137)rpsM (PRK05179)→ |
| env-372 | scaffold71124_1_V1.CD-8 | - | 108 | 57 | **RNA**→ |
| env-373 | scaffold75223_1_V1.UC-9 | - | 362 | 311 | **RNA**→ RpsM (COG0099)rpsM (PRK05179)→ |
| env-374 | 4491417.3_NODE_67088 | - | 675 | 624 | **RNA**→ RpsM (COG0099)rpsM (PRK05179)→ |
| env-375 | DLM027_scaffold24464_1 | - | 728 | 677 | **RNA**→ RpsM (COG0099)rpsM (PRK05179)→ PRK05309 (PRK05309)RpsK (COG0100)→ |
| env-376 | SRS016095_WUGC_scaffold_30576 | + | 143 | 194 | **RNA**→ RpsM (COG0099)rpsM (PRK05179)→ |
| env-377 | SRS012273_C3777396 | + | 2744 | 2795 | **RNA**→ |

| | | | | | |
|---|---|---|---|---|---|
| env-378 | SRS014313_C2660173 | + | 568 | 619 | **RNA**→ |
| env-379 | SRS014979_C2995935 | + | 209 | 260 | **RNA**→ |
| env-380 | SRS015217_C2217440 | + | 360 | 411 | **RNA**→ |
| env-381 | SRS017433_C1676010 | - | 110 | 59 | **RNA**→ hypo→ |
| env-382 | SRS051882_C2517682 | - | 110 | 59 | **RNA**→ |
| env-383 | SRS018817_C2062093 | + | 582 | 633 | **RNA**→ |
| env-384 | SRS050422_LANL_scaffold_21144 | - | 743 | 692 | **RNA**→ RpsM (COG0099)rpsM (PRK05179)→ PRK05309 (PRK05309)RpsK (COG0100)→ |
| env-385 | scaffold1165_1_V1.CD-4 | - | 107 | 56 | **RNA**→ |
| env-386 | DLM009_scaffold18031_8 | + | 36254 | 36305 | **RNA**→ RpsM (COG0099)rpsM (PRK05179)→ PRK05309 (PRK05309)RpsK (COG0100)→ RpsD (COG0522)rpsD (PRK05327)→ RpoA (COG0202)PRK05182 (PRK05182)→ |
| env-387 | DLM028_scaffold14333_2 | - | 459 | 408 | **RNA**→ RpsM (COG0099)rpsM (PRK05179)→ |
| env-388 | NLM027_scaffold1108_2 | - | 1341 | 1290 | **RNA**→ RpsM (COG0099)rpsM (PRK05179)→ PRK05309 (PRK05309)RpsK (COG0100)→ S4 (smart00363)RpsD (COG0522)→ |
| env-389 | SRS015782_C3477872 | + | 2144 | 2195 | **RNA**→ |
| env-390 | DOM013_scaffold7638_5 | + | 299 | 350 | **RNA**→ RpsM (COG0099)rpsM (PRK05179)→ PRK05309 (PRK05309)RpsK (COG0100)→ RpsD (COG0522)rpsD (PRK05327)→ PRK05182 (PRK05182)RNA_pol_L (pfam01193)→ |
| env-391 | scaffold52941_2_MH0011 | - | 294 | 243 | **RNA**→ rps13 (CHL00137)rpsM (PRK05179)→ |
| env-392 | scaffold841_6_MH0037 | + | 2857 | 2908 | **RNA**→ |
| env-393 | scaffold93665_1_MH0043 | - | 356 | 305 | **RNA**→ RpsM (COG0099)rpsM (PRK05179)→ |
| env-394 | scaffold44089_3_MH0060 | + | 396 | 447 | **RNA**→ RpsM (COG0099)rpsM (PRK05179)→ |
| env-395 | scaffold9918_1_MH0068 | - | 106 | 55 | **RNA**→ |
| env-396 | scaffold61996_1_MH0082 | - | 326 | 275 | **RNA**→ RpsM (COG0099)rpsM (PRK05179)→ |
| env-397 | scaffold49998_1_V1.CD-14 | - | 295 | 244 | **RNA**→ rps13 (CHL00137)rpsM (PRK05179)→ |
| env-398 | scaffold12031_1_V1.UC-15 | - | 266 | 215 | **RNA**→ rps13 (CHL00137)rpsM (PRK05179)→ |
| env-399 | 4491410.3_NODE_7014 | + | 1319 | 1370 | **RNA**→ RpsM (COG0099)rpsM (PRK05179)→ PRK05309 (PRK05309)RpsK (COG0100)→ RpsD (COG0522)rpsD (PRK05327)→ RpoA (COG0202)PRK05182 (PRK05182)→ |
| env-400 | 4491413.3_NODE_68514 | - | 12266 | 12215 | **RNA**→ RpsM (COG0099)rpsM (PRK05179)→ PRK05309 (PRK05309)RpsK (COG0100)→ |
| env-401 | DOF008_scaffold23304_2 | - | 4645 | 4594 | **RNA**→ RpsM (COG0099)rpsM (PRK05179)→ PRK05309 (PRK05309)RpsK (COG0100)→ RpsD (COG0522)rpsD (PRK05327)→ RpoA (COG0202)PRK05182 (PRK05182)→ |
| env-402 | DOM018_scaffold30267_2 | - | 9330 | 9279 | **RNA**→ RpsM (COG0099)rpsM (PRK05179)→ PRK05309 (PRK05309)RpsK (COG0100)→ RpsD (COG0522)rpsD (PRK05327)→ RpoA (COG0202)PRK05182 (PRK05182)→ |
| env-403 | NLM032_scaffold1492_2 | - | 4705 | 4654 | **RNA**→ RpsM (COG0099)rpsM (PRK05179)→ PRK05309 (PRK05309)RpsK (COG0100)→ RpsD (COG0522)rpsD (PRK05327)→ RpoA (COG0202)PRK05182 (PRK05182)→ |
| env-404 | ADJS01016619.1 | + | 2886 | 2937 | **RNA**→ RpsM (COG0099)rpsM (PRK05179)→ PRK05309 (PRK05309)RpsK (COG0100)→ RpsD (COG0522)rpsD (PRK05327)→ RpoA (COG0202)PRK05182 (PRK05182)→ rplQ (PRK05591)Ribosomal_L17 (pfam01196)→ |
| env-405 | DLF014_scaffold5097_1 | - | 28650 | 28599 | **RNA**→ RpsM (COG0099)rpsM (PRK05179)→ |
| env-406 | DLM016_scaffold2418_6 | - | 2957 | 2906 | **RNA**→ RpsM (COG0099)rpsM (PRK05179)→ PRK05309 (PRK05309)RpsK (COG0100)→ RpsD (COG0522)rpsD (PRK05327)→ RpoA (COG0202)PRK05182 (PRK05182)→ |
| env-407 | DOM026_scaffold10557_10 | + | 15236 | 15287 | **RNA**→ RpsM (COG0099)rpsM (PRK05179)→ PRK05309 (PRK05309)RpsK (COG0100)→ RpsD (COG0522)rpsD (PRK05327)→ RpoA (COG0202)PRK05182 (PRK05182)→ |
| env-408 | NOF012_scaffold6209_1 | + | 350 | 401 | **RNA**→ rps13 (CHL00137)rpsM (PRK05179)→ |
| env-409 | SRS014235_WUGC_scaffold_106228 | - | 130 | 79 | **RNA**→ |
| env-410 | SRS022071_C2808073 | - | 121 | 70 | **RNA**→ hypo→ |
| env-411 | SRS064276_LANL_scaffold_18813 | - | 435 | 384 | **RNA**→ RpsM (COG0099)rpsM (PRK05179)→ |
| env-412 | 4491401.3_NODE_45574 | + | 56 | 107 | **RNA**→ rps13 (CHL00137)rpsM (PRK05179)→ RpsM (COG0099)rpsM (PRK05179)→ PRK05309 (PRK05309)RpsK (COG0100)→ |
| env-413 | DLM018_scaffold30918_1 | + | 2 | 53 | **RNA**→ RpsM (COG0099)rpsM (PRK05179)→ PRK05309 (PRK05309)RpsK (COG0100)→ |
| env-414 | DOF003_scaffold5613_4 | + | 314 | 365 | **RNA**→ RpsM (COG0099)rpsM (PRK05179)→ PRK05309 (PRK05309)RpsK (COG0100)→ hypo→ |
| env-415 | DOF012_scaffold4673_1 | + | 384 | 435 | **RNA**→ RpsM (COG0099)rpsM (PRK05179)→ PRK05309 (PRK05309)RpsK (COG0100)→ RpsD (COG0522)rpsD (PRK05327)→ RpoA (COG0202)PRK05182 (PRK05182)→ |
| env-416 | NOF002_scaffold2669_5 | - | 615 | 564 | **RNA**→ RpsM (COG0099)rpsM (PRK05179)→ PRK05309 (PRK05309)RpsK (COG0100)→ |
| env-417 | NOM009_scaffold2007_2 | - | 2031 | 1980 | **RNA**→ RpsM (COG0099)rpsM (PRK05179)→ PRK05309 (PRK05309)RpsK (COG0100)→ RpsD (COG0522)rpsD (PRK05327)→ RNAP_alpha_NTD (cd06928)PRK05182 (PRK05182)→ |
| env-418 | NOM001_scaffold9282_1 | - | 151 | 100 | **RNA**→ rps13 (CHL00137)rpsM (PRK05179)→ |

| | | | | | |
|---|---|---|---|---|---|
| env-419 | SRS052697_LANL_scaffold_51469 | + | 80 | 131 | **RNA**→ |
| env-420 | scaffold85473_2_MH0025 | + | 479 | 530 | **RNA**→ |
| env-421 | scaffold52584_3_MH0045 | + | 1009 | 1060 | **RNA**→ |
| env-422 | scaffold37252_4_MH0062 | + | 528 | 579 | **RNA**→ |
| env-423 | scaffold96444_3_MH0069 | + | 741 | 792 | **RNA**→ |
| env-424 | scaffold2364_1_MH0071 | - | 108 | 57 | **RNA**→ |
| env-425 | scaffold88060_1_MH0083 | - | 108 | 57 | **RNA**→ |
| env-426 | scaffold91363_1_V1.UC-10 | + | 544 | 595 | **RNA**→ |
| env-427 | scaffold83915_1_V1.UC-8 | - | 108 | 57 | **RNA**→ hypo→ |
| env-428 | SRS020233_C3818898 | + | 153 | 205 | **RNA**→ |
| env-429 | SRS024132_C3632946 | - | 167 | 115 | **RNA**→ hypo→ |
| env-430 | SRS050422_LANL_scaffold_88482 | + | 593 | 645 | **RNA**→ |
| env-431 | SRS064276_C2411397 | - | 185 | 133 | **RNA**→ hypo→ |
| env-432 | UnmappedStool_Broad_scaffold_502667 | - | 347 | 295 | **RNA**→ rps13 (CHL00137)rpsM (PRK05179)→ |
| env-433 | AUXO010373404.1 | + | 958 | 1008 | **RNA**→ rps13 (CHL00137)rpsM (PRK05179)→ |
| env-434 | UnmappedStool_Broad_C251835313 | + | 227 | 284 | **RNA**→ |
| Bhy-1-1 | NZ_GG657686.1 | + | 284181 | 284238 | **RNA**→ RpsM (COG0099)rpsM (PRK05179)→ |
| env-435 | AUXO015141686.1 | - | 737 | 687 | **RNA**→ ←hypo |
| env-436 | AUXO012457309.1 | + | 505 | 561 | **RNA**→ RpsM (COG0099)rpsM (PRK05179)→ PRK05309 (PRK05309)RpsK (COG0100)→ |
| env-437 | AUXO017593349.1 | - | 353 | 299 | **RNA**→ RpsM (COG0099)rpsM (PRK05179)→ |
| env-438 | AUXO011871861.1 | + | 122 | 174 | **RNA**→ rps13 (CHL00137)rpsM (PRK05179)→ |
| Bsp-7-1 | NZ_AUIX01000015.1 | + | 102241 | 102293 | **RNA**→ RpsM (COG0099)rpsM (PRK05179)→ PRK05309 (PRK05309)RpsK (COG0100)→ |
| Bsp-9-1 | NZ_AUJO01000032.1 | + | 17808 | 17860 | **RNA**→ RpsM (COG0099)rpsM (PRK05179)→ PRK05309 (PRK05309)RpsK (COG0100)→ |
| Bsp-5-1 | NZ_KE384159.1 | - | 74816 | 74764 | **RNA**→ RpsM (COG0099)rpsM (PRK05179)→ PRK05309 (PRK05309)RpsK (COG0100)→ |
| Bsp-4-1 | NZ_KE384144.1 | + | 253197 | 253249 | **RNA**→ RpsM (COG0099)rpsM (PRK05179)→ PRK05309 (PRK05309)RpsK (COG0100)→ |
| Bsp-2-1 | NZ_ATVV01000009.1 | - | 41473 | 41421 | **RNA**→ RpsM (COG0099)rpsM (PRK05179)→ PRK05309 (PRK05309)RpsK (COG0100)→ |
| Bsp-8-1 | NZ_KE384200.1 | + | 31158 | 31210 | **RNA**→ RpsM (COG0099)rpsM (PRK05179)→ PRK05309 (PRK05309)RpsK (COG0100)→ |
| Bsp-3-1 | NZ_JNLQ01000002.1 | + | 492386 | 492438 | **RNA**→ RpsM (COG0099)rpsM (PRK05179)→ PRK05309 (PRK05309)RpsK (COG0100)→ |
| Bpr-2-1 | NZ_KB892647.1 | + | 780 | 851 | **RNA**→ RpsM (COG0099)rpsM (PRK05179)→ PRK05309 (PRK05309)RpsK (COG0100)→<br>RpsD (COG0522)rpsD (PRK05327)→ RpoA (COG0202)PRK05182 (PRK05182)→<br>rplQ (PRK05591)Ribosomal_L17 (pfam01196)→ |

#### 16.3 Conserved domains

Conserved domains found in protein-coding genes listed in Section 16.2 are shown below, with the first sentence in their description from the Conserved Domain Database (if any). Conserved domains associated with more than one S4-Clostridia RNA are assigned a color, while others are shown in gray. The number in parentheses after the colored domain name is the number of occurrences in Section 16.2.

cd00207 (5) 2Fe-2S iron-sulfur cluster binding domain.
cd06928 (3) N-terminal domain of the Alpha subunit of Bacterial RNA polymerase.
CHL00041 (2) ribosomal protein S11
CHL00137 (46) ribosomal protein S13; Validated
COG0099 (345) Ribosomal protein S13 [Translation, ribosomal structure and biogenesis]
COG0100 (189) Ribosomal protein S11 [Translation, ribosomal structure and biogenesis]
COG0202 (99) DNA-directed RNA polymerase, alpha subunit/40 kD subunit [Transcription]
COG0493 (5) NADPH-dependent glutamate synthase beta chain or related oxidoreductase [Amino acid transport and metabolism, General function prediction only]
COG0522 (132) Ribosomal protein S4 or related protein [Translation, ribosomal structure and biogenesis]
COG1146 (5) NAD-dependent dihydropyrimidine dehydrogenase, PreA subunit [Nucleotide transport and metabolism]
COG1597 (4) Diacylglycerol kinase family enzyme [Lipid transport and metabolism, General function prediction only]
COG1894 (5) NADH:ubiquinone oxidoreductase, NADH-binding 51 kD subunit (chain F) [Energy production and conversion]
COG1971 (4) Putative Mn2+ efflux pump MntP [Inorganic ion transport and metabolism]
pfam00163 (1) Ribosomal protein S4/S9 N-terminal domain.
pfam01193 (2) RNA polymerase Rpb3/Rpb11 dimerisation domain.
pfam01196 (44) Ribosomal protein L17.
pfam02256 (5) Iron hydrogenase small subunit.

pfam02906 (5) Iron only hydrogenase large subunit, C-terminal domain.
pfam10588 (5) NADH-ubiquinone oxidoreductase-G iron-sulfur binding region.
pfam10589 (5) NADH-ubiquinone oxidoreductase-F iron-sulfur binding region.
pfam12838 (5) 4Fe-4S dicluster domain.
pfam13450 (5) NAD(P)-binding Rossmann-like domain.
pfam13510 (5) 2Fe-2S iron-sulfur cluster binding domain.
PRK05179 (392) 30S ribosomal protein S13; Validated
PRK05182 (104) DNA-directed RNA polymerase subunit alpha; Provisional
PRK05309 (192) 30S ribosomal protein S11; Validated
PRK05327 (130) 30S ribosomal protein S4; Validated
PRK05591 (44) 50S ribosomal protein L17; Validated
PRK13337 (4) putative lipid kinase; Reviewed
smart00363 (3) S4 RNA-binding domain.
smart00902 (5) Iron hydrogenase small subunit. [4Fe-4S] iron-sulphur clusters. In FeFe hydrogenases, as in NiFe hydrogenases, the set of iron-sulphur clusters is dispersed regularly between the dinuclear Fe-Fe centre and the molecular surface. These clusters are distant by about 1.2 nm from each other but the [4Fe-4S] cluster closest to the dinuclear centre is covalently bound to one of the iron atoms though a thiolate bridging ligand. The moiety including the dinuclear centre, the thiolate bridging ligand, and the proximal [4Fe-4S]
TIGR02840 (4) putative sporulation protein YtaF. [Cellular processes, Sporulation and germination]
TIGR03631 (6) ribosomal protein uS13, bacterial form. [Protein synthesis, Ribosomal proteins: synthesis and modification]

#### 16.4 Multiple-sequence alignment

Each S4-Clostridia RNA is denoted by an abbreviation (like "Eco-1-1") that refers to a taxonomy given in Section 16.1. The alignment may include sequences containing the RNA motif, as well as flanking sequence. The S4-Clostridia RNA itself is denoted by the line underneath marked 5′ and 3′ on either end. Nucleotides in flanking sequences (i.e., not part of the motif) are written in gray letters. Stems of predicted rho-independent transcription terminators, if any, are shaded yellow. (Note: terminator predictions have not been analyzed manually, and many are likely to be false positives. Terminator predictions are those of the RNIE software.) Nucleotides predicted to function (as DNA) as transcription-factor binding sites, if any, are shaded in green. (But note: these predictions are manually annotated, so they might be under-predicted.) Annotated start codons, if any, are shaded green. (Note: start codons are frequently misannotated, especially in environmental samples.) Nucleotides proposed to basepair as part of the consensus structure are shaded in color when they comprise Watson-Crick or G-U pairs. Otherwise they are shaded gray. Conserved stems are also indicated at the bottom of the alignment by angle brackets, where matching $<$ and $>$ denote base-paired columns. Below these angle brackets, the symbol "2" denotes base pairs exhibiting covariation according to the statistically well-founded R-scape method. "1" denotes base pairs exhibiting covariation according to R2R's simplistic method. "0" denotes base pairs that are not observed to mutate and "?" denotes base pairs that have a significant frequency of non-canonical nucleotides for Watson-Crick or G-U pairs ($> 5\%$). Below these base pair annotation is the consensus sequence: "R" = "A" or "G", "Y" = "C" or "U", red nucleotides: nucleotide identity conserved more than 97% of the time, black nucleotides: 90%, gray nucleotides: 75%, red circle (●): nucleotide is present 97% of the time, black circle (●): 90%, gray circle (●): 75%, white circle (○): 50%. All percentages of sequences just described (e.g. 97% conserved) assume that sequences have been weighted by the GSC algorithm implemented by the Infernal software package.

Duplicate sequences: the following putative homologs are not shown in the alignment because their sequences are identical to a homolog already shown: Bha-1-1, Bsp-4-1, Bsp-8-1, Cba-2-1, Cne-1-1, Csc-1-1, Csp-1-1, Csp-2-1, Dfo-1-1, Dfo-2-1, Dlo-2-1, Lba-1-1, Lba-3-1, Lba-5-1, Lba-8-1, Lba-9-1, Rgn-2-1, Rgn-3-1, Rob-1-1, Rsp-2-1, Rto-1-1, env-100, env-104, env-105, env-107, env-108, env-109, env-110, env-111, env-112, env-113, env-114, env-115, env-121, env-122, env-125, env-126, env-131, env-132, env-133, env-134, env-135, env-141, env-142, env-143, env-144, env-145, env-146, env-147, env-148, env-15, env-157, env-158, env-166, env-173, env-174, env-180, env-193, env-196, env-197, env-198, env-199, env-203, env-204, env-205, env-206, env-207, env-208, env-209, env-210, env-213, env-214, env-215, env-224, env-225, env-233, env-247, env-248, env-249, env-250, env-251, env-253, env-257, env-26, env-269, env-27, env-270, env-274, env-297, env-31, env-312, env-316, env-330, env-333, env-334, env-345, env-347, env-376, env-378, env-379, env-380, env-381, env-382, env-40, env-400, env-401, env-402, env-403, env-404, env-406, env-407, env-415, env-416, env-417, env-421, env-422, env-423, env-424, env-425, env-426, env-427, env-431, env-44, env-56, env-57, env-58, env-61, env-65, env-69, env-70, env-71, env-72, env-73, env-74, env-84, env-85, env-86, env-98, env-99

alignment positions 1 · · · 180

```
env-1   UGAGAAUCCAAAGCAUAAACAGCGUCAGGGUUAAUGUACCGUCAUUCAUUAACAGACUGAACUUUUAUGCGGCAAGAAAAUAUUAUAAAGUGAGAUCUGACUGGCUGCGUCAGGAAUCCUCUAUAAUAUUUGCUUAAUCCCCGAAUCGAUCAUGAGGAGGGAAAGGCUGUGC..GGGA..
env-2   ..AGAAGUUAUACCGUCAAGAAAUAUAUGCUAUUAAAUAUAUGACAGAUAUAAUUAGAAUUUGUCGUGUGUUUACUUCUUGAUACCGAGAAAAGUUUCGCAGGAAACACGGUGGGAAAAAUGAAGACUGCCCACCUACAUCUGGAUAACCAGAUGCCGAUGUAGUAUCGGA...CAAAUC

        ....................................................................................................................................................................<<<<<<<<<<......
        .....................................................................................................................................................................222222.........
        ●●●●●●●●●●●●●●●●●●●●●●●●●●●●●●●●●●●●●●●●●●●●●●●●●●●●●●●●●●●●●●●●●●●●●●●●●●●●●●●●●●●●●●●●●●●●●●●●●●●●●●●●●●●●●●●●●●●●●●●●●●●●●●●●●●●●●●●●●●●●●●●●●●●●●YYGGAAAGGYYGY●○--●●●●○○
                                                                                                                                                                            5′
```

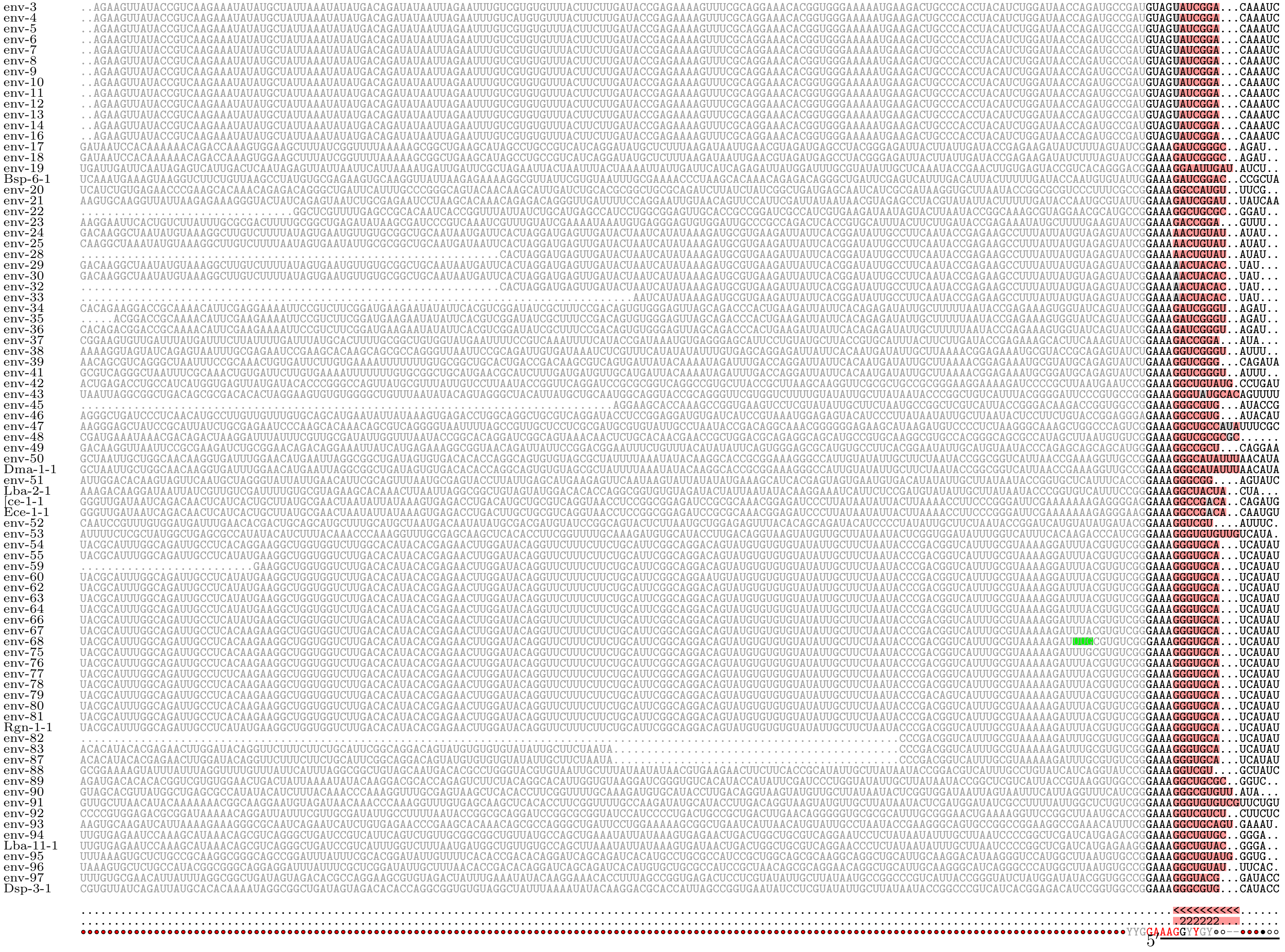
env-3
env-4
env-5
env-6
env-7
env-8
env-9
env-10
env-11
env-12
env-13
env-14
env-16
env-17
env-18
env-19
Bsp-6-1
env-20
env-21
env-22
env-23
env-24
env-25
env-28
env-29
env-30
env-32
env-33
env-34
env-35
env-36
env-37
env-38
env-39
env-41
env-42
env-43
env-45
env-46
env-47
env-48
env-49
env-50
Dma-1-1
env-51
Lba-2-1
[ce-1-1
Ece-1-1
env-52
env-53
env-54
env-55
env-59
env-60
env-62
env-63
env-64
env-66
env-67
env-68
env-75
env-76
env-77
env-78
env-79
env-80
env-81
Rgn-1-1
env-82
env-83
env-87
env-88
env-89
env-90
env-91
env-92
env-93
env-94
Lba-11-1
env-95
env-96
env-97
Dsp-3-1
5′

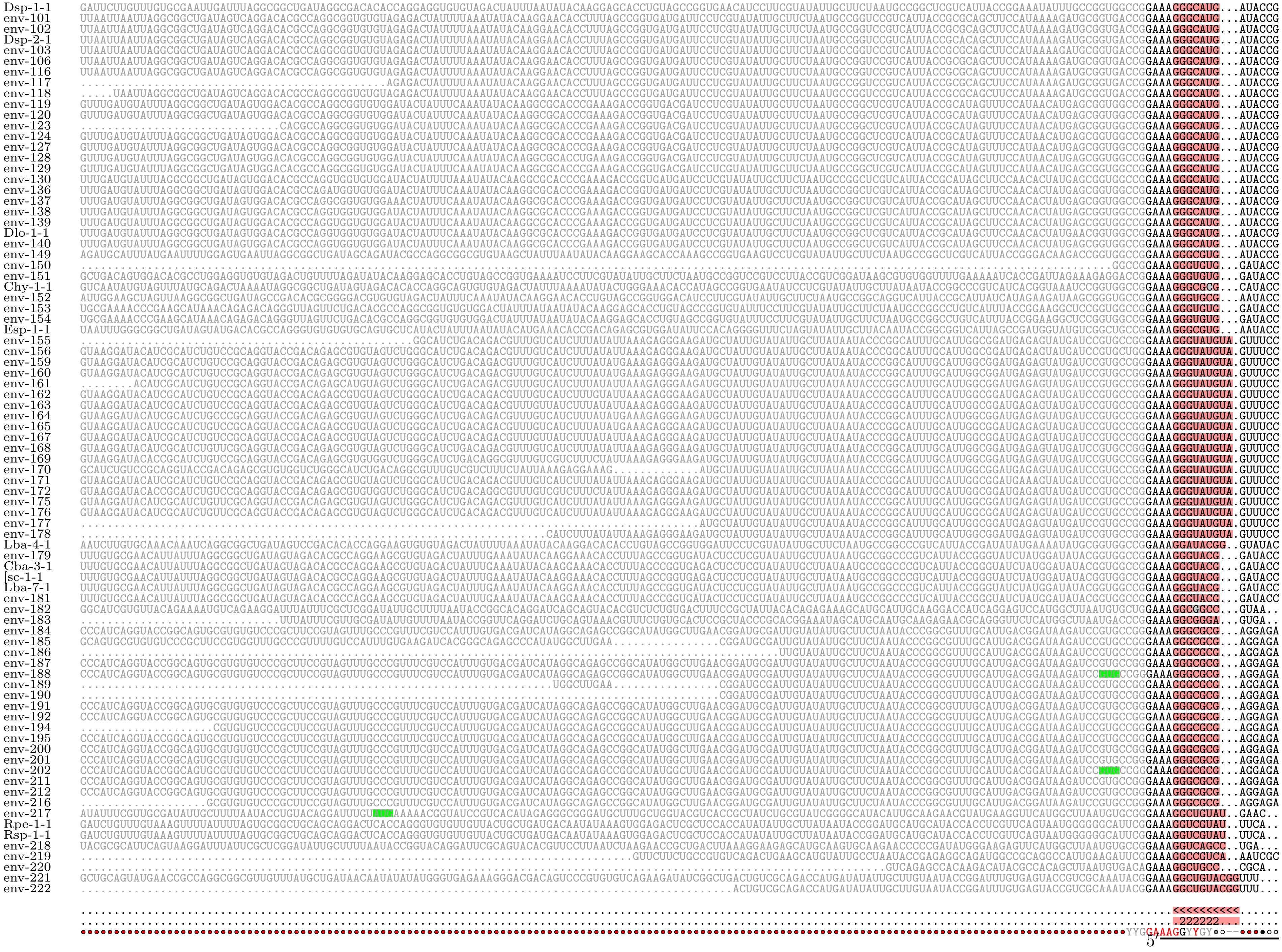
Dsp-1-1 GAAAGGGCAUG...AUACCG
env-101 GAAAGGGCAUG...AUACCG
env-102 GAAAGGGCAUG...AUACCG
Dsp-2-1 GAAAGGGCAUG...AUACCG
env-103 GAAAGGGCAUG...AUACCG
env-106 GAAAGGGCAUG...AUACCG
env-116 GAAAGGGCAUG...AUACCG
env-117 GAAAGGGCAUG...AUACCG
env-118 GAAAGGGCAUG...AUACCG
env-119 GAAAGGGCAUG...AUACCG
env-120 GAAAGGGCAUG...AUACCG
env-123 GAAAGGGCAUG...AUACCG
env-124 GAAAGGGCAUG...AUACCG
env-127 GAAAGGGCAUG...AUACCG
env-128 GAAAGGGCAUG...AUACCG
env-129 GAAAGGGCAUG...AUACCG
env-130 GAAAGGGCAUG...AUACCG
env-136 GAAAGGGCAUG...AUACCG
env-137 GAAAGGGCAUG...AUACCG
env-138 GAAAGGGCAUG...AUACCG
env-139 GAAAGGGCAUG...AUACCG
Dlo-1-1 GAAAGGGCAUG...AUACCG
env-140 GAAAGGGCAUG...AUACCG
env-149 GAAAGGGCGUG...AUACCG
env-150 GAAAGGGUGUG...GAUACC
env-151 GAAAGGGUGUG...GAUACC
Chy-1-1 GAAAGGGCGCG...CAUACC
env-152 GAAAGGGUGCG...AAUACC
env-153 GAAAGGGUGUG...GAUACC
env-154 GAAAGGGUGUG...GAUACC
Esp-1-1 GAAAGGGCGUG...AAUACC
env-155 GAAAGGGUAUGUA.GUUUCC
env-156 GAAAGGGUAUGUA.GUUUCC
env-159 GAAAGGGUAUGUA.GUUUCC
env-160 GAAAGGGUAUGUA.GUUUCC
env-161 GAAAGGGUAUGUA.GUUUCC
env-162 GAAAGGGUAUGUA.GUUUCC
env-163 GAAAGGGUAUGUA.GUUUCC
env-164 GAAAGGGUAUGUA.GUUUCC
env-165 GAAAGGGUAUGUA.GUUUCC
env-167 GAAAGGGUAUGUA.GUUUCC
env-168 GAAAGGGUAUGUA.GUUUCC
env-169 GAAAGGGUAUGUA.GUUUCC
env-170 GAAAGGGUAUGUA.GUUUCC
env-171 GAAAGGGUAUGUA.GUUUCC
env-172 GAAAGGGUAUGUA.GUUUCC
env-175 GAAAGGGUAUGUA.GUUUCC
env-176 GAAAGGGUAUGUA.GUUUCC
env-177 GAAAGGGUAUGUA.GUUUCC
env-178 GAAAGGGUAUGUA.GUUUCC
Lba-4-1 GAAAGGAUACGG..GUAUAC
env-179 GAAAGGGUACG...GAUACC
Cba-3-1 GAAAGGGUACG...GAUACC
[sc-1-1 GAAAGGGUACG...GAUACC
Lba-7-1 GAAAGGGUACG...GAUACC
env-181 GAAAGGGUACG...GAUACC
env-182 GAAAGGCGGCC...GUAA..
env-183 GAAAGGCGGGA...GUGA..
env-184 GAAAGGGCGCG...AGGAGA
env-185 GAAAGGGCGCG...AGGAGA
env-186 GAAAGGGCGCG...AGGAGA
env-187 GAAAGGGCGCG...AGGAGA
env-188 GAAAGGGCGCG...AGGAGA
env-189 GAAAGGGCGCG...AGGAGA
env-190 GAAAGGGCGCG...AGGAGA
env-191 GAAAGGGCGCG...AGGAGA
env-192 GAAAGGGCGCG...AGGAGA
env-194 GAAAGGGCGCG...AGGAGA
env-195 GAAAGGGCGCG...AGGAGA
env-200 GAAAGGGCGCG...AGGAGA
env-201 GAAAGGGCGCG...AGGAGA
env-202 GAAAGGGCGCG...AGGAGA
env-211 GAAAGGGCGCG...AGGAGA
env-212 GAAAGGGCGCG...AGGAGA
env-216 GAAAGGGCGCG...AGGAGA
env-217 GAAAGGCUGUAU..GAAC..
Rpe-1-1 GAAAGGUCGUAU..UUCA..
Rsp-1-1 GAAAGGUCGUAU..UUCA..
env-218 GAAAGGUCAGCC..UGA...
env-219 GAAAGGCCGUCA..AAUCGC
env-220 GAAAGGCUGCC...CGCA..
env-221 GAAAGGCUGUACGGUUU...
env-222 GAAAGGCUGUACGGUUU...
<<<<<<<<<<
.222222...
YYGGAAAGGYYGY
5′

```
env-223    GCUGCAGUAUGAACCGCCAGGCGGCGUUGUUUAUGCUGAUAACAAUAUAUAUGGGUGAGAAAGGGACGACAGUCCCGUGUGUCAGAAGAUAUCGGCUGACUGUCGCAGACCAUGAUAUAUUGCUUGUAAUACCGGAUUUGUGAGUACCGUCGCAAAUACGGAAAGGCUGUACGGUUU...
env-226    GCUGCAGUAUGAACCGCCAGGCGGCGUUGUUUAUGCUGAUAACAAUAUAUAUGGGUGAGAAAGGGACGACAGUCCCGUGUGUCAGAAGAUAUCGGCUGACUGUCGCAGACCAUGAUAUAUUGCUUGUAAUACCGGAUUUGUGAGUACCGUCGCAAAUACGGAAAGGCUGUACGGUUU...
env-227    GCUGCAGUAUGAACCGCCAGGCGGCGUUGUUUAUGCUGAUAACAAUAUAUAUGGGUGAGAAAGGGACGACAGUCCCGUGUGUCAGAAGAUAUCGGCUGACUGUCGCAGACCAUGAUAUAUUGCUUGUAAUACCGGAUUUGUGAGUACCGUCGCAAAUACGGAAAGGCUGUACGGUUU...
env-228    GCUGCAGUAUGAACCGCCAGGCGGCGUUGUUUAUGCUGAUAACAAUAUAUAUGGGUGAGAAAGGGACGACAGUCCCGUGUGUCAGAAGAUAUCGGCUGACUGUCGCAGACCAUGAUAUAUUGCUUGUAAUACCGGAUUUGUGAGUACCGUCGCAAAUACGGAAAGGCUGUACGGUUU...
env-229    GCUGCAGUAUGAACCGCCAGGCGGCGUUGUUUAUGCUGAUAACAAUAUAUAUGGGUGAGAAAGGGACGACAGUCCCGUGUGUCAGAAGAUAUCGGCUGACUGUCGCAGACCAUGAUAUAUUGCUUGUAAUACCGGAUUUGUGAGUACCGUCGCAAAUACGGAAAGGCUGUACGGUUU...
env-230    GCUGCAGUAUGAACCGCCAGGCGGCGUUGUUUAUGCUGAUAACAAUAUAUAUGGGUGAGAAAGGGACGACAGUCCCGUGUGUCAGAAGAUAUCGGCUGACUGUCGCAGACCAUGAUAUAUUGCUUGUAAUACCGGAUUUGUGAGUACCGUCGCAAAUACGGAAAGGCUGUACGGUUU...
env-231    NNNNNNNNNNNNNNNNNNNNNNNNNNNNNUUAUGCUGAUAACAAUAUAUAUGGGUGAGAAAGGGACGACAGUCCCGUGUGUCAGAAGAUAUCGGCUGACUGUCGCAGACCAUGAUAUAUUGCUUGUAAUACCGGAUUUGUGAGUACCGUCGCAAAUACGGAAAGGCUGUACGGUUU...
env-232    GCUGCAGUAUGAACCGCCAGGCGGCGUUGUUUAUGCUGAUAACAAUAUAUAUGGGUGAGAAAGGGACGACAGUCCCGUGUGUCAGAAGAUAUCGGCUGACUGUCGCAGACCAUGAUAUAUUGCUUGUAAUACCGGAUUUGUGAGUACCGUCGCAAAUACGGAAAGGCUGUACGGUUU...
env-234    GCUGCAGUAUGAACCGCCAGGCGGCGUUGUUUAUGCUGAUAACAAUAUAUAUGGGUGAGAAAGGGACGACAGUCCCGUGUGUCAGAAGAUAUCGGCUGACUGUCGCAGACCAUGAUAUAUUGCUUGUAAUACCGGAUUUGUGAGUACCGUCGCAAAUACGGAAAGGCUGUACGGUUU...
env-235    GCUGCAGUAUGAACCGCCAGGCGGCGUUGUUUAUGCUGAUAACAAUAUAUAUGGGUGAGAAAGGGACGACAGUCCCGUGUGUCAGAAGAUAUCGGCUGACUGUCGCAGACCAUGAUAUAUUGCUUGUAAUACCGGAUUUGUGAGUACCGUCGCAAAUACGGAAAGGCUGUACGGUUU...
env-236    GCUGCAGUAUGAACCGCCAGGCGGCGUUGUUUAUGCUGAUAACAAUAUAUAUGGGUGAGAAAAGGACGACAGUCCCGUGUGUCAGAAGAUAUCGGCUGACUGUCGCAGACCAUGAUAUAUUGCUUGUAAUACCGGAUUUGUGAGUACCGUCGCAAAUACGGAAAGGCUGUACGGUUU...
env-237    GCUGCAGUAUGAACCGCCAGGCGGCGUUGUUUAUGCUGAUAACAAUAUAUAUGGGUGAGAAAGGGACGACAGUCCCGUGUGUCAGAAGAUAUCGGCUGACUGUCGCAGACCAUGAUAUAUUGCUUGUAAUACCGGAUUUGUGAGUACCGUCGCAAAUACGGAAAGGCUGUACGGUUU...
env-238    GCUGCAGUAUGAACCGCCAGGCGGCGUUGUUUAUGCUGAUAACAAUAUAUAUGGGUGAGAAAGGGACGACAGUCCCGUGUGUCAGAAGAUAUCGGCUGACUGUCGCAGACCAUGAUAUAUUGCUUGUAAUACCGGAUUUGUGAGUACCGUCGCAAAUACGGAAAGGCUGUACGGUUU...
env-239    GCUGCAGUAUGAACCGCCAGGCGGCGUUGUUUAUGCUGAUAACAAUAUAUAUGGGUGAGAAAGGGACGACAGUCCCGUGUGUCAGAAGAUAUCGGCUGACUGUCGCAGACCAUGAUAUAUUGCUUGUAAUACCGGAUUUGUGAGUACCGUCGCAAAUACGGAAAGGCUGUACGGUUU...
env-240    ....................................................................................................................................................CCGUCGCAAAUACGGAAAGGCUGUACGGUUU...
env-241    AAAGUAACCUGCUGACCCUUUAUCGAAGAUAGAAGGAUUUGUUUCGCUCGUAUAUUGCUUUUAAUACCGGCCACAGGAUUGGCGGAACCGUGCUGCAUUAAUCCGCUAUUUGUGCAGGAAAACGCUUUUGCAAGGAAAAUGUCCAUGGCUUAAUGGGCCGGAAAGGCUGCAG..GCGG..
env-242    AAAGACAAGGAUAAUUAUCGUUGUCGAUUUUGUGCGUGAGAGCACAAACAUUAUUUAGGCGGCUGUAGUGUGACACGCCAGGCGGUGUGGAAGCACUAUUAAUAUACAGGGAAACUAUUCCUCGAUGUAUAUUGCUUAUAAUACCCGGUGUCAUUUCCGGGAAAGGCUGU....AGUAUC
env-243    AAAGACAAGGAUAAUUAUCGUUGUCGAUUUUGUGCGUGAGAGCACAAACAUUAUUUAGGCGGCUGUAGUGUGACACGCCAGGCGGUGUGGAAACACUAUUAAUAUACAGGGAAACUAUUCCUCGAUGUAUAUUGCUUAUAAUACCCGGUGUCAUUUCCGGGAAAGGCUGU....AGUAUC
env-244    AAAGACAAGGAUAAUUAUCGUUGUCGAUUUUGUGCGUGAGAGCACAAACAUUAUUUAGGCGGCUGUAGUGUGACACGCCAGGCGGUGUGGAAACACUAUUAAUAUACAGGGAAACUAUUCCUCGAUGUAUAUUGCUUAUAAUACCCGGUGUCAUUUCCGGGAAAGGCUGU....AGUAUC
env-245    AAAGACAAGGAUAAUUAUCGUUGUCGAUUUUGUGCGUGAGAGCACAAACAUUAUUUAGGCGGCUGUAGUGUGACACGCCAGGCGGUGUGGAAACACUAUUAAUAUACAGGGAAACUAUUCCUCGAUGUAUAUUGCUUAUAAUACCCGGUGUCAUUUCCGGGAAAGGCUGU....AGUAUC
env-246    AAAGACAAGGAUAAUUAUCGUUGUCGAUUUUGUGCGUGAGAGCACAAACAUUAUUUAGGCGGCUGUAGUGUGACACGCCAGGCGGUGUGGAAACACUAUUAAUAUACAGGGAAACUAUUCCUCGAUGUAUAUUGCUUAUAAUACCCGGUGUCAUUUCCGGGAAAGGCUGU....AGUAUC
env-252    AAAGACAAGGAUAAUUAUCGUUGUCGAUUUUGUGCGUGAGAGCACAAACAUUAUUUAGGCGGCUGUAGUGUGACACGCCAGGCGGUGUGGAAACACUAUUAAUAUACAGGGAAACUAUUCCUCGAUGUAUAUUGCUUAUAAUACCCGGUGUCAUUUCCGGGAAAGGCUGU....AGUAUC
env-254    ACAAACAAAGACAAGGAUAAUAAGCGUCGUUGUCGAAUUUGUGCGUAAAGCACAAAGAUUUAUUUAGGCGGCUGUAACAUGACACACCAGGGUGUGUGGUGUUAUUAAAAUAUACAAGAACAUACCGAUUGUAUAUUGCUUAUAAUACUGGCUCCGGCCAGAAAGGUCAC....GGGAAA
Lba-6-1    ACAAACAAAGACAAGGAUAAUAAGCGUCGUUGUCGAAUUUGUGCGUAAAGCACAAAGAUUUAUUUAGGCGGCUGUAACAUGACACACCAGGGUGUGUGGUGUUAUUAAAAUAUACAAGAACAUACCGAUUGUAUAUUGCUUAUAAUACUGGCUCCGGCCAGAAAGGUCACGG..GAAA..
env-255    .....................................................................................AUGACACACCAGGGUGUGUGGUGUUAUUAAAUAUACAAGAACAUACCGAUUGUAUAUUGCUUAUAAUACUGGCUCCGGCCAGAAAGGUUACAG..GAAA..
Lba-10-1   ACAAACAAAGACAAGGAUAAUAAGCGUCGUUGUCGAAUUUGUGCGUAAAGCACAAAAAUUUAUUUAGGCGGCUGUAACAUGACACACCAGGGUGUGUGGUGUUAUUAAAAUAUACAAGAACAUACCGAUUGUAUAUUGCUUAUAAUACUGGCUCCGGCCAGAAAGGUUACAG..GAAA..
env-256    ACAAACAAAGACAAGGAUAAUAAGCGUCGUUGUCGAAUUUGUGCGUAAAGCACAAAGAUUUAUUUAGGCGGCUGUAACAUGACACACCAGGGUGUGUGGUGUUAUUAAAAUAUACAAGAACAUACCGAUUGUAUAUUGCUUAUAAUACUGGCUCCGGCCAGAAAGGUCACAG..GAAA..
env-258    ACAAACAAAGACAAGGAUAAUAAGCGUCGUUGUCGAAUUUGUGCGUAAAGCACAAAAAUUUAUUUAGGCGGCUGUAACAUGACACACCAGGGUGUGUGGUGUUAUUAAAAUAUACAAGAACAUACCGAUUGUAUAUUGCUUAUAAUACUGGUUCCGGCCAGAAAGGUCACAG..GAAA..
env-259    ACAAACAAAGACAAGGAUAAUAAGCGUCGUUGUCGAAUUUGUGCGUAAAGCACAAAGAUUUAUUUAGGCGGCUGUAACAUGACACACCAGGGUGUGUGGUGUUAUUAAAAUAUACAAGAACAUACCGAUUGUAUAUUGCUUAUAAUACUGGCUCCGGCCAGAAAGGUCACAG..GAAA..
env-260    UCUGGCUUUUCUUAACGGGUUCAUUUCGUUGCGAUAUUGAUUUAAUACCGGAACAGGAUCGGCAGUACACGUAUCUGUGAUUUCCGCUAUUGCACAGAUAAAGCAUGUCAUUGCAGGAACAAGGGGCCGGCGGCCCGCAGGUUCAUGGCUUAAUGUCCCGGAAAGGCUGCAU..UUCC..
env-261    ACAUUUUAAAGUGACCUUCUUCAGGAUUAGGAUUAUUUCGUGCGAUAUUGCUUUUAAUACCGGCACAGGAUUGACAGUAUAACGUUUCUGCGUUUAUCCGUCUAUGGCGCAGAAAAAGCAUGUCCUUGCAAGAACACAGGUUCAUGGCUUAAUGUGCCGGAAAGGCUGCGUAU..UUUC..
env-262    UUUAGAAUUAACUUGAUUUUUGUGCAAACAAAUUAGGCGGCUGAUAGUUGACACACCAGGAAGUGUGUGAGACUAUUAAUAUAUAUUGAGCUGUGAGCUUCAUGUUAUAUUGCUUAUAAUACCGGCCCGUCAUUGCUGAGUAUGUACCGUCAGCGGCCGGAAAGGGCGUGC..AAAUGC
env-263    AUUGAAUUAAGAAAGUUGGAUUUUAUUUCGUUGCGAUAUUGGCUUUUAAUACCGACCCAGGAUUGAGACAGUAAACGGCUUCGCGUGAAACCGCUGUUCGCGAAGGCAGAAUGUCCUGCAAGAAUGAUCAUUUAUGAGCGAUUCACAGCUUAAUGGGUUGGAAAGGCCGCAG..GUGUG.
Bfo-1-1    GACAAUAUACGAACGGGACAUCCCAAUCCGCGGCAGGAAGCCGGACAUAUGAUCCGGAUGUGACGGUGCACCGCAUAUAAGGCGUGCGCAGGAUCCUUGGGCAGGAUGUCUCACAUAUAUUGCUUCUAAUACCGCAUUUCCGGGUACCGGCGGAUUUGCGGAAAGGUCGUCAG.U.....
Bpr-1-1    UUAAACGAUGUGAUUUGUGUGAAUUAUUUUAGUGCGGCUGCAGUAUGAAACGCGCGGUUGCGUUGUUCGUACUGAUUAAAUAUAGUAAUAUAUCACGUUUGUCGUCCUGCCGAGAGGGCGAUUACAGAAUAUUGCUUAAUACCGUCGCUCUGCCGUGACGGAAAGGCUGCGUAAAAUUUU
Cba-1-1    UGAUAGUUGAUGAGAUUUGUGUAAUAUUUUAGUGCGGCUGCAGUAUAAAACGCAAGGUUGCGUUGUUCAUACUGAUCAAAUAUAGUAAUAUAUCACGUUUGUCGUCCUGCCGAGAGGGCGAUUGCAGAAUAUUGCUUAAUACCGUCGCUCUGCCGUGACGGAAAGGCUGCGUAAAAUUUU
env-264    ............................................AUCAGAAUCAUCUGUGAAAAUCCGAAGCACAAACAGCGUCAGGGCUGAUCGCGGCUGACAACCUGACAGUACAUGUAUUGCCUAAUACCGAGAGGCAAUCGUGGCGUAAGCCACUAUCGGAAAGGCCGCG...UACAUU
env-265    CUGCGAGAAUCCGAAGCAUAAACAGCGUCAGGGAUAAUCAACAGGCUGUUUUUGCAGAAUGCGGCUGCAGCAAAAAGCCCCGUCAUGGCGUUUUGCUGUUACAUGAUUAACAUGUAUUGCCUAAUACCGAAGGCAUUGUGCAUUGCACAGAAGAAUCGGAAAGGCCGUGGAUCUUAC.
env-266    UUAUCUGUGAGAAUCCGAAGCACAAACAGCGCCAGGGCUGAUCGCCGGCGGUUGCUUCGAAAAUGCGGCUGCGGCUUGGGCCGCCGUCAUGGCCCGGGCUGUAUCAUGAAAAACAUGUAUUGCCUAAUACCGAAGGCAGCGCAGAUGCGCAAAUUUCGGAAAGGCUGCGG..AAGUUA
env-267    ..............................................................UUGAUACACCCGGAAGGGUUGUUCGAACUGUUUUAAUAUCUUAUUAAAGAAGCAUGUAUUGCCUAAUACCGAGAGGCAACAGCGUCCGUGAGGACACAAUUUCGGAAAGGUUGC....UUUACG
env-268    AAAACACAAACAGCGUCAGGGUUAAUUGACAAGUUUGUGAAACGUGCGGCUGCAGUUUGAUACACCCGGAAGGGUUGUUCGAACUGUUUUAAUAUCUUAUUAAAGAAGCAUGUAUUGCCUAAUACCGAGAGGCAACAGCGUCCGUGAGGACACAAUUUCGGAAAGGUCGC....UUUACG
env-271    AAAACACAAACAGCGUCAGGGUUAAUUGACAAGUUUGUGAAACGUGCGGCUGCAGUUUGAUACACCCGGAAGGGUUGUUCGAACUGUUUUAAUAUCUUAUUAAAGAAGCAUGUAUUGCCUAAUACCGAGAGGCAACAGCGUCCGUGAGGACACAAUUUCGGAAAGGUCGC....UUUACG
env-272    .................................AGUUUGUGAAACGUGCGGCUGCAGUUUGAUACACCNCGGAAGGGUUGUUCGAACUGUUUUAAUAUCUUAUUAAAGAAGCAUGUAUUGCCUAAUACCGAGAGGCAACAGCGUCCGUGAGGACACAAUUUCGGAAAGGUCGC....UUUACG
env-273    AAAACACAAACAGCGUCAGGGUUAAUUGACGGACAGUUUGUGAAACGUGCGGCUGCAGUUUGAUACACCCGGAAGGGUUGUUCGAACUGUUUUAAUAUCUUAUUAAAGAAGCAUGUAUUGCCUAAUACCGAGAGGCAACAGCGUCCGUGAGGACACAAUUUCGGAAAGGUCGUUCUUCGCC.
env-275    AAUCUGUGAAAAUCCAAAACACAAACAGCGUCAGGGUUGAUUGACUGAUUUGUGAAACGUGCGGCUGCAGUUUGAUACACCCGGAAGGGUUGUUCGGACUGUUUUAAUAUUUUAUUAAAGAAGCAUGUAUUGCCUAAUACCGAGAGGCAACAAAAUUUCGGAAAGGUCGCUUUUCGCC..
env-276    .........................................................................................ACCCGGAAGGGUUGUUCGGACUGUUUUAAUAUCUUAUUAAAGAAGCAUGUAUUGCCUAAUACCGAGAGGCAACAAAUUUCGGAAAGGUCGCUUU.UCGCCA
env-277    AAACACAAACAGCGCCAGGGUUAAUGACGGCAGUUUGUGAAACGUGCGGCUGCAGUUUGAUACACCCGGAAGGGUUGUUCGGACUGUUUGUAUAAUUGAUUUAUCAAAGCAUGUAUUGCCUAAUACCGAGAGGCAACAGCAGCAUUUAGCUGCAAUUUCGGAAAGGCUGUCUU.ACGCCA
env-278    AAACACAAACAGCGCCAGGGUUAAUGACGGCAGUUUGUGAAACGUGCGGCUGCAGUUUGAUACACCCGGAAGGGUUGUUCGGACUGUUUGUAUAAUUGAUUUAUCAAAGCAUGUAUUGCCUAAUACCGAGAGGCAACAGCAGCAUUUAGCUGCAAUUUCGGAAAGGCUGUCUU.ACGCCA
env-279    CCAAAAACACAAACAGCGCCAGGGUUGAUUCCGGCAGUUUGUGAAACGUGCGGCUGCAGUUUGAUACACCCGGAAGGGUUGUUCGGACUGUUUGAAUUUAAAAAAAGCAUGUAUUGCCUAAUACCGAGAGGCAACAGCGGUUAUUGCUGCAAUUUCGGAAAGGCUGUCU..UACGCC
env-280    AACAGCGUCAGGGUUGAUUUCCAUAUCGGAACUUCUUUGGAAUGUUUGUGAAACGUGCGGCUGCAGUAUGAAACACCCGGAAGGGUUGUUCUACUGAGUGAAUAAUAAGCAUGUAUUGCCUAAUACCGAGAGGCAACAGCGGUUAAGAUCCGCAAUUUCGGAAAGGCUGUGA..UUCGCC
env-281    AACAGCGUCAGGGUUGAUUUCCAUAUCGGAACUUCUUUGGAAUGUUUGUGAAACGUGCGGCUGCAGUAUGAAACACCCGGCAGGGUUGUUCUACUGAACAGAUAAUAAGCAUGUAUUGCCUAAUACCGAGAGGCAACAGCGGUUAAGAUCCGCAAUUUCGGAAAGGCUGU....GAUUCG
env-282    AACAGCGUCAGGGUUGAUUUUCAUAUCGGAACUUCUUUGGAAUGUUUGUGAAACGUGCGGCUGCAGUAUGAAACACCCGGCAGGGUUGUUCUACUGAACAGAUAAUAAGCAUGUAUUGCCUAAUACCGAGAGGCAACAGCGGUUAAGAUCCGCAAUUUCGGAAAGGCUGU....GAUUCG
Bsp-1-1    AACAGCGUCAGGGUUGAUUUCCAUAUCGGAACUUCUUUGGAAUGUUUGUGAAACGUGCGGCUGCAGUAUGAAACACCCGGUAGGGUUGUUCUACUGAACAGAUAAUAAGCAUGUAUUGCCUAAUACCGAGAGGCAACAGCGGUUAAGAUCCGCAAUUUCGGAAAGGCUGU....GAUUCG
env-283    AACAGCGUCAGGGUUGAUUUCCAUAUCGGAACUUCUUUGGAAUGUUUGUGAAACGUGCGGCUGCAGUAUGAAACACCCGGCAGGGUUGUUCUACUGAACAGAUAAUAAGCAUGUAUUGCCUAAUACCGAGAGGCAACAGCGGUUAAGAUCCGCAAUUUCGGAAAGGCUGU....GAUUCG
env-284    AACAGCGUCAGGGUUGAUUUCCAUAUCGGAACUUCUUUGGAAUGUUUGUGAAACGUGCGGCUGCAGUAUGAAACACCCGGUAGGGUUGUUCUGCUGAACAGAUAAUAAGCAUGUAUUGCCUAAUACCGAGAGGCAACAGCGGUUAAGAUCCGCAAUUUCGGAAAGGCUGU....GAUUCG
env-285    AACAGCGUCAGGGUUGAUUUCCAUAUCGGAACUUCUUUGGAAUGUUUGUGAAACGUGCGGCUGCAGUAUGAAACACCCGGUAGGGUUGUUCUACUGAACAGAUAAUAAGCAUGUAUUGCCUAAUACCGAGAGGCAACAGCGGUUAAGAUCCGCAAUUUCGGAAAGGCUGU....GAUUCG
env-286    AACAGCGUCAGGGUUGAUUUCCAUAUCGGAACUUCUUUGGAAUGUUUGUGAAACGUGCGGCUGCAGUAUGAAACACCCGGCAGGGUUGUUCUGCUGAACAGAUAAUAAGCAUGUAUUGCCUAAUACCGAGAGGCAACAGCGGUUAAGAUCCGCAAUUUCGGAAAGGCUGU....GAUUCG
env-287    AACAGCGUCAGGGUUGAUUUCCAUAUCGGAACUUCUUUGGAAUGUUUGUGAAACGUGCGGCUGCAGUAUGAAACACCCGGCAGGGUUGUUCUACUGAACAGAUAAUAAGCAUGUAUUGCCUAAUACCGAGAGGCAACAGCGGUUAAGAUCCGCAAUUUCGGAAAGGCUGU....GAUUCG
env-288    AACAGCGUCAGGGUUGAUUUCCAUAUCGGAACUUCUUUGGAAUGUUUGUGAAACGUGCGGCUGCAGUAUGAAACACCCGGUAGGGUUGUUCUACUGAACAGAUAAUAAGCAUGUAUUGCCUAAUACCGAGAGGCAACAGCGGUUAAGAUCCGCAAUUUCGGAAAGGCUGU....GAUUCG
env-289    AAACACAAACAGCGUCAGGGUUAAUUGACAGUUUAGAGUUUGUGUAACGUGCGGCUGCAGUGUGAGACACCCGGAAGGGUUGUUCAGACUGUUAAGAUAGUUUUAGCAUGUAUUGCCUAAUACCGAGAGGCAACGUUGGCCCGUUGGCCGCAAUAAUUCGGAAAGGCUGC....UUUUCG
env-290    AAACACAAACAGCGUCAGGGUUAAUUGACAGUUUAGAGUUUGUGUAACGUGCGGCUGCAGUGUGAGACACCCGGAAGGGUUGUUCAGACUGUUAAGAUAGUUUUAGCAUGUAUUGCCUAAUACCGAGAGGCAACGUUGGCCCGUUGGCCGCAAUAAUUCGGAAAGGCUGC....UUUUCG
env-291    AAACACAAACAGCGUCAGGGUUAAUUGACAGUUUAGAGUUUGUGUAACGUGCGGCUGCAGUGUGAGACACCCGGAAGGGUUGUUCAGACUGUUAAGAUAGUUUUAGCAUGUAUUGCCUAAUACCGAGAGGCAACGUUGGCCCGUUGGCCGCAAUAAUUCGGAAAGGCUGC....UUUUCG
env-292    AAACACAAACAGCGUCAGGGUUAAUUGACAGUUUAGAGUUUGUGUAACGUGCGGCUGCAGUGUGAGACACCCGGAAGGGUUGUUCAGACUGUUAAGAUAGUUUUAGCAUGUAUUGCCUAAUACCGAGAGGCAACGUUGGCCCGUUGGCCGCAAUAAUUCGGAAAGGCUGC....UUUUCG
env-293    AAACACAAACAGCGUCAGGGUUGAUUGACGGUUUAAAGUUUGUGUAACGUGCGGCUGCAGUGUGAGACGCCCGGAAGGGUUGUUCAGACUGUUAAGAUAGUUUUAGCAUGUAUUGCCUAAUACCGAGAGGCAACGUUGGCCCGUUGGCCGCAGUAAUUCGGAAAGGCUGC....UUUUCG
env-294    AAACACAAACAGCGUCAGGGUUGAUUGACGGUUUAAAGUUUGUGUAACGUGCGGCUGCAGUGUGAGACGCCCGGAAGGGUUGUUCAGACUGUUAAGAUAGUUUUAGCAUGUAUUGCCUAAUACCGAGAGGCAACGUUGGCCCGUUGGCCGCAGUAAUUCGGAAAGGCUGC....UUUUCG
env-295    AAACACAAACAGCGUCAGGGUUGAUUGACGGUUUAAAGUUUGUGUAACGUGCGGCUGCAGUGUGAGACGCCCGGAAGGGUUGUUCAGACUGUUAAGAUAGUUUUAGCAUGUAUUGCCUAAUACCGAGAGGCAACGUUGGCCCGUUGGCCGCAGUAAUUCGGAAAGGCUGC....UUUUCG
env-296    AUUGGCGGUUUAAAGUUUGUGUAACGUGCGGCUGCAGUGUGAGACGCCCGGAAGGGUUGUUCAGACUGUUAAGAUAGUUU......................UAGCAUGUAUUGCCUAAUACCGAGAGGCAACGUUGGCCCGUUGGCCGCAGUAAUUCGGAAAGGCUGC....UUUUCG
env-298    AUUGACGGUUUAAAGUUUGUGUAACGUGCGGCUGCAGUGUGAGACGCCCGGAAGGGUUGUUCAGACUGUUAAGAUAGUUU......................UAGCAUGUAUUGCCUAAUACCGAGAGGCAACGUUGGCCCGUUGGCCGCAGUAAUUCGGAAAGGCUGC....UUUUCG
env-299    ..............................UGCAGUGUGAGACGCCCGGAAGGGUUGUUCAGACUGUUAAGAUAGUUU......................UAGCAUGUAUUGCCUAAUACCGAGAGGCAACGUUGGCCCGUUGGCCGCAGUAAUUCGGAAAGGCUGC....UUUUCG
env-300    AUUGACAGUUUAAAGUUUGUGUAACGUGCGGCUGCAGUGUGAGACGCCCGGAAGGGUUGUUCAGACUGUUAAGAUAGUUU......................UAGCAUGUAUUGCCUAAUACCGAGAGGCAACGUUGGCCCGUUGGCCGCAGUAAUUCGGAAAGGCUGC....UUUUCG
env-301    AUUGACGGUUUAAAGUUUGUGUAACGUGCGGCUGCAGUGUGAGACGCCCGGAAGGGUUGUUCAGACUGUUAAGAUAGUUU......................UAGCAUGUAUUGCCUAAUACCGAGAGGCAACGUUGGCCCGUUGGCCGCAGUAAUUCGGAAAGGCUGC....UUUUCG
env-302    AUUGACAGUUUAAAGUUUGUGUAACGUGCGGCUGCAGUGUGAGACGCCCGGAAGGGUUGUUCAGACUGUUAAGAUAGUUU......................UAGCAUGUAUUGCCUAAUACCGAGAGGCAACGUUGGCCCGUUGGCCGCAGUAAUUCGGAAAGGCUGC....UUUUCG
env-303    AUUGACGGUUUAAAGUUUGUGUAACGUGCGGCUGCAGUGUGAGACGCCCGGAAGGGUUGUUCAGACUGUUAAGAUAGUUU......................UAGCAUGUAUUGCCUAAUACCGAGAGGCAACGUUGGCCCGUUGGCCGCAGUAAUUCGGAAAGGCUGC....UUUUCG
env-304    ...............UUGUGUAACGUGCGGCUGCAGUGUGAGACGCCCGGAAGGGUUGUUCAGACUGUUAAGAUAGUUU......................UAGCAUGUAUUGCCUAAUACCGAGAGGCAACGUUGGCCCGUUGGCCGCAGUAAUUCGGAAAGGCUGC....UUUUCG
env-305    AUUGACGGUUUAAAGUUUGUGUAACGUGCGGCUGCAGUGUGAGACGCCCGGAAGGGUUGUUCAGACUGUUAAGAUAGUUU......................UAGCAUGUAUUGCCUAAUACCGAGAGGCAACGUUGGCCCGUUGGCCGCAGUAAUUCGGAAAGGCUGC....UUUUCG
Bwe-2-1    AUUGACGGUUUAAAGUUUGUAUAACGUGCGGCUGCAGUGUGAGACGCCCGGAAGGGUUGUUCAGACUGUUAAGAUAGUUU......................UAGCAUGUAUUGCCUAAUACCGAGAGGCAACGUUGGCCCGUUGGCCGCAGUAAUUCGGAAAGGCUGC....UUUUCG
env-306    ...............................................................CGAGAGGCAACGU...................................................UGGCCCGUUGGCCGCAGUAAUUCGGAAAGGCUGC....UUUUCG
env-307    GAUUGACAGUUUAAAGUUUGUGUAACGUGCGGCUGCAGUGUGAGACACCCGGAAGGGUUGUUCAGACUGUUAAAAUAGUUU......................UAGCAUGUAUUGCCUAAUACCGAGAGGCAACGUUGGCCCGUUGGCCGCAGUAAUUCGGAAAGGCUGC....UUUUCG
env-308    ......................ACGUGCGGCUGCAGUGUGAGACGCCCGGAAGGGUUGUUCAGACUGUUAAGAUAGUUU......................UAGCAUGUAUUGCCUAAUACCGAGAGGCAACGUUGGCCCGUUGGCCGCAGUAAUUCGGAAAGGCUGC....UUUUCG
env-309    ..........................................................................................UAGCAUGUAUUGCCUAAUACCGAGAGGCAACGUUGGCCCGUUGGCCGCAGUAAUUCGGAAAGGCUGC....UUUUCG
env-310    AUUGACGGUUUAAAGUUUGUGUAACGUGCGGCUGCAGUGUGAGACGCCCGGAAGGGUUGUUCAGACUGUUAAGAUAGUUU......................UAGCAUGUAUUGCCUAAUACCGAGAGGCAACGUUGGCCCGUUGGCCGCAGUAAUUCGGAAAGGCUGC....UUUUCG
env-311    AUUGACAGUUUAAAGUUUGUGUAACGUGCGGCUGCAGUGUGAGACGCCCGGAAGGGUUGUUCAGACUGUUAAGAUAGUUU......................UAGCAUGUAUUGCCUAAUACCGAGAGGCAACGUUGGCCCGUUGGCCGCAGUAAUUCGGAAAGGCUGC....UUUUCG
env-313    AUUGACGGUUUAAAGUUUGUGUAACGUGCGGCUGCAGUGUGAGACGCCCGGAAGGGUUGUUCAGACUGUUAAGAUAGUUU......................UAGCAUGUAUUGCCUAAUACCGAGAGGCAACGUUGGCCCGUUGGCCGCAGUAAUUCGGAAAGGCUGC....UUUUCG

           ............................................................................................................................................................<<<<<<<<<<.....
           .............................................................................................................................................................222222........
           YYGGAAAGGYYGY
           5'
```

```
env-314  ......................................................................AAGAUAGUUU.......................UAGCAUGUAUUGCCUAAUACCGAGAGGCAACGUUGGCCCGUUGGCCGCAGUAAUUCGGAAAGGCUGC....UUUUCG
env-315  AUUGACGGUUUAAAGUUUGUAUAACGUGCGGCUGCAGUGUGAGACGCCCGGAAGGGUUGUUCAGACUGUUAAGAUAGUUU.......................UAGCAUGUAUUGCCUAAUACCGAGAGGCAACGUUGGCCCGUUGGCCGCAGUAAUUCGGAAAGGCUGC....UUUUCG
env-317  AUUGACAGUUUAAAGUUUGUGUAACGUGCGGCUGCAGUGUGAGACGCCCGGAAGGGUUGUUCAGACUGUUAAGAUAGUUU.......................UAGCAUGUAUUGCCUAAUACCGAGAGGCAACGUUGGCCCGUUGGCCGCAGUAAUUCGGAAAGGCUGC....UUUUCG
env-318  AUUGGCGGUUUAAAGUUUGUGUAACGUGCGGCUGCAGUGUGAGACGCCCGGAAGGGUUGUUCAGACUGUUAAGAUAGUUU.......................UAGCAUGUAUUGCCUAAUACCGAGAGGCAACGUUGGCCCGUUGGCCGCAGUAAUUCGGAAAGGCUGC....UUUUCG
Bwe-1-1  AUUGACAGUUUAAAGUUUGUGUAACGUGCGGCUGCAGUAUGAGACACCCGGAGAGGUUGUCCAGACUGUUAAGAUAGUUU.......................UAGCAUGUAUUGCCUAAUACCGAGAGGCAACGUUGGCCCGUUGGCCGCAGUAAUUCGGAAAGGCUGC....UUUUCG
env-319  ........................................................................................................................................UGGCCCGUUGGCCGCAGUAAUUCGGAAAGGCUGC....UUUUCG
env-320  AAACACAAACAGCGUCAGGGUUGAUUGACAGUUUAAAGUUUGUGUAACGUGCGGCUGCAGUGUGAGACGCCCGGAAGGGUUGUUCAGACUGUUAAGAUAGUUUAGCAUGUAUUGCCUAAUACCGAGAGGCAACGUUGGCCCGUUGGCCGCAAUAAUUCGGAAAGGCUGC....UUUUCG
env-321  AUUGACAGUUUAAAGUUUGUGUAACGUGCGGCUGCAGUGUGAGACGCCCGGAAGGGUUGUUCAGACUGUUAAGAUAGUUU.......................UAGCAUGUAUUGCCUAAUACCGAGAGGCAACGUUGGCCCGUUGGCCGCAGUAAUUCGGAAAGGCUGC....UUUUCG
env-322  .......................................................UGAUACACCCGGAAGGGUUGUUCGAACUGUUUUAAUAUCUUAUUAAAGAAGCAUGUAUUGCCUAAUACCGAGAGGCAACGUUGGCCCGUUGGCCGCAGUAAUUCGGAAAGGCUGC....UUUUCG
env-323  AAACACAAACAGCGUCAGGGUUGAUUGACGGUUUAAAGUUUGUAUAACGUGCGGCUGCAGUGUGAGACGCCCGGAAGGGUUGUUCAGACUGUUAAGAUAGUUUAGCAUGUAUUGCCUAAUACCGAGAGGCAACGUUGGCCCGUUGGCCGCAGUAAUUCGGAAAGGCUGC....UUUUCG
env-324  AAACACAAACAGCGUCAGGGUUGAUUGACAGUUUAAAGUUUGUGUAACGUGCGGCUGCAGUGUGAGACGCCCGGAAGGGUUGUUCAGACUGUUAAGAUAGUUUAGCAUGUAUUGCCUAAUACCGAGAGGCAACGUUGGCCCGUUGGCCGCAGUAAUUCGGAAAGGCUGC....UUUUCG
env-325  AAACACAAACAGCGUCAGGGUUGAUUGACAGUUUAAAGUUUGUGUAACGUGCGGCUGCAGUGUGAGACGCCCGGAAGGGUUGUUCAGACUGUUAAGAUAGUUUAGCAUGUAUUGCCUAAUACCGAGAGGCAACGUUGGCCCGUUGGCCGCAGUAAUUCGGAAAGGCUGC....UUUUCG
env-326  CCAAAACACAAACAGCGUCAGGGUUGAUUGACGGUUUAAAGUUUGUAUAACGUGCGGCUGCAGUGUGAGACGCCCGGAAGGGUUGUUCAGACUGUUAAGAUAGUUUUAGCAUGUAUUGCCUAAUACCGAGAGGCAACAGCGGUUAAGAUCCGCAAUUUCGGAAAGGCUGU....GAUUCG
env-327  ...............................................................................................UGUUUUAAAGUUUUAGCAUGUAUUGCCUAAUACCGAGAGGCAAGGCUGCAAAAGCAGCAUUUUCGGAAAGGCCGU....UAUUCG
env-328  .....AAACACAAACAGCGUCAGGGUUGAUGCCUGGAUUGUGUGGUAUGCGGCUGCAGUUGAGACACCCGGACAGGUUGUCAGGACUGAUGAAAUAAUGAACUAUUCAUGUAUUGCCUAAUACCGAGAGGCGACGUUGGCCCGUUGGCCGCAACAAAUCGGAAAGGCCGC....AACACG
env-329  AUCCAAAACACAAACAGCGUCAGGGUUGAUGAAUCCGAUUAGUUUGUGAAUGCGGCUGUAGAUGGAACACCCGGACGGGUUGUAACAUCUGACAAAUAAUGAACUAAGCAUGUAUUGCCUAAUACCGGAGGCAACGUUGGCCCGUUGGCCGCAAAUUUCGGAAAGGCUGC....GAAUCG
env-331  AUCCAAAACACAAACAGCGUCAGGGUUGAUGAAUCCGAUUAGUUUGUGAAUGCGGCUGUAGAUGGAACACCCGGACGGGUUGUAACAUCUGACAAAUAAUGAACUAAGCAUGUAUUGCCUAAUACCGGAGGCAACGUUGGCCCGUUGGCCGCAAAUUUCGGAAAGGCUGC....GAAUCG
env-332  GAAUCCAAAGCACAAACAGCGUCAGGGUUGAUUGAUCCAUUGUUUGUGAAUGCGGCUGCAGGUGAAACACCCGGAUGGGUUGUAUCAUCUGACAUAUAAUGAACUAAGCAUGUAUUGCCUAAUACCGGAGGCAACGUUGGCCCGUUGGCCGCAAAUUUCGGAAAGGCUGC....GAAUCG
env-335  GAAUCCAAAGCACAAACAGCGUCAGGGUUGAUUGAUCCAUUGUUUGUGAAUGCGGCUGCAGGUGAAACACCCGGACGGGUUGUAUCAUCUGACAUAUAAUGAACUAAGCAUGUAUUGCCUAAUACCGGAGGCAACGUUGGCCCGUUGGCCGCAAAUUUCGGAAAGGCUGC....GAAUCG
env-336  GAAUCCAAAGCACAAACAGCGUCAGGGUUGAUUGAUCCAUUGUUUGUGAAUGCGGCUGUAGGUGAAACACCCGGAUGGGUUGUAUCAUCUGACAUAUAAUGAACUAAGCAUGUAUUGCCUAAUACCGGAGGCAACGUUGGCCCGUUGGCCGCAAAUUUCGGAAAGGCUGC....GAAUCG
env-337  GAAUCCAAAGCACAAACAGCGUCAGGGUUGAUUGAUCCAUUGUUUGUGAAUGCGGCUGUAGGUGAAACACCCGGAUGGGUUGUAUCAUCUGACAUAUAAUGAACUAAGCAUGUAUUGCCUAAUACCGGAGGCAACGUUGGCCCGUUGGCCGCAAAUUUCGGAAAGGCUGC....GAAUCG
env-338  GAAUCCAAAGCACAAACAGCGUCAGGGUUGAUUGAUCCAUUGUUUGUGAAUGCGGCUGCAGGUGAAACACCCGGACGGGUUGUAUCAUCUGACAUAUAAUGAACUAAGCAUGUAUUGCCUAAUACCGGAGGCAACGUUGGCCCGUUGGCCGCAAAUUUCGGAAAGGCUGC....GAAUCG
env-339  GAAUCCAAAGCACAAACAGCGUCAGGGUUGAUUGAUCCAUUGUUUGUGAAUGCGGCUGCAGGUGAAACACCCGGAUGGGUUGUAUCAUCUGACAUAUAAUGAACUAAGCAUGUAUUGCCUAAUACCGGAGGCAACGUUGGCCCGUUGGCCGCAAAUUUCGGAAAGGCUGC....GAAUCG
env-340  ....CCAAAGCACAAACAGCGUCAGGGUUGAUUGAUCCAUUGUUUGUGAAUGCGGCUGCAGGUGAAACACCCGGAUGGGUUGUAUCAUCUGACAUAUAAUGAACUAAGCAUGUAUUGCCUAAUACCGGAGGCAACGUUGGCCCGUUGGCCGCAAAUUUCGGAAAGGCUGC....GAAUCG
env-341  ................................................................................................................................AGGCAACGUUGGCCCGUUGGCCGCAAAUUUCGGAAAGGCUGC....GAAUCG
env-342  .................................GAUCCAUUGUUUGUGAAUGCGGCUGCAGGUGAAACACCCGGAUGGGUUGUAUCAUCUGACAUAUAAUGAACUAAGCAUGUAUUGCCUAAUACCGGAGGCAACGUUGGCCCGUUGGCCGCAAAUUUCGGAAAGGCUGC....GAAUCG
Bsc-1-1  ACACAAACAGCGUCAGGGCUAAAAUCGGACUACAUUGUUUGUGGAGACGUGCGGCUGCAGUGUGAAACGCCCGGGAGGGUUGUUCAGACUGUUGAAAUAUUUUAAGCAUGUAUUGCCUAAUACCGAGAGGCAACGCCGGCCGGAUGGCCGCAUCAUUUCGGAAAGGCUGC....GGUUCG
Rsp-3-1  AAAACACAAACAGCGUCAGGGUUGAUUCACUGAUUCAGGUUUGUGAAACGUGCGGCUGCAGUGUGAGACACCCGGAAGGGUUGUUCAGACUGUUUAAAUAGUUUAGCAUGUAUUGCCUAAUACCGAGAGGCAACGUUGGCCCGUUGGCCGCAACAUUCGGAAAGGCUGC....UAUUCG
env-343  UUUAUGCGCCCGAUAUUAUUAUGUAAGACCCUGACUGGGCUGCGUCAGGAAAUCUGCUAAUAAUAUUACUUAAUACUCUCCGGCGCCUGCCCCGGGAGGCAGAGAGAUAACAGGAUCCGCAGGUCGAUUUGAGCGUCUGUGCGGGUUGCGUCUGUGAGAGAAAGGCCGC..U.GUCGUC
env-344  AAAUAUAAGACGAUUAGUUUGUGAAAAUUUUAGUGCGGCUGCAGUAUGAACCGCCAGGGUGCGUUGUUCGUACUGAUUAAAUAUAGUAAUAUUAUCACGUUAACUGUUUUACCGAGAACAGGAUAAAAUAUUGCUUAAUACCGGUGUCCGCCAAAUACCGGAAAGGCUGCGUAAAAUCUA
env-346  AAAUAUAAGACGAUUAGUUUGUGAAAAUUUUAGUGCGGCUGCAGUAUGAACCGCUAGGGUGCGUUGUUCGUACUGAUUAAAUAUAGUAAUAUUAUCACGUUAACUGUUUUACCGAGAACAGGAUAAAAUAUUGCUUAAUACCGGUGUCCGCCAAAUACCGGAAAGGCUGCGUAAAAUCUA
env-348  AAAUAUAAGACGAUUAGUUUGUGAAAAUUUUAGUGCGGCUGCAGUAUGAACCGCUAGGGUGCGUUGUUCGUACUGAUUAAAUAUAGUAAUAUUAUCACGUUAACUGUUUUACCGAGAACAGGAUAAAAUAUUGCUUAAUACCGGUGUCCGCCAAAUACCGGAAAGGCUGCGUAAAAUCUA
env-349  AAAUAUAAGACGAUUAGUUUGUGAAAAUUUUAGUGCGGCUGCAGUAUGAACCGCCAGGGUGCGUUGUUCGUACUGAUUAAAUAUAGUAAUAUUAUCACGUUAACUGUUUUACCGAGAACAGGAUAAAAUAUUGCUUAAUACCGGUGUCCGCCAAAUACCGGAAAGGCUGCGUAAAAUCUA
env-350  AAAUAUAAGACGAUUAGUUUGUGAAAAUUUUAGUGCGGCUGCAGUAUGAACCGCUAGGGUGCGUUGUUCGUACUGAUUAAAUAUAGUAAUAUUAUCACGUUAACUGUUUUACCGAGAACAGGAUAAAAUAUUGCUUAAUACCGGUGUCCGCCAAAUACCGGAAAGGCUGCGUAAAAUCUA
env-351  .............................................AUGAACCGCUAGGGUGCGUUGUUCGUACUGAUUAAAUAUAGUAAUAUUAUCACGUUAACUGUUUUACCGAGAACAGGAUAAAAUAUUGCUUAAUACCGGUGUCCGCCAAAUACCGGAAAGGCUGCGUAAAAUCUA
env-352  ............................................................................................................................................................CCGGAAAAGGCUGCGUAAAAUUAU
env-353  AACCUCAGAUGCAAGUGUAGUGAAUUUAGACGUACAUUUGCACUUUUAACGUGCGGCUGCAGCUUUGAACCACCCGGAAGUGUUGUUCAAGCUGUUACAAAUAUUAUUGUGAGAGUAUCACGUAUAAUAUUGCUUAAUACCGGUGCCGCCGUCGGUUCCGGAAAGGUCAUGC..CUUUAG
Rga-1-1  AUUCGAACUUCAGAUGCAAGUGUAAAAUGAAACUACAUUUGCACUUUUAACGUGCGGCUGCAGUUUGGAACCACCCGGAAGAGUUGUUCAGGCUGUUACUAAUAUUAUUGUGAGAGUAUCACGUAUAAUAUUGCUUAAUACCGGUGCCCCGUCGGCUCCGGAAAGGUCAUGC..GUUUAG
env-354  UAUUAUAAAGUGGGCCCCUGACUGGUGGCUGCGUCAGGAUGCCUUGGUAAAUGUGCAUCAGCACAUUCCGAAACUUUGUAAUGCUUCAGGCAUUAAAACGAUUGCAUUUUACUGCAUUCGCUUAAUACUCCUUCCUGAACCGUCACUCUUAAAGGGAGGAGAAAGGCCGUAA..GACAC.
env-355  AUAUUAGAUGAGAAUGUGGAAUAUUAGUAGAGAAUCAAGUGGGAGCGAAGCGAACAUUUGAUUUAAUCUGAUGUUUCAACGAAAGACCAAAGGAAUUAAGUUUUUUAAAAGGGUCUUUGUUUUUCAAUAAUAUUGCUUAAUACUGACAUCACCGAUGUCAGAAAGGUUGUGU..AAA...
env-356  AUAUUAGAUGAGAAUGUGGAAUAUUAGUAGAGAAUCAAGUGGGAGCGAAGCGAACAUUUGAUUUAAUCUGAUGUUUCAACGAAAGACCAAAGGAAUUAAGUUUUUUAAAAGGGUCUUUGUUUUUCAAUAAUAUUGCUUAAUACUGACAUCACCGAUGUCAGAAAGGUUGUGU..AAA...
env-357  UAUUAAAAUAUUAUAGAGUAUUAGCAUAAAUCAAGCACGAACAAAGUGAGUAUUUGGUUUUACUGAUAUUUAAACGAAAGACUGAAACGACUAAGUUUUAUAAAAUAGGUCUUUACGUUAUAUAAUAUUGCUUAAUACCGACAUCUUUAUUGAAGAGUCGGAAAGGUCAUG...GUAAAA
env-358  UAUUAAAAUAUUAUAGAGUAUUAGCAUAAAUCAAGCACGAACAAAGUGAGUAUUUGGUUUUACUGAUAUUUAAACGAAAGACCGAAACGAUUAAGUUUUAUAAAAUAGGUCUUUGCGUUAUAUAAUAUUGCUUAAUACCGACAUCUUUAUUGAAGAGUCGGAAAGGUCAUG...GUAAAA
env-359  UAUUAAAAUAUUAUAGAGUAUUAGCAUAAAUCAAGCACGAACAAAGUGAGUAUUUGGUUUUACUGAUAUUUAAACGAAAGACUGAAACGACUAAGUUUUAUAAAAUAGGUCUUUACGUUAUAUAAUAUUGCUUAAUACCGACAUCUUUAUUGAAGAGUCGGAAAGGUCAUG...GUAAAA
env-360  .....................................................................................................................................................UCUAAGGAACGGAAAGGUUGUGU..UUUAGC
env-361  AGGAUUCUAAUGAAAGUAAGAUCAUCUGUUAAACCUAUUUGCGAGAAAUGCAAAAUUAUUAAGAGAAAAGGAAGUAUCAGAGUAAUCUGUGAGAAUCCAAAGCAUAAACAACGUCAAGGUUAGACAUGGCGUUGCUUAAUACCGCCAUAUCUAAGCGGCGGAAAGGUUGUGU..UUUAGC
env-362  AGGAUUCUAAUGAAAGUAAGAUCAUCUGUUAAACCUAUUUGCGAGAAAUGCAAAAUUAUUAAGAGAAAAGGAAGUAUCAGAGUAAUCUGUGAGAAUCCAAAGCAUAAACAACGUCAAGGUUAGACAUGACGUUGCUUAAUACCGCCAUAUCUAAGCGGCGGAAAGGUUGUGU..UUUAGC
env-363  GAUUCUAAUGAAAGUAAGAUCAUCUGUUAAACCUAUUUGCGAAAAAUGCAAAGUCAUCAAGAGAAAAGGAAGUAUCAGAGUAAUCUGUGAGAAUCCAAAGCAUAAACAACGCCAAGGUUAAAUAACGGCGUUGCUUAAUACCGCCUAAUAAUGCAAGGCGGAAAGGUUGUGU..UUUAGC
env-364  AUUCGUCAUGAAAGUAAGAUCAUCUGUUAAACCUAUUUGCGAAAAAUGCAAGAUUAUCAAGAGAAAAGGAAGUAUCAGAGUAAUCUGUGAGAAUCCAAAGCAUAAACAACGUCAAGGUUAAAUAACGACGUUGCUUAAUACCGCCUAAUAAUGCGAGGCGGAAAGGUUGUGU..UUUAGC
env-365  .......................................................................................................................................UAAUACCGCCUAAUAAAGCGGGGCGGAAAGGUUGUGU..UUUAGC
env-366  GAUUCUAAUGAAAGUAAGAUCAUCUGUUAAACCUAUUUGCGAAAAAUGCAAAGUCAUCAAGAGAAAAGGAAGUAUCAGAGUAAUCUGUGAGAAUCCAAAGCAUAAACAACGUCAAGGUUAAAUAACGGCGUUGCUUAAUACCGCCUAAUAAUGCAAGGCGGAAAGGUUGUGU..UUUAGC
env-367  UGCAGCGAGGACGCGCGUGAAUGCGUUGUUCUGCUGUUAAUAUUAUAAAGUGAGACACCUGGCAAGUUGCAGCCAGGACACCUCCGGCGGGUGGACGCGCGGAGCGCAGACAUCCCUUAUAAUAUUGCUUAAUACUCCGAUCUGUCAGGUGGCAGAAGGAGAAAGGUCGUG...CGAA..
env-368  UGCAGCGAGGACGCGCGUGAAUGCGUUGUUCUGCUGUUAAUAUUAUAAAGUGAGACACCUGGCAAGUUGCAGCCAGGACACCUCCGGCGGGUGGACGCGCGGAGCGCAGACAUCCCUUAUAAUAUUGCUUAAUACUCCGAUCUGUCAGGUGGCAGAAGGAGAAAGGUCGUG...CGAA..
env-369  UGCAGCGAGGACGCGCGUGAAUGCGUUGUUCUGCUGUUAAUAUUAUAAAGUGAGACACCUGGCAAGUUGCAGCCAGGACACCUCCGGCGGGUGGACGCGCGGAGCGCAGACAUCCCUUAUAAUAUUGCUUAAUACUCCGAUCUGUCAGGUGGCAGAAGGAGAAAGGUCGUG...CGAA..
env-370  UGCAGCGAGGACGCGCGUGAAUGCGUUGUUCUGCUGUUAAUAUUAUAAAGUGAGACACCUGGCAAGUUGCAGCCAGGACACCUCCGGCGGGUGGACGCGCGGAGCGCAGACAUCCCUUAUAAUAUUGCUUAAUACUCCGAUCUGUCAGGUGGCAGAAGGAGAAAGGUCGUG...CGAA..
env-371  UGCAGCGAGGACGCGCGUGAAUGCGUUGUUCUGCUGUUAAUAUUAUAAAGUGAGACACCUGGCAAGUUGCAGCCAGGACACCUCCGGCGGGUGGACGCGCGGAGCGCAGACAUCCCUUAUAAUAUUGCUUAAUACUCCGAUCUGUCAGGUGGCAGAAGGAGAAAGGUCGUG...CGAA..
env-372  UGCAGCGAGGACGCGCGUGAAUGCGUUGUUCUGCUGUUAAUAUUAUAAAGUGAGACACCGGGCAAGUUGCAGCCAGGACACCUCCGGCGGGUGGACGCGCGGAGCGCAGACAUCCCUUAUAAUAUUGCUUAAUACUCCGAUCUGUCAGGUGGCAGAAGGAGAAAGGUCGUG...CGAA..
env-373  UGCAGCGAGGACGCGCGUGAAUGCGUUGUUCUGCUGUUAAUAUUAUAAAGUGAGACACCUGGCAAGUUGCAGCCAGGACACCUCCGGCGGGUGGACGCGCGGAGCGCAGACAUCCCUUAUAAUAUUGCUUAAUACUCCGAUCUGUCAGGUGGCAGAAGGAGAAAGGUCGUG...CGAA..
env-374  UGCAGCGAGGACGCGCGUGAAUGCGUUGUUCUGCUGUUAAUAUUAUAAAGUGAGACACCUGGCAAGUUGCAGCCAGGACACCUCCGGCGGGUGGACGCGCGGAGCGCAGACAUCCCUUAUAAUAUUGCUUAAUACUCCGAUCUGUCAGGUGGCAGAAGGAGAAAGGUCGUG...CGAA..
env-375  ..................GAAUGCGUUGUUCUGCUGUUAAUAUUAUAAAGUGAGACACCUGGCAAGUUGCAGCCAGGACACCUCCGGCGGGUGGACGCGCGGAGCGCAGACAUCCCUUAUAAUAUUGCUUAAUACUCCGAUCUGUCAGGUGGCAGAAGGAGAAAGGUCGUG...CGAA..
env-377  UGCAGCGAGGACGCGCGUGAAUGCGUUGUUCUGCUGUUAAUAUUAUAAAGUGAGACACCUGGCAAGUUGCAGCCAGGACACCUCCGGCGGGUGGACGCGCGGAGCGCAGACAUCCCUUAUAAUAUUGCUUAAUACUCCGAUCUGUCAGGUGGCAGAAGGAGAAAGGUCGUG...CGAA..
env-383  UGCAGCGAGGACGCGCGUGAAUGCGUUGUUCUGCUGUUAAUAUUAUAAAGUGAGACACCUGGCAAGUUGCAGCCAGGACACCUCCGGCGGGUGGACGCGCGGAGCGCAGACAUCCCUUAUAAUAUUGCUUAAUACUCCGAUCUGUCAGGUGGCAGAAGGAGAAAGGUCGUG...CGAA..
env-384  UGCAGCGAGGACGCGCGUGAAUGCGUUGUUCUGCUGUUAAUAUUAUAAAGUGAGACACCUGGCAAGUUGCAGCCAGGACACCUCCGGCGGGUGGACGCGCGGAGCGCAGACAUCCCUUAUAAUAUUGCUUAAUACUCCGAUCUGUCAGGUGGCAGAAGGAGAAAGGUCGUG...CGAA..
env-385  ACCUCCGGCGGGUGGACGCGCGGUGCGCAGACAUCCCUUAUAAUAUUGCUUAAUACUCCGAUCUGUCAGGUGGCAGAAGG...............................................................................AGAAAGGUCGUG...CGAA..
env-386  ACCUCCGGCGGAUGGACGCGCGGAGCGCAGACAUCCCUUAUAAUAUUGCUUAAUACUCCGAUCUGUCAGGUGGCAGAAGG...............................................................................AGAAAGGUCGUG...CGAA..
env-387  ..................CGCGGAGCGCAGACAUCCCUUAUAAUAUUGCUUAAUACUCCGAUCUGUCAGGUGGCAGAAGG...............................................................................AGAAAGGUCGUG...CGAA..
env-388  ..................CGCGGAGCGCAGACAUCCCUUAUAAUAUUGCUUAAUACUCCGAUCUGUCAGGUGGCAGAAGG...............................................................................AGAAAGGUCGUG...CGAA..
env-389  ACCUCCGGCGGAUGGACGCGCGGAGCGCAGACAUCCCUUAUAAUAUUGCUUAAUACUCCCAUCUGUCAGGUGGCAGAAGG...............................................................................AGAAAGGUCGUG...CGAA..
env-390  AGUGCGUUGUUUUGCUGUUAAUAUUAUAAAGUGAGACACCUGGCAGACAAAGCAGCCAGGACACCUCCGGCGGAUGGACG.................CGCGGAGCGCAGACAUCCCUUAUAAUAUUGCUUAAUACUCCGAUCUGUCAGGUGGCAGAAGGAGAAAGGUCGUG...CGAA..
env-391  UGCAGCGAGGACGCGCGUGAAUGCGUUGUUCUGCUGUUAAUAUUAUAAAGUGAGACACCUGGCAAGUUGCAGCCAGGACACCUCCGGCGGGUGGACGCGCGGAGCGCAGACAUCCCUUAUAAUAUUGCUUAAUACUCCGAUCUGUCAGGUGGCAGAAGGAGAAAGGUCGUG...CGAA..
env-392  UGCAGCGAGGACGCGCGUGAAUGCGUUGUUCUGCUGUUAAUAUUAUAAAGUGAGACACCUGGCAAGUUGCAGCCAGGACACCUCCGGCGGGUGGACGCGCGGAGCGCAGACAUCCCUUAUAAUAUUGCUUAAUACUCCGAUCUGUCAGGUGGCAGAAGGAGAAAGGUCGUG...CGAA..
env-393  UGCAGCGAGGACGCGCGUGAAUGCGUUGUUCUGCUGUUAAUAUUAUAAAGUGAGACACCUGGCAAGUUGCAGCCAGGACACCUCCGGCGGGUGGACGCGCGGAGCGCAGACAUCCCUUAUAAUAUUGCUUAAUACUCCGAUCUGUCAGGUGGCAGAAGGAGAAAGGUCGUG...CGAA..
env-394  UGCAGCGAGGACGCGCGUGAAUGCGUUGUUCUGCUGUUAAUAUUAUAAAGUGAGACCCCUGGCAAGUUGCAGCCAGGACACCUCCGGCGGGUGGACGCGCGGAGCGCAGACAUCCCUUAUAAUAUUGCUUAAUACUCCGAUCUGUCAGGUGGCAGAAGGAGAAAGGUCGUG...CGAA..
env-395  UGCAGCGAGGACGCGCGUGAAUGCGUUGUUCUGCUGUUAAUAUUAUAAAGUGAGACACCUGGCAAGUUGCAGCCAGGACACCUCCGGCGGGUGGACGCGCGGAGCGCAGACAUCCCUUAUAAUAUUGCUUAAUACUCCGAUCUGUCAGGUGGCAGAAGGAGAAAGGUCGUG...CGAA..
env-396  UGCAGCGAGGACGCGCGUGAAUGCGUUGUUCUGCUGUUAAUAUUAUAAAGUGAGACACCUGGCAAGUUGCAGCCAGGACACCUCCGGCGGGUGGACGCGCGGAGCGCAGACAUCCCUUAUAAUAUUGCUUAAUACUCCGAUCUGUCAGGUGGCAGAAGGAGAAAGGUCGUG...CGAA..
env-397  UGCAGCGAGGACGCGCGUGAAUGCGUUGUUCUGCUGUUAAUAUUAUAAAGUGAGACACCUGGCAAGUUGCAGCCAGGACACCUCCGGCGGGUGGACGCGCGGAGCGCAGACAUCCCUUAUAAUAUUGCUUAAUACUCCGAUCUGUCAGGUGGCAGAAGGAGAAAGGUCGUG...CGAA..
env-398  UGCAGCGAGGACGCGCGUGAAUGCGUUGUUCUGCUGUUAAUAUUAUAAAGUGAGACACCUGGCAAGUUGCAGCCAGGACACCUCCGGCGGGUGGACGCGCGGAGCGCAGACAUCCCUUAUAAUAUUGCUUAAUACUCCGAUCUGUCAGGUGGCAGAAGGAGAAAGGUCGUG...CGAA..
env-399  UGCAGCGAGGACGCGCGUGAAUGCGUUGUUCUGCUGUUAAUAUUAUAAAGUGAGACACCUGGCAAGUUGCAGCCAGGACACCUCCGGCGGGUGGACGCGCGGAGCGCAGACAUCCCUUAUAAUAUUGCUUAAUACUCCGAUCUGUCAGGUGGCAGAAGGAGAAAGGUCGUG...CGAA..
env-405  UGCAGCGAGGACGCGCGUGAAUGCGUUGUUCUGCUGUUAAUAUUAUAAAGUGAGACACCUGGCAAGUUGCAGCCAGGACACCUCCGGCGGGUGGACGCGCGGAGCGCAGACAUCCCUUAUAAUAUUGCUUAAUACUCCGAUCUGUCAGGUGGCAGAAGGAGAAAGGUCGUG...CGAA..
env-408  UGGAGCGAGGGCGCGCGUGAAUGCGUUGUUCUGCUGUUAAUAUUAUAAAGUGAGACACCUGGCAAGUUGCAGCCAGGACACCUCCGGCGGGUGGACGCGCGGAGCGCAGACAUCCCUUAUAAUAUUGCUUAAUACUCCGAUCUGUCAGGUGGCAGAAGGAGAAAGGUCGUG...CGAA..
env-409  UGCAGCGAGGACGCGCGUGAAUGCGUUGUUCUGCUGUUAAUAUUAUAAAGUGAGACACCUGGCAAGUUGCAGCCAGGACACCUCCGGCGGGUGGACGCGCGGAGCGCAGACAUCCCUUAUAAUAUUGCUUAAUACUCCGAUCUGUCAGGUGGCAGAAGGAGAAAGGUCGUG...CGAA..
env-410  UGCAGCGAGGACGCGCGUGAAUGCGUUGUUCUGCUGUUAAUAUUAUAAAGUGAGACACCUGGCAAGUUGCAGCCAGGACACCUCCGGCGGGUGGACGCGCGGAGCGCAGACAUCCCUUAUAAUAUUGCUUAAUACUCCGAUCUGUCAGGUGGCAGAAGGAGAAAGGUCGUG...CGAA..
env-411  UGCAGCGAGGACGCGCGUGAAUGCGUUGUUCUGCUGUUAAUAUUAUAAAGUGAGACACCUGGCAAGUUGCAGCCAGGACACCUCCGGCGGGUGGACGCGCGGAGCGCAGACAUCCCUUAUAAUAUUGCUUAAUACUCCGAUCUGUCAGGUGGCAGAAGGAGAAAGGUCGUG...CGAA..

         ....................................................................................................................................................................<<<<<<<<<<......
         ......................................................................................................................................................................222222.........
         ●●●●●●●●●●●●●●●●●●●●●●●●●●●●●●●●●●●●●●●●●●●●●●●●●●●●●●●●●●●●●●●●●●●●●●●●●●●●●●●●●●●●●●●●●●●●●●●●●●●●●●●●●●●●●●●●●●●●●●●●●●●●●●●●●●●●●●●●●●●●●●●●●●●●●YYGGAAAGGYYGY○○--●●●●○○
                                                                                                                                                                            5′
```

```
env-412   ..........................GCAGACAUCCCUUAUAAUAUUGCUUAAUACUCCGAUCUGUCAGGUGGCAGAAGG...............................................................................AGAAAGGUCGUG...CGAA..
env-413   ...............................................................................................................................................................AGAAAGGUCGUG...CGAA..
env-414   ACCUCCGGCGGAUGGACGCGCGGAGCGCAGACAUCCCUUAUAAUAUUGCUUAAUACUCCCAUCUGUCAGGUGGCAGAAGG...............................................................................AGAAAGGUCGUG...CGAA..
env-418   ACCUCCGGCGGAUGGACGCGCGGAGCGCAGACAUCCCUUAUAAUAUUGCUUAAUACUCCGAUCUGUCAGGUGGCAGAAGG...............................................................................AGAAAGGUCGUG...CGAA..
env-419   ..CUCCGGCGGGUGGACGCGCGGAGCGCAGACAUCCCUUAUAAUAUUGCUUAAUACUCCGAUCUGUCAGGCGGCAGAAGG...............................................................................AGAAAGGUCGUG...CGAA..
env-420   UGCAGCGAGGACGCGCGUGAAUGCGUUGUUCUGCUGUUAAUAUUAUAAAGUGAGACACCUGGCAAGUUGCAGCCAGGACACCUCCGGCGGGUGGACGCGCGGAGCGCAGACAUCCCUUAUAAUAUUGCUUAAUACUCCGAUCUGUCAGGUGGCAGAAGGAGAAAGGUCGUG...CGAA..
env-428   ........GUGCGGCUGCAGUAGGACACAUCUGGAUGUGUUGUACUGUUUAGAUAUUAUAAUGAGAGUGCUCCGGAGUGGUGUACCGGAGAGCCUCCGGCGGAUGAACGCCGUAGAGAGCGUUGCAUCCCUUAUAAUAUUGCUUAAUACCGGCUAAGCCGGAAAGGUCAUG...GAGA..
env-429   AAAAUUAAGUGCGGCUGCAGUAGGACACAUCUGGAUGUGUUGUACUGUUUAGAUAUUAUAAUGAGAGUGCUCCGGAGUGGUGUACCGGAGAGCCUCCGGCGGAUGAACGCCGUAGAGAGCGUUGCAUCCCUUAUAAUAUUGCUUAAUACCGGCUAAGCCGGAAAGGUCAUG...GAGA..
env-430   AAAAUUAAGUGCGGCUGCAGUAGGACACAUCUGGAUGUGUUGUACUGUUUAGAUAUUAUAAUGAGAGUGCUCCGGAGUGGUGUACCGGAGAGCCUCCGGCGGAUGAACGCCGUAGAGAGCGUUGCAUCCCUUAUAAUAUUGCUUAAUACCGGCUAAGCCGGAAAGGUCAUG...GAGA..
env-432   .....UAAGUGCGGCUGCAGUAGGACACAUCUGGAUGUGUUGUACUGUUUAGAUAUUAUAAUGAGAGUGCUCCGGAGUGGUGUACCGGAGAGCCUCCGGCGGAUGAACGCCGUAGAGAGCGUUGCAUCCCUUAUAAUAUUGCUUAAUACCGGCUAAGCCGGAAAGGUCAUG...GAGA..
env-433   CUGGUUUCAGGCUGCUGCGCCGGAAUGGUUUCAGAAUUAUUUCGAUCGAUAUUGGCUUUUAAUACCGGCACAGGAUCGGCAGUACACGUCUGCGCACGAAAGCUGACAAGCGCAGAAAGCAUGCUUUUGCAAGGACAAAGGUCCAUGGCUUAAUGUACCGGAAAGGCUGCUU..AAA...
env-434   CUGUUUAAAUAAUAUUGUGAGAGGUCACAAUCUCGCAGCUUUAUUUCCGAGGCAGAAACGUCACGAUUUCUAUCCUGUCUCGGGCGAAGUAUUGCCUAAUACCGAGAGGCUAUUUCGGAGGUCCUUUUCAGGACACGACGGGCGUUAAGCCGGUCAAUCGGAAAGGUUGUGU..UGCAAU
Bhy-1-1   CUGUUUAAAUAAUAUUGUGAGAGGUCACAAUCUCGCAGCUUUAUUUCCGAGGCAGAAACGUCACGAUUUCUAUCCUGUCUCGGGCGAAGUAUUGCCUAAUACCGAGAGGCUAUUUCGGAGGUCCUUUUCAGGACACGACGGGCGUUAAGCCGGUCAAUCGGAAAGGUUGUGU..UGCAAU
env-435   UCCCUCAACGAUCCCUAUUGCAGUUUAUGCGAACUAAUAUUAUAAAGUGAGACCUGACUGGCUGCGUCAGGAUGCCUCCGGCGGAUCAUACGCAGGAGCGUAAGAUCCCUUAUAAUAUUACUUAAAACUCUCAUGUACGGCUGCCCUGCCGGAUGUGGGAGAAAGGCCGU....UCAUCG
env-436   UGCUUAUGUCGGGCAAUUCCUCAGGAACUGUCCGGAAAGUCGAUUCCGCAGUUUGCGGUAUCGCAGAAUAUUAUAAUGUAAGAAACCUGACUGGGCUGCGUCAGGAUACCUUCUUUAUAAUAUUACUUAAUACUUCCGCCGCCCGGUCGGGUUUACGGAAGAAAGGCCGC....UGUCGU
env-437   UGAAGGUCAGAUCAUCCGUGAAACCGAUGUGUGAAAAGUGCAAGGUUAUCAAGAGAAAAGGCAGCAUCCGUAUUAUUUGUGAGAAUCCAAAACACAAACAGCGUCAGGGCUGAUCCCUCAUAGCCUUAUCGCUCGAUCCUCUCUGUUAGGCUGACAGAGAGAAAGGCCGUAU..UCGUCU
env-438   ........................................UGCAAGGUUAUCAAGAGAAAAGGCAGUAUUCGUAUCAUUUGUGAGAAUCCAAAACACAAACAGCGCCAGGGCUGAUCCCUCACAGCUUUGGACUUAAUCCUCUCUGUUAGGCUGACAGAGAGAAAGGCCAUAU..UCGUCU
Bsp-7-1   CAAAUGAAAGUUAGGUCUUCUGUUAAGCCUAUGUGCGAAAAGUGCAAGGUUAUCAAGAGAAAAGGCGUUAUUCGUGUAAUUUGCGAAAACCCUAAGCACAAGCAGAGACAGGGCUGAUUCGAGUCAUUUUGACAUUACUUUUUGAUACCAAUGCGUAUUGGAAAGAUCGGAU..UAACUA
Bsp-9-1   ACAAAUGAAAGUUAGGUCUUCUGUUAAGCCUAUGUGCGAAAAGUGCAAGGUUAUUAAGAGAAAAGGCGUUAUUCGUGUAAUUUGCGAAAACCCUAAGCACAAACAGAGACAGGGCUGAUUCGAGUCAUUUGACAUUACUUUUUGAUACCAAUGUGUAUUGGAAAGAUCGGAU..UAACUU
Bsp-5-1   ACAAAUGAAAGUUAGGUCUUCUGUUAAGCCUAUGUGCGAAAAGUGCAAGGUUAUUAAGAGAAAAGGCGUUAUUCGUGUAAUUUGCGAAAACCCUAAGCACAAACAGAGACAGGGCUGAUUCGAGUCAUUUGACAUUACUUUUUGAUACCAAUGUGUAUUGGAAAGAUCGGAU..UAACUU
Bsp-2-1   CAAAUGAAAGUUAGGUCUUCUGUUAAGCCUAUGUGCGAAAAGUGCAAGGUUAUCAAGAGAAAAGGCGUUAUUCGUGUAAUUUGCGAAAACCCUAAGCACAAACAGAGACAGGGCUAAUUCGAGUCAUUUUGACAUUACUUUUUGAUACCAAUGUGUAUUGGAAAGAUCGGAU..UAACUA
Bsp-3-1   CAAAUGAAAGUUAGGUCUUCUGUUAAGCCUAUGUGCGAAAAGUGCAAGGUUAUUAAGAGAAAAGGCGUUAUUCGUGUAAUUUGCGAAAACCCUAAGCACAAACAGAGACAGGGCUGAUUCGAGUCAUUUUGACAUUACUUUUUGAUACCAAUGUGUAUUGGAAAGAUCGGAU..UAAGUA
Bpr-2-1   UGAUAGUUGAUGAUAUUUGUGUAAUAUUUUAGUGCGGCUGCAGUAUGAAACGCAAGGUUGCGUUGUUCGUACUGAUCAAAUAUAGUAAUAUAUCACGUUUGUCGUCCUGCCGAGAGGGCGAUUGCAGAAUAUUGCUUAAUACCGUCGCUCUGCCGUGACGGAAAGGCUGCGUAAAAUUUU
          ....................................................................................................................................................................<<<<<<<<<<......
          ....................................................................................................................................................................222222.........
          ●●●●●●●●●●●●●●●●●●●●●●●●●●●●●●●●●●●●●●●●●●●●●●●●●●●●●●●●●●●●●●●●●●●●●●●●●●●●●●●●●●●●●●●●●●●●●●●●●●●●●●●●●●●●●●●●●●●●●●●●●●●●●●●●●●●●●●●●●●●●●●●●●●●●●●●●YYGGAAAGGYYGY○○--●●●●○○
                                                                                                                                                                           5′
```

alignment positions 181 · · · 360

```
env-1     ...................................................GCG.CAGCUGGAUGC..................................GUCAUACCG.......................................................................
env-2     GGAUUGUUA...........................................UC.CGAUCGGGUAU..................................UGU.............................................................................
env-3     GGAUUGUUA...........................................UC.CGAUCGGGUAU..................................UGU.............................................................................
env-4     GGAUUGUUA...........................................UC.CGAUCGGGUAU..................................UGU.............................................................................
env-5     GGAUUGUUA...........................................UC.CGAUCGGGUAU..................................UGU.............................................................................
env-6     GGAUUGUUA...........................................UC.CGAUCGGGUAU..................................UGU.............................................................................
env-7     GGAUUGUUA...........................................UC.CGAUCGGGUAU..................................UGU.............................................................................
env-8     GGAUUGUUA...........................................UC.CGAUCGGGUAU..................................UGU.............................................................................
env-9     GGAUUGUUA...........................................UC.CGAUCGGGUAU..................................UGU.............................................................................
env-10    GGAUUGUUA...........................................UC.CGAUCGGGUAU..................................UGU.............................................................................
env-11    GGAUUGUUA...........................................UC.CGAUCGGGUAU..................................UGU.............................................................................
env-12    GGAUUGUUA...........................................UC.CGAUCGGGUAU..................................UGU.............................................................................
env-13    GGAUUGUUA...........................................UC.CGAUCGGGUAU..................................UGU.............................................................................
env-14    GGAUUGUUA...........................................UC.CGAUCGGGUAU..................................UGU.............................................................................
env-16    GGAUUGUUA...........................................UC.CGAUCGGGUAU..................................UGU.............................................................................
env-17    ...................................................GCC.CGGUUGGGGAU..................................AUUUAAA.........................................................................
env-18    ...................................................GCC.CGGUUGGGGAU..................................AUUUAAA.........................................................................
env-19    ..................................................AUCA.AUUCUGGAAGCUG................................GAUAA...........................................................................
Bsp-6-1   CU.................................................GUC.CGAUUGGGC....................................AAGGAAAUAAUA....................................................................
env-20    ...................................................GCA.UGGCUGGGUACGG................................UACUU...........................................................................
env-21    CAACU..............................................GUC.CGAUUGGGCGU..................................AUAAG...........................................................................
env-22    ...................................................GCG.CAGCUGGAGGGCAG.........AGGAUAU.......UG......ACC..........................................................................CGAU
env-23    ....................................................UC.CGGUUGGGUUU..................................UAUUC...........................................................................
env-24    ...................................................AUG.CAGUUGGGUAUUUU........UCUUUGUACGUCAUUGAACAGG.AAUU..................................................................UCC.GUUUUG
env-25    ...................................................AUG.CAGUUGGGUAUUUU........UCUUUGUACGUCAUUGAACAGG.AAUU..................................................................UCC.GUUUUG
env-28    ...................................................AUG.CAGUUGGGUAUUUU........UCUUUGUACGUCAUUGAACAGG.AAUU..................................................................UCC.GUUUUG
env-29    ...................................................GUG.UAGUCGGGUAUUU..........CUUUGUACGUCAUU...AAGG.AAUU..................................................................UCCUGCUUUG
env-30    ...................................................GUG.UAGUCGGGUAUUU..........CUUUGUACGUCAUU.AGUAGG.AAUU..................................................................UCCUGCUUUG
env-32    ...................................................GUG.UAGUCGGGUAUUU..........CUUUGUACGUCAUU.AGUAGG.AAUU..................................................................UCCUGCUUUG
env-33    ...................................................GUG.UAGUCGGGUAUUU..........CUUUGUACGUCAUU.AGUAGG.AAUU..................................................................UCCUGCUUUG
env-34    ...................................................ACC.CGGUUGGGCGUCC................................ACU.............................................................................
env-35    ...................................................ACC.CGGUUGGGCGUCC................................ACU.............................................................................
env-36    ...................................................ACC.CGGUUGGGCGUCC................................ACU.............................................................................
env-37    ...................................................UUC.CGGUUGGGUU...................................AGAAAU..........................................................................
env-38    ...................................................ACC.CGGCUGGAUGCG.................................AACAC...........................................................................
env-39    ....................................................CC.CGGCUGGACGCGCG...............................AACU............................................................................
env-41    ...................................................AUC.CGGCUGGAUGUACC...............................GAAUGU..........................................................................
env-42    GAGGA.............................................CGUA.CAGCUGGAUAUGCA...............................CCUUCG..........................................................................
env-43    UA...............................................GUGCA.GGCCUGGAUGUUU.GGAUGAUGGCAGAA..........GCACCUUUCU...................................................................GGCAGC....
env-45    AA..................................................CA.CGCCUGGGAAC..................................CAGGA...........................................................................
env-46    ACCG................................................CA.CGGCUGGAUAC..................................CGGGC...........................................................................
env-47    CA...............................................UCCGG.CAGCUGGGUG...................................CAUACCGAGA......................................................................
env-48    .................................................GGGCG.CGGCUGGGUG...................................ACUAUGCG........................................................................
env-49    ACAGC...............................................GG.CGGCUGGACACUGC...CAUGA...GGCG................UACCGCAU......................................................................CG
env-50    CC...............................................GGAUA.UGCCUGGGCGCAUC...............................AGUAUUCCGUCAAAUAC...............................................................
Dma-1-1   CC...............................................GGAUA.UGCCUGGGCGCAUC...............................AGUAUUCCGUCAAAUAC...............................................................
env-51    UA...................................................C.UGCCUGGAUGUAUU.......................GAGAAAUGAAACAC................................................................CACUACUC..
Lba-2-1   ...................................................UAG.UAGCUGGAAGUA.................................GAGA............................................................................
[ce-1-1   U..................................................UGG.CGGCUGGAUGCA.................................UAUACC..........................................................................
Ece-1-1   ...................................................UGG.CGGCUGGAUGCA.................................UAUACC..........................................................................

          .........<<<<<<<<<<<......>>>>>>>>>>>.............>>>>>.>>>>><<<<<<<<<.........<<<<<<<.......<<<<.<<<......................................................................>>>.>>>>>>
          .........…10…0.1......1.0…01...............22.2222..0222222..........111111.......111...1......................................................................1...11111
          ○-------------------------------------------------○YR-CRRCUGGRYR●○-------------------------------●●●●○○--------------------------------------------------------------------------
```

```
env-52     .....................................................G.CGGCUGGGUGCUGG...............................AACAAUU..........................................................................
env-53     .................................................UGACA.UGCCUGGGCUG.GGAUAUCGGUGU.AUUU.CAUCGG....UAGGACAAAGCU................................................................UC.AA....G
env-54     U...................................................UG.CACCUG.GUGUGAU...............................UCGGCAAAGUCU.....................................................................
env-55     U...................................................UG.CACCUG.GUGUGAU...............................UCGGCAAAGUCU.....................................................................
env-59     U...................................................UG.CACCUG.GUGUGAU...............................UCGGCAAAGUCU.....................................................................
env-60     U...................................................UG.CACCUG.GUGUGAU...............................UCGGCAAAGUCU.....................................................................
env-62     U...................................................UG.CACCUG.GUGUGAU...............................UCGGCAAAGUCU.....................................................................
env-63     U...................................................UG.CACCUG.GUGUGAU...............................UCGGCAAAGUCU.....................................................................
env-64     U...................................................UG.CACCUG.GUGUGAU...............................UCGGCAAAGUCU.....................................................................
env-66     U...................................................UG.CACCUG.GUGUGAU...............................UCGGCAAAGUCU.....................................................................
env-67     U...................................................UG.CACCUG.GUGUGAU...............................UCAGCAAAGUCU.....................................................................
env-68     U...................................................UG.CACCUG.GUGUGAU...............................UCAGCAAAGUCU.....................................................................
env-75     U...................................................UG.CACCUG.GUGUGAU...............................UCAGCAAAGUCU.....................................................................
env-76     U...................................................UG.CACCUG.GUGUGAU...............................UCAGCAAAGUCU.....................................................................
env-77     U...................................................UG.CACCUG.GUGUGAU...............................UCAGCAAAGUCU.....................................................................
env-78     U...................................................UG.CACCUG.GUGUGAU...............................UCAGCAAAGUCU.....................................................................
env-79     U...................................................UG.CACCUG.GUGUGAU...............................UCAGCAAAGUCU.....................................................................
env-80     U...................................................UG.CACCUG.GUGUGAU...............................UCAGCAAAGUCU.....................................................................
env-81     U...................................................UG.CACCUG.GUGUGAU...............................UCAGCAAAGUCU.....................................................................
Rgn-1-1    U...................................................UG.CACCUG.GUGUGAU...............................UCAGCAAAGUCU.....................................................................
env-82     U...................................................UG.CACCUGG.UGUGAU...............................UCAGCGAAGUCU.....................................................................
env-83     U...................................................UG.CACCUGG.UGUGAU...............................UCAGCGAAGUCU.....................................................................
env-87     U...................................................UG.CACCUGG.UGUGAU...............................UCAGCGAAGUCU.....................................................................
env-88     UA...................................................A.CGGCUGGUUCUGA................................CUGGU............................................................................
env-89     ...................................................GCG.CGGCUGGGAGUGA............AAAA................AAC.............................................................................U
env-90     ..................................................AACA.UGCCUGGGCUG..............GGAU................AUCGGUGUAUUUUCGCCGGUAGGACAAGGUGGAAGC............................................G
env-91     GGAUAU..G........................................UGACA.UGCCUGGGCUG...GGAUAUCGG.UGGAU................UUUCACCGGUAGGAUAUAAUAACCGACAACGCAGUUGCUUGGUCCGUCAUUUUAUUGGCUAAU.................G
env-92     ACA......UUUCC........GACG......GGAGGCGGGG........AGGA.CGGCUGGAUGCAUG...............................UCCGG............................................................................
env-93     ........AUUCGCAGGUUCGGAAA.GUGAUCGUCACACUUUGCCCACAGACUG.CAGCUGGGCGUU.................................ACCGA............................................................................
env-94     ...................................................GCA.CAGCUGGAUGC..................................GUCAUACCG........................................................................
Lba-11-1   ...................................................GUA.CAGCUGGAUGC..................................GUCAUACCG........................................................................
env-95     ..................................................CAUA.CAGCUGGGUGU..................................CUAUGC...........................................................................
env-96     ...................................................AUA.CAGCUGGGUGU..................................CCAUGUU..........................................................................
env-97     GAAA................................................CG.UACCUGGGC....................................GAGCAAAGCGCAA....................................................................
Dsp-3-1    GA..................................................CA.CGCCUGGGAGC..................................AGCCAAA..........................................................................
Dsp-1-1    AA..................................................CA.UGCCUGGAC....................................GAAA.............................................................................
env-101    AA..................................................CA.UGCCUGGGACAUG................................GAGA.............................................................................
env-102    AA..................................................CA.UGCCUGGGACAUG................................GAGA.............................................................................
Dsp-2-1    AA..................................................CA.UGCCUGGGACAUG................................GAGA.............................................................................
env-103    AA..................................................CA.UGCCUGGGACAUG................................GAGA.............................................................................
env-106    AA..................................................CA.UGCCUGGGACAUG................................GAGA.............................................................................
env-116    AA..................................................CA.UGCCUGGGACAUG................................GAGA.............................................................................
env-117    AA..................................................CA.UGCCUGGGACAUG................................GAGA.............................................................................
env-118    AA..................................................CA.UGCCUGGGACAUG................................GAGA.............................................................................
env-119    AA..................................................CA.UGCCUGGGACA..................................AAAUCA...........................................................................
env-120    AA..................................................CA.UGCCUGGGACA..................................AAAUCA...........................................................................
env-123    AA..................................................CA.UGCCUGGGACA..................................AAAUCA...........................................................................
env-124    AA..................................................CA.UGCCUGGGACA..................................AAAUCA...........................................................................
env-127    AA..................................................CA.UGCCUGGGACA..................................AAAUCA...........................................................................
env-128    AA..................................................CA.UGCCUGGGACA..................................AAAUCA...........................................................................
env-129    AA..................................................CA.UGCCUGGGACA..................................AAAUCA...........................................................................
env-130    AA..................................................CA.UGCCUGGGAC...................................AAAAAUACAG.......................................................................
env-136    AA..................................................CA.UGCCUGGGAC...................................AAAAAUACAG.......................................................................
env-137    AA..................................................CA.UGCCUGGGAC...................................AAAAAUACAG.......................................................................
env-138    AA..................................................CA.UGCCUGGGAC...................................AAAAAUACAG.......................................................................
env-139    AA..................................................CA.UGCCUGGGAC...................................AAAAAUACAG.......................................................................
Dlo-1-1    AA..................................................CA.UGCCUGGGAC...................................AAAAAUACAG.......................................................................
env-140    AA..................................................CA.UGCCUGGGAC...................................AAAAAUACAG.......................................................................
env-149    AA..................................................CA.CGCCUGGGAAC..................................CAGGA............................................................................
env-150    GAAA................................................CG.CACCUGGGAUU..................................UAAA.............................................................................
env-151    GAAA................................................CA.CACCUGGAUU...................................UAAA.............................................................................
Chy-1-1    GA..................................................CA.CGCCUGGGAGA..................................CCAACC...........................................................................
env-152    GA..................................................CG.CACCUGGGUGGCGC...............................GAA..............................................................................
env-153    GA..................................................CA.CACCUGGAUGGC.................................GGCACA...........................................................................
env-154    GA..................................................CA.CACCUGGAUGGC.................................GGCACA...........................................................................
Esp-1-1    GA..................................................CA.CGCCUGGGCUG.GG........AUAUCGGA.......AC......AUUACUGC.....................................................................GUUC
env-155    .................................................GUACA.UACCUGGGCUGGGA........UAUCG..........GAAG....GAUACAUUCCGGUAGGAUGCAGCAGUAAAUGAGUUACACCAGACUUCAUUUAUAAAAAGAAGAGACUCA....UUUC..
env-156    .................................................GUACA.UACCUGGGCUGGGA........UAUCG..........GAAG....GAUACAUUCCGGUAGGAUGCAGCAGUAAAUGAGUUACACCAGACUUCAUUUAUAAAAAGAAGAGACUCA....UUUC..
env-159    .................................................GUACA.UACCUGGGCUGGGA........UAUCG..........GAAG....GAUACAUUCCGGUAGGAUGCAGCAGUAAAUGAGUUACACCAGACUUCAUUUAUAAAAAGAAGAGACUCA....UUUC..
env-160    .................................................GUACA.UACCUGGGCUGGGA........UAUCG..........GAAG....GAUACAUUCCGGUAGGAUGCAGCAGUAAAUGAGUUACACCAGACUUCAUUUAUAAAAAGAAGAGACUCA....UUUC..
env-161    .................................................GUACA.UACCUGGGCUGGGA........UAUCG..........GAAG....GAUACAUUCCGGUAGGAUGCAGCAGUAAAUGAGUUACACCAGACUUCAUUUAUAAAAAGAAGAGACUCA....UUUC..
env-162    .................................................GUACA.UACCUGGGCUGGGA........UAUCG..........GAAG....GAUACAUUCCGGUAGGAUGCAGCAGUAAAUGAGUUACACCAGACUUCAUUUAUAAAAAGAAGAGACUCA....UUUC..
env-163    .................................................GUACA.UACCUGGGCUGGGA........UAUCG..........GAAG....GAUACAUUCCGGUAGGAUGCAGCAGUAAAUGAGUUACACCAGACUUCAUUUAUAAAAAGAAGAGACUCA....UUUC..
env-164    .................................................GUACA.UACCUGGGCUGGGA........UAUCG..........GAAG....GAUACAUUCCGGUAGGAUGCAGCAGUAAAUGAGUUACACCAGACUUCAUUUAUAAAAAGAAGAGACUCA....UUUC..
env-165    .................................................GUACA.UACCUGGGCUGGGA........UAUCG..........GAAG....GAUACAUUCCGGUAGGAUGCAGCAGUAAAUGAGUUACACCAGACUUCAUUUAUAAAAAGAAGAGACUCA....UUUC..
env-167    .................................................GUACA.UACCUGGGCUGGGA........UAUCG..........GAAG....GAUACAUUCCGGUAGGAUGCAGCAGUAAAUGAGUUACACCAGACUUCAUUUAUAAAAAGAAGAGACUCA....UUUC..
env-168    G.................................................UACA.UGCCUGGGCUGGGA........UAUCG..........GAAG....GAUACAUUCCGGUAGGAUGCAGCAGUAAAUGAGUUACACCAGGCUUCAUUUAUAAAUAGAAGAUACUCA....UUUC..
env-169    G.................................................UACA.UGCCUGGGCUGGGA........UAUCG..........GAAG....GAUACAUUCCGGUAGGAUGCAGCAGUAAAUGAGUUACACCAGACUUCAUUUAUAAAUAGAAGAUACUCA....UUUC..
env-170    G.................................................UACA.UGCCUGGGCUGGGA........UAUCG..........GAAG....GAUACAUUCCGGUAGGAUGCAGCAGUAAAUGAGUUACACCAGACUUCAUUUAUAAAUAGAAGAUACUCA....UUUC..
env-171    G.................................................UACA.UGCCUGGGCUGGGA........UAUCG..........GAAG....GAUACAUUCCGGUAGGAUGCAGCAGUAAAUGAGUUACACCAGACUUCAUUUAUAAAUAGAAGAUACUCA....UUUC..
env-172    G.................................................UACA.UGCCUGGGCUGGGA........UAUCG..........GAAG....GAUACAUUCCGGUAGGAUGCAGCAGUAAAUGAGUUACACCAGACUUCAUUUAUAAAUAGAAGAUACUCA....UUUC..
env-175    G.................................................UACA.UGCCUGGGCUGGGA........UAUCG..........GAAG....GAUACAUUCCGGUAGGAUGCAGCAGUAAAUGAGUUACACCAGACUUCAUUUAUAAAUAGAAGAUACUCA....UUUC..
env-176    G.................................................UACA.UGCCUGGGCUGGGA........UAUCG..........GAAG....GAUACAUUCCGGUAGGAUGCAGCAGUAAAUGAGUUACACCAGACUUCAUUUAUAAAUAGAAGAUACUCA....UUUC..

           .........<<<<<<<<<<<......>>>>>>>>>>>............>>>>>.>>>>><<<<<<<<<.........<<<<<<<.......<<<<.<<<.......................................................................>>>.>>>>>>
           ............10...0.1......1.0...01..................22.2222..0222222..........111111.........111..1.........................................................................1..11111
           ○-------------------------------------------------○YR-CRRCUGGRY○○------------------------------------●●●○○○-------------------------------------------------------------------------
```

```
env-177   G.................................................UACA.UGCCUGGGCUGGGA........UAUCG..........GAAG....GAUACAUUCCGGUAGGAUGCAGCAGUAAAUGAGUUACACCAGACUUCAUUUAUAAAUAGAAGAUACUCA.....UUUC..
env-178   G.................................................UACA.UGCCUGGGCUGGGA........UAUCG..........GAAG....GAUACAUUCCGGUAGGAUGCAGCAGUAAAUGAGUUACACCAGACUUCAUUUAUAAAUAGAAGAUACUCA.....UUUC..
Lba-4-1   CGAA...............................................CCG.UAUCUGGAC....................................GAAAUCA.........................................................................
env-179   GAAA................................................CG.UACCUGGGC....................................GAGCAAAGCGCAA...................................................................
Cba-3-1   GAAA................................................CG.UACCUGGGC....................................GAGCAAAGCGCAA...................................................................
[sc-1-1   GAAA................................................CG.UACCUGG.....CG...............................AGCAAAG.........................................................................
Lba-7-1   GAAA................................................CG.UACCUGGGC....................................GAGCAAAGCGCAA...................................................................
env-181   GAAA................................................CG.UACCUGGGC....................................GAGCAAAGCGCAA...................................................................
env-182   ....................................................GG.CAGCUGGGUGU..................................CUAUGC..........................................................................
env-183   ....................................................UC.CUGCUGGGC....................................GUCGAAAGA.......................................................................
env-184   A...................................................CG.CGUCUGGGCUG............GGAU..................AUCGGCAAAAGGCCGGUAGGAUGCGGUAUUAUAAUAAGCAGAGC....................................
env-185   A...................................................CG.CGUCUGGGCUG............GGAU..................AUCGGCAAAAGGCCGGUAGGAUGCGGCAUUAUAAUAAGCAGAGC....................................
env-186   A...................................................CG.CGUCUGGGCUG............GGAU..................AUCGGCAAAAGGCCGGUAGGAUGCGGCAUUAUAAUAAGCAGAGC....................................
env-187   A...................................................CG.CGUCUGGGCUG............GGAU..................AUCGGCAAAAGGCCGGUAGGAUGCGGCAUUAUAAUAAGCAGAGC....................................
env-188   A...................................................CG.CGUCUGGGCUG............GGAU..................AUCGGCAAAAGGCCGGUAGGAUGCGGCAUUAUAAUAAGCAGAGC....................................
env-189   A...................................................CG.CGUCUGGGCUG............GGAU..................AUCGGCAAAAGGCCGGUAGGAUGCGGCAUUAUAAUAAGCAGAGC....................................
env-190   A...................................................CG.CGUCUGGGCUG............GGAU..................AUCGGCAAAAGGCCGGUAGGAUGCGGCAUUAUAAUAAGCAGAGC....................................
env-191   A...................................................CG.CGUCUGGGCUG............GGAU..................AUCGGCAAAAGGCCGGUAGGAUGCGGCAUUAUAAUAAGCAGAGC....................................
env-192   A...................................................CG.CGUCUGGGCUG............GGAU..................AUCGGCAAAAGGCCGGUAGGAUGCGGCAUUAUAAUAAGCAGAGC....................................
env-194   A...................................................CG.CGUCUGGGCUG............GGAU..................AUCGGCAAAAGGCCGGUAGGAUGCGGCAUUAUAAUAAGCAGAGC....................................
env-195   A...................................................CG.CGUCUGGGCUG............GGAU..................AUCGGCAAAAGGCCGGUAGGAUGCGGCAUUAUAAUAAGCAGAGC....................................
env-200   A...................................................CG.CGUCUGGGCUG............GGAU..................AUCGGCAAAAGGCCGGUAGGAUGCGGCAUUAUAAUAAGCAGAGC....................................
env-201   A...................................................CG.CGUCUGGGCUG............GGAU..................AUCGGCAAAAGGCCGGUAGGAUGCGGCAUUAUAAUAAGCAGAGC....................................
env-202   A...................................................CG.CGUCUGGGCUG............GGAU..................AUCGGCAAAAGGCCGGUAGGAUGCGGCAUUAUAAUAAGCAGAGC....................................
env-211   A...................................................CG.CGUCUGGGCUG............GGAU..................AUCGGCAAAAGGCCGGUAGGAUGCGGCAUUAUAAUAAGCAGAGC....................................
env-212   A...................................................CG.CGUCUGGGCUG............GGAU..................AUCGGCAAAAGGCCGGUAGGAUGCGGCAUUAUAAUAAGCAGAGC....................................
env-216   A...................................................CG.CGUCUGGGCUG............GGAU..................AUCGGCAAAAGGCCGGUAGGAUGCGGCAUUAUAAUAAGCAGAGC....................................
env-217   ...................................................AUA.CAGCUGGGUGUCA................................UGCACU..........................................................................
Rpe-1-1   ...................................................AUG.CGGCUGGGCA................................GGACCAUCUA...............................................................UCCG......
Rsp-1-1   ...................................................AUG.CGGCUGGGCA................................GGACCAUCUA...............................................................UCCG......
env-218   ...................................................GGC.UGGCUGGGC....................................AGCACGAAGGUG....................................................................
env-219   CGG................................................UGA.CGGCUGGGUGU..................................GACCGAG.........................................................................
env-220   ....................................................GG.CAGCUGGACAG..................................CUAUGU..........................................................................
env-221   .................................................UUGUG.CGGCUGGGC..........GG....................ACGGUUUUG.................................................................CCG.......
env-222   .................................................UUGUG.CGGCUGGGC..........GG....................ACGGUUUUG.................................................................CCG.......
env-223   .................................................UUGUG.CGGCUGGGC..........GG....................ACGGUUUUG.................................................................CCG.......
env-226   .................................................UUGUG.CGGCUGGGC..........GG....................ACGGUUUUG.................................................................CCG.......
env-227   .................................................UUGUG.CGGCUGGGC..........GG....................ACGGUUUUG.................................................................CCG.......
env-228   .................................................UUGUG.CGGCUGGGC..........GG....................ACGGUUUUG.................................................................CCG.......
env-229   .................................................UUGUG.CGGCUGGGC..........GG....................ACGGUUUUG.................................................................CCG.......
env-230   .................................................UUGUG.CGGCUGGGC..........GG....................ACGGUUUUG.................................................................CCG.......
env-231   .................................................UUGUG.CGGCUGGGC..........GG....................ACGGUUUUG.................................................................CCG.......
env-232   .................................................UUGUG.CGGCUGGGC..........GG....................ACGGUUUUG.................................................................CCG.......
env-234   .................................................UUGUG.CGGCUGGGC..........GG....................ACGGUUUUG.................................................................CCG.......
env-235   .................................................UUGUG.CGGCUGGGC..........GG....................ACGGUUUUG.................................................................CCG.......
env-236   .................................................UUGUG.CGGCUGGGC..........GG....................ACGGUUUUG.................................................................CCG.......
env-237   .................................................UUGUG.CGGCUGGGC..........GG....................ACGGUUUUG.................................................................CCG.......
env-238   .................................................UUGUG.CGGCUGGGC..........GG....................ACGGUUUUG.................................................................CCG.......
env-239   .................................................UUGUG.CGGCUGGGC..........GG....................ACGGUUUUG.................................................................CCG.......
env-240   .................................................UUGUG.CGGCUGGGC..........GG....................ACGGUUUUG.................................................................CCG.......
env-241   ...................................................CUG.CAGCUGGGCGU..................................UUGU............................................................................
env-242   UA...................................................G.CAGCUGGGUACGU................................UAUG............................................................................
env-243   UA...................................................G.CAGCUGGGUACGU................................UAUG............................................................................
env-244   UA...................................................G.CAGCUGGGUACGU................................UAUG............................................................................
env-245   UA...................................................G.CAGCUGGGUACGU................................UAUG............................................................................
env-246   UA...................................................G.CAGCUGGGUACGU................................UAUG............................................................................
env-252   UA...................................................G.CAGCUGGGUACGU................................UAUG............................................................................
env-254   UU...................................................G.UGGCUGGACGCAG...UGAAGAA..GGAU................GAAG...........................................................................A
Lba-6-1   ...................................................CUG.UGGCUGGACGCAG...UGAAGAA..GGAU................GAAG...........................................................................A
env-255   ...................................................CUG.UGGCUGGACGCAG...CGAAGAA..GGAU................GAAG...........................................................................A
Lba-10-1  ...................................................CUG.UGGCUGGACGCAG...CGAAGAA..GGAU................GAAG...........................................................................A
env-256   ...................................................CUG.UGGCUGGACGCAG...UGAAGAA..GGAU................GAAG...........................................................................A
env-258   ...................................................CUG.UGGCUGGACGCAG...UGAAGAA..GGAU................GAAG...........................................................................A
env-259   ...................................................CUG.UGGCUGGACGCAG...UGAAGAA..GGAU................GAAG...........................................................................A
env-260   ...................................................AUG.CAGCUGGGCAUA.................................CUGGUC..........................................................................
env-261   ...................................................AUA.CAGCUGGGUACC.................................ACAC............................................................................
env-262   AAA................................................GCACCGCCUGGGCGCA..GUCUAAAAGAACAA.................GACGUAGACAAGGAAAAG..............................................................
env-263   ...................................................CUG.CGGCUGGGCAG..................................CGCUUA..........................................................................
Bfo-1-1   .........AUUGCCGUUUGCUUAU.CAGACGGCAGUCAUUU........UUUG.CGGCUGGGU............GGUGGAAA................AGA............................................................................U
Bpr-1-1   A................................................UUUUG.CGGCUGGAUGCAUA...............................GAGACCCGUCAUUUU.................................................................
Cba-1-1   A................................................UUUUG.CGGCUGGAUGCAUA...............................GAGACCCGUCAUUUU.................................................................
env-264   CGCCGUCGU...........................................CG.CGGCUGGGUGUU.................................ACCGA...........................................................................
env-265   .................................................GGCCG.CGGCUGGGUGU..................................GAUACCGAG.......................................................................
env-266   UCUGAC.............................................CCG.CGGCUGGGUGUG.................................AAUACCGA........................................................................
env-267   CCAAAA...............................................G.CGACUGGGUGUUUU...............................ACC.............................................................................
env-268   CCAAAA...............................................G.CGGCUGGGUGUUUU...............................ACC.............................................................................
env-271   CCAAAA...............................................G.CGGCUGGGUGUUUU...............................ACC.............................................................................
env-272   CCAAAA...............................................G.CGGCUGGGUGUUUU...............................ACC.............................................................................
env-273   .................................................AAAGA.CGACUGGGUGUUUU...............................ACC.............................................................................
env-275   .................................................AAAGG.CGGCUGGGUGUUUU...............................ACC.............................................................................
env-276   ..................................................AAGG.CGGCUGGGUGUUUU...............................ACC.............................................................................
env-277   ..................................................AAGA.CGGCUGGGUGUUUU...............................ACC.............................................................................
env-278   ..................................................AAGA.CGGCUGGGUGUUUU...............................ACC.............................................................................

          .........<<<<<<<<<<<......>>>>>>>>>>>............>>>>>.>>>>><<<<<<<<<.........<<<<<<<.......<<<<.<<<......................................................................>>>.>>>>>>
          ............10...0.1......1.0...01..................22.2222...0222222.........111111........111...1.......................................................................1...11111
          ○---------------------------------------------------○YR-CRRCUGGRYR○○-------------------------------●●●○○○------------------------------------------------------------------------
```

```
env-279  AA.................................................AGA.CGGCUGGGUGUUUU...............................ACC..............................................................................
env-280  AA.................................................UUA.CAGCUGGGUGUUUU...............................ACC..............................................................................
env-281  CCAAUU...............................................A.CAGCUGGGUGUUUU...............................ACC..............................................................................
env-282  CCAAUU...............................................A.CAGCUGGGUGUUUU...............................ACC..............................................................................
Bsp-1-1  CCAAUU...............................................A.CAGCUGGGUGUUUU...............................ACC..............................................................................
env-283  CCAAUU...............................................A.CAGCUGGGUGUUUU...............................ACC..............................................................................
env-284  CCAAUU...............................................A.CAGCUGGGUGUUUU...............................ACC..............................................................................
env-285  CCAAUU...............................................A.CAGCUGGGUGUUUU...............................ACC..............................................................................
env-286  CCAAUU...............................................A.CAGCUGGGUGUUUU...............................ACC..............................................................................
env-287  CCAAUU...............................................A.CAGCUGGGUGUUUU...............................ACC..............................................................................
env-288  CCAAUU...............................................A.CAGCUGGGUGUUUU...............................ACC..............................................................................
env-289  CCAAUU...............................................G.CAGCUGGGUGUUUU...............................ACC..............................................................................
env-290  CCAAUU...............................................G.CAGCUGGGUGUUUU...............................ACC..............................................................................
env-291  CCAAUU...............................................G.CAGCUGGGUGUUUU...............................ACC..............................................................................
env-292  CCAAUU...............................................G.CAGCUGGGUGUUUU...............................ACC..............................................................................
env-293  CCAAUU...............................................G.CAGCUGGGUGUUUU...............................ACC..............................................................................
env-294  CCAAUU...............................................G.CAGCUGGGUGUUUU...............................ACC..............................................................................
env-295  CCAAUU...............................................G.CAGCUGGGUGUUUU...............................ACC..............................................................................
env-296  CCAAUU...............................................G.CAGCUGGGUGUUUU...............................ACC..............................................................................
env-298  CCAAUU...............................................G.CAGCUGGGUGUUUU...............................ACC..............................................................................
env-299  CCAAUU...............................................G.CAGCUGGGUGUUUU...............................ACC..............................................................................
env-300  CCAAUU...............................................G.CAGCUGGGUGUUUU...............................ACC..............................................................................
env-301  CCAAUU...............................................G.CAGCUGGGUGUUUU...............................ACC..............................................................................
env-302  CCAAUU...............................................G.CAGCUGGGUGUUUU...............................ACC..............................................................................
env-303  CCAAUU...............................................G.CAGCUGGGUGUUUU...............................ACC..............................................................................
env-304  CCAAUU...............................................G.CAGCUGGGUGUUUU...............................ACC..............................................................................
env-305  CCAAUU...............................................G.CAGCUGGGUGUUUU...............................ACC..............................................................................
Bwe-2-1  CCAAUU...............................................G.CAGCUGGGUGUUUU...............................ACC..............................................................................
env-306  CCAAUU...............................................G.CAGCUGGGUGUUUU...............................ACC..............................................................................
env-307  CCAAUU...............................................G.CAGCUGGGUGUUUU...............................ACC..............................................................................
env-308  CCAAUU...............................................G.CAGCUGGGUGUUUU...............................ACC..............................................................................
env-309  CCAAUU...............................................G.CAGCUGGGUGUUUU...............................ACC..............................................................................
env-310  CCAAUU...............................................G.CAGCUGGGUGUUUU...............................ACC..............................................................................
env-311  CCAAUU...............................................G.CAGCUGGGUGUUUU...............................ACC..............................................................................
env-313  CCAAUU...............................................G.CAGCUGGGUGUUUU...............................ACC..............................................................................
env-314  CCAAUU...............................................G.CAGCUGGGUGUUUU...............................ACC..............................................................................
env-315  CCAAUU...............................................G.CAGCUGGGUGUUUU...............................ACC..............................................................................
env-317  CCAAUU...............................................G.CAGCUGGGUGUUUU...............................ACC..............................................................................
env-318  CCAAUU...............................................G.CAGCUGGGUGUUUU...............................ACC..............................................................................
Bwe-1-1  CCAAUU...............................................G.CAGCUGGGUGUUUU...............................ACC..............................................................................
env-319  CCAAUU...............................................G.CAGCUGGGUGUUUU...............................ACC..............................................................................
env-320  CCAAUU...............................................G.CAGCUGGGUGUUUU...............................ACC..............................................................................
env-321  CCAAUU...............................................G.CAGCUGGGUGUUUU...............................ACC..............................................................................
env-322  CCAAUU...............................................G.CAGCUGGGUGUUUU...............................ACC..............................................................................
env-323  CCAAUU...............................................G.CAGCUGGGUGUUUU...............................ACC..............................................................................
env-324  CCAAUU...............................................G.CAGCUGGGUGUUUU...............................ACC..............................................................................
env-325  CCAAUU...............................................G.CAGCUGGGUGUUUU...............................ACC..............................................................................
env-326  CCAAUU...............................................A.CAGCUGGGUGUUUU...............................ACC..............................................................................
env-327  CCAGAA...............................................G.CGGCUGGGUGUUU................................ACC..............................................................................
env-328  CCAAA................................................G.CGGCUGGGUGUAUU...............................ACC..............................................................................
env-329  CCAAAA...............................................G.CAGCUGGGUGUUUU...............................ACC..............................................................................
env-331  CCAAAA...............................................G.CAGCUGGGUGUUUU...............................ACC..............................................................................
env-332  CCAAAA...............................................G.CAGCUGGGUGUUAU...............................ACC..............................................................................
env-335  CCAAAA...............................................G.CAGCUGGGUGUUAU...............................ACC..............................................................................
env-336  CCAAAA...............................................G.CAGCUGGGUGUUAU...............................ACC..............................................................................
env-337  CCAAAA...............................................G.CAGCUGGGUGUUAU...............................ACC..............................................................................
env-338  CCAAAA...............................................G.CAGCUGGGUGUUAU...............................ACC..............................................................................
env-339  CCAAAA...............................................G.CAGCUGGGUGUUAU...............................ACC..............................................................................
env-340  CCAAAA...............................................G.CAGCUGGGUGUUAU...............................ACC..............................................................................
env-341  CCAAAA...............................................G.CAGCUGGGUGUUAU...............................ACC..............................................................................
env-342  CCAAAA...............................................G.CAGCUGGGUGUUAU...............................ACC..............................................................................
Bsc-1-1  CCAAAU...............................................G.CAGCUGGGUGUUUU...............................ACC..............................................................................
Rsp-3-1  CCAAUU...............................................G.CAGCUGGGUGUUUU...............................ACC.
env-343  GUCAC.............................................G.UG.CGGCUGGAUGCU.................................UCAUACC..........................................................................
env-344  UAAUUA............................................UUUG.CGGCUGGAUGUAUA............GAAC...............CCCGUCA........................................................................GU
env-346  UAAUUA............................................UUUG.CGGCUGGAUGUAUA............GAAC...............CCCGUCA........................................................................GU
env-348  UAAUUA............................................UUUG.CGGCUGGAUGUAUA............GAAC...............CCCGUCA........................................................................GU
env-349  UAAUUA............................................UUUG.CGGCUGGAUGUAUA............GAAC...............CCCGUCA........................................................................GU
env-350  UAAUUA............................................UUUG.CGGCUGGAUGUAUA............GAAC...............CCCGUCA........................................................................GU
env-351  UAAUUA............................................UUUG.CGGCUGGAUGUAUA............GAAC...............CCCGUCA........................................................................GU
env-352  .................................................UUUUG.CGGCUGGAUAUGUA............AAAC...............CCCGUCA........................................................................GU
env-353  CA.................................................GCA.UGGCUGGAUGCUUG...............................CAUA.
Rga-1-1  CG.................................................GCA.UGGCUGGAUGCUUG...............................CAUA.
env-354  ...................................................UUG.CGGCUGGAUGUU.................................UUCAUACCG........................................................................
env-355  ...................................................ACA.CAACUGGAUGC..................................ACAACAUACCG......................................................................
env-356  ...................................................ACA.CAACUGGAUGC..................................ACAACAUACCG......................................................................
env-357  GAGA................................................CA.UGACUGGAUGC..................................ACAUACCG.........................................................................
env-358  GAGA................................................CA.UGACUGGAUGC..................................ACAUACCG.........................................................................
env-359  GAGA................................................CA.UGACUGGAUGC..................................ACAUACCG.........................................................................
env-360  ...................................................GCA.CAACUGGAUGCGU................................AACAUACC.........................................................................
env-361  ...................................................GCA.CAACUGGAUGC..................................GUACAUACCG.......................................................................
env-362  ...................................................GCA.CAACUGGAUGC..................................GUACAUACCG.......................................................................
env-363  ...................................................GCA.CAACUGGAUGC..................................GUUCAUACCGA......................................................................

         .........<<<<<<<<<<<......>>>>>>>>>>>.............>>>>>.>>>>><<<<<<<<<.........<<<<<<<.......<<<<.<<<......................................................................>>>.>>>>>>
         ............10...0.1......1.0...01..................22.2222..0222222..........111111.......111..1.......................................................................1...11111
         o-----------------------------------------------oYR-CRRCUGGRYRooo------------------------------------●●●ooo----------------------------------------------------------------------------
```

```
env-364  ..................................................GCA.CAACUGGAUGC................................GUACAUACCGA..................................................
env-365  ..................................................GCA.CAACUGGAUGCGUU.............................CAUACC.......................................................
env-366  ..................................................GCA.UGACUGGAUGCGUU.............................CAUACC.......................................................
env-367  ...................................................CA.CGGCUGGAUACA...............................CGCAUACC.....................................................
env-368  ...................................................CA.CGGCUGGAUACA...............................CGCAUACC.....................................................
env-369  ...................................................CA.CGGCUGGAUACA...............................CGCAUACC.....................................................
env-370  ...................................................CA.CGGCUGGAUACA...............................CGCAUACC.....................................................
env-371  ...................................................CA.CGGCUGGAUACA...............................CGCAUACC.....................................................
env-372  ...................................................CA.CGGCUGGAUACA...............................CGCAUACC.....................................................
env-373  ...................................................CA.CGGCUGGAUACA...............................CGCAUACC.....................................................
env-374  ...................................................CA.CGGCUGGAUACA...............................CGCAUACC.....................................................
env-375  ...................................................CA.CGGCUGGAUACA...............................CGCAUACC.....................................................
env-377  ...................................................CA.CGGCUGGAUACA...............................CGCAUACC.....................................................
env-383  ...................................................CA.CGGCUGGAUACA...............................CGCAUACC.....................................................
env-384  ...................................................CA.CGGCUGGAUACA...............................CGCAUACC.....................................................
env-385  ...................................................CA.CGGCUGGAUACA...............................CGCAUACC.....................................................
env-386  ...................................................CA.CGGCUGGAUACA...............................CGCAUACC.....................................................
env-387  ...................................................CA.CGGCUGGAUACA...............................CGCAUACC.....................................................
env-388  ...................................................CA.CGGCUGGAUACA...............................CGCAUACC.....................................................
env-389  ...................................................CA.CGGCUGGAUACA...............................CGCAUACC.....................................................
env-390  ...................................................CA.CGGCUGGAUACA...............................CGCAUACC.....................................................
env-391  ...................................................CA.CGGCUGGAUACA...............................CGCAUACC.....................................................
env-392  ...................................................CA.CGGCUGGAUACA...............................CGCAUACC.....................................................
env-393  ...................................................CA.CGGCUGGAUACA...............................CGCAUACC.....................................................
env-394  ...................................................CA.CGGCUGGAUACA...............................CGCAUACC.....................................................
env-395  ...................................................CA.CGGCUGGAUACA...............................CGCAUACC.....................................................
env-396  ...................................................CA.CGGCUGGAUACA...............................CGCAUACC.....................................................
env-397  ...................................................CA.CGGCUGGAUACA...............................CGCAUACC.....................................................
env-398  ...................................................CA.CGGCUGGAUACA...............................CGCAUACC.....................................................
env-399  ...................................................CA.CGGCUGGAUACA...............................CGCAUACC.....................................................
env-405  ...................................................CA.CGGCUGGAUACA...............................CGCAUACC.....................................................
env-408  ...................................................CA.CGGCUGGAUACA...............................CGCAUACC.....................................................
env-409  ...................................................CA.CGGCUGGAUACA...............................CGCAUACC.....................................................
env-410  ...................................................CA.CGGCUGGAUACA...............................CGCAUACC.....................................................
env-411  ...................................................CA.CGGCUGGAUACA...............................CGCAUACC.....................................................
env-412  ...................................................CA.CGGCUGGAUACA...............................CGCAUACC.....................................................
env-413  ...................................................CA.CGGCUGGAUACA...............................CGCAUACC.....................................................
env-414  ...................................................CA.CGGCUGGAUACA...............................CGCAUACC.....................................................
env-418  ...................................................CA.CGGCUGGAUACA...............................CGCAUACC.....................................................
env-419  ...................................................CA.CGGCUGGAUACA...............................CGCAUACC.....................................................
env-420  ...................................................CA.CGGCUGGAUACA...............................CGCAUACC.....................................................
env-428  ...................................................CA.UGGCUGGAUGC................................UUCACAUACCG..................................................
env-429  ...................................................CA.UGGCUGGAUGCU...............................UCACAUACC....................................................
env-430  ...................................................CA.UGGCUGGAUGCU...............................UCACAUACC....................................................
env-432  ...................................................CA.UGGCUGGAUGCU...............................UCACAUACC....................................................
env-433  ..................................................AGG.CAGCUGGAUGUA...............................AACCAA.......................................................
env-434  UGC...............................................ACG.CAACUGGGUGUAUU.............................ACC..........................................................
Bhy-1-1  UGC...............................................ACG.CAACUGGGUGUAUU.............................ACC..........................................................
env-435  UCGU...............................................G.CGGCUGGAUGC..................................AGAAA........................................................
env-436  CUAUAU.............................................G.CGGCUGGAUGC..................................AUCAUACCG....................................................
env-437  U.................................................AUA.CGGCUGGAUGC................................GACAGAAA.....................................................
env-438  ..................................................AUA.UGGCUGGAUGC................................GUCAGAA......................................................
Bsp-7-1  CU................................................GUC.CGAUUGGGC..................................UGAGUAUUC....................................................
Bsp-9-1  CU................................................GUC.CGAUUGGGC..................................UGAUUAUU.....................................................
Bsp-5-1  CU................................................GUC.CGAUUGGGC..................................UGAUCA.......................................................
Bsp-2-1  CU................................................GUC.CGAUUGGGC..................................UGAAUAAUC....................................................
Bsp-3-1  C................................................UGUC.CGAUUGGGCAUG...............................UUA..........................................................
Bpr-2-1  A...............................................UUUUG.CGGCUGGAUGCAUA.............................GAGACCCGUCAUUUU..............................................
         .........<<<<<<<<<<<.....>>>>>>>>>>>............>>>>>.>>>>><<<<<<<<<.........<<<<<<<......<<<<.<<<.......................................>>>.>>>>>>
         .........…10…0.1.....1.0…01…............…22.2222..0222222.........111111......111..1.......................................1..11111
         o-------------------------------------------------oYR-CRRCUGGRYRoo-------------------------------------●●●ooo-------------------------------------------
```

alignment positions 361 · · · 540

```
env-1    ...............................GCAUCCCAU....UUAGGUUAUUUAAAAGGA.....................................................................GGAACAAACAACAUGGCUCGUAUUUCUGGUGUAGAUUUACCGAGAGA
env-2    ...............................AUACCCCAAUCAAUUAGUUUACUAUUAAUAUA.....UGAUG...........................................................GAAUUUAUAUUU................................
env-3    ...............................AUACCCCAAUCAAUUAGUUUACUAUUAAUAUA.....UGAUG...........................................................GAAUUUAUAUUU................................
env-4    ...............................AUACCCCAAUCAAUUAGUUUACUAUUAAUAUA.....UGAUG...........................................................GAAUUUAUAUUU................................
env-5    ...............................AUACCCCAAUCAAUUAGUUUACUAUUAAUAUA.....UGAUG...........................................................GAAUUUAUAUUU................................
env-6    ...............................AUACCCCAAUCAAUUAGUUUACUAUUAAUAUA.....UGAUG...........................................................GAAUUUAUAUUU................................
env-7    ...............................AUACCCCAAUCAAUUAGUUUACUAUUAAUAUA.....UGAUG...........................................................GAAUUUAUAUUU................................
env-8    ...............................AUACCCCAAUCAAUUAGUUUACUAUUAAUAUA.....UGAUG...........................................................GAAUUUAUAUUU................................
env-9    ...............................AUACCCCAAUCAAUUAGUUUACUAUUAAUAUA.....UGAUG...........................................................GAAUUUAUAUUU................................
env-10   ...............................AUACCCCAAUCAAUUAGUUUACUAUUAAUAUA.....UGAUG...........................................................GAAUUUAUAUUU................................
env-11   ...............................AUACCCCAAUCAAUUAGUUUACUAUUAAUAUA.....UGAUG...........................................................GAAUUUAUAUUU................................
env-12   ...............................AUACCCCAAUCAAUUAGUUUACUAUUAAUAUA.....UGAUG...........................................................GAAUUUAUAUUU................................
env-13   ...............................AUACCCCAAUCAAUUAGUUUACUAUUAAUAUA.....UGAUG...........................................................GAAUUUAUAUUU................................
env-14   ...............................AUACCCCAAUCAAUUAGUUUACUAUUAAUAUA.....UGAUG...........................................................GAAUUUAUAUUU................................
env-16   ...............................AUACCCCAAUCAAUUAGUUUACUAUUAAUAUA.....UGAUG...........................................................GAAUUUAUAUUU................................

         >>>>>.......................>>>>>>>>>............................<<<<<.............<<<<<<.............<<<<<.....................<<<<<..........................................
         1111........................2222220..............................22222.............1111..............11111.....................11111..........................................
         ------------------------------oRYRYCCCAR----UUARRRA●R●●●●●●oooo------ooooo------------------------------ooo-----------------------●●●●ooo----------------------------------
                                                      3′
```

```
env-17   ................................AUCCCCCAAUCAAUUAGUAAC..................................................................................GGACUGCGGUUGCCAUGAGCAAUUGCGGCA.................
env-18   ................................AUCCCCCAAUCAAUUAGUAAC..................................................................................GGACUGCGGUUGCCAUGAGCAAUUGCGGCA.................
env-19   ...............................CAGUUUCUCAA...UUAGGGACAAUAUCAAUAA......GACAGAAGAAG......CGUACA..CAUUUCAGG.CACUU.........................UCCACCCGGGG....................................
Bsp-6-1  ...................................GCCCCAA...UCA.AAUAAGUAACAUU........AGUA.CAUACGAU....UUUAUC.GUAUAACUU..CCUGA.........................UAACGAGAAA.....................................
env-20   ...............................CCGCACCCCAA...UUAGGAAACCGUAAACAAUACA....................................................................GCAAAGAGUAACA..................................
env-21   ................................AUGCCCCAAUCAAUUAGUAACUGUAUGGUACAAA....GAUAU............................................................CAUACCUAUUU....................................
env-22   AUCCG.........................CAGCCGUCCCAG...UUAGGAAUACUGUAAGAUA.......................CCCGGA..........................................GCGGUAAGACCGG..................................
env-23   ...............................AAAACCCCAAUCAACUAGUAUCUGUUAAUUAU.......GUGC.............................................................AAAGGAUCAUCGUUUGGGUAUACGAUAGUUUUAUCCG..........
env-24   CAAAG.........................AACAUACCUCAA...UUUAGGACAAUAUAAUAA.......CAGUAAUUU........UAUACAAACAUUUCCU.UACAU..........................AUCAAGU........................................
env-25   CAAAG.........................AACAUACCUCAA...UUUAGGACAAUAUAAUAA.......CAGUAAUUU........UAUACAAACAUUUCCU.UACAU..........................AUCAAGU........................................
env-28   CAAAG.........................AACAUACCUCAA...UUUAGGACAAUAUAAUAA.......CAGUAAUUU........UAUACAAACAUUUCCU.UACAU..........................AUCAAGU........................................
env-29   CAAAGA.........................AGAUAUCUCAG...UUUAGGACAAUAUAAUAA.......UAGUAAUUU........UAUACAAACAUUUCCU.UACAU..........................AAUUAGU........................................
env-30   CAAAG.........................AAGAUAUCUCAGU..UUAGGA.CAAUAUAAUAA.......UAGUAAUUU........UAUACAAACAUUUCCU.UACAU..........................AAUUAGU........................................
env-32   CAAAG.........................AAGAUAUCUCAGU..UUAGGA.CAAUAUAAUAA.......UAGUAAUUU........UAUACAAACAUUUCCU.UACAU..........................AAUUAGU........................................
env-33   CAAAG.........................AAGAUAUCUCAGU..UUAGGA.CAAUAUAAUAA.......UAGUAAUUU........UAUACAAACAUUUCCU.UACAU..........................AAUUAGU........................................
env-34   ...............................GGAUGCCUCAA...UUAGGAACAAUAUGACG........CGAAGCUUUCAAAGUG.CGCACA..CAUUUCAGG.CACUG.........................CCCCGCAAG......................................
env-35   ...............................GGAUGCCUCAA...UUAGGAACAAUAUGACG........CGAAGCUUUCAAAGUG.CGCACA..CAUUUCAGG.CACUG.........................CCCCGCAAG......................................
env-36   ...............................GGAUGCCUCAA...UUAGGAACAAUAUGACG........CGAAGCUUUCAAAGUG.CGCACA..CAUUUCAGG.CACUG.........................CCCCGCAAG......................................
env-37   .................................AACCCCAAUCAAUUAGUAACUGUUAAUCAU.......GUGUGACAAUCAUUC....................UGCAU.........................UGUAAACAGGAACGCAGGCAG..........................
env-38   ................................CGCAUCCCAU...UUAGGAAAGUGUCAU.............................................CGGGC.........................ACCUAUUAAAAAAC.................................
env-39   ..............................UGCGUGUCUCAU...UUAGGAAAGUGUCAG.............................................CGGGC.........................ACCUAUUAAAAAAC.................................
env-41   ...............................GUAUAUCCCAU...UUAGGAAAGUGUCAG.............................................CGGGC.........................ACCUAUUAAAAAAC.................................
env-42   .............................UGUAUAUCCCACU...UUAAAGACAUUAUUUUUCAGGAG..GUCCCCAAAUGAUG...CGCAGC..........................................AUCAUUUUUGCAUGGGACCGACGACCUGCCGCCGAAGCUUUG.....
env-43   ..CUGA........................GGAACAUCCCAG...UUAGGGACAAUAUGAA.........................................................................................................................
env-45   .................................GUUCCCCUA...UUAAAGACAAUAUAAA..........................................................................GGGCGAGCAGUUCCUGCCGAUGAUGCAGAGCAACCGCAGCCUUAACG
env-46   ..................................GUAUCCCAA...UUAGGAAAAUAUUGAUUUAGAAAA.UGGAGGAAAUGUAAA..UGGCUC.........GUAUUUCU.........................GGUGUAGAUUUACCGAGAGACA.........................
env-47   ..................................CACCCCAA...UUAGGAGCGAAAAGAG.........AUGAGCAGAUUCUGA..UGCAUC.CCAUUUCAC..CUUGC.........................AGGUCCU........................................
env-48   ..................................CACCCCAG...UUAGGCACAAUAACC..........GAGUGCGG.........................................................AUCCCG.........................................
env-49   CC............................GAAGUGUCCCAA...UUAGUGACAAUAUGGA..........................CGGAGC..........................................UUAA...........................................
env-50   ..............................GGUACGCCCCAG...UUAAAGACAAUAUGAAACAA......................................................................AACAUGUA.......................................
Dma-1-1  ..............................GGUACGCCCCAG...UUAAAGACAAUAUGAAACAA......................................................................AACAUGUA.......................................
env-51   ...............................AAUAC.UCCCAA...UUAAAGACGAUAUUAAACAA......................................................................AAAACA.........................................
Lba-2-1  ................................UACUUCCCAAU..UUGAGA.CAAUAUUAACCAA.....UGGGAAUAGGUAUA...UACACA.ACAUUUCAGA.CAGUG.........................CAGAG..........................................
[ce-1-1  ................................UGCAUCCCAU...UUAAGAUGGUAUUAAAUUG......AUCGG............................................................CGAAAAAAGCAGCACUUUCAGA.........................
Ece-1-1  ................................UGCAUCCCAU...UUAAGAUGGUAUUAAAUU.......GAUCG............................................................GCGAAAAAAGCAGCACUUUCAGA........................
env-52   ..............................CCAGAACCCCAA...UUAAAG.ACAAUAU...........AGCUGAGCACCGGGCAGCGCACA.ACAUUUCAGA.CGUC..........................UGGCUA.........................................
env-53   GGU...........................CCGCAGCCUCAA...UUAGGGACAACGUAAACAAAUA...GAAUAUUAAUUCUAUUAUAGAAU.........AUUAAU...........................UCUAUUACAGAAU..................................
env-54   ...............................GACACAC.CCAG...UUAAAGACAAUAUUAAA.........................................................................UUUAGAUUCA.....................................
env-55   ...............................GACACAC.CCAG...UUAAAGACAAUAUUAAA.........................................................................UUUAGAUUCA.....................................
env-59   ...............................GACACAC.CCAG...UUAAAGACAAUAUUAAA.........................................................................UUUAGAUUCA.....................................
env-60   ...............................GACACAC.CCAG...UUAAAGACAAUAUUAAA.........................................................................UUUAGAUUCA.....................................
env-62   ...............................GACACAC.CCAG...UUAAAGACAAUAUUAAA.........................................................................UUUAGAUUCA.....................................
env-63   ...............................GACACAC.CCAG...UUAAAGACAAUAUUAAA.........................................................................UUUAGAUUCA.....................................
env-64   ...............................GACACAC.CCAG...UUAAAGACAAUAUUAAA.........................................................................UUUAGAUUCA.....................................
env-66   ...............................GACACAC.CCAG...UUAAAGACAAUAUUAAA.........................................................................UUUAGAUUCA.....................................
env-67   ...............................GACACAC.CCAG...UUNAAGACAAUAUUAAA.........................................................................UUUAGAUUCA.....................................
env-68   ...............................GACACAC.CCAG...UUAAAGACAAUAUUAAA.........................................................................UUUAGAUUCA.....................................
env-75   ...............................GACACAC.CCAG...UUAAAGACAAUAUUAAA.........................................................................UUUAGAUUCA.....................................
env-76   ...............................GACACAC.CCAG...UUAAAGACAAUAUUAAA.........................................................................UUUAGAUUCA.....................................
env-77   ...............................GACACAC.CCAG...UUAAAGACAAUAUUAAA.........................................................................UUUAGAUUCA.....................................
env-78   ...............................GACACAC.CCAG...UUAAAGACAAUAUUAAA.........................................................................UUUAGAUUCA.....................................
env-79   ...............................GACACAC.CCAG...UUAAAGACAAUAUUAAA.........................................................................UUUAGAUUCA.....................................
env-80   ...............................GACACAC.CCAG...UUAAAGACAAUAUUAAA.........................................................................UUUAGAUUCA.....................................
env-81   ...............................GACACAC.CCAG...UUAAAGACAAUAUUAAA.........................................................................UUUAGAUUCA.....................................
Rgn-1-1  ...............................GACACAC.CCAG...UUAAAGACAAUAUUAAA.........................................................................UUUAGAUUCA.....................................
env-82   ...............................GACACA.CCCAG...UUAAAGACAAUAUUAAA.........................................................................UUUAGAUUCA.....................................
env-83   ...............................GACACA.CCCAG...UUAAAGACAAUAUUAAA.........................................................................UUUAGAUUCA.....................................
env-87   ...............................GACACA.CCCAG...UUAAAGACAAUAUUAAA.........................................................................UUUAGAUUCA.....................................
env-88   ...............................UCAGGGCCCAA...UUUGAGACAAUAUUAACUAGA....GGGA.AUAUGAUUUG..UGC.......UGAAUGCUUCAG..........................AUAUU..........................................
env-89   UUU...........................CUUACUCCCCAA...UUUGAGACAAUAUUAAUAAGAGCA.GUCAUCA..........CAUACA..CAUUUCAGG.CUUUG.........................GCGA...........................................
env-90   UCCG.............................CAGCCUCAA...UUAGAGACAACGUAAACAC......................................................................................................................
env-91   UCCG.............................CAGCCUCAA...UUAGGGACAACGUAAACAGAUA...GAAAGAUUUUGAUUCU.................................................AUCACCA........................................
env-92   ..............................CAUGUAUCCCAA...UUAGAGACAAUACGGA............................................UGCGC.........................GGUGUGU........................................
env-93   ................................GGCGCCUCAA...UUAGGAAUCA...............GAUGAGCUGCGUGUGGAGGCAUC.CCAUUUCAC..CUUGC.........................AGAUCU.........................................
env-94   .................................GCAUCCCAU...UUAGGUUAUUUUUAAA.........GGAGGAACAAGUA....ACAUC.GCUCGUA.....UUUCU.....................GGU.GUAGAUU........................................
Lba-11-1 .................................GCAUCCCAU...UUAGGUUAUUUAAA...........GGAGGAACAAGUA....ACAUC.GCUCGUA.....UUUCU.....................GGU.GUAGAUU........................................
env-95   .................................ACACCCCAG...UCAGAAACAAUACACG..........................GAUAUC................C.........................UGAUC..........................................
env-96   .................................ACACCCCAG...UUAGAAACAAUACAC..........GGA................................UAUCC.........................CAACCA.........................................
env-97   ...................................GCCCCUA...UUAAAGACGAUAU................GUAAA........CUU........UUAAGA.CGUGC.........................UCGAAUCUUACGCG.................................
Dsp-3-1  .................................GCUCCCCAG...UUAGAGACGAUA.............UGCA.............................................................ACUAAAUGAGACGUGCACAGAUUUCACAAUUGGCUUUUGCAGGCUUG
Dsp-1-1  ...................................GUCCCUA...UUAAAGACAGUA..............................UGUACA............CUUGA.........................AGCAGGUUGGUUCUUAAUA............................
env-101  ...............................UGUGUCCCCUA...UUAAAGACGAUAACAAACAAAGA..................................................................................................................
env-102  ...............................UGUGUCCCCUA...UUAAAGACGAUAACAAACAAAGA..................................................................................................................
Dsp-2-1  ...............................UGUGUCCCCUA...UUAAAGACGAUAACAAACAAAGA..................................................................................................................
env-103  ...............................UGUGUCCCCUA...UUAAAGACGAUAACAAACAAAGA..................................................................................................................
env-106  ...............................UGUGUCCCCUA...UUAAAGACGAUAACAAACAAAGA..................................................................................................................
env-116  ...............................UGUGUCCCCUA...UUAAAGACGAUAACAAACAAAGA..................................................................................................................
env-117  ...............................UGUGUCCCCUA...UUAAAGACGAUAACAAACAAAGA..................................................................................................................
env-118  ...............................UGUGUCCCCUA...UUAAAGACGAUAACAAACAAAGA..................................................................................................................
env-119  .................................UGUCCCCUA...UUAAAGACAAUUAACAAUGAAAGA.................................................................................................................
env-120  .................................UGUCCCCUA...UUAAAGACAAUUAACAAUGAAAGA.................................................................................................................
env-123  .................................UGUCCCCUA...UUAAAGACAAUUAACAAUGAAAGA.................................................................................................................
env-124  .................................UGUCCCCUA...UUAAAGACAAUUAACAAUGAAAGA.................................................................................................................

         >>>>>.......................>>>>>>>>>.................................<<<<<............<<<<<<............<<<<<.....................<<<<<..............................................
         1111........................ .2222220..................................22222............1111..............11111.....................11111..............................................
         ------------------------------○RYRYCCAR---UUARRA●R●●●●●●○○○○------------○○○○○---------------------------○○○---------------------------●●●●○○○----------------------------------------------
                                                  3′
```

```
env-127  ...UGUCCCCUA...UUAAAGACAAUUAACAAUGAAAGA...
env-128  ...UGUCCCCUA...UUAAAGACAAUUAACAAUGAAAGA...
env-129  ...UGUCCCCUA...UUAAAGACAAUUAACAAUGAAAGA...
env-130  ...GUCCCCUA...UUAAAGACAAUUAACAA...UGAAAGA...
env-136  ...GUCCCCUA...UUAAAGACAAUUAACAA...UGAAAGA...
env-137  ...GUCCCCUA...UUAAAGACAAUUAACAA...UGAAAGA...
env-138  ...GUCCCCUA...UUAAAGACAAUUAACAA...UGAAAGA...
env-139  ...GUCCCCUA...UUAAAGACAAUUAACAA...UGAAAGA...
Dlo-1-1  ...GUCCCCUA...UUAAAGACAAUUAACAA...UGAAAGA...
env-140  ...GUCCCCUA...UUAAAGACAAUUAACAA...UGAAAGA...
env-149  ...GUUCCCCUA...UUAAAGACAAUAUGAAACAACA...GGGAGAA...AUGCUGAGUCCGAGGAAGACUGACGUUGCAAGCUUGCUUGCAAAACG
env-150  ...GAUCCCCUA...UUAAAGACGAUGUGCGCAAG...CUGA...AAAAGUGUUGUUUU...
env-151  ...GAUCCCCUA...UUAAAGACGAUGUGCGCAAG...CUGA...AAAAGUGUUGUUUU...
Chy-1-1  ...UCUCCCCAG...UUAGAGACAAUAUGAAA...CCAAUUGAGA.CGUGC...AAAGGAUUCAGAACUGGCUUUUUCAAGCUUGCUUGAUAAAAGACAGG
env-152  ...GCGCAACCCCUA...UUAAAGACGAUGUGCGUAA...GCU..GAAAAGUGCAUAGGUUGCACAUCGACUAUGUGAGUUUACUCACAACA
env-153  ...GCCAUCCCUA...UUAAAGACAGUAUCAAACAA...AAAAACAAAGAA...
env-154  ...GCCAUCCCUA...UUAAAGACAGUAUCAAAC...AAAAAAACAAAGAA...
Esp-1-1  CGGUAGGGAACAGGAAAAAAUCCAGUGCCGCAGCCUCAA...UUUAGGACAAUAUUAAAC...AACAAAGAUC...
env-155  .GCAU...UCGCAGCCUCAA...UUAGGGACAAUUAACAAG...AAAACCAGAAAAGA...
env-156  .GCAU...UCGCAGCCUCAA...UUAGGGACAAUUAACAAG...AAAACCAGAAAAGA...
env-159  .GCAU...UCGCAGCCUCAA...UUAGGGACAAUUAACAAG...AAAACCAGAAAAGA...
env-160  .GCAU...UCGCAGCCUCAA...UUAGGGACAAUUAACAAG...AAAACCAGAAAAGA...
env-161  .GCAU...UCGCAGCCUCAA...UUAGGGACAAUUAACAAG...AAAACCAGAAAAGA...
env-162  .GCAU...UCGCAGCCUCAA...UUAGGGACAAUUAACAAG...AAAACCAGAAAAGA...
env-163  .GCAU...UCGCAGCCUCAA...UUAGGGACAAUUAACAAG...AAAACCAGAAAAGA...
env-164  .GCAU...UCGCAGCCUCAA...UUAGGGACAAUUAACAAG...AAAACCAGAAAAGA...
env-165  .GCAU...UCGCAGCCUCAA...UUAGGGACAAUUAACAAG...AAAACCAGAAAAGA...
env-167  .GCAU...UCGCAGCCUCAA...UUAGGGACAAUUAACAAG...AAAACCAGAAAAGA...
env-168  .GCAU...UCGCAGCCUCAA...UUAGGGACAAUGAACAAG...AAAACCAGAAAAGA...
env-169  .GCAU...UCGCAGCCUCAA...UUAGGGACAAUGAACAAG...AAAACCAGAAAAGA...
env-170  .GCAU...UCGCAGCCUCAA...UUAGGGACAAUGAACAAG...AAAACCAGAAAAGA...
env-171  .GCAU...UCGCAGCCUCAA...UUAGGGACAAUGAACAAG...AAAACCAGAAAAGA...
env-172  .GCAU...UCGCAGCCUCAA...UUAGGGACAAUGAACAAG...AAAACCAGAAAAGA...
env-175  .GCAU...UCGCAGCCUCAA...UUAGGGACAAUGAACAAG...AAAACCAGAAAAGA...
env-176  .GCAU...UCGCAGCCUCAA...UUAGGGACAAUGAACAAG...AAAACCAGAAAAGA...
env-177  .GCAU...UCGCAGCCUCAA...UUAGGGACAAUGAACAAG...AAAACCAGAAAAGA...
env-178  .GCAU...UCGCAGCCUCAA...UUAGGGACAAUGAACAAG...AAAACCAGAAAAGA...
Lba-4-1  ...GUAGUCCCUA...UUAAAGUCAGUAUGAAA...AACAAAGA...
env-179  ...GCCCCUA...UUAAAGACGAUAUGUAAA...CUU..UUAA...[illegible]...CGCG...AAUCUUA...
Cba-3-1  ...GCCCCUA...UUAAAGACGAUAUGUAAA...CUU..UUAA...[illegible]...CGCG...AAUCUUA...
[sc-1-1  ...CGCAAGCCCCUA...UUAAAGACGAUAUGUAAA...CUUUUAA...[illegible]...CGCG...AAUCUU...
Lba-7-1  ...GCCCCUA...UUAAAGACGAUAUGUAAACU...UUUAA...[illegible]...CGCG...AAUCUU...
env-181  ...GCCCCUA...UUAAAGACGAUAUGUAAACU...UUUAA...[illegible]...CGCG...AAUCUU...
env-182  ...ACACCCCAA...UUAGAAACAAUACAA...UGAUGU...CCCCC...
env-183  ...GCCCCAA...UUAGAGACAAUACAAUGGCGA...GGC..AGGGA...[illegible]...GUGCCGA...
env-184  .GUCCG...CAGCCUCAA...UUAAAGACAGUAUAAACAAAGA...
env-185  .GUCCG...CAGCCUCAA...UUAAAGACAGUAUAAACAAAGA...
env-186  .GUCCG...CAGCCUCAA...UUAAAGACAGUAUAAACAAAGA...
env-187  .GUCCG...CAGCCUCAA...UUAAAGACAGUAUAAACAAAGA...
env-188  .GUCCG...CAGCCUCAA...UUAAAGACAGUAUAAACAAAGA...
env-189  .GUCCG...CAGCCUCAA...UUAAAGACAGUAUAAACAAAGA...
env-190  .GUCCG...CAGCCUCAA...UUAAAGACAGUAUAAACAAAGA...
env-191  .GUCCG...CAGCCUCAA...UUAAAGACAGUAUAAACAAAGA...
env-192  .GUCCG...CAGCCUCAA...UUAAAGACAGUAUAAACAAAGA...
env-194  .GUCCG...CAGCCUCAA...UUAAAGACAGUAUAAACAAAGA...
env-195  .GUCCG...CAGCCUCAA...UUAAAGACAGUAUAAACAAAGA...
env-200  .GUCCG...CAGCCUCAA...UUAAAGACAGUAUAAACAAAGA...
env-201  .GUCCG...CAGCCUCAA...UUAAAGACAGUAUAAACAAAGA...
env-202  .GUCCG...CAGCCUCAA...UUAAAGACAGUAUAAACAAAGA...
env-211  .GUCCG...CAGCCUCAA...UUAAAGACAGUAUAAACAAAGA...
env-212  .GUCCG...CAGCCUCAA...UUAAAGACAGUAUAAACAAAGA...
env-216  .GUCCG...CAGCCUCAA...UUAAAGACAGUAUAAACAAAGA...
env-217  ...UGACGCCCCAG...UUAGAGACAAUAGAG...UACUGCCUCA...[illegible]...GAUGUA...
Rpe-1-1  ...UGCCUCAA...UUAAGGACAAUAUAAU...UAAGGUUUCGAAAGG..[illegible]CA..CAUUUCAGA.CAUGC...AAAAG...
Rsp-1-1  ...UGCCUCAA...UUAAGGACAAUAUAAU...UAAGGUUUCGAAAGG..[illegible]CA..CAUUUCAGA.CAUGC...AAAAG...
env-218  ...CGCCCCAA...UUAGAGACAAUAGUC...GGAUGCAUGGG...
env-219  ...ACACCCCAG...UUAGGA.CAG...GAAAGGGUCCGCAUGA.[illegible].CCAUUUCAC..CUUAC...AAUAUUUU...
env-220  ...CUGUCCCAA...UUAGAGACAAUAU...GCAGC...[illegible]...CUUA...
env-221  ...UAAGCCUCAA...UUAGGGACAAUAUAAGGAA...GAAUGAAAAUG...[illegible]CA.UCAUUUCAGA.CGUUA...UCAUACCCGGA...
env-222  ...UAAGCCUCAA...UUAGGGACAAUAUAAGGAA...GAAUGAAAAUG...[illegible]CA.UCAUUUCAGA.CGUUA...UCAUACCCGGA...
env-223  ...UAAGCCUCAA...UUAGGGACAAUAUAAGGAA...GAAUGAAAAUG...[illegible]CA.UCAUUUCAGA.CGUUA...UCAUACCCGGA...
env-226  ...UAAGCCUCAA...UUAGGGACAAUAUAAGGAA...GAAUGAAAAUG...[illegible]CA.UCAUUUCAGA.CGUUA...UCAUACCCGGA...
env-227  ...UAAGCCUCAA...UUAGGGACAAUAUAAGGAA...GAAUGAAAAUG...[illegible]CA.UCAUUUCAGA.CGUUA...UCAUACCCGGA...
env-228  ...UAAGCCUCAA...UUAGGGACAAUAUAAGGAA...GAAUGAAAAUG...[illegible]CA.UCAUUUCAGA.CGUUA...UCAUACCCGGA...
env-229  ...UAAGCCUCAA...UUAGGGACAAUAUAAGGAA...GAAUGAAAAUG...[illegible]CA.UCAUUUCAGA.CGUUA...UCAUACCCGGA...
env-230  ...UAAGCCUCAA...UUAGGGACAAUAUAAGGAA...GAAUGAAAAUG...[illegible]CA.UCAUUUCAGA.CGUUA...UCAUACCCGGA...
env-231  ...UAAGCCUCAA...UUAGGGACAAUAUAAGGAA...GAAUGAAAAUG...[illegible]CA.UCAUUUCAGA.CGUUA...UCAUACCCGGA...
env-232  ...UAAGCCUCAA...UUAGGGACAAUAUAAGGAA...GAAUGAAAAUG...[illegible]CA.UCAUUUCAGA.CGUUA...UCAUACCCGGA...
env-234  ...UAAGCCUCAA...UUAGGGACAAUAUAAGGAA...GAAUGAAAAUG...[illegible]CA.UCAUUUCAGA.CGUUA...UCAUACCCGGA...
env-235  ...UAAGCCUCAA...UUAGGGACAAUAUAAGGAA...GAAUGAAAAUG...[illegible]CA.UCAUUUCAGA.CGUUA...UCAUACCCGGA...
env-236  ...UAAGCCUCAA...UUAGGGACAAUAUAAGGAA...GAAUGAAAAUG...[illegible]CA.UCAUUUCAGA.CGUUA...UCAUACCCGGA...
env-237  ...UAAGCCUCAA...UUAGGGACAAUAUAAGGAA...GAAUGAAAAUG...[illegible]CA.UCAUUUCAGA.CGUUA...UCAUACCCGGA...
env-238  ...UAAGCCUCAA...UUAGGGACAAUAUAAGGAA...GAAUGAAAAUG...[illegible]CA.UCAUUUCAGA.CGUUA...UCAUACCCGGA...

         >>>>>...>>>>>>>>>...<<<<<...<<<<<<...<<<<<...<<<<<...
         1111....2222220....22222...1111.....11111...11111...
         ---○RYRYCCCAR---UUARRRA●R●●●●○○○○○---○○○○○---○○○---○○○○○○○---
```

```
env-239    .............................UAAGCCUCAA....UUAGGGACAAUAUAAGGAA......GAAUGAAAAUG......UACACA.UCAUUUCAGA.CGUUA..........................UCAUACCCGGA.................................
env-240    .............................UAAGCCUCAA....UUAGGGACAAUAUAAGGAA......GAAUGAAAAUG......UACACA.UCAUUUCAGA.CGUUA..........................UCAUACCCGGA.................................
env-241    ..............................GCGCCCCAG....UUAGAGACAAUACGA.........UGA..............UACGGC...........................................ACAG........................................
env-242    ............................AUGUACCCCAA....UUUGAGACAAUAUUAACCAAUG...GGAAUAAAUAUC.....UGCACA..CAUUUCAGA.CGAUA..........................UUU.........................................
env-243    ............................AUGUACCCCAA....UUUGAGACAAUAUUAACCAAUG...GGAAUAAAUAUC.....UGCACA..CAUUUCAGG.CGAUA..........................UUU.........................................
env-244    ............................AUGUACCCCAA....UUUGAGACAAUAUUAACCAAUG...GGAAUAAAUAUC.....UGCACA..CAUUUCAGG.CGAUA..........................UUU.........................................
env-245    ............................AUGUACCCCAA....UUUGAGACAAUAUUAACCAAUG...GGAAUAAAUAUC.....UGCACA..CAUUUCAGG.CGAUA..........................UUU.........................................
env-246    ............................AUGUACCCCAA....UUUGAGACAAUAUUAACCAAUG...GGAAUAAAUAUC.....UGCACA..CAUUUCAGG.CGAUA..........................UUU.........................................
env-252    ............................AUGUAUCCCAAU...UUGAGA.CAAUAUUAACCAAUGGGAAUAAAUAUC........UGCACA..CAUUUCAGG.CGAUA..........................UUU.........................................
env-254    UCC.........................CGGCGUCCCAA....UUUGAGACAAUAUAAACUAC....GGCAGAAGAGGGUG...UAUACU.ACAUUUCAGA.CCUG...........................UAACC.......................................
Lba-6-1    UCC.........................CGGCGUCCCAA....UUUGAGACAAUAUAAACUAC....GGCAGAAGAGGGUG...UAUACU.ACAUUUCAGA.CCUG...........................UAACC.......................................
env-255    UCC.........................CGGCGUCCCAA....UUUGAGACAAUAUAAACUAC....GGCAGAAGAGGGUG...UAUACU.ACAUUUCAGA.CCUG...........................UAACC.......................................
Lba-10-1   UCC.........................CGGCGUCCCAA....UUUGAGACAAUAUAAACUAC....GGCAGAAGAGGGUG...UAUACU.ACAUUUCAGA.CCUG...........................UAACC.......................................
env-256    UCC.........................CGGCGUCCCAA....UUUGAGACAAUAUAAACUAC....GGCAGAAGAGGGUG...UAUACU.ACAUUUCAGA.CCUG...........................UAACC.......................................
env-258    UCC.........................CGGCGUCCCAA....UUUGAGACAAUAUAAACUAC....GGCAGAAGAGGGUG...UAUACU.ACAUUUCAGA.CCUG...........................UAACC.......................................
env-259    UCC.........................CGGCGUCCCAA....UUUGAGACAAUAUAAACUAC....GGCAGAAGAGGGUG...UAUACU.ACAUUUCAGA.CCUG...........................UAACC.......................................
env-260    .............................UAUGCCCCAG....UUAGAAUCAAUGAAC.........GGACGCAGGAG.............................................................................................
env-261    .............................GGUGCCCCAG....UUAGAGACAAUGGUAG........GAUGA............................CUGCU..........................UACAGAU.....................................
env-262    UUGUCG.....................C..GCGCCCCAG....UUAGGGACAAUAUGAAUGUAAUUAGA.....................................................................................................
env-263    ..............................CUGCCCCAG....UUAGAGACAAUAAAGA..........................................................................................................
Bfo-1-1    UUCCAA.....................A....ACCCCAA....UUAAAGACAAUAUAG.........CGAAGCACCGGGCUG..CGCACA.ACAUUUCAGA.CGUCU..........................UGCUAU......................................
Bpr-1-1    ...........................UAUGUAUCCCAA....UUAGGAACAAUAUUUUUUAAGAA.................................................................................................
Cba-1-1    ...........................UAUGUAUCCCAA....UUAGGAACAAUAUUUAUUAAGAA.................................................................................................
env-264    .............................GGCACCCCAA....UUAGGAAUAUUUA...........AAAAGCACCGACAGA..UGCACC.CCAUUUCAC..CUUGC..........................GGGA........................................
env-265    ..............................ACACCCCAA....UUAGGAAAAC..............GAGAGCAUUUGACAGA.GGCACC.CCAUUUCAC..CUUGC..........................GUCA........................................
env-266    .............................CACACCUCAA....UUAGGAAACGAAAGUAUCACA...GGCAGA...........UGCACC.CCAUUUCAC..CUUGC..........................CCCG........................................
env-267    ...........................GAGGCACCCCAA....UUAGGAAACUAGAAAAUCA.....GGAAGCUUGUGAAGA..UAUACC.CCAUUUCAC..CUUGC..........................GAGACA......................................
env-268    ...........................GAGGCACCCCAA....UUAGGAAACUAGAAAAUCA.....GGAAGCUUGUGAAGA..UAUACC.CCAUUUCAC..CUUGC..........................GAGACA......................................
env-271    ...........................GAGGCACCCCAA....UUAGGAAACUAGAAAAUCA.....GGAAGCUUGUGAAGA..UAUACC.CCAUUUCAC..CUUGC..........................GAGACA......................................
env-272    ...........................GAGGCACCCCAA....UUAGGAAACUAGAAAAUCA.....GGAAGCUUGUGAAGA..UAUACC.CCAUUUCAC..CUUGC..........................GAGACA......................................
env-273    ...........................GAGGCACCCCAA....UUAGGAAACUAGAAAAUCA.....GAAAGGUUUUGAAGA..ACUGCC.CCAUUUCAC..CUUGC..........................GAGACU......................................
env-275    ...........................GAGGCACCCCAA....UUAGGAAACUAGAAAAAUC.....GAGAGUUUUUGAAGA..UGUACC.CUAUUUCAC..CUUGC..........................GAGAUU......................................
env-276    ...........................GAGGCACCCCAA....UUAGGAAACUAGAAAACA......GAAAGUAUUGAAGA...UGUACC.CCAUUUCAC..CUUGC..........................GAAACU......................................
env-277    ...........................GAGGCACCCCAA....UUAGGAAACUAAAAACA.......GAAAGCAGACUUUCAGGUAUGCC.CCAUUUCAC..CUUGC..........................CAUCCG......................................
env-278    ...........................GAGGCACCCCAA....UUAGGAAACUAAAAACA.......GAAAGCAGACUUUCAGGUAUGCC.CCAUUUCAC..CUUGC..........................CAUCCG......................................
env-279    ...........................GAGGCACCCCAA....UUAGGAAACUAGAAACA.......GAAAGCAGGACUUAAGAUACACC.CCAUUUCAC..CUUGC..........................CUCAG.......................................
env-280    ...........................GAGGCACCCCAA....UUAGGAAACUAGAAAA........GAAAGCGGCAGGUUGA.UAUGUC.CCAUUUCAC..CUUGC..........................GGGA........................................
env-281    ...........................GAGGCACCCCAA....UUAGGAAACUAGAAAA........GAAAGAGGCAGGUCGA.UAUGUC.CCAUUUCAC..CUUGC..........................GAGA........................................
env-282    ...........................GAGGCACCCCAA....UUAGGAAACUAGAAAA........GAAAGAGGCAGGUCGA.UAUGUC.CCAUUUCAC..CUUGC..........................GAGA........................................
Bsp-1-1    ...........................GAGGCACCCCAA....UUAGGAAACUAGAAAA........GAAGGCGGCAGGUCGA.UAUGUC.CCAUUUCAC..CUUGC..........................GAGA........................................
env-283    ...........................GAGGCACCCCAA....UUAGGAAACUAGAAAA........GAAAGCGGCAGGUCGA.UAUGUC.CCAUUUCAC..CUUGC..........................GAGA........................................
env-284    ...........................GAGGCACCCCAA....UUAGGAAACUAGAAAA........GAAAGCGGCAGGUCGA.UAUGUC.CCAUUUCAC..CUUGC..........................GAGA........................................
env-285    ...........................GAGGCACCCCAA....UUAGGAAACUAGAAAA........GAAAGCGGCAGGUCGA.UAUGUC.CCAUUUCAC..CUUGC..........................GAGA........................................
env-286    ...........................GAGGCACCCCAA....UUAGGAAACUAGAAAA........GAAAGCGGCAGGUCGA.UAUGUC.CCAUUUCAC..CUUGC..........................GAGA........................................
env-287    ...........................GAGGCACCCCAA....UUAGGAAACUAGAAAA........GAAAGCGGCAGGUCGA.UAUGUC.CCAUUUCAC..CUUGC..........................GAGA........................................
env-288    ...........................GAGGCACCCCAA....UUAGGAAACUAGAAAA........GAAAGCGGCAGGUCGA.UAUGUC.CCAUUUCAC..CUUGC..........................GAGA........................................
env-289    ...........................GAGGCACCCCAA....UUAGGAACUAAGAAAUAA......GAAAGUGAAGUGACAGAUACAUC.CCAUUUCAC..CUUACAGGA................GUG..UAUUU.......................................
env-290    ...........................GAGGCACCCCAA....UUAGGAACUAAGAAAUAA......GAAAGUGAAGUGACAGAUACAUC.CCAUUUCAC..CUUACAGGA................GUG..UAUUU.......................................
env-291    ...........................GAGGCACCCCAA....UUAGGAACUAAGAAAUAA......GAAAGUGAAGUGACAGAUACAUC.CCAUUUCAC..CUUACAGGA................GUG..UAUUU.......................................
env-292    ...........................GAGGCACCCCAA....UUAGGAACUAAGAAAUAA......GAAAGUGAAGUGACAGAUACAUC.CCAUUUCAC..CUUACAGGA................GUG..UAUUU.......................................
env-293    ...........................GAGGCACCCCAA....UUAGGAACUGAGAAAUAA......GAAAGUGAUGUGUCCGAUACAUC.CCAUUUCAC..CUUACUUGA................GUGAGAGAA.........................................
env-294    ...........................GAGGCACCCCAA....UUAGGAACUGAGAAAUAA......GAAAGUGAUGUGUCCGAUACAUC.CCAUUUCAC..CUUACUUGA................GUGAGAGAA.........................................
env-295    ...........................GAGGCACCCCAA....UUAGGAACUGAGAAAUAA......GAAAGUGAUGUGUCCGAUACAUC.CCAUUUCAC..CUUACUUGA................GUGAGAGAA.........................................
env-296    ...........................GAGGCACCCCAA....UUAGGAACUGAGAAAUAA......GGAAGUGAUGUGUCCGAUACAUC.CCAUUUCAC..CUUACUUGA................GUGAGAGAA.........................................
env-298    ...........................GAGGCACCCCAA....UUAGGAACUGAGAAAUAA......GGAAGUGAUGUGUCCGAUACAUC.CCAUUUCAC..CUUACUUGA................GUGAGAGAA.........................................
env-299    ...........................GAGGCACCCCAA....UUAGGAACUGAGAAAUAA......GGAAGUGAUGUGUCCGAUACAUC.CCAUUUCAC..CUUACUUGA................GUGAGAGAA.........................................
env-300    ...........................GAGGCACCCCAA....UUAGGAACUGAGAAAUAA......GGAAGUGAUGUGUCCGAUACAUC.CCAUUUCAC..CUUACUUGA................GUGAGAGAA.........................................
env-301    ...........................GAGGCACCCCAA....UUAGGAACUGAGAAAUAA......GGAAGUGAUGUGUCCGAUACAUC.CCAUUUCAC..CUUACUUGA................GUGAGAGAA.........................................
env-302    ...........................GAGGCACCCCAA....UUAGGAACUGAGAAAUAA......GGAAGUGAUGUGUCCGAUACAUC.CCAUUUCAC..CUUACUUGA................GUGAGAGAA.........................................
env-303    ...........................GAGGCACCCCAA....UUAGGAACUGAGAAAUAA......GGAAGUGAUGUGUCCGAUACAUC.CCAUUUCAC..CUUACUUGA................GUGAGAGAA.........................................
env-304    ...........................GAGGCACCCCAA....UUAGGAACUGAGAAAUAA......GGAAGUGAUGUGUCCGAUACAUC.CCAUUUCAC..CUUACUUGA................GUGAGAGAA.........................................
env-305    ...........................GAGGCACCCCAA....UUAGGAACUGAGAAAUAA......GGAAGUGAUGUGUCCGAUACAUC.CCAUUUCAC..CUUACUUGA................GUGAGAGAA.........................................
Bwe-2-1    ...........................GAGGCACCCCAA....UUAGGAACUGAGAAAUAA......GGAAGUGAUGUGUCCGAUACAUC.CCAUUUCAC..CUUACUUGA................GUGAGAGAA.........................................
env-306    ...........................GAGGCACCCCAA....UUAGGAACUGAGAAAUAA......GGAAGUGAUGUGUCCGAUACAUC.CCAUUUCAC..CUUACUUGA................GUGAGAGAA.........................................
env-307    ...........................GAGGCACCCCAA....UUAGGAACUGAGAAAUAA......GGAAGUGAUGUGUCCGAUACAUC.CCAUUUCAC..CUUACUUGA................GUGAGAGAA.........................................
env-308    ...........................GAGGCACCCCAA....UUAGGAACUGAGAAAUAA......GGAAGUGAUGUGUCCGAUACAUC.CCAUUUCAC..CUUACUUGA................GUGAGAGAA.........................................
env-309    ...........................GAGGCACCCCAA....UUAGGAACUGAGAAAUAA......GGAAGUGAUGUGUCCGAUACAUC.CCAUUUCAC..CUUACUUGA................GUGAGAGAA.........................................
env-310    ...........................GAGGCACCCCAA....UUAGGAACUGAGAAAUAA......GGAAGUGAUGUGUCCGAUACAUC.CCAUUUCAC..CUUACUUGA................GUGAGAGAA.........................................
env-311    ...........................GAGGCACCCCAA....UUAGGAACUGAGAAAUAA......GGAAGUGAUGUGUCCGAUACAUC.CCAUUUCAC..CUUACUUGA................GUGAGAGAA.........................................
env-313    ...........................GAGGCACCCCAA....UUAGGAACUGAGAAAUAA......GGAAGUGAUGUGUCCGAUACAUC.CCAUUUCAC..CUUACUUGA................GUGAGAGAA.........................................
env-314    ...........................GAGGCACCCCAA....UUAGGAACUGAGAAAUAA......GGAAGUGAUGUGUCCGAUACAUC.CCAUUUCAC..CUUACUUGA................GUGAGAGAA.........................................
env-315    ...........................GAGGCACCCCAA....UUAGGAACUGAGAAAUAA......GGAAGUGAUGUGUCCGAUACAUC.CCAUUUCAC..CUUACUUGA................GUGAGAGAA.........................................
env-317    ...........................GAGGCACCCCAA....UUAGGAACUGAGAAAUAA......GGAAGUGAUGUGUCCGAUACAUC.CCAUUUCAC..CUUACUUGA................GUGAGAGAA.........................................
env-318    ...........................GAGGCACCCCAA....UUAGGAACUGAGAAAUAA......GGAAGUGAUGUGUCCGAUACAUC.CCAUUUCAC..CUUACUUGA................GUGAGAGAA.........................................
Bwe-1-1    ...........................GAGGCACCCCAA....UUAGGAACUGAGAAAUAA......GGAAGUGAUGUGUCCGAUACAUC.CCAUUUCAC..CUUACUUGA................GUGAGAGAA.........................................
env-319    ...........................GAGGCACCCCAA....UUAGGAACUGAGAAAUAA......GGAAGUGAUGUGUCCGAUACAUC.CCAUUUCAC..CUUACUUGA................GUGAGAGAA.........................................
env-320    ...........................GAGGCACCCCAA....UUAGGAACUGAGAAAUAA......GGAAGUGAUGUGUCCGAUACAUC.CCAUUUCAC..CUUACUUGA................GUGAGAGAA.........................................
env-321    ...........................GAGGCACCCCAA....UUAGGAACUGAGAAAUAA......GGAAGUGAUGUGUCCGAUACAUC.CCAUUUCAC..CUUACUUGA................GUGAGAGAA.........................................
env-322    ...........................GAGGCACCCCAA....UUAGGAACUGAGAAAUAA......GGAAGUGAUGUGUCCGAUACAUC.CCAUUUCAC..CUUACUUGA................GUGAGAGAA.........................................
env-323    ...........................GAGGCACCCCAA....UUAGGAACUGAGAAAUAA......GGAAGUGAUGUGUCCGAUACAUC.CCAUUUCAC..CUUACUUGA................GUGAGAGAA.........................................
env-324    ...........................GAGGCACCCCAA....UUAGGAACUGAGAAAUAA......GGAAGUGAUGUGUCCGAUACAUC.CCAUUUCAC..CUUACUUGA................GUGAGAGAA.........................................
env-325    ...........................GAGGCACCCCAA....UUAGGAACUGAGAAAUAA......GGAAGUGAUGUGUCCGAUACAUC.CCAUUUCAC..CUUACUUGA................GUGAGAGAA.........................................
env-326    ...........................GAGGCACCCCAA....UUAGGAACUGAGAAAUAA......GGAAGUGAUGUGUCCGAUACAUC.CCAUUUCAC..CUUACUUGA................GUGAGAGAA.........................................

           >>>>>.....................>>>>>>>>>..............................<<<<<............<<<<<<............<<<<<.................<<<<<...........................................
           1111......................2222220...............................22222............1111..............11111.................11111...........................................
           -------------------------oRYRYCCCAR---UUARRRA•R••••ooooo-------ooooo--------------------------------ooo-------------------------ooooooo-------------------------------------
                                                    3'
```

```
env-327   ............................GAGGCACCCCAA....UUAGGAACUAAAAGA..........GAAAGCAGCGUACAGA.CACAUG.CCAUUUCAC..CUUGU........................GCAAUAUGU.......................................
env-328   ............................GAGGCACCCCAA....UUAGGA.AUGAAGGA..........GAAGGCAGCGUACAGA.UGCACC.CCAUUUCAC..CUUGU........................AGAAAU..........................................
env-329   ............................GAGGCACCCCAA....UUAGGAACUAAAAAAGAA......GAAAGAGUGCGUUAAGAUGCACC.CCAUUUCAC..CUUGU...................AU...CUG.............................................
env-331   ............................GAGGCACCCCAA....UUAGGAACUAAAAAAGAA......GAAAGAGUGCGUUAAGAUGCACC.CCAUUUCAC..CUUGU...................AU...CUG.............................................
env-332   ............................GAGGCACCCCAA....UUAGGAACUAAGAAUUAA.......GAAAGAGUGCGUAUGGAUGCAUG.CCAUUUCAC..CUUGU...................AU...CUG.............................................
env-335   ............................GAGGCACCCCAA....UUAGGAACUAAGAAUUAA.......GAAAGAGUGCGUAUGGAUGCAUG.CCAUUUCAC..CUUGU...................AU...CUG.............................................
env-336   ............................GAGGCACCCCAA....UUAGGAACUAAGAAUUAA.......GAAAGAGUGCGUAUGGAUGCAUG.CCAUUUCAC..CUUGU...................AU...CUG.............................................
env-337   ............................GAGGCACCCCAA....UUAGGAACUAAGAAUUAA.......GAAAGAGUGCGUAUGGAUGCAUG.CCAUUUCAC..CUUGU...................AU...CUG.............................................
env-338   ............................GAGGCACCCCAA....UUAGGAACUAAGAAUUAA.......GAAAGAGUGCGUAUGGAUGCAUG.CCAUUUCAC..CUUGU...................AU...CUG.............................................
env-339   ............................GAGGCACCCCAA....UUAGGAACUAAGAAUUAA.......GAAAGAGUGCGUAUGGAUGCAUG.CCAUUUCAC..CUUGU...................AU...CUG.............................................
env-340   ............................GAGGCACCCCAA....UUAGGAACUAAGAAUUAA.......GAAAGAGUGCGUAUGGAUGCAUG.CCAUUUCAC..CUUGU...................AU...CUG.............................................
env-341   ............................GAGGCACCCCAA....UUAGGAACUAAGAAUUAA.......GAAAGAGUGCGUAUGGAUGCAUG.CCAUUUCAC..CUUGU...................AU...CUG.............................................
env-342   ............................GAGGCACCCCAA....UUAGGAACUAAGAAUUAA.......GAAAGAGUGCGUAUGGAUGCAUG.CCAUUUCAC..CUUGU...................AU...CUG.............................................
Bsc-1-1   ............................GAGGCACCCCAA....UUAGGAACUGGAAAUA.........GAGAGCGACGGACAGA.UACACC.CCAUUUCAC..CUUGU........................GCGUAUGU........................................
Rsp-3-1   ............................GAGGCACCCCAA....UUAGGAACUAAGAAAUAAUG.....AAAGCAGGUGACAGA..UGCACC.CCAUUUCAC..CUUGC........................GGAUCAAAUGGAUCA.................................
env-343   ..............................GGCAUCCCAA....UUAAGAUGAUAUUGAACA.......GAUCG.............GCCU.........................................AUCAUCGACUCAAAUUUCA.............................
env-344   UU..........................UAUAUAUCUCAA....UUAGGAACAAUAUUAACAGAA..........................................................................................................................
env-346   UU..........................UAUAUAUCUCAA....UUAGGAACAAUAUUAACAGAA..........................................................................................................................
env-348   UU..........................UAUAUAUCUCAA....UUAGGAACAAUAUUAACAGAA..........................................................................................................................
env-349   UU..........................UAUAUAUCUCAA....UUAGGAACAAUAUUAACAGAA..........................................................................................................................
env-350   UU..........................UAUAUAUCUCAA....UUAGGAACAAUAUUAACAGAA..........................................................................................................................
env-351   UU..........................UAUAUAUCUCAA....UUAGGAACAAUAUUAACAGAA..........................................................................................................................
env-352   UU..........................UAUAUAUCUCAA....UUAGGAACAUAUUACAGA..............................................................................................................................
env-353   ............................CCGGCAUCCCAG....UUAGGAAAAUAUUAACGAG......GAUUCAUGGGGUUG...CGCACA..CAUUUCAGG.CCUGU........................CAU.............................................
Rga-1-1   ............................CCGGCAUCCCAG....UUAGGAAAAUAUUAACGAG......GAUUCAUGGGGUUG...CGCACA..CAUUUCAGG.CCUGU........................UUU.............................................
env-354   ..............................GACAUCCCAA....UUAGGAAAAUUUUU...................................................................................................................................
env-355   ...............................GCAUCCCAA....UUAGGAAAAUAUUAACCAA......GAUUUAUGAAGGUG...UACACA..CAUUUCAGA.CUUGCAAGAUUUGUCACACU.AAAUACGAGAUACC.........................................
env-356   ...............................GCAUCCCAA....UUAGGAAAAUAUUAACCAA......GAUUUAUGAAGGUG...UACACA..CAUUUCAGA.CUUGCAAGAUUUGUCACACU.AAAUACGAGAUACC.........................................
env-357   ...............................GCAUCCCAA....UUAGGAAAAUAUUAACAAC......GAUUUAUAAAGA.....UACACACACAUUUCAGA.CCUA.........................GCAUUAUGG.......................................
env-358   ...............................GCAUCCCAA....UUAGGAAAAUAUUAACAAC......GAUUUAUAAAGA.....UACACACACAUUUCAGA.CCUA.........................GCAUUAUGG.......................................
env-359   ...............................GCAUCCCAA....UUAGGAAAAUAUUAACAAC......GAUUUAUAAAGA.....UACACACACAUUUCAGA.CCUA.........................GCAUUAUGG.......................................
env-360   .............................GAGCAUCCCAA....UUAGGUAAAUAUUAACCAA......GAUUUAUGAUUGG....CACACA..CAUUUCAGA.CCUGC........................UUGGAAGUUAAGAACAAAAA............................
env-361   ...............................GCAUCCCAA....UUAGGUAAAUAUUAACCAA......GAUUUAUGAUGAUG...UACACA..CAUUUCAGA.CCUGCCAA...................AGCAUGAUUAA.....................................
env-362   ...............................GCAUCCCAA....UUAGGUAAAUAUUAACCAA......GAUUUAUGAUGAUG...UACACA..CAUUUCAGA.CCUGCCAA...................AGCAUGAUUAA.....................................
env-363   ...............................GCAUCCCAA....UUAGGUAAAUAUUAACCAA......GAUUUAUAAACGAUG..UACACA..CAUUUCAGG.UCUGC........................UGUAAAA.........................................
env-364   ...............................GCAUCCCAA....UUAGGUAAAUAUUAACCAA......GAUUUAUAAACGA....UGUACACACAUUUCAGA.C............................UCACC...........................................
env-365   ............................GA.GCAUCCCAA....UUAGGUAAAUAUUAACCAA......GAUUUAUAAAUGA....UGUACACACAUUUCAGG.CUCAC........................UAGUAAGU........................................
env-366   ............................GA.GCAUCCCAA....UUAGGUAAAUAUUAACCAA......GAUUUAUAAAUGA....UGUACACACAUUUCAGG.CUCAC........................UAGUAAGU........................................
env-367   ..............................UGUAUCCCAA....UUAGGAAAAUAUUAAU.........................................................................GUUUUAGAA.......................................
env-368   ..............................UGUAUCCCAA....UUAGGAAAAUAUUAAU.........................................................................GUUUUAGAA.......................................
env-369   ..............................UGUAUCCCAA....UUAGGAAAAUAUUAAU.........................................................................GUUUUAGAA.......................................
env-370   ..............................UGUAUCCCAA....UUAGGAAAAUAUUAAU.........................................................................GUUUUAGAA.......................................
env-371   ..............................UGUAUCCCAA....UUAGGAAAAUAUUAAU.........................................................................GUUUUAGAA.......................................
env-372   ..............................UGUAUCCCAA....UUAGGAAAAUAUUAAU.........................................................................GUUUUAGAA.......................................
env-373   ..............................UGUAUCCCAA....UUAGGAAAAUAUUAAU.........................................................................GUUUUAGAA.......................................
env-374   ..............................UGUAUCCCAA....UUAGGAAAAUAUUAAU.........................................................................GUUUUAGAA.......................................
env-375   ..............................UGUAUCCCAA....UUAGGAAAAUAUUAAU.........................................................................GUUUUAGAA.......................................
env-377   ..............................UGUAUCCCAA....UUAGGAAAAUAUUAAU.........................................................................GUUUUAGAA.......................................
env-383   ..............................UGUAUCCCAA....UUAGGAAAAUAUUAAU.........................................................................GUUUUAGAA.......................................
env-384   ..............................UGUAUCCCAA....UUAGGAAAAUAUUAAU.........................................................................GUUUUAGAA.......................................
env-385   ..............................UGUAUCCCAA....UUAGGAAAAUAUUAAU.........................................................................GUUUUAGAA.......................................
env-386   ..............................UGUAUCCCAA....UUAGGAAAAUAUUAAU.........................................................................GUUUUAGAA.......................................
env-387   ..............................UGUAUCCCAA....UUAGGAAAAUAUUAAU.........................................................................GUUUUAGAA.......................................
env-388   ..............................UGUAUCCCAA....UUAGGAAAAUAUUAAU.........................................................................GUUUUAGAA.......................................
env-389   ..............................UGUAUCCCAA....UUAGGAAAAUAUUAAU.........................................................................GUUUUAGAA.......................................
env-390   ..............................UGUAUCCCAA....UUAGGAAAAUAUUAAU.........................................................................GUUUUAGAA.......................................
env-391   ..............................UGUAUCCCAA....UUAGGAAAAUAUUAAU.........................................................................GUUUUAGAA.......................................
env-392   ..............................UGUAUCCCAA....UUAGGAAAAUAUUAAU.........................................................................GUUUUAGAA.......................................
env-393   ..............................UGUAUCCCAA....UUAGGAAAAUAUUAAU.........................................................................GUUUUAGAA.......................................
env-394   ..............................UGUAUCCCAA....UUAGGAAAAUAUUAAU.........................................................................GUUUUAGAA.......................................
env-395   ..............................UGUAUCCCAA....UUAGGAAAAUAUUAAU.........................................................................GUUUUAGAA.......................................
env-396   ..............................UGUAUCCCAA....UUAGGAAAAUAUUAAU.........................................................................GUUUUAGAA.......................................
env-397   ..............................UGUAUCCCAA....UUAGGAAAAUAUUAAU.........................................................................GUUUUAGAA.......................................
env-398   ..............................UGUAUCCCAA....UUAGGAAAAUAUUAAU.........................................................................GUUUUAGAA.......................................
env-399   ..............................UGUAUCCCAA....UUAGGAAAAUAUUAAU.........................................................................GUUUUAGAA.......................................
env-405   ..............................UGUAUCCCAA....UUAGGAAAAUAUUAAU.........................................................................GUUUUAGAA.......................................
env-408   ..............................UGUAUCCCAA....UUAGGAAAAUAUUAAU.........................................................................GUUUUAGAA.......................................
env-409   ..............................UGUAUCCCAA....UUAGGAAAAUAUUAAU.........................................................................GUUUUAGAA.......................................
env-410   ..............................UGUAUCCCAA....UUAGGAAAAUAUUAAU.........................................................................GUUUUAGAA.......................................
env-411   ..............................UGUAUCCCAA....UUAGGAAAAUAUUAAU.........................................................................GUUUUAGAA.......................................
env-412   ..............................UGUAUCCCAA....UUAGGAAAAUAUUAAU.........................................................................GUUUUAGAA.......................................
env-413   ..............................UGUAUCCCAA....UUAGGAAAAUAUUAAU.........................................................................GUUUUAGAA.......................................
env-414   ..............................UGUAUCCCAA....UUAGGAAAAUAUUAAU.........................................................................GUUUUAGAA.......................................
env-418   ..............................UGUAUCCCAA....UUAGGAAAAUAUUAAU.........................................................................GUUUUAGAA.......................................
env-419   ..............................UGUAUCCCAA....UUAGGAAAAUAUUAAU.........................................................................GUUUUAGAA.......................................
env-420   ..............................UGUAUCCCAA....UUAGGAAAAUAUUAAU.........................................................................GUUUUAGAA.......................................
env-428   ...............................GCAUCCCAA....UUAGGAAAAUAUUAAUAUG.....ACCGGAUGA........GACACA..CAUUUCAGA.CAUAU........................GGUAAC..........................................
env-429   ..............................GGCAUCCCAA....UUAGGAAAAUAUUAAUAUG.....ACCGGAUAG.........GACACA..CAUUUCAGA.CAUAU........................GGUGAC..........................................
env-430   ..............................GGCAUCCCAA....UUAGGAAAAUAUUAAUAUG.....ACCGGAUAG.........GACACA..CAUUUCAGA.CAUAU........................GGUGAC..........................................
env-432   ..............................GGCAUCCCAA....UUAGGAAAAUAUUAAUAUG.....ACCGGAUAG.........GACACA..CAUUUCAGA.CAUAU........................GGUGAC..........................................
env-433   ..............................UACAUCCCAG....UUAAGCACAAUAAAC..........GGCAG............................................................CGAAAG..........................................
env-434   ............................GAGACACCCCAA....UUAGGAACAAUAUUAACU......GAGAGCAGUGAAAGA..UACGUC.CCAUUUCAC..CGUAU........................GGCUAC..........................................
Bhy-1-1   ............................GAGACACCCCAA....UUAGGAACAAUAUUAACU......GAGAGCAGUGAAAGA..UACGUC.CCAUUUCAC..CGUAU........................GGCUAC..........................................

          >>>>>.......................>>>>>>>>>...........................<<<<<............<<<<<<............<<<<<....................<<<<<................................................
          1111........................2222220............................22222............1111..............11111....................11111................................................
          ----------------------------oRYRYCCAR----UUARRRA•R•••••ooooo------ooooo-------------------------ooo----------------------ooooooo------------------------------------------------
                                                  3'
```

```
env-435   ........GCAUCCCAU....UUAAGCUAGUAUCAAAAU.......UGAUC..........CG.......................ACAUUCCAGGGACUUGUA.........
env-436   ........GCAUCCCAA....UUAAGAUGAUAUUUCA........GACAGAU........CAGCG.....................UAUCAUCGAAGAAAAACU.........
env-437   ........GCAUCCCAU....UCAAGU..............................................................AGGUAUUUUC..................
env-438   ........GCAUCCCAU....UUAAG...............................................................UUGUAGAUUUUU................
Bsp-7-1   ..........GCCCCAA....UCAAACAAGUAACAUUAAAUU...CGGAA...........................................ACAUUUGAU.............
Bsp-9-1   .........CGCCCCAA....UCAAACAAGUAACAUUAAUA....UGGAA...........................................AUGUAGUAU.............
Bsp-5-1   .......UUCGCCCCAA....UCAAACAAGUAACAUUAAUA....CGGAA...........................................AUAAAGUAU.............
Bsp-2-1   ..........GCCCCAA....UCAAACAAGUAACAUUUAAUUAU..GGAAACAAAAGG......UUUCC.................................................
Bsp-3-1   .......UACGCCCCAA....UCAAACAAGUAACAAU........AAGUUAAAUAGAAAU..CUCUGA....UUUCUGAAAUUAAC...................................
Bpr-2-1   ......UAUGUAUCCCAA....UUAGGAACAAUAUUUAUUAAGAA.............................................................................
          >>>>>.............>>>>>>>>>.......................<<<<<...........<<<<<<...........<<<<<..................<<<<<.............
          1111..............2222220.........................22222...........1111.............11111..................11111.............
          --------------------RYRYCCCAR---UUARRRA•R•••••◦◦◦◦-------◦◦◦◦◦-----------------------------◦◦◦----------------◦◦◦◦◦◦◦----------
                                           3'
```

alignment positions 541 · · · 720

```
env-1     CAAACGCGUAGAGAUAGGUCUUACCUAUAUCU..............................................................................
env-2     ..............................................................................................................
env-3     ..............................................................................................................
env-4     ..............................................................................................................
env-5     ..............................................................................................................
env-6     ..............................................................................................................
env-7     ..............................................................................................................
env-8     ..............................................................................................................
env-9     ..............................................................................................................
env-10    ..............................................................................................................
env-11    ..............................................................................................................
env-12    ..............................................................................................................
env-13    ..............................................................................................................
env-14    ..............................................................................................................
env-16    ..............................................................................................................
env-17    ..............................................................................................................
env-18    ..............................................................................................................
env-19    ..............................................................................................................
Bsp-6-1   ..............................................................................................................
env-20    ..............................................................................................................
env-21    ..............................................................................................................
env-22    ..............................................................................................................
env-23    ..............................................................................................................
env-24    ..............................................................................................................
env-25    ..............................................................................................................
env-28    ..............................................................................................................
env-29    ..............................................................................................................
env-30    ..............................................................................................................
env-32    ..............................................................................................................
env-33    ..............................................................................................................
env-34    ..............................................................................................................
env-35    ..............................................................................................................
env-36    ..............................................................................................................
env-37    ..............................................................................................................
env-38    ..............................................................................................................
env-39    ..............................................................................................................
env-41    ..............................................................................................................
env-42    ..............................................................................................................
env-43    ..............................................................................................................
env-45     CGUUUUGUA....................................................................................................
env-46     .............................................................................................................
env-47     .............................................................................................................
env-48     .............................................................................................................
env-49     .............................................................................................................
env-50     .............................................................................................................
Dma-1-1    .............................................................................................................
env-51     .............................................................................................................
Lba-2-1    .............................................................................................................
[ce-1-1    .............................................................................................................
Ece-1-1    .............................................................................................................
env-52     .............................................................................................................
env-53     .............................................................................................................
env-54     .............................................................................................................
env-55     .............................................................................................................
env-59     .............................................................................................................
env-60     .............................................................................................................
env-62     .............................................................................................................
env-63     .............................................................................................................
env-64     .............................................................................................................
env-66     .............................................................................................................
env-67     .............................................................................................................
env-68     .............................................................................................................
env-75     .............................................................................................................
env-76     .............................................................................................................

          ..........................................................................................................>>>>
          ..........................................................................................................1111
          --------------------------------------------------------------------------------------------------------------
```

```
env-77      ...
env-78      ...
env-79      ...
env-80      ...
env-81      ...
Rgn-1-1     ...
env-82      ...
env-83      ...
env-87      ...
env-88      ...
env-89      ...
env-90      ...
env-91      ...
env-92      ...
env-93      ...
env-94      ...U...AC
Lba-11-1    ...U...AC
env-95      ...
env-96      ...
env-97      ...
Dsp-3-1     CCUGCGUAAAGACAGAUUGGGAAAGAUGUAUAGGGCGAAAGCCCGCAUCGAGA...
Dsp-1-1     ...
env-101     ...
env-102     ...
Dsp-2-1     ...
env-103     ...
env-106     ...
env-116     ...
env-117     ...
env-118     ...
env-119     ...
env-120     ...
env-123     ...
env-124     ...
env-127     ...
env-128     ...
env-129     ...
env-130     ...
env-136     ...
env-137     ...
env-138     ...
env-139     ...
Dlo-1-1     ...
env-140     ...
env-149     GCAGAAUUCGAGGACGAAAUAGGGCGCCAGCCCAUCAUCGAGCCGCAGCAAAGCGGAGGCGAGAUGUGCAUUUCGA...
env-150     ...
env-151     ...
Chy-1-1     UUGUGAAGACUUUAUAGGGCGAUAGCCCGCAGCGAGAUGGAGCAAAGCGGAAUCGAGCUGGCACGGCGUACCCAUGUCGAAAGGACACGGCAAAAGCCGGGCAAAGAUAAGCGAAGCGAAAUCAAGCCGGCACAGCGUUCCGCGCCGAAAAGACGCGGCAAAAGCCGGGCAGGACA...
env-152     GAUGAAUGUGCAAGAUAUAUAGGGCGAAGCCCGACAACGAGAUGAAGCGAUAGCGAAAUCGAGUUGCUUUUC...AG
env-153     ...
env-154     ...
Esp-1-1     ...
env-155     ...
env-156     ...
env-159     ...
env-160     ...
env-161     ...
env-162     ...
env-163     ...
env-164     ...
env-165     ...
env-167     ...
env-168     ...
env-169     ...
env-170     ...
env-171     ...
env-172     ...
env-175     ...
env-176     ...
env-177     ...
env-178     ...
Lba-4-1     ...
env-179     ...
Cba-3-1     ...
[sc-1-1     ...
Lba-7-1     ...
env-181     ...
env-182     ...
env-183     ...
env-184     ...
env-185     ...
env-186     ...
env-187     ...
env-188     ...

            ...>>>>
            ...1111
            -------
```

```
env-189    ........................................................................................................................................................................................
env-190    ........................................................................................................................................................................................
env-191    ........................................................................................................................................................................................
env-192    ........................................................................................................................................................................................
env-194    ........................................................................................................................................................................................
env-195    ........................................................................................................................................................................................
env-200    ........................................................................................................................................................................................
env-201    ........................................................................................................................................................................................
env-202    ........................................................................................................................................................................................
env-211    ........................................................................................................................................................................................
env-212    ........................................................................................................................................................................................
env-216    ........................................................................................................................................................................................
env-217    ........................................................................................................................................................................................
Rpe-1-1    ........................................................................................................................................................................................
Rsp-1-1    ........................................................................................................................................................................................
env-218    ........................................................................................................................................................................................
env-219    ........................................................................................................................................................................................
env-220    ........................................................................................................................................................................................
env-221    ........................................................................................................................................................................................
env-222    ........................................................................................................................................................................................
env-223    ........................................................................................................................................................................................
env-226    ........................................................................................................................................................................................
env-227    ........................................................................................................................................................................................
env-228    ........................................................................................................................................................................................
env-229    ........................................................................................................................................................................................
env-230    ........................................................................................................................................................................................
env-231    ........................................................................................................................................................................................
env-232    ........................................................................................................................................................................................
env-234    ........................................................................................................................................................................................
env-235    ........................................................................................................................................................................................
env-236    ........................................................................................................................................................................................
env-237    ........................................................................................................................................................................................
env-238    ........................................................................................................................................................................................
env-239    ........................................................................................................................................................................................
env-240    ........................................................................................................................................................................................
env-241    ........................................................................................................................................................................................
env-242    ........................................................................................................................................................................................
env-243    ........................................................................................................................................................................................
env-244    ........................................................................................................................................................................................
env-245    ........................................................................................................................................................................................
env-246    ........................................................................................................................................................................................
env-252    ........................................................................................................................................................................................
env-254    ........................................................................................................................................................................................
Lba-6-1    ........................................................................................................................................................................................
env-255    ........................................................................................................................................................................................
Lba-10-1   ........................................................................................................................................................................................
env-256    ........................................................................................................................................................................................
env-258    ........................................................................................................................................................................................
env-259    ........................................................................................................................................................................................
env-260    ........................................................................................................................................................................................
env-261    ........................................................................................................................................................................................
env-262    ........................................................................................................................................................................................
env-263    ........................................................................................................................................................................................
Bfo-1-1    ........................................................................................................................................................................................
Bpr-1-1    ........................................................................................................................................................................................
Cba-1-1    ........................................................................................................................................................................................
env-264    ........................................................................................................................................................................................
env-265    ........................................................................................................................................................................................
env-266    ........................................................................................................................................................................................
env-267    ........................................................................................................................................................................................
env-268    ........................................................................................................................................................................................
env-271    ........................................................................................................................................................................................
env-272    ........................................................................................................................................................................................
env-273    ........................................................................................................................................................................................
env-275    ........................................................................................................................................................................................
env-276    ........................................................................................................................................................................................
env-277    ........................................................................................................................................................................................
env-278    ........................................................................................................................................................................................
env-279    ........................................................................................................................................................................................
env-280    ........................................................................................................................................................................................
env-281    ........................................................................................................................................................................................
env-282    ........................................................................................................................................................................................
Bsp-1-1    ........................................................................................................................................................................................
env-283    ........................................................................................................................................................................................
env-284    ........................................................................................................................................................................................
env-285    ........................................................................................................................................................................................
env-286    ........................................................................................................................................................................................
env-287    ........................................................................................................................................................................................
env-288    ........................................................................................................................................................................................
env-289    ......................................................................................................................................................................................UA
env-290    ......................................................................................................................................................................................UA
env-291    ......................................................................................................................................................................................UA
env-292    ......................................................................................................................................................................................UA
env-293    ....................................................................................................................................................................................UUCG

           ....................................................................................................................................................................................>>>>
           ....................................................................................................................................................................................1111
           ----------------------------------------------------------------------------------------------------------------------------------------------------------------------------------------
```

```
env-294   ......UUCG
env-295   ......UUCG
env-296   ......UUCG
env-298   ......UUCG
env-299   ......UUCG
env-300   ......UUCG
env-301   ......UUCG
env-302   ......UUCG
env-303   ......UUCG
env-304   ......UUCG
env-305   ......UUCG
Bwe-2-1   ......UUCG
env-306   ......UUCG
env-307   ......UUCG
env-308   ......UUCG
env-309   ......UUCG
env-310   ......UUCG
env-311   ......UUCG
env-313   ......UUCG
env-314   ......UUCG
env-315   ......UUCG
env-317   ......UUCG
env-318   ......UUCG
Bwe-1-1   ......UUCG
env-319   ......UUCG
env-320   ......UUCG
env-321   ......UUCG
env-322   ......UUUG
env-323   ......UUUG
env-324   ......UUUG
env-325   ......UUUG
env-326   ......UUCG
env-327   ......
env-328   ......
env-329   ......A
env-331   ......A
env-332   ......A
env-335   ......A
env-336   ......A
env-337   ......A
env-338   ......A
env-339   ......A
env-340   ......A
env-341   ......A
env-342   ......A
Bsc-1-1   ......
Rsp-3-1   ......
env-343   ......
env-344   ......
env-346   ......
env-348   ......
env-349   ......
env-350   ......
env-351   ......
env-352   ......
env-353   ......
Rga-1-1   ......
env-354   ......
env-355   ......UCGU
env-356   ......UCGU
env-357   ......
env-358   ......
env-359   ......
env-360   ......
env-361   ......GUGC
env-362   ......GUGC
env-363   ......
env-364   ......
env-365   ......
env-366   ......
env-367   ......
env-368   ......
env-369   ......
env-370   ......
env-371   ......
env-372   ......
env-373   ......
env-374   ......
env-375   ......
env-377   ......
env-383   ......
env-384   ......
env-385   ......
env-386   ......

          ......>>>>
          ......1111
```

```
env-387   ....................................................................................................................................................................................
env-388   ....................................................................................................................................................................................
env-389   ....................................................................................................................................................................................
env-390   ....................................................................................................................................................................................
env-391   ....................................................................................................................................................................................
env-392   ....................................................................................................................................................................................
env-393   ....................................................................................................................................................................................
env-394   ....................................................................................................................................................................................
env-395   ....................................................................................................................................................................................
env-396   ....................................................................................................................................................................................
env-397   ....................................................................................................................................................................................
env-398   ....................................................................................................................................................................................
env-399   ....................................................................................................................................................................................
env-405   ....................................................................................................................................................................................
env-408   ....................................................................................................................................................................................
env-409   ....................................................................................................................................................................................
env-410   ....................................................................................................................................................................................
env-411   ....................................................................................................................................................................................
env-412   ....................................................................................................................................................................................
env-413   ....................................................................................................................................................................................
env-414   ....................................................................................................................................................................................
env-418   ....................................................................................................................................................................................
env-419   ....................................................................................................................................................................................
env-420   ....................................................................................................................................................................................
env-428   ....................................................................................................................................................................................
env-429   ....................................................................................................................................................................................
env-430   ....................................................................................................................................................................................
env-432   ....................................................................................................................................................................................
env-433   ....................................................................................................................................................................................
env-434   ....................................................................................................................................................................................
Bhy-1-1   ....................................................................................................................................................................................
env-435   ....................................................................................................................................................................................
env-436   ....................................................................................................................................................................................
env-437   ....................................................................................................................................................................................
env-438   ....................................................................................................................................................................................
Bsp-7-1   ....................................................................................................................................................................................
Bsp-9-1   ....................................................................................................................................................................................
Bsp-5-1   ....................................................................................................................................................................................
Bsp-2-1   ....................................................................................................................................................................................
Bsp-3-1   ....................................................................................................................................................................................
Bpr-2-1   ....................................................................................................................................................................................
          ................................................................................................................................................................................>>>>
          ................................................................................................................................................................................1111
          ------------------------------------------------------------------------------------------------------------------------------------------------------------------------------------
```

alignment positions 721 · · · 900

```
env-1     ..............................................................................................AUGGUAUUGGAAGAGCAUCUUCCAACAGAAUCCUUGCGCAGGCAGGUGUCAACCCGGAUACCCGUGUCAGAGAUCUGACGGAUGAU
env-2     ...........................................CAUCAUGAAGCA...................AAAUGGAGGAA.U.UUUAACAUGGCUCGUAUUGCUGGUGUUGACUUACCAAGAGAAAAACGUGUAGAGA.....................................
env-3     ...........................................CAUCAUGAAGCA...................AAAUGGAGGAA.U.UUUAACAUGGCUCGUAUUGCUGGUGUUGACUUACCAAGAGAAAAACGUGU..........................................
env-4     ...........................................CAUCAUGAAGCA...................AAAUGGAGGAA.U.UUUAACAUGGCUCGUAUUGCUGGUGUUGACUUACCAAGAGAAAAACGUGUAGAGAUUGGCUUAACUUAUAUCUACGGAAUUGGUAGAGUAAG
env-5     ...........................................CAUCAUGAAGCA...................AAAUGGAGGAA.U.UUUAACAUGGCUCGUAUUGCUGGUGUUGACUUACCAAGAGAAAAACGUGUAGAGAUUGGCUUAACUUAUAUCUACGGAAUUGGUAGAGUAAG
env-6     ...........................................CAUCAUGAAGCA...................AAAUGGAGGAA.U.UUUAACAUGGCUCGUAUUGCUGGUGUUGACUUACCAAGAGAAAAACGUGUAGA.......................................
env-7     ...........................................CAUCAUGAAGCA...................AAAUGGAGGAA.U.UUUAACAUGGCUCGUAUUGCUGGUGUUGACUUACCAAGAGAAAAACGUGUAGAGAUCGGCUUAACUUAUAUCUACGGAAUUGGUAGAGUAAG
env-8     ...........................................CAUCAUGAAGCA...................AAAUGGAGGAA.U.UUUAACAUGGCUCGUAUUGCUGGUGUUGACUUACCAAGAGAAAAACGUGUAGAGAUUGGCUUAACUUAUAUCUACGGAAUUGGUAGAGUAAG
env-9     ...........................................CAUCAUGAAGCA...................AAAUGGAGGAA.U.UUUAACAUGGCUCGUAUUGCUGGUGUUGACUUACCAAGAGAAAAACGUGUAGAGAUCGGCUUAACUUAUAUCUACGGAAUUGGUAGAGUAAG
env-10    ...........................................CAUCAUGAAGCA...................AAAUGGAGGAA.U.UUUAACAUGGCUCGUAUUGCUGGUGUUGACUUACCAAGAGAAAAACGUGUAGAGAUUGGCUUAACUUAUAUCUACGGAAUUGGUAGAGUAAG
env-11    ...........................................CAUCAUGAAGCA...................AAAUGGAGGAA.U.UUUAACAUGGCUCGUAUUGCUGGUGUUGACUUACCAAGAGAAAAACGUGUAGAGAUUGGCUUAACUUAUAUCUACGGAAUUGGUAGAGUAAG
env-12    ...........................................CAUCAUGAAGCA...................AAAUGGAGGAA.U.UUUAACAUGGCUCGUAUUGCUGGUGUUGACUUACCAAGAGAAAAACGUGUAGAGAUUGGCUUAACUUAUAUCUACGGAAUUGGUAGAGUAAG
env-13    ...........................................CAUCAUGAAGCA...................AAAUGGAGGAA.U.UUUAACAUGGCUCGUAUUGCUGGUGUUGACUUACCAAGAGAAAAACGUGUAGAGAUCGGCUUAACUUAUAUCUACGGAAUUGGUAGAGUAAG
env-14    ...........................................CAUCAUGAAGCA...................AAAUGGAGGAA.U.UUUAACAUGGCUCGUAUUGCUGGUGUUGACUUACCAAGAGAAAAACGUGUAG........................................
env-16    ...........................................CAUCAUGAAGCA...................AAAUGGAGGAA.U.UUUAACAUGGCUCGUAUUGCUGGUGUUGACUUACCAAGAGAAAAACGUGUAGAGAUUGGCUUAACUUAUAUCUACGGAAUUGGUAGAGUAAG
env-17    .................................................AUUAAAC..................AAAUGGAGGAA.UUUUUAAAAUGGCUCGUAUUGCAGGUGUUGACUUACCAAGAGACAAACGUGUUGAAAUCGGUCUUACUUACAUCUAUGGAAUCGGCAGAGUAAG
env-18    .................................................AUUAAAC..................AAAUGGAGGAA.UUUUUAAAAUGGCUCGUAUUGCAGGUGUUGACUUACCAAGAGACAAACGUGUUGAAAUCGGUCUUACUUACAUCUAUGGAAUCGGCAGAGUAAG
env-19    ...............GAGUGA..GCUGCGUAGAG.........GUGUCUUAAUAAGA.................AAAUGGAGGAA.A.AUCA.CAUGGCUCGUAUUGCUGGUGUAGACUUACCAAGAGAAAAACGUGUAGAAAUUGGUCUAACUUAUAUAUAUGGAAUCGGCAGAGUUAG
Bsp-6-1   ...............UCAGGA..GAGACA...............AUCAAUUUAGUUGA................AAAUGGAGGAA.UUAGUAAAAUGGCUCGUAUUGCAGGUGUAGACUUACCGAGAGAUAAGCGAGUUGAGAUUGGUCUUACAUAUAUCUAUGGUAUUGGUAGAACAAG
env-20    ..........................................................................GGAUGGAGGAA....UUGCGAUGGCUCGUUUAGCCGGAGUAGAUUUACCGAGAGACAAACGGAUUGAAAUCGGCCUUACCUAUAUUUACGGUAUCGGCAGAAGCCA
env-21    ...........................................AUAUUCAUUU.....................AAUUGGAGGAA.UUAUUAAAAUGGCUCGUAUUGCAGGUGUAGACUUACCAAGAGAUAAGCGAGUAGAGAUCGGUUUGACUUACAUCUACGGAAUCGGUAGAACAAG
env-22    .......................UCCGGGCAGCAUGGCGC........AAAGGCGUCAUGAAAG..........CGAUGGAGGAA...AGAA.CAUGGCUCGUUUAGCAGGUGUUGAUCUGCCGAGGGACAAAAGAGUCGAGAUC...................................
env-23    ............................................GCGCCAAAAA...................GAAAGGAAGGA..AAUUAAACAUGGCUCGUAUUGCUGGUGUUGAUUUACCAAGAGACAAGCGUGUAGAGAUUGGUUUGACCAGUAUCUACGGAAUCGGAAGAGUAAG
env-24    ...............AUGUA...UGUGUAUAAAAUGAAAAA..CACUGAAAAAGA...................AGAUGGAGGAA.UAAUCA.CAUGGCUCGUAUCGCUGGUGUAGAUUUACCAAGAGAAAAACGUGUAGAAAUCGGAUUAACUUAUAUCUACGGAAUAGG.........
env-25    ...............AUGUA...UGUGUAUAAAAUGAAAAA..CACUGAAAAAGA...................AGAUGGAGGAA.UAAUCA.CAUGGCUCGUAUCGCUGGUGUAGAUUUACCAAGAGAAAAACGUGUAGAAAUCGGAUUAACUUAUAUCUACGGAAUAGGUAGAGUGAG
env-28    ...............AUGUA...UGUGUAUAAAAUGAAAAA..CACUGAAAAAGA...................AGAUGGAGGAA.UAAUCA.CAUGGCUCGUAUCGCUGGUGUAGAUUUACCAAGAGAAAAACGUGUAGAAAUCGGAUUAACUUAUAUCUACGGAAUAGGUAGAGUGAG
env-29    ...............AUGUA...UGUGUAUAAAAUGAAAAAC.CACUGAAAAAGA...................AGAUGGAGGAA.UAAUCA.CAUGGCUCGUAUCGCUGGUGUAGAUUUACCAAGAGAAAAACGUGUAGAAAUUGGAUUGACUUAUAUCUAUGGAAUAG..........
env-30    ...............AUGUA...UGUGUAUAAGAUGAAAAAC.CACUGAAAAAGA...................AGAUGGAGGAA.UAAUCA.CAUGGCUCGUAUCGCUGGUGUAGAUUUACCAAGAGAAAAACGUGUAGAAAUUGGAUUGACUUAUAUCUAUGGAAUAGGUAGAGUAAG
env-32    ...............AUGUA...UGUGUAUAAGAUGAAAAAC.CACUGAAAAAGA...................AGAUGGAGGAA.UAAUCA.CAUGGCUCGUAUCGCUGGUGUAGAUUUACCAAGAGAAAAACGUGUAGAAAUUGGAUUGACUUAUAUCUAUGGAAUAGGUAGAGUAAG
env-33    ...............AUGUA...UGUGUAUAAGAUGAAAAAC.CACUGAAAAAGA...................AGAUGGAGGAA.UAAUCA.CAUGGCUCGUAUCGCUGGUGUAGAUUUACCAAGAGAAAAACGUGUAGAAAUUGGAUUGACUUAUAUCUAUGGAAUAGGUAGAGUAAG
env-34    ...............CAGUGA..GCUGCGUGAUGGAGAUGG..CUUAG..A.......................AAAUGGAGGAA.UAAUCA.AAUGGCU................................................................................
env-35    ...............CAGUGA..GCUGCGUGAUGGAGAUGG..CUUAG..A.......................AAAUGGAGGAA.UAAUCA.AAUGGCUCGUAUUGCUGGUGUAGACUUACCAAGAGAAAAACGUGCAGAGAUUGGACUGACUUACAUCUACGGAAUCGGAAGAGUAAG
env-36    ...............CAGUGA..GCUGCGUGAUGGAGAUGG..CUUAG..A.......................AAAUGGAGGAA.UAAUCA.AAUGGCUCGUAUUGCUGGUGUAGACUUACCAA.......................................................
env-37    ...............AAGCAU......................CACACAAAAAG....................AAAUGGAGGAA.AU.UUAAAAUGGCUCGUAUUGCUGGUGUUGACUUGCCGAGAGACAAAAGAGUAGAAAUUGGUUUAACCAGUAUCUACGGAAU............

          >..............>>>>>...>>>>>>..............>>>>>....................................................................................................................................
          1..............11111.....1111..............22222....................................................................................................................................
          ---------------○○○○------------------------○○○○○○○○○○○○------------------AAAUGGAGGAA-○○●●YA-CAUGGCUCGUAU●GC●GGUGUAGAYUUACCAAGAGA●AAACG●RU●GARAUYGG●YU●ACYUAYAUYUAYGG●AUYGG●AGARYRAR
```

```
env-38    ...GCCCG...AACCCAAUUUUUUAUU...AAAUGGAGGAA.UAAUCA.AAUGGCUCGUAUUGCUGGUGUAGA...
env-39    ...GCCCGG...AACCCAAUUCUUUAUUU...AAAUGGAGGAA.UAAUCA.AAUGGCUCGUAUUUCUGGUGUAGACUUACCAAGAGAAAA...
env-41    ...GCCCGG...AACCCAAUUUUUU...AUUAGGAGGAA.UAAUCACAAUGGCUCGUAUUGCUGGUGUAGAUUUACC...
env-42    ...GCUGAG...GGGGCGACACUUA...AGCAGGAGGAA.UAAUUA.CAUGGCUCGUAUAGCUGGUGUAGACUUACCAAGAGACAAGCGAGUUGAGAUCGGCCUUACCUAUAUCUUCGGAAUCGGCCGUACCAC
env-43    ...AACAGAUUUG...AAAUGGAGGAA.A.GACA.CAUGGCUCGUAUUGCAGGUGUAGACUUACCAAGAGACAAACGUGUAGAAAUUGGCUUGACUUAUAUUUAUGGAAUUGGAAGAGUAAG
env-45    ...AAAAAAGG...AUGGAAAUUUAUAAAAAGCGGAUGCUUCGGAGACGGAAAGGAAAAUCAUCAUAACGGAGACAACGGAAAAUCACAUAGCUUUGA
env-46    ...AAAGAGUAGAGAUAGGACUUACCU...AUAUCUACGG...AAUUGGAAGAA...CAUCUUCAAACAGGAUCCUUGCCGAGGCAGGCGUAAAUCCGGAUACUCGUGUAAGAGAUCUUACAGACGACGAAGUUAAGAGG
env-47    ...GCAAGU..GAUGUACAUGAAUACCGG..CUCGUUUCACACA...UGAUUUAAGAA.UCCCUA....CAGAGAGGGAGUAUUAAGGAUCCCCGCAGAGGGAUAGUAUUAAGGAUCCCCGUAGAGGGACAGCAUUAAGAAUCCCCUGUAGA
env-48    ...CACUCAAAAAC...GAACGGAGGAA.AUAUU..CAUGGCUCGUAUUGCUGGUGUAGACUUACCAAGAGAGAAACGUAUCGAGAUCGGUCUUACCUAUAUCUAUGGGAUCGGCAGAACAAG
env-49    ...GCUCCGCAGGAA...CAAC...AGAUGGAGGAA.U..GAA.CAUGGCUCGUAUUGCAGGUGUGGAUUUACCAAGGGACAAGCGUGUCGAGAUCGGACUGACAUACAUUUAUGGCAUCGGGCUUCCCAG
env-50    ...AAAUGGAGGAAACAAG...UAUGGCUCGUAUUGCAGGUGUUGACU...
Dma-1-1   ...AAAUGGAGGAAACAAG...UAUGGCUCGUAUUGCAGGUGUUGACUUACCAAGAGACAAACGUGUAGAGAUUGGUCUGACUUAUAUCUACGGAAUCGGUAGAGUAAG
env-51    ...AAAUGGAGGAA.A.UUCA.CAUGGCUCGUAUUUCAGGUGUAGAUUUACCAAGAGACAAACGUGUUGAGAUUGGUUUGACUUAUAUUUAUGGAAUCGGAAGAACAAG
Lba-2-1   ...CAUUGG..GCUGUAUAUAUUUAUU...UACUAAGAUUCA...AAAUGGAGGAA.A.AUUA.CAUGGCUCGUAUUGCUGGUGUAGACUUACCAAGAGACAAACGUGUAGAAAUCGGAUUAACUUACAUUUACGGAAUCGGUAGAGUAAG
[ce-1-1   ...CCGAUCGCAAC...AAAAGGAGGAA.AACGCA.CAUGGCUCGUAUAGCUGGUGUAGAUUUACCAAGAGACAAACGUGUAGAGAUAGGUCUUACCUAUAUCUACGGAAUCGGAAGAACAAC
Ece-1-1   ...CUGAUCGCAAC...AAAAGGAGGAA.AACGCA.CAUGGCUCGUAUAGCUGGUGUAGAUUUACCAAGAGACAAACGUGUAGAAAUAGGUCUUACCUAUAUCUACGGAAUCGGAAGAACAAC
env-52    ...UGGCGG..GCUGCGUGUGAACGG...UAGCUUAUGAGA...AAAUGGAGGAA.AAAUUA.CAUGGCUCGUAUUGCUGGUGUAGACUUACCAAGAGAGAAAAGAAUUGAAAUCGGUUUGACUUACAUCUACGGUAUUGGUAGAGUAAG
env-53    ...AUUC..AUUCUAUUA...AAUUCAAGCAAGAUUUCA...AGAUGGAGGAA.AAAUUA.CAUGGCUCGUAUUUCAGGGGUAGACUUACCAAGAGGCAAGCGUGUAGAGAUUGGUUUGACUUACAUUUAUGGGAUUGGAAGAGCAAG
env-54    ...AAAUGGAGGAA.A.GACA.CAUGGC...
env-55    ...AAAUGGAGGAA.A.GACA.CAUGGCUCGUAUUGCAGGUGUAGACUUACCAAGAGACAAACGUGUAGAAAUCGGACUUACUUAUAUUUACGGAAUCGGAAGAACAUC
env-59    ...AAAUGGAGGAA.A.GACA.CAUGGCUCGUAUUGCAGGUGUAGACUUACCAAGAGACAAACGUGUAGAAAUCGGACUUACUUAUAUUUACGGAAUCGGAAGAACAUC
env-60    ...AAAUGGAGGAA.A.GACA.CAUGGCUCGUAUUGCAGGUGUAGACUUACCAAGAGACAAACGUGUAGAAAUCGGACUUACUUAUAUUUACGGAAUCGGAAGAACAUC
env-62    ...AAAUGGAGGAA.A.GACA.CAUGGCUCGUAUUGCAGGUG...
env-63    ...AAAUGGAGGAA.A.GACA.CAUGGCUCGUAUUGCAGUUGUAGACUUACCAAUAGACAAACGUGUAGAGAUCGGCUUGACUUAUAUUUACGGAAUCGGAAGAACAUC
env-64    ...AAAUGGAGGAA.A.GACA.CAUGGCUCGUAUUGCAGGUGUAGACUUACCAAGAGACAAAC...
env-66    ...AAAUGGAGGAA.A.GACA.CAUGGCUC...
env-67    ...AAAUGGAGGAA.A.GACA.CAUGGCUCGUAUUGCAGGUGUAGACUUACCAAGAGACAAACGUGUAGAAAUCGGACUUACUUAUAUUUACGGAAUCGGAAAAACAUC
env-68    ...AAAUGGAGGAA.A.GACA.CAUGGCUCGUAUUGCAGGUGUAGACUUACCAAGAGACAAACGUGUAGAAAUCGGACUUACUUAUAUUUACGGAAUCGGAAAAACAUC
env-75    ...AAAUGGAGGAA.A.GACA.CAUGGCUCGUAUUGCAGGUGUAGACUUACCAAGAGACAAACGUGUAGAAAUCGGACUUACUUAUAUUUACGGAAUCGGAAAAACAUC
env-76    ...AAAUGGAGGAA.A.GACA.CAUGGCUCGUAUUGCAGGUGUAGACUUACCAAGAGACAAACGUGUAGAAAUCGGACUUACUUAUAUUUACGGAAUCGGAAAAACAUC
env-77    ...AAAUGGAGGAA.A.GACA.CAUGGCUCGUAUUGCAGGUGUAGACUUACCAAGAGACAAACGUGUAGAAAUCGGACUUACUUAUAUUUACGGAAUCGGAAAAACAUC
env-78    ...AAAUGGAGGAA.A.GACA.CAUGGCUCGUAUUGCAGGUGUAGACUUACCAAGAGACAAACGUGUAGAAAUCGGACUUACUUAUAUUUACGGAAUCGGAA.......
env-79    ...AAAUGGAGGAA.A.GACA.CAUGGCUCGUAUUGCAGGUGUAGACUUACCAAGAGACAAACGUGUAGAAAUCGGACUUACUUAUAUUUACGGAAUCGGAAAAACAUC
env-80    ...AAAUGGAGGAA.A.GACA.CAUGGCUCGUAUUGCAGGUGUAGACUUACCAAGAGACAAACGUGUAGAAAUCGGACUUACUUAUAUUUACGGAAUCGGAAAAACAUC
env-81    ...AAAUGGAGGAA.A.GACA.CAUGGCUC...
Rgn-1-1   ...AAAUGGAGGAA.A.GACA.CAUGGCUCGUAUUGCAGGUGUAGACUUACCAAGAGACAAACGUGUAGAAAUCGGACUUACUUAUAUUUACGGAAUCGGAAAAACAUC
env-82    ...AAAUGGAGGAA.A.GACA.CAUGGCUCGUAUUGCAGGUGUAGACUUACCAAGAGACAAACGUGUAGAAAUCGGACUUACUUAUAUUUACGGAAUCGGAAAAACAUC
env-83    ...AAAUGGAGGAA.A.GACA.CAUGGCUCGUAUUGCAGGUGUAGACUUACCAAGAGACAAACGUGUAGAAAUCGGACUUACUUAUAUUUACGGAAUCGGAAAAACAUC
env-87    ...AAAUGGAGGAA.A.GACA.CAUGGCUCGUAUUGCAGGUGUAGACUUACCAAGAGACAAACGUGUAGAAAUCGGACUUACUUAUAUUUACGGAAUCGGAAAAACAUC
env-88    ...CUGAGCCGAUGCAUGAUGAUAACA..GUCCCAGAUUCA...AAAUGGAGGAA.A.UUCA.CAUGGCUCGUAUUGCUGGUGUAGACUUACCAAGAGACAAACGUGUAGAAAUCGGUCUUACUUAUAUCUAUGGUAUUGGUAGAACAAG
env-89    ...CAAAGA..GCUGUGUGAGCGGA...CUGACAGAAUUCA...AAAUGGAGGAA.U.UUUACAAUGGCUCGUAUUGCUGGUGUAGACUUACCAAGAGACAAACGUGUAGAAAUCGGACUUACUUAUAUUUAUGGAAUCGGAAGAGUAAG
env-90    ...AAGAUUUCA...AGAUGGAGGAA.AGAGUA.CAUGGCUCGUAUUUCAGGGGUAGACUUACCAAGAGGCAAGCGUGUAGAGAUUGGUCUGACUUACAUUUACGGGAUUGGAAGAGCAAG
env-91    ...GAUUCAAGCAAGAUUUCA...AAAUGGAGGAA.AAGAUA.CAUGGCUCGUAUUUCAGGGGUAGACUUACCAAGAGGCAAGCGUGUAGAGAUUGGAUUAACUUACAUUUAUGGGAUAGGAAGAGCAAG
env-92    ...GCGCAU...UCAGGAGAA...AGAUGGAGGAA.U.AACACAAUGGCUCGUAUAGCUGGUGUAGACUUACCAA...
env-93    ...GCAAGU..GAUGCCGCACCCGGAAGAU.UCAUCUCAAAAGA...AAAUGGAGGAA.ACAGUA.CAUGGCUCGUAUAGCUGGUAUCGACU...
env-94    CG...AGAGACAAACGUGUGGAAAUAGGUC..UUACCUAUAUCUACGG...UAUUGGAAGAG.CAUCUUCCAUGAGAAUCCUUGCGCAGGCAGGUGUCAACCCGGAUACCCGCGUCAGAGAUCUGACAGAUGAUGAGGUCAGAAAGAUCGCAGGCGU
Lba-11-1  CG...AGAGACAAACGUGUGGAAAUAGGUC..UUACCUAUAUCUACGG...AAUUGGAAGAG.CAUCUUCCAUGAAAAUCCUUGCGCAGGCAGGUGUCAACCCGGAUACCCGCGUCAGAGAUCUGACAGAUGAUGAAGUCAGAAAGAUCGCAGGCGU
env-95    ...G...GAUAUC...CAGACAGA...AAAUGGAGGAA.U.AACA.CAUGGCUCGUAUUGCUGGUGUAGACUUACCGAGAGAAAAACGUAUCGAGAUCGGUCUUACCUAUAUUUAUGGAAUUGGAAGAACCAG
env-96    ...GGAUA...UCCCCAAAAGA...AAAUGGAGGAA.AUUUCA.CAUGGCUCGUAUUGCUGGUGUAGACUUACCGAGAGAGAAACGUAUC...
env-97    ...GCGCGUCU...CAGCGCAACA...ACCAAGAAAACAGA...AAGUGGAGGAA.A.AUCA.CAUGGCUCGUAUUGCAGGUGUAGACUUACCAAGAGACAAACGUGUUGAAAUCGGAUUAACUUAUAUUUAUGGAAUUGGCAGAACAAG
Dsp-3-1   ...UGCA...GCGGAGCGGAA..UCGAGAUGGCACGGCGGGGCAGAAUGAAGCCAGCCGGCACAGGGCGAAAGCCCGCAUCAAGAUGCAGCGAAGCGGAAUCGAGAUGGCACGG
Dsp-1-1   ...ACUGGAUCAGGCAGGGCUUCGCAGCAAAAUU...UUUAAUCUUAUGAA...AAUUGGAGGAA.A.UUCA.CAUGGCACGUAUUGCAGGUGUAGAUUUACCAAGAGACAAACGCGUUGAGAUCGGACUGACUUAUAUCUAUGGGAUCGGCAGGCCGAG
env-101   ...AAGUGGAGGAA.CAGUCA.CAUGGCACGUAUUGCAGGUGUAGAUUUACCAAGAGACAAAC...
env-102   ...AAGUGGAGGAA.CAGUCA.CAUGGCACGUAUUGCAGGUGUAGAUUUACCAAGAGACAAACGAGUUGAGAUCGGAUUAACUUAUAUCUACGGAAUCGGUA.......
Dsp-2-1   ...AAGUGGAGGAA.CAGUCA.CAUGGCACGUAUUGCAGGUGUAGAUUUACCAAGAGACAAACGAGUUGAGAUCGGAUUAACUUAUAUCUACGGAAUCGGUAGAACAAG
env-103   ...AAGUGGAGGAA.CAGUCA.CAUGGCACGUAUUGCAGGUGUAGAUUUACCAA...
env-106   ...AAGUGGAGGAA.CAGUCA.CAUGGCACGUAUUGCAGGUGUAGAUUUACCAAGAGACAAACGAGUUGAGAUCGGAUUAACUUAUAUCUACGGAAUCGGUAGAACAAG
env-116   ...AAGUGGAGGAA.CAGUCA.CAUGGCACGUAUUGCAGGUGUA...
env-117   ...AAGUGGAGGAA.CAGUCA.CAUGGCACGUAUUGCAGGUGUAGAUUUACCAAGAGACAAACGAGUUGAGAUCGGAUUAACUUAUAUCUACGGAAUCGGUAGAACAAG
env-118   ...AAGUGGAGGAA.CAGUCA.CAUGGCACGUAUUGCAGGUGUAGAUUUACCAAGAGACAAACGAGUUGAGAUCGGAUUAACUUAUAUCUACGGAAUCGGUAGAACAAG
env-119   ...AAAUGGAGGAA.A.UUCA.CAUGGCUCGUAUAGCAGGUGUAGACUUACCAAGAGACAAACGUGUUGAGAUCGGAUUAACUUAUAUCUACGGAAUCGGUAGAACAAG
env-120   ...AAAUGGAGGAA.A.UUCA.CAUGGCUCGUAUAGCAGGUGUAGACUUACCAAGAGACA...
env-123   ...AAAUGGAGGAA.A.UUCA.CAUGGCUCGUAUAGCAGGUGUAGACUUACCAAGAGACAAACGUGUUGAGAUCGGAUUA...
env-124   ...AAAUGGAGGAA.A.UUCA.CAUGGCUCGUAUAGCAGGUGUAGACUUACCAAGAGACAAACGUGUUGAGAUCGGAUUAACUUAUAUCUACGGAAUCGGUAGAACAAG
env-127   ...AAAUGGAGGAA.A.UUCA.CAUGGCUCGUAUAGCAGGUGUAGACUUACCAAGAGACAAACGUGUUGAGAUCGGAUUAACUUAUAUCUACGGAAUCGGUAGAACAAG
env-128   ...AAAUGGAGGAA.A.UUCA.CAUGGCUCGUAUAGCAGGUGUAGACUUACCAAGAGACA...
env-129   ...AAAUGGAGGAA.A.UUCA.CAUGGCUCGUAUAGCAGGUGUAGACUUACCAAGAGACAAACGUGUUGAGAUUGGUCUGACCUAUAUCUACGGAAUCGGUAGAACAAG
env-130   ...AAAUGGAGGAA.A.UUCA.CAUGGCUCGUAUAGCAGGUGUAGACUUACCAAGAGACAAACGUGUUGAGAUCGGACUGACUUAUAUCUACGGAAUCGGUAGAACAAG
env-136   ...AAAUGGAGGAA.A.UUCA.CAUGGCUCGUAUAGCAGGUGUAGACUUACCAAGAGACAAACGUGUUGAGAUCGGACUGACUUAUAUCUACGGAAUCGGUAGAACAAG
env-137   ...AAAUGGAGGAA.A.UUCA.CAUGGCUCGUAUAGCAGGUGUAGACUUACCAAGAGACAAACGUGUUGAGAUCGGACUGACUUAUAUCUACGGAAUCGGUAGAACAAG
env-138   ...AAAUGGAGGAA.A.UUCA.CAUGGCUCGUAUAGCAGGUGUAGACUUACCAAGAGACAAACGUGUUGAGAUCGGACUGACUUAUAUCUACGGAAUCGGUAGAACAAG
env-139   ...AAAUGGAGGAA.A.UUCA.CAUGGCUCGUAUAGCAGGUGGAGACUUACCAAGAGACAAACGUGUUGAGAUCGGACUGACUUAUAUCUACGGAAUCGGUAGAACAAG
Dlo-1-1   ...AAAUGGAGGAA.A.UUCA.CAUGGCUCGUAUAGCAGGUGUAGACUUACCAAGAGACAAACGUGUUGAGAUCGGACUGACUUAUAUCUACGGAAUCGGUAGAACAAG
env-140   ...AAAUGGAGGAA.A.UUCA.CAUGGCUCGUAUAGCAGGUGUAGACUUACCAAGAGACAAACGUGUUGAGAUCGGACUGACUUAUAUCUACGGAAUCGGUAGAACAAG
env-149   ...CAACCAAAACAACAAGGAA...AAAUGGAGGAA.AAAUCA.CAUGGCUCGUAUUGCAGGUGUAGACUUACCAAGAGACAAACGUGUUGAGAUCGGAUUGACUUAUAUCUACGGAAUUGGUAGAACAAG
env-150   ...UCAGCG...AAACAAGAAAAAACAAGGA...AAAUGGAGGAA.AAUUCA.CAUGGCACGUAUUGCAGGUGUAGAUUUACCAAGAGACAAACGAGUAGAGAUCGGGCUUACUUAUAUCUACGGAAUCGGUAGAGCAAG
env-151   ...UCAGCG...AAACAAGAAAAAACAAGGA...AAAUGGAGGAA.AAUUCA.CAUGGCACGUAUUGCAGGUGUA...
Chy-1-1   ...GCACGUCAGAA...CAACAAAAAAAAGA...AAAUGGAGGAA.A.UUCA.CAUGGCUCGUAUAGCAGGUGUAGACUUACCAAGAGACAAACGUGUAGAGAUCGGACUUACUUAUAUUUAUGGGAUCGGAAGGACGAG
env-152   CAAGAACAAAAA...AUCAAAAAAACAGAA...AAAUGGAGGAA.A.UUCACAAUGGCUCGUAUAGCAGGUGUAGACUUACCAAGAGACAAACGUGUUGAGAUCGGAUUGACUUAUAUCUACGGAAUUGGUAGAACAAG
env-153   ...AUUUGGAGGAA.A.UUCA.CAUGGCUCGUAUUGCAGGUGUAGAUUUACCAAGAGACAAACGUGUUGAAAUCGGACUUACUUAUAUCUAUGGAAUCGGAAGAGCAAG
env-154   ...AUUUGGAGGAA.A.UUCA.CAUGGCUCGUAUUGCAGGUGUAG...
Esp-1-1   ...AAAUGGAGGAA.A.AUCA.CAUGGCUCGUAUUGCAGGUGUAGACUUACCAAGAGACAAACGUGUAGAAAUCGGACUGACCUAUAUUUACGGUAUCGGAAGAACGAG
env-155   ...AAAUGGAGGAA.A.GACA.CAUGGCUCGUAUUGCAGGUGUAGACUUACCAAGAGACAAACGUGUAGAAAUCGGUCUUACUUAUAUCUACGGAAUCGGUAGAACAAG
env-156   ...AAAUGGAGGAA.A.GACA.CAUGGCUCGUAUUGCAGGUGUAGACUUACCAAGAGACAAACGUGUAGAAAUCGGUCUUACUUAUAUCUACGGAAUCGGUAGAACAAG

          >...>>>>>...>>>>>>...>>>>>...
          1...11111.....1111...22222...
          ----oooo----ooooooooooo----AAAUGGAGGAA-◦◦••YA-CAUGGCUCGUAU•GC•GGUGUAGAYUUACCAAGAGA•AAACG•RU•GARAUYGG•YU•ACYUAYAUYUAYGG•AUYGG•AGARYRAR
```

```
env-159   ..........................................................................AAAUGGAGGAA.A.GACA.CAUGG..................................................................................
env-160   ..........................................................................AAAUGGAGGAA.A.GACA.CAUGGCUCGUAUUGCAGGUGUAGACUUACCAAGAGACAAACGUGUAGAAAUCGGUCUUACUUAUAUCUACGGAAUCGGUAGAACAAG
env-161   ..........................................................................AAAUGGAGGAA.A.GACA.CAUGGCUCGUAUUGCAGGUGUAGACUUACCAAGAGACAAACGUGUAGAAAUCGGUCUUACUUAUAUCUACGGAAUCGGUAGAACAAG
env-162   ..........................................................................AAAUGGAGGAA.A.GACA.CAUGGCUCGUAUUGCAGGUGUAGACUUACCAAGAGACAAACGUGUAGAAAUCGGUCUUACUUAUAUCUACGGAAUCGGUAGAACAAG
env-163   ..........................................................................AAAUGGAGGAA.A.GACA.CAUGGCUCGUAUUGCAGGUGUA.................................................................
env-164   ..........................................................................AAAUGGAGGAA.A.GACA.CAUGGCUC..............................................................................
env-165   ..........................................................................AAAUGGAGGAA.A.GACA.CAUGGCUCGUAUUGCAGGUGUAGACUUACCAAGAGACAAACGUGUAGAAAUCGGUCUUACUUAUAUCUACGGAAUCGGUAGAACAAG
env-167   ..........................................................................AAAUGGAGGAA.A.GACA.CAUG..................................................................................
env-168   ..........................................................................AAAUGGAGGAA.A.GACA.CAUGGCUCGUAUUGCAGGUGUAGACUUACCAAGAGACAAACGUGUAGAAAUCGGUCUUACUUAUAUCUACGGAAUCGGUAGAACAAG
env-169   ..........................................................................AAAUGGAGGAA.A.GACA.CAUGGCUCGUAUUGCAGGUGUAGACUUACCAAGAGACAAACGUGUAGAAAUCGGUCUUACUUAUAUCUACGGAAUCGGUAGAACAAG
env-170   ..........................................................................AAAUGGAGGAA.A.GACA.CAUGGCUCGUAUUGCAGGUGUAGACUUACCAAGAGACAAACGUGUAGAAAUCGGUCUUACUUAUAUCUACGGAAUCGGUAGAACAAG
env-171   ..........................................................................AAAUGGAGGAA.A.GACA.CAUGGCUCGUAUUGCAGGUGUAGACUUACCAAGAGACAAACGUGUAGAAAUCGGUCUUACUUAUAUCUACGGAAUCGGUAGAACAAG
env-172   ..........................................................................AAAUGGAGGAA.A.GACA.CAUGGCUCGUAUUGCAGGUGUAGACUUACCAAGAGACAAACGUGUAGAAAUCGGUCUUACUUAUAUCUACGGAAUCGGUAGAACAAG
env-175   ..........................................................................AAAUGGAGGAA.A.GACA.CAUGGCUCGUAUUGCAGGUGUAGACUUACCAAGAGACAAACGUGUAGAAAUCGGUCUUACUUAUAUCUACGGAAUCGGUAGAACAAG
env-176   ..........................................................................AAAUGGAGGAA.A.GACA.CAUGGCUCGUAUUGCAGGUGUAGACUUACCAAGAGACAAACGUGUAGAAAUCGGUCUUACUUAUAUCUACGGAAUCGGUAGAACAAG
env-177   ..........................................................................AAAUGGAGGAA.A.GACA.CAUGGCUCGUAUUGCAGGUGUAGACUUACCAAGAGACAAACGUGUAGAAAUCGGUCUUACUUAUAUCUACGGAAUCGGUAGAACAAG
env-178   ..........................................................................AAAUGGAGGAA.A.GACA.CAUGGCUCGUAUUGCAGGUGUAGACUUACCAAGAGACAAACGUGUAGAAAUCGGUCUUACUUAUAUCUACGGAAUCGGUAGAACAAG
Lba-4-1   ..........................................................................AAACGGAGGAA.A.AUUA.CAUGGCUCGUAUAGCAGGUGUAGACUUACCAAGAGACAAACGUGUUGAAAUCGGAUUAACUUAUAUCUACGGAAUCGGUAGAACAAG
env-179   ................CGCGG..CGCGUCUGAGCGCAACAA..CUAAGAAAACAGA..................AAGUGGAGGAA.A.AUCA.CAUGGCUCGUAUUGCAGGUGUAGACUUACCAAGAGACAAACGUGUUGAAAUCGGAUUAACUUAUAUUUAUGGAAUUGGCAGAACAAG
Cba-3-1   ................CGCGG..CGCGUCUGAGCGCAACAA..CCAAGAAAACAGA..................AAGUGGAGGAA.A.AUCA.CAUGGCUCGUAUUGCAGGUGUAGACUUACCAAGAGACAAACGUGUUGAAAUCGGAUUAACUUAUAUUUAUGGAAUUGGCAGAACAAG
[sc-1-1   ...............ACGCGG..CGCGUCUGAGCGCAACAA..CCAAGAAAACAGA..................AAGUGGAGGAA.A.AUCA.CAUGGCUCGUAUUGCAGGUGUAGACUUACCAAGAGACAAACGUGUUGAAAUCGGAUUAACUUAUAUUUAUGGAAUUGGCAGAACAAG
Lba-7-1   ...............ACGCGG..CGCGUCUGAGCGCAACAA..CUAAGAAAACAGG..................AAGUGGAGGAA.A.AUCA.CAUGGCUCGUAUUGCAGGUGUAGACUUACCAAGAGACAAACGUGUUGAAAUCGGAUUAACUUAUAUUUAUGGAAUUGGCAGAACAAG
env-181   ...............ACGCGG..CGCGUCUGAGCGCAACAA..CUAAGAAAACAGG..................AAGUGGAGGAA.A.AUCA.CAUGGCUCGUAUUGCAGGUGUAGACUUACCAAGAGACAAACGUGUAGAGAUCGGCUUGACUUAUAUUUAUGGAAUCGGAAGAA....
env-182   ...............GUGGG.......................CGUCACAAAA.....................AAACGGAGGAA.A.AAUA.CAUGGCUCGUAUAGCUGGUGUAGACUUACCAAGAGAGAAACGUGUAGAA......................................
env-183   .........................GUGCCGA.............GCCGAAAAGA...................AAAUGGAGGAA.A.AUCA.CAUGGCUCGUAUUGCUGGUGUAGACUUACCAAGAGAGAAGCGAAUCGAGAUCGGUCUGACUUACAUCUACGGAAUUGGCCGUACAAG
env-184   ..........................................................................AAAUGGAGGAA.A.GACA.CAUGGCUCGUAUUGCAGGUGUAGACUUACCAAGAGACAAACGUGUAGAGAUCGGCUUGACUUAUAUUUAUGGAAUCGGAAGAACAAG
env-185   ..........................................................................AAAUGGAGGAA.A.GACA.CAUGGCUCGUAUUGCAGUUGUAGACUUACCAAUAGACAAACGUGUAGAGAUCGGCUUGACUUAUAUUUACGGAAUCGGAAGAACAAG
env-186   ..........................................................................AAAUGGAGGAA.A.GACA.CAUGGCUCGUAUUGCAGGUGUAGACUUACCAAGAGACAAACGUGUAGAGAUCGGCUUGACUUAUAUUUAUGGAAUCGGAAGAACAAG
env-187   ..........................................................................AAAUGGAGGAA.A.GACA.CAUGGCUCGUAUUGCAGGUGUAGACUUACCAAGAGACAAAC.............................................
env-188   ..........................................................................AAAUGGAGGAA.A.GACA.CAUGGCUC..............................................................................
env-189   ..........................................................................AAAUGGAGGAA.A.GACA.CAUGGCUCGUAUUGCAGGUGUAGACUUACCAAGAGACAAACGUGUAGAGAUCGGCUUGACUUAUAUUUAUGGAAUCGGAAGAACAAG
env-190   ..........................................................................AAAUGGAGGAA.A.GACA.CAUGGCUCGUAUUGCAGGUGUAGACUUACCAAGAGACAAACGUGUAGAGAUCGGCUUGACUUAUAUUUAUGGAAUCGGAAGAACAAG
env-191   ..........................................................................AAAUGGAGGAA.A.GACA.CAUGGCUCGUAUUGCAGGUG...................................................................
env-192   ..........................................................................AAAUGGAGGAA.A.GACA.CAUGGCUCGUAUUGCAGGUGUAGACUUACCAAGAGACAAACGUGUAGAGAUCGGCUUGACUUAUAUUUAUGGAAUCGGAAGAACAAG
env-194   ..........................................................................AAAUGGAGGAA.A.GACA.CAUGGCUCGUAUUGCAGGUGUAGACUUACCAAGAGACAAACGUGUAGAGAUCGGCUUGACUUAUAUUUAUGGAAUCGGAAGAACAAG
env-195   ..........................................................................AAAUGGAGGAA.A.GACA.CAUGGCUCGUAUUGCAGGUGUAGACUUACCAAGAGACAAACGUGUAGAGAUCGGCUUGACUUAUAUUUAUGGAAUCGGAAGAACAAG
env-200   ..........................................................................AAAUGGAGGAA.A.GACA.CAUGGCUCGUAUUGCAGGUGUAGACUUACCAAGAGACAAACGUGUAGAGAUCGGCUUGACUUAUAUUUAUGGAAUCGGAAGAACAAG
env-201   ..........................................................................AAAUGGAGGAA.A.GACA.CAUGGCUCGUAUUGCAGGUGUAGACUUACCAAGAGACAAACGUGUAGA......................................
env-202   ..........................................................................AAAUGGAGGAA.A.GACA.CAUG..................................................................................
env-211   ..........................................................................AAAUGGAGGAA.A.GACA.CAUGGCUCGUAUUGCAG....................................................................
env-212   ..........................................................................AAAUGGAGGAA.A.GACA.CAUGGCUCGUAUUGCAGGUGUA.................................................................
env-216   ..........................................................................AAAUGGAGGAA.A.GACA.CAUGGCUCGUAUUGCAGGUGUAGACUUACCAAGAGACAAACGUGUAGAGAUCGGCUUGACUUAUAUUUAUGGAAUCGGAAGAACAAG
env-217   ........................CCGGAGUGUGUUUUGCGG.CAGUAUGAGUGAGA.................AAACGGAGGAA.UAUUA.CAUGGCUCGUAUAGCUGGUGUAGACUUACCAAGAGAAAAACGGAUCGAAAUCGGUCUGACCUACAUUUAUGGCAUUGGCAGAACAAC
Rpe-1-1   ...............GCAUGG..GCUGCGCGGCAUCGUAA...UCUUACAAGA.....................AAAUGGAGGAA.AUAUUA.CAUGGCUCGUAUUGCUGGUGUAGACUUACCAAGAGAAAAACGUGUAGAAAUUGGAUUAACUUACAUCUACGGAAUUGGUAGAGUAAG
Rsp-1-1   ...............GCAUGG..GCUGCGCGGCAUCGUAA...UCUUACAAGA.....................AAAUGGAGGAA.AUAUUA.CAUGGCUCGUAUUGCUGGUGUAGACUUACCAAGAGAAAAACGUGUAGAAAUUGGAUUAACUUACAUCUACGGAAUUGGUAGAGUAAG
env-218   ...........................................CAUCCAAAA......................GAAUGGAGGAA.A.AUCA.CAUGGCUCGUAUAGCUGGUGUAGACUUACCAAGAGAAAAGCGUGUCGAGAUCGGCCUGACCUACAUCUACGGAAUCGGCCGCACGAC
env-219   ...............GUAAGU..GAUGUACGGCGGAAAG....CUUUCAGAAAUUAUCAGAAUUAUCCAGUUUGGAAUGGAGGAAAACAGUA.CAUGGCUCGUAUAGCUGGUGUAGACUUACCAAGAGAUAAGAGAAUUGAGAUCGGACUUACCUAUAUUUACGGAAUCGGCAGAACAAG
env-220   .........................UGCG..............GCUGCGCACAGAGA.................AAAUGGAGGAA.CAAUUA.CAUGGCUCGUAUUGCUGGUGUAGACUUACCAAGAGAGAAACGUGCAGAGAUCGGUCUGACUUACAUUUACGGCAUUGGCCGUACAAG
env-221   ...............UGGCGG..GCUGUACAUAA.........CAUUCUCAAAGA...................AAAUGGAGGAA.UAAUUA.CAUGGCACGUAUUGCUGGUGUAGACUUACCAAGAGAAAAACGUAUCGAAAUUGGUCUGACUUACAUCUACGGUAUCGGUAGAGUAAG
env-222   ...............UGGCGG..GCUGUACAUAA.........CAUUCUCAAAGA...................AAAUGGAGGAA.UAAUUA.CAUGGCACGUAUUGCUGGUGUAGACUUACCAAGAGAAAAACGUAUCGAAAUUGGUCUGACUUACAUCUACGGUAUCGGUAGAGUAAG
env-223   ...............UGGCGG..GCUGUACAUAA.........CAUUCUCAAAGA...................AAAUGGAGGAA.UAAUUA.CAUGGCACGUAUUGCUGGUGUAGACUUACCAAGAGAAA..........................................
env-226   ...............UGGCGG..GCUGUACAUAA.........CAUUCUCAAAGA...................AAAUGGAGGAA.UAAUUA.CAUGGCACGUAUUGCUGGUGUAGACUUACCAAGAGAAAAACGUAUCGAAAU...............................
env-227   ...............UGGCGG..GCUGUACAUAA.........CAUUCUCAAAGA...................AAAUGGAGGAA.UAAUUA.CAUGGCACGUAUUGCUGGUGUAGACUUACCAAGAGAAAAACGUAUCGAAAUUGGUCUGACUUACAUCUACGGUAUCGGUAGAGUAAG
env-228   ...............UGGCGG..GCUGUACAUAA.........CAUUCUCAAAGA...................AAAUGGAGGAA.UAAUUA.CAUGGCACGUAUUGCUGGUGUAGACUUACCAAGAGAAAAACGUAUCGAAAUUGGUCUGACUUACAUCUACGGUAUCGGUAGAGUAAG
env-229   ...............UGGCGG..GCUGUACAUAA.........CAUUCUCAAAGA...................AAAUGGAGGAA.UAAUUA.CAUGGCACGUAUUGCUGGUGUAGACUUACCAAGAGA............................................
env-230   ...............UGGCGG..GCUGUACAUAA.........CAUUCUCAAAGA...................AAAUGGAGGAA.UAAUUA.CAUGGCACGUAUUGCUGGUGUAGACUUACCAAGAGAAAAACGUAUCGAAAUUGGUCUGACUUACAUCUACGGUAUCGGUAGAGUAAG
env-231   ...............UGGCGG..GCUGUACAUAA.........CAUUCUCAAAGA...................AAAUGGAGGAA.UAAUUA.CAUGGCACGUAUUGCUGGUGUAGACUUACCAAGAGAAAAACGUAUCGAAAUUGGUCUGACUUACAUCUACGGU..............
env-232   ...............UGGCGG..GCUGUACAUAA.........CAUUCUCAAAGA...................AAAUGGAGGAA.UAAUUA.CAUGGCACGUAUUGCUGGUGUAG................................................................
env-234   ...............UGGCGG..GCUGUACAUAA.........CAUUCUCAAAGA...................AAAUGGAGGAA.UAAUUA.CAUGGCACGUAUUGCUGG....................................................................
env-235   ...............UGGCGG..GCUGUACAUAA.........CAUUCUCAAAGA...................AAAUGGAGGAA.UAAUUA.CAUGGCACGUAUUGCUGGUGUAGACUUACCAAGAGAAAAACGUAUCGAAAUUGGUCUGACUUACAUCUACGGUAUCGGUAGAGUAAG
env-236   ...............UGGCGG..GCUGUACAUAA.........CAUUCUCAAAGA...................AAAUGGAGGAA.UAAUUA.CAUGGCACGUAUUGCUGGUGUAGACUUACCAAGAGAAAAACGUAUCGAAAUUGGUCUGACUUACAUCUACGGUAUCGGUAGAGUAAG
env-237   ...............UGGCGG..GCUGUACAUAA.........CAUUCUCAAAGA...................AAAUGGAGGAA.UAAUUA.CAUGGCACGUAUUGCUGGUGUAGACUUACCAAGAGAAAAACGUAUCGAAAUUGGUCUGACUUACAUCUACGGUAUCGGUAGAGUAAG
env-238   ...............UGGCGG..GCUGUACAUAA.........CAUUCUCAAAGA...................AAAUGGAGGAA.UAAUUA.CAUGGCACGUAUUGCUGGUG..................................................................
env-239   ...............UGGCGG..GCUGUACAUAA.........CAUUCUCAAAGA...................AAAUGGAGGAA.UAAUUA.CAUGGCACGUAUUGCUGGUGUAGACUUACCAAGAGAAAAACGUAUCGAAAUUGGUCUGACUUACAUCUACGGUAUCGGUAGAGUAAG
env-240   ...............UGGCGG..GCUGUACAUAA.........CAUUCUCAAAGA...................AAAUGGAGGAA.UAAUUA.CAUGGCACGUAUUGCUGGUGUAGACUUACCAAGAGAAAAACGUAUCGAAAUUGGUCUGACUUACAUCUACGGUAUCGGUAGAGUAAG
env-241   .......................GCUGUA................UCAAAAAGA....................AAACGGAGGAA.A.AACA.CAUGGCUCGUAUUGCUGGUGUAGACUUACCCAGAGAAAAACGCAUCGAAAUCGGUCUGACCUACAUUUUUGGUAUCGGCAGAACCAC
env-242   ...............UAUCGG..GCUGCAGUUAUUUA......UUUACUAAAAUUCA.................AGAUGGAGGAA.A.AUUA.CAUGGCUCGUAUUGCUGGUGUAGACUUACCAA.......................................................
env-243   ...............UAUCGG..GCUGCAGUUAUUUA......UUUACUAAAAUUCA.................AGAUGGAGGAA.A.AUUA.CAUGGCUCGUAUUGCUGGUGUAGACUUACCAAGAGACAAACGUGUAGAAAUCGGUUUGACUUACAUCUACGGAAUCGGUAGAGUAAG
env-244   ...............UAUCGG..GCUGCAGUUAUUUA......UUUACUAAAAUUCA.................AGAUGGAGGAA.A.AUUA.CAUGGCUCGUAUUGCUGGUGUAGACUUACCAAGAGA............................................
env-245   ...............UAUCGG..GCUGCAGUUAUUUA......UUUACUAAAAUUCA.................AGAUGGAGGAA.A.AUUA.CAUGGCUCGUA.........................................................................
env-246   ...............UAUCGG..GCUGCAGUUAUUUA......UUUACUAAAAUUCA.................AGAUGGAGGAA.A.AUUA.CAUGGCUCGUAUUGCUGGUGUAGACUUACCAAGAGACAAACGUGUAGAAAUCGGUUUGACUUACAUCUACGGAAUCGGUAGAGUAAG
env-252   ...............UAUCGG..GCUGCAGUUAUUUA......UUUACUAAAAUUCA.................AGAUGGAGGAA.A.AUUA.CAUGGCUCGUAUUGCUGGUGUAGACUUACCAAGAGACAAACGUGUAGAAAUCGGUUUGACUUACAUCUACGGAAUCGGUAGAGUAAG
env-254   ................CAGGG..GCUAUGCAUACUUCUUG...CUGACCAAGAUUCA.................AAAUGGAGGAA.A.AUCA.CAUGGCUCGUAUUGCUGGUGUAGACUUACCAAGAGACAAACGUGUAGAAAUCGGAUUAACUUACAUCUACGGAAUUGGUAGAGUAAG
Lba-6-1   ................CAGGG..GCUAUGCAUACUUCUUG...CUGACCAAGAUUCA.................AAAUGGAGGAA.A.AUCA.CAUGGCUCGUAUUGCUGGUGUAGACUUACCAAGAGACAAACGUGUAGAAAUCGGAUUAACUUACAUCUACGGAAUUGGUAGAGUAAG
env-255   ................CAGGG..GCUAUGCAUACUUCUUG...CUGACCAAGAUUCA.................AAAUGGAGGAA.A.AUCA.CAUGGCUCGUAUUGCUGGUGUAGACUUACCAAGAGACAAACGUGUAGAAAUCGGAUUAACUUACAUCUACGGAAUCGGUAGAGUAAG
Lba-10-1  ................CAGGG..GCUAUGCAUACUUCUUG...CUGACCAAGAUUCA.................AAAUGGAGGAA.A.AUCA.CAUGGCUCGUAUUGCUGGUGUAGACUUACCAAGAGACAAACGUGUAGAAAUCGGAUUAACUUACAUCUACGGAAUCGGUAGAGUAAG
env-256   ................CAGGG..GCUAUGCAUACUUCUUG...CUGACCAAGAUUCA.................AAAUGGAGGAA.A.AUCA.CAUGGCUCGUAUUGCUGGUGUAGACUUACCAAGAGACAAACGUGUAGAAAUCGGAUUAACUUACAUCUACGGAAUUGGUAGAGUAAG
env-258   ................CAGGG..GCUAUGCAUACUUCUUG...CUGACCAAGAUUCA.................AAAUGGAGGAA.A.AUCA.CAUGGCUCGUAUUGCUGGUGUAGACUUACCAAGAGACAAACGUGUAGAAAUCGGAUUAACUUACAU......................
env-259   ................CAGGG..GCUAUGCAUACUUCUUG...CUGACCAAGAUUCA.................AAAUGGAGGAA.A.AUCA.CAUGGCUCGUAUUGCUGGUGUAGACUUACCAAGAGACAAACGUGUAGAAAUCGGAUUAACUUACAUCUACGGAAUCGGUAGAGUAAG
env-260   ...........................................CGUCCAAACCAG...................AAACGGAGGAA.AAAU...AAUGGCUCGUAUAGCUGGUGUAGACUUACCGAGAG.....................................................
env-261   ...............GGCAGA......................UUAUCCGGAAGA...................AGAUGGAGGAA.UAAAUA.CAUGGCUCGUAUAGCUGGUGUAGACUUACCAAGAGAAAAACGUG........................................
env-262   ..........................................................................AAAUGGAGGAA.A.AACA.CAUGGCUCGUAU........................................................................
env-263   ..........................................................................AAAUGGAGGAA.CAAUUA.CAUGGCUCGUAUUGCUGGUGUAGACUUACCAAGAGAAAAACGUGCAGAAAUCGGUUUGACCUAUAUCUACGGAAUCGGUAGAACAAG
Bfo-1-1   ...............GGACGG..GCUGCGUGAUGACGGGAG..CUUAGAGA........................AAAUGGAGGAA.AAAUUA.CAUGGCUCGUAUUGCUGGUGUAGACUUACCAAGAGAGAAAAGAAUCGAAAUCGGUCUGACUUACAUUUACGGGAUUGGCAGAACAAG

          >..............>>>>>...>>>>>>..............>>>>>..........................................................................................................................
          1..............11111.....1111..............22222..........................................................................................................................
          ---------------○○○○----------------------○○○○○○○●○○○○------------------AAAUGGAGGAA-○○●●YA-CAUGGCUCGUAU●GC●GGUGUAGAYUUACCAAGAGA●AAACG●RU●GARAUYGG●YU●ACYUAYAUYUAYGG●AUYGG●AGARYRAR
```

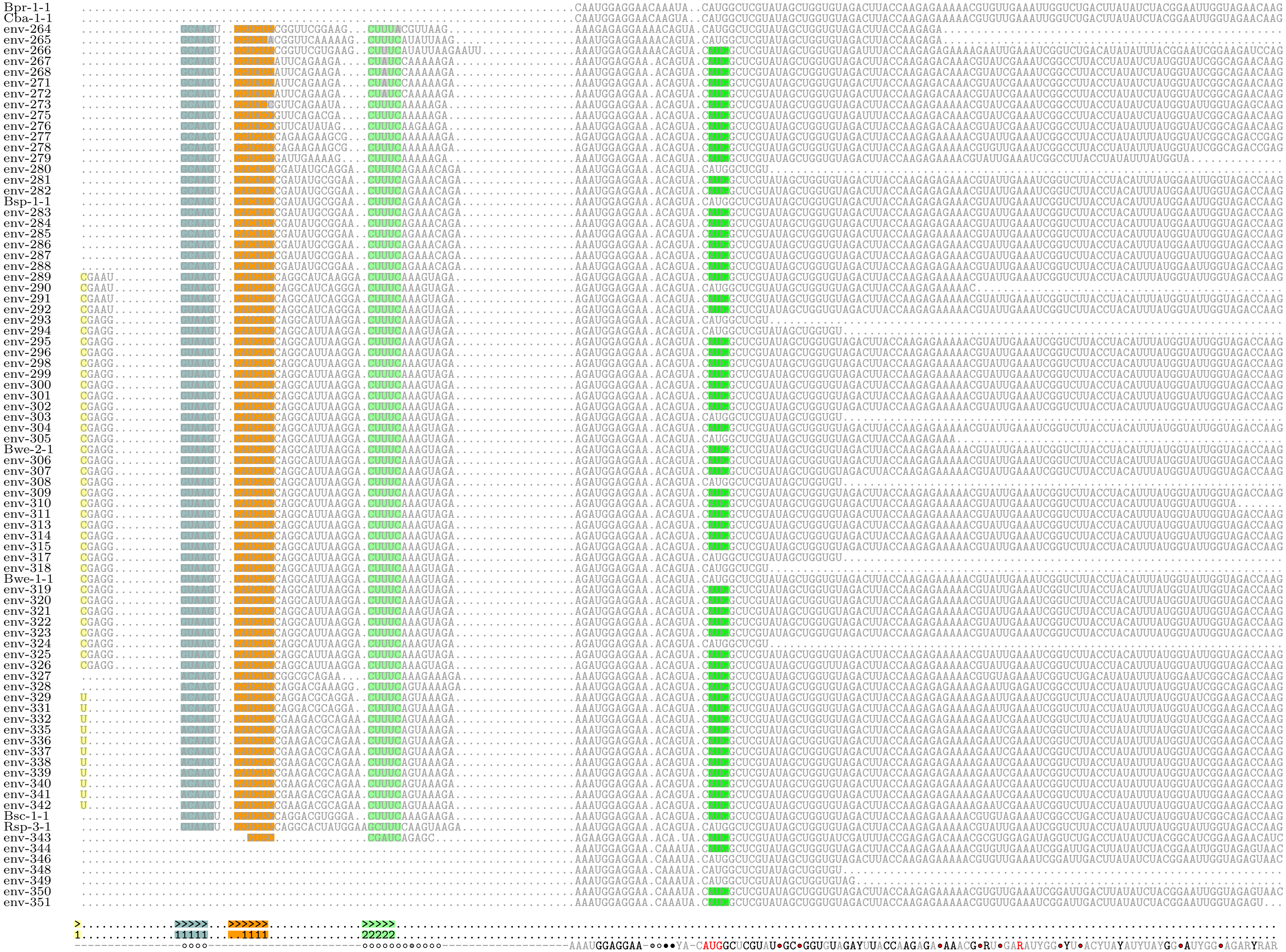
Bpr-1-1
Cba-1-1
env-264
env-265
env-266
env-267
env-268
env-271
env-272
env-273
env-275
env-276
env-277
env-278
env-279
env-280
env-281
env-282
Bsp-1-1
env-283
env-284
env-285
env-286
env-287
env-288
env-289
env-290
env-291
env-292
env-293
env-294
env-295
env-296
env-298
env-299
env-300
env-301
env-302
env-303
env-304
env-305
Bwe-2-1
env-306
env-307
env-308
env-309
env-310
env-311
env-313
env-314
env-315
env-317
env-318
Bwe-1-1
env-319
env-320
env-321
env-322
env-323
env-324
env-325
env-326
env-327
env-328
env-329
env-331
env-332
env-335
env-336
env-337
env-338
env-339
env-340
env-341
env-342
Bsc-1-1
Rsp-3-1
env-343
env-344
env-346
env-348
env-349
env-350
env-351

```
env-352   ...........................................................AAAUGGAGGAA.CAAAUA.CAUGGCUCGUAUAGCUGGUGUAGACUUACCAAGAGAAAACGUGUUGAAAUCGGMUUGACWUAUAUCUACGGAAUUGGUAGAGYAAGC
env-353   ...............GCAGGA..GCUGCGCAGAUUUUAU...GGAUCGGAAACAGA..................AAAUGGAGGAA.U.AAUA.CAUGGCUCGUAUAGCUGGUGUAGACUUACCAAGAGACAAACGUGUGGAAAUCGGAUUAACUUAUAUUUAUGGAAUCGGCAGAACAAG
Rga-1-1   ...............GCAGGA..GCUGCGUGUAUUUUAU...GGAUCGGAAACAGA..................AAAUGGAGGAA.U.AAUA.CAUGGCUCGUAUAGCUGGUGUAGACUUACCAAGAGACAAACGUGUGGAAAUCGGAUUAACUUAUAUCUAUGGAAUCGGCAGAACAAG
env-354   ..............................................UGA.......................AAGUGGAGGAA.AAAAUA.CAUGGCUCGUAUAGCUGGUGUAGAUUUACCGAGAGACAAAAGAGUAGAAAUAGGACUGACCUAUAUCUACGGC..............
env-355   UAAGUGAUUACAGUAGCAGGG..GCUGUGCAUUUUAGU....GAAUCAGCAGA.....................AAAUGGAGGAA...AUUA.CAUGGCUCGUAUAGCUGGUGUAGACUUACCAAGAGACAAACGUGUAGAGAUUGGAUUGACUUAUAUAUACGGAAUUGGCCUAGCAAG
env-356   UAAGUGAUUACAGUAGCAGGG..GCUGUGCAUUUUAGU....GAAUCAGCAGA.....................AAAUGGAGGAA...AUUA.CAUGGCUCGUAUAGCUGGUGUAGACUUACCAAGAGACAAACGUGUAGAGAUUGGAUUGACUUAUAUAUACGGAAUUGGCCUAGCAAG
env-357   ................UGGGGGCUGUGUAUGGUUAU......GAAUCAGCAG......................AAAUGGAGGAA...AUUA.CAUGGCUCGUAUAGCUGGUGUAGAUUUACCAAGAGACAAACGUGUAGAAAUCGGAUUAACUUACAUUUACGGAAUUGGCCUAUCAAG
env-358   ................UGGGGGCUGUGUAUGGUUAU......GAAUCAGCAG......................AAAUGGAGGAA...AUUA.CAUGGCUCGUAUAGCUGGUGUAGAUUUACCAAGAGACAAACGUGUAGAAAUCGGAUUAACUUACAUUUACGGAGUUGGCCUAUCAAG
env-359   ................UGGGGGCUGUGUAUGGUUAU......GAAUCAGCAG......................AAAUGGAGGAA...AUUA.CAUGGCUCGUAUAGCUGGUGUAGAUUUACCAAGAGACAAACGUGUAGAAAUCGGAUUAACUUACAUUUACGGAAUUGGCCUAUCAAG
env-360   ...............GCAGGG..GCUGUGCAUUUAUUAU...UAAUCAACAAA.....................AGAUGGAGGAA...AUUA.CAUGGCUCGUAUAGCUGGUGUAGACUUACCAAGAGACAAACGUGUAGAGUAUGGAUUAACUUACAUAUACGGAAUAGGCAUUUCAAG
env-361   UCAAA..........GCAGGG..GCUGUACAGAUAUUAU...UAAUCGGCUUG.....................AAACGGAGGAA...AUUA.CAUGGCUCGUAUAGCUGGUGUAGACUUACCAAGAGACAAACGUGUAGAAUAUGGAUUAACUUACAUUUACGGAAUAGGCAUUUCAAG
env-362   UCAAA..........GCAGGG..GCUGUACAGAUAUUAU...UAAUCGGCUUG.....................AAACGGAGGAA...AUUA.CAUGGCUCGUAUAGCUGGUGUAGACUUACCAAGAGACAAACGUGUAGAAUAUGGAUUAACUUACAUUUACGGAAUAGGCAUUUCAAG
env-363   ...............GCAGAA..GCUGUGCAUUUAUAU....GAAUCAGCAG......................AAACGGAGGAA...AUUA.CAUGGCUCGUAUAGCUGGUGUAGACUUACCAAG.........................................................
env-364   ...................GC..UGUGCAUUUAUAU......GAAUCAGCAG......................AAAUGGAGGAA...AUUA.CAUGGCUCGUAUAGCUGGUGUAGACUUACCAAGAGACAAACGCGUAGAGUAUGGAUUAACUUACAUUUACGGAAUUGGCAUUUCAAG
env-365   ...............GGGAGC..UGUGCAUUUAUAU......GAAUCAGCAG......................AAAUGGAGGAA...AUUA.CAUGGCUCGUAUAGCUGGUGUAGACUUACCAAGAGACAAACGCGUAGAGUAUGGAUUAACUUAUAUUUACGGAAUUGGCAUUUCAAG
env-366   ...............GGGAGC..UGUGCAUUUAUAU......GAAUCAGCAG......................AAAUGGAGGAA...AUUA.CAUGGCUCGUAUAGCUGGUGUAGACUUACCAAGAGACAAACGCGUAGAGUAUGGAUUAACUUACAUUUACGGAAUUGGCAUUUCAAG
env-367   ...........................................................AAAUGGAGGAC.AAAGUA.CAUGGCUCGUAUAGCUGGUGUAGACUUACCAAGAGACAAACGUGUUGAGAUUGGUCUGACCUAUAUCUACGGAAUCGGUAGAACAAG
env-368   ...........................................................AAAUGGAGGAC.AAAGUA.CAUGGCUCGUAUAGCUGGUGUAGACUUACCAAGAGACAAACGUGUUGAGAUUGGUCGGAGCUGUAUCUACGGAAUCGGUAGAACAAG
env-369   ...........................................................AAAUGGAGGAC.AAAGUA.CAUGGCUCGUAUAGCUGGGGUAGACUUACCAAGAGACAAACGUGUUGAGAUUGGUCUGACCUAUAUCUACGGAAUCGGUAGAACAAG
env-370   ...........................................................AAAUGGAGGAC.AAAGUA.CAUGGCUCGUAUAGCUGGUGUAGACUUACCAAGAGACAAACGUUUUGAGAUUGGUCUGACCUAUAUCUACGGAAUCGGUAGAACAAG
env-371   ...........................................................AAAUGGAGGAC.AAAGUA.CAUGGCUCGUAUAGCUGGUGUAGACUUACCAAGAGACAAACGUGUUGAGAUUGGUCUGACCUAUAUCUACGGAAUCGGUAGAACAAG
env-372   ...........................................................AAAUGGAGGAC.AAAGUA.CAUGGCUCGUAUAGCUGGU.........................................................................
env-373   ...........................................................AAAUGGAGGAC.AAAGUA.CAUGGCUCGUAUAGCUGGUGUAGACUUACCAAGAGACAAACGUGUUGAGAUUGGUCUGACCUAUAUCUACGGAAUCGGUAGAACAAG
env-374   ...........................................................AAAUGGAGGAC.AAAGUA.CAUGGCUCGUAUAGCUGGUGUAGACUUACCAAGAGACAAACGUGUUGAGAUUGGUCUGACCUAUAUCUACGGAAUCGGUAGAACAAG
env-375   ...........................................................AAAUGGAGGAC.AAAGUA.CAUGGCUCGUAUAGCUGGUGUAGACUUACCAAGAGACAAACGUGUUGAGAUUGGUCUGACCUAUAUCUACGGAAUCGGUAGAACAAG
env-377   ...........................................................AAAUGGAGGAC.AAAGUA.CAUGGCUCGUAUAGCUGGUGU.......................................................................
env-383   ...........................................................AAAUGGAGGAC.AAAGUA.CAUGGCUCGUAUAGCUGGUGUAGACUUACCAAGAGACAAACGUGUUGAGAUUGGUCUGACCUAUAUCUACGGAAUCGGUAGAACAA.
env-384   ...........................................................AAAUGGAGGAC.AAAGUA.CAUGGCUCGUAUAGCUGGUGUAGACUUACCAAGAGACAAACGUGUUGAGAUUGGUCUGACCUAUAUCUACGGAAUCGGUAGAACAAG
env-385   ...........................................................AAAUGGAGGAC.AAAGUA.CAUGGCUCGUAUAGCUGG..........................................................................
env-386   ...........................................................AAAUGGAGGAC.AAAGUA.CAUGGCUCGUAUAGCUGGUGUAGACUUACCAAGAGACAAACGUGUUGAGAUUGGUCUGACCUAUAUCUACGGAAUCGGUAGAACAAG
env-387   ...........................................................AAAUGGAGGAC.AAAGUA.CAUGGCUCGUAUAGCUGGUGUAGACUUACCAAGAGACAAACGUGUUGAGAUUGGUCUGACCUAUAUCUACGGAAUCGGUAGAACAAG
env-388   ...........................................................AAAUGGAGGAC.AAAGUA.CAUGGCUCGUAUAGCUGGUGUAGACUUACCAAGAGACAAACGUGUUGAGAUUGGUCUGACCUAUAUCUACGGAAUCGGUAGAACAAG
env-389   ...........................................................AAAUGGAGGAC.AAAGUA.CAUGGCUCGUAUAGCUGGUG........................................................................
env-390   ...........................................................AAAUGGAGGAC.AAAGUA.CAUGGCUCGUAUAGCUGGUGUAGACUUACCAAGAGACAAACGUGUUGAGAUUGGUCUGACCUAUAUCUACGGAAUCGGUAGAACAAG
env-391   ...........................................................AAAUGGAGGAC.AAAGUA.CAUGGCUCGUAUAGCUGGUGUAGACUUACCAAGAGACAAACGUGUUGAGAUUGGUCUGACCUACAUCUACGGAAUCGGUAGAACAAG
env-392   ...........................................................AAAUGGAGGAC.AAAGUA.CAUGGCUCGUAUAGCUGG..........................................................................
env-393   ...........................................................AAAUGGAGGAC.AAAGUA.CAUGGCUCGUAUAGCUGGUGUAGACUUACCAAGAGACAAACGUGUUGAGAUUGGUCUGACCUAUAUCUACGGAAUCGGUAGAACAAG
env-394   ...........................................................AAAUGGAGGAC.AAAGUA.CAUGGCUCGUAUAGCUGGUGUAGACUUACCAAGAGACAAACGUGUUGAGAUUGGUCUGACCUAUAUCUACGGAAUCGGUAGAACAAG
env-395   ...........................................................AAAUGGAGGAC.AAAGUA.CAUGGCCCGUAUAGCUG...........................................................................
env-396   ...........................................................AAAUGGAGGAC.AAAGUA.CAUGGCUCGUAUAGCUGGUGUAGACUUACCAAGAGACAAACGUGUUGAGAUUGGUCUGACCUAUAUCUACGGGAUCGGUAGAACAAG
env-397   ...........................................................AAAUGGAGGAC.AAAGUA.CAUGGCUCGUAUAGCUGGUGUAGACUUACCAAGAGACAAACGUGUUGAGAUUGGUCCGACCUAUAUCUACGGAAUCGGUAGAACAAG
env-398   ...........................................................AAAUGGAGGAC.AAAGUA.CAUGGCUCGUAUAGCUGGUGUAGACUUACCAAGAGACAAACGUGUUGAGAUUGGUCUGACCUAUAUCUACGGAAUCGGUAGAACAAG
env-399   ...........................................................AAAUGGAGGAC.AAAGUA.CAUGGCUCGUAUAGCUGGUGUAGACUUACCAAGAGACAAACGUGUUGAGAUUGGUCUGACCUAUAUCUACGGAAUCGGUAGAACAAG
env-405   ...........................................................AAAUGGAGGAC.AAAGUA.CAUGGCUCGUAUAGCUGGUGUAGACUUACCAAGAGACAAACGUGUUGAGAUUGGUCUGACCUAUAUCUACGGAAUCGGUAGAACAAG
env-408   ...........................................................AAAUGGAGGAC.AAAGUA.CAUGGCUCGUAUAGCUGGGGUAGACUUACCAAGAGACAAACGUGUUGAGAUUGGUCUGACCUAUAUCUACGGAAUCGGUAGAACAAG
env-409   ...........................................................AAAUGGAGGAC.AAAGUA.CAUGGCUCGUAUAGCUGGUGUAGACUUACCAAGAGACAAAC.....................................................
env-410   ...........................................................AAAUGGAGGAC.AAAGUA.CAUGGCUCGUAUAGCUGGUGUAGACUUACCAA..............................................................
env-411   ...........................................................AAAUGGAGGAC.AAAGUA.CAUGGCUCGUAUAGCUGGUGUAGACUUACCAAGAGACAAACGUGUUGAGAUUGGUCUGACCUAUAUCUACGGAAUCGGUAGAACAAG
env-412   ...........................................................AAAUGGAGGAC.AAAGUA.CAUGGCUCGUAUAGCUGGUGUAGACUUACCAAGAGACAAACGUGUUGAGAUUGGUCUGACCUAUAUCUACGGAAUCGGUAGAACAAG
env-413   ...........................................................AAAUGGAGGAC.AAAGUA.CAUGGCUCGUAUAGCUGGUGUAGACUUACCAAGAGACAAACGUGUUGAGAUUGGUCUGACCUAUAUCUACGGAAUCGGUAGAACAAG
env-414   ...........................................................AAAUGGAGGAC.AAAGUA.CAUGGCUCGUAUAGCUGGUGUAGACUUACCAAGAGACAAACGUGUUGAGAUUGGUCUGACCUAUAUCUACGGAAUCGGUAGAACAAG
env-418   ...........................................................AAAUGGAGGAC.AAAGUA.CAUGGCUCGUAUAGCUGGUGUAGACUUACCAAGAGACAAACGUGUUGAGAUUGGUCUGACCU...........................
env-419   ...........................................................AAAUGGAGGAC.AAAGUA.CAUGGCUCGUAUAGCUGGUGUNNNNUAGAACAAGUGCAGAUCGUAUCUUAAAAGAAGCUGGAGUAAAUCCGGACACACGCUGCAGAG
env-420   ...........................................................AAAUGGAGGAC.AAAGUA.CAUGGCUCGUAUAGCUGGU.........................................................................
env-428   ...............AUAUGG..GCUGUCUGGA.........CUGGUCAAGUAGA...................AGAUGGAGGAA.A.AUUA.CAUGGCACGUAUUGCUGGUGU.......................................................................
env-429   ...............AUAUGG..GCUGCCCCGA.........CUGGUCAAGUAGA...................AGAUGGAGGAA.A.AUUA.CAUGGCACGUAUUGCUGGUG........................................................................
env-430   ...............AUAUGG..GCUGCCCCGA.........CUGGUCAAGUAGA...................AGAUGGAGGAA.A.AUUA.CAUGGCACGUAUUGCUGGUGUAGACUUACCAAGAGAAA..........................................................
env-432   ...............AUAUGG..GCUGCCCCGA.........CUGGUCAAGUAGA...................AGAUGGAGGAA.A.AUUA.CAUGGCACGUAUUGCUGGUGUAGACUUACCAAGAGAAAAACGUAUUGAAAUCGGAUUAACUUAUAUCUACGGAAUCGGUAGAACAAG
env-433   ..........................................CUGUCAGAAGUA....................AAACGGAGGAA.A.AAUA.CAUGGCUCGUAUAGCUGGUGUAGACUUACCAAGAGAAAAACGUGUCGAAAUCGGUCUGACCUACAUUUACGGUAUCGGCAGAACAAC
env-434   ...............AUACGU..GACGUACGACACUGA....CUUUCCGUAGAA....................AAAUGGAGGAA.ACAUUA.CAUGGCUCGUAUAGCUGGUGUAG.....................................................................
Bhy-1-1   ...............AUACGU..GACGUACGACACUGA....CUUUCCGUAGAA....................AAAUGGAGGAA.ACAUUA.CAUGGCUCGUAUAGCUGGUGUAGACUUACCAAGAGACAAACGUGUAGAAAUCGGUCUGACCUAUAUUUAUGGUAUUGGAAGACCAAG
env-435   ..........................CG..............GAUCGCAAC.......................AGAAGGAGGAA.A.UGCA.CAUGGCUCGUAUAGCUGGUGUAGAUUUACCAAGAGACAAACGAGUAGAGAUAGGUCUGACCUAUAUCUAUGGCAUUGGAAGAACUUC
env-436   .......................UGCUG..............CUGGUCAGGA......................AAAAGGAGGAA.A.AGCA.CAUGGCUCGUAUAGCUGGUGUAGACUUACCAAGAGACAAGAGGGUAGAAAUAGGGCUUACCUAUAUCUACGGCAUUGGAAGAGCAUC
env-437   ...........................................................AGAAGGAGGAA.AAGGUA.CAUGGCUCGUAUAGCUGGUGUAGACUUACCAAGAGACAAACGCGUAGAAAUAGGCCUGACCUAUAUCUACGGCAUCGGAAGAACCUC
env-438   ...........................................................AAAAGGAGGAA.A.GAUA.CAUGGCUCGUAUAGCUGGUGUAGACUUACCAAGAGACAAACGCGUGGAGAUAGGUCUUACCUAUAUCUACGGAAUCGGAAGAACGUC
Bsp-7-1   ..........................................UUCCGCAAACAAAUCUUUUUUUAUU......AAAUGGAGGAA.U.UUUAAAAUGGCUCGUAUUGCAGGUGUAGACUUACCUAGAGAUAAGCGAGUUGAGAUUGGUCUUACAUACAUCUAUGGUAUUGGUAGAACAAG
Bsp-9-1   ..........................................UUCCGCGAACAAAUUUUUUUUAUU.......UAAUGGAGGAA.A.AUUAAAAUGGCUCGUAUUGCAGGUGUUGAUUUACCUAGAGAUAAGCGAGUUGAGAUUGGUCUUACAUACAUUUAUGGAAUUGGAAGAACAAG
Bsp-5-1   ..........................................UUCCGCAAAAACAAAUCUUUUUUAUU.....UAAUGGAGGAA.A.AUUAAAAUGGCUCGUAUUGCAGGUGUUGAUUUACCUAGAGAUAAGCGAGUUGAGAUUGGUCUUACAUACAUCUAUGGAAUUGGAAGAACAAG
Bsp-2-1   .......................GCAAACAAUUU........UUUUC..........................AAAUGGAGGAA.UAUUAAAAAUGGCUCGUAUUGCAGGUGUAGACUUACCCAGAGAUAAGCGAGUUGAGAUUGGUCUCACAUAUAUCUAUGGUAUUGGUAGAACAAG
Bsp-3-1   ...............UUAGUUUU...GCGGAAAAUGCGCAA..AACUUCACAUUAUUUUUAAUU........AAAUGGAGGAA.UUUUUAAAAUGGCUCGUAUUGCAGGUGUAGACUUACCUAGAGAUAAGCGAGUUGAGAUUGGUCUCACAUAUAUCUAUGGUAUUGGAAGAACAAG
Bpr-2-1   ...........................................................CAAUGGAGGAACAAGUA..CAUGGCUCGUAUAGCUGGUGUAGACUUACCAAGAGAAAAACGUGUUGAAAUUGGUCUGACUUAUAUCUACGGAAUUGGUAGAACAAG
          >..............>>>>>...>>>>>>..............>>>>>.......................................................................................................................
          1..............11111.....1111..............22222.......................................................................................................................
          ---------------oooo--------------------oooooooooooo-----------------AAAUGGAGGAA-oo••YA-CAUGGCUCGUAU•GC•GGUGUAGAYUUACCAAGAGA•AAACG•RU•GARAUYGG•YU•ACYUAYAUYUAYGG•AUYGG•AGARYRAR
```

alignment positions 901 · · · 1080

```
env-1     GAAGUCAGAAAGAUCGCAGGCGUGAUCGACGAGACACAGAUGGUAGAAGGUGAUCUGAGACGUGAGAUCGCGAUGAACAUCAAGCGUC...........................................................................................
env-2     ...........................................................................................................................................................................
env-3     ...........................................................................................................................................................................
env-4     UGCUGAUAAGAUCUUGGCAAAAGCAGGUGUAAAUCCUGACAUACGUGUAAGAGACUUAACAGAUGAUGAUGUUAAGAAGAUUAGUGCAGUUAUCGAUGAGGAGUAUAUGGUAGAAGGUGACCUCAGAAGAGAAAUCGCACUGAACAUUAAGAGAUUACAGGAAAU.............

          ...........................................................................................................................................................................
          ...........................................................................................................................................................................
          Y•C•R•Y•R•AUYYU•R•RR•RGCRRR•GU•RAYCC•RAYAY•CG•••YAGRGAYYU•AC•GAYGA•GARGU•RA•ARRAUY•RYR••GY•AUYGA•GAR•Y••••RY•RURGAAGR•GAY•U•••••G•GRRR•YRY••Y•A••A•YARR•R•YY•Y•R•ARR••RR••GCURY•R•RG
```

```
env-5     UGCUGAUAAGAUCUUGGCAAAAGCAGGUGUAAAUCCUGACAUACGUGUAAGAGACUUAACAGAUGAUGAUGUUAAGAAGAUUAGUGCAGUUAUCGAUGAGGAGUAUAUGGUAGAAGGUGACCUCAGAAGAGAAAUCGCACUGAACAUUAAGAGAUUACAGGAAAUCGGAUG.........
env-6     ...........................................................................................................................................................
env-7     UGCUGAUAAGAUCUUGGCAAAAGCAGGUGUAAAUCCUGACAUACGUGUAAGAGACUUAACAGAUGAUGAUGUUAAGAAGAUUAGUGCAGUUAUCGAUGAGGAGUAUAUGGUAGAAGGUGACCUCAGAAGA.....................................
env-8     UGCUGAUAAGAUCUUGGCAAAAGCAGGUGUAAAUCCUGACAUACGUGUAAGAGACUUAACAGAUGAUGAUGUUAAGAAGAUUAGUGCAGUUAUCGAUGAGGAGUAUAUGGUAGAAGGUGACCUCAGAAGAGA...................................
env-9     UGCUGAUAAGAUCUUGGCAAAAGCAGGUGUAAAUCCUGACAUACGUGUAAGAGACUUAACAGAUGAUGAUGUUAAGAAGAUUAGUGCAGUUAUCGAUGAGGAGUAUAUGGUAGAAGGUGACCUCAGAAGAGA...................................
env-10    UGCUGAUAAGAUCUUGGCAAAAGCAGGUGUAAAUCCUGACAUACGUGUAAGAGACUUAACAGAUGAUGAUGUUAAGAAGAUUAGUGCAGUUAUCGAUGAGGAGUAUAUGGUAGAAGGUGACCUCAGAAGAGAAAUCGCACUGAACAUUAAGAGAUUACAGGAAAUCGGAUGCUAUCGUGG
env-11    UGCUGAUAAGAUCUUGGCAAAAGCAGGUGUAAAUCCUGACAUACGUGUAAGAGACUUAACAGAUGAUGAUGUUAAGAAGAUUAGUGCAGUUAUCGAUGAGGAGUAUAUGGUAGAAGGUGACCUCAGAAGAGAAAUCGCACUGAACAUUAAGAGAUUACAGGAAAUCG.............
env-12    UGCUGAUAAGAUCUUGGCAAAAGCAGGUGUAAAUCCUGACAUACGUGUAAGAGACUUAACAGAUGAUGAUGUUAAGAAGAUUAGUGCAGUUAUCGAUGAGGAGUAUAUGGUAGAAGGUGACCUCAGAAGAGAAAUCGCACUGAACAUUAAGAGAUUACAGGAAAUCGGAUGCUAUCGUGG
env-13    UGCUGAUAAGAUCUUGGCAAAAGCAGGUGUAAAUCCUGACAUACGUGUAAGAGACUUAACAGAUGAUGAUGUUAAGAAGAUUAGUGCAGUUAUCGAUGAGGAGUAUAUGGUAGAAGGUGACCUCAGAAGAGAAAUCGCACUGAACAUUAAGAGAUUACAGGAAAUCGGAUGCUAUCGUGG
env-14    ...........................................................................................................................................................
env-16    UGCUGAUAAGAUCUUGGCAAAAGCAGGUGUAAAUCCUGACAUACGUGUAAGAGACUUAACAGAUGAUGAUGUUAAGAAGAUUAGUGCAGUUAUCGAUGAGGAGUAUAUGGUAGAAGGUGACCUCAGAAGAGAAAUCGCACUGAACAUUAAGAGAUUACAGGAAAUCGGAUGCU.......
env-17    UUCUAACCGUAUUUUAGCAGAAGCAAAAGUAAAUCCCGAUACACGUUGCAGGGAUUUAACAGAUGAAGAAGUAAAGAGAAUCAGUGAAGUAAUCGAUGCAACCCAGACUGUUGAAGGUGAUUUAAGAAGAGAAAUCGCUUUAAACAUUAAGAGAUUGCAGGAAAUCGGAUGCUAUAGAGG
env-18    UUCUAACCGUAUUUUAGCAGAAGCAAAAGUAAAUCCCGAUACACGUUGCAGGGAUUUAACAGAUGAAGAAGUAAAGAGAAUCAGUGAAGUAAUCGAUGCAACCCAGACUGUUGAAGGUGAUUUAAGAAGAGA...................................
env-19    UGCAACUAAAAUUUUAGAGGCAGCUAACGUGAAUGUAGAUACCAGAGUUAGAGACUUAACAGAUGAAGAAGUAUCAAGAAUCCGUGAUGUCAUGGAUGCAGAUUAUAAUGUAGAGGGUGAUUUAAGAAGAGAAAUCGCACUUAACAUCAAACGUCUUCAAGAAAUUGGGUGUUAUAGAGG
Bsp-6-1   CUCAAACAAGAUUCUUGAGGCUGCAAAGGUUAAUCCGGACACUCGUGUUCGUGAUCUCACAGAUGAUGAAGUAAAGAAAAUUGCUGAGAUAAUCGGUAAUGAUUAUGAAGUAGAAGGUGAUCUUAGAAGAGAAAUCGCUAUGAACAUCAAGCGUCUUACUGAGAUUGGUUGCUACAGAGG
env-20    GUCCAACAGGAUUCUCAAAGAAGCAAAAGUUGAUCCCAACACCCGCGUCCGCGAUCUGACGGAUGAGGAAGCGAACAGGAUCAAGGACGUCAUGGAUGCGUCCGUUACCGUGGAAGGCGAUCUCCGCCGUGAAGUUGCCAUGAAUAUCAAACGACUUCAGGAAAUCGGCUGCUACAGAGG
env-21    CUCAAACAAGAUUCUUGAGGCUGCAAAAGUUAAUCCAGAUACACGUGUUCGUGAUCUGACAGAUGAUGAAGUUAAGAAGAUUGCUGAGAUAAUCGGUGAUGAGUAUUUAGUUGAAGGUGACCUCAGAAGAGAAGUUGCCAUGAACAUCAAGAGACUUACAGAAAUCGGAUGCUUCAGAGG
env-22    ...........................................................................................................................................................
env-23    UUCCGGCAAGAUUUUGGAGGCAGCAGGAGUUAAUCCUGAUACCCGUUGCAGAGACUUAACGGACGAUGAAGUUAAGAAGAUCGGUGAAGUGAUUGCCGAUACUUACGUAGUGGAAGGUGAUUUAAAGAGAGAAGUUGCUCUUAACAUUAAGAGACUUCAGGAAAUCGGCUGCUAUCGUGG
env-24    ...........................................................................................................................................................
env-25    UUCAGGAAAGAUUCUUGAGCAGGCAAAAGUAAGCCCAGAUACACGUAUCAGAGAUUUGACAGACGAAGAAGUAAGCAGAAUCGUGAUGUUAUUGAAGAAGUACAGACUGUAGAAGGUGAUUUAAGAAGAGAAAUCGCUCUUAAUAUCAAAAGACUACAAGAGAUUGGAUGCUAUAGAGG
env-28    UUCAGGAAAGAUUCUUGAGCAGGCAAAAGUAAGCCCAGAUACACGUAUCAGAGAUUUGACAGACGAAGAAGUAAGCAGAAUCGUGAUGUUAUUGAAGAAGUACAGACUGUAGAAGGUGAUUUAAGAAGAGAAAUCGCUCUUAAUAUCAAAAGACUACAAGAGAUUGGAUGCUAUAGAGG
env-29    ...........................................................................................................................................................
env-30    UUCAAAUAAAAUUCUUGAGCAAGCAAAAGUAAGCCCAGAUACACGUGUUAGAGAUUUAACAGACGAAGAAGUAAGUAGAAUCCGUGAUGUUAUUGAUGAAGUACAAACUGUAGAAGGUGAUUUAAGAAGAGAAAUAGCUCUUAAUAUCAAAAGACUACAAGAAAUUGGAUGCUAUAGAGG
env-32    UUCAAAUAAAAUUCUUGAGCAAGCAAAAGUAAGCCCAGAUACACGUGUUAGAGAUUUAACAGACGAAGAAGUAAGUAGAAUCCGUGAUGUUAUUGAUGAAGUACAAACUGUAGAAGGUGAUUUAAGAAGAGAAAUAGCUCUUAAUAUCAAAAGACUACAAGAAAUUGGAUGCUAUAGAGG
env-33    UUCAAAUAAAAUUCUUGAGCAAGCAAAAGUAAGCCCAGAUACACGUGUUAGAGAUUUAACAGACGAAGAAGUAAGUAGAAUCCGUGAUGUUAUUGAUGAAGUACAAACUGUAGAAGGUGAUUUAAGAAGAGAAAUAGCUCUUAAUAUCAAAAGACUACAAGAAAUUGGAUGCUAUAGAGG
env-34    ...........................................................................................................................................................
env-35    CGCCAACAAAAUUUUAGCAGCAGCGAAUGUUAAUCCGGACACUCGUGUCAGAGACCUGACUGACGAAGAGGUUAAGAGACUGAGCGCUGUUAUCGAUGAACAGUACAUGGUAGAGGGCGACUUAAGAAGAGAUGUUGCCAUGAACAUCAAGAGACUUCAGGAGA...............
env-36    ...........................................................................................................................................................
env-37    ...........................................................................................................................................................
env-38    ...........................................................................................................................................................
env-39    ...........................................................................................................................................................
env-41    ...........................................................................................................................................................
env-42    CUCAACAAAGAUUCUCAGCAAGGCUGGAAUCGAUCCGAGUACCAGAGUGAGAGAUCUCACGGACGAGGAAGUCGAUAAGAUCA.......................................................................
env-43    UGCAGGCCGUAUCUUAGCAGCGGCAGGAGUAAACCCUGAUACCCGCGUAAGAGAUCUGACAAGUGAAGAAGUAAAGAAAAUCAGCGAAGUGAUUGAUGAGACACAGGUAGUAGAAGGUGAUCUCAGAAGAGAGAUCCAGCUGAAUAUUAAGAGACUGAAAGAAAUCGGCUGUUAUCGUGG
env-45    UGUCCGAAAGCGGAGACUAUACAUUGAUUUUAACUAAUGAUUCCGAUAAGCCAAGGUAUUAUGAUGUCUGCGUCAGGAAUAUAGAAAAAUGGGAUGAAGAGCUUCACUGCAUUGAAACCAGAGGACCGGAGGCAGGCGGCAGGUUUGAUUCGUCAUGGUUUAGGGUUAUAGAUAAAAUAA
env-46    AUUUCUACAGUAAUCGACGAGACGCAGACUGUAGAGGGUGAUCUGAGACGUGAGAUCGCUAUGAACAUCAAACGUCUUCAGGAGAUCGGAUGCUACAGAGGUAUCCGUCAUAGAAAAGGACUUCCUGUGCGUGGUCAGAAGACCAAAACAAACGCAAGAACUCGUAAGGGUCCGAGAAGA
env-47    GGGACAGUAAUGGAGGAAAACAGUAC[illegible]GCUCGUAUUGCAGGUAUUGACUUACCGAGAAAAACGCAUAGAGAUCGGUCUUACUUACAUUUAUGGAAUCGGCCGUUCCAGCUCCAACAAGAUCCUUGCCCAGGCAGGCGUAAAUCCUGAUAUCCGCGUCCGCGACCUUUCGGAUGAUG
env-48    CGCGGACAGAAUUCUUGCGGCAGCGAAUGUAAAUCCUGAUACCCGUGUCAGAGAUCUGACAGACGAAGAAGUUGACAAGAUCAGGGUCGCCAUUGAAGAACUCGGCAUCAUGGUAGAAGGUGAUCUUCGCCGUGACACCGCUAUGAACAUCAAGAGACUGCAGGAGAUUGGCUGCUACAG
env-49    UUCGCGCAGGAUCUGAAAGAUGCAGGGGUAAAUCCGGAUACACGUGUCAGAGAUCUUACAAGUGAAGAAGUCAAGAAUAUCAGUACUGUUAUUGAGAAUACGCAGACGGUUGAAGGUGUGUUAAGAACGGAAAUCGCCAUCAAUAUCAAACGUCUUCAGGAAAUUGGCUGCUAUCGCGG
env-50    ...........................................................................................................................................................
Dma-1-1   CGCAAACCGUAUUUUAACAGAAGCAGGUGUAAAUCCUGACACGAGAUGCAGAGACUUAACUGAUGAAGAAGUUAAAAAGAUCAGCUCUGUUAUCGAUGAAACUCAGGUUGUAGAAGGUGACCUGAGAAGAGAGAUCGCACUGAAUAUCAAGAGAUUACAGGAGAUCGGAUGCUACAGAGG
env-51    CGCAACACGCAUCUUAAAAGAAUCGGGUGUAGACCCAGACAUUCGCUGUAGAGAUUUAACAGACGACGAUGUAAAAGCAAUCAGUAAAGUAAUCGAAGAAUCACAAACAGUAGAAGGUGAACUUAGAAGAGAAAUCGCACUAAACAUCAAGAGAUUACAAGAAAUCGGAUGUUAUAGAGG
Lba-2-1   UGCAAAUAACAUUUUAGAGAAAGCUGGAGUAAAUCCGGAUACUCGUGUUAGAGAUUUAACAGACGAAGAAGUAAAUGAAUCAGUGCUGUAAUCGAUGAAAACUAUCAGGUAGAAGGUGAUUUACGUAGAGAAAUCGCUUUAAACAUUAAGAGAUUACAGGAAAUCGGUUGCUAUCGUGG
[ce-1-1   UUCCACAAAGCUUCUUGCACAGGCUGGAGUAAAUCCGGAUACACGUGUUAGAGACCUCACAGAUGACGAAGUAAAGAGCCUCACAAACGCAAUCGAGGAGCUUCAGGUUACUGUAGAGGGUGAUCUCAGACGUGAGAUUGCUAUGAACAUCAAACGUCUUCAGGAGAUCGGCUGCUACCG
Ece-1-1   UUCCACAAAGCUUCUUGCACAGGCUGGAGUAAAUCCGGAUACACGUGUUAGAGACCUCACAGAUGACGAAGUAAAGAGCCUCACAAACGCAAUCGAGGAGCUUCAGGUUACUGUAGAGGGUGAUCUCAGACGUGAGAUUGCUAUGAACAUCAAACGUCUUCAGGAGAUCGGCUGCUACCG
env-52    CGCAAACAAGAUCUUGGAAGCAGCAAAUGUAAAUCCUGAUACUCGUGUCAGAGAUCUUACAGAUGAAGAAGUUUCCAGAAUCCGUGACGUAAUGGAUGAACAUUAUACAGUAGAAGGUGACCUGAGAAGAGAAAUCGCUCUGAAUAUCAAGAGACUUCAGGAAAUCGGAUGUUACAGAGG
env-53    CUCAAACCGCAUCCUGGCAGAUGCGAAAGUAGAUCCAAACACCCGGGUUAGAGAUUUGACCCAAGAAGAUGUGAAGAAAAUCAGUGAUGUUAUUGAAGAAACGCAAAUGGUAGAAGGGGAUCUUCGCAGAGAGAUCCAAUUGAAUAUUAAGCGAUUGCAAGAGAUUGGCUGCUAUCGUGG
env-54    ...........................................................................................................................................................
env-55    CGCUAUCCGUAUCCUUGCAGAUGCAGGAGUAGAUCCGGAUACACGUGUUAGAGAUUUGACACAGGACGACGUUAAGAAAAUCAGUGAAGUGAUUGAUGAGACUCAGGUUGUAGAAGGUGAUCUCAGAAGAGAAAUCCAGUUAAACAUCAAGAGACUUAAAGAAAUCGGAUGCUACCGUGG
env-59    CGCUAUCCGUAUCCUUGCAGAUGCAGGAGUAGAUCCGGAUACACGUGUUAGAGAUUUGACACAGGACGACGUUAAGAAAAUCAGUGAAGUGAUUGAUGAGACUCAGGUUGUAGAAGGUGAUCUCAGAAGAGAAAUCCAGUUAAACAUCAAGAGACUUAAAGAAAUCGGAUGCUACCGUGG
env-60    CGCUAUCCGUAUCCUUGCAGAUGCAGGAGUAGAUCCGGAUACACGUGUUAGAGAUUUGACACAGGACGACGUUAAGAAAAUCAGUGAAGUGAUUGAUGAGACUCAGGUUGUAGAAGGUGAUCUCAGAAGAGAAAUCCAGUUAAACAUCAAGAGACUUAAAGAAAUCGGAUGCUACCGUGG
env-62    ...........................................................................................................................................................
env-63    CGCUAUCCGUAUCCUUGCAGAUGCAGGAGUAGAUCCGGAUACACGUGUUAGAGAUUUGACACAGGACGACGUUAAGAAAAUCAGUGAAGUGAUUGAUGAGACUCAGGUUGUAGAAGGUGAUCUCAGAAGAGAAAUCCAGUUAAACAUCAAGAGACUUAAAGAAAUCGGAUGCUACCGUGG
env-64    ...........................................................................................................................................................
env-66    ...........................................................................................................................................................
env-67    CGCUAUCCGUAUCCUUGCAGAUGCAGGAGUAGAUCCAGAUACACGUGUUAGAGAUUUGACACAGGACGACGUUNAGAAAAUCAGUGAAGUGAUCGAUGAGACUCAGGUUGUAGAAGGUGAUCUCAGAAGAGAAAUCCAGUUAAACAUCAAGAGACUUAAAGAAAUCGGAUGCUACCGUGG
env-68    CGCUAUCCGUAUCCUUGCAGAUGCAGGAGUAGAUCCAGAUACACGUGUUAGAGAUUUGACACAGGACGACGUUAAGAAAAUCAGUGAAGUGAUCGAUGAGACUCAGGUUGUAGAAGGUGAUCUCAGAAGAGAAAUCCAGUUAAACAUCAAGAGACUUAAAGAAAUCGGAUGCUACCGUGG
env-75    CGCUAUCCGUAUCCUUGCAGAUGCAGGAGUAGAUCCAGAUACACGUGUUAGAGAUUUGACACAGGACGACGUUAAGAAAAUCAGUGAAGUGAUCGAUGAGACUCAGGUUGUAGAAGGUGAUCUCAGAAGAGAAAUCCAGUUAAACAUCAAGAGACUUAAAGAAAUCGGAUGCUACCGUGG
env-76    CGCUAUCCGUAUCCUUGCAGAUGCAGGAGUAGAUCCAGAUACACGUGUUAGAGAUUUGACACAGGACGACGUUAAGAAAAUCAGUGAAGUGAUCGAUGAGACUCAGGUUGUAGAAGGUGAUCUCAGAAGAGAAAUCCAGUUAAACAUCAAGAGACUUAAAGAAAUCGGAUGCUACCGUGG
env-77    CGCUAUCCGUAUCCUUGCAGAUGCAGGAGUAGAUCCAGAUACACGUGUUAGAGAUUUGACACAGGACGACGUUAAGAAAAUCAGUGAAGUGAUCGAUGAGACUCAGGUUGUAGAAGGUGAUCUCAGAAGAGAAAUCCAGUUAAACAUCAAGAGACUUAAAGAAAUCGGAUGCUACCGUGG
env-78    ...........................................................................................................................................................
env-79    CGCUAUCCGUAUCCUUGCAGAUGCAGGAGUAGAUCCAGAUACACGUGUUAGAGAUUUGACACAGGACGACGUUAAGAAAAUCAGUGAAGUGAUCGAUGAGACUCAGGUUGUAGAAGGUGAUCUCAGAAGAGAAAUCCAGUUAAACAUCAAGAGACUUAAAGAAAUCGGAUGCUACCGUGG
env-80    CGCUAUCCGUAUCCUUGCAGAUGCAGGAGUAGAUCCAGAUACACGUGUUAGAGAUUUGACACAGGACGACGUUAAGAAAAUCAGUGAAGUGAUCGAUGAGACUCAGGUUGUAGAAGGUGAUCUCAGAAGAGAAAUCCAGUUAAAC.....................................
env-81    ...........................................................................................................................................................
Rgn-1-1   CGCUAUCCGUAUCCUUGCAGAUGCAGGAGUAGAUCCAGAUACACGUGUUAGAGAUUUGACACAGGACGACGUUAAGAAAAUCAGUGAAGUGAUCGAUGAGACUCAGGUUGUAGAAGGUGAUCUCAGAAGAGAAAUCCAGUUAAACAUCAAGAGACUUAAAGAAAUCGGAUGCUACCGUGG
env-82    CGCUAUCCGUAUCCUUGCAGAUGCAGGAGUAGAUCCAGAUACACGUGUUAGAGAUUUGACACAGGACGACGUUAAGAAAAUCAGUGAAGUGAUCGAUGAGACUCAGGUUGUAGAAGGUGAUCUCAGAAGAGAAAUCCAGUUAAACAUCAAGAGACUUAAAGAAAUCGGAUGCUACCGUGG
env-83    CGCUAUCCGUAUCCUUGCAGAUGCAGGAGUAGAUCCAGAUACACGUGUUAGAGAUUUGACACAGGACGACGUUAAGAAAAUCAGUGAAGUGAUCGAUGAGACUCAGGUUGUAGAAGGUGAUCUCAGAAGAGAAAUCCAGUUAAACAUCAAGAGACUUAAAGAAAUCGGAUGCUACCGUGG
env-87    CGCUAUCCGUAUCCUUGCAGAUGCAGGAGUAGAUCCAGAUACACGUGUUAGAGAUUUGACACAGGACGACGUUAAGAAAAUCAGUGAAGUGAUCGAUGAGACUCAGGUUGUA.....................................................................
env-88    UGCUGACCGUAUUUUGAAAGAAGCAGGGGUUAAUCCUGACACUCGCUGCAGAGAUUUAACCGAUGAAGAAGUUAAAAAAAUCAGUGCAGUUAUCGACGAAACACAGACUGUAGAAGGUGAUCUUCGUAGAGAAAUCGCUCUUAAUAUUAAGAGAUUACAGGAAAUCGGAUGCUACAGAGG
env-89    [illegible]CGAACCGUAUUUUUAACAGAGGCUGGUGUAAAUCCAGAUACUCGUGUAAGAGACUUAACAGAUGAUGAAGUUAAGAAAAUCAGUGCUGUAAUUGAUGAAACACAGAUGGUAGAAGGUGAUCUUCGUAGAGAAAUCGCUUUAAACAUCAAGAGAUUACAGGAGAUUGGAUGCUACAGAG
env-90    UUCAAACCGUAUCCUGGCAGAUGCAGGUGUGGAUCCAAACACUCGUGUCAGAGAUUUGACCCAAGAAGAUGUUAAGAAAAUCAGUGACGUUAUUGAAGAGACGCAAAUGGUAGAAGGGGAUCUUCGUAGAGAGAUUCAAUUGAAUAUUAAGCGAUUGCAAGAGAUUGGCUGCUAUCGUGG
env-91    CUCAAAUCGUAUCCUGGCAGAUGCGGGAGUAGAUCCAAACACCCGGGUCAGAGAUUUGACUCAAGAAGAUGUUAAGAAAAUCAGUGAUGUGAUUGAAGAAACGCAAAUGGUAGAAGGGGAUCUACGUAGAGAGAUUCAAUUGAAUAUUAAGCGUUUACAAGAGAUUGGCUGCUAUCGUGG
env-92    ...........................................................................................................................................................
env-93    ...........................................................................................................................................................
env-94    CAUCGAUGAGACGCAGAUGGUAGAAGGUGAUCUGAGACGUGAAAUCGCUAUGAACAUCAAACGUCUGCAGGAGAUCGGCUGCUAUCGCGGCAUCCGGCAUAGAAAAGGACUUCCGGUUCGUGGUCAGAAGACCAAGACAAAUGCAAGAACCUGUAAGGGUCCGAGAAGAACCGUUGCCAA
Lba-11-1  UAUCGACGAGACACAGAUGGUCGAGGGUGAUCUGAGACGUGAGAUCGCUAUGAAUAUCAAGCGUCUGCAGGAAAUCGGCUGCUAUCGCGGCAUCCGGCAUAGAAAAGGACUUCCGGUUCGUGGUCAGAAGACAAAGACGAAUGCAAGAACCUGUAAGGGUCCGAGAAGAACCGUUGCCAA
env-95    CUCGAACAAGAUUCUGGAAGCAGCGAACGUAAAUCCCGAUACCCGUGUCAGGGAUCUGACGGAUGAAGAAGUGAUAAGAUCCGCGUUUCUAUGGAAGAGCUCGGCAUCCUGGUAGAGGGUGACCUUCGCCGUGAGAUCGCGAUGAACAUCAAGAGACUUCAGGAAAUUGGUUGCUACAG
env-96    ...........................................................................................................................................................
env-97    CGCAACCCGUAUUUUGACAGAGGCAGGAGUUAAUCCUGACAUUCGCUGCAGAGAUCUUACAGAUGAAGAUGUAAAGAAAAUCAGCGCAGUAAUUGAUGAGACCCAGACGGUAGAAGGUGAUCUUAGAAGAGAGAUCGCUCUGAACAUCAAGAGAUUACAGGAAAUCGGAUGCUAUAGAGG
Dsp-3-1   CGGAAUCAAUCAAAAGAAAAUGGAGGAAAUUCAC[illegible]GCUCGUAUAGCAGGUGUAGACUUACCAAGAGACAAACGUGUAGAAAUCGGACUUACUUACAUUUAUGGAAUCGGAAGGACGAGUGCGACACGUAUCCUUAGCGAGGCAGGAGUUAAUCCUGACAUUCGCUGCAGAGACCUU
Dsp-1-1   CGCAACACGUAUCUUAUCAGAGGCAGGAGUUGAUCCUGAUAUCCGCUGCAGAGAUCUUACGGAUGAAGAUGUUAAGAAGAUCAGCGCAGUGAUCGAUGAGACACAGAUGGUAGAAGGUGAUCUUAGAAGAGAGAUCGCUCUGAACAUCAAGAGAUUACAGGAGAUCGGAUGCUACAGAGG
env-101   ...........................................................................................................................................................
env-102   ...........................................................................................................................................................

          ...........................................................................................................................................................
          Y•C•R•Y•R•AUYYU•R•RR•RGCRR•GU•RAYCC•RAYAY•CG•••YAGRGAYYU•AC•GAYGA•GARGU•RA•ARRAUY•RYR••GY•AUYGA•GAR•Y••••RY•RURGAAGR•GAY•U•••••G•GRRR•YRY••Y•A••A•YARR•R•YY•Y•R•ARR••RR••GCURY•R•RG
```

| | |
|---|---|
| Dsp-2-1 | CGCAACCCGUAUCUUAGCACAGGCAGGAGUUAAUCCUGAUAUUCGUUGUAGAGAUCUUACAGACGACGAUGUAAAGAAAAUCAGUGCAGUAAUCGACGAGACUCAGACUGUAGAAGGAGAUCUUAGAAGAGAGAUCGCUCUGAACAUCAAGAGAUUACAGGAGAUCGGAUGCUAUAGAGG |
| env-103 | ......................................................................................................................................................................................... |
| env-106 | CGCAACCCGUAUCUUAGCACAGGCAGGAGUUAAUCCUGAUAUUCGUUGUAGAGAUCUUACAGACGACGAUGUAAAGAAAAUCAGUGCAGUAAUCGACGAGACUCAGACUGUAGAAGGAGAUCUUAGAAGAGAGAUCGCUCUGAACAUCAAGAGAUUACAGGAGAUCGGAUGCUAUAGAGG |
| env-116 | ......................................................................................................................................................................................... |
| env-117 | CGCAACCCGUAUCUUAGCACAGGCAGGAGUUAAUCCUGAUAUUCGUUGUAGAGAUCUUACAGACGACGAUGUAAAGAAAAUCAGUGCAGUAAUCGACGAGACUCAGACUGUAGAAGGAGAUCUUAGAAGAGAGAUCGCUCUGAACAUCAAGAGAUUACAGGAGAUCGGAUGCUAUAGAGG |
| env-118 | CGCAACCCGUAUCUUAGCACAGGCAGGAGUUAAUCCUGAUAUUCGUUGUAGAGAUCUUACAGACGACGAUGUAAAGAAAAUCAGUGCAGUAAUCGACGAGACUCAGACUGUAGAAGGAGAUCUUAGAAGAGAGAUCGCUCUGAACAUCAAGAGAUUACAGGAGAUCGGAUGCUAUAGAGG |
| env-119 | UGCAACUCGUAUCUUAACAGAAGCAGGAGUUAAUCCUGAUAUUCGUUGUAGAGAUCUUACAGACGACGAUGUAAAGAAAAUCAGUGCUGUAAUCGAUGAGACUCAGACAGUAGAAGGUGAUCUUCGUAGAGAAAUCGCCCUCAACAUUAAGAGAUUACAGGAAAUCGGAUGUUAUAGAGG |
| env-120 | ......................................................................................................................................................................................... |
| env-123 | ......................................................................................................................................................................................... |
| env-124 | UGCAACUCGUAUCUUAACAGAAGCAGGAGUUAAUCCUGAUAUUCGUUGUAGAGAUCUUACAGACGACGAUGUAAAGAAAAUCAGUGCUGUAAUCGAUGAGACUCAGACAGUAGAAGGUGAUCUUCGUAGAGAAAUCGCCCUCAACAUUAAGAGAUUACAGGAAAUCGGAUGUUAUAGAGG |
| env-127 | UGCAACUCGUAUCUUAACAGAAGCAGGAGUUAAUCCUGAUAUUCGUUGUAGAGAUCUUACAGACGACGAUGUAAAGAAAAUCAGUGCUGUAAUCGAUGAGACUCAGACAGUAGAAGGUGAUCUUCGUAGAGAAAUCGCCCUCAACAUUAAGAGAUUACAGGAAAUCGGAUGCUAUAGAGG |
| env-128 | ......................................................................................................................................................................................... |
| env-129 | UGCAACUCGUAUCUUAACAGAAGCAGGAGUUAAUCCUGAUAUUCGUUGUAGAGAUCUUACAGACGACGAUGUAAAGAAAAUCAGUGCUGUAAUCGAUGAGACUCAGACAGUAGAAGGUGAUCUUCGUAGAGAAAUCGCCCUCAACAUUAAGAGAUUACAGGAAAUCGGAUGUUAUAGAGG |
| env-130 | CGCAACUCGUAUCUUGACAGAGGCAGGAGUUAAUCCUGAUAUUCGUUGUAGAGAUCUUACAGACGAUGAUGUAAAGAAAAUCAGUGCUGUAAUCGAUGAGACACAGACAGUAGAAGGUGAUCUUCGUAGAGAAAUCGCUCUCAACAUCAAGAGAUUACAGGAAAUCGGAUGCUAUAGAGG |
| env-136 | CGCAACUCGUAUCUUGACAGAGGCAGGAGUUAAUCCUGAUAUUCGUUGUAGAGAUCUUACAGACGAUGAUGUAAAGAAAAUCAGUGCUGUAAUCGAUGAGACACAGACAGUAGAAGGUGAUCUUCGUAGAGAAAUCGCUCUCAACAUCAAGAGAUUACAGGAAAUCGGAU.......... |
| env-137 | CGCAACUCGUAUCUUGACAGAGGCAGGAGUUAAUCCUGAUAUUCGUUGUAGAGAUCUUACAGACGAUGAUGUAAAGAAAAUCAGUGCUGUAAUCGAUGAGACACAGACAGUAGAAGGUGAUCUUCGUAGAGAAAUCGCUCUCAACAUCAAGAGAUUACAGGAAAUCGGAUGCUAUAGAGG |
| env-138 | CGCAACUCG................................................................................................................................................................................ |
| env-139 | CGCAACUCGUAUCUUGACAGAGGCAGGAGUUAAUCCUGAUAUUCGUUGUAGAGAUCUUACAGACGAUGAUGUAAAGAAAAUCAGUGCUGUAAUCGAUGAGACACAGACAGUAGAAGGUGAUCUUCGUAGAGAAAUCGCUCUCAACAUCAAGAGAUUACAGGAAAUCGGAUGCUAUAGAGG |
| Dlo-1-1 | CGCAACUCGUAUCUUGACAGAGGCAGGAGUUAAUCCUGAUAUUCGUUGUAGAGAUCUUACAGACGAUGAUGUAAAGAAAAUCAGUGCUGUAAUCGAUGAGACACAGACAGUAGAAGGUGAUCUUCGUAGAGAAAUCGCUCUCAACAUCAAGAGAUUACAGGAAAUCGGAUGCUAUAGAGG |
| env-140 | CGCAACUCGUAUCUUGACAGAGGCAGGAGUUAAUCCUGAUAUUCGUUGUAGAGAUCUUACAGACGAUGAUGUAAAGAAAAUCAGUGCUGUAAUCGAUGAGACACAGACAGUAGAAGGUGAUCUUCGUAGAGAAAUCGCUCUCAACAUCAAGAGAUUACAGGAAAUCGGAUGCUAUAGAGG |
| env-149 | UGCAACUCGUAUUUUGACAGAAGCAGGAGUUAAUCCUGACAUUCGCUGCAGAGAUCUGACAGACGAAGACGUUAAGAAGAUCAGUGCAGUAAUCGAUGAAACUCAGGUUGUAGAAGGUGAUCUUCGUAGAGAGAUCGCUCUGAACAUCAAGAGAUUACAGGAGAUCGGAUGCUACAGAGG |
| env-150 | UGCAACCCGUAUUCUGACAGAGGCAGGAGUAAAUCCUGACAUUCGCUGCAGAGAUCUGACAGACGAAGACGUUAAGAAGAUCAGCGCAGUUAUCGACGAGACACAGGUAGUAGAAGGUGACUUGAGAAGAGAGAUCGCUCUUAACAUUAAGAGACUUCAGGAAAUCGGAUGCUA...... |
| env-151 | ......................................................................................................................................................................................... |
| Chy-1-1 | UGCGACACGUAUUCUCAGCGAGGCAGGCGUUAGCCCUGACAUUCGCUGCAGAGACCUUACAGACGAAGACGUAAAGAAGAUCAGUGCUGUAAUCGAUGAAACUCAGACAGUAGAAGGUGAUCUCAGAAGAGAGAUCGCUCUGAACAUAAAGAGAUUACAGGAGAUCGGAUGCUACAGAGG |
| env-152 | C........................................................................................................................................................................................ |
| env-153 | CGCUACCCGUAUUCUGGCUGAUGCAGGAGUAGAUCCGGACAUCCGCUGCAGAGAUCUGACAGACGAAGA.................................................................................................................... |
| env-154 | ......................................................................................................................................................................................... |
| Esp-1-1 | CGCAACACGUAUUCUGAAAGAAGCAGGAGUUGAUCCGGACAUCCGCUGCAGAGAUCUGACAGACGACGAUGUAAAGAAGAUCAGCGCAGUCAUCGAUGAGACACAGAUGGUAGAAGGUGAUCUGAGAAGAGAGAUCGCGCUGAACAUCAAGAGAUUACAGGAGAUCGGAUGCUACAGAGG |
| env-155 | UGCAGAACGUAUCUUAAAGGAUGCAGGAGUUAAUCCUGACACUCGUGUAAGAGAUCUUACAAGUGAAGAAGUAAAGAAAAUCAGCGCAGUGAUCGAUGAGACUCAGGUUGUAGAAGGUGAUCUUAGAAGAGAGAUUCAGCUCAACAUCAAGAGACUUAAAGAGAUCGGAUGCUACCGUGG |
| env-156 | UGCAGAACGUAUCUUAAAGGAUGCAGGAGUUAAUCCUGAUACUCGUGUAAGAGAUCUUACAAGUGAAGAAGUAAAGAAAAUCAGCGCAGUGAUCGAUGAGACUCAGGUUGUAGAAGGUGAUCUCAGAAGAGAAAUCCAGCUCAACAUCAAGAGACUUAAAGAGAUCGGAUGCUACCGUGG |
| env-159 | ......................................................................................................................................................................................... |
| env-160 | UGCAGAACGUAUCUUAAAGGAUGCAGGAGUUAAUCCUGAUACUCGUGUAAGAGAUCUUACAAGUGAAGAAGUAAAGAAAAUCAGCGCAGUGAUCGAUGAGACUCAGGUUGUAGAAGGUGAUCUCAGAAGAGAAAUCCAGCUCAACAUCAAGAGACUUAAAGAGAUCGGAUGCUACCGUGG |
| env-161 | UGCAGAACGUAUCUUAAAGGAUGCAGGAGUUAAUCCUGACACUCGUGUAAGAGAUCUUACAAGUGAAGAAGUAAAGAAAAUCAGCGCAGUGAUCGAUGAGACUCAGGUUGUAGAAGGUGAUCUUAGAAGAGAGAUCCAGCUCAACAUCAAGAGACUUAAAGAGAUCGGAUGCUACCGUGG |
| env-162 | UGCAGAACGUAUCUUAAAGGAUGCAGGAGUUAAUCCUGAUACUCGUGUAAGAGAUCUUACAAGUGAAGAAGUAAAGAAAAUCAGCGCAGUGAUCGAUGAGACUCAGGUUGUAGAAGGUGAUCUCAGAAGAGAAAUCCAGCUCAACAUCAAGAGACUUAAAGAGAUCGGAUGCUACCGUGG |
| env-163 | ......................................................................................................................................................................................... |
| env-164 | ......................................................................................................................................................................................... |
| env-165 | UGCAGAACGUAUCUUAAAGGAUGCAGGAGUUAAUCCUGAUACUCGUGUAAGAGAUCUUACAAGUGAAGAAGUAAAGAAAAUCAGCGCAGUGAUCGAUGAGACUCAGGUUGUAGAAGGUGAUCUCAGAAGAGAAAUCCAGCUCAACAUCAAGAGACUUAAAGAGAUCGGAUGCUACCGUGG |
| env-167 | ......................................................................................................................................................................................... |
| env-168 | UGCAGAACGUAUCUUAAAGGAUGCAGGAGUUAAUCCUGACACUCGUGUAAGAGAUCUUACAAGUGAAGAAGUAAAGAAAAUCAGCGCAGUGAUCGAUGAGACUCAGGUUGUAGAAGGUGAUCUUAGAAGAGAGAUCCAGCUCAACAUCAAGAGACUUAAAGAGAUCGGAUGCUACCGUGG |
| env-169 | UGCAGAACGUAUCUUAAAGGAUGCAGGAGUUAAUCCUGACACUCGUGUAAGAGAUCUUACAAGUGAAGAAGUAAAGAAAAUCAGCGCAGUGAUCGAUGAGACUCAGGUUGUAGAAGGUGAUCUUAGAAGAGAGAUCCAGCUCAACAUCAAGAGACUUAAAGAGAUCGGAUGCUACCGUGG |
| env-170 | UGCAGAACGUAUCUUAAAGGAUGCAGGAGUUAAUCCUGACACUCGUGUAAGAGAUCUUACAAGUGAAGAAGUAAAGAAAAUCAGCGCAGUGAUCGAUGAGACUCAGGUUGUAGAAGGUGAUCUUAGAAGAGAGAUCCAGCUCAACAUCAAGAGACUUAAAGAGAUCGGAUGCUACCGUGG |
| env-171 | UGCAGAACGUAUCUUAAAGGAUGCAGGAGUUAAUCCUGAUACUCGUGUAAGAGAUCUUACAAGUGAAGAAGUAAAGAAAAUCAGCGCAGUGAUCGAUGAGACUCAGGUUGUAGAAGGUGAUCUUAGAAGAGAGAUCCAGCUCAACAUCAAGAGACUUAAAGAGAUCGGAUGCUACCGUGG |
| env-172 | UGCAGAACGUAUCUUAAAGGAUGCAGGAGUUAAUCCUGACACUCGUGUAAGAGAUCUUACAAGUGAAGAAGUAAAGAAAAUCAGCGCAGUGAUCGAUGAGACUCAGGUUGUAGAAGGUGAUCUUAGAAGAGAGAUCCAGCUCAACAUCAAGAGACUUAAAGAGAUCGGAUGCUACCGUGG |
| env-175 | UGCAGAACGUAUCUUAAAGGAUGCAGGAGUUAAUCCUGACACUCGUGUAAGAGAUCUUACAAGUGAAGAAGUAAAGAAAAUCAGCGCAGUGAUCGAUGAGACUCAGGUUGUAGAAGGUGAUCUCAGAAGAGAGAUUCAGCUCAACAUCAAGAGACUUAAAGAGAUCGGAUGCUACCGUGG |
| env-176 | UGCAGAACGUAUCUUAAAGGAUGCAGGAGUUAAUCCUGACACUCGUGUAAGAGAUCUUACAAGUGAAGAAGUAAAGAAAAUCAGCGCAGUGAUCGAUGAGACUCAGGUU........................................................................... |
| env-177 | UGCAGAACGUAUCUUAAAGGAUGCAGGAGUUAAUCCUGACACUCGUGUAAGAGAUCUUACAAGUGAAGAAGUAAAGAAAAUCAGCGCAGUGAUCGAUGAGACUCAGGUUGUAGAAGGUGAUCUUAGAAGAGAGAUCCAGCUCAACAUCAAGAGACUUAAAGAGAUCGGAUGCUACCGUGG |
| env-178 | UGCAGAACGUAUCUUAAAGGAUGCAGGAGUUAAUCCUGACACUCGUGUAAGAGAUCUUACAAGUGAAGAAGUAAAGAAAAUCAGCGCAGUGAUCGAUGAGACUCAGGUUGUAGAAGGUGAUCUUAGAAGAGAGAUCCAGCUCAACAUCAAGAGACUUAAAGAGAUCG............. |
| Lba-4-1 | UUCAAAUCGUAUUUUAGCAGAAGCAGGAGUUAGAUCCUGACAUCCGUUGCAGAGAUCUCACGGAUGAAGAUGUAAAGAAGAUCAGCACAGUAAUCGACGAGACACAGAUGGUAGAAGGUGAUCUUCGUAGAGAAGAUUGCUCUUAACAUCAAGAGAUUACAGGAAAUCGGAUGCUACAGAGG |
| env-179 | CGCAACCCGUAUUUUGACAGAGGCAGGAGUUAAUCCUGAUAUUCGCUGCAGAGAUCUUACAGAUGAAGAUGUAAAGAAAAUCAGCGCGGUAAUUGAUGAGACUCAGACGGUAGAAGGUGAUCUCAGAAGAGAGAUCGCUAUGAACAUCAAGAGAUUACAGGAAAUCGGAUGCUAUAGAGG |
| Cba-3-1 | CGCAACCCGUAUUUUGACAGAGGCAGGAGUUAAUCCUGACAUUCGCUGCAGAGAUCUUACAGAUGAAGAUGUAAAGAAAAUCAGCGCAGUAAUUGAUGAGACCCAGACGGUAGAAGGUGAUCUUAGAAGAGAGAUCGCUCUGAACAUCAAGAGAUUACAGGAAAUCGGAUGCUAUAGAGG |
| [sc-1-1 | CGCAACCCGUAUUUUGACAGAGGCAGGAGUUAAUCCUGACAUUCGCUGCAGAGAUCUUACAGAUGAAGAUGUAAAGAAAAUCAGCGCAGUAAUUGAUGAGACCCAGACGGUAGAAGGUGAUCUUAGAAGAGAGAUCGCUCUGAACAUCAAGAGAUUACAGGAAAUCGGAUGCUAUAGAGG |
| Lba-7-1 | CGCAACCCGUAUUUUGACAGAGGCAGGAGUUAAUCCUGAUAUUCGCUGCAGAGAUCUUACAGAUGAAGAUGUAAAGAAAAUCAGCGCGGUAAUUGAUGAGACUCAGACGGUAGAAGGUGAUCUCAGAAGAGAGAUCGCUAUGAACAUCAAGAGAUUACAGGAAAUCGGAUGCUAUAGAGG |
| env-181 | ......................................................................................................................................................................................... |
| env-182 | ......................................................................................................................................................................................... |
| env-183 | CGCAGACAGAAUUCUUGAAGCAGCGAAUGUAAAUCCCGACACCCGUGUCAGAGAUCUGACAAAUGAAGAAGUUGAUAAGAUCAAAGUUGCUAUUGAUGAACUUGGUAUCAUGGUCGAAGGUGACCUUCGCAGAUCUGUAGCAAUGGACAUCAAGACUCUUCAGGAAAUUGGCUGUUACAG |
| env-184 | CGCAAGCCGUAUUUUGGCAGAUGCAGGAGUAGAUCCGAAUACUCGUGUAAG...................................................................................................................................... |
| env-185 | CGCAAGCCGUAUUUUGGCAGAUGCAGGAGUAGAUCCGAAUACUCGUGUAAGAGAUCUUACAAGUGAAGAAGUAAAAAAGAUCAGCGCAGUGAUCGAUGAAACUCAGGUUGUAGAAGGUGAUCUUAGAAGAGA............................................... |
| env-186 | CGCAAGCCGUAUUUUGGCAGAUGCAGGAGUAGAUCCGAAUACUCGUGUAAGAGAUCUUACAAGUGAAGAAGUAAAAAAGAUCAGCGCAGUGAUCAAUGAAACUCAGGUUGUAGAAGGUGAUCUUAGAAGAGAGAUCCAGCUUAACAUCAAGAGACUGAAAGAAAUCGGAUGCUACCGUGG |
| env-187 | ......................................................................................................................................................................................... |
| env-188 | ......................................................................................................................................................................................... |
| env-189 | CGCAAGCCGUAUUUUGGCAGAUGCAGGAGUAGAUCCGAAUACUCGUGUAAGAGAUCUUACAAGUGAAGAAGUAAAAAAGAUCAGCGCAGUGAUCGAUGAAACUCAGGUUGUAGAAGGUGAUCUUAGAAGAGAGAUCCAGCUUAACAUCAAGAGACUGAAAGAAAUCGGAUGCUACCGUGG |
| env-190 | CGCAAGCCGUAUUUUGGCAGAUGCAGGAGUAGAUCCGAAUACUCGUGUAAGAGAUCUUACAAGUGAAGAAGUAAAAAAGAUCAGCGCAGUGAUCAAUGAAACUCAGGUUGUAGAAGGUGAUCUUAGAAGAGAGAUCCAGCUUAACAUCAAGAGACUGAAAGAAAUCGGAUGCUACCGUGG |
| env-191 | ......................................................................................................................................................................................... |
| env-192 | CGCAAGCCGUAUUUUGGCAGAUGCAGGAGUAGAUCCGAAUACUCGUGUAAGAGAUCUUACAAGUGAAGAAGUAAAAAAGAUCAGCGCAGUGAUCAAUGAAACUCAGGUUGUAGAAGGUGAUCUUAGAAGAGAGAUCCAGCUUAACAUCAAGAGACUGAAAGAAAUCGGAUGCUACCGUGG |
| env-194 | CGCAAGCCGUAUUUUGGCAGAUGCAGGAGUAGAUCCGAAUACUCGUGUAAGAGAUCUUACAAGUGAAGAAGUAAAAAAGAUCAGCGCAGUGAUCGAUGAAACUCAGGUUGUAGAAGGUGAUCUUAGAAGAGAGAUCCAGCUUAACAUCAAGAGACUGAAAGAAAUCGGAUGCUACCGUGG |
| env-195 | CGCAAGCCGUAUUUUGGCAGAUGCAGGAGUAGAUCCGAAUACUCGUGUAAGAGAUCUUACAAGUGAAGAAGUAAAAAAGAUCAGCGCAGUGAUCGAUGAAACUCAGGUUGUAGAAGGUGAUCUUAGAAGAGAGAUCCAGCUUAACAUCAAGAGACUGAAAGAAAUCGGAUGCUACCGUGG |
| env-200 | CGCAAGCCGUAUUUUGGCAGAUGCAGGAGUAGAUCCGAAUACUCGUGUAAGAGAUCUUACAAG.......................................................................................................................... |
| env-201 | ......................................................................................................................................................................................... |
| env-202 | ......................................................................................................................................................................................... |
| env-211 | ......................................................................................................................................................................................... |
| env-212 | ......................................................................................................................................................................................... |
| env-216 | CGCAAGCCGUAUUUUGGCAGAUGCAGGAGUAGAUCCGAAUACUCGUGUAAGAGAUCUUACAAGUGAAGAAGUAAAAAAGAUCAGCGCAGUGAUCGAUGAAACUCAGGUUGUAGAAGGUGAUCUUAGAAGAGAGAUCCAGCUUAACAUCAAGAGACUGAAAGAAAUCGGAUGCUACCGUGG |
| env-217 | AUCCAACAAGAUCCUUGCAGCGGCCGGCGUUGACCCGGACAUCCGUGUAAGAGACCUUACAGAUGAAGAUGUGGACAAGAUCCGUGUGAGCAUGGAAGAGCUUGGCGUAAUGGUGGAAGGUGAUCUUCGCCGUGAUAUCGCCAAUGAAUAUCAAGCGCCUUCAGGAGAUCGGCUGCUACAG |
| Rpe-1-1 | UGCAACAAAAUUCUGGAAUCUGCAAAGGUUAAUCCGGACACUCGUGUCAGAGAUUUAACUGAUGAAGAAGUAGGAAGAAUUCGUGACGUAAUUGAUGAAACACAGACUGUAGAAGGUGAUUUAAGAAGAGAAAUCGCUCUUAAUAUCAAGAGAUUACAGGAAAUCGGAUGCUAUCGUGG |
| Rsp-1-1 | UGCAACAAAAUUCUGGAAUCUGCAAAGGUUAAUCCGGACACUCGCGUCAGGGAUUUAACUGAUGAAGAAGUAGGAAGAAUUCGUGACGUAAUUGAUGAAACACAGACUGUAGAAGGUGAUUUAAGAAGAGAAAUCGCUCUUAAUAUCAAGAGAUUACAGGAAAUCGGAUGCUAUCGUGG |
| env-218 | AUCGAAUAAGAUCCUUGCCGCAGCGAAUGUUGACCCCAAUACGCGCGUCAGAGACCUUACGGAUGAAGAAGUUGAUAAGAUCCGCAGCUCGAUGGAAGAGCUGAAUGUUAUGGUGGAAGGUGACCUUCGCCGUGACACCGCUAUGAACAUCAAACGUCUGCAGGAGAUCGGCUGCUAUCG |
| env-219 | CAGCGACAAGAUCCUGGCUGCAGCCGGUGUAAACCCGGAUACCAGAGUAAGAGAUCUUACGGAUGAAGAAGUCAAGAAAAUCAGCACCGUUAUCGAAGAGACGAUGAUGGUUGAAGGUGAUCUUAGAAGAGAAGUCGCUCUGAACAUCAAGAGACUGCAGGAAAUCGGCUGCUACAGAGG |
| env-220 | CGCAAACAAGAUCCUUGCCGCGGCGAACGUGAAUCCCGACACACGUGUCAGAGAUUUAACCGAUGAAGAAGUAGCCC............................................................................................................ |
| env-221 | CGCUAACAAAAUCCUUGCAGAUGCAAAGGUUAAUCCGGAUACUCGCGUAAGAGAUCUGACUGACGAAGAAGUAA............................................................................................................... |
| env-222 | CGCUAACAAAAUCCUUGCAGAUGCAAAGGUUAAUCCGGAUACUCGCGUAAGAGAUCUGACUGACGAAGAAGUAAGCAGAAUCCGUGACGUAAUGGACGAGCAUUAUACAGUAGAAGGUGAUCUGAGAAGAGAGAUCGCUCUGAACAUCAAGAGACUUCAGGAGAUCGGAUGCUACAGAGG |
| env-223 | ......................................................................................................................................................................................... |
| env-226 | ......................................................................................................................................................................................... |
| env-227 | CGCUAACAAAAUCCUUGCAGAUGCAAAGGUUAAUCCGGAUACUCGCGUAAGAGAUCUGACUGACGAAGAAGUAAGCAGAAUCCGUGACGUAAUGGACGAGCAUUAUACAGUAGAAGGUGAUUUAAGAAGAGAG............................................. |
| env-228 | CGCUAACAAAAUCCUUGCAGAUGCAAAGGUUAAUCCGGAUACUCGCGUAAGAGAUCUGACUGACGAAGAAGUAAGCAGAAUCCGUGACGUAAUGGACGAGCAUUAUACAGUAGAAGGUGAUCUGAGAAGAGAGAUCGCUCUGAACAUCA............................... |

.........................................................................................................................................................................................

.........................................................................................................................................................................................

Y•C•R•Y•R•AUYYU•R•RR•RGCRRR•GU•RAYCC•RAYAY•CG•••YAGRGAYYU•AC•GAYGA•GARGU•RA•ARRAUY•RYR••GY•AUYGA•GAR•Y••••RY•RURGAAGR•GAY•U•••••G•GRRR•YRY••Y•A••A•YARR•R•YY•Y•R•ARR••RR••GCURY•R•RG

```
env-229   ..............................................................................................................................................................................................
env-230   CGCUAACAAAAUCCUUGCAGAUGCAAAGGUGAAUCCGGAUACUCGCGUAAGAGAUCUGACUGACGAAGAAGU........................................................................................................................
env-231   ..............................................................................................................................................................................................
env-232   ..............................................................................................................................................................................................
env-234   ..............................................................................................................................................................................................
env-235   CGCUAACAAAAUCCUUGCAGAUGCAAAGGUUAAUCCGGAUACUCGCGUAAGAGAUCUGACUGACGAAGAAGUAAGCAGAAUCCGUGACGUAAUGGACGAGCAUUAUACAGUAGAAGGUGAUCUGAGAAGAGA...........................................................
env-236   CGCUAACAAAAUCCUUGCAGAUGCAAAGGUUAAUCCGGAUACUCGCGUAAGAGAUCUGACUGACGAAGAAGUAAGCAGAAUCCGUGACGUAAUGGACGAGCAUUAUACAGUAGAAGGUGAUCUGAGAAGAGAGAUCGCUCUGAACAUCAAGAGACUUCAGGAGAUCGGAUGCUACAGAGG
env-237   CGCUAACAAAAUCCUUGCAGAUGCAAAGGUUAAUCCGGAUACUCGCGUAAGAGAUCUGACUGACGAAGAAGUAAGCAGAAUCCGUGACGUAAUGGACGAGCAUUAUACAGUAGAAGGUGAUCUGAGAAGAGNNNNNNNNNNNNNNNNNNNNNNNNNNNNNNNNNNNNNNNNNNNNNNNNACCUG
env-238   ..............................................................................................................................................................................................
env-239   CGCUAACAAAAUCCUUGCAGAUGCAAAGGUUAAUCCGGAUACUCGCGUAAGAGAUCUGACUGACGAAGAAGUAAGCAGAAUCCGUGACGUAAUGGACGAGCAUUAUACAGUAGAAGGUGAUCUGAGAAGAGAGA.........................................................
env-240   CGCUAACAAAAUCCUUGCAGAUGCAAAGGUUAAUCCGGAUACUCGCGUAAGAGAUCUGACUGACGAAGAAGUAAGCAGAAUCCGUGACGUAAUGGACGAGCAUUAUACAGUAGAAGGUGAUCUGAGAAGAGAGAUCGCUCUGAACAUCAAGAGACUUCAGGAGAUCGGAUGCUACAGAGG
env-241   UUCGAACAAAAUUCUUGAAGCAGCGAACGUAAAUCCUGAUACUCGUGUCAGAGAUCUUACAGAUGAAGAGGUAGACAAGAUCCGUAACUCUAUGGAAGAACUUGGCGUAAUGGUGGAAGGUGACCUUCGCCGCGACAUCGCCAUGAACAUCAAACGACUUCAGGAAAUCGGCUGCUACAG
env-242   ..............................................................................................................................................................................................
env-243   CGCAACAAAAAUUUUGGAGGCAGCUGGAGUAAAUCCAGAUACUCGUUGUAGAGACUUAACAGACGAAGAAGUAAAACAG.................................................................................................................
env-244   ..............................................................................................................................................................................................
env-245   ..............................................................................................................................................................................................
env-246   CGCAACAAAAAUUUUGGAGGCAGCUGGAGUAAAUCCAGAUACUCGUUGUAGAGACUUAACAGACGAAGAAGUAAAACAGAUCAGUGCUGUUAUCGAUGAAAGCUAUCAGGUAGAAGGUGAUCUUCGUAGAGAGAUUGCUUUAAACAUCAAGAGAUUACAGGAAAUCGGAUGCUACCGUGG
env-252   CGCAACAAAAAUUUUGGAGGCAGCUGGAGUAAAUCCAGAUACUCGUUGUAGAGACUUAACAGACGAAGAAGUAAAACAGAUCAGUGCUGUUAUCGAUGAAAGCUAUCAGGUAGAAGGUGAUCUUCGUAGAGAGAUUGCUUUAAACAUCAAGAGAUUACAGGAAAUCGGAUGCUACCGUGG
env-254   CUCUAAUCGUAUCUUAGAAGAGGCAGGAGUAAAUCCUGAUACUCGUGUCAGAGAUUUAACUGACGAAGAAGUAAAGAAAAUCAGUGCUGUAAUCGAUGAAACUCAGUUAGUAGAAGGUGAUCUUAGAAGAGAGAUCGCUCUUAACAUCAAGAGAUUACAGGAAAUCGGAUGCUAUAGAGG
Lba-6-1   CUCUAAUCGUAUCUUAGAAGAGGCAGGAGUAAAUCCUGAUACUCGUGUCAGAGAUUUAACUGACGAAGAAGUAAAGAAAAUCAGUGCUGUAAUCGAUGAAACUCAGUUAGUAGAAGGUGAUCUUAGAAGAGAGAUCGCUCUUAACAUCAAGAGAUUACAGGAAAUCGGAUGCUAUAGAGG
env-255   CUCUAAUCGUAUCUUAGAAGAGGCAGGAGUAAAUCCUGAUACUCGUGUCAGAGAUUUAACUGACGAAGAAGUAAAGAAAAUCAG............................................................................................................
Lba-10-1  CUCUAAUCGUAUCUUAGAAGAGGCAGGAGUAAAUCCUGAUACUCGUGUCAGAGAUUUAACUGACGAAGAAGUAAAGAAAAUCAGUGCUGUAAUCGAUGAAACUCAGUUAGUAGAAGGUGAUCUUAGAAGAGAGAUCGCUCUUAAUAUCAAGAGAUUACAGGAAAUCGGAUGCUAUAGAGG
env-256   CUCUAAUCGUAUCUUAGAAGAGGCAGGAGUAAAUCCUGAUACUCGUGUCAGAGAUUUAACUGACGAAGAAGUAAAGAAAAUCAGUGCUGUAAUCGAUGAAACUCAGUUAGUAGAAGGUGAUCUUAGAAGAGAGAUCGCUCUUAACAUCAAGAGAUUACAGGAAAUCGGAUGCUAUAGAGG
env-258   ..............................................................................................................................................................................................
env-259   CUCUAAUCGUAUCUUAGAAGAGGCAGGAGUAAAUCCUGAUACUCGUGUCAGAGAUUUAACUGACGAAGAAGUAAAGAAAAUCAGUGCUGUAAUCGAUGAAACUCAGUUAGUAGAAGGUGAUCUUAGAAGAGAGAUCGCUCUUAACAUCAAGAGAUUACAGGAAAUCGGAUGCUAUAGAGG
env-260   ..............................................................................................................................................................................................
env-261   ..............................................................................................................................................................................................
env-262   ..............................................................................................................................................................................................
env-263   CUCAAACAAGAUUCUGGCUGCAGCGAAUGUAAAUCCGGACACACGUGUCAGAGAUUUAACAGAUGAGGAAGUUGCGAGAAUCCGUGACAUCAUCGAUGAGCAGUUCACAGUAGAAGGUGAUCUCAGAAGAGAGACAGCCCUCAACAUCAAGAGACUGCAGGAGAUCGGAUGCUACAGAGG
Bfo-1-1   CUCGAACAAGAUUCUGGAGGCGGCAGGCGUAAAUCCGGAUACCCGUGUCAGAGAUCUGACGGACGAAGAGGUUUCCAAAAUCCGUGACGUAAUGGAUGAACAUUACACAGUGGAAGGUGACCUGAGAAGAGAAAUCGCUCUGAAUAUCAAGAGACUUCAGGAAAUCGGAUGCUACAGAGG
Bpr-1-1   UGCUGACCGUAUCUUAGUAGAGGCAGGAGUAAACCCUGACACUCGCUGCAGGGAUUUAACUGACGAAGAAGUAGGAAAGAUUCGUGAUGCAAUCGAUGAACUGAAUAUCGCUGUAGAGGGUGAUCUGAGAAGAGAAAUCGCUCUGAACAUCAAGAGAUUACAGGAAAUCGGAUGCUACAG
Cba-1-1   CGCUGACCGUAUCUUAGUAGAGGCAGGUGUAAACCCUGACACUCGCUGCAGGGAUUUAACUGACGAAGAAGUAGGAAAAAUCCGUGAUGCAAUCGAUGAACUGAAUAUCGCUGUAGAAGGUGAUCUGAGAAGAGAAAUCGCUCUGAACAUCAAGAGAUUACAGGAAAUCGGAUGCUACAG
env-264   ..............................................................................................................................................................................................
env-265   ..............................................................................................................................................................................................
env-266   CGCCACAAAGAUCGUGGAGCAGGCAGGAGUUAAUCCCGACACCCGUGUCAGAGAUCUUACAGAUGAAGAGGUCAAAAAACUGAGCAGCGUCAUCGAAGAGACAAUGAUGGUCGAAGGUGAUCUCAGAAGGGAAAUCGCCCUUAAUAUUAAGAGACUGCAGGAAAUCGGAUGCUACAGAGG
env-267   CGCUGACAAAAUUCUUGCUCAGGCAGGUGUAAAUCCUGACAUUCGUGUCAGAGACCUGACAGAUGAUGAUGUUAAGAAAAUCAGUGCUGUAAUUGAUGAAACAAUGACAGUUGAAGGUGAUCUUCGUCGUGAAAUCGCUCUGAAUAUCAAGAGACUGCAGGAAAUCGGAUGCUACAGAGG
env-268   CGCUGACAAAAUUCUUGCUCAGGCAGGUGUAAAUCCUGACAUUCGUGUCAGAGACCUGACAGAUGAUGAUGUUAAGAAAAUCAGUGCUGUAAUUGAUGAAACAAUGACAGUUGAAGGUGAUCUUCGUCGUGAAAUCGCUCUGAAUAUCAAGAGACUGCAGGAAAUCGGAUGCUACAGAGG
env-271   CGCUGACAAAAUUCUUGCUCAGGCAGGUGUAAAUCCUGACAUUCGUGUCAGAGACCUGACAGAUGAUGAUGUUAAGAAAAUCAGUGCUGUAAUUGAUGAAACAAUGACAGUUGAAGGUGAUCUUCGUCGUGAAAUCGCUCUGAAUAUC.................................
env-272   CGCUGACAAAAUUCUUGCUCAGGCAGGUGUAAAUCCUGACAUUCGUGUCAGAGACCUGACAGAUGAUGAUGUUNAGAAAAUCAGUGCUGUAAUUGAUGAAACAAUGACAGUUGAAGGUGAUCUUCGUCGUGAAAUCGCUCUGAAUAUCAAGAGACUGCAGGAAAUCGGAUGCUACAGAGG
env-273   CGCAGACAAAAUCCUUGCUCAGGCAGGCGUAAAUCCUGACAUUCGUGUCAGAGACCUGACAGACGAUGAUGUAAAGAAAAUCAGUGCUGUCAUUGAUGAGACAAUGAUGGUAGAAGGUGAUCUUCGUCGUGAAAUCGCUCUGAAUAUCAAGAGACUGCAGGAGAUCGGAUGCUACAGAGG
env-275   UGCAGACAAGAUUCUUGCACAGGCAGGAGUAAAUCCUGACAUCCGUGUCAGAGAUCUGACAGACGAUGAUGUAAAGAAAAUCAGUGCUGUUAUUGAUGAAACAAUGACAGUUGAAGGUGAUCUUCGUCGUGAAAUCGCUAUGAAUAUCAAGAGACUGCAGGAAAUCGGAUGCUACCGUGG
env-276   UGCAGACAAGAUUCUUGCACAGGCAGGAGUAAAUCCUGACAUCCGUGUCAGAGAUUUGACAGACGAUGAUGUAAAGAAAAUCAGUGCUGUAAUUGAUGAAACAAUGACAGUAGAAGGUGAUCUUCGUCGUGAAAUCGCUAUGAAUAUCAAGAGACUGCAGGAAAUCGGAUGCUACCGUGG
env-277   CGCAGACAAAAUUCUUGCGCAGGCAAAUGUAA................................................................................................................................................................
env-278   CGCAGACAAAAUUCUUGCGCAGGCAAAUGUAAAUCCUGACACACGUGUCAGAGAUCUGACAGAUGAAGAAGUAAAGAGAAUCAGCGCUGUCAUUGAUGAGACAAUGAUGGUAGAAGGUGAUCUUCGUCGUGAAAUCGCUCUUAAUAUCAAGAGACUGCAGGAAAUCGGAUGCUACAGAGG
env-279   ..............................................................................................................................................................................................
env-280   ..............................................................................................................................................................................................
env-281   CGCAGACAAAAUUCUUGCUAAAGCAGAGGUUAAUCCUGACACACGUGUCAGAGAUCUGACUGAUGACGAGGUUAAGAGACUGAGUGCUGUUAUUGAUGAGACAAUGACAGUAGAAGGUGAUCUGAGAAGGGAAAUCGCCCUUAACAUUAAGAGACUGCAGGAGAUUGGAUGCUACAGAGG
env-282   CGCAGACAAAAUUCUUGCUAAAGCAGAGGUUAAUCCUGACACACGUGUCAGAGAUCUGACUGAUGACGAGGUUAAGAGACUGAGUGCUGUUAUUGAUGAGACAAUGACAGUAGAAGGUGAUCUGAGAAGGGAAAUCGCCCUUAACAUCAAGAGACUGCAGGAGAUUGGAUGCUACAGAGG
Bsp-1-1   CGCAGACAAAAUUCUUGCUAAAGCAGAGGUUAAUCCUGACACACGUGUCAGAGAUCUGACUGAUGACGAGGUUAAGAGACUGAGUGCUGUUAUUGAUGAGACAAUGACAGUAGAAGGUGAUCUGAGAAGGGAAAUCGCCCUUAACAUCAAGAGACUGCAGGAGAUUGGAUGCUACAGAGG
env-283   CGCAGACAAAAUUCUUGCUAAAGCAGAGGUUAAUCCUGACACACGUGUCAGAGAUCUGACUGAUGACGAGGUUAAGAGACUGAGUGCUGUUAUUGAUGAGACAAUGACAGUAGAAGGUGAUCUGAGAAGGGAAAUCGCCCUUAACAUUAAGAGACUGCAGGAGAUUGGAUGCUAC.....
env-284   CGCAGACAAAAUUCUUGCUAAAGCAGAGGUUAAUCCUGACACACGUGUCAGAGAUCUGACUGAUGAUGAGGUUAAGAGACUGAGCGCUGUUAUUGAUGAGACAAUGACAGUAGAAGGUGAUCUGAGAAGGGAAAUCGCCCUUAACAUCAAGAGACUGCAGGAGAUUGGAUGCUACAGAGG
env-285   CGCAGACAAAAUUCUUGCUAAAGCAGAGGUUAAUCCUGACACACGUGUCAGAGAUCUGACUGAUGAUGAGGUUAAGAGACUGAGCGCUGUUAUUGAUGAGACAAUGACAGUAGAAGGUGAUCUGAGAAGGGAAAUCGCCCUUAACAUCAAGAGACUGCAGGAGAUUGGAUGCUACAGAGG
env-286   CGCAGACAAAAUUCUUGCUAAAGCAGAGGUUAAUCCUGACACACGUGUCAGAGAUCUGACUGAUGAUGAGGUUAAGAGACUGAGCGCUGUUAUUGAUGAGACAAUGACAGUAGAAGGUGAUCUGAGAAGGGAAAUCGCCCUUAACAUCAAGAGACUGCAGGAGAUUGGAUGCUACAGAGG
env-287   CGCAGACAAAAUCCUUGCAAAGGCAGAAGUUAAUCCUGACACACGUGUCAGAGAUCUGACUGAUGACGAAGUAAAGAGACUCAGCGCAGUUAUUGAUGAAACAAUGACAGUAGAAGGUGACCUUCGUAGAGAAAUCGCCCUCAACAUUAAGAGACUGCAGGAAAUUGGAUGCUACAGAGG
env-288   CGCAGACAAAAUUCUUGCUAAAGCAGAGGUUAAUCCUGACACACGUGUCAGAGAUCUGACUGAUGACGAGGUUAAGAGACUGAGUGCUGUUAUUGAUGAGACAAUGACAGUAGAAGGUGAUCUGAGAAGGGAAAUCGCCCUUAACAUCAAGAGACUGCAGGAGAUUGGAUGCUACAGAGG
env-289   CGCAGACAAGAUCCUUGCAAAGGCAGAAGUUAAUCCUGACACACGUGUCAGAGAUCUGACUGAUGACGAAGUUAAGAGACUUAGCGCAGUUAUUGAUGAAACAAUGACAGUAGAAGGUGACCUUCGUAGAGAAAUCGCCCUCAACAUUAAGAGACUGCAGGAAAUCGGAUGCUACAGAGG
env-290   ..............................................................................................................................................................................................
env-291   CGCAGACAAGAUCCUUGCAAAGGCAGAAGUUAAUCCUGACACACGUGUCAGAGAUCUGACUGAUGACGAAGUUAAGAGACUUAGCGCAGUUAUUGAUGAAACAAUGACAGUAGAAGGUGACCUUCGUAGAGAAAUCGCCCUCAACAUUAAGAGACUGCAGGAAAUCGGAUGCUACAGAGG
env-292   CGCAGAUAAGAUCCUUGCAAAGGCAGAAGUUAAUCCUGACACACGUGUCAGAGAUCUGACUGAUGACGAAGUUAAGAGACUCAGCGCAGUUAUUGAUGAAACAAUGACAGUAGAAGGUGACCUUCGUAGAGAAAUCGCCCUCAACAUUAAGAGACUGCAGGAAAUCGGAUGCUACAGAGG
env-293   ..............................................................................................................................................................................................
env-294   ..............................................................................................................................................................................................
env-295   CGCAGACAAAAUCCUUGCAAAGGCAGAAGUUAAUCCUGACACACGUGUCAGAG...........................................................................................................................................
env-296   CGCAGACAAAAUCCUUGCAAAGGCAGAAGUUAAUCCUGACACACGUGUCAGAGAUCUGACUGAUGACGAAGUAAAGAGACUCAGCGCAGUUAUUGAUGAAACAAUGACAGUAGAAGGUGACCUUCGUAGAGAAAUCGCCCUCAACAUUAAGAGACUGCAGGAAAUCGGAUGCUACAGAGG
env-298   CGCAGACAAGAUCCUUGCAAAGGCAGAAGUUAAUCCUGACACACGUGUCAGAGAUCUGACUGAUGACGAAGUAAAGAGACUCAGCGCAGUUAUUGAUGAAACAAUGACAGUAGAAGGUGACCUUCGUAGAGAAAUCGCCCUCAACAUUAAGAGACUGCAGGAAAUCGGAUGCUACAGAGG
env-299   CGCAGACAAAAUCCUUGCAAAGGCAGAAGUUAAUCCUGACACACGUGUCAGAGAUCUGACUGAUGACGAAGUAAAGAGACUCAGCGCAGUUAUUGAUGAAACAAUGACAGUAGAAGGUGACCUUCGUAGAGAAAUCGCCCUCAACAUUAAGAGACUGCAGGAAAUCGGAUGCUACAGAGG
env-300   CGCAGACAAGAUCCUUGCAAAGGCAGAAGUUAAUCCUGACACACGUGUCAGAGAUCUGACUGAUGACGAAGUAAAGAGACUCAGCGCAGUUAUUGAUGAAACAAUGACAGUAGAAGGUGACCUUCGUAGAGAAAUCGCCCUCAACAUUAAGAGACUGCAGGAAAUCGGAUGCUACAGAGG
env-301   CGCAGACAAAAUCCUUGCAAAGGCAGAGGUUAAUCCUGACACACGUGUCAGAGAUCUGACUGAUGACGAAGUAAAGAGACUCAGCGCAGUUAUUGAUGAAACAAUGACAGUAGAAGGUGACCUUCGUAGAGAAAUCGCCCUCAACAUUAAGAGACUGCAGGAAAUCGGAUGCUACAGAGG
env-302   CGCAGACAAGAUCCUUGCAAAGGCAGAAGUUAAUCCUGACACACGUGUCAGAGAUCUGACUGAUGACGAAGUAAAGAGACUCAGCGCAGUUAUUGAUGAAACAAUGACAGUAGAAGGUGACCUUCGUAGAGAAAUCGCCCUCAACAUUAAGAGACUGCAGGAAAUUGGAUGCUACAGAGG
env-303   ..............................................................................................................................................................................................
env-304   CGCAGACAAAAUCCUUGCAAAGGCAGAAGUUAAUCCUGACACACGUGUCAGAG...........................................................................................................................................
env-305   ..............................................................................................................................................................................................
Bwe-2-1   CGCAGACAAAAUCCUUGCAAAGGCAGAGGUUAAUCCUGACACACGUGUCAGAGAUCUGACUGAUGACGAAGUAAAGAGACUCAGCGCAGUUAUUGAUGAAACAAUGACAGUAGAAGGUGACCUUCGUAGAGAAAUCGCCCUCAACAUUAAGAGACUGCAGGAAAUCGGAUGCUACAGAGG
env-306   CGCAGACAAGAUCCUUGCAAAGGCAGAAGUUAAUCCUGACACACGUGUCAGAGAUCUGACUGAUGACGAAGUAAAGAGACUCAGCGCAGUUAUUGAUGAAACAAUGACAGUAGAAGGUGACCUUCGUAGAGAAAUCGCCCUCAACAUUAAGAGACUGCAGGAAAUUGGAUGCUACAGAGG
env-307   CGCAGACAAAAUCCUUGCAAAGGCAGAAGUUAAUCCUGACACACGUGUCAGAGAUUUGACUGAUGACGAAGUAAAGAGACUCAGCGCAGUUAUUGAUGAAACAAUGACAGUAGAAGGUGACCUUCGUAGAGAAAUCGCCCUCAACAUUAAGAGACUGCAGGAAAUCGGAUGCUACAGAGG
env-308   ..............................................................................................................................................................................................
env-309   CGCAGACAAAAUCCUUGCAAAGGCAGAAGUUAAUCCUGACACACGUGUCAGAGAUCUGACUGAUGACGAAGUAAAGAGACUCAGCGCAGUUAUUGAUGAAACAAUGACAGUAGAAGGUGACCUUCGUAGAGAAAUCGCCCUCAACAUUAAGAGACUGCAGGAAAUCGGAUGCUACAGAGG
env-310   ..............................................................................................................................................................................................
env-311   CGCAGACAAAAUCCUUGCAAAGGCAGAGGUUAAUCCUGACACACGUGUCAGAGAUCUGACUGAUGACGAAGUAAAGAGACUCAGCGCAGUUAUUGAUGAAACAAUGACAGUAGAAGGUGACCUUCGUAGAGAAAUCGCCCUCAACAUUAAGAGACUGCAGGAAAUCGGAUGCUACAGAGG
env-313   CGCAGACAAGAUCCUUGCAAAGGCAGAGGUUAAUCCUGACACACGUGUCAGAGAUCUGACUGAUGACGAAGUAAAGAGACUCAGCGCAGUUAUUGAUGAAACAAUGACAGUAGAAGGUGACCUUCGUAGAGAAAUCGCCCUCAACAUUAAGAGACUGCAGGAAAUCGGAUGCUACAGAGG
env-314   CGCAGACAAGAUCCUUGCAAAGGCAGAAGUUAAUCCUGACACACGUGUCAGAGAUCUGACUGAUGACGAAGUAAAGAGACUCAGCGCAGUUAUUGAUGAAACAAUGACAGUAGAAGGUGACCUUCGUAGAGAAAUCGCCCUCAACAUUAAGAGACUGCAGGAAAUCGGAUGCUACAGAGG
env-315   CGCAGACAAAAUCCUUGCAAAGGCAGAAGUUAAUCCUGACACACGUGUCAGAGAUCUGACUGAUGACGAAGUAAAGAGACUCAGCGCAGUUAUUGAUGAAACAAUGACAGUAGAAGGUGACCUUCGUAGAGAAAUCGCCCUCAACAUUAAGAGACUGCAGGAAAUCGGAUGCUACAGAGG
env-317   ..............................................................................................................................................................................................
env-318   ..............................................................................................................................................................................................
Bwe-1-1   CGCAGACAAGAUCCUUGCAAAGGCAGAAGUUAAUCCUGACACACGUGUCAGAGAUCUGACUGAUGACGAAGUAAAGAGACUCAGCGCAGUUAUUGAUGAAACAAUGACAGUAGAAGGUGACCUUCGUAGAGAAAUCGCCCUCAACAUUAAGAGACUGCAGGAAAUCGGAUGCUACAGAGG

          ..............................................................................................................................................................................................
          ..............................................................................................................................................................................................
          Y•C•R•Y•R•AUYYU•R•RR•RGCRRR•GU•RAYCC•RAYAY•CG•••YAGRGAYYU•AC•GAYGA•GARGU•RA•ARRAUY•RYR••GY•AUYGA•GAR•Y••••RY•RURGAAGR•GAY•U•••••G•GRRR•YRY••Y•A••A•YARR•R•YY•Y•R•ARR••RR••GCURY•R•RG
```

| | |
|---|---|
| env-319 | CGCAGACAAAAUCCUUGCAAAGGCAGAGGUUAAUCCUGACACACGUGUCAGAGAUCUGACUGAUGACGAAGUAAAGAGACUCAGCGCAGUUAUUGAUGAAACAAUGACAGUAGAAGGUGACCUUCGUAGAGAAAUCGCCCUCAACAUUAAGAGACUGCAGGAAAUCGGAUGCUACAGAGG |
| env-320 | CGCAGACAAAAUCCUUGCAAAGGCAGAAGUUAAUCCUGACACACGUGUCAGAGAUCUGACUGAUGACGAAGUAAAGAGACUCAGCGCAGUUAUUGAUGAAACAAUGACAGUAGAAGGUGACCUUCGUAGAGAAAUCGCCCUCAACAUUAAGAGACUGCAGGAAAUCGGAUGCUACAGAGG |
| env-321 | CGCAGACAAAAUCCUUGCAAAGGCAGAAGUUAAUCCUGACACACGUGUCAGAGAU... |
| env-322 | CGCAGACAAGAUCCUUGCAAAGGCAGAAGUUAAUCCUGACACACGUGUCAGAGAUCUGACUGAUGACGAAGUAAAGAGACUCAGCGCAGUUAUUGAUGAAACAAUGACAGUAGAAGGUGACCUUCGUAGAGAAAUCGCCC... |
| env-323 | CGCAGACAAGAUCCUUGCAAAGGCAGAAGUUAAUCCUGACACACGUGUCAGAGAUCUGACUGAUGACGAAGUAAAGAGACUCAGCGCAGUUAUUGAUGAAACAAUGACAGUAGAAGGUGACCUUCGUAGAGAAAUCGCCCUCAACAUUAAGAGACUGCAGGAAAUUGGAUGCUACAGAGG |
| env-324 | ... |
| env-325 | CGCAGACAAAAUCCUUGCAAAGGCAGAAGUUAAUCCUGACACACGUGUCAGAGAUCUGACUGAUGACGAAGUAAAGAGACUCAGCGCAGUUAUUGAUGAAACAAUGACAGUAGAAGGUGACCUUCGUAGAGAAAUCGCCCUCAACAUUAAGAGACUGCAGGAAAUUGGAUGCUACAGAGG |
| env-326 | CGCAGACAAAAUCCUUGCAAAGGCAGAGGUUAAUCCUGACACACGUGUCAGAGAUCUGACUGAUGACGAAGUAAAGAGACUCAGCGCAGUUAUUGAUGAAACAAUGACAGUAGAAGGUGACCUUCGUAGAGAAAUCGCCCUCAACAUUAAGAGACUGCAGGAAAUCUUAUGCUACAGAGG |
| env-327 | CGCAACUCAGAUCCUUGAG... |
| env-328 | UGCAGACAAGAUCCUUGCAC... |
| env-329 | UGCAGACAAAAUCCUUGCUCAGGCAAAUGUAAAUCCUGACACACGUGUCAGAGACCUGACUGAUGAUGAAGUUAAGAGACUUCAGGCUGUCAUUGAUGAGACAAUGAUGGUAGAAGGUGAUCUUCGUAGAGAAAUCGCCCUGAACAUUAAGAGACUGCAGGAAAUCGGAUGCUACAGAGG |
| env-331 | UGCAGACAAAAUCCUUGCUCAGGCAAAUGUAAAUCCUGACACACGUGUCAGAGACCUGACUGAUGAUGAAGUUAAGAGACUUCAGGCUGUCAUUGAUGAGACAAUGAUGGUAGAAGGUGAUCUUCGUAGAGAAAUCGCCCUGAACAUUAAGAGACUGCAGGAAAUCGGAUGCUACAG... |
| env-332 | UGCAGACAAAAUCCUUGCUCAGGCAAAUGUAAAUCCUGACACACGUGUCAGAGACCUGACUGAUGAUGAAGUUAAGAGACUUCAGGCUGUCAUUGAUGAGACAAUGAUGGUAGAAGGUGAUCUUCGUAGAGAAAUCGCCCUGAACAUUAAGAGACUGCAGGAAAUCGGAUGCUACAGAGG |
| env-335 | UGCAGACAAAAUCCUUGCUCAGGCAAAUGUAAAUCCUGACACACGUGUCAGAGACCUGACUGAUGAUGAAGUUAAGAGACUUCAGGCUGUCAUUGAUGAGACAAUGAUGGUAGAAGGUGAU... |
| env-336 | UGCAGACAAAAUCCUUGCUCAGGCAAAUGUAAAUCCUGACACACGUGUCAGAGACCUGACUGAUGAUGAAGUUAAGAGACUUCAGGCUGUCAUUGAUGAGACAAUGAUGGUAGAAGGUGAUCUUCGUAGAGAAA... |
| env-337 | UGCAGACAAAAUCCUUGCUCAGGCAAAUGUAAAUCCUGACACACGUGUCAGAGACCUGACUGAUGAUGAAGUUAAGAGACUUCAGGCUGUCAUUGAUGAGACAAUGAUGGUAGAAGGUGAUCUUCGUAGAGAAAUCGCCCUGAACAUUAAGAGACUGCAGGAAAUCGGAUGCUACAGAGG |
| env-338 | UGCAGACAAAAUCCUUGCUCAGGCAAAUGUAAAUCCUGACACACGUGUCAGAGACCUGACUGAUGAUGAAGUUAAGAGACUUCAGGCUGUCAUUGAUGAGACAAUGAUGGUAGAAGGUGAUCUUCGUAGAGAAAUCGCCCUGAACAUUAAGAGACUGCAGGAAAUCGGAUGCUACAGAGG |
| env-339 | UGCAGACAAAAUCCUUGCUCAGGCAAAUGUAAAUCCUGACACACGUGUCAGAGACCUGACUGAUGAUGAAGUUAAGAGACUUCAGGCUGUCAUUGAUGAGACAAUGAUGGUAGAAGGUGAUCUUCGUAGAGAAAUCGCCCUGAACAUUAAGAGACUGCAGGAAAUCGGAUGCUACAGAGG |
| env-340 | UGCAGACAAAAUCCUUGCUCAGGCAAAUGUAAAUCCUGACACACGUGUCAGAGACCUGACUGAUGAUGAAGUUAAGAGACUUCAGGCUGUCAUUGAUGAGACAAUGAUGGUAGAAGGUGAUCUUCGUAGAGAAAUCGCUCUCAACAUCAAGAGAUUACAGGAAAUCGGAU... |
| env-341 | UGCAGACAAAAUCCUUGCUCAGGCAAAUGUAAAUCCUGACACACGUGUCAGAGACCUGACUGAUGAUGAAGUUAAGAGACUUCAGGCUGUCAUUGAUGAGACAAUGAUGGUAGAAGGUGAUCUUCGUAGAGAAAUCGCCCUGAACAUUAAGAGACUGCAGGAAAUCG... |
| env-342 | UGCAGACAAAAUCCUUGCUCAGGCAAAUGUAAAUCCUGACACACGUGUCAGAGACCUGACUGAUGAUGAAGUUAAGAGACUUCAGGCUGUCAUUGAUGAGACAAUGAUGGUAGAAGGUGAUCUUCGUAGAGAAAUCGCCCUGAACAUUAAGAGACUGCAGGAAAUCGGAUGCUACAGAGG |
| Bsc-1-1 | UGCAACAAAAAUUCUUGCUCAGGCAAAUGUAAAUCCUGACACACGUGUCAGAGAUUUAACCGACGAAGAAGUAAACAGAAUCCGUGAGGUUAUUGACGAUUCAAUGACCGUAGAAGGUGAUCUUCGUAGAGAAAUCGCUCUGAACAUUAAGAGACUGCAGGAAAUCGGAUGCUACAGAGG |
| Rsp-3-1 | CGCUGACAAAAUUCUUGCAAAGGCUGAGGUUAAUCCUGACACACGUGUCAGAGAUCUGACUGAUGACGAAGUAAAGAGACUGAGCGCAGUAAUUGAUGAAACAAUGACAGUAGAAGGUGACCUUCGUCGUGAAAUCGCCCUCAACAUUAAGAGACUGCAGGAAAUCGGAUGCUACAGAGG |
| env-343 | UUCCAACCGCAUCCUUGCGCAGGCCGGCGUAAACCCGGAUACCCGUGUCAGAGACCUGACAGACGAUGAAGUCAAGAAGAUCGCGUCUGUUAUCGAUGAGACACAGGUGGUGGAAGGCGAUCUGAGACGUGAGAUUGCCAUGAAUAUCAAACGUCUCCAGGAGAUUGGCUGCUACCGCGG |
| env-344 | UUCAAACCGUAUUCUUGAAAAGGCAGGCGUAAAUCCUGACACUCGUGUCAGAGACGUUACAGACGAAGAAGUAGGAAAAAUCCGUGAUGCUAUCGAAGAAUUACAGAUUCCGGUAGAAGGUGAUUUAAGAAGAGAAAUCGCUUUAAACAUCAAGAGACUUCAGGAAAUCGGAUGUUAUAG |
| env-346 | UUCAAACCGUAUUCUUGAAAAGGCAGGCGUAAAUCCUGACACUCGUGUCAGAGACGUUACAG... |
| env-348 | ... |
| env-349 | ... |
| env-350 | UUCAAACCGUAUUCUUGAAAAGGCAGGCGUAAAUCCUGACACUCGUGUCAGAGACGUUACAGACGAAGAAGUAGGAAAAAUCCGUGAUGCUAUCGAAGAAUUACAGAUUCCGGUAGAAGGUGAUUUAAGAAGAGAAAUCGCUUUAAACAUCAAGAGACUUCAGGAAAUCGGAUGUUAUAG |
| env-351 | ... |
| env-352 | UUCAAACCGUAUUCUUAGAAAAGGCAGGCGUAAAUCCUGACACUCGUGUCAGAGAYGUUACWGACGAAGAAGUAGGAAAAAUCCGUGAYGCWAUYGAAGARCUUACAGAUUCCKGUAGAAGGUGAUUUGAGAAGAGAAAUCGCUCUGAACAUCAAAAGACUUCAGGAAAUCGGCUGCUA |
| env-353 | UGCUGACCGCAUCCUCAAAGAAGCAAAUGUGAAUCCGGAUACUCGAGUAAGAGAUUUGACUGACGAAGAAGUAAAAAGAAUCAGCGUAGUAAUCGAAGAGACACAGACGGUGGAAGGUGAUCUGAGAAGAGAGAUCGCACUGAACAUCAAACGUCUGCAGGAAAUCGGAUGCUACAGAGG |
| Rga-1-1 | UGCUGACCGCAUCCUGAAAGAAGCAAAUGUGAAUCCGGAUACUCGAGUAAGAGAUUUAACUGACGAAGAAGUAAAGAGAAUCAGCGUAGUAAUCGAAGAGACCCAGACAGUGGAAGGUGAUCUGAGAAGAGAGAUCGCUCUGAAUAUCAAACGUCUGCAGGAAAUCGGAUGCUACAGAGG |
| env-354 | ... |
| env-355 | UGCUAAACGUAUUGUUGCAGAUGCAAAAGUAAAUCCUGAUACAAGGGUCAGAGAUUUGACCGAUGACGAAGUUAAGAGACUUGCAGGAGUAAUUGACGAAACACAACAAGUAGAAGGUGAUUUAAGAAGAGAAAUUGCGAUGAACAUCAAGCGUCUUCAAGAAAUCGGAUGCUACCGUGG |
| env-356 | UGC... |
| env-357 | UGCUAAACGUAUUAUUGCAGAUGCAGGUGUUAAUCCAGAUACUAGAGUUAGAGACUUAACUGAUGAUGAAGUUAAGAAGCUAGCAGGAGUAAUUGACGAAACCCAACAAGUAGAGGGUGAUUUAAGAAGAGAAAUCGCAAUGAACAUUAAGCGUCUUCAAGAAAUCGGAUGCUACCGUGG |
| env-358 | UGCUAAACGUAUUAUUGCAGAUGCAGGUGUUAAUCCAGAUACUAGAGUUAGAGACUUAACUGAUGAUGAAGUUAAGAAGCUAGCAGGAGUAAUUGACGAAACCCAACAAGUAGAGGGUGAUUUAAGAAGAGAAAUCGCAAUGAACAUUAAGCGUCUUCAAGAAAUCGGAUGCUACCGUGG |
| env-359 | UGCUAAACGUAUUAUUGCAGAUGCAGGUGUUAAUCCAGAUACUAGAGUUAGAGACUUAACUGAUGAUGAAGUUAAGAAGCUAGCAGGAGUAAUUGACGAAACUCAACAAGUAGAGGGUGAUUUAAGAAGAGAAAUCGCAAUGAACAUUAAGCGUCUUCAAGAAAUCGGAUGCUACCGUGG |
| env-360 | UGCUAGACGUAUUGUAGCAGAAGCUAAAGUAAAUCCAGAUACUAGAGUAAGAGAUUUAACUGAUGAUGAAGUUAAAAGACUUGCGGGAGUAAUUGACGAAACACAACAAGUAGAAGGUGAUUUAAGAAGAGAAAUCGCAAUGAAUAUCAAGCGACUUCGUGAAAUCGGAUGCUACAGAGG |
| env-361 | UGCUAGACGUAUUGUAGCAGAAGCUAAAGUAAAUCCAGAUACUAGAAUAAGAGACUUAACUGAUGAUGAAGUUAAGAAACUUGCUGACGUAAUUGACGAAACACAACAAGUAGAAGGUGAUUUAAGAAGAGAAAUCGCGAUGAACAUCAAGCGACUUCGUGAAAUCGGUUGCUACCGUGG |
| env-362 | UGCUAGACGUAUUGUAGCAGAAGCUAAAGUAAAUCCAGAUACUAGAAUAAGAGACUUAACUGAUGAUGAAGUUAAGAAACUUGCUGACGUAAUUGACGAAACACAACAAGUAGAAGGUGAUUUAAGAAGAGAAAUCGCGAUGAACAUCAAGCGACUUCGUGAAAUCGGUUGCUACCGUGG |
| env-363 | ... |
| env-364 | UGCUAAACGUAUCGUAGCAGAAGCUAAAGUAGAUCCAGAUACUAGAGUAAGAGACUUAACUGAUGAUGAAGUUAAGAGACUUGCAGGCGUAAUUGACGAAACCCAACAAGUAGAAGGUGAUUUACGAAGAGAAAUCGCGAUGAAUAUCAAGCGUCUUCGUGAAAUCGGAUGCUACCGUGG |
| env-365 | UGCUAAACGUAUUGUAGCAGAAGCUAAAGUGAAUCCAGAUACUAGAGUAAGAGACUUGACUGAUGAUGAAGUUAAGAGACUUG... |
| env-366 | UGCUAAACGUAUCGUAGCAGAAGCUAAAGUAGAUCCAGAUACUAGAGUAAGAGACUUAACUGAUGAUGAAGUUAAGAGACUUGCAGGCGUAAUUGACGAAACCCAACAAGUAGAAGGUGAUUUACGAAGAGAAAUCGCGAUGAAUAUCAAGCGUCUUCGUGAAAUCGGAUGCUACCGUGG |
| env-367 | UGCAGAUCGUAUCUUAAAAGAAGCUGGAGUAAAUCCGGACACACGCUGCAGAGAUUUGACAGACGAUGAAGUAAAGAGACUCAGUGCAGUUAUCGAUGAGAGUCAGGUUGUAGAAGGAGAUCUUCGUAGAGAAAUCGCUAUGAACAUCAAACGU... |
| env-368 | UGCAGAUCGUAUCUUAAAAGAGGCUGGAGUAAAUCCGGACACACGCUGCAGAGAUUUGACAGACGAUGAAGUAAAGAGACUCAGCGCAGUUAUCGAUGAGAGUCAGGUUGUAGAAGGAGAUCUUCGUAGAGAAAUCGCUAUGAACAUCAAACGUCU... |
| env-369 | UGCAGAUCGUAUCUUAAAAGAAGCUGGAGUAAAUCCGGACACACGCUGCAGAGAUUUGACAGACGAUGAAGUAAAGAGACUCAGUGCAGUUAUCGAUGAGAGUCAGGUUGUAGAAGGAGAUCUUCGUAGAGAAAUCGCUAUGAACAU... |
| env-370 | UGCAGAUCGUAUCUUAAAAGAAGCUGGAGUAAAUCCGGACACACGCUGCAGAGAUUUGACAGACGAUGAAGUAAAGAGACUCAGUGCAGUUAUCGAUGAGAGUCAGGUUGUAGAAGGAGAUCUUCGUAGAGAAAUCGCUAUGAACAU... |
| env-371 | UGCAGAUCGUAUCUUAAAAGAAGCUGGAGUAAAUCCGGACACACGCUGCAGAGAUUUGACAGACGAUGAAGUAAAGAGACUCAGCGCAGU... |
| env-372 | ... |
| env-373 | UGCAGAUCGUAUCUUAAAAGAAGCUGGAGUAAAUCCGGACACACGCUGCAGAGAUUUGACAGACGAUGAAGUAAAGAGACUCAGCGCAGUUAUCGAUGAGAGUCAGGUUGUAGAAGGAGAUCUUCGUAGAGAAAUCGCUAUGAACAUCAAACGUCUGCAGGAGAUCGGAUGCUACAGAGG |
| env-374 | UGCAGAUCGUAUCUUAAAAGAAGCUGGAGUAAAUCCGGACACACGCUGCAGAGAUUUGACAGACNNNNNNNNNNNNNNNNNNNNNNNNNNNNNNNNNNNNNNNNNNNNNNNNNNNNNNNNNNNNNNNNNNNNNNNNNNNNNNNNNNNNNNNNNNNNNNNNNNNNNNNNNNNNNNNNNNNNNNNNNNNNNNNNNNN |
| env-375 | UGCAGAUCGUAUCUUAAAAGAAGCUGGAGUAAAUCCGGACACACGCUGCAGAGAUUUGACAGACGAUGAAGUAAAGAGACUCAGCGCAGUUAUCGAUGAGAGUCAGGUUGUAGAAGGAGAUCUUCGUAGAGAAAUCGCUAUGAACAUCAAACGUCUGCAGGAGAUCGGAUGCUACAGAGG |
| env-377 | ... |
| env-383 | ... |
| env-384 | UGCAGAUCGUAUCUUAAAAGAAGCUGGAGUAAAUCCGGACACACGCUGCAGAGAUUUGACAGACGAUGAAGUAAAGAGACUCAGCGCAGUUAUCGAUGAGAGUCAGGUUGUAGAAGGAGAUCUUCGUAGAGAANNNNNNNNNNNNNNNNNNNNNNNNNNNNNNNNNNNNNNNNNNNNNAA |
| env-385 | ... |
| env-386 | UGCAGAUCGUAUCUUAAAAGAAGCUGGAGUAAAUCCGGACACACGCUGCAGAGAUUUGACAGACGAUGAAGUAAAGAGACUCAGCGCAGUUAUCGAUGAGAGUCAGGUUGUAGAAGGAGAUCUUCGUAGAGAAAUCGCUAUGAACAUCAAACGUCUGCAGGAGAUCGGAUGCUACAGAGG |
| env-387 | UGCAGAUCGUAUCUUAAAAGAAGCUGGAGUAAAUCCGGACACACGCUGCAGAGAUUUGACAGACGAUGAAGUAAAGAGACUCAGCGCAGUUAUCGAUGAGAGUCAGGUUGUAGAAGGAGAUCUUCGUAGAGAAAUCGCUAUGAACAUCAAACGUCUGCAGGAGAUCGGAUGCUACAGAGG |
| env-388 | UGCAGAUCGUAUCUUAAAAGAAGCUGGAGUAAAUCCGGACACACGCUGCAGAGAUUUGACAGACGAUGAAGUAAAGAGACUCAGCGCAGUUAUCGAUGAGAGUCAGGUUGUAGAAGGAGAUCUUCGUAGAGAAAUCGCUAUGAACAUCAAACGUCUGCAGGAGAUCGGAUGCUACAGAGG |
| env-389 | ... |
| env-390 | UGCAGAUCGUAUCUUAAAAGAAGCUGGAGUAAAUCCGGACACACGCUGCAGAGAUUUGACAGACGAUGAAGUAAAGAGACUCAGCGCAGUUAUCGAUGAGAGUCAGGUUGUAGAAGGAGAUCUUCGUAGAGAAAUCGCUAUGAACAUCAAACGUCUGCAGGAGAUCGGAUGCUACAGAGG |
| env-391 | UGCAGAUCGUAUCUUAAAAGAAGCUGGAGUAAAUCCGGACACACGCUGCAGAGAUUUGACAGACGAUGAAGUAAAGAGACUCAGCGCAGUUAUCGAUGAGAGUCAGGUUGUAGAAGGA... |
| env-392 | ... |
| env-393 | UGCAGAUCGUAUCUUAAAAGAAGCUGGAGUAAAUCCGGACACACGCUGCAGAGAUUUGACAGACGAUGAAGUAAAGAGACUCAGUGCAGUUAUCGAUGAGAGUCAGGUUGUAGAAGGAGAUCUUCGUAGAGAAAUCGCUAUGAACAUCAAACGUCUGCAGGAGAUCGGAUGCUACAGAGG |
| env-394 | UGCAGAUCGUAUCUUAAAAGAAGCUGGAGUAAAUCCGGACACACGCUGCAGAGAUUUGACAGACGAUGAAGUAAAGAGACUCAGUGCAGUUAUCGAUGAGAGUCAGGUUGUAGAAGGAGAUCUUCGUAGAGAAAUCGCUAUGAACAUCAA... |
| env-395 | ... |
| env-396 | UGCAGAUCGUAUCUUAAAAGAAGCUGGAGUAAAUCCGGACACACGCUGCAGAGAUUUGACAGACGAUGAAGUAAAGAGACUCAGUGCAGUUAUCGAUGAGAGUCAGGUUGUAGAAGGAGAUCUUCGUAGAGAAAUCGCUAUGAACAUCAA... |
| env-397 | UGCAGAUCGUAUCUUAAAAGAAGCUGGAGUAAAUCCGGACACACGCUGCAGAGAUUUGACAGACGAUGAAGUAAAGAGACUCAGUGCAGUUAUCGAUGAGAGUCAGGUUGUAGAAGGAG... |
| env-398 | UGCAGAUCGUAUCUUAAAAGAAGCUGGAGUAAAUCCGGACACACGCUGCAGAGAUUUGACAGACGAUGAAGUAAAGAGACUCAGUGCAGU... |
| env-399 | UGCAGAUCGUAUCUUAAAAGAAGCUGGAGUAAAUCCGGACACACGCUGCAGAGAUUUGACAGACGAUGAAGUAAAGAGACUCAGUGCAGUUAUCGAUGAGAGUCAGGUUGUAGAAGGAGAUCUUCGUAGAGAAAUCGCUAUGAACAUCAAACGUCUGCAGGAGAUCGGAUGCUACAGAGG |
| env-405 | UGCAGAUCGUAUCUUAAAAGAAGCUGGAGUAAAUCCGGACACACGCUGCAGAGAUUUGACAGACGAUGAAGUAAAGAGACUCAGCGCAGUUAUCGAUGAGAGUCAGGUUGUAGAAGGAGAUCUUCGUAGAGAAAUCGCUAUGAACAUCAAACGUCUGCAGGAGAUCGGAUGCUACAGAGG |
| env-408 | UGCAGAUCGUAUCUUAAAAGAGGCUGGAGUAAAUCCAGAUACACGCUG... |
| env-409 | ... |
| env-410 | ... |
| env-411 | UGCAGAUCGUAUCUUAAAAGAAGCUGGAGUAAAUCCGGACACACGCUGCAGAGAUUUGACAGACGAUGAAGUAAAGAGACUCAGCGCAGUUAUCGAUGAGAGUCAGGUUGUAGAAGGAGAUCUUCGUAGAGAANNNNAGAAAACAAAGACCAACGCAAGAACCCGUAAAGGUCCUAAGA |
| env-412 | UGCAGAUCGUAUCUUAAAAGAAGCUGGAGUAAAUCCGNNNNNNNNNNNNNNNNNNNNNNNNNNNNNNNNNNNNNNNNNNNNNNNNNNNNNNNNNNNNNNNNNNNNNNNNNNNNNNNNNUUUGACAGACGAUGAAGUAAAGAGACUCAGUGCAGUUAUCGAUGAGAGUCAGGUUGUAGAAG |
| env-413 | UGCAGAUCGUAUCUUAAAAGAAGCUGGAGUAAAUCCGGACACACGCUGCAGAGAUUUGACAGACGAUGAAGUAAAGAGACUCAGCGCAGUUAUCGAUGAGAGUCAGGUUGUAGAAGGAGAUCUUCGUAGAGAAAUCGCUAUGAACAUCAAACGUCUGCAGGAGAUCGGAUGCUACAGAGG |
| env-414 | UGCAGAUCGUAUCUUAAAAGAGGCUGGAGUAAAUCCAGAUACACGCUGCAGAGAUUUGACAGACGAUGAAGUAAAGAGACUCAGCGCAGUUAUCGAUGAGACUCAGGUUGUAGAGGGAGAUCUUCGUAGAGAAAUCGCUAUGAACAUCAAACGUCUGCAGGAGAUCGGAUGCUACAGAGG |
| env-418 | ... |
| env-419 | AUUUGACAGACGAUGAAGUAAAGAGACUCAGUGCAGUUAUCGAUGAGAGUCAGGUUGUAGAAGGAGAUCUUCGUAGAGAAAUCGCUAUGAACAUCAAACGUCUGC... |
| env-420 | ... |
| | ... |
| | ... |
| | Y•C•R•Y•R•AUYYU•R•RR•RGCRRR•GU•RAYCC•RAYAY•CG•••YAGRGAYYU•AC•GAYGA•GARGU•RA•ARRAUY•RYR••GY•AUYGA•GAR•Y••••RY•RURGAAGR•GAY•U•••••G•GRRR•YRY••Y•A••A•YARR•R•YY•Y•R•ARR••RR••GCURY•R•RG |

| | |
|---|---|
| env-428 | ........................................................................................................................................................................................................ |
| env-429 | ........................................................................................................................................................................................................ |
| env-430 | ........................................................................................................................................................................................................ |
| env-432 | CGCUACCCGUAUUCUGGAAGCAGCUAAUGUAGAUCCGAACACCCGUGUUAGAGAUAUCACGGACGAAGAAGUAAAAAGAAUCAGUGCUGUAAUCGAUGAGACUCAGACUGUAG.................................................................................... |
| env-433 | UUCCAACAAGA............................................................................................................................................................................................ |
| env-434 | ........................................................................................................................................................................................................ |
| Bhy-1-1 | UGCAACUCGCAUCCUGGCUGAUGCGAAAGUAAAUCCUGACACUCGUGUUAGAGAUUUGACAGACGAAGAAGUAAACAGAAUCCGUGAAGUGAUUGAUGAGACUAUGAUGGUGGAAGGUGAUCUGAGAAGAGAGAUCGCUCUGAAUAUCAAGAGACUUCAGGAGAUUGGAUGCUACAGAGG |
| env-435 | UUCCAAUCGUAUCCUUGCGCAGGCCGGCGUUAACCCGGAUACCCGUGUCAGAGAUCUGACCGAUGACGAGGUCAAGAAGAUCGCUGGCGUGAUCGAAGAGACCCAGGUCGUAGAAGGUGAUCUGAGACGUGAAGUUGCUAUGAACAUCAAGCGUCUUCAGGAGAUCGGCUGCUACCGCGG |
| env-436 | UUCCAAUCGUAUCCUUGCGCAGGCUGGAGUAAAUCCGGAUACACGUGUGAGAGAUCUGACAGAUGAAGAAGUAAAGAAAAUCGCCGGAGUAAUCGACGAGACUCAGACGGUAGAAGGUGAUCUGAGACGUCAGGUUGCCAUGGAUAUCAAACGUCUUCAGGAGAUCGGAUGCUAUCGUGG |
| env-437 | UUCCAACCGUAUUCUGGCUGACGCAGGUGUUGACCCGAAUACACGCGUUCGUGAUCUUACGGACGACGAAGUUAAAAAGAUCGCUUCCGUGAUCGAAGAGACACAGGUUGUAGAAGGUGAUCUUCGCAGACAGACCGCUAUGGAUAUCAAGCGUCUGAUGGAGAUCGGCUGCUACCGCGG |
| env-438 | UUCCAACAGGAUCCUCGCUGAUGCGGGAGUAGAUCCCAAUACCCGUGUCCGUGAUCUUACUGAUGAAGA.......................................................................................................................... |
| Bsp-7-1 | CUCAAACAAAAUACUUGCUGCUGCAAAUGUAAACCCCGACACACGUGUUCGUGAUCUUACAGAUGAUGAGGUUAAGAAGAUUGCUGAGAUCAUCGGUAACGAUUAUGACGUAGAAGGUGAUCUUAGAAGAGAGAUCGCUAUGAACAUCAAGCGUCUUUCAGAAAUCGGAUGCUACAGAGG |
| Bsp-9-1 | CUCAAACAAAAUUCUUGAGGCUGCAAAAGUAAACCCGGACACACGUGUUCGUGAUCUUACAGAUGAUGAAGUAAAGAGAAUCGCAGAGAUUAUCGGUAAUGAUUAUGAAGUAGAAGGUGAUCUUAGAAGAGAAAUCGCUAUGAACAUCAAGCGUCUUUCAGAGAUCGGAUGCUACAGAGG |
| Bsp-5-1 | CUCAAACAAAAUUCUUGAGGUUGCAAAAGUAAACCCGGACACACGUGUUCGUGAUCUUACAGAUGAUGAAGUAAAGAGAAUCGCAGAGAUUAUCGGUAAGGAUUAUGAAGUAGAAGGUGAUCUUAGAAGAGAAAUCGCUAUGAACAUCAAGCGUCUUUCAGAGAUCGGAUGCUACAGAGG |
| Bsp-2-1 | CUCAAACAAAAUUCUUGCUGCUGCAAAUGUAAAUCCCGACACACGUGUUCGUGAUCUCACAGAUGAUGAGGUUAAGAAGAUUGCUGAGAUUAUCGGUAAUGAUUAUGACGUAGAAGGUGAUCUUAGAAGAGAGAUCGCUAUGAACAUCAAGCGUCUUUCAGAGAUCGGAUGCUACAGAGG |
| Bsp-3-1 | CUCAAAUAAGAUACUUGCUGCUGCAAAUGUAAAUCCCGACACACGUGUUCGUAAUCUUACAGAUGAUGAAGUAAAGAAGAUUGCUGAGAUUAUCGGUAAUGACUAUGACGUAGAAGGUGAUCUUAGAAGAGAAAUCGCUAUGAACAUCAAGCGUCUUUCAGAAAUUGGAUGCUACAGAGG |
| Bpr-2-1 | CGCUGACCGUAUCUUAGUAGAGGCAGGUGUAAACCCUGACACUCGCUGCAGGGAUUUGACUGACGAAGAAGUAGGAAAAAUCCGUGAUGCAAUCGAUGAACUGAAUAUCGCUGUAGAAGGUGAUCUGAGAAGAGAAAUCGCUCUGAACAUCAAGAGAUUACAGGAAAUCGGAUGCUACAG |
| | ........................................................................................................................................................................................................ |
| | ........................................................................................................................................................................................................ |
| | Y•C•R•Y•R•AUYYU•R•RR•RGCRRR•GU•RAYCC•RAYAY•CG•••YAGRGAYYU•AC•GAYGA•GARGU•RA•ARRAUY•RYR••GY•AUYGA•GAR•Y••••RY•RURGAAGR•GAY•U•••••G•GRRR•YRY••Y•A••A•YARR•R•YY•Y•R•ARR••RR••GCURY•R•RG |

alignment positions 1081 · · · 1117

| | |
|---|---|
| env-1 | ..................................... |
| env-2 | ..................................... |
| env-3 | ..................................... |
| env-4 | ..................................... |
| env-5 | ..................................... |
| env-6 | ..................................... |
| env-7 | ..................................... |
| env-8 | ..................................... |
| env-9 | ..................................... |
| env-10 | UAUCCGUCAUAGAAAGGGUCUUCCUGUUCGUGGUCAG |
| env-11 | ..................................... |
| env-12 | UAUCCGUCAUAGAAAGGGUCUUCCUGUUCGUGGCCAG |
| env-13 | UAUCCGUCAUAGAAAGGGUCUUCCUGUUCGUGGUCAG |
| env-14 | ..................................... |
| env-16 | ..................................... |
| env-17 | AAUCCGUCACAGAAAAGGACUUCCGGUUCGCGGUCAG |
| env-18 | ..................................... |
| env-19 | AAUUCGUCAUAGAAAAGGACUUCCAGUUCGUGGUCAG |
| Bsp-6-1 | UAUUCGUCAUAGAAGAGGCCUUCCUGUUCGUGGUCAG |
| env-20 | CAUUCGUCACAGAAAAGGACUUCCGGUUCGCGGUCAG |
| env-21 | UGUUCGCCACAGAAGAGGACUUCCUGUUCGUGGACAG |
| env-22 | ..................................... |
| env-23 | UAUCCGUCACAGAAAAGGACUUCCUGUUCGCGGACAG |
| env-24 | ..................................... |
| env-25 | UAUCCGUCAUAGAAAAGGACUUCCAGUUCGUGGUCAA |
| env-28 | UAUCCGUCAUAGAAAAGGACUUCCAGUUCGUGGUCAA |
| env-29 | ..................................... |
| env-30 | UAUCCGUCAUAGAAAAGGACUUCCUGUUCGUGGUCAG |
| env-32 | UAUCCGUCAUAGAAAAGGACUUCCUGUUCGUGGUCAG |
| env-33 | UAUCCGUCAUAGAAAAGGACUUCCUGUUCGUGGUCAG |
| env-34 | ..................................... |
| env-35 | ..................................... |
| env-36 | ..................................... |
| env-37 | ..................................... |
| env-38 | ..................................... |
| env-39 | ..................................... |
| env-41 | ..................................... |
| env-42 | ..................................... |
| env-43 | AAUCCGCCACAGAAAAGGCCUUCCGGUGCGCGGCCAA |
| env-45 | UUCCGGCGAAUAAUAAAACGGGCUGCUUUUACAGACU |
| env-46 | ACAGUUGCCAAUAAGAAGAAAUAAUAAGCAAUAAUAA |
| env-47 | AUGUCCGCAAGAUCAGCACCGUCAUCGAGGAGACCAU |
| env-48 | AGGCAUCCGUCACAGGAAGGGCCUUCCGGUCCGCGGA |
| env-49 | UGUCCGCCACAGGAAAGGACUUCCUGUCCGCGGUCAG |
| env-50 | ..................................... |
| Dma-1-1 | AAUCCGUCAUAGAAAAGGACUUCCGGUUCGUGGACAG |
| env-51 | GAUUCGUCAUAGAAAAGGACUUCCUGUUCGCGGUCAG |
| Lba-2-1 | UAUCCGCCACAGAAGAGGACUUCCGGUUCGUGGACAG |
| [ce-1-1 | UGGCAAACGUCACAGAGUAGGACUUCCUGUUCGUGGA |
| Ece-1-1 | UGGCAAACGUCACAGAGUAGGACUUCCUGUUCGUGGA |
| env-52 | AAUCCGUCAUCGUAAAGGACUUCCUGUUCGUGGUCAG |
| env-53 | GAUCCGUCAUAGAAGAGGAUUGCCGGUUCGUGGUCAA |
| env-54 | ..................................... |
| env-55 | AAUCCGUCACAGAAAAGGACUUCCGGUACGUGGUCAG |
| env-59 | AAUCCGUCACAGAAAAGGACUUCCGGUACGUGGUCAG |
| env-60 | AAUCCGUCACAGAAAAGGACUUCCGGUACGUGGUCAG |
| env-62 | ..................................... |
| env-63 | ..................................... |
| | ..................................... |
| | ..................................... |
| | •R•Y•RYCAY•R•ARARGR•••CY•RU•CRYRG•YRR |

| | |
|---|---|
| env-64 | ..................................... |
| env-66 | ..................................... |
| env-67 | AAUCCGUCACAGAAAAGGACUUCCGGUACGUGGUCAG |
| env-68 | AAUCCGUCACAGAAAAGGACUUCCGGUACGUGGUCAG |
| env-75 | AA................................... |
| env-76 | AAUCCGUCACAGAAAAGGACUUCCGGUACGUGGUCAG |
| env-77 | AAUCCGUCACAGAAAAGGACUUCCGGUACGUGGUCAG |
| env-78 | ..................................... |
| env-79 | AAUCCGUCACAGAAAAGGACUUCCGGUACGUGGUCAG |
| env-80 | ..................................... |
| env-81 | ..................................... |
| Rgn-1-1 | AAUCCGUCACAGAAAAGGACUUCCGGUACGUGGUCAG |
| env-82 | AAUCCGUCACAGAAAAGGACUUCCGGUACGUGGUCAG |
| env-83 | AAUCCGUCACAGAAAAGGACUUCCGGUACGUGGUCAG |
| env-87 | ..................................... |
| env-88 | UAUCCGUCAUAGAAAAGGACUUCCGGUUCGUGGUCAG |
| env-89 | GUAUCCGUCAUAGAAAAGGACUUCCUGUUCGUGGACA |
| env-90 | AAUCCGUCAUAGAAGAGGAUUGCCGGUUCGUGGUCAA |
| env-91 | GAUCCGUCAUAGAAGAGGAUUGCCGGUUCGUGGUCAA |
| env-92 | ..................................... |
| env-93 | ..................................... |
| env-94 | CAAGAAGAAAUAAUAAUUUAUAUAAGUAGAAAAGGUG |
| Lba-11-1 | CAAGAAGAAAUAAUUAUAUUUAGAAGAAAAGGUGAAA |
| env-95 | AGGUAUCCGUCACAGAAGAGGACUUCCGGUCCGCGGA |
| env-96 | ..................................... |
| env-97 | AAUCCGUCACAGAAAAGGACUUCCGGUUCGUGGUCAG |
| Dsp-3-1 | ACAGAUGAAGACGUUAAGAAAAUCAGUGCUGUAAUUG |
| Dsp-1-1 | UAUCCGUCACAGAAAAGGACUUCCGGUCCGCGGUCAG |
| env-101 | ..................................... |
| env-102 | ..................................... |
| Dsp-2-1 | AAUCCGUCACAGAAAAGGACUUCCGGUUCGUGGUCAG |
| env-103 | ..................................... |
| env-106 | AAUCCGUCACAGAAAAGGACUUCCGGUUCGUGGUCAG |
| env-116 | ..................................... |
| env-117 | AAUCCGUCACAGAAAAGGACUUCCGG........... |
| env-118 | AAUCCGUCACAGAAAAGGACUUCCGGUUCGUGGUCAG |
| env-119 | AAUCCGUCACAGAAAAGGACUUCCAGUUC........ |
| env-120 | ..................................... |
| env-123 | ..................................... |
| env-124 | AAUCCGUCACAGAAAAGGACUUCCAGUUCGUGGUCAG |
| env-127 | AAUCCGUCAUAGAAAAGGACUUCCAGUUCGUGGUCAG |
| env-128 | ..................................... |
| env-129 | AAUCCGUCACAGAAA...................... |
| env-130 | AAUCCGUCACAGAAAAGGACUUCCGGUUCGUGGUCAG |
| env-136 | ..................................... |
| env-137 | AAUCCGUCACAGAAGAGGACUUCCGGUACGUGGUCAG |
| env-138 | ..................................... |
| env-139 | AAUCCGUCACAGAAAAGGACUUCCGGUUCGUGGUCAG |
| Dlo-1-1 | AAUCCGUCACAGAAAAGGACUUCCGGUUCGUGGUCAG |
| env-140 | AAUCCGUCACAGAAAAGGACUUCCGGUUCGUGGUCAG |
| env-149 | UAUCCGUCACAGAAAAGGACUUCCGGUUCGUGGUCAG |
| env-150 | ..................................... |
| env-151 | ..................................... |
| Chy-1-1 | UAUCCGUCACAGAAAAGGACUUCCGGUGCGCGGUCAG |
| env-152 | ..................................... |
| env-153 | ..................................... |
| env-154 | ..................................... |
| Esp-1-1 | UAUCCGUCACAGAAAAGGACUCCCGGUUCGUGGACAG |
| env-155 | AAUCCGUCACAGAAAAGGACUUCCGGUACGUGGACAG |
| env-156 | AAUCCGUCAUAGAAAAGGACUUCCAGUACGUGGACAG |
| env-159 | ..................................... |
| env-160 | AAUCCGUCAUAGAAAAGGACUUCCAGUACGUGGACAG |
| env-161 | AAUCCGUCACAGAAAAGGACUUCCGGUACGUGGACAG |
| env-162 | AAUCCGUCAUAGAAAAGGACUUCCGGUACGUGGACAG |
| env-163 | ..................................... |
| env-164 | ..................................... |
| env-165 | AAUCCGUCAUAGAAAAGGACUUCCGGUACGUGGACAG |
| env-167 | ..................................... |
| env-168 | AAUCCGUCAUAGAAAAGGACUUCCGGUACGUGGACAG |
| env-169 | AAUCCGUCAUAGAAAAGGACUUCCAGUACGUGGACAG |
| env-170 | AAUCCGUCAUAGAAAAGGACUUCCGGUACGUGGACAG |
| env-171 | AAUCCGUCACAGAAAAGGACUUCCGGUACGUGGACAG |
| env-172 | AAUCCGUCAUAGAAAAGGACUUCCGGUACGUGGACAG |
| env-175 | AAUCCGUCACAGAAAAGGACUUCCGGUACGUGGACAG |
| env-176 | ..................................... |
| env-177 | AAUCCGUCAUAGAAAAGGACUUCCGGUACGUGGACAG |
| env-178 | ..................................... |
| Lba-4-1 | AAUCCGUCACAGAAAAGGCCUUCCGGUUCGCGGUCAA |
| env-179 | AAUCCGUCACAGAAAAGGACUUCCGGUUCGUGGUCAG |
| Cba-3-1 | AAUCCGUCACAGAAAAGGACUUCCGGUUCGUGGUCAG |
| [sc-1-1 | AAUCCGUCACAGAAAAGGACUUCCGGUUCGUGGUCAG |
| Lba-7-1 | AAUCCGUCACAGAAAAGGACUUCCGGUUCGUGGUCAG |
| env-181 | ..................................... |
| env-182 | ..................................... |
| | ..................................... |
| | ..................................... |
| | ●R●Y●RYCAY●R●ARARGR●●●CY●RU●CRYRG●YRR |

| | |
|---|---|
| env-183 | AGGCAUCCGUCACAG...................... |
| env-184 | ..................................... |
| env-185 | ..................................... |
| env-186 | AAUCCGUCAUAGAAAAGGACUUCCGGUGCGUGGUCAG |
| env-187 | ..................................... |
| env-188 | ..................................... |
| env-189 | AAUCCGUCAUAGAAAAGGACUUCCGGUGCGUGGUCAG |
| env-190 | AAUCCGUCAUAGAAAAGGACUUCCGGUGCGUGGUCAG |
| env-191 | ..................................... |
| env-192 | AAUCCGUCAUAGAAAAGGACUUCCGGUGCGUGGUCAG |
| env-194 | ..................................... |
| env-195 | AAUCCGUCAUAGAAAAGGACUUCCGGUGCGUGGUCAG |
| env-200 | ..................................... |
| env-201 | ..................................... |
| env-202 | ..................................... |
| env-211 | ..................................... |
| env-212 | ..................................... |
| env-216 | AAUCC................................ |
| env-217 | AGGCAUCCGUCACAGAAGAGGCCUUCCGGUUCGCGG. |
| Rpe-1-1 | AAUCCGUCAUAGAAAAGGACUUCCGGUUCGUGGACAG |
| Rsp-1-1 | AAUCCGUCAUAGAAAAGGACUUCCGGUUCGUGGACAG |
| env-218 | CGGCAUCCGCCACAGAAAGGGCC.............. |
| env-219 | UAUUCGUCAUCGUAAAGGACUUCCGGUCCGCGGACAG |
| env-220 | ..................................... |
| env-221 | ..................................... |
| env-222 | AAUCCGUCACAGAAAAGGUCUUCCGGUUCGUGGUCAG |
| env-223 | ..................................... |
| env-226 | ..................................... |
| env-227 | ..................................... |
| env-228 | ..................................... |
| env-229 | ..................................... |
| env-230 | ..................................... |
| env-231 | ..................................... |
| env-232 | ..................................... |
| env-234 | ..................................... |
| env-235 | ..................................... |
| env-236 | AAUCCGUCACAGAAAAGGUCUUCCGGUUCGUGGUCAG |
| env-237 | AGCAGGAAAUUGCAAAGCAGAUCAAGUUAACUAUAGA |
| env-238 | ..................................... |
| env-239 | ..................................... |
| env-240 | AAUCCGUCACAGAAAAGGUCUUCCGGUUCGUGGUCAG |
| env-241 | AGGCAUCCGUCACAGAAAAGGACUUCCGGUCCGCGGA |
| env-242 | ..................................... |
| env-243 | ..................................... |
| env-244 | ..................................... |
| env-245 | ..................................... |
| env-246 | UAUUCGUCAUAGAAAAGGACUUCCUGUUCGUGGUCAG |
| env-252 | UAUUCGUCAUAGAAAAGGACUUCCUGUUCGUGGUCAG |
| env-254 | AAUCCGUCACAGAAGAGGACUUCCGGUACGUGGUCAG |
| Lba-6-1 | AAUCCGUCACAGAAGAGGACUUCCGGUACGUGGUCAG |
| env-255 | ..................................... |
| Lba-10-1 | AAUCCGUCACAGAAGAGGACUUCCGGUACGUGGUCAG |
| env-256 | AAUCCGUCACAGAAGAGGACUUCCGGUACGUGGUCAG |
| env-258 | ..................................... |
| env-259 | AAUCCGUCACAGAAGAGGACUUCCGGUACGUGGUCAG |
| env-260 | ..................................... |
| env-261 | ..................................... |
| env-262 | ..................................... |
| env-263 | UAUCCGUCACAGAAAAGGCCUUCCGGUUCGUGGUCAG |
| Bfo-1-1 | UAUCCGCCAUCGUAAAGGGCUUCCGGUUCGUGGUCAG |
| Bpr-1-1 | AGGAAUCCGCCACAGAAAAGGACUUCCGGUUCGCGGA |
| Cba-1-1 | AGGAAUCCGUCACAGAAAAGGACUUCCGGUUCGUGGA |
| env-264 | ..................................... |
| env-265 | ..................................... |
| env-266 | UAUCCGUCAUCGUAAAGGACUUCCGGU.......... |
| env-267 | UAUCCGUCAUCGUAAAGGACUUCCGGUUCGUGGCCAG |
| env-268 | UAUCCGUCAUCGUAAAGGACUUCCGGUUCGUGGCCAG |
| env-271 | ..................................... |
| env-272 | UAUCCGUCAUCGUAAAGGACUUCCGGUUCGUGGCCAG |
| env-273 | UAUCCGUCAUCGUAAAGGACUUCCGGUUCGUGGCCAG |
| env-275 | UAUCCGUCAUCGUAAAGGACUUCCGGUUCGUGGCCAG |
| env-276 | UAUCCGU.............................. |
| env-277 | ..................................... |
| env-278 | AAUCCGUCAUCGUAAAGGACUUCCGGUUCGUGGUCAG |
| env-279 | ..................................... |
| env-280 | ..................................... |
| env-281 | UAUCCGUCAUCGUAAAGGACUCCCGGUUCGUGGUCAG |
| env-282 | UAUCCGUCAUCGUAAAGGACUCCCGGUUCGUGGUCAG |
| Bsp-1-1 | UAUCCGUCAUCGUAAAGGACUCCCGGUUCGUGGUCAG |
| env-283 | ..................................... |
| env-284 | UAUCCGUCAUCGUAAAGGACUCCCGGUUCGUGGUCAG |
| env-285 | UAUCCGUCAUCGUAAAGGACUCCCGGUUCGUGGUCAG |
| env-286 | UAUCCGUCAUCGUAAAGGACUCCCGGUUCGUGGUCAG |
| env-287 | UAUCCGUCAUCGUAAAGGACUUCCGGUUCGUGGUCAG |
| | ..................................... |
| | ..................................... |
| | •R•Y•RYCAY•R•ARARGR•••CY•RU•CRYRG•YRR |

```
env-288    UAUCCGUCAUCGUAAAGGACUCCCGGUUCGUGGUCAG
env-289    UAUCCGUCAUCGUAAAGGACUUCCGGUUCGUGGUCAG
env-290    .....................................
env-291    UAUCCGUCAUCGUAAAGGACUUCCGGUUCGUGGUCAG
env-292    UAUCCGUCAUCGUAAAGGACUUCCGGUUCGUGGUCAG
env-293    .....................................
env-294    .....................................
env-295    .....................................
env-296    UAUCCGUCAUCGUAAAGGACUUCCGGUUCGUGGUCAG
env-298    UAUCCGUCAUCGUAAAGGACUUCCGGUUCGUGGUCAG
env-299    UAUCCGUCAUCGUAAAGGACUUCCGGUUCGUGGUCAG
env-300    UAUCCGUCAUCGUAAAGGACUUCCGGUUCGUGGUCAG
env-301    UAUCCGUCAUCGUAAAGGACUUCCGGUUCGUGGUCAG
env-302    UAUCCGUCAUCGUAAAGGACUUCCGGUUCGUGGUCAG
env-303    .....................................
env-304    .....................................
env-305    .....................................
Bwe-2-1    UAUCCGUCAUCGUAAAGGACUUCCGGUUCGUGGUCAG
env-306    UAUCCGUCAUCGUAAAGGACUUCCGGUUCGUGGUCAG
env-307    .....................................
env-308    .....................................
env-309    UAUCCGUCAUCGUAAAGGACUUCCGGUUCGUGGUCAG
env-310    .....................................
env-311    UAUCCGUCAUCGUAAAGGACUUCCGGUUCGUGGUCAG
env-313    UAUCCGUCAUCGUAAAGGACUUCCGGUUCGUGGUCAG
env-314    UAUCCGUCAUCGUAAAGGACUUCCGGUUCGUGGUCAG
env-315    UAUCCGUCAUCGUAAAGGACUUCCGGUUCGUGGUCAG
env-317    .....................................
env-318    .....................................
Bwe-1-1    UAUCCGUCAUCGUAAAGGACUUCCGGUUCGUGGUCAG
env-319    UAUCCGUCAUCGUAAAGGACUUCCGGUUCGUGGUCAG
env-320    UAUCCGUCAUCGUAAAGGACUUCCGGUUCGUGGUCAG
env-321    .....................................
env-322    .....................................
env-323    UAUCCGUCAUCGUAAAGGACUUCCGGUUCGUGGUCAG
env-324    .....................................
env-325    UAUCCGUCAUCGUAAAGGACUUCCGGUUCGUGGUCAG
env-326    UAUCCGUCAUCGUAAAGGACUUCCGGUUCGUGGUCAG
env-327    .....................................
env-328    .....................................
env-329    AAUCCGUCAUCGUAAAGGACUUCCGGUUCGUGGUCAG
env-331    .....................................
env-332    AAUCCGUCAUCGUAAAGGACUUCCGGUUCGUGGUCAG
env-335    .....................................
env-336    .....................................
env-337    AAUCCGUCAUCGUAAAGGACUUCCGGUUCGUGGUCAG
env-338    AAUCCGUCAUCGUAAAGGACUUCCGGUUCGUGGUCAG
env-339    AAUCCGUCAUCGGAAAGGACUUCCGGUUCGUGGUCAG
env-340    .....................................
env-341    .....................................
env-342    .....................................
Bsc-1-1    UAUCCGUCAUCGUAAAGGACUUCCGGUUCGUGGUCAG
Rsp-3-1    UAUCCGUCAUCGUAAAGGACUUCCGGUUCGUGGUCAG
env-343    UAUCCGUCAUAGAAAAGGACUUCCCGUCCGCGGACAG
env-344    AGGUAUCCGUCAUCGUAAAGGACUUCCUGUUCGUGGU
env-346    .....................................
env-348    .....................................
env-349    .....................................
env-350    AGGUAUCCGUCAUCGUAAAGGACUUCCUGUUCGUGGU
env-351    .....................................
env-352    CAGAGGAAUCCGUCAUCGUAAAGGACUUCCUGUUCGU
env-353    AAUUCGUCAUAGAAAAGGACUUCCUGUUCGUGGGCAG
Rga-1-1    UAUUCGUCAUAGAAAAGGACUUCCUGUUCGUGGACAG
env-354    .....................................
env-355    AAUUCGUCACAGAAAAGGACUUCCAGUUCGUGGACAA
env-356    .....................................
env-357    AAUUCGUCAUAGAAGAGGACUUCCUGUUCGUGGUCAA
env-358    AAUUCGUCAUAGAAGAGGACUUCCUGUUCGUGGUCAA
env-359    AAUUCGUCAUAGAAGAGGACUUCCUGUUCGUGGUCAA
env-360    AAUUCGUCACAGAAAAGGACUUCCUGUUCGUGGUCAA
env-361    AAUUCGUCACAGAAAAGGACUUCCUGUUCGUGGUCAA
env-362    AAUUCGUCACAGAAAAGGACUUCCUGUUCGUGGUCAA
env-363    .....................................
env-364    AAUCC................................
env-365    .....................................
env-366    AAUCC................................
env-367    .....................................
env-368    .....................................
env-369    .....................................
env-370    .....................................
env-371    .....................................
env-372    .....................................
env-373    UAUCCG...............................
env-374    NNGUCAGGUUGUAGAAGGAGAUCUUCGUAGAGAAAUC

           .....................................
           .....................................
           •R•Y•RYCAY•R•ARARGR•••CY•RU•CRYRG•YRR
```

```
env-375   UAUCCGUCAUCGUAAAGGCCUUCCGGUUCGUGGUCAG
env-377   .....................................
env-383   .....................................
env-384   GGUCCUAAGAGAACUGUUGCAAAUAAGAAGAAAUAAU
env-385   .....................................
env-386   UAUCCGUCAUCGUAAAGGCCUUCCGGUUCGUGGUCAG
env-387   GAUCCGUCAUCGUAAAGGCCUUCCGGUUCGUGGUCAG
env-388   UAUCCGUCAUCGUAAAGGCCUUCCGGUUCGUGGUCAG
env-389   .....................................
env-390   UAUCCGUCAUCGUAAAGGCCUUCCGGUUCGUGGUCAG
env-391   .....................................
env-392   .....................................
env-393   .....................................
env-394   .....................................
env-395   .....................................
env-396   .....................................
env-397   .....................................
env-398   .....................................
env-399   UAUCCGUCAUCGUAAAGGCCUUCCGGUUCGUGGUCAG
env-405   UAUCCGUCAUCGUAAAGGCCUUCCGGUUCGUGGUCAG
env-408   .....................................
env-409   .....................................
env-410   .....................................
env-411   GAACUGUUGCAAAUAAGAAGAAAUAAUUUAACCAGUA
env-412   GAGAUCUUCGUAGAGAAAUCGCUAUGAACAUCAAACG
env-413   UAUCCGUCAUCGUAAAGGCCUUCCGGUUCGUGGUCAG
env-414   UAUCCGUCAUCGUAAAGGCCUUCCGGUUCGUGGUCAG
env-418   .....................................
env-419   .....................................
env-420   .....................................
env-428   .....................................
env-429   .....................................
env-430   .....................................
env-432   .....................................
env-433   .....................................
env-434   .....................................
Bhy-1-1   UAUCCGUCAUAGAAAAGGACUGCCGGUUCGUGGUCAG
env-435   UAUCCGUCAUAGAAAAGGACUUCCUGUCCGUGGUCAG
env-436   AAUCCGUCAUAGAAAAGGGCUUCCUGUCCGUGGUCAG
env-437   CAU..................................
env-438   .....................................
Bsp-7-1   AAUCCGUCACAGAAGAGGACUUCCUGUUCGUGGUCAG
Bsp-9-1   AAUUCGUCACAGAAGAGGACUUCCUGUUCGUGGUCAG
Bsp-5-1   AAUCCGUCACAGAAGAGGACUUCCUGUUCGUGGUCAG
Bsp-2-1   UAUCCGUCACAGAAGAGGACUUCCUGUUCGUGGUCAG
Bsp-3-1   UAUUCGUCACAGAAGAGGACUUCCUGUUCGCGGUCAG
Bpr-2-1   AGGAAUCCGUCACAGAAAAGGACUUCCGGUUCGUGGA
          .....................................
          .....................................
          •R•Y•RYCAY•R•ARARGR•••CY•RU•CRYRG•YRR
```

### 17 S4-Fusobacteriales

#### 17.1 Taxa

The taxonomy of each organism containing a putative S4-Fusobacteriales RNA is listed, with abbreviations identifying each hit (e.g., "Eco-1-1" and "Eco-1-2" might hypothetically represent two distinct RNAs in *E. coli*). The abbreviations will be used to identify each individual S4-Fusobacteriales RNA in Sections 17.2 and 17.4:.

| **abbrev. of hits** | **taxonomy of species** |
|---|---|
| Cso-1-1 | Bacteria Fusobacteria Fusobacteriales Fusobacteriaceae *Cetobacterium somerae* ATCC BAA-474 |
| Csp-1-1 | Bacteria Fusobacteria Fusobacteriales Fusobacteriaceae *Cetobacterium sp.* ZOR0034 |
| Csp-2-1 | Bacteria Fusobacteria Fusobacteriales Fusobacteriaceae *Cetobacterium sp.* ZWU0022 |
| Fgo-1-1 | Bacteria Fusobacteria Fusobacteriales Fusobacteriaceae *Fusobacterium gonidiaformans* ATCC 25563 |
| Fhw-1-1 | Bacteria Fusobacteria Fusobacteriales Fusobacteriaceae *Fusobacterium hwasookii* ChDC F145 |
| Fhw-2-1 | Bacteria Fusobacteria Fusobacteriales Fusobacteriaceae *Fusobacterium hwasookii* ChDC F174 |
| Fhw-3-1 | Bacteria Fusobacteria Fusobacteriales Fusobacteriaceae *Fusobacterium hwasookii* ChDC F206 |
| Fhw-4-1 | Bacteria Fusobacteria Fusobacteriales Fusobacteriaceae *Fusobacterium hwasookii* ChDC F300 |
| Fmo-1-1 | Bacteria Fusobacteria Fusobacteriales Fusobacteriaceae *Fusobacterium mortiferum* ATCC 9817 |
| Fne-1-1 | Bacteria Fusobacteria Fusobacteriales Fusobacteriaceae *Fusobacterium necrophorum* BFTR-1 |

| | |
|---|---|
| Fne-2-1 | Bacteria Fusobacteria Fusobacteriales Fusobacteriaceae *Fusobacterium necrophorum* BFTR-2 |
| Fne-3-1 | Bacteria Fusobacteria Fusobacteriales Fusobacteriaceae *Fusobacterium necrophorum* BL |
| Fne-4-1 | Bacteria Fusobacteria Fusobacteriales Fusobacteriaceae *Fusobacterium necrophorum* D12 |
| Fne-5-1 | Bacteria Fusobacteria Fusobacteriales Fusobacteriaceae *Fusobacterium necrophorum* DAB |
| Fne-6-1 | Bacteria Fusobacteria Fusobacteriales Fusobacteriaceae *Fusobacterium necrophorum* DJ-1 |
| Fne-7-1 | Bacteria Fusobacteria Fusobacteriales Fusobacteriaceae *Fusobacterium necrophorum* DJ-2 |
| Fne-8-1 | Bacteria Fusobacteria Fusobacteriales Fusobacteriaceae *Fusobacterium necrophorum* HUN048 |
| Fne-9-1 | Bacteria Fusobacteria Fusobacteriales Fusobacteriaceae *Fusobacterium necrophorum subsp. funduliforme* ATCC 51357 |
| Fne-10-1 | Bacteria Fusobacteria Fusobacteriales Fusobacteriaceae *Fusobacterium necrophorum subsp. funduliforme* B35 |
| Fne-11-1 | Bacteria Fusobacteria Fusobacteriales Fusobacteriaceae *Fusobacterium necrophorum subsp. funduliforme* Fnf 1007 |
| Fnu-1-1 | Bacteria Fusobacteria Fusobacteriales Fusobacteriaceae *Fusobacterium nucleatum* 13_3C |
| Fnu-2-1 | Bacteria Fusobacteria Fusobacteriales Fusobacteriaceae *Fusobacterium nucleatum* CC53 |
| Fnu-9-1 | Bacteria Fusobacteria Fusobacteriales Fusobacteriaceae *Fusobacterium nucleatum* ChDC F128 |
| Fnu-3-1 | Bacteria Fusobacteria Fusobacteriales Fusobacteriaceae *Fusobacterium nucleatum* CTI-1 |
| Fnu-4-1 | Bacteria Fusobacteria Fusobacteriales Fusobacteriaceae *Fusobacterium nucleatum* CTI-2 |
| Fnu-5-1 | Bacteria Fusobacteria Fusobacteriales Fusobacteriaceae *Fusobacterium nucleatum* CTI-3 |
| Fnu-6-1 | Bacteria Fusobacteria Fusobacteriales Fusobacteriaceae *Fusobacterium nucleatum* CTI-5 |
| Fnu-7-1 | Bacteria Fusobacteria Fusobacteriales Fusobacteriaceae *Fusobacterium nucleatum* CTI-6 |
| Fnu-8-1 | Bacteria Fusobacteria Fusobacteriales Fusobacteriaceae *Fusobacterium nucleatum* CTI-7 |
| Fnu-11-1 | Bacteria Fusobacteria Fusobacteriales Fusobacteriaceae *Fusobacterium nucleatum subsp. animalis* 11_3_2 |
| Fnu-12-1 | Bacteria Fusobacteria Fusobacteriales Fusobacteriaceae *Fusobacterium nucleatum subsp. animalis* 21_1A |
| Fnu-13-1 | Bacteria Fusobacteria Fusobacteriales Fusobacteriaceae *Fusobacterium nucleatum subsp. animalis* 3_1_33 |
| Fnu-14-1 | Bacteria Fusobacteria Fusobacteriales Fusobacteriaceae *Fusobacterium nucleatum subsp. animalis* 4_8 |
| Fnu-15-1 | Bacteria Fusobacteria Fusobacteriales Fusobacteriaceae *Fusobacterium nucleatum subsp. animalis* 7_1 |
| Fnu-16-1 | Bacteria Fusobacteria Fusobacteriales Fusobacteriaceae *Fusobacterium nucleatum subsp. animalis* ChDC F324 |
| Fnu-17-1 | Bacteria Fusobacteria Fusobacteriales Fusobacteriaceae *Fusobacterium nucleatum subsp. animalis* F0419 |
| Fnu-18-1 | Bacteria Fusobacteria Fusobacteriales Fusobacteriaceae *Fusobacterium nucleatum subsp. fusiforme* ATCC 51190 |
| Fnu-19-1 | Bacteria Fusobacteria Fusobacteriales Fusobacteriaceae *Fusobacterium nucleatum subsp. nucleatum* ATCC 23726 |
| Fnu-20-1 | Bacteria Fusobacteria Fusobacteriales Fusobacteriaceae *Fusobacterium nucleatum subsp. nucleatum* ATCC 25586 |
| Fnu-21-1 | Bacteria Fusobacteria Fusobacteriales Fusobacteriaceae *Fusobacterium nucleatum subsp. nucleatum* ChDC F316 |
| Fnu-22-1 | Bacteria Fusobacteria Fusobacteriales Fusobacteriaceae *Fusobacterium nucleatum subsp. polymorphum* ATCC 10953 |
| Fnu-23-1 | Bacteria Fusobacteria Fusobacteriales Fusobacteriaceae *Fusobacterium nucleatum subsp. polymorphum* F0401 |
| Fnu-24-1 | Bacteria Fusobacteria Fusobacteriales Fusobacteriaceae *Fusobacterium nucleatum subsp. vincentii* 3_1_27 |
| Fnu-25-1 | Bacteria Fusobacteria Fusobacteriales Fusobacteriaceae *Fusobacterium nucleatum subsp. vincentii* 3_1_36A2 |
| Fnu-26-1 | Bacteria Fusobacteria Fusobacteriales Fusobacteriaceae *Fusobacterium nucleatum subsp. vincentii* 4_1_13 |
| Fnu-27-1 | Bacteria Fusobacteria Fusobacteriales Fusobacteriaceae *Fusobacterium nucleatum subsp. vincentii* ATCC 49256 |
| Fnu-28-1 | Bacteria Fusobacteria Fusobacteriales Fusobacteriaceae *Fusobacterium nucleatum subsp. vincentii* ChDC F8 |
| Fnu-10-1 | Bacteria Fusobacteria Fusobacteriales Fusobacteriaceae *Fusobacterium nucleatum* W1481 |
| Fpe-1-1 | Bacteria Fusobacteria Fusobacteriales Fusobacteriaceae *Fusobacterium perfoetens* ATCC 29250 |
| Fpe-2-1 | Bacteria Fusobacteria Fusobacteriales Fusobacteriaceae *Fusobacterium periodonticum* 2_1_31 |
| Fpe-3-1 | Bacteria Fusobacteria Fusobacteriales Fusobacteriaceae *Fusobacterium periodonticum* ATCC 33693 |
| Fpe-4-1 | Bacteria Fusobacteria Fusobacteriales Fusobacteriaceae *Fusobacterium periodonticum* D10 |
| Fru-1-1 | Bacteria Fusobacteria Fusobacteriales Fusobacteriaceae *Fusobacterium russii* ATCC 25533 |
| Fsp-1-1 | Bacteria Fusobacteria Fusobacteriales Fusobacteriaceae *Fusobacterium sp.* 1_1_41FAA |
| Fsp-2-1 | Bacteria Fusobacteria Fusobacteriales Fusobacteriaceae *Fusobacterium sp.* 3_1_5R |
| Fsp-3-1 | Bacteria Fusobacteria Fusobacteriales Fusobacteriaceae *Fusobacterium sp.* CM1 |
| Fsp-4-1 | Bacteria Fusobacteria Fusobacteriales Fusobacteriaceae *Fusobacterium sp.* CM21 |
| Fsp-5-1 | Bacteria Fusobacteria Fusobacteriales Fusobacteriaceae *Fusobacterium sp.* CM22 |
| Fsp-6-1 | Bacteria Fusobacteria Fusobacteriales Fusobacteriaceae *Fusobacterium sp.* OBRC1 |
| Fsp-7-1 | Bacteria Fusobacteria Fusobacteriales Fusobacteriaceae *Fusobacterium sp. oral taxon* 370 str. F0437 |
| Ful-1-1 | Bacteria Fusobacteria Fusobacteriales Fusobacteriaceae *Fusobacterium ulcerans* 12-1B |
| Ful-2-1 | Bacteria Fusobacteria Fusobacteriales Fusobacteriaceae *Fusobacterium ulcerans* ATCC 49185 |
| Fva-1-1 | Bacteria Fusobacteria Fusobacteriales Fusobacteriaceae *Fusobacterium varium* ATCC 27725 |
| Ipo-1-1 | Bacteria Fusobacteria Fusobacteriales Fusobacteriaceae *Ilyobacter polytropus* DSM 2926 |
| Lbu-1-1 | Bacteria Fusobacteria Fusobacteriales Fusobacteriaceae *Leptotrichia buccalis* DSM 1135 |
| Lgo-1-1 | Bacteria Fusobacteria Fusobacteriales Fusobacteriaceae *Leptotrichia goodfellowii* F0264 |
| Pat-1-1 | Bacteria Fusobacteria Fusobacteriales Fusobacteriaceae *Psychrilyobacter atlanticus* DSM 19335 |
| Ste-1-1 | Bacteria Fusobacteria Fusobacteriales Fusobacteriaceae *Sebaldella termitidis* ATCC 33386 |
| Smo-1-1 | Bacteria Fusobacteria Fusobacteriales Fusobacteriaceae *Streptobacillus moniliformis* DSM 12112 |
| Lgo-2-1 | Bacteria Fusobacteria Fusobacteriales Leptotrichiaceae *Leptotrichia goodfellowii* DSM 19756 |
| Lsh-1-1 | Bacteria Fusobacteria Fusobacteriales Leptotrichiaceae *Leptotrichia shahii* DSM 19757 |
| Lsp-1-1 | Bacteria Fusobacteria Fusobacteriales Leptotrichiaceae *Leptotrichia sp. oral taxon* 212 |
| Lsp-2-1 | Bacteria Fusobacteria Fusobacteriales Leptotrichiaceae *Leptotrichia sp. oral taxon* 215 str. W9775 |
| Lsp-3-1 | Bacteria Fusobacteria Fusobacteriales Leptotrichiaceae *Leptotrichia sp. oral taxon* 225 str. F0581 |
| Lsp-4-1 | Bacteria Fusobacteria Fusobacteriales Leptotrichiaceae *Leptotrichia sp. oral taxon* 879 str. F0557 |

| | |
|---|---|
| Ltr-1-1 | Bacteria Fusobacteria Fusobacteriales Leptotrichiaceae *Leptotrichia trevisanii* DSM 22070 |
| Lwa-1-1 | Bacteria Fusobacteria Fusobacteriales Leptotrichiaceae *Leptotrichia wadei* DSM 19758 |
| Lwa-2-1 | Bacteria Fusobacteria Fusobacteriales Leptotrichiaceae *Leptotrichia wadei* F0279 |
| Sam-1-1 | Bacteria Fusobacteria Fusobacteriales Leptotrichiaceae *Sneathia amnii* |
| env-1 to env-482 | environmental samples |

#### 17.2 Gene contexts

Each S4-Fusobacteriales RNA (indicated by "RNA→") is listed. For each hit, the downstream genes predicted to reside in a regulated operon are listed. If the nearest downstream gene is encoding in the opposite strand (and therefore presumed to not be a part of a regulated operon), then that gene is still depicted. Some environmental sequences and some RefSeq entries lack gene annotations, and so no genes are available for such sequences. The direction of each gene is indicated with an arrow (→), and each predicted conserved domain in the gene is named. Conserved domains associated with more than one S4-Fusobacteriales RNA are assigned a color; other domains are gray. Information about these conserved domains is given in Section 17.3. The accession of the sequence containing each S4-Fusobacteriales RNA is given in the column named "Seq. accession". Accessions beginning with "NC_", "NS_", "NW_" or "NZ_" are contained in RefSeq. Other accession refer to environmental samples. Nucleotide coordinates are given for the 5′ and 3′ boundaries of each S4-Fusobacteriales RNA. If the 5′ coordinate is greater than the 3′ coordinate, the RNA is present on the reverse-complement strand of the containing genomic DNA sequence. Each hit is denoted by an abbreviation (like "Eco-1-1") that refers to a taxonomy given in Section 17.1.

| abbrev. | Seq. accession | | 5′ at | 3′ at | genes |
|---|---|---|---|---|---|
| env-1 | 4448820.3_Contig_4724 | - | 1508 | 1444 | **RNA**→ RpsM (COG0099)rpsM (PRK05179)→ |
| env-2 | 4444165.3.02425 | - | 1233 | 1174 | **RNA**→ RpsM (COG0099)rpsM (PRK05179)→ PRK05309 (PRK05309)RpsK (COG0100)→ PRK05309 (PRK05309)RpsK (COG0100)→ rpsD (PRK05327)Ribosomal_S4 (pfam00163)→ |
| env-3 | 4448815.3_Contig29902 | - | 527 | 468 | **RNA**→ RpsM (COG0099)rpsM (PRK05179)→ |
| env-4 | SRS064449_LANL_scaffold_2211 | + | 364 | 426 | **RNA**→ RpsM (COG0099)rpsM (PRK05179)→ PRK05309 (PRK05309)RpsK (COG0100)→ |
| env-5 | SRS043239_WUGC_scaffold_38909 | - | 221 | 159 | **RNA**→ rps13 (CHL00137)rpsM (PRK05179)→ |
| env-6 | SRS024318_LANL_scaffold_10757 | - | 103 | 41 | **RNA**→ |
| env-7 | LWFCAn_GLO1YSU01C2O9I | + | 165 | 222 | **RNA**→ RpsM (COG0099)rpsM (PRK05179)→ |
| env-8 | SRS019124_C1397861 | + | 326 | 392 | **RNA**→ RpsM (COG0099)rpsM (PRK05179)→ PRK05309 (PRK05309)RpsK (COG0100)→ |
| env-9 | SRS019125_WUGC_scaffold_10350 | + | 92 | 158 | **RNA**→ RpsM (COG0099)rpsM (PRK05179)→ PRK05309 (PRK05309)RpsK (COG0100)→ RpsD (COG0522)rpsD (PRK05327)→ RpoA (COG0202)PRK05182 (PRK05182)→ rplQ (PRK05591)Ribosomal_L17 (pfam01196)→ |
| env-10 | SRS013506_C1110483 | - | 685 | 618 | **RNA**→ RpsM (COG0099)rpsM (PRK05179)→ PRK05309 (PRK05309)RpsK (COG0100)→ |
| env-11 | CAM_READ_0099908967 | - | 481 | 418 | **RNA**→ RpsM (COG0099)rpsM (PRK05179)→ |
| env-12 | CAM_READ_0100166989 | + | 129 | 189 | **RNA**→ RpsM (COG0099)rpsM (PRK05179)→ |
| env-13 | CAM_READ_0099986119 | - | 478 | 417 | **RNA**→ RpsM (COG0099)rpsM (PRK05179)→ |
| env-14 | CAM_READ_0099775883 | - | 476 | 416 | **RNA**→ RpsM (COG0099)rpsM (PRK05179)→ |
| env-15 | CAM_READ_0099873361 | - | 474 | 414 | **RNA**→ RpsM (COG0099)uS13_bact (TIGR03631)→ |
| env-16 | CAM_READ_0100181569 | + | 35 | 95 | **RNA**→ |
| env-17 | 4444165.3.00747 | - | 3020 | 2960 | **RNA**→ RpsM (COG0099)rpsM (PRK05179)→ PRK05309 (PRK05309)RpsK (COG0100)→ rpsD (PRK05327)→ S4 (smart00363)RpsD (COG0522)→ |
| env-18 | CAM_READ_0100128399 | - | 372 | 312 | **RNA**→ RpsM (COG0099)rpsM (PRK05179)→ |
| env-19 | CAM_READ_0100169721 | + | 35 | 95 | **RNA**→ RpsM (COG0099)rpsM (PRK05179)→ |
| env-20 | 4444164.3.01074 | + | 805 | 865 | **RNA**→ RpsM (COG0099)rpsM (PRK05179)→ PRK05309 (PRK05309)RpsK (COG0100)→ RpsD (COG0522)rpsD (PRK05327)→ PRK05182 (PRK05182)RNA_pol_L (pfam01193)→ |
| Fpe-1-1 | NZ_JHXW01000013.1 | + | 27498 | 27558 | **RNA**→ RpsM (COG0099)rpsM (PRK05179)→ PRK05309 (PRK05309)RpsK (COG0100)→ RpsD (COG0522)rpsD (PRK05327)→ RpoA (COG0202)PRK05182 (PRK05182)→ RplQ (COG0203)rplQ (PRK05591)→ hypo→ |
| env-21 | CAM_READ_0099877645 | + | 39 | 98 | **RNA**→ RpsM (COG0099)rpsM (PRK05179)→ |
| env-22 | 4444164.3.00319 | + | 39 | 98 | **RNA**→ RpsM (COG0099)rpsM (PRK05179)→ |
| env-23 | 4444164.3.51166 | + | 542 | 602 | **RNA**→ RpsM (COG0099)rpsM (PRK05179)→ |

| | | | | | |
|---|---|---|---|---|---|
| env-24 | CAM_READ_0099837051 | - | 423 | 363 | **RNA**→ RpsM (COG0099)rpsM (PRK05179)→ |
| env-25 | SRS023604_Baylor_scaffold_1939 | + | 279 | 366 | **RNA**→ RpsM (COG0099)rpsM (PRK05179)→ PRK05309 (PRK05309)RpsK (COG0100)→ |
| env-26 | SRS011111_C480900 | + | 8 | 95 | **RNA**→ RpsM (COG0099)rpsM (PRK05179)→ |
| env-27 | SRS023604_Baylor_scaffold_1904 | + | 280 | 370 | **RNA**→ RpsM (COG0099)rpsM (PRK05179)→ PRK05309 (PRK05309)RpsK (COG0100)→ RpsD (COG0522)rpsD (PRK05327)→ RpoA (COG0202)PRK05182 (PRK05182)→ RplQ (COG0203)rplQ (PRK05591)→ |
| env-28 | SRS017497_Baylor_scaffold_808 | + | 42316 | 42406 | **RNA**→ RpsM (COG0099)rpsM (PRK05179)→ PRK05309 (PRK05309)RpsK (COG0100)→ RpsD (COG0522)rpsD (PRK05327)→ RpoA (COG0202)PRK05182 (PRK05182)→ RplQ (COG0203)rplQ (PRK05591)→ Acetyltransf_1 (pfam00583)Acetyltransf_7 (pfam13508)→ |
| Sam-1-1 | NZ_CP011280.1 | + | 495384 | 495474 | **RNA**→ RpsM (COG0099)rpsM (PRK05179)→ PRK05309 (PRK05309)RpsK (COG0100)→ RpsD (COG0522)rpsD (PRK05327)→ RpoA (COG0202)PRK05182 (PRK05182)→ RplQ (COG0203)rplQ (PRK05591)→ Acetyltransf_1 (pfam00583)Acetyltransf_7 (pfam13508)→ |
| env-29 | SRS011111_Baylor_scaffold_6158 | + | 13925 | 14015 | **RNA**→ RpsM (COG0099)rpsM (PRK05179)→ PRK05309 (PRK05309)RpsK (COG0100)→ RpsD (COG0522)rpsD (PRK05327)→ RpoA (COG0202)PRK05182 (PRK05182)→ RplQ (COG0203)rplQ (PRK05591)→ Acetyltransf_1 (pfam00583)Acetyltransf_7 (pfam13508)→ PRK00326 (PRK00326)MraZ_N (cd16320)MraZ_C (cd16321)→ RmsH (COG0275)PRK00050 (PRK00050)→ hypo→ |
| Ste-1-1 | NC_013517.1 | - | 4215417 | 4215353 | **RNA**→ RpsM (COG0099)rpsM (PRK05179)→ PRK05309 (PRK05309)RpsK (COG0100)→ RpsD (COG0522)rpsD (PRK05327)→ RpoA (COG0202)PRK05182 (PRK05182)→ RplQ (COG0203)rplQ (PRK05591)→ |
| Smo-1-1 | NC_013515.1 | + | 399617 | 399698 | **RNA**→ RpsM (COG0099)rpsM (PRK05179)→ PRK05309 (PRK05309)RpsK (COG0100)→ RpsD (COG0522)rpsD (PRK05327)→ RpoA (COG0202)PRK05182 (PRK05182)→ RplQ (COG0203)rplQ (PRK05591)→ hypo→ hypo→ hypo→ hypo→ hypo→ |
| env-30 | SRS016319_C3906942 | - | 367 | 278 | **RNA**→ RpsM (COG0099)rpsM (PRK05179)→ |
| env-31 | SRS049389_C4596854 | + | 103 | 192 | **RNA**→ RpsM (COG0099)rpsM (PRK05179)→ |
| env-32 | SRS015762_C3360097 | + | 103 | 192 | **RNA**→ rps13 (CHL00137)rpsM (PRK05179)→ |
| env-33 | SRS016225_C2898664 | - | 373 | 284 | **RNA**→ RpsM (COG0099)rpsM (PRK05179)→ |
| env-34 | SRS017120_Baylor_scaffold_77919 | + | 569 | 658 | **RNA**→ RpsM (COG0099)rpsM (PRK05179)→ |
| env-35 | SRS019219_WUGC_scaffold_70719 | - | 373 | 284 | **RNA**→ RpsM (COG0099)rpsM (PRK05179)→ |
| env-36 | SRS023352_LANL_scaffold_52359 | + | 475 | 564 | **RNA**→ |
| env-37 | SRS016569_C2417077 | - | 376 | 287 | **RNA**→ RpsM (COG0099)rpsM (PRK05179)→ |
| env-38 | SRS019026_C1965129 | + | 158 | 247 | **RNA**→ RpsM (COG0099)rpsM (PRK05179)→ PRK05309 (PRK05309)RpsK (COG0100)→ |
| env-39 | SRS019027_C1950830 | + | 158 | 247 | **RNA**→ RpsM (COG0099)rpsM (PRK05179)→ |
| env-40 | SRS019028_WUGC_scaffold_55374 | - | 365 | 276 | **RNA**→ RpsM (COG0099)rpsM (PRK05179)→ |
| env-41 | SRS017076_C2662400 | + | 122 | 211 | **RNA**→ RpsM (COG0099)rpsM (PRK05179)→ |
| env-42 | SRS014271_WUGC_scaffold_3488 | - | 457 | 368 | **RNA**→ RpsM (COG0099)rpsM (PRK05179)→ |
| env-43 | SRS014470_WUGC_scaffold_45929 | - | 373 | 284 | **RNA**→ RpsM (COG0099)rpsM (PRK05179)→ |
| env-44 | SRS014573_WUGC_scaffold_53471 | - | 376 | 287 | **RNA**→ RpsM (COG0099)rpsM (PRK05179)→ |
| env-45 | SRS015057_WUGC_scaffold_36508 | - | 376 | 287 | **RNA**→ RpsM (COG0099)rpsM (PRK05179)→ |
| env-46 | SRS019607_WUGC_scaffold_48886 | - | 645 | 556 | **RNA**→ RpsM (COG0099)rpsM (PRK05179)→ PRK05309 (PRK05309)RpsK (COG0100)→ |
| env-47 | SRS021496_Baylor_scaffold_82145 | + | 106 | 195 | **RNA**→ RpsM (COG0099)rpsM (PRK05179)→ |
| env-48 | SRS044373_WUGC_scaffold_7895 | + | 106 | 195 | **RNA**→ RpsM (COG0099)rpsM (PRK05179)→ |
| env-49 | SRS047219_WUGC_scaffold_21972 | + | 933 | 1022 | **RNA**→ RpsM (COG0099)rpsM (PRK05179)→ |
| env-50 | SRS047824_WUGC_scaffold_46958 | - | 645 | 556 | **RNA**→ RpsM (COG0099)rpsM (PRK05179)→ PRK05309 (PRK05309)RpsK (COG0100)→ |
| env-51 | SRS057791_LANL_scaffold_64973 | + | 798 | 887 | **RNA**→ rps13 (CHL00137)rpsM (PRK05179)→ |
| env-52 | Buccal_mucosa_LANL_scaffold_69102 | + | 103 | 192 | **RNA**→ RpsM (COG0099)rpsM (PRK05179)→ |
| env-53 | SRS013705_Baylor_scaffold_48015 | - | 839 | 750 | **RNA**→ RpsM (COG0099)rpsM (PRK05179)→ PRK05309 (PRK05309)RpsK (COG0100)→ |
| env-54 | SRS015038_WUGC_scaffold_27606 | - | 597 | 508 | **RNA**→ RpsM (COG0099)rpsM (PRK05179)→ |
| env-55 | SRS015272_WUGC_scaffold_163 | + | 206 | 295 | **RNA**→ rps13 (CHL00137)rpsM (PRK05179)→ |
| env-56 | SRS018357_C3855302 | - | 376 | 287 | **RNA**→ RpsM (COG0099)rpsM (PRK05179)→ |
| env-57 | SRS019894_WUGC_scaffold_51429 | + | 155 | 244 | **RNA**→ rps13 (CHL00137)rpsM (PRK05179)→ |
| env-58 | SRS024081_LANL_scaffold_63456 | - | 645 | 556 | **RNA**→ RpsM (COG0099)rpsM (PRK05179)→ PRK05309 (PRK05309)RpsK (COG0100)→ |

| | | | | | |
|---|---|---|---|---|---|
| env-59 | SRS053854_LANL_scaffold_54237 | - | 11790 | 11701 | **RNA**→ RpsM (COG0099)rpsM (PRK05179)→ PRK05309 (PRK05309)RpsK (COG0100)→ RpsD (COG0522)rpsD (PRK05327)→ RpoA (COG0202)PRK05182 (PRK05182)→ RplQ (COG0203)rplQ (PRK05591)→ |
| env-60 | SRS057539_LANL_scaffold_53408 | - | 376 | 287 | **RNA**→ RpsM (COG0099)rpsM (PRK05179)→ |
| env-61 | SRS064774_LANL_scaffold_7571 | - | 671 | 582 | **RNA**→ RpsM (COG0099)rpsM (PRK05179)→ hypo→ rps11 (CHL00041)PRK05309 (PRK05309)→ |
| env-62 | SRS015899_WUGC_scaffold_4042 | - | 330 | 246 | **RNA**→ RpsM (COG0099)rpsM (PRK05179)→ |
| env-63 | SRS018975_C1798714 | - | 194 | 110 | **RNA**→ RpsM (COG0099)rpsM (PRK05179)→ |
| env-64 | SRS064449_LANL_scaffold_25989 | - | 365 | 281 | **RNA**→ RpsM (COG0099)rpsM (PRK05179)→ |
| env-65 | SRS019389_C2168600 | - | 376 | 311 | **RNA**→ RpsM (COG0099)rpsM (PRK05179)→ |
| env-66 | SRS017808_Baylor_scaffold_37000 | + | 6883 | 6949 | **RNA**→ |
| env-67 | SRS022530_LANL_scaffold_52023 | + | 955 | 1052 | **RNA**→ RpsM (COG0099)rpsM (PRK05179)→ PRK05309 (PRK05309)RpsK (COG0100)→ |
| env-68 | SRS013164_Baylor_scaffold_58877 | + | 45 | 110 | **RNA**→ RpsM (COG0099)rpsM (PRK05179)→ PRK05309 (PRK05309)RpsK (COG0100)→ RpsD (COG0522)rpsD (PRK05327)→ RpoA (COG0202)PRK05182 (PRK05182)→ RplQ (COG0203)rplQ (PRK05591)→ |
| env-69 | SRS017209_Baylor_scaffold_57193 | - | 372 | 307 | **RNA**→ RpsM (COG0099)rpsM (PRK05179)→ |
| env-70 | SRS053603_LANL_scaffold_21307 | - | 669 | 604 | **RNA**→ RpsM (COG0099)rpsM (PRK05179)→ PRK05309 (PRK05309)RpsK (COG0100)→ |
| env-71 | SRS019607_WUGC_scaffold_56406 | - | 626 | 561 | **RNA**→ RpsM (COG0099)rpsM (PRK05179)→ PRK05309 (PRK05309)RpsK (COG0100)→ |
| env-72 | SRS013502_C2176709 | - | 644 | 579 | **RNA**→ RpsM (COG0099)rpsM (PRK05179)→ PRK05309 (PRK05309)RpsK (COG0100)→ |
| env-73 | SRS013705_Baylor_scaffold_95676 | + | 104 | 169 | **RNA**→ RpsM (COG0099)rpsM (PRK05179)→ PRK05309 (PRK05309)RpsK (COG0100)→ |
| env-74 | SRS014684_C3315306 | + | 4413 | 4478 | **RNA**→ RpsM (COG0099)rpsM (PRK05179)→ PRK05309 (PRK05309)RpsK (COG0100)→ |
| env-75 | SRS015209_WUGC_scaffold_55827 | - | 1494 | 1429 | **RNA**→ RpsM (COG0099)rpsM (PRK05179)→ PRK05309 (PRK05309)RpsK (COG0100)→ RpsD (COG0522)rpsD (PRK05327)→ |
| env-76 | SRS015434_WUGC_scaffold_26309 | - | 522 | 457 | **RNA**→ rps11 (CHL00041)PRK05309 (PRK05309)RpsM (COG0099)rpsM (PRK05179)→ |
| env-77 | SRS015762_WUGC_scaffold_32980 | - | 477 | 412 | **RNA**→ RpsM (COG0099)rpsM (PRK05179)→ |
| env-78 | SRS015893_WUGC_scaffold_40459 | + | 3690 | 3755 | **RNA**→ RpsM (COG0099)rpsM (PRK05179)→ |
| env-79 | SRS015941_C3324156 | - | 210 | 145 | **RNA**→ rps13 (CHL00137)rpsM (PRK05179)→ |
| env-80 | SRS016002_C2598934 | + | 157 | 222 | **RNA**→ RpsM (COG0099)rpsM (PRK05179)→ PRK05309 (PRK05309)RpsK (COG0100)→ |
| env-81 | SRS016037_C2049273 | + | 483 | 548 | **RNA**→ RpsM (COG0099)rpsM (PRK05179)→ PRK05309 (PRK05309)RpsK (COG0100)→ RpsD (COG0522)rpsD (PRK05327)→ PRK05182 (PRK05182)RNA_pol_L (pfam01193)→ |
| env-82 | SRS016319_WUGC_scaffold_50629 | + | 485 | 550 | **RNA**→ RpsM (COG0099)rpsM (PRK05179)→ |
| env-83 | SRS016569_C2462808 | - | 626 | 561 | **RNA**→ RpsM (COG0099)rpsM (PRK05179)→ PRK05309 (PRK05309)RpsK (COG0100)→ |
| env-84 | SRS018300_Baylor_scaffold_20150 | - | 420 | 355 | **RNA**→ RpsM (COG0099)rpsM (PRK05179)→ |
| env-85 | SRS018357_C3909917 | + | 104 | 169 | **RNA**→ RpsM (COG0099)rpsM (PRK05179)→ PRK05309 (PRK05309)RpsK (COG0100)→ |
| env-86 | SRS019026_C1948435 | + | 142 | 207 | **RNA**→ RpsM (COG0099)rpsM (PRK05179)→ |
| env-87 | SRS019027_WUGC_scaffold_19534 | + | 142 | 207 | **RNA**→ RpsM (COG0099)rpsM (PRK05179)→ |
| env-88 | SRS019045_WUGC_scaffold_56945 | - | 150 | 85 | **RNA**→ |
| env-89 | SRS019122_C5915383 | + | 2424 | 2489 | **RNA**→ |
| env-90 | SRS019126_C2215224 | + | 4877 | 4942 | **RNA**→ RpsM (COG0099)rpsM (PRK05179)→ PRK05309 (PRK05309)RpsK (COG0100)→ |
| env-91 | SRS019127_C1662274 | + | 888 | 953 | **RNA**→ RpsM (COG0099)rpsM (PRK05179)→ PRK05309 (PRK05309)RpsK (COG0100)→ |
| env-92 | SRS019219_C2851913 | + | 104 | 169 | **RNA**→ RpsM (COG0099)rpsM (PRK05179)→ |
| env-93 | SRS021496_Baylor_scaffold_36018 | - | 782 | 717 | **RNA**→ RpsM (COG0099)rpsM (PRK05179)→ PRK05309 (PRK05309)RpsK (COG0100)→ |
| env-94 | SRS022719_LANL_scaffold_70435 | + | 875 | 940 | **RNA**→ RpsM (COG0099)rpsM (PRK05179)→ PRK05309 (PRK05309)RpsK (COG0100)→ |
| env-95 | SRS023617_Baylor_scaffold_50101 | + | 369 | 434 | **RNA**→ RpsM (COG0099)rpsM (PRK05179)→ PRK05309 (PRK05309)RpsK (COG0100)→ RpsD (COG0522)rpsD (PRK05327)→ RpoA (COG0202)PRK05182 (PRK05182)→ RplQ (COG0203)rplQ (PRK05591)→ |
| env-96 | SRS024138_C2767007 | + | 4689 | 4754 | **RNA**→ RpsM (COG0099)rpsM (PRK05179)→ PRK05309 (PRK05309)RpsK (COG0100)→ |
| env-97 | SRS024580_LANL_scaffold_8234 | - | 788 | 723 | **RNA**→ RpsM (COG0099)rpsM (PRK05179)→ PRK05309 (PRK05309)RpsK (COG0100)→ |
| env-98 | SRS045127_LANL_scaffold_37515 | - | 578 | 513 | **RNA**→ RpsM (COG0099)rpsM (PRK05179)→ |
| env-99 | SRS049389_WUGC_scaffold_31108 | + | 747 | 812 | **RNA**→ RpsM (COG0099)rpsM (PRK05179)→ PRK05309 (PRK05309)RpsK (COG0100)→ |
| env-100 | SRS052227_C2920811 | + | 623 | 688 | **RNA**→ RpsM (COG0099)rpsM (PRK05179)→ PRK05309 (PRK05309)RpsK (COG0100)→ |
| env-101 | SRS055426_C3273637 | - | 680 | 615 | **RNA**→ RpsM (COG0099)rpsM (PRK05179)→ PRK05309 (PRK05309)RpsK (COG0100)→ |
| env-102 | SRS056622_LANL_scaffold_41405 | + | 1617 | 1682 | **RNA**→ RpsM (COG0099)rpsM (PRK05179)→ PRK05309 (PRK05309)RpsK (COG0100)→ |
| env-103 | SRS057205_LANL_scaffold_37743 | - | 1126 | 1061 | **RNA**→ RpsM (COG0099)rpsM (PRK05179)→ PRK05309 (PRK05309)RpsK (COG0100)→ rpsD (PRK05327)Ribosomal_S4 (pfam00163)→ |

| | | | | | |
|---|---|---|---|---|---|
| env-104 | SRS058336_LANL_scaffold_63124 | + | 2680 | 2745 | **RNA**→ RpsM (COG0099)rpsM (PRK05179)→ |
| env-105 | SRS062544_C2743506 | - | 183 | 118 | **RNA**→ RpsM (COG0099)rpsM (PRK05179)→ |
| env-106 | SRS064423_LANL_scaffold_46316 | + | 11043 | 11108 | **RNA**→ RpsM (COG0099)rpsM (PRK05179)→ PRK05309 (PRK05309)RpsK (COG0100)→ |
| env-107 | SRS065278_LANL_scaffold_19746 | + | 11609 | 11674 | **RNA**→ RpsM (COG0099)rpsM (PRK05179)→ |
| env-108 | SRS053917_LANL_scaffold_32361 | + | 1392 | 1457 | **RNA**→ |
| env-109 | SRS020334_C2558817 | + | 142 | 207 | **RNA**→ rps13 (CHL00137)rpsM (PRK05179)→ |
| env-110 | SRS021477_Baylor_scaffold_19050 | - | 755 | 690 | **RNA**→ RpsM (COG0099)rpsM (PRK05179)→ PRK05309 (PRK05309)RpsK (COG0100)→ |
| env-111 | SRS022725_LANL_scaffold_49014 | + | 367 | 432 | **RNA**→ RpsM (COG0099)rpsM (PRK05179)→ |
| env-112 | SRS044662_LANL_scaffold_17885 | + | 1141 | 1206 | **RNA**→ RpsM (COG0099)rpsM (PRK05179)→ PRK05309 (PRK05309)RpsK (COG0100)→ |
| | | | | | RpsD (COG0522)rpsD (PRK05327)→ RNAP_alpha_NTD (cd06928)PRK05182 (PRK05182)→ |
| env-113 | SRS050669_LANL_scaffold_31912 | + | 237 | 302 | **RNA**→ RpsM (COG0099)rpsM (PRK05179)→ PRK05309 (PRK05309)RpsK (COG0100)→ rpsD (PRK05327)→ |
| | | | | | S4 (smart00363)RpsD (COG0522)→ RpoA (COG0202)PRK05182 (PRK05182)→ |
| env-114 | SRS063603_LANL_scaffold_21218 | + | 1447 | 1512 | **RNA**→ RpsM (COG0099)rpsM (PRK05179)→ |
| env-115 | SRS065310_LANL_scaffold_6764 | + | 673 | 738 | **RNA**→ RpsM (COG0099)rpsM (PRK05179)→ PRK05309 (PRK05309)RpsK (COG0100)→ |
| env-116 | SRS011306_Baylor_scaffold_86858 | + | 43 | 108 | **RNA**→ RpsM (COG0099)rpsM (PRK05179)→ PRK05309 (PRK05309)RpsK (COG0100)→ |
| | | | | | RpsD (COG0522)rpsD (PRK05327)→ RpoA (COG0202)PRK05182 (PRK05182)→ |
| | | | | | RplQ (COG0203)rplQ (PRK05591)→ |
| env-117 | SRS018145_Baylor_scaffold_24537 | - | 588 | 523 | **RNA**→ RpsM (COG0099)rpsM (PRK05179)→ |
| env-118 | SRS023926_C2758757 | + | 68 | 133 | **RNA**→ RpsM (COG0099)rpsM (PRK05179)→ PRK05309 (PRK05309)RpsK (COG0100)→ |
| env-119 | SRS024015_C2994174 | + | 50 | 115 | **RNA**→ RpsM (COG0099)uS13_bact (TIGR03631)→ |
| env-120 | SRS023964_Baylor_scaffold_21763 | - | 794 | 729 | **RNA**→ RpsM (COG0099)rpsM (PRK05179)→ PRK05309 (PRK05309)RpsK (COG0100)→ |
| Lgo-2-1 | NZ_KI911785.1 | - | 2023857 | 2023791 | **RNA**→ RpsM (COG0099)rpsM (PRK05179)→ PRK05309 (PRK05309)RpsK (COG0100)→ |
| | | | | | RpsD (COG0522)rpsD (PRK05327)→ RpoA (COG0202)PRK05182 (PRK05182)→ |
| | | | | | RplQ (COG0203)rplQ (PRK05591)→ |
| Lgo-1-1 | NZ_ADAD01000134.1 | + | 34253 | 34319 | **RNA**→ RpsM (COG0099)rpsM (PRK05179)→ RpsM (COG0099)rpsM (PRK05179)→ |
| | | | | | PRK05309 (PRK05309)RpsK (COG0100)→ PRK05309 (PRK05309)RpsK (COG0100)→ |
| | | | | | RpsD (COG0522)rpsD (PRK05327)→ RpsD (COG0522)rpsD (PRK05327)→ |
| | | | | | RpoA (COG0202)PRK05182 (PRK05182)→ RpoA (COG0202)PRK05182 (PRK05182)→ |
| | | | | | RplQ (COG0203)rplQ (PRK05591)→ RplQ (COG0203)rplQ (PRK05591)→ |
| env-121 | SRS016043_C2119167 | + | 74 | 140 | **RNA**→ RpsM (COG0099)rpsM (PRK05179)→ |
| env-122 | SRS016575_Baylor_scaffold_23962 | - | 369 | 303 | **RNA**→ RpsM (COG0099)rpsM (PRK05179)→ |
| env-123 | SRS017445_Baylor_scaffold_6239 | - | 2604 | 2538 | **RNA**→ RpsM (COG0099)rpsM (PRK05179)→ PRK05309 (PRK05309)RpsK (COG0100)→ |
| | | | | | RpsD (COG0522)rpsD (PRK05327)→ RpoA (COG0202)PRK05182 (PRK05182)→ |
| env-124 | SRS019073_C2416263 | - | 249 | 183 | **RNA**→ RpsM (COG0099)rpsM (PRK05179)→ |
| env-125 | SRS063932_C3692410 | + | 186 | 252 | **RNA**→ RpsM (COG0099)rpsM (PRK05179)→ |
| env-126 | SRS022536_C4389246 | - | 210 | 144 | **RNA**→ RpsM (COG0099)rpsM (PRK05179)→ |
| env-127 | SRS015057_WUGC_scaffold_17812 | - | 437 | 374 | **RNA**→ RpsM (COG0099)rpsM (PRK05179)→ |
| env-128 | SRS022143_WUGC_scaffold_54212 | - | 269 | 206 | **RNA**→ RpsM (COG0099)rpsM (PRK05179)→ |
| env-129 | SRS023352_LANL_scaffold_3112 | - | 145 | 82 | **RNA**→ hypo→ |
| env-130 | SRS045715_LANL_scaffold_13578 | - | 494 | 431 | **RNA**→ RpsM (COG0099)rpsM (PRK05179)→ |
| env-131 | SRS019894_C3767991 | - | 146 | 38 | **RNA**→ hypo→ |
| env-132 | SRS017076_Baylor_scaffold_22004 | - | 540 | 432 | **RNA**→ RpsM (COG0099)rpsM (PRK05179)→ |
| env-133 | SRS011140_Baylor_scaffold_53017 | + | 666 | 774 | **RNA**→ RpsM (COG0099)rpsM (PRK05179)→ |
| env-134 | SRS017080_C1183285 | + | 157 | 265 | **RNA**→ RpsM (COG0099)rpsM (PRK05179)→ |
| env-135 | SRS022530_LANL_scaffold_23336 | + | 1435 | 1543 | **RNA**→ RpsM (COG0099)rpsM (PRK05179)→ PRK05309 (PRK05309)RpsK (COG0100)→ |
| env-136 | SRS023352_LANL_scaffold_46408 | - | 540 | 432 | **RNA**→ RpsM (COG0099)rpsM (PRK05179)→ |
| env-137 | SRS047210_WUGC_scaffold_54471 | - | 631 | 523 | **RNA**→ RpsM (COG0099)rpsM (PRK05179)→ PRK05309 (PRK05309)RpsK (COG0100)→ |
| env-138 | SRS056323_C2531100 | + | 190 | 298 | **RNA**→ RpsM (COG0099)rpsM (PRK05179)→ |
| env-139 | SRS064774_LANL_scaffold_45162 | - | 363 | 255 | **RNA**→ RpsM (COG0099)rpsM (PRK05179)→ |
| env-140 | SRS017808_C3851216 | - | 631 | 523 | **RNA**→ RpsM (COG0099)rpsM (PRK05179)→ PRK05309 (PRK05309)RpsK (COG0100)→ |
| env-141 | SRS016740_Baylor_scaffold_63596 | - | 601 | 493 | **RNA**→ RpsM (COG0099)rpsM (PRK05179)→ PRK05309 (PRK05309)RpsK (COG0100)→ |
| env-142 | SRS017209_Baylor_scaffold_58872 | - | 631 | 566 | **RNA**→ RpsM (COG0099)rpsM (PRK05179)→ PRK05309 (PRK05309)RpsK (COG0100)→ |
| env-143 | SRS015209_WUGC_scaffold_55797 | + | 3195 | 3260 | **RNA**→ RpsM (COG0099)rpsM (PRK05179)→ PRK05309 (PRK05309)RpsK (COG0100)→ |
| | | | | | RpsD (COG0522)rpsD (PRK05327)→ RpoA (COG0202)PRK05182 (PRK05182)→ |

| | | | | | |
|---|---|---|---|---|---|
| env-144 | SRS015272_C3450279 | - | 233 | 168 | RNA→ RpsM (COG0099)rpsM (PRK05179)→ |
| env-145 | SRS017120_Baylor_scaffold_63857 | + | 413 | 478 | RNA→ RpsM (COG0099)rpsM (PRK05179)→ |
| env-146 | SRS018300_Baylor_scaffold_29179 | - | 540 | 475 | RNA→ RpsM (COG0099)rpsM (PRK05179)→ |
| env-147 | SRS018591_WUGC_scaffold_4237 | + | 157 | 222 | RNA→ RpsM (COG0099)rpsM (PRK05179)→ PRK05309 (PRK05309)RpsK (COG0100)→<br>RpsD (COG0522)rpsD (PRK05327)→ RpoA (COG0202)PRK05182 (PRK05182)→<br>RplQ (COG0203)rplQ (PRK05591)→ |
| env-148 | SRS044486_C2170011 | - | 316 | 251 | RNA→ RpsM (COG0099)rpsM (PRK05179)→ |
| env-149 | SRS047824_WUGC_scaffold_43077 | + | 137 | 202 | RNA→ RpsM (COG0099)rpsM (PRK05179)→ PRK05309 (PRK05309)RpsK (COG0100)→ |
| env-150 | SRS053603_LANL_scaffold_76618 | - | 120 | 55 | RNA→ |
| env-151 | SRS056622_LANL_scaffold_51943 | - | 691 | 626 | RNA→ RpsM (COG0099)rpsM (PRK05179)→ PRK05309 (PRK05309)RpsK (COG0100)→ |
| env-152 | SRS064329_LANL_scaffold_26192 | - | 540 | 475 | RNA→ RpsM (COG0099)rpsM (PRK05179)→ |
| env-153 | SRS024081_LANL_scaffold_6310 | - | 649 | 584 | RNA→ RpsM (COG0099)rpsM (PRK05179)→ PRK05309 (PRK05309)RpsK (COG0100)→ |
| env-154 | SRS016086_C2470362 | + | 190 | 255 | RNA→ RpsM (COG0099)rpsM (PRK05179)→ |
| env-155 | SRS018739_WUGC_scaffold_22546 | + | 1090 | 1155 | RNA→ RpsM (COG0099)rpsM (PRK05179)→ PRK05309 (PRK05309)RpsK (COG0100)→ |
| env-156 | SRS018791_C2157240 | - | 181 | 116 | RNA→ RpsM (COG0099)rpsM (PRK05179)→ |
| env-157 | SRS020220_Baylor_scaffold_722 | - | 146 | 81 | RNA→ hypo→ |
| env-158 | SRS051791_C1815930 | - | 691 | 626 | RNA→ RpsM (COG0099)rpsM (PRK05179)→ PRK05309 (PRK05309)RpsK (COG0100)→ |
| env-159 | SRS013705_Baylor_scaffold_86091 | - | 631 | 566 | RNA→ RpsM (COG0099)rpsM (PRK05179)→ PRK05309 (PRK05309)RpsK (COG0100)→ |
| env-160 | SRS014271_WUGC_scaffold_1617 | + | 128 | 193 | RNA→ RpsM (COG0099)rpsM (PRK05179)→ |
| env-161 | SRS023926_Baylor_scaffold_56329 | + | 157 | 222 | RNA→ RpsM (COG0099)rpsM (PRK05179)→ PRK05309 (PRK05309)RpsK (COG0100)→<br>RpsD (COG0522)rpsD (PRK05327)→ RpoA (COG0202)PRK05182 (PRK05182)→<br>RplQ (COG0203)rplQ (PRK05591)→ |
| env-162 | SRS047219_WUGC_scaffold_31331 | - | 478 | 413 | RNA→ RpsM (COG0099)rpsM (PRK05179)→ |
| env-163 | SRS065335_LANL_scaffold_24633 | + | 231 | 296 | RNA→ RpsM (COG0099)rpsM (PRK05179)→ |
| env-164 | SRS019219_WUGC_scaffold_52621 | - | 783 | 718 | RNA→ RpsM (COG0099)rpsM (PRK05179)→ PRK05309 (PRK05309)RpsK (COG0100)→ |
| env-165 | SRS015893_WUGC_scaffold_14977 | + | 11585 | 11650 | RNA→ RpsM (COG0099)rpsM (PRK05179)→ PRK05309 (PRK05309)RpsK (COG0100)→ |
| env-166 | SRS024138_Baylor_scaffold_8471 | + | 1547 | 1612 | RNA→ RpsM (COG0099)rpsM (PRK05179)→ PRK05309 (PRK05309)RpsK (COG0100)→<br>RpsD (COG0522)rpsD (PRK05327)→ RpoA (COG0202)PRK05182 (PRK05182)→<br>RplQ (COG0203)rplQ (PRK05591)→ |
| env-167 | SRS065278_LANL_scaffold_50816 | - | 540 | 475 | RNA→ RpsM (COG0099)rpsM (PRK05179)→ |
| env-168 | SRS019607_C2720344 | + | 602 | 667 | RNA→ RpsM (COG0099)rpsM (PRK05179)→ |
| env-169 | SRS064423_LANL_scaffold_7006 | - | 793 | 728 | RNA→ RpsM (COG0099)rpsM (PRK05179)→ PRK05309 (PRK05309)RpsK (COG0100)→ |
| env-170 | SRS050628_C1893763 | + | 408 | 473 | RNA→ RpsM (COG0099)rpsM (PRK05179)→ |
| env-171 | SRS015278_WUGC_scaffold_6421 | + | 97 | 162 | RNA→ RpsM (COG0099)rpsM (PRK05179)→ |
| env-172 | SRS024144_LANL_scaffold_20370 | - | 551 | 486 | RNA→ RpsM (COG0099)rpsM (PRK05179)→ |
| env-173 | SRS055450_LANL_scaffold_40160 | - | 148 | 83 | RNA→ hypo→ |
| Lwa-1-1 | NZ_KB891061.1 | + | 14728 | 14793 | RNA→ RpsM (COG0099)rpsM (PRK05179)→ PRK05309 (PRK05309)RpsK (COG0100)→<br>RpsD (COG0522)rpsD (PRK05327)→ RpoA (COG0202)PRK05182 (PRK05182)→<br>RplQ (COG0203)rplQ (PRK05591)→ |
| Lwa-2-1 | NZ_KI271401.1 | - | 87815 | 87750 | RNA→ RpsM (COG0099)rpsM (PRK05179)→ PRK05309 (PRK05309)RpsK (COG0100)→<br>RpsD (COG0522)rpsD (PRK05327)→ RpoA (COG0202)PRK05182 (PRK05182)→<br>RplQ (COG0203)rplQ (PRK05591)→ |
| env-174 | SRS022083_Baylor_scaffold_19178 | - | 217 | 152 | RNA→ RpsM (COG0099)rpsM (PRK05179)→ |
| env-175 | SRS014690_C1889275 | - | 403 | 338 | RNA→ RpsM (COG0099)uS13_bact (TIGR03631)→ |
| env-176 | SRS017445_C3034105 | + | 35 | 100 | RNA→ RpsM (COG0099)rpsM (PRK05179)→ |
| env-177 | SRS047113_LANL_scaffold_39994 | + | 35 | 100 | RNA→ RpsM (COG0099)rpsM (PRK05179)→ |
| env-178 | SRS022536_LANL_scaffold_18564 | - | 468 | 403 | RNA→ RpsM (COG0099)rpsM (PRK05179)→ |
| env-179 | SRS013723_Baylor_scaffold_46255 | + | 46 | 111 | RNA→ RpsM (COG0099)uS13_bact (TIGR03631)→ |
| env-180 | SRS016360_Baylor_scaffold_58948 | - | 577 | 512 | RNA→ RpsM (COG0099)rpsM (PRK05179)→ |
| env-181 | SRS014476_C3328552 | + | 91 | 156 | RNA→ RpsM (COG0099)rpsM (PRK05179)→ |
| env-182 | SRS015470_C3307239 | + | 122 | 187 | RNA→ RpsM (COG0099)rpsM (PRK05179)→ |
| env-183 | SRS047265_C1024260 | + | 85 | 150 | RNA→ RpsM (COG0099)rpsM (PRK05179)→ |
| env-184 | SRS065310_LANL_scaffold_16461 | + | 1324 | 1389 | RNA→ RpsM (COG0099)rpsM (PRK05179)→ |
| env-185 | SRS053917_LANL_scaffold_85561 | + | 405 | 470 | RNA→ RpsM (COG0099)rpsM (PRK05179)→ |

| | | | | | |
|---|---|---|---|---|---|
| env-186 | SRS011343_Baylor_scaffold_32578 | - | 1439 | 1374 | **RNA**→ RpsM (COG0099)rpsM (PRK05179)→ PRK05309 (PRK05309)RpsK (COG0100)→<br>RpsD (COG0522)rpsD (PRK05327)→ |
| Lsp-3-1 | NZ_KI272907.1 | - | 14521 | 14456 | **RNA**→ RpsM (COG0099)rpsM (PRK05179)→ PRK05309 (PRK05309)RpsK (COG0100)→<br>RpsD (COG0522)rpsD (PRK05327)→ RpoA (COG0202)PRK05182 (PRK05182)→<br>RplQ (COG0203)rplQ (PRK05591)→ |
| env-187 | SRS014578_WUGC_scaffold_46133 | + | 84 | 149 | **RNA**→ hypo→ RpsM (COG0099)rpsM (PRK05179)→ |
| env-188 | SRS021477_Baylor_scaffold_2106 | - | 788 | 723 | **RNA**→ RpsM (COG0099)rpsM (PRK05179)→ PRK05309 (PRK05309)RpsK (COG0100)→ |
| env-189 | SRS015044_WUGC_scaffold_24016 | + | 23 | 88 | **RNA**→ RpsM (COG0099)rpsM (PRK05179)→ |
| env-190 | SRS063603_LANL_scaffold_95971 | - | 467 | 402 | **RNA**→ RpsM (COG0099)rpsM (PRK05179)→ |
| env-191 | SRS055378_LANL_scaffold_65216 | + | 150 | 215 | **RNA**→ RpsM (COG0099)rpsM (PRK05179)→ |
| env-192 | SRS052604_C1779911 | - | 369 | 304 | **RNA**→ RpsM (COG0099)rpsM (PRK05179)→ |
| Lsp-4-1 | NZ_KI271324.1 | + | 15601 | 15666 | **RNA**→ RpsM (COG0099)rpsM (PRK05179)→ PRK05309 (PRK05309)RpsK (COG0100)→<br>RpsD (COG0522)rpsD (PRK05327)→ RpoA (COG0202)PRK05182 (PRK05182)→<br>RplQ (COG0203)rplQ (PRK05591)→ |
| env-193 | SRS016200_WUGC_scaffold_19120 | - | 324 | 259 | **RNA**→ RpsM (COG0099)rpsM (PRK05179)→ |
| env-194 | SRS049318_LANL_scaffold_31958 | - | 260 | 195 | **RNA**→ RpsM (COG0099)rpsM (PRK05179)→ |
| env-195 | SRS024447_LANL_scaffold_24021 | + | 353 | 418 | **RNA**→ RpsM (COG0099)rpsM (PRK05179)→ PRK05309 (PRK05309)RpsK (COG0100)→ |
| Ltr-1-1 | NZ_AXVL01000009.1 | - | 21488 | 21423 | **RNA**→ RpsM (COG0099)rpsM (PRK05179)→ PRK05309 (PRK05309)RpsK (COG0100)→<br>RpsD (COG0522)rpsD (PRK05327)→ RpoA (COG0202)PRK05182 (PRK05182)→<br>RplQ (COG0203)rplQ (PRK05591)→ |
| Lsh-1-1 | NZ_KB890273.1 | - | 105178 | 105113 | **RNA**→ RpsM (COG0099)rpsM (PRK05179)→ PRK05309 (PRK05309)RpsK (COG0100)→<br>RpsD (COG0522)rpsD (PRK05327)→ RpoA (COG0202)PRK05182 (PRK05182)→<br>RplQ (COG0203)rplQ (PRK05591)→ |
| Lbu-1-1 | NC_013192.1 | + | 2131217 | 2131282 | **RNA**→ RpsM (COG0099)rpsM (PRK05179)→ PRK05309 (PRK05309)RpsK (COG0100)→<br>RpsD (COG0522)rpsD (PRK05327)→ RpoA (COG0202)PRK05182 (PRK05182)→<br>RplQ (COG0203)rplQ (PRK05591)→ hypo→ hypo→ Aes (COG0657)→<br>HAMP (smart00304)HATPase_c (smart00387)HisKA (smart00388)BaeS (COG0642)→ |
| env-196 | SRS018591_WUGC_scaffold_43849 | + | 45065 | 45132 | **RNA**→ RpsM (COG0099)rpsM (PRK05179)→ PRK05309 (PRK05309)RpsK (COG0100)→<br>RpsD (COG0522)rpsD (PRK05327)→ RpoA (COG0202)PRK05182 (PRK05182)→<br>RplQ (COG0203)rplQ (PRK05591)→ |
| env-197 | SRS063999_LANL_scaffold_22653 | - | 339 | 274 | **RNA**→ RpsM (COG0099)rpsM (PRK05179)→ |
| env-198 | SRS075404_LANL_scaffold_62603 | - | 150 | 85 | **RNA**→ hypo→ |
| env-199 | SRS014573_WUGC_scaffold_24913 | + | 104 | 169 | **RNA**→ RpsM (COG0099)rpsM (PRK05179)→ PRK05309 (PRK05309)RpsK (COG0100)→ |
| env-200 | SRS017120_C2908388 | + | 19 | 84 | **RNA**→ RpsM (COG0099)rpsM (PRK05179)→ |
| env-201 | SRS017533_C4103099 | - | 869 | 804 | **RNA**→ RpsM (COG0099)rpsM (PRK05179)→ PRK05309 (PRK05309)RpsK (COG0100)→ |
| env-202 | SRS018439_Baylor_scaffold_46100 | + | 34 | 99 | **RNA**→ RpsM (COG0099)rpsM (PRK05179)→ PRK05309 (PRK05309)RpsK (COG0100)→<br>RpsD (COG0522)rpsD (PRK05327)→ |
| env-203 | SRS018739_C3600550 | + | 123 | 188 | **RNA**→ RpsM (COG0099)rpsM (PRK05179)→ |
| env-204 | SRS019327_WUGC_scaffold_19629 | - | 1708 | 1643 | **RNA**→ RpsM (COG0099)rpsM (PRK05179)→ PRK05309 (PRK05309)RpsK (COG0100)→<br>RpsD (COG0522)rpsD (PRK05327)→ PRK05182 (PRK05182)RNA_pol_L (pfam01193)→ |
| env-205 | SRS022077_Baylor_scaffold_16023 | - | 575 | 510 | **RNA**→ RpsM (COG0099)rpsM (PRK05179)→ |
| env-206 | SRS024081_LANL_scaffold_33076 | + | 104 | 169 | **RNA**→ RpsM (COG0099)rpsM (PRK05179)→ |
| env-207 | SRS042131_WUGC_scaffold_3782 | + | 14639 | 14704 | **RNA**→ RpsM (COG0099)rpsM (PRK05179)→ PRK05309 (PRK05309)RpsK (COG0100)→<br>RpsD (COG0522)rpsD (PRK05327)→ RpoA (COG0202)PRK05182 (PRK05182)→<br>RplQ (COG0203)rplQ (PRK05591)→ |
| env-208 | SRS042643_WUGC_scaffold_36196 | + | 924 | 989 | **RNA**→ RpsM (COG0099)rpsM (PRK05179)→ |
| env-209 | SRS042910_C5617440 | + | 723 | 788 | **RNA**→ RpsM (COG0099)rpsM (PRK05179)→ PRK05309 (PRK05309)RpsK (COG0100)→ |
| env-210 | SRS043663_LANL_scaffold_40516 | - | 4450 | 4385 | **RNA**→ RpsM (COG0099)rpsM (PRK05179)→ PRK05309 (PRK05309)RpsK (COG0100)→<br>RpsD (COG0522)rpsD (PRK05327)→ RpoA (COG0202)PRK05182 (PRK05182)→<br>RplQ (COG0203)rplQ (PRK05591)→ |
| env-211 | SRS044373_WUGC_scaffold_60101 | - | 526 | 461 | **RNA**→ RpsM (COG0099)rpsM (PRK05179)→ |
| env-212 | SRS049268_LANL_scaffold_41923 | - | 669 | 604 | **RNA**→ RpsM (COG0099)rpsM (PRK05179)→ PRK05309 (PRK05309)RpsK (COG0100)→ |
| env-213 | SRS057022_LANL_scaffold_28937 | + | 206 | 271 | **RNA**→ RpsM (COG0099)rpsM (PRK05179)→ |
| env-214 | SRS057539_LANL_scaffold_29275 | + | 3873 | 3938 | **RNA**→ RpsM (COG0099)rpsM (PRK05179)→ |
| env-215 | SRS063193_LANL_scaffold_60270 | - | 626 | 561 | **RNA**→ RpsM (COG0099)rpsM (PRK05179)→ PRK05309 (PRK05309)RpsK (COG0100)→ |

| | | | | | |
|---|---|---|---|---|---|
| env-216 | SRS064774_C2799284 | + | 104 | 169 | **RNA**→ RpsM (COG0099)rpsM (PRK05179)→ |
| env-217 | SRS077736_C1759087 | - | 289 | 224 | **RNA**→ RpsM (COG0099)rpsM (PRK05179)→ |
| Lsp-2-1 | NZ_KI272826.1 | + | 100033 | 100098 | **RNA**→ RpsM (COG0099)rpsM (PRK05179)→ PRK05309 (PRK05309)RpsK (COG0100)→ RpsD (COG0522)rpsD (PRK05327)→ RpoA (COG0202)PRK05182 (PRK05182)→ RplQ (COG0203)rplQ (PRK05591)→ |
| env-218 | SRS023835_C3783904 | - | 527 | 462 | **RNA**→ RpsM (COG0099)rpsM (PRK05179)→ |
| env-219 | SRS055378_LANL_scaffold_20064 | - | 467 | 401 | **RNA**→ RpsM (COG0099)rpsM (PRK05179)→ |
| env-220 | SRS016043_WUGC_scaffold_6899 | + | 458 | 524 | **RNA**→ RpsM (COG0099)rpsM (PRK05179)→ |
| env-221 | SRS019591_WUGC_scaffold_33061 | - | 2209 | 2143 | **RNA**→ RpsM (COG0099)rpsM (PRK05179)→ PRK05309 (PRK05309)RpsK (COG0100)→ RpsD (COG0522)rpsD (PRK05327)→ RNAP_alpha_NTD (cd06928)PRK05182 (PRK05182)→ |
| env-222 | SRS013170_Baylor_scaffold_84939 | - | 535 | 469 | **RNA**→ RpsM (COG0099)rpsM (PRK05179)→ |
| env-223 | SRS017139_Baylor_scaffold_75273 | - | 382 | 316 | **RNA**→ RpsM (COG0099)rpsM (PRK05179)→ |
| env-224 | SRS023938_Baylor_scaffold_26910 | - | 1104 | 1038 | **RNA**→ RpsM (COG0099)rpsM (PRK05179)→ PRK05309 (PRK05309)RpsK (COG0100)→ rpsD (PRK05327)Ribosomal_S4 (pfam00163)→ |
| env-225 | SRS051930_C3131142 | - | 369 | 303 | **RNA**→ RpsM (COG0099)rpsM (PRK05179)→ |
| env-226 | SRS023841_Baylor_scaffold_19242 | + | 7063 | 7129 | **RNA**→ RpsM (COG0099)rpsM (PRK05179)→ PRK05309 (PRK05309)RpsK (COG0100)→ rpsD (PRK05327)Ribosomal_S4 (pfam00163)→ |
| env-227 | SRS011098_Baylor_scaffold_14455 | + | 724 | 790 | **RNA**→ RpsM (COG0099)rpsM (PRK05179)→ PRK05309 (PRK05309)RpsK (COG0100)→ RpsD (COG0522)rpsD (PRK05327)→ RpoA (COG0202)PRK05182 (PRK05182)→ RplQ (COG0203)rplQ (PRK05591)→ |
| env-228 | SRS012285_Baylor_scaffold_13798 | + | 101 | 167 | **RNA**→ RpsM (COG0099)rpsM (PRK05179)→ |
| env-229 | SRS013533_PGA_scaffold_82441 | - | 660 | 594 | **RNA**→ RpsM (COG0099)rpsM (PRK05179)→ RpsM (COG0099)rpsM (PRK05179)→ |
| env-230 | SRS014578_WUGC_scaffold_68326 | - | 305 | 239 | **RNA**→ RpsM (COG0099)rpsM (PRK05179)→ |
| env-231 | SRS015470_C3336101 | - | 669 | 603 | **RNA**→ RpsM (COG0099)rpsM (PRK05179)→ PRK05309 (PRK05309)RpsK (COG0100)→ |
| env-232 | SRS016575_C3616229 | - | 112 | 46 | **RNA**→ hypo→ |
| env-233 | SRS017511_C3312624 | - | 195 | 129 | **RNA**→ RpsM (COG0099)rpsM (PRK05179)→ |
| env-234 | SRS018394_Baylor_scaffold_45763 | + | 120 | 186 | **RNA**→ RpsM (COG0099)rpsM (PRK05179)→ |
| env-235 | SRS018665_WUGC_scaffold_479 | - | 1029 | 963 | **RNA**→ RpsM (COG0099)rpsM (PRK05179)→ PRK05309 (PRK05309)RpsK (COG0100)→ rpsD (PRK05327)→ |
| env-236 | SRS018975_C1710475 | - | 195 | 129 | **RNA**→ RpsM (COG0099)rpsM (PRK05179)→ |
| env-237 | SRS022149_LANL_scaffold_16029 | - | 599 | 533 | **RNA**→ RpsM (COG0099)rpsM (PRK05179)→ |
| env-238 | SRS024649_C3588658 | - | 373 | 307 | **RNA**→ RpsM (COG0099)rpsM (PRK05179)→ |
| env-239 | SRS045197_C3310867 | + | 117 | 183 | **RNA**→ RpsM (COG0099)rpsM (PRK05179)→ |
| env-240 | SRS051378_C1836081 | + | 393 | 459 | **RNA**→ RpsM (COG0099)rpsM (PRK05179)→ |
| env-241 | SRS052604_C1775703 | - | 372 | 306 | **RNA**→ RpsM (COG0099)rpsM (PRK05179)→ |
| env-242 | SRS055450_C2060497 | + | 104 | 170 | **RNA**→ RpsM (COG0099)rpsM (PRK05179)→ |
| env-243 | SRS075410_LANL_scaffold_38102 | - | 535 | 469 | **RNA**→ RpsM (COG0099)rpsM (PRK05179)→ |
| env-244 | SRS014476_C3276433 | - | 262 | 196 | **RNA**→ RpsM (COG0099)rpsM (PRK05179)→ |
| env-245 | SRS015215_WUGC_scaffold_30194 | - | 2281 | 2215 | **RNA**→ RpsM (COG0099)rpsM (PRK05179)→ PRK05309 (PRK05309)RpsK (COG0100)→ RpsD (COG0522)rpsD (PRK05327)→ RNAP_alpha_NTD (cd06928)PRK05182 (PRK05182)→ |
| env-246 | SRS015899_C1863770 | - | 113 | 47 | **RNA**→ hypo→ |
| env-247 | SRS019073_C2423699 | + | 117 | 183 | **RNA**→ RpsM (COG0099)rpsM (PRK05179)→ |
| env-248 | SRS019077_C2628206 | + | 123 | 189 | **RNA**→ RpsM (COG0099)rpsM (PRK05179)→ |
| env-249 | SRS020226_Baylor_scaffold_50340 | - | 366 | 300 | **RNA**→ RpsM (COG0099)rpsM (PRK05179)→ |
| env-250 | SRS024087_LANL_scaffold_39994 | + | 1637 | 1703 | **RNA**→ RpsM (COG0099)rpsM (PRK05179)→ |
| env-251 | SRS024381_LANL_scaffold_49494 | + | 578 | 644 | **RNA**→ RpsM (COG0099)rpsM (PRK05179)→ PRK05309 (PRK05309)RpsK (COG0100)→ |
| env-252 | SRS047113_LANL_scaffold_2260 | + | 101 | 167 | **RNA**→ RpsM (COG0099)rpsM (PRK05179)→ |
| env-253 | SRS051244_LANL_scaffold_43648 | - | 535 | 469 | **RNA**→ RpsM (COG0099)rpsM (PRK05179)→ |
| env-254 | SRS051941_LANL_scaffold_55544 | - | 535 | 469 | **RNA**→ RpsM (COG0099)rpsM (PRK05179)→ |
| Lsp-1-1 | NZ_CP012410.1 | + | 259790 | 259856 | **RNA**→ RpsM (COG0099)rpsM (PRK05179)→ PRK05309 (PRK05309)RpsK (COG0100)→ RpsD (COG0522)rpsD (PRK05327)→ RpoA (COG0202)PRK05182 (PRK05182)→ RplQ (COG0203)rplQ (PRK05591)→ |
| env-255 | SRS013723_Baylor_scaffold_66969 | + | 1803 | 1869 | **RNA**→ RpsM (COG0099)rpsM (PRK05179)→ PRK05309 (PRK05309)RpsK (COG0100)→ |

| | | | | | |
|---|---|---|---|---|---|
| env-256 | JGI24712J26585_10001668 | + | 13781 | 13845 | **RNA**→ RpsM (COG0099)rpsM (PRK05179)→ PRK05309 (PRK05309)RpsK (COG0100)→<br>RpsD (COG0522)rpsD (PRK05327)→ RpoA (COG0202)PRK05182 (PRK05182)→<br>RplQ (COG0203)rplQ (PRK05591)→ |
| env-257 | JGI24711J26586_10001349 | + | 13781 | 13845 | **RNA**→ RpsM (COG0099)rpsM (PRK05179)→ PRK05309 (PRK05309)RpsK (COG0100)→<br>RpsD (COG0522)rpsD (PRK05327)→ RpoA (COG0202)PRK05182 (PRK05182)→<br>RplQ (COG0203)rplQ (PRK05591)→ |
| env-258 | JGI24502J29692_10003819 | + | 16692 | 16756 | **RNA**→ RpsM (COG0099)rpsM (PRK05179)→ PRK05309 (PRK05309)RpsK (COG0100)→<br>RpsD (COG0522)rpsD (PRK05327)→ RpoA (COG0202)PRK05182 (PRK05182)→<br>RplQ (COG0203)rplQ (PRK05591)→ |
| env-259 | JGI24503J29689_10006856 | + | 15447 | 15511 | **RNA**→ RpsM (COG0099)rpsM (PRK05179)→ PRK05309 (PRK05309)RpsK (COG0100)→<br>RpsD (COG0522)rpsD (PRK05327)→<br>RNAP_alpha_NTD (cd06928)PRK05182 (PRK05182)RNA_pol_A_CTD (pfam03118)→<br>RplQ (COG0203)rplQ (PRK05591)→ |
| Cso-1-1 | NZ_KI518192.1 | + | 13030 | 13084 | **RNA**→ RpsM (COG0099)rpsM (PRK05179)→ PRK05309 (PRK05309)RpsK (COG0100)→<br>RpsD (COG0522)rpsD (PRK05327)→ RpoA (COG0202)PRK05182 (PRK05182)→<br>RplQ (COG0203)rplQ (PRK05591)→ |
| Csp-2-1 | NZ_JRJZ01000165.1 | + | 13067 | 13121 | **RNA**→ RpsM (COG0099)rpsM (PRK05179)→ PRK05309 (PRK05309)RpsK (COG0100)→<br>RpsD (COG0522)rpsD (PRK05327)→ RpoA (COG0202)PRK05182 (PRK05182)→<br>RplQ (COG0203)rplQ (PRK05591)→ |
| Csp-1-1 | NZ_JTLI01000001.1 | + | 13083 | 13137 | **RNA**→ RpsM (COG0099)rpsM (PRK05179)→ PRK05309 (PRK05309)RpsK (COG0100)→<br>RpsD (COG0522)rpsD (PRK05327)→ RpoA (COG0202)PRK05182 (PRK05182)→<br>RplQ (COG0203)rplQ (PRK05591)→ |
| env-260 | JGI994J12354_10000051 | - | 8355 | 8297 | **RNA**→ RpsM (COG0099)rpsM (PRK05179)→ PRK05309 (PRK05309)RpsK (COG0100)→<br>RpsD (COG0522)rpsD (PRK05327)→ RpoA (COG0202)PRK05182 (PRK05182)→<br>RplQ (COG0203)rplQ (PRK05591)→ |
| env-261 | 2227390564 | - | 241 | 183 | **RNA**→ RpsM (COG0099)rpsM (PRK05179)→ |
| env-262 | IMNBL1_c0066592 | + | 328 | 386 | **RNA**→ RpsM (COG0099)rpsM (PRK05179)→ rps11 (CHL00041)PRK05309 (PRK05309)→ |
| env-263 | BMHBC_159667 | + | 548 | 606 | **RNA**→ RpsM (COG0099)rpsM (PRK05179)→ PRK05309 (PRK05309)RpsK (COG0100)→<br>RpsD (COG0522)rpsD (PRK05327)→ RNAP_alpha_NTD (cd06928)PRK05182 (PRK05182)→ |
| Ipo-1-1 | NC_014632.1 | - | 1917483 | 1917425 | **RNA**→ RpsM (COG0099)rpsM (PRK05179)→ PRK05309 (PRK05309)RpsK (COG0100)→<br>RpsD (COG0522)rpsD (PRK05327)→ RpoA (COG0202)PRK05182 (PRK05182)→<br>RplQ (COG0203)rplQ (PRK05591)→ |
| env-264 | contig93332 | + | 673 | 734 | **RNA**→ RpsM (COG0099)rpsM (PRK05179)→ |
| env-265 | scaffold18717_1_V1.CD-11 | + | 1496 | 1557 | **RNA**→ RpsM (COG0099)rpsM (PRK05179)→ |
| env-266 | DLF004_scaffold35352_1 | + | 13177 | 13238 | **RNA**→ RpsM (COG0099)rpsM (PRK05179)→ PRK05309 (PRK05309)RpsK (COG0100)→<br>RpsD (COG0522)rpsD (PRK05327)→ RpoA (COG0202)PRK05182 (PRK05182)→<br>RplQ (COG0203)rplQ (PRK05591)→ |
| env-267 | DLM010_scaffold45172_1 | - | 4988 | 4927 | **RNA**→ RpsM (COG0099)rpsM (PRK05179)→ PRK05309 (PRK05309)RpsK (COG0100)→<br>RpsD (COG0522)rpsD (PRK05327)→ RpoA (COG0202)PRK05182 (PRK05182)→<br>RplQ (COG0203)rplQ (PRK05591)→ |
| env-268 | DLM014_scaffold52950_1 | + | 13617 | 13678 | **RNA**→ RpsM (COG0099)rpsM (PRK05179)→ PRK05309 (PRK05309)RpsK (COG0100)→<br>RpsD (COG0522)rpsD (PRK05327)→ RpoA (COG0202)PRK05182 (PRK05182)→<br>RplQ (COG0203)rplQ (PRK05591)→ |
| env-269 | DOM010_scaffold24129_1 | + | 9239 | 9300 | **RNA**→ RpsM (COG0099)rpsM (PRK05179)→ PRK05309 (PRK05309)RpsK (COG0100)→<br>RpsD (COG0522)rpsD (PRK05327)→ RpoA (COG0202)PRK05182 (PRK05182)→<br>RplQ (COG0203)rplQ (PRK05591)→ |
| env-270 | DOM016_scaffold37026_2 | - | 10720 | 10659 | **RNA**→ RpsM (COG0099)rpsM (PRK05179)→ PRK05309 (PRK05309)RpsK (COG0100)→<br>RpsD (COG0522)rpsD (PRK05327)→ RpoA (COG0202)PRK05182 (PRK05182)→<br>RplQ (COG0203)rplQ (PRK05591)→ |
| env-271 | DOM017_scaffold7272_3 | - | 2891 | 2830 | **RNA**→ RpsM (COG0099)rpsM (PRK05179)→ PRK05309 (PRK05309)RpsK (COG0100)→<br>RpsD (COG0522)rpsD (PRK05327)→ RpoA (COG0202)PRK05182 (PRK05182)→<br>rplQ (PRK05591)Ribosomal_L17 (pfam01196)→ |
| env-272 | NLF014_scaffold15038_1 | + | 529 | 590 | **RNA**→ RpsM (COG0099)rpsM (PRK05179)→ |
| env-273 | NLF015_scaffold34189_4 | - | 5098 | 5037 | **RNA**→ RpsM (COG0099)rpsM (PRK05179)→ PRK05309 (PRK05309)RpsK (COG0100)→<br>RpsD (COG0522)rpsD (PRK05327)→ RpoA (COG0202)PRK05182 (PRK05182)→<br>RplQ (COG0203)rplQ (PRK05591)→ |
| env-274 | NLM006_scaffold51268_3 | - | 727 | 666 | **RNA**→ RpsM (COG0099)rpsM (PRK05179)→ rps11 (CHL00041)PRK05309 (PRK05309)→ |

| | | | | | |
|---|---|---|---|---|---|
| env-275 | NOM026_scaffold3_1 | - | 17438 | 17377 | **RNA**→ RpsM (COG0099)rpsM (PRK05179)→ PRK05309 (PRK05309)RpsK (COG0100)→ RpsD (COG0522)rpsD (PRK05327)→ RpoA (COG0202)PRK05182 (PRK05182)→ RplQ (COG0203)rplQ (PRK05591)→ |
| env-276 | SRS017433_C1836359 | - | 3003 | 2942 | **RNA**→ RpsM (COG0099)rpsM (PRK05179)→ PRK05309 (PRK05309)RpsK (COG0100)→ RpsD (COG0522)rpsD (PRK05327)→ RpoA (COG0202)PRK05182 (PRK05182)→ RplQ (COG0203)rplQ (PRK05591)→ |
| env-277 | SRS023176_C936722 | - | 3456 | 3395 | **RNA**→ RpsM (COG0099)rpsM (PRK05179)→ PRK05309 (PRK05309)RpsK (COG0100)→ RpsD (COG0522)rpsD (PRK05327)→ RpoA (COG0202)PRK05182 (PRK05182)→ RplQ (COG0203)rplQ (PRK05591)→ |
| Fmo-1-1 | NZ_GL987993.1 | + | 62917 | 62978 | **RNA**→ RpsM (COG0099)rpsM (PRK05179)→ PRK05309 (PRK05309)RpsK (COG0100)→ RpsD (COG0522)rpsD (PRK05327)→ RpoA (COG0202)PRK05182 (PRK05182)→ RplQ (COG0203)rplQ (PRK05591)→ |
| env-278 | NLM024_scaffold20695_1 | - | 5661 | 5600 | **RNA**→ RpsM (COG0099)rpsM (PRK05179)→ PRK05309 (PRK05309)RpsK (COG0100)→ RpsD (COG0522)rpsD (PRK05327)→ RpoA (COG0202)PRK05182 (PRK05182)→ RplQ (COG0203)rplQ (PRK05591)→ |
| env-279 | NLM023_scaffold45840_1 | - | 36496 | 36435 | **RNA**→ RpsM (COG0099)rpsM (PRK05179)→ PRK05309 (PRK05309)RpsK (COG0100)→ RpsD (COG0522)rpsD (PRK05327)→ RpoA (COG0202)PRK05182 (PRK05182)→ RplQ (COG0203)rplQ (PRK05591)→ |
| env-280 | DOM005_scaffold13463_2 | - | 25370 | 25308 | **RNA**→ RpsM (COG0099)rpsM (PRK05179)→ PRK05309 (PRK05309)RpsK (COG0100)→ RpsD (COG0522)rpsD (PRK05327)→ RpoA (COG0202)PRK05182 (PRK05182)→ RplQ (COG0203)rplQ (PRK05591)→ |
| env-281 | DOM010_scaffold83_2 | + | 2172 | 2233 | **RNA**→ RpsM (COG0099)rpsM (PRK05179)→ PRK05309 (PRK05309)RpsK (COG0100)→ RpsD (COG0522)rpsD (PRK05327)→ RpoA (COG0202)PRK05182 (PRK05182)→ RplQ (COG0203)rplQ (PRK05591)→ |
| env-282 | 4448815.3_Contig41831 | + | 346 | 407 | **RNA**→ |
| env-283 | DOM023_scaffold22784_3 | - | 323 | 262 | **RNA**→ RpsM (COG0099)rpsM (PRK05179)→ |
| env-284 | NLM027_scaffold31130_3 | - | 5574 | 5513 | **RNA**→ RpsM (COG0099)rpsM (PRK05179)→ PRK05309 (PRK05309)RpsK (COG0100)→ RpsD (COG0522)rpsD (PRK05327)→ RpoA (COG0202)PRK05182 (PRK05182)→ RplQ (COG0203)rplQ (PRK05591)→ |
| env-285 | NOM022_scaffold9197_2 | - | 8382 | 8321 | **RNA**→ RpsM (COG0099)rpsM (PRK05179)→ PRK05309 (PRK05309)RpsK (COG0100)→ RpsD (COG0522)rpsD (PRK05327)→ RpoA (COG0202)PRK05182 (PRK05182)→ RplQ (COG0203)rplQ (PRK05591)→ |
| env-286 | NOM023_scaffold2804_1 | + | 2069 | 2130 | **RNA**→ RpsM (COG0099)rpsM (PRK05179)→ PRK05309 (PRK05309)RpsK (COG0100)→ RpsD (COG0522)rpsD (PRK05327)→ RpoA (COG0202)PRK05182 (PRK05182)→ RplQ (COG0203)rplQ (PRK05591)→ |
| env-287 | 4444165.3.11744 | + | 3036 | 3097 | **RNA**→ RpsM (COG0099)rpsM (PRK05179)→ PRK05309 (PRK05309)RpsK (COG0100)→ |
| env-288 | NOF009_scaffold43353_2 | - | 13425 | 13364 | **RNA**→ RpsM (COG0099)rpsM (PRK05179)→ PRK05309 (PRK05309)RpsK (COG0100)→ RpsD (COG0522)rpsD (PRK05327)→ RpoA (COG0202)PRK05182 (PRK05182)→ RplQ (COG0203)rplQ (PRK05591)→ |
| env-289 | NLM017_scaffold13921_5 | - | 472 | 411 | **RNA**→ RpsM (COG0099)rpsM (PRK05179)→ |
| env-290 | NOM004_scaffold18127_3 | + | 1020 | 1081 | **RNA**→ RpsM (COG0099)rpsM (PRK05179)→ PRK05309 (PRK05309)RpsK (COG0100)→ RpsD (COG0522)rpsD (PRK05327)→ RpoA (COG0202)PRK05182 (PRK05182)→ RplQ (COG0203)rplQ (PRK05591)→ |
| env-291 | UnmappedStool_Broad_C252318921 | - | 272 | 211 | **RNA**→ rps13 (CHL00137)rpsM (PRK05179)→ |
| env-292 | DLF008_C336303_1 | + | 18232 | 18293 | **RNA**→ RpsM (COG0099)rpsM (PRK05179)→ PRK05309 (PRK05309)RpsK (COG0100)→ RpsD (COG0522)rpsD (PRK05327)→ RpoA (COG0202)PRK05182 (PRK05182)→ RplQ (COG0203)rplQ (PRK05591)→ |
| env-293 | DOF006_scaffold7307_3 | - | 10364 | 10303 | **RNA**→ RpsM (COG0099)rpsM (PRK05179)→ PRK05309 (PRK05309)RpsK (COG0100)→ RpsD (COG0522)rpsD (PRK05327)→ RpoA (COG0202)PRK05182 (PRK05182)→ RplQ (COG0203)rplQ (PRK05591)→ |
| env-294 | DOM005_scaffold6985_2 | + | 829 | 890 | **RNA**→ RpsM (COG0099)uS13_bact (TIGR03631)→ |
| Ful-2-1 | NZ_JH815234.1 | + | 58593 | 58654 | **RNA**→ RpsM (COG0099)rpsM (PRK05179)→ PRK05309 (PRK05309)RpsK (COG0100)→ RpsD (COG0522)rpsD (PRK05327)→ RpoA (COG0202)PRK05182 (PRK05182)→ RplQ (COG0203)rplQ (PRK05591)→ |
| Ful-1-1 | NZ_KE161010.1 | + | 184763 | 184824 | **RNA**→ RpsM (COG0099)rpsM (PRK05179)→ PRK05309 (PRK05309)RpsK (COG0100)→ RpsD (COG0522)rpsD (PRK05327)→ RpoA (COG0202)PRK05182 (PRK05182)→ RplQ (COG0203)rplQ (PRK05591)→ |

| | | | | | |
|---|---|---|---|---|---|
| env-295 | 4491479.3_NODE_279 | + | 17428 | 17489 | **RNA**→ RpsM (COG0099)rpsM (PRK05179)→ PRK05309 (PRK05309)RpsK (COG0100)→ RpsD (COG0522)rpsD (PRK05327)→ RpoA (COG0202)PRK05182 (PRK05182)→ RplQ (COG0203)rplQ (PRK05591)→ |
| env-296 | NOF002_scaffold13914_10 | - | 446 | 385 | **RNA**→ RpsM (COG0099)rpsM (PRK05179)→ |
| Fva-1-1 | NZ_GL987999.1 | + | 52065 | 52126 | **RNA**→ RpsM (COG0099)rpsM (PRK05179)→ PRK05309 (PRK05309)RpsK (COG0100)→ RpsD (COG0522)rpsD (PRK05327)→ RpoA (COG0202)PRK05182 (PRK05182)→ RplQ (COG0203)rplQ (PRK05591)→ |
| env-297 | scaffold80362_1_V1.UC-17 | + | 2137 | 2199 | **RNA**→ RpsM (COG0099)rpsM (PRK05179)→ PRK05309 (PRK05309)RpsK (COG0100)→ RpsD (COG0522)rpsD (PRK05327)→ RpoA (COG0202)PRK05182 (PRK05182)→ RplQ (COG0203)rplQ (PRK05591)→ |
| Fsp-2-1 | NZ_GG657975.1 | - | 48581 | 48519 | **RNA**→ RpsM (COG0099)rpsM (PRK05179)→ RpsM (COG0099)rpsM (PRK05179)→ PRK05309 (PRK05309)RpsK (COG0100)→ PRK05309 (PRK05309)RpsK (COG0100)→ RpsD (COG0522)rpsD (PRK05327)→ RpsD (COG0522)rpsD (PRK05327)→ RpoA (COG0202)PRK05182 (PRK05182)→ RpoA (COG0202)PRK05182 (PRK05182)→ RplQ (COG0203)rplQ (PRK05591)→ RplQ (COG0203)rplQ (PRK05591)→ |
| Fgo-1-1 | NZ_KQ235756.1 | + | 86258 | 86320 | **RNA**→ RpsM (COG0099)rpsM (PRK05179)→ PRK05309 (PRK05309)RpsK (COG0100)→ RpsD (COG0522)rpsD (PRK05327)→ RpoA (COG0202)PRK05182 (PRK05182)→ RplQ (COG0203)rplQ (PRK05591)→ |
| Fne-5-1 | NZ_JAAF01000042.1 | + | 1430 | 1492 | **RNA**→ RpsM (COG0099)rpsM (PRK05179)→ PRK05309 (PRK05309)RpsK (COG0100)→ RpsD (COG0522)rpsD (PRK05327)→ RpoA (COG0202)PRK05182 (PRK05182)→ RplQ (COG0203)rplQ (PRK05591)→ |
| Fne-3-1 | NZ_JAAC01000019.1 | + | 1569 | 1631 | **RNA**→ RpsM (COG0099)rpsM (PRK05179)→ PRK05309 (PRK05309)RpsK (COG0100)→ RpsD (COG0522)rpsD (PRK05327)→ RpoA (COG0202)PRK05182 (PRK05182)→ RplQ (COG0203)rplQ (PRK05591)→ |
| Fne-10-1 | NZ_AOJP01000008.1 | - | 52376 | 52314 | **RNA**→ RpsM (COG0099)rpsM (PRK05179)→ PRK05309 (PRK05309)RpsK (COG0100)→ RpsD (COG0522)rpsD (PRK05327)→ RpoA (COG0202)PRK05182 (PRK05182)→ RplQ (COG0203)rplQ (PRK05591)→ |
| Fne-9-1 | NZ_AJSY01000009.1 | + | 2132 | 2194 | **RNA**→ RpsM (COG0099)rpsM (PRK05179)→ PRK05309 (PRK05309)RpsK (COG0100)→ RpsD (COG0522)rpsD (PRK05327)→ RpoA (COG0202)PRK05182 (PRK05182)→ RplQ (COG0203)rplQ (PRK05591)→ |
| Fne-1-1 | NZ_JAAE01000021.1 | - | 16569 | 16507 | **RNA**→ RpsM (COG0099)rpsM (PRK05179)→ PRK05309 (PRK05309)RpsK (COG0100)→ RpsD (COG0522)rpsD (PRK05327)→ RpoA (COG0202)PRK05182 (PRK05182)→ RplQ (COG0203)rplQ (PRK05591)→ |
| Fne-7-1 | NZ_JAAH01000082.1 | - | 16650 | 16588 | **RNA**→ RpsM (COG0099)rpsM (PRK05179)→ PRK05309 (PRK05309)RpsK (COG0100)→ RpsD (COG0522)rpsD (PRK05327)→ RpoA (COG0202)PRK05182 (PRK05182)→ RplQ (COG0203)rplQ (PRK05591)→ |
| Fne-2-1 | NZ_JAAG01000077.1 | - | 16625 | 16563 | **RNA**→ RpsM (COG0099)rpsM (PRK05179)→ PRK05309 (PRK05309)RpsK (COG0100)→ RpsD (COG0522)rpsD (PRK05327)→ RpoA (COG0202)PRK05182 (PRK05182)→ RplQ (COG0203)rplQ (PRK05591)→ |
| Fne-6-1 | NZ_JAAD01000236.1 | + | 1428 | 1490 | **RNA**→ RpsM (COG0099)rpsM (PRK05179)→ PRK05309 (PRK05309)RpsK (COG0100)→ RpsD (COG0522)rpsD (PRK05327)→ RpoA (COG0202)PRK05182 (PRK05182)→ RplQ (COG0203)rplQ (PRK05591)→ |
| Fne-4-1 | NZ_GL988017.1 | + | 49167 | 49229 | **RNA**→ RpsM (COG0099)rpsM (PRK05179)→ PRK05309 (PRK05309)RpsK (COG0100)→ RpsD (COG0522)rpsD (PRK05327)→ RpoA (COG0202)PRK05182 (PRK05182)→ RplQ (COG0203)rplQ (PRK05591)→ |
| Fne-11-1 | NZ_ALKK01000035.1 | - | 50617 | 50555 | **RNA**→ RpsM (COG0099)rpsM (PRK05179)→ PRK05309 (PRK05309)RpsK (COG0100)→ RpsD (COG0522)rpsD (PRK05327)→ RpoA (COG0202)PRK05182 (PRK05182)→ RplQ (COG0203)rplQ (PRK05591)→ |
| Fne-8-1 | NZ_JHWT01000020.1 | - | 51252 | 51190 | **RNA**→ RpsM (COG0099)rpsM (PRK05179)→ PRK05309 (PRK05309)RpsK (COG0100)→ RpsD (COG0522)rpsD (PRK05327)→ RpoA (COG0202)PRK05182 (PRK05182)→ RplQ (COG0203)rplQ (PRK05591)→ |
| env-298 | SRS043422_LANL_scaffold_508 | - | 307 | 245 | **RNA**→ RpsM (COG0099)rpsM (PRK05179)→ |
| env-299 | SRS065099_LANL_scaffold_56436 | + | 892 | 954 | **RNA**→ RpsM (COG0099)rpsM (PRK05179)→ rps11 (CHL00041)PRK05309 (PRK05309)→ |
| env-300 | SRS016360_Baylor_scaffold_43492 | + | 448 | 510 | **RNA**→ RpsM (COG0099)rpsM (PRK05179)→ PRK05309 (PRK05309)RpsK (COG0100)→ RpsD (COG0522)rpsD (PRK05327)→ RpoA (COG0202)rpoA (TIGR02027)→ |
| env-301 | SRS021960_Baylor_scaffold_29549 | - | 372 | 310 | **RNA**→ RpsM (COG0099)rpsM (PRK05179)→ |
| env-302 | SRS052876_LANL_scaffold_1775 | + | 450 | 512 | **RNA**→ rps13 (CHL00137)rpsM (PRK05179)→ |

| | | | | | |
|---|---|---|---|---|---|
| env-303 | SRS015985_WUGC_scaffold_11687 | - | 350 | 288 | **RNA**→ RpsM (COG0099)rpsM (PRK05179)→ |
| env-304 | SRS020340_Baylor_scaffold_22156 | - | 1355 | 1293 | **RNA**→ RpsM (COG0099)rpsM (PRK05179)→ PRK05309 (PRK05309)RpsK (COG0100)→ |
| | | | | | S4 (smart00363)RpsD (COG0522)→ |
| env-305 | SRS017025_C1636822 | + | 100 | 162 | **RNA**→ RpsM (COG0099)rpsM (PRK05179)→ |
| env-306 | SRS013170_Baylor_scaffold_81941 | + | 481 | 543 | **RNA**→ RpsM (COG0099)rpsM (PRK05179)→ |
| env-307 | SRS017445_Baylor_scaffold_8312 | + | 454 | 516 | **RNA**→ RpsM (COG0099)rpsM (PRK05179)→ rps11 (CHL00041)PRK05309 (PRK05309)→ |
| | | | | | PRK05309 (PRK05309)RpsK (COG0100)→ RpsD (COG0522)rpsD (PRK05327)→ |
| | | | | | PRK05182 (PRK05182)RNA_pol_L (pfam01193)→ |
| env-308 | SRS019073_WUGC_scaffold_286 | + | 20 | 82 | **RNA**→ RpsM (COG0099)rpsM (PRK05179)→ |
| env-309 | SRS020226_Baylor_scaffold_20989 | + | 451 | 513 | **RNA**→ RpsM (COG0099)rpsM (PRK05179)→ |
| env-310 | SRS047265_C1093350 | + | 348 | 410 | **RNA**→ RpsM (COG0099)rpsM (PRK05179)→ PRK05309 (PRK05309)RpsK (COG0100)→ rpsD (PRK05327)→ |
| Fhw-4-1 | NZ_ATKG01000094.1 | - | 9304 | 9242 | **RNA**→ RpsM (COG0099)rpsM (PRK05179)→ PRK05309 (PRK05309)RpsK (COG0100)→ |
| | | | | | RpsD (COG0522)rpsD (PRK05327)→ RpoA (COG0202)rpoA (TIGR02027)→ |
| | | | | | RpoA (COG0202)rpoA (TIGR02027)→ RplQ (COG0203)rplQ (PRK05591)→ |
| Fhw-1-1 | NZ_ATKE01000061.1 | - | 12226 | 12164 | **RNA**→ RpsM (COG0099)rpsM (PRK05179)→ PRK05309 (PRK05309)RpsK (COG0100)→ |
| | | | | | RpsD (COG0522)rpsD (PRK05327)→ RpoA (COG0202)rpoA (TIGR02027)→ |
| | | | | | RpoA (COG0202)rpoA (TIGR02027)→ RplQ (COG0203)rplQ (PRK05591)→ |
| Fhw-3-1 | NZ_ATKH01000028.1 | - | 9304 | 9242 | **RNA**→ RpsM (COG0099)rpsM (PRK05179)→ PRK05309 (PRK05309)RpsK (COG0100)→ |
| | | | | | RpsD (COG0522)rpsD (PRK05327)→ RpoA (COG0202)rpoA (TIGR02027)→ |
| | | | | | RpoA (COG0202)rpoA (TIGR02027)→ RplQ (COG0203)rplQ (PRK05591)→ |
| Fhw-2-1 | NZ_ATKF01000105.1 | - | 24308 | 24246 | **RNA**→ RpsM (COG0099)rpsM (PRK05179)→ PRK05309 (PRK05309)RpsK (COG0100)→ |
| | | | | | RpsD (COG0522)rpsD (PRK05327)→ RpoA (COG0202)rpoA (TIGR02027)→ |
| | | | | | RpoA (COG0202)rpoA (TIGR02027)→ RplQ (COG0203)rplQ (PRK05591)→ |
| Fsp-6-1 | NZ_JANA01000050.1 | + | 15841 | 15903 | **RNA**→ RpsM (COG0099)rpsM (PRK05179)→ PRK05309 (PRK05309)RpsK (COG0100)→ |
| | | | | | RpsD (COG0522)rpsD (PRK05327)→ RpoA (COG0202)rpoA (TIGR02027)→ |
| | | | | | RpoA (COG0202)rpoA (TIGR02027)→ RplQ (COG0203)rplQ (PRK05591)→ |
| env-311 | SRS018157_Baylor_scaffold_655 | + | 18 | 80 | **RNA**→ RpsM (COG0099)rpsM (PRK05179)→ |
| env-312 | SRS057022_C1993424 | - | 642 | 580 | **RNA**→ RpsM (COG0099)rpsM (PRK05179)→ |
| | | | | | PRK05309 (PRK05309)RpsK (COG0100)rps11p (PRK09607)arch_S11P (TIGR03628)uS11_bact (TIGR03632)Ribosomal_ |
| Fnu-9-1 | NZ_ALVD01000005.1 | + | 72312 | 72374 | **RNA**→ RpsM (COG0099)rpsM (PRK05179)→ PRK05309 (PRK05309)RpsK (COG0100)→ |
| | | | | | RpsD (COG0522)rpsD (PRK05327)→ RpoA (COG0202)rpoA (TIGR02027)→ |
| | | | | | RpoA (COG0202)rpoA (TIGR02027)→ RplQ (COG0203)rplQ (PRK05591)→ |
| env-313 | SRS015470_WUGC_scaffold_62148 | - | 625 | 563 | **RNA**→ RpsM (COG0099)rpsM (PRK05179)→ PRK05309 (PRK05309)→ |
| env-314 | SRS063351_C704759 | - | 376 | 314 | **RNA**→ RpsM (COG0099)rpsM (PRK05179)→ |
| Fnu-3-1 | NZ_KI518461.1 | - | 1355250 | 1355188 | **RNA**→ RpsM (COG0099)rpsM (PRK05179)→ PRK05309 (PRK05309)RpsK (COG0100)→ |
| | | | | | RpsD (COG0522)rpsD (PRK05327)→ RpoA (COG0202)rpoA (TIGR02027)→ |
| | | | | | RplQ (COG0203)rplQ (PRK05591)→ |
| Fnu-16-1 | NZ_ATKD01000080.1 | - | 9661 | 9599 | **RNA**→ RpsM (COG0099)rpsM (PRK05179)→ PRK05309 (PRK05309)RpsK (COG0100)→ |
| | | | | | RpsD (COG0522)rpsD (PRK05327)→ RpoA (COG0202)rpoA (TIGR02027)→ |
| | | | | | RplQ (COG0203)rplQ (PRK05591)→ |
| Fsp-3-1 | NZ_JANB01000001.1 | + | 5756 | 5818 | **RNA**→ RpsM (COG0099)rpsM (PRK05179)→ PRK05309 (PRK05309)RpsK (COG0100)→ |
| | | | | | RpsD (COG0522)rpsD (PRK05327)→ RpoA (COG0202)rpoA (TIGR02027)→ |
| | | | | | RplQ (COG0203)rplQ (PRK05591)→ |
| Fnu-10-1 | NZ_AXUR01000028.1 | - | 25993 | 25931 | **RNA**→ RpsM (COG0099)rpsM (PRK05179)→ PRK05309 (PRK05309)RpsK (COG0100)→ |
| | | | | | RpsD (COG0522)rpsD (PRK05327)→ RpoA (COG0202)rpoA (TIGR02027)→ |
| | | | | | RplQ (COG0203)rplQ (PRK05591)→ |
| env-315 | SRS051930_LANL_scaffold_6191 | + | 3750 | 3812 | **RNA**→ RpsM (COG0099)rpsM (PRK05179)→ PRK05309 (PRK05309)RpsK (COG0100)→ |
| | | | | | S4 (smart00363)RpsD (COG0522)→ RpoA (COG0202)rpoA (TIGR02027)→ |
| env-316 | SRS016541_C1528741 | + | 21 | 83 | **RNA**→ RpsM (COG0099)rpsM (PRK05179)→ |
| env-317 | SRS011126_Baylor_scaffold_58649 | - | 598 | 536 | **RNA**→ RpsM (COG0099)rpsM (PRK05179)→ |
| env-318 | SRS050628_C1835320 | - | 390 | 328 | **RNA**→ RpsM (COG0099)rpsM (PRK05179)→ |
| env-319 | SRS012285_Baylor_scaffold_1782 | + | 425 | 487 | **RNA**→ RpsM (COG0099)rpsM (PRK05179)→ |
| env-320 | SRS017691_Baylor_scaffold_50060 | + | 518 | 580 | **RNA**→ RpsM (COG0099)rpsM (PRK05179)→ |
| env-321 | SRS018443_C3801805 | + | 470 | 532 | **RNA**→ |
| env-322 | SRS014578_WUGC_scaffold_7353 | - | 722 | 660 | **RNA**→ RpsM (COG0099)rpsM (PRK05179)→ PRK05309 (PRK05309)RpsK (COG0100)→ |
| env-323 | SRS014690_C1866550 | - | 347 | 285 | **RNA**→ RpsM (COG0099)rpsM (PRK05179)→ |

| | | | | | |
|---|---|---|---|---|---|
| env-324 | SRS015278_WUGC_scaffold_42089 | + | 2204 | 2266 | **RNA**→ RpsM (COG0099)rpsM (PRK05179)→ PRK05309 (PRK05309)RpsK (COG0100)→ RpsD (COG0522)rpsD (PRK05327)→ RpoA (COG0202)rpoA (TIGR02027)→ RplQ (COG0203)rplQ (PRK05591)→ |
| env-325 | SRS015989_C2166928 | + | 538 | 600 | **RNA**→ RpsM (COG0099)rpsM (PRK05179)→ PRK05309 (PRK05309)RpsK (COG0100)→ |
| env-326 | SRS016043_WUGC_scaffold_19941 | - | 541 | 479 | **RNA**→ RpsM (COG0099)rpsM (PRK05179)→ |
| env-327 | SRS023358_C3483802 | - | 630 | 568 | **RNA**→ RpsM (COG0099)rpsM (PRK05179)→ PRK05309 (PRK05309)→ |
| env-328 | SRS023938_Baylor_scaffold_16323 | + | 244 | 306 | **RNA**→ RpsM (COG0099)rpsM (PRK05179)→ rps11 (CHL00041)PRK05309 (PRK05309)→ |
| env-329 | SRS024021_C2019934 | - | 499 | 437 | **RNA**→ RpsM (COG0099)rpsM (PRK05179)→ |
| env-330 | SRS024381_LANL_scaffold_17523 | + | 558 | 620 | **RNA**→ RpsM (COG0099)rpsM (PRK05179)→ rps11 (CHL00041)PRK05309 (PRK05309)→ |
| env-331 | SRS051378_C1822879 | + | 210 | 272 | **RNA**→ RpsM (COG0099)rpsM (PRK05179)→ |
| env-332 | SRS052604_C1807758 | + | 212 | 274 | **RNA**→ RpsM (COG0099)rpsM (PRK05179)→ |
| env-333 | SRS056892_C1486315 | + | 188 | 250 | **RNA**→ RpsM (COG0099)rpsM (PRK05179)→ |
| env-334 | SRS058808_LANL_scaffold_23314 | + | 321 | 383 | **RNA**→ RpsM (COG0099)rpsM (PRK05179)→ |
| Fnu-1-1 | NZ_KI965381.1 | - | 1470032 | 1469970 | **RNA**→ RpsM (COG0099)rpsM (PRK05179)→ PRK05309 (PRK05309)RpsK (COG0100)→ RpsD (COG0522)rpsD (PRK05327)→ RpoA (COG0202)rpoA (TIGR02027)→ RpoA (COG0202)rpoA (TIGR02027)→ RplQ (COG0203)rplQ (PRK05591)→ |
| Fnu-7-1 | NZ_AXNV01000011.1 | + | 20769 | 20831 | **RNA**→ RpsM (COG0099)rpsM (PRK05179)→ PRK05309 (PRK05309)RpsK (COG0100)→ RpsD (COG0522)rpsD (PRK05327)→ RpoA (COG0202)rpoA (TIGR02027)→ RpoA (COG0202)rpoA (TIGR02027)→ RplQ (COG0203)rplQ (PRK05591)→ |
| Fsp-5-1 | NZ_JAQC01000039.1 | + | 6874 | 6936 | **RNA**→ RpsM (COG0099)rpsM (PRK05179)→ PRK05309 (PRK05309)RpsK (COG0100)→ RpsD (COG0522)rpsD (PRK05327)→ RpoA (COG0202)rpoA (TIGR02027)→ RpoA (COG0202)rpoA (TIGR02027)→ RplQ (COG0203)rplQ (PRK05591)→ |
| env-335 | SRS055378_LANL_scaffold_3069 | + | 20 | 84 | **RNA**→ RpsM (COG0099)rpsM (PRK05179)→ PRK05309 (PRK05309)RpsK (COG0100)→ |
| env-336 | SRS014476_C3341139 | + | 43 | 105 | **RNA**→ RpsM (COG0099)rpsM (PRK05179)→ |
| Fnu-22-1 | NZ_CM000440.1 | + | 860522 | 860584 | **RNA**→ RpsM (COG0099)rpsM (PRK05179)→ RpsM (COG0099)rpsM (PRK05179)→ PRK05309 (PRK05309)RpsK (COG0100)→ PRK05309 (PRK05309)RpsK (COG0100)→ RpsD (COG0522)rpsD (PRK05327)→ RpsD (COG0522)rpsD (PRK05327)→ RpoA (COG0202)rpoA (TIGR02027)→ RpoA (COG0202)rpoA (TIGR02027)→ RpoA (COG0202)rpoA (TIGR02027)→ RplQ (COG0203)rplQ (PRK05591)→ RplQ (COG0203)rplQ (PRK05591)→ |
| env-337 | SRS064329_LANL_scaffold_32934 | + | 20 | 82 | **RNA**→ RpsM (COG0099)rpsM (PRK05179)→ PRK05309 (PRK05309)RpsK (COG0100)rpsD (PRK05327)Ribosomal_S4 (pfam00163)→ |
| env-338 | Throat_LANL_C5780661 | - | 163 | 101 | **RNA**→ |
| env-339 | SRS014684_WUGC_scaffold_2752 | + | 1181 | 1243 | **RNA**→ RpsM (COG0099)rpsM (PRK05179)→ PRK05309 (PRK05309)RpsK (COG0100)→ RpsD (COG0522)rpsD (PRK05327)→ RpoA (COG0202)rpoA (TIGR02027)→ rplQ (PRK05591)Ribosomal_L17 (pfam01196)→ |
| env-340 | SRS018591_C1305506 | - | 1175 | 1113 | **RNA**→ RpsM (COG0099)rpsM (PRK05179)→ PRK05309 (PRK05309)RpsK (COG0100)→ rpsD (PRK05327)Ribosomal_S4 (pfam00163)→ |
| env-341 | SRS019022_WUGC_scaffold_38441 | - | 3789 | 3727 | **RNA**→ RpsM (COG0099)rpsM (PRK05179)→ PRK05309 (PRK05309)RpsK (COG0100)→ RpsD (COG0522)rpsD (PRK05327)→ RpoA (COG0202)rpoA (TIGR02027)→ RplQ (COG0203)rplQ (PRK05591)→ |
| env-342 | SRS019045_WUGC_scaffold_36434 | - | 9682 | 9620 | **RNA**→ RpsM (COG0099)rpsM (PRK05179)→ PRK05309 (PRK05309)RpsK (COG0100)→ RpsD (COG0522)rpsD (PRK05327)→ RpoA (COG0202)rpoA (TIGR02027)→ RplQ (COG0203)rplQ (PRK05591)→ |
| env-343 | SRS019219_WUGC_scaffold_39593 | - | 1300 | 1238 | **RNA**→ RpsM (COG0099)rpsM (PRK05179)→ PRK05309 (PRK05309)RpsK (COG0100)→ RpsD (COG0522)rpsD (PRK05327)→ |
| env-344 | SRS019389_C2241031 | - | 648 | 586 | **RNA**→ RpsM (COG0099)rpsM (PRK05179)→ PRK05309 (PRK05309)RpsK (COG0100)→ |
| env-345 | SRS020220_C3612140 | - | 287 | 225 | **RNA**→ RpsM (COG0099)rpsM (PRK05179)→ |
| env-346 | SRS051791_LANL_scaffold_1205 | - | 1302 | 1240 | **RNA**→ RpsM (COG0099)rpsM (PRK05179)→ PRK05309 (PRK05309)RpsK (COG0100)→ RpsD (COG0522)rpsD (PRK05327)→ |
| env-347 | SRS053603_C3303360 | + | 458 | 520 | **RNA**→ RpsM (COG0099)rpsM (PRK05179)→ PRK05309 (PRK05309)RpsK (COG0100)→ |
| env-348 | SRS054687_C3486456 | - | 734 | 672 | **RNA**→ RpsM (COG0099)rpsM (PRK05179)→ PRK05309 (PRK05309)RpsK (COG0100)→ |
| env-349 | SRS065335_LANL_scaffold_29301 | - | 3749 | 3687 | **RNA**→ RpsM (COG0099)rpsM (PRK05179)→ PRK05309 (PRK05309)RpsK (COG0100)→ RpsD (COG0522)rpsD (PRK05327)→ RpoA (COG0202)rpoA (TIGR02027)→ RplQ (COG0203)rplQ (PRK05591)→ |

| | | | | | |
|---|---|---|---|---|---|
| env-350 | SRS017120_Baylor_scaffold_81370 | + | 457 | 519 | **RNA**→ RpsM (COG0099)rpsM (PRK05179)→ PRK05309 (PRK05309)RpsK (COG0100)→ RpsD (COG0522)rpsD (PRK05327)→ RNAP_alpha_NTD (cd06928)PRK05182 (PRK05182)→ |
| env-351 | SRS014692_WUGC_scaffold_12525 | - | 1378 | 1316 | **RNA**→ RpsM (COG0099)rpsM (PRK05179)→ PRK05309 (PRK05309)RpsK (COG0100)→ S4 (smart00363)RpsD (COG0522)→ |
| env-352 | SRS047824_WUGC_scaffold_34774 | - | 578 | 516 | **RNA**→ RpsM (COG0099)rpsM (PRK05179)→ |
| env-353 | SRS023930_C2177563 | - | 496 | 434 | **RNA**→ RpsM (COG0099)rpsM (PRK05179)→ |
| env-354 | SRS018665_WUGC_scaffold_5673 | - | 903 | 841 | **RNA**→ RpsM (COG0099)rpsM (PRK05179)→ PRK05309 (PRK05309)RpsK (COG0100)→ |
| env-355 | SRS018975_WUGC_scaffold_48555 | - | 578 | 516 | **RNA**→ RpsM (COG0099)rpsM (PRK05179)→ |
| env-356 | SRS020858_C751293 | - | 163 | 101 | **RNA**→ hypo→ |
| env-357 | SRS023987_C1155340 | - | 165 | 103 | **RNA**→ hypo→ |
| env-358 | SRS052668_LANL_scaffold_2440 | + | 371 | 433 | **RNA**→ RpsM (COG0099)rpsM (PRK05179)→ |
| env-359 | SRS011086_C2705778 | + | 490 | 552 | **RNA**→ |
| env-360 | SRS018971_C747146 | + | 209 | 271 | **RNA**→ |
| env-361 | SRS019128_WUGC_scaffold_1597 | - | 240 | 178 | **RNA**→ rps13 (CHL00137)rpsM (PRK05179)→ |
| env-362 | SRS023595_Baylor_scaffold_12376 | + | 20 | 82 | **RNA**→ RpsM (COG0099)rpsM (PRK05179)→ PRK05309 (PRK05309)RpsK (COG0100)→ rpsD (PRK05327)Ribosomal_S4 (pfam00163)→ |
| Fsp-7-1 | NZ_JH378968.1 | - | 23656 | 23594 | **RNA**→ RpsM (COG0099)rpsM (PRK05179)→ PRK05309 (PRK05309)RpsK (COG0100)→ RpsD (COG0522)rpsD (PRK05327)→ RpoA (COG0202)rpoA (TIGR02027)→ RpoA (COG0202)rpoA (TIGR02027)→ RplQ (COG0203)rplQ (PRK05591)→ |
| env-363 | SRS016740_Baylor_scaffold_31792 | - | 2409 | 2347 | **RNA**→ RpsM (COG0099)rpsM (PRK05179)→ PRK05309 (PRK05309)RpsK (COG0100)→ RpsD (COG0522)rpsD (PRK05327)→ RNAP_alpha_NTD (cd06928)PRK05182 (PRK05182)→ |
| env-364 | SRS055495_C1108972 | - | 120 | 58 | **RNA**→ |
| env-365 | SRS062878_LANL_scaffold_63865 | + | 495 | 557 | **RNA**→ RpsM (COG0099)rpsM (PRK05179)→ |
| env-366 | SRS075404_LANL_scaffold_14797 | - | 890 | 828 | **RNA**→ RpsM (COG0099)rpsM (PRK05179)→ |
| env-367 | SRS011140_Baylor_scaffold_51504 | + | 350 | 412 | **RNA**→ RpsM (COG0099)rpsM (PRK05179)→ PRK05309 (PRK05309)RpsK (COG0100)→ |
| env-368 | SRS011243_Baylor_scaffold_10689 | - | 3419 | 3357 | **RNA**→ RpsM (COG0099)rpsM (PRK05179)→ PRK05309 (PRK05309)RpsK (COG0100)→ RpsD (COG0522)rpsD (PRK05327)→ RpoA (COG0202)rpoA (TIGR02027)→ RplQ (COG0203)rplQ (PRK05591)→ |
| env-369 | SRS011247_C1351757 | + | 468 | 530 | **RNA**→ |
| env-370 | SRS011306_Baylor_scaffold_3104 | - | 3003 | 2941 | **RNA**→ RpsM (COG0099)rpsM (PRK05179)→ PRK05309 (PRK05309)RpsK (COG0100)→ RpsD (COG0522)rpsD (PRK05327)→ RpoA (COG0202)rpoA (TIGR02027)→ RplQ (COG0203)rplQ (PRK05591)→ |
| env-371 | SRS013164_Baylor_scaffold_46971 | - | 920 | 858 | **RNA**→ RpsM (COG0099)rpsM (PRK05179)→ PRK05309 (PRK05309)RpsK (COG0100)→ |
| env-372 | SRS013502_C2191535 | - | 163 | 101 | **RNA**→ |
| env-373 | SRS013705_Baylor_scaffold_64513 | + | 487 | 549 | **RNA**→ RpsM (COG0099)rpsM (PRK05179)→ PRK05309 (PRK05309)RpsK (COG0100)→ |
| env-374 | SRS013818_Baylor_scaffold_41472 | + | 473 | 535 | **RNA**→ RpsM (COG0099)rpsM (PRK05179)→ PRK05309 (PRK05309)RpsK (COG0100)→ RpsD (COG0522)rpsD (PRK05327)→ RpoA (COG0202)PRK05182 (PRK05182)→ |
| env-375 | SRS013879_WUGC_scaffold_41089 | - | 934 | 872 | **RNA**→ RpsM (COG0099)rpsM (PRK05179)→ PRK05309 (PRK05309)RpsK (COG0100)→ |
| env-376 | SRS013945_WUGC_scaffold_13633 | - | 525 | 463 | **RNA**→ RpsM (COG0099)rpsM (PRK05179)→ |
| env-377 | SRS013947_WUGC_scaffold_13273 | - | 7521 | 7459 | **RNA**→ RpsM (COG0099)rpsM (PRK05179)→ PRK05309 (PRK05309)RpsK (COG0100)→ RpsD (COG0522)rpsD (PRK05327)→ RpoA (COG0202)rpoA (TIGR02027)→ RplQ (COG0203)rplQ (PRK05591)→ |
| env-378 | SRS013948_C1529892 | + | 458 | 520 | **RNA**→ RpsM (COG0099)rpsM (PRK05179)→ PRK05309 (PRK05309)RpsK (COG0100)→ |
| env-379 | SRS014124_WUGC_scaffold_58834 | - | 1359 | 1297 | **RNA**→ RpsM (COG0099)rpsM (PRK05179)→ PRK05309 (PRK05309)RpsK (COG0100)→ S4 (smart00363)RpsD (COG0522)→ |
| env-380 | SRS014271_WUGC_scaffold_4413 | + | 497 | 559 | **RNA**→ RpsM (COG0099)rpsM (PRK05179)→ PRK05309 (PRK05309)RpsK (COG0100)→ RpsD (COG0522)rpsD (PRK05327)→ hypo→ |
| env-381 | SRS014470_WUGC_scaffold_38766 | - | 2083 | 2021 | **RNA**→ RpsM (COG0099)rpsM (PRK05179)→ PRK05309 (PRK05309)RpsK (COG0100)→ RpsD (COG0522)rpsD (PRK05327)→ RNAP_alpha_NTD (cd06928)PRK05182 (PRK05182)→ |
| env-382 | SRS014573_WUGC_scaffold_16390 | - | 844 | 782 | **RNA**→ RpsM (COG0099)rpsM (PRK05179)→ PRK05309 (PRK05309)RpsK (COG0100)→ PRK05309 (PRK05309)RpsK (COG0100)→ |
| env-383 | SRS014689_WUGC_scaffold_25280 | + | 2043 | 2105 | **RNA**→ RpsM (COG0099)rpsM (PRK05179)→ PRK05309 (PRK05309)RpsK (COG0100)→ RpsD (COG0522)rpsD (PRK05327)→ RpoA (COG0202)rpoA (TIGR02027)→ RplQ (COG0203)rplQ (PRK05591)→ |

| | | | | | |
|---|---|---|---|---|---|
| env-384 | SRS014888_WUGC_scaffold_2290 | + | 801 | 863 | RNA→ RpsM (COG0099)rpsM (PRK05179)→ PRK05309 (PRK05309)RpsK (COG0100)→ RpsD (COG0522)rpsD (PRK05327)→ RpoA (COG0202)rpoA (TIGR02027)→ RplQ (COG0203)rplQ (PRK05591)→ |
| env-385 | SRS015057_WUGC_scaffold_68354 | - | 734 | 672 | RNA→ RpsM (COG0099)rpsM (PRK05179)→ PRK05309 (PRK05309)RpsK (COG0100)→ |
| env-386 | SRS015154_C747837 | + | 666 | 728 | RNA→ RpsM (COG0099)rpsM (PRK05179)→ |
| env-387 | SRS015644_WUGC_scaffold_11157 | + | 577 | 639 | RNA→ RpsM (COG0099)rpsM (PRK05179)→ PRK05309 (PRK05309)RpsK (COG0100)→ RpsD (COG0522)rpsD (PRK05327)→ hypo→ |
| env-388 | SRS015745_C1206036 | - | 637 | 575 | RNA→ RpsM (COG0099)rpsM (PRK05179)→ |
| env-389 | SRS015762_C3450973 | - | 163 | 101 | RNA→ hypo→ |
| env-390 | SRS015797_WUGC_scaffold_30177 | - | 734 | 672 | RNA→ RpsM (COG0099)rpsM (PRK05179)→ PRK05309 (PRK05309)RpsK (COG0100)→ |
| env-391 | SRS015893_WUGC_scaffold_5057 | + | 854 | 916 | RNA→ RpsM (COG0099)rpsM (PRK05179)→ PRK05309 (PRK05309)RpsK (COG0100)→ RpsD (COG0522)rpsD (PRK05327)→ RpoA (COG0202)rpoA (TIGR02027)→ RplQ (COG0203)rplQ (PRK05591)→ |
| env-392 | SRS015941_WUGC_scaffold_78015 | - | 1690 | 1628 | RNA→ RpsM (COG0099)rpsM (PRK05179)→ PRK05309 (PRK05309)RpsK (COG0100)→ RpsD (COG0522)rpsD (PRK05327)→ hypo→ |
| env-393 | SRS016002_WUGC_scaffold_50772 | - | 2403 | 2341 | RNA→ RpsM (COG0099)rpsM (PRK05179)→ PRK05309 (PRK05309)RpsK (COG0100)→ RpsD (COG0522)rpsD (PRK05327)→ RNAP_alpha_NTD (cd06928)PRK05182 (PRK05182)→ |
| env-394 | SRS016037_WUGC_scaffold_2228 | - | 3491 | 3429 | RNA→ RpsM (COG0099)rpsM (PRK05179)→ PRK05309 (PRK05309)RpsK (COG0100)→ RpsD (COG0522)rpsD (PRK05327)→ RNAP_alpha_NTD (cd06928)PRK05182 (PRK05182)→ RNA_pol_A_CTD (pfam03118)HHH_5 (pfam14520)→ |
| env-395 | SRS016086_C2496538 | - | 737 | 675 | RNA→ RpsM (COG0099)rpsM (PRK05179)→ PRK05309 (PRK05309)RpsK (COG0100)→ |
| env-396 | SRS016225_WUGC_scaffold_27311 | - | 2971 | 2909 | RNA→ RpsM (COG0099)rpsM (PRK05179)→ PRK05309 (PRK05309)RpsK (COG0100)→ RpsD (COG0522)rpsD (PRK05327)→ RpoA (COG0202)rpoA (TIGR02027)→ RplQ (COG0203)rplQ (PRK05591)→ |
| env-397 | SRS016319_WUGC_scaffold_10490 | + | 864 | 926 | RNA→ RpsM (COG0099)rpsM (PRK05179)→ PRK05309 (PRK05309)RpsK (COG0100)→ RpsD (COG0522)rpsD (PRK05327)→ RNAP_alpha_NTD (cd06928)PRK05182 (PRK05182)→ |
| env-398 | SRS016569_Baylor_scaffold_43023 | - | 332 | 270 | RNA→ RpsM (COG0099)rpsM (PRK05179)→ |
| env-399 | SRS017209_Baylor_scaffold_12537 | + | 765 | 827 | RNA→ RpsM (COG0099)rpsM (PRK05179)→ PRK05309 (PRK05309)RpsK (COG0100)→ RpsD (COG0522)rpsD (PRK05327)→ RNAP_alpha_NTD (cd06928)PRK05182 (PRK05182)→ |
| env-400 | SRS017215_Baylor_scaffold_8618 | + | 162 | 224 | RNA→ RpsM (COG0099)rpsM (PRK05179)→ |
| env-401 | SRS017439_Baylor_scaffold_16569 | - | 2843 | 2781 | RNA→ RpsM (COG0099)rpsM (PRK05179)→ PRK05309 (PRK05309)RpsK (COG0100)→ RpsD (COG0522)rpsD (PRK05327)→ RpoA (COG0202)rpoA (TIGR02027)→ rplQ (PRK05591)Ribosomal_L17 (pfam01196)→ |
| env-402 | SRS017533_Baylor_scaffold_4653 | + | 366 | 428 | RNA→ RpsM (COG0099)rpsM (PRK05179)→ |
| env-403 | SRS017808_Baylor_scaffold_22336 | - | 2749 | 2687 | RNA→ RpsM (COG0099)rpsM (PRK05179)→ PRK05309 (PRK05309)RpsK (COG0100)→ RpsD (COG0522)rpsD (PRK05327)→ RpoA (COG0202)rpoA (TIGR02027)→ rplQ (PRK05591)Ribosomal_L17 (pfam01196)→ |
| env-404 | SRS018149_Baylor_scaffold_14890 | + | 287 | 349 | RNA→ RpsM (COG0099)rpsM (PRK05179)→ |
| env-405 | SRS018357_Baylor_scaffold_12839 | + | 1146 | 1208 | RNA→ RpsM (COG0099)rpsM (PRK05179)→ PRK05309 (PRK05309)RpsK (COG0100)→ |
| env-406 | SRS018739_WUGC_scaffold_20843 | + | 819 | 881 | RNA→ RpsM (COG0099)rpsM (PRK05179)→ PRK05309 (PRK05309)RpsK (COG0100)→ S4 (smart00363)RpsD (COG0522)→ |
| env-407 | SRS018969_C2947476 | - | 741 | 679 | RNA→ RpsM (COG0099)rpsM (PRK05179)→ PRK05309 (PRK05309)RpsK (COG0100)→ |
| env-408 | SRS019026_WUGC_scaffold_24792 | - | 723 | 661 | RNA→ RpsM (COG0099)rpsM (PRK05179)→ PRK05309 (PRK05309)RpsK (COG0100)→ |
| env-409 | SRS019122_WUGC_scaffold_7425 | - | 151 | 89 | RNA→ hypo→ |
| env-410 | SRS019126_WUGC_scaffold_31303 | + | 1131 | 1193 | RNA→ RpsM (COG0099)rpsM (PRK05179)→ |
| env-411 | SRS019127_WUGC_scaffold_14132 | + | 458 | 520 | RNA→ RpsM (COG0099)rpsM (PRK05179)→ PRK05309 (PRK05309)RpsK (COG0100)→ RpsD (COG0522)rpsD (PRK05327)→ hypo→ |
| env-412 | SRS019327_WUGC_scaffold_41934 | + | 485 | 547 | RNA→ RpsM (COG0099)rpsM (PRK05179)→ PRK05309 (PRK05309)RpsK (COG0100)→ RpsD (COG0522)rpsD (PRK05327)→ RpoA (COG0202)rpoA (TIGR02027)→ rplQ (PRK05591)Ribosomal_L17 (pfam01196)→ |
| env-413 | SRS019587_C1052430 | + | 490 | 552 | RNA→ RpsM (COG0099)rpsM (PRK05179)→ |
| env-414 | SRS019607_WUGC_scaffold_24021 | - | 1359 | 1297 | RNA→ RpsM (COG0099)rpsM (PRK05179)→ PRK05309 (PRK05309)RpsK (COG0100)→ S4 (smart00363)RpsD (COG0522)→ |
| env-415 | SRS019894_C3929909 | + | 2203 | 2265 | RNA→ RpsM (COG0099)rpsM (PRK05179)→ PRK05309 (PRK05309)RpsK (COG0100)→ |
| env-416 | SRS019974_Baylor_scaffold_593 | - | 1777 | 1715 | RNA→ RpsM (COG0099)rpsM (PRK05179)→ PRK05309 (PRK05309)RpsK (COG0100)→ RpsD (COG0522)rpsD (PRK05327)→ PRK05182 (PRK05182)RNA_pol_L (pfam01193)→ |

| | | | | | |
|---|---|---|---|---|---|
| env-417 | SRS020856_Baylor_scaffold_33762 | - | 2739 | 2677 | RNA→ RpsM (COG0099)rpsM (PRK05179)→ PRK05309 (PRK05309)RpsK (COG0100)→ RpsD (COG0522)rpsD (PRK05327)→ RpoA (COG0202)rpoA (TIGR02027)→ RplQ (COG0203)rplQ (PRK05591)→ |
| env-418 | SRS021496_Baylor_scaffold_62694 | - | 926 | 864 | RNA→ RpsM (COG0099)rpsM (PRK05179)→ PRK05309 (PRK05309)RpsK (COG0100)→ |
| env-419 | SRS021954_C3037904 | - | 734 | 672 | RNA→ RpsM (COG0099)rpsM (PRK05179)→ PRK05309 (PRK05309)RpsK (COG0100)→ |
| env-420 | SRS022143_WUGC_scaffold_83705 | - | 996 | 934 | RNA→ RpsM (COG0099)rpsM (PRK05179)→ PRK05309 (PRK05309)RpsK (COG0100)→ |
| env-421 | SRS022530_LANL_scaffold_49521 | - | 926 | 864 | RNA→ RpsM (COG0099)rpsM (PRK05179)→ PRK05309 (PRK05309)RpsK (COG0100)→ |
| env-422 | SRS022621_Baylor_scaffold_80815 | + | 486 | 548 | RNA→ RpsM (COG0099)rpsM (PRK05179)→ PRK05309 (PRK05309)RpsK (COG0100)→ |
| env-423 | SRS023352_LANL_scaffold_56800 | - | 3748 | 3686 | RNA→ RpsM (COG0099)rpsM (PRK05179)→ PRK05309 (PRK05309)RpsK (COG0100)→ RpsD (COG0522)rpsD (PRK05327)→ RpoA (COG0202)rpoA (TIGR02027)→ RplQ (COG0203)rplQ (PRK05591)→ |
| env-424 | SRS023835_Baylor_scaffold_7147 | + | 1830 | 1892 | RNA→ RpsM (COG0099)rpsM (PRK05179)→ PRK05309 (PRK05309)RpsK (COG0100)→ |
| env-425 | SRS023926_Baylor_scaffold_58416 | + | 523 | 585 | RNA→ |
| env-426 | SRS023958_Baylor_scaffold_28668 | + | 1028 | 1090 | RNA→ RpsM (COG0099)rpsM (PRK05179)→ PRK05309 (PRK05309)RpsK (COG0100)→ RpsD (COG0522)rpsD (PRK05327)→ RpoA (COG0202)rpoA (TIGR02027)→ RplQ (COG0203)rplQ (PRK05591)→ |
| env-427 | SRS024015_Baylor_scaffold_50485 | + | 758 | 820 | RNA→ RpsM (COG0099)rpsM (PRK05179)→ PRK05309 (PRK05309)RpsK (COG0100)→ |
| env-428 | SRS024138_Baylor_scaffold_11348 | + | 624 | 686 | RNA→ RpsM (COG0099)rpsM (PRK05179)→ PRK05309 (PRK05309)RpsK (COG0100)→ rpsD (PRK05327)Ribosomal_S4 (pfam00163)→ |
| env-429 | SRS024277_LANL_scaffold_17092 | - | 1783 | 1721 | RNA→ RpsM (COG0099)rpsM (PRK05179)→ PRK05309 (PRK05309)RpsK (COG0100)→ RpsD (COG0522)rpsD (PRK05327)→ PRK05182 (PRK05182)RNA_pol_L (pfam01193)→ |
| env-430 | SRS024375_LANL_scaffold_43939 | + | 472 | 534 | RNA→ RpsM (COG0099)rpsM (PRK05179)→ PRK05309 (PRK05309)RpsK (COG0100)→ RpsD (COG0522)rpsD (PRK05327)→ RpoA (COG0202)rpoA (TIGR02027)→ RplQ (COG0203)rplQ (PRK05591)→ |
| env-431 | SRS024441_LANL_scaffold_45032 | + | 350 | 412 | RNA→ RpsM (COG0099)rpsM (PRK05179)→ PRK05309 (PRK05309)RpsK (COG0100)→ |
| env-432 | SRS042131_WUGC_scaffold_44855 | - | 11940 | 11878 | RNA→ RpsM (COG0099)rpsM (PRK05179)→ PRK05309 (PRK05309)RpsK (COG0100)→ RpsD (COG0522)rpsD (PRK05327)→ RpoA (COG0202)rpoA (TIGR02027)→ RplQ (COG0203)rplQ (PRK05591)→ |
| env-433 | SRS042910_WUGC_scaffold_4026 | - | 142 | 80 | RNA→ |
| env-434 | SRS043663_LANL_scaffold_25082 | + | 1266 | 1328 | RNA→ RpsM (COG0099)rpsM (PRK05179)→ PRK05309 (PRK05309)RpsK (COG0100)→ RpsD (COG0522)rpsD (PRK05327)→ |
| env-435 | SRS044373_C2894119 | - | 163 | 101 | RNA→ |
| env-436 | SRS044662_LANL_scaffold_30324 | + | 1063 | 1125 | RNA→ RpsM (COG0099)rpsM (PRK05179)→ PRK05309 (PRK05309)RpsK (COG0100)→ RpsD (COG0522)rpsD (PRK05327)→ RNAP_alpha_NTD (cd06928)PRK05182 (PRK05182)→ |
| env-437 | SRS045127_LANL_scaffold_33107 | - | 934 | 872 | RNA→ RpsM (COG0099)rpsM (PRK05179)→ PRK05309 (PRK05309)RpsK (COG0100)→ |
| env-438 | SRS045978_C998448 | + | 162 | 224 | RNA→ RpsM (COG0099)rpsM (PRK05179)→ |
| env-439 | SRS047210_WUGC_scaffold_50675 | + | 485 | 547 | RNA→ RpsM (COG0099)rpsM (PRK05179)→ PRK05309 (PRK05309)RpsK (COG0100)→ RpsD (COG0522)rpsD (PRK05327)→ RpoA (COG0202)rpoA (TIGR02027)→ RplQ (COG0203)rplQ (PRK05591)→ |
| env-440 | SRS047219_C2250559 | + | 513 | 575 | RNA→ |
| env-441 | SRS048411_C6948333 | - | 287 | 225 | RNA→ RpsM (COG0099)rpsM (PRK05179)→ |
| env-442 | SRS048791_LANL_scaffold_14386 | - | 1359 | 1297 | RNA→ RpsM (COG0099)rpsM (PRK05179)→ PRK05309 (PRK05309)RpsK (COG0100)→ S4 (smart00363)RpsD (COG0522)→ |
| env-443 | SRS049147_LANL_scaffold_32903 | + | 345 | 407 | RNA→ RpsM (COG0099)rpsM (PRK05179)→ PRK05309 (PRK05309)RpsK (COG0100)→ |
| env-444 | SRS049389_C4447558 | - | 165 | 103 | RNA→ hypo→ |
| env-445 | SRS050029_C712147 | - | 1341 | 1279 | RNA→ RpsM (COG0099)rpsM (PRK05179)→ PRK05309 (PRK05309)RpsK (COG0100)→ S4 (smart00363)RpsD (COG0522)→ |
| env-446 | SRS052227_LANL_scaffold_45775 | + | 496 | 558 | RNA→ RpsM (COG0099)rpsM (PRK05179)→ PRK05309 (PRK05309)RpsK (COG0100)→ S4 (smart00363)RpsD (COG0522)→ |
| env-447 | SRS052874_LANL_scaffold_15177 | - | 214 | 152 | RNA→ rps13 (CHL00137)rpsM (PRK05179)→ |
| env-448 | SRS053854_LANL_scaffold_52950 | - | 165 | 103 | RNA→ |
| env-449 | SRS054569_C1459708 | + | 270 | 332 | RNA→ |
| env-450 | SRS055426_LANL_scaffold_39788 | - | 367 | 305 | RNA→ RpsM (COG0099)rpsM (PRK05179)→ |
| env-451 | SRS056323_C2439953 | - | 144 | 82 | RNA→ hypo→ |
| env-452 | SRS057205_LANL_scaffold_18775 | - | 538 | 476 | RNA→ RpsM (COG0099)rpsM (PRK05179)→ PRK05309 (PRK05309)RpsK (COG0100)→ |

| | | | | | |
|---|---|---|---|---|---|
| env-453 | SRS057355_LANL_scaffold_42476 | + | 233 | 295 | **RNA**→ RpsM (COG0099)rpsM (PRK05179)→ |
| env-454 | SRS057692_LANL_scaffold_15429 | - | 850 | 788 | **RNA**→ RpsM (COG0099)rpsM (PRK05179)→ PRK05309 (PRK05309)RpsK (COG0100)→ |
| env-455 | SRS057791_C3662634 | + | 525 | 587 | **RNA**→ RpsM (COG0099)rpsM (PRK05179)→ PRK05309 (PRK05309)RpsK (COG0100)→ |
| env-456 | SRS058336_LANL_scaffold_23007 | + | 495 | 557 | **RNA**→ RpsM (COG0099)rpsM (PRK05179)→ PRK05309 (PRK05309)RpsK (COG0100)→ RpsD (COG0522)rpsD (PRK05327)→ RpoA (COG0202)rpoA (TIGR02027)→ RplQ (COG0203)rplQ (PRK05591)→ |
| env-457 | SRS062544_LANL_scaffold_18883 | + | 811 | 873 | **RNA**→ RpsM (COG0099)rpsM (PRK05179)→ PRK05309 (PRK05309)RpsK (COG0100)→ RpsD (COG0522)rpsD (PRK05327)→ RNAP_alpha_NTD (cd06928)PRK05182 (PRK05182)→ |
| env-458 | SRS063193_LANL_scaffold_13819 | + | 549 | 611 | **RNA**→ RpsM (COG0099)rpsM (PRK05179)→ PRK05309 (PRK05309)RpsK (COG0100)→ RpsD (COG0522)rpsD (PRK05327)→ RpoA (COG0202)rpoA (TIGR02027)→ RplQ (COG0203)rplQ (PRK05591)→ |
| env-459 | SRS063288_LANL_scaffold_60004 | - | 1777 | 1715 | **RNA**→ RpsM (COG0099)rpsM (PRK05179)→ PRK05309 (PRK05309)RpsK (COG0100)→ RpsD (COG0522)rpsD (PRK05327)→ PRK05182 (PRK05182)RNA_pol_L (pfam01193)→ |
| env-460 | SRS064423_LANL_scaffold_34417 | + | 577 | 639 | **RNA**→ RpsM (COG0099)rpsM (PRK05179)→ PRK05309 (PRK05309)RpsK (COG0100)→ |
| env-461 | SRS064774_LANL_scaffold_4377 | + | 577 | 639 | **RNA**→ RpsM (COG0099)rpsM (PRK05179)→ PRK05309 (PRK05309)RpsK (COG0100)→ RpsD (COG0522)rpsD (PRK05327)→ RpoA (COG0202)rpoA (TIGR02027)→ rplQ (PRK05591)Ribosomal_L17 (pfam01196)→ |
| env-462 | SRS065278_LANL_scaffold_17949 | - | 734 | 672 | **RNA**→ RpsM (COG0099)rpsM (PRK05179)→ PRK05309 (PRK05309)RpsK (COG0100)→ |
| env-463 | SRS077736_LANL_scaffold_5525 | + | 6802 | 6864 | **RNA**→ RpsM (COG0099)rpsM (PRK05179)→ PRK05309 (PRK05309)RpsK (COG0100)→ RpsD (COG0522)rpsD (PRK05327)→ RpoA (COG0202)rpoA (TIGR02027)→ RplQ (COG0203)rplQ (PRK05591)→ |
| Fsp-1-1 | NZ_GG770381.1 | + | 968393 | 968455 | **RNA**→ RpsM (COG0099)rpsM (PRK05179)→ RpsM (COG0099)rpsM (PRK05179)→ PRK05309 (PRK05309)RpsK (COG0100)→ PRK05309 (PRK05309)RpsK (COG0100)→ RpsD (COG0522)rpsD (PRK05327)→ RpsD (COG0522)rpsD (PRK05327)→ RpoA (COG0202)rpoA (TIGR02027)→ RpoA (COG0202)rpoA (TIGR02027)→ RpoA (COG0202)rpoA (TIGR02027)→ RplQ (COG0203)rplQ (PRK05591)→ RplQ (COG0203)rplQ (PRK05591)→ |
| Fpe-4-1 | NZ_JH815417.1 | - | 11807 | 11745 | **RNA**→ RpsM (COG0099)rpsM (PRK05179)→ PRK05309 (PRK05309)RpsK (COG0100)→ RpsD (COG0522)rpsD (PRK05327)→ RpoA (COG0202)rpoA (TIGR02027)→ RplQ (COG0203)rplQ (PRK05591)→ |
| Fpe-2-1 | NZ_KN173677.1 | - | 26799 | 26737 | **RNA**→ RpsM (COG0099)rpsM (PRK05179)→ PRK05309 (PRK05309)RpsK (COG0100)→ RpsD (COG0522)rpsD (PRK05327)→ RpoA (COG0202)rpoA (TIGR02027)→ RplQ (COG0203)rplQ (PRK05591)→ |
| Fpe-3-1 | NZ_GG665896.1 | - | 54181 | 54119 | **RNA**→ RpsM (COG0099)rpsM (PRK05179)→ PRK05309 (PRK05309)RpsK (COG0100)→ RpsD (COG0522)rpsD (PRK05327)→ RpoA (COG0202)rpoA (TIGR02027)→ RpoA (COG0202)rpoA (TIGR02027)→ RplQ (COG0203)rplQ (PRK05591)→ |
| Fnu-13-1 | NZ_KI391961.1 | - | 1115280 | 1115218 | **RNA**→ RpsM (COG0099)rpsM (PRK05179)→ PRK05309 (PRK05309)RpsK (COG0100)→ RpsD (COG0522)rpsD (PRK05327)→ RpoA (COG0202)rpoA (TIGR02027)→ RplQ (COG0203)rplQ (PRK05591)→ |
| env-464 | SRS014107_C1350717 | - | 463 | 401 | **RNA**→ RpsM (COG0099)rpsM (PRK05179)→ |
| Fnu-20-1 | NC_003454.1 | - | 1944307 | 1944245 | **RNA**→ RpsM (COG0099)rpsM (PRK05179)→ PRK05309 (PRK05309)RpsK (COG0100)→ RpsD (COG0522)rpsD (PRK05327)→ RpoA (COG0202)rpoA (TIGR02027)→ RplQ (COG0203)rplQ (PRK05591)→ |
| Fnu-19-1 | NZ_ADVK01000050.1 | + | 10509 | 10571 | **RNA**→ RpsM (COG0099)rpsM (PRK05179)→ RpsM (COG0099)rpsM (PRK05179)→ PRK05309 (PRK05309)RpsK (COG0100)→ PRK05309 (PRK05309)RpsK (COG0100)→ RpsD (COG0522)rpsD (PRK05327)→ RpsD (COG0522)rpsD (PRK05327)→ RpoA (COG0202)rpoA (TIGR02027)→ RpoA (COG0202)rpoA (TIGR02027)→ RpoA (COG0202)rpoA (TIGR02027)→ RplQ (COG0203)rplQ (PRK05591)→ RplQ (COG0203)rplQ (PRK05591)→ |
| Fnu-4-1 | NZ_KI518439.1 | - | 69791 | 69729 | **RNA**→ RpsM (COG0099)rpsM (PRK05179)→ PRK05309 (PRK05309)RpsK (COG0100)→ RpsD (COG0522)rpsD (PRK05327)→ RpoA (COG0202)rpoA (TIGR02027)→ RpoA (COG0202)rpoA (TIGR02027)→ RplQ (COG0203)rplQ (PRK05591)→ |
| Fnu-21-1 | NZ_ATKC01000049.1 | + | 44411 | 44473 | **RNA**→ RpsM (COG0099)rpsM (PRK05179)→ PRK05309 (PRK05309)RpsK (COG0100)→ RpsD (COG0522)rpsD (PRK05327)→ RpoA (COG0202)rpoA (TIGR02027)→ RpoA (COG0202)rpoA (TIGR02027)→ RplQ (COG0203)rplQ (PRK05591)→ |
| env-465 | SRS023538_C1261540 | + | 17 | 79 | **RNA**→ RpsM (COG0099)rpsM (PRK05179)→ |

| | | | | | |
|---|---|---|---|---|---|
| env-466 | SRS024355_LANL_scaffold_106543 | + | 483 | 545 | **RNA**→ |
| env-467 | SRS014477_C1639124 | - | 1216 | 1154 | **RNA**→ RpsM (COG0099)rpsM (PRK05179)→ PRK05309 (PRK05309)RpsK (COG0100)→ rpsD (PRK05327)Ribosomal_S4 (pfam00163)→ |
| env-468 | SRS015062_WUGC_scaffold_604 | + | 27 | 89 | **RNA**→ RpsM (COG0099)rpsM (PRK05179)→ PRK05309 (PRK05309)RpsK (COG0100)→ RpsD (COG0522)rpsD (PRK05327)→ RpoA (COG0202)rpoA (TIGR02027)→ |
| Fnu-27-1 | NZ_AABF02000152.1 | - | 665 | 603 | **RNA**→ RpsM (COG0099)rpsM (PRK05179)→ RpsM (COG0099)rpsM (PRK05179)→ RpsM (COG0099)rpsM (PRK05179)→ |
| Fnu-18-1 | NZ_AKXI01000109.1 | - | 659 | 597 | **RNA**→ RpsM (COG0099)rpsM (PRK05179)→ PRK05309 (PRK05309)RpsK (COG0100)→ PRK05309 (PRK05309)RpsK (COG0100)→ |
| Fnu-8-1 | NZ_KI518350.1 | - | 29268 | 29206 | **RNA**→ RpsM (COG0099)rpsM (PRK05179)→ PRK05309 (PRK05309)RpsK (COG0100)→ RpsD (COG0522)rpsD (PRK05327)→ RpoA (COG0202)rpoA (TIGR02027)→ RplQ (COG0203)rplQ (PRK05591)→ |
| Fsp-4-1 | NZ_AZYU01000357.1 | + | 163603 | 163665 | **RNA**→ RpsM (COG0099)rpsM (PRK05179)→ PRK05309 (PRK05309)RpsK (COG0100)→ RpsD (COG0522)rpsD (PRK05327)→ RpoA (COG0202)rpoA (TIGR02027)→ RplQ (COG0203)rplQ (PRK05591)→ |
| Fnu-14-1 | NC_021281.1 | + | 757767 | 757829 | **RNA**→ RpsM (COG0099)rpsM (PRK05179)→ PRK05309 (PRK05309)RpsK (COG0100)→ RpsD (COG0522)rpsD (PRK05327)→ RpoA (COG0202)rpoA (TIGR02027)→ RpoA (COG0202)rpoA (TIGR02027)→ RplQ (COG0203)rplQ (PRK05591)→ |
| Fnu-11-1 | NZ_GL945391.1 | - | 1036500 | 1036438 | **RNA**→ RpsM (COG0099)rpsM (PRK05179)→ PRK05309 (PRK05309)RpsK (COG0100)→ RpsD (COG0522)rpsD (PRK05327)→ RpoA (COG0202)rpoA (TIGR02027)→ RplQ (COG0203)rplQ (PRK05591)→ |
| Fnu-17-1 | NZ_JH594451.1 | - | 821484 | 821422 | **RNA**→ RpsM (COG0099)rpsM (PRK05179)→ PRK05309 (PRK05309)RpsK (COG0100)→ RpsD (COG0522)rpsD (PRK05327)→ RpoA (COG0202)rpoA (TIGR02027)→ RplQ (COG0203)rplQ (PRK05591)→ |
| Fnu-5-1 | NZ_KI518408.1 | - | 1034316 | 1034254 | **RNA**→ RpsM (COG0099)rpsM (PRK05179)→ PRK05309 (PRK05309)RpsK (COG0100)→ RpsD (COG0522)rpsD (PRK05327)→ RpoA (COG0202)rpoA (TIGR02027)→ RplQ (COG0203)rplQ (PRK05591)→ |
| Fnu-6-1 | NZ_KI518379.1 | - | 4358 | 4296 | **RNA**→ RpsM (COG0099)rpsM (PRK05179)→ PRK05309 (PRK05309)RpsK (COG0100)→ RpsD (COG0522)rpsD (PRK05327)→ RpoA (COG0202)rpoA (TIGR02027)→ RplQ (COG0203)rplQ (PRK05591)→ |
| Fnu-15-1 | NZ_CP007062.1 | - | 1460036 | 1459974 | **RNA**→ RpsM (COG0099)rpsM (PRK05179)→ PRK05309 (PRK05309)RpsK (COG0100)→ RpsD (COG0522)rpsD (PRK05327)→ RpoA (COG0202)rpoA (TIGR02027)→ RplQ (COG0203)rplQ (PRK05591)→ |
| Fnu-12-1 | NZ_CM002368.1 | - | 1637804 | 1637742 | **RNA**→ RpsM (COG0099)rpsM (PRK05179)→ PRK05309 (PRK05309)RpsK (COG0100)→ RpsD (COG0522)rpsD (PRK05327)→ RpoA (COG0202)rpoA (TIGR02027)→ RplQ (COG0203)rplQ (PRK05591)→ |
| Fnu-23-1 | NZ_ADDB02000001.1 | - | 1457971 | 1457909 | **RNA**→ RpsM (COG0099)rpsM (PRK05179)→ PRK05309 (PRK05309)RpsK (COG0100)→ RpsD (COG0522)rpsD (PRK05327)→ RpoA (COG0202)rpoA (TIGR02027)→ RplQ (COG0203)rplQ (PRK05591)→ |
| env-469 | SRS011255_Baylor_scaffold_49939 | - | 316 | 254 | **RNA**→ RpsM (COG0099)rpsM (PRK05179)→ |
| env-470 | SRS017304_C1979295 | - | 442 | 380 | **RNA**→ RpsM (COG0099)rpsM (PRK05179)→ |
| env-471 | SRS024289_LANL_scaffold_19647 | - | 348 | 286 | **RNA**→ RpsM (COG0099)rpsM (PRK05179)→ |
| env-472 | SRS017076_C2692026 | + | 484 | 546 | **RNA**→ RpsM (COG0099)rpsM (PRK05179)→ |
| env-473 | SRS015061_WUGC_scaffold_2363 | + | 922 | 984 | **RNA**→ |
| env-474 | SRS063603_LANL_scaffold_101633 | + | 89 | 151 | **RNA**→ RpsM (COG0099)rpsM (PRK05179)→ rps11 (CHL00041)PRK05309 (PRK05309)→ |
| Fnu-25-1 | NC_022196.1 | - | 1518281 | 1518219 | **RNA**→ RpsM (COG0099)rpsM (PRK05179)→ PRK05309 (PRK05309)RpsK (COG0100)→ RpsD (COG0522)rpsD (PRK05327)→ RpoA (COG0202)rpoA (TIGR02027)→ RplQ (COG0203)rplQ (PRK05591)→ |
| Fnu-24-1 | NZ_CP007064.1 | + | 716475 | 716537 | **RNA**→ RpsM (COG0099)rpsM (PRK05179)→ PRK05309 (PRK05309)RpsK (COG0100)→ RpsD (COG0522)rpsD (PRK05327)→ RpoA (COG0202)rpoA (TIGR02027)→ RplQ (COG0203)rplQ (PRK05591)→ |
| Fnu-26-1 | NZ_KQ235737.1 | + | 623579 | 623641 | **RNA**→ RpsM (COG0099)rpsM (PRK05179)→ PRK05309 (PRK05309)RpsK (COG0100)→ RpsD (COG0522)rpsD (PRK05327)→ RpoA (COG0202)rpoA (TIGR02027)→ RplQ (COG0203)rplQ (PRK05591)→ |
| Fnu-2-1 | NZ_AOUL01000031.1 | + | 497 | 559 | **RNA**→ RpsM (COG0099)rpsM (PRK05179)→ PRK05309 (PRK05309)RpsK (COG0100)→ RpsD (COG0522)rpsD (PRK05327)→ RpoA (COG0202)rpoA (TIGR02027)→ RplQ (COG0203)rplQ (PRK05591)→ |

| | | | | | |
|---|---|---|---|---|---|
| Fnu-28-1 | NZ_ATKB01000107.1 | + | 34703 | 34765 | **RNA**→ RpsM (COG0099)rpsM (PRK05179)→ PRK05309 (PRK05309)RpsK (COG0100)→ RpsD (COG0522)rpsD (PRK05327)→ RpoA (COG0202)rpoA (TIGR02027)→ RplQ (COG0203)rplQ (PRK05591)→ |
| env-475 | SRS013946_WUGC_scaffold_10067 | - | 3701 | 3639 | **RNA**→ RpsM (COG0099)rpsM (PRK05179)→ RpsM (COG0099)rpsM (PRK05179)→ PRK05309 (PRK05309)RpsK (COG0100)→ RpsD (COG0522)rpsD (PRK05327)→ RpoA (COG0202)PRK05182 (PRK05182)→ RplQ (COG0203)rplQ (PRK05591)→ |
| env-476 | SRS014473_C1106123 | - | 312 | 250 | **RNA**→ RpsM (COG0099)rpsM (PRK05179)→ |
| Fru-1-1 | NZ_KB906917.1 | + | 50468 | 50530 | **RNA**→ RpsM (COG0099)rpsM (PRK05179)→ PRK05309 (PRK05309)RpsK (COG0100)→ RpsD (COG0522)rpsD (PRK05327)→ RpoA (COG0202)PRK05182 (PRK05182)→ RplQ (COG0203)rplQ (PRK05591)→ |
| Pat-1-1 | NZ_AUFS01000005.1 | + | 43017 | 43092 | **RNA**→ RpsM (COG0099)rpsM (PRK05179)→ PRK05309 (PRK05309)RpsK (COG0100)→ RpsD (COG0522)rpsD (PRK05327)→ RpoA (COG0202)PRK05182 (PRK05182)→ RplQ (COG0203)rplQ (PRK05591)→ |
| env-477 | ASRL01000232.1 | + | 4597 | 4673 | **RNA**→ RpsM (COG0099)rpsM (PRK05179)→ PRK05309 (PRK05309)RpsK (COG0100)→ RpsD (COG0522)rpsD (PRK05327)→ RpoA (COG0202)PRK05182 (PRK05182)→ RplQ (COG0203)rplQ (PRK05591)→ |
| env-478 | ASRO01002109.1 | - | 745 | 669 | **RNA**→ RpsM (COG0099)rpsM (PRK05179)→ rps11 (CHL00041)PRK05309 (PRK05309)→ |
| env-479 | ASRP01009088.1 | - | 232 | 156 | **RNA**→ RpsM (COG0099)rpsM (PRK05179)→ |
| env-480 | AYRG01000382.1 | - | 2353 | 2277 | **RNA**→ RpsM (COG0099)rpsM (PRK05179)→ PRK05309 (PRK05309)RpsK (COG0100)→ RpsD (COG0522)rpsD (PRK05327)→ RNAP_alpha_NTD (cd06928)PRK05182 (PRK05182)→ |
| env-481 | ASRM01008875.1 | - | 376 | 300 | **RNA**→ RpsM (COG0099)rpsM (PRK05179)→ |
| env-482 | KGI_S1_ANT01_95m_c10046776 | + | 726 | 802 | **RNA**→ RpsM (COG0099)rpsM (PRK05179)→ rps11 (CHL00041)PRK05309 (PRK05309)→ |

#### 17.3 Conserved domains

Conserved domains found in protein-coding genes listed in Section 17.2 are shown below, with the first sentence in their description from the Conserved Domain Database (if any). Conserved domains associated with more than one S4-Fusobacteriales RNA are assigned a color, while others are shown in gray. The number in parentheses after the colored domain name is the number of occurrences in Section 17.2.

cd06928 (15) N-terminal domain of the Alpha subunit of Bacterial RNA polymerase.
cd16320 (1) N-terminal subdomain of transcriptional regulator MraZ.
cd16321 (1) C-terminal subdomain of transcriptional regulator MraZ.
CHL00041 (11) ribosomal protein S11
CHL00137 (11) ribosomal protein S13; Validated
COG0099 (522) Ribosomal protein S13 [Translation, ribosomal structure and biogenesis]
COG0100 (308) Ribosomal protein S11 [Translation, ribosomal structure and biogenesis]
COG0202 (175) DNA-directed RNA polymerase, alpha subunit/40 kD subunit [Transcription]
COG0203 (145) Ribosomal protein L17 [Translation, ribosomal structure and biogenesis]
COG0275 (1) 16S rRNA C1402 N4-methylase RsmH [Translation, ribosomal structure and biogenesis]
COG0522 (199) Ribosomal protein S4 or related protein [Translation, ribosomal structure and biogenesis]
COG0642 (1) Signal transduction histidine kinase [Signal transduction mechanisms]
COG0657 (1) Acetyl esterase/lipase [Lipid transport and metabolism]
pfam00163 (9) Ribosomal protein S4/S9 N-terminal domain.
pfam00411 (1) Ribosomal protein S11.
pfam00583 (3) Acetyltransferase (GNAT) family.
pfam01193 (7) RNA polymerase Rpb3/Rpb11 dimerisation domain.
pfam01196 (7) Ribosomal protein L17.
pfam03118 (2) Bacterial RNA polymerase, alpha chain C terminal domain.
pfam13508 (3) Acetyltransferase (GNAT) domain.
pfam14520 (1) Helix-hairpin-helix domain.
PRK00050 (1) 16S rRNA m(4)C1402 methyltranserfase; Provisional
PRK00326 (1) cell division protein MraZ; Reviewed
PRK05179 (528) 30S ribosomal protein S13; Validated
PRK05182 (110) DNA-directed RNA polymerase subunit alpha; Provisional
PRK05309 (321) 30S ribosomal protein S11; Validated
PRK05327 (201) 30S ribosomal protein S4; Validated
PRK05591 (152) 50S ribosomal protein L17; Validated
PRK09607 (1) 30S ribosomal protein S11P; Reviewed
smart00304 (1) HAMP (Histidine kinases, Adenylyl cyclases, Methyl binding proteins, Phosphatases) domain.
smart00363 (11) S4 RNA-binding domain.
smart00387 (1) Histidine kinase-like ATPases.
smart00388 (1) His Kinase A (phosphoacceptor) domain.
TIGR02027 (87) DNA-directed RNA polymerase, alpha subunit, bacterial and chloroplast-type. [Transcription, DNA-dependent RNA polymerase]
TIGR03628 (1) ribosomal protein uS11P, archaeal form. [Protein synthesis, Ribosomal proteins: synthesis and modification]
TIGR03631 (5) ribosomal protein uS13, bacterial form. [Protein synthesis, Ribosomal proteins: synthesis and modification]
TIGR03632 (1) ribosomal protein uS11, bacterial form. [Protein synthesis, Ribosomal proteins: synthesis and modification]

#### 17.4 Multiple-sequence alignment

Each S4-Fusobacteriales RNA is denoted by an abbreviation (like "Eco-1-1") that refers to a taxonomy given in Section 17.1. The alignment may include sequences containing the RNA motif, as well as flanking sequence. The S4-Fusobacteriales RNA itself is denoted by the line underneath marked 5′ and 3′ on either end. Nucleotides in flanking sequences (i.e., not part of the motif) are written in gray letters. Stems of predicted rho-independent transcription terminators, if any, are shaded yellow. (Note: terminator predictions have not been analyzed manually, and many are likely to be false positives. Terminator predictions are those of the RNIE software.) Nucleotides predicted to function (as DNA) as transcription-factor binding sites, if any, are shaded in green. (But note: these predictions are manually annotated, so they might be under-predicted.) Annotated start codons, if any, are shaded green. (Note: start codons are frequently misannotated, especially in environmental samples.) Nucleotides proposed to basepair as part of the consensus structure are shaded in color when they comprise Watson-Crick or G-U pairs. Otherwise they are shaded gray. Conserved stems are also indicated at the bottom of the alignment by angle brackets, where matching < and > denote base-paired columns. Below these angle brackets, the symbol "2" denotes base pairs exhibiting covariation according to the statistically well-founded R-scape method. "1" denotes base pairs exhibiting covariation according to R2R's simplistic method. "0" denotes base pairs that are not observed to mutate and "?" denotes base pairs that have a significant frequency of non-canonical nucleotides for Watson-Crick or G-U pairs (> 5%). Below these base pair annotation is the consensus sequence: "R" = "A" or "G", "Y" = "C" or "U", red nucleotides: nucleotide identity conserved more than 97% of the time, black nucleotides: 90%, gray nucleotides: 75%, red circle (•): nucleotide is present 97% of the time, black circle (•): 90%, gray circle (•): 75%, white circle (◦): 50%. All percentages of sequences just described (e.g. 97% conserved) assume that sequences have been weighted by the GSC algorithm implemented by the Infernal software package.

Duplicate sequences: the following putative homologs are not shown in the alignment because their sequences are identical to a homolog already shown: Csp-1-1, Fgo-1-1, Fhw-1-1, Fhw-2-1, Fhw-3-1, Fmo-1-1, Fne-11-1, Fne-2-1, Fne-6-1, Fne-7-1, Fne-8-1, Fne-9-1, Fnu-1-1, Fnu-12-1, Fnu-15-1, Fnu-16-1, Fnu-17-1, Fnu-18-1, Fnu-19-1, Fnu-2-1, Fnu-21-1, Fnu-23-1, Fnu-24-1, Fnu-25-1, Fnu-26-1, Fnu-28-1, Fnu-4-1, Fnu-5-1, Fnu-6-1, Fnu-7-1, Fnu-8-1, Fpe-2-1, Fpe-4-1, Fsp-1-1, Fsp-2-1, Fsp-3-1, Fsp-4-1, Fsp-5-1, Fsp-6-1, Ful-1-1, Ful-2-1, Fva-1-1, Lsp-1-1, Lsp-2-1, Lsp-3-1, Ltr-1-1, Lwa-1-1, Lwa-2-1, Sam-1-1, env-100, env-101, env-102, env-103, env-104, env-105, env-106, env-107, env-109, env-110, env-111, env-112, env-113, env-114, env-115, env-117, env-118, env-119, env-122, env-123, env-124, env-125, env-128, env-129, env-130, env-133, env-134, env-135, env-136, env-137, env-138, env-139, env-144, env-145, env-146, env-147, env-148, env-149, env-15, env-150, env-151, env-153, env-155, env-156, env-157, env-158, env-16, env-160, env-161, env-162, env-163, env-17, env-172, env-173, env-177, env-183, env-184, env-186, env-190, env-199, env-200, env-201, env-202, env-203, env-204, env-205, env-206, env-207, env-208, env-209, env-210, env-211, env-212, env-213, env-214, env-215, env-216, env-217, env-22, env-221, env-223, env-224, env-225, env-227, env-228, env-229, env-230, env-231, env-232, env-233, env-234, env-235, env-236, env-237, env-238, env-239, env-240, env-241, env-242, env-243, env-245, env-246, env-247, env-248, env-249, env-250, env-251, env-252, env-253, env-254, env-257, env-258, env-259, env-262, env-265, env-266, env-267, env-268, env-269, env-270, env-271, env-272, env-273, env-274, env-275, env-276, env-277, env-28, env-283, env-284, env-285, env-286, env-287, env-290, env-292, env-293, env-294, env-296, env-301, env-302, env-307, env-308, env-309, env-31, env-310, env-312, env-317, env-318, env-322, env-323, env-324, env-325, env-326, env-327, env-328, env-329, env-33, env-330, env-331, env-332, env-333, env-334, env-339, env-34, env-340, env-341, env-342, env-343, env-344, env-345, env-346, env-347, env-348, env-349, env-35, env-352, env-354, env-355, env-356, env-357, env-358, env-36, env-360, env-362, env-364, env-365, env-366, env-367, env-368, env-369, env-370, env-371, env-372, env-373, env-374, env-375, env-376, env-377, env-378, env-379, env-38, env-380, env-381, env-382, env-383, env-384, env-385, env-386, env-387, env-388, env-389, env-39, env-390, env-391, env-392, env-393, env-394, env-395, env-396, env-397, env-398, env-399, env-40, env-400, env-401, env-402, env-403, env-404, env-405, env-406, env-407, env-408, env-409, env-410, env-411, env-412, env-413, env-414, env-415, env-416, env-417, env-418, env-419, env-42, env-420, env-421, env-422, env-423, env-424, env-425, env-426, env-427, env-428, env-429, env-43, env-430, env-431, env-432, env-433, env-434, env-435, env-436, env-437, env-438, env-439, env-44, env-440, env-441, env-442, env-443, env-444, env-445, env-446, env-447, env-448, env-449, env-45, env-450, env-451, env-452, env-453, env-454, env-455, env-456, env-457, env-458, env-459, env-46, env-460, env-461, env-462, env-463, env-47, env-473, env-474, env-476, env-478, env-479, env-48, env-480, env-49, env-50, env-51, env-53, env-54, env-55, env-56, env-57, env-58, env-59, env-60, env-63, env-64, env-73, env-74, env-75, env-76, env-77, env-78, env-79, env-80, env-81, env-82, env-83, env-84, env-85, env-86, env-87, env-88, env-89, env-9, env-90, env-91, env-92, env-93, env-94, env-95, env-96, env-97, env-98, env-99

alignment positions 1 · · · 159

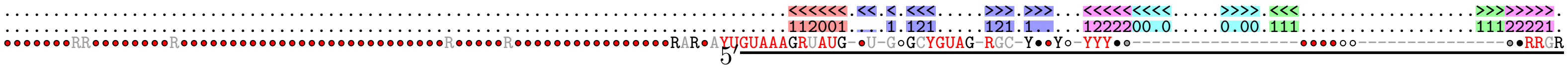

```
env-1      AAACCCUAAACACAAACAAGUACAAGGGUAAGAUUAUUUUACUGGAACAAAUUUAGUACUGACAUUGGAAGUACUGUAAAGACAUG.CA.GAGCUGUAG.AGC.CUGUU.CUUAC..............A..AUAUUAA............UGUAAGG
env-2      CCUAAACACAAACAAGUACAAGGAUAGUUUGGUUAUUUUACUGGAACAAAUUUAGUACUGACAUUGGAAGUACUGUAAAGACAUG.CA.GAGCUGUAG.AGC.CUGUU.CUUAC.................AAAAY...............GUAAGG
env-3      CUAAAACACAAACAAGUACAAGGAUAGUUUGGUUAUUUUACUGGAACAAAUUUAGUACUGACAUUGGAAGUACUGUAAAGACAUG.CA.GAGCUGUAG.AGC.CUGUU.CUUAC.................AAAAC...............GUAAGG
env-4      AUAAACAAGUUCAAGGAUAAUGUGUAAGAUCAAAAGUACAGCAUGAUUUUAAUAUAAUUAAGAUAAUGAAGUACUGUAAAGACAUG.GU.GAGUCGUAG.GAC.CAUC..UCCAU.................AAUUGCAGA...........AUGGGA
env-5      AUAAUGUGUAAGAUCAAAAGUACAGCAUAANAAAAGUACAGCAUGAUUUUAAUAUAAUUAAGAUAAUGAAGUACUGUAAAGACAUG.GU.GAGUCGUAG.GAC.CAUC..UCCAU.................AAUUGCAGA...........AUGGGA
env-6      CUGUGAAAAUCCUAAACAUAAACAAGUUCAAGGAUAAUGUGUAAGAUCAAAAGUACAGCAUAAUUUNGAAGUACUGUAAAGACAUG.GU.GAGUCGUAG.GAC.CAUC..UCCAU.................AAUUGCAGA...........AUGGGA
env-7      UGAAACGCCGAAACACAAGCAAAAGCAGGGUUAAAACGAUUUGUUAUUUGAGUUAUAAAACUUAACGUAGCGAUUGUAAAGGUAUA.UUUA.GCCGUAAGGGC.UUGUA.CUUC..................AAAA..................GAAGG
env-8      AUAUGCGAAAACCCUAAACACAAACAAGUACAAGGUUAAUAUCUAAAAUUUUUAUAUAUACUUUAUGUAAAAAUUGUAAAGAUAUG.UU.GAGUUGUAG.GACUCUAA..CUUUA.............CUAAAAUGUU...........UAGAAAAGG
env-10     AUAUGCGAAAACCCUAAACAUAAACAAGUACAAGGUUAAUAUCUAAAAUUUUUAUAUAUACUUUAUGUAAAAAUUGUAAAGAUAUG.UU.GAGUUGUAG.GACUCUAA..CUUUA.............CUAAAAUGUU...........UAGAAAAGG
env-11     ..........................................................................CUGUUAAAGAUAUGUUUG.GCUGUAG.GGU.CUAUU.CUUAG.............G..GAAA................CCUAAGG
env-12     AAGAUAAAAUUAUUUUAAAGAAAACUAGAUUAUUUUACUAACAGGAACAAAUUUAGUACUGACAUUGGAAGUACUGUAAAGAUAUG.UUUG.GCUGUAG.GGU.CUAUU.CUUAG.............G..GAAA................CCUGAGG
env-13     .......................................................................AAGUACUGUAAAGAUAUG.UUUGGUCUGUAG.GG.UCUAUU.CUUAG.............G..GAAA................CCUAAGG
env-14     ...................................................................GGAAGUACUGUAAAGAUAUG.UUUG.GCUGUAG.GGU.CUAUU.CUUAG.............G..GAAA................CCUAAGG
env-18     AAGAUAAAAUUAUUUUAAAGAAAACUAGAUUAUUUUACUAACAGGAACAAAUUUAGUACUGACAUUGGAAGUACUGUAAAGAUAUG.UUUG.GCUGUAG.GGU.CUAUU.CUUAG.............G..GAAA................CCUAAGG
env-19     ...............................................AACAGGAACAAAUUUAGUACUGACAUUGGAAGUACUGUAAAGAUAUG.UUUG.GCUGUAG.GGU.CUAUU.CUUAG.............G..GAAA................CCUAAGG
env-20     AAGAUAAAAUUAUUUUAAAGAAAACUAGAUUAUUUUACUAACAGGAACAAAUUUAGUACUGACAUUGGAAGUACUGUAAAGAUAUG.UUUG.GCUGUAG.GGU.CUAUU.CUUAG.............G..GAAA................CCUAAGG
Fpe-1-1    AUAAUUAAUAAACUUAUUUGGAAACUAGAUUAUUUUACUAACAGGAACAAAUUUAGUACUGACAUUGGAAGUACUGUAAAGAUAUGUGU.G.GCUGUAG.GGU.CUUUU.CUUAG.............G..UUAU................UCUAAGG
env-21     ..........................................AUUUUACUGGAACAAAUUUAGUACUGACAUUGGAAGUACUGUAAAGACAUG.CA.GAGCUGUAG.AGC.CUGUU.CUUAC.................AUAAA...............GUAAGG
env-23     CCCUAAACACAAACAAGUACAAGGAUAGUUUGGUUAUUUUACUGGAACAAAUUUAGUACUGACAUUGGAAGUACUGUAAAGACAUG.CA.GAGCUGUAG.AGC.CUGUU.CUUAC..............A..AAAA...............UGUAAGG
env-24     .CCCUAAACACAAACAAGUACAAGGAUAGUUUGGUUAUUUUACUGGAACAAAUUUAGUACUGACAUUGGAAGUACUGUAAAGACAUG.CA.GAGCUGUAG.AGC.CUGUU.CUUAC..............A..AAAA...............UGUAAGG
env-25     AUAAUCAAAGACAAGGAUAAGAAAAAUAGAUUAAACAUUAUUAAUAUUUUUAUACAUACCUAGAUAUAAAAAUUGUAAAGAUAUG.UUUA.GCUGUAG.GGC.UUGUU.CUUAU.............CA.AUAAAAUUAUG........UGGUAAGG
env-26     ........................................................................AUAAAAAUUGUAAAGAUAUG.UUUA.GCUGUAG.GGC.UUGUU.CUUAU.............CA.AUAAAAAUUAUG.......UGGUAAGG
env-27     UAAUCAAAAACAAGGAUAGUAAUAUUAAAGAACAUUAUUGAUAAUUUUUAUACAUACCUAGUAGAUAUAAAAAUUGUAAAGAUAUG.UUUG.GCCGUAG.AGC.UUAUU.CUUAU.................CGCUGAUUAUUUAAGGUA...AUAAGG
env-29     UAAUCAAAAACAAGGAUAGUAAUAUUAAAAAACAUUAUUGAUAAUUUUUAUACAUACCUAGUAGAUAUAAAAAUUGUAAAGAUAUG.UUUG.GCCGUAG.AGC.UUAUU.CUUAU.................CGCUGAUUAUUUAAGGUA...AUAAGG
Ste-1-1    AUCUGUGAAAAUCCGAAACAUAAACAGGUUCAAGGAUAAUAUGAGUUUAACUGUCUUUAUAUAUAUAUGAAAAUUGUAAAGGUAUG.UUGG.GCUGUAG.AGC.UUAUU.CUUAU.............GU.UUUAUC.............ACAUAAGG
Smo-1-1    AACCCUAAACACAAACAAGUACAAGGAUAAACAUCUAAAAUUUUUAUAUAUAUUUACUUAAAUUAUGUAAAAAUUGUAAAAAUAUG.UUUA.GCUGUAG.AGC.UUGUA.CUUUA.............U..GUUUGG.............GUAAAGG
env-30     GAUUGACAAAUAAACAAACUAUAAAGAUUUUUAUUAGAUAUACUUAUAUUAUAGAAGAUAUUGAAGGUAAAAAUUGUAAAAGUAUG.UUUG.GCUGUAG.AGC.UUAUU.CCUUA.............UA.AAACUUA...........UGUAAGGG
env-32     ...UGACAAAUAAACAAACUAUAAAGAUUUUUAUUAGAUAUACUUAUUAUUAUAAAAGAUAUUGAAGGUAAAAAUUGUAAAAGUAUG.UUUG.GCUGUAG.AGC.UUAUU.CCUUA.............UA.AAACUUA...........UGUAAGGG
env-37     UGACAAAUAAACAAACUAUAAAGAUUUUUAUUAGAUAUACUUAUAUUAUAGAAGAUAUUGAAGGUAAAAAUUGUAAAAGUAUG.UUUG.GCUGUAG.AGC.UUAUU.CCUUA.............UA.AAACUUA...........UGUAAGGG
env-41     UGACAAAUAAACAAACUAUAAAGAUUUUUAUUAGAUAUACUUAUUAUUAUAAAAGAUAUUGAAGGUAAAAAUUGUAAAAGUAUG.UUUG.GCUGUAG.AGC.UUAUU.CCUUA.............UA.AAACUUA...........UGUAAGGG
env-52     GAUUGACAAAUAAACAAACUAUAAAGAUUUUUAUUAGAUAUACUUAUAUUAUAGAAGAUAUUGAAGGUAAAAAUUGUAAAAGUAUG.UUUG.GCUGUAG.AGC.UUAUU.CCUUA.............UA.AAACUUA...........UGUAAGGG
env-61     GAUUGACAAAUAAACAAACUAUAAAGAUUUUUAUUAGAUAUACUUAUGUUAUAAAAGAUAUUGAAGGUAAAAAUUGUAAAAGUAUG.UUUG.GCUGUAG.AGC.UUAUU.CCUUA.............UA.AAACUUA...........UGUAAGGG
env-62     CAAGGAUAAGAAAACAAAUAAGCAGUCUUGAAAAAAUUUUUAUUUAAUUAUAUAUAGAUAUAACUGAUAAAAAUUGUAAAGAUAUGUU.UG.GCUGUAG.GGC.UUAUU.CCUUA.............UGUUUUUU............AUGUAAGGG
env-65     ..................................................GAUGGAAAUUAAGAUUACUAUGGAUAUUGAAAUAAAGAUUGUAAAGGUAUGUU.UG.GCUGUAG.AGC.UUAUU.CCUUA.............UA.AAACUUA...........UGUAAGGG
env-66     AAACAAAUAAGCAAUCUGAAAAAAUUUUUAUUUAAGUUAUAUAUAAUUCGAGAGAUAUGAUAUAAAUAAAAAUUGUAAAGAUAUG.UUUG.GCUGUAG.GGC.UUAUU.CCUUA.................GGUUUUUUUAUG........UAAGGG
env-67     ACAAAUAAGCAAUCUGAAAAAAUUUUUAUUUAAGUUAUAUAUAAUUCAAGAGAGAUAUGAUAUAAAUAAAAAUUGUAAAGAUAUG.UUUG.GCUGUAG.GGC.UUAUU.CCUUA.................GGUUUUUUUAUG........UAAGGG
env-68     ........................................UUAAAUUAUAUAUAAUUCGAGAGAGAUAUGAUAUGAAAUAAAAAUUGUAAAGAUAUG.UUUG.GCUGUAG.GGC.UUAUU.CCUUA.............UAUUUUUU............AUGUAAGGG
env-69     AACAAAUAAGCAAUCUGAAAAAAUUUUUAUUUAAAUUAUAUAUAAUUCGAGAGAGAUAUGAUAUGAAAUAAAAAUUGUAAAGAUAUG.UUUG.GCUGUAG.GGC.UUAUU.CCUUA.............UAUUUUUU............AUGUAAGGG
env-70     ACAAAUAAGCAAUCUGAAAAAAUUUUUAUUUAAGUUAUAUAUAAUUCGAGAGAGAUAUGAUAUAAAUAAAAAUUGUAAAGAUAUG.UUUG.GCUGUAG.GGC.UUAUU.CCUUA.............UAUUUUUU............AUGUAAGGG
env-71     ACAAAUAAGCAAUCUGAAAAAAUUUUUAUUUAAGUUAUAUAUAAUUCAAGAGAGAUAUGAUAUAAAUAAAAAUUGUAAAGAUAUG.UUUG.GCUGUAG.GGC.UUAUU.CCUUA.............UAUUUUUU............AUGUAAGGG
env-72     ACAAAUAAGCAAUCUGAAAAAAUUUUUAUUUAAAUUAUAUAUAAUUCGAGAGAGAUAUGAUAUGAAAUAAAAAUUGUAAAGAUAUG.UUUG.GCUGUAG.GGC.UUAUU.CCUUA.............UAUUUUUU............AUGUAAGGG
env-108    ACAAAUAAGCAAUCUGAAAAAAUUUUUAUUUAAGUUAUAUAUAAUUCAAGAGAGAUAUGAUAUAAAUAAAAAUUGUAAAGAUAUG.UUUG.GCUGUAG.GGC.UUAUU.CCUUA.............UAUUUUUU............AUGUAAGGG
env-116    .........................................AAAUUAUAUAUAAUUCGAGAGAGAUAUGAUAUGAAAUAAAAAUUGUAAAGAUAUG.UUUG.GCUGUAG.GGC.UUAUU.CCUUA.............UAUUUUUU............AUGUAAGGG
env-120    ..........AAUCUGAAAAAAUUUUUAUUUAAGUUAUAUAUAAUUCGAGAGAGAUAUGAUAUGAAAUAAAAAUUGUAAAGAUAUG.UUUG.GCUGUAG.GGC.UUAUU.CCUUA.............UAUUUUUU............AUGUAAGGG
Lgo-2-1    AGAUACAAAAAAGCAAUCUUGAAAAAAUUUUUAUUUAAUUAUAUACGAAAUAACAGAUAUAAAUAAUAAAAAUUGUAAAGAUAUG.UUUG.GCUGUAG.GGC.UUAUU.CCUUA.............UAUUUUAUU...........AUGUGAGGG
Lgo-1-1    AGAUCACAAAAAAGCAAUCUGAAAAAAUUUUUAUUUGAUUAUAUACGAAAUAACAGAUAUAAAUAAUAAAAAUUGUAAAGAUAUG.UUUG.GCUGUAG.GGC.UUAUU.CCUUA.............UAUUUUAUU...........AUGUGAGGG
env-121    CAAGGAUAAGAAAACAAAUAAGCAGUCUUGAAAAAAUUUUUAUUUAAUUAUAUAUAGAUAUAACUGAUAAAAAUUGUAAAGAUAUG.UUUG.GCUGUAG.GGC.UUAUU.CCUUA.............UGUUUUUUU...........AUGUAAGGG
env-126    AGGAUAAGAAAACAAAUAAGCAGUCUUGAAAAAAUUUUUAUUUAAUUAUAUAUAGAUAUAACUGAUAAAAAUUGUAAAGAUAUG.UUUG.GCUGUAG.GGC.UUAUU.CCUUA.............UGUUUUUUU...........AUGUAAGGG
env-127    AGGCGAAAAAAGUUAAAAACAAACUGAUUAAAAAAUUUUUAUUAUAAGACUUUUAAAGAUAUUUAAAUAAAAAUUGUAAAAGUAUGUU.UA.GCUGUAG.GGC.UUAUU.CCUUA.............UAGCAU..............UUGUGAGGG
env-131    CCUGAAAAAAAUUUUUAUUAAAAUAUAUAUAGACUGAAGGAAAUUGAGAUUACUAUGGAUAUUGAAAUAAAGAUUGUAAAGGUAUG.UUUG.GCUGUAG.AGC.UUAUU.CCUUA.............UAUUGCAU............AUGUGAGGG
env-132    CCUGAAAAAAAUUUUUAUUAAAAUAUAUAUAGACUGAAGGAAAUUAAGAUUACUAUGGAUAUUGAAAUAAAGAUUGUAAAGGUAUG.UUUG.GCUGUAG.AGC.UUAUU.CCUUA.............UAUUGCAU............AUGUGAGGG
env-140    .................................................AAUAUAUUAAGAUUACUAUGGAUAUUGAAAUAAAGAUUGUAAAGGUAUG.UUUG.GCUGUAG.AGC.UUAUU.CCUUA.............UAUUGCAU............AUGUGAGGG
env-141    ................AUUAAAAUAUAUAUAGACUGAAGGAAAUUAAGAUUACUAUGGAUAUUGAAAUAAAGAUUGUAAAGGUAUG.UUUG.GCUGUAG.AGC.UUAUU.CCUUA.............UAUUGCAU............AUGUGAGGG
env-142    ...............UAUUAAAAUAUAUAUAGACUGAAGGAAAUUAAGAUUACUAUGGAUAUUGAAAUAAAGAUUGUAAAGGUAUG.UUUG.GCUGUAG.AGC.UUAUU.CCUUA.............UAUUGUGU............AUGUGAGGG
env-143    CCUGAAAAAAAUUUUUAUUAAAAUAUAUAUAGACUGAAGGAAAUUAAGAUUACUAUGGAUAUUGAAAUAAAGAUUGUAAAGGUAUG.UUUG.GCUGUAG.AGC.UUAUU.CCUUA.............UAUUGUGU............AUGUGAGGG
env-152    CCUGAAAAAAAUUUUUAUUAAAAUAUAUAUAGACUGAAGGAAAUUAAGAUUGCUAUGGAUAUUGAAAUAAAGAUUGUAAAGGUAUG.UUUG.GCUGUAG.AGC.UUAUU.CCUUA.............UAUUGCGU............AUGUGAGGG
env-154    CCUGAAAAAAAUUUUUAUUAAAAUAUAUAUAGACUGAAGGAAAUUAAGAUUACUAUGGAUAUUGAAAUAAAGAUUGUAAAGGUAUG.UUUG.GCUGUAG.AGC.UUAUU.CCUUA.............UAUUGCGU............AUGUGAGGG
env-159    CCUGAAAAAGAUUUUUAUUAAAAUAUAUAUAGACUGAAGGAAAUUGAGAUUACUAUGGAUAUUGAAAUAAAGAUUGUAAAGGUAUG.UUUG.GCUGUAG.AGC.UUAUU.CCUUA.............UAUUGCGU............AUGUGAGGG
env-164    CCUGAAAAAAAUUUUUAUUAAAAUAUAUAUAGACUGAAGGAAAUUGAGAUUACUAUGGAUAUUGAAAUAAAGAUUGUAAAGGUAUG.UUUG.GCUGUAG.AGC.UUAUU.CCUUA.............UAUUGCGU............AUGUGAGGG
env-165    CCUGAAAAAAAUUUUUAUUAAAAUAUAUAUAGACUGAAAGAAAUUAAGAUUACUAUGGAUAUUGAAAUAAAGAUUGUAAAGGUAUG.UUUG.GCUGUAG.AGC.UUAUU.CCUUA.............UAUUGCGU............AUGUGAGGG
env-166    CCUGAAAAAAAUUUUUAUUAAAAUAUAUAUAGACUGAAGAAAAUUAAGAUUACUAUGGAUAUUGAAAUAAAGAUUGUAAAGGUAUG.UUUG.GCUGUAG.GGC.UUAUU.CCUUA.............UAUUGCGU............AUGUGAGGG
env-167    CCUGAAAAAAAUUUUUAUUAAAAUAUAUAUAGACUGAAGGAAAUUGAGAUUACUAUGGAUAUUGAAAUAAAGAUUGUAAAGGUAUG.UUUG.GCUGUAG.GGC.UUAUU.CCUUA.............UAUUGCGU............AUGUGAGGG
env-168    CCUGAAAAAGAUUUUUAUUAAAAUAUAUAUAGACUGAAGGAAAUUGAGAUUACUAUGGAUAUUGAAAUAAAGAUUGUAAAGGUAUG.UUUG.GCUGUAG.GGC.UUAUU.CCUUA.............UAUUGCGU............AUGUGAGGG
env-169    CCUGAAAAAAAUUUUUAUUAAAAUAUAUAUAGACUGAAAGAAAUUAAGAUUACUAUGGAUAUUGAAAUAAAGAUUGUAAAGGUAUG.UUUG.GCUGUAG.GGC.UUAUU.CCUUA.............UAUUGCGU............AUGUGAGGG
env-170    CUGAAAAAGAUUUUUAUUAAAAUAUAUAUAUAGAUUAAAGGAAAUAUAAGAUUACUAUGGAUAUUGAAAUAAAGAUUGUAAAGGUAUG.UUUG.GCUGUAG.GGC.UUAUU.CCUUA.............UAUUGCGU............AUGUGAGGG
env-171    CUGAAAAAGAUUUUUAUUAAAAUAUAUAUAUAGAUUAAAAGGAAAUAUAAGAUUACUAUGGAUAUUGAAAUAAAGAUUGUAAAGGUAUG.UUUG.GCUGUAG.GGC.UUAUU.CCUUA.............UAUUAUGU............AUGUGAGGG
env-174    AACCUAAAAACAUUUUUAUUAAAAUAUAUAUAGAUAAAAUGAGAUAAUGAUACUAUGGAUAUUUAAAUAAAGAUUGUAAAGGUAUG.UUUA.GCUGUAG.AGC.UUAUU.CCUUA.............UAUUGUAU............AUGUGAGGG
env-175    UAAACCUAAAAACAUUUUUAUUAAAAUAUAUUGUGAGAUAAAAAACAAAAAUACUAUGGAUAUUGAAAUAAAGAUUGUAAAGGUAUG.UUUA.GCUGUAG.AGC.UUAUU.CCUUA.............UAUUGUAU............AUGUGAGGG
env-176    ..................................................AGAUACAAAGACACUAUGGAUAUUUAAAUAAAGAUUGUAAAGGUAUG.UUUA.GCUGUAG.AGC.UUAUU.CCUUA.............UAUUGUGU............AUGUGAGGG
env-178    ................................................................UACUAUGGAUAUUUAAAUAAAGAUUGUAAAGGUAUG.UUUA.GCUGUAG.AGC.UUAUU.CCUUA.............UAUUGUGU............AUGUGAGGG
env-179    ....................................UAGAUUAAAGGAAAUAUAAGAUUACUAUGGAUAUUGAAAUAAAGAUUGUAAAGGUAUG.UUUA.GCUGUAG.AGC.UUAUU.CCUUA.............UAUUGUGU............AUGUGAGGG
env-180    ..............................................GAUGAAAAAACAAAAUACUAUGGAUAUUGAAAUAAAGAUUGUAAAGGUAUG.UUUA.GCUGUAG.AGC.UUAUU.CCUUA.............UAUUGUGU............AUGUGAGGG
env-181    AACCUAAAAACAUUUUUAUUAAAAUAUAUAUAGAUAAAAUGAGAUAAUGAUACUAUGGAUAUUUAAAUAAAGAUUGUAAAGGUAUG.UUUA.GCUGUAG.AGC.UUAUU.CCUUA.............UAUUGUGU............AUGUGAGGG
env-182    AAACCUAAAAACAUUUUUAUUAAAAUAUAUAUUGUGAGAUAAAAAACAAAAAUACUAUGGAUAUUUAAAUAAAGAUUGUAAAGGUAUG.UUUA.GCUGUAG.AGC.UUAUU.CCUUA.............UAUUGUGU............AUGUGAGGG
env-185    ...............UUAAAAUAUAUAUAGAUGAAAUUUAAGAUACAAGAUACUAUGGAUAUUGAAAUAAAGAUUGUAAAGGUAUG.UUUA.GCUGUAG.AGC.UUAUU.CCUUA.............UAUUGUGU............AUGUGAGGG
env-187    UAAACCUAAAAACAUUUUUAUUAAAAUAUAUUGUGAGAUAAAAAACAAAAAUACUAUGGAUAUUUAAAUAAAGAUUGUAAAGGUAUG.UUUA.GCUGUAG.AGC.UUAUU.CCUUA.............UAUUGUGU............AUGUGAGGG
env-188    CUAAAAACAUUUUUAUUAAAAUAUAUAUAGAUUAAAUUAAGAUACAAAGACACUAUGGAUAUUUAAAUAAAGAUUGUAAAGGUAUG.UUUA.GCUGUAG.AGC.UUAUU.CCUUA.............UAUUGUGU............AUGUGAGGG
env-189    ...............................................................UGAUAAAUAAAGAUUGUAAAGGUAUG.UUUA.GCUGUAG.AGC.UUAUU.CCUUA.............UAUUAUGU............AUGUGAGGG
env-191    UAAACCUAAAAACAUUUUUAUUAAAAUAUGUUAUAGAUAAAAAAACAAAAUACUAUGGAUAUUGAAAUAAAGAUUGUAAAGGUAUG.UUUA.GCUGUAG.GGC.UUAUU.CCUUA.............UAUUAUGU............AUGUGAGGG
env-192    CUAAAAACAUUUUUAUUAAAAUAUAUAUAGAUUAAAUUAAGAUACAAAGACACUAUGGAUAUUUAAAUAAAGAUUGUAAAGGUAUG.UUUA.GCUGUAG.GGC.UUAUU.CCUUA.............UAUUAUGU............AUGUGAGGG
Lsp-4-1    UAAACCUAAAAACAUUUUUAUUAAAAUAUUGUGAGAUAAAAAAACAAAAAUACUAUGGAUAUUGAAAUAAAGAUUGUAAAGGUAUG.UUUA.GCUGUAG.GGC.UUAUU.CCUUA.............UAUUGUAU............AUGUGAGGG
env-193    AAACCUAAAAACAUUUUUAUUAAAAUAUAUUGUGAGAUAAAAAAACAAAAAUACUAUGGAUAUUUAAAUAAAGAUUGUAAAGGUAUG.UUUA.GCUGUAG.GGC.UUAUU.CCUUA.............UAUUGUAU............AUGUGAGGG
env-194    ......UAAACCUAAAAACAUUUUUAUUAAAAUAUAUAGAUAAAAAAAAUUACUAUGGAUAUUGAAAUAAAGAUUGUAAAGGUAUG.UUUA.GCUGUAG.GGC.UUAUU.CCUUA.............UAUUGUGU............AUGUGAGGG

           ............................................................................<<<<<<.<<.<.<<<.....>>>.>>>...<<<<<<<<<.....>>>>.<<<.................>>>>>>>>.
           ............................................................................112001.....1.121.....121.1.....1222200.0.....0.00.111.................11122221.
           •••••••RR•••••••••R•••••••••••••••••••••••••R•••••R••••••••••••••••RAR•AYUGUAAAGRUAUG-•U-G•GCYGUAG-RGC-Y••Y•-YYY••--------------••••••--------------••RRGR
                                                                              5'
```

```
env-195   AACCUAAAAACAUUUUUAUUAAAAUAUAUAUAGAUUAAAUAAAAACAAAUUACUAUGGAUAUUGAAAUAAAGAUUGUAAAGGUAUG.UUUA.GCUGUAG.GGC.UUAUU.CCUUA..............UAUUGUGU.............AUGUGAGGG
Lsh-1-1   AGAUUAAUAAACCUAAAAACAUUUUUAUUAAAUAUAUAUAGAUAAAAAAAUUACUAUGGAUAUUGAAAUAAGAUUGUAAAGGUAUG.UUUA.GCUGUAG.GGC.UUAUU.CCUUA..............UAUUGUGU.............AUGUGAGGG
Lbu-1-1   AUAAACCUAAAAACAUUUUUAUUAAAAUAUGUUAUAGAUAAAAAACAAAAUACUAUGGAUAUUGAAAUAAAGAUUGUAAAGGUAUG.UUUA.GCUGUAG.GGC.UUAUU.CCUUA..............UAUUGUUU.............AUGUGAGGG
env-196   AAACAAAUAAGCAAUCUGAAAAAAUUUUUAUUUAAGUUAUAUAUAAUUCGAGAGAUAUGAUAUAAAAUAAAAAUUGUAAAGAUAUGUU.UG.GCUGUAG.GGC.UUAUU.CCUUA.................GGUUUUUUUUAUG........UAAGGG
env-197   ACAAAUAAGCAAUCUGAAAAAAAUUUUUAUUUAAGUUAUAUAUAAUUCGAGAGAGAUAUGAUAUAAAAUAAAAAUUGUAAAGAUAUGUU.UG.GCUGUAG.GGC.UUAUU.CCUUA.................GGUUUUUUAUG..........UAAGGG
env-198   .......................................................UAUGAUAUAAAAUAAAAAUUGUAAAGAUAUGUU.UG.GCUGUAG.GGC.UUAUU.CCUUA.................GGUUUUUUAUG..........UAAGGG
env-218   ....................................................AGAUAUGAUAUGAAAUAAAAAUUGUAAAGAUAUGUU.UG.GCUGUAG.GGC.UUAUU.CCUUA.................GGUUUUUUAUG..........UAAGGG
env-219   ACAAAUAAGCAAUCUGAAAAAAUUUUUAUUUAAGUUAUAUAUAAUUCGGGAGAGAUAUGAUAUAAAAUAAAAAUUGUAAAGAUAUGUU.UA.GCUGUAG.GGC.UUAUU.CCUUA..............UAUUUUUUU............GUGUAAGGG
env-220   ACAAAUAAGCAAUCUGAAAAAAUUUUUAUUUAAGUUAUAUAUAAUUCGAGAGAGAUAUGAUAUAAAAUAAAAAUUGUAAAGAUAUGUU.UG.GCUGUAG.GGC.UUAUU.CCUUA..............UAUUUUUUU............GUGUAAGGG
env-222   ....................................AUAUAUAAUUCGGGAGAGAUAUGAUAUAAAAUAAAAAUUGUAAAGAUAUGUU.UG.GCUGUAG.GGC.UUAUU.CCUUA..............UAUUUUUUU............GUGUAAGGG
env-226   ACAAAUAAGCAAUCUGAAAAAAUUUUUAUUUAAGUUAUAUAUAAUUCGAGAGAUAUGAUAUAAAAUAAAAAUUGUAAAGAUAUGUU.UG.GCUGUAG.GGC.UUAUU.CCUUA..............UAUUUUUUU............GUGUAAGGG
env-244   ACAAAUAAGCAAUCUGAAAAAAUUUUUAUUUAAGUUAUAUAUAAUUCGGGAGAGAUAUGAUAUAAAAUAAAAAUUGUAAAGAUAUGUU.UG.GCUGUAG.GGC.UUAUU.CCUUA..............UAUUUUUUU............GUGUAAGGG
env-255   ACAAAUAAGCAAUCUGAAAAAAUUUUUAUUUGAGUUAUAUAUAAUUCGGGAGAGAUAUGAUAUAAAAUAAAAAUUGUAAAGAUAUGUU.UG.GCUGUAG.GGC.UUAUU.CCUUA..............UAUUUUUUU............GUGUAAGGG
env-256   AUAAAAAGAAAAGGUAAAAUCAGAGUUAUAUGCGAAAAUGUUAAACACAAACAAGUACAAGGGUAAAGAAGAGCUGUAAAGGUAUAUU..UUGCUGUAG.GGC.A.AUA.UCCUC..............UUUUUAUC.............AAAGGGGAG
Cso-1-1   UGAAAACCCUAAGCACAAACAAGUACAAGGAUAACUUGAAAAGUUUUGCAAACUAGUACUGACAUGGGAAGUACUGUAAAGGUAUG.UU.AGGCUGUAG.AGC.UAGUA.CUC...................GAAA...................GAGG
Csp-2-1   UGAAAACCCUAAGCACAAACAAGUACAAGGAUAACUUGAAAAGUUUUGCAAACUAGUACUGACAUGGGAAGUACUGUAAAGGUAUG.UU.AGGCUGUAG.AGC.UAGUA.CUC...................GAAA...................GAGG
env-260   AAACAAGUGCAAGGGUAAUUUUUUUGAUUGAAAAUCAUUUGAUUAAAAUUUAAUUAUCAUGAAUUAUUUAGUACUGUAAAGAUAUGUAGAGAGCUGUAG.AGC.CUUAA.UCCC..................AAAU..................GGGGU
env-261   AAACAAGUGCAAGGGUAAUUUUUUUGAUUGAAAAUCAUUUGAUUAAAAUUUAAUUAUCAUGAAUUAUUUAGUACUGUAAAGAUAUGUAGAGAGCUGUAG.AGC.CGUAA.UCCC..................AAAU..................GGGGU
env-263   UUAUCAAAAGACAUGGAAAAAUCAGAGUAAUCUGCGAAAACCCAAAACACAAACAAGUACAAGGAUAAUCGUUCUGUAAAGAUAUG.UU.GAGCCGUAG.AGC.C.GCGACCUUU.................UACA.................AAAGGA
Ipo-1-1   CAUAAACAAGUUCAAGGAUAAUUUGUUGUUUGAUUGAAUUUUUUUAAGGUAAUUAUAGUACUGAUAUCAGGUACUGUAAAGGCAUG.UU.GAGCCGUAG.GGC.CACGA.CCUGA.................UUUU.................UCAGGG
env-264   UAAACACAAACAAGUUCAAGGAUAAUUUUGUAAAGAGAGUUUGUUAAAAGUACGACAAUUAUGACAUGUAGUACUGUAAAGAUAUG.UU.G.GCUGUAG.AGC.CAUC..UCCG..................AAUUUAGGAAA...........CGGAA
env-278   UAAACAUAAACAAGUUCAAGGAUAAUUUUGUAAAGAGAGUUUGUUAAAAGUACGACAAUUAUGACAUGUAGUACUGUAAAGAUAUG.UU.G.GCUGUAG.AGC.CAUC..UCCG..................AAUUUAGGAAA...........CGGAA
env-279   AACAUAAACAAGUUCAAGGAUAAUUUUGAAAGAGAGUUUGUUAAAAAGUACGACAAUUAUAAGAGAUGGAGUACUGUAAAGAUAUG.UU.G.GCUGUAG.AGC.CAUC..UCCG..................AAUUUAGGAAA...........CGGAA
env-280   AAACAUAAACAAGUUCAAGGGUAAUUUUGGAUUGAGAGUUUGUUAAAUAGUACGACAAUUAAGACGUGGAGUACUGUAAAGAUAUG.UU.G.GCCGUAG.GGC.CAUC..UCCG..................AGUUUUAGGAAG..........CGGAA
env-281   AACCCAAAACAUAAACAAGUUCAAGGAUAAUUUUACUAAAGAGAGUUUGUUAAAAAGUACUGAUAAUGGAGUGCUGUAAAGAUAUG.GU.G.GCUGUAG.AGU.CAUC..CCCAU.................AAUCUGGAA............AUGGGC
env-282   AACCCAAAACAUAAACAAGUUCAAGGAUAAUUUUACUAAAGAGAGUUUGUUAAAAAGUACUGAUAAUGGAGUGCUGUAAAGAUAUG.GU.G.GCUGUAG.AGU.CAUC..CCCAU.................AAUCUGGAA............AUGGGC
env-288   AAACACAAACAGGUUCAAGGGUAAUUUUGUAAUUUAAGAGAGUUUGUUAAAAAAAGUACUGAUAAUGGAGUGCUGUAAAGAUAUG.GU.GAGCUGUAG.AGC.CAUC..CCCAA.................AAUAGGAA.............UUGGGA
env-289   AAACACAAACAAGUUCAAGGGUAAUUUUGUAAUUUAAGAGAGUUUGUUAAAAAAAGUACUGAUAAUGGAGUGCUGUAAAGAUAUG.GU.GAGCUGUAG.AGC.CAUC..CCCAA.................AAUAGGAA.............UUGGGA
env-291   UAAACACAAACAAGUUCAAGGGUAAUUUUGUAAAAAGAAGAGUUUGUUAAAAAAAGUACUGAUAAUGGAGUGCUGUAAAGAUAUG.GU.GAGCUGUAG.AGC.CAUC..CCCAA.................AAUAGGAA.............UUGGGA
env-295   AAACACAAACAGGUUCAAGGGUAAUUUUGUAAUUUAAGAGAGUUUGUUAAAAAAAGUACUGAUAAUGGAGUGCUGUAAAGAUAUG.GU.GAGCUGUAG.AGC.CAUC..CCCAA.................AAUAGGAA.............UUGGGA
env-297   UGCAAGGAUAGUCUUUUGAAGAAGAGAUUUCAAAAAGUACUUAAAAAUUUUAAGUGAUUAAGAUUAUGGAGUACUGUAAAGAUAUG.GU.GAGUCGUAG.GAC.CAUC..CCCAU.................AAUCGCAGA............AUGGGA
Fne-5-1   GUGCAAGGAUAGUCUUUUGAGGAAGAGGUUUCAAAAAGUACUUAAAAUUUUAGAGGAUUAAGAUUGUGGAGUACUGUAAAGAUAUG.GU.GAGCUGUAG.AGC.CAUC..CCCAU.................AAUCGCAGA............AUGGGA
Fne-3-1   GUGCAAGGAUAGUCUUUUAAGGAAGAGGUUUCAAAAAGUACUUAAAAUUUUAGAGGAUUAAGAUUGUGGAGUACUGUAAAGAUAUG.GU.GAGCUGUAG.AGC.CAUC..CCCAU.................AAUCGCAGA............AUGGGA
Fne-10-1  GUGCAAGGAUAGUCUUUUGAGGAAGAGGUUUCAAAAAGUACUUAAAAUUUUAAAUCAUUAAGAUUGUGGAGUACUGUAAAGAUAUG.GU.GAGCUGUAG.AGC.CAUC..CCCAU.................AAUCGCAGA............AUGGGA
Fne-1-1   GUGCAAGGAUAGUCUUUUGAGGAAGAGGUUUCAAAAAGUACUUAAAAUUUUAGAGGAUUAAGAUUGUGGAGUACUGUAAAGAUAUG.GU.GAGCUGUAG.AGC.CAUC..CCCAU.................AAUCGCAGA............AUGGGA
Fne-4-1   GUGCAAGGAUAGUUUUUUGAGGAAGAGGUUUCAAAAAGUACUUAAAAUUUUAAAUCAUUAAGAUUGUGGAGUACUGUAAAGAUAUG.GU.GAGCUGUAG.AGC.CAUC..CCCAU.................AAUCGCAGA............AUGGGA
env-298   AACAAGUUCAAGGAUAAGAAGUAGUAAGAUCAAAAGUACAGCAUGAUUUUAAUAUUAUUAAGAUAAUGACGUACUGUAAAGACAUG.GU.GAGUCGUAG.GAC.CAUC..CCCAU.................AAUUGCAGA............AUGGGA
env-299   AUAAACAAGUUCAAGGAUAAUGUGUAAGAUCAAAAGUACAGCAUGAUUUUAAUAUAAUUAAGAUAAUGAAGUACUGUAAAGACAUG.GU.GAGUCGUAG.GAU.CAUC..UCCAU.................AAUUGCAGA............AUGGGA
env-300   AUAAACAAGUUCAAGGAUAAUGUGUAAGAUCAAAAGUACAGCAUGAUUUUAAUAUAAUUAAGAUAAUGAAGUACUGUAAAGACAUG.GU.GAGUCGUAG.GAC.CAUC..UCCAU.................AAUUGCAGA............AUGGGA
env-303   AUAAACAAGUUCAAGGAUAAUGUGUAAGAUCAAAAGUACAGCAUGAUUUUAAUAUAAUUAAGAUAAUGAAGUACUGUAAAGACAUG.GU.GAGUCGUAG.GAC.CAUC..UCCAU.................AAUUGCAGA............AUGGGA
env-304   ...................AUGUGUAAGAUCAAAAGUACAGCAUGAUUUUAAUAUAAUUAAGAUAAUGAAGUACUGUAAAGACAUG.GU.GAGUCGUAG.GAC.CAUC..UCCAU.................AAUUGUAGA............AUGGGA
env-305   AUAAACAAGUUCAAGGAUAAUGUGUAAGAUCAAAAGUACAGCAUGAUUUUAAUAUAAUUAAGAUAAUGAAGUACUGUAAAGACAUG.GU.GAGUCGUAG.GAC.CAUC..UCCAU.................AAUUGCAGA............AUGGGA
env-306   ......................................................UAAUUAAGAUAAUGAAGUACUGUAAAGACAUG.GU.GAGUCGUAG.GAC.CAUC..UCCAU.................AAUUGCAGA............AUGGGA
Fhw-4-1   AUAAACAAGUUCAAGGAUAAUGUGUAAGAUCAAAAGUACAGCAUGAUUUUAAUAUAAUUAAGAUAAUGAAGUACUGUAAAGACAUG.GU.GAGUCGUAG.GAC.CAUC..UCCAU.................AAUUGCAGA............AUGGGA
env-311   ........................................................AUUAAGAUAAUGAAGUACUGUAAAGACAUG.GU.GAGUCGUAG.GAC.CAUC..UCCAU.................AAUUGCAGA............AUGGGA
Fnu-9-1   AUAAACAAGUUCAAGGAUAAUGUGUAAGAUCAAAAGUACAGCAUGAUUUUAAUAUAUUUAAGAUAAUGAAGUACUGUAAAGACAUG.GU.GAGUCGUAG.GAC.CAUC..UCCAU.................AAUUGCAGA............AUGGGA
env-313   ......................................................UAAUUAAGAUAAUGAAGUACUGUAAAGACAUG.GU.GAGUCGUAG.GAC.CAUC..UCCAU.................AAUUGCAGA............AUGGGA
env-314   ......................................................UAAUUAAGAUAAUGAAGUACUGUAAAGACAUG.GU.GAGUCGUAG.GAC.CAUC..UCCAU.................AAUUGCAGA............AUGGAA
Fnu-3-1   AUAAACAAGUUCAAGGAUAAUAUGCAAGAUCAGAAGUACAGCAUAAUUUUAAUAUAAUUAAGAUAAUGAAGUACUGUAAAGACAUG.GU.GAGUCGUAG.GAC.CAUC..UCCAU.................AAUUGCAGA............AUGGAA
Fnu-10-1  AUAAACAAGUUCAAGGAUAAUAUGUAAGAUCAAAAGUACAGCAUGAUUUUAAUAUAAUUAAGAUAAUGAAGUACUGUAAAGACAUG.GU.GAGUCGUAG.GAC.CAUC..UCCAU.................AAUUGCAGA............AUGGAA
env-315   CAAGGAUAAUUUUUUAUUUUGUAGUAAGAGAAAAAGUACAGCAUGAUUUUAAUAUGAUUAAGAUAAUGAAGUACUGUAAAGACAUG.GU.GAGUCGUAG.GAC.CAUC..UCCAU.................AAUUACAGA............AUGGGA
env-316   ......................................................UAAUUAAGAUAAUGAAGUACUGUAAAGACAUG.GU.GAGUCGUAG.GAC.CAUC..UCCAU.................AAUUGCAGA............AUGGGA
env-319   AUAAACAAGUUCAAGGAUAAUGUGUAAGAUCAAAAGUACAGCAUGAUUUUAAUAUAAUUAAGAUAAUGAAGUACUGUAAAGACAUG.GU.GAGUCGUAG.GAC.CAUC..UCCAU.................AAUUGCAGA............AUGGGA
env-320   AUAAACAAGUUCAAGGAUAAUGUGUAAGAUCAAAAGUACAGCAUGAUUUUAAUAUAAUUAAGAUAAUGAAGUACUGUAAAGACAUG.GU.GAGUCGUAG.GAC.CAUC..UCCAU.................AAUUGCAGA............AUGGGA
env-321   AUAAACAAGUUCAAGGAUAAUGUGUAAGAUCAAAAGUACAGCAUGAUUUUAAUAUAAUUAAGAUAAUGAAGUACUGUAAAGACAUG.GU.GAGUCGUAG.GAC.CAUC..UCCAU.................AAUUGCAGA............AUGGGA
env-335   ......................................................UAAUUAAGAUGAUGAAGUACUGUAAAGACAUG.GU.GAGUCGUAG.GAC.CAUC..UCCAU.................AAUUGCAGA............AUGGGA
env-336   ...............................AAAAGUACAGCAUGAUUUUAAUAUAAUUAAGAUAAUGAAGUACUGUAAAGACAUG.GU.GAGUCGUAG.GAC.CAUC..UCCAU.................AAUUGCAGA............AUGGGA
Fnu-22-1  AACAAGUUCAAGGAUAAUGUGUAAGAUCAAAAGUACAGCAUAAUUUUAAUAAUUUAAUUAAGAUAAUGAAGUACUGUAAAGACAUG.GU.GAGUCGUAG.GAC.CAUC..UCCAU.................AAUUGCAGA............AUGGGA
env-337   ......................................................UAAUUAAGAUAAUGAAGUACUGUAAAGACAUG.GU.GAGUCGUAG.GAC.CAUC..UCCAU.................AAUUGCAGA............AUGGGA
env-338   AACAAGUUCAAGGAUAAUGUGUAAGAUCAAAAGUACAGCAUAAUUUUAAUAAUUUAAUUAAGAUAAUGAAGUACUGUAAAGACAUG.GU.GAGUCGUAG.GAC.CAUC..UCCAU.................AAUUGCAGA............AUGGGA
env-350   AACAAGUUCAAGGAUAAUGUGUAAGAUCAAAAGUACAGCAUAAUUUUAAUAAUUUAAUUAAGAUAAUGAAGUACUGUAAAGACAUG.GU.GAGUCGUAG.GAC.CAUC..UCCAU.................AAUUGCAGA............AUGGGA
env-351   ......................................................UAAUUAAGAUAAUGAAGUACUGUAAAGACAUG.GU.GAGUCGUAG.GAC.CAUC..UCCAU.................AAUUGCAGA............AUGGGA
env-353   AUAAACAAGUUCAAGGAUAAUGUGUAAGAUCAAAAGUACAGCAUGAUUUUAAUAUAAUUAAGAUAAUGAAGUACUGUAAAGACAUG.GU.GAGUCGUAG.GAC.CAUC..UCCAU.................AAUUGCAGA............AUGGGA
env-359   AACAAGUUCAAGGAUAAUGUGUAAGAUCAAAAGUACAGCAUAAUUUUAAUAAUUUAAUUAAGAUAAUGAAGUACUGUAAAGACAUG.GU.GAGUCGUAG.GAC.CAUC..UCCAU.................AAUUGCAGA............AUGGGA
env-361   ......................................................UAAUUAAGAUAAUGAAGUACUGUAAAGACAUG.GU.GAGUCGUAG.GAC.CAUC..UCCAU.................AAUUGCAGA............AUGGGA
Fsp-7-1   AACAAGUUCAAGGAUAAUGUGUAAGAUCAAAAGUACAGCAUAAUUUUAAUAAUUUAAUUAAGAUAAUGAAGUACUGUAAAGACAUG.GU.GAGUCGUAG.GAC.CAUC..UCCAU.................AAUUGCAGA............AUGGGA
env-363   AACAAGUUCAAGGAUAAUGUGUAAGAUCAAAAGUACAGCAUAAUUUUAAUAAUUUAAUUAAGAUAAUGAAGUACUGUAAAGACAUG.GU.GAGUCGUAG.GAC.CAUC..UCCAU.................AAUUGCAGA............AUGGGA
Fpe-3-1   AACAAGUUCAAGGAUAAUGUGUAAGAUCAAAAGUACAGCAUAAUUUUAAUAAUUUAAUUAAGAUGAUGAAGUACUGUAAAGACAUG.GU.GAGUCGUAG.GAC.CAUC..UCCAU.................AAUUGCAGA............AUGGGA
Fnu-13-1  AUAAACAAGUUCAAGGAUAAUAUGCAAGAUCAGAAGUACAGCAUAAUUUUAAUAUAAUUAAGAUAAUGAAGUACUGUAAAGACAUG.GU.GAGUCGUAG.GAC.CAUC..UCCAU.................AAUUGCAGA............AUGGAA
env-464   ...........................................UAAUUUUAAUAUAAUUAAGAUAAUGAAGUACUGUAAAGACAUG.GU.GAGUCGUAG.GAC.CAUC..UCCAU.................AAUUGCAGA............AUGGAA
Fnu-20-1  CAAGGAUAAUUUUUUAUUUUGUAGCAAGAGAAAAAGUACAGCAUGAUUUUAAUAUAAUUAAGAUAAUGAAGUACUGUAAAGACAUG.GU.GAGUCGUAG.GAC.CAUC..UCCAU.................AAUUGCAGA............AUGGGA
env-465   .........................................................UUAAGAUAAUGAAGUACUGUAAAGACAUG.GU.GAGUCGUAG.GAC.CAUC..UCCAU.................AAUUGCAGA............AUGGGA
env-466   AUAAACAAGUUCAAGGAUAAUGUGUAAGAUUAAAAGUACAGCAUGAUUUUAAUAUAAUUAAGAUAAUGAAGUACUGUAAAGACAUG.GU.GAGUCGUAG.GAC.CAUC..UCCAU.................AAUUGCAGA............AUGGGA
env-467   ........................................................AUUAAGAUAAUGAAGUACUGUAAAGACAUG.GU.GAGUCGUAG.GAC.CAUC..UCCAU.................AAUUGCAGA............AUGGAA
env-468   ...............................................UUUAAUAUGAUUAAGAUAAUGAAGUACUGUAAAGACAUG.GU.GAGUCGUAG.GAC.CAUC..UCCAU.................AAUUGCAGA............AUGGAA
Fnu-27-1  CAAGGAUAAUUUUUUAUUUUGUAGUAAGAGAAAAAGUACAGCAUGAUUUUAAUAUGAUUAAGAUAAUGAAGUACUGUAAAGACAUG.GU.GAGUCGUAG.GAC.CAUC..UCCAU.................AAUUACAGA............AUGGAA
Fnu-14-1  AUAAACAAGUUCAAGGAUAAUAUGCAAGAUUAGAAGUACAGCAUAAUUUUAAUAUAAUUAAGAUAAUGAAGUACUGUAAAGACAUG.GU.GAGUCGUAG.GAC.CAUC..UCCAU.................AAUUGCAGA............AUGGAA
Fnu-11-1  AUAAACAAGUUCAAGGAUAAUAUGCAAGAUCAGAAGUACAGCAUAAUUUUAAUAUAAUUAAGAUAAUGAAGUACUGUAAAGACAUG.GU.GAGUCGUAG.GAC.CAUC..UCCAU.................AAUUGCAGA............AUGGAA
env-469   ...............................AAAAGUACAGCAUGAUUUUAAUAUAAUUAAGAUAAUGAAGUACUGUAAAGACAUG.GU.GAGUCGUAG.GAC.CAUC..UCCAU.................AAUUGCAGA............AUGGAA
env-470   AUAAACAAGUUCAAGGAUAAUGUGUAAGAUCAAAAGUACAGCAUGAUUUUAAUAUAAUUAAGAUAAUGAAGUACUGUAAAGACAUG.GU.GAGUCGUAG.GAC.CAUC..UCCAU.................AAUUGCAGA............AUGGAA
env-471   AUAAACAAGUUCAAGGAUAAUAUGCAAGAUCAGAAGUACAGCAUAAUUUUAAUAUAAUUAAGAUAAUGAAGUACUGUAAAGACAUG.GU.GAGUCGUAG.GAC.CAUC..UCCAU.................AAUUGCAGA............AUGGAA
env-472   CAAGGAUAAUUUUUUAUUUUGUAGUAAGAGAAAAAGUACAGCAUGAUUUUAAUAUGAUUAAGAUAAUGAAGUACUGUAAAGACAUG.GU.GAGUCGUAG.GAC.CAUC..UCCAU.................AAUUGCAGA............AUGGAA
env-475   UAAACAAGUUCAAGGAUAGUUUGUAGUAAGAGAAAGUACAGAAGGAUUUUAAUUUUAUUAAGAUAAUGAAGUACUGUAAAGACAUG.GU.GAGUCGUAG.GAC.CAUC..UCCAU.................AAUUGCAGA............AUGGGA

          ................................................................................<<<<<<.<<.<.<<<.....>>>.>>>...<<<<<<<<<.....>>>>.<<<..................>>>>>>>>.
          ................................................................................112001....1.121.....121.1.....1222200.0.....0.00.111..................11122221.
          •••••••RR••••••••R•••••••••••••••••••••••••••R•••••R••••••••••••••••RAR•AYUGUAAAGRUAUG-•U-G○GCYGUAG-RGC-Y••Y○-YYY•○-----------------••••○○---------------○•RRGR
                                                                                  5′
```

```
Fru-1-1   CAAGGAUAGUCUUUAUUAAGAGAGUAUUAAAAAAGUACAACAUAAAUUUUAAAUUAUUUAAGAUGAUGAAGUACUGUAAAGACAUG.GU.GAGUCGUAG.GAC.CAUC..UCCAU.................AAUAGCAGA............AUGGGA
Pat-1-1   AUCUGUAAAUCUAAUCCUAAGCACAAGCAAGUUCAAGGGUAAUUGAAAGCAACACACAUUUGAAAAACGCGUACUGUAAAGAUAUG.UU.GACCCGUAG.GGG.CGGUA.CUUUUGGGUGAAA.AUUCGGA.GUAA..............UCAAAAGG
env-477   UGUAAAUCUAAUCCUAAGCACAAGCAAGUUCAAGGGUAAUUGUUGAAAGCAACACACAUUUGGAAAAUGCGUACUGUAAAGAUAUG.UU.GACCUGUAG.AGG.CGAUA.CUUUUGGAUGAAAAAUUCGGA.GUAA..............UCAAAAGG
env-481   UGUAAAUCUAAGCCUAAGCACAAGCAAGUUCAAGGGUAAUUGUUGAAAGCAACACACAUUUGGAAAAUGCGUACUGUAAAGAUAUG.UU.GACCUGUAG.AGG.CGAUA.CUUUUGGAUGAAAAAUUCGGA.GUAA..............UCAAAAGG
env-482   AUCUGUAAAUCUAAUCCUAAGCACAAGCAAGUUCAAGGGUAAUUGAAAGCAACACACAUUUGAAAAAUGCGUACUGUAAAGAUAUG.UU.GACCUGUAG.AGG.CGGUA.CUUUUGGAUGAAAAAUUCGAA.GUAA..............UCAAGAGG
          .............................................................................<<<<<<.<<.<.<<<.....>>>.>>>...<<<<<<<<<.....>>>>.<<<..................>>>>>>>>.
          .............................................................................112001....1.121.....121.1.....1222200.0.....0.00.111..................11122221.
          •••••••RR•••••••••R•••••••••••••••••••••••••••R•••••R•••••••••••••••••RAR•AYUGUAAAGRUAUG-•U-G○GCYGUAG-RGC-Y••Y○-YYY••------------------••••○○---------------○•RRGR
                                                                             5′
```

alignment positions 160 · · · 273

```
env-1     CAU.GUC..GAAG.AAAAGUGUUG..GCUGACCACAGAU.....GUC.AGC...GAAUAUAUAAAASUUUUGUGAA...............GAGGAGGAAAYAAU.....UUG
env-2     CAU.GUC..GAAG..AAAGUGUUG..GCUGACCACAGAU.....GUC.AGC...GAAUAUAUAAAAAUUUUUGUGAA..............GAGGAGGAAACAAU.....UUG
env-3     CAU.GUC..GAAG..AAAGUGUUG..GCUGACCACAGAU.....GUC.AGC...GAAUAUANAAAAAUUUUUUGUGAA.............GAGGAGGAAACAAU.....UUG
env-4     CAU.GUC..GAUG..AAAGUAUUA...GCCACUUAUAAANU...GUG.GUG...UAAUAUAUAAUUGUAACCUUUAAAAUUUUAUUUUA...AAAGAGGAGGAAAAAUU..UUG
env-5     CAU.GUC..GAUG..AAAGUAUUA...GCCACUAAAAGAAU...GUG.GUG...UAAUAUAUAAUAAUAACCUUUAAAAUUUUAUUUUAAAAGAGGAGGAAAAAUU.....UUG
env-6     CAU.GUC..GAUG..AAAGUAUUA...GCCACUAGAAU......GUG.GUG...UAAUAUAUAAUUGUAA.............................................
env-7     UAU.ACC..GAGG..AAGAUUAUAG...AAGGUACCUAGGUG..CCU.GAA...UAUAUUUAAAAUAAUUGA...................GAGGAGGAAACAAAAG...UUG
env-8     CAU.AUU..GAAG..AAGGUAAA.........AAA.....................UUUAUUU........................AAGGAGGAAAGGAU.....UUG
env-10    CAU.AUU..GAAG..AAGGUAAA.........AAA.....................UUUAUUU........................AAGGAGGAAAGGAU.....UUG
env-11    CAU.AUC..GAGGGAAAAGUGUUUG.GCUAACCUAAGAU.....GUU.AGC...GAAUAUAUAAAAAUUUUUGUGAAA..............UAGGAGGAAUCAGAU....UUG
env-12    CAU.AUC..GAGG..AAAGUGUUG..GCUAACCUAAGAU.....GUU.AGC...GAAUAUAUAAAAAUUUUUGUGAAA..............UAGGAGGAAUCAGAU....UUG
env-13    CAU.AUC..GAGG..AAAGUGUUG..GCUAACCUAAGAU.....GUU.AGC...GAAUAUAUAAAAAUUUUUGUGAAA..............UAGGAGGAAUCAGAU....UUG
env-14    CAU.AUC..GAGG..AAAGUGUUG..GCUAACCUAAGAU.....GUU.AGC...GAAUAUAUAAAAAUUUUUGUGAAA..............UAGGAGGAAUCAGAU....UUG
env-18    CAU.AUC..GAGG..AAAGUGUUG..GCUAACCUAAGAU.....GUU.AGC...GAAUAUAUAAAAAUUUUUUGUGAAA.............UAGGAGGAAUCAGAU....UUG
env-19    CAU.AUC..GAGG..AAAGUGUUG..GCUAACCUAAGAU.....GUU.AGC...GAAUAUAUAAAAAUUUUUGUGAAA..............UAGGAGGAAUCAGAU....UUG
env-20    CAU.AUC..GAGG..AAAGUGUUG..GCUAACCUAAGAU.....GUU.AGC...GAAUAUAUAAAAAUUUUUGUGAAA..............UAGGAGGAAUCAGAU....UUG
Fpe-1-1   CAU.AUC..GAGG..AAAGUGUUG..GCUAACCACAGAU.....GUU.AGC...GAAUAUAUAAAAAAUUUUUGUGAA..............UAGGAGGAAUCAGAU....UUG
env-21    CAU.GUC..GAAG..AAAGUGUUG..GCUGACCACAGAU.....GUC.AGC...GAAUAUAUAAAAAAUUUUUGUGAA..............GAGGAGGAAACAAU.....UUG
env-23    CAU.GUC..GAAG..AAAGUGUUG..GCUGACCACAGAU.....GUC.AGC...GAAUAUAUAAAAAAUUUUUGUGAA..............GAGGAGGAAACAAU.....UUG
env-24    CAU.GUC..GAAG..AAAGUGUUG..GCUGACCACAGAU.....GUC.AGC...GAAUAUAUAAAAAAUUUUUGUGAA..............GAGGAGGAAACAAU.....UUG
env-25    CAU.AUU..GAUG..AAGGUAAUA........UCAA....................AUUUA...........................AAGGAGGAAAAAAU___..UUG
env-26    CAU.AUU..GAUG..AAGGUAAUA........UCAA....................AUUUA...........................AAGGAGGAAAAAAU.....UUG
env-27    CAU.AUU..GAAG..AAGGUAAUA........UCGAA...................AUUAA............................AGGAGGAAAAAAU.....UUG
env-29    CAU.AUU..GAAG..AAGGUAAUA........UCGAA...................AUUAA............................AGGAGGAAAAAAU.....UUG
Ste-1-1   CAU.AUU..GAUG..AAGGUU...................................AAUAAAAUUAAGAAAAAAUU...........AUUAAGGAGGAAAAGAU.....UUG
Smo-1-1   CAU.AUU..GAUG..AAGGUA...........AAAAAAU.................UAUUUA...........................AGGAGGAAAGGAA.....UUG
env-30    CAU.AUU..GAAG..AAGAUUUAAAAAUAUCAAAAAAAUGAGG.AGG.AACGAAUUUGGCUAGGA...........................UAGCAGGAG..........UUG
env-32    CAU.AUU..GAAG..AAGAUUUAAAAAUAUCAAAAAAAUGAGG.AGG.AACGAAUUUGGCUAGGA...........................UAGCAGGAG..........UUG
env-37    CAU.AUU..GAAG..AAGAUUUAAAAAUAUCAAAAAAAUGAGG.AGG.AACGAAUUUGGCUAGGA...........................UAGCAGGAG..........UUG
env-41    CAU.AUU..GAAG..AAGAUUUAAAAAUAUCAAAAAAAUGAGG.AGG.AACGAAUUUGGCUAGAA...........................UAGCAGGAG..........UUG
env-52    CAU.AUU..GAAG..AAGAUUUAAAAAUAUCAAAAAAAUGAGG.AGG.AACGAAUUUGGCUAGAA...........................UAGCAGGAG..........UUG
env-61    CAU.AUU..GAAG..AAGAUUUAAAAAUAUCAAAAAAAUGAGG.AGG.AACGAAUUUGGCUAGGA...........................UAGCAGGAG..........UUG
env-62    CAU.AUU..GAAG..AAGAUU.UA........AAAA.................UAUCAAAAAC...........................AAGGAGGAAAAGAU___..UUG
env-65    CAU.AUU..GAAG..AAGAUU.UA........AAAA.................UAUCAAAAA............................AAUGAGGAGGAACGAAU..UUG
env-66    CAU.AUU..GAAG..AAGGUU...........AAAAAUAUCAAA.............AACA............................GAGGAGGAAAAAA..........
env-67    CAU.AUU..GAAG..AAGGUU...........AAAAAUAUCAAA.............AACA............................GAGGAGGAAAAAAU.....UUG
env-68    CAU.AUU..GAAG..AAGGUU...........AAAAAUAUCAAA.............AACA............................GAGGAGGAAAAAAAU....UUG
env-69    CAU.AUU..GAAG..AAGGUU...........AAAAAUAUCAAA.............AACA............................GAGGAGGAAAAAAAU....UUG
env-70    CAU.AUU..GAAG..AAGGUU...........AAAAAUAUCAAA.............AACA............................GAGGAGGAAAAAAAU....UUG
env-71    CAU.AUU..GAAG..AAGGUU...........AAAAAUAUCAAA.............AACA............................GAGGAGGAAAAAAAU....UUG
env-72    CAU.AUU..GAAG..AAGGUU...........AAAAAUAUCAAA.............AACA............................GAGGAGGAAAAAAAU....UUG
env-108   CAU.AUU..GAAG..AAGGUU...........AAAAAUAUCAAA.............AACA............................GAGGAGGAAAAAAAU....UUG
env-116   CAU.AUU..GAAG..AAGGUU...........AAAAAUAUCAAA.............AACA............................GAGGAGGAAAAAAAU....UUG
env-120   CAU.AUU..GAAG..AAGGUU...........AAAAAUAUCAAA.............AACA............................GAGGAGGAAAAAAAU....UUG
Lgo-2-1   CAU.AUU..GAAG..AAGGUUU..........AAA......................AAAUAUCAAAAAC......................AAGGAGGAAAAGAU.....UUG
Lgo-1-1   CAU.AUU..GAAG..AAGGUUU..........AAA......................AAAUAUCAAAAAC......................AAGGAGGAAAAGAU.....UUG
env-121   CAU.AUU..GAAG..AA.GAU...........UUAAAAAU..................AUCAAAAAC........................AAGGAGGAAAAGAU.....UUG
env-126   CAU.AUU..GAAG..AA.GAU...........UUAAAAAU..................AUCAAAAAC........................AAGGAGGAAAAGAU.....UUG
env-127   CAU.AUU..GAAG..AA.GAU...........UAAAAAU...................AUCAAA............................AACGAGGAGGAAACAAU..UUG
env-131   CAU.AUU..GAAG..AA.GAU...........UAAAAU....................AUCAAAAAU.........................GAGGAGGAAAAAAU.....UUG
env-132   CAU.AUU..GAAG..AA.GAU............UAAAAU...................AUCAAAAAU.........................GAGGAGGAAAAAAU.....UUG
env-140   CAU.AUU..GAAG..AA.GAU............UAAAAU...................AUCAAAAAU.........................GAGGAGGAAAAAAU.....UUG
env-141   CAU.AUU..GAAG..AA.GAU............UAAAAU...................AUCAAAAAU.........................GAGGAGGAAAAAAU.....UUG
env-142   CAU.AUU..GAAG..AA.GAU...........UAAAAU....................AUCAAAAAU.........................GAGGAGGAAAAAAU.....UUG
env-143   CAU.AUU..GAAG..AA.GAU...........UAAAAU....................AUCAAAAAU.........................GAGGAGGAAAAAAU.....UUG
env-152   CAU.AUU..GAAG..AA.GAU...........UAAAAU....................AUCAAAAAU.........................GAGGAGGAAAAAAU.....UUG
env-154   CAU.AUU..GAAG..AA.GAU...........UAAAAU....................AUCAAAAAU.........................GAGGAGGAAAAAAU.....UUG
env-159   CAU.AUU..GAAG..AA.GAU...........UAAAAU....................AUCAAAAAU.........................GAGGAGGAAAAAAU.....UUG
env-164   CAU.AUU..GAAG..AA.GAU...........UAAAAU....................AUCAAAAAU.........................GAGGAGGAAAAAAU.....UUG
env-165   CAU.AUU..GAAG..AA.GAU...........UAAAAU....................AUCAAAAAU.........................GAGGAGGAAAAAAU.....UUG
env-166   CAU.AUU..GAAG..AA.GAU...........UAAAAAU...................AUCAAAAAU.........................GAGGAGGAAAAAAU.....UUG
env-167   CAU.AUU..GAAG..AA.GAU...........UAAAAU....................AUCAAAAAU.........................GAGGAGGAAAAAAU.....UUG
env-168   CAU.AUU..GAAG..AA.GAU...........UAAAAU....................AUCAAAAAU.........................GAGGAGGAAAAAAU.....UUG
env-169   CAU.AUU..GAAG..AA.GAU...........UAAAAU....................AUCAAAAAU.........................GAGGAGGAAAAAAU.....UUG
env-170   CAU.AUU..GAAG..AA.GAU...........UAAAAU....................AUCAAAAAU.........................GAGGAGGAAAAAAU.....UUG
env-171   CAU.AUU..GAAG..AA.GAU...........UAAAAAU...................AUCAAAUAAC........................GAGGAGGAAACAAU.....UUG
env-174   CAU.AUU..GAAG..AA.GAU...........UAAAAAU...................AUCAAAAC..........................GAGGAGGAAAAAAU.....UUG
env-175   CAU.AUU..GAAG..AA.GAU...........UAAAAAU...................AUCAAAAC..........................GAGGAGGAAAAAAU.....UUG

          >>>.>>>...........<<<<<<..<<<<<<...........>>>.>>>...>>>>>>.................................................
          100.211...........2211....222..............222.......1122..................................................
          CAU-RUY--GA•G--AARGU••••--○○○○○○Y•••R○○-----○○○-○○○---○○○•AYA•AR○○○○○○○○------------------RAGGAGGAAR•RR•-----UUG
                          3′
```

```
env-176    CAU.AUU..GAAG..AA.GAU..........UAAAAAU..................AUCAAAAC..........................GAGGAGGAAAAAAU.....UUG
env-178    CAU.AUU..GAAG..AA.GAU..........UAAAAAU..................AUCAAAAC..........................GAGGAGGAAAAAAAU....UUG
env-179    CAU.AUU..GAAG..AA.GAU..........UAAAAAU..................AUCAAAAC..........................GAGGAGGAAAAAAU.....UUG
env-180    CAU.AUU..GAAG..AA.GAU..........UAAAAAU..................AUCAAAAC..........................GAGGAGGAAAAAAU.....UUG
env-181    CAU.AUU..GAAG..AA.GAU..........UAAAAAU..................AUCAAAAC..........................GAGGAGGAAAAAAU.....UUG
env-182    CAU.AUU..GAAG..AA.GAU..........UAAAAAU..................AUCAAAAC..........................GAGGAGGAAAAAAU.....UUG
env-185    CAU.AUU..GAAG..AA.GAU..........UAAAAAU..................AUCAAAAC..........................GAGGAGGAAAAAAU.....UUG
env-187    CAU.AUU..GAAG..AA.GAU..........UAAAAAU..................AUCAAAAC..........................GAGGAGGAAAAAAU.....UUG
env-188    CAU.AUU..GAAG..AA.GAU..........UAAAAAU..................AUCAAAAC..........................GAGGAGGAAAAAAAU....UUG
env-189    CAU.AUU..GAAG..AA.GAU..........UAAAAAU..................AUCAAAAC..........................GAGGAGGAAAAAAU.....UUG
env-191    CAU.AUU..GAAG..AA.GAU..........UAAAAAU..................AUCAAAAC..........................GAGGAGGAAAAAAU.....UUG
env-192    CAU.AUU..GAAG..AA.GAU..........UAAAAAU..................AUCAAAAC..........................GAGGAGGAAAAAAU.....UUG
Lsp-4-1    CAU.AUU..GAAG..AA.GAU..........UAAAAAU..................AUCAAAAC..........................GAGGAGGAAAAAAAU....UUG
env-193    CAU.AUU..GAAG..AA.GAU..........UAAAAAU..................AUCAAAAC..........................GAGGAGGAAAAAAU.....UUG
env-194    CAU.AUU..GAAG..AA.GAU..........UAAAAAU..................AUCAAAAC..........................GAGGAGGAAAAAAU.....UUG
env-195    CAU.AUU..GAAG..AA.GAU..........UAAAAAU..................AUCAAAAC..........................GAGGAGGAAAAAAU.....UUG
Lsh-1-1    CAU.AUU..GAAG..AA.GAU..........UAAAAAU..................AUCAAAAC..........................GAGGAGGAAAAAAU.....UUG
Lbu-1-1    CAU.AUU..GAAG..AA.GAU..........UAAAAAU..................AUCAAAAC..........................GAGGAGGAAAAAAU.....UUG
env-196    CAU.AUU..GAAG..AA.GGU..........UAAAAAU..................AUCAAAAACA........................GAGGAGGAAAAAAUU....UUG
env-197    CAU.AUU..GAAG..AA.GGU..........UAAAAAU..................AUCAAAAACA........................GAGGAGGAAAAAAU.....UUG
env-198    CAU.AUU..GAAG..AA.GGU..........UAAAAAU..................AUCAAAAACA........................GAGGAGGAAAAAAAU....UUG
env-218    CAU.AUU..GAAG..AA.GGU..........UAAAAAU..................AUCAAAAACA........................GAGGAGGAAAAAAAU....UUG
env-219    CAU.AUU..GAAG..AA.GGU..........UAAAAAU..................AUCAAAAACA........................GAGGAGGAAAAAAU.....UUG
env-220    CAU.AUU..GAAG..AA.GGU..........UAAAAAU..................AUCAAAAACA........................GAGGAGGAAAAAAAU....UUG
env-222    CAU.AUU..GAAG..AA.GGU..........UAAAAAU..................AUCAAAAACA........................GAGGAGGAAAAAAU.....UUG
env-226    CAU.AUU..GAAG..AA.GGU..........UAAAAAU..................AUCAAAAACA........................GAGGAGGAAAAAAU.....UUG
env-244    CAU.AUU..GAAG..AA.GGU..........UAAAAAU..................AUCAAAAACA........................GAGGAGGAAAAAAU.....UUG
env-255    CAU.AUU..GAAG..AA.GGU..........UAAAAAU..................AUCAAAAACA........................GAGGAGGAAAAAAU.....UUG
env-256    UAU.ACU..GAAG..AGAGUAUUU.AGAUAGUUCAUACU.....ACU.AUCGA.AAAUAUAUAUUAAUUUGAAAUUAACUUUUAAAAAAC..UGGGAGGAGGAAAACUAA.UUG
Cso-1-1    CAU.ACC..GAAG..AAAGUCUUA..GUCAGUUAAUAU.....ACU.GAC...GGAUAUAUAAAAUUUUCGGAA.................GAGGAGGAAAAAUU.....UUG
Csp-2-1    CAU.ACC..GAAG..AAAGUCUUA..GUCAGUUUAUAU.....ACU.GAC...GGAUAUAUAAAAUUUUCGGAA.................GAGGAGGAAGAAUU.....UUG
env-260    CAU.AUC..GAGG..AAAGUAUUU.AGCUAAUCUUUGUAGU...GUU.AGC...GAAUAUAUAAAUUUCUAAUAUAUU..............GAGGAGGAAAUAAUU....UUG
env-261    CAU.AUC..GAGG..AAAGUAUUU.AGCUAAUCUUUGUAGU...GUU.AGC...GAAUAUAUAAAUUUCUAAUAUAUU..............GAGGAGGAAAUAAUU....UUG
env-263    CAU.AUC..GAUG..AAAGUGUUUAGCGUGGUUAGUAU.....ACC.AAG...UAAUAUAUAUAUUUAUUU..................AAGGAGGAGCAAAG....UUG
Ipo-1-1    CAU.GCC..GAAG..AAAGUGUUU.AGUUGAUUGUUAU.....AUC.AAC...AAAUAUAUAAAUUUUCGG.................AAUAGGAGGAAGAAACG....UUG
env-264    CAU.AUC..GACG..AAAGUAUUG.AGCUGGUUUAUAUU....ACU.AGC.A.AAAUAUAUAAAAUUUUCGAU................AGAGGAGGAAACAAUU....UUG
env-278    CAU.AUC..GACG..AAAGUAUUG.AGCUGGUUUAUAUU....ACU.AGC.A.AAAUAUAUAAAAUUUUCGAU................AGAGGAGGAAACAAUU....UUG
env-279    CAU.AUC..GACG..AAAGUAUUG.AGCUAGUUCAUAUU....ACU.AGC.A.AAAUAUAUAAAAUUUUCGAU................AGAGGAGGAAACAAUU....UUG
env-280    CAU.AUC..GAUG..AAAGUAUUU.AGCUGGUUAUUAUC....ACC.GGC.AAAAAUAUAAAAAAUUUUCGAC................AGAGGAGGAAACAAUU....UUG
env-281    CAU.AUC..GAGG..AAAGUAUUU.AGCUGGUUAUUAU.....ACU.AGC.A.AAAUAUAUAAAUUUUCG.................AAAGAGGAGGAAACAAUU....UUG
env-282    CAU.AUC..GAGG..AAAGUAUUU.AGCUGGUUAUUAU.....ACU.AGC.A.AAAUAUAUAAAUUUUCG.................AAAGAGGAGGAAACAAUUUU.....
env-288    CAU.AUC..GAGG..AAAGUAUUU.AGUUGGUUAAUAU.....ACC.AGC.A.AAAUAUAUAAAUUUUCG.................AAAGAGGAGGAAAUAAUU....UUG
env-289    CAU.AUC..GAGG..AAAGUAUUU.AGUUGGUUAAUAU.....ACC.AGC.A.AAAUAUAUAAAUUUUCG.................AAAGAGGAGGAAAUAAUU....UUG
env-291    CAU.AUC..GAGG..AAAGUAUUU.AGUUGGUUAAUAU.....ACC.AGC.A.AAAUAUAUAAAUUUUCG.................AAAGAGGAGGAAAUAAUU....UUG
env-295    CAU.AUC..GAGG..AAAGUAUUU.AGUUGGUUAAUAU.....ACC.AGC.A.AAAUAUAUAAAUUUUCGG.................AAGAGGAGGAAAUAAUU....UUG
env-297    CAU.AUC..GAGG..AAAGUAUUU.AGCCGACUACGUU.....GUC.GAC.G.AAAUAUAAAAAACAAAAUUUGG..........AAAAGAGGAGGAAUAAUU....UUG
Fne-5-1    CAU.AUC..GAGG..AAAGUAUUU.AGCCGACUGUGUU.....GUC.GAC.G.AAAUAUAAAAAAACAAAAUUUGG..........AAAAGAGGAGGAAUAAUU.....UUG
Fne-3-1    CAU.AUC..GAGG..AAAGUAUUU.AGCCGACUGCGUU.....GUC.GAC.G.AAAUAUAAAAAACAAAAUUUGG..........AAAAGAGGAGGAAUAAUU.....UUG
Fne-10-1   CAU.AUC..GAGG..AAAGUAUUU.AGCCGACUGCGUU.....GUC.GAC.G.AAAUAUAAAAAAACAAAAUUUGG..........AAAAGAGGAGGAAUAAUU.....UUG
Fne-1-1    CAU.AUC..GAGG..AAAGUAUUU.AGCCGACUGCGUU.....GUC.GAC.G.AAAUAUAAAAAAACAAAAUUUGG..........AAAAGAGGAGGAAUAAUU.....UUG
Fne-4-1    CAU.AUC..GAGG..AAAGUAUUU.AGCCGACUGCGUU.....GUC.GAC.G.AAAUAUAAAAAAACAAAAUUUGG..........AAAAGAGGAGGAAUAAUU.....UUG
env-298    CAU.GUC..GAAG..AAAGUAUUA...GCCAACAAAAGA.....UUG.GCG...UAAUAUAUAAUAACCUUUAAAAUUUAUUUU....AAAAGAGGAGGAAAAAUU.....UUG
env-299    CAU.GUC..GAUG..AAAGUAUUA...GCCACUUAAUAAAGAAUGUG.GUG...UAAUAUAUAAUAAUAACCUUUAAAAUUUUAUUUU...AAAAGAGGAGGAAAAAUU..UUG
env-300    CAU.GUC..GAUG..AAAGUAUUA...GCCACUUAAUAAAGAAUGUG.GUG...UAAUAUAUAAUAAUAACCUUUAAAAUUUUAUUUU...AAAAGAGGAGGAAAAAUU..UUG
env-303    .AUAGUC..GAUG..AAAGUAUUA...GCCACUUAAUAAAGAAUGUG.GUG...UAAUAUAUAAUAAUAACCUUUAAAAUUUUAUUUU...AAAAGAGGAGGAAAAAUU..UUG
env-304    CAU.GUC..GAUG..AAAGUAUUA...GCCACUUAUGAAGAAU.GUG.GUG...UAAUAUAUAAUAAUAACCUUUAAAAUUUUAUUUU...AAAAGAGGAGGAAAAAUU..UUG
env-305    CAU.GUC..GAUG..AAAGUAUUA...GCCACUUAUGAAGAAU.GUG.GUG...UAAUAUAUAAUAAUAACCUUUAAAAUUUUAUUUU...AAAAGAGGAGGAAAAAUU..UUG
env-306    CAU.GUC..GAUG..AAAGUAUUA...GCCACUUAUGAAGAAU.GUG.GUG...UAAUAUAUAAUAAUAACCUUUAAAAUUUUAUUUU...AAAAGAGGAGGAAAAAUU..UUG
Fhw-4-1    CAU.GUC..GAUG..AAAGUAUUA...GCCACUUAUAAAGAAU.GUG.GUG...UAAUAUAUAAUAAUAACCUUUAAAAUUUUAUUUU...AAAAGAGGAGGAAAAAUU..UUG
env-311    CAU.GUC..GAUG..AAAGUAUUA...GCCACUUAUAAAGAAU.GUG.GUG...UAAUAUAUAAUAAUAACCUUUAAAAUUUUAUUUU...AAAAGAGGAGGAAAAAUU..UUG
Fnu-9-1    CAU.GUC..GAUG..AAAGUAUUA...GCCACUUAUAAAGAAU.GUG.GUG...UAAUAUAUAAUAAUAACCUUUAAAAUUUUAUUUU...AAAAGAGGAGGAAAAAUU..UUG
env-313    UAU.GUC..GAUG..AAAGUAUUA...GCCACUUAUAAAGAAU.GUG.GUG...UAAUAUAUAAUAAUAACCUUUAAAAUUUUAUUUU...AAAAGAGGAGGAAAAAUU..UUG
env-314    CAU.GUC..GAUG..AAAGUAUUA...GCCACUAAAGAAU....GUG.GUG...UAAUAUAUAAUAACCUUUAAAAUUUAAUUUU...AAAAGAGGAGGAAAAAUU.....UUG
Fnu-3-1    CAU.GUC..GAUG..AAAGUAUUA...GCCACUAAAGAAU....GUG.GUG...UAAUAUAUAAUAACCUUUAAAAUUUAAUUUU...AAAAGAGGAGGAAAAGUU.....UUG
Fnu-10-1   CAU.GUC..GAUG..AAAGUAUUA...GCCACUAAAAGAAU...GUG.GUG...UAAUAUAUGAAAUAACCUUUAAAAUUUAAUUUUAAAA.GAGGAGGAAAAAUU.....UUG
env-315    CAU.GUC..GAUG..AAAGUAUUA...GCCACUAAAAGAAU...GUG.GUG...UAAUAUAUAAUAAUAACCUUUAAAAUUUUAUUUUAAAAGAGGAGGAAAAAUU.....UUG
env-316    CAU.GUC..GAUG..AAAGUAUUA...GCCACUAAAAGAAU...GUG.GUG...UAAUAUAUAAUAAUAACCUUUAAAAUUUUAUUUUAAAAGAGGAGGAAAAAUU.....UUG
env-319    CAU.GUC..GAUG..AAAGUAUUA...GCCACUAAAAGAAU...GUG.GUG...UAAUAUAUAAUAAUAACCUUUAAAAUUUUAUUUUAAAAGAGGAGGAAAAGUU.....UUG
env-320    CAU.GUC..GAUG..AAAGUAUUA...GCCACUAAAAGAAU...GUG.GUG...UAAUAUAUAAUAAUAACCUUUAAAAUUUAAUUUUAAAAGAGGAGGAAAAAUU.....UUG
env-321    CAU.GUC..GAUG..AAAGUAUUA...GCCACUAAAAGAAU...GUG.GUG...UAAUAUAUAAUAAUAACCUUUAAAAUUUUAUUUUAAAAGAGGAGGAAAAAUU.....UUG
env-335    CAU.GUCUCGAUG..AAAGUAUUA...GCCACUAAAAGAAU...GUG.GUG...UAAUAUAUAAUAAUAACCUUUAAAAUUUUAUUUUAAAAGAGGAGGAAAAAUU.....UUG
env-336    CAU.GUC..GAUG..AAAGUAUUA...GCCACUAAAAGAAU...GUG.GUG...UAAUAUAUAAUAAUAACCUUUAAAAUUUAAUUUUAAAAGAGGAGGAAAAAUU.....UUG
Fnu-22-1   CAU.GUC..GAUG..AAAGUAUUA...GCCACUAAAAGAAU...GUG.GUG...UAAUAUAUAAUAAUAACCUUUAAAAUUUUAUUUUAAAAGAGGAGGAAAAAUU.....UUG
env-337    CAU.GUC..GAUG..AAAGUAUUA...GCCACUGGAAU......GUG.GUG...UAAUAUAUAAUUAUAACCUUUAAAAUUUUAUUUUAAAAGAGGAGGAAAAAUU.....UUG
env-338    CAU.GUC..GAUG..AAAGUAUUA...GCCACUAGAAU......GUG.GUG...UAAUAUAUAAUUAUAACCUUUAAAAUUUUAUUUUAAAAGAGGAGGAAAAAUU.....UUG
env-350    CAU.GUC..GAUG..AAAGUAUUA...GCCACUAGAAU......GUG.GUG...UAAUAUAUAAUUGUAACCUUUAAAAUUUUAUUUUAAAAGAGGAGGAAAAAUU.....UUG
env-351    CAU.GUC..GAUG..AAAGUAUUA...GCCACUAGAAU......GUG.GUG...UAAUAUAUAAUUAUAACCUUUAAAAUUUUAUUUUAAAAGAGGAGGAAAAAUU.....UUG
env-353    CAU.GUC..GAUG..AAAGUAUUA...GCCACUAAAAU......GUG.GUG...UAAUAUAUAAUUGUAACCUUUAAAAUUUUAUUUUAAAAGAGGAGGAAAAAUU.....UUG
env-359    CAU.GUC..GAUG..AAAGUAUUA...GCCACUAAAAU......GUG.GUG...UAAUAUAUAAUUGUAACCU..........................................
env-361    CAU.GUC..GAUG..AAAGUAUUA...GCCACUAAAAU......GUG.GUG...UAAUAUAUAAUAACCUUUAAAAUUUUAUUUU...AAAAGAGGAGGAAAAAUU.....UUG
Fsp-7-1    CAU.GUC..GAUG..AAAGUAUUA...GCCACUAAAAU......GUG.GUG...UAAUAUAUAAUGUAACCUUUAAAAUUUAUUUU..AAAAGAGGAGGAAAAAUU.....UUG
env-363    CAU.GUC..GAUG..AAAGUAUUA...GCCACUAAAAU......GUG.GUG...UAAUAUAUAAUUGUAACCUUUAAAAUUUUAUUU...........................
Fpe-3-1    CAU.GUC..GAUG..AAAGUAUUA...GCCACUAAAGU......GUG.GUG...UAAUAUAUAAUUGUAACCUUUAAAAUUUUAUUUU...AAAAGAGGAGGAAAAAUU..UUG

           >>>.>>>..........<<<<<<..<<<<<<.................>>>.>>>...>>>>>>.......................................................
           100.211...........2211.......222.................222.........1122.......................................................
           CAU-RUY--GA•G--AARGU••••--○○○○○○Y•••R••-----○○○-○○○---○○○○•AYA•AR••○○○○○○○-------------------RAGGAGGAAR•RR•-----UUG
                         3′
```

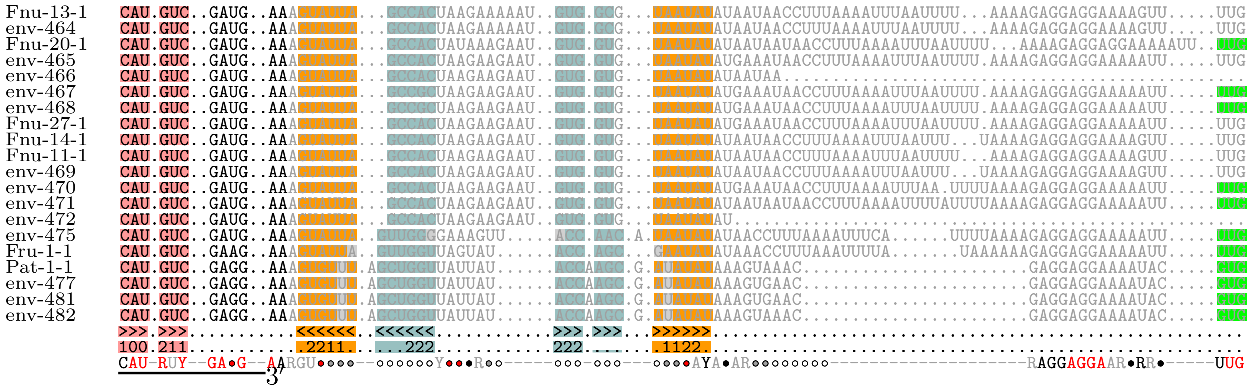

### 18 S6-S18-Chlorobi

#### 18.1 Taxa

The taxonomy of each organism containing a putative S6-S18-Chlorobi RNA is listed, with abbreviations identifying each hit (e.g., "Eco-1-1" and "Eco-1-2" might hypothetically represent two distinct RNAs in *E. coli*). The abbreviations will be used to identify each individual S6-S18-Chlorobi RNA in Sections 18.2 and 18.4:.

| abbrev. of hits | taxonomy of species |
|---|---|
| Cpa-1-1 | Bacteria Chlorobi Chlorobia Chlorobiales Chlorobiaceae *Chlorobaculum parvum* NCIB 8327 |
| Cte-1-1 | Bacteria Chlorobi Chlorobia Chlorobiales Chlorobiaceae Chlorobaculum *Chlorobium tepidum* TLS |
| Cch-1-1 | Bacteria Chlorobi Chlorobia Chlorobiales Chlorobiaceae Chlorobium/Pelodictyon group *Chlorobium chlorochromatii* CaD3 |
| Cfe-1-1 | Bacteria Chlorobi Chlorobia Chlorobiales Chlorobiaceae Chlorobium/Pelodictyon group *Chlorobium ferrooxidans* DSM 13031 |
| Cli-1-1 | Bacteria Chlorobi Chlorobia Chlorobiales Chlorobiaceae Chlorobium/Pelodictyon group *Chlorobium limicola* DSM 245 |
| Cph-1-1 | Bacteria Chlorobi Chlorobia Chlorobiales Chlorobiaceae Chlorobium/Pelodictyon group *Chlorobium phaeobacteroides* BS1 |
| Cph-2-1 | Bacteria Chlorobi Chlorobia Chlorobiales Chlorobiaceae Chlorobium/Pelodictyon group *Chlorobium phaeobacteroides* DSM 266 |
| Cph-3-1 | Bacteria Chlorobi Chlorobia Chlorobiales Chlorobiaceae Chlorobium/Pelodictyon group *Chlorobium phaeovibrioides* DSM 265 (Prosthecochloris vibrioformis |
| Clu-1-1 | Bacteria Chlorobi Chlorobia Chlorobiales Chlorobiaceae Chlorobium/Pelodictyon group Pelodictyon *Chlorobium luteolum* DSM 273 |
| Pph-1-1 | Bacteria Chlorobi Chlorobia Chlorobiales Chlorobiaceae Chlorobium/Pelodictyon group *Pelodictyon phaeoclathratiforme* BU-1 |
| Pae-1-1 | Bacteria Chlorobi Chlorobia Chlorobiales Chlorobiaceae *Prosthecochloris aestuarii* DSM 271 |
| env-1 to env-56 | environmental samples |

#### 18.2 Gene contexts

Each S6-S18-Chlorobi RNA (indicated by "RNA→") is listed. For each hit, the downstream genes predicted to reside in a regulated operon are listed. If the nearest downstream gene is encoding in the opposite strand (and therefore presumed to not be a part of a regulated operon), then that gene is still depicted. Some environmental sequences and some RefSeq entries lack gene annotations, and so no genes are available for such sequences. The direction of each gene is indicated with an arrow (→), and each predicted conserved domain in the gene is named. Conserved domains associated with more than one S6-S18-Chlorobi RNA are assigned a color; other domains are gray. Information about these conserved domains is given in Section 18.3. The accession of the sequence containing each S6-S18-Chlorobi RNA is given in the column named "Seq. accession". Accessions beginning with "NC_", "NS_", "NW_" or "NZ_" are contained in RefSeq. Other accession refer to environmental samples. Nucleotide coordinates are given for the 5′ and 3′ boundaries of each S6-S18-Chlorobi RNA. If the 5′ coordinate is greater than the 3′ coordinate, the RNA is present on the reverse-complement strand of the containing genomic DNA sequence. Each hit is denoted by an abbreviation (like "Eco-1-1") that refers to a taxonomy given in Section 18.1.

| abbrev. | Seq. accession | | 5′ at | 3′ at | genes |
|---|---|---|---|---|---|
| env-1 | F3OLC3J01EL6Z5 | - | 180 | 92 | **RNA**→ hypo→ |
| env-2 | F3OLC3J02I9N18 | + | 405 | 486 | **RNA**→ hypo→ |
| env-3 | F3OLC3J01B4FHU | - | 100 | 16 | **RNA**→ ←hypo |
| Cli-1-1 | NC_010803.1 | + | 167079 | 167163 | **RNA**→ rpsF (PRK00453)bS6 (cd00473)→ ssb (TIGR00621)SSB (pfam00436)→ rpsR (PRK00391)Ribosomal_S18 (pfam01084)→ RplI (COG0359)rplI (PRK00137)→ |
| env-4 | F3OLC3J01AOTFF | + | 81 | 168 | **RNA**→ hypo→ hypo→ |
| env-5 | F3OLC3J01ER1WJ | + | 100 | 184 | **RNA**→ rpsF (PRK00453)Ribosomal_S6 (pfam01250)→ |
| env-6 | F3OLC3J02FQPPB | + | 240 | 324 | **RNA**→ hypo→ |
| env-7 | F3OLC3J02HKSSD | + | 197 | 281 | **RNA**→ hypo→ |
| env-8 | F3OLC3J01EFF4C | + | 240 | 324 | **RNA**→ |
| env-9 | TB_FS08_3DRAFT_1000662 | - | 15648 | 15564 | **RNA**→ rpsF (PRK00453)Ribosomal_S6 (pfam01250)→ ssb (TIGR00621)SSB (pfam00436)→ rpsR (PRK00391)Ribosomal_S18 (pfam01084)→ RplI (COG0359)rplI (PRK00137)→ |
| env-10 | F3OLC3J01EZCCM | + | 89 | 173 | **RNA**→ rpsF (PRK00453)Ribosomal_S6 (pfam01250)→ |
| env-11 | F3OLC3J01BGKJE | + | 100 | 184 | **RNA**→ rpsF (PRK00453)Ribosomal_S6 (pfam01250)→ |
| env-12 | F3OLC3J01BIBU0 | - | 124 | 40 | **RNA**→ hypo→ |
| env-13 | F3OLC3J01B90M6 | + | 80 | 164 | **RNA**→ rpsF (PRK00453)Ribosomal_S6 (pfam01250)→ |
| env-14 | F3OLC3J01CB09B | + | 308 | 390 | **RNA**→ hypo→ |
| Cch-1-1 | NC_007514.1 | + | 109831 | 109904 | **RNA**→ rpsF (PRK00453)Ribosomal_S6 (pfam01250)→ ssb (TIGR00621)SSB (pfam00436)→ rpsR (PRK00391)Ribosomal_S18 (pfam01084)→ RplI (COG0359)rplI (PRK00137)→ |
| Cph-1-1 | NC_010831.1 | + | 160217 | 160291 | **RNA**→ rpsF (PRK00453)bS6 (cd00473)→ ssb (TIGR00621)SSB (pfam00436)→ rpsR (PRK00391)Ribosomal_S18 (pfam01084)→ RplI (COG0359)rplI (PRK00137)→ |
| env-15 | M750CN_1000441 | - | 2463 | 2386 | **RNA**→ rpsF (PRK00453)bS6 (cd00473)→ ssb (TIGR00621)SSB (pfam00436)→ rpsR (PRK00391)Ribosomal_S18 (pfam01084)→ RplI (COG0359)rplI (PRK00137)→ |
| env-16 | M740_1006987 | - | 2342 | 2265 | **RNA**→ rpsF (PRK00453)bS6 (cd00473)→ ssb (TIGR00621)SSB (pfam00436)→ rpsR (PRK00391)Ribosomal_S18 (pfam01084)→ RplI (COG0359)rplI (PRK00137)→ |
| Pae-1-1 | NC_011059.1 | + | 168156 | 168233 | **RNA**→ rpsF (PRK00453)bS6 (cd00473)→ ssb (TIGR00621)SSB (pfam00436)→ rpsR (PRK00391)Ribosomal_S18 (pfam01084)→ RplI (COG0359)rplI (PRK00137)→ |
| env-17 | M590M2_1034735 | - | 176 | 99 | **RNA**→ hypo→ |
| env-18 | M590M1_1003288 | + | 1442 | 1517 | **RNA**→ rpsF (PRK00453)bS6 (cd00473)→ ssb (TIGR00621)SSB (pfam00436)→ rpsR (PRK00391)Ribosomal_S18 (pfam01084)→ RplI (COG0359)rplI (PRK00137)→ |
| env-19 | M590M2_1009672 | - | 1755 | 1680 | **RNA**→ rpsF (PRK00453)bS6 (cd00473)→ ssb (TIGR00621)SSB (pfam00436)→ rpsR (PRK00391)Ribosomal_S18 (pfam01084)→ RplI (COG0359)rplI (PRK00137)→ |
| env-20 | GooDraft_1000020 | - | 11816 | 11741 | **RNA**→ rpsF (PRK00453)bS6 (cd00473)→ ssb (TIGR00621)SSB (pfam00436)→ rpsR (PRK00391)Ribosomal_S18 (pfam01084)→ RplI (COG0359)rplI (PRK00137)→ |
| env-21 | M740_1012895 | - | 3033 | 2959 | **RNA**→ rpsF (PRK00453)bS6 (cd00473)→ ssb (TIGR00621)SSB (pfam00436)→ rpsR (PRK00391)Ribosomal_S18 (pfam01084)→ RplI (COG0359)rplI (PRK00137)→ |
| env-22 | M750CN_1002294 | + | 1537 | 1611 | **RNA**→ rpsF (PRK00453)bS6 (cd00473)→ ssb (TIGR00621)SSB (pfam00436)→ rpsR (PRK00391)Ribosomal_S18 (pfam01084)→ RplI (COG0359)rplI (PRK00137)→ |
| Cpa-1-1 | NC_011027.1 | + | 129607 | 129681 | **RNA**→ rpsF (PRK00453)bS6 (cd00473)→ ssb (TIGR00621)SSB (pfam00436)→ rpsR (PRK00391)Ribosomal_S18 (pfam01084)→ RplI (COG0359)rplI (PRK00137)→ |
| env-23 | M590M1_1000641 | - | 1709 | 1635 | **RNA**→ rpsF (PRK00453)bS6 (cd00473)→ ssb (TIGR00621)SSB (pfam00436)→ rpsR (PRK00391)Ribosomal_S18 (pfam01084)→ RplI (COG0359)rplI (PRK00137)→ |
| env-24 | M590M2_1003423 | - | 1681 | 1607 | **RNA**→ rpsF (PRK00453)bS6 (cd00473)→ ssb (TIGR00621)SSB (pfam00436)→ rpsR (PRK00391)Ribosomal_S18 (pfam01084)→ rplI (PRK00137)Ribosomal_L9_N (pfam01281)→ |
| Cte-1-1 | NC_002932.3 | - | 2020240 | 2020165 | **RNA**→ rpsF (PRK00453)bS6 (cd00473)→ ssb (TIGR00621)SSB (pfam00436)→ |
| Cph-2-1 | NC_008639.1 | - | 2877490 | 2877409 | **RNA**→ rpsF (PRK00453)bS6 (cd00473)→ ssb (TIGR00621)SSB (pfam00436)→ rpsR (PRK00391)Ribosomal_S18 (pfam01084)→ RplI (COG0359)rplI (PRK00137)→ |
| env-25 | NCBI_READ_1113216244664 | - | 966 | 897 | **RNA**→ rpsF (PRK00453)Ribosomal_S6 (pfam01250)→ SSB_OBF (cd04496)ssb (TIGR00621)→ |
| env-26 | NCBI_READ_1113298190517 | - | 98 | 23 | **RNA**→ |
| env-27 | NCBI_READ_1113316091948 | + | 650 | 726 | **RNA**→ |
| env-28 | NCBI_READ_1113316246897 | + | 712 | 788 | **RNA**→ |
| env-29 | NCBI_READ_1112280131988 | + | 738 | 813 | **RNA**→ heptsyl_trn_II (TIGR02195)GT1_LPS_heptosyltransferase (cd03789)→ |
| env-30 | NCBI_READ_1113316182926 | + | 890 | 964 | **RNA**→ |
| env-31 | NCBI_READ_1112280190525 | + | 291 | 367 | **RNA**→ rpsF (PRK00453)bS6 (cd00473)→ |
| env-32 | NCBI_READ_1113235400465 | - | 346 | 270 | **RNA**→ rpsF (PRK00453)Ribosomal_S6 (pfam01250)→ |

| | | | | | |
|---|---|---|---|---|---|
| env-33 | NCBI_READ_1112280138898 | - | 302 | 226 | **RNA**→ rpsF (PRK00453)Ribosomal_S6 (pfam01250)→ |
| env-34 | NCBI_READ_1113315684505 | - | 349 | 273 | **RNA**→ rpsF (PRK00453)Ribosomal_S6 (pfam01250)→ |
| env-35 | NCBI_READ_1112280178452 | - | 455 | 379 | **RNA**→ rpsF (PRK00453)bS6 (cd00473)→ |
| env-36 | NCBI_READ_1113298125219 | - | 372 | 296 | **RNA**→ rpsF (PRK00453)Ribosomal_S6 (pfam01250)→ |
| env-37 | NCBI_READ_1113316249537 | + | 181 | 257 | **RNA**→ rpsF (PRK00453)bS6 (cd00473)→ |
| env-38 | NCBI_READ_1113316395352 | - | 471 | 395 | **RNA**→ rpsF (PRK00453)bS6 (cd00473)→ |
| env-39 | NCBI_READ_1113308556153 | - | 520 | 444 | **RNA**→ rpsF (PRK00453)bS6 (cd00473)→ |
| env-40 | NCBI_READ_1113316249529 | - | 226 | 150 | **RNA**→ rpsF (PRK00453)→ |
| env-41 | NCBI_READ_1113215706340 | + | 274 | 350 | **RNA**→ rpsF (PRK00453)Ribosomal_S6 (pfam01250)→ |
| env-42 | NCBI_READ_1113315555704 | - | 506 | 430 | **RNA**→ rpsF (PRK00453)bS6 (cd00473)→ |
| env-43 | NCBI_READ_1113316238233 | - | 471 | 395 | **RNA**→ rpsF (PRK00453)bS6 (cd00473)→ |
| env-44 | NCBI_READ_1113216208246 | - | 506 | 430 | **RNA**→ rpsF (PRK00453)bS6 (cd00473)→ |
| env-45 | NCBI_READ_1113316081654 | - | 811 | 735 | **RNA**→ rpsF (PRK00453)bS6 (cd00473)→ SSB_OBF (cd04496)ssb (TIGR00621)→ |
| env-46 | NCBI_READ_1113021287689 | - | 859 | 784 | **RNA**→ rpsF (PRK00453)bS6 (cd00473)→ |
| env-47 | NCBI_READ_1113289126331 | - | 828 | 752 | **RNA**→ rpsF (PRK00453)bS6 (cd00473)→ Ssb (COG0629)ssb (TIGR00621)→ |
| env-48 | NCBI_READ_1113215581360 | - | 190 | 114 | **RNA**→ |
| env-49 | NCBI_READ_1113235389909 | - | 644 | 568 | **RNA**→ rpsF (PRK00453)bS6 (cd00473)→ |
| Cph-3-1 | NC_009337.1 | + | 202059 | 202135 | **RNA**→ rpsF (PRK00453)Ribosomal_S6 (pfam01250)→ SSB_OBF (cd04496)ssb (TIGR00621)→<br>rpsR (PRK00391)Ribosomal_S18 (pfam01084)→ RplI (COG0359)rplI (PRK00137)→ |
| env-50 | NCBI_READ_1112308883715 | - | 917 | 846 | **RNA**→ rpsF (PRK00453)bS6 (cd00473)→ SSB_OBF (cd04496)ssb (TIGR00621)→ |
| env-51 | NCBI_READ_1112280218812 | - | 990 | 917 | **RNA**→ rpsF (PRK00453)Ribosomal_S6 (pfam01250)→ SSB_OBF (cd04496)ssb (TIGR00621)→ |
| env-52 | NCBI_READ_1113316170875 | - | 893 | 819 | **RNA**→ rpsF (PRK00453)bS6 (cd00473)→ |
| env-53 | TB_FS08_3DRAFT_1000257 | + | 16727 | 16803 | **RNA**→ rpsF (PRK00453)bS6 (cd00473)→ ssb (TIGR00621)SSB (pfam00436)→<br>rpsR (PRK00391)Ribosomal_S18 (pfam01084)→ RplI (COG0359)rplI (PRK00137)→ |
| Cfe-1-1 | NZ_AASE01000042.1 | - | 4661 | 4585 | **RNA**→ rpsF (PRK00453)bS6 (cd00473)→ rpsF (PRK00453)bS6 (cd00473)→ ssb (TIGR00621)SSB (pfam00436)→<br>ssb (TIGR00621)SSB (pfam00436)→ rpsR (PRK00391)Ribosomal_S18 (pfam01084)→<br>rpsR (PRK00391)Ribosomal_S18 (pfam01084)→ RplI (COG0359)rplI (PRK00137)→<br>RplI (COG0359)rplI (PRK00137)→ |
| Pph-1-1 | NC_011060.1 | + | 122430 | 122507 | **RNA**→ rpsF (PRK00453)bS6 (cd00473)→ ssb (TIGR00621)SSB (pfam00436)→<br>rpsR (PRK00391)Ribosomal_S18 (pfam01084)→ RplI (COG0359)rplI (PRK00137)→ |
| env-54 | TB03JUN2009H_c1 | + | 2274 | 2351 | **RNA**→ rpsF (PRK00453)bS6 (cd00473)→ ssb (TIGR00621)SSB (pfam00436)→<br>rpsR (PRK00391)Ribosomal_S18 (pfam01084)→ RplI (COG0359)rplI (PRK00137)→ |
| env-55 | TB18AUG2009E_c1 | + | 2274 | 2351 | **RNA**→ rpsF (PRK00453)bS6 (cd00473)→ ssb (TIGR00621)SSB (pfam00436)→<br>rpsR (PRK00391)Ribosomal_S18 (pfam01084)→ RplI (COG0359)rplI (PRK00137)→ |
| env-56 | TB18AUG2009H_c2 | + | 2274 | 2351 | **RNA**→ rpsF (PRK00453)bS6 (cd00473)→ ssb (TIGR00621)SSB (pfam00436)→<br>rpsR (PRK00391)Ribosomal_S18 (pfam01084)→ RplI (COG0359)rplI (PRK00137)→ |
| Clu-1-1 | NC_007512.1 | + | 128893 | 128969 | **RNA**→ rpsF (PRK00453)Ribosomal_S6 (pfam01250)→ ssb (TIGR00621)SSB (pfam00436)→<br>rpsR (PRK00391)Ribosomal_S18 (pfam01084)→ RplI (COG0359)rplI (PRK00137)→ |

#### 18.3 Conserved domains

Conserved domains found in protein-coding genes listed in Section 18.2 are shown below, with the first sentence in their description from the Conserved Domain Database (if any). Conserved domains associated with more than one S6-S18-Chlorobi RNA are assigned a color, while others are shown in gray. The number in parentheses after the colored domain name is the number of occurrences in Section 18.2.

cd00473 (36) Bacterial ribosomal protein S6.
cd03789 (1) Lipopolysaccharide heptosyltransferase is involved in the biosynthesis of lipooligosaccharide (LOS).
cd04496 (5) SSB_OBF: A subfamily of OB folds similar to the OB fold of ssDNA-binding protein (SSB).
COG0359 (24) Ribosomal protein L9 [Translation, ribosomal structure and biogenesis]
COG0629 (1) Single-stranded DNA-binding protein [Replication, recombination and repair]
pfam00436 (25) Single-strand binding protein family.
pfam01084 (25) Ribosomal protein S18.
pfam01250 (15) Ribosomal protein S6.
pfam01281 (1) Ribosomal protein L9, N-terminal domain.
PRK00137 (25) 50S ribosomal protein L9; Reviewed

PRK00391 (25) 30S ribosomal protein S18; Reviewed
PRK00453 (52) 30S ribosomal protein S6; Reviewed
TIGR00621 (31) single stranded DNA-binding protein (ssb). [DNA metabolism, DNA replication, recombination, and repair]
TIGR02195 (1) lipopolysaccharide heptosyltransferase II. [Cell envelope, Biosynthesis and degradation of surface polysaccharides and lipopolysaccharides]

#### 18.4 Multiple-sequence alignment

Each S6-S18-Chlorobi RNA is denoted by an abbreviation (like "Eco-1-1") that refers to a taxonomy given in Section 18.1. The alignment may include sequences containing the RNA motif, as well as flanking sequence. The S6-S18-Chlorobi RNA itself is denoted by the line underneath marked 5′ and 3′ on either end. Nucleotides in flanking sequences (i.e., not part of the motif) are written in gray letters. Stems of predicted rho-independent transcription terminators, if any, are shaded yellow. (Note: terminator predictions have not been analyzed manually, and many are likely to be false positives. Terminator predictions are those of the RNIE software.) Nucleotides predicted to function (as DNA) as transcription-factor binding sites, if any, are shaded in green. (But note: these predictions are manually annotated, so they might be under-predicted.) Annotated start codons, if any, are shaded green. (Note: start codons are frequently misannotated, especially in environmental samples.) Nucleotides proposed to basepair as part of the consensus structure are shaded in color when they comprise Watson-Crick or G-U pairs. Otherwise they are shaded gray. Conserved stems are also indicated at the bottom of the alignment by angle brackets, where matching < and > denote base-paired columns. Below these angle brackets, the symbol "2" denotes base pairs exhibiting covariation according to the statistically well-founded R-scape method. "1" denotes base pairs exhibiting covariation according to R2R's simplistic method. "0" denotes base pairs that are not observed to mutate and "?" denotes base pairs that have a significant frequency of non-canonical nucleotides for Watson-Crick or G-U pairs (> 5%). Below these base pair annotation is the consensus sequence: "R" = "A" or "G", "Y" = "C" or "U", red nucleotides: nucleotide identity conserved more than 97% of the time, black nucleotides: 90%, gray nucleotides: 75%, red circle (•): nucleotide is present 97% of the time, black circle (•): 90%, gray circle (•): 75%, white circle (◦): 50%. All percentages of sequences just described (e.g. 97% conserved) assume that sequences have been weighted by the GSC algorithm implemented by the Infernal software package.

Duplicate sequences: the following putative homologs are not shown in the alignment because their sequences are identical to a homolog already shown: Pae-1-1, env-10, env-11, env-12, env-19, env-22, env-24, env-28, env-32, env-33, env-34, env-35, env-36, env-37, env-38, env-39, env-40, env-41, env-42, env-43, env-44, env-55, env-56, env-8

alignment positions 1 · · · 180

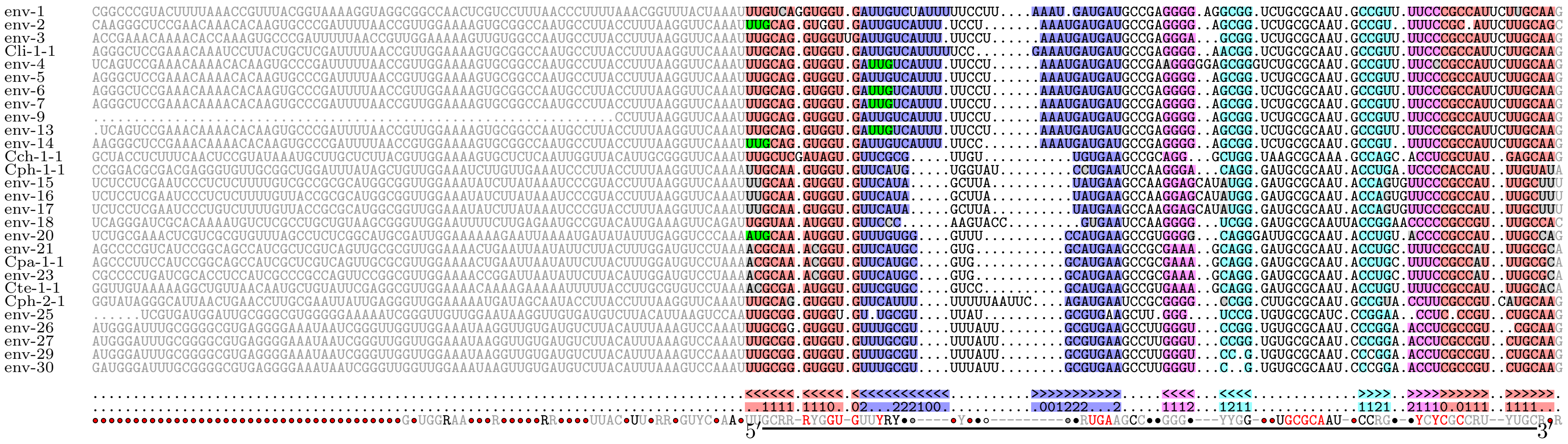

```
env-31    AUGGGAUUUGCGGGGCGUGAGGGGAAAUAAUCGGGUUGGUUGGAAAUAAGGUUGUGAUGUCUUACAUUUAAAGUCCAAAUUUGCGG.GUGGU.GUUUGCGU....UUUAUU........GCGUGAAGCCUUGGGU...CCGG.UGUGCGCAAU.CCCGGA.ACCUCGCCGU..CUGCAAG
env-45    GAUGGAUUUGCGGGGCGUGAGGGGAAAUAAUCGGGUUGGUUGGAAAUAAGGUUGUGAUGUCUUACAUUUAAAGUCCAAAUUUGCGG.GUGGU.GUUUGCGU....UUUAUU........GCGUGAAGCCUUGGGU...CCGG.UGUGCGCAAU.CCCGGA.ACCUCGCCGU..CUGCAAG
env-46    GACUUUUCGUGAUGGAUUGCGGGCGUGAGGGAAUAUCGGGUGGUGAAAUAAGGUUGUGAUGUCUACAUUAAAGUCCAAAUUUGCGG.GUGGU.GUUUGCGU....UUUAUU........GCGUGAAGC.UUGGGU...CCGG.UGUGCGCAAU.CCCGGA.ACCUCGCCGU..CUGCAAG
env-47    CGUGAUGGAUUGCGGGCGUGAGGGAAAUAAUCGGGUUGGUUGGAAAUAAGGUUGUGAUGUCUUACAUUUAAAGUCCAAAUUUGCGG.GUGGU.GUUUGCGU....UUUAUU........GCGUGAAGCUUUGGGU...CCGG.UGUGCGCAAU.CCCGGA.ACCUCGCCGU..CUGCAAG
env-48    AUGGGAUUUGCGGGGCGUGAGGGGAAAUAAUCGGGUUGGUUGGAAAUAAGGUUGUGAUGUCUUACAUUUAAAGUCCAAAUUUGCGG.GUGGU.GUUUGCGU....UUUAUU........GCGUGAAGCCUUGGGU...CCGG.UGUGCGCAAU.CUCGGA.ACCUCGCCGU..CUGCAAG
env-49    UGAGAUUUGCGGGGCGUGAGGGGAAAUAAUCGGGUUGGUUGGAAAAUAAGGUUGUGAUGUCUUACAUUUAAAGUCCAAAUUUGCGG.GUGGU.GUUUGCGU....UUUAUU........GCGUGAAGCUUUGGGU...CCGG.UGUGCGCAAU.CCGGGA.ACCUCGCUGU..CUGCAAG
Cph-3-1   UGGUGGUGAUUGUUGCGUGAGGGGAAAUCAGCGGUAAGGUUGGAAAUGAGAUUGUGAUGUCUUACAUUUAAGGUCCAAAUUUGCGG.GCGGU.GUUUGCGCU...UUCU.........GGCGUGAAGCCGUGGGU...CCGG.UGUGCGCAAU.CCCGGA.ACCUCGCCGU..CUGCAAG
env-50    UUCAGACCUUUCGUGAUGGAUUGCGGGCGUGAGGGAAUAUCGGGUGGUGAAUAAGGUUGUGAUGUCUACAUUAAGUCCAAUUUGCG.GGGGU.GU.UGCGU....UUAUU.........GCGUGAAGC..UUGGU...CCGG.UGUGCGCA.U.CCCGGA.ACCUCGCGUC..UGCAAGC
env-51    CAUUUCAGAACUUUCGGAUGGAUUGCGGGCGUGAGGGAAUAAUCGGUUGUGAAUAAGUUGUGAGUCUACAUUAAAGUCCAAAUUGC.GGUGU.GUUUGCGU....UUAU..........GCGUGAAGC.UUGGGU...CC.G.UGUGCGCAAU.CCUGGA.ACCUCGCGUC.UGCAAGCA
env-52    UUUCAGACUUUUCGUGAUGGAUUGCGGGCGGAGGGAAUAAUCGGGUGGUGAAUAGGUGUGAUGUCUACAUUAAAGUCCAAAUUUGC.GGUGU.GU.UGCGU....UUAUU.........GCGUGAAG.CUUGGGU...CCGG.UGUGCGCAAU.CCCGGA.ACCUCGCGUC.UGCAAGCA
env-53    AUGAACGAUGAUGCUCAUGGUUUUCUCUUUUUGCUAUGGUUGGAAAAAACGGGUAAAUAGAUUACUUUGAAUGUUCAAAUUUGCAG.AUGGU.GUUCC.CC....UUUAAU........GGUUGAAGCCGUGGGG...CUGA.UUAGCGCAAU.GUCAGA.CCCUCACCAU.AUUGCAAG
Cfe-1-1   UCCCGGGAAAACCUUCUUUUUAUGGUUUUAUCCGGCAGGUUGGAAAACUGGUGUAAAUAGAUUACCUUUAAUGUUCAAAUUUGCAG.AUGGU.GUUCAACGG...GAAA.........CCG.UGAAGCCGUGGGG...CUGG.UCUGCGCAAU.ACCGGA.CCCUCGCCGU.UUUGCAAG
Pph-1-1   CGUGACGAAAAAGCCAAUUCGAGGCAUUUUCACUCACGGUUGGAAAAAUGAUGCAAAUAGGUUACCUUUGAUGUUCAAAUUUGCAG.AUGGU.GUUCACAGG...GAAA.........CCUUUGAAGCCGUGGGG...CUGG.UCUGCGCAAU.GCCAGA.CCCUCGCCAU.AUUGCAAG
env-54    CUUGUCGGUUAAUCCGGGUUCAAGGUUUUUUCCAUGAGGUUGGAAAAAUGGUGCAAAUAGUUUACCUUUAAUGUUCAAAUUUGCAG.AUGGU.GUUCACAGG...GAGA.........CCUUUGAAGCCGUGGGG...CUGG.UCUGCGCAAU.ACCGGA.CCCUCGCCGU.UUUGCAAG
Clu-1-1   AUGAGGAUACCCGGUCAUUUUCUCGGUGAGUUGCGGCGGUUGGAAAAAAGAAUGCAAUUCCUUACAUUUAAGGUCCAAAUUUGCAC.GCGGU.GUUCACGU....CUGAAC........GCGUGAAGCCGUGGGG...CCGG.UAUGCGCAAU.UCCGGA.CCCUCGCCGC..AUGCAAG
          ................................................................................<<<<<<.<<<<<.<<<<<<<<<<<<..........>>>>>>>>>>>.....<<<<...<<<<.............>>>>..>>>>>>>>>>..>>>>>>.
          ..................................................................................1111.1110..02...222100...........001222...2.....1112...1211.............1121..21110.0111..1111...
          ••••••••••••••••••••••••••••••••••••••G•UGGRAA•••R•••••RR••••UUAC•UU•RR•GUYC•AA•UUGCRR-RYGGU-GUUYRY•∘----•Y••∘---------∘•RUGAAGCC••GGG•---YYGG-••UGCGCAAU-•CCRG•-•YCYCGCCRU--YUGCR•R
                                                                                5′——————————————————————————————————————————————————————————————————————————————————————————————————————3′
```

alignment positions 181 · · · 214

```
env-1     CAACUGAAGAGCUAACAUUAUCAGACU....UUG
env-2     AACUGAAGAGCUAACAUUANCAGUAC........
env-3     CAACUGAAGACUGA....................
Cli-1-1   CAACUGAAGAGCUAACAUUAUCAGACU....UUG
env-4     CAACUGAAGAGCUUAACAUUUAUCAGACU..UUG
env-5     CAACUGAAGAGCUAACAUUAUCAGACU....UUG
env-6     CAACUGAAGAGCUAACAUUUAUCAGACU...UUG
env-7     CAACUGAAGAGCUAAACAUUAUCAGACU...UUG
env-9     CAACUGAAGAGCUAACAUUAUCAGACU....UUG
env-13    CAACUGAAGAGCUAACAUUAUCAGACU....UUG
env-14    CAACUGAAAGAGCUAAACAUUAUCAGACU..UUG
Cch-1-1   CUAUUGAUCGAAAACACGAAAGCAUGA....UUG
Cph-1-1   AGCAAACAUAAUCUUUUUAAG..........UUG
env-15    UGUUUAAUUCAAUAUAAUGG...........UUG
env-16    UGUUUAAUUCAAUAUAAUGG...........UUG
env-17    UGUUUAAUUCAAUAUAAUGG...........UUG
env-18    UGUUCAGCAUAAUUAUUUAAGUUGCA.....UUG
env-20    UCUCUUAUCAUAUCAGAACAA..........UUG
env-21    GUUUGACAUCACUCAAGAACCA.........UUG
Cpa-1-1   GUUUGACAUCACUCAAGAACCA.........UUG
env-23    GUUUGACAUCACUCAAGAACCA.........UUG
Cte-1-1   GUAUGACUUCACUCAAAAACAA.........UUG
Cph-2-1   CAACUGAGGAAAGAACAAUGUUAUCAGGUAGUUG
env-25    CAACUGAGGACACUCAAUCAUUAUGCAA...UUG
env-26    CACCCAGCUUCCCCCCCCCG..............
env-27    CAACUGAGACACUCAAUCAUUAUUGCAA...UUG
env-29    CAACUGAGACACUCAAUCAUUAUUGCAA...UUG
env-30    CACUGAGACACUCAUCAUAUGCA........UUG
env-31    CAACUGAGGACACUCAAUCAUUAUUGCAA..UUG
env-45    CAACUGAGGACACUCAAUCAUUAUUGCAA..UUG
env-46    CAACUGAGGACACUCAAUCAUUAUUGCAA..UUG
env-47    CAACUGAGGACACUCAAUCAUUAUUGCAA..UUG
env-48    CAACUGAGGACGCUCAAUCAUUAUUGCAA..UUG
env-49    CAACUGAGGACACUCAAUCAUUAUUGCAA..UUG
Cph-3-1   CAACUGAGGACACUCAAUCAUUAUUGCAU..UUG
env-50    AACUGAGGACACUCAAUCAUUAUUGCAA...UUG
env-51    ACUGAGGACACUCAAUCAUUAUUGCAA....UUG
env-52    ACUGAGGACACUCAAUCAUUAUUGCAA....UUG
env-53    CAACUGAGGAGUAUUAAUUAUUACAAACAU.UUG
Cfe-1-1   CAACUGAGGAGUAAUAAUGAUUAUCAGCAAUUUG
Pph-1-1   CAACUGAGGACGAAUAAUACUAUCAGAUU..UUG
env-54    CAACUGAGGAGUAAUAAUGAUUAUCAGCUAUUUG
Clu-1-1   CAGCUGAGUACUCUCAAUCAUUAUUUACA..UUG
          ..................................
          ..................................
          •••Y•RRR••••••••••••••∘∘∘∘∘----UUG
```

### 19 S10-Clostridia

#### 19.1 Taxa

The taxonomy of each organism containing a putative S10-Clostridia RNA is listed, with abbreviations identifying each hit (e.g., "Eco-1-1" and "Eco-1-2" might hypothetically represent two distinct RNAs in *E. coli*). The abbreviations will be used to identify each individual S10-Clostridia RNA in Sections 19.2 and 19.4:.

| **abbrev. of hits** | **taxonomy of species** |
|---|---|
| Bhy-1-1 | Bacteria Firmicutes Clostridia Clostridiales *Blautia hydrogenotrophica* DSM 10507 (Ruminococcus |
| Cba-1-1 | Bacteria Firmicutes Clostridia Clostridiales *Clostridiaceae bacterium* MS3 |
| Cbo-1-1 | Bacteria Firmicutes Clostridia Clostridiales Clostridiaceae *Clostridium bolteae* 90A5 |
| Cbo-2-1 | Bacteria Firmicutes Clostridia Clostridiales Clostridiaceae *Clostridium bolteae* 90A9 |
| Cbo-3-1 | Bacteria Firmicutes Clostridia Clostridiales Clostridiaceae *Clostridium bolteae* 90B3 |
| Cbo-4-1 | Bacteria Firmicutes Clostridia Clostridiales Clostridiaceae *Clostridium bolteae* 90B7 |
| Cbo-5-1 | Bacteria Firmicutes Clostridia Clostridiales Clostridiaceae *Clostridium bolteae* 90B8 |
| Cbo-6-1 | Bacteria Firmicutes Clostridia Clostridiales Clostridiaceae *Clostridium bolteae* ATCC BAA-613 |
| Cci-1-1 | Bacteria Firmicutes Clostridia Clostridiales Clostridiaceae *Clostridium citroniae* WAL-17108 |
| Ccl-1-1 | Bacteria Firmicutes Clostridia Clostridiales Clostridiaceae *Clostridium clostridioforme* 2_1_49FAA |
| Ccl-2-1 | Bacteria Firmicutes Clostridia Clostridiales Clostridiaceae *Clostridium clostridioforme* 90A1 |
| Ccl-3-1 | Bacteria Firmicutes Clostridia Clostridiales Clostridiaceae *Clostridium clostridioforme* 90A3 |
| Ccl-4-1 | Bacteria Firmicutes Clostridia Clostridiales Clostridiaceae *Clostridium clostridioforme* 90A4 |
| Ccl-5-1 | Bacteria Firmicutes Clostridia Clostridiales Clostridiaceae *Clostridium clostridioforme* 90A6 |
| Ccl-6-1 | Bacteria Firmicutes Clostridia Clostridiales Clostridiaceae *Clostridium clostridioforme* 90A7 |
| Ccl-7-1 | Bacteria Firmicutes Clostridia Clostridiales Clostridiaceae *Clostridium clostridioforme* 90A8 |
| Ccl-8-1 | Bacteria Firmicutes Clostridia Clostridiales Clostridiaceae *Clostridium clostridioforme* 90B1 |
| Ccl-9-1 | Bacteria Firmicutes Clostridia Clostridiales Clostridiaceae *Clostridium clostridioforme* CM201 |
| Cha-1-1 | Bacteria Firmicutes Clostridia Clostridiales Clostridiaceae *Clostridium hathewayi* 12489931 |
| Cha-2-1 | Bacteria Firmicutes Clostridia Clostridiales Clostridiaceae *Clostridium hathewayi* WAL-18680 |
| Cne-1-1 | Bacteria Firmicutes Clostridia Clostridiales Clostridiaceae *Clostridium nexile* DSM 1787 |
| Cph-1-1 | Bacteria Firmicutes Clostridia Clostridiales Clostridiaceae *Clostridium phytofermentans* ISDg |
| Csa-1-1 | Bacteria Firmicutes Clostridia Clostridiales Clostridiaceae *Clostridium saccharolyticum* WM1 |
| Csc-1-1 | Bacteria Firmicutes Clostridia Clostridiales Clostridiaceae *Clostridium scindens* ATCC 35704 |
| Csp-1-1 | Bacteria Firmicutes Clostridia Clostridiales Clostridiaceae *Clostridium sp.* 7_3_54FAA |
| Csp-2-1 | Bacteria Firmicutes Clostridia Clostridiales Clostridiaceae *Clostridium sp.* ASF356 |
| Csp-3-1 | Bacteria Firmicutes Clostridia Clostridiales Clostridiaceae *Clostridium sp.* ASF502 |
| Csp-4-1 | Bacteria Firmicutes Clostridia Clostridiales Clostridiaceae *Clostridium sp.* D5 |
| Csp-5-1 | Bacteria Firmicutes Clostridia Clostridiales Clostridiaceae *Clostridium sp.* FS41 |
| Csp-6-1 | Bacteria Firmicutes Clostridia Clostridiales Clostridiaceae *Clostridium sp.* L2-50 |
| Csp-7-1 | Bacteria Firmicutes Clostridia Clostridiales Clostridiaceae *Clostridium sp.* M62/1 |
| Csy-1-1 | Bacteria Firmicutes Clostridia Clostridiales Clostridiaceae *Clostridium symbiosum* ATCC 14940 |
| Csy-2-1 | Bacteria Firmicutes Clostridia Clostridiales Clostridiaceae *Clostridium symbiosum* WAL-14163 |
| Csy-3-1 | Bacteria Firmicutes Clostridia Clostridiales Clostridiaceae *Clostridium symbiosum* WAL-14673 |
| Hha-1-1 | Bacteria Firmicutes Clostridia Clostridiales Clostridiaceae *Hungatella hathewayi* VE202-04 |
| Hha-2-1 | Bacteria Firmicutes Clostridia Clostridiales Clostridiaceae *Hungatella hathewayi* VE202-11 |
| Cba-2-1 | Bacteria Firmicutes Clostridia *Clostridiales bacterium* 1_7_47FAA |
| Cba-3-1 | Bacteria Firmicutes Clostridia *Clostridiales bacterium* OBRC5-5 |
| Cba-4-1 | Bacteria Firmicutes Clostridia *Clostridiales bacterium* VE202-07 |
| Cba-5-1 | Bacteria Firmicutes Clostridia *Clostridiales bacterium* VE202-16 |
| Cba-6-1 | Bacteria Firmicutes Clostridia *Clostridiales bacterium* VE202-26 |
| Cba-7-1 | Bacteria Firmicutes Clostridia *Clostridiales bacterium* VE202-28 |
| Cba-8-1 | Bacteria Firmicutes Clostridia *Clostridiales bacterium* VE202-29 |
| Ece-1-1 | Bacteria Firmicutes Clostridia Clostridiales Eubacteriaceae *Eubacterium cellulosolvens* 6 |
| Eha-1-1 | Bacteria Firmicutes Clostridia Clostridiales Eubacteriaceae *Eubacterium hallii* DSM 3353 |
| Ere-1-1 | Bacteria Firmicutes Clostridia Clostridiales Eubacteriaceae *Eubacterium rectale* ATCC 33656 |
| Esp-1-1 | Bacteria Firmicutes Clostridia Clostridiales Eubacteriaceae *Eubacterium sp.* 14-2 |
| [bo-1-1 | Bacteria Firmicutes Clostridia Clostridiales Lachnospiraceae [Clostridium] bolteae WAL-14578 |
| [ci-1-1 | Bacteria Firmicutes Clostridia Clostridiales Lachnospiraceae [Clostridium] citroniae WAL-19142 |
| [cl-1-1 | Bacteria Firmicutes Clostridia Clostridiales Lachnospiraceae [Clostridium] clostridioforme WAL-7855 |
| [sc-1-1 | Bacteria Firmicutes Clostridia Clostridiales Lachnospiraceae [Clostridium] scindens VE202-05 |
| Rgn-1-1 | Bacteria Firmicutes Clostridia Clostridiales Lachnospiraceae Blautia *Ruminococcus gnavus* AGR2154 |
| Rgn-2-1 | Bacteria Firmicutes Clostridia Clostridiales Lachnospiraceae Blautia *Ruminococcus gnavus* CC55_001C |
| Bfo-1-1 | Bacteria Firmicutes Clostridia Clostridiales Lachnospiraceae *Bryantella formatexigens* DSM 14469 |
| Bcr-1-1 | Bacteria Firmicutes Clostridia Clostridiales Lachnospiraceae *Butyrivibrio crossotus* DSM 2876 |
| Bfi-1-1 | Bacteria Firmicutes Clostridia Clostridiales Lachnospiraceae *Butyrivibrio fibrisolvens* AB2020 |
| Bfi-2-1 | Bacteria Firmicutes Clostridia Clostridiales Lachnospiraceae *Butyrivibrio fibrisolvens* FE2007 |
| Bfi-3-1 | Bacteria Firmicutes Clostridia Clostridiales Lachnospiraceae *Butyrivibrio fibrisolvens* ND3005 |
| Bfi-4-1 | Bacteria Firmicutes Clostridia Clostridiales Lachnospiraceae *Butyrivibrio fibrisolvens* WTE3004 |

| | |
|---|---|
| Bsp-1-1 | Bacteria Firmicutes Clostridia Clostridiales Lachnospiraceae *Butyrivibrio sp.* AD3002 |
| Bsp-2-1 | Bacteria Firmicutes Clostridia Clostridiales Lachnospiraceae *Butyrivibrio sp.* AE2015 |
| Bsp-3-1 | Bacteria Firmicutes Clostridia Clostridiales Lachnospiraceae *Butyrivibrio sp.* AE3009 |
| Bsp-4-1 | Bacteria Firmicutes Clostridia Clostridiales Lachnospiraceae *Butyrivibrio sp.* FCS006 |
| Bsp-5-1 | Bacteria Firmicutes Clostridia Clostridiales Lachnospiraceae *Butyrivibrio sp.* LB2008 |
| Bsp-6-1 | Bacteria Firmicutes Clostridia Clostridiales Lachnospiraceae *Butyrivibrio sp.* MC2021 |
| Bsp-7-1 | Bacteria Firmicutes Clostridia Clostridiales Lachnospiraceae *Butyrivibrio sp.* NC2007 |
| Bsp-8-1 | Bacteria Firmicutes Clostridia Clostridiales Lachnospiraceae *Butyrivibrio sp.* VCB2001 |
| Bsp-9-1 | Bacteria Firmicutes Clostridia Clostridiales Lachnospiraceae *Butyrivibrio sp.* WCD2001 |
| Bsp-10-1 | Bacteria Firmicutes Clostridia Clostridiales Lachnospiraceae *Butyrivibrio sp.* XPD2006 |
| Cmo-1-1 | Bacteria Firmicutes Clostridia Clostridiales Lachnospiraceae *Catonella morbi* ATCC 51271 |
| Cle-1-1 | Bacteria Firmicutes Clostridia Clostridiales Lachnospiraceae Cellulosilyticum *Clostridium lentocellum* DSM 5427 |
| Cco-1-1 | Bacteria Firmicutes Clostridia Clostridiales Lachnospiraceae *Coprococcus comes* ATCC 27758 |
| Ceu-1-1 | Bacteria Firmicutes Clostridia Clostridiales Lachnospiraceae *Coprococcus eutactus* ATCC 27759 |
| Csp-8-1 | Bacteria Firmicutes Clostridia Clostridiales Lachnospiraceae *Coprococcus sp.* HPP0048 |
| Csp-9-1 | Bacteria Firmicutes Clostridia Clostridiales Lachnospiraceae *Coprococcus sp.* HPP0074 |
| Dfo-1-1 | Bacteria Firmicutes Clostridia Clostridiales Lachnospiraceae *Dorea formicigenerans* 4_6_53AFAA |
| Dfo-2-1 | Bacteria Firmicutes Clostridia Clostridiales Lachnospiraceae *Dorea formicigenerans* ATCC 27755 |
| Dlo-1-1 | Bacteria Firmicutes Clostridia Clostridiales Lachnospiraceae *Dorea longicatena* AGR2136 |
| Dlo-2-1 | Bacteria Firmicutes Clostridia Clostridiales Lachnospiraceae *Dorea longicatena* DSM 13814 |
| Dsp-1-1 | Bacteria Firmicutes Clostridia Clostridiales Lachnospiraceae *Dorea sp.* 5-2 |
| Jig-1-1 | Bacteria Firmicutes Clostridia Clostridiales Lachnospiraceae *Johnsonella ignava* ATCC 51276 |
| Lsp-1-1 | Bacteria Firmicutes Clostridia Clostridiales Lachnospiraceae *Lachnoanaerobaculum sp.* MSX33 |
| Lbo-1-1 | Bacteria Firmicutes Clostridia Clostridiales Lachnospiraceae *Lachnobacterium bovis* |
| Lbo-2-1 | Bacteria Firmicutes Clostridia Clostridiales Lachnospiraceae *Lachnobacterium bovis* C6A12 |
| Lbo-3-1 | Bacteria Firmicutes Clostridia Clostridiales Lachnospiraceae *Lachnobacterium bovis* NK4B19 |
| Lba-1-1 | Bacteria Firmicutes Clostridia Clostridiales *Lachnospiraceae bacterium* 1_1_57FAA |
| Lba-2-1 | Bacteria Firmicutes Clostridia Clostridiales *Lachnospiraceae bacterium* 1_4_56FAA |
| Lba-3-1 | Bacteria Firmicutes Clostridia Clostridiales *Lachnospiraceae bacterium* 2_1_46FAA |
| Lba-4-1 | Bacteria Firmicutes Clostridia Clostridiales *Lachnospiraceae bacterium* 2_1_58FAA |
| Lba-5-1 | Bacteria Firmicutes Clostridia Clostridiales *Lachnospiraceae bacterium* 3-1 |
| Lba-6-1 | Bacteria Firmicutes Clostridia Clostridiales *Lachnospiraceae bacterium* 3-2 |
| Lba-7-1 | Bacteria Firmicutes Clostridia Clostridiales *Lachnospiraceae bacterium* 3_1_46FAA |
| Lba-8-1 | Bacteria Firmicutes Clostridia Clostridiales *Lachnospiraceae bacterium* 3_1_57FAA_CT1 |
| Lba-9-1 | Bacteria Firmicutes Clostridia Clostridiales *Lachnospiraceae bacterium* 4_1_37FAA |
| Lba-10-1 | Bacteria Firmicutes Clostridia Clostridiales *Lachnospiraceae bacterium* 5_1_57FAA |
| Lba-11-1 | Bacteria Firmicutes Clostridia Clostridiales *Lachnospiraceae bacterium* 8_1_57FAA |
| Lba-12-1 | Bacteria Firmicutes Clostridia Clostridiales *Lachnospiraceae bacterium* 9_1_43BFAA |
| Lba-13-1 | Bacteria Firmicutes Clostridia Clostridiales *Lachnospiraceae bacterium* A2 |
| Lba-14-1 | Bacteria Firmicutes Clostridia Clostridiales *Lachnospiraceae bacterium* A4 |
| Lba-15-1 | Bacteria Firmicutes Clostridia Clostridiales *Lachnospiraceae bacterium* ICM7 |
| Lba-16-1 | Bacteria Firmicutes Clostridia Clostridiales *Lachnospiraceae bacterium* M18-1 |
| Lba-17-1 | Bacteria Firmicutes Clostridia Clostridiales *Lachnospiraceae bacterium* NC2008 |
| Lba-18-1 | Bacteria Firmicutes Clostridia Clostridiales *Lachnospiraceae bacterium* NK4A136 |
| Lba-19-1 | Bacteria Firmicutes Clostridia Clostridiales *Lachnospiraceae bacterium* NK4A179 |
| Lba-22-1 | Bacteria Firmicutes Clostridia Clostridiales *Lachnospiraceae bacterium oral taxon* 082 str. F0431 |
| Lba-20-1 | Bacteria Firmicutes Clostridia Clostridiales *Lachnospiraceae bacterium* VE202-12 |
| Lba-21-1 | Bacteria Firmicutes Clostridia Clostridiales *Lachnospiraceae bacterium* VE202-23 |
| Lor-1-1 | Bacteria Firmicutes Clostridia Clostridiales *Lachnospiraceae oral taxon* 107 str. F0167 |
| Rho-1-1 | Bacteria Firmicutes Clostridia Clostridiales Lachnospiraceae *Roseburia hominis* A2-183 |
| Rin-1-1 | Bacteria Firmicutes Clostridia Clostridiales Lachnospiraceae *Roseburia intestinalis* L1-82 |
| Rin-2-1 | Bacteria Firmicutes Clostridia Clostridiales Lachnospiraceae *Roseburia inulinivorans* DSM 16841 |
| Slo-1-1 | Bacteria Firmicutes Clostridia Clostridiales Lachnospiraceae *Stomatobaculum longum* |
| Pba-1-1 | Bacteria Firmicutes Clostridia Clostridiales *Peptostreptococcaceae bacterium oral taxon* 113 str. W5053 |
| Rfl-1-1 | Bacteria Firmicutes Clostridia Clostridiales Ruminococcaceae *Ruminococcus flavefaciens* FD-1 |
| Rgn-3-1 | Bacteria Firmicutes Clostridia Clostridiales Ruminococcaceae *Ruminococcus gnavus* ATCC 29149 |
| Rla-1-1 | Bacteria Firmicutes Clostridia Clostridiales Ruminococcaceae *Ruminococcus lactaris* CC59_002D |
| Rsp-1-1 | Bacteria Firmicutes Clostridia Clostridiales Ruminococcaceae *Ruminococcus sp.* JC304 |
| Rto-1-1 | Bacteria Firmicutes Clostridia Clostridiales Ruminococcaceae *Ruminococcus torques* ATCC 27756 |
| env-1 to env-1176 | environmental samples |

#### 19.2 Gene contexts

Each S10-Clostridia RNA (indicated by "RNA→") is listed. For each hit, the downstream genes predicted to reside in a regulated operon are listed. If the nearest downstream gene is encoding in the opposite strand (and therefore presumed to not be a part of a regulated operon), then that gene is still depicted. Some environmental sequences and some RefSeq entries lack gene annotations, and so no genes are available for such sequences. The direction of each gene is indicated with an arrow (→), and each predicted conserved domain in the gene is named. Conserved domains associated with more than one S10-Clostridia RNA are assigned a color; other domains are gray. Information about these conserved domains is given in Section 19.3. The accession of the sequence containing each S10-Clostridia RNA is given in the column named "Seq. accession". Accessions beginning with "NC_", "NS_", "NW_" or "NZ_" are contained in RefSeq. Other accession refer to environmental samples. Nucleotide coordinates are given for the 5′ and 3′ boundaries of each S10-Clostridia RNA. If the 5′ coordinate is greater than the 3′ coordinate, the RNA is present on the reverse-complement strand of the containing genomic DNA sequence. Each hit is denoted by an abbreviation (like "Eco-1-1") that refers to a taxonomy given in Section 19.1.

| abbrev. | Seq. accession | | 5′ at | 3′ at | genes |
|---|---|---|---|---|---|
| env-1 | 2226990398 | - | 1238 | 1158 | RNA→ rpsJ (PRK00596)RpsJ (COG0051)→ |
| env-2 | IMNBL1_c0000333 | - | 31361 | 31281 | RNA→ rpsJ (PRK00596)RpsJ (COG0051)→ |
| env-3 | JGI994J12354_10007324 | + | 4642 | 4722 | RNA→ rpsJ (PRK00596)RpsJ (COG0051)→ |
| env-4 | 2227142228 | - | 1210 | 1130 | RNA→ rpsJ (PRK00596)RpsJ (COG0051)→ |
| env-5 | 2226961550 | - | 118 | 38 | RNA→ ←hypo |
| env-6 | 4491409.3_NODE_35665 | - | 1609 | 1535 | RNA→ rpsJ (PRK00596)RpsJ (COG0051)→ |
| env-7 | DLF009_scaffold18474_3 | + | 2148 | 2217 | RNA→ rpsJ (PRK00596)RpsJ (COG0051)→ |
| env-8 | NOF011_scaffold7296_4 | - | 966 | 897 | RNA→ rpsJ (PRK00596)RpsJ (COG0051)→ |
| env-9 | scaffold16313_1_O2.UC-19 | + | 689 | 758 | RNA→ |
| env-10 | SRS013687_Baylor_scaffold_43281 | + | 185 | 253 | RNA→ rpsJ (PRK00596)RpsJ (COG0051)→ |
| env-11 | SRS024625_C2443234 | - | 144 | 76 | RNA→ |
| env-12 | AUXO018559537.1 | - | 326 | 258 | RNA→ rpsJ (PRK00596)RpsJ (COG0051)→ |
| env-13 | SRS011302_Baylor_scaffold_3477 | + | 2312 | 2380 | RNA→ rpsJ (PRK00596)Ribosomal_S10 (pfam00338)→ |
| env-14 | 4491406.3_NODE_8170 | - | 1590 | 1522 | RNA→ rpsJ (PRK00596)RpsJ (COG0051)→ |
| env-15 | 4491409.3_NODE_14115 | + | 1704 | 1772 | RNA→ rpsJ (PRK00596)RpsJ (COG0051)→ |
| env-16 | 2013434593 | + | 25 | 93 | RNA→ |
| env-17 | scaffold41095_3_MH0053 | + | 367 | 435 | RNA→ |
| env-18 | SRS052697_LANL_scaffold_57795 | - | 276 | 208 | RNA→ hypo→ |
| env-19 | 4491407.3_NODE_19021 | + | 122 | 190 | RNA→ rpsJ (PRK00596)Ribosomal_S10 (pfam00338)→ RplC (COG0087)rplC (PRK00001)→ |
| env-20 | scaffold4401_13_MH0057 | + | 404 | 472 | RNA→ rpsJ (PRK00596)Ribosomal_S10 (pfam00338)→ |
| env-21 | 4491401.3_NODE_82882 | - | 1562 | 1494 | RNA→ rpsJ (PRK00596)RpsJ (COG0051)→ |
| env-22 | 4491487.3_NODE_33289 | - | 2671 | 2603 | RNA→ rpsJ (PRK00596)RpsJ (COG0051)→ |
| env-23 | scaffold21085_1_MH0037 | - | 150 | 82 | RNA→ |
| env-24 | SRS011061_C4645920 | + | 238 | 306 | RNA→ |
| env-25 | DLM001_scaffold9250_9 | + | 1391 | 1459 | RNA→ rpsJ (PRK00596)RpsJ (COG0051)→ |
| env-26 | NLM031_scaffold272_21 | - | 6329 | 6261 | RNA→ rpsJ (PRK00596)RpsJ (COG0051)→ |
| env-27 | SRS013521_LANL_scaffold_2326 | - | 159 | 91 | RNA→ |
| env-28 | DOM019_scaffold38828_2 | + | 759 | 827 | RNA→ hypo→ rpsJ (PRK00596)Ribosomal_S10 (pfam00338)→ |
| env-29 | SRS015663_WUGC_scaffold_1713 | - | 288 | 220 | RNA→ rpsJ (PRK00596)→ |
| env-30 | scaffold25433_1_O2.UC-19 | - | 264 | 196 | RNA→ hypo→ |
| env-31 | SRS016095_WUGC_scaffold_35063 | - | 1318 | 1250 | RNA→ rpsJ (PRK00596)RpsJ (COG0051)→ |
| env-32 | SRS017191_C1692823 | + | 852 | 920 | RNA→ hypo→ |
| env-33 | SRS017821_Baylor_scaffold_18138 | - | 237 | 169 | RNA→ |
| env-34 | SRS019030_Baylor_scaffold_35826 | - | 460 | 392 | RNA→ rpsJ (PRK00596)RpsJ (COG0051)→ |
| env-35 | SRS019161_WUGC_scaffold_56262 | + | 245 | 313 | RNA→ hypo→ rpsJ (PRK00596)Ribosomal_S10 (pfam00338)→ |
| env-36 | SRS022609_Baylor_scaffold_55492 | + | 119 | 187 | RNA→ hypo→ rpsJ (PRK00596)RpsJ (COG0051)→ |
| env-37 | SRS049712_C15422763 | - | 95 | 27 | RNA→ hypo→ |
| env-38 | SRS050422_LANL_scaffold_77388 | - | 273 | 205 | RNA→ hypo→ |
| env-39 | SRS051031_LANL_scaffold_19825 | + | 1380 | 1448 | RNA→ rpsJ (PRK00596)RpsJ (COG0051)→ |

| | | | | | |
|---|---|---|---|---|---|
| env-40 | SRS055982_C1608927 | + | 905 | 973 | **RNA**→ rpsJ (PRK00596)RpsJ (COG0051)→ |
| env-41 | SRS056259_LANL_scaffold_34991 | - | 207 | 139 | **RNA**→ |
| env-42 | SRS065504_LANL_scaffold_12366 | - | 375 | 307 | **RNA**→ rpsJ (PRK00596)Ribosomal_S10 (pfam00338)→ |
| env-43 | 4491417.3_NODE_11963 | - | 1588 | 1520 | **RNA**→ rpsJ (PRK00596)RpsJ (COG0051)→ |
| env-44 | SRS048870_WUGC_scaffold_9375 | - | 159 | 91 | **RNA**→ |
| env-45 | SRS049959_WUGC_scaffold_6371 | - | 438 | 370 | **RNA**→ rpsJ (PRK00596)Ribosomal_S10 (pfam00338)→ |
| env-46 | 4491477.3_NODE_36868 | + | 368 | 436 | **RNA**→ |
| env-47 | DLF013_scaffold3458_3 | + | 3322 | 3390 | **RNA**→ hypo→ rpsJ (PRK00596)RpsJ (COG0051)→ |
| env-48 | SRS049712_C15557807 | + | 731 | 810 | **RNA**→ rpsJ (PRK00596)Ribosomal_S10 (pfam00338)→ |
| env-49 | scaffold7779_5_O2.UC-12 | - | 667 | 588 | **RNA**→ rpsJ (PRK00596)RpsJ (COG0051)→ |
| env-50 | scaffold8248_1_O2.UC-13 | + | 480 | 559 | **RNA**→ rpsJ (PRK00596)RpsJ (COG0051)→ RplC (COG0087)rplC (PRK00001)→ rplD (PRK05319)Ribosomal_L4 (pfam00573)→ rplW (PRK05738)Ribosomal_L23 (pfam00276)→ |
| env-51 | scaffold6367_32_V1.CD-4 | + | 1972 | 2051 | **RNA**→ rpsJ (PRK00596)RpsJ (COG0051)→ RplC (COG0087)rplC (PRK00001)→ rplD (PRK05319)Ribosomal_L4 (pfam00573)→ rplW (PRK05738)Ribosomal_L23 (pfam00276)→ |
| env-52 | SRS049959_WUGC_scaffold_42217 | + | 15227 | 15306 | **RNA**→ rpsJ (PRK00596)RpsJ (COG0051)→ RplC (COG0087)rplC (PRK00001)→ |
| env-53 | SRS063040_Baylor_scaffold_53437 | - | 465 | 386 | **RNA**→ rpsJ (PRK00596)Ribosomal_S10 (pfam00338)→ |
| env-54 | DOM014_scaffold41515_2 | - | 624 | 545 | **RNA**→ rpsJ (PRK00596)RpsJ (COG0051)→ |
| env-55 | NLF007_scaffold12381_15 | + | 444 | 523 | **RNA**→ rpsJ (PRK00596)RpsJ (COG0051)→ RplC (COG0087)rplC (PRK00001)→ RplD (COG0088)rplD (PRK05319)→ |
| env-56 | NLM025_scaffold9675_4 | - | 702 | 623 | **RNA**→ rpsJ (PRK00596)RpsJ (COG0051)→ |
| env-57 | NOM018_scaffold2595_2 | + | 22357 | 22436 | **RNA**→ rpsJ (PRK00596)RpsJ (COG0051)→ RplC (COG0087)rplC (PRK00001)→ rplD (PRK05319)Ribosomal_L4 (pfam00573)→ rplW (PRK05738)Ribosomal_L23 (pfam00276)→ RplB (COG0090)rplB (PRK09374)→ rpsS (PRK00357)Ribosomal_S19 (pfam00203)→ rplV (PRK00565)Ribosomal_L22 (pfam00237)→ RpsC (COG0092)rpsC (PRK00310)→ rpl16 (CHL00044)rplP (PRK09203)→ PRK00306 (PRK00306)Ribosomal_L29 (pfam00831)→ rpsQ (PRK05610)Ribosomal_S17 (pfam00366)→ rplN (PRK05483)Ribosomal_L14 (pfam00238)→ RplX (COG0198)rplX (PRK00004)→ |
| env-58 | SRS015217_WUGC_scaffold_16052 | - | 586 | 507 | **RNA**→ rpsJ (PRK00596)RpsJ (COG0051)→ |
| env-59 | DOM015_scaffold18234_5 | + | 4775 | 4854 | **RNA**→ rpsJ (PRK00596)RpsJ (COG0051)→ RplC (COG0087)rplC (PRK00001)→ rplD (PRK05319)Ribosomal_L4 (pfam00573)→ rplW (PRK05738)Ribosomal_L23 (pfam00276)→ RplB (COG0090)rplB (PRK09374)→ rpsS (PRK00357)Ribosomal_S19 (pfam00203)→ rplV (PRK00565)Ribosomal_L22 (pfam00237)→ RpsC (COG0092)rpsC (PRK00310)→ rpl16 (CHL00044)rplP (PRK09203)→ PRK00306 (PRK00306)Ribosomal_L29 (pfam00831)→ rpsQ (PRK05610)Ribosomal_S17 (pfam00366)→ rplN (PRK05483)Ribosomal_L14 (pfam00238)→ RplX (COG0198)rplX (PRK00004)→ rplE (PRK00010)RplE (COG0094)→ rpsN (PRK08061)RpsN (COG0199)→ rpsH (PRK00136)Ribosomal_S8 (pfam00410)→ RplF (COG0097)rplF (PRK05498)→ rplR (PRK05593)Ribosomal_L18p (pfam00861)→ RpsE (COG0098)rpsE (PRK00550)→ Ribosomal_L30 (cd01658)rpmD (PRK05611)→ RplO (COG0200)rplO (PRK05592)→ SecY (COG0201)secY (PRK09204)→ adk (PRK00279)ADK (pfam00406)→ PRK05716 (PRK05716)MetAP1 (cd01086)→ infA (PRK00276)S1_IF1 (cd04451)→ rpmJ (PRK00465)Ribosomal_L36 (pfam00444)→ RpsM (COG0099)rpsM (PRK05179)→ PRK05309 (PRK05309)RpsK (COG0100)→ RpsD (COG0522)rpsD (PRK05327)→ RpoA (COG0202)PRK05182 (PRK05182)→ rplQ (PRK05591)Ribosomal_L17 (pfam01196)→ |
| env-60 | NOM029_scaffold14304_23 | - | 471 | 392 | **RNA**→ rpsJ (PRK00596)Ribosomal_S10 (pfam00338)→ |
| env-61 | scaffold9930_2_MH0009 | - | 424 | 345 | **RNA**→ rpsJ (PRK00596)Ribosomal_S10 (pfam00338)→ |
| env-62 | scaffold24332_11_MH0045 | - | 2664 | 2585 | **RNA**→ rpsJ (PRK00596)RpsJ (COG0051)→ RplC (COG0087)rplC (PRK00001)→ rplD (PRK05319)Ribosomal_L4 (pfam00573)→ rplW (PRK05738)Ribosomal_L23 (pfam00276)→ |
| env-63 | scaffold16949_1_MH0048 | + | 212 | 291 | **RNA**→ rpsJ (PRK00596)RpsJ (COG0051)→ RplC (COG0087)rplC (PRK00001)→ rplD (PRK05319)Ribosomal_L4 (pfam00573)→ rplW (PRK05738)Ribosomal_L23 (pfam00276)→ |
| env-64 | 4491484.3_NODE_8515 | - | 3592 | 3513 | **RNA**→ rpsJ (PRK00596)RpsJ (COG0051)→ RplC (COG0087)rplC (PRK00001)→ rplD (PRK05319)Ribosomal_L4 (pfam00573)→ rplW (PRK05738)Ribosomal_L23 (pfam00276)→ rplB (PRK09374)Ribosomal_L2 (pfam00181)→ rplB (PRK09374)Ribosomal_L2_C (pfam03947)→ rpsS (PRK00357)Ribosomal_S19 (pfam00203)→ |

| | | | | | |
|---|---|---|---|---|---|
| env-65 | NLM026_scaffold29289_1 | + | 154 | 233 | RNA→ rpsJ (PRK00596)RpsJ (COG0051)→ RplC (COG0087)rplC (PRK00001)→ rplD (PRK05319)Ribosomal_L4 (pfam00573)→ rplW (PRK05738)Ribosomal_L23 (pfam00276)→ RplB (COG0090)rplB (PRK09374)→ rpsS (PRK00357)Ribosomal_S19 (pfam00203)→ rplV (PRK00565)Ribosomal_L22 (pfam00237)→ RpsC (COG0092)rpsC (PRK00310)→ rpl16 (CHL00044)rplP (PRK09203)→ PRK00306 (PRK00306)Ribosomal_L29 (pfam00831)→ rpsQ (PRK05610)Ribosomal_S17 (pfam00366)→ rplN (PRK05483)Ribosomal_L14 (pfam00238)→ RplX (COG0198)rplX (PRK00004)→ rplE (PRK00010)RplE (COG0094)→ rpsN (PRK08061)RpsN (COG0199)→ rpsH (PRK00136)Ribosomal_S8 (pfam00410)→ RplF (COG0097)rplF (PRK05498)→ rplR (PRK05593)Ribosomal_L18p (pfam00861)→ RpsE (COG0098)rpsE (PRK00550)→ Ribosomal_L30 (cd01658)rpmD (PRK05611)→ RplO (COG0200)rplO (PRK05592)→ SecY (COG0201)secY (PRK09204)→ ADK (cd01428)AAA_17 (pfam13207)→ |
| env-66 | NLM031_scaffold32591_5 | - | 104111 | 104032 | RNA→ rpsJ (PRK00596)RpsJ (COG0051)→ RplC (COG0087)rplC (PRK00001)→ rplD (PRK05319)Ribosomal_L4 (pfam00573)→ rplW (PRK05738)Ribosomal_L23 (pfam00276)→ RplB (COG0090)rplB (PRK09374)→ rpsS (PRK00357)Ribosomal_S19 (pfam00203)→ rplV (PRK00565)Ribosomal_L22 (pfam00237)→ RpsC (COG0092)rpsC (PRK00310)→ rpl16 (CHL00044)rplP (PRK09203)→ PRK00306 (PRK00306)Ribosomal_L29 (pfam00831)→ rpsQ (PRK05610)Ribosomal_S17 (pfam00366)→ rplN (PRK05483)Ribosomal_L14 (pfam00238)→ RplX (COG0198)rplX (PRK00004)→ rplE (PRK00010)RplE (COG0094)→ rpsN (PRK08061)RpsN (COG0199)→ rpsH (PRK00136)Ribosomal_S8 (pfam00410)→ RplF (COG0097)rplF (PRK05498)→ rplR (PRK05593)Ribosomal_L18p (pfam00861)→ RpsE (COG0098)rpsE (PRK00550)→ Ribosomal_L30 (cd01658)rpmD (PRK05611)→ RplO (COG0200)rplO (PRK05592)→ SecY (COG0201)secY (PRK09204)→ |
| env-67 | NOM019_scaffold27930_10 | + | 83526 | 83605 | RNA→ rpsJ (PRK00596)RpsJ (COG0051)→ RplC (COG0087)rplC (PRK00001)→ rplD (PRK05319)Ribosomal_L4 (pfam00573)→ rplW (PRK05738)Ribosomal_L23 (pfam00276)→ RplB (COG0090)rplB (PRK09374)→ rpsS (PRK00357)RpsS (COG0185)→ |
| env-68 | SRS011302_Baylor_scaffold_21352 | - | 474 | 395 | RNA→ rpsJ (PRK00596)Ribosomal_S10 (pfam00338)→ |
| env-69 | SRS012273_Baylor_scaffold_14966 | - | 16160 | 16081 | RNA→ rpsJ (PRK00596)RpsJ (COG0051)→ RplC (COG0087)rplC (PRK00001)→ rplD (PRK05319)Ribosomal_L4 (pfam00573)→ rplW (PRK05738)Ribosomal_L23 (pfam00276)→ RplB (COG0090)rplB (PRK09374)→ rpsS (PRK00357)Ribosomal_S19 (pfam00203)→ rplV (PRK00565)Ribosomal_L22 (pfam00237)→ RpsC (COG0092)rpsC (PRK00310)→ rpl16 (CHL00044)rplP (PRK09203)→ PRK00306 (PRK00306)Ribosomal_L29 (pfam00831)→ rpsQ (PRK05610)Ribosomal_S17 (pfam00366)→ rplN (PRK05483)Ribosomal_L14 (pfam00238)→ RplX (COG0198)rplX (PRK00004)→ rplE (PRK00010)RplE (COG0094)→ rpsN (PRK08061)RpsN (COG0199)→ rpsH (PRK00136)Ribosomal_S8 (pfam00410)→ RplF (COG0097)rplF (PRK05498)→ rplR (PRK05593)Ribosomal_L18p (pfam00861)→ RpsE (COG0098)rpsE (PRK00550)→ Ribosomal_L30 (cd01658)rpmD (PRK05611)→ RplO (COG0200)rplO (PRK05592)→ SecY (COG0201)secY (PRK09204)→ |
| env-70 | SRS015133_WUGC_scaffold_36610 | - | 2755 | 2676 | RNA→ rpsJ (PRK00596)RpsJ (COG0051)→ RplC (COG0087)rplC (PRK00001)→ rplD (PRK05319)Ribosomal_L4 (pfam00573)→ rplW (PRK05738)Ribosomal_L23 (pfam00276)→ rplB (PRK09374)Ribosomal_L2 (pfam00181)→ |
| env-71 | SRS048870_WUGC_scaffold_45340 | - | 5622 | 5543 | RNA→ rpsJ (PRK00596)RpsJ (COG0051)→ RplC (COG0087)rplC (PRK00001)→ rplD (PRK05319)Ribosomal_L4 (pfam00573)→ rplW (PRK05738)Ribosomal_L23 (pfam00276)→ RplB (COG0090)rplB (PRK09374)→ rpsS (PRK00357)Ribosomal_S19 (pfam00203)→ rplV (PRK00565)Ribosomal_L22 (pfam00237)→ RpsC (COG0092)rpsC (PRK00310)→ rpl16 (CHL00044)rplP (PRK09203)→ PRK00306 (PRK00306)Ribosomal_L29 (pfam00831)→ rpsQ (PRK05610)Ribosomal_S17 (pfam00366)→ rplN (PRK05483)Ribosomal_L14 (pfam00238)→ |
| env-72 | SRS051031_LANL_scaffold_20990 | - | 7045 | 6966 | RNA→ rpsJ (PRK00596)RpsJ (COG0051)→ RplC (COG0087)rplC (PRK00001)→ rplD (PRK05319)Ribosomal_L4 (pfam00573)→ rplW (PRK05738)Ribosomal_L23 (pfam00276)→ RplB (COG0090)rplB (PRK09374)→ rpsS (PRK00357)Ribosomal_S19 (pfam00203)→ rplV (PRK00565)Ribosomal_L22 (pfam00237)→ RpsC (COG0092)rpsC (PRK00310)→ rpl16 (CHL00044)rplP (PRK09203)→ PRK00306 (PRK00306)Ribosomal_L29 (pfam00831)→ rpsQ (PRK05610)Ribosomal_S17 (pfam00366)→ rplN (PRK05483)Ribosomal_L14 (pfam00238)→ RplX (COG0198)rplX (PRK00004)→ rplE (PRK00010)RplE (COG0094)→ rpsN (PRK08061)RpsN (COG0199)→ |

| | | | | | |
|---|---|---|---|---|---|
| env-73 | SRS063985_LANL_scaffold_52558 | + | 6006 | 6085 | **RNA**→ rpsJ (PRK00596)RpsJ (COG0051)→ RplC (COG0087)rplC (PRK00001)→ rplD (PRK05319)Ribosomal_L4 (pfam00573)→ rplW (PRK05738)Ribosomal_L23 (pfam00276)→ RplB (COG0090)rplB (PRK09374)→ rpsS (PRK00357)Ribosomal_S19 (pfam00203)→ rplV (PRK00565)Ribosomal_L22 (pfam00237)→ |
| env-74 | SRS064276_LANL_scaffold_65717 | + | 3203 | 3282 | **RNA**→ rpsJ (PRK00596)RpsJ (COG0051)→ RplC (COG0087)rplC (PRK00001)→ rplD (PRK05319)Ribosomal_L4 (pfam00573)→ rplW (PRK05738)Ribosomal_L23 (pfam00276)→ RplB (COG0090)rplB (PRK09374)→ |
| env-75 | SRS065504_LANL_scaffold_36299 | + | 111 | 190 | **RNA**→ rpsJ (PRK00596)RpsJ (COG0051)→ rpl3 (CHL00143)rplC (PRK00001)→ |
| env-76 | NLM016_scaffold9518_14 | - | 10030 | 9953 | **RNA**→ rpsJ (PRK00596)RpsJ (COG0051)→ RplC (COG0087)rplC (PRK00001)→ rplD (PRK05319)Ribosomal_L4 (pfam00573)→ rplW (PRK05738)Ribosomal_L23 (pfam00276)→ RplB (COG0090)rplB (PRK09374)→ rpsS (PRK00357)Ribosomal_S19 (pfam00203)→ rplV (PRK00565)Ribosomal_L22 (pfam00237)→ RpsC (COG0092)rpsC (PRK00310)→ rpl16 (CHL00044)rplP (PRK09203)→ PRK00306 (PRK00306)Ribosomal_L29 (pfam00831)→ rpsQ (PRK05610)Ribosomal_S17 (pfam00366)→ rplN (PRK05483)Ribosomal_L14 (pfam00238)→ RplX (COG0198)rplX (PRK00004)→ rplE (PRK00010)RplE (COG0094)→ rpsN (PRK08061)RpsN (COG0199)→ rpsH (PRK00136)Ribosomal_S8 (pfam00410)→ RplF (COG0097)rplF (PRK05498)→ rplR (PRK05593)Ribosomal_L18p (pfam00861)→ RpsE (COG0098)rpsE (PRK00550)→ Ribosomal_L30 (cd01658)rpmD (PRK05611)→ RplO (COG0200)rplO (PRK05592)→ SecY (COG0201)secY (PRK09204)→ |
| env-77 | scaffold5973_1_MH0070 | + | 109 | 186 | **RNA**→ rpsJ (PRK00596)RpsJ (COG0051)→ RplC (COG0087)rplC (PRK00001)→ rplD (PRK05319)Ribosomal_L4 (pfam00573)→ rplW (PRK05738)Ribosomal_L23 (pfam00276)→ |
| env-78 | DLF010_scaffold363_8 | - | 13570 | 13493 | **RNA**→ rpsJ (PRK00596)RpsJ (COG0051)→ RplC (COG0087)rplC (PRK00001)→ rplD (PRK05319)Ribosomal_L4 (pfam00573)→ rplW (PRK05738)Ribosomal_L23 (pfam00276)→ RplB (COG0090)rplB (PRK09374)→ rpsS (PRK00357)Ribosomal_S19 (pfam00203)→ rplV (PRK00565)Ribosomal_L22 (pfam00237)→ RpsC (COG0092)rpsC (PRK00310)→ rpl16 (CHL00044)rplP (PRK09203)→ PRK00306 (PRK00306)Ribosomal_L29 (pfam00831)→ rpsQ (PRK05610)Ribosomal_S17 (pfam00366)→ rplN (PRK05483)Ribosomal_L14 (pfam00238)→ RplX (COG0198)rplX (PRK00004)→ rplE (PRK00010)RplE (COG0094)→ rpsN (PRK08061)RpsN (COG0199)→ rpsH (PRK00136)Ribosomal_S8 (pfam00410)→ RplF (COG0097)rplF (PRK05498)→ rplR (PRK05593)Ribosomal_L18p (pfam00861)→ RpsE (COG0098)rpsE (PRK00550)→ Ribosomal_L30 (cd01658)rpmD (PRK05611)→ RplO (COG0200)rplO (PRK05592)→ SecY (COG0201)secY (PRK09204)→ |
| env-79 | NOF005_scaffold4009_2 | + | 389 | 466 | **RNA**→ rpsJ (PRK00596)RpsJ (COG0051)→ RplC (COG0087)rplC (PRK00001)→ rplD (PRK05319)Ribosomal_L4 (pfam00573)→ rplW (PRK05738)Ribosomal_L23 (pfam00276)→ RplB (COG0090)rplB (PRK09374)→ |
| env-80 | 4491414.3_NODE_8789 | + | 2679 | 2756 | **RNA**→ rpsJ (PRK00596)RpsJ (COG0051)→ RplC (COG0087)rplC (PRK00001)→ rplD (PRK05319)Ribosomal_L4 (pfam00573)→ rplW (PRK05738)Ribosomal_L23 (pfam00276)→ RplB (COG0090)rplB (PRK09374)→ rpsS (PRK00357)Ribosomal_S19 (pfam00203)→ rplV (PRK00565)Ribosomal_L22 (pfam00237)→ RpsC (COG0092)rpsC (PRK00310)→ rpl16 (CHL00044)rplP (PRK09203)→ PRK00306 (PRK00306)Ribosomal_L29 (pfam00831)→ rpsQ (PRK05610)Ribosomal_S17 (pfam00366)→ rplN (PRK05483)Ribosomal_L14 (pfam00238)→ RplX (COG0198)rplX (PRK00004)→ rplE (PRK00010)RplE (COG0094)→ rpsN (PRK08061)RpsN (COG0199)→ rpsH (PRK00136)Ribosomal_S8 (pfam00410)→ RplF (COG0097)rplF (PRK05498)→ rplR (PRK05593)Ribosomal_L18p (pfam00861)→ RpsE (COG0098)rpsE (PRK00550)→ Ribosomal_L30 (cd01658)rpmD (PRK05611)→ RplO (COG0200)rplO (PRK05592)→ SecY (COG0201)secY (PRK09204)→ adk (PRK00279)ADK (pfam00406)→ PRK05716 (PRK05716)MetAP1 (cd01086)→ infA (PRK00276)S1_IF1 (cd04451)→ rpmJ (PRK00465)Ribosomal_L36 (pfam00444)→ RpsM (COG0099)rpsM (PRK05179)→ PRK05309 (PRK05309)RpsK (COG0100)→ |

| | | | | | |
|---|---|---|---|---|---|
| env-81 | DOM017_scaffold3301_19 | + | 250 | 327 | **RNA**→ rpsJ (PRK00596)RpsJ (COG0051)→ RplC (COG0087)rplC (PRK00001)→<br>rplD (PRK05319)Ribosomal_L4 (pfam00573)→ rplW (PRK05738)Ribosomal_L23 (pfam00276)→<br>RplB (COG0090)rplB (PRK09374)→ rpsS (PRK00357)Ribosomal_S19 (pfam00203)→<br>rplV (PRK00565)Ribosomal_L22 (pfam00237)→ RpsC (COG0092)rpsC (PRK00310)→<br>rpl16 (CHL00044)rplP (PRK09203)→ PRK00306 (PRK00306)Ribosomal_L29 (pfam00831)→<br>rpsQ (PRK05610)Ribosomal_S17 (pfam00366)→ rplN (PRK05483)Ribosomal_L14 (pfam00238)→<br>RplX (COG0198)rplX (PRK00004)→ rplE (PRK00010)RplE (COG0094)→ rpsN (PRK08061)RpsN (COG0199)→<br>rpsH (PRK00136)Ribosomal_S8 (pfam00410)→ RplF (COG0097)rplF (PRK05498)→<br>rplR (PRK05593)Ribosomal_L18p (pfam00861)→ RpsE (COG0098)rpsE (PRK00550)→<br>Ribosomal_L30 (cd01658)rpmD (PRK05611)→ RplO (COG0200)rplO (PRK05592)→<br>SecY (COG0201)secY (PRK09204)→ adk (PRK00279)ADK (pfam00406)→<br>PRK05716 (PRK05716)MetAP1 (cd01086)→ infA (PRK00276)S1_IF1 (cd04451)→<br>rpmJ (PRK00465)Ribosomal_L36 (pfam00444)→ RpsM (COG0099)rpsM (PRK05179)→<br>PRK05309 (PRK05309)RpsK (COG0100)→ hypo→ |
| env-82 | scaffold7442_2_MH0060 | + | 810 | 887 | **RNA**→ rpsJ (PRK00596)Ribosomal_S10 (pfam00338)→ |
| env-83 | DLM001_scaffold58196_1 | - | 43327 | 43250 | **RNA**→ rpsJ (PRK00596)RpsJ (COG0051)→ RplC (COG0087)rplC (PRK00001)→<br>rplD (PRK05319)Ribosomal_L4 (pfam00573)→ rplW (PRK05738)Ribosomal_L23 (pfam00276)→<br>RplB (COG0090)rplB (PRK09374)→ rpsS (PRK00357)Ribosomal_S19 (pfam00203)→<br>rplV (PRK00565)Ribosomal_L22 (pfam00237)→ RpsC (COG0092)rpsC (PRK00310)→<br>rpl16 (CHL00044)rplP (PRK09203)→ PRK00306 (PRK00306)Ribosomal_L29 (pfam00831)→<br>rpsQ (PRK05610)Ribosomal_S17 (pfam00366)→ rplN (PRK05483)Ribosomal_L14 (pfam00238)→<br>RplX (COG0198)rplX (PRK00004)→ rplE (PRK00010)RplE (COG0094)→ rpsN (PRK08061)RpsN (COG0199)→<br>rpsH (PRK00136)Ribosomal_S8 (pfam00410)→ RplF (COG0097)rplF (PRK05498)→<br>rplR (PRK05593)Ribosomal_L18p (pfam00861)→ RpsE (COG0098)rpsE (PRK00550)→<br>Ribosomal_L30 (cd01658)rpmD (PRK05611)→ RplO (COG0200)rplO (PRK05592)→<br>SecY (COG0201)secY (PRK09204)→ |
| env-84 | NLM015_scaffold954_6 | + | 5548 | 5625 | **RNA**→ rpsJ (PRK00596)RpsJ (COG0051)→ |
| env-85 | 4491401.3_NODE_15713 | + | 3028 | 3095 | **RNA**→ rpsJ (PRK00596)RpsJ (COG0051)→ |
| env-86 | SRS024435_LANL_scaffold_25731 | - | 322 | 255 | **RNA**→ rpsJ (PRK00596)RpsJ (COG0051)→ |
| env-87 | 4491482.3_NODE_33602 | + | 1071 | 1138 | **RNA**→ rpsJ (PRK00596)RpsJ (COG0051)→ |
| env-88 | UnmappedStool_Broad_scaffold_1102440 | + | 454 | 521 | **RNA**→ |
| env-89 | SRS050925_C2029917 | - | 176 | 109 | **RNA**→ |
| Pba-1-1 | NZ_KI259856.1 | - | 47470 | 47393 | **RNA**→ hypo→ rpsJ (PRK00596)RpsJ (COG0051)→ RplC (COG0087)rplC (PRK00001)→<br>rplD (PRK05319)Ribosomal_L4 (pfam00573)→ rplW (PRK05738)Ribosomal_L23 (pfam00276)→<br>RplB (COG0090)rplB (PRK09374)→ rps19 (CHL00050)rpsS (PRK00357)→<br>rplV (PRK00565)Ribosomal_L22 (pfam00237)→ RpsC (COG0092)rpsC (PRK00310)→<br>rpl16 (CHL00044)rplP (PRK09203)→ rpl16 (CHL00044)rplP (PRK09203)→<br>PRK00306 (PRK00306)Ribosomal_L29 (pfam00831)→ rpsQ (PRK05610)Ribosomal_S17 (pfam00366)→<br>rplN (PRK05483)Ribosomal_L14 (pfam00238)→ RplX (COG0198)rplX (PRK00004)→<br>rplE (PRK00010)RplE (COG0094)→ rplE (PRK00010)RplE (COG0094)→ rpsN (PRK08061)RpsN (COG0199)→<br>rpsH (PRK00136)Ribosomal_S8 (pfam00410)→ RplF (COG0097)rplF (PRK05498)→<br>rplR (PRK05593)Ribosomal_L18p (pfam00861)→ RpsE (COG0098)rpsE (PRK00550)→<br>Ribosomal_L30 (cd01658)rpmD (PRK05611)→ rplO (PRK05592)Ribosomal_L27A (pfam00828)→<br>SecY (COG0201)secY (PRK09204)→ |
| env-90 | ADJS01004188.1 | - | 54515 | 54436 | **RNA**→ rpsJ (PRK00596)RpsJ (COG0051)→ RplC (COG0087)rplC (PRK00001)→<br>rplD (PRK05319)Ribosomal_L4 (pfam00573)→ rplW (PRK05738)Ribosomal_L23 (pfam00276)→<br>RplB (COG0090)rplB (PRK09374)→ rpsS (PRK00357)Ribosomal_S19 (pfam00203)→<br>rplV (PRK00565)Ribosomal_L22 (pfam00237)→ RpsC (COG0092)rpsC (PRK00310)→<br>rpl16 (CHL00044)rplP (PRK09203)→ PRK00306 (PRK00306)Ribosomal_L29 (pfam00831)→<br>rpsQ (PRK05610)Ribosomal_S17 (pfam00366)→ rplN (PRK05483)Ribosomal_L14 (pfam00238)→<br>RplX (COG0198)rplX (PRK00004)→ rplE (PRK00010)RplE (COG0094)→ rpsN (PRK08061)RpsN (COG0199)→<br>rpsH (PRK00136)Ribosomal_S8 (pfam00410)→ RplF (COG0097)rplF (PRK05498)→<br>rplR (PRK05593)Ribosomal_L18p (pfam00861)→ RpsE (COG0098)rpsE (PRK00550)→<br>Ribosomal_L30 (cd01658)rpmD (PRK05611)→ RplO (COG0200)rplO (PRK05592)→<br>SecY (COG0201)secY (PRK09204)→ |

| | | | | | |
|---|---|---|---|---|---|
| env-91 | scaffold34758_2_V1.CD-9 | - | 3281 | 3202 | **RNA**→ rpsJ (PRK00596)RpsJ (COG0051)→ RplC (COG0087)rplC (PRK00001)→ rplD (PRK05319)Ribosomal_L4 (pfam00573)→ rplW (PRK05738)Ribosomal_L23 (pfam00276)→ RplB (COG0090)rplB (PRK09374)→ rpsS (PRK00357)Ribosomal_S19 (pfam00203)→ |
| env-92 | SRS017103_C2723434 | + | 281 | 357 | **RNA**→ rpsJ (PRK00596)RpsJ (COG0051)→ RplC (COG0087)rplC (PRK00001)→ rplD (PRK05319)Ribosomal_L4 (pfam00573)→ rplW (PRK05738)Ribosomal_L23 (pfam00276)→ RplB (COG0090)rplB (PRK09374)→ |
| env-93 | SRS024331_LANL_scaffold_32721 | - | 2639 | 2563 | **RNA**→ rpsJ (PRK00596)RpsJ (COG0051)→ RplC (COG0087)rplC (PRK00001)→ rplD (PRK05319)Ribosomal_L4 (pfam00573)→ rplW (PRK05738)Ribosomal_L23 (pfam00276)→ |
| env-94 | scaffold19892_1_MH0040 | + | 1899 | 1975 | **RNA**→ rpsJ (PRK00596)RpsJ (COG0051)→ RplC (COG0087)rplC (PRK00001)→ rplD (PRK05319)Ribosomal_L4 (pfam00573)→ rplW (PRK05738)Ribosomal_L23 (pfam00276)→ rplB (PRK09374)Ribosomal_L2 (pfam00181)→ |
| env-95 | scaffold1228_27_MH0072 | - | 10164 | 10088 | **RNA**→ rpsJ (PRK00596)RpsJ (COG0051)→ RplC (COG0087)rplC (PRK00001)→ rplD (PRK05319)Ribosomal_L4 (pfam00573)→ rplW (PRK05738)Ribosomal_L23 (pfam00276)→ RplB (COG0090)rplB (PRK09374)→ rpsS (PRK00357)RpsS (COG0185)→ rplV (PRK00565)Ribosomal_L22 (pfam00237)→ RpsC (COG0092)rpsC (PRK00310)→ rpl16 (CHL00044)rplP (PRK09203)→ PRK00306 (PRK00306)Ribosomal_L29 (pfam00831)→ rpsQ (PRK05610)Ribosomal_S17 (pfam00366)→ rplN (PRK05483)Ribosomal_L14 (pfam00238)→ RplX (COG0198)rplX (PRK00004)→ rplE (PRK00010)RplE (COG0094)→ |
| env-96 | scaffold14350_6_MH0053 | - | 1036 | 960 | **RNA**→ rpsJ (PRK00596)RpsJ (COG0051)→ RplC (COG0087)rplC (PRK00001)→ |
| env-97 | scaffold60961_3_MH0030 | - | 1485 | 1409 | **RNA**→ rpsJ (PRK00596)RpsJ (COG0051)→ |
| env-98 | scaffold51359_1_O2.UC-24 | - | 257 | 181 | **RNA**→ |
| env-99 | SRS016335_C5109370 | - | 393 | 317 | **RNA**→ rpsJ (PRK00596)Ribosomal_S10 (pfam00338)→ |
| env-100 | SRS018817_C2118834 | + | 952 | 1028 | **RNA**→ rpsJ (PRK00596)RpsJ (COG0051)→ rpl3 (CHL00143)rplC (PRK00001)→ |
| env-101 | SRS050752_LANL_scaffold_50853 | - | 2468 | 2392 | **RNA**→ rpsJ (PRK00596)RpsJ (COG0051)→ RplC (COG0087)rplC (PRK00001)→ rplD (PRK05319)Ribosomal_L4 (pfam00573)→ rplW (PRK05738)Ribosomal_L23 (pfam00276)→ RplB (COG0090)rplB (PRK09374)→ |
| env-102 | 4491405.3_NODE_1553 | - | 26834 | 26758 | **RNA**→ rpsJ (PRK00596)RpsJ (COG0051)→ |
| env-103 | DLM022_scaffold43434_1 | + | 1026 | 1102 | **RNA**→ rpsJ (PRK00596)RpsJ (COG0051)→ |
| env-104 | NLM021_scaffold24247_3 | - | 25270 | 25194 | **RNA**→ rpsJ (PRK00596)RpsJ (COG0051)→ |
| env-105 | SRS014235_WUGC_scaffold_122205 | - | 2336 | 2260 | **RNA**→ rpsJ (PRK00596)RpsJ (COG0051)→ |
| env-106 | SRS019601_C2046325 | - | 1588 | 1512 | **RNA**→ rpsJ (PRK00596)RpsJ (COG0051)→ |
| env-107 | DLF006_scaffold12563_13 | - | 870 | 794 | **RNA**→ rpsJ (PRK00596)RpsJ (COG0051)→ |
| env-108 | SRS024435_C3230964 | - | 2635 | 2559 | **RNA**→ rpsJ (PRK00596)RpsJ (COG0051)→ |
| env-109 | SRS017701_C1702668 | + | 150 | 226 | **RNA**→ |
| env-110 | scaffold12346_3_MH0031 | - | 1221 | 1145 | **RNA**→ rpsJ (PRK00596)RpsJ (COG0051)→ |
| env-111 | DLM019_scaffold63082_2 | - | 34456 | 34380 | **RNA**→ rpsJ (PRK00596)RpsJ (COG0051)→ |
| env-112 | scaffold8414_7_V1.CD-8 | - | 2764 | 2688 | **RNA**→ rpsJ (PRK00596)RpsJ (COG0051)→ |
| env-113 | 4491409.3_NODE_23640 | + | 28884 | 28960 | **RNA**→ rpsJ (PRK00596)RpsJ (COG0051)→ |
| env-114 | scaffold6976_8_MH0016 | + | 328 | 403 | **RNA**→ rpsJ (PRK00596)RpsJ (COG0051)→ RplC (COG0087)rplC (PRK00001)→ rplD (PRK05319)Ribosomal_L4 (pfam00573)→ rplW (PRK05738)Ribosomal_L23 (pfam00276)→ RplB (COG0090)rplB (PRK09374)→ |
| env-115 | scaffold81869_1_MH0033 | - | 212 | 137 | **RNA**→ |
| env-116 | 4491405.3_NODE_19252 | + | 230 | 305 | **RNA**→ rpsJ (PRK00596)Ribosomal_S10 (pfam00338)→ |

| | | | | | |
|---|---|---|---|---|---|
| env-117 | 4491410.3_NODE_2064 | + | 423 | 498 | **RNA**→ rpsJ (PRK00596)RpsJ (COG0051)→ RplC (COG0087)rplC (PRK00001)→ rplD (PRK05319)Ribosomal_L4 (pfam00573)→ rplW (PRK05738)Ribosomal_L23 (pfam00276)→ RplB (COG0090)rplB (PRK09374)→ rpsS (PRK00357)Ribosomal_S19 (pfam00203)→ rplV (PRK00565)Ribosomal_L22 (pfam00237)→ RpsC (COG0092)rpsC (PRK00310)→ rpl16 (CHL00044)rplP (PRK09203)→ PRK00306 (PRK00306)Ribosomal_L29 (pfam00831)→ rpsQ (PRK05610)Ribosomal_S17 (pfam00366)→ rplN (PRK05483)Ribosomal_L14 (pfam00238)→ RplX (COG0198)rplX (PRK00004)→ rplE (PRK00010)RplE (COG0094)→ rpsN (PRK08061)RpsN (COG0199)→ rpsH (PRK00136)Ribosomal_S8 (pfam00410)→ RplF (COG0097)rplF (PRK05498)→ rplR (PRK05593)Ribosomal_L18p (pfam00861)→ RpsE (COG0098)rpsE (PRK00550)→ Ribosomal_L30 (cd01658)rpmD (PRK05611)→ RplO (COG0200)rplO (PRK05592)→ SecY (COG0201)secY (PRK09204)→ adk (PRK00279)ADK (pfam00406)→ PRK05716 (PRK05716)MetAP1 (cd01086)→ KOW_RPL14 (cd06088)→ infA (PRK00276)S1_IF1 (cd04451)→ rpmJ (PRK00465)Ribosomal_L36 (pfam00444)→ RpsM (COG0099)rpsM (PRK05179)→ PRK05309 (PRK05309)RpsK (COG0100)→ RpsD (COG0522)rpsD (PRK05327)→ RpoA (COG0202)PRK05182 (PRK05182)→ rplQ (PRK05591)Ribosomal_L17 (pfam01196)→ |
| env-118 | 4491412.3_NODE_68334 | + | 16503 | 16578 | **RNA**→ rpsJ (PRK00596)RpsJ (COG0051)→ RplC (COG0087)rplC (PRK00001)→ rplD (PRK05319)Ribosomal_L4 (pfam00573)→ rplW (PRK05738)Ribosomal_L23 (pfam00276)→ |
| env-119 | NOF004_scaffold6915_3 | - | 18884 | 18809 | **RNA**→ rpsJ (PRK00596)RpsJ (COG0051)→ RplC (COG0087)rplC (PRK00001)→ rplD (PRK05319)Ribosomal_L4 (pfam00573)→ rplW (PRK05738)Ribosomal_L23 (pfam00276)→ RplB (COG0090)rplB (PRK09374)→ rpsS (PRK00357)Ribosomal_S19 (pfam00203)→ rplV (PRK00565)Ribosomal_L22 (pfam00237)→ RpsC (COG0092)rpsC (PRK00310)→ rpl16 (CHL00044)rplP (PRK09203)→ PRK00306 (PRK00306)Ribosomal_L29 (pfam00831)→ rpsQ (PRK05610)Ribosomal_S17 (pfam00366)→ rplN (PRK05483)Ribosomal_L14 (pfam00238)→ RplX (COG0198)rplX (PRK00004)→ rplE (PRK00010)RplE (COG0094)→ rpsN (PRK08061)RpsN (COG0199)→ rpsH (PRK00136)Ribosomal_S8 (pfam00410)→ RplF (COG0097)rplF (PRK05498)→ rplR (PRK05593)Ribosomal_L18p (pfam00861)→ RpsE (COG0098)rpsE (PRK00550)→ Ribosomal_L30 (cd01658)rpmD (PRK05611)→ RplO (COG0200)rplO (PRK05592)→ SecY (COG0201)secY (PRK09204)→ |
| env-120 | SRS052697_LANL_scaffold_30806 | + | 25704 | 25779 | **RNA**→ rpsJ (PRK00596)Ribosomal_S10 (pfam00338)→ |
| env-121 | SRS024435_C2906239 | - | 134 | 59 | **RNA**→ hypo→ |
| env-122 | SRS047014_WUGC_scaffold_72164 | - | 438 | 365 | **RNA**→ rpsJ (PRK00596)Ribosomal_S10 (pfam00338)→ |
| env-123 | scaffold29511_2_MH0002 | - | 2951 | 2878 | **RNA**→ rpsJ (PRK00596)RpsJ (COG0051)→ RplC (COG0087)rplC (PRK00001)→ rplD (PRK05319)Ribosomal_L4 (pfam00573)→ rplW (PRK05738)Ribosomal_L23 (pfam00276)→ RplB (COG0090)rplB (PRK09374)→ |
| env-124 | scaffold7276_3_MH0061 | - | 2984 | 2911 | **RNA**→ rpsJ (PRK00596)RpsJ (COG0051)→ RplC (COG0087)rplC (PRK00001)→ rplD (PRK05319)Ribosomal_L4 (pfam00573)→ rplW (PRK05738)Ribosomal_L23 (pfam00276)→ RplB (COG0090)rplB (PRK09374)→ |
| env-125 | scaffold78335_2_V1.UC-9 | + | 5667 | 5740 | **RNA**→ rpsJ (PRK00596)Ribosomal_S10 (pfam00338)→ |
| env-126 | SRS022524_Baylor_scaffold_45135 | + | 2738 | 2811 | **RNA**→ rpsJ (PRK00596)RpsJ (COG0051)→ RplC (COG0087)rplC (PRK00001)→ rplD (PRK05319)Ribosomal_L4 (pfam00573)→ rplW (PRK05738)Ribosomal_L23 (pfam00276)→ RplB (COG0090)rplB (PRK09374)→ |
| env-127 | SRS063040_Baylor_scaffold_52905 | - | 1532 | 1459 | **RNA**→ rpsJ (PRK00596)RpsJ (COG0051)→ RplC (COG0087)rplC (PRK00001)→ rplD (PRK05319)Ribosomal_L4 (pfam00573)→ |
| env-128 | 4491421.3_NODE_12916 | + | 171 | 244 | **RNA**→ rpsJ (PRK00596)RpsJ (COG0051)→ |
| Rfl-1-1 | NZ_ACOK01000089.1 | - | 23727 | 23654 | **RNA**→ rpsJ (PRK00596)RpsJ (COG0051)→ rpsJ (PRK00596)RpsJ (COG0051)→ |
| env-129 | AUXO012404992.1 | + | 861 | 934 | **RNA**→ |
| env-130 | scaffold4266_1_O2.UC-20 | - | 2230 | 2134 | **RNA**→ rpsJ (PRK00596)RpsJ (COG0051)→ |
| env-131 | SRS011306_Baylor_scaffold_82937 | - | 355 | 251 | **RNA**→ rpsJ (PRK00596)RpsJ (COG0051)→ |
| env-132 | SRS050244_C4490640 | + | 496 | 600 | **RNA**→ |
| env-133 | Throat_LANL_C5678524 | + | 67 | 171 | **RNA**→ hypo→ |
| env-134 | SRS022530_LANL_scaffold_32319 | - | 61485 | 61381 | **RNA**→ rpsJ (PRK00596)RpsJ (COG0051)→ |
| env-135 | SRS023617_C3177372 | + | 418 | 522 | **RNA**→ rpsJ (PRK00596)RpsJ (COG0051)→ |
| env-136 | SRS024015_Baylor_scaffold_39044 | + | 136 | 240 | **RNA**→ rpsJ (PRK00596)RpsJ (COG0051)→ |
| env-137 | SRS063288_LANL_scaffold_41293 | - | 688 | 584 | **RNA**→ rpsJ (PRK00596)RpsJ (COG0051)→ |
| env-138 | SRS048791_C3235272 | + | 564 | 668 | **RNA**→ |

| | | | | | |
|---|---|---|---|---|---|
| env-139 | SRS015057_C3265888 | + | 116 | 220 | **RNA**→ |
| env-140 | SRS055426_C3279481 | + | 716 | 820 | **RNA**→ |
| env-141 | SRS023557_C1764443 | + | 1478 | 1582 | **RNA**→ rpsJ (PRK00596)RpsJ (COG0051)→ |
| env-142 | SRS014888_C3060382 | + | 418 | 522 | **RNA**→ |
| env-143 | SRS024580_LANL_scaffold_44876 | - | 1561 | 1457 | **RNA**→ rpsJ (PRK00596)RpsJ (COG0051)→ |
| env-144 | SRS014124_C2621890 | - | 1301 | 1197 | **RNA**→ rpsJ (PRK00596)RpsJ (COG0051)→ |
| env-145 | SRS015893_WUGC_scaffold_31047 | + | 1959 | 2063 | **RNA**→ rpsJ (PRK00596)Ribosomal_S10 (pfam00338)→ |
| env-146 | SRS018591_WUGC_scaffold_38862 | - | 55571 | 55467 | **RNA**→ rpsJ (PRK00596)RpsJ (COG0051)→ |
| env-147 | SRS051791_LANL_scaffold_46426 | - | 75884 | 75780 | **RNA**→ rpsJ (PRK00596)RpsJ (COG0051)→ |
| env-148 | SRS011140_Baylor_scaffold_51353 | + | 94 | 198 | **RNA**→ rpsJ (PRK00596)RpsJ (COG0051)→ |
| env-149 | SRS012279_Baylor_scaffold_56090 | + | 690 | 794 | **RNA**→ |
| env-150 | SRS014271_C2921876 | - | 218 | 114 | **RNA**→ |
| env-151 | SRS014684_WUGC_scaffold_40865 | + | 1287 | 1391 | **RNA**→ rpsJ (PRK00596)RpsJ (COG0051)→ |
| env-152 | SRS014689_WUGC_scaffold_1936 | - | 994 | 890 | **RNA**→ rpsJ (PRK00596)RpsJ (COG0051)→ |
| env-153 | SRS019027_C1823783 | - | 185 | 81 | **RNA**→ hypo→ |
| env-154 | SRS024081_C2636285 | - | 218 | 114 | **RNA**→ |
| env-155 | SRS042910_C5605950 | + | 684 | 788 | **RNA**→ |
| env-156 | SRS062544_LANL_scaffold_74780 | + | 5551 | 5655 | **RNA**→ |
| env-157 | SRS064423_LANL_scaffold_33619 | + | 10606 | 10710 | **RNA**→ rpsJ (PRK00596)RpsJ (COG0051)→ |
| env-158 | SRS019122_C5850384 | + | 204 | 308 | **RNA**→ rpsJ (PRK00596)Ribosomal_S10 (pfam00338)→ |
| env-159 | SRS065278_LANL_scaffold_1918 | + | 684 | 788 | **RNA**→ rpsJ (PRK00596)RpsJ (COG0051)→ |
| env-160 | SRS014692_C1424509 | + | 220 | 324 | **RNA**→ |
| env-161 | SRS017713_C3272311 | + | 140 | 245 | **RNA**→ |
| Cmo-1-1 | NZ_KI535366.1 | - | 700966 | 700861 | **RNA**→ rpsJ (PRK00596)RpsJ (COG0051)→ |
| env-162 | SRS022725_C5792968 | - | 517 | 412 | **RNA**→ rpsJ (PRK00596)RpsJ (COG0051)→ |
| env-163 | SRS022602_Baylor_scaffold_110235 | + | 21 | 126 | **RNA**→ rpsJ (PRK00596)RpsJ (COG0051)→ |
| Ece-1-1 | NZ_CM001487.1 | + | 2574980 | 2575059 | **RNA**→ rpsJ (PRK00596)RpsJ (COG0051)→ rpsJ (PRK00596)RpsJ (COG0051)→ |
| env-164 | AUXO017143398.1 | + | 3141 | 3243 | **RNA**→ rpsJ (PRK00596)RpsJ (COG0051)→ |
| Bfi-3-1 | NZ_ATVY01000026.1 | - | 26672 | 26570 | **RNA**→ rpsJ (PRK00596)RpsJ (COG0051)→ |
| Bfi-4-1 | NZ_AUJV01000009.1 | + | 23517 | 23619 | **RNA**→ rpsJ (PRK00596)RpsJ (COG0051)→ |
| Bfi-1-1 | NZ_ATVZ01000006.1 | - | 67223 | 67121 | **RNA**→ rpsJ (PRK00596)RpsJ (COG0051)→ |
| Bfi-2-1 | NZ_AUJW01000021.1 | - | 71753 | 71651 | **RNA**→ rpsJ (PRK00596)RpsJ (COG0051)→ |
| env-165 | SRS013252_Baylor_scaffold_21948 | - | 7197 | 7095 | **RNA**→ rpsJ (PRK00596)RpsJ (COG0051)→ RplC (COG0087)rplC (PRK00001)→ |
| | | | | | rplD (PRK05319)Ribosomal_L4 (pfam00573)→ rplW (PRK05738)Ribosomal_L23 (pfam00276)→ |
| | | | | | RplB (COG0090)rplB (PRK09374)→ rpsS (PRK00357)RpsS (COG0185)→ |
| | | | | | rplV (PRK00565)Ribosomal_L22 (pfam00237)→ RpsC (COG0092)rpsC (PRK00310)→ |
| | | | | | rpl16 (CHL00044)rplP (PRK09203)→ PRK00306 (PRK00306)Ribosomal_L29 (pfam00831)→ |
| | | | | | rpsQ (PRK05610)Ribosomal_S17 (pfam00366)→ rplN (PRK05483)Ribosomal_L14 (pfam00238)→ |
| | | | | | RplX (COG0198)rplX (PRK00004)→ rplE (PRK00010)RplE (COG0094)→ rpsN (PRK08061)RpsN (COG0199)→ |
| env-166 | SRS011126_C4671534 | - | 242 | 140 | **RNA**→ |
| env-167 | SRS013533_C5425190 | + | 425 | 527 | **RNA**→ |
| env-168 | SRS045197_C3557001 | + | 691 | 793 | **RNA**→ |
| env-169 | SRS049318_LANL_scaffold_176197 | + | 2049 | 2151 | **RNA**→ rpsJ (PRK00596)RpsJ (COG0051)→ RplC (COG0087)rplC (PRK00001)→ |
| | | | | | rplD (PRK05319)Ribosomal_L4 (pfam00573)→ rplW (PRK05738)Ribosomal_L23 (pfam00276)→ |
| | | | | | rplB (PRK09374)Ribosomal_L2 (pfam00181)→ |
| env-170 | SRS013170_Baylor_scaffold_134367 | - | 1061 | 959 | **RNA**→ rpsJ (PRK00596)RpsJ (COG0051)→ RplC (COG0087)rplC (PRK00001)→ |
| env-171 | SRS011152_C4188878 | - | 220 | 118 | **RNA**→ hypo→ |
| env-172 | SRS022602_Baylor_scaffold_37959 | - | 16760 | 16658 | **RNA**→ rpsJ (PRK00596)RpsJ (COG0051)→ RplC (COG0087)rplC (PRK00001)→ |
| | | | | | rplD (PRK05319)Ribosomal_L4 (pfam00573)→ rplW (PRK05738)Ribosomal_L23 (pfam00276)→ |
| | | | | | RplB (COG0090)rplB (PRK09374)→ rpsS (PRK00357)RpsS (COG0185)→ |
| | | | | | rplV (PRK00565)Ribosomal_L22 (pfam00237)→ RpsC (COG0092)rpsC (PRK00310)→ |
| | | | | | rpl16 (CHL00044)rplP (PRK09203)→ PRK00306 (PRK00306)Ribosomal_L29 (pfam00831)→ |
| | | | | | rpsQ (PRK05610)Ribosomal_S17 (pfam00366)→ rplN (PRK05483)Ribosomal_L14 (pfam00238)→ |
| | | | | | RplX (COG0198)rplX (PRK00004)→ rplE (PRK00010)RplE (COG0094)→ rpsN (PRK08061)RpsN (COG0199)→ |

| | | | | | |
|---|---|---|---|---|---|
| env-173 | DLM028_scaffold27170_5 | - | 1772 | 1666 | **RNA**→ rpsJ (PRK00596)RpsJ (COG0051)→ |
| env-174 | DLF010_scaffold24_15 | - | 1235 | 1129 | **RNA**→ rpsJ (PRK00596)RpsJ (COG0051)→ |
| env-175 | scaffold958_4_MH0086 | - | 199 | 94 | **RNA**→ |
| env-176 | UnmappedStool_Broad_scaffold_346780 | + | 64 | 169 | **RNA**→ ←hypo |
| env-177 | scaffold29314_3_MH0080 | - | 202 | 97 | **RNA**→ |
| env-178 | 4491414.3_NODE_953 | - | 28628 | 28523 | **RNA**→ rpsJ (PRK00596)RpsJ (COG0051)→ |
| env-179 | scaffold20541_3_MH0083 | - | 207 | 102 | **RNA**→ |
| env-180 | SRS011061_C4862922 | - | 895 | 792 | **RNA**→ rpsJ (PRK00596)RpsJ (COG0051)→ RplC (COG0087)rplC (PRK00001)→ |
| env-181 | SRS016989_C1563263 | - | 492 | 389 | **RNA**→ rpsJ (PRK00596)RpsJ (COG0051)→ |
| env-182 | SRS019267_C3574650 | + | 121 | 224 | **RNA**→ rpsJ (PRK00596)RpsJ (COG0051)→ |
| env-183 | scaffold1958_7_MH0014 | - | 1410 | 1307 | **RNA**→ rpsJ (PRK00596)RpsJ (COG0051)→ |
| env-184 | scaffold37912_1_V1.CD-4 | + | 65 | 168 | **RNA**→ rpsJ (PRK00596)RpsJ (COG0051)→ |
| env-185 | scaffold70413_1_MH0035 | - | 229 | 126 | **RNA**→ |
| env-186 | NLM021_scaffold9243_17 | + | 2752 | 2855 | **RNA**→ rpsJ (PRK00596)RpsJ (COG0051)→ |
| env-187 | NLM031_scaffold29725_6 | + | 96 | 199 | **RNA**→ rpsJ (PRK00596)Ribosomal_S10 (pfam00338)→ |
| env-188 | SRS052697_LANL_scaffold_50890 | + | 162 | 265 | **RNA**→ rpsJ (PRK00596)RpsJ (COG0051)→ RplC (COG0087)rplC (PRK00001)→ RplD (COG0088)rplD (PRK05319)→ |
| env-189 | scaffold33193_1_V1.UC-14 | + | 65 | 168 | **RNA**→ rpsJ (PRK00596)RpsJ (COG0051)→ |
| env-190 | 4491490.3_NODE_5470 | + | 114 | 217 | **RNA**→ rpsJ (PRK00596)RpsJ (COG0051)→ |
| env-191 | 4491412.3_NODE_89684 | + | 7189 | 7292 | **RNA**→ rpsJ (PRK00596)RpsJ (COG0051)→ |
| env-192 | NLF005_scaffold9646_4 | - | 866 | 763 | **RNA**→ rpsJ (PRK00596)RpsJ (COG0051)→ |
| env-193 | scaffold8051_7_MH0055 | - | 726 | 623 | **RNA**→ rpsJ (PRK00596)RpsJ (COG0051)→ |
| env-194 | NOF008_scaffold890_10 | - | 417 | 314 | **RNA**→ rpsJ (PRK00596)Ribosomal_S10 (pfam00338)→ |
| env-195 | scaffold31333_4_MH0028 | - | 751 | 648 | **RNA**→ rpsJ (PRK00596)RpsJ (COG0051)→ |
| env-196 | scaffold6771_3_MH0033 | - | 1483 | 1380 | **RNA**→ rpsJ (PRK00596)RpsJ (COG0051)→ |
| env-197 | scaffold30659_3_V1.UC-10 | + | 703 | 806 | **RNA**→ rpsJ (PRK00596)Ribosomal_S10 (pfam00338)→ |
| env-198 | 4491487.3_NODE_19593 | + | 2229 | 2332 | **RNA**→ rpsJ (PRK00596)RpsJ (COG0051)→ |
| env-199 | SRS011239_C2472698 | + | 137 | 240 | **RNA**→ |
| env-200 | SRS013687_Baylor_scaffold_53562 | + | 167 | 270 | **RNA**→ rpsJ (PRK00596)RpsJ (COG0051)→ RplC (COG0087)rplC (PRK00001)→ rplD (PRK05319)Ribosomal_L4 (pfam00573)→ |
| env-201 | SRS014979_WUGC_scaffold_29225 | - | 1526 | 1423 | **RNA**→ rpsJ (PRK00596)RpsJ (COG0051)→ RplC (COG0087)rplC (PRK00001)→ RplD (COG0088)rplD (PRK05319)→ |
| env-202 | SRS015133_WUGC_scaffold_33347 | - | 754 | 651 | **RNA**→ rpsJ (PRK00596)RpsJ (COG0051)→ rplC (PRK00001)→ |
| env-203 | SRS015854_WUGC_scaffold_34494 | - | 1175 | 1072 | **RNA**→ rpsJ (PRK00596)RpsJ (COG0051)→ RplC (COG0087)rplC (PRK00001)→ |
| env-204 | SRS021948_C2578511 | + | 115 | 218 | **RNA**→ |
| env-205 | SRS051031_LANL_scaffold_55309 | + | 2873 | 2976 | **RNA**→ rpsJ (PRK00596)RpsJ (COG0051)→ |
| env-206 | SRS063985_LANL_scaffold_39954 | - | 2666 | 2563 | **RNA**→ rpsJ (PRK00596)RpsJ (COG0051)→ RplC (COG0087)rplC (PRK00001)→ RplD (COG0088)rplD (PRK05319)→ rplD (PRK05319)Ribosomal_L4 (pfam00573)→ rplW (PRK05738)Ribosomal_L23 (pfam00276)→ rplB (PRK09374)Ribosomal_L2 (pfam00181)→ |
| env-207 | SRS065504_LANL_scaffold_46409 | - | 492 | 389 | **RNA**→ rpsJ (PRK00596)RpsJ (COG0051)→ |
| env-208 | SRS014613_WUGC_scaffold_17595 | - | 3925 | 3822 | **RNA**→ rpsJ (PRK00596)RpsJ (COG0051)→ RplC (COG0087)rplC (PRK00001)→ rplD (PRK05319)Ribosomal_L4 (pfam00573)→ rplW (PRK05738)Ribosomal_L23 (pfam00276)→ RplB (COG0090)rplB (PRK09374)→ rpsS (PRK00357)Ribosomal_S19 (pfam00203)→ rplV (PRK00565)Ribosomal_L22 (pfam00237)→ |
| env-209 | SRS019161_WUGC_scaffold_54925 | + | 161 | 264 | **RNA**→ rpsJ (PRK00596)RpsJ (COG0051)→ RplC (COG0087)rplC (PRK00001)→ |
| env-210 | SRS053335_C2500585 | - | 1526 | 1423 | **RNA**→ rpsJ (PRK00596)RpsJ (COG0051)→ RplC (COG0087)rplC (PRK00001)→ RplD (COG0088)rplD (PRK05319)→ |
| env-211 | SRS064276_LANL_scaffold_32780 | - | 2864 | 2761 | **RNA**→ rpsJ (PRK00596)RpsJ (COG0051)→ RplC (COG0087)rplC (PRK00001)→ rplD (PRK05319)Ribosomal_L4 (pfam00573)→ rplW (PRK05738)Ribosomal_L23 (pfam00276)→ RplB (COG0090)rplB (PRK09374)→ |
| Lba-19-1 | NZ_ATWC01000002.1 | - | 171387 | 171260 | **RNA**→ rpsJ (PRK00596)RpsJ (COG0051)→ |
| Lba-17-1 | NZ_JHWS01000005.1 | + | 279183 | 279268 | **RNA**→ rpsJ (PRK00596)RpsJ (COG0051)→ |
| env-212 | AUXO018772739.1 | - | 475 | 390 | **RNA**→ rpsJ (PRK00596)RpsJ (COG0051)→ |
| Lba-18-1 | NZ_ATVW01000015.1 | + | 20018 | 20103 | **RNA**→ rpsJ (PRK00596)RpsJ (COG0051)→ |

| | | | | | |
|---|---|---|---|---|---|
| env-213 | 08_07_XOS_DNA_scaffold44549_1 | - | 5717 | 5619 | **RNA**→ rpsJ (PRK00596)RpsJ (COG0051)→ |
| env-214 | SRS011529_WUGC_scaffold_31649 | + | 2893 | 2977 | **RNA**→ |
| env-215 | scaffold20711_4_V1.CD-13 | - | 153 | 69 | **RNA**→ |
| env-216 | scaffold537_5_V1.UC-14 | + | 2694 | 2778 | **RNA**→ rpsJ (PRK00596)RpsJ (COG0051)→ |
| env-217 | 4491423.3_NODE_119558 | - | 2026 | 1942 | **RNA**→ rpsJ (PRK00596)Ribosomal_S10 (pfam00338)→ |
| env-218 | SRS019582_WUGC_scaffold_26056 | - | 3718 | 3634 | **RNA**→ RplC (COG0087)rplC (PRK00001)→ rplD (PRK05319)Ribosomal_L4 (pfam00573)→ rplW (PRK05738)Ribosomal_L23 (pfam00276)→ RplB (COG0090)rplB (PRK09374)→ rpsS (PRK00357)RpsS (COG0185)→ rplV (PRK00565)Ribosomal_L22 (pfam00237)→ |
| env-219 | scaffold4496_1_MH0063 | + | 3017 | 3101 | **RNA**→ rpsJ (PRK00596)Ribosomal_S10 (pfam00338)→ |
| env-220 | scaffold10033_5_MH0081 | - | 153 | 69 | **RNA**→ |
| env-221 | NLF007_scaffold23667_30 | - | 6135 | 6051 | **RNA**→ rpsJ (PRK00596)RpsJ (COG0051)→ |
| env-222 | NLM021_scaffold31892_2 | + | 6861 | 6945 | **RNA**→ rpsJ (PRK00596)RpsJ (COG0051)→ |
| env-223 | NLF013_scaffold4173_33 | - | 4292 | 4208 | **RNA**→ rpsJ (PRK00596)RpsJ (COG0051)→ |
| env-224 | SRS020328_Baylor_scaffold_12580 | - | 6998 | 6914 | **RNA**→ rpsJ (PRK00596)RpsJ (COG0051)→ |
| env-225 | scaffold27081_10_MH0017 | + | 123 | 207 | **RNA**→ rpsJ (PRK00596)RpsJ (COG0051)→ |
| env-226 | scaffold2277_2_MH0077 | - | 340 | 256 | **RNA**→ rpsJ (PRK00596)Ribosomal_S10 (pfam00338)→ |
| env-227 | DOM012_scaffold9226_17 | - | 1293 | 1209 | **RNA**→ rpsJ (PRK00596)RpsJ (COG0051)→ |
| env-228 | SRS015217_WUGC_scaffold_10150 | + | 19846 | 19930 | **RNA**→ rpsJ (PRK00596)RpsJ (COG0051)→ |
| Csp-6-1 | NZ_DS480326.1 | - | 124987 | 124903 | **RNA**→ rpsJ (PRK00596)RpsJ (COG0051)→ |
| env-229 | 4491421.3_NODE_4750 | + | 252 | 336 | **RNA**→ rpsJ (PRK00596)RpsJ (COG0051)→ |
| env-230 | 4491477.3_NODE_51506 | + | 30052 | 30136 | **RNA**→ rpsJ (PRK00596)RpsJ (COG0051)→ |
| env-231 | 4491487.3_NODE_4692 | + | 19908 | 19992 | **RNA**→ rpsJ (PRK00596)Ribosomal_S10 (pfam00338)→ |
| env-232 | SRS013951_WUGC_scaffold_4945 | - | 9240 | 9156 | **RNA**→ rpsJ (PRK00596)RpsJ (COG0051)→ |
| env-233 | SRS019161_WUGC_scaffold_49940 | - | 431 | 347 | **RNA**→ rpsJ (PRK00596)RpsJ (COG0051)→ |
| env-234 | scaffold7691_3_MH0009 | + | 6985 | 7069 | **RNA**→ |
| env-235 | scaffold13176_2_O2.UC-16 | - | 193 | 109 | **RNA**→ |
| env-236 | DLM018_scaffold17866_11 | - | 6083 | 5999 | **RNA**→ rpsJ (PRK00596)RpsJ (COG0051)→ |
| env-237 | DLF010_scaffold22271_21 | - | 10166 | 10082 | **RNA**→ rpsJ (PRK00596)RpsJ (COG0051)→ |
| env-238 | NOM012_scaffold41665_2 | + | 16835 | 16919 | **RNA**→ rpsJ (PRK00596)RpsJ (COG0051)→ |
| env-239 | NOM016_scaffold4013_10 | + | 667 | 751 | **RNA**→ rpsJ (PRK00596)Ribosomal_S10 (pfam00338)→ |
| env-240 | NOM017_scaffold8338_6 | - | 739 | 655 | **RNA**→ rpsJ (PRK00596)RpsJ (COG0051)→ |
| env-241 | SRS023914_Baylor_scaffold_6440 | + | 34973 | 35057 | **RNA**→ rpsJ (PRK00596)RpsJ (COG0051)→ |
| env-242 | ASRL01000150.1 | - | 7972 | 7887 | **RNA**→ rpsJ (PRK00596)RpsJ (COG0051)→ RplC (COG0087)rplC (PRK00001)→ rplD (PRK05319)Ribosomal_L4 (pfam00573)→ rplW (PRK05738)Ribosomal_L23 (pfam00276)→ RplB (COG0090)rplB (PRK09374)→ rpsS (PRK00357)RpsS (COG0185)→ rplV (PRK00565)Ribosomal_L22 (pfam00237)→ RpsC (COG0092)rpsC (PRK00310)→ rpl16 (CHL00044)rplP (PRK09203)→ PRK00306 (PRK00306)Ribosomal_L29 (pfam00831)→ rpsQ (PRK05610)Ribosomal_S17 (pfam00366)→ rplN (PRK05483)Ribosomal_L14 (pfam00238)→ rplX (PRK00004)KOW_RPL26 (cd06089)→ rplE (PRK00010)RplE (COG0094)→ rpsN (PRK08061)RpsN (COG0199)→ rpsH (PRK00136)Ribosomal_S8 (pfam00410)→ rpl6 (CHL00140)rplF (PRK05498)→ |
| env-243 | ASRM01003264.1 | - | 570 | 485 | **RNA**→ rpsJ (PRK00596)RpsJ (COG0051)→ |
| env-244 | ASRK01000096.1 | + | 8848 | 8933 | **RNA**→ rpsJ (PRK00596)RpsJ (COG0051)→ RplC (COG0087)rplC (PRK00001)→ rplD (PRK05319)Ribosomal_L4 (pfam00573)→ rplW (PRK05738)Ribosomal_L23 (pfam00276)→ RplB (COG0090)rplB (PRK09374)→ rpsS (PRK00357)RpsS (COG0185)→ rplV (PRK00565)Ribosomal_L22 (pfam00237)→ RpsC (COG0092)rpsC (PRK00310)→ rpl16 (CHL00044)rplP (PRK09203)→ PRK00306 (PRK00306)Ribosomal_L29 (pfam00831)→ rpsQ (PRK05610)Ribosomal_S17 (pfam00366)→ rplN (PRK05483)Ribosomal_L14 (pfam00238)→ rplX (PRK00004)KOW_RPL26 (cd06089)→ rplE (PRK00010)RplE (COG0094)→ rpsN (PRK08061)RpsN (COG0199)→ rpsH (PRK00136)Ribosomal_S8 (pfam00410)→ rpl6 (CHL00140)rplF (PRK05498)→ rplR (PRK05593)Ribosomal_L18p (pfam00861)→ RpsE (COG0098)rpsE (PRK00550)→ |
| env-245 | ASRN01000847.1 | - | 99 | 14 | **RNA**→ |
| env-246 | ASRP01009196.1 | - | 99 | 14 | **RNA**→ |

| | | | | | |
|---|---|---|---|---|---|
| env-247 | NLF011_scaffold23581_7 | - | 12837 | 12749 | **RNA**→ rpsJ (PRK00596)RpsJ (COG0051)→ |
| env-248 | NLM010_scaffold2128_3 | + | 5986 | 6074 | **RNA**→ rpsJ (PRK00596)RpsJ (COG0051)→ |
| env-249 | scaffold18342_7_V1.UC-21 | - | 2390 | 2302 | **RNA**→ rpsJ (PRK00596)RpsJ (COG0051)→ |
| env-250 | NLM029_scaffold31899_2 | + | 10319 | 10407 | **RNA**→ rpsJ (PRK00596)RpsJ (COG0051)→ |
| env-251 | SRS011271_WUGC_scaffold_26754 | + | 5636 | 5724 | **RNA**→ rpsJ (PRK00596)RpsJ (COG0051)→ |
| env-252 | scaffold20388_1_O2.UC-13 | + | 2191 | 2279 | **RNA**→ |
| env-253 | SRS014613_WUGC_scaffold_26039 | - | 294 | 206 | **RNA**→ |
| env-254 | SRS016095_WUGC_scaffold_35198 | - | 490 | 402 | **RNA**→ rpsJ (PRK00596)Ribosomal_S10 (pfam00338)→ |
| env-255 | SRS054956_LANL_scaffold_18901 | - | 202 | 114 | **RNA**→ |
| env-256 | SRS064276_LANL_scaffold_57665 | - | 294 | 206 | **RNA**→ hypo→ |
| env-257 | DLM028_scaffold41246_2 | + | 204 | 292 | **RNA**→ rpsJ (PRK00596)RpsJ (COG0051)→ |
| env-258 | DOF004_scaffold21058_13 | + | 2014 | 2102 | **RNA**→ rpsJ (PRK00596)RpsJ (COG0051)→ |
| env-259 | NOF008_scaffold41486_5 | + | 1075 | 1163 | **RNA**→ rpsJ (PRK00596)RpsJ (COG0051)→ |
| env-260 | NOF012_scaffold7935_4 | + | 3601 | 3689 | **RNA**→ rpsJ (PRK00596)RpsJ (COG0051)→ |
| env-261 | NOM007_scaffold1962_10 | + | 1752 | 1840 | **RNA**→ rpsJ (PRK00596)Ribosomal_S10 (pfam00338)→ |
| env-262 | NOF013_scaffold28708_1 | + | 3005 | 3093 | **RNA**→ rpsJ (PRK00596)RpsJ (COG0051)→ |
| env-263 | NOM010_scaffold39587_3 | - | 29192 | 29104 | **RNA**→ rpsJ (PRK00596)RpsJ (COG0051)→ |
| env-264 | SRS051031_LANL_scaffold_52982 | + | 5566 | 5654 | **RNA**→ rpsJ (PRK00596)RpsJ (COG0051)→ |
| env-265 | scaffold4680_1_V1.CD-14 | - | 1154 | 1066 | **RNA**→ rpsJ (PRK00596)RpsJ (COG0051)→ |
| env-266 | scaffold16707_1_MH0043 | - | 163 | 82 | **RNA**→ |
| env-267 | scaffold992_1_V1.CD-12 | + | 593 | 674 | **RNA**→ |
| env-268 | scaffold8291_2_V1.CD-3 | + | 6188 | 6269 | **RNA**→ |
| env-269 | scaffold15225_3_V1.UC-14 | + | 341 | 422 | **RNA**→ |
| env-270 | SRS022524_Baylor_scaffold_28649 | - | 181 | 100 | **RNA**→ |
| env-271 | SRS014287_WUGC_scaffold_7770 | + | 492 | 573 | **RNA**→ |
| env-272 | SRS015065_WUGC_scaffold_7960 | - | 167 | 86 | **RNA**→ |
| env-273 | SRS015264_WUGC_scaffold_19803 | + | 17439 | 17520 | **RNA**→ rpsJ (PRK00596)Ribosomal_S10 (pfam00338)→ |
| env-274 | SRS017701_Baylor_scaffold_14318 | + | 838 | 919 | **RNA**→ rpsJ (PRK00596)RpsJ (COG0051)→ |
| env-275 | SRS019968_Baylor_scaffold_7511 | - | 182 | 101 | **RNA**→ |
| env-276 | SRS048870_WUGC_scaffold_19339 | - | 180 | 99 | **RNA**→ |
| env-277 | SRS052697_LANL_scaffold_12986 | - | 165 | 84 | **RNA**→ |
| env-278 | SRS056519_LANL_scaffold_13514 | - | 181 | 100 | **RNA**→ |
| env-279 | SRS062427_LANL_scaffold_6177 | - | 181 | 100 | **RNA**→ |
| env-280 | scaffold9779_1_MH0068 | - | 164 | 83 | **RNA**→ |
| env-281 | scaffold17921_1_MH0044 | - | 167 | 86 | **RNA**→ |
| env-282 | 4491413.3_NODE_441 | - | 240 | 159 | **RNA**→ |
| env-283 | scaffold2177_1_V1.CD-9 | + | 18517 | 18598 | **RNA**→ rpsJ (PRK00596)RpsJ (COG0051)→ |
| env-284 | DLM018_scaffold20549_34 | - | 471 | 390 | **RNA**→ rpsJ (PRK00596)RpsJ (COG0051)→ |
| env-285 | NLF001_scaffold2_2 | - | 42911 | 42830 | **RNA**→ rpsJ (PRK00596)RpsJ (COG0051)→ |
| env-286 | NOF005_scaffold12064_3 | + | 6160 | 6241 | **RNA**→ rpsJ (PRK00596)RpsJ (COG0051)→ |
| env-287 | NOM015_scaffold4921_10 | + | 5913 | 5994 | **RNA**→ rpsJ (PRK00596)RpsJ (COG0051)→ |
| env-288 | DOM026_scaffold32809_1 | + | 1451 | 1532 | **RNA**→ rpsJ (PRK00596)RpsJ (COG0051)→ |
| env-289 | NLM029_scaffold10880_16 | + | 1008 | 1089 | **RNA**→ rpsJ (PRK00596)RpsJ (COG0051)→ |
| env-290 | SRS011271_WUGC_scaffold_252 | - | 2128 | 2047 | **RNA**→ RplC (COG0087)rplC (PRK00001)→ rplD (PRK05319)rplW (PRK05738)Ribosomal_L23 (pfam00276)Ribosomal_L4 (pfam00573)→ |
| env-291 | scaffold20849_1_MH0039 | - | 167 | 86 | **RNA**→ |
| env-292 | scaffold13214_3_MH0063 | + | 341 | 422 | **RNA**→ |
| env-293 | scaffold22108_1_V1.CD-8 | - | 167 | 86 | **RNA**→ |
| env-294 | SRS015217_WUGC_scaffold_8594 | - | 165 | 84 | **RNA**→ |
| env-295 | SRS019601_WUGC_scaffold_8958 | - | 167 | 86 | **RNA**→ |
| env-296 | SRS019685_WUGC_scaffold_21426 | - | 181 | 100 | **RNA**→ |
| env-297 | SRS042628_WUGC_scaffold_8232 | + | 494 | 575 | **RNA**→ |

| | | | | | |
|---|---|---|---|---|---|
| env-298 | SRS043701_LANL_scaffold_11145 | - | 181 | 100 | **RNA**→ |
| env-299 | SRS053335_LANL_scaffold_10264 | - | 181 | 100 | **RNA**→ hypo→ |
| env-300 | scaffold3402_4_MH0042 | + | 4470 | 4551 | **RNA**→ |
| env-301 | scaffold5536_2_MH0066 | - | 142 | 61 | **RNA**→ |
| env-302 | SRS017247_C943587 | - | 122 | 41 | **RNA**→ hypo→ |
| env-303 | SRS023526_Baylor_scaffold_5213 | - | 167 | 86 | **RNA**→ |
| env-304 | SRS043411_WUGC_scaffold_5719 | - | 1184 | 1103 | **RNA**→ rpsJ (PRK00596)Ribosomal_S10 (pfam00338)→ hypo→ |
| env-305 | SRS058723_Baylor_scaffold_3070 | - | 335 | 254 | **RNA**→ rpsJ (PRK00596)Ribosomal_S10 (pfam00338)→ |
| env-306 | BABB01001370.1 | - | 4795 | 4714 | **RNA**→ rpsJ (PRK00596)RpsJ (COG0051)→ |
| env-307 | scaffold23001_1_MH0033 | - | 167 | 86 | **RNA**→ |
| env-308 | SRS049164_WUGC_scaffold_27429 | - | 180 | 99 | **RNA**→ |
| env-309 | DLM023_scaffold28443_2 | - | 4447 | 4366 | **RNA**→ rpsJ (PRK00596)RpsJ (COG0051)→ |
| env-310 | NLF007_scaffold22991_5 | - | 331 | 250 | **RNA**→ rpsJ (PRK00596)Ribosomal_S10 (pfam00338)→ |
| env-311 | NOM001_scaffold36725_1 | - | 25726 | 25645 | **RNA**→ rpsJ (PRK00596)RpsJ (COG0051)→ |
| env-312 | ADJS01009405.1 | + | 3228 | 3309 | **RNA**→ rpsJ (PRK00596)RpsJ (COG0051)→ |
| env-313 | SRS015854_WUGC_scaffold_8959 | - | 679 | 598 | **RNA**→ RplC (COG0087)rplC (PRK00001)→ |
| env-314 | SRS024075_LANL_scaffold_4405 | + | 2401 | 2482 | **RNA**→ rpsJ (PRK00596)RpsJ (COG0051)→ |
| env-315 | SRS063040_Baylor_scaffold_7602 | - | 1030 | 949 | **RNA**→ RplC (COG0087)rplC (PRK00001)→ |
| env-316 | scaffold4160_1_MH0021 | - | 165 | 84 | **RNA**→ |
| env-317 | scaffold8722_1_MH0027 | - | 177 | 96 | **RNA**→ |
| env-318 | scaffold17060_5_MH0028 | + | 6414 | 6495 | **RNA**→ |
| env-319 | scaffold8943_6_MH0053 | + | 593 | 674 | **RNA**→ |
| env-320 | scaffold51963_3_MH0057 | + | 6208 | 6289 | **RNA**→ |
| env-321 | scaffold8695_1_MH0084 | - | 165 | 84 | **RNA**→ |
| env-322 | scaffold6602_1_O2.UC-14 | - | 165 | 84 | **RNA**→ |
| env-323 | scaffold5800_2_O2.UC-16 | + | 6220 | 6301 | **RNA**→ |
| env-324 | scaffold14590_1_V1.CD-4 | - | 163 | 82 | **RNA**→ |
| env-325 | scaffold3608_3_V1.UC-7 | + | 6221 | 6302 | **RNA**→ |
| env-326 | SRS015190_WUGC_scaffold_6257 | - | 167 | 86 | **RNA**→ |
| env-327 | SRS015782_WUGC_scaffold_7626 | - | 167 | 86 | **RNA**→ |
| env-328 | SRS042284_WUGC_scaffold_5200 | - | 167 | 86 | **RNA**→ hypo→ |
| env-329 | SRS047044_WUGC_scaffold_9146 | - | 181 | 100 | **RNA**→ hypo→ |
| env-330 | SRS053398_LANL_scaffold_13945 | + | 18555 | 18636 | **RNA**→ rpsJ (PRK00596)Ribosomal_S10 (pfam00338)→ |
| env-331 | scaffold18582_3_MH0040 | - | 322 | 241 | **RNA**→ rpsJ (PRK00596)RpsJ (COG0051)→ |
| env-332 | scaffold13759_1_MH0046 | - | 162 | 81 | **RNA**→ |
| env-333 | scaffold16705_1_O2.UC-22 | - | 165 | 84 | **RNA**→ |
| env-334 | 4491401.3_NODE_53304 | + | 618 | 699 | **RNA**→ rpsJ (PRK00596)Ribosomal_S10 (pfam00338)→ |
| env-335 | 4491417.3_NODE_3394 | + | 16865 | 16946 | **RNA**→ rpsJ (PRK00596)RpsJ (COG0051)→ |
| env-336 | DLF005_scaffold17120_4 | - | 40417 | 40336 | **RNA**→ rpsJ (PRK00596)RpsJ (COG0051)→ |
| env-337 | DLM008_scaffold2771_2 | + | 6173 | 6254 | **RNA**→ rpsJ (PRK00596)RpsJ (COG0051)→ |
| env-338 | DOF003_scaffold17942_4 | + | 360 | 441 | **RNA**→ rpsJ (PRK00596)RpsJ (COG0051)→ |
| env-339 | NLM004_scaffold3966_5 | + | 12804 | 12885 | **RNA**→ rpsJ (PRK00596)RpsJ (COG0051)→ |
| env-340 | NOM005_scaffold3277_7 | - | 7989 | 7908 | **RNA**→ rpsJ (PRK00596)RpsJ (COG0051)→ |
| env-341 | 4491409.3_NODE_22629 | - | 14580 | 14499 | **RNA**→ rpsJ (PRK00596)RpsJ (COG0051)→ |
| env-342 | 4491420.3_NODE_5117 | + | 55138 | 55219 | **RNA**→ rpsJ (PRK00596)RpsJ (COG0051)→ |
| env-343 | DOM017_scaffold32436_1 | - | 804 | 723 | **RNA**→ rpsJ (PRK00596)RpsJ (COG0051)→ |
| env-344 | NLF006_scaffold12057_2 | - | 969 | 888 | **RNA**→ rpsJ (PRK00596)RpsJ (COG0051)→ |
| env-345 | NLM006_scaffold7171_11 | - | 3023 | 2942 | **RNA**→ rpsJ (PRK00596)RpsJ (COG0051)→ |
| env-346 | SRS014683_WUGC_scaffold_18380 | + | 33 | 114 | **RNA**→ rpsJ (PRK00596)RpsJ (COG0051)→ |
| Ere-1-1 | NC_012781.1 | + | 383105 | 383186 | **RNA**→ rpsJ (PRK00596)RpsJ (COG0051)→ |
| env-347 | SRS015578_WUGC_scaffold_8463 | - | 165 | 84 | **RNA**→ |
| env-348 | SRS019161_WUGC_scaffold_21627 | - | 180 | 99 | **RNA**→ hypo→ |

| | | | | | |
|---|---|---|---|---|---|
| env-349 | SRS024132_Baylor_scaffold_5045 | - | 165 | 84 | **RNA**→ |
| env-350 | SRS048164_WUGC_scaffold_7310 | + | 1960 | 2041 | **RNA**→ rpsJ (PRK00596)RpsJ (COG0051)→ |
| env-351 | SRS078176_LANL_scaffold_9297 | - | 182 | 101 | **RNA**→ |
| env-352 | scaffold19886_8_MH0004 | + | 1468 | 1549 | **RNA**→ |
| env-353 | 4491414.3_NODE_10232 | + | 18886 | 18967 | **RNA**→ rpsJ (PRK00596)RpsJ (COG0051)→ |
| env-354 | DLM002_scaffold19292_4 | - | 1385 | 1304 | **RNA**→ rpsJ (PRK00596)RpsJ (COG0051)→ |
| env-355 | DOM013_scaffold53913_3 | - | 42788 | 42707 | **RNA**→ rpsJ (PRK00596)RpsJ (COG0051)→ |
| env-356 | NLM010_scaffold33432_2 | + | 26633 | 26714 | **RNA**→ rpsJ (PRK00596)RpsJ (COG0051)→ |
| env-357 | NLM032_scaffold5645_33 | - | 3767 | 3686 | **RNA**→ rpsJ (PRK00596)RpsJ (COG0051)→ |
| env-358 | NOM009_scaffold4493_3 | - | 42879 | 42798 | **RNA**→ rpsJ (PRK00596)RpsJ (COG0051)→ |
| env-359 | NLM022_scaffold12984_2 | + | 6167 | 6248 | **RNA**→ rpsJ (PRK00596)RpsJ (COG0051)→ |
| env-360 | NOM018_scaffold8271_1 | + | 75 | 156 | **RNA**→ rpsJ (PRK00596)RpsJ (COG0051)→ |
| env-361 | SRS014459_WUGC_scaffold_14109 | + | 10605 | 10686 | **RNA**→ rpsJ (PRK00596)RpsJ (COG0051)→ |
| env-362 | SRS019030_Baylor_scaffold_218 | + | 268 | 349 | **RNA**→ rpsJ (PRK00596)RpsJ (COG0051)→ |
| env-363 | SRS054956_LANL_scaffold_7970 | - | 167 | 86 | **RNA**→ |
| env-364 | BABG01016216.1 | - | 115 | 34 | **RNA**→ |
| env-365 | scaffold3598_2_MH0037 | + | 1682 | 1763 | **RNA**→ rpsJ (PRK00596)Ribosomal_S10 (pfam00338)→ |
| env-366 | DLM027_scaffold716_3 | - | 3549 | 3468 | **RNA**→ rpsJ (PRK00596)RpsJ (COG0051)→ |
| env-367 | NOF010_scaffold33104_25 | - | 4436 | 4355 | **RNA**→ rpsJ (PRK00596)RpsJ (COG0051)→ |
| env-368 | DLM028_scaffold6833_1 | - | 574 | 493 | **RNA**→ rpsJ (PRK00596)RpsJ (COG0051)→ |
| env-369 | NLF011_scaffold1666_1 | - | 295 | 214 | **RNA**→ rpsJ (PRK00596)RpsJ (COG0051)→ |
| env-370 | NLM008_scaffold17066_1 | - | 5551 | 5470 | **RNA**→ rpsJ (PRK00596)RpsJ (COG0051)→ |
| env-371 | NLM028_scaffold4257_4 | - | 40205 | 40124 | **RNA**→ rpsJ (PRK00596)RpsJ (COG0051)→ |
| env-372 | DLF003_scaffold1033_10 | + | 1103 | 1184 | **RNA**→ rpsJ (PRK00596)Ribosomal_S10 (pfam00338)→ |
| env-373 | DLM006_scaffold1766_11 | + | 1075 | 1156 | **RNA**→ rpsJ (PRK00596)RpsJ (COG0051)→ |
| env-374 | NLM021_scaffold5173_1 | - | 5388 | 5307 | **RNA**→ rpsJ (PRK00596)RpsJ (COG0051)→ |
| env-375 | scaffold16666_5_V1.CD-11 | - | 140 | 59 | **RNA**→ |
| env-376 | RUMENNODE_1856948_1 | + | 5532 | 5635 | **RNA**→ rpsJ (PRK00596)RpsJ (COG0051)→ |
| Bsp-5-1 | NZ_KK211308.1 | - | 207215 | 207112 | **RNA**→ rpsJ (PRK00596)RpsJ (COG0051)→ |
| env-377 | SRS017227_Baylor_scaffold_139101 | - | 221 | 145 | **RNA**→ |
| env-378 | SRS042984_LANL_scaffold_21113 | + | 2212 | 2288 | **RNA**→ |
| env-379 | SRS018443_Baylor_scaffold_52281 | - | 249 | 173 | **RNA**→ |
| env-380 | SRS018157_C4833961 | - | 217 | 141 | **RNA**→ |
| env-381 | SRS053917_LANL_scaffold_98590 | + | 169 | 245 | **RNA**→ rpsJ (PRK00596)RpsJ (COG0051)→ RplC (COG0087)rplC (PRK00001)→ |
| env-382 | SRS017139_Baylor_scaffold_54237 | + | 946 | 1022 | **RNA**→ rpsJ (PRK00596)RpsJ (COG0051)→ |
| env-383 | SRS024355_LANL_scaffold_90040 | + | 571 | 647 | **RNA**→ rpsJ (PRK00596)RpsJ (COG0051)→ |
| env-384 | SRS018394_Baylor_scaffold_71324 | + | 824 | 900 | **RNA**→ rpsJ (PRK00596)RpsJ (COG0051)→ RplC (COG0087)rplC (PRK00001)→ rplD (PRK05319)Ribosomal_L4 (pfam00573)→ |
| Lor-1-1 | NZ_GL890598.1 | + | 12752 | 12828 | **RNA**→ rpsJ (PRK00596)RpsJ (COG0051)→ RplC (COG0087)rplC (PRK00001)→ rplD (PRK05319)Ribosomal_L4 (pfam00573)→ rplW (PRK05738)Ribosomal_L23 (pfam00276)→ RplB (COG0090)rplB (PRK09374)→ rpsS (PRK00357)RpsS (COG0185)→ rplV (PRK00565)Ribosomal_L22 (pfam00237)→ RpsC (COG0092)rpsC (PRK00310)→ rpl16 (CHL00044)rplP (PRK09203)→ PRK00306 (PRK00306)Ribosomal_L29 (pfam00831)→ rpsQ (PRK05610)Ribosomal_S17 (pfam00366)→ rplN (PRK05483)Ribosomal_L14 (pfam00238)→ RplX (COG0198)rplX (PRK00004)→ rplE (PRK00010)RplE (COG0094)→ rpsN (PRK08061)RpsN (COG0199)→ hypo→ |
| env-385 | SRS014690_C1764201 | - | 155 | 79 | **RNA**→ |
| env-386 | SRS047634_LANL_scaffold_48885 | + | 6495 | 6571 | **RNA**→ rpsJ (PRK00596)RpsJ (COG0051)→ RplC (COG0087)rplC (PRK00001)→ rplD (PRK05319)Ribosomal_L4 (pfam00573)→ rplW (PRK05738)Ribosomal_L23 (pfam00276)→ RplB (COG0090)rplB (PRK09374)→ rpsS (PRK00357)RpsS (COG0185)→ rplV (PRK00565)Ribosomal_L22 (pfam00237)→ RpsC (COG0092)rpsC (PRK00310)→ rpl16 (CHL00044)rplP (PRK09203)→ PRK00306 (PRK00306)Ribosomal_L29 (pfam00831)→ rpsQ (PRK05610)Ribosomal_S17 (pfam00366)→ rplN (PRK05483)Ribosomal_L14 (pfam00238)→ RplX (COG0198)rplX (PRK00004)→ rplE (PRK00010)RplE (COG0094)→ rpsN (PRK08061)RpsN (COG0199)→ |

| | | | | | |
|---|---|---|---|---|---|
| env-387 | SRS017304_C2019822 | + | 4855 | 4931 | **RNA**→ rpsJ (PRK00596)RpsJ (COG0051)→ RplC (COG0087)rplC (PRK00001)→ rplD (PRK05319)Ribosomal_L4 (pfam00573)→ rplW (PRK05738)Ribosomal_L23 (pfam00276)→ RplB (COG0090)rplB (PRK09374)→ rpsS (PRK00357)RpsS (COG0185)→ rplV (PRK00565)Ribosomal_L22 (pfam00237)→ RpsC (COG0092)rpsC (PRK00310)→ rpl16 (CHL00044)rplP (PRK09203)→ PRK00306 (PRK00306)Ribosomal_L29 (pfam00831)→ rpsQ (PRK05610)Ribosomal_S17 (pfam00366)→ rplN (PRK05483)Ribosomal_L14 (pfam00238)→ RplX (COG0198)rplX (PRK00004)→ rplE (PRK00010)RplE (COG0094)→ rpsN (PRK08061)RpsN (COG0199)→ rpsH (PRK00136)Ribosomal_S8 (pfam00410)→ |
| env-388 | SRS047824_WUGC_scaffold_310 | + | 15740 | 15815 | **RNA**→ |
| Lba-15-1 | NZ_ALJL01000022.1 | + | 187340 | 187415 | **RNA**→ rpsJ (PRK00596)RpsJ (COG0051)→ rpsJ (PRK00596)RpsJ (COG0051)→ RplC (COG0087)rplC (PRK00001)→ RplC (COG0087)rplC (PRK00001)→ rplD (PRK05319)Ribosomal_L4 (pfam00573)→ rplD (PRK05319)Ribosomal_L4 (pfam00573)→ rplW (PRK05738)Ribosomal_L23 (pfam00276)→ rplW (PRK05738)Ribosomal_L23 (pfam00276)→ RplB (COG0090)rplB (PRK09374)→ RplB (COG0090)rplB (PRK09374)→ rpsS (PRK00357)RpsS (COG0185)→ rpsS (PRK00357)RpsS (COG0185)→ rplV (PRK00565)Ribosomal_L22 (pfam00237)→ rplV (PRK00565)Ribosomal_L22 (pfam00237)→ RpsC (COG0092)rpsC (PRK00310)→ RpsC (COG0092)rpsC (PRK00310)→ rpl16 (CHL00044)rplP (PRK09203)→ rpl16 (CHL00044)rplP (PRK09203)→ PRK00306 (PRK00306)Ribosomal_L29 (pfam00831)→ PRK00306 (PRK00306)Ribosomal_L29 (pfam00831)→ rpsQ (PRK05610)Ribosomal_S17 (pfam00366)→ rpsQ (PRK05610)Ribosomal_S17 (pfam00366)→ rplN (PRK05483)Ribosomal_L14 (pfam00238)→ rplN (PRK05483)Ribosomal_L14 (pfam00238)→ RplX (COG0198)rplX (PRK00004)→ RplX (COG0198)rplX (PRK00004)→ rplE (PRK00010)RplE (COG0094)→ rplE (PRK00010)RplE (COG0094)→ rpsN (PRK08061)RpsN (COG0199)→ rpsN (PRK08061)RpsN (COG0199)→ rpsH (PRK00136)Ribosomal_S8 (pfam00410)→ rpsH (PRK00136)Ribosomal_S8 (pfam00410)→ |
| env-389 | SRS063288_C2574221 | + | 262 | 337 | **RNA**→ rpsJ (PRK00596)RpsJ (COG0051)→ |
| env-390 | SRS013879_C2821049 | - | 196 | 121 | **RNA**→ |
| env-391 | SRS022530_LANL_scaffold_1983 | - | 1053 | 978 | **RNA**→ rpsJ (PRK00596)RpsJ (COG0051)→ RplC (COG0087)rplC (PRK00001)→ |
| env-392 | SRS024381_LANL_scaffold_11259 | + | 59 | 134 | **RNA**→ hypo→ rpsJ (PRK00596)RpsJ (COG0051)→ |
| env-393 | SRS055426_C3279857 | + | 722 | 797 | **RNA**→ |
| env-394 | SRS063193_LANL_scaffold_16021 | + | 86 | 161 | **RNA**→ |
| env-395 | SRS013502_C2153529 | + | 97 | 172 | **RNA**→ rpsJ (PRK00596)RpsJ (COG0051)→ |
| env-396 | SRS018969_WUGC_scaffold_55533 | + | 372 | 447 | **RNA**→ |
| env-397 | SRS023352_LANL_scaffold_14387 | - | 2374 | 2299 | **RNA**→ rpsJ (PRK00596)RpsJ (COG0051)→ RplC (COG0087)rplC (PRK00001)→ rplD (PRK05319)Ribosomal_L4 (pfam00573)→ rplW (PRK05738)Ribosomal_L23 (pfam00276)→ RplB (COG0090)rplB (PRK09374)→ |
| env-398 | SRS024580_C3068412 | - | 734 | 659 | **RNA**→ rpsJ (PRK00596)RpsJ (COG0051)→ rpl3 (CHL00143)rplC (PRK00001)→ |
| env-399 | SRS057692_LANL_scaffold_52939 | - | 1542 | 1467 | **RNA**→ rpsJ (PRK00596)RpsJ (COG0051)→ RplC (COG0087)rplC (PRK00001)→ rplD (PRK05319)Ribosomal_L4 (pfam00573)→ |
| env-400 | SRS013164_C2331459 | - | 765 | 690 | **RNA**→ rpsJ (PRK00596)RpsJ (COG0051)→ rpl3 (CHL00143)rplC (PRK00001)→ |
| env-401 | SRS018357_C3826668 | + | 11 | 86 | **RNA**→ hypo→ rpsJ (PRK00596)RpsJ (COG0051)→ |
| Lsp-1-1 | NZ_AZXX01000062.1 | - | 191602 | 191527 | **RNA**→ rpsJ (PRK00596)RpsJ (COG0051)→ RplC (COG0087)rplC (PRK00001)→ rplD (PRK05319)Ribosomal_L4 (pfam00573)→ rplW (PRK05738)Ribosomal_L23 (pfam00276)→ RplB (COG0090)rplB (PRK09374)→ rpsS (PRK00357)RpsS (COG0185)→ rplV (PRK00565)Ribosomal_L22 (pfam00237)→ RpsC (COG0092)rpsC (PRK00310)→ rpl16 (CHL00044)rplP (PRK09203)→ PRK00306 (PRK00306)Ribosomal_L29 (pfam00831)→ rpsQ (PRK05610)Ribosomal_S17 (pfam00366)→ rplN (PRK05483)Ribosomal_L14 (pfam00238)→ RplX (COG0198)rplX (PRK00004)→ rplE (PRK00010)RplE (COG0094)→ rpsN (PRK08061)RpsN (COG0199)→ |
| env-402 | SRS019026_C1857454 | + | 146 | 221 | **RNA**→ |
| env-403 | SRS024277_C2778072 | - | 387 | 312 | **RNA**→ rpsJ (PRK00596)Ribosomal_S10 (pfam00338)→ |
| env-404 | SRS050628_LANL_scaffold_2158 | - | 310 | 235 | **RNA**→ hypo→ rpsJ (PRK00596)Ribosomal_S10 (pfam00338)→ |
| env-405 | SRS057539_LANL_scaffold_17399 | + | 419 | 494 | **RNA**→ rpsJ (PRK00596)RpsJ (COG0051)→ |
| env-406 | SRS013234_C2990282 | - | 236 | 161 | **RNA**→ |

| | | | | | |
|---|---|---|---|---|---|
| Lba-22-1 | NZ_JH591133.1 | + | 12060 | 12135 | **RNA**→ rpsJ (PRK00596)RpsJ (COG0051)→ RplC (COG0087)rplC (PRK00001)→ hypo→ hypo→ rplD (PRK05319)Ribosomal_L4 (pfam00573)→ rplD (PRK05319)Ribosomal_L4 (pfam00573)→ rplW (PRK05738)Ribosomal_L23 (pfam00276)→ RplB (COG0090)rplB (PRK09374)→ rpsS (PRK00357)RpsS (COG0185)→ rplV (PRK00565)Ribosomal_L22 (pfam00237)→ RpsC (COG0092)rpsC (PRK00310)→ rpl16 (CHL00044)rplP (PRK09203)→ PRK00306 (PRK00306)Ribosomal_L29 (pfam00831)→ rpsQ (PRK05610)Ribosomal_S17 (pfam00366)→ rplN (PRK05483)Ribosomal_L14 (pfam00238)→ RplX (COG0198)rplX (PRK00004)→ rplE (PRK00010)RplE (COG0094)→ rpsN (PRK08061)RpsN (COG0199)→ rpsN (PRK08061)RpsN (COG0199)→ hypo→ |
| env-407 | SRS015893_C2284730 | + | 70 | 145 | **RNA**→ rpsJ (PRK00596)RpsJ (COG0051)→ |
| env-408 | SRS020856_C2635639 | + | 618 | 693 | **RNA**→ rpsJ (PRK00596)RpsJ (COG0051)→ RplC (COG0087)rplC (PRK00001)→ |
| env-409 | SRS024318_LANL_scaffold_69413 | + | 130 | 205 | **RNA**→ |
| env-410 | SRS053603_LANL_scaffold_68274 | + | 71 | 146 | **RNA**→ rpsJ (PRK00596)RpsJ (COG0051)→ RplC (COG0087)rplC (PRK00001)→ rplD (PRK05319)Ribosomal_L4 (pfam00573)→ rplW (PRK05738)Ribosomal_L23 (pfam00276)→ RplB (COG0090)rplB (PRK09374)→ |
| Cba-3-1 | NZ_JH815185.1 | - | 1384295 | 1384220 | **RNA**→ rpsJ (PRK00596)RpsJ (COG0051)→ RplC (COG0087)rplC (PRK00001)→ rplD (PRK05319)Ribosomal_L4 (pfam00573)→ rplW (PRK05738)Ribosomal_L23 (pfam00276)→ RplB (COG0090)rplB (PRK09374)→ rpsS (PRK00357)RpsS (COG0185)→ rplV (PRK00565)Ribosomal_L22 (pfam00237)→ RpsC (COG0092)rpsC (PRK00310)→ rpl16 (CHL00044)rplP (PRK09203)→ PRK00306 (PRK00306)Ribosomal_L29 (pfam00831)→ rpsQ (PRK05610)Ribosomal_S17 (pfam00366)→ rplN (PRK05483)Ribosomal_L14 (pfam00238)→ RplX (COG0198)rplX (PRK00004)→ rplE (PRK00010)RplE (COG0094)→ rpsN (PRK08061)RpsN (COG0199)→ |
| env-411 | SRS014684_C3308960 | - | 1490 | 1415 | **RNA**→ rpsJ (PRK00596)RpsJ (COG0051)→ RplC (COG0087)rplC (PRK00001)→ RplD (COG0088)rplD (PRK05319)→ |
| env-412 | SRS019027_C1949428 | + | 180 | 255 | **RNA**→ rpsJ (PRK00596)RpsJ (COG0051)→ |
| env-413 | SRS019045_C2675630 | + | 109 | 184 | **RNA**→ rpsJ (PRK00596)RpsJ (COG0051)→ |
| env-414 | SRS014573_WUGC_scaffold_30532 | - | 229 | 140 | **RNA**→ hypo→ |
| env-415 | SRS019071_C5206406 | + | 1242 | 1331 | **RNA**→ |
| env-416 | SRS018300_Baylor_scaffold_8057 | - | 2973 | 2884 | **RNA**→ rpsJ (PRK00596)RpsJ (COG0051)→ RplC (COG0087)rplC (PRK00001)→ rplD (PRK05319)Ribosomal_L4 (pfam00573)→ rplW (PRK05738)Ribosomal_L23 (pfam00276)→ RplB (COG0090)rplB (PRK09374)→ |
| env-417 | SRS016225_C2883210 | + | 201 | 290 | **RNA**→ |
| env-418 | SRS021496_C2575557 | - | 99 | 10 | **RNA**→ hypo→ |
| env-419 | SRS024441_C2869459 | - | 99 | 10 | **RNA**→ hypo→ |
| env-420 | SRS012279_Baylor_scaffold_34972 | - | 551 | 462 | **RNA**→ rpsJ (PRK00596)RpsJ (COG0051)→ |
| env-421 | SRS015762_C3488345 | - | 99 | 10 | **RNA**→ |
| env-422 | SRS017209_Baylor_scaffold_18507 | - | 18184 | 18095 | **RNA**→ rpsJ (PRK00596)RpsJ (COG0051)→ RplC (COG0087)rplC (PRK00001)→ rplD (PRK05319)Ribosomal_L4 (pfam00573)→ rplW (PRK05738)Ribosomal_L23 (pfam00276)→ RplB (COG0090)rplB (PRK09374)→ rpsS (PRK00357)RpsS (COG0185)→ rplV (PRK00565)Ribosomal_L22 (pfam00237)→ RpsC (COG0092)rpsC (PRK00310)→ rpl16 (CHL00044)rplP (PRK09203)→ |
| env-423 | SRS022077_C2576020 | + | 2624 | 2713 | **RNA**→ |
| env-424 | SRS050669_LANL_scaffold_22401 | + | 2618 | 2707 | **RNA**→ rpsJ (PRK00596)RpsJ (COG0051)→ RplC (COG0087)rplC (PRK00001)→ rplD (PRK05319)Ribosomal_L4 (pfam00573)→ rplW (PRK05738)Ribosomal_L23 (pfam00276)→ RplB (COG0090)rplB (PRK09374)→ |
| env-425 | SRS057205_C3496056 | - | 99 | 10 | **RNA**→ |
| env-426 | SRS057692_LANL_scaffold_28820 | - | 18094 | 18005 | **RNA**→ rpsJ (PRK00596)RpsJ (COG0051)→ RplC (COG0087)rplC (PRK00001)→ rplD (PRK05319)Ribosomal_L4 (pfam00573)→ rplW (PRK05738)Ribosomal_L23 (pfam00276)→ RplB (COG0090)rplB (PRK09374)→ rpsS (PRK00357)RpsS (COG0185)→ rplV (PRK00565)Ribosomal_L22 (pfam00237)→ RpsC (COG0092)rpsC (PRK00310)→ rpl16 (CHL00044)rplP (PRK09203)→ |
| env-427 | SRS015797_C2672144 | + | 11515 | 11604 | **RNA**→ |
| env-428 | SRS014683_WUGC_scaffold_48805 | + | 91 | 186 | **RNA**→ rpsJ (PRK00596)RpsJ (COG0051)→ |
| env-429 | scaffold12439_3_MH0067 | - | 209 | 114 | **RNA**→ |
| env-430 | NLM032_scaffold38249_2 | - | 401 | 306 | **RNA**→ rpsJ (PRK00596)Ribosomal_S10 (pfam00338)→ |

| | | | | | |
|---|---|---|---|---|---|
| env-431 | 4491413.3_NODE_14619 | - | 17813 | 17718 | **RNA**→ rpsJ (PRK00596)RpsJ (COG0051)→ |
| env-432 | SRS011271_C2970480 | + | 148 | 243 | **RNA**→ |
| env-433 | SRS023526_Baylor_scaffold_8422 | - | 1165 | 1070 | **RNA**→ rpsJ (PRK00596)RpsJ (COG0051)→ |
| env-434 | 4491406.3_NODE_11036 | - | 1072 | 977 | **RNA**→ rpsJ (PRK00596)RpsJ (COG0051)→ |
| env-435 | scaffold7196_1_MH0055 | - | 1151 | 1056 | **RNA**→ rpsJ (PRK00596)RpsJ (COG0051)→ |
| env-436 | scaffold9574_3_MH0014 | - | 209 | 114 | **RNA**→ hypo→ |
| env-437 | scaffold46288_2_MH0086 | + | 2014 | 2109 | **RNA**→ |
| env-438 | SRS011302_C3223053 | + | 118 | 213 | **RNA**→ |
| env-439 | SRS014459_WUGC_scaffold_42270 | - | 1165 | 1070 | **RNA**→ rpsJ (PRK00596)RpsJ (COG0051)→ |
| env-440 | SRS015133_C3303308 | - | 273 | 178 | **RNA**→ |
| env-441 | SRS015578_WUGC_scaffold_16888 | - | 1170 | 1075 | **RNA**→ rpsJ (PRK00596)RpsJ (COG0051)→ |
| env-442 | SRS015663_C2978496 | - | 273 | 178 | **RNA**→ |
| env-443 | SRS015782_C3242805 | - | 273 | 178 | **RNA**→ hypo→ |
| env-444 | SRS016018_C1196800 | - | 273 | 178 | **RNA**→ hypo→ |
| env-445 | SRS017103_C2464388 | - | 273 | 178 | **RNA**→ |
| env-446 | SRS018817_C1930195 | + | 96 | 191 | **RNA**→ |
| env-447 | SRS019968_C1951730 | - | 273 | 178 | **RNA**→ |
| env-448 | SRS042628_WUGC_scaffold_32234 | + | 9421 | 9516 | **RNA**→ rpsJ (PRK00596)RpsJ (COG0051)→ |
| env-449 | SRS048164_WUGC_scaffold_13083 | - | 3276 | 3181 | **RNA**→ rpsJ (PRK00596)RpsJ (COG0051)→ |
| env-450 | SRS050925_LANL_scaffold_67534 | + | 108 | 203 | **RNA**→ rpsJ (PRK00596)RpsJ (COG0051)→ |
| env-451 | SRS051031_C2444620 | + | 152 | 247 | **RNA**→ |
| env-452 | SRS053335_LANL_scaffold_29138 | - | 760 | 665 | **RNA**→ rpsJ (PRK00596)RpsJ (COG0051)→ |
| env-453 | SRS056259_LANL_scaffold_47056 | - | 1427 | 1332 | **RNA**→ rpsJ (PRK00596)RpsJ (COG0051)→ |
| env-454 | scaffold31147_1_MH0058 | - | 109 | 14 | **RNA**→ |
| env-455 | SRS015190_WUGC_scaffold_27972 | - | 760 | 665 | **RNA**→ rpsJ (PRK00596)RpsJ (COG0051)→ |
| env-456 | DLM028_scaffold21444_3 | + | 2563 | 2658 | **RNA**→ rpsJ (PRK00596)RpsJ (COG0051)→ |
| env-457 | scaffold5136_1_MH0064 | + | 124 | 219 | **RNA**→ rpsJ (PRK00596)RpsJ (COG0051)→ |
| env-458 | SRS045004_LANL_scaffold_32382 | - | 760 | 665 | **RNA**→ rpsJ (PRK00596)RpsJ (COG0051)→ |
| env-459 | UnmappedStool_Broad_scaffold_717639 | + | 562 | 657 | **RNA**→ |
| env-460 | SRS011586_C3298925 | - | 179 | 84 | **RNA**→ |
| env-461 | SRS019685_C8263105 | - | 253 | 158 | **RNA**→ |
| env-462 | SRS020233_C4054417 | + | 326 | 421 | **RNA**→ |
| env-463 | SRS049712_C15431011 | - | 273 | 178 | **RNA**→ hypo→ |
| env-464 | SRS017521_C2498973 | - | 169 | 74 | **RNA**→ hypo→ |
| env-465 | SRS019910_WUGC_scaffold_74754 | + | 108 | 203 | **RNA**→ rpsJ (PRK00596)RpsJ (COG0051)→ |
| env-466 | SRS053398_LANL_scaffold_39694 | + | 2463 | 2558 | **RNA**→ rpsJ (PRK00596)RpsJ (COG0051)→ |
| env-467 | 4491410.3_NODE_67969 | + | 66 | 161 | **RNA**→ rpsJ (PRK00596)Ribosomal_S10 (pfam00338)→ |
| env-468 | ADJS01016428.1 | + | 3391 | 3486 | **RNA**→ |
| env-469 | SRS016954_C2721358 | - | 273 | 178 | **RNA**→ hypo→ |
| env-470 | scaffold47201_1_O2.UC-22 | - | 1074 | 979 | **RNA**→ rpsJ (PRK00596)RpsJ (COG0051)→ |
| env-471 | scaffold32854_6_V1.UC-21 | + | 2000 | 2095 | **RNA**→ rpsJ (PRK00596)RpsJ (COG0051)→ |
| env-472 | NLM026_scaffold284_3 | - | 861 | 766 | **RNA**→ rpsJ (PRK00596)RpsJ (COG0051)→ |
| env-473 | scaffold3368_2_MH0011 | - | 209 | 114 | **RNA**→ |
| env-474 | scaffold23333_3_MH0069 | - | 209 | 114 | **RNA**→ |
| env-475 | scaffold52193_1_MH0075 | - | 232 | 137 | **RNA**→ |
| env-476 | scaffold40929_1_V1.UC-15 | - | 231 | 136 | **RNA**→ |
| env-477 | scaffold25089_3_V1.UC-18 | + | 908 | 1003 | **RNA**→ |
| env-478 | SRS016335_C5051729 | - | 108 | 13 | **RNA**→ |
| env-479 | SRS052697_LANL_scaffold_44674 | + | 797 | 892 | **RNA**→ |
| env-480 | SRS058770_C1525436 | - | 131 | 36 | **RNA**→ |
| env-481 | SRS065504_C3209247 | - | 273 | 178 | **RNA**→ |
| env-482 | SRS019582_WUGC_scaffold_83695 | + | 77 | 172 | **RNA**→ rpsJ (PRK00596)RpsJ (COG0051)→ |

| | | | | | |
|---|---|---|---|---|---|
| env-483 | SRS043411_WUGC_scaffold_47208 | + | 82 | 177 | **RNA**→ rpsJ (PRK00596)RpsJ (COG0051)→ |
| env-484 | scaffold74289_5_MH0035 | + | 677 | 772 | **RNA**→ |
| env-485 | DLM007_scaffold38_46 | + | 2742 | 2837 | **RNA**→ rpsJ (PRK00596)RpsJ (COG0051)→ |
| env-486 | SRS016095_C2550551 | - | 273 | 178 | **RNA**→ |
| env-487 | scaffold16164_3_MH0028 | + | 501 | 596 | **RNA**→ rpsJ (PRK00596)Ribosomal_S10 (pfam00338)→ |
| env-488 | scaffold14432_2_MH0039 | - | 208 | 113 | **RNA**→ |
| env-489 | scaffold7811_8_MH0072 | + | 11840 | 11919 | **RNA**→ rpsJ (PRK00596)RpsJ (COG0051)→ |
| env-490 | NLF001_scaffold1614_22 | + | 1732 | 1811 | **RNA**→ rpsJ (PRK00596)RpsJ (COG0051)→ |
| env-491 | NLM023_scaffold1764_27 | - | 4212 | 4133 | **RNA**→ rpsJ (PRK00596)RpsJ (COG0051)→ |
| env-492 | NOM009_scaffold20333_9 | + | 194 | 273 | **RNA**→ rpsJ (PRK00596)RpsJ (COG0051)→ |
| env-493 | SRS015578_WUGC_scaffold_19979 | - | 174 | 95 | **RNA**→ |
| env-494 | DLF007_scaffold11184_3 | + | 706 | 785 | **RNA**→ rpsJ (PRK00596)RpsJ (COG0051)→ |
| env-495 | DOM013_C728173_1 | + | 213 | 292 | **RNA**→ rpsJ (PRK00596)RpsJ (COG0051)→ |
| env-496 | SRS016335_WUGC_scaffold_19721 | - | 664 | 585 | **RNA**→ rpsJ (PRK00596)RpsJ (COG0051)→ |
| env-497 | DOM010_scaffold44278_1 | + | 145859 | 145938 | **RNA**→ rpsJ (PRK00596)RpsJ (COG0051)→ |
| env-498 | NLF005_scaffold29876_2 | - | 619 | 540 | **RNA**→ rpsJ (PRK00596)RpsJ (COG0051)→ |
| env-499 | SRS063040_Baylor_scaffold_18425 | - | 174 | 95 | **RNA**→ hypo→ |
| env-500 | DOF013_scaffold497_1 | - | 750 | 671 | **RNA**→ rpsJ (PRK00596)RpsJ (COG0051)→ |
| env-501 | NOM026_scaffold9794_3 | + | 191 | 270 | **RNA**→ rpsJ (PRK00596)RpsJ (COG0051)→ |
| env-502 | UnmappedStool_Broad_scaffold_88209 | - | 174 | 95 | **RNA**→ |
| env-503 | DOF004_scaffold20309_1 | - | 817 | 738 | **RNA**→ rpsJ (PRK00596)RpsJ (COG0051)→ |
| env-504 | DOM015_scaffold13120_2 | + | 30031 | 30110 | **RNA**→ rpsJ (PRK00596)RpsJ (COG0051)→ |
| env-505 | DOM016_scaffold32429_1 | - | 806 | 727 | **RNA**→ rpsJ (PRK00596)RpsJ (COG0051)→ |
| env-506 | NLF012_scaffold128_1 | - | 817 | 738 | **RNA**→ rpsJ (PRK00596)RpsJ (COG0051)→ |
| env-507 | NLM007_scaffold9577_1 | + | 38620 | 38699 | **RNA**→ rpsJ (PRK00596)RpsJ (COG0051)→ |
| env-508 | NLM022_scaffold16572_1 | + | 39462 | 39541 | **RNA**→ rpsJ (PRK00596)RpsJ (COG0051)→ |
| env-509 | NOM004_scaffold7882_14 | + | 3208 | 3287 | **RNA**→ rpsJ (PRK00596)RpsJ (COG0051)→ |
| env-510 | NOM014_scaffold27246_1 | - | 435 | 356 | **RNA**→ rpsJ (PRK00596)RpsJ (COG0051)→ |
| env-511 | NOM020_scaffold37871_1 | - | 399 | 320 | **RNA**→ rpsJ (PRK00596)Ribosomal_S10 (pfam00338)→ |
| env-512 | scaffold1071_13_V1.UC-21 | - | 1569 | 1489 | **RNA**→ RplC (COG0087)rplC (PRK00001)→ rplD (PRK05319)Ribosomal_L4 (pfam00573)→ |
| env-513 | scaffold63164_1_MH0035 | - | 406 | 326 | **RNA**→ rpsJ (PRK00596)Ribosomal_S10 (pfam00338)→ |
| env-514 | scaffold6429_5_MH0064 | - | 1119 | 1039 | **RNA**→ RplC (COG0087)rplC (PRK00001)→ |
| env-515 | scaffold4634_3_V1.UC-17 | - | 1536 | 1456 | **RNA**→ RplC (COG0087)rplC (PRK00001)→ RplD (COG0088)rplD (PRK05319)→ |
| env-516 | SRS014235_WUGC_scaffold_50709 | + | 24455 | 24535 | **RNA**→ rpsJ (PRK00596)RpsJ (COG0051)→ |
| env-517 | scaffold16691_6_MH0060 | - | 1548 | 1468 | **RNA**→ RplC (COG0087)rplC (PRK00001)→ RplD (COG0088)rplD (PRK05319)→ |
| env-518 | scaffold10811_1_MH0002 | - | 1137 | 1057 | **RNA**→ RplC (COG0087)rplC (PRK00001)→ |
| env-519 | scaffold66919_2_MH0012 | + | 21859 | 21939 | **RNA**→ rpsJ (PRK00596)Ribosomal_S10 (pfam00338)→ |
| env-520 | scaffold3200_2_MH0026 | + | 11311 | 11391 | **RNA**→ rpsJ (PRK00596)Ribosomal_S10 (pfam00338)→ |
| env-521 | scaffold3869_3_MH0028 | - | 1569 | 1489 | **RNA**→ RplC (COG0087)rplC (PRK00001)→ rplD (PRK05319)Ribosomal_L4 (pfam00573)→ |
| env-522 | scaffold64021_1_MH0050 | - | 1005 | 925 | **RNA**→ rpsJ (PRK00596)RpsJ (COG0051)→ |
| env-523 | scaffold41845_1_MH0051 | - | 684 | 604 | **RNA**→ rpsJ (PRK00596)RpsJ (COG0051)→ |
| env-524 | scaffold25986_1_MH0083 | + | 7731 | 7811 | **RNA**→ |
| env-525 | scaffold4557_5_O2.UC-24 | - | 329 | 249 | **RNA**→ rpsJ (PRK00596)RpsJ (COG0051)→ |
| env-526 | 4491414.3_NODE_889 | - | 1611 | 1531 | **RNA**→ rpsJ (PRK00596)RpsJ (COG0051)→ |
| env-527 | 4491423.3_NODE_15108 | + | 887 | 967 | **RNA**→ rpsJ (PRK00596)Ribosomal_S10 (pfam00338)→ |
| env-528 | 4491484.3_NODE_8519 | + | 26186 | 26266 | **RNA**→ rpsJ (PRK00596)RpsJ (COG0051)→ |
| env-529 | 4491487.3_NODE_1076 | - | 407 | 327 | **RNA**→ rpsJ (PRK00596)Ribosomal_S10 (pfam00338)→ |
| env-530 | DLF003_scaffold10360_5 | + | 291 | 371 | **RNA**→ |
| env-531 | DLF005_C790551_1 | - | 263 | 183 | **RNA**→ |
| env-532 | DLM028_scaffold47970_1 | + | 28193 | 28273 | **RNA**→ rpsJ (PRK00596)RpsJ (COG0051)→ |
| env-533 | NLF006_scaffold8323_3 | - | 902 | 822 | **RNA**→ rpsJ (PRK00596)RpsJ (COG0051)→ |
| env-534 | NLF007_scaffold181_2 | - | 911 | 831 | **RNA**→ rpsJ (PRK00596)RpsJ (COG0051)→ |

| | | | | | |
|---|---|---|---|---|---|
| env-535 | NLF011_scaffold2872_3 | + | 6086 | 6166 | **RNA**→ rpsJ (PRK00596)RpsJ (COG0051)→ |
| env-536 | NLF013_scaffold25874_1 | + | 29048 | 29128 | **RNA**→ rpsJ (PRK00596)RpsJ (COG0051)→ |
| env-537 | NLM001_scaffold144_5 | - | 903 | 823 | **RNA**→ rpsJ (PRK00596)RpsJ (COG0051)→ |
| env-538 | NLM004_scaffold47788_3 | - | 900 | 820 | **RNA**→ rpsJ (PRK00596)RpsJ (COG0051)→ |
| env-539 | NLM025_scaffold70_10 | - | 912 | 832 | **RNA**→ rpsJ (PRK00596)RpsJ (COG0051)→ |
| env-540 | NLM029_scaffold41578_1 | + | 28952 | 29032 | **RNA**→ rpsJ (PRK00596)RpsJ (COG0051)→ |
| env-541 | NOF004_scaffold164_8 | - | 869 | 789 | **RNA**→ rpsJ (PRK00596)RpsJ (COG0051)→ |
| env-542 | NOF014_scaffold249_4 | - | 49901 | 49821 | **RNA**→ rpsJ (PRK00596)RpsJ (COG0051)→ |
| env-543 | NOM010_scaffold40130_2 | + | 24785 | 24865 | **RNA**→ rpsJ (PRK00596)RpsJ (COG0051)→ |
| env-544 | NOM013_scaffold112_2 | - | 911 | 831 | **RNA**→ rpsJ (PRK00596)RpsJ (COG0051)→ |
| env-545 | NOM019_scaffold39671_3 | + | 26855 | 26935 | **RNA**→ rpsJ (PRK00596)RpsJ (COG0051)→ |
| env-546 | SRS011271_WUGC_scaffold_4629 | - | 677 | 597 | **RNA**→ rpsJ (PRK00596)RpsJ (COG0051)→ |
| env-547 | SRS013521_C1519763 | + | 436 | 516 | **RNA**→ |
| env-548 | SRS014459_WUGC_scaffold_70819 | + | 921 | 1001 | **RNA**→ rpsJ (PRK00596)RpsJ (COG0051)→ |
| env-549 | SRS015065_WUGC_scaffold_12019 | - | 3797 | 3717 | **RNA**→ rpsJ (PRK00596)RpsJ (COG0051)→ |
| env-550 | SRS015782_WUGC_scaffold_23904 | - | 233 | 153 | **RNA**→ |
| env-551 | SRS017307_Baylor_scaffold_21108 | + | 4913 | 4993 | **RNA**→ rpsJ (PRK00596)RpsJ (COG0051)→ |
| env-552 | SRS019582_WUGC_scaffold_43199 | - | 686 | 606 | **RNA**→ rpsJ (PRK00596)RpsJ (COG0051)→ |
| env-553 | SRS019685_WUGC_scaffold_38431 | - | 441 | 361 | **RNA**→ rpsJ (PRK00596)RpsJ (COG0051)→ |
| env-554 | SRS024132_Baylor_scaffold_10696 | - | 378 | 298 | **RNA**→ rpsJ (PRK00596)Ribosomal_S10 (pfam00338)→ |
| env-555 | SRS051031_LANL_scaffold_22555 | + | 19118 | 19198 | **RNA**→ rpsJ (PRK00596)RpsJ (COG0051)→ |
| env-556 | SRS011239_C2445216 | + | 87 | 167 | **RNA**→ hypo→ |
| env-557 | scaffold46369_1_MH0040 | + | 9444 | 9524 | **RNA**→ RplC (COG0087)rplC (PRK00001)→ |
| env-558 | scaffold1151_7_O2.UC-14 | + | 3292 | 3372 | **RNA**→ rpsJ (PRK00596)RpsJ (COG0051)→ |
| env-559 | NLM010_scaffold13542_1 | + | 29117 | 29197 | **RNA**→ rpsJ (PRK00596)RpsJ (COG0051)→ |
| env-560 | SRS016335_C5035038 | + | 205 | 285 | **RNA**→ |
| env-561 | scaffold5557_2_MH0036 | + | 17078 | 17158 | **RNA**→ rpsJ (PRK00596)Ribosomal_S10 (pfam00338)→ |
| env-562 | JGI994J12354_10001975 | + | 15910 | 16006 | **RNA**→ rpsJ (PRK00596)RpsJ (COG0051)→ RplC (COG0087)rplC (PRK00001)→<br>rplD (PRK05319)Ribosomal_L4 (pfam00573)→ rplW (PRK05738)Ribosomal_L23 (pfam00276)→ |
| env-563 | 2227187752 | - | 1209 | 1113 | **RNA**→ rpsJ (PRK00596)RpsJ (COG0051)→ RplC (COG0087)rplC (PRK00001)→ |
| env-564 | JGI994J12354_10004218 | - | 3107 | 3011 | **RNA**→ rpsJ (PRK00596)RpsJ (COG0051)→ RplC (COG0087)rplC (PRK00001)→<br>rplD (PRK05319)Ribosomal_L4 (pfam00573)→ rplW (PRK05738)Ribosomal_L23 (pfam00276)→ |
| env-565 | 2227464663 | - | 1219 | 1123 | **RNA**→ rpsJ (PRK00596)RpsJ (COG0051)→ RplC (COG0087)rplC (PRK00001)→ |
| Bsp-4-1 | NZ_KE384203.1 | - | 447202 | 447101 | **RNA**→ rpsJ (PRK00596)RpsJ (COG0051)→ |
| Bsp-7-1 | NZ_ATWY01000018.1 | - | 18605 | 18504 | **RNA**→ rpsJ (PRK00596)RpsJ (COG0051)→ |
| Bsp-8-1 | NZ_AUIZ01000002.1 | - | 353991 | 353890 | **RNA**→ rpsJ (PRK00596)RpsJ (COG0051)→ |
| Bsp-10-1 | NZ_ATVT01000003.1 | + | 3764 | 3865 | **RNA**→ rpsJ (PRK00596)RpsJ (COG0051)→ |
| env-566 | RUMENNODE_3010558_1 | + | 335 | 436 | **RNA**→ rpsJ (PRK00596)RpsJ (COG0051)→ |
| Bsp-2-1 | NZ_ATVR01000004.1 | + | 28829 | 28932 | **RNA**→ rpsJ (PRK00596)RpsJ (COG0051)→ |
| Bsp-3-1 | NZ_ATVS01000010.1 | + | 72924 | 73025 | **RNA**→ rpsJ (PRK00596)RpsJ (COG0051)→ |
| Bsp-6-1 | NZ_JHXX01000001.1 | - | 1304332 | 1304231 | **RNA**→ rpsJ (PRK00596)RpsJ (COG0051)→ |
| env-567 | 4491415.3_NODE_18881 | + | 4525 | 4614 | **RNA**→ rpsJ (PRK00596)RpsJ (COG0051)→ |
| env-568 | scaffold6327_2_MH0061 | + | 8496 | 8585 | **RNA**→ rpsJ (PRK00596)Ribosomal_S10 (pfam00338)→ |
| env-569 | NLF011_scaffold13765_1 | + | 307 | 396 | **RNA**→ rpsJ (PRK00596)RpsJ (COG0051)→ |
| env-570 | scaffold3636_3_O2.UC-1 | + | 564 | 653 | **RNA**→ rpsJ (PRK00596)Ribosomal_S10 (pfam00338)→ |
| env-571 | 4491409.3_NODE_31461 | - | 26153 | 26064 | **RNA**→ rpsJ (PRK00596)RpsJ (COG0051)→ |
| env-572 | 4491413.3_NODE_8050 | - | 15619 | 15530 | **RNA**→ rpsJ (PRK00596)RpsJ (COG0051)→ |
| env-573 | 4491477.3_NODE_7209 | + | 2699 | 2788 | **RNA**→ rpsJ (PRK00596)RpsJ (COG0051)→ |
| env-574 | NLM026_scaffold55560_5 | + | 459 | 548 | **RNA**→ rpsJ (PRK00596)RpsJ (COG0051)→ |
| env-575 | NOF004_scaffold311_1 | - | 5383 | 5294 | **RNA**→ rpsJ (PRK00596)RpsJ (COG0051)→ |
| env-576 | SRS014459_WUGC_scaffold_12493 | + | 297 | 386 | **RNA**→ rpsJ (PRK00596)RpsJ (COG0051)→ |
| env-577 | SRS017307_Baylor_scaffold_11892 | - | 793 | 704 | **RNA**→ rpsJ (PRK00596)RpsJ (COG0051)→ rplC (PRK00001)→ |
| env-578 | DLF004_scaffold35964_6 | - | 21511 | 21422 | **RNA**→ rpsJ (PRK00596)RpsJ (COG0051)→ |

| | | | | | |
|---|---|---|---|---|---|
| env-579 | DLF010_scaffold23_3 | - | 21633 | 21544 | **RNA**→ rpsJ (PRK00596)RpsJ (COG0051)→ |
| env-580 | DLM028_scaffold54551_3 | + | 250 | 339 | **RNA**→ rpsJ (PRK00596)RpsJ (COG0051)→ |
| env-581 | DLM014_scaffold42817_10 | + | 4633 | 4722 | **RNA**→ rpsJ (PRK00596)RpsJ (COG0051)→ |
| env-582 | DOM015_scaffold8595_1 | + | 338 | 427 | **RNA**→ rpsJ (PRK00596)RpsJ (COG0051)→ |
| env-583 | NLF002_scaffold9596_6 | + | 236 | 325 | **RNA**→ rpsJ (PRK00596)RpsJ (COG0051)→ |
| env-584 | NLF007_scaffold40019_1 | + | 4162 | 4251 | **RNA**→ rpsJ (PRK00596)RpsJ (COG0051)→ |
| env-585 | SRS011405_WUGC_scaffold_20040 | - | 262 | 173 | **RNA**→ |
| env-586 | SRS013951_WUGC_scaffold_50787 | - | 472 | 383 | **RNA**→ rpsJ (PRK00596)RpsJ (COG0051)→ |
| env-587 | SRS023914_Baylor_scaffold_17099 | + | 141 | 230 | **RNA**→ rpsJ (PRK00596)RpsJ (COG0051)→ |
| env-588 | NLM015_scaffold23209_1 | + | 303 | 392 | **RNA**→ rpsJ (PRK00596)RpsJ (COG0051)→ |
| env-589 | scaffold17594_1_MH0026 | - | 1172 | 1083 | **RNA**→ rpsJ (PRK00596)RpsJ (COG0051)→ |
| env-590 | scaffold24884_1_O2.UC-17 | + | 242 | 331 | **RNA**→ rpsJ (PRK00596)RpsJ (COG0051)→ |
| env-591 | 4491404.3_NODE_83831 | + | 1313 | 1402 | **RNA**→ hypo→ |
| env-592 | 4491421.3_NODE_88475 | - | 10256 | 10167 | **RNA**→ rpsJ (PRK00596)RpsJ (COG0051)→ |
| env-593 | 4491484.3_NODE_135 | + | 301 | 390 | **RNA**→ rpsJ (PRK00596)RpsJ (COG0051)→ |
| env-594 | DLF006_scaffold44611_3 | - | 3121 | 3032 | **RNA**→ rpsJ (PRK00596)RpsJ (COG0051)→ |
| env-595 | DLM005_scaffold21393_11 | - | 3900 | 3811 | **RNA**→ rpsJ (PRK00596)RpsJ (COG0051)→ |
| env-596 | DLM023_scaffold10572_11 | - | 2355 | 2266 | **RNA**→ rpsJ (PRK00596)RpsJ (COG0051)→ |
| env-597 | DOM017_scaffold16014_2 | + | 273 | 362 | **RNA**→ rpsJ (PRK00596)RpsJ (COG0051)→ |
| env-598 | NLF005_scaffold12_6 | + | 428 | 517 | **RNA**→ rpsJ (PRK00596)RpsJ (COG0051)→ |
| env-599 | NLM004_scaffold4711_1 | + | 328 | 417 | **RNA**→ rpsJ (PRK00596)RpsJ (COG0051)→ |
| env-600 | SRS013687_Baylor_scaffold_4489 | - | 566 | 477 | **RNA**→ rpsJ (PRK00596)RpsJ (COG0051)→ |
| env-601 | SRS015663_WUGC_scaffold_32361 | - | 261 | 172 | **RNA**→ |
| env-602 | NOM001_scaffold9206_1 | + | 179 | 268 | **RNA**→ rpsJ (PRK00596)Ribosomal_S10 (pfam00338)→ |
| env-603 | MA55A_contig00037 | + | 31742 | 31825 | **RNA**→ rpsJ (PRK00596)RpsJ (COG0051)→ |
| env-604 | SRS015038_C1830604 | - | 183 | 101 | **RNA**→ hypo→ |
| env-605 | SRS024318_LANL_scaffold_48420 | - | 9783 | 9701 | **RNA**→ rpsJ (PRK00596)RpsJ (COG0051)→ RplC (COG0087)rplC (PRK00001)→ rplD (PRK05319)Ribosomal_L4 (pfam00573)→ rplW (PRK05738)Ribosomal_L23 (pfam00276)→ RplB (COG0090)rplB (PRK09374)→ rpsS (PRK00357)RpsS (COG0185)→ rplV (PRK00565)Ribosomal_L22 (pfam00237)→ RpsC (COG0092)rpsC (PRK00310)→ rpl16 (CHL00044)rplP (PRK09203)→ PRK00306 (PRK00306)Ribosomal_L29 (pfam00831)→ rpsQ (PRK05610)Ribosomal_S17 (pfam00366)→ rplN (PRK05483)Ribosomal_L14 (pfam00238)→ RplX (COG0198)rplX (PRK00004)→ rplE (PRK00010)RplE (COG0094)→ rpsN (PRK08061)RpsN (COG0199)→ rpsH (PRK00136)Ribosomal_S8 (pfam00410)→ rplF (PRK05498)Ribosomal_L6 (pfam00347)→ |
| env-606 | SRS019389_C2174016 | + | 64 | 146 | **RNA**→ rpsJ (PRK00596)Ribosomal_S10 (pfam00338)→ |
| env-607 | SRS021496_Baylor_scaffold_61403 | - | 25384 | 25302 | **RNA**→ rpsJ (PRK00596)RpsJ (COG0051)→ RplC (COG0087)rplC (PRK00001)→ rplD (PRK05319)Ribosomal_L4 (pfam00573)→ rplW (PRK05738)Ribosomal_L23 (pfam00276)→ RplB (COG0090)rplB (PRK09374)→ rpsS (PRK00357)RpsS (COG0185)→ rplV (PRK00565)Ribosomal_L22 (pfam00237)→ RpsC (COG0092)rpsC (PRK00310)→ rpl16 (CHL00044)rplP (PRK09203)→ PRK00306 (PRK00306)Ribosomal_L29 (pfam00831)→ rpsQ (PRK05610)Ribosomal_S17 (pfam00366)→ rplN (PRK05483)Ribosomal_L14 (pfam00238)→ RplX (COG0198)rplX (PRK00004)→ rplE (PRK00010)RplE (COG0094)→ rpsN (PRK08061)RpsN (COG0199)→ rpsH (PRK00136)Ribosomal_S8 (pfam00410)→ rpl6 (CHL00140)rplF (PRK05498)→ |
| env-608 | SRS013705_Baylor_scaffold_10956 | - | 13141 | 13059 | **RNA**→ rpsJ (PRK00596)RpsJ (COG0051)→ RplC (COG0087)rplC (PRK00001)→ rplD (PRK05319)Ribosomal_L4 (pfam00573)→ rplW (PRK05738)Ribosomal_L23 (pfam00276)→ RplB (COG0090)rplB (PRK09374)→ rpsS (PRK00357)RpsS (COG0185)→ rplV (PRK00565)Ribosomal_L22 (pfam00237)→ RpsC (COG0092)rpsC (PRK00310)→ rpl16 (CHL00044)rplP (PRK09203)→ PRK00306 (PRK00306)Ribosomal_L29 (pfam00831)→ rpsQ (PRK05610)Ribosomal_S17 (pfam00366)→ rplN (PRK05483)Ribosomal_L14 (pfam00238)→ RplX (COG0198)rplX (PRK00004)→ rplE (PRK00010)RplE (COG0094)→ rpsN (PRK08061)RpsN (COG0199)→ rpsH (PRK00136)Ribosomal_S8 (pfam00410)→ rpl6 (CHL00140)rplF (PRK05498)→ |
| env-609 | SRS023926_Baylor_scaffold_57257 | - | 419 | 337 | **RNA**→ rpsJ (PRK00596)RpsJ (COG0051)→ |

| | | | | | |
|---|---|---|---|---|---|
| env-610 | SRS024081_LANL_scaffold_55960 | - | 2978 | 2896 | **RNA**→ rpsJ (PRK00596)RpsJ (COG0051)→ RplC (COG0087)rplC (PRK00001)→ rplD (PRK05319)Ribosomal_L4 (pfam00573)→ rplW (PRK05738)Ribosomal_L23 (pfam00276)→ RplB (COG0090)rplB (PRK09374)→ |
| env-611 | SRS024355_C4308570 | - | 2975 | 2893 | **RNA**→ rpsJ (PRK00596)RpsJ (COG0051)→ RplC (COG0087)rplC (PRK00001)→ rplD (PRK05319)Ribosomal_L4 (pfam00573)→ rplW (PRK05738)Ribosomal_L23 (pfam00276)→ RplB (COG0090)rplB (PRK09374)→ |
| env-612 | SRS051791_LANL_scaffold_39766 | - | 72601 | 72519 | **RNA**→ rpsJ (PRK00596)RpsJ (COG0051)→ RplC (COG0087)rplC (PRK00001)→ rplD (PRK05319)Ribosomal_L4 (pfam00573)→ rplW (PRK05738)Ribosomal_L23 (pfam00276)→ RplB (COG0090)rplB (PRK09374)→ rpsS (PRK00357)RpsS (COG0185)→ rplV (PRK00565)Ribosomal_L22 (pfam00237)→ RpsC (COG0092)rpsC (PRK00310)→ rpl16 (CHL00044)rplP (PRK09203)→ PRK00306 (PRK00306)Ribosomal_L29 (pfam00831)→ rpsQ (PRK05610)Ribosomal_S17 (pfam00366)→ rplN (PRK05483)Ribosomal_L14 (pfam00238)→ RplX (COG0198)rplX (PRK00004)→ rplE (PRK00010)RplE (COG0094)→ rpsN (PRK08061)RpsN (COG0199)→ rpsH (PRK00136)Ribosomal_S8 (pfam00410)→ rpl6 (CHL00140)rplF (PRK05498)→ |
| env-613 | SRS064774_LANL_scaffold_50591 | - | 419 | 337 | **RNA**→ rpsJ (PRK00596)RpsJ (COG0051)→ |
| env-614 | SRS057539_LANL_scaffold_48920 | - | 2989 | 2907 | **RNA**→ rpsJ (PRK00596)RpsJ (COG0051)→ RplC (COG0087)rplC (PRK00001)→ rplD (PRK05319)Ribosomal_L4 (pfam00573)→ rplW (PRK05738)Ribosomal_L23 (pfam00276)→ RplB (COG0090)rplB (PRK09374)→ |
| env-615 | Buccal_mucosa_LANL_C64980361 | + | 214 | 296 | **RNA**→ |
| env-616 | SRS014573_WUGC_scaffold_39733 | - | 183 | 101 | **RNA**→ |
| env-617 | SRS015272_WUGC_scaffold_20985 | - | 183 | 101 | **RNA**→ |
| env-618 | SRS019219_C2770408 | + | 150 | 232 | **RNA**→ |
| env-619 | SRS042131_C2238632 | - | 183 | 101 | **RNA**→ |
| env-620 | SRS047210_C3290105 | - | 183 | 101 | **RNA**→ |
| env-621 | SRS050244_C4357047 | + | 214 | 296 | **RNA**→ |
| env-622 | SRS063193_C2703828 | + | 165 | 247 | **RNA**→ |
| env-623 | SRS064329_C2690719 | + | 645 | 727 | **RNA**→ |
| env-624 | SRS019607_C2625282 | - | 183 | 101 | **RNA**→ hypo→ |
| env-625 | SRS022143_WUGC_scaffold_66638 | + | 646 | 728 | **RNA**→ rpsJ (PRK00596)RpsJ (COG0051)→ RplC (COG0087)rplC (PRK00001)→ |
| env-626 | Palatine_Tonsils_LANL_C6217726 | - | 183 | 101 | **RNA**→ |
| env-627 | SRS011140_C5100171 | + | 457 | 539 | **RNA**→ |
| env-628 | SRS014470_C3659642 | + | 443 | 525 | **RNA**→ |
| env-629 | SRS017120_Baylor_scaffold_61418 | - | 5322 | 5240 | **RNA**→ rpsJ (PRK00596)RpsJ (COG0051)→ RplC (COG0087)rplC (PRK00001)→ rplD (PRK05319)Ribosomal_L4 (pfam00573)→ rplW (PRK05738)Ribosomal_L23 (pfam00276)→ RplB (COG0090)rplB (PRK09374)→ rpsS (PRK00357)RpsS (COG0185)→ rplV (PRK00565)Ribosomal_L22 (pfam00237)→ RpsC (COG0092)rpsC (PRK00310)→ rpl16 (CHL00044)rplP (PRK09203)→ PRK00306 (PRK00306)Ribosomal_L29 (pfam00831)→ rpsQ (PRK05610)Ribosomal_S17 (pfam00366)→ |
| env-630 | SRS018357_C3905527 | + | 512 | 594 | **RNA**→ |
| env-631 | SRS053854_LANL_scaffold_37572 | + | 22349 | 22431 | **RNA**→ rpsJ (PRK00596)RpsJ (COG0051)→ RplC (COG0087)rplC (PRK00001)→ rplD (PRK05319)Ribosomal_L4 (pfam00573)→ rplW (PRK05738)Ribosomal_L23 (pfam00276)→ RplB (COG0090)rplB (PRK09374)→ rpsS (PRK00357)RpsS (COG0185)→ rplV (PRK00565)Ribosomal_L22 (pfam00237)→ RpsC (COG0092)rpsC (PRK00310)→ rpl16 (CHL00044)rplP (PRK09203)→ PRK00306 (PRK00306)Ribosomal_L29 (pfam00831)→ rpsQ (PRK05610)Ribosomal_S17 (pfam00366)→ rplN (PRK05483)Ribosomal_L14 (pfam00238)→ RplX (COG0198)rplX (PRK00004)→ rplE (PRK00010)RplE (COG0094)→ rpsN (PRK08061)RpsN (COG0199)→ rpsH (PRK00136)Ribosomal_S8 (pfam00410)→ rpl6 (CHL00140)rplF (PRK05498)→ |
| env-632 | SRS013836_Baylor_scaffold_6533 | + | 97 | 179 | **RNA**→ rpsJ (PRK00596)RpsJ (COG0051)→ RplC (COG0087)rplC (PRK00001)→ rplD (PRK05319)Ribosomal_L4 (pfam00573)→ rplW (PRK05738)Ribosomal_L23 (pfam00276)→ RplB (COG0090)rplB (PRK09374)→ rpsS (PRK00357)RpsS (COG0185)→ |
| env-633 | SRS016319_WUGC_scaffold_11309 | - | 183 | 101 | **RNA**→ hypo→ |
| env-634 | SRS019974_Baylor_scaffold_45226 | - | 1407 | 1325 | **RNA**→ rpsJ (PRK00596)RpsJ (COG0051)→ RplC (COG0087)rplC (PRK00001)→ RplD (COG0088)rplD (PRK05319)→ |

| | | | | | |
|---|---|---|---|---|---|
| env-635 | SRS023352_LANL_scaffold_27433 | + | 419 | 501 | **RNA**→ rpsJ (PRK00596)RpsJ (COG0051)→ RplC (COG0087)rplC (PRK00001)→ rplD (PRK05319)Ribosomal_L4 (pfam00573)→ rplW (PRK05738)Ribosomal_L23 (pfam00276)→ RplB (COG0090)rplB (PRK09374)→ rpsS (PRK00357)RpsS (COG0185)→ rplV (PRK00565)Ribosomal_L22 (pfam00237)→ RpsC (COG0092)rpsC (PRK00310)→ rpl16 (CHL00044)rplP (PRK09203)→ PRK00306 (PRK00306)Ribosomal_L29 (pfam00831)→ rpsQ (PRK05610)Ribosomal_S17 (pfam00366)→ rplN (PRK05483)Ribosomal_L14 (pfam00238)→ RplX (COG0198)rplX (PRK00004)→ rplE (PRK00010)RplE (COG0094)→ rpsN (PRK08061)RpsN (COG0199)→ rpsH (PRK00136)Ribosomal_S8 (pfam00410)→ rpl6 (CHL00140)rplF (PRK05498)→ |
| env-636 | SRS024447_LANL_scaffold_44990 | - | 725 | 643 | **RNA**→ rpsJ (PRK00596)RpsJ (COG0051)→ rpl3 (CHL00143)rplC (PRK00001)→ |
| env-637 | SRS014271_WUGC_scaffold_40283 | - | 1462 | 1380 | **RNA**→ rpsJ (PRK00596)RpsJ (COG0051)→ RplC (COG0087)rplC (PRK00001)→ RplD (COG0088)rplD (PRK05319)→ |
| env-638 | SRS044486_C2103388 | - | 215 | 133 | **RNA**→ hypo→ |
| env-639 | SRS062544_LANL_scaffold_1604 | - | 9172 | 9090 | **RNA**→ rpsJ (PRK00596)RpsJ (COG0051)→ RplC (COG0087)rplC (PRK00001)→ rplD (PRK05319)Ribosomal_L4 (pfam00573)→ rplW (PRK05738)Ribosomal_L23 (pfam00276)→ RplB (COG0090)rplB (PRK09374)→ rpsS (PRK00357)RpsS (COG0185)→ rplV (PRK00565)Ribosomal_L22 (pfam00237)→ RpsC (COG0092)rpsC (PRK00310)→ rpl16 (CHL00044)rplP (PRK09203)→ PRK00306 (PRK00306)Ribosomal_L29 (pfam00831)→ rpsQ (PRK05610)Ribosomal_S17 (pfam00366)→ rplN (PRK05483)Ribosomal_L14 (pfam00238)→ RplX (COG0198)rplX (PRK00004)→ rplE (PRK00010)RplE (COG0094)→ rpsN (PRK08061)RpsN (COG0199)→ rpsH (PRK00136)Ribosomal_S8 (pfam00410)→ rpl6 (CHL00140)rplF (PRK05498)→ |
| env-640 | SRS016740_Baylor_scaffold_57697 | - | 4537 | 4455 | **RNA**→ rpsJ (PRK00596)RpsJ (COG0051)→ RplC (COG0087)rplC (PRK00001)→ rplD (PRK05319)Ribosomal_L4 (pfam00573)→ rplW (PRK05738)Ribosomal_L23 (pfam00276)→ RplB (COG0090)rplB (PRK09374)→ rpsS (PRK00357)RpsS (COG0185)→ rplV (PRK00565)Ribosomal_L22 (pfam00237)→ RpsC (COG0092)rpsC (PRK00310)→ |
| env-641 | SRS015941_WUGC_scaffold_60199 | + | 239 | 321 | **RNA**→ rpsJ (PRK00596)RpsJ (COG0051)→ RplC (COG0087)rplC (PRK00001)→ rplD (PRK05319)Ribosomal_L4 (pfam00573)→ rplW (PRK05738)Ribosomal_L23 (pfam00276)→ RplB (COG0090)rplB (PRK09374)→ rpsS (PRK00357)RpsS (COG0185)→ rplV (PRK00565)Ribosomal_L22 (pfam00237)→ RpsC (COG0092)rpsC (PRK00310)→ rpl16 (CHL00044)rplP (PRK09203)→ PRK00306 (PRK00306)Ribosomal_L29 (pfam00831)→ rpsQ (PRK05610)Ribosomal_S17 (pfam00366)→ rplN (PRK05483)Ribosomal_L14 (pfam00238)→ RplX (COG0198)rplX (PRK00004)→ rplE (PRK00010)RplE (COG0094)→ rpsN (PRK08061)RpsN (COG0199)→ rpsH (PRK00136)Ribosomal_S8 (pfam00410)→ rplF (PRK05498)Ribosomal_L6 (pfam00347)→ |
| env-642 | SRS020220_Baylor_scaffold_40425 | - | 4359 | 4277 | **RNA**→ rpsJ (PRK00596)RpsJ (COG0051)→ RplC (COG0087)rplC (PRK00001)→ rplD (PRK05319)Ribosomal_L4 (pfam00573)→ rplW (PRK05738)Ribosomal_L23 (pfam00276)→ RplB (COG0090)rplB (PRK09374)→ rpsS (PRK00357)RpsS (COG0185)→ rplV (PRK00565)Ribosomal_L22 (pfam00237)→ RpsC (COG0092)rpsC (PRK00310)→ |
| env-643 | SRS047219_WUGC_scaffold_10994 | - | 4404 | 4322 | **RNA**→ rpsJ (PRK00596)RpsJ (COG0051)→ RplC (COG0087)rplC (PRK00001)→ rplD (PRK05319)Ribosomal_L4 (pfam00573)→ rplW (PRK05738)Ribosomal_L23 (pfam00276)→ RplB (COG0090)rplB (PRK09374)→ rpsS (PRK00357)RpsS (COG0185)→ rplV (PRK00565)Ribosomal_L22 (pfam00237)→ RpsC (COG0092)rpsC (PRK00310)→ |
| env-644 | SRS017076_Baylor_scaffold_27856 | + | 239 | 321 | **RNA**→ rpsJ (PRK00596)RpsJ (COG0051)→ RplC (COG0087)rplC (PRK00001)→ rplD (PRK05319)Ribosomal_L4 (pfam00573)→ rplW (PRK05738)Ribosomal_L23 (pfam00276)→ RplB (COG0090)rplB (PRK09374)→ rpsS (PRK00357)RpsS (COG0185)→ hypo→ |
| env-645 | SRS017080_C1208674 | + | 239 | 321 | **RNA**→ rpsJ (PRK00596)RpsJ (COG0051)→ |
| Jig-1-1 | NZ_JH378829.1 | - | 294163 | 294077 | **RNA**→ rpsJ (PRK00596)RpsJ (COG0051)→ RplC (COG0087)rplC (PRK00001)→ rplD (PRK05319)Ribosomal_L4 (pfam00573)→ rplW (PRK05738)Ribosomal_L23 (pfam00276)→ |
| env-646 | DLF013_scaffold30742_5 | + | 295 | 384 | **RNA**→ rpsJ (PRK00596)Ribosomal_S10 (pfam00338)→ |
| env-647 | ADJS01008642.1 | - | 6451 | 6362 | **RNA**→ rpsJ (PRK00596)RpsJ (COG0051)→ |
| Eha-1-1 | NZ_ACEP01000080.1 | + | 41279 | 41368 | **RNA**→ rpsJ (PRK00596)RpsJ (COG0051)→ rpsJ (PRK00596)RpsJ (COG0051)→ |
| env-648 | SRS017521_Baylor_scaffold_8153 | - | 190 | 101 | **RNA**→ hypo→ |
| env-649 | DLM013_scaffold7335_7 | - | 2319 | 2230 | **RNA**→ rpsJ (PRK00596)RpsJ (COG0051)→ |
| env-650 | scaffold2225_2_MH0044 | - | 1084 | 1004 | **RNA**→ rpsJ (PRK00596)RpsJ (COG0051)→ |
| env-651 | NLM026_scaffold21820_1 | + | 21714 | 21794 | **RNA**→ rpsJ (PRK00596)RpsJ (COG0051)→ |
| env-652 | NOF008_scaffold24903_1 | + | 21716 | 21796 | **RNA**→ rpsJ (PRK00596)RpsJ (COG0051)→ |
| env-653 | DLF003_scaffold30405_2 | + | 3989 | 4069 | **RNA**→ rpsJ (PRK00596)RpsJ (COG0051)→ |

| | | | | | |
|---|---|---|---|---|---|
| env-654 | SRS022609_Baylor_scaffold_47809 | + | 2933 | 3028 | **RNA**→ rpsJ (PRK00596)Ribosomal_S10 (pfam00338)→ |
| env-655 | scaffold48720_1_MH0045 | - | 185 | 90 | **RNA**→ |
| env-656 | DLM014_scaffold11994_1 | - | 368 | 273 | **RNA**→ rpsJ (PRK00596)Ribosomal_S10 (pfam00338)→ |
| env-657 | SRS052697_LANL_scaffold_39196 | - | 189 | 94 | **RNA**→ |
| env-658 | DLF005_scaffold27951_2 | - | 1013 | 918 | **RNA**→ rpsJ (PRK00596)RpsJ (COG0051)→ |
| env-659 | DOM024_scaffold33725_3 | + | 258 | 353 | **RNA**→ rpsJ (PRK00596)RpsJ (COG0051)→ |
| env-660 | NLF013_scaffold5635_8 | + | 41902 | 41997 | **RNA**→ rpsJ (PRK00596)RpsJ (COG0051)→ |
| env-661 | NLM029_scaffold14839_9 | + | 4602 | 4697 | **RNA**→ rpsJ (PRK00596)RpsJ (COG0051)→ |
| env-662 | NOM005_scaffold28840_1 | - | 381 | 286 | **RNA**→ rpsJ (PRK00596)Ribosomal_S10 (pfam00338)→ |
| env-663 | SRS063985_LANL_scaffold_18654 | - | 187 | 92 | **RNA**→ |
| env-664 | scaffold9404_4_V1.CD-11 | + | 127 | 222 | **RNA**→ rpsJ (PRK00596)Ribosomal_S10 (pfam00338)→ |
| env-665 | scaffold53060_1_V1.UC-6 | - | 185 | 90 | **RNA**→ |
| env-666 | NOF008_scaffold7151_1 | - | 255 | 160 | **RNA**→ |
| env-667 | SRS015578_WUGC_scaffold_23407 | - | 187 | 92 | **RNA**→ |
| env-668 | NLF005_scaffold32939_3 | - | 346 | 251 | **RNA**→ rpsJ (PRK00596)Ribosomal_S10 (pfam00338)→ |
| Lba-14-1 | NZ_KE159563.1 | - | 23368 | 23278 | **RNA**→ rpsJ (PRK00596)RpsJ (COG0051)→ RplC (COG0087)rplC (PRK00001)→ |
| | | | | | rplD (PRK05319)Ribosomal_L4 (pfam00573)→ rplW (PRK05738)Ribosomal_L23 (pfam00276)→ |
| | | | | | RplB (COG0090)rplB (PRK09374)→ rpsS (PRK00357)RpsS (COG0185)→ |
| | | | | | rplV (PRK00565)Ribosomal_L22 (pfam00237)→ RpsC (COG0092)rpsC (PRK00310)→ |
| | | | | | rpl16 (CHL00044)rplP (PRK09203)→ PRK00306 (PRK00306)Ribosomal_L29 (pfam00831)→ |
| | | | | | rpsQ (PRK05610)Ribosomal_S17 (pfam00366)→ rplN (PRK05483)Ribosomal_L14 (pfam00238)→ |
| | | | | | RplX (COG0198)rplX (PRK00004)→ rplE (PRK00010)RplE (COG0094)→ rpsN (PRK08061)RpsN (COG0199)→ |
| Bsp-1-1 | NZ_ATVV01000009.1 | - | 54629 | 54531 | **RNA**→ rpsJ (PRK00596)RpsJ (COG0051)→ |
| Bsp-9-1 | NZ_KE384200.1 | + | 18003 | 18101 | **RNA**→ rpsJ (PRK00596)RpsJ (COG0051)→ |
| env-669 | DLM009_scaffold18031_8 | + | 20467 | 20550 | **RNA**→ rpsJ (PRK00596)RpsJ (COG0051)→ |
| env-670 | NOM005_scaffold39343_1 | - | 364 | 281 | **RNA**→ rpsJ (PRK00596)Ribosomal_S10 (pfam00338)→ |
| env-671 | 4491417.3_NODE_55002 | - | 8825 | 8742 | **RNA**→ rpsJ (PRK00596)RpsJ (COG0051)→ |
| env-672 | 4491410.3_NODE_36037 | - | 7413 | 7330 | **RNA**→ rpsJ (PRK00596)RpsJ (COG0051)→ |
| env-673 | DLF014_scaffold5097_1 | - | 44435 | 44352 | **RNA**→ rpsJ (PRK00596)RpsJ (COG0051)→ |
| env-674 | DLM016_scaffold2775_3 | + | 1842 | 1925 | **RNA**→ rpsJ (PRK00596)RpsJ (COG0051)→ |
| env-675 | DLM018_scaffold25796_1 | + | 353 | 436 | **RNA**→ rpsJ (PRK00596)Ribosomal_S10 (pfam00338)→ |
| env-676 | DOF008_scaffold2592_2 | - | 581 | 498 | **RNA**→ rpsJ (PRK00596)RpsJ (COG0051)→ |
| env-677 | DOF012_scaffold5047_1 | - | 2352 | 2269 | **RNA**→ rpsJ (PRK00596)RpsJ (COG0051)→ |
| env-678 | DOM013_scaffold30571_6 | - | 470 | 387 | **RNA**→ rpsJ (PRK00596)RpsJ (COG0051)→ |
| env-679 | DOM026_scaffold10557_9 | + | 388 | 471 | **RNA**→ rpsJ (PRK00596)RpsJ (COG0051)→ |
| env-680 | NLM032_scaffold1492_2 | - | 20482 | 20399 | **RNA**→ rpsJ (PRK00596)RpsJ (COG0051)→ |
| env-681 | DOM018_scaffold445_3 | + | 1261 | 1344 | **RNA**→ rpsJ (PRK00596)RpsJ (COG0051)→ |
| env-682 | NOM009_scaffold4295_7 | + | 387 | 470 | **RNA**→ rpsJ (PRK00596)RpsJ (COG0051)→ |
| env-683 | 4491413.3_NODE_68514 | - | 29126 | 29043 | **RNA**→ rpsJ (PRK00596)RpsJ (COG0051)→ |
| env-684 | scaffold18238_14_MH0043 | + | 1123 | 1206 | **RNA**→ |
| env-685 | UnmappedStool_Broad_scaffold_262642 | - | 940 | 857 | **RNA**→ ←hypo |
| env-686 | DOF008_scaffold62_1 | - | 1739 | 1645 | **RNA**→ rpsJ (PRK00596)RpsJ (COG0051)→ RplC (COG0087)rplC (PRK00001)→ |
| | | | | | rplD (PRK05319)Ribosomal_L4 (pfam00573)→ |
| env-687 | DLM027_scaffold22199_1 | - | 5689 | 5595 | **RNA**→ rpsJ (PRK00596)RpsJ (COG0051)→ RplC (COG0087)rplC (PRK00001)→ |
| | | | | | rplD (PRK05319)Ribosomal_L4 (pfam00573)→ rplW (PRK05738)Ribosomal_L23 (pfam00276)→ |
| | | | | | RplB (COG0090)rplB (PRK09374)→ rpsS (PRK00357)Ribosomal_S19 (pfam00203)→ |
| | | | | | rplV (PRK00565)Ribosomal_L22 (pfam00237)→ RpsC (COG0092)rpsC (PRK00310)→ |
| | | | | | rpl16 (CHL00044)rplP (PRK09203)→ PRK00306 (PRK00306)Ribosomal_L29 (pfam00831)→ |
| | | | | | rpsQ (PRK05610)Ribosomal_S17 (pfam00366)→ rplN (PRK05483)Ribosomal_L14 (pfam00238)→ |

| | | | | | |
|---|---|---|---|---|---|
| env-688 | NLM026_scaffold16599_8 | + | 3734 | 3828 | **RNA**→ rpsJ (PRK00596)RpsJ (COG0051)→ RplC (COG0087)rplC (PRK00001)→<br>rplD (PRK05319)Ribosomal_L4 (pfam00573)→ rplW (PRK05738)Ribosomal_L23 (pfam00276)→<br>RplB (COG0090)rplB (PRK09374)→ rpsS (PRK00357)Ribosomal_S19 (pfam00203)→<br>rplV (PRK00565)Ribosomal_L22 (pfam00237)→ RpsC (COG0092)rpsC (PRK00310)→<br>rpl16 (CHL00044)rplP (PRK09203)→ PRK00306 (PRK00306)Ribosomal_L29 (pfam00831)→<br>rpsQ (PRK05610)Ribosomal_S17 (pfam00366)→ rplN (PRK05483)Ribosomal_L14 (pfam00238)→<br>RplX (COG0198)rplX (PRK00004)→ rplE (PRK00010)RplE (COG0094)→ |
| env-689 | NLM028_scaffold65_1 | - | 197 | 103 | **RNA**→ |
| env-690 | NOF010_scaffold14875_3 | - | 4414 | 4320 | **RNA**→ rpsJ (PRK00596)RpsJ (COG0051)→ RplC (COG0087)rplC (PRK00001)→<br>rplD (PRK05319)Ribosomal_L4 (pfam00573)→ rplW (PRK05738)Ribosomal_L23 (pfam00276)→<br>RplB (COG0090)rplB (PRK09374)→ rpsS (PRK00357)Ribosomal_S19 (pfam00203)→<br>rplV (PRK00565)Ribosomal_L22 (pfam00237)→ RpsC (COG0092)rpsC (PRK00310)→ |
| env-691 | SRS019161_WUGC_scaffold_28654 | - | 219 | 125 | **RNA**→ |
| env-692 | DOM003_scaffold5836_16 | + | 259 | 353 | **RNA**→ rpsJ (PRK00596)RpsJ (COG0051)→ |
| env-693 | NOM001_scaffold34192_1 | - | 968 | 874 | **RNA**→ rpsJ (PRK00596)RpsJ (COG0051)→ RplC (COG0087)rplC (PRK00001)→ |
| env-694 | longitudinal_441_49 | + | 131726 | 131820 | **RNA**→ rpsJ (PRK00596)RpsJ (COG0051)→ RplC (COG0087)rplC (PRK00001)→<br>rplD (PRK05319)Ribosomal_L4 (pfam00573)→ rplW (PRK05738)Ribosomal_L23 (pfam00276)→<br>RplB (COG0090)rplB (PRK09374)→ rpsS (PRK00357)RpsS (COG0185)→<br>rplV (PRK00565)Ribosomal_L22 (pfam00237)→ RpsC (COG0092)rpsC (PRK00310)→<br>rpl16 (CHL00044)rplP (PRK09203)→ PRK00306 (PRK00306)Ribosomal_L29 (pfam00831)→<br>rpsQ (PRK05610)Ribosomal_S17 (pfam00366)→ rplN (PRK05483)Ribosomal_L14 (pfam00238)→<br>RplX (COG0198)rplX (PRK00004)→ rplE (PRK00010)RplE (COG0094)→ rpsN (PRK08061)RpsN (COG0199)→<br>rpsH (PRK00136)Ribosomal_S8 (pfam00410)→ |
| env-695 | DOF009_scaffold11200_9 | + | 419 | 513 | **RNA**→ rpsJ (PRK00596)RpsJ (COG0051)→ |
| env-696 | longitudinal_21_17 | - | 22273 | 22179 | **RNA**→ rpsJ (PRK00596)RpsJ (COG0051)→ RplC (COG0087)rplC (PRK00001)→<br>rplD (PRK05319)Ribosomal_L4 (pfam00573)→ rplW (PRK05738)Ribosomal_L23 (pfam00276)→<br>RplB (COG0090)rplB (PRK09374)→ rpsS (PRK00357)RpsS (COG0185)→<br>rplV (PRK00565)Ribosomal_L22 (pfam00237)→ RpsC (COG0092)rpsC (PRK00310)→<br>rpl16 (CHL00044)rplP (PRK09203)→ PRK00306 (PRK00306)Ribosomal_L29 (pfam00831)→<br>rpsQ (PRK05610)Ribosomal_S17 (pfam00366)→ rplN (PRK05483)Ribosomal_L14 (pfam00238)→<br>RplX (COG0198)rplX (PRK00004)→ rplE (PRK00010)RplE (COG0094)→ rpsN (PRK08061)RpsN (COG0199)→<br>rpsH (PRK00136)Ribosomal_S8 (pfam00410)→ |
| env-697 | longitudinal_22_137 | - | 2800 | 2706 | **RNA**→ rpsJ (PRK00596)RpsJ (COG0051)→ RplC (COG0087)rplC (PRK00001)→<br>rplD (PRK05319)Ribosomal_L4 (pfam00573)→ rplW (PRK05738)Ribosomal_L23 (pfam00276)→<br>rplB (PRK09374)Ribosomal_L2 (pfam00181)→ |
| env-698 | longitudinal_23_385 | - | 119360 | 119266 | **RNA**→ rpsJ (PRK00596)RpsJ (COG0051)→ RplC (COG0087)rplC (PRK00001)→<br>rplD (PRK05319)Ribosomal_L4 (pfam00573)→ rplW (PRK05738)Ribosomal_L23 (pfam00276)→<br>RplB (COG0090)rplB (PRK09374)→ rpsS (PRK00357)RpsS (COG0185)→<br>rplV (PRK00565)Ribosomal_L22 (pfam00237)→ RpsC (COG0092)rpsC (PRK00310)→<br>rpl16 (CHL00044)rplP (PRK09203)→ PRK00306 (PRK00306)Ribosomal_L29 (pfam00831)→<br>rpsQ (PRK05610)Ribosomal_S17 (pfam00366)→ rplN (PRK05483)Ribosomal_L14 (pfam00238)→<br>RplX (COG0198)rplX (PRK00004)→ rplE (PRK00010)RplE (COG0094)→ rpsN (PRK08061)RpsN (COG0199)→<br>rpsH (PRK00136)Ribosomal_S8 (pfam00410)→ |
| env-699 | NOF014_scaffold54976_9 | + | 413 | 507 | **RNA**→ rpsJ (PRK00596)RpsJ (COG0051)→ RplC (COG0087)rplC (PRK00001)→<br>rplD (PRK05319)Ribosomal_L4 (pfam00573)→ rplW (PRK05738)Ribosomal_L23 (pfam00276)→<br>RplB (COG0090)rplB (PRK09374)→ rpsS (PRK00357)RpsS (COG0185)→<br>rplV (PRK00565)Ribosomal_L22 (pfam00237)→ RpsC (COG0092)rpsC (PRK00310)→<br>rpl16 (CHL00044)rplP (PRK09203)→ |
| env-700 | scaffold46017_3_MH0080 | - | 728 | 647 | **RNA**→ rpsJ (PRK00596)RpsJ (COG0051)→ |
| env-701 | scaffold248_6_O2.UC-4 | + | 3853 | 3934 | **RNA**→ rpsJ (PRK00596)Ribosomal_S10 (pfam00338)→ |
| env-702 | SRS014923_C3294507 | - | 94 | 13 | **RNA**→ |
| env-703 | SRS015854_WUGC_scaffold_39634 | - | 93 | 12 | **RNA**→ |
| env-704 | SRS050925_LANL_scaffold_33465 | + | 1412 | 1493 | **RNA**→ rpsJ (PRK00596)RpsJ (COG0051)→ |
| Ceu-1-1 | NZ_DS483526.1 | + | 154917 | 154998 | **RNA**→ rpsJ (PRK00596)RpsJ (COG0051)→ |
| env-705 | scaffold4547_6_MH0003 | + | 12959 | 13044 | **RNA**→ rpsJ (PRK00596)RpsJ (COG0051)→ |

| | | | | | |
|---|---|---|---|---|---|
| env-706 | scaffold24806_1_MH0065 | - | 95 | 10 | **RNA**→ |
| env-707 | scaffold15202_12_MH0075 | + | 1002 | 1087 | **RNA**→ rpsJ (PRK00596)Ribosomal_S10 (pfam00338)→ |
| env-708 | NLM004_scaffold50332_5 | + | 702 | 787 | **RNA**→ rpsJ (PRK00596)RpsJ (COG0051)→ |
| env-709 | scaffold7483_10_MH0050 | + | 16390 | 16475 | **RNA**→ rpsJ (PRK00596)RpsJ (COG0051)→ |
| env-710 | scaffold3174_3_MH0086 | - | 410 | 325 | **RNA**→ rpsJ (PRK00596)Ribosomal_S10 (pfam00338)→ |
| env-711 | SRS052697_C3345764 | - | 98 | 13 | **RNA**→ |
| env-712 | scaffold516_22_MH0034 | + | 1362 | 1447 | **RNA**→ rpsJ (PRK00596)RpsJ (COG0051)→ |
| env-713 | scaffold16883_4_MH0015 | - | 338 | 253 | **RNA**→ rpsJ (PRK00596)Ribosomal_S10 (pfam00338)→ |
| env-714 | scaffold3437_5_MH0042 | + | 12155 | 12240 | **RNA**→ |
| env-715 | scaffold32152_1_MH0052 | + | 799 | 884 | **RNA**→ hypo→ |
| env-716 | scaffold22830_6_MH0070 | + | 761 | 846 | **RNA**→ rpsJ (PRK00596)RpsJ (COG0051)→ |
| env-717 | scaffold48922_1_V1.CD-8 | - | 2029 | 1944 | **RNA**→ rpsJ (PRK00596)RpsJ (COG0051)→ |
| env-718 | 4491403.3_NODE_590 | - | 17129 | 17044 | **RNA**→ rpsJ (PRK00596)RpsJ (COG0051)→ |
| env-719 | SRS049164_C2339078 | + | 236 | 321 | **RNA**→ hypo→ |
| env-720 | NOM013_scaffold4188_8 | + | 111 | 196 | **RNA**→ rpsJ (PRK00596)RpsJ (COG0051)→ |
| env-721 | SRS012273_Baylor_scaffold_27311 | - | 447 | 362 | **RNA**→ rpsJ (PRK00596)RpsJ (COG0051)→ |
| env-722 | SRS014459_WUGC_scaffold_9463 | - | 179 | 94 | **RNA**→ |
| env-723 | SRS015065_WUGC_scaffold_19845 | - | 225 | 140 | **RNA**→ |
| env-724 | SRS047014_WUGC_scaffold_16737 | + | 1370 | 1455 | **RNA**→ |
| env-725 | SRS063040_Baylor_scaffold_3708 | - | 1563 | 1478 | **RNA**→ rpsJ (PRK00596)RpsJ (COG0051)→ |
| env-726 | scaffold19191_4_V1.UC-19 | - | 1168 | 1083 | **RNA**→ rpsJ (PRK00596)RpsJ (COG0051)→ |
| env-727 | SRS017103_C2720120 | + | 2366 | 2451 | **RNA**→ |
| env-728 | SRS024331_LANL_scaffold_46092 | - | 97 | 12 | **RNA**→ |
| env-729 | scaffold4163_12_MH0063 | + | 2063 | 2148 | **RNA**→ rpsJ (PRK00596)RpsJ (COG0051)→ |
| env-730 | DLM002_scaffold32671_6 | + | 53527 | 53612 | **RNA**→ rpsJ (PRK00596)RpsJ (COG0051)→ |
| env-731 | scaffold492_4_MH0046 | + | 3087 | 3172 | **RNA**→ rpsJ (PRK00596)RpsJ (COG0051)→ |
| Rsp-1-1 | NZ_HE978651.1 | - | 2542550 | 2542467 | **RNA**→ rpsJ (PRK00596)RpsJ (COG0051)→ RplC (COG0087)rplC (PRK00001)→ |
| | | | | | rplD (PRK05319)Ribosomal_L4 (pfam00573)→ rplW (PRK05738)Ribosomal_L23 (pfam00276)→ |
| | | | | | RplB (COG0090)rplB (PRK09374)→ rpsS (PRK00357)Ribosomal_S19 (pfam00203)→ |
| | | | | | rplV (PRK00565)Ribosomal_L22 (pfam00237)→ RpsC (COG0092)rpsC (PRK00310)→ |
| | | | | | rpl16 (CHL00044)rplP (PRK09203)→ PRK00306 (PRK00306)Ribosomal_L29 (pfam00831)→ |
| | | | | | rpsQ (PRK05610)Ribosomal_S17 (pfam00366)→ rplN (PRK05483)Ribosomal_L14 (pfam00238)→ |
| | | | | | RplX (COG0198)rplX (PRK00004)→ rplE (PRK00010)RplE (COG0094)→ rpsN (PRK08061)RpsN (COG0199)→ |
| env-732 | 4491479.3_NODE_8774 | + | 192531 | 192622 | **RNA**→ GTP_HydF (TIGR03918)→ |
| Rgn-1-1 | NZ_JAGQ01000004.1 | + | 1148637 | 1148728 | **RNA**→ rpsJ (PRK00596)RpsJ (COG0051)→ |
| env-733 | ADJT01004649.1 | + | 81 | 172 | **RNA**→ rpsJ (PRK00596)RpsJ (COG0051)→ |
| env-734 | DLM009_scaffold11270_2 | + | 1293 | 1384 | **RNA**→ rpsJ (PRK00596)RpsJ (COG0051)→ |
| env-735 | DOF012_scaffold7759_1 | + | 2947 | 3038 | **RNA**→ rpsJ (PRK00596)RpsJ (COG0051)→ |
| env-736 | longitudinal_230_114 | - | 55598 | 55507 | **RNA**→ rpsJ (PRK00596)RpsJ (COG0051)→ |
| env-737 | longitudinal_231_54 | - | 203125 | 203034 | **RNA**→ rpsJ (PRK00596)RpsJ (COG0051)→ |
| env-738 | longitudinal_232_15 | - | 49512 | 49421 | **RNA**→ rpsJ (PRK00596)RpsJ (COG0051)→ |
| env-739 | 4491482.3_NODE_3522 | + | 6776 | 6867 | **RNA**→ |
| env-740 | NOM002_scaffold8624_14 | - | 161 | 70 | **RNA**→ |
| env-741 | SRS042284_WUGC_scaffold_18018 | - | 166 | 75 | **RNA**→ hypo→ |
| env-742 | NOM026_scaffold565_8 | + | 1020 | 1111 | **RNA**→ rpsJ (PRK00596)RpsJ (COG0051)→ |
| env-743 | longitudinal_525_50 | + | 31964 | 32055 | **RNA**→ rpsJ (PRK00596)RpsJ (COG0051)→ |
| Rgn-2-1 | NZ_KI669415.1 | - | 308806 | 308715 | **RNA**→ rpsJ (PRK00596)RpsJ (COG0051)→ |
| Lba-4-1 | NZ_GL945339.1 | - | 18152 | 18061 | **RNA**→ rpsJ (PRK00596)RpsJ (COG0051)→ |
| env-744 | 4491488.3_NODE_2135 | - | 11126 | 11035 | **RNA**→ rpsJ (PRK00596)RpsJ (COG0051)→ |
| env-745 | DLM024_scaffold1675_1 | + | 6682 | 6773 | **RNA**→ rpsJ (PRK00596)RpsJ (COG0051)→ |
| env-746 | DOM016_scaffold28471_1 | - | 579 | 488 | **RNA**→ rpsJ (PRK00596)RpsJ (COG0051)→ |
| env-747 | NLM007_scaffold2834_3 | + | 4947 | 5038 | **RNA**→ rpsJ (PRK00596)RpsJ (COG0051)→ |
| env-748 | NLM027_scaffold10047_1 | - | 61541 | 61450 | **RNA**→ rpsJ (PRK00596)RpsJ (COG0051)→ |

| | | | | | |
|---|---|---|---|---|---|
| Rgn-3-1 | NZ_AAYG02000020.1 | - | 131008 | 130917 | **RNA**→ rpsJ (PRK00596)RpsJ (COG0051)→ rpsJ (PRK00596)RpsJ (COG0051)→<br>RplC (COG0087)rplC (PRK00001)→ RplC (COG0087)rplC (PRK00001)→<br>rplD (PRK05319)Ribosomal_L4 (pfam00573)→ rplD (PRK05319)Ribosomal_L4 (pfam00573)→<br>rplW (PRK05738)Ribosomal_L23 (pfam00276)→ rplW (PRK05738)Ribosomal_L23 (pfam00276)→<br>RplB (COG0090)rplB (PRK09374)→ RplB (COG0090)rplB (PRK09374)→ rpsS (PRK00357)RpsS (COG0185)→<br>rpsS (PRK00357)RpsS (COG0185)→ rplV (PRK00565)Ribosomal_L22 (pfam00237)→<br>rplV (PRK00565)Ribosomal_L22 (pfam00237)→ RpsC (COG0092)rpsC (PRK00310)→<br>RpsC (COG0092)rpsC (PRK00310)→ rpl16 (CHL00044)rplP (PRK09203)→ rpl16 (CHL00044)rplP (PRK09203)→<br>PRK00306 (PRK00306)Ribosomal_L29 (pfam00831)→ PRK00306 (PRK00306)Ribosomal_L29 (pfam00831)→<br>rpsQ (PRK05610)Ribosomal_S17 (pfam00366)→ rpsQ (PRK05610)Ribosomal_S17 (pfam00366)→<br>rplN (PRK05483)Ribosomal_L14 (pfam00238)→ rplN (PRK05483)Ribosomal_L14 (pfam00238)→<br>RplX (COG0198)rplX (PRK00004)→ RplX (COG0198)rplX (PRK00004)→ rplE (PRK00010)RplE (COG0094)→<br>rplE (PRK00010)RplE (COG0094)→ rpsN (PRK08061)RpsN (COG0199)→ rpsN (PRK08061)RpsN (COG0199)→<br>rpsH (PRK00136)Ribosomal_S8 (pfam00410)→ rpsH (PRK00136)Ribosomal_S8 (pfam00410)→ |
| env-749 | longitudinal_220_105 | + | 11476 | 11569 | **RNA**→ rpsJ (PRK00596)RpsJ (COG0051)→ RplC (COG0087)rplC (PRK00001)→<br>rplD (PRK05319)Ribosomal_L4 (pfam00573)→ rplW (PRK05738)Ribosomal_L23 (pfam00276)→ |
| env-750 | longitudinal_221_152 | + | 5020 | 5113 | **RNA**→ rpsJ (PRK00596)RpsJ (COG0051)→ RplC (COG0087)rplC (PRK00001)→<br>rplD (PRK05319)Ribosomal_L4 (pfam00573)→ rplW (PRK05738)Ribosomal_L23 (pfam00276)→ |
| env-751 | JGI994J12354_10010598 | - | 5140 | 5057 | **RNA**→ rpsJ (PRK00596)RpsJ (COG0051)→ |
| Cph-1-1 | NC_010001.1 | - | 4506701 | 4506618 | **RNA**→ rpsJ (PRK00596)RpsJ (COG0051)→ |
| Bfo-1-1 | NZ_ACCL02000013.1 | + | 96014 | 96105 | **RNA**→ rpsJ (PRK00596)RpsJ (COG0051)→ rpsJ (PRK00596)RpsJ (COG0051)→ |
| env-752 | RUMENNODE_3921064_1 | + | 21350 | 21432 | **RNA**→ rpsJ (PRK00596)RpsJ (COG0051)→ RplC (COG0087)rplC (PRK00001)→<br>rplD (PRK05319)Ribosomal_L4 (pfam00573)→ rplW (PRK05738)Ribosomal_L23 (pfam00276)→ |
| env-753 | AUXO012300854.1 | - | 288 | 206 | **RNA**→ |
| Lbo-3-1 | NZ_ATWB01000017.1 | - | 17560 | 17479 | **RNA**→ rpsJ (PRK00596)RpsJ (COG0051)→ |
| Lbo-2-1 | NZ_JHWP01000008.1 | - | 105109 | 105028 | **RNA**→ rpsJ (PRK00596)RpsJ (COG0051)→ |
| Lbo-1-1 | NZ_JHWB01000013.1 | + | 35396 | 35477 | **RNA**→ rpsJ (PRK00596)RpsJ (COG0051)→ |
| env-754 | 2227507959 | - | 19299 | 19219 | **RNA**→ rpsJ (PRK00596)RpsJ (COG0051)→ |
| env-755 | JGI994J12354_10001688 | + | 17679 | 17759 | **RNA**→ rpsJ (PRK00596)RpsJ (COG0051)→ |
| Csp-4-1 | NZ_GL870809.1 | + | 559679 | 559772 | **RNA**→ rpsJ (PRK00596)RpsJ (COG0051)→ rpsJ (PRK00596)RpsJ (COG0051)→<br>RplC (COG0087)rplC (PRK00001)→ RplC (COG0087)rplC (PRK00001)→<br>rplD (PRK05319)Ribosomal_L4 (pfam00573)→ rplD (PRK05319)Ribosomal_L4 (pfam00573)→<br>rplW (PRK05738)Ribosomal_L23 (pfam00276)→ rplW (PRK05738)Ribosomal_L23 (pfam00276)→<br>RplB (COG0090)rplB (PRK09374)→ RplB (COG0090)rplB (PRK09374)→ rpsS (PRK00357)RpsS (COG0185)→<br>rpsS (PRK00357)RpsS (COG0185)→ rplV (PRK00565)Ribosomal_L22 (pfam00237)→<br>rplV (PRK00565)Ribosomal_L22 (pfam00237)→ RpsC (COG0092)rpsC (PRK00310)→<br>RpsC (COG0092)rpsC (PRK00310)→ rpl16 (CHL00044)rplP (PRK09203)→ rpl16 (CHL00044)rplP (PRK09203)→<br>PRK00306 (PRK00306)Ribosomal_L29 (pfam00831)→ PRK00306 (PRK00306)Ribosomal_L29 (pfam00831)→<br>rpsQ (PRK05610)Ribosomal_S17 (pfam00366)→ rpsQ (PRK05610)Ribosomal_S17 (pfam00366)→<br>rplN (PRK05483)Ribosomal_L14 (pfam00238)→ rplN (PRK05483)Ribosomal_L14 (pfam00238)→<br>RplX (COG0198)rplX (PRK00004)→ RplX (COG0198)rplX (PRK00004)→ rplE (PRK00010)RplE (COG0094)→<br>rplE (PRK00010)RplE (COG0094)→ rpsN (PRK08061)RpsN (COG0199)→ rpsN (PRK08061)RpsN (COG0199)→ |
| env-756 | 4491477.3_NODE_8819 | - | 4476 | 4384 | **RNA**→ rpsJ (PRK00596)RpsJ (COG0051)→ |
| env-757 | NOF014_scaffold4940_42 | + | 2191 | 2283 | **RNA**→ rpsJ (PRK00596)RpsJ (COG0051)→ |
| env-758 | NLM008_scaffold14561_16 | + | 1445 | 1541 | **RNA**→ rpsJ (PRK00596)RpsJ (COG0051)→ RplC (COG0087)rplC (PRK00001)→ |
| Rla-1-1 | NZ_KI669408.1 | + | 599372 | 599468 | **RNA**→ rpsJ (PRK00596)RpsJ (COG0051)→ RplC (COG0087)rplC (PRK00001)→<br>rplD (PRK05319)Ribosomal_L4 (pfam00573)→ rplW (PRK05738)Ribosomal_L23 (pfam00276)→ |
| env-759 | 2227186343 | - | 2802 | 2722 | **RNA**→ rpsJ (PRK00596)RpsJ (COG0051)→ |
| env-760 | IMNBL1_c0024798 | + | 1559 | 1639 | **RNA**→ rpsJ (PRK00596)RpsJ (COG0051)→ |
| env-761 | JGI994J12354_10000212 | + | 20164 | 20244 | **RNA**→ rpsJ (PRK00596)RpsJ (COG0051)→ |
| env-762 | 2227058723 | + | 598 | 678 | **RNA**→ |
| env-763 | longitudinal_51_247 | - | 49840 | 49760 | **RNA**→ rpsJ (PRK00596)RpsJ (COG0051)→ |
| env-764 | longitudinal_411_28 | + | 32215 | 32297 | **RNA**→ rpsJ (PRK00596)RpsJ (COG0051)→ |
| Lba-6-1 | NZ_KE159795.1 | - | 1385114 | 1385002 | **RNA**→ rpsJ (PRK00596)RpsJ (COG0051)→ RplC (COG0087)rplC (PRK00001)→<br>rplD (PRK05319)Ribosomal_L4 (pfam00573)→ rplW (PRK05738)Ribosomal_L23 (pfam00276)→ |

| | | | | | |
|---|---|---|---|---|---|
| Dsp-1-1 | NZ_KE159794.1 | - | 173058 | 172976 | **RNA**→ rpsJ (PRK00596)RpsJ (COG0051)→ RplC (COG0087)rplC (PRK00001)→<br>rplD (PRK05319)Ribosomal_L4 (pfam00573)→ rplW (PRK05738)Ribosomal_L23 (pfam00276)→<br>RplB (COG0090)rplB (PRK09374)→ rpsS (PRK00357)RpsS (COG0185)→<br>rplV (PRK00565)Ribosomal_L22 (pfam00237)→ RpsC (COG0092)rpsC (PRK00310)→<br>rpl16 (CHL00044)rplP (PRK09203)→ PRK00306 (PRK00306)Ribosomal_L29 (pfam00831)→<br>rpsQ (PRK05610)Ribosomal_S17 (pfam00366)→ rplN (PRK05483)Ribosomal_L14 (pfam00238)→<br>RplX (COG0198)rplX (PRK00004)→ rplE (PRK00010)RplE (COG0094)→ rpsN (PRK08061)RpsN (COG0199)→ |
| env-765 | scaffold8116_4_MH0058 | + | 475 | 558 | **RNA**→ |
| env-766 | UnmappedStool_Broad_scaffold_811588 | + | 1503 | 1586 | **RNA**→ |
| env-767 | NOM005_scaffold16235_2 | + | 536 | 619 | **RNA**→ rpsJ (PRK00596)RpsJ (COG0051)→ |
| env-768 | scaffold879_1_MH0020 | - | 181 | 98 | **RNA**→ |
| env-769 | DOF003_scaffold12608_5 | - | 851 | 768 | **RNA**→ rpsJ (PRK00596)RpsJ (COG0051)→ |
| env-770 | DLF014_scaffold3019_1 | + | 38181 | 38264 | **RNA**→ rpsJ (PRK00596)RpsJ (COG0051)→ |
| env-771 | DLM004_scaffold2544_24 | + | 1613 | 1696 | **RNA**→ rpsJ (PRK00596)RpsJ (COG0051)→ |
| env-772 | NLM002_scaffold11552_20 | - | 6524 | 6441 | **RNA**→ rpsJ (PRK00596)RpsJ (COG0051)→ |
| env-773 | NLM007_scaffold8527_7 | + | 212 | 295 | **RNA**→ rpsJ (PRK00596)RpsJ (COG0051)→ |
| env-774 | NOM009_scaffold43909_1 | + | 265 | 348 | **RNA**→ rpsJ (PRK00596)RpsJ (COG0051)→ |
| env-775 | NOM028_scaffold32438_1 | - | 438 | 355 | **RNA**→ rpsJ (PRK00596)RpsJ (COG0051)→ |
| env-776 | SRS019397_WUGC_scaffold_4193 | - | 413 | 330 | **RNA**→ rpsJ (PRK00596)Ribosomal_S10 (pfam00338)→ |
| env-777 | DOM010_scaffold1445_8 | + | 1478 | 1561 | **RNA**→ rpsJ (PRK00596)RpsJ (COG0051)→ |
| env-778 | longitudinal_13_538 | - | 50782 | 50698 | **RNA**→ rpsJ (PRK00596)RpsJ (COG0051)→ RplC (COG0087)rplC (PRK00001)→<br>rplD (PRK05319)Ribosomal_L4 (pfam00573)→ rplW (PRK05738)Ribosomal_L23 (pfam00276)→<br>RplB (COG0090)rplB (PRK09374)→ rpsS (PRK00357)RpsS (COG0185)→<br>rplV (PRK00565)Ribosomal_L22 (pfam00237)→ RpsC (COG0092)rpsC (PRK00310)→<br>rpl16 (CHL00044)rplP (PRK09203)→ PRK00306 (PRK00306)Ribosomal_L29 (pfam00831)→<br>rpsQ (PRK05610)Ribosomal_S17 (pfam00366)→ rplN (PRK05483)Ribosomal_L14 (pfam00238)→<br>RplX (COG0198)rplX (PRK00004)→ rplE (PRK00010)RplE (COG0094)→ rpsN (PRK08061)RpsN (COG0199)→<br>rpsH (PRK00136)Ribosomal_S8 (pfam00410)→ |
| env-779 | longitudinal_14_60 | - | 50782 | 50698 | **RNA**→ rpsJ (PRK00596)RpsJ (COG0051)→ RplC (COG0087)rplC (PRK00001)→<br>rplD (PRK05319)Ribosomal_L4 (pfam00573)→ rplW (PRK05738)Ribosomal_L23 (pfam00276)→<br>RplB (COG0090)rplB (PRK09374)→ rpsS (PRK00357)RpsS (COG0185)→<br>rplV (PRK00565)Ribosomal_L22 (pfam00237)→ RpsC (COG0092)rpsC (PRK00310)→<br>rpl16 (CHL00044)rplP (PRK09203)→ PRK00306 (PRK00306)Ribosomal_L29 (pfam00831)→<br>rpsQ (PRK05610)Ribosomal_S17 (pfam00366)→ rplN (PRK05483)Ribosomal_L14 (pfam00238)→<br>RplX (COG0198)rplX (PRK00004)→ rplE (PRK00010)RplE (COG0094)→ rpsN (PRK08061)RpsN (COG0199)→<br>rpsH (PRK00136)Ribosomal_S8 (pfam00410)→ |
| env-780 | longitudinal_15_97 | - | 50782 | 50698 | **RNA**→ rpsJ (PRK00596)RpsJ (COG0051)→ RplC (COG0087)rplC (PRK00001)→<br>rplD (PRK05319)Ribosomal_L4 (pfam00573)→ rplW (PRK05738)Ribosomal_L23 (pfam00276)→<br>RplB (COG0090)rplB (PRK09374)→ rpsS (PRK00357)RpsS (COG0185)→<br>rplV (PRK00565)Ribosomal_L22 (pfam00237)→ RpsC (COG0092)rpsC (PRK00310)→<br>rpl16 (CHL00044)rplP (PRK09203)→ PRK00306 (PRK00306)Ribosomal_L29 (pfam00831)→<br>rpsQ (PRK05610)Ribosomal_S17 (pfam00366)→ rplN (PRK05483)Ribosomal_L14 (pfam00238)→<br>RplX (COG0198)rplX (PRK00004)→ rplE (PRK00010)RplE (COG0094)→ rpsN (PRK08061)RpsN (COG0199)→<br>rpsH (PRK00136)Ribosomal_S8 (pfam00410)→ |
| env-781 | longitudinal_40_24 | - | 50781 | 50697 | **RNA**→ rpsJ (PRK00596)RpsJ (COG0051)→ RplC (COG0087)rplC (PRK00001)→<br>rplD (PRK05319)Ribosomal_L4 (pfam00573)→ rplW (PRK05738)Ribosomal_L23 (pfam00276)→<br>RplB (COG0090)rplB (PRK09374)→ rpsS (PRK00357)RpsS (COG0185)→<br>rplV (PRK00565)Ribosomal_L22 (pfam00237)→ RpsC (COG0092)rpsC (PRK00310)→<br>rpl16 (CHL00044)rplP (PRK09203)→ PRK00306 (PRK00306)Ribosomal_L29 (pfam00831)→<br>rpsQ (PRK05610)Ribosomal_S17 (pfam00366)→ rplN (PRK05483)Ribosomal_L14 (pfam00238)→<br>RplX (COG0198)rplX (PRK00004)→ rplE (PRK00010)RplE (COG0094)→ rpsN (PRK08061)RpsN (COG0199)→<br>rpsH (PRK00136)Ribosomal_S8 (pfam00410)→ |

| | | | | | |
|---|---|---|---|---|---|
| env-782 | longitudinal_41_262 | + | 82107 | 82191 | **RNA**→ rpsJ (PRK00596)RpsJ (COG0051)→ RplC (COG0087)rplC (PRK00001)→ rplD (PRK05319)Ribosomal_L4 (pfam00573)→ rplW (PRK05738)Ribosomal_L23 (pfam00276)→ RplB (COG0090)rplB (PRK09374)→ rpsS (PRK00357)RpsS (COG0185)→ rplV (PRK00565)Ribosomal_L22 (pfam00237)→ RpsC (COG0092)rpsC (PRK00310)→ rpl16 (CHL00044)rplP (PRK09203)→ PRK00306 (PRK00306)Ribosomal_L29 (pfam00831)→ rpsQ (PRK05610)Ribosomal_S17 (pfam00366)→ rplN (PRK05483)Ribosomal_L14 (pfam00238)→ RplX (COG0198)rplX (PRK00004)→ rplE (PRK00010)RplE (COG0094)→ rpsN (PRK08061)RpsN (COG0199)→ rpsH (PRK00136)Ribosomal_S8 (pfam00410)→ |
| env-783 | longitudinal_42_626 | - | 50782 | 50698 | **RNA**→ rpsJ (PRK00596)RpsJ (COG0051)→ RplC (COG0087)rplC (PRK00001)→ rplD (PRK05319)Ribosomal_L4 (pfam00573)→ rplW (PRK05738)Ribosomal_L23 (pfam00276)→ RplB (COG0090)rplB (PRK09374)→ rpsS (PRK00357)RpsS (COG0185)→ rplV (PRK00565)Ribosomal_L22 (pfam00237)→ RpsC (COG0092)rpsC (PRK00310)→ rpl16 (CHL00044)rplP (PRK09203)→ PRK00306 (PRK00306)Ribosomal_L29 (pfam00831)→ rpsQ (PRK05610)Ribosomal_S17 (pfam00366)→ rplN (PRK05483)Ribosomal_L14 (pfam00238)→ RplX (COG0198)rplX (PRK00004)→ rplE (PRK00010)RplE (COG0094)→ rpsN (PRK08061)RpsN (COG0199)→ rpsH (PRK00136)Ribosomal_S8 (pfam00410)→ |
| env-784 | longitudinal_43_95 | + | 82106 | 82190 | **RNA**→ rpsJ (PRK00596)RpsJ (COG0051)→ RplC (COG0087)rplC (PRK00001)→ rplD (PRK05319)Ribosomal_L4 (pfam00573)→ rplW (PRK05738)Ribosomal_L23 (pfam00276)→ RplB (COG0090)rplB (PRK09374)→ rpsS (PRK00357)RpsS (COG0185)→ rplV (PRK00565)Ribosomal_L22 (pfam00237)→ RpsC (COG0092)rpsC (PRK00310)→ rpl16 (CHL00044)rplP (PRK09203)→ PRK00306 (PRK00306)Ribosomal_L29 (pfam00831)→ rpsQ (PRK05610)Ribosomal_S17 (pfam00366)→ rplN (PRK05483)Ribosomal_L14 (pfam00238)→ RplX (COG0198)rplX (PRK00004)→ rplE (PRK00010)RplE (COG0094)→ rpsN (PRK08061)RpsN (COG0199)→ rpsH (PRK00136)Ribosomal_S8 (pfam00410)→ |
| env-785 | longitudinal_44_5 | + | 13247 | 13331 | **RNA**→ rpsJ (PRK00596)RpsJ (COG0051)→ RplC (COG0087)rplC (PRK00001)→ rplD (PRK05319)Ribosomal_L4 (pfam00573)→ rplW (PRK05738)Ribosomal_L23 (pfam00276)→ RplB (COG0090)rplB (PRK09374)→ rpsS (PRK00357)RpsS (COG0185)→ rplV (PRK00565)Ribosomal_L22 (pfam00237)→ RpsC (COG0092)rpsC (PRK00310)→ rpl16 (CHL00044)rplP (PRK09203)→ PRK00306 (PRK00306)Ribosomal_L29 (pfam00831)→ rpsQ (PRK05610)Ribosomal_S17 (pfam00366)→ rplN (PRK05483)Ribosomal_L14 (pfam00238)→ RplX (COG0198)rplX (PRK00004)→ rplE (PRK00010)RplE (COG0094)→ rpsN (PRK08061)RpsN (COG0199)→ rpsH (PRK00136)Ribosomal_S8 (pfam00410)→ |
| env-786 | longitudinal_45_167 | - | 50782 | 50698 | **RNA**→ rpsJ (PRK00596)RpsJ (COG0051)→ RplC (COG0087)rplC (PRK00001)→ rplD (PRK05319)Ribosomal_L4 (pfam00573)→ rplW (PRK05738)Ribosomal_L23 (pfam00276)→ RplB (COG0090)rplB (PRK09374)→ rpsS (PRK00357)RpsS (COG0185)→ rplV (PRK00565)Ribosomal_L22 (pfam00237)→ RpsC (COG0092)rpsC (PRK00310)→ rpl16 (CHL00044)rplP (PRK09203)→ PRK00306 (PRK00306)Ribosomal_L29 (pfam00831)→ rpsQ (PRK05610)Ribosomal_S17 (pfam00366)→ rplN (PRK05483)Ribosomal_L14 (pfam00238)→ RplX (COG0198)rplX (PRK00004)→ rplE (PRK00010)RplE (COG0094)→ rpsN (PRK08061)RpsN (COG0199)→ rpsH (PRK00136)Ribosomal_S8 (pfam00410)→ |
| env-787 | longitudinal_46_248 | - | 50783 | 50699 | **RNA**→ rpsJ (PRK00596)RpsJ (COG0051)→ RplC (COG0087)rplC (PRK00001)→ rplD (PRK05319)Ribosomal_L4 (pfam00573)→ rplW (PRK05738)Ribosomal_L23 (pfam00276)→ RplB (COG0090)rplB (PRK09374)→ rpsS (PRK00357)RpsS (COG0185)→ rplV (PRK00565)Ribosomal_L22 (pfam00237)→ RpsC (COG0092)rpsC (PRK00310)→ rpl16 (CHL00044)rplP (PRK09203)→ PRK00306 (PRK00306)Ribosomal_L29 (pfam00831)→ rpsQ (PRK05610)Ribosomal_S17 (pfam00366)→ rplN (PRK05483)Ribosomal_L14 (pfam00238)→ RplX (COG0198)rplX (PRK00004)→ rplE (PRK00010)RplE (COG0094)→ rpsN (PRK08061)RpsN (COG0199)→ rpsH (PRK00136)Ribosomal_S8 (pfam00410)→ |
| env-788 | longitudinal_47_2 | - | 50781 | 50697 | **RNA**→ rpsJ (PRK00596)RpsJ (COG0051)→ RplC (COG0087)rplC (PRK00001)→ rplD (PRK05319)Ribosomal_L4 (pfam00573)→ rplW (PRK05738)Ribosomal_L23 (pfam00276)→ RplB (COG0090)rplB (PRK09374)→ rpsS (PRK00357)RpsS (COG0185)→ rplV (PRK00565)Ribosomal_L22 (pfam00237)→ RpsC (COG0092)rpsC (PRK00310)→ rpl16 (CHL00044)rplP (PRK09203)→ PRK00306 (PRK00306)Ribosomal_L29 (pfam00831)→ rpsQ (PRK05610)Ribosomal_S17 (pfam00366)→ rplN (PRK05483)Ribosomal_L14 (pfam00238)→ RplX (COG0198)rplX (PRK00004)→ rplE (PRK00010)RplE (COG0094)→ rpsN (PRK08061)RpsN (COG0199)→ rpsH (PRK00136)Ribosomal_S8 (pfam00410)→ |

| | | | | | |
|---|---|---|---|---|---|
| env-789 | longitudinal_48_317 | - | 50782 | 50698 | **RNA**→ rpsJ (PRK00596)RpsJ (COG0051)→ RplC (COG0087)rplC (PRK00001)→ rplD (PRK05319)Ribosomal_L4 (pfam00573)→ rplW (PRK05738)Ribosomal_L23 (pfam00276)→ RplB (COG0090)rplB (PRK09374)→ rpsS (PRK00357)RpsS (COG0185)→ rplV (PRK00565)Ribosomal_L22 (pfam00237)→ RpsC (COG0092)rpsC (PRK00310)→ rpl16 (CHL00044)rplP (PRK09203)→ PRK00306 (PRK00306)Ribosomal_L29 (pfam00831)→ rpsQ (PRK05610)Ribosomal_S17 (pfam00366)→ rplN (PRK05483)Ribosomal_L14 (pfam00238)→ RplX (COG0198)rplX (PRK00004)→ rplE (PRK00010)RplE (COG0094)→ rpsN (PRK08061)RpsN (COG0199)→ rpsH (PRK00136)Ribosomal_S8 (pfam00410)→ |
| env-790 | longitudinal_49_17 | + | 13247 | 13331 | **RNA**→ rpsJ (PRK00596)RpsJ (COG0051)→ RplC (COG0087)rplC (PRK00001)→ rplD (PRK05319)Ribosomal_L4 (pfam00573)→ rplW (PRK05738)Ribosomal_L23 (pfam00276)→ RplB (COG0090)rplB (PRK09374)→ rpsS (PRK00357)RpsS (COG0185)→ rplV (PRK00565)Ribosomal_L22 (pfam00237)→ RpsC (COG0092)rpsC (PRK00310)→ rpl16 (CHL00044)rplP (PRK09203)→ PRK00306 (PRK00306)Ribosomal_L29 (pfam00831)→ rpsQ (PRK05610)Ribosomal_S17 (pfam00366)→ rplN (PRK05483)Ribosomal_L14 (pfam00238)→ RplX (COG0198)rplX (PRK00004)→ rplE (PRK00010)RplE (COG0094)→ rpsN (PRK08061)RpsN (COG0199)→ rpsH (PRK00136)Ribosomal_S8 (pfam00410)→ |
| env-791 | longitudinal_50_78 | - | 50782 | 50698 | **RNA**→ rpsJ (PRK00596)RpsJ (COG0051)→ RplC (COG0087)rplC (PRK00001)→ rplD (PRK05319)Ribosomal_L4 (pfam00573)→ rplW (PRK05738)Ribosomal_L23 (pfam00276)→ RplB (COG0090)rplB (PRK09374)→ rpsS (PRK00357)RpsS (COG0185)→ rplV (PRK00565)Ribosomal_L22 (pfam00237)→ RpsC (COG0092)rpsC (PRK00310)→ rpl16 (CHL00044)rplP (PRK09203)→ PRK00306 (PRK00306)Ribosomal_L29 (pfam00831)→ rpsQ (PRK05610)Ribosomal_S17 (pfam00366)→ rplN (PRK05483)Ribosomal_L14 (pfam00238)→ RplX (COG0198)rplX (PRK00004)→ rplE (PRK00010)RplE (COG0094)→ rpsN (PRK08061)RpsN (COG0199)→ rpsH (PRK00136)Ribosomal_S8 (pfam00410)→ |
| env-792 | longitudinal_101_1 | + | 13247 | 13331 | **RNA**→ rpsJ (PRK00596)RpsJ (COG0051)→ RplC (COG0087)rplC (PRK00001)→ rplD (PRK05319)Ribosomal_L4 (pfam00573)→ rplW (PRK05738)Ribosomal_L23 (pfam00276)→ RplB (COG0090)rplB (PRK09374)→ rpsS (PRK00357)RpsS (COG0185)→ rplV (PRK00565)Ribosomal_L22 (pfam00237)→ RpsC (COG0092)rpsC (PRK00310)→ rpl16 (CHL00044)rplP (PRK09203)→ PRK00306 (PRK00306)Ribosomal_L29 (pfam00831)→ rpsQ (PRK05610)Ribosomal_S17 (pfam00366)→ rplN (PRK05483)Ribosomal_L14 (pfam00238)→ RplX (COG0198)rplX (PRK00004)→ rplE (PRK00010)RplE (COG0094)→ rpsN (PRK08061)RpsN (COG0199)→ rpsH (PRK00136)Ribosomal_S8 (pfam00410)→ |
| env-793 | longitudinal_102_26 | - | 50782 | 50698 | **RNA**→ rpsJ (PRK00596)RpsJ (COG0051)→ RplC (COG0087)rplC (PRK00001)→ rplD (PRK05319)Ribosomal_L4 (pfam00573)→ rplW (PRK05738)Ribosomal_L23 (pfam00276)→ RplB (COG0090)rplB (PRK09374)→ rpsS (PRK00357)RpsS (COG0185)→ rplV (PRK00565)Ribosomal_L22 (pfam00237)→ RpsC (COG0092)rpsC (PRK00310)→ rpl16 (CHL00044)rplP (PRK09203)→ PRK00306 (PRK00306)Ribosomal_L29 (pfam00831)→ rpsQ (PRK05610)Ribosomal_S17 (pfam00366)→ rplN (PRK05483)Ribosomal_L14 (pfam00238)→ RplX (COG0198)rplX (PRK00004)→ rplE (PRK00010)RplE (COG0094)→ rpsN (PRK08061)RpsN (COG0199)→ rpsH (PRK00136)Ribosomal_S8 (pfam00410)→ |
| env-794 | longitudinal_103_338 | - | 50782 | 50698 | **RNA**→ rpsJ (PRK00596)RpsJ (COG0051)→ RplC (COG0087)rplC (PRK00001)→ rplD (PRK05319)Ribosomal_L4 (pfam00573)→ rplW (PRK05738)Ribosomal_L23 (pfam00276)→ RplB (COG0090)rplB (PRK09374)→ rpsS (PRK00357)RpsS (COG0185)→ rplV (PRK00565)Ribosomal_L22 (pfam00237)→ RpsC (COG0092)rpsC (PRK00310)→ rpl16 (CHL00044)rplP (PRK09203)→ PRK00306 (PRK00306)Ribosomal_L29 (pfam00831)→ rpsQ (PRK05610)Ribosomal_S17 (pfam00366)→ rplN (PRK05483)Ribosomal_L14 (pfam00238)→ RplX (COG0198)rplX (PRK00004)→ rplE (PRK00010)RplE (COG0094)→ rpsN (PRK08061)RpsN (COG0199)→ rpsH (PRK00136)Ribosomal_S8 (pfam00410)→ |
| env-795 | longitudinal_104_5 | - | 50782 | 50698 | **RNA**→ rpsJ (PRK00596)RpsJ (COG0051)→ RplC (COG0087)rplC (PRK00001)→ rplD (PRK05319)Ribosomal_L4 (pfam00573)→ rplW (PRK05738)Ribosomal_L23 (pfam00276)→ RplB (COG0090)rplB (PRK09374)→ rpsS (PRK00357)RpsS (COG0185)→ rplV (PRK00565)Ribosomal_L22 (pfam00237)→ RpsC (COG0092)rpsC (PRK00310)→ rpl16 (CHL00044)rplP (PRK09203)→ PRK00306 (PRK00306)Ribosomal_L29 (pfam00831)→ rpsQ (PRK05610)Ribosomal_S17 (pfam00366)→ rplN (PRK05483)Ribosomal_L14 (pfam00238)→ RplX (COG0198)rplX (PRK00004)→ rplE (PRK00010)RplE (COG0094)→ rpsN (PRK08061)RpsN (COG0199)→ rpsH (PRK00136)Ribosomal_S8 (pfam00410)→ |

| | | | | | |
|---|---|---|---|---|---|
| env-796 | longitudinal_105_72 | + | 82106 | 82190 | **RNA**→ rpsJ (PRK00596)RpsJ (COG0051)→ RplC (COG0087)rplC (PRK00001)→ rplD (PRK05319)Ribosomal_L4 (pfam00573)→ rplW (PRK05738)Ribosomal_L23 (pfam00276)→ RplB (COG0090)rplB (PRK09374)→ rpsS (PRK00357)RpsS (COG0185)→ rplV (PRK00565)Ribosomal_L22 (pfam00237)→ RpsC (COG0092)rpsC (PRK00310)→ rpl16 (CHL00044)rplP (PRK09203)→ PRK00306 (PRK00306)Ribosomal_L29 (pfam00831)→ rpsQ (PRK05610)Ribosomal_S17 (pfam00366)→ rplN (PRK05483)Ribosomal_L14 (pfam00238)→ RplX (COG0198)rplX (PRK00004)→ rplE (PRK00010)RplE (COG0094)→ rpsN (PRK08061)RpsN (COG0199)→ rpsH (PRK00136)Ribosomal_S8 (pfam00410)→ |
| env-797 | longitudinal_106_357 | - | 50784 | 50700 | **RNA**→ rpsJ (PRK00596)RpsJ (COG0051)→ RplC (COG0087)rplC (PRK00001)→ rplD (PRK05319)Ribosomal_L4 (pfam00573)→ rplW (PRK05738)Ribosomal_L23 (pfam00276)→ RplB (COG0090)rplB (PRK09374)→ rps19 (CHL00050)rpsS (PRK00357)→ rplV (PRK00565)Ribosomal_L22 (pfam00237)→ RpsC (COG0092)rpsC (PRK00310)→ rpl16 (CHL00044)rplP (PRK09203)→ PRK00306 (PRK00306)Ribosomal_L29 (pfam00831)→ rpsQ (PRK05610)Ribosomal_S17 (pfam00366)→ rplN (PRK05483)Ribosomal_L14 (pfam00238)→ RplX (COG0198)rplX (PRK00004)→ rplE (PRK00010)RplE (COG0094)→ rpsN (PRK08061)RpsN (COG0199)→ rpsH (PRK00136)Ribosomal_S8 (pfam00410)→ |
| env-798 | longitudinal_107_17 | + | 13248 | 13332 | **RNA**→ rpsJ (PRK00596)RpsJ (COG0051)→ RplC (COG0087)rplC (PRK00001)→ rplD (PRK05319)Ribosomal_L4 (pfam00573)→ rplW (PRK05738)Ribosomal_L23 (pfam00276)→ RplB (COG0090)rplB (PRK09374)→ rpsS (PRK00357)RpsS (COG0185)→ rplV (PRK00565)Ribosomal_L22 (pfam00237)→ RpsC (COG0092)rpsC (PRK00310)→ rpl16 (CHL00044)rplP (PRK09203)→ PRK00306 (PRK00306)Ribosomal_L29 (pfam00831)→ rpsQ (PRK05610)Ribosomal_S17 (pfam00366)→ rplN (PRK05483)Ribosomal_L14 (pfam00238)→ RplX (COG0198)rplX (PRK00004)→ rplE (PRK00010)RplE (COG0094)→ rpsN (PRK08061)RpsN (COG0199)→ rpsH (PRK00136)Ribosomal_S8 (pfam00410)→ |
| env-799 | longitudinal_108_5 | + | 13247 | 13331 | **RNA**→ rpsJ (PRK00596)RpsJ (COG0051)→ RplC (COG0087)rplC (PRK00001)→ rplD (PRK05319)Ribosomal_L4 (pfam00573)→ rplW (PRK05738)Ribosomal_L23 (pfam00276)→ RplB (COG0090)rplB (PRK09374)→ rpsS (PRK00357)RpsS (COG0185)→ rplV (PRK00565)Ribosomal_L22 (pfam00237)→ RpsC (COG0092)rpsC (PRK00310)→ rpl16 (CHL00044)rplP (PRK09203)→ PRK00306 (PRK00306)Ribosomal_L29 (pfam00831)→ rpsQ (PRK05610)Ribosomal_S17 (pfam00366)→ rplN (PRK05483)Ribosomal_L14 (pfam00238)→ RplX (COG0198)rplX (PRK00004)→ rplE (PRK00010)RplE (COG0094)→ rpsN (PRK08061)RpsN (COG0199)→ rpsH (PRK00136)Ribosomal_S8 (pfam00410)→ |
| env-800 | longitudinal_109_7 | + | 13246 | 13330 | **RNA**→ rpsJ (PRK00596)RpsJ (COG0051)→ RplC (COG0087)rplC (PRK00001)→ rplD (PRK05319)Ribosomal_L4 (pfam00573)→ rplW (PRK05738)Ribosomal_L23 (pfam00276)→ RplB (COG0090)rplB (PRK09374)→ rpsS (PRK00357)RpsS (COG0185)→ rplV (PRK00565)Ribosomal_L22 (pfam00237)→ RpsC (COG0092)rpsC (PRK00310)→ rpl16 (CHL00044)rplP (PRK09203)→ PRK00306 (PRK00306)Ribosomal_L29 (pfam00831)→ rpsQ (PRK05610)Ribosomal_S17 (pfam00366)→ rplN (PRK05483)Ribosomal_L14 (pfam00238)→ RplX (COG0198)rplX (PRK00004)→ rplE (PRK00010)RplE (COG0094)→ rpsN (PRK08061)RpsN (COG0199)→ rpsH (PRK00136)Ribosomal_S8 (pfam00410)→ |
| env-801 | longitudinal_110_299 | + | 82113 | 82197 | **RNA**→ rpsJ (PRK00596)RpsJ (COG0051)→ RplC (COG0087)rplC (PRK00001)→ rplD (PRK05319)Ribosomal_L4 (pfam00573)→ rplW (PRK05738)Ribosomal_L23 (pfam00276)→ RplB (COG0090)rplB (PRK09374)→ rpsS (PRK00357)RpsS (COG0185)→ rplV (PRK00565)Ribosomal_L22 (pfam00237)→ RpsC (COG0092)rpsC (PRK00310)→ rpl16 (CHL00044)rplP (PRK09203)→ PRK00306 (PRK00306)Ribosomal_L29 (pfam00831)→ rpsQ (PRK05610)Ribosomal_S17 (pfam00366)→ rplN (PRK05483)Ribosomal_L14 (pfam00238)→ RplX (COG0198)rplX (PRK00004)→ rplE (PRK00010)RplE (COG0094)→ rpsN (PRK08061)RpsN (COG0199)→ rpsH (PRK00136)Ribosomal_S8 (pfam00410)→ |
| env-802 | NOF005_scaffold1985_8 | + | 1796 | 1880 | **RNA**→ rpsJ (PRK00596)RpsJ (COG0051)→ RplC (COG0087)rplC (PRK00001)→ rplD (PRK05319)Ribosomal_L4 (pfam00573)→ rplW (PRK05738)Ribosomal_L23 (pfam00276)→ |
| env-803 | NLM031_scaffold14783_14 | - | 386 | 302 | **RNA**→ rpsJ (PRK00596)Ribosomal_S10 (pfam00338)→ |
| env-804 | NOM009_scaffold30082_1 | - | 211 | 127 | **RNA**→ |
| env-805 | NOM026_scaffold8944_8 | + | 436 | 520 | **RNA**→ |

| | | | | | |
|---|---|---|---|---|---|
| Cco-1-1 | NZ_GG662012.1 | + | 43704 | 43788 | **RNA**→ rpsJ (PRK00596)RpsJ (COG0051)→ RplC (COG0087)rplC (PRK00001)→ rplD (PRK05319)Ribosomal_L4 (pfam00573)→ rplW (PRK05738)Ribosomal_L23 (pfam00276)→ RplB (COG0090)rplB (PRK09374)→ rpsS (PRK00357)RpsS (COG0185)→ rplV (PRK00565)Ribosomal_L22 (pfam00237)→ RpsC (COG0092)rpsC (PRK00310)→ rpl16 (CHL00044)rplP (PRK09203)→ PRK00306 (PRK00306)Ribosomal_L29 (pfam00831)→ hypo→ rpsQ (PRK05610)Ribosomal_S17 (pfam00366)→ rpsQ (PRK05610)Ribosomal_S17 (pfam00366)→ rplN (PRK05483)Ribosomal_L14 (pfam00238)→ hypo→ hypo→ RplX (COG0198)ribosomal_L24 (pfam17136)→ rplE (PRK00010)RplE (COG0094)→ rpsN (PRK08061)RpsN (COG0199)→ rpsH (PRK00136)Ribosomal_S8 (pfam00410)→ |
| env-806 | longitudinal_291_432 | - | 4369 | 4285 | **RNA**→ rpsJ (PRK00596)RpsJ (COG0051)→ RplC (COG0087)rplC (PRK00001)→ rplD (PRK05319)Ribosomal_L4 (pfam00573)→ rplW (PRK05738)Ribosomal_L23 (pfam00276)→ RplB (COG0090)rplB (PRK09374)→ rpsS (PRK00357)RpsS (COG0185)→ rplV (PRK00565)Ribosomal_L22 (pfam00237)→ RpsC (COG0092)rpsC (PRK00310)→ |
| env-807 | longitudinal_292_65 | - | 59834 | 59750 | **RNA**→ rpsJ (PRK00596)RpsJ (COG0051)→ RplC (COG0087)rplC (PRK00001)→ rplD (PRK05319)Ribosomal_L4 (pfam00573)→ rplW (PRK05738)Ribosomal_L23 (pfam00276)→ RplB (COG0090)rplB (PRK09374)→ rpsS (PRK00357)RpsS (COG0185)→ rplV (PRK00565)Ribosomal_L22 (pfam00237)→ RpsC (COG0092)rpsC (PRK00310)→ rpl16 (CHL00044)rplP (PRK09203)→ PRK00306 (PRK00306)Ribosomal_L29 (pfam00831)→ rpsQ (PRK05610)Ribosomal_S17 (pfam00366)→ rplN (PRK05483)Ribosomal_L14 (pfam00238)→ RplX (COG0198)rplX (PRK00004)→ rplE (PRK00010)RplE (COG0094)→ rpsN (PRK08061)RpsN (COG0199)→ rpsH (PRK00136)Ribosomal_S8 (pfam00410)→ |
| env-808 | longitudinal_293_123 | + | 13738 | 13822 | **RNA**→ rpsJ (PRK00596)RpsJ (COG0051)→ RplC (COG0087)rplC (PRK00001)→ rplD (PRK05319)Ribosomal_L4 (pfam00573)→ rplW (PRK05738)Ribosomal_L23 (pfam00276)→ RplB (COG0090)rplB (PRK09374)→ rpsS (PRK00357)RpsS (COG0185)→ rplV (PRK00565)Ribosomal_L22 (pfam00237)→ RpsC (COG0092)rpsC (PRK00310)→ rpl16 (CHL00044)rplP (PRK09203)→ PRK00306 (PRK00306)Ribosomal_L29 (pfam00831)→ rpsQ (PRK05610)Ribosomal_S17 (pfam00366)→ rplN (PRK05483)Ribosomal_L14 (pfam00238)→ RplX (COG0198)rplX (PRK00004)→ rplE (PRK00010)RplE (COG0094)→ rpsN (PRK08061)RpsN (COG0199)→ rpsH (PRK00136)Ribosomal_S8 (pfam00410)→ |
| env-809 | longitudinal_294_16 | - | 140925 | 140841 | **RNA**→ rpsJ (PRK00596)RpsJ (COG0051)→ RplC (COG0087)rplC (PRK00001)→ rplD (PRK05319)Ribosomal_L4 (pfam00573)→ rplW (PRK05738)Ribosomal_L23 (pfam00276)→ RplB (COG0090)rplB (PRK09374)→ rpsS (PRK00357)RpsS (COG0185)→ rplV (PRK00565)Ribosomal_L22 (pfam00237)→ RpsC (COG0092)rpsC (PRK00310)→ rpl16 (CHL00044)rplP (PRK09203)→ PRK00306 (PRK00306)Ribosomal_L29 (pfam00831)→ rpsQ (PRK05610)Ribosomal_S17 (pfam00366)→ rplN (PRK05483)Ribosomal_L14 (pfam00238)→ RplX (COG0198)rplX (PRK00004)→ rplE (PRK00010)RplE (COG0094)→ rpsN (PRK08061)RpsN (COG0199)→ rpsH (PRK00136)Ribosomal_S8 (pfam00410)→ |
| env-810 | longitudinal_344_234 | + | 686 | 770 | **RNA**→ rpsJ (PRK00596)RpsJ (COG0051)→ RplC (COG0087)rplC (PRK00001)→ rplD (PRK05319)Ribosomal_L4 (pfam00573)→ rplW (PRK05738)Ribosomal_L23 (pfam00276)→ RplB (COG0090)rplB (PRK09374)→ RplB (COG0090)rplB (PRK09374)→ rplB (PRK09374)Ribosomal_L2_C (pfam03947)→ rpsS (PRK00357)RpsS (COG0185)→ rplV (PRK00565)Ribosomal_L22 (pfam00237)→ RpsC (COG0092)rpsC (PRK00310)→ rpl16 (CHL00044)rplP (PRK09203)→ PRK00306 (PRK00306)Ribosomal_L29 (pfam00831)→ rpsQ (PRK05610)Ribosomal_S17 (pfam00366)→ rplN (PRK05483)Ribosomal_L14 (pfam00238)→ RplX (COG0198)rplX (PRK00004)→ rplE (PRK00010)RplE (COG0094)→ rpsN (PRK08061)RpsN (COG0199)→ rpsH (PRK00136)Ribosomal_S8 (pfam00410)→ |
| env-811 | longitudinal_345_116 | - | 4378 | 4294 | **RNA**→ rpsJ (PRK00596)RpsJ (COG0051)→ RplC (COG0087)rplC (PRK00001)→ rplD (PRK05319)Ribosomal_L4 (pfam00573)→ rplW (PRK05738)Ribosomal_L23 (pfam00276)→ RplB (COG0090)rplB (PRK09374)→ rpsS (PRK00357)RpsS (COG0185)→ rplV (PRK00565)Ribosomal_L22 (pfam00237)→ RpsC (COG0092)rpsC (PRK00310)→ |

| | | | | | |
|---|---|---|---|---|---|
| env-812 | longitudinal_133_22 | + | 48325 | 48409 | **RNA**→ rpsJ (PRK00596)RpsJ (COG0051)→ RplC (COG0087)rplC (PRK00001)→ rplD (PRK05319)Ribosomal_L4 (pfam00573)→ rplW (PRK05738)Ribosomal_L23 (pfam00276)→ RplB (COG0090)rplB (PRK09374)→ rpsS (PRK00357)RpsS (COG0185)→ rplV (PRK00565)Ribosomal_L22 (pfam00237)→ RpsC (COG0092)rpsC (PRK00310)→ rpl16 (CHL00044)rplP (PRK09203)→ PRK00306 (PRK00306)Ribosomal_L29 (pfam00831)→ rpsQ (PRK05610)Ribosomal_S17 (pfam00366)→ rplN (PRK05483)Ribosomal_L14 (pfam00238)→ RplX (COG0198)rplX (PRK00004)→ rplE (PRK00010)RplE (COG0094)→ rpsN (PRK08061)RpsN (COG0199)→ rpsH (PRK00136)Ribosomal_S8 (pfam00410)→ |
| env-813 | longitudinal_186_6 | + | 48325 | 48409 | **RNA**→ rpsJ (PRK00596)RpsJ (COG0051)→ RplC (COG0087)rplC (PRK00001)→ rplD (PRK05319)Ribosomal_L4 (pfam00573)→ rplW (PRK05738)Ribosomal_L23 (pfam00276)→ RplB (COG0090)rplB (PRK09374)→ rpsS (PRK00357)RpsS (COG0185)→ rplV (PRK00565)Ribosomal_L22 (pfam00237)→ rplV (PRK00565)Ribosomal_L22 (pfam00237)→ RpsC (COG0092)rpsC (PRK00310)→ rpl16 (CHL00044)rplP (PRK09203)→ PRK00306 (PRK00306)Ribosomal_L29 (pfam00831)→ rpsQ (PRK05610)Ribosomal_S17 (pfam00366)→ rplN (PRK05483)Ribosomal_L14 (pfam00238)→ RplX (COG0198)rplX (PRK00004)→ rplE (PRK00010)RplE (COG0094)→ rpsN (PRK08061)RpsN (COG0199)→ rpsH (PRK00136)Ribosomal_S8 (pfam00410)→ |
| env-814 | longitudinal_295_1 | - | 44521 | 44440 | **RNA**→ rpsJ (PRK00596)RpsJ (COG0051)→ RplC (COG0087)rplC (PRK00001)→ rplD (PRK05319)Ribosomal_L4 (pfam00573)→ rplW (PRK05738)Ribosomal_L23 (pfam00276)→ RplB (COG0090)rplB (PRK09374)→ rpsS (PRK00357)RpsS (COG0185)→ rplV (PRK00565)Ribosomal_L22 (pfam00237)→ RpsC (COG0092)rpsC (PRK00310)→ rpl16 (CHL00044)rplP (PRK09203)→ PRK00306 (PRK00306)Ribosomal_L29 (pfam00831)→ rpsQ (PRK05610)Ribosomal_S17 (pfam00366)→ rplN (PRK05483)Ribosomal_L14 (pfam00238)→ RplX (COG0198)rplX (PRK00004)→ rplE (PRK00010)RplE (COG0094)→ rpsN (PRK08061)RpsN (COG0199)→ |
| env-815 | longitudinal_296_205 | - | 4377 | 4296 | **RNA**→ rpsJ (PRK00596)RpsJ (COG0051)→ RplC (COG0087)rplC (PRK00001)→ rplD (PRK05319)Ribosomal_L4 (pfam00573)→ rplW (PRK05738)Ribosomal_L23 (pfam00276)→ RplB (COG0090)rplB (PRK09374)→ rpsS (PRK00357)RpsS (COG0185)→ rplV (PRK00565)Ribosomal_L22 (pfam00237)→ KH (smart00322)RpsC (COG0092)→ |
| env-816 | longitudinal_297_58 | - | 44521 | 44440 | **RNA**→ rpsJ (PRK00596)RpsJ (COG0051)→ RplC (COG0087)rplC (PRK00001)→ rplD (PRK05319)Ribosomal_L4 (pfam00573)→ rplW (PRK05738)Ribosomal_L23 (pfam00276)→ RplB (COG0090)rplB (PRK09374)→ rpsS (PRK00357)RpsS (COG0185)→ rplV (PRK00565)Ribosomal_L22 (pfam00237)→ RpsC (COG0092)rpsC (PRK00310)→ rpl16 (CHL00044)rplP (PRK09203)→ PRK00306 (PRK00306)Ribosomal_L29 (pfam00831)→ rpsQ (PRK05610)Ribosomal_S17 (pfam00366)→ rplN (PRK05483)Ribosomal_L14 (pfam00238)→ RplX (COG0198)rplX (PRK00004)→ rplE (PRK00010)RplE (COG0094)→ rpsN (PRK08061)RpsN (COG0199)→ |
| env-817 | longitudinal_298_873 | + | 4316 | 4397 | **RNA**→ rpsJ (PRK00596)RpsJ (COG0051)→ RplC (COG0087)rplC (PRK00001)→ rplD (PRK05319)Ribosomal_L4 (pfam00573)→ rplW (PRK05738)Ribosomal_L23 (pfam00276)→ RplB (COG0090)rplB (PRK09374)→ |
| Dfo-2-1 | NZ_AAXA02000012.1 | - | 139959 | 139878 | **RNA**→ rpsJ (PRK00596)RpsJ (COG0051)→ rpsJ (PRK00596)RpsJ (COG0051)→ RplC (COG0087)rplC (PRK00001)→ RplC (COG0087)rplC (PRK00001)→ rplD (PRK05319)Ribosomal_L4 (pfam00573)→ rplD (PRK05319)Ribosomal_L4 (pfam00573)→ rplW (PRK05738)Ribosomal_L23 (pfam00276)→ rplW (PRK05738)Ribosomal_L23 (pfam00276)→ |
| Dfo-1-1 | NZ_JH126494.1 | - | 105382 | 105301 | **RNA**→ rpsJ (PRK00596)RpsJ (COG0051)→ RplC (COG0087)rplC (PRK00001)→ rplD (PRK05319)Ribosomal_L4 (pfam00573)→ rplW (PRK05738)Ribosomal_L23 (pfam00276)→ RplB (COG0090)rplB (PRK09374)→ rpsS (PRK00357)RpsS (COG0185)→ rplV (PRK00565)Ribosomal_L22 (pfam00237)→ RpsC (COG0092)rpsC (PRK00310)→ rpl16 (CHL00044)rplP (PRK09203)→ PRK00306 (PRK00306)Ribosomal_L29 (pfam00831)→ rpsQ (PRK05610)Ribosomal_S17 (pfam00366)→ rplN (PRK05483)Ribosomal_L14 (pfam00238)→ RplX (COG0198)rplX (PRK00004)→ rplE (PRK00010)RplE (COG0094)→ rpsN (PRK08061)RpsN (COG0199)→ |
| env-818 | longitudinal_421_71 | + | 42399 | 42480 | **RNA**→ rpsJ (PRK00596)RpsJ (COG0051)→ RplC (COG0087)rplC (PRK00001)→ rplD (PRK05319)Ribosomal_L4 (pfam00573)→ rplW (PRK05738)Ribosomal_L23 (pfam00276)→ RplB (COG0090)rplB (PRK09374)→ rpsS (PRK00357)RpsS (COG0185)→ rplV (PRK00565)Ribosomal_L22 (pfam00237)→ RpsC (COG0092)rpsC (PRK00310)→ rpl16 (CHL00044)rplP (PRK09203)→ PRK00306 (PRK00306)Ribosomal_L29 (pfam00831)→ rpsQ (PRK05610)Ribosomal_S17 (pfam00366)→ rplN (PRK05483)Ribosomal_L14 (pfam00238)→ RplX (COG0198)rplX (PRK00004)→ rplE (PRK00010)RplE (COG0094)→ rpsN (PRK08061)RpsN (COG0199)→ |

| | | | | | |
|---|---|---|---|---|---|
| env-819 | longitudinal_187_175 | + | 27495 | 27576 | RNA→ rpsJ (PRK00596)RpsJ (COG0051)→ RplC (COG0087)rplC (PRK00001)→ rplD (PRK05319)Ribosomal_L4 (pfam00573)→ rplW (PRK05738)Ribosomal_L23 (pfam00276)→ RplB (COG0090)rplB (PRK09374)→ rpsS (PRK00357)RpsS (COG0185)→ rplV (PRK00565)Ribosomal_L22 (pfam00237)→ RpsC (COG0092)rpsC (PRK00310)→ rpl16 (CHL00044)rplP (PRK09203)→ PRK00306 (PRK00306)Ribosomal_L29 (pfam00831)→ rpsQ (PRK05610)Ribosomal_S17 (pfam00366)→ rplN (PRK05483)Ribosomal_L14 (pfam00238)→ RplX (COG0198)rplX (PRK00004)→ rplE (PRK00010)RplE (COG0094)→ rpsN (PRK08061)RpsN (COG0199)→ |
| env-820 | longitudinal_519_34 | + | 75799 | 75884 | RNA→ rpsJ (PRK00596)RpsJ (COG0051)→ |
| Cbo-6-1 | NZ_DS480701.1 | - | 29789 | 29704 | RNA→ rpsJ (PRK00596)RpsJ (COG0051)→ |
| Cbo-1-1 | NZ_KB851188.1 | + | 671984 | 672069 | RNA→ rpsJ (PRK00596)RpsJ (COG0051)→ |
| Cbo-5-1 | NZ_KB851140.1 | - | 31903 | 31818 | RNA→ rpsJ (PRK00596)RpsJ (COG0051)→ |
| Cbo-4-1 | NZ_KB851170.1 | - | 436154 | 436069 | RNA→ rpsJ (PRK00596)RpsJ (COG0051)→ |
| [bo-1-1 | NZ_KQ235847.1 | + | 638432 | 638517 | RNA→ rpsJ (PRK00596)RpsJ (COG0051)→ |
| [ci-1-1 | NZ_KQ235875.1 | - | 58399 | 58314 | RNA→ rpsJ (PRK00596)RpsJ (COG0051)→ |
| Csp-5-1 | NZ_JYHN01000073.1 | + | 12472 | 12557 | RNA→ rpsJ (PRK00596)RpsJ (COG0051)→ |
| Cci-1-1 | NZ_JH376430.1 | - | 58181 | 58096 | RNA→ rpsJ (PRK00596)RpsJ (COG0051)→ |
| env-821 | DOM008_scaffold5856_4 | - | 1719 | 1634 | RNA→ rpsJ (PRK00596)RpsJ (COG0051)→ |
| env-822 | DLM024_scaffold3474_1 | + | 221 | 306 | RNA→ rpsJ (PRK00596)RpsJ (COG0051)→ |
| env-823 | DOM022_scaffold7182_3 | - | 11175 | 11090 | RNA→ rpsJ (PRK00596)RpsJ (COG0051)→ |
| Cbo-2-1 | NZ_KB851182.1 | - | 3727490 | 3727405 | RNA→ rpsJ (PRK00596)RpsJ (COG0051)→ |
| Cbo-3-1 | NZ_KB851179.1 | - | 40006 | 39921 | RNA→ rpsJ (PRK00596)RpsJ (COG0051)→ |
| Cba-4-1 | NZ_BAHU02000028.1 | + | 24715 | 24800 | RNA→ rpsJ (PRK00596)RpsJ (COG0051)→ |
| Cba-7-1 | NZ_BAIK02000095.1 | - | 31314 | 31229 | RNA→ rpsJ (PRK00596)RpsJ (COG0051)→ |
| env-824 | DLF007_scaffold1906_9 | - | 8119 | 8034 | RNA→ rpsJ (PRK00596)RpsJ (COG0051)→ |
| env-825 | DOM005_scaffold2526_23 | + | 748 | 833 | RNA→ rpsJ (PRK00596)RpsJ (COG0051)→ |
| Cba-2-1 | NZ_DS990264.1 | + | 173299 | 173384 | RNA→ rpsJ (PRK00596)RpsJ (COG0051)→ rpsJ (PRK00596)RpsJ (COG0051)→ |
| env-826 | DOF010_scaffold27370_9 | - | 556 | 471 | RNA→ rpsJ (PRK00596)RpsJ (COG0051)→ |
| Ccl-6-1 | NZ_KB851048.1 | - | 152029 | 151944 | RNA→ rpsJ (PRK00596)RpsJ (COG0051)→ |
| env-827 | BABA01014305.1 | - | 333 | 248 | RNA→ |
| Ccl-3-1 | NZ_KB851058.1 | - | 172004 | 171919 | RNA→ hypo→ rpsJ (PRK00596)RpsJ (COG0051)→ |
| Ccl-9-1 | NZ_KB850936.1 | + | 2207674 | 2207759 | RNA→ hypo→ rpsJ (PRK00596)RpsJ (COG0051)→ |
| Ccl-4-1 | NZ_KB851076.1 | + | 62132 | 62217 | RNA→ hypo→ rpsJ (PRK00596)RpsJ (COG0051)→ |
| Ccl-8-1 | NZ_KB850966.1 | + | 1570893 | 1570978 | RNA→ hypo→ rpsJ (PRK00596)RpsJ (COG0051)→ |
| Ccl-5-1 | NZ_KB851039.1 | + | 308067 | 308152 | RNA→ hypo→ rpsJ (PRK00596)RpsJ (COG0051)→ |
| Ccl-2-1 | NZ_KB851121.1 | - | 453818 | 453733 | RNA→ hypo→ rpsJ (PRK00596)RpsJ (COG0051)→ |
| Ccl-7-1 | NZ_KB850990.1 | - | 33581 | 33496 | RNA→ hypo→ rpsJ (PRK00596)RpsJ (COG0051)→ |
| [cl-1-1 | NZ_KQ235916.1 | - | 65391 | 65306 | RNA→ hypo→ rpsJ (PRK00596)RpsJ (COG0051)→ |
| Ccl-1-1 | NZ_JH376872.1 | + | 684262 | 684347 | RNA→ hypo→ rpsJ (PRK00596)RpsJ (COG0051)→ |
| Lba-16-1 | NZ_KE159611.1 | - | 333481 | 333384 | RNA→ rpsJ (PRK00596)RpsJ (COG0051)→ |
| Csp-3-1 | NZ_KB822452.1 | + | 182613 | 182710 | RNA→ rpsJ (PRK00596)RpsJ (COG0051)→ |
| env-828 | 4491488.3_NODE_13653 | - | 1210 | 1116 | RNA→ rpsJ (PRK00596)RpsJ (COG0051)→ RplC (COG0087)rplC (PRK00001)→ |
| env-829 | DLM023_scaffold19875_8 | + | 194 | 288 | RNA→ rpsJ (PRK00596)RpsJ (COG0051)→ RplC (COG0087)rplC (PRK00001)→ |
| env-830 | NOF007_scaffold11968_1 | + | 314 | 408 | RNA→ hypo→ |
| env-831 | SRS017521_Baylor_scaffold_53704 | - | 166 | 72 | RNA→ |
| env-832 | NLM015_scaffold7043_13 | + | 1676 | 1770 | RNA→ hypo→ |
| env-833 | 4491420.3_NODE_1923 | - | 275 | 181 | RNA→ |
| env-834 | DOM022_scaffold13753_3 | - | 1709 | 1615 | RNA→ rpsJ (PRK00596)RpsJ (COG0051)→ RplC (COG0087)rplC (PRK00001)→ rplD (PRK05319)Ribosomal_L4 (pfam00573)→ |
| env-835 | NLM006_scaffold4964_7 | + | 3718 | 3812 | RNA→ rpsJ (PRK00596)RpsJ (COG0051)→ RplC (COG0087)rplC (PRK00001)→ rplD (PRK05319)Ribosomal_L4 (pfam00573)→ rplW (PRK05738)Ribosomal_L23 (pfam00276)→ RplB (COG0090)rplB (PRK09374)→ |
| env-836 | NLM031_scaffold4829_11 | + | 921 | 1015 | RNA→ rpsJ (PRK00596)RpsJ (COG0051)→ RplC (COG0087)rplC (PRK00001)→ RplD (COG0088)rplD (PRK05319)→ |

| | | | | | |
|---|---|---|---|---|---|
| Rto-1-1 | NZ_DS264365.1 | - | 34936 | 34842 | **RNA**→ hypo→ |
| env-837 | 4491479.3_NODE_8774 | + | 203259 | 203353 | **RNA**→ rpsJ (PRK00596)RpsJ (COG0051)→ RplC (COG0087)rplC (PRK00001)→ rplD (PRK05319)Ribosomal_L4 (pfam00573)→ rplW (PRK05738)Ribosomal_L23 (pfam00276)→ RplB (COG0090)rplB (PRK09374)→ rpsS (PRK00357)RpsS (COG0185)→ rplV (PRK00565)Ribosomal_L22 (pfam00237)→ rpsC (PRK00310)30S_S3_KH (cd02412)→ |
| env-838 | SRS022713_LANL_scaffold_9359 | - | 1239 | 1145 | **RNA**→ rpsJ (PRK00596)Ribosomal_S10 (pfam00338)→ |
| env-839 | DLM021_scaffold9512_3 | + | 55 | 149 | **RNA**→ rpsJ (PRK00596)RpsJ (COG0051)→ |
| env-840 | SRS014923_WUGC_scaffold_52186 | + | 3683 | 3777 | **RNA**→ |
| Lba-1-1 | NZ_GL945299.1 | - | 131629 | 131535 | **RNA**→ rpsJ (PRK00596)RpsJ (COG0051)→ RplC (COG0087)rplC (PRK00001)→ rplD (PRK05319)Ribosomal_L4 (pfam00573)→ rplW (PRK05738)Ribosomal_L23 (pfam00276)→ RplB (COG0090)rplB (PRK09374)→ rpsS (PRK00357)RpsS (COG0185)→ rplV (PRK00565)Ribosomal_L22 (pfam00237)→ RpsC (COG0092)rpsC (PRK00310)→ rpl16 (CHL00044)rplP (PRK09203)→ PRK00306 (PRK00306)Ribosomal_L29 (pfam00831)→ rpsQ (PRK05610)Ribosomal_S17 (pfam00366)→ rplN (PRK05483)Ribosomal_L14 (pfam00238)→ RplX (COG0198)rplX (PRK00004)→ rplE (PRK00010)RplE (COG0094)→ rpsN (PRK08061)RpsN (COG0199)→ rpsH (PRK00136)Ribosomal_S8 (pfam00410)→ |
| Lba-11-1 | NZ_GL622455.1 | + | 63318 | 63412 | **RNA**→ rpsJ (PRK00596)RpsJ (COG0051)→ RplC (COG0087)rplC (PRK00001)→ rplD (PRK05319)Ribosomal_L4 (pfam00573)→ rplW (PRK05738)Ribosomal_L23 (pfam00276)→ RplB (COG0090)rplB (PRK09374)→ rpsS (PRK00357)RpsS (COG0185)→ rplV (PRK00565)Ribosomal_L22 (pfam00237)→ RpsC (COG0092)rpsC (PRK00310)→ rpl16 (CHL00044)rplP (PRK09203)→ PRK00306 (PRK00306)Ribosomal_L29 (pfam00831)→ rpsQ (PRK05610)Ribosomal_S17 (pfam00366)→ rplN (PRK05483)Ribosomal_L14 (pfam00238)→ RplX (COG0198)rplX (PRK00004)→ rplE (PRK00010)RplE (COG0094)→ rpsN (PRK08061)RpsN (COG0199)→ rpsH (PRK00136)Ribosomal_S8 (pfam00410)→ |
| env-841 | DOM008_scaffold11099_2 | + | 379 | 473 | **RNA**→ rpsJ (PRK00596)RpsJ (COG0051)→ |
| Lba-7-1 | NZ_GL890521.1 | - | 333856 | 333762 | **RNA**→ rpsJ (PRK00596)RpsJ (COG0051)→ RplC (COG0087)rplC (PRK00001)→ rplD (PRK05319)Ribosomal_L4 (pfam00573)→ rplW (PRK05738)Ribosomal_L23 (pfam00276)→ RplB (COG0090)rplB (PRK09374)→ rpsS (PRK00357)RpsS (COG0185)→ rplV (PRK00565)Ribosomal_L22 (pfam00237)→ RpsC (COG0092)rpsC (PRK00310)→ rpl16 (CHL00044)rplP (PRK09203)→ PRK00306 (PRK00306)Ribosomal_L29 (pfam00831)→ rpsQ (PRK05610)Ribosomal_S17 (pfam00366)→ rplN (PRK05483)Ribosomal_L14 (pfam00238)→ RplX (COG0198)rplX (PRK00004)→ rplE (PRK00010)RplE (COG0094)→ rpsN (PRK08061)RpsN (COG0199)→ rpsH (PRK00136)Ribosomal_S8 (pfam00410)→ |
| env-842 | 4448814.3_Contig31048 | + | 343 | 437 | **RNA**→ hypo→ |
| env-843 | DLF009_scaffold139_11 | - | 490 | 383 | **RNA**→ rpsJ (PRK00596)RpsJ (COG0051)→ |
| env-844 | UnmappedStool_Broad_scaffold_696739 | - | 332 | 252 | **RNA**→ hypo→ |
| env-845 | 4491413.3_NODE_28238 | - | 28876 | 28796 | **RNA**→ rpsJ (PRK00596)RpsJ (COG0051)→ |
| env-846 | 4491423.3_NODE_19525 | - | 4056 | 3976 | **RNA**→ rpsJ (PRK00596)RpsJ (COG0051)→ |
| env-847 | SRS052697_LANL_scaffold_18998 | - | 390 | 310 | **RNA**→ rpsJ (PRK00596)Ribosomal_S10 (pfam00338)→ |
| env-848 | NLF013_scaffold9702_5 | - | 1227 | 1147 | **RNA**→ rpsJ (PRK00596)RpsJ (COG0051)→ |
| env-849 | NLF009_scaffold47128_2 | + | 415 | 495 | **RNA**→ |
| env-850 | scaffold72408_1_MH0058 | - | 228 | 148 | **RNA**→ hypo→ |
| env-851 | scaffold14894_1_MH0014 | - | 249 | 168 | **RNA**→ |
| env-852 | SRS015217_WUGC_scaffold_12868 | + | 2064 | 2145 | **RNA**→ |
| env-853 | DLM008_scaffold14278_10 | - | 4516 | 4435 | **RNA**→ rpsJ (PRK00596)RpsJ (COG0051)→ |
| env-854 | UnmappedStool_Broad_scaffold_412435 | - | 349 | 268 | **RNA**→ rpsJ (PRK00596)RpsJ (COG0051)→ |
| env-855 | DOM019_C726307_1 | - | 271 | 190 | **RNA**→ |
| env-856 | 4491417.3_NODE_36334 | + | 887 | 968 | **RNA**→ |
| env-857 | DLM028_scaffold37057_1_2 | - | 716 | 635 | **RNA**→ rpsJ (PRK00596)RpsJ (COG0051)→ |
| env-858 | NLM031_scaffold55673_5 | + | 29666 | 29747 | **RNA**→ rpsJ (PRK00596)RpsJ (COG0051)→ |
| env-859 | NOF013_scaffold6414_30 | + | 2033 | 2114 | **RNA**→ rpsJ (PRK00596)RpsJ (COG0051)→ |
| env-860 | longitudinal_438_77 | - | 42376 | 42295 | **RNA**→ rpsJ (PRK00596)RpsJ (COG0051)→ |
| Rho-1-1 | NC_015977.1 | - | 3143074 | 3142993 | **RNA**→ rpsJ (PRK00596)RpsJ (COG0051)→ |
| env-861 | NLM026_scaffold36927_9 | - | 437 | 356 | **RNA**→ rpsJ (PRK00596)Ribosomal_S10 (pfam00338)→ |

| | | | | | |
|---|---|---|---|---|---|
| env-862 | SRS049164_WUGC_scaffold_63181 | - | 252 | 171 | **RNA**→ ←hypo |
| env-863 | DLF012_scaffold4039_3 | + | 4877 | 4957 | **RNA**→ rpsJ (PRK00596)RpsJ (COG0051)→ |
| env-864 | SRS050422_LANL_scaffold_15527 | - | 181 | 101 | **RNA**→ |
| env-865 | 4491485.3_NODE_8452 | - | 8128 | 8047 | **RNA**→ hypo→ rpsJ (PRK00596)RpsJ (COG0051)→ |
| env-866 | DOM010_scaffold114_1 | + | 76895 | 76976 | **RNA**→ hypo→ rpsJ (PRK00596)RpsJ (COG0051)→ |
| env-867 | NLF010_scaffold280_62 | + | 221 | 302 | **RNA**→ rpsJ (PRK00596)RpsJ (COG0051)→ |
| env-868 | scaffold6333_2_O2.UC-11 | + | 2568 | 2649 | **RNA**→ rpsJ (PRK00596)Ribosomal_S10 (pfam00338)→ |
| env-869 | SRS014923_WUGC_scaffold_21701 | - | 206 | 125 | **RNA**→ |
| env-870 | SRS048164_WUGC_scaffold_6926 | - | 364 | 283 | **RNA**→ rpsJ (PRK00596)Ribosomal_S10 (pfam00338)→ |
| env-871 | SRS053398_LANL_scaffold_19345 | - | 207 | 126 | **RNA**→ hypo→ |
| env-872 | SRS024331_LANL_scaffold_6342 | - | 361 | 280 | **RNA**→ rpsJ (PRK00596)Ribosomal_S10 (pfam00338)→ |
| env-873 | SRS042628_WUGC_scaffold_13655 | - | 207 | 126 | **RNA**→ |
| env-874 | 4491413.3_NODE_36320 | + | 36281 | 36362 | **RNA**→ hypo→ rpsJ (PRK00596)RpsJ (COG0051)→ |
| env-875 | SRS049164_WUGC_scaffold_48687 | + | 416 | 497 | **RNA**→ hypo→ |
| env-876 | SRS047014_WUGC_scaffold_33266 | - | 207 | 126 | **RNA**→ |
| env-877 | DOF014_scaffold33897_7 | + | 141 | 222 | **RNA**→ rpsJ (PRK00596)Ribosomal_S10 (pfam00338)→ |
| env-878 | DOM024_scaffold8082_9 | + | 242 | 323 | **RNA**→ rpsJ (PRK00596)RpsJ (COG0051)→ |
| env-879 | SRS058770_LANL_scaffold_8927 | - | 361 | 280 | **RNA**→ rpsJ (PRK00596)Ribosomal_S10 (pfam00338)→ |
| env-880 | 4491403.3_NODE_43191 | - | 1209 | 1128 | **RNA**→ rpsJ (PRK00596)Ribosomal_S10 (pfam00338)→ |
| env-881 | 4491404.3_NODE_39204 | + | 16 | 97 | **RNA**→ rpsJ (PRK00596)RpsJ (COG0051)→ |
| env-882 | DLM015_scaffold18244_10 | + | 1214 | 1295 | **RNA**→ hypo→ rpsJ (PRK00596)RpsJ (COG0051)→ |
| env-883 | DLM021_scaffold23002_4 | - | 9326 | 9245 | **RNA**→ hypo→ rpsJ (PRK00596)RpsJ (COG0051)→ |
| env-884 | DOF011_scaffold5789_15 | + | 252 | 333 | **RNA**→ rpsJ (PRK00596)Ribosomal_S10 (pfam00338)→ |
| env-885 | DOM021_scaffold13341_7 | + | 164 | 245 | **RNA**→ rpsJ (PRK00596)RpsJ (COG0051)→ |
| env-886 | NLM008_scaffold17065_3 | - | 3947 | 3866 | **RNA**→ hypo→ rpsJ (PRK00596)RpsJ (COG0051)→ |
| env-887 | NLM009_scaffold5386_5 | + | 2122 | 2203 | **RNA**→ hypo→ rpsJ (PRK00596)RpsJ (COG0051)→ |
| env-888 | NLM021_scaffold13120_8 | + | 3360 | 3441 | **RNA**→ hypo→ rpsJ (PRK00596)RpsJ (COG0051)→ |
| env-889 | NOM020_scaffold2140_8 | - | 6244 | 6163 | **RNA**→ hypo→ rpsJ (PRK00596)RpsJ (COG0051)→ |
| env-890 | NOM025_scaffold13467_1 | - | 49759 | 49678 | **RNA**→ hypo→ rpsJ (PRK00596)RpsJ (COG0051)→ |
| env-891 | NOM029_scaffold63_2 | - | 3400 | 3319 | **RNA**→ hypo→ rpsJ (PRK00596)RpsJ (COG0051)→ |
| env-892 | 4491419.3_NODE_20619 | + | 4027 | 4108 | **RNA**→ hypo→ rpsJ (PRK00596)RpsJ (COG0051)→ |
| env-893 | DLM028_scaffold2088_3 | - | 11513 | 11432 | **RNA**→ hypo→ rpsJ (PRK00596)RpsJ (COG0051)→ |
| env-894 | NLM004_scaffold4826_13 | + | 1805 | 1886 | **RNA**→ hypo→ rpsJ (PRK00596)RpsJ (COG0051)→ |
| env-895 | NLM032_scaffold6862_1 | + | 3653 | 3734 | **RNA**→ hypo→ rpsJ (PRK00596)RpsJ (COG0051)→ |
| env-896 | NOF006_scaffold7748_15 | + | 408 | 489 | **RNA**→ hypo→ rpsJ (PRK00596)RpsJ (COG0051)→ |
| env-897 | NOF007_scaffold4503_1 | + | 37612 | 37693 | **RNA**→ hypo→ rpsJ (PRK00596)RpsJ (COG0051)→ |
| env-898 | DOM022_scaffold52_3 | - | 38941 | 38860 | **RNA**→ hypo→ rpsJ (PRK00596)RpsJ (COG0051)→ |
| env-899 | NOM002_scaffold8686_2 | + | 209 | 290 | **RNA**→ rpsJ (PRK00596)RpsJ (COG0051)→ |
| env-900 | SRS015264_WUGC_scaffold_25881 | - | 207 | 126 | **RNA**→ |
| env-901 | scaffold2311_1_V1.UC-21 | - | 204 | 123 | **RNA**→ |
| env-902 | 4491420.3_NODE_8720 | + | 93 | 174 | **RNA**→ |
| env-903 | DLM020_scaffold4730_1 | - | 107388 | 107307 | **RNA**→ hypo→ rpsJ (PRK00596)RpsJ (COG0051)→ |
| env-904 | DOM018_scaffold13457_5 | - | 1831 | 1750 | **RNA**→ rpsJ (PRK00596)RpsJ (COG0051)→ |
| env-905 | NOM007_scaffold35894_2 | - | 1318 | 1237 | **RNA**→ hypo→ rpsJ (PRK00596)RpsJ (COG0051)→ |
| env-906 | NOM022_scaffold18782_6 | + | 4188 | 4269 | **RNA**→ hypo→ rpsJ (PRK00596)RpsJ (COG0051)→ |
| Rin-2-1 | NZ_ACFY01000086.1 | + | 87520 | 87601 | **RNA**→ rpsJ (PRK00596)RpsJ (COG0051)→ rpsJ (PRK00596)RpsJ (COG0051)→ |
| env-907 | scaffold2088_3_MH0067 | + | 748 | 829 | **RNA**→ rpsJ (PRK00596)Ribosomal_S10 (pfam00338)→ |
| env-908 | scaffold3368_5_V1.UC-7 | - | 182 | 101 | **RNA**→ |
| env-909 | DLM002_scaffold8503_15 | + | 532 | 613 | **RNA**→ hypo→ rpsJ (PRK00596)RpsJ (COG0051)→ |
| env-910 | DOM017_scaffold4879_3 | + | 2589 | 2670 | **RNA**→ hypo→ rpsJ (PRK00596)RpsJ (COG0051)→ |
| env-911 | NOM014_scaffold11459_3 | - | 4961 | 4880 | **RNA**→ hypo→ rpsJ (PRK00596)RpsJ (COG0051)→ |
| env-912 | SRS064276_LANL_scaffold_12898 | - | 359 | 278 | **RNA**→ rpsJ (PRK00596)Ribosomal_S10 (pfam00338)→ |

| | | | | | |
|---|---|---|---|---|---|
| env-913 | NLF013_scaffold11235_2 | + | 33240 | 33321 | **RNA**→ rpsJ (PRK00596)RpsJ (COG0051)→ |
| env-914 | NOM026_scaffold13266_5 | - | 2455 | 2374 | **RNA**→ rpsJ (PRK00596)RpsJ (COG0051)→ |
| env-915 | NOM018_C815838_1 | - | 251 | 170 | **RNA**→ |
| env-916 | NLF002_scaffold7830_8 | + | 14588 | 14669 | **RNA**→ rpsJ (PRK00596)RpsJ (COG0051)→ |
| env-917 | NOF014_scaffold631_2 | - | 1554 | 1473 | **RNA**→ rpsJ (PRK00596)RpsJ (COG0051)→ |
| env-918 | longitudinal_461_56 | + | 96014 | 96096 | **RNA**→ rpsJ (PRK00596)RpsJ (COG0051)→ |
| env-919 | longitudinal_462_123 | + | 96021 | 96103 | **RNA**→ rpsJ (PRK00596)RpsJ (COG0051)→ |
| env-920 | longitudinal_463_7 | + | 96041 | 96123 | **RNA**→ rpsJ (PRK00596)RpsJ (COG0051)→ |
| Lba-10-1 | NZ_GL945245.1 | - | 74466 | 74384 | **RNA**→ rpsJ (PRK00596)RpsJ (COG0051)→ RplC (COG0087)rplC (PRK00001)→ rplD (PRK05319)Ribosomal_L4 (pfam00573)→ rplW (PRK05738)Ribosomal_L23 (pfam00276)→ RplB (COG0090)rplB (PRK09374)→ rpsS (PRK00357)RpsS (COG0185)→ rplV (PRK00565)Ribosomal_L22 (pfam00237)→ RpsC (COG0092)rpsC (PRK00310)→ rpl16 (CHL00044)rplP (PRK09203)→ PRK00306 (PRK00306)Ribosomal_L29 (pfam00831)→ rpsQ (PRK05610)Ribosomal_S17 (pfam00366)→ rplN (PRK05483)Ribosomal_L14 (pfam00238)→ RplX (COG0198)rplX (PRK00004)→ rplE (PRK00010)RplE (COG0094)→ rpsN (PRK08061)RpsN (COG0199)→ rpsH (PRK00136)Ribosomal_S8 (pfam00410)→ rpl6 (CHL00140)rplF (PRK05498)→ rplR (PRK05593)Ribosomal_L18p (pfam00861)→ RpsE (COG0098)rpsE (PRK00550)→ Ribosomal_L30 (cd01658)rpmD (PRK05611)→ RplO (COG0200)rplO (PRK05592)→ SecY (COG0201)secY (PRK09204)→ |
| [sc-1-1 | NZ_BAHS01000033.1 | + | 7765 | 7847 | **RNA**→ rpsJ (PRK00596)RpsJ (COG0051)→ |
| env-921 | DLF012_scaffold283_15 | + | 2596 | 2678 | **RNA**→ rpsJ (PRK00596)RpsJ (COG0051)→ |
| env-922 | longitudinal_337_87 | + | 26835 | 26917 | **RNA**→ rpsJ (PRK00596)RpsJ (COG0051)→ RplC (COG0087)rplC (PRK00001)→ rplD (PRK05319)Ribosomal_L4 (pfam00573)→ rplW (PRK05738)Ribosomal_L23 (pfam00276)→ RplB (COG0090)rplB (PRK09374)→ rpsS (PRK00357)RpsS (COG0185)→ rplV (PRK00565)Ribosomal_L22 (pfam00237)→ RpsC (COG0092)rpsC (PRK00310)→ rpl16 (CHL00044)rplP (PRK09203)→ PRK00306 (PRK00306)Ribosomal_L29 (pfam00831)→ rpsQ (PRK05610)Ribosomal_S17 (pfam00366)→ rplN (PRK05483)Ribosomal_L14 (pfam00238)→ RplX (COG0198)rplX (PRK00004)→ rplE (PRK00010)RplE (COG0094)→ rpsN (PRK08061)RpsN (COG0199)→ rpsH (PRK00136)Ribosomal_S8 (pfam00410)→ rpl6 (CHL00140)rplF (PRK05498)→ rplR (PRK05593)Ribosomal_L18p (pfam00861)→ RpsE (COG0098)rpsE (PRK00550)→ Ribosomal_L30 (cd01658)rpmD (PRK05611)→ RplO (COG0200)rplO (PRK05592)→ SecY (COG0201)secY (PRK09204)→ |
| env-923 | longitudinal_522_1406 | + | 359 | 441 | **RNA**→ rpsJ (PRK00596)RpsJ (COG0051)→ |
| Csc-1-1 | NZ_DS499705.1 | - | 73976 | 73894 | **RNA**→ rpsJ (PRK00596)RpsJ (COG0051)→ RplC (COG0087)rplC (PRK00001)→ rplD (PRK05319)Ribosomal_L4 (pfam00573)→ rplW (PRK05738)Ribosomal_L23 (pfam00276)→ RplB (COG0090)rplB (PRK09374)→ rpsS (PRK00357)RpsS (COG0185)→ rplV (PRK00565)Ribosomal_L22 (pfam00237)→ RpsC (COG0092)rpsC (PRK00310)→ rpl16 (CHL00044)rplP (PRK09203)→ PRK00306 (PRK00306)Ribosomal_L29 (pfam00831)→ rpsQ (PRK05610)Ribosomal_S17 (pfam00366)→ rplN (PRK05483)Ribosomal_L14 (pfam00238)→ RplX (COG0198)rplX (PRK00004)→ rplE (PRK00010)RplE (COG0094)→ rpsN (PRK08061)RpsN (COG0199)→ rpsH (PRK00136)Ribosomal_S8 (pfam00410)→ rpl6 (CHL00140)rplF (PRK05498)→ rplR (PRK05593)Ribosomal_L18p (pfam00861)→ rplR (PRK05593)Ribosomal_L18_L5e (cd00432)→ RpsE (COG0098)rpsE (PRK00550)→ Ribosomal_L30 (cd01658)rpmD (PRK05611)→ RplO (COG0200)rplO (PRK05592)→ SecY (COG0201)secY (PRK09204)→ |
| Cba-6-1 | NZ_BAII02000008.1 | + | 178284 | 178366 | **RNA**→ rpsJ (PRK00596)RpsJ (COG0051)→ |
| Bhy-1-1 | NZ_GG657686.1 | + | 266696 | 266777 | **RNA**→ rpsJ (PRK00596)RpsJ (COG0051)→ rpsJ (PRK00596)RpsJ (COG0051)→ RplC (COG0087)rplC (PRK00001)→ rplD (PRK05319)Ribosomal_L4 (pfam00573)→ rplW (PRK05738)Ribosomal_L23 (pfam00276)→ |
| env-924 | 4491487.3_NODE_119164 | + | 6041 | 6130 | **RNA**→ rpsJ (PRK00596)RpsJ (COG0051)→ |
| env-925 | NOM010_scaffold888_13 | - | 262 | 173 | **RNA**→ |
| env-926 | NLF007_scaffold25078_17 | - | 2019 | 1930 | **RNA**→ rpsJ (PRK00596)RpsJ (COG0051)→ |
| env-927 | NLM021_scaffold5854_5 | + | 3094 | 3183 | **RNA**→ rpsJ (PRK00596)RpsJ (COG0051)→ |
| Bcr-1-1 | NZ_GG663523.1 | - | 84988 | 84909 | **RNA**→ rpsJ (PRK00596)RpsJ (COG0051)→ rpsJ (PRK00596)RpsJ (COG0051)→ |
| env-928 | 4491405.3_NODE_49 | + | 11944 | 12023 | **RNA**→ rpsJ (PRK00596)RpsJ (COG0051)→ |

| | | | | | |
|---|---|---|---|---|---|
| env-929 | SRS019582_WUGC_scaffold_37983 | + | 23645 | 23724 | **RNA**→ rpsJ (PRK00596)RpsJ (COG0051)→ RplC (COG0087)rplC (PRK00001)→ rplD (PRK05319)Ribosomal_L4 (pfam00573)→ rplW (PRK05738)Ribosomal_L23 (pfam00276)→ |
| env-930 | 4491410.3_NODE_58360 | + | 14037 | 14116 | **RNA**→ rpsJ (PRK00596)RpsJ (COG0051)→ |
| env-931 | 4491484.3_NODE_3245 | - | 8920 | 8841 | **RNA**→ rpsJ (PRK00596)RpsJ (COG0051)→ |
| env-932 | scaffold20211_2_MH0025 | - | 397 | 318 | **RNA**→ rpsJ (PRK00596)Ribosomal_S10 (pfam00338)→ |
| env-933 | 4491414.3_NODE_1471 | - | 1688 | 1609 | **RNA**→ rpsJ (PRK00596)RpsJ (COG0051)→ |
| env-934 | NLM022_scaffold29854_2 | + | 26081 | 26161 | **RNA**→ |
| env-935 | 4491402.3_NODE_2732 | - | 220150 | 220070 | **RNA**→ rpsJ (PRK00596)RpsJ (COG0051)→ |
| env-936 | 4491419.3_NODE_5150 | - | 3952 | 3872 | **RNA**→ rpsJ (PRK00596)RpsJ (COG0051)→ |
| env-937 | NLM010_scaffold20898_13 | - | 1810 | 1730 | **RNA**→ rpsJ (PRK00596)RpsJ (COG0051)→ |
| env-938 | SRS013158_Baylor_scaffold_11109 | - | 287 | 207 | **RNA**→ hypo→ |
| env-939 | SRS015264_WUGC_scaffold_37105 | + | 430 | 510 | **RNA**→ rpsJ (PRK00596)RpsJ (COG0051)→ |
| env-940 | NLM004_scaffold7074_5 | + | 549 | 629 | **RNA**→ rpsJ (PRK00596)RpsJ (COG0051)→ |
| env-941 | SRS043701_LANL_scaffold_8835 | + | 329 | 409 | **RNA**→ rpsJ (PRK00596)RpsJ (COG0051)→ |
| env-942 | scaffold16680_3_MH0011 | + | 648 | 728 | **RNA**→ hypo→ |
| env-943 | NLF005_scaffold18267_1 | + | 154 | 234 | **RNA**→ rpsJ (PRK00596)RpsJ (COG0051)→ |
| env-944 | NOM009_scaffold7774_6 | - | 20905 | 20825 | **RNA**→ rpsJ (PRK00596)RpsJ (COG0051)→ |
| env-945 | DOM003_scaffold19611_1 | + | 136 | 216 | **RNA**→ rpsJ (PRK00596)RpsJ (COG0051)→ |
| env-946 | NLF013_scaffold11475_6 | + | 661 | 741 | **RNA**→ |
| env-947 | SRS015960_WUGC_scaffold_7480 | + | 329 | 409 | **RNA**→ |
| env-948 | SRS056519_LANL_scaffold_9865 | - | 288 | 208 | **RNA**→ |
| env-949 | NOM002_scaffold7458_1 | + | 147 | 227 | **RNA**→ rpsJ (PRK00596)RpsJ (COG0051)→ |
| env-950 | SRS022137_WUGC_scaffold_40873 | - | 287 | 207 | **RNA**→ hypo→ |
| Rin-1-1 | NZ_GG692728.1 | - | 64665 | 64585 | **RNA**→ rpsJ (PRK00596)RpsJ (COG0051)→ rpsJ (PRK00596)RpsJ (COG0051)→ |
| env-951 | 4491485.3_NODE_16391 | + | 24197 | 24277 | **RNA**→ rpsJ (PRK00596)RpsJ (COG0051)→ |
| env-952 | SRS019397_WUGC_scaffold_6745 | + | 328 | 408 | **RNA**→ rpsJ (PRK00596)RpsJ (COG0051)→ |
| env-953 | NOF005_scaffold5963_8 | - | 1692 | 1612 | **RNA**→ rpsJ (PRK00596)RpsJ (COG0051)→ |
| env-954 | SRS043411_WUGC_scaffold_12982 | - | 288 | 208 | **RNA**→ hypo→ |
| env-955 | 4491421.3_NODE_11491 | + | 13278 | 13358 | **RNA**→ rpsJ (PRK00596)RpsJ (COG0051)→ |
| env-956 | NOM022_scaffold14723_28 | - | 3683 | 3603 | **RNA**→ rpsJ (PRK00596)RpsJ (COG0051)→ |
| env-957 | NOM027_scaffold11125_1 | - | 446 | 366 | **RNA**→ rpsJ (PRK00596)Ribosomal_S10 (pfam00338)→ |
| env-958 | SRS014287_WUGC_scaffold_12600 | - | 819 | 739 | **RNA**→ rpsJ (PRK00596)RpsJ (COG0051)→ |
| env-959 | NOM002_scaffold929_13 | + | 1149 | 1229 | **RNA**→ ←hypo |
| env-960 | RUMENNODE_4338772_1 | - | 19974 | 19892 | **RNA**→ rpsJ (PRK00596)RpsJ (COG0051)→ RplC (COG0087)rplC (PRK00001)→ RplD (COG0088)rplD (PRK05319)→ rplD (PRK05319)Ribosomal_L4 (pfam00573)→ |
| env-961 | RUMENNODE_2488056_1 | - | 20859 | 20777 | **RNA**→ rpsJ (PRK00596)RpsJ (COG0051)→ RplC (COG0087)rplC (PRK00001)→ rplD (PRK05319)Ribosomal_L4 (pfam00573)→ |
| env-962 | SRS014923_WUGC_scaffold_57720 | - | 163 | 82 | **RNA**→ |
| env-963 | scaffold7909_1_MH0081 | - | 174 | 93 | **RNA**→ |
| env-964 | 4491408.3_NODE_8011 | - | 1352 | 1271 | **RNA**→ rpsJ (PRK00596)RpsJ (COG0051)→ RplC (COG0087)rplC (PRK00001)→ RplC (COG0087)rplC (PRK00001)→ |
| env-965 | IMNBL1_c0000852 | + | 20500 | 20581 | **RNA**→ rpsJ (PRK00596)RpsJ (COG0051)→ |
| env-966 | JGI994J12354_10001434 | - | 3364 | 3283 | **RNA**→ rpsJ (PRK00596)RpsJ (COG0051)→ |
| env-967 | 2227244666 | + | 3836 | 3917 | **RNA**→ rpsJ (PRK00596)RpsJ (COG0051)→ |
| env-968 | BABG01010268.1 | - | 446 | 363 | **RNA**→ rpsJ (PRK00596)RpsJ (COG0051)→ |
| env-969 | scaffold33008_3_MH0030 | + | 784 | 867 | **RNA**→ |
| env-970 | scaffold7906_1_V1.CD-1 | - | 885 | 802 | **RNA**→ rpsJ (PRK00596)RpsJ (COG0051)→ |
| env-971 | DLM023_scaffold1754_17 | - | 606 | 523 | **RNA**→ rpsJ (PRK00596)RpsJ (COG0051)→ |
| env-972 | 4491421.3_NODE_79900 | + | 381 | 464 | **RNA**→ |
| env-973 | scaffold17933_1_MH0058 | - | 182 | 99 | **RNA**→ |
| env-974 | scaffold9942_8_MH0073 | + | 2179 | 2262 | **RNA**→ |
| env-975 | scaffold7106_1_MH0074 | - | 181 | 98 | **RNA**→ |
| env-976 | scaffold6425_1_V1.UC-7 | - | 182 | 99 | **RNA**→ |

| | | | | | |
|---|---|---|---|---|---|
| env-977 | DOF003_scaffold16512_6 | + | 311 | 394 | **RNA**→ rpsJ (PRK00596)RpsJ (COG0051)→ |
| env-978 | DOM016_scaffold3915_5 | + | 242 | 325 | **RNA**→ rpsJ (PRK00596)Ribosomal_S10 (pfam00338)→ |
| env-979 | NOM002_scaffold75_3 | + | 434 | 517 | **RNA**→ rpsJ (PRK00596)RpsJ (COG0051)→ |
| Lba-3-1 | NZ_KE150017.1 | - | 1738715 | 1738634 | **RNA**→ rpsJ (PRK00596)RpsJ (COG0051)→ RplC (COG0087)rplC (PRK00001)→ |
| | | | | | rplD (PRK05319)Ribosomal_L4 (pfam00573)→ rplW (PRK05738)Ribosomal_L23 (pfam00276)→ |
| | | | | | RplB (COG0090)rplB (PRK09374)→ rpsS (PRK00357)RpsS (COG0185)→ |
| | | | | | rplV (PRK00565)Ribosomal_L22 (pfam00237)→ RpsC (COG0092)rpsC (PRK00310)→ |
| | | | | | rpl16 (CHL00044)rplP (PRK09203)→ PRK00306 (PRK00306)Ribosomal_L29 (pfam00831)→ |
| | | | | | rpsQ (PRK05610)Ribosomal_S17 (pfam00366)→ rplN (PRK05483)Ribosomal_L14 (pfam00238)→ |
| | | | | | RplX (COG0198)rplX (PRK00004)→ rplE (PRK00010)RplE (COG0094)→ rpsN (PRK08061)RpsN (COG0199)→ |
| env-980 | SRS057717_LANL_scaffold_6546 | - | 2301 | 2220 | **RNA**→ rpsJ (PRK00596)RpsJ (COG0051)→ |
| env-981 | 4491482.3_NODE_12049 | - | 97 | 16 | **RNA**→ |
| env-982 | NLM027_scaffold19893_2 | - | 681 | 600 | **RNA**→ rpsJ (PRK00596)RpsJ (COG0051)→ |
| Cne-1-1 | NZ_DS995339.4 | + | 431934 | 432015 | **RNA**→ rpsJ (PRK00596)RpsJ (COG0051)→ RplC (COG0087)rplC (PRK00001)→ |
| | | | | | rplD (PRK05319)Ribosomal_L4 (pfam00573)→ rplW (PRK05738)Ribosomal_L23 (pfam00276)→ |
| | | | | | RplB (COG0090)rplB (PRK09374)→ rpsS (PRK00357)RpsS (COG0185)→ |
| | | | | | rplV (PRK00565)Ribosomal_L22 (pfam00237)→ RpsC (COG0092)rpsC (PRK00310)→ |
| | | | | | rpl16 (CHL00044)rplP (PRK09203)→ PRK00306 (PRK00306)Ribosomal_L29 (pfam00831)→ |
| | | | | | rpsQ (PRK05610)Ribosomal_S17 (pfam00366)→ rplN (PRK05483)Ribosomal_L14 (pfam00238)→ |
| | | | | | RplX (COG0198)rplX (PRK00004)→ rplE (PRK00010)RplE (COG0094)→ rpsN (PRK08061)RpsN (COG0199)→ |
| env-983 | DOM008_scaffold9546_12 | - | 4435 | 4354 | **RNA**→ rpsJ (PRK00596)RpsJ (COG0051)→ RplC (COG0087)rplC (PRK00001)→ |
| | | | | | rplD (PRK05319)Ribosomal_L4 (pfam00573)→ rplW (PRK05738)Ribosomal_L23 (pfam00276)→ |
| | | | | | RplB (COG0090)rplB (PRK09374)→ rpsS (PRK00357)RpsS (COG0185)→ |
| | | | | | rplV (PRK00565)Ribosomal_L22 (pfam00237)→ RpsC (COG0092)rpsC (PRK00310)→ |
| env-984 | DOF006_scaffold10319_1 | - | 1987 | 1906 | **RNA**→ rpsJ (PRK00596)RpsJ (COG0051)→ RplC (COG0087)rplC (PRK00001)→ |
| | | | | | rplD (PRK05319)Ribosomal_L4 (pfam00573)→ |
| env-985 | DOF006_scaffold1788_9 | + | 437 | 518 | **RNA**→ rpsJ (PRK00596)RpsJ (COG0051)→ RplC (COG0087)rplC (PRK00001)→ |
| Csp-8-1 | NZ_KE150443.1 | + | 282818 | 282899 | **RNA**→ rpsJ (PRK00596)RpsJ (COG0051)→ RplC (COG0087)rplC (PRK00001)→ |
| | | | | | rplD (PRK05319)Ribosomal_L4 (pfam00573)→ rplW (PRK05738)Ribosomal_L23 (pfam00276)→ |
| | | | | | RplB (COG0090)rplB (PRK09374)→ rpsS (PRK00357)RpsS (COG0185)→ |
| | | | | | rplV (PRK00565)Ribosomal_L22 (pfam00237)→ RpsC (COG0092)rpsC (PRK00310)→ |
| | | | | | rpl16 (CHL00044)rplP (PRK09203)→ PRK00306 (PRK00306)Ribosomal_L29 (pfam00831)→ |
| | | | | | rpsQ (PRK05610)Ribosomal_S17 (pfam00366)→ rplN (PRK05483)Ribosomal_L14 (pfam00238)→ |
| | | | | | RplX (COG0198)rplX (PRK00004)→ rplE (PRK00010)RplE (COG0094)→ rpsN (PRK08061)RpsN (COG0199)→ |
| env-986 | DLF012_scaffold17148_1 | - | 413 | 332 | **RNA**→ rpsJ (PRK00596)RpsJ (COG0051)→ |
| Csp-9-1 | NZ_KE150436.1 | + | 356160 | 356241 | **RNA**→ rpsJ (PRK00596)RpsJ (COG0051)→ RplC (COG0087)rplC (PRK00001)→ |
| | | | | | rplD (PRK05319)Ribosomal_L4 (pfam00573)→ rplW (PRK05738)Ribosomal_L23 (pfam00276)→ |
| | | | | | RplB (COG0090)rplB (PRK09374)→ rpsS (PRK00357)RpsS (COG0185)→ |
| | | | | | rplV (PRK00565)Ribosomal_L22 (pfam00237)→ RpsC (COG0092)rpsC (PRK00310)→ |
| | | | | | rpl16 (CHL00044)rplP (PRK09203)→ PRK00306 (PRK00306)Ribosomal_L29 (pfam00831)→ |
| | | | | | rpsQ (PRK05610)Ribosomal_S17 (pfam00366)→ rplN (PRK05483)Ribosomal_L14 (pfam00238)→ |
| | | | | | RplX (COG0198)rplX (PRK00004)→ rplE (PRK00010)RplE (COG0094)→ rpsN (PRK08061)RpsN (COG0199)→ |
| env-987 | NLM027_scaffold221_5 | + | 1755 | 1836 | **RNA**→ rpsJ (PRK00596)RpsJ (COG0051)→ RplC (COG0087)rplC (PRK00001)→ |
| | | | | | rplD (PRK05319)Ribosomal_L4 (pfam00573)→ rplW (PRK05738)Ribosomal_L23 (pfam00276)→ |
| | | | | | RplB (COG0090)rplB (PRK09374)→ |
| env-988 | NOM028_scaffold23473_1 | - | 499 | 418 | **RNA**→ rpsJ (PRK00596)RpsJ (COG0051)→ |
| env-989 | longitudinal_520_20 | - | 28469 | 28388 | **RNA**→ rpsJ (PRK00596)RpsJ (COG0051)→ RplC (COG0087)rplC (PRK00001)→ |
| | | | | | rplD (PRK05319)Ribosomal_L4 (pfam00573)→ rplW (PRK05738)Ribosomal_L23 (pfam00276)→ |
| | | | | | RplB (COG0090)rplB (PRK09374)→ rpsS (PRK00357)RpsS (COG0185)→ |
| | | | | | rplV (PRK00565)Ribosomal_L22 (pfam00237)→ RpsC (COG0092)rpsC (PRK00310)→ |
| | | | | | rpl16 (CHL00044)rplP (PRK09203)→ PRK00306 (PRK00306)Ribosomal_L29 (pfam00831)→ |
| | | | | | rpsQ (PRK05610)Ribosomal_S17 (pfam00366)→ rplN (PRK05483)Ribosomal_L14 (pfam00238)→ |
| | | | | | RplX (COG0198)rplX (PRK00004)→ rplE (PRK00010)RplE (COG0094)→ rpsN (PRK08061)RpsN (COG0199)→ |

| | | | | | |
|---|---|---|---|---|---|
| env-990 | longitudinal_521_5 | - | 28425 | 28344 | **RNA**→ rpsJ (PRK00596)RpsJ (COG0051)→ RplC (COG0087)rplC (PRK00001)→ rplD (PRK05319)Ribosomal_L4 (pfam00573)→ rplW (PRK05738)Ribosomal_L23 (pfam00276)→ RplB (COG0090)rplB (PRK09374)→ rpsS (PRK00357)RpsS (COG0185)→ rplV (PRK00565)Ribosomal_L22 (pfam00237)→ RpsC (COG0092)rpsC (PRK00310)→ rpl16 (CHL00044)rplP (PRK09203)→ PRK00306 (PRK00306)Ribosomal_L29 (pfam00831)→ rpsQ (PRK05610)Ribosomal_S17 (pfam00366)→ rplN (PRK05483)Ribosomal_L14 (pfam00238)→ RplX (COG0198)rplX (PRK00004)→ rplE (PRK00010)RplE (COG0094)→ rpsN (PRK08061)RpsN (COG0199)→ |
| Lba-9-1 | NZ_GL872351.1 | - | 28391 | 28310 | **RNA**→ rpsJ (PRK00596)RpsJ (COG0051)→ RplC (COG0087)rplC (PRK00001)→ rplD (PRK05319)Ribosomal_L4 (pfam00573)→ rplW (PRK05738)Ribosomal_L23 (pfam00276)→ RplB (COG0090)rplB (PRK09374)→ rpsS (PRK00357)RpsS (COG0185)→ rplV (PRK00565)Ribosomal_L22 (pfam00237)→ RpsC (COG0092)rpsC (PRK00310)→ rpl16 (CHL00044)rplP (PRK09203)→ PRK00306 (PRK00306)Ribosomal_L29 (pfam00831)→ rpsQ (PRK05610)Ribosomal_S17 (pfam00366)→ rplN (PRK05483)Ribosomal_L14 (pfam00238)→ RplX (COG0198)rplX (PRK00004)→ rplE (PRK00010)RplE (COG0094)→ rpsN (PRK08061)RpsN (COG0199)→ |
| Lba-12-1 | NZ_GL890574.1 | - | 28756 | 28675 | **RNA**→ rpsJ (PRK00596)RpsJ (COG0051)→ RplC (COG0087)rplC (PRK00001)→ rplD (PRK05319)Ribosomal_L4 (pfam00573)→ rplW (PRK05738)Ribosomal_L23 (pfam00276)→ RplB (COG0090)rplB (PRK09374)→ rpsS (PRK00357)RpsS (COG0185)→ rplV (PRK00565)Ribosomal_L22 (pfam00237)→ RpsC (COG0092)rpsC (PRK00310)→ rpl16 (CHL00044)rplP (PRK09203)→ PRK00306 (PRK00306)Ribosomal_L29 (pfam00831)→ rpsQ (PRK05610)Ribosomal_S17 (pfam00366)→ rplN (PRK05483)Ribosomal_L14 (pfam00238)→ RplX (COG0198)rplX (PRK00004)→ rplE (PRK00010)RplE (COG0094)→ rpsN (PRK08061)RpsN (COG0199)→ |
| env-991 | longitudinal_114_32 | - | 33692 | 33610 | **RNA**→ rpsJ (PRK00596)RpsJ (COG0051)→ RplC (COG0087)rplC (PRK00001)→ rplD (PRK05319)Ribosomal_L4 (pfam00573)→ rplW (PRK05738)Ribosomal_L23 (pfam00276)→ RplB (COG0090)rplB (PRK09374)→ rpsS (PRK00357)RpsS (COG0185)→ rplV (PRK00565)Ribosomal_L22 (pfam00237)→ RpsC (COG0092)rpsC (PRK00310)→ rpl16 (CHL00044)rplP (PRK09203)→ PRK00306 (PRK00306)Ribosomal_L29 (pfam00831)→ rpsQ (PRK05610)Ribosomal_S17 (pfam00366)→ rplN (PRK05483)Ribosomal_L14 (pfam00238)→ RplX (COG0198)rplX (PRK00004)→ rplE (PRK00010)RplE (COG0094)→ rpsN (PRK08061)RpsN (COG0199)→ rpsH (PRK00136)Ribosomal_S8 (pfam00410)→ |
| env-992 | longitudinal_113_28 | - | 46667 | 46585 | **RNA**→ rpsJ (PRK00596)RpsJ (COG0051)→ RplC (COG0087)rplC (PRK00001)→ rplD (PRK05319)Ribosomal_L4 (pfam00573)→ rplW (PRK05738)Ribosomal_L23 (pfam00276)→ RplB (COG0090)rplB (PRK09374)→ rpsS (PRK00357)RpsS (COG0185)→ rplV (PRK00565)Ribosomal_L22 (pfam00237)→ RpsC (COG0092)rpsC (PRK00310)→ rpl16 (CHL00044)rplP (PRK09203)→ PRK00306 (PRK00306)Ribosomal_L29 (pfam00831)→ rpsQ (PRK05610)Ribosomal_S17 (pfam00366)→ rplN (PRK05483)Ribosomal_L14 (pfam00238)→ RplX (COG0198)rplX (PRK00004)→ rplE (PRK00010)RplE (COG0094)→ rpsN (PRK08061)RpsN (COG0199)→ rpsH (PRK00136)Ribosomal_S8 (pfam00410)→ |
| env-993 | SRS015782_WUGC_scaffold_40849 | + | 1056 | 1138 | **RNA**→ |
| Dlo-2-1 | NZ_DS264399.1 | + | 26108 | 26190 | **RNA**→ rpsJ (PRK00596)RpsJ (COG0051)→ RplC (COG0087)rplC (PRK00001)→ rplD (PRK05319)Ribosomal_L4 (pfam00573)→ rplW (PRK05738)Ribosomal_L23 (pfam00276)→ RplB (COG0090)rplB (PRK09374)→ rpsS (PRK00357)RpsS (COG0185)→ rplV (PRK00565)Ribosomal_L22 (pfam00237)→ RpsC (COG0092)rpsC (PRK00310)→ rpl16 (CHL00044)rplP (PRK09203)→ PRK00306 (PRK00306)Ribosomal_L29 (pfam00831)→ rpsQ (PRK05610)Ribosomal_S17 (pfam00366)→ rpsQ (PRK05610)Ribosomal_S17 (pfam00366)→ rplN (PRK05483)Ribosomal_L14 (pfam00238)→ RplX (COG0198)rplX (PRK00004)→ rplE (PRK00010)RplE (COG0094)→ rpsN (PRK08061)RpsN (COG0199)→ rpsH (PRK00136)Ribosomal_S8 (pfam00410)→ |
| env-994 | NOM009_scaffold172_2 | + | 629 | 711 | **RNA**→ rpsJ (PRK00596)RpsJ (COG0051)→ RplC (COG0087)rplC (PRK00001)→ |
| env-995 | 4491421.3_NODE_115793 | + | 3592 | 3674 | **RNA**→ rpsJ (PRK00596)RpsJ (COG0051)→ RplC (COG0087)rplC (PRK00001)→ rplD (PRK05319)Ribosomal_L4 (pfam00573)→ rplW (PRK05738)Ribosomal_L23 (pfam00276)→ RplB (COG0090)rplB (PRK09374)→ rpl2 (CHL00052)rplB (PRK09374)→ rpsS (PRK00357)RpsS (COG0185)→ rplV (PRK00565)Ribosomal_L22 (pfam00237)→ rpsC (PRK00310)30S_S3_KH (cd02412)→ |
| env-996 | 4491417.3_NODE_2013 | - | 1426 | 1344 | **RNA**→ rpsJ (PRK00596)RpsJ (COG0051)→ RplC (COG0087)rplC (PRK00001)→ RplD (COG0088)rplD (PRK05319)→ |

| | | | | | |
|---|---|---|---|---|---|
| env-997 | longitudinal_68_67 | + | 35973 | 36055 | **RNA**→ rpsJ (PRK00596)RpsJ (COG0051)→ RplC (COG0087)rplC (PRK00001)→ rplD (PRK05319)Ribosomal_L4 (pfam00573)→ rplW (PRK05738)Ribosomal_L23 (pfam00276)→ RplB (COG0090)rplB (PRK09374)→ rpsS (PRK00357)RpsS (COG0185)→ rplV (PRK00565)Ribosomal_L22 (pfam00237)→ RpsC (COG0092)rpsC (PRK00310)→ rpl16 (CHL00044)rplP (PRK09203)→ PRK00306 (PRK00306)Ribosomal_L29 (pfam00831)→ rpsQ (PRK05610)Ribosomal_S17 (pfam00366)→ rplN (PRK05483)Ribosomal_L14 (pfam00238)→ RplX (COG0198)rplX (PRK00004)→ rplE (PRK00010)RplE (COG0094)→ rpsN (PRK08061)RpsN (COG0199)→ rpsH (PRK00136)Ribosomal_S8 (pfam00410)→ |
| env-998 | longitudinal_299_23 | - | 39606 | 39524 | **RNA**→ rpsJ (PRK00596)RpsJ (COG0051)→ RplC (COG0087)rplC (PRK00001)→ rplD (PRK05319)Ribosomal_L4 (pfam00573)→ rplW (PRK05738)Ribosomal_L23 (pfam00276)→ RplB (COG0090)rplB (PRK09374)→ rpsS (PRK00357)RpsS (COG0185)→ rplV (PRK00565)Ribosomal_L22 (pfam00237)→ RpsC (COG0092)rpsC (PRK00310)→ |
| env-999 | longitudinal_300_84 | - | 39927 | 39845 | **RNA**→ rpsJ (PRK00596)RpsJ (COG0051)→ RplC (COG0087)rplC (PRK00001)→ rplD (PRK05319)Ribosomal_L4 (pfam00573)→ rplW (PRK05738)Ribosomal_L23 (pfam00276)→ RplB (COG0090)rplB (PRK09374)→ rpsS (PRK00357)RpsS (COG0185)→ rplV (PRK00565)Ribosomal_L22 (pfam00237)→ RpsC (COG0092)rpsC (PRK00310)→ rpl16 (CHL00044)rplP (PRK09203)→ PRK00306 (PRK00306)Ribosomal_L29 (pfam00831)→ rpsQ (PRK05610)Ribosomal_S17 (pfam00366)→ rplN (PRK05483)Ribosomal_L14 (pfam00238)→ RplX (COG0198)rplX (PRK00004)→ rplE (PRK00010)RplE (COG0094)→ rpsN (PRK08061)RpsN (COG0199)→ rpsH (PRK00136)Ribosomal_S8 (pfam00410)→ |
| env-1000 | longitudinal_301_117 | - | 4335 | 4253 | **RNA**→ rpsJ (PRK00596)RpsJ (COG0051)→ RplC (COG0087)rplC (PRK00001)→ rplD (PRK05319)Ribosomal_L4 (pfam00573)→ rplW (PRK05738)Ribosomal_L23 (pfam00276)→ RplB (COG0090)rplB (PRK09374)→ rpsS (PRK00357)RpsS (COG0185)→ rplV (PRK00565)Ribosomal_L22 (pfam00237)→ KH (smart00322)RpsC (COG0092)→ |
| env-1001 | longitudinal_302_50 | + | 39748 | 39830 | **RNA**→ rpsJ (PRK00596)RpsJ (COG0051)→ RplC (COG0087)rplC (PRK00001)→ rplD (PRK05319)Ribosomal_L4 (pfam00573)→ rplW (PRK05738)Ribosomal_L23 (pfam00276)→ RplB (COG0090)rplB (PRK09374)→ rpsS (PRK00357)RpsS (COG0185)→ rplV (PRK00565)Ribosomal_L22 (pfam00237)→ RpsC (COG0092)rpsC (PRK00310)→ rpl16 (CHL00044)rplP (PRK09203)→ PRK00306 (PRK00306)Ribosomal_L29 (pfam00831)→ rpsQ (PRK05610)Ribosomal_S17 (pfam00366)→ rplN (PRK05483)Ribosomal_L14 (pfam00238)→ RplX (COG0198)rplX (PRK00004)→ rplE (PRK00010)RplE (COG0094)→ rpsN (PRK08061)RpsN (COG0199)→ rpsH (PRK00136)Ribosomal_S8 (pfam00410)→ |
| env-1002 | longitudinal_346_79 | + | 7051 | 7133 | **RNA**→ rpsJ (PRK00596)RpsJ (COG0051)→ RplC (COG0087)rplC (PRK00001)→ rplD (PRK05319)Ribosomal_L4 (pfam00573)→ rplW (PRK05738)Ribosomal_L23 (pfam00276)→ RplB (COG0090)rplB (PRK09374)→ rpsS (PRK00357)RpsS (COG0185)→ rplV (PRK00565)Ribosomal_L22 (pfam00237)→ RpsC (COG0092)rpsC (PRK00310)→ rpl16 (CHL00044)rplP (PRK09203)→ PRK00306 (PRK00306)Ribosomal_L29 (pfam00831)→ rpsQ (PRK05610)Ribosomal_S17 (pfam00366)→ rplN (PRK05483)Ribosomal_L14 (pfam00238)→ RplX (COG0198)rplX (PRK00004)→ rplE (PRK00010)RplE (COG0094)→ rpsN (PRK08061)RpsN (COG0199)→ rpsH (PRK00136)Ribosomal_S8 (pfam00410)→ |
| env-1003 | longitudinal_347_101 | - | 67359 | 67277 | **RNA**→ rpsJ (PRK00596)RpsJ (COG0051)→ RplC (COG0087)rplC (PRK00001)→ rplD (PRK05319)Ribosomal_L4 (pfam00573)→ rplW (PRK05738)Ribosomal_L23 (pfam00276)→ RplB (COG0090)rplB (PRK09374)→ rpsS (PRK00357)RpsS (COG0185)→ rplV (PRK00565)Ribosomal_L22 (pfam00237)→ RpsC (COG0092)rpsC (PRK00310)→ rpl16 (CHL00044)rplP (PRK09203)→ PRK00306 (PRK00306)Ribosomal_L29 (pfam00831)→ rpsQ (PRK05610)Ribosomal_S17 (pfam00366)→ rplN (PRK05483)Ribosomal_L14 (pfam00238)→ RplX (COG0198)rplX (PRK00004)→ rplE (PRK00010)RplE (COG0094)→ rpsN (PRK08061)RpsN (COG0199)→ rpsH (PRK00136)Ribosomal_S8 (pfam00410)→ |
| env-1004 | AUXO010955118.1 | + | 38874 | 38956 | **RNA**→ |
| env-1005 | ADJS01006814.1 | + | 10307 | 10389 | **RNA**→ rpsJ (PRK00596)RpsJ (COG0051)→ RplC (COG0087)rplC (PRK00001)→ rplD (PRK05319)Ribosomal_L4 (pfam00573)→ rplW (PRK05738)Ribosomal_L23 (pfam00276)→ RplB (COG0090)rplB (PRK09374)→ |
| env-1006 | NOM029_scaffold8422_1 | - | 172 | 90 | **RNA**→ hypo→ |

| | | | | | |
|---|---|---|---|---|---|
| env-1007 | NLM027_scaffold13626_5 | - | 7795 | 7713 | **RNA**→ rpsJ (PRK00596)RpsJ (COG0051)→ RplC (COG0087)rplC (PRK00001)→ rplD (PRK05319)Ribosomal_L4 (pfam00573)→ rplW (PRK05738)Ribosomal_L23 (pfam00276)→ RplB (COG0090)rplB (PRK09374)→ rpsS (PRK00357)RpsS (COG0185)→ rplV (PRK00565)Ribosomal_L22 (pfam00237)→ RpsC (COG0092)rpsC (PRK00310)→ rpl16 (CHL00044)rplP (PRK09203)→ PRK00306 (PRK00306)Ribosomal_L29 (pfam00831)→ rpsQ (PRK05610)Ribosomal_S17 (pfam00366)→ rplN (PRK05483)Ribosomal_L14 (pfam00238)→ RplX (COG0198)rplX (PRK00004)→ rplE (PRK00010)RplE (COG0094)→ rpsN (PRK08061)RpsN (COG0199)→ rpsH (PRK00136)Ribosomal_S8 (pfam00410)→ |
| env-1008 | DLF013_scaffold33304_1 | - | 365 | 283 | **RNA**→ rpsJ (PRK00596)Ribosomal_S10 (pfam00338)→ |
| env-1009 | DLM009_scaffold5025_4 | + | 9759 | 9841 | **RNA**→ hypo→ |
| env-1010 | longitudinal_134_7 | + | 55196 | 55278 | **RNA**→ rpsJ (PRK00596)RpsJ (COG0051)→ RplC (COG0087)rplC (PRK00001)→ rplD (PRK05319)Ribosomal_L4 (pfam00573)→ rplW (PRK05738)Ribosomal_L23 (pfam00276)→ RplB (COG0090)rplB (PRK09374)→ rpsS (PRK00357)RpsS (COG0185)→ rplV (PRK00565)Ribosomal_L22 (pfam00237)→ RpsC (COG0092)rpsC (PRK00310)→ rpl16 (CHL00044)rplP (PRK09203)→ PRK00306 (PRK00306)Ribosomal_L29 (pfam00831)→ rpsQ (PRK05610)Ribosomal_S17 (pfam00366)→ rplN (PRK05483)Ribosomal_L14 (pfam00238)→ RplX (COG0198)rplX (PRK00004)→ rplE (PRK00010)RplE (COG0094)→ rpsN (PRK08061)RpsN (COG0199)→ rpsH (PRK00136)Ribosomal_S8 (pfam00410)→ |
| env-1011 | longitudinal_135_8 | - | 44719 | 44637 | **RNA**→ rpsJ (PRK00596)RpsJ (COG0051)→ RplC (COG0087)rplC (PRK00001)→ rplD (PRK05319)Ribosomal_L4 (pfam00573)→ rplW (PRK05738)Ribosomal_L23 (pfam00276)→ RplB (COG0090)rplB (PRK09374)→ rpsS (PRK00357)RpsS (COG0185)→ rplV (PRK00565)Ribosomal_L22 (pfam00237)→ RpsC (COG0092)rpsC (PRK00310)→ rpl16 (CHL00044)rplP (PRK09203)→ PRK00306 (PRK00306)Ribosomal_L29 (pfam00831)→ rpsQ (PRK05610)Ribosomal_S17 (pfam00366)→ rplN (PRK05483)Ribosomal_L14 (pfam00238)→ RplX (COG0198)rplX (PRK00004)→ rplE (PRK00010)RplE (COG0094)→ rpsN (PRK08061)RpsN (COG0199)→ rpsH (PRK00136)Ribosomal_S8 (pfam00410)→ |
| env-1012 | longitudinal_188_28 | + | 55196 | 55278 | **RNA**→ rpsJ (PRK00596)RpsJ (COG0051)→ RplC (COG0087)rplC (PRK00001)→ rplD (PRK05319)Ribosomal_L4 (pfam00573)→ rplW (PRK05738)Ribosomal_L23 (pfam00276)→ RplB (COG0090)rplB (PRK09374)→ rpsS (PRK00357)RpsS (COG0185)→ rplV (PRK00565)Ribosomal_L22 (pfam00237)→ RpsC (COG0092)rpsC (PRK00310)→ rpl16 (CHL00044)rplP (PRK09203)→ PRK00306 (PRK00306)Ribosomal_L29 (pfam00831)→ rpsQ (PRK05610)Ribosomal_S17 (pfam00366)→ rplN (PRK05483)Ribosomal_L14 (pfam00238)→ RplX (COG0198)rplX (PRK00004)→ rplE (PRK00010)RplE (COG0094)→ rpsN (PRK08061)RpsN (COG0199)→ rpsH (PRK00136)Ribosomal_S8 (pfam00410)→ |
| env-1013 | longitudinal_189_5 | - | 44722 | 44640 | **RNA**→ rpsJ (PRK00596)RpsJ (COG0051)→ RplC (COG0087)rplC (PRK00001)→ rplD (PRK05319)Ribosomal_L4 (pfam00573)→ rplW (PRK05738)Ribosomal_L23 (pfam00276)→ RplB (COG0090)rplB (PRK09374)→ rpsS (PRK00357)RpsS (COG0185)→ rplV (PRK00565)Ribosomal_L22 (pfam00237)→ RpsC (COG0092)rpsC (PRK00310)→ rpl16 (CHL00044)rplP (PRK09203)→ PRK00306 (PRK00306)Ribosomal_L29 (pfam00831)→ rpsQ (PRK05610)Ribosomal_S17 (pfam00366)→ rplN (PRK05483)Ribosomal_L14 (pfam00238)→ RplX (COG0198)rplX (PRK00004)→ rplE (PRK00010)RplE (COG0094)→ rpsN (PRK08061)RpsN (COG0199)→ rpsH (PRK00136)Ribosomal_S8 (pfam00410)→ |
| env-1014 | longitudinal_190_130 | + | 55195 | 55277 | **RNA**→ rpsJ (PRK00596)RpsJ (COG0051)→ RplC (COG0087)rplC (PRK00001)→ rplD (PRK05319)Ribosomal_L4 (pfam00573)→ rplW (PRK05738)Ribosomal_L23 (pfam00276)→ RplB (COG0090)rplB (PRK09374)→ rpsS (PRK00357)RpsS (COG0185)→ rplV (PRK00565)Ribosomal_L22 (pfam00237)→ RpsC (COG0092)rpsC (PRK00310)→ rpl16 (CHL00044)rplP (PRK09203)→ PRK00306 (PRK00306)Ribosomal_L29 (pfam00831)→ rpsQ (PRK05610)Ribosomal_S17 (pfam00366)→ rplN (PRK05483)Ribosomal_L14 (pfam00238)→ RplX (COG0198)rplX (PRK00004)→ rplE (PRK00010)RplE (COG0094)→ rpsN (PRK08061)RpsN (COG0199)→ rpsH (PRK00136)Ribosomal_S8 (pfam00410)→ |

| | | | | | |
|---|---|---|---|---|---|
| env-1015 | longitudinal_465_52 | + | 39026 | 39108 | **RNA**→ rpsJ (PRK00596)RpsJ (COG0051)→ RplC (COG0087)rplC (PRK00001)→ rplD (PRK05319)Ribosomal_L4 (pfam00573)→ rplW (PRK05738)Ribosomal_L23 (pfam00276)→ RplB (COG0090)rplB (PRK09374)→ rpsS (PRK00357)RpsS (COG0185)→ rplV (PRK00565)Ribosomal_L22 (pfam00237)→ RpsC (COG0092)rpsC (PRK00310)→ rpl16 (CHL00044)rplP (PRK09203)→ PRK00306 (PRK00306)Ribosomal_L29 (pfam00831)→ rpsQ (PRK05610)Ribosomal_S17 (pfam00366)→ rplN (PRK05483)Ribosomal_L14 (pfam00238)→ RplX (COG0198)rplX (PRK00004)→ rplE (PRK00010)RplE (COG0094)→ rpsN (PRK08061)RpsN (COG0199)→ rpsH (PRK00136)Ribosomal_S8 (pfam00410)→ |
| env-1016 | longitudinal_466_3 | - | 45264 | 45182 | **RNA**→ rpsJ (PRK00596)RpsJ (COG0051)→ RplC (COG0087)rplC (PRK00001)→ hypo→ RplC (COG0087)rplC (PRK00001)→ rplD (PRK05319)Ribosomal_L4 (pfam00573)→ rplW (PRK05738)Ribosomal_L23 (pfam00276)→ RplB (COG0090)rplB (PRK09374)→ rpsS (PRK00357)RpsS (COG0185)→ rplV (PRK00565)Ribosomal_L22 (pfam00237)→ RpsC (COG0092)rpsC (PRK00310)→ rpl16 (CHL00044)rplP (PRK09203)→ PRK00306 (PRK00306)Ribosomal_L29 (pfam00831)→ rpsQ (PRK05610)Ribosomal_S17 (pfam00366)→ rplN (PRK05483)Ribosomal_L14 (pfam00238)→ RplX (COG0198)rplX (PRK00004)→ rplE (PRK00010)RplE (COG0094)→ rpsN (PRK08061)RpsN (COG0199)→ rpsH (PRK00136)Ribosomal_S8 (pfam00410)→ |
| env-1017 | longitudinal_467_77 | + | 39024 | 39106 | **RNA**→ rpsJ (PRK00596)RpsJ (COG0051)→ RplC (COG0087)rplC (PRK00001)→ rplD (PRK05319)Ribosomal_L4 (pfam00573)→ rplW (PRK05738)Ribosomal_L23 (pfam00276)→ RplB (COG0090)rplB (PRK09374)→ rpsS (PRK00357)RpsS (COG0185)→ rplV (PRK00565)Ribosomal_L22 (pfam00237)→ RpsC (COG0092)rpsC (PRK00310)→ rpl16 (CHL00044)rplP (PRK09203)→ PRK00306 (PRK00306)Ribosomal_L29 (pfam00831)→ rpsQ (PRK05610)Ribosomal_S17 (pfam00366)→ rplN (PRK05483)Ribosomal_L14 (pfam00238)→ RplX (COG0198)rplX (PRK00004)→ rplE (PRK00010)RplE (COG0094)→ rpsN (PRK08061)RpsN (COG0199)→ rpsH (PRK00136)Ribosomal_S8 (pfam00410)→ |
| env-1018 | DLM012_scaffold1652_10 | + | 412 | 494 | **RNA**→ rpsJ (PRK00596)RpsJ (COG0051)→ RplC (COG0087)rplC (PRK00001)→ |
| env-1019 | longitudinal_464_28 | + | 39026 | 39108 | **RNA**→ rpsJ (PRK00596)RpsJ (COG0051)→ RplC (COG0087)rplC (PRK00001)→ rplD (PRK05319)Ribosomal_L4 (pfam00573)→ rplW (PRK05738)Ribosomal_L23 (pfam00276)→ RplB (COG0090)rplB (PRK09374)→ rpsS (PRK00357)RpsS (COG0185)→ rplV (PRK00565)Ribosomal_L22 (pfam00237)→ RpsC (COG0092)rpsC (PRK00310)→ rpl16 (CHL00044)rplP (PRK09203)→ PRK00306 (PRK00306)Ribosomal_L29 (pfam00831)→ rpsQ (PRK05610)Ribosomal_S17 (pfam00366)→ rplN (PRK05483)Ribosomal_L14 (pfam00238)→ RplX (COG0198)rplX (PRK00004)→ rplE (PRK00010)RplE (COG0094)→ rpsN (PRK08061)RpsN (COG0199)→ rpsH (PRK00136)Ribosomal_S8 (pfam00410)→ |
| Dlo-1-1 | NZ_KE384171.1 | - | 53064 | 52982 | **RNA**→ rpsJ (PRK00596)RpsJ (COG0051)→ RplC (COG0087)rplC (PRK00001)→ rplD (PRK05319)Ribosomal_L4 (pfam00573)→ rplW (PRK05738)Ribosomal_L23 (pfam00276)→ RplB (COG0090)rplB (PRK09374)→ rpsS (PRK00357)RpsS (COG0185)→ rplV (PRK00565)Ribosomal_L22 (pfam00237)→ RpsC (COG0092)rpsC (PRK00310)→ rpl16 (CHL00044)rplP (PRK09203)→ PRK00306 (PRK00306)Ribosomal_L29 (pfam00831)→ rpsQ (PRK05610)Ribosomal_S17 (pfam00366)→ rplN (PRK05483)Ribosomal_L14 (pfam00238)→ RplX (COG0198)rplX (PRK00004)→ rplE (PRK00010)RplE (COG0094)→ rpsN (PRK08061)RpsN (COG0199)→ rpsH (PRK00136)Ribosomal_S8 (pfam00410)→ |
| env-1020 | DLM008_scaffold98_16 | - | 4788 | 4708 | **RNA**→ rpsJ (PRK00596)RpsJ (COG0051)→ |
| env-1021 | 4491403.3_NODE_90116 | - | 1091 | 1011 | **RNA**→ rpsJ (PRK00596)RpsJ (COG0051)→ |
| env-1022 | 4491482.3_NODE_33763 | + | 29992 | 30072 | **RNA**→ rpsJ (PRK00596)RpsJ (COG0051)→ |
| Lba-2-1 | NZ_GL945165.1 | - | 181718 | 181624 | **RNA**→ hypo→ rpsJ (PRK00596)RpsJ (COG0051)→ |
| env-1023 | NLF005_scaffold8054_23 | - | 240 | 156 | **RNA**→ |
| env-1024 | scaffold48677_1_MH0016 | - | 226 | 142 | **RNA**→ hypo→ |
| env-1025 | SRS052697_LANL_scaffold_40762 | - | 228 | 144 | **RNA**→ |
| env-1026 | NOF014_scaffold21218_7 | + | 6081 | 6165 | **RNA**→ rpsJ (PRK00596)RpsJ (COG0051)→ RplC (COG0087)rplC (PRK00001)→ RplD (COG0088)rplD (PRK05319)→ |
| env-1027 | UnmappedStool_Broad_scaffold_56426 | + | 442 | 526 | **RNA**→ rpsJ (PRK00596)RpsJ (COG0051)→ rpl3 (CHL00143)rplC (PRK00001)→ |

| | | | | | |
|---|---|---|---|---|---|
| env-1028 | NOM007_scaffold46220_1 | + | 6092 | 6176 | **RNA**→ rpsJ (PRK00596)RpsJ (COG0051)→ RplC (COG0087)rplC (PRK00001)→ rplD (PRK05319)Ribosomal_L4 (pfam00573)→ rplW (PRK05738)Ribosomal_L23 (pfam00276)→ RplB (COG0090)rplB (PRK09374)→ rpsS (PRK00357)RpsS (COG0185)→ rplV (PRK00565)Ribosomal_L22 (pfam00237)→ RpsC (COG0092)rpsC (PRK00310)→ rpl16 (CHL00044)rplP (PRK09203)→ PRK00306 (PRK00306)Ribosomal_L29 (pfam00831)→ rpsQ (PRK05610)Ribosomal_S17 (pfam00366)→ rplN (PRK05483)Ribosomal_L14 (pfam00238)→ RplX (COG0198)rplX (PRK00004)→ rplE (PRK00010)RplE (COG0094)→ rpsN (PRK08061)RpsN (COG0199)→ |
| env-1029 | SRS015065_WUGC_scaffold_21436 | - | 228 | 144 | **RNA**→ |
| env-1030 | scaffold8950_1_MH0020 | - | 182 | 100 | **RNA**→ |
| env-1031 | scaffold13706_1_MH0082 | - | 182 | 100 | **RNA**→ |
| env-1032 | NLM023_scaffold21939_3 | + | 42 | 124 | **RNA**→ rpsJ (PRK00596)RpsJ (COG0051)→ |
| env-1033 | NOF005_scaffold41945_3 | - | 3030 | 2948 | **RNA**→ rpsJ (PRK00596)RpsJ (COG0051)→ |
| env-1034 | scaffold14387_1_O2.UC-20 | - | 183 | 101 | **RNA**→ |
| env-1035 | NLM005_scaffold29532_3 | - | 939 | 859 | **RNA**→ rpsJ (PRK00596)RpsJ (COG0051)→ |
| env-1036 | SRS014979_WUGC_scaffold_17094 | - | 219 | 139 | **RNA**→ |
| env-1037 | SRS015854_WUGC_scaffold_21352 | - | 218 | 138 | **RNA**→ hypo→ |
| env-1038 | scaffold11729_1_MH0046 | - | 176 | 96 | **RNA**→ |
| env-1039 | 4491404.3_NODE_80547 | + | 4100 | 4180 | **RNA**→ rpsJ (PRK00596)RpsJ (COG0051)→ |
| env-1040 | NLF012_scaffold34222_1 | + | 9882 | 9962 | **RNA**→ rpsJ (PRK00596)RpsJ (COG0051)→ |
| env-1041 | SRS014613_WUGC_scaffold_19279 | - | 219 | 139 | **RNA**→ |
| env-1042 | SRS015663_WUGC_scaffold_14486 | - | 205 | 125 | **RNA**→ |
| env-1043 | scaffold3356_1_MH0020 | - | 178 | 98 | **RNA**→ |
| env-1044 | scaffold41365_1_MH0048 | - | 215 | 135 | **RNA**→ hypo→ |
| env-1045 | DOM018_scaffold286_1 | - | 8103 | 8023 | **RNA**→ rpsJ (PRK00596)RpsJ (COG0051)→ |
| env-1046 | scaffold5676_2_MH0030 | + | 2969 | 3049 | **RNA**→ |
| env-1047 | NOM018_scaffold537_4 | + | 242 | 322 | **RNA**→ |
| env-1048 | ADJS01018011.1 | - | 102 | 22 | **RNA**→ |
| env-1049 | SRS012273_Baylor_scaffold_16633 | - | 217 | 137 | **RNA**→ |
| env-1050 | SRS051031_LANL_scaffold_29286 | - | 205 | 125 | **RNA**→ |
| env-1051 | SRS052027_LANL_scaffold_5017 | - | 180 | 100 | **RNA**→ |
| env-1052 | SRS015578_WUGC_scaffold_20977 | - | 205 | 125 | **RNA**→ |
| env-1053 | SRS058723_Baylor_scaffold_8787 | - | 220 | 140 | **RNA**→ |
| env-1054 | scaffold5589_1_V1.CD-15 | - | 176 | 96 | **RNA**→ |
| env-1055 | DLF006_scaffold6164_6 | - | 332 | 252 | **RNA**→ rpsJ (PRK00596)RpsJ (COG0051)→ |
| env-1056 | SRS011271_WUGC_scaffold_13792 | - | 203 | 123 | **RNA**→ |
| env-1057 | SRS015133_WUGC_scaffold_22071 | - | 217 | 137 | **RNA**→ |
| env-1058 | SRS050925_LANL_scaffold_18487 | - | 178 | 98 | **RNA**→ |
| env-1059 | 4491423.3_NODE_31871 | + | 887 | 967 | **RNA**→ rpsJ (PRK00596)RpsJ (COG0051)→ |
| env-1060 | DLM007_scaffold22614_6 | + | 347 | 427 | **RNA**→ rpsJ (PRK00596)RpsJ (COG0051)→ |
| env-1061 | DLM013_scaffold20638_2 | - | 589 | 509 | **RNA**→ rpsJ (PRK00596)RpsJ (COG0051)→ |
| env-1062 | DOM026_scaffold2783_5 | - | 1389 | 1309 | **RNA**→ rpsJ (PRK00596)RpsJ (COG0051)→ |
| env-1063 | NLM029_scaffold10879_6 | + | 10181 | 10261 | **RNA**→ rpsJ (PRK00596)RpsJ (COG0051)→ |
| env-1064 | NOM029_scaffold77_3 | - | 6632 | 6552 | **RNA**→ rpsJ (PRK00596)RpsJ (COG0051)→ |
| Cha-1-1 | NZ_KB850952.1 | - | 266489 | 266409 | **RNA**→ rpsJ (PRK00596)RpsJ (COG0051)→ |
| env-1065 | BABF01000827.1 | + | 5585 | 5665 | **RNA**→ rpsJ (PRK00596)RpsJ (COG0051)→ |
| Hha-2-1 | NZ_BAHY01000041.1 | + | 46269 | 46349 | **RNA**→ rpsJ (PRK00596)RpsJ (COG0051)→ |
| Hha-1-1 | NZ_BAHR02000176.1 | + | 17953 | 18033 | **RNA**→ rpsJ (PRK00596)RpsJ (COG0051)→ |
| env-1066 | DLM024_scaffold447_12 | - | 10332 | 10252 | **RNA**→ rpsJ (PRK00596)RpsJ (COG0051)→ |
| Cba-1-1 | NZ_CCYG01000030.1 | - | 17670 | 17589 | **RNA**→ rpsJ (PRK00596)RpsJ (COG0051)→ |
| Cba-8-1 | NZ_BAIL02000037.1 | + | 5849 | 5930 | **RNA**→ rpsJ (PRK00596)RpsJ (COG0051)→ |
| env-1067 | DLM003_scaffold37465_3 | - | 449 | 368 | **RNA**→ rpsJ (PRK00596)Ribosomal_S10 (pfam00338)→ |
| Lba-21-1 | NZ_BAIG01000037.1 | - | 23850 | 23769 | **RNA**→ rpsJ (PRK00596)RpsJ (COG0051)→ |
| Lba-20-1 | NZ_BAHZ01000017.1 | + | 31717 | 31798 | **RNA**→ rpsJ (PRK00596)RpsJ (COG0051)→ |

| | | | | | |
|---|---|---|---|---|---|
| Lba-8-1 | NZ_KE150406.1 | - | 356926 | 356845 | **RNA**→ rpsJ (PRK00596)RpsJ (COG0051)→ |
| env-1068 | DLM022_scaffold65033_2 | - | 21810 | 21729 | **RNA**→ rpsJ (PRK00596)RpsJ (COG0051)→ |
| Lba-13-1 | NZ_KE159636.1 | + | 315368 | 315449 | **RNA**→ rpsJ (PRK00596)RpsJ (COG0051)→ |
| Csp-7-1 | NZ_GG730310.1 | - | 511953 | 511870 | **RNA**→ rpsJ (PRK00596)RpsJ (COG0051)→ |
| env-1069 | NOM014_scaffold12257_6 | + | 214 | 297 | **RNA**→ rpsJ (PRK00596)RpsJ (COG0051)→ |
| env-1070 | 4448815.3_Contig39759 | - | 335 | 252 | **RNA**→ ←hypo |
| env-1071 | NOM004_scaffold1071_29 | + | 1697 | 1780 | **RNA**→ rpsJ (PRK00596)RpsJ (COG0051)→ |
| env-1072 | 4448817.3_Contig41879 | + | 547 | 630 | **RNA**→ |
| Esp-1-1 | NZ_KE159567.1 | + | 201686 | 201769 | **RNA**→ rpsJ (PRK00596)RpsJ (COG0051)→ |
| env-1073 | SRS014459_WUGC_scaffold_29318 | - | 183 | 101 | **RNA**→ |
| env-1074 | scaffold8733_1_MH0068 | - | 181 | 99 | **RNA**→ |
| env-1075 | 4448812.3_Contig6350 | + | 105 | 187 | **RNA**→ rpsJ (PRK00596)Ribosomal_S10 (pfam00338)→ |
| Csa-1-1 | NC_014376.1 | - | 4065824 | 4065743 | **RNA**→ rpsJ (PRK00596)RpsJ (COG0051)→ |
| env-1076 | 4491413.3_NODE_8993 | - | 365 | 282 | **RNA**→ |
| env-1077 | 4491487.3_NODE_55596 | - | 1190 | 1107 | **RNA**→ rpsJ (PRK00596)RpsJ (COG0051)→ |
| env-1078 | DLF014_scaffold222_7 | + | 23771 | 23854 | **RNA**→ rpsJ (PRK00596)RpsJ (COG0051)→ |
| env-1079 | NOF002_scaffold1633_2 | - | 30591 | 30508 | **RNA**→ rpsJ (PRK00596)RpsJ (COG0051)→ |
| env-1080 | NOM002_scaffold7212_2 | - | 50398 | 50315 | **RNA**→ rpsJ (PRK00596)RpsJ (COG0051)→ |
| env-1081 | NOM016_scaffold10985_7 | + | 832 | 915 | **RNA**→ |
| env-1082 | SRS049164_WUGC_scaffold_55041 | - | 222 | 139 | **RNA**→ |
| env-1083 | DOM012_scaffold51_17 | - | 3319 | 3236 | **RNA**→ rpsJ (PRK00596)RpsJ (COG0051)→ |
| env-1084 | DLM002_scaffold16134_2_1 | - | 304 | 221 | **RNA**→ |
| env-1085 | NOM017_scaffold25392_1 | + | 61 | 144 | **RNA**→ rpsJ (PRK00596)RpsJ (COG0051)→ |
| env-1086 | BABD01036607.1 | - | 431 | 348 | **RNA**→ rpsJ (PRK00596)Ribosomal_S10 (pfam00338)→ |
| Cha-2-1 | NZ_JH379029.1 | - | 49852 | 49771 | **RNA**→ rpsJ (PRK00596)RpsJ (COG0051)→ |
| Lba-5-1 | NZ_KE159571.1 | + | 661657 | 661740 | **RNA**→ rpsJ (PRK00596)RpsJ (COG0051)→ |
| Csp-1-1 | NZ_JH376516.1 | - | 458908 | 458824 | **RNA**→ rpsJ (PRK00596)RpsJ (COG0051)→ |
| env-1087 | DLM024_scaffold3420_12 | - | 30123 | 30039 | **RNA**→ rpsJ (PRK00596)RpsJ (COG0051)→ |
| Csy-3-1 | NZ_GL834357.1 | + | 437513 | 437597 | **RNA**→ rpsJ (PRK00596)RpsJ (COG0051)→ |
| Csy-2-1 | NZ_GL834305.1 | - | 526475 | 526391 | **RNA**→ rpsJ (PRK00596)RpsJ (COG0051)→ |
| Csy-1-1 | NZ_KE992799.1 | + | 25413 | 25497 | **RNA**→ rpsJ (PRK00596)RpsJ (COG0051)→ |
| Cba-5-1 | NZ_BAID02000107.1 | - | 26962 | 26878 | **RNA**→ rpsJ (PRK00596)RpsJ (COG0051)→ |
| env-1088 | RUMENNODE_4202514_70000 | - | 73558 | 73476 | **RNA**→ rpsJ (PRK00596)RpsJ (COG0051)→ RplC (COG0087)rplC (PRK00001)→ rplD (PRK05319)Ribosomal_L4 (pfam00573)→ rplW (PRK05738)Ribosomal_L23 (pfam00276)→ |
| env-1089 | SRS013818_Baylor_scaffold_51548 | + | 1200 | 1282 | **RNA**→ rpsJ (PRK00596)RpsJ (COG0051)→ rpl3 (CHL00143)rplC (PRK00001)→ |
| env-1090 | SRS024580_C3065196 | + | 214 | 296 | **RNA**→ rpsJ (PRK00596)RpsJ (COG0051)→ rpl3 (CHL00143)rplC (PRK00001)→ |
| env-1091 | SRS049389_WUGC_scaffold_44677 | + | 1748 | 1830 | **RNA**→ rpsJ (PRK00596)RpsJ (COG0051)→ RplC (COG0087)rplC (PRK00001)→ rplD (PRK05319)Ribosomal_L4 (pfam00573)→ rplW (PRK05738)Ribosomal_L23 (pfam00276)→ RplB (COG0090)rplB (PRK09374)→ |
| env-1092 | SRS051791_LANL_scaffold_6364 | - | 2179 | 2097 | **RNA**→ rpsJ (PRK00596)RpsJ (COG0051)→ RplC (COG0087)rplC (PRK00001)→ rplD (PRK05319)Ribosomal_L4 (pfam00573)→ rplW (PRK05738)Ribosomal_L23 (pfam00276)→ |
| env-1093 | SRS014689_WUGC_scaffold_23083 | + | 95 | 177 | **RNA**→ rpsJ (PRK00596)RpsJ (COG0051)→ RplC (COG0087)rplC (PRK00001)→ |
| env-1094 | SRS015434_C5946977 | + | 1744 | 1826 | **RNA**→ |
| env-1095 | SRS016342_Baylor_scaffold_32993 | + | 182 | 264 | **RNA**→ |

| | | | | | |
|---|---|---|---|---|---|
| env-1096 | SRS019022_WUGC_scaffold_7226 | - | 36273 | 36191 | **RNA**→ rpsJ (PRK00596)RpsJ (COG0051)→ RplC (COG0087)rplC (PRK00001)→ rplD (PRK05319)Ribosomal_L4 (pfam00573)→ rplW (PRK05738)Ribosomal_L23 (pfam00276)→ RplB (COG0090)rplB (PRK09374)→ rpsS (PRK00357)RpsS (COG0185)→ rplV (PRK00565)Ribosomal_L22 (pfam00237)→ RpsC (COG0092)rpsC (PRK00310)→ rpl16 (CHL00044)rplP (PRK09203)→ PRK00306 (PRK00306)Ribosomal_L29 (pfam00831)→ rpsQ (PRK05610)Ribosomal_S17 (pfam00366)→ rplN (PRK05483)Ribosomal_L14 (pfam00238)→ RplX (COG0198)rplX (PRK00004)→ rplE (PRK00010)RplE (COG0094)→ rpsN (PRK08061)RpsN (COG0199)→ rpsH (PRK00136)Ribosomal_S8 (pfam00410)→ RplF (COG0097)rplF (PRK05498)→ rplR (PRK05593)Ribosomal_L18p (pfam00861)→ RpsE (COG0098)rpsE (PRK00550)→ Ribosomal_L30 (cd01658)rpmD (PRK05611)→ rplO (PRK05592)Ribosomal_L27A (pfam00828)→ SecY (COG0201)secY (PRK09204)→ |
| env-1097 | SRS019894_C3850908 | - | 182 | 100 | **RNA**→ |
| env-1098 | SRS064774_C2746908 | - | 188 | 106 | **RNA**→ |
| env-1099 | SRS022719_LANL_scaffold_10343 | + | 154 | 236 | **RNA**→ rpsJ (PRK00596)Ribosomal_S10 (pfam00338)→ |
| env-1100 | SRS057692_C2605374 | - | 221 | 139 | **RNA**→ hypo→ |
| Slo-1-1 | NZ_JH590862.1 | - | 265923 | 265841 | **RNA**→ rpsJ (PRK00596)RpsJ (COG0051)→ RplC (COG0087)rplC (PRK00001)→ rplD (PRK05319)Ribosomal_L4 (pfam00573)→ rplW (PRK05738)Ribosomal_L23 (pfam00276)→ RplB (COG0090)rplB (PRK09374)→ rpsS (PRK00357)RpsS (COG0185)→ rplV (PRK00565)Ribosomal_L22 (pfam00237)→ RpsC (COG0092)rpsC (PRK00310)→ rpl16 (CHL00044)rplP (PRK09203)→ PRK00306 (PRK00306)Ribosomal_L29 (pfam00831)→ rpsQ (PRK05610)Ribosomal_S17 (pfam00366)→ rplN (PRK05483)Ribosomal_L14 (pfam00238)→ RplX (COG0198)rplX (PRK00004)→ rplE (PRK00010)RplE (COG0094)→ rplE (PRK00010)RplE (COG0094)→ rpsN (PRK08061)RpsN (COG0199)→ rpsH (PRK00136)Ribosomal_S8 (pfam00410)→ RplF (COG0097)rplF (PRK05498)→ rplR (PRK05593)Ribosomal_L18p (pfam00861)→ RpsE (COG0098)rpsE (PRK00550)→ Ribosomal_L30 (cd01658)rpmD (PRK05611)→ rplO (PRK05592)Ribosomal_L27A (pfam00828)→ SecY (COG0201)secY (PRK09204)→ |
| env-1101 | SRS018443_C3867499 | + | 1133 | 1215 | **RNA**→ hypo→ |
| env-1102 | SRS013879_C2852050 | - | 221 | 139 | **RNA**→ |
| env-1103 | SRS015644_WUGC_scaffold_48522 | - | 920 | 838 | **RNA**→ rpsJ (PRK00596)RpsJ (COG0051)→ rpl3 (CHL00143)rplC (PRK00001)→ |
| env-1104 | SRS019219_WUGC_scaffold_28960 | - | 350 | 268 | **RNA**→ rpsJ (PRK00596)Ribosomal_S10 (pfam00338)→ |
| env-1105 | SRS019974_C3184997 | + | 119 | 201 | **RNA**→ |
| env-1106 | SRS053854_LANL_scaffold_31660 | + | 1073 | 1155 | **RNA**→ rpsJ (PRK00596)RpsJ (COG0051)→ RplC (COG0087)rplC (PRK00001)→ rplD (PRK05319)Ribosomal_L4 (pfam00573)→ rplW (PRK05738)Ribosomal_L23 (pfam00276)→ |
| env-1107 | Palatine_Tonsils_LANL_scaffold_37420 | + | 240 | 322 | **RNA**→ |
| env-1108 | SRS016740_Baylor_scaffold_70786 | + | 1203 | 1285 | **RNA**→ rpsJ (PRK00596)RpsJ (COG0051)→ RplC (COG0087)rplC (PRK00001)→ rplD (PRK05319)Ribosomal_L4 (pfam00573)→ rplW (PRK05738)Ribosomal_L23 (pfam00276)→ RplB (COG0090)rplB (PRK09374)→ |
| env-1109 | SRS011306_Baylor_scaffold_89066 | + | 1041 | 1123 | **RNA**→ rpsJ (PRK00596)RpsJ (COG0051)→ RplC (COG0087)rplC (PRK00001)→ rplD (PRK05319)Ribosomal_L4 (pfam00573)→ rplW (PRK05738)Ribosomal_L23 (pfam00276)→ RplB (COG0090)rplB (PRK09374)→ |
| env-1110 | SRS015209_WUGC_scaffold_4051 | - | 58388 | 58306 | **RNA**→ rpsJ (PRK00596)RpsJ (COG0051)→ RplC (COG0087)rplC (PRK00001)→ rplD (PRK05319)Ribosomal_L4 (pfam00573)→ rplW (PRK05738)Ribosomal_L23 (pfam00276)→ RplB (COG0090)rplB (PRK09374)→ rpsS (PRK00357)RpsS (COG0185)→ rplV (PRK00565)Ribosomal_L22 (pfam00237)→ RpsC (COG0092)rpsC (PRK00310)→ rpl16 (CHL00044)rplP (PRK09203)→ PRK00306 (PRK00306)Ribosomal_L29 (pfam00831)→ rpsQ (PRK05610)Ribosomal_S17 (pfam00366)→ rplN (PRK05483)Ribosomal_L14 (pfam00238)→ RplX (COG0198)rplX (PRK00004)→ rplE (PRK00010)RplE (COG0094)→ rpsN (PRK08061)RpsN (COG0199)→ rpsH (PRK00136)Ribosomal_S8 (pfam00410)→ RplF (COG0097)rplF (PRK05498)→ rplR (PRK05593)Ribosomal_L18p (pfam00861)→ RpsE (COG0098)rpsE (PRK00550)→ Ribosomal_L30 (cd01658)rpmD (PRK05611)→ rplO (PRK05592)Ribosomal_L27A (pfam00828)→ SecY (COG0201)secY (PRK09204)→ |

| | | | | | |
|---|---|---|---|---|---|
| env-1111 | SRS016037_WUGC_scaffold_42187 | + | 1374 | 1456 | **RNA**→ rpsJ (PRK00596)RpsJ (COG0051)→ RplC (COG0087)rplC (PRK00001)→ rplD (PRK05319)Ribosomal_L4 (pfam00573)→ rplW (PRK05738)Ribosomal_L23 (pfam00276)→ RplB (COG0090)rplB (PRK09374)→ rpsS (PRK00357)RpsS (COG0185)→ rplV (PRK00565)Ribosomal_L22 (pfam00237)→ RpsC (COG0092)rpsC (PRK00310)→ rpl16 (CHL00044)rplP (PRK09203)→ PRK00306 (PRK00306)Ribosomal_L29 (pfam00831)→ rpsQ (PRK05610)Ribosomal_S17 (pfam00366)→ rplN (PRK05483)Ribosomal_L14 (pfam00238)→ RplX (COG0198)rplX (PRK00004)→ rplE (PRK00010)RplE (COG0094)→ rpsN (PRK08061)RpsN (COG0199)→ rpsH (PRK00136)Ribosomal_S8 (pfam00410)→ RplF (COG0097)rplF (PRK05498)→ rplR (PRK05593)Ribosomal_L18p (pfam00861)→ RpsE (COG0098)rpsE (PRK00550)→ Ribosomal_L30 (cd01658)rpmD (PRK05611)→ rplO (PRK05592)Ribosomal_L27A (pfam00828)→ SecY (COG0201)secY (PRK09204)→ adk (PRK00279)ADK (cd01428)→ PRK05716 (PRK05716)MetAP1 (cd01086)→ |
| env-1112 | SRS019127_C1647535 | - | 881 | 799 | **RNA**→ rpsJ (PRK00596)RpsJ (COG0051)→ rpl3 (CHL00143)rplC (PRK00001)→ |
| env-1113 | SRS021496_Baylor_scaffold_1948 | - | 18954 | 18872 | **RNA**→ rpsJ (PRK00596)RpsJ (COG0051)→ RplC (COG0087)rplC (PRK00001)→ rplD (PRK05319)Ribosomal_L4 (pfam00573)→ rplW (PRK05738)Ribosomal_L23 (pfam00276)→ RplB (COG0090)rplB (PRK09374)→ rpsS (PRK00357)RpsS (COG0185)→ rplV (PRK00565)Ribosomal_L22 (pfam00237)→ RpsC (COG0092)rpsC (PRK00310)→ rpl16 (CHL00044)rplP (PRK09203)→ PRK00306 (PRK00306)Ribosomal_L29 (pfam00831)→ rpsQ (PRK05610)Ribosomal_S17 (pfam00366)→ rplN (PRK05483)Ribosomal_L14 (pfam00238)→ RplX (COG0198)rplX (PRK00004)→ rplE (PRK00010)RplE (COG0094)→ rpsN (PRK08061)RpsN (COG0199)→ rpsH (PRK00136)Ribosomal_S8 (pfam00410)→ RplF (COG0097)rplF (PRK05498)→ rplR (PRK05593)Ribosomal_L18p (pfam00861)→ RpsE (COG0098)rpsE (PRK00550)→ Ribosomal_L30 (cd01658)rpmD (PRK05611)→ rplO (PRK05592)Ribosomal_L27A (pfam00828)→ SecY (COG0201)secY (PRK09204)→ |
| env-1114 | SRS056622_LANL_scaffold_50730 | - | 38763 | 38681 | **RNA**→ rpsJ (PRK00596)RpsJ (COG0051)→ RplC (COG0087)rplC (PRK00001)→ rplD (PRK05319)Ribosomal_L4 (pfam00573)→ rplW (PRK05738)Ribosomal_L23 (pfam00276)→ RplB (COG0090)rplB (PRK09374)→ rpsS (PRK00357)RpsS (COG0185)→ rplV (PRK00565)Ribosomal_L22 (pfam00237)→ RpsC (COG0092)rpsC (PRK00310)→ rpl16 (CHL00044)rplP (PRK09203)→ PRK00306 (PRK00306)Ribosomal_L29 (pfam00831)→ rpsQ (PRK05610)Ribosomal_S17 (pfam00366)→ rplN (PRK05483)Ribosomal_L14 (pfam00238)→ RplX (COG0198)rplX (PRK00004)→ rplE (PRK00010)RplE (COG0094)→ rpsN (PRK08061)RpsN (COG0199)→ rpsH (PRK00136)Ribosomal_S8 (pfam00410)→ RplF (COG0097)rplF (PRK05498)→ rplR (PRK05593)Ribosomal_L18p (pfam00861)→ RpsE (COG0098)rpsE (PRK00550)→ Ribosomal_L30 (cd01658)rpmD (PRK05611)→ rplO (PRK05592)Ribosomal_L27A (pfam00828)→ SecY (COG0201)secY (PRK09204)→ |
| env-1115 | SRS012279_Baylor_scaffold_39424 | + | 2148 | 2230 | **RNA**→ |
| env-1116 | SRS022077_C2487593 | + | 355 | 437 | **RNA**→ |
| env-1117 | SRS044662_C1603794 | - | 231 | 149 | **RNA**→ hypo→ |
| env-1118 | SRS014124_C2616663 | - | 366 | 284 | **RNA**→ rpsJ (PRK00596)Ribosomal_S10 (pfam00338)→ |
| env-1119 | SRS019045_WUGC_scaffold_27984 | - | 12270 | 12188 | **RNA**→ rpsJ (PRK00596)RpsJ (COG0051)→ RplC (COG0087)rplC (PRK00001)→ rplD (PRK05319)Ribosomal_L4 (pfam00573)→ rplW (PRK05738)Ribosomal_L23 (pfam00276)→ RplB (COG0090)rplB (PRK09374)→ rpsS (PRK00357)RpsS (COG0185)→ rplV (PRK00565)Ribosomal_L22 (pfam00237)→ RpsC (COG0092)rpsC (PRK00310)→ rpl16 (CHL00044)rplP (PRK09203)→ PRK00306 (PRK00306)Ribosomal_L29 (pfam00831)→ rpsQ (PRK05610)Ribosomal_S17 (pfam00366)→ rplN (PRK05483)Ribosomal_L14 (pfam00238)→ RplX (COG0198)rplX (PRK00004)→ rplE (PRK00010)RplE (COG0094)→ rpsN (PRK08061)RpsN (COG0199)→ rpsH (PRK00136)Ribosomal_S8 (pfam00410)→ RplF (COG0097)rplF (PRK05498)→ rplR (PRK05593)Ribosomal_L18p (pfam00861)→ RpsE (COG0098)rpsE (PRK00550)→ Ribosomal_L30 (cd01658)rpmD (PRK05611)→ rplO (PRK05592)Ribosomal_L27A (pfam00828)→ SecY (COG0201)secY (PRK09204)→ |
| env-1120 | SRS022621_Baylor_scaffold_71993 | - | 649 | 567 | **RNA**→ rpsJ (PRK00596)RpsJ (COG0051)→ |
| env-1121 | SRS023617_Baylor_scaffold_75006 | + | 4288 | 4370 | **RNA**→ rpsJ (PRK00596)RpsJ (COG0051)→ RplC (COG0087)rplC (PRK00001)→ rplD (PRK05319)Ribosomal_L4 (pfam00573)→ rplW (PRK05738)Ribosomal_L23 (pfam00276)→ RplB (COG0090)rplB (PRK09374)→ |
| env-1122 | SRS055426_LANL_scaffold_56046 | + | 434 | 516 | **RNA**→ rpsJ (PRK00596)RpsJ (COG0051)→ RplC (COG0087)rplC (PRK00001)→ rplD (PRK05319)Ribosomal_L4 (pfam00573)→ rplW (PRK05738)Ribosomal_L23 (pfam00276)→ |

| | | | | | |
|---|---|---|---|---|---|
| env-1123 | SRS015038_WUGC_scaffold_27865 | - | 2521 | 2439 | **RNA**→ rpsJ (PRK00596)RpsJ (COG0051)→ RplC (COG0087)rplC (PRK00001)→ rplD (PRK05319)Ribosomal_L4 (pfam00573)→ rplW (PRK05738)Ribosomal_L23 (pfam00276)→ RplB (COG0090)rplB (PRK09374)→ |
| env-1124 | Buccal_mucosa_LANL_scaffold_1634 | + | 150 | 232 | **RNA**→ rpsJ (PRK00596)RpsJ (COG0051)→ |
| env-1125 | SRS014888_C3099341 | - | 199 | 117 | **RNA**→ |
| env-1126 | SRS015762_WUGC_scaffold_42481 | + | 1202 | 1284 | **RNA**→ |
| env-1127 | SRS015797_C2554923 | + | 273 | 355 | **RNA**→ |
| env-1128 | SRS016569_C2502716 | + | 1756 | 1838 | **RNA**→ |
| env-1129 | SRS017808_C3738277 | + | 135 | 217 | **RNA**→ |
| env-1130 | SRS018300_Baylor_scaffold_24096 | + | 192 | 274 | **RNA**→ |
| env-1131 | SRS018439_C2357059 | + | 501 | 583 | **RNA**→ |
| env-1132 | SRS018739_C3682509 | + | 1199 | 1281 | **RNA**→ |
| env-1133 | SRS019122_C5774298 | - | 198 | 116 | **RNA**→ ←hypo |
| env-1134 | SRS019126_C2113536 | - | 241 | 159 | **RNA**→ |
| env-1135 | SRS019327_WUGC_scaffold_22335 | - | 366 | 284 | **RNA**→ rpsJ (PRK00596)Ribosomal_S10 (pfam00338)→ |
| env-1136 | SRS020856_C2575490 | + | 380 | 462 | **RNA**→ |
| env-1137 | SRS022530_C2080540 | + | 216 | 298 | **RNA**→ |
| env-1138 | SRS023926_Baylor_scaffold_581 | - | 11958 | 11876 | **RNA**→ rpsJ (PRK00596)RpsJ (COG0051)→ RplC (COG0087)rplC (PRK00001)→ rplD (PRK05319)Ribosomal_L4 (pfam00573)→ rplW (PRK05738)Ribosomal_L23 (pfam00276)→ RplB (COG0090)rplB (PRK09374)→ rpsS (PRK00357)RpsS (COG0185)→ rplV (PRK00565)Ribosomal_L22 (pfam00237)→ RpsC (COG0092)rpsC (PRK00310)→ rpl16 (CHL00044)rplP (PRK09203)→ PRK00306 (PRK00306)Ribosomal_L29 (pfam00831)→ rpsQ (PRK05610)Ribosomal_S17 (pfam00366)→ rplN (PRK05483)Ribosomal_L14 (pfam00238)→ RplX (COG0198)rplX (PRK00004)→ rplE (PRK00010)RplE (COG0094)→ rpsN (PRK08061)RpsN (COG0199)→ rpsH (PRK00136)Ribosomal_S8 (pfam00410)→ RplF (COG0097)rplF (PRK05498)→ rplR (PRK05593)Ribosomal_L18p (pfam00861)→ RpsE (COG0098)rpsE (PRK00550)→ Ribosomal_L30 (cd01658)rpmD (PRK05611)→ rplO (PRK05592)Ribosomal_L27A (pfam00828)→ SecY (COG0201)secY (PRK09204)→ |
| env-1139 | SRS024081_C2509738 | + | 250 | 332 | **RNA**→ |
| env-1140 | SRS024138_C2606537 | + | 134 | 216 | **RNA**→ |
| env-1141 | SRS024441_LANL_scaffold_49596 | + | 1744 | 1826 | **RNA**→ |
| env-1142 | SRS050244_LANL_scaffold_51697 | - | 366 | 284 | **RNA**→ rpsJ (PRK00596)Ribosomal_S10 (pfam00338)→ |
| env-1143 | SRS053603_LANL_scaffold_24802 | + | 1364 | 1446 | **RNA**→ |
| env-1144 | SRS057355_C2349863 | + | 1129 | 1211 | **RNA**→ |
| env-1145 | SRS017439_C3682793 | - | 198 | 116 | **RNA**→ hypo→ |
| env-1146 | SRS013705_Baylor_scaffold_81679 | - | 1216 | 1134 | **RNA**→ rpsJ (PRK00596)RpsJ (COG0051)→ RplC (COG0087)rplC (PRK00001)→ |
| env-1147 | SRS020220_C3614164 | + | 875 | 957 | **RNA**→ |
| env-1148 | SRS050669_LANL_scaffold_67916 | - | 1577 | 1495 | **RNA**→ rpsJ (PRK00596)RpsJ (COG0051)→ RplC (COG0087)rplC (PRK00001)→ rplD (PRK05319)Ribosomal_L4 (pfam00573)→ |
| env-1149 | SRS058336_LANL_scaffold_62543 | + | 1752 | 1834 | **RNA**→ |
| env-1150 | SRS011086_WUGC_scaffold_38837 | + | 190 | 272 | **RNA**→ rpsJ (PRK00596)RpsJ (COG0051)→ rpl3 (CHL00143)rplC (PRK00001)→ |
| env-1151 | SRS011140_C5153003 | + | 1201 | 1283 | **RNA**→ |
| env-1152 | SRS057539_C2728969 | + | 1202 | 1284 | **RNA**→ |
| env-1153 | SRS063288_C2538166 | - | 197 | 115 | **RNA**→ |
| env-1154 | SRS015395_C2397760 | + | 1066 | 1148 | **RNA**→ |
| env-1155 | SRS016002_WUGC_scaffold_44691 | - | 188 | 106 | **RNA**→ |
| env-1156 | SRS019607_C2614074 | - | 188 | 106 | **RNA**→ |
| env-1157 | SRS022536_LANL_scaffold_76935 | + | 51 | 133 | **RNA**→ rpsJ (PRK00596)Ribosomal_S10 (pfam00338)→ |
| env-1158 | SRS057791_LANL_scaffold_81273 | - | 188 | 106 | **RNA**→ |
| env-1159 | SRS043239_WUGC_scaffold_35420 | + | 57 | 139 | **RNA**→ rpsJ (PRK00596)Ribosomal_S10 (pfam00338)→ |
| Csp-2-1 | NZ_KB822565.1 | - | 883766 | 883673 | **RNA**→ rpsJ (PRK00596)RpsJ (COG0051)→ |
| env-1160 | DLM023_scaffold38722_2 | + | 555 | 632 | **RNA**→ rpsJ (PRK00596)RpsJ (COG0051)→ |
| env-1161 | NOM002_scaffold32230_2 | - | 52228 | 52151 | **RNA**→ rpsJ (PRK00596)RpsJ (COG0051)→ |

| | | | | | |
|---|---|---|---|---|---|
| env-1162 | DLF014_scaffold7481_2 | - | 97485 | 97408 | **RNA**→ rpsJ (PRK00596)RpsJ (COG0051)→ |
| env-1163 | scaffold1445_7_MH0020 | - | 5388 | 5311 | **RNA**→ rpsJ (PRK00596)RpsJ (COG0051)→ |
| env-1164 | C6522713_1_V1.UC-10 | - | 407 | 330 | **RNA**→ rpsJ (PRK00596)RpsJ (COG0051)→ |
| env-1165 | SRS024331_C2252267 | - | 502 | 425 | **RNA**→ rpsJ (PRK00596)RpsJ (COG0051)→ |
| env-1166 | scaffold50700_6_MH0006 | + | 13941 | 14018 | **RNA**→ rpsJ (PRK00596)RpsJ (COG0051)→ |
| env-1167 | JGI994J12354_10000791 | + | 2007 | 2104 | **RNA**→ rpsJ (PRK00596)RpsJ (COG0051)→ RplC (COG0087)rplC (PRK00001)→ |
| | | | | | rplD (PRK05319)Ribosomal_L4 (pfam00573)→ rplW (PRK05738)Ribosomal_L23 (pfam00276)→ |
| | | | | | RplB (COG0090)rplB (PRK09374)→ rpsS (PRK00357)RpsS (COG0185)→ |
| | | | | | rplV (PRK00565)Ribosomal_L22 (pfam00237)→ RpsC (COG0092)rpsC (PRK00310)→ |
| | | | | | rpl16 (CHL00044)rplP (PRK09203)→ PRK00306 (PRK00306)Ribosomal_L29 (pfam00831)→ |
| | | | | | rpsQ (PRK05610)Ribosomal_S17 (pfam00366)→ rplN (PRK05483)Ribosomal_L14 (pfam00238)→ |
| env-1168 | IMNBL1_c0000388 | + | 10662 | 10755 | **RNA**→ rpsJ (PRK00596)RpsJ (COG0051)→ RplC (COG0087)rplC (PRK00001)→ |
| | | | | | rplD (PRK05319)Ribosomal_L4 (pfam00573)→ rplW (PRK05738)Ribosomal_L23 (pfam00276)→ |
| | | | | | RplB (COG0090)rplB (PRK09374)→ rpsS (PRK00357)RpsS (COG0185)→ |
| | | | | | rplV (PRK00565)Ribosomal_L22 (pfam00237)→ RpsC (COG0092)rpsC (PRK00310)→ |
| | | | | | rpl16 (CHL00044)rplP (PRK09203)→ PRK00306 (PRK00306)Ribosomal_L29 (pfam00831)→ |
| | | | | | rpsQ (PRK05610)Ribosomal_S17 (pfam00366)→ rplN (PRK05483)Ribosomal_L14 (pfam00238)→ |
| env-1169 | JGI994J12354_10000720 | - | 21163 | 21067 | **RNA**→ rpsJ (PRK00596)RpsJ (COG0051)→ |
| env-1170 | 2227476835 | - | 442 | 346 | **RNA**→ rpsJ (PRK00596)RpsJ (COG0051)→ |
| env-1171 | IMNBL1_c0000272 | + | 15388 | 15484 | **RNA**→ rpsJ (PRK00596)RpsJ (COG0051)→ |
| env-1172 | SRS014459_WUGC_scaffold_61552 | - | 3023 | 2929 | **RNA**→ rpsJ (PRK00596)RpsJ (COG0051)→ |
| env-1173 | scaffold26742_2_MH0080 | - | 219 | 125 | **RNA**→ |
| env-1174 | DLM023_scaffold12761_3 | + | 988 | 1082 | **RNA**→ |
| env-1175 | SRS015217_WUGC_scaffold_38672 | - | 2002 | 1908 | **RNA**→ rpsJ (PRK00596)RpsJ (COG0051)→ |
| env-1176 | SRS015264_WUGC_scaffold_39672 | - | 16092 | 15998 | **RNA**→ rpsJ (PRK00596)RpsJ (COG0051)→ |
| Cle-1-1 | NC_015275.1 | - | 3861759 | 3861678 | **RNA**→ rpsJ (PRK00596)RpsJ (COG0051)→ RplC (COG0087)rplC (PRK00001)→ |
| | | | | | rplD (PRK05319)Ribosomal_L4 (pfam00573)→ rplW (PRK05738)Ribosomal_L23 (pfam00276)→ |
| | | | | | RplB (COG0090)rplB (PRK09374)→ rpsS (PRK00357)Ribosomal_S19 (pfam00203)→ |
| | | | | | rplV (PRK00565)Ribosomal_L22 (pfam00237)→ RpsC (COG0092)rpsC (PRK00310)→ |
| | | | | | rpl16 (CHL00044)rplP (PRK09203)→ PRK00306 (PRK00306)Ribosomal_L29 (pfam00831)→ |
| | | | | | RpsQ (COG0186)rpsQ (PRK05610)→ rplN (PRK05483)Ribosomal_L14 (pfam00238)→ |
| | | | | | rplX (PRK00004)KOW_RPL26 (cd06089)→ rplE (PRK00010)RplE (COG0094)→ |
| | | | | | rpsN (PRK08061)RpsN (COG0199)→ rpsH (PRK00136)Ribosomal_S8 (pfam00410)→ |
| | | | | | RplF (COG0097)rplF (PRK05498)→ rplR (PRK05593)Ribosomal_L18p (pfam00861)→ |
| | | | | | RpsE (COG0098)rpsE (PRK00550)→ Ribosomal_L30 (cd01658)rpmD (PRK05611)→ |
| | | | | | rplO (PRK05592)Ribosomal_L27A (pfam00828)→ SecY (COG0201)secY (PRK09204)→ |

#### 19.3 Conserved domains

Conserved domains found in protein-coding genes listed in Section 19.2 are shown below, with the first sentence in their description from the Conserved Domain Database (if any). Conserved domains associated with more than one S10-Clostridia RNA are assigned a color, while others are shown in gray. The number in parentheses after the colored domain name is the number of occurrences in Section 19.2.

cd00432 (1) Ribosomal L18/L5e: L18 (L5e) is a ribosomal protein found in the central protuberance (CP) of the large subunit.
cd01086 (5) Methionine Aminopeptidase 1.
cd01428 (2) Adenylate kinase (ADK) catalyzes the reversible phosphoryl transfer from adenosine triphosphates (ATP) to adenosine monophosphates (AMP) and to yield adenosine diphosphates (ADP).
cd01658 (25) Ribosomal protein L30, which is found in eukaryotes and prokaryotes but not in archaea, is one of the smallest ribosomal proteins with a molecular mass of about 7kDa.
cd02412 (2) K homology RNA-binding (KH) domain of the prokaryotic 30S small ribosomal subunit protein S3.
cd04451 (4) S1_IF1: Translation Initiation Factor IF1, S1-like RNA-binding domain.
cd06088 (1) KOW motif of Ribosomal Protein L14.
cd06089 (3) KOW motif of Ribosomal Protein L26.
CHL00044 (135) ribosomal protein L16
CHL00050 (2) ribosomal protein S19

CHL00052 (1) ribosomal protein L2
CHL00140 (11) ribosomal protein L6; Validated
CHL00143 (11) ribosomal protein L3; Validated
COG0051 (841) Ribosomal protein S10 [Translation, ribosomal structure and biogenesis]
COG0087 (263) Ribosomal protein L3 [Translation, ribosomal structure and biogenesis]
COG0088 (14) Ribosomal protein L4 [Translation, ribosomal structure and biogenesis]
COG0090 (179) Ribosomal protein L2 [Translation, ribosomal structure and biogenesis]
COG0092 (144) Ribosomal protein S3 [Translation, ribosomal structure and biogenesis]
COG0094 (127) Ribosomal protein L5 [Translation, ribosomal structure and biogenesis]
COG0097 (22) Ribosomal protein L6P/L9E [Translation, ribosomal structure and biogenesis]
COG0098 (26) Ribosomal protein S5 [Translation, ribosomal structure and biogenesis]
COG0099 (4) Ribosomal protein S13 [Translation, ribosomal structure and biogenesis]
COG0100 (4) Ribosomal protein S11 [Translation, ribosomal structure and biogenesis]
COG0185 (128) Ribosomal protein S19 [Translation, ribosomal structure and biogenesis]
COG0186 (1) Ribosomal protein S17 [Translation, ribosomal structure and biogenesis]
COG0198 (123) Ribosomal protein L24 [Translation, ribosomal structure and biogenesis]
COG0199 (124) Ribosomal protein S14 [Translation, ribosomal structure and biogenesis]
COG0200 (15) Ribosomal protein L15 [Translation, ribosomal structure and biogenesis]
COG0201 (25) Preprotein translocase subunit SecY [Intracellular trafficking, secretion, and vesicular transport]
COG0202 (2) DNA-directed RNA polymerase, alpha subunit/40 kD subunit [Transcription]
COG0522 (2) Ribosomal protein S4 or related protein [Translation, ribosomal structure and biogenesis]
pfam00181 (6) Ribosomal Proteins L2, RNA binding domain.
pfam00203 (24) Ribosomal protein S19.
pfam00237 (150) Ribosomal protein L22p/L17e.
pfam00238 (130) Ribosomal protein L14p/L23e.
pfam00276 (209) Ribosomal protein L23.
pfam00338 (95) Ribosomal protein S10p/S20e.
pfam00347 (2) Ribosomal protein L6.
pfam00366 (132) Ribosomal protein S17.
pfam00406 (4) Adenylate kinase.
pfam00410 (96) Ribosomal protein S8.
pfam00444 (4) Ribosomal protein L36.
pfam00573 (222) Ribosomal protein L4/L1 family.
pfam00828 (10) Ribosomal proteins 50S-L15, 50S-L18e, 60S-L27A.
pfam00831 (131) Ribosomal L29 protein.
pfam00861 (26) Ribosomal L18 of archaea, bacteria, mitoch.
pfam01196 (2) Ribosomal protein L17.
pfam03947 (2) Ribosomal Proteins L2, C-terminal domain.
pfam13207 (1) AAA domain.
pfam17136 (1) Ribosomal proteins 50S L24/mitochondrial 39S L24.
PRK00001 (276) 50S ribosomal protein L3; Validated
PRK00004 (125) 50S ribosomal protein L24; Reviewed
PRK00010 (127) 50S ribosomal protein L5; Validated
PRK00136 (96) 30S ribosomal protein S8; Validated
PRK00276 (4) translation initiation factor IF-1; Validated
PRK00279 (5) adenylate kinase; Reviewed
PRK00306 (131) 50S ribosomal protein L29; Reviewed
PRK00310 (144) 30S ribosomal protein S3; Reviewed
PRK00357 (154) 30S ribosomal protein S19; Reviewed
PRK00465 (4) 50S ribosomal protein L36; Reviewed
PRK00550 (26) 30S ribosomal protein S5; Validated
PRK00565 (150) 50S ribosomal protein L22; Reviewed
PRK00596 (937) 30S ribosomal protein S10; Reviewed
PRK05179 (4) 30S ribosomal protein S13; Validated
PRK05182 (2) DNA-directed RNA polymerase subunit alpha; Provisional
PRK05309 (4) 30S ribosomal protein S11; Validated
PRK05319 (236) 50S ribosomal protein L4; Provisional
PRK05327 (2) 30S ribosomal protein S4; Validated
PRK05483 (130) 50S ribosomal protein L14; Validated
PRK05498 (35) 50S ribosomal protein L6; Validated
PRK05591 (2) 50S ribosomal protein L17; Validated
PRK05592 (25) 50S ribosomal protein L15; Reviewed
PRK05593 (27) 50S ribosomal protein L18; Reviewed
PRK05610 (133) 30S ribosomal protein S17; Reviewed
PRK05611 (25) 50S ribosomal protein L30; Reviewed
PRK05716 (5) methionine aminopeptidase; Validated
PRK05738 (209) 50S ribosomal protein L23; Reviewed
PRK08061 (124) 30S ribosomal protein S14; Reviewed
PRK09203 (135) 50S ribosomal protein L16; Reviewed
PRK09204 (25) preprotein translocase subunit SecY; Reviewed
PRK09374 (188) 50S ribosomal protein L2; Validated
smart00322 (2) K homology RNA-binding domain.
TIGR03918 (1) [FeFe] hydrogenase H-cluster maturation GTPase HydF. [FeFe] hydrogenase H-cluster maturation GTPase HydF. This model describes the family of the [Fe] hydrogenase maturation protein HypF as characterized in Chlamydomonas reinhardtii and found, in an operon with radical SAM proteins HydE and HydG, in numerous bacteria. It has GTPase activity, can bind an 4Fe-4S cluster, and is essential for hydrogenase activity. [Protein fate, Protein modification and repair]

#### 19.4 Multiple-sequence alignment

Each S10-Clostridia RNA is denoted by an abbreviation (like "Eco-1-1") that refers to a taxonomy given in Section 19.1. The alignment may include sequences containing the RNA motif, as well as flanking sequence. The S10-Clostridia RNA itself is denoted by the line underneath marked 5′ and 3′ on either end. Nucleotides in flanking sequences (i.e., not part of the motif) are written in gray letters. Stems of predicted rho-independent transcription terminators, if any, are shaded yellow. (Note: terminator predictions have not been analyzed manually, and many are likely to be false positives. Terminator predictions are those of the RNIE software.) Nucleotides predicted to function (as DNA) as transcription-factor binding sites, if any, are shaded in green. (But note: these predictions are manually annotated, so they might be under-predicted.) Annotated start codons, if any, are shaded green. (Note: start codons are frequently misannotated, especially in environmental samples.) Nucleotides proposed to basepair as part of the consensus structure are shaded in color when they comprise Watson-Crick or G-U pairs. Otherwise they are shaded gray. Conserved stems are also indicated at the bottom of the alignment by angle brackets, where matching $<$ and $>$ denote base-paired columns. Below these angle brackets, the symbol "2" denotes base pairs exhibiting covariation according to the statistically well-founded R-scape method. "1" denotes base pairs exhibiting covariation according to R2R's simplistic method. "0" denotes base pairs that are not observed to mutate and "?" denotes base pairs that have a significant frequency of non-canonical nucleotides for Watson-Crick or G-U pairs ($>$ 5%). Below these base pair annotation is the consensus sequence: "R" = "A" or "G", "Y" = "C" or "U", red nucleotides: nucleotide identity conserved more than 97% of the time, black nucleotides: 90%, gray nucleotides: 75%, red circle (•): nucleotide is present 97% of the time, black circle (•): 90%, gray circle (•): 75%, white circle (◦): 50%. All percentages of sequences just described (e.g. 97% conserved) assume that sequences have been weighted by the GSC algorithm implemented by the Infernal software package.

Duplicate sequences: the following putative homologs are not shown in the alignment because their sequences are identical to a homolog already shown: Bfi-3-1, Bfi-4-1, Bsp-8-1, Cba-4-1, Cba-5-1, Cbo-1-1, Cbo-3-1, Cbo-4-1, Cbo-5-1, Cbo-6-1, Cci-1-1, Ccl-2-1, Ccl-4-1, Ccl-5-1, Ccl-7-1, Ccl-8-1, Ccl-9-1, Ceu-1-1, Cne-1-1, Csp-6-1, Csy-1-1, Csy-2-1, Csy-3-1, Ere-1-1, Hha-1-1, Hha-2-1, Lba-1-1, Lba-11-1, Lba-12-1, Lba-18-1, Lba-21-1, Lba-22-1, Lba-9-1, Lbo-1-1, Lbo-2-1, Lsp-1-1, Rgn-2-1, Rgn-3-1, Rho-1-1, Rla-1-1, Rto-1-1, [bo-1-1, [cl-1-1, env-100, env-1000, env-1001, env-1002, env-1003, env-1004, env-1005, env-101, env-1010, env-1011, env-1012, env-1013, env-1014, env-1015, env-1016, env-1017, env-1025, env-103, env-1031, env-104, env-1047, env-1048, env-1049, env-105, env-1050, env-1051, env-106, env-1060, env-1061, env-1062, env-1063, env-1064, env-1079, env-108, env-1080, env-1090, env-1091, env-1092, env-1098, env-1109, env-1110, env-1111, env-1112, env-1113, env-1114, env-1116, env-1117, env-1119, env-1120, env-1121, env-1122, env-1125, env-1126, env-1127, env-1128, env-1129, env-1130, env-1131, env-1132, env-1133, env-1134, env-1135, env-1136, env-1137, env-1138, env-1139, env-1140, env-1141, env-1142, env-1143, env-1144, env-115, env-1152, env-1153, env-116, env-1161, env-1164, env-1165, env-117, env-118, env-119, env-120, env-124, env-125, env-126, env-127, env-135, env-136, env-137, env-145, env-146, env-147, env-149, env-150, env-151, env-152, env-153, env-154, env-155, env-156, env-157, env-167, env-168, env-169, env-181, env-182, env-187, env-196, env-197, env-198, env-199, env-2, env-200, env-201, env-202, env-203, env-204, env-205, env-206, env-207, env-209, env-210, env-211, env-221, env-222, env-226, env-227, env-228, env-230, env-232, env-233, env-238, env-239, env-24, env-240, env-241, env-243, env-245, env-246, env-253, env-254, env-255, env-256, env-258, env-260, env-261, env-263, env-267, env-268, env-269, env-270, env-271, env-272, env-273, env-274, env-275, env-276, env-277, env-278, env-279, env-285, env-286, env-287, env-292, env-293, env-294, env-295, env-296, env-297, env-298, env-299, env-3, env-301, env-302, env-31, env-318, env-319, env-32, env-320, env-321, env-322, env-323, env-324, env-325, env-326, env-327, env-328, env-329, env-33, env-330, env-337, env-34, env-342, env-343, env-344, env-345, env-346, env-35, env-356, env-357, env-358, env-36, env-367, env-369, env-37, env-370, env-371, env-378, env-38, env-383, env-39, env-40, env-401, env-41, env-412, env-413, env-418, env-419, env-42, env-421, env-422, env-423, env-424, env-425, env-437, env-438, env-439, env-44, env-440, env-441, env-442, env-443, env-444, env-445, env-446, env-447, env-448, env-449, env-450, env-451, env-452, env-453, env-458, env-461, env-462, env-463, env-474, env-475, env-476, env-477, env-478, env-479, env-480, env-481, env-490, env-491, env-492, env-493, env-50, env-504, env-505, env-506, env-507, env-508, env-509, env-51, env-510, env-511, env-519, env-52, env-520, env-521, env-522, env-523, env-524, env-525, env-526, env-527, env-528, env-529, env-53, env-530, env-531, env-532, env-533, env-534, env-535, env-536, env-537, env-538, env-539, env-540, env-541, env-542, env-543, env-544, env-545, env-546, env-547, env-548, env-549, env-550, env-551, env-552, env-553, env-554, env-555, env-56, env-569, env-572, env-574, env-575, env-576, env-577, env-580, env-605, env-609, env-610, env-611, env-612, env-613, env-616, env-617, env-618, env-619, env-62, env-620, env-621, env-622, env-627, env-628, env-629, env-63, env-630, env-631, env-636, env-638, env-64, env-645, env-65, env-651, env-652, env-659, env-66, env-660, env-661, env-665, env-666, env-67, env-670, env-673, env-674, env-675, env-676, env-677, env-678, env-679, env-68, env-680, env-682, env-688, env-689, env-69, env-690, env-693, env-697, env-698, env-70, env-701, env-702, env-703, env-704, env-706, env-707, env-71, env-714, env-715, env-716, env-717, env-718, env-719, env-72, env-720, env-721, env-722, env-723, env-724, env-725, env-728, env-73, env-736, env-737, env-738, env-74, env-740, env-741, env-743, env-745, env-746, env-747, env-748, env-75, env-750, env-757, env-761, env-766, env-771, env-772, env-773, env-774, env-775, env-776, env-779, env-780, env-781, env-782, env-783, env-784, env-785, env-786, env-787, env-788, env-789, env-79, env-790, env-791, env-792, env-793, env-794, env-795, env-796, env-797, env-798, env-799, env-800, env-801, env-804, env-805, env-807, env-808, env-809, env-810, env-811, env-813, env-815, env-816, env-817, env-825, env-834, env-835, env-836, env-84, env-846, env-847, env-852, env-857, env-858, env-859, env-860, env-869, env-870, env-871, env-873, env-876, env-881, env-882, env-883, env-884, env-885, env-886, env-887, env-888, env-889, env-890, env-891, env-894, env-895, env-896, env-897, env-905, env-908, env-909, env-910, env-911, env-912, env-917, env-919, env-920, env-922, env-925, env-927, env-929, env-93, env-931, env-936, env-937, env-938, env-939, env-941, env-947, env-948, env-952, env-956, env-957, env-958, env-966, env-969, env-974, env-975, env-976, env-977, env-978, env-979, env-98, env-982, env-988, env-989, env-990, env-997, env-998, env-999

alignment positions 1 · · · 180

```
env-1     GAAAAUUAUCCUUGACAAGAGAGGGGAACUUUGUUAUCCUAAUAGGGCGCGUUAGGGUAAGUAUGCCCUAAAACGCAAAA..................AACUGCGAUGAAGCGGG.AGAUUGCGCC.AAAAA................................................
env-4     GAAAAUUAUCCUUGACAAGAGAGGGGAACUUUGUUAUUCUAAUAGGGCGCGUUAGGGUAAGUAUGCCCUAAAACGCAAAA..................AACUGCGAUGAAGCGGG.AGAUUGCGCC.AAAAA................................................
env-5     GAAAAUUAUCCUUGACAAGAGAGGGGAACUUUGUUAUCCUAAUAGGGCGCGUUAGGGUAAGUAUGCCCUAAAACGCAAAA..................AACUGCGAUGAAGCGGG.AGAUUGCGCC.AAAAA................................................
env-6     AAAAUAUUCAAAACCCCCUUGACACCGAGGGGGUUUUAUUAUAUGAUACUAAGGCGCGCGAAUGGGGGUAGCAACGCUACCA.GGCGU........GUCAAUGCGAUGAUGCAGG.AGAUUGCGUCCU....................................................
env-7     UCCCCUUGACACCAGCUGUGUUUUCCUAUACAAUAUACCGGUACGCGCGCAAUGGGCACAACACGCCCGGAGGCGUUUAC..................AAAUGCGAUGAACCGGG.AGAUUGCUCC.......................................GAAA...........
env-8     UACCCUUGACACCAGCUGUGUUUUCCUAUACAAUAUACCGGUACGCGCGCAAUGGGCACAACACGCCCGGAGGCGUUUAC..................AAAUGCGAUGAACCGGG.AGAUUGCUCC.......................................GAAA...........
env-9     UCCCCUUGACACCAGCUGUGUUUUCCUAUACAAUAUACCGGUACGCGCGCAAUGGGCACAACACGCCCGGAGGCGUUUAC..................AAAUGCGAUGAACCGGG.AGAUUGCUCC.......................................GAAA...........
env-10    GAGAAAAUCCUCUUGACAAUUAGAAGGGGAUCUUUUAUUAUAAUUAGGCGCGCGAUUGGGGUGUGUCUGCCCGUGCGCGC...................ACUGCGAUGAACCGGG.AGAUUGCUCC.GCA..................................................
env-11    GAGAAAAAGGUCUUGACAUUCCACUGAAUAUCCGAUAUGAUAAUCAAGUCCGCGUGGUGGAAGCCUUUCCGCGCGCAGAC...................ACUGCGAUGAACCGGG.AGAUUGCUCC.GCA..................................................
env-12    ...................................................................................................ACUGCGAUGAACCGGG.AGAUUGCUCC.GCA..................................................
env-13    CCCCUUGACACAAGGGGAUUUUUCCUUUAUUAUAUGUGGGUACGCGCGCGAAGGGUGCUGCACACCCGGAGGCGUUUGCA...................ACUGCGAUGAACCGGG.AGAUUGCUCC.GCA..................................................
env-14    CCCCUUGACACAAGGGGAUUUUUCCUUUAUUAUAUGUGGGUACGCGCGUGAAGGGUACUACACACCCGGAGGCGUUUGCA...................ACUGCGAUGAACCGGG.AGAUUGCUCC.GCA..................................................
env-15    CCCCUUGACACAAGGGGAUUUUUCCUUUAUUAUAUGUGGGUACGCGCGUGAAGGGUACUACACACCCGGAGGCGUUUGCA...................ACUGCGAUGAACCGGG.AGAUUGCUCC.GCA..................................................
env-16    ........................................................GAGGUAUGCCCGGGAGCGUUUACA...................ACUGCGAUGAACCGGG.AGAUUGCUCC.GCA..................................................
env-17    CCCCUUGACACAAGGGGAUUUUUCCUUUAUUAUAUGUGGGUACGCGCGUGAAGGGUGCUACACACCCGGAGGCGUUUGCA...................ACUGCGAUGAACCGGG.AGAUUGCUCC.GCA..................................................
env-18    GAGAAAAAGGUCUUGACAUUCCGCUGAAUAUCCGAUAUGAUAAUCAAGUCCGCGUGGUGGAAGCCUUUCUGCGCGCAGAC...................ACUGCGAUGAACCGGG.AGAUUGCUCC.GCA..................................................
env-19    CCCCUUGACACAAGGGGAUUUUUCCUUUAUUAUAUGUGGGUACGCGCGCGAAGGGUGCUGCACACCCGGAGGCGUUUGCA...................ACUGCGAUGAACCGGG.AGAUUGCUCC.GCA..................................................
env-20    CCCCUUGACACAAGGGGAUUUUUCCUUUAUUAUAUGUGGGUACGCGCGCGAAGGGUGCUGCACACCCGGAGGCGUUUGCA...................ACUGCGAUGAACCGGG.AGAUUGCUCC.GCA..................................................
env-21    UCCCCUUGACACAAGGGGAUUUUUCCUUUAUUAUAUGUGGGUACGCGCGUAAGGGGUGGUACACACCCGGAGGCGUUUAC...................ACUGCGAUGAACCGGG.AGAUUGCUCC.GCA..................................................
env-22    CCCCUUGACACAAGGGGAUUUUUCCUUUAUUAUAUAUGGGUACGCGCGUGAAGGGUGCUACACACCCGGAGGCGUUUGCA...................ACUGCGAUGAACCGGG.AGAUUGCUCC.GCA..................................................
env-23    CCCCUUGACAUAAGGGGAUUUUUCCUUUAUUAUAUGUGGGUACGCGCGCGAAGGGUGCUGCACACCCGGAGGCGUUUGCA...................ACUGCGAUGAACCGGG.AGAUUGCUCC.GCA..................................................
env-25    CCCCUUGACACAAGGGGAUUUUUCCUUUAUUAUAUGUGGGUACGCGCGCGAAGGGUGCUGCACACCCGGAGGCGUUUGCA...................ACUGCGAUGAACCGGG.AGAUUGCUCC.GCA..................................................
env-26    CCCCUUGACAUAAGGGGAUUUUUCCUUUAUUAUAUGUGGGUACGCGCGCGAAGGGUGCUGCACACCCGGAGGCGUUUGCA...................ACUGCGAUGAACCGGG.AGAUUGCUCC.GCA..................................................
env-27    CCCCUUGACACAAGGGGAUUUUUCCUUUAUUAUAUGUGGGUACGCGCGUGAAGGGUGCUACACACCCGGAGGCGUUUGCA...................ACUGCGAUGAACCGGG.AGAUUGCUCC.GCA..................................................
env-28    GAGAAAAAGGUCUUGACAUUCCGCUGAAUAUCCGAUAUGAUAAUCAAGUCCGCGUGGUGGAAGCCUUUCUGCGCGCAGAC...................ACUGCGAUGAACCGGG.AGAUUGCUCC.GCA..................................................
env-29    CCCCUUGACAUAAGGGGAUUUUUCCUUUAUUAUAUGUGGGUACGCGCGCGAAGGGUGCUGCACACCCGGAGGCGUUUGCA...................ACUGCGAUGAACCGGG.AGAUUGCUCC.GCA..................................................
env-30    GAGAAAAAGGUCUUGACAUUCCGCUGAAUAUCCGAUAUGAUAAUCAAGUCCGCGUGGUGGAAGCCUUUCUGCGCGCAGAC...................ACUGCGAUGAACCGGG.AGAUUGCUCC.GCA..................................................
env-43    CCCCUUGACACAAGGGGAUUUUUCCUUUAUUAUAUGUGGGUACGCGCGCGAAGGGUGCUACACACCCGGAGGCGUUUGCA...................ACUGCGAUGAACCGGG.AGAUUGCUCC.GCA..................................................
env-45    CCCCUUGACACAAGGGGAUUUUUCCUUUAUUAUAUGUGGGUACGCGCGUGAAGGGUGCUACACACCCGGAGGCGUUUGCA...................ACUGCGAUGAACCGGG.AGAUUGCUCC.GCA..................................................
env-46    CCUCUUGACAUAAGGGGAUUUUUCCUUUAUUAUAUAUGGGUACGCGCGUGAAGGGUGCUACACACCCGGAGGCGUUUGCA...................ACUGCGAUGAACCGGG.AGAUUGCUCC.GCA..................................................
env-47    GAGAAAAAGGUCUUGACAUUCCGCUGAAUAUCCGAUAUGAUAAUCAAGUCCGCGUGGUGGAAGCCUUUCUGCGCGCAGAC...................ACUGCGAUGAACCGGG.AGAUUGCUCC.GCA..................................................
env-48    GACAACAGGGGCUUUAUAAUAUAUAAUAAACGAGCGUGUGAACGAGGUAUAGUAUAUAAAUAUAUAUAUUUAUAUUUAUGCUUCACCAU........AGAUAUGCGAUGAUGUAUG.AGGUUGCGCCAUAACCCA..............................................
env-49    GACAACAGGGGCUUUAUAAUAUAUAAUAGACGAGCGUGUGAACGAGGUAUAGUAUAUAAAUAUAUAUAUUUAUAUUUAUGCUUCACCAU........AGAUAUGCGAUGAUGUAUG.AGGUUGCGCCAUAACCCA..............................................
env-54    ..........................................................................UUAUGCUUCACCAU........AGAUAUGCGAUGAUGUAUG.AGGUUGCGCCAUAACCCA..............................................
env-55    GACAACAGGGGCUUUAUAAUAUAUAAUAGACGAGCGUGUGAACGAGGUAUAGUAUAUAAAUAUAUAUAUUUAUAUUUAUGCUUCACCAU........AGAUAUGCGAUGAUGUAUG.AGGUUGCGCCAUAACCCA..............................................
env-57    UUGACAACAGGGGCUUUAUAAUAUAUAAUAGACGAGCGUGUGAACGAGGUAUAGUAUAUAAAUAUAUAUAUUUAUAUUUAUGCUUCACCAU........AGAUAUGCGAUGAUGUAUG.AGGUUGCGCCAUAACCCA..............................................
env-58    GACAACAGGGGCUUUAUAAUAUAUAAUAGACGAGCGUGUGAACGAGGUAUAGUAUAUAAAUAUAUAUAUUUAUAUUUAUGCUUCACCAU........AGAUAUGCGAUGAUGUAUG.AGGUUGCGCCAUAACCCA..............................................
env-59    GACAACAGGGGCUUUAUAAUAUAUAAUAGACGAGCGUGUGAACGAGGUAUAGUAUAUAAAUAUAUAUAUUUAUAUUUAUGCUUCACCAU........AGAUAUGCGAUGAUGUAUG.AGGUUGCGCCAUAACCCA..............................................
env-60    GACAACAGGGGCUUUAUAAUAUAUAAUAAACGAGCGUGUGAACGAGGUAUAGUAUAUAAAUAUAUAUAUUUAUAUUUAUGCUUCACCAU........AGAUAUGCGAUGAUGUAUG.AGGUUGCGCCAUAACCCA..............................................
env-61    ......UUGACAAGCAGGGCUUUAUAAUAUAUAAUAAACGAGCGUGUGAACGGGGUGUAUUAUAUAAAUAUAUAAUAUAUCA.GUGAU.UCACCAUAGAUAUGCGAUGAUGUAUG.AGGUUGCGUCAUAACCCA..............................................
env-76    GGCUUUAUAAUAUAUAAUAAACGAGCGUGUGGACGGGGUGUAUUAUAUAAAUAUAUAAUAUAUCAGUGAUUCACCAUAGA...................UAUGCGAUGAUGUAUG.AGGUUGCGUCAUAACCCA..............................................
env-77    GGCUUUAUAAUAUAUAAUAAACGAGCGUGUGAACGGGGUGUAUUAUAUAAAUAUAUAAUAUAUCAGUGAUUCACCAUAGA...................UAUGCGAUGAUGUAUG.AGGUUGCGUCAUAACCCA..............................................
env-78    GGCUUUAUAAUAUAUAAUAAACGAGCGUGUGAACAGGGUGUAUUAUAUAAAUAUAUAAUAUAUCAGUGAUUCACCAUAGA...................UAUGCGAUGAUGUAUG.AGGUUGCGUCAUAACCCA..............................................
env-80    GGCUUUAUAAUAUAUAAUAAACGAGCGUGUGAACGGGGUGUAUUAUAUAAAUAUAUAAUAUAUCAGUGAUUCACCAUAGA...................UAUGCGAUGAUGUAUG.AGGUUGCGUCAUAACCCA..............................................
env-81    GGCUUUAUAAUAUAUAAUAAACGAGCGUGUGAACGGGGUGUAUUAUAUAAAUAUAUAAUAUAUCAGUGAUUCACCAUAGA...................UAUGCGAUGAUGUAUG.AGGUUGCGUCAUAACCCA..............................................
env-82    GGCUUUAUAAUAUAUAAUAAACGAGCGUGUGAACGGGGUGUAUUAUAUAAAUAUAUAAUAUAUCAGUGAUUCACCAUAGA...................UAUGCGAUGAUGUAUG.AGGUUGCGUCAUAACCCA..............................................
env-83    GGCUUUAUAAUAUAUAAUAAACGAGCGUGUGAACAGGGUGUAUUAUAUAAAUAUAUAAUAUAUCAGUGAUUCACCAUAGA...................UAUGCGAUGAUGUAUG.AGGUUGCGUCAUAACCCA..............................................
env-85    AAAACCCCCUUGACACUGAGGGGGUUUUAUUAUAUGAUACUAAGGCGCGCGAAUGGGGGCAGCAACGCUACCACGCGUGU..................CAAUGCGAUGAUGCAGG.AGAUUGCGUCUU....................................................
env-86    AAAACCCCCUUGACACUGAGGGGGUUUUAUUAUAUGAUACUAAGGCGCACGAAUGGGGGUAGCAACGCUACCACGCGUGU..................CAAUGCGAUGAUGCAGG.AGAUUGCGUCUU....................................................
env-87    AAAACCCCCUUGACACCGAGGGGGUUUUAUUAUAUGAUACUAAGGCGCGCGAAUGGAAGUAGCAACGCUACCACGCGUGU..................CAAUGCGAUGAUGCAGG.AGAUUGCGUCUU....................................................
env-88    AUCCCUCUUGACAGCAAGAGGGAUUUAUUGUAGUAUAUAAAGGCGCGCGUAGGGAGCGGAAACAACGUGACCGUGCGUGU..................CAAUGCGAUGAUGCAGG.AGAUUGCGUCUU....................................................
env-89    AUCCCUCUUGACAGCAAGAGGGAUUUAUUGUAGUAUAUAAAGGCACGCGUAGGGAGCGGAAACAACGUGACCGUGCGUGU..................CAAUGCGAUGAUGCAGG.AGAUUGCGUCUU....................................................
Pba-1-1   AUUAACGAAAAAAAUAUCUAAAAAAUGAUUUUUUCUCUAAAAAGGUCUUGCUUUUUUUUGAAGGCUAAUGUAUUAUGUUAGG.GUGCU..UUAGGAUACACCGCGAUGAAACGGG.AGGUUGCCGCCAUUUU................................................
env-90    GAAAAUUCGUGAUUCCGCCGAAAUUUUUAUGAAAUAAUUGUUGCUUUUUCCAAAUCGGUGUGGUAUCAUAUUCAAGCAUGCU.GUGUA.GGUGUGCAGAUAUGCGAUGAUGCAGA.AGGUGGCCGCCCCUUGA...............................................
env-91    GAAAAUUCGUGAUUCCGCCGAAAUUUUUAUGAAAUAAUUGUUGCUUUUUCCAAAUCGGUGUGGUAUCAUAUUCAAGCAUGCU.GUGUA.GGUGUGCAGAUAUGCGAUGAUGCAGA.AGGUGGCCGCCCCUUGA...............................................
env-92    AUACAAGGAUUUGUGUGGUAUCAUUAUCAGGCAUGAUUGUGGGAAGUGGGGCAAGUAUGCCCUAAACACAAUCAGCUGAU....................AUGCGAUGAUGCA.GAAGGUGGCCGUCCUUUGA...............................................
env-94    AUACAAGGAUUUGUGUGGUAUCAUUAUCAGGCAUGAUUGUGGGAAGUGGGGCAAGUAUGCCCGAAACACAAUCUGCUGAU....................AUGCGAUGAUGCA.GAAGGUGGCCGUCCUUUGA...............................................
env-95    AUACAAGGAUUUGUGUGGUAUCAUUAUCAGGCAUGAUUGUGGGAAGUGGGGCAAGUAUGCCCGAAACACAAUCUGCUGAU....................AUGCGAUGAUGCA.GAAGGUGGCCGUCCUUUGA...............................................
env-96    AUACAAGGAUUUGUGUGGUAUCAUUAUCAGGCAUGAUUGUGGGAAGUGGGGCAAGUAUGCCCGAAACACAAUCUGCUGAU....................AUGCGAUGAUGCA.GAAGGUGGCCGUCCUUUGA...............................................
env-97    AUACAAGGAUUUGUGUGGUAUCAUUAUCAGGCAUGAUUGUGGGAAGUGGGGCAAGUAUGCCCGAAACACAAUCUGCUGAU....................AUGCGAUGAUGCA.GAAGGUGGCCGUCCUUUGA...............................................
env-99    AUACAAGGAUUUGUGUGGUAUCAUUAUCAGGCAUGAUUGUGGGAAGUGGGGUAAGUAUGCCCGAAACACAAUCUGCUGAU....................AUGCGAUGAUGCA.GAAGGUGGCCGUCCUUUGA...............................................
env-102   AUACAAGGCUUUGUGUGGUAUUAUUAUCAGGCAUGAUUGUGGGAAGUGGGGCAAGUACGCCCUAAACACAAUCAGCUGAU....................AUGCGAUGAUGCA.GAAGGUGGCCGUCCUUUGA...............................................
env-107   AUACAAGGCUUUGUGUGGUAUUAUUAUCAGGCAUGAUUGUGGGAAGUGGGGCAAGUACGCCCUAAACACAAUCAGCUGAU....................AUGCGAUGAUGCA.GAAGGUGGCCGUCCUUUGA...............................................
env-109   AUACAAGGAUUUGUGUGGUAUCAUUAUCAGGCAUGAUUGUGGGAAGUGGGGCAAGUACGCCCUAAACACAAUCAGCUGAU....................AUGCGAUGAUGCA.GAAGGUGGCCGUCCUUUGA...............................................
env-110   AUACAAGGAUUUGUGUGGUAUCAUUAUCAGGCAUGAUUGUGGGAAGUGGGGCAAGUAUGCCCGAAACACAAUCAGCUGAU....................AUGCGAUGAUGCA.GAAGGUGGCCGUCCUUUGA...............................................
env-111   AUACAAGGCUUUGUGUGGUAUUAUUAUCAGGCAUGAUUGUGGGAAGUGGGGCAAGUACGCCCUAAACACAAUCAGCUGAU....................AUGCGAUGAUGCA.GAAGGUGGCCGUCCUUUGA...............................................
env-112   AUACAAGGCUUUGUGUGGUAUUAUUAUCAGGCAUGAUUGUGGGAAGUGGGGCAAGUACGCCCUAAACACAAUCAGCUGAU....................AUGCGAUGAUGCA.GAAGGUGGCCGUCCUUUGA...............................................
env-113   AUACAAGGCUUUGUGUGGUAUUAUUAUCAGGCAUGAUUGUGGGAAGUGGGGCAAGUACGCCCUAAACACAAUCAGCUGAU....................AUGCGAUGAUGCA.GAAGGUGGCCGUCCUUUGA...............................................
env-114   AUAAAAAAUAUAUGCGAAUUUCUACGAAAAUCACAAGAAAAAAGCUUGCAAAAUGCAGGUCGGUAUGGUAUUAUAAUUGAGC.GUGAC..UGCGACAGAUAUGCGAUGAAGCGGG.AGGUUGCGACCAUAG.................................................
env-121   AUAAAAAAUAUAUGCGAAUUUCUACGAAAAUCACAAGAAAAAAGCUUGCAAAAUGCAGAUCGGUAUGGUAUUAUAAUUGAGC.GUGAC..UGCGACAGAUAUGCGAUGAAGCGGG.AGGUUGCGACCAUAG.................................................
env-122   UUAAAAUAUUUAUAAACUUCUAAUGAUAUAAUUGUGUAAAAACCUUGAAAAAGCCCAUUAAAUCUAGUAUAAUAAACCUUGC.GUGAC..UGCGAUAGAUAUGCGAAGAAGCGGG.AGGUUGCGAC.AGUUA................................................
env-123   UUAAAAUAUUUAUAAACUUCUAAUGAUAUAAUUGUGUAAAAACCUUGAAAAAGCCCAUUAAAUCUAGUAUAAUAAGCCUUGC.GUGAC..UGCGAUAGAUAUGCGAAGAAGCGGG.AGGUUGCGAC.AGUUA................................................
env-128   UUAAAAUAUUUAUAAACUUCUAAUGAUAUAAUUGUGUAAAAACCUUGAAAAAGCCCAUUAAAUCUAGUAUAAUAAACCUUGC.GUGAC..UGCGAUAGAUAUGCGAAGAAGCGGG.AGGUUGCGAC.AGUUA................................................
Rfl-1-1   AAUGGCUUGAUUUAUAUAAUAUAUUGUGAUAUAAUAAUUUACGUUGAGUGCGCUGAAUGCGUACUGACUGCAAUUAAGAU....................AUGCGUUGAAACG.GAAGGUUGCUGACGGUUU................................................
env-129   AACCGCUUGAUUUAUAUAAUAUAUUGUGAUAUAAUAAUUUACGUUGAGUGCGCUGAAUGCGUACUGACUGCAAUUAAGAU....................AUGCGUUGAAACG.GAAGGUUGCUGACGGUUU................................................
env-130   UAUCGGAAAUCAAAGAAAAAUUCUGAAAAAAUGCAAAAAAACACUUGCAAUCCCUGUCUUUAUGCCUUAUAAUAAUAGCUGUUGUGAC.........AUAGAGUGUUGAUACGGA.AGGUUGCCGAAGCGGAUGAAUCAAGGGCU...................................
env-131   ...............................UAUUAAAUAAAUGCUUGAUAAUACGCCAUCUUUGGCGUAUAAUAGCAAUGUUGUGGC........AUGAUAGCUAAGAAGCG.G.AGGUUGCCGAGAA.....................UACU..........GCAAA...........
env-132   UAAUUGUAGAAAAUAACUAAAAUUAUAUUUUUAUUAAAUAAAAGCUUGAUAAUACGUUGAUAUUGGCGUAUAAUGGCAAUGUUGUGGC........AUGAUAGCUAAGAAGCG.G.AGGUUGCCGAGAA.....................UACU..........GCAAA...........
env-133   .......................UAAAAAUUUGUGUAAAUUAUACUUGAAAAUAUAAGCAAAUUAGCAUAUAAUAGUACUGUUGUGGC........AUGAUAGCUAAGAAGCG.G.AGGUUGCCGAGAA.....................UACU..........GCAAA...........
env-134   UAAUUGUAGAAAAUAACUAAAAUUAUGCUUUUAUUAAAUAAAUGCUUGAUAAUACGCCAUCUUUGGCGUAUAAUAGCAAUGUUGUGGC........AUGAUAGCUAAGAAGCG.G.AGGUUGCCGAGAA.....................UACU..........GCAAA...........

          ................................................................................<<.<<<<<......................<<<.<.<<<...<<<<<.......................<<<<<<<<<<<<<<............>>>>
          ................................................................................21..1111......................222.2.012...12222.......................1111111.10..00............00..
          •••••••••••••••R••••••••••R••••••••RAAAAR•••YU••••••••••••••••••••••••••••••••••GYUGUGRC--------∘UGAYAGC••UGAAGCG∘G-AGGUUGCYRY••∘○○○-----------------------------------------------
                                                                                                5′
```

```
env-138   UAAUUGUAGAAAAUAACUAAAAUUAUGUUUUUAUUAAAUAAAUGCUUGAUAAUACGCCAUCUUUGGCGUAUAAUAGCAAUGUUGUGGC........AUGAUAGCUAAGAAGCG.G.AGGUUGCCGAGAA....................UACU..........GCAAA...........
env-139   UAAUUGUGGAAAAUAACUAAAAUUAUGCUUUUAUUAAAUAAAUGCUUGAUAAUACGCCAUCUUUGGCGUAUAAUAGCAAUGUUGUGGC........AUGAUAGCUAAGAAGCG.G.AGGUUGCCGAGAA....................UACU..........GCAAA...........
env-140   UAAUUGUAGAAAAUAACUAAAAUUAUGCUUUUAUUAAAUAAAUGCUUGAUAAUACGCCAUCUUUGGCGUAUAAUAACAAUGUUGUGGC........AUGAUAGCUAAGAAGCG.G.AGGUUGCCGAGAA....................UACU..........GCAAA...........
env-141   UAAUUGUGGAAAAUAACUAAAAUUAUGCUUUUAUUAAAUAAAUGCUUGAUAAUACGCCAUCUUUGGCGUAUAAUAGCAAUGUUGUGGC........AUGAUAGCUAAGAAGCG.G.AGGUUGCCGAGAA....................UACU..........GCAAA...........
env-142   UAAUUGUAGAAAAUAACUAAAAUUAUAUUUUUAUUAAAUAAAAGCUUGAUAAUACGUUGAUAUUGGCGUAUAAUGGCAAUGUUGUGGC........AUGAUAGCUAAGAAGCG.G.AGGUUGCCGAGAAUA..................CUGC..........AAAA............
env-143   ...........................................................................GCAAUGUUGUGGC........AUGAUAGCUAAGAAGCG.G.AGGUUGCCGAGAAUA..................CUGC..........AAAA............
env-144   UAAUUGUAGAAAAUAACUAAAAUUAUAUUUUUAUUAAAUAAAAGCUUGAUAAUACGUUGAUAUUGGCGUAUAAUGGCAAUGUUGUGGC........AUGAUAGCUAAGAAGCG.G.AGGUUGCCGAGAAUA..................CUGC..........AAAA............
env-148   UAAUUGUAGAAAAUAACUAAAAUUAUAUUUUUAUUAAAUAAAAGCUUGAUAAUACGUUGAUAUUGGCGUAUAAUGGCAAUGUUGUGGC........AUGAUAGCUAAGAAGCG.G.AGGUUGCCGAGAAUA..................CUGC..........AAAA............
env-158   UAAUUGUAGAAAAUAACUAAAAUUAUAUUUUUAUUAAAUAAAAGCUUGAUAAUACGUUGAUAUUGGCGUAUAAUGGCAAUGUUGUGGC........AUGAUAGCUAAGAAGCG.G.AGGUUGCCGAGAAUA..................CUGC..........AAAA............
env-159   UAAUUGUAGAAAAUAACUAAAAUUAUAUUUUUAUUAAAUAAAAGCUUGAUAAUACGCCAUCUUUGGCGUAUAAUGGCAAUGUUGUGGC........AUGAUAGCUAAGAAGCG.G.AGGUUGCCGAGAAUA..................CUGC..........AAAA............
env-160   UAAUUGUAGAAAAUAACUAAAAUUAUAUUUUUAUUAAAUAAAAGCUUGAUAAUACGUUGAUAUUGGCGUAUAAUGGCAAUGUUGUGGC........AUGAUAGCUAAGAAGCG.G.AGGUUGCCGAGAAUA..................CUGC..........AAAA............
env-161   UAAUUGUAGAAAAUAACUAAAAUUAUAUUUUUAUUAAAUAAAAGCUUGAUAAUACGUUGAUAUUGGCGUAUAAUGGCAAUGUUGUGGC........AUGAUAGCUAAGAAGCG.G.AGGUUGCCGAGAAAA..................CUG...........UGAAUU..........
Cmo-1-1   GGUCUUGUGCAGAAUAACUAAAAUGAUUUUUUAUAUAAAUAAUGCUUGAAAAAAGAUAAUAAAUAGCGUAUAAUAACAACGUUGUGGC........AUGAUAGCUAAGAAGCG.G.AGGUUGCCGAGAAAA..................CUG...........UGAAUU..........
env-162   GUCUUGUGCAAAACAACUAAAAUGAUUUUUUUCUGUAAAUAAUGCUUGAAAAGAGAAAAAUAAUAGCGUAUAAUAACAAUGUUGUGGC........AUGAUAGCUAAGAAGCG.G.AGGUUGCCGAGAAAA..................CUG...........UGAAUU..........
env-163   .....................................................................AUAAUAACAAUGUUGUGGC........AUGAUAGCUAAGAAGCG.G.AGGUUGCCGAGAAAA..................CUG...........UGAAUU..........
Ece-1-1   GGUUUUUCGUAUAAAAAAAUCUUUACAAAAAAGUUUUACAAAGUUCUUGCAUUUUCUACAAUGCGUGACUAUAAUGUAACUGUUGUGAC........AUGAAAGCUAUGAAGCGAG.AGAUUGCAGC.GAAGA...............................................
env-164   ACAAUCCUCUUUGUUUUAUUUUGACGAAAAAUCAUAUAAAAGGCUUGCAAAACAGAAAAAAAAGUAAUAUAGUAGCUAUUGCUGUGAC........AUGAUAGCGUUGAAGCGUG.AGGUUGCCACAUUAUCCAGUGA.........................................
Bfi-1-1   ACAAUCCUCUUUGUUUUAUUUUGACGAAAAAUCAUAUAAAAGGCUUGCAAAACAGAAAAAAAAGUAAUAUAGUAGCUAUUGCUGUGAC........AUGAUAGCGUUGAAGCGUG.AGGUUGCCACAUUAUCCAGUGA.........................................
Bfi-2-1   ACAAUCCUCUUUGCUUUAUUUUGACGAAAAAUCAUAUAAAAGGCUUGCAAAACAGAAAAAAAAGUAAUAUAGUAGCUAUUGCUGUGAC........AUGAUAGCGUUGAAGCGUG.AGGUUGCCACAUUAUCCAGUGA.........................................
env-165   AAAAGUUUGUAAUAUAUUUUUAGGAAAAUAUUCAAAAUAAAAUAUUGAUUAUUUUUCUAAAAGGUAGUAAUAUAUAGAAUGCUGUGGC.......AUUGAUAGCUGUGAAGUGGG.AGGUUGCUACUUUC...................AUUUUU........AAUUUA..........
env-166   AAAAGUUUGUAAUAUAUUUUUAGGAAAAUAUUCAAAAUAAAAUAUUGAUUAUUUUUCUAAAAGGUAGUAAUAUAUAGAAUGCUGUGGC.......AUUGAUAGCUGUGAAGUGGG.AGGUUGCUACUUUC...................AUUUUU........AAUUUA..........
env-170   ..................................................AUUUUUCUAAAAGGUAGUAAUAUAUAGAAUGCUGUGGC.......AUUGAUAGCUGUGAAGUGGG.AGGUUGCUACUUUC...................AUUUUU........AAUUUA..........
env-171   AAAACUUUGUGAUAUAUUUUUAUGAAAUUAUUUUAAAUAAGAUGUUGAUUAUUUUUGCGAAAGGUAGUAGUAUAUAGGAUACCGUGGC.......AUUGAUAGCUGUGAAGUGGG.AGGUUGCUACUU.....................UC.AUUUUU.....AAUUUA..........
env-172   ................UUUUUAUGAAAUUAUUUUAAAUAAGAUGUUGAUUAUUUUUGCGAAAGGUAGUAGUAUAUAGGAUACCGUGGC.......AUUGAUAGCUGUGAAGUGGG.AGGUUGCUACUU.....................UC.AUUUUU.....AAUUUA..........
env-173   AGAAAAUAAAAAGAAAAAAUACACAAAAUUUUUAAAAAAAUUUCUUGCAUAUUUCCCAAACAUCAUGUAUACUAACAAUUGCUGUGGC........AUGAUAGCGUUGAAGCGUG.AGGUUGCUGCAGG....................GCC...........GAAA............
env-174   AGAAAAUAAAAAGAAAAAAUACACAAAAUUUUUAAAAAAAUUUCUUGCAUAUUUCCCAAACAUCAUGUAUACUAACAAUUGCUGUGGC........AUGAUAGCGUUGAAGCGUG.AGGUUGCUGCAGG....................GCC...........GAAA............
env-175   UGUGGUUGUAUUGUUGAAACAAUGAUAAAAAAAUAAAAAAGUACUUGCAUUCCCCAAAAACUUAGUGUAUAAUAGCAAUUGCUGUGAC........AUGAUAGCGAAGAAGCGAG.AGGUUGCCAUCCGAUG...............................................
env-176   ..........................AAAAAAUAAAAAAAGUACUUGCAUUCCCCAAAAACUUAGUGUAUAAUAGCAAUUGCUGUGAC........AUGAUAGCGAAGAAGCGUG.AGGUUGCCAUCCGAUG...............................................
env-177   CUGAAAACGAUGAAAAAUAGCUUGCAAAAAAAUAAAAAAAGUACUUGCAUUCCCCAAAAACUUAGUGUAUAAUAGCAAUUGCUGUGAC........AUGAUAGCGAAGAAGCGUG.AGGUUGCCAUCCGAUG...............................................
env-178   CUGAAAACGAUGAAAAAUAGCUUGCAAAAAAAUAAAAAAAGUACUUGCAUUCCCCAAAAACUUAGUGUAUAAUAGCAAUUGCUGUGAC........AUGAUAGCGAAGAAGCGUG.AGGUUGCCAUCCGAUG...............................................
env-179   CUGAAAACGAUGAAAAAUAGCUUGCAAAAAAAUAAAAAAAGUACUUGCAUUCCCCAAAAACUUAGUGUAUAAUAGCAAUUGCUGUGAC........AUGAUAGCGAAGAAGCGUG.AGGUUGCCAUCCGAUG...............................................
env-180   UUUUACCUUUUUUGCAUGGCUUGUAGUUUUCGUGAAAAAAAGUGCUUGCAUUAUGAAAAAAAGGUAUGGUAGACUAAUAUUUGCUGUGAC........AUGAUAGCGAUGAAGCGUG.AGGUUGCCACCAUCC................................................
env-183   UUUUUCCUUUUUUGCAUGGCUUGUAGUUUUCGUGAAAAAAAGUGCUUGCAUUAUGAAAAAAAGGUAUGGUAGACUAAUAUUUGCUGUGAC........AUGAUAGCGAUGAAGCGUG.AGGUUGCCACCAUCC................................................
env-184   .......................GUUUUCGUGAAAAAAAGUGCUUGCAUUAUGAAAAAAAGGUAUGGUAGACUAAUAUUUGCUGUGAC........AUGAUAGCGAUGAAGCGUG.AGGUUGCCACCAUCC................................................
env-185   UUUUACCUUUUUUGCAUGGCUUGUAGUUUUCGUGAAAAAAAGUGCUUGCAUUAUGAAAAAAAGGUAUGGUAGACUAAUAUUUGCUGUGAC........AUGAUAGCGAUGAAGCGUG.AGGUUGCCACCAUCC................................................
env-186   UUUACCUUUUUUGCAUGGCUUGUAGUUUUCGUGAAAAAAAAGUGCUUGCAUUAUGGAAAAAGGUAUGGUAGACUAAUAUUUGCUGUGAC........AUGAUAGCGAUGAAGCGUG.AGGUUGCCACCAUCC................................................
env-188   UUUAUCUUUUUUGCAUGGCUUGUAGUUUUCGUGAAAAAAAAGUGCUUGCAUUAUGGAAAAAGGUAUGGUAGACUAAUAUUUGCUGUGAC........AUGAUAGCGAUGAAGCGUG.AGGUUGCCACCAUCC................................................
env-189   .......................GUUUUCGUGAAAAAAAGUGCUUGCAUUAUGAAAAAAAGGUAUGGUAGACUAAUAUUUGCUGUGAC........AUGAUAGCGAUGAAGCGUG.AGGUUGCCACCAUCC................................................
env-190   AAAGUGCCAACGAACACUUGACACAAACUUCGUGAAAAAAAGUGCUUGCAUUAUGGAAAAAGGUAUGGUAGACUAAUAUUUGCUGUGAC........AUGAUAGCGAUGAAGCGUG.AGGUUGCCACCAUCC................................................
env-191   UUUUACCUUUUUUGCAUGGCUUGUAGUUUUCGUGAAAAAAAGUGCUUGCAUUAUGGAAAAAGGUAUGGUAGACUAAUAUUUGCUGUGAC........AUGAUAGCGAUGAAGCGUG.AGGUUGCCACCAUCC................................................
env-192   UUUUACCUUUUUUGCAUGGCUUGUAGUUUUCGUGAAAAAAAGUGCUUGCAUUAUGGAAAAAGGUAUGGUAGACUAAUAUUUGCUGUGAC........AUGAUAGCGAUGAAGCGUG.AGGUUGCCACCAUCC................................................
env-193   ......CAACGAACACUUGACACAAACUUCGUGAAAAAAAGUUCUUGCAUUAUGGAAAAAGGUAUGGUAGACUAAUAUUUGCUGUGAC........AUGAUAGCGAUGAAGCGUG.AGGUUGCCACCAUCC................................................
env-194   UUUUACCUUUUUUGCAUGGCUUGUAGUUUUCGUGAAAAAAAGUGCUUGCAUUAUGGAAAAAGGUAUGGUAGACUAAUAUUUGCUGUGAC........AUGAUAGCGAUGAAGCGUG.AGGUUGCCACCAUCC................................................
env-195   UUUUACCUUUUUUGCAUGGCUUGUAGUUUUCGUGAAAAAAAGUGCUUGCAUUAUGGAAAAAGGUAUGGUAGACUAAUAUUUGCUGUGAC........AUGAUAGCGAUGAAGCGUG.AGGUUGCCACCAUCC................................................
env-208   UUUUACCUUUUUUGCAUGGCUUGUAGUUUUCGUGAAAAAAAGUGCUUGCAUUAUGAAAAAAAGGUAUGGUAGACUAAUAUUUGCUGUGAC........AUGAUAGCGAUGAAGCGUG.AGGUUGCCACCAUCC................................................
Lba-19-1  AGGGCUGUGAAUAGUAAUUAUUUGAAAAAUUGUACAAAAAGUCCUUGCAAUUUUCUGCAGACUAAUCUAUAAUCGGUCUUGCUGUGAC........AUGAUAGCUAUGAAGCGUG.AGGUUGCUGCCGCAACAAAAUAUAUAUAAGAAAC.............................
Lba-17-1  CAAAAAAUAAUUGAAAAAUUUAUUAAAAAAAAUAGAAAAAAGUACUUGCAUUCCUUAAAUCAAACAUUUAUACUAAUCAUUGCUGUGAC........AUGAUAGCUAUGAAGCGUG.AGGUUGCUAUUA.....................UGGC..........GAAA............
env-212   CAAAAAAUAAUUGCAAAAUUUAUUAAAAAAAAUAGAAAAAAGUACUUGCAUUCCUUAAAUCAAACAUUUAUACUAAUCAUUGCUGUGAC........AUGAUAGCUAUGAAGCGUG.AGGUUGCUAUUA.....................UGGC..........GAAA............
env-213   CUGCUCUUUAAAUCAAUUUUUUUGCAAAGAUUCAAAAAAAGUGCUUGCAUUUUGAAAAGUUAUCAUAUAUACUAUCUAUUGCUGUGGC........AUGAUAGCUAUGAAGCGUG.AGGUUGCUGCACAUGU...............................................
env-214   GGGAAAUAUAAUAAAAAAAUUUCUUAAAAAAAUAUAGAAAAAAUGCUUGCAUUCCGCAAAAACGUGUUAUAUAAUCAAACGUGCUGUGAC........AUGAUAGCGUUGAAACG.G.AGGUUGCUGUAUCGUAUAGA...........................................
env-215   GGGAAAUAUAAUAAAAAAAUUUCUUAAAAAAAUAUAGAAAAAAUGCUUGCAUUCCGCAAAAACGUGUUAUAUAAUCAAACGUGCUGUGAC........AUGAUAGCGUUGAAACG.G.AGGUUGCUGUAUCGUAUAGA...........................................
env-216   GGGAAAUAUAAUAAAAAAAUUUCUUAAAAAAAUAUAGAAAAAAUGCUUGCAUUCCGCAAAAACGUGUUAUAUAAUCAAACGUGCUGUGAC........AUGAUAGCGUUGAAACG.G.AGGUUGCUGUAUCGCAUAGA...........................................
env-217   ......................................................................AAUCAAACGUGCUGUGAC........AUGAUAGCGUUGAAACG.G.AGGUUGCUGUAUCGCAUAGA...........................................
env-218   GGGGAAUAUAAUAAAAAAAUUUCUUGAAAAAUAUAGAAAAAAUGCUUGCAUUCCGCAAAAACGUGUUAUAUAAUCAAACGUGCUGUGAC........AUGAUAGCGUUGAAACG.G.AGGUUGCUGUAUCGCAUAGA...........................................
env-219   GGGGAAUAUAAUAAAAAAAUUUCUUAAAAAAAUAUAGAAAAAAUGCUUGCAUUCCGCAAAAACGUGUUAUAUAAUCAAACGUGCUGUGAC........AUGAUAGCGUUGAAACG.G.AGGUUGCUGUAUCGCAUAGA...........................................
env-220   GGGAAAUAUAAUAAAAAAAUUUCUUAAAAAAAUAUAGAAAAAAUGCUUGCAUUCCGCAAAAACGUGUUAUAUAAUCAAACGUGCUGUGAC........AUGAUAGCGUUGAAACG.G.AGGUUGCUGUAUCGCAUAGA...........................................
env-223   GGGGAAUAUAAUAAAAAAAUUUCUUAAAAAAAUAUAGAAAAAAUGCUUGCAUUCCGCAAAAACGUGUUAUAUAAUCAAACGUGCUGUGAC........AUGAUAGCGUUGAAACG.G.AGGUUGCUGUAUCGCAUAGA...........................................
env-224   GGGAAAUAUAAUAAAAAAAAUUCUUAAAAAAAUAUAGAAAAAAUGCUUGCAUUCCGCAAAAACGUGUUAUAUAAUCAAACUUGCUGUGAC........AUGAUAGCGUUGAAACG.G.AGGUUGCUGUAUCGCAUAGA...........................................
env-225   GGGGAAUAUAAUAAAAAAAUUUCUUGAAAAAUAUAGAAAAAAUGCUUGCAUUCCGCAAAAACGUGUUAUAUAAUCAAACGUGCUGUGAC........AUGAUAGCGUUGAAACG.G.AGGUUGCUGUAUCGCAUAGA...........................................
env-229   GGGAAAUAUAAUAAAAAAAUUUCUUAAAAAAAUAUAGAAAAAAUGCUUGCAUUCCGCAAAAACGUGUUAUAUAAUCAAACGUGCUGUGAC........AUGAUAGCGUUGAAACG.G.AGGUUGCUGUAUCGCAUAGA...........................................
env-231   GGGGAAUAUAAUAAAAAAAUUUCUUAAAAAAAUAUAGAAAAAAUGCUUGCAUUCCGCAAAAACGUGUUAUAUAAUCAAACGUGCUGUGAC........AUGAUAGCGUUGAAACG.G.AGGUUGCUGUAUCGCAUAGA...........................................
env-234   GGGGAAUAUAAUAAAAAAAUUUCUUAAAAAAAUAUAGAAAAAAUGCUUGCAUUCCGCAAAAACGUGUUAUAUAAUCAAACGUGCUGUGAC........AUGAUAGCGUUGAAACG.G.AGGUUGCUGUAUCGCAUAGA...........................................
env-235   GGGAAAUAUAAUAAAAAAAUUUCUUAAAAAAAUAUAGAAAAAAUGCUUGCAUUCCGCAAAAACGUGUUAUAUAAUCAAACGUGCUGUGAC........AUGAUAGCGUUGAAACG.G.AGGUUGCUGUAUCGCAUAGA...........................................
env-236   GGGAAAUAUAAUAAAAAAAUUUCUUAAAAAAAUAUAGAAAAAAUGCUUGCAUUCCGCAAAAACGUGUUAUAUAAUCAAACGUGCUGUGAC........AUGAUAGCGUUGAAACG.G.AGGUUGCUGUAUCGCAUAGA...........................................
env-237   GGGAAAUAUAAUAAAAAAAAUUCUUAAAAAAAUAUAGAAAAAAUGCUUGCAUUCCGCAAAAACGUGUUAUAUAAUCAAACGUGCUGUGAC........AUGAUAGCGUUGAAACG.G.AGGUUGCUGUAUCGUAUAGA...........................................
env-242   AAUAUUCAUGAUUUUCAAGUGUAAAAAAAAGUUGAAAAUAAUACUUGCAUUGUAAGUGAAAAUGGUGUAUAAUUGCAAAUGCUGUGAC........AUGAGAGCGUUGAAGUAAG.AGGUUGCUACUUGAAAAGUA...........................................
env-244   AAUAUUCAUGAUUUUCAAGUGUAAAAAAAAGUUGAAAAUAAUACUUGCAUUGUAAGUGAAAAUGGUGUAUAAUUGCAAAUGCUGUGAC........AUGAGAGCGUUGAAGUAAG.AGGUUGCUACUUGAAAAGUA...........................................
env-247   UGAAACUGUGCUUUAGAAAAAAUUAAAAAAAAUUCAAAAAAAUGCUUGCAAACACUUGAAAAGUAUAGUAUAGUAAUAAAUGCUGUGGC........AUGAUAGCGUAGAAGCGUG.AGGUUGCUCUUAC....................CCG...........GCAA............
env-248   UGAAACUGUGCUUUAGAAAAAAUUAAAAAAAAUUCAAAAAAAUGCUUGCAAACCCUUGAAAAAUAUAGUAUAGUAAUAAAUGCUGUGAC........AUGAUAGCGUAGAAGCGCG.AGGUUGCUCUUAC....................CUG...........GUAA............
env-249   UUGAAACUGUGCUUGAAAAAAAAUUAAAAAAAAUGUAAAAAAAAUGCUUGCAAACCCUUGAAAAAUAUAGUAUAGUAAUAAAUGCUGUGAC........AUGAUAGCGUAGAAGCGUG.AGGUUGCUCUUAC....................CUG...........GUAA............
env-250   UGAAACUGUGCUUGAAAAAAAAUUAAAAAAAAUGUUAAAAAAAAUGCUUGCAAACCCUUGAAAAAUAUAGUAUAGUAAUAAAUGCUGUGAC........AUGAUAGCGUAGAAGCGUG.AGGUUGCUCUUAC....................CUG...........GUAA............
env-251   UGAAACUGUGCUUGAAAAAAAUUUAAAAAAAAUGUUAAAAAAAAUGCUUGCAAACCCUUGAAAAAUAUAGUAUAGUAAUAAAUGCUGUGAC........AUGAUAGCGUAGAAGCGUG.AGGUUGCUCUUAC....................CUG...........GUAA............
env-252   UUGAAACUGUGCUUGAAAAAAAAUUAAAAAAAAUGUAAAAAAAAUGCUUGCAAACCCUUGAAAAAUAUAGUAUAGUAAUAAAUGCUGUGAC........AUGAUAGCGUAGAAGCGUG.AGGUUGCUCUUAC....................CUG...........GUAA............
env-257   UGAAACUGUGCUUGAGAAAAAAAUUAAAAAAAAUGUAAAAAAAAUGCUUGCAAACCCUUGAAAAAUAUAGUAUAGUAAUAAAUGCUGUGAC........AUGAUAGCGUAGAAGCGUG.AGGUUGCUCUUAC....................CUG...........GUAA............
env-259   UGAAACUGUGCUUGAAAAAAAAUUAAAAAAAAUGUUAAAAAAAAUGCUUGCAAACCCUUGAAAAAUAUAGUAUAGUAAUAAAUGCUGUGAC........AUGAUAGCGUAGAAGCGUG.AGGUUGCUCUUAC....................CUG...........GUAA............
env-262   UUGAAACUGUGCUUGAAAAAAAAUUAAAAAAAAUGUUAAAAAAAAUGCUUGCAAACCCUUGAAAAAUAUAGUAUAGUAAUAAAUGCUGUGAC........AUGAUAGCGUAGAAGCGUG.AGGUUGCUCUUAC....................CUG...........GUAA............
env-264   UGAAACUGUGCUUGAGAAAAAAAUUAAAAAAAAUGUAAAAAAAAUGCUUGCAAACCCUUGAAAAAUAUAGUAUAGUAAUAAAUGCUGUGAC........AUGAUAGCGUAGAAGCGUG.AGGUUGCUCUUAC....................CUG...........GUAA............
env-265   UGAAACUGUGCUUGAAAAAAAUUUAAAAAAAAUGUUAAAAAAAAUGCUUGCAAACCCUUGAAAAAUAUAGUAUAGUAAUAAAUGCUGUGAC........AUGAUAGCGUAGAAGCGUG.AGGUUGCUCUUAC....................CUG...........GUAA............
env-266   AAUAUUACCCGAAAUAAAAAAAAUAAAUUUUUUUUGAAAAAAGUAUUGCAAUUAUUGGAAACGUCAUGUAUAAUAUCACUUGCUGUGGC........AUGAUAGCUAUGAAGUUUG.AAGUUACUGUCCAUGA...............................................
env-280   AAUAUUAUCCGAAAUAAAAAAAAUAAAUUUUUUUUGAAAAAAGUAUUGCAAUUAUUGUAAACGUCAUGUAUAAUAUCACUUGCUGUGGC........AUGAUAGCUAUGAAGUUUG.AAGUUACUGUCCAUGA...............................................
env-281   AAUAUUACCCGAAAUAAAAAAAAUAAAUUUUUUUUGAAAAAAGUAUUGCAAUUAUUGGAAACGUCAUGUAUAAUAUCACUUGCUGUGGC........AUGAUAGCUAUGAAGUUUG.AAGUUACUGUCCAUGA...............................................
env-282   AAUAUUAUCCGAAAUAAAAAAAAUAAAUUUUUUUUGAAAAAAGUAUUGCAAUUAUUGUAAACGUCAUGUAUAAUAUCACUUGCUGUGGC........AUGAUAGCUAUGAAGUUUG.AAGUUACUGUCCAUGA...............................................
env-283   AAUAUUACCCGAAAUAAAAAAAAUAAAUUUUUUUUGAAAAAAGUAUUGCAAUUAUUGGAAACGUCAUGUAUAAUAUCACUUGCUGUGGC........AUGAUAGCUAUGAAGUUUG.AAGUUACUGUCCAUGA...............................................

          ................................................................................<<.<<<<<......................<<<.<.<<<...<<<<<......................<<<<<<<<<<<<<<............>>>>
          ................................................................................21..1111......................222.2.012...12222......................1111111.10..00............00..
          ●●●●●●●●●●●●●●R●●●●●●●●●●R●●●●●●●●RAAAAR●●●YU●●●●●●●●●●●●●●●●●●●●●●●●●●●●●●●●●●●GYUGUGRC-------∘UGAYAGC●●UGAAGCG∘G-AGGUUGCYRY●●∘∘∘∘--------------------------------------------
                                                                                                 5′
```

```
env-284  AAUAUUACCCGAAAUAAAAAAAUAAAUUUUUUUUGAAAAAAGUAUUGCAAUUAUUGUAAACGUCAUGUAUAAUAUCACUUGCUGUGGC.......AUGAUAGCUAUGAAGUUUG.AAGUUACUGUCCAUGA.......................................................
env-288  AAUAUUACCCGAAAUAAAAAAAUAAAUUUUUUUUGAAAAAAGUAUUGCAAUUAUUGUAAACGUCAUGUAUAAUAUCACUUGCUGUGGC.......AUGAUAGCUAUGAAGUUUG.AAGUUACUGUCCAUGA.......................................................
env-289  AAUAUUAUCCGAAAUAAAAAAAUAAAUUUUUUUUGAAAAAAGUAUUGCAAUUAUUGGAAACGUCAUGUAUAAUAUCACUUGCUGUGGC.......AUGAUAGCUAUGAAGUUUG.AAGUUACUGUCCAUGA.......................................................
env-290  AAUAUUACCCGAAAUAAAAAAAUUAAUUUUUUUUGAAAAAAGUAUUGCAAUUAUUGUAAACGUCAUGUAUAAUAUCACUUGCUGUGGC.......AUGAUAGCUAUGAAGUUUG.AAGUUACUGUCCAUGA.......................................................
env-291  AAUAUUAUCCGAAAUAAAAAAAUAAAUUUUUUUUGAAAAAAGUAUUGCAAUUAUUGUAAACGUCAUGUAUAAUAUCACUUGCUGUGGC.......AUGAUAGCUAUGAAGUUUG.AAGUUACUGUCCAUGA.......................................................
env-300  AAUAUUACCCGAAAUAAAAAAAUAAAUUUUUUUUGAAAAAAGUAUUGCAAUUAUUGGAAACGUCAUGUAUAAUAUCACUUGCUGUGGC.......AUGAUAGCUAUGAAGUUUG.AAGUUACUGUCCAUGA.......................................................
env-303  AAUAUUACCCGAAAUAAAAAAAUUAAUUUUUUUUGAAAAAAGUAUUGCAAUUAUUGUAAACGUCAUGUAUAAUAUCACUUGCUGUGGC.......AUGAUAGCUAUGAAGUUUG.AAGUUACUGUCCAUGA.......................................................
env-304  AAUAUUACCCGAAAUAAAAAAAUAAAUUUUUUUUGAAAAAAGUAUUGCAAUUAUUGUAAACGUCAUGUAUAAUAUCACUUGCUGUGGC.......AUGAUAGCUAUGAAGUUUG.AAGUUACUGUCCAUGA.......................................................
env-305  AAUAUUAUCCGAAAUAAAAAAAUAAAUUUUUUUUGAAAAAAGUAUUGCAAUUAUUGUAAACGUCAUGUAUAAUAUCACUUGCUGUGGC.......AUGAUAGCUAUGAAGUUUG.AAGUUACUGUCCAUGA.......................................................
env-306  AAUAUUAUCCGAAAUAAAAAAAUAAAUUUUUUUUGAAAAAAGUAUUGCACUUAUUGUAAACGUCAUGUAUAAUAUCACUUGCUGUGGC.......AUGAUAGCUAUGAAGUUUG.AAGUUACUGUCCAUGA.......................................................
env-307  AAUAUUACCCGAAAUAAAAAAAUAAAUUUUUUUUGAAAAAAGUAUUGCAAUUAUUGUAAACGUCAUGUAUAAUAUCACUUGCUGUGGC.......AUGAUAGCUAUGAAGUUUG.AAGUUACUGUCCAUGA.......................................................
env-308  AAUAUUACCCGAAAUAAAAAAAUAAAUUUUUUUUGAAAAAAGUAUUGCAAUUAUUGGAAACGUCAUGUAUAAUAUCACUUGCUGUGGC.......AUGAUAGCUAUGAAGUUUG.AAGUUACUGUCCAUGA.......................................................
env-309  AAUAUUACCCUAAAUAAAAAAAUAAAUUUUUUUUGAAAAAAGUAUUGCAAUUAUUGGAAACGUCAUGUAUAAUAUCACUUGCUGUGGC.......AUGAUAGCUAUGAAGUUUG.AAGUUACUGUCCAUGA.......................................................
env-310  AAUAUUAUCCGAAAUAAAAAAAUAAAUUUUUUUUGAAAAAAGUAUUGCAAUUAUUGGGAACGUCAUGUAUAAUAUCACUUGCUGUGGC.......AUGAUAGCUAUGAAGUUUG.AAGUUACUGUCCAUGA.......................................................
env-311  AAUAUUACCCGAAAUAAAAAAAUAAAUUUUUUUUGAAAAAAGUAUUGCAAUUAUUGUAAACGUCAUGUAUAAUAUCACUUGCUGUGGC.......AUGAUAGCUAUGAAGUUUG.AAGUUACUGUCCAUGA.......................................................
env-312  GGAAUAUUACCCGAAAUAAAAAAAUAAAUUUUUUUUGAAAAAAGUAUUGCAAUUAUUGGAAACGUCAUGUAUAAUAUCACUUGCUGUGGC.......AUGAUAGCUAUGAAGUUUG.AAGUUACUGUCCAUGA.......................................................
env-313  AAUAUUACCCGAAAUAAAAAAAUAAAUUUUUUUUGAAAAAAGUAUUGCAAUUAUUGGAAACGUCAUGUAUAAUAUCACUUGCUGUGGC.......AUGAUAGCUAUGAAGUUUG.AAGUUACUGUCCAUGA.......................................................
env-314  AAUAUUACCCGAAAUAAAAAAAUAAAUUUUUUUUGAAAAAAGUAUUGCAAUUAUUGGAAACGUCAUGUAUAAUAUCACUUGCUGUGGC.......AUGAUAGCUAUGAAGUUUG.AAGUUACUGUCCAUGA.......................................................
env-315  AAUAUUACCCGAAAUAAAAAAAUAAAUUUUUUUUGAAAAAAGUAUUGCAAUUAUUGGAAACGUCAUGUAUAAUAUCACUUGCUGUGGC.......AUGAUAGCUAUGAAGUUUG.AAGUUACUGUCCAUGA.......................................................
env-316  AAUAUUACCCGAAAUAAAAAAAUAAAUUUUUUUUGAAAAAAGUAUUGCAAUUAUUGGGAACGUCAUGUAUAAUAUCACUUGCUGUGGC.......AUGAUAGCUAUGAAGUUUG.AAGUUACUGUCCAUGA.......................................................
env-317  AAUAUUACCCGAAAUAAAAAAAUAAAUUUUUUUUGAAAAAAGUAUUGCAAUUAUUGUAAACGUCAUGUAUAAUAUCACUUGCUGUGGC.......AUGAUAGCUAUGAAGUUUG.AAGUUACUGUCCAUGA.......................................................
env-331  AAUAUUAUCCGAAAUAAAAAAAUAAAUUUUUUUUGAAAAAAGUAUUGCAAUUAUUGUAAACGUCAUGUAUAAUAUCACUUGCUGUGGC.......AUGAUAGCUAUGAAGUUUG.AAGUUACUGUCCAUGA.......................................................
env-332  AAUAUUACCCGAAAUAAAAAAAUAAAUUUUUUUUGAAAAAAGUAUUGCAAUUAUUGUAAACGUCAUGUAUAAUAUCACUUGCUGUGGC.......AUGAUAGCUAUGAAGUUUG.AAGUUACUGUCCAUGA.......................................................
env-333  AAUAUUACCCGAAAUAAAAAAAUAAAUUUUUUUUGAAAAAAGUAUUGCAAUUAUUGGAAACGUCAUGUAUAAUAUCACUUGCUGUGGC.......AUGAUAGCUAUGAAGUUUG.AAGUUACUGUCCAUGA.......................................................
env-334  AAUAUUAUCCGAAAUAAAAAAAUAAAUUUUUUUUGAAAAAAGUAUUGCAAUUAUUGUAAACGUCAUGUAUAAUAUCACUUGCUGUGGC.......AUGAUAGCUAUGAAGUUUG.AAGUUACUGUCCAUGA.......................................................
env-335  AAUAUUACCCGAAAUAAAAAAAUAAAUUUUUUUUGAAAAAAGUAUUGCAAUUAUUGGAAACGUCAUGUAUAAUAUCACUUGCUGUGGC.......AUGAUAGCUAUGAAGUUUG.AAGUUACUGUCCAUGA.......................................................
env-336  AAUAUUAUCCGAAAUAAAAAAAUAAAUUUUUUUUGAAAAAAGUAUUGCAAUUAUUGGAAACGUCAUGUAUAAUAUCACUUGCUGUGGC.......AUGAUAGCUAUGAAGUUUG.AAGUUACUGUCCAUGA.......................................................
env-338  AAUAUUAUCCGAAAUAAAAAAAUAAAUUUUUUUUGAAAAAAGUAUUGCAAUUAUUGGAAACGUCAUGUAUAAUAUCACUUGCUGUGGC.......AUGAUAGCUAUGAAGUUUG.AAGUUACUGUCCAUGA.......................................................
env-339  AAUAUUACCCUAAAUAAAAAAAUAAAUUUUUUUUGAAAAAAGUAUUGCAAUUAUUGGAAACGUCAUGUAUAAUAUCACUUGCUGUGGC.......AUGAUAGCUAUGAAGUUUG.AAGUUACUGUCCAUGA.......................................................
env-340  AAUAUUAUCCGAAAUAAAAAAAUAAAUUUUUUUUGAAAAAAGUAUUGCAAUUAUUGUAAACGUCAUGUAUAAUAUCACUUGCUGUGGC.......AUGAUAGCUAUGAAGUUUG.AAGUUACUGUCCAUGA.......................................................
env-341  .........................................................AAACGUCAUGUAUAAUAUCACUUGCUGUGGC.......AUGAUAGCUAUGAAGUUUG.AAGUUACUGUCCAUGA.......................................................
env-347  AAUAUUACCCGAAAUAAAAAAAUAAAUUUUUUUUGAAAAAAGUAUUGCAAUUAUUGGAAAUGUCAUGUAUAAUAUCACUUGCUGUGGC.......AUGAUAGCUAUGAAGUUUG.AAGUUACUGUCCAUGA.......................................................
env-348  AAUAUUACCCGAAAUAAAAAAAAUAAAUUUUUUUGAAAAAAGUAUUGCAAUUAUUGGAAACGUCAUGUAUAAUAUCACUUGCUGUGGC.......AUGAUAGCUAUGAAGUUUG.AAGUUACUGUCCAUGA.......................................................
env-349  AAUAUUACCCGAAAUAAAAAAAUAAAUUUUUUUUGAAAAAAGUAUUGCAAUUAUUGGAAACGUCAUGUAUAAUAUCACUUGCUGUGGC.......AUGAUAGCUAUGAAGUUUG.AAGUUACUGUCCAUGA.......................................................
env-350  AAUAUUACCCGAAAUAAAAAAAUAAAUUUUUUUUGAAAAAAGUAUUGCAAUUAUUGGAAACGUCAUGUAUAAUAUCACUUGCUGUGGC.......AUGAUAGCUAUGAAGUUUG.AAGUUACUGUCCAUGA.......................................................
env-351  AAUAUUAUCUGAAAUAAAAAAAUAAAUUUUUUUUAAAAAAGGUAUUGCAAUUAUUGCAAACGUCAUGUAUAAUAUCACUUGCUGUGGC.......AUGAUAGCUAUGAAGUUUG.AAGUUACUGUCCAUGA.......................................................
env-352  AAUAUUAUCCGAAAUAAAAAAAUAAAUUUUUUUUGAAAAAAGUAUUGCAAUUAUUGGAAACGUCAUGUAUAAUAUCACUUGCUGUGGC.......AUGAUAGCUAUGAAGUUUG.AAGUUACUGUCCAUGA.......................................................
env-353  AAUAUUACCCGAAAUAAAAAAAUAAAUUUUUUUUAAAAAAAGUAUUGCAAUUAUUGAAAACGUCAUGUAUAAUAUCACUUGCUGUGGC.......AUGAUAGCUAUGAAGUUUG.AAGUUACUGUCCAUGA.......................................................
env-354  AAUAUUAUCCGAAAUAAAAAAAUAAAUUUUUUUUGAAAAAAGUAUUGCAAUUAUUGGAAACGUCAUGUAUAAUAUCACUUGCUGUGGC.......AUGAUAGCUAUGAAGUUUG.AAGUUACUGUCCAUGA.......................................................
env-355  AAUAUUAUCCGAAAUAAAAAAAUAAAUUUUUUUUGAAAAAAGUAUUGCAAUUAUUGGAAACGUCAUGUAUAAUAUCACUUGCUGUGGC.......AUGAUAGCUAUGAAGUUUG.AAGUUACUGUCCAUGA.......................................................
env-359  AAUAUUAUCCGAAAUAAAAAAAUAAAUUUUUUUUGAAAAAAGUAUUGCAAUUAUUGGGAACGUCAUGUAUAAUAUCACUUGCUGUGGC.......AUGAUAGCUAUGAAGUUUG.AAGUUACUGUCCAUGA.......................................................
env-360  ...............AAAAAAAUAAAUUUUUUUUGAAAAAAGUAUUGCAAUUAUUGUAAACGUCAUGUAUAAUAUCACUUGCUGUGGC.......AUGAUAGCUAUGAAGUUUG.AAGUUACUGUCCAUGA.......................................................
env-361  AAUAUUACCCGAAAUAAAAAAAUAAAUUUUUUUUGAAAAAAGUAUUGCAAUUAUUGGAAACGUCAUGUAUAAUAUCACUUGCUGUGGC.......AUGAUAGCUAUGAAGUUUG.AAGUUACUGUCCAUGA.......................................................
env-362  AAUAUUAUCCGAAAUAAAAAAAUAAAUUUUUUUUGAAAAAAGUAUUGCAAUUAUUGUAAACGUCAUGUAUAAUAUCACUUGCUGUGGC.......AUGAUAGCUAUGAAGUUUG.AAGUUACUGUCCAUGA.......................................................
env-363  AAUAUUACCCGAAAUAAAAAAAUAAAUUUUUUUUGAAAAAAGUAUUGCAAUUAUUGGAAACGUCAUGUAUAAUAUCACUUGCUGUGGC.......AUGAUAGCUAUGAAGUUUG.AAGUUACUGUCCAUGA.......................................................
env-364  AAUAUUAUCCGAAAUAAAAAAAUAAAUUUUUUUUGAAAAAAGUAUUGCAAUUAUUGGAAACGUCAUGUAUAAUAUCACUUGCUGUGGC.......AUGAUAGCUAUGAAGUUUG.AAGUUACUGUCCAUGA.......................................................
env-365  AAUAUUACCCGAAAUAAAAAAAUAAAUUUUUUUUGAAAAAAGUAUUGCAAUUAUUGUAAACGUCAUGUAUAAUAUCACUUGCUGUGGC.......AUGAUAGCUAUGAAGUUUG.AAGUUACUGUCCAUGA.......................................................
env-366  AAUAUUAUCCGAAAUAAAAAAAUAAAUUUUUUUUGAAAAAAGUAUUGCAAUUAUUGGAAACGUCAUGUAUAAUAUCACUUGCUGUGGC.......AUGAUAGCUAUGAAGUUUG.AAGUUACUGUCCGUGA.......................................................
env-368  AAUAUUAUCCGAAAUAAAAAAAUAAAUUUUUUUUGAAAAAAGUAUUGCAAUUAUUGGAAACGUCAUGUAUAAUAUCACUUGCUGUGGC.......AUGAUAGCUAUGAAGUUUG.AAGUUACUGUCCGUGA.......................................................
env-372  AAUAUUAUCCGAAAUAAAAAAAUAAAUUUUUUUUGAAAAAAGUAUUGCAAUUAUUGGAAACGUCAUGUAUAAUAUCACUUGCUGUGGC.......AUGAUAGCUAUGAAGUUUG.AAGUUACUGUCCGUGA.......................................................
env-373  AAUAUUAUCCGAAAUAAAAAAAUAAAUUUUUUUUGAAAAAAGUAUUGCAAUUAUUGGAAACGUCAUGUAUAAUAUCACUUGCUGUGGC.......AUGAUAGCUAUGAAGUUUG.AAGUUACUGUCCGUGA.......................................................
env-374  AAUAUUAUCCGAAAUAAAAAAAUAAAUUUUUUUUGAAAAAAGUAUUGCAAUUAUUGGAAACGUCAUGUAUAAUAUCAUUUGCUGUGGC.......AUGAUAGCUAUGAAGUUUG.AAGUUACUGUCCGUGA.......................................................
env-375  AAUAUUAUCCGAAAUAAAAAAAUAAAUUUUUUUUGAAAAAAGUAUUGCAAUUAUUGUAAACGUCAUGUAUAAUAUCACUUGCUGUGGC.......AUGAUAGCUAUGAAGUUUG.AAGUUACUGUCCGUGA.......................................................
env-376  AAACACUGAUUUUUGCGUAAAAGUCAUUAAUUUAUAAAAAGUAGUUGCAUAUACUUUUAAAAUGUUUUACAAUCAUAAACGCUGUGAC.......AUGAUAGCUAUGAAGCGUG.AGGUUGCUGCUUUCAUCCAGUUA..................................................
Bsp-5-1  AAACACUGAUUUUUGCGUAAAAGUCAUUAAUUUAUAAAAAGUGGUUGCAUAUACUUUUAAAAUGUUUUACAAUCAUAAACGCUGUGAC.......AUGAUAGCUAUGAAGCGUG.AGGUUGCUGCUUUCAUCCAGUUA..................................................
env-377  GUAUAAACAGAGCUUGAAAUAGUUUAAAAUUUUUUAAAAAAGUGCUUGCAUUCGUUUUUUUAAUAUGAUAUCCUAGUAUGGCUGUGAC......AUUGAUAGCUAUGAAGCA.U.AGGUUGCCGUUU.........................................................
env-379  UUUUAGAAGACUAUAGGCAGAAAAUAAAAUAGAAUAAAAAAGUGCUUGCAUUUGAUUUUCGUAUGUGAUACUAUGGAACAGCUGUGAC......AUUGAUAGCUAUGAAGCA.U.AGGUUGCCGUUU.........................................................
env-380  GUAUAAACAGAGCUUGAAAUAGUUUAAAAUUUUUUAAAAAAGUGCUUGCAUUCGUUUUUUUUAUAUGAUAUUCUAGUAUGGCUGUGAC......AUUGAUAGCUAUGAAGCA.U.AGGUUGCCGUUU.........................................................
env-381  GUAUAAACAGAGCUUGAAAUAGUUUAAAAUUUUUUAAAAAAGUGCUUGCAUUCGUUUUUUUAAUAUGAUAUCCUAGUAUGGCUGUGAC......AUUGAUAGCUAUGAAGCA.U.AGGUUGCCGUUU.........................................................
env-382  UUUUAGAAGACUAUAGGCAGAAAAUAAAAUAGAAUAAAAAAGUGCUUGCAUUUGAUUUUCGUAUGUGAUACUAUGGAACAGCUGUGAC......AUUGAUAGCUAUGAAGCA.U.AGGUUGCCGUUU.........................................................
env-384  AGCUUGAAAUAGCUUGAAAUAGUUUAAAAUUUUUUAAAAAAGUGCUUGCAUUCGUUUUUUUAAUAUGAUAUUCUAGUAUGGCUGUGAC......AUUGAUAGCUAUGAAGCA.U.AGGUUGCCGUUU.........................................................
Lor-1-1  GUGUAAACAGAGCUUGAAAUAGUUUAAAAUUUUUUAAAAAAGUGCUUGCAUUCGUUUUUUUAAUAUGAUAUCCUAGUAUGGCUGUGAC......AUUGAUAGCUAUGAAGCA.U.AGGUUGCCGUUU.........................................................
env-385  UUUUAGAAGACUACAGGCAGAAAAUAAAAUAGAAUAAAAAAGUGCUUGCAUUUGAUUUUCGUAUGUGAUACUAUGGAACAGCUGUGAC......AUUGAUAGCUAUGAAGCA.U.AGGUUGCCGUUU.........................................................
env-386  GUAUAAACAGAGCUUGAAAUAGUUUAAAAUUUUUUAAAAAAGUGCUUGCAUUCGUUUUUUUUAUAUGAUAUUCUAGUAUGGCUGUGAC......AUUGAUAGCUAUGAAGCA.U.AGGUUGCCGUUU.........................................................
env-387  UUUUAGAAGACUAUAGGCAGAAAAUAAAAUAGAAUAAAAAAGUGCUUGCAUUUGAUUUUCGUAUGUGAUACUAUGGAACAGCUGUGAC......AUUGAUAGCUAUGAAGCA.U.AGGUUGCCGUUU.........................................................
env-388  UACUAUAAAAGCUGUGAAAUAAAAAUAAAAAAAAAUAAAAAAGUGCUUGCAAUCGCUUUUCAUCUAUGCUAUCCUAGUAUGGCUGUGAC......AUUGAUAGCUGAGAAGCA.U.AGGUUGCCGUU..........................................................
Lba-15-1 UACUAUAAAAGCUGUGAAAUAAAAAUAAAAAAAAAUAAAAAAGUGCUUGCAAUCGCUUUUUAUCUAUGCUAUCCUAGUAUGGCUGUGAC......AUUGAUAGCUGAGAAGCA.U.AGGUUGCCGUU..........................................................
env-389  UACUAUAAAAGCUGUGAAAUAAAAAUAAAAAAAAAUAAAAAAGUGCUUGCAAUCGCUUUUCAUCUAUGCUAUCCUAGUAUGGCUGUGAC......AUUGAUAGCUGAGAAGCA.U.AGGUUGCCGUU..........................................................
env-390  UACUAUAAAAGCUGUGAAAUAAAAAUAAAAAAAAAUAAAAAAGUGCUUGCAAUCGCUUUUUAUCUAUGCUAUCCUAGUAUGGCUGUGAC......AUUGAUAGCUGAGAAGCA.U.AGGUUGCCGUU..........................................................
env-391  UAUAAAAGCUGUGAAAUAAAAAUAAAAAUAAUUUUAAAAAAAGUGCUUGCAAUCGGUUUUUGUUUGUGCUAUCCUAGUGUGGCUGUGAC......AUUGAUAGCUAAGAAGCA.U.AGGUUGCCGUU..........................................................
env-392  ................................UUUAAAAAAGUGCUUGCAAUCGGUUUUCAUCUGUGCUAUCCUAGUAUGGCUGUGAC......AUUGAUAGCUAAGAAGCA.U.AGGUUGCCGUU..........................................................
env-393  UAUAAAAGCUGUGAAAUAAAAAUAAAAAUAAUUUUAAAAAAAGUGCUUGCAAUCGGUUUUUGUUUGUGCUAUCCUAGUAUGGCUGUGAC......AUUGAUAGCUAAGAAGCA.U.AGGUUGCCGUU..........................................................
env-394  .....GAACAGUACAAACAAAAAAUAAAAUAAAAUAAAAAAAGUGCUUGCAAUCGGAUUCAAUCUGUGCUAUACUAGAACAGCUGUGAC......AUUGAUAGCUAAGAAGCA.U.AGGUUGCCGUU..........................................................
env-395  UACAAAAGCUGUGAAAUAAAAAUAAAAAUAAUUUUAAAAAAAGUGCUUGCAAUCAGUUUUUGUUUGUGCUAUCCUAGUAUGGCUGUGAC......AUUGAUAGCUAAGAAGCA.U.AGGUUGCCGUU..........................................................
env-396  UAUCAUAUACGGCUUAUAAAAAAAUAAAAUAAAAUAAAAAAGUGCUUGCAAUCGCUUUUCAUCUAUGCUAUCCUAGUAUGGCUGUGAC......AUUGAUAGCUAAGAAGCA.U.AGGUUGCCGUU..........................................................
env-397  ....................AAAAUAAAAUAAAAUAAAAAAAGUGCUUGCAAUCGGAUUCAAUCUGUGCUAUCCUAGAACAGCUGUGAC......AUUGAUAGCUAAGAAGCA.U.AGGUUGCCGUU..........................................................
env-398  UUUUGGAGCAGUACAAACAAAAAAUAAAAUAAAAUAAAAAAAGUGCUUGCAAUCGACUUUGAUAUGUGCUAUCCUAGAACAGCUGUGAC......AUUGAUAGCUAAGAAGCA.U.AGGUUGCCGUU..........................................................
env-399  UAUAGAAGCUGUGAAAUAAAAAUAAAAAUAAUUUUAAAAAAAGUGCUUGCAAUCGUUUUUUGUUUGUGCUAUCCUAGUAUGGCUGUGAC......AUUGAUAGCUAAGAAGCA.U.AGGUUGCCGUU..........................................................
env-400  ................................................................................GCUGUGAC......AUUGAUAGCUAAGAAGCA.U.AGGUUGCCGUU..........................................................
env-402  UUUUGGAGCAGUACAAACAAAAAACAAAAUAAAAUAAAAAAAGUGCUUGCAAUCGGAUUUAAUCUGUGCUAUCCUAGAACAGCUGUGAC......AUUGAUAGCUAAGAAGCA.U.AGGUUGCCGUU..........................................................
env-403  UUUUGGAGCAGUACAAACAAAAAAUAAAAUAAAAUAAAAAAAGUGCUUGCAAUCAGAUUCAAUCUGUGCUAUCCUAGAACAGCUGUGAC......AUUGAUAGCUAAGAAGCA.U.AGGUUGCCGUU..........................................................
env-404  AAUACUAUAAAAGCUGUGAAAUAAAAAUAAUUUUAAAAAAAGUGCUUGCAAUCGGUUUUUGUUUGUGCUAUCCUAGUAUGGCUGUGAC......AUUGAUAGCUAAGAAGCA.U.AGGUUGCCGUU..........................................................
env-405  UAUAAAAGCUGUGAAAUAAAAAUAAAAAUAAUUUUAAAAAAAGUGCUUGCAAUCGGUUUUUGUUUGUGCUAUCCUAGUAUGGCUGUGAC......AUUGAUAGCUAAGAAGCA.U.AGGUUGCCGUU..........................................................

         ................................................................................<<.<<<<<.......................<<<.<.<<<...<<<<<.........................<<<<<<<<<<<<<<.............>>>>
         ................................................................................21..1111.......................222.2.012...12222.........................1111111.10..00.............00..
         •••••••••••••••R••••••••••R••••••••RAAAAR•••YU••••••••••••••••••••••••••••••••••••GYUGUGRC--------•UGAYAGC••UGAAGCG•G-AGGUUGCYRY••••○○○---------------------------------------------------
                                                                                                 5′
```

```
env-406   UUUUGGAGCAGUACAAACAAAAAAUAAAAUAAAAUAAAAAAGUGCUUGCAAUCGACUUUGAUAUGUGCUAUCCUAGAACAGCUGUGAC.......AUUGAUAGCUAAGAAGCA.U.AGGUUGCCGUU.....................................................
env-407   .....................AAAUAAAAUAAAAUAAAAAAGUGCUUGCAAUCGCUUUUCAUCUAUGCUAUCCUAGUAUGGCUGUGAC.......AUUGAUAGCUAAGAAGCA.U.AGGUUGCCGUU.....................................................
env-408   UUUUGGAGCAGUACAAACAAAAAAUAAAAUAAAAUAAAAAAGUGCUUGCAAUCGCUUUUCAUCUAUGCUAUCCUAGUAUGGCUGUGAC.......AUUGAUAGCUAAGAAGCA.U.AGGUUGCCGUU.....................................................
env-409   UUUUGGAACAGUACAAACAAAAAAUAAAAUAAAAUAAAAAAGUGCUUGCAAUCGGAUUUAAUCUGUGCUAUCCUAGAACAGCUGUGAC.......AUUGAUAGCUAAGAAGCA.U.AGGUUGCCGUU.....................................................
env-410   ....................AAAAUAAAAUAAAAUAAAAAAGUGCUUGCAAUCGGAUUUGAUAUGUGCUAUCCUAGAACAGCUGUGAC.......AUUGAUAGCUAAGAAGCA.U.AGGUUGCCGUU.....................................................
Cba-3-1   AAAUACUAUAAAAGCUGUGAAAUAAAAAUAAUUUUAAAAAAGUGCUUGCAAUCGGUUUUUGUUUGUGCUAUCCUAGUAUGGCUGUGAC.......AUUGAUAGCUAAGAAGCA.U.AGGUUGCCGUU.....................................................
env-411   UUUUGGAGCAGUACAAACAAAAAAUAAAAUAAAAUAAAAAAGUGCUUGCAAUCGGAUUCAAUCUGUGCUAUCCUAGAACAGCUGUGAC.......AUUGAUAGCUAAGAAGCA.U.AGGUUGCCGUU.....................................................
env-414   CAGGUGGUCGAUCAAGGAAAAAAACAAAAAAAAUUUAAAACCCCCCUUGCACUCGGUUUAAAAUUAUGAUAAUAUAGUCGGGCUGUGAC.......AUUGAUAGCUAUGAAACA.C.AGGUUGCUAUAC......................CCC...........AUAAAAA.........
env-415   CAGGUGGUCGAUCAAGGAAAAAACCAAAAAAAAUUUUAAAUCCCCCUUGCACUCGGUUUAAAAUUAUGAUAAUAUAGUCGGGCUGUGAC.......AUUGAUAGCUAUGAAACA.C.AGGUUGCUAUAC......................CCC...........AUAAAAA.........
env-416   CAGGUGGUCGAUCAAGGAAAAAAACAAAAAAAAUUUAAAACCCCCCUUGCACUCGGUUUAAAAUUAUGAUAAUAUAGUCGGGCUGUGAC.......AUUGAUAGCUAUGAAACA.C.AGGUUGCUAUAC......................CCC...........AUAAAAA.........
env-417   CAGGUGGUCGAUCAAGGAAAAAACCAAAAAAAAUUUUAAAUCCCCCUUGCACUCGGUUUAAAAUUAUGAUAAUAUAGUCGGGCUGUGAC.......AUUGAUAGCUAUGAAACA.C.AGGUUGCUAUAC......................CCC...........AUAAAAA.........
env-420   CAGGUGGUCGAUCUAGGAAAAAACUAAAAAAAAUUUAAAACCCCCCUUGCACUCGGUUUAAAAUUAUGAUAAUAUAGUCGGGCUGUGAC.......AUUGAUAGCUAUGAAACA.C.AGGUUGCUAUAC......................CCC...........AUAAAAA.........
env-426   AGGUGGUCGAUCAAGGAAAAAAACAAAAAAAAAUUUAAAACCCCCCUUGCACUCGGUUUAAAAUUAUGAUAAUAUAGUCGGGCUGUGAC.......AUUGAUAGCUAUGAAACA.C.AGGUUGCUAUAC......................CCC...........AUAAAAA.........
env-427   CAGGUGGUCGAUCAAGGAAAAAACUAAAAAAAAUUUAAAACACCCCUUGCACUCGGUUUAAAAUUAUGAUAAUAUAGUCGGGCUGUGAC.......AUUGAUAGCUAUGAAACA.C.AGGUUGCUAUAC......................CCC...........AUAAAAA.........
env-428   UCGAAAAAUGUACAAAAAAAGAGAAAAGUUUUUAAAAAAAAGCCUUGCAAUUUUGCAGGGCUUACUUUAUACUACUAAGUGCUGUGAC........AUGAUAGCGAAGAAGCAUA.AGGUUGCUGCAAC.....................GGA...........AAAUACAUAA......
env-429   UCGAAAAAUGUACAAAAAAAGAGAAAAGUUUUUGAAAAAAAGCCUUGCAAUUUUGUAGGGCUUACUUUAUACUACUAAGUGCUGUGAC........AUGAUAGCGAAGAAGCAUA.AGGUUGCUGCAAC.....................GGA...........AAAUACAUAA......
env-430   UCGAAAAAUGUACAAUAAAAGAGAAAAGUUUUUGAAAAAAAGCCUUGCAAUUUUGCAGGGCUUACUUUAUACUACUAAGUGCUGUGAC........AUGAUAGCGAAGAAGCAUA.AGGUUGCUGCAAC.....................GGA...........AAAUACAUAA......
env-431   UCGAAAAAUGUACAAAAAAAGAGAAAAGUUUUUGAAAAAAAGCCUUGCAAUUUUGCAGGGCUUACUUUAUACUACUAAGUGCUGUGAC........AUGAUAGCGAAGAAGCAUA.AGGUUGCUGCAAC.....................GGA...........AAAUACAUAA......
env-432   UCGAAAAAUGUACAAAAAAAGAGAAAAGUUUUUGAAAAAAAGCCUUGCAAUUUUGCAGGGCUUACUUUAUACUACUAAGUGCUGUGAC........AUGAUAGCGAAGAAGCAUA.AGGUUGCUGCAAC.....................GGA...........AAAUACAUAA......
env-433   UCGAAAAAUGUACAAAAAAAGAGAAAAGUUUUUGAAAAAAAGCCUUGCAAUUUUGCAGGGCUUACUUUAUACUACUAAGUGCUGUGAC........AUGAUAGCGAAGAAGCAUA.AGGUUGCUGCAAC.....................GGA...........AAAUACAUAA......
env-434   UCGAAAAAUGUACAAAAAAAGAGAAAAGUUUUUGAAAAAAAGCCUUGCAAUUUUGCAGGGCUUACUUUAUACUACUAAGUGCUGUGAC........AUGAUAGCGAAGAAGCAUA.AGGUUGCUGCAAC.....................GGA...........AAAUACAUAA......
env-435   UCGAAAAAUGUACAAUAAAAGAGAAAAGUUUUUGAAAAAAAGCCUUGCAAUUUUGCAGGGCUUACUUUAUACUACUAAGUGCUGUGAC........AUGAUAGCGAAGAAGCAUA.AGGUUGCUGCAAC.....................GGA...........AAAUACAUAA......
env-436   UCGAAAAAUGUACAAAAAAAGAGAAAAGUUUUUGAAAAAAAGCCUUGCAAUUUUGCAGGGCUUACUUUAUACUACUAAGUGCUGUGAC........AUGAUAGCGAAGAAGCAUA.AGGUUGCUGCAAC.....................GGA...........AAAUACAUAA......
env-454   UCGAAAAAUGUACAAAAAAAGAGAAAAGUUUUUGAAAAAAAGCCUUGCAAUUUUGCAGGGCUUACUUUAUACUACUAAGUGCUGUGAC........AUGAUAGCAAAGAAGCAUA.AGGUUGCUGCAAC.....................GGA...........AAAUACAUAA......
env-455   UCGAAAAAUGUACAAAAAAAGAGAAAAGUUUUUGAAAAAAAGCCUUGCAAUUUUGCAGGGCUUACUUUAUACUACUAAGUGCUGUGAC........AUGAUAGCAAAGAAGCAUA.AGGUUGCUGCAAC.....................GGA...........AAAUACAUAA......
env-456   UCGAAAAAUGUACAAAAAAAGAGAAAAGUUUUUGAAAAAAAGCCUUGCAAUUUUGCAGGGCUUACUUUAUACUACUAAGUGCUGUGAC........AUGAUAGCAAAGAAGCAUA.AGGUUGCUGCAAC.....................GGA...........AAAUACAUAA......
env-457   UCGAAAAAUGUACAAAAAAAGAGAAAAGUUUUUGAAAAAAAGCCUUGCAAUUUUGCAGGGCUUACUUUAUACUACUAAGUGCUGUGAC........AUGAUAGCGAAGAAGCAUA.AGGUUGCUGCAAC.....................GGA...........AAAUACAUAA......
env-459   NNNNNNNAUGUACAAAAAAAGAGAAAAGUUUUUGAAAAAAAGCCUUGCAAUUUUGCAGGGCUUACUUUAUACUACUAAGUGCUGUGAC........AUGAUAGCGAAGAAGCAUA.AGGUUGCUGCAAC.....................GGA...........AAAUACAUAA......
env-460   UCGAAAAAUGUACAAAAAAAGAGAAAAGUUUUUGAAAAAAAGCCUUGCAAUUUUGCAGGGCUUACUUUAUACUACUAAGUGCUGUGAC........AUGAUAGCGAAGAAGCAUA.AGGUUGCUGCAAC.....................GGA...........AAAUACAUAA......
env-464   UCGAAAAAUGUACAAAAAAAGAGAAAAGUUUUUGAAAAAAAGCCUUGCAAUUUUGCAGGGCUUACUUUAUACUACUAAGUGCUGUGAC........AUGAUAGCGAAGAAGCAUA.AGGUUGCUGCAAC.....................GGA...........AAAUACAUAA......
env-465   UCGAAAAAUGUACAAAAAAAGAGAAAAGUUUUUGAAAAAAAGCCUUGCAAUUUUGCAGGGCUUACUUUAUACUACUAAGUGCUGUGAC........AUGAUAGCGAAGAAGCAUA.AGGUUGCUGCAAC.....................GGA...........AAAUACAUAA......
env-466   UCGAAAAAUGUACAAAAAAAGAGAAAAGUUUUUGAAAAAAAGCCUUGCAAUUUUGCAGGGCUUACUUUAUACUACUAAGUGCUGUGAC........AUGAUAGCGAAGAAGCAUA.AGGUUGCUGCAAC.....................GGA...........AAAUACAUAA......
env-467   ........................AAAGUUUUUGAAAAAAAGCCUUGCAAUUUUGCAGGGCUUACUUUAUACUACUAAGUGCUGUGAC........AUGAUAGCGAAGAAGCAUA.AGGUUGCUGCAAC.....................GGA...........AAAUACAUAA......
env-468   UCGAAAAAUGUACAAAAAAAGAGAAAAGUUUUUGAAAAAAAGCCUUGCAAUUUUGCAGGGCUUACUUUAUACUACUAAGUGCUGUGAC........AUGAUAGCGAAGAAGCAUA.AGGUUGCUGCAAC.....................GGA...........AAAUACAUAA......
env-469   UCGAAAAAUGUACAAAAAAAGAGAAAAGUUUUUGAAAAAAAGCCUUGCAAUUUUGCAGGGCUUACUUUAUACUACUAAGUGCUGUGAC........AUGAUAGCGAAGAAGCAUA.AGGUUGCUGCAAC.....................GGA...........AAAUACAUAA......
env-470   UCGAAAAAUGUACAAAAAAAGAGAAAAGUUUUUGAAAAAAAGCCUUGCAAUUUUGCAGGGCUUACUUUAUACUACUAAGUGCUGUGAC........AUGAUAGCGAAGAAGCAUA.AGGUUGCUGCAAC.....................GGA...........AAAUACAUAA......
env-471   UCGAAAAAUGUACAAAAAAAGAGAAAAGUUUUUGAAAAAAAGCCUUGCAAUUUUGCAGGGCUUACUUUAUACUACUAAGUGCUGUGAC........AUGAUAGCGAAGAAGCAUA.AGGUUGCUGCAAC.....................GGA...........AAAUACAUAA......
env-472   UCGAAAAAUGUACAAAAAAAGAGAAAAGUUUUUGAAAAAAAGCCUUGCAAUUUUGCAGGGCUUACUUUAUACUACUAAGUGCUGUGAC........AUGAUAGCGAAGAAGCAUA.AGGUUGCUGCAAC.....................GGA...........AAAUACAUAA......
env-473   UCGAAAAAUGUACAAAAAAAGAGAAAAGUUUUUGAAAAAAAGCCUUGCAAUUUUGCAGGGCUUACUUUAUACUACUAAGUGCUGUGAC........AUGAUAGCGAAGAAGCAUA.AGGUUGCUGCAAC.....................GGA...........AAAUACAUAA......
env-482   .............AAAAAAAGAGAAAAGUUUUUGAAAAAAAGCCUUGCAAUUUUGCAGGGCUUACUUUAUACUACUAAGUGCUGUGAC........AUGAUAGCGAAGAAGCAUA.AGGUUGCUGCAAC.....................GGA...........AAAUACAUAA......
env-483   ........UGUACAAAAAAAGAGAAAAGUUUUUGAAAAAAAGCCUUGCAAUUUUGCAGGGCUUACUUUAUACUACUAAGUGCUGUGAC........AUGAUAGCGAAGAAGCAUA.AGGUUGCUGCAAC.....................GGA...........AAAUACAUAA......
env-484   UCGAAAAAUGUACAAAAAAAGAGAAAAGUUUUUGAAAAAAAGCCUUGCAAUUUUGCAGGGCUUACUUUAUACUACUAAGUGCUGUGAC........AUGAUAGCGAAGAAGCAUA.AGGUUGCUGCAAC.....................GGA...........AAAUACAUAA......
env-485   UCGAAAAAUGUACAAUAAAAGAGAAAAGUUUUUGAAAAAAAGCCUUGCAAUUUUGCAGGGCUUACUUUAUACUACUAAGUGCUGUGAC........AUGAUAGCGAAGAAGCAUA.AGGUUGCUGCAAC.....................GGA...........AAAUACAUAA......
env-486   UCGAAAAAUGUACAAAAAAAGAGAAAAGUUUUUGAAAAAAAGCCUUGCAAUUUUGCAGGGCUUACUUUAUACUACUAAGUGCUGUGAC........AUGAUAGCGAAGAAGCAUA.AGGUUGCUGCAAC.....................GGA...........AAAUACAUAA......
env-487   UCGAAAAAUGUACAAAAAAAGAGAAAAGUUUUUGAAAAAAAGCCUUGCAAUUUUGCAGGGCUUACUUUAUACUAAUAAGUGCUGUGAC........AUGAUAGCGAAGAAGCAUA.AGGUUGCUGCAAC.....................GGA...........AAAUACAUAA......
env-488   UCGAAAAAUGUACAAAAAAAGAGGAAAGUUUUUGAAAAAAAGCCUUGCAAUUUUGCAGGGCUUACUUUAUACUACUAAGUGCUGUGAC........AUGAUAGCGAAGAAGCAUA.AGGUUGCUGCAAC.....................GGA...........AAAUACAUAA......
env-489   AAUGACAACAGGGUGAUUGUUCUUAAAAAAGAACAAUUAUCCUGUUGAUUUUAUUAGUAUAGUCGUGUAUACUAGCAAUUGCUGUGAC.......AUUGAUAGCUGUGAAGCGUG.AGGUUGCUGCGAUACAAU...............................................
env-494   AAUGACAACAGGGUGAUUGUUCUUAAAAAAGAACAAUUAUCCUGUUGAUUUUAUUAGUAUAGUCGUGUAUACUAGCAAUUGCUGUGAC.......AUUGAUAGCUGUGAAGCGUG.AGGUUGCUGCGAUACAAU...............................................
env-495   AAUGACAACAGGGUGAUUGUUCUUAAAAAAGAACAAUUAUCCUGUUGAUUUUAUUAGUAUAGUCGUGUAUACUAGCAAUUGCUGUGAC.......AUUGAUAGCUGUGAAGCGUG.AGGUUGCUGCGAUACAAU...............................................
env-496   AAUGACAACAGGGUGAUUGUUCUUAAAAAAGAACAAUUAUCCUGUUGAUUUUAUUAGUAUAGUCGUGUAUACUAGCAAUUGCUGUGAC.......AUUGAUAGCUGUGAAGCGUG.AGGUUGCUGCGAUACAAU...............................................
env-497   AAUAACAACAGGGUGAUUGUUCUUAAAAAAGAACAAUUAUCCUGUUGAUUUUAUUAGUAUAGUCGUGUAUACUAGCAAUUGCUGUGAC.......AUUGAUAGCUGUGAAGCGUG.AGGUUGCUGCGAUACAAU...............................................
env-498   AAUGACAACAGGGUGAUUGUUCUUAAAAAAGAACAAUUAUCCUGUUGAUUUUAUUAGUAUAGUCGUGUAUACUAGCAAUUGCUGUGAC.......AUUGAUAGCUGUGAAGCGUG.AGGUUGCUGCGAUACAAU...............................................
env-499   AAUAACAACAGGGUGAUUGUUCUUAAAAAAGAACAAUUAUCCUGUUGAUUUUAUUAGUAUAGUCGUGUAUACUAGCAAUUGCUGUGAC.......AUUGAUAGCUGUGAAGCGUG.AGGUUGCUGCGAUACAAU...............................................
env-500   AAUGACAACAGGGUGAUUGUUCUUAAAAAAGAACAAUUAUCCUGUUGAUUUUAUUAGUAUAGUCGUGUAUACUAGCAAUUGCUGUGAC.......AUUGAUAGCUGUGAAGCGUG.AGGUUGCUGCGAUACAAU...............................................
env-501   AAUGACAACAGGGUGAUUGUUCUUAAAAAAGAACAAUUAUCCUGUUGAUUUUAUUAGUAUAGUCGUGUAUACUAGCAAUUGCUGUGAC.......AUUGAUAGCUGUGAAGCGUG.AGGUUGCUGCGAUACAAU...............................................
env-502   AAUGACAACAGGGUGAUUGUUCUUAAAAAAGAACAAUUAUCCUGUUGAUUUUAUUAGUAUAGUCGUGUAUACUAGCAAUUGCUGUGAC.......AUUGAUAGCUGUGAAGCGUG.AGGUUGCUGCGAUACAAU...............................................
env-503   AAUGACAACAGGGUGAUUGUUCUUAAAAAAGAACAAUUAUCCUGUUGAUUUUAUUAGUAUAGUCGUGUAUACUAGCAAUUGCUGUGAC.......AUUGAUAGCUGUGAAGCGUG.AGGUUGCUGCGAUACAAU...............................................
env-512   AAUACCAGCGUAUCGAAAAAAAUACAAAAAAAUAAAAAAAAACUCUUGCAUUUACAGGUGGUAUUGAGUAUAAUAACCCUUGCUGUGAC.......AUUGAUAGCGUAGAAGCG.A.AGGUUGCUAUGCAUGA.................................................
env-513   AAUACCAGCGUAUCGAAAAAAAUACAAAAAAAUAAAAAAAAACUCUUGCAUUUACAGGUGGUAUUGAGUAUAAUAACCCUUGCUGUGAC.......AUUGAUAGCGUAGAAGCG.A.AGGUUGCUAUGCAUGA.................................................
env-514   AAUACCAGCGUAUCGAAAAAAAUACAAAAAAAUAAAAAAAAACUCUUGCAUUUACAGGUGGUAUUGAGUAUAAUAACCCUUGCUGUGAC.......AUUGAUAGCGUAGAAGCG.A.AGGUUGCUAUGCAUGA.................................................
env-515   AAUACCAGCGUAUCGAAAAAAAUACAAAAAAAUAAAAAAAAACUCUUGCAUUUACAGGUGGUAUUGAGUAUAAUAACCCUUGCUGUGAC.......AUUGAUAGCGUAGAAGCG.A.AGGUUGCUAUGCAUGA.................................................
env-516   NNNNNNNNNNNNNNNNNNNNNNNNNNNNNNNNNNNNNNNNNNNNNNNNNNUUUACAGGUGGUAUUGAGUAUAAUAACCCUUGCUGUGAC.......AUUGAUAGCGUAGAAGCG.A.AGGUUGCUAUGCAUGA.................................................
env-517   AAUACCAGCGUAUCGAAAAAAAUACAAAAAAAUAAAAAAAAACUCUUGCAUUUACAGGUGGUAUUGAGUAUAAUAACCCUUGCUGUGAC.......AUUGAUAGCGUAGAAGCG.A.AGGUUGCUAUGCAUGA.................................................
env-518   AAUACCAGCGUAUCGAAAAAAAUACAAAAAAAUAAAAAAAAACUCUUGCAUUUACAGGUGGUAUUGAGUAUAAUAACCCUUGCUGUGAC.......AUUGAUAGCGUAGAAGCG.A.AGGUUGCUAUGCAUGA.................................................
env-556   ....CGAUUGUAUACAAAAAAAUACAAUUUUUCUGAAAAAAGUAUUGCAAAAGCGAAACCUGUCGUGUAUACUAAUCCUUGCUGUGAC.......AUUGAUAGCGUAGAAGCG.A.AGGUUGCUAUGCAUGA.................................................
env-557   AAUACCAGCGUAUCGAAAAAAAUACAAAAAAAUAAAAAAAAACUCUUGCAUUUACAGGUGGUAUUGAGUAUAAUAACCCUUGCUGUGAC.......AUUGAUAGCGUAGAAGCG.A.AGGUUGCUAUGCAUGA.................................................
env-558   AAUACCAGCGUAUCGAAAAAAAUACAAAAAAAUAAAAAAAAACUCUUGCAUUUACAGGUGGUAUUGAGUAUAAUAACCCUUGCUGUGAC.......AUUGAUAGCGUAGAAGCG.A.AGGUUGCUAUGCAUGA.................................................
env-559   AAUACCAGCGUAUCGAAAAAAAUACAAAAAAAUAAAAAAAAAUACUUGCAUUUACAGAUGGUAUUGAGUAUAAUAACCCUUGCUGUGAC.......AUUGAUAGCGUAGAAGCG.A.AGGUUGCUAUGCAUGA.................................................
env-560   AAAGAAACCCGCAAAAAAAUAUGAAAGAAAAAUGGAAAAAAUACUUGCAUUUCACAAAAUUAUCGUGUAUACUAAGCAAUGCUGUGAC.......AUUGAUAGCGUAGAAGCG.A.AGGUUGCUAUGCAUGA.................................................
env-561   AAUACCAGCGUAUCGAAAAAAAUACAAAAAAAUAAAAAAAAACUCUUGCAUUUACAGGUGGUAUUGAGUAUAAUAACCCUUGCUGUGAC.......AUUGAUAGCGUAGAAGCG.A.AGGUUGCUAUGCAUGA.................................................
env-562   CUAAUCGUUUAACACAAGAUAAAGUAAGAAAAUAGUCAAAAACACUUGCAAACACAGUGUUUAUCGAGUAUACUAAUAGAGCUGUGAC.........AAAUAGCAAAGAAGCGCG.AGGUUGCUGUAU......................ACCC..........GAAUACAUAA......
env-563   UUAGGGAGUGGGUUUUGAACAAUUUGAAAAAUAUUAGAAAAACACUUGCAAACCCCACGUUUAUCAAGUAUACUAAUAGAGCUGUGAC.........AAAUAGCAAAGAAGCGCG.AGGUUGCUGUAU......................AUCC..........GAAUACAUAA......
env-564   CCUUUUUUGAACGCAAAAACAAUUUGAAAAAUAUUAGAAAAACACUUGCAAACCCCACGUUUAUCAAGUAUACUAAUAGAGCUGUGAC.........AAAUAGCAAAGAAGCGCG.AGGUUGCUGUAU......................AUCC..........GAAUACAUAA......
env-565   AUAAAUGAAAAAAUUAACAUAAAGUAAGAAAAUAGUCAAAAACACUUGCAAAUACAGUGUUUAUCAAGUAUACUAAUAGAGCUGUGAC.........AAAUAGCAAAGAAGUGUG.AGGUUGCUGUAU......................ACCC..........GAAUACAUAA......
Bsp-4-1   AAAGGCUGAUUUUGUAUUAAAAGUGAUUAAUUUUCAAAAACUUGUUGCAUAUGCACAAAAAAUGUUUUAUACUAACAAACGCUGUGAC........AUGAUAGCUAUGAAGCGUG.AGGUUGCUGCUUAUCCAGUUG............................................
Bsp-7-1   AAAGGCUGAUUUUAUAUUAAAAGUGAUUAAUUUUCAAAAACUUGUUGCAUAUGCACAAAAAAUGUUUUAUACUAACAAACGCUGUGAC........AUGAUAGCUAUGAAGCGUG.AGGUUGCUGCUUAUCCAGUUG............................................
Bsp-10-1  AAAGGCUGAUUUUGUAUUAAAAGUGAUUAAUUUUCAAAAACUUGUUGCAUAUGCACAAAAAAUGUUUUAUACUAACAAACGCUGUGAC........AUGAUAGCUAUGAAGCGUG.AGGUUGCUGCUUAUCCAGUUG............................................
env-566   UUCCAGGGUGCUAUUAAGAAGAAUGGUCAGCACAGNNNNNNNNNNUGCAUAUGCACAAAAAAUGUUUUAUACUAACAAACGCUGUGAC........AUGAUAGCUAUGAAGCGUG.AGGUUGCUGCUUAUCCAGUUG............................................
Bsp-2-1   AAAGGCUGAUUUCACGAUAAAAGUGAUUAAUUUUAAAAAAGUGGUUGCAUUAUAAAAUAAAAUGCUUUAUAGUAACAUAUGCUGUGAC........AUGAUAGCUAUGAAGCGUG.AGGUUGCUGCUUUAUCCAGUUA...........................................
Bsp-3-1   AAACACCUAAUAUGCGAAAAAAGAGAUUAAUUUUAAAAAAGGUGUUGCAUUAAAUACCAAAAUGUUUUAUACUCAUAAACGCUGUGAC........AUGAUAGCUAUGAAGCGUG.AGGUUGCUGCUUUAUCCAGUUA...........................................
Bsp-6-1   AAACACUGAUUUUGCGAAAAAACUGAUUAAUUUACAAAAAGGUGUUGCAUAUCAGAUAAAAAUGAUUUAUACUCAACCAUGCUGUGAC........AUGAUAGCUAUGAAGCGUG.AGGUUGCUGCUUUAUCCAGUUA...........................................
env-567   AAAUAAAUUGUUUUAAAAAAAAAUACAAUUUUUUGAAAAAAAGUCUUGCAAUGCCCAUAAAUUCAAUGUAUACUAAUCCUUGCUGUGGC........AUGAUAGCUGUGAAGCGUG.AGGUUGUCGGUUC.....................CCA...........GUAA............
env-568   AAAAGAAUUGUAUCUAAAAAAAAUACAAUUUUUUUAAAAAAAGUCUUGCAAUGCCCGUAAAUUCAAUGUAUACUAAUCCUUGCUGUGGC........AUGAUAGCUGUGAAGCGUG.AGGUUGCCGGUUC.....................CCA...........GUAA............

          ................................................................................<<.<<<<<......................<<<.<.<<<...<<<<<.......................<<<<<<<<<<<<<<............>>>>
          ................................................................................21..1111......................222.2.012...12222.......................1111111.10..00............00..
          ●●●●●●●●●●●●●●R●●●●●●●●●●R●●●●●●●●RAAAAR●●●YU●●●●●●●●●●●●●●●●●●●●●●●●●●●●●●●●●●●GYUGUGRC-------∘UGAYAGC••UGAAGCG∘G-AGGUUGCYRY••∘∘∘∘---------------------------------------------------
                                                                                                 5′
```

env-570 AAAGAAAUUGUAUCAAAAAAAAUACAAUUUUUUUAAAAAAAGUCUUGCAAUGCCCAUAAAUUCAAUGUAUACUAAUCCUUGGUGUGGC.......AUGAUAGCUGUGAAGCGUG.AGGUUGCCGGUUC.....................CCA..........GUAA............
env-571 AAAGAAAUUGUAUCAAAAAAAAUACAAUUUUUUUAAAAAAAGUCUUGCAAUGCCCGUAAAUUCAAUGUAUACUAAUCCUUGGUGUGGC.......AUGAUAGCUGUGAAGCGUG.AGGUUGCCGGUUC.....................CCA..........GUAA............
env-573 AAAGAAAUUGUAUCAAAAAAAAUACAAUUUUUUUAAAAAAAGUCUUGCAAUGCCCAUAAAUUCAAUGUAUACUAAUCCUUGGUGUGGC.......AUGAUAGCUGUGAAGCGUG.AGGUUGCCGGUUC.....................CCA..........GUAA............
env-578 AAAGAAAUUGUAUCAAAAAAAAUACAUUUUUUUUAAAAAAAGUCUUGCAAUGCCCGUAAAUUCAAUGUAUACUAAUCCUUGGUGUGGC.......AUGAUAGCUGUGAAGCGUG.AGGUUGCCGGUUC.....................CCA..........GUAA............
env-579 AAAGAAAUUGUAUCAAAAAAAAUACAAUUUUUUUAAAAAAAGUCUUGCAAUGCCCAUAAAUUCAAUGUAUACUAAUCCUUGGUGUGGC.......AUGAUAGCUGUGAAGCGUG.AGGUUGCCGGUUC.....................CCA..........GUAA............
env-581 AAUAAGAUUGUAUCAAAAAAAAUACAAUAUUUUGAAAAAAAGUCUUGCAAUGCCCAUAAAUUCAAUGUAUACUAAUCCUUGGUGUGGC.......AUGAUAGCUGUGAAGCGUG.AGGUUGCCGGUUC.....................CCA..........GUAA............
env-582 AAAGAAUUGUAUCUAAAAAAAAUACAAUUUUUUGAAAAAAAGUCUUGCAAUGCCCGUAAAUUCAAUGUAUACUAAUCCUUGGUGUGGC.......AUGAUAGCUGUGAAGCGUG.AGGUUGCCGGUUC.....................CCA..........GUAA............
env-583 AAAAGAAUUGUAUCUAAAAAAAUGCAAUUUUUUGAAAAAAAGUCUUGCAAUGCCCGUAAAUUCAAUGUAUACUAAUCCUUGGUGUGGC.......AUGAUAGCUGUGAAGCGUG.AGGUUGCCGGUUC.....................CCA..........GUAA............
env-584 AAAUAAAUUGUUUUAAAAAAAAUACAAUUUUUUUAAAAAAAGUCUUGCAAUGCCCAUAAAUUCAAUGUAUACUAAUCCUUGGUGUGGC.......AUGAUAGCUGUGAAGCGUG.AGGUUGCCGGUUC.....................CCA..........GUAA............
env-585 AAAGAAAUUGUAUCAAAAAAAAUACAAUUUUUUUAAAAAAAGUCUUGCAAUGCCCGUAAAUUCAAUGUAUACUAAUCCUUGGUGUGGC.......AUGAUAGCUGUGAAGCGUG.AGGUUGCCGGUUC.....................CCA..........GUAA............
env-586 ........................CAAUUUUUUUAAAAAAAGUCUUGCAAUGCCCAUAAAUUCAAUGUAUACUAAUCCUUGGUGUGGC.......AUGAUAGCUGUGAAGCGUG.AGGUUGCCGGUUC.....................CCA..........GUAA............
env-587 AAAAGAAUUGUAUCUAAAAAAAUACAAUUUUUUGAAAAAAAGUCUUGCAAUGCCCAUAAAUUCAAUGUAUACUAAUCCUUGGUGUGGC.......AUGAUAGCUGUGAAGCGUG.AGGUUGCCGGUUC.....................CCA..........GUAA............
env-588 AAAAGAAUUGUAUCUAAAAAAAUACAAUUUUUUUAAAAAAAGUCUUGCAAUGCCCGUAAAUUCAAUGUAUACUAAUCCUUGGUGUGGC.......AUGAUAGCUGUGAAGCGUG.AGGUUGCCGGUUC.....................CCA..........GUAA............
env-589 AAAAGAAUUGUAUCUAAAAAAAUACAAUUUUUUGAAAAAAAGUCUUGCAAUGCCCGUAAAUUCAAUGUAUACUAAUCCUUGGUGUGGC.......AUGAUAGCUGUGAAGCGUG.AGGUUGCCGGUUC.....................CCA..........GUAA............
env-590 AAAUAAAUUGUUUUAAAAAAAAUACAAUUUUUUGAAAAAAAGUCUUGCAAUGCCCAUAAAUUCAAUGUAUACUAAUCCUUGGUGUGGC.......AUGAUAGCUGUGAAGCGUG.AGGUUGCCGGUUC.....................CCA..........GUAA............
env-591 AAAGAAAUUGUAUCAAAAAAAAUACAUUUUUUUUAAAAAAAGUCUUGCAAUGCCCAUAAAUUCAAUGUAUACUAAUCCUUGGUGUGGC.......AUGAUAGCUGUGAAGCGUG.AGGUUGCCGGUUC.....................CCA..........GUAA............
env-592 AAAAGAAUUGUAUCUAAAAAAAUACAAUUUUUUGAAAAAAAGUCUUGCAAUGCCCGUAAAUUCAAUGUAUACUAAUCCUUGGUGUGGC.......AUGAUAGCUGUGAAGCGUG.AGGUUGCCGGUUC.....................CCA..........GUAA............
env-593 AAAGAAAUUGUAUCAAAAAAAAUACAUUUUUUUUAAAAAAAGUCUUGCAAUGCCCAUAAAUUCAAUGUAUACUAAUCUUUGGUGUGGC.......AUGAUAGCUGUGAAGCGUG.AGGUUGCCGGUUC.....................CCA..........GUAA............
env-594 ......................................................CAUAAAUUCAAUGUAUACUAAUCCUUGGUGUGGC.......AUGAUAGCUGUGAAGCGUG.AGGUUGCCGGUUC.....................CCA..........GUAA............
env-595 AAUAAGAUUGUAUCAAAAAAAAUACAAUAUUUUGAAAAAAAGUCUUGCAAUGCCCAUAAAUUCAAUGUAUACUAAUCCUUGGUGUGGC.......AUGAUAGCUGUGAAGCGUG.AGGUUGCCGGUUC.....................CCA..........GUAA............
env-596 AAAGAAUUGUAUCUAAAAAAAAUGCAAUUUUUUGAAAAAAAGUCUUGCAAUGCCCGUAAAUUCAAUGUAUACUAAUCCUUGGUGUGGC.......AUGAUAGCUGUGAAGCGUG.AGGUUGCCGGUUC.....................CCA..........GUAA............
env-597 AAAGAAAUUGUAUCAAAAAAAAUACAAUUUUUUUAAAAAAAGUCUUGCAAUGCCCAUAAAUUCAAUGUAUACUAAUCCUUGGUGUGGC.......AUGAUAGCUGUGAAGCGUG.AGGUUGCCGGUUC.....................CCA..........GUAA............
env-598 AAAGAAAUUGUAUCAAAAAAAAUACAAUUUUUUUAAAAAAAGUCUUGCAAUGCACGUAAAUUCAAUGUAUACUAAUCCUUGGUGUGGC.......AUGAUAGCUGUGAAGCGUG.AGGUUGCCGGUUC.....................CCA..........GUAA............
env-599 AAAAGAAUUGUAUCUAAAAAAAUACAAUUUUUUGAAAAAAAGUCUUGCAAUGCCCGUAAAUUCAAUGUAUACUAAUCCUUGGUGUGGC.......AUGAUAGCUGUGAAGCGUG.AGGUUGCCGGUUC.....................CCA..........GUAA............
env-600 AAAAGAAUUGUAUCUAAAAAAAUACAAUUUUUUUAAAAAAAGUCUUGCAAUGCCCAUAAAUUCAAUGUAUACUAAUCCUUGGUGUGGC.......AUGAUAGCUGUGAAGCGUG.AGGUUGCCGGUUC.....................CCA..........GUAA............
env-601 AAAGGAAUUGUUUUAAAAAAAAUACAAUUUUUUGAAAAAAAGUCUUGCAAUGCCCGUAAAUUCAAUGUAUACUAAUCCUUGGUGUGGC.......AUGAUAGCUGUGAAGCGUG.AGGUUGCCGGUUC.....................CCA..........GUAA............
env-602 AAAAGAAUUGUAUCUAAAAAAAUACAAUUUUUUGAAAAAAAGUCUUGCAAUGCCCACAAAUUCAAUGUAUACUAAUCCUUGGUGUGGC.......AUGAUAGCUGUGAAGCGUG.AGGUUGCCGGUUC.....................CCA..........GUAA............
env-603 AGACAAAAAUUUGUAAAAAUCUUGGGUUAACUGAAAAAAAGUGCUUGCAAAUAGCAUUUGCGUAGUGUAAUAUCAACUGUUGUGAG......GUUGAUAGCGUUGAUGCGUA.AGGUUGCUACUGGAUUUC...............................................
env-604 GUAAUAAUGUUUUUUAGAAAAAAACAAAAAAACCGAAAAAAGCCCUUGUCAAGCACACUUGAGUGUGCUAUCAUAAACGAGGUGUGAG......AUUGAUAGCUGUGAAGCGUG.AGGUUGCCGUCUCAAGAG.............................................
env-606 ..........................AAAAACCGAAAAAAGCCCUUGUCAAGCACACUUGAGUGUGCUAUCAUAAACGAGGUGUGAG......AUUGAUAGCUGUGAAGCGUG.AGGUUGCCGUCUCAAGAG.............................................
env-607 GUAAUAAUGUUUUUUAGAAAAAAACAAAAAAACCGAAAAAAGCCCUUGUCAAGCACACUUGAGUGUGCUAUCAUAAACGAGGUGUGAG......AUUGAUAGCUGUGAAGCGUG.AGGUUGCCGUCUCAAGAG.............................................
env-608 .........UUUUUUAGAAAAAAAUAAAAAAACCGAAAAAAGCCCUUGUCAAGCACACUUGAGUGUGCUAUCAUAAACGAGGUGUGAG......AUUGAUAGCUGUGAAGCGUG.AGGUUGCCGUCUCAAGAG.............................................
env-614 AUAAUAAUGUUUUUUAGAAAAAAAUAAAAAAACCGAAAAAAGCCCUUGUCAAGCACACUUGAGUGUGCUAUCAUAAACGAGGUGUGAG......AUUGAUAGCUGUGAAGCGUG.AGGUUGCCGUCUCAAGAG.............................................
env-615 AUAAUAAUGUUUUUUAGAAAAAAAUAAAAAAACCGAAAAAAGCCCUUGUCAAGCACACUUGAGUGUGCUAUCAUAAACGAGGUGUGAG......AUUGAUAGCUGUGAAGCGUG.AGGUUGCCGUCUCAAGAG.............................................
env-623 AUAAUAAUGUUUUUUAGAAAAAAAUAAAAAAACCGAAAAAAGCCCUUGUCAAGCACACUUGAGUGUGCUAUCAUAAACGAGGUGUGAG......AUUGAUAGCUGUGAAGCGUG.AGGUUGCCGUCUCAAGAG.............................................
env-624 AUAAUAAUGUUUUUUAGAAAAAAAUAAAAAAACCGAAAAAAGCCCUUGUCAAGCACACUUGAGUGUGCUAUCAUAAACGAGGUGUGAG......AUUGAUAGCUGUGAAGCGUG.AGGUUGCCGUCUCAAGAG.............................................
env-625 AUAAUAAUGUUUUUUAGAAAAAAAUAAAAAAACCGAAAAAAGCCCUUGUCAAGCACACUUGAGUGUGCUAUCAUAAACGAGGUGUGAG......AUUGAUAGCUGUGAAGCGUG.AGGUUGCCGUCUCAAGAG.............................................
env-626 AUAAUAAUGUUUUUUAGAAAAAAACAAAAAAACCGAAAAAAGCCCUUGUCAAGCACACUUGAGUGUGCUAUCAUAAACGAGGUGUGAG......AUUGAUAGCUGUGAAGCGUG.AGGUUGCCGUCUCAAGAG.............................................
env-632 UUUAAGACACACCCUAGAAAAAAAUAAAAAAACCGAAAAAAGCCCUUGUCAAGCACACUUGAGUGUGCUAUCAUAAACAAGGUGUGAG......AUUGAUAGCUGUGAAGCGUG.AGGUUGCCGUCUCAAGAG.............................................
env-633 AUAAUAAUAUUUUUUAGAAAAAAAUAAAAAAACCGAAAAAAGCCCUUGUCAAGCACACUUGAGUGUGCUAUCAUAAACGAGGUGUGAG......AUUGAUAGCUGUGAAGCGUG.AGGUUGCCGUCUCAAGAG.............................................
env-634 AUAAUAAUGUUUUUUAGAAAAAAAUAAAAAAACCGAAAAAAGCCCUUGUCAAGCACACUUGAGUGUGCUAUCAUAAACGAGGUGUGAG......AUUGAUAGCUGUGAAGCGUG.AGGUUGCCGUCUCAAGAG.............................................
env-635 GUAAUAAUGUUUUUUAGAAAAAAAUAAAAAAACCGAAAAAAGCCCUUGUCAAGCACACUUGAGUGUGCUAUCAUAAACGAGGUGUGAG......AUUGAUAGCUGUGAAGCGUG.AGGUUGCCGUCUCAAGAG.............................................
env-637 ....UAAUGUUUUUUAGAAAAAAAUAAAAAAACCGAAAAAAGCCCUUGUCAAGCACACUUGAGUGUGCUAUCAUAAACGAGGUGUGAG......AUUGAUAGCUGUGAAGCGUG.AGGUUGCCGUCUCAAGAG.............................................
env-639 AUAAUAAUGUUUUUUAGAAAAAAAUAAAAAAACCGAAAAAAGCCCUUGUCAAGCACACUUGAGUGUGCUAUCAUAAACGAGGUGUGAG......AUUGAUAGCUGUGAAGCGUG.AGGUUGCCGUCUCAAGAG.............................................
env-640 AUAAUAAUGUUUUUUAGAAAAAAACAAAAAAACCGAAAAAAGCCCUUGUCAAGCACACUUGAGUGUGCUAUCAUAAACGAGGUGUGAG......AUUGAUAGCUGUGAAGCGUG.AGGUUGCCGUCUCAAGAG.............................................
env-641 AUAAUAAUGUUUUUUAGAAAAAAAUAAAAAAACCGAAAAAAGCCCUUGUCAAGCACACUUGAGUGUGCUAUCAUAAACGAGGUGUGAG......AUUGAUAGCUGUGAAGCGUG.AGGUUGCCGUCUCAAGAG.............................................
env-642 GUAAUAAUGUUUUUUAGAAAAAAAUAAAAAAACCGAAAAAAGCCCUUGUCAAGCACACUUGAGUGUGCUAUCAUAAACGAGGUGUGAG......AUUGAUAGCUGUGAAGCGUG.AGGUUGCCGUCUCAAGAG.............................................
env-643 AUAAUAAUGUUUUUUAGAAAAAAAUAAAAAAACCGAAAAAAGCCCUUGUCAAGCACACUUGAGUGUGCUAUCAUAAACGAGGUGUGAG......AUUGAUAGCUGUGAAGCGUG.AGGUUGCCGUCUCAAGAG.............................................
env-644 AUAAUAAUGUUUUUUAGAAAAAAAUAAAAAAACCGAAAAAAGCCCUUGUCAAGCACACUUGAGUGUGCUAUCAUAAACGAGGUGUGAG......AUUGAUAGCUGUGAAGCGUG.AGGUUGCCGUCUCAAGAG.............................................
Jig-1-1 ACAGCGAGAAAUUCCAGGGUGGACGAAAGCCAAAGUAAAAAAUACUUGCAAUCCGUUUAAAUAUAUGUUAUUCUUACGGAGGUGUGGC......AUUGAUAGCUGUGAAGCGUG.AGGUUGCCGUUCCUGUUGG............................................
env-646 AAAACUUCUUGAUAUAACUUCAAAAGUUUUUUUAAAAAAGUACUUGAUUUCUCGGGAAAAUCGUGUAUAAUAAAAAGCGUUGU.GUGAG.......UGAUAGCGAUGAAGCGAG.AGGUUGCUGUG.......................AUGGA.........AAAA............
env-647 AAACUUUCUUGAUAUUCCUUUAAAACUUUUUGAAAAAAACACUUGAUUUAUCCUAGAAUCGUGUAUAAUAGAAAGCGUUGU.GUGAG.......UGAUAGCAAAGAAGCGCG.AGGUUGCUGUG.......................AUGGA.........AAAA............
Eha-1-1 AAACUUUCUUGAUAUUCCUUUAAAACUUUUUGAAAAAAACACUUGAUUUAUCCUAGAAUCGUGUAUAAUAGAAAGCGUUGU.GUGAG.......UGAUAGCAAAGAAGCGCG.AGGUUGCUGUG.......................AUGGA.........AAAA............
env-648 AAACUUUCUUGAUAUUCCUUUAAAACUUUUUGAAAAAAACACUUGAUUUAUUCCUGGAAUCGUGUAUAAUAAAAAGCGUUGU.GUGAG.......UGAUAGCAAAGAAGCGCG.AGGUUGCUGUG.......................AUGGA.........AAAA............
env-649 AAACUUUCUUGAUAUUCCUUUAAAACUUUUUGAAAAAAACACUUGAUUUAUCCUGGAAUCGUGUAUAAUAGAAAGCGUUGU.GUGAG.......UGAUAGCAAAGAAGCGCG.AGGUUGCUGUG.......................AUGGA.........AAAA............
env-650 UAAAAUUUACCAAAAAAAAUUGUGGAAAUUAUGAAUAAAAAAGUCUUGCAUAACUGGAAACACUGGUAUAUAAUCAUUCUUGUGUGAG......AUUGAUAGCGUAGAAGCGUG.AGGUUGCUGUAUAUAA...............................................
env-653 AUAAAAUUUACCAAAAAAAUUGUGAAAAUUAUGAAUAAAAAAGCUUGCAUAACUGGAAACACUGGUAUAUAAUCAUUCUUGUGUGAG......AUUGAUAGCGUAGAAGCG.G.AGGUUGCUGUAUAUAA...............................................
env-654 AAAAUUAUAUUAAGAUUGUACUAAAAAAAGAACAAUAAAAUUUAUUGAUUUUAUCAAACUUAUCAUGUAUACUAUCACUUGUGUGAG......AUUGAUAGCGUAGAAGCGUG.AGGUUGCUGCCGCAA...............................................
env-655 AAAAUUAUAUUAAGAUUGUACUAAAAAAAGAACAAUAAAAUUUAUUGAUUUUAUCAAACUUAUCAUGUAUACUAUCACUUGUGUGAG......AUUGAUAGCGUAGAAGCGUG.AGGUUGCUGCCGCAA...............................................
env-656 AAAAUUAUAUUAAUAUUGUACUAAAAAAAGAACAAUAAAAUUUAUUGAUUUUAUCAAACUUAUCAUGUAUACUAUCACUUGUGUGAG......AUUGAUAGCGUAGAAGCGUG.AGGUUGCUGCCGCAA...............................................
env-657 AAAAUUAUAUUAAGAUUGUACUAAAAAAAGAACAAUAAAAUUUAUUGAUUUUAUCAAACUUAUCAUGUAUACUAUCACUUGUGUGAG......AUUGAUAGCGUAGAAGCGUG.AGGUUGCUGCCGCAA...............................................
env-658 AAAAUUAUAUUAAUAUUGUACUAAAAAAAGAACAAUAAAAUUUAUUGAUUUUAUCAAACUUAUCAUGUAUACUAUCACUUGUGUGAG......AUUGAUAGCGUAGAAGCGUG.AGGUUGCUGCCGCAA...............................................
env-662 AAAAUUAUAUUAAUAUUGUACUAAAAAAAGAACAAUAAAAUUUAUUGAUUUUAUCAAACUUAUCAUGUAUACUAUCACUUGUGUGAG......AUUGAUAGCGUAGAAGCGUG.AGGUUGCUGCCGCAA...............................................
env-663 AAAAUCAUAUUAAGAUUGUACUAAAAAAAGAACAAUAAAAUUUAUUGAUUUUAUUAAACUUAUCAUGUAUACUAUCACUUGUGUGAG......AUUGAUAGCGUAGAAGCGUG.AGGUUGCUGCCGCAA...............................................
env-664 AAAAUUAUAUUAAUAUUGUACUAAAAAAAGAACAAUAAAAUUUAUUGAUUUUAUCAAACUUAUCAUGUAUACUAUCACUUGUGUGAG......AUUGAUAGCGUAGAAGCGUG.AGGUUGCUGCCGCAA...............................................
env-667 AAAAUUAUAUUAAGAUUGUACUAAAAAAAGAACAAUAAAAUUUAUUGAUUUUAUCAAACUUAUCAUGUAUACUAUCACUUGUGUGAG......AUUGAUAGCGUAGAAGCGUG.AGGUUGCUGCCGCAA...............................................
env-668 AAAAUUAUAUUAAUAUUGUACUAAAAAAAGAACAAUAAAAUUUAUUGAUUUUAUCAAACUUAUCAUGUAUACUAUCACUUGUGUGAG......AUUGAUAGCGUAGAAGCGUG.AGGUUGCUGCCGCAA...............................................
Lba-14-1 AUUACUUUGUAUGGAAAAAAAUACAAUAAAAAAACAAAAAAUACUUGCGAAAGCCAAAAGGUUACGAUAUACUACUAAUUGGUGUGGC.......UGAUAGCGAUGAAACGUG.AGGUUGCUGC.......................AUAUG.........GAGAAA..........
Bsp-1-1 AUUGCCUAUUUUUAUUAAAAAGUGAUUAAUUUUGAAAAUAACGCUUGCUGAAAUGAAAAAUGGGAUAUAGUUAUCCAUGGUGUGAG.......UGAUAGCUGUGAAGCGUG.AGGUUGCUGUCUUAUCCAGUGA.........................................
Bsp-9-1 AUUGCCUAUUAUUAUUAAAAAGUGAUUAAUUUUGAAAAUAACGCUUGCUGAAAUGAAAAAUGGGAUAUAGUUAUCCAUGGUGUGAG.......UGAUAGCUGUGAAGCGUG.AGGUUGCUGUCUUAUCCAGUGA.........................................
env-669 UUUUUGCAUAAAAACGGAUAAAUCGAAAAAAAUAAAAAAAACCUUGAUUUUUCGUGGAGUUUCUUGUAUACUAACAACUGUGUGAG......AUUGAUAGCGUUGAAGCGCG.AGGUUGCCGCUGAGAAAC.............................................
env-671 UUUUUGCAUAAAAACGGAUAAAUCGAAAAAAAUAAAAAAAACCUUGAUUUUUCGCGUGCUUUCUUGUAUACUAACAACUGUGUGAG......AUUGAUAGCGUUGAAGCGCG.AGGUUGCCGCUGAGAAAC.............................................
env-672 UUUUUGCAUAAAAACGGAUAAAUCGAAAAAAAUAAAAAAAACCUUGAUUUUUCGCGUGCUUUCUUGUAUACUAACAACUGUGUGAG......AUUGAUAGCGUUGAAGCGCG.AGGUUGCCGCUGAGAAAC.............................................
env-681 UUUUUGCAUAAAAACGGAUAAAUCGAAAAAAAUAAAAAAAACCUUGAUUUUUCGUGGAGUUUCUUGUAUACUAACAACUGUGUGAG......AUUGAUAGCGUUGAAGCGCG.AGGUUGCCGCUGAGAAAC.............................................
env-683 NNNNNNNNNNNNNNNNNNNNNNNNNNNNNNNNNNNNNNNNNAAACCUUGAUUUUUCGCGUGCUUUCUUGUAUACUAACAACUGUGUGAG......AUUGAUAGCGUUGAAGCGCG.AGGUUGCCGCUGAGAAAC.............................................
env-684 UUUUUGCAUAAAAACGGAUAAAUCGAAAAAAAUAAAAAAAACCUUGAUUUUUCGCGUGCUUUCUUGUAUACUAACAACUGUGUGAG......AUUGAUAGCGUUGAAGCGCG.AGGUUGCCGCUGAGAAAC.............................................
env-685 UUUUUGCAUAAAAACGGAUAAAUCGAAAAAAAUAAAAAAAACCUUGAUUUUUCGCGUGCUUUCUUGUAUACUAACAACUGUGUGAG......AUUGAUAGCGUUGAAGCGCG.AGGUUGCCGCUGAGAAAC.............................................
env-686 UAAAUAGAAAAAAGAUUGUACUAAAAAAAAACAAAGAAAGUACUUGCAAUUUUCGGUGCUCUCGGAUAUACUAAGAGAGUGUGAG.......UGAUAGCAAUGAAGCGCG.AGGUUGCUGC.......................CCGGC.........AAAUGAAUAA......
env-687 AUAAAUAGAAAAAAGAUUGUACUUAAAAAAAAACAAAGAAAGUACUUGCAAUUUUCGGUACUCUCGGAUAUACUAAGAGAGUGUGAG.......UGAUAGCAAUGAAGCGCG.AGGUUGCUGC.......................CCGGC.........AAAUGAAUAA......
env-691 AUAAAGAAGGAAAAGAUUGUACUCGAAAAAAAACAAAGAAAGUACUUGCAAUUUUGCACUCUCUCGGAUAUACUAAGAGAGUGUGAG.......UGAUAGCAAUGAAGCGCG.AGGUUGCUGC.......................CCGGC.........AAAUGAAUAA......
env-692 AUAAAUAGAAAAAAGAUUGUACUAAAAAAAAACAAAGAAAGUACUUGCAAUUUUCGGUGCUCUCGGAUAUACUAAGAGAGUGUGAG.......UGAUAGCAAUGAAGCGCG.AGGUUGCUGC.......................CCGGC.........AAAUGAAUAA......
env-694 AAUAAAGAAGAAAAAAUUGUACUCAAAAAAAAACAAAAAAAGUACUUGCAAUUUUGCACUCUCUCGGAUAUACUAAGAGAGUGUGAG.......UGAUAGCAAUGAAGCGCG.AGGUUGCUGC.......................CCGGC.........AAAUGAAUAA......
env-695 AAUAAAGAAGAAAAAGAUUGUACUUGAAAAAAAACAAAAAAAGUACUUGCAAUUUUGCACUCUCUCGGAUAUACUAAGAGAGUGUGAG.......UGAUAGCAAUGAAGCGCG.AGGUUGCUGC.......................CCGGC.........AAAUGAAUAA......

..................................................................................<<.<<<<<......................<<<.<.<<<...<<<<<.......................<<<<<<<<<<<<<<............>>>>
..................................................................................21..1111......................222.2.012...12222.......................1111111.10..00............00..
••••••••••••••R••••••••••R••••••••RAAAAR•••YU•••••••••••••••••••••••••••••••••••GYUGUGRC-------∘UGAYAGC••UGAAGCG∘G-AGGUUGCYRY••∘∘∘∘------------------------------------------------
5′

```
env-696   AUAAAGAAGAAAAAGAUUGUACUUGAAAAAAAACAAAAAAAGUACUUGCAAUUUUACACUCUCUCAGAUAUACUAAGAGAGCUGUGAC........AUGAUAGCAAUGAAGCGCG.AGGUUGCUGC.......................CCGGC.........AAAUGAAUAA......
env-699   AUAAAGAAGAAAAAGAUUGUACUUGAAAAAAAACAAAAAAAGUACUUGCAAUUUUGCGCUCUCUCGGAUAUACUAAGAGAGCUGUGAC........AUGAUAGCAAUGAAGCGCG.AGGUUGCUGC.......................CCGGC.........AAAUGAAUAA......
env-700   UAAAAUAAUAAUUAAAAAAACCAUAAAAAAAAUUAAAAAAACCUCUUGAUUUAAAGAAAUCCCUAGUAUAUAAUCAGAUUUGCUGUGAC........AUGAUAGCUAUGAAGCG.G.AGGUUGCUGCGUACAUAU.............................................
env-705   CGUUAAAAAAAAUUCAAAAAAACAUAAAAAAAAUUAAAAAAAACUCUUGAUUUAAAGAAAUCCCUAGUAUAUAAUCAAACGUGCUGUGGC........AUGAUAGCUGAGAAGCG.G.AGGUUGCUGCGUACAUAUAUGA.........................................
env-708   UUAAAAAAUAAUUCAAAAAAACAUAAAAAAAAUUAAAAAAAACUCUUGAUUUAAAGAAAUCCCUAGUAUAUAAUCAAACGUGCUGUGGC........AUGAUAGCUGAGAAGCG.G.AGGUUGCUGCGUACAUAUAUGA.........................................
env-709   CGUUAAAAAAAAUUCAAAAAAACAUAAAAAAAAUUAAAAAAAACCUCUUGAUUUAAAGAAAUCCCUAGUAUAUAAUCAAACGUGCUGUGGC........AUGAUAGCUGAGAAGCG.G.AGGUUGCUGCGUACAUAUAUGA.........................................
env-710   GUUAAAAAAUAAUUCAAAAAAACAUAAAAAAAAUUAAAAAAAACUCUUGAUUUGAAGAAAUCCCUAGUAUAUAAUCAAACGUGCUGUGGC........AUGAUAGCUGAGAAGCG.G.AGGUUGCUGCGUACAUAUAUGA.........................................
env-711   GUUAAAAAAAAUUCAAAAAAACAUAAAAAAAAUUAAAAAAAACCUCUUGAUUUAAAGAAAUCCCUAGUAUAUAAUCAAACGUGCUGUGGC........AUGAUAGCUGAGAAGCG.G.AGGUUGCUGCGUACAUAUAUGA.........................................
env-712   CGUUAAAAAAAAUUCAAAAAAACAUAAAAAAAAUUAAAAAAAACUCUUGAUUUAAAGAAAUCCCUAGUAUAUAAUCAAACGUGCUGUGGC........AUGAUAGCUGAGAAGCG.G.AGGUUGCUGCGUACAUAUAUGA.........................................
env-713   CGUUAAAAAAAAUUCAAAAAAACAUAAAAAAAAUUAAAAAAAACCUCUUGAUUUAAAGAAAUCCCUAGUAUAUAAUCAAACGUGCUGUGGC........AUGAUAGCUGAGAAGCG.G.AGGUUGCUGCGUACAUAUAUGA.........................................
env-726   CGUUAAAAAAAAUUCAAAAAAACAUAAAAAAAAUUAAAAAAAACCUCUUGAUUUAAAGAAACCCCUAGUAUAUAAUCAAACGUGCUGUGGC........AUGAUAGCUGAGAAGCG.G.AGGUUGCUGCGUACAUAUAUGA.........................................
env-727   CCGUUAAAAAAAAUUCAAAAAAACAUAAAAAAAAUUAAAAAAAACCUCUUGAUUUAAAGAAAUCCCUAGUAUAUAAUCAAACGUGCUGUGGC........AUGAUAGCUGAGAAGCG.G.AGGUUGCUGCGUACAUAUAUGA.........................................
env-729   CCGUUAAAAAAAUUCAAAAAAACAUAAAAAAAAUUAAAAAAAACCUCUUGAUUUAAAGAAAUCCCUAGUAUAUAAUCAAACGUGCUGUGGC........AUGAUAGCUGAGAAGCG.G.AGGUUGCUGCGUACAUAUAUGA.........................................
env-730   UUAAAAAAUAAUUCAAAAAAACAUAAAAAAAAUUAAAAAAAACCUCUUGAUUUAAAGAAAUCCCUAGUAUAUAAUCAAACGUGCUGUGGC........AUGAUAGCUGAGAAGCG.G.AGGUUGCUGCGUACAUAUAUGA.........................................
env-731   CGUUAAAAAAAAUUCAAAAAAACAUAAAAAAAAUUAAAAAAAACUCUUGAUUUAAAGAAAUCCCUAGUAUAUAAUCAAACGUGCUGUGGC........AUGAUAGCUGAGAAGCG.G.AGGUUGCUGCGUACAUAUAUGA.........................................
Rsp-1-1   UCACAGGGUAGAAUAAACAAAAAAUAAAAAAAAUAAAAAAAAAUACUUGCAAUUUCCCAAAAGGUCGUGUAUACUCCUAUUUGUUGUGAC.......AUUGAUAGCAAUGAAGCGCG.AGGUUGCUGCCACAAAGCU..............................................
env-732   CGGCUGUAUUGUAUUAAAAAAAAAUACAAAAAAAUAAAAAAAAGUACUUGAAAAACGAGAAUAAGUUCGUUAUACUAAGAGAGCUGUGGC........AUGAUAGCGUUGAAGCGUG.AGGUUGCCGCUUA....................GUGA..........GAAA............
Rgn-1-1   UGGCUGUAUUGUAUUAAAAAAAAAUACAAAAAAAUAAAAAAAAGUACUUGAAAAAUGAGAAUAAGUUCGUUAUACUAAGAGAGCUGUGGC........AUGAUAGCGUUGAAGCGUG.AGGUUGCCGCUUA....................GUGA..........GAAA............
env-733   .........UGUAUUAAAAAAAAAUACAAAAAAAUAAAAAAAAGUACUUGAAAAAUGAGAAUAAGUUCGUUAUACUAAGAGAGCUGUGGC........AUGAUAGCGUUGAAGCGUG.AGGUUGCCGCUUA....................GUGA..........GAAA............
env-734   CGGCUGUAUUGUAUUAAAAAAAAAUACAAAAAAAUAAAAAAAAGUACUUGAAAAAUGAGAAUAAGUUCGUUAUACUAAGAGAGCUGUGGC........AUGAUAGCGUUGAAGCGUG.AGGUUGCCGCUUA....................GUGA..........GAAA............
env-735   CGGCUGUAUUGUAUUAAAAAAAAAUACAAAAAAAUAAAAAAAAGUACUUGAAAAAUGAGAAUAAGUUCGUUAUACUAAGAGAGCUGUGGC........AUGAUAGCGUUGAAGCGUG.AGGUUGCCGCUUA....................GUGA..........GAAA............
env-739   CGGCUGUAUUGUAUUAAAAAAAAAUACAAAAAAAUAAAAAAAAGUACUUGAAAAAUGAGAAUAAGUUCGUUAUACUAAGAGAGCUGUGGC........AUGAUAGCGUUGAAGCGUG.AGGUUGCCGCUUA....................GUGA..........GAAA............
env-742   CGGCUGUAUUGUAUUAAAAAAAAAUACAAAAAAAUAAAAAAAAGUACUUGAAAAACGAGAAUAAGUUCGUUAUACUAAGAGAGCUGUGGC........AUGAUAGCGUUGAAGCGUG.AGGUUGCCGCUUA....................GUGA..........GAAA............
Lba-4-1   GAGCGGCUGUAUUGUAUUAAAAAAAUACAAAAAAUAAAAAAAGUACUUGAAAAAUGAGAAUAAGUUCGUUAUACUAAGAGAGCUGUGGC........AUGAUAGCGUUGAAGCGUG.AGGUUGCCGCUUA....................GUGA..........GAAA............
env-744   CGGCUGUAUUGUAUUAAAAAAAAAUACAAAAAAAUAAAAAAAAGUACUUGAAAAAUGAGAAUAAGUUCGUUAUACUAAGAGAGCUGUGGC........AUGAUAGCGUUGAAGCGUG.AGGUUGCCGCUUA....................GUGA..........GAAA............
env-749   AAGAAAAAUUGUUCUGGAAAAAAUACAAUUAUUCGGAAAAAGCUCUUGAAAUAUAGAGGUGAAUCCGCUAUACUAAUAAAGCUGUGGC........AUGAUAGCGAUGAAGCGAG.AGGUUGCUGUUUA....................GUG...........UAAAA...........
env-751   AAAACAAAUUGUAUUGAAAAAAAAACAAUAUUUCUUGAAAAACCCUUGCAAACGCAGGGGUUUUCCUAUAUAAUAGCGAUGUUGUGAC.......CUUGAUAGCGUAGAAACGUG.AGGUUGCUGUGGAACUUAC..............................................
Cph-1-1   AAAUAAAGGAAAUAAUUUACUAGAAAGAAAAAAUAAAAAAAAUACUUGCAAAUAUUUUUGUUCUAUUAUAUAAUAACUACUGUUGUGAC.......GAUGAUAGCGUAGAAACGUG.AGGUUGCUGUGUCAAAUGA..............................................
Bfo-1-1   AAACAGGAUUGUACGAAAAAAAAAUACAAUAUUUUUAAAAAACUCCUUGCAUAUGGUUAAAAUUUCAUAUAUAAUGCAGAUGUUGUGAC.......CUUGAUAGCGUUGAAGCGUG.AGGUUGCUGCGGGAAAAGGAAAAAC.........................................
env-752   ACUAAAAAAAAAUACAAUCUUUUUUGCAAAAUCUUUAAAAAAGACUUGCAAGGCCCAAAACUAUGGGUUAUACUAUUUUUUGCUGUGAC........AUGAUAGCUAUGAAGCGCG.AGGUUGCGGCCUUUAUAU.............................................
env-753   ACUAAAAAAAAAUACAAUCUUUUUUGCAAAAUCUUUAAAAAAGACUUGCAAGGCCCAAAACUAUGGGUUAUACUAUCUUUUGCUGUGAC........AUGAUAGCUAUGAAGCGCG.AGGUUGCGGCCUUUAUAU.............................................
Lbo-3-1   GUAAAUAUUGGUGAUAAAUUUUAAAAAAUAAAAACGGAAAAUAUGUUGCAAAAAAGAUGAUAUUAGGUUAUAAUAUGAAUUGCUGUGAC.......AUUGAUAGCGAUGAAAUGUG.AGGUUGCUACCCGAUGU...............................................
env-754   AAAAAAGGUUUUUUUAUAUUAUAAAUUGAUAAUAUAAAAAAAAUGCUUGCAUUCAUGUGAAUAUUGUUAUAUACUAUUAAGGCUGUGGC........AUGAUAGCUGUGAAACGUG.AGGUUGUUACCUUUAU...............................................
env-755   GAAAAUGGGUUGUUUGAUAUUAUAAUUUAUAAUAUAAAAAAAACACUUGCAUUCAUUUGAAUAUUGUUAUAUAAUAUUAAGGCUGUGGC........AUGAUAGCUGUGAAACGUG.AGGUUGUUACCUUUAU...............................................
Csp-4-1   AAAUUGGAUUGUAUCAAAAAAAAAUACAAUAAUUCAAAAAAACCUCUUGAAAUAUAUGGAUGAAUCCGCUAUACUAAGAGAGCUGUGGC........AUGAUAGCGUGGAAGCGUG.AGGUUGCUGCCGG.....................UUGC..........GAAGA...........
env-756   UUUCAAAAAAGUACAAUUUAAAUACAAUUUUUUUGAAAAAGGACUUGCUUUUUUUAGAGGAGUAAUGUAAAAUACUCCUUGCUGUGGC........AUGAUAGCGAUGAAGCGUG.AGGUUGCUGCUGA....................UGG...........GUGAUG..........
env-758   AAUCUGCAAUGUACGAAAAAAAAAUACAAUAAAUAAAAAAAAAGACUUGAAAUAAAGGAAAGUAUCAUUUAUACUAAGAGAGCUGUGAC........AUGAUAGCAAUGAAGCGUG.AGGUUGCUGUCC......................UGC...........AAAAAAAGAUAA....
env-759   UAUAGUCCCCAUGAAUAAUUAAUAAAAUUUUUUAUAUUUAAGUCUUGCAAAUACCCAAGACAUACGAUAUACUUAGUAAUGUUGUGAU.......AUUGAUAGCGUAGAAGCGUG.AGGUUGCUACAAGAUU................................................
env-760   UAAAAUCUAAAAUAGGUAAAAAUAAAAAUUUUUUAUAUUUAAGUCUUGCAAAUACCAAAGACAUACGAUAUACUUAGUAAUGUUGUGAU.......AUUGAUAGCGUAGAAGCGUG.AGGUUGCUACAAGAUU................................................
env-762   UAAAAUCUAAAAUAGGUAAAAAUAAAAAUUUUUUAUAUUUAAGUCUUGCAAAUACUCAAGACAUACGAUAUACUUAGUAAUGUUGUGAU.......AUUGAUAGCGUAGAAGCGUG.AGGUUGCUACAAGAUU................................................
env-763   CUGAGAUAAAUUGAGAAAAAAAAUCAAAAAAAUAAAAAAACCCCUUGCAUUUGCGAGGGGUUUCGUUUAUAAUCAUUCAUGUUGUGAC.......AUUGAUAGCUGUGAAGCGCG.AGGUUGCUGCAGAAAC................................................
env-764   GAAAAAUAGUGUAUAAGAAAAAAAUACAAAAAAAAAUCAAAAAACCCUUGCAAAAUAAGAAAAUAUCAUUUAUAAUUCAAUCGUUGUGAC.......CUUGAUAGCGUUGAAGCGUG.AGGUUGCUGUAGAAAUUC..............................................
Lba-6-1   AACACUGAUUGUAUGAAAAAAAAAUACAAUAAUUCGAUAAAAGGGCUUGAAAUCAGUUCUUUCAUCAUAUAUACUAAAAAAGCUGUGAC........AUGAUAGCAAUGAAGCGUG.AGGUUGCUGCCGGGAC..................UUCUGAAAC....AACAGCA.........
Dsp-1-1   AUUUCUGAUUGUACGAAAAAAAAAUACAAUAUUUGAAAGAAAAGGCUUGAAAAAUCUUUCUCUGUCGGAUAUACUGAAAAAGCUGUGAC........AUGAUAGCGUCGAAGCGAG.AGGUUGCUGCCCGUAAGU.............................................
env-765   AUAAAACCCCUUAUAAAAAUUAUUUAGAAAAAUACAAAAAAAAUGCUUGCAUUUCCCUGGAAUAUAUGUUAUCUUAGAACAGCUGUGAC........AUGAUAGCAAAGAAACGUG.AGGUUGCUGCCUUACAAAA............................................
env-767   AUAAAACCCCUUAUAAAAAUUAUUUAGAAAAAUACAAAAAAAAUGCUUGCAUUUCUCUGGAAUAUAUGUUAUCUUAGAACAGCUGUGAC........AUGAUAGCAAAGAAACGUG.AGGUUGCUGCCUUACAAAA............................................
env-768   AUAAAACCCCUUAUAAAAAUUAUUUAGAAAAAUACAAAAAAAAUGCUUGCAUUUCUCUGGAAUAUAUGUUAUCUUAGAACAGCUGUGAC........AUGAUAGCAAAGAAACGUG.AGGUUGCUGCCUUACAAAA............................................
env-769   ................AAAUUAUUUAGAAAAAUACAAAAAAAAUGCUUGCAUUUCCCUGGAAUAUAUGUUAUCUUAGAACAGCUGUGAC........AUGAUAGCAAAGAAACGUG.AGGUUGCUGCCUUACAAAA............................................
env-770   AUAAAACCCCUUAUAAAAAUUAUUUAGAAAAAUACAAAAAAAAUGCUUGCAUUUCCCUGGAAUAUAUGUUAUCUUAGAACAGCUGUGAC........AUGAUAGCAAAGAAACGUG.AGGUUGCUGCCUUACAAAA............................................
env-777   CUGUUUCCCCGUCUGGGGGAAAUUUAAAAAAAUUUAAAAAAGUACUUGCAUUUCCUGUAAACAUAUGCUAUCCUUGAAAAGCUGUGAC........AUGAUAGCAAAGAAGCGAG.AGGUUGCUGCCCACCAUAA............................................
env-778   AGAAAAUUUUGUAUGAAAAAAAAGUACAAAAAAAUAAAAUAAAAUGCUUGCAAAAUGCAGAAACAUCGGGUAUACUAAAAAAGCUGUGAC........AUGAUAGCAAAGAAGCGCG.AGGUUGCUGCCGAGAAGAAU...........................................
env-802   AGAAAAUUUUGUAUGAAAAAAAAGUACAAAAAAAUAAAAUAAAAUGCUUGCAAAAUGCAGAAACAUCGGGUAUACUAAAAAAGCUGUGAC........AUGAUAGCAAAGAAGCGCG.AGGUUGCUGCCGAGAAGAAU...........................................
env-803   AGAAAAUUUUGUAUGAAAAAAAAGUACAAAAAAAUAAAAUAAAAUGCUUGCAAAAUGCAGAAACAUCGGGUAUACUAAAAAAGCUGUGAC........AUGAUAGCAAAGAAGCGCG.AGGUUGCUGCCGAGAAGAAU...........................................
Cco-1-1   AGAAAAUUUUGUAUGAAAAAAAAGUACAAAAAAAUAAAAUAAAAUGCUUGCAAAAUGCAGAAACAUCGGGUAUACUAAAAAAGCUGUGAC........AUGAUAGCAAAGAAGCGCG.AGGUUGCUGCCGAGAAGAAU...........................................
env-806   AGAAAAUUUUGUAUGAAAAAAAAGUACAAAAAAAUAAAAUAAAAUGCUUGCAAAAUAUAGAAACAUCGGGUAUACUAAAAAAGCUGUGAC........AUGAUAGCAAAGAAGCGCG.AGGUUGCUGCCGAGAAGAAU...........................................
env-812   GAAAAAUUUUGUAUGAAAAAAAAGUACAAAAAAAUAAAAUAAAAUGCUUGCAAAAUAUAGAAACAUCGGGUAUACUAAAAAAGCUGUGAC........AUGAUAGCAAAGAAGCGCG.AGGUUGCUGCCGAGAAGAAU...........................................
env-814   GUUUUGGAUUGUACGUGAAAAAAAUACAAAAAAAAUAAAAAAAAGACUUGCAUUAUUGCCUUGGAUUAGAUAUACUAAGAGAGCUGUGAC........AUGAUAGCAAUGAAGCGCG.AGGUUGCCGUCAAGAAU.............................................
Dfo-2-1   UUUUUAGAUUGUACUGGAAAAAGUACAAAAAAAAUAAAAAAAAGACUUGCAUUAUUGCCUUGGAUUAGAUAUACUAAGAGAGCUGUGAC........AUGAUAGCAAUGAAGCGCG.AGGUUGCCGUCGAGAAU.............................................
Dfo-1-1   GUUUUGGAUUGUACGUGAAAAAAAUACAAAAAAAAUAAAAAAAAGAACUUGCAUUAUUGCCGCGGAUUGGAUAUACUAAGAGAGCUGUGAC........AUGAUAGCAAUGAAGCGCG.AGGUUGCCGUCGAGAAU.............................................
env-818   CUUUUAGAUUGUACUGGAAAAAGUACAAAAAAAAUAAAAAAAAGACUUGCAUUAUUGCCUUGGAUUAGAUAUACUAAGAGAGCUGUGAC........AUGAUAGCAAUGAAGCGCG.AGGUUGCCGUCGAGAAU.............................................
env-819   CUCUUGGAUUGUACGGAAAAAAGUACAAAAAAAAUAAAAAAAAGACUUGCAUUAUUGCCUUGGAUUAGAUAUACUAAGAGAGCUGUGAC........AUGAUAGCAAUGAAGCGCG.AGGUUGCCGUCGAGAAU.............................................
env-820   GGCAAAGGCUGACGAAGCACUGACGAAGAAAAUUGAAAAAAAAUACUUGCAUUUAUUUGGAAUGUGUGCUAUCAUACAAAGGCUGUGGC........AUGAUAGCGUUGAAGCGAG.AGGUUGCUGCC......................CAC...........AAA.............
[ci-1-1   UAAAUAGCAAUGCAGAUAACUAUGCAGAAAAAAAAUAAAAAAGUACUUGCAUUUAAUGGAGAUGUAUGCUAUCAUAACAAGGCUGUGGC........AUGAUAGCGUUGAAGCGAG.AGGUUGCCGCC......................CAC...........AAA.............
Csp-5-1   UAAAUAGCAAUGCAGAUAACUAUGCAGAAAAAAAAUAAAAAAGUACUUGCAUUUAAUGGAGAUGUAUGCUAUCAUAACAAGGCUGUGGC........AUGAUAGCGUUGAAGCGAG.AGGUUGCCGCC......................CAC...........AAA.............
env-821   GGCAAAGGCUGACAAAGCACUGACGAAGAAAAUUGAAAAAAAAUACUUGCAUUUAUUUGGAAUGUGUGCUAUCAUACAAAGGCUGUGGC........AUGAUAGCGUUGAAGCGAG.AGGUUGCUGCC......................CAC...........AAA.............
env-822   GGCAAAGGCUGACAAAGCAUUGACGAAGAAAAUUGAAAAAAAAUACUUGCAUUUAUUUGGAAUGUGUGCUAUCAUACAAAGGCUGUGGC........AUGAUAGCGUUGAAGCGAG.AGGUUGCUGCC......................CAC...........AAA.............
env-823   GGCAAAGGCUGACAAAGCACUGACGAAGAAAAUUGAAAAAAAAUACUUGCAUUUAUUUGGAAUGUGUGCUAUCAUACAAAGGCUGUGGC........AUGAUAGCGUUGAAGCGAG.AGGUUGCUGCC......................CAC...........AAA.............
Cbo-2-1   GGCAAAGGCUGACGAAGCACUGACGAAGAAAAUUGAAAAAAAAUACUUGCAUUUAUUUGGAAUGUGUGCUAUCAUACAAAGGCUGUGGC........AUGAUAGCGUUGAAGCGAG.AGGUUGCUGCC......................CAC...........AAA.............
Cba-7-1   AAAAAAGGUUAUAAAAAAAAUGCAUGAAUAAAAAAAUAAAAAAGUACUUGCAUUUAUAAAAGAAAUGUGCUAAUAUAACAAGGCUGUGGC........AUGAUAGCGUUGAAGCGAG.AGGUUGCUGCC......................CAC...........AAA.............
env-824   GGCAAAGGCUGACAAAGCACUGACGAAGAAAAUUGAAAAAAAAUACUUGCAUUUAUUUGGAAUGUGUGCUAUCAUACAAAGGCUGUGGC........AUGAUAGCGUUGAAGCGAG.AGGUUGCUGCC......................CAC...........AAA.............
Cba-2-1   AAAAAGGGUUAUAAAAAAAAUGCAUGAAUAAAAAAAUAAAAAAGUACUUGCAUUUAUAAAAGAAAUGUGCUAAUAUAACAAGGCUGUGGC........AUGAUAGCGUUGAAGCGAG.AGGUUGCUGCC......................CAC...........AAA.............
env-826   ACAUAACUCCUAACUAAGGCCGUAAAGGAAAAUUGAAAAAAAAUACUUGCAUUUAUUUGGAAUGUGUGCUAUCAUACAAAGGCUGUGGC........AUGAUAGCGUUGAAGCGAG.AGGUUGCUGCC......................CAC...........AAA.............
Ccl-6-1   GGCAAAGGCUGACGAAGCACUGACGAAGAAAAUUGAAAAAAAAUACUUGCAUUUAUUUGGAAUGUGUGCUAUCAUACAAAGGCUGUGGC........AUGAUAGCGUUGAAGCGAG.AGGUUGCUGCC......................CAC...........AAA.............
env-827   GGCAAAGGCUGACGAAGCACUGACGAAGAAAAUUGAAAAAAAAUACUUGCAUUUAUUUGGAAUGUGUGCUAUCAUACAAAGGCUGUGGC........AUGAUAGCGUUGAAGCGAG.AGGUUGCUGCC......................CAC...........AAA.............
Ccl-3-1   ACAUAACUCCUAACUAAGGCCGUAAAGGAAAAUUGAAAAAAAAUACUUGCAUUUAUUUGGAAUGUGUGCUAUCAUACAAAGGCUGUGGC........AUGAUAGCGUUGAAGCGAG.AGGUUGCUGCC......................CAC...........AAG.............
Ccl-1-1   CAUAACUCCUAACUAAGGCCGUAAAGGAAAAUUGAAAAAAAAAUACUUGCAUUUAUUUGGAAUGUGUGCUAUCAUACAAAGGCUGUGGC........AUGAUAGCGUUGAAGCGAG.AGGUUGCUGCC......................CAC...........AAG.............
Lba-16-1  CUAACAGAUCAAGGAGAAAAAAUAACAAAAAUUUUGAAAAACUACUUGAAUUUUCCGGAGCUAUUCGUUAUACUAAAAAAGCUGUGGC........AUGAUAGCGAUGAAGCGUG.AGGUUGCUGCCAAG...................UGCAUG........AAC.............
Csp-3-1   UUUUCCGGGUUUGUGAAAAAAAAUAUGAAAAAAUCGAAAAAACCACUUGAAUUUUCCAGAACUAUUCGUUAUACUAAAAAAGCUGUGAC........AUGAUAGCGAUGAAGCGUG.AGGUUGCUGCCAAG...................UGCAUG........AAC.............
env-828   AAAUGAUAUUGUAUUGAAAAAAAAUACAAUUUUUCAUAAAAAAUAGCUUGAAAUACAAUGUUAUUUCCUAUAUACUAAAAAAGCUGUGAC........AUGAUAGCUGUGAAGCGUG.AGGUUGCUGCC......................CGGC..........AAAAAGAUAA......
env-829   AAAUGAUAUUGUAUUGAAAAAAAAUACAAUUUUUCAUAAAAAAUAGCUUGAAAUACAAUGUUAUUUCCUAUAUACUAAAAAAGCUGUGAC........AUGAUAGCUGUGAAGCGUG.AGGUUGCUGCC......................CGGC..........AAAAAGAUAA......
env-830   AAAUGAUAUUGUAUUGAAAAAAAAUGCAAUUUUUCAUAAAAAAUAGCUUGAAAUACAAUGUUAUUUCCUAUAUACUAAAAAAGCUGUGAC........AUGAUAGCUGUGAAGCGUG.AGGUUGCUGCC......................CGGC..........AAAAAGAUAA......
env-831   CUAUCUGAUUGUACAUAAAAAAAAUACAAUUUUUCAUAAAAAAUAGCUUGAAAUACAAUGUUAUUUCUUAUAUACUAAAAAAGCUGUGAC........AUGAUAGCUGUGAAGCGUG.AGGUUGCUGCC......................CGGC..........AAAAAGAUAA......
env-832   AAAUGAUAUUGUAUUGAAAAAAAAUACAAUUUUUCAUAAAAAAUAGCUUGAAAUACAAUGUUAUUUCCUAUAUACUAAAAAAGCUGUGAC........AUGAUAGCUGUGAAGCGUG.AGGUUGCUGCC......................CGGC..........AAAAAGAUAA......

          ................................................................................<<.<<<<<.......................<<<.<.<<<...<<<<<.......................<<<<<<<<<<<<<<............>>>>
          ................................................................................21..1111.......................222.2.012...12222.......................1111111.10..00............00..
          •••••••••••••••R••••••••••R••••••••RAAAAR•••YU••••••••••••••••••••••••••••••••••GYUGUGRC-------∘UGAYAGC••UGAAGCG∘G-AGGUUGCYRY••∘∘∘∘∘----------------------------------------------
                                                                                                   5′
```

```
env-833   AAAUGAUAUUGUAUUGAAAAAAAUACAAUUUUUCAUAAAAAAUAGCUUGAAAUACAAUGUUAUUUCCUAUAUACUAAAAAAGCUGUGAC........AUGAUAGCUGUGAAGCGUG.AGGUUGCUGCC......................CGGC..........AAAAAGAUAA......
env-837   CUAUCUGAUUGUACAUAAAAAAAAUACAAUUUUUCAUAAAAAAUAGCUUGAAAUACAAUGUUAUUUCCUAUAUACUAAAAAAGCUGUGAC........AUGAUAGCUGUGAAGCGUG.AGGUUGCUGCC......................CGGC..........AAAAAGAUAA......
env-838   CUAUCUGAUUGUACAUAAAAAAAAUACAAUUUUUCAUAAAAAAUAGCUUGAAAUACAAUGUUAUUUCCUAUAUACUAAAAAAGCUGUGAC........AUGAUAGCUGUGAAGCGUG.AGGUUGCUGCC......................CGGC..........AAAAAGAUAA......
env-839   ....................................UAAAAAAUAGCUUGAAAUACAAUGUUAUUUCCUAUAUACUAAAAAAGCUGUGAC........AUGAUAGCUGUGAAGCGUG.AGGUUGCUGCC......................CGGC..........AAAAAGAUAA......
env-840   CUAUCUGAUUGUACAUAAAAAAAAUACAAUUUUUCAUAAAAAAUAGCUUGAAAUACAAUGUUAUUUCUUAUAUACUAAAAAAGCUGUGAC........AUGAUAGCUGUGAAGCGUG.AGGUUGCUGCC......................CGGC..........AAAAAGAUAA......
env-841   AAAUGAUAUUGUAUUGAAAAAAAUACAAUUUUUCAUAAAAAAUAGCUUGAAAUACAAUGUUAUUUCCUAUAUACUAAAAAAGCUGUGAC........AUGAUAGCUGUGAAGCGUG.AGGUUGCUGCC......................CGGC..........AAAAAGAUAA......
Lba-7-1   CUAUCUGAUUGUACAUAAAAAAAAUACAAUUUUUCAUAAAAAAUAGCUUGAAAUACAAUGUUAUUUCUUAUAUACUAAAAAAGCUGUGAC........AUGAUAGCUGUGAAGCGUG.AGGUUGCUGCC......................CGGC..........AAAAAGAUAA......
env-842   AUGAUAUUGUAUUGAAAAAAAUACAAUUUUUCAUAAAAAAUAGCUUGAAAUACAAUGUUAUUUCCUAUAUACUAAAAAAGCUGUGAC........AUGAUAGCUGUGAAGCGUG.AGGUUGCUGCC......................CGGC..........AAAAAGAUAA......
env-843   GAUGGGAAAAAAUCAAAAAAAAUUAUAAAAUAAAGAAAAAAAUGCUUGCAAUGUAAAAAAAAGUGUGCUAUUCUGUAAAGGCUGUGGC........AUGAUAGCUGUGAAGCGUG.AGGUUGCUGUC......................UGAAUUCACCGAACGAAA........GUGA
env-844   UAUAACCAAGCCGCAAAAAUAUUUUAAAAAAUUUUAAAAAGUACUUGCAUUUUCCUAAGAUAUAUAGUAACAUAUCAUUUGCUGUGAC.......AUUGAUAGCGUAGAAGCGUG.AGGUUGCUGCAUUGAU..................................
env-845   UAUAACCAAGCCGCAAAAAUAUUUUAAAAAAUUUUAAAAAGUACUUGCAUUUUCCUAAAAUAUAUAGUAACAUAUCAUUUGCUGUGAC.......AUUGAUAGCGUAGAAGCGUG.AGGUUGCUGCAUUGAU..................................
env-848   UAUAACCAAGCCGCAAAAAUAUUUUAAAAAAUUUUAAAAAGUACUUGCAUUUUCCUAAAAUAUAUAGUAACAUAUCAUUUGCUGUGAC.......AUUGAUAGCGUAGAAGCGUG.AGGUUGCUGCAUUGAU..................................
env-849   UAUAACCAAGCCGCAAAAAUAUUUUAAAAAAUUUUAAAAAGUACUUGCAUUUUCCUAAGAUAUAUAGUAACAUAUCAUUUGCUGUGAC.......AUUGAUAGCGUAGAAGCGUG.AGGUUGCUGCAUUGAU..................................
env-850   UAUAACCAAGCCGCAAAAAUAUUUUAAAAAAUUUUAAAAAGUACUUGCAUUUUCCUAAGAUAUAUAGUAACAUAUCAUUUGCUGUGAC.......AUUGAUAGCGUAGAAGCGUG.AGGUUGCUGCAUUGAU..................................
env-851   GUGGAUGGGGCGGCGCGAAAAUGCAAAAAACUGAAAAAAAAUCCUUGCAAUUCGCGGUUUCAUCGUGUAUACUAUGCAUUGCUGUGAC.......AUUGAUAGCGUUGAAGCGUG.AGGUUGCUGCAGAGCAU.................................
env-853   AGUGGAUGGGGCGGCGCGAAAAUGCAAAAAACUGAAAAAAAAUCCUUGCAAUUCGCGGUUUCAUCGUGUAUACUAUGCAUUGCUGUGAC.......AUUGAUAGCGUUGAAGCGUG.AGGUUGCUGCAGAGCAU.................................
env-854   AGUGGAUGGGGCGGCGCGAAAAUGCAAAAAACUGAAAAAAAAUCCUUGCAAUUCGCGGUUUCAUCGUGUAUACUAUGCAUUGCUGUGAC.......AUUGAUAGCGUUGAAGCGUG.AGGUUGCUGCAGAGCAU.................................
env-855   AGCGGAUGGGGCGGCGCGAAAAUGCAAAAAACUGAAAAAAAAUCCUUGCAAUUCGCGGUUUCAUCGUGUAUACUAUGCAUUGCUGUGAC.......AUUGAUAGCGUUGAAGCGUG.AGGUUGCUGCAGAGCAU.................................
env-856   AGUGGAUGGGGCGGCGCGAAAAUGCAAAAAACUGAAAAAAAAUCCUUGCAAUUCGCGGUUUCAUCGUGUAUACUAUGCAUUGCUGUGAC.......AUUGAUAGCGUUGAAGCGUG.AGGUUGCUGCAGAGCAU.................................
env-861   AGUGGAUGGGGCGGCGCGAAAAUGCAAAAAACUGAAAAAAAAUCCUUGCAAUUCGCGGUUUCAUCGUGUAUACUAUGCAUUGCUGUGAC.......AUUGAUAGCGUUGAAGCGUG.AGGUUGCUGCAGAGCAU.................................
env-862   GUGGAUGGGGCGGCGCGAAAAUGCAAAAAACUGAAAAAAAAAUCCUUGCAAUUCGCGGUUUCAUCGUGUAUACUAUGCAUUGCUGUGAC.......AUUGAUAGCGUUGAAGCGUG.AGGUUGCUGCAGAGCAU.................................
env-863   UUCCGAUAUCCUGUAAAAAAAAAUGAAUUUUUUUUCAGAAACCCCUUGCUUUUGCCAGGGGUUUCGUUUAUAAUAACUCAUGUUGUGAC.......AUUGAUAGCGAUGAAGCGUG.AGGUUGCUGCAGAGAU..................................
env-864   AAAAUAAGAAAAUAAAAAUUUUUGAAAAAAAUUUGAAAAACCCCUUGCAUUUGCGAGGGGUUUCGUUUAUAAUAACCCAUGUUGUGAC.......AUUGAUAGCGAUGAAGCGCG.AGGUUGCUGCACUAAU..................................
env-865   GAGCGGAUUAUGCAAAAAAAAAUUAAAAAAAAUUGAAAAAAACACUUGCAAUAUAAAGGAACAUCAUGUAAAAUAGCAACUGUUGUGAC.......AUUGAUAGCGUAGAAACGCG.AGGUUGCUGCAUGAAAA.................................
env-866   AGAGCGGAUUAUGCAAAAAAAAAAUAAAAAAAAUUAAAAAAAAACACUUGCAAUAUAAAGGAACAUCAUGUAAAAUAGCAACUGUUGUGAC.......AUUGAUAGCGUAGAAACGCG.AGGUUGCUGCAUGAAAA.................................
env-867   GAGCGGAUUAUGCAAAAAAAAAUUAAAAAAAAUUAAAAAAAAACACUUGCAAUAUAAAGGAACAUCAUGUAAAAUAGCAACUGUUGUGAC.......AUUGAUAGCGUAGAAACGCG.AGGUUGCUGCAUGAAAA.................................
env-868   GAGCGGAUUAUGCAAAAAAAAAUUAAAAAAAAAUUGAAAAAAACACUUGCAAUAUAAAGGAACAUCAUGUAAAAUAGCAACUGUUGUGAC.......AUUGAUAGCGUAGAAACGCG.AGGUUGCUGCAUGAAAA.................................
env-872   AGAGCGGAUUAUGCAAAAAAAAAUUAAAAAAAAUUGAAAAAAACACUUGCAAUAUAAAGGAACAUCAUGUAAAAUAGCAACUGUUGUGAC.......AUUGAUAGCGUAGAAACGCG.AGGUUGCUGCAUGAAAA.................................
env-874   AGAGCGGAUUAUGCAAAAAAAAAUUAAAAAAAAUUGAAAAAAACACUUGCAAAAUAAAGGAACAUCAUGUAAAAUAGCAACUGUUGUGAC.......AUUGAUAGCGUAGAAACGCG.AGGUUGCUGCAUGAAAA.................................
env-875   GAGAGCGGAUUAUGCAAAAAAAAAUUAAAAAAAAUUGAAAAAAACACUUGCAAUAUAAAGGAACAUCAUGUAAAAUAGCAACUGUUGUGAC.......AUUGAUAGCGUAGAAACGCG.AGGUUGCUGCAUGAAAA.................................
env-877   GAGCGGAUUAUGCAAAAAAAAAUUAAAAAAAAAUUGAAAAAAACACUUGCAAUAUAAAGGAACAUCAUGUAAAAUAGCAACUGUUGUGAC.......AUUGAUAGCGUAGAAACGCG.AGGUUGCUGCAUGAAAA.................................
env-878   AGAGCGGAUUAUGCAAAAAAAAAUUAAAAAAAAUUAAAAAAAAACACUUGCAAUAUAAAGGAACAUCAUGUAAAAUAGCAACUGUUGUGAC.......AUUGAUAGCGUAGAAACGCG.AGGUUGCUGCAUGAAAA.................................
env-879   AGAGCGGAUUAUGCAAAAAAAAAUUAAAAAAAAUUGAAAAAAACACUUGCAAUAUAAAGGAACAUCAUGUAAAAUAGCAAGUGUUGUGAC.......AUUGAUAGCGUAGAAACGCG.AGGUUGCUGCAUGAAAA.................................
env-880   ..............................................................................CAACUGUUGUGAC.......AUUGAUAGCGUAGAAACGCG.AGGUUGCUGCAUGAAAA.................................
env-892   AGAGCGGAUUAUGCAAAAAAAAAUUAAAAAAAAUUGAAAAAAACACUUGCAAUAUAAAGGAACAUCAUGUAAAAUAGCAACUGUUGUGAC.......AUUGAUAGCGUAGAAACGCG.AGGUUGCUGCAUGAAAA.................................
env-893   GAGCGGAUUAUGCAAAAAAAAAUUAAAAAAAAAUUGAAAAAAACACUUGCAAUAUAAAGGAACAUCAUGUAAAAUAGCAACUGUUGUGAC.......AUUGAUAGCGUAGAAACGCG.AGGUUGCUGCAUGAAAA.................................
env-898   AGCGGAUUAUGCAAAAAAAAAAUUAAAAAAAAAUUAAAAAAAAACACUUGCAAUAUAAAGGAACAUCAUGUAAAAUAGCAACUGUUGUGAC.......AUUGAUAGCGUAGAAACGCG.AGGUUGCUGCAUGAAAA.................................
env-899   GAGCGGAUUAUGCAAAAAAAAAUUAAAAAAAAAUUGAAAAAAACACUUGCAAUAUAAAGGAACAUCAUGUAAAAUAGCAACUGUUGUGAC.......AUUGAUAGCGUAGAAACGCG.AGGUUGCUGCAUGAAAA.................................
env-900   GAGCGGAUUAUGCAAAAAAAAAUUAAAAAAAAAUUGAAAAAAACACUUGCAAUAUAAAGGAACAUCAUGUAAAAUAGCAACUGUUGUGAC.......AUUGAUAGCGUAGAAACGCG.AGGUUGCUGCAUGAAAA.................................
env-901   AGAGCGGAUUAUGCAAAAAAAAAUUAAAAAAAAUUGAAAAAAACACUUGCAAUAUAAAGGAACAUCAUGUAAAAUAGCAACUGUUGUGAC.......AUUGAUAGCGUAGAAACGCG.AGGUUGCUGCAUGAAAA.................................
env-902   GAGCGGAUUAUGCAAAAAAAAAUUAAAAAAAAAUUGAAAAAAACACUUGCAAUAUAAAGGAACAUCAUGUAAAAUAGCAACUGUUGUGAC.......AUUGAUAGCGUAGAAACGCG.AGGUUGCUGCAUGAAAA.................................
env-903   AGAGCGGAUUAUGCAAAAAAAAAUUAAAAAAAAUUGAAAAAAACACUUGCAAUAUAAAGGAACAUCAUGUAAAAUAGCAACUGUUGUGAC.......AUUGAUAGCGUAGAAACGCG.AGGUUGCUGCAUGAAAA.................................
env-904   AGAGCGGAUUAUGCAAAAAAAAAUUAAAAAAAAUUGAAAAAAACACUUGCAAUAUAAAGGAACAUCAUGUAAAAUAGCAACUGUUGUGAC.......AUUGAUAGCGUAGAAACGCG.AGGUUGCUGCAUGAAAA.................................
env-906   AGAGAGGAUUAUGCAAAAAAAAAUUAAAAAAAAUUGAAAAAAACACUUGCAAUAUAAAGGAACAUCAUGUAAAAUAGCAACUGUUGUGAC.......AUUGAUAGCGUAGAAACGCG.AGGUUGCUGCAUGAAAA.................................
Rin-2-1   GAGAGAGGAUUAUGCAAAAAAAAAUUAAAAAAAUUGAAAAAAACACUUGCAAUAUAAAGGAACAUCAUGUAAAAUAGCAACUGUUGUGAC.......AUUGAUAGCGUAGAAACGCG.AGGUUGCUGCAUGAAAA.................................
env-907   AGAGAGGAUUAUGCAAAAAAAAAUUAAAAAAAAUUGAAAAAAACACUUGCAAUAUAAAGGAACAUCAUGUAAAAUAGCAACUGUUGUGAC.......AUUGAUAGCGUAGAAACGCG.AGGUUGCUGCAUGAAAA.................................
env-913   GAGCGGAUUAUGUGAAAAAAAAUUAAAAAAAAAUUGAAAAAAACACUUGCAAUAUAAAGGAACAUCAUGUAAAAUAGCUACUGUUGUGAC.......AUUGAUAGCGUAGAAACGUG.AGGUUGCUGCAUUAGAU.................................
env-914   GAGCGGAUUAUGCAAAAAAAAAUUAAAAAAAAUUAAAAAAAAACACUUGCAAUAUAAAGGAACAUCAUGUAAAAUAGCAACUGUUGUGAC.......AUUGAUAGCGUAGAAACGUG.AGGUUGCUGCAUUAGAU.................................
env-915   GAGCGGAUUAUGUGAAAAAAAAUUAAAAAAAAAUUGAAAAAAACACUUGCAAUAUAAAGGAACAUCAUGUAAAAUAGCUAAUGUUGUGAC.......AUUGAUAGCGUAGAAACGUG.AGGUUGCUGCAUUAGAU.................................
env-916   GAGCGGAUUAUGUGAAAAAAAAUUAAAAAAAAAUUGAAAAAAACACUUGCAAUAUAAAGGAACAUCAUGUAAAAUAGCUACUGUUGUGAC.......AUUGAUAGCGUAGAAACGUG.AGGUUGCUGCAUUAGAU.................................
env-918   GACUGAUAUUGUAUGGAAAAAAAUACAAUAUUUCCCCAAAAAGGCUUGCAAAAUGCUUUUCCAUCAGAUAUACUAAGAGGGCUGUGAC.......AUUGAUAGCGUAGAAGCGCG.AGGUUGCUACUUAUAUAU................................
Lba-10-1  GACUGAUAUUGUAUGGAAAAAAAUACAAUAUUUCCCCGAAAAGGCUUGCAAAAUACUUUCCCAUCAGAUAUACUAAGAGGGCUGUGAC.......AUUGAUAGCGUAGAAGCGCG.AGGUUGCUACUUAUAUAU................................
[sc-1-1   GACUGAUAUUGUAUGGAAAAAAAUACAAUAUUUCCCCGAAAAGGCUUGCAAAAUGCUUUUCCAUCAGAUAUACUAAGAGGGCUGUGAC.......AUUGAUAGCGUAGAAGCGCG.AGGUUGCUACUUAUAUAU................................
env-921   GACUGAUAUUGUAUGAAAAAAAAUACAAUAUUUCCCCGAAAAGGCUUGCAAAAUACUUUCCCAUCAGAUAUACUAAGAGGGCUGUGAC.......AUUGAUAGCGUAGAAGCGCG.AGGUUGCUACUUAUAUAU................................
env-923   GACUGAUAUUGUAUGGAAAAAAAUACAAUAUUUCCCCAAAAAGGCUUGCAAAAUGCUUUUCCAUCAGAUAUACUAAGAGGGCUGUGAC.......AUUGAUAGCGUAGAAGCGCG.AGGUUGCUACUUAUAUAU................................
Csc-1-1   GACUGAUAUUGUAUGAAAAAAAAUACAAUAUUUCCCCGAAAAGGCUUGCAAAAUACUUUCCCAUCAGAUAUACUAAGAGGGCUGUGAC.......AUUGAUAGCGUAGAAGCGCG.AGGUUGCUACUUAUAUAU................................
Cba-6-1   GACUGAUAUUGUAUGGAAAAAAAUACAAUAUUUCCCCGAAAAGGCUUGCAAAAUGCUUUUCCAUCAGAUAUACUAAGAGGGCUGUGAC.......AUUGAUAGCGUAGAAGCGCG.AGGUUGCUACUUAUAUAU................................
Bhy-1-1   UGUAUCCUUGCUUUUUCUGCUCAUGAAAAAAAACAAAAAAAAGCUUGCAUUUUUUAGAAACGUCCUGUAUACUCAUAGAUGCUGUGAC........AUGAUAGCUGUGAAGCGUG.AGGUUGCUGUCCACUGU..................................
env-924   AAACGUAGAAAAAGUAAAAAAUACAAAAAAAAUAAAAAAAGUCCUUGAAAAGCCUGUAAAAUCUUGAUACAAUAGCAAAUGCUGUGAC.......AUUGAUAGCUGUGAAGCGUG.AGGUUGCUGUCAUCUGGUAACGAAU.........................
env-926   AAUCGUAGAAAAAUAAAAAAAGAUGAAAAAAACGAAAAAAAACCUUGAAAAGCCUGUAAAAUCUUGCUACAAUAGCAAAUGCUGUGAC.......AUUGAUAGCUGUGAAGCGCG.AGGUUGCUGUCAUCUGGUAACGAAU.........................
Bcr-1-1   CAUUAUUUUUUUCUAUUGUAUUAAAAAAAAGUGCAAUAAAAAAUAAUUGAUUUUUUUCUGUAUCUACUAUAUACUAAUCAUUGCUGUGAC........AUGAUAGCUGUGAAGCGUG.AGGUUGCUGUCGUAA.....................................
env-928   ACAAUAUUUUUUCUAUUGUAUUAAAAAAAAGUGCAAUAAAAAAUAAUUGAUUUUUUUCUGUAUCUACUAUAUACUAAUCAUUGCUGUGAC........AUGAUAGCUGUGAAGCGUG.AGGUUGCUGUCGUAA.....................................
env-930   ACAAUAUUUUUUCUAUUGUAUUAAAAAAAAGUGCAAUAAAAAAUAAUUGAUUUUUUUCUGUAUCUACUAUAUACUAAUCAUUGCUGUGAC........AUGAUAGCUGUGAAGCGUG.AGGUUGCUGUCGUAA.....................................
env-932   ACAAUAUUUUUUCUAUUGUAUUAAAAAAAAGUGCAAUAAAAAAUAAUUGAUUUUUUUCUGCAUCUACUAUAUACUAAUCAUUGCUGUGAC........AUGAUAGCUGUGAAGCGUG.AGGUUGCUGUCGUAA.....................................
env-933   ACAAUAUUUUUUCUAUUGUAUUAAAAAAAAAUGCAAUAAAAAAUAAUUGAUUUUUUUCUGUAUCUACUAUAUACUAAUCAUUGCUGUGAC........AUGAUAGCUGUGAAGCGUG.AGGUUGCUGUCGUAA.....................................
env-934   AAGAAACCCGCAAAAAAAAUAUGAAAGAAAAAUGGAAAAAAUACUUGCAUUUCACAAAAUUAUCGUAUAUACUAAGCAGUGCUGUGAC.......AUUGAUAGCGUAGAAGCGUG.AGGUUGCUGCCGAUAU..................................
env-935   AAAGAAACCCGCAAAAAAAUAUGAAAGAAAAAUGGAAAAAAUACUUGCAUUUCACAAAAUUAUCGUGUAUACUAAGCAAUGCUGUGAC.......AUUGAUAGCGUAGAAGCGUG.AGGUUGCUGCCGAUAU..................................
env-940   AAAGAAACCCGCAAAAAAAUAUGAAAGAAAAAUGGAAAAAAUACUUGCAUUUCACAAAAUUAUCGUGUAUACUAAGCAAUGCUGUGAC.......AUUGAUAGCGUAGAAGCGUG.AGGUUGCUGCCGAUAU..................................
env-942   AAAGAAACCCGCAAAAAAAUAUGAAAGAAAAAUGGAAAAAAUACUUGCAUUUCUCAAAAUUAUCGUAUAUACUAAGCAGUGCUGUGAC.......AUUGAUAGCGUAGAAGCGUG.AGGUUGCUGCCGAUAU..................................
env-943   AGUAUCUUCUAUAAUAGAAGAAACACAAAAAGUGAAAAAAGUACUUGCAUUUCUGCAAAUCAUCGUGUAUACUAAGAAUUGCUGUGAC.......AUUGAUAGCGUAGAAGCGUG.AGGUUGCUGCCGAUAU..................................
env-944   AGUAUCUUCUAUAAUAUAAGAAAUACAAAAAGUGAAAAAAGUACUUGCAUUUCUGCAAAUCAUCGUGUAUACUAAGCAUUGCUGUGAC.......AUUGAUAGCGUAGAAGCGUG.AGGUUGCUGCCGAUAU..................................
env-945   AGUAUCUUCUAUAAUAGAAGAAACACAAAAAGUGAAAAAAGUACUUGCAUUUCUGCAGAUCAUCGUGUAUACUAAGCAUUGCUGUGAC.......AUUGAUAGCGUAGAAGCGUG.AGGUUGCUGCCGAUAU..................................
env-946   AAAGAAACCCGCAAAAAAAUAUGAAAGAAAAAUGGAAAAAAUACUUGCAUUUCUCAAAAUUAUCGUAUAUACUAAGCAGUGCUGUGAC.......AUUGAUAGCGUAGAAGCGUG.AGGUUGCUGCCGAUAU..................................
env-949   AGUAUCUUCUAUAAUAGAAGAAACACAAAAAGUGAAAAAAGUACUUGCAUUUCUGUAAAUCAUCGUGUAUACUAAGCAUUGCUGUGAC.......AUUGAUAGCGUAGAAGCGUG.AGGUUGCUGCCGAUAU..................................
env-950   AAAGAAACCCGCAAAAAAAUAUGAAAGAAAAAUGGAAAAAAUACUUGCAUUUCUCAAAAUUAUCGUAUAUACUAAGCAGUGCUGUGAC.......AUUGAUAGCGUAGAAGCGUG.AGGUUGCUGCCGAUAU..................................
Rin-1-1   AAAGAAACCCGCAAAAAAAUAUGAAAGAAAAAUGGAAAAAAUACUUGCAUUUCUCAAAAUUAUCGUAUAUACUAAGCAGUGCUGUGAC.......AUUGAUAGCGUAGAAGCGUG.AGGUUGCUGCCGAUAU..................................
env-951   CAAAGAAACCCGCAAAAAAAUAUGAAAGAAAAAUGGAAAAAAUACUUGCAUUUCACAAAAUUAUCGUGUAUACUAAGCAGUGCUGUGAC.......AUUGAUAGCGUAGAAGCGUG.AGGUUGCUGCCGAUAU..................................
env-953   AAAGAAACCCGCAAAAAAAUAUGAAAGAAAAAUGGAAAAAAUACUUGCAUUUCUCAAAAUUAUCGUAUAUACUAAGCAGUGCUGUGAC.......AUUGAUAGCGUAGAAGCGUG.AGGUUGCUGCCGAUAU..................................
env-954   AAAGAAACACGCAAAAAAAUAUGAAAGAAAAAUGGAAAAAAUACUUGCAUUUCUCAAAAUUAUCGUAUAUACUAAGCAGUGCUGUGAC.......AUUGAUAGCGUAGAAGCGUG.AGGUUGCUGCCGAUAU..................................
env-955   AAAGAAACCCGCAAAAAAAUAUGAAAGAAAAAUGGAAAAAAUACUUGCAUUUCACAAAAUUAUCGUAUAUACUAAGCAGUGCUGUGAC.......AUUGAUAGCGUAGAAGCGUG.AGGUUGCUGCCGAUAU..................................
env-959   UUUUGUAGAAAAAUGAAAAAAGUUGUGAAAAACUGAAAAAAGGCUUGCAAAACCUUUUGAGUCGGUGUAUAAUAGCAACUGCUGUGAC.......AUUGAUAGCUGUGAAGCGUG.AGGUUGCUGCCCUGAU..................................
env-960   AAGAAUCUUAAGAAAAAACAAAAGAAAAUAAAGCGAUAUUUCGCUUGCAAAGAUGCUUUUCCUGUGCUAAAUUAUCCCUUGCUGUGAC.......AUUGAUAGCGGUGAAGCGUG.AGGUUGCUGCCCGCCCGU................................
env-961   AAGAAUAUUAAGGAAAAACAAAAGAAAAUUAAGUGAUAUUUCGCUUGCAAUGCCGGGGCGGCUGUGCUAAAUUAUCCCUUGCUGUGAC.......AUUGAUAGCGGUGAAGCGUG.AGGUUGCUGCCCGCCCGU................................

          ..................................................................................<<.<<<<<.....................<<<.<.<<<...<<<<<......................<<<<<<<<<<<<<<............>>>>
          ..................................................................................21..1111.....................222.2.012...12222......................1111111.10..00............00..
          ●●●●●●●●●●●●●●R●●●●●●●●●●R●●●●●●●●RAAAAR●●●YU●●●●●●●●●●●●●●●●●●●●●●●●●●●●●●●●●●GYUGUGRC-------○UGAYAGC●●UGAAGCG○G-AGGUUGCYRY●●○○○○---------------------------------------
                                                                                                         5′
```

```
env-962   AAAUGGAUUGUAUACAAAAAAAAUACAAUUUUUCUGAAAAAAGUAUUGCAAAAGCGUAACCUGUCGUGUAUACUAAUCCUUGCUGUGAC......AUUGAUAGCGUAGAAGCGUG.AGGUUGCUGCCCGCUGU.............................................
env-963   AAAUGGAUUGUAUACAAAAAAAAUACAAUUUUUCUGAAAAAAGUAUUGCAAAAGCGAAACCUGUCGUGUAUACUAAUCCUUGCUGUGAC......AUUGAUAGCGUAGAAGCGUG.AGGUUGCUGCCCGCUGU.............................................
env-964   AAAUGGAUUGUAUACAAAAAAAAUACAAUUUUUCUGAAAAAAGUAUUGCAAAAGCGAAACCUGUCGUGUAUACUAAUCCUUGCUGUGAC......AUUGAUAGCGUAGAAGCGUG.AGGUUGCUGCCCGCUGU.............................................
env-965   CAUUUUAUAAAAACACAAAAAUACAUAAAAAAUAUUUAGAAACCCCUUGCGUUUGCAAGGGGUUUCGUGUAUAAUAGCACUGUUGUGAC......AUUGAUAGCGUUGAAGCGUG.AGGUUGCUACAUAUUAA.............................................
env-967   CAUUUUAUAAAAAUACAAAAAUACAUAAAAAAUAUUUAGAAACCCCUUGCGUUUGCAAGGGGUUUCGUGUAUAAUAGCACUGUUGUGAC......AUUGAUAGCAUUGAAGCGUG.AGGUUGCUACAUAUUAG.............................................
env-968   UCCUGUUUUUUCUUGAAAAAUAUAGAUUUUUUCUUAAAAAAACACUUGCAUUUCCCGGAAAGCUGUGUUAUUAUAACGAAGCUGUGGC.......AUGAUAGCGAUGAAACGAG.AGGUUGCUGCUUAUGAUA............................................
env-970   UCCUGUUUUUUCUUGAAAAAUAUAGAUUUUUUCUUAAAAAAACACUUGCAUUUCCCAGAAAGCUGUGUUAUUAUAACGAAGCUGUGGC.......AUGAUAGCGAUGAAACGAG.AGGUUGCUGCUUAUGAUA............................................
env-971   UCCUGUUUUUUCUUGAAAAAUAUAGAUUUUUUCUUAAAAAAACACUUGCAUUUCCCAGAAAGCUGUGUUAUUAUAACGAAGCUGUGGC.......AUGAUAGCGAUGAAACGAG.AGGUUGCUGCUUAUGAUA............................................
env-972   NNNNNNNNNNNNNNNNNNNNNNNNNNNNNNNNNNNNNNNNNNNNNNNNNNNNNNNNNNNNNNNNNNNNNNNNNNNNNNNNGCUGUGGC.......AUGAUAGCGAUGAAACGAG.AGGUUGCUGCUUAUGAUA............................................
env-973   UCCUGUUUUUUCUUGAAAAAUAUAGAUUUUUUCUUAAAAAAACACUUGCAUUUCCCAGAAAGCUGUGUUAUUAUAACGAAGCUGUGGC.......AUGAUAGCGAUGAAACGAG.AGGUUGCUGCUUAUGAUA............................................
Lba-3-1   AAAAUAAAUUGUAUUGAAAAAAGUACAAUAUUUUUCAAAAAAGACUUGAAAAAAGCAGAAUGGUUUAGUAUACUUAAAAAGCUGUGAC.......AUGAUAGCGAUGAAACGUG.AGGUUGCUACUUAUAAU.............................................
env-980   AUUUUCUCUUGUAAACGAACUGGCAGAAAAAAAUAAAAAAAAGUCUUGAAAAAUGAAAAGAAAUCAUGUAUACUAAAAAAGCUGUGAC.......AUGAUAGCGUUGAAGCGUG.AGGUUGCUACUUAUAAU.............................................
env-981   AUUUUCUCUUGUAAACGAACUGGCAGAAAAAAAUAAAAAAAAGUCUUGAAAAAUGAAAAGAAAUCAUGUAUACUAAAAAAGCUGUGAC.......AUGAUAGCGUUGAAGCGUG.AGGUUGCUACUUAUAAU.............................................
env-983   AUUUUCUCUUGUAAACGAACUGGCAGAAAAAAAUAAAAAAAAGUCUUGAAAAAUGAAAAGAAAUCAUGUAUACUAAAAAAGCUGUGAC.......AUGAUAGCGUUGAAGCGUG.AGGUUGCUACUUAUAAU.............................................
env-984   AAUUUUCUCUUGUAAACGAACUGGCAGAAAAAAAUAAAAAAAGUCUUGAAAAAUGAAAAGAAAUCAUGUAUAAUAAAAAAGCUGUGAC.......AUGAUAGCGUUGAAGCGUG.AGGUUGCUACUUAUAAU.............................................
env-985   ACAUUAAAAAAUACAAUAAAAAAAUCGAAAAAAAUGUAAAAAAUAGUUGCAAAACGAAAAAGUAUCUUGUAUACUAAGAGAGCUGUGAC.......AUGAUAGCGAUGAAGCGUG.AGGUUGCUGCUUAACAU.............................................
Csp-8-1   ACAUUAAAAAAUACAAUGAAAAAUCGAAAAAAAUGUAAAAAAUAGUUGCAAAACUAAAAAGUAUCUUGUAUACUAAGAGAGCUGUGAC.......AUGAUAGCGAUGAAGCGUG.AGGUUGCUGCUUAACAU.............................................
env-986   ACAUUAAAAAAUACAAUAAAAAAAUCGAAAAAAAUGUAAAAAAUAGUUGCAAAACUAAAAAGUAUCUUGUAUACUAAGAGAGCUGUGAC.......AUGAUAGCGAUGAAGCGUG.AGGUUGCUGCUUAACAU.............................................
Csp-9-1   ACAUUAAAAAAUACAAUAAGAAAUCGAAAAAAAUGUAAAAAAUAGUUGCAAAACUAAAAAGUAUCUUGUAUACUAAGAGAGCUGUGAC.......AUGAUAGCGAUGAAGCGUG.AGGUUGCUGCUUAACAU.............................................
env-987   ACAUUAAAAAAUACAAUAAAAAAAUCGAAAAAAAUGUAAAAAAUAGUUGCAAAACUAAAAAGUAUCUUGUAUACUAAGAGAGCUGUGAC.......AUGAUAGCGAUGAAGCGUG.AGGUUGCUGCUUAACAU.............................................
env-991   GAAAGUACAAAAUAAAUUGUAUGAAUAAAAAAACAAAAAAAGUACUUGCAAAAUGUAUGGGAAUCGGAUAUACUAAGAGAGCUGUGAC.......AUGAUAGCGUUGAAGCGUG.AGGUUGCUGCGUGGAAUU............................................
env-992   GAAAGUACAAAAUAAAUUGUAUGAAUAAAAAAACAAAAAAAGUACUUGCAAAAUGUAUGGGAAUCGGAUAUACUAAGAGAGCUGUGAC.......AUGAUAGCGUUGAAGCGUG.AGGUUGCUGCUAGGAAUU............................................
env-993   GAAAGUACAAAAUAAAUUGUAUGAAUAAAAAAACAAAAAAAGUACUUGCAAAAUGUAUGGAAAUCGGAUAUACUAAGAGAGCUGUGAC.......AUGAUAGCGUUGAAGCGUG.AGGUUGCUGCUAGGAAUU............................................
Dlo-2-1   GGAAAGUACAAAAUAAAUUGUAUGAAUAAAAAAACAAAAAAGUACUUGCAAAAUGUAUGGGAAUCGGAUAUACUAAGAGAGCUGUGAC.......AUGAUAGCGUUGAAGCGUG.AGGUUGCUGCUAGGAAUU............................................
env-994   GAAAGUACAAAAUAAAUUGUAUGAAUAAAAAAACAAAAAAAGUACUUGCAAAAUGUAUGGGAAUCGGAUAUACUAAGAGAGCUGUGAC.......AUGAUAGCGUUGAAGCGUG.AGGUUGCUGCUAGGAAUU............................................
env-995   GAAAGUACAAAAUAAAUUGUAUGAAUAAAAAAACAAAAAAAGUACUUGCAAAAUGUAUGGGAAUCGGAUAUACUAAGAGAGCUGUGAC.......AUGAUAGCGUUGAAGCGUG.AGGUUGCUGCUAGGAAUU............................................
env-996   GAAAGUACAAAAUAAAUUGUAUGAAUAAAAAAACAAAAAAAGUACUUGCAAAAUGUAUGGGAAUCGGAUAUACUAAGAGAGCUGUGAC.......AUGAUAGCGUUGAAGCGUG.AGGUUGCUGCUAGGAAUU............................................
env-1006  AAGAAAAUCUAAAAGAUUGUACGGUAAAAAAAAACAAAAAUAGUGCUUGCAUUAUCUGCCGGAUUGGGAUAUACUAAGAGAGCUGUGAC.......AUGAUAGCGUUGAAGCGUG.AGGUUGCUGCUGGAGAUC............................................
env-1007  AAGAGAAUCUGAAAGAUUGUACGGUAAAAAAAAACAAAAAUAGUACUUGCAUUAUCUGCCGGAUUGGGAUAUACUAAGAGAGCUGUGAC.......AUGAUAGCGUUGAAGCGUG.AGGUUGCUGCUGGAGAUC............................................
env-1008  AAGAGAAUCUGAAAGAUUGUACGGUAAAAAAAAACAAAAAUAGUACUUGCAUUAUCUGCCGGAUUGGGAUAUACUAAGAGAGCUGUGAC.......AUGAUAGCGUUGAAGCGUG.AGGUUGCUGCUGGAGAUC............................................
env-1009  AAGAGAAUCUGAAAGAUUGUACGGUAAAAAAAAACAAAAAUAGUACUUGCAUUAUCUGCCGGAUUGGGAUAUACUAAGAGAGCUGUGAC.......AUGAUAGCGUUGAAGCGUG.AGGUUGCUGCUGGAGAUC............................................
env-1018  AAGAGAAUCUGAAAGAUUGUACGGUAAAAAAAAACAAAAAUAGUACUUGCAUUAUCUGCCGGAUUGGGAUAUACUAAGAGAGCUGUGAC.......AUGAUAGCGUUGAAGCGUG.AGGUUGCUGCUGGAGAUC............................................
env-1019  AAGAGAAUCUGAAAGAUUGUACGGUAAAAAAAAACAAAAAUAGUACUUGCAUUAUCUGCCGGAUUGGGAUAUACUAAGAGAGCUGUGAC.......AUGAUAGCGUUGAAGCGUG.AGGUUGCUGCUGGAGAUC............................................
Dlo-1-1   AAAAGAAUCUGAAAGAUUGUACGGUAAAAAAAAACAAAAAUAGUACUUGCAUUAUCUGCCAGAUUGGGAUAUACUAAGAGAGCUGUGAC.......AUGAUAGCGUUGAAGCGUG.AGGUUGCUGCUGGAGAUC............................................
env-1020  ACGAAAAAAAAAUACAAAAAAAAAUACAAUUUCGCAAAAAAAAAUACUUGCAAUCCCAUAGACCCUGGUUUAUACUCAAUAUUGCUGUGGC.......AUGAUAGCUAUGAAGCGUG.AGGUUGCUGCAUUCUA..............................................
env-1021  ACGAAAAAAAAAUACAAAAAAAAAUACAAUUUCGCAAAAAAAAAUACUUGCAAUCCCAUAGACCCUGGUUUAUACUCAAUAUUGCUGUGGC.......AUGAUAGCUAUGAAGCGUG.AGGUUGCUGCAUUCUA..............................................
env-1022  ACGAAAAAAAAAUACAAAAAAAAAUACAAUUUCGCAAAAAAAAAUACUUGCAAUCCCAUAGACCCUGGUUUAUACUCAAUAUUGCUGUGGC.......AUGAUAGCUAUGAAGCGUG.AGGUUGCUGCAUUCUA..............................................
Lba-2-1   UGUCAAAAAAAUCGGUUAAAAAAUUUUUGAAAAAUUAAAAAACUACUUGCAAAAUCCGGUGUGAUUCGCUAUACUAAGAGGGCUGUGAC.......AUGAUAGCUGUGAAGCGUG.AGGUUGCUGC.CGA..................GACGA........AGAA............
env-1023  AAAUAAUAAGUAAAAGAAAAAAUAAAAAAAAAUUAUAAAAAACACUUGCAAAAUAUAGAAACAUACGAUAUACUAUCAAUUGCUGUGAC.......AUGAUAGCGAUGAAACGUG.AGGUUGCUGCUUCAGAAUGA..........................................
env-1024  AAAUAAUAAGUAAAAGAAAAAAUAAAAAAAAAUUAUAAAAAACACUUGCAAAAUAUAGAAACAUACGAUAUACUAUCAAUUGCUGUGAC.......AUGAUAGCGAUGAAACGUG.AGGUUGCUGCUUCAGAAUGA..........................................
env-1026  CAGAAAAAAGUAAAAUAAAAAAUAAAAAAAAAUUAUAAAAAACACUUGCAAAAUAUAGAAACAUACGAUAUACUAUCAAUUGCUGUGAC.......AUGAUAGCGAUGAAACGUG.AGGUUGCUGCUUCAGAAUGA..........................................
env-1027  AAAUAAUAAGUAAAAGAAAAAAUAAAAAAAAAUUAUAAAAAACACUUGCAAAAUAUAGAAACAUACGAUAUACUAUCAAUUGCUGUGAC.......AUGAUAGCGAUGAAACGUG.AGGUUGCUGCUUCAGAAUGA..........................................
env-1028  AAAUAAUAAGUAAAAGAAAAAAUAAAAAAAAAUUAUAAAAAACACUUGCAAAAUAUAGAAACAUACGAUAUACUAUCAAUUGCUGUGAC.......AUGAUAGCGAUGAAACGUG.AGGUUGCUGCUUCAGAAUGA..........................................
env-1029  CAGAAAAAAGUAAAAUAAAAAAUAAAAAAAAAUUAUAAAAAACACUUGCAAAAUAUAGAAACAUACGAUAUACUAUCAAUUGCUGUGAC.......AUGAUAGCGAUGAAACGUG.AGGUUGCUGCUUCAGAAUGA..........................................
env-1030  UUAUCCACAAAUUGAAAAAACCCUUGAAAAAUAAGCAAAAAAAGCUUGAUUUUUCGAUUUCGAUGUGUUAUUCUUAUCUGGCUGUGAC.......AUGAUAGCGUUGAAGCGAG.AGGUUGCUGCCUGAGAAU............................................
env-1032  ..............................................AUUUUUCGAUUUCGAUGUGUUAUUCUUAUCUGGCUGUGAC.......AUGAUAGCGUUGAAACGAG.AGGUUGCUGCCUGAGAAU............................................
env-1033  UUAUCCACAAAUUGAAAAAACCCUUGAAAAAUAAGCAAAAAAAGCUUGAUUUUUCGAUUUCGAUGUGUUAUUCUUAUCUGGCUGUGAC.......AUGAUAGCGUUGAAACGAG.AGGUUGCUGCCUGAGAAU............................................
env-1034  UAUCCACAAAUUGAAAAAAACCCUUGAAAAAUAAGCAAAAAAAGCUUGAUUUUUCGAUUUCGAUGUGUUAUUCUUAUCUGGCUGUGAC.......AUGAUAGCGUUGAAACGAG.AGGUUGCUGCCUGAGAAU............................................
env-1035  AAUUUUGGGGCGAAAAAACGUAAAAGAAAAAAUAAAAAAAGUUCUUGCUUUUUUGGGAAAGACCAUGUAUAAUAUCAAUUGCUGUGGC.......AUGAUAGCUGUGAAACGUG.AGGUUGCUGCCUAUGU..............................................
env-1036  UCCUUUUCAUUUUUCUCUUUUUUGACAGAAAAUCAAAAAAGUUCUUGCUUUUUUGGAAAGACCAUGUAUAAUAUCAAUUGCUGUGGC.......AUGAUAGCUGUGAAACGUG.AGGUUGCUGCCUAUGU..............................................
env-1037  AAAUUUUGGGGCGAAAAAACGUAAAAGAAAAAAUAAAAAAAGUUCUUGCUUUUUUGGGAAAGACCAUGUAUAAUAUCAAUUGCUGUGGC.......AUGAUAGCUGUGAAACGUG.AGGUUGCUGCCUAUGU..............................................
env-1038  UCCUUUUCAUUUUUCUCUUUUUUGACAGAAAAUCAAAAAAGUUCUUGCUUUUUUGGGAAAGACCAUGUAUAAUAUCAAUUGCUGUGGC.......AUGAUAGCUGUGAAACGUG.AGGUUGCUGCCUAUGU..............................................
env-1039  UCCUUUUUAUUUUUCUCUUUUUUGACAGAAAAUCAAAAAAGUUCUUGCUUUUUUGGGAAAGACCAUGUAUAAUAUCAAUUGCUGUGGC.......AUGAUAGCUGUGAAACGUG.AGGUUGCUGCCUAUGU..............................................
env-1040  AAUUUUGGGGCGAAAAAACGUAAAAGAAAAAAUAAAAAAAGUUCUUGCUUUUUUGAAUAAGACCAUGUAUAAUAUCAAUUGCUGUGGC.......AUGAUAGCUGUGAAACGUG.AGGUUGCUGCCUAUGU..............................................
env-1041  UCCUUUUUAUUUUUCUCUUUUUUGACAGAAAAUCAAAAAAGUUCUUGCUUUUUUGGGAAAGACCAUGUAUAAUAUCAAUUGCUGUGGC.......AUGAUAGCUGUGAAACGUG.AGGUUGCUGCCUAUGU..............................................
env-1042  AAUUUUGGGGCGAAAAAACGUAAAAGAAAAAAUAAAAAAAGUUCUUGCUUUUUUGGAAAGACCAUGUAUAAUAUCAAUUGCUGUGGC.......AUGAUAGCUGUGAAACGUG.AGGUUGCUGCCUAUGU..............................................
env-1043  AAUUUUGGGGCGAAAAAACGUAAAAGAAAAAAUAAAAAAAGUUCUUGCUUUUUUGAAUAAGACCAUGUAUAAUAUCAAUUGCUGUGGC.......AUGAUAGCUGUGAAACGUG.AGGUUGCUGCCUAUGU..............................................
env-1044  UCCUUUUCAUUUUUCUCUUUUUUGACAGAAAAUCAAAAAAGUUCUUGCUUUUUUGGGAAAGACCAUGUAUAAUAUCAAUUGCUGUGGC.......AUGAUAGCUGUGAAACGUG.AGGUUGCUGCCUAUGU..............................................
env-1045  AAUUUUGGGGGCGAAAAAACGUAAAAGAAAAAAUAAAAAAGUUCUUGCUUUUUUGGGAAAGACCAUGUAUAAUAUCAAUUGCUGUGGC.......AUGAUAGCUGUGAAACGUG.AGGUUGCUGCCUAUGU..............................................
env-1046  AAUUUUGGGGCGAAAAAACGUAAAAGAAAAAAUAAAAAAAGUUCUUGCUUUUUUGGGAAAGACCAUGUAUAAUAUCAAUUGCUGUGGC.......AUGAUAGCUGUGAAACGUG.AGGUUGCUGCCUAUGU..............................................
env-1052  UCCUUUUCAUUUUUCUCUUUUUUGACAGAAAAUCAAAAAAGUUCUUGCUUUUUUGGGAAAGACCAUGUAUAAUAUCAAUUGCUGUGGC.......AUGAUAGCUGUGAAACGUG.AGGUUGCUGCCUAUGU..............................................
env-1053  UCCUUUUCAUUUUUCUCUUUUUUGACAGAAAAUCAAAAAAGUUCUUGCUUUUUUGGAAAGACCAUGUAUAAUAUCAAUUGCUGUGGC.......AUGAUAGCUGUGAAACGUG.AGGUUGCUGCCUAUGU..............................................
env-1054  AAUUUUGGGGCGAAAAAACGUAAAAGAAAAAAUAAAAAAAGUUCUUGCUUUUUUGGGAAAGACCAUGUAUAAUAUCAAUUGCUGUGGC.......AUGAUAGCUGUGAAACGUG.AGGUUGCUGCCUAUGU..............................................
env-1055  AAUUUUGGGGGCGAAAAACGUAAAAGAAAAAAUAAAAAAAGUUCUUGCUUUUUUGAAUAAGACCAUGUAUAAUAUCAAUUGCUGUGGC.......AUGAUAGCUGUGAAACGUG.AGGUUGCUGCCUAUGU..............................................
env-1056  AAUUUUGGGGCGAAAAAACGUAAAAGAAAAAAUAAAAAAAGUUCUUGCUUUUUUGGGAAAGACCAUGUAUAAUAUCAAUUGCUGUGGC.......AUGAUAGCUGUGAAACGUG.AGGUUGCUGCCUAUGU..............................................
env-1057  AAUUUUGGGGGCGAAAAACGUAAAAGAAAAAAUCAAAAAAGUUCUUGCUUUUUUGGGAAAGACCAUGUAUAAUAUCAAUUGCUGUGGC.......AUGAUAGCUGUGAAACGUG.AGGUUGCUGCCUAUGU..............................................
env-1058  UCCUUUUCAUUUUUCUCCUUUUUGACAGAAAAUCAAAAAAGUUCUUGCUUUUUUGGGAAAGACCAUGUAUAAUAUCAAUUGCUGUGGC.......AUGAUAGCUGUGAAACGUG.AGGUUGCUGCCUAUGU..............................................
env-1059  AAUUUUGGGGCGAAAAAACGUAAAAGAAAAAAUAAAAAAAGUUCUUGCUUUUUUGGGAAAGACCAUGUAUAAUAUCAAUUGCUGUGGC.......AUGAUAGCUGUGAAACGUG.AGGUUGCUGCCUAUGU..............................................
Cha-1-1   AUCAGCUAUCGAAUAAAAAAAAAUUAAGAAAUAAAGAAAAAAAAUGCUUGCAAUGUAAAAAAAAGUGUGCUAUCCUGUAAAGGCUGUGGC.......AUGAUAGCUGUGAAGCGUG.AGGUUGCUGCCCAAGA..............................................
env-1065  AUCAGCCAUCGAAUAAAAAAAAAUUAAGAAAUAAAGAAAAAAAAUGCUUGCAAUGUAAAAAAAAGUGUGCUAUCCUGUAAAGGCUGUGGC.......AUGAUAGCUGUGAAGCGUG.AGGUUGCUGCCCAAGA..............................................
env-1066  AUCAGCCAUCAAAGAAAAAAAAAUUAAGAAAUAAAGAAAAAAAAUGCUUGCAAUGUAAAAAAAAGUGUGCUAUCCUGUAAAGGCUGUGGC.......AUGAUAGCUGUGAAGCGUG.AGGUUGCUGCCCAAGA..............................................
Cba-1-1   UAUAGAAAAAAAAAUAAAAAAAAUUUUAUUUUUUAAGAAAAAAAGCUUGCAACUUCAGGAAAAUACCAGUAUAAUAACAAAUGCUGUGAC.......AUGAUAGCGAUGAAGCGUG.AGGUUGCUGCCUGAAAG.............................................
Cba-8-1   AUAAAAAAAAAAAUUAAAAAAAAUUUUAUUUUUUAAGAAAAAAAGCUUGCAACUUCAGGAAAAUACCAGUAUAAUAACAAAUGCUGUGAC.......AUGAUAGCGAUGAAGCGUG.AGGUUGCUGCCUGAAAG.............................................
env-1067  ...................................AAAAAGCUUGCAACUUCAGGAAAAUACCAGUAUAAUAACAAAUGCUGUGAC.......AUGAUAGCGAUGAAGCGUG.AGGUUGCUGCCUGAAAG.............................................
Lba-20-1  UUAUAAAAAAAAAUUAAAAAAAAUUUUAUUUUUUAAGAAAAAAAGCUUGCAACUUCAGGAAAAUACCAGUAUAAUAACAAAUGCUGUGAC.......AUGAUAGCGAUGAAGCGUG.AGGUUGCUGCCUGAAAG.............................................
Lba-8-1   UAUAAAAAAAAAAUUAAAAAAAAUUUUAUUUUUUAAGAAAAAAAGCUUGCAACUUCAGGAAAAUACCAGUAUAAUAACAAAUGCUGUGAC.......AUGAUAGCGAUGAAGCGUG.AGGUUGCUGCCUGAAAG.............................................
env-1068  UAUAGAAAAAAAAUUAAAAAAAAUUUUAUUUUUUAAGAAAAAAAGCUUGCAACUUCAGGAAAAUACCAGUAUAAUAACAAAUGCUGUGAC.......AUGAUAGCGAUGAAGCGUG.AGGUUGCUGCCUGAAAG.............................................
Lba-13-1  UCUGGCCAGUCCCAUUUACCGUCCAAAAAAAAUUGAAUUUACCCCUUGCAAUCUUUGAACUUUUGCAUUAUAAUAGUACCUGCUGUGGC.......AUGAUAGCAAUGAAGCGCG.AGGUUGCUGCCUGAUAU.............................................
Csp-7-1   CGGAAAUUUCUUUGAAAAAAAUGAGGAAAAACCCUGAAAAAUCCCUUGCAAUUCAAGGAAACUUAUGAUAUGAUACAGAGGCUGUGGC.......AUGAUAGCGAUGAAGCGUG.AGGUUGCUGCCUGAGAGAU...........................................
env-1069  CGCGAUUUUCUAAGGAAAAAUCGGCGAAAAAAGUAGAAAAAUCCCUUGCAAUUACAGGAAACAUAUGAUAUGAUACAGAGGCUGUGGC.......AUGAUAGCAAUGAAGCGCG.AGGUUGCUGCCUGAGAGAU...........................................
env-1070  .........CUAAGGAAAAUCGGCGAAAAAAGUAGAAAAAUCCCUUGCAAUUACAGGAAACAUAUGAUAUGAUACAGAGGCUGUGGC.......AUGAUAGCAAUGAAGCGCG.AGGUUGCUGCCUGAGAGAU...........................................
env-1071  CGCGAUUUUCUAAGGAAAAAUCGGCGAAAAAAGUAGAAAAAUCCCUUGCAAUUACAGGAAACAUAUGAUAUGAUACAGAGGCUGUGGC.......AUGAUAGCAAUGAAGCGCG.AGGUUGCUGCCUGAGAGAU...........................................
env-1072  CGCGAUUUUCUAAGGAAAAAUCGGCGAAAAAAGUGAAAAAAUCCCUUGCAAUUACAGGAAACAUAUGAUAUGAUACAGAGGCUGUGGC.......AUGAUAGCAAUGAAGCGCG.AGGUUGCUGCCUGAGAGAU...........................................

          ..............................................................................<<.<<<<<.....................<<<.<.<<<...<<<<<......................<<<<<<<<<<<<<<............>>>>
          ..............................................................................21..1111.....................222.2.012...12222......................1111111.10..00............00..
          ●●●●●●●●●●●●●●R●●●●●●●●●●R●●●●●●●●RAAAAR●●●YU●●●●●●●●●●●●●●●●●●●●●●●●●●●●●●●●●●●●GYUGUGRC-------∘UGAYAGC●●UGAAGCG∘G-AGGUUGCYRY●●∘∘∘∘--------------------------------------------
                                                                                                 5′
```

```
Esp-1-1    GAAUCACCCUUAUCACCCCUUUACAAAAUACAGGAAAAAAAUACUUGCAAUAUGGAUAAAUUUGUUUUAUAAUAAUACGUGCUGUGCG.......AUGAUAGCGAUGAAGCGUG.AGGUUGCUGCCCGAAGAAA.............................................
env-1073   AGAAAAACAGGAUAUGCACAGAUUGUAAAAAUUGCGGAAAAGUGCUUGCAUUUUAAAAGAUUUUAUGUUAUGCUGUCAGAGCUGUGCG.......AUGAUAGCUAAGAAGCGUG.AGGUUGCUGCCCUGUAAAU..............................................
env-1074   AGAAAAACAGGAUAUGCACAGAUUGUAAAAAUUGCGGAAAAGUGCUUGCAUUUUAAAAGAUUUUAUGUUAUGCUGUCAGAGCUGUGCG.......AUGAUAGCUAAGAAGCGUG.AGGUUGCUGCCCUGUAAAU..............................................
env-1075   GAAAACAGGAUAUGCACAGUAUUGUAAAAUUGCGGAAAAGUUGCUUGCUAUUUUAAAAGAUUUUAUGUUAUGCUGUCAGAGCUGUGCG.......AUGAUAGCUAAGAAGCGUG.AGGUUGCUGCCCUGUAAAU..............................................
Csa-1-1    AAACUGGGAAAAUCAGAUUUUGCGCCAUUUUUUUCAAAAAAAUGCUUGCAAUGUAAAAAAAAAUAUGCUAUCAUACAAAGGCUGUGCG.......AUGAUAGCUGUGAAGCGUG.AGGUUGCUGCUUGUAUA...............................................
env-1076   GAAAUGGAAGGGCUGCAAGUAAAUAAAAAAUAUUAAAAAAAUACUUGCAUUUUUGAAAAUACCAUAUAUAAUAGCAAGUGCUGUGCG......AUUGAUAGCUGUGAAGCGUG.AGGUUGCUGCUUACAAAUU.............................................
env-1077   GAAAUGGAAGGGCUGCAAGUAAAUAAAAAAUAUUAAAAAAAUACUUGCAUUUUUGAAAAUACCAUAUAUAAUAGCAAGUGCUGUGCG......AUUGAUAGCUGUGAAGCGUG.AGGUUGCUGCUUACAAAUU.............................................
env-1078   AAUUUUGGGGCGAAAAACGUAAAAGAAAAAAUAAAAAAAGUUCUUGCUUUUUUGAAUAAGACCAUGUAUAAUAUCAAUUGCUGUGCG.......AUGAUAGCUGUGAAGCGCG.AGGUUGCUGCUUAUAAAUU..............................................
env-1081   AAUUUGGGGGGCGAAAAACGUAAAAGAAAAAAUAAAAAAAGUUCUUGCUUUUUUGAAUAAGACCAUGUAUAAUAUCAAUUGCUGUGCG.......AUGAUAGCUGUGAAGCGCG.AGGUUGCUGCUUAUAAAUU..............................................
env-1082   AAUUUUGGGGCGAAAAAACGUAAAAGAAAAAAUAAAAAAAGUUCUUGCUUUUUUGAAUAAGACCAUGUAUAAUAUCAAUUGCUGUGCG.......AUGAUAGCUGUGAAGCGCG.AGGUUGCUGCUUAUAAAUU..............................................
env-1083   AAUUUUGGGGCGAAAAAACGUAAAAGAAAAAAUAAAAAAAGUUCUUGCUUUUUUGAAUAAGACCAUGUAUAAUAUCAAUUGCUGUGCG.......AUGAUAGCUGUGAAGCGCG.AGGUUGCUGCUUAUAAAUU..............................................
env-1084   AAUUUUGGGGCGAAAAACGUAAAAGAAAAAAUAAAAAAAGUUCUUGCUUUUUUGAAUAAGACCAUGUAUAAUAUCAAUUGCUGUGCG.......AUGAUAGCUGUGAAGCGCG.AGGUUGCUGCUUAUAAAUU..............................................
env-1085   ..............................AAAAUAAAAAAAGUUCUUGCUUUUUUGAAUAAGACCAUGUAUAAUAUCAAUUGCUGUGCG.......AUGAUAGCUGUGAAGCGCG.AGGUUGCUGCUUAUAAAUU..............................................
env-1086   AAUUUUGGGGGCGAAAAACGUAAAAGAAAAAAUAAAAAAAGUUCUUGCUUUUUUGAAUAAGACCAUGUAUAAUAUCAAUUGCUGUGCG.......AUGAUAGCUGUGAAGCGCG.AGGUUGCUGCUUAUAAAUU..............................................
Cha-2-1    GAGUGUUUUCUUUUUGCCAAGGCCAAAAUAUUUUUGAAAAAGUUCUUGCAAUUAAAAAAACUUUGUGAUAAAAAUAACAAGGCUGUGAG.......AUGAUAGCGUUGAAGCGUG.AGGUUGCCGUCCCUUGU..............................................
Lba-5-1    AAGAUGAGUUGUUAAAAGUUAUUGAAAAAAAUACAAUUUAGUACUUGCAUUAUACAAAAUUUUGUUUUAUAAUAACAAAUGCUGUGAG.......AUGAUAGCAACGAAGCGAG.AGGUUGCUGCCCAAUGGUU..............................................
Csp-1-1    UGCAUAAUUAAAACCCAAUAAUACAAAAAAAUAUGAAAAAAAUCCUUGCAAUCCCUGUAAAUGUAUGAUAUGCUAUCAAAGCUGUGCG.......AUGAUAGCUAUGAAGCGUG.AGGUUGCUGCCCUGAAACAG.............................................
env-1087   UGCAUAAUUAAAUCCCAAUAAUACAAAAAAAUAUGAAAAAAAUCCUUGCAAUCCCUGUAAAUGUAUGAUAUGCUAUCAAAGCUGUGCG.......AUGAUAGCUAUGAAGCGUG.AGGUUGCUGCCCUGAAACAG.............................................
env-1088   UUAACAAUCUUAAGAAAAAUCCUUUAAAAUCAAGAAAACAAAAACUUGCACAAAUGCAGGCCAGUUGUAUUUUUAGAGGGCUGUGAG......AUUGAUAGCGUAGAUCUGUG.AGGUUGCUGUCGUGUGU..............................................
env-1089   GGACACUGAGUCGAAUUGUUUCUAAAAAAGAGCUGUGCUUUUACUUGCGCAGCUCUUUUUUUUGUGUUAAGCUAUAUGGCGCUGUGAG.......AUGAUAGCUGUGAAGCGAG.AGGUUGCUGUCUCGAAAG..............................................
env-1093   GUUCAUCUUACAAUCAUCCAGCUAAAAAAGAGCUGCGACACUACUUGCGCAGCUCUUUUUUUUGUGGUUAAGCUAUAUGGCGCUGUGAG.......AUGAUAGCUGUGAAGCGAG.AGGUUGCUGUCUCGAAAG..............................................
env-1094   GGACAUUGAGUCGAAUUGUUUCUAAAAAAGAGCUGUGCUUUUACUUGCGCAGCUCUUUUUUUUGUGUUAAGCUAUAUGGCGCUGUGAG.......AUGAUAGCUGUGAAGCGAG.AGGUUGCUGUCUCGAAAG..............................................
env-1095   AUGCCGGAGGAGUCGGAAUCCGCUCACUAAAGCCAAAAAAAAGAGNNNNNNNNNNNNNNNNNNNNNNNNNNNNNNNNNNNNNNGCUGUGAG.......AUGAUAGCUGUGAAGCGAG.AGGUUGCUGUCUCGAAAG..............................................
env-1096   GGACACUGAGUCGAAUUGUUUCUAAAAAAGAGCUGUGCUUUUGCUUGCACAGCUCUUUCUUUUGUGUUAAGCUAUAUGGCGCUGUGAG.......AUGAUAGCUGUGAAGCGAG.AGGUUGCUGUCUCGAAAG..............................................
env-1097   UCGGAAUCCGCUCACUAAAGCUAAAAAAGAGCUGCGACACUACUUGCGCAGCUCUUUUUUUUGUGGUUAAGCUAUAUGGCGCUGUGAG.......AUGAUAGCUGUGAAGCGAG.AGGUUGCUGUCUCGAAAG..............................................
env-1099   GGACACUGAGUCGAAUUGUUUCUAAAAAAGAGCUGUGCUUUUACUUGCACAGCUCUUUUUUUUGUGUUAAGCUAUAUGGCGCUGUGAG.......AUGAUAGCUGUGAAGCGAG.AGGUUGCUGUCUCGAAAG..............................................
env-1100   GGACACUGAGUCGAAUUGUUUCUAAAAAAGAGCUGUGUUUUUACUUGCGCAGCUCUUUUUUUUGUGUUAAGCUAUAUGGCGCUGUGAG.......AUGAUAGCUGUGAAGCGAG.AGGUUGCUGUCUCGAAAG..............................................
Slo-1-1    UCGGAAUCCGCUCACUAAAGCUAAAAAAGAGCUGCGACACUACUUGCGCAGCUCUUUUUUUUGUGGUUAAGCUAUAUGGCGCUGUGAG.......AUGAUAGCUGUGAAGCGAG.AGGUUGCUGUCUCGAAAG..............................................
env-1101   CGGAAUCCGCUCACUAAAGCUAAAAAAAGAGCUGCGACACUACUUGCGCAGCUCUUUUUUUUGUGGUUAAGCUAUAUGGCGCUGUGAG.......AUGAUAGCUGUGAAGCGAG.AGGUUGCUGUCUCGAAAG..............................................
env-1102   GGCCACUGAGUCGAAUUGUUUCUAAAAAAGAGCUGUGCUUUUACUUGCGCAGCUCUUUUUUUUGUGUUAAGCUAUAUGGCGCUGUGAG.......AUGAUAGCUGUGAAGCGAG.AGGUUGCUGUCUCGAAAG..............................................
env-1103   GGCCACUGAGUCGAAUUGUUUCUAAAAAAGAGCUGUGCUUUUACUUGCGCAGCUCUUUUUUUUGUGUUAAGCUAUAUGGCGCUGUGAG.......AUGAUAGCUGUGAAGCGAG.AGGUUGCUGUCUCGAAAG..............................................
env-1104   ........................................................................UAAGCUAUAUGGCGCUGUGAG.......AUGAUAGCUGUGAAGCGAG.AGGUUGCUGUCUCGAAAG..............................................
env-1105   GGACACUGAGUCGAAUUGUUUCUAAAAAAGAGCUGUGAUUUUACUUGCGCAGCUCUUUUUUUUGUGUUAAGCUAUAUGGCGCUGUGAG.......AUGAUAGCUGUGAAGCGAG.AGGUUGCUGUCUCGAAAG..............................................
env-1106   UCGGAAUCCGCUCACUAAAGCUAAAAAAGAGCUGCGACACUACUUGCGCAGCUCUUUUUUUUGUGGUUAAGCUAUAUGGCGCUGUGAG.......AUGAUAGCUGUGAAGCGAG.AGGUUGCUGUCUCGAAAG..............................................
env-1107   GGACACUGAGUCGAAUUGUUUCUAAAAAAGAGCUGUGCUUUUGCUUGCACAGCUCUUUCUUUUGUGUUAAGCUAUAUGGCGCUGUGAG.......AUGAUAGCUGUGAAGCGAG.AGGUUGCUGUCUCGAAAG..............................................
env-1108   GGACACUGAGUCGAAUUGUUUCUAAAAAAGAGCUGUGCUUUUACUUGCGCAGCUCUUUUUUUUGUGUUAAGCUAUAUGGCGCUGUGAG.......AUGAUAGCUGUGAAGCGAG.AGGUUGCUGUCUCGAAAG..............................................
env-1115   GGACACUGAGUCGAAUUGUUUCUAAAAAAGAGCUGUGCUUUUACUUGCACAGCUCUUUUUUUUGUGUUAAGCUAUAUGGCGCUGUGAG.......AUGAUAGCUGUGAAGCGAG.AGGUUGCUGUCUCGAAAG..............................................
env-1118   GGACACUGAGUCGAAUUGUUUCUAAAAAAGAGCUGUGCUUUUGCUUGCACAGCUCUUUCUUUUGUGUUAAGCUAUAUGGCGCUGUGAG.......AUGAUAGCUGUGAAGCGAG.AGGUUGCUGUCUCGAAAG..............................................
env-1123   ...................................CGACACUACUUGCGCAGCUCUUUUUUUUGUGGUUAAGCUAUAUGGCGCUGUGAG.......AUGAUAGCUGUGAAGCGAG.AGGUUGCUGUCUCGAAAG..............................................
env-1124   GGACACUGAGUCGAAUUGUUUCUAAAAAAGAGCUGUGCUUUUACUUGCGCAGCUCUUUUUUUUGUGUUAAGCUAUAUGGCGCUGUGAG.......AUGAUAGCUGUGAAGCGAG.AGGUUGCUGUCUCGAAAG..............................................
env-1145   GGACACUGAGUCGAAUUGUUUCUAAAAAAGAGCUGUGCUUUUACUUGCGCAGCUCUUUUUUUUGUGUUAAGCUAUAUGGCGCUGUGAG.......AUGAUAGCUGUGAAGCGAG.AGGUUGCUGUCUCGAAAG..............................................
env-1146   GGACACUGAGUCGAAUUGUUUCUAAAAAAGAGCUGUGCUUUUACUUGCGCAGCUCUUUUUUUUGUGUUAAGCUAUAUGGCGCUGUGAG.......AUGAUAGCUGUGAAGCGAG.AGGUUGCUGUCUCGAAAG..............................................
env-1147   CGGAAUCCGCUCACUAAAGCCAAAAAAAGAGCUGCGACACUACUUGCGCAGCUCUUUUUUUUGUGGUUAAGCUAUAUGGCGCUGUGAG.......AUGAUAGCUGUGAAGCGAG.AGGUUGCUGUCUCGAAAG..............................................
env-1148   GGACACUGAGUCGAAUUGUUUCUAAAAAAGAGCUGUGCUUUUACUUGCGCAGCUCUUUUUUUUGUGUUAAGCUAUAUGGCGCUGUGAG.......AUGAUAGCUGUGAAGCGAG.AGGUUGCUGUCUCGAAAG..............................................
env-1149   GGACACUGAGUCGAAUUGUUUCUAAAAAAGAGCUGUGCUUUUACUUGCGCAGCUCUCCUUUUUGUGUUAAGCUAUAUGGCGCUGUGAG.......AUGAUAGCUGUGAAGCGAG.AGGUUGCUGUCUCGAAAG..............................................
env-1150   GACAUUGAGUCGAAUUGUUUCUAAAAAAGAGCUGUGCUUUUACUUGCGCAGCUCUUUUUUUUGUGUUAAGCUAUAUGGCGCUGUGAG.......AUGAUAGCUGUGAAGCGAG.AGGUUGCUGUCUCGAAAG..............................................
env-1151   GGACACUGAGUCGAAUUGUUUCUAAAAAAGAGCUGUGCUUUUGCUUGCACAGCUCUUUCUUUUGUGUUAAGCUAUAUGGCGCUGUGAG.......AUGAUAGCUGUGAAGCGAG.AGGUUGCUGUCUCGAAAG..............................................
env-1154   UCGGAAUCCGCUCACUAAAGCUAAAAAAGAGCUGCGACACUACUUGCGCAGCUCUUUUUUUUGUGGUUAAGCUAUAUGGCGCUGUGAG.......AUGAUAGCUGUGAAGCGAG.AGGUUGCUGUCUCGAAAG..............................................
env-1155   GGACACUGAGUCGAAUUGUUUCUAAAAAAGAGCUGUGCUUUUACUUGCGCAGCUCUCCUUUUUGUGUUAAGCUAUAUGGCGCUGUGAG.......AUGAUAGCUGUGAAGCGAG.AGGUUGCUGUCUCGAAAG..............................................
env-1156   UCGGAAUCCGCUCACUAAAGCUAAAAAAGAGCUGCGACACUACUUGCGCAGCUCUUUUUUUUGUGGUUAAGCUAUAUGGCGCUGUGAG.......AUGAUAGCUGUGAAGCGAG.AGGUUGCUGUCUCGAAAG..............................................
env-1157   ........................................ACUACUUGCGCAGCUCUUUUUUUUGUGGUUAAGCUAUAUGGCGCUGUGAG.......AUGAUAGCUGUGAAGCGAG.AGGUUGCUGUCUCGAAAG..............................................
env-1158   CGGAAUCCGCUCACUAAAGCCAAAAAAAGAGCUGCGACACUACUUGCGCAGCUCUUUUUUUUGUGGUUAAGCUAUAUGGCGCUGUGAG.......AUGAUAGCUGUGAAGCGAG.AGGUUGCUGUCUCGAAAG..............................................
env-1159   ...............................UGUGCUUUUACUUGCGCAGCUCUUUUUUUUGUGUUAAGCUAUAUGGCGCUGUGAG.......AUGAUAGCUGUGAAGCGAG.AGGUUGCUGUCUCGAAAG..............................................
Csp-2-1    UGAAAAUUAUUACAAUAUUUUUGUAACAAAGUCAAAAAAAAUACUUGCAAACAAAUGAAUUUUAAUGUAUACUGAAUUACGUUGUGAG.......AUACGAGCGAUGAUACAGA.AGGUUGCGGCA......................UACAA........AAUGAG..........
env-1160   GUGAAAAAUUGUAGAUUUAUUUUGUAAAAAAAAUAAAAAAAUGCUUGCACUCUGGCUUAAACUAUUGUAUAAUACAUAUUGUUGUGAG.......AUAGGAGCGAUGAUGCA.G.AGGUUGCCUA.GAGA...............................................
env-1162   GUGAAAAAUUGUAGAUUUAUUUUGUAAAAAAAAUAAAAAAAUGCUUGCACUCUGGCUUAAACUAUUGUAUAAUAUAUAUUGUUGUGAG.......AUAGGAGCGAUGAUGCA.G.AGGUUGCCUA.GAGA...............................................
env-1163   ..................................AAAUAAAAAAAUGCUUGCACUCUGGCUUAAACUAUUGUAUAAUACAUAUUGUUGUGAG.......AUAGGAGCGAUGAUGCA.G.AGGUUGCCUA.GAGA...............................................
env-1166   GUGAAAAAUUGUAGAUUUAUUUUGUAAAAAAAAUAAAAAAAUGCUUGCACUCUGGCUUAAACUAUUGUAUAAUACAUAUUGUUGUGAG.......AUAGGAGCGAUGAUGCA.G.AGGUUGCCUA.GAGA...............................................
env-1167   AAAAUUUUUUCGAUAAAAAAAUGAAAAAAAAUCUUAUAAAACGCUUGCACUGAAUGAAAAUCUAUAGUAAGAUAUAAAUGUUGUGAG........AGAGAGCGUUGAUUUGAA.AGGUUGCUGUG......................AUUUCUU.......AAUUAA..........
env-1168   AAAAUUUUUUCGAUAAAAAAAUGAAAAAAAAUCUUAUAAAACGCUUGCACUGAAUGAAAAUCUAUAGUAAGAUAUAAAUGUUGUGAG........AGAGAGCGUUGAUUUGAA.AGGUUGCUUUG......................AUUUCU........UAAA...........
env-1169   UAUGAAAAACAGAAGAAAAUUUUUCGAAAAACAGAUAGAAAAGCUUGCAAUCAUAGGGUUUUUCCUGUAUACUAAUUGAUGUUGUGAG........AUAGAGUGUUGAGUCGAA.AGGUUGCCGAAA....................ACCUU.........AAAAAUCA........
env-1170   UAUGAAAAACAGAAGAAAAUUUUUCGAAAAACAGAAGAAAAGCUUGCAAUCAUAGGGUUUUUCCUGUAUACUAAUUGAUGUUGUGAG........AUAGAGUGUUGAGUCGAA.AGGUUGCCGAAA....................ACCUU.........AAAAAUCA........
env-1171   UAUGAAAAACAGAAGAAAAUUUUUCGAAAAACAGAUGGAAAAGCUUGCAAUCAUAGGGUUUUUCCUGUAUACUAAUUGAUGUUGUGAG........AUAGAGUGUUGAGUCGAA.AGGUUGCCGAAA....................ACCUU.........AAAAAUCA........
env-1172   GUUAAAAUUCUAAGAAAUUUUUUUAAAAAAAUACAAAAAAACCCUUGCCAAAUCAGGAAAAGUCGUGUAUACUUGUUUUGUUGUGAG........AUAGAGUGUUGAUACGGA.AGGUUGC....UGA....................GAGCCG........AAUUGAG.........
env-1173   GUUAAAAUUCUAAGAAAUUUUUUUGAAAAAAUACAAAAAAACUCUUGCCAAAUCAGGAAAAGUCGUGUAUACUUGUUUAUGUUGUGAG........AUAGAGUGUUGAUACGGA.AGGUUGC....UGA....................GAGCCG........AAUUGAG.........
env-1174   GUUAAAAUUCAAAGAAAUUUUUUUAAAAAAAUACAAAAAAACUCUUGCCAAAUCAGGAAAAGUCGUGUAUACUUGUUUUGUUGUGAG........AUAGAGUGUUGAUACGGA.AGGUUGC....UGA....................GAGCCG........AAUUGAG.........
env-1175   GUUAAAAUUCUAAGAAAUUUUUUUGAAAAAAUACAAAAAAACUCUUGUCAAAUCAGGAAAAGUCGUGUAUACUUAUUUGUGUUGUGAG........AUAGAGUGUUGAUACGGA.AGGUUGCUGAU.....................GGCCG.........AAUUGAG.........
env-1176   UUAAAAUUCUAAGAAAUUUUUUUAGAAAAAAUACAAAAAAACUCUUGCCAAAUCAGGAAAAGUCGUGUAUACUUGUUUUGUUGUGAG........AUAGAGUGUUGAUACGGA.AGGUUGCUGAG.....................AGCCG.........AAUUGAG.........
Cle-1-1    UGCAUAAAUUACCAGAAUAAAAUUGUAAAAAAAAUAAAAAAAUCCUUGUCAUGUACAAGAAGAUAGUAUAUAAUAAAUACGUUGUGAG......AUUGAGAACUAUGAUCCAGG.AGGUUACGCAUCGCA...............................................
           ..........................................................................<<.<<<<<.......................<<<.<.<<<...<<<<<......................<<<<<<<<<<<<<<...........>>>>
           ..........................................................................21..1111.......................222.2.012...12222......................1111111.10..00...........00..
           •••••••••••••R••••••••••R••••••••RAAAAR•••YU•••••••••••••••••••••••••••••••GYUGUGRC-------•UGAYAGC••UGAAGCG•G-AGGUUGCYRY••••○○○-------------------------------------------
                                                                                                 5′
```

alignment positions 181 · · · 360

```
env-1      .................GGCG.GUAAC.UUU...CGUGGAGC..AUGUCCGUCGUCU..GU..UUGUUU..............GCA.............GACGAGU.........AAUCGGGCGGCUGAGGUUUUCCUGGGGGCGUGGUCGAAAGGCCGUAUAUCGUCUGGACGGGA
env-4      .................GGCG.GUAAC.UUU...CGUGGAGC..AUGUCCGUCGUCU..GU..UUGUUU..............GCA.............GACGAGU.........AAUCGGGCGGCUGAGGUUUUCCUGGGGGCGUGGUCGAAAGGCCGUAUAUCGUCUGGACGGGA
env-5      .................GGCG.GUAAC.UUU...CGUGGAGC..AUGUCCGUCGUCU..GU..UUGUUU..............GCA.............GACGAGU.........AAUCGGGCGGCUGAGGUUUUCCUGGGGGCGUGGUC..........................
env-6      ................GGACG.GUAAC.UUC...UGCGGAGUAUGUCCG.......G..UC..AUCGGGC.............GGC.............UGGG.A..GCUAUCAGCGUAUAUCGCCGCAGUGGAACAAGGCCGACAACGUCGGUCUUUUUCAUGGUGCGGAAU
           >>>>>>>>>>.......>>>>>.....>.>>>...>>>.......<<<<.......<..<<<....................>>>>.............>>>>...>.>>>>..>>..................................................
           01.1111111.......22221.....2.102...222.......111.......2..212....................2122.............111...1.111..12..................................................
           ----------------•GYRG-GU--U-UUC---CGYGGAGCGAAUGUC•R-----G--•Y•R•RRR•--------------•G•C-------------GRCAA--G-UCACUGUAC•RR•••••••••••••••••••••••••••••••••••••••••
                                                                                                     3′
```

```
env-7     .................GGAG.GUAAC.UUC...CGGGGAGU..AUGUC.......C..G..AUAAU.................CG..............GGCGGC..........UGAGAGAUUUCUGUUCGUCGGCGUAGAUCGCCGCGCCAGAAGGACACCGGCCCAAAGCCGGU
env-8     .................GGAG.GUAAC.UUC...CGGGGAGU..AUGUC.......C..G..AUAAU.................CG..............GGCGGC..........UGAGAGAUUUCUGUUCGUCGGCGUAGAUCGCCGCGCCAGAAGGACACCGGCCCAAAGCCGGU
env-9     .................GGAG.GUAAC.UUC...CGGGGAGU..AUGUC.......C..G..AUAAU.................CG..............GGCGGC..........UGAGAGAUUUCUGUUCGUCGGCGUAGAUCGCCGCGCCAGAAGGACACCGGCCCAAAGCCGGU
env-10    ................AGGAG.GUAAC.UUC...CGGGGAGU..AUGUC.......C..G..AUAAU.................CG..............GGCGGC..........UGAGAAGAAUCUGUACGUUGCGUAGAUCGCUGCGCCAUGACAGGAUACCGGCCGUAGACAAC
env-11    ................AGGAG.GUAAC.UUC...CGGGGAGU..AUGUC.......C..G..AUAAU.................CG..............GGCGGC..........UGAGAAGGUCCUGCACGCAGCGUAGAUCGCCGUGCCAUGGACGAGUACCGGCCCUGUGCCGG
env-12    ................AGGAG.GUAAC.UUC...CGGGGAGU..AUGUC.......C..G..AUAAU.................CG..............GGCGGC..........UGAGGAUGUUCUGUUCGCCGGCGUAGAUCGCCGCGCCAGAGCCACACCGGCCACGUCUGCGU
env-13    ................AGGAG.GUAAC.UUC...CGGGGAGU..AUGUC.......C..G..AUAAU.................CG..............GGCGGC..........UGAGAGAUCUCUGUACGUCGGCGUAGAUCGCCGCGCCAGAAGGACACCGGCCUGAUCGCCGG
env-14    ................AGGAG.GUAAC.UUC...CGGGGAGU..AUGUC.......C..G..AUAAU.................CG..............GGCGGC..........UGAGAGAUCUCUGUACGUCGGCGUAGAUCGCCGCGCCAGAAGGACACCGGCCUGGUCGCCGG
env-15    ................AGGAG.GUAAC.UUC...CGGGGAGU..AUGUC.......C..G..AUAAU.................CG..............GGCGGC..........UGAGAGAUCUCUGUACGUCGGCGUAGAUCGCCGCGCCAGAAGGACACCGGCCUGUUGCCGGU
env-16    ................AGGAG.GUAAC.UUC...CGGGGAGU..AUGUC.......C..G..AUAAU.................CG..............GGCGGC..........UGAGAGGAUUCUGUGCGUCAGCGG........................................
env-17    ................AGGAG.GUAAC.UUC...CGGGGAGU..AUGUC.......C..G..AUAAU.................CG..............GGCGGC..........UGAGAGAUCUCUGUACGUCGGCGUAGAUCGCCGCGCCAGAAGGACACCGGCCUGGUCGCCGG
env-18    ................AGGAG.GUAAC.UUC...CGGGGAGU..AUGUC.......C..G..AUAAU.................CG..............GGCGGC..........UGAGAAGUCCUGUAUGCGGCGUAGAUCGCCAUACCAUGGGACAAGUACCGGCCACAGUGCCG
env-19    ................AGGAG.GUAAC.UUC...CGGGGAGU..AUGUC.......C..G..AUAAU.................CG..............GGCGGC..........UGAGAGAUCUCUGUACGUCGGCGUAGAUCGCCGCGCCAGAAGGACACCGGCCUGGUCGCCGG
env-20    ................AGGAG.GUAAC.UUC...CGGGGAGU..AUGUC.......C..G..AUAAU.................CG..............GGCGGC..........UGAGAGAUUUCUGUACGUCGGCGUAGAUCGCCGCGCCAGAAGGACACCGGCCUGGUCGCCGG
env-21    ................AGGAG.GUAAC.UUC...CGGGGAGU..AUGUC.......C..G..AUAAU.................CG..............GGCGGC..........UGAGAGAUUUCUGUUCGCCGGCGUAGAUCGCCGCGCCAGAAGGACACCGGCCUGUUGCCGGU
env-22    ................AGGAG.GUAAC.UUC...CGGGGAGU..AUGUC.......C..G..AUAAU.................CG..............GGCGGC..........UGAGAGAUCUCUGUACGUCGGCGUAGAUCGCCGCGCCAGAAGGACACCGGCCUGAUCGCCGG
env-23    ................AGGAG.GUAAC.UUC...CGGGGAGU..AUGUC.......C..G..AUAAU.................CG..............GGCGGC..............................................................................
env-25    ................AGGAG.GUAAC.UUC...CGGGGAGU..AUGUC.......C..G..AUAAU.................CG..............GGCGGC..........UGAGAGAUCUCUGUACGUCGGCGUAGAUCGCCGCGCCAGAAGGACACCGGCCUGGUCGCCGG
env-26    ................AGGAG.GUAAC.UUC...CGGGGAGU..AUGUC.......C..G..AUAAU.................CG..............GGCGGC..........UGAGAGAUCUCUGUACGUCGGCGUAGAUCGCCGCGCCAGAAGGACACCGGCCUGCUGCCGGU
env-27    ................AGGAG.GUAAC.UUC...CGGGGAGU..AUGUC.......C..G..AUAAU.................CG..............GGCGGC..........UGAGAGAUCUCUGUACGUCGGCGUAGAUCGCCGCGCCAGAAGGACACCGGCCUGAUCGCCGG
env-28    ................AGGAG.GUAAC.UUC...CGGGGAGU..AUGUC.......C..G..AUAAU.................CG..............GGCGGC..........UGAGAAGUCCUGUAUGCGGCGUAGAUCGCCAUACCAUGGGACAAGUACCGGCCACAGUGCCG
env-29    ................AGGAG.GUAAC.UUC...CGGGGAGU..AUGUC.......C..G..AUAAU.................CG..............GGCGGC..........UGAGAGAUUUCUGUUCGUCGGCGUAGAUCGCCGCGCCAGAAGGACACCGGCCUGAUCGCCGG
env-30    ................AGGAG.GUAAC.UUC...CGGGGAGU..AUGUC.......C..G..AUAAU.................CG..............GGCGGC..........UGAGAAG...............................................
env-43    ................AGGAG.GUAAC.UUC...CGGGGAGU..AUGUC.......C..G..AUAAU.................CG..............GGCGGC..........UGAGAGAUUUCUGUACGUCGGCGUAGAUCGCCGCGCCAGAAGGACACCGGCCUGGUCGCCGG
env-45    ................AGGAG.GUAAC.UUC...CGGGGAGU..AUGUC.......C..G..AUAAU.................CG..............GGCGGC..........UGAGAGAUCUCUGUACGUCGGCGUAGAUCGCCGCGCCAGAAGGACACCGGCCUGGUCGCCGG
env-46    ................AGGAG.GUAAC.UUC...CGGGGAGU..AUGUC.......C..G..AUAAU.................CG..............GGCGGC..........UGAGA.................................................
env-47    ................AGGAG.GUAAC.UUC...CGGGGAGU..AUGUC.......C..G..AUAAU.................CG..............GGCGGC..........UGAGAAGUCCUGUAUGCGGCGUAGAUCGCCAUACCAUGGGACAAGUACCGGCCACAGUGCCG
env-48    ................CGGCG.GU..AAUUCA..UACGGAGU..AUGUC.......C..G..AUUUUAAAC.............CG..............GGCGAA.A.CGAA.GAUGAUUUGUAAUCAUCUGAACCGAUAGGAAACUACAAGCUGUGACUUGCGAACGCAGAUUUUC
env-49    ................CGGCG.GU..AAUUCA..UACGGAGU..AUGUC.......C..G..AUUUUAAAC.............CG..............GGCGAA.A.CGAA.GAUGAUUUGUAAUCAUCUGAACCGAUAGGAAACUACAAGCUGUGACUUGCGAACGCAGAUUUUC
env-54    ................CGGCG.GU..AAUUCA..UACGGAGU..AUGUC.......C..G..AUUUUAAAC.............CG..............GGCGAA.A.CGAA.GAUGAUUUGUAAUCAUCUGAACCGAUAGGAAACUACAAGCUGUGACUUGCGAACGCAGAUUUUC
env-55    ................CGGCG.GU..AAUUCA..UACGGAGU..AUGUC.......C..G..AUUUUAAAC.............CG..............GGCGAA.A.CGAA.GAUGAUUUGUAAUCAUCUGAACCGAUAGGAAACUACAAGCUGUGACUUGCGAACGCAGAUUUUC
env-57    ................CGGCG.GU..AAUUCA..UACGGAGU..AUGUC.......C..G..AUUUUAAAC.............CG..............GGCGAA.A.CGAA.GAUGAUUUGUAAUCAUCUGAACCGAUAGGAAACUACAAGCUGUGACUUGCGAACGCAGAUUUUC
env-58    ................CGGCG.GU..AAUUCA..UACGGAGU..AUGUC.......C..G..AUUUUAAAC.............CG..............GGCGAA.A.CGAA.GAUGAUUUGUAAUCAUCUGAACCGAUAGGAAACUACAAGCUGUGACUUGCGAACGCAGAUUUUC
env-59    ................CGGCG.GU..AAUUCA..UACGGAGU..AUGUC.......C..G..AUUUUAAAC.............CG..............GGCGAA.A.CGAA.GAUGAUUUGUAAUCAUCUGAACCGAUAGGAAACUACAAGCUGUGACUUGCGAACGCAGAUUUUC
env-60    ................CGGCG.GU..AAUUCA..UACGGAGU..AUGUC.......C..G..AUUUUAAAC.............CG..............GGCGAA.A.CGAA.GAUGAUUUGUAAUCAUCUGAACCGAUAGGAAACUACAAGCUGUGACUUGCGAACGCAGAUUUUC
env-61    ................CGACG.GU..AAUUCA..UACGGAGU..AUGUC.......C..G..AUUUUAAAC.............CG..............GGCGA..A.ACGA.AGAUGAUUUGUAAUCAUCUGAACCGAUAGGAAACUACAAGCUGUGACUUGCGAACGCAGAUUUU
env-76    ................CGACG.GU..AAUUCA..UACGGAGU..AUGUC.......C..G..AUUUUAAAC.............CG..............GGCGAAA.........CGAAGAUGAUUUGUAAUCAUCUGAACCGAUAGGAAACUACAAGCUGUGACUUGCGAACGCAG
env-77    ................CGACG.GU..AAUUCA..UACGGAGU..AUGUC.......C..G..AUUUUAAAC.............CG..............GGCGAAA.........CGAAGAUGAUUUUUAAUCAUCUGAACCGAUAGGAAACUACAAGCUGUGACUUGCGAACGCAG
env-78    ................CGACG.GU..AAUUCA..UACGGAGU..AUGUC.......C..G..AUUUUAAAC.............CG..............GGCGAAA.........CGAAGAUGAUUUGUAAUCAUCUGAACCGAUAGGAAACUACAAGCUGUGACUUGCGAACGCAG
env-80    ................CGACG.GU..AAUUCA..UACGGAGU..AUGUC.......C..G..AUUUUAAAC.............CG..............GGCGAAA.........CGAAGAUGAUUUGUAAUCAUCUGAACCGAUAGGAAACUACAAGCUGUGACUUGCGAACGCAG
env-81    ................CGACG.GU..AAUUCA..UACGGAGU..AUGUC.......C..G..AUUUUAAAC.............CG..............GGCGAAA.........CGAAGAUGAUUUGUAAUCAUCUGAACCGAUAGGAAACUACAAGCUGUGACUUGCGAACGCAG
env-82    ................CGACG.GU..AAUUCA..UACGGAGU..AUGUC.......C..G..AUUUUAAAC.............CG..............GGCGAAA.........CGAAGAUGAUUUGUAAUCAUCUGAACCGAUAGGAAACUACAAGCUGUGACUUGCGAACGCAG
env-83    ................CGACG.GU..AAUUCA..UACGGAGU..AUGUC.......C..G..AUUUUAAAC.............CG..............GGCGAAA.........CGAAGAUGAUUUGUAAUCAUCUGAACCGAUAGGAAACUACAAGCUGUGACUUGCGAACGCAG
env-85    ................GGACG.GUAAC.UUC...UGCGGAGU..AUGUC.......C..G..GUCAU.................CG..............GGCG...........GCUGGGAGUCCUAUGCAGCGUAUAUCGCCGCAGUGGAACAAGGCCGACAACGUCGGUCUUUU
env-86    ................GGACG.GUAAC.UUC...UGCGGAGU..AUGUC.......C..G..GUCAU.................CG..............GGCG...........GCUGGGAGUCCUAUGCAGCGUAUAUCGCCGCAGUGGAACAAGGCCGACAACGUCGGUCUUUU
env-87    ................GGACG.GUAAC.UUC...UGCGGAGU..AUGUC.......C..G..GUCAU.................CG..............GGCG...........GCUGGGAGUCCUAUGCAGCGUAUAUCGCCGCAGUGGAACAAGGCCGACAACGUCGGUCUUUU
env-88    ................GGACG.GUAAC.UUC...UGCGGAGU..AUGUC.......C..G..GUCAU.................CG..............GGCG...........GCUGAGGAUCACGAUACGGCGUAUAUCGCCGCAGUGAACAGGCCGACAACGUCGG.......
env-89    ................GGACG.GUAAC.UUC...UGCGGAGU..AUGUC.......C..G..GUCAU.................CG..............GGCG...........GCUGAGGAUCACAAUACAGCGUAUAUCGCUGCAGUGAUGAAAGGCCGACAACGUCGGUCUUU
Pba-1-1   ................GGCAG.GAAAU.UUC...CGUGGAGU..AUGUC.......C..AAACUUGGA...............UUUG..............GGCG...UGGAGA.GCUGUAUUUUCAAUGACCGUAGGUCAUUUUUUUAAAGGAAAAGCACAACACACCCGAGAGUGUG
env-90    ................GGCGG.GGAAU.UUC...UGCGGAGU..AUGUC.......C..G..AUUUACAAAC............CG..............GGCGAA.AGAAUACUGGCUCGCUUCGGCCGGCCGCAGUUUGUGCUGCGGCGGUUUUCUGCGGCCAGGCGCGAACAAU
env-91    ................GGCGG.GGAAU.UUC...UGCGGAGU..AUGUC.......C..G..AUUUACAAAC............CG..............GGCGAA.AGAAUACUGGCUCGCUUCGGCCGGCCGCAGUUUGUGCUGCGGCGGUUUUCUGCGGCCAGGCGCGAACCAU
env-92    ................GACGG.GGAAU.UUC...UGCGGAGU..AUGUC.......C..G..AUUUUAAAAC............CG..............GGCGA..........AACAGGUCUGACUGCCGCACGGGGUGCGUUUCCCUCGCACCUUCGCUUGAGGCAGCAGUAAU
env-94    ................GACGG.GGAAU.UUC...UGCGGAGU..AUGUC.......C..G..AUUUUAAAAC............CG..............GGCGA..........AACAGGUCUGACUGCCGCACGGGGUGCGUUUCCCUCGCACCUUCGCUUGAGGCAGCAGUAAU
env-95    ................GACGG.GGAAU.UUC...UGCGGAGU..AUGUC.......C..G..AUUUUAAAAC............CG..............GGCGA..........AACAGGUCUGACUGCCGCACGGGGUGCGUUUCCCUCGCACCUUCGCUUGAGGCAGCAGUAAU
env-96    ................GACGG.GGAAU.UUC...UGCGGAGU..AUGUC.......C..G..AUUUUAAAAC............CG..............GGCGA..........AACAGGUCUGACUGCCGCACGGGGUGCGUUUCCCUCGCACCUUCGCUUGAGGCAGCAGUAAU
env-97    ................GACGG.GGAAU.UUC...UGCGGAGU..AUGUC.......C..G..AUUUUAAAAC............CG..............GGCGA..........AACAGGUCUGACUGCCGCACGGGGUGCGUUUCCCUCGCACCUUCGCUUGAGGCAGCAGUAAU
env-99    ................GACGG.GGAAU.UUC...UGCGGAGU..AUGUC.......C..G..AUUUUAAAAC............CG..............GGCGA..........AACAGGUCUGACUGCCGCACGGGGUGCGUUUCCCUCGCACCUUCGCUUGAGGCAGCAGUAAU
env-102   ................GACGG.GGAAU.UUC...UGCGGAGU..AUGUC.......C..G..AUUUUAAAAC............CG..............GGCGA..........AACAGGUCUGACUGCCGCACGGGGUGCGUUUCCCUCGCACCUUCGCUUGAGGCAGCAGUAAU
env-107   ................GACGG.GGAAU.UUC...UGCGGAGU..AUGUC.......C..G..AUUUUAAAAC............CG..............GGCGA..........AACAGGUCUGACUGCCGCACGGGGUGCGUUUCCCUCGCACCUUCGCUUGAGGCAGCAGUAAU
env-109   ................GACGG.GGAAU.UUC...UGCGGAGU..AUGUC.......C..G..AUUUUAAAAC............CG..............GGCGA..........AACAGGUCUGACUGCCGCACGGGGUGCGUUUCCCUCGCACCUUCGCUUGAGGCAGCAGUAAU
env-110   ................GACGG.GGAAU.UUC...UGCGGAGU..AUGUC.......C..G..AUUUUAAAAC............CG..............GGCGA..........AACAGGUCUGACUGCCGCACGGGGUGCGUUUCCCUCGCACCUUCGCUUGAGGCAGCAGUAAU
env-111   ................GACGG.GGAAU.UUC...UGCGGAGU..AUGUC.......C..G..AUUUUAAAAC............CG..............GGCGA..........AACAGGUCUGACUGCCGCACGGGGUGUGUUUCCCUCGCACCUUCGCUUGAGGCAGCAGUAAU
env-112   ................GACGG.GGAAU.UUC...UGCGGAGU..AUGUC.......C..G..AUUUUAAAAC............CG..............GGCGA..........AACAGGUCUGACUGCCGCACGGGGUGCGUUUCCCUCGCACCUUCGCUUGAGGCAGCAGUAAU
env-113   ................GACGG.GGAAU.UUC...UGCGGAGU..AUGUC.......C..G..AUUUUAAAAC............CG..............GGCGA..........AACAGGUCUGACUGCCGCACGGGGUGCGUUUCCCUCGCACCUUCGCUUGAGGCAGCAGUAAU
env-114   ................GGUCG.GU..UUUUC...CGUGGAGU..AUGUC.......C..G..AUUUUUAAAC............CG..............GGCG...A.AAGA.AUAAAGAAGGAAGUUAAGAGAUACUGAGGGUUGCUGUGUCUCAGAACUACUACGACCUUUUUCU
env-121   ................GGUCG.GU..UUUUC...CGUGGAGU..AUGUC.......C..G..AUUUUUAAAC............CG..............GGCG...A.AAGA.AUAAAGAAGGAAGUUAAGAGAUACUGAGGGUUGCUGUGUCUCAGAACUACUAC..........
env-122   .................GUCG.GU..UUUUC...CGUGGAGU..AUGUC.......C..G..AUUUUAAAC.............CG..............GGCG...A.UUAA.GUUAAGAUUGAGCAAAUAUUACCCAAUAGGGUAACACUAUGCUCUUAAUGCGCUUUGAACUGCC
env-123   .................GUCG.GU..UUUUC...CGUGGAGU..AUGUC.......C..G..AUUUUAAAC.............CG..............GGCG...A.UUAA.GUUAAGAUUGAGCAAAUAUUACCCAAUAGGGUAACACUAUGCUCUUAAUGCGCUUUGAACUGCC
env-128   .................GUCG.GU..UUUUC...CGUGGAGU..AUGUC.......C..G..AUUUUAAAC.............CG..............GGCG...A.UUAA.GUUAAGAUUGAGCAAAUAUUACCCAAUAGGGUAACACUAUGCUCUUAAUGCGCUUUGAACUGCC
Rfl-1-1   ................GUCAG.GUAAU.UUC...CGUGGAGU..AUGUC.......C..G..AUUUUAAAC.............CG..............GGCG...........AUUGGAAUAGCGAACACAGUUUGUGCUUUUAUUCUCACGCAGCUUGCGUAUGCGUGAACUAU
env-129   ................GUCAG.GUAAU.UUC...CGUGGAGU..AUGUC.......C..G..AUUUUAAAC.............CG..............GGCG...........AUUGGAAUA.............................................
env-130   ................UACGG.GU..U.UUUC..CGAGGAGCG.AUGUCCA.....G..UUCAAAGAAACC...........GGGC..............GACAA..GGUCACUGUACUGAAUAUUUUGGGUUUGACUUACCGACAGGGUAAGUUUGCGUAUCUGACGUUGCUGGAUA
env-131   ......AGUGGGU....UUUGG.GU..U.UUU...CGUGGAGCGAAUGUCAAGA...G..GCCGAA.A...............GGUCUGUAAACCUGGC..GACAA..G.UCACUGUACAGAAAACAACUAAUAUGUUCUACAGAGUAGGGCGUGGUGUGAUUAUCAGUAUGUUUAUGA
env-132   ......AGUGGGU....UUUGG.GU..U.UUU...CGUGGAGCGAAUGUCAAGA...G..GCCGAA.A...............GGUCUGUAAACCUGGC..GACAA..G.UCACUGUACAGAAAACAACUAAUAUGUUCUACAGAGUAGGGCGUGGUGUGAUUAUCAGUAUGUUUAUGA
env-133   ......AGUGGGU....UUUGG.GU..U.UUU...CGUGGAGCGAAUGUCAAGA...G..GCCGAA.A...............GGUCUGUAAACCUGGC..GACAA..G.UCACUGUACAGAAAACAACUAAUAUGUUCUACAGAGUAGGGCGUGGUGUGAUUAUCAGUAUGUUUAUGA
env-134   ......AGUGGGU....UUUGG.GU..U.UUU...CGUGGAGCGAAUGUCAAGA...G..GCCGAA.A...............GGUCUGUAAACCUGGC..GACAA..G.UCACUGUACAGAAAACAACUAAUAUGUUCUACAGAGUAGGGCGUGGUGUGAUUAUCAGUAUGUUUAUGA
env-138   ......AGUGGGU....UUUGG.GU..U.UUU...CGUGGAGCGAAUGUCAAGA...G..GCCGAA.A...............GGUCUGUAAACCUGGC..GACAA..G.UCACUGUACAGAAAACAACUAAUAUGUUCUACAGAGUAGGGCGUGGUGUGAUUAUCAGUAUGUUUAUGA
env-139   ......AGUGGGU....UUUGG.GU..U.UUU...CGUGGAGCGAAUGUCAAGA...G..GCCGAA.A...............GGUCUGUAAACCUGGC..GACAA..G.UCACUGUACAGAAAACAACUAAUAUGUUCUACAGAGUAGGGCGUGGUGUGAUUAUCAGUAUGUUUAUGA
env-140   ......AGUGGGU....UUUGG.GU..U.UUU...CGUGGAGCGAAUGUCAAGA...G..GCCGAA.A...............GGUCUGUAAACCUGGC..GACAA..G.UCACUGUACAGAAAACAACUAAUAUGUUCUACAGAGUAGGGCGUGGUGUGAUUAUCAGUAUGUUUAUGA
env-141   ......AGUGGGU....UUUGG.GU..U.UUU...CGUGGAGCGAAUGUCAAGA...G..GCCGAA.A...............GGUCUGUAAACCUGGC..GACAA..G.UCACUGUACAGAAAACAACUAAUAUGUUCUACAGAGUAGGGCGUGGUGUGAUUAUCAGUAUGUUUAUGA
env-142   ......GCAGGU.....UUUGG.GU..U.UUU...CGUGGAGCGAAUGUCAAGA...G..ACCGAA.A...............GGUCUGUAAACCUGGC..GACAA..G.UCACUGUACAGAAAACAACUAAUAUGUUCUACAGAGUAGGGCGUGGUGUGAUUAUCAGUAUGUUUAUGA
env-143   ......GCAGGU.....UUUGG.GU..U.UUU...CGUGGAGCGAAUGUCAAGA...G..ACCGAA.A...............GGUCUGUAAACCUGGC..GACAA..G.UCACUGUACAGAAAACAACUAAUAUGUUCUACAAAGUAGGGCGUGGUGUGAUUAUCAGUAUGUUUAUGA
env-144   ......GCAGGU.....UUUGG.GU..U.UUU...CGUGGAGCGAAUGUCAAGA...G..ACCGAA.A...............GGUCUGUAAACCUGGC..GACAA..G.UCACUGUACAGAAAACAACUAAUAUGUUCUACAGAGUAGGGCGUGGUGUGAUUAUCAGUAUGUUUAUGA

          >>>>>>>>>>.......>>>>>.....>.>>>...>>>........<<<<.......<..<<<...................>>>>..............>>>>...>.>>>>.>>..........................................................
          01.1111111.......22221.....2.102...222.........111.......2..212...................2122..............111....1.111..12..........................................................
          ----------------•GYRG-GU--U-UUC---CGYGGAGCGAAUGUC•R-----G--•Y•R•RRR•---------------•G•C-------------GRCAA--G-UCACUGUAC•RR••••••••••••••••••••••••••••••••••••••••••••••••••••••
                                                                                                              3′
```

```
env-148   ......GCAGGU.....UUUGG.GU..U.UUU...CGUGGAGCGAAUGUCAAGA...G..ACCGAA.A...............GGUCUGUAAACCUGGC..GACAA..G.UCACUGUACAGAAAACAACUAAUAUGUUCUACAGAGUAGGGCGUGGUGUGAUUAUCAGUAUGUUUAUGA
env-158   ......GCAGGU.....UUUGG.GU..U.UUU...CGUGGAGCGAAUGUCAAGA...G..ACCGAA.A...............GGUCUGUAAACCUGGC..GACAA..G.UCACUGUACAGAAAACAACUAAUAUGUUCUACAGAGUAGGGCGUGGUGUGAUUAUCAGUAUGUUUAUGA
env-159   ......GCAGGU.....UUUGG.GU..U.UUU...CGUGGAGCGAAUGUCAAGA...G..ACCGAA.A...............GGUCUGUAAACCUGGC..GACAA..G.UCACUGUACAGAAAACAACUAAUAUGUUCUACAGAGUAGGGCGUGGUGUGAUUAUCAGUAUGUUUAUGA
env-160   ......GCAGGU.....UUUGG.GU..U.UUU...CGUGGAGCGAAUGUCAAGA...G..ACCGAA.A...............GGUCUGUAAACCUGGC..GACAA..G.UCACUGUACAGAA..........................................................
env-161   .......CAGGUU....UUUGG.GU..U.UUU...CGUGGAGCGAAUGUCAAGA...G..ACCGAA.A...............GGUCUGUAAACCUGGC..GACAA..G.UCACUGUACAGAAAACAACUAAUAUGUUCUACAGAGUAGGGCGUGGUGUGAUUAUCAGUAUGUUUAUGA
Cmo-1-1   .......CAGGUU....UUUGG.GU..U.UUU...CGUGGAGCGAAUGUCAAGA...G..ACCGAA.A...............GGUCUGUAAACCUGGC..GACAA..G.UCACUGUACAGAAAACAACUAAUAUGUUCUACAAAGUAGGGCGUGGUGUGAGUAAAAAUAUUUGCGUUU
env-162   .......CAGGUU....UUUGG.GU..U.UUU...CGUGGAGCGAAUGUCAAGA...G..ACCGAA.A...............GGUCUGUAAACCUGGC..GACAA..G.UCACUGUACAGAAAACAACUAAUAUGUUCUACAAGGUAGGGCGUGGUGUGAGUAAAAAUAGUUUAUGUU
env-163   .......CAGGUU....UUUGG.GU..U.UUU...CGUGGAGCGAAUGUCAAGA...G..ACCGAA.A...............GGUCUGUAAACCUGGC..GACAA..G.UCACUGUACAGAAAACAACUAAUAUGUUCUACAAGGUAGGGCGUGGUGUGAGUAAAAAUAAUUUAUGUU
Ece-1-1   ..................GCUG.GU..AACUC...CGUGGAGCAAAUGUCUA.....G..UC.AAGAUACA...............GGC.............GAAAC..G.UCACUGUACAACUGAAUGGUCCACCACAGUGUGGAAACAACGUGUGCGUUUUAGGACACACACGGAAA
env-164   .................UGUGG.GU..U.UUC...CGUGGAGCGAAUGUCAGUAG..G..CUGCAUAG.................CGGCCACAAAAUCAUGACGACAA..G.UCACUGUACAAGCAAAUGAUGUCUGCUGCAAAUCCCAAAGGAUUGAUGGGCAGAAACGACGUGAAGUG
Bfi-1-1   .................UGUGG.GU..U.UUC...CGUGGAGCGAAUGUCAGUAG..G..CUGCAUAG.................CGGCCACAAAAUCAUGACGACAA..G.UCACUGUACAAGCAAAUGAUGUCUGUUGCAAAUCCCAAAGGAUUGAUGGGCAGAAACGACGUGAAGUG
Bfi-2-1   .................UGUGG.GU..U.UUC...CGUGGAGCGAAUGUCAGUAG..G..CUGCAUAG.................CGGCCACAAAAUCAUGACGACAA..G.UCACUGUACAAGCAAAUGAUGUCUGCUGCAAAUCCCAAAGGAUUGAUGGGCAGAAACGACGUGAAGUG
env-165   ....AAAGAU...AGAAAGUAG.GU..U.UUCA..CACUGAGCGAAGGUCAU.....G..UUACUAAAAC..............UGGC.............GACAA..G.UCACUGUACAAUUUACAGAAGUUUCUAAGUGGAAAUGAUGAGGUGUGUCAUUUUUAUGACACGCACGGA
env-166   ....AAAGAU...AGAAAGUAG.GU..U.UUCA..CACUGAGCGAAGGUCAU.....G..UUACUAAAAC..............UGGC.............GACAA..G.UCACUGUACAAU..........................................................
env-170   ....AAAGAU...AGAAAGUAG.GU..U.UUCA..CACUGAGCGAAGGUCAU.....G..UUACUAAAAC..............UGGC.............GACAA..G.UCACUGUACAAUUUACAGAAGUUUCUAAGUGGAAAUGAUGAGGUGUGUCAUUUUUAUGACACGCACGGA
env-171   .AAAGAUAGAA......AGUAG.GU..U.UUCA..CACUGAGCGAAGGUCAU.....G..UUACUAAAAC..............UGGC.............GACAA..G.UCACUGUACAAUUUACAGAAGUUUCUAAGUGGAAAUGAUGAGGUGUGUCAUUUUUAUGACACGCACGGA
env-172   .AAAGAUAGAA......AGUAG.GU..U.UUCA..CACUGAGCGAAGGUCAU.....G..UUACUAAAAC..............UGGC.............GACAA..G.UCACUGUACAAUUUACAGAAGUUUCUAAGUGGAAAUGAUGAGGUGUGUCAUUUUUAUGACACGCACGGA
env-173   .......GGCAC.....UGCAG.GU..U.UUC...CGUGGAGCGAAUGUCAAGA...G..GCCGCGUAAGAU............GGCCAAAGAACCUGAC..GGCAA..G.UCACUGUACAAACAAUUUGUCUGCAAGAUUGCAGAGACACGGGCGUUAAGCGUACCGGACAUAAGAGCG
env-174   .......GGCAC.....UGCAG.GU..U.UUC...CGUGGAGCGAAUGUCAAGA...G..GCCGCGUAAGAU............GGCCAAAGAACCUGAC..GGCAA..G.UCACUGUACAAACAAUUUGUCUGCAAGAUUGCAGAGACACGGGCGUUAAGCGUACCGGCAUAAGAGUGC
env-175   .................GAUGG.GU..U.UUC...CGUGGAGCGAAUGUCAAGA...G..ACCGUAGAUCAGCAUAUCAC....GGUCGGUAAACCUGAC..GGCAA..G.UCACUGUACGUAACAUAGUUUCUGCCAGAUAGAGAGCAGAUAAACGUGUGAGAAAGUAUGCCACGAUGU
env-176   .................GAUGG.GU..U.UUC...CGUGGAGCGAAUGUCAAGA...G..ACCGUAGAUCAGCAUAUCAC....GGUCAGUAAACCUGAC..GGCAA..G.UCACUGUACGUAACAUAGUUUCUGCCAGAUAGAGAGCAGAUAAACGUGUGAGAAAGUAUGCCACGAUGU
env-177   .................GAUGG.GU..U.UUC...CGUGGAGCGAAUGUCAAGA...G..ACCGUAGAUCAGCAUAUCAC....GGUCAGUAAACCUGAC..GGCAA..G.UCACUGUACGUAACAUAGUUUCUGCCAGAUAGAGAGCAGAUAAACGUGUGAGAAAGUAUGCCACGAUGU
env-178   .................GAUGG.GU..U.UUC...CGUGGAGCGAAUGUCAAGA...G..ACCGUAGAUCAGCAUAUCAC....GGUCAGUAAACCUGAC..GGCAA..G.UCACUGUACGUAACAUAGUUUCUGCCAGAUAGAGAGCAGAUAAACGUGUGAGAAAGUAUGCCACGAUGU
env-179   .................GAUGG.GU..U.UUC...CGUGGAGCGAAUGUCAAGA...G..ACCGUAGAUCAGCAUAUCAC....GGUCAGUAAACCUGAC..GGCAA..G.UCACUGUACGUAACAUAGUUUCUGCCAGAUAGAGAGCAGAUAAACGUGUGAGAAAGUAUGCCACGAUGU
env-180   .................GGUGG.GU..UUUUC...CGUGGAGCGAAUGUCAAGA...G..GUCGAGAUCAGCAAAUUC......GGCCGGGAACCCUGAC..GGCAA..G.UCACUGUACAGUAACAUAUUUUCUGCCAAGGAGCAGAUAAACGUGUGAGUCAAGCCUCCAUAGGAGGA
env-183   .................GGUGG.GU..UUUUC...CGUGGAGCGAAUGUCAAGA...G..GUCGAGAUCAGCAAAUUC......GGCCGGGAACCCUGAC..GGCAA..G.UCACUGUACAGUAACAUAUUUUCUGCCAAGGAGCAGAUAAACGUGUGAGUCAAGCCUCCGUAGGAGGA
env-184   .................GGUGG.GU..UUUUC...CGUGGAGCGAAUGUCAAGA...G..GUCGAGAUCAGCAAAUUC......GGCCGGGAACCCUGAC..GGCAA..G.UCACUGUACAGUAACAUAUUUUCUGCCAAGGAGCAGAUAAACGUGUGAGUCAAGCCUCCAUAGGAGGA
env-185   .................GGUGG.GU..UUUUC...CGUGGAGCGAAUGUCAAGA...G..GUCGAGAUCAGCAAAUUC......GGCCGGGAACCCUGAC..GGCAA..G.UCACUGUACAGUAACAUAUUUUCUGCCAAGGAGCAGAUAAACGUGUGAGUCAAGCCUCCAUAGGAGGA
env-186   .................GGUGG.GU..UUUUC...CGUGGAGCGAAUGUCAAGA...G..GCCGAGAUCAGCAAAUUC......GGCCGGGAACCCUGAC..GGCAA..G.UCACUGUACAGUAACAUAUUUUCUGCCAAGGAGCAGAUAAACGUGUGAGUCAAGCCUCCAUAGGAGGA
env-188   .................GGUGG.GU..UUUUC...CGUGGAGCGAAUGUCAAGA...G..GCCGAGAUCAGCAAAUUC......GGCCGGGAACCCUGAC..GGCAA..G.UCACUGUACAGUAACAUAUUUUCUGCCAAGGAGCAGAUAAACGUGUGAGUCAAGCCUCUGUAGGAGGA
env-189   .................GGUGG.GU..UUUUC...CGUGGAGCGAAUGUCAAGA...G..GCCGAGAUCAGCAAAUUC......GGCCGGGAACCCUGAC..GGCAA..G.UCACUGUACAGUAACAUAUUUUCUGCCAAGGAGCAGAUAAACGUGUGAGUCAAGCCUCCAUAGGAGGA
env-190   .................GGUGG.GU..UUUUC...CGUGGAGCGAAUGUCAAGA...G..GCCGAGAUCAGCAAAUUC......GGCCGGGAACCCUGAC..GGCAA..G.UCACUGUACAGUAACAUAUUUUCUGCCAAGGAGCAGAUAAACGUGUGAGUCAAGCCUCCAUAGGAGGA
env-191   .................GGUGG.GU..UUUUC...CGUGGAGCGAAUGUCAAGA...G..GCCGAGAUCAGCAAAUUC......GGCCGGGAACCCUGAC..GGCAA..G.UCACUGUACANNNNNNNNNNNNNNNNNNNNNNNNNNNNNNNNNNNNNNNNNNNNNNNNNNNNNNNNNNN
env-192   .................GGUGG.GU..UUUUC...CGUGGAGCGAAUGUCAAGA...G..GCCGAGAUCAGCAAAUUC......GGCCGGGAACCCUGAC..GGCAA..G.UCACUGUACAGUAACAUAUUUUCUGCCAAGGAGCAGAUAAACGUGUGAGUCAAGCCUCCAUAGGAGGA
env-193   .................GGUGG.GU..UUUUC...CGUGGAGCGAAUGUCAAGA...G..GCCGAGAUCAGCAAAUUC......GGCCGGGAACCCUGAC..GGCAA..G.UCACUGUACAGUAACAUAUUUUCUGCCAAGGAGCAGAUAAACGUGUGAGUCAAGCCUCCAUAGGAGGA
env-194   .................GGUGG.GU..UUUUC...CGUGGAGCGAAUGUCAAGA...G..GCCGAGAUCAGCAAAUUC......GGCCGGGAACCCUGAC..GGCAA..G.UCACUGUACAGUAACAUAUUUUCUGCCAAGGAGCAGAUAAACGUGUGAGUCAAGCCUCCGUAGGAGGA
env-195   .................GGUGG.GU..UUUUC...CGUGGAGCGAAUGUCAAGA...G..GCCGAGAUCAGCAAAUUC......GGCCGGGAACCCUGAC..GGCAA..G.UCACUGUACAGUAACAUAUUUUCUGCCAAGGAGCAGAUAAACGUGUGAGUCAAGCCUCCAUAGGAGGA
env-208   .................GGUGG.GU..UUUUC...CGUGGAGCGAAUGUCAAGA...G..GUCGAGAUCAGCAAAUUC......GGCC.GGAAACCCUGAC.GGCAA..G.UCACUGUACAGUAACAUAUUUUCUGCCAAGGAGCAGAUAAACGUGUGAGUCAAGCCUCCAUAGGAGGA
Lba-19-1  .................GGCAG.GU..U.UUC...CGCGGAGCGAAUGUUGAUG...G..CGGGUAAUACAGAUACGAUGUAUUCCGCAAGAAUAUUAAC..GACAA..G.UCACUGUACAGAUGGGAACCGUCCGGGAAACCGGAAACGACGCGUGGUCCUGCGUCUUUUUAAAGGA
Lba-17-1  ......GCUGU......AAUAG.GU..U.UUC...CGUGGAGCGAAUGUCA......G..AUAAUA..................UGAC.............GGCAA..G.UCACUGUACAAAGCUUAAACAGUCUACAAUAUAGUAGAAACGACGCGUGAGGACGCGCUUUUAAAAUGA
env-212   ......GCUGU......AAUAG.GU..U.UUC...CGUGGAGCGAAUGUCA......G..AUAAUA..................UGAC.............GGCAA..G.UCACUGUACAAAGCUUAAACAGUCUACAAUAUAGUAGAAACGACGCGUGAGGACGCGCUUUUAAAAUGA
env-213   .................UGCAG.GU..U.UUC...CGUGGAGCGAAUGUCAGU....G..GCGGAUAUAUGUUC..........CGCCGAUAAUAUUGAC..GACAA..G.UCACUGUACAAUGCUUAGACAGUCUGCAUGAAAGCAGAAACGACGCGUGUGAAUCAUGGCUCUGAAAGUUC
env-214   .................UACAG.GU..U.UUU...CGUGGAGCGAAUGUCA......A..UAAAGAAUUAC.............UGAC.............GACAA..G.UCACUGUACCAAUAACUGUCUAUCAAAAGGAUAGAAACGACGUGUUGAUUCUGAUUUAAUUCAGGAUAC
env-215   .................UACAG.GU..U.UUU...CGUGGAGCGAAUGUCA......A..UAAAGAAUUAC.............UGAC.............GACAA..G.UCACUGUACCAAUAACUGUCUAUCAAAAGGAUAGAAACGACGUGUUGAUUCUGAUUUAAUUCAGGAU..
env-216   .................UACAG.GU..U.UUU...CGUGGAGCGAAUGUCAAU........AAAGAAUUAC.............UGAC.............GACAA..G.UCACUGUACCAAUAACUGUCUAUCAAAAGGAUAGAAACGACGUGUUGAUUCUGAUUUAAUUCAGGAUAC
env-217   .................UACAG.GU..U.UUU...CGUGGAGCGAAUGUCAAU........AAAGAAUUAC.............UGAC.............GACAA..G.UCACUGUACCAAUAACUGUCUAUCAAAAGGAUAGAAACGACGUGUUGAUUCUGAUUUAAUUCAGGAUAC
env-218   .................UACAG.GU..U.UUU...CGUGGAGCGAAUGUCAAU........AAAGAAUUAC.............UGAC.............GACAA..G.UCACUGUACCAAUAACUGUCUAUCAAAAGGAUAGAAACGACGUGUUGAUUCUGAUUUAAUUCAGGAUAC
env-219   .................UACAG.GU..U.UUU...CGUGGAGCGAAUGUCAAU........AAAGAAUUAC.............UGAC.............GACAA..G.UCACUGUACCAAUAACUGUCUAUCAAAAGGAUAGAAACGACGUGUUGGUUCUGAUUUAAUUCAGGAUAC
env-220   .................UACAG.GU..U.UUU...CGUGGAGCGAAUGUCAAU........AAAGAAUUAC.............UGAC.............GACAA..G.UCACUGUACCAAUAACUGUCUAUCAAAAGGAUAGAAACGACGUGUUGAUUCUGAUUUAAUUCAGGAU..
env-223   .................UACAG.GU..U.UUU...CGUGGAGCGAAUGUCAAU........AAAGAAUUAC.............UGAC.............GACAA..G.UCACUGUACCAAUAACUGUCUAUCAAAAGGAUAGAAACGACGUGUUGAUUCUGAUUUAAUUCAGGAUAC
env-224   .................UACAG.GU..U.UUU...CGUGGAGCGAAUGUCAAU........AAAGAAUUAC.............UGAC.............GACAA..G.UCACUGUACCAAUAACUGUCUAUCAAAAGGAUAGAAACGACGUGUUGAUUCUGAUUUAAUUCAGGAUAC
env-225   .................UACAG.GU..U.UUU...CGUGGAGCGAAUGUCAAU........AAAGAAUUAC.............UGAC.............GACAA..G.UCACUGUACCAAUAACUGUCUAUCAAAAGGAUAGAAACGACGUGUUGAUUCUGAUUUAAUUCAGGAUAC
env-229   .................UACAG.GU..U.UUU...CGUGGAGCGAAUGUCAAU........AAAGAAUUAC.............UGAC.............GACAA..G.UCACUGUACCAAUAACUGUCUAUCAAAAGGAUAGAAACGACGUGUUGAUUCUGAUUUAAUUCAGGAUAC
env-231   .................UACAG.GU..U.UUU...CGUGGAGCGAAUGUCAAU........AAAGAAUUAC.............UGAC.............GACAA..G.UCACUGUACCAAUAACUGUCUAUCAAAAGGAUAGAAACGACGUGUUGGUUCUGAUUUAAUUCAGGAUAC
env-234   .................UACAG.GU..U.UUU...CGUGGAGCGAAUGUCAAU........AAAGAAUUAC.............UGAC.............GACAA..G.UCACUGUACCA..........................................................
env-235   .................UACAG.GU..U.UUU...CGUGGAGCGAAUGUCAAU........AAAGAAUUAC.............UGAC.............GACAA..G.UCACUGUACCAAUAACUGUCUAUCAAAAGGAUAGAAACGACGUGUUGAUUCUGAUUUAAUUCAGGAUAC
env-236   .................UACAG.GU..U.UUU...CGUGGAGCGAAUGUCAAU........AAAGAAUUAC.............UGAC.............GACAA..G.UCACUGUACCAAUAACUGUCUAUCAAAAGGAUAGAAACGACGUGUUGAUUCUGAUUUAAUUCAGGAUAC
env-237   .................UAUAG.GU..U.UUU...CGUGGAGCGAAUGUCAA.......UAAAAAA..................UUACUGAC..........GACAA..G.UCACUGUACCAAUAACUGUCUAUCAAAAGGAUAGAAACGACGUGUUGAUUCUGAUUUAAUUCAGGAUAC
env-242   .................AGUAG.GU..U.UUCU..UAUGGAGCG.AUGUCUA.....GU.UCAAGAAAC..............UGAC.............GACAA..AUCACUGUACAAAUAUCUUAAAGAUUUAGCCAUGUGAACCGGAUGAUACACGUGGAAAGUGUACAGUCA
env-244   .................AGUAG.GU..U.UUCU..UAUGGAGCG.AUGUCUA.....GU.UCAAGAAAC..............UGAC.............GACAA..AUCACUGUACAA..........................................................
env-247   .......CGGAU.....AAGAG.GU..U.UUC...CGUGGAGCGAAUGUCAA.....G..UUGAGAAU................UGGC.............GACAA..G.UCACUGUACGAAAAUCCAAUAUUCUGUAGAGCAUAUUUGUAAGAAUGUGCCCGGAAAACACGGAAGAG
env-248   .......CGGAU.....AAGAG.GU..U.UUC...CGUGGAGCGAAUGUCAA.....G..UUGAGAAU................UGGC.............GACAA..G.UCACUGUACGAAAAUCCAAUAUUCUGUAGAGCAUAUUUGUAAGAAUGUGCCCGGAAAACACGGAAGAG
env-249   .......CGGAU.....AAGAG.GU..U.UUC...CGUGGAGCGAAUGUCAA.....G..UUGAGAAU................UGGC.............GACAA..G.UCACUGUACGAAAAUCCAAUAUUCUGUAGAGCAUAUUUGUAAGAAUAUGCCCGGAAAACACGGAAGAG
env-250   .......CGGAU.....AAGAG.GU..U.UUC...CGUGGAGCGAAUGUCAA.....G..UUGAGAAU................UGGC.............GACAA..G.UCACUGUACGAAAAUCCAAUAUUCUGUAGAGCAUAUUUGUAAGAAUAUGCCCGGAAAACACGGAAGAG
env-251   .......CGGAU.....AAGAG.GU..U.UUC...CGUGGAGCGAAUGUCAA.....G..UUGAGAAU................UGGC.............GACAA..G.UCACUGUACGAAAAUCCAAUAUUCUGUAGAGCAUAUUUGUAAGAAUAUGCCCGGAAAACACGGAAGAG
env-252   .......CGGAU.....AAGAG.GU..U.UUC...CGUGGAGCGAAUGUCAA.....G..UUGAGAAU................UGGC.............GACAA..G.UCACUGUACGAAAAUCCAAUAUUCUGUAGAGCAUAUUUGUAAGAAUAUGCCCGGAAAACACGGAAGAG
env-257   .......CGGAU.....AAGAG.GU..U.UUC...CGUGGAGCGAAUGUCAA.....G..UUGAGAAU................UGGC.............GACAA..G.UCACUGUACGAAAAUCCAAUAUUCUGUAGAGCAUAUUUGUAAGAAUAUGCCCGGAAAACACGGAAGAG
env-259   .......CGGAU.....AAGAG.GU..U.UUC...CGUGGAGCGAAUGUCAA.....G..UUGAGAAU................UGGC.............GACAA..G.UCACUGUACGAAAAUCCAAUAUUCUGUAGAGCAUAUUUGUAAGAAUAUGCCCGGAAAACACGGAAGAG
env-262   .......CGGAU.....AAGAG.GU..U.UUC...CGUGGAGCGAAUGUCAA.....G..UUGAGAAU................UGGC.............GACAA..G.UCACUGUACGAAAAUCCAAUAUUCUGUAGAGCAUAUUUGUAAGAAUAUGCCCGGAAAACACGGAAGAG
env-264   .......CGGAU.....AAGAG.GU..U.UUC...CGUGGAGCGAAUGUCAA.....G..UUGAGAAU................UGGC.............GACAA..G.UCACUGUACGAAAAUCCAAUAUUCUGUAGAGCAUAUUUGUAAGAAUAUGCCCGGAAAACACGGAAGAG
env-265   .......CGGAU.....AAGAG.GU..U.UUC...CGUGGAGCGAAUGUCAA.....G..UUGAGAAU................UGGC.............GACAA..G.UCACUGUACGAAAAUCCAAUAUUCUGUAGAGCAUAUUUGUAUGAAUAUGCCCGGAAAACACGGGAAGAG
env-266   .................GACAGAGA...A.UUC...GGCGGAGCGAAUGUCAA.....G..UUAGGAAAC...............UGAC.............GGCAA..G.UCACUGUACAGAACAGAUAUGGUCUACAAAAUGUGAAACAACGUGUGAAAGUCCACAGGACACACACG
env-280   .................GACAGAGA...A.UUC...GGCGGAGCGAAUGUCAA.....G..UUAGGAAAC...............UGAC.............GGCAA..G.UCACUGUACAGAACAGAUAUGGUCUACAAAAUGUGAAACAACGUGUGAAAGUCCACACGACACACACG
env-281   .................GACAGAGA...A.UUC...GGCGGAGCGAAUGUCAA.....G..UUAGGAAAC...............UGAC.............GGCAA..G.UCACUGUACAGAACAGAUAUGGUCUACAAAAUGUGAAACAACGUGUGAAAGUCCACAGGACACACACG
env-282   .................GACAGAGA...A.UUC...GGCGGAGCGAAUGUCAA.....G..UUAGGAAAC...............UGAC.............GGCAA..G.UCACUGUACAGAACAGAUAUGGUCUACAAAAUGUGAAACAACGUGUGAAAGUCCACAGGACACACACG
env-283   .................GACAGAGA...A.UUC...GGCGGAGCGAAUGUCAA.....G..UUAGGAAAC...............UGAC.............GGCAA..G.UCACUGUACAGAACAGAUAUGGUCUACAAAAUGUGAAACAACGUGUGAAAGUCCACAGGACACACACG
env-284   .................GACAGAGA...A.UUC...GGCGGAGCGAAUGUCAA.....G..UUAGGAAAC...............UGAC.............GGCAA..G.UCACUGUACAGAACAGAUAUGGUCUACAAAAUGUGAAACAACGUGUGAAAGCCCACAGGACACACACG
env-288   .................GACAGAGA...A.UUC...GGCGGAGCGAAUGUCAA.....G..UUAGGAAAC...............UGAC.............GGCAA..G.UCACUGUACAGAACAGAUAUGGUCUACAAAAUGUGAAACAACGUGUGAAAGUCCACAGGACACACACG
env-289   .................GACAGAGA...A.UUC...GGCGGAGCGAAUGUCAA.....G..UUAGGAAAC...............UGAC.............GGCAA..G.UCACUGUACAGAACAGAUAUGGUCUACAAUAUGUGAAACAACGUGUGAAAGUCCACAGGACACACACG
env-290   .................GACAGAGA...A.UUC...GGCGGAGCGAAUGUCAA.....G..UUAGGAAAC...............UGAC.............GGCAA..G.UCACUGUACAGAACAGAUAUGGUCUACAAAAUGUGAAACAACGUGUGAAAGUCCACAGNNNNNNNNNN
env-291   .................GACAGAGA...A.UUC...GGCGGAGCGAAUGUCAA.....G..UUAGGAAAC...............UGAC.............GGCAA..G.UCACUGUACAGAACAGAUAUGGUCUACAAAAUGUGAAACAACGUGUGAAAGUCCACAGGACACACACG
env-300   .................GACAGAGA...A.UUC...GGCGGAGCGAAUGUCAA.....G..UUAGGAAAC...............UGAC.............GGCAA..G.UCACUGUACAGAACAGAUAUGGUCUACAAAAUGUGAAA.............................
env-303   .................GACAGAGA...A.UUC...GGCGGAGCGAAUGUCAA.....G..UUAGGAAAC...............UGAC.............GGCAA..G.UCACUGUACAGAACAGAUAUGGUCUACAAAAUGUGAAACAACGUGUGAAAGUCCACAGGACACACACG

          >>>>>>>>>>.......>>>>>.....>.>>>...>>>.........<<<<.......<..<<<......................>>>>.............>>>>...>.>>>>.>>.................................................................
          01.1111111.......22221.....2.102...222.........111.......2..212.......................2122.............111...1.111..12..................................................................
          ----------------•GYRG-GU--U-UUC---CGYGGAGCGAAUGUC•R-----G--•Y•R•RRR•---------------•G•C-------------GRCAA--G-UCACUGUAC•RR•••••••••••••••••••••••••••••••••••••••••••••••••••••
                                                                                                                   3′
```

```
env-304    .................GACAGAGA..A.UUC...GGCGGAGCGAAUGUCAA.....G..UUAGGAAAC...............UGAC..............GGCAA..G.UCACUGUACAGAACAGAUAUGGUCUACAAAAUGUAGAAACAACGUGUGAAAGUCCACAGGACACACACG
env-305    .................GACAGAGA..A.UUC...GGCGGAGCGAAUGUCAA.....G..UUAGGAAAC...............UGAC..............GGCAA..G.UCACUGUACAGAACAGAUAUGGUCUACAAAAUGUAGAAACAACGUGUGAAAGUCCACAGGACACACACG
env-306    .................GACAGAGA..A.UUC...GGCGGAGCGAAUGUCAA.....G..UUAGGAAAC...............UGAC..............GGCAA..G.UCACUGUACAGAACAGAUAUGGUCUACAAAAUGUAGAAACAACGUGUGAAAGUCCGCAGGACACACACG
env-307    .................GACAGAGA..A.UUC...GGCGGAGCGAAUGUCAA.....G..UUAGGAAAC...............UGAC..............GGCAA..G.UCACUGUACAGAACAGAUAUGGUCUACAAUAUGUAGAAACAACGUGUGAAAGUCCACAGGACACACACG
env-308    .................GACAGAGA..A.UUC...GGCGGAGCGAAUGUCAA.....G..UUAGGAAAC...............UGAC..............GGCAA..G.UCACUGUACAGAACAGAUAUGGUCUACAAAAUGUAGAAACAACGUGUGAAAGUCUACAGGACACACACG
env-309    .................GACAGAGA..A.UUC...GGCGGAGCGAAUGUCAA.....G..UUAGGAAAC...............UGAC..............GGCAA..G.UCACUGUACAGAACAGAUAUGGUCUACAAAAUGUAGAAACAACGUGUGAAAGUCCACAGGACACACACG
env-310    .................GACAGAGA..A.UUC...GGCGGAGCGAAUGUCAA.....G..UUAGGAAAC...............UGAC..............GGCAA..G.UCACUGUACAGAACAGAUAUGGUCUACAAUAUGUAGAAACAACGUGUGAAAGUCCACAGGACACACACG
env-311    .................GACAGAGA..A.UUC...GGCGGAGCGAAUGUCAA.....G..UUAGGAAAC...............UGAC..............GGCAA..G.UCACUGUACAGAACAGAUAUGGUCUACAAAAUGUAGAAACAACGUGUGAAAGUCCACAGGACACACACG
env-312    .................GACAGAGA..A.UUC...GGCGGAGCGAAUGUCAA.....G..UUAGGAAAC...............UGAC..............GGCAA..G.UCACUGUACAGAACAGAUAUGGUCUACAAAAUGUAGAAACAACGUGUGAAAGUCCACAGGACACACACG
env-313    .................GACAGAGA..A.UUC...GGCGGAGCGAAUGUCAA.....G..UUAGGAAAC...............UGAC..............GGCAA..G.UCACUGUACAGAACAGAUAUGGUCUACAAAAUGUAGAAACAACGUGUGAAAGUCCACAGGACACACACG
env-314    .................GACAGAGA..A.UUC...GGCGGAGCGAAUGUCAA.....G..UUAGGAAAC...............UGAC..............GGCAA..G.UCACUGUACAGAACAGAUAUGGUCUACAAAAUGUAGAAACAACGUGUGAAAGUCCACAGGACACACACG
env-315    .................GACAGAGA..A.UUC...GGCGGAGCGAAUGUCAA.....G..UUAGGAAAC...............UGAC..............GGCAA..G.UCACUGUACAGAACAGAUAUGGUCUACAAAAUGUAGAAACAACGUGUGAAAGUCCACAGGACACACACG
env-316    .................GACAGAGA..A.UUC...GGCGGAGCGAAUGUCAA.....G..UUAGGAAAC...............UGAC..............GGCAA..G.UCACUGUACAGAACAGAUAUGGUCUACAAAAUGUAGAAACAACGUGUGAAAGUCCACAGGACACACACG
env-317    .................GACAGAGA..A.UUC...GGCGGAGCGAAUGUCAA.....G..UUAGGAAAC...............UGAC..............GGCAA..G.UCACUGUACAGAACAGAUAUGGUCUACAAAAUGUAGAAACAACGUGUGAAAGUCCACAGGACACACACG
env-331    .................GACAGAGA..A.UUC...GGCGGAGCGAAUGUCAA.....G..UUAGGAAAC...............UGAC..............GGCAA..G.UCACUGUACAGAACAGAUAUGGUCUACAAAAUGUAGAAACAACGUGUGAAAGUCCACAGGACACACACG
env-332    .................GACAGAGA..A.UUC...GGCGGAGCGAAUGUCAA.....G..UUAGGAAAC...............UGAC..............GGCAA..G.UCACUGUACAGAACAGAUAUGGUCUACAAAAUGUAGAAACAACGUGUGAAAGUCCACAGGACACACACG
env-333    .................GACAGAGA..A.UUC...GGCGGAGCGAAUGUCAA.....G..UUAGGAAAC...............UGAC..............GGCAA..G.UCACUGUACAGAGCAGAUAUGGUCUACAAAAUGUAGAAACAACGUGUGAAAGUCCACAGGACACACACG
env-334    .................GACAGAGA..A.UUC...GGCGGAGCGAAUGUCAA.....G..UUAGGAAAC...............UGAC..............GGCAA..G.UCACUGUACAGAACAGAUAUGGUCUACAAAAUGUAGAAACAACGUGUGAAAGUCCACAGGACACACACG
env-335    .................GACAGAGA..A.UUC...GGCGGAGCGAAUGUCAA.....G..UUAGGAAAC...............UGAC..............GGCAA..G.UCACUGUACAGAACAGAUAUGGUCUACAAAAUGUAGAAACAACGUGUGAAAGUCCACAGGACACACACG
env-336    .................GACAGAGA..A.UUC...GGCGGAGCGAAUGUCAA.....G..UUAGGAAAC...............UGAC..............GGCAA..G.UCACUGUACAGAACAGAUAUGGUCUACAAUAUGUAGAAACAACGUGUGAAAGUCCACAGGACACACACG
env-338    .................GACAGAGA..A.UUC...GGCGGAGCGAAUGUCAA.....G..UUAGGAAAC...............UGAC..............GGCAA..G.UCACUGUACAGAACAGAUAUGGUCUACGAAAUGUAGAAACAACGUGUGAAAGUCCACAGGACACACACG
env-339    .................GACAGAGA..A.UUC...GGCGGAGCGAAUGUCAA.....G..UUAGGAAAC...............UGAC..............GGCAA..G.UCACUGUACAGAACAGAUAUGGUCUACAAAAUGUAGAAACAACGUGUGAAAGUCCACAGGACACACACG
env-340    .................GACAGAGA..A.UUC...GGCGGAGCGAAUGUCAA.....G..UUAGGAAAC...............UGAC..............GGCAA..G.UCACUGUACAGAACAGAUAUGGUCUACAAUAUGUAGAAACAACGUGUGAAAGUCCACAGGACACACACG
env-341    .................GACAGAGA..A.UUC...GGCGGAGCGAAUGUCAA.....G..UUAGGAAAC...............UGAC..............GGCAA..G.UCACUGUACAGAACAGAUAUGGUCUACAAAAUGUAGAAACAACGUGUGAAAGUCCACAGGACACACACG
env-347    .................GACAGAGA..A.UUC...GGCGGAGCGAAUGUCAA.....G..UUAGGAAAC...............UGAC..............GGCAA..G.UCACUGUACAGAACAGAUAUGGUCUACAAAAUGUAGAAACAACGUGUGAAAGUCCACAGGACACACACG
env-348    .................GACAGAGA..A.UUC...GGCGGAGCGAAUGUCAA.....G..UUAGGAAAC...............UGAC..............GGCAA..G.UCACUGUACAGAACAGAUAUGGUCUACAAAAUGUAGAAACAACGUGUGAAAGUCCACAGGACACACACG
env-349    .................GACAGAGA..A.UUC...GGCGGAGCGAAUGUCAA.....G..UUAGGAAAC...............UGAC..............GGCAA..G.UCACUGUACAGAACAGAUAUGGUCUACAAAAUGUAGAAACAACGUGUGAAAGUCAACAGGACACACACG
env-350    .................GACAGAGA..A.UUC...GGCGGAGCGAAUGUCAA.....G..UUAGGAAAC...............UGAC..............GGCAA..G.UCACUGUACAGAACAGAUAUGGUCUACAAAAUGUAGAAACAACGUGUGAAAGUCUACAGGACACACACG
env-351    .................GACAGAGA..A.UUC...GGCGGAGCGAAUGUCAA.....G..UUAGGAAAC...............UGAC..............GGCAA..G.UCACUGUACAGAACAGAUAUGGUCUACAAAAUGUAGAAACAACGUGUGAAAGUCCACAGGACACACACG
env-352    .................GACAGAGA..A.UUC...GGCGGAGCGAAUGUCAA.....G..UUAGGAAAC...............UGAC..............GGCAA..G.UCACUGUACAGAACAGAUAUGGUCUACAAAAUGUAGAAACAACGUGUGAAAGUCCACAGGACACACACG
env-353    .................GACAGAGA..A.UUC...GGCGGAGCGAAUGUCAA.....G..UUAGGAAAC...............UGAC..............GGCAA..G.UCACUGUACAGAACAGAUAUGGUCUACAAUAUGUAGAAACAACGUGUGAAAGUCCACAGGACACACACG
env-354    .................GACAGAGA..A.UUC...GGCGGAGCGAAUGUCAA.....G..UUAGGAAAC...............UGAC..............GGCAA..G.UCACUGUACAGAACAGAUAUGGUCUACAAUAUGUAGAAACAACGUGUGAAAGUCCACAGGACACACACG
env-355    .................GACAGAGA..A.UUC...GGCGGAGCGAAUGUCAA.....G..UUAGGAAAC...............UGAC..............GGCAA..G.UCACUGUACAGAACAGAUAUGGUCUACAAUAUGUAGAAACAACGUGUGAAAGUCUACAGGACACACACG
env-359    .................GACAGAGA..A.UUC...GGCGGAGCGAAUGUCAA.....G..UUAGGAAAC...............UGAC..............GGCAA..G.UCACUGUACAGAACAGAUAUGGUCUACAAUAUGUAGAAACAACGUGUGAAAGUCCACAGGACACACACG
env-360    .................GACAGAGA..A.UUC...GGCGGAGCGAAUGUCAA.....G..UUAGGAAAC...............UGAC..............GGCAA..G.UCACUGUACAGAACAGAUAUGGUCUACAAUAUGUAGAAACAACGUGUGAAAGUCCGCAGGACACACACG
env-361    .................GACAGAGA..A.UUC...GGCGGAGCGAAUGUCAA.....G..UUAGGAAAC...............UGAC..............GGCAA..G.UCACUGUACAGAACAGAUAUGGUCUACAAAAUGUAGAAACAACGUGUGAAAGUCCACAGGACACACACG
env-362    .................GACAGAGA..A.UUC...GGCGGAGCGAAUGUCAA.....G..UUAGGAAAC...............UGAC..............GGCAA..G.UCACUGUACAGAACAGAUAUGGUCUACAAAAUGUAGAAACAACGUGUGAAAGUCCACAGGACACACACG
env-363    .................GACAGAGA..A.UUC...GGCGGAGCGAAUGUCAA.....G..UUAGGAAAC...............UGAC..............GGCAA..G.UCACUGUACAGAACAGAUAUGGUCUACAAUAUGUAGAAACAACGUGUGAAAGUCCACAGGACACACACG
env-364    .................GACAGAGA..A.UUC...GGCGGAGCGAAUGUCAA.....G..UUAGGAAAC...............UGAC..............GGCAA..G.UCACUGUACAGAACAGAUAUGUUUACAAUAUG.....................................
env-365    .................GACAGAGA..A.UUC...GGCGGAGCGAAUGUCAA.....G..UUAGGAAAC...............UGAC..............GGCAA..G.UCACUGUACAGAACAGAUAUGGUCUACAAAAUGUAGAAACAACGUGUGAAAGUCCACAGGACACACACG
env-366    .................GACAGAGA..A.UUC...GGCGGAGCGAAUGUCAA.....G..UUAGGAAAC...............UGAC..............GGCAA..G.UCACUGUACAGAACAGAUAUGGUCUACAAUAUGUAGAAACAACGUGUGAAAGUCCACAGGACACACACG
env-368    .................GACAGAGA..A.UUC...GGCGGAGCGAAUGUCAA.....G..UUAGGAAAC...............UGAC..............GGCAA..G.UCACUGUACAGAACAGAUAUGGUCUACAAUAUGUAGAAACAACGUGUGAAAGUCUACAGGACACACACG
env-372    .................GACAGAGA..A.UUC...GGCGGAGCGAAUGUCAA.....G..UUAGGAAAC...............UGAC..............GGCAA..G.UCACUGUACAGAACAGAUAUGGUCUACAAAAUGUAGAAACAACGUGUGAAAGUCCACAGGACACACACG
env-373    .................GACAGAGA..A.UUC...GGCGGAGCGAAUGUCAA.....G..UUAGGAAAC...............UGAC..............GGCAA..G.UCACUGUACAGAACAGAUAUGGUCUACAAAAUGUAGAAACAACGUGUGAAAGUCCACAGGACACACACG
env-374    .................GACAGAGA..A.UUC...GGCGGAGCGAAUGUCAA.....G..UUAGGAAAC...............UGAC..............GGCAA..G.UCACUGUACAGAACAGAUAUGGUCUACAAUAUGUAGAAACAACGUGUGAAAGUCUACAGGACACACACG
env-375    .................GACAGAGA..A.UUC...GGCGGAGCGAAUGUCAA.....G..UUAGGAAAC...............UGAC..............GGCAA..G.UCACUGUACAGAACAGAUAUGGUCUACAAAAUGUAGAAACAACGUGUGAAAGUCCAC............
env-376    .................AGCAG.GU..U.UUC...CGUGGAGCGAAUGUCUAGAG..A..CCACUAAUG...............UGGUCAAACAAACUGAC.GACAA..G.UCACUGUACAGACAAUUAAGUCUGCCGACUGACCCAAGGCAGAAACGACGUGUAGGGAUUCUAUGGCCU
Bsp-5-1    .................AGCAG.GU..U.UUC...CGUGGAGCGAAUGUCUAGAG..A..CCACUAAUG...............UGGUCAAACAAACUGAC.GACAA..G.UCACUGUACAGACAAUUAAGUCUGCCGACUAACCCAAGGCAGAAACGACGUGUAGGGAUUCUAUGGCCU
env-377    .................AUCGG.GU..U.UUA...UGUGGAGCGAAGGUCAU.....G..UUGCAAAAAC..............UGGC..............GACAA..G.UCACUGUACAAACAGCAUCUCAUAUUGAGUAAUUUCGUGGGUUAAUUUUUAUGACACGCACGGAAGAGU
env-379    .................AUCGG.GU..U.UUA...UGUGGAGCGAAGGUCAU.....G..UUGCAAAAAC..............UGGC..............GACAA..G.UCACUGUACAAACAGCAUCUCAUAUUGAGUAAUUUCGUGCUUUUAUUUUUAUGACACGCACGGAAGAGU
env-380    .................AUCGG.GU..U.UUA...UGUGGAGCGAAGGUCAU.....G..UUGCAAAAAC..............UGGC..............GACAA..G.UCACUGUACAAUUUACAGAAGUUUCUAAGUGGAAAUGAUGAGGUGUGUCAUUUUUAUGACACGCACGGA
env-381    .................AUCGG.GU..U.UUA...UGUGGAGCGAAGGUCAU.....G..UUGCAAAAAC..............UGGC..............GACAA..G.UCACUGUACAAACAGCAUCUCAUAUUGAGUAAUUUCGUGGGUUUAUUUUUAUGACACGCACGGAAGAGU
env-382    .................AUCGG.GU..U.UUA...UGUGGAGCGAAGGUCAU.....G..UUGCAAAAAC..............UGGC..............GACAA..G.UCACUGUACAAACAGCAUCUCAUAUUGAGUAAUUUCGUGCUUUUAUUUUUAUGACACGCACGGAAGAGU
env-384    .................AUCGG.GU..U.UUA...UGUGGAGCGAAGGUCAU.....G..UUGCAAAAAC..............UGGC..............GACAA..G.UCACUGUACAAACAGCAUCUCAUAUUGAGUAAUUUCGUGUGUUUAUUUUUAUGACACGCACGGAAGAGU
Lor-1-1    .................AUCGG.GU..U.UUA...UGUGGAGCGAAGGUCAU.....G..UUGCAAAAAC..............UGGC..............GACAA..G.UCACUGUACAAACAGCAUCUCAUAUUGAGUAAUUUCGUGGGUUUAUUUUUAUGACACGCACGGAAGAGU
env-385    .................AUCGG.GU..U.UUA...UGUGGAGCGAAGGUCAU.....G..UUGCAAAAAC..............UGGC..............GACAA..G.UCACUGUACAAACAGCAUCUCAUAUUGAGUAAUUUCGUGCUUUUAUUUUUAUGACACGCACGGAAGAGU
env-386    .................AUCGG.GU..U.UUA...UGUGGAGCGAAGGUCAU.....G..UUGCAAAAAC..............UGGC..............GACAA..G.UCACUGUACAAACAGCAUCUCAUAUUGAGUAAUUUCGUGUGUUUAUUUUUAUGACACGCACGGAAGAGU
env-387    .................AUCGG.GU..U.UUA...UGUGGAGCGAAGGUCAU.....G..UUGCAAAAAC..............UGGC..............GACAA..G.UCACUGUACAAACAGCAUCUCAUAUUGAGUAGUUUCGUGCUUUUAUUUUUAUGACACGCACGGAAGAGU
env-388    .................UACGG.GU..U.UUA...UGUGGAGCGAAGGUCAU.....G..UUGCAAAAAC..............UGGC..............GACAA..G.UCACUGUACAAACAGCAUCUCAUAUUGAGUAUUUUCGUGGGUUAGUCACUAUGACACGCACGGAAGAGU
Lba-15-1   .................UACGG.GU..U.UUA...UGUGGAGCGAAGGUCAU.....G..UUGCAAAAAC..............UGGC..............GACAA..G.UCACUGUACAAACAGCAUCUCAUAUUGAGUAUUUUCGUGGGUUAGUCACUAUGACACGCACGGAAGAGU
env-389    .................UACGG.GU..U.UUA...UGUGGAGCGAAGGUCAU.....G..UUGCAAAAAC..............UGGC..............GACAA..G.UCACUGUACAAACAGCAUCUCAUAUUGAGUAUUUUCGUGGGUUAGUCACUAUGACACGCACGGAAGAGU
env-390    .................UACGG.GU..U.UUA...UGUGGAGCGAAGGUCAU.....G..UUGCAAAAAC..............UGGC..............GACAA..G.UCACUGUACAAACAGCAUCUCAUAUUGAGUAUUUUCGUGGGUUAGUCACUAUGACACGCACGGAAGAGU
env-391    .................UACGG.GU..U.UUA...UGUGGAGCGAAGGUCAU.....G..UUGCAAAAAC..............UGGC..............GACAA..G.UCACUGUACAAACAGCAUCUCAUAUUGAGUAUUUUCGUGGGUUAGUCACUAUGACACGCACGGAAGAGU
env-392    .................UACGG.GU..U.UUA...UGUGGAGCGAAGGUCAU.....G..UUGCAAAAAC..............UGGC..............GACAA..G.UCACUGUACAAACAGCAUCUCAUAUUGAGUAUUUUCGUGGGUUGGUCACUAUGACACGCACGGAAGAGU
env-393    .................UACGG.GU..U.UUA...UGUGGAGCGAAGGUCAU.....G..UUGCAAAAAC..............UGGC..............GACAA..G.UCACUGUACAAACAGCAUCUCAUAUUGAGUAUUUUCGUGGGUUAGUCACUAUGACACGCACGGAAGAGU
env-394    .................UACGG.GU..U.UUA...UGUGGAGCGAAGGUCAU.....G..UUGCAAAAAC..............UGGC..............GACAA..G.UCACUGUACAAACAGCAUCUCAUAUUGAGUAUUUUCGUGGGUUAGUCACUAUGACACGCACGGAAGAGU
env-395    .................UACGG.GU..U.UUA...UGUGGAGCGAAGGUCAU.....G..UUGCAAAAAC..............UGGC..............GACAA..G.UCACUGUACAAACAGCAUCUCAUAUUGAGUAUUUUCGUGGGUUGGUCACUAUGACACGCACGGAAGAGU
env-396    .................UACGG.GU..U.UUA...UGUGGAGCGAAGGUCAU.....G..UUGCAAAAAC..............UGGC..............GACAA..G.UCACUGUACAAACAGCAUCUCAUAUUGAGU.......................................
env-397    .................UACGG.GU..U.UUA...UGUGGAGCGAAGGUCAU.....G..UUGCAAAAAC..............UGGC..............GACAA..G.UCACUGUACAAACAGCAUCUCAUAUUGAGUAUUUCGUGUGGUUAAGUUUUACGACACGCACGGAAGAGU
env-398    .................UACGG.GU..U.UUA...UGUGGAGCGAAGGUCAU.....G..UUGCAAAAAC..............UGGC..............GACAA..G.UCACUGUACAAACAGCAUCUCAUAUUGAGUAUUUCGUGUAGUUAAGUUUUACGACACGCACGGAAGAGU
env-399    .................UACGG.GU..U.UUA...UGUGGAGCGAAGGUCAU.....G..UUGCAAAAAC..............UGGC..............GACAA..G.UCACUGUACAAACAGCAUCUCAUAUUGAGUAUUUUCGUGGGUUAGUCACUAUGACACGCACGGAAGAGU
env-400    .................UACGG.GU..U.UUA...UGUGGAGCGAAGGUCAU.....G..UUGCAAAAAC..............UGGC..............GACAA..G.UCACUGUACAAACAGCAUCUCAUAUUGAGUAUUUUCGUGGGUUAGUCACUAUGACACGCACGGAAGAGU
env-402    .................UACGG.GU..U.UUA...UGUGGAGCGAAGGUCAU.....G..UUGCAAAAAC..............UGGC..............GACAA..G.UCACUGUACAAACAGCAUCUCAUAUUGAGUAUUUCGUGUGGUCGAGUUUAUGACACGCACGGAAGAGUG
env-403    .................UACGG.GU..U.UUA...UGUGGAGCGAAGGUCAU.....G..UUGCAAAAAC..............UGGC..............GACAA..G.UCACUGUACAAACAGCAUCUCAUAUUGAGUAUUUUCGUGGGUUGGUCACCAUGACACGCACGGAAGAGU
env-404    .................UACGG.GU..U.UUA...UGUGGAGCGAAGGUCAU.....G..UUGCAAAAAC..............UGGC..............GACAA..G.UCACUGUACAAACAGCAUCUCAUAUUGAGUAUUUUCGUGGGUUAGUCACUAUGACACGCACGGAAGAGU
env-405    .................UACGG.GU..U.UUA...UGUGGAGCGAAGGUCAU.....G..UUGCAAAAAC..............UGGC..............GACAA..G.UCACUGUACAAACAGCAUCUCAUAUUGAGUAUUUUCGUGGGUUAGUCACUAUGACACGCACGGAAGAGU
env-406    .................UACGG.GU..U.UUA...UGUGGAGCGAAGGUCAU.....G..UUGCAAAAAC..............UGGC..............GACAA..G.UCACUGUACAAACAGCAUCUCAUAUUGAGUAUUUCGUGUAGUUAAGUUUUACGACACGCACGGAAGAGU
env-407    .................UACGG.GU..U.UUA...UGUGGAGCGAAGGUCAU.....G..UUGCAAAAAC..............UGGC..............GACAA..G.UCACUGUACAAACAGCAUCUCAUAUUGAGUAUUUCGUGUGGUUAAGUUUUACGACACGCACGGAAGAGU
env-408    .................UACGG.GU..U.UUA...UGUGGAGCGAAGGUCAU.....G..UUGCAAAAAC..............UGGC..............GACAA..G.UCACUGUACAAACAGCAUCUCAUAUUGAGUAUUUCGUGUGGUUAAGUUUUACGACACGCACGGAAGAGU
env-409    .................UACGG.GU..U.UUA...UGUGGAGCGAAGGUCAU.....G..UUGCAAAAAC..............UGGC..............GACAA..G.UCACUGUACAAACAGCAUCUCAUAUUGAGUAUUUCGUGUGGUUAAGUUUUACGACACGCACGGAAGAGU
env-410    .................UACGG.GU..U.UUA...UGUGGAGCGAAGGUCAU.....G..UUGCAAAAAC..............UGGC..............GACAA..G.UCACUGUACAAACAGCAUCUCAUAUUGAGUAUUUCGUGUGGUUAAGUUUUACGACACGCACGGAAGAGU
Cba-3-1    .................UACGG.GU..U.UUA...UGUGGAGCGAAGGUCAU.....G..UUGCAAAAAC..............UGGC..............GACAA..G.UCACUGUACAAACAGCAUCUCAUAUUGAGUAUUUUCGUGGGUUAGUCACUAUGACACGCACGGAAGAGU
env-411    .................UACGG.GU..U.UUA...UGUGGAGCGAAGGUCAU.....G..UUGCAAAAAC..............UGGC..............GACAA..G.UCACUGUACAAACAGCAUCUCAUAUUGAGUAUUUCGUGUGGUCGAGUUUAUGACACGCACGGAAGAGUG

           >>>>>>>>>>.......>>>>>.....>.>>>...>>>........<<<<.......<..<<<.....................>>>>..............>>>>...>.>>>>.>>..............................................................
           01.1111111.......22221.....2.102...222.........111.......2..212.....................2122..............111....1.111..12..............................................................
           ----------------•GYRG-GU--U-UUC---CGYGGAGCGAAUGUC∘R-----G--•Y∘R•RRR∘---------------∘G•C-------------GRCAA--G-UCACUGUAC•RR••••••••••••••••••••••••••••••••••••••••••••••••••••••••
                                                                                                                3′
```

```
env-414   .......GGG.......CAUAG.GU..U.UUG...UGUGGAGCGAAGGUCAU.....G..UUACAAAAAC..............UGGC..............GACAA..G.UCACUGUACAAACAACAGCAAAUAAUACCACUAUGUGGUAUUUCCGUGUGAAUUAAAGAUGACACACAC
env-415   .......GGG.......CAUAG.GU..U.UUG...UGUGGAGCGAAGGUCAU.....G..UUACAAAAAC..............UGGC..............GACAA..G.UCACUGUACAAACAACAGCAAAUAAUACCACUAUGUGGUAUUUCCGUGUGAAUUAAAGAUGACACACAC
env-416   .......GGG.......CAUAG.GU..U.UUG...UGUGGAGCGAAGGUCAU.....G..UUACAAAAAC..............UGGC..............GACAA..G.UCACUGUACAAACAACAGCAAAUAAUACCACUAUGUGGUAUUUCCGUGUGAAUUAAAGAUGACACACAC
env-417   .......GGG.......CAUAG.GU..U.UUG...UGUGGAGCGAAGGUCAU.....G..UUACAAAAAC..............UGGC..............GACAA..G.UCACUGUA.............................................................
env-420   .......GGG.......CAUAG.GU..U.UUG...UGUGGAGCGAAGGUCAU.....G..UUACAAAAAC..............UGGC..............GACAA..G.UCACUGUA.............................................................
env-426   .......GGG.......CAUAG.GU..U.UUG...UGUGGAGCGAAGGUCAU.....G..UUACAAAAAC..............UGGC..............GACAA..G.UCACUGUACAAACAACAGCAAAUAAUACCACUAUGUGGUAUUUCCGUGUGAAUUAAAGAUGACACACAC
env-427   .......GGG.......CAUAG.GU..U.UUG...UGUGGAGCGAAGGUCAU.....G..UUACAAAAAC..............UGGC..............GACAA..G.UCACUGUACAAACAACAGCAAAUAAUACCACUAUGUGGUAUUUCCGUGUGAAUUAAAGAUGACACACAC
env-428   .......UCCC......UGCAG.GU..U.UUUA..UGCGGAGCGAAUGUCAA.....G..UUAGUAAAC...............UGAC..............GACAA..G.UCACUGUACCAGAUAACAGCACCUGUCACGAGACAGGGGAAACGACGUGGAGUCCACGAUAAGAGAAAG
env-429   .......UCCC......UGCAG.GU..U.UUUA..UGCGGAGCGAAUGUCAA.....G..UUAGUAAAC...............UGAC..............GACAA..G.UCACUGUACCAGAUAACAGCACCUGUCACAAGACAGGGGAAACGACGUGGAGUCCACGAUAAGAGAAAG
env-430   .......UCCC......UGCAG.GU..U.UUUA..UGCGGAGCGAAUGUCAA.....G..UUAGUAAAC...............UGAC..............GACAA..G.UCACUGUACCAGAUAACAGCACCUGUCAUAAGACAGGGGAAACGACGUGGAGUCCACGAUAAGAGAAAG
env-431   .......UCCC......UGCAG.GU..U.UUUA..UGCGGAGCGAAUGUCAA.....G..UUAGUAAAC...............UGAC..............GACAA..G.UCACUGUACCAGAUAACAGCACCUGUCACGAGACAGGGGAAACGACGUGGAGUCCACGAUAAGAGAAAG
env-432   .......UCCC......UGCAG.GU..U.UUUA..UGCGGAGCGAAUGUCAA.....G..UUAGUAAAC...............UGAC..............GACAA..G.UCACUGUACCAGAUAACAGCACCUGUCACAAGACAGGGGAAACGACGUGGAGUCCACGAUAAGAGAAAG
env-433   .......UCCC......UGCAG.GU..U.UUUA..UGCGGAGCGAAUGUCAA.....G..UUAGUAAAC...............UGAC..............GACAA..G.UCACUGUACCAGAUAACAGCACCUGUCACAAGACAGGGGAAACGACGUGGAGUCCACGAUAAGAGAAAG
env-434   .......UCCC......UGCAG.GU..U.UUUA..UGCGGAGCGAAUGUCAA.....G..UUAGUAAAC...............UGAC..............GACAA..G.UCACUGUACCAGAUAACAGCACCUGUCACGAGACAGGGGAAACGACGUGGAGUCCACGAUAAGAGAAAG
env-435   .......UCCC......UGCAG.GU..U.UUUA..UGCGGAGCGAAUGUCAA.....G..UUAGUAAAC...............UGAC..............GACAA..G.UCACUGUACCAGAUAACAGCACCUGUCACGAGACAGGGGAAACGACGUGGAGUCCACGAUAAGAGAAAG
env-436   .......UCCC......UGCAG.GU..U.UUUA..UGCGGAGCGAAUGUCAA.....G..UUAGUAAAC...............UGAC..............GACAA..G.UCACUGUAC............................................................
env-454   .......UCCC......UGCAG.GU..U.UUUA..UGUGGAGCGAAUGUCAA.....G..UUAGUAAAC...............UGAC..............GACAA..G.UCACUGUACCAA.........................................................
env-455   .......UCCC......UGCAG.GU..U.UUUA..UGUGGAGCGAAUGUCAA.....G..UUAGUAAAC...............UGAC..............GACAA..G.UCACUGUACCAGAUAACAGCACCUGUCACGAGACAGGGGAAACAACGUGGAGUCCAUGAUAAGAGAAAG
env-456   .......UCCC......UGCAG.GU..U.UUUA..UGUGGAGCGAAUGUCAA.....G..UUAGUAAAC...............UGAC..............GACAA..G.UCACUGUACCAGAUAACAGCACCUGUCACGAGACAGGGGAAACGACGUGGAGUCCACGAUAAGAGAAAG
env-457   .......UCCC......UGCAG.GU..U.UUUA..UGUGGAGCGAAUGUCAA.....G..UUAGUAAAC...............UGAC..............GACAA..G.UCACUGUACCAGAUAACAGCACCUGUCACGAGACAGGGGAAACGACGUGGAGUCCACGAUAAGAGAAAG
env-459   .......UCCC......UGCAG.GU..U.UUUA..UGUGGAGCGAAUGUCAA.....G..UUAGUAAAC...............UGAC..............GACAA..G.UCACUGUACCAGAUAACAGCACCUGUCACGAGACAGGGGAAACGACGUGGAGUCCACGAUAAGAGAAAG
env-460   .......UCCC......UGCAG.GU..U.UUUA..UGUGGAGCGAAUGUCAA.....G..UUAGUAAAC...............UGAC..............GACAA..G.UCACUGUACCAGAUAACAGCACCUGUCACGAGACAGGGGAAACAACGUGGAGUCCAUGAUAAGAGAAAG
env-464   .......UCCC......UGCAG.GU..U.UUUA..UGUGGAGCGAAUGUCAA.....G..UUAGUAAAC...............UGAC..............GACAA..G.UCACUGUACCAGAUAACAGCACCUGUCACAAGACAGGGGAAACGACGUGGAGUCCACGAUAAGAGAAAG
env-465   .......UCCC......UGCAG.GU..U.UUUA..UGUGGAGCGAAUGUCAA.....G..UUAGUAAAC...............UGAC..............GACAA..G.UCACUGUACCAGAUAACAGCACCUGUCACGAGACAGGGGAAACAACGUGGAGUCCACGAUAAGAGAAAG
env-466   .......UCCC......UGCAG.GU..U.UUUA..UGUGGAGCGAAUGUCAA.....G..UUAGUAAAC...............UGAC..............GACAA..G.UCACUGUACCAGAUAACAGCACCUGUCACAAGACAGGGGAAACGACGUGGAGUCCACGAUAAGAGAAAG
env-467   .......UCCC......UGCAG.GU..U.UUUA..UGUGGAGCGAAUGUCAA.....G..UUAGUAAAC...............UGAC..............GACAA..G.UCACUGUACCAGAUNNNNNNNNNNNNNNNNNNNNNNNNNNNNNNNNNNNNNNNNNNNNNNNNNNNNNNN
env-468   .......UCCC......UGCAG.GU..U.UUUA..UGUGGAGCGAAUGUCAA.....G..UUAGUAAAC...............UGAC..............GACAA..G.UCACUGUACCAGAUAACAGCACCUGUCACAAGACAGGGGAAACGACGUGGAGUCCACGAUAAGAGAAGA
env-469   .......UCCC......UGCAG.GU..U.UUUA..UGUGGAGCGAAUGUCAA.....G..UUAGUAAAC...............UGAC..............GACAA..G.UCACUGUACCAGAUAACAGCACCUGUCAUGAGACAGGGGAAACGACGUGGAGUCCACGAUAAGAGAAAG
env-470   .......UCCC......UGCAG.GU..U.UUUA..UGUGGAGCGAAUGUCAA.....G..UUAGUAAAC...............UGAC..............GACAA..G.UCACUGUACCAGAUAACAGCACCUGUCAUAAGACAGGGGAAACGACGUGGAGUCCACGAUAAGAGAAAG
env-471   .......UCCC......UGCAG.GU..U.UUUA..UGUGGAGCGAAUGUCAA.....G..UUAGUAAAC...............UGAC..............GACAA..G.UCACUGUACCAGAUAACAGCACCUGUCACGAGACAGGGGAAACAACGUGGAGUCCACGAUAAGAGAAAG
env-472   .......UCCC......UGCAG.GU..U.UUUA..UGUGGAGCGAAUGUCAA.....G..UUAGUAAAC...............UGAC..............GACAA..G.UCACUGUACCAGAUAACAGCACCUGUCACGAGACAGGGGAAACGACGUGGAGUCCACGAUAAGAGAAAG
env-473   .......UCCC......UGCAG.GU..U.UUUA..UGUGGAGCGAAUGUCAA.....G..UUAGUAAAC...............UGAC..............GACAA..G.UCACUGUACCA..........................................................
env-482   .......UCCC......UGCAG.GU..U.UUUA..UGUGGAGCGAAUGUCAA.....G..UUAGUAAAC...............UGAC..............GACAA..G.UCACUGUACCAGAUAACAGCACCUGUCACGAGACAGGGGAAACAACGUGGAGUCCACGAUAAGAGAAAG
env-483   .......UCCC......UGCAG.GU..U.UUUA..UGUGGAGCGAAUGUCAA.....G..UUAGUAAAC...............UGAC..............GACAA..G.UCACUGUACCAGAUAACAGCACCUGUCACGAGACAGGGGAAACGACGUGGAGUCCACGAUAAGAGAAAG
env-484   .......UCCC......UGCAG.GU..U.UUUA..UGUGGAGCGAAUGUCAA.....G..UUAGUAAAC...............UGAC..............GACAA..G.UCACUGUACCAGAUAACAGCACCUGUCACGAGACAGGGGAAACAACGUGGAGUCCACGAUAAGAGAAAG
env-485   .......UCCC......UGCAG.GU..U.UUUA..UGUGGAGCGAAUGUCAA.....G..UUAGUAAAC...............UGAC..............GACAA..G.UCACUGUACCAGAUAACAGCACCUGUCACAAGACAGGGGAAACGACGUGGAGUCCACGAUAAGAGAAAG
env-486   .......UCCC......UGCAG.GU..U.UUUA..UGUGGAGCGAAUGUCAA.....G..UUAGUAAAC...............UGAC..............GACAA..G.UCACUGUACCAGAUAACAGCACCUGUCACGAGACAGGGGAAACAACGUGGAGUCCAUGAUAAGAGAAAG
env-487   .......UCCC......UGCAG.GU..U.UUUA..UGUGGAGCGAAUGUCAA.....G..UUAGUAAAC...............UGAC..............GACAA..G.UCACUGUACCAGAUAACAGCACCUGUCACAAGACAGGGGAAACGACGUGGAGUCCACGAUAAGAGAAAG
env-488   .......UCCC......UGCAG.GU..U.UUUA..UGUGGAGCGAAUGUCAA.....G..UUAGUAAAC...............UGAC..............GACAA..G.UCACUGUACCAGAUAACAGCACCUGUCACGAGACAGGGGAAACGACGUGGAGUCCACGAUAAGAGAAAG
env-489   .................CGCAG.GU..U.UUC...CGUGGAGCGAAUGUCAA.....A..UAAUUC..................UGGC..............GACAA..G.UCACUGUACAAUAUCAUCUGUAAAUGAAUAAGAUGACCCGUAGGUUCACAACGUCCUAAGAGAUAUUGU
env-494   .................CGCAG.GU..U.UUC...CGUGGAGCGAAUGUCAA.....A..UAAUUC..................UGGC..............GACAA..G.UCACUGUACAAUAUCAUCUGUAAAUGAAUAAGAUGACCCGUAGGUUCACAAUGUCCUAAGAGAUACUGU
env-495   .................CGCAG.GU..U.UUC...CGUGGAGCGAAUGUCAA.....A..UAAUUC..................UGGC..............GACAA..G.UCACUGUACAAUAUCAUCUGUAAAUGAAUAAGAUGACCCGUAGAUUCACAAUGUCCUAAGAGAUACUGU
env-496   .................CGCAG.GU..U.UUC...CGUGGAGCGAAUGUCAA.....A..UAAUUC..................UGGC..............GACAA..G.UCACUGUACAAUAUCAUCUGUAAAUGAAUAAGAUGACCCGUAGGUUCACAAUGCCCUAAGAGAUAUUGU
env-497   .................CGCAG.GU..U.UUC...CGUGGAGCGAAUGUCAA.....A..UAAUUC..................UGGC..............GACAA..G.UCACUGUACAAUAUCAUCUGUAAAUGAAUAAGAUGACCCGUAGGUUCACAAUGUCCUAAGAGAUGCUGU
env-498   .................CGCAG.GU..U.UUC...CGUGGAGCGAAUGUCAA.....A..UAAUUC..................UGGC..............GACAA..G.UCACUGUACAAUAUCAUCUGUAAAUGAAUAAGAUGACCCGUAGGUUCACAAUGUCCUAAGAGAUACUGU
env-499   .................CGCAG.GU..U.UUC...CGUGGAGCGAAUGUCAA.....A..UAAUUC..................UGGC..............GACAA..G.UCACUGUACAAUAUCAUCUGUAAAUGAAUAAGAUGACCCGUAGGUUCACAAUGCCCUAAGAGAUAUUGU
env-500   .................CGCAG.GU..U.UUC...CGUGGAGCGAAUGUCAA.....A..UAAUUC..................UGGC..............GACAA..G.UCACUGUACAAUAUCAUCUGUAAAUGAAUAAGAUGACCCGUAGAUUCACAAUGUCCUAAGAGAUACUGU
env-501   .................CGCAG.GU..U.UUC...CGUGGAGCGAAUGUCAA.....A..UAAUUC..................UGGC..............GACAA..G.UCACUGUACAAUAUCAUCUGUAAAUGAAUAAGAUGACCCGUAGGCUCACAAUGUCCUAAGAGAUACUGU
env-502   .................CGCAG.GU..U.UUC...CGUGGAGCGAAUGUCAA.....A..UAAUUC..................UGGC..............GACAA..G.UCACUGUACAAUAUCAUCUGUAAAUGAAUAAGAUGACCCGUAGGUUCACAAUGUCCUAAGAGAUACUGU
env-503   .................CGCAG.GU..U.UUC...CGUGGAGCGAAUGUCAA.....A..UAAUUC..................UGGC..............GACAA..G.UCACUGUACAAUAUCAUCUGUAAAUGAAUAAGAUGACCCGUAGGUUCACAAUGUCCUAAGAGAUACUGU
env-512   .................CAUAG.GU..U.UUU...CGUGGAGCGAAUGUCAA.....U..UAACGAUAAC..............UGGC..............GACAA..G.UCACUGUACUCAAUAACCAUCUGGCAAUGCUAGAGAUGACGUGUGAAGAUAUCUGAAACACCUUGGUGU
env-513   .................CAUAG.GU..U.UUU...CGUGGAGCGAAUGUCAA.....U..UAACGAUAAC..............UGGC..............GACAA..G.UCACUGUACUCAAUAACCAUCUGGCAAUGCCAGAGAUGACGUGUGAAGAUAUCUGAAACACCUUGGUGU
env-514   .................CAUAG.GU..U.UUU...CGUGGAGCGAAUGUCAA.....U..UAACGAUAAC..............UGGC..............GACAA..G.UCACUGUACUCAAUAACCAUCUGGCAAUGCCAGAGAUGACGUGUGAAGAUAUCUGAAACACCUUGGUGU
env-515   .................CAUAG.GU..U.UUU...CGUGGAGCGAAUGUCAA.....U..UAACGAUAAC..............UGGC..............GACAA..G.UCACUGUACUCAAUAACCAUCUGGCAAUGCCAGAGAUGACGUGUGAAGAUAUCUGAAACACCUUGGUGU
env-516   .................CAUAG.GU..U.UUU...CGUGGAGCGAAUGUCAA.....U..UAACGAUAAC..............UGGC..............GACAA..G.UCACUGUACUCAAUAACCAUCUGGCAAUGCCAGAGAUGACGUGUGAAGAUAUCUGAAACACCUUGGUGU
env-517   .................CAUAG.GU..U.UUU...CGUGGAGCGAAUGUCAA.....U..UAACGAUAAC..............UGGC..............GACAA..G.UCACUGUACUCAAUAACCAUCUGGCAAUGCCAGAGAUGACGUGUGAAGAUAUCUGAAACACCUUGGUGU
env-518   .................CAUAG.GU..U.UUU...CGUGGAGCGAAUGUCAA.....U..UAACGAUAAC..............UGGC..............GACAA..G.UCACUGUACUCAAUAACCAUCUGGCAAUGCCAGAGAUGACGUGUGAAGAUAUCUGAAACACCUUGGUGU
env-556   .................CAUAG.GU..U.UUU...CGUGGAGCGAAUGUCAA.....U..UAACGAUAAC..............UGGC..............GACAA..G.UCACUGUACUCAAUAACCAUCUGGCAAUGCCAGAGAUGACGUGUGAAGAUAUCUGAAACACCUUGGUGU
env-557   .................CAUAG.GU..U.UUU...CGUGGAGCGAAUGUCAA.....U..UAACGAUAAC..............UGGC..............GACAA..G.UCACUGUACCCAAUAACCAUCUGGCAAUGCCAGAGAUGACGUGUGAAGAUAUCUGAAACACCUUGGUGU
env-558   .................CAUAG.GU..U.UUU...CGUGGAGCGAAUGUCAA.....U..UAACGAUAAC..............UGGC..............GACAA..G.UCACUGUACUCAAUAACCAUCUGGCAAUGCCAGAGAUGACGUGUGAAGAUAUCUGAAACACCUUGGUGU
env-559   .................CAUAG.GU..U.UUU...CGUGGAGCGAAUGUCAA.....U..UAACGAUAAC..............UGGC..............GACAA..G.UCACUGUACUCAAUAACCAUCUGGCAAUGCCAGAGAUGACGUGUGAAGAUAUCUGAAACACCUUGGUGU
env-560   .................CAUAG.GU..U.UUU...CGUGGAGCGAAUGUCAA.....U..UAACGAUAAC..............UGGC..............GACAA..G.UCACUGUACUCAAUAACCAUCUGGCAAUGCCAGAGAUGACGUGUGAAGAUAUCUGAAACACCUUGGUGU
env-561   .................CAUAG.GU..U.UUU...CGUGGAGCGAAUGUCAA.....U..UAACGAUAAC..............UGGC..............GACAA..G.UCACUGUACUCAAUAACCAUCUGGCAAUGCCAGAGAUGACGUGUGAAGAUAUCUGAAACACCUUGGUGU
env-562   ......GGGUU......UGCAG.GU..U.UUCA..CGCAGAGCGAAUGUCAA.....G..UUAGGAAAC...............UGGC..............GACAA..G.UCACUGUACGAAUCAGGAGUCCACGAAACGAGUGGAAACGGCGUGUGCGUAUUUCACGACACACACGGC
env-563   ......GGGUU......UGCAG.GU..U.UUCA..CGUGGAGCGAAUGUCAA.....G..UUAGGAAAC...............UGGC..............GACAA..G.UCACUGUACGAAUCAGAAGUCCACGAAUAGGUGGAAACGGCGUGUGGGUAUUUCACGACACACACGGCA
env-564   ......GGGUU......UGCAG.GU..U.UUCA..CGUGGAGCGAAUGUCAA.....G..UUAGGAAAC...............UGGC..............GACAA..G.UCACUGUACGAAUCAGAAGUCCACGAAUAGGUGGAAACGGCGUGUGGGUAUUUCACGACACACACGGCA
env-565   ......GGGUU......UACAG.GU..U.UUCA..CGCGGAGCGAAUGUCAA.....G..UUAGGAAAC...............UGGC..............GACAA..G.UCACUGUACGAAUCAGGAGUCCACGAAACGAGUGGAAACGGCGUGUGCGUAUUUCACGACACACACGGC
Bsp-4-1   .................AGCAG.GU..U.UUC...CGUGGAGCGAAUGUCUAGA...G..GCCGGAAUAAU.............GGCCAAACAAACUGAC..GACAA..G.UCACUGUACAGACAAUUGUAAGUCUGCCUACUAACCCAAAGGCAGAAACGACGUGUAGAUAGUUCCAAG
Bsp-7-1   .................AGCAG.GU..U.UUC...CGUGGAGCGAAUGUCUAGA...G..GCCGGAAUAAU.............GGCCAAACAAACUGAC..GACAA..G.UCACUGUACAGACAAUUGUAAGUCUGCCUACUAACCCAAAGGCAGAAACGACGUGUAGAUAGUUCCAAG
Bsp-10-1  .................AGCAG.GU..U.UUC...CGUGGAGCGAAUGUCUAGA...G..GCCGGAAUAAU.............GGCCAAACAAACUGAC..GACAA..G.UCACUGUACAGACAAUUGUAAGUCUGCCUACUAACCCAAAGGCAGAAACGACGUGUAGAGAGUUCAAAG
env-566   .................AGCAG.GU..U.UUC...CGUGGAGCGAAUGUCUAGA...G..GCCGGAAUAAU.............GGCCAAACAAACUGAC..GACAA..G.UCACUGUACAGACAAUUGUAAGUCUGCCUACUAACCCAAAGGCAGAAACGACGUGUAGAUAGUUCCAAG
Bsp-2-1   .................AGCAG.GU..U.UUC...CGUGGAGCGAAUGUCUAGA...G..GCCAGUUGCUUU............GGCCAACCAAACUGAC..GACAA..G.UCACUGUACAGUUAAUUCAAUGUCUGCCGACUAACCCAAGGCAGAAACGACGUGUAGAGAUCCAGGACC
Bsp-3-1   .................AGCAG.GU..U.UUC...CGUGGAGCGAAUGUCAAG....GA.CCGAGUGAU...............CGGCCAAACAAACUGAC.GACAA..G.UCACUGUACAGUUAAUUUAGGUCUGCCGACUAACCCAAGGCAGAAACGACGUGUAGGAUAAGUUCGGGA
Bsp-6-1   .................AGCAG.GU..U.UUC...CGUGGAGCGAAUGUCAAG....GA.CCGAGUGAU...............CGGCCAAACAAACUGAC.GACAA..G.UCACUGUACAGUUAAUUUAGGUCUGCCGACUAACCCAAGGCAGAAACGACGUGUAGGAUAAGUUCGGGA
env-567   .......UGGAG.....AUCGG.GU..U.UUC...CGUGGAGCGAAUGUCAA.....G..UUAGAAAAC...............UGAC..............GGCAA..G.UCACUGUACCAAUUAACAUCUGCCUUGAGCAGUAACAACGUGUAGGUUUAUGACCGGAGAGAAGAAAGC
env-568   .......UGGAG.....AUCGG.GU..U.UUC...CGUGGAGCGAAUGUCAA.....G..UUAGAAAAC...............UGAC..............GGCAA..G.UCACUGUACCAAUUAACAUCUGCCUUGAGCAGUAACAACGUGUAAGUUUAUGACCGGAGAGAAGAAAGC
env-570   .......UGGAG.....AUCGG.GU..U.UUC...CGUGGAGCGAAUGUCAA.....G..UUAGAAAAC...............UGAC..............GGCAA..G.UCACUGUACCAAUUAACAUCUGCCUUGAGCAGUAACAACGUGUAAGUUUAUGGCCGGAGAGAAGAAAGC
env-571   .......UGGAG.....AUCGG.GU..U.UUC...CGUGGAGCGAAUGUCAA.....G..UUAGAAAAC...............UGAC..............GGCAA..G.UCACUGUACCAAUUAACAUCUGCCUUGAGCAGUAACAACGUGUAAGUUUAUGGCCGGAGAGAAGAAAGC
env-573   .......UGGAG.....AUCGG.GU..U.UUC...CGUGGAGCGAAUGUCAA.....G..UUAGAAAAC...............UGAC..............GGCAA..G.UCACUGUACCAAUUAACAUCUGCCUUGAGCAGUAACAACGUGUAAGUUUAUGGCCGGAGAGAAGAAAGC
env-578   .......UGGAG.....AUCGG.GU..U.UUC...CGUGGAGCGAAUGUCAA.....G..UUAGAAAAC...............UGAC..............GGCAA..G.UCACUGUACCAAUUAACAUCUGCCUUGAGCAGUAACAACGUGUAAGUUUAUGGCCGGAGAGAAGAAAGC
env-579   .......UGGAG.....AUCGG.GU..U.UUC...CGUGGAGCGAAUGUCAA.....G..UUAGAAAAC...............UGAC..............GGCAA..G.UCACUGUACCAAUUAACAUCUGCCUUGAGCAGUAACAACGUGUAAGUUUAUGACCGGAGAGAAGAAAGC
env-581   .......UGGAG.....AUCGG.GU..U.UUC...CGUGGAGCGAAUGUCAA.....G..UUAGAAAAC...............UGAC..............GGCAA..G.UCACUGUACCAAUUAACAUCUGCCUUGAGCAGUAACAACGUGUAAGUUUAUGACCGGAGAGAAGAAAGC
env-582   .......UGGAG.....AUCGG.GU..U.UUC...CGUGGAGCGAAUGUCAA.....G..UUAGAAAAC...............UGAC..............GGCAA..G.UCACUGUACCAAUUAACAUCUGCCUUGAGCAGUAACAACGUGUAAGUUUAUGACCGGAGAGAAGAAAGC

          >>>>>>>>>>.......>>>>>.....>.>>>...>>>........<<<<.......<..<<<.....................>>>>..............>>>>.....>.>>>>.>>..............................................................
          01.1111111.......22221.....2.102...222.........111.......2..212.....................2122..............111......1.111..12..............................................................
          ----------------•GYRG-GU--U-UUC---CGYGGAGCGAAUGUC•R-----G--•Y•R•RRR•--------------•G•C-------------GRCAA--G-UCACUGUAC•RR••••••••••••••••••••••••••••••••••••••••••••••••••••••
                                                                                                                    3′
```

```
env-583  .......UGGAG.....AUCGG.GU..U.UUC...CGUGGAGCGAAUGUCAA.....G..UUAGAAAAC...............UGAC..............GGCAA..G.UCACUGUACCAAUUAACAUCUGCCUUGAGCAGUAACAACGUGUAAGUUUAUGACCGGAGAGAAGAAAGC
env-584  .......UGGAG.....AUCGG.GU..U.UUC...CGUGGAGCGAAUGUCAA.....G..UUAGAAAAC...............UGAC..............GGCAA..G.UCACUGUACCAAUUAACAUCUGCCUUGAGCAGUAACAACGUGUAAGUUUAUGGCCGGAGAGAAGAAAGC
env-585  .......UGGAG.....AUCGG.GU..U.UUC...CGUGGAGCGAAUGUCAA.....G..UUAGAAAAC...............UGAC..............GGCAA..G.UCACUGUACCAAUUAACAUCUGCCUUGAGCAGUAACAACGUGUAAGUUUAUGACCGGAGAGAAGAAAGC
env-586  .......UGGAG.....AUCGG.GU..U.UUC...CGUGGAGCGAAUGUCAA.....G..UUAGAAAAC...............UGAC..............GGCAA..G.UCACUGUACCAAUUAACAUCUGCCUUGAGCAGUAACAACGUGUAAGUUUAUGACCGGAGAGAAGAAAGC
env-587  .......UGGAG.....AUCGG.GU..U.UUC...CGUGGAGCGAAUGUCAA.....G..UUAGAAAAC...............UGAC..............GGCAA..G.UCACUGUACCAAUUAACAUCUGCCUUGAGCAGUAACAACGUGUAAGUUUAUGACCGGAGAGAAGAAAGC
env-588  .......UGGAG.....AUCGG.GU..U.UUC...CGUGGAGCGAAUGUCAA.....G..UUAGAAAAC...............UGAC..............GGCAA..G.UCACUGUACCAAUUAACAUCUGCCUUGAGCAGUAACAACGUGUAAGUUUAUGACCGGAGAGAAGAAAGC
env-589  .......UGGAG.....AUCGG.GU..U.UUC...CGUGGAGCGAAUGUCAA.....G..UUAGAAAAC...............UGAC..............GGCAA..G.UCACUGUACCAAUUAACAUCUGCCUUGAGCAGUAACAACGUGUAAGUUUAUGACCGGAGAGAAGAAAGC
env-590  .......UGGAG.....AUCGG.GU..U.UUC...CGUGGAGCGAAUGUCAA.....G..UUAGAAAAC...............UGAC..............GGCAA..G.UCACUGUACCAAUUAACAUCUGCCUUGAGCAGUAACAACGUGUAGGUUUAUGACCGGAGAGAAGAAAGC
env-591  .......UGGAG.....AUCGG.GU..U.UUC...CGUGGAGCGAAUGUCAA.....G..UUAGAAAAC...............UGAC..............GGCAA..G.UCACUGUACCAAUUAACAUCUGCCUUGAGCAGUAACAACGUGUAAGUUUAUGNNNNNNNNNNNNNNNNN
env-592  .......UGGAG.....AUCGG.GU..U.UUC...CGUGGAGCGAAUGUCAA.....G..UUAGAAAAC...............UGAC..............GGCAA..G.UCACUGUACCAAUUAACAUCUGCCUUGAGCAGUAACAACGUGUAAGUUUAUGACCGGAGAGAAGAAAGC
env-593  .......UGGAG.....AUCGG.GU..U.UUC...CGUGGAGCGAAUGUCAA.....G..UUAGAAAAC...............UGAC..............GGCAA..G.UCACUGUACCAAUUAACAUCUGCCUUGAGCAGUAACAACGUGUAAGUUUAUGGCCGGAGAGAAGAAAGC
env-594  .......UGGAG.....AUCGG.GU..U.UUC...CGUGGAGCGAAUGUCAA.....G..UUAGAAAAC...............UGAC..............GGCAA..G.UCACUGUACCAAUUAACAUCUGCCUUGAGCAGUAACAACGUGUAAGUUUAUGACCGGAGAGAAGAAAGC
env-595  .......UGGAG.....AUCGG.GU..U.UUC...CGUGGAGCGAAUGUCAA.....G..UUAGAAAAC...............UGAC..............GGCAA..G.UCACUGUACCAAUUAACAUCUGCCUUGAGCAGUAACAACGUGUAAGUUUAUGACCGGAGAGAAGAAAGC
env-596  .......UGGAG.....AUCGG.GU..U.UUC...CGUGGAGCGAAUGUCAA.....G..UUAGAAAAC...............UGAC..............GGCAA..G.UCACUGUACCAAUUAACAUCUGCCUUGAGCAGUAACAACGUGUAAGUUUAUGACCGGAGAGAAGAAAGC
env-597  .......UGGAG.....AUCGG.GU..U.UUC...CGUGGAGCGAAUGUCAA.....G..UUAGAAAAC...............UGAC..............GGCAA..G.UCACUGUACCAAUUAACAUCUGCCUUGAGCAGUAACAACGUGUAAGUUUAUGGCCGGAGAGAAGAAAGC
env-598  .......UGGAG.....AUCGG.GU..U.UUC...CGUGGAGCGAAUGUCAA.....G..UUAGAAAAC...............UGAC..............GGCAA..G.UCACUGUACCAAUUAACAUCUGCCUUGAGCAGUAACAACGUGUAAGUUUAUGACCGGAGAGAAGAAAGC
env-599  .......UGGAG.....AUCGG.GU..U.UUC...CGUGGAGCGAAUGUCAA.....G..UUAGAAAAC...............UGAC..............GGCAA..G.UCACUGUACCAAUUAACAUCUGCCUUGAGCAGUAACAACGUGUAAGUUUAUGACCGGAGAGAAGAAAGC
env-600  .......UGGAG.....AUCGG.GU..U.UUC...CGUGGAGCGAAUGUCAA.....G..UUAGAAAAC...............UGAC..............GGCAA..G.UCACUGUACCAAUUAACAUCUGCCUUGAGCAGUAACAACGUGUAAGUUUAUGACCGGAGAGAAGAAAGC
env-601  .......UGGAG.....AUCGG.GU..U.UUC...CGUGGAGCGAAUGUCAA.....G..UUAGAAAAC...............UGAC..............GGCAA..G.UCACUGUACCAAUUAACAUCUGCCUUGAGCAGUAACAACGUGUAAGUUUAUGACCGGAGAGAAGAAAGC
env-602  .......UGGAG.....AUCGG.GU..U.UUC...CGUGGAGCGAAUGUCAA.....G..UUAGAAAAC...............UGAC..............GGCAA..G.UCACUGUACCAAUUAACAUCUGCCUUGAGCAGUAACAACGUGUAAGUUUAUGACCGGAGAGAAGAAAGC
env-603  .................AGUAG.GU..U.UUUA..CGCGGAGCGAAUGUCAA.....G..UUAUAAAAC...............UGGC..............GACAA..G.UCACUGUACAAGAACAUUCUGCUUUUGGCAGAUAAACGUGUGAUAGUUCAAUAACAUAGGCAUCCGGAG
env-604  .................GAUGG.GU..U.UUC...CGUGGAGCGAAAGUCAU.....G..UUACUAAAC...............UGGC..............GACAA..G.UCACUGUACAAGUUAACAGCAACCCAACAGGGAGGUAGUUUUUUCGGGUAGAAUAGAGAAGACACGCCC
env-606  .................GAUGG.GU..U.UUC...CGUGGAGCGAAAGUCAU.....G..UUACUAAAC...............UGGC..............GACAA..G.UCACUGUACAAGUUAACAGCAACCCAACAGGGAGGUAGUUUUUUCGGGUAAAAUAGAGAAGACACACCC
env-607  .................GAUGG.GU..U.UUC...CGUGGAGCGAAAGUCAU.....G..UUACUAAAC...............UGGC..............GACAA..G.UCACUGUACAAGUUAACAGCAACCCAACAGGGAGGUAGUUUUUUCGGGUAGAAUAGAGAAGACACGCCC
env-608  .................GAUGG.GU..U.UUC...CGUGGAGCGAAAGUCAU.....G..UUACUAAAC...............UGGC..............GACAA..G.UCACUGUACAAGUUAACAGCAACCCAACAGGGAGGUAGUUUUUUCGGGUAGAAUAGAGAAGACACGCCC
env-614  .................GAUGG.GU..U.UUC...CGUGGAGCGAAAGUCAU.....G..UUACUAAAC...............UGGC..............GACAA..G.UCACUGUACAAGUUAACAGCAACCCAACAGGGAGGUAGUUUUUUCGGGUAAAAUAGAGAAGACACACCC
env-615  .................GAUGG.GU..U.UUC...CGUGGAGCGAAAGUCAU.....G..UUACUAAAC...............UGGC..............GACAA..G.UCACUGUACAAGUUAACAGCAACCCAACAGGGAGGUAGUUUUUUCGGGUAGAAUAGAGAAGACACGCCC
env-623  .................GAUGG.GU..U.UUC...CGUGGAGCGAAAGUCAU.....G..UUACUAAAC...............UGGC..............GACAA..G.UCACUGUACAAGUUAACAGCAACCCAACAGGGAGGUAGUUUUUUCGGGUAGAAUAGAGAAGACACGCCC
env-624  .................GAUGG.GU..U.UUC...CGUGGAGCGAAAGUCAU.....G..UUACUAAAC...............UGGC..............GACAA..G.UCACUGUACAAGUUAACAGCAACCCAACAGGGAGGUAGUUUUUUCGGGUAAAAUAGAGAAGACACACCC
env-625  .................GAUGG.GU..U.UUC...CGUGGAGCGAAAGUCAU.....G..UUACUAAAC...............UGGC..............GACAA..G.UCACUGUACAAGUUAACAGCAACCCAACAGGGAGGUAGUUUUUUCGGGUAGAAUAGAGAAGACACGCCC
env-626  .................GAUGG.GU..U.UUC...CGUGGAGCGAAAGUCAU.....G..UUACUAAAC...............UGGC..............GACAA..G.UCACUGUACAA..........................................................
env-632  .................GAUGG.GU..U.UUC...CGUGGAGCGAAAGUCAU.....G..UUACUAAAC...............UGGC..............GACAA..G.UCACUGUACAAGUUAACAGCAACCCAACAGGGAGGUAGUUUUUUCGGGUAGAAUAGAGAAGACACGCCC
env-633  .................GAUGG.GU..U.UUC...CGUGGAGCGAAAGUCAU.....G..UUACUAAAC...............UGGC..............GACAA..G.UCACUGUACAAGUUAACAGCAACCCAACAGGGAGGUAGUUUUUUCGGGUAAAAUAGAGAAGACACGCCC
env-634  .................GAUGG.GU..U.UUC...CGUGGAGCGAAAGUCAU.....G..UUACUAAAC...............UGGC..............GACAA..G.UCACUGUACAAGUUAACAGCAACCCAACAGGGAGGUAGUUUUUUCGGGUAGAAUAGAGAAGACACGCCC
env-635  .................GAUGG.GU..U.UUC...CGUGGAGCGAAAGUCAU.....G..UUACUAAAC...............UGGC..............GACAA..G.UCACUGUACAAGUUAACAGCAACCCAACAGGGAGGUAGUUUUUUCGGGUAGAAUAGAGAAGACACGCCC
env-637  .................GAUGG.GU..U.UUC...CGUGGAGCGAAAGUCAU.....G..UUACUAAAC...............UGGC..............GACAA..G.UCACUGUACAAGUUAACAGCAACCCAACAGGGAGGUAGUUUUUUCGGGUAGAAUAGAGAAGACACGCCC
env-639  .................GAUGG.GU..U.UUC...CGUGGAGCGAAAGUCAU.....G..UUACUAAAC...............UGGC..............GACAA..G.UCACUGUACAAGUUAACAGCAACCCAACAGGGAGGUAGUUUUUUCGGGUAGAAUAGAGAAGACACGCCC
env-640  .................GAUGG.GU..U.UUC...CGUGGAGCGAAAGUCAU.....G..UUACUAAAC...............UGGC..............GACAA..G.UCACUGUACAAGUUAACAGCAACCCAACAGGGAGGUAGUUUUUUCGGGUAGAAUAGAGAAGACACGCCC
env-641  .................GAUGG.GU..U.UUC...CGUGGAGCGAAAGUCAU.....G..UUACUAAAC...............UGGC..............GACAA..G.UCACUGUACAAGUUAACAGCAACCCAACAGGGAGGUAGUUUUUUCGGGUAGAAUAGAGAAGACACGCCC
env-642  .................GAUGG.GU..U.UUC...CGUGGAGCGAAAGUCAU.....G..UUACUAAAC...............UGGC..............GACAA..G.UCACUGUACAAGUUAACAGCAACCCAACAGGGAGGUAGUUUUUUCGGGUAGAAUAGAGAAGACACGCCC
env-643  .................GAUGG.GU..U.UUC...CGUGGAGCGAAAGUCAU.....G..UUACUAAAC...............UGGC..............GACAA..G.UCACUGUACAAGUUAACAGCAACCCAACAGGGAGGUAGUUUUUUCGGGUAGAAUAGAGAAGACACGCCC
env-644  .................GAUGG.GU..U.UUC...CGUGGAGCGAAAGUCAU.....G..UUACUAAAC...............UGGC..............GACAA..G.UCACUGUACAAGUUAACAGCAACCCAACAGGGAGGUAGUUUUUUCGGGUAAAAUAGAGAAGACACACCC
Jig-1-1  .................GAUGGGGU..U.UUC...CGUGGAGCGAAAGUCAU.....G..UUACUAUAAAC.............UGGC..............GACAA..G.UCACUGUACCUGAAAAGAUAGGGCUCCGUAAGGAGCAUAGUGUGUUUAGUCCAAAUAUCGACACACACG
env-646  .....UCCAU.......UGCAG.GU..U.UUC...CGUGGAGCGAAUGUCAA.....G..UUUAGAAAC...............UGGC..............GACAA..G.UCACUGUACAAUAUUUACAAUCUACAAAUUCAGCGUAGAGACGCAGUGUGAUAUCUUAGAUGAUACACA
env-647  .....UCCAU.......UGCAG.GU..U.UUC...CGUGGAGCGAAUGUCAA.....G..UUUAGAAAC...............UGGC..............GACAA..G.UCACUGUACAAUAUUUACAAUCUACAAAUUCAGCGUAGAGACGCAGUGUGGUAUCAUAGAUGAUACACA
Eha-1-1  .....UCCAU.......UGCAG.GU..U.UUC...CGUGGAGCGAAUGUCAA.....G..UUUAGAAAC...............UGGC..............GACAA..G.UCACUGUACAAUAUUUACAAUCUACAAAUUCAGCGUAGAGACGCAGUGUGGUAUCAUAGAUGAUACACA
env-648  .....UCCAU.......UGCAG.GU..U.UUC...CGUGGAGCGAAUGUCAA.....G..UUUAGAAAC...............UGGC..............GACAA..G.UCACUGUACAAUAUUUACAAUCUACAAAUUCAGCGUAGAGACGCAGUGUGGUAUCAUAGAUGAUACACA
env-649  .....UCCAU.......UGCAG.GU..U.UUC...CGUGGAGCGAAUGUCAA.....G..UUUAGAAAC...............UGGC..............GACAA..G.UCACUGUACAAUAUUUACAAUCUACAAAUUCAGCGUAGAGACGCAGUGUGGUAUCAUAGAUGAUACACA
env-650  .................UACAG.GU..UAUUU...CGUGGAGCGAAUGUCAA.....U..UUACAAAAC...............UGAC..............GACAA..G.UCACUGUACAAAAGAACCGUCUGCCAGAGAGCAGAGACGACGUGUAUGUUGUGAUAAUAAAUACGACAC
env-653  .................UACAG.GU..UAUUU...CGUGGAGCGAAUGUCAA.....U..UUACAAAAC...............UGAC..............GACAA..G.UCACUGUACAAAAGAACCGUCUGCCAGAGAGCAGAGACGACGUGUAUGUUGCGAUAAUAAAUACGACAC
env-654  .................GGCAG.GU..U.UUC...CGUGGAGCGAAUGUCAAG....G..CCGGUAUUAAC.............CGGCAGAUGAACCUGGC.GACAA..G.UCACUGUACAAAUAUUAAUCAUCUGCAUGAGCAGAGAUGACACGCAGUUAUUGAAAUGAGAAUUUUCGA
env-655  .................GGCAG.GU..U.UUC...CGUGGAGCGAAUGUCAAG....G..CCGGUAUUAAC.............CGGCAGAUGAACCUGGC.GACAA..G.UCACUGUACAAAUAUUAAUCAUCUGCAUGAGCAGAGAUGACACGCAGUUAUUGAGAUGAGAAUUUUCGA
env-656  .................GGCAG.GU..U.UUC...CGUGGAGCGAAUGUCAAG....G..CCGGUAUUAAC.............CGGCAGAUGAACCUGGC.GACAA..G.UCACUGUACAAAUAUUAAUCAUCUGCAUGAGCAGAGAUGACACGCAAUUAUUGAGAUGAGAAUUUUCGA
env-657  .................GGCAG.GU..U.UUC...CGUGGAGCGAAUGUCAAG....G..CCGGUAUUAAC.............CGGCAGAUGAACCUGGC.GACAA..G.UCACUGUACAAAUAUUAAUCAUCUGCAUGAGCAGAGAUGACACGCAGUAAUUGAGAUGAGAAUUUUCGA
env-658  .................GGCAG.GU..U.UUC...CGUGGAGCGAAUGUCAAG....G..CCGGUAUUAAC.............CGGCAGAUGAACCUGGC.GACAA..G.UCACUGUACAAAUAUUAAUCAUCUGCAUGAGCAGAGAUGACACGCAGUUAUUGAGAUGAGAAUUUUCGA
env-662  .................GGCAG.GU..U.UUC...CGUGGAGCGAAUGUCAAG....G..CCGGUAUUAAC.............CGGCAGAUGAACCUGGC.GACAA..G.UCACUGUACAAAUAUUAAUCAUCUGCAUGAGCAGAGAUGACACGCAGUUAUUGAGAUGAGAAUUUUCGA
env-663  .................GGCAG.GU..U.UUC...CGUGGAGCGAAUGUCAAG....G..CCGGUAUUAAC.............CGGCAGAUGAACCUGGC.GACAA..G.UCACUGUACAAAUAUUAAUCAUCUGCAUGAGCAGAGAUGACACGCAGUUAUUGAGAUGAGAAUUUUCGA
env-664  .................GGCAG.GU..U.UUC...CGUGGAGCGAAUGUCAAG....G..CCGGUAUUAAC.............CGGCAGAUGAACCUGGC.GACAA..G.UCACUGUACAAAUAUUAAUCAUCUGCAUGAGCAGAGAUGACACGCAGUUAUUGAGAUGAGAAUUUUCGA
env-667  .................GGCAG.GU..U.UUC...CGUGGAGCGAAUGUCAAG....G..CCGGUAUUAAC.............CGGCAGAUGAACCUGGC.GACAA..G.UCACUGUACAAAUAUUAAUCAUCUGCAUGAGCAGAGAUGACACGCAGUUAUUGAGAUGAGAAUUUUCGA
env-668  .................GGCAG.GU..U.UUC...CGUGGAGCGAAUGUCAAG....G..CCGGUAUUAAC.............CGGCAGAUGAACCUGGC.GACAA..G.UCACUGUACAAAUAUUAAUCAUCUGCAUGAGCAGAGAUGACACGCAGUUAUUGAGAUGAGAAUUUUCGA
Lba-14-1 .....CAUAUG.......GCAG.GU..U.UUC...CGCGGAGCGAAUGUCAA.....G..UUAGGAAAC...............UGGC..............GACAA..G.UCACUGUACAAAGGUAUGACCCCAGGGUCAGCCAUACUGUGAAGCAACAGACAAUCACAGUAAAGAACG
Bsp-1-1  .................GACAG.GU..U.UUC...CGUGGAGCGAAUGUCAAGA...G..ACC.GCAA................GGUCAGACAAUCUGAC..GACAA..G.UCACUGUACAAACAAUUUGUCUGCGAAGCUAUGCUAGCAGAAACGACGUGUGAGAACAGGAUGAGAAUG
Bsp-9-1  .................GACAG.GU..U.UUC...CGUGGAGCGAAUGUCAAGA...G..ACC.GCAA................GGUCAGACAAUCUGAC..GACAA..G.UCACUGUACAAACAAUUUGUCUGCGAAGCUAUGCUAGCAGAAACGACGUGUGAGAACAGGAUGAGAAUG
env-669  .................AGCGG.GU..U.UUC...CGUGGAGCGAAUGUCAA.....G..UUAAAGAAAC..............UGGC..............GACAA..G.UCACUGUACCCACAAUUAAGUCCACCUGUAAGAGUGGAAACGACCGUGUGCAAACAUCAGGACACACAC
env-671  .................AGCGG.GU..U.UUC...CGUGGAGCGAAUGUCAA.....G..UUAAAGAAAC..............UGGC..............GACAA..G.UCACUGUACCCACAAUUAAGUCCACCUGUAAGAGUGGAAACGACCGUGUGCAAACAUCAGGACACACAC
env-672  .................AGCGG.GU..U.UUC...CGUGGAGCGAAUGUCAA.....G..UUAAAGAAAC..............UGGC..............GACAA..G.UCACUGUACCCACAAUUAAGUCCACCUGUAAGAGUGGAAACGACCGUGUGCAAACAUCAGGACACACAC
env-681  .................AGCGG.GU..U.UUC...CGUGGAGCGAAUGUCAA.....G..UUAAAGAAAC..............UGGC..............GACAA..G.UCACUGUACCCACAAUUAAGUCCACCUGUAAGAGUGGAAACGACCGUGUGCAAACAUUAGGACACACAC
env-683  .................AGCGG.GU..U.UUC...CGUGGAGCGAAUGUCAA.....G..UUAAAGAAAC..............UGGC..............GACAA..G.UCACUGUACCCACAAUUAAGUCCACCUGUAAGAGUGGAAACGACCGUGUGCAAACAUCAGGACACACAC
env-684  .................AGCGG.GU..U.UUC...CGUGGAGCGAAUGUCAA.....G..UUAAAGAAAC..............UGGC..............GACAA..G.UCACUGUACCCACAAUUAAGUCCACCUGUAAGAGUGGAAACGACCGUGUGCAAACAUUAGGACACACAC
env-685  .................AGCGG.GU..U.UUC...CGUGGAGCGAAUGUCAA.....G..UUAAAGAAAC..............UGGC..............GACAA..G.UCACUGUACCCACAAUUAAGUCCACCUGUAAGAGUGGAAACGACCGUNNNNNNNNNNNNNNNNNNNNN
env-686  .....GCUGGU.......GCAG.GU..U.UUC...CGUGGAGCGAAUGUCAA.....G..UUAGGAAAC...............UGGC..............GACAA..G.UCACUGUACGAAUCAAAAACUGAUUACUAUUUGCCCCAAUGGGAUAACCAGGGUAGGAGUAUGAAACAA
env-687  .....GCUGGU.......GCAG.GU..U.UUC...CGUGGAGCGAAUGUCAA.....G..UUAGGAAAC...............UGGC..............GACAA..G.UCACUGUACGAAUCAAAAACUGAUUACUAUUUGCCCCAAUGGGAUAACCAGGGUAGGAGUAUGAAACAA
env-691  .....GCUGGU.......GCAG.GU..U.UUC...CGUGGAGCGAAUGUCAA.....G..UUAGGAAAC...............UGGC..............GACAA..G.UCACUGUACGAAUCAAAAACUGAUUACUAUUUGCCCCAAUGGGAUAACCAGGGUAGGAGUAUGAAACAA
env-692  .....GCUGGU.......GCAG.GU..U.UUC...CGUGGAGCGAAUGUCAA.....G..UUAGGAAAC...............UGGC..............GACAA..G.UCACUGUACGAAUCAAAAACUGAUUACUAUUUGCCCCAAUGGGAUAACCAGGGUAGGAGUAUGAAACAA
env-694  .....GCUGGU.......GCAG.GU..U.UUC...CGUGGAGCGAAUGUCAA.....G..UUAGGAAAC...............UGGC..............GACAA..G.UCACUGUACGAAUCAAAAACUGAUUACUAUUUGCCCCAAUGGGAUAACCAGGGUAGGAGUAUGAAACAA
env-695  .....GCUGGU.......GCAG.GU..U.UUC...CGUGGAGCGAAUGUCAA.....G..UUAGGAAAC...............UGGC..............GACAA..G.UCACUGUACGAAUCAAAAACUGAUUACUAUUUGCCCCAAUGGGAUAACCAGGGUAGGAGUAUGAAACAA
env-696  .....GCUGGU.......GCAG.GU..U.UUC...CGUGGAGCGAAUGUCAA.....G..UUAGGAAAC...............UGGC..............GACAA..G.UCACUGUACGAAUCAAAAACUGAUUACUAUUUGCCCCAAUGGGAUAACCAGGGUAGGAGUAUGAAACAA
env-699  .....GCUGGU.......GCAG.GU..U.UUC...CGUGGAGCGAAUGUCAA.....G..UUAGGAAAC...............UGGC..............GACAA..G.UCACUGUACGAAUCAAAAACUGAUUACUAUUUGCCCCAAUGGGAUAACCAGGGUAGGAGUAUGAAACAA
env-700  .................CGCAG.GU..U.UUU...CGUGGAGCGAAUGUCAA.....U..UUACAAAAC...............UGGC..............GACAA..G.UCACUGUACA...........................................................
env-705  .................CGCAG.GU..U.UUU...CGUGGAGCGAAUGUCAA.....U..UUACAAAAC...............UGGC..............GACAA..G.UCACUGUA.............................................................
env-708  .................CGCAG.GU..U.UUU...CGUGGAGCGAAUGUCAA.....U..UUACAAAAC...............UGGC..............GACAA..G.UCACUGUACAAUAUAACCGUCUGUCAAGAGGACAGAAACGACGUGAUGGUUCUGGGUUUUAACUUAGGA
env-709  .................CGCAG.GU..U.UUU...CGUGGAGCGAAUGUCAA.....U..UUACAAAAC...............UGGC..............GACAA..G.UCACUGUACAAUAUAACCGUCUGUCAAGAGGACAGAAACGACGUGAUGGUUCUGGGUUUUAACUUAGGA
env-710  .................CGCAG.GU..U.UUU...CGUGGAGCGAAUGUCAA.....U..UUACAAAAC...............UGGC..............GACAA..G.UCACUGUACAAUAUAACCGUCUGUCAAGAGGACAGAAACGACGUGAUGGUUCUGGGUUUUAACUUAGGA

         >>>>>>>>>>.......>>>>>.....>.>>>...>>>........<<<<.......<..<<<.....................>>>>..............>>>>...>.>>>>.>>..............................................................
         01.1111111.......22221.....2.102...222.........111.......2..212.....................2122..............111....1.111..12..............................................................
         -----------------•GYRG-GU--U-UUC---CGYGGAGCGAAUGUC∘R-----G--•Y∘R•RRR∘---------------∘G•C--------------GRCAA--G-UCACUGUAC•RR•••••••••••••••••••••••••••••••••••••••••••••••••••••••••
                                                                                                                3′
```

```
env-711   .................CGCAG.GU..U.UUU...CGUGGAGCGAAUGUCAA......U..UUACAAAAC................UGGC..............GACAA..G.UCACUGUACAA..........................................................
env-712   .................CGCAG.GU..U.UUU...CGUGGAGCGAAUGUCAA......U..UUACAAAAC................UGGC..............GACAA..G.UCACUGUACAAUAUAACCGUCUGUCAAGAGGACAGAAACGACGUGAUGGUUCUGGGUUUUAACUUAGGA
env-713   .................CGCAG.GU..U.UUU...CGUGGAGCGAAUGUCAA......U..UUACAAAAC................UGGC..............GACAA..G.UCACUGUACAAUAUAACCGUCUGUCAAGAGGACAGAAACGACGUGAUGGUUCUGGGUUUUAACUUAGGA
env-726   .................CGCAG.GU..U.UUU...CGUGGAGCGAAUGUCAA......U..UUACAAAAC................UGGC..............GACAA..G.UCACUGUACAAUAUAACCGUCUGUCAAGAGGACAGAAACGACGUGAUGGUUCUGGGUUUUAACUUAGGA
env-727   .................CGCAG.GU..U.UUU...CGUGGAGCGAAUGUCAA......U..UUACAAAAC................UGGC..............GACAA..G.UCACUGUACA...........................................................
env-729   .................CGCAG.GU..U.UUU...CGUGGAGCGAAUGUCAA......U..UUACAAAAC................UGGC..............GACAA..G.UCACUGUACAAUAUAACCGUCUGUCAAGAGGACAGAAACGACGUGAUGGUUCUGGGUUUUAACUUAGGA
env-730   .................CGCAG.GU..U.UUU...CGUGGAGCGAAUGUCAA......U..UUACAAAAC................UGGC..............GACAA..G.UCACUGUACAAUAUAACCGUCUGUCAAGAGGACAGAAACGACGUGAUGGUUCUGGGUUUUAACUUAGGA
env-731   .................CGCAG.GU..U.UUU...CGUGGAGCGAAUGUCAA......U..UUACAAAAC................UGGC..............GACAA..G.UCACUGUACAAUAUAACCGUCUGUCAAGAGGACAGAAACGACGUGAUGGUUCUGGGUUUUAACUUAGGA
Rsp-1-1   .................GGCAG.GU..U.UUC...CGUGGAGCGAAUGUCAA......G..UUUGAAAAC................UGAC..............GACAA..G.UCACUGUACAAGUUCAGAAAGACCAUCAUGGGAUGGGCGUGUGUGGAAAAAUUUGAUGACACACACGGA
env-732   ......UCACAU.....AGCGG.GU..U.UUC...CGUGGAGCGAAUGUCAA......G..UUAGGAAAC................UGGC..............GACAA..G.UCACUGUACGAAUCACAACUGAUUACUGCUUACCGAUAAUGGGUAAAGGUAUGAAACAACGUGUGAGUU
Rgn-1-1   ......UCACAU.....AGCGG.GU..U.UUC...CGUGGAGCGAAUGUCAA......G..UUAGGAAAC................UGGC..............GACAA..G.UCACUGUACGAAUCACAACUGAUUACUGCUUACCGAUAAUGGGUAAAGGUAUGAAACAACGUGUGAGUU
env-733   ......UCACAU.....AGCGG.GU..U.UUC...CGUGGAGCGAAUGUCAA......G..UUAGGAAAC................UGGC..............GACAA..G.UCACUGUACGAAUCACAACUGAUUACUGCUUACCGAUAAUGGGUAAAGGUAUGAAACAACGUGUGAGUU
env-734   ......UCACAU.....AGCGG.GU..U.UUC...CGUGGAGCGAAUGUCAA......G..UUAGGAAAC................UGGC..............GACAA..G.UCACUGUACGAAUCACAACUGAUUACUGCUUACCGAUAAUGGGUAAAGGUAUGAAACAACGUGUGAGUU
env-735   ......UCACAU.....AGCGG.GU..U.UUC...CGUGGAGCGAAUGUCAA......G..UUAGGAAAC................UGGC..............GACAA..G.UCACUGUACGAAUCACAACUGAUUACUGCUUACCGAUAAUGGGUAAAGGUAUGAAACAACGUGUGAGUU
env-739   ......UCACAU.....AGCGG.GU..U.UUC...CGUGGAGCGAAUGUCAA......G..UUAGGAAAC................UGGC..............GACAA..G.UCACUGUACGAAUC.......................................................
env-742   ......UCACAU.....AGCGG.GU..U.UUC...CGUGGAGCGAAUGUCAA......G..UUAGGAAAC................UGGC..............GACAA..G.UCACUGUACGAAUCACAACUGAUUACUGCUUACCGAUAAUGGGUAAAGGUAUGAAACAACGUGUGAGUU
Lba-4-1   ......UCACAU.....AGCGG.GU..U.UUC...CGUGGAGCGAAUGUCAA......G..UUAGGAAAC................UGGC..............GACAA..G.UCACUGUACGAAUCACAACUGAUUACUGCUUACCGAUAAUGGGUAAAGGUAUGAAACAACGUGUGAGUU
env-744   ......UCACAU.....AGCGG.GU..U.UUC...CGUGGAGCGAAUGUCAA......G..UUAGGAAAC................UGGC..............GACAA..G.UCACUGUACGAAUCACAACUGAUUACUGCUUACCGAUAAUGGGUAAAGGUAUGAAACAACGUGUGAGUU
env-749   .......CAUAUGCU..GGCAG.GU..U.UUC...CGUGGAGCGAAUGUCAA......G..UUAGGAAAC................UGGC..............GACAA..G.UCACUGUACGAAUCACAAUGGUAACUCGUUAAAUGUAUGACCGCAAAUUAGAGCUUUAAAAGAUGAAAG
env-751   .................CGCAG.GU..U.UUC...CGUGGAGCGAAUGUCAA......G..UUAAGAAAC................UGAC..............GACAA..G.UCACUGUACAAUUAAGAAUGACCAUGAAAGUCUUGGUUCUAAGACCGAGCAAAAGAGCAUGGAACUGUG
Cph-1-1   .................UACAG.GU..U.UUC...CGUGGAGCGAAUGUCAU......G..UUAUGAAAC................UGGC..............GACAA..G.UCACUGUACAAGAAUAUAGUUCUGCUACGGGCAGAUAAACCGUGUUGAAACGAAGGCUUUUCGGGAUGA
Bfo-1-1   .................CGCAG.GU..U.UUC...CGUGGAGCGAAUGUCAA......GUUCUGAGAAAC................UGGC..............GACAA..G.UCACUGUACAAGAACAACUGUCCGGCAAAGACCGGAAACAACGUGUGGAUUAUUCAGGAUACACACGGG
env-752   .................GGCCG.GU..U.UUC...CGUGGAGCGAAUGUCAA......G..UUAGGAAAC................UGAC..............GACAG..G.UCACUGUACCAAAGAAAUGUCUACAAAACAGUAGAAACGACGUGUGAAAAUCCUUUGAAAACAGGAUGA
env-753   .................GGCCG.GU..U.UUC...CGUGGAGCGAAUGUCAA......G..UUAGGAAAC................UGAC..............GACAG..G.UCACUGUACCAAAGAAAUGUCUACAAAACAGUAGAAACGACGUGUGAAAAUCCUCUUGAAUUUACAGGA
Lbo-3-1   .................GGUAG.GU..U.UUC...CAUGGAGCGAAUGUCAA......G..UUAGGAAAC................UGAC..............GACAA..G.UCACUGUACAAACAAUAUGGACUACCAAAGUGUAGGGAUAACGUGUGUUCAAAUAGAUGACACACACGG
env-754   .................GGUAA.GU..A.UUC...CGUGGAGCGAAUGUCAA......G..UUAGGAAAC................UGGC..............GACAA..G.UCACUGUACGAACGAAAUAUCCACUUUAUGUGGAAACUUGUGUAUAAAGUUUUAUGAUACACACGGGAA
env-755   .................GGUAA.GU..A.UUC...CGUGGAGCGAAUGUCAA......G..UUAGGAAAC................UGGC..............GACAA..G.UCACUGUACGAACGAAAUAUCCAUUUAAUGUGGAAACUUGUGUAUAUAGCUUUAUGAUACACACGGGAA
Csp-4-1   ......GCAGGAU....GGCAG.GU..U.UUC...CGUGGAGCGAAUGUCAA......G..UUAGGAAAC................UGGC..............GACAA..G.UCACUGUACGAAUCAAAACUGAUUACUGAUUAUCAAAGGAUAAGAAGUAUGAAACAACGUGUGAGUUUC
env-756   .......CCGGAU....AGCAG.GU..U.UUC...CGUGGAGCGAAUGUCAA......G..UUAGGAAAC................UGAC..............GACAA..G.UCACUGUACAGACAAUCAGGUCUGCCGAUAGGCAGAAACAACCGUGAGAAACGAAGCAGUUCAAAUACA
env-758   .......GCAAG.....GGCAG.GU..U.UUC...CGUGGAGCGAAUGUCAA......G..UUAGGAAAC................UGGC..............GACAA..G.UCACUGUACGAAUCACAACUGAUUACUGUUUACCAUACCAUAAGGUAGUAGUAUGAAACAACGUGUGAG
env-759   .................UGUAG.GU..A.UUC...CGUGGAGCGAAUGUCAA......G..UUAUGAAAC................UGGC..............GACAA..G.UCACUGUACAAACAUAUAACGUCCAUCAAAGAAUGGAAACAACGUGUGAGAUUUUAAAGAUACACACGG
env-760   .................UGUAG.GU..A.UUC...CGUGGAGCGAAUGUCAA......G..UUAUGAAAC................UGGC..............GACAA..G.UCACUGUACAAACAUAUAACGUCCAUCAAAGAAUGGAAACAACGUGUGAGAUUUUAAAGAUACACACGG
env-762   .................UGUAG.GU..A.UUC...CGUGGAGCGAAUGUCAA......G..UUAUGAAAC................UGGC..............GACAA..G.UCACUGUACAAACAUAUAACGUCCAUCAAAGAAUGGAAAUAACGUGUGAGAUUUUAAAGAUACACACGG
env-763   .................UGCAG.GU..U.UUC...CGUGGAGCGAAUGUCAA......G..UUAGAAAAC................UGGC..............GACAA..G.UCACUGUACAAGUUAAAAAAGUCCAUCGUAAAGAGUGGAAUGCCGCGUGUGAAAUUAGGACACACCCGG
env-764   .................UACAG.GU..U.UUC...CGUGGAGCGAAUGUCAA......G..UUAGAAAAC................UGGC..............GACAA..G.UCACUGUACAAAUUAACUGUCCGGCUAUGACCGGAAACAACGUGUGGAGAAACGGAGUAUGCUCUGAGA
Lba-6-1   .GUUUGAGAAAAGAUAUGGCAG.GU..U.UUC...CGUGGAGCGAAUGUCAA......G..UUAGGAAAC................UGGC..............GACAA..G.UCACUGUAUCACAUAUAGCCCGUGUAAGGUUCUCACGACACACACGGCAGAGUGUACAGUCACCGCUUG
Dsp-1-1   .................GGCAG.GU..U.UUC...CGUGGAGCGAAUGUCAA......G..UUAAGAAAC................UGGC..............GACAA..G.UCACUGUAACACUGUAAUCGUUGGCACACAUGUGCCGCAUGUAAGCAGUUAGAAAAAAAAACUGCGAAA
env-765   .................GGCAG.GU..U.UUC...CGUGGAGCGAAUGUCAA......G..UUAGGAAAC................UGGC..............GACAA..G.UCACUGUACCAAUUAAAGAAACUGCAAUUUAGGGCAGAAUCAGUGUGUAGUUCUGUGAGGACACACACG
env-767   .................GGCAG.GU..U.UUC...CGUGGAGCGAAUGUCAA......G..UUAGGAAAC................UGGC..............GACAA..G.UCACUGUACCAAUUAAAGAAACUGCAAUUUAGGGCAGAAUCAGUGUGUAGUUCUGUGAGGACACACACG
env-768   .................GGCAG.GU..U.UUC...CGUGGAGCGAAUGUCAA......G..UUAGGAAAC................UGGC..............GACAA..G.UCACUGUACCAAUUAAAGAAACUGCAAUUUAGGGCAGAAUCAGUGUGUAGUUCUGUGAGGACACACACG
env-769   .................GGCAG.GU..U.UUC...CGUGGAGCGAAUGUCAA......G..UUAGGAAAC................UGGC..............GACAA..G.UCACUGUACCAAUUAAAGAAUCUGCAAACAAGAGCAGGAUCAGUGUGGAGUUCUAUGAGGACACACACG
env-770   .................GGCAG.GU..U.UUC...CGUGGAGCGAAUGUCAA......G..UUAGGAAAC................UGGC..............GACAA..G.UCACUGUACCAAUUAAAGAAACUGCAAUUUAGGGCAGAAUCAGUGUGUAGUUCUGUGAGGACACACACG
env-777   .................GGCAG.GU..U.UUC...CGUGGAGCGAAUGUCAA......G..UUAGGAAAC................UGGC..............GACAA..G.UCACUGUACCAAUUAAAGAAACUGCAGCAAAGAGAGCAGAAUCAGUGUGGGGUUCCGUGAGGACACACA
env-778   .................GGCAG.GU..U.UUC...CGUGGAGCGAAUGUCAA......G..UUAGGAAAC................UGGC..............GACAA..G.UCACUGUACGAAAUCAUAACAGCUGCUGAGAUGCAGUAACAACGUGUGAGUUUCUCACGACACACACGG
env-802   .................GGCAG.GU..U.UUC...CGUGGAGCGAAUGUCAA......G..UUAGGAAAC................UGGC..............GACAA..G.UCACUGUACGAAAUCAUAACAGCUGCUGAAAUGCAGUAACAACGUGUGAGUUUCUCACGACACACACGG
env-803   .................GGCAG.GU..U.UUC...CGUGGAGCGAAUGUCAA......G..UUAGGAAAC................UGGC..............GACAA..G.UCACUGUACGAAAUCAUAACAGCUGCUGAGAUGCAGUAACAACGUGUGAGUUUCUCACGACACACACGG
Cco-1-1   .................GGCAG.GU..U.UUC...CGUGGAGCGAAUGUCAA......G..UUAGGAAAC................UGGC..............GACAA..G.UCACUGUACGAAAUCAUAACAGCUGCUGAGAUGCAGUAACAACGUGUGAGUUUCUCACGACACACACGG
env-806   .................GGCAG.GU..U.UUC...CGUGGAGCGAAUGUCAA......G..UUAGGAAAC................UGGC..............GACAA..G.UCACUGUACGAAAUCAUAACAGCUGCUGAGAUGCAGUAACAACGUGUGAGUUUCUCACGACACACACGG
env-812   .................GGCAG.GU..U.UUC...CGUGGAGCGAAUGUCAA......G..UUAGGAAAC................UGGC..............GACAA..G.UCACUGUACGAAAUCAUAACAGCUGCUGAGAUGCAGUAACAACGUGUGAGUUUCUCACGACACACACGG
env-814   .................GACGG.GU..U.UUC...CGUGGAGCGAAUGUCAA......G..UUAGGAAAC................UGGC..............GACAA..G.UCACUGUACGAAUCAAAAACUGAUUACUGAUUACCAUUAGGUAUAAGUAUGAAACAACGUGUGAGUUUC
Dfo-2-1   .................GACGG.GU..U.UUC...CGUGGAGCGAAUGUCAA......G..UUAGGAAAC................UGGC..............GACAA..G.UCACUGUACGAAUCAAAAACUGAUUACUGAUUACCAUUAGGUAUAAGUAUGAAACAACGUGUGAGUUUC
Dfo-1-1   .................GACGG.GU..U.UUC...CGUGGAGCGAAUGUCAA......G..UUAGGAAAC................UGGC..............GACAA..G.UCACUGUACGAAUCAAAAACUGAUUACUGAUUACCAUUAGGUAUAAGUAUGAAACAACGUGUGAGUUUC
env-818   .................GACGG.GU..U.UUC...CGUGGAGCGAAUGUCAA......G..UUAGGAAAC................UGGC..............GACAA..G.UCACUGUACGAAUCAAAAACUGAUUACUGAUUACCAUUAGGUAUAAGUAUGAAACAACGUGUGAGUUUC
env-819   .................GACGG.GU..U.UUC...CGUGGAGCGAAUGUCAA......G..UUAGGAAAC................UGGC..............GACAA..G.UCACUGUACGAAUCAAAAACUGAUUACUGAUUACCAUUAGGUAUAAGUAUGAAACAACGUGUGAGUUUC
env-820   .......GUGA......GGCGG.GU..U.UUC...CGUGGAGCGAAUGUCAA......G..UUAGGAAAC................UGGC..............GACAA..G.UCACUGUACCAAUUAAAGAAACUGCAAUGUUAGAACGCAGAAUCAGUGUGGAGUUCUGUUAGGACACAC
[ci-1-1   .......GUGA......GGCGG.GU..U.UUC...CGUGGAGCGAAUGUCAA......G..UUAGGAAAC................UGGC..............GACAA..G.UCACUGUACCAAUUAAAGAAACUGCAGGAUAAAUGGCAGAAUCAGUGUGGAGUUCCGUUGGGACACACA
Csp-5-1   .......GUGA......GGCGG.GU..U.UUC...CGUGGAGCGAAUGUCAA......G..UUAGGAAAC................UGGC..............GACAA..G.UCACUGUACCAAUUAAAGAAACUGCAGGAUAAAUGGCAGAAUCAGUGUGGAGUUCCGUUGGGACACACA
env-821   .......GUGA......GGCAG.GU..U.UUC...CGUGGAGCGAAUGUCAA......G..UUAGGAAAC................UGGC..............GACAA..G.UCACUGUACCAAUUAAAGAAACUGCAAUGUUAGAACGCAGAAUCAGUGUGGAGUUCUGUUAGGACACAC
env-822   .......GUGA......GGCAG.GU..U.UUC...CGUGGAGCGAAUGUCAA......G..UUAGGAAAC................UGGC..............GACAA..G.UCACUGUACCAAUUAAAGAAACUGCAAUGUUAGAACGCAGAAUCAGUGUGGAGUUCUGUUAGGACACAC
env-823   .......GUGA......GGCAG.GU..U.UUC...CGUGGAGCGAAUGUCAA......G..UUAGGAAAC................UGGC..............GACAA..G.UCACUGUACCAAUUAAAGAAACUGCAAUGUUAGAACGCAGAAUCAGUGUGGAGUUCUGUUAGGACACAC
Cbo-2-1   .......GUGA......GGCAG.GU..U.UUC...CGUGGAGCGAAUGUCAA......G..UUAGGAAAC................UGGC..............GACAA..G.UCACUGUACCAAUUAAAGAAACUGCAAUGUUAGAACGCAGAAUCAGUGUGGAGUUCUGUUAGGACACAC
Cba-7-1   .......GUGA......GGCAG.GU..U.UUC...CGUGGAGCGAAUGUCAA......G..UUAGGAAAC................UGGC..............GACAA..G.UCACUGUACCAAUUAAAGAAACUGCAGGAUAAUCAGCAGAAUCAGUGUGGAGUUCUUUUAGGACACACA
env-824   .......GUGA......GGCAG.GU..U.UUC...CGUGGAGCGAAUGUCAA......G..UUAGGAAAC................UGGC..............GACAA..G.UCACUGUACCAAUUAAAGAAACUGCAAUGUUAGAACGCAGAAUCAGUGUGGAGUUCUGUUAGGACACAC
Cba-2-1   .......GUGA......GGCAG.GU..U.UUC...CGUGGAGCGAAUGUCAA......G..UUAGGAAAC................UGGC..............GACAA..G.UCACUGUACCAAUUAAAGAAACUGCAGGAUAAUCAGCAGAAUCAGUGUGGAGUUCUUUUAGGACACACA
env-826   .......GUGA......GGCAG.GU..U.UUC...CGUGGAGCGAAUGUCAA......G..UUAGGAAAC................UGGC..............GACAA..G.UCACUGUACCAAUUAAAGAAACUGCAAUGUUAGAACGCAGAAUCAGUGUGGAGUUCUGUUAGGACACAC
Ccl-6-1   .......GUGA......GGCAG.GU..U.UUC...CGUGGAGCGAAUGUCAA......G..UUAGGAAAC................UGGC..............GACAA..G.UCACUGUACCAAUUAAAGAAACUGCAAUGUUAGAACGCAGAAUCAGUGUGGAGUUCUGUUAGGACACAC
env-827   .......GUGA......GGCAG.GU..U.UUC...CGUGGAGCGAAUGUCAA......G..UUAGGAAAC................UGGC..............GACAA..G.UCACUGUACCAAUUAAAGAAACUGCAAUGUUAGAACGCAGAAUCAGUGUGGAGUUCUGUUAGGACACAC
Ccl-3-1   .......GUGA......GGCAG.GU..U.UUC...CGUGGAGCGAAUGUCAA......G..UUAGGAAAC................UGGC..............GACAA..G.UCACUGUACCAAUUAAAGAAACUGCAAUGUUAGAUGGCAGAAUCAGUGUGGAGUUCCGUUGGGACACAC
Ccl-1-1   .......GUGA......GGCAG.GU..U.UUC...CGUGGAGCGAAUGUCAA......G..UUAGGAAAC................UGGC..............GACAA..G.UCACUGUACCAAUUAAAGAAACUGCAAUGUUAGAUGGCAGAAUCAGUGUGGAGUUCCGUUGGGACACAC
Lba-16-1  ....CAUGCAGGAG...GGCAG.GU..U.UUC...CGUGGAGCGAAUGUCAA......G..UUAGGAAAC................UGGC..............GACAA..G.UCACUGUACGAAUCACAACAGACUGCUGGUCUGCCGUAAGGCGCAGUAAGAAGUAACGUGUAUGUUUCU
Csp-3-1   ....CAUGCAGGAG...GGCAG.GU..U.UUC...CGUGGAGCGAAUGUCAA......G..UUAGGAAAC................UGGC..............GACAA..G.UCACUGUACGAAUCACAACAGACUACUGGUCUGCCGUAAGGCGCAGUAAGAAGUAACGUGUAUGUUUCU
env-828   ......GCUGA......GGCAG.GU..U.UUC...CGUGGAGCGAAUGUCAA......G..UUAGGAAAC................UGGC..............GACAA..G.UCACUGUACGAAUCACAACUGAUUACUGCUUACCGAUAAUGGGUAAAGGUAUGAAACAACGUGUGAGUU
env-829   ......GCUGA......GGCAG.GU..U.UUC...CGUGGAGCGAAUGUCAA......G..UUAGGAAAC................UGGC..............GACAA..G.UCACUGUACGAAUCACAACUGAUUACUGAUUACCACAAGGGAAAAGUAUGAAACAACGUGUGAGUUUCU
env-830   ......GCUGA......GGCAG.GU..U.UUC...CGUGGAGCGAAUGUCAA......G..UUAGGAAAC................UGGC..............GACAA..G.UCACUGUACGAAUCACAACUGAUUACUGAUUACCACAAGGGAAAAGUAUGAAACAACGUGUGAGUUUCU
env-831   ......GCUGA......GGCAG.GU..U.UUC...CGUGGAGCGAAUGUCAA......G..UUAGGAAAC................UGGC..............GACAA..G.UCACUGUACGAAUCACAACUGAUUACUGAUUACCACAAGGGAAAAGUAUGAAACAACGUGUGAGUUUCU
env-832   ......GCUGA......GGCAG.GU..U.UUC...CGUGGAGCGAAUGUCAA......G..UUAGGAAAC................UGGC..............GACAA..G.UCACUGUACGAAUCACAACUGAUUACUGAUUACCACAAGGGAAAAGUAUGAAACAACGUGUGAGUUUCU
env-833   ......GCUGA......GGCAG.GU..U.UUC...CGUGGAGCGAAUGUCAA......G..UUAGGAAAC................UGGC..............GACAA..G.UCACUGUACGAAUCACAACUGAUUACUGAUUACCACAAGGGAAAAGUAUGAAACAACGUGUGAGUUUCU
env-837   ......GCUGA......GGCAG.GU..U.UUC...CGUGGAGCGAAUGUCAA......G..UUAGGAAAC................UGGC..............GACAA..G.UCACUGUACGAAUCACAACUGAUUACUGAUUACCACAAGGGAAAAGU[illegible]AAACAACGUGUGAGUUUCU
env-838   ......GCUGA......GGCAG.GU..U.UUC...CGUGGAGCGAAUGUCAA......G..UUAGGAAAC................UGGC..............GACAA..G.UCACUGUACGAAUCACAACUGAUUACUGAUUACCACAAGGGAAAAGUAUGAAACAACGUGUGAGUUUCU
env-839   ......GCUGA......GGCAG.GU..U.UUC...CGUGGAGCGAAUGUCAA......G..UUAGGAAAC................UGGC..............GACAA..G.UCACUGUACAAAACAUAAGUCCACCAUAAGAGUGGAAACACCGAGUUAAGGUUGCAAGAGAUUGCGACA
env-840   ......GCUGA......GGCAG.GU..U.UUC...CGUGGAGCGAAUGUCAA......G..UUAGGAAAC................UGGC..............GACAA..G.UCACUGUACGAAUCACAACUGAUUACUGAUUACCACAAGGGAAAAGUAUGAAACAACGUGUGAGUUUC.
env-841   ......GCUGA......GGCAG.GU..U.UUC...CGUGGAGCGAAUGUCAA......G..UUAGGAAAC................UGGC..............GACAA..G.UCACUGUACGAAUCACAACUGAUUACUGAUUACCACAAGGGAAAAGUAUGAAACAACGUGUGAGUUUCU
Lba-7-1   ......GCUGA......GGCAG.GU..U.UUC...CGUGGAGCGAAUGUCAA......G..UUAGGAAAC................UGGC..............GACAA..G.UCACUGUACGAAUCACAACUGAUUACUGAUUACCACAAGGGAAAAGUAUGAAACAACGUGUGAGUUUCU

          >>>>>>>>>>.......>>>>>.....>.>>>...>>>.........<<<<.......<..<<<......................>>>>..............>>>>...>.>>>>.>>................................................................
          01.1111111.......22221.....2.102...222..........111.......2..212......................2122..............111....1.111..12................................................................
          -----------------•GYRG-GU--U-UUC---CGYGGAGCGAAUGUC•R-----G--•Y•R•RRR•---------------•G•C-------------GRCAA--G-UCACUGUAC•RR••••••••••••••••••••••••••••••••••••••••••••••••••••••••••••••
                                                                                                            3′
```

```
env-842   ......GCUGA.....GGCAG.GU..U.UUC...CGUGGAGCGAAUGUCAA.....G..UUAGGAAAC..............UGGC..............GACAA..G.UCACUGUACGAAUCACAACUGAUUACUGAUUACCACAAGGGGAAAAGUAUGAAACAACGUGUGAGUUUC
env-843   GG.GAAG.CAAA....GGCAG.GU..U.UUC...CGUGGAGCGAAUGUCAA.....G..UUAGGAAAC..............UGGC..............GACAA..G.UCACUGUACAAUUAAAGAAUUCUGUUUCGAAAAAACGGAAAGUCAGUGUGGGGUCCUGGAGGACACACA
env-844   ................UGCAG.GU..G.UUC...CGUGGAGCGAAUGUCAA.....G..UUAGGAAAC..............UGGC..............GACAA..G.UCACUGUACAAGUUAAGAAUCUGUCGAGCGAAAGCUUUGAAGACAGAGAGAAACGCGAAGAAGCUUACU
env-845   ................UGCAG.GU..G.UUC...CGUGGAGCGAAUGUCAA.....G..UUAGGAAAC..............UGGC..............GACAA..G.UCACUGUACAAGUUAAGAAUCUGUCGAGCGAAAGCUUUGAAGACAGAGAGAAACGCGAAGAAGCUUACU
env-848   ................UGCAG.GU..G.UUC...CGUGGAGCGAAUGUCAA.....G..UUAGGAAAC..............UGGC..............GACAA..G.UCACUGUACAAGUUAAGAAUCUGUCGAGCGAAAGCUUUGAAGACAGAGAGAAACGCGAAGAAGCUUACU
env-849   ................UGCAG.GU..G.UUC...CGUGGAGCGAAUGUCAA.....G..UUAGGAAAC..............UGGC..............GACAA..G.UCACUGUACAAGUUAAGAAUCUGUCGAGCGAAAGCUUUGAAGACAGAGAGAAACGCGAAGAAGCUUACU
env-850   ................UGCAG.GU..G.UUC...CGUGGAGCGAAUGUCAA.....G..UUAGGAAAC..............UGGC..............GACAA..G.UCACUGUACAAGUUAAGAAUCUGUCGAGCGAAAGCUUUGAAGACAGAGAGAAACGCGAAGAAGCUUACU
env-851   ................UGCAG.GU..U.UUC...CGUGGAGCGAAUGUCAA.....G..UUAGGAAAC..............UGGC..............GACAA..G.UCACUGUAUCACGAAUAGAAUUUGUCCGAAGGACAGAGGGAUGGCGUGUAGGAAGCGCGGGCUUCUUUG
env-853   ................UGCAG.GU..U.UUC...CGUGGAGCGAAUGUCAA.....G..UUAGGAAAC..............UGGC..............GACAA..G.UCACUGUAUCACGAAUAGAAUUUGUCCGAAGGACAGAGGGAUGGCGUGUAGGAAGCGCGGGCUUCUUUG
env-854   ................UGCAG.GU..U.UUC...CGUGGAGCGAAUGUCAA.....G..UUAGGAAAC..............UGGC..............GACAA..G.UCACUGUAUCACGAAUAGAAUUUGUCCGAAGGACAGAGGGAUGGCGUGUAGGAAGCGCGGGCUUCUUUG
env-855   ................UGCAG.GU..U.UUC...CGUGGAGCGAAUGUCAA.....G..UUAGGAAAC..............UGGC..............GACAA..G.UCACUGUAUCACGAAUAGAAUUUGUCCGAAGGACAGAGGGAUGGCGUGUAGGAAGCGCGGGCUUCUUUG
env-856   ................UGCAG.GU..U.UUC...CGUGGAGCGAAUGUCAA.....G..UUAGGAAAC..............UGGC..............GACAA..G.UCACUGUAUCACGAAUAGAAUUUGUCCGAAGGACAGAGGGAUGGCGUGUAGGAAGCGCGGGCUUCUUUG
env-861   ................UGCAG.GU..U.UUC...CGUGGAGCGAAUGUCAA.....G..UUAGGAAAC..............UGGC..............GACAA..G.UCACUGUAUCACGAAUAGAAUUUGUCCGAAGGACAGAGGGAUGGCGUGUAGGAAGCGCGGGCUUCUUUG
env-862   ................UGCAG.GU..U.UUC...CGUGGAGCGAAUGUCAA.....G..UUAGGAAAC..............UGGC..............GACAA..G.UCACUGUAUCACGAAUAGAAUUUGUCCGAAGGACAGAGGGAUGGCGUGUAGGAAGCGCGGGCUUCUUUG
env-863   ................UGCAG.GU..U.UUC...CGUGGAGCGAAUGUCAA.....G..UUAAGAAAC..............UGGC..............GACAA..G.UCACUGUAUACAAGUCAAGGAAGUCCGCUGAAAAGAGCGGAGCCGUGUGAAGAUCAGCAAGACACACAC
env-864   ................UGCAG.GU..U.UUC...CGUGGAGCGAAUGUCAA.....G..UUAAGAAAC..............UGGC..............GACAA..G.UCACUGUAUACAUGUUAAAUACGUCCACCAGAAGUGUGGAAAACCGCGUGAAAGUCCGGAGGACACACA
env-865   ................UGCAG.GU..U.UUC...CGUGGAGCGAAUGUCAA.....G..UUAAGAAAC..............UGGC..............GACAA..G.UCACUGUAUCACGAAUAACGUCUACAAAAGGUAGAAACAACGUGCCAAAUCAUCAACGGGAAGAUGUUU
env-866   ................UGCAG.GU..U.UUC...CGUGGAGCGAAUGUCAA.....G..UUAAGAAAC..............UGGC..............GACAA..G.UCACUGUAUCACGAAUAACGUCUACAAAAGGUAGAAACAACGUGCCAAAUCAUCAACGGGAAGAUGUUU
env-867   ................UGCAG.GU..U.UUC...CGUGGAGCGAAUGUCAA.....G..UUAAGAAAC..............UGGC..............GACAA..G.UCACUGUAUCACGAAUAACGUCUACAAAAGGUAGAAACAACGUGCCAAAUCAUCAACGGGAAGAUGUUU
env-868   ................UGCAG.GU..U.UUC...CGUGGAGCGAAUGUCAA.....G..UUAAGAAAC..............UGGC..............GACAA..G.UCACUGUAUCACGAAUAACGUCUACAAAAGGUAGAAACAACGUGCCAAAUCAUCAACGGGAAGAUGUUU
env-872   ................UGCAG.GU..U.UUC...CGUGGAGCGAAUGUCAA.....G..UUAAGAAAC..............UGGC..............GACAA..G.UCACUGUAUCACUAAUAACGUCUACAAAAGGUAGAAACAACGUGCCAAAUCAUCAACGGGAAGAUGUUU
env-874   ................UGCAG.GU..U.UUC...CGUGGAGCGAAUGUCAA.....G..UUAAGAAAC..............UGGC..............GACAA..G.UCACUGUAUCACGAAUAACGUCUACAAAAGGUAGAAACAACGUGCCAAAUCAUCAACGGGAAGAUGUUU
env-875   ................UGCAG.GU..U.UUC...CGUGGAGCGAAUGUCAA.....G..UUAAGAAAC..............UGGC..............GACAA..G.UCACUGUAUCACGAAUAACGUCUACAAAAGGUAGAAACAACGUGCCAAAUCAUCAACGGGAAGAUGUUU
env-877   ................UGCAG.GU..U.UUC...CGUGGAGCGAAUGUCAA.....G..UUAAGAAAC..............UGGC..............GACAA..G.UCACUGUAUCACGAAUAACGUCUACAAAAGGUAGAAACAACGUGCCAAAUCAUCAACGGGAAGAUGUUU
env-878   ................UGCAG.GU..U.UUC...CGUGGAGCGAAUGUCAA.....G..UUAAGAAAC..............UGGC..............GACAA..G.UCACUGUAUCACGAAUAACGUCUACAAAAGGUAGAAACAACGUGCCAAAUCAUCAACGGGAAGAUGUUU
env-879   ................UGCAG.GU..U.UUC...CGUGGAGCGAAUGUCAA.....G..UUAAGAAAC..............UGGC..............GACAA..G.UCACUGUAUCACGAAUAACGUCUACAAAAGGUAGAAACAACGUGCCAAAUCAUCAACGGGAAGAUGUUU
env-880   ................UGCAG.GU..U.UUC...CGUGGAGCGAAUGUCAA.....G..UUAAGAAAC..............UGGC..............GACAA..G.UCACUGUAUCACGAAUAACGUCUACAAAAGGUAGAAACAACGUGCCAAAUCAUCAACGGGAAGAUGUUU
env-892   ................UGCAG.GU..U.UUC...CGUGGAGCGAAUGUCAA.....G..UUAAGAAAC..............UGGC..............GACAA..G.UCACUGUAUCACGAAUAACGUCUACAAAAGGUAGAAACAACGUGCCAAAUCAUCAACGGGAAGAUGUUU
env-893   ................UGCAG.GU..U.UUC...CGUGGAGCGAAUGUCAA.....G..UUAAGAAAC..............UGGC..............GACAA..G.UCACUGUAUCACGAAUAACGUCUACAAAAGGUAGAAACAACGUGCCAAAUCAUCAACGGGAAGAUGUUU
env-898   ................UGCAG.GU..U.UUC...CGUGGAGCGAAUGUCAA.....G..UUAAGAAAC..............UGGC..............GACAA..G.UCACUGUAUCACGAAUAACGUCUACAAAAGGUAGAAACAACGUGCCAAAUCAUCAACGGGAAGAUGUUU
env-899   ................UGCAG.GU..U.UUC...CGUGGAGCGAAUGUCAA.....G..UUAAGAAAC..............UGGC..............GACAA..G.UCACUGUAUCACGAAUAACGUCUACAAAAGGUAGAAACAACGUGCCAAAUCAUCAACGGGAAGAUGUUU
env-900   ................UGCAG.GU..U.UUC...CGUGGAGCGAAUGUCAA.....G..UUAAGAAAC..............UGGC..............GACAA..G.UCACUGUAUCACGAAUAACGUCUACAAAAGGUAGAAACAACGUGCCAAAUCAUCAACGGGAAGAUGCUU
env-901   ................UGCAG.GU..U.UUC...CGUGGAGCGAAUGUCAA.....G..UUAAGAAAC..............UGGC..............GACAA..G.UCACUGUAUCACGAAUAACGUCUACAAAAGGUAGAAACAACGUGCCAAAUCAUCAACGGGAAGAUGUUU
env-902   ................UGCAG.GU..U.UUC...CGUGGAGCGAAUGUCAA.....G..UUAAGAAAC..............UGGC..............GACAA..G.UCACUGUAUCACGAAUAACGUCUACAAAAGGUAGAAACAACGUGCCAAAUCAUCAACGGGAAGAUGUUU
env-903   ................UGCAG.GU..U.UUC...CGUGGAGCGAAUGUCAA.....G..UUAAGAAAC..............UGGC..............GACAA..G.UCACUGUAUCACGAAUAACGUCUACAAAAGGUAGAAACAACGUGCCAAAUCAUCAACGGGAAGAUGUUU
env-904   ................UGCAG.GU..U.UUC...CGUGGAGCGAAUGUCAA.....G..UUAAGAAAC..............UGGC..............GACAA..G.UCACUGUAUCACGAAUAACGUCUACAAAAGGUAGAAACAACGUGCCAAAUCAUCAACGGGAAGAUGUUU
env-906   ................UGCAG.GU..U.UUC...CGUGGAGCGAAUGUCAA.....G..UUAAGAAAC..............UGGC..............GACAA..G.UCACUGUAUCACGAAUAACGUCUACAAAAGGUAGAAACAACGUGCCAAAUCAUCAACGGGAAGAUGUUU
Rin-2-1   ................UGCAG.GU..U.UUC...CGUGGAGCGAAUGUCAA.....G..UUAAGAAAC..............UGGC..............GACAA..G.UCACUGUAUCACGAAUAACGUCUACAAAAGGUAGAAACAACGUGCCAAAUCAUCAACGGGAAGAUGUUU
env-907   ................UGCAG.GU..U.UUC...CGUGGAGCGAAUGUCAA.....G..UUAAGAAAC..............UGGC..............GACAA..G.UCACUGUAUCACGAAUAACGUCUACAAAAGGUAGAAACAACGUGCCAAAUCAUCAACGGGAAGAUGUUU
env-913   ................UGCAG.GU..U.UUC...CGUGGAGCGAAUGUCAA.....G..UUAAGAAAC..............UGGC..............GACAA..G.UCACUGUAUCACGAAUAACGUCUACAAAAGGUAGAAACAACGUGCCAAAUCAUCAACGGGAAGAUGUUU
env-914   ................UGCAG.GU..U.UUC...CGUGGAGCGAAUGUCAA.....G..UUAAGAAAC..............UGGC..............GACAA..G.UCACUGUAUCACGAAUAACGUCUACAAAAGGUAGAAACAACGUGCCAAAUCAUCAACGGGAAGAUGUUU
env-915   ................UGCAG.GU..U.UUC...CGUGGAGCGAAUGUCAA.....G..UUAAGAAAC..............UGGC..............GACAA..G.UCACUGUAUCACGAAUAACGUCUACAAAAGGUAGAAACAACGUGCCAAAUCAUCAACGGGAAGAUGUUU
env-916   ................UGCAG.GU..U.UUC...CGUGGAGCGAAUGUCAA.....G..UUAAGAAAC..............UGGC..............GACAA..G.UCACUGUAUCACGAAUAACGUCUACAAAAGGUAGAAACAACGUGCCAAAUCAUCAACGGGAAGAUGUUU
env-918   ................AGUAG.GU..U.UUC...CGUGGAGCGAAUGUCAA.....G..UUAGGAAAC..............UGGC..............GACAA..G.UCACUGUACGAAACUAUAACAGCCACUGAGAUGUGGUAACAACGUGUGAGUUUCUCACGACACACACGG
Lba-10-1  ................AGUAG.GU..U.UUC...CGUGGAGCGAAUGUCAA.....G..UUAGGAAAC..............UGGC..............GACAA..G.UCACUGUACGAAACUAUAACAGCCACUGAGAUGUGGUAACAACGUGUGAGUUUCUCACGACACACACGG
[sc-1-1   ................AGUAG.GU..U.UUC...CGUGGAGCGAAUGUCAA.....G..UUAGGAAAC..............UGGC..............GACAA..G.UCACUGUACGAAACUAUAACAGCCACUGAGAUGUGGUAACAACGUGUGAGUUUCUCACGACACACACGG
env-921   ................AGUAG.GU..U.UUC...CGUGGAGCGAAUGUCAA.....G..UUAGGAAAC..............UGGC..............GACAA..G.UCACUGUACGAAACUAUAACAGCCACUGAGAUGUGGUAACAACGUGUGAGUUUCUCACGACACACACGG
env-923   ................AGUAG.GU..U.UUC...CGUGGAGCGAAUGUCAA.....G..UUAGGAAAC..............UGGC..............GACAA..G.UCACUGUACGAAACUAUAACAGCCACUGAGAUGUGGUAACAACGUGUGAGUUUCUCACGACACACACGG
Csc-1-1   ................AGUAG.GU..U.UUC...CGUGGAGCGAAUGUCAA.....G..UUAGGAAAC..............UGGC..............GACAA..G.UCACUGUACGAAACUAUAACAGCCACUGAGAUGUGGUAACAACGUGUGAGUUUCUCACGACACACACGG
Cba-6-1   ................AGUAG.GU..U.UUC...CGUGGAGCGAAUGUCAA.....G..UUAGGAAAC..............UGGC..............GACAA..G.UCACUGUACGAAACUAUAACAGCCACUGAGAUGUGGUAACAACGUGUGAGUUUCUCACGACACACACGG
Bhy-1-1   ................GGCAG.GU..U.UUC...CGUGGAGCGAAUGUCAA.....G..UUAGGAAAC..............UGAC..............GACAA..G.UCACUGUACAAGUUCAGAUAUCCACAGAAAGUGUGGAAAUAGUGUGUGUGAAAGUUAGGACACACACGG
env-924   ................GACAG.GU..U.UUC...CGUGGAGCGAAUGUCAA.....G..UUAGGAAAC..............UGGC..............GACAA..G.UCACUGUACCAAUUUAACAUUCUGCGAAAGCAGAUAGAGACAGAAGGGAUGUCCUGCGUUACCCGAUGU
env-926   ................GACAG.GU..U.UUC...CGUGGAGCGAAUGUCAA.....G..UUAGGAAAC..............UGGC..............GACAA..G.UCACUGUACCAAUUUAACAAUCCGUGUGAACGGAUAAGAACAGAAGGGAAGUCCUGCGUUACCCGAUGU
Bcr-1-1   ................GACAG.GU..U.UUC...CGUGGAGCGAAUGUCAA.....G..UUAGGAAAC..............UGAC..............GACAA..G.UCACUGUACAACUUACCAUCUGCACAUGGCAGAGAUGACCGUUUGAGGUUGCAUUAUAGAAGAAGGUUU
env-928   ................GACAG.GU..U.UUC...CGUGGAGCGAAUGUCAA.....G..UUAGGAAAC..............UGAC..............GACAA..G.UCACUGUACAACUUACCAUCUGCACAUGGCAGAGAUGACCGUAUGAGGUUGCAUUAUAGAAGAAGGUUU
env-930   ................GACAG.GU..U.UUC...CGUGGAGCGAAUGUCAA.....G..UUAGGAAAC..............UGAC..............GACAA..G.UCACUGUACAACUUACCAUCUGCACAUGGCAGAGAUGACCGUUUGAGGUUGCAUUAUAGAAGAAGGUUU
env-932   ................GACAG.GU..U.UUC...CGUGGAGCGAAUGUCAA.....G..UUAGGAAAC..............UGAC..............GACAA..G.UCACUGUACAACUUACCAUCUGCACAUGGCAGAGAUGACCGUUUGAGGUUGCAUUAUAGAAGAAGGUUU
env-933   ................GACAG.GU..U.UUC...CGUGGAGCGAAUGUCAA.....G..UUAGGAAAC..............UGAC..............GACAA..G.UCACUGUACAACUUACCAUCUGCACAUGGCAGAGAUGACCGUUUGAGGUUGCAUUAUAGAAGAAGGUUU
env-934   ................GGCAG.GU..U.UUC...CGUGGAGCGAAUGUCAA.....G..UUAGGAAAC..............UGGC..............GACAA..G.UCACUGUAUACGAGUAACAUCUACAAAGUAGAAACAACGUGCGGGCAGACCGGAAGGCUGUCAGAAGAA
env-935   ................GGCAG.GU..U.UUC...CGUGGAGCGAAUGUCAA.....G..UUAGGAAAC..............UGGC..............GACAA..G.UCACUGUAUACGAGUAACAUCUACAAAGUAGAAACAACGUGCGGGCAGACCGGAAGGCUGUCAGAAGAA
env-940   ................GGCAG.GU..U.UUC...CGUGGAGCGAAUGUCAA.....G..UUAGGAAAC..............UGGC..............GACAA..G.UCACUGUAUACGAGUAACAUCUACAAAGUAGAAACAACGUGCGGGCAGACCGGAAGGCUGUCAGAAGAA
env-942   ................GGCAG.GU..U.UUC...CGUGGAGCGAAUGUCAA.....G..UUAGGAAAC..............UGGC..............GACAA..G.UCACUGUAUACGAGUAACAUCUACAAAGUAGAAACAACGUGCGGGCAGACCGGAAGGCUGUCAGAAGAA
env-943   ................GGCAG.GU..U.UUC...CGUGGAGCGAAUGUCAA.....G..UUAGGAAAC..............UGGC..............GACAA..G.UCACUGUAUACGAGUAACAUCUACAAAGUAGAAACAACGUGCGGGCAGACCGGAAGGCUGUCAGAAGAA
env-944   ................GGCAG.GU..U.UUC...CGUGGAGCGAAUGUCAA.....G..UUAGGAAAC..............UGGC..............GACAA..G.UCACUGUAUACGAGUAACAUCUACAAAGUAGAAACAACGUGCGGGCAGACCGGAAGGCUGUCAGAAGAA
env-945   ................GGCAG.GU..U.UUC...CGUGGAGCGAAUGUCAA.....G..UUAGGAAAC..............UGGC..............GACAA..G.UCACUGUAUACGAGUAACAUCUACAAAGUAGAAACAACGUGCGGGCAGACCGGAAGGCUGUCAGAAGAA
env-946   ................GGCAG.GU..U.UUC...CGUGGAGCGAAUGUCAA.....G..UUAGGAAAC..............UGGC..............GACAA..G.UCACUGUAUACGAGUAACAUCUACAAAGUAGAAACAACGUGCGGGCAGACCGGAAGGCUGUCAGAAGAA
env-949   ................GGCAG.GU..U.UUC...CGUGGAGCGAAUGUCAA.....G..UUAGGAAAC..............UGGC..............GACAA..G.UCACUGUAUACGAGUAACAUCUACAAAGUAGAAACAACGUGCGGGCAGACCGGAAGGCUGUCAGAAGAA
env-950   ................GGCAG.GU..U.UUC...CGUGGAGCGAAUGUCAA.....G..UUAGGAAAC..............UGGC..............GACAA..G.UCACUGUAUACGAGUAACAUCUACAAAGUAGAAACAACGUGCGGGCAGACCGGAAGGCUGUCAGAAGAA
Rin-1-1   ................GGCAG.GU..U.UUC...CGUGGAGCGAAUGUCAA.....G..UUAGGAAAC..............UGGC..............GACAA..G.UCACUGUAUACGAGUAACAUCUACAAAGUAGAAACAACGUGCGGGCAGACCGGAAGGCUGUCAGAAGAA
env-951   ................GGCAG.GU..U.UUC...CGUGGAGCGAAUGUCAA.....G..UUAGGAAAC..............UGGC..............GACAA..G.UCACUGUAUACGAGUAACAUCUACAAAGUAGAAACAACGUGCGGGCAGACCGGAAGGCUGUCAGAAGAA
env-953   ................GGCAG.GU..U.UUC...CGUGGAGCGAAUGUCAA.....G..UUAGGAAAC..............UGGC..............GACAA..G.UCACUGUAUACGAGUAACAUCUACAAAGUAGAAACAACGUGCGGGCAGACCGGAAGGCUGUCAGAAGAA
env-954   ................GGCAG.GU..U.UUC...CGUGGAGCGAAUGUCAA.....G..UUAGGAAAC..............UGGC..............GACAA..G.UCACUGUAUACGAGUAACAUCUACAAAGUAGAAACAACGUGCGGGCAGACCGGAAGGCUGUCAGAAGAA
env-955   ................GGCAG.GU..U.UUC...CGUGGAGCGAAUGUCAA.....G..UUAGGAAAC..............UGGC..............GACAA..G.UCACUGUAUACGAGUAACAUCUACAAAGUAGAAACAACGUGCGGGCAGACCGGAAGGCUGUCAGAAGAA
env-959   ................GGCAG.GU..U.UUC...CGUGGAGCGAAUGUCAA.....G..UUAGGAAAC..............UGGC..............GACAA..G.UCACUGUACAACAAUUUGAUAUCUUAUAUGUAACAUUAGGAAACAUAUUUAGAUAAGGCGGAUUUUCU
env-960   ................GGCAG.GU..U.UUC...CGUGGAGCGAAUGUCAA.....G..UUAGGAAAC..............UGGC..............GACAA..G.UCACUGUACCAAGACAAUCUGUGAAAAACAGAGAUAACGUGUGAACAGUCUCAGGACACACACGGAGGC
env-961   ................GGCAG.GU..U.UUC...CGUGGAGCGAAUGUCAA.....G..UUAGGAAAC..............UGGC..............GACAA..G.UCACUGUACCAGACAAUCUGUGAAAAACAGAGAUAACGUGUGAACAGUCUCAGGACACACACGGAGGCG
env-962   ................GGCAG.GU..U.UUC...CGUGGAGCGAAUGUCAA.....G..UUAGGAAAC..............UGGC..............GACAA..G.UCACUGUAUCACGAAUAUGGUCUACAGAGGUAGAAACAACGUGUGUAAAGCUUAUGACACACACGGAGA
env-963   ................GGCAG.GU..U.UUC...CGUGGAGCGAAUGUCAA.....G..UUAGGAAAC..............UGGC..............GACAA..G.UCACUGUAUCACGAAUAUGGUCUACAGAGGUAGAAACAACGUGUGUAAAGCUUAUGACACACACGGAGA
env-964   ................GGCAG.GU..U.UUC...CGUGGAGCGAAUGUCAA.....G..UUAGGAAAC..............UGGC..............GACAA..G.UCACUGUAUCACGAAUAUGGUCUACAGAGGUAGAAACAACGUGUGUAAAGCUU[illegible]ACACACGGAGA
env-965   ................UGUAG.GU..G.UUC...CGUGGAGCGAAUGUCAA.....G..UUAGGAAAC..............UGGC..............GACAA..G.UCACUGUACUAAAAAUUAUAUGUGUGAAGCUUUAUGACACACACGGUAAUGUGUACAGUCACCGCUUGU
env-967   ................UGUAG.GU..U.UUC...CGUGGAGCGAAUGUCAA.....G..UUAGGAAAC..............UGGC..............GACAA..G.UCACUGUACUAAAAAUUAUAUGUGUGAAGCUUUAUGACACACACGGUAAUGUGUACAGUCACCGCUUGU
env-968   ................AGCAG.GU..UAUUC...CGCGGAGCGAAUGUCAA.....G..UUAGGAAAC..............UGGC..............GACAA..G.UCACUGUACAAGAAUAGAGUAUCCUGAAGACAAAUGACGGAUUUGGUGUGUGGACUGCGAGGACACACA
env-970   ................AGCAG.GU..UAUUC...CGCGGAGCGAAUGUCAA.....G..UUAGGAAAC..............UGGC..............GACAA..G.UCACUGUACAAGAAUAGAGUAUCCUGAAGACAAAUGACGGAUUUGGUGUGUGGACUGCGAGGACACACA

          >>>>>>>>>>.......>>>>>.....>.>>>...>>>........<<<<......<..<<<..................>>>>..............>>>>...>.>>>>.>>.......................................................
          01.1111111.......22221.....2.102...222........111.......2..212..................2122..............111....1.111..12.......................................................
          ---------------•GYRG-GU--U-UUC---CGYGGAGCGAAUGUC•R-----G--•Y•R•RRR•--------------•G•C-------------GRCAA--G-UCACUGUAC•RR•••••••••••••••••••••••••••••••••••••••••••••••••••••
                                                                                                         3′
```

```
env-971   ................AGCAG.GU..UAUUC...CGCGGAGCGAAUGUCAA.....G..UUAGGAAAC..............UGGC.............GACAA..G.UCACUGUACAAGAAUAGAGUAUCCUGAAGACAAAUGACGGAUUUGGUGUGUGGACUGCGAGGACACACA
env-972   ................AGCAG.GU..UAUUC...CGCGGAGCGAAUGUCAA.....G..UUAGGAAAC..............UGGC.............GACAA..G.UCACUGUACAAGAAUAGAGUAUCCUGAAGACAAAUGACGGAUUUGGUGUGUGGACUGCGAGGACACACA
env-973   ................AGCAG.GU..UAUUC...CGCGGAGCGAAUGUCAA.....G..UUAGGAAAC..............UGGC.............GACAA..G.UCACUGUACAAGAAUAGAGUAUCCUGAAGACAAAUGACGGAUUUGGUGUGUGGACUGCGAGGACACACA
Lba-3-1   ................AGUAG.GU..G.UUC...CGCGGAGCGAAUGUCAA.....G..UUAGGAAAC..............UGGC.............GACAA..G.UCACUGUACGAAUCAAUAACGGCCACUGAAACGUGGUAACAACGUGUGAAGAGAUUAGGACACACACGG
env-980   ................AGUAG.GU..G.UUC...CGUGGAGCGAAUGUCAA.....G..UUAGGAAAC..............UGGC.............GACAA..G.UCACUGUACGAAUCAAUAACAACCGAUGAAAAACGGUAACAACGUGUGAGUUCUCACGACACACACGGG
env-981   ................AGUAG.GU..G.UUC...CGUGGAGCGAAUGUCAA.....G..UUAGGAAAC..............UGGC.............GACAA..G.UCACUGUACGAAUC.......................................................
env-983   ................AGUAG.GU..G.UUC...CGUGGAGCGAAUGUCAA.....G..UUAGGAAAC..............UGGC.............GACAA..G.UCACUGUACGAAUCAAUAACAACCGAUGAAAAACGGUAACAACGUGUGAGUUCUCACGACACACACGGG
env-984   ................AGUAG.GU..G.UUC...CGUGGAGCGAAUGUCAA.....G..UUAGGAAAC..............UGGC.............GACAA..G.UCACUGUACGAAUCAAUAACAACCGAUGAAAAACGGUAACAACGUGUGAGUUCUCACGACACACACGGG
env-985   ................AGCAG.GU..G.UUC...CGUGGAGCGAAUGUCAA.....G..UUAGGAAAC..............UGGC.............GACAA..G.UCACUGUACGAAUCAAUAACAGCCGAUGAAAAACGGUAACAACGUGUGAGAGUUUUAUGACACACACGG
Csp-8-1   ................AGCAG.GU..G.UUC...CGUGGAGCGAAUGUCAA.....G..UUAGGAAAC..............UGGC.............GACAA..G.UCACUGUACGAAUCAAUAACAGCCGAUGAAAAACGGUAACAACGUGUGAGAGUUUUAUGACACACACGG
env-986   ................AGCAG.GU..G.UUC...CGUGGAGCGAAUGUCAA.....G..UUAGGAAAC..............UGGC.............GACAA..G.UCACUGUACGAAACUAUAACAGCCACUGAGAUGUGGUAACAACGUGUGAGUUUCUCACGACACACACGG
Csp-9-1   ................AGCAG.GU..G.UUC...CGUGGAGCGAAUGUCAA.....G..UUAGGAAAC..............UGGC.............GACAA..G.UCACUGUACGAAUCAAUAACAGCCGAUGAAAAACGGUAACAACGUGUGAGAGUUUUAUGACACACACGG
env-987   ................AGCAG.GU..G.UUC...CGUGGAGCGAAUGUCAA.....G..UUAGGAAAC..............UGGC.............GACAA..G.UCACUGUACGAAUCAAUAACAGCCGAUGAAAAACGGUAACAACGUGUGAGAGUUUUAUGACACACACGG
env-991   ................AGCAG.GU..U.UUC...CGUGGAGCGAAUGUCAA.....G..UUAGGAAAC..............UGGC.............GACAA..G.UCACUGUACGAAACUACUAUAACGGCCACUGCGAUGUGGUAACAACGUGUGGGUUCUCACGACACACAC
env-992   ................AGCAG.GU..U.UUC...CGUGGAGCGAAUGUCAA.....G..UUAGGAAAC..............UGGC.............GAAAA..G.UCACUGUACGAAACUACUAUAACGGCCACUGCGAUGUGGUAACAACGUGUGGGUUCUCACGACACACAC
env-993   ................AGCAG.GU..U.UUC...CGUGGAGCGAAUGUCAA.....G..UUAGGAAAC..............UGGC.............GACAA..G.UCACUGUACGAAACUACUAUAACGGCCACUGCGAUGUGGUAACAACGUGUGGGUUCUCACGACACACAC
Dlo-2-1   ................AGCAG.GU..U.UUC...CGUGGAGCGAAUGUCAA.....G..UUAGGAAAC..............UGGC.............GACAA..G.UCACUGUACGAAACUACUAUAACGGCCACUGCGAUGUGGUAACAACGUGUGGGUUCUCACGACACACAC
env-994   ................AGCAG.GU..U.UUC...CGUGGAGCGAAUGUCAA.....G..UUAGGAAAC..............UGGC.............GACAA..G.UCACUGUACGAAACUACUAUAACGGCCACUGCGAUGUGGUAACAACGUGUGGGUUCUCACGACACACAC
env-995   ................AGCAG.GU..U.UUC...CGUGGAGCGAAUGUCAA.....G..UUAGGAAAC..............UGGC.............GACAA..G.UCACUGUNNNNNNNNNNNNNNNNNNNNNNNNNNNNNNNNNNNNNNNNNNNNNNNNNNNNNNNNNNNNNN
env-996   ................AGCAG.GU..U.UUC...CGUGGAGCGAAUGUCAA.....G..UUAGGAAAC..............UGGC.............GACAA..G.UCACUGUA.............................................................
env-1006  ................AGCAG.GU..U.UUC...CGUGGAGCGAAUGUCAA.....G..UUAGGAAAC..............UGGC.............GACAA..G.UCACUGUACGAAACUACUAUAACAGCCACUGCGAUGUGGUAACAACGUGUGGGUUCUCACGACACACAC
env-1007  ................AGCAG.GU..U.UUC...CGUGGAGCGAAUGUCAA.....G..UUAGGAAAC..............UGGC.............GACAA..G.UCACUGUACGAAACUACUAUAACAGCCACUGCGAUGUGGUAACAACGUGUGGGUUCUCACGACACACAC
env-1008  ................AGCAG.GU..U.UUC...CGUGGAGCGAAUGUCAA.....G..UUAGGAAAC..............UGGC.............GACAA..G.UCACUGUACGAAUCACAACUGAUUACUGAUUACCACAAGGGAAAAGUAUGAAACAACGUGUGAGUUUCU
env-1009  ................AGCAG.GU..U.UUC...CGUGGAGCGAAUGUCAA.....G..UUAGGAAAC..............UGGC.............GACAA..G.UCACUGUACGAAACUACUAUAACAGCCACUGCGAUGUGGUAACAACGUGUGGGUUCUCACGACACACAC
env-1018  ................AGCAG.GU..U.UUC...CGUGGAGCGAAUGUCAA.....G..UUAGGAAAC..............UGGC.............GACAA..G.UCACUGUACGAAACUACUAUAACAGCCACUGCGAUGUGGUAACAACGUGUGGGUUCUCACGACACACAC
env-1019  ................AGCAG.GU..U.UUC...CGUGGAGCGAAUGUCAA.....G..UUAGGAAAC..............UGGC.............GACAA..G.UCACUGUACGAAACUACUAUAACAGCCACUGCGAUGUGGUAACAACGUGUGGGUUCUCACGACACACAC
Dlo-1-1   ................AGCAG.GU..U.UUC...CGUGGAGCGAAUGUCAA.....G..UUAGGAAAC..............UGGC.............GACAA..G.UCACUGUACGAAACUACUAUAACAGCCACUGCGAUGUGGUAACAACGUGUGGGUUCUCACGACACACAC
env-1020  ................UGCAG.GU..U.UUC...CGUGGAGCGAAUGUCAA.....G..UUAGGAAAC..............UGAC.............GACAA..G.UCACUGUACCAAAUAUGGAGUCUGCCAAAGAGCAGAAACGACGUGUGAGUCUAGGACACACACGGGAGA
env-1021  ................UGCAG.GU..U.UUC...CGUGGAGCGAAUGUCAA.....G..UUAGGAAAC..............UGAC.............GACAA..G.UCACUGUACCAAAUAUGGAGUCUGCCAAAGAGCAGAAACGACGUGUGAGUUCAGGACACACACGGGAGA
env-1022  ................UGCAG.GU..U.UUC...CGUGGAGCGAAUGUCAA.....G..UUAGGAAAC..............UGAC.............GACAA..G.UCACUGUACCAAAUAUGGAGUCUGCCAAAGAGCAGAAACGACGUGUGAGUCCAGGACACACACGGGAGA
Lba-2-1   ....UCGUCGAUA....GCAG.GU..U.UUC...CGUGGAGCGAAUGUCAA.....G..UUAGGAAAC..............UGGC.............GACAA..G.UCACUGUACGAAUCACAACUGAUUACUUAUUACCAAAAGGGAAGAGUAUGAAACAACGUGUGAGUUUCU
env-1023  ................GGCAG.GU..U.UUC...CGCGGAGCGAAUGUCAA.....G..UUAGGAAAC..............UGGC.............GACAA..G.UCACUGUACAAGGUUAGAAUGCCUACCGAACAUAUUGGUAGAGACGAUGUGAAGAAUGCAAACUUCUAG
env-1024  ................GGCAG.GU..U.UUC...CGCGGAGCGAAUGUCAA.....G..UUAGGAAAC..............UGGC.............GACAA..G.UCACUGUACAAGGUUAGAAUGCCUACCGAACAUAUUGGUAGAGACGAUGUGAAGAAUGCAAACUUCUGG
env-1026  ................GGCAG.GU..U.UUC...CGCGGAGCGAAUGUCAA.....G..UUAGGAAAC..............UGGC.............GACAA..G.UCACUGUACAAGGUUAGAAUGCCUACCGAACAUAUUGGUAGAGACGAUGUGAAGAAUGCAAACUUCUAG
env-1027  ................GGCAG.GU..U.UUC...CGCGGAGCGAAUGUCAA.....G..UUAGGAAAC..............UGGC.............GACAA..G.UCACUGUACAAGGUUAGAAUGCCUACCGAACAUAUUGGUAGAGACGAUGUGAAGAAUGCAAACUUCUGG
env-1028  ................GGCAG.GU..U.UUC...CGCGGAGCGAAUGUCAA.....G..UUAGGAAAC..............UGGC.............GACAA..G.UCACUGUACAAGGUUAGAAUGCCUACCGAACAUAUUGGUAGAGACGAUGUGAAGAAUGCAAACUUCUAG
env-1029  ................GGCAG.GU..U.UUC...CGCGGAGCGAAUGUCAA.....G..UUAGGAAAC..............UGGC.............GACAA..G.UCACUGUACAAGGUUAGAAUGCCUACCGAACAUAUUGGUAGAGACGAUGUGAAGAAUGCAAACUUCUGA
env-1030  ................GGCAG.GU..G.UUC...CGUGGAGCGAAUGUCAA.....G..UUAGGAAAC..............UGGC.............GACAA..G.UCACUGUACCAAAUUAACAGCUGCUCAGGGUAAGGGGCAGUAACAACGUGUGAAUCGGCAUGAUACACA
env-1032  ................GGCAG.GU..G.UUC...CGUGGAGCGAAUGUCAA.....G..UUAGGAAAC..............UGGC.............GACAA..G.UCACUGUACCAAAUUAACAGCUGCUCAGGGUAAGGGGCAGUAACAACGUGUGAAUCGGCAUGAUACACA
env-1033  ................GGCAG.GU..G.UUC...CGUGGAGCGAAUGUCAA.....G..UUAGGAAAC..............UGGC.............GACAA..G.UCACUGUACCAAAUUAACAGCUGCUCAGGGUAAGGGGCAGUAACAACGUGUGAAUCGGCAUGAUACACA
env-1034  ................GGCAG.GU..G.UUC...CGUGGAGCGAAUGUCAA.....G..UUAGGAAAC..............UGGC.............GACAA..G.UCACUGUACCAAAUUAACAGCUGCUCAGGGUAAGGGGCAGUAACAACGUGUGAAUCGGCAUGAUACACA
env-1035  ................GGCAG.GU..U.UUC...CGUGGAGCGAAUGUCAA.....G..UUAGGAAAC..............UGAC.............GACAA..G.UCACUGUACAAACAAUAUGGUCUAUCAAAGAUAGAAACAACGUGUGAAAAAAUGCGAAUCGCCUGACGA
env-1036  ................GGCAG.GU..U.UUC...CGUGGAGCGAAUGUCAA.....G..UUAGGAAAC..............UGAC.............GACAA..G.UCACUGUACAAACAAUAUGGUCUAUCAAAGAUAGAAACAACGUGUGAAAAAAAUGCGAAACGCCUGACA
env-1037  ................GGCAG.GU..U.UUC...CGUGGAGCGAAUGUCAA.....G..UUAGGAAAC..............UGGC.............GACAA..G.UCACUGUACAAACAAUAUGGUCUAUCAAAGAUAGAAACAACGUGUGAAAAAAUGCGAAUCGCCUGACGA
env-1038  ................GGCAG.GU..U.UUC...CGUGGAGCGAAUGUCAA.....G..UUAGGAAAC..............UGAC.............GACAA..G.UCACUGUACAAACAAUAUGGUCUAUCAAAGAUAGAAACAACGUGUGAAAAAAUGCGAAUCGCCUGACAA
env-1039  ................GGCAG.GU..U.UUC...CGUGGAGCGAAUGUCAA.....G..UUAGGAAAC..............UGAC.............GACAA..G.UCACUGUACAAACAAUAUGGUCUAUCAAAGAUAGAAACAACGUGUGAAAAAAUGCGAAUCGCCUGACAA
env-1040  ................GGCAG.GU..U.UUC...CGUGGAGCGAAUGUCAA.....G..UUAGGAAAC..............UGAC.............GACAA..G.UCACUGUACAAACAAUAUGGUCUAUCAAAGAUAGAAACAACGUGUGAAAAAAUGCGAAUCGCCUGACGA
env-1041  ................GGCAG.GU..U.UUC...CGUGGAGCGAAUGUCAA.....G..UUAGGAAAC..............UGAC.............GACAA..G.UCACUGUACAAACAAUAUGGUCUAUCAAAGAUAGAAACAACGUGUGAAAAAAUGCGAAUCGCCUGAUAA
env-1042  ................GGCAG.GU..U.UUC...CGUGGAGCGAAUGUCAA.....G..UUAGGAAAC..............UGAC.............GACAA..G.UCACUGUACAAACAAUAUGGUCUAUCAAAGAUAGAAACAACGUGUGAAAAAAUGCGAAUCGCCUGACGA
env-1043  ................GGCAG.GU..U.UUC...CGUGGAGCGAAUGUCAA.....G..UUAGGAAAC..............UGAC.............GACAA..G.UCACUGUACAAACAAUAUGGUCUAUCAAAGAUAGAAACAACGUGUGAAAAAAUGCGAAUCGCCUGACGA
env-1044  ................GGCAG.GU..U.UUC...CGUGGAGCGAAUGUCAA.....G..UUAGGAAAC..............UGGC.............GACAA..G.UCACUGUACAAACAAUAUGGUCUAUCAAAGAUAGAAACAACGUGUGAAAAAAUGCGAAUCGCCUGACAA
env-1045  ................GGCAG.GU..U.UUC...CGUGGAGCGAAUGUCAA.....G..UUAGGAAAC..............UGAC.............GACAA..G.UCACUGUACAAACAAUAUGGUCUAUCAAAGAUAGAAACAACGUGUGAAAAAAUGCGAAUCGCCUGACGA
env-1046  ................GGCAG.GU..U.UUC...CGUGGAGCGAAUGUCAA.....G..UUAGGAAAC..............UGAC.............GACAA..G.UCACUGUACAAACAAUAUGG.................................................
env-1052  ................GGCAG.GU..U.UUC...CGUGGAGCGAAUGUCAA.....G..UUAGGAAAC..............UGAC.............GACAA..G.UCACUGUACAAACAAUAUGGUCUAUCAAAGAUAGAAACAACGUGUGAAAAAAUGCGAAUCGCCUGACAA
env-1053  ................GGCAG.GU..U.UUC...CGUGGAGCGAAUGUCAA.....G..UUAGGAAAC..............UGAC.............GACAA..G.UCACUGUACAAACAAUAUGGUCUAUCAAAGAUAGAAACAACGUGUGAAAAAAAUGCGAAACGCCUGACA
env-1054  ................GGCAG.GU..U.UUC...CGUGGAGCGAAUGUCAA.....G..UUAGGAAAC..............UGAC.............GACAA..G.UCACUGUACAAACAAUAUGGUCUAUCAAAGAUAGAAACAACGUGUGAAAAAAUGCGAAUCGCCUGACAA
env-1055  ................GGCAG.GU..U.UUC...CGUGGAGCGAAUGUCAA.....G..UUAGGAAAC..............UGAC.............GACAA..G.UCACUGUACAAACAAUAUGGUCUAUCAAAGAUAGAAACAACGUGUGAAAAAAUGCGAAUCGCCUGACGA
env-1056  ................GGCAG.GU..U.UUC...CGUGGAGCGAAUGUCAA.....G..UUAGGAAAC..............UGAC.............GACAA..G.UCACUGUACAAACAAUAUGGUCUAUCAAAGAUAGAAACAACGUGUGAAAAAAUGCGAAUCGCCUGACAA
env-1057  ................GGCAG.GU..U.UUC...CGUGGAGCGAAUGUCAA.....G..UUAGGAAAC..............UGAC.............GACAA..G.UCACUGUACAAACAAUAUGGUCUAUCAAAGAUAGAAACAACGUGUGAAAAAAUGCGAAUCGCCUGACAA
env-1058  ................GGCAG.GU..U.UUC...CGUGGAGCGAAUGUCAA.....G..UUAGGAAAC..............UGAC.............GACAA..G.UCACUGUACAAACAAUAUGGUCUAUCAAAGAUAGAAACAACGUGUGAAAAAAUGCGAAUCGCCUGACAA
env-1059  ................GGCAG.GU..U.UUC...CGUGGAGCGAAUGUCAA.....G..UUAGGAAAC..............UGAC.............GACAA..G.UCACUGUACAAACAAUAUGGUCUAUCAAAGAUAGAAACAACGUGUGAAAAAAUGCGAAUCGCCUGACGA
Cha-1-1   ................GGCAG.GU..U.UUC...CGUGGAGCGAAUGUCAA.....G..UUAGGAAAC..............UGGC.............GACAA..G.UCACUGUACAAUUAAAGAAUUCUGUUUCUAAAAAACGGAAAAUCAGUGUGGGGUCCUGGAGGACACACA
env-1065  ................GGCAG.GU..U.UUC...CGUGGAGCGAAUGUCAA.....G..UUAGGAAAC..............UGGC.............GACAA..G.UCACUGUACAAUUAAAGAAUUCUGUUUCAAAAAAACGGAAAAUCAGUGUGGGGUCUUGGAGGACACACA
env-1066  ................GGCAG.GU..U.UUC...CGUGGAGCGAAUGUCAA.....G..UUAGGAAAC..............UGGC.............GACAA..G.UCACUGUACAAUUAAAGAAUUCUGUUUCAAAAAAACGGAAAAUCAGUGUGGGGUCCUGGAGGACACACA
Cba-1-1   ................GGCGG.GU..U.UUC...CGUGGAGCGAAUGUCAA.....G..UUAGGAAAC..............UGAC.............GACAA..G.UCACUGUACAAGCAAUGAAGUCUGUUAUUAAACAGAAACAACCGCGUGAAAAGUCCUGAAGGAACGGGA
Cba-8-1   ................GGCGG.GU..U.UUC...CGUGGAGCGAAUGUCAA.....G..UUAGGAAAC..............UGAC.............GACAA..G.UCACUGUACAAGCAAUGAAGUCUGUUAUUAAACAGAAACAACCGCGUGAAAAGUCCUGAAGGAACGGGA
env-1067  ................GGCGG.GU..U.UUC...CGUGGAGCGAAUGUCAA.....G..UUAGGAAAC..............UGAC.............GACAA..G.UCACUGUACAAGCAAUGAAGUCUGUUAUUAAACAGAAACAACCGCGUGAAAAGUCCUGAAGGAACGGGA
Lba-20-1  ................GGCGG.GU..U.UUC...CGUGGAGCGAAUGUCAA.....G..UUAGGAAAC..............UGAC.............GACAA..G.UCACUGUACAAGCAAUGAAGUCUGUUAUUAAACAGAAACAACCGCGUGAAAAGUCCUGAAGGAACGGGA
Lba-8-1   ................GGCGG.GU..U.UUC...CGUGGAGCGAAUGUCAA.....G..UUAGGAAAC..............UGAC.............GACAA..G.UCACUGUACAAGCAAUGAAGUCUGUUAUUAAACAGAAACAACCGCGUGAAAAGUCCUGAAGGAACGGGA
env-1068  ................GGCGG.GU..U.UUC...CGUGGAGCGAAUGUCAA.....G..UUAGGAAAC..............UGAC.............GACAA..G.UCACUGUACAAGCAAUGAAGUCUGUUGUUAAACAGAAACAACCGCGUGAAAAGUCCUGAAGGAACGGGA
Lba-13-1  ................GGCAG.GU..U.UUC...CGUGGAGCGAAUGUCAA.....G..UUAAGAAAC..............UGGC.............GACAA..G.UCACUGUAUCACAACAAGGAAUCUGCAAGGGGCAGACCCGUGUGGGGUAUAUACGACACACACGGAGAA
Csp-7-1   ................GGCAG.GU..U.UUC...CGUGGAGCGAAUGUCAA.....G..UUAGGAAAC..............UGGC.............GACAA..G.UCACUGUACAACUUUAGAGAAUACCUGAUUAUAUGGAGGAUUCAGUGUGGAGUUUUGUAUGACACACAC
env-1069  ................GGCAG.GU..U.UUC...CGUGGAGCGAAUGUCAA.....G..UUAGGAAAC..............UGGC.............GACAA..G.UCACUGUACAACUUUAGAGAAUACCUAAUAUAUGGAGGAUUCAGUGUGGAGUUUUGUAUGACACACACG
env-1070  ................GGCAG.GU..U.UUC...CGUGGAGCGAAUGUCAA.....G..UUAGGAAAC..............UGGC.............GACAA..G.UCACUGUACAACUUUAGAGAAUACCUAAUAUAUGGAGGAUUCAGUGUGGAGUUUUGUAUGACACACACG
env-1071  ................GGCAG.GU..U.UUC...CGUGGAGCGAAUGUCAA.....G..UUAGGAAAC..............UGGC.............GACAA..G.UCACUGUACAACUUUAGAGAAUACCUAAUAUAUGGAGGAUUCAGUGUGGAGUUUUGUAUGACACACACG
env-1072  ................GGCAG.GU..U.UUC...CGUGGAGCGAAUGUCAA.....G..UUAGGAAAC..............UGGC.............GACAA..G.UCACUGUACAACUUUAGAGAAUACCUAAUAUAUGGAGGAUUCAGUGUGGAGUUUUGUAUGACACACACG
Esp-1-1   ................GGCAG.GU..U.UUC...CGUGGAGCGAAUGUCAA.....G..UUAGGAAAC..............UGGC.............GACAA..G.UCACUGUAUCACAAUAAAAAAUCUGCAGGAACGCAGAAGACGUGUGUAGAAGUUAUGACACACACGGAG
env-1073  ................GGCAG.GU..U.UUC...CGUGGAGCGAAUGUCAA.....G..UUAGGAAAC..............UGGC.............GACAA..G.UCACUGUACCAAAUUAACAGCUGCUCUUGGUAAGGGGCAGUAACAACGUGUGGAUCGGAUGAUACACAC
env-1074  ................GGCAG.GU..U.UUC...CGUGGAGCGAAUGUCAA.....G..UUAGGAAAC..............UGGC.............GACAA..G.UCACUGUACCAAAUUAACAGCUGCUCUUGGUAAGGGGCAGUAACAACGUGUGGAUCGGAUGAUACACAC
env-1075  ................GGCAG.GU..U.UUC...CGUGGAGCGAAUGUCAA.....G..UUAGGAAAC..............UGGC.............GACAA..G.UCACUGUACCAAAUUAACAGCUGCUCUUGGUAAGGGGCAGUAACAACGUGUGGAUCGGAUGAUACACAC
Csa-1-1   ................AGCAG.GU..U.UUC...CGUGGAGCGAAUGUCAA.....G..UUAGGAAAC..............UGGC.............GACAA..G.UCACUGUACAAUUUAAAAUUCUGUUUAUAUAAGCAGAAAGUCAGUGUGUGAUCCUGAAGGACACACACG
env-1076  ................AGCAG.GU..U.UUC...CGUGGAGCGAAUGUCAA.....G..UUAGGAAAC..............UGAC.............GACAA..G.UCACUGUACAAACAAUAUGGUCUACAGUAAAGUAGAAUGAUGUGUGAAGAAUGCUUAAGGAUGGAGCA
env-1077  ................AGCAG.GU..U.UUC...CGUGGAGCGAAUGUCAA.....G..UUAGGAAAC..............UGAC.............GACAA..G.UCACUGUACAACUUACCAUCUGCACAUGGCAGAGAUGACCGUAUGAGGUUGCAUUAUAGAAGAAGGUUU

          >>>>>>>>>>......>>>>>.....>.>>>...>>>.........<<<<.......<..<<<..............>>>>.............>>>>...>.>>>>.>>..............................................................
          01.1111111......22221.....2.102...222..........111.......2..212..............2122.............111....1.111..12..............................................................
          ----------------•GYRG-GU--U-UUC---CGYGGAGCGAAUGUC•R-----G--•Y•R•RRR•---------------•G•C-------------GRCAA--G-UCACUGUAC•RR•••••••••••••••••••••••••••••••••••••••••••••••••••••
                                                                                                                3′
```

```
env-1078  ................AGCAG.GU..U.UUC...CGUGGAGCGAAUGUCAA.....G..UUAGGAAAC...............UGAC..............GACAA..G.UCAGUGUACAAACAAUAUGGUCUAUCAAAGAUAGAAACAACGUGUGAAAAAAUGCGAAUCGCCUGACGA
env-1081  ................AGCAG.GU..U.UUC...CGUGGAGCGAAUGUCAA.....G..UUAGGAAAC...............UGAC..............GACAA..G.UCAGUGUACAAACAAUAUGGUCUAUCAAAGAUAGAAACAACGUGUGAAAAAAUGCGAAUCGCCUGACGA
env-1082  ................AGCAG.GU..U.UUC...CGUGGAGCGAAUGUCAA.....G..UUAGGAAAC...............UGAC..............GACAA..G.UCAGUGUACAAACAAUAUGGUCUAUCAAAGAUAGAAACAACGUGUGAAAAAAUGCGAAUCGCCUGACGAU
env-1083  ................AGCAG.GU..U.UUC...CGUGGAGCGAAUGUCAA.....G..UUAGGAAAC...............UGAC..............GACAA..G.UCAGUGUACAAACAAUAUGGUCUAUCAAAGAUAGAAACAACGUGUGAAAAAAUGCGAAUCGCCUGACGA
env-1084  ................AGCAG.GU..U.UUC...CGUGGAGCGAAUGUCAA.....G..UUAGGAAAC...............UGAC..............GACAA..G.UCAGUGUACAAACAAUAUGGUCUAUCAAAGAUAGAAACAACGUGUGAAAAAAUGCGAAUCGCCUGACGA
env-1085  ................AGCAG.GU..U.UUC...CGUGGAGCGAAUGUCAA.....G..UUAGGAAAC...............UGAC..............GACAA..G.UCAGUGUACAAACAAUAUGGUCUAUCAAAGAUAGAAACAACGUGUGAAAAAAUGCGAAUCGCCUGACAA
env-1086  ................AGCAG.GU..U.UUC...CGUGGAGCGAAUGUCAA.....G..UUAGGAAAC...............UGAC..............GACAA..G.UCAGUGUACAAACAAUAUGGUCUAUCAAAGAUAGAAACAACGUGUGAAAAAAUGCGAAUCGCCUGACGA
Cha-2-1   ................GGCGG.GU..U.UUC...CGUGGAGCGAAUGUCAA.....G..UUAGGAAAC...............UGGC..............GACAA..G.UCAGUGUACAAACAAAUAUUCUGCAGCUAUGUAGAUUAGAAGGAUGUGAACCAUCAGGACACACCCGGA
Lba-5-1   ................GGCAG.GU..UUUUC...CGUGGAGCGAAUGUCAA.....G..UUAGGAAAC...............UGGC..............GACAA..G.UCAGUGUAUCACAAUCAUAUGUCUGCAAGAGCAGCGCGUGUGUAAAACAUGCGACACACGGAGAAG
Csp-1-1   ................GGCAG.GU..U.UUC...CGUGGAGCGAAUGUCAA.....G..UCGAAGAAAC..............UGGC..............GACAA..G.UCAGUGUACAACUACAGAAAUAUCCCGAAAAUAAAGAAGGAUUCCGUGUGAGGUCUAAGAUGACACC
env-1087  ................GGCAG.GU..U.UUC...CGUGGAGCGAAUGUCAA.....G..UCGAAGAAAC..............UGGC..............GACAA..G.UCAGUGUACAACUACAGAAAUAUCCCGAAAAUAAAGAAGGAUUCCGUGUGAGGUCUAAGAUGACACC
env-1088  ................GACAG.GU..U.UUCA..CAGGGAGUAAAUGUCAA.....G..UUAGGAAAC...............UGGC..............GACAA..G.UCAGUGUACUGAAUACGAUCUGCAUAGCGCAGAGAUGACGUGUGAGACCGAUCAGGAAACACACGGCA
env-1089  ................AGCAG.GA..G.UUC...CGCGGAGCGAAUGUCAA.....G..UCUCGAUAC...............UGAC..............GACAA..G.UCAGUGUACAAACGAAGAGCGCCGUAUUUCGAAAGAGAGAAGGCCUGCCAUGGGUGUUUCGCAAGGACA
env-1093  ................AGCAG.GA..G.UUC...CGCGGAGCGAAUGUCAA.....G..UCUCGAUAC...............UGAC..............GACAA..G.UCAGUGUACAAACGAAGAGCGCCGNNNNNNNNNNNNNNNNNNNNNNNNNNNNNNNNNNNNNNNNNACUGUGU
env-1094  ................AGCAG.GA..G.UUC...CGCGGAGCGAAUGUCAA.....G..UCUCGAUAC...............UGAC..............GACAA..G.UCAGUGUACAAACGAAGAGCGCCGUAUUUCGAAAGAGAGAAGGCCUGCCAUGGGUGUUUCGCAAGGACA
env-1095  ................AGCAG.GA..G.UUC...CGCGGAGCGAAUGUCAA.....G..UCUCGAUAC...............UGAC..............GACAA..G.UCAGUGUACAAACGAAGAGCGCCGGAUUUCAAAGUGAAAGAUGGCCUGCUGUGAGGUUUCGAGAGGAC
env-1096  ................AGCAG.GA..G.UUC...CGCGGAGCGAAUGUCAA.....G..UCUCGAUAC...............UGAC..............GACAA..G.UCAGUGUACAAACGAAGAGCGCCGUAUUUCGAAAGAGAGAAGGCCUGCCUUGGGUGUUUCGAAGGACAC
env-1097  ................AGCAG.GA..G.UUC...CGCGGAGCGAAUGUCAA.....G..UCUCGAUAC...............UGAC..............GACAA..G.UCAGUGUACAAACGAAGAGCGCCGACUUUCGAAAGAAGGAAGGCCUGCCGUUGGGUUUCCGGGAGGACA
env-1099  ................AGCAG.GA..G.UUC...CGCGGAGCGAAUGUCAA.....G..UCUCGAUAC...............UGAC..............GACAA..G.UCAGUGUACAAACGAAGAGCGCCGUAUUUCGAAAGAGAGAAGGCCUGCCAUGGGUGUUUCGCAAGGACA
env-1100  ................AGCAG.GA..G.UUC...CGCGGAGCGAAUGUCAA.....G..UCUCGAUAC...............UGAC..............GACAA..G.UCAGUGUACAAACGAAGAGCGCCGUAUUUCGAAAGAGAGAAGGCCUGCCAUGGGUGUUUCACAAGGACA
Slo-1-1   ................AGCAG.GA..G.UUC...CGCGGAGCGAAUGUCAA.....G..UCUCGAUAC...............UGAC..............GACAA..G.UCAGUGUACAAACGAAGAGCGCCGGAUUUCAAAACGAAAGAUGGCCUGCUGUGGGGUUUCCAAGAGGAC
env-1101  ................AGCAG.GA..G.UUC...CGCGGAGCGAAUGUCAA.....G..UCUCGAUAC...............UGAC..............GACAA..G.UCAGUGUACAAACGAAGAGCGCCGGAUUUCAAAACGAAAGAUGGCCUGCUGUGGGGUUUCCGAGAGGAC
env-1102  ................AGCAG.GA..G.UUC...CGCGGAGCGAAUGUCAA.....G..UCUCGAUAC...............UGAC..............GACAA..G.UCAGUGUACAAACGAAGAGCGCCGUAUUUCGAAAGAGAGAAGGCCUGCCAUGGGUGUUUCGCAAGGACA
env-1103  ................AGCAG.GA..G.UUC...CGCGGAGCGAAUGUCAA.....G..UCUCGAUAC...............UGAC..............GACAA..G.UCAGUGUACAAACGAAGAGCGCCGUAUUUCGAAAGAGAGAAGGCCUGCCAUGGGUGUUUCGCAAGGACA
env-1104  ................AGCAG.GA..G.UUC...CGCGGAGCGAAUGUCAA.....G..UCUCGAUAC...............UGAC..............GACAA..G.UCAGUGUACAAACGAAGAGCGCCGACUUUCGAAAGAAGGAAGGCCUGCCGUUGGGUUUCCGGGAGGACA
env-1105  ................AGCAG.GA..G.UUC...CGCGGAGCGAAUGUCAA.....G..UCUCGAUAC...............UGAC..............GACAA..G.UCAGUGUACAAACGAAGAGCGCCGACUUUCGAAAGAAGGAAGGCCUGCCGUUGGGUUUCCGGGAGGACA
env-1106  ................AGCAG.GA..G.UUC...CGCGGAGCGAAUGUCAA.....G..UCUCGAUAC...............UGAC..............GACAA..G.UCAGUGUACAAACGAAGAGCGCCGACUUUCGAAAGAAGGAAGGCCUGCCGUUGGGUUUCCGGGAGGACA
env-1107  ................AGCAG.GA..G.UUC...CGCGGAGCGAAUGUCAA.....G..UCUCGAUAC...............UGAC..............GACAA..G.UCAGUGUACAAACGAAGAGCGCCGUAUUUCGAAAGAGAGAAGGCCUGCCAUGGGUGUUUCGAAGGACAC
env-1108  ................AGCAG.GA..G.UUC...CGCGGAGCGAAUGUCAA.....G..UCUCGAUAC...............UGAC..............GACAA..G.UCAGUGUACAAACGAAGAGCGCCGUAUUUCGAAAGAGAGAAGGCCUGCCAUGGGUGUUUCGCAAGGACA
env-1115  ................AGCAG.GA..G.UUC...CGCGGAGCGAAUGUCAA.....G..UCUCGAUAC...............UGAC..............GACAA..G.UCAGUGUACAAACGAAGAGCGCCGUAUUUCGAAAGAGAGAAGGCCUGCCAUGGGUGUUUCGCAAGGACA
env-1118  ................AGCAG.GA..G.UUC...CGCGGAGCGAAUGUCAA.....G..UCUCGAUAC...............UGAC..............GACAA..G.UCAGUGUACAAACGAAGAGCGCCGUAUUUCGAAAGAGAGAAGGCCUGCCAUGGGUGUUUCGAAGGACAC
env-1123  ................AGCAG.GA..G.UUC...CGCGGAGCGAAUGUCAA.....G..UCUCGAUAC...............UGAC..............GACAA..G.UCAGUGUACAAACGAAGAGCGCCGACUUUCGAAAGAAGGAAGGCCUGCCGUUGGGUUUCCGGGAGGACA
env-1124  ................AGCAG.GA..G.UUC...CGCGGAGCGAAUGUCAA.....G..UCUCGAUAC...............UGAC..............GACAA..G.UCAGUGUACAAACGAAGAGCGCCGUAUUUCGAAAGAGAGAAGGCCUGCCAUGGGUGUUUCGCAAGGACA
env-1145  ................AGCAG.GA..G.UUC...CGCGGAGCGAAUGUCAA.....G..UCUCGAUAC...............UGAC..............GACAA..G.UCAGUGUACAAACGAAGAGCGCCGUAUUUCGAAAGAGAGAAGGCCUGCCAUGGGUGUUUCACAAGGACA
env-1146  ................AGCAG.GA..G.UUC...CGCGGAGCGAAUGUCAA.....G..UCUCGAUAC...............UGAC..............GACAA..G.UCAGUGUACAAACGAAGAGCGCCGUAUUUCGAAAGAGAGAAGGCCUGCCAUGGGUGUUUCGCAAGGACA
env-1147  ................AGCAG.GA..G.UUC...CGCGGAGCGAAUGUCAA.....G..UCUCGAUAC...............UGAC..............GACAA..G.UCAGUGUACAAACGAAGAGCGCCG..........................................
env-1148  ................AGCAG.GA..G.UUC...CGCGGAGCGAAUGUCAA.....G..UCUCGAUAC...............UGAC..............GACAA..G.UCAGUGUACAAACGAAGAGCGCCGUAUUUCGAAAGAGAGAAGGCCUGCCAUGGGUGUUUCGCAAGGACA
env-1149  ................AGCAG.GA..G.UUC...CGCGGAGCGAAUGUCAA.....G..UCUCGAUAC...............UGAC..............GACAA..G.UCAGUGUACAAACGAAGAGCGCCGUAUUUCGAAAGAGAGAAGGCCUGCCAUGGGUGUUUCGCAAGGACA
env-1150  ................AGCAG.GA..G.UUC...CGCGGAGCGAAUGUCAA.....G..UCUCGAUAC...............UGAC..............GACAA..G.UCAGUGUACAAACGAAGAGCGCCGNNNNNNNNNNNNNNNNNNNNNNNNNNNNNNNNNNNNNNNNNNNNNNNN
env-1151  ................AGCAG.GA..G.UUC...CGCGGAGCGAAUGUCAA.....G..UCUCGAUAC...............UGAC..............GACAA..G.UCAGUGUACAAACGAAGAGCGCCGUAUUUCGAAAGAGAGAAGGCCUGCCAUGGGUGUUUCGAAGGACAC
env-1154  ................AGCAG.GA..G.UUC...CGCGGAGCGAAUGUCAA.....G..UCUCGAUAC...............UGAC..............GACAA..G.UCAGUGUACAAACGAAGAGUGCCGUAUUUCGAAAGAGAGAAGGCCUGCCAUGGGUGUUUCGCAAG....
env-1155  ................AGCAG.GA..G.UUC...CGCGGAGCGAAUGUCAA.....G..UCUCGAUAC...............UGAC..............GACAA..G.UCAGUGUACAAACGAAGAGCGCCGUAUUUCGAAAGAGAGAAGGCCUGCCAUGGGUGUUUCGCAAGGACA
env-1156  ................AGCAG.GA..G.UUC...CGCGGAGCGAAUGUCAA.....G..UCUCGAUAC...............UGAC..............GACAA..G.UCAGUGUACAAACGAAGAGCGCCGGAUUUCGAAAGAGAGAAGGCCUGCCGUGGGGUUUCCGUGAGGACA
env-1157  ................AGCAG.GA..G.UUC...CGCGGAGCGAAUGUCAA.....G..UCUCGAUAC...............UGAC..............GACAA..G.UCAGUGUACAAACGAAGAGCGCCGGAUUUCAAAACGAAAGAUGGCCUGCUGUGGGGUUUCCGAGAGGAC
env-1158  ................AGCAG.GA..G.UUC...CGCGGAGCGAAUGUCAA.....G..UCUCGAUAC...............UGAC..............GACAA..G.UCAGUGUACAAACGAAGAGCGCCGACUUUCGAAAGAAGGAAGGCCUGCCGUUGGGUUUCCGGGAGGACA
env-1159  ................AGCAG.GA..G.UUC...CGCGGAGCGAAUGUCAA.....G..UCUCGAUAC...............UGAC..............GACAA..G.UCAGUGUACAAACGAAGAGCGCCGUAUUUCGAAAGAGAGAAGGCCUGCCUUGGGUNNNNNNNNNNNNNNNN
Csp-2-1   .....UUGUG.......UGCAG.GU..U.UUUC..UGAGGAGCG.AUGUCCA....G..UUCAAGAAACC.............GGGC..............GACAU..GUCAGUGUACUGAAUAUAGGAAAAGACAUUCUCAAAAAAAGGGAAUAGUUCUGUGUAUCUUUACGUUGCG
env-1160  ..................UAGG.GU..U.UUC...UGCGGAGCG.AUGUCCA....G..UUCAAGAAACC.............GGGC..............GACAAG..G.UCAGAGUGCUAAAUAAUGAAUUGUGAGUUUCAUAUGGGAUAUGUGCCCGCGUUGCGAUGAAACACGCA
env-1162  ..................UAGG.GU..U.UUC...UGCGGAGCG.AUGUCCA....G..UUCAAGAAACC.............GGGC..............GACAAG..G.UCAGAGUGCUAAAUAAUGAAUUGUGAGUUUCAUACGGGAUAGUGCCCGCGUUGCGAUGAAACACGCA
env-1163  ..................UAGG.GU..U.UUC...UGCGGAGCG.AUGUCCA....G..UUCAAGAAACC.............GGGC..............GACAAG..G.UCAGAGUGCUAAAUAAUGAAUUGUGAGUUUCAUACGGGAUAGUGCCCGCGUUGCGAUGGAACACGCA
env-1166  ..................UAGG.GU..U.UUC...UGCGGAGCG.AUGUCCA....G..UUCAAGAAACC.............GGGC..............GACAAG..G.UCAGAGUGCUAAAUAAUGAAUUGUGAGUUUCAUAUGGGAUAUUAUGCCCACGUUGCGAUGAAACACGCA
env-1167  ...AAGGAAU.......CAAAG.GU..U.UUUU..CAAGGAGCG.AUGUCCA....G..UUCAUGAAAUG.............GGAC..............GACAAA..G.UCAGUGUAUUAAAUACUUUUGGUGUAAUUAUGAAACAUCACCAGACCGAAGUUUAAAAAAUUUAAACAC
env-1168  ....AGGAAU.......CAAAG.GU..U.UUUU..CAAGGAGCG.AUGUCCA....G..UUCAUGAAAUG.............GGAC..............GACAAA..G.UCAGUGUAUUAAAUAUUUUGGUGUAAUUAUGAAACAUCACCAGACCGAAGUUUAAAAAAUUUAAACAC
env-1169  .....AGGGU.......UUCGG.GU..UUUUU...CGAGGAGCG.AUGUCCA....G..UUCAUGAAACC.............GGAC..............GGCAA.GG.UCAGUGUACUUAAUAUUUAGGGGGAGUUUGCGUGAGAAAGCGCAGCUCUGGAAUCCGACGUUGCAGGAU
env-1170  .....AGGGU.......UUCGG.GU..UUUUU...CGAGGAGCG.AUGUCCA....G..UUCAUGAAACC.............GGAC..............GGCAA.GG.UCAGUGUACUUAAUAUUUAGGGGGAGUUUGCGUGAGAAAGUGCAGCUCUGGAAUCCGACGUUGCAGGAU
env-1171  .....AGGGU.......UUCGG.GU..UUUUU...CGAGGAGCG.AUGUCCA....G..UUCAUGAAACC.............GGAC..............GGCAA.GG.UCAGUGUACUUAAUAUUUAGGAGAGUUUGCGUGAGAAAGCGCAGCUCUGGAAUCCGACGUUGCAGGAUA
env-1172  ....CGGCUUGAU........G.GU..U.UUUC..CGAGGAGCG.AUGUCCA....G..UUCAAGAAACC.............GGGC..............GACAAG..G.UCAGUGUACUGAAUAUUUAGGGUAAGUUAUCUCGACAGGAUAACUUUGCGUAUCUGACGUUGCAGGAUA
env-1173  ....CGGCUUGAU........G.GU..U.UUUC..CGAGGAGCG.AUGUCCA....G..UUCAAGAAACC.............GGGC..............GACAAG..G.UCAGUGUACUGAAUAUUUAGGGUAAGUUAUCUCGACAGGAUAACUUUGCGUAUCUGACGUUGCAGGAUA
env-1174  ....CGGCUUGAU........G.GU..U.UUUC..CGAGGAGCG.AUGUCCA....G..UUCAAGAAACC.............GGGC..............GACAAG..G.UCAGUGUACUGAAUAUUUAGGGUAAGUUAUCUCGACAGGAUAACUUUGCGUAUCUUUCGUUGCAGGAUA
env-1175  .....CGGCUUG.....AU..G.GU..U.UUUC..CGAGGAGCG.AUGUCCA....G..UUCAAGAAACC.............GGGC..............GACAA.GG.UCAGUGUACUGAAUAUUUAGGGUAAGUUAUCUCGACAGGAUAACUUUGCGUAUCUGACGUUGCAGGAUA
env-1176  .....CGGCU.......UGAAG.GU..U.UUUC..CGAGGAGCG.AUGUCCA....G..UUCAAGAAACC.............GGGC..............GACAA.GG.UCAGUGUACUGAAUAUUUAGGGUAAGUUAUCUCGACAGGAUAACUUUGCGUAUCUUUCGUUGCAGGAUA
Cle-1-1   ................AGAUG.GA..U.UUC...UGGGGAGCGAAUGUCAG.....G..UU.CAAGAAACCA............GGC..............GGCAG..G.UCAGUGUACUAUAUUGAAUAGGUGUUAGGCGUGUCCAUAGUUAUAUGGAAAUGCUUGAUAUAAGUUUGC
          >>>>>>>>>>>.......>>>>>.....>.>>>....>>>.........<<<<.......<..<<<....................>>>>..............>>>>....>.>>>>..>>...........................................
          01.1111111.......22221.....2.102....222.........111.......2..212....................2122..............111.....1.111..12...........................................
          ---------------•GYRG-GU--U-UUC---CGYGGAGCGAAUGUC•R-----G--•Y•R•RRR•---------------•G•C-------------GRCAA--G-UCACUGUAC•RR•••••••••••••••••••••••••••••••••••••••••••••••
                                                                                                                    3'
```

alignment positions 361 · · · 540

```
env-1   UAAAACCGGCCUCUUUGUGCCGGUUUGUUUUUCGGUCAGAGUCUGAUACGCACGGGAACGUGUCUGGAAAUCUCAACCGUACGAGCAGACGGUCCCAAUGCCGUCAAAAUUAUCGUACGAUUUGGAGGAAUUACAUUUGCAGCAGCAAACAAAAAAAUGAUUCGCAUCAGACUAAAAGCC
env-4   UAAAACCGGCCUCUUUGUGCCGGUUUGUUUUUCGGUCAGAGUCUGAUACGCACGGGAACGUGUCUGGAAAUCUCAACCGUACGAGCAGACGGUCCCAAUGCCGUCAAAAUUAUCGUACGAUUUGGAGGAAUUACAUUUGCAGCAGCAAACAAAAAAAUGAUUCGCAUCAGACUAAAAGCC
env-5   .............................................................................................................................................................................
env-6   GAUACGCACGGCUUAGCGUAUACGGAAUCCCGACCGUACCCAGAGACACAGAAAACUGAGUACGUGAUUCAAGGAGGAACACAAACUUUGCAAAACAGGAAAUGAUCAGAAUUCGCCUGAAGGCUUAUGAUCAUCAGCUCAUCGACGCGAGUGCAGAGAAGAUCGUCGAGACCGCCAAGC
env-7   GUUUCUUAUGAGCCGGAAUGAUACGCACGGAACGGUGAACAAGAAAUUUCCCGCCGUACGGCAGUAGGUCACGCAGAAAAAAACCGUACGAAAACAUGGAGGAAAAGACCAAUUUGCAAAGAAAUGAUUCGCAUCCGUCUGAAGGCCUAUGACCAUCAGGUCAUCGACCAGUCCGCAGAG
env-8   GUUUCUUAUGAGCCGGAAUGAUACGCACGGAACGGUGAACAAGAAAUUUCCCGCCGUACGGCAGUAGGUCACGCAGAAAAAAACCGUACGAAAACAUGGAGGAAAAGACCAAUUUGCAAAAGAAAUGAUUCGCAUCCGUCUGAAGGCCUAUGACCAUCAGGUCAUCGACCAGUCCGCAGAG
env-9   GUUUCUUAUGAGCCGGAAUGAUACGCACGGAACGGUGAACAAGAAAUUUCCCGCCGUACGGCAGUAGGUCACGCAGAAAAAAACCGUACGAAAACAUGGAGGAAAAGACCAAAUGGCAAAAGAAAUGAUU.........................................
env-10  UGCGCCGGUAUUUUUCGUGAGCCGGAAUGAUACGCACGGUUAAGCGUAUAGAAAAAUUCUCUCGCCGUAGGUCGUCGAGACUUCGUGAAAACUACCUACGAAAUCAGGAGGAAAGAACCUUUGCAGCACAGAAAGAAAUGAUCGCAUUCGCCUGAAGGCCUAUGACCACCAGGUGAUCG
env-11  UAUUUUUCAUGA.................................................................................................................................................................
env-12  GCCGGUGUUUCUUAUGAGCCGGAAUGAUACGCACGGAACGGCGAUAGAGAGACCUCACCGUACGGAAUUGGUCAAAGCAAACAAAACCGUACAUUGAAGUAAGGAGUAAACAAGAAUUUGCAAAGGAAAUGAUUCGCAUUCGUCUGAAGGCCUAUGACCACCAGGUCAUCGACCAGAGC
env-13  UGUUUCUUAUGAGCCGGAAUGAUACGCACGGAACGGCGUAUAAGAAAUUUCCCGCCGUACGGCAUUGGUCAAAGCAACAAAAACCGUACGAAUUUUAUGGAGGAACAAACCAAUUUGCAAAAGAAAUGAUCCGCAUUCGUCUGAAGGCCUAUGACCAUCAGGUCAUCGACCAGUCCGCAG
env-14  UGUUUCUUAUGAGCCGGAAUGAUACGCACGGAACGGCGUAUAAGAAAUUUCCCGCCGUACGGCAUUGGUCAAAGCAACAAAAACCGUACGAAUUUUAUGGAGGAACAAACCAAUUUGCAAAAGAAAUGAUCCGCAUUCGUCUGAAGGCCUAUGACCAUCAGGUCAUCGACCAGUCCGCAG

        .............................................................................................................................................................................
        .............................................................................................................................................................................
        ••••••••••••••••••••••••••••••••••••••••••••••••••••••••••••••••••••••••••••••••••••••••••••••••••••••••••••••••••••••••••••••••••••••••••••••••••••••••••••••R••••••••••••••••••••
```

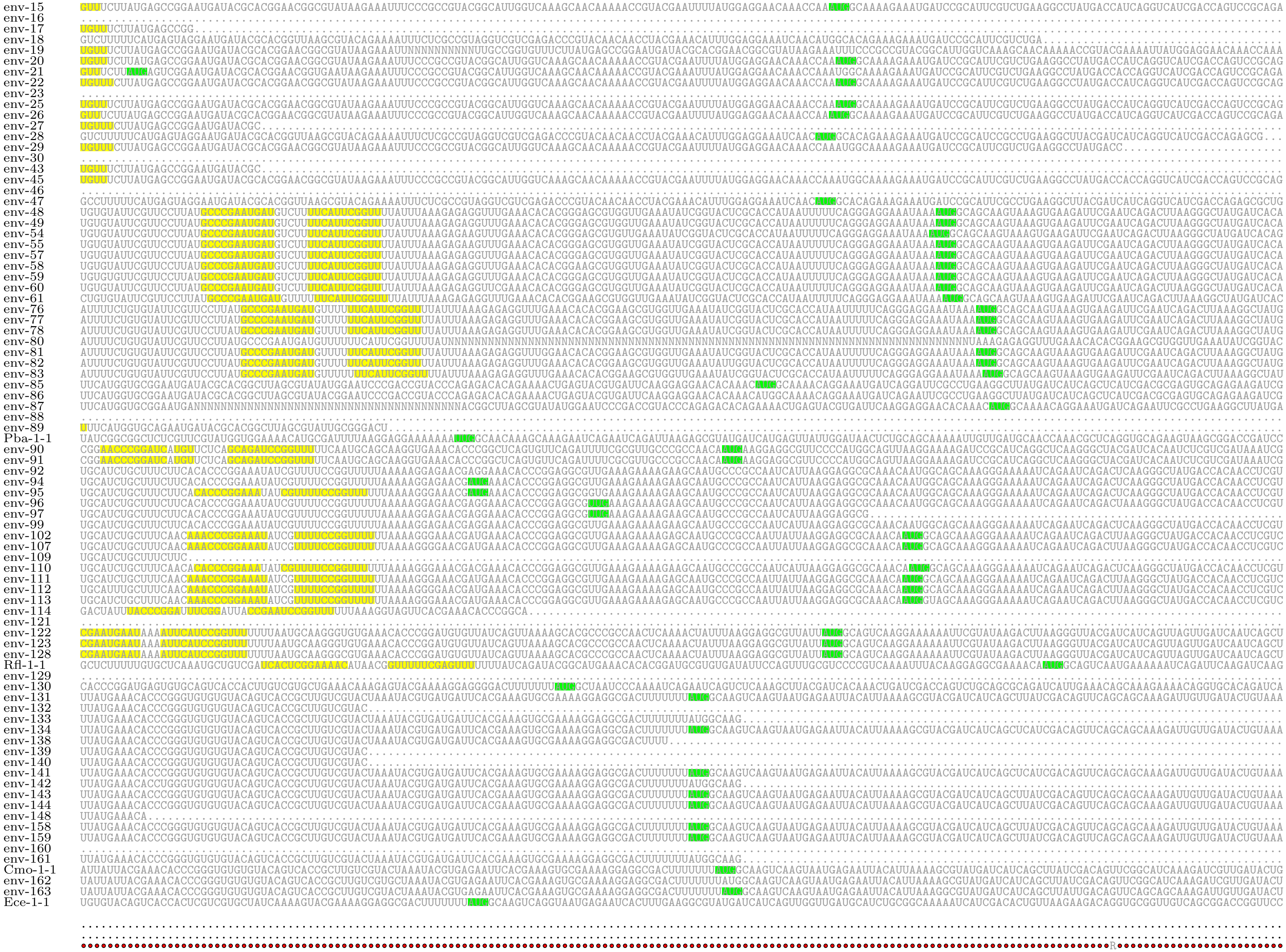

env-15
env-16
env-17
env-18
env-19
env-20
env-21
env-22
env-23
env-25
env-26
env-27
env-28
env-29
env-30
env-43
env-45
env-46
env-47
env-48
env-49
env-54
env-55
env-57
env-58
env-59
env-60
env-61
env-76
env-77
env-78
env-80
env-81
env-82
env-83
env-85
env-86
env-87
env-88
env-89
Pba-1-1
env-90
env-91
env-92
env-94
env-95
env-96
env-97
env-99
env-102
env-107
env-109
env-110
env-111
env-112
env-113
env-114
env-121
env-122
env-123
env-128
Rfl-1-1
env-129
env-130
env-131
env-132
env-133
env-134
env-138
env-139
env-140
env-141
env-142
env-143
env-144
env-148
env-158
env-159
env-160
env-161
Cmo-1-1
env-162
env-163
Ece-1-1

```
env-164  AAAAUCUGAGUUGUAAUUUGGGUUGCGAAUCAGGACACACACGGGAGAAUGUACAGUCACCGCUUGUCGUACUUAAACAAAAGUGCGAAAAGGAGGCGACUUUUUUUAUGGCAAAUCAGGUAAUGAGGAUUACUCUUAAAGCAUAUGAUCAUCAGUUAGUUGAUGCAUCUGCUAAGAAGA
Bfi-1-1  AAAAUCUGAGUUGUAAUUUGGGUUGCGAAUCAGGACACACACGGGAGAAUGUACAGUCACCGCUUGUCGUACUUAAACAAAAGUGCGAAAAGGAGGCGACUUUUUUUAUGGCAAAUCAGGUAAUGAGGAUUACUCUUAAAGCAUAUGAUCAUCAGUUAGUUGAUGCAUCUGCUAAGAAGA
Bfi-2-1  AAAAUCUGAGUUGUAAUUUGGGUUGCGAAUCAGGACACACACGGGAGAAUGUACAGUCACCGCUUGUCGUACUUAAACAAAAGUGCGAAAAGGAGGCGACUUUUUUUAUGGCAAAUCAGGUAAUGAGGAUUACUCUUAAAGCAUAUGAUCAUCAGUUAGUUGAUGCAUCUGCUAAGAAGA
env-165  GAUGUGUACAGUCACCGCUUGUCGUACUUAUAUUUUAGAAAGUGCGAAAAGGAGGCGACUUUUUUUAUGGCAAAUCAAGUUAUGAGAAUUACAUUAAAAGCUUAUGACCACCAGUUAGUAGACAGUUCUGCAAGUAAGAUUGUAGAGUCUGUUAAGAAAACAGGUGCUAAGGUUUCAGGA
env-166  ....................................................................................................................................................................................
env-170  GAUGUGUACAGUCACCGCUUGUCGUACUUAUAUUUUAGAAAGUGCGAAAAGGAGGCGACUUUUUUUAUGGCAAAUCAAGUUAUGAGAAUUACAUUAAAAGCUUAUGACCACCAGUUAGUAGACAGUUCUGCAAGUAAGAUUGUAGAGUCUGUUAAGAAAACAGGUGCUAAGGUUUCAGGA
env-171  GAUGUGUACAGUCACCGCUUGUCGUACUAAAUACGUGAGAAUUCACG.....................................................................................................................................
env-172  GAUGUGUACAGUCACCGCUUGUCGUACUUAUAUUUUAGAAAGUGCGAAAAGGAGGCGACUUUUUUUAUGGCAAAUCAAGUUAUGAGAAUUACAUUAAAAGCUUAUGACCACCAGUUAGUAGACAGUUCUGCAAGUAAGAUUGUAGAGUCUGUUAAGAAAACAGGUGCUAAGGUUUCAGGA
env-173  CGGAAAUCUGCGCAUUGGCUUAUGAAAAGGCAGGACACGCACGGAGGAGUGUACAGUCACCGCUUGUCGUACUUCUGAGGAAUAUCAUUCCUAUUAUUAAAAAGUGCGAAAAGGAGGCGACUUUUUUUAUGGCAAGUCAAGUAAUGAGAAUCACACUCAAGGCUUAUGAUCAUCAGUUAG
env-174  GGAUGAAACCGCACAUCGUCUUAUGAAAAGGUAGGACACGCACGGAGGAGUGUACAGUCACCGCUUGUCGUACUUCUAAGGAAUAUCAUUCCUAUUAUUAAAAAGUGCGAAAAGGAGGCGACUUUUUUUAUGGCAAGUCAAGUAAUGAGAAUCACACUCAAGGCUUAUGAUCAUCAGUUA
env-175  GGCAUUUUCGUGUGAACCGGCAC.............................................................................................................................................................
env-176  GGCAUUUUCGUGUGAACCGGCACGANNNNNNNNNNNNNNNNNNNNNNNNNNNNNNNNNNNNNNNNNNNNNNNNNNNNNNNNNNNNNNNNNNNNNNNNNNNNNNNNNNNNNNNNNNNNNNNNNNNNNNNNNNNGAACUAUUAUUAAUUCAGUACGAAAAGGAGGCGGCUUUUUU....
env-177  GGCAUUUUCGUGUGAACCGGCACGAU..........................................................................................................................................................
env-178  GGCAUUUUCGUGUGAACCGGCACGAUACACACGGAUGAGUGUACAGUCACCACUUGUCGUACGAACUAUUAUUAAUUCAGUACGAAAAGGAGGCGGCUUUUUUUAUGGCAAGUCAGCAGACAAUGAGAAUCAUUUUAAAAGCAUACGAUCAUCAGUUAAUCGAUAUGGCAGCAGCCAAAC
env-179  GGCAUUUUCGUGUGAACCGGCACGAUACACA.....................................................................................................................................................
env-180  UUUUUGAAUGAACCGAUAUGACACACACGGAUGAGUGUACAGUCACCACUUGUCGUACUAUUACCAAAUAAAGUACGAAAAGGAGGCGACUAUUUUUUAUGGCAAGUAAAAAAAGUCAGAAAAUGAGAAUUAUCCUGAAGGCAUACGAUCAUCAGUUGAUCGACUUUGCAGCAUCUAAGC
env-183  UUUUUGAAUGAACCGAUAUGACACACACGGAUGAGUGUACAGUCACCACUUGUCGUACUAUUACCAAAUAAAGUACGAAAAGGAGGCGACUAUUUUUUAUGGCAAGUAAAAAAAGUCAGAAAAUGAGAAUUAUCCUGAAGGCAUACGAUCAUCAGUUGAUCGACUUUGCAGCAUCUAAGC
env-184  UUUUUGAAUGAACCGAUAUGACACACACGGAUGAGUGUACAGUCACCACUUGUCGUACUAUUACCAAAUAAAGUACGAAAAGGAGGCGACUAUUUUUUAUGGCAAGUAAAAAAAGUCAGAAAAUGAGAAUUAUCCUGAAGGCAUACGAUCAUCAGUUGAUCGACUUUGCAGCAUCUAAGC
env-185  UUUUUGAAUGAACCGAUAUGACACACACGGAUGAGUGUACAGUCACCACUUGUCG.............................................................................................................................
env-186  UUUUUGAAUGAACCGAUAUGACACACACGGAUGAGUGUACAGUCACCACUUGUCGUACUAUUACCAAAUAAAGUACGAAAAGGAGGCGACUAUUUUUUAUGGCAAGUAAAAAAAGUCAGAAAAUGAGAAUUAUCCUGAAGGCAUACGAUCAUCAGUUGAUCGACUUUGCAGCAUCUAAGC
env-188  UUUUUGAAUGAACCGAUAUGACACACACGGAUGAGUGUACAGUCACCACUUGUCGUACUAUUACCAAAUAAAGUACGAAAAGGAGGCGACUAUUUUUUAUGGCAAGUAAAAAAAGUCAGAAAAUGAGAAUUAUCCUGAAGGCAUACGAUCAUCAGUUGAUCGACUUUGCAGCAUCUAAGU
env-189  UUUUUGAAUGAACCGAUAUGACACACACGGAUGAGUGUACAGUCACCACUUGUCGUACUAUUACCAAAUAAAGUACGAAAAGGAGGCGACUAUUUUUUAUGGCAAGUAAAAAAAGUCAGAAAAUGAGAAUUAUCCUGAAGGCAUACGAUCAUCAGUUGAUCGACUUUGCAGCAUCUAAGU
env-190  UUUUUGAAUGAACCGAUAUGACACACACGGAUGAGUGUACAGUCACCACUUGUCGUACUAUUACCAAAUAAAGUACGAAAAGGAGGCGACUAUUUUUUAUGGCAAGUAAAAAAAGUCAGAAAAUGAGAAUUAUCCUGAAGGCAUACGAUCAUCAGUUGAUCGACUUUGCAGCAUCUAAGU
env-191  NNNNNNNNNNNNNNNNNNAUUUUCUGCCAAGGAGCAGAUAAACGUGUGAGUCAAGCCUCCAUAGGGAGGAUUUUUGAAUGAACCGAUAUGACACACACGGAUGAGUGUACAGUCACCACUUGUCGUACUAUUACCAAAUAAAGUACGAAAAGGAGGCGACUAUUUUUUAUGGCAAGUAAA
env-192  UUUUUGAAUGAACCGAUAUGACACACACGGAUGAGUGUACAGUCACCACUUGUCGUACUAUUACCAAAUAAAGUACGAAAAGGAGGCGACUAUUUUUUAUGGCAAGUAAAAAAAGUCAGAAAAUGAGAAUUAUCCUGAAGGCAUACGAUCAUCAGUUGAUCGACUUUGCAGCAUCUAAGC
env-193  UUUUUGAAUGAACCGAUAUGACACACACGGAUGAGUGUACAGUCACCACUUGUCGUACUAUUACCAAAUAAAGUACGAAAAGGAGGCGACUAUUUUUUAUGGCAAGUAAAAAAAGUCAGAAAAUGAGAAUUAUCCUGAAGGCAUACGAUCAUCAGUUGAUCGACUUUGCAGCAUCUAAGC
env-194  UUUUUGAAUGAACCGAUAUGACACACACGGAUGAGUGUACAGUCACCACUUGUCGUACUAUUACCAAAUAAAGUACGAAAAGGAGGCGACUAUUUUUUAUGGCAAGUAAAAAAAGUCAGAAAAUGAGAAUUAUCCUGAAGGCAUACGAUCAUCAGUUGAUCGACUUUGCAGCAUCUAAGC
env-195  UUUUUGAAUGAACCGAUAUGACACACACGGAUGAGUGUACAGUCACCACUUGU...............................................................................................................................
env-208  UUUUUGAAUGAACCGAUAUGACACACACGGAUGAGUGUACAGUCACCACUUGUCGUACUAUUACCAAAUAAAGUACGAAAAGGAGGCGACUAUUUUUUAUGGCAAGUAAAAAAAGUCAGAAAAUGAGAAUUAUCCUGAAGGCAUACGAUCAUCAGUUGAUCGACUUUGCAGCAUCUAAGC
Lba-19-1 AAUCAGGACACACACGGGAGAGUGGGAAGGACAAAGAUCCUGACCGAUGUGUACAGUCGCCGCUUGUAAAGAACAGAGGAGACUUUUUUUAUGCAGGCAAACCAGAGAAUGAGAGUAAUCUUAAAGGCAUACGAUCAUCAGCUGAUCGAUGCAGCUGCCAAGAAGAUCGUUGACAGCG
Lba-17-1 AAGCGAUGGACACACACGGGAAAGUGGUACAGUCACCAGCUUGUCGUACUUACAAAAAAUAAUAAGUGCGAAAAGGAGGCGACUUUUUUUAUGGCAAGUCAGAAAAUGAGAGUCACAUUAAAAGCGUAUGAUCAUCAGCUAGUAGAUGCAUCAGCAAAGAAGAUUAUUGACACAGUUAAG
env-212  AAGCGAUGGACACACACGGGAAAGUGGUACAGUCACCAGCUUGUCGUACUUACAAAAAAUAAUAAGUGCGAAAAGGAGGCGACUUUUUUUAUGGCAAGUCAGAAAAUGAGAGUCACAUUAAAAGCGUAUGAUCAUCAGCUAGUAGAUGCAUCAGCAAAGAAGAUUAUUGACACAGUUAAG
env-213  AGGACACACACGGGAAAGUGUACAGUCACCAAGCUUGUCGUGCUUGAAAACAAAGUACGAAAAGGAGGCGACUUUUUUUAUGGCAAGUCAGAAAAUGAGAGUCACAUUAAAAGCGUAUGAUCAUCAGUUGGUUGAUGCAUCAGCAAAGAAGAUCAUUGAGGCAGUGAGGAAGAACGGUGG
env-214  ACACGGGAGAGUGUACAGUCACCACUUGUUGUACUAUUAAAAAGUGCGAAAAGGAGGCGGCUUUUUUUAUG.............................................................................................................
env-215  ....................................................................................................................................................................................
env-216  ACACGGGAGAGUGUACAGUCACCACCUGUUGUACUAUUAAAAAGUGCGAAAAGGAGGCGGCUUUUUUUAUGGCAAAACAGAUUAUGAGAAUCACAUUGAAGGCGUACGACCACAAACUUAUCGAUCAGGCUGCUGGACAGAUCAUCGAUACUGUUAAAAAGACAGGAGCUAAAGUCAGCG
env-217  ACACGGGAGAGUGUACAGUCACCACUUGUUGUACUAUUAAAAAGUGCGAAAAGGAGGCGGCUUUUUUUAUGGCAAAUCAGAUUAUGAGAAUCACAUUGAAGGCGUACGACCACAAACUUAUCGAUCAGGCUGCUGGACAGAUCAUCGAUACUGUUAAAAAGACAGGAGCUAAAGUCAGCG
env-218  ACACGGGAGAGUGUACAGUCACCACUUGUUGUACUAUUAAAAAGUGCGAAAAGGAGGCGGCUUUUUUUAUGGCAAAUCAGAUUAUGAGAAUCACAUUGAAGGCGUACGACCACAAACUUAUCGAUCAGGCUGCUGGACAGAUCAUCGAUACUGUUAAAAAGACAGGAGCUAAAGUCAGCG
env-219  ACACGGGAGAGUGUACAGUCACCACUUGUUGUACUUUUAAAAAGUGCGAAAAGGAGGCGGCUUUUUUUAUGGCAAAUCAGAUUAUGAGAAUCACAUUGAAGGCGUACGACCACAAACUUAUCGAUCAGGCUGCUGGACAGAUCAUCGAUACUGUUAAAAAGACAGGAGCUAAAGUCAGCG
env-220  ....................................................................................................................................................................................
env-223  ACACGGGAGAGUGUACAGUCACCACUUGUUGUACUAUUAAAAAGUGCGAAAAGGAGGCGGCUUUUUUUAUGGCAAAUCAGAUUAUGAGAAUCACAUUGAAGGCGUACGACCACAAACUUAUCGAUCAGGCUGCUGGACAGAUCAUCGAUACUGUUAAAAAGACAGGAGCUAAAGUCAGCG
env-224  ACACGGGAGAGUGUACAGUCACCACUUGUUGUACUAUUAAAAAGUGCGAAAAGGAGGCGGCUUUUUUUAUGGCAAAACAGAUUAUGAGAAUCACAUUGAAGGCGUACGACCACAAACUUAUCGAUCAGGCUGCUGGACAGAUCAUCGAUACUGUUAAAAAGACAGGAGCUAAAGUCAGCG
env-225  ACACGGGAGAGUGUACAGUCACCACUUGUUGUACUAUUAAAAAGUGCGAAAAGGAGGCGGCUUUUUUUAUGGCAAAUCAGAUUAUGAGAAUCACAUUGAAGGCGUACGACCACAAACUUAUCGAUCAGGCUGCUGGACAGAUCAUCGAUACUGUUAAAAAGACAGGAGCUAAAGUCAGCG
env-229  ACACGGGAGAGUGUACAGUCACCACUUGUUGUACUAUUAAAAAGUGCGAAAAGGAGGCGGCUUUUUUUAUGGCAAAUCAGAUUAUGAGAAUCACAUUGAAGGCGUACGACCACAAACUUAUCGAUCAGGCUGCUGGACAGAUCAUCGAUACUGUUAAAAAGACAGGAGCUAAAGUCAGCG
env-231  ACACGGGAGAGUGUACAGUCACCACUUGUUGUACUAUUAAAAAGUGCGAAAAGGAGGCGGCUUUUUUUAUGGCAAAUCAGAUUAUGAGAAUCACAUUGAAGGCGUACGACCACAAACUUAUCGAUCAGGCUGCUGGACAGAUCAUCGAUACUGUUAAAAAGACAGGAGCUAAAGUCAGCG
env-234  ....................................................................................................................................................................................
env-235  ACACGGGAGAGUGUACAGUCACCACUUGUUGUACUAUU..............................................................................................................................................
env-236  ACACGGGAGAGUGUACAGUCACCACUUGUUGUACUAUUAAAAAGUGCGAAAAGGAGGCGGCUUUUUUUAUGGCAAAACAGAUUAUGAGAAUCACAUUGAAGGCGUACGACCACAAACUUAUCGAUCAGGCUGCUGGACAGAUCAUCGAUACUGUUAAAAAGACAGGAGCUAAAGUCAGCG
env-237  ACACGGGAGAGUGUACAGUCACCACUUGUUGUACUAUUAAAAAGUGCGAAAAGGAGGCGGCUUUUUUUAUGGCAAAACAGAUUAUGAGAAUCACAUUGAAGGCGUACGACCACAAACUUAUCGAUCAGGCUGCUGGACAGAUCAUCGAUACUGUUAAAAAGACAGGAGCUAAAGUCAGCG
env-242  CCACUUGUCGUACAACAAGUACGAAAAGGAGGCGGCUUUUUUUAUGGCAACACAAAAAAUGCGAAUUAGCUUAAAGGCAUACGAUCACCAAUUAAUCGAUCAAUCAUCACAAAAGAUUAUUGAAACAGCAAAGAAAACUGGAGCUAAGGUAAGUGGACCGAUUCCAUUACCAACUAAGAA
env-244  ....................................................................................................................................................................................
env-247  UUCUGUAUAAUUUGUAAACCGACAGGACACACCCGGAUGAGUGUACAGUCACCACUUGUUGCGCUGAAACAGGCGGGCAUGGCGAAGAGCCAAGAUCAGUAAUGUCCGUGAAAGUGCGAAAAGGAGGCGGCUUUUUUUAUGGCAACAAAGGUAAUGAGAAUUACACUUAAGGCGUAUGAC
env-248  UUCUGUAUAUUUUGUAAACCGACAGGACACACCCGGAUGAGUGUACAGUCACCACUUGUUGCGCUGAAACAGGCGGACAUGGCGAAGAGCCAAGAUCAGUAAUGUCCGUGAAAGUGCGAAAAGGAGGCGGCUUUUUUUAUGGCAACAAAGGUAAUGAGAAUUACACUUAAAGCGUAUGAC
env-249  UUCUGUAUAAUUUGUAAACCGACAGGACACACCCGGAUGAGUGUACAGUCACCACUUGUUGCGCUGAAACAGGCGGACAUGGCGAAGAGCCAAGAUCAGUAAUGUCCGUGAAAGUGCGAAAAGGAGGCGGCUUUUUUUAUGGCAACAAAGGUAAUGAGAAUUACACUUAAGGCGUAUGAC
env-250  UUCUGUAUAAUUUGUAAACCGACAGGACACACCCGGAUGAGUGUACAGUCACCACUUGUUGCGCUGAAACAGGCGGGCAUGGCGAAGAGCCAAGAUCAGUAAUGUCCGUGAAAGUGCGAAAAGGAGGCGGCUUUUUUUAUGGCAACAAAGGUAAUGAGAAUUACACUUAAGGCGUAUGAC
env-251  UUCUGUAUAAUUUGUAAACCGACAGGACACACCCGGAUGAGUGUACAGUCACCACUUGUUGCGCUGAAACAGGCGGGCAUGGCGAAGAGCCAAGAUCAGUAAUGUCCGUGAAAGUGCGAAAAGGAGGCGGCUUUUUUUAUGGCAACAAAGGUAAUGAGAAUUACACUUAAGGCGUAUGAC
env-252  UUCUGUAUAAUUUGUAAACCGACAGGACACACCCGGAUGAGUG.........................................................................................................................................
env-257  UUCUGUAUAAUUUGUAAACCGACAGGACACACCCGGAUGAGUGUACAGUCACCACUUGUUGCGCUGAAACAGGCGGGCAUGGCGAAGAGCCAAGAUCAGUAAUGUCCGUGAAAGUGCGAAAAGGAGGCGGCUUUUUUUAUGGCAACAAAGGUAAUGAGAAUUACACUUAAGGCGUAUGAC
env-259  UUCUGUAUAAUUUGUAAACCGACAGGACACACCCGGAUGAGUGUACAGUCACCACUUGUUGCGCUGAAACAGGCGGACAUGGCGAAGAGCCAAGAUCAGUAAUGUCCGUGAAAGUGCGAAAAGGAGGCGGCUUUUUUUAUGGCAACAAAGGUAAUGAGAAUUACACUUAAAGCGUAUGAC
env-262  UUCUGUAUAAUUUGUAAACCGACAGGACACACCCGGAUGAGUGUACAGUCACCACUUGUUGCGCUGAAACAGGCGGGCAUGGCGAAGAGCCAAGAUCAGUAAUGUCCGCGAAAGUGCGAAAAGGAGGCGGCUUUUUUUAUGGCAACAAAGGUAAUGAGAAUUACACUUAAGGCGUAUGAC
env-264  UUCUGUAUAAUUUGUAAACCGACAGGACACACCCGGAUGAGUGUACAGUCACCACUUGUUGCGCUGAAACAGGCGGACAUGGCGAAGAGCCAAGAUCAGUAAUGUCCGUGAAAGUGCGAAAAGGAGGCGGCUUUUUUUAUGGCAACAAAGGUAAUGAGAAUUACACUUAAGGCGUAUGAC
env-265  UUCUGUAUAAUUUGUAAACCGACAGGACACACCCGGAUGAGUGUACAGUCACCACUUGUUGCGCUGAAACAGGCGGACAUGGCGAAGAGCCAAGAUCAGUAAUGUCCGUGAAAGUGCGAAAAGGAGGCGGCUUUUUUUAUGGCAACAAAGGUAAUGAGAAUUACACUUAAAGCGUAUGAC
env-266  GAGAUGUGUAC.........................................................................................................................................................................
env-280  GAGAUGUGUACA........................................................................................................................................................................
env-281  GAGAUGUGUACAG.......................................................................................................................................................................
env-282  GAGAUGUGUACAGUCACCGCUUGUCGUACUGAAGUUAAAAGUGCGAAAAGGAGGCGACUUUUUUUAUGACAAGUCAAGUAAUGAGAAU............................................................................................
env-283  GAGAUGUGUACAGUCACCGCUUGUCGUAC.......................................................................................................................................................
env-284  GAGAUGUGUACAGUCACCGCUUGUCGUACUGAAGUUAAAAGUGCGAAAAGGAGGCGACUUUUUUUAUGGCAAGUCAAGUAAUGAGAAUUACUCUCAAGGCGUAUGAUCACGAGUUAGUAGAUUCAAGCGCAAAGAAAAUCAUCGAGACUGUAAAGAAGAAUGGAUCACAGGUGAGCGGAC
env-288  GAGAUGUGUACAGUCACCGCUUGUCGUACUGAAGUUAAAAGUGCGAAAAGGAGGCGACUUUUUUUAUGGCAAGUCAAGUAAUGAGAAUUACUCUCAAGGCGUAUGAUCACGAGUUAGUAGAUUCAAGCGCAAAGAAAAUCAUCGAGACUGUAAAGAAGAAUGGAUCACAGGUGAGCGGAC
env-289  GAGAUGUGUACAGUCACCGCUUGUCGUACUGAAGUUAAAAGUGCGAAAAGGAGGCGACUUUUUUUAUGGCAAGUCAAGUAAUGAGAAUUACUCUCAAGGCGUAUGAUCACGAGUUAGUAGAUUCAAGCGCAAAGAAAAUCAUCGAGACUGUAAAGAAGAAUGGAUCACAGGUGAGCGGAC
env-290  NNNNNNNNNNNCAAGUCAAGUAAUGAGAAUCACUCUCAAGGCGUAUGAUCACGAGUUAGUAGAUUCAAGCGCAANNNNNNNNNNNNNNNNNNNNNCUCAUAAGAGACUUAUCGAUAUCAUCAAUCCAACACAGAAGACAACAGAGUCUUUAUCAAGGAUUGAAAUGCCAGCUGGUGUUUA
env-291  GAGAUGUGUACAG.......................................................................................................................................................................
env-300  ....................................................................................................................................................................................
env-303  GAGAUGUGUACAGUC.....................................................................................................................................................................
env-304  GAGAUGUGUACAGUCACCGCUUGUCGUACUGAAGUUAAAAGUGCGAAAAGGAGGCGACUUUUUUUAUGGCAAGUCAAGUAAUGAGAAUUACUCUCAAGGCGUAUGAUCACGAGUUAGUAGAUUCAAGCGCAAAGAAAAUCAUCGAGACUGUAAAGAAGAAUGGAUCACAGGUGAGCGGAC
env-305  GAGAUGUGUACAGUCACCGCUUGUCGUACUGAAGUUAAAAGUGCGAAAAGGAGGCGACUUUUUUUAUGGCAAGUCAAGUAAUGAGAAUCACUCUCAAGGCGUAUGAUCACGAGUUAGUAGAUUCAAGCGCAAAGAAAAUCAUCGAGACUGUAAAGAAGAAUGGAUCACAGGUGAGCGGAC
env-306  GAGAUGUGUACAGUCACCGCUUGUCGUACUGAAGUUAAAAGUGCGAAAAGGAGGCGACUUUUUUUAUGGCAAGUCAAGUAAUGAGAAUUACUCUCAAGGCGUAUGAUCACGAGUUAGUAGAUUCAAGCGCAAAGAAAAUCAUCGAGACUGUAAAGAAGAAUGGAUCACAGGUGAGCGGAC
env-307  GAGAUGUGUACAGUC.....................................................................................................................................................................
env-308  GAGAUGUGUACAGUCACCGCUUGUCGUA........................................................................................................................................................
env-309  GAGAUGUGUACAGUCACCGCUUGUCGUACUGAAGUUAAAAGUGCGAAAAGGAGGCGACUUUUUUUAUGGCAAGUCAAGUAAUGAGAAUUACUCUCAAGGCGUAUGAUCACGAGUUAGUAGAUUCAAGCGCAAAGAAAAUCAUCGAGACUGUAAAGAAGAAUGGAUCACAGGUGAGCGGAC
env-310  GAGAUGUGUACAGUCACCGCUUGUCGUACUGAAGUUAAAAGUGCGAAAAGGAGGCGACUUUUUUUAUGGCAAGUCAAGUAAUGAGAAUUACUCUCAAGGCGUAUGAUCACGAGUUAGUAGAUUCAAGCGCAAAGAAAAUCAUCGAGACUGUAAAGAAGAAUGGAUCACAGGUGAGCGGA.
env-311  GAGAUGUGUACAGUCACCGCUUGUCGUACUGAAGUUAAAAGUGCGAAAAGGAGGCGACUUUUUUUAUGGCAAGUCAAGUAAUGAGAAUUACUCUCAAAGCGUAUGAUCACGAGUUAGUAGAUUCAAGCGCAAAGAAAAUCAUCGAGACUGUAAAGAAGAAUGGAUCACAGGUGAGCGGAC
env-312  GAGAUGUGUACAGUCACCGCUUGUCGUACUGAAGUUAAAAGUGCGAAAAGGAGGCGACUUUUUUUAUGGCAAGUCAAGUAAUGAGAAUUACUCUCAAGGCGUAUGAUCACGAGUUAGUAGAUUCAAGCGCAAAGAAAAUCAUCGAGACUGUAAAGAAGAAUGGAUCACAGGUGAGCGGAC
env-313  GAGNNNNNCAAGUCAAGUAAUGAGAAUCACUCUCAAGGCGUAUGAUCACGAGUUAGUAGAUUCAAGCGCAAAGNNNNNNNNNNNNNNNGAACUCAUAAGAGACUUAUCGAUAUCAUCAAUCCAACACAGAAGACAACAGAGUCUUUAUCAAGGAUUGAAAUGCCAGCUGGUGUUUACAUU

         ....................................................................................................................................................................................
         ....................................................................................................................................................................................
         ..........................................................................................................................................................R.........................
```

| | |
|---|---|
| env-314 | GAGAUGUGUACAGUCNNNNNNNNNNNNNNNNUACUCUCAAGGCGUAUGAUCACGAGUUAGUAGAUUCAAGCGCAAAGAAAAUCAUCGAGACUGUAAAGAAGAAUGGAUCACAGGUGAGCGGACCGGUACCUCUUCCAACAAAGAAGGAAGUAGUUACAAUUCUUAGAGCUACUCACAAG |
| env-315 | GAGAUGUGUACAGUCACCGCUUGUCGUACUGAAGUUAAAAGUGCGAAAAGGAGGCGACUUUUUUUAUGGCAAGUCAAGUAAUGAGAAUUACUCUCAAGGCGUAUGAUCACGAGUUAGUAGAUUCAAGCGCAAAGNNNNNNNNNNNNNNNNNNNNNNNNNNNNNNNNNNNNNNNNNNNG |
| env-316 | GAGAUGUGUACAG… |
| env-317 | GAGAUGUGUAC… |
| env-331 | GAGAUGUGUACAGUCACCGCUUGUCGUACUGAAGUUAAAAGUGCCAAAAGGAGGCGACUUUUUUUAUGGCAAGUCAAGUAAUGAGAAUUACUCUCAAGGCGUAUGAUCACGAGUUAGUAGAUUCAAGCGCAAAGAAAAUCAUCGAGACUGUAAAGAAGAAUGGAUCACAG… |
| env-332 | GAGAAGUGUA… |
| env-333 | GAGAUGUGUACAG… |
| env-334 | GAGAUGUGUACAGUCACCGCUUGUCGUACUGAAGUUAAAAGUGCGAAAAGGAGGCGACUUUUUUUAUGGCAAGUCAAGUAAUGAGAAUUACUCUCAAGGCGUAUGAUCACGAGUUAGUAGAUUCAAGCGCAAAGAAAAUCAUCGAGACUGUAAAGAAGAAUGGAUCACAGGUGAGCGGAC |
| env-335 | GAGAUGUGUACAGUCACCGCUUGUCGUACUGAAGUUAAAAGUGCGAAAAGGAGGCGACUUUUUUUAUGGCAAGUCAAGUAAUGAGAAUUACUCUCAAGGCGUAUGAUCACGAGUUAGUAGAUUCAAGCGCAAAGAAAAUCAUCGAGACUGUAAAGAAGAAUGGAUCACAGGUGAGCGGAC |
| env-336 | GAGAUGUGUACAGUCACCGCUUGUCGUACUGAAGUUGAAAGUGCGAAAAGGAGGCGACUUUUUUUAUGGCAAGUCAAGUAAUGAGAAUUACUCUCAAGGCGUAUGAUCACGAGUUAGUAGAUUCAAGCGCAAAGAAAAUCAUCGAGACUGUAAAGAAGAAUGGAUCACAGGUGAGCGGAC |
| env-338 | GAGAUGUGUACAGUCACCGCUUGUCGUACUGAAGUUAAAAGUGCGAAAAGGAGGCGACUUUUUUUAUGGCAAGUCAAGUAAUGAGAAUUACUCUCAAGGCGUAUGAUCACGAGUUAGUAGAUUCAAGCGCAAAGAAAAUCAUCGAGACUGUAAAGAAGAAUGGAUCACAGGUGAGCGGAC |
| env-339 | GAGAUGUGUACAGUCACCGCUUGUCGUACUGAAGUUAAAAGUGCGAAAAGGAGGCGGCUUUUUUUAUGGCAAGUCAAGUAAUGAGAAUUACUCUCAAGGCGUAUGAUCACGAGUUAGUAGAUUCAAGCGCAAAGAAAAUCAUCGAGACUGUAAAGAAGAAUGGAUCACAGGUGAGCGGAC |
| env-340 | GAGAUGUGUACAGUCACCGCUUGUCGUACUGAAGUUAAAAGUGCGAAAAGGAGGCGACUUUUUUUAUGGCAAGUCAAGUAAUGAGAAUUACUCUCAAGGCGUAUGAUCACGAGUUAGUAGAUUCAAGCGCAAAGAAAAUCAUCGAGACUGUAAAGAAGAAUGGAUCACAGGUGAGCGGAC |
| env-341 | GAGAUGUGUACAGUCACCGCUUGUCGUACUGAAGUUAAAAGUGCGAAAAGGAGGCGACUUUUUUUAUGGCAAGUCAAGUAAUGAGAAUUACUCUCAAGGCGUAUGAUCACGAGUUAGUAGAUUCAAGCGCAAAGAAAAUCAUCGAGACUGUAAAGAAGAAUGGAUCACAGGUGAGCGGAC |
| env-347 | GAGAUGUGUACAG… |
| env-348 | GAGAUGUGUACAGUCACCGCUUGUCGUA… |
| env-349 | GAGAUGUGUACAG… |
| env-350 | GAGAUGUGUACAGUCACCGCUUGUCGUACUGAAGUUAAAAGUGCGAAAAGGAGGCGACUUUUUUUAUGGCAAGUCAAGUAAUGAGAAUUACUCUCAAGGCGUAUGAUCAUGAGUUAGUAGAUUCAAGCGCAAAGAAAAUCAUCGAGACUGUAAAGAAGA… |
| env-351 | GAGAUGUGUACAGUCACCGCUUGUCGUACU… |
| env-352 | GAGAUGUGUACAGUC… |
| env-353 | GAGAUGUGUACAGUCACCGCUUGUCGUACUGAAGUUAAAAGUGCGAAAAGGAGGCGACUUUUUUUAUGGCAAGUCAAGUAAUGAGAAUUACUCUCAAGGCGUAUGAUCACGAGUUAGUAGAUUCAAGCGCAAAGAAAAUCAUCGAGACUGUAAAGAAGAAUGGAUCACAGGUGAGCGGAC |
| env-354 | GAGAUGUGUACAGUCACCGCUUGUCGUACUGAAGUUAAAAGUGCGAAAAGGAGGCGACUUUUUUUAUGGCAAGUCAAGUAAUGAGAAUUACUCUCAAGGCGUAUGAUCACGAGUUAGUAGAUUCAAGCGCAAAGAAAAUCAUCGAGACUGUAAAGAAGAAUGGAUCACAGGUGAGCGGAC |
| env-355 | GAGAUGUGUACAGUCACCGCUUGUCGUACUGAAGUUAAAAGUGCGAAAAGGAGGCGACUUUUUUUAUGGCAAGUCAAGUAAUGAGAAUUACUCUCAAGGCGUAUGAUCACGAGUUAGUAGAUUCAAGCGCAAAGAAAAUCAUCGAGACUGUAAAGAAGAAUGGAUCACAGGUGAGCGGAC |
| env-359 | GAGAUGUGUACAGUCACCGCUUGUCGUACUGAAGUUAAAAGUGCGAAAAGGAGGCGACUUUUUUUAUGGCAAGUCAAGUAAUGAGAAUUACUCUCAAGGCGUAUGAUCACGAGUUAGUAGAUUCAAGCGCAAAGAAAAUCAUCGAGACUGUAAAGAAGAAUGGAUCACAGGUGAGCGGAC |
| env-360 | GAGAUGUGUACAGUCACCGCUUGUCGUACUGAAGUUAAAAGUGCGAAAAGGAGGCGACUUUUUUUAUGGCAAGUCAAGUAAUGAGAAUUACUCUCAAGGCGUAUGAUCACGAGUUAGUAGAUUCAAGCGCAAAGAAAAUCAUCGAGACUGUAAAGAAGAAUGGAUCACAGGUGAGCGGAC |
| env-361 | GAGAUGUGUACAGUCACCGCUUGUCGUACUGAAGUUAAAAGUGCGAAAAGGAGGCGACUUUUUUUAUGGCAAGUCAAGUAAUGAGAAUUACUCUCAAGGCGUAUGAUCACGAGUUAGUAGAUUCAAGCGCAAAGAAAAUCAUCGAGACUGUAAAGAAGAAUGGAUCACAGGUGAGCGGAC |
| env-362 | GAGAUGUGUACAGUCNNNNNNNNNNNNNNNNUACUCUCAAGGCGUAUGAUCACGAGUUAGUAGAUUCAAGCGCAAAGAAAAUCAUCGAGACUGUAAAGAAGAAUGGAUCACAGGUGAGCGGACCGGUACCUCUUCCAACAAAGAAGGAAGUAGUUACAAUUCUUAGAGCUACUCAC |
| env-363 | GAGAUGUGUACAGUC… |
| env-364 | … |
| env-365 | GAGAUGUGUACAGUCACCGCUUGUCGUACUGAAGUUAAAAGUGCGAAAAGGAGGCGACUUUUUUUAUGGCAAGUCAAGUAAUGAGAAUUACUCUCAAGGCGUAUGAUCACGAGUUAGUAGAUUCAAGCGCAAAGAAAAUCAUCGAGACUGUAAAGAAGAAUGGAUCACAGGUGAGCGGAC |
| env-366 | GAGAUGUGUACAGUCACCGCUUGUCGUACUGAAGUUAAAAGUGCGAAAAGGAGGCGACUUUUUUUAUGGCAAGUCAAGUAAUGAGAAUUACUCUCAAGGCGUAUGAUCACGAGUUAGUAGAUUCAAGCGCAAAGAAAAUCAUCGAGACUGUAAAGAAGAAUGGAUCACAGGUGAGCGGAC |
| env-368 | GAGAUGUGUACAGUCACCGCUUGUCGUACUGAAGUUAAAAGUGCGAAAAGGAGGCGACUUUUUUUAUGGCAAGUCAAGUAAUGAGAAUUACUCUCAAGGCGUAUGAUCACGAGUUAGUAGAUUCAAGCGCAAAGAAAAUCAUC… |
| env-372 | GAGAUGUGUACAGUCACCGCUUGUCGUACUGAAGUCAAAAGUACGAAAAGGAGGCGACUUUUUUUAUGGCAAGUCAAGUAAUGAGAAUUACUCUCAAGGCGUAUGAUCACGAGUUAGUAGAUUCAAGCGCAAAGAAAAUCAUCGAGACUGUAAAGAAGAAUGGAUCACAGGUGAGCGGAC |
| env-373 | GAGAUGUGUACAGUCACCGCUUGUCGUACUGAAGUUAAAAGUGCGAAAAGGAGGCGACUUUUUUUAUGGCAAGUCAAGUAAUGAGAAUUACUCUCAAGGCGUAUGAUCACGAGUUAGUAGAUUCAAGCGCAAAGAAAAUCAUCGAGACUGUAAAGAAGAAUGGAUCACAGGUGAGCGGAC |
| env-374 | GAGAUGUGUACAGUCACCGCUUGUCGUACUGGAGUUAAAAGUGCGAAAAGGAGGCGACUUUUUUUAUGGCAAGUCAAGUAAUGAGAAUUACUCUCAAGGCGUAUGAUCACGAGUUAGUAGAUUCAAGCGCAAAGAAAAUCAUCGAGACUGUAAAGAAGAAUGGAUCACAGGUGAGCGGAC |
| env-375 | … |
| env-376 | UAAUUUGUGUUAAGACGUAGGACACACACGGGAGAGUGUACAGUCACCGCUUGUCGUACUUGUAUAUAAAAGUGCGAAAAGGAGGCGACUUUUUUUAUGGCAAAUCAGGUAAUGAGAAUUACUCUUAAGGCUUAUGAUCAUCAGCUUGUUGAUGCAUCAGCCAAGAAGAUUAUCGAGACA |
| Bsp-5-1 | UAAUUUGUGUUAAGACGUAGGACACACACGGGAGAGUGUACAGUCACCGCUUGUCGUACUUGUAUAUAAAAGUGCGAAAAGGAGGCGACUUUUUUUAUGGCAAAUCAGGUAAUGAGAAUUACACUUAAGGCUUACGAUCAUCAGCUUGUUGAUGCAUCAGCCAAGAAGAUUAUCGAGACA |
| env-377 | GUACAGUCACCGCUUGUCAAUUUCUUUUAUAA… |
| env-379 | GUACAGUCACCGCUUGUCAAUUUCUUUUAUAAAGAUUGAAAAGGAGGCGGCUUUUUUUAUGGCAAAUCAGGUAAUGAGAAUCACAUUAAAAGCUUAUGAUCA… |
| env-380 | GAUGUGUACAGUCACCGCUUGUCGAACUUUGUAUAAAAGUACGAAAAGGAGGCGACUUUUUUUAUGGCAA… |
| env-381 | GUACAGUCACCGCUUGUCAAUUUCUUUUAUAAGAUUGAAAAGGAGGCGGCUUUUUUUAUGGCAAAUCAGGUAAUGAGAAUCACAUUAAAAGCUUAUGAUCAUCAAUUGGUUGACAGCUGUGCUGCAAGAAUUGUUGAUACUGUAAAGAAGACAGGAGCAGCUGUUUCAGGACCUGUCCA |
| env-382 | GUACAGUCACCGCUUGUCAAUUUCUUUUAUAAAGAUUGAAAAGGAGGCGGCUUUUUUUAUGGCAAAUCAGGUAAUGAGAAUCACAUUAAAAGCUUAUGAUCACCAAUUGGUUGACAGCUGUGCUGCAAGAAUUGUUGAUACUGUAAAGAAGACAGGAGCAGCUGUUUCAGGACCUGUUCC |
| env-384 | GUACAGUCACCGCUUGUCAAUUUCUUUUAUAAGAUUGAAAAGGAGGCGGCUUUUUUUAUGGCAAAUCAGGUAAUGAGAAUCACAUUAAAAGCUUAUGAUCACCAAUUGGUUGACAGCUGUGCUGCAAGAAUUGUUGAUACUGUAAAGAAGACAGGAGCAGCUGUUUCAGGACCUGUCCA |
| Lor-1-1 | GUACAGUCACCGCUUGUCAAUUUCUUUUAUAAGAUUGAAAAGGAGGCGGCUUUUUUUAUGGCAAAUCAGGUAAUGAGAAUCACAUUAAAAGCUUAUGAUCACCAAUUGGUUGACAGCUGUGCUGCAAGAAUUGUUGAUACUGUAAAGAAGACAGGAGCAGCUGUUUCAGGACCUGUCCA |
| env-385 | GUACAGUC… |
| env-386 | GUACAGUCACCGCUUGUCAAUUUCUUUUAUAAGAUUGAAAAGGAGGCGGCUUUUUUUAUGGCAAAUCAGGUAAUGAGAAUCACAUUAAAAGCUUAUGAUCACCAAUUGGUUGACAGCUGUGCUGCAAGAAUUGUUGAUACUGUAAAGAAGACAGGAGCAGCUGUUUCAGGACCUGUCCA |
| env-387 | GUACAGUCACCGCUUGUCAAUUUCUUUUAUAAAGAUUGAAAAGGAGGCGGCUUUUUUUAUGGCAAAUCAGGUAAUGAGAAUCACAUUAAAAGCUUAUGAUCACCAAUUGGUUGACAGCUGUGCUGCAAGAAUUGUUGAUACUGUAAAGAAGACAGGAGCAGCUGUUUCAGGACCUGUUCC |
| env-388 | GUACAGUC… |
| Lba-15-1 | GUACAGUCACCGCUUGUCAAUUUCUUUUAUAAAGAUUGAAAAGGAGGCGGCUUUUUUUUAUGGCAAAUCAGGUAAUGAGAAUCACAUUAAAAGCUUAUGAUCAUCAAUUAGUUGACAGCUGUGCUGCAAGAAUUGUUGAUACUGUAAAGAAGACAGGAGCAGCUGUUUCAGGACCUGUUCC |
| env-389 | GUACAGUCACCGCUUGUCAAUUUCUUUUAUAAAGAUUGAAAAGGAGGCGGCUUUUUUUUAUGGCAAAUCAGGUAAUGAGAAUCACAUUAAAAGCUUAUGAUCAUCAAUUAGUUGACAGCUGUGCUGCAAGAAUUGUUGAUACUGUAAAGAAGACAGGAGCAGCUGUUUCAGGACCUGUUCC |
| env-390 | GUACAGUCACCGCUUGUCAAUUUCUUUUAUAAAGAUUGAAAAGGAGGCGG… |
| env-391 | GUACAGUCACCACUUGUCAAUUUCUUUUAUAAAGAUUGAAAAGGAGGCGGCUUUUUUUUAUGGCAAAUCAGGUAAUGAGAAUCACAUUAAAAGCUUAUGAUCAUCAAUUAGUUGACAGCUGUGCUGCAAGAAUUGUUGAUACUGUAAAGAAGACAGGAGCAGCUGUUUCAGGACCUGUUCC |
| env-392 | GUACAGUCACCACUUGUCAAUUUCUUUUAUAAAGAUUGAAAAGGAGGCGGCUUUUUUUUAUGGCAAAUCAGGUAAUGAGAAUCACAUUAAAAGCUUAUGAUCAUCAAUUAGUUGACAGCUGUGCUGCAAGAAUUG… |
| env-393 | GUACAGUCACCACUUGUCAAUUUCUUUUAUAAAGAUUGAAAAGGAGGCGGCUUUUUUUUAUGGCAAAUCAGGUAAU… |
| env-394 | GUACAGUCACCGCUUGUCAAUUUCUUUUAUAAAGAUUGAAAAGGAGGCGGCUUUUUUUUAUGGCAAAUCAGGUAAU… |
| env-395 | GUACAGUCACCACUUGUCAAUUUCUUUUAUAAAGAUUGAAAAGGAGGCGGCUUUUUUUUAUGGCAAAUCAGGUAAUGAGAAUCACAUUAAAAGCUUAUGAUCAUCAAUUAGUUGACAGCUGUGCUGCAAGAAUUGUUGAUACUGUAAAGAAGACAGGAGCAGCUGUUUCAGGACCUGUUCC |
| env-396 | … |
| env-397 | GUACAGUCACCGCUUGUCAAUUUCUUUUAUAAAGAUUGAAAAGGAGGCGGCUUUUUUUUAUGGCAAAUCAGGUAAUGAGAAUCACAUUAAAAGCUUAUGAUCACCAAUUGGUUGACAGCUGUGCUGCAAGAAUUGUUGAUACUGUAAAGAAGACAGGAGCAGCUGUUUCAGGACCUGUUCC |
| env-398 | GUACAGUCACCGCUUGUCAAUUUCUUUUAUAAAGAUUGAAAAGGAGGCGGCUUUUUUUUAUGGCAAAUCAGGUAAUGAGAAUCACAUUAAAAGCUUAUGAUCACCAAUUGGUUGACAGCUGUGCUGCAAGAAUUGUUGAUACUGUAAAGAAGACAGGAGCAGCUGUUUCAGGACCUGUUCC |
| env-399 | GUACAGUCACCACUUGUCAAUUUCUUUUAUAAAGAUUGAAAAGGAGGCGGCUUUUUUUUAUGGCAAAUCAGGUAAUGAGAAUCACAUUAAAAGCUUAUGAUCAUCAAUUAGUUGACAGCUGUGCUGCAAGAAUUGUUGAUACUGUAAAGAAGACAGGAGCAGCUGUUUCAGGACCUGUUCC |
| env-400 | GUACAGUCACCACUUGUCAAUUUCUUUUAUAAAGAUUGAAAAGGAGGCGGCUUUUUUUUAUGGCAAAUCAGGUAAUGAGAAUCACAUUAAAAGCUUAUGAUCAUCAAUUAGUUGACAGCUGUGCUGCAAGAAUUGUUGAUACUGUAAAGAAGACAGGAGCAGCUGUUUCAGGACCUGUUCC |
| env-402 | UACAGUCACCGCUUGUCAAUUUCUUUUAUAAAGAUUGAAAAGGAGGCGGCUUUUUUUUAUGGCAAAUCAGGUAAU… |
| env-403 | GUACAGUCACCACUUGUCAAUUUCUUUUAUAAAGAUUGAAAAGGAGGCGGCUUUUUUUUAUGGCAAAUCAGGUAAUGAGAAUCACAUUAAAAGCUUAUGAUCAUCAAUUAGUUGACAGCUGUGCUGCAAGAAUUGUUGAUACUGUAAAGAAGACAGGAGCAGCUGUUUCAGGACCUGUUCC |
| env-404 | GUACAGUCACCACUUGUCAAUUUCUUUUAUAAAGAUUGAAAAGGAGGCGGCUUUUUUUUAUGGCAAAUCAGGUAAUGAGAAUCACAUUAAAAGCUUAUGAUCAUCAAUUAGUUGACAGCUGUGCUGCAAGAAUUGUUGAUACUGUAAAGAAGACAGGAGCAGCUG… |
| env-405 | GUACAGUCACCACUUGUCAAUUUCUUUUAUAAAGAUUGAAAAGGAGGCGGCUUUUUUUUAUGGCAAAUCAGGUAAUGAGAAUCACAUUAAAAGCUUAUGAUCAUCAAUUAGUUGACAGCUGUGCUGCAAGAAUUGUUGAUACUGUAAAGAAGACAGGAGCAGCUGUUUCAGGACCUGUUCC |
| env-406 | GUACAGUCACCGCUUGUCAAUUUCUUUUAUAAAGAUUGAAAAGGAGGCGGCUUUUUUUUAUGGCAAAUCAGGUAAUGAGAAUCACAUUAAA… |
| env-407 | GUACAGUCACCGCUUGUCAAUUUCUUUUAUAAAGAUUGAAAAGGAGGCGGCUUUUUUUUAUGGCAAAUCAGGUAAUGAGAAUCACAUUAAAAGCUUAUGAUCACCAAUUGGUUGACAGCUGUGCUGCAAGAAUUGUUGAUACUGUAAAGAAGACAGGAGCAGCUGUUUCAGGACCUGUUCC |
| env-408 | GUACAGUCACCGCUUGUCAAUUUCUUUUAUAAAGAUUGAAAAGGAGGCGGCUUUUUUUUAUGGCAAAUCAGGUAAUGAGAAUCACAUUAAAAGCUUAUGAUCAUCAAUUAGUUGACAGCUGUGCUGCAAGAAUUGUUGAUACUGUAAAGAAGACAGGAGCAGCUGUUUCAGGACCUGUUCC |
| env-409 | GUACAGUCACCGCUUGUCAAUUUCUUUUAUAAAGAUUGAAAAGGAGGCGGCUUUUUUUUAUGGCAAAUCAGGUAAU… |
| env-410 | GUACAGUCACCGCUUGUCAAUUUCUUUUAUAAAGAUUGAAAAGGAGGCGGCUUUUUUUUAUGGCAAAUCAGGUAAUGAGAAUCACAUUAAAAGCUUAUGAUCACCAAUUGGUUGACAGCUGUGCUGCAAGAAUUGUUGAUACUGUAAAGAAGACAGGAGCAGCUGUUUCAGGACCUGUUCC |
| Cba-3-1 | GUACAGUCACCACUUGUCAAUUUCUUUUAUAAAGAUUGAAAAGGAGGCGGCUUUUUUUUAUGGCAAAUCAGGUAAUGAGAAUCACAUUAAAAGCUUAUGAUCAUCAAUUAGUUGACAGCUGUGCUGCAAGAAUUGUUGAUACUGUAAAGAAGACAGGAGCAGCUGUUUCAGGACCUGUUCC |
| env-411 | UACAGUCACCGCUUGUCAAUUUCUUUUAUAAAGAUUGAAAAGGAGGCGGCUUUUUUUUAUGGCAAAUCAGGUAAUGAGAAUCACAUUAAAAGCUUAUGAUCACCAAUUGGUUGACAGCUGUGCUGCAAGAAUUGUUGAUACUGUAAAGAAGACAGGAGCAGCUGUUUCAGGACCUGUCCA |
| env-414 | GGAGAUGUGUACAGUCACCGCUUGUCGAACUUUGUUUAAAAGUACGAAAAGGAGGCGACUUUUUUUUAUG… |
| env-415 | GGAGAUGUGUACAGUCACCGCUUGUCGAACUUUGUUUAAAAGUACGAAAAGGAGGCGACUUUUUUUUAUG… |
| env-416 | GGAGAUGUGUACAGUCACCGCUUGUCGAACUUUGUUUAAAAGUACGAAAAGGAGGCGACUUUUUUUUAUGGCAAGUCAAGUUAUGAGAAUUAUACUAAAGGCGUACGACCACCAACUGCUUGAUAGCUGUGCUCUUAAGAUAGUUGACACAGUUAAGAAGGCGGGUGCUAAGGUUUCAGGU |
| env-417 | … |
| env-420 | … |
| env-426 | GGAGAUGUGUACAGUCACCGCUUGUCGAACUUUGUUUAAAAGUACGAAAAGGAGGCGACUUUUUUUUAUGGCAAGUCAAGUUAUGAGAAUUAUACUAAAGGCGUACGACCACCAACUGCUUGAUAGCUGUGCUCUUAAGAUAGUUGACACAGUUAAGAAGGCGGGUGCUAAGGUUUCAGGU |
| env-427 | GGAGAUGUGUACAGUCACCGCUUGUCGAACUUUGUUUAAAAGUACGAAAAGGAGGCGACUUUUUUUUAUG… |
| env-428 | AAAGGAUCUUUCAGGGUUGCUUAUCAAGAGGACACACACGGGAACGUGUACAGUCACCGCUUGUCGCACUUAUCUCAAAAGCGCGAAAAGGAGGCGACUUUUUUUAUGGCAAAUCAGGUAAUGAGGAUUACACUGAAAGCUUACGAUCAUCAGUUAGUUGAUGCGUCUGCGAAGAAAAUC |
| env-429 | AAAGAAUCUUUCAGGGUUGCUUAUCAAGAGGACACACACGGGA… |
| env-430 | AAAGGAUCUUUCAGGGUUGCUUAUCAAGAGGACACACACGGGAACGUGUACAGUCACCGCUUGUCGCACUUAUCUCAAAAGCGCGAAAAGGAGGCGACUUUUUUUAUGGCAAAUCAGGUAAUGAGGAUUACACUGAAAGCUUACGAUCAUCAGUUAGUUGAUGCGUCUGCGAAGAAAAUC |
| env-431 | AAAGGAUCUUUCAGGGUUGCUUAUCAAGAGGACACACACGGGAACGUGUACAGUCACCGCUUGUCGCACUUAUCUCAAAAGCGCGAAAAGGAGGCGACUUUUUUUAUGGCAAAUCAGGUAAUGAGGAUUACACUGAAAGCUUACGAUCAUCAGUUAGUUGAUGCGUCUGCGAAGAAAAUC |
| | … |
| | … |
| | …R… |

| | |
|---|---|
| env-432 | AAAGGAUCUUUCAGGGUUGCUUAUCAAGAGGACACACACGGGAACGUGUACAGUCACCGCUUGUCGCACUUAUCUCAAAAGCGCGAAAAGGAGGCGACUUUUUUUUAU.......................................................................... |
| env-433 | AAAGGAUCUUUCAGGGUUGCUUAUCAAGAGGACACACACGGGAACGUGUACAGUCACCGCUUGUCGCACUUAUCUCAAAAGCGCGAAAAGGAGGCGACUUUUUUUUAUGGCAAAUCAGGUAAUGAGGAUUACACUGAAAGCUUACGAUCAUCAGUUAGUUGAUGCGUCUGCGAAGAAAAUC |
| env-434 | AAAGGAUCUUUCAGGGUUGCUUAUCAAGAGGACACACACGGGAACGUGUACAGUCACCGCUUGUCGCACUUAUCUCAAAAGCGCGAAAAGGAGGCGACUUUUUUUUAUGGCAAAUCAGGUAAUGAGGAUUACACUGAAAGCUUACGAUCAUCAGUUAGUUGAUNNNNNNNNNNNNNNNNNN |
| env-435 | AAAGGAUCUUUCAGGGUUGCUUAUCAAGAGGACACACACGGGAACGUGUACAGUCACCGCUUGUCGCACUUAUCUCAAAAGCGCGAAAAGGAGGCGACUUUUUUUUAUGGCAAAUCAGGUAAUGAGGAUUACACUGAAAGCUUACGAUCAUCAGUUAGUUGAUGCGUCUGCGAAGAAAAUC |
| env-436 | ........................................................................................................................................................................................ |
| env-454 | ........................................................................................................................................................................................ |
| env-455 | AAAAGAUCUUUCAGGGUUGCUUAUCAAGAGGACACACACGGGAACGUGUACAGUCACCGCUUGUCGCACUUAUCUCAAAAGCGCGAAAAGGAGGCGACUUUUUUUUAUGGCAAAUCAGGUAAUGAGGAUUACACUGAAAGCUUACGAUCAUCAGUUAGUUGAUGCGUCUGCGAAGAAAAUC |
| env-456 | AAAGGAUCUUUCAGGGUUGCUUAUCAAGAGGACACACACGGGAACGUGUACAGUCACCGCUUGUCGCACUUAUCUCAAAAGCGCGAAAAGGAGGCGACUUUUUUUUAUGGCAAAUCAGGUAAUGAGGAUUACACUGAAAGCUUACGAUCAUCAGUUAGUUGAUGCGUCUGCGAAGAAAAUC |
| env-457 | AAAGGAUCUUUCAGGGUUGCUUAUCAAGAGGACACACACGGGAACGUGUACAGUCACCGCUUGUCGCACUUAUCUCAAAAGCGCGAAAAGGAGGCGACUUUUUUUUAUGGCAAAUCAGGUAAUGAGGAUUACACUGAAAGCUUACGAUCAUCAGUUAGUUGAUGCGUCUGCGAAGAAAAUC |
| env-459 | AAA..................................................................................................................................................................................... |
| env-460 | AAAAGAUCUUUCA........................................................................................................................................................................... |
| env-464 | AAA..................................................................................................................................................................................... |
| env-465 | AAAGGAUCUUUCAGGGUUGCUUAUCAAGAGGACACACACGGGAACGUGUACAGUCACCGCUUGUCGCACUUAUCUCAAAAGCGCGAAAAGGAGGCGACUUUUUUUUAUGGCAAAUCAGGUAAUGAGGAUUACACUGAAAGCUUACGAUCAUCAGUUAGUUGAUGCGUCUGCGAAGAAAAUC |
| env-466 | AAAGGAUCUUUCAGGGUUGCUUAUCAAGAGGACACACACGGGAACGUGUACAGUCACCGCUUGUCGCACUUAUCUCAAAAGCGCGAAAAGGAGGCGACUUUUUUUUAUGGCAAAUCAGGUAAUGAGGAUUACACUGAAAGCUUACGAUCAUCAGUUAGUUGAUGCGUCUGCGAAGAAAAUC |
| env-467 | NNNNNNNNNNNNNNNNNNNNNNNNNNNNNNNNNNNNNNNNNNNNNNNNNNNNNNNNNNNNNNNNNNNNNNNNNNNNNNNNNNNNNNNNAAGAGGACACACACGGGAACGUGUACAGUCACCGCUUGUCGCACUUAUCUCAAAAGCGCGAAAAGGAGGCGACUUUUUUUUAUGGCAAA |
| env-468 | AAGGAUCUUUCAGGGUUGCUUAUCAA.............................................................................................................................................................. |
| env-469 | AAAGGAUCUUUCAGGGUUGCUUAUCAAGAGGACACACACGGGAACGUGUACAGUCACCGCUUGUCGCACUUAUCUCAAAAGCGCGAAAAGGAGGCGACUUUUUUUUAU.......................................................................... |
| env-470 | AAAGGAUCUUUCAGGGUUGCUUAUCAAGAGGACACACACGGGAACGUGUACAGUCACCGCUUGUCGCACUUAUCUCAAAAGCGCGAAAAGGAGGCGACUUUUUUUUAUGGCAAAUCAGGUAAUGAGGAUUACACUGAAAGCUUACGAUCAUCAGUUAGUUGAUGCGUCUGCGAAGAAAAUC |
| env-471 | AAAGGAUCUUUCAGGGUUGCUUAUCAAGAGGACACACACGGGAACGUGUACAGUCACCGCUUGUCGCACUUAUCUCAAAAGCGCGAAAAGGAGGCGACUUUUUUUUAUGGCAAAUCAGGUAAUGAGGAUUACACUGAAAGCUUACGAUCAUCAGUUAGUUGAUGCGUCUGCGAAGAAAAUC |
| env-472 | AAAGGAUCUUUCAGGGUUGCUUAUCAAGAGGACACACACGGGAACGUGUACAGUCACCGCUUGUCGCACUUAUCUCAAAAGCGCGAAAAGGAGGCGACUUUUUUUUAUGGCAAAUCAGGUAAUGAGGAUUACACUGAAAGCUUACGAUCAUCAGUUAGUUGAUGCGUCUGCGAAGAAAAUC |
| env-473 | ........................................................................................................................................................................................ |
| env-482 | AAAGGAUCUUUCAGGGUUGCUUAUCAAGAGGACACACACGGGAACGUGUACAGUCACCGCUUGUCGCACUUAUCUCAAAAGCGCGAAAAGGAGGCGACUUUUUUUUAUGGCAAAUCAGGUAAUGAGGAUUACACUGAAAGCUUACGAUCAUCAGUUAGUUGAUGCGUCUGCGAAGAAAAUC |
| env-483 | AAAGGAUCUUUCAGGGUUGCUUAUCAAGAGGACACACACGGGAACGUGUACAGUCACCGCUUGUCGCACUUAUCUCAAAAGCGCGAAAAGGAGGCGACUUUUUUUUAUGGCAAAUCAGGUAAUGAGGAUUACACUGAAAGCUUACGAUCAUCAGUUAGUUGAUGCGUCUGCGAAGAAAAUC |
| env-484 | AAAGGAUCUUUCAGGGUUGCUUAUCAAGAGGACACACACGGGAACGUGUACAGUCACCGCUUGUCGCACUUAUCUCAAAAGCGCGAAAAGGAGGCGAC.................................................................................... |
| env-485 | AAAGGAUCUUUCAGGGUUGCUUAUCAAGAGGACACACACGGGAACGUGUACAGUCACCGCUUGUCGCACUUAUCUCAAAAGCGCGAAAAGGAGGCGACUUUUUUUUAUGGCAAAUCAGGUAAUGAGGAUUACACUGAAAGCUUACGAUCAUCAGUUAGUUGAUGCGUCUGCGAAGAAAAUC |
| env-486 | AAAAGAUCUUUCAGGGUUGCUUAUCAAGAGGACACACACGGGAACGUGUACAGUCACCGCUUGUCGCACUUAUCUCAAAAGCGCGAAAAGGAGGCGACUUUUUUUUAU.......................................................................... |
| env-487 | AAAGGAUCUUUCAGGGUUGCUUAUCAAGAGGACACACACGGGAACGUGUACAGUCACCGCUUGUCGCACUUAUCUCAAAAGCGCGAAAAGGAGGCGACUUUUUUUUAUGGCAAAUCAGGUAAUGAUGAUUACACUGAAAGCUUACGAUCAUCAGUUAGUUGAUGCGUCUGCGAAGAAAAUC |
| env-488 | AAAGGAUCUUUCAGGGUUGCUUAUCAAGAGGACACACACGGG.............................................................................................................................................. |
| env-489 | GAUACGCGUGGGAGUGUGUACAGU................................................................................................................................................................ |
| env-494 | GAUACGCGUGGGAGUGUGUACAGUCACCGCUUGUCGUACUUAAACAAUAAAGUGCGAAGGAGGCGACUUUUUUUAUGGCAAAUCAGGUAAUUAGAAUUACACUGAAGGCUUAUGAUCAUCAGUUAAUCGAUCAGUCAGCAAAGAAAAUCAUUGAAACUGCUAAGAAAACAGGAGCACAGG |
| env-495 | GAUACGCGUGGGAGUGUGUACAGUCACCGCUUGUCGUACUUAAAUCUAGAAAGUGCGAAGGAGGCGACUUUUUUUUAUGGCAAAUCAGGUAAUUAGAAUUACACUGAAGGCUUAUGAUCAUCAGUUAAUCGAUCAGUCAGCAAAGAAA.................................. |
| env-496 | GAUACGCGUGGGAGUGUGUACAGUCACCGCUUGUCGUACUUAAAUCUAGAAAGUGCGAAGGAGGCGACUUUUUUUUAUGGCAAAUCAGGUAAUUAGAAUUACACUGAAGGCUUAUGAUCAUCAGUUAAUCGAUCAGUCAGCAAAGAAAAUCAUUGAAACUGCUAAGAAAACAGGAGCACAG |
| env-497 | GAUACGCGUGGGAGUGUGUACAGUCACCGCUUGUCGUACUUAAAUCUAGAAAGUGCGAAGGAGGCGACUUUUUUUUAUGGCAAAUCAGGUAAUUAGAAUUACACUGAAGGCUUAUGAUCAUCAGUUAAUCGAUCAGUCAGCAAAGAAAAUCAUUGAAACUGCUAAGAAAACAGGAGCACAG |
| env-498 | GAUACGCGUGGGAGUGUGUACAGUCACCGCUUGUCGUGCUUAAACAAUAAAGUGCGAAGGAGGCGACUUUUUUUAUGGCAAAUCAGGUAAUUAGAAUUACACUGAAAGCUUAUGAUCAUCAGUUAAUCGAUCAGUCAGCAAAGAAAAUCAUUGAAACUGCUAAGAAAACAGGAGCACAGG |
| env-499 | GAUACGCGUGGGAGUGUGUACAGU................................................................................................................................................................ |
| env-500 | GAUACGCGUGGGAGUGUGUACAGUCACCGCUUGUCGUACUUAAACAAUAAAGUGCGAAGGAGGCGACUUUUUUUAUGGCAAAUCAGGUAAUUAGAAUUACACUGAAGGCUUAUGAUCAUCAGUUAAUCGAUCAGUCAGCAAAGAAAAUCAUUGAAACUGCUAAGAAAACAGGAGCACAGG |
| env-501 | GAUACGCGUGGGAGUGUGUACAGUCACCGCUUGUCGUACUUAAAUCUAGAAAGUGCGAAGGAGGCGACUUUUUUUAUGGCAAAUCAGGUAAUUAGAAUUACACUGAAGGCUUAUGAUCAUCAGUUAAUCGAUCAGUCAGCAAAGAAAAUCAUUGAAACUGCUAAGAAAACAGGAGCACAGG |
| env-502 | GAUACGCGUGGGAGUGUGUACAGU................................................................................................................................................................ |
| env-503 | GAUACGCGUGGGAGUGUGUACAGUCACCGCUUGUCGUACUUAAAUCUAGAAAGUGCGAAGGAGGCGACUUUUUUUUAUGGCAAAUCAGGUAAUUAGAAUUACACUGAAGGCUUAUGAUCAUCAGUUAAUCGAUCAGUCAGCAAAGAAAAUCAUUGAAACUGCUAAGAAAACAGGAGCACAG |
| env-512 | GAGAUAAGAUAAGAAACACACGGGAAAGUUGUACAGUCACCACUUGUCGUACUACAAAAAGUGCGAAAAGGAGGCGGCUUUUUUUUAUGGCAAGUCAAGUAAUGAGAAUCACAUUGAAGGCGUACGAUCACAAGUUAAUCGAUCAGGCUGCAAAGCAGAUCAUCGAGACGAUCAAGAGUAC |
| env-513 | GAGAUAAGAUAAGAAACACACGGGAAAGUUGUACAGUCACCACUUGUCGUACUACAAAAAGUGCGAAAAGGAGGCGGCUUUUUUUUAUGGCAAGUCAAGUAAUGAGAAUCACAUUGAAGGCGUACGAUCACAAGUUAAUCGAUCAGGCUGCAAAGCAGAUCAUCGAGACGAUCAAGAGUAC |
| env-514 | GAGAUAAGAUAAGAAACACACGGGAAAGUUGUACAGUCACCACUUGUCGUACUACAAAAGGUGCGAAAAGGAGGCGGCUUUUUUUUAUGGCAAGUCAAGUAAUGAGAAUCACAUUGAAGGCGUACGAUCACAAGUUAAUCGAUCAGGCUGCAAAGCAGAUCAUCGAGACGAUCAAGAGUAC |
| env-515 | GAGAUAAGAUAAGAAACACACGGGAAAGUUGUACAGUCACCACUUGUCGUACUACAAAAAGUGCGAAAAGGAGGCGGCUUUUUUUUAUGGCAAAGCAAGUAAUGAGAAUCACAUUGAAGGCGUACGAUCACAAGUUAAUCGAUCAGGCUGCAAAGCAGAUCAUCGAGACGAUCAAGAGUAC |
| env-516 | GAGAUAAGAUAAGAAACACACGGGAAAGUUGUACAGUCACCACUUGUCGUACUACAAAAAGUGCGAAAAGGAGGCGGCUUUUUUUAUGGCAAGUCAAGUAAUGAGAAUCACAUUGAAGGCGUACGAUCACAAGUUAAUCGAUCAGGCUGCAAAGCAGAUCAUCGAGACGAUCAAGAGUAC |
| env-517 | GAGAUAAGAUAAGAAACACACGGGAAAGUUGUACAGUCACCACUUGUCGUACUACAAAAAGUGCGAAGAGGAGUCGGCUUUUUUUUAUGGCAAGUCAAGUAAUGAGAAUCACAUUGAAGGCGUACGAUCACAAGUUAAUCGAUCAGGCUGCAAAGCAGAUCAUCGAGACGAUCAAGAGUAC |
| env-518 | GAGAUAAGAUAAGAAACACACGGGAAAGUUGUACAGUCACCACUUGUCGUAC.................................................................................................................................... |
| env-556 | GAGAUAAGAUAAGAAACACACGGGAAAGUUGUACAGUCACCACUUGUCGUACUACAAAAAGUGCGAAAAGGAGGCGGCUUUUUUUUAUGGCAAGUCAAGUAAU................................................................................ |
| env-557 | GAGAUAAGAUAAGAAACACACGGGAAAGUUGUACAGUCACCACUUGUCGUACUACAAAAAGUGCGAAAAGGAGGCGGCUUUUUUUUAUGGCAAGUCAAGUAAUGAGAAUCACAUUGAAGGCGUACGAUCACAAGUUAAUCGAUCAGGCUGCAAAGCAGAUCAUCGAGACGAUCAAGAGUAC |
| env-558 | GAGAUAAGAUAAGAAACACACGGGAAAGUUGUACAGUCACCACUUGUCGUACUAAAAAAAGAGCGAAAAGGAGGCGGCUUUUUUUAUGGCAAGUCAAGUAAUGAGAAUCACAUUGAAGGCGUACGAUCACAAGUUAAUCGAUCAGGCUGCAAAGCAGAUCAUCGAGACGAUCAAGAGUAC |
| env-559 | GAGAUAAGAUAAGAAACACACGGGAAAGUUGUACAGUCACCACUUGUCGUACUACAAAAAGUGCGAAAAGGAGGCGGCUUUUUUUAUGGCAAGUCAAGUAAUGAGAAUCACAUUGAAGGCGUACGAUCACAAGUUAAUCGAUCAGGCUGCAAAGCAGAUCAUCGAGACGAUCAAGAGUAC |
| env-560 | GAGAUAAGAUAAGAAACACACGGGAAAGUUGUACAGUCACCACUUGUCGUACUACAAAAAGUGCGAAAAGGAGGCGGCUUU....................................................................................................... |
| env-561 | GAGAUAAGAUAAGAAACACACGGGAAAGUUGUACAGUCACCACUUGUCGUACUACAAAAAGUGUGAGAAAAGGAGGCGGCUUUUUUUAUGGCAAGUCAAGUAAUGAGAAUCACAUUGAAGGCGUACGAUCACAAGUUAAUCGAUCAGGCUGCAAAGCAGAUCAUCGAGACGAUCAAGAGUAC |
| env-562 | AGAGUGUACAGUCACCGCUUGUCGUACUGGAAAUUAGAAAGUACGAAAAGGAGGCGACUUUUUUUAUGGCAAGUCAAGUAAUGAGAAUCACAUUAAAAGCAUAUGAACAUCAAUUAGUGAGUGCAUCUGCGAAGAAGAUUAUCGAAACUGUAAAGAAGAACGGAGCACAAGUGAGUGGGC |
| env-563 | GAGUGUACAGUCACCGCUUGUCGUACUAGAAAUUUAGAAAGUACGAAAAGGAGGCGACUUUUUUUAUGGCAAGUCAAGUAAUGAGAAUCACAUUGAAAGCAUAUGAACAUCAAUUAGUGAUGCAUCUGCACAAAAAAUCAUCGAAACUGUAAAGAAAAACGGAUCACAAGUGAGUGGAC |
| env-564 | GAGUGUACAGUCACCGCUUGUCGUACUAGAAAUUUAGAAAGUACGAAAAGGAGGCGACUUUUUUUAUGGCAAGUCAAGUAAUGAGAAUCACAUUGAAAGCAUAUGAACAUCAAUUAGUGAUGCAUCUGCACAAAAAAUCAUCGAAACUGUAAAGAAAAACGGAUCACAAGUGAGUGGAC |
| env-565 | AGAGUGUACAGUCACCGCUUGUCGUACUGGAAAUUAGAAAGUACGAAAAGGAGGCGACUUUUUUUAUGGCAAGUCAAGUAAUGAGAAUCACAUUAAAAGCAUAUGAACAUCAAUUAGUGAGUGCAUCUGCGAAGAAGAUUAUCGAAACUGUAAAGAAGAACGGAGCACAAGUGAGUGGGC |
| Bsp-4-1 | GCCUUACUUUGUGAUAAGGGUCAGGACACACACGGGAGAGUGUACAGUCACCGCUUGUCGUACUUAGAUCAAAAAGUGCGAAAAGGAGGCGACUUUUUUUUAUGGCAAAUCAGGUAAUGAGAAUUACUCUUAAGGCUUAUGAUCAUCAGCUUGUUGAUGCAUCAGCCAAGAAGAUCAUCGA |
| Bsp-7-1 | GCCUUACUUUGUGAUAAGGGUCAGGACACACACGGGAGAGUGUACAGUCACCGCUUGUCGUACUUAGAUCAAAAAGUGCGAAAAGGAGGCGACUUUUUUUAUGGCAAAUCAGGUAAUGAGAAUUACUCUUAAGGCUUAUGAUCAUCAGCUUGUUGAUGCAUCAGCCAAGAAGAUCAUCGA |
| Bsp-10-1 | GCCUUACUUUGUGUUAAGGGUCAGGACACACACGGGAGAGUGUACAGUCACCGCUUGUCGUACUUAGAUCAAAAAGUGCGAAAAGGAGGCGACUUUUUUUUAUGGCAAAUCAGGUAAUGAGAAUUACUCUUAAGGCUUAUGAUCAUCAGCUCGUUGAUGCAUCAGCCAAGAAGAUCAUCGA |
| env-566 | GCCUUACUUUGUGAUAAGGGUCAGGACACACACGGGAGAGUGUACAGUCACCGCUUGUCGUACUUAGAUCAAAAAGUGCGAAAAGGAGGCGACUUUUUUUAUGGCAAAUCAGGUAAUGAGAAUUACUCUUAAGGCUUAUGAUCAUCAGCUUGUUGAUGCAUCAGCCAAGAAGAUCAUCGA |
| Bsp-2-1 | UUAUUAUGUAUUUGAGGUUAGGACACACACGGGAGAGUGUACAGUCACCGCUUGUCGUACUUCAUUAAAAGUGCGAAAAGGAGGCGACUUUUUUUAUGGCAAAUCAGGUAAUGAGAAUUACACUUAAGGCAUAUGAUCAUCAGCUCGUUGAUGCAUCUGCCAAGAAGAUUAUCGAGACAG |
| Bsp-3-1 | UCUUAUUUUGCGAUGAGGUUCUGGACACACACGGGAGAGUGUACAGUCACCGCUUGUCGUACUUAUUCAUAAAAGUGCGAAAAGGAGGCGACUUUUUUUUAUGGCAAAUCAGGUAAUGAGAAUUACACUUAAGGCUUAUGAUCAUCAGCUUGUUGAUGCAUCUGCCAAGAAGAUUAUCGAG |
| Bsp-6-1 | UCUUAUUUUGCGAUGAGGUUCUGGACACACACGGGAGAGUGUACAGUCACCGCUUGUCGUACUUAUUCAUAAAAAGUGCGAAAAGGAGGCGACUUUUUUUUAUGGCAAAUCAGGUAAUGAGAAUUACACUUAAGGCUUAUGAUCAUCAGCUUGUUGAUGCAUCUGCCAAGAAGAUUAUCGAG |
| env-567 | GUUGCACAAAGGUCGCAGACACACACGGGAAAGUGUACAGUCACCGCUUGUCCGUACAAAAAGUAAGUAUUAACUUACAAAAGUGCGAAAAGGAGGCGACUUUUUUUAUGGCAAAUCAGGUAAUGAGAAUUACAUUGAAAGCUUAUGAACAUCAGCUUGUUGAUGCAUCUGCCAAAAAAA |
| env-568 | GUUGCACAAAGGUCGCAGACACACACGGGAAAGUGUACAGUCACCGCUUGUCCGUACAAAAAGUAAGUAUCUACUUACAAAAGUGCGAAAAGGAGGCGACUUUUUUUUAUGGCAAAUCAGGUAAUGAGAAUUACAUUGAAAGCUUAUGAACAUCAGCUUGUUGAUGCAUCUGCCAAAAAAA |
| env-570 | GUUGCACAAAGGUCGCAGACACACACGGGAAAGUGUACAGUCACCGCUUGUCCGUACAAAAAGUAAGUAUUUACUUACAAAAGUGCGAAAAGGAGGCGACUUUUUUUAUGGCAAGUCAAGUAAUGAGAAUUACAUUGAAAGCUUAUGAACAUCAGCUUGUUGAUGCAUCUGCCAAAAAAA |
| env-571 | GUUGCACAAAGGUCGCAGACACACACGGGAAAGUGUACAGUCACCGCUUGUCCGUACAAAAAGUAAGUAUUUACUUACAAAAGUGCGAAAAGGAGGCGACUUUUUUUAUGGCAAAUCAGGUAAUGAGAAUUACAUUGAAAGCUUAUGAACAUCAGCUUGUUGAUGCAUCUGCCAAAAAAA |
| env-573 | GUUGCACAAAGGUCGCAGACACACACGGGAAAGUGUACAGUCACCGCUUGUCCGUACAAAAAGUAAGUAUUUACUUACAAAAGUGCGAAAAGGAGGCGACUUUUUUUAUGGCAAAUCAGGUAAUGAGAAUUACAUUGAAAGCUUAUGAACAUCAGCUUGUUGAUGCAUCUGCCAAAAAAA |
| env-578 | GUUGCACAAAGGUCGCAGACACACACGGGAAAGUGUACAGUCACCGCUUGUCCGUACAAAAAGUAAGUAUUUACUUACAAAAGUGCGAAAAGGAGGCGACUUUUUUUAUGGCAAAUCAGGUAAUGAGAAUUACAUUGAAAGCUUAUGAACAUCAGCUUGUUGAUGCAUCUGCCAAAAAAA |
| env-579 | GUUGCACAAAGGUCGCAGACACACACGGGAAAGUGUACAGUCACCGCUUGUCCGUACAAAAAGUAAGUAUUAACUUACAAAAGUGCGAAAAGGAGGCGACUUUUUUUAUGGCAAAUCAGGUAAUGAGAAUUACAUUGAAAGCUUAUGAACAUCAGCUUGUUGAUGCAUCUGCCAAAAAAA |
| env-581 | GUUGCACAAAGGUCGCAGACACACACGGGAAAGUGUACAGUCACCGCUUGUCCGUACAAAAAGUAAGUAUUUACUUACAAAAGUGCGAAAAGGAGGCGACUUUUUUUAUGGCAAAUCAGGAUAAGAGAAUUACAUUGAAAGCUUAUGAACAUCAGCUUGUUGAUGCAUCUGCCAAAAAAA |
| env-582 | GUUGCACAAAGGUCGCAGACACACACGGGAAAGUGUACAGUCACCGCUUGUCCGUACAAAAAGUAAGUAUUAACUUACAAAAGUGCGAAAAGGAGGCGACUUUUUUUAUGGCAAAUCAGGUAAUGAGAAUUACAUUGAAAGCUUAUGAACAUCAGCUUGUUGAUGCAUCUGCCAAAAAAA |
| env-583 | GUUGCACAAAGGUCGCAGACACACACGGGAAAGUGUACAGUCACCGCUUGUCCGUACAAAAAGUAAGUAUUUACUUACAAAAGUGCGAAAAGGAGGCGACUUUUUUUAUGGCAAAUCAGGUAAUGAGAAUUACAUUGAAAGCUUAUGAACAUCAGCUUGUUGAUGCAUCUGCCAAAAAAA |
| env-584 | GUUGCACAAAGGUCGCAGACACACACGGGAAAGUGUACAGUCACCGCUUGUCCGUACAAAAAGUAAGUAUUUACUUACAAAAGUGCGAAAAGGAGGCGACUUUUUUUAUGGCAAAUCAGGUAAUGAGAAUUACAUUGAAAGCUUAUGAACAUCAGCUUGUUGAUGCAUCUGCCAAAAAAA |
| env-585 | GUUGCACAAAGGUCGCAGACACACACGGGAAAGUGUACAGUCACCGCUUGUCCGUACAAAAAGUAAGUAUUAACUUACAAAAGUGCGAAAAGGAGGCGACUU................................................................................ |
| env-586 | GUUGCACAAAGGUCGCAGACACACACGGGAAAGUGUACAGUCACCGCUUGUCCGUACAAAAAGUAAGUAUCUACUUACAAAAGUGCGAAAAGGAGGCGACUUUUUUUUAUGGCAAAUCAGGUAAUGAGAAUUACAUUGAAAGCUUAUGAACAUCAGCUUGUUGAUGCAUCUGCCAAAAAAA |
| env-587 | GUUGCACAAAGGUCGCAGACACACACGGGAAAGUGUACAGUCACCGCUUGUCCGUACAAAAAGUAAGUAUCUACUUACAAAAGUGCGAAAAGGAGGCGACUUUUUUUAUGGCAAAUCAGGUAAUGAGAAUUACAUUGAAAGCUUAUGAACAUCAGCUUGUUGAUGCAUCUGCCAAAAAAA |
| env-588 | GUUGCACAAAGGUCGCAGACACACACGGGAAAGUGUACAGUCACCGCUUGUCCGUACAAAAAGUAAGUAUUAACUUACAAAAGUGCGAAAAGGAGGCGACUUUUUUUAUGGCAAAUCAGGUAAUGAGAAUUACAUUGAAAGCUUAUGAACAUCAGCUUGUUGAUGCAUCUGCCAAAAAAA |
| env-589 | GUUGCACAAAGGUCGCAGACACACACGGGAAAGUGUACAGUCACCGCUUGUCCGUACAAAAAGUAAGUAUCUACUUACAAAAGUGCGAAAAGGAGGCGACUUUUUUUAUGGCAAAUCAGGUAAUGAGAAUUACAUUGAAAGCUUAUGAACAUCAGCUUGUUGAUGCAUCUGCCAAAAAAA |
| env-590 | GUUGCACAAAGGUCGCAGACACACACGGGAAAGUGUACAGUCACCGCUUGUCCGUACAAAAAGUAAGUAUUAACUUACAAAAGUGCGAAAAGGAGGCGACUUUUUUUUAUGGCAAGUCAAGUAAUGAGAAUUACAUUGAAAGCUUAUGAACAUCAGCUUGUUGAUGCAUCUGCCAAAAAA |
| env-591 | NNNNNNNNNNNNNNNNNNNNNNNNNNNNNNNNNNNNNNNNNNNNNNNNNNNNNNNNNNNNNNNNNNNNNNNNNNNNCGGAGAGAAGAAAGCGUUGCACAAAGGUCGCAGACACACACGGGAAAGUGUACAGUCACCGCUUGUCCGUACAAAAAGUAAGUAUUUACUUACAAAAGU |
| env-592 | GUUGCACAAAGGUCGCAGACACACACGGGAAAGUGUACAGUCACCGCUUGUCCGUACAAAAAGUAAGUAUCUACUUACAAAAGUGCGAAAAGGAGGCGACUUUUUUUUAUGGCAAAUCAGGUAAUGAGAAUUACAUUGAAAGCUUAUGAACAUCANNNNNNNNNNNNNNNNNNNNNNNNNNNNN |
| env-593 | GUUGCACAAAGGUCGCAGACACACACGGGAAAGUGUACAGUCACCGCUUGUCCGUACAAAAAGUAAGUAUUUACUUACAAAAGUGCGAAAAGGAGGCGACUUUUUUUAUGGCAAAUCAGGUAAUGAGAAUUACAUUGAAAGCUUAUGAACAUCAGCUUGUUGAUGCAUCUGCCAAAAAAA |
| env-594 | GUUGCACAAAGGUCGCAGACACACACGGGAAAGUGUACAGUCACCGCUUGUCCGUACAAAAAGUAAGUAUUUACUUACAAAAGUGCGAAAAGGAGGCGACUUUUUUUAUGGCAAAUCAGGUAAUGAGAAUUACAUUGAAAGCUUAUGAACAUCAGCUUGUUGAUGCAUCUGCCAAAAAAA |
| | ........................................................................................................................................................................................ |
| | ........................................................................................................................................................................................ |
| | ....................................................................................................................................................................R................... |

```
env-595   GUUGCACAAAGGUCGCAGACACACACGGGAAAGUGUACAGUCACCGCUUGUCCGUACAAAAAGUAAGUAUUUACUUACAAAAGUGCGAAAAGGAGGCGACUUUUUUUAUGGCAAAUCAGGUAAUGAGAAUUACAUUGAAAGCUUAUGAACAUCAGCUUGUUGAUGCAUCUGCCAAAAAAA
env-596   GUUGCACAAAGGUCGCAGACACACACGGGAAAGUGUACAGUCACCGCUUGUCCGUACAAAAAGUAAGUAUCUACUUACAAAAGUGCGAAAAGGAGGCGACUUUUUUUAUGGCAAAUCAGGUAAUGAGAAUUACAUUGAAAGCUUAUGAACAUCAGCUUGUUGAUGCAUCUGCCAAAAAAA
env-597   GUUGCACAAAGGUCGCAGACACACACGGGAAAGUGUACAGUCACCGCUUGUCCGUACAAAAAGUAAGUAUUUACUUACAAAAGUGCGAAAAGGAGGCGACUUUUUUUAUGGCAAAUCAGGUAAUGAGAAUUACAUUGAAAGCUUAUGAACAUCAGCUUGUUGAUGCAUCUGCCAAAAAAA
env-598   GUUGCACAAAGGUCGCAGACACACACGGGAAAGUGUACAGUCACCGCUUGUCCGUACAAAAAGUAAGUAUUAACUUACAAAAGUGCGAAAAGGAGGCGACUUUUUUUAUGGCAAAUCAGGUAAUGAGAAUUACAUUGAAAGCUUAUGAACAUCAGCUUGUUGAUGCAUCUGCCAAAAAAA
env-599   GUUGCACAAAGGUCGCAGACACACACGGGAAAGUGUACAGUCACCGCUUGUCCGUACAAAAAGUAAGUAUUAACUUACAAAAGUGCGAAAAGGAGGCGACUUUUUUUAUGGCAAAUCAGGUAAUGAGAAUUACAUUGAAAGCUUAUGAACAUCAGCUUGUUGAUGCAUCUGCCAAAAAAA
env-600   GUUGCACAAAGGUCGCAGACACACACGGGAAAGUGUACAGUCACCGCUUGUCCGUACAAAAAGUAAGUAUUUACUUACAAAAGUGCGAAAAGGAGGCGACUUUUUUUAUGGCAAAUCAGGUAAUGAGAAUUACAUUGAAAGCUUAUGAACAUCAGCUUGUUGAUGCAUCUGCCAAAAAAA
env-601   GUUGCACAAAGGUCGCAGACACACACGGGAAAGUGUACAGUCACCGCUUGUCCGUACAAAAAGUAAGUAUCUACUUACAAAAGUGCGAAAAGGAGGCGACU.................................................................................
env-602   GUUGCACAAAGGUCGCAGACACACACGGGAAAGUGUACAGUCACCGCUUGUCCGUACAAAAAGUAAGUAUUAACUUACAAAAGUGCGAAAAGGAGGCGACUUUUUUUAUGGCAAAUCAGGUAAUGAGAAUUACAUUGAAAGCUUAUGAACAUCAGCUUGUUGAUGCAUCUGCCAAAAAAA
env-603   CAUACGAUUAUUGACAGUUCUGACAGACAGCGCUUAUCAUAUAAUCCUAGGAGACCGGAUUAUACGUAUGGAAAACGAAGAUACACACGGAUGAGUGUACAGUCACCACUUGUCGUACUGAUAUUAAAGUGCGAAAAGGAGGCGGCUUUUUUUUAUGGCAAAUCAAAUUAUGAGAAUCAC
env-604   GGAAAUGUGUACAGUCACCGCUUGUCGUAC...........................................................................................................................................................
env-606   GGAAAUGUGUACAGUCACCGCUUGUCGUACUUAAAUCAAAAAGUGCGAAAAGGAGGCGACUUUUUUUAUGGCAAAUCAGGUAAUGAGAAUCACAUUAAAGGCUUAUGACCAUCAAUUGGUAGACAAUUCAGCUAGUAAAAUUGUCGAAACUGUAAAGAAGACAGGAGCAAAGGUGUCAGG
env-607   GGAAAUGUGUACAGUCACCGCUUGUCGUACUUAAAUCAAAAAGUGCGAAAAGGAGGCGACUUUUUUUAUGGCAAAUCAGGUAAUGAGAAUCACAUUAAAGGCUUAUGACCAUCAAUUGGUAGACAAUUCAGCUAGUAAAAUUGUCGAAACUGUAAAGAAGACAGGAGCAAAGGUGUCAGG
env-608   GGAAAUGUGUACAGUCACCGCUUGUCGUACUUAAAUCAAAAAGUGCGAAAAGGAGGCGACUUUUUUUAUGGCAAAUCAGGUAAUGAGAAUCACAUUAAAGGCUUAUGACCAUCAAUUGGUAGACAAUUCAGCUAGUAAAAUUGUCGAAACUGUAAAGAAGACAGGAGCAAAGGUGUCAGG
env-614   GGAAAUGUGUACAGUCACCGCUUGUCGUACUUAAAUCAAAAAGUGCGAAAAGGAGGCGACUUUUUUUAUGGCAAAUCAGGUAAUGAGAAUCACAUUAAAGGCUUAUGACCAUCAAUUGGUAGACAAUUCAGCUAGUAAAAUUGUCGAAACUGUAAAGAAGACAGGAGCAAAGGUGUCAGG
env-615   GGAAAUGUGUACAGUCACCGCUUGUCGUA............................................................................................................................................................
env-623   GGAAAUGUGUACAGUCACCGCUUGUCGUAC...........................................................................................................................................................
env-624   GGAAAUGUGUACAGUCACCGCUUGUCGUAC...........................................................................................................................................................
env-625   GGAAAUGUGUACAGUCACCGCUUGUCGUACNNNNNNNNNNNNNNNNNNGGCUUAUGACCAUCAAUUGGUAGACAAUUCAGCUAGUAAAAUUGUCGAAACUGUAAAGAAGACAGGAGCAAAGGUGUCAGGACCAGUUCCACUUCCAACAAAGAAGGAAGUAGUAACAAUCUUGAGAGCGGUU
env-626   .........................................................................................................................................................................................
env-632   GGAAAUGUGUACAGUCACCGCUUGUCGUACUUAAAUCAAAAAGUGCGAAAAGGAGGCGACUUUUUUUAUGGCAAAUCAGGUAAUGAGAAUCACAUUAAAGGCUUAUGACCAUCAAUUGGUAGACAAUUCAGCUAGUAAAAUUGUCGAAACUGUAAAGAAGACAGGAGCAAAGGUGUCAGG
env-633   GGAAAUGUGUACAGUCACCGCUUGUCGUAC...........................................................................................................................................................
env-634   GGAAANNNNNNNNNNNNNNNAUCAGGUAAUGAGAAUCACAUUAAAGGCUUAUGACCAUCAAUUGGUAGACAAUUCAGCUAGUAAAAUUGUCGAAACUGUAAAGAAGACAGGAGCAAAGGUGUCAGGACCAGUUCCACUUCCAACAAAGAAGGAAGUAGUAACAAUCUUGAGAGCGGUUCAU
env-635   GGAAAUGUGUACAGUCACCGCUUGUCGUACUUAAAUCAAAAAGUGCGAAAAGGAGGCGACUUUUUUUAUGGCAAAUCAGGUAAUGAGAAUCACAUUAAAGGCUUAUGACCAUCAAUUGGUAGACAAUUCAGCUAGUAAAAUUGUCGAAACUGUAAAGAAGACAGGAGCAAAGGUGUCAGG
env-637   GGAAAUGUGUACAGUCACCGCUUGUCGUACUUAAAUCAAAAAGUGCGAAAAGGAGGCGACUU...........................................................................................................................
env-639   GGAAANNNNNNNNNNNNNNNNNNNAUCAGGUAAUGAGAAUCACAUUAAAGGCUUAUGACCAUCAAUUGGUAGACAAUUCAGCUAGUAAAAUUGUCGAAACUGUAAAGAAGACAGGAGCAAAGGUGUCAGGACCAGUUCCACUUCCAACAAAGAAGGAAGUAGUAACAAUCUUGAGAGCGGUU
env-640   GGAAAUGUGUACAGUCACCGCUUGUCGUACUUAAAUCAAAAAGUGCGAAAAGGAGGCGACUUUUUUUAUGGCAAAUCAGGUAAUGAGAAUCACAUUAAAGGCUUAUGACCAUCAAUUGGUAGACAAUUCAGCUAGUAAAAUUGUCGAAACUGUAAAGAAGACAGGAGCAAAGGUGUCAGG
env-641   GGAAAUGUGUACAGUCACCGCUUGUCGUACNNNNGGCUUAUGACCAUCAAUUGGUAGACAAUUCAGCUAGUAAAAUUGUCGAAACUGUAAAGAAGACAGGAGCAAAGGUGUCAGGACCAGUUCCACUUCCAACAAAGAAGGAAGUAGUAACAAUCUUGAGAGCGGUUCAUAAGUAUAAGG
env-642   GGAAANNNNNNNNNNNNNNNNNNNAUCAGGUAAUGAGAAUCACAUUAAAGGCUUAUGACCAUCAAUUGGUAGACAAUUCAGCUAGUAAAAUUGUCGAAACUGUAAAGAAGACAGGAGCAAAGGUGUCAGGACCAGUUCCACUUCCAACAAAGAAGGAAGUAGUAACAAUCUUGAGAGCGGUU
env-643   GGAAANNNNNNNNAUCAGGUAAUGAGAAUCACAUUAAAGGCUUAUGACCAUCAAUUGGUAGACAAUUCAGCUAGUAAAAUUGUCGAAACUGUAAAGAAGACAGGAGCAAAGGUGUCAGGACCAGUUCCACUUCCAACAAAGAAGGAAGUAGUAACAAUCUUGAGAGCGGUUCAUAAGUA
env-644   GGAAAUGUGUACAGUCACCGCUUGUCGUACUUAAAUCAAAAAGUGCGAAAAGGAGGCGACUUUUUUUAUGGCAAAUCAGGUAAUGAGAAUCACAUUAAAGGCUUAUGACCAUCAAUUGGUAGACAAUUCAGCUAGUAAAAUUGUCGAAACUGUAAAGAAGACAGGAGCAAAGGUAUCAGG
Jig-1-1   GCAGAGUGUACAGUCACCGCUUGUCGAACUUUUUUAAAAGUGCGAAAAGGAGGCGACUUUUUUUAUGGCAACUGAAGUAAUGAGGAUCAUAUUAAAGGCUUAUGACCAUCAGCUGGUGGACAGCUCGGCAGCCAAGAUAGUUGAUACUGUUAAAAGGAGCGGAGCAAAGGUUUCAGGACC
env-646   CGGAGGAGUGUACAGUCACCGCUUGUCGUACUAGAUUAAGUGCGAAAAGGAGGCGACUUUUUUUAUGGCAAGUCAAAUUAUGAGAAUCACGUUGAAGGCGUAUGAUCAUCAGUUAAUCGAUGCAUCCGCAAAGAGCAUUAUCGAAACUGUAAAGAAAACAGGAUCUAAGGUGAGCGGUCC
env-647   CGGAGGAGUGUACAGUCACCGCUUGUCGUACUAGAUUAAGUGCGAAAAGGAGGCGACUUUUUUUAUGGCAAGUCAAAUUAUGAGAAUCACGUUGAAGGCGUAUGAUCAUCAGUUAAUCGAUGCAUCCGCAAAGAGCAUCAUCGAAACUGUAAAGAAAACAGGAUCUAAGGUGAGCGGUCC
Eha-1-1   CGGAGGAGUGUACAGUCACCGCUUGUCGUACUAGAUUAAGUGCGAAAAGGAGGCGACUUUUUUUAUGGCAAGUCAAAUUAUGAGAAUCACGUUGAAAGCGUAUGAUCAUCAGUUAAUCGAUGCAUCCGCAAAGAGCAUCAUCGAAACUGUAAAGAAAACAGGAUCUAAGGUGAGCGGUCC
env-648   CGGAGGAGUGUACAGUCACCGCUUGUCGUA...........................................................................................................................................................
env-649   CGGAGGAGUGUACAGUCACCGCUUGUCGUACUAGAUUAAGUGCGAAAAGGAGGCGACUUUUUUUAUGGCAAGUCAAAUUAUGAGAAUCACGUUGAAAGCGUAUGAUCAUCAGUUAAUCGAUGCAUCCGCAAAGAGCAUCAUCGAAACUGUAAAGAAAACAGGAUCUAAGGUGAGCGGUCC
env-650   ACACGGGAGAGUGUACAGUCACCACUUGUUGUACUAAUCAAAAGUGCGAAAAGGAGGCGGCUUUUUUUAUGGCAAAUCAGAUUAUGAGAAUCACACUUAAGGCAUAUGACCACAAGCUUAUCGAUCAGGCUGCAAGCCAGAUCAUCGAUACUGUUAAGAAGACAGGUGCUAAGGUUAGCG
env-653   ACACGGGAGAGUGUACAGUCACCACUUGUUGUACUAAUCAAAAGUGCGAAAAGGAGGCGGCUUUUUUUAUGGCAAAUCAGAUUAUGAGAAUCACACUUAAGGCAUAUGACCACAAGCUUAUCGAUCAGGCUGCAAGCCAGAUCAUCGAUACUGUUAAGAAGACAGGUGCUAAGGUUAGCG
env-654   UACGCGUGGGAGUGUGUACAGUCACCGCUUGUCGUACUUACAUUAUAAAGUGCGAAAAGGAGGCGACUUUUUUUAUGGCAAAUCAGGUAAUAAGAAUUACACUGAAAGCUUACGAUCAUCAGUUAGUUGAUCAGUCUGCGAAGAAAAUCAUCGAAACAGUUAAGAAAACAGGAGCUCAGG
env-655   UACGCGUGGGAGUGUGUAC......................................................................................................................................................................
env-656   UACGCGUGGGAGUGUGUACAGUCACCGCUUGUCGUACUUACAUUAUAAAGUGCGAAAAGGAGGCGACUUUUUUUAUGGCAAAUCAGGUAAUAAGAAUUACACUGAAAGCUUACGAUCAUCAGUUAGUUGAUCAGUCUGCGAAGAAAAUCAUCGAAACAGUUAAGAAAACAGGAGCUCAGG
env-657   UACGCGUGGGAGUGUGUACAGUC..................................................................................................................................................................
env-658   UACGCGUGGGAGUGUGUACAGUCACCGCUUGUCGUACUUACAUUAUAAAGUGCGAAAAGGAGGCGACUUUUUUUAUGGCAAAUCAGGUAAUAAGAAUUACACUGAAAGCUUACGAUCAUCAGUUAGUUGAUCAGUCUGCGAAGAAAAUCAUCGAAACAGUUAAGAAAACAGGAGCUCAGG
env-662   UACGCGUGGGAGUGUGUACAGUCACCGCUUGUCGUACUUAAACAAUAAAGUGCGAAGGAGGCGACUUUUUUUAUGGCAAAUCAGGUAAUAAGAAUUACACUGAAAGCUUACGAUCAUCAGUUAGUUGAUCAGUCUGCGAAGAAAAUCAUCGAAACAGUUAAGAAAACAGGAGCUCAGGUG
env-663   UACGCGUGGGAGUGUGUACAG....................................................................................................................................................................
env-664   UACGCGUGGGAGUGUGUAC......................................................................................................................................................................
env-667   UACGCGUGGGAGUGUGUACAG....................................................................................................................................................................
env-668   UACGCGUGGGAGUGUGUACAGUCACCGCUUGUCGUGCUUAAACAAUAAAGUGCGAAGGAGGCGACUUUUUUUAUGGCAAAUCAGGUAAUAAGAAUUACACUGAAAGCUUACGAUCAUCAGUUAGUUGAUCAGUCUGCGAAGAAAAUCAUCGAAACAGUUAAGAAAACAGGAGCUCAGGUG
Lba-14-1  AUGUGUGAAGCUGUAAUGAGAGCGCAGAGAACUUCUGCCGGGGAGUACAGCAUGACACACACGGAGGAGUGUACAGUCACCGCUUGUCGUACUUAUGUAUAAAAAGUGCGAAAAGGAGGCGACUUUUUUUAUGGCAAGUCAGGUAAUGAGAAUUACAUUAAAAGCUUAUGAUCAUCAGUU
Bsp-1-1   GUUUGAGUCCAUGACACACACGGGAGAGUGUACAGUCACCGCUUGUCGUACUUAUCUAAUUAAAGUGCGAAAAGGAGGCGACUUUUUUUAUGGCAAAUCAGGUAAUGAGAAUUACACUUAAGGCGUAUGAUCAUCAGCUCGUAGAUGCAUCUGCAAAGAAGAUCAUCGAAACAGUAAAGA
Bsp-9-1   GUUUGAGUCCAUGACACACACGGGAGAGUGUACAGUCACCGCUUGUCGUACUUAUCUAAUUAAAGUGCGAAAAGGAGGCGACUUUUUUUAUGGCAAAUCAGGUAAUGAGAAUUACACUUAAGGCGUAUGAUCAUCAGCUCGUAGAUGCAUCUGCAAAGAAGAUCAUCGAAACAGUAAAGA
env-669   GGGUAAGUGUACAGUCACCGCUUGUCGUACUCGUAAAAGUGCGAAAAGGAGGCGACUUUUUUUAUGGCAAGUCAAGUAAUGAGAAUUACACUGAAAGCGUAUGAUCAUCAGUUAGUAGAUGCAUCUGCACAGAAAAUUAUCGAAACUGUAAAGAAGAACGGAGCAACGGUGAGCGGACCA
env-671   GGGUAAGUGUACAGUCACCGCUUGUCGUACUCGUAAAAGUGCGAAAAGGAGGCGACUUUUUUUAUGGCAAGUCAAGUAAUGAGAAUUACACUGAAAGCGUAUGAUCAUCAGUUAGUAGAUGCAUCUGCACAGAAAAUUAUCGAAACUGUAAAGAAGAACGGAGCAACGGUGAGCGGACCA
env-672   GGGUAAGUGUACAGUCACCGCUUGUCGUACUCGUAAAAGUGCGAAAAGGAGGCGACUUUUUUUAUGGCAAGUCAAGUAAUGAGAAUUACACUGAAAGCGUAUGAUCAUCAGUUAGUAGAUGCAUCUGCACAGAAAAUUAUCGAAACUGUAAAGAAGAACGGAGCAACGGUGAGCGGACCA
env-681   GGGUAAGUGUACAGUCACCGCUUGUCGUACUCGUAAAAGUGCGAAAAGGAGGCGACUUUUUUUAUGGCAAGUCAAGUAAUGAGAAUUACACUGAAAGCGUAUGAUCAUCAGUUAGUAGAUGCAUCUGCACAGAAAAUUAUUGAAACUGUAAAGAAGAACGGAGCAACGGUGAGCGGACCA
env-683   GGGUAAGUGUACAGUCACCGCUUGUCGUACUCGUAAAAGUGCGAAAAGGAGGCGACUUUUUUUAUGGCAAGUCAAGUAAUGAGAAUUACACUGAAAGCGUAUGAUCAUCAGUUAGUAGAUGCAUCUGCACAGAAAAUUAUCGAAACUGUAAAGAAGAACGGAGCAACGGUGAGCGGACCA
env-684   GGGUAAGUGUACAGUCACCGCUUGUCG..............................................................................................................................................................
env-685   NNNNNNNNNNNNNNNNNNNNNNNNNNNNNNNNNNNNNNNNNNNNNNNNNNNNNNNNNNNNNNNNNNNCGAAACUGUAAAGAAGAACGGAGCAACGGUGAGCGGACCAGUACCGCUUCCAACGAAGAAAGAGGNNNNNNNNNNNNNNNNNNNNNNNNNNNNNNNNNNNNNNNNNNNNNN
env-686   CGUGUGAGUUUCUCACGACACACACGGGAGAGUGUACAGUCACCGCUUGUCGUACUGAUGUGAAAAGUACGAAAAGGAGGCGACUUUUUUUAUGGCAAGUCAAGUAAUGAGAAUCACAUUGAAGGCGUAUGAUCAUCAGCUUGUUGAUGCAUCCGCAAAGAAAAUCAUCGAAACUGUAAA
env-687   CGUGUGAGUUUCUCACGACACACACGGGAGAG.........................................................................................................................................................
env-691   CGUGUGAGUUUCUCACGACACACACGGGAGAGUGUACAGUCACCGCUUGUCGUA...................................................................................................................................
env-692   CGUGUGAGUUUCUCACGACACACACGGGAGAGUGUACAGUCACCGCUUGUCGUACUGAUGUGAAAAGUACGAAAAGGAGGCGACUUUUUUUAUGGCAAGUCAAGUAAUGAGAAUCACAUUGAAGGCGUAUGAUCAUCAGCUUGUUGAUGCAUCCGCAAAGAAAAUCAUCGAAACUGUAAA
env-694   CGUGUGAGUUUCUCACGACACACACGGGAGAGUGUACAGUCACCGCUUGUCGUACUGAUGUGAAAAGUACGAAAAGGAGGCGACUUUUUUUAUGGCAAGUCAAGUAAUGAGAAUCACAUUGAAGGCGUAUGAUCAUCAGCUUGUUGAUGCAUCCGCAAAGAAAAUCAUCGAAACUGUAAA
env-695   CGUGUGAGUUUCUCACGACACACACGGGAGAGUGUACAGUCACCGCUUGUCGUACUGAUGUGAAAAGUACGAAAAGGAGGCGACUUUUUUUAUGGCAAGUCAAGUAAUGAGAAUCACAUUGAAGGCGUAUGAUCAUCAGCUUGUUGAUGCAUCCGCAAAGAAAAUCAUCGAAACUGUAAA
env-696   CGUGUGAGUUUCUCACGACACACACGGGAGAGUGUACAGUCACCGCUUGUCGUACUGAUGUGAAAAGUACGAAAAGGAGGCGACUUUUUUUAUGGCAAGUCAAGUAAUGAGAAUCACAUUGAAGGCGUAUGAUCAUCAGCUUGUUGAUGCAUCCGCAAAGAAAAUCAUCGAAACUGUAAA
env-699   CGUGUGAGUUUCUCACGACACACACGGGAGAGUGUACAGUCACCGCUUGUCGUACUGAUGUGAAAAGUACGAAAAGGAGGCGACUUUUUUUAUGGCAAGUCAAGUAAUGAGAAUCACAUUGAAGGCGUAUGAUCAUCAGCUUGUUGAUGCAUCCGCAAAGAAAAUCAUCGAAACUGUAAA
env-700   .........................................................................................................................................................................................
env-705   .........................................................................................................................................................................................
env-708   CACACACGGGAGAGUGUACAGUCACCGCUUGUUGUACUAAGAUUAAGUGCGAAAAGGAGGCGGCUUUUUUUAUGGCAAAUCAGAUUAUGAGAAUUACACUUAAGGCUUAUGACCACAAACUUAUCGACAAGGCAGCAGCACAGAUUGUUGAUACAGUUAAGAAGACUGGCGCAAAGGUAA
env-709   CACACACGGGAGAGUGUACAGUCACCGCUUGUUGUACUAAGAUUAAGUGCGAAAAGGAGGCGGCUUUUUUUAUGGCAAAUCAGAUUAUGAGAAUUACACUUAAGGCUUAUGACCACAAACUUAUCGACAAGGCAGCAGCACAGAUUGUUGAUACAGUUAAGAAGACUGGCGCAAAGGUAA
env-710   CACACACGGGAGAGUGUACAGUCACCGCCUGUUGUACUAAGAUUAAGCGCGAAAAGGAGGCGGCUUUUUUUAUGGCAAAUCAGAUUAUGAGAAUUACACUUAAGGCUUAUGACCACAAACUUAUCGACAAGGCAGCAGCACAGAUUGUUGAUACAGUUAAGAAGACUGGCGCAAAGGUAA
env-711   .........................................................................................................................................................................................
env-712   CACACACGGGAGAGUGGACAGUCACCGCUUGUUGUACUAAGAUUAAGUGCGAAAAGGAGGCGGCUUUUUUUAUGGCAAAUCAGAUUAUGAGAAUUACACUUAAGGCUUAUGACCACAAACUUAUCGACAAGG...............................................
env-713   CACACACGGGAGAGUGUA.......................................................................................................................................................................
env-726   CACACACGGGAGAGUGUACAGUCACCGCUUGUUGUACUAAGAUUAAGUGCGAAAAGGAGGCGGCUUUUUUUAUGGCAAAUCAGAUUAUGAGAAUUACACUUAAGGCUUAUGACCACAAACUUAUCGACAAGGCAGCAGCACAGAUUGUUGAUACAGUUAAGAAGACUGGCGCAAAGGUAA
env-727   .........................................................................................................................................................................................
env-729   CACACACGGGAGAGUGUACAGUCACCGCUUGUUGUAUUAAGGUUAAGUGCGAAAAGGAGGCGGCUUUUUUUAUGGCAAAUCAGAUUAUGAGAAUUACACUUAAGGCUUAUGACCACAAACUUAUCGACAAGGCAGCAGCACAGAUUGUUGAUACAGUUAAGAAGACUGGCGCAAAGGUAA
env-730   CACACACGGGAGAGUGUACAGUCACCGCUUGUUGUACUAAGAUUAAGUGCGAAAAGGAGGCGGCUUUUUUUAUGGCAAAUCAGAUUAUGAGAAUUACACUUAAGGCUUAUGACCACAAACUUAUCGACAAGGCAGCAGCACAGAUUGUUGAUACAGUUAAGAAGACUGGCGCAAAGGUAA
env-731   CACACACGGGAGAGUGUACAGUCACCGAUUGUUGUACUAAGAUUAAGUGCGAAAAGGAGGCGGCUUUUUUUAUGGCAAAUCAGAUUAUGAGAAUUACACUUAAGGCUUAUGACCACAAACUUAUCGACAAGGCAGCAGCACAGAUUGUUGAUACAGUUAAGAAGACUGGCGCAAAGGUAA
Rsp-1-1   GAUGUGUACAGUCGCCGCUUGUCGUACUGCUUAAAAAGUACGAAUAGGAGGAGACUUUUUUUAUGGCAAAUCAGGUAAUGAGAAUCACAUUAAAGGCAUAUGAUCAUCAGUUAGUAGAUGCAUCUGCUGCAAAAAUCAUCGAAACUGUUAAAAAGAACGGGGCAAUGGUAAGCGGACCGG
env-732   UCUCACGACACACACGGGAGAGUGUACAGUCACCGCUUGUCGUACUGAUGUGAAAAGUACGAAAAGGAGGCGACUUUUUUUAUGGCAAGUCAAGUAAUGAGAAUCACAUUGAAAGCNNNNNNNNNNAUUCACUGCGGCGGCUGUAUGCUGAAUGAGCGGGAGAUGAAGUACCGUAUGAGU
Rgn-1-1   UCUCACGACACACACGGGAGAGUGUACAGUCACCGCUUGUCGUACUGAUGUGAAAAGUACGAAAAGGAGGCGACUUUUUUUAUGGCAAGUCAAGUAAUGAGAAUCACAUUGAAAGCAUAUGAUCACCAGCUGGUUGAUGCAUCUGCAAAAAAAAUCAUCGAAACUGUAAAGAAAAACGGA
env-733   UCUCACGACACACACGGGAGAGUGUACAGUCACCGCUUGUCGUACUGAUGUGAAAAGUACGAAAAGGAGGCGACUUUUUUUAUGGCAAGUCAAGUAAUGAGAAUCACAUUGAAAGCAUAUGAUCACCAGCUGGUUGAUGCAUCUGCAAAAAAAAUCAUCGAACUGUAAAGAAAAACGGAAG
env-734   UCUCACGACACACACGGGAGAGUGUACAGUCACCGCUUGUCGUACUCGAUUAUAAGUACGAAAAGGAGGCGACUUUUUUUAUGGCAAGUCAAGUAAUGAGAAUCACAUUGAAAGCAUAUGAUCACCAGCUGGUUGAUGCAUCUGCAAAAAAAAUCAUCGAAACUGUAAAGAAAAACGGAG

          .........................................................................................................................................................................................
          .........................................................................................................................................................................................
          ..........................................................................................................................................................R..............................
```

```
env-735   UCUCACGACACACACGGGAGAGUGUACAGUCACCGCUUGUCGUACUGAAGUGAAAAGUACGAAAAGGAGGCGACUUUUUUUAUGGCAAGUCAAGUAAUGAGAAUCACAUUGAAAGCAUAUGAUCACCAGUUGGUUGAUGCAUCUGCAAAAAAAAUCAUCGAAACUGUAAAAAAGAACGGA
env-739   ....................................................................................................................................................................................
env-742   UCUCACGACACACACGGGAGAGUGUACAGUCACCGCUUGUCGUACUGAUGUGAAAAGUACGAAAAGGAGGCGACUUUUUUUAUGGCAAGUCAAGUAAUGAGAAUCACAUUGAAAGCAUAUGAUCACCAGCUGGUUGAUGCAUCUGCAAAAAAAAUCAUCGAAACUGUAAAGAAAAACGGA
Lba-4-1   UCUCACGACACACACGGGAGAGUGUACAGUCACCGCUUGUCGUACUGAUGUGAAAAGUACGAAAAGGAGGCGACUUUUUUUAUGGCAAGUCAAGUAAUGAGAAUCACAUUGAAAGCAUAUGAUCACCAGCUGGUUGAUGCAUCUGCAAAAAAAAUCAUCGAAACUGUAAAGAAAAACGGA
env-744   UCUCACGACACACACGGGAGAGUGUACAGUCACCGCUUGUCGUACUGAUGUGAAAAGUACGAAAAGGAGGCGACUUUUUUUAUGGCAAGUCAAGUAAUGAGAAUCACAUUGAAAGCAUAUGAUCACCAGCUGGUUGAUGCAUCUGCAAAAAAAAUCAUCGAAACUGUAAAGAAAAACGGA
env-749   CCAAUAGCGGUAACUGGCAUGUAACCAGCCAAAAGACAGCUGAGGAGUGCUUAUGACGAGAAACAACGUGUGAGUUUCUCACGACACACACGGGAGAGUGUACAGUCACCGCUUGUCGUACUGAGGUUAAUAGUACGAAAAGGAGGCGACUUUUUUUAUGGCAAGUCAAGUAAUGAGAAU
env-751   UGUUUACAUCUAUGAUACACACGGGAGAGUGUACAGUCACCGCUUGUCGUACUAAUUAAAGUGCGAAAAGGAGGCGACUUUUUUUAUGGCAAGUCAAGUAAUGAGAAUUACGCUUAAGGCGUACGAUCAUCAGUUAGUUGAUGCAUCGGCAAAAAAAAUCAUCGAAACUGUAAAGAAGAA
Cph-1-1   AGCGAUUUUUAUUGACACCUAGGACGGGUUGCGUCUUAGGUUAUAUAUUUCCGACAAGUAUGAAACACACGGGUGAGUGUACAGUCACCACUUGUCGUACUGAAAUGAAGUGCGAAAAGGAGGCGGCUUUUUUUAUGGCAAAUCAAGUUAUGAGAAUUACUCUCAAGGCGUAUGAUCAUC
Bfo-1-1   AGCGUGUACAGUCACCGCUUGUCGUACUGAUGUUAAGUGCGAAAAGGAGGCGACUUUUUUUAUGGCAAGUCAAGUAAUGAGAAUCACACUGAAAGCGUAUGACCAUCAGCUGAUUGAUGCAUCCGCAAAGAAAAUCAUCGAAACUGUAAAAAAGAACGGAUCACAGGUGAGCGGACCGGU
env-752   AAUCCGGGAUUUCGGAAUGUGAAUAAAUAGGACACACACGGGAGAGUGUACAGUCACCGCUUGUCGUACUAAGUAACAAAAGUGCGAAAAGGAGGCGACUUUUUUUAUGGCAACUCAGGUAAUGAGAAUCACAUUAAAGGCGUAUGAUCAUCAGUUAGUUGAUGCAUCUGCAAAGAAAAU
env-753   UGAAAUCCGGGAUUUCGGAAUGUGAAUAAAUAGGACACACACGGGAGAGUGUACAGUCACCGCUUGUCGUACUAAGUAACAAAAGUGCGAAAAGGAGGCGACUUUUUUUAUGGCAACUCAGGUAAUGAGAAUCAC............................................
Lbo-3-1   GGAAGUGUACAGUCACCGCUUGUCGUACUGAAGUAAAAGUGCGAAAAGGAGGCGACUUUUUUUAUGGCAAGUCAAGUAAUGAGAAUUACUCUUAAAGCUUAUGAUCACGAAUUAGUUGAUUCAUCAGCAAAAAAAAUUAUCGAAACUGUAAAGAAGAACGGAUCACAGGUGAGCGGUCCA
env-754   CGUGUACAGUCACCGCUUGUCGUACUUCGAAUAAUAAGUGCGAAAAGGAGGCGACUUUUUUUAUGGCAAGUCAAGUAAUGAGAAUCACAUUAAAAGCUUAUGAUCAUCAAUUAGUGGAUCAAUCAGCGAGCAAGAUUAUUGAUACUGCUAAGAAGACAGGAUCACAGGUGAGCGGACCAG
env-755   CGUGUACAGUCACCGCUUGUCGUACUUCGAAUAAUAAGUGCGAAAAGGAGGCGACUUUUUUUAUGGCAAGUCAAGUAAUGAGAAUCACAUUAAAAGCUUAUGAUCAUCAAUUAGUGGAUCAGUCAGCAAGCAAGAUUAUUGAUACUGCUAAGAAGACAGGAUCACAGGUGAGCGGACCAG
Csp-4-1   UCACGACACACACGGGAGAGUGUACAGUCACCGCUUGUCGUACUGAUGGAAUAGUACGAAAAGGAGGCGACUUUUUUUAUGGCAAGUCAAGUAAUGAGAAUCACAUUGAAGGCAUAUGAUCACCAGCUGGUAGAUGCAUCUGCAAAGAAAAUCAUCGAAACUGUAAAAAAGAACGGAUCA
env-756   UUGUGUUUGACUGUAGGACACACACGGGAGAGUGUACAGUCACCGCUUGUCGUACUUCAAUUAAAAGUGCGAAAAGGAGGCGACUUUUUUUAUGGCAACGAGUCAAGUAAUGAGAAUUACACUGAAAGCGUACGAUCAUCAGUUAGUAGAUGCAUCUGCCAAAAAAAAUCAUCGAAACAGU
env-758   UUUCUCACGACACACACGGGAGAGUGUACAGUCACCGCUUGUCGUACUGAAGUCAAAAGUACGAAAAGGAGGCGACUUUUUUUAUGGCAAGUCAAGUAAUGAGAAUCACAUUAAAGGCGUAUGAUCAUCAGCUCGUUGAUGCAUCCGCAAAGAAAAUCAUCGAAACUGUAAAAAAGAACG
env-759   GAGAGUGUACAGUCACCGCUUGUCGUACUGAAAUUAAGUGCGAAAAGGAGGCGACUUUUUUUAUGGCAAGUCAAGUAAUGAGAAUAACAUUAAAGGCGUAUGAUCAUCAAUUAGUUGAUGCAUCAUCAAAGAAAAUCAUCGAAACUGUAAAGAAAAAUGGACUACAGGUGAGUGGACCAG
env-760   GAGAGUGUACAGUCACCGCUUGUCGUACUGAAAUUAAGUGCGAAAAGGAGGCGACUUUUUUUAUGGCAAGUCAAGUAAUGAGAAUAACAUUAAAGGCGUAUGAUCAUCAAUUAGUUGAUGCAUCAUCAAAGAAAAUCAUCGAAACUGUAAAGAAAAAUGGACUACAGGUGAGUGGACCAG
env-762   GAGAGUGUA...........................................................................................................................................................................
env-763   AGAUGUGUACAGUCACCGCUUGUCGUACUCGAAAAGUGCGAAAAGGAGGCGACUUUUUUUAUGGCAAGUCAAGUAAUGAGAAUCACAUUAAAAGCUUAUGAUCAUCAGCUGGUUGAUGCAUCCGCAAGCAAAAUUAUCGAAACUGUAAAAAAGAAUGGAGCAGUAGUGAGUGGACCGGUC
env-764   UACACACGGGAGCGUGUACAGUCACCGCUUGUCGUACUUAGCAAUAAAAGUGCGAAAAGGAGGCGACUUUUUUUAUGGCAAGUCAAGUAAUGAGAAUCACAUUAAAGGCAUAUGAUCAUCAGCUGGUUGAUGCAUCUGCUAAGAAAAUCAUCGAAACUGUAAAAAAGAAUGGAGCUCAGG
Lba-6-1   UCGUACUGGGGUUUAUAGUACGAAAAGGAGGCGACUUUUUUUAUGGCAAGUCAAGUAAUGAGAAUCACUUUGAAAGCGUAUGAUCAUCAACUGGUUGAUGCAUCCGCGAAGAAAAUCAUCGAAACUGUUAAGAAGAAUGGGGCAAAGGUGAGUGGCCCGGUACCUCUUCCGACUAAAAAG
Dsp-1-1   UGCACAUAACGGCGUGUGAGGUUCUCGCGACACACACGGAAGAGUGUACAGUCACCGCUUGUCGUACUGAAGUUAGAAGUACGAAAAGGAGGCGACUUUUUUUAUGGCAAGUCAAGUAAUGAGAAUCACUUUGAAAGCGUAUGAUCAUCAGAUGGUAGAUGCAUCCGCUAAGAAAAUCAU
env-765   GAAACGUGUA..........................................................................................................................................................................
env-767   GAAACGUGUACAGUCACCGCUUGUCGUACUUGGUUUAUAAAGUGCGAAAAGGAGGCGACUUUUUUUAUGGCAAGUCAAGUAAUGAGAAUCACAUUAAAAGCUUAUGAUCAUCAGUUAGUAGAUCAGUCCGCUGGAAAGAUCAUCGAGACAGUAAAAAAGACAGGAUCUCAGGUGAGCGGA
env-768   GAAACGUGUACAGUCACCGCUUGUCGU.........................................................................................................................................................
env-769   GAAACGUGUACAGUCACCGCUUGUCGUACUUGGUUUAUAAAGUGCGAAAAGGAGGCGACUUUUUUUAUGGCAAGUCAAGUAAUGAGAAUCACAUUAAAAGCUUAUGAUCAUCAGUUAGUAGAUCAGUCCGCUGGAAAGAUCAUCGAGACAGUAAAAAAGACAGGAUCUCAGGUGAGCGGA
env-770   GAAACGUGUACAGUCACCGCUUGUCGUACUUGGUUUAUAAAGUGCGAAAAGGAGGCGACUUUUUUUAUGGCAAGUCAAGUAAUGAGAAUCACAUUAAAAGCUUAUGAUCAUCAGUUAGUAGAUCAGUCCGCUGGAAAGAUCAUCGAGACAGUAAAAAAGACAGGAUCUCAGGUGAGCGGA
env-777   CGGAAACGUGUACAGUCACCGCUUGUCGUACUUGGUUUAUAAAGUGCGAAAAGGAGGCGACUUUUUUUAUGGCAAGUCAAGUAAUGAGAAUCACAUUAAAAGCUUAUGAUCAUCAGUUAGUAGAUCAGUCCGCUGGAAAGAUCAUCGAGACAGUAAAAAAGACAGGAUCUCAGGUGAGCG
env-778   GGGAGUGUACAGUCACCGCUUGUCGUACUGAGGUUUAAGUACGAAAAGGAGGCGACUUUUUUUAUGGCAAGUCAAGUAAUGAGAAUCACAUUGAAGGCAUAUGAUCAUCAGCUUGUAGAUGCAUCUGCCAAGAAAAUUAUCGAAACUGUAAAGAAAAAUGGAGCAAAGGUUAGCGGACCG
env-802   GGAAGUGUACAGUCACCGCUUGUCGUACUGAGGUUUAAGUACGAAAAGGAGGCGACUUUUUUUAUGGCAAGUCAAGUAAUGAGAAUCACAUUGAAGGCAUAUGAUCAUCAGCUUGUAGAUGCAUCUGCCAAGAAAAUUAUCGAAACUGUAAAGAAAAAUGGAGCAAAGGUUAGCGGACCG
env-803   GGAAGUGUACAGUCACCGCUUGUCGUACUGAGG...................................................................................................................................................
Cco-1-1   GGAAGUGUACAGUCACCGCUUGUCGUACUGAGGUUUAAGUACGAAAAGGAGGCGACUUUUUUUAUGGCAAGUCAAGUAAUGAGAAUCACAUUGAAGGCAUAUGAUCAUCAGCUUGUAGAUGCAUCUGCCAAGAAAAUUAUCGAAACUGUAAAGAAAAAUGGAGCAAAGGUUAGCGGACCG
env-806   GGAAGUGUACAGUCACCGCUUGUCGUACUGAGGUUUAAGUACGAAAAGGAGGCGACUUUUUUUAUGGCAAGUCAAGUAAUGAGAAUCACAUUGAAGGCAUAUGAUCAUCAGCUUGUAGAUGCAUCUGCCAAGAAAAUUAUCGAAACUGUAAAGAAAAAUGGAGCAAAGGUUAGCGGACCG
env-812   GGAAGUGUACAGUCACCGCUUGUCGUACUGAGGUUUAAGUACGAAAAGGAGGCGACUUUUUUUAUGGCAAGUCAAGUAAUGAGAAUCACAUUGAAGGCAUAUGAUCAUCAGCUUGUAGAUGCAUCUGCCAAGAAAAUUAUCGAAACUGUAAAGAAAAAUGGAGCAAAGGUUAGCGGACCG
env-814   UCACGACACACACGGGAGAGUGUACAGUCACCGCUUGUCGUACUGAAGUUAAAAGUACGAAAAGGAGGCGACUUUUUUUAUGGCAAGUCAAGUAAUGAGAAUCACAUUAAAAGCGUAUGAUCAUCAGUUAGUAGAUGCAUCCGCAAAAAAAAUCAUCGAAACUGUAAAGAAGAACGGAGC
Dfo-2-1   UCACGACACACACGGGAGAGUGUACAGUCACCGCUUGUCGUACUGAAGUUAAAAGUACGAAAAGGAGGCGACUUUUUUUAUGGCAAGUCAAGUAAUGAGAAUCACAUUAAAAGCGUAUGAUCAUCAGUUAGUAGAUGCAUCCGCAAAAAAAAUCAUCGAAACUGUAAAGAAGAACGGAGC
Dfo-1-1   UCACGACACACACGGGAGAGUGUACAGUCACCGCUUGUCGUACUGAAGUUAAAAGUACGAAAAGGAGGCGACUUUUUUUAUGGCAAGUCAAGUAAUGAGAAUCACAUUAAAAGCGUAUGAUCAUCAGUUAGUAGAUGCAUCCGCAAAAAAAAUCAUCGAAACUGUAAAGAAGAACGGAGC
env-818   UCACGACACACACGGGAGAGUGUACAGUCACCGCUUGUCGUACUGAAGUUAAAAGUACGAAAAGGAGGCGACUUUUUUUAUGGCAAGUCAAGUAAUGAGAAUCACAUUAAAAGCGUAUGAUCAUCAGUUAGUAGAUGCAUCCGCAAAAAAAAUCAUCGAAACUGUAAAGAAGAACGGAGC
env-819   UCACGACACACACGGGAGAGUGUACAGUCACCGCUUGUCGUACUGAAGUUAAAAGUACGAAAAGGAGGCGACUUUUUUUAUGGCAAGUCAAGUAAUGAGAAUCACAUUAAAAGCGUAUGAUCAUCAGUUAGUAGAUGCAUCCGCAAAAAAAAUCAUCGAAACUGUAAAGAAGAACGGAGC
env-820   ACGGAAACGUGUACAGUCACCGCUUGUCGUACUUGGUUUGAAAGUGCGAAAAGGAGGCGACUUUUUUUAUGGCAAGUCAAGUAAUGAGAAUCACUUUAAAGGCUUAUGAUCAUCAGUUGGUAGAUCAGUCCGCCGGAAAAAUCAUCGAGACUGUAAAGAAGACAGGAUCACAGGUGAGCG
[ci-1-1   CGGAAACGUGUACAGUCACCGCUUGUCGUACUUGGUUUAGAAGUGCGAAAAGGAGGCGACUUUUUUUAUGGCAAGUCAAGUAAUGAGAAUCACUUUAAAGGCUUAUGAUCAUCAGUUAGUAGAUCAGUCCGCCGGCAAGAUCAUCGAGACUGUAAAGAAGACAGGAUCACAGGUGAGCGG
Csp-5-1   CGGAAACGUGUACAGUCACCGCUUGUCGUACUUGGUUUAGAAGUGCGAAAAGGAGGCGACUUUUUUUAUGGCAAGUCAAGUAAUGAGAAUCACUUUAAAGGCUUAUGAUCAUCAGUUAGUAGAUCAGUCCGCCGGCAAAAUCAUCGAGACUGUAAAGAAGACAGGAUCACAGGUGAGCGG
env-821   ACGGAAACGUGUACAGUCACCGCUUGUCGUACUUGGUUUAUAAAGUGCGAAAAGGAGGCGACUUUUUUUAUGGCAAGUCAAGUAAUGAGAAUCACUUUAAAGGCUUAUGAUCAUCAGUUGGUAGAUCAGUCCGCCGGAAAAAUCAUCGAGACUGUAAAGAAGACAGGAUCACAGGUGAGC
env-822   ACGGAAACGUGUACAGUCACCGCUUGUCGUACUUGGUUUGAAAGUGCGAAAAGGAGGCGACUUUUUUUAUGGCAAGUCAAGUAAUGAGAAUCACUUUAAAGGCUUAUGAUCAUCAGUUGGUAGAUCAGUCCGCCGGAAAAAUCAUCGAGACUGUAAAGAAGACAGGAUCACAGGUGAGCG
env-823   ACGGAAACGUGUACAGUCACCGCUUGUCGUACUUGGUUUGAAAGUGCGAAAAGGAGGCGACUUUUUUUAUGGCAAGUCAAGUAAUGAGAAUCACUUUAAAGGCUUAUGAUCAUCAGUUAGUAGAUCAGUCCGCCGGAAAAAUCAUCGAGACUGUAAAGAAGACAGGAUCACAGGUGAGCG
Cbo-2-1   ACGGAAACGUGUACAGUCACCGCUUGUCGUACUUGGUUUGAAAGUGCGAAAAGGAGGCGACUUUUUUUAUGGCAAGUCAAGUAAUGAGAAUCACUUUAAAGGCUUAUGAUCAUCAGUUGGUAGAUCAGUCCGCCGGAAAAAUCAUCGAGACUGUAAAGAAGACAGGAUCACAGGUGAGCG
Cba-7-1   CGGAAACGUGUACAGUCACCGCUUGUCGUACUUGGUUUAAAAGUGCGAAAAGGAGGCGACUUUUUUUAUGGCAAGUCAAGUAAUGAGAAUCACUUUAAAGGCUUAUGAUCAUCAGUUAGUAGAUCAGUCUGCCGGCAAAAUCAUCGAGACUGUAAAGAAGACAGGAUCACAGGUGAGCGG
env-824   ACGGAAACGUGUACAGUCACCGCUUGUCGUACUUGGUUUGAAAGUGCGAAAAGGAGGCGACUUUUUUUAUGGCAAGUCAAGUAAUGAGAAUCACUUUAAAGGCUUAUGAUCAUCAGUUGGUAGAUCAGUCCGCCGGAAAAAUCAUCGAGACUGUAAAGAAGACAGGAUCACAGGUGAGCG
Cba-2-1   CGGAAACGUGUACAGUCACCGCUUGUCGUACUUGGUUUAAAAGUGCGAAAAGGAGGCGACUUUUUUUAUGGCAAGUCAAGUAAUGAGAAUCACUUUAAAGGCUUAUGAUCAUCAGUUAGUAGAUCAGUCUGCCGGCAAAAUCAUCGAGACUGUAAAGAAGACAGGAUCACAGGUGAGCGG
env-826   ACGGAAACGUGUACAGUCACCGCUUGUCGUACUUGGUUUGAAAGUGCGAAAAGGAGGCGACUUUUUUUAUGGCAAGUCAAGUAAUGAGAAUCACUUUAAAGGCUUAUGAUCAUCAGUUGGUAGAUCAGUCCGCCGGAAAAAUCAUCGAGACUGUAAAGAAGACAGGAUCACAGGUGAGCG
Ccl-6-1   ACGGAAACGUGUACAGUCACCGCUUGUCGUACUUGGUUUGAAAGUGCGAAAAGGAGGCGACUUUUUUUAUGGCAAGUCAAGUAAUGAGAAUCACUUUAAAGGCUUAUGAUCAUCAGUUAGUAGAUCAGUCCGCCGGAAAAAUCAUCGAGACUGUAAAGAAGACAGGAUCACAGGUGAGCG
env-827   ACGGAAACGUGUACAGUCACCGCUUGUCGUACUUGGUUUGAAAGUGCGAAAAGGAGGCGACUUUUUUUAUGGCAAGUCAAGUAAUGAGAAUCACUUUAAAGGCUUAUGAUCAUCAGUUAGUAGAUCAGUCCGCCGNGAAAAAUCAUCGAGACUGUAAAGAAGACAGACGAUUCACCC...
Ccl-3-1   ACGGAAACGUGUACAGUCACCGCUUGUCGUACUUGGUUUGAAAGUGCGAAAAGGAGGCGACUUUUUUUAUGGCAAGUCAAGUAAUGAGAAUCACUUUAAAGGCUUAUGAUCAUCAGUUGGUAGAUCAGUCCGCCGGAAAAAUCAUCGAGACUGUAAAGAAGACAGGAUCACAGGUGAGCG
Ccl-1-1   ACGGAAACGUGUACAGUCACCGCUUGUCGUACUUGGUUUGAAAGUGCGAAAAGGAGGCGACUUUUUUUAUGGCAAGUCAAGUAAUGAGAAUCACUUUAAAGGCUUAUGAUCAUCAGUUGGUAGAUCAGUCCGCCGGAAAAAUCAUCGAGACUGUAAAGAAGACAGGAUCACAGGUGAGCG
Lba-16-1  CACGACACACACGGGAGAGUGUACAGUCACCGCUUGUCGUACUGAAGUGAAUAGUACGAAAAGGAGGCGACUUUUUUUAUGGCAAGUCAAGUAAUGAGAAUCACAUUGAAGGCAUACGAUCAUCAGCUUGUUGAUGCAUCUGCCAGAAAAAUCAUCGAAACUGUAAAGAAAAACGGAUCU
Csp-3-1   CACGACACACACGGGAGAGUGUACAGUCACCGCUUGUCGUACUGAAGUGAAUAGUACGAAAAGGAGGCGACUUUUUUUAUGGCAAGUCAAGUAAUGAGAAUCACAUUGAAGGCAUACGAUCAUCAGCUUGUUGAUGCAUCUGCCAGAAAAAUCAUCGAAACUGUAAAGAAAAACGGAUCU
env-828   UCUCACGACACACACGGGAGAGUGUACAGUCACCGCUUGUCGUACUGAUGUGAAAAGUACGAAAAGGAGGCGACUUUUUUUAUGGCAAGUCAAGUAAUGAGAAUCACAUUGAAAGCGUAUGAUCAUCAGCUCGUUGAUGCAUCCGCAAAGAAAAUCAUCGAAACUGUAAAAAAGAAUGGG
env-829   CACGACACACACGGGAGAGUGUACAGUCACCGCUUGUCGUACUGAAGUCAAAAGUACGAAAAGGAGGCGACUUUUUUUAUGGCAAGUCAAGUAAUGAGAAUCACAUUGAAAGCGUAUGAUCAUCAGCUCGUUGAUGCAUCCGCAAAGAAAAUCAUCGAAACUGUAAAAAAGAAUGGGUCA
env-830   CACGACACACACGGGAGAGUGUACAGUCACCGCUUGUCGUACU.........................................................................................................................................
env-831   C...................................................................................................................................................................................
env-832   CACGACACACACGGGAGAGUGUACAGUCACCGCUUGUCGUACUGAUGUGAAAAGUACGAAAAGGAGGCGACUUUUU.......................................................................................................
env-833   CACGACACACACGGGAGAGUGUACAGUCACCGCUUGUCGUACUGAAGUCAAAAGUACGAAAAGGAGGCGACUUUUUUUAUGGCAAGUCAAGUAAUGAGAAUCACAUUGAA...................................................................
env-837   CACGACACACACGGGAGAGUGUACAGUCACNNNNNNNNNNNNNNNNNNNNNNNNNNNNNNNNNNNNNNNNNNNNNNNNNNNNNNNNNNNNNNNNNNNNNNNNNNNNNNNNNNNNNNNNNNNNNNNNNNNNNNNNNNNNNNNNNNNNNNNNNNNNNNNNNNNNNNNNNNNNNNNNNNNNNNNNN
env-838   CACGACACACACGGGAGAGUGUACAGUCACCGCUUGUCGUACUGAAGUCAAAAGUACGAAAAGGAGGCGACUUUUUUUAUGGCAAGUCAAGUAAUGAGAAUCACAUUGAAAGCGUAUGAUCAUCAGCUCGUUGAUGCAUCCGCAAAGAAAAUCAUCGAAACUGUAAAAAAGAAUGGGUCA
env-839   CGCCCGGAUAGGUUGUACGGUCACCGCUUGUCGUGCUAAAUAGUACGAAAAGGAGGCGGCUUUUUUUAUGGCACAAGUAAUGAGAAUUACACUUAAAGCUUACGAUCACAAAUUAGUAGAUCAGUCAGCUAAGAAAAUUGUUGAAACUGCAAAGAACUCAGGAUCAGAAGUUAGCGGUCC
env-840   ....................................................................................................................................................................................
env-841   CACGACACACACGGGAGAGUGUACAGUCACCGCUUGUCGUACUUGAGUUUAAUAGUACGAAAAGGAGGCGACUUUUUUUAUGGCAAGUCAAGUAAUGAGAAUCACAUUGAAAGCGUAUGAUCAUCAGCUCGUUGAUGCAUCCGCAAAGAAAAUCAUCGAAACUGUAAAAAAGAAUGGGUC
Lba-7-1   CACGACACACACGGGAGAGUGUACAGUCACCGCUUGUCGUACUGAAGUCAAAAGUACGAAAAGGAGGCGACUUUUUUUAUGGCAAGUCAAGUAAUGAGAAUCACAUUGAAAGCGUAUGAUCAUCAGCUCGUUGAUGCAUCCGCAAAGAAAAUCAUCGAAACUGUAAAAAAGAAUGGGUCA
env-842   UCACGACACACACGGGAGAGUGUACAGUCACCGCUUGUCGUACUGAAGUCAAAAGUACGAAAAGGAGGCGACUUUUUUUANGGCAAGUCAAGUAAUGAGAAUCACAUUGAAAGCGUAUGAUCAUCAGCUCGUUGAUGCAUCCGCAAAGAAAUCAUCGAAACUGUAAAAAAGAAUGGGUCA
env-843   CGGGAACGUGUACAGUCACCGCUUGUCGUACUUCUAUUAAGUGCGAAAAGGAGGCGACUUUUUUUAUGGCAAGUCAAGUAAUGAGAAUCACAUUAAAGGCGUACGAUCAUCAGUUAGUUGAUCAGUCUGCAAGCAAAAUCAUCGACACUGUGAAAAAAACAGGAUCACAGGUGAGCGGAC
env-844   GCACGAAAAGUAUGCACAUUUGUGUGCGAAUGAGCAGAGGAUACGCGCGGGCAUNNNNNNNNNNNNNNNNNNNNNNNNNNNNNNNNNNNNNNNNNNNAUCAUCAGUUAGUUGAUCAGUCAGCCAAGAAGAUUAUCGAGAGUGCUAAGAAGACAGGGGCUCAGGUGAGCGGACCGGUACCU
env-845   GCACGAAAAGUAUGCACAUUUGUGUGCAAAUGAGCAGAGGAUACGCGCGGGCAUGUGUACAGUCACCACUUGUCGUACUGAUUUAUAGUGCGAAAAGGAGGCGGCUUUUUUUAUGGCAAGUCAAGUAAUGAGAAUCACAUUGAAGGCUUACGAUCAUCAGUUAGUUGAUCAGUCAGCCAA
env-848   GCACGAAAAGUAUGCACAUUUGUGUGCGAAUGAGCAGAGGAUACGCGCGGGCAUGUGUACAGUCACCACUUGUCGUACUGAUUUAUAGUGCGAAAAGGAGGCGGCUUUUUUUAUGGCAAGUCAAGUAAUGAGAAUCACAUUGAAGGCUUACGAUCAUCAGUUAGUUGAUCAGUCAGCCAA
env-849   GCACGAAAAGUAUGCACAUUUGUGUGCGAAUGAGCAGAGGAUACGCGCGGGCAUGUGUACAGUCACCACUUGUCGUACUGAUUUAUAGU...........................................................................................
env-850   GCACGAAAAGUAUGCACAUUUGUGUGCGAAUGAGCAGAGGAUACGCGCGGGCAUGUGUACAGUCACCACUUGUCGUA.......................................................................................................
env-851   UGGGCGUGAGUCCAUGAUACACGCGGGUAUGUGUGCAGUCACCGCUUGUCGUACAAAAAGCGAUUACAUCGCUUGUAAAAGUGCGAAAAGGAGGCGA..................................................................................
env-853   UGGGCGUGAGUCCAUGAUACACGCGGGUAUGUGUGCAGUCACCGCUUGUCGUACAAAAAGCGAUUACAUCGCUUGUAAAAGUGCGAAAAGGAGGCGACUUUUUUUAUGGCAAGUCAAGUAAUGAGAAUCACUCUGAAGGCUUAUGAUCACAAGUUAGUUGAUUCAUCAGCACAGAAAAUC
env-854   UGGGCGUGAGUCCAUGAUACACGCGGGGAUGUGUGCAGUCACCGCUUGUCGUACAAAAAGCGAUUACAUCGCUUGUAAAAGUGCGAAAAGGAGGCGACUUUUUUUAUGGCAAGUCAAGUAAUGAGAAUCACUCUGAAGGCUUAUGAUCACAAGUUAGUUGAUUCAUCAGCACAGAAAAUC
env-855   UGGGCGUGAGUCCAUGAUACACGCGGGUAUGUGUGCAGUCACCGCUUGUCGUACAAAAAGCGAUUACAUCGCUUGUAAAAGUGCGAAAAGGAGGCGACUUUUUUUAUGGCAAGUCAAGU.............................................................
env-856   UGGGCGUGAGUCCAUGAUACACGCGGGUAUGUGUGCAGUCACCGCUUGUCGUACAAAAAGCGAUU...................................................................................................................
env-861   UGGGCGUGAGUCCAUGAUACACGCGGGUAUGUGUGCAGUCACCGCUUGUCGUACAAAAAGCGAUUACAUCGCUUGUAAAAGUGCGAAAAGGAGGCGACUUUUUUUAUGGCAAGUCAAGUAAUGAGAAUCACUCUGAAGGCUUAUGAUCACAAGUUAGUUGAUUCAUCAGCGCAGAAAAUC
env-862   UGGGCGUGAGUCCAUGAUACACGCGGGUAUGUGUGCAGUCACCGCUUGUCGUACAAAAAGCGAUUACAUCGCUUGUAAAAGUGCGAAAAGGAGGCGACUU................................................................................

          ....................................................................................................................................................................................
          ....................................................................................................................................................................................
          ••••••••••••••••••••••••••••••••••••••••••••••••••••••••••••••••••••••••••••••••••••••••••••••••••••••••••••••••••••••••••••••••••••••••••••••••••••••••••••••R•••••••••••••••••••••
```

```
env-863    GGGUAAGUGUACAGUCACCGCUUGUCGUACUCGUAAAGUGCGAAAAGGAGGCGACUUUUUUUAUGGCAAGUCAAGUAAUGAGAAUCACAUUAAAAGCUUACGAUCAUCAGCUGGUUGAUGCAUCUGCAAGCAAAAUCAUCGAAACUGUAAAAAAGAACGGAUCUGUAGUGAGUGGACCUG
env-864    CGGGAAAGUGUACAGUCACCGCUUGUCGUA..................................................................................................................................................
env-865    UGAAAUCGAAAGUGGAUGACACGCACGGGAAUGUGUACAGUCACCGCUUGUCGUACUGAUUUAUAAGUGCGAAAAGGAGGCGACUUUUUUUAUGGCAAGUCAAGUAAUGAGAAUCACAUUGAAAGCUUAUGAUCAUGAAUUAGUUGAUGCUUCAGCUAAGAAGAUCAUCGAGACUGUAAA
env-866    UGAAAUCGAAAGUGGAUGACACGCACGGGAAUGUGUACAGUCACCGCUUGUCGUACUGAUUUAUAAGUGCGAAAAGGAGGCGACUUUUUUUAUGGCAAGUCAAGUAAUGAGAAUCACAUUGAAAGCUUAUGAUCAUGAAUUAGUUGAUGCUUCAGCUAAGAAGAUCAUCGAGACUGUAAA
env-867    UGAAAUCGAAAGUGGAUGACACGCACGGGAAUGUGUACAGUCACCGCUUGUCGUACUGAUUUAUAAGUGCGAAAAGGAGGCGACUUUUUUUAUGGCAAGUCAAGUAAUGAGAAUCACAUUGAAAGCUUAUGAUCAUGAAUUAGUUGAUGCUUCAGCUAAGAAGAUCAUCGAGACUGUAAA
env-868    UGAAAUCGAAAGUGGAUGACACGCACGGGAAUGUGUACAGUCACCGCUUGUCGU..........................................................................................................................
env-872    UGAAAUCGAAAGUGGAUGACACGCACGGGAAUGUGUACAGUCACCGCUUGUCGUA.........................................................................................................................
env-874    UGAAAUCGAAAGUGGAUGACACGCACGGGAAUGUGUACAGUCACCGCUUGUCGUACUGAUUUAUAAGUGCGAAAAGGAGGCGACUUUUUUUAUGGCAAGUCAAGUAAUGAGAAUCACAUUGAAAGCUUAUGAUCAUGAAUUAGUUGAUGCUUCAGCUAAGAAGAUCAUCGAGACUGUAAA
env-875    UGAAAUCGAAAGUGGAUGACACGCACGGGAAUGUGUACAGUCACCGCUUGUCGUA.........................................................................................................................
env-877    UGAAAUCGAAAGUGGAUGACACGCACGGGAAUGUGUACAGUCACCGCUUGUCGUACUGAUUUAUAAGUGCGAAAAGGAGGCGACUUUUUUUAUGGCAAGUCAAGUAAUGAGAAUCACAUUAAAAGCUUAUGAUCACGAAUUAGUAGAUUCAUCAGCAAAAAAAAUCAUCGAGACUGUAAA
env-878    UGAAAUCGAAAGUGGAUGACACGCACGGGAAUGUGUACAGUCACCGCUUGUCGUACUGAUUUAUAAGUGCGAAAAGGAGGCGACUUUUUUUAUGGCAAGUCAAGUAAUGAGAAUCACAUUGAAAGCUUAUGAUCAUGAAUUAGUUGAUGCUUCAGCUAAGAAGAUCAUCGAGACUGUAAA
env-879    UGAAAUCGAAAGUGGAUGACACGCACGGGAAUGUGUACAGUCACCGCUUGUCGUACUGAUUUAUAAGUGCGAAAAGGAGGCGACUUUUUUUAUGGCAAGUCAAGUAAUGAGAAUCACAUUGAAAGCUUAUGAUCAUGAAUUAGUUGAUGCUUCAGCUAAGAAGAUCAUCGAGACUGUAAA
env-880    UGAAAUCGAAAGUGGAUGACACGCACGGGAAUGUGUACAGUCACCGCUUGUCGUACUGAUUUAUAAGUGCGAAAAGGAGGCGACUUUUUUUAUGGCAAGUCAAGUAAUGAGAAUCACAUUGAAAGCUUAUGAUCAUGAAUUAGUUGAUGCUUCAGCUAAGAAGAUCAUCGAGACUGUAAA
env-892    UGAAAUCGAAAGUGGAUGACACGCACGGGAAUGUGUACAGUCACCGCUUGUCGUACUGAUUUAUAAGUGCGAAAAGGAGGCGACUUUUUUUAUGGCAAGUCAAGUAAUGAGAAUCACAUUGAAAGCUUAUGAUCAUGAAUUAGUUGAUGCUUCAGCUAAGAAGAUCAUCGAGACUGUAAA
env-893    UGAAAUCGAAAGUGGAUGACACGCACGGGAAUGUGUACAGUCACCGCUUGUCGUACUGAUUUAUAAGUGCGAAAAGGAGGCGACUUUUUUUAUGGCAAGUCAAGUAAUGAGAAUCACAUUGAAAGCUUAUGAUCAUGAAUUAGUUGAUGCUUCAGCUAAGAAGAUCAUCGAGACUGUAAA
env-898    UGAAAUCGAAAGUGGAUGACACGCACGGGAAUGUGUACAGUCACCGCUUGUCGUACUGAUUUAUAAGUGCGAAAAGGAGGCGACUUUUUUUAUGGCAAGUCAAGUAAUGAGAAUCACAUUGAAAGCUUAUGAUCAUGAAUUAGUUGAUGCUUCAGCUAAGAAGAUCAUCGAGACUGUAAA
env-899    UGAAAUCGAAAGUGGAUGACACGCACGGGAAUGUGUACAGUCACCGCUUGUCGUACUGAUUUAUAAGUGCGAAAAGGAGGCGACUUUUUUUAUGGCAAGUCAAGUAAUGAGAAUCACAUUGAAAGCUUAUGAUCAUGAAUUAGUUGAUGCUUCAGCUAAGAAGAUCAUCGAGACUGUAAA
env-900    UGAAAUCGAAAGUGGAUGACACGCACGGGAAUGUGUACAGUCACCGCUUGUCGUA.........................................................................................................................
env-901    UGAAAUCGGAAGUGGAUGACACGCACGGGAAUGUGUACAGUCACCGCUUGUC............................................................................................................................
env-902    UGAAAUCGAAAGUGGAUGACACGCACGGGAAUGUGUACAGUCACCGCUUGUCGUACUGAUUUAUAAGUGCGAAAAGGAGGCGACUUUUUUUAUGGCAAGUCAAGUAAUGAGAAUCACAUUGAA...................................................................
env-903    UGAAAUCGAAAGUGGGUGACACGCACGGGAAUGUGUACAGUCACCGCUUGUCGUACUGAUUUAUAAGUGCGAAAAGGAGGCGACUUUUUUUAUGGCAAGUCAAGUAAUGAGAAUCACAUUGAAAGCUUAUGAUCAUGAAUUAGUUGAUGCUUCAGCUAAGAAGAUCAUCGAGACUGUAAA
env-904    UGAAAUCGAAAGUGGAUGACACGCACGGGAAUGUGUACAGUCACCGCUUGUCGUACUGAUUUAUAAGUGCGAAAAGGAGGCGACUUUUUUUAUGGCAAGUCAAGUAAUGAGAAUCACAUUGAAAGCUUAUGAUCAUGAAUUAGUUGAUGCUUCAGCUAAGAAGAUCAUCGAGACUGUAAA
env-906    UGAAAUCGAAAGUGGAUGACACGCACGGGAAUGUGUACAGUCACCGCUUGUCGUACUGAUUUAUAAGUGCGAAAAGGAGGCGACUUUUUUUAUGGCAAGUCAAGUAAUGAGAAUCACAUUGAAAGCUUAUGAUCAUGAAUUAGUUGAUGCUUCAGCUAAGAAGAUCAUCGAGACUGUAAA
Rin-2-1    UGAAAUCGAAAGUGGAUGACACGCACGGGAAUGUGUACAGUCACCGCUUGUCGUACUGAUUUAUAAGUGCGAAAAGGAGGCGACUUUUUUUUAUGGCAAGUCAAGUAAUGAGAAUCACAUUGAAAGCUUAUGAUCAUGAAUUAGUUGAUGCUUCAGCUAAGAAGAUCAUCGAGACUGUAA
env-907    UGAAAUCGAAAGUGGAUGACACGCACGGGA..................................................................................................................................................
env-913    UGAAAUCGAAAGUGGAUGACACGCACGGGAAUGUGUACAGUCACCGCUUGUCGUACUGAUUUAUAAGUGCGAAAAGGAGGCGACUUUUUUUAUGGCAAGUCAAGUAAUGAGAAUCACAUUGAAAGCUUAUGAUCAUGAAUUAGUUGAUGCUUCAGCUAAAAAGAUCAUCGAGACUGUAAA
env-914    UGAAAUCGAAAGUGGAUGACACGCACGGGAAUGUGUACAGUCACCGCUUGUCGUACUGAUUUAUAAGUGCGAAAAGGAGGCGACUUUUUUUAUGGCAAGUCAAGUAAUGAGAAUCACAUUGAAAGCUUAUGAUCAUGAAUUAGUUGAUGCUUCAGCUAAAAAGAUCAUCGAGACUGUAAA
env-915    UGAAAUCGAAAGUGGAUGACACGCACGGGAAUGUGUACAGUCACCGCUUGUCGUACUGAUUUAUAAGUGCGAAAAGGAGGCGACUUUUUUUAUGGCAAG............................................................................
env-916    UGAAAUCGAAAGUGGAUGACACGCACGGGAAUGUGUACAGUCACCGCUUGUCGUACUGAUUUAUAAGUGCGAAAAGGAGGCGACUUUUUUUAUGGCAAGUCAAGUAAUGAGAAUCACAUUGAAAGCUUAUGAUCAUGAAUUAGUUGAUGCUUCAGCUAAAAAGAUCAUCGAGACUGUAAA
env-918    GAGAGUGUACAGUCACCGCUUGUCGUACUGAAGUUAAAAGUACGAAAAGGAGGCGACUUUUUUUAUGGCAAGUCAAGUAAUGAGAAUCACUUUGAAAGCGUAUGAUCAUCAGCUGGUUGAUGCAUCCGCAAAGAAAAUCAUCGAAACUGUAAAAAAGAAUGGAUCACAGGUGAGCGGACC
Lba-10-1   GAGAGUGUACAGUCACCGCUUGUCGUACUGAAGUUAAAAGUACGAAAAGGAGGCGACUUUUUUUAUGGCAAGUCAAGUAAUGAGAAUCACUUUGAAAGCGUAUGAUCAUCAGUUGGUUGAUGCAUCCGCAAAGAAAAUCAUCGAAACUGUAAAAAAGAAUGGAUCACAGGUGAGCGGGCC
[sc-1-1    GAGAGUGUACAGUCACCGCUUGUCGUACUGAAGUUAAAAGUACGAAAAGGAGGCGACUUUUUUAUGGCAAGUCAAGUAAUGAGAAUCACUUUGAAAGCGUAUGAUCAUCAGUUGGUUGAUGCAUCCGCAAAGAAAAUCAUCGAAACUGUAAAAAAGAAUGGAUCACAGGUGAGCGGACCG
env-921    GAGAGUGUACAGUCACCGCUUGUCGUACUGAAGUUAAAAGUACGAAAAGGAGGCGACUUUUUUUAUGGCAAGUCAAGUAAUGAGAAUCACUUUGAAAGCGUAUGAUCAUCAGUUGGUUGAUGCAUCCGCAAAGAAAAUCAUCGAAACUGUAAAAAAGAAUGGAUCACAGGUGAGCGGGCC
env-923    GAGAGUGUACAGUCACCGCUUGUCGUACUGAAGUUAAAAGUACGAAAAGGAGGCGACUUUUUUUAUGGCAAGUCAAGUAAUGAGAAUCACUUUGAAAGCGUAUGAUCAUCAGCUGGUUGAUGCAUCCGCAAAGAAAAUCAUCGAAACUGUAAAAAAGAAUGGAUCACAGGUGAGCGGACC
Csc-1-1    GAGAGUGUACAGUCACCGCUUGUCGUACUGAAGUUAAAAGUACGAAAAGGAGGCGACUUUUUUUAUGGCAAGUCAAGUAAUGAGAAUCACUUUGAAAGCGUAUGAUCAUCAGUUGGUUGAUGCAUCCGCAAAGAAAAUCAUCGAAACUGUAAAAAAGAAUGGAUCACAGGUGAGCGGGCC
Cba-6-1    GAGAGUGUACAGUCACCGCUUGUCGUACUGAAGUUAAAAGUACGAAAAGGAGGCGACUUUUUUUAUGGCAAGUCAAGUAAUGAGAAUCACUUUGAAAGCGUAUGAUCAUCAGUUGGUUGAUGCAUCCGCAAAGAAAAUCAUCGAAACUGUAAAAAAGAAUGGAUCACAGGUGAGCGGACC
Bhy-1-1    AGAUGUGUACAGUCACCGCUUGUCGUACUGAUAAUAAGUGCGAAAAGGAGGCGACUUUUUUUAUGGCAAGUCAAGUAAUGAGAAUCACAUUAAAGGCAUAUGAUCAUCAGUUGGUAGAUGCAUCUGCGAAGAAAAUUAUCGACACUGUAAAGAAAAAUGGAGCAAUGGUGAGCGGUCCGG
env-924    GAUUCAACAGGACACACCCGGAUAAGUGUACAGUCACCACUUGUUGCGCUAUUACAACGAGCAUGAUUCGAUGCCAAUCAGAUCAGUUUGCUCGCAAAAGUG..........................................................................
env-926    GAUUCAACAGGACACACCCGGAUAAGUGUACAGUCACCACUUGUUGCGCUAUUACAACGAGCAUGAUUCGAUGCCAAUCAGAUCAGUUUGCUCGCAAAAGUGCGAAAAGGAGGCGGCUUUUUUUAUGGCUAACGUAAUGAGAAUUACACUGAAGGCGUAUGACCACAAUCAGAUUGAUCA
Bcr-1-1    UUCCACUUCUUUUUUGAUCAGUCAGAUGCGACACACGCGGGGAUGUGUACAGUCACCACUUGUUGUACUGAGAUACGAGAUAACAACGUAUCAUUUUCAAAAGUACGAAAAGGAGGCGGCUUUUUUUAUGGCAAGUCAAGUAAUGAGAAUUACACUUAAAGCUUACGAUCAUCAGAUGGU
env-928    UUCCACUUCUUUUUUGAUCAGUCAGAUGCGACACACGCGGGGAUGUGUACAGUCACCACUUGUUGUACUGAGAUACGAGAUAACAACGUAUCAUUUUCAAAAGUACGAAAAGGAGGCGGCUUUUUUUAUGGCAAGUCAAGUAAUGAGAAUUACACUUAAAGCUUACGAUCAUCAGAUGGU
env-930    UUCCACUUCUUUUUUGAUCAGUCAGAUGCGACACACGCGGGGAUGUGUACAGUCACCACUUGUUGUACUGAGAUACGAGAUAACAACGUAUCAUUUUCAAAAGUACGAAAAGGAGGCGGCUUUUUUUAUGGCAAGUCAAGUAAUGAGAAUUACACUUAAAGCUUACGAUCAUCAGAUGGU
env-932    UUCCACUUCUUUUUUGAUCAGUCAGAUGCGACACACGCGGGGAUGUGUACAGUCACCACUUGUUGUACUGAGAUACGAGAUAACAACGUAUCAUUUUCAAAAGUACGAAAAGGAGGCGGCUUUUUUUAUGGCACAAGUAAUGAGAAUUACACUUAAAGCUUACGAUCAUCAGAUGGUUGA
env-933    UUCCACUUCUUUUUUGAUCAGUCAGAUGCGACACACGCGGGGAUGUGUACAGUCACCACUUGUUGUACUGAGAUACGAGAUAACAACGUAUCAUUUUCAAAAGUACGAAAAGGAGGCGGCUUUUUUUAUGGCAAGUCAAGUAAUGAGAAUUACACUUAAAGCUUACGAUCAUCAGAUGGU
env-934    CUCCUAGCAUUCGAUGGUGAGGAGAAGUUUCUUUUUAUGGGUAAAAACUCAUGAUACGCACGGGAAUGUGUGCAGUCACCGCUUGUCGUACCAAAGCGAGUUUAUCGCUUGCAAAAGUGCGAAAAGGAGGCGACUUUUUUUAUGGCAAGUCAAGU........................
env-935    CUCCUAGCAUUCGAUGGUGCGGAGAAGUUUCUUUUUAUGGGUAAAAGCUCAUGAUACGCACGGGAAUGUGUGCAGUCACCGCUUGUCGUACCAAAGCGAGUUUAUCGCUUGUAAAAGUGCGAAAAGGAGGCGACUU.........................................
env-940    CUCCUAGCAUUCGAUGGUGCGGAGAAGUUUCUUUUUAUGGGUAAAAGCUCAUGAUACGCACGGGAAUGUGUGCAGUCACCGCUUGUCGUACCAAAGCGAGUUUAUCGCUUGCAAAAGUGCGAAAAGGAGGCGACUUUUUUUAUGGCAAGUCAAGUAAUGAGAAUCACAUUAAAAGCUUAU
env-942    CUCCUAGCAUUCGAUGGUGCGGAGAAGUUUCUUUUUAUGGGUAAAAGCUCAUGAUACGCACGGGAAUGUGUGCAGUCACCGCUUGUCGUACCAAAGCGAGUUUAUCGCUUGCAAAAGUGCGAAAAGGAGGCGAC...........................................
env-943    CUUUAAGCAUUCGAUGGUGCAAAGAAGUUUCUUUUUAUGGGUAAAAGCUCAUGAUACGCACGGGAAUGUGUGCAGUCACCGCUUGUCGUACCAAAGCGAGUUUAUCGCUUGCAAAAGUGCGAAAAGGAGGCGACUUUUUUUAUGGCAAGUCAAGUAAUGAGAAUCACAUUAAAAGCUUAU
env-944    CUUUAAGCAUUCGAUGGUGCAAAGAAGUUUCUUUUUAUGGGUAAAAGCUCAUGAUACGCACGGGAAUGUGUGCAGUCACCGCUUGUCGUACCAAAGCGAGUUUAUCGCUUGCAAAAGUGCGAAAAGGAGGCGACUUUUUUUAUGGCAAGUCAAGUAAUGAGAAUCACAUUAAAAGCUUAU
env-945    CUUUAAGCAUUCGAUGGUGCAAAGAAGUUUCUUUUUAUGGGUAAAAGCUCAUGAUACGCACGGGAAUGUGUGCAGUCACCGCUUGUCGUACCAAAGCGAGUUUAUCGCUUGCAAAAGUGCGAAAAGGAGGCGACUUUUUUUAUGGCAAGUCAAGUAAUGAGAAUCACAUUAAAAGCUUAU
env-946    CUCCUAGCAUUCGAUGGUGCGAAGAAGUUUCUUUUUAUGGGUAAAAGCUCAUGAUACGCACGGGAAUGUGU........................................................................................................
env-949    CUUUAAGCAUUCGAUGGUGCAAAGAAGUUUCUUUUUAUGGGUAAAAGCUCAUGAUACGCACGGGAAUGUGUGCAGUCACCGCUUGUCGUACCAAAGCGAGUUUAUCGCUUGCAAAAGUGCGAAAAGGAGGCGACUUUUUUUAUGGCAAGUCAAGUAAUGAGAAUCACAUUAAAAGCUUAU
env-950    CUCCUAGCAUUCGAUGGUGCGGAGAAGUUUCUUUUUAUGGGUAAAAGCUCAUGAUACGCACGGGAAUGUGUGCAGUCACCGCUUGUCGUACCAAAGCGAGUUUAUCGCUUGCAAAAGUGCGAAAAGGAGGCGACUU.........................................
Rin-1-1    CUCCUAGCAUUCGAUGGUGCGGAGAAGUUUCUUUUUAUGGGUAAAAGCUCAUGAUACGCACGGGAAUGUGUGCAGUCACCGCUUGUCGUACCAAAGCGAGUUUAUCGCUUGCAAAAGUGCGAAAAGGAGGCGACUUUUUUUAUGGCAAGUCAAGUAAUGAGAAUCACAUUAAAAGCUUAU
env-951    CUCCUAGCAUUCGAUGGUGCGAAGAAGUUUCUUUUUAUGGGUAAAAGCUCAUGAUACGCACGGGAAUGUGUGCAGUCACCGCUUGUCGUACCAAAGCGAGUUUAUCGCUUGCAAAAGUGCGAAAAGGAGGCGACUUUUUUUAUGGCAAGUCAAGUAAUGAGAAUCACAUUAAAAGCUUAU
env-953    CUCCUAGCAUUCGAUGGUGCGAAGAAGUUUCUUUUUAUGGGUAAAAGCUCAUGAUACGCACGGGAAUGUGUGCAGUCACCGCUUGUCGUACCAAAGCGAGUUUAUCGCUUGCAAAAGUGCGAAAAGGAGGCGACUUUUUUUAUGGCAAGUCAAGUAAUGAGAAUCACUCUCAAGGCUUAU
env-954    CUCCUAGCAUUCGAUGGUGCGAAGAAGUUUCUUUUUAUGGGUAAAAGCUCAUGAUACGCACGGGAAUGUGUGCAGUCACCGCUUGUCGUACCAAAGCGAGUUUAUCGCUUGCAAAAGUGCGAAAAGGAGGCGACUUU........................................
env-955    CUCCUAGCAUUCGAUGGUGCGGAGAAGUUUCUUUUUAUGGGUAAAAGCUCAUGAUACGCACGGGAAUGUGUGCAGUCACCGCUUGUCGUACCAAAGCGAGUUUAUCGCUUGCAAAAGUGCGAAAAGGAGGCGACUUUUUUUAUGGCAAGUCAAGUAAUGAGAAUCACAUUAAAAGCUUAU
env-959    GCGGCUUAAUUGUAAACCAUAGCAGGAAACACCCGGAUAAGUGUACAGUCACCACUUGUUCCGCUGAAAUUAGCAGACUUGGCAAAAAGCCAGAUCAGUAUGUCUGCAAAAAGUGCGAAAAGGAGGCGGCUUUUUUUAUGGCAACAAAUGUAAUGAGAAUUACACUUAAAGCGUAUGACC
env-960    GUGUACAGUCACCGCUUGUCGUACUUCCAAAAAAGUACGAAUAAGGAGGCGACUUUUUUUAUGGCAAACCAGAUCAUGAGAAUCACUUUGAAGGCUUACGAACACCAGUCCGUAGACCGCAGCGCGGCGGAGAUCGUCGAAGCGGUCAAGAGGACCGGCGCCCAGGUCAGCGGACCCGUC
env-961    UGUACAGUCACCGCUUGUCGUACUUCCACAAGUACGAAUAAGGAGGCGACUAUUUUUUAUGGCAAACCAAAUCAUGAGAAUCACCCUGAAGGCUUAUGAACACCAGUCGGUGGACCGCAGCGCGGCGGAGAUCGUCGAAGCGGUCAAGAAGACCGGAGCCCAGGUCAGCGGCCCCGUCCC
env-962    AGUGUACAGUC.....................................................................................................................................................................
env-963    AGUGUACAGUCACCGCUUGUCG..........................................................................................................................................................
env-964    AGUGUACAGUCACCGCUUGUCGUACUGAAUUAAAGUGCGAAAAGGAGGCGACUUUUUUUAUGGCAAGUCAAGUAAUGAGAAUCACAUUGAAAGCUUAUGAUCACAAAUUAGUAGAUUCAUCAGCAAAAAAAAUCAUCGAGACUGUAAAGAAAAACGGAUCACAGGUGAGCGGACCGGUGC
env-965    CGUACUAUUUAAGAGUGCGAAAAGGAGGCGACUUUUUUUAUGGCAAAUCAAGUAAUGAGAAUCACAUUGAAAGCUUACGAACAUCAAUUAGUUGAUGCUUCAGCAAGCAAAAUCAUCGAAACUGUAAAGAAAAACGGAGCAAUCGUGAGUGGGCCGGUACCACUUCCAACUAAGAAGGAA
env-967    CGUACUAUUUAAGAGUGCGAAAAGGAGGCGACUUUUUUUAUGGCAAAUCAAGUAAUGAGAAUCACAUUGAAAGCUUAUGAACAUCAAUUAGUUGAUGCUUCAGCAAGCAAAAUCAUCGAAACUGUAAAGAAAAACGGAGCAAUCGUGAGUGGGCCGGUACCACUUCCAACUAAGAAGGAA
env-968    CGGGUAUGUGUACAGUCA..............................................................................................................................................................
env-970    CGGGUAUGCGUACAGACACCGCUUGUCGUGCUUAAUAGAACGAAAAGGAGGCGACUUUUUUUAUGGCAAGUCAAGUAAUGAGAAUCACACUGAAAGCUUAUGAUCAUCAGCUGGUAGAUACAUCCGCUGCAAAAAUCAUUGAGACUGUAAAGAAAAACGGAUCAAAGGUGAGUGGACCGG
env-971    CGGGUAUGUGUACAGUCACCGCUUGUCGUGCUUAAUAGUACGAAAAGGAGGCGACUUUUUUUAUGGCAAGUCAAGUAAUGAGAAUCACACUGAAAGCUUAUGAUCAUCAGCUGGUAGAUACAUCCGCUGCAAAAAUCAUUGAGACUGUAAAGAAAAACGGAUCAAAGGUGAGUGGACCGG
env-972    CGGGUAUGUGUA....................................................................................................................................................................
env-973    CGGGUAUGUGUACAGUCACCGCUUGUC.....................................................................................................................................................
Lba-3-1    GAGAGUGUACAGUCACCGCUUGUCGUACUGAAGUUAAAAGUACGAAAAGGAGGCGACUUUUUUUAUGGCAAGUCAAGUAAUGAGAAUUACACUUAAAGCAUAUGAUCAUCAGUUAGUUGAUUCAUCUGCGAAAAAAAUUAUCGAAACUGUAAAGAAAAAUGGAGCACAGGUGAGUGGACC
env-980    AGAGUGUACAGUCACCGCUUGUCGUACUUGAGUUUAAUAGUACGAAAAGGAGGCGACUUUUUUUAUGGCAAGUCAAGUAAUGAGAAUUACACUGAAAGCAUAUGAUCACCAAUUGGUUGAUGCAUCUGCAAAGAAAAUUAUCGAAACUGUAAAGAAAAAUGGAUCACAGGUGNUCAUAAA
env-981    ................................................................................................................................................................................
env-983    AGAGUGUACAGUCACCGCUUGUCGUACUUGAGUUUAAUAGUACGAAAAGGAGGCGACUUUUUUUAUGGCAAGUCAAGUAAUGAGAAUUACACUGAAAGCAUAUGAUCACCAAUUGGUUGAUGCAUCUGCAAAGAAAAUUAUCGAAACUGUAAAGAAAAAUGGAUCACAGGUGAGCGGACC
env-984    AGAGUGUACAGUCACCGCUUGUCGUACUUGAGUUUAAUAGUACGAAAAGGAGGCGACUUUUUUUAUGGCAAGUCAAGUAAUGAGAAUUACACUGAAAGCAUAUGAUCACCAAUUGGUUGAUGCAUCUGCAAAGAAAAUUAUCGAAACUGUAAAGAAAAAUGGAUCACAGGUGAGCGGACC
env-985    GAGAGUGUACAGUCACCGCUUGUCGUACUUAAGUUGAAUAGUACGAAAAGGAGGCGACUUUUUUUUAUGGCAAGUCAAGUAAUGAGAAUCACAUUAAAAGCAUAUGAUCAUCAGUUAGUAGAUGCAUCAGCAAAGAAAAUUAUCGAAACUGUAAAGAAAAAUGGAGCACAGGUGAGUGGAC
Csp-8-1    GAGAGUGUACAGUCACCGCUUGUCGUACUUAAGUUGAAUAGUACGAAAAGGAGGCGACUUUUUUUUAUGGCAAGUCAAGUAAUGAGAAUCACAUUAAAAGCAUAUGAUCAUCAGUUAGUAGAUGCAUCAGCAAAGAAAAUUAUCGAAACUGUAAAGAAAAAUGGAGCACAGGUGAGUGGAC
env-986    GAGAGUGUACAGUCACCGCUUGUCGUACUGAAGUUAAAAGUACGAAAAGGAGGCGACUUUUUUUAUGGCAAGUCAAGUAAUGAGAAUCACAUUAAAAGCAUAUGAUCAUCAGUUAGUAGAUGCAUCAGCAAAGAAAAUUAUCGAAACUGUAAAGAAAAAUGGAGCACAGGUGAGUGGACC
Csp-9-1    GAGAGUGUACAGUCACCGCUUGUCGUACUUAAGUUGAAUAGUACGAAAAGGAGGCGACUUUUUUUUAUGGCAAGUCAAGUAAUGAGAAUCACAUUAAAAGCAUAUGAUCAUCAGUUAGUAGAUGCAUCAGCAAAGAAAAUUAUCGAAACUGUAAAGAAAAAUGGAGCACAGGUGAGUGGAC
env-987    GAGAGUGUACAGUCACCGCUUGUCGUACUUAAGUUGAAUAGUACGAAAAGGAGGCGACUUUUUUUUAUGGCAAGUCAAGUAAUGAGAAUCACAUUAAAAGCAUAUGAUCAUCAGUUAGUAGAUGCAUCAGCAAAGAAAAUUAUCGAAACUGUAAAGAAAAAUGGAGCACAGGUGAGUGGAC
env-991    GGGAGAGUGUACAGUCACCGCUUGUCGUACUCGAUUAUAAGUACGAAAAGGAGGCGACUUUUUUUUAUGGCAAGUCAAGUAAUGAGAAUCACUUUGAAAGCGUAUGAUCAUCAGUUAGUAGAUGCAUCCGCUAAGAAAAUCAUCGAAACUGUAAAGAAAAACGGAUCUAAGGUGAGUGGAC
env-992    GGGAGAGUGUACAGUCACCGCUUGUCGUACUCGAUUAUAAGUACGAAAAGGAGGCGACUUUUUUUUAUGGCAAGUCAAGUAAUGAGAAUCACUUUGAAAGCGUAUGAUCAUCAGUUAGUAGAUGCAUCCGCUAAGAAAAUCAUCGAAACUGUAAAGAAAAACGGAUCUAAGGUGAGUGGAC

           ................................................................................................................................................................................
           ................................................................................................................................................................................
           ........................................................................................................................................................R.......................
```

```
env-993   GGGAGAGUG..........................................................................................................................................................
Dlo-2-1   GGGAGAGUGUACAGUCACCGCUUGUCGUACUCGAUUAUAAGUACGAAAAGGAGGCGACUUUUUUUAUGGCAAGUCAAGUAAUGAGAAUCACUUUGAAAGCGUAUGAUCAUCAGUUAGUAGAUGCAUCCGCUAAGAAAAUCAUCGAAACUGUAAAGAAAAACGGAUCUAAGGUGAGUGGAC
env-994   GGGAGAGUGUACAGUCACCGCUUGUCGUACUGAUGUGAAAAGUACGAAAAGGAGGCGACUUUUUUUAUGGCAAGUCAAGUAAUGAGAAUCACAUUGAAAGCGUAUGAUCAUCAGUUAGUAGAUGCAUCCGCUAAGAAAAUCAUCGAAACUGUAAAGAAAAAUGGAUCUAAGGUGAGUGGA
env-995   NNNNNNNNNNNNNNNNNNNNNNNNNNNNNNNNNNNNNNNNNNNNNNNNNNNNNNNNNNNNNNNNNNNNNNNNNNNNNNNNNNNNNNNNNNNNNNNNNNNNNNNNNNNNNNNNNNNNNNNNNNNNNNNNNNNNNNNNNNNNNNNNGAUGGAUCAUCAGUUA
env-996   .......................................................................................................................................................................
env-1006  GGGAGAGUGUACAGUCACC....................................................................................................................................................
env-1007  GGGAGAGUGUACAGUCACCGCUUGUCGUACUCGAUUAUAAGUACGAAAAGGAGGCGACUUUUUUUAUGGCAAGUCAAGUAAUGAGAAUCACUUUAAAGGCGUAUGAUCAUCAGUUAGUAGAUGCAUCCGCUAAGAAAAUCAUCGAAACUGUAAAGAAAAAUGGAUCUAAGGUGAGUGGAC
env-1008  CACGACACACACGGGAGAGUGUACAGUCACCGCUUGUCGUACUGAAGUCAAAAGUACGAAAAGGAGGCGACUUUUUUUAUGGCAAGUCAAGUAAUGAGAAUCACUUUAAAGGCGUAUGAUCAUCAGUUAGUAGAUGCAUCCGCUAAGAAAAUCAUCGAAACUGUAAAGAAAAAUGGAUCU
env-1009  GGGAGAGUGUACAGUCACC....................................................................................................................................................
env-1018  GGGAGAGUGUACAGUCACCGCUUGUCGUACUGAAGUGAAAAGUACGAAAAGGAGGCGACUUUUUUUAUGGCAAGUCAAGUAAUGAGAAUCACUUUAAAGGCGUAUGAUCAUCAGUUAGUAGAUGCAUCCGCUAAGAAAAUCAUCGAAACUGUAAAGAAAAAUGGAUCUAAGGUGAGUGGA
env-1019  GGGAGAGUGUACAGUCACCGCUUGUCGUACUCGAUUAUAAGUACGAAAAGGAGGCGACUUUUUUUAUGGCAAGUCAAGUAAUGAGAAUCACUUUAAAGGCGUAUGAUCAUCAGUUAGUAGAUGCAUCCGCUAAGAAAAUCAUCGAAACUGUAAAGAAAAACGGAUCUAAGGUGAGUGGAC
Dlo-1-1   GGGAGAGUGUACAGUCACCGCUUGUCGUACUCGAUUAUAAGUACGAAAAGGAGGCGACUUUUUUUAUGGCAAGUCAAGUAAUGAGAAUCACUUUAAAGGCGUAUGAUCAUCAGUUAGUAGAUGCAUCCGCUAAGAAAAUCAUCGAAACUGUAAAGAAAAAUGGAUCUAAGGUGAGUGGAC
env-1020  GUGUACAGUCACCGCUUGUCGUACUUAAGAAACAAAAGUGCGAAAAGGAGGCGACUUUUUUUAUGGCAAAUCAACAGGUAAUGAGAAUUACAUUGAAAGCCUACGAUCAUCAAUUAGUUGAUGCAUCUGCAAAAAAAAUCAUCGAAACAGUUAAGAAGAACGGAUCUCAGGUAAGUGGACC
env-1021  GUGUACAGUCACCGCUUGUCGUACUUAAGAAACAAAAGUGCGAAAAGGAGGCGACUUUUUUUAUGGCAAAUCAACAGGUAAUGAGAAUUACAUUGAAAGCCUACGAUCAUCAAUUAGUUGAUGCAUCUGCAAAAAAAAUCAUCGAAACAGUUAAGAAGAACGGAUCUCAGGUAAGUGGAC
env-1022  GUGUACAGUCACCGCUUGUCGUACUUAAGAAACAAAAGUGCGAAAAGGAGGCGACUUUUUUUAUGGCAAAUCAACAGGUAAUGAGAAUUACAUUGAAAGCCUACGAUCAUCAAUUAGUUGAUGCAUCUGCAAAAAAAAUCAUCGAAACAGUUAAGAAGAACGGAUCUCAGGUAAGUGGAC
Lba-2-1   CACGACACACACGGGAGAGUGUACAGUCACCGCUUGUCGUACUGAGUUUAAAAGUACGAAAAGGAGGCGACUUUUUUUAUGGCAAGUCAAGUAAUGAGAAUCACAUUGAAAGCGUAUGAUCACCAGCUGGUAGAUGCAUCCGCAAAGAAAAUCAUCGAAACUGUAAAAAAGAACGGAUCA
env-1023  AUAAAAGGAGUGCGCCUUUGGUGGUGUGAAAUGCAUGACACACACGGCAGAGUGUACAGUCACCGCUUGUCGUACUUAGAAAUGG..................................................................................
env-1024  AUAAAAGGAGUGCGCCUUUGGUGGUGUGAAAUGCAUGACACACACGGCAGAGUGUACAGUCACCGCUUGUC................................................................................................
env-1026  AUAAAAGGAGUGCGCCUUUGGUGGUGUGAAAUGCAUGACACACACGGCAGAGUGUACAGUCACCGCUUGUCGUACUUAGAAAUGGAAAGUGCGAAAAGGAGGCGACUUUUUUUAUGGCAAGUCAGGUAAUGAGAAUUACAUUAAAAGCUUACGAUCAUCAGUUAGUUGAUGCAUCUGCCA
env-1027  AUAAAAGGAGUGCGCCUUUGGUGGUGUGAAAUGCAUGACACACACGGCAGAGUGUACAGUCACCGCUUGUCGUACUUAGAAAUGGAAAGUGCGAAAAGGAGGCGACUUUUUUUAUGGCAAGUCAGGUAAUGAGAAUUACAUUAAAAGCUUACGAUCAUCAGUUAGUUGAUGCAUCUGCCA
env-1028  AUAAAAGGAGUGCGCCUUUGGUGGUGUGAAAUGCAUGACACACACGGCAGAGUGUACAGUCACCGCUUGUCGUACUUAGAAAUGGAAAGUGCGAAAAGGAGGCGACUUUUUUUAUGGCAAGUCAGGUAAUGAGAAUUACAUUAAAAGCUUACGAUCAUCAGUUAGUUGAUGCAUCUGCCA
env-1029  AUAAAAGGAGUGCGCCUUUGGUGGUGUGAAAUGCAUGACACACACGGCAGAGUGUACAGUCACCGCUUGUCGU..............................................................................................
env-1030  CGAGACCGUGUACAGUCACCGCUUGUCGU..........................................................................................................................................
env-1032  CGAGACCGUGUACAGUCACCGCUUGUCGUACUUCGAUUAAAAGUGCGAAAAGGAGGCGACUUUUUUUAUGGCAAGUCAGGUUAUGAGAAUCACAUUAAAGGCUUACGACCAUCAGCUGAUCGACCAGUCCGCUGGAAAGAUCAUCGAGACAGUAAAGAAAACAGGAGCACAGGUGAGUGG
env-1033  CGAGACCGUGUACAGUCACCGCUUGUCGUACUUCGAUUAAAAGUGCGAAAAGGAGGCGACUUUUUUUAUGGCAAGUCAGGUUAUGAGAAUCACAUUAAAGGCUUACGACCAUCAGCUGAUCGACCAGUCCGCUGGAAAGAUCAUCGAGACAGUAAAGAAAACAGGAGCACAGGUGAGUGG
env-1034  CGAGACCGUGUACAGUCACCGCUUGUCGUG.........................................................................................................................................
env-1035  UGGGGCGAUACAUCAGAGAUGUACCGCAGGACACACACGGAGAAGUGUACAGUCACCGCUUGUCGUACUAUUUACAAAGUGCGAAAAGGAGGCGACUUUUUUUAUGCAAGUCAAGUAAUGAGAAUCACUCUCAAGGCUUAUGAUCACGAAUUAGUUGAUUCAUCAGCAAAGAAAAUCAUC
env-1036  AUGGGGCGAUACAUCAGAGAUGUACAGCAGGACACACACGGAGAAGUGUACAGUCACCGCUUGUCGUA...................................................................................................
env-1037  UGGGGCGAUACAUCAGAGAUGUACCGCAGGACACACACGGAGAAGUGUACAGUCACCGCUUGUCGUA....................................................................................................
env-1038  UGGGGCGAUACAUCGGAGAUGUACC..............................................................................................................................................
env-1039  UGGGGCGAUACAUCAGAGAUGUACCGCAGGACACACACGGAGAAGUGUACAGUCACCGCUUGUCGUACUAUUUACAAAGUGCGAAAAGGAGGCGACUUUUUUUAUGGCAAGUCAAGUAAUGAGAAUCACUCUCAAGGCUUAUGAUCACGAAUUAGUUGAUUCAUCAGCAAAGAAAAUCAU
env-1040  UGGGGCGAUACAUCAGAGGUGUACCGCAGGACACACACGGAGAAGUGUACAGUCACCGCUUGUCGUACUAUUUACAAAGUGCGAAAAGGAGGCGACUUUUUUUAUGGCAAGUCAAGUAAUGAGAAUCACUCUCAAGGCUUAUGAUCACGAAUUAGUUGAUUCAUCAGCAAAGAAAAUCAU
env-1041  UGGGGCGAUACAUCAGAGAUGUACCGCAGGACACACACGGAGAAGUGUACAGUCACCGCUUGUCGUAC...................................................................................................
env-1042  UGAGGCGAUACAUCAGAGAUGUACCGCAGGACACACACGGAGAAGUGUACAGUC.................................................................................................................
env-1043  UGGGGCGAUACAUCAGAGAUGUACCGC............................................................................................................................................
env-1044  UGGGGCGAUACAUCAAAGAUGUACCGCAGGACACACACGGAGAAGUGUACAGUCACCGCUUGUC.......................................................................................................
env-1045  UGGGGCGAUACAUCAGAGAUGUACCGCAGGACACACACGGAGAAGUGUACAGUCACCGCUUGUCGUACUAUUUACAAAGUGCGAAAAGGAGGCGACUUUUUUUAUGGCAAGUCAAGUAAUGAGAAUCACUCUCAAGGCUUAUGAUCACGAAUUAGUUGAUUCAUCAGCAAAGAAAAUCAU
env-1046  .......................................................................................................................................................................
env-1052  UGGGGCGAUACAUCAGAGAUGUACCGCAGGACACACACGGAGAAGUGUACAGUC.................................................................................................................
env-1053  AUGGGGCGAUACAUCAGAGAUGUACAGCAGGACACACACGGAGAAGUGUACAGUCACCGCUUGUCGUAC..................................................................................................
env-1054  UGGGGCGAUACAUCGGAGAUGUACC..............................................................................................................................................
env-1055  UGGGGCGAUACAUCAGAGAUGUACCGCAGGACACACACGGAGAAGUGUACAGUCACCGCUUGUCGUACUAUUUACAAAGUGCGAAAAGGAGGCGACUUUUUUUAUGGCAAGUCAAGUAAUGAGAAUCACUCUCAAGGCUUAUGAUCACGAAUUAGUUGAUUCAUCAGCAAAGAAAAUCAU
env-1056  UGGGGCGAUACAUCAGAGAUGUACCGCAGGACACACACGGAGAAGUGUACAG...................................................................................................................
env-1057  UGGGGCGAUACAUCAGAGAUGUACCGCAGGACACACACGGAGAAGUGUACAGUCACCGCUUGUCGU.....................................................................................................
env-1058  UGGGGCGAUACAUCAAAGAUGUACCGC............................................................................................................................................
env-1059  UGGGGCGAUACAUCAGAGAUGUACCGCAGGACACACACGGAGAAGUGUACAGUCACCGCUUGUCGUACUAUUUACAAAGUGCGAAAAGGAGGCGACUUUUUUUAUGGCAAGUCAAGUAAUGAGAAUCACUCUCAAGGCUUAUGAUCACGAAUUAGUUGAUUCAUCAGCAAAGAAAAUCAU
Cha-1-1   CGGGAACGUGUACAGUCACCGCUUGUCGUACUUCUACUAAGUGCGAAAAGGAGGCGACUUUUUUUAUGGCAAGUCAAGUAAUGAGAAUCACAUUAAAGGCGUACGAUCAUCAGUUAGUUGAUCAGUCUGCAAGCAAAAUCAUCGACACUGUGAAAAAAACAGGAUCACAGGUGAGCGGAC
env-1065  CGGGAACGUGUACAGUCACCGCUUGUCGUACUUCUACUAAGUGCGAAAAGGAGGCGACUUUUUUUAUGGCAAGUCAAGUAAUGAGAAUCACAUUAAAGGCGUACGAUCAUCAGUUAGUUGAUCAGUCUGCAAGCAAAAUCAUCGACACUGUGAAAAAAACAGGAUCACAGGUGAGCGGAC
env-1066  CGGGAACGUGUACAGUCACCGCUUGUCGUACUUCUACUAAGUGCGAAAAGGAGGCGACUUUUUUUAUGGCAAGUCAAGUAAUGAGAAUCACAUUAAAGGCGUACGAUCAUCAGUUAGUUGAUCAGUCUGCAAGCAAAAUCAUCGACACUGUGAAAAAAACAGGAUCACAGGUGAGCGGAC
Cba-1-1   UGAGGUUCCAACAGUCGCAGACGUAUUGAUACGUCAAAAUUUAGGACACACACGGGAGAGUGUACAGUCACCGCUUGUCGUACUUCAACAAAAAGUGCGAAAAGGAGGCGACUUUUUUUAUGGCAAAUCAGGUAAUGAGAAUUACAUUGAAAGCUUAUGACCAUCAGCUUGUUGAUGCGU
Cba-8-1   UGAGGUUCCAACAGUCGCAGACGUAUUGAUACGUCAAAAUUUAGGACACACACGGGAGAGUGUACAGUCACCGCUUGUCGUACUUCAACAAAAAGUGCGAAAAGGAGGCGACUUUUUUUAUGGCAAAUCAGGUAAUGAGAAUUACAUUGAAAGCUUAUGACCAUCAGCUUGUUGAUGCGU
env-1067  UGAGGUUCCAACAGUCGCAGACGUAUUGAUACGUCAAAAUUUAGGACACACACGGGAGAGUGUACAGUCACCGCUUGUCGUACUUCAACAAAAAGUGCGAAAAGGAGGCGACUUUUUUUAUGGCAAAUCAGGUAAUGAGAAUUACAUUGAAAGCUUAUGACCAUCAGCUUGUUGAUGCGU
Lba-20-1  UGAGGUUCCAACAGUCGCAGACGUAUUGAUACGUCAAAAUUUAGGACACACACGGGAGAGUGUACAGUCACCGCUUGUCGUACUUCAACAAAAAGUGCGAAAAGGAGGCGACUUUUUUUAUGGCAAAUCAGGUAAUGAGAAUUACAUUGAAAGCUUAUGACCAUCAGCUUGUUGAUGCGU
Lba-8-1   UGAGGUUCCAACAGUCGCAGACGUAUUGAUACGUCAAAAUUUAGGACACACACGGGAGAGUGUACAGUCACCGCUUGUCGUACUUCAACAAAAAGUGCGAAAAGGAGGCGACUUUUUUUAUGGCAAAUCAGGUAAUGAGAAUUACAUUGAAAGCUUAUGACCAUCAGCUUGUUGAUGCGU
env-1068  UGGGGUUCCAACAGUCGCAGACGUAUUGAUACGUCAAAAUUUAGGACACACACGGGAGAGUGUACAGUCACCGCUUGUCGUACUUCAACAAAAAGUGCGAAAAGGAGGCGACUUUUUUUAUGGCAAAUCAGGUAAUGAGAAUUACAUUGAAAGCUUAUGACCAUCAGCUUGUUGAUGCGU
Lba-13-1  GUGUACAGUCACCGCUUGUCGUACUGAAGUAAAGUGCGAAAAGGAGGCGGCUUUUUUUAUGGCAAGUCAAGUAAUGAGAAUCACACUGAAGGCAUACGACCAUCAGUUGGUGGAUGCAUCUGCAAAGAAAAUUAUCGAAACUGUGAAAAAGAAUGGCUCACAGGUGAGCGGCCCUGUGCC
Csp-7-1   GGAUAUGUGUACAGUCACCGCUUGUCGUGCUUCAAAUAGUACGAAAAGGAGGCGACUUUUUUUAUGGCAAGUCAAGUAAUGAGAAUCACAUUGAAAGCUUAUGAUCAUCAGCUGGUUGACCAGUCUGCUGCAAAAAUCAUCGAGACUGUAAAGAAGAACGGAUCUAAGGUGAGCGGACCG
env-1069  GAUAUGUGUACAGUCACCGCUUGUCGUGCUUCAAAUAGUACGAAAAGGAGGCGACUUUUUUUAUGGCAAGUCAAGUAAUGAGAAUCACAUUAAAAGCUUAUGAUCAUCAGUUAGUAGAUCAGUCCGCUGGAAAGAUCAUCGAGACAGUAAAAAAGACAGGAUCUCAGGUGAGCGGACCAG
env-1070  GAUAUGUGUACAGUCACCGCUUGUCGUGCUUCAAAUAGUACGAAAAGGAGGCGACUUUUUUUAUGGCAAGUCAAGUAAUGAGAAUCACACUUAAAGCUUAUGAUCAUCAGCUAGGUUGACCAGUCUGCUGCAAAAAUCAUUGAGACUGUAAAGAAGAACGGAUCAAAGGUGAGCGGACCG
env-1071  GAUAUGUGUACAGUCACCGCUUGUCGUGCUUCAAAUAGUACGAAAAGGAGGCGACUUUUUUUAUGGCAAGUCAAGUAAUGAGAAUCACACUUAAAGCUUAUGAUCAUCAGCUGGUUGACCAGUCUGCUGCAAAAAUCAUUGAGACUGUAAAGAAGAACGGAUCAAAGGUGAGCGGACCGG
env-1072  GAUAUGUGUA.............................................................................................................................................................
Esp-1-1   AAGUGUACAGUCACCGCUUGUCGUACUGAGGUAAAGUGCGAAAAGGAGGCGACUUUUUUUAUGGCAAGUCAGGUAAUGAGAAUCACACUGAAGGCAUAUGACCAUCAGUUAGUGGAUGCAUCUGCGAAGAAAAUCAUCGAAACUGUAAAGAAAAACGGAUCACAGGUGAGCGGCCCGGUG
env-1073  GAGACCGUGUACAGUCACCGCUUGUCGUAC.........................................................................................................................................
env-1074  GAGACCGUGUACAGUCACCGCUUGUCGU...........................................................................................................................................
env-1075  GAGACCGUGUACAGUCACCGCUUGUCGUACUUCGUUAAAAGUGCGAAAAGGAGGCGACUUUUUUUAUGGCAAGUCAGGUUAUGAGAAUCACAUUAAAGGCGUACGACCAUCAGCUGGUUGAUCAGUCUGCCGCUGGAAAAAUCAUCGAGACAGUAAAARAGCAGGAUCUAAGGUGAGUGGACC
Csa-1-1   GGAACGUGUACAGUCACCGCUUGUCGUACUUCUACUAAGUGCGAAAAGGAGGCGACUUUUUUUAUGGCAAGUCAAGUAAUGAGAAUCACAUUAAAAGCGUAUGAUCAUCAGUUGGUUGAUCAGUCUGCAGGCAAAAUCAUCGACACUGUAAAAAAGACAGGAUCAAAGUGAGCGGACCG
env-1076  UGACAGAUAAACGACUGUAAGGUUGCGAAGAGAUUAGCUGUAAGCGACAAUCCUUUGGAAGGACGACGGAAAUCGGUUAUAUGGAAUCCGCCGCAGACAACCGCCCUUAUGACAGGAAACACACGGGUAUGUGUACAGUCACCGCUUGUCGUACUAUUUAAAAGUGCGAAAAGGAGGCGA
env-1077  UUCCACUUCUUUUUUGAUCAGUCAGAUGCGACACACGCGGGGAUGUGUACAGUCACCACUUGUUGUACUGAGAUACGAGAUAACAACGUAUCAUUUUCAAAAGUACGAAAAGGAGGCGGCUUUUUUUAUGGCAAGUCAAGUAAUGAGAAUUACACUUAAAGCUUACGAUCAUCAGAUGGU
env-1078  UGGGGCGAUACAUCAGAGAUGUACCGCAGGACACACACGGAGAAGUGUACAGUCACCGCUUGUCGUACUAUUUACAAAGUGCGAAAAGGAGGCGACUUUUUUUAUGGCAAGUCAAGUAAUGAGAAUCACUCUCAAGGCUUAUGAUCACGAAUUAGUUGAUUCAUCAGCAAAGAAAAUCAU
env-1081  UGGGGCGACACAUCAGAGAUGUACCGCAGGACACACACGGAGAAGUGUACAGUCACCGCUUGUCGUACUAUUUACAAAGUGCGAAAAGGAGGCGACUUUUUUUAUGGCAAGUCAAGUA...............................................
env-1082  GGGGCGAUACAUCAGAGAUGUACCGCAGGACACACACGGAGAAGUGUACAGUCACCGCUUGUCGUACU...................................................................................................
env-1083  UGGGGCGAUACAUCAGAGAUGUACCGCAGGACACACACGGAGAAGUGUACAGUCACCGCUUGUCGUACUAUUUACAAAGUGCGAAAAGGAGGCGACUUUUUUUAUGGCAAGUCAAGUAAUGAGAAUCACUCUCAAGGCUUAUGAUCACGAAUUAGUUGAUUCAUCAGCAAAGAAAAUCAU
env-1084  UGGGGCGAUACAUCAGAGAUGUACCGCAGGACACACACGGAGAAGUGUACAGUCACCGCUUGUCGUACUAUUUACAAAGUGCGAAAAGGAGGCGACUUUUUUUAUGGCAAGUCAAGUAAUGAGAAUCACAUUGAAAGCUUAUGAUCAUGA.................
env-1085  UGGGGCGAUACAUCGGAGAUGUACCGCAGGACACACACGGAGAAGUGUACAGUCACCGCUUGUCGUACUAUUUACAAAGUGCGAAAAGGAGGCGACUUUUUUUAUGGCAAGUCAAGUAAUGAGAAUCACAUUAAAAGCUUAUGAUCACGAAUUAGUAGAUUCAUCAGCAAAAAAAAUCAU
env-1086  UGGGGCGAUACAUCAGAGAUGUACCGCAGGACACACACGGAGAAGUGUACAGUCACCGCUUGUCGUACUAUUUACAAAGUGCGAAAAGGAGGCGACUUUUUUUAUGGCAAGUCAAGUAAUGAGAAUCACUCUCAAGGCUUAUGAUCACGAAUUAGUUGAUUCAUCAGCAAAGAAAAUCAU
Cha-2-1   AACGUGUACAGUCACCGCUUGUCGUACUUCAAUAAUAGUGCGAAAAGGAGGCGACUUUUUUUAUGGCAAGUCAAGUAAUGAGAAUCACAUUGAAGGCUUAUGAUCAUCAGUUAGUAGAUCAGUCUGCUGGAAAAAUCAUCGAAACUGUAAAGAAGACAGGGGCGCAGGUGAGCGGACCA
Lba-5-1   UGUACAGUCACCGCUUGUCGUACUGAGGUAAAGUGCGAAAAGGAGGCGGCUUUUUUUAUGGCAAGUCAAGUAAUGAGAAUCACACUGAAGGCAUAUGACCAUCAGUUAGUGGAUGCAUCUGCGAAGAAAAUUAUCGAAACUGUAAAGAAAAAUGGAUCACAGGUGAGCGGACCCGUACCA
Csp-1-1   CGGGCAUGUGUACAGUCACCGCUUGUCGUGCUUAAAUAGUACGAAAAGGAGGCGACUUUUUUUAUGGCAAGUCAAGUAAUGAGAAUCACAUUAAAAGCUUAUGAUCAUCAGCUGGUUGACAUCCGCUGCAAAAAUCAUCGAGACUGUAAAGAAAAACGGAUCAAAGGUGAGCGGACCG
env-1087  CGGGCAUGUGUACAGUCACCGCUUGUCGUGCUUAAAUAGUACGAAAAGGAGGCGACUUUUUUUAUGGCAAGUCAAGUAAUGAGAAUCACAUUAAAAGCUUAUGAUCAUCAGCUGGUUGACACAUCCGCUGCAAAAAUCAUCGAGACUGUAAAGAAAAACGGAUCAAAGGUGAGCGGACCG
env-1088  AGUGUACAGUCACCGCUUGUCGUACUUCUUCCAAAAGAAAGUGCGAAAAGGAGGCGACUUUUUUUAUGGCAAACCAGGUAAUGAGAAUCACUUUGAAGGCUUAUGAACACCAGCUGGUGGACACGAGCGCAGCCCGCAUCGUUGACACGGUCAAGAAGACCGGCGCUGCGGUGAGCGGACC
env-1089  CGCCCGGCACUGUGUACAGUCACCGCUUGUCGUACUUUCAAAAAAGUGCGAAAAGGAGGCGACUUUUUUUAUGGCAAAUCAGGUAAUGAGAAUCACGUUGAAGGCGUAUGAUCACAAGCUGGUUGAUCAGUCUGCCGGAAAGAUUGUCGACACGGUGAAGAGAACGGGAGCAAAGGUUCUC
env-1093  ACAGUCACCGCUUGUCGUACUUUAAAAAAGUGCGAAAAGGAGGCGACUUUUUUUAUGGCAAAUCAGGUAAUGAGAAUCACGUUGAAGGCGUAUGAUCACAAGCUGGUUGAUCAGUCUGCCGGAAAGAUUGUCGACACGGUGAAGAGAACGGGAGCUAAGGUUUCCGGCCCGAUUCCGCUG
env-1094  CGCCCGGCACUGUGUACAGUCACCGCUUGUCGUAC....................................................................................................................................
env-1095  ACACACGGCACUGUGUACAGUCACCGCUUGUCGUACUUUAAAAAAGUGCGAAAAGGAGGCGACUUUUUUUAUGGCAAAUCAGGUAAU...............................................................................

          .......................................................................................................................................................................
          .......................................................................................................................................................................
          ...................................................................................................................................................R...................
```

```
env-1096  GCCCGGCACUGUGUACAGUCACCGCUUGUCGUACUUUCAAAAAAAGUGCGAAAAGGAGGCGACUUUUUUUAUGGCAAAUCAGGUAAUGAGAAUCACGUUGAAGGCGUAUGAUCACAAGCUGGUUGAUCAGUCUGCCGGAAAGAUUGUCGACACGGUGAAGAGAACGGGAGCAAAGGUCUC
env-1097  CACACGGCACUGUGUACAGUCACCGCUUG........................................................................................................................................................
env-1099  CGCCCGGCACUGUGUACAGUCACCGCUUGUCGUACUUUCAAAAAAAGUGCGAAAAGGAGGCGACUUUUUUUAUGGCAAAUCAGGUANNNNNNNNNNNNNNNNNNNNNNNNNNNNNNNNNNNNNNNNNNNNNNNNNNNNNNNNNNNNNNNNNNNNNNNNNNNAGGAAGUAGUUACAAUUCUCCGU
env-1100  CGCCCGGCACUGUGUACAGUCACCGCUUGUCGUACUUUCAAAAAAAGUGCGAAAAGGAGGCGACUUUU.................................................................................................................
Slo-1-1   ACACACGGCACUGUGUACAGUCACCGCUUGUCGUACUUUAAAAAAAGUGCGAAAAGGAGGCGACUUUUUUUAUGGCAAAUCAGGUAAUGAGAAUCACGUUGAAGGCGUAUGAUCACAAGCUGGUUGAUCAGUCUGCCGGAAAGAUUGUCGACACGGUGAAGAGAACGGGAGCUAAGGUUUC
env-1101  ACACACGGCACUGUGUACAGUCACCGCUUGUCGUACUUUAAAAAAAGUGCGAAAAGGAGGCGACUUUU.................................................................................................................
env-1102  CGCCCGGCACUGUGUACAGUCACCGCUUGUCGUACUUUCAAAAAAAGUGCGAAAAGGAGGCGACUUUU.................................................................................................................
env-1103  CGCCCGGCACUGUGUACAGUCACCGCUUGUCGUACUUUCAAAAAAAGUGCGAAAAGGAGGCGACUUUUUUUAUGGCAAAUCAGGUAAUGAGAAUCACGUUGAAGGCGUAUGAUCACAAGCUGGUUGAUCAGUCUGCCGGAAAGAUUGUCGACACGGUGAAGAGAACGGGAGCAAAGGUCU
env-1104  CACANNNNNNNNNNNNNNNNNNNNNNNNNNNNNNNNNNNNNNNNNNNNNNNNNNNNNNNNNUCGACACGGUGAAGAGAACGGGAGCUAAGGUUUCCGGCCCGAUUCCGCUGCCGACCAAGAAAGAGGUAGUUACAAUUCUUCGUGCGGUCCACAAGUAUAAGGACAGCCGCGAGCAGUU
env-1105  CACACGGCACUGUGUACAGUCACCGCUUG........................................................................................................................................................
env-1106  CACACGGCACUGUGUACAGUCACCGCUUGUCGUACUUUAAAAAAAGUGCGAAAAGGAGGCGACUUUUUUUAUGGCAAAUCAGGUAAUGAGAAUCACGUUGAAGGCGUAUGAUCACAAGCUGGUUGAUCAGUCUGCCGGAAAGAUUGUCGACACGGUGAAGAGAACGGGAGCUAAGGUUUCC
env-1107  GCCCGGCACUGUGUACAGUCACCGCUUG.........................................................................................................................................................
env-1108  CGCCCGGCACUGUGUACAGUCACCGCUUGUCGUACUUUCAAAAAAAGUGCGAAAAGGAGGCGACUUUUUUUAUGGCAAAUCAGGUAAUGAGAAUCACGUUGAAGGCGUAUGAUCACAAGCUGGUUGAUCAGUCUGCCGGAAAGAUUGUCGACACGGUGAAGAGAACGGGAGCAAAGGUCU
env-1115  CGCCCGGCACUGUGUACAGUCACCGCUUGUCGUAC..................................................................................................................................................
env-1118  GCCCGGCACUGUGUACAGUCACCGCUUGUCGUACUUUCAAAAAAAGUGCGAAAAGGAGGCGACUUUUUUUAUGGCAAAUCAGGUAAUGAGAAUCACGUUGAAGGCGUAUGAUCACAAGCUGGUUGAUCAGUCUGCCGGAAAGAUUGUCGACACGGUGAAGAGAACGGGAGCAAAGGUCUC
env-1123  CACACGGCACUGUGUACAGUCACCGCUUGUCGUACUUUAAAAAAAGUGCGAAAAGGAGGCGACUUUUUUUAUGGCAAAUCAGGUAAUGAGAAUCACGUUGAAGGCGUAUGAUCACAAGCUGGUUGAUCAGUCUGCCGGAAAGAUUGUCGACACGGUGAAGAGAACGGGAGCUAAGGUUUCC
env-1124  CGCCCGGCACUGUGUACAGUCACCGCUUGUCGUAC..................................................................................................................................................
env-1145  CGCCCGGCACUGUGUACAGUCACCGCUUGUCGUACUUUCAAAAAA........................................................................................................................................
env-1146  CGCCCGGCACUGUGUACAGUCACCGCUUGUCGUACUUUCAAAAAAAGUGCGAAAAGGAGGCGACUUUUUUUAUGGCAAAUCAGGUAAUGAGAAUCACGUUGAAGGCGUAUGAUCACAAGCUGGUUGAUCAGUCUGCCGGAAAGAUUGUCGACACGGUGAAGAGAACGGGAGCAAAGGUCUC
env-1147  .....................................................................................................................................................................................
env-1148  CGCCCGGCACUGUGUACAGUCACCGCUUGUCGUACUUUCAAAAAAAGUGCGAAAAGGAGGCGACUUUUUUUAUGGCAAAUCAGGUAAUGAGAAUCACGUUGAAGGCGUAUGAUCACAAGCUGGUUGAUCAGUCUGCCGGAAAGAUUGUCGACACGGUGAAGAGAACGGGAGCAAAGGUCUC
env-1149  CGCCCGGCACUGUGUACAGUCACCGCUUGUCGUACUUUCAAAAAAA.......................................................................................................................................
env-1150  NNNNNNNNNNNNNNNNNNNNNNNNNNNNNNNNNNNNNNNNNNNNNNNNNNNNNNNNNNNNNNNNNNACUUUCGAAAGAAGGAAGGCCUGCUGUUGGGUUUCCGGGAGGACACACACGGCACUGUGUACAGUCACCGCUUGUCGUACUUUCAAAAAAAGUGCGAAAAGGAGGCGACUUUUU
env-1151  GCCCGGCACUGUGUACAGUCACCGCUUGUCGUAC...................................................................................................................................................
env-1154  .....................................................................................................................................................................................
env-1155  CGCCCGGCACUGUGUACAGUCACCGCUUGUCGUAC..................................................................................................................................................
env-1156  CACACGGCACUGUGUACAGUCACCGCUUGUCGUAC..................................................................................................................................................
env-1157  ACACACGGCACUGUGUACAGUCACCGCUUGUCGUACUUUAAAAAAAGUGCGAAAAGGAGGCGACUUUUUUUAUGGCAAAUCAGGUAAUGAGAAUCACGUUGAAGGCGUAUGAUCACAAGCUGGUUGAUCAGUCUGCCGGAAAGAUUGUCGACACGGUGAAGAGAACGGGAGCUANNNNNNN
env-1158  CACACGGCACUGUGUACAGUCACCGCUUGUCGUAC..................................................................................................................................................
env-1159  NNNNNNNNNNNUUUUUUUAUGGCAAAUCAGGUAAUGAGAAUCACGUUGAAGGCGUAUGAUCACAAGCUGGUUGAUCAGUCUGCCGGAAAGAUUGUCGACACGGUGAAGAGAACGGGAGCAAAGGUCUCCGGUCCGAUUCCGAUGCCGACCAAGAAGGAAGUGGUUACAAUUCUCCGUGC
Csp-2-1   AGAUACGUCCGAAUGUGUGUGCAGUCACCGCUUGUCGUGCUGAAGUUCAAUAGUGCGAAAAGGAGGGGACUUUUUUUAUGGCUAACCCAAAAAUCAGAAUCAGUCUAAAAGCGUACGAUCAUAAAUUGAUCGACCAGUCCGCAGCUCAAAUUAUAGAGACUGCAAAGAAAACAGGUGCUC
env-1160  GAUGAGUUGUGCAGUCACCACUUGUCGUACUAGAAAAUCUAAAAAGUGCGAAAAGGAGGGGACUUUUUUUAUGGCUAAUCAGAAGAUUAGAAUCAGUCUUAAGUCUUUCGAUCACAAAUUGAUCGACCAGUCAGCUUCUCAGAUUAUUGAGACUGCUAAGAAAACAGGUGCUAAGGUUAG
env-1162  GAUGAGUUGUGCAGUCACCACUUGUCGUACUAGAAAAUCUAAAAAGUGCGAAAAGGAGGGGACUUUUUUUAUGGCUAAUCAGAAGAUUAGAAUCAGUCUUAAGUCUUUCGAUCACAAAUUGAUCGACCAGUCAGCUUCUCAGAUUAUUGAGACUGCUAAGAAAACAGGUGCUAAGGUUAG
env-1163  GAUGAGUUGUGCAGUCACCACUUGUCGUACUAGAAAAUCUAAAAAGUGCGAAAAGGAGGGGACUUUUUUUAUGGCUAAUCAGAAGAUUAGAAUCAGUCUUAAGUCUUUCGAUCACAAAUUGAUCGACCAGUCAGCUUCUCAGAUUAUUGAGACUGCUAAGAAAACAGGUGCUAAGGUUAG
env-1166  GAUGAGUUGUGCAGUCACCACUUGUCGUACUAGAAAAUCUAAAAAGUGCGAAAAGGAGGGGACUUUUUUUAUGGCUAAUCAGAAGAUUAGAAUCAGUCUUAAGUCUUUCGAUCACAAAUUGAUCGACCAGUCAGCUUCUCAGAUUAUUGAGACUGCUAAGAAAACAGGUGCUAAGGUUAG
env-1167  UCGGGUGUGUUGUACAGUCGCUACUUGUCGUGCUAGGAAAUUGUGCGAAAAGGAGGGGCUUUUUUUAUGGCUAACGAAAAAAUUAGAAUAAGCUUAAAAGCUUAUGACCACAAACUUAUCGACCAGUCCGUUAAACAAAUUAUUGAAGCGGCAAAAAAGACCGGCGCAAAAGUUUCAGGU
env-1168  UCGGGUGUGUUGUACAGUCGCUACUUGUCGUGCUAGGAAAUUGUGCGAAAAGGAGGGGCUUUUUUUAUGGCUAACGAAAAAAUUAGAAUAAGCUUAAAAGCUUAUGACCACAAACUUAUCGACCAGUCCGUUAAACAAAUUAUUGAAGCGGCAAAAAAGACUGGCGCAAAAGUUUCAGGU
env-1169  ACACCCGGAUAAGUGUGCAGUCGCUACUUGUCGUGCUGAAAUUUAAAAGUACGAAAAGGAGGGGACUUUUUUUAUGGCUAACCCAAAAAUCAGAAUCAGUCUCAAAGCUUACGAUCACAAGUUGAUCGACCAAUCUGCAGCGCAGAUCAUUGAAACAGCGAAGAAAACAGGAGCGCAAAU
env-1170  ACACCCGGAUAAGUGUGCAGUCGCUACUUGUCGUGCUGAAAUUUAAAAGUACGAAAAGGAGGGGACUUUUUUUAUGGCUAACCCAAAAAUCAGAAUCAGUCUCAAAGCUUACGAUCACAAGUUGAUCGACCAAUCUGCAGCGCAGAUCAUUGAAACAGCGAAGAAAACAGGAGCGCAAAU
env-1171  CACCCGGAUAAGUGUGCAGUCGCUACUUGUCGUGCUGAAAUUUAAAAGUACGAAAAGGAGGGGACUUUUUUUAUGGCUAACCCAAAAAUCAGAAUCAGUCUCAAAGCUUACGAUCACAAGUUGAUCGACCAAUCUGCAGCGCAGAUCAUUGAAACAGCGAAGAAAACAGGAGCGCAAAUC
env-1172  CACCCGGAUAAGUGUGCAGUCACCACUUGUCGUGCUGAAAGUUAAAGUACGAAAAGGAGGGGACUUUUUUUAUGGCUAACCCAAAAAUCAGAAUCAGUCUCAAAGCUUACGAUCACAAGCUGAUCGAUCAGUCUGCAGCGCAGAUCAUUGAAACAGCAAAGAAAACAGGAGCGCAGAUCA
env-1173  CACCCGGAUAAGUGUGCAGUCACCACUUGUCGUGCUGAAAGUUAAAGUACGAA................................................................................................................................
env-1174  .....................................................................................................................................................................................
env-1175  CACCCGGAUAAGUGUGCAGUCACCACUUGUCGUGCUGAAAGUUAAAGUACGAAAAGGAGGGGACUUUUUUUAUGGCUAACCCAAAAAUCAGAAUCAGUCUCAAAGCUUACGAUCACAAGCUGAUCGAUCAGUCUGCAGCGCAGAUCAUUGAAACAGCAAAGAAAACAGGAGCGCAGAUCA
env-1176  CACCCGGAUAAGUGUGCAGUCACCACUUGUCGUGCUGAAAGUUAAAGUACGAAAAGGAGGGGACUUUUUUUAUGGCUAACCCAAAAAUCAGAAUCAGUCUCAAAGCUUACGAUCACAAGCUGAUCGAUCAGUCUGCAGCGCAGAUCAUUGAAACAGCAAAGAAAACAGGAGCGCAGAUCA
Cle-1-1   GCUUAUUAACACCAGCAGAGUGUACAGUCACCACUAGUCGUACUAGGUUCAAAAAAGUACGAAAAGGAGGGGACUUUUUUUAUGGCAAGUCAAAAAAUCCGUAUUACAUUAAAAGCUUAUGAUCACAAAUUAAUCGAUCAAACAGCUGCACAAAUCGUAGAGAUCGCUAAAAAAACUGGC
          .....................................................................................................................................................................................
          .....................................................................................................................................................................................
          ●●●●●●●●●●●●●●●●●●●●●●●●●●●●●●●●●●●●●●●●●●●●●●●●●●●●●●●●●●●●●●●●●●●●●●●●●●●●●●●●●●●●●●●●●●●●●●●●●●●●●●●●●●●●●●●●●●●●●●●●●●●●●●●●●●●●●●●●●●●●●●●●●●●●●●●●R●●●●●●●●●●●●●●●●●●●●●●●●●
```

alignment positions 541 · · · 598

```
env-1     UAUGACCACCAGCUUAUCGACAAGGCUUCCGAGACUAUCGUGGCAACUGCAAAGCGCA
env-4     UAUGACCACCAGCUUAUCGACAAGGCUUCCGAGACUAUCGUGGCAACUGCAAAGCGCA
env-5     ..........................................................
env-6     GCAACGGCGCUUCCGUCUCCGGCCCCAUCCCCCUGCCCACCAAGAAGGAAGUCGUCAC
env-7     AAGAUCGUCGAGACCGCCAAGCGCACCGGCGCCACCGUCUCCGGCCCCAUCCCUCUGC
env-8     AAGAUCGUCGAGACCGCCAAGCGCACCGGCGCCACCGUCUCCGGCCCCAUCCCUCUGC
env-9     ..........................................................
env-10    ACCAGAGUGCAGAGAAGAUCGUGGAGACCGCCAAGCGCACCGGCGCCGAGGUUUCCGG
env-11    ..........................................................
env-12    GCAGAGAAGAUCGU............................................
env-13    AGAAGAUCGUCGAGACGGCCAAGCGCACCGGCGCUACCGUGUCCGGGCCCAUCCCUCU
env-14    AGAAGAUCGUCGAGACCGCCAAGCGCACCGGCGCUACCGUGUCCGGGCCCAUCCCUCU
env-15    GAAGAUCGUCGAGACGGCUAAGCGCACCGGCGCUACCGUGUCCGGGCCCAUCCCUCUG
env-16    ..........................................................
env-17    ..........................................................
env-18    ..........................................................
env-19    UGGCAAAAGAAAUGAUCCGCAUUCGUCUGAAGGCCUAUGACCAUCAGGUCAUCGACCA
env-20    AGAAGAUCGUCGAGACGGCCAAGCGCACCGGCGCUACCGUGUCUGGGCCCAUCCCUCU
env-21    GAAGAUCGUCGAGACCGCCAAGNNNNNNNNNNNNNNNNNNNNNNNNNNNNNNNNNNNN
env-22    AGAAGAUCGUCGAGACGGCCAAGCGCACCGGCGCUACCGUGUCCGGGCCCAUCCCUCU
env-23    ..........................................................
env-25    AGAAGAUCGUCGAGACCGCCAAGCGCACCGGCGCUUCCGUCUCCGGCCCCAUCCCUCU
env-26    GAAGAUCGUCGAGACCGCUAAGCGCACCGGCGCUUCCGUGUCCGGCCCCAUCCCUCUG
env-27    ..........................................................
env-28    ..........................................................
env-29    ..........................................................
env-30    ..........................................................
env-43    ..........................................................
env-45    AGAAGAUCGUCGAGACGGCCAAGCGCACCGGCGCUACCGUGUCCGGGCCCAUCCCUCU

          ..........................................................
          ..........................................................
          ●●●●●●●●●●●●●●●●●●●●●●●●●●●●●●●●●●●●●●●●●●●●●●●●●●●●●●●●●●
```

```
env-46     ..........................................................
env-47     AGAAGAUCGUAGAGACCGCCAAGCGCACCGGCGCCCAGGUGUCCGGCCCCAUCCCCCU
env-48     GCUUAGUUGAUCAGGCUGCUGAGAAGAUUGUUGACACUGCAAAGAGAACAGGCGCUCA
env-49     GCUUAGUUGAUCAGGCUGCUGAGAAGAUUGUUGACACUGCAAAGAGAACAGGCGCUCA
env-54     CUUAGUUGAUCAGGCUGCUGAGAAGAUUGUUGACACUGCAAAGAGAACAGGCGCUCAG
env-55     GCUUAGUUGAUCAGGCUGCUGAGAAGAUUGUUGACACUGCAAAGAGAACAGGCGCUCA
env-57     GCUUAGUUGAUCAGGCUGCUGAGAAGAUUGUUGACACUGCAAAGAGAACAGGCGCUCA
env-58     GCUUAGUUGAUCAGGCUGCUGAGAAGAUUGUUGACACUGCAAAGAGAACAGGCGCUCA
env-59     GCUUAGUUGAUCAGGCUGCUGAGAAGAUUGUUGACACUGCAAAGAGAACAGGCGCUCA
env-60     GCUUAGUUGAUCAGGCUGCUGAGAAGAUUGUUGACACUGCAAAGAGAACAGGCGCUCA
env-61     AGUUUAGUUGAUCAGGCUGCUGAGAAGAUUGUUGACACUGCAAAGAGAACAGGCGCAC
env-76     AUCACAGUUUAGUUGAUCAGGCUGCUGAGAAGAUUGUUGACACUGCAAAGAGAACAGG
env-77     AUCACAGUUUAGUUGAUCAGGCUGCUGAGAAGAUUGUUGACACUGCAAAGAGAACAGG
env-78     AUCACAGUUUAGUUGAUCAGGCUGCUGAGAAGAUUGUUGACACUGCAAAGAGAACAGG
env-80     UCGCACCAUAAUUUUUCAGGGAGGAAAUAAAAUGGCAGCAAGUAAAGUGAAGAUUCGA
env-81     AUCACAGUUUAGUUGAUCAGGCUGCUGAGAAGAUUGUUGACACUGCAAAGAGAACAGG
env-82     AUCACAGUUUAGUUGAUCAGGCUGCUGAGAAGAUUGUUGACACUGCAAAGAGAACAGG
env-83     GAUCACAGUUUAGUUGAUCAGGCUGCUGAGAAGAUUGUUGACACUGCAAAGAGAACAG
env-85     UCGAGACCGCCAAGCGCAACGGCGCUUCCGUCUCCGGCCCCAUCCCCCUGCCCACCAA
env-86     UCGAGACCGCCA..............................................
env-87     UCAUCAGCUCAUNNNNNNNNNNNNNNNNNNNNNNNNNNNNNNNNNNNNNNNNNNNNNN
env-88     ..........................................................
env-89     ..........................................................
Pba-1-1    CUCUUCCCACAGAGAAGGAGAUUAUCACGAUCCUACGUAGUGUUCAUAAGCAUAAGGA
env-90     GCGGAGAAGAUUGUUGAGACGGCCAGAAACACAGGCGCGCAGGUCUCCGGUCCCGUGC
env-91     GCGGAGAAGAUUGUUGAGACGGCCAGAAACACAGGCGCGCAGGUCUCCGGUCCCGUGC
env-92     CGACAAGGCGGCGGAGAAGAUCGUAGAGACCGCAAAGAGAACCGAUGCUCAGGUCUCA
env-94     CGACAAGGCGGCGGAGAAGAUCGUAGAGACCGCAAAGAGAACCGAUGCUCAGGUCUCA
env-95     CGACAAGGCGGCGGAGAAGAUCGUAGAGACCGCAAAGAGACCCGAUGCUCAGGUCUCA
env-96     CGACAAGGCGGCGGAGAAGAUCGUAGAGACCGCAAAGAGAACCGAUGCUCAGGUCUCA
env-97     ..........................................................
env-99     CGACAAGGCGGCGGAGAAGAUCGUAGAGACCGCAAAGAGAACCGAUGCUCAGGUCUCA
env-102    GACAAGGCGGCGGAGAAGAUCGUAGAGACCGCAAAGAGAACCGAUGCUCAGGUUUCCG
env-107    GACAAGGCGGCGGAGAAGAUCGUAGAGACCGCAAAGAGAACCGAUGCUCAGGUUUCCG
env-109    ..........................................................
env-110    CGACAAGGCGGCGGAGAAGAUCGUAGAGACCGAAAAGAGAACCGAUGCUCAGGUUUCC
env-111    GACAAGGCGGCGGAGAAGAUCGUAGAGACCGCAAAGAGAACCGAUGCUCAGGUUUCCG
env-112    GACAAGGCGGCGGAGAAGAUCGUAGAGACCGCAAAGAGAACCGAUGCUCAGGUUUCCG
env-113    GACAAGGCGGCGGAGAAGAUCGUAGAGACCGCAAAGAGAACCGAUGCUCAGGUUUCCG
env-114    ..........................................................
env-121    ..........................................................
env-122    GAAAGAAUCGUUGCAACAGCCAAGAGAACAGGUGCUAGAGUUUCAGGUCCUGUACCUC
env-123    GAAAGAAUCGUUGCAACAGCCAAGAGAACAGGUGCUAGAGUUUCAGGUCCUGUACCUC
env-128    GAAAGAAUCGUUGCAACAGCCAAGAGAACAGGUGCUAGAGUUUCAGGUCCUGUACCUC
Rfl-1-1    GGUUACGAUCACGCUACUGUUGAUAUCGCAGCAGCUAAGAUCGUUGAGGCAGCUAAGA
env-129    ..........................................................
env-130    GCGGCCCUAUCCCUCUUCCCACAAAGAAGGAAGUUGUAACAAUCAUCAGAGCGGUGCA
env-131    ..........................................................
env-132    ..........................................................
env-133    ..........................................................
env-134    GAAGGCUGGAGCAAAGGUUAGUGGUCCUGUACCUCUUCCAACAAAGAAAGAGAUAGUU
env-138    ..........................................................
env-139    ..........................................................
env-140    ..........................................................
env-141    GAAGGCUGGAGCAAAGGUUAGUGGUCCUGUACCUCUUCCAACAAAGAAAGAGAUAGUU
env-142    ..........................................................
env-143    GAAGGCUGGGGCAAAGGUAAGUGGUCCUGUACCUCUUCCAACAAAGAAAGAGAUAGUU
env-144    GAAGGCUGGGGCAAAGGUAAGUGGUCCUGUACCUCUUCCAACAAAGAAGGAGAUAGUU
env-148    ..........................................................
env-158    GAAGGCUGGAGCAAAGGUUAGUGGUCCUGUACCUCUUCCAACAAAGAAAGAGAUAGUU
env-159    GAAGGCUGGGGCAAAGGUAAGUGGUCCUGUACCUCUUCCAACAAAGAAAGAGAUAGUU
env-160    ..........................................................
env-161    ..........................................................
Cmo-1-1    UAAAGAAGGCUGGAGCAAAGGUUAGUGGUCCUGUACCUCUUCCAACAAAGAAAGAGAU
env-162    GUAAAGAAGGCUGGAGCAAAGGUUAGUGGUCCUGUACCUCUUCCAACAAAGAAAGAGA
env-163    GUAAAGAAGGCUGGAGCAAAGGUUAGUGGUCCUGUACCUCUUCCAACAAAGAAAGAAA
Ece-1-1    GCUUCCUACAAAGAAGGAAGUUGUAACUAUUCUGCGUGCGGUACAUAAGUACAAAGAC
env-164    UCAUCGAGACAGUCAAGAAGAACGGAUCUCAGGUUUCAGGACCAGUUCCGCUUCCUAC
Bfi-1-1    UCAUCGAGACAGUCAAGAAGAACGGAUCUCAGGUUUCAGGACCGGUUCCGCUUCCUAC
Bfi-2-1    UCAUCGAGACAGUCAAGAAGAACGGAUCUCAGGUUUCAGGACCAGUUCCGCUUCCUAC
env-165    CCUGUUCCACUUCCAACUAAAAAGGAAGUAGUUACAAUAUUAAGAGCUGUACAUAAGU
env-166    ..........................................................
env-170    CCUGUUCCACUUCCAACUAAAAAGGAAGUAGUUACAAUAUUAAGAGCUGUACAUAAGU
env-171    ..........................................................
env-172    CCUGUUCCACUUCCAACUAAAAAGGAAGUAGUUACAAUAUUAAGAGCUGUACAUAAGU
env-173    UAGAUGCAUCUGCCAAGAAAAUCAUCGAAACAGUCAAGAAGAACGGAUCCCAGGUAAG
env-174    GUAGAUGCAUCUGCCAAGAAAAUCAUCGAAACAGUCAAGAAGAACGGAUCUCAGGUAA
env-175    ..........................................................
env-176    ..........................................................
env-177    ..........................................................
env-178    UGAUUGAUACAAUUAAGAAAACAGGAGCAAAAGUAAGCGGUCCGGUACCUCUUCCUAC
env-179    ..........................................................
env-180    UGAUUGAUACUGUAAAGAAAACGGGAGCAAAGGUGAGUGGGCCUGUGCCACUUCCUAC
env-183    UGAUUGAUACUGUAAAGAAAACGGGAGCAAAGGUGAGUGGGCCUGUGCCACUUCCUAC

           ..........................................................
           ..........................................................
           ●●●●●●●●●●●●●●●●●●●●●●●●●●●●●●●●●●●●●●●●●●●●●●●●●●●●●●●●●●
```

| | |
|---|---|
| env-184 | UGAUUGAUACUGUAAAGAAAACGGGAGCAAAGGUGAGUGGGCCUGUGCCACUUCCUAC |
| env-185 | .......................................................... |
| env-186 | UGAUUGAUACUGUAAAGAAAACGGGAGCAAAGGUGAGUGGGCCUGUGCCACUUCCUAC |
| env-188 | UGAUUGAUACUGUAAAGAAAACGGGAGCAAAGGUGAGUGGGCCUGUGCCACUUCCUAC |
| env-189 | UGAUUGAUACUGUAAAGAAAACGGGAGCAAAGGUGAGUGGGCCUGUGCCACUUCCUAC |
| env-190 | UGAUUGAUACUGUAAAGAAAACGGGAGCAAAGGUGAGUGGGCCUGUGCCACUUCCUAC |
| env-191 | AAAAGUCAGAAAAUGAGAAUUAUCCUGAAGGCAUACGAUCAUNNNNNNNNNNNNNNNN |
| env-192 | UGAUUGAUACUGUAAAGAAAACGGGAGCAAAGGUGAGUGGGCCUGUGCCACUUCCUAC |
| env-193 | UGAUUGAUACUGUAAAGAAAACGGGAGCAAAGGUGAGUGGGCCUGUGCCACUUCCUAC |
| env-194 | UGAUUGAUACUGUAAAGAAAACGGGAGCAAAGGUGAGUGGGCCUGUGCCACUUCCUAC |
| env-195 | .......................................................... |
| env-208 | UGAUUGAUACUGUAAAGAAAACGGGAGCAAAGGUGAGUGGGCCUGUGCCACUUCCUAC |
| Lba-19-1 | UUAAGAAGAACGGUGGAGCAGUAAGCGGUCCCGUUCCCCUUCCUACAAAGAGGGAAGU |
| Lba-17-1 | AAGAAUGGUGGAGAGACAUCUGGUCCAGUACCUCUUCCAACAAAGAAGGAAGUAGUUA |
| env-212 | AAGAAUGGUGGAGAGACAUCUGGUCCAGUACCUCUUCCAACAAAGAAGGAAGUAGUUA |
| env-213 | GGAGACAAGCGGUCCGGUUCCUCUUCCUACAAAGAAGGAAGUGGUUACGAUCCUGCGU |
| env-214 | .......................................................... |
| env-215 | .......................................................... |
| env-216 | GACCGGUUCCAAUGCCUACAAAGGUUGAGAAGAUUACAAUCCUCAGAGCUGUUCAUAA |
| env-217 | GACCGNNNNNNNNNNNNNNNNNNNNNNNNNNNNNNNNNNNNNNNNNNNNNNNNNNNNN |
| env-218 | GACCGNNNNNNNNNNNNNNNNNGAACUCAUAAGAGAUUAAUCGACGUAAUUGCACCAA |
| env-219 | GACCGGUUCCAAUGCCUACAAAGGCUGAGAAGAUCACAAUCCUCAGAGCUGUUCAUAA |
| env-220 | .......................................................... |
| env-223 | GACCGGUUCCAAUGCCUACAAAGGUUGAGAAGAUCACAAUCCUCAGAGCUGUUCAUAA |
| env-224 | GACCGGUUCCAAUGCCUACAAAGGUUGAGAAGAUUACAAUCCUCAGAGCUGUUCAUAA |
| env-225 | GACCG..................................................... |
| env-229 | GACCGGUUCCAAUGCCUACAAAGGUUGAGAAGAUCACAAUCCUCAGAGCUGUUCAUAA |
| env-231 | GACCGGUUCCAAUGCCUACAAAGGUUGAGAAGAUCACAAUCCUCAGAGCUGUUCAUAA |
| env-234 | .......................................................... |
| env-235 | .......................................................... |
| env-236 | GACCGGUUCCAAUGCCUACAAAGGUUGAGAAGAUUACAAUCCUCAGAGCUGUUCAUAA |
| env-237 | GACCGGUUCCAAUGCCUACAA..................................... |
| env-242 | AGAAAUCGUAACAAUUCUUAGAGCAGUACAUAAGUACAAAGAUUCCAGAGAGCAAUUC |
| env-244 | .......................................................... |
| env-247 | CACAAUCAGAUCGAUCAUGCAGCUGGUAAAAUUAUUGAUACGGUAAAGAAGACAGGAG |
| env-248 | CACAAUCAGAUCGAUCAUGCAGCUGGUAAAAUUAUUGAUACGGUAAAGAAGACAGGAG |
| env-249 | CACAAUCAGAUCGAUCAUGCAGCUGGUAAAAUUAUUGAUACGGUAAAGAAGACAGGAG |
| env-250 | CACAAUCAGAUCGAUCAUGCAGCUGGUAAAAUUAUUGAUACGGUAAAGAAGACAGGAG |
| env-251 | CACAAUCAGAUCGAUCAUGCAGCUGGUAAAAUUAUUGAUACGGUAAAGAAGACAGGAG |
| env-252 | .......................................................... |
| env-257 | CACAAUCAGAUCGAUCAUGCAGCUGGUAAAAUUAUUGAUACGGUAAAGAAGACAGGAG |
| env-259 | CACAAUCAGAUCGAUCAUGCAGCUGGUAAAAUUAUUGAUACGGUAAAGAAGACAGGAG |
| env-262 | CACAAUCAGAUCGAUCAUGCAGCUGGUAAAAUUAUUGAUACGGUAAAGAAGACAGGAG |
| env-264 | CACAAUCAGAUCGAUCAUGCAGCUGGUAAAAUUAUUGAUACGGUAAAGAAGACAGGAG |
| env-265 | CACAAUCAGAUCGAUCAUGCAGCUGGUAAAAUUAUUGAUACGGUAAAGAAGACAGGAG |
| env-266 | .......................................................... |
| env-280 | .......................................................... |
| env-281 | .......................................................... |
| env-282 | .......................................................... |
| env-283 | .......................................................... |
| env-284 | CGGUACCUCUUCCAACAAAGAAGGAAGUAGUUACAAUUCUUAGAGCUACUCACAAGUA |
| env-288 | CGGUACCUCUUCCAACAAAGAAGGAAGUAGUUACAAUUCUUAGAGCUACUCACAAGUA |
| env-289 | CGGUACCUCUUCCAACAAAGAAGGAAGUAGUUACAAUUCUUAGAGCUACUCACAAGGA |
| env-290 | CAUUGAUAUCAAAAUGAAGCAGAAAUAAGCUUCAACAAUUAAUUAGUAUAUCCCCGAA |
| env-291 | .......................................................... |
| env-300 | .......................................................... |
| env-303 | .......................................................... |
| env-304 | CGGUACCUCUUCCAACAAAGAAGGAAGUAGUUACAAUUCUUAGAGCUACUCACAAGUA |
| env-305 | CGG....................................................... |
| env-306 | CGGUACCUCUUCCAACAAAGAAGGAAGUAGUUACAAUUCUUAGAGCUACUCACAAGUA |
| env-307 | .......................................................... |
| env-308 | .......................................................... |
| env-309 | CGGUACCUCUUCCAACAAAGAAGGAAGUAGUUACAAUUCUUAGAGCUACUCACAAGUA |
| env-310 | .......................................................... |
| env-311 | CGGUACCUCUUCCAACAAAGAAGGAAGUAGUUACAAUUCUUAGAGCUACUCACAAGUA |
| env-312 | CGGUACCUCUUCCAACAAAGAAGGAAGUAGUUACAAUUCUUAGAGCUACUCACAAGUA |
| env-313 | GAUAUCAAAAUGAAGCAGAAAUAAGCUUCAACAAUUAAUUAGUAUAUCCCCGAAUGGU |
| env-314 | UACAAGGAUUCUCGUGAACAGUUCGAGCAGAGAACUCAUAAGAGACUUAUCGAUAUCA |
| env-315 | AGAACUCAUAAGAGACUUAUCGAUAUCAUCAAUCCAACACAGAAGACAACAGAGUCUU |
| env-316 | .......................................................... |
| env-317 | .......................................................... |
| env-331 | .......................................................... |
| env-332 | .......................................................... |
| env-333 | .......................................................... |
| env-334 | CGGUACCUCUUCCAACAAAGAAGGAAGUAGUUANNNNNNNNNNNNNNNNNNNNNNNNN |
| env-335 | CGGUACCUCUUCCAACAAAGAAGGAAGUAGUUACAAUUCUUAGAGCUACUCACAAGUA |
| env-336 | CGGUACCUCUUCCAACAAAGAAGGAAGUAGUUACAAUUCUUAGAGCUACUCACAAGUA |
| env-338 | CGGUACCUCUUCCAACAAAGAAGGAAGUAGUUACAAUUCUUAGAGCUACUCACAAGUA |
| env-339 | CGGUACCUCUUCCAACAAAGAAGGAAGUAGUUACAAUUCUUAGAGCUACUCACAAGUA |
| env-340 | CGGUACCUCUUCCAACAAAGAAGGAAGUAGUUACAAUUCUUAGAGCUACUCACAAGUA |
| env-341 | CGGUACCUCUUCCAACAAAGAAGGAAGUAGUUACAAUUCUUAGAGCUACUCACAAGUA |
| env-347 | .......................................................... |
| env-348 | .......................................................... |
| env-349 | .......................................................... |
| | .......................................................... |
| | .......................................................... |
| | ●●●●●●●●●●●●●●●●●●●●●●●●●●●●●●●●●●●●●●●●●●●●●●●●●●●●●●●●●● |

| | |
|---|---|
| env-350 | .......................................................... |
| env-351 | .......................................................... |
| env-352 | .......................................................... |
| env-353 | CGGUACCUCUUCCAACAAAGAAGGAAGUAGUUACAAUUCUUAGAGCUACUCACAAGUA |
| env-354 | CGGUACCUCUUCCAACAAAGAAGGAAGUAGUUACAAUUCUUAGAGCUACUCACAAGUA |
| env-355 | CGGUACCUCUUCCAACAAAGAAGGAAGUAGUUACAAUUCUUAGAGCUACUCACAAGUA |
| env-359 | CGGUACCUCUUCCGACAAAGAAGGAAGUAGUUACAAUUCUUAGAGCUACUCACAAGUA |
| env-360 | CGGUACCUCUUCCAACAAAGAAGGAAGUAGUUACAAUUCUUAGAGCUACUCACAAGUA |
| env-361 | CGGUACCUCUUCCAACAAAGAAGGAAGUAGUUACAAUUCUUAGAGCUACUCACAAGUA |
| env-362 | AAGUACAAGGAUUCUCGUGAACAGUUCGAGCAGAGAACUCAUAAGAGACUUAUCGAUA |
| env-363 | .......................................................... |
| env-364 | .......................................................... |
| env-365 | CGGUACCUCUUCCAACAAAGAA.................................... |
| env-366 | CGGUACCUCUUCCAACAAAGAAGGAAGUAGUUACAAUUCUUAGAGCUACUCACAAGUA |
| env-368 | .......................................................... |
| env-372 | CGGUACCUCUUCCAACAAAGAAGGAAGUAGUUAC........................ |
| env-373 | CGGUACCUCUUCCAACAAAGAAGGAAGUAGUUACAAUUCUUAGAGCUACUCACAAGUA |
| env-374 | CGGUACCUCUUCCAACAAAGAAGGAAGUAGUUACAAUUCUUAGAGCUACUCACAAGUA |
| env-375 | .......................................................... |
| env-376 | GUCAAGAAGAACGGAUCUCAGGUUAGCGGACCGGUACCCCUUCCUACUAAGAAGGAAG |
| Bsp-5-1 | GUCAAGAAGAACGGAUCUCAGGUUAGCGGACCGGUACCCCUUCCUACUAAGAAGGAAG |
| env-377 | .......................................................... |
| env-379 | .......................................................... |
| env-380 | .......................................................... |
| env-381 | CUACCAACAAAGAAGGAAGUGGUUACUAUCCUUAGAGCUGUACAUAAGUACAAAGACU |
| env-382 | ACUACCGACAAAGAAGGAAGUGGUAACUAUCCUUAGAGCUGUACAUAAGUACAAAGAC |
| env-384 | CUACCAACAAAGAAGGAAGUGGUUACUAUCCUUAGAGCUGUACAUAAGUACAAAGACU |
| Lor-1-1 | CUACCAACAAAGAAGGAAGUGGUUACUAUCCUUAGAGCUGUACAUAAGUACAAAGACU |
| env-385 | .......................................................... |
| env-386 | CUACCAACAAAGAAGGAAGUGGUUACUAUCCUUAGAGCUGUACAUAAGUACAAAGACU |
| env-387 | ACUACCGACAAAGAAGGAAGUGGUAACUAUCCUUAGAGCUGUACAUAAGUACAAAGAC |
| env-388 | .......................................................... |
| Lba-15-1 | ACUACCAACAAAGAAGGAAGUGGUUACUAUCCUUAGAGCUGUACAUAAGUACAAAGAC |
| env-389 | ACUACCAACAAAGAAGGAAGUGGUUACUAUCCUUAGAGCUGUACAUAAGUACAAAGAC |
| env-390 | .......................................................... |
| env-391 | ACUACCAACAAAGAAGGAAGUGGUUACUAUCCUUAGAGCUGUACAUAAGUACAAAGAC |
| env-392 | .......................................................... |
| env-393 | .......................................................... |
| env-394 | .......................................................... |
| env-395 | ACUACCAACAAAGAAGGAAGUGGUUACUAUCCUUAGAGCUGUACAUAAGUACAAAGAC |
| env-396 | .......................................................... |
| env-397 | ACUACCAACAAAGAAGGAAGUGGUAACUAUCCUUAGAGCUGUACAUAAGUACAAAGAC |
| env-398 | ACUACCAACAAAGAAGGAAGUGGUAACUAUCCUUAGAGCUGUACAUAAGUACAAAGAC |
| env-399 | ACUACCAACAAAGAAGGAAGUGGUUACUAUCCUUAGAGCUGUACAUAAGUACAAAGAC |
| env-400 | ACUACCAACAAAGAAGGAAGUGGUUACUAUCCUUAGAGCUGUACAUAAGUACAAAGAC |
| env-402 | .......................................................... |
| env-403 | ACUACCAACAAAGAAGGAAGUGGUUACUAUCCUUAGAGCUGUACAUAAGUACAAAGAC |
| env-404 | .......................................................... |
| env-405 | ACUACCAACAAAGAAGGAAGUGGUUACUAUCCUUAGAGCUGUACAUAAGUACAAAGAC |
| env-406 | .......................................................... |
| env-407 | ACUACCAACAAAGAAGGAAGUGGUAACUAUCCUUAGAGCUGUACAUAAGUACAAAGAC |
| env-408 | ACUACCAACAAAGAAGGAAGUGGUAACUAUCCUUAGAGCUGUACAUAAGUACAAAGAC |
| env-409 | .......................................................... |
| env-410 | ACUACCAACAAAGAAGGAAGUGGUAACUAUCCUUAGAGCUGUACAUAAGUACAAAGAC |
| Cba-3-1 | ACUACCAACAAAGAAGGAAGUGGUUACUAUCCUUAGAGCUGUACAUAAGUACAAAGAC |
| env-411 | CUACCAACAAAGAAGGAAGUGGUAACUAUCCUUAGAGCUGUACAUAAGUACAAAGACU |
| env-414 | .......................................................... |
| env-415 | .......................................................... |
| env-416 | CCUGUGCCGCUGCCUACCAAGAAAGAGGUAGUUACAAUACUCCGUGCAGUUCACAAAU |
| env-417 | .......................................................... |
| env-420 | .......................................................... |
| env-426 | CCUGUGCCGCUGCCUACUAAGAAAGAGGUAGUUACAAUACUCCGUGCAGUUCACAAAU |
| env-427 | .......................................................... |
| env-428 | AUCGAAACAGUCAAGAAAAAUGGAGCACAGGUGAGCGGACCGGUACCGCUUCCCACUA |
| env-429 | .......................................................... |
| env-430 | AUCGAAACAGUCAAGAAAAAUGGAGCACAGGUGAGCGGACCGGUACCGCUUCCUA... |
| env-431 | AUCGAAACAGUCAAGAAAAAUGGAGCACAGGUGAGCGGACCGGUACCGCUUCCCACUA |
| env-432 | .......................................................... |
| env-433 | AUCGAAACAGUCAAGAAAAAUGGAGCACAGGUGAGCGGACCGGUACCGCUUCCCACUA |
| env-434 | NNNNNNNNNNNNNNNNNNNNNNNNNNNNNNNNNNNNNNNNNNNNNNNNNNNNNNNNNN |
| env-435 | AUCGAAACAGUCAAGAAAAAUGGAGCACAGGUGAGCGGACCGGUACCGCUUCCUACUA |
| env-436 | .......................................................... |
| env-454 | .......................................................... |
| env-455 | AUCGAAACAGUCAAGAAAAAUGGAGCACAGGUGAGCGGACCGGUACCGCUUCCCACUA |
| env-456 | AUCGAAACAGUCAAGAAAAAUGGAGCACAGGUGAGCGGACCGGUACCGCUUCCUACUA |
| env-457 | AUCGAAACAGUCAAGAAAAAUGGAGCACAGGUGAGCGGACCGGUACCGCUUCCCACUA |
| env-459 | .......................................................... |
| env-460 | .......................................................... |
| env-464 | .......................................................... |
| env-465 | AUCGAAACAGUCAAGAAAAAUGGAGCACAGGUGAGCGGACCGGUACCGCUUCCCACUA |
| env-466 | AUCGAAACAGUCAAGAAAAAUGGAGCACAGGUGAGCGGACCGGUACCGCUUCCCACUA |
| env-467 | UCAGGUAAUGAGGAUUACACUGAAAGCUUACGAUCAUCAGUUAGUUGAUGCGUCUGCG |
| env-468 | .......................................................... |
| env-469 | .......................................................... |
| | .......................................................... |
| | .......................................................... |

```
env-470    AUCGAAACAGUCAAGAAAAAUGGAGCACAGGUGAGCGGACCGGUACCGCUUCCCACUA
env-471    AUCGAAACAGUCAAGAAAAAUGGAGCACAGGUGAGCGGACCGGUACCGCUUCCCACUA
env-472    AUCGAAACAGUCAAGAAAAAUGGAGCACAGGUGAGCGGACCGGUACCGCUUCCCACUA
env-473    ..........................................................
env-482    AUCGAAACAGUCAAGAAAAAUGGAGCACAGGUGAGCGGACCGGUACCGCUUCCCACUA
env-483    AUCGAAACAGUCAAGAAAAAUGGAGCACAGGUGAGCGGACCGGUACCGCUUCCUACUA
env-484    ..........................................................
env-485    AUCGAAACAGUCAAGAAAAAUGGAGCACAGGUGAGCGGACCGGUACCGCUUCCCACUA
env-486    ..........................................................
env-487    AUCGAAACAGUCAAGAAAAAUGGAGCACAGGUGAGCGGACCGGUACCGCUUCCUACUA
env-488    ..........................................................
env-489    ..........................................................
env-494    UAAGUGGUCCGGUACCUCUUCCAACAAAGAAGGAAGUUGUUACAAUUCUUAGAGCUGU
env-495    ..........................................................
env-496    GUAAGUGGUCCGGUACCUCUUCCAACAAAGAAGGAAGUUGUUACAAUUCUUAGAGCUG
env-497    GUAAGUGGUCCGGUACCUCUUCCAACAAAGAAGGAAGUUGUUACAAUUCUUAGAGCUG
env-498    UAAGUGGUCCGGUACCUCUUCCAACAAAGAAGGAAGUUGUUACAAUUCUUAGAGCUGU
env-499    ..........................................................
env-500    UAAGUGGUCCGGUACCUCUUCCAACAAAGAAGGAAGUUGUUACUAUUCUUAGAGCUGU
env-501    UAAGUGGUCCGGUACCUCUUCCAACAAAGAAGGAAGUUGUUACAAUUCUUAGAGCUGU
env-502    ..........................................................
env-503    GUAAGUGGUCCGGUACCUCUUCCAACAAAGAAGGAAGUUGUUACAAUUCUUAGAGCUG
env-512    AGGAGCAAAGGUUUCAGGUCCUGUUCCAAUGCCUACAAAGGUUGAGAAGAUCACAAUC
env-513    AGGAGCAAAGGUUUCAGGUCCUGUUCCAAUGCCUACAAAGGUUGAGAAGAUCACAAUC
env-514    AGGAGCAAAGGUUUCAGGUCCUGUUCCAAUGCCUACAAAGGUUGAGAAGAUCACAAUC
env-515    AGGAGCAAAGGUUUCAGGUCCUGUUCCAAUGCCUACAAAGGUUGAGAAGAUCACAAUC
env-516    AGGAGCAAAGGUUUCAGGUCCUGUUCCAAUGCCUACAAAGGUUGAGAAGAUCACAAUC
env-517    AGGAGCAAAGGUUUCAGGUCCUGUUCCAAUGCCUACAAAGGUUGAGAAGAUCACAAUC
env-518    ..........................................................
env-556    ..........................................................
env-557    AGGAGCAAAGGUUUCAGGUCCUGUUCCAAUGCCUACAAAGGUUGAGAAGAUCACAAUC
env-558    AGGAGCAAAGGUUUCAGGUCCUGUUCCAAUGCCUACAAAGGUUGAGAAGAUCACAAUC
env-559    AGGAGCAAAGGUUUCAGGUCCUGUUCCAAUGCCUACAAAGGUUGAGAAGAUCACAAUC
env-560    ..........................................................
env-561    AGGAGCAAAGGUUUCAGGUCCUGUUCCAAUGCCUACAAAGGUUGA.............
env-562    CCGUGCCACUUCCAACGAAGAAGGAAGUAGUAACAAUCUUAAGAGCGGUACAUAAGUA
env-563    CCGUGCCACUUCCAACAAAAAAGGAAGUAGUAACAAUCUUAAGAGCUGUGCAUAAGUA
env-564    CCGUGCCACUUCCAACAAAAAAGGAAGUAGUAACAAUCUUAAGAGCUGUGCAUAAGUA
env-565    CCGUGCCACUUCCAACGAAGAAGGAAGUAGUAACAAUCUUAAGAGCGGUACAUAAGUA
Bsp-4-1    GACAGUCAAGAAGAACGGAUCUCAGGUUAGUGGACCAGUGCCUCUUCCUACUAAGAAG
Bsp-7-1    GACAGUCAAGAAGAACGGAUCUCAGGUUAGUGGACCAGUGCCUCUUCCUACUAAGAAG
Bsp-10-1   GACAGUCAAGAAGAACGGAUCUCAGGUUAGUGGACCAGUGCCUCUUCCUACUAAAAAG
env-566    GACAGUCAAGAAGAACGGAUCUCAGGUUAGUGGACCAGUGCCUCUUCCUACUAAGAAG
Bsp-2-1    UCAAGAAGAACGGUUCUCAGGUAAGUGGACCAGUACCUCUUCCUACAAAGAAGGAAGU
Bsp-3-1    ACAGUCAAGAAGAAUGGAUCUCAGGUUUCCGGACCUGUACCUCUUCCUACUAAGAAGG
Bsp-6-1    ACAGUCAAGAAGAACGGAUCUCAGGUUUCCGGACCUGUACCUCUUCCUACUAAGAAGG
env-567    UCAUCGAAACUGUUAAGAAGACCGGAGCUCAGGUGAGUGGUCCCGUGCCUCUUCCUAC
env-568    UCAUCGAAACUGUUAAGAAGACCGGAGCUCAGGUGAGCGGUCCCGUACCUCUUCCUAC
env-570    UCAUCGAAACUGUUAAGAAGACCGGAUCUCAGGUAAGUGGUCCCGUGCCUCUUCCUAC
env-571    UCAUCGAAACUGUUAAGAAGACCGGAUCUCAGGUAAGUGGUCCCGUGCCUCUUCCUAC
env-573    UCAUCGAAACUGUUAAGAAGACCGGAUCUCAGGUAAGUGGUCCCGUGCCUCUUCCUAC
env-578    UCAUCGAAACUGUUAAGAAGACCGGAUCUCAGGUAAGUGGUCCCGUGCCUCUUCCUAC
env-579    UCAUCGAAACUGUUAAGAAGACCGGAUCUCAGGUAAGUGGUCCCGUGCCUCUUCCUAC
env-581    UCAUCGAAACUGUUAAGAAGACCGGAUCUCAGGUAAGUGGUCCUGUGCCUCUUCCUAC
env-582    UCAUCGAAACUGUUAAGAAGACCGGAGCUCAGGUGAGCGGUCCCGUGCCUCUUCCUAC
env-583    UCAUCGAAACUGUUAAGAAGACCGGAGCUCAGGUGAGCGGUCCCGUACCUCUUCCUAC
env-584    UCAUCGAAACUGUUAAGAAGACCGGAUCUCAGGUAAGUGGUCCCGUGCCUCUUCCUAC
env-585    ..........................................................
env-586    UCAUCGAAACUGUUAAGAAGACCGGAGCUCAGGUGAGCGGUCCCGUACCUCUUCCUAC
env-587    UCAUCGAAACUGUUAAGAAGACCGGAGCUCAGGUGAGCGGUCCCGUACCUCUUCCUAC
env-588    UCAUCGAAACUGUUAAGAAGACCGGAGCUCAGGUGAGUGGUCCCGUACCUCUUCCUAC
env-589    UCAUCGAAACUGUUAAGAAGACCGGAGCUCAGGUGAGCGGUCCCGUACCUCUUCCUAC
env-590    AUCAUCGAAACUGUUAAGAAGACCGGAGCUCAGGUGAGUGGUCCCGUGCCUCUUCCUA
env-591    GCGAAAAGGAGGCGACUUUUUUUUAUGGCANNNNNNNNNNNNNNNNNNNNNNNNNNNN
env-592    NNNNNNNNNNNNNNNNNNNNNNNNNNNNNNNNNNNNNNNNNNNNNNNNNNNNNNNNNN
env-593    UCAUCGAAACUGUUAAGAAGACCGGAUCUCAGGUAAGUGGUCCCGUGCCUCUUCCUAC
env-594    UCAUCGAAACUGUUAAGAAGACCGGAUCUCAGGUAAGUGGUCCCGUGCCUCUUCCUAC
env-595    UCAUCGAAACUGUUAAGAAGACCGGAUCUCAGGUAAGUGGUCCUGUGCCUCUUCCUAC
env-596    UCAUCGAAACUGUUAAGAAGACCGGAGCUCAGGUGAGCGGUCCCGUACCUCUUCCUAC
env-597    UCAUCGAAACUGUUAAGAAGACCGGAUCUCAGGUAAGUGGUCCCGUGCCUCUUCCUAC
env-598    UCAUCGAAACUGUUAAGAAGACCGGAUCUCAGGUAAGUGGUCCCGUGCCUCUUCCUAC
env-599    UCAUCGAAACUGUUAAGAAGACCGGAGCUCAGGUGAGUGGUCCCGUGCCUCUUCCUAC
env-600    UCAUCGAAACUGUUAAGAAGACCGGAGCUCAGGUGAGCGGUCCCGUACCUCUUCCUAC
env-601    ..........................................................
env-602    UCAUCGAAACUGUUAAGAAGACCGGAGCUCAGGUGAGCGGUCCCGUGCCUCUUCCUAC
env-603    ACUGAAGGCAUAUGAUCAUCAGCUUAUCGAUCAAUCUGCCGGAAAGAUUAUUGAAACU
env-604    ..........................................................
env-606    ACCAGUUCCACUUCCAACAAAGAAGGAAGUAG..........................
env-607    ACCAGUUCCACUUCCAACAAAGAAGGAAGUAGUAACAAUCUUGAGAGCGGUUCAUAAG
env-608    ACCAGUUCCACUUCCAACAAAGAAGGAAGUAGUAACAAUCUUGAGAGCGGUUCAUAAG
env-614    ACCAGUUCCACUUCCAACAAAGAAGGAAGUAGUAACAAUCUUGAGAGCGGUUCAUAAG
env-615    ..........................................................
env-623    ..........................................................
env-624    ..........................................................

           ..........................................................
           ..........................................................
           ..........................................................
```

```
env-625    CAUAAGUAUAAGGACUCUCGUGAGCAGUUCGAGCAGAGAACACAUAAGAGAUUGAUUG
env-626    ..........................................................
env-632    ACCAGUUCCACUUCCAACAAAGAAGGAAGUAGUAACAAUCUUGAGAGCGGUUCAUAAG
env-633    ..........................................................
env-634    AAGUAUAAGGACUCUCGUGAGCAGUUCGAGCAGAGAACACAUAAGAGAUUGAUUGAUG
env-635    ACCAGUUCCACUUCCAACAAAGAAGGAAGUAGUAACAAUCUUGAGAGCGGUUCAUAAG
env-637    ..........................................................
env-639    CAUAAGUAUAAGGACUCUCGUGAGCAGUUCGAGCAGAGAACACAUAAGAGAUUGAUUG
env-640    ACCAGUUCCACUUCCAACAAAGAAGGAAGUAGUAACAAUCUUGAGAGCGGUUCAUAAG
env-641    ACUCUCGUGAGCAGUUCGAGCAGAGAACACAUAAGAGAUUGAUUGAUGUUGUUGCACC
env-642    CAUAAGUAUAAGGACUCUCGUGAGCAGUUCGAGCAGAGAACACAUAAGAGAUUGAUUG
env-643    UAAGGACUCUCGUGAGCAGUUCGAGCAGAGAACACAUAAGAGAUUGAUUGAUGUUGUU
env-644    ACCAGUUCCACUUCCAACAAAGAAGGAAGUAGUAACAAUCUUGAGAGCGGUUCAUAAG
Jig-1-1    GGUGCCGCUGCCUACUAAAAAGGAAGUGGUAACUAUACUUAGAGCGGUUCAUAAGUAC
env-646    AGUACCGAUGCCAACUAAGAAAGAGGUAGUUACAAUUCUUAGAGCGGUUCACAAAUAC
env-647    AGUACCGAUGCCAACUAAGAAAGAGGUAGUUACAAUUCUUAGAGCGGUUCACAAAUAU
Eha-1-1    AGUACCGAUGCCAACUAAGAAAGAGGUAGUUACAAUUCUUAGAGCGGUUCACAAAUAU
env-648    ..........................................................
env-649    AGUACCGAUGCCAACUAAGAAAGAGGUAGUUACAAUUCUUAGAGCGGUUCACAAAUAU
env-650    GACCAGUUCCAAUGCCAACAAAGGUAGAAAAGGUGACAAUCUUAAGAGCUGUUCAUAA
env-653    GACCAGUUCCAAUGCCAACAAAGGUAGAAAAGGUGACAAUCUUAAGAGCUGUUCAUAA
env-654    UGAGUGGUCCGGUACCUCU.......................................
env-655    ..........................................................
env-656    UGAGUGGUCCGGUACCUCUUCC....................................
env-657    ..........................................................
env-658    UGAGUGGUCCGGUACCUCUUCCAACAAAGAAGGAAGUUGUUACAAUUCUUAGAGCUGU
env-662    AGUGGUCCGGUACCUCUUCCAACAAAGAAGGAAGU.......................
env-663    ..........................................................
env-664    ..........................................................
env-667    ..........................................................
env-668    ..........................................................
Lba-14-1   GGUUGAUGCAUCUGCCAAAAAAAUAAUUGAAACAGUUAAGAAAAAUGGAUCACAGGUG
Bsp-1-1    AGAACGGAUCUCAGGUUAGCGGACCUGUACCGCUUCCUACUAAGAAGGAAGUAGUUAC
Bsp-9-1    AGAACGGAUCUCAGGUUAGCGGACCUGUACCGCUUCCGACUAAGAAGGAAGUAGUUAC
env-669    GUACCGCUUCCAACGAAGAAAGAGGUAGUA............................
env-671    GUACCGCUUCCAACGAAGAAAGAGGUAGUAACAAUCCUUCGUGCAGUUCACAAGUACA
env-672    GUACCGCUUCCAACGAAGAAAG....................................
env-681    GUACCGCUUCCAACGAAGAAAGAGGUAGUAACAAUCCUUCGUGCCGUUCAUAAGUACA
env-683    GUACCGCUUCCAACGAAGAAAGAGGUAGUAACAAUCCUUCGUGCAGUUCACAAGUACA
env-684    ..........................................................
env-685    NNNNNNNNNNNNNNNNNNNNNNNNNNNNNNNNNNNNNNNNAUCAAGAUGAAAAACAAAU
env-686    AAAGAACGGGUCACAGGUGAGUGGACCGGUACCACUUCCGACUAAGAAGGAAGUUGUA
env-687    ..........................................................
env-691    ..........................................................
env-692    AAAGAACGGGUCACAGGUGAGUGGACCGGUACCACUUCCGACUAAGAAGGAAGUUGUA
env-694    AAAGAACGGGUCACAGGUGAGUGGACCGGUACCACUUCCGACUAAGAAGGAAGUUGUA
env-695    AAAGAACGGGUCACAGGUGAGUGGACCGGUACCACUUCCGACUAAGAAGGAAGUUGUA
env-696    AAAGAACGGGUCACAGGUGAGUGGACCGGUACCACUUCCGACUAAGAAGGAAGUUGUA
env-699    AAAGAACGGGUCACAGGUGAGUGGACCGGUACCACUUCCGACUAAGAAGGAAGUUGUA
env-700    ..........................................................
env-705    ..........................................................
env-708    GCGGACCAGUUCCAAUGCCUACUAAGAUUGAGAAGAUCACAAUCUUAAGAGCAGUUCA
env-709    GCGGACCAGUUCCAAUGCCUACUAAGAUUGAGAAGAUCACAAUCUUAAGAGCAGUUCA
env-710    GCGGACCAGUUCCAAUGCCUACUAAGAUUGAGAAGAUCACAAUCUUAAGAGCAGUUCA
env-711    ..........................................................
env-712    ..........................................................
env-713    ..........................................................
env-726    GCGGACCAGUUCCAAUGCCUACUAAGAUUGAGAAGAUCACAAUCUUAAGAGCAGUUCA
env-727    ..........................................................
env-729    GCGGACCAGUUCCAAUGCCUACUAAGAUUGAGAAGAUCACAAUCUUAAGAGCAGUUCA
env-730    GCGGACCAGUUCCAAUGCCUACUAAGAUUGAGAAGAUCACAAUCUUAAGAGCAGUUCA
env-731    GCGGACCAGUUCCAAUGCCUACUAAGAUUGAGAAGAUCACAAUCUUAAGAGCAGUUCA
Rsp-1-1    UUCCACUUCCUACAAAGAAAGAGGUUGUUACAAUCUUAAGAGCCGUUCACAAAUACAA
env-732    UGCGCCACUGAUCAGAAAACUACAAUGACGAAUUACGGAAUCGCAAUCGCCUAUAUGA
Rgn-1-1    GCACAGGUAAGCGGGCCGGUACCACUUCCAACUAAGAAGGAAGUUGUUACAAUCUUAA
env-733    CACAGGUAAGCGGACCGGUACCACUUCCAACUAAGAAGGAAGUUGUUACAAUCUUAAG
env-734    CACAGGUAAGCGGACCGGUACCACUUCCAACUAAGAAGGAAGUUGUUACAAUCUUAAG
env-735    GCACAGGUAAGCGGACCGGUACCACUUCCAACUAAGAAGGAAGUUGUAACAAUCUUAA
env-739    ..........................................................
env-742    GCACAGGUAAGCGGACCGGUACCACUUCCAACUAAGAAGGAAGUUGUUACAAUCUUAA
Lba-4-1    GCACAGGUAAGCGGACCGGUACCACUUCCAACUAAGAAGGAAGUUGUUACAAUCUUAA
env-744    GCACAGGUAAGCGGACCGGUACCACUUCCAACUAAGAAGGAAGUUGUUACAAUCUUAA
env-749    CACAUUGAAAGCAUAUGAUCACCAGUUGGUUGAUGCAUCUGCUAAGAAAAUCAUCGAA
env-751    UGGAUCACAGGUGAGCGGACCAGUGCCGUUACCAACAAAAAAGGAAGUAAUCACCAUU
Cph-1-1    AUUUAAUCGAUGCAUCUGCAGGAAAAAUUAUCGAAACUGUAAAGAAGACAGGAUCGAA
Bfo-1-1    GCCCCUUCCGACAAAGAAGGAAGUAGUUACCAUUCUCCGUGCUGUACACAAGUACAAA
env-752    CAUCGAGACUGUAAAGAAGAACGGAUCACAGGUAAGCGGUCCCGUACCCCUUCCUACU
env-753    ..........................................................
Lbo-3-1    GUACCUCUUCCAACAAAGAAGGAAGUUGUUACUAUCUUAAGAGCUGUUCACAAAUAUA
env-754    UUCCGUUGCCAACUAAGAAAGAAAUUGUUACAAUUCUUAGAGCGGUUCAUAAGUACAA
env-755    UUCCGUUGCCAACUAAGAAAGAAAUUGUUACAAUUCUUAGAGCGGUUCAUAAGUACAA
Csp-4-1    CAAGUGAGUGGACCAGUGCCACUUCCAACUAAGAAGGAAGUAGUAACAAUCUUAAGAG
env-756    CAAGAAGAAUGGAGCUCAGGUAAGCGGACCGGUACCCCUUCCUACUAAAAAGGAAGUA
env-758    GGUCACAGGUGAGUGGACCGGUACCACUUCCGACUAAGAAGGAAGUUGUAACAAUUCU

           ..........................................................
           ..........................................................
           ●●●●●●●●●●●●●●●●●●●●●●●●●●●●●●●●●●●●●●●●●●●●●●●●●●●●●●●●●●
```

```
env-759    UACCACUACCAACGAAAAAAGAGGUAGUUACAAUUCUUAGAGCGGUACACAAGUACAA
env-760    UACCACUACCAACGAAAAAAGAGGUAGUUACAAUUCUUAGAGCGGUACACAAGUACAA
env-762    ..........................................................
env-763    CCACUUCCAACUAAGAAAGAGGUAGUUACAAUUCUUAGAGCGGUUCAUAAGUACAAAG
env-764    UAAGUGGACCGGUACCUCUUCCUACUAAGAAGGAAGUUGUUACAAUUCUUCGUGCCGU
Lba-6-1    GAAGUUGUAACUAUCUUAAGAGCUGUUCACAAAUAUAAAGAUUCCAGAGAGCAGUUUG
Dsp-1-1    CGAAACUGUUAAGAAGAAUGGAUCAAAGGUGAGCGGACCGGUACCGCUUCCGACUAAG
env-765    ..........................................................
env-767    CCAGUGCCGUUACCAACUAAGAAAGAGGUUGUAACAAUUCUGCGUGCGGUUCAUAAGU
env-768    ..........................................................
env-769    CCAGUGCCGUUACCAACUAAGAAAGAGGUUGUAACAAUUCUGCGUGCGGUUCAUAAGU
env-770    CCAGUGCCGUUACCAACUAAGAAAGAGGUUGUAACAAUUCUGCGUGCGGUUCAUAAGU
env-777    GACCGGUGCCGUUACCAACUAAGAAAGAGGUUGUAACCAUUCUGCGUGCGGUUCAUAA
env-778    GUACCGCUUCCGACAAAGAAGGAAGUUGUAACAAUUUUAAGAGCGGUUCACAAAUACA
env-802    GUACCGCUUCCGACAAAGAAGGAAGUUGUAACAAUCUUAAGAGCGGUUCACAAAUACA
env-803    ..........................................................
Cco-1-1    GUACCGCUUCCGACAAAGAAGGAAGUUGUAACAAUUUUAAGAGCGGUUCACAAAUACA
env-806    GUACCGCUUCCGACAAAGAAGGAAGUUGUAACAAUCUUAAGAGCGGUUCACAAAUACA
env-812    GUACCGCUUCCGACAAAGAAGGAAGUUGUAACAAUCUUAAGAGCGGUUCAUAAAUACA
env-814    ACAGGUGAGCGGACCGGUUCCACUUCCAACUAAGAAGGAAGUUGUAACAAUCUUAAGA
Dfo-2-1    ACAGGUGAGCGGACCGGUUCCACUUCCAACUAAGAAGGAAGUUGUAACAAUCUUAAGA
Dfo-1-1    ACAGGUGAGCGGACCGGUUCCACUUCCAACUAAGAAGGAAGUUGUAACAAUCUUAAGA
env-818    ACAGGUGAGCGGACCGGUUCCACUUCCAACUAAGAAGGAAGUUGUAACAAUCUUAAGA
env-819    ACAGGUGAGCGGACCGGUUCCACUUCCAACUAAGAAGGAAGUUGUAACAAUCUUAAGA
env-820    GACCGGUGCCGUUACCAACCAAGAAAGAGGUAGUAACUAUUCUGCGUGCGGUUCAUAA
[ci-1-1    ACCGGUACCGUUACCAACCAAGAAAGAGGUAGUAACCAUUCUGCGUGCUGUUCAUAAG
Csp-5-1    ACCGGUACCGUUACCAACCAAGAAAGAGGUAGUAACCAUUCUGCGUGCUGUUCAUAAG
env-821    GGACCGGUGCCGUUACCAACCAAGAAAGAGGUAGUAACUAUUCUGCGUGCGGUUCAUA
env-822    GACCGGUGCCGUUACCAACCAAGAAAGAGGUAGUAACUAUUCUGCGUGCGGUUCAUAA
env-823    GACCGGUGCCGUUACCAACCAAGAAAGAGGUAGUAACUAUUCUGCGUGCGGUUCAUAA
Cbo-2-1    GACCGGUGCCGUUACCAACCAAGAAAGAGGUAGUAACUAUUCUGCGUGCGGUUCAUAA
Cba-7-1    ACCGGUGCCGUUACCAACCAAGAAAGAGGUAGUAACCAUUCUGCGUGCUGUUCAUAAG
env-824    GACCGGUGCCGUUACCAACCAAGAAAGAGGUAGUAACUAUUCUGCGUGCGGUUCAUAA
Cba-2-1    ACCGGUGCCGUUACCAACCAAGAAAGAGGUAGUAACUAUUCUGCGUGCUGUUCAUAAG
env-826    GACCGGUGCCGUUACCAACCAAGAAAGAGGUAGUAACUAUUCUGCGUGCGGUUCAUAA
Ccl-6-1    GACCGGUGCCGUUACCAACCAAGAAAGAGGUAGUAACUAUUCUGCGUGCGGUUCAUAA
env-827    ..........................................................
Ccl-3-1    GACCGGUGCCGUUACCAACCAAGAAAGAGGUAGUAACUAUUCUGCGUGCGGUUCAUAA
Ccl-1-1    GACCGGUGCCGUUACCAACCAAGAAAGAGGUAGUAACUAUUCUGCGUGCGGUUCAUAA
Lba-16-1   CAGGUGAGCGGUCCGGUACCACUUCCGACCAAGAAGGAAGUAGUUACGAUCCUGCGGG
Csp-3-1    CAGGUGAGCGGUCCGGUACCACUUCCGACCAAGAAGGAAGUAGUUACAAUCCUGCGGG
env-828    UCACAGGUGAGCGGACCGGUGCCGCUGCCGACAAAGAAAGAGGUAGUGACAAUUCUUA
env-829    CAGGUGAGCGGACCGGUGCCGCUGCCGACAAAGAAAGAGGUAGUGACAAUUCUUAGAG
env-830    ..........................................................
env-831    ..........................................................
env-832    ..........................................................
env-833    ..........................................................
env-837    NNNNNNNNNNNNNNNNGUAUGAUCAUCAGCUCGUUGAUGCAUCCGCAAAGAAAAUCAU
env-838    CAGGUGAGCGGACCGGUGCCGCUGCCGACAAAGAAAGAGGUAGUGACAAUUCUUAGAG
env-839    UGUACCUCUUCCAACAAAGAAGGAAGUUGUAACAAUUCUUAGAGCUGUACACAAGUAC
env-840    ..........................................................
env-841    ACAGGUGAGCGGACCGGUGCCGCUGCCGACAAAGAAAGAGGUAGUGACAAUUCUUAGA
Lba-7-1    CAGGUGAGCGGACCGGUGCCGCUGCCGACAAAGAAAGAGGUAGUGACAAUUCUUAGAG
env-842    CAGGUGAGCG................................................
env-843    CGGUGCCGUUACCAACGAAGAAGGAAGUGGUAACCAUCUUACGUGCAGUUCACAAGUA
env-844    C.........................................................
env-845    GAAGAUUAUCGAGAGUGCUAAGAAGACAGGGGCUCAGGUGAGCGGACCGGUACCUCUU
env-848    GAAGAUUAUCGAGAGUGCUAAGAAGACAGGGGCUCAGGUGAGCGGACCGGUACCUCUU
env-849    ..........................................................
env-850    ..........................................................
env-851    ..........................................................
env-853    AUCGAAACUGUAAAGAAGAACGGAUCACAGGUGAGCGGACCGGUACCUCUUCCUACAA
env-854    AUCGAAACUGUAAAGAA.........................................
env-855    ..........................................................
env-856    ..........................................................
env-861    AUCGAAACUGUAAAGAAGAACGGAUCACAGGUGAGCGGACCGGUACCUCUUCCUACAA
env-862    ..........................................................
env-863    UACCACUUCCCACUAAGAAAGAAGUAGUAACUAUCUUACGUGCGGUUCAUAAAUACAA
env-864    ..........................................................
env-865    GAAAAACGGAUCACAGGUGAGCGGACCGGUACCACUUCCAACUAAGAGAGAAGUUGUU
env-866    GAAAAACGGAUCACAGGUGAGCGGACCGGUACCACUUCCAACUAAGAGAGAAGUUGUU
env-867    GAAAAACGGAUCACAGGUGAGCGGACCGGUACCACUUCCAACUAAGAGAGAAGUUGUU
env-868    ..........................................................
env-872    ..........................................................
env-874    GAAAAACGGAUCACAGGUGAGCGGACCGGUACCACUUCCAACUAAGAGAGAAGUUGUU
env-875    ..........................................................
env-877    GAAGAACGGAUCACAGGUGAGCGGACCGGUACCUCUUCCUACUAAGAAGGAAGUAGUU
env-878    GAAAAACGGAUCACAGGUGAGCGGACCGGUACCACUUCCAACUAAGAGAGAAGUUGUU
env-879    GAAAAACGGAUCACAGGUGAGCGGACCGG.............................
env-880    GAAAAACGGAUCACAGGUGAGCGGACCGGUACCACUUCCAACUAAGAGAGAAGUUGUU
env-892    GAAAAACGGAUCACAGGUGAGCGGACCGGUACCACUUCCAACUAAGAGAGAAGUUGUU
env-893    GAAAAACGGAUCACAGGUGAGCGGACCGGUACCACUUCCGACUAAGAGAGAAGUUGUU
env-898    GAAAAACGGAUCACAGGUGAGCGGACCGGUACCACUUCCAACUAAGAGAGAAGUUGUU
env-899    GAAAAACGGAUCACAGGUGAGCGGACCGGUACCACUUCCAACUAAGAGAGAAGUUGUU

           ..........................................................
           ..........................................................
           ●●●●●●●●●●●●●●●●●●●●●●●●●●●●●●●●●●●●●●●●●●●●●●●●●●●●●●●●●●
```

```
env-900     ..........................................................
env-901     ..........................................................
env-902     ..........................................................
env-903     GAAAAACGGAUCACAGGUGAGCGGACCGGUACCACUUCCAACUAAGAGAGAAGUUGUU
env-904     GAAAAACGGAUCACAGGUGAGCGGACCGGUACCACUUCCGACUAAGAGAGAAGUUGUU
env-906     GAAAAACGGAUCACAGGUGAGCGGACCGGUACCACUUCCGACUAAGAGAGAAGUUGUU
Rin-2-1     AGAAAAACGGAUCACAGGUGAGCGGACCGGUACCACUUCCAACUAAGAGAGAAGUUGU
env-907     ..........................................................
env-913     GAAAAACGGAUCACAGGUGAGCGGACCGGUACCACUUCCUACUAAGAGAGAAGUUGUU
env-914     GAAAAACGGAUCACAGGUGAGCGGACCAGUACCACUUCCUACUAAGAGAGAAGUUGUU
env-915     ..........................................................
env-916     GAAAAACGGAUCACAGGUGAGCGGACCAGUACCACUUCCUACUAAGAGAGAAGUUGUU
env-918     GGUACCACUUCCAACUAAGAAGGAAGUAGUAACAAUCUUAAGAGCGGUUCACAAAUAC
Lba-10-1    GGUGCCACUUCCAACUAAGAAGGAAGUAGUGACAAUCUUAAGGGCGGUUCACAAAUAC
[sc-1-1     GUACCACUUCCAACUAAGAAGGAAGUAGUAACAAUCUUAAGAGCGGUUCACAAAUACA
env-921     GGUGCCACUUCCAACUAAGAAGGAAGUAGUGACAAUCUUAAGGGCGGUUCACAAAUAC
env-923     GGUACCACUUCCAACUAAGAAGGAAGUAGUAACAAUCUUAAGAGCGGUUCACAGAUAC
Csc-1-1     GGUGCCACUUCCAACUAAAAAGGAAGUAGUGACAAUCUUAAGGGCGGUUCACAAAUAC
Cba-6-1     GGUACCACUUCCAACUAAGAAGGAAGUAGUAACAAUCUUAAGAGCGGUUCACAAAUAC
Bhy-1-1     UACCUCUUCCGACAAAGAAGGAAGUAGUUACCAUCCUGAGAGCGGUUCACAAGUAUAA
env-924     ..........................................................
env-926     UGCUGCUGGGAAGAUUAUUGAGACGGCAAAGAAGACUGGCGCUAAUGUAAGCGGUCCG
Bcr-1-1     UGAUCAGUCAGCUAAGAAAAUUAUCGAGACUGUUAAGAAGACAGGAUCUCAGGUGAGU
env-928     UGAUCAGUCAGCUAAGAAAAUUAUCGAGACUGUUAAGAAGACAGGAUCUCAGGUGAGU
env-930     UGAUCAGUCAGCUAAGAAAAUUAUCGAGACUGUUAAGAAGACAGGAUCUCAGGUGAGU
env-932     UCAGUCAGCUAAGAAAAUUAUCGAGACUGUUAAGAAGACAGGAUCUCAGGUGAGUGGU
env-933     UGAUCAGUCAGCUAAGAAAAUUAUCGAGACUGUUAAGAAGACAGGAUCUCAGGUGAGU
env-934     ..........................................................
env-935     ..........................................................
env-940     GAUCACGAAUUAGUAGAUUCAUCAGCAAAAAAAAUCAUCGAGACUGUAAAGAAGA...
env-942     ..........................................................
env-943     GAUCACGAAUUAGUAGAUUCAUCAGCAAAAAAAAUCAUCGAGACUGUAAAGAAGAACG
env-944     GAUCACGAAUUAGUAGAUUCAUCAGCAAAAAAAAUCAUCGAGACUGUAAAGAAGAACG
env-945     GAUCACGAAUUAGUAGAUUCAUCAGCAAAAAAAAUCAUCGAGACUGUAAAGAAGAACG
env-946     ..........................................................
env-949     GAUCACGAAUUAGUAGAUUCAUCAGCAAAAAAAAUCAUCGAGACUGUAAAGAAGAACG
env-950     ..........................................................
Rin-1-1     GAUCACGAAUUAGUAGAUUCAUCAGCAAAAAAAAUCAUCGAGACUGUAAAGAAGAACG
env-951     GAUCACGAAUUAGUAGAUUCAUCAGCAAAAAAAAUCAUCGAGACUGUAAAGAAGAACG
env-953     GAUCACGAAUUAGUUGAUUCAUCAGCAAAGAAAAUCAUCGAGACUGUAAAGAAGAACG
env-954     ..........................................................
env-955     GAUCACGAAUUAGUAGAUUCAUCAGCAAAAAAAAUCAUCGAGACUGUAAAGAAGAACG
env-959     ACAAUCAGAUCGAUCAUGCUGCUGGAAAGAUUAUUGAGACAUUGAAGAAAAACGGGGC
env-960     CCGCUUCCCACGAAGCGUGAGGUCGUGACCAUCCUGCGUGCCGUUCAUAAGUACAAAG
env-961     GCUUCCCACGAAGCGCGAGAUCGUGACCAUCCUGCGCGCGGUGCAUAAGUACAAAGAU
env-962     ..........................................................
env-963     ..........................................................
env-964     CGCUGCCAACUAAGAAAGAGGUAGUUACUAUUCUUAGAGCAGUACACAAGUACAAAGA
env-965     GUAGUAACAAUUCUAAGAGCGGUACAUAAGUACAAAGACUCUAGAGAACAGUUUGAAC
env-967     GUAGUAACAAUUCUAAGAGCGGUACAUAAGUACAAAGACUCUAGAGAACAGUUUGAAC
env-968     ..........................................................
env-970     UGCCGCUGCCAACUAAGAAAGAGGUAGUAACCAUUCUGCGAGCAGUUCAUAAGUACAA
env-971     UGCCGCUGCCAACUAAGAAAGAGGUAGUAACCAUUCUGCGAGCAGUUCAUAAGUACAA
env-972     ..........................................................
env-973     ..........................................................
Lba-3-1     GGUACCACUUCCAACUAAGAAGGAAGUUGUAACAAUCUUAAGAGCGGUUCACAAAUAC
env-980     AGACUGAUCGAUAUCAUCACACCAACACAGAAAACUGUUGAUGCAUUAUCAAGAUUAG
env-981     ..........................................................
env-983     GGUACCACUUCCAACUAAAAAAGAAGUAGUAACAAUCUUAAGAGCCGUACACAAAUAC
env-984     GGUACCACUUCCAACUAAAAAAGAAGUAGUAACAAUCUUAAGAGCCGUACACAAAUAC
env-985     CGGUACCACUUCCAACUAAGAAGGAAGUUGUAACAAUCUUAAGAGCGGUACACAAAUA
Csp-8-1     CGGUACCACUUCCAACUAAGAAGGAAGUUGUAACAAUCUUAAGAGCGGUACACAAAUA
env-986     GGUACCACUUCCAACUAAGAAGGAAGUUGUAACAAUCUUAAGAGCGGUACACAAAUAC
Csp-9-1     CGGUACCACUUCCAACUAAGAAGGAAGUUGUAACAAUCUUAAGAGCGGUGCACAAAUA
env-987     CGGUACCACUUCCAACUAAGAAGGAAGUUGUAACAAUCUUAAGAGCGGUACACAAAUA
env-991     CUGUACCACUUCCGACAAAGAAGGAAGUAGUUACAAUCUUAAGAGCUGUUCACAAAUA
env-992     CUGUACCACUUCCGACAAAGAAGGAAGUAGUUACAAUCUUAAGAGCUGUUCACAAAUA
env-993     ..........................................................
Dlo-2-1     CUGUACCACUUCCGACAAAGAAGGAAGUAGUUACAAUCUUAAGAGCUGUUCACAAAUA
env-994     CCUGUACCACUUCCGACAAAGAAGGAAGUAGUUACAAUCUUAAGAGCUGUUCACAAAU
env-995     GUAGAUGCAUCCGCUAAGAAAAUCAUCGAAACUGUAAAGAAAAACGGAUCUAAGGUGA
env-996     ..........................................................
env-1006    ..........................................................
env-1007    CUGUACCACUUCCGACAAAGAAGGAAGUAGUUACAAUCUUAAGAGCUGUUCACAAAUA
env-1008    AAGGUGAGUGGACCUGUACCACUUCCGACAAA..........................
env-1009    ..........................................................
env-1018    CCUGUACCACUUCCGACAAAGAAGGAAGUAGUUACAAUCUUAAGAGCUGUUCACAAAU
env-1019    CUGUACCACUUCCGACAAAGAAGGAAGUAGUUACAAUCUUAAGAGCUGUUCACAAAUA
Dlo-1-1     CUGUACCACUUCCGACAAAGAAGGAAGUAGUUACAAUCUUAAGAGCUGUUCACAAAUA
env-1020    UGUACCGCUUCCUACUAAAAAGGAAGUAGUUACAAUCCUUCGUGCAGUUCACAAGUAC
env-1021    CUGUACCGCUUCCUACUAAAAAGGAAGUAGUUACAAUCCUUCGUGCAGUUCACAAGUA
env-1022    CUGUACCGCUUCCUACUAAAAAGGAAGUAGUUACAAUCCUUCGUGCAGUUCACAAGUA
Lba-2-1     CAGGUGAGCGGUCCAGUGCCACUUCCGACUAAGAAGGAAGUUGUAACAAUUCUGAGAG
env-1023    ..........................................................

            ..........................................................
            ..........................................................
            ●●●●●●●●●●●●●●●●●●●●●●●●●●●●●●●●●●●●●●●●●●●●●●●●●●●●●●●●●●
```

```
env-1024   ..........................................................
env-1026   AAAAAAUUAUCGAAACAGUAAAGAAAAAUGGUUCACAGGUGAGUGGACCGGUACCUCU
env-1027   AAAAAAUUAUCGAAACAGUAAAGAAAAAUGGUUCACAGGUGAGUGGACCGGUACCUCU
env-1028   AAAAAAUUAUCGAAACAGUAAAGAAAAAUGGUUCACAGGUGAGUGGACCGGUACCUCU
env-1029   ..........................................................
env-1030   ..........................................................
env-1032   UCCGGUUCCGCUGCCGACAAAGAAAGAGGUUGUAACAAUCUUAAGAGCUGUUCAUAAG
env-1033   UCCGGUUCCGCUGCCGACAAAGAAAGAGGUUGUAACAAUCUUAAGAGCUGUUCAUAAG
env-1034   ..........................................................
env-1035   GAGACUGUAAAGAAGAACGGAUCACAGGUGAGCGGACCGGUACCUCUUCCUACUAAGA
env-1036   ..........................................................
env-1037   ..........................................................
env-1038   ..........................................................
env-1039   CGAGACUGUAAAGAAGAACGGAUCACAGGUGAGCGGACCGGUACCUCUUCCUACUAAG
env-1040   CGAGACUGUAAAGAAGAACGGAUCACAGGUGAGCGGACCGGUACCUCUUCCUACUAAG
env-1041   ..........................................................
env-1042   ..........................................................
env-1043   ..........................................................
env-1044   ..........................................................
env-1045   CGAGACUGUAAAGAAGAACGGAUCACAGGUGAGCGGACCGGUACCUCUUCCUACUAAG
env-1046   ..........................................................
env-1052   ..........................................................
env-1053   ..........................................................
env-1054   ..........................................................
env-1055   C.........................................................
env-1056   ..........................................................
env-1057   ..........................................................
env-1058   ..........................................................
env-1059   CGAGACUGUAAAGAAGAACGGAUCACAGGUGAGCGGACCGGUACCUCUUCCUACUAAG
Cha-1-1    CGGUGCCGUUACCAACGAAGAAAGAAGUGGUAACUAUCUUACGUGCAGUUCACAAGUA
env-1065   CGGUGCCGUUACCAACGAAGAAAGAAGUGGUAACUAUCUUACGUGCAGUUCACAAGUA
env-1066   CGGUGCCGUUACCAACGAAGAAAGAAGUGGUAACUAUCUUACGUGCAGUUCACAAGUA
Cba-1-1    CUGCCAAGAAAAUCAUCGAAACAGUUAAGAAGAACGGAGCACAGGUGAGCGGACCGGU
Cba-8-1    CUGCCAAGAAAAUCAUCGAAACAGUUAAGAAGAACGGAGCACAGGUGAGCGGACCGGU
env-1067   CUGCCAAGAAAAUCAUCGAAACAGUUAAGAAGAACGGAGCACAGGUGAGCGGACCGGU
Lba-20-1   CUGCCAAGAAAAUCAUCGAAACAGUUAAGAAGAACGGAGCACAGGUGAGCGGACCGGU
Lba-8-1    CUGCCAAGAAAAUCAUCGAAACAGUUAAGAAGAACGGAGCACAGGUGAGCGGACCGGU
env-1068   CUGCCAAGAAAAUCAUCGAAACAGUUAAGAAGAACGGAGCACAGGUGAGCGGACCGGU
Lba-13-1   ACUCCCAACCAAAAGGGAAGUAGUGACAAUCUUAAGGGCAGUCCACAAGUAUAAGGAC
Csp-7-1    GUACCGUUACCAACAAAGAAAGAGGUUGUUACAAUCUUAAGAGCUGUUCAUAAGUACA
env-1069   UGCCGUUACCAACUAAGAAAGAGGUUGUAACAAUCUUAAGAGCUGUUCAUAAGUACAA
env-1070   G.........................................................
env-1071   UGCCGUUACCAACUAAGAAAGAGGUUGUAACAAUCUUAAGAGCUGUUCAUAAGUACAA
env-1072   ..........................................................
Esp-1-1    CCCCUUCCCACCAAACGGGAAGUAGUAACCAUUUUGAGAGCGGUUCACAAAUACAAAG
env-1073   ..........................................................
env-1074   ..........................................................
env-1075   GGUUCCGCUGCCGACCAAGAAAGAGGUUGUAACGAUCUUAAGAGCUGUUCAUAAGUAC
Csa-1-1    GUGCCGUUACCAACCAAGAAGGAAGUGGUAACCAUCUUAAGGGCGGUUCACAAGUACA
env-1076   CUUUUUUUAUGGCAAGUCAAGUAAUGAGAAU...........................
env-1077   UGAUCAGUCAGCUAAGAAAAUUAUCGAGACUGUUAAGAAGACAGGAUCUCAGGUGAGU
env-1078   CGAGACUGUAAAGAAGAACGGAUCACAGGUGAGCGGACCGGUACCUCUUCCUACUAAG
env-1081   ..........................................................
env-1082   ..........................................................
env-1083   CGAGACUGUAAAGAAGAACGGAUCACAGGUGAGCGGACCGGUACCUCUUCCUACUAAG
env-1084   ..........................................................
env-1085   CGAGACUGUAAAGAAGAACGGAUCACAGGUGAGCGGACCGGUACCUCUUCCUACUAAG
env-1086   CGAGACUGUAAAGAAGAACGGAUCACGGGUGAGCGGACCGGUACCUCUUCCUACUAAG
Cha-2-1    GUUCCGUUACCAACAAAGAAAGAAGUAGUUACUAUCUUACGAGCAGUUCACAAGUACA
Lba-5-1    CUUCCCACCAAGAAGGAAGUAGUUACAAUUUUGCGUGCAGUUCACAAAUAUAAGGAUU
Csp-1-1    GUGCCGUUACCGACAAAGAAGGAAGUAGUAACAAUUUUAAGGGCUGUUCAUAAGUACA
env-1087   GUGCCGUUACCGACAAAGAAGGAAGUAGUAACAAUUUUAAGGGCUGUUCAUAAGUACA
env-1088   GGUUCCGCUUCCUACUAAAAAGGAAGUUGUGACCAUCCUGCGUGCGGUACAUAAGUAC
env-1089   CGGUCCGAUUCCGAUGCCGACCAAGAAGGAAGUAGUUACAAUUCUCCGUGCGGUUCAC
env-1093   CCGACCAAGAAAGAGGUAGUUACAAUUCUUCGUGCGGUCCACAAGUAUAAGGACAGCC
env-1094   ..........................................................
env-1095   ..........................................................
env-1096   CGGUCCGAUUCCGAUGCCGACCAAGAAGGAAGUGGUUACAAUUCUCCGUGCGGUUCAC
env-1097   ..........................................................
env-1099   GCGGUUCACAAGUAUAAGGACAGCCGUGAGCAGUUCGAGCAGAGAACGCAUAAGAGAC
env-1100   ..........................................................
Slo-1-1    CGGUCCGAUUCCGCUGCCGACCAAGAAAGAGGUAGUUACAAUUCUUCGUGCGGUCCAC
env-1101   ..........................................................
env-1102   ..........................................................
env-1103   CCGGUCCGAUUCCGAUGCCGACCAAGAAGGAAGUAGUUACAAUUCUCCGUGCGGUUCA
env-1104   CGAGCAGAGAACUCACA.........................................
env-1105   ..........................................................
env-1106   GGCCCGAUUCCGCUGCCGACCAAGAAAGAGGUAGUUACAAUUCUUCGUGCGGUCCACA
env-1107   ..........................................................
env-1108   CCGGUCCGAUUCCGAUGCCGACCAAGAAGGAAGUAGUUACAAUUCUCCGUGCGGUUCA
env-1115   ..........................................................
env-1118   CGGUCCGAUUCCGAUGCCGACCAAGAAGGAAGU.........................
env-1123   GGCCCGAUUCCGCUGCCGACCAAGAAAGAGGUAGUUACAAUUCUUCGUGCGGUCCACA
env-1124   ..........................................................

           ..........................................................
           ..........................................................
           ●●●●●●●●●●●●●●●●●●●●●●●●●●●●●●●●●●●●●●●●●●●●●●●●●●●●●●●●●●
```

```
env-1145   ..........................................................
env-1146   CGGUCCGAUUCCGAUGCCGACCAAGAAGGAAGU.........................
env-1147   ..........................................................
env-1148   CGGUCCGAUUCCGAUGCCGACCAAGAAGGAAGUAGUUACAAUUCUCCGUGCGGUCCAC
env-1149   ..........................................................
env-1150   UUAUGGCAAAUCAGGUAAUGAGAAUCACGUUGAAGGCGUAUGAUCACAAGCUGGUUGA
env-1151   ..........................................................
env-1154   ..........................................................
env-1155   ..........................................................
env-1156   ..........................................................
env-1157   NNNNNNNNNNNNNNNNNNNNNNNNNNNNNNNNNNNNNNNNNNNNNNNNNNNNNNNNNN
env-1158   ..........................................................
env-1159   GGUUCACAAGUA..............................................
Csp-2-1    AGGUAAGUGGACCUAUUCCACUUCCAACAAAAAAAGAAGUUGUAACAAUAUUAAGAGC
env-1160   CGGACCUAUUCCACUUCCUACUAAGAAGGAAGUUGUAACAAUCCUUAGAGCUGUACAC
env-1162   CGGACCUAUUCCACUUCCUACUAAGAAGGAAGUUGUAACAAUCCUUAGAGCUGUACAC
env-1163   CGGACCUAUUCCACUUCCUACUAAGAAGGAAGUUGUAACAAUCCUUAGAGCUGUACAC
env-1166   CGGACCUAUUCCACUUCCUACUAAGAAGGAAGUUGUAACAAUCCUUAGAGCUGUACAC
env-1167   CCAAUUCCAUUACCUACGAAAAGAGAAGUAGUUACAAUCAUCAGAGCUGUCCAUAAAU
env-1168   CCAAUUCCAUUACCUACGAAAAGAGAGGUAGUUACAAUCAUCAGAGCUGUCCAUAAAU
env-1169   CAGUGGUCCAAUCCCUCUUCCCACAAAGAAGGAAAUUGUGACUAUUAUUAGAGCGGUA
env-1170   CAGUGGUCCAAUCCCUCUUCCCACAAAGAAGGAAAUUGUGACUAUUAUUAGAGCGGUA
env-1171   AGUGGUCCAAUCCCUCUUCCCACAAAGAAGGAAAUUGUGACUAUUAUUAGAGCGGUAC
env-1172   GUGGUCCUAUUCCACUUCCCACUAAGAAGGAAGUUGUGACCAUCAUCAGAGCAACACA
env-1173   ..........................................................
env-1174   ..........................................................
env-1175   GUGGUCCUAUUCCACUUCCCACUAAGAAGGAAGUUGUGACCAUCAUCAGAGCAACACA
env-1176   GUGGUCCUAUUCCACUUCCCACUAAGAAGGAAGUUGUGACCAUCAUCAGAGCAACACA
Cle-1-1    GCUCAAGUAAGUGGACCAAUCCCACUUCCAACUAAAAAAGAAGUUGUAACAAUUCUAC
           ..........................................................
           ..........................................................
           ●●●●●●●●●●●●●●●●●●●●●●●●●●●●●●●●●●●●●●●●●●●●●●●●●●●●●●●●●●
```

### 20 S15-Halobacteria

#### 20.1 Taxa

The taxonomy of each organism containing a putative S15-Halobacteria RNA is listed, with abbreviations identifying each hit (e.g., "Eco-1-1" and "Eco-1-2" might hypothetically represent two distinct RNAs in *E. coli*). The abbreviations will be used to identify each individual S15-Halobacteria RNA in Sections 20.2 and 20.4:.

| **abbrev. of hits** | **taxonomy of species** |
|---|---|
| h3A-1-1 | Archaea Euryarchaeota Halobacteria Halobacteriales haloarchaeon 3A1_DGR |
| Hsa-1-1 | Archaea Euryarchaeota Halobacteria Halobacteriales Haloarculaceae *Halapricum salinum* |
| CHa-1-1 | Archaea Euryarchaeota Halobacteria Halobacteriales Halobacteriaceae *Candidatus* Halobonum tyrrellensis G22 |
| Hci-1-1 | Archaea Euryarchaeota Halobacteria Halobacteriales Halobacteriaceae *Haladaptatus cibarius* D43 |
| Hpa-1-1 | Archaea Euryarchaeota Halobacteria Halobacteriales Halobacteriaceae *Haladaptatus paucihalophilus* DX253 |
| Hje-1-1 | Archaea Euryarchaeota Halobacteria Halobacteriales Halobacteriaceae *Halalkalicoccus jeotgali* B3 |
| Hsu-1-1 | Archaea Euryarchaeota Halobacteria Halobacteriales Halobacteriaceae *Halanaeroarchaeum sulfurireducens* |
| Hac-1-1 | Archaea Euryarchaeota Halobacteria Halobacteriales Halobacteriaceae *Halarchaeum acidiphilum* MH1-52-1 |
| Ham-1-1 | Archaea Euryarchaeota Halobacteria Halobacteriales Halobacteriaceae *Haloarcula amylolytica* JCM 13557 |
| Har-1-1 | Archaea Euryarchaeota Halobacteria Halobacteriales Halobacteriaceae *Haloarcula argentinensis* DSM 12282 |
| Hca-1-1 | Archaea Euryarchaeota Halobacteria Halobacteriales Halobacteriaceae *Haloarcula californiae* ATCC 33799 |
| Hja-1-1 | Archaea Euryarchaeota Halobacteria Halobacteriales Halobacteriaceae *Haloarcula japonica* DSM 6131 |
| Hma-1-1 | Archaea Euryarchaeota Halobacteria Halobacteriales Halobacteriaceae *Haloarcula marismortui* ATCC 43049 |
| Hsa-2-1 | Archaea Euryarchaeota Halobacteria Halobacteriales Halobacteriaceae *Haloarcula salaria* |
| Hsi-1-1 | Archaea Euryarchaeota Halobacteria Halobacteriales Halobacteriaceae *Haloarcula sinaiiensis* ATCC 33800 |
| Hva-1-1 | Archaea Euryarchaeota Halobacteria Halobacteriales Halobacteriaceae *Haloarcula vallismortis* ATCC 29715 |
| Hsp-1-1 | Archaea Euryarchaeota Halobacteria Halobacteriales Halobacteriaceae *Halobacterium sp.* NRC-1 |
| Hla-1-1 | Archaea Euryarchaeota Halobacteria Halobacteriales Halobacteriaceae *Halobiforma lacisalsi* AJ5 |
| Hni-1-1 | Archaea Euryarchaeota Halobacteria Halobacteriales Halobacteriaceae *Halobiforma nitratireducens* JCM 10879 |
| Hha-1-1 | Archaea Euryarchaeota Halobacteria Halobacteriales Halobacteriaceae *Halococcus hamelinensis* 100A6 |
| Hmo-1-1 | Archaea Euryarchaeota Halobacteria Halobacteriales Halobacteriaceae *Halococcus morrhuae* DSM 1307 |
| Hsa-3-1 | Archaea Euryarchaeota Halobacteria Halobacteriales Halobacteriaceae *Halococcus saccharolyticus* DSM 5350 |
| Hsa-4-1 | Archaea Euryarchaeota Halobacteria Halobacteriales Halobacteriaceae *Halococcus salifodinae* DSM 8989 |
| Hth-1-1 | Archaea Euryarchaeota Halobacteria Halobacteriales Halobacteriaceae *Halococcus thailandensis* JCM 13552 |

| | |
|---|---|
| Hde-1-1 | Archaea Euryarchaeota Halobacteria Halobacteriales Halobacteriaceae *Haloferax denitrificans* ATCC 35960 |
| Hel-1-1 | Archaea Euryarchaeota Halobacteria Halobacteriales Halobacteriaceae *Haloferax elongans* ATCC BAA-1513 |
| Hgi-1-1 | Archaea Euryarchaeota Halobacteria Halobacteriales Halobacteriaceae *Haloferax gibbonsii* ATCC 33959 |
| Hla-2-1 | Archaea Euryarchaeota Halobacteria Halobacteriales Halobacteriaceae *Haloferax larsenii* JCM 13917 |
| Hme-1-1 | Archaea Euryarchaeota Halobacteria Halobacteriales Halobacteriaceae *Haloferax mediterranei* ATCC 33500 |
| Hmu-1-1 | Archaea Euryarchaeota Halobacteria Halobacteriales Halobacteriaceae *Haloferax mucosum* ATCC BAA-1512 |
| Hpr-1-1 | Archaea Euryarchaeota Halobacteria Halobacteriales Halobacteriaceae *Haloferax prahovense* DSM 18310 |
| Hsp-2-1 | Archaea Euryarchaeota Halobacteria Halobacteriales Halobacteriaceae *Haloferax sp.* ATCC BAA-646 |
| Hsu-2-1 | Archaea Euryarchaeota Halobacteria Halobacteriales Halobacteriaceae *Haloferax sulfurifontis* ATCC BAA-897 |
| Hvo-1-1 | Archaea Euryarchaeota Halobacteria Halobacteriales Halobacteriaceae *Haloferax volcanii* DS2 |
| Hbo-1-1 | Archaea Euryarchaeota Halobacteria Halobacteriales Halobacteriaceae *Halogeometricum borinquense* DSM 11551 |
| Hsa-5-1 | Archaea Euryarchaeota Halobacteria Halobacteriales Halobacteriaceae *Halogranum salarium* B-1 |
| Hka-1-1 | Archaea Euryarchaeota Halobacteria Halobacteriales Halobacteriaceae *Halomicrobium katesii* DSM 19301 |
| Hmu-2-1 | Archaea Euryarchaeota Halobacteria Halobacteriales Halobacteriaceae *Halomicrobium mukohataei* DSM 12286 |
| Hxa-1-1 | Archaea Euryarchaeota Halobacteria Halobacteriales Halobacteriaceae *Halopiger xanaduensis* SH-6 |
| Hti-1-1 | Archaea Euryarchaeota Halobacteria Halobacteriales Halobacteriaceae *Halorhabdus tiamatea* SARL4B |
| Hut-1-1 | Archaea Euryarchaeota Halobacteria Halobacteriales Halobacteriaceae *Halorhabdus utahensis* DSM 12940 |
| Hai-1-1 | Archaea Euryarchaeota Halobacteria Halobacteriales Halobacteriaceae *Halorubrum aidingense* JCM 13560 |
| Har-2-1 | Archaea Euryarchaeota Halobacteria Halobacteriales Halobacteriaceae *Halorubrum arcis* JCM 13916 |
| Hca-2-1 | Archaea Euryarchaeota Halobacteria Halobacteriales Halobacteriaceae *Halorubrum californiensis* DSM 19288 |
| Hco-1-1 | Archaea Euryarchaeota Halobacteria Halobacteriales Halobacteriaceae *Halorubrum coriense* DSM 10284 |
| Hdi-1-1 | Archaea Euryarchaeota Halobacteria Halobacteriales Halobacteriaceae *Halorubrum distributum* JCM 10118 |
| Hho-1-1 | Archaea Euryarchaeota Halobacteria Halobacteriales Halobacteriaceae *Halorubrum hochstenium* ATCC 700873 |
| Hko-1-1 | Archaea Euryarchaeota Halobacteria Halobacteriales Halobacteriaceae *Halorubrum kocurii* JCM 14978 |
| Hli-1-1 | Archaea Euryarchaeota Halobacteria Halobacteriales Halobacteriaceae *Halorubrum lipolyticum* DSM 21995 |
| Hli-2-1 | Archaea Euryarchaeota Halobacteria Halobacteriales Halobacteriaceae *Halorubrum litoreum* JCM 13561 |
| Hsa-6-1 | Archaea Euryarchaeota Halobacteria Halobacteriales Halobacteriaceae *Halorubrum saccharovorum* DSM 1137 |
| Hsp-3-1 | Archaea Euryarchaeota Halobacteria Halobacteriales Halobacteriaceae *Halorubrum sp.* T3 |
| Hte-1-1 | Archaea Euryarchaeota Halobacteria Halobacteriales Halobacteriaceae *Halorubrum tebenquichense* DSM 14210 |
| Hpa-2-1 | Archaea Euryarchaeota Halobacteria Halobacteriales Halobacteriaceae *Halosarcina pallida* JCM 14848 |
| Hca-3-1 | Archaea Euryarchaeota Halobacteria Halobacteriales Halobacteriaceae *Halosimplex carlsbadense* 2-9-1 |
| Hli-3-1 | Archaea Euryarchaeota Halobacteria Halobacteriales Halobacteriaceae *Haloterrigena limicola* JCM 13563 |
| Hsa-7-1 | Archaea Euryarchaeota Halobacteria Halobacteriales Halobacteriaceae *Haloterrigena salina* JCM 13891 |
| Hth-2-1 | Archaea Euryarchaeota Halobacteria Halobacteriales Halobacteriaceae *Haloterrigena thermotolerans* DSM 11522 |
| Htu-1-1 | Archaea Euryarchaeota Halobacteria Halobacteriales Halobacteriaceae *Haloterrigena turkmenica* DSM 5511 |
| Has-1-1 | Archaea Euryarchaeota Halobacteria Halobacteriales Halobacteriaceae *Halovivax asiaticus* JCM 14624 |
| Nae-1-1 | Archaea Euryarchaeota Halobacteria Halobacteriales Halobacteriaceae *Natrialba aegyptia* DSM 13077 |
| Nas-1-1 | Archaea Euryarchaeota Halobacteria Halobacteriales Halobacteriaceae *Natrialba asiatica* DSM 12278 |
| Nch-1-1 | Archaea Euryarchaeota Halobacteria Halobacteriales Halobacteriaceae *Natrialba chahannaoensis* JCM 10990 |
| Nhu-1-1 | Archaea Euryarchaeota Halobacteria Halobacteriales Halobacteriaceae *Natrialba hulunbeirensis* JCM 10989 |
| Nma-1-1 | Archaea Euryarchaeota Halobacteria Halobacteriales Halobacteriaceae *Natrialba magadii* ATCC 43099 |
| Nta-1-1 | Archaea Euryarchaeota Halobacteria Halobacteriales Halobacteriaceae *Natrialba taiwanensis* DSM 12281 |
| Nga-1-1 | Archaea Euryarchaeota Halobacteria Halobacteriales Halobacteriaceae *Natrinema gari* JCM 14663 |
| Npa-1-1 | Archaea Euryarchaeota Halobacteria Halobacteriales Halobacteriaceae *Natrinema pallidum* DSM 3751 |
| Npe-1-1 | Archaea Euryarchaeota Halobacteria Halobacteriales Halobacteriaceae *Natrinema pellirubrum* DSM 15624 |
| Nve-1-1 | Archaea Euryarchaeota Halobacteria Halobacteriales Halobacteriaceae *Natrinema versiforme* JCM 10478 |
| Ngr-1-1 | Archaea Euryarchaeota Halobacteria Halobacteriales Halobacteriaceae *Natronobacterium gregoryi* SP2 |
| Nam-1-1 | Archaea Euryarchaeota Halobacteria Halobacteriales Halobacteriaceae *Natronococcus amylolyticus* DSM 10524 |
| Nje-1-1 | Archaea Euryarchaeota Halobacteria Halobacteriales Halobacteriaceae *Natronococcus jeotgali* DSM 18795 |
| Noc-1-1 | Archaea Euryarchaeota Halobacteria Halobacteriales Halobacteriaceae *Natronococcus occultus* SP4 |
| Nin-1-1 | Archaea Euryarchaeota Halobacteria Halobacteriales Halobacteriaceae *Natronolimnobius innermongolicus* JCM 12255 |
| Nmo-1-1 | Archaea Euryarchaeota Halobacteria Halobacteriales Halobacteriaceae *Natronomonas moolapensis* 8.8.11 |
| Nph-1-1 | Archaea Euryarchaeota Halobacteria Halobacteriales Halobacteriaceae *Natronomonas pharaonis* DSM 2160 |
| Nba-1-1 | Archaea Euryarchaeota Halobacteria Halobacteriales Halobacteriaceae *Natronorubrum bangense* JCM 10635 |
| Nsu-1-1 | Archaea Euryarchaeota Halobacteria Halobacteriales Halobacteriaceae *Natronorubrum sulfidifaciens* JCM 14089 |
| Nti-1-1 | Archaea Euryarchaeota Halobacteria Halobacteriales Halobacteriaceae *Natronorubrum tibetense* GA33 |
| Ssp-1-1 | Archaea Euryarchaeota Halobacteria Halobacteriales Halobacteriaceae *Salinarchaeum sp.* Harcht-Bsk1 |
| Hag-1-1 | Archaea Euryarchaeota Halobacteria Halobacteriales Halococcaceae *Halococcus agarilyticus* |
| Hse-1-1 | Archaea Euryarchaeota Halobacteria Halobacteriales Halococcaceae *Halococcus sediminicola* |
| Hru-1-1 | Archaea Euryarchaeota Halobacteria Haloferacales Haloferacaceae *Halobellus rufus* |
| Hal-1-1 | Archaea Euryarchaeota Halobacteria Haloferacales Haloferacaceae *Haloferax alexandrinus* |
| Hgi-2-1 | Archaea Euryarchaeota Halobacteria Haloferacales Haloferacaceae *Haloferax gibbonsii* |
| Hsp-4-1 | Archaea Euryarchaeota Halobacteria Haloferacales Haloferacaceae *Haloferax sp.* ATB1 |
| Hna-1-1 | Archaea Euryarchaeota Halobacteria Haloferacales Haloferacaceae *Haloplanus natans* DSM 17983 |
| Hru-2-1 | Archaea Euryarchaeota Halobacteria Haloferacales Halorubraceae *Halolamina rubra* |

| | |
|---|---|
| Hha-2-1 | Archaea Euryarchaeota Halobacteria Haloferacales Halorubraceae *Halorubrum halophilum* |
| Hsa-8-1 | Archaea Euryarchaeota Halobacteria Haloferacales Halorubraceae *Halorubrum saccharovorum* |
| Hsp-5-1 | Archaea Euryarchaeota Halobacteria Haloferacales Halorubraceae *Halorubrum sp.* BV1 |
| Hdj-1-1 | Archaea Euryarchaeota Halobacteria Natrialbales Natrialbaceae *Halopiger djelfimassiliensis* |
| Hgo-1-1 | Archaea Euryarchaeota Halobacteria Natrialbales Natrialbaceae *Halopiger goleimassiliensis* |
| Hsa-9-1 | Archaea Euryarchaeota Halobacteria Natrialbales Natrialbaceae *Halopiger salifodinae* |
| Hla-3-1 | Archaea Euryarchaeota Halobacteria Natrialbales Natrialbaceae *Halostagnicola larsenii* XH-48 |
| Hsp-6-1 | Archaea Euryarchaeota Halobacteria Natrialbales Natrialbaceae *Halostagnicola sp.* A56 |
| Hje-2-1 | Archaea Euryarchaeota Halobacteria Natrialbales Natrialbaceae *Haloterrigena jeotgali* A29 |
| Nal-1-1 | Archaea Euryarchaeota Halobacteria Natrialbales Natrialbaceae *Natrinema altunense* |
| Nsp-1-1 | Archaea Euryarchaeota Halobacteria Natrialbales Natrialbaceae *Natrinema sp.* J7-1 |
| env-1 to env-185 | environmental samples |

#### 20.2 Gene contexts

Each S15-Halobacteria RNA (indicated by "RNA→") is listed. For each hit, the downstream genes predicted to reside in a regulated operon are listed. If the nearest downstream gene is encoding in the opposite strand (and therefore presumed to not be a part of a regulated operon), then that gene is still depicted. Some environmental sequences and some RefSeq entries lack gene annotations, and so no genes are available for such sequences. The direction of each gene is indicated with an arrow (→), and each predicted conserved domain in the gene is named. Conserved domains associated with more than one S15-Halobacteria RNA are assigned a color; other domains are gray. Information about these conserved domains is given in Section 20.3. The accession of the sequence containing each S15-Halobacteria RNA is given in the column named "Seq. accession". Accessions beginning with "NC_", "NS_", "NW_" or "NZ_" are contained in RefSeq. Other accession refer to environmental samples. Nucleotide coordinates are given for the 5′ and 3′ boundaries of each S15-Halobacteria RNA. If the 5′ coordinate is greater than the 3′ coordinate, the RNA is present on the reverse-complement strand of the containing genomic DNA sequence. Each hit is denoted by an abbreviation (like "Eco-1-1") that refers to a taxonomy given in Section 20.1.

| abbrev. | Seq. accession | | 5′ at | 3′ at | genes |
|---|---|---|---|---|---|
| env-1 | CETH01443816.1 | - | 36475 | 36410 | **RNA**→ rps15p (PRK08561)Ribosomal_S15 (pfam00312)→ PRK14888 (PRK14888)→ RPS3A (COG1890)PRK04057 (PRK04057)→ |
| env-2 | CEWR01249401.1 | - | 634 | 569 | **RNA**→ rps15p (PRK08561)Ribosomal_S15 (pfam00312)→ |
| env-3 | CESY01224593.1 | + | 483 | 548 | **RNA**→ |
| env-4 | CEQN01077410.1 | - | 89 | 24 | **RNA**→ |
| env-5 | CEUZ01094134.1 | + | 294 | 359 | **RNA**→ Ribosomal_S15 (pfam00312)Ribosomal_S13_N (pfam08069)→ |
| env-6 | CEVQ01090649.1 | - | 366 | 301 | **RNA**→ Ribosomal_S13_N (pfam08069)→ |
| env-7 | JXWU01104845.1 | + | 1 | 97 | **RNA**→ S15_NS1_EPRS_RNA-bind (cd00677)Ribosomal_S13_N (pfam08069)→ |
| Hut-1-1 | NC_013158.1 | - | 714527 | 714434 | **RNA**→ rps15p (PRK08561)Ribosomal_S15p_S13e (cd00353)→ hypo→ PRK14888 (PRK14888)→ RPS3A (COG1890)PRK04057 (PRK04057)→ |
| env-8 | JXWS01091643.1 | - | 837 | 774 | **RNA**→ rps15p (PRK08561)Ribosomal_S15 (pfam00312)→ PRK07409 (PRK07409)→ |
| env-9 | GQKUMWE02J03Y4 | - | 278 | 180 | **RNA**→ rps15p (PRK08561)→ |
| Hti-1-1 | NZ_AFNT02000006.1 | - | 58722 | 58627 | **RNA**→ RpsO (COG0184)rps15p (PRK08561)→ hypo→ PRK14888 (PRK14888)→ RPS3A (COG1890)PRK04057 (PRK04057)→ |
| env-10 | GSLNARP_contig10642 | + | 226 | 286 | **RNA**→ rps15p (PRK08561)Ribosomal_S15p_S13e (cd00353)→ PHA03247 (PHA03247)→ |
| env-11 | GSLNARP_GFPJP1N02I2Z83 | - | 234 | 174 | **RNA**→ rps15p (PRK08561)→ |
| env-12 | SS_3KL_010_SOIL_10126881 | - | 211 | 126 | **RNA**→ ←hypo |
| env-13 | 2140918017-39245 | - | 377 | 279 | **RNA**→ Ribosomal_S13_N (pfam08069)→ |
| env-14 | GSLNAS_contig01475 | - | 166 | 104 | **RNA**→ rps15p (PRK08561)→ |
| env-15 | APHM01016997.1 | + | 513 | 574 | **RNA**→ RpsO (COG0184)rps15p (PRK08561)→ hypo→ |
| Hac-1-1 | NZ_BATA01000023.1 | + | 18137 | 18202 | **RNA**→ RpsO (COG0184)rps15p (PRK08561)→ hypo→ Pcc1 (COG2892)PRK14888 (PRK14888)→ RPS3A (COG1890)PRK04057 (PRK04057)→ |
| env-16 | GSLNARP_contig03059 | + | 263 | 328 | **RNA**→ rps15p (PRK08561)→ |
| Hsa-5-1 | NZ_ALJD01000003.1 | - | 168876 | 168813 | **RNA**→ ←hypo |
| env-17 | P_A23_Liq_1_Fmt_1021563 | + | 164 | 234 | **RNA**→ hypo→ |

| | | | | | |
|---|---|---|---|---|---|
| env-18 | P_A23_Liq_3_Fmt_1005281 | - | 131 | 61 | **RNA**→ |
| env-19 | P_1C_Liq_3_UnCty_1085606 | - | 92 | 22 | **RNA**→ ←hypo |
| env-20 | P_1C_Sed_2_UnCty_1004024 | + | 2712 | 2782 | **RNA**→ rps15p (PRK08561)Ribosomal_S15p_S13e (cd00353)→ hypo→ |
| Har-2-1 | NZ_AOJJ01000049.1 | - | 118618 | 118548 | **RNA**→ rps15p (PRK08561)Ribosomal_S15p_S13e (cd00353)→ hypo→ PRK14888 (PRK14888)→ RPS3A (COG1890)PRK04057 (PRK04057)→ |
| Hsp-3-1 | NZ_JH815329.1 | + | 386490 | 386560 | **RNA**→ rps15p (PRK08561)Ribosomal_S15p_S13e (cd00353)→ hypo→ PRK14888 (PRK14888)→ RPS3A (COG1890)PRK04057 (PRK04057)→ |
| env-21 | P_1C_Liq_1_UnCty_1154732 | + | 171 | 241 | **RNA**→ |
| env-22 | P_1C_Liq_1_UnCty_1093462 | + | 241 | 311 | **RNA**→ ←hypo |
| env-23 | P_A23_Liq_1_Fmt_1001349 | + | 3246 | 3316 | **RNA**→ rps15p (PRK08561)Ribosomal_S15p_S13e (cd00353)→ |
| Hdi-1-1 | NZ_AOJN01000047.1 | + | 134657 | 134727 | **RNA**→ rps15p (PRK08561)Ribosomal_S15p_S13e (cd00353)→ hypo→ PRK14888 (PRK14888)→ RPS3A (COG1890)PRK04057 (PRK04057)→ |
| Hli-2-1 | NZ_AOJF01000030.1 | - | 133338 | 133268 | **RNA**→ rps15p (PRK08561)Ribosomal_S15p_S13e (cd00353)→ PRK13800 (PRK13800)→ PRK14888 (PRK14888)→ RPS3A (COG1890)PRK04057 (PRK04057)→ |
| Hsa-8-1 | NZ_JNFH02000131.1 | + | 4096 | 4166 | **RNA**→ rps15p (PRK08561)Ribosomal_S15p_S13e (cd00353)→ |
| env-24 | P_1C_Liq_1_UnCty_1068401 | - | 82 | 12 | **RNA**→ ←hypo |
| Hha-2-1 | NZ_BBJP01000023.1 | + | 160344 | 160414 | **RNA**→ rps15p (PRK08561)Ribosomal_S15p_S13e (cd00353)→ |
| Hko-1-1 | NZ_AOJH01000076.1 | + | 51508 | 51578 | **RNA**→ rps15p (PRK08561)Ribosomal_S15p_S13e (cd00353)→ |
| env-25 | SL_6KL_011_BRINE_1017551 | + | 2402 | 2473 | **RNA**→ ←hypo |
| env-26 | ADL20m3uS_contig00641 | - | 3847 | 3777 | **RNA**→ rps15p (PRK08561)Ribosomal_S15p_S13e (cd00353)→ |
| env-27 | SL_6KL_011_BRINE_1076960 | - | 86 | 16 | **RNA**→ |
| env-28 | SL_6KL_011_BRINE_1017271 | + | 2449 | 2519 | **RNA**→ ←hypo |
| env-29 | SL_6KL_011_BRINE_1090146 | - | 309 | 239 | **RNA**→ rps15p (PRK08561)→ |
| env-30 | SL_6KL_011_BRINE_1023965 | - | 103 | 33 | **RNA**→ ←cad (TIGR00779)CadD (COG4300) |
| env-31 | SL_6KL_011_BRINE_1037136 | - | 103 | 33 | **RNA**→ |
| env-32 | SL_6KL_011_BRINE_1181093 | - | 313 | 245 | **RNA**→ ←hypo |
| env-33 | SL_6KL_011_BRINE_1011159 | - | 349 | 278 | **RNA**→ rps15p (PRK08561)→ |
| env-34 | SL_5KL_010_SED_10000937 | + | 4177 | 4247 | **RNA**→ rps15p (PRK08561)Ribosomal_S15p_S13e (cd00353)→ hypo→ PRK14888 (PRK14888)→ PRK04057 (PRK04057)Ribosomal_S3Ae (pfam01015)→ |
| env-35 | SL_6KL_011_BRINE_1004176 | - | 2444 | 2374 | **RNA**→ rps15p (PRK08561)Ribosomal_S15p_S13e (cd00353)→ hypo→ PRK14888 (PRK14888)→ |
| env-36 | SL_5KL_010_BRINE_10006748 | - | 203 | 133 | **RNA**→ |
| env-37 | SL_5KL_010_BRINE_10374007 | + | 150 | 220 | **RNA**→ ←hypo |
| env-38 | SL_4KL_010_BRINE_10048044 | - | 170 | 100 | **RNA**→ |
| env-39 | P_1C_Liq_3_UnCty_1004643 | + | 1043 | 1114 | **RNA**→ |
| env-40 | P_1C_Liq_1_UnCty_1024121 | - | 147 | 77 | **RNA**→ |
| env-41 | JXWS01080544.1 | - | 204 | 134 | **RNA**→ |
| Hsp-5-1 | NZ_KN050825.1 | + | 8179 | 8249 | **RNA**→ rps15p (PRK08561)Ribosomal_S15p_S13e (cd00353)→ hypo→ PRK14888 (PRK14888)→ RPS3A (COG1890)PRK04057 (PRK04057)→ |
| Hca-2-1 | NZ_AOJK01000009.1 | - | 96124 | 96054 | **RNA**→ rps15p (PRK08561)Ribosomal_S15p_S13e (cd00353)→ hypo→ PRK14888 (PRK14888)→ RPS3A (COG1890)PRK04057 (PRK04057)→ |
| Hli-1-1 | NZ_AOJG01000013.1 | - | 3341 | 3271 | **RNA**→ rps15p (PRK08561)Ribosomal_S15p_S13e (cd00353)→ |
| Hte-1-1 | NZ_AOJD01000053.1 | - | 79083 | 79012 | **RNA**→ rps15p (PRK08561)Ribosomal_S15p_S13e (cd00353)→ hypo→ PRK14888 (PRK14888)→ RPS3A (COG1890)PRK04057 (PRK04057)→ |
| Hho-1-1 | NZ_AOJO01000016.1 | - | 18126 | 18055 | **RNA**→ rps15p (PRK08561)Ribosomal_S15p_S13e (cd00353)→ hypo→ PRK14888 (PRK14888)→ RPS3A (COG1890)PRK04057 (PRK04057)→ |
| Hsa-6-1 | NZ_AOJE01000062.1 | + | 16777 | 16847 | **RNA**→ rps15p (PRK08561)Ribosomal_S15p_S13e (cd00353)→ hypo→ hypo→ PRK14888 (PRK14888)→ RPS3A (COG1890)PRK04057 (PRK04057)→ |
| Hco-1-1 | NZ_AOJL01000060.1 | + | 22697 | 22767 | **RNA**→ rps15p (PRK08561)Ribosomal_S15p_S13e (cd00353)→ A-tRNA_syn_arch (TIGR03683)→ PRK14888 (PRK14888)→ RPS3A (COG1890)PRK04057 (PRK04057)→ |
| Hai-1-1 | NZ_AOJI01000013.1 | + | 26726 | 26797 | **RNA**→ rps15p (PRK08561)Ribosomal_S15p_S13e (cd00353)→ hypo→ PRK14888 (PRK14888)→ RPS3A (COG1890)PRK04057 (PRK04057)→ |
| env-42 | P_1C_Sed_2_UnCty_1023573 | + | 227 | 298 | **RNA**→ rps15p (PRK08561)Ribosomal_S15p_S13e (cd00353)→ hypo→ |
| env-43 | GSLNARP_contig15969 | - | 992 | 921 | **RNA**→ rps15p (PRK08561)Ribosomal_S15p_S13e (cd00353)→ hypo→ |
| env-44 | SL_6KL_011_BRINE_1115284 | + | 186 | 256 | **RNA**→ rps15p (PRK08561)→ |
| env-45 | SL_6KL_011_BRINE_1080748 | + | 491 | 561 | **RNA**→ hypo→ |

| | | | | | |
|---|---|---|---|---|---|
| env-46 | GSLNARP_GFPJP1N02H805S | - | 330 | 265 | **RNA**→ Ribosomal_S13_N (pfam08069)→ |
| Hpa-1-1 | NZ_AEMG01000012.1 | + | 19787 | 19853 | **RNA**→ rps15p (PRK08561)Ribosomal_S15p_S13e (cd00353)→ rps15p (PRK08561)Ribosomal_S15p_S13e (cd00353)→ hypo→ hypo→ PRK14888 (PRK14888)→ PRK14888 (PRK14888)→ RPS3A (COG1890)PRK04057 (PRK04057)→ RPS3A (COG1890)PRK04057 (PRK04057)→ hypo→ |
| Hci-1-1 | NZ_JDTH01000002.1 | - | 369624 | 369515 | **RNA**→ RpsO (COG0184)rps15p (PRK08561)→ hypo→ PRK14888 (PRK14888)→ RPS3A (COG1890)PRK04057 (PRK04057)→ |
| env-47 | P_1C_Sed_2_UnCty_1119332 | - | 170 | 102 | **RNA**→ |
| env-48 | GSLSAAI_contig12508 | - | 55982 | 55920 | **RNA**→ rps15p (PRK08561)Ribosomal_S15p_S13e (cd00353)→ hypo→ PRK14888 (PRK14888)→ RPS3A (COG1890)PRK04057 (PRK04057)→ |
| env-49 | ABOG01057646.1 | - | 101 | 46 | **RNA**→ ←hypo |
| env-50 | ABOG01040231.1 | + | 23 | 85 | **RNA**→ hypo→ |
| env-51 | PR_CR_10_Liq_1_inCR_1012431 | + | 1250 | 1312 | **RNA**→ rps15p (PRK08561)Ribosomal_S15 (pfam00312)→ hypo→ |
| env-52 | GSLSAAI_contig09376 | + | 9954 | 10016 | **RNA**→ rps15p (PRK08561)S15_NS1_EPRS_RNA-bind (cd00677)→ hypo→ PRK14888 (PRK14888)→ PRK04057 (PRK04057)Ribosomal_S3Ae (pfam01015)→ |
| env-53 | GSLSAAI_contig12665 | - | 1260 | 1198 | **RNA**→ rps15p (PRK08561)Ribosomal_S15p_S13e (cd00353)→ hypo→ |
| env-54 | GSLSAAI_contig00663 | - | 9050 | 8988 | **RNA**→ rps15p (PRK08561)Ribosomal_S15p_S13e (cd00353)→ hypo→ PRK14888 (PRK14888)→ PRK04057 (PRK04057)Ribosomal_S3Ae (pfam01015)→ |
| env-55 | GSLSAAI_contig05944 | + | 161 | 223 | **RNA**→ rps15p (PRK08561)Ribosomal_S15 (pfam00312)→ hypo→ PRK14888 (PRK14888)→ PRK04057 (PRK04057)Ribosomal_S3Ae (pfam01015)→ |
| env-56 | PR_CR_10_Liq_1_inCR_1198478 | + | 83 | 145 | **RNA**→ rps15p (PRK08561)Ribosomal_S13_N (pfam08069)→ |
| env-57 | GSLSAAI_contig11173 | - | 1195 | 1133 | **RNA**→ rps15p (PRK08561)Ribosomal_S15p_S13e (cd00353)→ hypo→ |
| env-58 | GSLSAAI_contig11466 | + | 1789 | 1851 | **RNA**→ Ribosomal_S13_N (pfam08069)→ |
| Hla-2-1 | NZ_AOLI01000007.1 | - | 53069 | 53009 | **RNA**→ rps15p (PRK08561)Ribosomal_S15p_S13e (cd00353)→ PRK13800 (PRK13800)→ PRK14888 (PRK14888)→ PRK04057 (PRK04057)Ribosomal_S3Ae (pfam01015)→ |
| Hsp-4-1 | NZ_JPES01000120.1 | + | 18134 | 18194 | **RNA**→ rps15p (PRK08561)Ribosomal_S15p_S13e (cd00353)→ hypo→ hypo→ |
| Hsp-2-1 | NZ_AOLD01000018.1 | + | 222134 | 222194 | **RNA**→ rps15p (PRK08561)Ribosomal_S15p_S13e (cd00353)→ hypo→ PRK14888 (PRK14888)→ hypo→ hypo→ PRK04057 (PRK04057)Ribosomal_S3Ae (pfam01015)→ RPS3A (COG1890)PRK04057 (PRK04057)→ |
| Hme-1-1 | NZ_AOLO01000006.1 | + | 119016 | 119079 | **RNA**→ rps15p (PRK08561)Ribosomal_S15p_S13e (cd00353)→ hypo→ Sema_3G (cd11255)PRK14888 (PRK14888)→ PRK04057 (PRK04057)Ribosomal_S3Ae (pfam01015)→ |
| Hel-1-1 | NZ_AOLK01000015.1 | - | 258454 | 258394 | **RNA**→ rps15p (PRK08561)Ribosomal_S15p_S13e (cd00353)→ PRK13800 (PRK13800)→ PRK14888 (PRK14888)→ PRK04057 (PRK04057)Ribosomal_S3Ae (pfam01015)→ |
| Hvo-1-1 | NZ_AOHU01000091.1 | + | 222642 | 222702 | **RNA**→ rps15p (PRK08561)Ribosomal_S15p_S13e (cd00353)→ hypo→ PRK14888 (PRK14888)→ hypo→ hypo→ PRK04057 (PRK04057)Ribosomal_S3Ae (pfam01015)→ |
| Hsu-2-1 | NZ_AOLM01000003.1 | - | 127653 | 127593 | **RNA**→ rps15p (PRK08561)Ribosomal_S15p_S13e (cd00353)→ hypo→ PRK14888 (PRK14888)→ PRK04057 (PRK04057)Ribosomal_S3Ae (pfam01015)→ |
| Hde-1-1 | NZ_AOLP01000019.1 | + | 71205 | 71265 | **RNA**→ rps15p (PRK08561)Ribosomal_S15p_S13e (cd00353)→ hypo→ PRK14888 (PRK14888)→ PRK04057 (PRK04057)Ribosomal_S3Ae (pfam01015)→ |
| Hpr-1-1 | NZ_AOLG01000012.1 | - | 2307 | 2247 | **RNA**→ rps15p (PRK08561)Ribosomal_S15p_S13e (cd00353)→ hypo→ PRK14888 (PRK14888)→ hypo→ PRK04057 (PRK04057)Ribosomal_S3Ae (pfam01015)→ |
| Hgi-1-1 | NZ_AOLJ01000025.1 | + | 12076 | 12136 | **RNA**→ rps15p (PRK08561)Ribosomal_S15p_S13e (cd00353)→ hypo→ PRK14888 (PRK14888)→ PRK04057 (PRK04057)Ribosomal_S3Ae (pfam01015)→ |
| Hal-1-1 | NZ_LK053000.1 | - | 2441666 | 2441606 | **RNA**→ rps15p (PRK08561)Ribosomal_S15p_S13e (cd00353)→ hypo→ PRK14888 (PRK14888)→ PRK04057 (PRK04057)Ribosomal_S3Ae (pfam01015)→ |
| Hgi-2-1 | NZ_CP011947.1 | - | 1061334 | 1061274 | **RNA**→ rps15p (PRK08561)Ribosomal_S15p_S13e (cd00353)→ hypo→ PRK14888 (PRK14888)→ PRK04057 (PRK04057)Ribosomal_S3Ae (pfam01015)→ |
| Hmu-1-1 | NZ_AOLN01000018.1 | + | 214475 | 214538 | **RNA**→ rps15p (PRK08561)Ribosomal_S15p_S13e (cd00353)→ hypo→ PRK14888 (PRK14888)→ PRK04057 (PRK04057)Ribosomal_S3Ae (pfam01015)→ |
| env-59 | JXWS01117778.1 | - | 479 | 415 | **RNA**→ Ribosomal_S15 (pfam00312)Ribosomal_S13_N (pfam08069)→ |
| env-60 | SL_4KL_010_BRINE_10436734 | - | 180 | 109 | **RNA**→ |
| env-61 | SL_4KL_010_BRINE_10606398 | - | 98 | 13 | **RNA**→ ←hypo |
| env-62 | GSLNAS_F7I09SK02FQQIZ | - | 461 | 396 | **RNA**→ Ribosomal_S13_N (pfam08069)→ |
| env-63 | SS_3KL_010_SOIL_10006438 | - | 147 | 81 | **RNA**→ |
| env-64 | SS_3KL_010_SOIL_10046089 | - | 149 | 81 | **RNA**→ ←hypo |
| Has-1-1 | NZ_AOIQ01000008.1 | + | 125599 | 125690 | **RNA**→ hypo→ rps15p (PRK08561)→ hypo→ rps15p (PRK08561)Ribosomal_S15p_S13e (cd00353)→ hypo→ PRK14888 (PRK14888)→ RPS3A (COG1890)PRK04057 (PRK04057)→ |

| | | | | | |
|---|---|---|---|---|---|
| env-65 | JXWU01059374.1 | + | 232 | 306 | **RNA**→ |
| h3A-1-1 | NZ_KK033115.1 | - | 376523 | 376455 | **RNA**→ RpsO (COG0184)rps15p (PRK08561)→ hypo→ PRK14888 (PRK14888)→ PRK04057 (PRK04057)Ribosomal_S3Ae (pfam01015)→ RNAP_L (cd06927)PRK01146 (PRK01146)→ |
| env-66 | ADL20m3uS_contig11732 | + | 2953 | 3015 | **RNA**→ rps15p (PRK08561)Ribosomal_S15p_S13e (cd00353)→ hypo→ PRK14888 (PRK14888)→ RPS3A (COG1890)PRK04057 (PRK04057)→ |
| env-67 | GSLNARP_contig20163 | + | 786 | 851 | **RNA**→ rps15p (PRK08561)Ribosomal_S15p_S13e (cd00353)→ hypo→ |
| Hka-1-1 | NZ_KB905379.1 | + | 563781 | 563847 | **RNA**→ rps15p (PRK08561)Ribosomal_S15p_S13e (cd00353)→ hypo→ PRK14888 (PRK14888)→ RPS3A (COG1890)PRK04057 (PRK04057)→ PRK04057 (PRK04057)Ribosomal_S3Ae (pfam01015)→ |
| Hmu-2-1 | NC_013202.1 | - | 2042594 | 2042528 | **RNA**→ rps15p (PRK08561)Ribosomal_S15p_S13e (cd00353)→ hypo→ PRK14888 (PRK14888)→ PRK04057 (PRK04057)Ribosomal_S3Ae (pfam01015)→ |
| env-68 | P_A23_Liq_1_Fmt_1018204 | - | 215 | 150 | **RNA**→ |
| Ham-1-1 | NZ_AOLW01000050.1 | - | 65561 | 65496 | **RNA**→ RpsO (COG0184)rps15p (PRK08561)→ hypo→ PRK14888 (PRK14888)→ PRK04057 (PRK04057)Ribosomal_S3Ae (pfam01015)→ |
| Hva-1-1 | NZ_AOLQ01000067.1 | - | 102654 | 102589 | **RNA**→ rps15p (PRK08561)Ribosomal_S15p_S13e (cd00353)→ hypo→ PRK14888 (PRK14888)→ PRK04057 (PRK04057)Ribosomal_S3Ae (pfam01015)→ |
| Hja-1-1 | NZ_AOLY01000009.1 | - | 228112 | 228047 | **RNA**→ RpsO (COG0184)rps15p (PRK08561)→ hypo→ PRK14888 (PRK14888)→ PRK04057 (PRK04057)Ribosomal_S3Ae (pfam01015)→ |
| Hma-1-1 | NC_006396.1 | + | 1271347 | 1271412 | **RNA**→ RpsO (COG0184)rps15p (PRK08561)→ hypo→ PRK14888 (PRK14888)→ RPS3A (COG1890)PRK04057 (PRK04057)→ |
| Hsa-2-1 | NZ_JXAO01000006.1 | + | 33018 | 33083 | **RNA**→ RpsO (COG0184)rps15p (PRK08561)→ hypo→ PRK14888 (PRK14888)→ PRK04057 (PRK04057)Ribosomal_S3Ae (pfam01015)→ |
| Hca-1-1 | NZ_AOLS01000008.1 | + | 111762 | 111827 | **RNA**→ RpsO (COG0184)rps15p (PRK08561)→ hypo→ hypo→ |
| Hsi-1-1 | NZ_AOLR01000002.1 | + | 110713 | 110778 | **RNA**→ RpsO (COG0184)rps15p (PRK08561)→ hypo→ PRK14888 (PRK14888)→ RPS3A (COG1890)PRK04057 (PRK04057)→ |
| Har-1-1 | NZ_AOLX01000027.1 | - | 130959 | 130894 | **RNA**→ RpsO (COG0184)rps15p (PRK08561)→ hypo→ PRK14888 (PRK14888)→ PRK04057 (PRK04057)Ribosomal_S3Ae (pfam01015)→ |
| env-69 | P_1C_Sed_2_UnCty_1113621 | + | 256 | 321 | **RNA**→ |
| env-70 | ADL5mRS1u_GM034OG01ABXFC | - | 278 | 212 | **RNA**→ rps15p (PRK08561)→ |
| env-71 | APHM01021510.1 | + | 41573 | 41636 | **RNA**→ rps15p (PRK08561)Ribosomal_S15p_S13e (cd00353)→ |
| env-72 | P_A23_Liq_2_Fmt_1000692 | - | 6203 | 6140 | **RNA**→ rps15p (PRK08561)Ribosomal_S15p_S13e (cd00353)→ |
| env-73 | P_A23_Liq_1_Fmt_1001408 | + | 1210 | 1273 | **RNA**→ rps15p (PRK08561)Ribosomal_S15p_S13e (cd00353)→ |
| env-74 | GSLNARP_contig28817 | - | 4621 | 4558 | **RNA**→ rps15p (PRK08561)Ribosomal_S15p_S13e (cd00353)→ hypo→ PRK14888 (PRK14888)→ PRK04057 (PRK04057)Ribosomal_S3Ae (pfam01015)→ |
| env-75 | APHM01021421.1 | - | 191186 | 191123 | **RNA**→ rps15p (PRK08561)Ribosomal_S15p_S13e (cd00353)→ |
| env-76 | SL_4KL_010_BRINE_10225816 | - | 172 | 88 | **RNA**→ ←hypo |
| Hca-3-1 | NZ_AOIU01000035.1 | + | 133178 | 133243 | **RNA**→ RpsO (COG0184)rps15p (PRK08561)→ hypo→ PRK14888 (PRK14888)→ RPS3A (COG1890)PRK04057 (PRK04057)→ |
| env-77 | SL_5KL_010_BRINE_10001052 | + | 13389 | 13455 | **RNA**→ |
| env-78 | SL_4KL_010_BRINE_10019393 | - | 210 | 144 | **RNA**→ |
| env-79 | SL_6KL_011_BRINE_1125381 | + | 228 | 294 | **RNA**→ ←hypo |
| env-80 | SL_4KL_010_BRINE_10007098 | - | 1494 | 1429 | **RNA**→ ←hypo |
| env-81 | SL_4KL_010_BRINE_10047307 | + | 2075 | 2140 | **RNA**→ |
| Hsa-1-1 | NZ_BBMO01000001.1 | + | 1186894 | 1186960 | **RNA**→ RpsO (COG0184)rps15p (PRK08561)→ hypo→ PRK14888 (PRK14888)→ RPS3A (COG1890)PRK04057 (PRK04057)→ |
| env-82 | GSLNARP_GFPJP1N02JIM46 | + | 6 | 71 | **RNA**→ rps15p (PRK08561)→ |
| env-83 | SL_4KL_010_BRINE_10142426 | - | 703 | 638 | **RNA**→ rps15p (PRK08561)Ribosomal_S15p_S13e (cd00353)→ |
| env-84 | GSLNAS_F7I09SK02IMHTW | - | 401 | 336 | **RNA**→ Ribosomal_S13_N (pfam08069)→ |
| env-85 | SL_4KL_010_BRINE_10262119 | + | 294 | 359 | **RNA**→ hypo→ |
| env-86 | SL_4KL_010_BRINE_10060909 | + | 1536 | 1601 | **RNA**→ ←hypo |
| env-87 | SL_6KL_011_BRINE_1008478 | - | 2259 | 2195 | **RNA**→ ←MarR (COG1846)HTH_27 (pfam13463) |
| env-88 | APHM01021551.1 | + | 1361 | 1425 | **RNA**→ RpsO (COG0184)rps15p (PRK08561)→ hypo→ PRK14888 (PRK14888)→ RPS3A (COG1890)PRK04057 (PRK04057)→ |
| env-89 | GSLSAAI_GFP0ZZ301BID9D | - | 389 | 325 | **RNA**→ Ribosomal_S13_N (pfam08069)→ |
| env-90 | GSLNAS_F7I09SK02H8PXO | + | 23 | 89 | **RNA**→ Ribosomal_S13_N (pfam08069)→ |
| env-91 | GSLNAS_F7I09SK02JLDI6 | - | 289 | 223 | **RNA**→ rps15p (PRK08561)→ |

| | | | | | |
|---|---|---|---|---|---|
| env-92 | SL_6KL_011_BRINE_1005455 | - | 398 | 338 | **RNA**→ Ribosomal_S13_N (pfam08069)→ |
| env-93 | SL_5KL_010_BRINE_10006403 | - | 4853 | 4793 | **RNA**→ rps15p (PRK08561)Ribosomal_S15p_S13e (cd00353)→ hypo→ PRK14888 (PRK14888)→ RPS3A (COG1890)PRK04057 (PRK04057)→ |
| env-94 | GRCP39E01BOGZV | - | 496 | 438 | **RNA**→ Ribosomal_S13_N (pfam08069)→ |
| env-95 | P_A23_Liq_1_Fmt_1014328 | - | 338 | 279 | **RNA**→ Ribosomal_S13_N (pfam08069)→ |
| env-96 | APHM01010841.1 | - | 1489 | 1430 | **RNA**→ rps15p (PRK08561)Ribosomal_S15p_S13e (cd00353)→ hypo→ |
| env-97 | P_A23_Liq_3_Fmt_1003525 | + | 2170 | 2229 | **RNA**→ Ribosomal_S13_N (pfam08069)→ |
| env-98 | APHM01021204.1 | - | 7840 | 7781 | **RNA**→ rps15p (PRK08561)Ribosomal_S15p_S13e (cd00353)→ A-tRNA_syn_arch (TIGR03683)→ PRK14888 (PRK14888)→ PRK04057 (PRK04057)Ribosomal_S3Ae (pfam01015)→ |
| env-99 | P_A23_Liq_1_Fmt_1003446 | + | 2004 | 2063 | **RNA**→ ←hypo |
| env-100 | ADL20m3uS_contig11784 | + | 13949 | 14008 | **RNA**→ Ribosomal_S15 (pfam00312)Ribosomal_S13_N (pfam08069)→ |
| env-101 | ADL20m3uS_contig11574 | - | 35601 | 35542 | **RNA**→ rps15p (PRK08561)Ribosomal_S15p_S13e (cd00353)→ hypo→ PRK14888 (PRK14888)→ RPS3A (COG1890)PRK04057 (PRK04057)→ |
| env-102 | SL_4KL_010_BRINE_10059133 | - | 223 | 164 | **RNA**→ |
| env-103 | SL_6KL_011_BRINE_1000187 | + | 3664 | 3723 | **RNA**→ rps15p (PRK08561)Ribosomal_S15p_S13e (cd00353)→ hypo→ PRK14888 (PRK14888)→ PRK04057 (PRK04057)Ribosomal_S3Ae (pfam01015)→ |
| env-104 | ADL5mRS1u_GM034OG02GJ0EH | - | 169 | 110 | **RNA**→ hypo→ |
| env-105 | P_1C_Sed_2_UnCty_1025083 | + | 307 | 366 | **RNA**→ rps15p (PRK08561)Ribosomal_S15 (pfam00312)→ hypo→ |
| env-106 | SL_4KL_010_BRINE_10300408 | + | 236 | 296 | **RNA**→ ←hypo |
| env-107 | SL_4KL_010_BRINE_10350117 | + | 300 | 360 | **RNA**→ |
| env-108 | SL_4KL_010_BRINE_10637802 | - | 76 | 16 | **RNA**→ |
| env-109 | SL_4KL_010_BRINE_10714498 | + | 107 | 167 | **RNA**→ ←hypo |
| env-110 | SL_5KL_010_BRINE_10197968 | + | 189 | 249 | **RNA**→ ←hypo |
| env-111 | SL_4KL_010_BRINE_10023425 | + | 3324 | 3384 | **RNA**→ ←hypo |
| env-112 | SL_4KL_010_BRINE_10204576 | + | 115 | 175 | **RNA**→ rps15p (PRK08561)→ |
| env-113 | SL_5KL_010_SED_10041201 | - | 980 | 920 | **RNA**→ rps15p (PRK08561)Ribosomal_S15p_S13e (cd00353)→ hypo→ |
| env-114 | SL_6KL_011_BRINE_1114368 | + | 262 | 322 | **RNA**→ ←hypo |
| env-115 | SL_4KL_010_BRINE_10020685 | - | 82 | 22 | **RNA**→ |
| env-116 | SL_6KL_011_BRINE_1011655 | - | 284 | 224 | **RNA**→ rps15p (PRK08561)→ |
| env-117 | SL_5KL_010_BRINE_10209515 | + | 284 | 344 | **RNA**→ |
| env-118 | SL_6KL_011_BRINE_1001544 | - | 2246 | 2186 | **RNA**→ rps15p (PRK08561)Ribosomal_S15p_S13e (cd00353)→ alaS (PRK13902)→ PRK14888 (PRK14888)→ RPS3A (COG1890)PRK04057 (PRK04057)→ |
| env-119 | SL_6KL_011_BRINE_1024707 | - | 74 | 14 | **RNA**→ |
| env-120 | SL_5KL_010_BRINE_10059954 | + | 1069 | 1129 | **RNA**→ ←hypo |
| env-121 | SL_4KL_010_BRINE_10154812 | - | 369 | 309 | **RNA**→ Ribosomal_S13_N (pfam08069)→ |
| env-122 | SL_4KL_010_BRINE_10414697 | + | 252 | 312 | **RNA**→ |
| env-123 | SL_5KL_010_BRINE_10109794 | - | 494 | 434 | **RNA**→ Ribosomal_S13_N (pfam08069)→ |
| env-124 | SL_5KL_010_BRINE_10069484 | - | 177 | 117 | **RNA**→ ←hypo |
| env-125 | SL_4KL_010_BRINE_10162859 | - | 206 | 146 | **RNA**→ ←hypo |
| env-126 | SL_4KL_010_BRINE_10275765 | - | 177 | 117 | **RNA**→ ←hypo |
| env-127 | GSLSAAI_GFP0ZZ301EKX2T | - | 305 | 245 | **RNA**→ Ribosomal_S13_N (pfam08069)→ |
| env-128 | SL_4KL_010_BRINE_10097307 | - | 214 | 154 | **RNA**→ ←hypo |
| env-129 | SL_4KL_010_BRINE_10048125 | + | 2101 | 2161 | **RNA**→ |
| env-130 | SL_4KL_010_BRINE_10328064 | + | 187 | 247 | **RNA**→ |
| env-131 | SL_6KL_011_BRINE_1018831 | - | 500 | 440 | **RNA**→ Ribosomal_S13_N (pfam08069)→ |
| env-132 | SL_4KL_010_BRINE_10013651 | - | 210 | 150 | **RNA**→ ←hypo |
| env-133 | P_A23_Liq_1_Fmt_1006894 | - | 1048 | 989 | **RNA**→ rps15p (PRK08561)Ribosomal_S15p_S13e (cd00353)→ hypo→ |
| env-134 | P_1C_Liq_3_UnCty_1011064 | + | 95 | 154 | **RNA**→ rps15p (PRK08561)Ribosomal_S15p_S13e (cd00353)→ |
| env-135 | P_A23_Liq_2_Fmt_1004041 | - | 190 | 131 | **RNA**→ |
| env-136 | APHM01021593.1 | + | 13468 | 13527 | **RNA**→ rps15p (PRK08561)Ribosomal_S15p_S13e (cd00353)→ hypo→ |
| env-137 | P_A23_Liq_1_Fmt_1011742 | + | 225 | 284 | **RNA**→ rps15p (PRK08561)Ribosomal_S15 (pfam00312)→ |
| env-138 | P_A23_Liq_3_Fmt_1033915 | - | 244 | 185 | **RNA**→ rps15p (PRK08561)→ |
| env-139 | SL_5KL_010_BRINE_10002441 | - | 7903 | 7844 | **RNA**→ rps15p (PRK08561)Ribosomal_S15 (pfam00312)→ hypo→ PRK14888 (PRK14888)→ RPS3A (COG1890)PRK04057 (PRK04057)→ |

| | | | | | |
|---|---|---|---|---|---|
| env-140 | P_A23_Liq_1_Fmt_1003155 | - | 239 | 180 | **RNA**→ ←hypo |
| CHa-1-1 | NZ_ASGZ01000049.1 | + | 3617 | 3683 | **RNA**→ rps15p (PRK08561)Ribosomal_S15p_S13e (cd00353)→ PRK13800 (PRK13800)PBP2_HisK (cd13704)→ |
| env-141 | SL_4KL_010_BRINE_10085604 | - | 150 | 84 | **RNA**→ |
| env-142 | SL_6KL_011_BRINE_1044936 | - | 158 | 90 | **RNA**→ ←hypo |
| Nsu-1-1 | NZ_AOHX01000042.1 | - | 103712 | 103642 | **RNA**→ RpsO (COG0184)rps15p (PRK08561)→ PRK13800 (PRK13800)→ PRK14888 (PRK14888)→ RPS3A (COG1890)PRK04057 (PRK04057)→ |
| Hla-1-1 | NZ_AGFZ01000124.1 | + | 17222 | 17292 | **RNA**→ RpsO (COG0184)rps15p (PRK08561)→ RpsO (COG0184)rps15p (PRK08561)→ hypo→ hypo→ |
| env-143 | SL_5KL_010_BRINE_10055562 | - | 160 | 95 | **RNA**→ ←hypo |
| env-144 | SL_5KL_010_BRINE_10106739 | + | 525 | 590 | **RNA**→ |
| env-145 | SL_5KL_010_BRINE_10001172 | + | 12650 | 12715 | **RNA**→ |
| Hxa-1-1 | NC_015666.1 | + | 2172566 | 2172639 | **RNA**→ RpsO (COG0184)rps15p (PRK08561)→ hypo→ PRK14888 (PRK14888)→ PRK04057 (PRK04057)Ribosomal_S3Ae (pfam01015)→ |
| env-146 | SL_6KL_011_BRINE_1235537 | - | 113 | 48 | **RNA**→ |
| env-147 | SL_4KL_010_BRINE_10453247 | - | 143 | 76 | **RNA**→ ←hypo |
| Ngr-1-1 | NZ_AOIC01000118.1 | + | 22540 | 22604 | **RNA**→ RpsO (COG0184)rps15p (PRK08561)→ hypo→ PRK14888 (PRK14888)→ RPS3A (COG1890)PRK04057 (PRK04057)→ |
| env-148 | SL_5KL_010_BRINE_10000150 | + | 26232 | 26305 | **RNA**→ rps15p (PRK08561)Ribosomal_S15p_S13e (cd00353)→ hypo→ PRK14888 (PRK14888)→ RPS3A (COG1890)PRK04057 (PRK04057)→ |
| env-149 | SL_8KL_010_SED_10418808 | - | 198 | 125 | **RNA**→ rps15p (PRK08561)→ |
| env-150 | SL_5KL_010_BRINE_10011173 | - | 233 | 168 | **RNA**→ hypo→ |
| env-151 | SL_4KL_010_BRINE_10233198 | - | 160 | 95 | **RNA**→ ←hypo |
| Hdj-1-1 | NZ_HG315685.1 | - | 946456 | 946386 | **RNA**→ RpsO (COG0184)rps15p (PRK08561)→ hypo→ PRK14888 (PRK14888)→ RPS3A (COG1890)PRK04057 (PRK04057)→ |
| env-152 | ADL20m3uS_GLR497S01CYFB9 | - | 217 | 152 | **RNA**→ hypo→ |
| env-153 | 2140918017-18446 | - | 216 | 151 | **RNA**→ |
| env-154 | SL_8KL_010_SED_10420993 | + | 127 | 192 | **RNA**→ ←hypo |
| env-155 | SL_4KL_010_BRINE_10209625 | - | 174 | 109 | **RNA**→ |
| env-156 | SL_4KL_010_BRINE_10009987 | - | 1525 | 1460 | **RNA**→ rps15p (PRK08561)Ribosomal_S15p_S13e (cd00353)→ RecJ (COG0608)→ |
| env-157 | SL_6KL_011_BRINE_1000505 | - | 8160 | 8095 | **RNA**→ rps15p (PRK08561)Ribosomal_S15p_S13e (cd00353)→ hypo→ PRK14888 (PRK14888)→ RPS3A (COG1890)PRK04057 (PRK04057)→ |
| env-158 | SL_6KL_011_BRINE_1064947 | + | 350 | 413 | **RNA**→ Ribosomal_S13_N (pfam08069)→ |
| env-159 | SL_5KL_010_BRINE_10001414 | + | 3775 | 3841 | **RNA**→ RpsO (COG0184)rps15p (PRK08561)→ hypo→ PRK14888 (PRK14888)→ RPS3A (COG1890)PRK04057 (PRK04057)→ |
| env-160 | SL_4KL_010_BRINE_10410649 | + | 75 | 141 | **RNA**→ rps15p (PRK08561)→ |
| Hgo-1-1 | NZ_HG315690.1 | + | 210551 | 210620 | **RNA**→ RpsO (COG0184)rps15p (PRK08561)→ minC (PRK00030)→ PRK14888 (PRK14888)→ RPS3A (COG1890)PRK04057 (PRK04057)→ |
| Npa-1-1 | NZ_AOII01000045.1 | + | 2929 | 2999 | **RNA**→ rps15p (PRK08561)Ribosomal_S15p_S13e (cd00353)→ hypo→ PRK14888 (PRK14888)→ RPS3A (COG1890)PRK04057 (PRK04057)→ |
| Nga-1-1 | NZ_AOIJ01000082.1 | + | 8646 | 8716 | **RNA**→ rps15p (PRK08561)Ribosomal_S15p_S13e (cd00353)→ hypo→ PRK14888 (PRK14888)→ RPS3A (COG1890)PRK04057 (PRK04057)→ |
| Nsp-1-1 | NZ_AJVG01000015.1 | + | 33476 | 33546 | **RNA**→ rps15p (PRK08561)Ribosomal_S15p_S13e (cd00353)→ hypo→ PRK14888 (PRK14888)→ RPS3A (COG1890)PRK04057 (PRK04057)→ |
| Nal-1-1 | NZ_JNCS01000002.1 | - | 209831 | 209761 | **RNA**→ rps15p (PRK08561)Ribosomal_S15p_S13e (cd00353)→ hypo→ PRK14888 (PRK14888)→ RPS3A (COG1890)PRK04057 (PRK04057)→ |
| env-161 | ADL20m3uS_GLR497S01D4APN | + | 16 | 86 | **RNA**→ Ribosomal_S13_N (pfam08069)→ |
| Hli-3-1 | NZ_AOIT01000016.1 | + | 177551 | 177622 | **RNA**→ RpsO (COG0184)rps15p (PRK08561)→ hypo→ PRK14888 (PRK14888)→ RPS3A (COG1890)PRK04057 (PRK04057)→ |
| Nve-1-1 | NZ_AOID01000012.1 | - | 23261 | 23191 | **RNA**→ rps15p (PRK08561)Ribosomal_S15p_S13e (cd00353)→ hypo→ PRK14888 (PRK14888)→ RPS3A (COG1890)PRK04057 (PRK04057)→ |
| Npe-1-1 | NZ_AOIE01000076.1 | + | 117112 | 117182 | **RNA**→ RpsO (COG0184)rps15p (PRK08561)→ hypo→ PRK14888 (PRK14888)→ RPS3A (COG1890)PRK04057 (PRK04057)→ |
| Hje-2-1 | NZ_JDTG01000005.1 | + | 36819 | 36889 | **RNA**→ RpsO (COG0184)rps15p (PRK08561)→ hypo→ PRK14888 (PRK14888)→ RPS3A (COG1890)PRK04057 (PRK04057)→ |
| Hth-2-1 | NZ_AOIR01000045.1 | + | 2606 | 2676 | **RNA**→ RpsO (COG0184)rps15p (PRK08561)→ hypo→ hypo→ |
| Nje-1-1 | NZ_AOIA01000160.1 | + | 12251 | 12321 | **RNA**→ rps15p (PRK08561)Ribosomal_S15p_S13e (cd00353)→ hypo→ PRK14888 (PRK14888)→ PRK04057 (PRK04057)Ribosomal_S3Ae (pfam01015)→ |

| | | | | | |
|---|---|---|---|---|---|
| env-162 | SS_3KL_010_SOIL_10055065 | + | 1009 | 1078 | **RNA**→ ←hypo |
| Noc-1-1 | NC_019974.1 | - | 817241 | 817172 | **RNA**→ rps15p (PRK08561)Ribosomal_S15p_S13e (cd00353)→ hypo→ PRK14888 (PRK14888)→ RPS3A (COG1890)PRK04057 (PRK04057)→ |
| Nam-1-1 | NZ_AOIB01000013.1 | + | 141311 | 141380 | **RNA**→ rps15p (PRK08561)Ribosomal_S15p_S13e (cd00353)→ PRK13029 (PRK13029)→ PRK14888 (PRK14888)→ RPS3A (COG1890)PRK04057 (PRK04057)→ |
| Nhu-1-1 | NZ_AOIM01000039.1 | - | 10511 | 10445 | **RNA**→ RpsO (COG0184)rps15p (PRK08561)→ hypo→ PRK14888 (PRK14888)→ RPS3A (COG1890)PRK04057 (PRK04057)→ |
| Nma-1-1 | NZ_AOHS01000030.1 | - | 82382 | 82316 | **RNA**→ RpsO (COG0184)rps15p (PRK08561)→ |
| Nta-1-1 | NZ_AOIL01000028.1 | + | 66155 | 66221 | **RNA**→ RpsO (COG0184)rps15p (PRK08561)→ hypo→ PRK14888 (PRK14888)→ RPS3A (COG1890)PRK04057 (PRK04057)→ |
| Nas-1-1 | NZ_AOIO01000033.1 | - | 16936 | 16870 | **RNA**→ RpsO (COG0184)rps15p (PRK08561)→ hypo→ PRK14888 (PRK14888)→ RPS3A (COG1890)PRK04057 (PRK04057)→ |
| Nch-1-1 | NZ_AOIN01000092.1 | + | 72402 | 72468 | **RNA**→ RpsO (COG0184)rps15p (PRK08561)→ |
| Nae-1-1 | NZ_AOIP01000015.1 | - | 106319 | 106249 | **RNA**→ RpsO (COG0184)rps15p (PRK08561)→ hypo→ PRK14888 (PRK14888)→ RPS3A (COG1890)PRK04057 (PRK04057)→ |
| Nba-1-1 | NZ_AOHY01000002.1 | - | 121091 | 121021 | **RNA**→ RpsO (COG0184)rps15p (PRK08561)→ PRK13800 (PRK13800)→ PRK14888 (PRK14888)→ PRK04057 (PRK04057)Ribosomal_S3Ae (pfam01015)→ |
| Nin-1-1 | NZ_AOHZ01000012.1 | + | 7343 | 7413 | **RNA**→ RpsO (COG0184)rps15p (PRK08561)→ hypo→ PRK14888 (PRK14888)→ PRK04057 (PRK04057)Ribosomal_S3Ae (pfam01015)→ |
| Hni-1-1 | NZ_AOMA01000110.1 | + | 847 | 917 | **RNA**→ RpsO (COG0184)rps15p (PRK08561)→ hypo→ hypo→ |
| Hsa-7-1 | NZ_AOIS01000067.1 | - | 104689 | 104619 | **RNA**→ RpsO (COG0184)rps15p (PRK08561)→ hypo→ PRK14888 (PRK14888)→ PRK04057 (PRK04057)Ribosomal_S3Ae (pfam01015)→ |
| Htu-1-1 | NC_013743.1 | - | 1454550 | 1454480 | **RNA**→ RpsO (COG0184)rps15p (PRK08561)→ hypo→ PRK14888 (PRK14888)→ PRK04057 (PRK04057)Ribosomal_S3Ae (pfam01015)→ |
| Hsa-9-1 | NZ_JROF01000003.1 | + | 151229 | 151299 | **RNA**→ RpsO (COG0184)rps15p (PRK08561)→ hypo→ PRK14888 (PRK14888)→ RPS3A (COG1890)PRK04057 (PRK04057)→ |
| Nti-1-1 | NZ_AOHW01000044.1 | + | 85110 | 85180 | **RNA**→ rps15p (PRK08561)Ribosomal_S15p_S13e (cd00353)→ hypo→ hypo→ hypo→ hypo→ hypo→ PRK14888 (PRK14888)→ RPS3A (COG1890)PRK04057 (PRK04057)→ |
| env-163 | SL_5KL_010_BRINE_10001436 | + | 10905 | 10968 | **RNA**→ rps15p (PRK08561)Ribosomal_S15p_S13e (cd00353)→ hypo→ |
| env-164 | GSLNARP_GFPJP1N02G46KO | + | 349 | 411 | **RNA**→ hypo→ |
| env-165 | GSLNARP_GFPJP1N02HKHM0 | + | 321 | 383 | **RNA**→ ←hypo |
| env-166 | GSLNARP_GFPJP1N02HIBH2 | + | 301 | 362 | **RNA**→ hypo→ |
| Hpa-2-1 | NZ_AOIV01000045.1 | + | 239288 | 239352 | **RNA**→ rps15p (PRK08561)Ribosomal_S15p_S13e (cd00353)→ hypo→ PRK14888 (PRK14888)→ PRK04057 (PRK04057)Ribosomal_S3Ae (pfam01015)→ |
| Hru-1-1 | NZ_BBJO01000031.1 | + | 112826 | 112888 | **RNA**→ rps15p (PRK08561)Ribosomal_S15p_S13e (cd00353)→ hypo→ hypo→ |
| Hbo-1-1 | NZ_AOHT01000051.1 | - | 203113 | 203048 | **RNA**→ RpsO (COG0184)rps15p (PRK08561)→ hypo→ PRK14888 (PRK14888)→ PRK04057 (PRK04057)Ribosomal_S3Ae (pfam01015)→ |
| Ssp-1-1 | NC_021313.1 | - | 2955315 | 2955225 | **RNA**→ rps15p (PRK08561)Ribosomal_S15p_S13e (cd00353)→ hypo→ hypo→ PRK14888 (PRK14888)→ RPS3A (COG1890)PRK04057 (PRK04057)→ DUF2127 (pfam09900)→ |
| Hje-1-1 | NZ_AXZD01000009.1 | - | 4784 | 4656 | **RNA**→ rps15p (PRK08561)Ribosomal_S15p_S13e (cd00353)→ hypo→ PRK14888 (PRK14888)→ PRK04057 (PRK04057)Ribosomal_S3Ae (pfam01015)→ |
| env-167 | P_1C_Sed_2_UnCty_1161793 | + | 207 | 266 | **RNA**→ hypo→ |
| env-168 | P_1C_Sed_2_UnCty_1391468 | + | 115 | 172 | **RNA**→ hypo→ |
| Hna-1-1 | NZ_KE386573.1 | + | 1370676 | 1370733 | **RNA**→ rps15p (PRK08561)Ribosomal_S15p_S13e (cd00353)→ hypo→ PRK14888 (PRK14888)→ RPS3A (COG1890)PRK04057 (PRK04057)→ HATPase_c (smart00387)BaeS (COG0642)PAS (pfam00989)PAS_9 (pfam13426)→ |
| env-169 | GSLNARP_GFPJP1N02I1Q94 | - | 238 | 180 | **RNA**→ rps15p (PRK08561)→ |
| env-170 | SL_6KL_011_BRINE_1044879 | + | 455 | 520 | **RNA**→ rps15p (PRK08561)Ribosomal_S15p_S13e (cd00353)→ hypo→ |
| env-171 | SL_4KL_010_BRINE_10071208 | + | 486 | 551 | **RNA**→ rps15p (PRK08561)Ribosomal_S15p_S13e (cd00353)→ hypo→ |
| Nph-1-1 | NC_007426.1 | + | 1546623 | 1546688 | **RNA**→ rps15p (PRK08561)Ribosomal_S15p_S13e (cd00353)→ hypo→ PRK14888 (PRK14888)→ RPS3A (COG1890)PRK04057 (PRK04057)→ |
| Nmo-1-1 | NC_020388.1 | + | 1095664 | 1095729 | **RNA**→ rps15p (PRK08561)Ribosomal_S15p_S13e (cd00353)→ hypo→ PRK14888 (PRK14888)→ RPS3A (COG1890)PRK04057 (PRK04057)→ |
| env-172 | GSLNARP_contig01185 | + | 384 | 449 | **RNA**→ Ribosomal_S13_N (pfam08069)→ |
| env-173 | SL_6KL_011_BRINE_1143269 | + | 264 | 329 | **RNA**→ |
| env-174 | SL_4KL_010_BRINE_10124144 | - | 762 | 697 | **RNA**→ rps15p (PRK08561)Ribosomal_S15p_S13e (cd00353)→ hypo→ |

| | | | | | |
|---|---|---|---|---|---|
| env-175 | SL_6KL_011_BRINE_1002809 | + | 7175 | 7240 | **RNA**→ Ribosomal_S13_N (pfam08069)→ rps15p (PRK08561)Ribosomal_S15 (pfam00312)→ hypo→ |
| env-176 | SL_4KL_010_BRINE_10155541 | + | 78 | 143 | **RNA**→ ←hypo |
| env-177 | SL_4KL_010_BRINE_10020079 | + | 3724 | 3789 | **RNA**→ |
| env-178 | SL_4KL_010_BRINE_10196029 | + | 155 | 220 | **RNA**→ Ribosomal_S13_N (pfam08069)→ |
| Hsp-6-1 | NZ_JMIP02000018.1 | + | 49610 | 49681 | **RNA**→ rps15p (PRK08561)Ribosomal_S15p_S13e (cd00353)→ hypo→ PRK14888 (PRK14888)→ RPS3A (COG1890)PRK04057 (PRK04057)→ |
| Hla-3-1 | NZ_CP007055.1 | + | 931368 | 931439 | **RNA**→ RpsO (COG0184)rps15p (PRK08561)→ hypo→ PRK14888 (PRK14888)→ RPS3A (COG1890)PRK04057 (PRK04057)→ |
| env-179 | P_A23_Liq_1_Fmt_1002256 | - | 2002 | 1926 | **RNA**→ RpsO (COG0184)rps15p (PRK08561)→ hypo→ hypo→ |
| env-180 | JXWS01021119.1 | - | 732 | 656 | **RNA**→ RpsO (COG0184)rps15p (PRK08561)→ |
| env-181 | APHM01021963.1 | - | 6541 | 6476 | **RNA**→ rps15p (PRK08561)Ribosomal_S15p_S13e (cd00353)→ |
| env-182 | GSLNARP_contig29640 | + | 334 | 399 | **RNA**→ rps15p (PRK08561)Ribosomal_S15p_S13e (cd00353)→ hypo→ |
| env-183 | GSLNARP_contig29701 | + | 717 | 782 | **RNA**→ rps15p (PRK08561)Ribosomal_S15p_S13e (cd00353)→ hypo→ |
| Hsu-1-1 | NZ_CP008874.1 | + | 1124695 | 1124758 | **RNA**→ rps15p (PRK08561)Ribosomal_S15p_S13e (cd00353)→ hypo→ PRK14888 (PRK14888)→ RPS3A (COG1890)PRK04057 (PRK04057)→ archae_ser_T (TIGR04217)CDP-OH_P_transf (pfam01066)→ HEAT (COG1413)prfB (PRK00578)HEAT_2 (pfam13646)→ |
| Hha-1-1 | NZ_AJRK01000105.1 | - | 627 | 565 | **RNA**→ rps15p (PRK08561)Ribosomal_S15 (pfam00312)→ rps15p (PRK08561)Ribosomal_S15 (pfam00312)→ |
| Hru-2-1 | NZ_BBJN01000039.1 | - | 65806 | 65742 | **RNA**→ rps15p (PRK08561)Ribosomal_S15p_S13e (cd00353)→ hypo→ |
| env-184 | 2140918017-23842 | + | 118 | 178 | **RNA**→ rps15p (PRK08561)→ |
| Hsa-4-1 | NZ_AOME01000070.1 | + | 267801 | 267861 | **RNA**→ rps15p (PRK08561)Ribosomal_S15 (pfam00312)→ hypo→ PRK14888 (PRK14888)→ RPS3A (COG1890)PRK04057 (PRK04057)→ |
| Hag-1-1 | NZ_BAFM01000018.1 | - | 8492 | 8432 | **RNA**→ rps15p (PRK08561)Ribosomal_S15p_S13e (cd00353)→ hypo→ PRK14888 (PRK14888)→ RPS3A (COG1890)PRK04057 (PRK04057)→ |
| Hsa-3-1 | NZ_AOMD01000016.1 | + | 129258 | 129318 | **RNA**→ rps15p (PRK08561)Ribosomal_S15p_S13e (cd00353)→ hypo→ PRK14888 (PRK14888)→ RPS3A (COG1890)PRK04057 (PRK04057)→ |
| Hse-1-1 | NZ_BBMP01000004.1 | + | 37396 | 37456 | **RNA**→ rps15p (PRK08561)Ribosomal_S15 (pfam00312)→ hypo→ PRK14888 (PRK14888)→ RPS3A (COG1890)PRK04057 (PRK04057)→ |
| Hth-1-1 | NZ_AOMF01000159.1 | - | 66211 | 66151 | **RNA**→ rps15p (PRK08561)Ribosomal_S15 (pfam00312)→ hypo→ PRK14888 (PRK14888)→ RPS3A (COG1890)PRK04057 (PRK04057)→ |
| Hmo-1-1 | NZ_AOMC01000070.1 | + | 55554 | 55614 | **RNA**→ rps15p (PRK08561)Ribosomal_S15p_S13e (cd00353)→ hypo→ PRK14888 (PRK14888)→ RPS3A (COG1890)PRK04057 (PRK04057)→ |
| env-185 | APHM01010805.1 | + | 863 | 937 | **RNA**→ Ribosomal_S15 (pfam00312)Ribosomal_S13_N (pfam08069)→ |
| Hsp-1-1 | NC_002607.1 | - | 593897 | 593832 | **RNA**→ RpsO (COG0184)rps15p (PRK08561)→ hypo→ PRK14888 (PRK14888)→ PRK14888 (PRK14888)→ RPS3A (COG1890)PRK04057 (PRK04057)→ |

#### 20.3 Conserved domains

Conserved domains found in protein-coding genes listed in Section 20.2 are shown below, with the first sentence in their description from the Conserved Domain Database (if any). Conserved domains associated with more than one S15-Halobacteria RNA are assigned a color, while others are shown in gray. The number in parentheses after the colored domain name is the number of occurrences in Section 20.2.

cd00353 (98) Ribosomal protein S15 (prokaryotic)_S13 (eukaryotic) binds the central domain of 16S rRNA and is required for assembly of the small ribosomal subunit and for intersubunit association, thus representing a key element in the assembly of the whole ribosome.
cd00677 (2) S15/NS1/EPRS_RNA-binding domain.
cd06927 (1) L subunit of Archaeal RNA polymerase.
cd11255 (1) The Sema domain, a protein interacting module, of semaphorin 3G (Sema3G).
cd13704 (1) The periplasmic sensor domain of histidine kinase receptors; the type 2 periplasmic binding fold protein.
COG0184 (44) Ribosomal protein S15P/S13E [Translation, ribosomal structure and biogenesis]
COG0608 (1) Single-stranded DNA-specific exonuclease, DHH superfamily, may be involved in archaeal DNA replication intiation [Replication, recombination and repair]
COG0642 (1) Signal transduction histidine kinase [Signal transduction mechanisms]
COG1413 (1) HEAT repeat [General function prediction only]
COG1846 (1) DNA-binding transcriptional regulator, MarR family [Transcription]
COG1890 (69) Ribosomal protein S3AE [Translation, ribosomal structure and biogenesis]
COG2892 (1) tRNA threonylcarbamoyladenosine modification (KEOPS) complex, Pcc1 subunit [Translation, ribosomal structure and biogenesis]
COG4300 (1) Cadmium resistance protein CadD, predicted permease [Inorganic ion transport and metabolism]
pfam00312 (18) Ribosomal protein S15.

pfam00989 (1) PAS fold.
pfam01015 (36) Ribosomal S3Ae family.
pfam01066 (1) CDP-alcohol phosphatidyltransferase.
pfam08069 (27) Ribosomal S13/S15 N-terminal domain.
pfam09900 (1) Predicted membrane protein (DUF2127).
pfam13426 (1) PAS domain.
pfam13463 (1) Winged helix DNA-binding domain.
pfam13646 (1) HEAT repeats.
PHA03247 (1) large tegument protein UL36; Provisional
PRK00030 (1) septum formation inhibitor; Provisional
PRK00578 (1) peptide chain release factor 2; Validated
PRK01146 (1) DNA-directed RNA polymerase subunit L; Provisional
PRK04057 (105) 30S ribosomal protein S3Ae; Validated
PRK07409 (1) threonine synthase; Validated
PRK08561 (176) 30S ribosomal protein S15P; Reviewed
PRK13029 (1) 2-oxoacid ferredoxin oxidoreductase; Provisional
PRK13800 (6) putative oxidoreductase/HEAT repeat-containing protein; Provisional
PRK13902 (1) alanyl-tRNA synthetase; Provisional
PRK14888 (105) KEOPS complex Pcc1-like subunit; Provisional
smart00387 (1) Histidine kinase-like ATPases.
TIGR00779 (1) cadmium resistance transporter (or sequestration) family protein. [Transport and binding proteins, Amino acids, peptides and amines]
TIGR03683 (2) alanyl-tRNA synthetase.
TIGR04217 (1) archaetidylserine synthase.

#### 20.4 Multiple-sequence alignment

Each S15-Halobacteria RNA is denoted by an abbreviation (like "Eco-1-1") that refers to a taxonomy given in Section 20.1. The alignment may include sequences containing the RNA motif, as well as flanking sequence. The S15-Halobacteria RNA itself is denoted by the line underneath marked 5′ and 3′ on either end. Nucleotides in flanking sequences (i.e., not part of the motif) are written in gray letters. Stems of predicted rho-independent transcription terminators, if any, are shaded yellow. (Note: terminator predictions have not been analyzed manually, and many are likely to be false positives. Terminator predictions are those of the RNIE software.) Nucleotides predicted to function (as DNA) as transcription-factor binding sites, if any, are shaded in green. (But note: these predictions are manually annotated, so they might be under-predicted.) Annotated start codons, if any, are shaded green. (Note: start codons are frequently misannotated, especially in environmental samples.) Nucleotides proposed to basepair as part of the consensus structure are shaded in color when they comprise Watson-Crick or G-U pairs. Otherwise they are shaded gray. Conserved stems are also indicated at the bottom of the alignment by angle brackets, where matching < and > denote base-paired columns. Below these angle brackets, the symbol "2" denotes base pairs exhibiting covariation according to the statistically well-founded R-scape method. "1" denotes base pairs exhibiting covariation according to R2R's simplistic method. "0" denotes base pairs that are not observed to mutate and "?" denotes base pairs that have a significant frequency of non-canonical nucleotides for Watson-Crick or G-U pairs (> 5%). Below these base pair annotation is the consensus sequence: "R" = "A" or "G", "Y" = "C" or "U", red nucleotides: nucleotide identity conserved more than 97% of the time, black nucleotides: 90%, gray nucleotides: 75%, red circle (•): nucleotide is present 97% of the time, black circle (•): 90%, gray circle (•): 75%, white circle (∘): 50%. All percentages of sequences just described (e.g. 97% conserved) assume that sequences have been weighted by the GSC algorithm implemented by the Infernal software package.

alignment positions 1 · · · 180

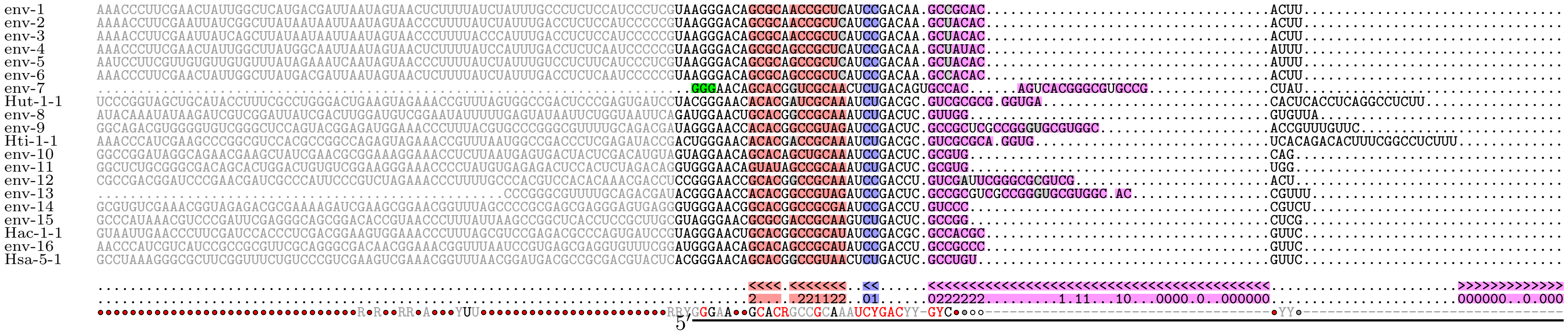

```
env-17   AAACCCUCUCGUCUCAGCUCCGCGGCGCGUCCACACGAACACGCUUAUCAACCACGACCGCGACCGUUCGNACGGGAACAGCACAGCUGCAGAUCCGACGC.GCCG.GGGCG.................................CUCU.................................
env-18   AAACCCUCUCGUCUCAGCUCCGCGGCGCGUCCACACGAACACGCUUAUCAACCACGACCGCGACCGUUCGAACGGGAACAGCACAGCUGCAGAUCCGACGC.GCCG.GGGCG.................................CUCU.................................
env-19   CGGGCGGGGUUCUCCCGGCCGGGCCGCGGACCACAAGAACACGCUUAAGAACCCCGACGGCGACUUCCGGAACGGGAACAGCACAGCUGCAGAUCCGACUC.GCCG.GGGC..................................CUCUUC...............................
env-20   GCGUUCGCCAGACGGAGUCGCCUCACGCCGGGACAAGAACACGCUUAAUACCCCCGACGGCGACCGUCCCCACGGGAACAGCACAGCUGCAGAUCCGACUC.GCCG.GGGC..................................CCUUUC...............................
Har-2-1  CGGAAGCGAGGACGCCUCCGCCGCUCCGAUCCACAAGAACACGCUUAAGAACCCGGACGGCGACCCCGAACACGGGAACAGCACAGCUGCAGAUCCGACUC.GCCG.GGGC..................................CCUUUC...............................
Hsp-3-1  AGAGCGCUGGAACGCGCCCGCCACUCCGCCCCACAAGAACACGCUUAAGAACCCGGACGGCGACUCCGCAGACGGGAACAGCACAGCUGCAGAUCCGACUC.GCCG.GGGC..................................CCUUUC...............................
env-21   GCGCUCCCCCGACGGCGUCGCCCCGCCGCGGGACAAGAACACGCUUAAGACCCCCGACGGCGACCGUCCUGACGGGAACAGCACAGCUGCAGAUCCGACUC.GCCG.GGGC..................................CCUUUC...............................
env-22   UGUGUGGCCCGAAACGGGCGGCGUCCGGACCGACAAGAACACCCUUAAGACCCCCGACGGCGACCGUCGGAACGGGAACAGCACAGCUGCAGAUCCGACUC.GCCG.GGGC..................................CCUUUC...............................
env-23   GCGUUCGCCCGACGGAAUCGCCUCACGCGGGGACAAGAACACGCUUAAGACCCCCGACGGCGACCGUCUUGACGGGAACAGCACAGCUGCAGAUCCGACUC.GCCG.GGGC..................................CCUUUC...............................
Hdi-1-1  CGGAAGCGAGGACGCCUCCGCCGCUCCGACCCACAAGAACACGCUUAAGAACCCUGACGGCGACUCCGAACACGGGAACAGCACAGCUGCAGAUCCGACUC.GCCG.GGGC..................................CCUUUC...............................
Hli-2-1  CGGAAGCGAGGACGCCUCCGCCGCUCCGACCCACAAGAACACGCUUAAGAACCCGGACGGCGACUCCGAAUACGGGAACAGCACAGCUGCAGAUCCGACUC.GCCG.GGGC..................................CCUUUC...............................
Hsa-8-1  CUCCCAAUGUCCCGCACUCGCUCGAGCGGACGACAAGAACACGCUUAACAACCCGGACGACGACGCCUCAUACGGGAACAGCACAGCUGCAGAUCCGACGC.GCCG.GGGU..................................CUUUUC...............................
env-24   ACGCAAUCCCACCGGAACCGACCGACCGGGCCACACGAACACGCUUAAGAACCCCCACGCCGACCGUUCGAACGGGAACAGCACAGCUGCAGAUCCGACGC.GCCG.GGGU..................................CUUUUC...............................
Hha-2-1  CUCUAGAUGUCCCGCACUCGCCCGAGCGGACGACAAGAACACGCUUAACAACCCGGACGACGACGCCUCAUACGGGAACAGCACAGCUGCAGAUCCGACGC.GCCG.GGGU..................................CUUUUC...............................
Hko-1-1  CCGAUCUGCCCGCCGCGCUCGCCCGGCAGGCCACAAGAACACUCUUAAGAACCCCGACACCGACCGUUCACACGGGAACAGCACAGCUGCAGAUCCGACGC.GCCG.GGGU..................................CUUUUC...............................
env-25   GCUACUUAAGUGAACGACGUCAGGGCGGACGCACAAGAACACGCUUAAGAAGGGUGACGACGACGGUGAAAACGGGAACAGCACAGCUACAGAUCCGACGC.GCCG.GGGC..................................CUCUUUC..............................
env-26   UUCCUCGAUCUCCGUAUCCAUCCGAGCGCCCCACAAGAACACGCUUAAGAACCCCGACGACGACCCCUCACACGGGAACAGCACAGCUGCAGAUCCGACGC.GCCG.GGGC..................................CUCUUC...............................
env-27   GGAAGNAAAACGGUUCGGACGCCCGCGACCAGACAAGAACACGCUUAAGAGGGGUGACGGCGAGAUAUUUGAUGGGAACAGCACAGCUGCGGAUCCGACGC.GCCG.GGGU..................................CUGUUC...............................
env-28   GGUCGAAGGCGAGUGCUUCACGCGGCGUACGGACAAGAACACGCUUAAGAGGGGCGACGGCGAGGUGUUCGAUGGGAACAGCACAGCUGCGGAUCCGACGC.GCCG.GGGU..................................CUAUUC...............................
env-29   ACGGGAACUGUCGCCGUCGGCGGCGUCGAUCCACAAGAACACCCUUAAGAGGAGCGACGGUGAGAUGUUCGAUGGGAACAGCACAGCUGCGGAUCCGACGC.GCCG.GGGC..................................CUCUUC...............................
env-30   UGUCGGCGCCGAUGAGUCCGCCCGUCGGACACACAAGAACACGCUUAAGAGGCGUGGCGGCGAUCUCUGCGGUGGGAACAGCACAGCUGCGGAUCCGACGC.GCCG.GGGC..................................CUGUUC...............................
env-31   GGAAGCAAAACGGUUCGGACGCCUGCGACCGGACAAGAACACGCUUAAGAGGGGUGACGGCGAGGCGAUAAACGGGAACAGCACAGCUGCGGAUCCGACGC.GCCG.GGGC..................................CUGUUC...............................
env-32   ..........................................................................GGGAACAGCACAGCUGCGGAUCCGACGC.GCCG.GGGC..................................CUGUUC...............................
env-33   GAGGCGCCGGUCGCCGGGGAGCGACCGACCGCACAAGAACACGCUUAAGAAGGGUGACGACGACGUUGACGGUGGGAACAGCACAGCUACAGAUCCGACGC.GCCG.GGGC..................................UUGUUUC..............................
env-34   UAACUCACUCCCCAAGUCCGGAUAUCGGGACGAAGAGAACACGCUUAAGAGGGCCGACGGCGACCGUCGGGAUGGGAACAGCACAGCUAUAGAUCCGACGC.GCCG.GGGC..................................CUUUUC...............................
env-35   UAACUCACUCCCCAAGUCCGGAUAUCGGGACGAAGAGAACACGCUUAAGAGGGCCGACGGCGACCGUNGGGAUGGGAACAGCACAGCUAUAGAUCCGACGC.GCCG.GGGC..................................CUUUUC...............................
env-36   UAACUCACUCCCCAAGUCCGGAUAUCGGGACGAAGAGAACACGCUUAAGAGGGCCGACGGCGACCGUGGGGAUGGGAACAGCACAGCUAUAGAUCCGACGC.GCCG.GGGC..................................CUUUUC...............................
env-37   CGAGAGACCACAAUCCGGGACGCGACGCACGCACAAGAACACCCUUAAGAGGACCGACGGCGACCGUCGGUAUGGGAACAGCACAGCUAUAGAUCCGACGC.GCCG.GGGC..................................CUUUUC...............................
env-38   ACUCUCGUCGGCGUUUCACCCCCGCCGAACGCACAAGAACACCCUUAAGAAGGGUGACGACGACGGUGAUGAUGGGAACAGCACAGCUACAGAUCCGACGC.GCCG.GGGC..................................UUUUUC...............................
env-39   GGCCCUGUGCCCGCUCGAUCGGAGCCGACCCGACACGGACACCCUUAAGAACCCGUGCGGCGUCGCUCACGAUGGGAACAGCACAGCUGCGGAUCCGACGC.GCCGGGG....................................AUUGUUUCGC...........................
env-40   GGCUCUAUGCCAGCUCGAUCGGGGUCGGCCCGACACGGACACGCUUAAGAACCCGUGCGGCGUCGCUCACGAUGGGAACAGCACAGCUGCGGAUCCGACGC.GCCGGGG....................................AUCAGUUGC............................
env-41   GGCUCUCUGCCCGCUCGACCAGGGUCGGCCCGACACGGACACGCUUAAGAACCCGUGCGGCGUCGCUCACGAUGGGAACAGCACAGCUGCGGAUCCGACGC.GCCGGGG....................................AUCAGUUGC............................
Hsp-5-1  CUAAAUCCUCGUUUCUGUGCCGCGGCCGAGCCACACGAACACGCUUAAGAACCUCGGCGGCGACCGGUCGGACGGGAACAGCACAGCCGCAGAUCCGACUC.GCCG.GGGU..................................AUUUUC...............................
Hca-2-1  GGGCUCCCCCGACGGAGCCGCCUCGCCGCGGGACAAGAACACGCUUAAGAACCCCGACGGCGACCCCUCACACGGGAACAGCACAGCUGCAGAUCCGACUC.GCCG.GGGC..................................UCUUUC...............................
Hli-1-1  AGUUCAUCCCGCCGCGCUCGCCAGCGCAGCCCACAAGAACACGCUUAAGAACCCCGACGCCGACCCUCGACAUGGGAACAGCACAGCUGCAGAUCCGACGC.GCCG.GGGU..................................CUUUUC...............................
Hte-1-1  ACCGAGGGGGCGGACGCACGGCGCUCGUCCCCACAAGAACACGCUUAAGAACCCCGACGGCGAGGUCCCGAACGGGAACAGCACAGCUGCAGAUCCGACGU.GCCGGGGC..................................CUGUUC...............................
Hho-1-1  GACCGAGGGGCGGACGCACGGCGCUCGUCCCCACAAGAACACGCUUAAGAACCCCGACGGCGAGGUCUCGAACGGGAACAGCACAGCUGCAGAUCCGACGU.GCCGGGGC..................................CUGUUC...............................
Hsa-6-1  GUUCACCCCCGCCGCGCCCGCUCCAGCGACUCACAAGAACACGCUUAAGAACCCCGGCGACGACCCCUCACACGGGAACAGCACAGCUGCAGAUCCGACGC.GCCG.GGGC..................................CUUUUC...............................
Hco-1-1  GCGUUCGCCCGACGGAGUCGCCUCGGCGAGGGACAAGAACACGCUUAAGACCCCCGACGGCGACCCUCCUGACGGGAACAGCACAGCUGCAGAUCCGACGC.GCCG.GGGC..................................CCUUUC...............................
Hai-1-1  GCGGUUUCCGUCCGCGAUCCCGCGCGAGCGCCACAAGAACACCCUUAAGAACCCCGACGGCGACGUGACGAACGGGAACAGCACAGCUGCAGAUCCGACGC.GCCGGGGC..................................CUUUUC...............................
env-42   CGGGCGGGUCUCUCCCGGCCGGGCCGCGGACCACAAGAACACGCUUAAGAACCCCGACGGCGACUCCCGGAACGGGAACAGCACAGCUGCAGAUCCGACGC.GCCGGGGC..................................CUCUUC...............................
env-43   ACCGACGGAGACGACAAGCGGGCCGCCGGCACACAAGAACACGCUUAUCACUCCCGGCAACGACCGCUUUGACGGGAACAGCACAGCUGCGGAUCCGACGC.GCCG.GGGU..................................UUGUUUU..............................
env-44   GCCGAGGAGUGAUCGGUAGGGCCGUCAGCUGCACAAGAACACCCUUAAGAGGGGCGACGGCGACCGAUCGAACGGGAACAGCACAGCUGCAAAUCCGACCC.GCCGGGG....................................ACGUUUCGC............................
env-45   AUCGGGCUGGCACGGGGUCGUCGGGUCGAUCAACAAGAACACGCUUAAGAGGGGAGCGGACCUACGUUCCGGUGGGAACAGCACAGCUGCAGAUCCGACGC.GCCGGGG....................................ACCUUUCGC............................
env-46   CGGUUCUUAACUCCCUCGGAUCGGGGCGUCCGUAUGGCAACCCUUUAGUCCGCGAGAGCAGUUCGAAUCGGUAGGGAACGGCACAGCCGCAGAUCCGACGC.GCCGUGU....................................GUUU.................................
Hpa-1-1  CUCCAUCCCCGUCGCCGCCGAAACCGUACUCGAAUGGAAACCCUUUAACCCGCGAGAACGGAACGCGUCUGUAGGGAACGGCACAGCCGCAAAUCCGACUC.GUCGUGA....................................GAUU.................................
Hci-1-1  UGGGCGAGGUGGACUGUAGCACGGCGGCCUGAAUACCAAUCUUUUAACUGCGUCCACGAAUUGCUACUCACUAGGGAACGGCACAGCCGCAAAUCCGACUC.GUUGCGUGUGUUCGCGGGCACCUGCCC...............GCUCUCUUUCG..........................
env-47   ACGACAGCCGUUCGCUCCUCACGCCGCGGGAUGUCGUAACCGUUUAGUUCGGCGCGAGGAGGAAGAUAGGUAGGGAACCAGCACGGCCGCAAAUCUGACUC.GCUCGGCG...................................UCGA.................................
env-48   AGCCGGUGAUUAAGCACUUCGACCCGUGUCGGUGAUGCCAACGGUUAUAGCGACCGAUGGAGUGGCCACAACCGGGAAGAACACGGCCGCAAGUCCGACGC.GUCGGC.....................................AUC..................................
env-49   ...............................................................................ACACACGGCCGCAAGUCCGACAU.GUCGGC.....................................GAU..................................
env-50   ..................................................CCGGCUGCUGGAGUGGUUUGUGUCGGGAAGCACACGGCCGCAAGUCCGACAU.GUCGGC.....................................GAU..................................
env-51   AGUCCGUGAUUAAGCCCUUCGACCCACAUCGAUGAUGCCAACGGUUAAACGGGCUGCUGGAGUGGCUUAUAUCGGGAAGCACACGGCCGCAAGUCCGACAU.GUCGGC.....................................GAU..................................
env-52   AGUCCGUGAUUAAGCCCUUCGACCCACAUCGAUGAUGCCAACGGUUAAACGGGCUGCUGGAGUGAUUUAUCUCGGGAAGCACACGGCCGCAAGUCCGACAU.GUCGGC.....................................GAU..................................
env-53   AGACUGUGAUUAAGCGCUUCGACCCACGGGGGUGAUGCCAACGGUUAUACCGCCCGAUAGUGACCUCACAGUCGGGAAACACACGGCCGCAACUCCGACGC.GUCGGC.....................................GCU..................................
env-54   GAGUCGGUGAUUAAGAGCUUCGACACUCAUCGGUGAUGCCAACGGUUAUACCGGACAGUGGAGUGUGAUAUGCGGGAAGCACACGGCCGCAACUCCGACGC.GUCGGC.....................................GGU..................................
env-55   AGACCGUGAUUAAGCGCUUCGACCCACGUCGGUGAUGCCAACGGUUAUACCGGCCAACGGAGUGAUUCCAGCCGGGAAGCACACGGCCGCAACUCCGACGC.GUCGGC.....................................GGU..................................
env-56   AGUCGGUGAUUAAGAGCUUCGACACUCAUCGGUGAUGCCAACGGUUAUACCGGACAAUGGAGUGGUGAUAUUCGGGAAGCACACGGCCGCAACUCCGACGC.GUCGGC.....................................GGU..................................
env-57   .....................................................................ACAGCCGGGAAAACACGGCCGCAACUCCGACGU.GUCGGC.....................................GCU..................................
env-58   AGACCGUGAUUAAGCGCUUCGACCCACGGCGGUGAUGCCAACGGUUAUACGGGCCGAUGGAGUCCUCACAGCCGGGAAAAACACGGCCGCAACUCCGACGU.GUCGGC.....................................GCU..................................
Hla-2-1  UUUAGAAGCCCUUCGAUUCGGUCGCGCAGCGAGUCGAAACCGUUUAAGUCUCGGCACGAGGAGUGACGCAUGGGAAUCCAGCACGGCCGUAAAUCUGACUC.GCCCG......................................AUU..................................
Hsp-4-1  CUUCGAUUCCGACCGGGACGAGGCGGGGCCGAGUCGAAACGGUUUAACCCGGCGCGCGUGGACAACCCGAUGGGAAUCCAGCACGGCCGCAAAUCUGACUUCGCCC.......................................GUUU.................................
Hsp-2-1  CUUCGAUUCCGACCGGGGCGAGUCGGGCCCGAGUCGAAACGGUUUAACCCGGCGCGCGUGGACAACCCGAUGGGAAUCCAGCACGGCCGCAAAUCUGACUUCGCCC.......................................GUUU.................................
Hme-1-1  UUUAGAAGCCCUUCGAUUCUCAUGCGCCGCGAGUCGAAACGGUUUAACCCGGCGCGCGUGCAGAUGUCGAUGGGAAUCUAGCACGGCCGCAAAUCUGACGU.GCCCGC.....................................GUUU.................................
Hel-1-1  UUUAGAAGCCCUUCGAUUCGACCGCGCAGCGAGUCGAAACCGUUUAAGUCUCGGCACGAGGAGUGACGCAUGGGAAUCCAGCACGGCCGUAAAUCUGACUC.GCCCA......................................AUU..................................
Hvo-1-1  ACCGGACCGAGUCGCGGUCAAUUCGGAGCCGAGUCGAAACGGUUUAACCCGGCGCGCGUGGACAACCCGAUGGGAAUCCAGCACGGCCGCAAAUCUGACUC.GCCCA......................................GUU..................................
Hsu-2-1  CUUCGAUUCCGACCGGGUCGAGUCGGGACCGAGUCGAAACGGUUUAACCCGGCGCGCGUGGACAACCCGAUGGGAAUCCAGCACGGCCGCAAAUCUGACUC.GCCCA......................................GUU..................................
Hde-1-1  CUUCGAUUCCGACCGGGUCGAGUCGGGACCGAGUCGAAACGGUUUAACCCGGCGCGCGUGGACAACCCGAUGGGAAUCCAGCACGGCCGCAAAUCUGACUC.GCCCG......................................GUU..................................
Hpr-1-1  CCCGAGGCGAGUUGAGGCCGAGUCGAAUCCGAGUCGAAACGGUUUAACCCGGCGCGCGUGGACAACACGAUGGGAAUCCAGCACGGCCGCAAAUCUGACUC.GCCCG......................................GUU..................................
Hgi-1-1  ACCGAGGCGAGUUGAGGCCGAGUCGAAUCCGAGUCGAAACGGUUUAACCCGGCGCGCGUGGACAACCCGAUGGGAAUCCAGCACGGCCGCAAAUCUGACUC.GCCCG......................................GUU..................................
Hal-1-1  UCCGAGGCGAGUUGAGGCCGAGUCGAAUCCGAGUCGAAACGGUUUAACCCGGCGCGCGUGGACAACACGAUGGGAAUCCAGCACGGCCGCAAAUCUGACUC.GCCCG......................................GUU..................................
Hgi-2-1  ACCGAGGCGAGUUGAGACCGAGUCGAAUCCGAGUCGAAACGGUUUAACCCGGCGCGCGUGGACAACCCGAUGGGAAUCCAGCACGGCCGCAAAUCUGACUC.GCCCG......................................GUU..................................
Hmu-1-1  UUUAGAAGCCCUUCGAUUCUGACGCGCGUCGAGUCGAAACGGUUUAACCCGCCGCGUGUGCAGAUGUCGAUGGGAAUCUAGCACGGCCGCAAAUCUGACUC.GCCCGC.....................................GUUU.................................
env-59   .........................AGAGACGGGAGUAACAACCCAUAAGUACGCGAGCGGGAGAAUACGUGGUAGGAACGGCACAGCCGCAAAUCCGACCU.GCCAC.C....................................GAUU.................................
env-60   CCCAUCAGACGCCCGGUCGCCAGCCCGACCGCGAUCAGUAACGCUUACCUUGCUGACUCCCAAACCAUGCUUCGGGAAGCGCACAGCCGCAAAUCUGACUGUGUCACC.CGA..................................UUCG.................................
env-61   ACCCCUCGAAUCGUCGCGGCGUAUGCGAGUCGGUUACGGAACGUUUUUCAGCGCCUCCCGACCACACACGACUGGGAAGCGCACGGCCGCAAAUCUGACCC.GUCGUCCAGACGGCGAC..........................CCCU.................................
env-62   GAGGGGAGUAGACGGCAUGACGGCCCGUGGUGAGAUGAAAACGGUUUAGUCGCGUGACCGUCGACUUCACAACGGGAACUGCACGACCGCAAAUCUGACUC.GCCGU......................................GAAUUUCA.............................
env-63   UACCCGUGGCCUCGGCAUAAGCGCUUCGAACGGAGGUAGUAAGACUUACCUGCGAGAGCGCCCCGUCUCCGGUGGGAACGGCACGGCUGCAAAUCUGACCC.GUCGCGCG...................................GUU..................................
env-64   UAUCCUCGGCUGGGUGAUAAGCGCUUCGAACGUCGGUAGUAAGACUUACUUGCCCGAGCGGCCGUCAUUCGGUGGGAACGGCACGGCUGCAAAUCUGACCC.GUCGUGCG...................................GCUU.................................
Has-1-1  CGUGUCCUGCCACCCGAUUUCCCGUCGUCCCGUUCUAGAAACCCUUUUGCCCCACAGCCCACAACGACGCGUUGGGAAUCGCACGGCCGCAAAUCUGACCC.GUCGCGC......CUCGCGUCC....................ACCUC................................
env-65   CCCCGCAGGGAGCGCGUGAGUGGGGUGAACGCGAACGACAACGGAUAAGUGCGCGAAGGCACUCCUCACGGCUGGGAACAGCACAGCCGCAUAUCCGACUUCGUCGCGCAGGC................................CUCGC................................
h3A-1-1  UUCUCCGCGUCAUCGUCGACGCCCGUUGCACCGAAGAGAACACCCUUAAGAGGCUCGGCGGGAACUCAUUCGUGGGAACAGCACAGCCGCAAGUCUGACUC.GCCG.GGGC..................................GCUC.................................
env-66   AGCAAUUCACACUCCUCGACGGCGCUCGCGGCCGAUGGAAAGCCAUUAAGCCCCUCGACUGCCACCUUCCACUGGGAACAGCACGGCCGCAAAUCUGACUUUGCCCA......................................UCG..................................
env-67   GGGUUCCGACGGCCGUCCGGCUCGCCCCCUCCGCGGAUAGAAACUGUUUUCAGCGCGAGCCGACAGAUUCGAGUGGGAACACACGGCCGCAAAUCCGACGC.GCCGCUC....................................GCUU.................................

         ................................................................................<<<<.<<<<<<<..<<......<<<<<<<<<<<<<<<<<<<<<<<<<<<<<<<<<<<<<<<<<<<.......................>>>>>>>>>>>>>
         ................................................................................2....221122..01......0222222.........1.11...10...0000.0..000000.......................000000..0.000
         ●●●●●●●●●●●●●●●●●●●●●●●●●●●●●●●●R●R●●RR●A●●●YUU●●●●●●●●●●●●●●●●●●●●●●●●RRYGGAA●●GCACRGCCGCAAAUCYGACYY-GYC●●○○-----------------------------------●YY●---------------------------
                                                                              5′
```

```
Hka-1-1   GCCCGUCACCGUUCACGGACAGCCACCGGCCGCAGUAGAAACGGUUUUACGCGCCGAGUGGGGAGAGUACGUCGGGAACAGCACGACCGCAAAUCUGACCC.GCCACGC...................................GGUU................................
Hmu-2-1   GCCGGUCACCGUUCACCGACAGUCACCGGCCGCAGUAGAAACGGUUUUACGCGCCGAGUGGGGAGAGUACGUCGGGAACAGCACGACCGCAAAUCUGACCC.GCCACGC...................................GGUU................................
env-68    ACGGCGGUAUGGCCGUCCCGGAACGCGAUCGGGAAGCGGAACGGUUUAGUCGCGCCCUCUCAGACAGGGAGAUGGGAACAGCACGACCGCAAAUCUGACUC.GCCGUGG...................................GUUU................................
Ham-1-1   ACGACUGUGUCGCCGUUUCGAGACGCGACCGGGAAGCGGAACGGUUUAGUCGCGCCCUCUCAGAAAGGGAGAUGGGAACAGCACGACCGCAAAUCUGACUC.GCUGUGA...................................AUUU................................
Hva-1-1   GAAGCGGUAUCGCCGUUUCGAGACCCGGCCGGGAAGCGGAACGGUUUAGUCGCGCCCUCUCAGACAGACAGUUGGGAACAGCACGACCGCAAAUCUGACUC.GCUGUGG...................................ACUU................................
Hja-1-1   ACGACGAUGUCGUCGGUUAGAGACGCGGCCGGGAAACGGAACGGUUUAGUCGCGCCCUCUCAGACAGGGAGAUGGGAACAGCACGACCGCAAAUCUGACUC.GCUGUGG...................................ACUU................................
Hma-1-1   ACGACGAUGUCGGCACUUCCAGACGCGACCGGGAAGCGGAACGGUUUAGUCGCGCCCUCUCAGACAGGGAGAUGGGAACAGCACGACCGCAAAUCUGACUC.GCUGUGG...................................ACUU................................
Hsa-2-1   UCGCGGCUAUCCUCGUUUCGAGACGCGACCGGGAAGCGGAACGGUUUAGUCGCGCCCUCUCAGACAGGGAGAUGGGAACAGCACGACCGCAAAUCUGACUC.GCUGUGG...................................ACUU................................
Hca-1-1   UGGACGGUGUCGUCGUUUCGAGACUCGGCUGGGAAGCGGAACGGUUUAGUCGCGCCCUCUCAGACAGGAAGAUGGGAACAGCACGACCGCAAAUCUGACUC.GCUGUGG...................................ACUU................................
Hsi-1-1   UGGACGGUGUCGUCGUUUCAAGACUCGGCCGGGAAGCGGAACGGUUUAGUCGCGCCCCCUCAGACAGGAAGAUGGGAACAGCACGACCGCAAAUCUGACUC.GCUGUGG...................................ACUU................................
Har-1-1   GAGACGGUGUCCUCGUUUCGAGACGCGGCCGGGAAGCGGAACGGUUUAGUCGCGCCCUCUCAGAUAGGGAAACGGGAACAGCACGACCGCAAAUCUGACUC.GCUAUGG...................................ACUU................................
env-69    CGCGGGAGGCAGUAGCGUGGUGGACGCUCCGGAGAUGGAAACGGUUUAGUUCGGUGACUGAUGACCAUACAUCGGGAACAGCACGACCGCAAAUCUGACAC.GCUGUGG...................................GUUU................................
env-70    GUUCGCGCGCGAGCGACCGGUUUCGGACGGACAGAUGGAAACCGUUUAGUCGGGCGCUCGUCGUUGUAGAGACGGGAACAGCACGACCGCAAAUCUGACUC.GCCGUCC...................................GUUU................................
env-71    UCGACUCAUACAUUAGUAUUUUCCACAGGUAUCGGUCGCAAUGUUUUUAAUAUACUCAACCGCUAACUAUGUUGGGAACAGCACGGCCGCAAGUCUGACUC.GCCGU.....................................AUUUUG..............................
env-72    CAUAUAUCAGUACCAAUAUCAUCGGUAUCUGUCAGUCGCAAUGUUUUUUAUACGCUCAACCGAUAAUUUAGUUGGGAACAGCACGGCCGCAAAUCUGACUU.GCCGUC....................................UCUA................................
env-73    CAUAUAUCAGUACCAAUAUCAUCGGUAUCUGUCAGUCGCAAUGUUUUUUAUACACUCAACCGAUAAUUUAGUUGGGAACAGCACGGCCGCAAAUCUGACUU.GCCGUC....................................UCUA................................
env-74    CAUAUAUCAGUACCAAUAUCAUCGGUAUCUGUCAGUCGCAAUGUUUUUUAUACGCUCAACCGAUAAUUUAGCUGGGAACAGCACGGCCGCAAAUCUGACUU.GCCGUC....................................UCUA................................
env-75    CGACUCAUACGUCAAGUUGCGCCGGUAUCCGUCGGUCGCAAUGUUUUUUAUACGCUCAACUGGUAAUUCAACUGGGAACAGCACGGCCGCAAAUCUGACUU.GCCGUC....................................UCUA................................
env-76    GACCGCGGUCUCGGCAGCAGGAUGCCGAAACCAGGUAGAAACCCUUUUGACCGCCCCAAUACACCGAUGACCUGGGAAGCGCACGGCCGCAAAUCUGACUC.GUCGUCCAC..GCCACC.........................CACUC...............................
Hca-3-1   AGAUUCGUCGAACGACCGCUGGGUCGGCCGGGUCCGAUGGAAACUGUUUAGUGUCGCAAAGGGGACUCUCCGGUGGGAACGCACGACCGCAAAUCUGACUC.GCUGUGC...................................GCUU................................
env-77    GGUUUUCUGCAUUCUACGUGUUUCUAUCAACCGUAACCGAGAGGUUUAUCAUGCGCAGUCGUAUUUUCACGAGUGGGAACACACGGCCGCAAAUCUGACAUCGCCGUCC...................................GCUU................................
env-78    CCUCCGAGACUGCGACAAGGCGCUUAGCACCCACAACCGAGACGUUUAUACCGCACAGUCCAGUUGCUACGAGUGGGAACGCACGGCCGCAAAUCUGACAUCGCCGUCC...................................GCUU................................
env-79    GAUUUUCCCCCUUCUACCCGGUUGUUUCGACCGUAACCGAGAGGUUUAUGCCGCAUAGUCGUAUUGCUAUGAGUGGGAACGCACGGCCGCAAAUCUGACAUCGCCGUCC...................................GCUU................................
env-80    UUUUUGGCCAAGGCUCGACCGUCCAGAAACGUCGAACAGUAACGGUUAACAGUGCUGCUCCGUUUUCCAACAGUGGGAACGCACGACCGCAAAUCUGACUC.GCCGUCC...................................GUUU................................
env-81    CUCGGACGAUCACGUUCUCCCUGCCCUGGCUGCGAACAGUAACCGUUAACAGUGGUGGUCCGUUUUGCUCGAACGGGAACGCACGACCGCAAAUCUGACUC.GCCGUCC...................................GCUU................................
Hsa-1-1   CUAUUGGCCGUUUCUCAGCGAUGAGUCUCGGCCCGGAUAGAAACCGUCAAGUGGCCCAGCGAGAAGAGGACAGUGGGAACACACGACCGCAAAUCUGACUC.GCCGUCC...................................GAUU................................
env-82    ..................................................................AAGCGGGUGGGAACGCACGACCGCAAAUCUGACUC.GCCGCAC...................................GAUU................................
env-83    ....UGCGAAAUUACGGUAUUCGUUCCCGACGCGAAAAGUAACCGUUAACAGUCCUGGCCGACAGGGUUCGCAUGGGAACGCACGGCCGCAAAUCUGACUC.GUCGUCC...................................ACUU................................
env-84    GAUCCGUGUCGGUUCGGCGUAGACGGGCAGCCAGGACAGUAACAGUUAACAGUAGCCGGACGCUCCGUAGAGGUGGGAACGCACGACCGCAAAUCUGACUC.GUCGCCC...................................GCUU................................
env-85    UGGAAAAAAUCAGUUUUGGAGCCGGUGAGCACCGAAAAGUAACCGUUAACAGUGCUGGCCUGUUUCGUCCGGGUGGGAACGCACGACCGCAAAUCUGACUC.GCCGUCU...................................ACUU................................
env-86    CGGAGAAGUGGGUGACGACUAGACAGUGGGCGAGAACAGUAACGGUUAACAGUGCUGACCCGCUGGCUUUCGAUGGGAACGCACGACCGCAAAUCUGACUC.GCCGUCU...................................GUUU................................
env-87    UGAGUCCCCCGCUGCGUGUACGGCGACAGAAGGGAGCCGUAACCGUUAACACACUCGAACGGCUGUUCUCGGGUGGGAACGCACGGCCGCAAAUCUGACUC.GCCGUCG...................................ACG.................................
env-88    GCCGUGGUGCGGGCACACAGACACGCCCGUCAGAAUGAAAACGAUUAACGGCCGUGAUGAAUUACCUUCUCUUUGGGAACACACGGCCGCAAAUCUGACUACGCCGCC....................................GAUU................................
env-89    AUCGGGCGCGGACCGAGCCGAGAGCGGGCGUCCGGAAGGAAACGGUUAACAGCCGAUCGGCCGAAACUCAGGUGGGAACGCACGGCCGCAACUCUGACAACGCCGCC....................................GAUU................................
env-90    .................................................AGCGCGCCUCGCGGAGGUGUUCGUGGGAACGGCACGGUCGCAAAUCUGACUC.GCCGCCCG..................................UUC.................................
env-91    GGCGUGUGGUCGGGUGCCGUGUUCUGUGAGUCGUUUGGAAACCCUUUUGAGCGCGCCUCGCGGAGGUGUUGGUGGGAACGGCACGGUCGCAAAUCUGACUC.GCUGCCCG..................................GUU.................................
env-92    GACGAACUCGGAUCGCAGCGAGCGAAGCCAGCGAAUAGAAAACGCUUAACCGCCGCGUCGACCACCUCCCGGUGGGAACAGCACAGUCGUAACUCUGACUC.GCC.......................................CCGUUGN.............................
env-93    AGCGGAUCGACAAAGACUCACACAGCGACUGGCGAUGGAAAACCAUUAAGCCUCGAAGCCAGACAGUAUCAGUGGGAACAGCACAGUCGUAACUCUGACUC.GCCCA.....................................UGU.................................
env-94    ...........................................................CCGUGGAGACACCAUUGGGAACAGCACAGUCGUAACUCUGACUC.GCCC......................................UCG.................................
env-95    CGGGGAACUGGGUACUGUUUCGGCAGUCGCGCCGAAUAGAAAGCCAUUAACGCCCACAAUGGAGACUACCGGUGGGAACAGCACAGUCGUAACUCUGACUC.GCCU......................................UCAC................................
env-96    GGCCUUCGACUCGGGGUAAUGAACCCGUCAGCAAAUAGAAAGCCCUUAACGCCCACAGCAGGGACAAAUCGGUGGGAACAGCACAGUCGUAACUCUGACUC.GCCC......................................AUAC................................
env-97    GGCCUUCGACUCGGGGUAAUGAACCCAUCAGCAAAUAGAAAGCCAUUAACGCCCACAACGGAGACAGAUCAGUGGGAACAGCACAGUCGUAACUCUGACUC.GCCC......................................ACAC................................
env-98    GGGGAAAUGGGUGGCUAUCUCGGCCGUCGUACCGAAUAGAAAGCUCUUAACGCCCACAAUAGAGACUACCGGUGGGAACAGCACAGUCGUAACUCUGACUC.GCCC......................................ACAC................................
env-99    AGGCCUUCGACUCGGGGUAAUGAACCCAUCAGCAAAUAGAAAGCCCUUAACGCCCACAACGGAGACAAUCAGUGGGAACAGCACAGUCGUAACUCUGACUC.GCCC......................................ACAC................................
env-100   ......................................................GCGUCGACCAGACACCGAUGGGAACAGCACAGUCGUAACUCUGACUC.GCCC......................................CUCG................................
env-101   GACACUGAGUCAGUCGACGGCACACCCAUACGGAAUAGAAAACCCUUAACGCCCACACCGUGGAGACACCAUUGGGAACAGCACAGUCGUAACUCUGACUC.GCCC......................................CUCG................................
env-102   ACGGAUGUCAACACCAAACACCCAGCGCGCACGAAUAGAAAACCCUUAACGCCCGCGUGGUAGAAGAUCCGAUGGGAACAGCACAGUCGUAACUCUGACUC.GCCC......................................UUCG................................
env-103   UCGGCUGUCAACACCGAACACCCAGCGCGCACGAAUAGAAAACUCUUAACGCCCGCGUGGUAGAAGAUCAGGUGGGAACAGCACAGUCGUAACUCUGACUC.GCCC......................................UUCG................................
env-104   GACACUGAGUCAGUCGACGGCACACCCAUACGGAAUAGAAAACCCUUAACGCCCACACCGUGGAGACACCAGUGGGAACAGCACAGUCGUAACUCUGACUC.GCCC......................................CUCG................................
env-105   CGGCCAGACAACCGUCGAUCAACUGCCGACCAGAAUAGAAAACCCUUAACGCCCACACCGUUCACACACCAGUGGGAACAGCACAGUCGUAACUCUGACUC.GCCC......................................CUCG................................
env-106   AUCACUUGACAUCGUCCAGCGACUCCGUCUACAAAUAGAAAACGCUUAACCGACGCGUCGACCACUUCCCGAUGGGAACAGCACAGUCGUAACUCUGACUC.GCCCC.....................................GUU.................................
env-107   CGACUCGACAGCAGCUGGCGGCUGUCAGCCACGAAUGGAAAACCCUUAACGCACGCGUCGACGACCCUCCGAUGGGAACAGCACAGUCGUAACUCUGACUC.GCCCC.....................................GUU.................................
env-108   UCGAUUUUUCGUUCGGCCCCUCGAUAAGAGCCGAAUAGAAAACGCUUAAGCGCCGCGUCGACCACUCCCCGAUGGGAACAGCACAGUCGUAACUCUGACUC.GCCCC.....................................GUU.................................
env-109   CUCGCUUUUACAUUCGGCCCCUCGACAAGAGCUAAAUAGAAAACGCUUAACCGCCGCGUCGACCACUCCCGAUGGGAACAGCACAGUCGUAACUCUGACUC.GCCCC.....................................GUU.................................
env-110   UCGCUUUUUCGUUCGGCCCCUCGAUAAGAGCCGAAUAGAAAACGCUUAAGCGCCGCGUCGACCACCUUCCGAUGGGAACAGCACAGUCGUAACUCUGACUC.GCCCC.....................................GUU.................................
env-111   GAGCGGUUAGGGUGGUUUUGCCCUAACGACUGAAAUAGAAAACUCUUAACGCCCACGUCGACCACCACCUGAUGGGAACAGCACAGUCGUAACUCUGACUC.GCCCC.....................................GUU.................................
env-112   CGACUCGACAGCAGCUGGCGGCUGUCAGCCACGAAUGGAAAACCCUUAACGCACGCGUCGACNACCCUCCGAUGGGAACAGCACAGUCGUAACUCUGACUC.GCCCC.....................................GUU.................................
env-113   UCGCUUUUUCGUUCGGCCCCUCGAUAAGAGCCGAAUAGAAAACGCUUAACCGACGCGUCGACCACCCUCCGAUGGGAACAGCACAGUCGUAACUCUGACUC.GCCCC.....................................GUU.................................
env-114   AUCACUUGACAUCGUCCAGCGACUCCGUCUACAAAUAGAAAACGCUUAACCGACACGUCGACCACUUCCCGAUGGGAACAGCACAGUCGUAACUCUGACUC.GCCCC.....................................GUU.................................
env-115   GCUUAUCGAGAGAACAGCACGCAGAGAGAGACGAAUAGAAAACCCUUAACCGACGCGUCGACCACCUCCCGAUGGGAACAGCACAGUCGUAACUCUGACUC.GCCCC.....................................GUU.................................
env-116   UCUUGACGCUGAGCGUUCGCCAGUCGUAGGUGGAAUAGAAAACGCUUAACCGACGCGUCGACCACCCACCGAUGGGAACAGCACAGUCGUAACUCUGACUC.GCCCC.....................................GUU.................................
env-117   CGACUCGACAGCAGCUGGCGGCUGUCAGCCACGAAUGGAAAACCCUUAACGCACGCGUCGACCACCCUCCGAUGGGAACAGCACAGUCGUAACUCUGACUC.GCCCC.....................................GUU.................................
env-118   GAGCNGUGAGGCUGNCUUUGUNCCAGCNACUGAAAUAGAAAAGCCUUAACGCCCACGUCGACCACCGUCCGAUGGGAACAGCACAGUCGUAACUCUGACUC.GCCCC.....................................GUU.................................
env-119   AUCACUUGACAUCGUCCAGCGACUCCGUCGGCAAAUAGAAAACCCUUAACCGACGCGUCGACCACCCACCGAUGGGAACAGCACAGUCGUAACUCUGACUC.GCCCC.....................................GUU.................................
env-120   UCACUCGGUAGCAGCCAACGGCUGUCAGCGACGAAUGGAAAACUCUUAACGCACGCGUCGACGACCUUCCGAUGGGAACAGCACAGUCGUAACUCUGACUC.GCCCC.....................................GUU.................................
env-121   CGCCCGGUUAGAAUAGAUGAGACGCUCGAUUGGAAUAGAAAACGCUUAACCGACGCGUCGACCACCGUUCGAUGGGAACAGCACAGUCGUAACUCUGACUC.GCCCC.....................................GUU.................................
env-122   UCUCGACGCCGAGCGUUCGCCGGUCGUAAGGAGAAUAGAAAACGCUUAACCGACGCGUCGACGACCCUCCGAUGGGAACAGCACAGUCGUAACUCUGACUC.GCCCC.....................................GUU.................................
env-123   UCGCUUUUACAUUCGGCCCCUCGACAAGAGCUAAAUAGAAAACGCUUAACCGCCGCGUCGACCACUCACCGAUGGGAACAGCACAGUCGUAACUCUGACUC.GCCCC.....................................GUU.................................
env-124   AAUACAUAGUUGACGGCAAACAGCGUCGGCUGAAAUAGAAAACCCUUAACGUUCAGGUCGACCACCACCCGAUGGGAACAGCACAGUCGUAACUCUGACUC.GCCCC.....................................GUU.................................
env-125   GAGCAGUGAGGCUGGCUUUGUUCCAGCCACUGAAAUAGAAAAGCCUUAACGCCCACGUCGACCACCGNCCNAUGGGAACAGCACAGUCGUAACUCUGACUC.GCCCC.....................................GUU.................................
env-126   GAGCGGUGAGGCUGUCUUUGUCCCAGCGACUGAAAUAGAAAACUCUUAACGCCCACGUCGACCACCGUCCGUUGGGAACAGCACAGUCGUAACUCUGACUC.GCCCC.....................................GUU.................................
env-127   GACACUGAGUCAGUCGACGGCACACCACCGCCGAAUAGAAAACCAUUAACGCCCGCGUCGACCACUCCCCGUUGGGAACAGCACAGUCGUAACUCUGACUC.GCCCC.....................................GUU.................................
env-128   AUCACUUGACAUCGUCCAGCGACUCCGUCGGCAAAUAGAAAACGCUUAAGCGCCGCGUCGACCACCUCCCGGUGGGAACAGCACAGUCGUAACUCUGACUC.GCCCC.....................................GUU.................................
env-129   CAUUCGACAUCGUUUAGCGACUCCGUCGAAGUGAAAUAGAAAACGCUUAACCGACGCGUCGACCACCCUCCGUGGGAACAGCACAGUCGUAACUCUGACUC.GCCCC.....................................GUU.................................
env-130   GACGAACUCGGAUCGCAGCGAGCGAAGCCAGCGAAUAGAAAACGCUUAACCGCCGCGUCGACCACCUCCCGGUGGGAACAGCACAGUCGUAACUCUGACUC.GCCCC.....................................GUU.................................
env-131   AUCACUUGACAUCGUCCAGCGACUCCGUCGGCAAAUAGAAAACGCUUAACCGCCGCGUCGACCACCUCCCGGUGGGAACAGCACAGUCGUAACUCUGACUC.GCCCC.....................................GUU.................................
env-132   GCGUCGAGCACGGGCCGGUAGCUCGACGACUGAAAUAGAAAACCCUUAACGCCCACGCCGAGCAGUGAUCAGUGGGAACAGCACAGUCGUAACUCUGACUC.GCCCC.....................................GUU.................................
env-133   GCAGUCACAACCCGGUUUGCGGUGUCGCCAACAAAUAGAAAGCUCUUAACACCCACAGCGGGCACAGUUUGAUGGGAACAGCACAGUCGUAACUCUGACUC.GCCC......................................GUUU................................
env-134   GGCCUUCGACUCGGGGUAAUGAGCCCGUCGGCGAAUAGAAAGCCCUUAACGCCCACAGCGGCGAGCAAUCGCUGGGAACAGCACAGUCGUAACUCUGACUC.GCCC......................................GUUU................................
env-135   GCAGUCACAACCCGGUUUGCGGUGUCGCCAUCAAAUAGAAAGCUCUUAACACCCACAGCGAGCACGGUUUGGUGGGAACAGCACAGUCGUAACUCUGACUC.GCCC......................................GUCU................................
env-136   GUAGUCACGACCGUGUUUGCGGUGACGCCAACAAAUAGAAACCUCUUAACACCCACAGCGGACAUGAUCUGGUGGGAACAGCACAGUCGUAACUCUGACUC.GCCC......................................GUCU................................
env-137   GGCCUUCGACUCGGGGUUAUGCACCCGUCAGCAAAUAGAAAGCCAUUAACGCCCACAGCAGCGAUCGACUGGUGGGAACAGCACAGUCGUAACUCUGACUC.GCCC......................................AUCU................................
env-138   GGCCUUCGACUCGGGGUUAUGCACCCGUCAGCAAAUAGAAAGCCCUUAACGCCCACAGCGGCGACCGACUGGUGGGAACAGCACAGUCGUAACUCUGACUC.GCCC......................................AUCU................................

          ................................................................................<<<<.<<<<<<<..<<......<<<<<<<<<<<<<<<<<<<<<<<<<<<<<<<<<<<<<<<<<<.......................>>>>>>>>>>>>>
          ................................................................................2....221122..01......0222222.........1.11...10...0000.0..000000.......................000000..0.000
          ●●●●●●●●●●●●●●●●●●●●●●●●●●●●●●●●R●R●●RR●A●●●YUU●●●●●●●●●●●●●●●●●●●●●●●RRYGGAA●●GCACRGCCGCAAAUCYGACYY-GYC●∘∘∘--------------------------------------●YY∘-------------------------------
                                                                             5′
```

```
env-139   CAGCUACGCCGGGAGCGACGUGUCGACAACACAAAUAGAAAACCCUUAACGCUCGCGUGGCUGAAGAGUCAGUGGGAACAGCACAGUCGUAACUCUGACUC.GCCC......................................AUCU................................
env-140   GCAGUCACAACCCGGUUUGCGGUGUCGCCAUCAAAUAGAAAGCUCUUAACACCCACAGCGAGCACGGUUUGGUGGGAACAGCACAGUCGUAACUCUGACUC.GCCC......................................GUUU................................
CHa-1-1   GAAAACCGGCCGGGCCGACGGCGGGUCGGACGGAUGGAAAGCCAUUAACCGACGGACACCCUCGGUGGCGACGGGAACGGCACGGCCGCAAAUCUGACUC.GCCUCGC...................................ACCC................................
env-141   CGUGGGAACCGAUCGCGUGAUCGGUCGAUCGCGAUCACGCAACGUUUUAGUCGCUGCUCACGAUAGCACAGGUGGGAAUCGCACGGCCGCAAAUCUGACCC.GCCGCCG...................................UUUC................................
env-142   GGCCGCGAUCACCGUACGACCCGCUCGAAAGACAGGUCGCAACCCUUUUAGGCCAUCGAGCCAGUCUCAACUGGGAAGCGCACGGCCGCAAAUCUGACUC.GUCGCC....................................CGAUUCGUU...........................
Nsu-1-1   GCUCUCGAGUCUCCCGUCGGAGCCCCGAAUCCGGUCUAGAAACCCUUUAGUCCGUCCGACGGAAAGCGAUGGUGGGAAGCGCGCGGCCGCAAAUCCGACUGUGUCAACCG..................................CUGUUU..............................
Hla-1-1   GAGUGGGUUUCGUUCCGGUCCCGACGCGGAACGGUUACGCAACCCUUUAGUCGGUGCUCACGAUACGUCCGGUGGGAAGCGCACGGCCGCAAAUCCGACUC.GUCACCCG..................................UCGCUCU.............................
env-143   CUCGAGAAGCGAACGCUCGCACUCGAGACAGAGAGGUAGAAACCCUUUAGUCGCUUACUCCAGAAGAAAGGGUGGGAAGCGCACGGCCGCAAAUCUGACCU.GUCGUCG...................................GUUU................................
env-144   ACACCCUCUGAGAGAAACUCGAGUCCGAGACGGAGGUAGAAACCCUUUAGUCGCUCAGACGGAAAGUACGGCUGGGAAGCGCACGGCCGCAAAUCUGACCU.GUCGCCG...................................UUUC................................
env-145   GCUGUCGAGCACCGGAAGCGAUCGCACGGCGCCGUGACGCAACCCUUUAGUUGCUACUCGCGAUAGCUCAGAUGGGAACCGCACGGCCGCAAAUCUGACUC.GCCGCCG...................................UUUC................................
Hxa-1-1   GUCAAUCACUGGCACGGCCCGCUGCGCGCGCUGACCUAGAAACGCUUUAGUCCAUCCCGGCGAAAGCCCCGGUGGGAAGCGCACGGCCGCAAAUCUGACUGCGUCACC....................................GACUGCUAUUCGA.......................
env-146   GACACGGUGUAUCAGUGAUACUCUCCGAAACGGAGGUAGAAACCCUUUAGUCGCUCACCCCGGAAGGAGGGGUGGGAAGCGCACGGCCGCAAAUCUGACUUCGUCACC....................................GAUUG...............................
env-147   CGAUCGACAGCGCCGAUCUGCGUCCGAAACCGAUGACGCAAGUCAUUUAGUCGCUGGCCGAUAUGUAACGGCUGGGAAGCGCACGGCCGCAAAUCUGACUUCGUCACGC...................................CAAUU...............................
Ngr-1-1   GAGUCUCUGUCACGCCAGCACGGUCACCAGCCGGUUACGCAACGUUUUAGUUCGUCGUAGUGAUAGGCUCAGUGGGAAGCGCACGGCCGCAAAUCUGACUC.GUCACC....................................CUUGC...............................
env-148   GUCAGCCGACAGUCGUCCCUCGAGUGGACACGCAGGUCGCAACGCUUUUGUCUCACACGGGGGAACGAAGGGUGGGAAGCGCACGGCCGCAAAUCUGACUC.GUCACCCGU.................................CGAUUUUC............................
env-149   GUCAGCCGACGGUUGUCCCUCGAGUGGACACACAGGUCGCAACGCUUUUGUCUCACACGGGGGAACGAAGGGUGGGAAGCGCACGGCCGCAAAUCUGACUC.GUCACCCGU.................................CGAUUUUC............................
env-150   ACCGUCGAGGACCGAAAGCGAUCGCACGGAGCCGUGACGCAACCCUUUAGUCGCUGCUCACGAUAGCUCCGGUGGGAACCGCACGGCCGCAAAUCUGACUU.GCCAUCG...................................GUUC................................
env-151   GGCGGUGGGGAGUCUCUCACAUACUCGAGUGGGUUACGGAACGUUUUAGUCUGGGCCUGCGGUAAGGCUGUGUGGGAAGCGCACGGCCGCAAAUCUGACUU.GUCAUCC...................................GGUU................................
Hdj-1-1   AAGCGAUGUCUCGAGGAGUCUGACAGCGAGCCGUUCUAGAAACCCUUUAGUCCGUUCCGCUGGAAGCGAGUGUGGGAAGCGCACGGCCGCAAAUCCGACGUUGUCAUCCGG.................................GCUU................................
env-152   ......................CCGGCGAGGGGAGGUCGCAAUGCUUUUGAGCACCACGAGCAAGGGUUAGAUGGGAACAACACGGCCGCAAAUCUGACUC.GCCGUCC...................................GCUU................................
env-153   CUCCGUGUGACGCCCUGACGCCCCGGCGAGGGGAGGUCGCAAUGCUUUUGAGCACCACGAGCAAGGGUUAGAUGGGAACAACACGGCCGCAAAUCUGACUC.GCCGUCC...................................GCUU................................
env-154   UCGUUCCGUGAGCAGUAGUGAGCGUUCGAGUGGAGGUCGCAACGCUUUUGAGCACCACGAGCAAGAUACAAAUGGGAACAACACGGCCGCAAAUCUGACUC.GCCGUCC...................................GUUC................................
env-155   UCGUUCCGUGAGCAGUAGUGAGCGNUCGAGUGGAGGUCGCAACGCUUUUGAGCACCACGAGCAAGAUNCAAAUGGGAACAACACGGCCGCAAAUCUGACUC.GCCGUCC...................................GUUC................................
env-156   UGUGCUGUGACGCGGCACUGCGCGAUCGGUCGAAGGUCGCAACGCUUUUGAGCACCACGAGCAAGAUGCAAGUGGGAACAACACGGCCGCAAAUCUGACUC.GCCGUCC...................................ACAC................................
env-157   UAUCGAAGUUCCACGUACAGAGCGCUCGAACGGAGGUCGCAAUGCUUUUUACCAUCACGGGCGAGAUGCAAGUGGGAACAACACGGCCGCAAAUCUGACUC.GCCGUCC...................................ACUC................................
env-158   UCCCACCGAUGGGAUCGAGGGGCGUCAGGUCGGUUACGCAACGCUUUUGUCCGCGACGGCGUUACGCCCCGUCGGGAAGCGCACGGCCGCAAAUCCGACGUCGCCGUC....................................GUU.................................
env-159   GUCCUUGAGGGAGCGGUCCGUGGUCGCCGGACAGGUACGCAACCCUUUAGUCGGUGCUCACGAUACCACAGGUGGGAAGCGCACGGCCGCAAAUCCGACAUCGCUGUCC...................................GCUC................................
env-160   GGAGAACGGUGUGGUUUCUCUUCCGGUCGAACGGGUACGCAACGCUUUAGUCGGUGCUCGCGAUAGCACAGGUGGGAAGCGCACGGCCGCAAAUCUGACAUCGCUGUCC...................................GCUU................................
Hgo-1-1   GAGCGGACCGCCGAGCGAGGAGAAGUCCGCUGUAGUCCGAUACCCUUUUACUGGUACUCCCCGGAAUGCCGAUGGGAAGCGCACGGCCGCAAAUCCGACUC.GCCACCC...................................GCUAUUCG............................
Npa-1-1   CCGAACCGGCCGACCGUGCGCGGUGGGAUGGCGAUGUAGAAACCCUUUAGUCCCCGACGCGGGAAGUGAGCAUGGGAAGCGCACGGCCGCAAAUCCGACUGCGUCACCC...................................CCUCUUUC............................
Nga-1-1   CGGAGAUCGGCCGACCGCGCGUGUGAGAUGGCGAUGUAGAAACCCUUUAGUCCCCGACGCGGGAAGUGAGCGUGGGAAGCGCACGGCCGCAAAUCCGACUGUGUCACCC...................................ACUCUUCC............................
Nsp-1-1   CGGAGAUCGGCCGACCGCGCGUGUGAGAUGGCGAUGUAGAAACCCUUUAGUCCCCGACGCGGGAAAUGAGCGUGGGAAGCGCACGGCCGCAAAUCCGACUGUGUCACCC...................................ACUCUUCC............................
Nal-1-1   CUGGCCUGGCCGACCGCGUGCGGUGGGAUGGCGAUGUAGAAACCCUUUAGUCCCCGACGCGGGAAGUGAGCGUGGGAAGCGCACGGCCGCAAAUCCGACUGCGUCACCC...................................CCUCUUCC............................
env-161   ..........................................................GACCGGGAAGCGGUAGUGGGAAGCGUACGGCCGCAAAUCCGACUGCGUCACCC...................................CCUCUUUC............................
Hli-3-1   CGAGUGCGGGCCCGACCGUGUCGGCGUCGCAGCGAGUAGAAACCCUUUAGUCUGCAACGACGGUACCGACGGUGGGAAGCGCACGGCCGCAAAUCCGACUGUGUCACCC...................................CCCUUUUCC...........................
Nve-1-1   UCGGUACUCGAGCGAGAAUCGGCGGUGCCGGCGAUGUAGAAACCCUUUAGUCCCCGACGAGGGAAGGACCGGUGGGAAGCGCACGGCCGCAAAUCCGACUGCGCCAUCC...................................GCUCUUUG............................
Npe-1-1   CGAGACAGCGAACCGUCCGAGGCGUUGACCGGCGAUGAGAAACCCUUUAGUCUCCGACCCCGGAAGCGGUAGUGGGAAGCGCACGGCCGCAAAUCCGACUCCGCCGCCC...................................GCUCUUUG............................
Hje-2-1   UCUUCACGGUCGUCCUGUAGGGCGCUGACCGGCGAUGAGAAACCCUUUAGUCCUCGACGGCGGAAGCGGUAGUGGGAAGCGCACGGCCGCAAAUCCGACUCCGUCGCCC...................................GCUCUUUG............................
Hth-2-1   UCUUCACGGUCAUCCUGUAGGUCGCUGACCGGCGAUGAGAAACCCUUUAGUCCUCGACGGCGGAAGCGGUAGUGGGAAGCGCACGGCCGCAAAUCCGACUCCGUCGCCC...................................GCUCUUUG............................
Nje-1-1   ACGAGGAGUCGAUGGUCGGGCAGGGCUGACGGCAGGACACAACUCUUUAGUCGCUGCUCCCGGUAGCGACGGUGGGAAGCGCACGGCCGCAAAUCCGACACCGCCGUCC...................................ACCCUUUC............................
env-162   GUCACGACGUCGUGCGAUCCGCCGUCGCGGCCGAUGACGCAACCCUUUAGUCGCUGCUCCCGGUAGCGACGAUGGGAAGCGCACGGCCGCAAAUCCGACUC.GCUGUCC...................................ACCCUUUC............................
Noc-1-1   ACGCUUCCUGCGAGCGCGGCCGAUGGCCGCCUCAGGACGCAACCCUUUAGUCGUUGCUCCCGGUAGCGACGGUGGGAAGCGCACGGCCGCAAAUCCGACUC.GCUGUCC...................................ACCCUUUC............................
Nam-1-1   ACGAGGAGUCGAUGGUCGGACAGGGCUGACGGCAGGACGCAACCCUUUAGUCGCUGCUCCCGGUAGCGACGGUGGGAAGCGCACGGCCGCAAAUCCGACUC.GCUGUCC...................................ACCCUUUC............................
Nhu-1-1   UCCGGUUUACAACCCACACCGACUGUGACCGCUGUCUAGAAACCCUUUAGUCCGUGCCGCGGGAAGCGACGGUGGGAAGCGCACGGCCGCAAAUCUGACAGUGCCACUC...................................GCUU................................
Nma-1-1   UCGGGCUCACAACCCAUACCGACUGUGGCCGCUGUCUAGAAACCCUUUAGUCCGUGCCGCGGGAAGCGACGGUGGGAAGCGCACGGCCGCAAAUCUGACAGUGCCACUC...................................GCUU................................
Nta-1-1   GCAUAUUGAAGACUGCCGCGACGGUCGCUCUAGAUCUAGAAACCCUUUAGUCCGUCCCGCUGAAAGCGACGGUGGGAAGCGCACGGCCGCAAAUCUGACUGUGCCACUC...................................GCUU................................
Nas-1-1   GCGUAUUGAAGACUGCCGCGACGGUCGCUCUAGAUCUAGAAACCCUUUAGUCCGUCCCGCUGAAAGCGACGGUGGGAAGCGCACGGCCGCAAAUCUGACUGUGCCACUC...................................GCUU................................
Nch-1-1   UCCGGUUCACGACGUGCACCGACUGUGACCGCUGUCUAGAAACCCUUUAGUCCGUGCCGCGGGAAGCGACGAUGGGAAGCGCACGGCCGCAAAUCUGACAGUGCCACUC...................................ACUU................................
Nae-1-1   GCGUAUUGGUGGCUGUCACGGCGGUCGCUCUAGAUCUAGAAACCCUUUAGUUCGUUCCGCUGAAAGCGACGGUGGGAAGCGCACGGCCGCAAAUCUGACUGCGCCACUC...................................ACUCUUUC............................
Nba-1-1   GCUGUCGACGGCAGAGUCGUGGUCGCGAGCACGGUCUAGAAACCCUUUAGUCCGUUCGACGGAAAGCAGGCAUGGGAAGCGCACGGCCGCAAAUCCGACUGUGCCAUCC...................................ACUGUUUC............................
Nin-1-1   CGAGCACCCACCCGAUACGACUCGAGGAGCCCAGUCUAGAAACUCUUUAGUCCGUCCGAAAGAAGACAUCGGUGGGAAGCGCACGGCCGCAAAUCUGACUGCGCCGUCC...................................ACUAUUCG............................
Hni-1-1   GCCCGAUCAUCCAACGCUCGAGUUCCCCGGGUCGUUACGCAACCCUUUAGUCGGUGCUCACGAUACGUCCGGUGGGAAGCGCACGGCCGCAAAUCCGACUGUGCCGUCC...................................ACUCUUUC............................
Hsa-7-1   UGCGUAUUCUAUAGAGGCGACUCGAGCAACCCAGUCUAGAAACCCUUUAGUCCGUCCGAGCGAAGACACUGGUGGGAAGCGCACGGCCGCAAAUCUGACUGCGCCGUCC...................................ACUGUUUC............................
Htu-1-1   CGAGCACCGACCAGCUACGGUUCGAGGAAGCCAGUCUAGAAACCCUUUAGUCCGUCCGAGCGAAGACACUGGUGGGAAGCGCACGGCCGCAAAUCUGACUGCGCCGUCC...................................ACUGUUUC............................
Hsa-9-1   UCGACCGGCGUCCGAGCAGCGGCCACGGUGCCGGUCUAGAAACCCUUUAGUUCGUCCGAACGAAGAGAUGAGUGGGAAGCGCACGGCCGCAAAUCUGACUCCGCCGUCC...................................ACUGUUCC............................
Nti-1-1   CCCACCAGAGCCACGACUCCCGAGCCGGUUCCGGUCUAGAAACCCUUUAGUCCGUCCGAGCGAAAGCGCUGGUGGGAAGCGCACGGCCGCAAAUCUGACUGCGCCGUCC...................................ACUCUUUC............................
env-163   GGUUCCCCGUUCUAACUUAAACCCGCGGACGGCCGAACACAACGUUUUGUUCGCGACGACCGACUCCCGAGCGGGAAGCGCACGGCCGCAAAUCCGACAUCGCCGUC....................................CCC.................................
env-164   CUUAAACGCUUCGACUCGAUCACUCGUGGUCAGGGUCGCAACCCUUUUUAUCCAGUCGCCGGUGACUACGGAUGGGAACAGCACGGCCGCAAGUCCGACUC.GCCGCA....................................GCU.................................
env-165   UUAAGCGCUUCGACUCGAUCGGACGCGGUCGCGGGUCGCAACCCUUUUGCCGCCAUCGCCGGUGAAAUCAGAUGGGAACAGCACGGCCGCAAGUCCGACUC.GCCGC.....................................CGCGC...............................
env-166   UUAAGCGCUUCGACUCGACCGGACGCAACCGCGGGUCGCAAUGCUUUAGUCGCCGUCUACAGUGAUAUCGGAUGGGAACAGCACGGCCGCAAGUCCGACUC.GCCGC.....................................CCGU................................
Hpa-2-1   AUUAAGGCCUUCGACUCGGAGUUCCCGACCCCGUGUCGCAACGCUUUUCAGUCGGUCAGGGCAGGGUACACACGGGAACAGCACGGCCGCAAAUCUGACUC.GCCGCGC...................................CUC.................................
Hru-1-1   UUAAGGGCUUCGACUUGCCGAACGUCGACCGCGGGUCGGAAUCCUUUUGUCGACCACGGAGGAGAGUACAGACGGGAACAGCACGGCCGCAAAUCUGACUC.GCCGC.....................................CUCGC...............................
Hbo-1-1   UUAAGGCCUUCGACUCGGAUUUCAUACGGCCAGUGUCGCAACCCUUUUCAAUCGGUCAAGGGUGGGUAUACACGGGAACAGCACGGCCGCAAAUCUGACUC.GCCGCGC...................................GGUU................................
Ssp-1-1   GGUCGUGCGCACCGAAGAUUAGCCCUGUCACCGAGUAGAAACCCUUUAGAACGGUCCACGGGUACGGAUCGUUGGGAACAGCACGGCCGCAAGUCUGACUC.GCCGCCGCGUGUCUUCGCGGAG....................CGUCC...............................
Hje-1-1   GGAGUGAUAAACGCUUCGAACGCCCGUCGUGGCGAGUCGCAAUCCUUUAGACCGCAGACGGACGAAUCACGGUGGGAACAGCACGGCCGCAAAUCCGACUC.GCCACG.AGCCGUCG...AUGUCUC..CGCGA.CUCGCGGGCUAC....................GCCCGCUCGUUCG
env-167   CCGUAUGCGGGCCGGUGACACCGCCCUCGACGCCUUCGAAACGGUUUAGUGCUGGAUUCCCGAACUGACGACCGGGAACGGCGCGACCGCAAAUCUGACUC.GCCC......................................AUCU................................
env-168   UCGGUAUGCGGGGCAGUCACACCGGUCUCGUGCCUUCGAAACGGUUUAGUGCCGAAUCCCCGAACGUACCCUCGGGAACGGCGCGACCGCAAAUCCGACUC.GCC.......................................UUCG................................
Hna-1-1   CCGUAUGCGGGGCGGUCACACCGGCCUCGAGGCCUUCGAAACGGUUUAGUGCUCGAUUCCCGAACGGUUCGACGGGAACGGCGCGACCGCAAAUCCGACUC.GCC.......................................UUCG................................
env-169   UUCGGAUCGGGCGCCGCGGGCAGUGACACCCCCGAUUCGAAACGACUUAAUGUAUGAACGGGAAACGUUCAAUGGGAACGGCACGACCGCAAAUCCGACUC.GCC.......................................UUCG................................
env-170   UUCUGAGCGGAACCGACGGCUGAGCCGAACCCGGAGUAGAAACCGUUUAAUGUCCGAACCCACGAGUACUACUGGGAACCGCACGAUCGCAAAUCCGACUC.GCCGGUC...................................GAUU................................
env-171   UUUUGAGCGAGAUCGACGGCUGAGCCGAACCGGGAGUAGAAACCGUUUAAUGUCCGAACCCACGAGUACCACUGGGAACCGCACGAUCGCAAAUCCGACUC.GCCGGUC...................................GAUU................................
Nph-1-1   GGGCGACCGGGGCCAGCGCCGCUUCAGGUGGCGAGUAGAAACGGUUUUGUGUCCGACCGCUUGAGGUACACCCGGGAACCGCACGACCGCAAAUCCGACUC.GCCGGCC...................................GCUU................................
Nmo-1-1   UCUGGACGGGACGUAGCAGGCAUCCUCUCGCGGAGUAGAAACCCUUUUGCGUGCGACCGCGCGAACUACUGAUGGGAACCGCACGACCGCAAAUCCGACUC.GCCGGCC...................................GGUU................................
env-172   CGCGGACGCGAGCGUUCCUGGACGGGAGCACCGAGUAGAAACCGUUUUGUGUGCGACCGCGCGAGAUACCGGUGGGAACCGCACGACCGUAAAUCCGACUC.GCCGGCC...................................GGUU................................
env-173   CGCGCGUUCGGGCGUUCCUCGGGGAAAACGCCGAGUAGAAACCGUUUUGUGUGCGAGUACACGAGGUAGAGACGGGAACCGCACGACCGCAAAUCCGACUC.GCCGGCC...................................GAUU................................
env-174   GCGUCCGUAGACGACGCUUCGGGCAAACGACCGAGUAGAAACCGUUUUGUGUCCGAGCACACGAUGUACGCCCGGGAACCGCACGACCGCAAAUCCGACUC.GCCGGCC...................................GAUU................................
env-175   GCGUCCGUAGACGACGCUUCGGGCAAACGACCGAGUAGAAACCGUUUUGUGUCCGAGCACACGAAGUACGACCGGGAACCGCACGACCGCAAAUCCGACUC.GCCGGCC...................................GAUU................................
env-176   UAGACGUUCUGUAACGAGCCGAUGGCACGUCGGAGUAGAAACCGUUUUGUGUCCGUACCGUCGACGAACGUGUGGGAACAGCACGACCGCAAAUCCGACUC.GUCGGCC...................................GAUU................................
env-177   UAGACGUUCUGUAACGAGCCGGUGGCACGUCGGAGUAGAAACCGUUUUGUGUCCGUGCCGUCGACUUACCCACGGGAACAGCACGACCGCAAAUCCGACUC.GUCGGCC...................................GAUU................................
env-178   GAGCGGGUGAAGCUCCCGAAAAUCGACAGCCGAGUAGAAACCGUUUAAUGUUCGAACGCACGACGUACGCCUGGGAACCUGCACGACCGCAAAUCCGACUC.GCCGGCC...................................GAUU................................
Hsp-6-1   ACCCAGACCGACGAACACAGCGCACGCGAUCGGAGGUAGAAACGCUUUAGUCGCUGACCGAAUAGGUACCAGUGGGAACCGCACGGCCGCAAAUCUGACCCCGUCGCCCG..................................CUCUCUU.............................
Hla-3-1   CCUCUCUACCGAGACCCAUCAUUGCAGAGGCGGAGGUAGAAACCCUUUAGUCGCUCACCGAAUAGAUACCAGUGGGAACCGCACGGCCGCAAAUCUGACCCCGUCGCCCG..................................CUCUCUU.............................
env-179   GCCGAGCAGUGGAGCGACAGCGGCGUCGACAGAGACCGAAAACGGUUACACCCCGCGGCCGCCAGCAACAGACGGGAACAGCACGGCCGCAAAUCUGACUC.GUCCCGCAGGGGA.............................CUCG................................

          ................................................................................<<<<.<<<<<<<..<<......<<<<<<<<<<<<<<<<<<<<<<<<<<<<<<<<<<<<<<<<<<.......................>>>>>>>>>>>>>
          ................................................................................2.....221122..01......0222222.........1.11...10...0000.0..000000.......................000000..0.000
          ••••••••••••••••••••••••••••••••R•R••RR•A•••YUU•••••••••••••••••••••RRYGGAA••GCACRGCCGCAAAUCYGACYY-GYC•∘∘∘------------------------------------•YY∘--------------------------------
                                                                                5′
```

```
env-180   GACCCAGCGGCCGCCACGGUCGGGGUCGACACAGACCGAAAACGGUUACACCCUACGGCCGCCAGCGACAGACGGGAACAGCACGGCCGCAAAUCUGACUC.GUCCCGCAGGGGA.............................CUCG................................
env-181   GUCCGGAUGCUACUCCCCGGGACCGGAACCGUCGAAUGGAAACGGUUAACAGCGGGACAGACCAUACACAGACGGGAACGUACGGCCGCAACUCCGACGU.GCCGCCC...................................GCUU................................
env-182   CGAACUCGGCCCGACAGAGUCUGGGUGGUCACCGAAGGGAAACGAUUAACAGCGCAGACAGACUCACCUAGAGUGGGAACGCACGGUCGUAACUCUGACUC.GCCGUUC...................................GCUU................................
env-183   AAGUACGACAGGGCCAGAGUUCGGACGGUGGCCGAAGGGAAACGGUUAACAGCACAGACAGACUUCCUUCUGAUGGGAACGCACGGUCGUAACUCCGACUC.GCCGUCC...................................GCUU................................
Hsu-1-1   CCAGGGAAUUGUGACCGUCGAGAAACGGUCCGGAUAGUAAGGCUUUACCCUGCGGCCCCAGUUUCUCUCGAUAGGGAACCGCACGACCGCAAAUCUGACUC.GUCCAC....................................GCUC................................
Hha-1-1   GGUCCGUCGGUUAAACCCAUCGAACCCCUCUCGAAGUCGAAACGGUUUAACCCCGCGCACGGUUCCCGUCGAUGGGAACAGCACGGCCACGAAUCCGACUC.GUCCG.....................................GCGG................................
Hru-2-1   CUGCGUCGACGCCACAUCGACGUCGACGCCGCGGAUGGAAAGCCAUUAAGCCCCUCGCCGGUCACUCACCGUUGGGAACAGCACGGCCGCAAAUCUGACUC.GCUCGG....................................ACCGC...............................
env-184   ...................................................................AUGGAUGGGAACAGCACGGCCGCAGAUCUGACUC.GCCC......................................ACGC................................
Hsa-4-1   CGUUUUCAUAAACCCAUCGAACCCGACGCGCCGAGUCGAAACGGUUUAAUCCGGCCCACGGGUGUGGGAGGAUGGGAACAGCACGGCCGCAAAUCUGACAC.GCCC......................................UCGC................................
Hag-1-1   CGAUCGCAUAAACCCAUCGAACCCGACGCACCGAGUCGAAACGGUUUAACCCGGCCCACGGAUGAGUGAGGAUGGGAACAGCACGGCCGCAAAUCUGACUC.GCCC......................................ACGU................................
Hsa-3-1   CCGUCGCAUAAACCCAUCGAACCCGGCGCGUCGAGUCGAAACGGUUUACUGCGCCCCACGGAUGAGUGGGGAUGGGAACAGCACGGCCGCAAAUCUGACUC.GCCC......................................GCAC................................
Hse-1-1   UAUCGACGGCGAUUGGAUAAACCCAUCGUAUCGAGUCGAAACGGUUUACUUCGGUGCGGAGAGUGAUGGAGAUGGGAACAGCACGGCCGCAAAUCCGACUC.GUCU......................................ACGU................................
Hth-1-1   AUCGCCGUCGAUCCCAUAAACCCAUCGUCAUCGAGUCGAAACGGUUUACUGCGGUGCGGGAAGUGUUGGGAGUGGGAACAGCACGGCCGCAAAUCCGACUC.GUCC......................................ACGU................................
Hmo-1-1   AUCGCCGUCUAUCUCAUAAACCCAUCGUCAUCGAGUCGAAACGGUUUACUUCGGUGCGGGAAGUGUUGGGAGUGGGAACAGCACGGCCGCAAAUCCGACUC.GUCC......................................ACGU................................
env-185   GCCCAGUAGGGGAGCGGGUCGCCGGUCGACAGAGACCGAAAACGGUUACACCCGCGGCCGCCAGCAGUAGAUGGGAACAGCACGGCCGCAAAUCUGACUC.GUCCC.GCGGGG..............................GACU................................
Hsp-1-1   UUGAUAAACCCCGCGAUUCGAGUGCCUCGGCGGGAUGGAAACGUUUUACCAGGCGGACCGUCUCCGACGGUGUAGGGAACGCACGGCCGUAGAUCCGACAC.GCCCGCC...................................GGUU................................
          ................................................................................<<<<.<<<<<<<..<<......<<<<<<<<<<<<<<<<<<<<<<<<<<<<<<<<<<<<<<<<<<.......................>>>>>>>>>>>>>
          ................................................................................2.....221122..01......0222222........1.11...10...0000.0..000000.......................000000..0.000
          ••••••••••••••••••••••••••••••••R•R••RR•A•••YUU•••••••••••••••••••••••RRYGGGAA••GCACRGCCGCAAAUCYGACYY-GYC•∘∘∘---------------------------------------•YY∘-------------------------------
                                                                                5′
```

alignment positions 181 · · · 281

```
env-1     ......................GUGUUGC.U.A.GGG.AAGGUGGUAGUGCUGGCAAUAUGCCCACUAUAUAUCAGCUGACGUGUUGUUCCACACAUACUA
env-2     ......................GUGUUGC.U.A.GGG.AAGGUGGUAGUGCUGGCAAUAUGCCCACUAUAUCGGCUGACGUGUUGUUCCACAAAUAGCUAA
env-3     ......................GUGUUGC.U.A.GGG.AAGGUGGUAGUGCUGGCAAUAUGCCCACUAUAUCGGCUGACGUGUUGUUCCACACAUACCAAC
env-4     ......................GUAUUGC.U.A.GGG.AAGGUGGUAGUGCUGGCAAUAUGCCCACUAUAUAUC...........................
env-5     ......................GUGUUGC.U.A.GGG.AAGGUGGUAGUGCUGGCAAUAUGCCCACUAUAUAUCAGCUGACGUGUUGUUCCACACAUACUA
env-6     ......................GUGUUGC.U.A.GGG.AAGGUGGUAGUGCUGGCAAUAUGCCCACUAUAUAUCAGCUGACGUGUUGUUCCACACAUACUA
env-7     ..CGGU.CGUCCGUG.CUCGUUACGUGGC.U.C.GGG.AUUGCGAAAGUGUUUCGGCGGACUGCCGACCAGGCCGGUGGCCCGGUAACGGCCAGCGGUCGG
Hut-1-1   ...............UCGCCGUGCGCGAC.U.U.GGG.AUUGCGGACGUGUCCGGCCGGGAUCGGCGACCCGACCAGUCGGUCGCGUCGGUCGAUCCGGCC
env-8     ........................CCAAC.U.U.GGG.AUUGCGGAAGUGCGGCAGGUCUUCAACUGCCAAGAUAACUUCUGCGGUCAGUGCUGUUCCAUC
env-9     ........GCCACGCUGCCGG.G.GCGGC.U.U.GGG.ACUGCGGUAGUGUCGUGGCACCGUCGGGUGCCGCCAGCGUUUCGUGUAGUUCCAAUCCGAACA
Hti-1-1   ................CGCCGUGCGCGAC.U.U.GGG.AUUGCGGACGUGUCCGGCAGGGAUCGUCGGCCCGACCAGUGGGUCGCGUCGCUCGAUCCGUGC
env-10    ........................UGCGC.U.U.GGG.AUUGCGGCAGUGCAUCCGUCCUGUUUUGCAGAGACGGACGAUAUCACGCGUUCUGCGGUUGUG
env-11    ........................UGCGC.U.U.GGG.AUUGCGGUAGUGCGUUCGGCUGUGCGCUCGUGAUCGAGGGGACAGCCGCUAUCACACAUUCUG
env-12    ...........CGACCCGCCCGG.ACGAC.U.U.GGG.AUUGCGGAAGUGCACCCGGCCACGGGGCCGGACGACCGAGCGGUCAGUGCGAUUCCUAUCCUC
env-13    ....GUCGCCACGCUGCCGG.G..GCGGC.U.U.GGG.ACUGCGGUAGUGUCGUGGCACCGUCGGUGCCGCCAGCGUUUCGUGUAGUUCCAAUCCGAACAA
env-14    ........................GGGGC.U.U.GGG.AUCGCGGCAGUGCGCCGAGAGCGGCGCUUCUGCGGUCGUGCGUUCCAACCACCAAUGGCACGA
env-15    ........................CCGGC.U.U.GGG.AUUGCGGUAGUGCUCGACACACAUGUCGAACUAUCCUGCGGCUCGUGUCGUUCCAAUCACCUA
Hac-1-1   ......................GCGUGGC.U.U.GGG.AGUGUGGCAGUGCUCGGGCGCACCCGCGCCCAGCGCCGCACAGCGGUUCGUGUCGUUCCAAUU
env-16    ......................GGGUGGC.U.U.GGG.AGUGCGGCGGUGCAGGCCGACAGCGGCGCGACGCCGCACCGGCCACGGCAUCGCGGCCCUCGC
Hsa-5-1   .......................GCAGGC.U.U.GGG.AUUACGGAGGUGCGCCGAGAGCCGUUCGUACCGAAUACGCGGCUCUCGGACACGCUUUCUGCG
env-17    ...................CGCCCACGGC.U.U.GGG.AAUGUAGCGGUGCGCCGUCGUCCCCGAUGCGCCGGGCGCGGCCCGUCGGUCGCGCCGAGACGA
env-18    ...................CGCCCACGGC.U.U.GGG.AAUGUAGCGGUGCGCCGUCGUCCCCGAUGCGCCGGGCGCGGCCCGUCGGUCGCGCCGAGACGA
env-19    ....................GCCCGCGGC.U.U.GGG.AAUGUAGCAGUGCACCGACGCCUCCGGCGUCGCG.............................
env-20    ....................GCCCGCGGC.U.U.GGG.AAUGUAGCAGUGCGCCGACGCCUCCGGCGUCGCGGUCGGGCGGUCCGCGCGAGCGGGCCGUCC
Har-2-1   ....................GCCCGCGGC.U.U.GGG.AAUGUAGCAGUGCGCCGACGCCUCCGGCGUCGCGGUCGGGCGGGCCGCUCGCGCGGCCCGUCC
Hsp-3-1   ....................GCCCGCGGC.U.U.GGG.AAUGUAGCAGUGCGCCGACGCCUCCGGCGUCGCGGUCGGGCGGGCCGCUCGCGCGGCCCGUCC
env-21    ....................GCCCGCGGC.U.U.GGG.AAUGUAGCAGUGCGCCGACGCCUCCGG....................................
env-22    ....................GCCCGCGGC.U.U.GGG.AAUGUAGCAGUGCGCCGACGCCUCCGG....................................
env-23    ....................GCCCGCGGC.U.U.GGG.AAUGUAGCAGUGCGCCGACGCCUCCGGCGUCGCGGUCGGGCGGCCCGCUCGCGCGGACCGUCC
Hdi-1-1   ....................GCCCGCGGC.U.U.GGG.AAUGUAGCAGUGCGCCGACGCCUCCGGCGUCGCGGUCGGGCGGGCCGCUCGCGCGGCCCGUCC
Hli-2-1   ....................GCCCGCGGC.U.U.GGG.AAUGUAGCAGUGCGCCGACGCCUCCGGCGUCGCGGUCGGGCGGGCCGCUCGCGCGGCCCGUCC
Hsa-8-1   ....................GCCCACGGC.U.U.GGG.AAUGUAGCAGUGCGCCGACGCCUCCGGCGUCGCGGUCGGACCGUCCGCUUGCGCGGGCCGUCC
env-24    ....................GCCCACGGC.U.U.GGG.AAUGUAGCAGUGCGCCGACGCCUC.......................................
Hha-2-1   ....................GCCCACGGC.U.U.GGG.AAUGUAGCAGUGCGCCGACGCCUCCGGCGUCGCGGUCGGACCGUCCGCUUGCGCGGGCCGUCC
Hko-1-1   ....................GCCCACGGC.U.U.GGG.AAUGUAGCAGUGCGCCGACGCCUCCGGCGUCGCGGUCGGGCCGUCCGCUCGCGCGGACCGUCC
env-25    ....................GCCCACGGC.U.U.GGG.AAUGUAGCAGUGCACCGACGCCUCCGGCGUCGCGGCCGGUCGAUCGUUCACCCGCACGGGGGA
env-26    ....................GCCCACGGC.U.U.GGG.AAUGUAGCAGUGCGCCGACGCCUCCGGCGUCGCGGUCGGACCGUCCGCUCACGCGGGUUGUCC
env-27    ....................GCCCACGGC.U.U.GGG.AACGUAGCAGUGCACCGCCGUCUCCGAC...................................
env-28    ....................GCCCACGGC.U.U.GGG.AACGUAGCAGUGCACCGUCGUCUCCGACGGCGCGACCACGCUGGCCCGCAACGCGCGGACCGA
env-29    ....................GCCCACGGC.U.U.GGG.AACGUAGCAGUGCACCGCCGCCUCCGACGACGGGGUCGAGGCGUCCGCUCGCGCGGAUCGCCU
env-30    ....................GCCCACGGC.U.U.GGG.AACGUAGCAGUGCGCCGCCGUCUCCGACGGCGUGGCUCGAGUCGA..................
env-31    ....................GCCCACGGC.U.U.GGG.AACGUAGCAGUGCGCCGCCGUCUCCGACGGCGUGGCUCGAGUCGA..................
env-32    ....................GCCCACGGC.U.U.GGG.AACGUAGCAGUGCGCCGCCGUCUCCGACGGCGUGGCUCGAGUCGAACACCGUCCACGCGGACC
env-33    ....................GCCCACGGC.U.U.GGG.AAUGUAGCAGUGCACCGACGCCUCCGACGUCGCGGCCGGUCGAACACUCGCCGAACGGGCGAC
env-34    ....................GCCCACGGC.U.U.GGG.AAUGUAGCAGUGCACCGCCGUCUCCGGCGCGUGGCCGGAUCGCUGGUCCGCCCACGGGAACGA
env-35    ....................GCCCACGGC.U.U.GGG.AAUGUAGCAGUGCACCGCCGUCUCCGGCGCGUGGCCGGAUCGCGGGUCCGCCCACGGGAACGA
env-36    ....................GCCCACGGC.U.U.GGG.AAUGUAGCAGUGCACCGCCGUCUCCGGCGCGUGGCCGGAUCGCGGGUCCGCCCACGGGAACGA
env-37    ....................GCCCACGGC.U.U.GGG.AAUGUAGCAGUGCACCGCCGUCUCCGGCGCGUGGC............................
env-38    ....................GCCCACGGC.U.U.GGG.AAUGUAGCAGUGCACCGACGCCUCCGGCGUCGCGGCCGGUCGAUCGUUCACCCGAACGGGGGA
env-39    ......................CCACGGC.U.U.GGG.AACGUAGCAGUGCGCCGCCGUCGCCGCCCCUGCGGCCACCGC.....................
env-40    ......................CCACGGC.U.U.GGG.AACGUAGCAGUGCGCCGCCGUCGCCGCCUCACAGCCCGUGUGGCUCGUAUCGACGAGCCGUCU
env-41    ......................CCACGGC.U.U.GGG.AACGUAGCAGUGCGCCGCCGUCGCCGCCUCACAGCCCGUGUGGCUCGUAUCGACGAUCCGUCU
Hsp-5-1   ....................GCCCGCGGC.U.U.GGG.AAUGUGGCAGUGCGCCGACGCCUCCGGCGUCGCGGCCGGGAUGGGUCCGCGCUCGCGAACCCG
Hca-2-1   ....................GCCCGCGGC.U.U.GGG.AAUGUAGCAGUGCGCCGACGCCUCCGGCGUCGCGGUCGGGCGGUCCGCUCGCGCGGAUCGGCC
Hli-1-1   ....................GCCCGCGGC.U.U.GGG.AAUGUAGCAGUGCGCCGACGCCUCCGGCGUCGCGGUCGGACCGUCCGCUCGCGCGGAUCGUCC
Hte-1-1   ....................GCCCGCGGC.U.U.GGG.AAUGUAGCAGUGCGCCGACGCCUCCGGCGUCGCGGUCGGGCGGUCCGCUCGCGCGGGCCGGCC
Hho-1-1   ....................GCCCGCGGC.U.U.GGG.AAUGUAGCAGUGCGCCGACGCCUCCGGCGUCGCGGUCGGGCGGUCCGCUCGCGCGGGCCGUCC

          >>>>>>>>>>>>>>>>>>>>>>>>>>>>>......>>..>>>>>>>.>>>>..................................................
          0...01...11.1.........2222220......10..221122.....2..................................................
          ----------------------∘∘••GGC-U-U-GGG-AUUGCGG•AGUGY••••••••••••••••••••••••••••••••••••••••••••••••••
                                                   3′
```

```
Hsa-6-1   ....................GCCCACGGC.U.U.GGG.AAUGUAGCAGUGCGCCGACGCCUCCGGCGUCGCGGUCGGAGCGCCCGCUCGCACGGCCGUCC
Hco-1-1   ....................GCCCGCGGC.U.U.GGG.AAUGUAGCAGUGCGCCGACGCCUCCGGCGUCGCGGUCGGGUGGUCCGCUCGCGCGGACCGUCC
Hai-1-1   ....................GCCCGCGGC.U.U.GGG.AAUGUAGCAGUGCACCGACGCCUCCGGCGUCGCGGCCCGGUCUGUCCGCUCGCGCGGACCGAC
env-42    ....................GCCCGCGGC.U.U.GGG.AAUGUAGCAGUGCACCGACGCCUCCGGCGUCGCGGCUUCGGACGGUCCGCUCGCGCGGGCCGU
env-43    ....................GCCCACGGC.U.U.GGG.AACGUAGCAGUGCGCCGCCGUCUCCGGCGUCGCGGCCGGAGCCGCUCGCGUUCGCGAGCCGCU
env-44    ......................CCACGGC.U.U.GGG.AUUGUAGCAGUGCGCCGACGUCUCCCGCUACGCGCUCGCGAUGCCGGCGUUCGCCGGCGUCUC
env-45    ......................CCACGGC.U.U.GGG.AAUGUAGCAGUGCACCGACGCCUCCGGCGUCGUGGUUGGGCGGUCCGCUCGGGCGGGUCUCCC
env-46    ......................GCACGGC.U.U.GGG.AAUGCGGCAGUGCAGUCGGCGCCGAGACGGCGCCACGCAUCAGCGGCUCGUGUUCGUUCCAAU
Hpa-1-1   ......................UCACGGC.UCU.GGG.AUUGCGGACGUGCAACGUGACUCAUGUCACGACUGGCGGAACCCACCGCCAGCACAGCGGCUC
Hci-1-1   ..GGGCAUUUGCCCUCU...UUGCGCAAC.U.U.GGG.AUUGCGGACGUGCACCGUGGCAAUCGCCACGCAGGCGGAAUUUCCGCCGACAACACUCUAGCG
env-47    .....................CGUCGGGC.UUC.GAG.AUUGCGGAAGUGCGUCGGUAGAUUCGACGCUCUCCUGCGGUCUGUGGGUUCCAAUUCAACCAA
env-48    .......................GCCGAC.U.U.GGG.AUUGCGGAUGUGUCCACCUGCGGAGGGUUGGGAACAUCACACCGCGGUCAGUCGAUCCCGCUC
env-49    .......................GCCGAC.U.G.GGG.AUUGCGGAUGUGUCCACCUGUGGAGGGUUAGGACCAUCACAACCGCGGUCAGUCGAUC.....
env-50    .......................GCCGAC.U.G.GGG.AUUGCGGAUGUGUCCACCUGUGGAGGGUUAGG...............................
env-51    .......................GCCGAC.U.G.GGG.AUUGCGGAUGUGUCCACCUGUAGAGGGCUAGGACCAUCACAACCGCGGUCAGUCGAUCCCGCU
env-52    .......................GCCGAC.U.G.GGG.AUUGCGGAUGUGUCCACCUGUGGAGGGUUAGGACCAUCACAACCGCGGUCAGUCGAUCCCGCU
env-53    .......................GCCGAC.U.U.GGG.AUUGCGGAUGUGUCCACCUGCGAAGGCUUGGGAACAUCGCAACGCGGUCCGUCGAUCCCGCUC
env-54    .......................GCCGAC.U.U.GGG.AUUGCGGAUGUGUCCACCUGCGGAGGGUUGGGAACAUCGCAACGCGGUCAGUCGAUCCCGCUC
env-55    .......................GCCGAC.U.U.GGG.AUUGCGGAUGUGUCCACCUGCGGAGGGUUGGGAACAUCGCAACGCGGUCAGUCGAUCCCGCUC
env-56    .......................GCCGAC.U.U.GGG.AUUGCGGAUGUGUCCACCUGCGGAGGGUUGGGAACAUCGCAACGCGGUCAGUCGAUCCCGCUC
env-57    .......................GCCGAC.U.U.GGG.AUUGCGGAUGUGUCCACCUGCGAAGGGUUGGGAACAUCGCAACGCGGUCCGUCGAUCCCGCUC
env-58    .......................GCCGAC.U.U.GGG.AUUGCGGAUGUGUCCACCUGCGAAGGGUUGGGAACAUCGCAACGCGGUCCGUCGAUCCCGCUC
Hla-2-1   ........................UGGGC.U.U.GGG.AUUGCGGUAGUGCCCCAGCAGUCGCGAGUGCGACUGCGGGGUACUCGUUUCUGCGGUCAGUGC
Hsp-4-1   .........................GGGC.U.U.GGG.AUUGCGGUAGUGCCCCGGCAGUCGGUGACACGACUGCGGGGUACUCGUUUCUGCGGUCAGUGC
Hsp-2-1   .........................GGGC.U.U.GGG.AUUGCGGUAGUGCCCCGGUAGUCGGUGAUACGACUGCGGGGUACUCGUUUCUGCGGUCAGUGC
Hme-1-1   .......................GCGGGC.U.U.GGG.AUUGCGGUAGUGCCCCAGUAGUCGUCAAUGCGACUACGGGGUACUCGUUUCUGCGGUCAGUGC
Hel-1-1   ........................UGGGC.U.U.GGG.AUUGCGGUAGUGCCCCAGCAGUCGCGAGUGCGACUGCGGGGUACUCGUUUCUGCGGUCAGUGC
Hvo-1-1   ........................UGGGC.U.U.GGG.AUUGCGGUAGUGCCCCGGUAGUCGGUGAUACGACUGCGGGGUACUCGUUUCUGCGGUCAGUGC
Hsu-2-1   ........................UGGGC.U.U.GGG.AUUGCGGUAGUGCCCCGGUAGUCGGUGAUACGACUGCGGGGUACUCGUUUCUGCGGUCAGUGC
Hde-1-1   ........................CGGGC.U.U.GGG.AUUGCGGUAGUGCCCCGGUAGUCGGUGAUACGACUGCGGGGUACUCGUUUCUGCGGUCAGUGC
Hpr-1-1   ........................CGGGC.U.U.GGG.AUUGCGGUAGUGCCCCAGCAGUCGGUGAUACGACUGCGGGGUACUCGUUUCUGCGGUCAGUGC
Hgi-1-1   ........................CGGGC.U.U.GGG.AUUGCGGUAGUGCCCCAGCAGUCGGUGAGACGACUGCGGGGUACUCGUUUCUGCGGUCAGUGC
Hal-1-1   ........................CGGGC.U.U.GGG.AUUGCGGUAGUGCCCCAGCAGUCGGUGAUACGACUGCGGGGUACUCGUUUCUGCGGUCAGUGC
Hgi-2-1   ........................CGGGC.U.U.GGG.AUUGCGGUAGUGCCCCAGCAGUCGGUGAGACGACUGCGGGGUACUCGUUUCUGCGGUCAGUGC
Hmu-1-1   .......................GUGGGC.U.U.GGG.AUUGCGGUAGUGCCCCAGUAGUCGUGUAUGCGACUACGGGGUACUCGUUUCUGCGGUCAGUGC
env-59    ......................GUGUGGC.U.U.GGG.AUUGCGGACGUGCGAGUAGACCUCGCUGUAGCGGCUCGUGUUCGUUCAAUCAAUCUAUGGCAC
env-60    ...................UCGUGGUGAC.U.C.GAG.AUUGCGGACGUGCCGUGGCGAGAGCCACGUUCAUACGACGCGGUGCGUGCGGUUCCUAUCCUC
env-61    ............GUCGCCGUCUGGACGAC.U.C.GAG.AUUGCGGAAGUGCACCGGCAGCACC......................................
env-62    ........................ACGGC.U.U.GGG.AUUGCGGCCGUGUUCGGCGCACCAGCGCCAACACCGCAUCGAAACCUGUUGCAGCACAGCGGU
env-63    .....................CGCGCGAC.U.C.GAG.AUUGCAGUAGUGCGAGUGCUCUACUGCGGUCUGUGCGGUUCCUAUCCUCACUAUGGCACGAAU
env-64    .....................CGCACGAC.UCC.GAG.AUUGCAGUAGUGCGAGUGCUCUACUGCGGUCUGUGCGGUUCCUAUCCUCACUAUGGCACGAAU
Has-1-1   ......GGGCGUGAUCUAUCUCGCGCGAC.U.C.GAG.AUUGCGGAAGUGCGCCCGGCACGGAGCCGGAUCUUCGCGGUCAGUGUGGUUCCUAUCCUCAGC
env-65    ..................GCCU.CGCGGC.U.U.GGG.AGUGCGGACGUGCAGCCGGCAGGUACGCCUGCCGUUUCGUUUCGCGGGCCAGUGGCCCGUACG
h3A-1-1   ....................GCCCACGGC.U.C.GGG.AUUGCGGCAGUGCACCGCGAGUCCCCGAUGGGGGGCCCGCGGUCACGCUUUUGCGGUUGUGCC
env-66    ........................UGGGC.UUU.GAG.AUUGCGGAAGUGUACCGACAGAGCCGUCGGUCUUUUCGCGGUCAGUGCCGUUCCAACUCAGAU
env-67    ......................GAGCAGC.U.U.GGG.AUUGCGGUGGUGUCCGCGCCACCAGGCGCAACGCCGGCGGUCCGUAGUUCCAACCUAUCAUGG
Hka-1-1   ......................GCGUGGC.UUC.GAG.AUUGCGGCCGUGUCCGGUGUUUCGACACCACACGCUCGCGCUGCGAGUAGCACGUAGCGGUCU
Hmu-2-1   ......................GCGUGGC.UUC.GAG.AUUGCGGCCGUGUCCGGUGUUUCGACACCACACGCUCGCGCUGCGAGUAGCACGUAGCGGUCU
env-68    ......................CCAUGGC.U.C.GGG.AUUGCGGCCGUGUUCGGCACUGUACGGUGCCAAUACGCCUCGUAACGAGAGCGGUUCGUGCGG
Ham-1-1   ......................UCACAGC.U.C.GGG.AUUGCGGCCGUGUUCGGCACUGUACAGUGCCAAUACGCCUCGUAACGAGAGCGGUUCGUGCGG
Hva-1-1   ......................UCACAGC.U.C.GGG.AUUGCGGCCGUGUUCGGCACUGUACAGUGCCAAUACGCCUCGUAACGAGAGCGGUUCGUGCGG
Hja-1-1   ......................UCACAGC.U.C.GGG.AUUGCGGCCGUGUUCGGCACUGUACAGUGCCAAUACGCCUCGUAACGAGAGCGGUUCGUGCGG
Hma-1-1   ......................UCACAGC.U.C.GGG.AUUGCGGCCGUGUUCGGCACUGUACAGUGCCAAUACGCCUCGUAACGAGAGCGGUUCGUGCGG
Hsa-2-1   ......................UCACAGC.U.C.GGG.AUUGCGGCCGUGUUCGGCACUGUACAGUGCCAAUACGCCUCGUAACGAGAGCGGUUCGUGCGG
Hca-1-1   ......................UCACAGC.U.C.GGG.AUUGCGGCCGUGUUCGGCACUGUACAGUGCCAAUACGCCUCGUAACGAGAGCGGUUCGUGCGG
Hsi-1-1   ......................UCACAGC.U.C.GGG.AUUGCGGCCGUGUUCGGCACUGUACAGUGCCAAUACGCCUCGUAACGAGAGCGGUUCGUGCGG
Har-1-1   ......................UCAUAGC.U.C.GGG.AUUGCGGCCGUGUUCGGCACUGUACAGUGCCAAUACGCCUCGUAACGAGAGCGGUUCGUGCGG
env-69    ......................CCACAGC.U.U.GAG.AUUGCGGCCGUGUUCGGCUCCGUAGCCGAUACGCCUCGCUACACGAGCGGUUCGUGCGGUUCC
env-70    ......................GGGCGGC.UCC.GAG.AUUGCGGCCGUGUUCGGCGACGCACUGGAUGCGACAGAUCCAGGCGUUCGCCAAUCACGCCAG
env-71    ........................ACGGC.U.U.GGG.AUUGCGGAAGUGUGAUACUAGAGUCAAUACUCGUCUCUUAUUUCACCGCUUUUCUGCGGUCUG
env-72    .......................GACGGC.U.U.GGG.AUUGCGGAAGUGUGACACAAGAGCCGUAUCUGAUACUCGUCUCUUAUUUCACCGCUUUUCUGC
env-73    .......................GACGGC.U.U.GGG.AUUGCGGAAGUGUGACACAAGAGCCGUAUCUGAUACUCGUCUCUUAUUUCACCGCUUUUCUGC
env-74    .......................GACGGC.U.U.GGG.AUUGCGGAAGUGUGACACAAGAGCCGUAUCUGAUACUCGUCUCUUAUUUCACCGCUUUUCUGC
env-75    .......................GACGGC.U.U.GGG.AUUGCGGAAGUGUGACACAAGAGCCGUAUCUGACACUCGUCUCUUAUUUCACCGCUUUUCUGC
env-76    ............GGUGUCGGUCGGGCGAC.U.C.GAG.AUUGCGGAAGUGCAGGGCCAGCAGACACUGCGAGGCCGUGGCGGUCAGUGCGAUUCCUAUCCU
Hca-3-1   ......................GCACAGC.U.C.GGG.AUUGCGGCCGUGUCCGCGUCGAUCGGGCUUCGCAUCGCGAAGCGCGCGGCGCACACACGCACG
env-77    ......................GGGCGGC.U.U.GGG.AUUGCGGAAGUGUUCAGCUCAGCCGGAGCCACGAUGGCGCGGCUGAGACGUGACUGAUAUCAU
env-78    ......................GGACGGC.U.U.GGG.AUUGCGGAAGUGUACAACCAGCCGCUGCCGCGGUGGUGCGGCUGAAUCGCGACUUUGCUGACA
env-79    ......................GGGCGGC.U.U.GGG.AUUGCGGAAGUGUUCAAUCCAGCCGCAGCUACUGUAGCGCGGCUGGAACGUGACUGAUAUCAU
env-80    ......................GGGCGGC.U.U.GGG.AUUGUGGUAGUGUCGGCCGACAGCCCGUGUUCCUCACGGAGGGCGGGCGAGGCGGCCACAUCG
env-81    ......................GGGCGGC.U.U.GGG.AUUGUGGGAGUGUCGGCCGACAGCCGCUGUCCCGGGAGACCGGUACAGCCGGAAACACGGCCA
Hsa-1-1   ......................GGGCGGC.UCC.GGG.AUUGCGGACGUGUCCGGCGUCACGCGCCGACCGCAGCCCUGCUGCAGACGCGCGAAUGAUAGU
env-82    ......................GUGCGGC.U.C.GGG.AUUGCGGCCGUGUUCGCGCCACCGCGCGCAUCGCGGCGCCGCCGUCUGGCGGACAGCGGUACG
env-83    ......................GGGCGGC.U.C.GGG.AUUGCGGUAGUGUCACCCGCCGACCGUUGGCUGUCUCUGGGUUCUCGCCCAGCGCCAGCAGUC
env-84    ......................GGGCGGC.U.C.GGG.AUUGCGGUGGUGUCGACCGUCGACCGGAGUCCCACCCCACGUGGACCGGGACUCUCGCGGUCG
env-85    ......................GGGCGGC.U.U.GGG.AUUGCGGGAGUGUCGGCCAAUCGCCAGUCGUCCGUGCGUGGCUUCCACGCCCUGGACGACCGG
env-86    ......................GGAUGGC.U.U.GGG.AUUGCGGGAGUGUCGGCCGACGCUUUCGUCCGCUCUCGGGAUUGCCGGGUUCGGGACGAAAGU
env-87    ......................CGACGGC.U.C.GGG.AUUGCGGUAGUGUCGACCGCUGACCGCCGACGUCCGGCAGAGACACCUGUCUCUGGAGGCGUC
env-88    .......................GGCGGC.U.U.GAG.AUUGCGGUGGUGUCACACACACCGUGUGGUGACGUCCGCCCGGACGUCGAGUAGUCACCAGUG
env-89    .......................GGCGGC.U.C.GGG.AUUGCGGUGGUGUCGCGCGAGAGCGCGGUGGCGUCCGCCUGGACGCCGAUCACCGAGCGGUCC
env-90    .....................UGGGUGGC.U.U.GGG.AUUGCGAUAGUGCGGCCCGACCGCGAGCCCCGCCUGGCGGGUGGGGUGGCGGUCGGAAGACUA
env-91    .....................CGGGUGGC.U.U.GGG.AUUGCGAUAGUGCGGGCCCGACCGCGAGCCCUGUCUGCAGGAUGGGGUGGCGGUCGGGGAGCU
env-92    ..........................GGC.U.U.GGG.AUUGCGACAGUGCGCCACCAGCCGCGGCAGACGCACGGUCGACGCUGUCGACCCCGUCCGUGC

          >>>>>>>>>>>>>>>>>>>>>>>>>>>>>......>>..>>>>>>>.>>>>..................................................
          0...01...11.1.........2222220......10..221122....2..................................................
          --------------------○○●●GGC-U-U-GGG-AUUGCGG●AGUGY●●●●●●●●●●●●●●●●●●●●●●●●●●●●●●●●●●●●●●●●●●●●●●●●●●
                                                        3′
```

```
env-93    ........................UGGGC.U.U.GGG.AUUGCGAUAGUGCGCCGGUACCCCGUGAGCUGAUCGAUCGUAGUUCGAUCGUGAGCCUCGGGA
env-94    .........................GGGC.U.U.GGG.AUUGCGACAGUGCGCUGCCAGCCCCCGGUGGCGGUCGACCGACGGUCGACCCUGCCGCGGGAC
env-95    .........................GGGC.U.U.GGG.AUUGCGACAGUGCGCCACCAGCCGGACGGGGGGUGUGGCCCCCGGGCCAGCCGUCCGUUUCGG
env-96    .........................GGGC.U.U.GGG.AUUGCGACAGUGCGCCACCAGCCCGCGACGACCAGUUGGUCGUCUCUGGUCUGUGUGGACACG
env-97    .........................GGGC.U.U.GGG.AUUGCGACAGUGCGCCACCAGCCCGCGACGAUCAGUUGGUCGUCUCUGGGCUGUGUGGACACG
env-98    .........................GGGC.U.U.GGG.AUUGCGACAGUGCGCCACCAGCCGGACGGGGGGUGUGGCCCCCGGGCCAGCCGCCCCUCUCGG
env-99    .........................GGGC.U.U.GGG.AUUGCGACAGUGCGCCACCAGCCCGCGACGAUCAGUUGGUCGUCUCUGGGCUGUGUGGACACG
env-100   .........................GGGC.U.U.GGG.AUUGCGACAGUGCGCUGCCAGCCCCCGGUGGCGGUCGACUGCUGUCGACCCUGCUGUGGGACU
env-101   .........................GGGC.U.U.GGG.AUUGCGACAGUGCGCUGCCAGCCCCCGGUGGCGGUCGACCCGACGGUCGACCCUGCCGCGGGA
env-102   .........................GGGC.U.U.GGG.AUUGCGACAGUGCGCCACCAGCCCCAGCAGGCGGACCAUCGGUCCGUGAGACAGCAGUGCUGU
env-103   .........................GGGC.U.U.GGG.AUUGCGACAGUGCGCCACCAGCCCCAGCAGGCGGACCCUCGGUCCGUAAGACAGCUGUGCCGU
env-104   .........................GGGC.U.U.GGG.AUUGCGACAGUGCGCUGCCAGCCCCCGGUGGCGGUCGACACCGCAGUCGACCCUGCCGCGGGA
env-105   .........................GGGC.U.U.GGG.AUUGCGACAGUGCGCUGCCAGCCCCCGGUAGCGGUCGACACAGCAGUCGACCCUGCCGCGGGA
env-106   ........................GGGGC.U.U.GGG.AUUGCGACAGUGCGCCGCCAGCCGCAGCAGGCGUGGGUCGACGCUGUCGACCACCUGUGGGGC
env-107   ........................GGGGC.U.U.GGG.AUUGCGACAGUGCG.................................................
env-108   ........................GGGGC.U.U.GGG.AUUGCGACAGUGCGCCGCCAGCCGCAGC...................................
env-109   ........................GGGGC.U.U.GGG.AUUGCGACAGUGCGCCGCCAGCCGCAGCAGGCGUGGGUCGACACUGUCGA.............
env-110   ........................GGGGC.U.U.GGG.AUUGCGACAGUGCGCCGCCAGCUGCAGCAGGCGUGCGGUCGAGACUGUCGACCACCUGUGCGG
env-111   ........................GGGGC.U.U.GGG.AUUGCGACAGUGCGCCGCCAGCCGCAGCAGGCGUUGGGUCGAUGCUGUCGAUCCCGUCCGUGC
env-112   ........................GGGGC.U.U.GGG.AUUGCGACAGUGCGCCGCCAGCCGCAGCAGGCGUGGGUCGACGACUGUCGACCACCUGUGCGG
env-113   ........................GGGGC.U.U.GGG.AUUGCGACAGUGCGCCGCCAGCCGCAGCAGGCGUGGGUCGACGCUGUCGACCACCUGUGCGGC
env-114   ........................GGGGC.U.U.GGG.AUUGCGACAGUGCGCCGCCAGCCGCAGCAGGCGUGCGGUCGAGACUGUCGACCACCUGUGCGG
env-115   ........................GGGGC.U.U.GGG.AUUGCGACAGUGCGCUGUCAGCACCCGGUGUCGG.............................
env-116   ........................GGGGC.U.U.GGG.AUUGCGACAGUGCGCCGCCAGCCGCAGCAGGCGUGGGUCGACGACUGUCGACCACCUGUGCGG
env-117   ........................GGGGC.U.U.GGG.AUUGCGACAGUGCGCCGCCAGCCGCAGCAG.................................
env-118   ........................GGGGC.U.U.GGG.AUUGCGACAGUGCGCUCACAGCGCCCGGUGGCGGUCGAGCAGUCGACCGUCUCACGGGACUGC
env-119   ........................GGGGC.U.U.GGG.AUUGCGACAGUGCGCCACCAGCCGCA.....................................
env-120   ........................GGGGC.U.U.GGG.AUUGCGACAGUGCGCCGCCAGCCGCAGCAGGCGUGGG..........................
env-121   ........................GGGGC.U.U.GGG.AUUGCGACAGUGCGCUGUCAGCCUCUGGUCGCGCUCGACCCGCCAGUCGACCGACCCGGGAGC
env-122   ........................GGGGC.U.U.GGG.AUUGCGACAGUGCGCUGUUAGCCUCC.....................................
env-123   ........................GGGGC.U.U.GGG.AUUGCGACAGUGCGCCGCCAGCCGCAGCAGGCGUGGGUCGACACUGUCGACCACCUGUGCGGC
env-124   ........................GGGGC.U.U.GGG.AUUGCGACAGUGCGCCGCCAGCCGCAGCAGGUGUGGGUCGACGCUGUCGACCACCUGUGCGGC
env-125   ........................GGGGC.U.U.GGG.AUUGCGACAGUGCGCCACCAGCCGCGGCAGACGCACGGUCGACGCUGUCGACCGCGUCCGUCU
env-126   ........................GGGGC.U.U.GGG.AUUGCGACAGUGCGCCACCAGCCGCGGCAGACGCACGGUCGACGCUGUCGACCACCUGUGCGG
env-127   ........................GGGGC.U.U.GGG.AUUGCGACAGUGCGCUGCCAGCCCCCGGUGGCGGUCGACACGGGAGUCGACCCUGCCGCGGGA
env-128   ........................GGGGC.U.U.GGG.AUUGCGACAGUGCGCUCACAGCCCCCGGUGGCGGUCGACCCACCAGUCGGCCGUCCCGCGGGA
env-129   ........................GGGGC.U.U.GGG.AUUGCGACAGUGCGCCGCCAGCCGCAGCAG.................................
env-130   ........................GGGGC.U.U.GGG.AUUGCGACAGUGCGCCACCAGCCGCGGCAGACGCACGGUCGACGCUGUCGACCCCGUCCGUGC
env-131   ........................GGGGC.U.U.GGG.AUUGCGACAGUGCGCUCACCAGCCNCNGNNGNCGGUCGACCCACCAGUUGGCCGUCUCGCGGG
env-132   ........................GGGGC.U.U.GGG.AUUGCGACAGUGCGCUGUGAGCCCUUGGUCGAGUACUCGACCAGCCGAUGGGCUCGCAGACAC
env-133   .........................GGGC.U.U.GGG.AUUGCGACAGUGCGCCGCCAGCCCGCGGCGACCAGUCUGUCGUCUCGGGCUGUGUGGACACGU
env-134   .........................GGGC.U.U.GGG.AUUGCGACAGUGCGCCACCAGCCCGUGGCGACCAGUCUGUCGUCUCUGGGCUGUGUGGACACG
env-135   .........................GGGC.U.U.GGG.AUUGCGACAGUGCGCCACCAGCCCGCAGCGACUAGUUGGUCGUCUCUGGGCGGUGUGGACACG
env-136   .........................GGGC.U.U.GGG.AUUGCGACAGUGCGCCGCCAGCCCGCGGCGACCAGUCUGUCGUCUCGGGCUGUGUGGACACGU
env-137   .........................GGGC.U.U.GGG.AUUGCGACAGUGCGCCACCAGCCCGCGGCGACUAGUUGGUCGUCUCUGGGCUGUGUGGACACG
env-138   .........................GGGC.U.U.GGG.AUUGCGACAGUGCGCCACCAGCCCGCGGCGACUAGUUGGUCGUCUCUGGGCUGUGUGGACACG
env-139   .........................GGGC.U.U.GGG.AUUGCGACAGUGCGCCACCAGCCGUUGGCGUCUCGCCGAGUCGGUGGACGCGACGGCCGCGUG
env-140   .........................GGGC.U.U.GGG.AUUGCGACAGUGCGCCACCAGCCCGCAGCGACCAGUUAGUCGUUUUUGGGCGGUGUGGACACG
CHa-1-1   ......................GCGAGGCUU.C.GGG.AUUGCGGAGGUGCCCCCGUCGGCCAGCCGCGGGACGCCGUACGGCGUCACAGCGGCGGUACGG
env-141   ......................UGGUGGC.UUC.GAG.AUUGCGGCAGUGCACGGUGAGAGCCGGAUCUGCGAUGCGGUCAGUGCGUUUCCUAUCCUCAUA
env-142   .......................GGCGAC.U.U.GGG.AUUGCGGAAGUGCACCCGGCACAGUGCCGGAUACAUCACCUCGCGGUCAGUGCGGUUCCUAUC
Nsu-1-1   .....................CGGUUGGC.U.U.GGG.AUUGCGGAAGUGCACCGACAGUCGGUCGAGCCGACUGUUACGUACGGAGCGGUCAGUGCGGUU
Hla-1-1   .....................UGGGUGGC.U.U.GGG.AUUGCGGAAGUGCACCGCCAGUCGUCGGUCGAUUCCAACGACUGGCGCGCUCUCGUGCGGUCA
env-143   ......................UGGCGAC.U.U.GGG.AUUGCGGAAGUGCAAACCGGCUCGAGGAGGCCAAGACCGACUCGAGCCAGGACCACAGCCGGC
env-144   ......................UGGCGAC.U.U.GGG.AUUGCGGAAGUGCAUAUCGGCUCGAGGUGGCCACUAGGCUUACUCGAGCCAGGACCACGGCCG
env-145   ......................UGGCGGC.U.U.GGG.AUUGCGGAAGUGCACGGCACGCGCCGAUGCUGUCGGCUACCGGUUGUCCCGGCGAUCGCACCG
Hxa-1-1   .......................GGUGAC.U.U.GGG.AUUGCGGAAGUGCACCGGCAGUCGCCCGACCACAUCGGCGACUGUCGCGUCAUCGUGCGGUCA
env-146   .......................GGUGAC.U.C.GAG.AUUGCGGAAGUGCAAACCGGCUCGAGCAGGCUGAUCGACAUGCCGACUCGAGCCAGGACC...
env-147   ......................GCGUGAC.U.C.GGG.AUUGCGGAAGUGCAGUUCGACUCGACGGAACCACCUCGAGUCGGUUUCACGCCAGCGGUCAGU
Ngr-1-1   .......................GGUGAC.U.U.GGG.AUUGCGGAAGUGCAUUGUCAGUCGGUCGUUUCGGUCGACUGGCGCGCUAUCGUGCGGUCAGUG
env-148   ....................GCGGGUGAC.U.U.GGG.AUUGCGGAAGUGCAGCCGGCCAGUGCCGGAACUGUAAGCGUCGCGGUCAGUGCGAUUCCUAUC
env-149   ....................GCGGGUGAC.U.U.GGG.AUUGCGGAAGUGCAGCCGGCCAGUGCCGGAACUGUAAGCGUCGCGGUCAGUGCGAUUCCUAUC
env-150   ......................UGAUGGC.U.U.GGG.AUUGCGGAAGUGCACGGCAAGCGCCGAAUCUGUCGGCUGACAGUCGUCCCGUGGAUCGCGCUG
env-151   ......................GGAUGGC.U.U.GGG.AUUGCGGAAGUGCAUCGACAGCUGUGAGCUGUCUGGAGCGGUCAGUGCGAUUCCUAUCCUCAC
Hdj-1-1   ....................UCGGAUGGC.U.U.GGG.AUUGCGGAAGUGCACCGGCAGUCGGUCGUCCCGACUGUCGCACAUCGAGCGGUCAGUGCGGUU
env-152   ......................GGGCGGC.U.U.GGG.AUUGCGGAAGUGAACCCGGCCCACCGGCCGGAGUAUCGUUGAGCGGUCUGUGCGGUUCCUAUC
env-153   ......................GGGCGGC.U.U.GGG.AUUGCGGAAGUGAACCCGGCCCACCGGCCGGAGUAUCGUUGAGCGGUCUGUGCGGUUCCUAUC
env-154   ......................GGGCGGC.U.U.GGG.AUUGCGGAAGUGAACCCGGCCAGGUGGCCGGUCUAUCGUGUAGCGGUCUGUGCGGUUCCUAUC
env-155   ......................GGGCGGC.U.U.GGG.AUUGCGGAAGUGAACCCGGCCAGGUGGCCGGUCUAUCGUGUAGCGGUCUGUGCGGUUCCUAUC
env-156   ......................GGGCGGC.U.U.GGG.AUUGCGGAAGUGAACCCGGCCCAGCGGCCGGAGUAUUGAAUAUAGCGGUCUGUGCGGUUCCUA
env-157   ......................GGGCGGC.U.U.GGG.AUUGCGGAAGUGAACCCGGCCAGGUGGCCGGUCUAUCGUUUAGCGGUCUGUGCGGUUCCUAUC
env-158   .......................GACGGC.U.U.GGG.AUUGCGGAGGUGCGACCGGCGAGAGCCGCGUUCACGCAGGGCGGUCGGUGCGAUUCCUAUCCU
env-159   ......................GGACGGC.U.U.GGG.AUUGCGGAAGUGCACCCGGCGAGAGCCGGAUUUACGCGACGCGGUCAGUGCGGUUCCUAUCCU
env-160   ......................GGGCAGC.U.U.GGG.AUUGCGGAAGUGCACCCGGCGAGAGCCGGAUUUACACGACGCGGUCAGUGCGAUUCCUAUCCU
Hgo-1-1   ......................GGGUGGC.U.U.GGG.AUUGCGGAAGUGCACCGACAGUCCCCCGUGGACUGUCGCACGUCUCGAGCGGUCAGUGCGGUU
Npa-1-1   ......................GGGUGAC.U.U.GGG.AUUGCGGAAGUGCACGGCGAGAGCCAGUACUGUCGUUCGACAGUACACUGAGCGGUCAGUGCG
Nga-1-1   ......................GGGUGGC.U.U.GGG.AUUGCGGAAGUGCAUGGCGAGAGCCAGUACUGUCGUUCGACAGUAUACUGAGCGGUCAGUGCG
Nsp-1-1   ......................GGGUGGC.U.U.GGG.AUUGCGGAAGUGCAUGGCGAGAGCCAGUACUGUCGUUCGACAGUAUACUGAGCGGUCAGUGCG
Nal-1-1   ......................GGGUGGC.U.U.GGG.AUUGCGGAAGUGCACGGCGAGAGCCAAUACUGUCGUUCGACAGUAUACUGAGCGGUCAGUGCG
env-161   ......................GGGUGGC.U.U.GGG.AUUGCGGAAGUGCACGGCGAGAGCCACGUACUGUCGGACAGCGCGACUCCCGACGCGGGAGCA
Hli-3-1   ......................GGGUGGC.U.U.GGG.AUUGCGGAAGUGCACGGCGAGAGCCGGCUGCUGUCGGCGGACACGCUGCCAUCGGUGCGCGUG
Nve-1-1   ......................GGGUGGC.U.U.GGG.AUUGCGGAAGUGCACGGCGAGAGCCAGAUGUUGUCUUCGACAGCAUCCCGAGCGGUCAGUGCG
Npe-1-1   ......................GGGCGGC.U.U.GGG.AUUGCGGAAGUGCAGACGAGAGCAGUACAACGCAGCGGUCAGUGCGAUUCCUAUCCUCAACAA

          >>>>>>>>>>>>>>>>>>>>>>>>>>>>>......>>..>>>>>>>.>>>>..................................................
          0...01...11.1.........2222220......10..221122.....2..................................................
          ---------------------○○●●GGC-U-U-GGG-AUUGCGG●AGUGY●●●●●●●●●●●●●●●●●●●●●●●●●●●●●●●●●●●●●●●●●●●●●●●●●●
                                                        3′
```

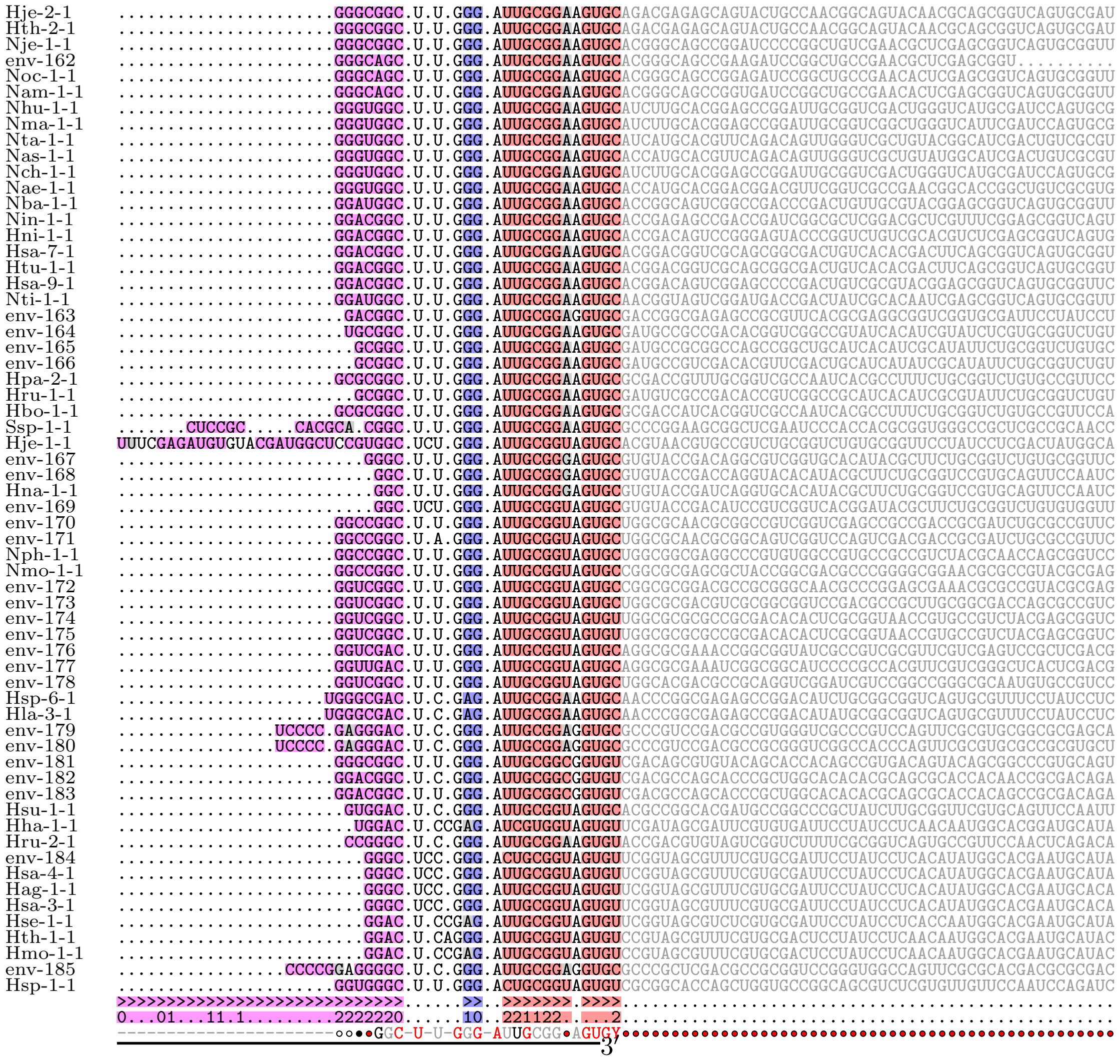

### 21 S15-Methanobacteria

#### 21.1 Taxa

The taxonomy of each organism containing a putative S15-Methanobacteria RNA is listed, with abbreviations identifying each hit (e.g., "Eco-1-1" and "Eco-1-2" might hypothetically represent two distinct RNAs in *E. coli*). The abbreviations will be used to identify each individual S15-Methanobacteria RNA in Sections 21.2 and 21.4:.

| abbrev. of hits | taxonomy of species |
|---|---|
| CMe-1-1 | Archaea Euryarchaeota Methanomicrobia Methanomicrobiales Generaincertae sedis *Candidatus* Methanoregula boonei 6A8 |
| Mpa-1-1 | Archaea Euryarchaeota Methanomicrobia Methanomicrobiales Generaincertae sedis *Methanosphaerula palustris* E1-9c |
| Mba-1-1 | Archaea Euryarchaeota Methanomicrobia Methanomicrobiales Methanocorpusculaceae *Methanocorpusculum bavaricum* DSM 4179 |
| Mla-1-1 | Archaea Euryarchaeota Methanomicrobia Methanomicrobiales Methanocorpusculaceae *Methanocorpusculum labreanum* Z |
| Mbo-1-1 | Archaea Euryarchaeota Methanomicrobia Methanomicrobiales Methanomicrobiaceae *Methanoculleus bourgensis* MS2 |
| Mma-1-1 | Archaea Euryarchaeota Methanomicrobia Methanomicrobiales Methanomicrobiaceae *Methanoculleus marisnigri* JR1 |
| Mse-1-1 | Archaea Euryarchaeota Methanomicrobia Methanomicrobiales Methanomicrobiaceae *Methanoculleus sediminis* |
| Msp-1-1 | Archaea Euryarchaeota Methanomicrobia Methanomicrobiales Methanomicrobiaceae *Methanoculleus sp.* MH98A |
| Mli-1-1 | Archaea Euryarchaeota Methanomicrobia Methanomicrobiales Methanomicrobiaceae *Methanofollis liminatans* DSM 4140 |
| Mpa-2-1 | Archaea Euryarchaeota Methanomicrobia Methanomicrobiales Methanomicrobiaceae *Methanolacinia paynteri* |
| Mmo-1-1 | Archaea Euryarchaeota Methanomicrobia Methanomicrobiales Methanomicrobiaceae *Methanomicrobium mobile* BP |
| Mli-2-1 | Archaea Euryarchaeota Methanomicrobia Methanomicrobiales Methanomicrobiaceae *Methanoplanus limicola* DSM 2279 |
| Mpe-1-1 | Archaea Euryarchaeota Methanomicrobia Methanomicrobiales Methanomicrobiaceae *Methanoplanus petrolearius* DSM 11571 |
| Mta-1-1 | Archaea Euryarchaeota Methanomicrobia Methanomicrobiales Methanoregulaceae *Methanolinea tarda* NOBI-1 |
| Mfo-1-1 | Archaea Euryarchaeota Methanomicrobia Methanomicrobiales Methanoregulaceae *Methanoregula formicica* SMSP |
| env-1 to env-179 | environmental samples |

#### 21.2 Gene contexts

Each S15-Methanobacteria RNA (indicated by "RNA→") is listed. For each hit, the downstream genes predicted to reside in a regulated operon are listed. If the nearest downstream gene is encoding in the opposite strand (and therefore presumed to not be a part of a regulated operon), then that gene is still depicted. Some environmental sequences and some RefSeq entries lack gene annotations, and so no genes are available for such sequences. The direction of each gene is indicated with an arrow (→), and each predicted conserved domain in the gene is named. Conserved domains associated with more than one S15-Methanobacteria RNA are assigned a color; other domains are gray. Information about these conserved domains is given in Section 21.3. The accession of the sequence containing each S15-Methanobacteria RNA is given in the column named "Seq. accession". Accessions beginning with "NC_", "NS_", "NW_" or "NZ_" are contained in RefSeq. Other accession refer to environmental samples. Nucleotide coordinates are given for the 5′ and 3′ boundaries of each S15-Methanobacteria RNA. If the 5′ coordinate is greater than the 3′ coordinate, the RNA is present on the reverse-complement strand of the containing genomic DNA sequence. Each hit is denoted by an abbreviation (like "Eco-1-1") that refers to a taxonomy given in Section 21.1.

| abbrev. | Seq. accession | | 5′ at | 3′ at | genes |
|---|---|---|---|---|---|
| env-1 | RUMENNODE_3956981_1 | - | 89863 | 89776 | **RNA**→ rps15p (PRK08561)Ribosomal_S15p_S13e (cd00353)→ RecJ (COG0608)recJ (TIGR00644)→ |
| Mmo-1-1 | NZ_JOMF01000007.1 | - | 133803 | 133716 | **RNA**→ rps15p (PRK08561)Ribosomal_S15p_S13e (cd00353)→ RecJ (COG0608)recJ (TIGR00644)→ PRK14888 (PRK14888)→ PRK04057 (PRK04057)Ribosomal_S3Ae (pfam01015)→ hypo→ |
| env-2 | JGI24731J21663_1004595 | + | 6197 | 6276 | **RNA**→ |
| env-3 | JGI24731J21663_1017363 | - | 1544 | 1465 | **RNA**→ rps15p (PRK08561)Ribosomal_S15p_S13e (cd00353)→ RecJ (COG0608)recJ (TIGR00644)→ |
| env-4 | JGI24732J26686_1057961 | - | 395 | 316 | **RNA**→ Ribosomal_S13_N (pfam08069)→ |
| env-5 | JGI24728J21555_1000004 | - | 136779 | 136704 | **RNA**→ ←hypo |
| env-6 | 3300000558_11395505 | + | 169 | 244 | **RNA**→ Ribosomal_S13_N (pfam08069)→ |
| env-7 | JGI24729J20445_1005959 | - | 2081 | 2005 | **RNA**→ RpsO (COG0184)rps15p (PRK08561)→ RecJ (COG0608)recJ (TIGR00644)→ PRK14888 (PRK14888)→ |
| env-8 | SL_5KL_010_SED_10017420 | - | 502 | 427 | **RNA**→ Ribosomal_S13_N (pfam08069)→ |
| env-9 | SL_3KL_010_SED_11192733 | + | 39 | 114 | **RNA**→ hypo→ |
| env-10 | SL_3KL_010_SED_10014053 | - | 606 | 533 | **RNA**→ RpsO (COG0184)rps15p (PRK08561)→ |
| env-11 | SL_3KL_010_SED_10006919 | - | 1903 | 1830 | **RNA**→ RpsO (COG0184)rps15p (PRK08561)→ RecJ (COG0608)A-tRNA_syn_arch (TIGR03683)→ |
| Mpa-1-1 | NC_011832.1 | - | 2788986 | 2788870 | **RNA**→ rps15p (PRK08561)Ribosomal_S15p_S13e (cd00353)→ RecJ (COG0608)recJ (TIGR00644)→ PhnD (COG3221)PRK14888 (PRK14888)→ PRK00960 (PRK00960)Gly_His_Pro_Ser_Thr_tRS_core (cd00670)→ PRK04057 (PRK04057)Ribosomal_S3Ae (pfam01015)→ |
| env-12 | DHTCA2_GJDNVXK02I5RIS | - | 305 | 187 | **RNA**→ hypo→ rps15p (PRK08561)→ |
| env-13 | BDMC2_contig04064 | - | 1051 | 974 | **RNA**→ RpsO (COG0184)rps15p (PRK08561)→ RecJ (COG0608)recJ (TIGR00644)→ hypo→ |
| env-14 | 2205091243 | + | 132 | 214 | **RNA**→ |
| env-15 | 2204637606 | + | 116 | 198 | **RNA**→ rps15p (PRK08561)→ |
| env-16 | JGI20214J14112_1457962 | + | 22 | 99 | **RNA**→ hypo→ rps15p (PRK08561)→ |

| | | | | | |
|---|---|---|---|---|---|
| env-17 | SakLake120mDRAFT_c0001602 | + | 6343 | 6421 | **RNA**→ rps15p (PRK08561)Ribosomal_S15p_S13e (cd00353)→ RecJ (COG0608)recJ (TIGR00644)→ PRK14888 (PRK14888)→ PRK00960 (PRK00960)Gly_His_Pro_Ser_Thr_tRS_core (cd00670)→ PRK04057 (PRK04057)Ribosomal_S3Ae (pfam01015)→ |
| env-18 | JGI24729J20445_1013682 | - | 766 | 687 | **RNA**→ RpsO (COG0184)rps15p (PRK08561)→ RecJ (COG0608)recJ (TIGR00644)→ |
| env-19 | JGI24730J26740_1000033 | + | 32652 | 32731 | **RNA**→ RpsO (COG0184)rps15p (PRK08561)→ RecJ (COG0608)recJ (TIGR00644)→ PRK14888 (PRK14888)→ PRK04057 (PRK04057)Ribosomal_S3Ae (pfam01015)→ |
| env-20 | 3300000558_10042858 | - | 212 | 133 | **RNA**→ ←hypo |
| env-21 | 3300001592_10223493 | + | 286 | 365 | **RNA**→ Ribosomal_S13_N (pfam08069)→ |
| env-22 | 3300000032_0002816 | - | 6001 | 5922 | **RNA**→ rps15p (PRK08561)Ribosomal_S15p_S13e (cd00353)→ RecJ (COG0608)recJ (TIGR00644)→ PRK14888 (PRK14888)→ PRK00960 (PRK00960)Gly_His_Pro_Ser_Thr_tRS_core (cd00670)→ PRK04057 (PRK04057)Ribosomal_S3Ae (pfam01015)→ iPGM_like (cd16011)PhosphMutase (pfam10143)→ hypo→ |
| env-23 | M940CN_1015758 | - | 1243 | 1152 | **RNA**→ rps15p (PRK08561)Ribosomal_S15p_S13e (cd00353)→ RecJ (COG0608)recJ (TIGR00644)→ |
| Mpe-1-1 | NC_014507.1 | + | 2024969 | 2025060 | **RNA**→ rps15p (PRK08561)Ribosomal_S15p_S13e (cd00353)→ RecJ (COG0608)recJ (TIGR00644)→ PRK14888 (PRK14888)→ PRK04057 (PRK04057)Ribosomal_S3Ae (pfam01015)→ hypo→ hypo→ BluB (TIGR02476)Nitroreductase (pfam00881)→ PRK03922 (PRK03922)DUF555 (pfam04475)→ ribokinase_group_A (cd01942)PTZ00247 (PTZ00247)→ COG3356 (COG3356)DUF2070 (pfam09843)→ |
| Mpa-2-1 | NZ_KN360943.1 | + | 84264 | 84354 | **RNA**→ rps15p (PRK08561)Ribosomal_S15p_S13e (cd00353)→ RecJ (COG0608)recJ (TIGR00644)→ PRK14888 (PRK14888)→ PRK04057 (PRK04057)Ribosomal_S3Ae (pfam01015)→ hypo→ hypo→ BluB (TIGR02476)Nitroreductase (pfam00881)→ PRK03922 (PRK03922)DUF555 (pfam04475)→ ribokinase_group_A (cd01942)D_ribokin_bact (TIGR02152)→ COG3356 (COG3356)DUF2070 (pfam09843)→ |
| env-24 | SakLake120mDRAFT_c0582226 | - | 239 | 158 | **RNA**→ Ribosomal_S13_N (pfam08069)→ |
| env-25 | 3300001567_10013045 | - | 512 | 429 | **RNA**→ Ribosomal_S13_N (pfam08069)→ |
| env-26 | BDMC2_GDQ3S3O02JCPBT_left | - | 164 | 85 | **RNA**→ |
| env-27 | WSSedA1Ba3_1085468 | - | 477 | 405 | **RNA**→ Ribosomal_S15 (pfam00312)Ribosomal_S13_N (pfam08069)→ |
| env-28 | JGI20214J14112_1152162 | + | 106 | 178 | **RNA**→ Ribosomal_S13_N (pfam08069)→ |
| env-29 | 3300001567_10002105 | + | 3402 | 3481 | **RNA**→ RpsO (COG0184)rps15p (PRK08561)→ RecJ (COG0608)recJ (TIGR00644)→ PRK14888 (PRK14888)→ PRK00960 (PRK00960)Gly_His_Pro_Ser_Thr_tRS_core (cd00670)→ PRK04057 (PRK04057)Ribosomal_S3Ae (pfam01015)→ |
| env-30 | JGI20214J14112_1280193 | + | 182 | 255 | **RNA**→ |
| env-31 | WSSedB1B2_c063428 | - | 318 | 245 | **RNA**→ Ribosomal_S13_N (pfam08069)→ |
| env-32 | SL_5KL_010_SED_10000187 | + | 25577 | 25650 | **RNA**→ rps15p (PRK08561)Ribosomal_S15p_S13e (cd00353)→ RecJ (COG0608)→ PRK14888 (PRK14888)→ PRK04057 (PRK04057)Ribosomal_S3Ae (pfam01015)→ apgM (TIGR00306)iPGM_like (cd16011)→ COG4801 (COG4801)glmU (TIGR01173)→ hypo→ |
| env-33 | JGI24713J26584_10001722 | - | 13168 | 13096 | **RNA**→ hypo→ rps15p (PRK08561)Ribosomal_S15p_S13e (cd00353)→ RecJ (COG0608)recJ (TIGR00644)DHHA1 (pfam02272)→ PRK14888 (PRK14888)→ PRK04057 (PRK04057)Ribosomal_S3Ae (pfam01015)→ |
| env-34 | JGI24714J26587_10001783 | - | 13168 | 13096 | **RNA**→ PRK11160 (PRK11160)→ rps15p (PRK08561)Ribosomal_S15p_S13e (cd00353)→ RecJ (COG0608)recJ (TIGR00644)DHHA1 (pfam02272)→ PRK14888 (PRK14888)→ PRK04057 (PRK04057)Ribosomal_S3Ae (pfam01015)→ |
| env-35 | JGI24709J26583_10009162 | + | 2538 | 2609 | **RNA**→ rps15p (PRK08561)Ribosomal_S15 (pfam00312)→ RecJ (COG0608)recJ (TIGR00644)→ PRK14888 (PRK14888)→ PRK04057 (PRK04057)Ribosomal_S3Ae (pfam01015)→ |
| env-36 | JGI24707J26582_10010674 | - | 1854 | 1783 | **RNA**→ rps15p (PRK08561)Ribosomal_S15p_S13e (cd00353)→ RecJ (COG0608)recJ (TIGR00644)→ |
| env-37 | JGI24711J26586_10015339 | - | 2787 | 2716 | **RNA**→ rps15p (PRK08561)Ribosomal_S15p_S13e (cd00353)→ RecJ (COG0608)recJ (TIGR00644)→ hypo→ PRK14888 (PRK14888)→ PRK04057 (PRK04057)Ribosomal_S3Ae (pfam01015)→ |
| env-38 | JGI24710J26742_10008399 | + | 5929 | 6000 | **RNA**→ rps15p (PRK08561)Ribosomal_S15p_S13e (cd00353)→ RecJ (COG0608)recJ (TIGR00644)→ |
| env-39 | JGI24712J26585_10018987 | + | 1068 | 1139 | **RNA**→ rps15p (PRK08561)Ribosomal_S15p_S13e (cd00353)→ RecJ (COG0608)recJ (TIGR00644)→ PRK14888 (PRK14888)→ PRK04057 (PRK04057)Ribosomal_S3Ae (pfam01015)→ |
| env-40 | JGI24708J26588_10009965 | - | 2892 | 2821 | **RNA**→ rps15p (PRK08561)Ribosomal_S15p_S13e (cd00353)→ RecJ (COG0608)recJ (TIGR00644)→ PRK14888 (PRK14888)→ PRK04057 (PRK04057)Ribosomal_S3Ae (pfam01015)→ |
| env-41 | JGI24712J26585_10013199 | - | 199 | 130 | **RNA**→ |
| env-42 | JGI24711J26586_10077988 | + | 748 | 817 | **RNA**→ rps15p (PRK08561)→ |
| env-43 | JGI24710J26742_10001551 | - | 12821 | 12752 | **RNA**→ rps15p (PRK08561)Ribosomal_S15p_S13e (cd00353)→ RecJ (COG0608)recJ (TIGR00644)→ PRK14888 (PRK14888)→ PRK04057 (PRK04057)Ribosomal_S3Ae (pfam01015)→ |
| env-44 | JGI24709J26583_10004510 | + | 8218 | 8287 | **RNA**→ rps15p (PRK08561)Ribosomal_S15p_S13e (cd00353)→ RecJ (COG0608)recJ (TIGR00644)→ PRK14888 (PRK14888)→ PRK04057 (PRK04057)Ribosomal_S3Ae (pfam01015)→ |
| env-45 | JGI24711J26586_10013429 | + | 3611 | 3680 | **RNA**→ |

| | | | | | |
|---|---|---|---|---|---|
| env-46 | JGI24712J26585_10007655 | + | 6829 | 6898 | **RNA**→ rps15p (PRK08561)Ribosomal_S15p_S13e (cd00353)→ |
| env-47 | JGI24712J26585_10045214 | - | 875 | 806 | **RNA**→ rps15p (PRK08561)Ribosomal_S15 (pfam00312)→ RecJ (COG0608)→ |
| env-48 | JGI24709J26583_10006535 | - | 7792 | 7723 | **RNA**→ Ribosomal_S13_N (pfam08069)→ RecJ (COG0608)recJ (TIGR00644)→ PRK14888 (PRK14888)→ PRK04057 (PRK04057)Ribosomal_S3Ae (pfam01015)→ |
| env-49 | JGI24707J26582_10115681 | + | 545 | 614 | **RNA**→ rps15p (PRK08561)→ |
| env-50 | JGI24711J26586_10231301 | - | 163 | 94 | **RNA**→ |
| env-51 | JGI24710J26742_10017231 | - | 2099 | 2030 | **RNA**→ rps15p (PRK08561)Ribosomal_S15p_S13e (cd00353)→ RecJ (COG0608)recJ (TIGR00644)→ PRK14888 (PRK14888)→ |
| env-52 | JGI24708J26588_10034739 | + | 663 | 732 | **RNA**→ rps15p (PRK08561)Ribosomal_S15p_S13e (cd00353)→ RecJ (COG0608)recJ (TIGR00644)→ |
| env-53 | JGI24707J26582_10002665 | - | 12607 | 12538 | **RNA**→ RpsO (COG0184)rps15p (PRK08561)→ RecJ (COG0608)→ PRK14888 (PRK14888)→ PRK04057 (PRK04057)Ribosomal_S3Ae (pfam01015)→ |
| env-54 | JGI24708J26588_10002365 | - | 13618 | 13549 | **RNA**→ RpsO (COG0184)rps15p (PRK08561)→ RecJ (COG0608)→ PRK14888 (PRK14888)→ PRK04057 (PRK04057)Ribosomal_S3Ae (pfam01015)→ |
| env-55 | JGI24713J26584_10003246 | + | 742 | 811 | **RNA**→ RpsO (COG0184)rps15p (PRK08561)→ RecJ (COG0608)→ PRK14888 (PRK14888)→ PRK04057 (PRK04057)Ribosomal_S3Ae (pfam01015)→ |
| env-56 | JGI24714J26587_10001431 | + | 3149 | 3218 | **RNA**→ RpsO (COG0184)rps15p (PRK08561)→ RecJ (COG0608)→ PRK14888 (PRK14888)→ PRK04057 (PRK04057)Ribosomal_S3Ae (pfam01015)→ |
| env-57 | JGI24712J26585_10004187 | - | 10439 | 10370 | **RNA**→ RpsO (COG0184)rps15p (PRK08561)→ RecJ (COG0608)→ PRK14888 (PRK14888)→ PRK04057 (PRK04057)Ribosomal_S3Ae (pfam01015)→ |
| env-58 | JGI24711J26586_10011349 | - | 3687 | 3618 | **RNA**→ RpsO (COG0184)rps15p (PRK08561)→ RecJ (COG0608)→ PRK14888 (PRK14888)→ PRK04057 (PRK04057)Ribosomal_S3Ae (pfam01015)→ |
| env-59 | JGI24709J26583_10004141 | - | 10439 | 10370 | **RNA**→ RpsO (COG0184)rps15p (PRK08561)→ RecJ (COG0608)→ PRK14888 (PRK14888)→ PRK04057 (PRK04057)Ribosomal_S3Ae (pfam01015)→ |
| env-60 | JGI24710J26742_10004693 | - | 10439 | 10370 | **RNA**→ RpsO (COG0184)rps15p (PRK08561)→ RecJ (COG0608)→ PRK14888 (PRK14888)→ PRK04057 (PRK04057)Ribosomal_S3Ae (pfam01015)→ |
| Mbo-1-1 | NC_018227.2 | + | 355435 | 355504 | **RNA**→ rps15p (PRK08561)Ribosomal_S15 (pfam00312)→ RecJ (COG0608)recJ (TIGR00644)→ PRK14888 (PRK14888)→ PRK04057 (PRK04057)Ribosomal_S3Ae (pfam01015)→ |
| env-61 | JGI24712J26585_10064911 | + | 1269 | 1338 | **RNA**→ |
| env-62 | JGI24711J26586_10108516 | + | 580 | 649 | **RNA**→ |
| env-63 | JGI24709J26583_10179352 | - | 163 | 94 | **RNA**→ |
| env-64 | 3300000558_10029409 | + | 126 | 197 | **RNA**→ rps15p (PRK08561)Ribosomal_S15p_S13e (cd00353)→ RecJ (COG0608)recJ (TIGR00644)→ |
| env-65 | 3300000568_10009510 | - | 2212 | 2141 | **RNA**→ rps15p (PRK08561)Ribosomal_S15p_S13e (cd00353)→ RecJ (COG0608)recJ (TIGR00644)→ PRK14888 (PRK14888)→ PRK04057 (PRK04057)Ribosomal_S3Ae (pfam01015)→ |
| env-66 | 3300000032_0014754 | + | 78 | 149 | **RNA**→ rps15p (PRK08561)Ribosomal_S15p_S13e (cd00353)→ RecJ (COG0608)recJ (TIGR00644)→ |
| env-67 | 3300000032_0066330 | + | 155 | 226 | **RNA**→ Ribosomal_S13_N (pfam08069)→ |
| env-68 | 3300001592_10784659 | + | 44 | 115 | **RNA**→ rps15p (PRK08561)→ |
| env-69 | 3300000507_100903 | - | 1882 | 1811 | **RNA**→ rps15p (PRK08561)Ribosomal_S15p_S13e (cd00353)→ RecJ (COG0608)recJ (TIGR00644)→ hypo→ |
| Msp-1-1 | NZ_JMIO01000090.1 | + | 12381 | 12453 | **RNA**→ hypo→ hypo→ |
| Mma-1-1 | NC_009051.1 | - | 416072 | 416000 | **RNA**→ rps15p (PRK08561)Ribosomal_S15p_S13e (cd00353)→ RecJ (COG0608)recJ (TIGR00644)→ hypo→ PRK14888 (PRK14888)→ PRK04057 (PRK04057)Ribosomal_S3Ae (pfam01015)→ hypo→ apgM (TIGR00306)iPGM_like (cd16011)→ lipid_A_lpxD (TIGR01853)Bactofilin (pfam04519)→ hypo→ |
| env-70 | JGI24712J26585_10323374 | - | 185 | 113 | **RNA**→ |
| Mse-1-1 | NZ_JXOJ01000001.1 | - | 229493 | 229421 | **RNA**→ rps15p (PRK08561)Ribosomal_S15p_S13e (cd00353)→ RecJ (COG0608)recJ (TIGR00644)→ PRK14888 (PRK14888)→ PRK04057 (PRK04057)Ribosomal_S3Ae (pfam01015)→ hypo→ ApgM (COG3635)apgM (TIGR00306)→ lipid_A_lpxD (TIGR01853)Bactofilin (pfam04519)→ hypo→ |
| env-71 | 3300000568_10009858 | - | 298 | 228 | **RNA**→ Ribosomal_S13_N (pfam08069)→ |
| env-72 | 3300001567_10003374 | - | 12269 | 12199 | **RNA**→ rps15p (PRK08561)Ribosomal_S15p_S13e (cd00353)→ RecJ (COG0608)recJ (TIGR00644)→ PRK14888 (PRK14888)→ PRK04057 (PRK04057)Ribosomal_S3Ae (pfam01015)→ |
| env-73 | 3300000032_0044310 | - | 13186 | 13116 | **RNA**→ rps15p (PRK08561)Ribosomal_S15p_S13e (cd00353)→ RecJ (COG0608)recJ (TIGR00644)→ PRK14888 (PRK14888)→ PRK04057 (PRK04057)Ribosomal_S3Ae (pfam01015)→ |
| env-74 | 3300000558_11565244 | + | 1079 | 1149 | **RNA**→ rps15p (PRK08561)Ribosomal_S15p_S13e (cd00353)→ RecJ (COG0608)→ |
| env-75 | 3300001592_10006172 | - | 8474 | 8404 | **RNA**→ rps15p (PRK08561)Ribosomal_S15p_S13e (cd00353)→ RecJ (COG0608)recJ (TIGR00644)→ PRK14888 (PRK14888)→ PRK04057 (PRK04057)Ribosomal_S3Ae (pfam01015)→ |
| env-76 | JGI26437J51864_10033120 | + | 500 | 575 | **RNA**→ rps15p (PRK08561)Ribosomal_S15p_S13e (cd00353)→ |

| | | | | | |
|---|---|---|---|---|---|
| env-77 | 3300001580_10010117 | - | 5601 | 5525 | **RNA**→ RpsO (COG0184)rps15p (PRK08561)→ RecJ (COG0608)recJ (TIGR00644)→ PRK14888 (PRK14888)→ PRK00960 (PRK00960)Gly_His_Pro_Ser_Thr_tRS_core (cd00670)→ PRK04057 (PRK04057)Ribosomal_S3Ae (pfam01015)→ apgM (TIGR00306)iPGM_like (cd16011)→ hypo→ |
| env-78 | 2232139342 | - | 237 | 160 | **RNA**→ rps15p (PRK08561)→ |
| env-79 | 3300001592_10034324 | + | 2742 | 2834 | **RNA**→ RpsO (COG0184)rps15p (PRK08561)→ |
| env-80 | 2013874838 | - | 1123 | 1033 | **RNA**→ rps15p (PRK08561)Ribosomal_S15 (pfam00312)→ RecJ (COG0608)recJ (TIGR00644)→ |
| env-81 | BMHB3a_c125774 | + | 423 | 513 | **RNA**→ Ribosomal_S13_N (pfam08069)→ |
| env-82 | JGI2065J20421_1144564 | + | 131 | 221 | **RNA**→ hypo→ |
| env-83 | JGI26339J46600_10075181 | - | 619 | 538 | **RNA**→ rps15p (PRK08561)Ribosomal_S15 (pfam00312)→ |
| env-84 | JGI20225J20221_1000010 | + | 326457 | 326544 | **RNA**→ rps15p (PRK08561)Ribosomal_S15 (pfam00312)→ RecJ (COG0608)recJ (TIGR00644)→ PRK14888 (PRK14888)→ PRK04057 (PRK04057)Ribosomal_S3Ae (pfam01015)→ |
| env-85 | PRSSGFe2_Sequence0000002097 | + | 2324 | 2413 | **RNA**→ rps15p (PRK08561)Ribosomal_S15 (pfam00312)Ribosomal_S13_N (pfam08069)→ RecJ (COG0608)recJ (TIGR00644)→ PRK14888 (PRK14888)→ PRK04057 (PRK04057)Ribosomal_S3Ae (pfam01015)→ apgM (TIGR00306)iPGM_like (cd16011)→ PRK15319 (PRK15319)Bactofilin (pfam04519)→ Hanta_G2 (pfam01561)→ |
| env-86 | WSSedL1Cam_c0002 | + | 96851 | 96938 | **RNA**→ rps15p (PRK08561)Ribosomal_S15 (pfam00312)→ RecJ (COG0608)recJ (TIGR00644)→ PRK14888 (PRK14888)→ PRK04057 (PRK04057)Ribosomal_S3Ae (pfam01015)→ |
| CMe-1-1 | NC_009712.1 | + | 2304410 | 2304492 | **RNA**→ rps15p (PRK08561)Ribosomal_S15 (pfam00312)→ RecJ (COG0608)→ PRK14888 (PRK14888)7tmA_CXCR1_2 (cd15178)→ PRK04057 (PRK04057)Ribosomal_S3Ae (pfam01015)→ |
| env-87 | JGI24729J20445_1138040 | - | 236 | 148 | **RNA**→ ←hypo |
| env-88 | JGI24730J26740_1000008 | + | 72470 | 72558 | **RNA**→ rps15p (PRK08561)Ribosomal_S15 (pfam00312)→ RecJ (COG0608)recJ (TIGR00644)→ PRK14888 (PRK14888)→ PRK04057 (PRK04057)Ribosomal_S3Ae (pfam01015)→ |
| Mli-1-1 | NZ_CM001555.1 | - | 2222769 | 2222698 | **RNA**→ RpsO (COG0184)rps15p (PRK08561)→ RpsO (COG0184)rps15p (PRK08561)→ RecJ (COG0608)→ RecJ (COG0608)→ PRK14888 (PRK14888)→ PRK14888 (PRK14888)→ PRK04057 (PRK04057)Ribosomal_S3Ae (pfam01015)→ PRK04057 (PRK04057)Ribosomal_S3Ae (pfam01015)→ |
| Mfo-1-1 | NC_019943.1 | - | 469269 | 469194 | **RNA**→ rps15p (PRK08561)Ribosomal_S15 (pfam00312)→ RecJ (COG0608)recJ (TIGR00644)→ PRK14888 (PRK14888)→ PRK04057 (PRK04057)Ribosomal_S3Ae (pfam01015)→ |
| env-89 | 3300000558_10035676 | + | 11455 | 11529 | **RNA**→ rps15p (PRK08561)→ |
| env-90 | 3300001605_10975353 | + | 94 | 168 | **RNA**→ Ribosomal_S13_N (pfam08069)→ |
| env-91 | JGI24730J26740_1005506 | - | 3828 | 3750 | **RNA**→ RpsO (COG0184)rps15p (PRK08561)→ RecJ (COG0608)→ PRK14888 (PRK14888)→ PRK04057 (PRK04057)Ribosomal_S3Ae (pfam01015)→ |
| env-92 | JGI24729J20445_1023293 | + | 517 | 593 | **RNA**→ rps15p (PRK08561)Ribosomal_S15p_S13e (cd00353)→ |
| env-93 | JGI24728J21555_1000106 | + | 20382 | 20458 | **RNA**→ rps15p (PRK08561)Ribosomal_S15p_S13e (cd00353)→ RecJ (COG0608)recJ (TIGR00644)DHHA1 (pfam02272)→ PRK14888 (PRK14888)→ PRK04057 (PRK04057)Ribosomal_S3Ae (pfam01015)→ |
| env-94 | JGI24732J26686_1000182 | - | 32050 | 31974 | **RNA**→ rps15p (PRK08561)Ribosomal_S15p_S13e (cd00353)→ RecJ (COG0608)recJ (TIGR00644)DHHA1 (pfam02272)→ PRK14888 (PRK14888)→ PRK04057 (PRK04057)Ribosomal_S3Ae (pfam01015)→ |
| env-95 | JGI24732J26686_1000319 | + | 14050 | 14126 | **RNA**→ rps15p (PRK08561)Ribosomal_S15p_S13e (cd00353)→ RecJ (COG0608)recJ (TIGR00644)DHHA1 (pfam02272)→ PRK14888 (PRK14888)→ PRK04057 (PRK04057)Ribosomal_S3Ae (pfam01015)→ |
| env-96 | JGI24728J21555_1038878 | + | 851 | 927 | **RNA**→ |
| env-97 | JGI24730J26740_1222110 | - | 264 | 188 | **RNA**→ rps15p (PRK08561)→ |
| env-98 | 3300001605_10027048 | - | 295 | 219 | **RNA**→ Ribosomal_S13_N (pfam08069)→ |
| env-99 | 3300000185_101747 | - | 1221 | 1145 | **RNA**→ rps15p (PRK08561)Ribosomal_S15 (pfam00312)→ RecJ (COG0608)recJ (TIGR00644)→ |
| env-100 | 3300000032_0027748 | - | 1766 | 1690 | **RNA**→ rps15p (PRK08561)Ribosomal_S15 (pfam00312)→ RecJ (COG0608)recJ (TIGR00644)→ |
| env-101 | 3300000558_11833810 | - | 1221 | 1145 | **RNA**→ rps15p (PRK08561)Ribosomal_S15 (pfam00312)→ RecJ (COG0608)recJ (TIGR00644)→ |
| env-102 | 3300000568_10087874 | + | 4955 | 5031 | **RNA**→ rps15p (PRK08561)Ribosomal_S15 (pfam00312)→ RecJ (COG0608)recJ (TIGR00644)→ |
| env-103 | 3300001580_10011015 | - | 3080 | 3002 | **RNA**→ rps15p (PRK08561)Ribosomal_S15p_S13e (cd00353)→ RecJ (COG0608)recJ (TIGR00644)DHHA1 (pfam02272)→ PRK14888 (PRK14888)→ PRK04057 (PRK04057)Ribosomal_S3Ae (pfam01015)→ MFS (cd06174)efflux_EmrB (TIGR00711)→ |
| env-104 | LWFCAn_GLO1YSU01E41BP | + | 16 | 91 | **RNA**→ |
| env-105 | 3300001592_10847636 | + | 30 | 107 | **RNA**→ rps15p (PRK08561)→ |
| env-106 | 3300001567_10049145 | - | 100 | 26 | **RNA**→ |
| env-107 | JGI26437J51864_10417374 | + | 109 | 183 | **RNA**→ hypo→ |

| | | | | | |
|---|---|---|---|---|---|
| env-108 | JGI20214J14112_1001966 | + | 2925 | 3001 | **RNA**→ rps15p (PRK08561)Ribosomal_S15 (pfam00312)→ |
| env-109 | SakLake120mDRAFT_c0148215 | - | 531 | 456 | **RNA**→ rps15p (PRK08561)Ribosomal_S15 (pfam00312)→ |
| env-110 | JGI20214J14112_1066793 | - | 466 | 393 | **RNA**→ Ribosomal_S13_N (pfam08069)→ |
| env-111 | 3300001567_10018396 | - | 99 | 26 | **RNA**→ ←hypo |
| env-112 | 3300000558_10022832 | - | 2400 | 2328 | **RNA**→ rps15p (PRK08561)Ribosomal_S15p_S13e (cd00353)→ RecJ (COG0608)recJ (TIGR00644)→ PRK14888 (PRK14888)→ PRK04057 (PRK04057)Ribosomal_S3Ae (pfam01015)→ |
| env-113 | 3300001580_10015327 | + | 3751 | 3823 | **RNA**→ rps15p (PRK08561)Ribosomal_S15p_S13e (cd00353)→ RecJ (COG0608)recJ (TIGR00644)→ PRK14888 (PRK14888)→ PRK04057 (PRK04057)Ribosomal_S3Ae (pfam01015)→ |
| env-114 | 3300001605_10381060 | - | 80 | 8 | **RNA**→ |
| env-115 | JGI24729J20445_1000035 | + | 33077 | 33154 | **RNA**→ rps15p (PRK08561)Ribosomal_S15 (pfam00312)→ RecJ (COG0608)recJ (TIGR00644)→ PRK14888 (PRK14888)→ PRK04057 (PRK04057)Ribosomal_S3Ae (pfam01015)→ |
| env-116 | JGI24732J26686_1000038 | + | 17911 | 17988 | **RNA**→ rps15p (PRK08561)Ribosomal_S15 (pfam00312)→ RecJ (COG0608)recJ (TIGR00644)→ PRK14888 (PRK14888)→ PRK04057 (PRK04057)Ribosomal_S3Ae (pfam01015)→ |
| env-117 | BDMC2_FXAG6RB01C8MB5 | - | 494 | 416 | **RNA**→ Ribosomal_S13_N (pfam08069)→ |
| env-118 | JGI24729J20445_1010075 | + | 1466 | 1542 | **RNA**→ rps15p (PRK08561)Ribosomal_S15 (pfam00312)→ RecJ (COG0608)→ |
| env-119 | JGI24728J21555_1016609 | - | 489 | 413 | **RNA**→ Ribosomal_S13_N (pfam08069)→ |
| env-120 | TB_GS10_10_10107963 | - | 629 | 548 | **RNA**→ rps15p (PRK08561)Ribosomal_S15 (pfam00312)→ |
| env-121 | TB_GS09_5_10262278 | - | 191 | 110 | **RNA**→ hypo→ |
| env-122 | JGI26437J51864_10012658 | + | 109 | 188 | **RNA**→ rps15p (PRK08561)Ribosomal_S15p_S13e (cd00353)→ RecJ (COG0608)recJ (TIGR00644)→ |
| env-123 | 3300000032_0017538 | + | 5307 | 5386 | **RNA**→ rps15p (PRK08561)Ribosomal_S15p_S13e (cd00353)→ RecJ (COG0608)recJ (TIGR00644)→ |
| env-124 | WSSedA2C_c014251 | - | 708 | 630 | **RNA**→ rps15p (PRK08561)Ribosomal_S15p_S13e (cd00353)→ hypo→ |
| env-125 | WSSedA1TJGI_c016288 | + | 290 | 368 | **RNA**→ rps15p (PRK08561)Ribosomal_S15p_S13e (cd00353)→ RecJ (COG0608)recJ (TIGR00644)→ |
| env-126 | WSSedA1Ba2_1007239 | + | 706 | 784 | **RNA**→ rps15p (PRK08561)Ribosomal_S15p_S13e (cd00353)→ RecJ (COG0608)→ |
| env-127 | WSSedA2Ca3_1014251 | - | 708 | 630 | **RNA**→ rps15p (PRK08561)Ribosomal_S15p_S13e (cd00353)→ hypo→ |
| env-128 | WSSedA1Ba3_1001463 | - | 6148 | 6070 | **RNA**→ rps15p (PRK08561)Ribosomal_S15p_S13e (cd00353)→ RecJ (COG0608)recJ (TIGR00644)→ PRK14888 (PRK14888)→ PRK04057 (PRK04057)Ribosomal_S3Ae (pfam01015)→ |
| env-129 | WSSedB1B2_c008138 | + | 171 | 249 | **RNA**→ rps15p (PRK08561)Ribosomal_S15p_S13e (cd00353)→ RecJ (COG0608)recJ (TIGR00644)→ |
| env-130 | WSSedB2T_c084959 | + | 143 | 221 | **RNA**→ ←hypo |
| env-131 | WSSedB2T_c089397 | - | 217 | 139 | **RNA**→ ←hypo |
| env-132 | JGI20213J14113_1035134 | + | 232 | 310 | **RNA**→ Ribosomal_S13_N (pfam08069)→ |
| env-133 | JGI20214J14112_1021896 | - | 626 | 548 | **RNA**→ rps15p (PRK08561)Ribosomal_S15p_S13e (cd00353)→ |
| env-134 | JGI26437J51864_10213950 | + | 73 | 151 | **RNA**→ Ribosomal_S13_N (pfam08069)→ |
| env-135 | WSSedA1Ba3_1000357 | - | 6727 | 6647 | **RNA**→ rps15p (PRK08561)Ribosomal_S15p_S13e (cd00353)→ RecJ (COG0608)recJ (TIGR00644)→ PRK14888 (PRK14888)→ PRK04057 (PRK04057)Ribosomal_S3Ae (pfam01015)→ |
| env-136 | WSSedA1Ba3_1272904 | + | 141 | 221 | **RNA**→ hypo→ |
| env-137 | WSSedA1Ba_1001070 | + | 2641 | 2721 | **RNA**→ rps15p (PRK08561)Ribosomal_S15p_S13e (cd00353)→ RecJ (COG0608)recJ (TIGR00644)→ PRK14888 (PRK14888)→ PRK04057 (PRK04057)Ribosomal_S3Ae (pfam01015)→ hypo→ PRK04024 (PRK04024)iPGM_like (cd16011)→ |
| env-138 | WSSedA2C_c018532 | - | 789 | 709 | **RNA**→ rps15p (PRK08561)Ribosomal_S15p_S13e (cd00353)→ RecJ (COG0608)recJ (TIGR00644)→ |
| env-139 | WSSedA1TJGI_c000079 | - | 10509 | 10429 | **RNA**→ rps15p (PRK08561)Ribosomal_S15p_S13e (cd00353)→ RecJ (COG0608)recJ (TIGR00644)→ PRK14888 (PRK14888)→ PRK04057 (PRK04057)Ribosomal_S3Ae (pfam01015)→ |
| env-140 | WSSedB1B2_c003916 | + | 734 | 814 | **RNA**→ rps15p (PRK08561)Ribosomal_S15p_S13e (cd00353)→ |
| env-141 | WSSedB2T_c002759 | + | 1013 | 1093 | **RNA**→ rps15p (PRK08561)Ribosomal_S15p_S13e (cd00353)→ RecJ (COG0608)→ |
| env-142 | WSSedA1Ba2_1000860 | + | 4312 | 4392 | **RNA**→ rps15p (PRK08561)Ribosomal_S15p_S13e (cd00353)→ RecJ (COG0608)recJ (TIGR00644)→ |
| env-143 | WSSedB2Ba_1002821 | - | 672 | 592 | **RNA**→ rps15p (PRK08561)Ribosomal_S15p_S13e (cd00353)→ |
| env-144 | WSSedB2Ca_1002616 | + | 792 | 872 | **RNA**→ rps15p (PRK08561)Ribosomal_S15p_S13e (cd00353)→ |
| env-145 | WSSedA2Ca3_1018532 | - | 789 | 709 | **RNA**→ rps15p (PRK08561)Ribosomal_S15p_S13e (cd00353)→ RecJ (COG0608)recJ (TIGR00644)→ |
| env-146 | JGI20214J14112_1000149 | - | 1161 | 1081 | **RNA**→ rps15p (PRK08561)Ribosomal_S15p_S13e (cd00353)→ RecJ (COG0608)recJ (TIGR00644)→ |
| env-147 | JGI20213J14113_1001057 | + | 1284 | 1364 | **RNA**→ rps15p (PRK08561)Ribosomal_S15p_S13e (cd00353)→ RecJ (COG0608)recJ (TIGR00644)→ PRK14888 (PRK14888)→ PRK04057 (PRK04057)Ribosomal_S3Ae (pfam01015)→ |
| env-148 | WSSedA1Ba3_1006166 | + | 2461 | 2541 | **RNA**→ rps15p (PRK08561)Ribosomal_S15p_S13e (cd00353)→ |
| env-149 | WSSedA1TJGI_c233835 | + | 122 | 202 | **RNA**→ |
| env-150 | JGI26437J51864_10004063 | - | 209 | 129 | **RNA**→ hypo→ |

| | | | | | |
|---|---|---|---|---|---|
| env-151 | 3300001567_10000108 | - | 84134 | 84057 | RNA→ hypo→ rps15p (PRK08561)Ribosomal_S15p_S13e (cd00353)→ RecJ (COG0608)→ PRK14888 (PRK14888)→ PRK04057 (PRK04057)Ribosomal_S3Ae (pfam01015)→ PRK04057 (PRK04057)Ribosomal_S3Ae (pfam01015)→ |
| env-152 | WSSedA1Ba3_1177753 | + | 108 | 186 | RNA→ Ribosomal_S13_N (pfam08069)→ |
| env-153 | WSSedA1Ba_1152331 | - | 288 | 210 | RNA→ Ribosomal_S13_N (pfam08069)→ |
| env-154 | B3_all_c_Cons177848 | - | 485 | 407 | RNA→ rps15p (PRK08561)→ |
| env-155 | 3300000568_10530120 | + | 366 | 444 | RNA→ hypo→ |
| env-156 | JGI24729J20445_1002298 | - | 5486 | 5409 | RNA→ rps15p (PRK08561)Ribosomal_S15 (pfam00312)→ RecJ (COG0608)recJ (TIGR00644)→ PRK14888 (PRK14888)→ PRK04057 (PRK04057)Ribosomal_S3Ae (pfam01015)→ |
| env-157 | JGI24729J20445_1020946 | - | 87 | 10 | RNA→ |
| env-158 | JGI24732J26686_1097519 | - | 87 | 10 | RNA→ |
| env-159 | JGI24732J26686_1003719 | - | 4772 | 4695 | RNA→ rps15p (PRK08561)Ribosomal_S15 (pfam00312)→ RecJ (COG0608)recJ (TIGR00644)→ PRK14888 (PRK14888)→ PRK04057 (PRK04057)Ribosomal_S3Ae (pfam01015)→ |
| env-160 | 3300001567_10000097 | + | 54256 | 54336 | RNA→ rps15p (PRK08561)Ribosomal_S15p_S13e (cd00353)→ RecJ (COG0608)recJ (TIGR00644)→ PRK14888 (PRK14888)→ PRK04057 (PRK04057)Ribosomal_S3Ae (pfam01015)→ |
| env-161 | 3300000568_10641437 | - | 280 | 200 | RNA→ rps15p (PRK08561)→ |
| env-162 | JGI20214J14112_1020774 | - | 594 | 514 | RNA→ rps15p (PRK08561)Ribosomal_S15p_S13e (cd00353)→ |
| env-163 | 3300001580_10805722 | + | 98 | 181 | RNA→ rps15p (PRK08561)→ |
| env-164 | taComm3_C4593 | - | 845 | 762 | RNA→ RpsO (COG0184)rps15p (PRK08561)→ RecJ (COG0608)recJ (TIGR00644)→ |
| env-165 | 3300001592_10767277 | + | 24 | 111 | RNA→ hypo→ rps15p (PRK08561)→ |
| env-166 | JGI12104J13512_1000881 | + | 4950 | 5036 | RNA→ RpsO (COG0184)rps15p (PRK08561)→ RecJ (COG0608)recJ (TIGR00644)→ PRK14888 (PRK14888)→ |
| env-167 | JGI11944J13513_1000579 | + | 5848 | 5934 | RNA→ RpsO (COG0184)rps15p (PRK08561)→ RecJ (COG0608)recJ (TIGR00644)→ PRK14888 (PRK14888)→ |
| env-168 | JGI12104J13512_1055609 | - | 126 | 41 | RNA→ |
| env-169 | 3300000032_0028051 | - | 6644 | 6556 | RNA→ RpsO (COG0184)rps15p (PRK08561)→ RecJ (COG0608)recJ (TIGR00644)→ PRK14888 (PRK14888)→ |
| env-170 | 3300000558_10008220 | - | 2159 | 2071 | RNA→ RpsO (COG0184)rps15p (PRK08561)→ RecJ (COG0608)recJ (TIGR00644)→ PRK14888 (PRK14888)→ hypo→ |
| env-171 | 3300000568_10642328 | + | 2264 | 2352 | RNA→ RpsO (COG0184)rps15p (PRK08561)→ RecJ (COG0608)recJ (TIGR00644)→ PRK14888 (PRK14888)→ |
| env-172 | 3300000568_10908886 | - | 124 | 36 | RNA→ |
| env-173 | 3300001580_10001748 | + | 3910 | 3998 | RNA→ RpsO (COG0184)rps15p (PRK08561)→ RecJ (COG0608)recJ (TIGR00644)→ PRK14888 (PRK14888)→ |
| Mta-1-1 | NZ_AGIY02000001.1 | - | 138618 | 138536 | RNA→ RpsO (COG0184)rps15p (PRK08561)→ RecJ (COG0608)recJ (TIGR00644)→ PRK14888 (PRK14888)→ PRK04057 (PRK04057)Ribosomal_S3Ae (pfam01015)→ |
| env-174 | taComm3_C6165 | - | 76136 | 76054 | RNA→ RpsO (COG0184)rps15p (PRK08561)→ RecJ (COG0608)recJ (TIGR00644)→ PRK14888 (PRK14888)→ |
| env-175 | JGI12104J13512_1000508 | + | 4260 | 4342 | RNA→ RpsO (COG0184)rps15p (PRK08561)→ RecJ (COG0608)recJ (TIGR00644)→ PRK14888 (PRK14888)→ PRK04057 (PRK04057)Ribosomal_S3Ae (pfam01015)→ |
| env-176 | JGI11944J13513_1004555 | + | 4268 | 4350 | RNA→ RpsO (COG0184)rps15p (PRK08561)→ RecJ (COG0608)recJ (TIGR00644)→ PRK14888 (PRK14888)→ PRK04057 (PRK04057)Ribosomal_S3Ae (pfam01015)→ |
| Mli-2-1 | NZ_CM001436.1 | - | 2802614 | 2802534 | RNA→ rps15p (PRK08561)Ribosomal_S15 (pfam00312)→ rps15p (PRK08561)Ribosomal_S15 (pfam00312)→ RecJ (COG0608)recJ (TIGR00644)→ RecJ (COG0608)recJ (TIGR00644)→ PRK14888 (PRK14888)→ PRK14888 (PRK14888)→ PRK04057 (PRK04057)Ribosomal_S3Ae (pfam01015)→ PRK04057 (PRK04057)Ribosomal_S3Ae (pfam01015)→ hypo→ hypo→ hypo→ hypo→ PRK10765 (PRK10765)Nitroreductase (pfam00881)→ PRK10765 (PRK10765)Nitroreductase (pfam00881)→ PRK03922 (PRK03922)DUF555 (pfam04475)→ PRK03922 (PRK03922)DUF555 (pfam04475)→ ribokinase_group_A (cd01942)PTZ00247 (PTZ00247)→ ribokinase_group_A (cd01942)PTZ00247 (PTZ00247)→ |
| Mba-1-1 | NZ_AUMX01000001.1 | + | 125785 | 125859 | RNA→ rps15p (PRK08561)Ribosomal_S15p_S13e (cd00353)→ RecJ (COG0608)recJ (TIGR00644)→ PRK14888 (PRK14888)→ PRK04057 (PRK04057)Ribosomal_S3Ae (pfam01015)→ |
| Mla-1-1 | NC_008942.1 | - | 271642 | 271569 | RNA→ rps15p (PRK08561)Ribosomal_S15p_S13e (cd00353)→ RecJ (COG0608)recJ (TIGR00644)→ PRK14888 (PRK14888)→ PRK04057 (PRK04057)Ribosomal_S3Ae (pfam01015)→ |
| env-177 | BDMC2_GGZN29857_b1 | - | 505 | 433 | RNA→ Ribosomal_S15 (pfam00312)Ribosomal_S13_N (pfam08069)→ |
| env-178 | BDMC2_FXAG6RB01AV6H8 | - | 77 | 5 | RNA→ |
| env-179 | JGI20163J15578_11006719 | + | 71 | 153 | RNA→ Ribosomal_S13_N (pfam08069)→ |

#### 21.3 Conserved domains

Conserved domains found in protein-coding genes listed in Section 21.2 are shown below, with the first sentence in their description from the Conserved Domain Database (if any). Conserved domains associated with more than one S15-Methanobacteria RNA are assigned a color, while others are shown in gray. The number in parentheses after the colored domain name is the number of occurrences in Section 21.2.

cd00353 (67) Ribosomal protein S15 (prokaryotic)_S13 (eukaryotic) binds the central domain of 16S rRNA and is required for assembly of the small ribosomal subunit and for intersubunit association, thus representing a key element in the assembly of the whole ribosome.
cd00670 (5) Gly_His_Pro_Ser_Thr_tRNA synthetase class II core domain.
cd01942 (4) Ribokinase-like subgroup A.
cd06174 (1) The Major Facilitator Superfamily (MFS) is a large and diverse group of secondary transporters that includes uniporters, symporters, and antiporters.
cd15178 (1) CXC chemokine receptor types 1 and 2, member of the class A family of seven-transmembrane G protein-coupled receptors.
cd16011 (6) uncharacterized subfamily of alkaline phosphatase, homologous to 2 3 bisphosphoglycerate independent phosphoglycerate mutase (iPGM) and bacterial phosphopentomutases.
COG0184 (31) Ribosomal protein S15P/S13E [Translation, ribosomal structure and biogenesis]
COG0608 (107) Single-stranded DNA-specific exonuclease, DHH superfamily, may be involved in archaeal DNA replication intiation [Replication, recombination and repair]
COG3221 (1) ABC-type phosphate/phosphonate transport system, periplasmic component [Inorganic ion transport and metabolism]
COG3356 (2) Predicted membrane-associated lipid hydrolase, neutral ceramidase superfamily [Lipid transport and metabolism]
COG3635 (1) 2,3-bisphosphoglycerate-independent phosphoglycerate mutase, archeal type [Carbohydrate transport and metabolism]
COG4801 (1) Predicted acyltransferase, contains DUF342 domain [General function prediction only]
pfam00312 (27) Ribosomal protein S15.
pfam00881 (4) Nitroreductase family.
pfam01015 (68) Ribosomal S3Ae family.
pfam01561 (1) Hantavirus glycoprotein G2.
pfam02272 (6) DHHA1 domain.
pfam04475 (4) Protein of unknown function (DUF555).
pfam04519 (3) Polymer-forming cytoskeletal.
pfam08069 (25) Ribosomal S13/S15 N-terminal domain.
pfam09843 (2) Predicted membrane protein (DUF2070).
pfam10143 (1) 2,3-bisphosphoglycerate-independent phosphoglycerate mutase. [2-phospho-D-glycerate + 2,3-diphosphoglycerate = 3-phospho-D-glycerate + 2,3-diphosphoglycerate]
PRK00960 (5) seryl-tRNA synthetase; Provisional
PRK03922 (4) hypothetical protein; Provisional
PRK04024 (1) cofactor-independent phosphoglycerate mutase; Provisional
PRK04057 (68) 30S ribosomal protein S3Ae; Validated
PRK08561 (137) 30S ribosomal protein S15P; Reviewed
PRK10765 (2) nitroreductase A; Provisional
PRK11160 (1) cysteine/glutathione ABC transporter membrane/ATP-binding component; Reviewed
PRK14888 (76) KEOPS complex Pcc1-like subunit; Provisional
PRK15319 (1) AIDA autotransporter-like protein ShdA; Provisional
PTZ00247 (3) adenosine kinase; Provisional
TIGR00306 (5) phosphoglycerate mutase (2,3-diphosphoglycerate-independent), archaeal form. [Energy metabolism, Glycolysis/gluconeogenesis]
TIGR00644 (87) single-stranded-DNA-specific exonuclease RecJ. [DNA metabolism, DNA replication, recombination, and repair]
TIGR00711 (1) drug resistance transporter, EmrB/QacA subfamily. [Cellular processes, Toxin production and resistance, Transport and binding proteins, Other]
TIGR01173 (1) UDP-N-acetylglucosamine diphosphorylase/glucosamine-1-phosphate N-acetyltransferase. [Cell envelope, Biosynthesis and degradation of murein sacculus and peptidoglycan, Cell envelope, Biosynthesis and degradation of surface polysaccharides and lipopolysaccharides, Central intermediary metabolism, Amino sugars]
TIGR01853 (2) UDP-3-O-[3-hydroxymyristoyl] glucosamine N-acyltransferase LpxD. [3-hydroxymyristoyl] glucosamine N-acyltransferase LpxD. This model describes LpxD, an enzyme for the biosynthesis of lipid A, a component oflipopolysaccharide (LPS) in the outer membrane outer leaflet of most Gram-negative bacteria. Some differences are found between lipid A of different species. This protein represents the third step from UDP-N-acetyl-D-glucosamine. The group added at this step generally is 14:0(3-OH) (myristate) but may vary; in Aquifex it appears to be 16:0(3-OH) (palmitate). [Cell envelope, Biosynthesis and degradation of surface polysaccharides and lipopolysaccharides]
TIGR02152 (1) ribokinase. [Energy metabolism, Sugars]
TIGR02476 (2) 5,6-dimethylbenzimidazole synthase. [Biosynthesis of cofactors, prosthetic groups, and carriers, Heme, porphyrin, and cobalamin]
TIGR03683 (1) alanyl-tRNA synthetase.

#### 21.4 Multiple-sequence alignment

Each S15-Methanobacteria RNA is denoted by an abbreviation (like "Eco-1-1") that refers to a taxonomy given in Section 21.1. The alignment may include sequences containing the RNA motif, as well as flanking sequence. The S15-Methanobacteria RNA itself is denoted by the line underneath marked 5′ and 3′ on either end. Nucleotides in flanking sequences (i.e., not part of the motif) are written in gray letters. Stems of predicted rho-independent transcription terminators, if any, are shaded yellow. (Note: terminator predictions have not been analyzed manually, and many are likely to be false positives. Terminator predictions are those of the RNIE software.) Nucleotides predicted to function (as DNA) as transcription-factor binding sites, if any, are shaded in green. (But note: these predictions are manually annotated, so they might be under-predicted.) Annotated start codons, if any, are shaded green. (Note: start codons are frequently misannotated, especially in environmental samples.) Nucleotides proposed to basepair as part of the consensus structure are shaded in color when they comprise Watson-Crick or G-U pairs. Otherwise they are shaded gray. Conserved stems are also indicated at the bottom of the alignment by angle brackets, where matching < and > denote base-paired columns. Below these angle brackets, the symbol "2" denotes base pairs exhibiting covariation according to the statistically well-founded R-scape method. "1" denotes base pairs exhibiting covariation according to R2R's simplistic method. "0" denotes base pairs that are not observed to mutate and "?" denotes base pairs that have a significant frequency of non-canonical nucleotides for Watson-Crick or G-U pairs (> 5%). Below these base pair annotation is the consensus sequence: "R" = "A" or "G", "Y" = "C" or "U", red nucleotides: nucleotide identity conserved more than 97% of the time, black nucleotides: 90%, gray nucleotides: 75%, red circle (•): nucleotide is present 97% of the time, black circle (•): 90%, gray circle (•): 75%, white circle (◦): 50%. All percentages of sequences just described (e.g. 97% conserved) assume that sequences have been weighted by the GSC algorithm implemented by the Infernal software package.

Duplicate sequences: the following putative homologs are not shown in the alignment because their sequences are identical to a homolog already shown: Mmo-1-1, env-100, env-101, env-102, env-113, env-116, env-119, env-121, env-125, env-126, env-127, env-130, env-139, env-140, env-141, env-142, env-143, env-144, env-145, env-146, env-147, env-158, env-167, env-170, env-171, env-172, env-173, env-175, env-176, env-19, env-37, env-38, env-43, env-50, env-54, env-55, env-56, env-57, env-58, env-59, env-60, env-62, env-63, env-74, env-75, env-9, env-93, env-94, env-95

alignment positions 1 · · · 180

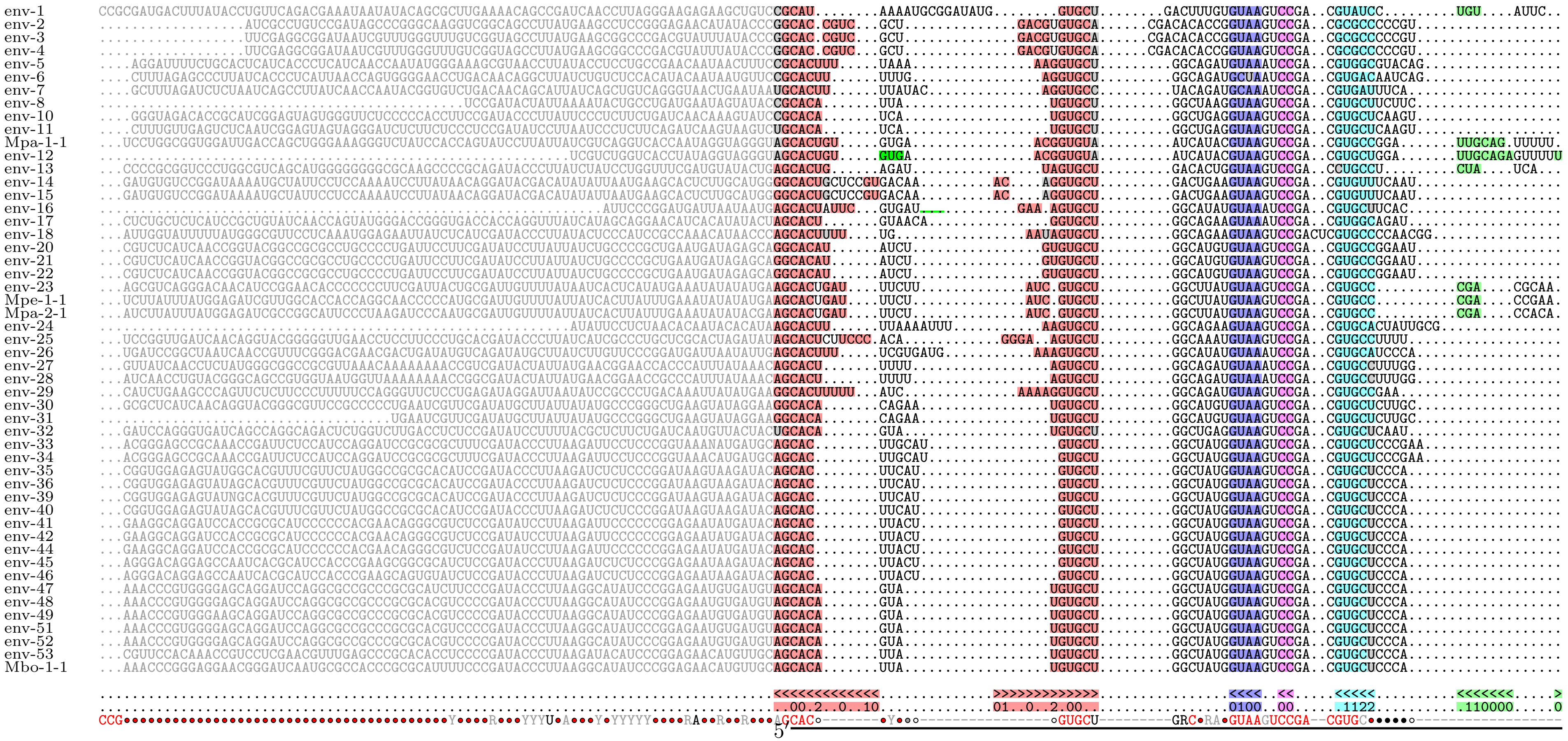

```
env-61    ...AAACCCGGGAGGAACGGGAUCAAUGCGCCACCCGCGCAUUUUCCCGAUACCCUUAAGGCAUAUCCCGGAGAACAUGUUGCAGCACA.......UUA..................UGUGCU.........GGCUAUGGUAAGUCCGA..CGUGCUCCCA..................
env-64    ...GGGUGGGGUCUUCCCGCUCCGCCCCUUCACCCCGUUGGAGGGUCCGAUACUCUUAAGAUUCUUCCCGGUAAAUGAAGUAGAGCAC........UGUAU.................GUGCU.........GGCUAUGGUAAGUCCGA..CGUGCUCCCGGA................
env-65    ...GCCCUCACUGGUUUGAUGGCCUCCCUUCACCCCGCUGGAGAGUCCGAUACUCUUAAGAUUCUUCCCGGUAAAUGAAGUAGAGCAC........UGUAU.................GUGCU.........GGCUAUGGUAAGUCCGA..CGUGCUCCCGGA................
env-66    ......UGGGGUCUUCCCGCUCCGCCCCUUCACCCCGUUGGAGGGUCCGAUACUCUUAAGAUUCUUCCCGGUAAAUGAAGUAGAGCAC........UGUAU.................GUGCU.........GGCUAUGGUAAGUCCGA..CGUGCUCCCGGA................
env-67    ...GCCCUCACUGGUUUGAUGGCCUCCCUUCACCCCGCUGGAGAGUCCGAUACUCUUAAGAUUCUUCCCGGUAAAUGAAGUAGAGCAC........UGUAU.................GUGCU.........GGCUAUGGUAAGUCCGA..CGUGCUCCCGGA................
env-68    ......................................GAGAGUCCGAUACUCUUAAGAUUCUUCCCGGUAAAUGAAGUAGAGCAC........UGUAU.................GUGCU.........GGCUAUGGUAAGUCCGA..CGUGCUCCCGGA................
env-69    ...AACCAUCCGCGCAACGUUCUUCCGCCGGGGGUUUUGCACCCGCCCGAUACCCUUAAGAUACCUCCCGGAGAAUAUGUUGCAGCACA.......AUA..................UGUGCU.........GGCUAUGGUAAGUCCGA..CGUGCUCCCGGA................
Msp-1-1   ...AACCUCCGUACAACGUAUUUCGCCCCCCGGCACGCGCGCCUGUCCGAUACCCUUAAGAUUCCUCCCGGAGAAUAUGUUGCAGCACG.......AUCA.................CGUGCU.........GACUGUGGUAAGUCCGA..CGUGCUCCCGGA................
Mma-1-1   ...AACCUCCGUACAACGUAUUUCGCUCCCCGGUACGCGCGCCUGUCCGAUACCCUUAAGAUUCCUCCCGGAGAAUAUGUUGCAGCACG.......AUCA.................CGUGCU.........GACUGUGGUAAGUCCGA..CGUGCUCCCGGA................
env-70    ...AAAACCCCAGGAAUUGCAUUUCCUGGGAAACCGGGCGCGCCUCCGAUAUCCUUAAGGUGUCCUGUAGCGAACGAUACUGCAGCACG.......AUAA.................CGUGCU.........GGCUAUGGUAAGUCCGA..CGUGCUCCCGGA................
Mse-1-1   ...CGCCCCCCUGUCGCCGGAAGGGGGCGCGGGCCGCGCAACCCAUACGAUACCCUUAAGAUUCCUCCCGGAGAAUAUGAUGCAGCACG.......AAUA.................CGUGCU.........GGCUAUGGUAAGUCCGA..CGUGCUCCCGGA................
env-71    ...ACUCCACGGAUCAAUCCUGCCACAGGUGCCGCAGCACACCCGCCCGAUACCCUUAAGACGCCUCCCGGUGAAUAUGAUGCAGCAC........CAUAU.................GUGCU.........GGCUAUGGUAAGUCCGA..CGUGCUCCCGG.................
env-72    ...ACUCCACGGAUCAAUCCUGCCACAGGUGCCGCAGCACACCCGCCCGAUACCCUUAAGACGCCUCCCGGUGAAUAUGAUGCAGCAC........CAUAU.................GUGCU.........GGCUAUGGUAAGUCCGA..CGUGCUCCCGG.................
env-73    ...ACUCCACGGAUCAAUCCUGCCACAGGUGCCGCAGCACACCCGCCCGAUACCCUUAAGACGCCUCCCGGUGAAUAUGAUGCAGCAC........CAUAU.................GUGCU.........GGCUAUGGUAAGUCCGA..CGUGCUCCCGG.................
env-76    ...UCAACCUCAACGGUCAGCCGUGCCCCUGACCCGUGCUCCUUCGAUACCCUUAUUAUCCCAACCCACUGAUAAAUACCACAAGCACG.......AAAU.................CGUGCU.........GGCAUAUGUAAGUCCGA..CGUGCCUUUU..................
env-77    ...UCAACCGCUACGGCAGGGAGCGGUGAGGGUCCCGUCAUCUUCGAUAUGCUUAUCAUCCUGCCGGCCUUAUAAAUGAGACAAGCAC........GAAAAU................GUGCU.........GGCAUAUGUAAGUCCGA..CGUGCCGUACC.................
env-78    ...UCAACCGCUACGGCAGGGAGCGGUGAGGGUCCCGUCAUCUUCGAUAUGCUUAUCAUCCUGCCGGCCUUAUAAAUGAGACAAGCACG.......AAAA.................UGUGCU.........GGCAUAUGUAAGUCCGA..CGUGCCGUACC.................
env-79    ...CUGAACAAGNGGUUUUUUCCCCAGUCCGUUUCUCUGGCAGGUACGAUAGCAUUAUUAUACAAUCUGCUGAUUUAGUAGGUAGCACUCUUUCAUUUUU..........AUGAA..AGUGCU.........GGCAUAUGUAAGUCCGA..CGUGCCGCUCCC................
env-80    ...GUUCCCGACUCUAAAGGCAGUUCAUGGUGAGGCAGCGAUCCUCAGAUACCCUUAAUAUUUGCCGGUGUCAAUAAUAGGACAGCACUU......UUUAAAAC............AGGUGCU.........GACAGAUGUAAGUCCGA..CGUGUCCGUUUCGGC.............
env-81    ...GUUCCCGACUCUAAAGGCAGUUCAUGGUGAGGCAGCGAUCCUCAGAUACCCUUAAUAUUUGCCGGUGUCAAUAAUAGGACAGCACUU......UUUAAAAC............AGGUGCU.........GACAGAUGUAAGUCCGA..CGUGUCCGUUUCGGC.............
env-82    ...GUUCCCGACUCUAAAGGCAGUUCAUGGUGAGGCAGCGAUCCUCAGAUACCCUUAAUAUUUGCCGGUGUUAAUAAUAGAACAGCACUU......UUUAAAAC............AGGUGCU.........GACAGAUGUAAGUCCGA..CGUGUCCGUUUCGGC.............
env-83    ...UCAGGUUAUCCCAGGCACACGGCCAUAAAUGCGGCCCGACCACAGAUACCCUUAAUAUUUGCAGGAGUUAAUAGUAGGACAGCACUUU.....UUAAC..............AGGGUGCU.........GACAGAUGUAAGUCCGA..CGUGCCGUUUU.................
env-84    ...UGCCGGAUCCCACCGACAGGCCCCGGUGACGGCGGCAGGUAUCAGAUACCCUUAAUAUCGCCAGCGGUCAAUAAUACAACAGCACUU......UUUAAAAC............AGGUGCU.........GACAGAUGUAAGUCCGA..CGUGCCCGUUUCAA..............
env-85    ...AGGUGCCGGACGCCAUGCGGUCCCGGUGACGGCGGCAGGGAUCAGAUACCCUUAAUAUCGGCAGCGGCCAAUAAUAGGACAGCACUU......UUUAAAAC............AGGUGCU.........GACAGAUGUAAGUCCGA..CGUGCCCGUUUCGAC.............
env-86    ...UGCCGGACGCCACCGGGCCUGCAUAACAGCGGGUACGCUCCUCAGAUACCCUUAAUAUUUGCCGGUGUCAAUAAUAGGACAGCACUU......UUUAAAAC............AGGUGCU.........GACAGAUGUAAGUCCGA..CGUGCCCGUUUCGAC.............
CMe-1-1   ...AGGAAAUGCCCGCUGAUCUCUUACAAAAGCGGCACCGGGACACAGAUACCAUUAAUAUUUGCAGCCGUCAAUAGUAGGGUAGCACUUU.....CUUAU.............AAGGUGCU.........GACAGAUGUAAGUCCGA..CGUGCCUUUCUU................
env-87    ............CAAAGGAAGUACACGGUAAGGCAACGAUCCUCAGAUACCCUUAAUAUUUGCCGAUGUUAAUAAUAGGACAGCACUU......UUUAAAAC............AGGUGCU.........GACAGAUGUAAGUCCGA..CGUGCCCGAUUAAA..............
env-88    ...GUUCCCGACUCCAAAGGAAGUACACGGUAAGGCAACGAUCCUCAGAUACCCUUAAUAUUUGCCGAUGUUAAUAAUAGGACAGCACUU......UUUAAAAC............AGGUGCU.........GACAGAUGUAAGUCCGA..CGUGCCCGAUUAAA..............
Mli-1-1   ...GCAUACUUUCGGGAGGGAGAUUAAAGGUUUCGCUUUUGUUCUGCCGUCAGCCUUAAGAAGAAGGAACGAGAUUUAUAUAUAGCAC........UUUUAU................GUGCU.........GGCCGAUGUAAGUCCGA..CGUGCCGUUCC.................
Mfo-1-1   ...CGCACCGGGCAGGACGCUCCUUUGUGAUGGGGGCUGACACUACGGUGCCUUUUAUUACCCCUCCCCGCCAAUAGUAAGCCAGCAC........UGUUUAAU..............GUGCU.........GACAGAUGUAAGUCCGA..CGUGCGACCAC.................
env-89    ...GCAAUUCUUCUCCUCUUCCAGGUCCUGACGGACCUGUCAGAUGCCGCAACCCUUAAUAUCCCUCCCGUCGAUUGAUGAUCAGCACU.......UUUAAC...............GGUGCU.........GACAGAUGUAAGUCCGA..CGUGCGACAC..................
env-90    ...GCAAUUCUUCUCCUCUUCCAGGUCCUGACGAACCUGUCAGAUGCCGCAACCCUUAAUAUCCCUCCCGUCGAUUGAUGAUCAGCACU.......UUUAAC...............GGUGCU.........GACAGAUGUAAGUCCGA..CGUGCGACAC..................
env-91    ...GAGGUUUGUGGAAGGGACAUCUCCCACCAGGCUAGAAUUCCCCCGUUCCCUUUAUUAUCCCUCCCCGCCAAUAGAUAUCUAGCACU.......AUUCACA..............GGUGCU.........GACAGAUGUAAGUCCGA..CGUGCGACAC..................
env-92    ...CUGCUAAGAAGGACGAUGAAUGAAAAACGGCGUACUUCUCUCCCGUACCCUUUAUUACCCCUCCCCACCAAUAGUAGGAUAGCAC........UUUACAU...............GUGCU.........GACAGAUGUAAGUCCGA..CGUGCGACAC..................
env-96    ...CAAACGGUACCCGGUGUCCGUGUGAGGCUACGGGAUUUUCUCCCGUACCCUUUAUUACCCCUCCCCACCAAUAGUAGAAUAGCAC........UUUAUAU...............GUGCU.........GACAGAUGUAAGUCCGA..CGUGCGACAC..................
env-97    ..................................GAUUUUCUCCCGUACCCUUUAUUACCCCUCCCCGCCAAUAGUAGGACAGCAC........UUUAUAU...............GUGCU.........GACAGAUGUAAGUCCGA..CGUGCGACAC..................
env-98    ...AUACAAUGGUACAGGGCAGACCGUGGCAGGUAUGGCAGGUCAUCCGAUACCCUUAUUAUACUACCCGCCGAUUAAUGCUCAGCAC........UUUUCAGAU.............GUGCU.........GACAGAUGUAAGUCCGA..CGUGCGACAC..................
env-99    ...AUACAAUGGUACAGGGCAGACCGUGGCAGGUAUGGCAGGUCAUCCGAUACCCUUAUUAUACUACCCGCCGAUUAAUGCUCAGCAC........UUUUCAGAU.............GUGCU.........GACAGAUGUAAGUCCGA..CGUGCGACAC..................
env-103   ...CCCGUUUUACCAUCAUGCACCCUUUUCGAGAUGACCAACCGGGACGCAACCCUUAUUAUACCUCCCGUCGAAUGAUGGUAAGCAC........UUUUACCAAAU...........GUGCU.........GACAGAUGUAAGUCCGA..CGUGCGAACCAA................
env-104   ...................................................................GGCAAAUAGUAGAUCAGCAC........UUUUCAUAU.............GUGCU.........GACAGAUGUAAGUCCGA..CGUGCUUAAC..................
env-105   ......................................................CUUAUUAUACCUCCCGUCGAAUGAUGAUAAGCAC........UUUUUAACAACU..........GUGCU.........GACAGAUGUAAGUCCGA..CGUGCGACAC..................
env-106   ...ACGUGAACGGAAUCAGCAGGUCAGGCGGGCAUAUCGGCCUUGUCCGAUACCCUUAUUAUUCCACCCGGCGAUUACUGGUAAGCAC........UUUUCAGAU.............GUGCU.........GACAGAUGUAAGUCCGA..CGUGCGACAC..................
env-107   ...CCGGAAACGGAUUUUUCAGCUCCGGCAGGUAUAUCGGCCUUGUCCGAUACCCUUAUUAUUCCACCCGGCGAUUACUGGUAAGCAC........UUUUCAGAU.............GUGCU.........GACAGAUGUAAGUCCGA..CGUGCGACAC..................
env-108   ...GGGGUACCUGGUUUCCGCAUCCAUCAACUCCUGACACACCCCUCCGCAAUGCUUAUCAUACCACCAGUCGAUUGAUGAUAAGCAC........UUUUCAUAUU............GUGCU.........GACAGAUGUAAGUCCGA..CGUGCGACAC..................
env-109   ......................UAGGAAUGGCGGGAGGGGGCAUCCGCCGCAACCCUUAAUAUCCCAUCCGUCGAAUGAUGGUAAGCAC........UUUUAACAAU............GUGCU.........GACAGAUGUAAGUCCGA..CGUGCGACAC..................
env-110   ...ACGACGAUACGGAAACAUGAUCCGUAUCCGGCCGGCCCUCCCGCCGCAACCCUUAAUAUCCCUCCCGUCGAUUGAUGAUCAGCAC........UUUUACCU..............GUGCU.........GACAGAUGUAAGUCCGA..CGUGCGACAC..................
env-111   ...GAGGAUAAACUAACAUACGAAUCUGUGAUGAGAACUGUCCCCUCAGAUAUCCUUAAUAUCCCUCCCGUCGAUUGAUGAUCAGCAC........UUUUAACU..............GUGCU.........GACAGAUGUAAGUCCGA..CGUGCGACAC..................
env-112   ...GAGGAUAAACUCACAUACGAAUCUGUGAUGAGAACGGUCCCCUCAGAUAUCCUUAAUAUCCCUCCCGUCGAUUGAUGAUCAGCAC........UUUUACU...............GUGCU.........GACAGAUGUAAGUCCGA..CGUGCGACAC..................
env-114   ...GAGGAUAAACUCACAUACGAAUCUGUGAUGAGAACGGUCCCCUCAGAUAUCCUUAAUAUCCCUCCCGUCGAUUGAUGAUCAGCAC........UUUUACU...............GUGCU.........GACAGAUGUAAGUCCGA..CGUGCGACAC..................
env-115   ...GGGUGUUCGAGCAAAUCCUUAAGCUGACCCAACUCAUGCAUCUGCGAUACCCUUAUUAUACCUCCCGUCGAAUGAUGGUAAGCAC........UUUUAACAAU............GUGCU.........GACAGAUGUAAGUCCGA..CGUGCGACAC..................
env-117   .............................................................AUACCUCCCGUCGAAUGAUGGUAAGCAC........UUUUUAACAAU...........GUGCU.........GACAGAUGUAAGUCCGA..CGUGCGACAC..................
env-118   ...AGGCUUUUAUCCGCGUCACCGUCCGGAUCAGGGUUGCCGCGGGGCGAUACCAUUAAUAGAGCUCCCAUCGAAUUAUGGUAAGCAC........UUUUCAUAU.............GUGCU.........GACAGAUGUAAGUCCGA..CGUGCGACAC..................
env-120   ...GAAUAGUGGGGAGGUCAUGGACGAAAAGCCGGGAAGUCCGGCUCAGUUACCCUUAUUAUACCUCACAUCGAAUGAUGGUAAGCAC........UUUUAAUAAAU...........GUGCU.........GGCAGAUGUAAGUCCGA..CGUGCGAUACAAA...............
env-122   ...GUGAAAUAAUCGAGCAAGAUGUUUCAAUCACCACUCCUUCCCAUCGCAACCCUUAUAAUUCCUCCCCCAAAUAGAGUCAAGCAC........UUUUAAUAAAU...........GUGCU.........GACAGAUGUAAGUCCGA..CGUGCUAUCAA.................
env-123   ...UAUUUGAAUUAGUCAUAAUACUCCCCCUUCCCCGGAAAAACCAUCGCAACCCUUAUAAUUCCUCCCCCAAAUACAGUCAAGCAC........UUUUAAAAAUU...........GUGCU.........GACAGAUGUAAGUCCGA..CGUGCUGUUAC.................
env-124   ...CUCAUCCCUUUCGUUCCUAACCUGUUAUUACCACAAAAAACCAUCGCAACCCUUAUUAUUCCUCCACCCAAAUACAGUCAAGCAC........UUUUACAAUU............GUGCU.........GACAGAUGUAAGUCCGA..CGUGCUUUUAA.................
env-128   ...CUCAUCCCUUUCGUUCCUAACCUGUUAUUACCACAAAAAACCAUCGCAACCCUUAUUAUUCCUCCACCCAAAUACAGUCAAGCAC........UUUUACAAUU............GUGCU.........GACAGAUGUAAGUCCGA..CGUGCUUUUAA.................
env-129   ...CUCAUCCCUUUCGUUCCUAACCUGUUAUUACCACAAAAAACCAUCGCAACCCUUAUUAUUCCUCCACCCAAAUACAGUCAAGCAC........UUUUACAAUU............GUGCU.........GACAGAUGUAAGUCCGA..CGUGCUUUUAA.................
env-131   ...CUCAUCCCUUUCGUUCCUAACCUGUUAUUACCACAAAAAACCAUCGCAACCCUUAUUAUUCCUCCACCCAAAUACAGUCAAGCAC........UUUUACAAUU............GUGCU.........GACAGAUGUAAGUCCGA..CGUGCUUUUAA.................
env-132   ...CUCAUCCCUUUCGUUCCUAACCUGUUAUUACCACAAAAAACCAUCGCAACCCUUAUUAUUCCUCCACCCAAAUACAGUCAAGCAC........UUUUACAAUU............GUGCU.........GACAGAUGUAAGUCCGA..CGUGCUUUUAA.................
env-133   ...CUCAUCCCUUUCGUUCCUAACCUGUUAUUACCACAAAAAACCAUCGCAACCCUUAUUAUUCCUCCACCCAAAUACAGUCAAGCAC........UUUUACAAUU............GUGCU.........GACAGAUGUAAGUCCGA..CGUGCUUUUAA.................
env-134   ..........GACGUUCACUGCCCGAAGAUCAUCAANNNANNCCAUCGCAACCCUUAUUAUUCCUCCACCCAAAUANNGUCAAGCAC........UUUUACAAUU............GUGCU.........GACAGAUGUAAGUCCGA..CGUGCUUUUAA.................
env-135   ...AAAUUCCGCAAAAAUUACAUUGAGAAAUUGGCGUGAAAAACCAUCGCAACCCUUAUAAUUGCUUAACCCAAAUAGAGUCAAGCAC........UUUUACAUAUU...........GUGCU.........GACAGAUGUAAGUCCGA..CGUGCUUUUUUG................
env-136   ...AAAUUCUGGAAAAAUGGCUUUGCAGACAUUGCCGUGAAAACCAUCGCAACCCUUAUAAUUCCUCCACCCAAAUACAGUCAAGCAC........UUUUACAUAUU...........GUGCU.........GACAGAUGUAAGUCCGA..CGUGCUUUUUUG................
env-137   ...AAAUUCCGCAAAAAUUGCAUUGAGAAAUUGNCNUGAAAAACCAUCGCAACCCUUAUAAUUGCUUAACCCAAAUAGAGUCAAGCAC........UUUUACAUAUU...........GUGCU.........GACAGAUGUAAGUCCGA..CGUGCUUUUUUG................
env-138   ...AAAUUCCGCAAAAAUUGCAUUGAGAAAUUGACUUGAAAAACCAUCGCAACCCUUAUAAUUGCUUAACCCAAAUAGAGUCAAGCAC........UUUUACAUAUU...........GUGCU.........GACAGAUGUAAGUCCGA..CGUGCUUUUUUG................
env-148   ...CAUAGGCCUUAGUAAGUCCCCCAGAUAUUCUGCCGGAAAACCAUCGCAACCCUUAUAAUUCCUCCACCCAAAUACAGUCAAGCAC........UUUUACAUAUU...........GUGCU.........GACAGAUGUAAGUCCGA..CGUGCUUUUUUA................
env-149   ...CAUAGGCCUUAGUAAGUCCCCCAGAUAUUCUGCCGGAAAACCAUCGCAACCCUUAUAAUUCCUCCACCCAAAUACAGUCAAGCAC........UUUUACAUAUU...........GUGCU.........GACAGAUGUAAGUCCGA..CGUGCUUUUUUA................
env-150   ...AAAUUCCGGAAAAAUUGCAUGAAGAAAUUGGCGUGAAAAACCAUCGCAACCCUUAUAAUUGCUCAACCCAAAUGUAGUCAGGCAC........UUUUACAUAUU...........GUGCU.........GACAGAUGUAAGUCCGA..CGUGCUUUUUUA................
env-151   ...GCUCCAGGAUGAAGGUCGUUGCAGAAAUCGGCAGGGAAAUCCAUCACAACCCUUAUAAUUUCUCCACCCAAAUAGAGUCAGGCAC........UUUUAGAAAU............GUGCU.........GACAGAUGUAAGUCCGA..CGUGCUUCAC..................
env-152   ...AAUUCCAAAAAAAGAUAGCGUUGCAAAGACCGAUGGAAAACCAUCGCAACCCUUAUAAUUCCUCCACCCAAAUACAGUCAGGCAC........UUUUACCAAUU...........GUGCU.........GACAGAUGUAAGUCCGA..CGUGCUUCAC..................
env-153   ...AAUUCCAAAAAAAAGAUAGCGUUGCAAAGACCGAUGGAAAACCAUCGCAACCCUUAUAAUUCCUCCACCCAAAUACAGUCAGGCAC........UUUUACCAAUU...........GUGCU.........GACAGAUGUAAGUCCGA..CGUGCUUCAC..................
env-154   ...UCUCGGCAUUCCUGCUGAACCGCUCUUCCGAUCUUCACCCCAUCACAACCCUUAUCAUUCCUCUCACCUCAUAAUAGUCAAGCAC........UUUUACAAAUU...........GUGCU.........GACAGAUGUAAGUCCGA..CGUGCUUCAC..................
env-155   ...AUGAAUCCAUCAGAAGUCUUGCAGAAAUUAGCAUGAAAAACCAUCACAACCCUUAUAAUUUCUCCACCCAAAUACAGUCAAGCAC........UUUUACAAAUU...........GUGCU.........GACAGAUGUAAGUCCGA..CGUGCUUCAC..................
env-156   ...CCUCGUUUCCGGCACCGUGAAUCCCUGAGGGCCCGGUCUCUCUCAGUUACCCUUAUUAUACCUCCAGUCAAAUGAUGGUAAGCAC........UUUUAACAAU............GUGCU.........GGCAGAUGUAAGUCCGA..CGUGCGACAA..................
env-157   ...ACCUGGUUUCCGGUACCGUAAAUCCUUGAGGAAGUAUCCGCCUCAGUUACCCUUAUUAUACCUCCAGUCGAAUGAUGGUAAGCAC........UUUUAACAAU............GUGCU.........GGCAGAUGUAAGUCCGA..CGUGCGACAA..................
env-159   ..........................................................................AAUGAUGGUAAGCAC........UUUUAACAAU............GUGCU.........GGCAGAUGUAAGUCCGA..CGUGCGACAA..................
env-160   ...GAUUUGAAUUAGUCAUAAUACUCCCCCUUCACCGGAAAAACCAUCGCAACCCUUAUAAUUCCUCCCCCAAAUAGAGUCAAGCACUUUU....UAAU...............AAAUGUGCU.........GACAGAUGUAAGUCCGA..CGUGCUGUUAC.................
env-161   ...AAUUCAGGAUAGAACUCAUUGUAAAAAUUCCCGCAAAAAACCAUCGCAACCCUUAUAAUUCACCACCCAAAUACAGUCAAGCACUUUU....AAUA...............AAUUGUGCU.........GACAGAUGUAAGUCCGA..CGUGCUUCACA.................
env-162   ...CAGAGAGAUUAAGUUUUGGCAGAUCCUUUUUUUGAAAAACCAUCACAACCCUUAAUAUUCCUCCGCCUAAAUAGAGUCAAGCACUUUU....ACCA...............AAAUGUGCU.........GACAGAUGUAAGUCCGA..CGUGUUUCACA.................
env-163   ...CCAGGUCAACCGCUACGGCAGCACCUGACCGCACCUGCCGCCCACGAUAGCAUUAAUAUUCCACCACCCAAAGAAAAUACAGCACU.......UCUGGAU..............AGUGCU.........GGCAGAUGUAAGUCCGA..CGUGCGGCAA..................
env-164   ...GCAGAUCAACCGCUACGGCAGCACCUGACCGCACCUGCCGCCCACGAUAGCAUUAAUAGUCCACCACCCAAAGGAGCUACAGCACU.......UCUGGAU..............AGUGCU.........GGCAGAUGUAAGUCCGA..CGUGCGGCAA..................
env-165   ...........................................................AUUCCCCCGCCCAAAUGGAUUGAAGCACU.......CUUGGAU..............AGUGCU.........GGCAGAUGUAAGUCCGA..CGUGCGGCAA..................

          .....................................................................................<<<<<<<<<<<<<..............>>>>>>>>>>>>>.................<<<<..<<.....<<<<<...........<<<<<<<.....>
          .......................................................................................00.2..0..10..............01..0..2.00..................0100..00......1122............110000.....0
          CCG••••••••••••••••••••••••••••••••••••••Y••••R•••YYYU•A•••Y•YYYYY••••RA••R••R•••AGCAC○-------•Y•••○---------------○GUGCU--------GRC•RA•GUAAGUCCGA--CGUGC•••••○------------------
                                                                                        5′
```

```
env-166   ...UGUAUUCAGGACGCACGGGGAAGCAGGUGACCCGGCCUCCCCGCGAUAUCAUUAAGAGAAUUCCCGUUCCACGGAUAUGCAGCACUG......UUUCUAC.............UGGUGCU.........GGCAGAUGUAAGUCCGA..CGUGCACCACGC.................
env-168   ...CGCAUUCAGGACGCACGGGGGAGCAGGUGAGCCGGACUCCCCGCGAUAUCAUUAAGAGAAUUCCCGCUCCAUGGAUAUGCAGCACUG......UUUCUAC.............UGGUGCU.........GGCAGAUGUAAGUCCGA..CGUGCACCACGC.................
env-169   ...GGCUCAUCAACCGGUACGGAAGCACCUGAUACCAGGUGCAUGCCCGAUAGCCUUAAUAUUCAUUCAAUCAAAUGAAAUCAGGCACUUUU....GAAA..............AAAUGUGCU.........GGCAGAUGUAAGUCCGA..CGUGCAUCACACA................
Mta-1-1   ...GAUCCGGCUCAUCAACCGCUACGGGCAGCCGCUGUAGCCCCCGCGAUAUCAUUAAGAGUACGCCUGCCAAAUGGAUAGACAGCACUU......AUAA................GGGUGCU.........GGCAGAUGUAAGUCCGA..CGUGCAGACCA..................
env-174   ...GAUCCGGCUCAUCAACCGCUACGGGGAGCCGCUGUAGCCCCCGCGAUAUCAUUAAGAGUACGCCUGCCAAAUGGAUAGACAGCACUU......AUAA................GGGUGCU.........GGCAGAUGUAAGUCCGA..CGUGCAGACCA..................
Mli-2-1   ...GAAGAGGUGUUAUGUAUUGCUGCCGCCCCGGAAUAAAUCCCUGCGAUAUUAUUAAUAUCUCAUCCAUCCUAGUUUUAUGAAGCACUUAUC...UUU..............GAU.AGUGCU.........GGCUUUUGUAAGUCCGA..CGUGUUCUGUACAACA.............
Mba-1-1   ...GAUCACAUCAUACUCACGCAUAUUCAUUGUAUGUACGUACGUCCCAAACCCUUAUUAUUGAUCCCCUCCAACAUAUACGAAGCAC........UUAG..................GUGCU.........GACUGAAGUAAGUCCGA..CGUGUCUGUCUACUAA.............
Mla-1-1   ...GAUCACAUCAUACUCAUGCAUACUCAUUGUAUGUACGUACGUCCCAAACCCUUAUUAUAAAUCCCCUCCAACAUAUACGAAGCAC........UUAG..................GUGCU.........GACUGAAGUAAGUCCGA..CGUGUCUGUCUACUAA.............
env-177   ...CCGGUUCACGACAACCACGUGAGGAGGUACGAAUCGUUCCCGGAGGAUAUCCUUAUCUCGUCUGCAUAAGAAGGUACUACAGCACUU......UUCG................AAGUGCU.........GGCUGAUGUAAGUCCGA..CGUGUCGUCC...................
env-178   ...CCGGUUCACGACAACCACGUGAGGAGGUACGAAUCGUUCCCGGAGGAUAUCCUUAUCUCGUCUGCAUAAGAAGGUACUACAGCACUU......UUCG................AAGUGCU.........GGCUGAUGUAAGUCCGA..CGUGUCGUCC...................
env-179   .............UUUUUUAUCGAUAUUCACCACGCCUCAAAUACAACACUACUUUUAAUAUAACACCCGUAGAAAGUAUAUUAGCACUCAU....AAC...............AUG.GUGCU.........GACAGAAGUACAUCCGA..CGUGACAUCAAU.................
          ...................................................................................<<<<<<<<<<<<<..............>>>>>>>>>>>>>................<<<<..<<....<<<<<...........<<<<<<<....>
          .....................................................................................00.2..0..10..............01..0..2.00..................0100..00.....1122...........110000....0
          CCG••••••••••••••••••••••••••••••••••••••••Y••••R•••YYYU•A•••Y•YYYYY••••RA••R••R•••AGCAC○-------•Y•◦○---------------○GUGCU--------GRC•RA•GUAAGUCCGA--CGUGC•••••○-----------------
                                                                                           5′
```

alignment positions 181 · · · 360

```
env-1     ...ACG......................GAUACUGCCA........GAUA................UGGCUUUGGG.AUUACAAAAAAGGUCAGUAAUGGCUCUUUUUGUGGACUUACAUGUAUCUCAUCAGAAAAAGAGAUACAUGGACAAAAAAGCGGAAGCCCCGCCGAUUUGUCCC
env-2     ............................GGCGC.GGCCU.......UCC................AGGCU..UGGG.AUUACGGAGCGUGCGUCUC...................................................................................
env-3     ............................GGCGC.GGCCU.......UCC................AGGCU..UGGG.AUUACGGAGCGUGCGUCUCAUCGCUGCUCUUCUCGGGACUAUCGGAAUCGGAGUAGAACUAUGGCACGAAUGCAUGCGCGACGGAGAGGCAAGUCGGGAUCUG
env-4     ............................GGCGC.GGCCU.......UCC................AGGCU..UGGG.AUUACGGAGCGUGCGUCUCAUCGCUGCUCUUCUCGGGACUAUCGGAAUCGGAGUAGAACUAUGGCACGAAUGCAUGCGCGACGGAGAGGCAAGUCGGGAUCUG
env-5     ............................GCCAC.GGCC........UUCA................GGCUU.UGGG.AUUGCAGGGAGUCAGCAGAAAUAGCUGUCACUCUUUUCCCGUAGGAUUUUAUACUCUGUUCAUGUAACGGAGGCUUUUAGAUAUGGCAAGAAUGCAUGCUCGC
env-6     ............................GUCACUGCCU........GUAA................AGGCUUUGGG.AUUGCAGAGAGUUGGCAGAUUAAUCACCUGACUCUUUCUCUCUCAGGAUUUUGCACUGUUUACAGGAGGUUAUAGAAAAUGGCACGAAUGCAUGCCCGCAGAA
env-7     ............................AUCAC.GGCCC.......GGUCA..............GGGCUU.UGGG.AUUGCAGAAAGUUGGCAGAAUUAAUUAUCUGGCUCUUUCUAUCGCAGGAUUUUGCUCUGUUUUCAGGAGGAUUUUGUUAUGGCAAGAAUGCAUGCCCGAAGAA
env-8     ............................AGCACUGCUCCUU.....UUG..............GGGGGGCU.UGGG.AUUACGGCGAGUCAGUUUCGGCUCUUUUCGUGACUUACAGUGAAUGGCAGCUGCCAGACACUCCUGCAGGAUUCUUCUGCACGGCAGGCUAUAAACAACAAUU
env-10    ............................AGCACUGCCC........UGUAUGU.............GGGCU.UGGG.AUUGCGGCGAGUCAGAUACGGCUCUUUUCGUGACUUACGGUGAACGGUAUGACCGGACACAUCUGCUGUAACCAGCAGGCAGCUAAAAAAUUUCGAGGUACCA
env-11    ............................AGCACUGCCCU.......GUAUG..............GGGGCU.UGGG.AUUGCGGCGAGUCAGAUACGGCUCUUUUCGUGACUUACGGUGAACGGUAUGACCGGACACACCUGCUAUAACCGGCAGGCAGCUAUAGAAUUUCGAGGUACCA
Mpa-1-1   CUGCAGAUUCCUGAACAAUCAGGG.GAUGGCACUGCCCAG......AGAU..............CUGGGCUCAGGG.AUUACAGAGCUGAUGCAACAGACGUACGCACAAAUACGUACAUGCUCUUUUCUGGACUUACUCUGAUAGAUUUGGAGGUGAAUUAUGGCACGAAUGCAUGCCC
env-12    CUGCAGAUCCCUGAACAAUCAGGG.GAAGGCACUGCCCG.......AAAAAC.............CGGGCUCAGGG.AUUACAGAGCUGAUGCAACAGACGUACGCACAAAUCCGUACAUGCUCUUUUCUGGACUUACUCUGAUAAAUUUGGAGGUGAAAUAUGGCACGAAUGCAUGCCC
env-13    ...UGG......................GGCAUCGCC.........GUAAU................GGCU.UGGG.AUUACAGAGAGUCAGUAAAUACGGACUCUUUUUCCAAGGACCCCAUCGUGUCCAUUCGGAGGUGAACGUAUGGCAAGAAUGCAUGCCCGAAGGAAAGGGAUCU
env-14    ............................GACACUGCC.........CUUGA................GGCUUUGGG.AUUACGUAUCGAGUCAGCUACAUUACAG.........................................................................
env-15    ............................GACACUGCC.........CUUGA................GGCUUUGGG.AUUACGUAUCGAGUCAGCUACAUUACAGCAAAUGCUAAGGCUGCUCUUUUCGUGACUUACACCCGCAUAUCCAACUACAAACGCGAUAAGGAUAUCCGGGCAG
env-16    ............................CGCAC.GGCC........UGCGU...............GGCU..GGGG.AUUACUGAGGGUCAGCAGUUAUGACUCUUUUUCCCGGGUUUACAUGAUUUAAGGAGGUGAAAGAUGGCACGAAUGCAUGCCCGGCGUCGGGGAAAGUCAGGAU
env-17    ............................GUCAC.GGCCUGA.....AUGAA............UCAGGCUC.AGGG.AUUACCGGGGUCAGUAUAAACAGGGAUACCUGUGGCUCUUUUCCCGUAUGGAUUAUUCUGUUACGAACACAGAUAGGAGGUAGUAUUCAUGGCACGUAUGCAU
env-18    ............................GGCACUGCC.........CUUUGU...............GGCU.GGGG.AUUACGGGAGUCAGUAUCUGGCCCCUUUUUUGGACUUACGUACGUCGUUUGGAGGAUUGUUCGAAUGGCAAGAAUGCAUACCCGCAGGAGAGGGAAGUCCAGC
env-20    ............................GGCAC.GGCCU.......UGCGGGAAUC.........AGGCU..CGGG.AUUACAGGAGAGUCAGCAGAGACGACUCUUUUUCCGGAUUUACAUGGUGCCAUCUGGAGGUGAACCAUGGCACGUAUACACGCACGCCGACGAGGCGAUUCAG
env-21    ............................GGCAC.GGCCU.......UGCUGGAAUC.........AGGCU..CGGG.AUUACAGGAGAGUCAGCAGAGACGACUCUUUUUCCGGAUUUACAUGGUGUCAUCUGGAGGUGAACCAUGGCACGUAUACACGCACGCCGGAGAGGCGAUUCAG
env-22    ............................GGCAC.GGCCU.......UGCUGGAAUC.........AGGCU..CGGG.AUUACAGGAGAGUCAGCAGAGACGACUCUUUUUCCGGAUUUACAUGGUGUCAUCUGGAGGUGAACCAUGGCACGAAUACACGCACGCCGACGAGGCGAUUCAG
env-23    ...UUG..................AGACGGCACUGCCC.........UGUUCAU.............GGGCUUUGGG.AUUACAAAAAGGUCAGUUAUGGCUCUUUUUGUGGACUUACAUGACGAAUGCGAAUCAUAUCUGUGAUCGCAAAAAAAGUAUCUGUUAUAAUAAAAUAGGUGGU
Mpe-1-1   ...UUG..................AGACGGCACUGCCCUG......AAUC..............CAGGGCUUUGGG.AUUACAAAAAGGUCAGUUAUGGCUCUUUUUGUGGACUUACAUGACAUUGCGGACAAUAUUUUCGUGUUCGCUGAAAUUAUCUUCAAAACACGCGUGAAAAGAU
Mpa-2-1   ...UUG..................AGACGGCACUGCCCGG......GAU...............CCGGGCUUUGGG.AUUACAAAAAGGUCAGUUAUGGCUCUUUUUGUGGACUUACAUGACAUUGCGGACUAUUUUUCUUGUGUCCGCAGGAAGUAUCUUAAAACACGCGUGAAAAGAU
env-24    ............................UGCACUGCCC........AACU................GGGCU.UGGG.AUUACCGGGAGUCAGCGACUGCUGCUCUUUUUGGACUUACAUAAUGUCGUAAGGAGGUUAUACUUUUGGCAAGAAUGCAUGCCCGCAGAAGGGGCAAAUCAUG
env-25    ............................GGCACUGGCCUGC.....CCU..............GCAGGCU..UGGG.AUUACAGAGAGUCAGCGGAAUUAUGACUCCUUUUCGGAUUUACAUGGGAUCCCCCGAUGAGGGGCUUCCCUUGCAGAAUACAUCUGGAAAUUUCACUGGAAUU
env-26    ............................UGCAC.GGCC........CGCAGU..............GGCU..GGGG.AUUACAGAGGGUCAGCAGUUAUGACUCUUUUUCCCGGGUUUACAUGAUUGUUAGGAGGUGAAUGAUGGCACGAAUGCAUGCCCGGCGUC..............
env-27    ............................UGCAC.GGCC.C......AUUC..............GUGGCU..CGGG.AUUACGGGGAGUCAGCAGUAGCGGCUCUUUUCCCGGAUUUACUUUUCGCACAGUAGAGGUGAACUAUGGCAAGAAUGUAUGCGCGAAGGAGAGGCCAGUCAGG
env-28    ............................UGCAC.GGCC.C......AAUC..............GUGGCU..CGGG.AUUACGGGGAGUCAGCAGUAGCGGCUCUUUUCCCGGAUUUACUUUUCGCACAGUAGAGGUGAACUAUGGCAAGAAUGUAUGCGCGAAGGAGAGGCCAGUCAGG
env-29    ............................GGCAC.GGCCUG......CAUUA.............CAGGCU..AGGG.AUUACAGGGAGUCAGCAGAAUCAGCGACUCUUUUCGGAUUUACAUAGAUAUUUAUGGAGGUGAAUUUAUGGCACGAAUGCAUGCACGCCGCAGGGGCGAUUCC
env-30    ............................AGCAC.GGCCUU......GCU...............AGGGCU..CGGG.AUUACAGGAGAGUCAGCAGAAGCUGCUCUUUUUCCGGAUUUACAUGGCGCCUUAA....................................................
env-31    ............................AGCAC.GGCCUU......GCU...............AGGGCU..CGGG.AUUACAGGAGAGUCAGCAGAAGCUGCUCUUUUUCCGGAUUUACAUGGCGCCUUAAGGAGGUGAACCAUGGCACGUAUACACGCACGCCGGAGAGGCGAUUCAG
env-32    ............................UGCACUGCCC........GAUUUUGG............GGGCU.UGGG.AUUACGAUGAGUCAGUUUCGGCUCUUUUCGUGACUUACAGUGACUGGCAGUUGCCAGACACACCUGCGGUACAAACUGCGGGCAGCUACAAUAAUUGACGAGG
env-33    ............................AGCACUGCCC........UCUC................GGGCU.UGGG.AUUACAGAGAUUAGUCAGUAAAAGACUCUUUUCUGGACUUAACAUAGCGUAUCUAACGGAGGUGUAUGAAUGGCAAGAAUGUAUGCUCGGCGCCGCGGAACCA
env-34    ............................AGCACUGCCC........UCUC................GGGCU.UGGG.AUUACAGAGAUUAGUCAGUAAAAGACUCUUUUCUGGACUUAACAUAGCGUAUCUAACGGAGGUGUAUGAAUGGCAAGAAUGUAUGCUCGGCGCCGCGGAACCA
env-35    ............................CGCACUGCCCC.......UGCA...............GGGGCU.CGGG.AUUACAGAGAAUAGUCAGUCUACGGACUUUUUUCUGGACUUAACAUAGUGUAUAAUCGGAGGUGUAUGAAUGGCAAGAAUGUACGCUCGGCGUCGCGGGACCA
env-36    ............................CGCACUGCCCC.......UGCA...............GGGGCU.CGGG.AUUACAGAGAAUAGUCAGUCUACGGACUUUUUUCUGGACUUAACAUAGUGUAUAAUCGGAGGUGUAUGAAUGGCAAGAAUGUACGCUCGGCGUCGCGGGACCA
env-39    ............................CGCACUGCCCC.......UGCA...............GGGGCU.CGGG.AUUACAGAGAAUAGUCAGUCUACGGACUUUUUUCUGGACUUAACAUAGUGUAUAAUCGGAGGUGUAUGAAUGGCAAGAAUGUACGCUCGGCGUCGCGGGACCA
env-40    ............................CGCACUGCCCC.......UGCA...............GGGGCU.CGGG.AUUACAGAGAAUAGUCAGUCUACGGACUUUUUUCUGGACUUAACAUAGUGUAUAAUCGGAGGUGUAUGAAUGGCAAGAAUGUACGCUCGGCGUCGCGGGACCA
env-41    ............................CGCACUGCCC........GCAA................GGGCU.UGGG.AUUACAGAGAUUAGUCAGUUUACGGACUCUUUUCUGGACUUACAUAGUGUAUAAUCGGAGGUGUAUUGAAUGGCAAGAAUGUAUGCCCGGCGUCGCGGAACCA
env-42    ............................CGCACUGCCC........GCAA................GGGCU.UGGG.AUUACAGAGAUUAGUCAGUUUACGGACUCUUUUCUGGACUUACAUAGUGUAUAAUCGGAGGUGUAUUGAAUGGCAAGAAUGUAUGCCCGGCGUCGCGGAACCA
env-44    ............................CGCACUGCCC........GCAA................GGGCU.UGGG.AUUACAGAGAUUAGUCAGUUUACGGACUCUUUUCUGGACUUACAUAGUGUAUAAUCGGAGGUGUAUUGAAUGGCAAGAAUGUAUGCCCGGCGUCGCGGAACCA
env-45    ............................CGCACUGCCC........GAAA................GGGCU.UGGG.AUUACAGAGAUUAGUCAGUUUACGGACUCUUUUCUGGACUUAACAUAGUAUAUAAUCGGAGGUGUAUUGAAUGGCAAGAAUGUAUGCCCGGCGUCGCGGAACC
env-46    ............................CGCACUGCCC........GAAA................GGGCU.UGGG.AUUACAGAGAUUAGUCAGUUUACGGACUCUUUUCUGGACUUAACAUAGUNUAUAAUCGGAGGUGUAUUGAAUGGCAAGAAUGUAUGCCCGGCGUCGCGGAACC
env-47    ............................UGCACUGCCC........GCAA................GGGCU.UGGG.AUUACAGAGAUUAGUCAGUUUACGGACUCUUUUCUGGACUUAACAUAGUGUAUCUAACGGAGGUGUAUGAAUGGCAAGAAUGUACGCUCGGCGUCGCGGAACC
env-48    ............................UGCACUGCCC........GCAA................GGGCU.UGGG.AUUACAGAGAUUAGUCAGUUUACGGACUCUUUUCUGGACUUAACAUAGUGUAUCUAACGGAGGUGUAUGAAUGGCAAGAAUGUACGCUCGGCGUCGCGGAACC
env-49    ............................UGCACUGCCC........GCAA................GGGCU.UGGG.AUUACAGAGAUUAGUCAGUUUACGGACUCUUUUCUGGACUUAACAUAGUGUAUCUAACGGAGGUGUAUGAAUGGCAAGAAUGUACGCUCGGCGUCGCG.....
env-51    ............................UGCACUGCCC........GCAA................GGGCU.UGGG.AUUACAGAGAUUAGUCAGUUUACGGACUCUUUUCUGGACUUAACAUAGUGUAUCUAACGGAGGUGUAUGAAUGGCAAGAAUGUACGCUCGGCGUCGCGGAACC
env-52    ............................UGCACUGCCC........GCAA................GGGCU.UGGG.AUUACAGAGAUUAGUCAGUUUACGGACUCUUUUCUGGACUUAACAUAGUGUAUCUAACGGAGGUGUAUGAAUGGCAAGAAUGUACGCUCGGCGUCGCGGAACC
env-53    ............................CGCACUGCCC........GCGA................GGGCU.CGGG.AUUACAGAGAAUAGUCAGUCAAAGGACUCUUUUCUGGACUUAACAUAGUGUAUCUAACGGAGGUGUAUAAAUGGCAAGAAUGUACGCUCGGCGUCGUGGAACC
Mbo-1-1   ............................UGCACUGCCC........GCAA................GGGCU.CGGG.AUUACAGAGAUUAGUCAGUUUACGGACUCUUUUCUGGACUUAACAUAGUGUAUCUAACGGAGGUGUAUGAAUGGCAAGAAUGUACGCUCGGCGUCGCGGAACC
env-61    ............................UGCACUGCCC........GCAA................GGGCU.CGGG.AUUACAGAGAUUAGUCAGUUUACGGACUCUUUUCUGGACUUAACAUAGUGUAUCUAACGGAGGUGUAUGAAUGGCAAGAAUGUACGCUCGGCGUCGCG.....
env-64    ............................AGCACUGCCC........GCAA................GGGCU.CGGG.AUUACAGAGAUUAGUCAGGAAACGGACUCUUUUCUGGACUUACAUAGUUAAUACCUAACGGAGGUGCAUGAAUGGCAAGAAUGUACGCUCGGCGCCGGGGAAC
env-65    ............................AGCACUGCCC........GCAA................GGGCU.CGGG.AUUACAGAGAUUAGUCAGGAAACGGACUCUUUUUCUGGACUUACAUAGUUAAUACCUAACGGAGGUGCAUGAAUGGCAAGAAUGUACGCUCGGCGCCGGGGAA
env-66    ............................AGCACUGCCC........GCAA................GGGCU.CGGG.AUUACAGAGAAUAGUCAGGAAACGGACUCUUUUCUGGACUUACAUAGUUAAUACCUAACGGAGGUGCAUGAAUGGCAAGAAUGUACGCUCGGCGCCGGGGAAC
env-67    ............................AGCACUGCCC........GCAA................GGGCU.CGGG.AUUACAGAGAUUAGUCAGGAAACGGACUCUUUUCUGGACUUACAUAGUUAAUACCUAACGGAGGUGCAUGAAUGGCAAGAAUGUACGCUCGGCGCCGGGGAAC
env-68    ............................AGCACUGCCC........GCAA................GGGCU.CGGG.AUUACAGAGAAUAGUCAGGAAACGGACUCUUUUCUGGACUUACAUAGUUAAUACCUAACGGAGGUGCAUGAAUGGCAAGAAUGUACGCUCGGCGCCGGGGAAC
env-69    ............................AGCACUGCCC........GUAA................GGGCU.CGGG.AUUACAGAGAUUAGUCAGUAAAAGGACUCUUUUCUGGACUUAACAUAGUACAUCUAACGGAGGUGCAUGAAUGGCAAGAAUGUACGCUCGGCGCCGCGGAACC
Msp-1-1   ............................AGCACUGCCC........GCAA................GGGCU.CGGG.AUUACAGAGAUUAGUCAGUUAAAGGACUCUUUUCUGGACUUAACAUAGUAUAUCUAUCGGAGGUGCAUGAAUGGCAAGAAUGUACGCUCGGCGUCGCGGAACA
Mma-1-1   ............................AGCACUGCCC........GCAA................GGGCU.CGGG.AUUACAGAGAUUAGUCAGUUAAAGGACUCUUUUCUGGACUUAACAUAGUAUAUCUAUCGGAGGUGCAUGAAUGGCAAGAAUGUACGCUCGGCGUCGCGGAACC
env-70    ............................AGCACUGCCC........GCAA................GGGCU.CGGG.AUUACAGAGAUUAGUCAGUAAAAGGACUCUUUUCUGGACUUAACAUAGUAUAUCUAUCGGAGGUGCAUGAAUGGCAAGAAUGUACGCUCGGCGUCGCGGAACC
Mse-1-1   ............................AGCACUGCCC........GCAA................GGGCU.CGGG.AUUACAGAGAUUAGUCAGUUAAAGGACUCUUUUCUGGACUUAACAUAGUAUAUCUAUCGGAGGUGCAUGAAUGGCAAGAAUGUACGCUCGGCGUCGCGGAACC

          >>>>>>..<<<<<<....>>>>>>....>>>>>.<<<<<<<<<<<<............>>>>>>>>>>>>....>>..>>>>..................................................................................................
          00011...00000....00000.....2211..1222211.1000............0001.1122221....00..0010..................................................................................................
          ----------------------------•GCAC○G•CY--------••••○---------------RG•YU-UGGG-AUUACRGRGARUYRRY•R••••••••••YYYYYYY•••••Y•••••••••••••••••••••••••••••••••••••••••••••••••R•••••
                                                                              3′
```

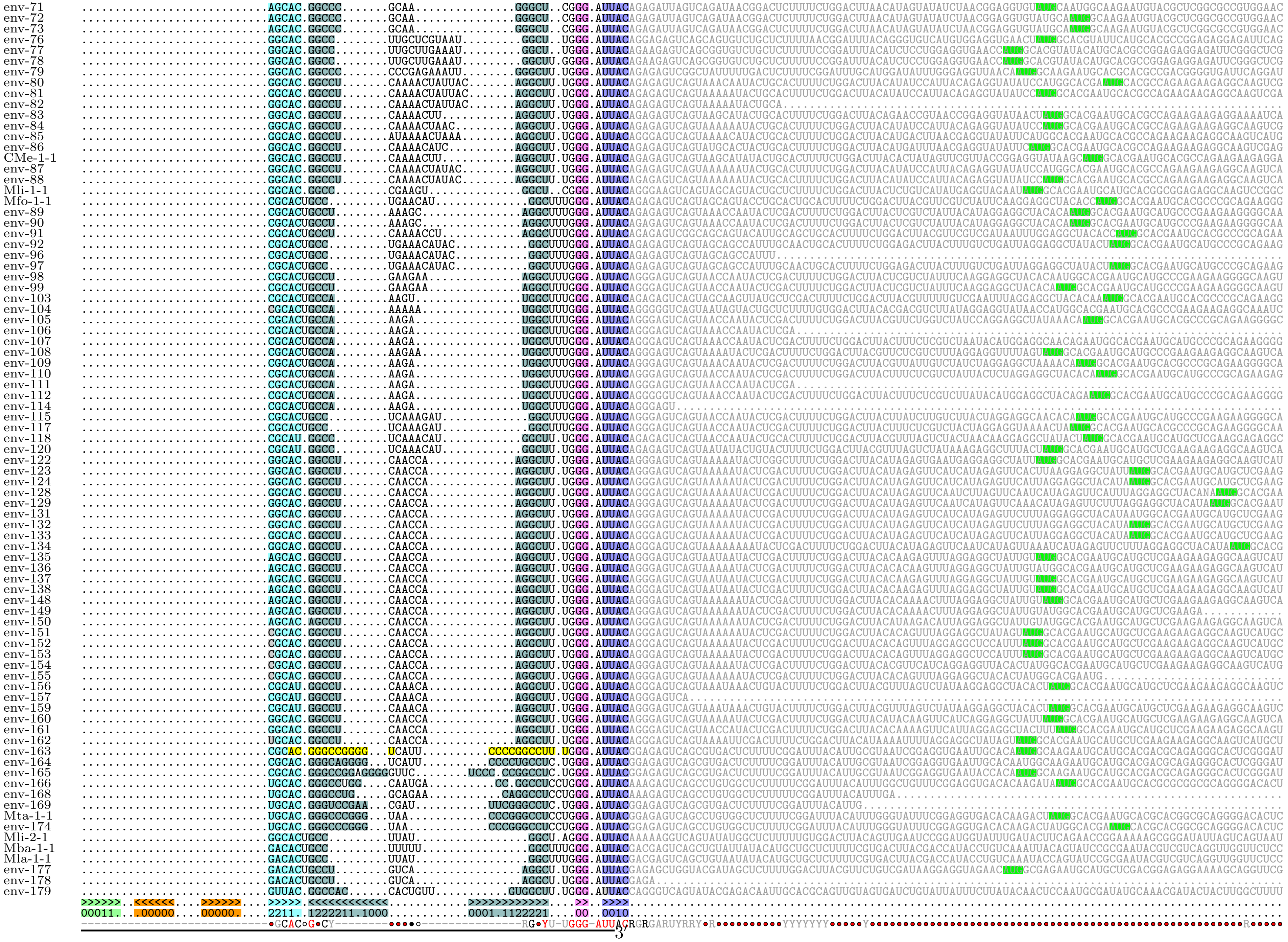

alignment positions 361 · · · 540

```
env-1     UUGUUUCUUCUUCGCAAUGAUGAAAACUAAUUAUCAACGGUGGUGAACUAAUGGCACGAAUUCAUGCCCGAAGGCGCGGAACGUCCGGUUCGGUACGUCCGCUUCGCACAGAGGCACCGGAAUGGUCAAAUAAAGACAAAGCAGAGAUCGAGAAAAUAAUCGUCGAACUCAGAAAGAACG
env-2     ....................................................................................................................................................................................
env-3     UCCGGCCACACCGGGCGGACCCGCCCGCGUGGUCGACGACCGACGCUGCCGAGAUCGAGAAGCUCAUCGUCGAGCUCCACAAAAAGGAGCUCUCGAGCGCCGCGAUCGGAACGAUCCUGCGCGACCAGUACGCAGUCCCGGACGUGAAGCUCGCGACCGGGAAGCGGGUCGGCGAGAUCC
env-4     UCCGGCCACACCGGGCGGACCCGCCCGCGUGGUCGACGACCGACGCUGCCGAGAUCGAGAAGCUCAUUGUCGAGCUCCACAAAAAGGAGCUCUCGAGCGCCGCGAUCGGAACGAUCCUGCGCGACCAGUACGCAGUCCCGGACGUGAAGCUCGCGACCGGGAAGCGGGUCGGCGAGAUCC
env-5     AGAAGAGGUAUCUCCCGUUCAGUACGGCCGUACCGCACCUCUGCGCCCGAAUGGUCAAAUACCGACAAGGAAACGAUCAUCAAGCAGGUCGUUGAUAUGAAGAAGGAGGGCCUUUCAAGUGCCCGGAUAGGUCUCGUCCUUCGUGACAAGUAUGGUGUACCCAGUGUCAAACUCGCAACA
env-6     GAGGUAUCUCACGCUCCGUAAGACCUUACCGGACAGAGGCACCGGAAUGGUCAAACAGAGACAAAGAUACCAUCAUAAAGCAGAUUGUUGACCUGCGCAGACAGGGAACAUCCACAGCCCAGAUCGGACUUAU..............................................
env-7     GAGGUAUCUCCCGUUCAGUCAGACCAUACCGGACAGAAGCACCGGAAUGGUCCAACAAGGAUAAAGAUACAAUUAUAAAACAGAUCAUCGAUCUGCGCAGGAAAGGUAUGUCGACAGCCGAGAUAGGGCUGGUAAUGCGCGACAAGUAUGGAAUUCCAAGAGUAAAACUGGUUACAGGAA
env-8     UCGAGGUAUACAUGGCACGAAUGCAUGCACGACGGAGAGGAUCAUCGGGUUCA...............................................................................................................................
env-10    UAUGGCACGAAUGUAUGCACGACGGAAGGGAUCGUCCGGCUCCGUCCGUCCUCACAGGAAGGAUGUUCCGGAAUGGUUCAAUACCGAUCUCGCUGAGAUCGAGAAGCUGGUCGUCGAUAUGAGAAAGGAAGGUCUUUCAACAGCCGUCAUCGGGACGAUUCUCCGUGAUAAAUACGGCGU
env-11    UAUGGCACGAAUGUAUGCACGACGGAAGGGAUCGUCCGGCUCCGUCCGUCCCCACAGGAAGGACGUUCCGGAAUGGUUCAAUACUGAUCUCGCUGAGAUCGAGAAGCUGGUCGUCGAUAUGAGAAAGGAAGGUCUCUCGACCGCCGUCAUUGGGACGGUUCUCCGUGAUAAGUACGGUGU
Mpa-1-1   GAUGCAGGGGUAAAUCCGGCUCAGUGAGGCCGCACAGGACAGAAGCGCCUGUCUGGUCCACGACUGACAGCACAGAGAUUGAGAAACUGAUCGUCGAACUCAAGAAGAACGAGAAGUCGACCAGUGAGAUCGGGUUGAUCCUCCGUGACCGGUACAGCGUCCCUGAUGUGAAGCUGGCGA
env-12    GAUGUAGGGGUAAAUCCGGUUCAGUGAGGCCGCACAGGACAGAAGCGCCUGCCUGGUCCACAACUGACAGCACAGAGAUUGAGAAACU...............................................................................................
env-13    CGCGAUCGGUACGACCCUACCGCAAGGAAGUCCCCUCCUGGUCGACUUCUGAUGCGAACGAGAUCGAGAAGACCAUUCUCGAUCUGCGGAAGCAGGGAAAGACCAUGAGCGAGAUCGGAUUGAUUCUGCGGGAUAAACACGGUGUCCCGAACGUGAAACUGGCCACCGGAAAAAGGAUAG
env-14    ....................................................................................................................................................................................
env-15    GUUUACUUGAUUGUACGACAGGUAAACGCAGUCAUUUAUCGCAAAGGUGAAUUAAAAUGGCACGAAUGCACGCAAGAAGACGGGGAAUUGCAGCCUCAGUCAGACCAUACAGAAAAGAAGUUC.......................................................
env-16    CGGUGCGGCCCCACCGCAAAGAAGCCCCUGAAUGGUCUAAUACUGACGUGAAGG..............................................................................................................................
env-17    GCCCGCCGCAGAGGUAUUUCAGGAUCAGUACGUCCGUACCGAACGAAAGCACCCGAGUGGUCAAAUACAGAUACAGAACAGAUCGAAACGUUAAUUAUCGAUUUAAAGAAACAGGGUGAAUCAAGCGCCCGGAUAGGCAUCAUCCUGAGAGACAAGUACGGCGUUCCAAACGUGAAACUG
env-18    UCGGUCCGUCCGUUCCGGAAAGAAGUUCCCGCAUGGUCGAACACCGAUGUCCAGAGCAUCGAGAAGAUCAUCGUGGACCUUCAGAAAGAGGGGACGUCCAUGGCGAAGAUCGGCCUCAUCCUGCGCGACAGGUACGGCGUACCCGACGUCAAGCUCCUGACCGGAAAGCGGAUCGGCGAG
env-20    GAUCGGUCAGACCGCACCGCAGCGAGGCCCCGGC..................................................................................................................................................
env-21    GAUCGGUCAGACCGCACCGCAGCGAGGCCCCGGCCUGGUCAAAUACCGACGUGAAAUCAAUCGAGAAAGUGAUUGUUGACCUGAGAAAAGAGGGGUACUCGAGCAGCCGGAUCGGCCUCGUCCUCCGUGACCAGUACGGGGUCCCUGACGUCAAACUGGUGAUGGGAAAACGGAUCGACC
env-22    GAUCGGUCAGACCGCACCGCAGCGAGGCCCCGGCCUGGUCAAAUACCGACGUGAAAUCAAUCGAGAAAGUGAUUGUUGACCUGAGAAAAGAGGGGUACUCGAGCAGCCGGAUCGGCCUCGUCCUCCGUGACCAGUACGGGGUCCCUGANGUCAAACUGGUGAUGGGAAAACGGAUCGACC
env-23    GAAAAAUGGCAAGAAUGCAUGCCCGUCGGCGCGGAAAAUCCGGAUCUGUCCGCCCGCUCCGUACCGAGGCACCCGAAUGGUCGAACAAAGACCCCGCAGAGAUCGAAAAAACCAUCAUUGAUCUUAGAAAGGAAGGAAUGUCCAGUGCAAAAAUUGGACUUGUAAUGCGUGACAAAUACG
Mpe-1-1   UCUUUCAUGCGAUGUUUGCCGGAAAUCACACAGUGAUUUCGAAAUAAAAAUAAGGUGGUGAAAAAUGGCAAGAAUGCAUGCCCGUCGGCGCGGAAAAUCCGGUUCGGUUCGUCCGCUCCGCACAGAGGCACCAGAAUGGUCAAACAAGGACCCCUCAGAGAUCGAAAAAACCAUCAUAGA
Mpa-2-1   UCUUUCAUGCGAUGUUUGCCGGAAAUCACACAGUGAUUUCGAAAUAAAAAUAAGGUGGUGAAAAAUGGCAAGAAUGCAUGCCCGUCGGCGCGGAAAAUCCGGUUCGGUUCGUCCGCUCCGCACAGAGGCACCAGAAUGGUCAAACAAGGACCCCUCAGAGAUCGAAAAAACCAUCAUUGA
env-24    UUCCGUCCGCCCCUACCGGAAGGAGGCGCCCUCGUGGUCCAACACGGACAUAAAGGGGA.........................................................................................................................
env-25    GGGAGGUGAACAGGACAUGGCACGAAUGCAUGCACGACGGAGGGGACAUUCAGGUUCCGUGCGCCCCUACCGCAAAGAGACGCCCUCUUGGGCCAACACCGAUCAAAAGGCCAUAGAAAAGGUCAUCGUGGACCUUCGCAAGGAAGGAAUGCCAAGCAGCCGGAUCGGGCUCAUUCUGCG
env-26    ....................................................................................................................................................................................
env-27    AUCGGUCAGGCCAUACCGGAAAGAAGCACCGGAAUGGUCCAACACCGAUGUGGCCGCGAUCGAGAAGGUCAUUGUUGACCUGAGUAAAGAGGGCGUAUCAACCGCCCGGAUCGGUCUGGUCCUCCGUGACCGCUAUGGAGUCCCUGAUGUGAAACUCGCUACAGGGAAGCGGAUAGGACA
env-28    AUCGGUCAGGCCAUACCGGAAAGAAGCACCGGAAUGGUCCAANACCGAUGUGGCCGCGAUCGAGAAGGUCAUUGUUGACCUGAGUAAAGAGGGCAUAUCAACCGCCCGGAUCGGCCU..........................................................
env-29    GGUUCAGUACGGCCAUACCGUAAGGAAAACCCGGAAUGGGCCAAUGCCGACGUGGCGGCGAUUGAGAAGAUCAUCGUCGAUUUGAAAAAGGAAGGAGUCUCUACCAGCCGUAUCGGCCUUAUCCUGAGAGACAAGCACGGAGUACCGGACGUCAAGCUUGCGACAGGCAAACGUAUCGGG
env-30    ....................................................................................................................................................................................
env-31    GAUCGGUACGCCCGCACCGCAGCGAGGCUCCCGCCUGGUCCAAUACCGACGUGAAAGCAAUCGAGAAAGUCAUUGUUGACUUGAAAAAAGAGGGUUACUCAAGCAGUAAAAUCGGCCUUGUCCUCCGUGAUCAGUACGGAGUCCCC...................................
env-32    UAGACAUGGCACGAAUGCAUGCAAGACGGAGAGGAUCAUCCGGAUCUGUCCGGCCGUACAGAAAGGACGCUCCUGAGUGGUCCAAUACCGAUCAACAAGAGAUCGAGAAGCAGAUUAUUGAUAUGAAGAAGGAAGGUCUUUCCUCCUCUGUUAUCGGAAUGGUGAUGCGCGACAAGUAUG
env-33    GCGGUUCCGUCCGACCCUACCGGAAGGAAACUCCCGAGUGGUCCAACACGGAUACAGCGGAGAUCGAGAAGAUCAUCAUCGAUCUUCGCAAAGACGGCCUCUCGGCCAGCCAGAUCGGCCUCGUGCUGCGGGACAGAUAUGCCGUCCCUGAUGUCAAACUCGCCACGGGCAAGCGGAUAA
env-34    GCGGUUCCGUCCGACCCUACCGGAAGGAAACUCCCGAGUGGUCCAACACGGAUACAGCGGAGAUCGAGAAGAUCAUCAUCGAUCUUCGCAAAGACGGCCUCUCGGCCAGCCAGAUCGGCCUCGUGCUGCGGGACAGAUAUGCCGUCCCUGAUGUCAAACUCGCCACGGGCAAGCGGAUAA
env-35    GCGGGUCCGUCCGACCCUAUCGGAAAGAGGCGCCCGAGUGGUCCAACACGGACGCAGCGGAGAUAGAGAAGAUCAUAGUUGAGCUCCGCAAAGAUGGUAUGUCGAGCAGCCAGAUCGGCCUUGCGCUGCGGGAUAAAUACGGUGUUCCUGAUGUCAAGCUCGCCACCGGCAAGCGCGUGA
env-36    GCGGGUCCGUCCGACCCUAUCGGAAAGAGGCGCCCGAGUGGUCCAACACGGACGCAGCGGAGAUAGAGAAGAUCAUAGUUGAGCUCCGCAAAGAUGGUAUGUCGAGCAGCCAGAUCGGCCUUGCGCUGCGGGAUAAAUACGGUGUUCCUGAUGUCAAGCUCGCCACCGGCAAGCGCGUGA
env-39    GCGGGUCNGUCCGACCCUAUCGGAAAGAGGCGCCCGAGUGGUCCAACACGGACGCAGCGGAGAUAGAGAAGAUCAUAGUUGAGCUCCGCAAAGAUGGUAUGUCGAGCAGCCAGAUCGGCCUUGCGCUGCGGGAUAAAUACGGUGUUCCUGAUGUCAAGCUCGCCACCGGCAAGCGCGUGA
env-40    GCGGGUCUGUCCGACCCUAUCGGAAAGAGGCGCCCGAGUGGUCCAACACGGACGCAGCGGAGAUAGAGAAGAUCAUAGUUGAGCUCCGCAAAGAUGGUAUGUCGAGCAGCCAGAUCGGCCUUGCGCUGCGGGAUAAAUACGGUGUUCCUGAUGUCAAGCUCGCCACCGGCAAGCGCGUGA
env-41    GUGGUUCCGUCAAACCCUACCGGAAGGAAGC.....................................................................................................................................................
env-42    GUGGUUCCGUCAAACCCUACCGGAAGGAAGCACCCGGGUGGUCCAAC.....................................................................................................................................
env-44    GUGGUUCCGUCAAACCCUACCGGAAGGAAGCACCCGGGUGGUCCAACACGGAUACGGCGGAGAUCGAGAAGAUCAUCGUUGAGCUCCGCAAAGACGGUAUGUCGAGCAGCCAGAUCGGCCUUGUGCUACGGGACAAGUAUGGUGUCCCCGACGUCAAGCUCGCUACCGGUAAACGCGUGA
env-45    AGUGGUUCCGUCAAACCCUACCGGAAGGAAGCACCCGGGUGGUC........................................................................................................................................
env-46    AGUGGUNCCGUCAAACCCUACCGGAAGGAAGCACCCGGGUGGUCCAACACGGAUACGGCGGAGAUCGAGAAGAUCAUCGUUGAGCUCCGCAAAGACGGCAUGUCGAGCAGCCAGAUUGGCCUUGUGCUCCGGGACAAGUACGGUGUCCCGGACGUCAAGCUCGCCACCGGCAAGCGCGUG
env-47    GCCGGUUCCGUCCGACCCUACCGGAAGGAAGCACCCGAGUGGUCCAACACGGACGCAGCGGAGAUCGAGAAGAUCGUCGUUGAGCUCCGCAAAGACGGCAUGUCGAGCAGCCAGAUUGGCCUUGCGUUGCGGGACAGGUACGGUGUCCCCGACGUCAAGCUCGCCACCGGCAAACGCGUG
env-48    GCCGGUUCCGUCCGACCCUACCGGAAGGAAGCACCCGAGUGGUCCAACACGGACGCAGCGGAGAUCGAGAAGAUCGUCGUUGAGCUCCGCAAAGACGGCAUGUCGAGCAGCCAGAUCGGCCUUGCGUUGCGGGACAGGUACGGUGUCCCCGACGUCAAGCUCGCCACCGGCAAGCGCGUG
env-49    ....................................................................................................................................................................................
env-51    GCCGGUUCCGUCCGACCCUACCGGAAGGAAGCACCCGAGUGGUCCAACACGGACNCAGCGGAGAUCGAGAAGAUCGUCGUUGAGCUCCGCAAAGACGGCAUGUCGAGCAGCCAGAUNGGCCUUGCGUUGCGGGACAGGUACGGUGUCCCCGACGUCAAGCUCGCCACCGGCAAGCGCGUG
env-52    GCCGGUUCCGUCCGACCCUACCGGAAGGAAGCACCCGAGUGGUCCAACACGGACNCAGCGGAGAUCGAGAAGAUCGUCGUUGAGCUCCGCAAAGACGGCAUGUCGAGCAGCCAGAUUGGCCUUGCGUUGCGGGACAGGUACGGUGUCCCCGACGUCAAGCUCGCCACCGGCAAGCGCGUG
env-53    AGCGGUUCCGUCCGACCCUACCGGAAGGAAGCACCCGAGUGGUCCAACACGGACGCGGCGGAGAUAGAGAAGAUCAUCAUUGGACUCCGCAAGGACGGCAUGUCGAGCAGCCAGAUCGGCCUUGUGCUGCGGGACAAAUACGGCGUCCCUGACGUCAAGCUCGCCACUGGCAAACGCGUG
Mbo-1-1   GCUGGUUCUGUCCGACCCUACCGGAAGGAGGCACCCGAGUGGUCCAACACGGACGCAGCGGAGAUCGAGAAGAUCAUCGUUGAGCUCCGCAAAGACGGCAUGUCGAGCAGCCAGAUUGGCCUUGCGUUGCGGGACAAGUACGGUGUCCCCGACGUCAAGCUCGCCACCGGCAAACGCGUG
env-61    ....................................................................................................................................................................................
env-64    CAGCGGUUCAGUCCGCCCCUAUCGUAAAGAGGCGCCCGAGUGGUCAAACACGGAUGCAGCGGAGAUCGAGAAGAUCAUCAUCGGCCUCCGCAAAGACGGCCUCUCGGCCAGCCAGAUCGGGCUUGUCCUGCGCGACAGGCACGCCGUCCCUGACGUCAGGCUCGCCACAGGCAAACGUAU
env-65    CCAGCGGUUCAGUCCGCCCCUAUCGUAAAGAGGCGCCCGAGUGGUCAAACACGGAUGCAGCGGAGAUCGAGAAGAUCAUCAUCGGCCUCCGCAAAGACGGCCUCUCGGCCAGCCAGAUCGGGCUUGUCCUGCGCGACAGGCACGCCGUCCCUGACGUCAGGCUCGCCACAGGCAAACGUA
env-66    CAGCGGUUCAGUCCGCCCCUAUCGUAAAGAGGCGCCCGAGUGGUCAAACACGGAUGCAGCGGAGAUCGAGAAGAUCAUCAUCGGCCUCCGCAAAGACGGCCUCUCGGCCAGCCAGAUCGGGCUUGUCCUGCGCGACAGGCACGCCGUCCCUGACGUCAGGCUCGCCACAGGCAAACGUAU
env-67    CAGCGGUUCAGUCCGCCCCUAUCGUAAAGAGGCGCCCGAGUGGUCAAACACGGAUGCAGCGGAGAUCGAGAAGAUCAUCAUCGGCCUCCGCAAAGACGGCCUCUCGGCCAGCCAGAUCGGGCUUGUCCUGCGCGACAGGCACGCCGUCCCUGACGUCAGGCUCGCCACAGGCAAACGUAU
env-68    CAGCGGUUCAGUCCGCCCCUAUCGUAAAGAGGCGCCCGAGUGGUCAAACACGGAUGCAGCGGAGAUCGAGAAGAUCAUCAUCG.................................................................................................
env-69    AGUGGUUCCGUCCGACCUUACCGGAAGGAAGCGCCCGAGUGGUCCAACACGGACACAGCUGAGAUCGAGAAGAUCAUCGUCGAGCUCCGUAAAGACGGCAUGUCGGCAAGCCAGAUCGGCCUUGCAUUGCGUGACAGGUACGCCGUCCCCGACGUCAAGCUCGCCACCGGCAAACGCGUG
Msp-1-1   AGCAGUUCCGUCCGACCCUACCGGAAGGAAGCACCCGAGUGGUCCAACACGGACGCAACAGAGAUCGAGAAGAUCGUCGUCGAUCUUCGCAAAGACGGCAUGUCGACCAGCCAGAUCGGCCUUGUGCUGCGUGACAGGUACGCCGUCCCCGA.............................
Mma-1-1   AGCAGUUCCGUCCGACCCUACCGAAAGGAGGCACCCGAGUGGUCCAAUACGGACGCAACAGAGAUCGAGAAGAUCGUCGUCGAUCUUCGCAAAGACGGCAUGUCGACCAGCCAGAUCGGCCUUGUGCUACGUGACAGGUACGCCGUCCCCGACGUCAAGCUCGCCACCGGCAAGCGCAUA
env-70    AGCAGUUCCGUCCG......................................................................................................................................................................
Mse-1-1   AGCAGUUCCGUCCGACCCUACCGGAAGGAGGCACCCGAGUGGUCCAACACGGACAUAGCAGAGAUCGAGAAGAUCAUCGUCGAUCUUCGCAAAGACGGCAUGUCGACCAGCCAGAUCGGCCUUGCGCUGCGUGACAGGUACGCCGUCCCCGACGUCAAGCUCGCCACCGGCAAGCGCAUA
env-71    CAGCGGUUCUGUCCGACCUUACCGGAAGGAAGCACCCGGGUAGGUCCAACACGGACGCACCGGAGAUCGAGAAGAUUAUCGUUGAUCUCCGCAAAGACGGCAUGUCGACCAGUCAGAUUGGUCCUUGCG................................................
env-72    CAGCGGUUCUGUCCGACCUUACCGGAAGGAAGCACCCGGGUGGUCCAACACGGACGCACCGGAGAUCGAGAAGAUUAUCGUUGAUCUCCGCAAAGACGGCAUGUCGACCAGUCAGAUUGGCCUUGCGCUGCGGGACAGUUACGCUGUCCCUGACGUCAAGCUCGCCACUGGCAAGCGCGU
env-73    CAGCGGUUCUGUCCGACCUUACCGGAAGGAAGCACCCGGGUGGUCCAACACGGACGCACCGGAGAUCGAGAAGAUUAUCGUUGAUCUCCGCAAAGACGGCAUGUCGACCAGUCAGAUUGGCCUUGCGUUGCGGGACAGUUACGCUGUCCCUGACGUCAAGCUCGCCACUGGCAAGCGCGU
env-76    GUUCGGUACGCCCCCACCGUACUGAGGCGCCGUCCUGGUCAAAUACCGACGUGGAAGCAAUUGAGAAAGUCAUCAUUACCCUGAAAAAGGAUGGGCAUUCAAGCAGCCGCAUCGGCCUCAUCCUUCGUGAUCGCUAUGGCGUUCCUGAUGUCAAACUGGUUAUGGGGAAAAGGAUCGAUC
env-77    GUCCGUCCUCACCGCACAGAGGCGCCGUCCUGGUCUAAUACCGACGUGAAAGCCAUCGAGAAAGUCAUUAUCGACCUCAAGAAAGAGGGAUACUCAAGCAGCCGGAUCGGGCUCGUCCUCCGUGACCGUUAUGGCGUACCUGACGUCAAGCUGGUUAUGGGGAAGCGGAUCGACCAGAUC
env-78    GUCCGUCCUCACCGCACAGAGGCGCCGUCCUGGUCUAAUACCGACGUGAAAGCCAUCGAGA.......................................................................................................................
env-79    CCGUCCGGCCAUACAGGAAAGAAUCCCCUGAAUGGGCCAAUACAGAUGUAGAGGCAAUCGAGAAGAUCAUCGUUGAUCUCAAGAAGGAAGGGGUCUCCAGCAGCCGGAUCGGGCUCAUUCUCCGUGACAAGUACGGUGUGCCUGAUGUGAAGCUGAUCACCGGAAAGCGUAUCGGCCAGA
env-80    AGUUCUGUGCGGCCGGCACGCAGUGAAGCGCCCGACUGGUCCAACACGGACAAGGCAGCGAUCGAGAAACUCAUCGUCGAUCUCCGCAAGGAAGGCACAUCGAGCAGCAUGAUUGGCCUGAUCCUCCGGGACCGCUACGGGGUUCCCGAUGUUAAGCUGGUCACCGGCAAGCGCAUCGGC
env-81    GUUCUGUGCGGCCGGCACGCAGCGAAGCGCCCGACUGGUCCAACACGGACAAGGCAGCGAUCGAGAAACUCAUCGUCGAUCUCCGCAAGGAAGGCACAUCGAGCAGCAUGAUUGGCCUGAUCCUCCGGGACCGCUACGGGGUUCCCGAUGUCAAGCUGGUCACCGGCAAGCGCAUCGGCG
env-82    ....................................................................................................................................................................................
env-83    AGCUCAGUGCGACCAGCGCGCAAAGAAGCGCCUGCCUGGUCCAACACGGACAAGGCAGCGAUCGAGAAACUUAUCGUCGAUCUCCGCAAGGAAGGCACAUCGGGCAGUAUGAUUGGCCUCAUCCUGCGGGACCGCUACGGCGUUCCCGAUGUCAAGAUGGUCACCGGCAAGCGUAUUGGC
env-84    AGUUCUGUCCGACCGGCACGCAGCGAAGCGCCCGCAUGGUCCAACACGGACAAGGCAGCGAUCGAGAAACUUAUCGUCGAUCUCCGCAAGGAAGGCACAUCGAGCAGCAUGAUCGGUCUCGUCCUGCGGGACCGCUACGGCGUUCCCGAUGUCAAGCUGGUCACCGGUAAGCGUAUCGGU
env-85    CUCUGUGCGGCCAGCACGGAACGAAGCUCCUGCAUGGUCCAACACGGACAAGGCAGCGAUCGAGAAGAUUAUUGUCGAUCUCCGUAAGGAAGGCACAUCGAGCAGCAUGAUCGGCCUGAUCCUGCGGGACCGCUACGGCGUCCCUGAUGUCAAGAUGGUCACCGGCAAAAGCGUCGGCGA
env-86    UUCUGUUAGACCGGCACGCAGCGAAGCGCCCGCAUGGUCCAACACGGACAAGGCAGCGAUCGAGAAACUUAUCGUCGAUCUCCGCAAGGAAGGCACAUCAAGCAGCAUGAUCGGUCUCGUUCUCCGGGACCGCUACGGCGUUCCCGAUGUCAAGCUGGUCACCGGCAAGCGUAUCGGCGA
CMe-1-1   AAGUCAAGCUCGGUGCGGCCGGCGCGUAAUGAAGCGCCCGCGUGGUCCAACACGGACAAGGCAGCAAUCGAGAAGCUCAUCGUCGAUCUCCGCAAGGAAGGCACAUCGGCCAGCAUGAUUGGCCUUGUCCUGCGGGACCGCUACGGUGUCCCCGAUGUCAAGAUGGUCACCGGAAAAAGU
env-87    AGUUCUGUGCGGCCGGCACGCAGCGAAGCGCCCGACUGGUCCAACACGG...................................................................................................................................
env-88    AGUUCUGUGCGGCCGGCACGCAGCGAAGCGCCCGACUGGUCCAACACGGACAAGGCAGCGAUCGAGAAACUCAUCGUCGAUCUCCGCAAGGAAGGCACAUCGAGCAGCAUGAUUGGCCUGAUCCUCCGGGACCGCUACGGGGUUCCCGAUGUCAAGCUGGUCACCGGCAAGCGCAUCGGC
Mli-1-1   UCAGUCGCACCGUACCGCACCGAGGCACCGGAAUGGACCGGGAUCUCGGCGGACGAGGCAGAGAAGGUUGUUGUCGAUCUCCGUAAGAAAGGCAAAUCCAGCAGCGAGAUCGGGUUCGUCCUGAGAGACAAGUACGGCGUUCCGGACGUGAAACUCGUGGCCGGGAAGAAGAUCGGCGAG
Mfo-1-1   GCAAGUCAUGCUCUGUCCGCCCCUACCGGAAACAGGCGCCCGCCUGGUCGAACACUGACCCCGCGGCGAUCACCAAGAUCAUCCUGGACCUCCGGAAGGAAGGCGCAUCGAGCGCAAAGAUCGGUCUGGUCCUGAGGGACCGGUACGGCGUACCCGACGUCAAGCUGGCAACCGGCAAGC

          ....................................................................................................................................................................................
          ....................................................................................................................................................................................
          ●●●●●●●●●●●●●●●●●●●●●●●●●●●●●●●●●●●●●●●●●●●●●●●●●●●●●●●●●●●●●●●●●●●●●●●●●●●●●●●●●●●●●●●●●●●●●●●●●●●●●●●●●●●●●●●●●●●●●●●●●●●●●●●●●●●●●●●●●●●●●●●●●●●●●●●●●●●●●●●●●●●●●●●●●●●●●●●●●●
```

```
env-89    GUCAUGCUCAGUCCGCCCCUACCGGAAACAGGCCCCCGCGUGGUCCAACACGGACGCAAAGGCGAUCGAGAAGAUC..........................................................................
env-90    GUCAUGCUCAGUCCGCCCCUACCGGAAACAGGCCCCCGCGUGGUCCAACACGGACGCAAAGGCGAUCGAGAAGAUCAUCCUGGAUCUGAGAAAGGAAGGCGCAUCGAGCGCGAAGAUCGGA.................................
env-91    GGGGCAAGUCAUGCUCAGUGCGCCCCUACCGGAACCAGGCGCCCGCGUGGUCGAAUGCUGACCCCGCCGCGAUCACCAAGAUCGUUCUGGACCUGAGGAAGGAUGGCGCAUCGAGCGCGAAGAUCGGUCUGGUCCUGCGGGACCGGUACGGCGUACCUGACGUCAAGCUUGCCACCGGCA
env-92    GGGCAAGUCAUGCUCUGUCCGCCCCUACCGGAACCAGGCCCCCGCGUGGUCAAAUGCUGACCCCGAGGCGAUCACCAAGAUCAUCCUGGAACUGAGAAAGGACGGCGCAUCGAGCGCAAAGAUCGGCCUGGUCCUCAGGGACCGGCACGGCGUACCUGACGUCAAGCUUGCCACCGGCAA
env-96    ....................................................................................................................................................
env-97    GGGCAAGUCAUGCUCUGUCCGCCCCUACCGGAACCAGGCCCCCGCGUGGUCAAAUGCUGACCCCGAGGCGAUCACCAAGAUCAUCCUGG.........................................................
env-98    CAUGCUCGGUCCGUCCCUACCGGAAACAGGCACCCGCGUGGUCCAACACGGACGCAAAGGCGAUCGAGAAGAUCAUCCUGGAUCUGAGAAAGGAAGGGCGCAUCGAGCGCGAAGAUCGGA...................................
env-99    CAUGCUCGGUCCGUCCCUACCGGAAACAGGCACCCGCGUGGUCCAACACGGACGCAAAGGCGAUCGAGAAGAUCAUCCUGGAUCUGAGAAAGGAAGGCGCAUCGAGCGCGAAGAUCGGACUGGUACUCCGGGACCGCCACGGUGUUCCCGACGUGAAACUUGCGACCGGCAAGCGCAUCG
env-103   GGCAAGUCAUGCUCCGUGCGCCCCUACCGGAAACAGGCACCCGCGUGGUCCAAUACCGACGCAAAAGCAAUCGAAAAGAUUGUCCUUGACCUGAGAAAGGAAGGCGCAUCGAGCGCAAAGAUCGGACUGGUACUCCGCGACAGCUAUGGCGUCCCGGACGUCAAGCUCGCCACCGGCAAG
env-104   AUGCUCAGUCCG........................................................................................................................................
env-105   AAGUCAUGCUCCGUGCGCCCUUACCGGAAACAGGCCCCCGCGUGGUCAAACACCGACGCAAAGGCAAUCGAGAAGG..........................................................................
env-106   ....................................................................................................................................................
env-107   CAAGUCAUGCUCAGUCCGCCCCUACCGGAAACAGGCUCCCGCGUGGUCCAACACGGACACAAAGG.....................................................................................
env-108   UGCUCCGUCCGCCCCUACCGGAAACAGGCGCCUGCAUGGUCCAAUACGGACGCAAAGGCGAUCGAGAAGAUCAUCCUGGAUCUCCGCAAGGAAGGCGCAUCGAGCGCGAAAAUCGGAUUGGUACUCCGGGACCGCUACGGCGUCCCCGACGUGAAACUCGCCAUGGGCAAGAGGAUCGGA
env-109   AGUCAUGCUCCGUGCGUCCCUACCGUAAAGAAGCCCCCGCGUGGUCCAACACCGACGCAAAGGCUAUCGAGAAGAUCGUUCUCGACUUACGGAAGGAAGGCGCAUCGAGCGCAAAGAUCGGACUGGUACUCCGGGACCGCUACGGCGUUCCUGAUGUUAAGCUCGCCACCGGCAAGCGUA
env-110   GCAAGUCAUGCUCAGUCCGCCCCUACCGGAAACAGGCACCCGCGUGGUCCAACACGGACACAAAGGCGAUCGAGAAGAUCAUCCUGGAUCUGAGAAAGGAAGGCGCAUCGAGCGCAAAGAUCGGACUGGUACUCCGGGACCGCUAUGGCGUCCCUGACGUGAAACUCGCGACCGGCAAGC
env-111   ....................................................................................................................................................
env-112   CAAGUCAUGCUCAGUCCGCCCCUACCGGAAACAGGCCCCCGCGUGGUCCAACACGGACACAAAGGCGAUCGAAAAGAUCAUCCUGGAUCUGAGAAAGGAAGGCGUGUCGAGCGCGAAGAUCGGACUGGUACUCCGGGACCGCUACGGUGUUCCCGACGUGAAACUUGCGACCGGCAAGCG
env-114   ....................................................................................................................................................
env-115   AGUCAUGCUCCGUCCGCCCCUACCGGAAACAGGCACCUGCAUGGUCCAACACGGACAAGGCAGCGAUCGAAAAGGUCAUCCUGGAUCUGAGAAAGGAAGGCGCAUCGAGCGCAAAGAUCGGACUCGUACUCAGGGACCGCUACGGUGUUCCCGAUGUGAAACUUGCCCUCGGCAAGCGUA
env-117   GUCAUGCUCCGUGCGCCCUACCGGAAACAGGCACCCGCGUGGUCCGUUGUGGACACAAAGGCGAUCGAGAAGAUCAUCCUGGAUCUGAGAAAGGAAGGCGCAUCGAGCGCAAAGAUCGGACUGGUUCUCCGGGACCGCUACGGCGUCCCCGACGUUAAGCUCGCUACCGGCAAGCGUGUC
env-118   AAAUCAUGCUCUGUCCGCCCCUACCGUAAGCAGGCACCUGCAUGGUCCAACACGGACGUUCAGGCAAUCGAGAAGAUCAUUCUCGAGCUCCGGAAGGAAGGCGUAUCGAGCAGCAAGAUCGGUCUGAUACUCCGUGACCGUCACGGAGUUCCGGACGUGAAGCUCGCGACCGGCAAGCGG
env-120   UGCUCAGUCCG.........................................................................................................................................
env-122   GCUCUGUCCGCCCCCACCGGAAAUCGGCACCUGCGUGGUCCAAUACGGACGCAAAGGCGAUCACCAAGGUGAUCCUGGAUCUCCGCAAGGAAGGCGCAUCGAGCAGCAGAAUUGGGCUUGUCCUCAGGGACCGCUAUGGCGUUCCUGACGUGAAACUUGCCCUUGGUAAGCGUAUCGGUG
env-123   AAGAGGCAAGUCAUGCUCUGUCCGCCCCCACCGGAAACAGGCACCUGCGUGGUCCAAUACGGACGCAAAGGCGAUCACCAAAGUGAUCCUGGAUCUCCGCAAGGAAGGCGCAUCGAGCAGCAAAAUUGGGCUUGUCCUCAGGGACCGCUAUGGCGUCCCUGACGUAAAACUUGCCCUCGG
env-124   AAGAGGCAAAUCAUGCUCAGUCCGCCCCUACCGGAAACAGGCACCCGCGUGGUCCAACACGGACGCAAAGGCAAUAACAAAAGUUAUCCUGGAACUCCGCAAGGAAGGCGCAUCGAGCAGCAGGAUUGGUCUGAUCCUCAGGGAUCGCCAUGGCGUCCCGGACGUGAAACUUGCAACAGG
env-128   UGCAUGCUCGAAGAAGAGGCAAAUCAUGCUCAGUCCGCCCCUACCGGAAACAGGCACCCGCGUGGUCCAACACGGACNCAAAGGCAAUAACAAAAGUUAUCCUGGAACUCCGCAAGGAAGGCGCAUCGAGCAGCANGAUUGGUCUGAUCCUCAGGGAUCGCCAUGGCGUCCCGGACGUGA
env-129   GCAUGCUCGAAGAAGAGGCAAAUCAUGCUCAGUCCGCCCCUACCGG......................................................................................................
env-131   AAGAGGCAAAUCAUGCUCAGUCCGCCCCUACCGGAAACAG............................................................................................................
env-132   AAGAGGCAAAUCAUGCUCAGUCCGCCCCUACCGGAAACAGGCACCCGCGUGGUCCAACACGGACGCAAAGGCAAUAACAAAAGUUAUCCUGGAACUCCGCAAGGAAGGCGCAUCGAGCAGCAGGAUUGGUCUGAUCCUCAGGGAUCGCCAUGGCGUCCCGGACGUGAAACUUGCAACAGG
env-133   AAGAGGCAAAUCAUGCUCAGUCCGCCCCUACCGGAAACAGGCACCCGCGUGGUCCAACACGGACACAAAGGCAAUAACAAAAGUUAUCCUGGAACUCCGCAAGGAAGGCGCAUCGAGCAGCAGGAUUGGUCUGAUCCUCAGGGAUCGCCAUGGCGUCCCGGACGUGAAACUUGCAACAGG
env-134   AAUGCAUGCUCGAAGAAGAGGCAAAUCAUGCUCAGUCCGCCCCUACCGGAAACAGGCACCUGCGUGGUCCAACAUGGACGCAAAGGCGAUCACCAAAGUGAUCCUGGAUCUCCGCAAGGAAGGCGCAUCGAGCAGCAAAAUUGGGCUUGUCCUGCGGGACAGCUACGGCGUCCCGGACGU
env-135   GCUCUGUCCGCCCCUACCGGAAACAGGCACCCGCGUGGUCCAAUAUGGACGCAAAGGCGAUCACCAAAGUGAUCCUGGAUCUCCGCAAGGAAGGCGCAUCGAGCAGCAAAAUUGGGCUUGUCCUGCGGGAUAGCUACGGCGUCCCUGACGUGAAACUUGCCCUAGGUAAGCGUAUCGGUG
env-136   GCUCUGUCCGCCCCUACC..................................................................................................................................
env-137   GCUCUGUCCGCCCCUACCGGAAACAGGCACCCGCGUGGUCCAAUAUGGACGCAAAGGCGAUCACCAAAGUGAUCCUGGAUCUCCGCAAGGAAGGCGCAUCGAGCAGCAAAAUUGGGCUUGUCCUGCGGGAUAGCUACGGCGUCCCUGACGUGAAACUUGCCCUAGGUAAGCGUAUCGGUG
env-138   GCUCUGUCCGCCCCUACCGGAAACAGGCACCCGCGUGGUCCAAUAUGGACGCAAAGGCGAUCACCAAAGUGAUCCUGGAUCUCCGCAAGGAAGGCGCAUCGAGCAGCAAAAUUGGGCUUGUCCUGCGGGAUAGCUACGGCGUCCCUGACGUGAAACUUGCCCUAGGUAAGCGUAUCGGUG
env-148   UGCUCUGUCCGCCCCUACCGGAAACAGGCACCUGCGUGGUCCAAUAUGGACGCAAAGGCGAUCACAAAAGUGAUCCUGGAUCUCCGCAAGGAAGGCGCAUCGAGCAGCAAAAUUGGGCUUGUCCUGCGGGACAGCUACGGCGUCCCGGACGUGAAACUUGCCCUCGGUAAGCGUAUCGGU
env-149   ....................................................................................................................................................
env-150   UGCUCUGUCCGCCCCUACCGGAAACAGGCA......................................................................................................................
env-151   UCUGUCCGCCCCUACCGGAAACAGGCACCUGCGUGGUCCAACACGGACGCAAAGGCCAUCACCAAAGUGAUCCUGGAUCUCCGCAAGGAAGGCGCAUCGGGCAGCAGGAUCGGACUGAUCCUGCGGGACCGCUACGGCGUCCCGGAUGUGAAGCUUGCUCUCGGUAAGCGUAUCGGAGAU
env-152   UCUGUCCGCCCCUACCGGAAACAGGCACCUGCGUGGUCCAACACGGACGCAAAGGCCAUCACAAAAGUAAUCCUGGAUCUCCGCAAGGAAGGCGCAUCGGGCAGCAGGAUCGGAUUGAUCCU.................................
env-153   UCUGUCCGCCCCUACCGGAAACAGGCACCUGCGUGGUCCAACACGGACGCAAAGGCCAUCACAAAAGUAAUCCUGGAUCUCCGCAAGGAAGGCGCAUCGGGCAGCAGGAUC...........................................
env-154   CUCUGUCCGCCCCUACCGGAAACAGGCACCAUCGUGGUCCAACGUGGACACGAAGGCUAUCACCAAGGUGAUCCUGGAGCUCCGGAAGGAAGGCGCGUCGAGCGCCAGGAUCGGACUGAUCCUGCGGGACCGCUAUGGCGUCCCGGAUGUGAAACUUGCCCUCGGUAAGCGUAUCGGGGA
env-155   ....................................................................................................................................................
env-156   AUGCUCAGUCCGCCCCUACCGCAAGGAAGUCCCCGCAUGGUCCAACAAGGACGUAGCAGGGAUUACAAAAGUUAUCCUUGAACUCCGGAAGGAAGGCGCGUCGAGCAGCAAGAUUGGACUGGUACUCAGGGACAGCUACGGAGUUCCCGAUAUUAAACUUGCCACCGGCAAGCGUCUCGG
env-157   ....................................................................................................................................................
env-159   AUGCUCAGUCCGCCCCUACCGCAAGGAAGUCCCCGCAUGGUCCAACAAGGACGUAGCAGGGAUNACAAAAGUUAUCCUUGAACUCCGGAAGGAAGGCGCGUCGAGCAGCAAGAUUGGACUGGUACUCAGGGACAGCUACGGAGUUCCCGAUAUUAAACUUGCCACCGGCAAGCGUCUCGG
env-160   UGCUCUGUCCGCCCCCACCGGAAACAGGCACCUGCGUGGUCCAAUACGGACGCAAAGGCGAUCACCAAAGUGAUCCUGGAUCUCCGCAAGGAAGGCGCAUCGAGCAGCAAAAUUGGGCUUGUCCUCAGGGACCGCUAUGGCGUCCCUGACGUGAAACUUGCCCUCGGUAAGCGUAUCGGG
env-161   CAUGCUCUGUCCGCCCCCUCCGGAAACAGGCACCUGAGUGGUCCAAUACGGACGCAAAGGCGAUCACCAAAGUGAUCCUGGAUCUCCGCAAGGAAGGCGCA.....................................................
env-162   CUGUCCGCCCCUACCGGAAACAGGCACCUGAGUGGUCCAAUACGGACGCAAAGGCGAUUACCAAGGUAAUCCUGGAUCUCCGCAAGGAAGGCGCAUCGAGCAGCAGGAUUGGACUUGUCCUGCGGGACCGUUACGGCGUCCCGGACGUGAAACUUGCCCUUGGUAAGCGAAUCGGUGAUG
env-163   CUGUGCGGCCCUUCCGCAAGGAGGCCCCGUCCUGGGCGAACACCGAUGUCGCGGCGAUCGAGAAGAUCAUCCUGGAC.........................................................................
env-164   CUGUGCGGCCCUUCCGCAAGGAGGCCCCGUCCUGGGCGAACACCGAUGUCGCGGCGAUCGAGAAGAUCAUCCUUGACCUGCGAAAGGAGGGAGUCUCCACCAGUAAAAUCGGGAUGAUCCUCCGCGACCGGCACGGCGUCCCGGACGUCAAGCUGGUCACCGGAAAGCGGAUCGGCCAGA
env-165   CCGUUCGACCCUUCCGCAAGGAGGCCCCUUCCUGGGCCAACACCGAUGUCGCGGCGAUCGAGAAGAUCAUCCUGGACCUGCGCAAGGAGGGC...........................................................
env-166   CAGGGUCGGUCCGUCCGUUCCGCAGGGAAGCCCCGUCAUGGGCAAACACGGAUGUCGCGGCCAUCGAGAAGAUCAUCGUCGACCUGCGCAGGGAAGGCCACUCCAUGAGCCACAUCGGCCAGAUCCUGCGGGACCGCCACGGAGUCCCUGACGUCAAGCUGAUCACGGGCAAGCGGAUCA
env-168   ....................................................................................................................................................
env-169   ....................................................................................................................................................
Mta-1-1   GGGAUCGGUCCGCCCGUUCCGCAAGGAAGCCCCGUCAUGGGCGAACACGGAUGUUGCGGCCAUCGAGAAGAUCAUCGUCGACCUGCGGAAGGAAGGCAUGUCCGCGAGCCGGAUCGGCCUGAUCCUGCGGGACCGCCACGGCGUCCCCGACGUGAAACUGGUGACGGGAAAACGGAUCAG
env-174   GGGAUCGGUCCGCCCGUUCCGCAAGGAAGCCCCGUCAUGGGCGAACACGGAUGUCGCGGCCAUCGAGAAGAUCAUCGUCGAUUUGCGGAAGGAAGGCAUGUCCGCGAGCCGGAUCGGCCUGAUCCUGCGGGACCGCCACGGCGUCCCCGACGUGAAACUGGUGACGGGAAAACGGAUCAG
Mli-2-1   UGUCCCUGGGCAACUCUGCUGAAUGCGCAAUUAAUCAUGCGCAAAGGACGCAGAUGCCCGUUAAAGUAUCUUUUAAAAUAAAGGUGGUGAAAAAUCGCACGAAUGCACGCCCGCAGGCGCGGUAAAUCCGGCUCAGUCCGUCCACUCCGCACAGAAGCACCGGAAUGGUCCAACAAGGAC
Mba-1-1   GCCCGGUACGGAUGGGCGGACAGUCUCUUUCGCGAGGUGAUAAAUUAUGGCACGAAUGCAUGCUAGAAGAAGAGGAAUUGCAUCCUCUGUGAGACCAUACAGAAAAGAAGUCCCUGCCUGGUCCAACUCCGAUGUCAAAGAGAUCGAGGGUAAAAUCGUCGAGCUGCGCAAAGCAGGUCU
Mla-1-1   GCCCGGUACGGAUGGCGGACAGUCUCAUUCGCGAGGUGAUAAAUUAUGGCACGAAUGCAUGCUAGAAGAAGAGGAAUUGCAUCCUCUGUCAGACCAUACAGAAAAGAAGUCCCUGCCUGGUCCAACUCCGAUGUCAAAGAGAUCGAGGGUAAAAUCGUCGAGCUGCGCAAAGCAGGUCUU
env-177   GUUCGUCCGUACCGGAAAGAGGCUCCCGAAUGGUCCAACAAGGAUCUCACCGAGAUCGAAAAGGUCAUUCUGGAGCUGCGCCGGCAGGGCAAAUCGAGCAGCGAGAUUGGUAUGAUACUCCGUGACAAGCACGGUGUACCGGAUAUCAAACUGGUUGCCGGAAAACGGAUCAAUCAGAUC
env-178   ....................................................................................................................................................
env-179   UUCCCGUUGGAUCACUACUGUGAUCAGGAGGUUAAUCACAUGGCACGAAUGUAUGCCCGACGUAGGGGCGUAUCAAGUUCAGUCAGACCACACCGGACAGAGGCACCGGAAUGGUCUAACCAGAACAAAGAACAGAUCGAGAAAUUAGUUGUUGACAUGAAAAAACAAGGUCUUCAAAGU
          ....................................................................................................................................................
          ....................................................................................................................................................
          ••••••••••••••••••••••••••••••••••••••••••••••••••••••••••••••••••••••••••••••••••••••••••••••••••••••••••••••••••••••••••••••••••••••••••••••••••••
```

alignment positions 541 · · · 562

```
env-1     GUAUGUCAAGCGCACAGAUUGG
env-2     ......................
env-3     UUGAGGCGAACGGGUUCAGACC
env-4     UUGAGUCGAACGGGUUCAGACC
env-5     GGAAAGCGUGUUGACGAGAUUC
env-6     ......................
env-7     AGCGAAUCACAGAAAUUCUCAA
env-8     ......................
env-10    UCCUGAUGUAAAACUUGCAACA
env-11    UCCUGACGUAAAACUUGCAACA
Mpa-1-1   CCGGAAAGCGGAUCAACGCGGU

          ......................
          ......................
          ••••••••••••••••••••••
```

| | |
|---|---|
| env-12 | ...................... |
| env-13 | GGCGGAUCCUUGCAUCGGCGGG |
| env-14 | ...................... |
| env-15 | ...................... |
| env-16 | ...................... |
| env-17 | GCAACCGGAAAAUCAAUAGGCC |
| env-18 | AUCCUGGAAUCCCAGGGUGUCA |
| env-20 | ...................... |
| env-21 | AGAUCAUCCGGGAUAACGGCCU |
| env-22 | AGAUCAUCCGGGAUAACGGCCU |
| env-23 | GAGUUCCAAGCGUCAAACUUGC |
| Mpe-1-1 | UCUUAGAAAGGAAGGAAUGUCA |
| Mpa-2-1 | UCUUAGAAAGGAAGGAAUGUCC |
| env-24 | ...................... |
| env-25 | GGACAGGUACGGAGUCCCCGAU |
| env-26 | ...................... |
| env-27 | GAUCGUCAAGGAAAAGGGACUG |
| env-28 | ...................... |
| env-29 | GAGAUCUUAAAAGAGCAGGGCA |
| env-30 | ...................... |
| env-31 | ...................... |
| env-32 | GUGUUCCCGACAUCAAACUCGC |
| env-33 | CGGAGAUCCUCCGCGAAAAAGG |
| env-34 | CGGAGAUCCUCCGCGAAAAAGG |
| env-35 | ACCAGAUCCUUCGCGAAAAUGA |
| env-36 | ACCAGAUCCUUCGCGAAAAUGA |
| env-39 | ACCAGAUCCUUCGCGAAAAUGA |
| env-40 | ACCAGAUCCUUCGCGAAAAUGA |
| env-41 | ...................... |
| env-42 | ...................... |
| env-44 | ACGAGAUCCUCCGCGAAAACGA |
| env-45 | ...................... |
| env-46 | AACGAGAUCCUCCGCGAAAAGG |
| env-47 | AACGAGAUCCUCCGCGAGAAGG |
| env-48 | AACGAGAUUCCUCCGCGAGAAG |
| env-49 | ...................... |
| env-51 | AACGAGAUCCUCCGCGAGAAGG |
| env-52 | AACGAGAUCCUCCGCGAGAAGG |
| env-53 | AAUCAGAUCCUUCGCGAGAACG |
| Mbo-1-1 | AACGAGAUCCUCCGCGAGAAGG |
| env-61 | ...................... |
| env-64 | AACGGAGAUUCUCCGUGAGAAC |
| env-65 | UAACGGAGAUUCUCCGUGAGAA |
| env-66 | AACGGAGAUUCUCCGUGAGAAC |
| env-67 | AACGGAGAUUCUCCGUGAGAAC |
| env-68 | ...................... |
| env-69 | AACGAAAUCCUCCGCGAGAAGG |
| Msp-1-1 | ...................... |
| Mma-1-1 | GGCGAGAUCCUCCGCGAGAAGG |
| env-70 | ...................... |
| Mse-1-1 | AGCGAGAUCCUCCGCGAGAAGG |
| env-71 | ...................... |
| env-72 | AAACGAGAUCCUCCGCGAGAAG |
| env-73 | AAACGAGAUCCUCCGCGAGAAG |
| env-76 | AGAUACUCAAGGAGAAGGGCCU |
| env-77 | CUGGAGGAGAACGGUCUGCAGU |
| env-78 | ...................... |
| env-79 | UCAUCAGGGAGAAAGGAAUGGA |
| env-80 | GACAUCUUAAAGGAGAACAAGG |
| env-81 | ACAUCUUAAAGGAGAACAAGGU |
| env-82 | ...................... |
| env-83 | GACAUCUUAAAAGAGAACAAGG |
| env-84 | GACAUCUUGAAAGAGAACAAGG |
| env-85 | UAUCUUAGUCGAGAACAAGAUC |
| env-86 | CAUCUUGAAAGAGAACCAGGUC |
| CMe-1-1 | AUCGGUGACAUCUUAACGGAGA |
| env-87 | ...................... |
| env-88 | GACAUCUUAAAGGAGAACAAGA |
| Mli-1-1 | AUCCUCAGCCAGAACGGGCUCG |
| Mfo-1-1 | GGAUCGGGACCAUUCUCCGCGA |
| env-89 | ...................... |
| env-90 | ...................... |
| env-91 | AGCGGAUCGGAACCAUCCUGAG |
| env-92 | GCGGAUCGGCACCAUCUUACGG |
| env-96 | ...................... |
| env-97 | ...................... |
| env-98 | ...................... |
| env-99 | GGGACAUCCUGAGAGCGAACAA |
| env-103 | CGUAUCGGCGAUAUCUUAAAAG |
| env-104 | ...................... |
| env-105 | ...................... |
| env-106 | ...................... |
| env-107 | ...................... |
| env-108 | GAUAUCCUGCGGGAGAACAAAG |
| | ...................... |
| | ...................... |
| | ●●●●●●●●●●●●●●●●●●●●●● |

```
env-109    UCGGAGACAUCUUAAAAGAGAA
env-110    GUAUCGGGGACAUCCUGAGAGC
env-111    ......................
env-112    CAUCGGGGAUAUCCUGAGAGCA
env-114    ......................
env-115    UCGGGGAGAUCUUAAGAGAGCA
env-117    GGGCAGAUCUUGCGCGAGAACA
env-118    AUCGGCGCAAUCUUACGCGAAC
env-120    ......................
env-122    AUAUCUUAAAAGAGAACAAGGU
env-123    UAAGCGUAUCGGGGAUGUUUUG
env-124    UAAGCGUAUCGGUGAUAUCUUA
env-128    AACUUGCAACAGGUAAGCGUAU
env-129    ......................
env-131    ......................
env-132    UAAGCGUAUCGGUGAUAUCUUA
env-133    UAAGCGUAUCGGUGAUAUCUUA
env-134    GAAACUUGCCCUCGGUAAGCGU
env-135    AUAUUUUAAAAGAGAACAAGGU
env-136    ......................
env-137    AUAUUUUAAAAGAGAACAAGGU
env-138    AUAUUUUAAAAGAGAACAAGGU
env-148    GAUAUUUUAAAAGAGAACAAGG
env-149    ......................
env-150    ......................
env-151    AUAUUAAAAGAGAACAAGGUUG
env-152    ......................
env-153    ......................
env-154    UAUUUUGAAAGAGAACAAGGUC
env-155    ......................
env-156    GGACAUCUUAAAGGAGAACAAG
env-157    ......................
env-159    GGACAUCUUAAAGGAGAACAAG
env-160    GAUGUUUUGAAAGAGAACAAGG
env-161    ......................
env-162    UCUUAAAAGAGAACAAGGUCGC
env-163    ......................
env-164    UACUCAAAGAGAAGGGCAUGGA
env-165    ......................
env-166    GCCAGAUCCUCAAGGAAAAGGG
env-168    ......................
env-169    ......................
Mta-1-1    CCAAAUCCUGAAAGAAAAGGGG
env-174    CCAAAUCCUGAAAGAAAAGGGG
Mli-2-1    GCAGCAGAGAUCAUAAAGACAG
Mba-1-1    UACCUGUGCACAGAUCGGUCUC
Mla-1-1    ACCUGUGCACAGAUCGGUCUCG
env-177    CUCGAAGAAAAAGGCGUCGAAU
env-178    ......................
env-179    GCGGAGAUCGGUCUU.......
           ......................
           ......................
           ......................
```

### 22 S15-Flavobacteria

#### 22.1 Taxa

The taxonomy of each organism containing a putative S15-Flavobacteria RNA is listed, with abbreviations identifying each hit (e.g., "Eco-1-1" and "Eco-1-2" might hypothetically represent two distinct RNAs in *E. coli*). The abbreviations will be used to identify each individual S15-Flavobacteria RNA in Sections 22.3 and 22.5:.

| abbrev. of hits | taxonomy of species |
| --- | --- |
| Fba-1-1 | Bacteria Bacteroidetes *Flavobacteria bacterium* BBFL7 |
| Fba-2-1 | Bacteria Bacteroidetes *Flavobacteria bacterium* MS024-2A |
| Fba-3-1 | Bacteria Bacteroidetes Flavobacteria Flavobacteriales *Flavobacteria bacterium* BAL38 |
| Cgi-1-1 | Bacteria Bacteroidetes Flavobacteria Flavobacteriales Flavobacteriaceae *Capnocytophaga gingivalis* ATCC 33624 |
| Coc-1-1 | Bacteria Bacteroidetes Flavobacteria Flavobacteriales Flavobacteriaceae *Capnocytophaga ochracea* DSM 7271 |
| Csp-1-1 | Bacteria Bacteroidetes Flavobacteria Flavobacteriales Flavobacteriaceae *Capnocytophaga sputigena* ATCC 33612 |
| Fba-4-1 | Bacteria Bacteroidetes Flavobacteria Flavobacteriales *Flavobacteriaceae bacterium* 3519-10 |
| Fjo-1-1 | Bacteria Bacteroidetes Flavobacteria Flavobacteriales Flavobacteriaceae *Flavobacterium johnsoniae* UW101 |

| | |
|---|---|
| Gfo-1-1 | Bacteria Bacteroidetes Flavobacteria Flavobacteriales Flavobacteriaceae *Gramella forsetii* KT0803 |
| Kal-1-1 | Bacteria Bacteroidetes Flavobacteria Flavobacteriales Flavobacteriaceae *Kordia algicida* OT-1 |
| Lbl-1-1 | Bacteria Bacteroidetes Flavobacteria Flavobacteriales Flavobacteriaceae *Leeuwenhoekiella blandensis* MED217 |
| Pir-1-1 | Bacteria Bacteroidetes Flavobacteria Flavobacteriales Flavobacteriaceae *Polaribacter irgensii* 23-P |
| Rbi-1-1 | Bacteria Bacteroidetes Flavobacteria Flavobacteriales Flavobacteriaceae *Robiginitalea biformata* HTCC2501 |
| Zpr-1-1 | Bacteria Bacteroidetes Flavobacteria Flavobacteriales Flavobacteriaceae *Zunongwangia profunda* SM-A87 |
| Fba-5-1 | Bacteria Bacteroidetes Flavobacteria *Flavobacteriales bacterium* ALC-1 |
| Fba-6-1 | Bacteria Bacteroidetes Flavobacteria *Flavobacteriales bacterium* HTCC2170 |
| Cca-1-1 | Bacteria Bacteroidetes Flavobacteriia Flavobacteriales Crocinitomicaceae *Crocinitomix catalasitica* ATCC 23190 |
| Fta-1-1 | Bacteria Bacteroidetes Flavobacteriia Flavobacteriales Cryomorphaceae *Fluviicola taffensis* DSM 16823 |
| Aca-1-1 | Bacteria Bacteroidetes Flavobacteriia Flavobacteriales Flavobacteriaceae *Aequorivita capsosiphonis* DSM 23843 |
| Asu-1-1 | Bacteria Bacteroidetes Flavobacteriia Flavobacteriales Flavobacteriaceae *Aequorivita sublithincola* DSM 14238 |
| Avl-1-1 | Bacteria Bacteroidetes Flavobacteriia Flavobacteriales Flavobacteriaceae *Aequorivita vladivostokensis* |
| Ale-1-1 to Ale-1-2 | Bacteria Bacteroidetes Flavobacteriia Flavobacteriales Flavobacteriaceae *Algibacter lectus* |
| Ale-2-1 | Bacteria Bacteroidetes Flavobacteriia Flavobacteriales Flavobacteriaceae *Altibacter lentus* |
| Aag-1-1 | Bacteria Bacteroidetes Flavobacteriia Flavobacteriales Flavobacteriaceae *Aquimarina agarilytica* ZC1 |
| Aat-1-1 | Bacteria Bacteroidetes Flavobacteriia Flavobacteriales Flavobacteriaceae *Aquimarina atlantica* |
| Ala-1-1 | Bacteria Bacteroidetes Flavobacteriia Flavobacteriales Flavobacteriaceae *Aquimarina latercula* DSM 2041 |
| Ama-1-1 | Bacteria Bacteroidetes Flavobacteriia Flavobacteriales Flavobacteriaceae *Aquimarina macrocephali* JAMB N27 |
| Ame-1-1 | Bacteria Bacteroidetes Flavobacteriia Flavobacteriales Flavobacteriaceae *Aquimarina megaterium* XH134 |
| Amu-1-1 | Bacteria Bacteroidetes Flavobacteriia Flavobacteriales Flavobacteriaceae *Aquimarina muelleri* DSM 19832 |
| Apa-1-1 | Bacteria Bacteroidetes Flavobacteriia Flavobacteriales Flavobacteriaceae *Aquimarina pacifica* |
| Aal-1-1 | Bacteria Bacteroidetes Flavobacteriia Flavobacteriales Flavobacteriaceae *Arenibacter algicola* |
| Ace-1-1 | Bacteria Bacteroidetes Flavobacteriia Flavobacteriales Flavobacteriaceae *Arenibacter certesii* DSM 19833 |
| Ala-2-1 | Bacteria Bacteroidetes Flavobacteriia Flavobacteriales Flavobacteriaceae *Arenibacter latericius* DSM 15913 |
| Bzo-1-1 | Bacteria Bacteroidetes Flavobacteriia Flavobacteriales Flavobacteriaceae *Bergeyella zoohelcum* ATCC 43767 |
| Bzo-2-1 | Bacteria Bacteroidetes Flavobacteriia Flavobacteriales Flavobacteriaceae *Bergeyella zoohelcum* CCUG 30536 |
| Bar-1-1 | Bacteria Bacteroidetes Flavobacteriia Flavobacteriales Flavobacteriaceae *Bizionia argentinensis* JUB59 |
| Cca-2-1 to Cca-2-3 | Bacteria Bacteroidetes Flavobacteriia Flavobacteriales Flavobacteriaceae *Capnocytophaga canimorsus* |
| Cca-3-1 | Bacteria Bacteroidetes Flavobacteriia Flavobacteriales Flavobacteriaceae *Capnocytophaga canimorsus* Cc5 |
| Cca-4-1 to Cca-4-3 | Bacteria Bacteroidetes Flavobacteriia Flavobacteriales Flavobacteriaceae *Capnocytophaga canis* |
| Ccy-1-1 to Ccy-1-3 | Bacteria Bacteroidetes Flavobacteriia Flavobacteriales Flavobacteriaceae *Capnocytophaga cynodegmi* |
| Ccy-2-1 | Bacteria Bacteroidetes Flavobacteriia Flavobacteriales Flavobacteriaceae *Capnocytophaga cynodegmi* DSM 19736 |
| Coc-2-1 | Bacteria Bacteroidetes Flavobacteriia Flavobacteriales Flavobacteriaceae *Capnocytophaga ochracea* |
| Coc-3-1 | Bacteria Bacteroidetes Flavobacteriia Flavobacteriales Flavobacteriaceae *Capnocytophaga ochracea* F0287 |
| Coc-4-1 | Bacteria Bacteroidetes Flavobacteriia Flavobacteriales Flavobacteriaceae *Capnocytophaga ochracea str.* Holt 25 |
| Csp-2-1 | Bacteria Bacteroidetes Flavobacteriia Flavobacteriales Flavobacteriaceae *Capnocytophaga sp. oral taxon* 324 str. F0483 |
| Csp-3-1 | Bacteria Bacteroidetes Flavobacteriia Flavobacteriales Flavobacteriaceae *Capnocytophaga sp. oral taxon* 326 str. F0382 |
| Csp-4-1 | Bacteria Bacteroidetes Flavobacteriia Flavobacteriales Flavobacteriaceae *Capnocytophaga sp. oral taxon* 329 str. F0087 |
| Csp-5-1 | Bacteria Bacteroidetes Flavobacteriia Flavobacteriales Flavobacteriaceae *Capnocytophaga sp. oral taxon* 332 str. F0381 |
| Csp-6-1 | Bacteria Bacteroidetes Flavobacteriia Flavobacteriales Flavobacteriaceae *Capnocytophaga sp. oral taxon* 335 str. F0486 |
| Csp-7-1 | Bacteria Bacteroidetes Flavobacteriia Flavobacteriales Flavobacteriaceae *Capnocytophaga sp. oral taxon* 336 str. F0502 |
| Csp-8-1 | Bacteria Bacteroidetes Flavobacteriia Flavobacteriales Flavobacteriaceae *Capnocytophaga sp. oral taxon* 338 str. F0234 |
| Csp-9-1 | Bacteria Bacteroidetes Flavobacteriia Flavobacteriales Flavobacteriaceae *Capnocytophaga sp. oral taxon* 380 str. F0488 |
| Csp-10-1 | Bacteria Bacteroidetes Flavobacteriia Flavobacteriales Flavobacteriaceae *Capnocytophaga sp. oral taxon* 412 str. F0487 |
| Cal-1-1 | Bacteria Bacteroidetes Flavobacteriia Flavobacteriales Flavobacteriaceae *Cellulophaga algicola* DSM 14237 |
| Cba-1-1 | Bacteria Bacteroidetes Flavobacteriia Flavobacteriales Flavobacteriaceae *Cellulophaga baltica* 13 |
| Cba-2-1 | Bacteria Bacteroidetes Flavobacteriia Flavobacteriales Flavobacteriaceae *Cellulophaga baltica* 18 |
| Cba-3-1 | Bacteria Bacteroidetes Flavobacteriia Flavobacteriales Flavobacteriaceae *Cellulophaga baltica* 4 |
| Cba-4-1 | Bacteria Bacteroidetes Flavobacteriia Flavobacteriales Flavobacteriaceae *Cellulophaga baltica* NN016038 |
| Cge-1-1 | Bacteria Bacteroidetes Flavobacteriia Flavobacteriales Flavobacteriaceae *Cellulophaga geojensis* KL-A |
| Cly-1-1 | Bacteria Bacteroidetes Flavobacteriia Flavobacteriales Flavobacteriaceae *Cellulophaga lytica* |
| Cly-2-1 | Bacteria Bacteroidetes Flavobacteriia Flavobacteriales Flavobacteriaceae *Cellulophaga lytica* DSM 7489 |
| Csp-11-1 | Bacteria Bacteroidetes Flavobacteriia Flavobacteriales Flavobacteriaceae *Cellulophaga sp.* E6(2014) |
| Csp-12-1 | Bacteria Bacteroidetes Flavobacteriia Flavobacteriales Flavobacteriaceae *Cellulophaga sp.* Hel_I_12 |
| Can-1-1 | Bacteria Bacteroidetes Flavobacteriia Flavobacteriales Flavobacteriaceae *Chryseobacterium angstadtii* |
| Cca-5-1 | Bacteria Bacteroidetes Flavobacteriia Flavobacteriales Flavobacteriaceae *Chryseobacterium caeni* DSM 17710 |
| Cda-1-1 | Bacteria Bacteroidetes Flavobacteriia Flavobacteriales Flavobacteriaceae *Chryseobacterium daeguense* DSM 19388 |
| Cfo-1-1 | Bacteria Bacteroidetes Flavobacteriia Flavobacteriales Flavobacteriaceae *Chryseobacterium formosense* |
| Cga-1-1 | Bacteria Bacteroidetes Flavobacteriia Flavobacteriales Flavobacteriaceae *Chryseobacterium gallinarum* |
| Cgl-1-1 | Bacteria Bacteroidetes Flavobacteriia Flavobacteriales Flavobacteriaceae *Chryseobacterium gleum* ATCC 35910 |
| Cha-1-1 | Bacteria Bacteroidetes Flavobacteriia Flavobacteriales Flavobacteriaceae *Chryseobacterium haifense* DSM 19056 |
| Cha-2-1 | Bacteria Bacteroidetes Flavobacteriia Flavobacteriales Flavobacteriaceae *Chryseobacterium halperniae* |
| Chi-1-1 | Bacteria Bacteroidetes Flavobacteriia Flavobacteriales Flavobacteriaceae *Chryseobacterium hispalense* DSM 25574 |
| Cin-1-1 | Bacteria Bacteroidetes Flavobacteriia Flavobacteriales Flavobacteriaceae *Chryseobacterium indologenes* |

| | |
|---|---|
| Cin-2-1 | Bacteria Bacteroidetes Flavobacteriia Flavobacteriales Flavobacteriaceae *Chryseobacterium indologenes* NBRC 14944 |
| Cje-1-1 | Bacteria Bacteroidetes Flavobacteriia Flavobacteriales Flavobacteriaceae *Chryseobacterium jeonii* |
| Cko-1-1 | Bacteria Bacteroidetes Flavobacteriia Flavobacteriales Flavobacteriaceae *Chryseobacterium koreense* CCUG 49689 |
| Clu-1-1 | Bacteria Bacteroidetes Flavobacteriia Flavobacteriales Flavobacteriaceae *Chryseobacterium luteum* |
| Cor-1-1 to Cor-1-2 | Bacteria Bacteroidetes Flavobacteriia Flavobacteriales Flavobacteriaceae *Chryseobacterium oranimense* G311 |
| Cpi-1-1 | Bacteria Bacteroidetes Flavobacteriia Flavobacteriales Flavobacteriaceae *Chryseobacterium piperi* |
| Cso-1-1 | Bacteria Bacteroidetes Flavobacteriia Flavobacteriales Flavobacteriaceae *Chryseobacterium soli* |
| Cso-2-1 | Bacteria Bacteroidetes Flavobacteriia Flavobacteriales Flavobacteriaceae *Chryseobacterium solincola* |
| Csp-13-1 | Bacteria Bacteroidetes Flavobacteriia Flavobacteriales Flavobacteriaceae *Chryseobacterium sp.* BLS98 |
| Csp-14-1 | Bacteria Bacteroidetes Flavobacteriia Flavobacteriales Flavobacteriaceae *Chryseobacterium sp.* CF365 |
| Csp-15-1 | Bacteria Bacteroidetes Flavobacteriia Flavobacteriales Flavobacteriaceae *Chryseobacterium sp.* FH1 |
| Csp-16-1 | Bacteria Bacteroidetes Flavobacteriia Flavobacteriales Flavobacteriaceae *Chryseobacterium sp.* FH2 |
| Csp-17-1 | Bacteria Bacteroidetes Flavobacteriia Flavobacteriales Flavobacteriaceae *Chryseobacterium sp.* JM1 |
| Csp-18-1 | Bacteria Bacteroidetes Flavobacteriia Flavobacteriales Flavobacteriaceae *Chryseobacterium sp.* OV259 |
| Csp-19-1 | Bacteria Bacteroidetes Flavobacteriia Flavobacteriales Flavobacteriaceae *Chryseobacterium sp.* OV705 |
| Csp-20-1 | Bacteria Bacteroidetes Flavobacteriia Flavobacteriales Flavobacteriaceae *Chryseobacterium sp.* OV715 |
| Csp-21-1 | Bacteria Bacteroidetes Flavobacteriia Flavobacteriales Flavobacteriaceae *Chryseobacterium sp.* P1-3 |
| Csp-22-1 | Bacteria Bacteroidetes Flavobacteriia Flavobacteriales Flavobacteriaceae *Chryseobacterium sp.* StRB126 |
| Csp-23-1 | Bacteria Bacteroidetes Flavobacteriia Flavobacteriales Flavobacteriaceae *Chryseobacterium sp.* UNC8MFCol |
| Csp-24-1 | Bacteria Bacteroidetes Flavobacteriia Flavobacteriales Flavobacteriaceae *Chryseobacterium sp.* YR459 |
| Csp-25-1 | Bacteria Bacteroidetes Flavobacteriia Flavobacteriales Flavobacteriaceae *Chryseobacterium sp.* YR460 |
| Csp-26-1 | Bacteria Bacteroidetes Flavobacteriia Flavobacteriales Flavobacteriaceae *Chryseobacterium sp.* YR477 |
| Csp-27-1 | Bacteria Bacteroidetes Flavobacteriia Flavobacteriales Flavobacteriaceae *Chryseobacterium sp.* YR480 |
| Csp-28-1 | Bacteria Bacteroidetes Flavobacteriia Flavobacteriales Flavobacteriaceae *Chryseobacterium sp.* YR485 |
| Csp-29-1 | Bacteria Bacteroidetes Flavobacteriia Flavobacteriales Flavobacteriaceae *Chryseobacterium sp.* YR561 |
| Cta-1-1 | Bacteria Bacteroidetes Flavobacteriia Flavobacteriales Flavobacteriaceae *Chryseobacterium taiwanense* |
| Cte-1-1 | Bacteria Bacteroidetes Flavobacteriia Flavobacteriales Flavobacteriaceae *Chryseobacterium tenax* DSM 16811 |
| Cvr-1-1 | Bacteria Bacteroidetes Flavobacteriia Flavobacteriales Flavobacteriaceae *Chryseobacterium vrystaatense* |
| Ddo-1-1 | Bacteria Bacteroidetes Flavobacteriia Flavobacteriales Flavobacteriaceae *Dokdonia donghaensis* DSW-1 |
| Dsp-1-1 | Bacteria Bacteroidetes Flavobacteriia Flavobacteriales Flavobacteriaceae *Dokdonia sp.* MED134 |
| Dsp-2-1 | Bacteria Bacteroidetes Flavobacteriia Flavobacteriales Flavobacteriaceae *Dokdonia sp.* PRO95 |
| Ean-1-1 to Ean-1-4 | Bacteria Bacteroidetes Flavobacteriia Flavobacteriales Flavobacteriaceae *Elizabethkingia anophelis* |
| Ean-2-1 | Bacteria Bacteroidetes Flavobacteriia Flavobacteriales Flavobacteriaceae *Elizabethkingia anophelis* Ag1 |
| Ean-3-1 | Bacteria Bacteroidetes Flavobacteriia Flavobacteriales Flavobacteriaceae *Elizabethkingia anophelis* FMS-007 |
| Ean-4-1 | Bacteria Bacteroidetes Flavobacteriia Flavobacteriales Flavobacteriaceae *Elizabethkingia anophelis* NUH1 |
| Ean-5-1 | Bacteria Bacteroidetes Flavobacteriia Flavobacteriales Flavobacteriaceae *Elizabethkingia anophelis* NUH11 |
| Ean-6-1 | Bacteria Bacteroidetes Flavobacteriia Flavobacteriales Flavobacteriaceae *Elizabethkingia anophelis* NUH4 |
| Ean-7-1 | Bacteria Bacteroidetes Flavobacteriia Flavobacteriales Flavobacteriaceae *Elizabethkingia anophelis* NUH6 |
| Ean-8-1 | Bacteria Bacteroidetes Flavobacteriia Flavobacteriales Flavobacteriaceae *Elizabethkingia anophelis* NUHP1 |
| Ean-9-1 | Bacteria Bacteroidetes Flavobacteriia Flavobacteriales Flavobacteriaceae *Elizabethkingia anophelis* NUHP2 |
| Ean-10-1 | Bacteria Bacteroidetes Flavobacteriia Flavobacteriales Flavobacteriaceae *Elizabethkingia anophelis* NUHP3 |
| Ean-11-1 | Bacteria Bacteroidetes Flavobacteriia Flavobacteriales Flavobacteriaceae *Elizabethkingia anophelis* PW2809 |
| Ean-12-1 | Bacteria Bacteroidetes Flavobacteriia Flavobacteriales Flavobacteriaceae *Elizabethkingia anophelis* R26 |
| Eme-1-1 | Bacteria Bacteroidetes Flavobacteriia Flavobacteriales Flavobacteriaceae *Elizabethkingia meningoseptica* 502 |
| Eme-2-1 to Eme-2-2 | Bacteria Bacteroidetes Flavobacteriia Flavobacteriales Flavobacteriaceae *Elizabethkingia meningoseptica* ATCC 13253 = NBRC 12535 |
| Emi-1-1 | Bacteria Bacteroidetes Flavobacteriia Flavobacteriales Flavobacteriaceae *Elizabethkingia miricola* |
| Ebr-1-1 | Bacteria Bacteroidetes Flavobacteriia Flavobacteriales Flavobacteriaceae *Empedobacter brevis* NBRC 14943 = ATCC 43319 |
| Efa-1-1 | Bacteria Bacteroidetes Flavobacteriia Flavobacteriales Flavobacteriaceae *Empedobacter falsenii* |
| Ead-1-1 | Bacteria Bacteroidetes Flavobacteriia Flavobacteriales Flavobacteriaceae *Eudoraea adriatica* DSM 19308 |
| Fic-1-1 | Bacteria Bacteroidetes Flavobacteriia Flavobacteriales Flavobacteriaceae *Flaviramulus ichthyoenteri* Th78 |
| Fba-7-1 | Bacteria Bacteroidetes Flavobacteriia Flavobacteriales *Flavobacteriaceae bacterium* HQM9 |
| Fba-8-1 | Bacteria Bacteroidetes Flavobacteriia Flavobacteriales *Flavobacteriaceae bacterium* P7-3-5 |
| Fba-9-1 | Bacteria Bacteroidetes Flavobacteriia Flavobacteriales *Flavobacteriaceae bacterium* S85 |
| Fan-1-1 | Bacteria Bacteroidetes Flavobacteriia Flavobacteriales Flavobacteriaceae *Flavobacterium antarcticum* DSM 19726 |
| Faq-1-1 | Bacteria Bacteroidetes Flavobacteriia Flavobacteriales Flavobacteriaceae *Flavobacterium aquatile* LMG 4008 = ATCC 11947 |
| Fbe-1-1 | Bacteria Bacteroidetes Flavobacteriia Flavobacteriales Flavobacteriaceae *Flavobacterium beibuense* F44-8 |
| Fbr-1-1 | Bacteria Bacteroidetes Flavobacteriia Flavobacteriales Flavobacteriaceae *Flavobacterium branchiophilum* FL-15 |
| Fca-1-1 to Fca-1-2 | Bacteria Bacteroidetes Flavobacteriia Flavobacteriales Flavobacteriaceae *Flavobacterium cauense* R2A-7 |
| Fch-1-1 | Bacteria Bacteroidetes Flavobacteriia Flavobacteriales Flavobacteriaceae *Flavobacterium chungangense* LMG 26729 |
| Fco-1-1 | Bacteria Bacteroidetes Flavobacteriia Flavobacteriales Flavobacteriaceae *Flavobacterium columnare* ATCC 49512 |
| Fda-1-1 | Bacteria Bacteroidetes Flavobacteriia Flavobacteriales Flavobacteriaceae *Flavobacterium daejeonense* DSM 17708 |
| Fde-1-1 | Bacteria Bacteroidetes Flavobacteriia Flavobacteriales Flavobacteriaceae *Flavobacterium denitrificans* DSM 15936 |
| Fen-1-1 to Fen-1-2 | Bacteria Bacteroidetes Flavobacteriia Flavobacteriales Flavobacteriaceae *Flavobacterium enshiense* DK69 |
| Ffi-1-1 | Bacteria Bacteroidetes Flavobacteriia Flavobacteriales Flavobacteriaceae *Flavobacterium filum* DSM 17961 |
| Ffr-1-1 | Bacteria Bacteroidetes Flavobacteriia Flavobacteriales Flavobacteriaceae *Flavobacterium frigidarium* DSM 17623 |

| | |
|---|---|
| Ffr-2-1 | Bacteria Bacteroidetes Flavobacteriia Flavobacteriales Flavobacteriaceae *Flavobacterium frigoris* PS1 |
| Fge-1-1 | Bacteria Bacteroidetes Flavobacteriia Flavobacteriales Flavobacteriaceae *Flavobacterium gelidilacus* DSM 15343 |
| Fgi-1-1 | Bacteria Bacteroidetes Flavobacteriia Flavobacteriales Flavobacteriaceae *Flavobacterium gilvum* |
| Fhi-1-1 | Bacteria Bacteroidetes Flavobacteriia Flavobacteriales Flavobacteriaceae *Flavobacterium hibernum* |
| Fhy-1-1 | Bacteria Bacteroidetes Flavobacteriia Flavobacteriales Flavobacteriaceae *Flavobacterium hydatis* |
| Fin-1-1 | Bacteria Bacteroidetes Flavobacteriia Flavobacteriales Flavobacteriaceae *Flavobacterium indicum* GPTSA100-9 |
| Fli-1-1 | Bacteria Bacteroidetes Flavobacteriia Flavobacteriales Flavobacteriaceae *Flavobacterium limnosediminis* JC2902 |
| Fps-1-1 to Fps-1-10 | Bacteria Bacteroidetes Flavobacteriia Flavobacteriales Flavobacteriaceae *Flavobacterium psychrophilum* |
| Fps-2-1 | Bacteria Bacteroidetes Flavobacteriia Flavobacteriales Flavobacteriaceae *Flavobacterium psychrophilum* FPG101 |
| Fps-3-1 | Bacteria Bacteroidetes Flavobacteriia Flavobacteriales Flavobacteriaceae *Flavobacterium psychrophilum* FPG3 |
| Fps-4-1 | Bacteria Bacteroidetes Flavobacteriia Flavobacteriales Flavobacteriaceae *Flavobacterium psychrophilum* JIP02/86 |
| Fri-1-1 | Bacteria Bacteroidetes Flavobacteriia Flavobacteriales Flavobacteriaceae *Flavobacterium rivuli* DSM 21788 |
| Fri-2-1 | Bacteria Bacteroidetes Flavobacteriia Flavobacteriales Flavobacteriaceae *Flavobacterium rivuli* WB 3.3-2 = DSM 21788 |
| Fsa-1-1 | Bacteria Bacteroidetes Flavobacteriia Flavobacteriales Flavobacteriaceae *Flavobacterium saliperosum* S13 |
| Fsa-2-1 | Bacteria Bacteroidetes Flavobacteriia Flavobacteriales Flavobacteriaceae *Flavobacterium sasangense* DSM 21067 |
| Fse-1-1 | Bacteria Bacteroidetes Flavobacteriia Flavobacteriales Flavobacteriaceae *Flavobacterium seoulense* |
| Fso-1-1 | Bacteria Bacteroidetes Flavobacteriia Flavobacteriales Flavobacteriaceae *Flavobacterium soli* DSM 19725 |
| Fsp-1-1 | Bacteria Bacteroidetes Flavobacteriia Flavobacteriales Flavobacteriaceae *Flavobacterium sp.* 316 |
| Fsp-2-1 | Bacteria Bacteroidetes Flavobacteriia Flavobacteriales Flavobacteriaceae *Flavobacterium sp.* 83 |
| Fsp-3-1 | Bacteria Bacteroidetes Flavobacteriia Flavobacteriales Flavobacteriaceae *Flavobacterium sp.* ABG |
| Fsp-4-1 | Bacteria Bacteroidetes Flavobacteriia Flavobacteriales Flavobacteriaceae *Flavobacterium sp.* ACAM 123 |
| Fsp-5-1 | Bacteria Bacteroidetes Flavobacteriia Flavobacteriales Flavobacteriaceae *Flavobacterium sp.* AED |
| Fsp-6-1 | Bacteria Bacteroidetes Flavobacteriia Flavobacteriales Flavobacteriaceae *Flavobacterium sp.* CF136 |
| Fsp-7-1 | Bacteria Bacteroidetes Flavobacteriia Flavobacteriales Flavobacteriaceae *Flavobacterium sp.* F52 |
| Fsp-8-1 | Bacteria Bacteroidetes Flavobacteriia Flavobacteriales Flavobacteriaceae *Flavobacterium sp.* JRM |
| Fsp-9-1 | Bacteria Bacteroidetes Flavobacteriia Flavobacteriales Flavobacteriaceae *Flavobacterium sp.* KJJ |
| Fsp-10-1 | Bacteria Bacteroidetes Flavobacteriia Flavobacteriales Flavobacteriaceae *Flavobacterium sp.* KMS |
| Fsp-11-1 | Bacteria Bacteroidetes Flavobacteriia Flavobacteriales Flavobacteriaceae *Flavobacterium sp.* MEB061 |
| Fsp-12-1 | Bacteria Bacteroidetes Flavobacteriia Flavobacteriales Flavobacteriaceae *Flavobacterium sp.* URHB0058 |
| Fsp-13-1 | Bacteria Bacteroidetes Flavobacteriia Flavobacteriales Flavobacteriaceae *Flavobacterium sp.* WG21 |
| Fsu-1-1 to Fsu-1-2 | Bacteria Bacteroidetes Flavobacteriia Flavobacteriales Flavobacteriaceae *Flavobacterium subsaxonicum* WB 4.1-42 = DSM 21790 |
| Fsu-2-1 | Bacteria Bacteroidetes Flavobacteriia Flavobacteriales Flavobacteriaceae *Flavobacterium succinicans* LMG 10402 |
| Fsu-3-1 to Fsu-3-2 | Bacteria Bacteroidetes Flavobacteriia Flavobacteriales Flavobacteriaceae *Flavobacterium suncheonense* GH29-5 = DSM 17707 |
| Fte-1-1 | Bacteria Bacteroidetes Flavobacteriia Flavobacteriales Flavobacteriaceae *Flavobacterium tegetincola* DSM 22377 |
| Fag-1-1 | Bacteria Bacteroidetes Flavobacteriia Flavobacteriales Flavobacteriaceae *Formosa agariphila* KMM 3901 |
| Fsp-14-1 | Bacteria Bacteroidetes Flavobacteriia Flavobacteriales Flavobacteriaceae *Formosa sp.* AK20 |
| Gsa-1-1 | Bacteria Bacteroidetes Flavobacteriia Flavobacteriales Flavobacteriaceae *Gaetbulibacter saemankumensis* DSM 17032 |
| Gsp-1-1 | Bacteria Bacteroidetes Flavobacteriia Flavobacteriales Flavobacteriaceae *Galbibacter sp. ck*-I2-15 |
| Gme-1-1 | Bacteria Bacteroidetes Flavobacteriia Flavobacteriales Flavobacteriaceae *Gelidibacter mesophilus* DSM 14095 |
| Gli-1-1 | Bacteria Bacteroidetes Flavobacteriia Flavobacteriales Flavobacteriaceae *Gillisia limnaea* DSM 15749 |
| Gsp-2-1 | Bacteria Bacteroidetes Flavobacteriia Flavobacteriales Flavobacteriaceae *Gillisia sp.* CAL575 |
| Gsp-3-1 | Bacteria Bacteroidetes Flavobacteriia Flavobacteriales Flavobacteriaceae *Gillisia sp.* CBA3202 |
| Gsp-4-1 | Bacteria Bacteroidetes Flavobacteriia Flavobacteriales Flavobacteriaceae *Gillisia sp.* Hel_I_29 |
| Gsp-5-1 | Bacteria Bacteroidetes Flavobacteriia Flavobacteriales Flavobacteriaceae *Gillisia sp.* JM1 |
| Gec-1-1 | Bacteria Bacteroidetes Flavobacteriia Flavobacteriales Flavobacteriaceae *Gramella echinicola* DSM 19838 |
| Gpo-1-1 | Bacteria Bacteroidetes Flavobacteriia Flavobacteriales Flavobacteriaceae *Gramella portivictoriae* DSM 23547 |
| Iha-1-1 | Bacteria Bacteroidetes Flavobacteriia Flavobacteriales Flavobacteriaceae *Imtechella halotolerans* K1 |
| Jpa-1-1 to Jpa-1-3 | Bacteria Bacteroidetes Flavobacteriia Flavobacteriales Flavobacteriaceae *Jejuia pallidilutea* |
| Jma-1-1 | Bacteria Bacteroidetes Flavobacteriia Flavobacteriales Flavobacteriaceae *Joostella marina* DSM 19592 |
| Kje-1-1 | Bacteria Bacteroidetes Flavobacteriia Flavobacteriales Flavobacteriaceae *Kordia jejudonensis* |
| Kzh-1-1 | Bacteria Bacteroidetes Flavobacteriia Flavobacteriales Flavobacteriaceae *Kordia zhangzhouensis* |
| Ksp-1-1 | Bacteria Bacteroidetes Flavobacteriia Flavobacteriales Flavobacteriaceae *Krokinobacter sp.* 4H-3-7-5 |
| Lja-1-1 | Bacteria Bacteroidetes Flavobacteriia Flavobacteriales Flavobacteriaceae *Lacinutrix jangbogonensis* |
| Lsp-1-1 | Bacteria Bacteroidetes Flavobacteriia Flavobacteriales Flavobacteriaceae *Lacinutrix sp.* 5H-3-7-4 |
| Lsp-2-1 | Bacteria Bacteroidetes Flavobacteriia Flavobacteriales Flavobacteriaceae *Lacinutrix sp.* Hel_I_90 |
| Lsp-3-1 | Bacteria Bacteroidetes Flavobacteriia Flavobacteriales Flavobacteriaceae *Leeuwenhoekiella sp.* Hel_I_48 |
| Lsp-4-1 | Bacteria Bacteroidetes Flavobacteriia Flavobacteriales Flavobacteriaceae *Leeuwenhoekiella sp.* MAR_2009_132 |
| Myu-1-1 | Bacteria Bacteroidetes Flavobacteriia Flavobacteriales Flavobacteriaceae *Mangrovimonas yunxiaonensis* |
| Man-1-1 | Bacteria Bacteroidetes Flavobacteriia Flavobacteriales Flavobacteriaceae *Maribacter antarcticus* DSM 21422 |
| Mfo-1-1 | Bacteria Bacteroidetes Flavobacteriia Flavobacteriales Flavobacteriaceae *Maribacter forsetii* DSM 18668 |
| Msp-1-1 | Bacteria Bacteroidetes Flavobacteriia Flavobacteriales Flavobacteriaceae *Maribacter sp.* Hel_I_7 |
| Mth-1-1 | Bacteria Bacteroidetes Flavobacteriia Flavobacteriales Flavobacteriaceae *Maribacter thermophilus* |
| Mze-1-1 | Bacteria Bacteroidetes Flavobacteriia Flavobacteriales Flavobacteriaceae *Mesoflavibacter zeaxanthinifaciens* DSM 18436 |
| Mze-2-1 | Bacteria Bacteroidetes Flavobacteriia Flavobacteriales Flavobacteriaceae *Mesoflavibacter zeaxanthinifaciens* S86 |
| Mmo-1-1 | Bacteria Bacteroidetes Flavobacteriia Flavobacteriales Flavobacteriaceae *Mesonia mobilis* DSM 19841 |

| | |
|---|---|
| Mru-1-1 | Bacteria Bacteroidetes Flavobacteriia Flavobacteriales Flavobacteriaceae *Muricauda ruestringensis* DSM 13258 |
| Msp-2-1 | Bacteria Bacteroidetes Flavobacteriia Flavobacteriales Flavobacteriaceae *Muricauda sp.* MAR_2010_75 |
| M[o-1-1 | Bacteria Bacteroidetes Flavobacteriia Flavobacteriales Flavobacteriaceae *Myroides [odoratimimus]* CIP 103059 |
| Min-1-1 | Bacteria Bacteroidetes Flavobacteriia Flavobacteriales Flavobacteriaceae *Myroides injenensis* M09-0166 |
| Mod-1-1 | Bacteria Bacteroidetes Flavobacteriia Flavobacteriales Flavobacteriaceae *Myroides odoratimimus* |
| Mod-2-1 | Bacteria Bacteroidetes Flavobacteriia Flavobacteriales Flavobacteriaceae *Myroides odoratimimus* CCUG 10230 |
| Mod-3-1 | Bacteria Bacteroidetes Flavobacteriia Flavobacteriales Flavobacteriaceae *Myroides odoratimimus* CCUG 12700 |
| Mod-4-1 | Bacteria Bacteroidetes Flavobacteriia Flavobacteriales Flavobacteriaceae *Myroides odoratimimus* CCUG 12901 |
| Mod-5-1 | Bacteria Bacteroidetes Flavobacteriia Flavobacteriales Flavobacteriaceae *Myroides odoratimimus* CCUG 3837 |
| Mod-6-1 | Bacteria Bacteroidetes Flavobacteriia Flavobacteriales Flavobacteriaceae *Myroides odoratimimus* CIP 101113 |
| Mod-7-1 | Bacteria Bacteroidetes Flavobacteriia Flavobacteriales Flavobacteriaceae *Myroides odoratimimus* H1bi |
| Mod-8-1 | Bacteria Bacteroidetes Flavobacteriia Flavobacteriales Flavobacteriaceae *Myroides odoratus* DSM 2801 |
| Mpr-1-1 | Bacteria Bacteroidetes Flavobacteriia Flavobacteriales Flavobacteriaceae *Myroides profundi* |
| Msp-3-1 | Bacteria Bacteroidetes Flavobacteriia Flavobacteriales Flavobacteriaceae *Myroides sp.* A21 |
| Nma-1-1 | Bacteria Bacteroidetes Flavobacteriia Flavobacteriales Flavobacteriaceae *Nonlabens marinus* S1-08 |
| Nul-1-1 to Nul-1-6 | Bacteria Bacteroidetes Flavobacteriia Flavobacteriales Flavobacteriaceae *Nonlabens ulvanivorans* |
| Oma-1-1 | Bacteria Bacteroidetes Flavobacteriia Flavobacteriales Flavobacteriaceae *Olleya marilimosa* CAM030 |
| Osp-1-1 | Bacteria Bacteroidetes Flavobacteriia Flavobacteriales Flavobacteriaceae *Olleya sp.* VCSA23 |
| Osp-2-1 | Bacteria Bacteroidetes Flavobacteriia Flavobacteriales Flavobacteriaceae *Olleya sp.* VCSM12 |
| Orh-1-1 | Bacteria Bacteroidetes Flavobacteriia Flavobacteriales Flavobacteriaceae *Ornithobacterium rhinotracheale* DSM 15997 |
| Orh-2-1 | Bacteria Bacteroidetes Flavobacteriia Flavobacteriales Flavobacteriaceae *Ornithobacterium rhinotracheale* H06-030791 |
| Orh-3-1 | Bacteria Bacteroidetes Flavobacteriia Flavobacteriales Flavobacteriaceae *Ornithobacterium rhinotracheale* ORT-UMN 88 |
| Psp-1-1 | Bacteria Bacteroidetes Flavobacteriia Flavobacteriales Flavobacteriaceae *Polaribacter sp.* Hel1_33_49 |
| Psp-2-1 | Bacteria Bacteroidetes Flavobacteriia Flavobacteriales Flavobacteriaceae *Polaribacter sp.* Hel1_85 |
| Psp-3-1 | Bacteria Bacteroidetes Flavobacteriia Flavobacteriales Flavobacteriaceae *Polaribacter sp.* Hel_I_88 |
| Psp-4-1 | Bacteria Bacteroidetes Flavobacteriia Flavobacteriales Flavobacteriaceae *Polaribacter sp.* MED152 |
| Pgo-1-1 | Bacteria Bacteroidetes Flavobacteriia Flavobacteriales Flavobacteriaceae *Psychroflexus gondwanensis* ACAM 44 |
| Pto-1-1 | Bacteria Bacteroidetes Flavobacteriia Flavobacteriales Flavobacteriaceae *Psychroflexus torquis* ATCC 700755 |
| Ptr-1-1 | Bacteria Bacteroidetes Flavobacteriia Flavobacteriales Flavobacteriaceae *Psychroflexus tropicus* DSM 15496 |
| Pbu-1-1 | Bacteria Bacteroidetes Flavobacteriia Flavobacteriales Flavobacteriaceae *Psychroserpens burtonensis* DSM 12212 |
| Pda-1-1 | Bacteria Bacteroidetes Flavobacteriia Flavobacteriales Flavobacteriaceae *Psychroserpens damuponensis* |
| Pja-1-1 | Bacteria Bacteroidetes Flavobacteriia Flavobacteriales Flavobacteriaceae *Psychroserpens jangbogonensis* |
| Pme-1-1 | Bacteria Bacteroidetes Flavobacteriia Flavobacteriales Flavobacteriaceae *Psychroserpens mesophilus* |
| Psp-5-1 | Bacteria Bacteroidetes Flavobacteriia Flavobacteriales Flavobacteriaceae *Psychroserpens sp.* Hel_I_66 |
| Ran-1-1 to Ran-1-2 | Bacteria Bacteroidetes Flavobacteriia Flavobacteriales Flavobacteriaceae *Riemerella anatipestifer* |
| Ran-2-1 to Ran-2-2 | Bacteria Bacteroidetes Flavobacteriia Flavobacteriales Flavobacteriaceae *Riemerella anatipestifer* ATCC 11845 = DSM 15868 |
| Ran-3-1 | Bacteria Bacteroidetes Flavobacteriia Flavobacteriales Flavobacteriaceae *Riemerella anatipestifer* RA-CH-2 |
| Ran-4-1 | Bacteria Bacteroidetes Flavobacteriia Flavobacteriales Flavobacteriaceae *Riemerella anatipestifer* RA-GD |
| Ran-5-1 | Bacteria Bacteroidetes Flavobacteriia Flavobacteriales Flavobacteriaceae *Riemerella anatipestifer* RA-SG |
| Ran-6-1 | Bacteria Bacteroidetes Flavobacteriia Flavobacteriales Flavobacteriaceae *Riemerella anatipestifer* RA-YM |
| Ran-7-1 | Bacteria Bacteroidetes Flavobacteriia Flavobacteriales Flavobacteriaceae *Riemerella anatipestifer* Yb2 |
| Ssp-1-1 | Bacteria Bacteroidetes Flavobacteriia Flavobacteriales Flavobacteriaceae *Salegentibacter sp.* Hel_I_6 |
| Ste-1-1 | Bacteria Bacteroidetes Flavobacteriia Flavobacteriales Flavobacteriaceae *Salinimicrobium terrae* DSM 17865 |
| Sxi-1-1 | Bacteria Bacteroidetes Flavobacteriia Flavobacteriales Flavobacteriaceae *Salinimicrobium xinjiangense* DSM 19287 |
| Ssp-2-1 | Bacteria Bacteroidetes Flavobacteriia Flavobacteriales Flavobacteriaceae *Sediminibacter sp.* Hel_I_10 |
| Sze-1-1 | Bacteria Bacteroidetes Flavobacteriia Flavobacteriales Flavobacteriaceae *Siansivirga zeaxanthinifaciens* CC-SAMT-1 |
| Tna-1-1 | Bacteria Bacteroidetes Flavobacteriia Flavobacteriales Flavobacteriaceae *Tamlana nanhaiensis* |
| Tse-1-1 | Bacteria Bacteroidetes Flavobacteriia Flavobacteriales Flavobacteriaceae *Tamlana sedimentorum* |
| Tma-1-1 | Bacteria Bacteroidetes Flavobacteriia Flavobacteriales Flavobacteriaceae *Tenacibaculum maritimum* NCIMB 2154 |
| Tme-1-1 | Bacteria Bacteroidetes Flavobacteriia Flavobacteriales Flavobacteriaceae *Tenacibaculum mesophilum* |
| Tov-1-1 | Bacteria Bacteroidetes Flavobacteriia Flavobacteriales Flavobacteriaceae *Tenacibaculum ovolyticum* DSM 18103 |
| Tsp-1-1 | Bacteria Bacteroidetes Flavobacteriia Flavobacteriales Flavobacteriaceae *Tenacibaculum sp.* 47A_GOM-205m |
| Wma-1-1 | Bacteria Bacteroidetes Flavobacteriia Flavobacteriales Flavobacteriaceae *Weeksella massiliensis* |
| Wvi-1-1 | Bacteria Bacteroidetes Flavobacteriia Flavobacteriales Flavobacteriaceae *Weeksella virosa* DSM 16922 |
| Wps-1-1 | Bacteria Bacteroidetes Flavobacteriia Flavobacteriales Flavobacteriaceae *Winogradskyella psychrotolerans* RS-3 |
| Wsp-1-1 | Bacteria Bacteroidetes Flavobacteriia Flavobacteriales Flavobacteriaceae *Winogradskyella sp.* PG-2 |
| Zam-1-1 | Bacteria Bacteroidetes Flavobacteriia Flavobacteriales Flavobacteriaceae *Zhouia amylolytica* AD3 |
| Zga-1-1 | Bacteria Bacteroidetes Flavobacteriia Flavobacteriales Flavobacteriaceae *Zobellia galactanivorans* |
| Zul-1-1 | Bacteria Bacteroidetes Flavobacteriia Flavobacteriales Flavobacteriaceae *Zobellia uliginosa* |
| Psp-6-1 | Bacteria Proteobacteria Gammaproteobacteria Alteromonadales Psychromonadaceae *Psychromonas sp.* SP041 |
| env-1 to env-2152 | environmental samples |

#### 22.2 Notes

Superscript numbers are used to annotate any S15-Flavobacteria RNA that has special characteristics (described below). These numeric annotations will be used in Sections 22.3 and 22.5:. The meaning of the numbers are as follows.

[1] No UAnnUUUG is apparent in the 100 nucleotides upstream of this motif instance.

#### 22.3 Gene contexts

Each S15-Flavobacteria RNA (indicated by "RNA→") is listed. For each hit, the downstream genes predicted to reside in a regulated operon are listed. If the nearest downstream gene is encoding in the opposite strand (and therefore presumed to not be a part of a regulated operon), then that gene is still depicted. Some environmental sequences and some RefSeq entries lack gene annotations, and so no genes are available for such sequences. The direction of each gene is indicated with an arrow (→), and each predicted conserved domain in the gene is named. Conserved domains associated with more than one S15-Flavobacteria RNA are assigned a color; other domains are gray. Information about these conserved domains is given in Section 22.4. The accession of the sequence containing each S15-Flavobacteria RNA is given in the column named "Seq. accession". Accessions beginning with "NC_", "NS_", "NW_" or "NZ_" are contained in RefSeq. Other accession refer to environmental samples. Nucleotide coordinates are given for the 5′ and 3′ boundaries of each S15-Flavobacteria RNA. If the 5′ coordinate is greater than the 3′ coordinate, the RNA is present on the reverse-complement strand of the containing genomic DNA sequence. Each hit is denoted by an abbreviation (like "Eco-1-1") that refers to a taxonomy given in Section 22.1. (Superscript numbers refer to annotations listed in Section 22.2)

| abbrev. | Seq. accession | | 5′ at | 3′ at | genes |
|---|---|---|---|---|---|
| Pgo-1-1 | NZ_APLF01000002.1 | - | 80877 | 80813 | RNA→ rpsO (PRK05626)Ribosomal_S15 (pfam00312)→ |
| env-1 | P_2C_Liq_1_UnCty_1066413 | - | 280 | 216 | RNA→ rpsO (PRK05626)Ribosomal_S15 (pfam00312)→ |
| env-2 | SL_9KL_010_SED_10540992 | + | 126 | 190 | RNA→ rpsO (PRK05626)Ribosomal_S15 (pfam00312)→ |
| env-3 | JGI11876J14442_10000003 | + | 205280 | 205344 | RNA→ rpsO (PRK05626)Ribosomal_S15 (pfam00312)→ |
| Pto-1-1 | NC_018721.1 | + | 147626 | 147690 | RNA→ rpsO (PRK05626)Ribosomal_S15 (pfam00312)→ |
| Ptr-1-1 | NZ_KB905343.1 | - | 600481 | 600417 | RNA→ rpsO (PRK05626)Ribosomal_S15 (pfam00312)→ |
| env-4 | JGI1221J11331_1000106 | - | 12367 | 12303 | RNA→ rpsO (PRK05626)Ribosomal_S15 (pfam00312)→ |
| env-5 | P_1C_Liq_1_UnCty_1089716 | - | 107 | 43 | RNA→ |
| env-6 | P_1C_Liq_3_UnCty_1064543 | - | 231 | 167 | RNA→ rpsO (PRK05626)Ribosomal_S15 (pfam00312)→ |
| env-7 | P_1C_Liq_1_UnCty_1008265 | + | 1415 | 1479 | RNA→ rpsO (PRK05626)Ribosomal_S15 (pfam00312)→ |
| env-8 | P_2C_Liq_1_UnCty_1170116 | + | 90 | 154 | RNA→ rpsO (PRK05626)Ribosomal_S15 (pfam00312)→ |
| Mod-8-1 | NZ_CM001437.1 | - | 1513261 | 1513195 | RNA→ rpsO (PRK05626)Ribosomal_S15 (pfam00312)→ rpsO (PRK05626)Ribosomal_S15 (pfam00312)→ |
| M[o-1-1 | NZ_JH815539.1 | + | 176108 | 176174 | RNA→ rpsO (PRK05626)Ribosomal_S15 (pfam00312)→ |
| env-9 | JGI20156J14371_10110659 | - | 680 | 612 | RNA→ rpsO (PRK05626)Ribosomal_S15 (pfam00312)→ |
| env-10 | JGI20159J14440_10000064 | - | 3008 | 2940 | RNA→ rpsO (PRK05626)Ribosomal_S15 (pfam00312)→ |
| env-11 | JGI20160J14292_10230840 | + | 25 | 93 | RNA→ rpsO (PRK05626)Ribosomal_S15 (pfam00312)→ |
| env-12 | JGI20155J14468_10405117 | - | 344 | 276 | RNA→ rpsO (PRK05626)Ribosomal_S15 (pfam00312)→ |
| env-13 | JGI20159J14440_10135058 | + | 106 | 174 | RNA→ rpsO (PRK05626)Ribosomal_S15 (pfam00312)→ |
| env-14 | JGI20155J14468_10023041 | - | 280 | 212 | RNA→ rpsO (PRK05626)Ribosomal_S15 (pfam00312)→ |
| env-15 | CAM_READ_0235006571 | - | 166 | 99 | RNA→ rpsO (PRK05626)RpsO (COG0184)→ |
| env-16 | 3300001592_10000361 | - | 46071 | 46015 | RNA→ rpsO (PRK05626)RpsO (COG0184)→ |
| env-17 | P_2C_Liq_1_UnCty_1027718 | + | 347 | 414 | RNA→ rpsO (PRK05626)Ribosomal_S15 (pfam00312)→ |
| Fbr-1-1 | NC_016001.1 | - | 1162758 | 1162692 | RNA→ rpsO (PRK05626)Ribosomal_S15 (pfam00312)→ |
| env-18 | JGI25922J50271_10000339 | - | 8516 | 8450 | RNA→ rpsO (PRK05626)Ribosomal_S15 (pfam00312)→ |
| env-19 | JGI25921J50272_10046219 | - | 307 | 241 | RNA→ rpsO (PRK05626)Ribosomal_S15 (pfam00312)→ |
| env-20 | soil_XZS65148_x1 | - | 304 | 236 | RNA→ rpsO (PRK05626)Ribosomal_S15 (pfam00312)→ |
| env-21 | BS_KBA_SWE21_205m_10189866 | + | 318 | 376 | RNA→ |
| env-22 | CEUB01052019.1 | + | 160 | 227 | RNA→ rpsO (PRK05626)RpsO (COG0184)→ |
| env-23 | CEOP01451304.1 | + | 607 | 674 | RNA→ rpsO (PRK05626)RpsO (COG0184)→ PRK11824 (PRK11824)RNase_PH (pfam01138)RNase_PH_C (pfam03725)→ |

| | | | | | |
|---|---|---|---|---|---|
| env-24 | CETG01141171.1 | + | 474 | 541 | **RNA**→ rpsO (PRK05626)Ribosomal_S15 (pfam00312)→ |
| env-25 | CETM01278025.1 | + | 477 | 544 | **RNA**→ |
| Fba-9-1 | NZ_AFPK01000014.1 | + | 150749 | 150814 | **RNA**→ rpsO (PRK05626)Ribosomal_S15 (pfam00312)→ rpsO (PRK05626)Ribosomal_S15 (pfam00312)→ |
| env-26 | CEOM01058151.1 | - | 607 | 540 | **RNA**→ rpsO (PRK05626)Ribosomal_S15 (pfam00312)→ |
| env-27 | JGI25913J50563_1244176 | + | 71 | 137 | **RNA**→ rpsO (PRK05626)Ribosomal_S15 (pfam00312)→ |
| env-28 | CETR01087310.1 | - | 18809 | 18745 | **RNA**→ rpsO (PRK05626)Ribosomal_S15 (pfam00312)→ |
| env-29 | CEVJ01009927.1 | + | 44140 | 44204 | **RNA**→ rpsO (PRK05626)Ribosomal_S15 (pfam00312)→ |
| env-30 | CEVK01027416.1 | - | 18585 | 18521 | **RNA**→ rpsO (PRK05626)Ribosomal_S15 (pfam00312)→ |
| env-31 | CEVM01013066.1 | + | 53334 | 53398 | **RNA**→ rpsO (PRK05626)Ribosomal_S15 (pfam00312)→ |
| env-32 | CEVO01125175.1 | + | 10666 | 10730 | **RNA**→ rpsO (PRK05626)Ribosomal_S15 (pfam00312)→ |
| env-33 | CEVQ01101129.1 | + | 770 | 834 | **RNA**→ rpsO (PRK05626)Ribosomal_S15 (pfam00312)→ |
| env-34 | JGI20154J14316_10054052 | - | 581 | 522 | **RNA**→ rpsO (PRK05626)RpsO (COG0184)→ |
| env-35 | JGI20157J14317_10003423 | - | 3710 | 3651 | **RNA**→ rpsO (PRK05626)RpsO (COG0184)→ |
| env-36 | DelMOSum2010_c10728123 | - | 223 | 164 | **RNA**→ rpsO (PRK05626)Ribosomal_S15 (pfam00312)→ |
| env-37 | JCVI_SCAF_1096627992251 | + | 1032 | 1089 | **RNA**→ rpsO (PRK05626)Ribosomal_S15 (pfam00312)→ |
| env-38 | 2236876001_130459 | + | 242 | 300 | **RNA**→ rpsO (PRK05626)Ribosomal_S15 (pfam00312)→ |
| env-39 | DelMOSum2010_c10011046 | - | 5729 | 5669 | **RNA**→ rpsO (PRK05626)RpsO (COG0184)→ |
| env-40 | DelMOSum2011_c10604815 | - | 74 | 14 | **RNA**→ |
| env-41 | DelMOSum2011_c10341564 | + | 308 | 368 | **RNA**→ |
| env-42 | JGI24723J26617_10012210 | - | 2791 | 2732 | **RNA**→ rpsO (PRK05626)RpsO (COG0184)→ |
| env-43 | JGI25925J51416_10027951 | + | 789 | 854 | **RNA**→ rpsO (PRK05626)Ribosomal_S15 (pfam00312)→ |
| Myu-1-1 | NZ_JPFK01000007.1 | - | 20817 | 20726 | **RNA**→ rpsO (PRK05626)RpsO (COG0184)→ hypo→ hypo→ |
| env-44 | CERM01235517.1 | + | 1791 | 1859 | **RNA**→ |
| env-45 | M3P_c10006697 | - | 1022 | 954 | **RNA**→ rpsO (PRK05626)Ribosomal_S15 (pfam00312)→ |
| env-46 | JGI25913J50563_1345866 | + | 57 | 125 | **RNA**→ rpsO (PRK05626)Ribosomal_S15 (pfam00312)→ |
| env-47 | _GVICOHK01AKHQ1 | + | 147 | 215 | **RNA**→ |
| env-48 | GXP7IEG01BCPE8 | + | 165 | 233 | **RNA**→ |
| env-49 | TFO_HY__Contig_16695 | - | 374 | 310 | **RNA**→ rpsO (PRK05626)Ribosomal_S15 (pfam00312)→ |
| env-50 | ATED1026_F5B15RW02JR6D3 | + | 134 | 198 | **RNA**→ rpsO (PRK05626)Ribosomal_S15 (pfam00312)→ |
| env-51 | JGI20157J14317_10000285 | + | 5024 | 5082 | **RNA**→ rpsO (PRK05626)RpsO (COG0184)→ |
| env-52 | SA_S1_NOR08_45m_c10119203 | - | 441 | 384 | **RNA**→ rpsO (PRK05626)Ribosomal_S15 (pfam00312)→ |
| env-53 | 20090826-Illumina_contig28958 | - | 458 | 392 | **RNA**→ rpsO (PRK05626)RpsO (COG0184)→ |
| Fsa-1-1 | NZ_AVFO01000002.1 | + | 264122 | 264190 | **RNA**→ rpsO (PRK05626)Ribosomal_S15 (pfam00312)→ |
| Fca-1-1 | NZ_AVBI01000015.1 | - | 13067 | 12999 | **RNA**→ rpsO (PRK05626)Ribosomal_S15 (pfam00312)→ |
| Fca-1-2 | NZ_JRLU01000001.1 | - | 375317 | 375249 | **RNA**→ rpsO (PRK05626)Ribosomal_S15 (pfam00312)→ |
| env-54 | CESA01099157.1 | + | 266 | 332 | **RNA**→ rpsO (PRK05626)Ribosomal_S15 (pfam00312)→ |
| env-55 | CESN01196738.1 | + | 1485 | 1551 | **RNA**→ rpsO (PRK05626)Ribosomal_S15 (pfam00312)→ |
| env-56 | CESP01155747.1 | + | 1403 | 1469 | **RNA**→ rpsO (PRK05626)Ribosomal_S15 (pfam00312)→ |
| env-57 | CEVS01034257.1 | - | 7704 | 7638 | **RNA**→ rpsO (PRK05626)Ribosomal_S15 (pfam00312)→ |
| env-58 | CEVU01127351.1 | + | 981 | 1047 | **RNA**→ rpsO (PRK05626)Ribosomal_S15 (pfam00312)→ |
| env-59 | LBBO01003853.1 | + | 546 | 600 | **RNA**→ rpsO (PRK05626)RpsO (COG0184)→ |
| env-60 | JGI20157J14317_10520585 | - | 115 | 48 | **RNA**→ |
| env-61 | JGI20152J14361_10102081 | + | 445 | 512 | **RNA**→ hypo→ |
| env-62 | JGI20154J14316_10389205 | + | 184 | 251 | **RNA**→ |
| env-63 | JGI20153J14295_10154293 | + | 431 | 498 | **RNA**→ hypo→ |
| env-64 | JGI20153J14318_10154293 | + | 431 | 498 | **RNA**→ hypo→ |
| env-65 | DelMOSum2010_c10048941 | - | 1015 | 948 | **RNA**→ rpsO (PRK05626)Ribosomal_S15 (pfam00312)→ |
| env-66 | JGI20152J14361_10018736 | + | 1929 | 1996 | **RNA**→ rpsO (PRK05626)Ribosomal_S15 (pfam00312)→ |
| env-67 | DelMOSum2011_c10028302 | - | 2345 | 2278 | **RNA**→ rpsO (PRK05626)Ribosomal_S15 (pfam00312)→ |
| env-68 | JGI20152J14361_10074584 | - | 336 | 269 | **RNA**→ rpsO (PRK05626)Ribosomal_S15 (pfam00312)→ |
| env-69 | JCVI_SCAF_1096626866200 | - | 228 | 161 | **RNA**→ rpsO (PRK05626)Ribosomal_S15 (pfam00312)→ |
| env-70 | 2236876010_0476296 | + | 34 | 101 | **RNA**→ rpsO (PRK05626)Ribosomal_S15 (pfam00312)→ |

| | | | | | |
|---|---|---|---|---|---|
| env-71 | DelMOSum2010_c10114672 | - | 927 | 860 | **RNA**→ rpsO (PRK05626)Ribosomal_S15 (pfam00312)→ |
| env-72 | JGI20156J14371_10028011 | + | 1695 | 1762 | **RNA**→ rpsO (PRK05626)Ribosomal_S15 (pfam00312)→ |
| env-73 | JGI20154J14316_10016589 | - | 2489 | 2422 | **RNA**→ rpsO (PRK05626)Ribosomal_S15 (pfam00312)→ |
| env-74 | JGI20160J14292_10093798 | + | 200 | 267 | **RNA**→ rpsO (PRK05626)Ribosomal_S15 (pfam00312)→ |
| env-75 | JGI20157J14317_10004874 | - | 1379 | 1312 | **RNA**→ rpsO (PRK05626)Ribosomal_S15 (pfam00312)→ |
| env-76 | JGI20152J14361_10042016 | + | 1197 | 1264 | **RNA**→ |
| env-77 | JGI20153J14295_10052012 | + | 1350 | 1417 | **RNA**→ |
| env-78 | JGI20153J14318_10052012 | + | 1350 | 1417 | **RNA**→ |
| env-79 | PML_READ_00443584 | - | 242 | 175 | **RNA**→ rpsO (PRK05626)Ribosomal_S15 (pfam00312)→ |
| env-80 | PML_READ_00298684 | + | 75 | 143 | **RNA**→ |
| env-81 | CESE01042213.1 | - | 314 | 247 | **RNA**→ rpsO (PRK05626)Ribosomal_S15 (pfam00312)→ |
| env-82 | CESJ01033522.1 | + | 13583 | 13650 | **RNA**→ rpsO (PRK05626)Ribosomal_S15 (pfam00312)→ |
| env-83 | CEST01015177.1 | + | 5814 | 5881 | **RNA**→ rpsO (PRK05626)Ribosomal_S15 (pfam00312)→ |
| env-84 | CESU01002158.1 | - | 2307 | 2240 | **RNA**→ rpsO (PRK05626)Ribosomal_S15 (pfam00312)→ |
| env-85 | JGI20152J14361_10001968 | - | 123 | 56 | **RNA**→ |
| env-86 | JGI20154J14316_10153966 | - | 465 | 398 | **RNA**→ rpsO (PRK05626)Ribosomal_S15 (pfam00312)→ |
| env-87 | JGI20153J14295_10006772 | - | 4012 | 3945 | **RNA**→ rpsO (PRK05626)Ribosomal_S15 (pfam00312)→ |
| env-88 | JGI20153J14318_10006772 | - | 4012 | 3945 | **RNA**→ rpsO (PRK05626)Ribosomal_S15 (pfam00312)→ |
| env-89 | CERM01030429.1 | - | 985 | 918 | **RNA**→ rpsO (PRK05626)Ribosomal_S15 (pfam00312)→ |
| env-90 | CESE01041678.1 | - | 313 | 247 | **RNA**→ rpsO (PRK05626)Ribosomal_S15 (pfam00312)→ |
| env-91 | CEST01086311.1 | + | 5957 | 6023 | **RNA**→ rpsO (PRK05626)Ribosomal_S15 (pfam00312)→ |
| env-92 | CEQA01115990.1 | + | 458 | 525 | **RNA**→ rpsO (PRK05626)Ribosomal_S15 (pfam00312)→ |
| env-93 | TahiMoana_1000156 | - | 8657 | 8590 | **RNA**→ rpsO (PRK05626)Ribosomal_S15 (pfam00312)→ |
| env-94 | JGI24723J26617_10000024 | - | 87111 | 87044 | **RNA**→ rpsO (PRK05626)Ribosomal_S15 (pfam00312)→ |
| env-95 | CENR01073469.1 | - | 3738 | 3671 | **RNA**→ rpsO (PRK05626)Ribosomal_S15 (pfam00312)→ |
| env-96 | CENX01105912.1 | - | 734 | 680 | **RNA**→ rpsO (PRK05626)Ribosomal_S15 (pfam00312)→ PRK11824 (PRK11824)RNase_PH (pfam01138)→ |
| env-97 | CEOP01457717.1 | + | 2903 | 2957 | **RNA**→ rpsO (PRK05626)Ribosomal_S15 (pfam00312)→ Pnp (COG1185)PRK11824 (PRK11824)→ |
| | | | | | RpoD (COG0568)sigma70-ECF (TIGR02937)→ |
| env-98 | CEOF01208035.1 | - | 315 | 261 | **RNA**→ rpsO (PRK05626)Ribosomal_S15 (pfam00312)→ |
| env-99 | P_1C_Liq_1_UnCty_1000027 | - | 17086 | 17028 | **RNA**→ rpsO (PRK05626)Ribosomal_S15 (pfam00312)→ Pnp (COG1185)PRK11824 (PRK11824)→ |
| | | | | | TonB (COG0810)tonB_Cterm (TIGR01352)→ |
| env-100 | P_2C_Liq_1_UnCty_1195289 | + | 93 | 151 | **RNA**→ rpsO (PRK05626)Ribosomal_S15 (pfam00312)→ |
| env-101 | JCVI_SCAF_1101667010888 | + | 552 | 612 | **RNA**→ rpsO (PRK05626)Ribosomal_S15 (pfam00312)→ |
| env-102 | CERI01036223.1 | - | 4293 | 4241 | **RNA**→ rpsO (PRK05626)Ribosomal_S15 (pfam00312)→ Pnp (COG1185)PRK11824 (PRK11824)→ |
| | | | | | RpoD (COG0568)sigma70-ECF (TIGR02937)→ PRK05581 (PRK05581)RPE (cd00429)→ |
| env-103 | CENP01014601.1 | + | 918 | 969 | **RNA**→ rpsO (PRK05626)Ribosomal_S15 (pfam00312)→ |
| env-104 | CENX01209556.1 | - | 1161 | 1110 | **RNA**→ rpsO (PRK05626)Ribosomal_S15 (pfam00312)→ |
| | | | | | PRK11824 (PRK11824)RNase_PH (pfam01138)RNase_PH_C (pfam03725)→ |
| env-105 | CEOF01432258.1 | + | 505 | 556 | **RNA**→ rpsO (PRK05626)Ribosomal_S15 (pfam00312)→ |
| | | | | | PRK11824 (PRK11824)RNase_PH (pfam01138)RNase_PH_C (pfam03725)→ |
| env-106 | CEOK01358130.1 | + | 1063 | 1114 | **RNA**→ rpsO (PRK05626)Ribosomal_S15 (pfam00312)→ |
| env-107 | CETK01306832.1 | - | 576 | 524 | **RNA**→ rpsO (PRK05626)Ribosomal_S15 (pfam00312)→ RNase_PH_PNPase_1 (cd11363)PRK11824 (PRK11824)– |
| env-108 | CESW01023746.1 | + | 683 | 736 | **RNA**→ rpsO (PRK05626)Ribosomal_S15 (pfam00312)→ Pnp (COG1185)PRK11824 (PRK11824)→ |
| env-109 | CESP01265619.1 | + | 546 | 598 | **RNA**→ rpsO (PRK05626)Ribosomal_S15 (pfam00312)→ |
| Min-1-1 | NZ_BAEX01000001.1 | + | 19959 | 20024 | **RNA**→ rpsO (PRK05626)Ribosomal_S15 (pfam00312)→ rpsO (PRK05626)Ribosomal_S15 (pfam00312)→ |
| env-110 | CESJ01281228.1 | - | 803 | 743 | **RNA**→ rpsO (PRK05626)Ribosomal_S15 (pfam00312)→ PRK11824 (PRK11824)RNase_PH (pfam01138)→ |
| env-111 | SA_S1_NOR05_45m_c10263701 | + | 212 | 278 | **RNA**→ rpsO (PRK05626)RpsO (COG0184)→ |
| env-112 | SA_S1_NOR08_45m_c10224370 | + | 426 | 492 | **RNA**→ |
| env-113 | SA_S1_NOR05_45m_c10350149 | + | 212 | 278 | **RNA**→ |
| env-114 | SA_S1_NOR08_45m_c10075071 | + | 209 | 275 | **RNA**→ rpsO (PRK05626)Ribosomal_S15 (pfam00312)→ |
| env-115 | SA_S2_NOR15_50m_c10328732 | + | 371 | 437 | **RNA**→ |
| env-116 | P_2C_Liq_1_UnCty_1015831 | + | 1190 | 1252 | **RNA**→ rpsO (PRK05626)Ribosomal_S15 (pfam00312)→ |
| Nul-1-1 | NZ_BBMM01000004.1 | - | 180573 | 180485 | **RNA**→ rpsO (PRK05626)Ribosomal_S15 (pfam00312)→ rpsO (PRK05626)Ribosomal_S15 (pfam00312)→ |

| | | | | | |
|---|---|---|---|---|---|
| Nul-1-2 | NZ_BBNT01000011.1 | - | 28548 | 28460 | **RNA**→ rpsO (PRK05626)Ribosomal_S15 (pfam00312)→ |
| Fba-1-1 | NZ_CH672374.1 | + | 1359719 | 1359807 | **RNA**→ rpsO (PRK05626)Ribosomal_S15 (pfam00312)→ rpsO (PRK05626)Ribosomal_S15 (pfam00312)→ |
| Nul-1-3 | NZ_JPJI01000032.1 | - | 1047570 | 1047482 | **RNA**→ rpsO (PRK05626)Ribosomal_S15 (pfam00312)→ |
| Nul-1-4 | NZ_BBMJ01000005.1 | - | 254727 | 254639 | **RNA**→ rpsO (PRK05626)Ribosomal_S15 (pfam00312)→ |
| Nul-1-5 | NZ_BBMK01000001.1 | + | 51054 | 51142 | **RNA**→ rpsO (PRK05626)Ribosomal_S15 (pfam00312)→ rpsO (PRK05626)Ribosomal_S15 (pfam00312)→ |
| Nul-1-6 | NZ_BBLG01000005.1 | - | 173187 | 173099 | **RNA**→ rpsO (PRK05626)Ribosomal_S15 (pfam00312)→ |
| env-117 | P_2C_Liq_1_UnCty_1000001 | + | 20247 | 20338 | **RNA**→ rpsO (PRK05626)Ribosomal_S15 (pfam00312)→ |
| env-118 | P_2C_Liq_2_UnCty_1019503 | + | 802 | 893 | **RNA**→ rpsO (PRK05626)Ribosomal_S15 (pfam00312)→ |
| env-119 | CEOM01017949.1 | - | 2624 | 2561 | **RNA**→ rpsO (PRK05626)Ribosomal_S15 (pfam00312)→ Pnp (COG1185)PRK11824 (PRK11824)→ |
| env-120 | CENX01076851.1 | - | 336 | 273 | **RNA**→ rpsO (PRK05626)Ribosomal_S15 (pfam00312)→ |
| env-121 | CEOF01146199.1 | + | 578 | 641 | **RNA**→ rpsO (PRK05626)Ribosomal_S15 (pfam00312)→ Pnp (COG1185)PRK11824 (PRK11824)→ |
| Fsu-1-1 | NZ_AUGP01000017.1 | + | 428051 | 428115 | **RNA**→ rpsO (PRK05626)Ribosomal_S15 (pfam00312)→ |
| Fsu-1-2 | NZ_JRLY01000002.1 | - | 230583 | 230519 | **RNA**→ rpsO (PRK05626)Ribosomal_S15 (pfam00312)→ |
| Gsp-1-1 | NZ_AMSG01000007.1 | + | 33255 | 33321 | **RNA**→ rpsO (PRK05626)Ribosomal_S15 (pfam00312)→ rpsO (PRK05626)Ribosomal_S15 (pfam00312)→ |
| env-122 | JGI11876J14442_11183631 | + | 151 | 208 | **RNA**→ |
| env-123 | JGI11876J14442_10280383 | - | 423 | 359 | **RNA**→ rpsO (PRK05626)RpsO (COG0184)→ |
| env-124 | CETR01192788.1 | - | 196 | 132 | **RNA**→ rpsO (PRK05626)Ribosomal_S15 (pfam00312)→ |
| env-125 | SI60aug11_200m_c1160499 | - | 131 | 66 | **RNA**→ rpsO (PRK05626)→ |
| env-126 | 2236876001_545617 | + | 56 | 124 | **RNA**→ rpsO (PRK05626)Ribosomal_S15 (pfam00312)→ |
| env-127 | 4491417.3_NODE_64483 | + | 5852 | 5910 | **RNA**→ rpsO (PRK05626)Ribosomal_S15 (pfam00312)→ |
| env-128 | SRS052697_C3073829 | + | 11 | 69 | **RNA**→ rpsO (PRK05626)Ribosomal_S15 (pfam00312)→ |
| env-129 | scaffold49704_5_MH0038 | + | 198 | 256 | **RNA**→ rpsO (PRK05626)RpsO (COG0184)→ |
| env-130 | SRS024435_C2966745 | + | 226 | 284 | **RNA**→ rpsO (PRK05626)Ribosomal_S15 (pfam00312)→ |
| env-131 | KGI_S1_ANT01_95m_c10061386 | + | 974 | 1038 | **RNA**→ rpsO (PRK05626)Ribosomal_S15 (pfam00312)→ |
| env-132 | KGI_S1_ANT02_95m_c10009452 | - | 2499 | 2435 | **RNA**→ rpsO (PRK05626)Ribosomal_S15 (pfam00312)→ |
| env-133 | CESB01280198.1 | - | 828 | 770 | **RNA**→ rpsO (PRK05626)Ribosomal_S15 (pfam00312)→ PRK11824 (PRK11824)RNase_PH (pfam01138)→ |
| env-134 | TB_GS09_5_10008900 | - | 6175 | 6110 | **RNA**→ rpsO (PRK05626)Ribosomal_S15 (pfam00312)→ |
| env-135 | JGI20154J14316_10009605 | - | 4721 | 4654 | **RNA**→ rpsO (PRK05626)Ribosomal_S15 (pfam00312)→ |
| env-136 | JGI20153J14295_10017197 | + | 805 | 872 | **RNA**→ rpsO (PRK05626)Ribosomal_S15 (pfam00312)→ |
| env-137 | JGI20153J14318_10017197 | + | 805 | 872 | **RNA**→ rpsO (PRK05626)Ribosomal_S15 (pfam00312)→ |
| env-138 | DelMOSum2010_c10000887 | - | 4525 | 4458 | **RNA**→ rpsO (PRK05626)Ribosomal_S15 (pfam00312)→ |
| env-139 | DelMOSum2011_c10406029 | - | 342 | 275 | **RNA**→ rpsO (PRK05626)Ribosomal_S15 (pfam00312)→ |
| env-140 | SI36aug09_100m_1134333 | - | 170 | 103 | **RNA**→ rpsO (PRK05626)RpsO (COG0184)→ |
| env-141 | JGI20154J14316_10002772 | + | 14364 | 14431 | **RNA**→ rpsO (PRK05626)→ |
| env-142 | JGI20160J14292_10003420 | + | 11666 | 11733 | **RNA**→ rpsO (PRK05626)Ribosomal_S15 (pfam00312)→ |
| env-143 | JGI20153J14295_10019149 | - | 130 | 63 | **RNA**→ rpsO (PRK05626)→ |
| env-144 | JGI20153J14318_10019149 | - | 130 | 63 | **RNA**→ rpsO (PRK05626)→ |
| env-145 | JGI20157J14317_10004310 | - | 467 | 400 | **RNA**→ rpsO (PRK05626)Ribosomal_S15 (pfam00312)→ |
| env-146 | JGI20160J14292_10025459 | - | 466 | 400 | **RNA**→ rpsO (PRK05626)Ribosomal_S15 (pfam00312)→ |
| env-147 | DelMOSum2011_c10000072 | + | 60905 | 60972 | **RNA**→ rpsO (PRK05626)Ribosomal_S15 (pfam00312)→ |
| env-148 | JGI20160J14292_10243857 | + | 387 | 454 | **RNA**→ hypo→ |
| env-149 | JGI20153J14295_10195907 | - | 246 | 179 | **RNA**→ rpsO (PRK05626)Ribosomal_S15 (pfam00312)→ |
| env-150 | JGI20153J14318_10195907 | - | 246 | 179 | **RNA**→ rpsO (PRK05626)Ribosomal_S15 (pfam00312)→ |
| env-151 | 2236876010_0615026 | - | 87 | 20 | **RNA**→ |
| env-152 | JGI20154J14316_10005663 | + | 965 | 1032 | **RNA**→ rpsO (PRK05626)Ribosomal_S15 (pfam00312)→ |
| env-153 | DelMOSum2010_c10007098 | - | 7581 | 7514 | **RNA**→ rpsO (PRK05626)Ribosomal_S15 (pfam00312)→ |
| env-154 | SI34jun09_10m_1109894 | + | 227 | 294 | **RNA**→ hypo→ |
| env-155 | DelMOSum2010_c10176236 | - | 322 | 255 | **RNA**→ rpsO (PRK05626)Ribosomal_S15 (pfam00312)→ |
| env-156 | DelMOSum2010_c10534266 | + | 94 | 161 | **RNA**→ rpsO (PRK05626)Ribosomal_S15 (pfam00312)→ |
| env-157 | DelMOSum2010_c10110255 | - | 223 | 156 | **RNA**→ rpsO (PRK05626)Ribosomal_S15 (pfam00312)→ |
| env-158 | LPaug08P2610m_c1011541 | + | 445 | 512 | **RNA**→ rpsO (PRK05626)Ribosomal_S15 (pfam00312)→ |
| env-159 | JGI20153J14295_10000898 | + | 23567 | 23634 | **RNA**→ rpsO (PRK05626)Ribosomal_S15 (pfam00312)→ |

| | | | | | |
|---|---|---|---|---|---|
| env-160 | JGI20153J14318_10000898 | + | 23567 | 23634 | **RNA**→ rpsO (PRK05626)Ribosomal_S15 (pfam00312)→ |
| env-161 | DelMOSum2011_c10006148 | + | 4842 | 4908 | **RNA**→ rpsO (PRK05626)Ribosomal_S15 (pfam00312)→ |
| env-162 | JGI20154J14316_10039500 | + | 1690 | 1757 | **RNA**→ rpsO (PRK05626)Ribosomal_S15 (pfam00312)→ |
| env-163 | JGI20157J14317_10001809 | + | 18612 | 18679 | **RNA**→ rpsO (PRK05626)Ribosomal_S15 (pfam00312)→ |
| env-164 | JGI20157J14317_10230393 | - | 112 | 45 | **RNA**→ |
| env-165 | CESU01027531.1 | - | 2628 | 2561 | **RNA**→ rpsO (PRK05626)Ribosomal_S15 (pfam00312)→ |
| env-166 | Antarctic1_1233742 | + | 29 | 96 | **RNA**→ rpsO (PRK05626)Ribosomal_S15 (pfam00312)→ |
| env-167 | JGI20154J14316_10002676 | - | 481 | 414 | **RNA**→ rpsO (PRK05626)Ribosomal_S15 (pfam00312)→ |
| env-168 | JGI20153J14295_10002037 | - | 495 | 428 | **RNA**→ rpsO (PRK05626)Ribosomal_S15 (pfam00312)→ |
| env-169 | JGI20153J14318_10002037 | - | 495 | 428 | **RNA**→ rpsO (PRK05626)Ribosomal_S15 (pfam00312)→ |
| env-170 | JGI20157J14317_10482150 | - | 149 | 82 | **RNA**→ rpsO (PRK05626)RpsO (COG0184)→ |
| env-171 | DelMOSum2010_c10084194 | + | 926 | 993 | **RNA**→ rpsO (PRK05626)Ribosomal_S15 (pfam00312)→ |
| env-172 | JGI20157J14317_10001466 | + | 8836 | 8903 | **RNA**→ rpsO (PRK05626)Ribosomal_S15 (pfam00312)→ |
| env-173 | DelMOSum2010_c10625437 | + | 182 | 249 | **RNA**→ |
| env-174 | CESI01018756.1 | - | 84 | 17 | **RNA**→ |
| env-175 | CESU01045154.1 | - | 572 | 505 | **RNA**→ rpsO (PRK05626)Ribosomal_S15 (pfam00312)→ |
| env-176 | JGI20154J14316_10053478 | - | 87 | 20 | **RNA**→ |
| env-177 | JGI20154J14316_10077664 | + | 1096 | 1163 | **RNA**→ |
| env-178 | JGI20157J14317_10014000 | + | 4902 | 4969 | **RNA**→ |
| env-179 | JGI20157J14317_10311427 | + | 322 | 389 | **RNA**→ |
| env-180 | DelMOSum2010_c10437430 | + | 283 | 350 | **RNA**→ ←hypo |
| env-181 | JGI20156J14371_10063188 | - | 538 | 471 | **RNA**→ rpsO (PRK05626)Ribosomal_S15 (pfam00312)→ |
| env-182 | JGI20160J14292_10007193 | + | 2729 | 2796 | **RNA**→ rpsO (PRK05626)Ribosomal_S15 (pfam00312)→ |
| env-183 | JGI20153J14295_10004189 | + | 1692 | 1759 | **RNA**→ rpsO (PRK05626)Ribosomal_S15 (pfam00312)→ |
| env-184 | JGI20153J14318_10004189 | + | 1692 | 1759 | **RNA**→ rpsO (PRK05626)Ribosomal_S15 (pfam00312)→ |
| env-185 | DelMOSum2010_c10470294 | - | 79 | 12 | **RNA**→ ←hypo |
| env-186 | DelMOSum2010_c10178987 | + | 509 | 576 | **RNA**→ rpsO (PRK05626)Ribosomal_S15 (pfam00312)→ |
| env-187 | JGI20153J14295_10013902 | + | 4069 | 4136 | **RNA**→ |
| env-188 | JGI20153J14318_10013902 | + | 4069 | 4136 | **RNA**→ |
| env-189 | JGI20153J14295_10133651 | - | 491 | 425 | **RNA**→ rpsO (PRK05626)Ribosomal_S15 (pfam00312)→ |
| env-190 | JGI20153J14318_10133651 | - | 491 | 425 | **RNA**→ rpsO (PRK05626)Ribosomal_S15 (pfam00312)→ |
| env-191 | JGI20154J14316_10101586 | - | 128 | 62 | **RNA**→ |
| env-192 | JGI20160J14292_10004452 | + | 816 | 882 | **RNA**→ rpsO (PRK05626)Ribosomal_S15 (pfam00312)→ |
| env-193 | JGI20153J14295_10014690 | + | 3555 | 3621 | **RNA**→ rpsO (PRK05626)Ribosomal_S15 (pfam00312)→ |
| env-194 | JGI20153J14318_10014690 | + | 3555 | 3621 | **RNA**→ rpsO (PRK05626)Ribosomal_S15 (pfam00312)→ |
| env-195 | JGI20154J14316_10305977 | + | 282 | 348 | **RNA**→ |
| env-196 | JGI20157J14317_10001080 | + | 26023 | 26090 | **RNA**→ rpsO (PRK05626)→ |
| env-197 | DelMOSum2010_c10351914 | - | 182 | 115 | **RNA**→ rpsO (PRK05626)Ribosomal_S15 (pfam00312)→ |
| env-198 | JGI20154J14316_10018926 | - | 1714 | 1647 | **RNA**→ rpsO (PRK05626)Ribosomal_S15 (pfam00312)→ |
| env-199 | JGI20160J14292_10001541 | - | 464 | 397 | **RNA**→ rpsO (PRK05626)Ribosomal_S15 (pfam00312)→ |
| env-200 | JGI20160J14292_10012905 | + | 4746 | 4813 | **RNA**→ rpsO (PRK05626)Ribosomal_S15 (pfam00312)→ |
| env-201 | CESI01128275.1 | - | 83 | 16 | **RNA**→ |
| env-202 | CESU01085162.1 | + | 222 | 289 | **RNA**→ rpsO (PRK05626)Ribosomal_S15 (pfam00312)→ |
| env-203 | CENJ01024989.1 | - | 40023 | 39970 | **RNA**→ rpsO (PRK05626)Ribosomal_S15 (pfam00312)→ Pnp (COG1185)PRK11824 (PRK11824)→ RpoD (COG0568)sigma70-ECF (TIGR02937)→ PRK05581 (PRK05581)RPE (cd00429)→ |
| env-204 | CEOK01177274.1 | - | 588 | 535 | **RNA**→ rpsO (PRK05626)Ribosomal_S15 (pfam00312)→ RNase_PH_PNPase_1 (cd11363)PRK11824 (PRK11824)– |
| env-205 | CEVN01281347.1 | + | 297 | 350 | **RNA**→ rpsO (PRK05626)RpsO (COG0184)→ |
| env-206 | CEVZ01405612.1 | - | 177 | 124 | **RNA**→ rpsO (PRK05626)Ribosomal_S15 (pfam00312)→ |
| env-207 | CETH01159915.1 | - | 862 | 809 | **RNA**→ rpsO (PRK05626)RpsO (COG0184)→ PRK11824 (PRK11824)RNase_PH (pfam01138)→ |
| env-208 | CETM01029131.1 | + | 33254 | 33307 | **RNA**→ rpsO (PRK05626)RpsO (COG0184)→ RNase_PH_PNPase_1 (cd11363)PRK11824 (PRK11824)→ |
| env-209 | CEUL01222522.1 | + | 158 | 211 | **RNA**→ rpsO (PRK05626)RpsO (COG0184)→ |
| env-210 | CEVL01135957.1 | + | 141 | 194 | **RNA**→ rpsO (PRK05626)RpsO (COG0184)→ |

| | | | | | |
|---|---|---|---|---|---|
| env-211 | CETL01011955.1 | - | 471 | 418 | **RNA**→ rpsO (PRK05626)Ribosomal_S15 (pfam00312)→ hypo→ |
| env-212 | CESZ01084417.1 | - | 85 | 17 | **RNA**→ |
| env-213 | LGVF01484185.1 | + | 293 | 357 | **RNA**→ rpsO (PRK05626)RpsO (COG0184)→ |
| env-214 | LGOV01011148.1 | + | 10676 | 10740 | **RNA**→ rpsO (PRK05626)RpsO (COG0184)→ hypo→ |
| env-215 | LGVD01117462.1 | + | 298 | 362 | **RNA**→ rpsO (PRK05626)RpsO (COG0184)→ |
| env-216 | 3300001239_10037134 | + | 37 | 106 | **RNA**→ rpsO (PRK05626)RpsO (COG0184)→ |
| env-217 | 3300001605_10046264 | - | 862 | 793 | **RNA**→ rpsO (PRK05626)RpsO (COG0184)→ |
| env-218 | CESI01059016.1 | + | 1093 | 1152 | **RNA**→ |
| env-219 | CESU01035381.1 | + | 515 | 574 | **RNA**→ |
| env-220 | LWAnNN_GHFF8UE02IP5ZP | - | 108 | 47 | **RNA**→ hypo→ |
| env-221 | JGI24768J34885_10063673 | - | 1068 | 1001 | **RNA**→ rpsO (PRK05626)Ribosomal_S15 (pfam00312)→ |
| env-222 | JGI24768J34885_10402764 | - | 304 | 237 | **RNA**→ rpsO (PRK05626)Ribosomal_S15p_S13e (cd00353)→ |
| Fli-1-1 | NZ_AVGG01000002.1 | + | 225714 | 225782 | **RNA**→ rpsO (PRK05626)Ribosomal_S15 (pfam00312)→ |
| Gme-1-1 | NZ_AUHD01000001.1 | - | 60972 | 60906 | **RNA**→ rpsO (PRK05626)Ribosomal_S15 (pfam00312)→ hypo→ hypo→ |
| env-223 | topACOD_F95O9CU01BVTXN | - | 187 | 122 | **RNA**→ rpsO (PRK05626)Ribosomal_S15 (pfam00312)→ |
| env-224 | JGI25914J50564_10243922 | + | 296 | 364 | **RNA**→ |
| env-225 | JGI25930J51415_1071170 | + | 134 | 202 | **RNA**→ rpsO (PRK05626)RpsO (COG0184)→ |
| env-226 | JGI12421J11937_10355000 | + | 222 | 290 | **RNA**→ |
| env-227 | JGI12421J11937_10090896 | + | 762 | 830 | **RNA**→ |
| env-228 | JGI12421J11937_10098907 | + | 706 | 774 | **RNA**→ |
| env-229 | JGI24768J34885_10985290 | + | 138 | 206 | **RNA**→ |
| env-230 | JGI25910J50241_10014301 | - | 1797 | 1729 | **RNA**→ rpsO (PRK05626)Ribosomal_S15 (pfam00312)→ |
| env-231 | JGI25920J50251_10021433 | + | 1008 | 1076 | **RNA**→ rpsO (PRK05626)Ribosomal_S15 (pfam00312)→ |
| env-232 | JGI25911J50253_10009747 | + | 2185 | 2253 | **RNA**→ rpsO (PRK05626)Ribosomal_S15 (pfam00312)→ |
| env-233 | JGI25912J50252_10017251 | + | 1608 | 1676 | **RNA**→ rpsO (PRK05626)Ribosomal_S15 (pfam00312)→ |
| env-234 | JGI25919J51413_1008186 | + | 196 | 264 | **RNA**→ rpsO (PRK05626)Ribosomal_S15 (pfam00312)→ |
| env-235 | JGI12421J11937_10068959 | - | 121 | 53 | **RNA**→ |
| env-236 | JGI25914J50564_10005894 | + | 1142 | 1210 | **RNA**→ rpsO (PRK05626)Ribosomal_S15 (pfam00312)→ |
| env-237 | JGI25925J51416_10005604 | + | 1735 | 1803 | **RNA**→ rpsO (PRK05626)Ribosomal_S15 (pfam00312)→ |
| env-238 | JGI25910J50241_10558763 | - | 157 | 89 | **RNA**→ hypo→ |
| env-239 | CEVJ01089541.1 | + | 746 | 812 | **RNA**→ rpsO (PRK05626)Ribosomal_S15 (pfam00312)→ |
| env-240 | CEVK01107692.1 | + | 697 | 763 | **RNA**→ rpsO (PRK05626)Ribosomal_S15 (pfam00312)→ |
| env-241 | CEVM01127682.1 | - | 8640 | 8574 | **RNA**→ rpsO (PRK05626)Ribosomal_S15 (pfam00312)→ |
| env-242 | CEVO01247303.1 | - | 757 | 691 | **RNA**→ rpsO (PRK05626)Ribosomal_S15 (pfam00312)→ |
| env-243 | CEVQ01258568.1 | + | 523 | 589 | **RNA**→ rpsO (PRK05626)RpsO (COG0184)→ |
| env-244 | P_1C_Liq_3_UnCty_1003320 | - | 752 | 689 | **RNA**→ rpsO (PRK05626)RpsO (COG0184)→ |
| env-245 | P_1C_Liq_1_UnCty_1001124 | + | 88 | 151 | **RNA**→ rpsO (PRK05626)RpsO (COG0184)→ |
| env-246 | P_1C_Liq_2_UnCty_1013913 | - | 483 | 420 | **RNA**→ rpsO (PRK05626)RpsO (COG0184)→ |
| env-247 | P_1C_Liq_2_UnCty_1115639 | - | 112 | 49 | **RNA**→ |
| env-248 | GSLSAS_GLMUJHB02IJCCC | - | 69 | 6 | **RNA**→ |
| env-249 | P_1C_Liq_1_UnCty_1006195 | - | 740 | 677 | **RNA**→ rpsO (PRK05626)Ribosomal_S15 (pfam00312)→ |
| env-250 | P_2C_Liq_1_UnCty_1185669 | + | 181 | 244 | **RNA**→ rpsO (PRK05626)RpsO (COG0184)→ |
| env-251 | CEQG01151576.1 | - | 14001 | 13934 | **RNA**→ rpsO (PRK05626)Ribosomal_S15 (pfam00312)→ |
| env-252 | CERQ01228901.1 | + | 560 | 627 | **RNA**→ rpsO (PRK05626)Ribosomal_S15 (pfam00312)→ |
| env-253 | CETB01276978.1 | + | 270 | 337 | **RNA**→ rpsO (PRK05626)Ribosomal_S15 (pfam00312)→ |
| env-254 | CETR01042174.1 | + | 13985 | 14052 | **RNA**→ rpsO (PRK05626)Ribosomal_S15 (pfam00312)→ |
| env-255 | CEQS01121120.1 | - | 530 | 463 | **RNA**→ rpsO (PRK05626)Ribosomal_S15 (pfam00312)→ |
| env-256 | CEVH01121698.1 | + | 336 | 403 | **RNA**→ rpsO (PRK05626)Ribosomal_S15 (pfam00312)→ |
| env-257 | JCVI_SCAF_1096627482312 | + | 478 | 545 | **RNA**→ rpsO (PRK05626)Ribosomal_S15 (pfam00312)→ |
| env-258 | CEQG01262115.1 | + | 449 | 514 | **RNA**→ rpsO (PRK05626)Ribosomal_S15 (pfam00312)→ |
| env-259 | CEVM01147883.1 | + | 152 | 217 | **RNA**→ rpsO (PRK05626)Ribosomal_S15 (pfam00312)→ |
| env-260 | CEVO01240128.1 | - | 371 | 306 | **RNA**→ rpsO (PRK05626)Ribosomal_S15 (pfam00312)→ |

| | | | | | |
|---|---|---|---|---|---|
| env-261 | CEVJ01132551.1 | + | 371 | 436 | **RNA**→ rpsO (PRK05626)Ribosomal_S15 (pfam00312)→ |
| Nma-1-1 | NZ_AP014548.1 | - | 1343589 | 1343499 | **RNA**→ rpsO (PRK05626)Ribosomal_S15 (pfam00312)→ |
| env-262 | topACOD_F95O9CU02GIL5N | - | 178 | 115 | **RNA**→ rpsO (PRK05626)Ribosomal_S15 (pfam00312)→ |
| env-263 | CEVX01088623.1 | - | 401 | 336 | **RNA**→ rpsO (PRK05626)Ribosomal_S15 (pfam00312)→ |
| env-264 | CENF01080549.1 | - | 982 | 917 | **RNA**→ rpsO (PRK05626)Ribosomal_S15 (pfam00312)→ |
| | | | | | PRK11824 (PRK11824)RNase_PH (pfam01138)RNase_PH_C (pfam03725)→ |
| env-265 | CEVP01232759.1 | + | 45 | 110 | **RNA**→ rpsO (PRK05626)Ribosomal_S15 (pfam00312)→ RNase_PH_PNPase_1 (cd11363)PRK11824 (PRK11824)– |
| env-266 | CEQE01251909.1 | + | 229 | 292 | **RNA**→ rpsO (PRK05626)Ribosomal_S15 (pfam00312)→ |
| env-267 | CERQ01096008.1 | - | 324 | 261 | **RNA**→ rpsO (PRK05626)Ribosomal_S15 (pfam00312)→ |
| env-268 | CEUY01328535.1 | + | 133 | 196 | **RNA**→ rpsO (PRK05626)Ribosomal_S15 (pfam00312)→ |
| env-269 | JGI11881J13070_1009257 | - | 658 | 595 | **RNA**→ rpsO (PRK05626)Ribosomal_S15 (pfam00312)→ |
| env-270 | JGI12148J13107_117495 | + | 102 | 165 | **RNA**→ rpsO (PRK05626)Ribosomal_S15 (pfam00312)→ |
| env-271 | CETU01138396.1 | + | 53 | 116 | **RNA**→ rpsO (PRK05626)Ribosomal_S15 (pfam00312)→ |
| env-272 | CEVI01059114.1 | - | 539 | 476 | **RNA**→ rpsO (PRK05626)Ribosomal_S15 (pfam00312)→ |
| Mmo-1-1 | NZ_KE384013.1 | - | 343223 | 343160 | **RNA**→ rpsO (PRK05626)Ribosomal_S15 (pfam00312)→ |
| env-273 | SRS016095_C2559685 | + | 1214 | 1274 | **RNA**→ rpsO (PRK05626)Ribosomal_S15 (pfam00312)→ |
| env-274 | SRS019968_C2017240 | - | 402 | 342 | **RNA**→ rpsO (PRK05626)Ribosomal_S15 (pfam00312)→ |
| env-275 | SRS064557_C1810798 | - | 387 | 327 | **RNA**→ rpsO (PRK05626)Ribosomal_S15 (pfam00312)→ |
| env-276 | CENF01212627.1 | + | 1179 | 1247 | **RNA**→ rpsO (PRK05626)RpsO (COG0184)→ hypo→ |
| env-277 | CERG01152242.1 | + | 414 | 482 | **RNA**→ rpsO (PRK05626)RpsO (COG0184)→ |
| env-278 | CERR01009272.1 | - | 2063 | 1995 | **RNA**→ rpsO (PRK05626)RpsO (COG0184)→ Pnp (COG1185)PRK11824 (PRK11824)→ |
| env-279 | CERV01011610.1 | + | 2857 | 2925 | **RNA**→ rpsO (PRK05626)RpsO (COG0184)→ PRK11824 (PRK11824)RNase_PH (pfam01138)→ |
| env-280 | CETB01333319.1 | - | 576 | 508 | **RNA**→ rpsO (PRK05626)RpsO (COG0184)→ RNase_PH_PNPase_1 (cd11363)PRK11824 (PRK11824)→ |
| env-281 | CERB01165146.1 | - | 781 | 714 | **RNA**→ rpsO (PRK05626)RpsO (COG0184)→ PRK11824 (PRK11824)RNase_PH (pfam01138)→ |
| env-282 | CEUN01041406.1 | + | 479 | 546 | **RNA**→ rpsO (PRK05626)Ribosomal_S15 (pfam00312)→ |
| env-283 | CEUQ01052428.1 | + | 476 | 543 | **RNA**→ rpsO (PRK05626)RpsO (COG0184)→ PRK11824 (PRK11824)RNase_PH (pfam01138)→ |
| env-284 | CEVZ01245147.1 | + | 1848 | 1915 | **RNA**→ rpsO (PRK05626)RpsO (COG0184)→ |
| env-285 | CEVA01049874.1 | - | 671 | 604 | **RNA**→ rpsO (PRK05626)RpsO (COG0184)→ PRK11824 (PRK11824)RNase_PH (pfam01138)→ |
| env-286 | CENY01189203.1 | + | 413 | 480 | **RNA**→ |
| env-287 | CEOQ01377130.1 | - | 582 | 515 | **RNA**→ rpsO (PRK05626)RpsO (COG0184)→ PRK11824 (PRK11824)RNase_PH (pfam01138)→ |
| env-288 | CETG01040803.1 | + | 479 | 546 | **RNA**→ rpsO (PRK05626)Ribosomal_S15 (pfam00312)→ |
| env-289 | CEPK01084254.1 | - | 717 | 650 | **RNA**→ rpsO (PRK05626)RpsO (COG0184)→ PRK11824 (PRK11824)RNase_PH (pfam01138)→ |
| env-290 | CETB01107343.1 | - | 567 | 500 | **RNA**→ rpsO (PRK05626)RpsO (COG0184)→ PRK11824 (PRK11824)RNase_PH (pfam01138)→ |
| env-291 | CETM01027306.1 | - | 537 | 470 | **RNA**→ rpsO (PRK05626)RpsO (COG0184)→ RNase_PH_PNPase_1 (cd11363)PRK11824 (PRK11824)→ |
| env-292 | CEUS01161241.1 | + | 657 | 724 | **RNA**→ rpsO (PRK05626)RpsO (COG0184)→ |
| env-293 | CEUZ01497389.1 | - | 394 | 327 | **RNA**→ rpsO (PRK05626)RpsO (COG0184)→ |
| env-294 | CETW01247773.1 | - | 486 | 419 | **RNA**→ rpsO (PRK05626)RpsO (COG0184)→ hypo→ |
| env-295 | CEUY01024920.1 | + | 693 | 760 | **RNA**→ rpsO (PRK05626)RpsO (COG0184)→ PRK11824 (PRK11824)RNase_PH (pfam01138)→ |
| env-296 | CEVN01169446.1 | - | 230 | 163 | **RNA**→ rpsO (PRK05626)Ribosomal_S15 (pfam00312)→ |
| env-297 | CERB01253177.1 | - | 256 | 189 | **RNA**→ rpsO (PRK05626)Ribosomal_S15 (pfam00312)→ |
| env-298 | CETP01182766.1 | + | 1115 | 1170 | **RNA**→ |
| env-299 | CEUW01218718.1 | - | 377 | 322 | **RNA**→ rpsO (PRK05626)Ribosomal_S15 (pfam00312)→ |
| env-300 | CEUF01031262.1 | - | 3989 | 3934 | **RNA**→ rpsO (PRK05626)Ribosomal_S15 (pfam00312)→ Pnp (COG1185)PRK11824 (PRK11824)→ |
| | | | | | RpoD (COG0568)sigma70-ECF (TIGR02937)→ PRK05581 (PRK05581)RPE (cd00429)→ |
| env-301 | CEUG01214040.1 | - | 119 | 64 | **RNA**→ hypo→ |
| env-302 | CETK01110687.1 | - | 61390 | 61335 | **RNA**→ rpsO (PRK05626)Ribosomal_S15 (pfam00312)→ Pnp (COG1185)PRK11824 (PRK11824)→ |
| | | | | | RpoD (COG0568)sigma70-ECF (TIGR02937)→ PRK05581 (PRK05581)RPE (cd00429)→ |
| env-303 | CETL01064018.1 | - | 538 | 483 | **RNA**→ rpsO (PRK05626)Ribosomal_S15 (pfam00312)→ RNase_PH_PNPase_1 (cd11363)PRK11824 (PRK11824)– |
| env-304 | CESW01089027.1 | + | 96 | 152 | **RNA**→ rpsO (PRK05626)Ribosomal_S15 (pfam00312)→ PRK11824 (PRK11824)RNase_PH (pfam01138)→ |
| env-305 | CAM_READ_0234091691 | - | 79 | 24 | **RNA**→ |
| env-306 | CESZ01040152.1 | + | 353 | 408 | **RNA**→ rpsO (PRK05626)RpsO (COG0184)→ |
| env-307 | CESA01198900.1 | + | 376 | 431 | **RNA**→ rpsO (PRK05626)RpsO (COG0184)→ |
| env-308 | CEQS01074332.1 | - | 162559 | 162500 | **RNA**→ rpsO (PRK05626)Ribosomal_S15 (pfam00312)→ Pnp (COG1185)PRK11824 (PRK11824)→ |

| | | | | | |
|---|---|---|---|---|---|
| Lsp-3-1 | NZ_JHZY01000002.1 | - | 172107 | 172041 | RNA→ rpsO (PRK05626)Ribosomal_S15 (pfam00312)→ |
| env-309 | CEVM01090294.1 | - | 285 | 219 | RNA→ rpsO (PRK05626)RpsO (COG0184)→ |
| env-310 | CETA01052158.1 | + | 5065 | 5131 | RNA→ rpsO (PRK05626)Ribosomal_S15 (pfam00312)→ |
| env-311 | CETB01058157.1 | + | 1085 | 1151 | RNA→ rpsO (PRK05626)Ribosomal_S15 (pfam00312)→ |
| env-312 | CEVJ01090841.1 | - | 927 | 861 | RNA→ rpsO (PRK05626)Ribosomal_S15 (pfam00312)→ |
| env-313 | CETU01159611.1 | - | 800 | 734 | RNA→ rpsO (PRK05626)Ribosomal_S15 (pfam00312)→ |
| env-314 | CETT01019050.1 | + | 500 | 566 | RNA→ rpsO (PRK05626)Ribosomal_S15 (pfam00312)→ |
| env-315 | CEVK01011741.1 | - | 496 | 430 | RNA→ rpsO (PRK05626)Ribosomal_S15 (pfam00312)→ |
| env-316 | CERQ01230915.1 | - | 1011 | 945 | RNA→ rpsO (PRK05626)Ribosomal_S15 (pfam00312)→ |
| env-317 | CEUY01464639.1 | - | 77 | 11 | RNA→ |
| env-318 | CEPD01093744.1 | - | 461 | 395 | RNA→ rpsO (PRK05626)Ribosomal_S15 (pfam00312)→ |
| env-319 | CERY01058984.1 | - | 449 | 383 | RNA→ rpsO (PRK05626)Ribosomal_S15 (pfam00312)→ |
| env-320 | CEPZ01179059.1 | + | 369 | 435 | RNA→ rpsO (PRK05626)Ribosomal_S15 (pfam00312)→ |
| env-321 | CERE01075129.1 | + | 1132 | 1198 | RNA→ rpsO (PRK05626)Ribosomal_S15 (pfam00312)→ |
| env-322 | CESC01013190.1 | + | 80317 | 80383 | RNA→ rpsO (PRK05626)Ribosomal_S15 (pfam00312)→ |
| env-323 | CEUN01016453.1 | + | 2805 | 2871 | RNA→ rpsO (PRK05626)Ribosomal_S15 (pfam00312)→ |
| Lbl-1-1 | NZ_CH672395.1 | - | 1099698 | 1099632 | RNA→ rpsO (PRK05626)Ribosomal_S15 (pfam00312)→ |
| env-324 | CETS01234391.1 | + | 379 | 445 | RNA→ rpsO (PRK05626)Ribosomal_S15 (pfam00312)→ |
| env-325 | CEWI01015492.1 | - | 496 | 430 | RNA→ rpsO (PRK05626)Ribosomal_S15 (pfam00312)→ |
| env-326 | CEOS01054411.1 | - | 238 | 172 | RNA→ rpsO (PRK05626)Ribosomal_S15 (pfam00312)→ |
| Fri-1-1 | NZ_KB899968.1 | - | 97102 | 97037 | RNA→ rpsO (PRK05626)Ribosomal_S15 (pfam00312)→ |
| Fri-2-1 | NZ_JRLX01000004.1 | + | 133005 | 133070 | RNA→ rpsO (PRK05626)Ribosomal_S15 (pfam00312)→ |
| env-327 | JCVI_SCAF_1096626847824 | + | 1625 | 1679 | RNA→ rpsO (PRK05626)Ribosomal_S15 (pfam00312)→ PRK11824 (PRK11824)RNase_PH (pfam01138)RNase_PH_C (pfam03725)→ |
| env-328 | CETY01060541.1 | - | 800 | 747 | RNA→ rpsO (PRK05626)Ribosomal_S15 (pfam00312)→ PRK11824 (PRK11824)RNase_PH (pfam01138)→ |
| env-329 | CESE01249388.1 | + | 30 | 85 | RNA→ rpsO (PRK05626)RpsO (COG0184)→ PRK11824 (PRK11824)RNase_PH (pfam01138)→ |
| env-330 | CESJ01046981.1 | + | 6198 | 6253 | RNA→ rpsO (PRK05626)RpsO (COG0184)→ PRK11824 (PRK11824)RNase_PH (pfam01138)→ |
| env-331 | CEVF01245326.1 | + | 262 | 318 | RNA→ rpsO (PRK05626)RpsO (COG0184)→ |
| env-332 | JCVI_SCAF_1096628209080 | - | 1575 | 1519 | RNA→ rpsO (PRK05626)RpsO (COG0184)→ PRK11824 (PRK11824)RNase_PH (pfam01138)→ |
| env-333 | CEUA01276370.1 | - | 476 | 420 | RNA→ rpsO (PRK05626)RpsO (COG0184)→ hypo→ |
| env-334 | CESJ01065949.1 | - | 404 | 345 | RNA→ rpsO (PRK05626)Ribosomal_S15 (pfam00312)→ |
| env-335 | GSLSAS_GBFV41Z02H1HZP | + | 224 | 282 | RNA→ |
| env-336 | GSLSAAI_GFP0ZZ301DBBHK | - | 358 | 298 | RNA→ rpsO (PRK05626)Ribosomal_S15 (pfam00312)→ |
| env-337 | JGI11876J14442_10048055 | + | 1753 | 1811 | RNA→ rpsO (PRK05626)Ribosomal_S15 (pfam00312)→ |
| env-338 | JCVI_READ_1102140320562 | + | 586 | 640 | RNA→ rpsO (PRK05626)Ribosomal_S15 (pfam00312)→ |
| env-339 | JCVI_READ_1108829504037 | + | 48 | 102 | RNA→ rpsO (PRK05626)Ribosomal_S15 (pfam00312)→ PRK11824 (PRK11824)RNase_PH (pfam01138)→ |
| env-340 | JCVI_READ_1101733741174 | - | 276 | 222 | RNA→ |
| env-341 | CEOO01040203.1 | + | 631 | 685 | RNA→ rpsO (PRK05626)→ |
| env-342 | CEWA01031860.1 | + | 385 | 439 | RNA→ rpsO (PRK05626)Ribosomal_S15 (pfam00312)→ Pnp (COG1185)PRK11824 (PRK11824)→ |
| env-343 | CEOV01172242.1 | + | 485 | 539 | RNA→ |
| env-344 | CEUR01130058.1 | - | 88 | 34 | RNA→ |
| env-345 | CETK01152287.1 | - | 721 | 667 | RNA→ rpsO (PRK05626)Ribosomal_S15 (pfam00312)→ PRK11824 (PRK11824)RNase_PH (pfam01138)→ |
| env-346 | CEUG01240482.1 | - | 938 | 884 | RNA→ rpsO (PRK05626)Ribosomal_S15 (pfam00312)→ PRK11824 (PRK11824)RNase_PH (pfam01138)→ |
| env-347 | CETE01019518.1 | - | 64087 | 64033 | RNA→ rpsO (PRK05626)Ribosomal_S15 (pfam00312)→ Pnp (COG1185)PRK11824 (PRK11824)→ RpoD (COG0568)sigma70-ECF (TIGR02937)→ PRK05581 (PRK05581)RPE (cd00429)→ |
| env-348 | CETQ01056045.1 | - | 3058 | 3004 | RNA→ rpsO (PRK05626)Ribosomal_S15 (pfam00312)→ Pnp (COG1185)PRK11824 (PRK11824)→ sigma70-ECF (TIGR02937)Sigma70_r1_2 (pfam00140)Sigma70_r2 (pfam04542)→ |
| env-349 | CEUH01289173.1 | - | 939 | 885 | RNA→ rpsO (PRK05626)Ribosomal_S15 (pfam00312)→ PRK11824 (PRK11824)RNase_PH (pfam01138)→ |
| env-350 | CEWG01025700.1 | + | 54379 | 54433 | RNA→ rpsO (PRK05626)Ribosomal_S15 (pfam00312)→ Pnp (COG1185)PRK11824 (PRK11824)→ RpoD (COG0568)sigma70-ECF (TIGR02937)→ PRK05581 (PRK05581)RPE (cd00429)→ |
| env-351 | CEWH01036275.1 | - | 2198 | 2144 | RNA→ rpsO (PRK05626)Ribosomal_S15 (pfam00312)→ Pnp (COG1185)PRK11824 (PRK11824)→ |
| env-352 | CEVG01088667.1 | + | 633 | 687 | RNA→ rpsO (PRK05626)→ |
| env-353 | CEWJ01239071.1 | - | 240 | 186 | RNA→ rpsO (PRK05626)Ribosomal_S15 (pfam00312)→ |

| | | | | | |
|---|---|---|---|---|---|
| env-354 | CEVH01159213.1 | + | 181 | 235 | **RNA**→ rpsO (PRK05626)Ribosomal_S15 (pfam00312)→ PRK11824 (PRK11824)RNase_PH (pfam01138)→ |
| env-355 | CEPT01275949.1 | - | 108 | 54 | **RNA**→ |
| env-356 | CEUM01290778.1 | + | 535 | 589 | **RNA**→ rpsO (PRK05626)Ribosomal_S15 (pfam00312)→ hypo→ |
| env-357 | PBR_1004362 | - | 413 | 348 | **RNA**→ rpsO (PRK05626)Ribosomal_S15 (pfam00312)→ |
| Fta-1-1 | NC_015321.1 | + | 2727109 | 2727174 | **RNA**→ rpsO (PRK05626)Ribosomal_S15 (pfam00312)→ |
| env-358 | KGI_S1_ANT02_95m_c10279369 | - | 384 | 320 | **RNA**→ rpsO (PRK05626)Ribosomal_S15 (pfam00312)→ |
| env-359 | P_1C_Liq_1_UnCty_1146961 | + | 178 | 240 | **RNA**→ |
| env-360 | P_2C_Liq_1_UnCty_1109790 | - | 320 | 258 | **RNA**→ rpsO (PRK05626)Ribosomal_S15 (pfam00312)→ |
| env-361 | SL_7KL_010_BRINE_10000173 | - | 3812 | 3748 | **RNA**→ rpsO (PRK05626)Ribosomal_S15 (pfam00312)→ |
| env-362 | SL_9KL_010_SED_10000238 | + | 12358 | 12422 | **RNA**→ rpsO (PRK05626)Ribosomal_S15 (pfam00312)→ |
| Mod-7-1 | NZ_AYMS01000053.1 | - | 7079 | 7009 | **RNA**→ rpsO (PRK05626)Ribosomal_S15 (pfam00312)→ |
| Mod-6-1 | NZ_JH590839.1 | + | 306818 | 306888 | **RNA**→ rpsO (PRK05626)Ribosomal_S15 (pfam00312)→ |
| Mod-5-1 | NZ_JH815537.1 | + | 340698 | 340768 | **RNA**→ rpsO (PRK05626)Ribosomal_S15 (pfam00312)→ |
| Mod-2-1 | NZ_KE161017.1 | + | 300297 | 300367 | **RNA**→ rpsO (PRK05626)Ribosomal_S15 (pfam00312)→ |
| Msp-3-1 | NZ_CP010327.1 | - | 814045 | 813975 | **RNA**→ rpsO (PRK05626)Ribosomal_S15 (pfam00312)→ |
| Mod-3-1 | NZ_KE340321.1 | + | 302906 | 302976 | **RNA**→ rpsO (PRK05626)Ribosomal_S15 (pfam00312)→ |
| Mpr-1-1 | NZ_CP010817.1 | - | 3674296 | 3674226 | **RNA**→ rpsO (PRK05626)Ribosomal_S15 (pfam00312)→ |
| Mod-4-1 | NZ_JH590835.1 | - | 245456 | 245386 | **RNA**→ rpsO (PRK05626)Ribosomal_S15 (pfam00312)→ |
| Mod-1-1 | NZ_JWJO01000006.1 | + | 753 | 821 | **RNA**→ rpsO (PRK05626)Ribosomal_S15 (pfam00312)→ |
| Fin-1-1 | NC_017025.1 | + | 2426652 | 2426716 | **RNA**→ rpsO (PRK05626)Ribosomal_S15 (pfam00312)→ |
| env-363 | JGI25910J50241_10022696 | + | 1815 | 1881 | **RNA**→ rpsO (PRK05626)Ribosomal_S15 (pfam00312)→ |
| env-364 | JGI25910J50241_10032142 | - | 1405 | 1339 | **RNA**→ rpsO (PRK05626)Ribosomal_S15 (pfam00312)→ |
| env-365 | JGI25911J50253_10054721 | + | 193 | 259 | **RNA**→ rpsO (PRK05626)Ribosomal_S15 (pfam00312)→ |
| env-366 | JGI25908J49247_10520374 | + | 33 | 99 | **RNA**→ rpsO (PRK05626)Ribosomal_S15 (pfam00312)→ |
| env-367 | JGI25920J50251_10045732 | + | 42 | 108 | **RNA**→ rpsO (PRK05626)Ribosomal_S15 (pfam00312)→ |
| env-368 | JGI25910J50241_10027611 | - | 1889 | 1823 | **RNA**→ rpsO (PRK05626)Ribosomal_S15 (pfam00312)→ |
| env-369 | P_1C_Liq_1_UnCty_1013506 | + | 698 | 774 | **RNA**→ rpsO (PRK05626)RpsO (COG0184)→ |
| env-370 | P_2C_Liq_1_UnCty_1023963 | + | 685 | 761 | **RNA**→ rpsO (PRK05626)RpsO (COG0184)→ PRK11824 (PRK11824)RNase_PH (pfam01138)→ |
| env-371 | P_2C_Liq_2_UnCty_1019436 | + | 188 | 264 | **RNA**→ rpsO (PRK05626)RpsO (COG0184)→ PRK11824 (PRK11824)RNase_PH (pfam01138)RNase_PH_C (pfam03725)→ |
| env-372 | P_1C_Liq_2_UnCty_1015786 | - | 536 | 460 | **RNA**→ rpsO (PRK05626)RpsO (COG0184)→ RNase_PH_PNPase_1 (cd11363)PRK11824 (PRK11824)→ |
| env-373 | CENP01234479.1 | + | 54048 | 54107 | **RNA**→ rpsO (PRK05626)Ribosomal_S15 (pfam00312)→ Pnp (COG1185)PRK11824 (PRK11824)→ RpoD (COG0568)sigma70-ECF (TIGR02937)→ PRK05581 (PRK05581)RPE (cd00429)→ |
| env-374 | CEOK01051128.1 | + | 11806 | 11865 | **RNA**→ rpsO (PRK05626)Ribosomal_S15 (pfam00312)→ Pnp (COG1185)PRK11824 (PRK11824)→ RpoD (COG0568)sigma70-ECF (TIGR02937)→ PRK05581 (PRK05581)RPE (cd00429)→ |
| env-375 | CEOF01428303.1 | + | 1460 | 1519 | **RNA**→ |
| env-376 | CERM01056288.1 | - | 643 | 576 | **RNA**→ rpsO (PRK05626)RpsO (COG0184)→ |
| env-377 | CERM01008643.1 | + | 3284 | 3351 | **RNA**→ rpsO (PRK05626)RpsO (COG0184)→ |
| env-378 | CEVX01053585.1 | + | 2992 | 3059 | **RNA**→ rpsO (PRK05626)RpsO (COG0184)→ |
| env-379 | CESF01002239.1 | - | 6290 | 6222 | **RNA**→ rpsO (PRK05626)Ribosomal_S15 (pfam00312)→ |
| env-380 | CESO01109506.1 | - | 5554 | 5486 | **RNA**→ rpsO (PRK05626)Ribosomal_S15 (pfam00312)→ |
| env-381 | CETY01366948.1 | + | 307 | 375 | **RNA**→ rpsO (PRK05626)Ribosomal_S15 (pfam00312)→ |
| env-382 | CESM01460641.1 | + | 62 | 130 | **RNA**→ rpsO (PRK05626)Ribosomal_S15 (pfam00312)→ |
| env-383 | CEVP01299765.1 | + | 326 | 394 | **RNA**→ rpsO (PRK05626)Ribosomal_S15 (pfam00312)→ |
| env-384 | CEWG01217676.1 | + | 305 | 373 | **RNA**→ rpsO (PRK05626)Ribosomal_S15 (pfam00312)→ |
| env-385 | CERK01085407.1 | + | 265 | 333 | **RNA**→ rpsO (PRK05626)Ribosomal_S15 (pfam00312)→ |
| env-386 | CERW01016535.1 | + | 3627 | 3695 | **RNA**→ rpsO (PRK05626)Ribosomal_S15 (pfam00312)→ |
| env-387 | CESA01134118.1 | + | 1636 | 1704 | **RNA**→ rpsO (PRK05626)Ribosomal_S15 (pfam00312)→ |
| env-388 | CEVX01008274.1 | + | 4128 | 4196 | **RNA**→ rpsO (PRK05626)Ribosomal_S15 (pfam00312)→ |
| env-389 | CERM01128748.1 | + | 189 | 257 | **RNA**→ rpsO (PRK05626)Ribosomal_S15 (pfam00312)→ |
| env-390 | JCVI_SCAF_1096627802799 | + | 1198 | 1266 | **RNA**→ rpsO (PRK05626)Ribosomal_S15 (pfam00312)→ |
| env-391 | CEOM01115757.1 | - | 1876 | 1808 | **RNA**→ rpsO (PRK05626)Ribosomal_S15 (pfam00312)→ |
| env-392 | CEWK01131420.1 | + | 392 | 460 | **RNA**→ rpsO (PRK05626)Ribosomal_S15 (pfam00312)→ |

| | | | | | |
|---|---|---|---|---|---|
| env-393 | CESW01060567.1 | - | 673 | 605 | **RNA**→ rpsO (PRK05626)Ribosomal_S15 (pfam00312)→ |
| env-394 | CEOM01357113.1 | + | 580 | 648 | **RNA**→ rpsO (PRK05626)Ribosomal_S15 (pfam00312)→ |
| env-395 | CESP01222974.1 | - | 266 | 198 | **RNA**→ rpsO (PRK05626)Ribosomal_S15 (pfam00312)→ |
| env-396 | CEST01081216.1 | - | 179 | 120 | **RNA**→ rpsO (PRK05626)Ribosomal_S15 (pfam00312)→ |
| env-397 | NCBI_READ_1112532322036 | + | 335 | 394 | **RNA**→ rpsO (PRK05626)Ribosomal_S15 (pfam00312)→ |
| env-398 | NCBI_READ_1112777362018 | + | 412 | 471 | **RNA**→ rpsO (PRK05626)Ribosomal_S15 (pfam00312)→ |
| env-399 | NCBI_READ_1112533192499 | - | 730 | 671 | **RNA**→ rpsO (PRK05626)Ribosomal_S15 (pfam00312)→ PRK11824 (PRK11824)RNase_PH (pfam01138)→ |
| env-400 | CESE01040916.1 | + | 2914 | 2973 | **RNA**→ rpsO (PRK05626)Ribosomal_S15 (pfam00312)→ |
| env-401 | CESJ01014804.1 | - | 376 | 317 | **RNA**→ rpsO (PRK05626)Ribosomal_S15 (pfam00312)→ |
| env-402 | CESN01055916.1 | - | 278 | 222 | **RNA**→ rpsO (PRK05626)Ribosomal_S15 (pfam00312)→ |
| env-403 | CESP01147131.1 | - | 157 | 101 | **RNA**→ rpsO (PRK05626)RpsO (COG0184)→ |
| env-404 | CESW01247893.1 | + | 149 | 205 | **RNA**→ rpsO (PRK05626)Ribosomal_S15 (pfam00312)→ PRK11824 (PRK11824)RNase_PH (pfam01138)→ |
| env-405 | CETY01067238.1 | - | 908 | 856 | **RNA**→ rpsO (PRK05626)Ribosomal_S15 (pfam00312)→ PRK11824 (PRK11824)RNase_PH (pfam01138)→ |
| env-406 | CEOP01095216.1 | - | 241 | 189 | **RNA**→ rpsO (PRK05626)Ribosomal_S15 (pfam00312)→ |
| env-407 | CEUR01327391.1 | + | 1009 | 1065 | **RNA**→ rpsO (PRK05626)Ribosomal_S15 (pfam00312)→ |
| | | | | | PRK11824 (PRK11824)RNase_PH (pfam01138)RNase_PH_C (pfam03725)→ |
| env-408 | CEWJ01231416.1 | - | 68 | 12 | **RNA**→ |
| env-409 | CAM_READ_0231777305 | + | 246 | 302 | **RNA**→ hypo→ |
| env-410 | CAM_READ_0233120627 | + | 246 | 302 | **RNA**→ |
| env-411 | CEOK01184894.1 | - | 9274 | 9218 | **RNA**→ rpsO (PRK05626)Ribosomal_S15 (pfam00312)→ Pnp (COG1185)PRK11824 (PRK11824)→ |
| | | | | | RpoD (COG0568)sigma70-ECF (TIGR02937)→ PRK05581 (PRK05581)RPE (cd00429)→ |
| env-412 | CESP01118836.1 | + | 165 | 221 | **RNA**→ rpsO (PRK05626)Ribosomal_S15 (pfam00312)→ |
| env-413 | JCVI_SCAF_1096628396118 | + | 1238 | 1294 | **RNA**→ rpsO (PRK05626)Ribosomal_S15 (pfam00312)→ |
| env-414 | CENP01152184.1 | - | 17026 | 16970 | **RNA**→ rpsO (PRK05626)Ribosomal_S15 (pfam00312)→ Pnp (COG1185)PRK11824 (PRK11824)→ |
| | | | | | RpoD (COG0568)sigma70-ECF (TIGR02937)→ PRK05581 (PRK05581)RPE (cd00429)→ |
| env-415 | CESI01270948.1 | + | 837 | 890 | **RNA**→ |
| env-416 | CESI01305757.1 | + | 1479 | 1532 | **RNA**→ rpsO (PRK05626)Ribosomal_S15 (pfam00312)→ |
| | | | | | PRK11824 (PRK11824)RNase_PH (pfam01138)RNase_PH_C (pfam03725)→ |
| env-417 | CESU01062348.1 | + | 413 | 466 | **RNA**→ rpsO (PRK05626)Ribosomal_S15 (pfam00312)→ |
| | | | | | PRK11824 (PRK11824)RNase_PH (pfam01138)RNase_PH_C (pfam03725)→ |
| env-418 | CESJ01175686.1 | - | 363 | 309 | **RNA**→ rpsO (PRK05626)Ribosomal_S15 (pfam00312)→ |
| Fbe-1-1 | NZ_JRLV01000023.1 | - | 16228 | 16166 | **RNA**→ rpsO (PRK05626)Ribosomal_S15 (pfam00312)→ |
| env-419 | CERZ01029826.1 | - | 671 | 607 | **RNA**→ rpsO (PRK05626)Ribosomal_S15 (pfam00312)→ |
| Ksp-1-1 | NC_015496.1 | + | 583327 | 583390 | **RNA**→ rpsO (PRK05626)Ribosomal_S15 (pfam00312)→ |
| Dsp-2-1 | NZ_CM001837.1 | + | 492542 | 492605 | **RNA**→ rpsO (PRK05626)Ribosomal_S15 (pfam00312)→ |
| Dsp-1-1 | NZ_CP009301.1 | + | 1271436 | 1271503 | **RNA**→ rpsO (PRK05626)Ribosomal_S15 (pfam00312)→ |
| Ddo-1-1 | NZ_JSAQ01000001.1 | + | 1202154 | 1202221 | **RNA**→ rpsO (PRK05626)Ribosomal_S15 (pfam00312)→ |
| Iha-1-1 | NZ_AJJU01000004.1 | + | 333125 | 333187 | **RNA**→ rpsO (PRK05626)Ribosomal_S15 (pfam00312)→ rpsO (PRK05626)Ribosomal_S15 (pfam00312)→ |
| env-420 | CENJ01055151.1 | - | 1153 | 1089 | **RNA**→ rpsO (PRK05626)Ribosomal_S15 (pfam00312)→ RNase_PH_PNPase_1 (cd11363)PRK11824 (PRK11824)– |
| env-421 | LPaug09P1610m_c1073129 | - | 145 | 81 | **RNA**→ rpsO (PRK05626)RpsO (COG0184)→ |
| env-422 | A09P0410a_Sequence0000045006 | + | 360 | 424 | **RNA**→ rpsO (PRK05626)Ribosomal_S15 (pfam00312)→ RNase_PH_PNPase_1 (cd11363)PRK11824 (PRK11824)– |
| env-423 | CEOM01146924.1 | - | 65 | 1 | **RNA**→ |
| env-424 | CENN01066147.1 | + | 1036 | 1100 | **RNA**→ rpsO (PRK05626)Ribosomal_S15 (pfam00312)→ Pnp (COG1185)PRK11824 (PRK11824)→ |
| env-425 | CENU01000611.1 | - | 5201 | 5137 | **RNA**→ rpsO (PRK05626)Ribosomal_S15 (pfam00312)→ Pnp (COG1185)PRK11824 (PRK11824)→ |
| env-426 | CEOK01054851.1 | - | 1174 | 1110 | **RNA**→ rpsO (PRK05626)Ribosomal_S15 (pfam00312)→ RNase_PH_PNPase_1 (cd11363)PRK11824 (PRK11824)– |
| env-427 | CENP01019191.1 | + | 2058 | 2122 | **RNA**→ rpsO (PRK05626)Ribosomal_S15 (pfam00312)→ Pnp (COG1185)PRK11824 (PRK11824)→ |
| env-428 | CENU01173310.1 | - | 367 | 303 | **RNA**→ rpsO (PRK05626)Ribosomal_S15 (pfam00312)→ |
| env-429 | CESS01087722.1 | - | 2995 | 2931 | **RNA**→ rpsO (PRK05626)Ribosomal_S15 (pfam00312)→ Pnp (COG1185)PRK11824 (PRK11824)→ |
| env-430 | JCVI_SCAF_1096627377572 | + | 2760 | 2824 | **RNA**→ rpsO (PRK05626)Ribosomal_S15 (pfam00312)→ |
| env-431 | CEOK01021460.1 | - | 1198 | 1134 | **RNA**→ rpsO (PRK05626)Ribosomal_S15 (pfam00312)→ RNase_PH_PNPase_1 (cd11363)PRK11824 (PRK11824)– |
| env-432 | CENJ01182900.1 | + | 248 | 312 | **RNA**→ rpsO (PRK05626)Ribosomal_S15 (pfam00312)→ |
| env-433 | CENU01105541.1 | + | 758 | 822 | **RNA**→ |
| env-434 | CEOM01022458.1 | - | 1511 | 1447 | **RNA**→ rpsO (PRK05626)Ribosomal_S15 (pfam00312)→ Pnp (COG1185)PRK11824 (PRK11824)→ |

| | | | | | |
|---|---|---|---|---|---|
| env-435 | JCVI_SCAF_1096627389939 | + | 60590 | 60654 | **RNA**→ rpsO (PRK05626)Ribosomal_S15 (pfam00312)→ Pnp (COG1185)PRK11824 (PRK11824)→ |
| env-436 | CENG01086864.1 | - | 453 | 389 | **RNA**→ rpsO (PRK05626)Ribosomal_S15 (pfam00312)→ |
| env-437 | CENN01059376.1 | - | 230 | 166 | **RNA**→ rpsO (PRK05626)Ribosomal_S15 (pfam00312)→ |
| env-438 | CESW01265115.1 | + | 247 | 311 | **RNA**→ rpsO (PRK05626)Ribosomal_S15 (pfam00312)→ Pnp (COG1185)PRK11824 (PRK11824)→ |
| env-439 | CEOM01082955.1 | + | 4838 | 4902 | **RNA**→ rpsO (PRK05626)Ribosomal_S15 (pfam00312)→ PRK11824 (PRK11824)RNase_PH (pfam01138)→ |
| env-440 | CENP01061027.1 | + | 20726 | 20790 | **RNA**→ rpsO (PRK05626)Ribosomal_S15 (pfam00312)→ |
| env-441 | CENU01097470.1 | - | 401 | 337 | **RNA**→ rpsO (PRK05626)Ribosomal_S15 (pfam00312)→ |
| env-442 | CENX01202860.1 | - | 8088 | 8024 | **RNA**→ rpsO (PRK05626)Ribosomal_S15 (pfam00312)→ Pnp (COG1185)PRK11824 (PRK11824)→ |
| env-443 | CEOF01213869.1 | + | 2775 | 2839 | **RNA**→ rpsO (PRK05626)Ribosomal_S15 (pfam00312)→ Pnp (COG1185)PRK11824 (PRK11824)→ |
| env-444 | CEOK01043507.1 | + | 75 | 139 | **RNA**→ rpsO (PRK05626)Ribosomal_S15 (pfam00312)→ RNase_PH_PNPase_1 (cd11363)PRK11824 (PRK11824)– |
| Kje-1-1 | NZ_LBMG01000021.1 | - | 84468 | 84404 | **RNA**→ rpsO (PRK05626)Ribosomal_S15 (pfam00312)→ |
| Kal-1-1 | NZ_DS544873.1 | - | 4807475 | 4807411 | **RNA**→ rpsO (PRK05626)Ribosomal_S15 (pfam00312)→ |
| Kzh-1-1 | NZ_LBMH01000003.1 | + | 156456 | 156520 | **RNA**→ rpsO (PRK05626)Ribosomal_S15 (pfam00312)→ |
| env-445 | JCVI_SCAF_1096627449837 | + | 1096 | 1163 | **RNA**→ |
| env-446 | CEQG01221456.1 | - | 120 | 62 | **RNA**→ rpsO (PRK05626)→ |
| env-447 | CEQY01168872.1 | + | 258 | 316 | **RNA**→ rpsO (PRK05626)Ribosomal_S15 (pfam00312)→ PRK11824 (PRK11824)RNase_PH (pfam01138)→ |
| env-448 | CERC01134264.1 | + | 7325 | 7383 | **RNA**→ rpsO (PRK05626)Ribosomal_S15 (pfam00312)→ Pnp (COG1185)PRK11824 (PRK11824)→ |
| env-449 | P_2C_Liq_1_UnCty_1172006 | + | 238 | 295 | **RNA**→ |
| env-450 | CEOO01008234.1 | + | 2508 | 2575 | **RNA**→ rpsO (PRK05626)Ribosomal_S15 (pfam00312)→ |
| env-451 | CESA01225116.1 | + | 184 | 251 | **RNA**→ rpsO (PRK05626)Ribosomal_S15 (pfam00312)→ |
| env-452 | CESZ01040124.1 | - | 244 | 177 | **RNA**→ rpsO (PRK05626)Ribosomal_S15 (pfam00312)→ |
| env-453 | CESZ01039937.1 | - | 243 | 177 | **RNA**→ rpsO (PRK05626)Ribosomal_S15 (pfam00312)→ |
| env-454 | CEOM01051750.1 | - | 464 | 397 | **RNA**→ rpsO (PRK05626)Ribosomal_S15 (pfam00312)→ |
| env-455 | CEVU01350716.1 | - | 629 | 562 | **RNA**→ rpsO (PRK05626)Ribosomal_S15 (pfam00312)→ |
| env-456 | P_2C_Liq_1_UnCty_1017278 | - | 872 | 816 | **RNA**→ rpsO (PRK05626)Ribosomal_S15 (pfam00312)→ PRK11824 (PRK11824)RNase_PH (pfam01138)→ |
| env-457 | P_2C_Liq_2_UnCty_1110420 | - | 101 | 45 | **RNA**→ |
| env-458 | GQKUMWE01EWSJM | + | 239 | 300 | **RNA**→ ←hypo |
| env-459 | CENT01012923.1 | - | 2675 | 2616 | **RNA**→ rpsO (PRK05626)Ribosomal_S15 (pfam00312)→ Pnp (COG1185)PRK11824 (PRK11824)→ |
| env-460 | CERW01249250.1 | + | 463 | 522 | **RNA**→ rpsO (PRK05626)Ribosomal_S15 (pfam00312)→ RNase_PH_PNPase_1 (cd11363)PRK11824 (PRK11824)– |
| env-461 | CEVS01053715.1 | - | 327 | 268 | **RNA**→ rpsO (PRK05626)Ribosomal_S15 (pfam00312)→ |
| env-462 | CEOP01008691.1 | - | 1501 | 1441 | **RNA**→ rpsO (PRK05626)Ribosomal_S15 (pfam00312)→ RNase_PH_PNPase_1 (cd11363)PRK11824 (PRK11824)– |
| env-463 | CEVP01023094.1 | + | 259 | 319 | **RNA**→ rpsO (PRK05626)Ribosomal_S15 (pfam00312)→ RNase_PH_PNPase_1 (cd11363)PRK11824 (PRK11824)– |
| env-464 | JCVI_READ_1104230391553 | + | 800 | 860 | **RNA**→ rpsO (PRK05626)RpsO (COG0184)→ |
| env-465 | JCVI_READ_1104230226763 | - | 664 | 604 | **RNA**→ rpsO (PRK05626)Ribosomal_S15 (pfam00312)→ PRK11824 (PRK11824)RNase_PH (pfam01138)→ |
| env-466 | CETQ01008005.1 | - | 1003 | 943 | **RNA**→ rpsO (PRK05626)Ribosomal_S15 (pfam00312)→ |
| | | | | | PRK11824 (PRK11824)RNase_PH (pfam01138)RNase_PH_C (pfam03725)→ |
| env-467 | CESF01056087.1 | + | 108 | 168 | **RNA**→ rpsO (PRK05626)Ribosomal_S15 (pfam00312)→ PRK11824 (PRK11824)RNase_PH (pfam01138)→ |
| env-468 | CESO01047575.1 | - | 1806 | 1746 | **RNA**→ rpsO (PRK05626)Ribosomal_S15 (pfam00312)→ Pnp (COG1185)PRK11824 (PRK11824)→ |
| env-469 | CEWG01134969.1 | + | 150 | 210 | **RNA**→ rpsO (PRK05626)Ribosomal_S15 (pfam00312)→ PRK11824 (PRK11824)RNase_PH (pfam01138)→ |
| env-470 | CESM01335079.1 | - | 449 | 389 | **RNA**→ rpsO (PRK05626)Ribosomal_S15 (pfam00312)→ |
| env-471 | JGI25920J50251_10101252 | - | 633 | 565 | **RNA**→ rpsO (PRK05626)RpsO (COG0184)→ |
| env-472 | 2236876001_042780 | - | 239 | 185 | **RNA**→ rpsO (PRK05626)Ribosomal_S15 (pfam00312)→ |
| env-473 | SI34jun09_10m_1003614 | - | 2745 | 2691 | **RNA**→ rpsO (PRK05626)RpsO (COG0184)→ Pnp (COG1185)PRK11824 (PRK11824)→ |
| | | | | | PRK09210 (PRK09210)Sigma70_r1_2 (pfam00140)→ |
| env-474 | 2236876001_503295 | + | 35 | 89 | **RNA**→ |
| env-475 | LAZR01010698.1 | - | 870 | 806 | **RNA**→ rpsO (PRK05626)RpsO (COG0184)→ |
| env-476 | JGI20160J14292_10000017 | - | 161918 | 161862 | **RNA**→ rpsO (PRK05626)Ribosomal_S15 (pfam00312)→ |
| env-477 | JGI20156J14371_10003440 | + | 5695 | 5749 | **RNA**→ rpsO (PRK05626)Ribosomal_S15 (pfam00312)→ |
| env-478 | JGI20157J14317_10056873 | + | 532 | 586 | **RNA**→ rpsO (PRK05626)Ribosomal_S15 (pfam00312)→ |
| env-479 | JGI20159J14440_10211892 | + | 222 | 276 | **RNA**→ rpsO (PRK05626)Ribosomal_S15 (pfam00312)→ |
| env-480 | CESJ01009412.1 | - | 295 | 238 | **RNA**→ rpsO (PRK05626)Ribosomal_S15 (pfam00312)→ |
| env-481 | DelMOSum2010_c10003273 | + | 6417 | 6474 | **RNA**→ rpsO (PRK05626)Ribosomal_S15 (pfam00312)→ |
| env-482 | JGI20154J14316_10013644 | + | 4557 | 4614 | **RNA**→ rpsO (PRK05626)RpsO (COG0184)→ |

| | | | | | |
|---|---|---|---|---|---|
| env-483 | JGI20157J14317_10026517 | + | 2621 | 2678 | **RNA**→ rpsO (PRK05626)RpsO (COG0184)→ |
| env-484 | JGI20153J14295_10010421 | - | 315 | 258 | **RNA**→ rpsO (PRK05626)Ribosomal_S15 (pfam00312)→ |
| env-485 | JGI20153J14318_10010421 | - | 315 | 258 | **RNA**→ rpsO (PRK05626)Ribosomal_S15 (pfam00312)→ |
| env-486 | 2236876001_158288 | - | 282 | 226 | **RNA**→ rpsO (PRK05626)Ribosomal_S15 (pfam00312)→ |
| env-487 | JGI20152J14361_10115381 | + | 161 | 218 | **RNA**→ rpsO (PRK05626)Ribosomal_S15 (pfam00312)→ |
| env-488 | JGI20160J14292_10003695 | - | 5908 | 5851 | **RNA**→ rpsO (PRK05626)Ribosomal_S15 (pfam00312)→ |
| env-489 | CESI01136105.1 | - | 853 | 796 | **RNA**→ rpsO (PRK05626)Ribosomal_S15 (pfam00312)→ |
| env-490 | DelMOSum2010_c10699725 | - | 59 | 2 | **RNA**→ |
| env-491 | JGI20152J14361_10001110 | + | 2581 | 2638 | **RNA**→ rpsO (PRK05626)RpsO (COG0184)→ |
| env-492 | JGI20156J14371_10001772 | + | 5804 | 5861 | **RNA**→ rpsO (PRK05626)RpsO (COG0184)→ |
| env-493 | JGI20154J14316_10024766 | + | 2744 | 2801 | **RNA**→ rpsO (PRK05626)RpsO (COG0184)→ |
| env-494 | JGI20160J14292_10090187 | - | 774 | 717 | **RNA**→ rpsO (PRK05626)RpsO (COG0184)→ |
| env-495 | JGI20157J14317_10011759 | - | 3922 | 3865 | **RNA**→ rpsO (PRK05626)RpsO (COG0184)→ |
| env-496 | JGI20153J14295_10003124 | + | 11087 | 11144 | **RNA**→ rpsO (PRK05626)Ribosomal_S15 (pfam00312)→ |
| env-497 | JGI20153J14318_10003124 | + | 11087 | 11144 | **RNA**→ rpsO (PRK05626)Ribosomal_S15 (pfam00312)→ |
| env-498 | JGI20153J14295_10020066 | - | 1717 | 1660 | **RNA**→ rpsO (PRK05626)RpsO (COG0184)→ |
| env-499 | JGI20153J14318_10020066 | - | 1717 | 1660 | **RNA**→ rpsO (PRK05626)RpsO (COG0184)→ |
| env-500 | JGI20159J14440_10267556 | + | 379 | 436 | **RNA**→ ←hypo |
| env-501 | JGI20154J14316_10180509 | + | 119 | 176 | **RNA**→ rpsO (PRK05626)RpsO (COG0184)→ |
| env-502 | CERI01279172.1 | + | 1860 | 1917 | **RNA**→ rpsO (PRK05626)Ribosomal_S15 (pfam00312)→ |
| env-503 | CEGC01022181.1 | + | 362 | 426 | **RNA**→ rpsO (PRK05626)Ribosomal_S15 (pfam00312)→ |
| env-504 | CEQG01070770.1 | - | 2992 | 2928 | **RNA**→ rpsO (PRK05626)Ribosomal_S15 (pfam00312)→ |
| env-505 | CEQQ01079414.1 | + | 374 | 438 | **RNA**→ rpsO (PRK05626)Ribosomal_S15 (pfam00312)→ |
| env-506 | CEPG01012475.1 | - | 616 | 548 | **RNA**→ rpsO (PRK05626)RpsO (COG0184)→ |
| env-507 | P_1C_Liq_2_UnCty_1052240 | - | 318 | 260 | **RNA**→ rpsO (PRK05626)Ribosomal_S15 (pfam00312)→ |
| env-508 | P_1C_Liq_1_UnCty_1125732 | - | 94 | 36 | **RNA**→ |
| env-509 | P_2C_Liq_1_UnCty_1007508 | + | 1871 | 1929 | **RNA**→ rpsO (PRK05626)RpsO (COG0184)→ PRK11824 (PRK11824)RNase_PH (pfam01138)RNase_PH_C (pfam03725)→ |
| env-510 | P_2C_Liq_2_UnCty_1069522 | - | 283 | 225 | **RNA**→ rpsO (PRK05626)RpsO (COG0184)→ |
| env-511 | ElkS_mat_MD6A_1029892 | - | 674 | 608 | **RNA**→ rpsO (PRK05626)Ribosomal_S15 (pfam00312)→ |
| env-512 | JGI11876J14442_10002542 | + | 13284 | 13350 | **RNA**→ rpsO (PRK05626)Ribosomal_S15 (pfam00312)→ |
| Fhi-1-1 | NZ_JPRK01000005.1 | + | 251976 | 252039 | **RNA**→ rpsO (PRK05626)Ribosomal_S15 (pfam00312)→ |
| Fsp-9-1 | NZ_JARP01000002.1 | - | 708129 | 708066 | **RNA**→ rpsO (PRK05626)Ribosomal_S15 (pfam00312)→ |
| Asu-1-1 | NC_018013.1 | + | 528300 | 528368 | **RNA**→ rpsO (PRK05626)Ribosomal_S15 (pfam00312)→ |
| Aca-1-1 | NZ_AUBG01000007.1 | - | 10769 | 10702 | **RNA**→ rpsO (PRK05626)Ribosomal_S15 (pfam00312)→ |
| env-513 | JGI24723J26617_10077130 | - | 289 | 222 | **RNA**→ rpsO (PRK05626)Ribosomal_S15 (pfam00312)→ |
| env-514 | JGI24723J26617_10048522 | + | 1324 | 1391 | **RNA**→ rpsO (PRK05626)Ribosomal_S15 (pfam00312)→ |
| env-515 | CERO01082754.1 | + | 566 | 633 | **RNA**→ rpsO (PRK05626)Ribosomal_S15 (pfam00312)→ |
| env-516 | CERP01000066.1 | - | 89558 | 89491 | **RNA**→ rpsO (PRK05626)Ribosomal_S15 (pfam00312)→ |
| env-517 | CERU01094717.1 | + | 740 | 807 | **RNA**→ rpsO (PRK05626)Ribosomal_S15 (pfam00312)→ |
| Avl-1-1 | NZ_JSVU01000005.1 | + | 248306 | 248373 | **RNA**→ rpsO (PRK05626)Ribosomal_S15 (pfam00312)→ |
| env-518 | CERY01108911.1 | + | 730 | 797 | **RNA**→ rpsO (PRK05626)Ribosomal_S15 (pfam00312)→ |
| env-519 | JGI24723J26617_10269373 | - | 259 | 192 | **RNA**→ rpsO (PRK05626)Ribosomal_S15 (pfam00312)→ |
| env-520 | JGI24723J26617_10077183 | - | 289 | 222 | **RNA**→ rpsO (PRK05626)Ribosomal_S15 (pfam00312)→ |
| env-521 | CEVN01317104.1 | + | 431 | 498 | **RNA**→ rpsO (PRK05626)Ribosomal_S15 (pfam00312)→ |
| env-522 | JGI24723J26617_10004896 | + | 8365 | 8432 | **RNA**→ rpsO (PRK05626)Ribosomal_S15 (pfam00312)→ |
| env-523 | CENR01030012.1 | + | 19705 | 19772 | **RNA**→ rpsO (PRK05626)Ribosomal_S15 (pfam00312)→ |
| env-524 | CEOG01036059.1 | + | 11314 | 11381 | **RNA**→ rpsO (PRK05626)Ribosomal_S15 (pfam00312)→ |
| env-525 | CERJ01077310.1 | + | 15535 | 15602 | **RNA**→ rpsO (PRK05626)Ribosomal_S15 (pfam00312)→ |
| env-526 | CERN01056527.1 | + | 206606 | 206673 | **RNA**→ rpsO (PRK05626)Ribosomal_S15 (pfam00312)→ |
| env-527 | CERO01015055.1 | - | 94 | 27 | **RNA**→ |
| env-528 | CETB01099781.1 | - | 934 | 867 | **RNA**→ rpsO (PRK05626)Ribosomal_S15 (pfam00312)→ |

| | | | | | |
|---|---|---|---|---|---|
| env-529 | CETR01093448.1 | - | 3881 | 3814 | **RNA**→ rpsO (PRK05626)Ribosomal_S15 (pfam00312)→ |
| env-530 | CEVM01236749.1 | - | 847 | 780 | **RNA**→ rpsO (PRK05626)Ribosomal_S15 (pfam00312)→ |
| env-531 | CEVO01146838.1 | - | 280 | 213 | **RNA**→ rpsO (PRK05626)Ribosomal_S15 (pfam00312)→ |
| env-532 | JGI24723J26617_10024902 | - | 107 | 40 | **RNA**→ |
| env-533 | JGI24723J26617_10051817 | - | 111 | 44 | **RNA**→ |
| env-534 | CENX01022829.1 | + | 1733 | 1788 | **RNA**→ rpsO (PRK05626)Ribosomal_S15 (pfam00312)→ Pnp (COG1185)PRK11824 (PRK11824)→ |
| | | | | | sigma70-ECF (TIGR02937)Sigma70_r1_2 (pfam00140)Sigma70_r2 (pfam04542)→ |
| env-535 | CEOF01084231.1 | + | 2913 | 2968 | **RNA**→ rpsO (PRK05626)Ribosomal_S15 (pfam00312)→ Pnp (COG1185)PRK11824 (PRK11824)→ |
| | | | | | RpoD (COG0568)sigma70-ECF (TIGR02937)→ PRK05581 (PRK05581)RPE (cd00429)→ |
| env-536 | CENP01067191.1 | - | 87 | 32 | **RNA**→ |
| env-537 | CENG01112007.1 | + | 4191 | 4246 | **RNA**→ rpsO (PRK05626)Ribosomal_S15 (pfam00312)→ Pnp (COG1185)PRK11824 (PRK11824)→ |
| env-538 | CENU01401332.1 | + | 472 | 527 | **RNA**→ rpsO (PRK05626)Ribosomal_S15 (pfam00312)→ |
| | | | | | PRK11824 (PRK11824)RNase_PH (pfam01138)RNase_PH_C (pfam03725)→ |
| env-539 | CENJ01312050.1 | - | 97 | 42 | **RNA**→ |
| env-540 | CENP01044677.1 | + | 77 | 132 | **RNA**→ rpsO (PRK05626)Ribosomal_S15 (pfam00312)→ PRK11824 (PRK11824)RNase_PH (pfam01138)→ |
| Apa-1-1 | NZ_JACC01000030.1 | - | 38381 | 38316 | **RNA**→ rpsO (PRK05626)RpsO (COG0184)→ |
| Ama-1-1 | NZ_JACA01000052.1 | - | 55849 | 55785 | **RNA**→ rpsO (PRK05626)Ribosomal_S15 (pfam00312)→ |
| Aat-1-1 | NZ_AQRA01000003.1 | - | 469802 | 469738 | **RNA**→ rpsO (PRK05626)Ribosomal_S15 (pfam00312)→ |
| Ame-1-1 | NZ_JACB01000015.1 | + | 60959 | 61023 | **RNA**→ rpsO (PRK05626)Ribosomal_S15 (pfam00312)→ |
| Ala-1-1 | NZ_KE387187.1 | + | 63041 | 63106 | **RNA**→ rpsO (PRK05626)RpsO (COG0184)→ |
| Amu-1-1 | NZ_AUML01000001.1 | - | 274510 | 274444 | **RNA**→ rpsO (PRK05626)RpsO (COG0184)→ |
| Fsu-3-1 | NZ_AUCZ01000003.1 | + | 320199 | 320263 | **RNA**→ rpsO (PRK05626)Ribosomal_S15 (pfam00312)→ |
| Fsu-3-2 | NZ_JRLW01000014.1 | + | 90546 | 90610 | **RNA**→ rpsO (PRK05626)Ribosomal_S15 (pfam00312)→ |
| Fen-1-1 | NZ_AVCS01000009.1 | - | 299934 | 299870 | **RNA**→ rpsO (PRK05626)Ribosomal_S15 (pfam00312)→ |
| Fen-1-2 | NZ_JRLZ01000004.1 | + | 212529 | 212593 | **RNA**→ rpsO (PRK05626)Ribosomal_S15 (pfam00312)→ |
| Fco-1-1 | NC_016510.2 | + | 875063 | 875127 | **RNA**→ rpsO (PRK05626)Ribosomal_S15 (pfam00312)→ |
| env-541 | JGI24770J26754_10043964 | + | 1074 | 1138 | **RNA**→ rpsO (PRK05626)Ribosomal_S15 (pfam00312)→ |
| env-542 | JGI25913J50563_1022430 | - | 297 | 233 | **RNA**→ rpsO (PRK05626)Ribosomal_S15 (pfam00312)→ |
| env-543 | LWAeNNiSIP_GDP3C2T02IBMEZ | - | 77 | 13 | **RNA**→ |
| env-544 | LWAnNN_GDP3C2T02IBMEZ | - | 77 | 13 | **RNA**→ |
| env-545 | JGI25920J50251_10102258 | + | 416 | 480 | **RNA**→ rpsO (PRK05626)Ribosomal_S15 (pfam00312)→ |
| env-546 | TDF_OR_ARG04_113m_c1238479 | - | 103 | 38 | **RNA**→ |
| Fps-1-1 | NZ_CP010278.1 | + | 2255345 | 2255410 | **RNA**→ rpsO (PRK05626)Ribosomal_S15 (pfam00312)→ |
| Fps-1-2 | NZ_CP010276.1 | + | 2300567 | 2300632 | **RNA**→ rpsO (PRK05626)Ribosomal_S15 (pfam00312)→ |
| Fps-1-3 | NZ_CP010274.1 | + | 2298136 | 2298201 | **RNA**→ rpsO (PRK05626)Ribosomal_S15 (pfam00312)→ |
| Jma-1-1 | NZ_JH651379.1 | - | 1502761 | 1502696 | **RNA**→ rpsO (PRK05626)Ribosomal_S15 (pfam00312)→ rpsO (PRK05626)Ribosomal_S15 (pfam00312)→ |
| env-547 | KVRMV2_100016903 | - | 110 | 57 | **RNA**→ |
| env-548 | CEQT01124934.1 | - | 606 | 550 | **RNA**→ rpsO (PRK05626)Ribosomal_S15 (pfam00312)→ PRK11824 (PRK11824)RNase_PH (pfam01138)→ |
| env-549 | CEVG01326867.1 | + | 310 | 366 | **RNA**→ rpsO (PRK05626)Ribosomal_S15 (pfam00312)→ PRK11824 (PRK11824)RNase_PH (pfam01138)→ |
| env-550 | CEQI01034781.1 | + | 143 | 199 | **RNA**→ rpsO (PRK05626)Ribosomal_S15 (pfam00312)→ Pnp (COG1185)PRK11824 (PRK11824)→ |
| env-551 | P_2C_Liq_1_UnCty_1000352 | + | 10687 | 10752 | **RNA**→ rpsO (PRK05626)Ribosomal_S15 (pfam00312)→ |
| env-552 | P_2C_Liq_2_UnCty_1001631 | - | 4100 | 4035 | **RNA**→ rpsO (PRK05626)Ribosomal_S15 (pfam00312)→ |
| env-553 | P_2C_Liq_3_UnCty_100392 | + | 7024 | 7089 | **RNA**→ rpsO (PRK05626)Ribosomal_S15 (pfam00312)→ |
| env-554 | JGI20153J14295_10009471 | - | 2396 | 2331 | **RNA**→ rpsO (PRK05626)Ribosomal_S15 (pfam00312)→ |
| env-555 | JGI20153J14318_10009471 | - | 2396 | 2331 | **RNA**→ rpsO (PRK05626)Ribosomal_S15 (pfam00312)→ |
| env-556 | JCVI_SCAF_1096626865111 | - | 109 | 44 | **RNA**→ |
| env-557 | JGI20154J14316_10022771 | + | 940 | 1005 | **RNA**→ rpsO (PRK05626)Ribosomal_S15 (pfam00312)→ |
| Ffi-1-1 | NZ_KE384381.1 | + | 65930 | 65995 | **RNA**→ rpsO (PRK05626)Ribosomal_S15 (pfam00312)→ |
| env-558 | JGI24768J34885_10768442 | + | 60 | 125 | **RNA**→ rpsO (PRK05626)Ribosomal_S15 (pfam00312)→ |
| env-559 | JGI20156J14371_10000091 | - | 48791 | 48726 | **RNA**→ rpsO (PRK05626)Ribosomal_S15 (pfam00312)→ |
| env-560 | JGI20154J14316_10038808 | - | 884 | 819 | **RNA**→ rpsO (PRK05626)Ribosomal_S15 (pfam00312)→ |
| env-561 | DelMOWin2010_c10012594 | - | 4166 | 4099 | **RNA**→ rpsO (PRK05626)Ribosomal_S15 (pfam00312)→ |
| env-562 | SI48aug10_10m_c1029846 | + | 215 | 282 | **RNA**→ rpsO (PRK05626)Ribosomal_S15 (pfam00312)→ |

| | | | | | |
|---|---|---|---|---|---|
| env-563 | JGI20160J14292_10046285 | - | 129 | 62 | **RNA**→ |
| env-564 | JGI20155J14468_10071765 | - | 979 | 912 | **RNA**→ rpsO (PRK05626)Ribosomal_S15 (pfam00312)→ |
| env-565 | DelMOSum2010_c10029733 | - | 95 | 28 | **RNA**→ |
| env-566 | JGI20160J14292_10025745 | + | 2542 | 2609 | **RNA**→ rpsO (PRK05626)Ribosomal_S15 (pfam00312)→ |
| env-567 | JGI20153J14295_10164677 | - | 300 | 233 | **RNA**→ rpsO (PRK05626)Ribosomal_S15 (pfam00312)→ |
| env-568 | JGI20153J14318_10164677 | - | 300 | 233 | **RNA**→ rpsO (PRK05626)Ribosomal_S15 (pfam00312)→ |
| env-569 | JGI20151J14362_10106988 | + | 760 | 827 | **RNA**→ rpsO (PRK05626)Ribosomal_S15 (pfam00312)→ |
| env-570 | JGI20155J14468_10226844 | - | 184 | 117 | **RNA**→ rpsO (PRK05626)Ribosomal_S15 (pfam00312)→ |
| env-571 | JGI20156J14371_10017225 | + | 4008 | 4075 | **RNA**→ |
| env-572 | JGI20157J14317_10097956 | - | 992 | 925 | **RNA**→ rpsO (PRK05626)Ribosomal_S15 (pfam00312)→ |
| env-573 | JGI20159J14440_10001664 | - | 3899 | 3832 | **RNA**→ rpsO (PRK05626)Ribosomal_S15 (pfam00312)→ |
| env-574 | JGI20151J14362_10553745 | + | 192 | 259 | **RNA**→ |
| env-575 | JGI20156J14371_10055795 | - | 129 | 62 | **RNA**→ |
| env-576 | JGI20156J14371_10169803 | + | 499 | 566 | **RNA**→ |
| env-577 | JGI20157J14317_10131865 | + | 746 | 813 | **RNA**→ |
| env-578 | 2236876007_0312397 | - | 171 | 104 | **RNA**→ rpsO (PRK05626)Ribosomal_S15 (pfam00312)→ |
| env-579 | DelMOSum2010_c10010162 | + | 1748 | 1815 | **RNA**→ rpsO (PRK05626)Ribosomal_S15 (pfam00312)→ |
| env-580 | LPaug09P1610m_c1025198 | - | 345 | 278 | **RNA**→ rpsO (PRK05626)Ribosomal_S15 (pfam00312)→ |
| env-581 | JGI20152J14361_10068744 | + | 344 | 411 | **RNA**→ rpsO (PRK05626)Ribosomal_S15 (pfam00312)→ |
| env-582 | JGI20151J14362_10030196 | + | 80 | 147 | **RNA**→ rpsO (PRK05626)Ribosomal_S15 (pfam00312)→ |
| env-583 | JGI20156J14371_10081935 | - | 1117 | 1050 | **RNA**→ rpsO (PRK05626)Ribosomal_S15 (pfam00312)→ |
| env-584 | JGI20160J14292_10175855 | + | 558 | 625 | **RNA**→ |
| env-585 | JGI20157J14317_10317670 | + | 324 | 391 | **RNA**→ |
| env-586 | JGI20155J14468_10022308 | + | 3140 | 3207 | **RNA**→ |
| env-587 | JGI20155J14468_10502791 | - | 79 | 12 | **RNA**→ |
| env-588 | JGI20158J14315_10014377 | - | 461 | 394 | **RNA**→ rpsO (PRK05626)Ribosomal_S15 (pfam00312)→ |
| env-589 | JGI20158J14315_10054385 | + | 304 | 371 | **RNA**→ rpsO (PRK05626)Ribosomal_S15 (pfam00312)→ |
| env-590 | LPjun09P1210m_c1021057 | - | 265 | 198 | **RNA**→ rpsO (PRK05626)Ribosomal_S15 (pfam00312)→ |
| env-591 | CESF01026423.1 | - | 426 | 359 | **RNA**→ rpsO (PRK05626)Ribosomal_S15 (pfam00312)→ |
| env-592 | LPaug08P2610m_c1011929 | - | 656 | 589 | **RNA**→ rpsO (PRK05626)Ribosomal_S15 (pfam00312)→ |
| env-593 | JGI20152J14361_10054897 | + | 879 | 946 | **RNA**→ |
| env-594 | JGI20153J14295_10031185 | + | 1241 | 1308 | **RNA**→ rpsO (PRK05626)Ribosomal_S15 (pfam00312)→ |
| env-595 | JGI20153J14318_10031185 | + | 1241 | 1308 | **RNA**→ rpsO (PRK05626)Ribosomal_S15 (pfam00312)→ |
| env-596 | JGI20156J14371_10287260 | + | 272 | 339 | **RNA**→ |
| env-597 | JGI20160J14292_10262699 | - | 129 | 62 | **RNA**→ |
| env-598 | JGI20154J14316_10027342 | - | 1569 | 1502 | **RNA**→ rpsO (PRK05626)Ribosomal_S15 (pfam00312)→ |
| env-599 | JGI20157J14317_10012354 | + | 3736 | 3803 | **RNA**→ rpsO (PRK05626)Ribosomal_S15 (pfam00312)→ |
| env-600 | JCVI_SCAF_1096626862538 | + | 155 | 222 | **RNA**→ rpsO (PRK05626)Ribosomal_S15 (pfam00312)→ PRK11824 (PRK11824)RNase_PH (pfam01138)RNase_PH_C (pfam03725)→ |
| env-601 | JGI20156J14371_10124702 | - | 670 | 604 | **RNA**→ rpsO (PRK05626)Ribosomal_S15 (pfam00312)→ PRK11824 (PRK11824)RNase_PH (pfam01138)→ |
| env-602 | CESO01123227.1 | - | 455 | 390 | **RNA**→ rpsO (PRK05626)Ribosomal_S15 (pfam00312)→ |
| env-603 | CETY01169678.1 | - | 8265 | 8200 | **RNA**→ rpsO (PRK05626)Ribosomal_S15 (pfam00312)→ Pnp (COG1185)PRK11824 (PRK11824)→ RpoD (COG0568)sigma70-ECF (TIGR02937)→ PRK05581 (PRK05581)RPE (cd00429)→ anmK (PRK09585)AnmK (pfam03702)→ |
| env-604 | Antartic2_1179192 | + | 146 | 211 | **RNA**→ rpsO (PRK05626)Ribosomal_S15 (pfam00312)→ |
| env-605 | CEWO01136473.1 | + | 175 | 241 | **RNA**→ rpsO (PRK05626)Ribosomal_S15 (pfam00312)→ |
| env-606 | NCBI_READ_1112493426512 | + | 347 | 411 | **RNA**→ rpsO (PRK05626)Ribosomal_S15 (pfam00312)→ |
| env-607 | CESJ01083807.1 | - | 353 | 289 | **RNA**→ rpsO (PRK05626)Ribosomal_S15 (pfam00312)→ |
| env-608 | LPaug08P2610m_c1089043 | + | 81 | 145 | **RNA**→ rpsO (PRK05626)Ribosomal_S15 (pfam00312)→ |
| env-609 | JGI24816J26688_1216386 | + | 208 | 272 | **RNA**→ |
| env-610 | JGI24817J26689_1164217 | + | 175 | 239 | **RNA**→ rpsO (PRK05626)RpsO (COG0184)→ |

| | | | | | |
|---|---|---|---|---|---|
| env-611 | CESL01010325.1 | + | 5005 | 5069 | **RNA**→ rpsO (PRK05626)Ribosomal_S15 (pfam00312)→ Pnp (COG1185)PRK11824 (PRK11824)→ RpoD (COG0568)sigma70-ECF (TIGR02937)→ PRK05581 (PRK05581)RPE (cd00429)→ anmK (PRK09585)AnmK (pfam03702)→ |
| env-612 | CETA01075676.1 | - | 752 | 688 | **RNA**→ rpsO (PRK05626)Ribosomal_S15 (pfam00312)→ PRK11824 (PRK11824)RNase_PH (pfam01138)→ |
| env-613 | SI47jul10_100m_c1098805 | - | 210 | 146 | **RNA**→ rpsO (PRK05626)Ribosomal_S15 (pfam00312)→ |
| env-614 | CEQF01068126.1 | + | 924 | 988 | **RNA**→ rpsO (PRK05626)Ribosomal_S15 (pfam00312)→ PRK11824 (PRK11824)RNase_PH (pfam01138)→ |
| env-615 | CERL01242589.1 | + | 2074 | 2138 | **RNA**→ rpsO (PRK05626)Ribosomal_S15 (pfam00312)→ |
| env-616 | JCVI_SCAF_1101667292350 | - | 651 | 587 | **RNA**→ rpsO (PRK05626)Ribosomal_S15 (pfam00312)→ PRK11824 (PRK11824)RNase_PH (pfam01138)→ |
| env-617 | CESM01026631.1 | + | 383 | 447 | **RNA**→ rpsO (PRK05626)Ribosomal_S15 (pfam00312)→ PRK11824 (PRK11824)RNase_PH (pfam01138)RNase_PH_C (pfam03725)→ |
| env-618 | CEUB01079894.1 | - | 176 | 112 | **RNA**→ rpsO (PRK05626)Ribosomal_S15 (pfam00312)→ |
| env-619 | CESU01087378.1 | + | 261 | 325 | **RNA**→ rpsO (PRK05626)Ribosomal_S15 (pfam00312)→ PRK11824 (PRK11824)RNase_PH (pfam01138)RNase_PH_C (pfam03725)→ |
| env-620 | CEVX01097219.1 | + | 420 | 484 | **RNA**→ rpsO (PRK05626)Ribosomal_S15 (pfam00312)→ PRK11824 (PRK11824)RNase_PH (pfam01138)→ |
| env-621 | CEWO01070284.1 | - | 70 | 6 | **RNA**→ |
| env-622 | CEOP01011972.1 | - | 955 | 891 | **RNA**→ rpsO (PRK05626)Ribosomal_S15 (pfam00312)→ PRK11824 (PRK11824)RNase_PH (pfam01138)→ |
| env-623 | CERW01092002.1 | - | 678 | 614 | **RNA**→ rpsO (PRK05626)Ribosomal_S15 (pfam00312)→ PRK11824 (PRK11824)RNase_PH (pfam01138)→ |
| env-624 | JCVI_SCAF_1096626863907 | - | 2148 | 2084 | **RNA**→ rpsO (PRK05626)Ribosomal_S15 (pfam00312)→ Pnp (COG1185)PRK11824 (PRK11824)→ |
| env-625 | DelMOSum2010_c10030564 | - | 2683 | 2619 | **RNA**→ rpsO (PRK05626)Ribosomal_S15 (pfam00312)→ Pnp (COG1185)PRK11824 (PRK11824)→ |
| env-626 | JGI20156J14371_10004976 | - | 5364 | 5300 | **RNA**→ rpsO (PRK05626)Ribosomal_S15 (pfam00312)→ Pnp (COG1185)PRK11824 (PRK11824)→ RpoD (COG0568)sigma70-ECF (TIGR02937)bSCRAP40 (bSCRAP40)bSCRAP106 (bSCRAP106)→ PRK05581 (PRK05581)RPE (cd00429)→ anmK (PRK09585)AnmK (pfam03702)→ |
| env-627 | JGI20160J14292_10560358 | - | 67 | 3 | **RNA**→ |
| env-628 | JGI20155J14468_10155867 | + | 228 | 292 | **RNA**→ rpsO (PRK05626)Ribosomal_S15 (pfam00312)→ RNase_PH_PNPase_1 (cd11363)PRK11824 (PRK11824)→ |
| env-629 | CERI01034968.1 | - | 2137 | 2073 | **RNA**→ rpsO (PRK05626)Ribosomal_S15 (pfam00312)→ Pnp (COG1185)PRK11824 (PRK11824)→ |
| env-630 | CERM01237716.1 | + | 160 | 224 | **RNA**→ rpsO (PRK05626)Ribosomal_S15 (pfam00312)→ Pnp (COG1185)PRK11824 (PRK11824)→ |
| env-631 | CESF01120804.1 | - | 1011 | 947 | **RNA**→ rpsO (PRK05626)Ribosomal_S15 (pfam00312)→ PRK11824 (PRK11824)RNase_PH (pfam01138)RNase_PH_C (pfam03725)→ |
| env-632 | CESO01082605.1 | + | 3598 | 3662 | **RNA**→ rpsO (PRK05626)Ribosomal_S15 (pfam00312)→ PRK11824 (PRK11824)RNase_PH (pfam01138)→ |
| env-633 | CETY01057294.1 | + | 812 | 876 | **RNA**→ rpsO (PRK05626)Ribosomal_S15 (pfam00312)→ |
| env-634 | CEWO01335636.1 | + | 832 | 896 | **RNA**→ |
| env-635 | CESJ01039048.1 | + | 2696 | 2760 | **RNA**→ |
| env-636 | LPjun09P1210m_c1043297 | - | 263 | 199 | **RNA**→ rpsO (PRK05626)Ribosomal_S15 (pfam00312)→ |
| env-637 | CENF01241446.1 | - | 475 | 411 | **RNA**→ rpsO (PRK05626)Ribosomal_S15 (pfam00312)→ hypo→ |
| env-638 | LPaug09P1610m_c1079724 | - | 152 | 88 | **RNA**→ rpsO (PRK05626)RpsO (COG0184)→ |
| env-639 | LPaug08P2610m_c1008313 | - | 509 | 445 | **RNA**→ rpsO (PRK05626)Ribosomal_S15 (pfam00312)→ RNase_PH_PNPase_1 (cd11363)PRK11824 (PRK11824)→ |
| env-640 | CESI01003936.1 | - | 2614 | 2550 | **RNA**→ rpsO (PRK05626)Ribosomal_S15 (pfam00312)→ Pnp (COG1185)PRK11824 (PRK11824)→ |
| env-641 | SI54feb11_100m_c1030638 | - | 504 | 440 | **RNA**→ rpsO (PRK05626)Ribosomal_S15 (pfam00312)→ RNase_PH_PNPase_1 (cd11363)PRK11824 (PRK11824)→ |
| env-642 | SI60aug11_100m_c1000018 | + | 42717 | 42781 | **RNA**→ rpsO (PRK05626)Ribosomal_S15 (pfam00312)→ Pnp (COG1185)PRK11824 (PRK11824)→ RpoD (COG0568)sigma70-ECF (TIGR02937)bSCRAP40 (bSCRAP40)bSCRAP106 (bSCRAP106)→ PRK05581 (PRK05581)RPE (cd00429)→ AnmK (COG2377)anmK (PRK09585)→ |
| env-643 | SI47jul10_120m_c1021072 | - | 725 | 661 | **RNA**→ rpsO (PRK05626)Ribosomal_S15 (pfam00312)→ PRK11824 (PRK11824)RNase_PH (pfam01138)→ |
| env-644 | SI36aug09_120m_1000015 | + | 9032 | 9096 | **RNA**→ rpsO (PRK05626)Ribosomal_S15 (pfam00312)→ Pnp (COG1185)PRK11824 (PRK11824)→ RpoD (COG0568)sigma70-ECF (TIGR02937)bSCRAP40 (bSCRAP40)bSCRAP106 (bSCRAP106)→ PRK05581 (PRK05581)RPE (cd00429)→ anmK (PRK09585)AnmK (pfam03702)→ |
| env-645 | A09P04500a_Sequence0000024128 | - | 490 | 426 | **RNA**→ rpsO (PRK05626)Ribosomal_S15 (pfam00312)→ RNase_PH_PNPase_1 (cd11363)PRK11824 (PRK11824)→ |
| env-646 | J09P20500a_Sequence0000027602 | - | 643 | 579 | **RNA**→ rpsO (PRK05626)Ribosomal_S15 (pfam00312)→ PRK11824 (PRK11824)RNase_PH (pfam01138)→ |
| env-647 | F10SI03120a_Sequence0000002163 | + | 5397 | 5461 | **RNA**→ rpsO (PRK05626)Ribosomal_S15 (pfam00312)→ |
| env-648 | SI39nov09_120m_c1156358 | + | 103 | 167 | **RNA**→ rpsO (PRK05626)Ribosomal_S15 (pfam00312)→ |
| env-649 | SI60aug11_100m_c1008765 | + | 593 | 657 | **RNA**→ rpsO (PRK05626)Ribosomal_S15 (pfam00312)→ Pnp (COG1185)PRK11824 (PRK11824)→ |
| env-650 | SI47jul10_100m_c1000490 | - | 5467 | 5403 | **RNA**→ rpsO (PRK05626)Ribosomal_S15 (pfam00312)→ Pnp (COG1185)PRK11824 (PRK11824)→ RpoD (COG0568)sigma70-ECF (TIGR02937)bSCRAP40 (bSCRAP40)bSCRAP106 (bSCRAP106)→ PRK05581 (PRK05581)RPE (cd00429)→ anmK (PRK09585)AnmK (pfam03702)→ |
| env-651 | SI39nov09_135m_c1093920 | + | 131 | 195 | **RNA**→ rpsO (PRK05626)Ribosomal_S15 (pfam00312)→ |

| | | | | | |
|---|---|---|---|---|---|
| env-652 | SI60aug11_200m_c1241338 | + | 85 | 149 | RNA→ rpsO (PRK05626)RpsO (COG0184)→ |
| env-653 | SI47jul10_135m_c1051027 | - | 459 | 395 | RNA→ rpsO (PRK05626)Ribosomal_S15 (pfam00312)→ hypo→ |
| env-654 | SI34jun09_120m_1000008 | - | 61442 | 61378 | RNA→ rpsO (PRK05626)Ribosomal_S15 (pfam00312)→ Pnp (COG1185)PRK11824 (PRK11824)→ RpoD (COG0568)sigma70-ECF (TIGR02937)bSCRAP40 (bSCRAP40)bSCRAP106 (bSCRAP106)→ PRK05581 (PRK05581)RPE (cd00429)→ anmK (PRK09585)AnmK (pfam03702)→ |
| env-655 | SI34jun09_135m_1000014 | - | 50759 | 50695 | RNA→ rpsO (PRK05626)Ribosomal_S15 (pfam00312)→ Pnp (COG1185)PRK11824 (PRK11824)→ RpoD (COG0568)sigma70-ECF (TIGR02937)bSCRAP40 (bSCRAP40)bSCRAP106 (bSCRAP106)→ PRK05581 (PRK05581)RPE (cd00429)→ anmK (PRK09585)AnmK (pfam03702)→ |
| env-656 | SI39nov09_100m_1072152 | - | 470 | 406 | RNA→ rpsO (PRK05626)Ribosomal_S15 (pfam00312)→ hypo→ |
| env-657 | SI36aug09_100m_1060339 | + | 326 | 390 | RNA→ |
| env-658 | SI34jun09_100m_1011554 | - | 1665 | 1601 | RNA→ rpsO (PRK05626)Ribosomal_S15 (pfam00312)→ Pnp (COG1185)PRK11824 (PRK11824)→ |
| env-659 | CEWR01151164.1 | - | 180 | 116 | RNA→ rpsO (PRK05626)Ribosomal_S15 (pfam00312)→ |
| env-660 | CESE01088802.1 | - | 353 | 289 | RNA→ rpsO (PRK05626)Ribosomal_S15 (pfam00312)→ |
| env-661 | CETV01112507.1 | + | 2433 | 2497 | RNA→ rpsO (PRK05626)Ribosomal_S15 (pfam00312)→ Pnp (COG1185)PRK11824 (PRK11824)→ RpoD (COG0568)sigma70-ECF (TIGR02937)→ PRK05581 (PRK05581)RPE (cd00429)→ anmK (PRK09585)AnmK (pfam03702)→ |
| env-662 | A09P041000a_Sequence0000001821 | + | 883 | 947 | RNA→ rpsO (PRK05626)Ribosomal_S15 (pfam00312)→ PRK11824 (PRK11824)RNase_PH (pfam01138)→ |
| env-663 | CESM01243490.1 | + | 238 | 303 | RNA→ rpsO (PRK05626)Ribosomal_S15 (pfam00312)→ |
| env-664 | BS_KBA_SWE12_21m_c10000131 | + | 18308 | 18374 | RNA→ rpsO (PRK05626)Ribosomal_S15 (pfam00312)→ |
| env-665 | BS_KBB_SWE26_205m_c1000040 | - | 13487 | 13421 | RNA→ rpsO (PRK05626)Ribosomal_S15 (pfam00312)→ |
| env-666 | BS_KBA_SWE07_21m_c1009901 | - | 943 | 877 | RNA→ rpsO (PRK05626)Ribosomal_S15 (pfam00312)→ |
| env-667 | BS_KBA_SWE21_205m_10000004 | + | 15225 | 15291 | RNA→ rpsO (PRK05626)Ribosomal_S15 (pfam00312)→ |
| env-668 | BS_KBA_SWE02_21m_10000008 | + | 18316 | 18382 | RNA→ rpsO (PRK05626)Ribosomal_S15 (pfam00312)→ |
| env-669 | KGI_S1_ANT02_95m_c10523594 | - | 194 | 129 | RNA→ rpsO (PRK05626)Ribosomal_S15 (pfam00312)→ |
| env-670 | SA_S1_NOR08_45m_c10397994 | - | 292 | 227 | RNA→ rpsO (PRK05626)Ribosomal_S15 (pfam00312)→ |
| env-671 | SA_S2_NOR15_50m_c10501265 | + | 214 | 279 | RNA→ rpsO (PRK05626)→ |
| env-672 | CEUH01124891.1 | + | 289 | 346 | RNA→ rpsO (PRK05626)Ribosomal_S15 (pfam00312)→ |
| env-673 | CESI01168467.1 | - | 345 | 287 | RNA→ rpsO (PRK05626)Ribosomal_S15 (pfam00312)→ |
| env-674 | CESU01168001.1 | - | 537 | 479 | RNA→ rpsO (PRK05626)Ribosomal_S15 (pfam00312)→ RNase_PH_PNPase_1 (cd11363)PRK11824 (PRK11824)→ |
| env-675 | CESM01131397.1 | - | 185 | 127 | RNA→ rpsO (PRK05626)Ribosomal_S15 (pfam00312)→ |
| env-676 | CESZ01043879.1 | - | 5070 | 5012 | RNA→ rpsO (PRK05626)Ribosomal_S15 (pfam00312)→ Pnp (COG1185)PRK11824 (PRK11824)→ |
| env-677 | CENG01063226.1 | - | 2602 | 2544 | RNA→ rpsO (PRK05626)Ribosomal_S15 (pfam00312)→ Pnp (COG1185)PRK11824 (PRK11824)→ |
| env-678 | CESA01043993.1 | + | 34 | 92 | RNA→ rpsO (PRK05626)Ribosomal_S15 (pfam00312)→ Pnp (COG1185)PRK11824 (PRK11824)→ |
| env-679 | CEQX01317913.1 | - | 490 | 432 | RNA→ rpsO (PRK05626)Ribosomal_S15 (pfam00312)→ PRK11824 (PRK11824)→ |
| env-680 | CESQ01364581.1 | + | 618 | 676 | RNA→ hypo→ |
| env-681 | CETE01036010.1 | - | 845 | 787 | RNA→ rpsO (PRK05626)Ribosomal_S15 (pfam00312)→ PRK11824 (PRK11824)RNase_PH (pfam01138)→ |
| env-682 | CETK01367368.1 | + | 610 | 668 | RNA→ rpsO (PRK05626)Ribosomal_S15 (pfam00312)→ PRK11824 (PRK11824)RNase_PH (pfam01138)→ |
| env-683 | CETQ01203907.1 | + | 444 | 502 | RNA→ rpsO (PRK05626)Ribosomal_S15 (pfam00312)→ |
| env-684 | CEUH01124635.1 | - | 299 | 241 | RNA→ rpsO (PRK05626)Ribosomal_S15 (pfam00312)→ |
| env-685 | CEUW01071784.1 | - | 737 | 679 | RNA→ rpsO (PRK05626)Ribosomal_S15 (pfam00312)→ PRK11824 (PRK11824)RNase_PH (pfam01138)→ |
| env-686 | CEWH01138841.1 | - | 766 | 708 | RNA→ rpsO (PRK05626)Ribosomal_S15 (pfam00312)→ PRK11824 (PRK11824)RNase_PH (pfam01138)→ |
| env-687 | CEUK01251936.1 | + | 511 | 569 | RNA→ |
| env-688 | CEUU01042451.1 | - | 497 | 439 | RNA→ rpsO (PRK05626)Ribosomal_S15 (pfam00312)→ RNase_PH_PNPase_1 (cd11363)PRK11824 (PRK11824)→ |
| env-689 | CEVF01030331.1 | - | 845 | 787 | RNA→ rpsO (PRK05626)Ribosomal_S15 (pfam00312)→ PRK11824 (PRK11824)RNase_PH (pfam01138)→ |
| env-690 | CEWJ01102386.1 | + | 420 | 478 | RNA→ rpsO (PRK05626)Ribosomal_S15 (pfam00312)→ PRK11824 (PRK11824)RNase_PH (pfam01138)→ |
| env-691 | JGI24820J26691_1081338 | - | 226 | 168 | RNA→ rpsO (PRK05626)Ribosomal_S15 (pfam00312)→ |
| env-692 | JCVI_SCAF_1096627810395 | + | 221 | 279 | RNA→ rpsO (PRK05626)Ribosomal_S15 (pfam00312)→ PRK11824 (PRK11824)RNase_PH (pfam01138)→ PRK11824 (PRK11824)RNase_PH (pfam01138)RNase_PH_C (pfam03725)→ |
| env-693 | CESM01131398.1 | + | 31 | 89 | RNA→ rpsO (PRK05626)Ribosomal_S15 (pfam00312)→ PRK11824 (PRK11824)RNase_PH (pfam01138)→ |
| env-694 | CERI01037614.1 | + | 3582 | 3642 | RNA→ rpsO (PRK05626)Ribosomal_S15 (pfam00312)→ Pnp (COG1185)PRK11824 (PRK11824)→ RpoD (COG0568)sigma70-ECF (TIGR02937)→ PRK05581 (PRK05581)RPE (cd00429)→ |
| env-695 | CESF01030271.1 | - | 5728 | 5668 | RNA→ rpsO (PRK05626)Ribosomal_S15 (pfam00312)→ Pnp (COG1185)PRK11824 (PRK11824)→ RpoD (COG0568)sigma70-ECF (TIGR02937)→ PRK05581 (PRK05581)RPE (cd00429)→ |
| env-696 | CESO01050962.1 | - | 2098 | 2038 | RNA→ rpsO (PRK05626)Ribosomal_S15 (pfam00312)→ Pnp (COG1185)PRK11824 (PRK11824)→ |

| | | | | | |
|---|---|---|---|---|---|
| env-697 | CETY01400875.1 | - | 635 | 575 | **RNA**→ rpsO (PRK05626)Ribosomal_S15 (pfam00312)→ PRK11824 (PRK11824)RNase_PH (pfam01138)→ |
| env-698 | CEVX01285939.1 | + | 370 | 430 | **RNA**→ rpsO (PRK05626)Ribosomal_S15 (pfam00312)→ |
| env-699 | CEOP01004610.1 | + | 2645 | 2705 | **RNA**→ rpsO (PRK05626)Ribosomal_S15 (pfam00312)→ Pnp (COG1185)PRK11824 (PRK11824)→ |
| env-700 | JCVI_SCAF_1096626851857 | - | 1063 | 1003 | **RNA**→ rpsO (PRK05626)Ribosomal_S15 (pfam00312)→ RNase_PH_PNPase_1 (cd11363)PRK11824 (PRK11824)– |
| env-701 | CENT01185471.1 | - | 1093 | 1032 | **RNA**→ rpsO (PRK05626)Ribosomal_S15 (pfam00312)→ RNase_PH_PNPase_1 (cd11363)PRK11824 (PRK11824)– |
| env-702 | LPaug09P1610m_c1022285 | - | 558 | 498 | **RNA**→ rpsO (PRK05626)Ribosomal_S15 (pfam00312)→ RNase_PH_PNPase_1 (cd11363)PRK11824 (PRK11824)– |
| env-703 | CENG01117863.1 | + | 933 | 989 | **RNA**→ |
| env-704 | CENJ01221902.1 | - | 62 | 6 | **RNA**→ ←hypo |
| env-705 | NCBI_BBAY_READ_1105609640893 | - | 179 | 122 | **RNA**→ |
| env-706 | CEOK01189823.1 | - | 83 | 26 | **RNA**→ |
| env-707 | JCVI_SCAF_1101668161477 | - | 498 | 441 | **RNA**→ rpsO (PRK05626)Ribosomal_S15 (pfam00312)→ |
| env-708 | JGI20157J14317_10026495 | - | 1195 | 1138 | **RNA**→ rpsO (PRK05626)Ribosomal_S15 (pfam00312)→ RNase_PH_PNPase_1 (cd11363)PRK11824 (PRK11824)– |
| env-709 | HBC_scaff_10024157 | + | 21661 | 21709 | **RNA**→ rpsO (PRK05626)Ribosomal_S15 (pfam00312)→ |
| env-710 | HBC_ctgs_1000293 | - | 1761 | 1713 | **RNA**→ rpsO (PRK05626)Ribosomal_S15 (pfam00312)→ |
| env-711 | JGI24023J19991_10000252 | - | 22396 | 22343 | **RNA**→ rpsO (PRK05626)Ribosomal_S15 (pfam00312)→ Pnp (COG1185)PRK11824 (PRK11824)→ |
| env-712 | CERI01113722.1 | - | 1102 | 1043 | **RNA**→ rpsO (PRK05626)Ribosomal_S15 (pfam00312)→ RNase_PH_PNPase_1 (cd11363)PRK11824 (PRK11824)– |
| Fba-2-1 | NZ_ABVV01000001.1 | - | 275587 | 275526 | **RNA**→ rpsO (PRK05626)Ribosomal_S15 (pfam00312)→ rpsO (PRK05626)Ribosomal_S15 (pfam00312)→ hypo→ Pnp (COG1185)PRK11824 (PRK11824)→ Pnp (COG1185)PRK11824 (PRK11824)→ RpoD (COG0568)sigma70-ECF (TIGR02937)bSCRAP40 (bSCRAP40)bSCRAP106 (bSCRAP106)→ RpoD (COG0568)sigma70-ECF (TIGR02937)→ PRK05581 (PRK05581)RPE (cd00429)→ PRK05581 (PRK05581)RPE (cd00429)→ |
| env-713 | DelMOSum2010_c10000077 | - | 70521 | 70460 | **RNA**→ rpsO (PRK05626)Ribosomal_S15 (pfam00312)→ Pnp (COG1185)PRK11824 (PRK11824)→ RpoD (COG0568)sigma70-ECF (TIGR02937)bSCRAP40 (bSCRAP40)bSCRAP106 (bSCRAP106)→ PRK05581 (PRK05581)RPE (cd00429)→ |
| env-714 | SI34jun09_10m_1001264 | - | 3237 | 3176 | **RNA**→ rpsO (PRK05626)Ribosomal_S15 (pfam00312)→ Pnp (COG1185)PRK11824 (PRK11824)→ RpoD (COG0568)sigma70-ECF (TIGR02937)→ |
| env-715 | JGI20152J14361_10000391 | - | 11554 | 11493 | **RNA**→ rpsO (PRK05626)Ribosomal_S15 (pfam00312)→ Pnp (COG1185)PRK11824 (PRK11824)→ RpoD (COG0568)sigma70-ECF (TIGR02937)bSCRAP40 (bSCRAP40)bSCRAP106 (bSCRAP106)→ PRK05581 (PRK05581)RPE (cd00429)→ |
| env-716 | JGI20160J14292_10000039 | + | 19294 | 19355 | **RNA**→ rpsO (PRK05626)Ribosomal_S15 (pfam00312)→ Pnp (COG1185)PRK11824 (PRK11824)→ RpoD (COG0568)sigma70-ECF (TIGR02937)bSCRAP40 (bSCRAP40)bSCRAP106 (bSCRAP106)→ PRK05581 (PRK05581)RPE (cd00429)→ |
| env-717 | JGI20157J14317_10000232 | + | 16872 | 16933 | **RNA**→ rpsO (PRK05626)Ribosomal_S15 (pfam00312)→ Pnp (COG1185)PRK11824 (PRK11824)→ RpoD (COG0568)sigma70-ECF (TIGR02937)bSCRAP40 (bSCRAP40)bSCRAP106 (bSCRAP106)→ PRK05581 (PRK05581)RPE (cd00429)→ |
| env-718 | DelMOSum2011_c10015687 | - | 2741 | 2680 | **RNA**→ rpsO (PRK05626)Ribosomal_S15 (pfam00312)→ Pnp (COG1185)PRK11824 (PRK11824)→ PRK09210 (PRK09210)Sigma70_r1_2 (pfam00140)→ |
| env-719 | JGI20151J14362_10000070 | - | 48313 | 48252 | **RNA**→ rpsO (PRK05626)Ribosomal_S15 (pfam00312)→ Pnp (COG1185)PRK11824 (PRK11824)→ RpoD (COG0568)sigma70-ECF (TIGR02937)bSCRAP40 (bSCRAP40)bSCRAP106 (bSCRAP106)→ PRK05581 (PRK05581)RPE (cd00429)→ |
| env-720 | JGI20156J14371_10000130 | + | 17278 | 17339 | **RNA**→ rpsO (PRK05626)Ribosomal_S15 (pfam00312)→ Pnp (COG1185)PRK11824 (PRK11824)→ RpoD (COG0568)sigma70-ECF (TIGR02937)bSCRAP40 (bSCRAP40)bSCRAP106 (bSCRAP106)→ PRK05581 (PRK05581)RPE (cd00429)→ |
| env-721 | JGI20154J14316_10000543 | - | 13329 | 13268 | **RNA**→ rpsO (PRK05626)Ribosomal_S15 (pfam00312)→ Pnp (COG1185)PRK11824 (PRK11824)→ RpoD (COG0568)sigma70-ECF (TIGR02937)bSCRAP40 (bSCRAP40)bSCRAP106 (bSCRAP106)→ PRK05581 (PRK05581)RPE (cd00429)→ |
| env-722 | JGI20153J14295_10000212 | - | 48142 | 48081 | **RNA**→ rpsO (PRK05626)Ribosomal_S15 (pfam00312)→ Pnp (COG1185)PRK11824 (PRK11824)→ RpoD (COG0568)sigma70-ECF (TIGR02937)bSCRAP40 (bSCRAP40)bSCRAP106 (bSCRAP106)→ PRK05581 (PRK05581)RPE (cd00429)→ |
| env-723 | JGI20153J14318_10000212 | - | 48142 | 48081 | **RNA**→ rpsO (PRK05626)Ribosomal_S15 (pfam00312)→ Pnp (COG1185)PRK11824 (PRK11824)→ RpoD (COG0568)sigma70-ECF (TIGR02937)bSCRAP40 (bSCRAP40)bSCRAP106 (bSCRAP106)→ PRK05581 (PRK05581)RPE (cd00429)→ |
| env-724 | JGI20159J14440_10000191 | - | 27139 | 27078 | **RNA**→ rpsO (PRK05626)Ribosomal_S15 (pfam00312)→ Pnp (COG1185)PRK11824 (PRK11824)→ RpoD (COG0568)sigma70-ECF (TIGR02937)bSCRAP40 (bSCRAP40)bSCRAP106 (bSCRAP106)→ PRK05581 (PRK05581)RPE (cd00429)→ |

| | | | | | |
|---|---|---|---|---|---|
| env-725 | JGI20155J14468_10000002 | - | 197670 | 197609 | **RNA**→ rpsO (PRK05626)Ribosomal_S15 (pfam00312)→ Pnp (COG1185)PRK11824 (PRK11824)→ RpoD (COG0568)sigma70-ECF (TIGR02937)bSCRAP40 (bSCRAP40)bSCRAP106 (bSCRAP106)→ PRK05581 (PRK05581)RPE (cd00429)→ |
| env-726 | JGI20158J14315_10001443 | - | 14132 | 14071 | **RNA**→ rpsO (PRK05626)Ribosomal_S15 (pfam00312)→ Pnp (COG1185)PRK11824 (PRK11824)→ RpoD (COG0568)sigma70-ECF (TIGR02937)bSCRAP40 (bSCRAP40)bSCRAP106 (bSCRAP106)→ PRK05581 (PRK05581)RPE (cd00429)→ |
| env-727 | SI60aug11_100m_c1210894 | - | 111 | 50 | **RNA**→ |
| env-728 | SI48aug10_10m_c1076796 | - | 113 | 52 | **RNA**→ |
| env-729 | SI48aug10_100m_1053294 | - | 541 | 480 | **RNA**→ rpsO (PRK05626)Ribosomal_S15 (pfam00312)→ RNase_PH_PNPase_1 (cd11363)PRK11824 (PRK11824)→ |
| env-730 | JGI20152J14361_10285356 | + | 41 | 101 | **RNA**→ rpsO (PRK05626)Ribosomal_S15 (pfam00312)→ |
| env-731 | JGI20151J14362_10001694 | + | 13999 | 14059 | **RNA**→ rpsO (PRK05626)Ribosomal_S15 (pfam00312)→ Pnp (COG1185)PRK11824 (PRK11824)→ |
| env-732 | JGI20156J14371_10036665 | + | 758 | 818 | **RNA**→ rpsO (PRK05626)Ribosomal_S15 (pfam00312)→ Pnp (COG1185)PRK11824 (PRK11824)→ |
| env-733 | JGI20159J14440_10045471 | + | 1257 | 1317 | **RNA**→ rpsO (PRK05626)Ribosomal_S15 (pfam00312)→ |
| env-734 | JGI20160J14292_10275000 | + | 128 | 188 | **RNA**→ rpsO (PRK05626)Ribosomal_S15 (pfam00312)→ |
| env-735 | JGI20155J14468_10035248 | + | 2056 | 2116 | **RNA**→ rpsO (PRK05626)Ribosomal_S15 (pfam00312)→ |
| env-736 | JGI20158J14315_10222416 | + | 279 | 339 | **RNA**→ rpsO (PRK05626)Ribosomal_S15 (pfam00312)→ |
| env-737 | CENJ01283541.1 | + | 787 | 847 | **RNA**→ |
| env-738 | JCVI_SCAF_1096626899896 | + | 1250 | 1310 | **RNA**→ |
| env-739 | CENG01040397.1 | - | 73 | 13 | **RNA**→ |
| env-740 | CESI01023085.1 | - | 644 | 584 | **RNA**→ rpsO (PRK05626)Ribosomal_S15 (pfam00312)→ PRK11824 (PRK11824)RNase_PH (pfam01138)→ |
| env-741 | CETK01101931.1 | - | 10939 | 10891 | **RNA**→ rpsO (PRK05626)Ribosomal_S15 (pfam00312)→ Pnp (COG1185)PRK11824 (PRK11824)→ RpoD (COG0568)sigma70-ECF (TIGR02937)→ PRK05581 (PRK05581)RPE (cd00429)→ |
| env-742 | CEVG01048401.1 | - | 452 | 404 | **RNA**→ rpsO (PRK05626)Ribosomal_S15 (pfam00312)→ |
| env-743 | CESS01088671.1 | + | 460 | 508 | **RNA**→ rpsO (PRK05626)RpsO (COG0184)→ PRK11824 (PRK11824)RNase_PH (pfam01138)RNase_PH_C (pfam03725)→ |
| env-744 | CENP01025453.1 | + | 853 | 901 | **RNA**→ |
| env-745 | CENU01400115.1 | + | 500 | 548 | **RNA**→ rpsO (PRK05626)RpsO (COG0184)→ PRK11824 (PRK11824)RNase_PH (pfam01138)→ |
| env-746 | JCVI_SCAF_1096627284330 | - | 131 | 83 | **RNA**→ |
| env-747 | CENN01195248.1 | + | 427 | 475 | **RNA**→ rpsO (PRK05626)RpsO (COG0184)→ Pnp (COG1185)PRK11824 (PRK11824)→ RpoD (COG0568)sigma70-ECF (TIGR02937)→ |
| env-748 | CEWA01144527.1 | + | 171 | 219 | **RNA**→ rpsO (PRK05626)Ribosomal_S15 (pfam00312)→ |
| env-749 | CEUH01264893.1 | - | 295 | 247 | **RNA**→ rpsO (PRK05626)Ribosomal_S15 (pfam00312)→ |
| env-750 | CEWG01086055.1 | + | 2874 | 2922 | **RNA**→ rpsO (PRK05626)Ribosomal_S15 (pfam00312)→ |
| env-751 | JCVI_SCAF_1101668541356 | - | 246 | 198 | **RNA**→ rpsO (PRK05626)Ribosomal_S15 (pfam00312)→ |
| env-752 | CEUB01094603.1 | + | 343 | 391 | **RNA**→ rpsO (PRK05626)Ribosomal_S15 (pfam00312)→ |
| env-753 | CENG01122428.1 | - | 281 | 233 | **RNA**→ rpsO (PRK05626)Ribosomal_S15 (pfam00312)→ |
| env-754 | CENJ01278422.1 | + | 82 | 130 | **RNA**→ rpsO (PRK05626)Ribosomal_S15 (pfam00312)→ PRK11824 (PRK11824)RNase_PH (pfam01138)→ |
| env-755 | CEWH01115234.1 | - | 417 | 368 | **RNA**→ rpsO (PRK05626)Ribosomal_S15 (pfam00312)→ |
| env-756 | CESF01312487.1 | + | 10933 | 10981 | **RNA**→ rpsO (PRK05626)Ribosomal_S15 (pfam00312)→ Pnp (COG1185)PRK11824 (PRK11824)→ |
| env-757 | CESO01048748.1 | + | 2685 | 2733 | **RNA**→ rpsO (PRK05626)Ribosomal_S15 (pfam00312)→ Pnp (COG1185)PRK11824 (PRK11824)→ RpoD (COG0568)sigma70-ECF (TIGR02937)→ PRK05581 (PRK05581)RPE (cd00429)→ |
| env-758 | CEVG01111268.1 | - | 830 | 782 | **RNA**→ rpsO (PRK05626)Ribosomal_S15 (pfam00312)→ PRK11824 (PRK11824)RNase_PH (pfam01138)→ |
| env-759 | CENG01071379.1 | + | 1616 | 1664 | **RNA**→ rpsO (PRK05626)RpsO (COG0184)→ Pnp (COG1185)PRK11824 (PRK11824)→ RpoD (COG0568)sigma70-ECF (TIGR02937)→ |
| env-760 | CENT01073065.1 | + | 574 | 622 | **RNA**→ rpsO (PRK05626)RpsO (COG0184)→ RNase_PH_PNPase_1 (cd11363)PRK11824 (PRK11824)→ |
| env-761 | CEQP01301392.1 | + | 997 | 1051 | **RNA**→ hypo→ |
| env-762 | CETB01210491.1 | + | 254 | 308 | **RNA**→ rpsO (PRK05626)Ribosomal_S15 (pfam00312)→ |
| env-763 | CESM01391969.1 | - | 199 | 145 | **RNA**→ rpsO (PRK05626)Ribosomal_S15 (pfam00312)→ |
| env-764 | CEWG01210985.1 | - | 2290 | 2236 | **RNA**→ rpsO (PRK05626)Ribosomal_S15 (pfam00312)→ Pnp (COG1185)PRK11824 (PRK11824)→ |
| env-765 | JCVI_SCAF_1096626961789 | - | 286 | 232 | **RNA**→ rpsO (PRK05626)Ribosomal_S15 (pfam00312)→ |
| env-766 | CEUF01056559.1 | + | 193 | 247 | **RNA**→ rpsO (PRK05626)Ribosomal_S15 (pfam00312)→ PRK11824 (PRK11824)RNase_PH (pfam01138)RNase_PH_C (pfam03725)→ |
| env-767 | CEOV01002341.1 | - | 270 | 216 | **RNA**→ rpsO (PRK05626)Ribosomal_S15 (pfam00312)→ |
| env-768 | CEOW01267461.1 | + | 1775 | 1829 | **RNA**→ rpsO (PRK05626)Ribosomal_S15 (pfam00312)→ |

| | | | | | |
|---|---|---|---|---|---|
| env-769 | CEPA01173849.1 | + | 414 | 468 | **RNA**→ hypo→ |
| env-770 | CEPT01173609.1 | + | 76 | 130 | **RNA**→ rpsO (PRK05626)Ribosomal_S15 (pfam00312)→ |
| env-771 | CEVG01159809.1 | - | 1097 | 1043 | **RNA**→ rpsO (PRK05626)Ribosomal_S15 (pfam00312)→ |
| | | | | | PRK11824 (PRK11824)RNase_PH (pfam01138)RNase_PH_C (pfam03725)→ |
| env-772 | CEWA01198601.1 | - | 261 | 207 | **RNA**→ rpsO (PRK05626)Ribosomal_S15 (pfam00312)→ |
| env-773 | CETK01338805.1 | + | 612 | 666 | **RNA**→ rpsO (PRK05626)Ribosomal_S15 (pfam00312)→ |
| env-774 | CEUH01280879.1 | + | 105 | 159 | **RNA**→ rpsO (PRK05626)Ribosomal_S15 (pfam00312)→ PRK11824 (PRK11824)RNase_PH (pfam01138)→ |
| env-775 | CEVH01186256.1 | + | 1905 | 1959 | **RNA**→ rpsO (PRK05626)Ribosomal_S15 (pfam00312)→ |
| env-776 | CEWH01028013.1 | + | 91 | 145 | **RNA**→ rpsO (PRK05626)Ribosomal_S15 (pfam00312)→ Pnp (COG1185)PRK11824 (PRK11824)→ |
| env-777 | CENJ01092577.1 | - | 1007 | 952 | **RNA**→ rpsO (PRK05626)Ribosomal_S15 (pfam00312)→ |
| | | | | | PRK11824 (PRK11824)RNase_PH (pfam01138)RNase_PH_C (pfam03725)→ |
| env-778 | CEOK01031026.1 | + | 658 | 713 | **RNA**→ rpsO (PRK05626)Ribosomal_S15 (pfam00312)→ |
| env-779 | CEVU01050680.1 | - | 328 | 273 | **RNA**→ rpsO (PRK05626)Ribosomal_S15 (pfam00312)→ |
| env-780 | CEVP01013039.1 | + | 315 | 369 | **RNA**→ rpsO (PRK05626)Ribosomal_S15 (pfam00312)→ PRK11824 (PRK11824)RNase_PH (pfam01138)→ |
| env-781 | CEQN01285656.1 | + | 461 | 515 | **RNA**→ rpsO (PRK05626)Ribosomal_S15 (pfam00312)→ |
| env-782 | CEQY01281147.1 | - | 162 | 108 | **RNA**→ rpsO (PRK05626)Ribosomal_S15 (pfam00312)→ |
| env-783 | CESB01256322.1 | - | 418 | 364 | **RNA**→ rpsO (PRK05626)Ribosomal_S15 (pfam00312)→ |
| env-784 | CENX01069896.1 | - | 443 | 388 | **RNA**→ rpsO (PRK05626)Ribosomal_S15 (pfam00312)→ |
| env-785 | CEOM01046697.1 | - | 328 | 273 | **RNA**→ rpsO (PRK05626)Ribosomal_S15 (pfam00312)→ |
| env-786 | CENP01062878.1 | + | 5461 | 5515 | **RNA**→ rpsO (PRK05626)Ribosomal_S15 (pfam00312)→ Pnp (COG1185)PRK11824 (PRK11824)→ |
| env-787 | CEOF01162626.1 | + | 613 | 667 | **RNA**→ rpsO (PRK05626)Ribosomal_S15 (pfam00312)→ |
| env-788 | CEOK01252908.1 | + | 73 | 127 | **RNA**→ rpsO (PRK05626)Ribosomal_S15 (pfam00312)→ RNase_PH_PNPase_1 (cd11363)PRK11824 (PRK11824)– |
| env-789 | SR_TTP_S3_1008655 | - | 875 | 808 | **RNA**→ rpsO (PRK05626)Ribosomal_S15 (pfam00312)→ |
| env-790 | SR_TP_S13sol_1003857 | + | 417 | 484 | **RNA**→ rpsO (PRK05626)Ribosomal_S15 (pfam00312)→ |
| env-791 | NCBI_READ_1112684383788 | - | 614 | 547 | **RNA**→ rpsO (PRK05626)Ribosomal_S15 (pfam00312)→ |
| env-792 | CEST01074805.1 | - | 430 | 363 | **RNA**→ rpsO (PRK05626)Ribosomal_S15 (pfam00312)→ |
| env-793 | CESU01086043.1 | - | 552 | 485 | **RNA**→ rpsO (PRK05626)Ribosomal_S15 (pfam00312)→ |
| env-794 | NCBI_READ_1112777362025 | + | 472 | 539 | **RNA**→ rpsO (PRK05626)Ribosomal_S15 (pfam00312)→ |
| env-795 | CESE01059308.1 | - | 72 | 5 | **RNA**→ |
| env-796 | NCBI_READ_1112684267655 | - | 819 | 752 | **RNA**→ hypo→ |
| env-797 | NCBI_READ_1112777442403 | - | 338 | 271 | **RNA**→ rpsO (PRK05626)Ribosomal_S15 (pfam00312)→ |
| env-798 | CESJ01108224.1 | - | 304 | 237 | **RNA**→ rpsO (PRK05626)Ribosomal_S15 (pfam00312)→ |
| env-799 | DelMOSum2010_c10152077 | + | 650 | 699 | **RNA**→ rpsO (PRK05626)Ribosomal_S15 (pfam00312)→ |
| Cca-4-1 | NZ_CDOL01000024.1 | - | 3056 | 3001 | **RNA**→ rpsO (PRK05626)RpsO (COG0184)→ |
| Cca-4-2 | NZ_CDOI01000148.1 | - | 32622 | 32567 | **RNA**→ rpsO (PRK05626)RpsO (COG0184)→ |
| Cca-4-3 | NZ_CDOH01000002.1 | + | 17823 | 17878 | **RNA**→ rpsO (PRK05626)RpsO (COG0184)→ |
| env-800 | JGI11876J14442_10315541 | - | 187 | 137 | **RNA**→ rpsO (PRK05626)Ribosomal_S15 (pfam00312)→ |
| env-801 | JCVI_SCAF_1101668147026 | + | 1128 | 1191 | **RNA**→ rpsO (PRK05626)Ribosomal_S15 (pfam00312)→ |
| env-802 | JCVI_SCAF_1101668641004 | - | 585 | 522 | **RNA**→ rpsO (PRK05626)RpsO (COG0184)→ |
| env-803 | KVRMV2_100003289 | + | 329 | 382 | **RNA**→ rpsO (PRK05626)RpsO (COG0184)→ |
| env-804 | KVRMV2_100403512 | + | 971 | 1024 | **RNA**→ rpsO (PRK05626)RpsO (COG0184)→ |
| env-805 | CERW01009190.1 | - | 617 | 563 | **RNA**→ rpsO (PRK05626)Ribosomal_S15 (pfam00312)→ PRK11824 (PRK11824)RNase_PH (pfam01138)→ |
| env-806 | CERK01116695.1 | + | 478 | 532 | **RNA**→ rpsO (PRK05626)Ribosomal_S15 (pfam00312)→ |
| env-807 | NCBI_BBAY_READ_1106105340888 | + | 777 | 830 | **RNA**→ |
| env-808 | CEVG01118472.1 | - | 579 | 526 | **RNA**→ rpsO (PRK05626)Ribosomal_S15 (pfam00312)→ RNase_PH_PNPase_1 (cd11363)PRK11824 (PRK11824)– |
| env-809 | CESM01310257.1 | - | 73 | 20 | **RNA**→ |
| Zga-1-1 | NC_015844.1 | + | 930103 | 930167 | **RNA**→ rpsO (PRK05626)RpsO (COG0184)→ |
| env-810 | JGI26524J50256_1000051 | + | 47029 | 47095 | **RNA**→ rpsO (PRK05626)Ribosomal_S15 (pfam00312)→ |
| env-811 | JGI26523J50269_1000086 | - | 81858 | 81792 | **RNA**→ rpsO (PRK05626)Ribosomal_S15 (pfam00312)→ |
| env-812 | JGI26534J51046_1001036 | + | 19681 | 19747 | **RNA**→ rpsO (PRK05626)Ribosomal_S15 (pfam00312)→ |
| env-813 | JGI25911J50253_10156824 | - | 177 | 112 | **RNA**→ |
| env-814 | JGI25910J50241_10411963 | + | 60 | 125 | **RNA**→ rpsO (PRK05626)Ribosomal_S15 (pfam00312)→ |
| env-815 | JGI25912J50252_10064814 | - | 743 | 678 | **RNA**→ rpsO (PRK05626)Ribosomal_S15 (pfam00312)→ |

| | | | | | |
|---|---|---|---|---|---|
| env-816 | JGI25908J49247_10047059 | - | 348 | 283 | RNA→ rpsO (PRK05626)Ribosomal_S15 (pfam00312)→ |
| env-817 | JGI25914J50564_10007880 | + | 233 | 298 | RNA→ rpsO (PRK05626)Ribosomal_S15 (pfam00312)→ |
| env-818 | JGI25920J50251_10000314 | - | 11641 | 11576 | RNA→ rpsO (PRK05626)Ribosomal_S15 (pfam00312)→ |
| env-819 | JGI25920J50251_10070705 | - | 70 | 5 | RNA→ |
| env-820 | JGI25914J50564_10014061 | - | 82 | 17 | RNA→ |
| env-821 | JCVI_SCAF_1101668133290 | + | 581 | 646 | RNA→ rpsO (PRK05626)Ribosomal_S15 (pfam00312)→ PRK11824 (PRK11824)RNase_PH (pfam01138)RNase_PH_C (pfam03725)→ |
| env-822 | JGI25920J50251_10005537 | + | 3465 | 3530 | RNA→ rpsO (PRK05626)Ribosomal_S15 (pfam00312)→ |
| env-823 | JGI12421J11937_10023408 | + | 486 | 551 | RNA→ rpsO (PRK05626)Ribosomal_S15 (pfam00312)→ |
| env-824 | CEWG01134755.1 | - | 547 | 492 | RNA→ rpsO (PRK05626)Ribosomal_S15 (pfam00312)→ PRK11824 (PRK11824)RNase_PH (pfam01138)→ |
| env-825 | DelMOSum2011_c10066067 | + | 20 | 85 | RNA→ rpsO (PRK05626)Ribosomal_S15 (pfam00312)→ PRK11824 (PRK11824)PNPase (pfam03726)RNase_PH (pfam01138)→ |
| env-826 | CESJ01016598.1 | + | 4458 | 4522 | RNA→ rpsO (PRK05626)Ribosomal_S15 (pfam00312)→ RNase_PH_PNPase_1 (cd11363)PRK11824 (PRK11824)– |
| env-827 | CESI01019537.1 | - | 314 | 249 | RNA→ rpsO (PRK05626)Ribosomal_S15 (pfam00312)→ |
| env-828 | JCVI_SCAF_1096627403573 | - | 332 | 267 | RNA→ rpsO (PRK05626)Ribosomal_S15 (pfam00312)→ |
| env-829 | PML_READ_01087893 | - | 217 | 151 | RNA→ rpsO (PRK05626)Ribosomal_S15 (pfam00312)→ |
| env-830 | PML_READ_00236953 | + | 109 | 174 | RNA→ rpsO (PRK05626)RpsO (COG0184)→ |
| env-831 | PML_READ_01045449 | - | 100 | 35 | RNA→ |
| env-832 | CESU01015641.1 | - | 388 | 323 | RNA→ rpsO (PRK05626)Ribosomal_S15 (pfam00312)→ |
| env-833 | LPaug08P2610m_c1025036 | - | 264 | 199 | RNA→ rpsO (PRK05626)Ribosomal_S15 (pfam00312)→ |
| env-834 | LPjun09P1210m_c1044378 | - | 232 | 167 | RNA→ rpsO (PRK05626)Ribosomal_S15 (pfam00312)→ |
| env-835 | DelMOSum2010_c10056809 | - | 1201 | 1136 | RNA→ rpsO (PRK05626)RpsO (COG0184)→ PRK11824 (PRK11824)RNase_PH (pfam01138)RNase_PH_C (pfam03725)→ |
| env-836 | SI60aug11_200m_c1181697 | + | 116 | 180 | RNA→ rpsO (PRK05626)Ribosomal_S15 (pfam00312)→ |
| env-837 | SI36aug09_120m_1049180 | + | 473 | 537 | RNA→ rpsO (PRK05626)Ribosomal_S15 (pfam00312)→ |
| env-838 | SI47jul10_100m_c1026668 | - | 558 | 494 | RNA→ rpsO (PRK05626)Ribosomal_S15 (pfam00312)→ RNase_PH_PNPase_1 (cd11363)PRK11824 (PRK11824)– |
| env-839 | SI60aug11_100m_c1000082 | + | 8859 | 8923 | RNA→ rpsO (PRK05626)Ribosomal_S15 (pfam00312)→ Pnp (COG1185)PRK11824 (PRK11824)→ RpoD (COG0568)sigma70-ECF (TIGR02937)bSCRAP40 (bSCRAP40)bSCRAP106 (bSCRAP106)→ PRK05581 (PRK05581)RPE (cd00429)→ anmK (PRK09585)AnmK (pfam03702)→ |
| env-840 | SI48aug10_10m_c1011319 | + | 396 | 460 | RNA→ rpsO (PRK05626)Ribosomal_S15 (pfam00312)→ |
| env-841 | SI34jun09_10m_1000012 | + | 25879 | 25943 | RNA→ rpsO (PRK05626)Ribosomal_S15 (pfam00312)→ Pnp (COG1185)PRK11824 (PRK11824)→ RpoD (COG0568)sigma70-ECF (TIGR02937)bSCRAP40 (bSCRAP40)bSCRAP106 (bSCRAP106)→ PRK05581 (PRK05581)RPE (cd00429)→ anmK (PRK09585)AnmK (pfam03702)→ |
| env-842 | SI34jun09_120m_1001580 | - | 5416 | 5352 | RNA→ rpsO (PRK05626)Ribosomal_S15 (pfam00312)→ Pnp (COG1185)PRK11824 (PRK11824)→ RpoD (COG0568)sigma70-ECF (TIGR02937)bSCRAP40 (bSCRAP40)bSCRAP106 (bSCRAP106)→ PRK05581 (PRK05581)RPE (cd00429)→ anmK (PRK09585)AnmK (pfam03702)→ |
| env-843 | SI34jun09_135m_1005514 | + | 3165 | 3229 | RNA→ rpsO (PRK05626)Ribosomal_S15 (pfam00312)→ Pnp (COG1185)PRK11824 (PRK11824)→ |
| env-844 | SI36aug09_100m_1014540 | - | 871 | 807 | RNA→ rpsO (PRK05626)Ribosomal_S15 (pfam00312)→ PRK11824 (PRK11824)RNase_PH (pfam01138)→ |
| env-845 | SI34jun09_100m_1000069 | - | 38756 | 38692 | RNA→ rpsO (PRK05626)Ribosomal_S15 (pfam00312)→ Pnp (COG1185)PRK11824 (PRK11824)→ RpoD (COG0568)sigma70-ECF (TIGR02937)bSCRAP40 (bSCRAP40)bSCRAP106 (bSCRAP106)→ PRK05581 (PRK05581)RPE (cd00429)→ anmK (PRK09585)AnmK (pfam03702)→ |
| env-846 | SI48aug10_100m_1002881 | - | 383 | 319 | RNA→ rpsO (PRK05626)Ribosomal_S15 (pfam00312)→ |
| env-847 | DelMOSum2011_c10015265 | - | 605 | 540 | RNA→ rpsO (PRK05626)Ribosomal_S15 (pfam00312)→ PRK11824 (PRK11824)RNase_PH (pfam01138)→ |
| env-848 | JGI20157J14317_10011885 | - | 386 | 321 | RNA→ rpsO (PRK05626)Ribosomal_S15 (pfam00312)→ |
| env-849 | DelMOSum2010_c10060176 | - | 479 | 414 | RNA→ rpsO (PRK05626)Ribosomal_S15 (pfam00312)→ hypo→ |
| env-850 | SI34jun09_100m_1160428 | + | 29 | 94 | RNA→ rpsO (PRK05626)Ribosomal_S15 (pfam00312)→ |
| env-851 | JGI20156J14371_10000759 | + | 27598 | 27663 | RNA→ rpsO (PRK05626)Ribosomal_S15 (pfam00312)→ |
| env-852 | JGI20157J14317_10012691 | - | 388 | 323 | RNA→ rpsO (PRK05626)Ribosomal_S15 (pfam00312)→ |
| env-853 | JGI20155J14468_10000146 | - | 524 | 459 | RNA→ rpsO (PRK05626)Ribosomal_S15 (pfam00312)→ RNase_PH_PNPase_1 (cd11363)PRK11824 (PRK11824)– |
| env-854 | CESO01040974.1 | + | 1915 | 1980 | RNA→ |
| env-855 | CEWO01311710.1 | + | 677 | 742 | RNA→ |
| env-856 | 2236876009_219883 | - | 133 | 67 | RNA→ rpsO (PRK05626)RpsO (COG0184)→ |
| env-857 | SI39nov09_10m_c1013699 | + | 177 | 242 | RNA→ rpsO (PRK05626)Ribosomal_S15 (pfam00312)→ RNase_PH_PNPase_1 (cd11363)PRK11824 (PRK11824)– |
| env-858 | SI48aug10_10m_c1001560 | - | 392 | 327 | RNA→ rpsO (PRK05626)Ribosomal_S15 (pfam00312)→ |

| | | | | | |
|---|---|---|---|---|---|
| env-859 | SI48aug10_100m_1003033 | + | 2938 | 3003 | **RNA**→ rpsO (PRK05626)Ribosomal_S15 (pfam00312)→ Pnp (COG1185)PRK11824 (PRK11824)→ |
| env-860 | JGI20152J14361_10014083 | - | 305 | 240 | **RNA**→ rpsO (PRK05626)Ribosomal_S15 (pfam00312)→ |
| env-861 | JGI20160J14292_10040223 | + | 2051 | 2116 | **RNA**→ rpsO (PRK05626)Ribosomal_S15 (pfam00312)→ |
| env-862 | JGI20159J14440_10514398 | - | 304 | 239 | **RNA**→ rpsO (PRK05626)Ribosomal_S15 (pfam00312)→ |
| env-863 | CESF01137326.1 | + | 1896 | 1961 | **RNA**→ |
| env-864 | CETY01149387.1 | - | 339 | 274 | **RNA**→ rpsO (PRK05626)Ribosomal_S15 (pfam00312)→ |
| env-865 | SI54feb11_100m_c1083680 | - | 282 | 217 | **RNA**→ rpsO (PRK05626)RpsO (COG0184)→ |
| env-866 | JGI20156J14371_10129797 | - | 113 | 48 | **RNA**→ |
| env-867 | JGI20155J14468_10151635 | + | 628 | 693 | **RNA**→ |
| env-868 | JGI20160J14292_10059861 | - | 181 | 116 | **RNA**→ rpsO (PRK05626)Ribosomal_S15 (pfam00312)→ |
| env-869 | JCVI_SCAF_1101668107239 | + | 814 | 879 | **RNA**→ rpsO (PRK05626)Ribosomal_S15 (pfam00312)→ |
| env-870 | JGI20154J14316_10048683 | - | 1152 | 1087 | **RNA**→ rpsO (PRK05626)Ribosomal_S15 (pfam00312)→ |
| | | | | | PRK11824 (PRK11824)RNase_PH (pfam01138)RNase_PH_C (pfam03725)→ |
| env-871 | JGI20153J14295_10057124 | - | 943 | 878 | **RNA**→ rpsO (PRK05626)Ribosomal_S15 (pfam00312)→ PRK11824 (PRK11824)RNase_PH (pfam01138)→ |
| env-872 | JGI20153J14318_10057124 | - | 943 | 878 | **RNA**→ rpsO (PRK05626)Ribosomal_S15 (pfam00312)→ PRK11824 (PRK11824)RNase_PH (pfam01138)→ |
| env-873 | JGI20159J14440_10010099 | + | 2098 | 2163 | **RNA**→ rpsO (PRK05626)Ribosomal_S15 (pfam00312)→ Pnp (COG1185)PRK11824 (PRK11824)→ |
| | | | | | sigma70-ECF (TIGR02937)Sigma70_r1_2 (pfam00140)Sigma70_r2 (pfam04542)→ |
| env-874 | JGI20158J14315_10004959 | - | 6347 | 6282 | **RNA**→ rpsO (PRK05626)Ribosomal_S15 (pfam00312)→ PRK11824 (PRK11824)RNase_PH (pfam01138)→ |
| | | | | | S1 (smart00316)KH (smart00322)Pnp (COG1185)→ |
| | | | | | RpoD (COG0568)sigma70-ECF (TIGR02937)bSCRAP40 (bSCRAP40)bSCRAP106 (bSCRAP106)→ |
| | | | | | PRK05581 (PRK05581)RPE (cd00429)→ anmK (PRK09585)AnmK (pfam03702)→ |
| env-875 | CEOP01365269.1 | - | 98 | 33 | **RNA**→ |
| env-876 | DelMOWin2010_c10528752 | + | 190 | 255 | **RNA**→ rpsO (PRK05626)→ |
| env-877 | JGI20156J14371_10036705 | + | 2103 | 2168 | **RNA**→ rpsO (PRK05626)Ribosomal_S15 (pfam00312)→ |
| env-878 | JGI20160J14292_10049262 | - | 1686 | 1621 | **RNA**→ rpsO (PRK05626)Ribosomal_S15 (pfam00312)→ PRK11824 (PRK11824)RNase_PH (pfam01138)→ |
| | | | | | PRK11824 (PRK11824)RNase_PH (pfam01138)→ |
| env-879 | JGI20157J14317_10057593 | - | 704 | 639 | **RNA**→ rpsO (PRK05626)Ribosomal_S15 (pfam00312)→ PRK11824 (PRK11824)RNase_PH (pfam01138)→ |
| env-880 | JGI20155J14468_10064733 | - | 1016 | 951 | **RNA**→ rpsO (PRK05626)Ribosomal_S15 (pfam00312)→ |
| | | | | | PRK11824 (PRK11824)RNase_PH (pfam01138)RNase_PH_C (pfam03725)→ |
| env-881 | JGI20152J14361_10000414 | - | 3697 | 3632 | **RNA**→ rpsO (PRK05626)Ribosomal_S15 (pfam00312)→ Pnp (COG1185)PRK11824 (PRK11824)→ |
| | | | | | RpoD (COG0568)sigma70-ECF (TIGR02937)bSCRAP40 (bSCRAP40)bSCRAP106 (bSCRAP106)→ |
| | | | | | PRK05581 (PRK05581)RPE (cd00429)→ |
| env-882 | JGI20151J14362_10002476 | + | 4075 | 4140 | **RNA**→ rpsO (PRK05626)Ribosomal_S15 (pfam00312)→ Pnp (COG1185)PRK11824 (PRK11824)→ |
| | | | | | RpoD (COG0568)sigma70-ECF (TIGR02937)bSCRAP40 (bSCRAP40)bSCRAP106 (bSCRAP106)→ |
| | | | | | PRK05581 (PRK05581)RPE (cd00429)→ anmK (PRK09585)AnmK (pfam03702)→ |
| env-883 | 2236876001_196002 | - | 290 | 236 | **RNA**→ rpsO (PRK05626)Ribosomal_S15 (pfam00312)→ |
| env-884 | JGI20155J14468_10058899 | + | 1110 | 1164 | **RNA**→ rpsO (PRK05626)Ribosomal_S15 (pfam00312)→ |
| env-885 | JGI20158J14315_10054641 | - | 1345 | 1291 | **RNA**→ rpsO (PRK05626)Ribosomal_S15 (pfam00312)→ |
| | | | | | PRK11824 (PRK11824)PNPase (pfam03726)RNase_PH (pfam01138)RNase_PH_C (pfam03725)→ |
| env-886 | CENT01010038.1 | + | 1338 | 1392 | **RNA**→ rpsO (PRK05626)Ribosomal_S15 (pfam00312)→ |
| env-887 | CESF01070596.1 | + | 472 | 526 | **RNA**→ rpsO (PRK05626)Ribosomal_S15 (pfam00312)→ |
| env-888 | CESO01000708.1 | - | 1675 | 1621 | **RNA**→ rpsO (PRK05626)Ribosomal_S15 (pfam00312)→ Pnp (COG1185)PRK11824 (PRK11824)→ |
| env-889 | CEVU01160017.1 | + | 113 | 167 | **RNA**→ rpsO (PRK05626)Ribosomal_S15 (pfam00312)→ PRK11824 (PRK11824)RNase_PH (pfam01138)→ |
| env-890 | CEWO01095888.1 | - | 470 | 416 | **RNA**→ rpsO (PRK05626)Ribosomal_S15 (pfam00312)→ hypo→ |
| env-891 | JCVI_SCAF_1096627428780 | - | 347 | 293 | **RNA**→ rpsO (PRK05626)Ribosomal_S15 (pfam00312)→ |
| env-892 | JCVI_SCAF_1096627340448 | - | 710 | 656 | **RNA**→ rpsO (PRK05626)Ribosomal_S15 (pfam00312)→ PRK11824 (PRK11824)RNase_PH (pfam01138)→ |
| env-893 | CEWG01034068.1 | - | 8107 | 8053 | **RNA**→ rpsO (PRK05626)Ribosomal_S15 (pfam00312)→ Pnp (COG1185)PRK11824 (PRK11824)→ |
| | | | | | RpoD (COG0568)sigma70-ECF (TIGR02937)→ PRK05581 (PRK05581)RPE (cd00429)→ |
| | | | | | AnmK (COG2377)anmK (PRK09585)→ |
| env-894 | DelMOWin2010_c10023670 | + | 1770 | 1824 | **RNA**→ rpsO (PRK05626)Ribosomal_S15 (pfam00312)→ |
| | | | | | PRK11824 (PRK11824)PNPase (pfam03726)RNase_PH (pfam01138)RNase_PH_C (pfam03725)→ |
| env-895 | CENF01414486.1 | + | 550 | 604 | **RNA**→ rpsO (PRK05626)Ribosomal_S15 (pfam00312)→ |
| | | | | | PRK11824 (PRK11824)PNPase (pfam03726)RNase_PH (pfam01138)RNase_PH_C (pfam03725)→ |
| env-896 | CEOP01028619.1 | - | 1417 | 1363 | **RNA**→ rpsO (PRK05626)Ribosomal_S15 (pfam00312)→ |
| | | | | | PRK11824 (PRK11824)RNase_PH (pfam01138)RNase_PH_C (pfam03725)→ |

| | | | | | |
|---|---|---|---|---|---|
| env-897 | CESM01030051.1 | - | 2678 | 2624 | **RNA**→ rpsO (PRK05626)Ribosomal_S15 (pfam00312)→ Pnp (COG1185)PRK11824 (PRK11824)→ |
| env-898 | CETY01071453.1 | - | 87 | 33 | **RNA**→ |
| env-899 | CEVX01269025.1 | - | 90 | 36 | **RNA**→ |
| env-900 | JCVI_SCAF_1096627909011 | + | 651 | 705 | **RNA**→ PRK11824 (PRK11824)RNase_PH (pfam01138)RNase_PH_C (pfam03725)→ |
| env-901 | LPaug09P1610m_c1031693 | - | 129 | 73 | **RNA**→ rpsO (PRK05626)→ |
| env-902 | LPjun09P1210m_c1013475 | - | 432 | 376 | **RNA**→ rpsO (PRK05626)Ribosomal_S15 (pfam00312)→ |
| env-903 | DelMOSum2010_c10000068 | + | 18253 | 18310 | **RNA**→ rpsO (PRK05626)Ribosomal_S15 (pfam00312)→ |
| env-904 | A09P0410a_Sequence0000021066 | - | 462 | 405 | **RNA**→ rpsO (PRK05626)Ribosomal_S15 (pfam00312)→ |
| env-905 | JGI20157J14317_10344599 | - | 183 | 126 | **RNA**→ rpsO (PRK05626)Ribosomal_S15 (pfam00312)→ |
| env-906 | JGI25920J50251_10011692 | - | 2092 | 2026 | **RNA**→ rpsO (PRK05626)Ribosomal_S15 (pfam00312)→ |
| env-907 | JGI25911J50253_10208029 | - | 174 | 108 | **RNA**→ |
| env-908 | JGI25910J50241_10133287 | - | 519 | 453 | **RNA**→ rpsO (PRK05626)Ribosomal_S15 (pfam00312)→ hypo→ |
| Fge-1-1 | NZ_KE383905.1 | - | 172236 | 172170 | **RNA**→ rpsO (PRK05626)Ribosomal_S15 (pfam00312)→ |
| Ale-2-1 | NZ_KN050846.1 | - | 2805051 | 2804985 | **RNA**→ rpsO (PRK05626)Ribosomal_S15 (pfam00312)→ |
| env-909 | PBR_1019653 | - | 5944 | 5878 | **RNA**→ rpsO (PRK05626)Ribosomal_S15 (pfam00312)→ |
| env-910 | JGI24723J26617_10002167 | - | 11651 | 11585 | **RNA**→ rpsO (PRK05626)Ribosomal_S15 (pfam00312)→ |
| env-911 | JCVI_SCAF_1101668691680 | + | 126 | 192 | **RNA**→ rpsO (PRK05626)Ribosomal_S15 (pfam00312)→ |
| env-912 | CEWG01179869.1 | + | 230 | 296 | **RNA**→ rpsO (PRK05626)Ribosomal_S15 (pfam00312)→ |
| env-913 | CETK01031226.1 | - | 238 | 171 | **RNA**→ rpsO (PRK05626)Ribosomal_S15 (pfam00312)→ |
| env-914 | CESF01019749.1 | - | 849 | 782 | **RNA**→ rpsO (PRK05626)Ribosomal_S15 (pfam00312)→ |
| env-915 | CESO01019946.1 | + | 842 | 909 | **RNA**→ rpsO (PRK05626)Ribosomal_S15 (pfam00312)→ |
| env-916 | CEUH01062216.1 | - | 338 | 272 | **RNA**→ rpsO (PRK05626)Ribosomal_S15 (pfam00312)→ |
| Man-1-1 | NZ_KK211215.1 | - | 65801 | 65738 | **RNA**→ rpsO (PRK05626)RpsO (COG0184)→ |
| env-917 | JGI20156J14371_10007079 | - | 147 | 90 | **RNA**→ rpsO (PRK05626)→ |
| env-918 | JGI20160J14292_10016935 | - | 147 | 90 | **RNA**→ rpsO (PRK05626)→ |
| env-919 | CEVW01069772.1 | - | 669 | 612 | **RNA**→ rpsO (PRK05626)Ribosomal_S15 (pfam00312)→ PRK11824 (PRK11824)RNase_PH (pfam01138)→ |
| env-920 | JGI20156J14371_10415361 | + | 33 | 90 | **RNA**→ rpsO (PRK05626)Ribosomal_S15 (pfam00312)→ |
| env-921 | JGI20160J14292_10453555 | - | 300 | 243 | **RNA**→ rpsO (PRK05626)Ribosomal_S15 (pfam00312)→ |
| env-922 | JGI20155J14468_10000239 | - | 6353 | 6296 | **RNA**→ rpsO (PRK05626)Ribosomal_S15 (pfam00312)→ Pnp (COG1185)PRK11824 (PRK11824)→ |
| env-923 | DelMOSum2010_c10011135 | - | 3298 | 3241 | **RNA**→ rpsO (PRK05626)Ribosomal_S15 (pfam00312)→ Pnp (COG1185)PRK11824 (PRK11824)→ |
| env-924 | DelMOSum2010_c10190970 | + | 126 | 183 | **RNA**→ rpsO (PRK05626)Ribosomal_S15 (pfam00312)→ RNase_PH_PNPase_1 (cd11363)PRK11824 (PRK11824)- |
| env-925 | JGI20156J14371_10100972 | + | 860 | 917 | **RNA**→ |
| env-926 | JGI20160J14292_10127509 | - | 69 | 12 | **RNA**→ |
| env-927 | JGI20159J14440_10000775 | - | 3153 | 3096 | **RNA**→ rpsO (PRK05626)Ribosomal_S15 (pfam00312)→ Pnp (COG1185)PRK11824 (PRK11824)→ |
| env-928 | CEUW01210784.1 | + | 690 | 747 | **RNA**→ |
| env-929 | CEUK01207791.1 | + | 22 | 79 | **RNA**→ rpsO (PRK05626)Ribosomal_S15 (pfam00312)→ RNase_PH_PNPase_1 (cd11363)PRK11824 (PRK11824)- |
| env-930 | CEUU01245541.1 | - | 1236 | 1179 | **RNA**→ rpsO (PRK05626)Ribosomal_S15 (pfam00312)→ RNase_PH_PNPase_1 (cd11363)PRK11824 (PRK11824)- |
| env-931 | CEVF01306501.1 | - | 4315 | 4258 | **RNA**→ rpsO (PRK05626)Ribosomal_S15 (pfam00312)→ Pnp (COG1185)PRK11824 (PRK11824)→ |
| env-932 | CEQD01093616.1 | + | 62 | 120 | **RNA**→ rpsO (PRK05626)Ribosomal_S15 (pfam00312)→ |
| | | | | | PRK11824 (PRK11824)RNase_PH (pfam01138)RNase_PH_C (pfam03725)→ |
| env-933 | Y607_1006247 | + | 254177 | 254245 | **RNA**→ rpsO (PRK05626)RpsO (COG0184)→ |
| env-934 | 3300000558_11729233 | - | 567 | 500 | **RNA**→ rpsO (PRK05626)RpsO (COG0184)→ |
| env-935 | 3300001605_10025997 | - | 1325 | 1258 | **RNA**→ rpsO (PRK05626)RpsO (COG0184)→ |
| env-936 | AYRH01021120.1 | - | 183 | 121 | **RNA**→ rpsO (PRK05626)Ribosomal_S15 (pfam00312)→ |
| env-937 | CEWO01187126.1 | - | 947 | 880 | **RNA**→ rpsO (PRK05626)Ribosomal_S15 (pfam00312)→ |
| env-938 | CEVS01341770.1 | - | 695 | 640 | **RNA**→ rpsO (PRK05626)Ribosomal_S15 (pfam00312)→ PRK11824 (PRK11824)RNase_PH (pfam01138)→ |
| env-939 | CEVW01355221.1 | - | 618 | 563 | **RNA**→ rpsO (PRK05626)Ribosomal_S15 (pfam00312)→ PRK11824 (PRK11824)RNase_PH (pfam01138)→ |
| env-940 | CEQM01182343.1 | + | 515 | 570 | **RNA**→ |
| env-941 | CEQN01033481.1 | - | 720 | 665 | **RNA**→ rpsO (PRK05626)Ribosomal_S15 (pfam00312)→ PRK11824 (PRK11824)RNase_PH (pfam01138)→ |
| env-942 | CESQ01380936.1 | - | 103 | 48 | **RNA**→ |
| env-943 | CEVP01035867.1 | + | 331 | 386 | **RNA**→ rpsO (PRK05626)Ribosomal_S15 (pfam00312)→ PRK11824 (PRK11824)RNase_PH (pfam01138)→ |
| env-944 | CEQO01283396.1 | + | 1289 | 1344 | **RNA**→ rpsO (PRK05626)Ribosomal_S15 (pfam00312)→ |

| | | | | | |
|---|---|---|---|---|---|
| env-945 | CERS01106140.1 | + | 701 | 756 | **RNA**→ rpsO (PRK05626)Ribosomal_S15 (pfam00312)→ |
| env-946 | CEUA01394830.1 | + | 729 | 784 | **RNA**→ |
| Tma-1-1 | NZ_BAUG01000053.1 | - | 3972 | 3908 | **RNA**→ rpsO (PRK05626)Ribosomal_S15 (pfam00312)→ |
| env-947 | JGI25920J50251_10315894 | + | 94 | 151 | **RNA**→ rpsO (PRK05626)Ribosomal_S15 (pfam00312)→ |
| Fsp-12-1 | NZ_AUEU01000012.1 | + | 243905 | 243971 | **RNA**→ rpsO (PRK05626)Ribosomal_S15 (pfam00312)→ |
| Fde-1-1 | NZ_AUDL01000013.1 | - | 39996 | 39930 | **RNA**→ rpsO (PRK05626)Ribosomal_S15 (pfam00312)→ |
| Fba-7-1 | NZ_AFPB01000144.1 | - | 4726 | 4662 | **RNA**→ rpsO (PRK05626)RpsO (COG0184)→ rpsO (PRK05626)RpsO (COG0184)→ |
| Ale-1-1 | NZ_BBNQ01000031.1 | - | 6559 | 6493 | **RNA**→ rpsO (PRK05626)Ribosomal_S15 (pfam00312)→ |
| Ale-1-2 | NZ_BBNU01000036.1 | + | 8531 | 8597 | **RNA**→ rpsO (PRK05626)Ribosomal_S15 (pfam00312)→ |
| Ffr-1-1 | NZ_AUDO01000008.1 | - | 223593 | 223528 | **RNA**→ rpsO (PRK05626)Ribosomal_S15 (pfam00312)→ |
| env-948 | JGI25921J50272_10193124 | - | 177 | 111 | **RNA**→ rpsO (PRK05626)RpsO (COG0184)→ |
| Fsp-1-1 | NZ_JYGZ01000002.1 | - | 102653 | 102588 | **RNA**→ rpsO (PRK05626)Ribosomal_S15 (pfam00312)→ |
| env-949 | 3300001592_10204136 | - | 110 | 44 | **RNA**→ |
| env-950 | M3P_c10003408 | - | 87 | 21 | **RNA**→ |
| Fsa-2-1 | NZ_JMLU01000001.1 | + | 65197 | 65263 | **RNA**→ rpsO (PRK05626)Ribosomal_S15 (pfam00312)→ |
| env-951 | 3300001605_10665184 | + | 444 | 510 | **RNA**→ |
| env-952 | 3300001605_10033828 | + | 4228 | 4294 | **RNA**→ rpsO (PRK05626)Ribosomal_S15 (pfam00312)→ |
| Fba-3-1 | NZ_AAXX01000002.1 | - | 363051 | 362985 | **RNA**→ rpsO (PRK05626)Ribosomal_S15 (pfam00312)→ rpsO (PRK05626)Ribosomal_S15 (pfam00312)→ |
| env-953 | TB_FS06_10DRAFT_1000175 | - | 6311 | 6245 | **RNA**→ rpsO (PRK05626)Ribosomal_S15 (pfam00312)→ |
| env-954 | 3300001605_10453220 | - | 110 | 44 | **RNA**→ |
| env-955 | CESI01049345.1 | + | 1523 | 1589 | **RNA**→ rpsO (PRK05626)Ribosomal_S15 (pfam00312)→ |
| env-956 | CEST01010500.1 | - | 551 | 485 | **RNA**→ rpsO (PRK05626)Ribosomal_S15 (pfam00312)→ |
| env-957 | NCBI_READ_1112788336194 | + | 594 | 660 | **RNA**→ |
| env-958 | JCVI_SCAF_1101667195802 | - | 126 | 70 | **RNA**→ hypo→ |
| env-959 | SRS018157_Baylor_scaffold_12805 | + | 66 | 120 | **RNA**→ rpsO (PRK05626)Ribosomal_S15 (pfam00312)→ |
| Csp-1-1 | NZ_ABZV01000001.1 | + | 56329 | 56384 | **RNA**→ rpsO (PRK05626)Ribosomal_S15 (pfam00312)→ rpsO (PRK05626)Ribosomal_S15 (pfam00312)→ |
| env-960 | SRS024289_LANL_scaffold_56301 | + | 344 | 399 | **RNA**→ rpsO (PRK05626)→ |
| env-961 | SRS051244_LANL_scaffold_25997 | + | 197 | 252 | **RNA**→ rpsO (PRK05626)Ribosomal_S15 (pfam00312)→ |
| Csp-3-1 | NZ_KB291394.1 | - | 29522 | 29467 | **RNA**→ rpsO (PRK05626)Ribosomal_S15 (pfam00312)→ |
| env-962 | SRS021496_Baylor_scaffold_73949 | - | 2767 | 2712 | **RNA**→ rpsO (PRK05626)Ribosomal_S15 (pfam00312)→ |
| env-963 | SRS022530_LANL_scaffold_20874 | - | 38233 | 38178 | **RNA**→ rpsO (PRK05626)Ribosomal_S15 (pfam00312)→ |
| env-964 | SRS063288_LANL_scaffold_3750 | - | 772 | 717 | **RNA**→ rpsO (PRK05626)Ribosomal_S15 (pfam00312)→ |
| env-965 | SRS011098_Baylor_scaffold_11988 | + | 19624 | 19680 | **RNA**→ rpsO (PRK05626)Ribosomal_S15 (pfam00312)→ |
| env-966 | SRS023938_Baylor_scaffold_45086 | + | 428 | 483 | **RNA**→ |
| env-967 | SRS014573_C3519028 | + | 210 | 265 | **RNA**→ rpsO (PRK05626)→ |
| env-968 | SRS015434_C5951484 | + | 1370 | 1425 | **RNA**→ rpsO (PRK05626)Ribosomal_S15 (pfam00312)→ |
| env-969 | SRS018439_Baylor_scaffold_4715 | + | 14608 | 14663 | **RNA**→ rpsO (PRK05626)Ribosomal_S15 (pfam00312)→ |
| env-970 | SRS022719_C2913748 | - | 436 | 381 | **RNA**→ rpsO (PRK05626)Ribosomal_S15 (pfam00312)→ |
| env-971 | SRS024441_LANL_scaffold_31186 | + | 18667 | 18722 | **RNA**→ rpsO (PRK05626)Ribosomal_S15 (pfam00312)→ |
| env-972 | SRS015470_WUGC_scaffold_17737 | + | 950 | 1005 | **RNA**→ rpsO (PRK05626)Ribosomal_S15 (pfam00312)→ |
| env-973 | SRS015947_WUGC_scaffold_20781 | - | 17604 | 17549 | **RNA**→ rpsO (PRK05626)Ribosomal_S15 (pfam00312)→ |
| env-974 | SRS016092_WUGC_scaffold_7783 | - | 4221 | 4166 | **RNA**→ rpsO (PRK05626)Ribosomal_S15 (pfam00312)→ |
| env-975 | SRS019591_WUGC_scaffold_41796 | + | 10308 | 10363 | **RNA**→ rpsO (PRK05626)Ribosomal_S15 (pfam00312)→ |
| env-976 | SRS017814_C2664019 | - | 331 | 276 | **RNA**→ rpsO (PRK05626)Ribosomal_S15 (pfam00312)→ |
| env-977 | SRS019029_C3201897 | + | 280 | 335 | **RNA**→ rpsO (PRK05626)Ribosomal_S15 (pfam00312)→ |
| env-978 | SRS019077_C2652316 | + | 764 | 819 | **RNA**→ rpsO (PRK05626)→ |
| env-979 | SRS051378_C1874895 | - | 114 | 59 | **RNA**→ rpsO (PRK05626)→ |
| env-980 | SRS019127_WUGC_scaffold_15951 | - | 88 | 33 | **RNA**→ |
| env-981 | SRS055450_LANL_scaffold_46997 | + | 520 | 575 | **RNA**→ |
| env-982 | SRS019387_C1738776 | + | 554 | 609 | **RNA**→ |
| env-983 | SRS020862_Baylor_scaffold_705 | + | 10311 | 10366 | **RNA**→ rpsO (PRK05626)Ribosomal_S15 (pfam00312)→ |
| env-984 | SRS024021_Baylor_scaffold_18404 | + | 1381 | 1436 | **RNA**→ rpsO (PRK05626)Ribosomal_S15 (pfam00312)→ |

| | | | | | |
|---|---|---|---|---|---|
| env-985 | SRS047100_C2096794 | - | 1382 | 1327 | **RNA**→ rpsO (PRK05626)Ribosomal_S15 (pfam00312)→ |
| env-986 | SRS050628_LANL_scaffold_20741 | + | 893 | 948 | **RNA**→ rpsO (PRK05626)Ribosomal_S15 (pfam00312)→ |
| env-987 | SRS065310_LANL_scaffold_14848 | - | 379 | 324 | **RNA**→ rpsO (PRK05626)Ribosomal_S15 (pfam00312)→ |
| env-988 | SRS063603_LANL_scaffold_3555 | - | 469 | 414 | **RNA**→ rpsO (PRK05626)Ribosomal_S15 (pfam00312)→ |
| env-989 | SRS015540_C1142881 | + | 29 | 84 | **RNA**→ rpsO (PRK05626)Ribosomal_S15 (pfam00312)→ |
| env-990 | SRS013879_WUGC_scaffold_44390 | - | 1424 | 1369 | **RNA**→ rpsO (PRK05626)Ribosomal_S15 (pfam00312)→ |
| env-991 | SRS015797_WUGC_scaffold_12357 | - | 4177 | 4122 | **RNA**→ rpsO (PRK05626)Ribosomal_S15 (pfam00312)→ |
| env-992 | SRS023926_Baylor_scaffold_18801 | - | 32149 | 32094 | **RNA**→ rpsO (PRK05626)Ribosomal_S15 (pfam00312)→ |
| env-993 | SRS014888_C3106033 | - | 1913 | 1858 | **RNA**→ rpsO (PRK05626)Ribosomal_S15 (pfam00312)→ |
| env-994 | SRS015440_WUGC_scaffold_28748 | + | 418 | 473 | **RNA**→ rpsO (PRK05626)Ribosomal_S15 (pfam00312)→ |
| env-995 | SRS015899_WUGC_scaffold_18506 | + | 1223 | 1278 | **RNA**→ rpsO (PRK05626)Ribosomal_S15 (pfam00312)→ |
| env-996 | SRS018145_Baylor_scaffold_48780 | - | 786 | 731 | **RNA**→ rpsO (PRK05626)Ribosomal_S15 (pfam00312)→ |
| env-997 | SRS019225_WUGC_scaffold_29613 | + | 4319 | 4374 | **RNA**→ rpsO (PRK05626)Ribosomal_S15 (pfam00312)→ |
| env-998 | SRS057692_LANL_scaffold_3535 | + | 4937 | 4992 | **RNA**→ rpsO (PRK05626)Ribosomal_S15 (pfam00312)→ |
| env-999 | SRS016037_WUGC_scaffold_44207 | + | 7031 | 7086 | **RNA**→ rpsO (PRK05626)Ribosomal_S15 (pfam00312)→ |
| env-1000 | SRS016225_C2977708 | + | 383 | 438 | **RNA**→ rpsO (PRK05626)Ribosomal_S15 (pfam00312)→ |
| env-1001 | SRS017511_C3344598 | + | 142 | 197 | **RNA**→ rpsO (PRK05626)Ribosomal_S15 (pfam00312)→ |
| env-1002 | SRS018573_WUGC_scaffold_30691 | + | 507 | 562 | **RNA**→ rpsO (PRK05626)RpsO (COG0184)→ |
| env-1003 | SRS019073_WUGC_scaffold_15301 | - | 330 | 275 | **RNA**→ rpsO (PRK05626)Ribosomal_S15 (pfam00312)→ |
| env-1004 | SRS019122_C5920199 | + | 1857 | 1912 | **RNA**→ rpsO (PRK05626)Ribosomal_S15 (pfam00312)→ |
| env-1005 | SRS019333_C1243671 | + | 173 | 228 | **RNA**→ rpsO (PRK05626)RpsO (COG0184)→ |
| env-1006 | SRS020340_C2878267 | + | 340 | 395 | **RNA**→ |
| env-1007 | SRS021477_Baylor_scaffold_40483 | + | 1827 | 1882 | **RNA**→ rpsO (PRK05626)Ribosomal_S15 (pfam00312)→ |
| env-1008 | SRS024561_LANL_scaffold_37452 | + | 268 | 323 | **RNA**→ rpsO (PRK05626)RpsO (COG0184)→ |
| env-1009 | SRS047265_WUGC_scaffold_19746 | - | 470 | 415 | **RNA**→ rpsO (PRK05626)Ribosomal_S15 (pfam00312)→ hypo→ |
| env-1010 | SRS055401_LANL_scaffold_34453 | + | 585 | 640 | **RNA**→ rpsO (PRK05626)→ |
| env-1011 | SRS053630_C928372 | - | 3451 | 3396 | **RNA**→ rpsO (PRK05626)Ribosomal_S15 (pfam00312)→ |
| env-1012 | CEOK01091424.1 | + | 379 | 434 | **RNA**→ rpsO (PRK05626)Ribosomal_S15 (pfam00312)→ |
| env-1013 | CENG01166087.1 | + | 243 | 298 | **RNA**→ rpsO (PRK05626)Ribosomal_S15 (pfam00312)→ |
| env-1014 | CENJ01054220.1 | + | 696 | 751 | **RNA**→ rpsO (PRK05626)Ribosomal_S15 (pfam00312)→ PRK11824 (PRK11824)RNase_PH (pfam01138)→ |
| env-1015 | CENX01011707.1 | + | 917 | 972 | **RNA**→ rpsO (PRK05626)Ribosomal_S15 (pfam00312)→ |
| env-1016 | CENP01029996.1 | + | 133 | 188 | **RNA**→ rpsO (PRK05626)Ribosomal_S15 (pfam00312)→ PRK11824 (PRK11824)RNase_PH (pfam01138)→ |
| env-1017 | CETL01077499.1 | + | 235 | 290 | **RNA**→ rpsO (PRK05626)Ribosomal_S15 (pfam00312)→<br>PRK11824 (PRK11824)RNase_PH (pfam01138)RNase_PH_C (pfam03725)→ |
| env-1018 | CESP01137952.1 | + | 2063 | 2118 | **RNA**→ rpsO (PRK05626)Ribosomal_S15 (pfam00312)→<br>PRK11824 (PRK11824)RNase_PH (pfam01138)RNase_PH_C (pfam03725)→ |
| env-1019 | CENX01186301.1 | - | 389 | 334 | **RNA**→ rpsO (PRK05626)Ribosomal_S15 (pfam00312)→ |
| env-1020 | CEOF01186277.1 | - | 389 | 334 | **RNA**→ rpsO (PRK05626)Ribosomal_S15 (pfam00312)→ |
| env-1021 | CESW01208627.1 | + | 457 | 513 | **RNA**→ rpsO (PRK05626)Ribosomal_S15 (pfam00312)→ |
| env-1022 | CEQX01005238.1 | - | 426 | 371 | **RNA**→ rpsO (PRK05626)Ribosomal_S15 (pfam00312)→ |
| env-1023 | CEPU01014169.1 | - | 422 | 367 | **RNA**→ rpsO (PRK05626)Ribosomal_S15 (pfam00312)→ |
| env-1024 | CEVH01129166.1 | - | 2144 | 2089 | **RNA**→ rpsO (PRK05626)Ribosomal_S15 (pfam00312)→ Pnp (COG1185)PRK11824 (PRK11824)→ |
| env-1025 | CEOW01121765.1 | - | 962 | 907 | **RNA**→ rpsO (PRK05626)RpsO (COG0184)→<br>PRK11824 (PRK11824)RNase_PH (pfam01138)RNase_PH_C (pfam03725)→ |
| env-1026 | CEVX01126697.1 | - | 281 | 226 | **RNA**→ rpsO (PRK05626)Ribosomal_S15 (pfam00312)→ |
| env-1027 | CEVS01109929.1 | - | 229 | 174 | **RNA**→ rpsO (PRK05626)Ribosomal_S15 (pfam00312)→ |
| env-1028 | CEVH01173857.1 | - | 230 | 174 | **RNA**→ rpsO (PRK05626)Ribosomal_S15 (pfam00312)→ |
| env-1029 | CEUW01249740.1 | - | 307 | 251 | **RNA**→ rpsO (PRK05626)Ribosomal_S15 (pfam00312)→ |
| env-1030 | CEUM01264125.1 | - | 71 | 15 | **RNA**→ |
| env-1031 | CEUV01191480.1 | - | 71 | 15 | **RNA**→ |
| env-1032 | CEUU01248441.1 | + | 1447 | 1503 | **RNA**→ rpsO (PRK05626)Ribosomal_S15 (pfam00312)→ |
| env-1033 | CEVF01023934.1 | + | 1550 | 1606 | **RNA**→ rpsO (PRK05626)Ribosomal_S15 (pfam00312)→ |
| env-1034 | CETP01104982.1 | + | 419 | 475 | **RNA**→ rpsO (PRK05626)Ribosomal_S15 (pfam00312)→ |

| | | | | | |
|---|---|---|---|---|---|
| env-1035 | CEWG01181614.1 | - | 71 | 15 | **RNA**→ |
| env-1036 | CAM_READ_0233394955 | - | 381 | 324 | **RNA**→ ←hypo |
| env-1037 | CETQ01034087.1 | - | 365 | 310 | **RNA**→ rpsO (PRK05626)Ribosomal_S15 (pfam00312)→ |
| env-1038 | JCVI_SCAF_1101668393340 | + | 580 | 636 | **RNA**→ rpsO (PRK05626)Ribosomal_S15 (pfam00312)→ |
| env-1039 | CEOX01118814.1 | + | 508 | 564 | **RNA**→ |
| env-1040 | CEOV01159916.1 | + | 492 | 548 | **RNA**→ |
| env-1041 | CEOK01026918.1 | - | 67 | 10 | **RNA**→ |
| env-1042 | CENJ01059679.1 | + | 390 | 447 | **RNA**→ rpsO (PRK05626)Ribosomal_S15 (pfam00312)→ PRK11824 (PRK11824)RNase_PH (pfam01138)→ |
| env-1043 | CENP01107856.1 | - | 726 | 669 | **RNA**→ rpsO (PRK05626)Ribosomal_S15 (pfam00312)→ PRK11824 (PRK11824)RNase_PH (pfam01138)→ |
| env-1044 | JCVI_SCAF_1101668367367 | + | 220 | 278 | **RNA**→ rpsO (PRK05626)Ribosomal_S15 (pfam00312)→ PRK11824 (PRK11824)RNase_PH (pfam01138)→ |
| env-1045 | JCVI_READ_1103359209347 | + | 374 | 430 | **RNA**→ rpsO (PRK05626)Ribosomal_S15 (pfam00312)→ PRK11824 (PRK11824)RNase_PH (pfam01138)→ |
| env-1046 | CENJ01271348.1 | + | 690 | 746 | **RNA**→ |
| env-1047 | CERR01108442.1 | - | 2061 | 2005 | **RNA**→ rpsO (PRK05626)Ribosomal_S15 (pfam00312)→ Pnp (COG1185)PRK11824 (PRK11824)→ |
| env-1048 | CEVW01297929.1 | - | 67 | 11 | **RNA**→ |
| env-1049 | CERK01007231.1 | + | 11630 | 11686 | **RNA**→ rpsO (PRK05626)Ribosomal_S15 (pfam00312)→ Pnp (COG1185)PRK11824 (PRK11824)→ |
| | | | | | RpoD (COG0568)sigma70-ECF (TIGR02937)→ PRK05581 (PRK05581)RPE (cd00429)→ |
| env-1050 | CERW01018950.1 | + | 11301 | 11357 | **RNA**→ rpsO (PRK05626)Ribosomal_S15 (pfam00312)→ Pnp (COG1185)PRK11824 (PRK11824)→ |
| | | | | | RpoD (COG0568)sigma70-ECF (TIGR02937)→ PRK05581 (PRK05581)RPE (cd00429)→ |
| env-1051 | CERV01210325.1 | + | 688 | 744 | **RNA**→ |
| env-1052 | CEWB01130717.1 | + | 504 | 560 | **RNA**→ |
| env-1053 | CEVX01029906.1 | - | 1046 | 990 | **RNA**→ rpsO (PRK05626)Ribosomal_S15 (pfam00312)→ PRK11824 (PRK11824)RNase_PH (pfam01138)→ |
| env-1054 | CEWO01080375.1 | + | 31704 | 31760 | **RNA**→ rpsO (PRK05626)Ribosomal_S15 (pfam00312)→ Pnp (COG1185)PRK11824 (PRK11824)→ |
| | | | | | RpoD (COG0568)sigma70-ECF (TIGR02937)→ PRK05581 (PRK05581)RPE (cd00429)→ |
| env-1055 | CERG01075223.1 | - | 301 | 245 | **RNA**→ rpsO (PRK05626)Ribosomal_S15 (pfam00312)→ |
| env-1056 | JCVI_SCAF_1101668453793 | - | 755 | 699 | **RNA**→ rpsO (PRK05626)RpsO (COG0184)→ PRK11824 (PRK11824)RNase_PH (pfam01138)→ |
| env-1057 | CETE01088223.1 | - | 254 | 198 | **RNA**→ rpsO (PRK05626)Ribosomal_S15 (pfam00312)→ |
| env-1058 | CEVG01324060.1 | + | 1087 | 1143 | **RNA**→ |
| env-1059 | CEQX01186058.1 | - | 92 | 36 | **RNA**→ |
| env-1060 | CETL01280057.1 | + | 711 | 767 | **RNA**→ |
| env-1061 | CEOP01144955.1 | - | 414 | 358 | **RNA**→ rpsO (PRK05626)Ribosomal_S15 (pfam00312)→ |
| env-1062 | CEOM01088735.1 | - | 235 | 179 | **RNA**→ rpsO (PRK05626)Ribosomal_S15 (pfam00312)→ |
| env-1063 | CEOK01164348.1 | + | 1258 | 1314 | **RNA**→ rpsO (PRK05626)Ribosomal_S15 (pfam00312)→ |
| | | | | | PRK11824 (PRK11824)RNase_PH (pfam01138)RNase_PH_C (pfam03725)→ |
| env-1064 | CEOO01015551.1 | + | 104 | 160 | **RNA**→ rpsO (PRK05626)Ribosomal_S15 (pfam00312)→ |
| env-1065 | CEPW01148047.1 | - | 367 | 311 | **RNA**→ rpsO (PRK05626)Ribosomal_S15 (pfam00312)→ |
| env-1066 | CESQ01107160.1 | + | 179 | 235 | **RNA**→ rpsO (PRK05626)Ribosomal_S15 (pfam00312)→ |
| env-1067 | CEVP01076511.1 | + | 372 | 428 | **RNA**→ rpsO (PRK05626)Ribosomal_S15 (pfam00312)→ |
| env-1068 | CEWO01041439.1 | + | 478 | 534 | **RNA**→ rpsO (PRK05626)Ribosomal_S15 (pfam00312)→ |
| env-1069 | JCVI_SCAF_1101668473583 | - | 731 | 675 | **RNA**→ rpsO (PRK05626)Ribosomal_S15 (pfam00312)→ PRK11824 (PRK11824)RNase_PH (pfam01138)→ |
| env-1070 | CENF01040883.1 | + | 158 | 214 | **RNA**→ rpsO (PRK05626)Ribosomal_S15 (pfam00312)→ |
| env-1071 | CERK01009463.1 | - | 541 | 485 | **RNA**→ rpsO (PRK05626)Ribosomal_S15 (pfam00312)→ RNase_PH_PNPase_1 (cd11363)PRK11824 (PRK11824)– |
| env-1072 | CERR01043387.1 | + | 2545 | 2601 | **RNA**→ rpsO (PRK05626)Ribosomal_S15 (pfam00312)→ |
| env-1073 | CERW01074256.1 | - | 390 | 334 | **RNA**→ rpsO (PRK05626)Ribosomal_S15 (pfam00312)→ |
| env-1074 | CETI01126305.1 | - | 941 | 885 | **RNA**→ rpsO (PRK05626)Ribosomal_S15 (pfam00312)→ PRK11824 (PRK11824)RNase_PH (pfam01138)→ |
| env-1075 | CEUB01187863.1 | + | 575 | 631 | **RNA**→ rpsO (PRK05626)Ribosomal_S15 (pfam00312)→ Pnp (COG1185)PRK11824 (PRK11824)→ |
| env-1076 | CEVX01089166.1 | + | 423 | 479 | **RNA**→ rpsO (PRK05626)Ribosomal_S15 (pfam00312)→ |
| env-1077 | CEOM01102445.1 | - | 218 | 162 | **RNA**→ rpsO (PRK05626)Ribosomal_S15 (pfam00312)→ |
| env-1078 | CEOP01072639.1 | - | 187 | 131 | **RNA**→ rpsO (PRK05626)Ribosomal_S15 (pfam00312)→ |
| env-1079 | CESM01057045.1 | - | 387 | 331 | **RNA**→ rpsO (PRK05626)Ribosomal_S15 (pfam00312)→ |
| env-1080 | CETE01063798.1 | + | 198 | 254 | **RNA**→ rpsO (PRK05626)Ribosomal_S15 (pfam00312)→ |
| env-1081 | CEVX01052949.1 | + | 126 | 182 | **RNA**→ rpsO (PRK05626)Ribosomal_S15 (pfam00312)→ |
| env-1082 | JCVI_SCAF_1096627072953 | + | 895 | 951 | **RNA**→ rpsO (PRK05626)Ribosomal_S15 (pfam00312)→ PRK11824 (PRK11824)RNase_PH (pfam01138)→ |
| env-1083 | CENT01157003.1 | - | 481 | 425 | **RNA**→ rpsO (PRK05626)Ribosomal_S15 (pfam00312)→ hypo→ |

| | | | | | |
|---|---|---|---|---|---|
| env-1084 | CEVU01001438.1 | - | 529 | 473 | **RNA**→ rpsO (PRK05626)Ribosomal_S15 (pfam00312)→ RNase_PH_PNPase_1 (cd11363)PRK11824 (PRK11824)– |
| env-1085 | JCVI_READ_1103180834378 | - | 500 | 444 | **RNA**→ rpsO (PRK05626)Ribosomal_S15 (pfam00312)→ hypo→ |
| env-1086 | JCVI_READ_1108830074671 | - | 624 | 568 | **RNA**→ rpsO (PRK05626)Ribosomal_S15 (pfam00312)→ |
| env-1087 | CEWG01092952.1 | - | 74 | 18 | **RNA**→ |
| env-1088 | CESW01098570.1 | - | 598 | 542 | **RNA**→ rpsO (PRK05626)Ribosomal_S15 (pfam00312)→ PRK11824 (PRK11824)RNase_PH (pfam01138)→ |
| env-1089 | CENG01044642.1 | - | 530 | 474 | **RNA**→ rpsO (PRK05626)Ribosomal_S15 (pfam00312)→ RNase_PH_PNPase_1 (cd11363)PRK11824 (PRK11824)– |
| env-1090 | CEOM01078862.1 | + | 107 | 163 | **RNA**→ rpsO (PRK05626)Ribosomal_S15 (pfam00312)→ RNase_PH_PNPase_1 (cd11363)PRK11824 (PRK11824)– |
| env-1091 | CEOY01062467.1 | + | 215 | 271 | **RNA**→ rpsO (PRK05626)Ribosomal_S15 (pfam00312)→ |
| env-1092 | CEPU01224898.1 | - | 334 | 278 | **RNA**→ rpsO (PRK05626)Ribosomal_S15 (pfam00312)→ |
| env-1093 | CESN01053220.1 | - | 69 | 13 | **RNA**→ |
| env-1094 | CETI01258919.1 | + | 443 | 499 | **RNA**→ |
| env-1095 | CEVZ01325749.1 | + | 257 | 313 | **RNA**→ rpsO (PRK05626)Ribosomal_S15 (pfam00312)→ |
| env-1096 | CAM_READ_0233245559 | + | 268 | 324 | **RNA**→ rpsO (PRK05626)Ribosomal_S15 (pfam00312)→ |
| env-1097 | JCVI_READ_1108830074672 | - | 627 | 571 | **RNA**→ rpsO (PRK05626)Ribosomal_S15 (pfam00312)→ PRK11824 (PRK11824)RNase_PH (pfam01138)→ |
| env-1098 | JCVI_SCAF_1096626319309 | + | 109 | 165 | **RNA**→ rpsO (PRK05626)Ribosomal_S15 (pfam00312)→ PRK11824 (PRK11824)RNase_PH (pfam01138)→ |
| env-1099 | 85126184 | + | 606 | 662 | **RNA**→ |
| env-1100 | JCVI_READ_1103180276409 | + | 606 | 662 | **RNA**→ |
| env-1101 | CENP01224387.1 | - | 579 | 523 | **RNA**→ rpsO (PRK05626)Ribosomal_S15 (pfam00312)→ PRK11824 (PRK11824)RNase_PH (pfam01138)→ |
| env-1102 | CEOO01058349.1 | + | 262 | 318 | **RNA**→ rpsO (PRK05626)Ribosomal_S15 (pfam00312)→ |
| env-1103 | CEPL01104959.1 | + | 227 | 283 | **RNA**→ rpsO (PRK05626)Ribosomal_S15 (pfam00312)→ |
| env-1104 | CEQN01113421.1 | - | 266 | 210 | **RNA**→ rpsO (PRK05626)Ribosomal_S15 (pfam00312)→ |
| env-1105 | CESZ01006198.1 | + | 105 | 161 | **RNA**→ rpsO (PRK05626)Ribosomal_S15 (pfam00312)→ PRK11824 (PRK11824)RNase_PH (pfam01138)→ |
| env-1106 | CEVX01162813.1 | + | 10219 | 10275 | **RNA**→ rpsO (PRK05626)Ribosomal_S15 (pfam00312)→ Pnp (COG1185)PRK11824 (PRK11824)→ |
| env-1107 | CAM_READ_0234149989 | - | 230 | 174 | **RNA**→ |
| env-1108 | JCVI_READ_1108830083326 | - | 137 | 81 | **RNA**→ rpsO (PRK05626)→ |
| env-1109 | JCVI_READ_1108799804924 | + | 237 | 293 | **RNA**→ rpsO (PRK05626)Ribosomal_S15 (pfam00312)→ PRK11824 (PRK11824)RNase_PH (pfam01138)→ |
| env-1110 | CEWB01036815.1 | - | 340 | 283 | **RNA**→ rpsO (PRK05626)Ribosomal_S15 (pfam00312)→ |
| env-1111 | JCVI_SCAF_1096628021305 | - | 1523 | 1467 | **RNA**→ rpsO (PRK05626)Ribosomal_S15 (pfam00312)→ PRK11824 (PRK11824)RNase_PH (pfam01138)→<br>PRK11824 (PRK11824)RNase_PH (pfam01138)RNase_PH_C (pfam03725)→ |
| env-1112 | CESN01190334.1 | + | 896 | 952 | **RNA**→ rpsO (PRK05626)Ribosomal_S15 (pfam00312)→ |
| env-1113 | CETE01015087.1 | - | 366 | 310 | **RNA**→ rpsO (PRK05626)Ribosomal_S15 (pfam00312)→ |
| env-1114 | CETK01092484.1 | + | 149 | 205 | **RNA**→ rpsO (PRK05626)Ribosomal_S15 (pfam00312)→ |
| env-1115 | CETJ01158801.1 | - | 614 | 558 | **RNA**→ rpsO (PRK05626)Ribosomal_S15 (pfam00312)→ PRK11824 (PRK11824)RNase_PH (pfam01138)→ |
| env-1116 | CETO01171402.1 | - | 663 | 607 | **RNA**→ rpsO (PRK05626)Ribosomal_S15 (pfam00312)→ PRK11824 (PRK11824)RNase_PH (pfam01138)→ |
| env-1117 | CEUJ01011635.1 | - | 1182 | 1126 | **RNA**→ rpsO (PRK05626)Ribosomal_S15 (pfam00312)→<br>PRK11824 (PRK11824)RNase_PH (pfam01138)RNase_PH_C (pfam03725)→ |
| env-1118 | CEUL01314205.1 | - | 583 | 527 | **RNA**→ rpsO (PRK05626)Ribosomal_S15 (pfam00312)→ PRK11824 (PRK11824)RNase_PH (pfam01138)→ |
| env-1119 | CEUQ01085079.1 | + | 281 | 337 | **RNA**→ rpsO (PRK05626)Ribosomal_S15 (pfam00312)→ |
| env-1120 | JGI24820J26691_1000397 | + | 11685 | 11741 | **RNA**→ rpsO (PRK05626)Ribosomal_S15 (pfam00312)→<br>PRK11824 (PRK11824)RNase_PH (pfam01138)RNase_PH_C (pfam03725)→ |
| env-1121 | CETT01075724.1 | - | 760 | 704 | **RNA**→ rpsO (PRK05626)Ribosomal_S15 (pfam00312)→ PRK11824 (PRK11824)RNase_PH (pfam01138)→ |
| env-1122 | CENX01010795.1 | + | 29698 | 29754 | **RNA**→ rpsO (PRK05626)RpsO (COG0184)→ Pnp (COG1185)PRK11824 (PRK11824)→<br>RpoD (COG0568)sigma70-ECF (TIGR02937)→ PRK05581 (PRK05581)RPE (cd00429)→ |
| env-1123 | CEPT01054511.1 | + | 99 | 155 | **RNA**→ rpsO (PRK05626)Ribosomal_S15 (pfam00312)→ PRK11824 (PRK11824)RNase_PH (pfam01138)→ |
| env-1124 | CESW01141929.1 | - | 852 | 796 | **RNA**→ rpsO (PRK05626)Ribosomal_S15 (pfam00312)→ PRK11824 (PRK11824)RNase_PH (pfam01138)→ |
| env-1125 | CETG01325760.1 | - | 690 | 634 | **RNA**→ rpsO (PRK05626)Ribosomal_S15 (pfam00312)→ PRK11824 (PRK11824)RNase_PH (pfam01138)→ |
| env-1126 | CETM01138654.1 | + | 38 | 94 | **RNA**→ rpsO (PRK05626)Ribosomal_S15 (pfam00312)→ PRK11824 (PRK11824)RNase_PH (pfam01138)→ |
| env-1127 | CETS01042552.1 | + | 1863 | 1919 | **RNA**→ rpsO (PRK05626)Ribosomal_S15 (pfam00312)→ PRK11824 (PRK11824)RNase_PH (pfam01138)→ |
| env-1128 | CEOF01214230.1 | - | 326 | 270 | **RNA**→ rpsO (PRK05626)RpsO (COG0184)→ |
| env-1129 | JGI25921J50272_10110947 | - | 112 | 45 | **RNA**→ hypo→ |
| env-1130 | JGI25921J50272_10064100 | + | 59 | 126 | **RNA**→ rpsO (PRK05626)Ribosomal_S15p_S13e (cd00353)→ |
| env-1131 | JGI12547J11936_1016467 | - | 880 | 813 | **RNA**→ rpsO (PRK05626)RpsO (COG0184)→ |
| env-1132 | JGI25912J50252_10000040 | + | 10957 | 11024 | **RNA**→ rpsO (PRK05626)RpsO (COG0184)→ |

| | | | | | |
|---|---|---|---|---|---|
| env-1133 | JGI25925J51416_10000721 | - | 2637 | 2570 | **RNA**→ rpsO (PRK05626)RpsO (COG0184)→ |
| env-1134 | JGI12421J11937_10092536 | - | 740 | 673 | **RNA**→ rpsO (PRK05626)RpsO (COG0184)→ |
| env-1135 | JGI25910J50241_10000033 | + | 10957 | 11024 | **RNA**→ rpsO (PRK05626)RpsO (COG0184)→ |
| env-1136 | JGI25911J50253_10000079 | - | 16190 | 16123 | **RNA**→ rpsO (PRK05626)RpsO (COG0184)→ |
| env-1137 | JGI25914J50564_10211613 | - | 350 | 283 | **RNA**→ rpsO (PRK05626)RpsO (COG0184)→ |
| env-1138 | M3P_c10368127 | - | 78 | 11 | **RNA**→ |
| env-1139 | JGI25921J50272_10001427 | - | 6248 | 6181 | **RNA**→ rpsO (PRK05626)RpsO (COG0184)→ Pnp (COG1185)PRK11824 (PRK11824)→ |
| env-1140 | JGI25913J50563_1000060 | - | 14177 | 14110 | **RNA**→ rpsO (PRK05626)Ribosomal_S15 (pfam00312)→ |
| env-1141 | CEOK01126743.1 | + | 2956 | 3010 | **RNA**→ rpsO (PRK05626)Ribosomal_S15 (pfam00312)→ |
| env-1142 | CENJ01063439.1 | - | 3516 | 3462 | **RNA**→ rpsO (PRK05626)Ribosomal_S15 (pfam00312)→ |
| env-1143 | CENP01230084.1 | + | 4337 | 4391 | **RNA**→ rpsO (PRK05626)Ribosomal_S15 (pfam00312)→ |
| Lsp-4-1 | NZ_JPOL01000002.1 | - | 3091910 | 3091846 | **RNA**→ rpsO (PRK05626)Ribosomal_S15 (pfam00312)→ |
| Cca-5-1 | NZ_AUFK01000009.1 | - | 843585 | 843537 | **RNA**→ rpsO (PRK05626)Ribosomal_S15 (pfam00312)→ |
| env-1144 | SwRhRL2b_contig_307599 | - | 287 | 239 | **RNA**→ rpsO (PRK05626)RpsO (COG0184)→ |
| Fan-1-1 | NZ_ATTM01000001.1 | - | 367999 | 367938 | **RNA**→ rpsO (PRK05626)Ribosomal_S15 (pfam00312)→ |
| Fte-1-1 | NZ_AUDN01000008.1 | + | 180924 | 180985 | **RNA**→ rpsO (PRK05626)Ribosomal_S15 (pfam00312)→ |
| env-1145 | JGI24723J26617_10000025 | - | 95199 | 95137 | **RNA**→ rpsO (PRK05626)Ribosomal_S15 (pfam00312)→ |
| env-1146 | JGI1221J11331_1065710 | + | 185 | 247 | **RNA**→ rpsO (PRK05626)Ribosomal_S15 (pfam00312)→ |
| env-1147 | CEOS01027552.1 | + | 66892 | 66954 | **RNA**→ rpsO (PRK05626)Ribosomal_S15 (pfam00312)→ |
| env-1148 | CEOX01194702.1 | - | 519 | 457 | **RNA**→ rpsO (PRK05626)Ribosomal_S15 (pfam00312)→ |
| env-1149 | CESL01150597.1 | - | 869 | 807 | **RNA**→ rpsO (PRK05626)Ribosomal_S15 (pfam00312)→ |
| env-1150 | CETA01146872.1 | - | 546 | 484 | **RNA**→ rpsO (PRK05626)Ribosomal_S15 (pfam00312)→ |
| env-1151 | CETB01171796.1 | + | 30062 | 30124 | **RNA**→ rpsO (PRK05626)Ribosomal_S15 (pfam00312)→ |
| env-1152 | CETR01014521.1 | + | 66892 | 66954 | **RNA**→ rpsO (PRK05626)Ribosomal_S15 (pfam00312)→ |
| env-1153 | CETS01016621.1 | + | 66892 | 66954 | **RNA**→ rpsO (PRK05626)Ribosomal_S15 (pfam00312)→ |
| env-1154 | CETT01201410.1 | + | 158 | 220 | **RNA**→ rpsO (PRK05626)Ribosomal_S15 (pfam00312)→ |
| env-1155 | CEVJ01093656.1 | - | 1367 | 1305 | **RNA**→ rpsO (PRK05626)Ribosomal_S15 (pfam00312)→ |
| env-1156 | CEVK01102272.1 | - | 1022 | 960 | **RNA**→ rpsO (PRK05626)Ribosomal_S15 (pfam00312)→ |
| env-1157 | CEVQ01351872.1 | + | 781 | 843 | **RNA**→ rpsO (PRK05626)Ribosomal_S15 (pfam00312)→ |
| env-1158 | CEWI01263175.1 | - | 378 | 316 | **RNA**→ rpsO (PRK05626)Ribosomal_S15 (pfam00312)→ |
| env-1159 | CERQ01119226.1 | - | 71 | 9 | **RNA**→ |
| env-1160 | CEVI01194451.1 | - | 71 | 9 | **RNA**→ |
| env-1161 | CETG01242168.1 | + | 199 | 261 | **RNA**→ rpsO (PRK05626)Ribosomal_S15 (pfam00312)→ |
| env-1162 | JGI24025J20009_10171038 | + | 263 | 325 | **RNA**→ rpsO (PRK05626)Ribosomal_S15 (pfam00312)→ |
| env-1163 | JGI20152J14361_10000034 | - | 29790 | 29725 | **RNA**→ rpsO (PRK05626)Ribosomal_S15 (pfam00312)→ |
| env-1164 | JGI20156J14371_10001636 | - | 2827 | 2762 | **RNA**→ rpsO (PRK05626)Ribosomal_S15 (pfam00312)→ |
| env-1165 | JGI20153J14295_10056361 | - | 492 | 427 | **RNA**→ rpsO (PRK05626)Ribosomal_S15 (pfam00312)→ |
| env-1166 | JGI20153J14318_10056361 | - | 492 | 427 | **RNA**→ rpsO (PRK05626)Ribosomal_S15 (pfam00312)→ |
| env-1167 | DelMOSum2011_c10000619 | + | 21340 | 21404 | **RNA**→ rpsO (PRK05626)Ribosomal_S15 (pfam00312)→ |
| env-1168 | 2236876010_0099679 | - | 354 | 287 | **RNA**→ |
| env-1169 | 2236876001_018781 | - | 282 | 215 | **RNA**→ hypo→ |
| Psp-6-1 | NZ_CBRF010000389.1 | - | 277 | 212 | **RNA**→ rpsO (PRK05626)Ribosomal_S15 (pfam00312)→ |
| env-1170 | CESU01016827.1 | - | 658 | 593 | **RNA**→ rpsO (PRK05626)Ribosomal_S15 (pfam00312)→ |
| env-1171 | CESI01053671.1 | - | 1054 | 989 | **RNA**→ rpsO (PRK05626)Ribosomal_S15 (pfam00312)→ |
| env-1172 | NCBI_READ_1112493756060 | + | 800 | 866 | **RNA**→ |
| env-1173 | NCBI_READ_1112290405280 | - | 757 | 690 | **RNA**→ rpsO (PRK05626)Ribosomal_S15 (pfam00312)→ |
| env-1174 | 2236876010_0188177 | + | 202 | 268 | **RNA**→ rpsO (PRK05626)RpsO (COG0184)→ |
| Tov-1-1 | NZ_AUMF01000013.1 | + | 158381 | 158445 | **RNA**→ rpsO (PRK05626)Ribosomal_S15 (pfam00312)→ |
| Pir-1-1 | NZ_CH724148.1 | + | 565143 | 565208 | **RNA**→ rpsO (PRK05626)Ribosomal_S15 (pfam00312)→ |
| env-1175 | NCBI_READ_1112290390846 | + | 257 | 324 | **RNA**→ rpsO (PRK05626)Ribosomal_S15 (pfam00312)→ |
| env-1176 | NCBI_READ_1112788434094 | + | 709 | 776 | **RNA**→ |
| env-1177 | NCBI_READ_1112777244532 | - | 379 | 312 | **RNA**→ rpsO (PRK05626)Ribosomal_S15 (pfam00312)→ |

| | | | | | |
|---|---|---|---|---|---|
| env-1178 | NCBI_READ_1112637244083 | + | 363 | 430 | **RNA**→ rpsO (PRK05626)Ribosomal_S15 (pfam00312)→ |
| env-1179 | NCBI_READ_1112637711645 | + | 430 | 497 | **RNA**→ rpsO (PRK05626)Ribosomal_S15 (pfam00312)→ |
| env-1180 | CESE01018107.1 | + | 12288 | 12355 | **RNA**→ rpsO (PRK05626)Ribosomal_S15 (pfam00312)→ |
| env-1181 | CEST01011319.1 | + | 1239 | 1306 | **RNA**→ |
| env-1182 | NCBI_READ_1112788335987 | - | 294 | 227 | **RNA**→ rpsO (PRK05626)Ribosomal_S15 (pfam00312)→ |
| env-1183 | wf3_AHAI5724_y1 | - | 298 | 233 | **RNA**→ rpsO (PRK05626)RpsO (COG0184)→ |
| env-1184 | DelMOWin2010_c10256912 | + | 1 | 66 | **RNA**→ rpsO (PRK05626)Ribosomal_S15 (pfam00312)→ |
| env-1185 | SI60aug11_100m_c1001628 | - | 875 | 810 | **RNA**→ rpsO (PRK05626)Ribosomal_S15 (pfam00312)→ |
| env-1186 | JGI20152J14361_10009221 | - | 790 | 725 | **RNA**→ rpsO (PRK05626)Ribosomal_S15 (pfam00312)→ |
| Psp-1-1 | NZ_JPDI01000001.1 | - | 301145 | 301080 | **RNA**→ rpsO (PRK05626)Ribosomal_S15 (pfam00312)→ |
| env-1187 | JGI20160J14292_10015031 | + | 1484 | 1549 | **RNA**→ rpsO (PRK05626)Ribosomal_S15 (pfam00312)→ |
| env-1188 | JGI20153J14295_10072278 | - | 131 | 66 | **RNA**→ |
| env-1189 | JGI20153J14318_10072278 | - | 131 | 66 | **RNA**→ |
| env-1190 | 2236876009_034639 | - | 316 | 251 | **RNA**→ rpsO (PRK05626)Ribosomal_S15 (pfam00312)→ rpsO (PRK05626)Ribosomal_S15 (pfam00312)→ |
| env-1191 | SI60aug11_200m_c1030663 | - | 361 | 296 | **RNA**→ rpsO (PRK05626)Ribosomal_S15 (pfam00312)→ |
| env-1192 | SI34jun09_10m_1045700 | + | 218 | 283 | **RNA**→ rpsO (PRK05626)Ribosomal_S15 (pfam00312)→ |
| env-1193 | JGI20154J14316_10009990 | + | 1427 | 1492 | **RNA**→ rpsO (PRK05626)Ribosomal_S15 (pfam00312)→ |
| env-1194 | JGI20153J14295_10000097 | - | 96 | 31 | **RNA**→ |
| env-1195 | JGI20153J14318_10000097 | - | 96 | 31 | **RNA**→ |
| env-1196 | DelMOSum2010_c10004701 | + | 2457 | 2522 | **RNA**→ rpsO (PRK05626)Ribosomal_S15 (pfam00312)→ |
| env-1197 | NCBI_READ_1112532466838 | + | 411 | 476 | **RNA**→ rpsO (PRK05626)Ribosomal_S15 (pfam00312)→ |
| env-1198 | JGI20154J14316_10069178 | + | 223 | 288 | **RNA**→ rpsO (PRK05626)Ribosomal_S15 (pfam00312)→ |
| env-1199 | CEWR01072923.1 | + | 9501 | 9566 | **RNA**→ |
| env-1200 | NCBI_READ_1112494129442 | - | 686 | 621 | **RNA**→ rpsO (PRK05626)Ribosomal_S15 (pfam00312)→ |
| env-1201 | NCBI_READ_1112493965127 | - | 731 | 666 | **RNA**→ rpsO (PRK05626)Ribosomal_S15 (pfam00312)→ |
| env-1202 | CEST01080561.1 | - | 520 | 455 | **RNA**→ rpsO (PRK05626)Ribosomal_S15 (pfam00312)→ |
| env-1203 | CESI01299916.1 | - | 89 | 24 | **RNA**→ |
| env-1204 | NCBI_READ_1112493618954 | - | 348 | 283 | **RNA**→ rpsO (PRK05626)Ribosomal_S15 (pfam00312)→ |
| env-1205 | P_2C_Liq_3_UnCty_100110 | - | 10558 | 10493 | **RNA**→ rpsO (PRK05626)Ribosomal_S15 (pfam00312)→ |
| env-1206 | P_2C_Liq_1_UnCty_1000155 | + | 17803 | 17868 | **RNA**→ rpsO (PRK05626)Ribosomal_S15 (pfam00312)→ |
| env-1207 | P_2C_Liq_2_UnCty_1014517 | - | 424 | 359 | **RNA**→ rpsO (PRK05626)Ribosomal_S15 (pfam00312)→ |
| Psp-3-1 | NZ_JHZZ01000001.1 | + | 843905 | 843970 | **RNA**→ rpsO (PRK05626)Ribosomal_S15 (pfam00312)→ hypo→ Pnp (COG1185)PRK11824 (PRK11824)→ |
| env-1208 | DelMOSum2011_c10451001 | - | 187 | 122 | **RNA**→ rpsO (PRK05626)Ribosomal_S15 (pfam00312)→ |
| env-1209 | JGI20156J14371_10087495 | + | 211 | 276 | **RNA**→ rpsO (PRK05626)Ribosomal_S15 (pfam00312)→ |
| env-1210 | JGI20159J14440_10134887 | - | 66 | 1 | **RNA**→ |
| env-1211 | JGI20152J14361_10000060 | + | 50242 | 50307 | **RNA**→ rpsO (PRK05626)Ribosomal_S15 (pfam00312)→ |
| env-1212 | P_2C_Liq_1_UnCty_1001158 | - | 2910 | 2845 | **RNA**→ rpsO (PRK05626)Ribosomal_S15 (pfam00312)→ |
| env-1213 | JGI20153J14295_10391622 | - | 175 | 109 | **RNA**→ rpsO (PRK05626)RpsO (COG0184)→ |
| env-1214 | JGI20153J14318_10391622 | - | 175 | 109 | **RNA**→ rpsO (PRK05626)RpsO (COG0184)→ |
| Psp-2-1 | NZ_JPDS01000003.1 | + | 146619 | 146684 | **RNA**→ rpsO (PRK05626)Ribosomal_S15 (pfam00312)→ |
| Psp-4-1 | NC_020830.1 | - | 2479424 | 2479359 | **RNA**→ rpsO (PRK05626)Ribosomal_S15 (pfam00312)→ |
| env-1215 | 2236876010_0335376 | + | 192 | 256 | **RNA**→ hypo→ |
| env-1216 | JCVI_SCAF_1096627466044 | + | 161 | 226 | **RNA**→ rpsO (PRK05626)Ribosomal_S15 (pfam00312)→ |
| env-1217 | 2236876010_0251574 | - | 232 | 167 | **RNA**→ rpsO (PRK05626)Ribosomal_S15 (pfam00312)→ |
| env-1218 | 2236876010_0458838 | - | 157 | 92 | **RNA**→ rpsO (PRK05626)RpsO (COG0184)→ |
| env-1219 | 2236876010_0341471 | - | 321 | 256 | **RNA**→ rpsO (PRK05626)Ribosomal_S15 (pfam00312)→ |
| env-1220 | DelMOSum2011_c10002449 | - | 3023 | 2958 | **RNA**→ rpsO (PRK05626)Ribosomal_S15 (pfam00312)→ |
| env-1221 | CERI01273154.1 | - | 366 | 301 | **RNA**→ rpsO (PRK05626)Ribosomal_S15 (pfam00312)→ |
| env-1222 | SI34jun09_10m_1064123 | + | 231 | 296 | **RNA**→ rpsO (PRK05626)Ribosomal_S15 (pfam00312)→ |
| env-1223 | JGI20151J14362_10003118 | + | 10963 | 11028 | **RNA**→ rpsO (PRK05626)Ribosomal_S15 (pfam00312)→ |
| env-1224 | JGI20159J14440_10001807 | - | 6646 | 6581 | **RNA**→ rpsO (PRK05626)Ribosomal_S15 (pfam00312)→ |
| env-1225 | JGI20158J14315_10040249 | - | 1575 | 1510 | **RNA**→ rpsO (PRK05626)Ribosomal_S15 (pfam00312)→ |

| | | | | | |
|---|---|---|---|---|---|
| env-1226 | JGI20154J14316_10029064 | - | 823 | 758 | RNA→ rpsO (PRK05626)Ribosomal_S15 (pfam00312)→ |
| env-1227 | JGI20153J14295_10003610 | - | 825 | 760 | RNA→ rpsO (PRK05626)Ribosomal_S15 (pfam00312)→ |
| env-1228 | JGI20153J14318_10003610 | - | 825 | 760 | RNA→ rpsO (PRK05626)Ribosomal_S15 (pfam00312)→ |
| env-1229 | JGI20155J14468_10055793 | - | 1584 | 1519 | RNA→ rpsO (PRK05626)Ribosomal_S15 (pfam00312)→ |
| env-1230 | 2236876010_0648566 | + | 166 | 232 | RNA→ rpsO (PRK05626)→ |
| env-1231 | 2236876010_0594514 | + | 218 | 283 | RNA→ |
| env-1232 | 2236876009_222657 | - | 142 | 77 | RNA→ rpsO (PRK05626)→ |
| env-1233 | 2236876010_0518603 | - | 123 | 58 | RNA→ hypo→ |
| env-1234 | 2236876010_0164712 | + | 288 | 353 | RNA→ |
| env-1235 | 2236876010_0454774 | + | 51 | 116 | RNA→ rpsO (PRK05626)RpsO (COG0184)→ |
| env-1236 | DelMOWin2010_c10635086 | + | 50 | 115 | RNA→ rpsO (PRK05626)Ribosomal_S15 (pfam00312)→ |
| env-1237 | JGI20152J14361_10030741 | - | 526 | 461 | RNA→ rpsO (PRK05626)Ribosomal_S15 (pfam00312)→ |
| env-1238 | JGI20156J14371_10002279 | - | 527 | 462 | RNA→ rpsO (PRK05626)Ribosomal_S15 (pfam00312)→ |
| env-1239 | JGI20160J14292_10001237 | - | 71 | 6 | RNA→ |
| env-1240 | JGI20157J14317_10001850 | + | 18454 | 18519 | RNA→ rpsO (PRK05626)Ribosomal_S15 (pfam00312)→ |
| env-1241 | 2236876010_0276371 | + | 118 | 182 | RNA→ |
| env-1242 | JGI20154J14316_10081544 | - | 110 | 45 | RNA→ hypo→ |
| env-1243 | JGI20153J14295_10059635 | - | 113 | 48 | RNA→ hypo→ |
| env-1244 | JGI20153J14318_10059635 | - | 113 | 48 | RNA→ hypo→ |
| env-1245 | PML_READ_00810429 | + | 39 | 104 | RNA→ rpsO (PRK05626)RpsO (COG0184)→ |
| env-1246 | PML_READ_00749665 | + | 164 | 229 | RNA→ hypo→ |
| env-1247 | JGI20156J14371_10251340 | - | 138 | 74 | RNA→ |
| env-1248 | JGI20152J14361_10042742 | + | 1189 | 1254 | RNA→ hypo→ |
| env-1249 | JGI20160J14292_10013292 | + | 5005 | 5070 | RNA→ |
| env-1250 | JGI20153J14295_10037233 | - | 1940 | 1875 | RNA→ rpsO (PRK05626)Ribosomal_S15 (pfam00312)→ |
| env-1251 | JGI20153J14318_10037233 | - | 1940 | 1875 | RNA→ rpsO (PRK05626)Ribosomal_S15 (pfam00312)→ |
| env-1252 | CESU01096896.1 | - | 500 | 435 | RNA→ rpsO (PRK05626)Ribosomal_S15 (pfam00312)→ |
| env-1253 | PML_READ_00817042 | + | 104 | 169 | RNA→ rpsO (PRK05626)Ribosomal_S15 (pfam00312)→ |
| env-1254 | DelMOSum2010_c10498216 | + | 248 | 313 | RNA→ |
| env-1255 | SI34jun09_100m_1156551 | + | 262 | 327 | RNA→ |
| env-1256 | JGI20154J14316_10022051 | + | 3363 | 3428 | RNA→ hypo→ |
| env-1257 | JGI20153J14295_10059402 | - | 113 | 48 | RNA→ hypo→ |
| env-1258 | JGI20153J14318_10059402 | - | 113 | 48 | RNA→ hypo→ |
| env-1259 | JGI20152J14361_10005596 | + | 1194 | 1259 | RNA→ rpsO (PRK05626)Ribosomal_S15 (pfam00312)→ |
| env-1260 | JGI20156J14371_10001740 | + | 17326 | 17391 | RNA→ rpsO (PRK05626)Ribosomal_S15 (pfam00312)→ hypo→ |
| env-1261 | JGI20157J14317_10050291 | - | 461 | 396 | RNA→ rpsO (PRK05626)Ribosomal_S15 (pfam00312)→ hypo→ |
| env-1262 | JGI20160J14292_10037220 | - | 1297 | 1232 | RNA→ rpsO (PRK05626)Ribosomal_S15 (pfam00312)→ |
| env-1263 | JGI20151J14362_10001103 | - | 1600 | 1535 | RNA→ rpsO (PRK05626)RpsO (COG0184)→ hypo→ Pnp (COG1185)PRK11824 (PRK11824)→ |
| env-1264 | SI34jun09_10m_1001615 | - | 5506 | 5441 | RNA→ rpsO (PRK05626)Ribosomal_S15 (pfam00312)→ hypo→ Pnp (COG1185)PRK11824 (PRK11824)→ |
| env-1265 | SI34jun09_10m_1017464 | + | 625 | 690 | RNA→ rpsO (PRK05626)Ribosomal_S15 (pfam00312)→ |
| env-1266 | SI34jun09_135m_1013108 | - | 2236 | 2171 | RNA→ rpsO (PRK05626)Ribosomal_S15 (pfam00312)→ |
| env-1267 | SI34jun09_100m_1007927 | + | 444 | 509 | RNA→ rpsO (PRK05626)Ribosomal_S15 (pfam00312)→ |
| env-1268 | SI48aug10_100m_1159875 | + | 93 | 158 | RNA→ rpsO (PRK05626)Ribosomal_S15 (pfam00312)→ |
| env-1269 | JGI20154J14316_10014588 | - | 110 | 45 | RNA→ hypo→ |
| env-1270 | CESF01290968.1 | - | 205 | 140 | RNA→ rpsO (PRK05626)Ribosomal_S15 (pfam00312)→ |
| env-1271 | JCVI_SCAF_1096627401693 | - | 1341 | 1276 | RNA→ rpsO (PRK05626)Ribosomal_S15 (pfam00312)→ |
| env-1272 | SI48aug10_10m_c1023266 | + | 218 | 283 | RNA→ rpsO (PRK05626)Ribosomal_S15 (pfam00312)→ |
| env-1273 | SI34jun09_120m_1022537 | + | 54 | 119 | RNA→ rpsO (PRK05626)Ribosomal_S15 (pfam00312)→ |
| env-1274 | JGI20156J14371_10172854 | + | 486 | 551 | RNA→ |
| env-1275 | JGI20153J14295_10026978 | - | 113 | 48 | RNA→ hypo→ |
| env-1276 | JGI20153J14318_10026978 | - | 113 | 48 | RNA→ hypo→ |
| env-1277 | 2236876009_230751 | - | 266 | 201 | RNA→ rpsO (PRK05626)Ribosomal_S15 (pfam00312)→ |

| | | | | | |
|---|---|---|---|---|---|
| env-1278 | CESE01261489.1 | + | 340 | 405 | **RNA**→ rpsO (PRK05626)Ribosomal_S15 (pfam00312)→ |
| env-1279 | CEST01084614.1 | - | 1635 | 1570 | **RNA**→ rpsO (PRK05626)Ribosomal_S15 (pfam00312)→ |
| Ccy-1-1 | NZ_CDOF01000013.1 | - | 34423 | 34369 | **RNA**→ rpsO (PRK05626)Ribosomal_S15 (pfam00312)→ |
| Ccy-1-2 | NZ_CDOG01000011.1 | - | 34424 | 34370 | **RNA**→ rpsO (PRK05626)Ribosomal_S15 (pfam00312)→ |
| Ccy-2-1 | NZ_KB900704.1 | - | 181426 | 181372 | **RNA**→ rpsO (PRK05626)Ribosomal_S15 (pfam00312)→ |
| Ccy-1-3 | NZ_CDOD01000067.1 | + | 17098 | 17152 | **RNA**→ rpsO (PRK05626)Ribosomal_S15 (pfam00312)→ |
| Tme-1-1 | NZ_LDOD01000002.1 | + | 281523 | 281586 | **RNA**→ rpsO (PRK05626)Ribosomal_S15 (pfam00312)→ |
| Tsp-1-1 | NZ_JADN01000008.1 | - | 275749 | 275686 | **RNA**→ rpsO (PRK05626)RpsO (COG0184)→ |
| env-1280 | JGI20154J14316_10212002 | + | 357 | 424 | **RNA**→ |
| env-1281 | DelMOSum2011_c10031224 | + | 1515 | 1582 | **RNA**→ rpsO (PRK05626)Ribosomal_S15 (pfam00312)→ |
| env-1282 | NCBI_READ_1112493817117 | + | 489 | 556 | **RNA**→ rpsO (PRK05626)Ribosomal_S15 (pfam00312)→ |
| env-1283 | NCBI_READ_1112532339470 | - | 139 | 72 | **RNA**→ |
| env-1284 | JGI20154J14316_10007880 | + | 7231 | 7298 | **RNA**→ |
| env-1285 | JGI20153J14295_10044777 | - | 1598 | 1531 | **RNA**→ rpsO (PRK05626)Ribosomal_S15 (pfam00312)→ |
| env-1286 | JGI20153J14318_10044777 | - | 1598 | 1531 | **RNA**→ rpsO (PRK05626)Ribosomal_S15 (pfam00312)→ |
| env-1287 | NCBI_READ_1112493996523 | - | 766 | 699 | **RNA**→ rpsO (PRK05626)Ribosomal_S15 (pfam00312)→ |
| env-1288 | NCBI_READ_1112493957189 | - | 592 | 525 | **RNA**→ rpsO (PRK05626)Ribosomal_S15 (pfam00312)→ |
| env-1289 | NCBI_READ_1112494170669 | - | 517 | 450 | **RNA**→ rpsO (PRK05626)Ribosomal_S15 (pfam00312)→ |
| env-1290 | CESE01136377.1 | + | 326 | 393 | **RNA**→ rpsO (PRK05626)Ribosomal_S15 (pfam00312)→ |
| env-1291 | CEST01080931.1 | + | 326 | 393 | **RNA**→ rpsO (PRK05626)Ribosomal_S15 (pfam00312)→ |
| env-1292 | CEWR01087712.1 | + | 3318 | 3385 | **RNA**→ rpsO (PRK05626)Ribosomal_S15 (pfam00312)→ |
| env-1293 | NCBI_READ_1112493907442 | + | 499 | 566 | **RNA**→ rpsO (PRK05626)Ribosomal_S15 (pfam00312)→ |
| env-1294 | NCBI_READ_1112494181464 | + | 794 | 861 | **RNA**→ |
| env-1295 | DelMOSum2010_c10001171 | + | 1998 | 2065 | **RNA**→ rpsO (PRK05626)Ribosomal_S15 (pfam00312)→ |
| env-1296 | JGI20154J14316_10023581 | - | 1717 | 1650 | **RNA**→ rpsO (PRK05626)Ribosomal_S15 (pfam00312)→ |
| env-1297 | JGI20160J14292_10181928 | - | 501 | 434 | **RNA**→ rpsO (PRK05626)Ribosomal_S15 (pfam00312)→ |
| env-1298 | JGI20157J14317_10011652 | + | 375 | 442 | **RNA**→ rpsO (PRK05626)Ribosomal_S15 (pfam00312)→ |
| env-1299 | NCBI_READ_1112493510178 | + | 737 | 804 | **RNA**→ |
| env-1300 | DelMOWin2010_c10091431 | - | 949 | 884 | **RNA**→ rpsO (PRK05626)Ribosomal_S15 (pfam00312)→ |
| env-1301 | 2236876010_0513203 | + | 165 | 230 | **RNA**→ rpsO (PRK05626)Ribosomal_S15 (pfam00312)→ |
| env-1302 | 2236876010_0390247 | + | 46 | 111 | **RNA**→ rpsO (PRK05626)Ribosomal_S15 (pfam00312)→ |
| env-1303 | 2236876010_0377328 | + | 221 | 286 | **RNA**→ ←hypo |
| env-1304 | CEPZ01057464.1 | + | 94811 | 94876 | **RNA**→ rpsO (PRK05626)Ribosomal_S15 (pfam00312)→ |
| env-1305 | SA_S2_NOR15_50m_c10278819 | + | 84 | 148 | **RNA**→ rpsO (PRK05626)Ribosomal_S15 (pfam00312)→ |
| env-1306 | SA_S1_NOR08_45m_c10000468 | + | 15998 | 16062 | **RNA**→ rpsO (PRK05626)Ribosomal_S15 (pfam00312)→ |
| env-1307 | SA_S1_NOR05_45m_c10094816 | + | 548 | 612 | **RNA**→ rpsO (PRK05626)→ |
| env-1308 | SA_S1_NOR08_45m_c10341122 | - | 105 | 42 | **RNA**→ |
| env-1309 | SA_S1_NOR08_45m_c10492149 | + | 268 | 332 | **RNA**→ |
| env-1310 | KGI_S1_ANT02_95m_c10084079 | - | 684 | 620 | **RNA**→ rpsO (PRK05626)RpsO (COG0184)→ |
| env-1311 | KGI_S2_ANT04_2345m_c1062656 | + | 34 | 98 | **RNA**→ rpsO (PRK05626)RpsO (COG0184)→ |
| Tna-1-1 | NZ_JTDV01000003.1 | - | 177687 | 177624 | **RNA**→ rpsO (PRK05626)RpsO (COG0184)→ |
| env-1312 | NCBI_READ_1112308404053 | - | 82 | 19 | **RNA**→ |
| Cly-1-1 | NZ_CP009239.1 | + | 671680 | 671743 | **RNA**→ rpsO (PRK05626)Ribosomal_S15 (pfam00312)→ |
| Cge-1-1 | NZ_ARZX01000025.1 | - | 10591 | 10528 | **RNA**→ rpsO (PRK05626)Ribosomal_S15 (pfam00312)→ |
| Cly-2-1 | NC_015167.1 | + | 719362 | 719425 | **RNA**→ rpsO (PRK05626)Ribosomal_S15 (pfam00312)→ |
| Sxi-1-1 | NZ_KE384055.1 | - | 51323 | 51260 | **RNA**→ rpsO (PRK05626)Ribosomal_S15 (pfam00312)→ |
| Ste-1-1 | NZ_KE384221.1 | - | 180772 | 180709 | **RNA**→ rpsO (PRK05626)Ribosomal_S15 (pfam00312)→ |
| Cal-1-1 | NC_014934.1 | - | 2557672 | 2557605 | **RNA**→ rpsO (PRK05626)Ribosomal_S15 (pfam00312)→ |
| env-1313 | G312J29652_10037410 | + | 360 | 426 | **RNA**→ rpsO (PRK05626)Ribosomal_S15 (pfam00312)→ |
| Mze-2-1 | NZ_AFOE01000038.1 | - | 92933 | 92869 | **RNA**→ rpsO (PRK05626)Ribosomal_S15 (pfam00312)→ rpsO (PRK05626)Ribosomal_S15 (pfam00312)→ |
| env-1314 | BS_KBB_SWE26_205m_c1009634 | + | 1084 | 1151 | **RNA**→ rpsO (PRK05626)Ribosomal_S15 (pfam00312)→ |
| env-1315 | BS_KBA_SWE21_205m_10014103 | + | 1629 | 1696 | **RNA**→ rpsO (PRK05626)Ribosomal_S15 (pfam00312)→ |

| | | | | | |
|---|---|---|---|---|---|
| env-1316 | BS_KBA_SWE12_21m_c10159732 | + | 214 | 281 | **RNA**→ rpsO (PRK05626)Ribosomal_S15 (pfam00312)→ |
| env-1317 | JGI24770J26754_10000514 | + | 13589 | 13655 | **RNA**→ rpsO (PRK05626)Ribosomal_S15 (pfam00312)→ |
| env-1318 | JGI24770J26754_10369233 | + | 63 | 130 | **RNA**→ rpsO (PRK05626)RpsO (COG0184)→ |
| Fgi-1-1 | NZ_JNCP01000044.1 | - | 44328 | 44263 | **RNA**→ rpsO (PRK05626)Ribosomal_S15 (pfam00312)→ |
| Fsu-2-1 | NZ_JATV01000015.1 | + | 31818 | 31885 | **RNA**→ rpsO (PRK05626)Ribosomal_S15 (pfam00312)→ |
| env-1319 | JGI24770J26754_10511844 | - | 322 | 256 | **RNA**→ rpsO (PRK05626)Ribosomal_S15 (pfam00312)→ |
| env-1320 | BS_KBB_SWE26_205m_c1206028 | + | 132 | 198 | **RNA**→ rpsO (PRK05626)Ribosomal_S15 (pfam00312)→ |
| env-1321 | BS_KBA_SWE12_21m_c10074375 | - | 497 | 431 | **RNA**→ rpsO (PRK05626)Ribosomal_S15 (pfam00312)→ |
| env-1322 | BS_KBA_SWE21_205m_10062507 | - | 450 | 384 | **RNA**→ rpsO (PRK05626)Ribosomal_S15 (pfam00312)→ |
| Fjo-1-1 | NC_009441.1 | - | 1659768 | 1659702 | **RNA**→ rpsO (PRK05626)Ribosomal_S15 (pfam00312)→ |
| Fsp-7-1 | NZ_AKZQ01000027.1 | - | 119685 | 119618 | **RNA**→ rpsO (PRK05626)Ribosomal_S15 (pfam00312)→ rpsO (PRK05626)Ribosomal_S15 (pfam00312)→ |
| Fsp-13-1 | NZ_AMYW01000001.1 | - | 340269 | 340204 | **RNA**→ rpsO (PRK05626)Ribosomal_S15 (pfam00312)→ |
| Fsp-6-1 | NZ_AKJZ01000010.1 | - | 81893 | 81828 | **RNA**→ rpsO (PRK05626)Ribosomal_S15 (pfam00312)→ rpsO (PRK05626)Ribosomal_S15 (pfam00312)→ |
| Fsp-8-1 | NZ_JSYO01000054.1 | + | 12825 | 12890 | **RNA**→ rpsO (PRK05626)Ribosomal_S15 (pfam00312)→ |
| Fsp-10-1 | NZ_JSYP01000001.1 | + | 564883 | 564948 | **RNA**→ rpsO (PRK05626)Ribosomal_S15 (pfam00312)→ |
| Fhy-1-1 | NZ_JPRM01000021.1 | + | 90628 | 90693 | **RNA**→ rpsO (PRK05626)Ribosomal_S15 (pfam00312)→ |
| env-1323 | ARWTCa_c258734 | + | 36 | 101 | **RNA**→ rpsO (PRK05626)Ribosomal_S15 (pfam00312)→ |
| env-1324 | PBR_1164806 | - | 6336 | 6271 | **RNA**→ rpsO (PRK05626)Ribosomal_S15 (pfam00312)→ |
| Fsp-11-1 | NZ_JXQR01000045.1 | - | 39476 | 39411 | **RNA**→ rpsO (PRK05626)Ribosomal_S15 (pfam00312)→ |
| Fch-1-1 | NZ_JASY01000002.1 | - | 513988 | 513923 | **RNA**→ rpsO (PRK05626)Ribosomal_S15 (pfam00312)→ |
| env-1325 | BS_KBA_SWE21_205m_10269192 | - | 314 | 250 | **RNA**→ rpsO (PRK05626)Ribosomal_S15 (pfam00312)→ |
| env-1326 | BS_KBA_SWE21_205m_10250829 | + | 230 | 294 | **RNA**→ rpsO (PRK05626)RpsO (COG0184)→ |
| env-1327 | BS_KBA_SWE02_21m_10049645 | - | 190 | 126 | **RNA**→ rpsO (PRK05626)Ribosomal_S15 (pfam00312)→ |
| env-1328 | BS_KBA_SWE12_21m_c10022633 | + | 520 | 584 | **RNA**→ rpsO (PRK05626)Ribosomal_S15 (pfam00312)→ |
| env-1329 | BS_KBB_SWE26_205m_c1001111 | - | 783 | 719 | **RNA**→ rpsO (PRK05626)Ribosomal_S15 (pfam00312)→ |
| env-1330 | JGI25914J50564_10000261 | - | 10238 | 10176 | **RNA**→ rpsO (PRK05626)Ribosomal_S15 (pfam00312)→ |
| env-1331 | JGI25917J50250_1000514 | + | 5135 | 5197 | **RNA**→ rpsO (PRK05626)Ribosomal_S15 (pfam00312)→ |
| env-1332 | TB_FS06_10DRAFT_1000014 | - | 38913 | 38850 | **RNA**→ rpsO (PRK05626)RpsO (COG0184)→ |
| env-1333 | TB_GS09_5_10013944 | + | 2931 | 2994 | **RNA**→ rpsO (PRK05626)Ribosomal_S15 (pfam00312)→ |
| Gsp-4-1 | NZ_JQLP01000005.1 | - | 753960 | 753897 | **RNA**→ rpsO (PRK05626)Ribosomal_S15 (pfam00312)→ |
| Gli-1-1 | NZ_JH594606.1 | + | 1080060 | 1080123 | **RNA**→ rpsO (PRK05626)Ribosomal_S15 (pfam00312)→ rpsO (PRK05626)Ribosomal_S15 (pfam00312)→ |
| env-1334 | ACOFG987_F36MELC02HNE7Q | - | 136 | 72 | **RNA**→ rpsO (PRK05626)RpsO (COG0184)→ |
| env-1335 | 3300001197_1006016 | - | 258 | 195 | **RNA**→ rpsO (PRK05626)Ribosomal_S15 (pfam00312)→ |
| env-1336 | KGI_S1_ANT02_95m_c10124631 | + | 34 | 97 | **RNA**→ rpsO (PRK05626)Ribosomal_S15 (pfam00312)→ |
| env-1337 | KGI_S1_ANT02_95m_c10029222 | + | 1799 | 1862 | **RNA**→ |
| env-1338 | KGI_S1_ANT01_95m_c10051775 | - | 89 | 26 | **RNA**→ |
| env-1339 | KGI_S1_ANT01_95m_c10120864 | + | 17 | 80 | **RNA**→ rpsO (PRK05626)RpsO (COG0184)→ |
| env-1340 | KGI_S2_ANT04_2345m_c1012199 | + | 2487 | 2550 | **RNA**→ ←hypo |
| env-1341 | KGI_S1_ANT02_95m_c10492692 | - | 191 | 128 | **RNA**→ rpsO (PRK05626)Ribosomal_S15 (pfam00312)→ |
| Gsp-5-1 | NZ_KE384218.1 | - | 925343 | 925280 | **RNA**→ rpsO (PRK05626)RpsO (COG0184)→ |
| env-1342 | KGI_S1_ANT01_95m_c10008468 | - | 78 | 15 | **RNA**→ ←hypo |
| env-1343 | KGI_S1_ANT01_95m_c10008586 | - | 85 | 22 | **RNA**→ |
| env-1344 | KGI_S1_ANT01_95m_c10068748 | - | 92 | 29 | **RNA**→ |
| env-1345 | KGI_S2_ANT04_2345m_c1047821 | + | 810 | 873 | **RNA**→ |
| env-1346 | KGI_S2_ANT04_2345m_c1020564 | - | 89 | 26 | **RNA**→ |
| Gsp-3-1 | NZ_AJLT01000036.1 | - | 4218 | 4155 | **RNA**→ rpsO (PRK05626)Ribosomal_S15 (pfam00312)→ rpsO (PRK05626)Ribosomal_S15 (pfam00312)→ |
| Gsp-2-1 | NZ_APHJ01000020.1 | - | 9198 | 9135 | **RNA**→ rpsO (PRK05626)RpsO (COG0184)→ |
| env-1347 | KGI_S1_ANT01_95m_c10037026 | - | 89 | 26 | **RNA**→ |
| env-1348 | KGI_S1_ANT02_95m_c10022116 | + | 2196 | 2259 | **RNA**→ |
| env-1349 | JGI1221J11331_1000234 | - | 4346 | 4283 | **RNA**→ rpsO (PRK05626)Ribosomal_S15 (pfam00312)→ |
| env-1350 | KGI_S1_ANT01_95m_c10095535 | - | 799 | 736 | **RNA**→ rpsO (PRK05626)Ribosomal_S15 (pfam00312)→ |
| Gec-1-1 | NZ_KE383973.1 | - | 8392 | 8330 | **RNA**→ rpsO (PRK05626)Ribosomal_S15 (pfam00312)→ |

| | | | | | |
|---|---|---|---|---|---|
| Gpo-1-1 | NZ_AUHF01000001.1 | - | 869907 | 869845 | RNA→ rpsO (PRK05626)Ribosomal_S15 (pfam00312)→ |
| env-1351 | CESL01230473.1 | + | 117 | 179 | RNA→ rpsO (PRK05626)Ribosomal_S15 (pfam00312)→ |
| env-1352 | CESF01213480.1 | + | 362 | 424 | RNA→ rpsO (PRK05626)RpsO (COG0184)→ |
| env-1353 | CETA01214501.1 | - | 215 | 153 | RNA→ rpsO (PRK05626)Ribosomal_S15 (pfam00312)→ |
| Gfo-1-1 | NC_008571.1 | - | 445156 | 445094 | RNA→ rpsO (PRK05626)Ribosomal_S15 (pfam00312)→ hypo→ Pnp (COG1185)PRK11824 (PRK11824)→ hypo→ hypo→ hypo→ |
| env-1354 | CEVK01150598.1 | + | 458 | 520 | RNA→ hypo→ |
| env-1355 | CEVO01219582.1 | + | 51 | 113 | RNA→ rpsO (PRK05626)Ribosomal_S15 (pfam00312)→ |
| env-1356 | CETX01274527.1 | - | 892 | 830 | RNA→ rpsO (PRK05626)Ribosomal_S15 (pfam00312)→ |
| env-1357 | CEUY01486981.1 | + | 1938 | 2000 | RNA→ rpsO (PRK05626)Ribosomal_S15 (pfam00312)→ |
| env-1358 | CETR01013089.1 | - | 4212 | 4150 | RNA→ rpsO (PRK05626)Ribosomal_S15 (pfam00312)→ |
| env-1359 | CETS01018170.1 | + | 146 | 208 | RNA→ rpsO (PRK05626)Ribosomal_S15 (pfam00312)→ |
| env-1360 | CEOS01036999.1 | + | 5728 | 5790 | RNA→ rpsO (PRK05626)Ribosomal_S15 (pfam00312)→ |
| env-1361 | CEPD01004855.1 | + | 10048 | 10110 | RNA→ rpsO (PRK05626)Ribosomal_S15 (pfam00312)→ |
| env-1362 | CERQ01017587.1 | - | 4212 | 4150 | RNA→ rpsO (PRK05626)Ribosomal_S15 (pfam00312)→ |
| env-1363 | CETA01098167.1 | - | 4212 | 4150 | RNA→ rpsO (PRK05626)Ribosomal_S15 (pfam00312)→ |
| env-1364 | CETG01119913.1 | - | 4015 | 3953 | RNA→ rpsO (PRK05626)Ribosomal_S15 (pfam00312)→ |
| env-1365 | CEUX01200289.1 | - | 611 | 549 | RNA→ rpsO (PRK05626)Ribosomal_S15 (pfam00312)→ |
| env-1366 | CEVI01039718.1 | + | 2211 | 2273 | RNA→ rpsO (PRK05626)Ribosomal_S15 (pfam00312)→ |
| Zpr-1-1 | NC_014041.1 | - | 1015479 | 1015417 | RNA→ rpsO (PRK05626)Ribosomal_S15 (pfam00312)→ |
| env-1367 | CEOX01230633.1 | + | 1223 | 1285 | RNA→ rpsO (PRK05626)Ribosomal_S15 (pfam00312)→ |
| env-1368 | CETU01337419.1 | - | 615 | 553 | RNA→ rpsO (PRK05626)Ribosomal_S15 (pfam00312)→ |
| env-1369 | CEVM01259718.1 | + | 226 | 288 | RNA→ rpsO (PRK05626)Ribosomal_S15 (pfam00312)→ |
| env-1370 | CETM01453121.1 | - | 2692 | 2630 | RNA→ rpsO (PRK05626)Ribosomal_S15 (pfam00312)→ |
| env-1371 | CESL01138425.1 | - | 5314 | 5252 | RNA→ rpsO (PRK05626)Ribosomal_S15 (pfam00312)→ |
| env-1372 | CEVA01504643.1 | - | 1690 | 1628 | RNA→ rpsO (PRK05626)Ribosomal_S15 (pfam00312)→ |
| env-1373 | CETB01101212.1 | - | 4212 | 4150 | RNA→ rpsO (PRK05626)Ribosomal_S15 (pfam00312)→ |
| env-1374 | CETO01054190.1 | - | 4212 | 4150 | RNA→ rpsO (PRK05626)Ribosomal_S15 (pfam00312)→ |
| env-1375 | CETT01093762.1 | - | 4212 | 4150 | RNA→ rpsO (PRK05626)Ribosomal_S15 (pfam00312)→ |
| env-1376 | CEVJ01161702.1 | + | 426 | 488 | RNA→ rpsO (PRK05626)Ribosomal_S15 (pfam00312)→ |
| env-1377 | CEWI01262926.1 | - | 4015 | 3953 | RNA→ rpsO (PRK05626)Ribosomal_S15 (pfam00312)→ |
| env-1378 | JGI1221J11331_1000144 | - | 6864 | 6801 | RNA→ rpsO (PRK05626)Ribosomal_S15 (pfam00312)→ |
| env-1379 | CEVI01097594.1 | - | 909 | 846 | RNA→ rpsO (PRK05626)Ribosomal_S15 (pfam00312)→ |
| env-1380 | CESF01293253.1 | - | 867 | 804 | RNA→ rpsO (PRK05626)Ribosomal_S15 (pfam00312)→ |
| env-1381 | CEVA01269270.1 | - | 286 | 223 | RNA→ rpsO (PRK05626)Ribosomal_S15 (pfam00312)→ |
| env-1382 | JGI11876J14442_10131815 | - | 547 | 484 | RNA→ rpsO (PRK05626)Ribosomal_S15 (pfam00312)→ |
| env-1383 | ElkS_mat_MD6A_1190057 | - | 183 | 120 | RNA→ rpsO (PRK05626)Ribosomal_S15 (pfam00312)→ |
| Ssp-1-1 | NZ_JQNQ01000001.1 | + | 899638 | 899701 | RNA→ rpsO (PRK05626)Ribosomal_S15 (pfam00312)→ |
| Tse-1-1 | NZ_JTDW01000005.1 | + | 421254 | 421317 | RNA→ rpsO (PRK05626)RpsO (COG0184)→ |
| Lsp-2-1 | NZ_JYNQ01000001.1 | - | 2806058 | 2805994 | RNA→ rpsO (PRK05626)Ribosomal_S15 (pfam00312)→ |
| Jpa-1-1 | NZ_BBNS01000015.1 | - | 28578 | 28513 | RNA→ rpsO (PRK05626)RpsO (COG0184)→ |
| Jpa-1-2 | NZ_BBNR01000016.1 | + | 55491 | 55556 | RNA→ rpsO (PRK05626)RpsO (COG0184)→ |
| Jpa-1-3 | NZ_BBNY01000095.1 | + | 87840 | 87905 | RNA→ rpsO (PRK05626)RpsO (COG0184)→ |
| Ssp-2-1 | NZ_JHZX01000001.1 | - | 2892930 | 2892865 | RNA→ rpsO (PRK05626)Ribosomal_S15 (pfam00312)→ |
| env-1384 | JGI24023J19991_10003832 | + | 3258 | 3322 | RNA→ rpsO (PRK05626)Ribosomal_S15 (pfam00312)→ |
| env-1385 | SA_S1_NOR08_45m_c10123497 | - | 596 | 531 | RNA→ rpsO (PRK05626)Ribosomal_S15 (pfam00312)→ |
| env-1386 | SA_S1_NOR05_45m_c10247099 | + | 238 | 303 | RNA→ rpsO (PRK05626)→ |
| env-1387 | KGI_S2_ANT05_2345m_c1162791 | + | 181 | 246 | RNA→ rpsO (PRK05626)Ribosomal_S15 (pfam00312)→ |
| env-1388 | KGI_S1_ANT03_95m_c1210231 | - | 114 | 49 | RNA→ hypo→ |
| env-1389 | KGI_S1_ANT01_95m_c10052137 | - | 700 | 635 | RNA→ rpsO (PRK05626)Ribosomal_S15 (pfam00312)→ |
| env-1390 | KGI_S1_ANT02_95m_c10092914 | + | 320 | 385 | RNA→ rpsO (PRK05626)Ribosomal_S15 (pfam00312)→ |
| env-1391 | SA_S2_NOR15_50m_c10267897 | + | 343 | 408 | RNA→ rpsO (PRK05626)Ribosomal_S15 (pfam00312)→ |
| env-1392 | CEVA01348977.1 | - | 69 | 4 | RNA→ |

| | | | | | |
|---|---|---|---|---|---|
| env-1393 | CEQX01182899.1 | + | 35 | 100 | **RNA**→ rpsO (PRK05626)RpsO (COG0184)→ |
| env-1394 | CEUN01355620.1 | + | 760 | 825 | **RNA**→ |
| env-1395 | CESF01312563.1 | - | 1354 | 1289 | **RNA**→ rpsO (PRK05626)RpsO (COG0184)→ |
| env-1396 | ElkS_mat_MD6A_1000645 | - | 997 | 932 | **RNA**→ rpsO (PRK05626)Ribosomal_S15 (pfam00312)→ |
| env-1397 | JGI11876J14442_10001894 | - | 4964 | 4899 | **RNA**→ rpsO (PRK05626)Ribosomal_S15 (pfam00312)→ |
| env-1398 | CGUI_GFCP3IQ02HOG5X | - | 70 | 5 | **RNA**→ |
| env-1399 | CEVA01011410.1 | - | 380 | 315 | **RNA**→ rpsO (PRK05626)RpsO (COG0184)→ |
| env-1400 | CEQH01190568.1 | - | 69 | 4 | **RNA**→ |
| env-1401 | CEUN01165117.1 | + | 490 | 555 | **RNA**→ rpsO (PRK05626)RpsO (COG0184)→ |
| env-1402 | CEUF01199729.1 | - | 259 | 194 | **RNA**→ rpsO (PRK05626)RpsO (COG0184)→ |
| env-1403 | JGI11876J14442_10807421 | - | 249 | 184 | **RNA**→ rpsO (PRK05626)RpsO (COG0184)→ |
| env-1404 | TDF_OR_ARG04_113m_c1142939 | + | 11 | 76 | **RNA**→ rpsO (PRK05626)Ribosomal_S15 (pfam00312)→ |
| env-1405 | KGI_S1_ANT03_95m_c1006422 | + | 12 | 77 | **RNA**→ rpsO (PRK05626)Ribosomal_S15 (pfam00312)→ |
| env-1406 | KGI_S1_ANT01_95m_c10028809 | - | 1106 | 1041 | **RNA**→ rpsO (PRK05626)Ribosomal_S15 (pfam00312)→ |
| env-1407 | KGI_S1_ANT01_95m_c10013761 | + | 2777 | 2842 | **RNA**→ rpsO (PRK05626)Ribosomal_S15 (pfam00312)→ |
| env-1408 | KGI_S2_ANT06_2345m_c1017384 | - | 817 | 752 | **RNA**→ rpsO (PRK05626)Ribosomal_S15 (pfam00312)→ |
| env-1409 | KGI_S2_ANT04_2345m_c1018151 | - | 817 | 752 | **RNA**→ rpsO (PRK05626)Ribosomal_S15 (pfam00312)→ |
| env-1410 | KGI_S1_ANT02_95m_c10015429 | - | 1648 | 1583 | **RNA**→ rpsO (PRK05626)Ribosomal_S15 (pfam00312)→ |
| env-1411 | SA_S1_NOR08_45m_c10818453 | + | 54 | 119 | **RNA**→ rpsO (PRK05626)Ribosomal_S15 (pfam00312)→ |
| Wsp-1-1 | NZ_AP014583.1 | - | 2199375 | 2199310 | **RNA**→ rpsO (PRK05626)Ribosomal_S15 (pfam00312)→ |
| env-1412 | KGI_S1_ANT01_95m_c10108529 | + | 255 | 320 | **RNA**→ rpsO (PRK05626)RpsO (COG0184)→ |
| env-1413 | SA_S1_NOR05_45m_c10222323 | - | 117 | 52 | **RNA**→ |
| env-1414 | KGI_S1_ANT02_95m_c10018235 | + | 1331 | 1396 | **RNA**→ rpsO (PRK05626)RpsO (COG0184)→ |
| env-1415 | KGI_S1_ANT01_95m_c10439590 | - | 216 | 151 | **RNA**→ rpsO (PRK05626)Ribosomal_S15 (pfam00312)→ |
| env-1416 | JGI24024J18818_10456388 | - | 203 | 138 | **RNA**→ rpsO (PRK05626)Ribosomal_S15 (pfam00312)→ |
| Bar-1-1 | NZ_AFXZ01000012.1 | + | 168728 | 168815 | **RNA**→ rpsO (PRK05626)RpsO (COG0184)→ rpsO (PRK05626)RpsO (COG0184)→ |
| env-1417 | JGI24724J26744_10000170 | - | 8375 | 8311 | **RNA**→ rpsO (PRK05626)RpsO (COG0184)→ |
| env-1418 | CEWQ01060895.1 | - | 188 | 124 | **RNA**→ rpsO (PRK05626)Ribosomal_S15 (pfam00312)→ |
| env-1419 | CESO01252919.1 | + | 473 | 537 | **RNA**→ rpsO (PRK05626)Ribosomal_S15 (pfam00312)→ |
| env-1420 | JGI12097J13213_1000798 | - | 3534 | 3470 | **RNA**→ rpsO (PRK05626)Ribosomal_S15 (pfam00312)→ |
| Pme-1-1 | NZ_JXJP01000001.1 | - | 451220 | 451155 | **RNA**→ rpsO (PRK05626)Ribosomal_S15 (pfam00312)→ |
| env-1421 | CEWR01222457.1 | - | 577 | 512 | **RNA**→ rpsO (PRK05626)Ribosomal_S15 (pfam00312)→ |
| Psp-5-1 | NZ_JUGU01000001.1 | - | 3801057 | 3800992 | **RNA**→ rpsO (PRK05626)Ribosomal_S15 (pfam00312)→ |
| Pbu-1-1 | NZ_AUDE01000023.1 | - | 4567 | 4502 | **RNA**→ rpsO (PRK05626)RpsO (COG0184)→ |
| env-1422 | JGI24025J20009_10010090 | + | 1129 | 1194 | **RNA**→ rpsO (PRK05626)Ribosomal_S15 (pfam00312)→ |
| env-1423 | JGI24724J26744_10030194 | - | 1651 | 1586 | **RNA**→ rpsO (PRK05626)Ribosomal_S15 (pfam00312)→ |
| env-1424 | JGI24723J26617_10025495 | - | 1546 | 1481 | **RNA**→ rpsO (PRK05626)Ribosomal_S15 (pfam00312)→ |
| env-1425 | SA_S1_NOR08_45m_c10367044 | + | 199 | 264 | **RNA**→ rpsO (PRK05626)Ribosomal_S15 (pfam00312)→ |
| Pja-1-1 | NZ_JSWG01000009.1 | - | 755527 | 755462 | **RNA**→ rpsO (PRK05626)Ribosomal_S15 (pfam00312)→ |
| env-1426 | JGI24724J26744_10068913 | - | 1143 | 1078 | **RNA**→ rpsO (PRK05626)Ribosomal_S15 (pfam00312)→ |
| env-1427 | JGI24723J26617_10012390 | + | 378 | 443 | **RNA**→ rpsO (PRK05626)Ribosomal_S15 (pfam00312)→ |
| env-1428 | JGI24723J26617_10058375 | - | 123 | 58 | **RNA**→ |
| env-1429 | JGI20151J14362_10049778 | + | 238 | 303 | **RNA**→ rpsO (PRK05626)Ribosomal_S15 (pfam00312)→ |
| env-1430 | JGI20156J14371_10001957 | - | 2149 | 2084 | **RNA**→ rpsO (PRK05626)Ribosomal_S15 (pfam00312)→ |
| env-1431 | JGI20158J14315_10011911 | + | 3564 | 3629 | **RNA**→ rpsO (PRK05626)Ribosomal_S15 (pfam00312)→ |
| env-1432 | CERM01194957.1 | + | 488 | 553 | **RNA**→ rpsO (PRK05626)Ribosomal_S15 (pfam00312)→ |
| env-1433 | 2236876011_209432 | + | 56 | 121 | **RNA**→ rpsO (PRK05626)Ribosomal_S15 (pfam00312)→ |
| env-1434 | JGI11876J14442_10001335 | - | 9660 | 9595 | **RNA**→ rpsO (PRK05626)RpsO (COG0184)→ hypo→ |
| env-1435 | ElkS_mat_MD6A_1062416 | + | 49 | 114 | **RNA**→ rpsO (PRK05626)RpsO (COG0184)→ |
| env-1436 | KGI_S2_ANT04_2345m_c1011179 | - | 642 | 578 | **RNA**→ rpsO (PRK05626)Ribosomal_S15 (pfam00312)→ |
| Oma-1-1 | NZ_KI912443.1 | - | 246097 | 246033 | **RNA**→ rpsO (PRK05626)Ribosomal_S15 (pfam00312)→ |
| env-1437 | wf3_AHAI12229_x1 | - | 290 | 226 | **RNA**→ rpsO (PRK05626)Ribosomal_S15 (pfam00312)→ |

| | | | | | |
|---|---|---|---|---|---|
| env-1438 | JGI24724J26744_10003514 | - | 4752 | 4688 | **RNA**→ rpsO (PRK05626)Ribosomal_S15 (pfam00312)→ |
| env-1439 | CESF01013150.1 | - | 383089 | 383025 | **RNA**→ rpsO (PRK05626)Ribosomal_S15 (pfam00312)→ |
| env-1440 | CESO01127605.1 | - | 775 | 711 | **RNA**→ rpsO (PRK05626)Ribosomal_S15 (pfam00312)→ |
| Sze-1-1 | NZ_CP007202.1 | - | 1742017 | 1741953 | **RNA**→ rpsO (PRK05626)Ribosomal_S15 (pfam00312)→ hypo→ hypo→ |
| Gsa-1-1 | NZ_AUDV01000006.1 | + | 383148 | 383212 | **RNA**→ rpsO (PRK05626)RpsO (COG0184)→ hypo→ hypo→ |
| env-1441 | KGI_S1_ANT01_95m_c10171289 | - | 192 | 128 | **RNA**→ rpsO (PRK05626)Ribosomal_S15 (pfam00312)→ |
| env-1442 | ElkS_mat_MD6A_1000561 | - | 14496 | 14432 | **RNA**→ rpsO (PRK05626)Ribosomal_S15 (pfam00312)→ |
| env-1443 | JGI11876J14442_10000603 | - | 26866 | 26802 | **RNA**→ rpsO (PRK05626)Ribosomal_S15 (pfam00312)→ |
| Mze-1-1 | NZ_KE383820.1 | - | 249635 | 249571 | **RNA**→ rpsO (PRK05626)Ribosomal_S15 (pfam00312)→ |
| env-1444 | CEUX01207121.1 | + | 1570 | 1634 | **RNA**→ rpsO (PRK05626)Ribosomal_S15 (pfam00312)→ |
| env-1445 | KGI_S2_ANT04_2345m_c1032766 | - | 578 | 514 | **RNA**→ rpsO (PRK05626)Ribosomal_S15 (pfam00312)→ |
| env-1446 | KGI_S1_ANT02_95m_c10040186 | + | 144 | 208 | **RNA**→ rpsO (PRK05626)Ribosomal_S15 (pfam00312)→ |
| env-1447 | SA_S1_NOR02_45m_c1045496 | - | 449 | 385 | **RNA**→ rpsO (PRK05626)Ribosomal_S15 (pfam00312)→ |
| env-1448 | SA_S2_NOR13_50m_c1120752 | + | 50 | 114 | **RNA**→ rpsO (PRK05626)Ribosomal_S15 (pfam00312)→ |
| env-1449 | SA_S2_NOR13_50m_c1031479 | - | 458 | 394 | **RNA**→ rpsO (PRK05626)Ribosomal_S15 (pfam00312)→ |
| env-1450 | SA_S1_NOR05_45m_c10015538 | + | 1854 | 1918 | **RNA**→ rpsO (PRK05626)Ribosomal_S15 (pfam00312)→ hypo→ |
| env-1451 | SA_S1_NOR08_45m_c10002201 | - | 821 | 757 | **RNA**→ rpsO (PRK05626)Ribosomal_S15 (pfam00312)→ hypo→ |
| env-1452 | SA_S2_NOR15_50m_c10001453 | + | 9897 | 9961 | **RNA**→ rpsO (PRK05626)Ribosomal_S15 (pfam00312)→ hypo→ |
| env-1453 | SA_S2_NOR18_50m_1054082 | + | 176 | 240 | **RNA**→ rpsO (PRK05626)Ribosomal_S15 (pfam00312)→ |
| env-1454 | KGI_S1_ANT02_95m_c10590781 | + | 23 | 87 | **RNA**→ rpsO (PRK05626)Ribosomal_S15 (pfam00312)→ |
| env-1455 | wf3_AHAI10363_y1 | - | 203 | 139 | **RNA**→ rpsO (PRK05626)→ |
| Wps-1-1 | NZ_ATMR01000007.1 | + | 31896 | 31960 | **RNA**→ rpsO (PRK05626)Ribosomal_S15 (pfam00312)→ |
| env-1456 | wf1_AGZO11054_b2 | + | 348 | 412 | **RNA**→ rpsO (PRK05626)RpsO (COG0184)→ |
| env-1457 | JGI24723J26617_10002165 | + | 14707 | 14771 | **RNA**→ rpsO (PRK05626)Ribosomal_S15 (pfam00312)→ |
| Osp-2-1 | NZ_JADT01000002.1 | + | 86800 | 86864 | **RNA**→ rpsO (PRK05626)Ribosomal_S15 (pfam00312)→ |
| Osp-1-1 | NZ_JADS01000007.1 | + | 86864 | 86928 | **RNA**→ rpsO (PRK05626)Ribosomal_S15 (pfam00312)→ |
| env-1458 | JGI24724J26744_10155390 | + | 492 | 556 | **RNA**→ rpsO (PRK05626)Ribosomal_S15 (pfam00312)→ |
| env-1459 | JGI24723J26617_10081657 | + | 261 | 325 | **RNA**→ rpsO (PRK05626)Ribosomal_S15 (pfam00312)→ |
| env-1460 | SA_S1_NOR08_45m_c10027555 | + | 957 | 1021 | **RNA**→ rpsO (PRK05626)Ribosomal_S15 (pfam00312)→ |
| env-1461 | JGI24724J26744_10015488 | + | 1261 | 1325 | **RNA**→ rpsO (PRK05626)Ribosomal_S15 (pfam00312)→ |
| env-1462 | JGI24723J26617_10001993 | + | 11787 | 11851 | **RNA**→ rpsO (PRK05626)Ribosomal_S15 (pfam00312)→ |
| env-1463 | KGI_S1_ANT02_95m_c10261135 | + | 309 | 373 | **RNA**→ hypo→ |
| Lsp-1-1 | NC_015638.1 | + | 582743 | 582807 | **RNA**→ rpsO (PRK05626)Ribosomal_S15 (pfam00312)→ |
| env-1464 | SA_S2_NOR15_50m_c10085505 | + | 209 | 273 | **RNA**→ rpsO (PRK05626)Ribosomal_S15 (pfam00312)→ |
| env-1465 | KGI_S1_ANT02_95m_c10028795 | + | 1129 | 1193 | **RNA**→ rpsO (PRK05626)Ribosomal_S15 (pfam00312)→ |
| env-1466 | KGI_S1_ANT01_95m_c10028466 | - | 325 | 261 | **RNA**→ rpsO (PRK05626)Ribosomal_S15 (pfam00312)→ |
| env-1467 | KGI_S2_ANT06_2345m_c1003951 | - | 969 | 905 | **RNA**→ rpsO (PRK05626)RpsO (COG0184)→ |
| env-1468 | KGI_S2_ANT05_2345m_c1024722 | + | 428 | 492 | **RNA**→ rpsO (PRK05626)Ribosomal_S15 (pfam00312)→ rpsO (PRK05626)Ribosomal_S15 (pfam00312)→ |
| env-1469 | KGI_S2_ANT06_2345m_c1002277 | - | 564 | 500 | **RNA**→ rpsO (PRK05626)Ribosomal_S15 (pfam00312)→ |
| env-1470 | KGI_S1_ANT03_95m_c1005571 | + | 822 | 886 | **RNA**→ rpsO (PRK05626)Ribosomal_S15 (pfam00312)→ |
| env-1471 | CEQG01149785.1 | - | 2866 | 2802 | **RNA**→ rpsO (PRK05626)RpsO (COG0184)→ |
| Fba-5-1 | NZ_ABHI01000002.1 | - | 642916 | 642852 | **RNA**→ rpsO (PRK05626)Ribosomal_S15 (pfam00312)→ rpsO (PRK05626)Ribosomal_S15 (pfam00312)→ |
| Fic-1-1 | NZ_KN525716.1 | + | 1154245 | 1154309 | **RNA**→ rpsO (PRK05626)Ribosomal_S15 (pfam00312)→ hypo→ hypo→ |
| env-1472 | JGI24025J20009_10267879 | - | 111 | 47 | **RNA**→ hypo→ |
| Lja-1-1 | NZ_JSWF01000036.1 | + | 51771 | 51835 | **RNA**→ rpsO (PRK05626)Ribosomal_S15 (pfam00312)→ |
| env-1473 | JGI24025J20009_10053124 | - | 1206 | 1142 | **RNA**→ rpsO (PRK05626)RpsO (COG0184)→ |
| Fba-8-1 | NZ_ALIH01000008.1 | + | 90157 | 90221 | **RNA**→ rpsO (PRK05626)RpsO (COG0184)→ hypo→ hypo→ |
| env-1474 | JGI24723J26617_10480515 | - | 213 | 149 | **RNA**→ rpsO (PRK05626)Ribosomal_S15 (pfam00312)→ |
| env-1475 | JGI11876J14442_10711006 | - | 246 | 182 | **RNA**→ rpsO (PRK05626)Ribosomal_S15 (pfam00312)→ |
| env-1476 | JGI24723J26617_10000349 | - | 8866 | 8802 | **RNA**→ rpsO (PRK05626)Ribosomal_S15 (pfam00312)→ |
| env-1477 | SA_S1_NOR05_45m_c10059526 | - | 797 | 733 | **RNA**→ rpsO (PRK05626)Ribosomal_S15 (pfam00312)→ |
| env-1478 | SA_S2_NOR15_50m_c10011596 | - | 1359 | 1295 | **RNA**→ rpsO (PRK05626)Ribosomal_S15 (pfam00312)→ |

| | | | | | |
|---|---|---|---|---|---|
| env-1479 | SA_S1_NOR08_45m_c10057202 | + | 230 | 294 | **RNA**→ rpsO (PRK05626)Ribosomal_S15 (pfam00312)→ |
| env-1480 | SA_S1_NOR08_45m_c10566095 | - | 249 | 185 | **RNA**→ rpsO (PRK05626)Ribosomal_S15 (pfam00312)→ |
| env-1481 | SA_S2_NOR15_50m_c10000178 | + | 21761 | 21825 | **RNA**→ rpsO (PRK05626)Ribosomal_S15 (pfam00312)→ |
| env-1482 | KGI_S1_ANT03_95m_c1016120 | - | 66 | 2 | **RNA**→ |
| env-1483 | CEUD01278901.1 | + | 657 | 721 | **RNA**→ rpsO (PRK05626)Ribosomal_S15 (pfam00312)→ |
| env-1484 | KGI_S1_ANT01_95m_c10027578 | - | 798 | 734 | **RNA**→ rpsO (PRK05626)Ribosomal_S15 (pfam00312)→ |
| env-1485 | SA_S1_NOR05_45m_c10358528 | + | 155 | 219 | **RNA**→ rpsO (PRK05626)RpsO (COG0184)→ |
| env-1486 | SA_S1_NOR08_45m_c10002212 | - | 620 | 556 | **RNA**→ rpsO (PRK05626)Ribosomal_S15 (pfam00312)→ |
| env-1487 | SA_S2_NOR15_50m_c10549841 | - | 196 | 132 | **RNA**→ rpsO (PRK05626)Ribosomal_S15 (pfam00312)→ |
| env-1488 | KGI_S1_ANT02_95m_c10003746 | - | 2134 | 2070 | **RNA**→ rpsO (PRK05626)Ribosomal_S15 (pfam00312)→ |
| env-1489 | SA_S1_NOR05_45m_c10195079 | + | 321 | 385 | **RNA**→ hypo→ |
| env-1490 | SA_S1_NOR02_45m_c1140729 | + | 236 | 300 | **RNA**→ ←hypo |
| env-1491 | JGI20152J14361_10000356 | + | 14689 | 14753 | **RNA**→ rpsO (PRK05626)Ribosomal_S15 (pfam00312)→ |
| env-1492 | JGI20154J14316_10013934 | - | 3274 | 3210 | **RNA**→ rpsO (PRK05626)Ribosomal_S15 (pfam00312)→ |
| env-1493 | JGI20160J14292_10006817 | - | 6691 | 6627 | **RNA**→ rpsO (PRK05626)Ribosomal_S15 (pfam00312)→ |
| env-1494 | JGI20153J14295_10001021 | + | 14651 | 14715 | **RNA**→ rpsO (PRK05626)Ribosomal_S15 (pfam00312)→ |
| env-1495 | JGI20153J14318_10001021 | + | 14651 | 14715 | **RNA**→ rpsO (PRK05626)Ribosomal_S15 (pfam00312)→ |
| env-1496 | A09P04_1300_Sequence0000005964 | - | 443 | 379 | **RNA**→ rpsO (PRK05626)Ribosomal_S15 (pfam00312)→ |
| env-1497 | SA_S1_NOR05_45m_c10061437 | + | 558 | 622 | **RNA**→ rpsO (PRK05626)Ribosomal_S15 (pfam00312)→ |
| env-1498 | SA_S2_NOR15_50m_c10172884 | - | 143 | 79 | **RNA**→ rpsO (PRK05626)→ |
| env-1499 | SA_S2_NOR18_50m_1220507 | - | 273 | 209 | **RNA**→ rpsO (PRK05626)Ribosomal_S15 (pfam00312)→ |
| env-1500 | SA_S1_NOR08_45m_c10021859 | - | 176 | 112 | **RNA**→ rpsO (PRK05626)Ribosomal_S15 (pfam00312)→ |
| env-1501 | SA_S2_NOR15_50m_c10038244 | - | 971 | 907 | **RNA**→ rpsO (PRK05626)Ribosomal_S15 (pfam00312)→ hypo→ |
| env-1502 | SA_S1_NOR05_45m_c10068838 | + | 336 | 400 | **RNA**→ rpsO (PRK05626)Ribosomal_S15 (pfam00312)→ |
| env-1503 | SA_S1_NOR08_45m_c10016610 | + | 2682 | 2746 | **RNA**→ rpsO (PRK05626)Ribosomal_S15 (pfam00312)→ hypo→ |
| env-1504 | SA_S1_NOR08_45m_c10435309 | + | 269 | 333 | **RNA**→ |
| env-1505 | SA_S1_NOR08_45m_c10144408 | - | 487 | 423 | **RNA**→ rpsO (PRK05626)Ribosomal_S15 (pfam00312)→ hypo→ |
| env-1506 | KGI_S2_ANT04_2345m_c1296240 | - | 281 | 217 | **RNA**→ rpsO (PRK05626)Ribosomal_S15 (pfam00312)→ |
| Pda-1-1 | NZ_JXJO01000023.1 | + | 45518 | 45582 | **RNA**→ rpsO (PRK05626)Ribosomal_S15 (pfam00312)→ |
| env-1507 | SA_S1_NOR08_45m_c10764472 | - | 134 | 71 | **RNA**→ rpsO (PRK05626)RpsO (COG0184)→ |
| env-1508 | BS_KBA_SWE12_21m_c10000055 | + | 27026 | 27089 | **RNA**→ rpsO (PRK05626)RpsO (COG0184)→ |
| env-1509 | BS_KBB_SWE26_205m_c1007908 | + | 294 | 357 | **RNA**→ rpsO (PRK05626)RpsO (COG0184)→ |
| env-1510 | BS_KBB_SWE26_205m_c1015589 | + | 493 | 556 | **RNA**→ rpsO (PRK05626)RpsO (COG0184)→ |
| env-1511 | BS_KBA_SWE07_21m_c1008693 | + | 159 | 222 | **RNA**→ rpsO (PRK05626)RpsO (COG0184)→ |
| env-1512 | BS_KBA_SWE21_205m_10000856 | + | 7869 | 7932 | **RNA**→ rpsO (PRK05626)RpsO (COG0184)→ |
| env-1513 | BS_KBA_SWE21_205m_10001614 | - | 7075 | 7012 | **RNA**→ rpsO (PRK05626)RpsO (COG0184)→ |
| env-1514 | BS_KBA_SWE02_21m_10000074 | + | 25016 | 25079 | **RNA**→ rpsO (PRK05626)RpsO (COG0184)→ |
| env-1515 | 3300001592_10093314 | - | 181 | 117 | **RNA**→ rpsO (PRK05626)Ribosomal_S15 (pfam00312)→ |
| env-1516 | KVRMV2_100601001 | + | 114 | 178 | **RNA**→ rpsO (PRK05626)RpsO (COG0184)→ |
| env-1517 | CEOS01128007.1 | - | 419 | 357 | **RNA**→ rpsO (PRK05626)Ribosomal_S15 (pfam00312)→ |
| env-1518 | CERQ01221307.1 | + | 319 | 381 | **RNA**→ rpsO (PRK05626)Ribosomal_S15 (pfam00312)→ |
| env-1519 | BS_KBA_SWE21_205m_10160982 | - | 121 | 58 | **RNA**→ hypo→ |
| env-1520 | BS_KBA_SWE02_21m_10189888 | - | 297 | 234 | **RNA**→ rpsO (PRK05626)Ribosomal_S15 (pfam00312)→ |
| env-1521 | BS_KBA_SWE12_21m_c10156796 | + | 253 | 316 | **RNA**→ rpsO (PRK05626)Ribosomal_S15 (pfam00312)→ |
| env-1522 | CEQZ01049836.1 | - | 1155 | 1068 | **RNA**→ rpsO (PRK05626)RpsO (COG0184)→ |
| env-1523 | CERU01038294.1 | - | 1201 | 1114 | **RNA**→ rpsO (PRK05626)RpsO (COG0184)→ |
| Fsp-14-1 | NZ_ANLA01000005.1 | - | 82448 | 82361 | **RNA**→ rpsO (PRK05626)RpsO (COG0184)→ |
| env-1524 | CERY01028704.1 | - | 371 | 284 | **RNA**→ rpsO (PRK05626)RpsO (COG0184)→ |
| env-1525 | KGI_S1_ANT01_95m_c10252634 | - | 205 | 141 | **RNA**→ rpsO (PRK05626)Ribosomal_S15 (pfam00312)→ |
| env-1526 | KGI_S1_ANT02_95m_c10380062 | - | 148 | 84 | **RNA**→ rpsO (PRK05626)→ |
| env-1527 | SA_S1_NOR08_45m_c10034326 | + | 179 | 243 | **RNA**→ rpsO (PRK05626)Ribosomal_S15 (pfam00312)→ |
| env-1528 | TB_PC08_66_10134032 | + | 487 | 551 | **RNA**→ rpsO (PRK05626)Ribosomal_S15 (pfam00312)→ |

| | | | | | |
|---|---|---|---|---|---|
| env-1529 | BS_KBB_SWE26_205m_c1003922 | + | 1790 | 1854 | **RNA**→ rpsO (PRK05626)RpsO (COG0184)→ |
| env-1530 | BS_KBA_SWE12_21m_c10003244 | - | 3098 | 3034 | **RNA**→ rpsO (PRK05626)RpsO (COG0184)→ |
| env-1531 | BS_KBB_SWE26_205m_c1000422 | + | 8956 | 9020 | **RNA**→ rpsO (PRK05626)RpsO (COG0184)→ |
| env-1532 | BS_KBA_SWE21_205m_10001432 | + | 5368 | 5432 | **RNA**→ rpsO (PRK05626)RpsO (COG0184)→ |
| env-1533 | BS_KBA_SWE02_21m_10001031 | - | 1511 | 1447 | **RNA**→ rpsO (PRK05626)RpsO (COG0184)→ |
| Fag-1-1 | NZ_HG315671.1 | - | 573237 | 573173 | **RNA**→ rpsO (PRK05626)RpsO (COG0184)→ |
| [1]env-1534 | KGI_S1_ANT01_95m_c10197802 | + | 270 | 324 | **RNA**→ rpsO (PRK05626)Ribosomal_S15 (pfam00312)→ rpsO (PRK05626)Ribosomal_S15 (pfam00312)→ |
| env-1535 | KGI_S1_ANT01_95m_c10417003 | + | 73 | 137 | **RNA**→ rpsO (PRK05626)Ribosomal_S15 (pfam00312)→ |
| env-1536 | KGI_S1_ANT01_95m_c10197802 | + | 146 | 210 | **RNA**→ rpsO (PRK05626)→ rpsO (PRK05626)→ rpsO (PRK05626)Ribosomal_S15 (pfam00312)→ rpsO (PRK05626)Ribosomal_S15 (pfam00312)→ |
| env-1537 | KGI_S2_ANT04_2345m_c1079454 | - | 304 | 240 | **RNA**→ rpsO (PRK05626)RpsO (COG0184)→ |
| env-1538 | SA_S1_NOR05_45m_c10195650 | - | 251 | 186 | **RNA**→ rpsO (PRK05626)Ribosomal_S15 (pfam00312)→ |
| env-1539 | SA_S2_NOR13_50m_c1186424 | + | 131 | 196 | **RNA**→ rpsO (PRK05626)→ |
| env-1540 | SA_S1_NOR08_45m_c10053497 | - | 624 | 559 | **RNA**→ rpsO (PRK05626)RpsO (COG0184)→ |
| env-1541 | KGI_S1_ANT02_95m_c10081391 | + | 477 | 542 | **RNA**→ rpsO (PRK05626)RpsO (COG0184)→ |
| env-1542 | KGI_S1_ANT01_95m_c10127393 | - | 466 | 401 | **RNA**→ rpsO (PRK05626)RpsO (COG0184)→ |
| env-1543 | SA_S1_NOR05_45m_c10021755 | + | 211 | 276 | **RNA**→ rpsO (PRK05626)RpsO (COG0184)→ |
| env-1544 | SA_S2_NOR15_50m_c10037074 | - | 1773 | 1708 | **RNA**→ rpsO (PRK05626)RpsO (COG0184)→ |
| env-1545 | SA_S2_NOR18_50m_1058205 | + | 140 | 205 | **RNA**→ rpsO (PRK05626)RpsO (COG0184)→ |
| env-1546 | SA_S1_NOR02_45m_c1038615 | + | 422 | 487 | **RNA**→ rpsO (PRK05626)→ |
| env-1547 | SA_S1_NOR05_45m_c10018119 | + | 1240 | 1305 | **RNA**→ rpsO (PRK05626)RpsO (COG0184)→ |
| env-1548 | JGI24024J18818_10009536 | - | 3130 | 3065 | **RNA**→ rpsO (PRK05626)Ribosomal_S15 (pfam00312)→ |
| env-1549 | SA_S1_NOR08_45m_c10538363 | + | 246 | 311 | **RNA**→ |
| env-1550 | JGI24024J18818_10055265 | + | 859 | 923 | **RNA**→ rpsO (PRK05626)Ribosomal_S15 (pfam00312)→ |
| env-1551 | JGI24023J19991_10458387 | - | 291 | 227 | **RNA**→ rpsO (PRK05626)Ribosomal_S15 (pfam00312)→ |
| env-1552 | SA_S2_NOR15_50m_c10835355 | - | 217 | 153 | **RNA**→ rpsO (PRK05626)Ribosomal_S15 (pfam00312)→ |
| env-1553 | SA_S1_NOR08_45m_c10167461 | - | 625 | 561 | **RNA**→ rpsO (PRK05626)Ribosomal_S15 (pfam00312)→ |
| env-1554 | SA_S1_NOR08_45m_c10626463 | + | 221 | 285 | **RNA**→ |
| env-1555 | JGI11876J14442_10165063 | + | 433 | 498 | **RNA**→ rpsO (PRK05626)Ribosomal_S15 (pfam00312)→ |
| env-1556 | ElkS_mat_MD6A_1051781 | - | 206 | 141 | **RNA**→ rpsO (PRK05626)Ribosomal_S15 (pfam00312)→ |
| env-1557 | ElkS_mat_MD6A_1006443 | + | 2218 | 2283 | **RNA**→ rpsO (PRK05626)Ribosomal_S15 (pfam00312)→ |
| env-1558 | JGI11876J14442_10009103 | - | 1440 | 1375 | **RNA**→ rpsO (PRK05626)Ribosomal_S15 (pfam00312)→ |
| env-1559 | SA_S1_NOR02_45m_c1138063 | - | 201 | 137 | **RNA**→ rpsO (PRK05626)Ribosomal_S15 (pfam00312)→ |
| env-1560 | SA_S1_NOR05_45m_c10066143 | + | 481 | 545 | **RNA**→ rpsO (PRK05626)Ribosomal_S15 (pfam00312)→ |
| env-1561 | SRS043422_C2682413 | + | 1246 | 1291 | **RNA**→ rpsO (PRK05626)Ribosomal_S15 (pfam00312)→ |
| env-1562 | SRS046686_C887109 | + | 19 | 64 | **RNA**→ rpsO (PRK05626)Ribosomal_S15 (pfam00312)→ |
| env-1563 | Buccal_mucosa_LANL_scaffold_1631 | + | 636 | 681 | **RNA**→ rpsO (PRK05626)Ribosomal_S15 (pfam00312)→ |
| env-1564 | SRS014687_C1256631 | - | 75 | 30 | **RNA**→ |
| env-1565 | SRS013506_C1077366 | + | 303 | 348 | **RNA**→ |
| env-1566 | TrichMG_c100003 | - | 114945 | 114878 | **RNA**→ rpsO (PRK05626)Ribosomal_S15 (pfam00312)→ |
| Zam-1-1 | NZ_AYXY01000022.1 | + | 24468 | 24536 | **RNA**→ rpsO (PRK05626)Ribosomal_S15 (pfam00312)→ |
| env-1567 | JGI24770J26754_10002254 | - | 8613 | 8546 | **RNA**→ rpsO (PRK05626)Ribosomal_S15 (pfam00312)→ |
| Ffr-2-1 | NZ_AHKF01000016.1 | - | 97801 | 97735 | **RNA**→ rpsO (PRK05626)Ribosomal_S15 (pfam00312)→ rpsO (PRK05626)Ribosomal_S15 (pfam00312)→ |
| env-1568 | KGI_S1_ANT01_95m_c10016611 | - | 1987 | 1921 | **RNA**→ rpsO (PRK05626)Ribosomal_S15 (pfam00312)→ |
| env-1569 | KGI_S2_ANT04_2345m_c1022905 | - | 1437 | 1371 | **RNA**→ rpsO (PRK05626)Ribosomal_S15 (pfam00312)→ |
| env-1570 | KGI_S1_ANT02_95m_c10016777 | - | 2633 | 2567 | **RNA**→ rpsO (PRK05626)Ribosomal_S15 (pfam00312)→ |
| Fsp-2-1 | NZ_JQMS01000001.1 | - | 1756958 | 1756892 | **RNA**→ rpsO (PRK05626)Ribosomal_S15 (pfam00312)→ |
| env-1571 | JGI24770J26754_10342540 | - | 429 | 363 | **RNA**→ rpsO (PRK05626)Ribosomal_S15 (pfam00312)→ |
| env-1572 | KGI_S1_ANT01_95m_c10332605 | - | 256 | 190 | **RNA**→ rpsO (PRK05626)Ribosomal_S15 (pfam00312)→ |
| env-1573 | KGI_S1_ANT01_95m_c10257100 | + | 199 | 265 | **RNA**→ rpsO (PRK05626)Ribosomal_S15 (pfam00312)→ |
| env-1574 | KGI_S1_ANT02_95m_c10031472 | + | 1706 | 1772 | **RNA**→ |
| env-1575 | KGI_S1_ANT01_95m_c10019254 | - | 1136 | 1070 | **RNA**→ rpsO (PRK05626)Ribosomal_S15 (pfam00312)→ |

| | | | | | |
|---|---|---|---|---|---|
| env-1576 | KGI_S2_ANT06_2345m_c1022245 | - | 116 | 50 | **RNA**→ |
| Fsp-5-1 | NZ_JSYM01000001.1 | - | 1738792 | 1738726 | **RNA**→ rpsO (PRK05626)Ribosomal_S15 (pfam00312)→ |
| Fsp-3-1 | NZ_LEKS01000001.1 | - | 40307 | 40241 | **RNA**→ rpsO (PRK05626)Ribosomal_S15 (pfam00312)→ |
| env-1577 | KGI_S2_ANT04_2345m_c1000215 | + | 4350 | 4416 | **RNA**→ rpsO (PRK05626)Ribosomal_S15 (pfam00312)→ |
| env-1578 | KGI_S1_ANT02_95m_c10005510 | - | 1136 | 1070 | **RNA**→ rpsO (PRK05626)Ribosomal_S15 (pfam00312)→ |
| env-1579 | KGI_S1_ANT01_95m_c10022295 | - | 134 | 68 | **RNA**→ hypo→ |
| Fsp-4-1 | NZ_AJXL01000050.1 | - | 10986 | 10920 | **RNA**→ rpsO (PRK05626)Ribosomal_S15 (pfam00312)→ |
| env-1580 | JGI26055J44792_1116768 | + | 87 | 134 | **RNA**→ rpsO (PRK05626)Ribosomal_S15 (pfam00312)→ |
| env-1581 | JGI25831J46370_119340 | - | 174 | 127 | **RNA**→ rpsO (PRK05626)Ribosomal_S15 (pfam00312)→ |
| Wma-1-1 | NZ_CCMH01000008.1 | - | 91520 | 91474 | **RNA**→ rpsO (PRK05626)Ribosomal_S15 (pfam00312)→ |
| Wvi-1-1 | NC_015144.1 | - | 1924258 | 1924211 | **RNA**→ rpsO (PRK05626)Ribosomal_S15 (pfam00312)→ |
| Orh-1-1 | NC_018016.1 | + | 365476 | 365528 | **RNA**→ rpsO (PRK05626)Ribosomal_S15 (pfam00312)→ |
| Orh-3-1 | NZ_CP006828.1 | + | 371699 | 371751 | **RNA**→ rpsO (PRK05626)Ribosomal_S15 (pfam00312)→ |
| Orh-2-1 | NZ_AXDE01000002.1 | + | 306778 | 306830 | **RNA**→ rpsO (PRK05626)Ribosomal_S15 (pfam00312)→ |
| env-1582 | 3300001197_1085176 | - | 1056 | 1010 | **RNA**→ rpsO (PRK05626)Ribosomal_S15 (pfam00312)→ |
| env-1583 | APMI01076955.1 | + | 2558 | 2605 | **RNA**→ rpsO (PRK05626)Ribosomal_S15 (pfam00312)→ |
| env-1584 | JGI25853J50183_17647 | - | 50 | 4 | **RNA**→ |
| env-1585 | 3300001592_10080689 | - | 473 | 427 | **RNA**→ rpsO (PRK05626)Ribosomal_S15 (pfam00312)→ |
| env-1586 | JGI26538J50259_1000062 | + | 73740 | 73786 | **RNA**→ rpsO (PRK05626)Ribosomal_S15 (pfam00312)→ |
| env-1587 | 3300001592_10000999 | + | 9714 | 9760 | **RNA**→ rpsO (PRK05626)Ribosomal_S15 (pfam00312)→ |
| env-1588 | 2210193624 | + | 353 | 399 | **RNA**→ |
| Ebr-1-1 | NZ_KB908301.1 | + | 70085 | 70131 | **RNA**→ rpsO (PRK05626)Ribosomal_S15 (pfam00312)→ |
| env-1589 | JGI26051J44788_1005993 | + | 1041 | 1087 | **RNA**→ rpsO (PRK05626)Ribosomal_S15 (pfam00312)→ |
| env-1590 | JGI26053J44789_1004649 | + | 636 | 682 | **RNA**→ rpsO (PRK05626)Ribosomal_S15 (pfam00312)→ |
| env-1591 | JGI26055J44792_1013313 | - | 518 | 472 | **RNA**→ rpsO (PRK05626)Ribosomal_S15 (pfam00312)→ |
| env-1592 | JGI26057J44791_1000035 | + | 25918 | 25964 | **RNA**→ rpsO (PRK05626)Ribosomal_S15 (pfam00312)→ |
| env-1593 | JGI26054J46599_1000016 | - | 39347 | 39301 | **RNA**→ rpsO (PRK05626)Ribosomal_S15 (pfam00312)→ |
| env-1594 | JGI26052J44787_1039558 | + | 277 | 323 | **RNA**→ rpsO (PRK05626)Ribosomal_S15 (pfam00312)→ |
| Efa-1-1 | NZ_JSYQ01000009.1 | + | 118305 | 118351 | **RNA**→ rpsO (PRK05626)Ribosomal_S15 (pfam00312)→ |
| env-1595 | JGI26050J44790_1040856 | - | 81 | 35 | **RNA**→ hypo→ |
| Bzo-1-1 | NZ_JH932293.1 | - | 2055058 | 2055010 | **RNA**→ rpsO (PRK05626)Ribosomal_S15 (pfam00312)→ |
| Bzo-2-1 | NZ_JH932282.1 | + | 1022343 | 1022391 | **RNA**→ rpsO (PRK05626)Ribosomal_S15 (pfam00312)→ |
| env-1596 | CEVG01031283.1 | + | 424 | 489 | **RNA**→ rpsO (PRK05626)RpsO (COG0184)→ PRK11824 (PRK11824)RNase_PH (pfam01138)→ |
| env-1597 | CEWK01195648.1 | - | 283 | 218 | **RNA**→ rpsO (PRK05626)RpsO (COG0184)→ |
| env-1598 | CEPJ01242679.1 | - | 79 | 14 | **RNA**→ |
| env-1599 | CEWH01147159.1 | + | 7783 | 7848 | **RNA**→ rpsO (PRK05626)Ribosomal_S15 (pfam00312)→ |
| env-1600 | JCVI_SCAF_1101667581186 | + | 571 | 636 | **RNA**→ rpsO (PRK05626)Ribosomal_S15 (pfam00312)→ |
| env-1601 | CEVH01171373.1 | + | 701 | 766 | **RNA**→ rpsO (PRK05626)Ribosomal_S15 (pfam00312)→ |
| env-1602 | CEUH01137174.1 | - | 914 | 849 | **RNA**→ rpsO (PRK05626)RpsO (COG0184)→ PRK11824 (PRK11824)RNase_PH (pfam01138)→ |
| env-1603 | CEVG01256217.1 | - | 217 | 152 | **RNA**→ rpsO (PRK05626)Ribosomal_S15 (pfam00312)→ |
| env-1604 | CEWK01191961.1 | + | 343 | 408 | **RNA**→ rpsO (PRK05626)RpsO (COG0184)→ |
| env-1605 | CEVS01311665.1 | + | 167 | 232 | **RNA**→ rpsO (PRK05626)RpsO (COG0184)→ RNase_PH_PNPase_1 (cd11363)PRK11824 (PRK11824)→ |
| env-1606 | CEOM01011442.1 | - | 239 | 174 | **RNA**→ rpsO (PRK05626)Ribosomal_S15 (pfam00312)→ |
| env-1607 | JCVI_READ_1104230051515 | - | 1009 | 945 | **RNA**→ rpsO (PRK05626)RpsO (COG0184)→ PRK11824 (PRK11824)RNase_PH (pfam01138)RNase_PH_C (pfam03725)→ |
| env-1608 | CENF01403979.1 | + | 590 | 654 | **RNA**→ rpsO (PRK05626)RpsO (COG0184)→ PRK11824 (PRK11824)RNase_PH (pfam01138)→ |
| env-1609 | CEOC01304602.1 | - | 743 | 678 | **RNA**→ rpsO (PRK05626)Ribosomal_S15 (pfam00312)→ PRK11824 (PRK11824)RNase_PH (pfam01138)→ |
| env-1610 | CETE01250240.1 | - | 394 | 329 | **RNA**→ rpsO (PRK05626)Ribosomal_S15 (pfam00312)→ |
| env-1611 | CEUR01041503.1 | - | 281 | 216 | **RNA**→ rpsO (PRK05626)Ribosomal_S15 (pfam00312)→ |
| env-1612 | CEVG01012585.1 | - | 211 | 146 | **RNA**→ rpsO (PRK05626)Ribosomal_S15 (pfam00312)→ |
| env-1613 | CEWG01074623.1 | + | 1532 | 1597 | **RNA**→ rpsO (PRK05626)Ribosomal_S15 (pfam00312)→ |
| env-1614 | CENG01192212.1 | - | 188 | 123 | **RNA**→ rpsO (PRK05626)Ribosomal_S15 (pfam00312)→ |

| | | | | | |
|---|---|---|---|---|---|
| env-1615 | CENO01053689.1 | + | 270 | 335 | **RNA**→ rpsO (PRK05626)Ribosomal_S15 (pfam00312)→ RNase_PH_PNPase_1 (cd11363)PRK11824 (PRK11824)→ |
| env-1616 | CEUM01074299.1 | + | 1019 | 1084 | **RNA**→ |
| env-1617 | CEUV01130266.1 | - | 5263 | 5198 | **RNA**→ rpsO (PRK05626)Ribosomal_S15 (pfam00312)→ Pnp (COG1185)PRK11824 (PRK11824)→ PLN02198 (PLN02198)Ggt (COG0405)→ PRK10115 (PRK10115)PtrB (COG1770)→ |
| env-1618 | CEVH01019235.1 | - | 182 | 117 | **RNA**→ rpsO (PRK05626)Ribosomal_S15 (pfam00312)→ |
| env-1619 | CEWH01078602.1 | - | 279 | 214 | **RNA**→ rpsO (PRK05626)Ribosomal_S15 (pfam00312)→ |
| env-1620 | JCVI_SCAF_1096627336466 | + | 264 | 329 | **RNA**→ rpsO (PRK05626)Ribosomal_S15 (pfam00312)→ |
| env-1621 | JCVI_SCAF_1096628013630 | - | 1884 | 1819 | **RNA**→ rpsO (PRK05626)RpsO (COG0184)→ |
| env-1622 | sludgePhrap_Contig10770 | + | 80 | 144 | **RNA**→ rpsO (PRK05626)RpsO (COG0184)→ |
| env-1623 | JGI25920J50251_10058504 | + | 359 | 425 | **RNA**→ rpsO (PRK05626)Ribosomal_S15 (pfam00312)→ |
| env-1624 | JGI25911J50253_10258121 | - | 271 | 205 | **RNA**→ rpsO (PRK05626)Ribosomal_S15 (pfam00312)→ |
| env-1625 | JGI25910J50241_10053410 | - | 957 | 891 | **RNA**→ rpsO (PRK05626)Ribosomal_S15 (pfam00312)→ |
| env-1626 | JGI25910J50241_10095017 | - | 415 | 349 | **RNA**→ rpsO (PRK05626)Ribosomal_S15 (pfam00312)→ |
| env-1627 | JGI25910J50241_10069387 | + | 906 | 972 | **RNA**→ |
| env-1628 | JGI12421J11937_10000034 | + | 36707 | 36773 | **RNA**→ rpsO (PRK05626)Ribosomal_S15 (pfam00312)→ |
| env-1629 | LM20APR2010E_c396 | + | 1869 | 1935 | **RNA**→ rpsO (PRK05626)Ribosomal_S15 (pfam00312)→ |
| env-1630 | TB_PC08_66_10060944 | - | 109 | 43 | **RNA**→ |
| env-1631 | TB_GS09_5_10171935 | - | 305 | 239 | **RNA**→ rpsO (PRK05626)Ribosomal_S15 (pfam00312)→ |
| env-1632 | TB_PC08_66_10061930 | + | 1206 | 1272 | **RNA**→ |
| env-1633 | AglaG_GDN60OX02H4ZEN | - | 102 | 35 | **RNA**→ |
| env-1634 | JGI24770J26754_10074152 | + | 350 | 417 | **RNA**→ rpsO (PRK05626)Ribosomal_S15 (pfam00312)→ |
| env-1635 | TB03JUN2009E__Contig_145064 | - | 74 | 7 | **RNA**→ |
| Faq-1-1 | NZ_JRHH01000001.1 | - | 229487 | 229420 | **RNA**→ rpsO (PRK05626)Ribosomal_S15 (pfam00312)→ |
| env-1636 | JFJP01044092.1 | - | 4887 | 4820 | **RNA**→ rpsO (PRK05626)Ribosomal_S15 (pfam00312)→ |
| env-1637 | JFJP01001225.1 | + | 19883 | 19950 | **RNA**→ rpsO (PRK05626)Ribosomal_S15 (pfam00312)→ |
| env-1638 | SRS045197_WUGC_scaffold_71844 | + | 1187 | 1244 | **RNA**→ rpsO (PRK05626)Ribosomal_S15 (pfam00312)→ |
| env-1639 | SRS053584_LANL_scaffold_46820 | + | 1187 | 1244 | **RNA**→ rpsO (PRK05626)Ribosomal_S15 (pfam00312)→ |
| env-1640 | SRS015803_WUGC_scaffold_1822 | - | 19092 | 19035 | **RNA**→ rpsO (PRK05626)Ribosomal_S15 (pfam00312)→ |
| env-1641 | SRS018157_Baylor_scaffold_79748 | + | 2305 | 2362 | **RNA**→ rpsO (PRK05626)Ribosomal_S15 (pfam00312)→ |
| env-1642 | SRS019077_WUGC_scaffold_63873 | - | 412 | 355 | **RNA**→ rpsO (PRK05626)Ribosomal_S15 (pfam00312)→ |
| Csp-6-1 | NZ_AKFR01000034.1 | + | 60045 | 60102 | **RNA**→ rpsO (PRK05626)Ribosomal_S15 (pfam00312)→ |
| env-1643 | SRS047113_C5238338 | - | 69 | 13 | **RNA**→ |
| env-1644 | SRS024144_C1787925 | + | 808 | 864 | **RNA**→ |
| env-1645 | SRS043772_WUGC_scaffold_9035 | + | 507 | 563 | **RNA**→ rpsO (PRK05626)→ |
| Coc-2-1 | NZ_JVFB01000069.1 | - | 264032 | 263976 | **RNA**→ rpsO (PRK05626)Ribosomal_S15 (pfam00312)→ |
| env-1646 | SRS014578_WUGC_scaffold_50948 | - | 289 | 233 | **RNA**→ |
| Csp-9-1 | NZ_KB291210.1 | + | 18472 | 18527 | **RNA**→ rpsO (PRK05626)Ribosomal_S15 (pfam00312)→ |
| env-1647 | SRS011343_Baylor_scaffold_28900 | + | 921 | 976 | **RNA**→ |
| env-1648 | SRS015215_WUGC_scaffold_40668 | - | 90 | 35 | **RNA**→ |
| env-1649 | SRS024087_C3535890 | - | 68 | 13 | **RNA**→ |
| env-1650 | SRS018337_Baylor_scaffold_6407 | + | 3 | 58 | **RNA**→ rpsO (PRK05626)Ribosomal_S15 (pfam00312)→ |
| env-1651 | SRS055378_LANL_scaffold_11295 | - | 202 | 147 | **RNA**→ rpsO (PRK05626)Ribosomal_S15 (pfam00312)→ |
| env-1652 | SRS011255_Baylor_scaffold_50639 | - | 90 | 35 | **RNA**→ |
| env-1653 | SRS014476_WUGC_scaffold_37616 | - | 90 | 35 | **RNA**→ |
| env-1654 | SRS016043_WUGC_scaffold_47910 | - | 90 | 35 | **RNA**→ |
| env-1655 | SRS021960_Baylor_scaffold_21591 | + | 91 | 146 | **RNA**→ rpsO (PRK05626)RpsO (COG0184)→ |
| env-1656 | SRS013252_Baylor_scaffold_38320 | - | 68 | 13 | **RNA**→ |
| env-1657 | SRS053917_LANL_scaffold_37634 | - | 1535 | 1481 | **RNA**→ rpsO (PRK05626)Ribosomal_S15 (pfam00312)→ |
| Coc-3-1 | NZ_GL573160.1 | + | 348550 | 348604 | **RNA**→ rpsO (PRK05626)Ribosomal_S15 (pfam00312)→ |
| Coc-4-1 | NZ_AKFV01000036.1 | + | 171648 | 171702 | **RNA**→ rpsO (PRK05626)Ribosomal_S15 (pfam00312)→ |
| Coc-1-1 | NC_013162.1 | + | 2020418 | 2020473 | **RNA**→ rpsO (PRK05626)Ribosomal_S15 (pfam00312)→ |
| Csp-7-1 | NZ_KE150489.1 | - | 245846 | 245793 | **RNA**→ rpsO (PRK05626)Ribosomal_S15 (pfam00312)→ |

| | | | | | |
|---|---|---|---|---|---|
| Csp-10-1 | NZ_AJZR01000054.1 | + | 24577 | 24630 | **RNA**→ rpsO (PRK05626)Ribosomal_S15 (pfam00312)→ rpsO (PRK05626)Ribosomal_S15 (pfam00312)→ |
| Csp-2-1 | NZ_KB291539.1 | + | 18156 | 18209 | **RNA**→ rpsO (PRK05626)Ribosomal_S15 (pfam00312)→ |
| env-1658 | G312J29652_10291633 | + | 142 | 208 | **RNA**→ rpsO (PRK05626)Ribosomal_S15 (pfam00312)→ |
| env-1659 | M3P_c10401073 | - | 92 | 26 | **RNA**→ hypo→ |
| env-1660 | JGI25913J50563_1004702 | - | 1156 | 1090 | **RNA**→ rpsO (PRK05626)Ribosomal_S15 (pfam00312)→ |
| env-1661 | 3300000052_045724 | + | 487 | 553 | **RNA**→ rpsO (PRK05626)Ribosomal_S15 (pfam00312)→ |
| env-1662 | JGI25913J50563_1032633 | + | 701 | 767 | **RNA**→ |
| env-1663 | JCVI_SCAF_1101668144518 | - | 828 | 762 | **RNA**→ rpsO (PRK05626)Ribosomal_S15 (pfam00312)→ |
| env-1664 | SRS043422_LANL_scaffold_27461 | + | 70 | 126 | **RNA**→ rpsO (PRK05626)Ribosomal_S15 (pfam00312)→ |
| env-1665 | SRS013533_PGA_scaffold_76303 | + | 331 | 387 | **RNA**→ rpsO (PRK05626)Ribosomal_S15 (pfam00312)→ |
| env-1666 | SRS014888_C3093205 | - | 468 | 412 | **RNA**→ rpsO (PRK05626)Ribosomal_S15 (pfam00312)→ |
| env-1667 | SRS050669_LANL_scaffold_37527 | - | 1248 | 1192 | **RNA**→ rpsO (PRK05626)Ribosomal_S15 (pfam00312)→ |
| env-1668 | SRS015063_C2120774 | + | 197 | 253 | **RNA**→ rpsO (PRK05626)Ribosomal_S15 (pfam00312)→ |
| env-1669 | SRS015762_WUGC_scaffold_51343 | - | 4350 | 4294 | **RNA**→ rpsO (PRK05626)Ribosomal_S15 (pfam00312)→ |
| env-1670 | SRS019029_WUGC_scaffold_45090 | + | 1155 | 1211 | **RNA**→ rpsO (PRK05626)Ribosomal_S15 (pfam00312)→ |
| env-1671 | SRS019122_C5910877 | - | 652 | 596 | **RNA**→ rpsO (PRK05626)Ribosomal_S15 (pfam00312)→ |
| env-1672 | SRS057205_LANL_scaffold_58800 | + | 12511 | 12567 | **RNA**→ rpsO (PRK05626)Ribosomal_S15 (pfam00312)→ |
| env-1673 | SRS016092_C1171386 | + | 397 | 453 | **RNA**→ rpsO (PRK05626)Ribosomal_S15 (pfam00312)→ |
| env-1674 | SRS017227_Baylor_scaffold_110356 | - | 740 | 684 | **RNA**→ rpsO (PRK05626)Ribosomal_S15 (pfam00312)→ |
| env-1675 | SRS019591_WUGC_scaffold_42767 | - | 4361 | 4305 | **RNA**→ rpsO (PRK05626)Ribosomal_S15 (pfam00312)→ |
| env-1676 | SRS019027_C1985251 | - | 2922 | 2866 | **RNA**→ rpsO (PRK05626)Ribosomal_S15 (pfam00312)→ |
| Csp-4-1 | NZ_KN389985.1 | - | 83839 | 83783 | **RNA**→ rpsO (PRK05626)Ribosomal_S15 (pfam00312)→ |
| env-1677 | SRS019077_C2547620 | - | 115 | 59 | **RNA**→ rpsO (PRK05626)→ |
| env-1678 | SRS063288_C2556324 | + | 441 | 497 | **RNA**→ rpsO (PRK05626)→ |
| env-1679 | SRS019906_WUGC_scaffold_52431 | - | 674 | 618 | **RNA**→ rpsO (PRK05626)Ribosomal_S15 (pfam00312)→ |
| env-1680 | SRS021960_C2067122 | + | 215 | 271 | **RNA**→ |
| env-1681 | SRS023538_Baylor_scaffold_38631 | + | 1459 | 1515 | **RNA**→ rpsO (PRK05626)Ribosomal_S15 (pfam00312)→ |
| env-1682 | SRS024289_C2070569 | - | 115 | 59 | **RNA**→ rpsO (PRK05626)→ |
| env-1683 | SRS045715_C4722109 | - | 89 | 33 | **RNA**→ |
| env-1684 | SRS050244_LANL_scaffold_99594 | + | 959 | 1015 | **RNA**→ rpsO (PRK05626)Ribosomal_S15 (pfam00312)→ |
| env-1685 | SRS051378_C1779297 | - | 115 | 59 | **RNA**→ rpsO (PRK05626)→ |
| env-1686 | SRS054653_C1990320 | + | 34 | 90 | **RNA**→ rpsO (PRK05626)Ribosomal_S15 (pfam00312)→ |
| env-1687 | SRS058336_C2081196 | + | 224 | 280 | **RNA**→ rpsO (PRK05626)Ribosomal_S15 (pfam00312)→ |
| env-1688 | Buccal_mucosa_LANL_scaffold_150368 | - | 636 | 580 | **RNA**→ rpsO (PRK05626)Ribosomal_S15 (pfam00312)→ |
| env-1689 | SRS023930_C2191794 | - | 1919 | 1863 | **RNA**→ rpsO (PRK05626)Ribosomal_S15 (pfam00312)→ |
| env-1690 | SRS018975_WUGC_scaffold_37944 | + | 424 | 480 | **RNA**→ rpsO (PRK05626)Ribosomal_S15 (pfam00312)→ |
| env-1691 | SRS012285_Baylor_scaffold_28847 | - | 817 | 761 | **RNA**→ rpsO (PRK05626)Ribosomal_S15 (pfam00312)→ |
| env-1692 | SRS014473_C1058379 | + | 108 | 164 | **RNA**→ rpsO (PRK05626)RpsO (COG0184)→ |
| env-1693 | SRS015044_WUGC_scaffold_70508 | + | 49 | 105 | **RNA**→ rpsO (PRK05626)Ribosomal_S15 (pfam00312)→ |
| env-1694 | SRS015434_C5932842 | + | 276 | 332 | **RNA**→ rpsO (PRK05626)Ribosomal_S15 (pfam00312)→ |
| env-1695 | SRS015899_WUGC_scaffold_29136 | + | 830 | 886 | **RNA**→ rpsO (PRK05626)Ribosomal_S15 (pfam00312)→ |
| env-1696 | SRS016297_C532059 | + | 45 | 101 | **RNA**→ rpsO (PRK05626)Ribosomal_S15 (pfam00312)→ |
| env-1697 | SRS018300_Baylor_scaffold_13428 | + | 11276 | 11332 | **RNA**→ rpsO (PRK05626)Ribosomal_S15 (pfam00312)→ |
| env-1698 | SRS019071_C5001202 | + | 275 | 331 | **RNA**→ |
| env-1699 | SRS019225_WUGC_scaffold_37789 | + | 737 | 793 | **RNA**→ rpsO (PRK05626)Ribosomal_S15 (pfam00312)→ |
| env-1700 | SRS021477_Baylor_scaffold_50396 | + | 770 | 826 | **RNA**→ rpsO (PRK05626)→ |
| env-1701 | SRS022725_LANL_scaffold_57376 | + | 10486 | 10542 | **RNA**→ rpsO (PRK05626)Ribosomal_S15 (pfam00312)→ |
| env-1702 | SRS023595_Baylor_scaffold_65314 | - | 1031 | 975 | **RNA**→ rpsO (PRK05626)Ribosomal_S15 (pfam00312)→ |
| env-1703 | SRS024355_LANL_scaffold_78151 | + | 324 | 380 | **RNA**→ rpsO (PRK05626)Ribosomal_S15 (pfam00312)→ |
| env-1704 | SRS065099_LANL_scaffold_43567 | - | 1245 | 1189 | **RNA**→ rpsO (PRK05626)Ribosomal_S15 (pfam00312)→ |
| env-1705 | SRS047634_LANL_scaffold_56906 | + | 930 | 986 | **RNA**→ rpsO (PRK05626)Ribosomal_S15 (pfam00312)→ |
| env-1706 | SRS050628_C1885263 | + | 48 | 104 | **RNA**→ rpsO (PRK05626)Ribosomal_S15 (pfam00312)→ |

| | | | | | |
|---|---|---|---|---|---|
| env-1707 | SRS051941_LANL_scaffold_40092 | - | 624 | 568 | **RNA**→ rpsO (PRK05626)Ribosomal_S15 (pfam00312)→ PRK11824 (PRK11824)RNase_PH (pfam01138)→ |
| env-1708 | SRS057022_C1966776 | - | 419 | 363 | **RNA**→ rpsO (PRK05626)Ribosomal_S15 (pfam00312)→ |
| env-1709 | SRS020222_C1001786 | + | 45 | 101 | **RNA**→ rpsO (PRK05626)Ribosomal_S15 (pfam00312)→ |
| env-1710 | SRS063215_LANL_scaffold_746 | + | 1746 | 1802 | **RNA**→ rpsO (PRK05626)Ribosomal_S15 (pfam00312)→ |
| Cko-1-1 | NZ_LFNG01000006.1 | - | 73410 | 73364 | **RNA**→ rpsO (PRK05626)Ribosomal_S15 (pfam00312)→ |
| env-1711 | SRS019128_WUGC_scaffold_40428 | - | 250 | 198 | **RNA**→ rpsO (PRK05626)RpsO (COG0184)→ |
| env-1712 | SRS024561_LANL_scaffold_27863 | - | 246 | 194 | **RNA**→ rpsO (PRK05626)RpsO (COG0184)→ |
| env-1713 | SRS063932_LANL_scaffold_85576 | - | 294 | 242 | **RNA**→ rpsO (PRK05626)Ribosomal_S15 (pfam00312)→ |
| env-1714 | SRS011126_Baylor_scaffold_15902 | - | 467 | 415 | **RNA**→ rpsO (PRK05626)Ribosomal_S15 (pfam00312)→ |
| env-1715 | SRS016360_Baylor_scaffold_10965 | + | 1870 | 1922 | **RNA**→ rpsO (PRK05626)Ribosomal_S15 (pfam00312)→ |
| env-1716 | SRS017227_Baylor_scaffold_133947 | + | 2861 | 2913 | **RNA**→ rpsO (PRK05626)Ribosomal_S15 (pfam00312)→ |
| env-1717 | SRS018665_WUGC_scaffold_9476 | - | 1439 | 1387 | **RNA**→ rpsO (PRK05626)Ribosomal_S15 (pfam00312)→ |
| env-1718 | SRS024087_C3338429 | + | 348 | 400 | **RNA**→ |
| env-1719 | SRS063999_C3578431 | - | 65 | 13 | **RNA**→ |
| env-1720 | SRS052876_LANL_scaffold_43042 | - | 2813 | 2761 | **RNA**→ rpsO (PRK05626)Ribosomal_S15 (pfam00312)→ |
| env-1721 | SRS063603_LANL_scaffold_104178 | + | 1536 | 1588 | **RNA**→ rpsO (PRK05626)Ribosomal_S15 (pfam00312)→ |
| env-1722 | SRS015803_WUGC_scaffold_46814 | + | 633 | 685 | **RNA**→ rpsO (PRK05626)Ribosomal_S15 (pfam00312)→ |
| Csp-5-1 | NZ_KB291273.1 | + | 7305 | 7357 | **RNA**→ rpsO (PRK05626)Ribosomal_S15 (pfam00312)→ |
| env-1723 | SRS047113_LANL_scaffold_40316 | - | 65 | 13 | **RNA**→ |
| env-1724 | BS_KBA_SWE02_21m_10002378 | - | 6637 | 6571 | **RNA**→ rpsO (PRK05626)Ribosomal_S15 (pfam00312)→ |
| env-1725 | BS_KBA_SWE12_21m_c10257970 | - | 316 | 250 | **RNA**→ rpsO (PRK05626)Ribosomal_S15 (pfam00312)→ |
| env-1726 | BS_KBA_SWE21_205m_10065151 | - | 791 | 725 | **RNA**→ rpsO (PRK05626)Ribosomal_S15 (pfam00312)→ |
| env-1727 | BS_KBA_SWE07_21m_c1084691 | + | 116 | 182 | **RNA**→ rpsO (PRK05626)Ribosomal_S15 (pfam00312)→ |
| Csp-8-1 | NZ_GL872413.1 | + | 1146487 | 1146540 | **RNA**→ rpsO (PRK05626)RpsO (COG0184)→ |
| env-1728 | SRS013950_C2010154 | + | 28 | 81 | **RNA**→ rpsO (PRK05626)Ribosomal_S15 (pfam00312)→ |
| env-1729 | SRS024381_LANL_scaffold_23096 | - | 964 | 909 | **RNA**→ rpsO (PRK05626)RpsO (COG0184)→ |
| env-1730 | SRS064329_C2708069 | + | 592 | 647 | **RNA**→ rpsO (PRK05626)RpsO (COG0184)→ |
| env-1731 | SRS012279_C2755382 | + | 1578 | 1633 | **RNA**→ rpsO (PRK05626)RpsO (COG0184)→ |
| env-1732 | SRS016319_C4022715 | - | 1914 | 1859 | **RNA**→ rpsO (PRK05626)RpsO (COG0184)→ |
| env-1733 | SRS017209_Baylor_scaffold_62146 | + | 1830 | 1885 | **RNA**→ rpsO (PRK05626)RpsO (COG0184)→ |
| env-1734 | SRS018300_Baylor_scaffold_54476 | - | 1573 | 1518 | **RNA**→ rpsO (PRK05626)RpsO (COG0184)→ |
| env-1735 | SRS019122_C5923314 | - | 6524 | 6469 | **RNA**→ rpsO (PRK05626)RpsO (COG0184)→ |
| env-1736 | SRS019127_C1675613 | + | 2142 | 2197 | **RNA**→ rpsO (PRK05626)RpsO (COG0184)→ |
| env-1737 | SRS021496_Baylor_scaffold_73479 | - | 8230 | 8175 | **RNA**→ rpsO (PRK05626)RpsO (COG0184)→ |
| env-1738 | SRS022077_Baylor_scaffold_57464 | + | 1502 | 1557 | **RNA**→ rpsO (PRK05626)RpsO (COG0184)→ |
| env-1739 | SRS022530_LANL_scaffold_23647 | - | 9977 | 9922 | **RNA**→ rpsO (PRK05626)RpsO (COG0184)→ |
| env-1740 | SRS024081_LANL_scaffold_27019 | - | 8253 | 8198 | **RNA**→ rpsO (PRK05626)RpsO (COG0184)→ |
| env-1741 | SRS050244_LANL_scaffold_22625 | + | 4127 | 4182 | **RNA**→ rpsO (PRK05626)RpsO (COG0184)→ |
| env-1742 | SRS050669_LANL_scaffold_21743 | - | 1678 | 1623 | **RNA**→ rpsO (PRK05626)RpsO (COG0184)→ |
| env-1743 | SRS063288_C2588099 | - | 397 | 342 | **RNA**→ rpsO (PRK05626)RpsO (COG0184)→ |
| env-1744 | SRS011126_Baylor_scaffold_9375 | + | 632 | 687 | **RNA**→ rpsO (PRK05626)RpsO (COG0184)→ |
| env-1745 | SRS013533_PGA_scaffold_2853 | + | 34 | 89 | **RNA**→ rpsO (PRK05626)RpsO (COG0184)→ |
| env-1746 | SRS015063_WUGC_scaffold_46448 | + | 1494 | 1549 | **RNA**→ |
| env-1747 | SRS018665_C3617939 | + | 376 | 431 | **RNA**→ |
| env-1748 | SRS018778_C1276842 | - | 1257 | 1202 | **RNA**→ rpsO (PRK05626)RpsO (COG0184)→ |
| env-1749 | SRS019980_Baylor_scaffold_28622 | + | 2149 | 2204 | **RNA**→ rpsO (PRK05626)RpsO (COG0184)→ |
| env-1750 | SRS020340_C2796995 | + | 244 | 299 | **RNA**→ |
| env-1751 | SRS043755_WUGC_scaffold_43382 | + | 2651 | 2706 | **RNA**→ |
| env-1752 | SRS023964_C2569817 | - | 445 | 390 | **RNA**→ rpsO (PRK05626)RpsO (COG0184)→ |
| env-1753 | SRS049268_C4218766 | - | 234 | 179 | **RNA**→ rpsO (PRK05626)Ribosomal_S15 (pfam00312)→ |
| env-1754 | SRS058336_C2129811 | - | 202 | 147 | **RNA**→ rpsO (PRK05626)Ribosomal_S15 (pfam00312)→ |
| env-1755 | SRS063215_C2276132 | - | 206 | 151 | **RNA**→ rpsO (PRK05626)Ribosomal_S15 (pfam00312)→ |

| | | | | | |
|---|---|---|---|---|---|
| Cgi-1-1 | NZ_ACLQ01000019.1 | - | 321176 | 321120 | **RNA**→ rpsO (PRK05626)RpsO (COG0184)→ rpsO (PRK05626)RpsO (COG0184)→ |
| env-1756 | SRS053630_C926066 | + | 2157 | 2212 | **RNA**→ |
| env-1757 | Tongue_dor_WUGI_scaffold_5146 | + | 1830 | 1885 | **RNA**→ |
| env-1758 | SRS013705_C3757970 | + | 419 | 474 | **RNA**→ |
| env-1759 | SRS014573_C3640759 | + | 448 | 503 | **RNA**→ |
| env-1760 | SRS015762_WUGC_scaffold_51227 | - | 69 | 14 | **RNA**→ |
| env-1761 | SRS016225_C2972871 | + | 722 | 777 | **RNA**→ |
| env-1762 | SRS019071_WUGC_scaffold_32009 | + | 372 | 427 | **RNA**→ |
| env-1763 | SRS042131_WUGC_scaffold_54007 | + | 6552 | 6607 | **RNA**→ |
| env-1764 | SRS057205_C3544919 | + | 1708 | 1763 | **RNA**→ |
| env-1765 | SRS012285_Baylor_scaffold_29593 | + | 360 | 415 | **RNA**→ |
| env-1766 | SRS014888_C3089015 | - | 58 | 3 | **RNA**→ |
| env-1767 | SRS018357_C3820809 | + | 107 | 162 | **RNA**→ rpsO (PRK05626)Ribosomal_S15 (pfam00312)→ |
| env-1768 | SRS018739_C3515427 | - | 340 | 285 | **RNA**→ rpsO (PRK05626)RpsO (COG0184)→ |
| env-1769 | SRS019327_C2770117 | + | 716 | 771 | **RNA**→ rpsO (PRK05626)RpsO (COG0184)→ |
| env-1770 | SRS020226_Baylor_scaffold_48761 | + | 121 | 176 | **RNA**→ rpsO (PRK05626)RpsO (COG0184)→ |
| env-1771 | SRS021477_Baylor_scaffold_53832 | - | 3224 | 3169 | **RNA**→ rpsO (PRK05626)RpsO (COG0184)→ |
| env-1772 | SRS024138_Baylor_scaffold_48685 | - | 5719 | 5664 | **RNA**→ rpsO (PRK05626)RpsO (COG0184)→ |
| env-1773 | SRS049389_C4680666 | + | 115 | 170 | **RNA**→ rpsO (PRK05626)RpsO (COG0184)→ |
| env-1774 | SRS062544_C3032576 | - | 2729 | 2674 | **RNA**→ rpsO (PRK05626)RpsO (COG0184)→ |
| env-1775 | SRS016541_Baylor_scaffold_10002 | - | 8245 | 8190 | **RNA**→ rpsO (PRK05626)RpsO (COG0184)→ |
| env-1776 | SRS011343_Baylor_scaffold_51832 | + | 619 | 674 | **RNA**→ rpsO (PRK05626)RpsO (COG0184)→ |
| env-1777 | SRS018573_WUGC_scaffold_37052 | - | 745 | 690 | **RNA**→ rpsO (PRK05626)RpsO (COG0184)→ |
| env-1778 | SRS055401_LANL_scaffold_16367 | - | 3521 | 3466 | **RNA**→ rpsO (PRK05626)RpsO (COG0184)→ |
| env-1779 | SRS018439_C2379934 | - | 315 | 259 | **RNA**→ rpsO (PRK05626)Ribosomal_S15 (pfam00312)→ |
| env-1780 | SRS011255_Baylor_scaffold_57867 | + | 143 | 199 | **RNA**→ rpsO (PRK05626)RpsO (COG0184)→ |
| env-1781 | SRS015755_WUGC_scaffold_31067 | + | 144 | 200 | **RNA**→ rpsO (PRK05626)RpsO (COG0184)→ |
| env-1782 | SRS017511_Baylor_scaffold_52784 | - | 1865 | 1809 | **RNA**→ rpsO (PRK05626)RpsO (COG0184)→ |
| env-1783 | SRS013949_C3427567 | - | 63 | 7 | **RNA**→ |
| env-1784 | SRS047265_C1018884 | - | 63 | 7 | **RNA**→ |
| env-1785 | SRS054430_C1991223 | - | 63 | 7 | **RNA**→ |
| env-1786 | SRS015215_C2602102 | + | 1914 | 1970 | **RNA**→ rpsO (PRK05626)RpsO (COG0184)→ |
| env-1787 | SRS018157_Baylor_scaffold_17243 | + | 5326 | 5382 | **RNA**→ rpsO (PRK05626)RpsO (COG0184)→ |
| env-1788 | SRS018394_Baylor_scaffold_16582 | + | 207 | 263 | **RNA**→ rpsO (PRK05626)RpsO (COG0184)→ |
| env-1789 | SRS018975_WUGC_scaffold_27725 | - | 2246 | 2190 | **RNA**→ rpsO (PRK05626)RpsO (COG0184)→ |
| env-1790 | SRS023538_C1297948 | + | 625 | 681 | **RNA**→ rpsO (PRK05626)RpsO (COG0184)→ |
| env-1791 | SRS024021_Baylor_scaffold_35694 | - | 3813 | 3757 | **RNA**→ rpsO (PRK05626)RpsO (COG0184)→ |
| env-1792 | SRS051930_LANL_scaffold_30198 | - | 1945 | 1889 | **RNA**→ rpsO (PRK05626)RpsO (COG0184)→ |
| env-1793 | SRS053584_LANL_scaffold_27427 | + | 1111 | 1167 | **RNA**→ rpsO (PRK05626)RpsO (COG0184)→ |
| env-1794 | SRS064449_LANL_scaffold_54476 | - | 821 | 765 | **RNA**→ rpsO (PRK05626)RpsO (COG0184)→ |
| env-1795 | SRS015797_WUGC_scaffold_40800 | - | 3734 | 3678 | **RNA**→ rpsO (PRK05626)RpsO (COG0184)→ |
| env-1796 | SRS049268_LANL_scaffold_48367 | - | 1235 | 1179 | **RNA**→ rpsO (PRK05626)RpsO (COG0184)→ |
| env-1797 | SRS016575_Baylor_scaffold_62429 | - | 468 | 412 | **RNA**→ rpsO (PRK05626)RpsO (COG0184)→ |
| env-1798 | SRS021960_Baylor_scaffold_32080 | + | 1206 | 1262 | **RNA**→ rpsO (PRK05626)RpsO (COG0184)→ |
| env-1799 | SRS043018_WUGC_scaffold_17072 | - | 1686 | 1630 | **RNA**→ rpsO (PRK05626)RpsO (COG0184)→ |
| env-1800 | SRS052604_C1741226 | + | 192 | 248 | **RNA**→ rpsO (PRK05626)Ribosomal_S15 (pfam00312)→ |
| env-1801 | SRS075406_C835402 | + | 1492 | 1548 | **RNA**→ rpsO (PRK05626)Ribosomal_S15 (pfam00312)→ |
| env-1802 | SRS015044_C2954296 | - | 71 | 14 | **RNA**→ |
| env-1803 | SRS017445_Baylor_scaffold_67615 | - | 71 | 14 | **RNA**→ |
| env-1804 | SRS063999_LANL_scaffold_69744 | - | 714 | 657 | **RNA**→ rpsO (PRK05626)RpsO (COG0184)→ |
| env-1805 | SRS018443_C3761441 | + | 440 | 496 | **RNA**→ |
| env-1806 | SRS023841_Baylor_scaffold_2255 | + | 17160 | 17216 | **RNA**→ |

| | | | | | |
|---|---|---|---|---|---|
| env-1807 | SRS013723_Baylor_scaffold_96492 | + | 1691 | 1747 | **RNA**→ |
| env-1808 | SRS015158_C2638461 | - | 70 | 14 | **RNA**→ |
| env-1809 | SRS015899_C1864030 | - | 70 | 14 | **RNA**→ |
| env-1810 | SRS015989_WUGC_scaffold_9962 | - | 70 | 14 | **RNA**→ |
| env-1811 | SRS016331_C4186867 | + | 1034 | 1090 | **RNA**→ |
| env-1812 | SRS017227_Baylor_scaffold_129048 | - | 70 | 14 | **RNA**→ |
| env-1813 | SRS023938_Baylor_scaffold_54567 | - | 70 | 14 | **RNA**→ |
| env-1814 | SRS043772_C1805334 | + | 959 | 1015 | **RNA**→ |
| env-1815 | SRS047634_LANL_scaffold_135361 | + | 1682 | 1738 | **RNA**→ |
| env-1816 | SRS051941_LANL_scaffold_63198 | - | 70 | 14 | **RNA**→ |
| env-1817 | SRS055450_LANL_scaffold_60723 | + | 443 | 499 | **RNA**→ |
| env-1818 | SRS011098_Baylor_scaffold_47355 | - | 447 | 391 | **RNA**→ rpsO (PRK05626)RpsO (COG0184)→ |
| env-1819 | SRS011152_Baylor_scaffold_56328 | - | 448 | 392 | **RNA**→ rpsO (PRK05626)RpsO (COG0184)→ |
| env-1820 | SRS017139_Baylor_scaffold_22700 | - | 713 | 657 | **RNA**→ rpsO (PRK05626)RpsO (COG0184)→ |
| env-1821 | SRS013252_Baylor_scaffold_570 | - | 3369 | 3313 | **RNA**→ rpsO (PRK05626)RpsO (COG0184)→ |
| env-1822 | SRS024561_LANL_scaffold_19776 | + | 521 | 577 | **RNA**→ rpsO (PRK05626)RpsO (COG0184)→ |
| env-1823 | SRS014894_WUGC_scaffold_44021 | + | 2091 | 2147 | **RNA**→ rpsO (PRK05626)RpsO (COG0184)→ |
| env-1824 | SRS015278_WUGC_scaffold_55991 | - | 313 | 257 | **RNA**→ rpsO (PRK05626)Ribosomal_S15 (pfam00312)→ |
| env-1825 | SRS047113_LANL_scaffold_63026 | + | 134 | 190 | **RNA**→ rpsO (PRK05626)Ribosomal_S15 (pfam00312)→ |
| env-1826 | SRS016043_WUGC_scaffold_15210 | + | 52 | 108 | **RNA**→ rpsO (PRK05626)RpsO (COG0184)→ |
| env-1827 | SRS017025_C1711249 | + | 439 | 495 | **RNA**→ rpsO (PRK05626)Ribosomal_S15 (pfam00312)→ |
| env-1828 | SRS022725_LANL_scaffold_39329 | - | 181 | 125 | **RNA**→ rpsO (PRK05626)Ribosomal_S15 (pfam00312)→ |
| env-1829 | SRS056622_C2506152 | - | 456 | 400 | **RNA**→ rpsO (PRK05626)Ribosomal_S15 (pfam00312)→ |
| env-1830 | SRS014578_WUGC_scaffold_10366 | + | 9436 | 9493 | **RNA**→ rpsO (PRK05626)RpsO (COG0184)→ |
| env-1831 | SRS016200_C2197115 | + | 122 | 179 | **RNA**→ rpsO (PRK05626)RpsO (COG0184)→ |
| env-1832 | SRS019225_WUGC_scaffold_3776 | + | 18738 | 18795 | **RNA**→ rpsO (PRK05626)RpsO (COG0184)→ |
| env-1833 | SRS053917_LANL_scaffold_17810 | + | 480 | 536 | **RNA**→ RNase_PH_PNPase_1 (cd11363)PRK11824 (PRK11824)→ |
| env-1834 | SRS014476_WUGC_scaffold_21840 | + | 108 | 161 | **RNA**→ rpsO (PRK05626)Ribosomal_S15 (pfam00312)→ |
| env-1835 | SRS015278_WUGC_scaffold_27115 | - | 448 | 395 | **RNA**→ rpsO (PRK05626)RpsO (COG0184)→ |
| env-1836 | SRS019073_C2403742 | - | 61 | 8 | **RNA**→ |
| env-1837 | SRS023938_C2280951 | + | 621 | 674 | **RNA**→ |
| env-1838 | SRS042984_LANL_scaffold_33434 | - | 1845 | 1792 | **RNA**→ rpsO (PRK05626)RpsO (COG0184)→ |
| env-1839 | SRS045197_WUGC_scaffold_57665 | + | 563 | 616 | **RNA**→ rpsO (PRK05626)Ribosomal_S15 (pfam00312)→ |
| env-1840 | SRS011126_Baylor_scaffold_28444 | - | 814 | 761 | **RNA**→ rpsO (PRK05626)RpsO (COG0184)→ |
| env-1841 | SRS013723_Baylor_scaffold_97854 | + | 595 | 648 | **RNA**→ rpsO (PRK05626)RpsO (COG0184)→ |
| env-1842 | SRS018157_Baylor_scaffold_39858 | + | 111 | 164 | **RNA**→ rpsO (PRK05626)RpsO (COG0184)→ |
| env-1843 | SRS022536_LANL_scaffold_17148 | + | 75494 | 75547 | **RNA**→ rpsO (PRK05626)RpsO (COG0184)→ |
| env-1844 | SRS049318_LANL_scaffold_66197 | - | 467 | 414 | **RNA**→ rpsO (PRK05626)RpsO (COG0184)→ |
| env-1845 | SRS065099_LANL_scaffold_25462 | - | 1601 | 1548 | **RNA**→ rpsO (PRK05626)RpsO (COG0184)→ |
| env-1846 | SRS011255_C2425047 | + | 575 | 628 | **RNA**→ |
| env-1847 | SRS014578_WUGC_scaffold_64330 | + | 620 | 673 | **RNA**→ rpsO (PRK05626)RpsO (COG0184)→ |
| env-1848 | SRS015899_C1850213 | + | 925 | 978 | **RNA**→ |
| env-1849 | SRS017445_C3104098 | - | 61 | 8 | **RNA**→ |
| env-1850 | SRS019980_C2969016 | - | 61 | 8 | **RNA**→ |
| env-1851 | SRS024144_C1802205 | + | 72 | 125 | **RNA**→ rpsO (PRK05626)RpsO (COG0184)→ |
| env-1852 | SRS043755_WUGC_scaffold_43461 | - | 467 | 414 | **RNA**→ rpsO (PRK05626)RpsO (COG0184)→ |
| env-1853 | SRS063932_LANL_scaffold_49231 | + | 1550 | 1603 | **RNA**→ rpsO (PRK05626)Ribosomal_S15 (pfam00312)→ |
| env-1854 | Buccal_mucosa_LANL_scaffold_38424 | + | 493 | 546 | **RNA**→ rpsO (PRK05626)Ribosomal_S15 (pfam00312)→ |
| env-1855 | SRS018665_WUGC_scaffold_60069 | - | 1584 | 1530 | **RNA**→ rpsO (PRK05626)Ribosomal_S15 (pfam00312)→ |
| env-1856 | SRS015989_WUGC_scaffold_7147 | - | 469 | 415 | **RNA**→ rpsO (PRK05626)Ribosomal_S15 (pfam00312)→ |
| env-1857 | SRS053917_LANL_scaffold_6591 | + | 5942 | 5996 | **RNA**→ rpsO (PRK05626)Ribosomal_S15 (pfam00312)→ |
| env-1858 | SRS022725_LANL_scaffold_59028 | + | 5381 | 5435 | **RNA**→ rpsO (PRK05626)Ribosomal_S15 (pfam00312)→ |

| | | | | | |
|---|---|---|---|---|---|
| env-1859 | SRS015899_WUGC_scaffold_38801 | - | 470 | 416 | **RNA**→ rpsO (PRK05626)Ribosomal_S15 (pfam00312)→ |
| env-1860 | SRS015158_WUGC_scaffold_8656 | - | 470 | 416 | **RNA**→ rpsO (PRK05626)Ribosomal_S15 (pfam00312)→ |
| env-1861 | SRS043018_WUGC_scaffold_62457 | + | 332 | 386 | **RNA**→ rpsO (PRK05626)Ribosomal_S15 (pfam00312)→ |
| env-1862 | SRS045197_WUGC_scaffold_57684 | - | 249 | 197 | **RNA**→ rpsO (PRK05626)RpsO (COG0184)→ |
| env-1863 | SRS016360_Baylor_scaffold_47393 | + | 1132 | 1185 | **RNA**→ |
| env-1864 | SRS014477_C1535248 | - | 232 | 179 | **RNA**→ rpsO (PRK05626)Ribosomal_S15 (pfam00312)→ |
| env-1865 | CEWR01254345.1 | + | 688 | 738 | **RNA**→ rpsO (PRK05626)Ribosomal_S15 (pfam00312)→ |
| Eme-2-1 | NZ_BARD01000020.1 | + | 39213 | 39259 | **RNA**→ rpsO (PRK05626)Ribosomal_S15 (pfam00312)→ rpsO (PRK05626)Ribosomal_S15 (pfam00312)→ hypo→ hypo→ |
| Eme-2-2 | NZ_ASAN01000003.1 | + | 99334 | 99380 | **RNA**→ rpsO (PRK05626)Ribosomal_S15 (pfam00312)→ hypo→ hypo→ |
| Cca-1-1 | NZ_JHXV01000017.1 | + | 23329 | 23378 | **RNA**→ rpsO (PRK05626)Ribosomal_S15 (pfam00312)→ |
| Emi-1-1 | NZ_CP011059.1 | + | 4046339 | 4046387 | **RNA**→ rpsO (PRK05626)Ribosomal_S15 (pfam00312)→ SMC_prok_A (TIGR02169)DUF1003 (pfam06210)→ |
| Cte-1-1 | NZ_AUAA01000028.1 | + | 23283 | 23330 | **RNA**→ rpsO (PRK05626)Ribosomal_S15 (pfam00312)→ |
| Csp-15-1 | NZ_JPLZ01000006.1 | + | 1840 | 1887 | **RNA**→ rpsO (PRK05626)Ribosomal_S15 (pfam00312)→ |
| Cha-2-1 | NZ_JPLY01000004.1 | + | 813621 | 813669 | **RNA**→ rpsO (PRK05626)Ribosomal_S15 (pfam00312)→ |
| Ean-3-1 | NZ_CP006576.1 | - | 2427035 | 2426987 | **RNA**→ rpsO (PRK05626)Ribosomal_S15 (pfam00312)→ SMC_prok_A (TIGR02169)DUF1003 (pfam06210)→ |
| Ean-5-1 | NZ_ASYK01000015.1 | + | 43042 | 43090 | **RNA**→ rpsO (PRK05626)Ribosomal_S15 (pfam00312)→ SMC_prok_A (TIGR02169)DUF1003 (pfam06210)→ |
| Ean-7-1 | NZ_ASYJ01000011.1 | - | 311125 | 311077 | **RNA**→ rpsO (PRK05626)Ribosomal_S15 (pfam00312)→ SMC_prok_A (TIGR02169)DUF1003 (pfam06210)→ |
| Ean-1-1 | NZ_CCAC010000078.1 | - | 65873 | 65825 | **RNA**→ rpsO (PRK05626)Ribosomal_S15 (pfam00312)→ SMC_prok_A (TIGR02169)DUF1003 (pfam06210)→ |
| Ean-1-2 | NZ_CCAB010000101.1 | - | 104723 | 104675 | **RNA**→ rpsO (PRK05626)Ribosomal_S15 (pfam00312)→ SMC_prok_A (TIGR02169)DUF1003 (pfam06210)→ |
| Ean-1-3 | NZ_LFKT01000009.1 | + | 119469 | 119517 | **RNA**→ rpsO (PRK05626)Ribosomal_S15 (pfam00312)→ SMC_prok_A (TIGR02169)DUF1003 (pfam06210)→ |
| Ean-11-1 | NZ_CBYE010000026.1 | + | 27074 | 27122 | **RNA**→ rpsO (PRK05626)Ribosomal_S15 (pfam00312)→ SMC_prok_A (TIGR02169)DUF1003 (pfam06210)→ |
| Ean-2-1 | NZ_AHHG01000045.1 | - | 182436 | 182388 | **RNA**→ rpsO (PRK05626)Ribosomal_S15 (pfam00312)→ rpsO (PRK05626)Ribosomal_S15 (pfam00312)→ SMC_prok_A (TIGR02169)DUF1003 (pfam06210)→ SMC_prok_A (TIGR02169)DUF1003 (pfam06210)→ |
| Ean-8-1 | NZ_CP007547.1 | - | 3067040 | 3066992 | **RNA**→ rpsO (PRK05626)Ribosomal_S15 (pfam00312)→ SMC_prok_A (TIGR02169)DUF1003 (pfam06210)→ |
| Eme-1-1 | NZ_AVCQ01000015.1 | - | 442021 | 441973 | **RNA**→ rpsO (PRK05626)Ribosomal_S15 (pfam00312)→ rpsO (PRK05626)Ribosomal_S15 (pfam00312)→ SMC_prok_A (TIGR02169)DUF1003 (pfam06210)→ |
| Ean-6-1 | NZ_ASYI01000016.1 | + | 45322 | 45370 | **RNA**→ rpsO (PRK05626)Ribosomal_S15 (pfam00312)→ SMC_prok_A (TIGR02169)DUF1003 (pfam06210)→ |
| Ean-10-1 | NZ_ASYG01000010.1 | + | 159214 | 159262 | **RNA**→ rpsO (PRK05626)Ribosomal_S15 (pfam00312)→ SMC_prok_A (TIGR02169)DUF1003 (pfam06210)→ |
| Ean-9-1 | NZ_ASYF01000003.1 | - | 448232 | 448184 | **RNA**→ rpsO (PRK05626)Ribosomal_S15 (pfam00312)→ SMC_prok_A (TIGR02169)DUF1003 (pfam06210)→ |
| Ean-4-1 | NZ_ASYH01000003.1 | - | 448359 | 448311 | **RNA**→ rpsO (PRK05626)Ribosomal_S15 (pfam00312)→ SMC_prok_A (TIGR02169)DUF1003 (pfam06210)→ |
| Ean-1-4 | NZ_JNCG01000021.1 | - | 185482 | 185434 | **RNA**→ rpsO (PRK05626)Ribosomal_S15 (pfam00312)→ SMC_prok_A (TIGR02169)DUF1003 (pfam06210)→ |
| Ean-12-1 | NZ_ANIW01000060.1 | - | 190333 | 190285 | **RNA**→ rpsO (PRK05626)Ribosomal_S15 (pfam00312)→ SMC_prok_A (TIGR02169)DUF1003 (pfam06210)→ |
| Csp-18-1 | NZ_KN050755.1 | - | 1438373 | 1438327 | **RNA**→ rpsO (PRK05626)Ribosomal_S15 (pfam00312)→ |
| Cje-1-1 | NZ_JSYL01000004.1 | + | 28177 | 28223 | **RNA**→ rpsO (PRK05626)Ribosomal_S15 (pfam00312)→ |
| env-1866 | JGI26523J50269_1016856 | - | 1306 | 1260 | **RNA**→ rpsO (PRK05626)Ribosomal_S15 (pfam00312)→ |
| env-1867 | JGI26527J51213_1032686 | - | 406 | 360 | **RNA**→ rpsO (PRK05626)Ribosomal_S15 (pfam00312)→ |
| env-1868 | JGI26531J51214_10008686 | - | 2483 | 2437 | **RNA**→ rpsO (PRK05626)Ribosomal_S15 (pfam00312)→ |
| Cha-1-1 | NZ_JASZ01000256.1 | + | 10759 | 10805 | **RNA**→ rpsO (PRK05626)Ribosomal_S15 (pfam00312)→ |
| env-1869 | JGI26530J50255_1106976 | + | 312 | 358 | **RNA**→ rpsO (PRK05626)Ribosomal_S15 (pfam00312)→ |
| env-1870 | topACOD_contig19283 | - | 304 | 259 | **RNA**→ rpsO (PRK05626)Ribosomal_S15 (pfam00312)→ |
| Chi-1-1 | NZ_JARQ01000003.1 | + | 453031 | 453077 | **RNA**→ rpsO (PRK05626)Ribosomal_S15 (pfam00312)→ |
| env-1871 | 3300001197_1001501 | + | 9230 | 9275 | **RNA**→ rpsO (PRK05626)Ribosomal_S15 (pfam00312)→ |
| Cso-2-1 | NZ_JSYK01000004.1 | - | 52246 | 52199 | **RNA**→ rpsO (PRK05626)Ribosomal_S15 (pfam00312)→ |
| Cda-1-1 | NZ_AUMT01000001.1 | + | 62762 | 62807 | **RNA**→ rpsO (PRK05626)Ribosomal_S15 (pfam00312)→ |
| Cpi-1-1 | NZ_JPRJ01000010.1 | + | 67169 | 67214 | **RNA**→ rpsO (PRK05626)Ribosomal_S15 (pfam00312)→ |
| env-1872 | 07_22_INULIN_DNA_scaffold18418_31 | - | 14224 | 14179 | **RNA**→ rpsO (PRK05626)Ribosomal_S15 (pfam00312)→ |
| Fba-4-1 | NC_013062.1 | - | 480533 | 480488 | **RNA**→ rpsO (PRK05626)Ribosomal_S15 (pfam00312)→ |
| Cfo-1-1 | NZ_JPRP01000005.1 | + | 14172 | 14217 | **RNA**→ rpsO (PRK05626)Ribosomal_S15 (pfam00312)→ |
| env-1873 | APMI01051208.1 | - | 1050 | 1005 | **RNA**→ rpsO (PRK05626)Ribosomal_S15 (pfam00312)→ |
| Clu-1-1 | NZ_JPRO01000006.1 | - | 65330 | 65285 | **RNA**→ rpsO (PRK05626)Ribosomal_S15 (pfam00312)→ |
| env-1874 | 2200753365 | - | 6123 | 6078 | **RNA**→ rpsO (PRK05626)Ribosomal_S15 (pfam00312)→ |
| Cta-1-1 | NZ_JWTA01000021.1 | - | 33380 | 33335 | **RNA**→ rpsO (PRK05626)Ribosomal_S15 (pfam00312)→ |

| | | | | | |
|---|---|---|---|---|---|
| Csp-29-1 | NZ_JUGT01000011.1 | - | 37505 | 37460 | RNA→ rpsO (PRK05626)Ribosomal_S15 (pfam00312)→ |
| Csp-24-1 | NZ_JUGI01000017.1 | - | 175541 | 175496 | RNA→ rpsO (PRK05626)Ribosomal_S15 (pfam00312)→ |
| Csp-19-1 | NZ_JUHD01000005.1 | + | 223343 | 223388 | RNA→ rpsO (PRK05626)Ribosomal_S15 (pfam00312)→ |
| Cor-1-1 | NZ_CCAV010000003.1 | + | 217492 | 217537 | RNA→ rpsO (PRK05626)Ribosomal_S15 (pfam00312)→ |
| Cor-1-2 | NZ_CDHM01000003.1 | + | 217492 | 217537 | RNA→ rpsO (PRK05626)Ribosomal_S15 (pfam00312)→ |
| Csp-16-1 | NZ_LFNE01000002.1 | + | 313537 | 313582 | RNA→ rpsO (PRK05626)Ribosomal_S15 (pfam00312)→ |
| Can-1-1 | NZ_LFND01000006.1 | - | 295584 | 295539 | RNA→ rpsO (PRK05626)Ribosomal_S15 (pfam00312)→ |
| Cgl-1-1 | NZ_GL379781.1 | - | 3418094 | 3418049 | RNA→ rpsO (PRK05626)Ribosomal_S15 (pfam00312)→ |
| Csp-26-1 | NZ_KN549099.1 | - | 1525773 | 1525728 | RNA→ rpsO (PRK05626)Ribosomal_S15 (pfam00312)→ |
| Csp-23-1 | NZ_KI535291.1 | + | 490120 | 490165 | RNA→ rpsO (PRK05626)Ribosomal_S15 (pfam00312)→ |
| Cin-2-1 | NZ_BAVL01000002.1 | + | 189412 | 189457 | RNA→ rpsO (PRK05626)Ribosomal_S15 (pfam00312)→ |
| Cso-1-1 | NZ_JPRH01000003.1 | + | 437436 | 437481 | RNA→ rpsO (PRK05626)RpsO (COG0184)→ |
| Csp-17-1 | NZ_JPRN01000005.1 | - | 444957 | 444912 | RNA→ rpsO (PRK05626)Ribosomal_S15 (pfam00312)→ |
| Csp-20-1 | NZ_JUGX01000001.1 | - | 451596 | 451551 | RNA→ rpsO (PRK05626)Ribosomal_S15 (pfam00312)→ |
| env-1875 | JGI24741J21665_1003225 | + | 547 | 592 | RNA→ rpsO (PRK05626)Ribosomal_S15 (pfam00312)→ |
| env-1876 | SwRhRL2b_contig_2736501 | - | 126 | 81 | RNA→ rpsO (PRK05626)RpsO (COG0184)→ |
| Csp-27-1 | NZ_JUGN01000014.1 | - | 542127 | 542082 | RNA→ rpsO (PRK05626)Ribosomal_S15 (pfam00312)→ |
| Csp-28-1 | NZ_JUGO01000011.1 | + | 158290 | 158335 | RNA→ rpsO (PRK05626)Ribosomal_S15 (pfam00312)→ |
| Cin-1-1 | NZ_LAZY01000038.1 | - | 61237 | 61192 | RNA→ rpsO (PRK05626)Ribosomal_S15 (pfam00312)→ |
| Csp-13-1 | NZ_LFNF01000003.1 | + | 556712 | 556757 | RNA→ rpsO (PRK05626)Ribosomal_S15 (pfam00312)→ |
| Csp-21-1 | NZ_JPEQ01000008.1 | - | 148879 | 148834 | RNA→ rpsO (PRK05626)Ribosomal_S15 (pfam00312)→ |
| Csp-25-1 | NZ_KN549083.1 | + | 153277 | 153322 | RNA→ rpsO (PRK05626)Ribosomal_S15 (pfam00312)→ |
| Csp-22-1 | NZ_AP014624.1 | + | 1694760 | 1694805 | RNA→ rpsO (PRK05626)Ribosomal_S15 (pfam00312)→ |
| Cga-1-1 | NZ_CP009928.1 | + | 3037125 | 3037170 | RNA→ rpsO (PRK05626)Ribosomal_S15 (pfam00312)→ |
| Cvr-1-1 | NZ_JPRI01000002.1 | - | 96400 | 96355 | RNA→ rpsO (PRK05626)Ribosomal_S15 (pfam00312)→ |
| Csp-14-1 | NZ_JQJM01000004.1 | + | 231770 | 231815 | RNA→ rpsO (PRK05626)Ribosomal_S15 (pfam00312)→ |
| env-1877 | JGI24735J21928_10576231 | - | 306 | 261 | RNA→ rpsO (PRK05626)Ribosomal_S15 (pfam00312)→ |
| env-1878 | SRS050628_LANL_scaffold_18598 | + | 464 | 518 | RNA→ rpsO (PRK05626)RpsO (COG0184)→ |
| env-1879 | SRS023595_C6923925 | - | 62 | 7 | RNA→ |
| env-1880 | JGI26529J51215_1088893 | - | 319 | 271 | RNA→ rpsO (PRK05626)Ribosomal_S15 (pfam00312)→ |
| env-1881 | JGI26530J50255_1057323 | - | 544 | 496 | RNA→ rpsO (PRK05626)Ribosomal_S15 (pfam00312)→ |
| env-1882 | JGI26527J51213_1015831 | - | 1014 | 966 | RNA→ rpsO (PRK05626)Ribosomal_S15 (pfam00312)→ |
| env-1883 | JGI25921J50272_10550746 | - | 155 | 89 | RNA→ |
| env-1884 | SRS011243_Baylor_scaffold_47946 | - | 2597 | 2550 | RNA→ rpsO (PRK05626)Ribosomal_S15 (pfam00312)→ |
| env-1885 | SRS013164_Baylor_scaffold_59591 | - | 4089 | 4042 | RNA→ rpsO (PRK05626)Ribosomal_S15 (pfam00312)→ |
| env-1886 | SRS014124_WUGC_scaffold_51556 | - | 1480 | 1433 | RNA→ rpsO (PRK05626)Ribosomal_S15 (pfam00312)→ |
| env-1887 | SRS018739_C3569612 | + | 103 | 150 | RNA→ rpsO (PRK05626)Ribosomal_S15 (pfam00312)→ |
| env-1888 | SRS019327_WUGC_scaffold_53864 | - | 2020 | 1973 | RNA→ rpsO (PRK05626)Ribosomal_S15 (pfam00312)→ |
| env-1889 | SRS022719_LANL_scaffold_27291 | + | 47554 | 47601 | RNA→ rpsO (PRK05626)Ribosomal_S15 (pfam00312)→ |
| env-1890 | SRS050669_LANL_scaffold_22387 | + | 10680 | 10727 | RNA→ rpsO (PRK05626)Ribosomal_S15 (pfam00312)→ |
| env-1891 | SRS077736_LANL_scaffold_38299 | + | 9035 | 9082 | RNA→ rpsO (PRK05626)Ribosomal_S15 (pfam00312)→ |
| env-1892 | SRS015060_C1545315 | + | 15 | 62 | RNA→ rpsO (PRK05626)Ribosomal_S15 (pfam00312)→ |
| env-1893 | SRS015434_C5634447 | - | 55 | 8 | RNA→ |
| env-1894 | SRS018300_Baylor_scaffold_41675 | + | 423 | 470 | RNA→ |
| env-1895 | SRS019071_C5124367 | + | 543 | 590 | RNA→ |
| env-1896 | SRS024441_C2974660 | + | 377 | 424 | RNA→ |
| env-1897 | SRS050244_LANL_scaffold_85939 | + | 1274 | 1321 | RNA→ |
| env-1898 | SRS015762_C3672449 | - | 55 | 8 | RNA→ |
| env-1899 | SRS054687_C3386725 | - | 55 | 8 | RNA→ |
| env-1900 | SRS015803_WUGC_scaffold_2285 | + | 78 | 125 | RNA→ rpsO (PRK05626)Ribosomal_S15 (pfam00312)→ |
| env-1901 | SRS016037_C2014584 | - | 305 | 258 | RNA→ rpsO (PRK05626)Ribosomal_S15 (pfam00312)→ |
| env-1902 | SRS016200_WUGC_scaffold_25492 | - | 60 | 13 | RNA→ |

| | | | | | |
|---|---|---|---|---|---|
| env-1903 | SRS016569_C2492960 | + | 724 | 771 | **RNA**→ rpsO (PRK05626)Ribosomal_S15 (pfam00312)→ |
| env-1904 | SRS018357_Baylor_scaffold_29493 | - | 767 | 720 | **RNA**→ rpsO (PRK05626)Ribosomal_S15 (pfam00312)→ |
| env-1905 | SRS019045_C2658954 | + | 176 | 223 | **RNA**→ rpsO (PRK05626)Ribosomal_S15 (pfam00312)→ |
| env-1906 | SRS019587_C1053840 | - | 327 | 280 | **RNA**→ rpsO (PRK05626)Ribosomal_S15 (pfam00312)→ |
| env-1907 | SRS021496_Baylor_scaffold_63711 | + | 53 | 100 | **RNA**→ rpsO (PRK05626)Ribosomal_S15 (pfam00312)→ |
| env-1908 | SRS022149_C2781104 | + | 1 | 48 | **RNA**→ rpsO (PRK05626)Ribosomal_S15 (pfam00312)→ |
| env-1909 | SRS023557_C1767695 | - | 247 | 200 | **RNA**→ rpsO (PRK05626)Ribosomal_S15 (pfam00312)→ |
| env-1910 | SRS042910_C5602692 | - | 383 | 336 | **RNA**→ rpsO (PRK05626)Ribosomal_S15 (pfam00312)→ |
| env-1911 | SRS044373_C2907430 | - | 742 | 695 | **RNA**→ rpsO (PRK05626)Ribosomal_S15 (pfam00312)→ |
| env-1912 | SRS048791_C3091469 | - | 313 | 266 | **RNA**→ rpsO (PRK05626)Ribosomal_S15 (pfam00312)→ |
| env-1913 | SRS051244_LANL_scaffold_30757 | + | 464 | 511 | **RNA**→ rpsO (PRK05626)Ribosomal_S15 (pfam00312)→ |
| env-1914 | SRS055450_LANL_scaffold_72003 | - | 364 | 317 | **RNA**→ rpsO (PRK05626)Ribosomal_S15 (pfam00312)→ |
| env-1915 | SRS057692_LANL_scaffold_6218 | + | 21 | 68 | **RNA**→ rpsO (PRK05626)Ribosomal_S15 (pfam00312)→ |
| env-1916 | SRS058336_C2042484 | - | 153 | 106 | **RNA**→ rpsO (PRK05626)Ribosomal_S15 (pfam00312)→ |
| env-1917 | SRS064423_LANL_scaffold_57424 | + | 419 | 466 | **RNA**→ rpsO (PRK05626)Ribosomal_S15 (pfam00312)→ |
| env-1918 | SRS043422_C2694041 | - | 1486 | 1439 | **RNA**→ rpsO (PRK05626)Ribosomal_S15 (pfam00312)→ |
| env-1919 | SRS014473_C1126565 | + | 640 | 687 | **RNA**→ rpsO (PRK05626)Ribosomal_S15 (pfam00312)→ |
| env-1920 | SRS019125_C625732 | + | 7434 | 7481 | **RNA**→ rpsO (PRK05626)Ribosomal_S15 (pfam00312)→ |
| env-1921 | SRS064329_C2689539 | - | 427 | 380 | **RNA**→ rpsO (PRK05626)Ribosomal_S15 (pfam00312)→ |
| env-1922 | SRS012279_Baylor_scaffold_66478 | + | 7846 | 7893 | **RNA**→ rpsO (PRK05626)Ribosomal_S15 (pfam00312)→ |
| env-1923 | SRS022077_Baylor_scaffold_42750 | + | 9400 | 9447 | **RNA**→ rpsO (PRK05626)Ribosomal_S15 (pfam00312)→ |
| env-1924 | SRS013947_C1568635 | + | 15 | 62 | **RNA**→ rpsO (PRK05626)Ribosomal_S15 (pfam00312)→ |
| env-1925 | SRS014573_WUGC_scaffold_22664 | + | 73 | 120 | **RNA**→ rpsO (PRK05626)Ribosomal_S15 (pfam00312)→ |
| env-1926 | SRS015158_WUGC_scaffold_53335 | + | 495 | 542 | **RNA**→ rpsO (PRK05626)Ribosomal_S15 (pfam00312)→ |
| env-1927 | SRS015644_WUGC_scaffold_47575 | + | 224 | 271 | **RNA**→ rpsO (PRK05626)Ribosomal_S15 (pfam00312)→ |
| env-1928 | SRS015797_C2629919 | + | 42 | 89 | **RNA**→ rpsO (PRK05626)Ribosomal_S15 (pfam00312)→ |
| env-1929 | SRS016002_C2634626 | + | 834 | 881 | **RNA**→ rpsO (PRK05626)Ribosomal_S15 (pfam00312)→ |
| env-1930 | SRS016086_C2525590 | - | 620 | 573 | **RNA**→ rpsO (PRK05626)Ribosomal_S15 (pfam00312)→ |
| env-1931 | SRS016225_C3015844 | - | 61 | 14 | **RNA**→ |
| env-1932 | SRS024081_LANL_scaffold_72737 | + | 2277 | 2324 | **RNA**→ |
| env-1933 | SRS042131_C2167664 | + | 340 | 387 | **RNA**→ |
| env-1934 | SRS049147_C4008101 | + | 291 | 338 | **RNA**→ |
| env-1935 | SRS018145_Baylor_scaffold_19145 | - | 1405 | 1358 | **RNA**→ rpsO (PRK05626)Ribosomal_S15 (pfam00312)→ |
| env-1936 | SRS019027_C1962878 | - | 130 | 83 | **RNA**→ rpsO (PRK05626)RpsO (COG0184)→ |
| env-1937 | SRS019122_C5834060 | + | 331 | 378 | **RNA**→ rpsO (PRK05626)Ribosomal_S15 (pfam00312)→ |
| env-1938 | SRS019591_WUGC_scaffold_17646 | - | 61 | 14 | **RNA**→ |
| env-1939 | SRS022083_C1886151 | - | 305 | 258 | **RNA**→ rpsO (PRK05626)Ribosomal_S15 (pfam00312)→ |
| env-1940 | SRS022530_LANL_scaffold_2248 | + | 50661 | 50708 | **RNA**→ rpsO (PRK05626)Ribosomal_S15 (pfam00312)→ |
| env-1941 | SRS024277_LANL_scaffold_38800 | - | 331 | 284 | **RNA**→ rpsO (PRK05626)Ribosomal_S15 (pfam00312)→ |
| env-1942 | SRS042984_LANL_scaffold_420 | + | 1488 | 1535 | **RNA**→ rpsO (PRK05626)Ribosomal_S15 (pfam00312)→ |
| env-1943 | SRS047824_C2273016 | - | 257 | 210 | **RNA**→ rpsO (PRK05626)Ribosomal_S15 (pfam00312)→ |
| env-1944 | SRS049389_WUGC_scaffold_29036 | + | 784 | 831 | **RNA**→ rpsO (PRK05626)Ribosomal_S15 (pfam00312)→ |
| env-1945 | SRS057539_LANL_scaffold_61878 | - | 651 | 604 | **RNA**→ rpsO (PRK05626)Ribosomal_S15 (pfam00312)→ |
| env-1946 | SRS057791_C3560696 | + | 282 | 329 | **RNA**→ rpsO (PRK05626)Ribosomal_S15 (pfam00312)→ |
| env-1947 | SRS063288_LANL_scaffold_13079 | + | 772 | 819 | **RNA**→ rpsO (PRK05626)Ribosomal_S15 (pfam00312)→ |
| env-1948 | SRS065278_C2443864 | + | 63 | 110 | **RNA**→ rpsO (PRK05626)Ribosomal_S15 (pfam00312)→ |
| env-1949 | Palatine_Tonsils_LANL_C6176060 | - | 63 | 16 | **RNA**→ |
| env-1950 | SRS045313_C1321695 | + | 55 | 102 | **RNA**→ rpsO (PRK05626)Ribosomal_S15 (pfam00312)→ |
| env-1951 | SRS018443_C3710842 | + | 364 | 411 | **RNA**→ |
| env-1952 | SRS023538_Baylor_scaffold_34561 | - | 542 | 495 | **RNA**→ rpsO (PRK05626)Ribosomal_S15 (pfam00312)→ |
| env-1953 | SRS051378_LANL_scaffold_19919 | + | 108 | 155 | **RNA**→ rpsO (PRK05626)Ribosomal_S15 (pfam00312)→ |
| env-1954 | SRS055401_LANL_scaffold_31063 | + | 824 | 871 | **RNA**→ |

| | | | | | |
|---|---|---|---|---|---|
| env-1955 | SRS015985_WUGC_scaffold_29053 | - | 1071 | 1024 | **RNA**→ rpsO (PRK05626)Ribosomal_S15 (pfam00312)→ |
| env-1956 | SRS015989_WUGC_scaffold_321 | - | 1263 | 1216 | **RNA**→ rpsO (PRK05626)Ribosomal_S15 (pfam00312)→ |
| env-1957 | SRS011140_C5162271 | - | 1086 | 1039 | **RNA**→ rpsO (PRK05626)Ribosomal_S15 (pfam00312)→ |
| env-1958 | SRS011343_Baylor_scaffold_58822 | - | 579 | 532 | **RNA**→ rpsO (PRK05626)Ribosomal_S15 (pfam00312)→ |
| env-1959 | SRS013950_C2176163 | - | 1345 | 1298 | **RNA**→ rpsO (PRK05626)Ribosomal_S15 (pfam00312)→ |
| env-1960 | SRS019129_WUGC_scaffold_25059 | + | 248 | 295 | **RNA**→ rpsO (PRK05626)Ribosomal_S15 (pfam00312)→ |
| env-1961 | SRS014894_WUGC_scaffold_48888 | + | 400 | 447 | **RNA**→ |
| env-1962 | SRS015278_WUGC_scaffold_56393 | + | 212 | 259 | **RNA**→ rpsO (PRK05626)Ribosomal_S15 (pfam00312)→ |
| env-1963 | SRS015470_WUGC_scaffold_46669 | + | 449 | 496 | **RNA**→ rpsO (PRK05626)Ribosomal_S15 (pfam00312)→ |
| env-1964 | SRS016043_C2059938 | - | 61 | 14 | **RNA**→ |
| env-1965 | SRS016746_Baylor_scaffold_53508 | + | 20 | 67 | **RNA**→ rpsO (PRK05626)Ribosomal_S15 (pfam00312)→ |
| env-1966 | SRS017439_C3749384 | + | 21 | 68 | **RNA**→ rpsO (PRK05626)Ribosomal_S15 (pfam00312)→ |
| env-1967 | SRS017691_C5063285 | + | 292 | 339 | **RNA**→ |
| env-1968 | SRS018394_Baylor_scaffold_1898 | - | 472 | 425 | **RNA**→ rpsO (PRK05626)Ribosomal_S15 (pfam00312)→ |
| env-1969 | SRS019128_WUGC_scaffold_19415 | + | 88 | 135 | **RNA**→ rpsO (PRK05626)Ribosomal_S15 (pfam00312)→ |
| env-1970 | SRS019906_WUGC_scaffold_36213 | - | 53 | 6 | **RNA**→ |
| env-1971 | SRS022536_LANL_scaffold_43917 | - | 900 | 853 | **RNA**→ rpsO (PRK05626)Ribosomal_S15 (pfam00312)→ |
| env-1972 | SRS023595_C6958305 | + | 381 | 428 | **RNA**→ |
| env-1973 | SRS024021_Baylor_scaffold_13409 | - | 8309 | 8262 | **RNA**→ rpsO (PRK05626)Ribosomal_S15 (pfam00312)→ |
| env-1974 | SRS024447_LANL_scaffold_52352 | - | 305 | 258 | **RNA**→ rpsO (PRK05626)Ribosomal_S15 (pfam00312)→ |
| env-1975 | SRS047113_LANL_scaffold_72955 | + | 244 | 291 | **RNA**→ rpsO (PRK05626)Ribosomal_S15 (pfam00312)→ |
| env-1976 | SRS047634_LANL_scaffold_135861 | - | 561 | 514 | **RNA**→ rpsO (PRK05626)Ribosomal_S15 (pfam00312)→ |
| env-1977 | SRS053584_LANL_scaffold_40362 | - | 1914 | 1867 | **RNA**→ rpsO (PRK05626)Ribosomal_S15 (pfam00312)→ |
| env-1978 | SRS058808_LANL_scaffold_50592 | - | 215 | 168 | **RNA**→ rpsO (PRK05626)Ribosomal_S15 (pfam00312)→ |
| env-1979 | SRS063932_LANL_scaffold_80836 | + | 169 | 216 | **RNA**→ rpsO (PRK05626)Ribosomal_S15 (pfam00312)→ |
| env-1980 | SRS023930_C2188236 | - | 1153 | 1106 | **RNA**→ rpsO (PRK05626)Ribosomal_S15 (pfam00312)→ |
| env-1981 | SRS011126_Baylor_scaffold_69832 | - | 338 | 291 | **RNA**→ rpsO (PRK05626)Ribosomal_S15 (pfam00312)→ |
| env-1982 | SRS011152_Baylor_scaffold_54314 | - | 364 | 317 | **RNA**→ rpsO (PRK05626)Ribosomal_S15 (pfam00312)→ |
| env-1983 | SRS013252_C3586002 | - | 55 | 8 | **RNA**→ |
| env-1984 | SRS014578_WUGC_scaffold_33113 | + | 891 | 938 | **RNA**→ rpsO (PRK05626)Ribosomal_S15 (pfam00312)→ |
| env-1985 | SRS015215_C2492460 | - | 441 | 394 | **RNA**→ rpsO (PRK05626)Ribosomal_S15 (pfam00312)→ |
| env-1986 | SRS015440_C2591831 | + | 803 | 850 | **RNA**→ |
| env-1987 | SRS015574_WUGC_scaffold_74608 | + | 333 | 380 | **RNA**→ rpsO (PRK05626)Ribosomal_S15 (pfam00312)→ |
| env-1988 | SRS016331_C3952241 | + | 270 | 317 | **RNA**→ |
| env-1989 | SRS052876_LANL_scaffold_31436 | + | 425 | 472 | **RNA**→ |
| env-1990 | SRS017025_Baylor_scaffold_2897 | - | 844 | 797 | **RNA**→ rpsO (PRK05626)Ribosomal_S15 (pfam00312)→ |
| env-1991 | SRS017511_C3486220 | + | 49 | 96 | **RNA**→ rpsO (PRK05626)Ribosomal_S15 (pfam00312)→ |
| env-1992 | SRS017814_C2648955 | - | 305 | 258 | **RNA**→ rpsO (PRK05626)Ribosomal_S15 (pfam00312)→ |
| env-1993 | SRS018971_C748940 | - | 283 | 236 | **RNA**→ rpsO (PRK05626)Ribosomal_S15 (pfam00312)→ |
| env-1994 | SRS019387_C1725966 | - | 63 | 16 | **RNA**→ |
| env-1995 | SRS019980_Baylor_scaffold_58741 | + | 1292 | 1339 | **RNA**→ |
| env-1996 | SRS022725_LANL_scaffold_113781 | + | 1278 | 1325 | **RNA**→ rpsO (PRK05626)Ribosomal_S15 (pfam00312)→ |
| env-1997 | SRS023964_Baylor_scaffold_44464 | - | 2165 | 2118 | **RNA**→ rpsO (PRK05626)Ribosomal_S15 (pfam00312)→ |
| env-1998 | SRS024381_LANL_scaffold_36143 | - | 1503 | 1456 | **RNA**→ rpsO (PRK05626)Ribosomal_S15 (pfam00312)→ |
| env-1999 | SRS043755_WUGC_scaffold_39570 | + | 517 | 564 | **RNA**→ rpsO (PRK05626)RpsO (COG0184)→ |
| env-2000 | SRS047265_C1087503 | - | 931 | 884 | **RNA**→ rpsO (PRK05626)Ribosomal_S15 (pfam00312)→ |
| env-2001 | SRS051930_C3123168 | + | 372 | 419 | **RNA**→ |
| env-2002 | SRS054430_LANL_scaffold_24266 | + | 1510 | 1557 | **RNA**→ |
| env-2003 | SRS063603_LANL_scaffold_97549 | - | 336 | 289 | **RNA**→ rpsO (PRK05626)Ribosomal_S15 (pfam00312)→ |
| env-2004 | SRS063999_LANL_scaffold_80667 | - | 1141 | 1094 | **RNA**→ rpsO (PRK05626)Ribosomal_S15 (pfam00312)→ |
| env-2005 | SRS016541_Baylor_scaffold_17430 | + | 71 | 118 | **RNA**→ rpsO (PRK05626)Ribosomal_S15 (pfam00312)→ |
| Ran-4-1 | NC_017569.1 | - | 1632510 | 1632464 | **RNA**→ rpsO (PRK05626)Ribosomal_S15 (pfam00312)→ |

| | | | | | |
|---|---|---|---|---|---|
| Ran-1-1 | NZ_CP007504.1 | + | 677153 | 677199 | **RNA**→ rpsO (PRK05626)Ribosomal_S15 (pfam00312)→ |
| Ran-1-2 | NZ_CP007503.1 | + | 665064 | 665110 | **RNA**→ rpsO (PRK05626)Ribosomal_S15 (pfam00312)→ |
| Ran-2-1 | NC_014738.1 | + | 675647 | 675693 | **RNA**→ rpsO (PRK05626)Ribosomal_S15 (pfam00312)→ |
| Ran-6-1 | NZ_AENH01000022.1 | + | 24212 | 24258 | **RNA**→ rpsO (PRK05626)Ribosomal_S15 (pfam00312)→ |
| Ran-5-1 | NZ_KB206037.1 | - | 211263 | 211217 | **RNA**→ rpsO (PRK05626)Ribosomal_S15 (pfam00312)→ |
| Ran-3-1 | NC_020125.1 | - | 1053939 | 1053893 | **RNA**→ rpsO (PRK05626)Ribosomal_S15 (pfam00312)→ |
| Ran-2-2 | NC_017045.1 | + | 837166 | 837212 | **RNA**→ rpsO (PRK05626)Ribosomal_S15 (pfam00312)→ |
| Ran-7-1 | NZ_CP007204.1 | + | 663670 | 663716 | **RNA**→ rpsO (PRK05626)Ribosomal_S15 (pfam00312)→ |
| Cca-2-1 | NZ_CDOE01000080.1 | - | 107055 | 107000 | **RNA**→ rpsO (PRK05626)Ribosomal_S15 (pfam00312)→ |
| Cca-2-2 | NZ_CDOK01000138.1 | + | 7933 | 7987 | **RNA**→ rpsO (PRK05626)Ribosomal_S15 (pfam00312)→ |
| Cca-2-3 | NZ_CDOJ01000102.1 | + | 7875 | 7929 | **RNA**→ rpsO (PRK05626)Ribosomal_S15 (pfam00312)→ |
| Cca-3-1 | NC_015846.1 | - | 2206651 | 2206597 | **RNA**→ rpsO (PRK05626)Ribosomal_S15 (pfam00312)→ |
| env-2006 | JGI12420J12819_1001506 | - | 9008 | 8962 | **RNA**→ rpsO (PRK05626)Ribosomal_S15 (pfam00312)→ |
| env-2007 | TFO_HY__Contig_14630 | - | 7035 | 6950 | **RNA**→ rpsO (PRK05626)RpsO (COG0184)→ |
| Mru-1-1 | NC_015945.1 | - | 2244219 | 2244134 | **RNA**→ rpsO (PRK05626)RpsO (COG0184)→ |
| env-2008 | JGI11876J14442_10002026 | + | 6244 | 6306 | **RNA**→ rpsO (PRK05626)RpsO (COG0184)→ |
| env-2009 | ElkS_mat_CD2A_1000757 | - | 3779 | 3717 | **RNA**→ rpsO (PRK05626)RpsO (COG0184)→ |
| env-2010 | CEOE01009065.1 | + | 2223 | 2285 | **RNA**→ |
| env-2011 | CEOE01041545.1 | + | 1214 | 1276 | **RNA**→ rpsO (PRK05626)RpsO (COG0184)→ |
| env-2012 | CEQQ01164659.1 | - | 411 | 349 | **RNA**→ rpsO (PRK05626)RpsO (COG0184)→ |
| env-2013 | CESC01055335.1 | + | 492 | 554 | **RNA**→ rpsO (PRK05626)RpsO (COG0184)→ |
| env-2014 | CEVF01145327.1 | + | 1277 | 1339 | **RNA**→ rpsO (PRK05626)Ribosomal_S15 (pfam00312)→ |
| env-2015 | CEOS01127385.1 | + | 891 | 953 | **RNA**→ |
| env-2016 | CEQH01250475.1 | - | 233 | 171 | **RNA**→ rpsO (PRK05626)Ribosomal_S15 (pfam00312)→ |
| env-2017 | CETR01116473.1 | - | 448 | 386 | **RNA**→ rpsO (PRK05626)RpsO (COG0184)→ |
| env-2018 | CEQG01236235.1 | - | 299 | 237 | **RNA**→ rpsO (PRK05626)RpsO (COG0184)→ |
| Msp-2-1 | NZ_JQNJ01000001.1 | - | 2554372 | 2554309 | **RNA**→ rpsO (PRK05626)RpsO (COG0184)→ |
| Cba-2-1 | NZ_CP009976.1 | - | 2003323 | 2003261 | **RNA**→ rpsO (PRK05626)Ribosomal_S15 (pfam00312)→ |
| Cba-3-1 | NZ_ATLG01000025.1 | + | 148886 | 148948 | **RNA**→ rpsO (PRK05626)Ribosomal_S15 (pfam00312)→ |
| Csp-11-1 | NZ_JQCT01000001.1 | + | 36796 | 36858 | **RNA**→ rpsO (PRK05626)Ribosomal_S15 (pfam00312)→ |
| Cba-4-1 | NZ_CP009887.1 | - | 1991741 | 1991679 | **RNA**→ rpsO (PRK05626)Ribosomal_S15 (pfam00312)→ |
| Cba-1-1 | NZ_ATLH01000029.1 | + | 35153 | 35215 | **RNA**→ rpsO (PRK05626)Ribosomal_S15 (pfam00312)→ |
| env-2019 | P_1C_Sed_3_UnCty_1112983 | + | 133 | 197 | **RNA**→ rpsO (PRK05626)RpsO (COG0184)→ |
| env-2020 | SA_S1_NOR08_45m_c10098966 | - | 656 | 592 | **RNA**→ rpsO (PRK05626)Ribosomal_S15 (pfam00312)→ |
| Csp-12-1 | NZ_JUHB01000001.1 | - | 3425850 | 3425786 | **RNA**→ rpsO (PRK05626)Ribosomal_S15 (pfam00312)→ |
| Zul-1-1 | NZ_JQMD01000002.1 | + | 2791513 | 2791577 | **RNA**→ rpsO (PRK05626)RpsO (COG0184)→ |
| env-2021 | CEQH01267320.1 | - | 701 | 637 | **RNA**→ rpsO (PRK05626)Ribosomal_S15 (pfam00312)→ |
| env-2022 | CEQR01110231.1 | + | 2717 | 2781 | **RNA**→ rpsO (PRK05626)Ribosomal_S15 (pfam00312)→ |
| env-2023 | CEQT01008943.1 | - | 946 | 882 | **RNA**→ rpsO (PRK05626)Ribosomal_S15 (pfam00312)→ |
| env-2024 | CERC01066844.1 | - | 30660 | 30596 | **RNA**→ rpsO (PRK05626)Ribosomal_S15 (pfam00312)→ |
| env-2025 | CERD01127367.1 | + | 35670 | 35734 | **RNA**→ rpsO (PRK05626)Ribosomal_S15 (pfam00312)→ |
| env-2026 | CERE01165140.1 | - | 692 | 628 | **RNA**→ rpsO (PRK05626)Ribosomal_S15 (pfam00312)→ |
| env-2027 | CERX01241151.1 | - | 248 | 184 | **RNA**→ rpsO (PRK05626)Ribosomal_S15 (pfam00312)→ |
| env-2028 | CENJ01132294.1 | - | 814 | 750 | **RNA**→ rpsO (PRK05626)Ribosomal_S15 (pfam00312)→ |
| env-2029 | JGI24723J26617_10000090 | - | 7488 | 7425 | **RNA**→ rpsO (PRK05626)Ribosomal_S15 (pfam00312)→ |
| Ace-1-1 | NZ_AUCB01000005.1 | + | 34830 | 34893 | **RNA**→ rpsO (PRK05626)Ribosomal_S15 (pfam00312)→ |
| Ala-2-1 | NZ_KE384290.1 | - | 34915 | 34852 | **RNA**→ rpsO (PRK05626)Ribosomal_S15 (pfam00312)→ |
| env-2030 | KGI_S1_ANT01_95m_c10011027 | + | 4052 | 4115 | **RNA**→ ←hypo |
| env-2031 | KGI_S2_ANT06_2345m_c1000769 | - | 806 | 743 | **RNA**→ rpsO (PRK05626)RpsO (COG0184)→ |
| env-2032 | SA_S1_NOR05_45m_c10101907 | - | 644 | 581 | **RNA**→ rpsO (PRK05626)RpsO (COG0184)→ |
| env-2033 | SA_S2_NOR15_50m_c10610413 | + | 126 | 189 | **RNA**→ rpsO (PRK05626)Ribosomal_S15 (pfam00312)→ |
| env-2034 | KGI_S2_ANT05_2345m_c1051480 | + | 342 | 405 | **RNA**→ rpsO (PRK05626)RpsO (COG0184)→ |

| | | | | | |
|---|---|---|---|---|---|
| env-2035 | KGI_S1_ANT02_95m_c10160552 | + | 8 | 71 | **RNA**→ rpsO (PRK05626)RpsO (COG0184)→ |
| env-2036 | JGI24025J20009_10098606 | + | 246 | 309 | **RNA**→ rpsO (PRK05626)RpsO (COG0184)→ |
| env-2037 | KGI_S1_ANT01_95m_c10032764 | + | 1919 | 1982 | **RNA**→ rpsO (PRK05626)→ |
| env-2038 | KGI_S1_ANT01_95m_c10238661 | - | 453 | 390 | **RNA**→ rpsO (PRK05626)RpsO (COG0184)→ |
| env-2039 | TDF_OR_ARG04_113m_c1031286 | + | 374 | 437 | **RNA**→ rpsO (PRK05626)Ribosomal_S15 (pfam00312)→ |
| env-2040 | TDF_MC_ARG01_113m_c1031732 | - | 233 | 170 | **RNA**→ rpsO (PRK05626)Ribosomal_S15 (pfam00312)→ |
| env-2041 | SA_S1_NOR08_45m_c10338978 | + | 208 | 271 | **RNA**→ rpsO (PRK05626)Ribosomal_S15 (pfam00312)→ |
| env-2042 | TDF_MC_ARG02_113m_c1027467 | + | 202 | 265 | **RNA**→ rpsO (PRK05626)Ribosomal_S15 (pfam00312)→ |
| env-2043 | KGI_S1_ANT03_95m_c1036695 | + | 376 | 439 | **RNA**→ rpsO (PRK05626)Ribosomal_S15 (pfam00312)→ |
| env-2044 | KGI_S1_ANT02_95m_c10041159 | + | 904 | 967 | **RNA**→ rpsO (PRK05626)RpsO (COG0184)→ |
| env-2045 | JGI24023J19991_10153635 | + | 471 | 534 | **RNA**→ rpsO (PRK05626)Ribosomal_S15 (pfam00312)→ |
| env-2046 | KGI_S1_ANT01_95m_c10009658 | - | 886 | 823 | **RNA**→ rpsO (PRK05626)RpsO (COG0184)→ |
| env-2047 | KGI_S2_ANT04_2345m_c1021074 | + | 852 | 915 | **RNA**→ rpsO (PRK05626)RpsO (COG0184)→ |
| env-2048 | KGI_S1_ANT02_95m_c10001832 | + | 7430 | 7493 | **RNA**→ rpsO (PRK05626)RpsO (COG0184)→ |
| env-2049 | P_2C_Liq_1_UnCty_1014332 | - | 1481 | 1417 | **RNA**→ rpsO (PRK05626)RpsO (COG0184)→ |
| env-2050 | PML_READ_00802233 | + | 187 | 251 | **RNA**→ |
| env-2051 | DelMOSum2011_c10009803 | - | 282 | 218 | **RNA**→ rpsO (PRK05626)Ribosomal_S15 (pfam00312)→ |
| env-2052 | DelMOWin2010_c10595920 | + | 80 | 144 | **RNA**→ rpsO (PRK05626)Ribosomal_S15 (pfam00312)→ |
| env-2053 | P_2C_Liq_2_UnCty_1013023 | - | 1456 | 1392 | **RNA**→ rpsO (PRK05626)RpsO (COG0184)→ |
| env-2054 | PML_READ_00935082 | + | 13 | 77 | **RNA**→ rpsO (PRK05626)Ribosomal_S15 (pfam00312)→ |
| env-2055 | DelMOSum2010_c10010410 | + | 322 | 386 | **RNA**→ rpsO (PRK05626)RpsO (COG0184)→ |
| env-2056 | JGI20152J14361_10001126 | - | 5003 | 4939 | **RNA**→ rpsO (PRK05626)RpsO (COG0184)→ |
| env-2057 | JGI20151J14362_10008756 | + | 270 | 334 | **RNA**→ rpsO (PRK05626)RpsO (COG0184)→ |
| env-2058 | JGI20156J14371_10000639 | - | 17377 | 17313 | **RNA**→ rpsO (PRK05626)RpsO (COG0184)→ |
| env-2059 | JGI20154J14316_10003245 | + | 8271 | 8335 | **RNA**→ rpsO (PRK05626)RpsO (COG0184)→ |
| env-2060 | JGI20160J14292_10001354 | + | 13475 | 13539 | **RNA**→ rpsO (PRK05626)RpsO (COG0184)→ |
| env-2061 | JGI20153J14295_10000494 | + | 27409 | 27473 | **RNA**→ rpsO (PRK05626)RpsO (COG0184)→ |
| env-2062 | JGI20153J14318_10000494 | + | 27409 | 27473 | **RNA**→ rpsO (PRK05626)RpsO (COG0184)→ |
| env-2063 | JGI20157J14317_10000832 | + | 24813 | 24877 | **RNA**→ rpsO (PRK05626)RpsO (COG0184)→ |
| env-2064 | JGI20159J14440_10001145 | - | 5009 | 4945 | **RNA**→ rpsO (PRK05626)RpsO (COG0184)→ |
| env-2065 | JGI20155J14468_10001234 | + | 14134 | 14198 | **RNA**→ rpsO (PRK05626)RpsO (COG0184)→ |
| env-2066 | JGI20158J14315_10000172 | + | 28789 | 28853 | **RNA**→ rpsO (PRK05626)RpsO (COG0184)→ |
| env-2067 | F10SI0310a_Sequence0000028922 | - | 721 | 657 | **RNA**→ rpsO (PRK05626)RpsO (COG0184)→ |
| Ead-1-1 | NZ_KB907552.1 | + | 40646 | 40709 | **RNA**→ rpsO (PRK05626)RpsO (COG0184)→ |
| env-2068 | TDF_MC_ARG02_113m_c1042706 | + | 63 | 126 | **RNA**→ rpsO (PRK05626)RpsO (COG0184)→ |
| env-2069 | TDF_OR_ARG04_113m_c1023153 | - | 698 | 635 | **RNA**→ rpsO (PRK05626)RpsO (COG0184)→ |
| env-2070 | TDF_MC_ARG01_113m_c1008162 | - | 795 | 732 | **RNA**→ rpsO (PRK05626)RpsO (COG0184)→ |
| env-2071 | JGI24024J18818_10072705 | + | 731 | 794 | **RNA**→ rpsO (PRK05626)RpsO (COG0184)→ |
| env-2072 | TDF_OR_ARG05_123m_c1037339 | - | 340 | 277 | **RNA**→ rpsO (PRK05626)RpsO (COG0184)→ |
| env-2073 | JGI24723J26617_10002068 | - | 2830 | 2767 | **RNA**→ rpsO (PRK05626)Ribosomal_S15 (pfam00312)→ |
| env-2074 | JGI24723J26617_10248799 | + | 233 | 296 | **RNA**→ rpsO (PRK05626)Ribosomal_S15 (pfam00312)→ |
| env-2075 | CENR01000826.1 | - | 4573 | 4510 | **RNA**→ rpsO (PRK05626)Ribosomal_S15 (pfam00312)→ |
| Aal-1-1 | NZ_JPOO01000001.1 | - | 1038671 | 1038608 | **RNA**→ rpsO (PRK05626)Ribosomal_S15 (pfam00312)→ |
| env-2076 | JGI24723J26617_10486333 | - | 145 | 82 | **RNA**→ rpsO (PRK05626)RpsO (COG0184)→ |
| env-2077 | JGI24724J26744_10000006 | - | 23136 | 23073 | **RNA**→ rpsO (PRK05626)Ribosomal_S15 (pfam00312)→ |
| env-2078 | JGI24724J26744_10987649 | + | 71 | 135 | **RNA**→ rpsO (PRK05626)Ribosomal_S15 (pfam00312)→ |
| env-2079 | JGI24723J26617_10013201 | + | 2526 | 2590 | **RNA**→ rpsO (PRK05626)Ribosomal_S15 (pfam00312)→ |
| Fba-6-1 | NC_014472.1 | + | 156641 | 156705 | **RNA**→ rpsO (PRK05626)RpsO (COG0184)→ |
| env-2080 | JGI24723J26617_10539426 | - | 214 | 150 | **RNA**→ rpsO (PRK05626)Ribosomal_S15 (pfam00312)→ |
| env-2081 | JGI25321J50212_10002132 | + | 13149 | 13213 | **RNA**→ rpsO (PRK05626)RpsO (COG0184)→ |
| env-2082 | JGI25321J39296_1282722 | + | 90 | 154 | **RNA**→ |
| env-2083 | KGI_S1_ANT01_95m_c10067499 | + | 739 | 804 | **RNA**→ rpsO (PRK05626)Ribosomal_S15 (pfam00312)→ |

| | | | | | |
|---|---|---|---|---|---|
| env-2084 | KGI_S2_ANT06_2345m_c1000581 | + | 4905 | 4970 | **RNA**→ rpsO (PRK05626)Ribosomal_S15 (pfam00312)→ |
| Msp-1-1 | NZ_JHZW01000003.1 | + | 2014167 | 2014232 | **RNA**→ rpsO (PRK05626)Ribosomal_S15 (pfam00312)→ |
| env-2085 | LAZR01000507.1 | + | 2889 | 2954 | **RNA**→ rpsO (PRK05626)RpsO (COG0184)→ |
| env-2086 | JGI11876J14442_10355929 | - | 340 | 276 | **RNA**→ rpsO (PRK05626)Ribosomal_S15 (pfam00312)→ |
| env-2087 | JGI11876J14442_10036760 | - | 2240 | 2176 | **RNA**→ rpsO (PRK05626)Ribosomal_S15 (pfam00312)→ |
| env-2088 | ElkS_mat_CD6A_1068972 | - | 237 | 173 | **RNA**→ rpsO (PRK05626)Ribosomal_S15 (pfam00312)→ |
| Mth-1-1 | NZ_LDAS01000001.1 | - | 4009227 | 4009163 | **RNA**→ rpsO (PRK05626)Ribosomal_S15 (pfam00312)→ |
| env-2089 | CEVJ01146598.1 | - | 214 | 150 | **RNA**→ rpsO (PRK05626)Ribosomal_S15 (pfam00312)→ |
| env-2090 | CETA01166376.1 | + | 440 | 504 | **RNA**→ |
| env-2091 | CETS01181212.1 | + | 283 | 347 | **RNA**→ rpsO (PRK05626)Ribosomal_S15 (pfam00312)→ |
| env-2092 | CETB01163303.1 | - | 1560 | 1496 | **RNA**→ rpsO (PRK05626)Ribosomal_S15 (pfam00312)→ |
| env-2093 | CETR01268141.1 | - | 1727 | 1663 | **RNA**→ rpsO (PRK05626)Ribosomal_S15 (pfam00312)→ |
| env-2094 | CEVI01056776.1 | - | 475 | 411 | **RNA**→ rpsO (PRK05626)Ribosomal_S15 (pfam00312)→ |
| env-2095 | CEVK01107237.1 | - | 796 | 732 | **RNA**→ rpsO (PRK05626)Ribosomal_S15 (pfam00312)→ |
| env-2096 | CEVM01115743.1 | + | 2313 | 2377 | **RNA**→ rpsO (PRK05626)Ribosomal_S15 (pfam00312)→ |
| env-2097 | CEVO01246946.1 | - | 1295 | 1231 | **RNA**→ rpsO (PRK05626)Ribosomal_S15 (pfam00312)→ |
| env-2098 | CEVQ01106901.1 | + | 43110 | 43174 | **RNA**→ rpsO (PRK05626)Ribosomal_S15 (pfam00312)→ |
| Rbi-1-1 | NC_013222.1 | + | 3406822 | 3406886 | **RNA**→ rpsO (PRK05626)RpsO (COG0184)→ |
| Mfo-1-1 | NZ_JQLH01000001.1 | + | 3616475 | 3616539 | **RNA**→ rpsO (PRK05626)Ribosomal_S15 (pfam00312)→ |
| env-2099 | SRS018149_C2886978 | - | 75 | 30 | **RNA**→ |
| env-2100 | 3300000558_10272742 | + | 581 | 647 | **RNA**→ rpsO (PRK05626)Ribosomal_S15 (pfam00312)→ |
| env-2101 | 3300001605_10046245 | + | 2444 | 2510 | **RNA**→ rpsO (PRK05626)Ribosomal_S15 (pfam00312)→ |
| Fse-1-1 | NZ_JNCA01000025.1 | - | 40213 | 40146 | **RNA**→ rpsO (PRK05626)Ribosomal_S15 (pfam00312)→ |
| Fda-1-1 | NZ_AUDK01000001.1 | + | 111173 | 111240 | **RNA**→ rpsO (PRK05626)Ribosomal_S15 (pfam00312)→ |
| env-2102 | KGI_S1_ANT02_95m_c10000014 | - | 20719 | 20655 | **RNA**→ rpsO (PRK05626)Ribosomal_S15 (pfam00312)→ |
| env-2103 | CEQG01122965.1 | + | 138 | 206 | **RNA**→ rpsO (PRK05626)Ribosomal_S15 (pfam00312)→ |
| env-2104 | CEQP01291574.1 | - | 688 | 620 | **RNA**→ rpsO (PRK05626)Ribosomal_S15 (pfam00312)→ |
| env-2105 | PBR_1135713 | - | 3768 | 3704 | **RNA**→ rpsO (PRK05626)Ribosomal_S15 (pfam00312)→ rpsO (PRK05626)Ribosomal_S15 (pfam00312)→ |
| Fso-1-1 | NZ_AUGO01000002.1 | - | 155840 | 155776 | **RNA**→ rpsO (PRK05626)Ribosomal_S15 (pfam00312)→ |
| env-2106 | PBDCA2_FISUTAU01D46T8 | + | 233 | 297 | **RNA**→ rpsO (PRK05626)Ribosomal_S15 (pfam00312)→ |
| env-2107 | SwRhRL2b_contig_129529 | + | 1053 | 1118 | **RNA**→ |
| env-2108 | SwRhRL2b_contig_2550456 | + | 642 | 706 | **RNA**→ rpsO (PRK05626)Ribosomal_S15 (pfam00312)→ |
| env-2109 | JGI26527J51213_1149640 | + | 13 | 79 | **RNA**→ rpsO (PRK05626)Ribosomal_S15 (pfam00312)→ |
| env-2110 | JGI26529J51215_1006318 | + | 1509 | 1575 | **RNA**→ rpsO (PRK05626)Ribosomal_S15 (pfam00312)→ |
| Aag-1-1 | NZ_JH621247.1 | - | 228487 | 228423 | **RNA**→ rpsO (PRK05626)RpsO (COG0184)→ |
| env-2111 | JGI24724J26744_10001654 | + | 4696 | 4763 | **RNA**→ rpsO (PRK05626)Ribosomal_S15 (pfam00312)→ |
| env-2112 | JGI24723J26617_10011881 | - | 2756 | 2690 | **RNA**→ rpsO (PRK05626)Ribosomal_S15 (pfam00312)→ |
| env-2113 | CEPR01003942.1 | + | 62986 | 63052 | **RNA**→ rpsO (PRK05626)Ribosomal_S15 (pfam00312)→ |
| env-2114 | CERJ01037600.1 | - | 198 | 132 | **RNA**→ rpsO (PRK05626)Ribosomal_S15 (pfam00312)→ |
| env-2115 | CETR01183773.1 | + | 156 | 222 | **RNA**→ rpsO (PRK05626)Ribosomal_S15 (pfam00312)→ |
| env-2116 | SI34jun09_100m_1197366 | - | 207 | 141 | **RNA**→ rpsO (PRK05626)Ribosomal_S15 (pfam00312)→ |
| env-2117 | CERM01080683.1 | + | 40389 | 40455 | **RNA**→ rpsO (PRK05626)Ribosomal_S15 (pfam00312)→ |
| env-2118 | DelMOSum2010_c10000067 | + | 83929 | 83995 | **RNA**→ rpsO (PRK05626)Ribosomal_S15 (pfam00312)→ |
| env-2119 | SI34jun09_10m_1052143 | - | 402 | 336 | **RNA**→ rpsO (PRK05626)Ribosomal_S15 (pfam00312)→ |
| env-2120 | JGI20154J14316_10000666 | - | 10629 | 10563 | **RNA**→ rpsO (PRK05626)Ribosomal_S15 (pfam00312)→ |
| env-2121 | DelMOSum2011_c10001660 | + | 6855 | 6921 | **RNA**→ rpsO (PRK05626)Ribosomal_S15 (pfam00312)→ |
| env-2122 | A09P04_1300_Sequence0000000013 | + | 12503 | 12569 | **RNA**→ rpsO (PRK05626)Ribosomal_S15 (pfam00312)→ |
| env-2123 | A09P04_1300_Sequence0000004592 | + | 1617 | 1683 | **RNA**→ rpsO (PRK05626)Ribosomal_S15 (pfam00312)→ |
| env-2124 | CESJ01250408.1 | + | 408 | 474 | **RNA**→ rpsO (PRK05626)Ribosomal_S15 (pfam00312)→ |
| env-2125 | SI39nov09_120m_c1053170 | + | 111 | 177 | **RNA**→ rpsO (PRK05626)Ribosomal_S15 (pfam00312)→ |
| env-2126 | CEWR01159061.1 | - | 91 | 25 | **RNA**→ |
| env-2127 | CESM01110142.1 | + | 653 | 720 | **RNA**→ rpsO (PRK05626)Ribosomal_S15 (pfam00312)→ |

| | | | | | |
|---|---|---|---|---|---|
| env-2128 | SA_S1_NOR08_45m_c10035062 | + | 255 | 321 | **RNA**→ rpsO (PRK05626)Ribosomal_S15 (pfam00312)→ |
| env-2129 | SA_S1_NOR08_45m_c10000452 | - | 838 | 771 | **RNA**→ rpsO (PRK05626)Ribosomal_S15 (pfam00312)→ |
| env-2130 | SA_S2_NOR15_50m_c10000459 | - | 223 | 156 | **RNA**→ rpsO (PRK05626)Ribosomal_S15 (pfam00312)→ |
| env-2131 | SA_S1_NOR05_45m_c10453048 | + | 105 | 172 | **RNA**→ rpsO (PRK05626)Ribosomal_S15 (pfam00312)→ |
| env-2132 | 2236876010_0472988 | - | 323 | 256 | **RNA**→ |
| env-2133 | JGI20158J14315_10190251 | + | 302 | 369 | **RNA**→ rpsO (PRK05626)Ribosomal_S15 (pfam00312)→ |
| env-2134 | SI34jun09_10m_1034686 | - | 415 | 348 | **RNA**→ rpsO (PRK05626)Ribosomal_S15 (pfam00312)→ |
| env-2135 | JGI20155J14468_10136840 | - | 350 | 283 | **RNA**→ rpsO (PRK05626)Ribosomal_S15 (pfam00312)→ |
| env-2136 | CESF01030834.1 | - | 322 | 255 | **RNA**→ rpsO (PRK05626)Ribosomal_S15 (pfam00312)→ |
| env-2137 | DelMOWin2010_c10000940 | + | 13266 | 13333 | **RNA**→ rpsO (PRK05626)Ribosomal_S15 (pfam00312)→ |
| env-2138 | 2236876010_0612317 | - | 99 | 31 | **RNA**→ |
| env-2139 | 2236876009_226267 | + | 99 | 166 | **RNA**→ rpsO (PRK05626)Ribosomal_S15 (pfam00312)→ |
| env-2140 | JGI24723J26617_10002855 | + | 10915 | 10980 | **RNA**→ rpsO (PRK05626)Ribosomal_S15 (pfam00312)→ |
| env-2141 | KVRMV2_105383168 | + | 103 | 169 | **RNA**→ |
| env-2142 | SA_S1_NOR08_45m_c10517261 | - | 204 | 138 | **RNA**→ rpsO (PRK05626)Ribosomal_S15 (pfam00312)→ |
| env-2143 | JGI24724J26744_10022380 | - | 2421 | 2353 | **RNA**→ rpsO (PRK05626)RpsO (COG0184)→ |
| env-2144 | KVRMV2_100000644 | - | 1868 | 1802 | **RNA**→ rpsO (PRK05626)Ribosomal_S15 (pfam00312)→ |
| env-2145 | 3300001605_10000882 | + | 723 | 789 | **RNA**→ rpsO (PRK05626)Ribosomal_S15 (pfam00312)→ |
| env-2146 | 3300001605_10008345 | - | 347 | 281 | **RNA**→ rpsO (PRK05626)Ribosomal_S15 (pfam00312)→ |
| env-2147 | 3300001605_10233542 | + | 724 | 790 | **RNA**→ rpsO (PRK05626)Ribosomal_S15 (pfam00312)→ |
| env-2148 | 2210106066 | - | 148 | 89 | **RNA**→ rpsO (PRK05626)RpsO (COG0184)→ |
| env-2149 | 2157111858 | + | 47 | 106 | **RNA**→ |
| env-2150 | JGI25908J49247_10464678 | - | 201 | 146 | **RNA**→ rpsO (PRK05626)Ribosomal_S15 (pfam00312)→ |
| env-2151 | JGI25911J50253_10473464 | + | 71 | 126 | **RNA**→ rpsO (PRK05626)Ribosomal_S15 (pfam00312)→ |
| env-2152 | M3P_1084921 | - | 420 | 362 | **RNA**→ rpsO (PRK05626)Ribosomal_S15 (pfam00312)→ |
| Fps-4-1 | NC_009613.3 | + | 1899222 | 1899287 | **RNA**→ rpsO (PRK05626)Ribosomal_S15 (pfam00312)→ |
| Fps-1-4 | NZ_CP008878.1 | + | 2162665 | 2162730 | **RNA**→ rpsO (PRK05626)Ribosomal_S15 (pfam00312)→ |
| Fps-1-5 | NZ_CP008883.1 | + | 2156137 | 2156202 | **RNA**→ rpsO (PRK05626)Ribosomal_S15 (pfam00312)→ |
| Fps-1-6 | NZ_CP008881.1 | + | 2153198 | 2153263 | **RNA**→ rpsO (PRK05626)Ribosomal_S15 (pfam00312)→ |
| Fps-1-7 | NZ_CP007627.1 | - | 849155 | 849090 | **RNA**→ rpsO (PRK05626)Ribosomal_S15 (pfam00312)→ |
| Fps-2-1 | NZ_CP007206.1 | - | 797073 | 797008 | **RNA**→ rpsO (PRK05626)Ribosomal_S15 (pfam00312)→ |
| Fps-3-1 | NZ_CP007207.1 | - | 1689453 | 1689388 | **RNA**→ rpsO (PRK05626)Ribosomal_S15 (pfam00312)→ |
| Fps-1-8 | NZ_CP008902.1 | + | 2199599 | 2199664 | **RNA**→ rpsO (PRK05626)Ribosomal_S15 (pfam00312)→ |
| Fps-1-9 | NZ_CP010275.1 | + | 2296737 | 2296802 | **RNA**→ rpsO (PRK05626)Ribosomal_S15 (pfam00312)→ |
| Fps-1-10 | NZ_CP010277.1 | + | 2258520 | 2258585 | **RNA**→ rpsO (PRK05626)Ribosomal_S15 (pfam00312)→ |

#### 22.4 Conserved domains

Conserved domains found in protein-coding genes listed in Section 22.3 are shown below, with the first sentence in their description from the Conserved Domain Database (if any). Conserved domains associated with more than one S15-Flavobacteria RNA are assigned a color, while others are shown in gray. The symbols 'd' and 'u' (if any) to the left of the colored domain name indicate whether the domain occurs upstream or downstream of the RNA motif. Domains marked with lower-case 'd' occur (at least once) present downstream of some S15-Flavobacteria RNA in the opposite orientation. Domains marked with capital 'D' occur downstream and in the same orientation (i.e., a potential *cis*-regulatory arrangement). Domains marked with a 'u' occur upstream of the RNA in either orientation. If the 'd'/'u' symbols are missing, then the domain is downstream and in the same strand. (This mode is used for motifs predicted as *cis*-regulatory.) The number in parentheses after the colored domain name is the number of occurrences in Section 22.3.

bSCRAP106 (26) Self-Cleaving Ribozyme Associated Protein number 106 (gi=2031276512 by jackhmmer)
bSCRAP40 (26) Self-Cleaving Ribozyme Associated Protein number 40 (gi=10605909721 by jackhmmer)
cd00353 (2) Ribosomal protein S15 (prokaryotic)_S13 (eukaryotic) binds the central domain of 16S rRNA and is required for assembly of the small ribosomal subunit and for intersubunit association, thus representing a key element in the assembly of the whole ribosome.
cd00429 (50) Ribulose-5-phosphate 3-epimerase (RPE).
cd11363 (45) Polyribonucleotide nucleotidyltransferase, repeat 1.
COG0184 (344) Ribosomal protein S15P/S13E [Translation, ribosomal structure and biogenesis]
COG0405 (1) Gamma-glutamyltranspeptidase [Amino acid transport and metabolism]
COG0568 (54) DNA-directed RNA polymerase, sigma subunit (sigma70/sigma32) [Transcription]
COG0810 (1) Periplasmic protein TonB, links inner and outer membranes [Cell wall/membrane/envelope biogenesis]
COG1185 (118) Polyribonucleotide nucleotidyltransferase (polynucleotide phosphorylase) [Translation, ribosomal structure and biogenesis]
COG1770 (1) Protease II [Amino acid transport and metabolism]
COG2377 (2) 1,6-Anhydro-N-acetylmuramate kinase [Cell wall/membrane/envelope biogenesis]
pfam00140 (5) Sigma-70 factor, region 1.
pfam00312 (1729) Ribosomal protein S15.
pfam01138 (150) 3' exoribonuclease family, domain 1.
pfam03702 (14) Anhydro-N-acetylmuramic acid kinase.
pfam03725 (39) 3' exoribonuclease family, domain 2.
pfam03726 (4) Polyribonucleotide nucleotidyltransferase, RNA binding domain.
pfam04542 (3) Sigma-70 region 2.
pfam06210 (18) Protein of unknown function (DUF1003).
PLN02198 (1) glutathione gamma-glutamylcysteinyltransferase
PRK05581 (50) ribulose-phosphate 3-epimerase; Validated
PRK05626 (2112) 30S ribosomal protein S15; Reviewed
PRK09210 (2) RNA polymerase sigma factor RpoD; Validated
PRK09585 (16) anhydro-N-acetylmuramic acid kinase; Reviewed
PRK10115 (1) protease 2; Provisional
PRK11824 (313) polynucleotide phosphorylase/polyadenylase; Provisional
smart00316 (1) Ribosomal protein S1-like RNA-binding domain.
smart00322 (1) K homology RNA-binding domain.
TIGR01352 (1) TonB family C-terminal domain. [Transport and binding proteins, Cations and iron carrying compounds]
TIGR02169 (18) chromosome segregation protein SMC, primarily archaeal type. [Cellular processes, Cell division, DNA metabolism, Chromosome-associated proteins]
TIGR02937 (57) RNA polymerase sigma factor, sigma-70 family.

#### 22.5 Multiple-sequence alignment

Each S15-Flavobacteria RNA is denoted by an abbreviation (like "Eco-1-1") that refers to a taxonomy given in Section 22.1. (Superscript numbers refer to annotations listed in Section 22.2) The alignment may include sequences containing the RNA motif, as well as flanking sequence. The S15-Flavobacteria RNA itself is denoted by the line underneath marked 5′ and 3′ on either end. Nucleotides in flanking sequences (i.e., not part of the motif) are written in gray letters. Stems of predicted rho-independent transcription terminators, if any, are shaded yellow. (Note: terminator predictions have not been analyzed manually, and many are likely to be false positives. Terminator predictions are those of the RNIE software.) Nucleotides predicted to function (as DNA) as transcription-factor binding sites, if any, are shaded in green. (But note: these predictions are manually annotated, so they might be under-predicted.) Annotated start codons, if any, are shaded green. (Note: start codons are frequently misannotated, especially in environmental samples.) Nucleotides proposed to basepair as part of the consensus structure are shaded in color when they comprise Watson-Crick or G-U pairs. Otherwise they are shaded gray. Conserved stems are also indicated at the bottom of the alignment by angle brackets, where matching $<$ and $>$ denote base-paired columns. Below these angle brackets, the symbol "2" denotes base pairs exhibiting covariation according to the statistically well-founded R-scape method. "1" denotes base pairs exhibiting covariation according to R2R's simplistic method. "0" denotes base pairs that are not observed to mutate and "?" denotes base pairs that have a significant frequency of non-canonical nucleotides for Watson-Crick or G-U pairs ($> 5\%$). Below these base pair annotation is the consensus sequence: "R" = "A" or "G", "Y" = "C" or "U", red nucleotides: nucleotide identity conserved more than 97% of the time, black nucleotides: 90%, gray nucleotides: 75%, red circle (•): nucleotide is present 97% of the time, black circle (•): 90%, gray circle (•): 75%, white circle (○): 50%. All percentages of sequences just described (e.g. 97% conserved) assume that sequences have been weighted by the GSC algorithm implemented by the Infernal software package.

Duplicate sequences: the following putative homologs are not shown in the alignment because their sequences are identical to a homolog already shown: Aal-1-1, Cba-1-1, Cba-3-1, Cca-2-3, Cca-3-1, Ccy-1-2, Ccy-2-1, Cin-2-1, Coc-4-1, Cor-1-2, Csp-4-1, Ean-1-1, Ean-1-2, Ean-1-4, Ean-10-1, Ean-11-1, Ean-12-1, Ean-2-1, Ean-4-1, Ean-5-1, Ean-6-1, Ean-7-1, Ean-8-1, Ean-9-1, Eme-1-1, Eme-2-2, Fca-1-2, Fen-1-2, Fps-1-10, Fps-1-2, Fps-1-3, Fps-1-4, Fps-1-5, Fps-1-6, Fps-1-7, Fps-1-8, Fps-1-9, Fps-2-1, Fps-3-1, Fri-2-1, Fsp-14-1, Fsu-1-2, Fsu-3-2, Jpa-1-2, Jpa-1-3, Lbl-1-1, M[o-1-1, Mod-2-1, Mod-5-1, Msp-3-1, Orh-2-1, Orh-3-1, Psp-1-1, Ran-1-1, Ran-1-2, Ran-2-2, Ran-3-1, Ran-5-1, Ran-6-1, Ran-7-1, Zpr-1-1, env-10, env-1000, env-1014, env-1020, env-1031, env-104, env-1043, env-105, env-1050, env-1052, env-1054, env-1071, env-1072, env-1073, env-1074, env-1075, env-1076, env-1083, env-1084, env-1100, env-1114, env-1117, env-1118, env-1119, env-1120, env-1132, env-1133, env-1136, env-1148, env-1149, env-1150, env-1151, env-1152, env-1153, env-1154, env-1155, env-1156, env-1157, env-1158, env-1160, env-1164, env-1166, env-1178, env-1179, env-1180, env-1189, env-1195, env-1201, env-1202, env-1206, env-1207, env-1214, env-1221, env-1224, env-1225, env-1228, env-1244, env-1249, env-1251, env-1258, env-1261, env-1265, env-1266, env-1267, env-1276, env-1279, env-1286, env-1288, env-1289, env-1290, env-1291, env-1292, env-1297, env-1315, env-1322, env-1329, env-136, env-1361, env-1362, env-1363, env-1364, env-1365, env-1366, env-1368, env-1369, env-137, env-1372, env-1390, env-1407, env-1408, env-1409, env-1410, env-1423, env-1424, env-1430, env-1431, env-1432, env-144, env-1440, env-1443, env-1446, env-1450, env-1451, env-1452, env-1453, env-1470, env-1492, env-1493, env-1494, env-1495, env-1496, env-150, env-1509, env-1510, env-1511, env-1512, env-1513, env-1514, env-1518, env-1523, env-1531, env-1532, env-1533, env-1544, env-1545, env-1558, env-1569, env-1570, env-1578, env-1590, env-1591, env-1592, env-1593, env-160, env-1648, env-1653, env-1654, env-1667, env-1670, env-1671, env-1672, env-1675, env-1678, env-168, env-169, env-1690, env-1704, env-1715, env-1716, env-1719, env-1721, env-1731, env-1732, env-1733, env-1734, env-1735, env-1736, env-1737, env-1738, env-1739, env-1740, env-1741, env-1742, env-1743, env-1751, env-1758, env-1759, env-1760, env-1761, env-1762, env-1763, env-1764, env-1776, env-1777, env-1778, env-178, env-1781, env-1782, env-1784, env-1785, env-1787, env-1788, env-1789, env-1790, env-1791, env-1792, env-1793, env-1794, env-1796, env-1799, env-1803, env-1806, env-1807, env-1808, env-1809, env-1810, env-1811, env-1812, env-1813, env-1814, env-1815, env-1816, env-1817, env-1819, env-182, env-1820, env-1822, env-1825, env-183, env-1831, env-184, env-1841, env-1842, env-1843, env-1844, env-1845, env-1849, env-1858, env-1861, env-1867, env-1868, env-188, env-1882, env-1887, env-1888, env-1889, env-1890, env-1891, env-1894, env-1895, env-1896, env-1897, env-1899, env-190, env-1914, env-1919, env-1920, env-1923, env-1932, env-1933, env-1934, env-194, env-1953, env-1956, env-1960, env-1989, env-199, env-200, env-2013, env-2022, env-2023, env-2024, env-2025, env-2026, env-204, env-2047, env-2048, env-2056, env-2057, env-2058, env-2059, env-2060, env-2061, env-2062, env-2063, env-2064, env-2065, env-2066, env-2067, env-208, env-2084, env-209, env-2093, env-2094, env-2095, env-2096, env-2097, env-2098, env-2101, env-2104, env-2122, env-2123, env-219, env-232, env-233, env-240, env-241, env-242, env-245, env-246, env-252, env-253, env-254, env-259, env-260, env-275, env-279, env-284, env-29, env-291, env-292, env-293, env-30, env-31, env-311, env-312, env-319, env-32, env-321, env-322, env-323, env-348, env-349, env-350, env-351, env-365, env-370, env-371, env-372, env-374, env-380, env-401, env-41, env-425, env-426, env-436, env-443, env-448, env-468, env-483, env-485, env-493, env-495, env-497, env-499, env-512, env-516, env-524, env-525, env-526, env-529, env-530, env-535, env-538, env-544, env-555, env-560, env-568, env-570, env-58, env-595, env-597, env-630, env-631, env-632, env-64, env-642, env-643, env-644, env-645, env-646, env-647, env-654, env-665, env-666, env-667, env-668, env-695, env-696, env-697, env-710, env-714, env-715, env-716, env-717, env-719, env-72, env-720, env-721, env-722, env-723, env-724, env-725, env-726, env-73, env-732, env-733, env-74, env-75, env-757, env-78, env-782, env-790, env-792, env-793, env-804, env-811, env-812, env-839, env-840, env-841, env-842, env-843, env-844, env-845, env-846, env-858, env-859, env-867, env-872, env-873, env-874, env-88, env-880, env-882, env-885, env-893, env-904, env-918, env-926, env-931, env-935, env-963, env-964, env-969, env-97, env-970, env-971, env-975, env-979, env-981, env-987, env-991, env-992, env-996, env-997, env-998

alignment positions 1 · · · 79

```
Pgo-1-1   UUAUACUUGGAUCUGAUCUUGAAUCAACCCGUAAGAGCUUAAUUUUUAUAAAUUUAAUAUUCAAUUUCAAUUUAAAAGC
env-1     UCUGUAUAUAGAUUUAAUAAUGAAUGCGCCUGUGAGAGCUUAAUUUUUAAAAGAAAUACGUUUAUUUCAUUUUAAAUAC
env-2     CCUUUACAUAGAUUUGAUUAUGAAUCAACCUGUGAGAGCAUAAUUUUUAAAUUAAUGUCAUCAAUUUAAAAUUAAUAUU
env-3     AUCUUUAUAUAGACCUUAUUACAAAUGAGCCUGUUAGAGCAUAAUCUCGUAAAAAAGGUUCCUAUUUCAAAUUAAAAGA
Pto-1-1   UUAUACUUGGAUCUGAUCUUGAAUCCACCAGUGAGGGCUUAUUUUUUAUAAAUUUGAUAUUUAGAUUCAAUAUAAAAGG
Ptr-1-1   UCUUUACAUAGACCUUAUUCUAAAUCAGCCAGUAAGAGCAUAAUUUUUUUAGAAGAGGUUGCUAUCUCCAUUUAAAAGA
env-4     UUAUACUUGGAUCUCAUCUUGAAUCAACCAGUGAGAGCUUAAUUUUUAUAAAUUUAAUAUUCAGAUUCAAUAUAAAAGG
env-5     AUUUAUACAUAGAUCUUAUUAGGAACCAGCAAGUAAGAGCAUAAAUUCACUGAAUUACUUUGCAAUUAAAUUUAAAAGA
env-6     ......................AAAUGAGCCUGUUAGGGCAUAAUCUUAUAAAAAAGACUCCUAUUUCAAAUUAAAAGA
env-7     AUCUUUAUAUAGACCUUAUUACAAAUGAGCCUGUUAGGGCAUAAUCUUAUAAAAAAGACUCCUAUUUCAAAUUAAAAGA
env-8     AUCUUUAUAUAGACCUUAUUACAAAUGAGCCUGUUAGGGCAUAAUCUUAUAAAAAAGACUCCUAUUUCAAAUUAAAAGA
Mod-8-1   UGUAAUAAGGUGUUAGCAAGGAACUUAUUAUUUUUUUGAAAAAAAAUAUAGAACUGUGAUGAUGGUUUGAAAAAUAAAA
env-9     CUUUGGCGCUUUCAUUGAAGCUUUUUAGAUCAUAAAAGUUAAAAAAAUAAUAAACAAGGCUUGUAAUAAGGAUAAAUUAC
env-11    ........................................................GGCUUGUAAUAAGGAUAAAUUAC
env-12    ..............................................UAAUAAACAAGGCUUGUAAUAAGGAUAAAUUAC
env-13    CCUUGGCGCUUUCAUUGAAGCUUUUUAGAUCAUAAAAGUUAAAAAAAUAAUAAACAAGGCUUGUAAUCAGGAUAAAAUAC
env-14    CCUUGGCGCUUUCAUUGAAGCUUUUUAGAUCAUAAAAGUUAAAAAAAUAAUAAACAAGGCUUGUAAUCAGGAUAAAAUAC
env-15    UCCGCGUCAGGAGCUGCGUGAGCGCGUUGCGCAAGCGUUGAAAAUGAUGGAAGUCCCUGCGUAAAACCAGAAAGAAAAC
env-16    AAAUGUGGCUAAACUCAAUUUAAUAAAAAGCUGAUUAUUAAGUUUUUUAACAUAGAGUAACAAGUUAACAGAAUAAUGC
env-17    AAUUGUGAAACAUCAGUUACUAGAUUUCAUAUUCAUUUUCGAUUUCUAUUUCUAUUUCAACUUCAAUUUCAAAAAUAGU
Fbr-1-1   AAAUUGCUAUUGUAAAAAUAAAUUCUAAAAUAUUCUUACUAUAAAUUUAACUAAAAUGUUGUAUGUUCCAAAAAAUUAA
env-18    ACUUAGAUGUAGAUUUGGUCUGUUCUCCUACGCAUUCUUGGAUUUCUUUUUAACGUCCUUGUGUUUUAAGAUUAAUUCA
env-19    ACUUAGAUGUAGAUUUGGUCUGUUCUCCUACGCAUUCUUGGAUUUCUUUUUAACGUCCUUGUGUUUUAAGAUUAAUUCA
env-20    GGUUUUAACAGGAUUUUGGUUUUUUCACCUGAUCCCCAAACUUUUUUCACCAACUUUACUCCCAUUCAAAAAUAAUUCC
env-21    GAAAGAAAAAACUUGAAAUCUAAAGUGUCACAAUUUAUUAGAAUGUUGAAUAAUUAAUUAGGAUGUGUAUCGAUUUUAA
env-22    ACCUUUAUCUUGAUCUUAUUCUUAAUAGAACCAUAAGAGAAUAAAUAUUCCUAAAUAAAUCGAAGUUUCCUUAUUUUUA
env-23    ACCUUUAUCUUGAUCUUAUUCUUAAUAGAACCAUAAGAGAAUAAAUAUUCCUAAAUAAAUCGAAGUUUCCUUAUUUUUA
env-24    ACCUUUAUCUUGAUCUUAUUCUUAAUAGAACCAUAAGAGAAUAAAUAUUCCUAAAUAAAUCGAAGUUUCCUUAUUUUUA
env-25    ACCUUUAUCUUGAUCUUAUUCUUAAUAGAACCAUAAGAGAAUAAAUAUUCCUAAAUAAAUCGAAGUUUCCUUAUUUUUA

          ...............................................................................
          ...............................................................................
          ●●●●●●●●●●●●●●●●●●●●●●●●●●●●●●●●●●●●●●●●●●●●●●●●●●●●●●●●●●●●●●●●●●Y●●●●RR●●●●●●
```

```
Fba-9-1   GUUUAAAAAAGAUUCUUAUCAAACCAGAAGGAAAGAAAGAAUUAAUCCAUCAAUAACAAUUGUAAAUCAAAUAAUUAGU
env-26    CCGUUACAUUGAUUUGAUUCUAAAUCAACCUCUAAGAGAAUAAUUUUUUUAAAGCAAAGCACAAUUUUCCUGAAAUUGU
env-27    ..........UUAUCGUAACAUGGUAAUCCUUUCGUAUUGAAUAGUUGUUAUAAAUACUUGCUAAUUAAAAAUCUUUAC
env-28    AGCAGAUUAGAAGUUAAUAAAGGCUUUAAAACCUUAAUUUUCGAAAAGUAAAAAAACACUUUACUUCUCAAAAUUAAUU
env-33    AGCAGAUUAGAAGUUAAUAAAGGCUUUAAAACCUUAAUUUUCGAAAAGUAAAAAAACACUUUACUUCUCAAAAUUAAUU
env-34    AAUUAACCUCUAUGUAGAUCUAAUUCUCAAUCAUCCCAUAAGAAAGUAAAGCCAAGCCUUUUUACUUAGAUAAAUUGGU
env-35    AAUUAACCUCUAUGUAGAUCUAAUUCUCAAUCAUCCCAUAAGAAAGUAAAGCCAAGCCUUUUUACUUAGAUAAAUUGGU
env-36    ............................................AGUAAAGCCAAGCCUUUUUACUUAGAUAAAUUGGU
env-37    NNNNNNNNNNNNNNNNNNNNNNNNNNNNNNNNNNNACCCAUAAGAAAGUAAGAUAAAGCAUCCGUCUUUCACAAAUUAGU
env-38    UAAAGUCAAUCUCUACAUUGAUCUUGUUUAAAUCAACCCAUAAGAAAUAAAAUAGGGUAAUAGACUUUGAUAAAUUAGU
env-39    ............CAUUGACUUAAUUUUAAAUCACCCUAUAAGAAAGUAAAGCGAGGCCUUUUUACUUAGAUAAAUUGGU
env-40    GAUCAACCUUUACGUCGACUUAAUUUUAAAUCACCCUAUAAGAAAGUAAAGCAAGGCCUUUUUACUUAGAUAAAUUGGU
env-42    CAAAUAAGUAUAGAAAUACCUAUAAAAGAAAAUGUAAAAGAAAAAAAUUAAGGUUAAAUGAUUUGGUAAUAAAUAAAUAC
env-43    GAAACCUUCCCAUGGAGCCAUUCACCUUCCUUGCGCCAAAAAAAUAAGACGUCCACUUUUGGUUUAUAGGGUUUUAUCC
Myu-1-1   UUUGUAUAUCGAUUUAAUUACAAAUCAGCCCUUAAGAGCAUAAUUUUAAGUCCAGCAUCUUGUUACUAACAAAUAAUUA
env-44    AAUACAUUGAUUUAAUUUUAAACUUACCGCUCAGAGCCUAAUUAAUAAUCACAAUAGGUUAAUAAUUCAAACAAAAUUA
env-45    AAAACGGUGCCUACGAACGCGACGGAUUGUUAUAAUUCAGCGUUUCUUUCAAUAAUAUUUUAUUUAUAACAAAGAAUUA
env-46    ........................GAUUGUUAUAAUUCAGCGUUUCUUUCAAUAAUAUUUUAUUUAUAACAAAGAAUUA
env-47    AAAACGGUGCCUACGAACGCGACGGAUUGUUAUAAUUCAGCGUUUCUUUCAAUAAUAUUUUAUUUAUAACAAAGAAUUA
env-48    AAAAUGGCGCUUACGAGCGCGAUGGAUUAUUAUAAUUCUACGUUUCUUUCAAUAAUAUUUUCUUUUAACAAAGAAUUA
env-49    CUACCAUCGCUAAUAAUCCGUAUAACAUGGUUUAAGUUUUAAUUUACCUUCGAAAUAUAAGGUUUUUAGAAAACUAGUG
env-50    GAAAGGUUUAAACCGUUCAGCUAAAGAAAUCAUUCUGACCAAAAUAGAUUAACAAAAAAUUGCGCAUCUAAAAAAAGGC
env-51    GAUCAAUCUCUAUAUUGAUCUUAUUCUAAAUCAACCCAUAAGAAAUUAAAUUUAGGACGUUAGUCUUACAUAAAUUGGU
env-52    AGAUUACGUUGAAUGUGGGAUUCAAUAAAUGAUUUUUAAUAAAGUUAGUUUAUUAAUUUCAUUUAUUAAUUAAGAUUAU
env-53    UUUUGGGUAAAGAUAAUCAACGGAAAAAAGCUCACCGCCAGUUUAGAUGUGUUUUUUUUGGUAGUGCGCAAAAAAGACC
Fsa-1-1   UUAGAGUUUGUAAAUCAGGAGAUUGCCAAACUUUUUUAGGGAAGUUUCCGUAAUUAUUUGCCUAUUCGGAAAAUAAAAA
Fca-1-1   CUUGGAGUUUGUAAACCAAGAAAUUGCAAAACUUUUUUAGGCAACUUUGACGUUUUAUUUGUCAAUUCAAAAAUAAAAU
env-54    ACCAAUAUAUUGAUUUAGUUCUCAACGAUGUACUGCGUAAAUAAAAUCUAUACUUAAAUCCACAUUUUAACGAAAAUGU
env-55    ACCAAUAUAUUGACUUAGUUCUCAACGAUGUAUUGCGUAAAUAAAAUCUAUACUUAAAUCCACAUUUUAACGAAAAUGU
env-56    ACCAAUAUAUUGACUUAGUUCUCAACGAUGCACUGCGUAAAUAAAAUCUAUACUUAAAUCCACAUUUUAACGAAAAUGU
env-57    ACCAAUAUAUUGAUUUAGUUCUCAACGAUGUAGUGCGUAAAUAAAAUCUAUACUUAAAUCCACAUUUUAACGAAAAUGU
env-59    UUCUAGACAGGAUUUGAAGGAAAAAAUUGCGCUUUCUAUCAAAUUGUUUAAUAAUUAAAGAAAUGUAGUUAUAUUUUAA
env-60    UGUAAGAGUAUUGAAAAAAGUAAGGUCUCAGGUUUUUUAAAUAAUGUGCUAUCAAACAAUGCCAUAUCAGUGAAUUUUA
env-61    UGUAAGAGUAUUGAAAAAAGUAAGGUCUCAGGUUUUUUAAAUAAUGUGCUAUCAAACAAUGCCAUAUCAGUGAAUUUUA
env-62    UGUAAGAGUAUUGAAAAAAGUAAGGUCUCAGGUUUUUUAAAUAAUGUGCUAUCAAACAAUGCCAUAUCAGUGAAUUUUA
env-63    UGUAAGAGUAUUGAAAAAAGUAAGGUCUCAGGUUUUUUAAAUAAUGUGCUAUCAAACAAUGCCAUAUCAGUGAAUUUUA
env-65    UGUAAGAGUAUUAAAAAAAGUAGGGUCUCAGGUUUUUUAAAUAAUAUGCUAUCAAACAAUGCCAUAUCAUUGAAUUUUA
env-66    UGUAAGAGUAUUAAAAAAAGUAGGGUCUCAGGUUUUUUAAAUAAUAUGCUAUCAAACAAUGCCAUAUCAUUGAAUUUUA
env-67    UGUAAGAGUAUUGAAAAAAGUAAGGUCUCAGGUUUUUUAAAUAAUAUGCUAUCAAACAAUGCCAUAUCAGUGAAUUUUA
env-68    UGAAAGAGUAUUGAAAAAAGUAAGGUCUCAAGUUUUUUAAAUAAUAUGCCAUCAAACAAUGCCAUAUCAUUGAAUUUUA
env-69    UGAAAGAGUAUUGAAAAAAGUAAGGUCUCAUGUUUUUUAAAUAAUAUAUCAUCAAACAAUGCCAUAUCAUUGAAUUUUA
env-70    ...............................................AUCAUCAAACAAUGCCAUAUCAUUGAAUUUUA
env-71    UGAAAGAGUAUUGAAAAAAGUAAGGUCUCAUGUUUUUUAAAUAAUAUAUCAUCAAACAAUGCCAUAUCAUUGAAUUUUA
env-76    UGAAAGAGUAUUGAAAAAAGUAAGGUCUCAUGUUUUUUAAAUAAUAUAUCAUCAAACAAUGCCAUAUCAUUGAAUUUUA
env-77    UGAAAGAGUAUUGAAAAAAGUAAGGUCUCAUGUUUUUUAAAUAAUAUAUCAUCAAACAAUGCCAUAUCAUUGAAUUUUA
env-79    .....................................................................UUGAAUUUUA
env-80    ......AGUAUUGAAAAAAGUAAGGUCUCAUGUUUUUUAAAUAAUAUAUCAUCAAACAAUGCCAUAUCAUUGAAUUUUA
env-81    AACACUUUGUAUUUUAAAAUACAAAGUGUUUUAUUUUUAAAUAAAGCACUACCAUACAAUGCCAUAUCAUUGAAUUUUA
env-82    AACACUUUGUAUUUUAAAAUACAAAGUGUUUUAUUUUUAAAUAAAGCACUACCAUACAAUGCCAUAUCAUUGAAUUUUA
env-83    AACACUUUGUAUUUUAAAAUACAAAGUGUUUUAUUUUUAAAUAAAGCACUACCAUACAAUGCCAUAUCAUUGAAUUUUA
env-84    AACACUUUGUAUUUUAAAAUACAAAGUGUUUUAUUUUUAAAUAAAGCACUACCAUACAAUGCCAUAUCAUUGAAUUUUA
env-85    AUACACCUUGUGUCUAAAUUCAAGGUGUAUUAUUUUUUAAAUAAAAUACUACCUUACAAUGCCAUAUCAUUGAAUUUUA
env-86    AUACACCUUGUGUCUAAAUUCAAGGUGUAUUAUUUUUUAAAUAAAAUACUACCUUACAAUGCCAUAUCAUUGAAUUUUA
env-87    AUACACCUUGUGUCUAAAUUCAAGGUGUAUUAUUUUUUAAAUAAAAUACUACCUUACAAUGCCAUAUCAUUGAAUUUUA
env-89    AUACACCUUGUGUCUAAAUUCAAGGUGUAUUAUUUUUUAAAUAAAAUACUACCUUACAAUGCCAUAUCAUUGAAUUUUA
env-90    UUUUAGUAUUUUAAAAUAGUAAAAGAAGUAACUUUUUAAAUAAUUAUCUACUCAAACAAUGCCAUAUCAUUGAAUUUCA
env-91    UUUUAGUAUUUUAAAAUAGUAAAAGAAGUAACUUUUUAAAUAAUUAUCUACUCAAACAAUGCCAUAUCAUUGAAUUUCA
env-92    CUCCACUUUUCACCCUUCACUCUUCACUCUUCCCUCCUCACAAAAUUAAAAAACACACCUUGCAACUCACAUAUUUUUA
env-93    CUCCACUUUUCACCUUUUACUCUUCACUCUUCCCUCCUCACAAAAUUAAAAAACACACCUUGCAACUCACAUAUUUUUA
env-94    AAAUUAUUUCGAAAUAGUUAUAAGGUUUGAGGUUUGAGGAAAAAUCUACGAACAUAAAUCAUUAAUCAAAAAUAAUUCA
env-95    CAGGGUUGAAAUUGAGAUUAUAUCAAGGUAAAGUCUAAAGUAAAAAAUAAAAAAUAAUGCUGUACAUCGUUAAUUUUUA
env-96    AAUCUUUAUCUGGACUUGAUUCUAAAUCAUCCCUUAAGAGAUUAAUCUAGAAACAGGUCAGUAUAUCAAAUAAUUACAU
env-98    AAUCUUUAUCUGGACUUGAUUCUAAAUCAUCCCUUAAGAGAUUAAUCUAGAAACAGGUCAGUAUAUCAAAUAAUUACAU
env-99    UUGUUUAGGGGGUAUAAGGCUACACCUCAUUUUUAUGGCAAAAAAAAAAUCCAAAAUACUGUCAAACAAAAUAUUUCAA
env-100   UUGUUUAGGGGGUAUAAGGCUACACCUCAUUUUUAUGGCAAAAAAAAAAUCCAAAAUACUGUCAAACAAAAUAUUUCAA
env-101   UAAUACUAAAUCAACCUGUAAGAAAAUAAUUACACCUAUUUAAUUCUAAAUAAAAUUAUUGUAGUGCACGGGUUUAGUC
env-102   CUAAAUCGCCCUUUGCGUAAAUAGCUUAUUAACUGUAAUUUAAUUAAUUAGGUAUUGGUUGUUUUAUAAAGGUUAAGUU
env-103   CUAAAUCGCCCUUUGCGUAAAUAGCUUAUUAACUGUAAUUUAAUUAAUUAGGGAUUGGUUGUUUUGUAAAGGUUAAGUU
env-106   CUAAAUCGCCCUUUGCGUAAAUAGCUUAUUAACUGUAAUUUAAUUAAUUAGGGAUUGGUUGUUUUGUAAAGGUUAAGUU
env-107   CUAAAUCGUCCUUUGCGUAAAUAGCUUAUUAACUGUAAUUUAAUUAAUUAGGGAUUGGUUGUUUUAUAAAGGUUAAGUU
env-108   CUAAAUCGCCCACUGCGUAAAUAGCUCAUUACCUGUAAUUUAAUUAAUUAGGUAUUGGUUGUUUUAUAAAGGUUAAGUU
env-109   CUAAAUCGCCCACUGCGUAAAUAGCUCAUUAACUGUAAUUUAAUUAAUUAGGUAUUGAUUGUUUUAUAAAGGUUAAGUU
Min-1-1   AUGCUUAACGUGAAGUGUAUAUUGUUCUUUGAAGUAGAAAAAAAAUAUUCAUUUAAUAGAUGGAAUUUGAAAAAUAAAA
env-110   AGAAAUUAGAGAACAAUUAUCACAGUUGUUGAGAAUGUUCAAAAACUAAGUGAAAAACAAUUGUCAUUCAAUUUAUUGA
env-111   CCCUGAUUCAUAAUUACUAAAACAAUUUUAGCUUUAGAUUAAAAAUCAGCGUAAAUGAUUGUUAUUCAAAAUAAUAAUU
env-112   CCCUGACUCAUACUUACUAAAACGAUUUUAGUAUUAGAUUAAAAAUCAGCGCAAAUGAUUGUUAUUCAAAAUAAUAAUU
env-113   CCCUGACUCAUAAUUACUAAAACAAUUUUAAUGUUAGAUUAAAAAUCAACGCAAAUGAUUGUUAUUCAAAAUAAUAAUU
env-114   CCCUGACUCAUAAUUACUAAAACAAUUUUAAUGUUAGAUUAAAAAUCAACGCAAAUGAUUGUUAUUCAAAAUAAUAAUU
env-115   CCCUGACUCAUAAUUACUAAAACAAUUUUAAUGUUAGAUUAAAAAUCAACGCAAAUGAUUGUUAUUCAAAAUAAUAAUU
env-116   AUAAAUACCUAGAUUUAAUAUUAAAUAGACCUUUACGAGCAUAAAAACAUUUUAAAACAUUGUAAAUUAAAAUUUUAUU
Nul-1-1   UAUAAAACGCCUGUAAUUUUAAAUCACAGGCGUUUUUAAUUUAAAGAUAUUUAUAGAGUAAUUGGAUUUAAUUAAAAGU
Nul-1-2   UAUAAAACGCCUGUAAUUUUAAAUCACAGGCGUUUUUAAUUUAAAGAUAUUUAUAGAGUAAUUGGAUUUAAUUAAAAGU
Fba-1-1   UAUAAAACGCCUGUAAUUUUAAUUUACAGGCGUUUUUAAUUUAAAGAUAUUUAUAGAGUAAUUGAAUUUAAUUAAAAGU
Nul-1-3   UAUAAAACGCCUGUAAUUUUAAAUCACAGGCGUUUUUAAUUUAAAGAUAUUUAUAGAGUAAUUGGAUUUAAUUAAAAGU

          ...............................................................................
          ...............................................................................
          ●●●●●●●●●●●●●●●●●●●●●●●●●●●●●●●●●●●●●●●●●●●●●●●●●●●●●●●●●●●●●●●●●●Y●●●●RR●●●●●●
```

```
Nul-1-4   UUAUAAAACGCCUCUGAAUUAAAAUCACAGGCGUUUUUAAUUAAAGAUAUUUAUAAGAUAACCGGAUUUAAUUAAAAGU
Nul-1-5   UUAUAAAACGCCUCUGAAUUAAAAUCACAGGCGUUUUUAAUUAAAGAUAUUUAUAAGAUAACCGGAUUUAAUUAAAAGU
Nul-1-6   UAUAAAACGCCUGUAAUUUUAAAUCACAGGCGUUUUUAAUUUAAAGAUAUUUAUAGAGUAAUUGGAUUUAAUUAAAAGU
env-117   UAUUAAGAGCCUGUACCUAUCAAGCUACAGGCUUUUUUUAUAAAAAAACCUGCAUUAAAACGUAGAUACGAUUAAAAGU
env-118   UAUUAAGAGCCUGUACCUAUCAAGCUACAGGCUUUUUUUAUAAAAAAACCUGCAUUAAAACGUAGAUACGAUUAAAAGU
env-119   GAUUAAUUUGUAUAUAGAUCUUAUACUAAACCGGAAAAUUCGCAAAAUUACUAAGAAUGGCUCAUUAUAAAAUUUAGUA
env-120   GAUUAAUUUGUAUAUAGAUCUUAUACUAAACCGGAAAAUUCGCAAAAUUACUAAGAAUGGCUCAUUAUAAAAUUUAGUA
env-121   GAUUAAUUUGUAUAUAGAUCUUAUACUAAACCGGAAAAUUCGCAAAAUUACUAAGAAUGGCUCAUUAUAAAAUUUAGUA
Fsu-1-1   AAGUCGAAAGUUUACGCACGUUACUGGACUUUAAGACUUUAAAAACUUUACGACUUUUAACGGUAAUUAAAAUUUUUGU
Gsp-1-1   AAUUAAGAUCUAACAAAAUAGGCUCUUUUACCUAAUAGGAUUAAUUAUUAAUUUUAUCGGUUUAAAAUUAAAAUAAUUA
env-122   CCAAAGAAUCUGAAGAAACAGCCGAAGUAAAAUAACAAUAAAAUAAAAUGGUGAAAGAUAGUGGUAUUCGAAAUUAGUG
env-123   AAACAAAAAGGUUUAAUUUGAACUAAAAUUUUCUGAAAAUCAAUCGCUACAAACUUUUAAAUCAUUUGAAUAAAAAGAA
env-124   ACUUAUAUAUUGAUCUUAUUACCAAUAAGCCCAUAAGAAACUAAUUGAUUCAUUGAAAUAGUUAUAUUUCAAAAAAUAU
env-125   AAAACAAAUUCUUAAUUGCUAUUACUUUUAAUUAAUCGUGUUUUUCUACAUUUUUAACUGAAUGAUAUUUAAAUUAAUA
env-126   .........................CCCCCUGAGAGCCUAAUUAAUAAUCACAAUAAAUCAUAAAUUCAAACAAAAUUA
env-127   AGGACGGGUGUUUUUCACAAAAUACUCAAGGGGUGAAAAAUAAUGCCGAAAAACAUUUUGGACGAACCGCAAACUAUCC
env-128   ......................................................................AAACUAUCC
env-129   AGGGCAGGGGUUUUUCACAAAAUACUCAAGGGGUGAAAAAUAAUGGCGAAAAACCUUUUGGACGAACCGCAAACUAUCC
env-130   AGGGCAGGGGUUUUUCACAAAAUACUCAAGGGGUGAAAAAUAAUGGCGAAAAACCUUUUGGACGAACCGCAAACUAUCC
env-131   ACUGCAUCAGGCGGAUUAACACAAUUAAUAACAUAGAAAAUCAAAAGCCAAUACAAGGUUUGGAUUCACUAAAAAAUCA
env-132   ACUGCAUCAGGCGGAUUAACACAAUUAAUAACAUAGAAAAUCAAAAGCCAAUACAAGGUUUGGAUUCACUAAAAAAUCA
env-133   UGAUCCUAAACCGUCCUAUCCGUAAUUAAAUUAUAAAUUAGUUAAUAUCCUUUUUUUAUUGUUUAACAAGAGUUUAAAC
env-134   AAAAUUUUUCGUUUACAGUCAAAAAUCAAAAUAUAUACAAAUAAUCUGAAAAAUAUAAUUCAUAAUCUAAAUUAAUUUU
env-135   AAUACAUUGAUUUAAUUUUAAACCAACCACUAAGACCCUAAUAAAUAGUCACAAUAGGUAACUAAUUCAAACAAAAUUA
env-138   AAUACAUUGAUUUAAUUUUAAACCAACCACUAAGACCCUAAUAAAUAGUCACAAUAGGUAACUAAUUCAAACAAAAUUA
env-139   ...............................................................................
env-140   AAUACAUUGAUUUAAUUUUAAACCAACCACUAAGAGCCUAAUAAAUAGUCACAAUAGGUAACUAAUUCAAACAAAAUUA
env-141   UUGGCUUGGUUUUGAAAUUUCAGAGAAUCCUCUUUCAAAAUAAAAUAGAUAACUAAACCCUUUAGAUUCCUAAAAAUCA
env-142   UUGGCUUGGUUUUGAAAUUUCAGAGAAUCCUCUUUCAAAAUAAAAUAGAUAACUAAACCCUUUAGAUUCCUAAAAAUCA
env-143   UUGGCUUGGUUUUGAAAUUUCAGAGAAUCCUCUUUCAAAAUAAAAUAGAUAACUAAACCCUUUAGAUUCCUAAAAAUCA
env-145   UUGGCUUGGUUUUGAAAUUUCAGAGAAUCCUCUUUCAAAAUAAAAUAGAUAACUAAACCCUUUAGAUUCCUAAAAAUCA
env-146   GUGCUUAAGCCUCAGUUUAAGGGAGACAUCCUAGUUCAAAAUAAAAUAAAAACUAGAGGCUUUAGGUUCCUGAAAAUCC
env-147   UUUUGAUUAAUUUUAAAAUUUUAGAAAAUCCUCUUUCAAAAUAAAAUAGAAAACUGAACCCUUUGGAUUACUAAAAUCA
env-148   UUUUGAUUAAUUUUAAAAUUUUAGAAAAUCCUCUUUCAAAAUAAAAUAGAAAACUAAACCCUUUGGAUUACUAAAAUCA
env-149   UUUUGAUUAAUUUUAAAAUUUUAGAAAAUCCUCUUUCAAAAUAAAAUAGAAAACUAAACCCUUUGGAUUACUAAAAUCA
env-151   CAGUACAUUGAUUUAAUUUUAAAUCAACCUAUUAGAGCCUAAUAGUAACCACAAUAGGUUUCUAAUUCAAACAAAAUUA
env-152   CAGUACAUUGAUUUAAUUUUAAAUCAACCUCUUAGAGCCUAAUAGUAACCACAAUAGGUUUCUAAUUCAAACAAAAUUA
env-153   .........GAUUUAAUUUUAAAUCAACCUCUUAGAGCCUAAUAGUAACCACAAUAGGUUUCUAAUUCAAACAAAAUUA
env-154   CAGUACAUUGAUUUAAUUUUAAAUCAACCUCUUAGAGCCUAAUAGUAACCACAAUAGGUUUCUAAUUCAAACAAAAUUA
env-155   CAGUACAUUGAUUUAAUUUUAAAUCAACCACUUAGAGCCUAAUAGUAACCACAUUAGGUUUCUAAUUCAAACAAAAUUA
env-156   CAGUACAUUGAUUUAAUUUUAAAUCAACCUCUUAGAGCCUAAUAGUAACCACAAUAGGUUUCUAAUUCAAACAAAAUUA
env-157   CAGUACAUUGAUUUAAUUUUAAAUCAACCUNUUAGAGCCUAAUAGUAACCACAAUAGGUUUCUAAUUCAAACAAAAUUA
env-158   CAGUACAUUGAUUUAAUUUUAAAUCAACCUCUUAGAGCCUAAUAGUAACCACAAUAGGUUUCUAAUUCAAACAAAAUUA
env-159   CAGUACAUUGAUUUAAUUUUAAAUCAACCUCUUAGAGCCUAAUAGUAACCACAAUAGGUUUCUAAUUCAAACAAAAUUA
env-161   GUAAUAACUGUUGCACUUAUUGUUUGAACUUAGUAAAACUAAAAUACUAUUCACUCUAGUGUCAUUAAUAAUAAUUAGU
env-162   GCGAUAAAACCGGUAACUAGAGCCUGUUUGAGCAACAAAAUAAAAAUAAAAACUACACGGCUUAACUAAUCGAAAAUCA
env-163   CAUAAUAUCUAAUAUUAUCAGGACAUUUGAUUGUUCUAAAUAAAAAUACCAACUUGAGGGGAUAAACAAUCAAAAAUCA
env-164   UAUUUUGAUGACUACAAUAAGAUAUCAUAUUAGUUUUGAAUAAAUUAAACAACGACAAGGCAUAAACAAUAGAAAAUCC
env-165   CAGUAUAUUGAUUUAAUUUUAAACCAACCUCUUAGAACCUAAUUCUUACCUCAAUAAGUUUCUAAUCCAAACAAAAUUA
env-166   ....................................................AAUAAGUUUCUAAUCCAAACAAAAUUA
env-167   GGAAAUGCUCAAUAUCUUGAAGAUGAUUGUUUGCUAUAAAUAAAACAGGAAACUUCAAGGCAUAAACAACUGGAAAUUA
env-170   ACUAUACUAAAUUUUCUAGAGAACGUUUUUUAGCAGUAAAUAAAAACAACAACUUCAGGUCAUAAACAAUCAAAAUUCA
env-171   CUAUACCGAAUUUAAUAGAGAACGUUUUUUUAGCUGUAAAUAAAAACAGCAACUUCAGGUCAUAAACAAUCAAAAUUCA
env-172   CUAUACCGAAUUUAAUAGAGAACGUUUUUUUAGCUGUAAAUAAAAACAGCAACUUCAGGUCAUAAACAAUCAAAAUUCA
env-173   GAUAUCAUUAAAAAAAUUGAACCCACUAAUUAACUUAAAAUAAAAUAGAUAACCACAAGGGUUCGCCAACAGAAAAUCA
env-174   UAAAACGAGCCUAUGGUUCGUUUUUUUGUAUCACGACUGAUAAAUUAGAUAACCACAAGGCAUAAGCAAUAGAAAAUCA
env-175   UAAAACGAGCCUAUGGUUCGUUUUUUUGUAUCACGACUGAUAAAUUAGAUAACCACAAGGCAUAAGCAAUAGAAAAUCA
env-176   UAAAACGAGCCUAUGGUUCGUUUUUUUGUAUCACGACUGAUAAAUUAGAUAACCACAAGGCAUAAGCAAUAGAAAAUCA
env-177   CUAUUAAAAUGAGCCAAUGGUUCGUUUUUUUUGCUGUAAAUAAAAUCGAGAACCUAAAGGUAUAAACAACAGAAAAUCA
env-179   CUUUAUAAUUAAUAAAUCAAACCCAUUAAUUGACUUAAAAUAAAAUAGAUAACCACAUGGGUUAACCAACAGAAAAUCA
env-180   UAAAACGAGCCUAUGGUUCGUUUUUUUGUAUCACGACUGAUAAAUUAGAUAACCACAAGGCAUAAGCAAUAGAAAAUCA
env-181   CUAUUAAAAUGAGCCAAUGGUUCGUUUUUUUUGCUGUAAAUAAAAUCGAGAACCUAAAGGUAUAAACAACAGAAAAUCA
env-185   CUAUUAAAAUGAGCCAAUGGUUCGUUUUUUUUGCUGUAAAUAAAAUCGAGAACCUAAAGGUAUAAACAACAGAAAAUCA
env-186   CUUUAUAAUUAAUAAAUCAAACCCAUUAAUUGACUUAAAAUAAAAUAGAUAACCACAAGGGUUAACCAACAGAAAAUCA
env-187   CUAUUAAAAUGAGCCAAUGGUUCGUUUUUUUUGCUGUAAAUAAAAUCGAGAACCUAAAGGUAUAAACAACAGAAAAUCA
env-189   UUUUUGUGCUAAGGUUUAUAGCUGAAUCUCGGUCCCCAAAUAAAACAGAAAACGACAAGGAAUAAAUAGUCGAAAAUCA
env-191   UUUUUUGUGUUAAGGUUUAUAACUGAAUCUCGCCCCUAAAUAAAACAGAAAACCACAAGGAAUAAAUAGUCGAAAAUCA
env-192   GGAUUAAAGUCACUAACUAAAGUCCAUUGAUGCACCGAAAUAAAAAAUAAAACGACAAGGCUUAAAUAAGCGAAAAUCA
env-193   GGAUUAAAGUCACUAACUAAAGUCCAUUGAUGCACCGAAAUAAAAAAUAAAACGAGAAGGCUUAAAUAAGCGAAAAUCA
env-195   UGGACUACUGAUAAGUAUGAAGCCAUUGGUUGGCUUUAAAUAAAAUAGAAAACGACAAGCGUUAAGCAAACGAAAAUCA
env-196   CAGUACAUUGAUUUAAUUUCAAAUCAACCCCUUAGAGCCUAAUAGUAUCAACAAUAGGUUUCUAAUUCAAACAAAAUUA
env-197   CAGUACAUUGAUUUAAUUUCAAAUCAACCCCUUAGAGCCUAAUAGUAUCAACAAUAGGUUUCUAAUUCAAACAAAAUUA
env-198   CAGUACAUUGAUUUAAUUUCAAAUCAACCCCUUAGAGCCUAAUAGUAUCAACAAUAGGUUUCUAAUUCAAACAAAAUUA
env-201   CAGUACAUUGAUUUAAUUUCAAAUCAACCUCUUAGAGCCUAAUAGUAUCAACAAUAGGUUUCUAAUUCAAACAAAAUUA
env-202   CAGUACAUUGAUUUAAUUUCAAAUCAACCUCUUAGAGCCUAAUAGUAUCAACAAUAGGUUUCUAAUUCAAACAAAAUUA
env-203   UGAAAUAAACAACUAUCUGGAUUUGGUUCUCAACCGCCAUCUUCGUUCUUAGUUUUACUUUGUUAUCAGACAAUUAAGU
env-205   AAUAUAUAGAUCUCUUACUUAAUCUUAAAUUAAGGCAAUAAUUAGUUCAAAUAUAACGUUAUAGAAAGGUUAAAUACUA
env-206   AAUAUAUAGAUCUCUUACUUAAUCUUAAAUUAAGGCAAUAAUUAGUUCAAAUAUGUCGUUAUAGAAAGGUUAAAUACUA
env-207   AAUAUAUAGAUCUCUUACUUAAUCUUAAAUUAAGGCAAUAAUUACUUCAAAUAUGACGUUAUAGAAACGCUAAAUACUA
env-210   AAUAUAUAGAUCUCUUACUUAAUCUUAAAUUAAGACAAUAAUUAGUUCAAAUAUGACGUUAUAGAAACGUUAAAUACUA
env-211   UUAACUUAUACCUCGACUUAAUUCUAAAUGAACCUAUAAGGUAUUAAUUUACAAAUAGAGUCGUGUAUCAAAAAAUUAC
env-212   UAAAUACAUUGAUUUAAUUUCAAACCAAGAGCUUAGGUUAUAAAAUAUCCAAAGCUUAGCUUGUUUUUAUCGAAAAUGU
env-213   UUAUUUUGAAAAGAAGCACUGAUUAACAUGAUUCGUUAUAAAUAACAAAUAGUAAUAAUUCUUAUAUUAAUUAAAAUCA
env-214   UUAUUUUGAAAAGAAGCACUGAUUAACAUGAUUCGUUAUAAAUAACAAAUAGUAAUAAUUCUUAUAUUAAUUAAAAUCA

          ...............................................................................
          ...............................................................................
          ●●●●●●●●●●●●●●●●●●●●●●●●●●●●●●●●●●●●●●●●●●●●●●●●●●●●●●●●●●●●●●●●●●Y●●●●RR●●●●●●
```

```
env-215   UUAUUUUGAAAAGAAGCACUGAUUAACAUGAUUCGUUAUAAAUAACAAAUAGUAAUAAUUCUUAUAUUAAUUAAAAUCA
env-216   .............................................UAUUCCGAAUUAACUUGUCAAACCCAAAAAACUUA
env-217   AUUUUACUUCUUUCUUAAAAUAAGAAACAUAAGCACCGGAGAAAUAUUCCGAAUUAACUUGUCAAACCCAAAAAACUUA
env-218   AAAUCAGCCCCUCAGAAAAUAAUUUAUUUAAAAAUUUAAUAUAACUUAAAGCAAGGCAUUGUUUACUUAAGCUUUAAUA
env-220   .GCUCAACGAUUACACAAUUUUACAUUCUCAAAUUUUAAAUAAAAACACCUCCAAUCGCUUGAUAAUUAAAAAAAUAAU
env-221   AAAUAAAGUCCCACCAAAUAAUAAAAAACCUCUUCGGAGGUUUUUUUAUGCAAUCACUUUGCGUAUUUCAAAAGAAUAA
env-222   AAAUAAAGUCCCACCAAAUAAUAAAAAACCUCUUCGGAGGUUUUUUUAUGCAAUCACUUUGCGUAUUUCAAAAGAAUAA
Fli-1-1   AAUCAAGAGGUUGCAAAACUUUUUUAAGUCGUUUUUUAAAAAAAUUCACAAUUUGUGAUUUAUAAUUAAUAAUUAUUUU
Gme-1-1   AAUAUUUAUAGUCAAUUUCCUUCAAAUCAUCGAUUAAAAGAAAUAUCAAUGGGCAGUAUUAUUCAUAUUAUAAUAAUUA
env-223   UUGAGAGUGUAAUAAUAAAACGAACCAAUAGGUUCGUUUUUUCUUUUCAAUAGAUUCCUUGUAAAUCAUUAAAAAAUUA
env-224   GACAAUAUUUUAGUUAUAUUGGUUUUAAAAACUCUUUUCAUGUUUUUUUGUAAUCUCUUUGUGUAUUUCAAAAGAAUAG
env-225   AGACAAUAUUUAGCUAUAUUGGUUUUAAAAACUCUUUUCAUGUUUUUUUGUAAUCUCUUUGUGUAUUUCAAAAGAAUAU
env-226   AGACAAUAUUUUGUUAUAUUGGUUUUAAAAACUCUUUUCAUGUUUUUUUGUAAUCUCUUUGUGUAUUUCAAAAGAAUAU
env-227   AAUAAGAAUAUCUAAUUUUAGUAUUUUUUCCUUAAUUUGAUUUUUUUUUAUCAUCUCUUUGUGUAUUUCAAAAGAAUAU
env-228   AAAAAGAAUCUCCAAUUUUAGUACUUCUUCCUUAAUUUGAUUUUUUUUUAUCAUCACUUUGCGUAUUUCAAAAGAAUAA
env-229   AGACAAUAUUUAGUUAUAUUGGUUUUAAAAACUCUUUUCAUGUUUUUUUGUAAUCUCUUUGUGUAUUUCAAAAGAAUAU
env-230   AGACAAUAUUUAGNUAUAUUGGUUUUAAAAACUCUUUUCAUGUUUUUUUGUAAUCUCUUUGUGUAUUUCAAAAGAAUAU
env-231   AGACAAUAUUUAGUUAUAUUGGUUUUAAAAACUCUUUUCAUGUUUUUUUGUAAUCUCUUUGUGUAUUUCAAAAGAAUAU
env-234   GACAAUAUUUUAGUUAUAUUGGUUUUAAAAACUCUUUUCAUGUUUUUUUGUAAUCUCUUUGUGUAUUUCAAAAGAAUAU
env-235   AGACAAUAUUUAGUUAUAUUGGUUUUAAAAACUCUUUUCAUGUUUUUUUGUAAUCUCUUUGUGUAUUUCAAAAGAAUAU
env-236   AGACAAUAUUUAGUUAUAUUGGUUUUAAAAACUCUUUUCAUGUUUUUUUGUAAUCUCUUUGUGUAUUUCAAAAGAAUAU
env-237   AGACAAUAUUUAGUUAUAUUGGUUUUAAAAACUCUUUUCAUGUUUUUUUGUAAUCUCUUUGUGUAUUUCAAAAGAAUAU
env-238   AUAACUUGUUAUCUUACAAUUAAAAGCCUCCUUCGGGAGGUUUUUUUAUGCAAUCACUUUGCGUAUUUCAAAACUAUAG
env-239   UUAAAACGACAAAACGUUUAAUCUAGUGCCUGAGGACAAAUAAAAUAGCAAUUCGUAAAACAGGUAUUAAACUAUUUUU
env-243   UUAAAACGACAAAACGUUUAAUCUAGUGCCUGAGGACAAAUAAAAUAGCAAUUCGUAAAACAGGUAUUAAACUAUUUUU
env-244   ACAAAAAUGGUUUUUAUCAGUAAAAAGUUAACCAGGAAGGGAAUUAUUCACCUGUAAUAAUAAAUUGCAAUUUAUUUUC
env-247   AAAAUUAGCUAAAUAUAAUUUAAUAAUUUUUGAUCCAUUUUUGAGUCUUACUUCUAAUAAUUGUUGCAAUUAAUUUUA
env-248   AUUGCUUAAUUAAAAGUCAGCUUUUACGAGUAAUAUUUUUUUAAUAGAAUCUUCUGAUAACAGUUGAAAUUAAUUUUUU
env-249   AAAAUUAGCUAAAUAUAAUUUAAUAAUUUUUAGAUCCAUUUUUGAGUCUUACUUCUAAUAAUUGUUGCAAUUAAUUUUA
env-250   AAAAUUAGCUAAAUAUAAUUUAAUAAUUUUUAGAUCCAUUUUUGAGUCUUACUUCUAAUAAUUGUUGCAAUUAAUUUUA
env-251   AAAAACGUCUCAUAGUUUGAUUACGUAUUUUCAGUAAUAAAUAUUUAUACAAUACUACCUUUUUUUUCAAUACGAUUUU
env-255   UAGAACGUAAAAUGGUUUAAUUCAGUAUUUUUCGAACUAAAUAUUUAUACAAUAUUAGGCUUUUUUUAAAUUGGAUUUU
env-256   CAUCCGCCUCGAUUAAGGCUUAGAAACAAGCUCACGCAGGAUUUUAUCACAAUUAUAAAGAUAAAAAUGAUUUAAUUUC
env-257   AAUGGUAAACAACCGACCCGCUUGUAAUGGGGAAUGAUUGAUUUUAUUACAAUUAUACAGAUAAUAAUAAUUCAAUUUU
env-258   AAACCUGUACCUGGACCUUAUAUUGAAUGAACCCAUACGCAAUUAAUUGAACUAGAGGUUUUGCAUAAUGAUAAUUUGA
env-261   AAACCUGUACCUGGACCUUAUAUUGAAUGAACCCAUACGCAAUUAAUUGAACUAGAGGUUUUGCAUAAUGAUAAUUUGA
Nma-1-1   GGCAAUCAUCAAUAGACCAUAUUGAAACUUGUAGCUUUAAAGUUUGAUUUUGUUUUUCAUGUUGUUUUUAUUUUAAACU
env-262   AACAAAAGUAGAUUCGACAAAACACCUUUACAUUUGGUCGAAAAUAACGAUCUUGCAUUUGCUUUUCCAAAAAAAAGGA
env-263   UAAAUUUAUAUUUUGAUUUAAUACUUAAUAGACCUGUUAGAACUAAAAACACUUAAUUCGAAAAUUCCAAAAUUUAAUC
env-264   UAAAUUUAUAUUUUGAUUUAAUACUUAAUAGACCUGUUAGAACUAAAAACACUUAAUUCGAAAAUUCCAAAAUUUAAUC
env-265   ....................................UUAGAACUAAAAACACUUAAUUCGAAAAUUCCAAAAUUUAAUC
env-266   AUAGAUUUAAUUCUAAAUCAGCCUAUUAGAGCGUAAUAAAAAUAAUUCGAGUAAAGAAAUUGUUUAUUAAAUAAUAAAU
env-267   AUAGAUUUAAUUUUAAAUCAGCCCAUUAGAGCGUAAUAAAAAUAAUUCGAGUAAAGAAAUUGUUUAUUAAAUAAUAAAU
env-268   AUAGAUUUAAUUCUAAAUCAGCCUAUUAGAGCGUAAUAAAAAUAAUUCGAGUAAAGAAAUUGUUUAUUAAAUAAUAAAU
env-269   AUAGAUUUAAUUUUAAAUCAGCCUAUUAGAGCGUAAUAAAAAUAAUUCGAGUAAAGAAAUUGUUUAUUAAAUAAUAAAU
env-270   AUAGAUUUAAUUUUAAAUCAGCCUAUUAGAGCGUAAUAAAAAUAAUUCGAGUAAAGAAAUUGUUUAUUAAAUAAUAAAU
env-271   ............................GAGCGUAAUAAAAAUAAUUCGAGUAAAGAAAUUGUUUAUUAAAUAAUAAAU
env-272   AUAGAUUUAAUUUUAAAUCAGCCUAUUAGAGCGUAAUAAAAAUAAUUCGAGUAAAGAAAUUGUUUAUUAAAUAAUAAAU
Mmo-1-1   AUAGAUUUAAUUUUAAAUCAGCCUAUUAGAGCGUAAUAAAAAUAAUUCGAGUAAAGAAAUUGUUUAUUAAAUAAUAAAU
env-273   AGAGUGUUGCAUGUAAGUAAUAAAAGCCCUUGAUGGAAUUAUAGUGUGCGAUUAUUCUUGGAAGAGUCGAAAACUAUCC
env-274   AAAGUGUUGCAUGUAAGUAAUAAAAGCCCUUGAUGGAAUUAUAGUGUGCGAUUAUUCUUGGAAGAGUCGAAAACUAUCC
env-276   ACCUUUAUCUUGAUCUUAUUCUUAAUAGAACCAUAAGAGAAUAAAUAUUCCUAAAUAAAUCGAAGUUUCCUUAUUUUUA
env-277   ACCUUUAUCUUGAUCUUAUUCUUAAUAGAACCAUAAGAGAAUAAAUAUUCCUAAAUAAAUCGAAGUUUCCUUAUUUUUA
env-278   ACCUUUAUCUUGAUCUUAUUCUUAAUAGAACCAUAAGAGAAUAAAUAUUCCUAAAUAAAUCGAAGUUUCCUUAUUUUUA
env-280   AUCUUUAUAUUGAUCUUAUCCUUAACAAACCUAUAAGAGAAUAAAAUAUUAGAUAUAAAUCUAAGUUUCCUUAUUUUUA
env-281   AUCUUUAUAUUGAUCUUAUCCUUAACAAACCUAUAAGAGAAUAAAAUAUUAGAUAUAAAUCUAAGUUUCCUUAUUUUUA
env-282   CAACCUUUAUCUUGAUCUUGUUCUAAACAGGCCUAUAAGAGAAAUAUCUGCUUAAAUAAUCUAUGUUUCCUUAUUUUUA
env-283   CAACCUUUAUCUAGAUCUUGUUCUAAACAGACCUAUAAGAGAAAUAUCUGCUUAAAUAAUCUAUGUUUCCUUAUUUUUA
env-285   CAACCUUUAUCUUGAUCUUGUUCUAAACAGACCUAUAAGAGAAAUAUCUGCUUAAAUAAUCUAUGUUUCCUUAUUUUUA
env-286   CAACCUUUAUCUAGAUCUUGUUCUAAACAGACCUAUAAGAGAAAUAUCUGCUUAAAUAAUCUAUGUUUCCUUAUUUUUA
env-287   .................................................................UUUCCUUAUUUUAUA
env-288   CAACCUUUAUCUUGAUCUUGUUCUAAACAGGCCUAUAAGAGAAAUAUCUGCUUAAAUAAUCUAUGUUUCCUUAUUUUUA
env-289   CAACCUUUACCUUGAUCUUGUUCUAAACAGACCUAUAAGAGAAAUAUCUGCUUAAAUAAUCUAUGUUUCCUUAUUUUUA
env-290   CAACCUUUAUCUUGAUCUUGUUCUAAACAGGCCUAUAAGAGAAAUAUCUGCUUAAAUAAUCUAUGUUUCCUUAUUUUUA
env-294   CAACCUUUAUCUUGAUCUUAUUCUAAACAGGCCUAUAAGAGAAAUAUCUGCUUAAAUAAUCUAUGUUUCCUUAUUUUUA
env-295   CAACCUUUAUCUUGAUCUUGUUCUAAACAGGCCUAUAAGAGAAAUAUCUGCUUAAAUAAUCUAUGUUUCCUUAUUUUUA
env-296   CAACCUUUACCUUGAUCUUGUUCUAAACAGGCCUAUAAGAGAAAUAUCUGCUUAAAUAAUCUAUGUUUCCUUAUUUUUA
env-297   CAACCUUUAUCUUGAUCUUGUUCUAAACAGGCCUAUAAGAGAAAUAUCUGCUUAAAUAAUCUAUGUUUCCUUAUUUUUA
env-298   AAUCGAGAGAUUAGAAGGUAAAUAAAUAUACCUUCUUAUUUUAUUGCCAAUAAAAUUAUUGCUUUAGUGUACAUUCCUU
env-299   AAUCGAGAGAUUAGAAGGUAAAUAAAUAUACCUUCUUAUUUUAUUGCCAAUAAAAUUAUUGCUUUAGUGUACAUUCCUU
env-300   AAUCGUGAGAUUAGAAGGUAAAUAAAUAUACCUUCUGAUUUUAUUGCCAAUAAAAUUAUUGCUUUAGUGUACAUUCCUU
env-301   AAUCGAGAGAUUAGAAGGUAAAUAAAUAUACCUUCUUAUUUUAUUGCCAAUAAAAUUAUUGCUUUAGUGUACAUUCCUU
env-302   AUCGAGAGAUCAGAAGGUAAAAUAAAUAUAUCUUCUUGUUUUAUUGCCAAUAAAAUUAUUGCUUUAAUGUACAUUCCUU
env-303   ..............................CCUUCUUAUUUUAUUGCCAACAAAAUUAUUGCUUUAAUAUACAUUCCUU
env-304   AUCAAGAAAUAAGGAUAUAAAACACGUAAAUAACUCAUUUUAAUUUGAUUAAAAUUGUUGGUUUAAGUUAGAUUACUC
env-305   AGAGAUUUAAAGAAAAAAAUUAAAUUUAUUUAUUGAUCUAAUUUCUAAUAGACCUGUGAGAGAAUAAUCUAUUUUUUU
env-306   GAAGAGAUUUAAAGAAAAAAAUAAAUUUAUUUAUCGAUCUAAUUUCUAAUAGACCUGUAAGAGAAUAAUCUAUUUUUUU
env-307   GAAGAGAUUUAAAGAAAAAAAUAAAUUUAUUUAUCGAUCUAAUUUCUAAUAGACCUGUAAGAGAAUAAUCUAUUUUUUU
env-308   ACAAGCUGAUAAGUGGUAGGGUUUGCAGUUUAACAGUUUAAAAAAGAAUCAGAUUAUUCUUGUAAAACGAAUAAUUCAU
Lsp-3-1   AAUAUAAAUCUCUAUAUAGACUUAAUUCUCAAUCAAGCUAUACGUAAUUAAAAAUAUCGUUUUUCUUUAAGAAUUAAUU
env-309   UAUCAACCUUUACAUAGACUUGAUUUUAAAUCAGCCUGUUCGCAAUUAAAAAUAACAGACCGGUAUUUAGAAAUUAAUU
env-310   UAUCAACCUUUACAUAGACUUGAUUUUAAAUCAGCCUGUUCGCAAUUAAAAAUAACAGACCGGUAUUUAGAAAUUAAUU
env-313   UAUCAACCUUUACAUAGACUUGAUUUUAAAUCAGCCUGUUCGCAAUUAAAAAUAACAGACCGGUAUUUAGAAAUUAAUU
env-314   UAUCAACCUUUACAUAGACUUGAUUUUAAAUCAGCCUGUUCGCAAUUAAAAAUAACAGACCGGUAUUUAGAAAUUAAUU

          ...............................................................................
          ...............................................................................
          ●●●●●●●●●●●●●●●●●●●●●●●●●●●●●●●●●●●●●●●●●●●●●●●●●●●●●●●●●●●●●●●●●Y●●●●RR●●●●●●
```

```
env-315   UAUCAACCUUUACAUAGACUUGAUUUUAAAUCAGCCUGUUCGCAAUUAAAAAUAACAGACCGGUACUUAGAAAUUAAUU
env-316   AUCAAUCUUUAUAUAGACUUGAUUCUCAAUCAGGCAAUUCGCAAUUAAAAAAAGUUAGGUUAAGAUUGAUAAAUUAAUU
env-317   AUCAAUCUUUAUAUAGACUUGAUUCUUAAUCAGGCAAUUCGCAAUUAAAAAAAGUUAGGUUAAGAUUGAUAAAUUAAUU
env-318   AUCAAUCUUUAUAUAGACUUGAUUCUCAAUCAGGCAAUUCGCAAUUAAAAAAAGUUAGGUUAAGAUUGAUAAAUUAAUU
env-320   AUCAAUCUUUAUAUAGACUUGAUUCUUAAUCAGGCAAUUCGCAAUUAAAAAAAGUUAGGUUAAGAUUGAUAAAUUAAUU
env-324   UAUCAACCUUUACAUAGACUUGAUUUUAAAUCAGCCUGUUCGCAAUUAAAAAUAACAGACCGGUACUUAGAAAUUAAUU
env-325   CAUCAAUCUUUAUAUAGACUUGAUUUUAAAUCAGCCUGUUCGCAAUUAAAAAUAACAGACCGGUACUUAGAAAUUAAUU
env-326   AUCAAUCUUUAUAUAGACUUGAUUCUUAAUCAGGCAAUUCGCAAUUAAAAAAAGUUAGGUUAAGAUUGAUAAAUUAAUU
Fri-1-1   CUUUGCAACUUUGCCACUAUGUAACUUUGCUGCUAAAAACUUUAUAACUUUUUUACCCUUGCAGGUGUAAAAAUAAUGU
env-327   AAUCUUUAUCUGGACUUGAUUCUAAAUCAUCCCUUAAGAGAUUAAUCUAGAAACAGGUCAGUAUAUCAAAUAAUUACAU
env-328   AAUCUUUAUCUGGACUUGAUUCUAAAUCAACCCUUAAGAGAUUAAUCCAGAAACAGGUCAGUAUAUCAAAUAAUUACAU
env-329   ....................................................AUUUAAAAGAUCUAAUUAUUAAAAAUAC
env-330   AAUAAAUACAUUGAUUUGAUACUAAAUCAAAAGAUUAGAGAAUAAUUUAGGAUUUAAAAGAUCUAAUUAUUAAAAAUAC
env-331   UAUUGAAUCAGCAUUAACAGUAAAAGAUGGGUUUAUAUUUAUGGAAUAAAAUAUUUUUUUAUAGUUGAAAUUUAAAUAU
env-332   UUGAAUCAGCAUUAACAGUAAAAGAUGGGUAUAUAUUUAUGGAAUAAAAUAUUUUUUUUAAUAGUUGAUAUUUAAAUAU
env-333   ...............AAGUAAAAGAUGGAUAUAUAUUUAUCGAAUAAAAUAUUUUUUUAAUAGUUGGAAUUUAAAUAU
env-334   ACCAGCCAAUCAGAAAGUAAUAACAAGAAUAUUCAAGGUGUAAAUAAACACAAAUUCGUUGUUUUGUAUAGGUUUAGUU
env-335   CGGGGAAACCCAGAAAGAGACUGGAAGAGCCACAAAAGUUGAGCCGUAAUAUUUGGUGUUGCUUUUCUUAAAUAAAGUU
env-336   ........CCCAGAAAGAGACUGGAAGAGCCACAAAAGUUGAGCCGUAAUAUUUGGUGUUGCUUUUCUUAAAUAAAGUU
env-337   AGAAAUUGGGAUAGCUGAAAAGACUAGCAUGAAGUGAUUUUAUUAUUUUAUUAUUCAAUUGCGGGAAUUAAAUAAUAAU
env-338   AUCUAAUUUUAAACAGACCAAUUAGAAGUUAAUUAUAUUCUCAUAUUCUUAAAGUUUGUUGUGUAAGUCUACAUUAAUU
env-339   .................................UAGCUGAUCUAUAUUCUUAAGUUUGUUGUGUAAGUUAACAUUAAUU
env-340   AUCUAAUUUUAAACAGACCAAUUAGAAGUUGAUUAUAUUCUCAUAUUCUUAAAGUUUGUUGUGAAAGCCAACAUUAAUU
env-341   AUCUAAUUUUAAACAGACCAAUUAGAAGUUGAUUAUAUUCUCAUAUUCUUAAAGUUUGUUGUGUAAGUUAACAUUAAUU
env-342   AUCUAAUUUUAAACAGACCAAUUAGAAGUUGAUUAUAUUCUCAUAUUCUUAAAGUUUGUUGUGUAAGUUAACAUUAAUU
env-343   AUCUAAUUUUAAACAGACCAAUUAGAAGUUGAUUAUAUUCUCAUAUUCUUAAAGUUUGUUGUGUAAGUCAACAUUAAUU
env-344   AUCUAAUUUUAAACAGACCAAUUAGAAGUUGAUUAUAUUCUCAUAUUCUUAAAGUUUGUUGUGUAAGUCAACAUUAAUU
env-345   AUCUAAUUUUAAACAGACCAAUUAGAAAUUAAUUAUACUCUCAUAUUCGUAAAGUUUGUUGUGAAAGUCUACAUUAAUU
env-346   ........UUAAACAGACCAAUUAGAAAUUAAUUAUACUCUCAUAUUCGUAAAGUUUGUUGUGAAAGUCUACAUUAAUU
env-347   AUCUAAUUUUAAACAGACCAAUUAGAAGUUGAUUAUAUUCUCAUAUUCUUAAAGUUUGUUGUGUAAGUCAACAUUAAUU
env-352   AUCUAAUUUUAAACAGACCAAUUAGAAGUUGAUUAUAUUCUCAUAUUCUUAAAGUUUGUUGUGUAAGUCAACAUUAAUU
env-353   AUCUAAUUUUAAACAGACCAAUUAGAAGUUGAUUAUAUUCUCAUAUUCUUAAAGUUUGUUGUGUAAGUCAACAUUAAUU
env-354   AUCUAAUUUUAAACAGACCAAUUAGAAGUUGAUUAUAUUCUCAUAUUCUUAAAGUUUGUUGUGUAAGUCAACAUUAAUU
env-355   AUCUAAUUUUAAACAGACCAAUUAGAAGUUGAUUAUAUUCUCAUAUUCUUAAAGUUUGUUGUGUAAGUCAACAUUAAUU
env-356   AUCUAAUUUUAAACAGACCAAUUAGAAGUUGAUUAUAUUCUCAUAUUCUUAAAGUUUGUUGUGUAAGUCAACAUUAAUU
env-357   CACUUGCUCAAUUAUGAAUUAUCAAUAGAUCGUCACUUGAUAAUUAUGCCAUUGAUAAUUGAUAAUUCGGAAUUAUAUA
Fta-1-1   UUUUCAAAGAUGGACUUUACGUUUCUGAUGGAGGACUGUAAUUAAUUCAUAAUUCACACUCGUAAUUCGUAAUUAGUUG
env-358   .............................................................................AAUUA
env-359   AUUUAUACCUAGAUUUAAUUCAGAAUCAGAAAAUAAGAGCAUAAAAACAAUCUUAAAUUAGGGAAUAUAAAAAAAAUAGU
env-360   AUUUAUACCUAGAUUUAAUUCAGAAUCAGAAAAUAAGAGCAUAAAAACAAUCUUAAAUUAGGGAAUAUAAAAAAAAUAGU
env-361   GUAUCCCUAUUACUAGAGAGCUUUUGCAAAAAAAAAUCUAUUUAUUUACAAGGAUUAUAGCCUAUUAUAAAAUUUAAUU
env-362   GUUUUAGCUGUUAGACUUUACUAUUUUUUUAUUUAUCUAAUUAAUUGAACGUAUUAGACUCAUAAGAAAAAAAUUAGUU
Mod-7-1   UUGGUAAUCAAUGUACUAGAGGGUAAAUAAGAACCUAAAAAAAAAGUGUAUUAUUUAUAGUUGUGUUUUGAAAAAUAAAA
Mod-6-1   UUGGUAAUCAAUGUACUAGAGGGUAAAUAAGAACCUAAAAAAAAAGUGUAUUAUUUAUAGUUGUAUUUUGAAAAAUAAAA
Mod-3-1   UUGGUAAUCAAUGUACUAGAGGGUAAAUAAGAACCUGAAAAAAAAGUGUAUUAUUUAUAGUUGUAUUUUGAAAAAUAAAA
Mpr-1-1   UUGGUAAUCAAUGUACUAGCGGGUAGAUAAGAACCUAAAAAAAAAGUAUAUUAUUUAUAGUUGUAUUUUGAAAAAUAAAA
Mod-4-1   UUGGUAAUCAAUGUACUAGAGGGUAAAUAAGAACCUAAAAAAAAAGUGUAUUAUUUAUAGUUGUAUUUUGAAAAAUAAAA
Mod-1-1   CUGAUAAUCAAAUAAGUAAGUUCAAUAUGAGACUUUUAAAAAAAGUCGAUUAUUUAUAGUUGUAAUUUCAAAAAUAAAA
Fin-1-1   CUUAGAAUUUGUAAAUGAACAAGUUGCAACACUUUUUUAAUCAAUAAACAAGAAAAGGUUUGGUAAUUAAAAAAUAAAG
env-363   UGUUCUCGUUUCCACAAUCUUUAAAUCAUUUAAUCUUUAAAUUUUUCAAUGUGAAUUCUUUUCCAUAUCAAAAUAAAUA
env-364   UGUUCUCGUUUCCACAAUCUUUAAAUCAUUUUAUCUUUAAAUUUUUCAAUGUGAAUUCUUUUCCAUCUCAAAAUAAAUA
env-366   ................................................AUGUGAAUUCUUUUCCAUCUCAAAAUAAAUA
env-367   .......................................AAUUUUUCAAUGUGAAUUCUUUUCCAUAUCAAAAUAAAUA
env-368   ....................UUAAAUCAUUUAAUCUUUAAAUUUUUCAAUGUGAAUUCUUUUCCAUAUCAAAAUAAAUA
env-369   UUUAACCCGGCAUAAAAUAAAAAGAGCGUCAAAUUCAUAAGCUUUUUUACCGGUGCGCACACUGUUCAACUAAAUCAUU
env-373   UGGUUUUAAAUCAUCCUAUACGCAUUUAGGUCAUAAAUAACUUAAUAUCCUUUUUUUAUUGUAUAUCAACAGUUUAAAC
env-375   UGGUUUUAAAUCAUCCUAUACGCAUUUAGGUCAUAAAUAACUUAAUAUCCUUUUUUUAUUGUAUAUCAACAGUUUAAAC
env-376   AUCAAUACAUUGAUUUAAUUUUAAAUCGGCCGUUAAGAGCAUAAUUUCUAUAAAAUUAUAUUUUUGUUAACCAAUUUUC
env-377   AUCAAUACAUCGAUUUGAUCUUGAAUCAACCGAUCAGAGCAUAAUUUCUUUAAAAUUAUGUUUAUAUCACACGAUUUUC
env-378   AUCAAUACAUUGAUUUGAUUCAAAACCAGGUUUUAAGAUUAUAAAAUUUUUCAAGGAAGUAUUGUUAUAUCUAAAAUGC
env-379   CAAAUACAUUGAUUUGAUCUCAAAUCAAGAGCUCAGGCUAUAAAAUAUCCAAAGCAAAGCUUCGUUUUAUCGAAAAUGU
env-381   CAAAUACAUUGAUUUGAUCUCAAAUCAAGAGCUCAGGCUAUAAAAUAUCCAAAGCAAAGCUUCGUUUUAUCGAAAAUGU
env-382   ...................UCAAAUCAAGAGCUCAGGCUAUAAAAUAUCCAAAGCAAAGCUUCGUUUUAUCGAAAAUGU
env-383   CAAAUACAUUGAUUUGAUUUCAAAUCAAAAGCUCAGGCCAUAAAAUAUCCAAAGCAAAGCUUCGUUUUAUCGAAAAUGU
env-384   CAAAUACAUUGAUUUGAUCUCAAAUCAAGAGCUCAGGCCAUAAAAUAUCCAAAGCAAAGCUUCGUUUUAUCGAAAAUGU
env-385   CAAAUACAUUGAUUUGAUCUCAAAUCAAGAGCUCAGGCUAUAAAAUAUCCAAAGCAAAGCUUCGUUUUAUCGAAAAUGU
env-386   CAAAUACAUUGAUUUGAUAUCAAAUCAAGAGCUCAGGCUAUAAAAUAUCCAAAGCAAAGCUUCGUUUUAUCGAAAAUGU
env-387   CAAAUACAUUGAUUUGAUCUCAAAUCAAGAGCUCAGGCUAUAAAAUAUCCAAAGCAAAGCUUCGUUUUAUCGAAAAUGU
env-388   CAAAUACAUUGAUUUGAUCUCAAAUCAAGAGCUCAGGCUAUAAAAUAUCCAAAGCAAAGCUUCGUUUUAUCGAAAAUGU
env-389   UAAAUACAUUGAUUUGAUCUCAAAUCAAGAGCUCAGGUUAUAAAAUAUCCAAAGCAAAGCUUCGUUUUAUCGAAAAUGU
env-390   CAAAUACAUUGAUUUGAUCUCAAAUCAAAAACUCAGGCCAUAAAAUAUUUAAAGCAAAGCUUCAUUUUACCGAAAAUGU
env-391   CAAAUACAUUGAUUUGAUCUCAAAUCAACAACUCAGGCCAUAAAAUAUUUAAAGCAAAGCUUCAUUUUACUGAAAAUGU
env-392   CAAAUACAUUGAUUUGAUCUCAAAUCAACAACUCAGGCCAUAAAAUAUUUAAAGCAAAGCUUCAUUUUACUGAAAAUGU
env-393   CCAAUACAUCGAUUUGAUUCUUAAUCAACCUCUAAGAGCAUAAUUUUUUUAAAGCAAAGCACAAUUUUGUUGAAAUUGU
env-394   CCAAUACAUCGAUUUGAUUCUAAAUCAACCUCUAAGAGCAUAAUUUUUUUAAAGCAAAGCGCAAUUUUGUUGAAAUUGU
env-395   CCAAUACAUCGAUUUGAUUCUAAAUCAACCUCUAAGAGCAUAAUUUUUUUAAAGCAAAGCGCAAUUUUGUUGAAAUUGU
env-396   AAUCAACCUCUUAGGAAAUAAUAUAUUCCUAAAAUAGAAUAUAACUCAUAGCAAGGCAUUGUUUACUUAAGCUUUAAUA
env-397   AAUCAACCUCUUAGGAAAUAAUAUAUUCCUAAAAUAGAAUAUAACUCAUAGCAAGGCAUUGUUUACUUAAGCUUUAAUA
env-398   AAUCAACCUCUUAGGAAAUAAUAUAUUCCUAAAAUAGAAUAUAACUCAUAGCAAGGCAUUGUUUACUUAAGCUUUAAUA
env-399   AAUCAGCCUCUCAGGAAAUAAUUUAUUCCUAAAAUAGAAUAUAACUCAUAGCAAGGCAUUGUUUACUUAAGCUUUAAUA
env-400   AAUCAACCUCUUAGGAAAUAAUAUAUUCCUAAAAUAGAAUAUAACUCAUAGCAAGGCAUUGUUUACUUAAGCUUUAAUA
env-402   AUCAAGAAAUAAGAAUAUAAAACACGUAAACAAUUCAUUUUAAUUUUGAUUAAAAUUGUUGGUUUAAAUUAGAUUACUC
env-403   AAUCAAGAAAUAAGAAUAUAAAACACGUAAACAAUUCAUUUUAAUUUUGAUUAAAAUUGUUGGUUUAAAUAGAUUACUC
env-404   AAUCAAGAAAUAAGAAUAUAAAACACGUAAACAAUUCAUUUUAAUUUUGAUUAAAAUUGUUGGUUUAAAUAGAUUACUC

          ................................................................................
          ................................................................................
          ●●●●●●●●●●●●●●●●●●●●●●●●●●●●●●●●●●●●●●●●●●●●●●●●●●●●●●●●●●●●●●●●●●●Y●●●●RR●●●●●●
```

```
env-405   UCCGUUGCGCAAGUAGUUCAUCGUUUCAAUAUAUAAUAUAUUAAAAAUUUGAUAUUGGUUGUUUUGUAGAGGUUAAGUU
env-406   UAAAUCGUCCGUUGCGCAAAUAGUUCAUCGUUUCAAUAUAUUAAAAAUUUGAAAUUGGUUGUUUUAUAGAGGUUAAGUU
env-407   AUCAAGAAAUAAGGAUAUAAAACACGUAAAUGAAUCAUUUUAAUUUUAAUUAAAAUUGUUGGUUUAAAUUAGAUUACUC
env-408   AUCAAGAAAUAAGGAUAUAAAACACGUAAAUGAAUCAUUUUAAUUUUAAUUAAAAUUGUUGGUUUAAAUUAGAUUACUC
env-409   AUCAAGAAAUAAGGAUAUAAAACACGUAAAUGAAUCAUUUUAAUUUUAAUUAAAAUUGUUGGUUUAAAUUAGAUUACUC
env-410   AUCAAGAAAUAAGGAUAUAAAACACGUAAAUGAAUCAUUUUAAUUUUAAUUAAAAUUGUUGGUUUAAAUUAGAUUACUC
env-411   UCAAGAAAUAAGGUGACAAAACCUAAAAAAAUAACCCCUAUUAAUUAUAAUUUAAAUUGUUGGUUUAAACUAGAUUACUC
env-412   UCAAGAAAUAAGGUGACAAAAUCUAAAAAAAUAACCCCUAUUAAUUAUAAUUUAAAUUGUUGGUUUAAACUAGAUUACUC
env-413   GAAUCAAGAAAUAAGGUGACAAAAUCUAAAAAAUAACCCCUAAUUAUAAUUUAAAUUGUUGGUUUAAACUAGAUUACUC
env-414   UCAAGAAAUAAGGUGACAAAAUCUAAAAAAAUAACCCCUAUUAAUUAUAAUUUAAAUUGUUGGUUUAAACUAGAUUACUC
env-415   UUAAAAAAAGAGAUUAAUCUUUAUAUUGAUUUGAUCCUUAAUCAACCCAUUAGAAAAUAACGAUUUAUAAAAAAUAAGC
env-416   UAAAUACAUUGAUUUAAUACUUAACCAACCUAUUAGAGAAUAAUUAAUUACAAUAGAAAGCACUUAUUCUCAAAAAUAA
env-417   UAAAUACAUUGAUUUAAUACUUAACCAACCUAUUAGAGAAUAAUUAAUUACAAUAGAAAGCACUUAUUCUCAAAAAUAA
env-418   AUAAAUACAUUGAUUUGAUCCUUAAUCAAGCUGUUAGACAAUAAUUAAUGAUAAUAAAAGGACUUCUUAUUAAAAAGAA
Fbe-1-1   UUCUAACUUACUGAUUACCUGACACACUACACACCUUUUAAAGUUAUUCAGCAAAGUAUUGCGGAUAUUAAAAUAUUGA
env-419   UCUAUUAAAACGCAAACGUUUAAUUUUAUUUCACAUUUUUUUAAUAAAAAUGAUCCCUAGGCACUUUUUAAAUAAAAUU
Ksp-1-1   AAAAACUAACACUAAAAGGUCUUAUUUACAGAAAUGGCUGUAAAUUAAGCGUCAACUGUUUGAUAAUACAAAGAAUAAU
Dsp-2-1   AAAAACUAGCGGUAAAGGCUUUAAUUUACAGAAAUGGCUGUAAAUUGAGCCUCAACAGUUUGAUAAUACAAAGAAUAAU
Dsp-1-1   UGUUAAGACUUUAUUAAAAUCAAUAGGUUUAAUCACAAACAAAUAUUUGUGCGAUUUAUUUUAUUUGAUAUUAUUAUUU
Ddo-1-1   UGUUAAGACUUUAUUAAAAUCAAUAGGUUUAAUCACAAACAAAUAUUUGUGCGAUUUAUUUUAUUUGAUAUUAUUAUUU
Iha-1-1   CAUUACAAAUAGAAUUAGACAGCUGAAAUACAAAGACAUAAUUAAAACAACAUCUCUAUUUAUCAAUUCAAAAAAAGUA
env-420   GAAAAACAUCAACCUAUAUCUUGAUCUAAUUUUAAAUCGUCCACUUUCAACAGAGGCGUAAUUUCAUUCAAUAAUUAUA
env-421   GAAGAACAUCAAUCUCUACCUCGAUUUAAUAUUAAAUCGACCGCUUUCAUCACAGCUGUAAUUUCAAUCAAUUAUUGUG
env-422   AAAGAACAUCAACCUAUACCUCGAUUUAAUAUUAAAUCGACCGCUUUCAGAACAGCUGUAAUUCUAAUCGAUUAAAGUG
env-423   AAAGAACAUUAAUCUCUACCUCGAUUUAAUAUUAAAUCGACCGCUUUCAAAAGAGGUGUGAUUUCAAUCAAUUAUUGUA
env-424   GAAAAACAUCAACCUAUAUCUUGAUCUAAUUUUAAAUCGUCCACUUUCAACAGAGGCGUAAUUUCAUUCAAUAAUUAUA
env-427   GAAAAACAUCAACCUAUAUCUUGAUCUAAUUUUAAAUCGUCCACUUUCAACAGAGGCGUAAUUUCAUUCAAUAAUUAUA
env-428   GAAAAACAUCAACCUAUACCUCGAUUUAAUUUUAAAUAGACCUCUUGCGUCAGAGGCGUAAUUUCAUUCAAUAAUUAUA
env-429   GAAAAACAUCAACCUAUAUCUUGAUCUAAUUUUAAAUCGUCCACUUUCAACAGAGGCGUAAUUUCAUUCAAUAAUUAUA
env-430   GAAGAACAUCAAUCUUUACCUCGACUUAAUAUUAAAUCGACCGCUUUCAAAUCAGCUGUAAUUUCAAUCAAUUAUUAUG
env-431   ....AACAUCAAUCUUUACCUCGAUUUAAUAUUAAAUCGACCGCUUUCAAAUCAGCUGUAAUUUCAAUCAAUUAUUAUG
env-432   GAAGAACAUCAAUCUUUACCUCGAUUUAAUAUUAAAUCGACCGCUUUCAGCUCAGCUGUAAUUUCAAUCAAUUAUUAUG
env-433   GAAGAACAUCAAUCUUUACCUCGAUUUAAUAUUAAAUCGACCGCUUUCAGCUCAGCUGUAAUUUCAAUCAAUUAUUAUG
env-434   GAAGAACAUCAAUCUUUACCUCGAUUUAAUAUUAAAUCGACCGCUUUCAGCUCAGCUGUAAUUUCAAUCAAUUAUUAUG
env-435   GAAGAACAUCAAUCUUUACCUCGAUUUAAUAUUAAAUCGACCGCUUUCAGCUCAGCUGUAAUUUCAAUCAAUUAUUAUG
env-437   GAAGAACAUCAAUCUUUACCUCGAUUUAAUAUUAAAUCGACCGCUUUCAGCUCAGCUGUAAUUUCAAUCAAUUAUUAUG
env-438   GAAGAAUAUCAAUCUCUACCUCGAUUUAAUAUUAAAUCGACCGCUUUCAACUCAGCUGUAAUUUCAAUCAAUUAUUGUG
env-439   GAAAAACAUCAAUCUUUACCUAGAUUUAAUAUUAAAUCGGCCGCUUUCAGCACAACUGUGAUUUCAAUCAAUUAUUAUG
env-440   GAAAAACAUCAAUCUUUACCUAGAUUUAAUAUUAAAUCGACCGCUUUCAGCACAACUGUGAUUUCAAUCAAUUAUUAUG
env-441   GAAAAACAUCAAUCUUUACCUAGAUUUAAUAUUAAAUCGGCCGCUUUCAGCACAACUGUGAUUUCAAUCAAUUAUUAUG
env-442   GAAAAACAUCAAUCUUUACCUAGAUUUAAUAUUAAAUCGACCGCUUUCAGCACAACUGUGAUUUCAAUCAAUUAUUAUG
env-444   ......CAUCAAUCUUUACCUAGAUUUAAUAUUAAAUCGACCGCUUUCAGCACAACUGUGAUUUCAAUCAAUUAUUAUG
Kje-1-1   ACAUCUUUUAACAUUUAUAAUCUUAAGACUUUACAGAAUCAAAAAGGUUUAAUAUUAAUUUGGAAUCUAAAAAAAUAGUA
Kal-1-1   GUUUUUUAACAUUUGUUGCACUUAGGACUUCAAGUAUUCAAAAAGGUUUAAUUAAACAUUUGGAUUCUAAAAAAAUAGUA
Kzh-1-1   CAUUUUUCAGCAUGUAUAACAUAAUAAACAACAUUUCUCAAAAAGGUUAAAUUAAUAAUUUGGAAUCUAAAAAAAUAGUA
env-445   NNNNNNNNNNNNNNNNNNNNCCAAUUAUUGAUAAAAAAAAAACAAAAUUUAUUUAUUUUGAAUGUUUCAAUAUUUUGUA
env-446   GAAAUCAGAGAGAAAAUAGCUCAAACGCUCCAACUCUUUAAAAAUUAAUCACAAAGACAAAGUUUUAUCAAUUUAAUUG
env-447   GAAAUCAGAGAGAAAAUAGCUCAAACGCUCCAACUCUUUAAAAAUUAAUCACAAAGACAAAGUUUUAUCAAUUUAAUUG
env-449   UUUUAAAGCUUUAGGAUUUGUAUUAAUGUAAUAUUCACAUUAAAUUUAAUUAUAAUUGUUACAUAAUCAAAUAAAAGCA
env-450   AAUCUGUAUAUUGAUUUAAUUCAAAACCUUCCUUUAAGGCCAUAAAACUUAAUUAAUAGUAUUUUUUUAUUUAAAAUAC
env-451   AAUCUGUAUAUUGAUUUAAUUCAAAACCUUCCUUUAAGAUCAUAAAACCUAUUUAAUAGUAUUUUUUUAUUUAAAAUAC
env-452   AAUCUGUAUAUUGAUUUAAUUCAAAACCUUCCUUUAAGAUCAUAAAACCUAUUUAAUAGUAUUUUUUUAUUUAAAAUAC
env-453   AAUCUAUAUAUUGAUUUAAUUCAAAACCUCCCUUUAAGAUCAUAAAAUCUAUUUAAUAGUAUUUUUUUAUUUAAAAUAC
env-454   AAUCUAUAUAUUGAUUUAAUUCAAAACCUUCCUUUAAGGUCAUAAAACCUAUUUAAUAGUAUUUUUUUAUUUAAAAUAC
env-455   AAUCUAUAUAUUGAUUUAAUUCAAAACCGUCCUUUAAGAUCAUAAAGCCUAUUUAAUAGUAUUUUUUUAUUUAAAAUAC
env-456   UAGAUAGACGCGAACUCAAAAAUAAAAUGGAUCAAGUUCUUAGGCUGUUCUCCAACUAGAUCUGUAUCGCAGAAUAGUU
env-457   UAGAUAGACGCGAACUCAAAAAUAAAAUGGAUCAAGUUCUUAGGCUGUUCUCCAACUAGAUCUGUAUCGCAGAAUAGUU
env-458   AAAAUUAAGUUUAACGGGAUUAGACUUUUAAAGGCUUUGGAACUAAGCUUUUUUUUCGGCAUAAUUUCUCAAAAUAAUC
env-459   UGAUAUUAAAUCAUCCUAUUCGGCCAUAAAUUUCCAACUUCAUAAGUUAUUAUUUUUAUUGUUUUACAUUGGUUAAAUU
env-460   UAAUAUUAAAUCAUCCUAUUCGGCCAUAAAUUUCCAAUUUCAUAAGUUAUUAUUUUUAUUGUUUUACAUUGGUUAAAUU
env-461   UGAUAUUAAAUCAUCCUAUUCGGCCAUAAAUUUCCAACUUCAUAAGUUAUUAUUUUUAUUGUUUUACAUUGGUUAAAUU
env-462   UAAUAUUAAAUCAUCCUAUUCGGCCAUAAAUUUCCAACUUCAUAAGUUAUUAUUUUUAUUGUUUUACAUUGGUUAAAUU
env-463   UAAUAUUAAAUCAUCCUAUUCGGCCAUAAAUUUCCAACUUCAUAAGUUAUUAUUUUUAUUGUUUUACAUUGGUUAAAUU
env-464   UAAUAUUAAAUCAUCCUAUUCGGCGAUAAAUUUCCAACUUCAUAAGUUAUUAUUUUUAUUGUUUUACAUUGGUUAAAUU
env-465   UAAUAUUAAAUCAUCCUAUUCGGCCAUAAAUUUCCAAUUUCAUAAGUUAUUAUUUUUAUUGUUUUACAUUGGUUAAAUU
env-466   UAAUAUUAAAUCAUACUAUUCGGCCAUAAAUUUCCAACUUCAUAAGUUAUUAUUUUUAUUGUUUUACAUUAGUUAAAUU
env-467   UAAUAUUAAAUCAUCCUAUUAGACAAUAAAUUCCGAGCUUCAUAAUUUAUUAUUUUUAUUGUUUUACAUUGGUUUAAUU
env-469   UUAAUAUUAAAUCAUCCUAUUAGACAAUAAAUUCCGAGCUUCAUAAUUUAUUUUUUUAUUGUUUUACAUUGGUUUAAUU
env-470   UAAUAUUAAAUCAUCCAGUUCGGCCAUAAUUUUCCAAUUUCAUAAGUAAUUAUUUUUAUUGUUUUACAUUGGUUUAAUU
env-471   ..................................................GAUUUUAUUUGUCAAUUAAAAAAGUAUCA
env-472   GAAAUCAAUGCCUAUCUCGAUUUGGUAUUGAACCGACACUACCGCGCUUAGGUUUUACUUUGUUAUCAGACAAUUAAGU
env-473   GAAAUCAAUGCCUAUCUCGAUUUGGUAUUGAACCGACACUACCGCGCUUAGGUUUUACUUUGUUAUCAGACAAUUAAGU
env-474   ..............................................CUUAGGUUUUACUUUGUUAUCAGACAAUUAAGU
env-475   GAUGCUACCGCGGUACUUAAUCACAGUACUUUCUCUCUUAAAAUAAUGCCCUUCAAUCACAGCAACUUACAAAAUAAUC
env-476   UUAAAAAAUAAAAUCAAUCUAUACAUUGAUUUAAUCCUUAAUCAACCCGUAAGAAACUAAGCUUUAAUCAAAAAAAAGU
env-477   CUAAAAAAAGAGAUUAAUCUUUAUAUUGAUUUGAUCCUCAAUCAGCCCGUUAGAAAAUAACGAUUUAUAAAAAAUAAGU
env-478   CUAAAAAAAGAGAUUAAUCUUUAUGUUGAUUUGAUCCUCAAUCAGCCCGUUAGAAAAUAACGAUUUAUAAAAAAUAAGU
env-479   CUAAAAAAAGAGAUGAAUCUUUAUAUUGAUUUGAUCCUCAAUCAACCCGUUAGAAAAUAACGAUUUAUAAAAAAUAAGU
env-480   UAUUAAAAAAGAGAUUAAUCUUUAUAUUGAUUUGAUCCUUAAUCAACCCAUUAGAAAAUAACGAUUUAAAAAAAUAAGU
env-481   CUAAAAAAAGAGAUUAAUCUUUAUAUUGAUUUGAUCCUCAAUCAACCCGUUAGAAAAUAACGAUUUAUAAAAAAUAAGU
env-482   CUAAAAAACAAGAUCAAUCUAUACAUUGACUUGAUCCUCAAUCAGCCGAUUAGAAAGUAAGCAUUAAUCAAAAAUAAGU
env-484   CUAAAAAACAAGAUUAAUCUAUACAUUGACUUGAUCCUCAAUCAGCCGAUUAGAAAGUAAGCAUUAAUCAAAAAUAAGU
env-486   CUAAAAAACAAGAUUAAUCUAUACAUUGACUUGAUCCUCAAUCAGCCGAUUAGAAAGUAAGCAUUAAUACAAAAUAAGU
env-487   CUAAAAAAAGAGAUGAAUCUUUAUAUUGAUUUGAUCCUCAAUCAACCCGUUAGAAAAUAACGAUUUAUAAAAAAUAAGU

          ...............................................................................
          ...............................................................................
          ●●●●●●●●●●●●●●●●●●●●●●●●●●●●●●●●●●●●●●●●●●●●●●●●●●●●●●●●●●●●●●●●●●Y●●●●RR●●●●●●
```

```
env-488   CUAAAAAAAGAGAUGAAUCUUUAUAUUGAUUUGAUCCUCAAUCAACCCGUUAGAAAAUAACGAUUUAUAAAAAAUAAGU
env-489   CUAAAAAAAGAGAUGAAUCUUUAUAUUGAUUUGAUCCUCAAUCAACCCGUUAGAAAAUAACGAUUUAUAAAAAAUAAGU
env-490   CUAAAAAACAAGAUCAAUCUAUACAUUGACUUGAUUCUCAAUCAGCCGAUUAGAAACUAAGCAUUAAUCAAAAAUAAGU
env-491   CUAAAAAACAAGAUCAAUCUAUACAUUGACUUGAUUCUCAAUCAGCCGAUUAGAAACUAAGCAUUAAUCAAAAAUAAGU
env-492   CUAAAAAACAAGAUCAAUCUAUACAUUGACUUGAUUCUCAAUCAGCCGAUUAGAAACUAAGCAUUAAUCAAAAAUAAGU
env-494   CUAAAAAACAAGAUCAAUCUAUACAUUGACUUGAUUCUCAAUCAGCCGAUUAGAAACUAAGCAUUAAUCAAAAAUAAGU
env-496   CUAAAAAACAAGAUCAAUCUAUACAUUGACUUGAUUCUCAAUCAGCCGAUUAGAAACUAAGCAUUAAUCAAAAAUAAGU
env-498   CUAAAAAACAAGAUCAAUCUAUACAUUGACUUGAUUCUCAAUCAGCCGAUUAGAAACUAAGCAUUAAUCAAAAAUAAGU
env-500   CUAAAAAACAAGAUCAAUCUAUACAUUGACUUGAUUCUCAAUCAGCCGAUUAGAAACUAAGCAUUAAUCAAAAAUAAGU
env-501   CUAAAAAACAAGAUCAAUCUAUACAUUGAUUUGAUCCUCAAUCAGCCGAUUAGAAAGUAAGCAUUAACCAAAAAUAAGU
env-502   CUUAAAAACAAAAUCAAUUUGUACAUUGAUUUAAUUCUUAAUCAACCUAUAAGAAACUAAGAAUUAAUCAAAAAUAUAU
env-503   UUAAAGCACAUAUAUUUUACACAAAAUUGUUAUAAAAAAGCAAAUACACUAUAAUAUAUUGCCGAUAUCAAAAAUAUAA
env-504   UUUAUAAGCCACUGAUUAUUAAACACAAACAUUAAACUUGAAACUUGAAACUUUACGCUUGCACGAACAGUUUUUUUGU
env-505   UUUAUAAGCCACUGAUUAUUAAACACAAACAUUAAACUUGAAACUUGAAACUUUACGCUUGCACGAACAGUUUUUUUGU
env-506   UUGGCAACUUUGUAAAAGUUUAAUUUUAUGUUAAUUAAAAUGAAAAUCCAUAGUUUCCUUGCAGGAAUUUUGAAAAGAA
env-507   ...............................CAACUCUUUAAAAAUUAACUUGAAAGCAAGUUUGUGUGAGUUAUUCUA
env-508   AGAGGCCAAGGCAAAAAUUGCUCAAACGCUACAACUCUUUAAAAAUUAACUUGAAAGCAAGUUUGUGUGAGUUAUUCUA
env-509   AGAGGCCAAGGCAAAAAUUGCUCAAACGCUACAACUCUUUAAAAAUUAACUUGAAAGCAAGUUUGUGUGAGUUAUUCUA
env-510   AGAGGCCAAGGCAAAAAUUGCUCAAACGCUACAACUCUUUAAAAAUUAACUUGAAAGCAAGUUUGUGUGAGUUAUUCUA
env-511   AAGCCACUUUAAAAAGGUUUAAUUUCAUUCUUAAUUUCCAAAAGAAAUACUAUUUAAUGACAUUAAUUUUCAUUUUUUU
Fhi-1-1   GUAAAUUACUCAAAAAACAAAAAUUAAUAUACUAUUUUAAGAUUCUUUUAUUUUUGUUUGAUUCUUUUGUUUUUUUAAG
Fsp-9-1   AAAAAAAGAUUAUUUUCAGCCUAAAUUGUUUAAUCUCUAAAUUUUUCAUUCUUUAAAUUUAUUUUUCAAAAAAAAGUA
Asu-1-1   GAAAAAGUUGAAAAAUCGGCUUGAUUUUCUAAAAUUUAAAAAAAACUGGCUAUAGAAGGCCGUAUACUGAUUACUUUUA
Aca-1-1   GAGCAAGUUGAUAAAUCCGCAUAAGAUUUUAAAGUUUUAAAAUAACUGGAUCCUGAGCACUGUAUACUGAAUACUUUUU
env-513   UCUAUAACGCUGAAAUAUCUCAUUUCAAUAUAUUUGAAUUAAUUUUUAUUUGAAACACAGCUAUUUCAAAAUCGAUUUU
env-514   UCUAUAACGCUGAAAAAGAAGAAAUAAAUAUAAUUUGAAUUAUUUUUAUUCGAAACACAUCUAUUUCAAAACUGAUUUU
env-515   CUUAUAACGCUGAAAAGCAUCUAUUUGAUAUAUUUUGAAUUAAUUUUAUUCGAAACACUUCAAUAUCAAAAUCGAUUUU
env-517   CUUAUAACGCUGAAAAGCAUCUAUUUGAUAUAUUUUGAAUUAAUUUUAUUCGAAACACUUCAAUAUCAAAAUCGAUUUU
Avl-1-1   CUUAUAACGCUGAAAAGCAUCUAUUUGAUAUAUUUUGAAUUAAUUUUAUUCGAAACACUUCAAUAUCAAAAUCGAUUUU
env-518   CUUAUAACGCUGAAAAGCAUCUAUUUGAUAUAUUUUGAAUUAAUUUUAUUCGAAACACUUCAAUAUCAAAAUCGAUUUU
env-519   UUAUAACGCUGAAAAAGCAUCUAUUUGAUAUAUUUCGAAUUAAUUUUAUUCGAAACACUUCAAUAUCAAAAUCGAUUUU
env-520   GUCUAUAACGCUGAAAACCUCAUUUCAAUAUAUUUGAAUUAAUUUUUAUUUGAAACACAGCUAUUUCAAAAUCGAUUUU
env-521   CCUAUAACGCUGAAAAACAUCAUUUCAAUAUAUUUGAAUUAAUUUUUAUUUGAAAUACAGCUAUUUCAAAAUCGAUUUU
env-522   CCCAUAACGCUGAAAAACAUCAUUUCAAUAUAUUUGAAUUAAUUUUUAUUUGAAAUACAGCUAUUUCAAAAUCGAUUUU
env-523   CCUAUAACGCUGAAAAACAUCAUUUCAAUAUAUUUGAAUUAAUUUUUAUUUGAAAUACAGCUAUUUCAAAAUCGAUUUU
env-527   CCUAUAACGCUGAAAAACAUCAUUUCAAUAUAUUUGAAUUAAUUUUUAUUUGAAAUACAGCUAUUUCAAAAUCGAUUUU
env-528   CUAUAACGCUGAGAAUGGUAAAUUCAAUAUAUUUAAUGUAAAUUUUUAUUUGAAAUACAGCUAUUUCAAAAUCGAUUUU
env-531   CUAUAACGCUGAGAAUGGUAAAUUCAAUAUAUUUAAUGUAAAUUUUUAUUUGAAAUACAGCUAUUUCAAAAUCGAUUUU
env-532   CCUAUAACGCUAAAAAAUUAAGAAUUUAAUAUAUCUAAAUUAAUUUCCCUCAAAACACACCAAUUUCAAAUUCGAUUUU
env-533   UAUAACGCUGAAAAUUGCAAAUCCAAUAUACUUUAAAAUUAUUUUUUAUUCGAAACACAUUAAUUUCAGAUUCGAUUUU
env-534   CUCUAUAUCGAUUUGAUUCAAAAUCAACAGAUAAGAGAAUAAAAUAGGGAGGCAAUUAAGUCAUAAAUCAAUAUAAAUC
env-536   UCUUUAUAUAGAUUUGAUUCAAAAUCAACAGAUAAGAGAAUAAAAUAGGGAAGCAAUAAGUUAUAAAUCAAUAUAAAUC
env-537   UCUUUAUAUCGAUUUGAUUCAAAAUCAACAGAUAAGAGAAUAAAAUAGGGAGGCAAUAAGUUAUAAAUCAAUAUAAAUC
env-539   UCUUUAUAUCGAUUUGAUUCAAAAUCAACAGAUAAGAGAAUAAAAUAGGGAGGCAAUAAGUUAUAAAUCAAUAUAAAUC
env-540   ....UAUAUCGAUUUGAUUCAAAAUCAACAGAUAAGAGAAUAAAAUAGGGAGGCAAUAAGUUAUAAAUCAAUAUAAAUC
Apa-1-1   CUAGAGAGGCAAUAAUGUACUAGUAACACUUUGUUCAUUUCAAUUUAUGAAAUAAACCAUUGUAAUGUUUUAAAAAUUA
Ama-1-1   UACUAAGACCGUAUAUUAUGCAAAACGUUUAAUUGCAUUAUGUUUUUUCAUAUAAAAUGCAUUUCUAAUCCAAAAAAUA
Aat-1-1   UACUAAGACCAUAUAUUAUGCAAAACGUUUAAUUGCAAUAUGUUUUUUCAUAUAAAAUGCAUUUCUAAUCCAAAAAAUA
Ame-1-1   UACUAAGACUAUAUAUUAUGUAAAACGUUUAAUUGCAUUAUGUUUUUUCAUAUAAAAUGCGUUUCUAAUUCAAAAAAUA
Ala-1-1   CCUCUAGGACUUAAUUAUUUACAAAAGGUUUAAUCAAACAUGUUUUUUUUCAUAAGAAGCAAUUCUAAUAUAAAAAAUA
Amu-1-1   AAUAAAAAAGAAAUAUAUUUACAAUAAAAUAUAGCUUUUAUGUUUUUUUAGACAAAAUAUAAUUAUAAAGCAAAAAACA
Fsu-3-1   CUGGAGUUUGUAAAUCAGGAAGUUGCAAAAUUAUUUUAAGGAAGUUUCGGCAAAUAAUUUGUGAUUCUGAAAAAUAAAG
Fen-1-1   UUAGAAUUUGUAAAUCAGGAAGUUGCAAAACUUUUUUAAGCAUGUUUCGGCAAUUUGUUUGUGAGUUUAAAAAAUAAAA
Fco-1-1   UUUAGAAUUUGUAAAUUCAGAAGUUGCUAAAAUAUUUUAAAAAUUUUUAUAAUUCCUGUUUUAUAAUUCAUAAUUAUUU
env-541   UUACGAUAAAAAAAGAGAGUUUAGAAACCUGUGGUUAAAAUAAAAAACCAGUAUAAACUUGUAACUUCAAAAAUAAUUU
env-542   GAUUAUGACAAAAAGAAAUCUUUUAGAAAGUUGUGGUUAAAAUAAUAAGAGCUAAAAGUUGCAUCUUCAAAAAUAAUUU
env-543   UUACGAUAAAAAAAUGGAGUUUAGAAAAUUGUGGUUGAAAUAAAAAACAAGUAUAAACUUGCAACUUCGGAAAUUAAUU
env-545   GAUUAUGACAAAAAUAAAUUGUUUAGAAAUUUGUGGUUGAAAUAAAAAGAGCUAAAAGUUGCAACUUCAAAAUUAAUUU
env-546   AAACUUCAAAAUAGUUUAAUUAGACACCCGAAAAAGAUUAAAACAUUUUAGAGAUAGUAUUGUAAAUCAUUUAAAAUCA
Fps-1-1   UUUUAUUUUUGAUGGAAUGAUCAAAAACAUUGCCAAGGCUAAAUAAUAAGUAAAAACAUUGCCACAUCAAAAAUAAAGU
Jma-1-1   UGUUUAUAAUUAAUAAAUUAGCUAUAUUUUAGCCGUUAAAUAAUCUACCAUAAAAAACGCUAGAUAAUAAAAUUUAUUG
env-547   UCUAAGACUUUGUCUUUUUAAAAUGGUUUAAAUUGAUAAACCUUUUUUUUAUAAUUGAUUGCUCUUCAAAAUAAUAAUU
env-548   AAUUUGUACCUAGACUUAAUUCUUAACAAUCCAAUCAGGGAAUAAAUUAGACAUUUACAGCCUAGUAUAUCUUUUUUCA
env-549   AAUCUAUACCUAGACUUAAUUCUCAAUAAUCCAAUCAGGGAAUAAAUUAAACAUUUUCAACCUAGUAUAUCUUUUUUCA
env-550   AAUUUGUACCUAGACUUAAUUCUCAAUAAUCCAAUCAGGGAAUAAAUUAAACAUUUACAACCUAGUAUAUCUUUUUUCA
env-551   GGUAUCUAAGUUUACUUAAAAACAGCAAUUUAGACUAUAAAAUAGUAAGCUAUUAAAGUUUUUUAAUAAAAAAAAUAAUC
env-552   GGUAUCUAAGUUUACUUAAAAACAGCAAUUUAGACUAUAAAAUAGUAAGCUAUUAAAGUUUUUUAAUAAAAAAAAUAAUC
env-553   GGUAUCUAAGUUUACUUAAAAACAGCAAUUUAGACUAUAAAAUAGUAAGCUAUUAAAGUUUUUUAAUAAAAAAAAUAAUC
env-554   AAGAAAAAAUUUAAAAAAGCAAGUAAACUUAUACAUCGAUUUAAUACAGAAUAUACCUGUAAGAAAUCAAUAAAUAAUA
env-556   AAGAAAAAAUUUAAAAAAGCAAGUAAACUUAUACAUCGAUUUAAUACAGAAUAUACCUGUAAGAAAUCAAUAAAUAAUA
env-557   AAGAAAAAAUUUAAAAAAGCAAGUAAACUUAUACAUCGAUUUAAUACAGAAUAUACCUGUAAGAAAUCAAUAAAUAAUA
Ffi-1-1   AGAUGAACGCUUUUUAGAAUUUGUAACCGAAGAAGUAGCAAAGUUGUUUUAGAAAAACUUGUUCAUUCAAAAAAUAAAU
env-558   .....................CCAAUUUUUAUUUUUAUAAAAUCUUUUUAUUUUUAAACUCCUUUUUUAAAAUUAAAUA
env-559   UAAUAUUUUAAAAGGUUUAAUUAGGUGUUUAAAAAAAGCUAAAUAAUUCUCUUUUAGAUUUGUAGAAGUUUAAUUUUUU
env-561   UUAAAUACUCUUUUUUUUAUCUCAAAUUUAAAGAAGAUUUCUAGAUUGUAAAUAAUUCUGAUUUUAAUCCAAUUUUUCA
env-562   UUAAAUACCUUUUUUUUUAUCUCAAAUUUAAAGAAGAUUUCUAGAUUGUAAAUAAAUCUGAAUUUAAUCUAAUUUUUCA
env-563   UUUAAAAGUAACUAAAUUCUUUCUUUUUUCAAAAAAAUGUUUUAGGUUAAAAUUAAUCAUUCUUUAAUCUAAUUUUUCA
env-564   UUUAAAAGUAACUAAAUUCUUUCUUUUUUCAAAAAAAUGUUUUAGGUUAAAAUUAAUCAUUCUUUAAUCUAAUUUUUCA
env-565   UUUAAAAAUAACUAAAUUCUUUCUUUUUUCAAAAAAAUGUUUAGGUUAAAAAUUAAUCAUUAUUUAAUCUAAUUUUUCA
env-566   UAUACACUCUUGUUUUUAGUUACAAAUUUAAAGAAGAGUUCUAGAUUAUAAAUAAUUCUAAUUUUAAUCUAAUUUUUCA
env-567   UAUACACUCUUGUUUUUAGUUACAAAUUUAAAGAAGAGUUCUAGAUUAUAAAUAAUUCUAAUUUUAAUCUAAUUUUUCA
env-569   UAUACACUCUUGUUUUUAGUUACAAAUUUAAAGAAGAGUUCUAGAUUAUAAAUAAUUCUAAUUUUAAUCUAAUUUUUCA
env-571   UAUACACUCUUGUUUUUAGUUACAAAUUUAAAGAAGAGUUCUAGAUUAUAAAUAAUUCUAAUUUUAAUCUAAUUUUUCA
env-572   ...ACACUCUUGUUUUUAGUUACAAAUUUAAAGAAGAGUUCUAGAUUAUAAAUAAUUCUAAUUUUAAUCUAAUUUUUCA

          ...............................................................................
          ...............................................................................
          ●●●●●●●●●●●●●●●●●●●●●●●●●●●●●●●●●●●●●●●●●●●●●●●●●●●●●●●●●●●●●●●●●●Y●●●●RR●●●●●●
```

```
env-573   UAUACACUCUUGUUUUUAGUUACAAAUUUAAAGAAGAGUUCUAGAUUAUAAAUAAUUCUAAUUUUAAUCUAAUUUUUCA
env-574   CUUAAAAAUAAAUAAAUUCUUUCUUUUUUCAAAAAAAUGUUUUAGGUUAAAAAUUAAUCAUUCUUUAAUCCAAUUUUUCA
env-575   UUAAAAGUAANUAAAUUCUUUCUUUUUUCAAAAAAAUGUUUUAGGUUAAAAAUUAAUCAUUCUUUAAUCUAAUUUUUCA
env-576   CUUAAAAAUAAAUAAAUUAUUUCUUUUUUCAAAAAAAUGUUUUAGGUUAAAAAUUAAUCAUUCUUUAAUCCAAUUUUUCA
env-577   UUUAAAAGUAAAUAAAUUCUUUCUUUUUUCAAAAAAAUGUUUUGGUUAAAAAUUAAUCAUUCUUUAAUCUAAUUUUUCA
env-578   UAAAUACUCGUUUUUUUAAUUAUAAAUUUAAAGAAGAGUUCUUGAUUACAAAUAAUUCUUAUUUUAAUCUAAUUUUUCA
env-579   UAAAUACUCGUUUUUUUAAUUAUAAAUUUAAAGAAGAGUUCUUGAUUACAAAUAAUUCUUAUUUUAAUCUAAUUUUUCA
env-580   UAAAUACUCGUUUUUUUAAUUAUAAAUUUAAAGAAGAGUUCUUGAUUACAAAUAAUUCUUAUUUUAAUCUAAUUUUUCA
env-581   UUUAAAAGUAACUAAAUUCUUUCUUUUUUCAAAAAAAUGUUUUAGGUUAAAAAUUAAUCAUUCUUUAAUCUAAUUUUUCA
env-582   .AUACACUCUUGUUUUUAGUUACAAAUUUAAAGAAGAGUUCUAGAUUAUAAAUAAUUCUAAUUUUAAUCUAAUUUUUCA
env-583   ......................................................................AAUUUUUCA
env-584   CUUAAAAAUAAAUAAAUUAUUUCUUUUUUCAAAAAAAUGUUUUAGGUUAAAAAUUAAUCAUUCUUUAAUCCAAUUUUUCA
env-585   CUUAAAAAUAAAUAAAUUAUUUCUUUUUUCAAAAAAAUGUUUUAGGUUAAAANUNAUCAUUCUUUAAUCCAAUUUUUCA
env-586   CUUAAAAAUAAAUAAAUUAUUUCUUUUUUCAAAAAAAUGUUUUAGGUUAAAAAUUAAUCAUUCUUUAAUCCAAUUUUUCA
env-587   UUAAAUACUCUUUUUUUUAUCUCAAAUUUAAAGAAGAUUUCUAGAUUGUAAAUAAUUCUGAUUUUAAUCCAAUUUUUCA
env-588   UAUACACUCUUGUUUUUAGUUACAAAUUUAAAGAAGAGUUCUAGAUUAUAAAUAAUUCUAAUUUUAAUCUAAUUUUUCA
env-589   UUUAAAAAUAAAUAAAUUCUUUCUUUUUUCAAAAAAAUGUUUUAGGUUAAAAAUUAAUCAUUCUUUAAUCCAAUUUUUCA
env-590   UAAAUACUCGUUUUUUUAAUUAUAAAUUUAAAGAAGAGUUCUUGAUUACAAAUAAUUCUUAUUUUAAUCUAAUUUUUCA
env-591   UUUAAAAAUAACUAAAUUCUUUCUUUUUUCAAAAAAAUGUUUUAGGUUAAAAAUUAAUCAUUCUUUAAUCUAAUUUUUCA
env-592   UAAAUACUCGUUUUUUUAAUUAUAAAUUUAAAGAAGAGUUCUUGAUUACAAAUAAUUCUUAUUUUAAUCUAAUUUUUCA
env-593   UAAAUACUCGUUUUUUUAAUUAUAAAUUUAAAGAAGAGUUCUUGAUUACAAAUAAUUCUUAUUUUAAUCUAAUUUUUCA
env-594   UAAAUACUCGUUUUUUUAAUUAUAAAUUUAAAGAAGAGUUCUUGAUUACAAAUAAUUCUUAUUUUAAUCUAAUUUUUCA
env-596   UAAAUACUCGUUUUUUUAAUUAUAAAUUUAAAGAAGAGUUCUUGAUUACAAAUAAUUCUUAUUUUAAUCUAAUUUUUCA
env-598   UAAAUACUCGUUUUUUUAAUUAUAAAUUUAAAGAAGAGUUCUUGAUUACAAAUAAUUCUUAUUUUAAUCUAAUUUUUCA
env-599   UAAAUACUCGUUUUUUUAAUUAUAAAUUUAAAGAAGAGUUCUUGAUUACAAAUAAUUCUUAUUUUAAUCUAAUUUUUCA
env-600   UAAACACUCUUGUUUUUAGUUACAAAUUUAAAGAAGAGUUCUAGAUUAUAAAUAAUUCUGAUUUUAAUCUAAUUUUUCA
env-601   UAAAUUUGUAUUUGGAUUUAAUACUCAACAGACCUCUCAGAUCAAAUAAUAUUUAAUUACCUCUUUUCGAAAUUUAAUC
env-602   UAAAUUUAUAUUUUGAUUUAAUACUUAAUAGACCUCUAAGAUCAAAAAAUAUUUAAUUACCUCUUUUCGAAAUUUAAUC
env-603   UAAAUUUAUAUUUUGAUUUAAUACUUAAUAGACCUCUAAGAUCAAAUAAUAUUUAAUUACCUCUUUUCGAAAUUUAAUC
env-604   UAAAUUUAUAUUUUGAUUUAAUACUAAACAGACCUCUCAGAUCAAAUAAUAUUUAAUUACCUCUUUUCGAAAUUUAAUC
env-605   UAAAUUUGUAUUUGGAUUUAAUACUCAAUAGACCUCUGAGAUCAAAUAAUAUUUAAUUGCCUCUUUUCGAAAUUUAAUC
env-606   UCAAUCUUUACUUUGAUUUAGUUCUAAAUAGACCCGUUAGGCAAAUCAAUAAAUAGUUGCUUAUUUCUAAAAUUUAAUC
env-607   UCAAUCUUUACUUUGAUUUAGUUCUAAAUAGACCCGUUAGGCAAAUCAAUAAAUAGUUGCUUAUUUCUAAAAUUUAAUC
env-608   UAAAUUUAUAUUUUGAUUUAAUACUUAACAGACCUAUAAGAUCAAAUAAUAUUUAAUUACCUCUUUUCGAAAUUUAAUC
env-609   UAAAUUUAUAUUUUGAUUUAAUACUUAAUAGACCUAUUAGAAAGAACAAUAUUUAAUUCUAUCAUUUCAAAAUUUAAUC
env-610   UAAAUUUAUAUUUUGAUUUAAUACUUAAUAGACCUAUUAGAAAGAACAAUAUUUAAUUCUAUCAUUUCAAAAUUUAAUC
env-611   UAAAUUUAUAUUUUGAUUUAAUACUUAAUAGACCUAUUAGAAAGAACAAUAUUUAAUUCUAUCAUUUCAAAAUUUAAUC
env-612   UAAAUUUAUAUUUUGAUUUAAUACUUAAUAGACCUAUUAGAAAGAACAAUAUUUAAUUCUAUCAUUUCAAAAUUUAAUC
env-613   UAAAUUUAUAUUUUGAUUUAAUACUUAAUAGACCUAUUAGAAAGAACAGUAUUUAAUUCUAUCAUUUCAAAAUUUAAUC
env-614   UAAAUUUAUAUUUUGAUUUAAUACUUAAUAGACCUAUUAGAAAGAACAAUAUUUAAUUCUAUCAUUUCAAAAUUUAAUC
env-615   UAAAUUUAUAUUUUGAUUUAAUACUUAAUAGACCUAUUAGAAAGAACAGUAUUUAAUUCUAUCAUUUCAAAAUUUAAUC
env-616   UUAAUUUAUAUUUUGAUUUAAUCCUUAACAGACCUGUUAGAACCAAUACUAUUUAAUUGAAACAUUUCAAAAUUUAAUC
env-617   UAAAUUUAUAUUUUGAUUUAAUCCUUAAUAGACCCGUUAGAACCAACAUUAUUUAAUUCGAUCAUUUCAAAAUUUAAUC
env-618   UAAAUUUAUAUUUUGAUUUAAUCCUUAAUAGACCCGUUAGAACCAACAUUAUUUAAUUCGAUCAUUUCAAAAUUUAAUC
env-619   AAAUUUAUAUUUUGACUUAAUACUUAAUAGGCCCAUUCGUUCUAAUAAAAUUUAAUUACUCCUUUUAUAAAAUUUAAUA
env-620   UAAAUUUAUAUUUUGAUUUAAUAUUAAAUAGGCCUGUUAGGUCUUCAAAUAUUUAAUUACUUCUUUUCAAAAUUUAAUC
env-621   UAAAUUUAUAUUUUGAUUUAAUAUUAAAUAGGCCUGUUAGGUCUUCAAAUAUUUAAUUACUUCUUUUCAAAAUUUAAUC
env-622   UAAAUUUAUAUUUUGAUUUAAUAUUAAAUAGGCCUGUUAGGUCUUCAAAUAUUUAAUUACUUCUUUUCAAAAUUUAAUC
env-623   UAAAUUUAUAUUUUGAUUUAAUAUUAAAUAGGCCUGUUAGGUCUUCAAAUAUUUAAUUACUUCUUUUCAAAAUUUAAUC
env-624   UUAAUCUAUAUUUUGAUUUGAUACUUAACAACCCUGUUAGAACAAAUAAUUUAUAAUUGUUUCUUUUCAAAAUUUAAUC
env-625   UUAAUCUAUAUUUUGAUUUGAUACUUAACAACCCUGUUAGAACAAAUAAUUUAUAAUUGUUUCUUUUCAAAAUUUAAUC
env-626   UUAAUCUAUAUUUUGAUUUGAUACUUAACAACCCUGUUAGAACAAAUAAUUUAUAAUUGUUUCUUUUCAAAAUUUAAUC
env-627   UUAAUCUAUAUUUUGAUUUGAUACUUAACAACCCUGUUAGAACAAAUAAUUUAUAAUUGUUUCUUUUCAAAAUUUAAUC
env-628   UUAAUCUAUAUUUUGAUUUGAUACUUAACAACCCUGUUAGAACAAAUAAUUUAUAAUUGUUUCUUUUCAAAAUUUAAUC
env-629   UUAAUCUAUAUUUUGAUUUGAUACUUAACAACCCUGUUAGAACAAAUAAUUUAUAAUUGUUUCUUUUCAAAAUUUAAUC
env-633   UUAAUCUAUAUUUUGAUUUGAUACUUAACAACCCUGUUAGAACAAAUAAUUUAUAAUUGUUUCUUUUCAAAAUUUAAUC
env-634   UUAAUCUAUAUUUUGAUUUGAUACUUAACAACCCUGUUAGAACAAAUAAUUUAUAAUUGUUUCUUUUCAAAAUUUAAUC
env-635   UAAAUUUGUAUUUAGAUUUAAUACUUAACAGAGCUGUUAGACCAAAUAACAUUUAAUUACUUCUUUUCGAAAUUUAAUC
env-636   AAAUUUAUAUUUUGAUUUAAUACUUAAUAGGCCCAUUCGUUCUAAUAAAAUUUAAUUACCUCUUUUAUAAAAUUUAAUA
env-637   ..................................UUUAAGGUCUUCAAAUAUUUAAUUACUUCUUUUCAAAAUUUAAUC
env-638   UAAAUUUAUAUUUAGAUUUAAUCUUAAAUAGACCUGUUAGAUCAAUUAAUAUUUAAUUGCCUCUUUUCGAAAUUUAAUC
env-639   UAUAUUUUGAUUUAAUACUUAAUAGGCCCAUUCUGUUNGAUCAAUNAAUAUUUAAUUACCUCUUUUNNAAAAUUUAAUN
env-640   UAAAUUUAUAUUUUGAUUUAAUACUAAACAGACCUCUCAGAUCAAAUAAUAUUUAAUUACCUCUUUUCGAAAUUUAAUC
env-641   UAAAUCUUUACUUUGAUUUGGUUCUGAACAGACCCAUUAGAUCAAUAAAUAAAUAGUUGCGAGUUAAUAAAAUUUAAUC
env-648   UAAAUCUUUACUUUGAUUUNGUUCUGAANAGACCCNUUAGAUCAAUAAAUAAAUAGUUGCGAGUUAAUAAAAUUUAAUC
env-649   UAAAUCUUUACUUUGAUUUGGUUCUGAACAGACCCAUUAGAUCAAUAAAUAAAUAGUUGCGAGUUAAUAAAAUUUAAUC
env-650   UAAAUCUUUACUUUGAUUUNGUUCUGAACAGACCCNUUAGAUCAAUAAAUAAAUAGUUGCGAGUUAAUAAAAUUUAAUC
env-651   UAAAUCUUUACUUUGAUUUGGUUCUGAACAGACCCAUUAGAUCAAUAAAUAAAUAGUUGCGAGUUAAUAAAAUUUAAUC
env-652   UAAAUCUUUACUUUGAUUUGGUUCUGAACAGACCCAUUAGAUCAAUAAAUAAAUAGUUGCGAGUUAAUAAAAUUUAAUC
env-653   UAAAUCUUUACUUUGAUUUGGUUCUGAACAGACCCAUUAGAUCAAUAAAUAAAUAGUUGCGAGUUAAUAAAAUUUAAUC
env-655   UAAAUCUUUACUUUGAUUUNGUUCUGAANAGACCCNUUAGAUCAAUAAAUAAAUAGUUGCGAGUUAAUAAAAUUUAAUC
env-656   ......................UCUGAACAGACCCAUUAGAUCAAUAAAUAAAUAGUUGCGAGUUAAUAAAAUUUAAUC
env-657   UAAAUCUUUACUUUGAUUUGGUUCUGAACAGACCCAUUAGAUCAAUAAAUAAAUAGUUGCGAGUUAAUAAAAUUUAAUC
env-658   UAAAUCUUUACUUUGAUUUAGUUCUGAANAGACCCCUUAGAUCAAUAAAUAAAUAGUUGCGAGUUAAUAAAAUUUAAUC
env-659   UAAAUCUUUACUUUGAUUUAGUUCUGAACAGACCCAUUAGAUCAAUAAAUAAAUAGUUGCGAGUUAAUAAAAUUUAAUC
env-660   AAAUCUUUACUUUGAUUUAAUCCUAAAUAGAGCUGUUAGGCAAUUGAACAAAUAGUUGCUAAAUUCUAAAAAUUUAAUC
env-661   UCAAUCUUUACUUUGAUUUAGUUCUAAACAGACCCAUUAGGCAAAUCAAUAAAUAGUUGCUAAUUUCUAAAAUAUAAUC
env-662   UCAAUCUUUACUUUGAUUUAGUUCUAAACAGACCCGUUAGGCAAAUCAAUAAAUAGUUGCUAAUUUCUAAAAUAUAAUC
env-663   UUAAUUUGUAUUUGGACUUGAUUCUUAACAGACCUGUGAGAUUCAAAAAUAAUUAAUUAUCUCUUUUCAAAAUUUAAUC
env-664   UAGAACGCAAAAAAAUGUAAGUCAGUAUAAUUUAUUGUUAAUUUAGUGUAAUUAAACAUCAAUAUGUUGUUAAGUAUUA
env-669   UAGAACGCAAAAAAGUUUAACCCAGUAUAAUUAAUUGUUAAUUAGGUCUAUCUAAUAUGUCAAAUAUGAAUCGAAAUUU
env-670   ...AACGCAAAAAAGUUUAACCCAGUAUAAUUAAUUGUUAAUUAGGUCUAUCUAAUAUGUCAAAUAUGAAUCGAAAUUU
env-671   UAGAACGCAAAAAAGUUUAACCCAGUAUAAUUAAUUGUUAAUUAGGUCUAUCUAAUAUGUCAAAUAUGAAUCGAAAUUU
env-672   AUUUUUUUCAUAAAAUGUUCUUUGGAUUUAACAAGCAGGUGAUUUCUUGGUGAUAAUCUUGUUUAUUGAAAAACUAUCC

          ...............................................................................
          ...............................................................................
          ●●●●●●●●●●●●●●●●●●●●●●●●●●●●●●●●●●●●●●●●●●●●●●●●●●●●●●●●●●●●●●●●●●Y●●●●RR●●●●●●
```

```
env-673   UUUUUUUUGUGUGUAUUUUGCUGGAGUAUGACUGAUUUCUUAUUUAUUCGGGAUAAUAUUGUUUAUUGAAAAACUAUCC
env-674   UUUUUUUUGUGUGUAUUUUGCUGGAGUAUGACUGAUUUCUUAUUUAUUCGGGAUAAUAUUGUUUAUUGAAAAACUAUCC
env-675   UUUUUUGCUUGAAUAAGUCCUUAGGAUGUAAUGAGUAGGUUAUUUCUUGGUGAUUAUCUUGUUUAUUGAAAAACUAUCC
env-676   CUUUUUUUUUUGUGGCUUCGGGUUAGGAUGAAGAGAAUGUUAUUUCUAUGUGAUAAUCUUGUUUAUUGAAAAACUAUCC
env-677   UUUUUUUUAUUCAGAAAUCAGGUGCGUGUGAAGAGAAUGUUAUUUCUAGGGGAUAAUCUUGUUUAUUGAAAAACUAUCC
env-678   ...............................................AUGUGAUAAUCUUGUUUAUUGAAAAACUAUCC
env-679   UUUUUUUUGCUGUGGCGUCAGGUUAGGAUGAAGAGAAUGUUAUUUCUAUGUGAUAAUCUUGUUUAUUGAAAAACUAUCC
env-680   UUUUUUUUGCUGUGGCGUCAGGUUAGGAUGAAGAGAAUGUUAUUUCUAUGUGAUAAUCUUGUUUAUUGAAAAACUAUCC
env-681   UUAUUUUUUGAUGAAGUAUCCUUUGUAUUUAUAAUCAGGUUAUUUCUUGGUGAUAAUCUUGUUUAUUGGAAAACUAUCC
env-682   AUUUUUUUUGAUGAAGUAUCCUUUGUAUUUAUAAUUAGGUUAUUUCUAGGUGAUAAUCUUGUUUAUUGGAAAACUAUCC
env-683   UUAUUUUUUGAUGAAGUAUCCUUUGUAUUUAUAAUCAGGUUAUUUCUUGGUGAUAAUCUUGUUUAUUGGAAAACUAUCC
env-684   AUUUUUUUUGAUGAAGUAUCCUUUGUAUUUAUAAUUAGGUUAUUUCUAGGUGAUAAUCUUGUUUAUUGGAAAACUAUCC
env-685   UAUUUUUUUGAUGAAGUAUCCUUUGUAUUUACAAUCAGGUUAUUUCUUGGUGAUAAUCUUGUUUAUUGGAAAACUAUCC
env-686   UUAUUUUUUGAUGAAGUAUCCUUUGUAUUUAUAAUCAGGUUAUUUCUUGGUGAUAAUCUUGUUUAUUGGAAAACUAUCC
env-687   AUUUUUUUUGAUGAAGUAUCCUUUGUAUUUAUAAUUAGGUUAUUUCUAGGUGAUAAUCUUGUUUAUUGGAAAACUAUCC
env-688   UAUUUUUUUGAUGAAGUAUCCUUUGUAUUUACAAUCAGGUUAUUUCUUGGUGAUAAUCUUGUUUAUUGGAAAACUAUCC
env-689   UAUUUUUUUGAUGAAGUAUCCUUUGAAUUUAUAAUCAGGUUAUUUCUUGGUGAUAAUCUUGUUUAUUGGAAAACUAUCC
env-690   AUUUUUUUUGAUGAAGUAUCCUUUGUAUUUAUAAUUAGGUUAUUUCUAGGUGAUAAUCUUGUUUAUUGGAAAACUAUCC
env-691   NNUUUUUUUGAUGAAGUAUCCUUUGUAUUUAUAAUUAGGUUAUUUCUAGGUGAUAAUCUUGUUUAUUGGAAAACUAUCC
env-692   UUUUUUGCUUGAAUAAGUCCUUAGGAUGUAAUGAGUAGGUUAUUUCUUGGUGAUUAUCUUGUUUAUUGAAAAACUAUCC
env-693   ..................................................UGAUUAUCUUGUUUAUUGAAAAACUAUCC
env-694   UGAUUUUAAAUCAUCCUCUUCGACAAUAAACUUAUAACUACUUAAUUAUAUUUUUAUAUUGUCUAACGUUUGUUUAAUU
env-698   UGAUUUUAAAUCAUCCUCUUCGACAAUAAACUUAUAACUACUUAAUUAUAUUUUUGUAUUGUCUAACGUUUGUUUAAUU
env-699   UGAUUUUAAAUCAUCCUCUUCGACAAUAAACUUAUAACUACUUAAUUAUAUUUUUGUAUUGUCUAACGUUUGUUUAAUU
env-700   UGAUUUUAAAUCAUCCUCUUCGACAAUAAACUUAUAAUCACUUAAUUAUAUUUUUAUAUUGUCUAACGUUUGUUUAAUU
env-701   ...................................................UUUUAUAUUGUCUAACGUUUGUUUAAUU
env-702   UGAUUUUAAAUCAUCCUCUUCGACAAUAAACUUAUAACAACUUAAUUAUAUUUUUAUAUUGUCUAACGUUUGUUUAAUU
env-703   UCAGCCUUUGAGAAAUUAGGCUGUCGGUGAGACACCCAUUUUAUUCAAUCCCAAUUGGUUGUUUAAUCCUUCCGAUGUU
env-704   UCAGCCUUUGAGAAAUUAGGCUGUCGGUGAGACACCCAUUUUAUUCAAUCCCAAUUGGUUGUUUAAUCCUUCCGAUGUU
env-705   CAACCUUUGAGAAACUGAGUCGUUGGUUAAACAUCUAUUUUUAUUCAAUCGCUAUUGAUUGUUUAAUCCUUUCGAAGUU
env-706   CAACCUUUGAGAAACUGAGUCGUUGGUUAUACAUCUAUUUUUAUUCAAUCGCUAUUGAUUGUUUAAUCCUUUCGAAGUU
env-707   AAUCAACCCUUACGCAACUAAGUUGCUGAUUUUCAUAACAUAAUUAAAUUUAAAAAUGUUGUAUAAUACAGCCGAUGUU
env-708   AAUCAACCCUUAAGGAAGUAAGUUGCUGAUUUUCAAAAUUUAAAUAAAAUUAAAAGUAUUGUAUAAUACGCCUUAAGUU
env-709   CCUCUCCGAGUGAGCAAAAAUAUAAAAAGAGUGAAUUUUUUGUAAAAUCACUCUUAUUUUCUUUUCAUCAAAAAAAGUU
env-711   GGUUUUUUAGGUUUUUAUAAAUCUUAAAAACUAUUUGGCAAUAUUCAAGAUUAAUGCCUUGUUAAUUAAAAUAAAGUUC
env-712   AAUCAGCCGAUAAGAAAAUAAUCUCAAAACUUAUUUAAUUUAAUUUGAGUCUAAUUUCUUUGUUUGCAAGGGUAAUAUU
Fba-2-1   AAUCAACCAAUUAGAAAGUGAUUUCACCAAAACACCUAUUUAAUUUAAAAUAAAUUUGUUGGUUUAAAUAGGUUAUAUU
env-713   AAUCAACCAAUUAGAAAGUGAUUUCACCAAAACACCUAUUUAAUUUAAAAUAAAUUUGUUGGUUUAAAUAGGUUAUAUU
env-718   AAUCAACCGAUUAGAAAGUGAUUUCACCAAAACAACUAUUUAAUUUAAAAUAAAUUUGUUGGUUUAAAUAGGUUAUAUU
env-727   AAUCAACCAAUUAGAAAGUGAUUUCACCAAAACACCUAUUUAAUUUAAAAUAAAUUUGUUGGUUUAAAUAGGUUAUAUU
env-728   AAUCAACCAAUUAGAAAGUGAUUUCACCAAAACACCUAUUUAAUUUAAAAUAAAUUUGUUGGUUUAAAUAGGUUAUAUU
env-729   ........AAUUAGAAAGUGAUUUCACCAAAACACCUAUUUAAUUUAAAAUAAAUUUGUUGGUUUAAAUAGGUUAUAUU
env-730   ........................................UAAUUUGGGUCUAAUUUGUUUGUUUACAAGGGUAAUAUU
env-731   AUUUAAUUCUAAAUCAGCCUGUUCGAAAAUAAUUUCAUCCUUAUUUGAUUGAAAUUAAUUGGUUUACAUUUCUUUAAUU
env-734   AUUUAAUUCUAAAUCAGCCUGUUCGAAAAUAAUUUCAUCCUUAUUUGAUUGAAAUUAAUUGGUUUACAUUUCUUUAAUU
env-735   AUUUAAUUCUAAAUCAGCCUGUUCGAAAAUAAUUUCAUCCUUAUUUGAUUGAAAUUAAUUGGUUUACAUUUCUUUAAUU
env-736   AUUUAAUUCUAAAUCAGCCUGUUCGAAAAUAAUUUCAUCCUUAUUUGAUUGAAAUUAAUUGGUUUACAUUUCUUUAAUU
env-737   AUUUAAUUCUAAAUCAGCCUGUUCGAAAAUAAUUUCAUCCUUAUUUGAUUGAAAUUAAUUGGUUUACAUUUCUUUAAUU
env-738   AUUUGAUCUUAAAUCAACCCAUUAGAGAUUAAUACUCUCCUUAAUUUUGUUGAAAUUAUUGUUUUACAAUCCUUUAAUU
env-739   AUUUAAUUCUAAAUCAGCCUGUUCGAAAAUAAUUUCAUCCUUAUUUGAUUGAAAUUAAUUGGUUUACAUUUCUUUAAUU
env-740   ............................................AUUAUAUUUUUUUAUUGUCUAGCGUUUGUUUAAUU
env-741   AUCUAUACUUGGAUUUGAUACUUAACCAACCUCUAAGAGAUUAAUUUUACCAAUAAGAUUGUAAUUCUGAUUAUUACAA
env-742   AUCUAUACUUGGAUUUGAUACUUAAUCAACCUCUAAGAGAUUAAUUAUACCAAUAAGAUUGUAAUUCUGAUUAUUACAA
env-743   AUUUGUAUUUGGAUUUAAUCAUGAACCAGCCCUUGAGAGAUUAAUUAUGCCAUUUCAAUUGUAGAUUUAAUAUUAACAA
env-744   AUUUGUAUUUGGAUUUAAUCAUGAACCAGCCCUUGAGAGAUUAAUUAUGCCAUUUCAAUUGUAGAUUUAAUAUUAACAA
env-745   AUUUGUAUUUGGAUUUAAUCAUGAACCAGCCCUUGAGAGAUUAAUUAUGCCAUUUCAAUUGUAGAUUUAAUAUUAACAA
env-746   AUUUGUAUUUGGAUUUAAUCAUGAACCAGCCCUUUAGAGAUUAAUUAUGCCAUUUCAAUUGUAGAUUUAAUAUUAACAA
env-747   AUUUGUAUUUGGAUUUAAUCAUGAACCAGCCCUUGAGAGAUUAAUUAUGCCAUUUCAAUUGUAGAUUUAAUAUUAACAA
env-748   AUCUAUACCUUGACUUGAUCCUCAACCAACCCUUGAGAGAUUAAUUAUUACAAUGUAGUUGUAAUUAUGAUUAUUACAU
env-749   AUCUAUACCUUGACUUGAUCCUCAACCAACCCCUGAGAGAUUAAUUAUUAUGAUCAAGUUGUAAUUAUGAAUAUUACAU
env-750   AUCUAUACCUUGACUUGAUCCUCAACCAACCCUUGAGAGAUUAAUUAUUAUGAUCAAGUUGUAAUUAUGAAUAUUACAU
env-751   AUCUGUACCUUGACUUGAUCCUCAACGAACCCUUGAGAGAUUAAUUAUUACGAUGUAGUUGUAAUUAUGAUUAUUACAU
env-752   AUCUGUACCUUGACUUGAUCCUCAACGAACCCUUGAGAGAUUAAUUAUUACGAUGUAGUUGUAAUUAUGAUUAUUACAU
env-753   ACCUUUAUUUUGAUUUGAUCCUCAAUCAACCCCUAAGGGAUUAAUUAGUACAAUAGGGUUGUAUUUCUGAUUAUUACAA
env-754   ACCUUUAUUUUGAUUUGAUCCUCAAUCAACCCCUAAGGGAUUAAUUAGUACAAUAGGGUUGUAUUUCUGAUUAUUACAA
env-755   AUCUAUACUUGGAUUUGAUACUUAACCAACCUCUUAGAGAUUAAUUAUACCAACAAGAUUGUAAUUCUGAUUAUUAGAA
env-756   AUCUUUACUUGGACUUGAUCCUUAACCAACCUCUUAGGGAUUAAUUAGAUUAUUAGGGUUGUAAUUCUAAUUAUUACAU
env-758   ACCUUUACUUAGACUUGAUCCUCAACCAACCCUUGAGAGAUUAAUUAGUGUAAUAGGGUUGUAAUUCUGAUUAUUACAA
env-759   AUUUGUAUUUGGAUUUAAUCAUGAACCAUCGCUUAAGAGAUUAAUUAUGCCAUUUCAAUUGUAGAUUUAAUAUUAACAA
env-760   AUUUGUAUUUGGAUUUAAUCAUGAACCAUCGCUUAAGAGAUUAAUUAUGCCAUUUCAAUUGUAGAUUUAAUAUUAACAA
env-761   AACCUAUAUCUUGACCUAAUUUCAAAUCAACCUGUCAGGGAAUAAUUUAUGAAUUGAGUUGUAAAUAAAAGAAUAACAU
env-762   AACCUAUAUCUUGACCUAAUUUCAAAUCAACCUGUCAGGGAAUAAUUUAUGAAUUGAGUUGUAAAUAAAAGAAUAACAU
env-763   AACUUAUAUAUUGACCUAAUUUCAAAUCAACCCGUAAGGGAAUAAUUUAUGAAUUGAGUUGUAAAUAAAAGAAUAACAU
env-764   AACUUAUAUAUUGACCUAAUUUCAAAUCAACCCGUAAGGGAAUAAUUUAUGAAUUGAGUUGUAAAUAAAAGAAUAACAU
env-765   AACUUAUAUAUUGACCUAAUUUCAAAUCAACCCAUAAGGGAAUAAUUUAUGAAUUGAGUUGUAAAUAAAAGAAUAACAU
env-766   AACUUAUAUAUUGACCUAAUUUCAAAUCAACCCGUAAGGGAAUAAUUUAUGAAUUGAGUUGUAAAUAAAAGAAUAACAU
env-767   AACUUAUAUAUUGACCUAAUUUCAAAUCAACCCGUGAGGGAAUAAUUUAUGAAUUGAGUUGUAAAUAAAAGAAUAACAU
env-768   AACUUAUAUAUUGACCUAAUUUCAAAUCAACCCGUGAGGGAAUAAUUUAUGAAUUGAGUUGUAAAUAAAAGAAUAACAU
env-769   AACUUAUAUAUUGACCUAAUUUCAAAUCGACCCGUAAGGGAAUAAUUUAUGAAUUGAGUUGUAAAUAAAAGAAUAACAU
env-770   .....AUAUAUUGACCUAAUUUCAAAUCAACCCGUAAGGGAAUAAUUUAUGAAUUGAGUUGUAAAUAAAAGAAUAACAU
env-771   AACUUAUAUAUUGACCUAAUUUCAAAUCAACCCGUGAGGGAAUAAUUUAUGAAUUGAGUUGUAAAUAAAAGAAUAACAU
env-772   AACUUAUAUAUUGACCUAAUUUCAAAUCAACCCGUGAGGGAAUAAUUUAUGAAUUGAGUUGUAAAUAAAAGAAUAACAU
env-773   AACUUAUAUAUUGACCUAAUUUCAAAUCAACCCGUAAGGGAAUAAUUUAUGAAUUAAGUUGUAAAUAAAAGAAUAACAU
env-774   AACUUAUAUAUUGACCUAAUUUCAAAUCAACCCGUAAGGGAAUAAUUUAUGAAUUGAGUUGUAAAUAAAAGAAUAACAU

          ...............................................................................
          ...............................................................................
          ••••••••••••••••••••••••••••••••••••••••••••••••••••••••••••••••••Y••••RR••••••
```

```
env-775   AACUUAUAUAUUGACCUAAUUUCAAAUCAACCCGUAAGGGAAUAAUUUAUGAAUUGAGUUGUAAAUAAAAGAAUAACAU
env-776   AACUUAUAUAUUGACCUAAUUUCAAAUCAACCCGUGAGGGAAUAAUUUAUGAAUUGAGUUGUAAAUAAAAGAAUAACAU
env-777   AACCUAUAUCUUGAUUUAAUUCUAAAUCAUCCCUUGAGGGAUUAAUUUACAAAUAGAGUUGUAGAUCAAAGAAUUACAU
env-778   AACCUAUAUCUUGAUUUAAUUCUAAAUCAUCCCUUGAGGGAUUAAUUUACAAAUAGAGUUGUAGAUCAAAGAAUUACAU
env-779   AACCUAUAUCUUGAUUUAAUUCUAAAUCAACCCUUAAGGGAAUAAUUUACAAAUAGAGUUGUAGAUCAAAGAAUUACAU
env-780   AACCUAUAUCUUGAUUUAAUUCUAAAUCAUCCCUUGAGGGAUUAAUUUACAAAUAGAGUUGUAGAUCAAAGAAUUACAU
env-781   AACCUAUAUCUUGAUUUAAUUCUAAAUCAUCCCUUGAGGGAUUAAUUUACAAAUAGAGUUGUAGAUCAAAGAAUUACAU
env-783   AACCUAUAUCUUGAUUUAAUUCUAAAUCAACCCUUAAGGGAAUAAUUUACAAAUAGAGUUGUAGAUCAAAGAAUUACAU
env-784   AACCUAUAUCUCGACUUAAUACUAAAUCAACCCUUGAGGGAUUAAUUUACAAAUAGAGUUGUAGAUCAAAGAAUUACAU
env-785   AACCUAUAUCUCGACUUAAUACUAAAUCAACCCUUAAGGGAUUAAUUUACAAAUAGAGUUGUAGAUCAAAGAAUUACAU
env-786   AACCUAUAUCUAGACUUAAUUCUAAAUCAACCCUUGAGGGAUUAAUUUACAAAAAGGGUUAUAGAUCAAAGAAUUACAU
env-787   AACCUAUAUCUAGACUUAAUUCUAAAUCAACCCUUGAGGGAUUAAUUUACAAAAAGGGUUAUAGAUCAAAGAAUUACAU
env-788   ........UCUAGACUUAAUUCUAAAUCAACCCUUGAGGGAUUAAUUUACAAAAAGGGUUAUAGAUCAAAGAAUUACAU
env-789   CAAAGAUAACUUAUCAGAAACGAAUAUUUCACUGAAACAAAUAAAAAUUAGCAAGCCCUAAAAAUUAGAAAAUAAAAUA
env-791   AAGAAAAAAUUUAAAAAAGCAAGUGAAUUUAUACAUAGAUUUAAUCCAGAAUUUACCUGUAAGAAAUCAAUAAAUAAUA
env-794   AGAAAAAAUUUAAAAAAGCAAGUGAAUUUAUACAUAGAUUUAAUCCAGAAUUUACCGGGGGGGAAGUCAAUAAAUAAUA
env-795   AAGAAAAAAUUUAAAAAAGCAAGUGAAUUUAUACAUAGAUUUAAUCCAGAAUUUACCUGUAAGAAAUCAAUAAAUAAUA
env-796   CGAAAGAAAAAUUUAAAAAAGCAAGUGAAUUUAUACAUAGAUUAAUCCAGAAUUUACUGUAAGAAAUCAAUAAAUAAUA
env-797   AAGAAAAAAUUUAAAAAAGCAAGUGAAUUUAUACAUAGAUUUAAUCCAGAAUUUACCUGUAAGAAAUCAAUAAAUAAUA
env-798   AAGAAAAAAUUUAAAAAAGCAAGUGAAUUUAUACAUAGAUUUAAUCCAGAAUUUACCUGUAAGAAAUCAAUAAAUAAUA
env-799   GCUUAAAAUUGAACAAGUAGAGGCGUAAUUCACAUUCUCAAUAAUUUUAAAUAAAAAAUGCGUAGAUAGAAAAUUAGUA
Cca-4-1   ACUGUUAUGCAUGGCUUCUCCUACAAUUCAGAACGAUGAAAAAUAAAAUAUAAAAAAUUUGGAGGUUACAUAUUUUUAU
Cca-4-2   ACUGUUAUGCAUGGCUUCUCCUACAAUUGAGAACGAUGAAAAAUAAAUUAUAAAAAAUUUGGAGGUUACAUAUUUUUAU
Cca-4-3   ACUGUUAUGCAUGGCUUCUCCUACAAUUCAGAACGAUGAAAAAUAAAUUAUAAAAAAUUUGGAGGUUACAUAUUUUUAU
env-800   CAGAAACCUGAAAGAAAACCUCGCCAAUAUUUUAAGAAUGUUGAAAAACUAGGUUUUAUUGGUUUUGAGAAAAAUAGAA
env-801   GAAAAGAAUUAAAAAACAAGCUUAAUCUCUACAUCGAUUUAAUUCAAAAUAAGCCUGUGCGUAAUUAGGAUAUUUUAUU
env-802   CUCUACAUCGAUUUGAUUUUGAAUCUUCCGCUUCGAGAAGAAAAACAAAAGGAAGAUCUAGAAUAACUGAGAAUAAAAC
env-803   UUCUAAGACUUUGUCUUUAUAAAAUGGUUUAAUUUCUAAACCUUUUUUUUAUAAUUGAUUGCGCUUCAAAAUAAUAAUU
env-805   AGAAAAUCAAUUUAUAUUUUGACCUUAUUUUAAAUAGACCAUUAAGAUCUAUAUAACAUAGUAUUUUAGAAAAAUAAUC
env-806   AGAAAAUCAAUUUAUAUUUUGACCUUAUUUUAAAUAGACCAUUAAGAUCUAUAUAACAUAGUAUUUUAGAAAAAUAAUC
env-807   AUAAAAUCAACUUAUAUUUUGAUUUAGUUCUAAACAGACCUUUAAGAUCAUAUUAAAGGUAUUUUUUUCAAAAUUAAUA
env-808   AUAAAAUCAACUUAUAUUUUGAUUUAGUUCUAAACAGACCUUUAAGAUCAUAUUAAAAGUAUUUUUUUCAAAAUUAAUA
env-809   AUAAAAUCAACUUAUAUUUUGAUUUAGUUCUAAACAGACCUUUAAGAUCAUAUUAAAAGUAUUUUUUUCAAAAUUAAUA
Zga-1-1   AAAGCAGACCCGCUUCCAUCAGGAUGCGAAUUUUAGUCCACAUAUUAAGAGGGCUCCCUUUAAAAUAUCAAUUUUAAUU
env-810   CUGGAGUUUGUAAAUCAGGAAGUUGCAAAACUUUUUUAACGAUGUUUGGUCAUUUUAUUUGUCAAUUAAAAAAAUAAAU
env-813   UGAUUAUGAUAAAAAUAAAGCUUUCAGAGAUUUAUGGUUAUAAUAAAGUAUAAAAUUCUUGUAGUUUAAAAAAUAAAUA
env-814   .....................CCUUUAGAAAUUUAUGGUUAGAAUAAUAUAUAAAGUAUUGCAUUUUAAAAAAUAAAUA
env-815   AUGAUUAUGACAAAAACAAGGCCUUUAGAAAUUUAUGGUUAGAAUAAUAUAUAAAGUAUUGCAUUUUAAAAAAUAAAUA
env-816   AUGAUUAUGACAAAAACAAGGCCUUUAGAAAUUUAUGGUUAGAAUAAUAUAUAAAGUAUUGCAUUUUAAAAAAUAAAUA
env-817   AAAAAUAUAUCCCCAAUUAUUAUUUCUUUAGCAGUUUUCAAAAUAAAGAAAUGUUUAUUUGGUAAUUAAAAAAUAAGUA
env-818   GAAUUUUAGCAACUGAUUAUCAGAUAAUUAUAAAUUAGCAAUAGAGAUUGAUUUUUAUUUGGUAAUUAAAAAAUAAGUA
env-819   AAAAAUAUAUCCCCAAUUAUUAUUUCUUUAGCAGUUUUCAAAAUAAAGAAAUGUUUAUUUGGUAAUUAAAAAAUAAGUA
env-820   AUGAUUAUGAUAAAAACAAGGCCUUUAGAAAUUUAUGGUUAGAAUAAUAAAUAAAGUAUUGCAUUUUAAAAAAUAAAUA
env-821   GAUUUUUAACAAUUGACUAUCAGAUAAUUAUAAAUUGGCAUUAGAGAUUGAUUUUUAUUUGGUAAUUAAAAAAUAAGUA
env-822   AUGAUUAUGAUAAAAACAAGGCCUUUAGAAAUUUAUGGUUAGAAUAAUAUAUAAAGUAUUGCAUUUUAAAAAAUAAAUA
env-823   AUGAUUAUGAUAAAAACAAGGCCUUUAGAAAUUUAUGGUUAGAAUAAUAUAUAAAGUAUUGCAUUUUAAAAAAUAAAUA
env-824   GAGAUUUAAAGAAAAAAAUAAAUUUAUUUAUUGAUCUAAUUUCUAAUAGACCUGUAAGAGAAUAAUAAUCUAUUUUUUU
env-825   .............................................................UAUUUACCAAAUUUAAUC
env-826   AAAAUUAAUCUUUAUUUUGAUUUAAUUUUAAACAGACCAUUAAGGUCAAUUUAAAAUUACAUAUUUCUAAAAUUUAAUC
env-827   AAAAUUAAUCUUUAUUUUGAUUUAAUUCUUAAUAGACCUUUAAGGUCAAUUUAAAAUUAUAUAUUUCUAAAAUUUAAUC
env-828   AAAAUUAAUCUUUAUUUUGAUUUGAUCCUCAAUAGACCAUUAAGGUCUAUUUAAAAUUACAUAUUUACCAAAUUUAAUC
env-829   AAUUAAAUCUUUAUUUUUGAUUUGAUCCUCAAUAGACCAUUAAGGUCUAUUUAAAAUUACAUAUUUACCAAAUUUAAUC
env-830   AAAUUAAUCUUUAUUUUGAUUUGAUCCCUUAAUAGACCAUUAAGGUCAAUUUAAAAUUACAUAUAUACCAAAUUUAAUC
env-831   AAAAUUAAUCUUUAUUUUGAUUUGAUCCUUAAUAGACCAUUAAGGUCAAUUUAAAAUUACAUAUAUACCAAAUUUAAUC
env-832   AAAAUUAAUCUUUAUUUUGAUUUGAUCCUUAAUAGACCAUUAAGGUCUAUUUAAAAUUACAUAUUUACCAAAUUUAAUC
env-833   AAAAUUAAUCUUUAUUUUGAUUUGAUCCUUAAUAGACCAUUAAGGUCUAUUUAAAAUUACAUAUUUACCAAAUUUAAUC
env-834   AAAAUUAAUCUUUAUUUUGAUUUGAUCCUUAAUAGACCAUUAAGGUCUAUUUAAAAUUACAUAUUUACCAAAUUUAAUC
env-835   AAAAUUAAUCUUUAUUUUGAUUUGAUCCUUAAUAGACCAUUAAGGUCUAUUUAAAAUUACAUAUUUACCAAAUUUAAUC
env-836   AAAAUCAAUUUAUAUUUUGAUUUAAUUCUUAACAGACCAUUAAGGUCAAUUUAAAAUUACUUAUUUCUCAAAUAUAAUC
env-837   AAAAUCAAUUUAUAUUUUGAUUUAAUUCUUAACAGACCAUUAAGGUCAAUUUAAAAUUACUUAUUUCUCAAAUAUAAUC
env-838   AAAAUCAAUUUAUAUUUUGAUUUAAUUCUUAACAGACCAUUAAGGUCAAUUUAAAAUUACUUAUUUCUCAAAUAUAAUC
env-847   AAAAUAAAUUUAUAUUUUGAUUUAAUUCUUAACAGACCAUUAAGGUCAAUUUAAAAUUACUUAUUUAUCAAAUAUAAUC
env-848   AAAAUAAAUUUAUAUUUUGAUUUAAUUCUUAACAGACCAUUAAGGUCAAUUUAAAAUUACUUAUUUAUCAAAUAUAAUC
env-849   AAAAUNAAUUUAUAUUUUGAUUUAAUUCUUAACAGACCAUUAAGGUCAAUUUAAAAUUACUUAUUUAUCAAAUAUAAUC
env-850   ....................................................AAAUUACAUAUUUAUCAAAAUAUAAUC
env-851   AAAUUAAUCUUUAUUUUGAUUUAGUCCUCAAUAGACCAAUAAGGUCAAUUUAAAAUUACAUAUUUAUCAAAAUAUAAUC
env-852   AAAUUAAUCUUUAUUUUGAUUUAGUCCUCAAUAGACCACUAAGGUCAAUUUAAAAUUACAUAUUUAUCAAAAUAUAAUC
env-853   AAAUUAAUCUUUAUUUUGAUUUAGUCCUCAAUAGACCAAUAAGGUCAAUUUAAAAUUACAUAUUUAUCAAAAUAUAAUC
env-854   AAAUUAAUCUAUAUUUUGAUUUAAUUCUCAAUAGACCAUUAAGGUCAAUUUAAAAUUACAUAUUUAUCAAAAUAUAAUC
env-855   AAAUUAAUCUUUAUUUUGAUUUAGUCCUCAAUAGACCAAUAAGGUCAAUUUAAAAUUACAUAUUUAUCAAAAUAUAAUC
env-856   AAUUAAUCUUUAUUUUUGAUUUAGUCCUCAAUAGACCAAUAAGGUCAAUUUAAAAUUACAUAUUUAUCAAAAUAUAAUC
env-857   AAAUUAAUCUUUAUUUCGAUUUAGUCCUCAAUAGACCACUAAGGUCAAUUUAAAAUUACAUAUUUAUCAAAAUAUAAUC
env-860   AAAUUAAUCUUUAUUUUGAUUUAGUCCUCAAUAGACCAAUAAGGUCAAUUUAAAAUUACAUAUUUAUCAAAAUAUAAUC
env-861   AAAUUAAUCUUUAUUUUGAUUUAGUCCUCAAUAGACCAAUAAGGUCAAUUUAAAAUUACAUAUUUAUCAAAAUAUAAUC
env-862   ...............................................................................
env-863   AAAUUAAUCUAUAUUUUGAUUUAAUUCUCAAUAGACCAUUAAGGUCAAUUUAAAAUUACAUAUUUAUCAAAAUAUAAUC
env-864   AAAUUAAUCUUUAUUUCGAUUUAGUCCUCAAUAGACCACUAAGGUCAAUUUAAAAUUACAUAUUUAUCAAAAUAUAAUC
env-865   AAAUUAAUCUUUAUUUNGAUUUAGUCCUCAAUAGACCANUAAGGUCAAUUUAAAAUUACAUAUUUAUCAAAAUAUAAUC
env-866   AAAAUUAAUCUGUAUUUUGAUUUAAUUCUCAACAGACCAUUAAGGUCAAUUUAAAAUUACAUUUUUAUCAAAUAUAAUC
env-868   AAAAUUAAUCUUUAUUUUGAUUUAAUUCUCAACAGACCAUUAAGGUCAAUUUAAAAUUACAUUUUUAUCAAAUAUAAUC
env-869   AAAAUUAAUCUUUAUUUUGAUUUAAUUCUCAACAGACCAUUAAGGUCAAUUUAAAAUUACAUUUUUAUCAAAUAUAAUC
env-870   AAAUUAAUCUUUAUUUUGAUUUAGUCCUCAAUAGACCAAUAAGGUCAAUUUAAAAUUACAUAUUUAUCAAAAUAUAAUC
env-871   AAAAUUAAUCUUUAUUUUGAUUUAAUUCUCAACAGACCAUUAAGGUCAAUUUAAAAUUACAUUUUUAUCAAAUAUAAUC
env-875   AAAAUUAAUCUUUAUUUUGAUUUAAUUCUCAACAGACCAUUAAGGUCAAUUUAAAAUUACAUUUUUAUCAAAUAUAAUC

          ...............................................................................
          ...............................................................................
          ●●●●●●●●●●●●●●●●●●●●●●●●●●●●●●●●●●●●●●●●●●●●●●●●●●●●●●●●●●●●●●●●●●Y●●●●RR●●●●●●
```

| | |
|---|---|
| env-876 | AAAAUUAAUCUUUAUUUUGAUUUAAUUCUCAACAGACCAUUAAGGUCAAUUUAAAAUUACAUUUUUAUCAAAUAUAAUC |
| env-877 | AAAAUUAAUCUUUAUUUUGAUUUAAUUCUCAACAGACCAUUAAGGUCAAUUUAAAAUUACAUUUUUAUCAAAUAUAAUC |
| env-878 | AAAAUUAAUCUGUAUUUUGAUUUAAUUCUCAACAGACCAUUAAGGUCAAUUUAAAAUUACAUUUUUAUCAAAUAUAAUC |
| env-879 | AAAAUUAAUCUUUAUUUUGAUUUAAUUCUCAACAGACCAUUAAGGUCAAUUUAAAAUUACAUUUUUAUCAAAUAUAAUC |
| env-881 | AAAAUUAAUCUUUAUUUUGAUUUAAUUCUCAACAGACCAUUAAGGUCAAUUUAAAAUUACAUUUUUAUCAAAUAUAAUC |
| env-883 | AAGAAAUUAACUUAUAUUUUGAUUUAGUACUUAAUAGACCAUUAAGAUCAGUUUAAACUACUUUUUUAUAAAAUUAAUA |
| env-884 | AGAAAAUUAACUUAUAUUUUGAUUUAGUACUUAAUAGACCAUUAAGAUCAGUUUAAACUACUUUUUUAUAAAAUUAAUA |
| env-886 | AGAAAAUUAACUUGUAUUUUGAUUUAGUACUUAAUAGACCAUUAAGAUCAGUUUAAACUACUUUUUUAUAAAAUUAAUA |
| env-887 | AGAAAAUUAACUUAUAUUUUGAUUUAGUGCUUAAUAGACCAUUAAGAUCAGUUUAAACUACUUUUUUAUAAAAUUAAUA |
| env-888 | AGAAAAUUAACUUAUAUUUUGAUUUAGUGCUUAAUAGACCAUUAAGAUCAGUUUAAACUACUUUUUUAUAAAAUUAAUA |
| env-889 | AUAAAAUUAACUUAUAUUUUGAUUUAGUACUUAAUAGACCAUUAAGAUCAGUUUAAACUACUUUUUUAUAAAAUUAAUA |
| env-890 | AGAAAAUUAACUUAUAUUUUGAUUUAGUACUUAAUAGACCAUUAAGAUCAGUUUAAACUACUUUUUUAUAAAAUUAAUA |
| env-891 | UACAACCGGNNNNNNNNNNNNNNNNNNNNNNNNNNNNNNNNNNNNNNNNNNNNNNNNNNNNNCUUUCUUAUAAAAUUAAUA |
| env-892 | AGAAAAUUAACUUAUAUUUUGAUUUAGUGCUUAAUAGACCAUUAAGAUCAGUUUAAACUACUUUUUUAUAAAAUUAAUA |
| env-894 | AGAAAAUUAACUUNUAUUUUGAUUUAGUACUUAAUAGACCAUUAAGAUCAGUUUAAACUACUUUUUUAUAAAAUUAAUA |
| env-895 | AGAAAAUUAACUUAUAUUUUGAUUUAGUACUUAAUAGACCAUUAAGAUCAGUUUAAACUACUUUUUUAUAAAAUUAAUA |
| env-896 | AGAAAAUUAACUUAUAUUUUGAUUUAGUACUUAAUAGACCAUUAAGAUCAGUUUAAACUACUUUUUUAUAAAAUUAAUA |
| env-897 | AGAAAAUUAACUUAUAUUUUGAUUUAGUGCUUAAUAGACCAUUAAGAUCAGUUUAAACUACUUUUUUAUAAAAUUAAUA |
| env-898 | AGAAAAUUAACUUAUAUUUUGAUUUAGUACUUAAUAGACCAUUAAGAUCAGUUUAAACUACUUUUUUAUAAAAUUAAUA |
| env-899 | AGAAAAUUAACUUAUAUUUUGAUUUAGUACUUAAUAGACCAUUAAGAUCAGUUUAAACUACUUUUUUAUAAAAUUAAUA |
| env-900 | AGAAAAUUAACUUAUAUUUUGAUUUAGUACUUAAUAGACCAUUAAGAUCAGUUUAAACUACUUUUUUAUAAAAUUAAUA |
| env-901 | AAUCACUACAUAGAUCUGAUCUUAAAUCAACCCCUUAGGGAUUAAUUUAUUACAAAGAUUGUAUAUGAUCUAAUUAACA |
| env-902 | AAUCACUACAUAGAUCUGAUCUUAAAUCAACCCCUUAGGGAUUAAUUUAUUACAAAGAUUGUAUAUGAUCUAAUUAACA |
| env-903 | AAUCACUAUCUCGAUCUGAUCUUAAAUCAACCCCUUAGGGAUUAAUUCAUUGCAAAGAUUGUAUAUCAUCUAAUUAACA |
| env-905 | AAUCACUAUCUCGAUCUGAUCUUAAAUCAACCCCUUAGGGAUUAAUUCAUUGCAAAGAUUGUAUAUCAUCUAAUUAACA |
| env-906 | CUUUUUUAUUGCCAAUUUAUAUUCAAAAUAAAAGGUAAAAUAAUUUCAUAUUUCAUAAUUUAUAAUCAAAAAAUAAAAU |
| env-907 | CUUUUUUAUUGCCAAUUUAUAUUCAAAAUAGAAGGUAAAAUAAUUUCAUAUUUCAUAAUUUAUAAUCAAAAAAUAAAAU |
| env-908 | CUUUUUUAUUGCCAAUUUAUAUUCAAAAUAGAAGGUAAAAUAAUUUCAUAUUUCAUAAUUUAUAAUCAAAAAAUAAAAU |
| Fge-1-1 | UGUAAAUGAGGAAGUUGCAAAAUUAUUUUAAAGCAACGAAAAUAAUUCAUAACUCAUAAUUCAUAAUUCAUAAUUAUGU |
| Ale-2-1 | UUAAAACGUAUAAAAGUUUAAUUUCGUACGCAAUUUGUAAGAAUUUGCGUAAGAACCCACUCUUUCUGAAAUAGAUUUU |
| env-909 | GCCGAACUGUUGGAGCGCAGCGGAAAAAAAACAUUUUCAAAAUAAUUCAUCAUUCCUAUUUCAUAACUGAGUUUUAUUU |
| env-910 | UAGAACGGAAAAAGGCUUAAUUCAGUAUAAUUUAUUGUUAAUUUAGAAGGAAAAGGAAUUAAUAUAUUAGUUAGAUUUU |
| env-911 | UCAGUACAUUGAUUUAAUUCAAAAUCACCCUCUGAGGUCAUAAAAUACAUUUAACUUAGUAUUCUUUUCAUCAAAUUGG |
| env-912 | UCAGUACAUUGAUUUAAUUCAAAAUCACCCUCUGAGGUCAUAAAAUACAUUUAACUUAGUAUUCUUUUCAUCAAAUUGG |
| env-913 | UCACUACAUUGAUUUAAUUCAAAAUCAACCUCUGAGGUAAUAAAACACAUUUAACUUAUUAUUCUUUUAAUCAAAUUGG |
| env-914 | UCAGUACAUUGAUUUAAUUCAAAAUCAACCUCUGAGGUCAUAAAACACAUUUAACUUAGUAUUCUUUUAAUCAAAUUGG |
| env-915 | UCAGUACAUUGAUUUAAUUCAAAAUCAGCCUCUGAGGUCAUAAAACACAUUUAACUUAGUAUUCUUUUAAUCAAAUUGG |
| env-916 | UCAGUACAUUGAUUUAAUUCAAAAUCAACCUCUGAGGUCAUAAAACUCAUUUAACUUAGUAUACUUUUAAUCAAAUUGG |
| Man-1-1 | AUUUUGACCAAGUGUAAAAUAGCUAUAAAGGCUAGCUUUAGAUUUUGGUUUUCAAAAAAUCAAAAUUUAAAAAAUAAUA |
| env-917 | GUGAAUAUUACGACGGAUUAUUUUGAAUGAUAAUUCUAAUUUAAAUCGCAACUAAAUAUUGUAUAUUAAAAAAGUAUCC |
| env-919 | GCAUUAAAUAAAGGACAAUAUCGAGCUUAUAUUUUAUGAUUUAAUCGUCUUUUAGUUAUUGUAUAUUAAAAAAGUAUCC |
| env-920 | ................................................CAACUAAAUAUUGUAUAUUAAAAAAGUAUCC |
| env-921 | ................................................CAACUAAAUAUUGUAUAUUAAAAAAGUAUCC |
| env-922 | GUGAAUAUUACGACGGAUUAUUUUGAAUGAUAAUUCUAAUUUAAAUCGCAACUAAAUAUUGUAUAUUAAAAAAGUAUCC |
| env-923 | GUGAAUAUUACGGGGGAUUAUUUUGAAUGAUAAUUCUAAUUUAAAUCGCAACUAAAUAUUGUAUAUUAAAAAAGUAUCC |
| env-924 | UUCACAGAACUAAGGAGUUUAGUUUAAUGUCAAUUUGUAUUUAAAUCACAACUAAAUAUUGGAUAUUAAAAAAGUAUCC |
| env-925 | UUCACAGUAUUAAUGAGUUUAGUUUAAUGUCAAUUUGUAUUUAAAUCACAACUAAAUAUUGUAUAUUAAAAAAGUAUCC |
| env-927 | GUGAAUAUUACGACGGAUUAUUUUGAAUGAUAAUUCUAAUUUAAAUCGCAACUAAAUAUUGUAUAUUAAAAAAGUAUCC |
| env-928 | UACUGAAAUACAGAAUGAUAUCUCUUGUAUAUUUUAUGAUUUAAUCACCUUUAAGUUAUGGUAUAUUAAAAAAGUAUCC |
| env-929 | ...........................................................GGUAUAUUAAAAAAGUAUCC |
| env-930 | UACUGAAAUACAGAAUGAUAUCUCUUGUAUAUUUUAUGAUUUAAUCACCUUUAAGUUAUGGUAUAUUAAAAAAGUAUCC |
| env-932 | ...................AUAGAUGACUUAAAUUAUUAGAUAAAUACCAAUUAAUUAUGUUUUAUUAAAAAAGUAUCC |
| env-933 | AGAUGAAAAAUUUCUAGAAUUUGUAAACGAAGAAGUUGCAAAGUUGUUUUAGAAAAACUUGUCCAUUAAAAAAAUAAAA |
| env-934 | GACGAACGUUUUUUGGAAUUUGUAAACAAGGAAGUUGCAAAGUUGUAUUAGAAAAAACUUGUCCAUCUAAAAAAAUAAAA |
| env-936 | CUUGAAUCCUGAAAAAUCAACUGUGAAUUAUCUAUUAUCAAUUAUUUUCUACAAUAAAUAGUAUAACAAAAUAAAACCG |
| env-937 | GUUUAAUUCAUAACAUAAAUAUUUAAAAAAAAAUGAAUUCUUAAGACACAAAAUAAAUCCUUAUUUAAUCUAUUUUUUA |
| env-938 | UAAUAAAUACAUUGAUUUAAUUCUUAAUCAAACUUUAAGAGAAUAAUUAGCAAUUCAAAGAUCUAAUUAUCAAAAUUAA |
| env-939 | UAAUAAAUACAUUGAUUUAAUUCUUAAUCAAACUUUAAGAGAAUAAUUAGCAAUUCGAAGAUCUAAUUAUCAAAAUUAA |
| env-940 | UAAUAAAUACAUUGAUUUAAUUCUUAAUCAAACUUUAAGAGAAUAAUUAGCAAUUCAAAGAUCUAAUUAUCAAAAUUAA |
| env-941 | CAAUAAAUACAUUGAUUUAAUUCUUAAUCAAACUUUAAGAGAAUAAUUAGCAAUUCAAAGAUCUAAUUAUCAAAAUUAA |
| env-942 | UAAUAAAUACAUUGAUUUAAUUCUUAAUCAACCUUUAAGAGAAUAAUUAGCAAUUCGAAGAUCUAAUUAUCAAAAUUAA |
| env-943 | UAAUAAAUACAUUGAUUUAAUUCUUAAUCAAACUUUAAGAGAAUAAUUAGCAAUUCGAAGAUCUAAUUAUCAAAAUUAA |
| env-944 | UAAUAAAUACAUUGAUUUAAUUCUUAAUCAAACUUUAAGAGAAUAAUUAGCAAUUCGAAGAUCUAAUUAUCAAAAUUAA |
| env-945 | UAAUAAAUACAUUGAUUUAAUUCUUAAUCAAACUUUAAGAGAAUAAUUAGCAAUUCGAAGAUCUAAUUAUCAAAAUUAA |
| env-946 | UAAUAAAUACAUUGAUUUAAUUCUUAAUCAAACUUUAAGAGAAUAAUUAGCAAUUCGAAGAUCUAAUUAUCAAAAUUAA |
| Tma-1-1 | AAGAUUUACGUAUCACAGACCACCUUCCUUUCCUUGCUAUUUAGACGAAAAAAAUCUGACUCUCAAUCAAAAAAUAAUU |
| env-947 | CUGUAAAAGAGGAUUUUAUUUUGAAUUGGCGAUGAGAUAUGUUUAAUUAAAAAAAGGUUUGUUAAUAUAAUAAUUAUGU |
| Fsp-12-1 | UCCUGUUCGGGUUCUUUAUUCUUUACCAAUAACAAUUACAAAAAAGGACUAAUAAAAUUUAUUUUUCGAAAAAAAAGUA |
| Fde-1-1 | CAAAAUUAUUCAAAAAUCAAAAAGCGAUAUUCUUUUUUUAGAUUCUUUUAUUUUUGUUUGAUUCUUUUGUUUUUUUACC |
| Fba-7-1 | AUUAUCAAAAGCUUUAAAAUAAGAAUUCCCUUAACUAAAAUUUUAGUAUCCCUAAAAUAGAUUAGUAUCAGAAAAAAGA |
| Ale-1-1 | CUUUUACAUAUUACACUAUAUUAAAACGUUAAAAGGACAUCAAAAAUUAGAUAAAAAAAUUAGAUAAAUGUAAAAAAGUC |
| Ale-1-2 | CUUUUACAUAUUACACUAUAUUAAAACGUUAAAAGGACAUCAAAAAUUAGAUAAAAAAAUUAGAUAAAUGUAAAAAAGUC |
| Ffr-1-1 | UUUGUUAAUGCAGAGGUUGCGAAGUUAUAUUAAGUAUAAAAUAAAUGAUGAUUAAUAAUUCAAAUUUGAAAUUUAUUAA |
| env-948 | AAUCGAUUGAAAAAAUAGUUUAAUUUUAAACUUUUUUGAAAAUAAUUCUGAAUUCAUAAUUCAAAAUUCUAAAAUAUCU |
| Fsp-1-1 | AUAAGUCUCAUUUAAAAAAAUAUGAAAAGCAAAUAUAAAAAAAUCAACACUAUAAUGAUUGUUAAUACUAAAAUUAAAC |
| env-949 | AAUCGAUUGAUUAAAUUGCUUAAGUUUUGAAUUUGGAAAAAAUAAUUCCUGAUUUAUAAUUCAUAAUUCACAAUUAUUU |
| env-950 | AAAUCGAUUGACAAAGCCCUCUAAUUUUAAUGUUUUUAAAAAUAUUUCGGCAAUCAUAAUUCAAAAUUCCAAAAUAUUU |
| Fsa-2-1 | AAAUCGAUUGAUAAAAUUGCUUAAGUUUUGAAUUUGAAAAAAUAAUUCCUGAUUUAUAAUUCAUAAUUCACAAUUAUUU |
| env-951 | GAUGAACGAUUUUUAGAAUAUGUGAAUCAAGAGAUUGAAAGAUUAUUUCUUUAAUUUCUUGUAAUUUUAAAAAAUAGAA |
| env-952 | GACGAACGUUUUUUAGAAUAUGUAAAUCAAGAGAUUGAAAGAUUAUUUCUUUAAUUUCUUGUNAUUUUAAAAAAUAGNA |
| Fba-3-1 | GACGAGCGCUUUUUAGAAUUUGUAAAUCAAGAAGUAGCAAAGUUAUUUUUAUAAUUACUUGUGAUUUUAAAAAAUAAAA |
| env-953 | AAAAAAGUAUUUAAUUUUUUUUAGACAAAUCUUUUUUUAAAUAAUUUAUAAUUUACAAUUCAUAAUUCACAAUUAUUA |
| env-954 | AUCGAUUGAUAAUUAAGAAGAAUAUCUUGCCGUAUUUUUAAAAAUAAAAAGGGAUUACUUGUAUUUUAAAAAAAUAAAA |
| env-955 | AAGAAUUACUACUGACGAACUUCUUAACAUUAACCAAGCUCUUAUGAUAAAAUAAAUCCUGUUUUAAUCUAUUUUUUUA |
| | ............................................................................... |
| | ............................................................................... |
| | ••••••••••••••••••••••••••••••••••••••••••••••••••••••••••••••••••Y••••RR•••••• |

```
env-956    AAGAAUUACUACAGACAAAAUUCUUAACAUUAGCCAAGCUCUUAUGAUAAAAUAAAUCCUGUUUUAAUCUAUUUUUUUA
env-957    AAGAAUUACUACUGACAAACUUCUUAACAUUAGCCAAGCUCUUAUGAUAAAAUAAAUCCUGUUUUAAUCUAUUUUUUUA
env-958    ........AUUUAGAUUUAAUACUUAUCAACCCGGUUAGAGAAUAAAUUAUCUUUUUUUAUUGAAUACCCCAUUCUUUA
env-959    ...............CACAAAUUUCGUGCGUUUGACCUAGAAUUUUUCAAGAUAAAUUUGCACAGCUAAAAAAAAGGUU
Csp-1-1    AGAACUCGUCCACUUAGAAGAUGUAAGUGAAUAAAAAAUAAAAAUAAGAGCUAUUUCUUUGGUAGUUAGAAAAAUUUUA
env-960    GAAGUCGUCCAUUUAGAAGAUGUAAGUGAAUAAAAAAAUAAAAAUAAGUGCCAUUUCUUUGGUAGUUAGAAAAUAUUUA
env-961    CGCGUGGUGUAUAAGGAAGGCUNNNNNNNNNNNNNNNNNNNNNNNNNNNNNNNNNNNCUUUGGUAGUUAGAAAAUAUUUA
Csp-3-1    GAAGUCGUCCAUUUAGAAGAUAUAAGUGAAUAAAAAAAUAAAAAUAAGUGCUAUUUCUUUGGUAGUUAGAAAAUAUUUA
env-962    AGAACUCGUCCACUUAGAAGAUGUAAGUGAAUAAAAAAUAAAAAUAAGAGCUAUUUCUUUGGUAGUUAGAAAAAUUUUA
env-965    AGAACUCGUCCACUUAGAAGAUGUAAGUGAAUAAAAAAUAAAAAUAAGUGCUAUUUCUUUGGUAGUUAGAAAAUAUUUA
env-966    AGAACUCGUCCACUUAGAAGAUGUAAGUGAAUAAAAAAUAAAAAUAAGUGCUAUUUCUUUGGUAGUUAGAAAAUAUUUA
env-967    AGAACUCGUCCACUUAGAAGAUGUAAGUGAAUAAAAAAUAAAAAUAAGUGCUAUUUCUUUGGUAGUUAGAAAUUUUUCA
env-968    AGAACUCGUCCACUUAGAAGAUGUAAGUGAAUAAAAAAUAAAAAUAAGUGCUAUUUCUUUGGUAGUUAGAAAAUUUUUA
env-972    AGAACUCGUCCACUUAGAAGAUGUAAGUGAAUAAAAAAUAAAAAUAAGUGCUAUUUCUUUGGUAGUUAGAAAAUAUUUA
env-973    AGAACUCGUCCACUUAGAAGAUGUAAGUGAAUAAAAAAUAAAAAUAAGUGCUAUUUCUUUGGUAGUUAGAAAAUUUUUA
env-974    AGAACUCGUCCACUUAGAAGAUGUAAGUGAAUAAAAAAUAAAAAUAAGUGCUAUUUCUUUGGUAGUUAGAAAAUAUUUA
env-976    ...................................................CUAUUUCUUUGGUAGUUAGAAAAUAUUUA
env-977    AGAACUCGUCCACUUAGAAGAUGUAAGUGAAUAAAAAAUAAAAAUAAGUGCUAUUUCUUUGGUAGUUAGAAAAUAUUUA
env-978    AGAACUCGUCCACUUAGAAGAUGUAAGUGAAUAAAAAAUAAAAAUAAGUGCUAUUUCUUUGGUAGUUAGAAAAUAUUUA
env-980    AGAACUCGUCCACUUAGAAGAUGUAAGUGAAUAAAAAAUAAAAAUAAGUGCUAUUUCUUUGGUAGUUAGAAAAUUUUUA
env-982    AGAACUCGUCCACUUAGAAGAUGUAAGUGAAUAAAAAAUAAAAAUAAGUGCUAUUUCUUUGGUAGUUAGAAAAUAUUUA
env-983    AGAACUCGUCCACUUAGAAGAUGUAAGUGAAUAAAAAAUAAAAAUAAGUGCUAUUUCUUUGGUAGUUAGAAAAUUUUUA
env-984    AGAACUCGUCCACUUAGAAGAUGUAAGUGAAUAAAAAAUAAAAAUAAGUGCUAUUUCUUUGGUAGUUAGAAAAUAUUUA
env-985    AGAACUCGUCCACUUAGAAGAUGUAAGUGAAUAAAAAAUAAAAAUAAGUGCUAUUUCUUUGGUAGUUAGAAAAUUUUUA
env-986    AGAACUCGUCCACUUAGAAGAUGUAAGUGAAUAAAAAAUAAAAAUAAGUGCUAUUUCUUUGGUAGUUAGAAAAUUUUUA
env-988    AGAACUCGUCCACUUAGAAGAUGUAAGUGAAUAAAAAAUAAAAAUAAGUGCUAUUUCUUUGGUAGUUAGAAAAUAUUUA
env-989    .....................................................AUUUCUUUGGUAGUUAGAAAAUAUUUA
env-990    AGAACUCGUCCACUUAGAAGAUGUAAGUGAAUAAAAAAUAAAAAUAAGUGCUAUUUCUUUGGUAGUUAGAAAAUUUUUA
env-993    AGAACUCGUCCACUUAGAAGAUGUAAGUGAAUAAAAAAUAAAAAUAAGUGCUAUUUCUUUGGUAGUUAGAAAUUUUUCA
env-994    AGAACUCGUCCACUUAGAAGAUGUAAGUGAAUAAAAAAUAAAAAUAAGUGCUAUUUCUUUGGUAGUUAGAAAAUAUUUA
env-995    AGAACUCGUCCACUUAGAAGAUGUAAGUGAAUAAAAAAUAAAAAUAAGUGCUAUUUCUUUGGUAGUUAGAAAAUAUUUA
env-999    AGAAGUCGUCCACUUAGAAGAUGUAAGUGAAUAAAAAAUAAAAAUAAGUGCUAUUUCUUUGGUAGUUAGAAAAUUUUUA
env-1001   AGAACUCGUCCACUUAGAAGAUGUAAGUGAAUAAAAAAUAAAAAUAAGUGCUAUUUCUUUGGUAGUUAGAAAAUAUUUA
env-1002   AGAACUCGUCCACUUAGAAGAUGUAAGUGAAUAAAAAAUAAAAAUAAGUGCUAUUUCUUUGGUAGUUAGAAAAUAUUUA
env-1003   AGAACUCGUCCACUUAGAAGAUGUAAGUGAAUAAAAAAUAAAAAUAAGUGCUAUUUCUUUGGUAGUUAGAAAUUUUUUA
env-1004   AGAACUCGUCCACUUAGAAGAUGUAAGUGAAUAAAAAAUAAAAAUAAGUGCUAUUUCUUUGGUAGUUAGAAAAUUUUUA
env-1005   AGAACUCGUCCACUUAGAAGAUGUAAGUGAAUAAAAAAUAAAAAUAAGUGCUAUUUCUUUGGUAGUUAGAAAAUAUUUA
env-1006   AGAACUCGUCCACUUAGAAGAUGUAAGUGAAUAAAAAAUAAAAAUAAGUGCUAUUUCUUUGGUAGUUAGAAAAUAUUUA
env-1007   AGAACUCGUCCACUUAGAAGAUGUAAGUGAAUAAAAAAUAAAAAUAAGUGCUAUUUCUUUGGUAGUUAGAAAAUAUUUA
env-1008   AGAACUCGUCCACUUAGAAGAUGUAAGUGAAUAAAAAAUAAAAAUAAGUGCUAUUUCUUUGGUAGUUAGAAAAUAUUUA
env-1009   AGAACUCGUCCACUUAGAAGAUGUAAGUGAAUAAAAAAUAAAAAUAAGUGCUAUUUCUUUGGUAGUUAGAAAAUAUUUA
env-1010   AGAACUCGUCCACUUAGAAGAUGUAAGUGAAUAAAAAAUAAAAAUAAGUGCUAUUUCUUUGGUAGUUAGAAAAUUUUUA
env-1011   AGAACUCGUCCACUUAGAAGAUGUAAGUGAAUAAAAAAUAAAAAUAAGUACUAUUUCUUUGGUAGUUAGAAAAUUUUUA
env-1012   CGUAGAGAUUUAAAGAAAAAAAUAAAUCUAUUUAUUGAUUUAAUUUCAAACAAACCUGUUAGGUAAUAAUCUAUUUUUU
env-1013   CGUAGAGAUUUAAAGAAAAAAAUAAAUCUAUUUAUUGAUUUAAUUUCAAACAAACCUGUUAGGUAAUAAUCUAUUUUUU
env-1015   CGAAGAGAUUUGAAGAAAAAAAUAAAUCUAUUUAUUGAUUUAAUUUCAAACAAACCUGUUAGGUAAUAAUCUAUUUUUU
env-1016   CGAAGAGAUUUGAAGAAAAAAAUAAAUCUAUUUAUUGAUUUAAUUUCAAAUAAACCUGUUAGGUAAUAAUCUAUUUUUU
env-1017   GAAGAGAUUUAAAGAAAAAAAUAAAUUUAUUUAUUGAUUUAAUUUCAAAUAAACCUGUUAGGGAAUAGUCUAAUUUUUU
env-1018   GAAGAGAUUUAAAGAAAAAAAUAAAUUUAUUUAUUGAUUUAAUUUCAAAUAAACCUGUAAGGGAAUAGUCUAAUUUUUU
env-1019   AGAAGAGAUUUAAAGAAAAAAAUAAAUCUAUUUAUUGAUUUAAUUUCAAAUAAAGCUGUAAGGGAAUAAUCUAUUUUUA
env-1021   ACUGAUUUGCAGCAAGCUUCGGCAAAUAUUAAUUGGAAAAAAACAGAAGAAAUUAAUAUUUGAGUUAUUAUUAUUAUUG
env-1022   GAAGAGAUUUAAAGAAAAAAAUAAAUCUGUUUAUUGAUUUAAUUUCAAAUAAACCUGUAAGGGAAUAAUCUAAUUUUUU
env-1023   GAAAAGAUUUGAAGAAAAAAAUAAAUUUGUUUAUUGAUUUAAUUUCAAAUAAACCUGUAAGGGAAUAAUCUAAUUUUUU
env-1024   GAAGGGAUUUGAAGAAAAAAAUAAAUUUGUUUAUUGAUUUAAUUUCAAAUAAACCUGUAAGGUAAUAAUCGAAUUUUUU
env-1025   GAAGGGAUUUGAAGAAAAAAAUAAAUUUGUUUAUUGAUUUAAUUUCAAAUAAACCUAUAAGGGAAUAAUCUAAUUUUUU
env-1026   GUAGGGAUUUAAAGAAAAAAAUAAAUCUGUUUAUCGACUUAAUUUCAAAUAAACCUGUAAGGGAAUAAUGAAUUUUUUU
env-1027   GAAGAGAUUUAAAGAAAAAAAUAAAUUUAUUUAUUGAUUUAAUUUCAAAUAAACCUGUUAGGGAAUAAUCUAUUUUUUU
env-1028   AAACCUAUAUUUAGAUUUAAUUCUUAAUCAGCCUAUUAGAGAAUAAAUUAUCUUUUUUCAUAUAAUAUACCAUUUUUUA
env-1029   GAUUUUACAGAUAUGCAGCAGGCUUCCUCAAAUAUAGAUUGGGAAAAAACAGAUAUAAUAAAUAUUUAAGUUAUUAUUA
env-1030   GAUUUUACAGAUAUGCAGCAGGCUUCCUCAAAUAUAGAUUGGGAAAAAACAGAUAUAAUAAAUAUUUAAGUUAUUAUUA
env-1032   GAUUUUACAGAUAUGCAGCAGGCUUCCUCAAAUAUAGAUUGGGAAAAAACAGAUAUAAUAAAUAUUUAAGUUAUUAUUA
env-1033   GAUUUUACAGAUAUGCAGCAGGCUUCCUCAAAUAUAGAUUGGGAAAAAACAGAUAUAAUAAAUAUUUAAGUUAUUAUUA
env-1034   GAUUUUACAGAUAUGCAGCAGGCUUCCUCAAAUAUAGAUUGGGAAAAAACAGAUAUAAUAAAUAUUUAAGUUAUUAUUA
env-1035   GAUUUUACAGAUAUGCAGCAGGCUUCCUCAAAUAUAGAUUGGGAAAAAACAGAUAUAAUAAAUAUUUAAGUUAUUAUUA
env-1036   ...................AAUACUCAACAACCCUGUAAGAGAAUAAAUUAUCUUUUUUAUUGAAUACCCAUUUUUUA
env-1037   AAACCUUUACAUAGAUUUGAUACUUAACAACCCUAUUAGAUAAUAAAUUAUCUUAUUUAAUCAAAUAUCUUAUUUUUUA
env-1038   CAAUCUUUAUUUGGAUUUAAUACUUAAUAAUCCUAUCAGAGAAUAAAUUAUCUUUUUUCUUCGUAUAUCCUAUUUUUUA
env-1039   CAAUCUUUAUUUAGAUUUAAUACUUAAUAAUCCUAUCAGAGAAUAAAUCAUCUUUUUUCUUCAUAUAAGCUAUUUUUUA
env-1040   CAAUCUUUAUUUAGAUUUAAUACUUAAUAAUCCUAUCAGAGAAUAAAUCAUCUUUUUUCUUCAUAUAAGCUAUUUUUUA
env-1041   AUAAAUCUGUAUUUGGAUUUAAUACUUAACAAGCCAAUUAGAUAAUAAAUAUUUUUUUCUACUGUAAAUCCUUUUUUUA
env-1042   AUAAAUCUGUAUUUGGAUUUAAUACUUAACAAGCCAAUUAGAUAAUAAAUAUUUUUUUCUACUGUAAAUCCUUUUUUUA
env-1044   UAAAUCUAUAUUUAGAUUUAAUACUUAACAAGCCAAUUAGAGAAUAAAUAUUUUUUUUCUACUGUAAACCCUUUUUUUA
env-1045   AAAUCUAUAUUUAGAUUUAAUACUUAACAAUCCGGUAAGAGAAUAAAUUAUCUUUUUUUAUUGAAUACCCUAUUUUUUA
env-1046   UCAGAUAUGCAACAGGCCUCCUCAAAUAUAGAUUGGAAAAAAACAGAAAUAAUAAAUAUUUAAGUUAUUAUUAUUAUUA
env-1047   AAUUUAUUUUUAGAUUUAAUACUAAAUAAACCUGUUAGACAAUAAAAAGGUUAUUUUACUCUGAAUAAUCGAUUUUUUA
env-1048   AAUUUAUUUUUAGAUUUAAUACUAAAUAAACCUGUUAGACAAUAAAAAGGUUAUUUUACUCUGAAUAAUCGAUUUUUUA
env-1049   AAUUUAUUUUUAGAUUUAAUACUAAAUAAACCUGUUAGACAAUAAAAAGGUUAUUUUACUCUGAAUAAUCGAUUUUUUA
env-1051   AAUUUAUUUUUAGAUUUAAUACUAAAUAAACCUGUUAGACAAUAAAAAGGUUAUUUUACUCUGAAUAAUCGAUUUUUUA
env-1053   AAUUUAUUUUUAGAUUUAAUACUAAAUAAACCUGUUAGACAAUAAAAAGGUUAUUUUACUCUGAAUAAUCGAUUUUUUA
env-1055   AAUUUAUUUUUAGAUUUAAUACUAAAUAAACCUGUUAGACAAUAAAAAGGUUAUUUUACUCUGAAUAAUCGAUUUUUUA
env-1056   AAACCUAUUUCUGGAUUUAAUACUUAACAAUCCAAUAAGAGAAUAAAUUAUCUUUUUUCUUCCUAUAACCCAUUUUUUA
env-1057   AAGAAAAAAAUAAACCUAUACUUAGAUUUAAUUCUAAAUAAUCCGAUAAGAAAUUAAUCUUCAUUUAAUCUUAUUUUUUA
env-1058   GAUUUUAUUGAUUUGCAGCAGGCUUCCUCAAAUAUUAAUUGGAAAAAAACAGAGAUAAUAAAUAUUUAAGUUAUUAUUA
env-1059   UAAAUUAUUAGGUAGACAGAAAAUAUUAAUAUUUUAAUUUACUCAGUUUAUCAAUUUCUUUAUUUAAUCUUAUUUUUUA

           ................................................................................
           ................................................................................
           ●●●●●●●●●●●●●●●●●●●●●●●●●●●●●●●●●●●●●●●●●●●●●●●●●●●●●●●●●●●●●●●●●●●Y●●●●RR●●●●●●
```

```
env-1060  AAAUCAAUAUAUUGAUUUAAUUCUAAAUCAACCUAUUAGAGUUUAAAUUUCGUUUUUUUUCUCUGUUAUUCAUUUUUUU
env-1061  GUUAAUACUACGUGAUACCAUGUUCCUUGAGAGUAUGUGUUAUCAGCUUAUAAAUUAAUCUUCAUUUAAUCUUAUUUUA
env-1062  AACCUAUACUUAGAUUUAAUUCUUAAUAAAACAAUCAGGGAAUAAAAACUUUAUUUUAUACCCAAUCCUCUAAUUUUUA
env-1063  AACCUUUACUUAGAUUUAAUUCUCAAUAAACCAAUCAGAGAAUAAAACUUUUAUUUUUAACCCAAUCAUCUAAUUUUUA
env-1064  AAAUCUAUAUUUAGAUUUAAUACUCAACAAUCCGGUGAGAGAAUAAAUUAUCUUUUUUCAUUGAAUACCCCAUUUUUUA
env-1065  UAAUCUUUAUUUAGAUUUAAUACUUAAUAAUCCUAUCAGAGAAUAAAUUAUCUUUUUUCUUCGUAUAUCCUAUUUUUUA
env-1066  AAAUCUUUAUCUUGAUUUAAUAUUAAACAAUCCUGUUAGAGAAUAAACUUACUUUUUUCUCUGUAUUUCUUAAUUUUUA
env-1067  AACCUUUAUUUAGAUUUAAUUCUCAAUAAUCCAAUCAGAGAAUAAAACUUUUAUUUUUCACCCAAUCAUCUAAUUUUUA
env-1068  AACCUUUAUUUAGAUUUAAUUCUCAAUAAUCCAAUCAGAGAAUAAAACUUUUAUUUUUCACCCAAUCAUCUAAUUUUUA
env-1069  AACCUUUAUUUAGAUUUAAUUCUGAAUAAUCCAAUCAGAGAAUAAAAAUUUUAUUUUUCACCCAAUCAUCUAAUUUUUA
env-1070  AAAUCUUUAUCUUGAUUUAAUAUUAAACAAUCCUGUUAGAGAAUAAACUUACUUUUUUCUCUGUAUUUCUUAAUUUUUA
env-1077  AACCUUUAUUUAGAUUUAAUUCUCAAUAAACCUAUCAGAGAAUAAAACUUAUAUUUUUCACCCAAUCAUCUAAUUUUUA
env-1078  AAAUCUUUAUCUUGAUUUAAUAUUAAACAAUCCUGUUAGAGAAUAAACUUACUUUUUUCUCUGUAUUUCUUAAUUUUUA
env-1079  AACCUUUAUUUAGAUUUAAUUCUCAAUAAUCCAAUCAGAGAAUAAAACUUUUAUUUUUCACCCAAUCAUCUAAUUUUUA
env-1080  AACCUUUAUUUAGAUUUAAUUCUCAAUAAUCCAAUCAGAGAAUAAAACUUUUAUUUUUCACCCAAUCAUCUAAUUUUUA
env-1081  AACCUUUAUUUAGAUUUAAUUCUCAAUAAACCAAUCAGAGAAUAAAACUUUUAUUUUUCACCCAAUCAUCUAAUUUUUA
env-1082  AACCUUUAUUUAGAUUUAAUUCUCAAUAAUCCAAUCAGAGAAUAAAACUUUUAUUUUUCACCCAAUCAUCUAAUUUUUA
env-1085  AAAUCUGUAUUUAGAUUUAAUACUCAACAAACCGGUGAGAGAAUAAAUUAUCUUUUUUUAUUGAAUACCUCAUUUUUUA
env-1086  AAAUCUAUAUUUAGAUUUAAUACUCAACAACCCUGUAAGAGAAUAAAUUAUCUUUUUUUAUUGAAUACCCCAUUUUUUA
env-1087  AAAUCUAUAUUUAGAUUUAAUACUCAAUAAACCUGUAAGAGAAUAAAUUAUCUUUUUUUAUUCAUUACCUCAUUUUUUA
env-1088  AUUAAUCUUUAUUUAGAUUUAAUACUUAAUAAUCCUAUCAGAGAAUAAAUUAUUUUUUCUUCGUAUUUCCUAUUUUUUA
env-1089  AAAUCUAUAUUUAGAUUUAAUACUCAACAAUCCGGUGAGAGAAUAAAUUAUCUUUUUUCAUUGAAUACCCCAUUUUUUA
env-1090  AAAUCUAUAUUUAGAUUUAAUACUCAACAAUCCGGUGAGAGAAUAAAUUAUCUUUUUUCAUUGAAUACCCCAUUUUUUA
env-1091  UAAUCUUUAUUUAGAUUUAAUACUUAAUAAUCCUAUCAGAGAAUAAAUUAUCUUUUUUCUUCGUAUAUCCUAUUUUUUA
env-1092  AAAUCUAUAUUUAGAUUUAAUACUCAACAACCCUGUAAGAGAAUAAAUUAUCUUUUUUUAUUGAAUACCCCAUUUUUUA
env-1093  CAAUCUUUAUUUAGAUUUAAUACUUAAUAAUCCUAUCAGAGAAUAAAUCAUCUUUUUUCUUCUUACGUCCUAUUUUUUA
env-1094  UAAUCUUUAUAUAGAUUUAAUACUUAAUAAUCCUAUCAGAGAAUAAAUUAUCUUUUUUCUUCGUAUAUCCUAUUUUUUA
env-1095  UAAUCUUUAUUUAGAUUUAAUACUUAAUAAUCCUAUCAGAGAAUAAAUUAUCUUUUUUCUUCGUAUAUCCUAUUUUUUA
env-1096  UAAUCUUUAUUUAGAUUUAAUACUUAAUAAUCCUAUCAGAGAAUAAAUUAUCUUUUUUCUUCGUAUAUCCUAUUUUUUA
env-1097  AAAUCUAUAUUUAGAUUUAAUACUCAACAACCCUGUAAGAGAAUAAAUUAUCUUUUUUUAUUGAAUACCCCAUUUUUUA
env-1098  AACGAAGCAAACAAAGCUAUAAAUUGGAGUAAGGUGCAAGAAUAUAAUAUUUAAAAUAUAUCUGGUUUCAUAUUAUUUG
env-1099  UAAUCUUUAUUUAGAUUUAAUACUUAAUAAUCCUAUCAGAGAAUAAAUUAUCUUUUUUCUUCGUAUAUCCUAUUUUUUA
env-1101  GAGUUUACGGAUUUGCAUAUGGCCUCCUCAAAUAUAGACUGGGAAAAAACAGAAAUAAUAAAUAUUUAAGUUAUUAUUG
env-1102  AACCUUUAUUUAGAUUUAAUUCUCAAUAAUCCAAUCAGAGAAUAAAACUUUUAUUUUUCACCCAAUCAUCUAAUUUUUA
env-1103  AAAUCUAUAUUUAGAUUUAAUACUUAACAAUCCGGUGAGAGAAUAAAUUAUCUUUUUUUAUUGAAUACCCUAUUUUUUA
env-1104  CAAUCUUUAUUUAGAUUUAAUACUUAAUAAUCCUAUCAGAGAAUAAAUCAUCUUUUUUCUUCGUAUAUCCUAUUUUUUA
env-1105  AACCUUUAUUUAGAUUUAAUUCUCAAUAAUCCAAUCAGAGAAUAAAACUUCUAUUUUUAACCCAAUCAUCUAAUUUUUA
env-1106  AAACCUUUACAUAGAUUUGAUACUUAACAGCCCUCUUAGAUAAUAAAUUAUCUUAUUUCAUCAAAUAUCUUAUUUUUUA
env-1107  UCUAUAUUUAGAUUUUAAUACUCAAACAAUUCCGGUGAGAGAAUAAAUUAUCUUUUUUUAUUGAAUACCCCAUUUUUUA
env-1108  UAAUCUUUAUUUAGAUUUAAUACUUAAUAAUCCUAUCAGAGAAUAAAUUAUCUUUUUUCUUCGUAUAUCCUAUUUUUUA
env-1109  UAAUCUUUAUUUAGAUUUAAUACUUAAUAAUCCUAUCAGAGAAUAAAUUAUCUUUUUUCUUCGUAUAUCCUAUUUUUUA
env-1110  AAAUCUUUAUCUUGAUUUAAUAUUAAACAAUCCUGUUAGAGAAUAAACUUACUUUUUUCUCUGUAUUUCUUAAUUUUUA
env-1111  AAACCUAUAUUUAGAUUUAAUACUUAACAAGCCGGUAAGAGAAUAAAUUAUCUUUUUUUAUUGAAUACCCUAUUUUUUA
env-1112  AAACCUUUAUUUAGAUUUAAUUCUUAAUAAUCCAAUCAGAGAAUAAAUUAUCUUUUUUCUUCGAAUAUCCUAUUUUUUA
env-1113  AAACCUUUAUUUAGAUUUAAUUCUUAAUAAUCCCAUCAGAGAAUAAAUUAUCUUUUUUCUUCGAAUAUCCUAUUUUUUA
env-1115  AAACCUUUAUUUAGAUUUAAUUCUUAAUAAUCCAAUCAGAGAAUAAAUUAUCUUUUUUCUUCGAAUAUCCUAUUUUUUA
env-1116  AAACCUUUAUUUAGAUUUAAUUCUUAAUAAUCCAAUCAGAGAAUAAAUUAUCUUUUUUCUUCGAAUAUCCUAUUUUUUA
env-1121  AAACCUUUAUUUAGAUUUAAUUCUUAAUAAUCCAAUCAGAGAAUAAAUUAUCUUUUUUCUUCGAAUAUCCUAUUUUUUA
env-1122  AAAUCAAUAUAUUGAUUUAAUUCUAAAUCAACCUAUUAGAGUUUAAAUUUCGUUUUUUUUCUCUGUUAUUCAUUUUUUU
env-1123  UAAUCUUUAUUUAGAUUUAAUACUUAAUAAUCCUAUCAGAGAAUAAAUUAUCUUUUUUCUUCGUAUAUCCUAUUUUUUA
env-1124  AAACCUUUAUUUAGAUUUAAUUCUUAAUAAUCCAAUCAGAGAAUAAAUUAUCUUUUUUCUUCGAAUAUACUAUUUUUUA
env-1125  ................................................................AAUAUCCUAUUUUUUA
env-1126  ............................................AUAAAUUAUUUUUUUCUUCUAAUAUCCUAUUUUUUA
env-1127  AAACCUUUAUUUAGAUUUAAUUCUUAAUAAUCCCAUCAGAGAAUAAAUUAUCUUUUUUCUUCGAAUAUCCUAUUUUUUA
env-1128  AAAUCAAUAUAUUGAUUUAAUUCUAAAUCAACCUAUUAGAGUUUAAAUUUCGUUUUUUUUCUCUGUUAUUCAUUUUUUU
env-1129  AAUCAAAAUAGAGGUCCGUGUUUAAAAAGUACGGUAAGUCCAAUAUAUUGCGGUUUUUUGUUCAAUUCAAAAAGAGUUC
env-1130  .......................AGAACGCGUCAAAGUUAAGUUUAAUUUUGGAUUCUAUUGUGAUUCAGAAAGAUUCAU
env-1131  CGACAAGCGGACACCCAAGAACGGUUCAAAGGUAACAAUAAAAACCACAUGCAAUUUUUGUUGGAAUCAAAAAGAGUUC
env-1134  CGACAAGCGGACACCCAAGAACGGUUCAAAGGUAACAAUAAAAACCACAUGCAAUUUUUGUUGGAAUCAAAAAGAGUUC
env-1135  CGACAAGCGGACACCCAAUAACGGUUCAAAGGUAACAAUAAAAACCACAUGCAAUUUUUGUUGGAAUCAAAAAGAGUUC
env-1137  CGACAAGCGGACACCCAAGAACGGUUCAAAGGUAACAAUAAAAACCACAUGCAAUUUUUGUUGGAAUCAAAAAGAGUUC
env-1138  UUCGACAAGCGAACACCUAAGAACACAGCAAAGAUAGCAAUAAAAACCGGGUUUGGUUUUGCGGGUAUCAAAAAGAGUUC
env-1139  CAAGAAUAAAUAUUUUUGUUUUGAUAUAGAGCCGCUUCGUUUUGGAGCGGCUUUUUUUUGCUCGUUUCGAAAAGAGUAC
env-1140  AAUUCCAAAGUGCUAAUGGCGGAAUUUUUUUAUCCUAUGGAAUUUGGAAUUUAAAAUAUUGGAAUUUAAAAAUUAAAUA
env-1141  AAUUUAUAUCUCGAUCUAAUCCUAAAUCAACCUCUAAGGGAUUAAUUCCCUAAUCGGGUUGUAUAUCAAAGAAUUACAA
env-1142  AAUUUAUAUCUCGAUCUAAUCCUAAAUCAACCUCUAAGGGAUUAAUUCCCUAAUCGGGUUGUAUAUCAAAGAAUUACAA
env-1143  AAUUUAUAUCUCGAUCUAAUCCUAAAUCAACCUCUAAGGGAUUAAUUCCCUAAUCGGGUUGUAUAUCAAAGAAUUACAA
Lsp-4-1   AGUAAACCUUUACCUCGACUUGAUUCAGAAUCAACCCGUUAGAAAAUAACUUUUCAUAACGCUGCACAGAUUAAAAAAU
Cca-5-1   CUUUCAACAUUUGCAAAAUAAAACUUCUGCCUUUGCGAUAAAAUAUAUUGAUUUACAGUUUCUAAUCCGGAAAUAAAUC
env-1144  GUUUGAAGCAAUGAAAAACUACCUGAUAGAGGAGAAGUGAUAAUUUUUUGAUUUACAGUUUCUAAUCCAGAAAUAAAUC
Fan-1-1   AUAAAGAGUGAUUUAUCAUGGAUUCGAAAACAUCUGUUAUUAAUUUUACCGAAUAGCAUUAUUUAAAUAAAAAAAUGUC
Fte-1-1   AACCUCAAUAUAUCAUUGCGUUAGACGGUAGAAAUAUAAUUUCUUUUCGAUCAAAGCAUUAUUUAAAUAAAAAAAUGUC
env-1145  AUAUGAAUCUAUCUUUCAAAUUAGGGUUUAGAACCAUAAAAAUUCUUCUACUAAAAAACAAAUUAUUAAAUUUUAAGCU
env-1146  ACUAUUUAAGACAUUGACCCCCUACUCUUUAAGCAUAAAAAAUAUGUAAGUCAAAAAAUAGGUUAUUCAUAAUUUAGCU
env-1147  AUUGAUCUACAAUCAUUUAGAAAAAAGGAAGGCUCAAAAAGGAAAUCGAAAAUUUCAAUAGACAAAUAAAAAAAAUAAAU
env-1159  AUUGAUCUACAAUCAUUUAGAAAAAAGGAAGGCUCAAAAAGGAAAUCGAAAAUUUCAAUAGACAAAUAAAAAAAAUAAAU
env-1161  AUUGAUCUACAAUCAUUUAGAAAAAAGGAAGGCUCAAAAAGGAAAUCGAAAAUUUCAAUAGACAAAUAAAAAAAAUAAAU
env-1162  GAUUUAAACCUUUUAAUCAGCUUAAUAAAUAGGUAACAAAUUGAAACUAAAAUUUUAUCUGAAAAAAUAAAUAAUAAGU
env-1163  AAGAAAAAAUUUAAAAAAGCACGUAAAUUUAUACAUCGAUUUAAUACAGAAUAUACCUGUAAGAAAUCAAUAAAUAAUA
env-1165  AAGAAAAAAUUUAAAAAAGCACGUAAAUUUAUACAUCGAUUUAAUACAGAAUAUACCUGUAAGAAAUCAAUAAAUAAUA
env-1167  AAGAAAAAAUUUAAAAAAGCAGGUAAAUUUAUACAUCGAUUUAAUACAGAAUAUACCUGUAAGAAAUCAAUAAAUAAUA
env-1168  .....................................................................UAAAAAUAAUU
env-1169  CUAGUAUUCAAUAGAUUACACCGUAGCUCGCACUAAAUUCAUACUAAACUUAUUUUAUAGGUUACCUUAUAAAAUAAUU
Psp-6-1   CCUUUAAACAAAUUAUUAUACAUAUUACAAAGGGCUAAAUAAUCUAUAUAUUUUACUAUCUCAAAUUAAAAAAAAUAAUC

          ................................................................................
          ................................................................................
          ●●●●●●●●●●●●●●●●●●●●●●●●●●●●●●●●●●●●●●●●●●●●●●●●●●●●●●●●●●●●●●●●●●Y●●●●RR●●●●●●
```

| | |
|---|---|
| env-1170 | AUUGAUUUAAUUCAGAAUUUACCUAUUAGAGCUUAAUAAAAAAGGGCUGUAUCUAAGGUUUAUAUUUAAAAAAAUAAUU |
| env-1171 | AUUGAUUUAAUUCAGAAUUUACCUAUAAGAGCUUAAUAAAAAAGGGCUGUAUCUAAGGUUUAUAUUUAAAAAAAUAAUU |
| env-1172 | UAAGGGAAUAAGCGCUUUGUAACAGCACUAAUUGUACACAAAUAUCUUGCUGCAAGAGUGAAUUAAAUAAAAAAUAGUA |
| env-1173 | GCUUAGAAAUUAGACUGAUAAACACCAUUAAUUGUACAGGAAUAUCUUUUCACAAGAAUGAUUUAAAUAAAAAAUAAUA |
| env-1174 | ACUAGUAAUCAAUAGAUUACAACGUAUCUCGCACUAAAUUCAUCUAAACUUAUUUUGUAGGUUACCUUAUAAAAUAAUU |
| Tov-1-1 | CCAGCAAAUGCUGGCUCUUUUGUUUUAUUUACACCAUAAAAAACACUGAUUUCUAUACAUUUCAAAUUAAAAAUUAUUU |
| Pir-1-1 | AAACACCUUAAAUGCUGUUUAACAGUUCUUAAUCGAACAUAAUAUCUCUGUAAAAUCAUUCGUAAAUUAGAAAAUAAUA |
| env-1175 | CACCACCUUAAAUGCUGCCUAACAGUUCUUAAUAGGGUAUAAUAUCUGAUUAAAAUCACAAGUAAAUUAGAAAAUAAUG |
| env-1176 | CACCCCCUUAAAUGCUGCCUAACAGUUCUUAAUAGGGUAUAAUAUCUGAUUAAAAUCACAAGUAAAUUAGAAAAUAAUG |
| env-1177 | CACCCCCUUAAAUGCUGCCUAACAGUUCUUAAUAGGGUAUAAUAUCUGAUUAAAAUCACAAGUAAAUUAGAAAAUAAUG |
| env-1181 | CACCCCCUUAAAUGCUGCCUAACAGUUCUUAAUAGGGUAUAAUAUCUGAUUAAAAUCACAAGUAAAUUAGAAAAUAAUG |
| env-1182 | CACCCCCUUAAAUGCUGCCUAACAGUUCUUAAUAGGGUAUAAUAUCUGAUUAAAAUCACAAGUAAAUUAGAAAAUAAUG |
| env-1183 | AAAAUAAUCUAUAAUUUAUAUCGUAUUCAAAACCAGCAAAAGAACUAUUUUUUUGUUGGUUUUCUUUUUAAAAAUAACA |
| env-1184 | ............................................................................... |
| env-1185 | AUUGAUUUAAUUCAGAAUCUACCAAUUAGAGCAUAAGUAAAAAGCACGCACUGUUGGGUUUCUAUUCAAAAAAAUAAUU |
| env-1186 | AUUGAUUUAAUUCAGAAUCUACCAAUUAGAGCAUAAGUAAAAAGCACGCACUGUUGGGUUUCUAUUCAAAAAAAUAAUU |
| env-1187 | AAUAAGAGCUUAAAGACAAAGUAGACAGCAAAAACCUUUAAAAGAGGUGGUUGGUAAACUUUUUUUUAAAAAAAUAAUC |
| env-1188 | AAUAAGAGCUUAAAGACAAAGUAGACAACAAAAACCUUUAAAAGAGGUGGUUUGUAAACUUUUCUUUAAAAAAAUAAUC |
| env-1190 | AAUAAGGGCUUAAAGACAAAGUAGACAACAAAAACCUUUAAAAGAGGUGGUUUGUAAACUUUUCUUUAAAAAAAUAAUC |
| env-1191 | AUUGAUUUAAUUCAGAAUCUACCAAUUAGAGCAUAAGUAAAAAGCACGCACUGUUGGGUUUCUAUUCAAAAAAAUAAUU |
| env-1192 | CAAUAAGAGCUUAAAGACAAAGUAGACAUCAAAAACCUUUAAAAGAGGUGGUUUGUAAACUUUUUUUAAAAAAAUAAUC |
| env-1193 | CAAUAAGGGCUUAAAGACAAAGUAGACAGCAAAAACCUUUAAAAGAGGUGGUUUGUAAACUUUUUUUAAAAAAAUAAUC |
| env-1194 | AUUGAUUUAAUUCAGAAUCUACCAAUUAGAGCAUAAGUAAAAAGCACGCACUGUUGGGUUUCUAUUCAAAAAAAUAAUU |
| env-1196 | CAAUAAGAGCUUAAAGACAAAGUAGACAGCAAAAACCUUUAAAAGAGGUGGUUUGUAAACUUUUUUUAAAAAAAUAAUC |
| env-1197 | AUUGAUUUAAUUCAGAAUUUACCAAUUAGAGCAUAACAAAAAAGCGCACGCUCUAAGGUUUCUAUUCAAGAAAAUAAUU |
| env-1198 | AGCAAAUUGCUGUCCAUGAUAUUUUUUAGGUAAAACCUUUAAAAGAAGAACUUUCAUUCUGUUUUUAUUCAAAAAAAUU |
| env-1199 | AUUGAUUUAAUUCAGAAUUUACCAAUUAGAGCAUAACAAAAAAGCGCACGCUCUAAGGUUUCUAUUCAAGAAAAUAAUU |
| env-1200 | AUUGAUUUAAUUCAGAAUUUACCAAUUAGAGCAUAACAAAAAAGCGCACGCUCUAAGGUUUCUAUUCAAGAAAAUAAUU |
| env-1203 | CAAUAAGAGCUUAAAGACAAAGUAGACGGCAAAAACCUUUAAAAGAGGUGGUUUGUAAACUUUUUUUAAAAAAAUAAUC |
| env-1204 | AUUGAUUUAAUUCAGAAUUUACCAAUUAGAGCAUAACAAAAAAGCGCACGCUCUAAGGUUUCUAUUCAAGAAAAUAAUU |
| env-1205 | GCUUAAGAACAUACUAUUUCCAAUUAAUUUAAAACCAAUAAAAGAUAUUCUUUUAUUGGUUUUCUUUUAAAAAAUAAUU |
| Psp-3-1 | AAAACCAUCCAUAAUUUAUAAACAAUUCAAAACCAACAAAAGAACUAUUUUUUUGUUGGUUUUCUUCUUAAAAAAUAACA |
| env-1208 | AAUUCUUUUAAUCAACAAUUUACCAUUAGCUAUAGAUUUCAAAAUCAUAAAGUAUUUAGGGUAAAAAAUAAAAAAUAGUU |
| env-1209 | AAGCUAAAAAAAAACAAUAAACCAUUGUUUUUAAGAUUAUUAAAUUCAGCUGGUAUUUUGCUGGAAAAAUAAAAAAUAGUU |
| env-1210 | AGCUAAAAAAAAACAAUAAACCAUUGUUUUUAAGAUUAUUAAAUUCAGCUGGUAUUUUGCUGGAAAAAUAAAAAAUAGUU |
| env-1211 | AGCUAAAAAAAAACAAUAAACCAUUGUUUUUAAGAUUAUUAAAUUCAGCUGGUAUUUUGCUGGAAAAAUAAAAAAUAGUU |
| env-1212 | GCCAGCAGUUGCUGGCUUUUUUAUUUCAAUGACUUUGUAAAAAGAAAGAGGUUACAUAUGUAGGUUAUGAAAAAUAAUC |
| env-1213 | UACCUUCUAUCAAGCCAUUAAACAACCUAAAAGAAAUACAUUUUGAAUACUUCAUUGCGCUUAAAAAAUAAAAAUUAGU |
| Psp-2-1 | GCAAUUGCUGGCUUUUUUAUUUUUAGAGUAAAACUACAAAAGAAGAACUUUGGUUCCGUUUUUAUUCAAAAAAAUAAUU |
| Psp-4-1 | UAUAGAUUUAAUUCAAAAUAUACCUGUAAGAGCCUAAAUAAAAAGGUUUAAGUUCAAUUAGUUAUUCUAAAAAAUAAGA |
| env-1215 | AGUAGUAAUCAAUAGAUUACAACGUAUCUCGCACUAAAUUCAUCUAAACUUAUUUUGUAGGUUACCUUAUAAAAUAAUU |
| env-1216 | ACUAGUAUUCAAUAGAUUACAACGUAUCUCGCACUAAAUUCAUCUAAACUUAUUUUGUAGGUUACCUUAUAAAAUAAUU |
| env-1217 | ACUAGUAAUCAAUAGAUUACAACGUAUCUCGCACUAAAUUCAUCUAAACUUAUUUUGUAGGUUACCUUAUAAAAUAAUU |
| env-1218 | CUAGUAAUCAAUAGAUUACAACGUAUCUCGCACUAAAUUCAUCUAAACUUAUUUUGUAGGUUACCUUAUAAAAUAAUU |
| env-1219 | AGUAGUAAUCAAUAGAUUACAACGUAUCUCGCACUAAAUUCAUCUAAACUUAUUUUGUAGGUUACCUUAUAAAAUAAUU |
| env-1220 | ACUAGUAUUCAAUAGAUUACAACGUAUCUCGCACUAAAUUCAUCUAAACUUAUUUUGUAGGUUACCUUAUAAAAUAAUU |
| env-1222 | ACUAGUAUUCAAUAGAUUACAACGUAUCUCGCACUAAAUUCAUCUAAACUUAUUUUAUAGGUUACCUUAUAAAAUAAUU |
| env-1223 | ACUAGUAUUCAAUAGAUUACAACGUAUCUCUCACUAAAUUCAUCUAAACUUAUUUUGUAGGUUACCUUAUAAAAUAAUU |
| env-1226 | ACUAGUAUUCAAUAGAUUACAACGUAUUUCUCACUAAAUUCAUCUAAACUUAUUUUGUAGGUUACCUUAUAAAAUAAUU |
| env-1227 | ACUAGUAUUCAAUAGAUUACAACGUAUCUCUCACUAAAUUCAUCUAAACUUAUUUUGUAGGUUACCUUAUAAAAUAAUU |
| env-1229 | ...............................CACUAAAUUCAUCUAAACUUAUUUUGUAGGUUACCUUAUAAAAUAAUU |
| env-1230 | ACUAGUAAUCAAUAGAUUACAACGUAUCUCGCACUAAAUUCAUCUAAACUUAUUUUGUAGGUUACCUUAUAAAAUAAUU |
| env-1231 | ACUAGUAUUCAAUAGAUUACAACGUAUCUCGCACUAAAUUCAUCUAAACUUAUUUUGUAGGUUACCUUAUAAAAUAAUU |
| env-1232 | ACUAGUAAUCAAUAGAUUACAACGUAUCUCGCACUAAAUUCAUCUAAACUUAUUUUGUAGGUUACCUUAUAAAAUAAUU |
| env-1233 | ACUAGUAAUCAAUAGAUUACAACGUAUCUCGCACUAAAUUCAUCUAAACUUAUUUUGUAGGUUACCUUAUAAAAUAAUU |
| env-1234 | ACUAGUAAUCAAUAGAUUACAAAGUAUCUCGCACUAAAUUCAUCUAAACUUAUUUUGUAGGUUACCUUAUAAAAUAAUU |
| env-1235 | ..............................GCACUAAAUUCAUCUAAACUUAUUUUGUAGGUUACCUUAUAAAAUAAUU |
| env-1236 | ...............................CACUAAAUUCAUCUAAACUUAUUUUGUAGGUUACCUUAUAAAAUAAUU |
| env-1237 | ACUAGUAUUCAAUAGAUUACAACGUAUCUCUCACUAAAUUCAUCUAAACUUAUUUUGUAGGUUACCUUAUAAAAUAAUU |
| env-1238 | ACUAGUAUUCAAUAGAUUACAAUGUAUCUCGCACUAAAUUCAUCUAAACUUAUUUUGUAGGUUACCUUAUAAAAUAAUU |
| env-1239 | ACUAGUAUUCAAUAGAUUACAACGUAUCUCUCACUAAAUUCAUCUAAACUUAUUUUGUAGGUUACCUUAUAAAAUAAUU |
| env-1240 | ACUAGUAUUCAAUAGAUUACAACGUAUUUCUCACUAAAUUCAUCUAAACUUAUUUUGUAGGUUACCUUAUAAAAUAAUU |
| env-1241 | GUAGUAAUCAAUAGAUUACAACGUAAUCUCGCACUAAAUUCAUCUAAACUUAUUUUGUAGGUUACCUUAUAAAAUAAUU |
| env-1242 | CCUCUAUUGUAAUAAGUUAUCCCGCUUCUUUACCUUUAUUUAUCACUAUAAAAUUUAUAGGUUAACAUAUAAAAUAAUU |
| env-1243 | CCUCUAUUGUAAUAAGUUAUCCCGCUUCUUUACCUUUAUUUAUCACUAUAAAAUUUAUAGGUUAACAUAUAAAAUAAUU |
| env-1245 | ..........................................UCACUAUAAAAUUUAUAGGUUAACAUAUAAAAUAAUU |
| env-1246 | CCUCUAUUGUAAUAAGUUAUCCCGCUUCUUUACCUUUAUUUAUCACUAUAAAAUUUAUAGGUUAACAUAUAAAAUAAUU |
| env-1247 | AGACCUUUAUAGCCUGAGAUAGUUUGUUUUUAUCAAGAUAAAAUAGCCAAUCCCAGUGAUUUAGAUUUAAAAAAUAAUU |
| env-1248 | CCUCUAUUGUAAUAAGUUAUCCCGCUUCUUUACCUUUAUUUAUCACUAUAAAAUUUAUAGGUUAACAUAUAAAAUAAUU |
| env-1250 | .........................................UAUCCUUUUAAUUUUAUAGGUUAGCAUAUAAAAUAAUU |
| env-1252 | CCUCUAUUGUAAUAAGUUAUCCCGCUUCUUUACCUUUAUUUAUCACUAUAAAAUUUAUAGGUUAACAUAUAAAAUAAUU |
| env-1253 | CCUCUAUUGUAAUAAGUUAUCCCGCUUCUUUACCUUUAUUUAUCACUAUAAAAUUUAUAGGUUAACAUAUAAAAUAAUU |
| env-1254 | CCUCUAUUGUAAUAAGUUAUCCCGCUUCUUUACCUUUAUUUAUCACUAUAAAAUUUAUAGGUUAACAUAUAAAAUAAUU |
| env-1255 | UCUCGCCUAGAACAAGUUACCCAUGAGCUAUGCUACUAUUUAUCGCUGUAAAAUUUAUAGGUUAGCAUAUAAAAUAAUU |
| env-1256 | GCUCUAUAGCAAUAAGUUAUCCCGCUUCUUCACCUUUAUUUAUCACUAUAAAAUUUAUAGGUUAACAUAUAAAAUAAUU |
| env-1257 | GCUCUAUAGCAAUAAGUUAUCCCGCUUCUUCACCUUUAUUUAUCACUAUAAAAUUUAUAGGUUAACAUAUAAAAUAAUU |
| env-1259 | GCUCUAUAGCAAUAAGUUAUCCCGCUUCUUCACCUUUAUUUAUCACUAUAAAAUUUAUAGGUUAACAUAUAAAAUAAUU |
| env-1260 | GCUCUAUAGCAAUAAGUUAUCCCGCUUCUUCACCUUUAUUUAUCACUAUAAAAUUUAUAGGUUAACAUAUAAAAUAAUU |
| env-1262 | GCUCUAUAGCAAUAAGUUAUCCCGCUUCUUCACCUUUAUUUAUCACUAUAAAAUUUAUAGGUUAACAUAUAAAAUAAUU |
| env-1263 | GCUCUAUAGCAAUAAGUUAUCCCGCUUCUUCACCUUUAUUUAUCACUAUAAAAUUUAUAGGUUAACAUAUAAAAUAAUU |
| env-1264 | UCUAAUCUAGAACAAGUUACCAUAGAACUUUGCUUCUAUACAUCACUACAAAAUUUAUAGGUUACCAUAUAAAAUAAUU |
| env-1268 | CUAAUCUAGAACAAGUUACCACNNAAUCUUUGCUUCUNUACAUCACUANAAAAUUUAUAGGUUACCAUAUAAAAUAAUU |
| env-1269 | UCUAGUCUAGAACAAGUUACCCUUAAUCUUUGCUUCUUUACAUCACUAUAAAAUUUAUAGGUUACCAUAUAAAAUAAUU |
| env-1270 | UCUAGUCUAGAACAAGUUACCAAAGAACUUUGCUUCUAUACAUCGCUACAAAAUUUAUAGGUUACCAUAUAAAAUAAUU |
| | ............................................................................... |
| | ............................................................................... |
| | ●●●●●●●●●●●●●●●●●●●●●●●●●●●●●●●●●●●●●●●●●●●●●●●●●●●●●●●●●●●●●●●●●●Y●●●●RR●●●●●● |

```
env-1271   UCUAGUCUAGAACAAGUUACCCUUAAUCUUUGCUUCUUUACAUCACUAUAAAAUUUAUAGGUUACCAUAUAAAAUAAUU
env-1272   UCUAAUCUAGAACAAGUUACCACAGAACUUUGCUUCUAUACAUCACUACAAAAUUUAUAGGUUACCAUAUAAAAUAAUU
env-1273   ...........................CUUUGCUUCUUUACAUCACUACAAAAUUUAUAGGUUACCAUAUAAAAUAAUU
env-1274   UCUAGUCUAGAACAAGUUACCCUUAAUCUUUGCUUCUUUACAUCACUAUAAAAUUUAUAGGUUACCAUAUAAAAUAAUU
env-1275   UCUAGUCUAGAACAAGUUACCCUUAAUCUUUGCUUCUUUACAUCACUAUAAAAUUUAUAGGUUACCAUAUAAAAUAAUU
env-1277   UCUAAUCUAGAACAAGUUACCAUAGAACUUUGCUUCUAUACAUCACUACAAAAUUUAUAGGUUACCAUAUAAAAUAAUU
env-1278   UCUCGCUUAUAACACUUUACACUAAGUAUUUUGUUCUAUUCAUCCCCGUGAAAUUUAUAGGUUAACAUAUAAAAUAAUU
Ccy-1-1    UAAUCAAAAUUUUAAUUGAAUUUCAGAAAGAAAAUAAAUUAAAAAAAUCACAUAUUAACUUGUGGAUUAAAUAUUUUUCC
Ccy-1-3    UAAUCAAAAUUUUAAUUGAAUUUCAGAAAGAAAAUAAAUUAAAAAAAUUACAUAUUAACUUGUGGAUUAAAUAUUUUUCC
Tme-1-1    GUAAGAGCAUAACAAAAAAGCUAGCGAUUGCUAGCUUUUUUGCUUUCUAACUUUUUCGUUACCAUUCAAAAAAAUAAUC
Tsp-1-1    AAAAAAGCUAGCGUUUGCUAGCUUUUUUGUUUUCUGUCAUUUUCAAUAUACAUGAAUAGUUUUCAUUCAAAAAAUAAUU
env-1280   AGACGCUUAAACACUAUUUGAUACCCCUUAAUAUCAAGGAAAUAUCUCUUUACUAGAAUGAAUUAAAUAAAAAAUAAUA
env-1281   AGACGCUUAAAGACUAUUUGAUACCCCUUAAUAUCAAGGAAAUAUCUCUUUACUAGAAUGAAUUAAAUAAAAAAUAAUA
env-1282   AGACGCUUAAACACUAUUUAAUAACCAUUAAUAUCAAGGAAAUAUCUCUUUACAAGAAUGAAUUAAAUAAAAAAUAAUA
env-1283   AGACGCUUAAACACUAUUUAAUAACCAUUAAUAUCAAGGAAAUAUCUCUUUACAAGAAUGAAUUAAAUAAAAAAUAAUA
env-1284   AGACGCUUAAACACUAUUUAAUAACCAUUAAUAUCAAGGAAAUAUCUCUUUACUAGAAUGAAUUAAAUAAAAAAUAAUA
env-1285   AGACGCUUAAACACUAUUUAAUAACCAUUAAUAUCAAGGAAAUAUCUCUUUACUAGAAUGAAUUAAAUAAAAAAUAAUA
env-1287   AGACGCUUAAACACUAUUUAAUAACCAUUAAUAUCAAGGAAAUAUCUCUUUACAAGAAUGAAUUAAAUAAAAAAUAAUA
env-1293   AGACGCUUAAACACUAUUUAAUAACCAUUAAUAUCAAGGAAAUAUCUCUUUACAAGAAUGAAUUAAAUAAAAAAUAAUA
env-1294   AGACGCUUAAACACUAUUUAAUAACCAUUAAUAUCAAGGAAAUAUCUCUUUACAAGAAUGAAUUAAAUAAAAAAUAAUA
env-1295   AGAGGCUUAAACACUAUGUGAUACCCCUUAAUAUCAAGGAAAUAUCUCUUUACUAGAAUGAAUUAAAUGAAAAAUAAUA
env-1296   AGACGCUUAAACACUAUUUAAUAACCAUUAAUAUCAAGGAAAUAUCUCUUUACUAGAAUGAAUUAAAUAAAAAAUAAUA
env-1298   AGACGCUUAAACAAUAUUUAAUACCCGUUAAUAUCAAGGAAAUAUCUCUUUACUAGAAUGAAUUAAAUAAAAAAUAAUA
env-1299   AGACGCUUAAACACUAUUUAAUAACCAUUAAUAUCAAGGAAAUAUCUCUUUACAAGAAUGAAUUAAAUAAAAAAUAAUA
env-1300   UCUCAACUGUAAUAAGUUAUCCAUUAUCUUUUCCUUUUUUUAUAGAAAUAAAAUUUAUAGGUUAACAUAUAAAAUAAUU
env-1301   UCUCUAUUGUAAUAAGUUAUCCCAUUUCUUCACCUUUAUUUAUCACUACAAAAUUUAUAGGUUAACAUAUAAAAUAAUU
env-1302   ...................................UUAUUUAUCACUACAAAAUUUAUAGGUUAACAUAUAAAAUAAUU
env-1303   AUUUGCCCGCUUUAUUAUUCUUAGAGUAAAACUCUAAAAAAGAAGAAGUUCAGCAAUGUUUCUAUUCAAAAAAUAAAUU
env-1304   AAUAUUGGAGCCUUUUAUGAGAACCCAGCCAUCUUCAAUUAAUAUUCCUCAUAAUCAAUCAGAUAUCUAGAAAAUAGUU
env-1305   AACAUUUAUGGGCAGUUGAAAAGUCUUGAUCAGCAAAAAAGAUCCAGUAAGUUAAUUAGGCAGAAUAGCAAAAAUAUUU
env-1306   AACAUUUAUGGGCAGUUGAAAAGUCUUGAUCAGCAAAAAAGAUCCAGUAAGUUAAUUAGGCAGAAUAGCAAAAAUAUUU
env-1307   AACAUUUAUGGGCAGUUGAAAAGUCUUGAUCAGCAAAAAAGAUCCAGUAAGUUAAUUAGGCAGAAUAGCAAAAAUAUUU
env-1308   AUUACAAUGCGCACUACUACGAGCAAAAAACCAUAGCAAAAAAUUUCUAUUUUAAGGCGUUUUACAUUCCAAAAUAAUA
env-1309   AACAUUUAUGGGCAGUUGAGAUGCCUUAAUCAACAAAAAAGAUCUAGCAAUUUAUUUAGGUAGAAAUGCGAAAAUAUUU
env-1310   AACAUUUACGGGGAGUCAGGAAGCAUUGACCAAUAAAAAAGAUCCGGUAAGUUAAUUGGGCUGAAUAGCAAAAAUAUUU
env-1311   ...............................................UAAGUUAAUUGGGCUGAAUAGCAAAAAUAUUU
Tna-1-1    AAAAAAUUAACACCACCAAUGUAAGAUUUUUAUGCUAAUUAAAAAAAACAAAUCAUUUUAGUUUUUUACUAAAAAAAACAG
env-1312   CAUUUGCCUAAGCCGCUUGCAAUAAAGCAUUUAGGCUUGUAAAAAAUAAAAAAAAGUAGUGUGUUUUCACUAACAUUUA
Cly-1-1    AACAUAUCCAAUUAAAGUAAAAAUAGUAUUCGCACAAAAAAAAUCUACACAGGUUAAGUUAAAGUGUAAUAAAUAAAUU
Cge-1-1    CAACAUAUCCAAUUAAAGUAAAAAUAGUAUUCGCACAAAAAAAUCUACACAGGUUAAGCUAAAGUGUAAUAAAUAAAUU
Cly-2-1    CAACAUAUCCAAUUAAAGUAAAAAUAGUAUUCGCACAAAAAAAUCUACACAGGUUAAGUUAAAGUGUAAUAAAUAAAUU
Sxi-1-1    CCAAUAAGAGCUUAAAACAGCAUUUUUAUCCUGAUUUCCAGAAAAGUAAAAAAUUACAUCGCUCAUAUUUUAUAUUAUA
Ste-1-1    CCAAUAAGAGCUUAAAAAUGUCAUUUCUGACUGACCUUCAGAAAAUUAAAAAUUUACACAGCUAAUAUUUUAUAAUAUC
Cal-1-1    UACAUGAUACUGAUCAAUUUAGAAAAAGAGGAAGAUUUAAAAUUUCUCUUUUAUAGAAUAAGAUUAUCAUAAAAUAAUU
env-1313   GGCGUUAUCAUUCACAACAACAUUUUUCAAAAAUACCGGCGAACAACUUCCAAAAAACUUGUUUUUCAAAAAAAUAAAU
Mze-2-1    UAUUGCGAUCAAUUAAUCCAUUUGAUUUUACGGAUUAUACCGAUAUUUAACUUUUUGAAAAUCUGUUUCAAAAAAAGCA
env-1314   GACGAACGCUUUUUAGAAUUUGUGAAUGAAGAAGUUGCAAAGUUGUUUUAGAAAAAACUUGUACUUUUAAAAAAUAAAA
env-1316   GACGAACGCUUUUUAGAAUUUGUGAAUGAAGAAGUUGCAAAGUUGUUUUAGAAAAAACUUGUACUUUUAAAAAAUAAAA
env-1317   GCACUUUUUGGAAUUUGUGAAUCGUGAAAUUGCGAAGUUGUAUUAACGCGCAAAUUACUUGGAACUUUAAAAAUAAAGU
env-1318   ..................UCAAGAAGUUGCAAAAUUAUUUUAAACUGUACUUUUUUAUUUGCCUAAUUAAAAAAGAAGA
Fgi-1-1    AUAAUUCACAAUUCAUAAUUGCUUCGCCGAUUCAAAAAAUAAAAAUAUUUUUUUUCGGGUCAUAAUUCAUAAUUAUUUU
Fsu-2-1    UUUUAAACAACUAACACCUAACCACUUGUCAACUCUAAAAAUAAAGAAACGUUUCUUUUGUUCUAGUCAAAAUUAUUUU
env-1319   ...............AUUGGGCNCAUUGACUCCUUUUUUUAGAUUUUNAUAAUAUUUUAGAUUUUUCAAAAAAUAAAUA
env-1320   CCAUAGUGGUCUAGAAGUUGUUUUUUAUUGAUUAGGUAAAAUAAUUCAUAAAGUUUAAUUCACAAUUCAAAAUUAUUAC
env-1321   CCAUAGUGGUCUAGAAGUUGUUUUUUAUUGAUUAGGUAAAAUAAUUCAUAAAGUUUAAUUCACAAUUCAAAAUUAUUAC
Fjo-1-1    UCUAAAUCAUUAAAAAAGAAUUUCUUAGGUCUUAUUUUUAGAUUCUUUUAUUUUUUUUUGAUUUUUCUGUUUUUUUAAU
Fsp-7-1    UCUAAAUAUUUAAAAACAAAAAUCUUAUAUCCAUUUUUUAGAUUCUUUUAUUUUUUUUUGAUUUUUCUGUUUUUUUAAU
Fsp-13-1   ACCUGCAAGGAUUCUAUACAAACUCCCCAUUACAAUUGUAAAAAACGGACGAUAAAAUUCGUUUUUCCAAAAAAAAGUU
Fsp-6-1    AAAAAAGGAUUAAUUUUAUCUAAAAUUGUUUAAUCUUUAAAUUUUUCAAUCUUUAAAUUUAUUUUUUCGAAAAAAAGAU
Fsp-8-1    UCCGAUAAGAGUUCUGUACACAUUACCUAUAACAAUUAGAAAAAAUACUCCAUAAAAUUUAUUUUUAGAGAAAAAAAGUA
Fsp-10-1   UCCAAGCCGAGUUAGCUAUUCAUUACCUAUAACAAUUAGAAAAAAUACUCAAUAAAAUUUAUUUUUAGAGAAAAAAAGUA
Fhy-1-1    UCCUAGGCGAGUUAGCUAUUCGUUACCUAUAACAAUUAGAAAAAAUAAUCAAUAAAAUUUAUUUUUAGAGAAAAAAAGUA
env-1323   ..............................................UCAAUCUUUAAAUUUAUUUUUCGAAAAAAAGAU
env-1324   AAAAAAGGAUUAUUUUUACCUGAAUUUGAUUAAUUAUUAAAAUUUUCAAAGUUUAAAUUUAUUUUUUCGGAAAAAAAGAU
Fsp-11-1   AAAUAUUGCUCAAAAAACAAAAUUUGGUAUUCUAUUUUAAGAUUCUCUUAUUUUUGUUUGAUUCUUUUGUUUUUUUACC
Fch-1-1    AAAAAGAAUUAUUUCCACCUGAAAUUGACUAAUUAUUAAAAUUUUCAAAGUUUAAAUUUAUUUUUUCAGAAAAAAAAGAU
env-1325   .............................GUGUUUAUAAAAUAAUUCAUAAUUCAUAAUUCAUAAUUGAUUAUUAUUUU
env-1326   GACUUUAAUUAAUUGUAUUUUGUAAUUUAGUGUUUAUAAAAUAAUUCAUAAUUCAUAAUUCAUAAUUGAUUAUUAUUUU
env-1327   GACUUUAAUUAAUUGUAUUUUGUAAUUUAGUGUUUAUAAAAUAAUUCAUAAUUCAUAAUUCAUAAUUGAUUAUUAUUUU
env-1328   GACUUUAAUUAAUUGUAUUUUGUAAUUUAGUGUUUAUAAAAUAAUUCAUAAUUCAUAAUUCAUAAUUGAUUAUUAUUUU
env-1330   UGACUAAUGCUCUUGUUUGAUUUGUUGAAAUUGCUCAAAAACUUUCCUUGAAAAAUAAUUUGUUUUUGAAUAUUAUUUU
env-1331   UGACUAAUGCUCUUGUUUGAUUUGUUGAAAUUGCUCAAAAACUUUCCUUGAAAAAUAAUUUGUUUUUGAAUAUUAUUUU
env-1332   UUUUUCCAUCAUAACAUUUUUUAGGGGAAAGUUAUGAUGGUUUUUUUUGCUUAAAAUUUGCUUACUCUAAAAAAUAUUC
env-1333   UUUUCCAUCAUAACAAUUUUUAGGGGGAAGUUAUGAUGGUUUUUUUUGCUUAAAAUUUGCUAACUCUAAAAAAAUAUUC
Gsp-4-1    AAUAAUAUAUCAGAAUGCUAUCUAUAGAAAGGCUGAAAUAUUAAAAUUUGACUUGAGAAUAAUUUAUAAGAAAAAUAUA
Gli-1-1    CCUGCCCGACGGUCAGGUGGAUUAAACCCUCAAUGAGGGAUUAAAAUUAAAUUUCAGAAUAAAUUUUAAUAAUAAUAUA
env-1334   ...............................AUGAGGGAGUUAAAAUUAAAUUUCAGAAUAAAUUUUAAUAAUAAUAUA
env-1335   UAUCUCGAUCUAAUUCUAAAUCAGCCUUUAAGGGCUUAAAAAGUUUUGAUCACGCCGGUACGAUAUUAUUAAAAAUAUA
env-1336   ...............................................UUUUUGUUGGAAUAAGUAUUCAUAAAAUAGUA
env-1337   CUUAAAACUUAAAGAACAAUCCCACAACUACUUAUCACAAUAAUUUCUUUUUGUUGGAAUAAGUAUUCAUAAAAUAGUA
env-1338   GCUAAAAGAAGGGUUUUAAAGAAUUUAUUUAGCCUGAUUAAUAAAAGAUUUCACACGUAUUACCUUUAAUAAAAAUAUA
env-1339   ................................................................UUUAAUAAAAAUAUA
env-1340   AGCUAAAAGAAGGGUUUUAAAGAAUUUAUUUAGCCUGAUAAUAAAAGAUUUCACACGUAUUACCUUUAAUAAAAAUAUA
env-1341   UUAAUUCCAGUUUAUUCAGCUAAUUGAUUUAGUUGCAUAAUAAAUGAAUUCUACUUCUAUAGCUUUAAAUAAAAAUAUA

           ...............................................................................
           ...............................................................................
           ●●●●●●●●●●●●●●●●●●●●●●●●●●●●●●●●●●●●●●●●●●●●●●●●●●●●●●●●●●●●●●●●●●Y●●●●RR●●●●●●
```

```
Gsp-5-1    GUUAAAUUAGGAUGUUUUAGUAAUUCCUUUUACUUGAUCAAUAAAAGAUUUCACUUCUAUAAGGUUUAAUAAAAAAUAUA
env-1342   AGCUAAAAGAAGGGUUUUAAAGAAUUUAUUUAGCCUGAUAAUAAAAGAUUUCACACGUAUUACCUUUAAUAAAAAAUAUA
env-1343   AGUUAAAUUUCAUUUCAUUUUAAUAGUUUUUGGUUUUUUAAUAAAAGAUUUUUCCUCUAUAACAUUUAAUAAAAAAUAUA
env-1344   UUAAUUCCAGUUUAUUCAGCUAAUUGAUUUAGUUGCAUAAUAAAUGAAUUCUACUUCUAUAGCUUUAAAUAAAAAAUAUA
env-1345   UUAAUUCCAGUUUAUUCAGCUAAUUGAUUUAGUUGCAUAAUAAAUGAAUUCUACUUCUAUAGCUUUAAAUAAAAAAUAUA
env-1346   GCUAAAAUUAGUACUUUCAGGAAUUUGGAUUGCUAGAUUAAUAAAUGAUUUUACAACUAUAACCUUUAAUAAAAAAUAUA
Gsp-3-1    UUUAACAUUUAAUUAUUUAGCAAUUUUAGUUCACUGAAUAAUAAAAGAAUUAACAUCUAAAUAUUUUAAGAAUAAUAUA
Gsp-2-1    AGCUAAAAGAAGGGUUUUAAAGAAUUUAUUUAGCCUGAUAAUAAAAGAUUUCACACGUAUUACCUUUAAUAAAAAAUAUA
env-1347   GCUAAAAUUAGUACUUUCAGGAAUUUGGAUUGCUAGAUUAAUAAAUGAUUUUACAACAAUAGCUUUUAAUAAAAAAUAUA
env-1348   CUUAAAACUUAAAGGACAAUCACACAACUACUUAUCACAAUAAUUUCUUUUUGUUGGAAUAAGUAUUCAUAAAAAUAGUA
env-1349   CUUAAAACUUAAAGGACAAUCACACAACUACUUAUCACAAUAAUUUCUUUUUGUUGGAAUAAGUAUUCAUAAAAAUAAUA
env-1350   CUUAAAACUUAAAGGACAAUCACACAACUACUUAUCACAAUAAUUUCUUUUUGUUGGAAUAAGUAUUCAUAAAAAUAGUA
Gec-1-1    UUAUUCAGAAUCAACCAGUAAGAGCAUAAAUACCUUGAAAUAAAAAAUUUAGGUUUAGAUAGGUAUUUUUUAAAUUAGUA
Gpo-1-1    AUUUUCACUUUGCAAAAAUAUGAAUUUUUGAAGAGUUUAUAAAUUUAAAAGCUUAUACAUACGGAAUAAAAAAUAAAUG
env-1351   AUUUUCACUUUGCAAAAAUACGACUUUUGAUUGAGUCUAUAAAUUUAAAACCUAAUACAUACGGAAUAAAAAAUAAAUG
env-1352   UUAUUCAGAAUCAACCGGUAAGAGCCUGAUAGCUUUCAAAUAAAAAAUUAGGUUUAGAUAGGUAAUUUUUAAAUUAAUA
env-1353   UUAUUCAGAAUCAACCGGUAAGAGCCUGAUAGCUUUCAAAUAAAAAAUUAGGUUUAGAUAGGUAAUUUUUAAAUUAAUA
Gfo-1-1    UUAUUCAGAAUCAACCGGUAAGAGCUUAAUUAGCACAAAAUAAUAAUAGCGGUUUAAAUAGGUAUUUUUUAAAUUAGUA
env-1354   UGAAUAGGAAAACAGUAUUAUAUCCCUUGCCCUGUGGCAAUAAAAUUCCAAGAAUAAAUAGCAUUUAUACAAAAAAGUA
env-1355   ..............................CCUGUGGCAAUAAAAUUCCAAGAAUAAAUAGCAUUUAUACAAAAAAGUA
env-1356   ................................................................................
env-1357   UGAAUAGGAAAACAGUAUUAUAUCCCUUGCCCUGUGGCAAUAAAAUUCCAAGAAUAAAUAGCAUUUAUACAAAAAAGUA
env-1358   UGAAUAGGAAAACAGUAUUAUAUCCCUUGCCCUGUGGUAAUAAAAUUCCAAUAAUAAAUAGCAUUUAUACAAAAAAAUA
env-1359   UGAAUAGGAAAACAGUAUUAUAUCCCUUGCCCUGUGGCAAUAAAAUUCCAAGAAUAAAUAGCAUUUAUACAAAAAAAUA
env-1360   UGAAUAGGAAAACAGUAUUAUAUCCCUUGCCCUGUGGCAAUAAAAUUCCAAGAAUAAAUAGCAUUUAUACAAAAAAGUA
env-1367   UGAAUAGGAAAACAGUAUUAUAUCCCUUGCCCUGUGGCAAUAAAAUUCCAAGAAUAAAUAGCAUUUAUACAAAAAAGUA
env-1370   ..............................................................................UA
env-1371   AGAAUAGGAAAACAGUAUUAUAUCCCUUGCCCUGUGGUAAUAAAAUUCCAAUAAUAAAUAGCAUUUAUACAAAAAAAUA
env-1373   AGAAUAGGAAAACAGUAUUAUAUCCCUUGCCCUGUGGUAAUAAAAUUCCAAUAAUAAAUAGCAUUUAUACAAAAAAAUA
env-1374   AGAAUAGGAAAACAGUAUUAUAUCCCUUGCCCUGUGGUAAUAAAAUUCCAAUAAUAAAUAGCAUUUAUACAAAAAAAUA
env-1375   .....................................................AAUAAAUAGCAUUUAUACAAAAAAGUA
env-1376   UGAAUAGGAAAACAGUAUUAUAUCCCUUGCCCUGUGGCAAUAAAAUUCCAAGAAUAAAUAGCAUUUAUACAAAAAAGUA
env-1377   .....................................................................ACAAAAAAGUA
env-1378   CUUAUUCAGAAUCAACCGGUAAGAGCCUAAAACAAAUUUCAAAAAGAUAAACCUGCAGUAGGUAAUUCUCAAGAAAGUA
env-1379   .............AUUCAAAAUCAACCGGUAAGAGCUUAAAAUAAAUUUUCCUUUAAGUAGGUAAUUAUAUAGAAAGUA
env-1380   AUAUUUGGAUCUUAUUCAAAAUCAACCGGUAAGAGCUUAAAAUAAAUUUUCCUUUAAGUAGGUAAUUAUAUAGAAAGUA
env-1381   GUAUUUGGAUCUUAUUCAAAAUCAACCGGUAAGAGCUUAAAAUAAAUUUUUCUUUAAGUAGGUAAUUAUAUAGAAAAUA
env-1382   GUAUUUGGAUCUUAUUCAAAAUCAACCGGUAAGAGCUUAAAAUAAAUUUUUCUUUAAGUAGGUAAUUAUAUAGAAAAUA
env-1383   GUAUUUGGAUCUUAUUCAAAAUCAACCGGUAAGAGCUUAAAAUAAAUUUUUCUUUAAGUAGGUAAUUAUAUAGAAAAUA
Ssp-1-1    GUAUUUGGAUCUUAUUCAGAAUCAACCAGUAAGGGCUUAAAAAAAAUAUUUUCUUUCCGUAGGUAAUUAUAUAGAAAGUA
Tse-1-1    UGAAAAAUUAACAACAUUAUUGUAAGAUUUUUACAUCAAAAAAAAUAAAUAGUUUUAACUAUUCAUAAAUAAAAUAUGU
Lsp-2-1    GGCUAUUAUGGGACUUUUUGUACCAAUAUAAAGCAAUACGCAUUUUGAAAUAAAAAUAUUUGAAAAUCAAUUAAAAAGC
Jpa-1-1    AUAGCUACUUUAGGGAGGGCAACUUUUUAAAAAUCAAUGUAAAAGUUACAUAAAAAAACAAUUAAAUACAAAAAAGAUA
Ssp-2-1    GAGUGAAGUCGUACCGUAUAAGCAAAAUUAUUACCCCAAAAGAAACUGAUUACGAGCAAUUAAAUUUACAAAUAAUUUA
env-1384   GAUGUGCUGAAUUAAAUGAAGUAAAUUUCAUUUUCAACAUCAAAAAGUUACGGAAAAAAUACCAAAUCACAAAAAAAUA
env-1385   AAGGUUAAUAGCCUCGUAAAAACCAUUCAUUUAAAAAUGAUUCUUUAGACUAAAAAAAUUAUGGAAUCAUUAAAAAUCA
env-1386   AAGGUUAAUAGCCUCGUAAAAACCAUUCAUUUAAAAAUGAUUCUUUAGACUAAAAAAAUUAUGGAAUCAUUAAAAAUCA
env-1387   CUAUUAGUCUAAGGAGAAUACUUAAUACGUAUAAGGACUAAAAGUUACAAAAAAAUUAAAUAUAAAUUGCGAAAAAGUU
env-1388   CUAUUAGUCUAAGGAGAAUACUUAAUACGUAUAAGGACUAAAAGUUACAAAAAAAUUAAAUAUAAAUUGCGAAAAAGUU
env-1389   CUAUUAGUCUAAGGAGAAUACUUAAUACGUAUAAGGACUAAAAGUUACAAAAAAAUUAAAUAUAAAUUGCGAAAAAGUU
env-1391   UAUUAGUAUAAGGGGGAUACUUAAUGUGUAUAAGGACUAAAAGUUACAAAAAAAAUUAAAUAUAAAUUGCGAAAAAGUU
env-1392   CCAGUCGAGGGGAGACUAUAUUAAAACACUGUAAUACAGAUUAAAAGUUACAAAAAAAUCCACAAAUCACAAAAAAGUA
env-1393   ...............................................GUUACAAAAAAAUCCACAAAUCACAAAAAAGUA
env-1394   CCAGUCGAGGGGAGACUAUAUUAAAACACUGUAAUACAGAUUAAAAGUUACAAAAAAAUCCACAAAUCACAAAAAAGUA
env-1395   CCAGUCGAGGGGAGACUAUAUUAAAACACUGUAAUACAGAUUAAAAGUUACAAAAAAAUCCACAAAUCACAAAAAAGUA
env-1396   AGCUAAUUGCAAAUGGUUUAAAAAAAUAAUUUCAAUAAGCAAAAGAAAAGAAGGCAGGUGGUUUAGUCAGCAAUUUUUU
env-1397   AGCUAAUUGCAAAUGGUUUAAAAAAAUAAUUUCAAUAAGCAAAAGAAAAGAAGGCAGGUGGUUUAGUCAGCAAUUUUUU
env-1398   AAGAGGGAUACAAUGGUUUAAUACCGUUUAUCGAAAAAAUAAUUUUUCAGCAUAACAACAGCGAAAUUACUUAUUUUAC
env-1399   AUUAGUCAGGGGAGACUUACUAAAACAUUGUUAUACAGAUUAAAAGUUACAAAAAAAACCACAAAUCACAAAGAAAGUA
env-1400   AUUAGUCAGGGGAGACUUACUAAAACAUUGUUAUACAGAUUAAAAGUUACAAAAAAAACCACAAAUCACAAAGAAAGUA
env-1401   AUUAGUCAGGGGAGACUUACUAAAACAUUGUUAUACAGAUUAAAAGUUACAAAAAAAACCACAAAUCACAAAGAAAGUA
env-1402   AUUAGUCAGGGGAGACUUACUAAAACAUUGUUAUACAGAUUAAAAGUUACAAAAAAAACCACAAAUCACAAAGAAAGUA
env-1403   ............................AUGUUAACGCUAAAGAAUUACGAUUUAUCAACAAGAUUUGUUAAUUAUUC
env-1404   .......................................................................UAAAAAUUA
env-1405   ......................................................................UAAAAAAGUU
env-1406   GAUAGACUUAUCUAAGGAAAAGACUGAAUGAAAAUAAGAAUUAAAAGUAACAAAAAAAAAAUAUAAAGGGUAAAAAAGUU
env-1411   ...........................UUGAAAAUAAGAAUUAAAAGUAACAAAAAAAAAAUAUAAAGGGUAAAAAAGUU
Wsp-1-1    UCAAUCGAAAAGUGAGAAAUUAAAACAAAGUUAUACUCUAAGAAAUGCAUCAAAAAAAAUAAGCAAAUCACAAAAAAGUA
env-1412   AAGGUUUAAAAUCACUAUAAUAAAUAUUUCAAAAGUAAAAACAAGAGUAUCCAAUUUGUUAUUAUUCAAAAUAAUAGUU
env-1413   AGGUUUAAAAACGUUAUCAAAAAUUAUUCUAUCUUAAAAAACUAGAGGAUUCAAUUUGUUGCUAAUCAAAAUAAUAGUU
env-1414   AAGGUUUAAAAUCACUAUAAUAAAUAUUUCAAAAGUAAAAACAAGAGUAUCCAAUUUGUUAUUAUUCAAAAUAAUAGUU
env-1415   AAGGUUUAAAAUCACUAUAAUAAAUAUUUCAAAAGUAAAAACAAGAGUAUCCAAUUUGUUAUUAUUCAAAAUAAUAGUU
env-1416   AAAAGUUUAAAAUUGCUUUAAUUUUUUUUCAACCUUAUAAACUAGAACAUACAAUUUAUUGUUAAUCAAAAUAAUAGUU
Bar-1-1    GCAAUUUUCUGUUAGAUAAAGCGCUACAAAACAGUGGUUGUCUUUUUUUAACAAAAAACGAUUGAAUAAUUAAAAUUUA
env-1417   GUUACUUAUUAGUAGCUAAUACGCUAAAAAACAGAGCGGGACUUUUUCAAACAAAAAACGGCUGAAUAAUUAAAAUUUA
env-1418   GCGCAUUUUUAAAGAAUCUAUUUUAUUUAAAACAGCGUUGUCUUUUUCAAACAAAAAAUGGCUAAAUAAUUAAAAUUUA
env-1419   AAAGGUUUAAUAUCAUUAUAAAAAAUUCGAUGAAUGACAUCAAAAAAAUAAGCUUUAGCUUGUAAAUCACUAAAAAUUA
env-1420   AAAGGUUUAAUAUCAUUAUAAAAAAUUCGAUGAAUGACAUCAAAAAAAUAAGCUUUAGCUUGUAAAUCACUAAAAAUUA
Pme-1-1    AAGUUGAAAGUCAUCAGGUUGUCAUUCAAUAAAAUAGAAAACUAUUAAGACGCUAAUAUUUGUUAAUCAUUAAAAAUUA
env-1421   AACAAAUAAACCAAACAAGUUAGAACCCGAUAUAAAAAUAAAACUAAGACAUCCUCUAUUUGUAAAUCAUUAAAAAUUA
Psp-5-1    UAAUAGCUAUUUUAUAAAAAAAACAUAAAAUAUUAACUCAAAAACUUUUUUGAAGAACUUUUAGAAUUGUAAAAAAAGA
Pbu-1-1    CCUAAAAUGAAAACGUUUAAUUUUCUGUGUAAAAAAAACAAAACCAACAUUGUAAAUAGAUACAAAUCAUUAAAAAUUA
env-1422   UAUAACGAGAUGGUUUAAAAAAAAUUAUUAGUUCAUAAUAAUAAUCUUUUUACAAAAGAUUGCAAUUCAUGAAAAAGUA
env-1425   ACAAAAUUGAAAACGUUUAAUUUUCGGACCAAAAAACACAAAACUAAAAUUGCGAACCCUUGUAAAUCAUUAAAAAUUA

           ................................................................................
           ................................................................................
           ●●●●●●●●●●●●●●●●●●●●●●●●●●●●●●●●●●●●●●●●●●●●●●●●●●●●●●●●●●●●●●●●●●●Y●●●●RR●●●●●●
```

| | |
|---|---|
| Pja-1-1 | AUCAAUUAGAAAAGGUUUAAUCUUUCCGUUCAAAAUGGAAAAAACAGUGAUUUCAAAAUUUGUAAAUCAUUAAAAAAUUA |
| env-1426 | UAUCAAUUAUAAAAGGUUUAAUCUUUCAAUUUAAAAUAGAAAAACAGUGGUUUGUAAAUUUGUAAAUCAUUAAAAAAUUA |
| env-1427 | CACUUUAUUUAUUCUUUCUCAUAAAUCAAAUUCAAAAUAAAAAACAACCUUCAGUAAUCUUGUAAAUCAUUAAAAAAUUA |
| env-1428 | CUAAGAGAUUUAUCAAGAAAUAACUUUUAAUUAGAUAUAAAAAACAAUGGCAUCUAAAUUUGCAAAUCAUUAAAAAAUUA |
| env-1429 | AAACCUUUACUCAAUGUCGAAGUUUAGAAUAAAAUUAUAUUCUUUCGAAACAAAAACAACAUCAAAUAACUUAAAAUCA |
| env-1433 | ..........................AGAAUAAAAUUAUAUUCUUUCGAAACAAAAACAACAUCAAAUAACUUAAAAUCA |
| env-1434 | UAAAAACUUACAAAAAAAAGAGAUUAAUGAAACGCUAAAAAAGAUUUUUUACUUAUUAUUAUGCACAUUAAAUUAUUCA |
| env-1435 | .................................CGCUAAAAAAGAUUUUUUACUUAUUAUUAUGCACAUUAAAUUAUUCA |
| env-1436 | AACACAAUGUUAACCACUAUCUUGAUCUCAUACAAAACCAUCCACUAAGAGAAUAAAUGAAGGCUAUUUUAUUCAAAUU |
| Oma-1-1 | CCUAAAAAUAGACCAUACUGCAUUUUUAAAAAAAUUUUAGAAGAAUUUGUUUUAAAAAUUACGAAAUCAUUAAAAAAUAA |
| env-1437 | CCUAAAAAUAGUUCAUACUACAUUUUUAAAAAAAUUUUAGAAGAAUUUGUUUUAAAAAUUACGAAAUCAUUAAAAAAUAA |
| env-1438 | CCUAAAAAUAGUUCAUACUGUAUUUUUAAAAAAAUUUUAGAAGAAUUUGUUUUAAAAAUUACGAAAUCAUUAAAAAAUAA |
| env-1439 | CCUAAAAAUAGUUCAUACUGUAUUUUUAAAAAAAUUUUAGAAGAAUUUGUUUUAAAAAUUACGAAAUCAUUAAAAAAUAA |
| Sze-1-1 | CCUUUAAGAGCCUAAGUAUUAACUUUAGUUUUAAGUGGAAAUAAAGCCAACAAAGUUUUCAACUUUCAACUUAAAAGCA |
| Gsa-1-1 | UAAGAGCAUAACAAUAAAAGCGACUAAAAUUAGUCGCUUUUUCUUUUUUAUUAUUAGGCUUUUAUAUCAGUAUAAAUUA |
| env-1441 | AUACAUGAAAAAGGUUUAAAAGAACACCAAAAACAAUUCGAAACAAUUAUAACUAUCACUUGCAAAUCAUUAAAAAAUUA |
| env-1442 | UAUAAAACGAUACAAAAGUUCAACACUUCUAAAAUAAUAUCAAAAAUCACAAAAAAAACCUCUCAAAAGCAAAAAAGCA |
| Mze-1-1 | GUUAUAAACUCAAAAAAAAUUAAAAAAAAUAUCUAAUUAGAUGUUUUAUACAUUUAAACAUAGAGUUUUAAUAAAAAUAC |
| env-1444 | GAUAUAAACUCAAAAAAAAUUAAAAAAAAUAUCUAAUUAGAUGUUUUAUACAUUUAAAUAUAUAGUUUUAAUAAAAAUAC |
| env-1445 | AUUAAAACGGAAAGGUUUAAACUUAAGUAUCUAAAUCUUUAAACAAUUUUCGCAAUUACUUAUAAAUCAUUAAAAAAUUA |
| env-1447 | .........................GCUUAGGGAUUAUAAGCAAAAGUAACAAAAAAAAAUGUUUCAAACCAAAAAAUUA |
| env-1448 | ................................GAUUAUUAGCUAAAGUAAUGAAAAAAAAUGUUUCAAACCAAAAAUUCA |
| env-1449 | AUAUUAUUAACUAUAAAUCUUAGGGCUUAGGGAUUAUAAGCAAAAGUAACAAAAAAAAAUGUUUCAAACCAAAAAAUUA |
| env-1454 | ...........................................................UAUCUAUAUAAAUAAAAAUAA |
| env-1455 | AAAAUCGUAAUUAGGGCAUAAAUCUAUUCUUUAGGAUAUGCCAAAAGUUACGAAAAAAAUUUCAAAUCACAAAAAAGUA |
| Wps-1-1 | GGAAAUUUUAGUUAGGACAUAAUUAUACGCCUUAGGAUAUCAAAAAGUUACGAAAAAAAUUCCAAAUCACAAAAAAGUA |
| env-1456 | GAAAAUCGUAAUUAGGGCAUAAAUCUAUUCUUUAGGAUAUCAAAAAGUUACGAAAAAAAUUUCAAAUCACAAAAAAGUA |
| env-1457 | CACACUAACAAACAAGGCCUUUAUUUUACAAUAAUAAAGGUCUUUUUUAUUUUUUAUAUAUCUAUAUAAAUAAAAAAUAA |
| Osp-2-1 | GCAACACUUAGAGUUUAAUAGCUACAAAUUAUACCUUAUUUAUAAAUUAAAUCAAAUGUUUGCCAUUUAAUAAAAAAUAA |
| Osp-1-1 | GCAACACUUAGAGUUUAAUAGCUACAAAUUAUACCUUAUUUAUAAAUUAAAUCAAAUGUUUGCCAUUUAAUAAAAAAUAA |
| env-1458 | GUUAACUAGGUUAAAUCAUUAUAAAUAAUAUUUUUGACAUCAAAAAGUUACGAAAAAAAAAUCAAAUCACAAAAAAAAUA |
| env-1459 | GUUAACUAGGUUAAAUCAUUAUAAAUAAUAUUUUUGACAUCAAAAAGUUACGAAAAAAAAAUCAAAUCACAAAAAAAAUA |
| env-1460 | GUCAAAGGGGUGACUUAAUUCAAACAUUGAAAUACUGAUUAAAAGUUACGAAAAAAAAUUAACAACUUGCAAAAAAGUG |
| env-1461 | GUUUGUUGCUUUAAAUUAUUUCUAAUCGUAAUUUCAACAUCAAAAAGUUACGAAAAAAAUACCAAAUCACAAAAAAAAUA |
| env-1462 | GUUUGUUGCUUUAAAUUAUUUCUAAUCGUAAUUUCAACAUCAAAAAGUUACGAAAAAAAUACCAAAUCACAAAAAAAAUA |
| env-1463 | GAAAAUCUUAGUUAGGACAUAAUUACACUCUUUAGGAUAUCAAAAAGUUACGAAAAAAAUUCCAAAUCACAAAAAAGUA |
| Lsp-1-1 | UAGUUUUACGAUAAAAUACAUGAGAAAAUAUUAAAAAAAAGGUUUUAGAUAUAUAAAUUUAAGAAUUCAAUAAAAAAUAA |
| env-1464 | GUCAAAGGGGUGACUUAAUUCAAACAUUGAAAUACUGAUUAAAAGUUACGAAAAAAAAUUAACAACUUGCAAAAAAGUG |
| env-1465 | GUCAAAGGGGUGACUUAAUUCAAACAUUGAAAUACUGAUUAAAAGUUACGAAAAAAAAUUAACAACUUGCAAAAAAGUG |
| env-1466 | GUCAAAGGGGUGACUUAAUUCAAACAUUGAAAUACUGAUUAAAAGUUACGAAAAAAAAUUAACAACUUGCAAAAAAGUG |
| env-1467 | GAUUGUUAUAUUAAAUCAUUCCUGAUUUAAAAUUCGACAUCAAAAAGUUACGAAAAAAAUAUCAAAUCACAAAAAAAUG |
| env-1468 | GUCAAAGGGGUGACUUAAUUCAAACAUUGAAAUACUGAUUAAAAGUUACGAAAAAAAAUUAACAACUUGCAAAAAAGUG |
| env-1469 | GUCAAAGGGGUGACUUAAUUCAAACAUUGAAAUACUGAUUAAAAGUUACGAAAAAAAAUUAACAACUUGCAAAAAAGUG |
| env-1471 | UGCUAUUGAUUUACAAAACCAUAAACUCAAAAAUCAAUUAUAAAAAGUUACGAAAAAAAUACUAAAUGACAAAAAAACA |
| Fba-5-1 | GUCUACAGAAUUAAAACAUUACUAUUUUAAUAUUAUAUCUCAAAAAGUUACGAAAAAAAUACCAAAUCACAAAAAAAACA |
| Fic-1-1 | AUUAAGAGCAUAACAUAAAAAGCGACUUUUCUAGUCGCUUUUCUUUUUUAUAGAUUUCCUAUAAAAUCAUUAAAAAAUUA |
| env-1472 | ACUGUUGAGACAACAAAUUACUUCAUUAAUAUUCAACACCAUAAAAGUUACGAAAAAUAUACCAAAUCACAAAAAAAAUA |
| Lja-1-1 | AUAUUAAACAGUUCAAAAAACUAUUAUUUUUGAACUGUUUUUAUUUUAUAUAAAGGCAUUUGCAUUUCAAUAAAAAAUAC |
| env-1473 | ACAAUAGACAAAAAGACCUUUUCUUCUUAAAUAGAAAAGGUCUUUUUUUGUUUUUAUAUAUUUAUGUAAAUAAAAAAUAC |
| Fba-8-1 | CCAUGGAUAUUAAGUAUUUUCAUGUUAAUUUAAUAAAAAUUUUACAGAAAAAGAAGUCUUGUUUAAUUAUAAAAAAAUAC |
| env-1474 | GAAUCAGCCUUUAAGAGCUUAAUUAAAAGUAACUACAAAAUUGUCUUUUCUUUUUAAGUUUGCUAAUCAUUAAAAAAUUA |
| env-1475 | ....CGAGCAAAAGGUUUAAUUGGGCGCGCUAAAAACCAAAAAGAAAUAAAUUCAUAGAUUGUAAAUAGCUAAAAAAUUA |
| env-1476 | AGCCUUUAAGAGAAAUAAAUGAAGGUUAAUUUAUUCAAUUCCUUUUUUCAUAAAACAUCUUGUAAAUAACUAAAAAAUUA |
| env-1477 | AACCCUUAAGAUAAUAAGGGAAAGUUAUUUUACUCAAAUUCCUUUUUUUAUAAAACAUCUUGUAAAUCAUCAAAAAAUCA |
| env-1478 | AACCCUUAAGAUAAUAAGGAAAAGUUAUUUUACUCAAAUUCCUUUUUUUAUAAAACAUCUUGUAAAUCAUCAAAAAAUCA |
| env-1479 | AACCCUUAAGAUAAUAAGGGAAAGUUAUUUUACUCAAAUUCCUUUUUUUAUAAAACAUCUUGUAAAUCAUCAAAAAAUCA |
| env-1480 | ....................UAGAGUUUAAAAACCUGAUAGGUCUUUUUAUUUAAAAGCCUUGUAAAUCAGCAAAAUUUA |
| env-1481 | AACCAUUAAGAGAAUAAAUGAGGGUUGUUUUAUUCAAAUUCCUUUUUUUAAAAAACAUCUUGUAAAUAACUAAAAAAUUA |
| env-1482 | AUCCACUAAGAGAAUAAAUGAAGGCUAUUUUAUUCAAAUUCCUUUUUUUAAAAAACAUCUUAUAAAUAACUAAAAAAUUA |
| env-1483 | AUAAAAAGGUAACGCUAUUUUUCAUCUUUAAUAAUUUAAACAAGUUUCUAAAUUAAUCAUUUGUUUUCAAUAAAAAAUAA |
| env-1484 | AUCCACUAAGAGAAUAAAUGAAGGCUAUUUUAUUCAAAUUCCUUUUUUUAAAAAACAUCUUAUAAAUAACUAAAAAAUUA |
| env-1485 | AAAAUAGAUUCAGAUUUGUUAGGGUUUAAAAACCUGAUAGGUCUUUUUAUUUAAAAGCCUUGUAAAUCAGCAAAAUUUA |
| env-1486 | AACCAUUAAGAGAAUAAAUGAGGGUUGUUUUAUUCAAAUUCCUUUUUUUAAAAAACAUCUUGUAAAUAACUAAAAAAUUA |
| env-1487 | AAAAUAGAUUCAGAUUUGUUAGAGUUUAAAAACCUGAUAGGUCUUUUUAUUUAAAAGCCUUGUAAAUCAGCAAAAUUUA |
| env-1488 | AUCCACUAAGAGAAUAAAUGAAGGCUAUUUUAUUCAAAUUCCUUUUUUUAAAAAACAUCUUAUAAAUAACUAAAAAAUUA |
| env-1489 | AACCAUUAAGAGAAUAAAUGAGGGUUGUUUUAUUCAAAUUCCUUUUUUUAAAAAACAUCUUGUAAAUAACUAAAAAAUUA |
| env-1490 | UUACUGAAACUAUAAAUUUAAUUCUUGGGAGAAAUAAUACUAAAAGUUACGCAAAAAAAUGUUUCAAACCAAAAAAGUA |
| env-1491 | UACUUAAACUAUAAAAUUCAAUUCUUGGGAGAAAUAAUAGUAAAAGUUACGCAAAAAAAUGUUUCAAACCAAAAAAGUA |
| env-1497 | UUACUGAAACUAUAAAUUUAAUUCUUGGGAGAAAUAAUACUAAAAGUUACGCAAAAAAAUGUUUCAAACCAAAAAAGUA |
| env-1498 | UUACUGAAACUAUAAAUUUAAUUCUUGGGAGAAAUAAUACUAAAAGUUACGCAAAAAAAUGUUUCAAACCAAAAAAGUA |
| env-1499 | ........................................................................AAAAAGUA |
| env-1500 | UUACUGAAACUAUAAAUUUAAUUCUUGGGAGAAAUAAUACUAAAAGUUACGCAAAAAAAUGUUUCAAACCAAAAAAGUA |
| env-1501 | UUACUCAAACUAUAAAUUUAAUUCUUGGGAUAAAUAAUAGUAAAAGUUACAUAAAAAAAACGUAAAACACCAAAAAAGUA |
| env-1502 | UUACUCAAACUAUAAAUUUAAUUCUUGGGAUAAAUAAUAGUAAAAGUUACAUAAAAAAAACGUAAAACACCAAAAAAGUA |
| env-1503 | UUACUCAAACUAUAAAUUUAAUUCUUGGGAUAAAUAAUAGUAAAAGUUACAUAAAAAAAACGUAAAACACCAAAAAAGUA |
| env-1504 | UAAUCAAACUAUAAAUUUUAAUUCUUGGGAGAAAUAAUAGUAAAAGUUACGUAAAAAAAUCGUAUAAAACCAAAAAAGUA |
| env-1505 | AAUACGAGCAUAAAUAUAAAAGCGACUAAAUAGUCGCUUUUUCUUUUUUAUACAAUUGCUUGAAAAUAAUUAAAAAAACA |
| env-1506 | ......................................................UUAUUUAAUCAAAUAAAUAAAAAUAA |
| Pda-1-1 | UAAUAAUUUUCUAUUCUAUUUAAAUUAAUAGACGACUAUUUAAUUUUAUAAAAAGCAAUUAGCUAUUAAAUAAAAAAUAA |
| env-1507 | ACCCUUAAGAGCUUAAAAUAUAGUUGAAAUUCAAUUCAGAUUUAUUUCUAAUGUAAUAUAUGUAAAUCAUUAAUUAUUA |
| env-1508 | ACCUUUAAGAGCUUAAAAUAUAAUUGACUUUCAAUUUACAUUUAUUUCUAAUCUAAUUUAUGUAAAUCAUUAAUUAUUA |
| env-1515 | UUAAUCACCAAUCAACCAUUACGAGCAUAAUUAAUAUAUAUUUAUUUUUAAUGAGGUAUUCGUAAAUCAUUAAUUAUUA |
| env-1516 | AGGGAAAUGGUUUAAUGUGAGGUAAAAAAAUAUAAAUCAUCUUUUUAGUACUUGAUUAAAGCCUAUCUAAUUAAAAUAC |
| env-1517 | CUUAUUCAGAAUCAACCGGUUAGAGCUUAGGCAUUUCCGAAUAAUUCUUUCCAAAAGAUAGGUUAUUCUGAAAUUAGUA |
| | ................................................................................ |
| | ................................................................................ |
| | ●●●●●●●●●●●●●●●●●●●●●●●●●●●●●●●●●●●●●●●●●●●●●●●●●●●●●●●●●●●●●●●●●●●Y●●●●RR●●●●●● |

```
env-1519  AACCAUUAAGAGUUUAAAAAGUGGCUUCUAAGUUACUUUUCCUUUUUUGAUAAAAUAACUUGUAAGUCAUUAAAAAUUA
env-1520  AACCAUUAAGAGUUUAAAAAGUGGCUUCUAAGUUACUUUUCCUUUUUUGAUAAAAUAACUUGUAAGUCAUUAAAAAUUA
env-1521  AACCAUUAAGAGUUUAAAAAGUGGCUUCUAAGUUACUUUUCCUUUUUUGAUAAAAUAACUUGUAAGUCAUUAAAAAUUA
env-1522  AAAACGUGAUUUUUGUUAAAAAAUUAUGAAUCCCCUCAAAAUAAAAUAUUUACAAAAGAUUGCAAUUAGUGAAAAAGUA
env-1524  AAAACGUGAUUUUUGUUAAAAAAUUAUGAAUCCCCUCAAAAUAAAAUAUUUACAAAAGAUUGCAAUUAGUGAAAAAGUA
env-1525  AAUCAACCUUUAAGAGUCUAGUUUUUCAUUGUUGCAUCAAAUAAUAUUUUUACAAAAGAUUACAAUUGGCGAAAAAGUA
env-1526  CAAUCAACCUUUAAGAGUGUGAUAUUUAAUGUAGCAUAAAAUAAUAUUUUUACAAGAGAUUACUAUUGGCGAAAAAGUA
env-1527  ACCUUUAAGAGUAUAAAAAACUGCCCAUUAAAUCACUUUUCCUUUUUUAACAAAAUAACUUGUAAGUCGUUAAAAAUUA
env-1528  ACCUUUAAGAGUAUAAUAAAAGCGCCUUGUAGGUCGUUUUCCUUUUUUUCUAAAAUAACUUGUAAGUCAUUAAAAAUUA
env-1529  CUUUAAGGGUAUGAUAAAAGCGACCUGUUGGUCGCUUUUCCUUUUUUAUACAAAAUAACUUGUAAGUCAUUAAAAAUUA
env-1530  CUUUAAGGGUAUGAUAAAAGCGACCUGUUGGUCGCUUUUCCUUUUUUAUACAAAAUAACUUGUAAGUCAUUAAAAAUUA
1 Fag-1-1 AAAAAGAGACGUAAUCGUCUCUUUUUUUUAUCCUCUUUAUAAAAGAUUGUCAAUAUAAUAUUUAAAUGAUAUAAUAUUA
env-1534  AUUUAAAUAAAUAUUGGAAUGUAUUUAACAAAAGAAGCAAAAGAAGAACUCUUCGCAAAACACGGUAAAGGAAAAAACG
env-1535  ........AAAGUUUCAUUUAAAUUACAUUUUGAUAACAAUAAUUAAUCAAAGUAAAACUUGUAAAUGACUAAAAAUUA
env-1536  CUAACGUGAAAGUUUAAUAUAAAUUACAUUUUGAUAACAAUAAUUAAUCAAAGUAAAACUUGUAAAUGACUAAAAAUUA
env-1537  CUAACGUGAAAGUUUAAUAUAAAUUACAUUUUGAUAACAAUAAUUAAUCAAAGUAAAACUUGUAAAUGACUAAAAAUUA
env-1538  AAAAAGUUUAAAAUUGAAUCAACUUUUUUCAAAAUAAUAAACUAGGGCAUACAAUUUAUUGCUAAUCAAAAUAAUAGUU
env-1539  AAAAAGUUUAAAAUUGAAUCAACUUUUUUCAAAAUAAUAAACUAGGGCAUACAAUUUAUUGCUAAUCAAAAUAAUAGUU
env-1540  AAAAGUUUAAAAUUGAAGCGAACUUUUUUCAAAAUAAUAAACUAGGACAUACAAUUUAUUGCUAAUCAAAAUAAUAGUU
env-1541  AAAAGUUUAAAAUUGAAGCGAACUUUUUUCAAAAUAAUAAACUAGGACAUACAAUUUAUUGCUAAUCAAAAUAAUAGUU
env-1542  AAAAGUUUAAAAUUGAAGCGAACUUUUUUCAAAAUAAUAAACUAGGACAUACAAUUUAUUGCUAAUCAAAAUAAUAGUU
env-1543  AAAAGUUUAAAAUUGAAGCGAACUUUUUUCAAAAUAAUAAACUAGGACAUACAAUUUAUUGCUAAUCAAAAUAAUAGUU
env-1546  AAAAGUUUAAAAUUGAAGCGAACUUUUUUCAAAAUAAUAAACUAGGACAUACAAUUUAUUGCUAAUCAAAAUAAUAGUU
env-1547  AAAAGUUUAAAAUUGAAGCGAACUUUUUUCAAAAUAAUAAACUAGGACAUACAAUUUAUUGCUAAUCAAAAUAAUAGUU
env-1548  AAAAAGUUUAAAUUGAAGUGAUUUUUUUUCAAAAUUAUAAUCUAGAGCAUACAAUUUAUUGCUAAUCAAAAUAAUAGUU
env-1549  UUACCUCGAUUUAGUUACUAAUCAACCGCUAAGAGCAUAAAAAAACAACAUUUUUUUGGUUUAUAUAUAGUUAAAAUUA
env-1550  UUAAAACGAAAAGGUUUAAAAAAAAUUAUGAUACAUAUUAAAACAUUAAUAGGUUUCAUAUGUAAAUCACUAAAAAUUA
env-1551  .........................................................AUAUGUAAAUCACUAAAAAUUA
env-1552  ....................AAAAAUUAUUUCUCAUAUCAAAACAUUUAUAGGUUUCAUAUGUAAAUCAUUAAAAAUAA
env-1553  .............................................................................AA
env-1554  UUAAAACGAAAAGGUUUAAAAAAAAUUAUUUCUCAUAUCAAAACAUUUAUAGGUUUCAUAUGUAAAUCAUUAAAAAUAA
env-1555  CCAAUCAUUUGGAGGAACCCUAUUCAGCUUGAAUUAAUAUGAUUUUAUAUAAAAUUUCUUAGCAUCUCAUCUAAUUUGG
env-1556  CCAAUCAUUUGGAGGAACCCUAUUCAGCUUGAAUUAAUAUGAUUUUAUAUAAAAUUUCUUAGCAUCUCAUCUAAUUUGG
env-1557  CCAUUAUAUUUGCGAUCCAUAUUUCAGUUUGAAGUAAUGUGAUUUUAUAUAUAAUUUAUUAGCAUCUCAUCUAAUUUGA
env-1559  AUAAAAAGGUUUAAAAUGCAUCUAAAAUUUUUCCAAAAAAACAACUUCAGCUAACCUAUUGCAAUUCAAAAUAAUAGUU
env-1560  AUAAAAAGGUUUAAAAUGCAUCUAAAAUUUUUCCAAAAAAACAACUUCAGCUAACCUAUUGCAAUUCAAAAUAAUAGUU
env-1561  CCAAAUAAUCAGAUUUAUAAAAUUUUGUUAAAGAAAACUGCAACGCACGGAAAAUAUUUGUUAGAUUCGGAGAAAAAGC
env-1562  ..............................................................AGAUUCGGAGAAAAAGC
env-1563  CCAAAUAAUCAGAUUUAUAAAAUUUUGUUAAAGAAAACUGCAACGCACGGAAAAUAUUUGUCAGAUUCGGAGAAAAAGC
env-1564  CCAAAUAAUCAGAUUUAUAAAAUUUUGUUAAAGAAAACUGCAACGCACGGAAAAUAUUUGUCAGAUUCGGAGAAAAAGC
env-1565  CCAAAUAAUCAGAUUUAUAAAAUUUUGUUAAAGAAAACUGCAACGCACGGAAAAUAUUUGUCAGAUUCGGAGAAAAAGC
env-1566  CUAUUAACAUAAAAUAGUUUAACGAGAUUACUUGAUUUCUUUAUUUUAACAAUUAUAGGCUUUUUUUCAAUUGAUUUUC
Zam-1-1   UCCUGAAUGCUAAAGCCCUAUUGCCUUUAAAUUGAAAAGAUUUAAUUUGAUUAAAACGGAUCAGUUAGCGAAAAUAUUU
env-1567  CUUUUUAGAAUAUGUAAAUCAGGAAGUUUCGAAAUUAUUCUAAACUACCUUUUUUUAUUUGCCUAAUCAAAAAAGAAGA
Ffr-2-1   AGUUCUCUACUCUCUUCCUAUUACAAUAGCAAAAUAACAUUAAUUAUUAAAUUUAUUUUAGAUUUCUCCAAAAUAAAUA
env-1568  AGUUCUCUACUCCCUUCCUAUCACAAUAGCAAAAUAACAUUAAUUAUUAAAUUUAUUUUAGAUUUCUCCAAAAUAAAUA
Fsp-2-1   UUGACAAUUCAUAAUUCAUAAUUGCUUCACCAGUUCACAAAAUAAGGAAGGUUUAUUUUGUUCGUGUCAUAAUUAUUUC
env-1571  ...................GAAGUUGUAUUAAGAUUUAAAAUAAUUCACAAUUCAUAAUUCAUAAUUCAUAAUUAUUUU
env-1572  AACGUUUACAGUUUAACAUCCUGAAAACUAUUGCAGUAAAUCUUAGAUAAAAUUUAUAUUGUCAUUUUGAAAAUAAAUA
env-1573  UAUAAGGUGCUUCAUACAAUACCACUUGUAUUAUGGUGAUCGAUUUAAAUAAUUAUACUUGUCUGAUAAAAAAUAAAUA
env-1574  AACUUUCACUGUUUAACACUCCGAAUACUGUAGCGGUAAAUCUUAGUUAAAAUUAAUAUUGUCAUUUAAAAAUUAAAUA
env-1575  AACUUUCACUGUUUAACACUCCGAAUACUGUAGCGGUAAAUCUUAGUUAAAAUUAAUAUUGUCAUUUAAAAAUUAAAUA
env-1576  AACUUUCACUGUUUAACACUCCGAAUACUGUAGCGGUAAAUCUUAGUUAAAAUUAAUAUUGUCAUUUAAAAAUUAAAUA
Fsp-5-1   UUGACAAUUCAUAAUUCAUAAUUGCUUCGCCAGUUCACAAAAUAAGGGAGGUUUAUUUUGUUCGUGUCAUAAUUAUUUU
Fsp-3-1   AUCAAAUGGUGCUUUUUCGAUUAAAAUAUUUAAUAUUUAAAUUUUUCAAUCUUUAAAUUUAUUUUUUCGAAAAAAAAAU
env-1577  UAAAAGAACCAACUCAAAAACGGGAAUUGAUUUAAACUUAAAAUAUUUAUAAAUUUAAAUAUUAUUUCAAAAAUAAAUA
env-1579  UAAAAGAACCAACUCAAAAACGGGAAUUGAUUUAAACUUAAAAUAUUUAUAAAUUUAAAUAUUAUUUCAAAAAUAAAUA
Fsp-4-1   AACGUUCACAGUGUAGUAAAUUACAUUUUAUUGCGGUCAAUCUUAGAUAAAAUUUAUAUUGCCGUUAUGAAAAUAAAUA
env-1580  UCAACCUGCCUACCUAAAAGAGACUAAGUAAAUAUUUAGUCUCUUUUUUUUAAAAAGGUUAAUUUAUUGUAAAAUAGUC
env-1581  ACUUUUUAACAUUGUAAAAUAAAACUCUGCUUUUGCGAUAAAAUAUAUUGACUUACAGUUUCUAAUCCGGAAAUAAAUC
Wma-1-1   UUAGUGAGAGAGAACGCUAGAAUUUUGAUGCGAUAGAAAAAUCUAUCGCAUUUUUUUAUUAUUAAUUGAAAAAUUAAUC
Wvi-1-1   AAUUCUAGAAAAGAACAUUAAGCCAUAUUACAGAAGAAAAAUAAAUUACAUUUUUUUAUUAAUUAUCCAAAAAAUAAUC
Orh-1-1   GUUUUUUGUUAAAAUAUUCAACUUGCUUUCAAGCAACUAAGAUUUUUCCAUAUAAAGUUUGGUAAUUCAAUAAAAAGGC
env-1582  UAUUAUUUAAGAUUUUUUUAACAAUUUUUCUACUAUAGUACUUGUUUUUAUCUAGAGUUGUGUGUUUUAUAAAUAAGUA
env-1583  AUUUAAGAAAAUUUUAACAAAUUUAUCUAUUAAGUAUCAUUAAUGUUUUCGAUAAUAGUUUGUAUUUCGAAAAAUUAUU
env-1584  AAAUUUAAGAAUUUGUUAACAUUAUUGCGGGAAUGUCUUUAGAAUCUAAGAAUAAAUUUUUCGGUUUAAAAAUAAAGUU
env-1585  AAAUUUAAGAAUUUGUUAACAUUAUUAUGGGAAUGUCUUUAGAAUCUAAGAAUAAAUUUUUCGGUUUAAAAAUAAAGUU
env-1586  AUUUAAGAAAUAUUUAACAUUUUUUCAAAUGGCUGUAUUUGUAAGAAUAAAAAUGCUUUUGCGGGUUCAUAUUAUUAUU
env-1587  AUUUAAGAAAUAUUUAACAUUUUUUCAAAUGGCUGUAUCUGUAAGAAUAAAAAUGCUUUUGCGGGUUCAUAUUAUUAUU
env-1588  UUUUUUAAGAAAAUUUUAACAUUUUAUGUGUUUCAGGUAUAAUUGUUGGUGAGUACGCUUUAUAAUUAAUAAAAAAGAU
Ebr-1-1   CAAAUUUAAAAGAGACCAAACCAACCAAAUGGUCUCUUUUUUUAUUUUAAUAAAUAAUCAAUUAAUUAUAAAAUUAGUU
env-1589  CAAAUUUUUAAAAAGUGACCGAAUUAACCAAUCGGUCACUUUUAUUUUUGUACAAAGUUUAUUAUUUCAUAUAAUAGUU
env-1594  CAAAUUUUUAAAAAGUGACCGAAUUAACCAAUCGGUCACUUUUAUUUUUGUACAAAGUUUAUUAUUUCAUAUAAUAGUU
Efa-1-1   CAAAUUUUUAAAAAGUGACCGAAUUAACCAAUCGGUCACUUUUAUUUUUGUACAAAGUUUAUUAUUUCAUAUAAUAGUU
env-1595  CAAAUUUUUAAAAAGUGACCGAAUUAACCAAUCGGUCACUUUUAUUUUUGUACAAAGUUUAUUAUUUCAUAUAAUAGUU
Bzo-1-1   CUACGGGAUGGGAUUUUUUGUUUGAAUAGAUUUACAAUAAAGAUUCUUAAAUUCUCUUUUGUUAUUCCGGAAAAAAACU
Bzo-2-1   CUAUGGGAUGGGAUUUUUUGUUUGAAUAGAUUUACAAUAAAGAUUUUUAAAUUCUCUUUUGUUAUUCCGGAAAAAAACU
env-1596  AAAGAGUAAGGUAAAUCUAUAUUUGGACCUAGUUCUCAACCGGCCAGUAGCUAAAGCAUAAGUUUUCACAUUUAAUCC
env-1597  AAAAAAGAAGGUAAACCUUUACUUAGACUUGGUUCUCAACCGAACAAUAAGCAAAUCAUAGCUUUUUCAUAUUAAAUCC
env-1598  AAAAAAGAAGGUAAACCUUUACUUAGACUUGGUUCUCAACCGAACAAUAAGCAAAUCAUAGCUUUUUCGUAUUAAAUCC
env-1599  AAAAAAGAAGGUAAACCUGUACUUAGACUUGGUUCUCAACCGAACGAUAAGCAAAUCAUAGCUUUUUCGUAUUAAAUCC
env-1600  AAAAAAGAAGGUGAACCUGUACUUAGACUUGGUUCUCAACCGAACGAUAAGCAAAUCAUAGCUUUUUCGUAUUAAAUUC
env-1601  AAAAAAGAAGGUAAACCUGUACUUAGACUUGGUUCUCAACCGAACAAUAAGCAAAUCAUAGCUUUUUCGUAUUAAAUUC
env-1602  AAAAAAGAAGGUAAACCUGUACUUAGACUUGGUUCUCAACCGAACGAUAAGCAAAUCAUAGCUUUUUCGUAUUAAAUCC

          ...............................................................................
          ...............................................................................
          ●●●●●●●●●●●●●●●●●●●●●●●●●●●●●●●●●●●●●●●●●●●●●●●●●●●●●●●●●●●●●●●●●●Y●●●●RR●●●●●●
```

```
env-1603   AAAGCAUAAAAUAAACCUUUAUUUGGAUCUUGUUCUGAACCGACCACUGGCGAAAGCAUAGAUUUUUCACAUUUAAUCC
env-1604   AAAGCGAAAAGUAAAUCUAUAUUUGGACUUGGUUCUCAACCGGCCAGUAGUGAAAGCAUAGGUUUUUCACAUUUAAUCC
env-1605   AAAGCGAAAAGUAAAUCUAUAUUUGGACUUGGUUCUCAACCGACCAGUGGUGAAAGCAUAGGUUUUUCACAUUUAAUCC
env-1606   AAAGCGAAAAGUAAAUCUAUAUUUGGACUUGGUUCUCAACCGACCAGUGGUGAAAGCAUAGGUUUUUCACAUUUAAUCC
env-1607   AAGAUUUAAAAGUAAAGUAAAUUAUAUUUGAACUGUCUCACCGACCAGUGUGAAAGCAUAGGUUUUUCACAUUUAAUCC
env-1608   AAAGCGAAAAGUAAAUCUAUAUUUGAACUUGGUUCUCAACCGACCAGUGGUGAAAGCAUAGGUUUUUCACAUUUAAUCC
env-1609   AAAAAAGAAGGUAAAUUUGUAUUUAGACUUGGUUCUAAACCGUCCCAUAAGCAAAACAUAGCUUUUUAGGAUUUAAUUC
env-1610   AAAAAAGAAGGUAAACUUGUAUUUAGACUUGGUUCUAAACCGUCCCAUAAGCAAAACAUAGCUUUUUAGGAUUUAAUUC
env-1611   AAAAAAGAAGGUAAACUUGUAUUUAGACUUGGUUCUAAACCGUCCCAUAAGCAAAACAUAGCUUUUUAGUAUUUAAUUC
env-1612   AAAAAAGAAGGUAAACUUGUAUUUAGACUUGGUUCUAAACCGUCCCAUAAGCAAAACAUAGCUUUUUAGGAUUUAAUUC
env-1613   AAAAAAGAAGGUAAACUUGUAUUUAGACUUGGUUCUAAACCGUCCCAUAAGCAAAACAUAGCUUUUUAGGAUUUAAUUC
env-1614   AAAAAGUAAAGUAAAUUUAUAUUUGGACUUGGUUCUCAACCGACCAGUGGUGAAAGCAUAGGUUUUUCACAUUUAAUCC
env-1615   AAAAAAGAAGGUAAACUUGUAUUUAGACUUGGUUCUAAACCGUCCCACAAGCAAAACAUAGCUUUUUAGGAUUUAAUUC
env-1616   AAAAAAGAAGGUAAACUUGUAUUUAGACUUGGUUCUAAACCGUCCCAUAAGCAAAACAUAGCUUUUUAGUAUUUAAUUC
env-1617   AAAAAAGAAGGUAAACUUGUAUUUAGACUUGGUUCUAAACCGUCCCAUAAGCAAAACAUAGCUUUUUAGUAUUUAAUUC
env-1618   AAAAAAGAAGGUAAACUUGUAUUUAGACUUGGUUCUAAACCGUCCCAUAAGCAAAACAUAGCUUUUUAGGAUUUAAUUC
env-1619   AAAAAAGAAGGUAAACUUGUAUUUAGACUUGGUUCUAAACCGUCCCAUAAGCAAAACAUAGCUUUUUAGGAUUUAAUUC
env-1620   AAAAAAGAAGGUAAACUUGUAUUUAGACUUGGUUCUAAACCGUCCUGUUAGCAAAACAUAGCUUUUUAGGAUUUAAUUC
env-1621   .AAGAGUAAGGUAAAUCUAUAUUUAGACCUAGUUCUUAACCGCCCUGUUGCAAAAGCAUAAGUUUUUCACAUUUAAUCC
env-1622   .UUUAUCCAAAGAAAUAAUAGGUUGAUUUUGUUCCUGAAAAAUAAUUCAAAAUUUAUAAUUCAAAAUUUAUAAUUAUUU
env-1623   UUAGAAUUUGUUAAUCAAGAAGUUGCCAAACUUUUUUAAGGUAAGAGUGUCUUUUUAUUUGCCUAUUAAAAAAAGAUAA
env-1624   AAAAAUAAUUCGUAAUUCGCAAUUGCUUCGCCAGUUCNCAAAAUAAAUUGGAUUUAUUUUGCUCGGGUCGUAAUUAUUU
env-1625   AAAAAUAAUUCGUAAUUCGCAAUUGCUUCGCCAGUUCCCAAAAUAAAUUGGAUUUAUUUUGCUCGGGUCGUAAUUAUUU
env-1626   AAAAAUAAUUCGUAAUUCGCAAUUGCUUCGCCAGUUCCCAAAAUAAAUUGGAUUUAUUUUGCUCGGGUCGUAAUUAUUU
env-1627   AAAAAUAAUUCGUAAUUCGCAAUUGCUUCGCCAGUUCCCAAAAUAAAUUGGAUUUAUUUUGCUCGGGUCGUAAUUAUUU
env-1628   AAUUUCUUUUUGCUAUUUGCUGCUUAAAGUGAUUCAUAAAAUAAUUCAUAAUUUACAAUUCACAAUUAAUAAUUAUUCU
env-1629   UUAAUGUAAAAUCAAAGCGAUAAAAGCUAUUCCGUUCCAAAUUAAUUUUUCUUACUCAUUGCAAAUCCAAAAAUAAAAA
env-1630   UUCUAUGCCAAUCGCAAUAACAACUCAAUAAAAUCAAUGUUUUUUAUUUUGCAAAACCUUGCAAAUCCAAAAAUAAAAA
env-1631   CAGUCAGAAAAUAAUUCAUAAUUACUUCACCAGUUCACAAAAUAAAAAAGGUUUAUUUUGUUCGUGUCAUAAUUAUUAU
env-1632   UCCAAUAACAUUUAAGUAAAUCUAAUCCAUCAAUUUAAAAAUAAUUCCUAUUUCAUAAUUCAUAAUUCAUAAUUAUUUA
env-1633   GUAAAUCAAGAAGUUGCUAAAAUUGUUUUAAUACAAACAAAAAUAAUUCAUAAUUCAUAAUUCAUAAUUCAUAAUUAUU
env-1634   CUUUUUAGAAUUUGUAAAUCAAGAAGUUGCAAAAUUAUUUUAAACUAUAACUUUUUAUUUGUCUAAUCAAAAAAGAAGA
env-1635   CCAAAUUGUUUUAGUGACAAAUUAUUUGCAAAAAAAUAUCAAUAAACAGCUAUUUUUCUUGUGUUUUUAAAAAAAAAGU
Faq-1-1    UCCGUUUACCUUAUCAAGUGAAUAGUAAAAAGCAAUUUCACAAAUUCCAAAAUAUCACUUGUUCUUUCGAAAAAUAAAA
env-1636   AUGUUCCUUUAGUAUACCUAUCUCAAUUCAACCAUCAAGAUAAUUAUACGUAAAAAUAUUGUGCUUUCAAAAAAUAAAA
env-1637   UGGAAGAUGGAAGAUGGAAGUAGGAGGUUGGAAGAUUUUUUUUUUGGUAUUUAAAAAAUUGGAAUUUUAAAAUUAAAUU
env-1638   UUUUUGAACAAAAUGAAUUUAGAAGGAAAAUAUUUUUAAAAAAAGUUCUAAUUAAAUUUGCAUAGAAAAAAAAAUAUAUU
env-1639   UUUUUGAACAAAAUGAAUUUAGAAGAAAAAUAUUUUUAAAAAAAGUUCUAAUUAAAUUUGCAUAGAAAAAAAAAUAUAUU
env-1640   UUUUUGAACAAAAUGAAUUUAGAAGGAAAAUAUUUUUAAAAAAAGUUCUAAUUAAAUUUGCAUAGAAAAAAAAAUAUAUU
env-1641   UUUUUGAACAAAAUGAAUUUAGAAGGAAAAUAUUUUAAAAAAAAGUUCUAAUUAAAUUUGCAUAGAAAAAAAAAUAUAUU
env-1642   UUUUUGAACAAAAUGAAUUUAGAAGGAAAAUAUUUUUAAAAAAAGUUCUAAUUAAAUUUGCAUAGAAAAAAAAAUAUAUU
Csp-6-1    UAUAAGCUUUGUAUAGGUUAAGGAUAAAUUGAGAAGAGGAAAAAUCUAUCUAUAAAUUUGCACAGCUUAUAAAAUUUUU
env-1643   UAUAAGCUCUGUAUAAGCUAAGCAUAAGUUGAGAAGAGGAAAAAUCUAUCUAUAAAUUUGCACAGCUUAUAAAAUCUUU
env-1644   UAUAAGCUCUGUAUAAGCUAAGCAUAAGUUGAGAAGAGGAAAAAUCUAUCUAUAAAUUUGCACAGCUUAUAAAAUCUUU
env-1645   UAUAAGCUUUGUAUAGGUUAAGGAUAAAUUGAGAAGAGGAAAAAUCUAUCUAUAAAUUUGCACAGCUUAUAAAAUCUUU
Coc-2-1    UAUAAGCUUUGUAUAGGUUAAGGAUAAAUUGAGAAGAGGAAAAAUCUAUCUAUAAAUUUGCACAGCUUAUAAAAUCUUU
env-1646   UAUAAGCUCUGUAUAAGCUAAGCAUAAGUUGAGAAGAGGAAAAAUCUAUCUAUAAAUUUGCACAGCUUAUAAAAUCUUU
Csp-9-1    UAUAAGCUUUGUAUAGGUUAAGGAUAAAUUGAGAAGAGGAAAAAUCUAUCUAUAAAUUUGCACAGCUUAUAAAAUCUUU
env-1647   UAUAAGCUUUGUAUAGGUUAAGGAUAAAUUGAGAAGAGGAAAAAUCUAUCUAUAAAUUUGCACAGCUUAUAAAAUCUUU
env-1649   UAUAAGCUCUGUAUAAGCUAAGCAUAAGUUGAGAAGAGGAAAAAUCUAUCUAUAAAUUUGCACAGCUUAUAAAAUCUUU
env-1650   ...............................................................................U
env-1651   UAUAAGCUUUGUAUAGGUUAAGGAUAAAUUGAGAAGAAGAAAAAUCUAUCUAUAAAUUUGCACAGCUUAUAAAAUCUUU
env-1652   UAUAAGCUUUGUAUAGGUUAAGGAUAAAUUGAGAAGAAGAAAAAUCUAUCUAUAAAUUUGCACAGCUUAUAAAAUCUUU
env-1655   UAUAAGCUUUGUAUAGGUUAAGGAUAAAUUGAGAAGAAGAAAAAUCUAUCUAUAAAUUUGCACAGCUUAUAAAAUCUUU
env-1656   UAUAAGCUUUGUAUAGGUUAAGGAUAAAUUGAGAAGAAGAAAAAUCUAUCUAUAAAUUUGCACAGCUUAUAAAAUCUUU
env-1657   UAUAAGCUUUGUAUAGGUUAAGGAUAAAUUGAGAAGAGGAAAAAUCUAUCUAUAAAUUUGCACAGCUUAUAAAAUCUUU
Coc-3-1    UAUAAGCUUUGUAUAGGUUAAGGAUAAAUUGAGAAGAGGAAAAAUCUAUCUAUAAAUUUGCACAGCUUAUAAAAUCUUU
Coc-1-1    UAUAAGCUUUGUAUAGGUUAAGGAUAAAUUGAGAAGAGGAAAAAUCUAUCUAUAAAUUUGCACAGCUUAUAAAAUCUUU
Csp-7-1    UAUAAGCUUUGUAUAGGUUAAGGAUAAAUUGAGAAGAGGAAAAAUCUAUCUAUAAAUUUGCACAGCUUAUAAAAUCUUU
Csp-10-1   UAUAAGCUCUGUAUAAGCUAAGCAUAAGUUGAGAAGAAGAAAAAUCUAUUUAUAAAUUUGCACAGCUUAUAAAAUCUUU
Csp-2-1    UAUAAGCUCUGUAUAAGCUAAGCAUAAGUUGAGAAGAAGAAAAAUCUAUUUAUAAAUUUGCACAGCUUAUAAAAUCUUU
env-1658   UUAGAAUUUGUAAAUCAGGAAGUUGCAAAAUUAUUUUCAAAAUAAUUUAUAACUCAUAAUCCAUAACUCAUAAUUAUUU
env-1659   UUCCUUACCUAUUACCAUUAAAUCCAAUUACUAAUUAAAAACUAAAUGUGAUUUCUUUUGAAUUUUUCAAUAUUUUAUU
env-1660   UUCCUUACCUAUUACCAUUAAAUCCAAUUACUAAUUAAAAACUAAAUGUGAUUUCUUUUGAAUUUUUCAAUAUUUUAUU
env-1661   UUCCUUACCUAUUACCAUUAAAUCCAAUUACUAAUUAAAAACUAAAUGUGAUUUCUUUUGAAUUUUUCAAUAUUUUAUU
env-1662   AUACCUAUUGAUCAAUCAUUUUACUUGCCUAUUACUUUAAAAUAAUUUGUUGUUUUGAUUUAUUUAUAAAAAAAUAAAUA
env-1663   AUACCUAUUGAUCAAUCAUUUUACUUGCCUAUUACUUUAAAAUAAUUUGUUGUUUUGAUUUAUUUAUAAAAAAAUAAAUA
env-1664   ...........UGCACUUGAUGGUGGAAUGGGCGCAAAAUAAUAUUUCAUCUAAAAUUUGCAUAUCUAAAAAAAAAGGUA
env-1665   NNNNNNNNNNNNNNNNNNNNNNNNNNNNNNNNNNNNNNNNNNNNNNNNNNNNNNNNUUUGCAUAUCUAAAAAAAAAGGUA
env-1666   CGAGAGUUUAGUUAGUAGUUAGUGCAAUGGGCGGCAAAAUAAUAUUUCAUCUAAAAUUUGCAUAUCUAAAAAAAAAGGUA
env-1668   CGAGAGUUUAGUUAGUAGUUAGUGCAAUGGGCGGCAAAAUAAUAUUUCAUCUAAAAUUUGCAUAUCUAAAAAAAAAGGUA
env-1669   CGAGAGUUUAGUUAGUAGUUAGUGCAAUAGGCGGCAAAAUAAUAUUUCAUCUAAAAUUUGCAUAUCUAAAAAAAAAGGUA
env-1673   ACGAGAGUUUAGUUAGUAGUUAGUGCAAUGGACGCAAAAUAAUAUUUCAUCUAAAAUUUGCAUAUCUAAAAAAAAAGGUA
env-1674   CGAGAGUUUAGUUAGUAGUUAGUGCAAUGGGCGGCAAAAUAAUAUUUCAUCUAAAAUUUGCAUAUCUAAAAAAAAAGGUA
env-1676   CGAGAGUUUAGUUAGUAGUUAGUGCAAUGGGCGGCAAAAUAAUAUUUCAUCUAAAAUUUGCAUAUCUAAAAAAAAAGGUA
env-1677   CGAGAGUUUAGUUAGUAGUUAGUGCAAUGGGCGGCAAAAUAAUAUUUCAUCUAAAAUUUGCAUAUCUAAAAAAAAAGGUA
env-1679   CGAGAGUUUAGUUAGUAGUUAGUGCAAUGGGCGGCAAAAUAAUAUUUCAUCUAAAAUUUGCAUAUCUAAAAAAAAAGGUA
env-1680   CGAGAGUUUAGUUAGUAGUUAGUGCAAUGGGCGGCAAAAUAAUAUUUCAUCUAAAAUUUGCAUAUCUAAAAAAAAAGGUA
env-1681   GCGAGAGUUUAGUUAGUAGUUAGUGCAAUGGGCGAAAAAUAAUAUUUCAUCUAAAAUUUGCAUAUCUAAAAAAAAAGGUA
env-1682   AGCGAGAGUUUAGUUAGUAGUUAGUGCAAUGGCGCAAAAUAAUAUUUCAUCUAAAAUUUGCAUAUCUAAAAAAAAAGGUA
env-1683   CGAGAGUUUAGUUAGUAAUUAGUGCAAUGGGCGGCAAAAUAAUAUUUCAUCUAAAAUUUGCAUAUCUAAAAAAAAAGGUA
env-1684   CGAGAGUUUAGUUAGUAGUUAGUGCAAUGGGCGGCAAAAUAAUAUUUCAUCUAAAAUUUGCAUAUCUAAAAAAAAAGGUA
env-1685   CGAGAGUUUAGUUAGUAAUUAGUGCAAUGGGCGGCAAAAUAAUAUUUCAUCUAAAAUUUGCAUAUCUAAAAAAAAAGGUA
env-1686   ...............................................CAUCUAAAAUUUGCAUAUCUAAAAAAAAAGGUA

           ................................................................................
           ................................................................................
           ●●●●●●●●●●●●●●●●●●●●●●●●●●●●●●●●●●●●●●●●●●●●●●●●●●●●●●●●●●●●●●●●●●●Y●●●●RR●●●●●●
```

```
env-1687  ACGAGAGUUUAGUUAGUAGUUAGUGCAAUGGGCGCAAAAUAAUAUUUCAUCUAAAAUUUGCAUAUCUAAAAAAAAAGGUA
env-1688  ............................GGCGCAAAAUAAUAUUUCAUCUAAAAAAUUUGCAUAUCUAAAAAAAAAGGUA
env-1689  ACGAGAGUUUAGUUAGUAGUUAGUGCAAUGGGCGCAAAAUAAUAUUUCAUCUAAAAUUUGCAUAUCUAAAAAAAAAGGUA
env-1691  CGAGAGUUUAGUUAGUAGAUAGUGCAAUGGGUGGCAAAAUAAUAUUUCAUCUAAAAUUUGCAUAUCUAAAAAAAAAGGUA
env-1692  CGAGAGUUUAGUUAGUAGUUAGUGCAAUGGGAGGCAAAAUAAUAUUUCAUCUAAAAUUUGCGUAUCUAAAAAAAAAGGUA
env-1693  ................................GGCAAAAUAAUAUUUCAUCUAAAAUUUGCAUAUCUAAAAAAAAAGGUA
env-1694  CGAGAGUUUAGUUAGUAGUUAGUGCAAUGGGCGGCAAAAUAAUAUUUCAUCUAAAAUUUGCAUAUCUAAAAAAAAAGGUA
env-1695  CGAGAGUUUAGUUAGUAGUUAGUGCAAUAGGCGGCAAAAUAAUAUUUCAUCUAAAAUUUGCAUAUCUAAAAAAAAAGGUA
env-1696  ....................................AAAUAAUAUUUCAUCUAAAAUUUGCAUAUCUAAAAAAAAAGGUA
env-1697  CGAGAGUUUAGUUAGUAGUUAGUGCAAUGGGCGGCAAAAUAAUAUUUCAUCUAAAAUUUGCAUAUCUAAAAAAAAAGGUA
env-1698  ACGAGAGUUUAGUUAGUAGUUAGUGCAAUGGGCGCAAAAUAAUAUUUCAUCUAAAAUUUGCAUAUCUAAAAAAAAAGGUA
env-1699  CGAGAGUUUAGUUAGUAGUUAGUGCAAUGGGCGGCAAAAUAAUAUUUCAUCUAAAAUUUGCAUAUCUAAAAAAAAAGGUA
env-1700  CGAGAGUUUAGUUAGUAGUUAGUGCAAUGGGCGGCAAAAUAAUAUUUCAUCUAAAAUUUGCAUAUCUAAAAAAAAAGGUA
env-1701  CGAGAGUUUAGUUAGUAGUUAGUGCAAUGGGCGGCAAAAUAAUAUUUCAUCUAAAAUUUGCAUAUCUAAAAAAAAAGGUA
env-1702  CGAGAGUUUAGUUAGUAGUUAGUGCAAUGGGCGGCAAAAUAAUAUUUCAUCUAAAAUUUGCAUAUCUAAAAAAAAAGGUA
env-1703  AACGAGAGUUUAGUUAGUAGUUAGCAAUGGGCGGCAAAAUAAUAUUUCAUCUAAAAUUUGCAUAUCUAAAAAAAAAGGUA
env-1705  CGAGAGUUUAGUUAGUAAUUAGUGCAAUGGGCGGCAAAAUAAUAUUUCAUCUAAAAUUUGCAUAUCUAAAAAAAAAGGUA
env-1706  .................................GCAAAAUAAUAUUUCAUCUAAAAUUUGCAUAUCUAAAAAAAAAGGUA
env-1707  CGAGAGUUUAGUUAGUAGUUAGUGCAAUGGGCGGCAAAAUAAUAUUUCAUCUAAAAUUUGCAUAUCUAAAAAAAAAGGUA
env-1708  ...............................................................................A
env-1709  ....................................AAAUAAUAUUUCAUCUAAAAUUUGCAUAUCUAAAAAAAAAGGUA
env-1710  CGAGAGUUUAGUUAGUAGUUAGUGCAAUGGGCGGCAAAAUAAUAUUUCAUCUAAAAUUUGCAUAUCUAAAAAAAAAGGUA
Cko-1-1   UGAGAUGGGGGAAAUAUAAAUUUCGUUCUUUUCUUUUUUCCCCGAUCCAAAUUCUACUUUGUGAAUUAAGAAAAUAGUU
env-1711  AAAAGGUACGAUUUUGCCCAAAGUUUAGGACUUCACCGAAAAUAUUCGUUAUAAAAUUUGCGUAUAUAAAAAAAAAGGUU
env-1712  AAAGGUACGAUUUUGCCCAAAGUUUAGGACUUCACCGAAAAAUAUUCGUUAUAAAAUUUGCGUAUAUAAAAAAAAAGGUU
env-1713  AAAAGGUACGAUUUUGCCCAAAGUUUAGGACUUCACCGAAAAUAUUCGUUAUAAAAUUUGCGUAUAUAAAAAAAAAGGUU
env-1714  AAAAGGUACGAUUUUGCCCAAAGUUUAGGACUUCACCGAAAAUAUUCGUUAUAAAAUUUGCGUAUAUAAAAAAAAAGGUU
env-1717  AAAAGGUAUGAUUUUAUCCAAAGUUUAGGACUUUACCGAAAAUAUUCGUUAUAAAAUUUGCGUAUAUAAAAAAAAAGGUU
env-1718  AAAAGGUACGAUUUUGCCCAAAGUUUAGGACUUCACCGAAAAUAUUCGUUAUAAAAUUUGCGUAUAUAAAAAAAAAGGUU
env-1720  AAAAGGUAUGGUUUUAUUCAAAGUUUAGGACUUUACCGAAAAUAUUCGUUAUAAAAUUUGCGUAUAUAAAAAAAAAGGUU
env-1722  UAAAAGGUACGAUUUUGCCAAAGUUUAGGACUUUACCGAAAAUAUUCGUUAUAAAAUUUGCGUAUAUAAAAAAAAAGGUU
Csp-5-1   AAAAGGUAUGAUUUUAUCCAAAGUUUAGGAUUUUACCGAAAAUAUUCGUUAUAAAAUUUGCGUAUAUAAAAAAAAAGGUU
env-1723  AAAAGGUAUGAUUUUAUCCAAAGUUUAGGAUUUUACCGAAAAUAUUCGUUAUAAAAUUUGCGUAUAUAAAAAAAAAGGUU
env-1724  CAAGAUAAAAUAAUUCAUAAUUGCUUCGCCAGUUCACAAAAUAAAGAAAUGAUUAUUUUGUUCUUGUCAUAAUUAUUGU
env-1725  CAAGAUAAAAUAAUUCAUAAUUGCUUCGCCAGUUCACAAAAUAAAGAAAUGAUUAUUUUGUUCUUGUCAUAAUUAUUGU
env-1726  ...................................................GAUUAUUUUGUUCUUGUCAUAAUUAUUGU
env-1727  CAAGAUAAAAUAAUUCAUAAUUGCUUCGCCAGUUCACAAAAUAAAGAAAUGAUUAUUUUGUUCUUGUCAUAAUUAUUGU
Csp-8-1   AAGUAAGAGAUAAUUAUUGAUCAUAAUUAGUGGAUAGUUUUUUUUUUCUCAUAAAUGUUGUGUGUAAAAAAAAAAAUAUA
env-1728  .....................................................AAUGUUGUGUGUAAAAAAAAAAAUAUA
env-1729  UCCUGCAAAUAGCAGGAGUAUUUCUUUUGAGGAAAAUAAAGAUUUUUUUAGAAAAAGUUGCGUAUAAAAAAAUAUUUGUA
env-1730  CUCCUGCAAUAGCAGGAGUAUUUCUUUUGAGGAAAAUAAAGAUUUUCUUAGAAAAAGUUGCGUAUAAAAAAAUAUUUGUA
env-1744  CGAAGAAGAUGAUGACGAUGAAGAUGAAGAGGNNNNNNNNNNNNNNNNNNNNNNNNNNNGCGUAUAAAAAAAUAUUUGUA
env-1745  ...............................................UUAGAAAAAGUUGCGUAUAAAAAAAUAUUUGUA
env-1746  UCCUGCAAAUAGCAGGAGUAUUUCUUUUGAGGAAAAUAAAGAUUUUUUUAGAAAAAGUUGCGUAUAAAAAAAUAUUUGUA
env-1747  UCCUGCAAAUAGCAGGAGUAUUUCUUUUGAGGAAAAUAAAGAUUUUUUUAGAAAAAGUUGCGUAUAAAAAAAUAUUUGUA
env-1748  CUCCUGCAAUAGCAGGAGUAUUUCUUUUGAGGGAAAUAAAGAUUUUCUUAGAAAAAGUUGCGUAUAAAAAAAUAUUUGUA
env-1749  CUCCUGCAAUAGCAGGAGUAUUUCUUUUGAGGAAAAUAAAGAUUUUUUUAGAAAAAGUUGCGUAUAAAAAAAUAUUUGUA
env-1750  CUCCUGCAAUAGCAGGAGUAUUUCUUUUGAGGAAAAUAAAGAUUUUUUUAGAAAAAGUUGCGUAUAAAAAAAUAUUUGUA
env-1752  UCCUGCAAAUAGCAGGAGUAUUUCUUUUGAGGAAAAUAAAGAUUUUUUUAGAAAAAGUUGCGUAUAAAAAAAUAUUUGUA
env-1753  UCCUGCAAAUAGCAGGAGUAUUUCUUUUGAGGAAAAUAAAGAUUUUUUUAGAAAAAGUUGCGUAUAAAAAAAUAUUUGUA
env-1754  CUCCUGCAAUAGCAGGAGUAUUUCUUUUGAGGAAAAUAAAGAUUUUCUUAGAAAAAGUUGCGUAUAAAAAAAUAUUUGUA
env-1755  CUCCUGCAAUAGCAGGAGUAUUUCUUUUGAGGAAAAUAAAGAUUUUUUUAGAAAAAGUUGCGUAUAAAAAAAUAUUUGUA
Cgi-1-1   CUCCUGCAAUAGCAGGAGUAUUUCUUUUGAGGAAAAUAAAGAUUUUUUUAGAAAAAGUUGCGUAUAAAAAAAUAUUUGUA
env-1756  UCCUGCAAAUAGCAGGAGUAUUUCUUUUGAGGAAAAUAAAGAUUUUUUUAGAAAAAGUUGCGUAUAAAAAAAUAUUUGUA
env-1757  CUCCUGCAAUAGCAGGAGUAUUUCUUUUGAGGAAAAUAAAGAUUUUCUUAGAAAAAGUUGCGUAUAAAAAAAUAUUUGUA
env-1765  CUCCUGCAAUAGCAGGAGUAUUUCUUUUGAGGAAAAUAAAGAUUUUUUUAGAAAAAGUUGCGUAUAAAAAAAUAUUUGUA
env-1766  CUCCUGCAAUAGCAGGAGUAUUUCUUUUGAGGAAAAUAAAGAUUUUCUUAGAAAAAGUUGCGUAUAAAAAAAUAUUUGUA
env-1767  UCCUGCAAUAGCAGGAGUAUUUCUUUUGAGGAAAAAUAAAGAUUUUCUUAGAAAAAGUUGCGUAUAAAAAAAUAUUUGUA
env-1768  .....GCAAUAGCAGGAGUAUUUCUUUUGAGGAAAAUAAAGAUUUUCUUAGAAAAAGUUGCGUAUAAAAAAAUAUUUGUA
env-1769  CUCCUGCAAUAGCAGGAGUAUUUCUUUUGAGGAAAAUAAAGAUUUUCUUAGAAAAAGUUGCGUAUAAAAAAAUAUUUGUA
env-1770  CUCCUGCAAUAGCAGGAGUAUUUCUUUUGAGGAAAAUAAAGAUUUUCUUAGAAAAAGUUGCGUAUAAAAAAAUAUUUGUA
env-1771  CUCCUGCAAUAGCAGGAGUAUUUCUUUUGAGGAAAAUAAAGAUUUUCUUAGAAAAAGUUGCGUAUAAAAAAAUAUUUGUA
env-1772  CUCCUGCUAUUGCAGGAGUAUUUCUUUUGAGGAAAAUAAAGAUUUUCUUAGAAAAAGUUGCGUAUAAAAAAAUAUUUGUA
env-1773  CUCCUGCAAUAGCAGGAGUAUUUCUUUUGAGGAAAAUAAAGAUUUUCUUAGAAAAAGUUGCGUAUAAAAAAAUAUUUGUA
env-1774  ................AGUAUUUCUUUUGAGGAAAAUAAAGAUUUUCUUAGAAAAAGUUGCGUAUAAAAAAAUAUUUGUA
env-1775  UCCUGCAAAUAGCAGGAGUAUUUCUUUUGAGGAAAAUAAAGAUUUUUUUAGAAAAAGUUGCGUAUAAAAAAAUAUUUGUA
env-1779  CUCCUGCAAUAGCAGGAGUAUUUCUUUUGAGGAAAAUAAAGAUUUUCUUAGAAAAAGUUGCGUAUAAAAAAAUAUUUGUA
env-1780  UCUGCAAUAGUAGGAGUAUUUCUUUUUUUGAGGGAAUAAAGAUUUUCUUAGAAAAAGUUGCGUAUAAAAAAAUAUUUACA
env-1783  UCUGCAAUAGCAGGAGUAUUUCUUUUUUCGAGGGAAUAAAGAUUUUCUUAGAAAAAGUUGUGUAUAAAAAAAUAUUUAUA
env-1786  UCUGCAAUAGCAGGAGUAUUUCUUUUUUCGAGGGAAUAAAGAUUUUCUUAGAAAAAGUUGUGUAUAAAAAAAUAUUUAUA
env-1795  UCUGCAAUAGCAGGAGUAUUUCUUUUUUCGAGGGAAUAAAGAUUUUCUUAGAAAAAGUUGUGUAUAAAAAAAUAUUUAUA
env-1797  UCUGCAAUAGCAGGAGUAUUUCUUUUUUCGAGGGAAUAAAGAUUUUCUUAGAAAAAGUUGUGUAUAAAAAAAUAUUUAUA
env-1798  UCUGCAAUAGCAGGAGUAUUUCUUUUUUCGAGGGAAUAAAGAUUUUCUUAGAAAAAGUUGUGUAUAAAAAAAUAUUUAUA
env-1800  UCUGCAAUAGCAGGAGUAUUUCUUUUUUCGAGGGAAUAAAGAUUUUCUUAGAAAAAGUUGUGUAUAAAAAAAUAUUUAUA
env-1801  UCUGCAAUAGCAGGAGUAUUUCUUUUUUCGAGGGAAUAAAGAUUUUCUUAGAAAAAGUUGUGUAUAAAAAAAUAUUUAUA
env-1802  UCUGCAAUAGCAGGAGUAUUUCUUUUUUCGAGGGAAUAAAGAUUUUUUUAGAAAAAGUUGUGUAUAAAAAAAUAUUUAUA
env-1804  ..............................................................................UA
env-1805  UCUGCAAUAGCAGGAGUAUUUCUUUUUUCGAGGGAAUAAAGAUUUUCUUAGAAAAAGUUGUGUAUAAAAAAAUAUUUAUA
env-1818  UCUGCAAUAGCAGGAGUAUUUCUUUUUUCGAGGGAAUAAAGAUUUUCUUAGAAAAAGUUGUGUAUAAAAAAAUAUUUAUA
env-1821  UCUGCAAUAGCAGGAGUAUUUCUUUUUUCGAGGGAAUAAAGAUUUUCUUAGAAAAAGUUGCGUAUAAAAAAAUAUUUAUA
env-1823  UCUGCAAUAGCAGGAGUAUUUCUUUUUUCGAGGGAAUAAAGAUUUUCUUAGAAAAAGUUGUGUAUAAAAAAAUAUUUAUA
env-1824  UCUGCAAUAGCAGGAGUAUUUCUUUUUUCGAGGGAAUAAAGAUUUUCUUAGAAAAAGUUGUGUAUAAAAAAAUAUUUAUA
env-1826  .............................GAGGGAAUAAAGAUUUUCUUAGAAAAAGUUGUGUAUAAAAAAAUAUUUAUA
env-1827  UCUGCAAUAGCAGGAGUAUUUCUUUUUUCGAGGGAAUAAAGAUUUUCUUAGAAAAAGUUGUGUAUAAAAAAAUAUUUAUA

          ................................................................................
          ................................................................................
          ●●●●●●●●●●●●●●●●●●●●●●●●●●●●●●●●●●●●●●●●●●●●●●●●●●●●●●●●●●●●●●●●●●●Y●●●●RR●●●●●●
```

```
env-1828   UCUGCAAUAGCAGGAGUAUUUCUUUUUUCGAGGGAAUAAAGAUUUUCUUAGAAAAAGUUGUGUAUAAAAAAUAUUUAUA
env-1829   UCUGCAAUAGCAGGAGUAUUUCUUUUUUCGAGGGAAUAAAGAUUUUCUUAGAAAAAGUUGUGUAUAAAAAAUAUUUAUA
env-1830   UCUGCAAUAGCAGGAGUAUUUCUUUUUUCGAGGGAAUAAAGAUUUUUUUAGAAAAAGUUGUGUAUAAAAAAUAUUUAUA
env-1832   UCUGCAAUAGCAGGAGUAUUUCUUUUUUCGAGGGAAUAAAGAUUUUCUUAGAAAAAGUUGUGUAUAAAAAAUAUUUAUA
env-1833   UCUGCAAUAGCAGGAGUAUUUCUUUUUUCGAGGGAAUAAAGAUUUUCUUAGAAAAAGUUGUGUAUAAAAAAUAUUUAUA
env-1834   GACAAAGAAGAUGAGGACGAUGAGGAUGACGAGGACUAAAUAAUUUUUCAAAAAAAGUUGUAUGUGAAGAAUAUUUAUA
env-1835   GACAAAGAAGAUGAGGACGAUGAGGAUGACGAGGACUAAAUAAUUUUUCAAAAAAAGUUGUAUGUGAAGAAUAUUUAUA
env-1836   GAGGACGAUGACAAAGAAGAUGAGGAUGACGAGGACUAAAUAAUUUUUCAAAAAAAGUUGUAUGUGAAGAAUAUUUAUA
env-1837   GACAAAGAAGAUGAGGACGAUGAGGAUGACGAGGACUAAAUAAUUUUUCAAAAAAAGUUGUAUGUGAAGAAUAUUUAUA
env-1838   GACAAAGAAGAUGAGGAAGAUGAGGAUGACGAGGACUAAAUAAUUUUUCAAAAAAAGUUGUAUGUGAAGAAUAUUUAUA
env-1839   GACAAAGAAGAUGAGGACGAUGAGGAUGACGAGGACUAAAUAAUUUUUCAAAAAAAGUUGUAUGUGAAGAAUAUUUAUA
env-1840   GACAAAGAAGAUGAGGACGAUGAGGAUGACGAGGACUAAAUAAUUUUUCAAAAAAAGUUGUAUGUGAAGAAUAUUUAUA
env-1846   GACAAAGAAGAUGAGGACGAUGAGGAUGACGAGUACUAAAUAAUUUUUCAAAAAAAGUUGUAUGUGAAGAAUAUUUAUA
env-1847   GAUGAGGACGAUGAGGACGAUGAGGAUGACGAGUACUAAAUAAUUUUUCAAAAAAAGUUGUAUGUGAAGAAUAUUUAUA
env-1848   GACAAAGAAGAUGAGGACGAUGAGGAUGACGAGGACUAAAUAAUUUUUCAAAAAAAGUUGUAUGUGAAGAAUAUUUAUA
env-1850   GACAAAGAAGAUGAGGACGAUGAGGAUGACGAGUACUAAAUAAUUUUUCAAAAAAAGUUGUAUGUGAAGAAUAUUUAUA
env-1851   .........GAUGAGGACGAUGAGGAUGACGAGGACUAAAUAAUUUUUCAAAAAAAGUUGUAUGUGAAGAAUAUUUAUA
env-1852   ...............GACGAUGAGGAUGACGAGGACUAAAUAAUUUUUCAAAAAAAGUUGUAUGUGAAGAAUAUUUAUA
env-1853   NNNNNNNNNNNNNNNNNNNNNNNNNNNNNNNNNNNNACUAAAUAAUUUUUCAAAAAAAGUUGUAUGUGAAGAAUAUUUAUA
env-1854   GACAAAGAAGAUGAGGACGAUGAGGAUGACGAGGACUAAAUAAUUUUUCAAAAAAAGUUGUAUGUGAAGAAUAUUUAUA
env-1855   AUGAAAGGUAAUAGGGGAGCUAGAAUAAAAAACAAGAAAAAAUUAACUAUAAUUUAUUUUUUAUAAUUAUUCUUUUUUU
env-1856   AUGAAAGGUAAUAGGAGAGCUAGAAUAAAAAACAAGAAAAAAUUAACUAUAAUUUAUUUUUUAUAAUUAUUCUUUUUUU
env-1857   AUGAAAGGUAAUAGGAGAGCUAGAAUAAAAAACAAGAAAAAAUUAACUAUAAUUUAUUUUUUAUAAUUAUUCUUUUUUU
env-1859   AUAAAAGGUAAUAGGAGAGCUAGAAUAAAAAACAAGAAAAAAUUAACUAUAAUUUAUUUUUUAUAAUUAUUCUUUUUUU
env-1860   UAAAAGGUAAUAGGAGAGCUAGAAUAAAAAACAAGAAAAAAUUAACUAUAAUUUAUUUUUUAUAAUUAUUCUUUUUUU
env-1862   GAGGAUGAUGAAGAGGAGGACGACGACGAGGAUGAUUAAAUAUUUUUUAAAAAAAGUUGUAUGUGAAAAAUAUUUAUA
env-1863   CGACCAAGACGAAGACGACGAUGAAGAAGAUGAGGAUUAAAUAUUUUAAAAAAAGGUUGUGUAUAAAAAAUAUUUAUA
env-1864   ...CCAAGACGAAGACGAUGAUGAAGAAGAUGAGGAUUAAAUAUUUUAAAAAAAGGUUGUGUAUAAAAAAUAUUUAUA
env-1865   AAUUAUGAUGGAACAUCAAUUACAAAGAGAAUCAUAAAAAUAUAAUAUUAAACAUAUUCGAGAUUACUAAAUAAAAAUA
Eme-2-1    UAAUAAAACAUAAACUUUUUUGAAUUAUUUCAAUUAGAUUUAAAAAUAAUAAUUCGUAUUGUAGAUGAAGAAAUUAGUU
Cca-1-1    UGAUUGAUAUGUUUGAAAAUUGAUUUCGCGUUCACUUGAAUAAAUAAUUUAAAAAAUAGAGGUAAUUAAUUUAUUACUA
Emi-1-1    UAAAAAAGUAUAAAGUUUUUUGAAUUAUUUGAAUUUGAAUUUUAAAACAGAAUCAGUAUUGUGAAUGAAGAAAUUAGUC
Cte-1-1    UUAUGGCAUAAUUAAAAUAAAUAUCUUGCGUCUUUGCGAUAAUUUUUUGAGUUAUAGGUUUCUAAUCCCAAAUAAAAUC
Csp-15-1   ACAAAAUCGUGUUUUAUUGAAUAGAAUUUAUUCCAAUAAAAAUACUUUUGAUUUUAAGUUCUUAAUCCCAAAUAAAAUC
Cha-2-1    ACAAAAAUGUGUUUUAUUGAAUAGAAUUUAUUCCAAUAAAAAUAGUUUUGAUUUUAAGUUUUUAAUCUAAAAUAAAAUC
Ean-3-1    UAAAAAAAUAUAAAGUUUUUUGAAUUAUAUGAAUUUUAAUUUUAAAACAGAAUCUAUAUUGUGAAUGAAGAAAUUAGUC
Ean-1-3    UAAAAAAAUAUAAAGUUUUUUGAAUUAUAUGAAUUUUAAUUUUAAAACAGAAUCUAUAUUGUGAAUGAAGAAAUUAGUC
Csp-18-1   UUAAGAUUUUUUUAACAUAUAUUUUAUGUUUAAAUUAUAAAUGAAAUCAAUAAUGUUUUGUUGAAUUUAGAAAAUAGUC
Cje-1-1    UUUUAAGAAAUUUUUAACACUUAUAAAUUUCCUUUUAAUUCAAAGUCUGCAUAACGUUUUAGAGAUUCAGAAAAUAGUU
env-1866   UUUAAGAUUUCUUUAACAUUUCUUUCUAUUUCGCUCCAUUCACAGGAUCAAUAACGUUUUCGGGAUUAAAAAAAUAGUU
Cha-1-1    UUUAAGGUUUUUUUAACAUUUAUUUUGAAUAAUUUAAUUCAUUAGAUUAAUAAUGUUUUUCAGGAAAGAAAAUAAUGUU
env-1869   UUUAAGAUUUCUUUAACAUUUCUUUCUAUUUCGCUCCAUUCACAGGAUCAAUAACGUUUUCGGGAUUAAAAAAAUAGUU
env-1870   .....................UUUUCUAUAAUUACCCAUUGAUGUUUUCCAUAAACAUUUUGCAGGGACAAAAAUUGUC
Chi-1-1    UUUUAAGAAAUGUUUAACAUUUUUUUCUAUCAAAAUCAAUUUAUGAAAUCAAUAAUGUUUGUGAAAUCAAAAAUUUGUC
env-1871   AAUUAAGAUUUUUUUAACAUUUUUCAGUCAACACAUCAGUCAUUAAUUCAAUAAUGUUUUCAUAGAAAAUAAAAUUGUC
Cso-2-1    AAACAAUAAAUUUUUAAGAUAUUUUAACAAACUAUAUAAAAUGGCGGUAAAUUAUGUUUCCGCGUACCAAAAAAAUAGUU
Cda-1-1    UUUUAAGAUAUUUUUAACAUUUUUUUCUAUCAAAUUUCAUUUACAGAAUCAAUAAUGUUUGCAGAAUAAAAAAUAAAGUC
Cpi-1-1    AUUAAGAAAUUUUUAACAUUUUUUUCUAUCAAUUUUGAUUUAUGAAAUCAAUAAUGUUUUGCUGACUUAAAAAAAUAGUU
env-1872   UUUAGGAAAAAUGUAGAGGCUUUCAAUAGGUGUCUCAUCAUUAAUUUAAACUACUUUUUUGGUGAUUAAAAAAAAUAGUC
Fba-4-1    UUUUAAGAAAAAUUUAACAGGAUUAAACAAUGCCUUGUUUUUAAUGCAAAUAAAUGAUUGCCAGAUACCUUGAUUUGUU
Cfo-1-1    AUUUUUUUUAAUUAAAAUUGAUCUUAAUGAUUGUAAUAUUUAAUUCCGACUUAAAUUUUGCGGAGUUAAAAAAAAUAGUC
env-1873   UUUUAAGAAAUUUUUAACAUUUUUUUCUAUCAAAAUCAAUAAAUGAAAUCAAUAAUGUUGGCAGAAUAAAAAAAAAUAGUC
Clu-1-1    AUUAAGAAUUUUUUAACAUUUUUUUCUAUCAAAUUUGAUAUAUGAAAUCAAUAAUGUUUUGCUGAAUUAAAAAAAUAGUU
env-1874   AUUAAGAAAUUUUUAACAUUUUUUUCUAUUAAUUUUAAUAAAUGAAAUCAAUAAUGUUUUGUGGGCUUGAAAAUUAGUU
Cta-1-1    UUUUAAGAAAUUUUUAACAUUUUUUUCUAUCAAAUUCCAUAUAUGAAAUCAAUAAUGUUUGCAGAAUCAAAAUAAAGUC
Csp-29-1   UUUUAAGAAAUUUUUAACAUUUUUUUCUAUCAAAUUCCAUAUAUGAAAUCAAUAAUGUUUGCAGAAUCAAAAAAAUAGUC
Csp-24-1   UAUUAAGAAAUUUUUAACAUUUUUUUCUAUCAAAUUUGAUUUAUGAAAUCAAUAAUGUUUGCAGAAUCAAAAAAAUAGUC
Csp-19-1   UAUUAAGAAAUUUUUAACAUUUUUUUCUAUCAAUUUUGAUUUAUGAAAUCAAUAAUGUUUGCAGAAUCAAAAAAAUAGUC
Cor-1-1    AUUAAGAAUUUUUUAACAUUUUUUUCUAUUAAUUUUGAUACACGAAAUCAAUAAUGUUUGGCUGAAUUAAAAAAAUAGUU
Csp-16-1   UUUUAAGAAAUUUUUAACAUUUUUUUCUAUUAAAUUUGAUUUAUGAAAUCAAUAAUGUUUGUGGAAUCAAAAAAAUAGUC
Can-1-1    UAUUAAGAAAUUUUUAACAUUUUUUUCUAUCAAUUUUGAUAUAUGAAAUCAAUAAUGUUUGCAGAAUCAAAAAAAUAGUC
Cgl-1-1    UAUUAAGAAUUUUUUAACAUUUUUUUCUAUCAAAUUUGAUUUAUGAAAUCAAUAAUGUUUGCAGAAUCAAAAAAAUAGUC
Csp-26-1   UGUUAAGAAAUUUUUAACAUUUUUUUCUAUCAAAUUUGAUAUAUGAAAUCAAUAAUGUUUGCAGAAUCAAAAAAAUAGUC
Csp-23-1   UUUUUAACAUUUUUUUCUAUCAAAUUUGAUUUAUGAAAUCAAUAAUGUGAGAAGAUGUUUGCAGAAUCAAAAAAAUAGUU
Cso-1-1    AUUAAGAAAUUUUUAACAUUUUUUUCUAUCAAUUUUGAUACAUGGAAUCAAUAAUGUUUUGCCGGGUUAGAAAAUAGUU
Csp-17-1   AUUAAGAAAUUUUUAACAUUUUUUUCUAUCAAUUUUGAUACAUGGAAUCAAUAAUGUUUUGCUGAAUUAAAAAAAUAGUU
Csp-20-1   AUUAAGAAAUUUUUAACAUUUUUUUCUAUCAAUUUUGAUACAUGGAAUCAAUAAUGUUUUGCUGAAUUAAAAAAAUAGUU
env-1875   UAUUAAGAAAUUUUUAACAUUUUUUUCUAUCAAAUUUGAUUUAUGAAAUCAAUAAUGUUUGCAGAAUCAAAAAAAUAGUC
env-1876   UAUUAAGAAAUUUUUAACAUUUUUUUCUAUCAAUUUUGAUUUAUGAAAUCAAUAAUGUUUGCAGAAUCAAAAAAAUAGUC
Csp-27-1   UAUUAAGAAAUUUUUAACAUUUUUUUCUAUCAAAUUUGAUUUAUGGAAUCAAUAAUGUUUGCAGAAUCAAAAAAAUAGUC
Csp-28-1   UAUUAAGAAAUUUUUAACAUUUUUUUCUAUCAAAUUUGAUUUAUGAAAUCAAUAAUGUUUGCAGAAUCAAAAAAAUAGUC
Cin-1-1    UAUUAAGAAAUUUUUAACAUUUUUUUCUAUCAAAUUUGAUUUAUGAAAUCAAUAAUGUUUGCAGAUUCAAAAAAAUAGUC
Csp-13-1   AUUAAGAAAUUUUUAACAUUUUUUUCUAUUAAUUUUGAUACACGAAAUCAAUAAUGUUUGGCCGAAUUAAAAAAAUAGUU
Csp-21-1   UAUUAAGAAAUUUUUAACAUUUUUUUCUAUCAAUUUUAAUUUAUGAAAUCAAUAAUGUUUGCAAAAUCAAAAAAAUAGUC
Csp-25-1   UUUUAAGAAAUAUUUAACAUUUUUUUCUAUCAAAAUUGAUAAAUGAAAUCGAUAAUGUUUGCAGAAUCAAAAAUAAGUC
Csp-22-1   UAUUAAGAAAUUUUUAACAUUUUUUUCUAUCAAAUUUGAUAUAUGAAAUCAAUAAUGUUUGCAGAAUCAAAAAAAUAGUC
Cga-1-1    UUAUUAAGAAUUUUUUAACAUUUUUUUCUAUCAAUUUUAAUUUAUGAAAUCAUAAUGUUUGCAAAAUCAAAAAAAUAGUC
Cvr-1-1    AUUAAGAAAUUUUUAACAUUUUUUUCUAUCAAUUUUGAUACAUGAAAUCAAUAAUGUUUUGCUGAAUUGAAAAAUAGUU
Csp-14-1   CAUUAAGAAAUUUUUAACAUUUUUUUCUAUCAAAAUUGAUAUAUGAAAUCAAUAAUGUUUGCAGAAUCAAAAAAAUAGUC
env-1877   .................................................AAUAAUGUUUGGCUGAAUUAAAAAAAUAGUU
env-1878   UCCUGCAAAUAGCAGGAGUAUUUCUUUUGAGGAAAAUAAAGAUUUUUUUAGAAAAAGUUGCGUAUAAAAAAUAUUUGUA
env-1879   UCUGCAAUAGCAGGAGUAUUUCUUUUUUCGAGGGAAUAAAGAUUUUCUUAGAAAAAGUUGUGUAUAAAAAAUAUUUAUA
env-1880   UUUUGAAUCUUUGAAAAGUUAUCUGAUGAAGGAGAAGUGAUAAUUUUUUGAUUUACAGUUUUUAAUCCGAAAAAUAAUC
env-1881   UUUUGAAUCUUUGAAAAGUUAUCUGAUGAAGGAGAAGUGAUAAUUUUUUGAUUUACAGUUUUUAAUCCGAAAAAUAAUC

           ...............................................................................
           ...............................................................................
           ••••••••••••••••••••••••••••••••••••••••••••••••••••••••••••••••••Y••••RR••••••
```

```
env-1883  AUUAGGUUGGAAGUUGGGCGUUGCAAGUAAAAAACACGCAACAAACUAUUUAAUUUCCUUGUAACUUCAAAAAUAAAGU
env-1884  .......................................................................AAAUAAUUU
env-1885  ........................................................UUUUUUAUUUAAAAAAAUAAUUU
env-1886  ACUUUUUUCCGAUUCAUAUUCCGCACAAAGAUACUUUUAAAAAUAUAUCAUAAAAAAAUUUUUUAUUUAAAAAAAUAAUUU
env-1892  .................................................................UAAAAAAUAAUUU
env-1893  ACUUUUUUCCGAUUCAUAUUCCGCACAAAGAUACUUUUAAAAAUAUAUCAUAAAAAAAUUUUUUAUUUAAAAAAAUAAUUU
env-1898  ACUUUUUUCCGAUUCAUAUUCUGCACAAAGAUACUUUUAAAAAUAUAUCAUAAAAAAAUUUUUUAUUUAAAAAAAUAAUUU
env-1900  ...UUCAUAUUCCGCACAAAGAUACUUUAAUUUAUUUACAAAAAUUAGCAUAAAAAAAUUUUUUAUUCUAAAAAAUAAUUU
env-1901  ACUUUUUUCCGAUUCAUAUUCUGCACAAAGAUACUUUUAAAAAUAUAUCAUAAAAAAAUUUUUUAUUUAAAAAAAUAAUUU
env-1902  CGAUUCAUAUUCCGCACAAAGAUACUUUAAUUUAUUUACAAAAAUUAGCAUAAAAAAAUUUUUUAUUCUAAAAAAUAAUUU
env-1903  ACUUUUUUCCGAUUCAUAUUCCGCACAAAGAUACUUUUAAAAAUAUAUCAUAAAAAAAUUUUUUAUUUAAAAAAAUAAUUU
env-1904  ACUUUUUUCCGAUUCAUAUUCCGCACAAAGAUACUUUUAAAAAUAUAUCAUAAAAAAAUUUUUUAUUUAAAAAAAUAAUUU
env-1905  ACUUUUUUCCGAUUCAUAUUCCGCACAAAGAUACUUUUAAAAAUAUAUCAUAAAAAAAUUUUUUAUUUAAAAAAAUAAUUU
env-1906  ACUUUUUUCCGAUUCAUAUUCCGCACAAAGAUACUUUUAAAAAUAUAUCAUAAAAAAAUUUUUUAUUUAAAAAAAUAAUUU
env-1907  ............................AGAUACUUUUAAAAAUAUAUCAUAAAAAAAUUUUUUAUUUAAAAAAAUAAUUU
env-1908  ................................................................................
env-1909  ACUUUUUUCCGAUUCAUAUUCCGCACAAAGAUACUUUUAAAAAUAUAUCAUAAAAAAAUUUUUUAUUUAAAAAAAUAAUUU
env-1910  ACUUUUUUCCGAUUCAUAUUCUGCACAAAGAUACUUUUAAAAAUAUAUCAUAAAAAAAUUUUUUAUUUAAAAAAAUAAUUU
env-1911  ACUUUUUUCCGAUUCAUAUUCUGCACAAAGAUACUUUUAAAAAUAUAUCAUAAAAAAAUUUUUUAUUUAAAAAAAUAAUUU
env-1912  ..................................................UAAAAAAUUUUUUAUUUAAAAAAAUAAUUU
env-1913  CGAUUCAUAUUCCGCACAAAGAUACUUUAAUUUAUUUACAAAAAUUAGCAUAAAAAAAUUUUUUAUUCUAAAAAAUAAUUU
env-1915  .............................................................UUUAUUUAAAAAAAUAAUUU
env-1916  ACUUUUUUCCGAUUCAUAUUCUGCACAAAGAUACUUUUAAAAAUAUAUCAUAAAAAAAUUUUUUAUUUAAAAAAAUAAUUU
env-1917  ACUUUUUUCCGAUUCAUAUUCUGCACAAAGAUACUUUUAAAAAUAUAUCAUAAAAAAAUUUUUUAUUUAAAAAAAUAAUUU
env-1918  ACUUUUUUCCGAUUCAUAUUCUGCACAAAGAUACUUUUAAAAAUAUAUCAUAAAAAAAUUUUUUAUUUAAAAAAAUAAUUU
env-1921  ACUUUUUUCCGAUUCAUAUUCUGCACAAAGAUACUUUUAAAAAUAUAUCAUAAAAAAAUUUUUUAUUUAAAAAAAUAAUUU
env-1922  ACUUUUUUCCGAUUCAUAUUCUGCACAAAGAUACUUUUAAAAAUAUAUCAUAAAAAAAUUUUUUAUUUAAAAAAAUAAUUU
env-1924  .................................................................UAAAAAAUAAUUU
env-1925  ........CCGAUUCAUAUUCCGCACAAAGAUACUUUUAAAAAUAUAUCAUAAAAAAAUUUUUUAUUUAAAAAAAUAAUUU
env-1926  CGAUUCAUAUUCCGCACAAAGAUACUUUAAUUUAUUUACAAAAAUUAGCAUAAAAAAAUUUUUUAUUCUAAAAAAUAAUUU
env-1927  ACUUUUUUCCGAUUCAUAUUCCGCACAAAGAUACUUUUAAAAAUAUAUCAUAAAAAAAUUUUUUAUUUAAAAAAAUAAUUU
env-1928  .......................................AAAAUAUAUCAUAAAAAAAUUUUUUAUUUAAAAAAAUAAUUU
env-1929  ACUUUUUUCCGAUUCAUAUUCUGCACAAAGAUACUUUUAAAAAUAUAUCAUAAAAAAAUUUUUUAUUUAAAAAAAUAAUUU
env-1930  ACUUUUUUCCGAUUCAUAUUCUGCACAAAGAUACUUUUAAAAAUAUAUCAUAAAAAAAUUUUUUAUUUAAAAAAAUAAUUU
env-1931  ACUUUUUUCCGAUUCAUAUUCUGCACAAAGAUACUUUUAAAAAUAUAUCAUAAAAAAAUUUUUUAUUUAAAAAAAUAAUUU
env-1935  ................................................................................
env-1936  ACUUUUUUCCGAUUCAUAUUCCGCACAAAGAUACUUUUAAAAAUAUAUCAUAAAAAAAUUUUUUAUUUAAAAAAAUAAUUU
env-1937  ACUUUUUUCCGAUUCAUAUUCCGCACAAAGAUACUUUUAAAAAUAUAUCAUAAAAAAAUUUUUUAUUUAAAAAAAUAAUUU
env-1938  ACUUUUUUCCGAUUCAUAUUCCGCACAAAGAUACUUUUAAAAAUAUAUCAUAAAAAAAUUUUUUAUUUAAAAAAAUAAUUU
env-1939  GAUUCAUAUUCCGCACAAAGAUACUUUAAUUUAUUUACUAAAAAUUAGCAUAAAAAAAUUUUUUAUUCUAAAAAAUAAUUU
env-1940  ACUUUUUUCCGAUUCAUAUUCCGCACAAAGAUACUUUUAAAAAUAUAUCAUAAAAAAAUUUUUUAUUUAAAAAAAUAAUUU
env-1941  ...................................................................UUAAAAAAUAAUUU
env-1942  GCAGCCUCAUACAUAAUAAUCAUUUAAAUNUUUAUUUACAAAAAUUAGCAUAAAAAAAUUUUUUAUUCUAAAAAAUAAUUU
env-1943  .....................UGCACAAAGAUACUUUUAAAAAUAUAUCAUAAAAAAAUUUUUUAUUUAAAAAAAUAAUUU
env-1944  ACUUUUUUCCGAUUCAUAUUCCGCACAAAGAUACUUUUAAAAAUAUAUCAUAAAAAAAUUUUUUAUUUAAAAAAAUAAUUU
env-1945  ACUUUUUUCCGAUUCAUAUUCUGCACAAAGAUACUUUUAAAAAUAUAUCAUAAAAAAAUUUUUUAUUUAAAAAAAUAAUUU
env-1946  ACUUUUUUCCGAUUCAUAUUCCGCACAAAGAUACUUUUAAAAAUAUAUCAUAAAAAAAUUUUUUAUUUAAAAAAAUAAUUU
env-1947  ACUUUUUUCCGAUUCAUAUUCCGCACAAAGAUACUUUUAAAAAUAUAUCAUAAAAAAAUUUUUUAUUUAAAAAAAUAAUUU
env-1948  ..................UUCUGCACAAAGAUACUUUUAAAAAUAUAUCAUAAAAAAAUUUUUUAUUUAAAAAAAUAAUUU
env-1949  ACUUUUUUCCGAUUCAUAUUCCGCACAAAGAUACUUUUAAAAAUAUAUCAUAAAAAAAUUUUUUAUUUAAAAAAAUAAUUU
env-1950  ..........................UAAUUUAUUUACUAAAAAUUCGCAUAAAAAAAUUUUUUAUUCUAAAAAAUAAUUU
env-1951  GAUUCAUAUUCCGCACAAAGAUACUUUAAUUUAUUUACUAAAAAUUAGCAUAAAAAAAUUUUUUAUUCUAAAAAAUAAUUU
env-1952  CGAUUCAUAUUCCGCACAAAGAUACUUUAAUUUAUUUACAAAAAUUAGCAUAAAAAAAUUUUUUAUUCUAAAAAAUAAUUU
env-1954  GAUUCAUAUUCCGCACAAAGAUAAUUUAAUUUAUUUACUAAAAAUUCGCAUAAAAAAAUUUUUUAUUCUAAAAAAUAAUUU
env-1955  GAUUCAUAUUCCGCACAAAGAUAAUUUAAUUUAUUUACUAAAAAUUCGCAUAAAAAAAUUUUUUAUUCUAAAAAAUAAUUU
env-1957  CGAUUCAUAUUCCGCACAAAGAUACUUUAAUUUAUUUACAAAAAUUAGCAUAAAAAAAUUUUUUAUUCUAAAAAAUAAUUU
env-1958  .........................................................UUUUUUAUUCUAAAAAAUAAUUU
env-1959  CGAUUCAUAUUCCGCACAAAGAUACUUUAAUUUAUUUACAAAAAUCAGCAUAAAAAAAUUUUUUAUUCUAAAAAAUAAUUU
env-1961  GAUUCAUAUUCCGCACAAAGAUAAUUUAAUUUAUUUACUAAAAAUUCGCAUAAAAAAAUUUUUUAUUCUAAAAAAUAAUUU
env-1962  CGAUUCAUAUUCCGUACAAAGAUACUUUAAUUUAUUUACAAAAAUUAACAUAAAAAAAUUUUUUAUUCUAAAAAAUAAUUU
env-1963  CGAUUCAUAUUCCGCACAAAGAUACUUUAAUUUAUUUACAAAAAUCAGCAUAAAAAAAUUUUUUAUUCUAAAAAAUAAUUU
env-1964  GAUUCAUAUUCCGCACAAAGAUACUUUAAUUUAUUUACUAAAAAUUAGCAUAAAAAAAUUUUUUAUUCUAAAAAAUAAUUU
env-1965  ..............................................................UUAUUCUAAAAAAUAAUUU
env-1966  .............................................................UUUAUUCUAAAAAAUAAUUU
env-1967  AUUCUCCCCAAUACUUUGAUAUUAUUGUAAAUACCUAUCAAAAAAGAUAGAAAAAACUUUUGUGAUUAAAAAAAUAAUUU
env-1968  GAUUCAUAUUCCGCACAAAGAUACUUUAAUUUAUUUACUAAAAAUUAGUAUAAAAAAAUUUUUUAUUCUAGAAAUAAUUU
env-1969  CGAUUCAUAUUCCGCACAAAGAUACUUUAAUUUAUUUACAAAAAUUAGCAUAAAAAAAUUUUUUAUUCUAAAAAAUAAUUU
env-1970  CGAUUCAUAUUCCGUACAAAGAUACUUUAAUUUAUUUACAAAAAUUAACAUAAAAAAAUUUUUUAUUCUAAAAAAUAAUUU
env-1971  GAUUCAUAUUCCGCACAAAGAUACUUUAAUUUAUUUACUAAAAAUUAGUAUAAAAAAAUUUUUUAUUCUAAAAAAUAAUUU
env-1972  CGAUUCAUAUUCCGCACAAAGAUACUUUAAUUUAUUUACAAAAAUUAGCAUAAAAAAAUUUUUUAUUCUAAAAAAUAAUUU
env-1973  CGAUUCAUAUUCCGCACAAAGAUACUUUAAUUUAUUUACAAAAAUUAGCAUAAAAAAAUUUUUUAUUCUAAAAAAUAAUUU
env-1974  GAUUCAUAUUCCGCACAAAGAUAAUUUAAUUUAUUUACUAAAAAUUAGUAUAAAAAAAUUUUUUAUUCUAAAAAAUAAUUU
env-1975  CGAUUCAUAUUCCGCACAAAGAUACUUUAAUUUAUUUACAAAAAUUAGCAUAAAAAAAUUUUUUAUUCUAAAAAAUAAUUU
env-1976  UUCUCCACAAUACUUUGAUAUUAUUGUAAAUACCUAUCAAAAAAGAUAGAAAAAACUUUUGUGAUUAAAAAAAUAAUUU
env-1977  CGAUUCAUAUUCCGCACAAAGAUACUUUAAUUUAUUUACAAAAAUUAGCAUAAAAAAAUUUUUUAUUCUAAAAAAUAAUUU
env-1978  UUCUCCCCAAUACUUUGAUAUUAUUGUAAAUACCUAUCAAAAAAGAUAGAAAAAAACUUUUGUGAUUAAAAAAAUAAUUU
env-1979  CGAUUCAUAUUCCGCACAAAGAUAAUUUAAUUUAUUUACAAAAAUUAGCAUAAAAAAAUUUUUUAUUCUAAAAAAUAAUUU
env-1980  GAUUCAUAUUCCGCACAAAGAUAAUUUAAUUUAUUUACUAAAAAUUAGCAUAAAAAAAUUUUUUAUUCUAAAAAAUAAUUU
env-1981  UUCUCCCCAAUACUUUGAUAUUAUUGUAAAUACCUAUCAAAAAAGAUAGAAAAAACUUUUGUGAUUAAAAAAAUAAUUU
env-1982  CGAUUCAUAUUCCGUACAAAGAUACUUUAAUUUAUUUACAAAAAUUAGCAUAAAAAAAUUUUUUAUUCUAAAAAAUAAUUU
env-1983  GAUUCAUAUUCCGCACAAAGAUAAUUUAAUUUAUUUACUAAAAAUUAGCAUAAAAAAAUUUUUUAUUCUAAAAAAUAAUUU
env-1984  ACUUUUUUCCGAUUCAUAUUCCGCACAAAGAUACUUUUAAAAAUAUAUCAUAAAAAAAUUUUUUAUUUAAAAAAAUAAUUU
env-1985  .....................AUACUUUAAUUUAUUUACAAAAAUUAGCAUAAAAAAAUUUUUUAUUCUAAAAAAUAAUUU
env-1986  CGAUUCAUAUUCCGCACAAAGAUACUUUAAUUUAUUUACAAAAAUCAGCAUAAAAAAAUUUUUUAUUCUAAAAAAUAAUUU

          ................................................................................
          ................................................................................
          ●●●●●●●●●●●●●●●●●●●●●●●●●●●●●●●●●●●●●●●●●●●●●●●●●●●●●●●●●●●●●●●●●Y●●●●RR●●●●●●
```

| | |
|---|---|
| env-1987 | UUCUCCCCAAUACUUUGAUAUUAUUGUAAAUACCUAUCAAAAAAGAUAGAAAAAACUUUUGUGAUUAAAAAAAUAAUUU |
| env-1988 | CGAUUCAUAUUCCGCACAAAGAUACUUUAAUUUAUUUACAAAAAUUAGCAUAAAAAAAUUUUUUAUUCUAAAAAAUAAUUU |
| env-1990 | ............................................UUAGUAUAAAAAAUUUUUUAUUCUAGAAAUAAUUU |
| env-1991 | ................................AUUUACUAAAAAUUCGCAUAAAAAAAUUUUUUAUUCUAAAAAAUAAUUU |
| env-1992 | .......................................................AAUUUUUUAUUCUAAAAAAUAAUUU |
| env-1993 | ..........................................AAUUAGUAUAAAAAAAUUUUUUAUUCUAAAAAAUAAUUU |
| env-1994 | CGAUUCAUAUUCCGCACAAAGAUACUUUAAUUUAUUUACAAAAAUUAGCAUAAAAAAAUUUUUUAUUCUAAAAAAUAAUUU |
| env-1995 | AUUCUCCCCAAUACUUUGAUAUUAUUGUAAAUACCUAUCAAAAAAGAUAGAAAAAACUUUUGUGAUUAAAAAAAUAAUUU |
| env-1996 | UUCUCCGCAAUAUUUUGAUAUUAUUGUAAAUACCUAUCAAAAAAGAUAGAAAAAACUUUUGUGAUUUAAAAAAAUAAUUU |
| env-1997 | .....................................CUAAAAAUUAGUAUAAAAAAAUUUUUUAUUCUAAAAAAUAAUUU |
| env-1998 | ....................................................AAAAAUUUUUUAUUCUAAAAAAUAAUUU |
| env-1999 | GAUUCAUAUUCCGCACAAAGAUACUUUAAUUUAUUUACUAAAAAUUAGUAUAAAAAAAUUUUUUAUUCUAAAAAAUAAUUU |
| env-2000 | .................................................AUAAAAAAUUUUUUAUUCUAAAAAAUAAUUU |
| env-2001 | GAUUCAUAUUCCGCACAAAGAUAAUUUAAUUUAUUUACUAAAAAUUAGUAUAAAAAAAUUUUUUAUUCUAAAAAAUAAUUU |
| env-2002 | GAUUCAUAUUCCGCACAAAGAUACUUUAAUUUAUUUACUAAAAAUUAGUAUAAAAAAAUUUUUUAUUCUAGAAAUAAUUU |
| env-2003 | ..........................UAAUUUAUUUACUAAAAAUUAGUAUAAAAAAAUUUUUUAUUCUAAAAAAUAAUUU |
| env-2004 | ....................................................................AAAAAUAAUUU |
| env-2005 | ..........CCGCACAAAGAUAAUUUAAUUUAUUUACUAAAAAUUAGCAUAAAAAAAUUUUUUAUUCUAAAAAAUAAUUU |
| Ran-4-1 | CAGAAAUAGGUCUAAAAAAGGAUUUUAAAAACUAAAAAGACUCUACUUCAAAUACUUCUUGUGCGAUACAAAAAAAAAUA |
| Ran-2-1 | AGAAAUAGGUCUAAAAAAGGAUUUUAAAAACUAAAAAGACUCUACUUCAAAUACUUCUUGUGCGAUACAAAAAAAAAAUA |
| Cca-2-1 | GAAUAUUGCAAAAACAACAAAAGUUUUGGUUAGUAAUGCAAUUUUGUUACUCUUAAGUUUGUAGAUUACAUAAUUUUAU |
| Cca-2-2 | GAAUAUUGCAAAAACAACAAAAGUUUUGGUUAGUAAUGCAAUUUUGUUACUCUUAAGUUUGUAGAUUACAUAAUUUUAU |
| env-2006 | UCUCACUGCUGCAAACUCAAAUUCACGGCCUUUAGCAGGCAGUUUUUUGCAAUAAAAUUUUCAGUUUUGUUAUUUAAUU |
| env-2007 | UUUAAAGGUUUGGCUCUUUGACCACCGCUUUUGCAAAAGGUUUAAUUUCUUGUUAAAAUUCGUAGUUUCAAAAAAAGGA |
| Mru-1-1 | UUUGUUUUACAAAAAGAAAGUCUGCUACAUACCGACCUUAAAAGUUUGAUCGAUAAGUCUUUUAAAUUCAAAAAAACAC |
| env-2008 | UAUUUUAUAAAGUGAAAGUCUGCCGCAUACUGCCCUUUAAAGUUUAUAUUAGAUAAGUCUUUAAAAUUCAAAAAAACGC |
| env-2009 | UAUUUUAUAAAGUGAAAGUCUGCCGCAUACUGCCCUUUAAAGUUUAUAUUAGAUAAGUCUUUAAAAUUCAAAAAAACGC |
| env-2010 | UUAUGGUUUUGCUUCUUUGACACACGGAUCCACAAAAGGUUUAAUUUCUUGUUAAAAUUCGUAGUUUAAAAAAAAAGAAC |
| env-2011 | UGUAUUAGAAAGAGAAAGUCUGCUGCGUACCAACCUUAAAAGUUUAUAUUCAAUAAGUAUUUUAAAUUCAAAAAAACGC |
| env-2012 | UGAUUUUAAUCGGGGGAUUUACAGGCCAUGCCAACAUGUAAAGAUUUUUCCAAUAAAUCUUUUGAUUUCAAAUAAAAGC |
| env-2014 | UAAUAGGUUUGCCCCUUUGACUCAAAUAGUUUGAAAAGGUUUAAUUUUUUGUUAGAAUUCGUAGAUUAGAAAAAAGAAC |
| env-2015 | UGAUUUUAAUCGGGGGAUUUACAGGCCAUGCCAACAUGUAAAGAUUUUUCCAAUAAAUCUUUUGAUUUCAAAUAAAAGC |
| env-2016 | UGAUUUUAAUCGGGGGAUUUACAGGCCAUGCCAACAUGUAAAGAUUUUUCCAAUAAAUCUUUUGAUUUCAAAUAAAAGC |
| env-2017 | AAAAAAGCCCCGAAAUUCGGGGCUUUUUUGUUUUACCAAUGGAAAAUCGGCAAGAAGUCUUUAAAAUUCAAAUAAAAGC |
| env-2018 | UGAUUUUAAUCGGGGGAUUUACAGGCCAUGCCAACAUGUAAAGAUUUUUCCAAUAAAUCUUUUGAUUUCAAAUAAAAGC |
| Msp-2-1 | UUUGUUUUAUAAAAUGUGCAAAUACAAUAUACCGGCCUCAAAAGUUUGGUUCAUUGCUCUUUUCAAUUCAAAAAAAGAC |
| Cba-2-1 | UUUUACAUGAUCUAUAUCUAGCAAUAAAUAGAAGCUUUAAAUUUUCUGUUUUCUAGAAUAUGAUUAUCAUAAAAAUAGUU |
| Csp-11-1 | UUUUACAUGAUCUAUAUCUGGGAAUAAAAAGAAGCUUUAAAUUUUCUGUUUUCUAGAUUAUGAUUAUCACAAAAUAGUU |
| Cba-4-1 | UUUUACAUGAUCUAUAUCUAACAAUAAAUAGAAGCUUUAAAUUUUCUGUUUUCUAGAAUAUGAUUAUCAUAAAAAUAGUU |
| env-2019 | UUUUUGGACCCCGGGAAACGCAAAAGGUUUAAAAAGCAGGCAAGCAAACCGGCAAAAUGCACUUUAUGGGAAAUAAUUG |
| env-2020 | AAUACAGAACAAUCCUGUCAGGUCAUAAUAAGGUCUAUGUUUUUCUUGAAUUGAAAUAGUUUCUUAUUUCAAAAAAAGA |
| Csp-12-1 | GAAUCCCCAAAAAAAUACAACGUGCUUUUCAAAAAAUGAGCACAAAGUUUAAUUCGUAUUGUUUUUUCUUAAAAAGUA |
| Zul-1-1 | GAAAACUGAACCACUACCUACGGGUUGUGCUUUUUAAUAAACAUAUUUAAAAGCUCGUUAAAAAAAUAUCAAUAAUAAUU |
| env-2021 | AUCUUAGAGUGCUCGCCCUAUCAAUUGAUUCGGCGAGCAUUUUUGAUUAUAAAAUAUACUGCACAUUUCAAUAAUAGUU |
| env-2027 | AUCUUAGAGUGCUCGCCCUAUCAAUUGAUUCGGCGAGCAUUUUUGAUUAUAAAAUAUACUGCACAUUUCAAUAAUAGUU |
| env-2028 | .......................................................AUACUGCACAUUUCAAUAAUAAUU |
| env-2029 | GCAGGGAUUUAAAAAAGAAAAUUAAUCUUUAUAUCGACUUAAUUCAGAACAAACCUGUUAGAUCGUAAUAAAAAAUAAUU |
| Ace-1-1 | UUUUUUUAUUUACCCCUUGCAGUUGCUAUUAAAUCUUUUGAAAAAUCAUCAAACAUGCUCGUUGGAUAUAAAAAAUAGC |
| Ala-2-1 | UUUUAGUUAUACACCUAGGCAAUAGGCAUUAUAACCAACGAAAAAAGAUUAUUAAUACCCGUUGGAUGUAAAAAAUAAC |
| env-2030 | UCUUUGACCAACAAAAGAUGAAAAGGUUUAAUAUUGAUUUAAAAAAAGAAUAGCUUGCUGCCCGAAACAAAAAAAUAUUC |
| env-2031 | UCUUUGACCAACAAAAGAUGAAAAGGUUUAAUAUUGAUUUAAAAAAAGAAUAGCUUGCUGCCCGAAACAAAAAAAUAUUC |
| env-2032 | ............................................................................UUC |
| env-2033 | AAUUUGACCAAGGAAUGAUCAAAAGGUUUAAUGUUGAUUUAAAAAAAGAAUACCUUGCUGCCCUAAACAAAAAAAUAUUC |
| env-2034 | UCUUUGACCAACAAAAAAUGAAAAGGUUUAAUAUUGAUUUAAAAAAAGAAUAGCUUGCUGCCCGAAACAAAAAAAUAUUC |
| env-2035 | .........................................................................AUAUUC |
| env-2036 | AAUUUGACCUAUGAAAACCGAAAAGGUUUAAUAUUGACUUAAAAAAAGAAUACCUUGCUUUCCGAAACAAAAAAAUAUUC |
| env-2037 | AAUUUGACCAAGGAAUGAUCAAAAGGUUUAAUGUUUAUUUAAAAAAAGAAUACCUUGCUGUCCUAAACAAAAAAAUAUUC |
| env-2038 | .....................................................................AAAAAUAUUC |
| env-2039 | AAUUUGACCAAGGAAAGAUCAAAAGGUUUAAUGUUGAUUUAAAAAAAGAAUACCUUGCUGCCCGAAACAAAAAAAUAUUC |
| env-2040 | AAUUUGACCAAGGAAAGAUCAAAAGGUUUAAUGUUGAUUUAAAAAAAGAAUACCUUGCUGCCCGAAACAAAAAAAUAUUC |
| env-2041 | UCUUUGACCAACAAAAGAUGAAAAGGUUUAAUAUUGAUUUAAAAAAAGAAUAGCUUGCUGCCCGAAACAAAAAAAUAUUC |
| env-2042 | AAUUUGACCAAGGAAAGAUCAAAAGGUUUAAUGUUGAUUUAAAAAAAGAAUACCUUGCUGCCCGAAACAAAAAAAUAUUC |
| env-2043 | UCUUUGACCAACAAAAGAUGAAAAGGUUUAAUAUUGAUUUAAAAAAAGAAUAGCUUGCUGCCCGAAACAAAAAAAUAUUC |
| env-2044 | AAUUUGACCAAGGAAUGAUCAAAAGGUUUAAUGUUUAUUUAAAAAAAGAAUACCUUGCUGUCCUAAACAAAAAAAUAUUC |
| env-2045 | AAUUGGACCUGGAAAAGAUCAAAAGGUUUAAUGCUGAUUGAAAAAAAUAAUACUGCGCUGUCCGAAACAAAAAAAUAUUC |
| env-2046 | UCUUUGACCAACAAAAAAUGAAAAGGUUUAAUAUUGAUUUAAAAAAAGAAUAGCUUGCUGCCCGAAACAAAAAAAUAUUC |
| env-2049 | CUGUAGCAUAAGUAAUUCAUAUAAAAGCAAUAACAACUCAUAAAAAAAUUGAAAAACAAAGUUAAUAUCAUUUAGUUUUU |
| env-2050 | UGUAUCCUAAGUGCUUCAUUUAAAAAGCAUUAGCAUUAAAUAAAAAAUUGAAAAAGAAAGUUAAUUUCAUUUAGUUUUU |
| env-2051 | UGUAUCCUAAGUGCUUCAUUUAAAAAGCAUUAGCAUUAAAUAAAAAAUUGAAAAAGAAAGUUAAUUUCAUUUAGUUUUU |
| env-2052 | .UAUAGUAUAAGUGCCUCAUAUAAAAGCCAUAGCAACUAAUAAAAAAUUAAAAAACAAAGUUAAUAUCAUUUAGUUUUU |
| env-2053 | CUGUAGCAUAAGUAAUUCAUAUAAAAGCAAUAACAACUCAUAAAAAAAUUGAAAAACAAAGUUAAUAUCAUUUAGUUUUU |
| env-2054 | ....................................................................AUUUAGUUUUU |
| env-2055 | UGUAUCCUAAGUGCUUCAUUUAAAAAGCAUUAGCAUUAAAUAAAAAAUUGAAAAAGAAAGUUAAUUUCAUUUAGUUUUU |
| Ead-1-1 | CCUUGGACCCGGGUUAUUCAUAAAGGUUUAAUCGGUGAUCAAAAAAAGUGCAUGACAUCUGCUUUAUUUAAAUUUAUUA |
| env-2068 | ..................CACAAUGGUUUAAUUGAAGUUCAAAAAAAAUAAAUGUCAUCCGCUUUAUAUAAAUUUAUUA |
| env-2069 | .................................................AAAUGUCAUCCGCUUUAUAUAAAUUUAUUA |
| env-2070 | .........................GUUUAAUUGAAGUUCAAAAAAAAUAAAUGUCAUCCGCUUUAUAUAAAUUUAUUA |
| env-2071 | CCUUUGACCCCAGGUUUUCAUAAAGGUUUAAUUGAAGUUCAAAAAAAAUAAAUGACAUCCGCUUUAUAUAAAUUUAUUA |
| env-2072 | ...AUGACCCCGGGUAUUCACAAUGGUUUAAUUGAAGUUCAAAAAAAAUAAAUGUCAUCCGCUUUAUAUAAAUUUAUUA |
| env-2073 | UUAGGACGAACUAUUUUAAAAAAAGUUUAAUCCACACUUCAAAAAACAUUGGGAUGGCUCUAGGUUUCUAAAAAAAAGU |
| env-2074 | AUUUGACGGUGAUUAAGUCAAAAAGUUUAAUCUACCUUUCAAAAAACAUUAUGUUUACUCAAGGUUUCUAAAAAAAAGU |
| env-2075 | AUAGGACGCCGGAUAAUUAAAAAAGUUUAAUACACCUUUCAAAAAAUAUUAGGAUUGCACAAGGUUUCUAAAAAAAAGU |
| env-2076 | UNUAUGACCGCGUUUAAAUAAAAAAGUUUAAUCCAUGAUUAAAAAAAAGCUGGAUUGGACGGGGAUUCUAAAAAAUAGU |
| env-2077 | AAUUUGACCCCAAUUAAUUAAAAAGUUUAAUUCAUCUAUCAAAAAACAUCAUGUUUAACCAAGGUUUCUAAAAAAAAGU |
| | ............................................................................... |
| | ............................................................................... |
| | ..................................................................Y....RR...... |

```
env-2078  ..........CUUUAAAUCAUGAAAAGGUUUAAUAUUAUCUAAAAAAAGAUACCUUGCCGUCUGAAACAAAAAUUAUUC
env-2079  AAACUAAGAACCACUGCCAACAGGUUGUACCUUUUAAUUCAAAUUAUUAAAAUAUAGAAUAUACUUUUCAAUAAUAAUU
Fba-6-1   AUUCUUUGACCAGCGAAAUUCAGAAAGGUUUAAUAUUGAUUAAAAAGAAAUACCUUGACGGCUUAAACAAAAAAAUAUUC
env-2080  AUUCAAAACAAUCCUGUUAGAUCGUAAAUCGGGCUGAUAUAUUUCUUGAUUUGAAAUAGUUACAUGUUUCAAAAAAAGA
env-2081  CCCCAUUACCGGAGGGAUGUGCGUUUUAAAUACACCCUAUUAAAAAUAUGGGCAAUCCGUUUCGUUUCUAAAAAUAUCC
env-2082  CCCCAUUACCGGAGGGAUGUGCGUUUUAAAUACACCCUAUUAAAAAUAUGGGCAAUCCGUUUCGUUUCUAAAAAUAUCC
env-2083  AAUUCAAAAUAAUCCAGUUAGAAAGUAAUACUAUUAGAUGCUUUCUAGAUUUGAAAUAGUUUUAUAUUUCAAAAAAAGA
Msp-1-1   AUUCAGAACAACCCUGUGAGAUCUAAAGAAGCCUAAAUUUAUUUCUUGAUUCGAUAUAGUUUCUUAUAUUAAAAAAAGA
env-2085  AUUCAGAACAACCCUGUGAGAUCAUAAAAAGCUCAAAUUUAUUUCUUGAUUUGAUAUAGUUUCUUAUUUCAAAAAAAGA
env-2086  CUCAUCCAGAAUAAUCCCAUUCGAAAAAUAGAGGCAGAGGCUUAAUUCGAAAAAUUCAUUAAAGAUUAUAAAAAAAGAU
env-2087  UAUACAGAACAAUCCCAUAAGGGCCUAGUUCCCUGACCAUUGAUUUUUAUGCAAGAGCGCAGAUAAUCGUAAAAUAAAU
env-2088  UAUACAGAACAAUCCCAUAAGGGCCUAGUUCCCUGACCAUUGAUUUUUAUGCAAGAGCGCAGAUAAUCGUAAAAUAAAU
Mth-1-1   CCUUAUUCAAAAUAAUCCGGUACGUAAAAUAGAAGAAACUGCUUAAUCACAAAAAAUUAGUUCAAAUUCUAAAAAAGAC
env-2089  GAAGAGAUUUGAAGAAAAAAGUAAAUCUGUAUUUAGAUUUAAUUCAGAACAAUCCUGUUAGGAGUUCAGAAAAUUAGUA
env-2090  GAAGAGAUUUGAAGAAAAAAGUAAAUCUGUAUUUAGAUUUAAUUCAGAACAAUCCUGUUAGGAGUUCAGAAAAUUAGUA
env-2091  GAAGAGAUUUGAAGAAAAAAGUAAAUCUGUAUUUAGAUUUAAUUCAGAACAAUCCUGUUAGGAGUUCAGAAAAUUAGUA
env-2092  GAAGAGAUUUGAAGAAAAAAGUAAAUCUGUAUUUAGAUUUAAUUCAGAACAAUCCUGUUAGGAGUUCAGAAAAUUAGUA
Rbi-1-1   AGUGCCGGAAGAAUAAGCGGGAAGCCAUUGGGACAUAAAAUUUCCCGGGACCGCCCGGAUAGAAAUCCUGAAAUUAUUC
Mfo-1-1   AACGUAGAACCAUUACCAAAUGGUUGUGCUUUUUGAUUUUAAAUUAUUAAAAUUUAAAAUAAACAUUUCAAAAAAUAUC
env-2099  CCAAAUAAUCAGAUUUAUAAAAUUUUGUUAAAGAAAACUGCAACGCACGGAAAAUAUUUGUCAGAUUCAGAGAAAAAGC
env-2100  GACGAACGCUUUCUGGAAUAUGUAAAUAAAGAAGUGGCAAAGUUGUAUUAGAAAAAACUUGUUCAUUAAAAAAAAUAAAA
Fse-1-1   AUAGAUUGAGAAUUAUUCGAAUAAAAUAAAAAAGAAGAAAAUAAUUGACUCAUCAUCAUUUAUAUUCUAUAAUAAUUUU
Fda-1-1   ACUCUAUCCCUAUUACAUUAAGAGGUAGCAAAUAAAGAAUCAAAACAUUUGAAUUGCAUUGCCAAUUCAAAAUAAAGUA
env-2102  UAAAAAAGAGAUUUUAUAAUUUAUAGUAAUUUACAUCUUUUAUAUACUAUUCCUAUUGUUUGCUAUUCAAUAAAAUUAA
env-2103  AAAAAGUAAAAAGGAUGUCGUUUCCAUCUGGGCUAUUAAAAAUAAAUACUAAAACAUAGUAAUUAAUAUAUUAUAAUUU
env-2105  ACAAUUCUGAAUGAUUUGUUUAAUUUUAAAUAAUCUUUAAAUUUUUCAAUCUUUAAAUUUUUCAAUCUCAAAAUAAUGG
Fso-1-1   ACAAUUUUUAGCGAUUUGUUUAAUUUUAAAUAAUCAUUAAAUUUUUCAACCUUUAAAUUUUUCAAUCCCAAAAUAAUGG
env-2106  ACUUUAAAUUUUUGUAAACAAGGCAAUUACAAAACCUAUUCACAAAAAUAAAAAACAAUUGCAGUUCUAAAAAUAAAUA
env-2107  GAAUUUAAAGAUUUUGCUUACGCUUUGAAACAAUUUUCAAAUUUUUCAAUCUUUAAAUUUUUCAAUCUAAAAAUAAAAG
env-2108  CCAUUAGUCUAGAAUUGUAUAGAAGAUUUUAUUUUUUUAAAUCUUUCAAUCUGAAAAUUAUUUUUUCUAAAAAUAAAUA
env-2109  ....................................................................GAAAAUAAAUA
env-2110  CCGUUUAAACAAAACAAACUCUACAAACCGUAUUUUUUAAAAUCUUUUUUCUGAAAAUUAUUUUUUCUGAAAAUAAAUA
Aag-1-1   UUCUAAUGUCCUAAAUUAAUAUGUAUUAAAAAACAUCCAUUAUAUUUUACAUAUUAAGCAAAAUAUAAUAAUAAAAAUG
env-2111  ACAAAAACGUAAAAAAGUUUAAAAUCAUACGUAAAACAAUAAAAAUUACUAUUAUUCUAAAUAUUAUUAAUAAGAUUUU
env-2112  AUCCAAAAACAGUUAAAAAAUUAGCUUUAGAUAAAAUAUAAAAACAAUGUCAUUAAAAGCAAAUUUAUUAUUUUUUAGC
env-2113  UUAGUUUCACGAUGCAAAUUUACUAAAUUUCAGCUAUUAAAAAACAAUAGCAUUAAAGGCUUAUUUAUUAUUUUUUAGC
env-2114  UUAGUUUCACGAUGCAAAUUUACUAAAUUUCAGCUAUUAAAAAACAAUAGCAUUAAAGGCUUAUUUAUUAUUUUUUAGC
env-2115  UUAGUUUCACGAUGCAAAUUUACUAAAUUUCAGCUAUUAAAAAACAAUAGCAUUAAAGGCUUAUUUAUUAUUUUUUAGC
env-2116  GAAUUUCAAUAAUAAAAAGAAAUACUUUUUGUUUAUUAAAAUAAAGCAGUAAUUCUAUUAGGUUUAUUAUAGAAAAGCA
env-2117  GACUUUCAAUAAUAAAUAGAAAUACUUUUUGUCUAUUAAAAUAAAGCAAUAAUUCUAUUAGGUUUAUUAUAGAAAAGCA
env-2118  GAAUUUCAAUAAUAAAAAGAAAUACUUUUUGUUUAUUAAAAUAAAGCAGUAAUUCUAUUAGGUUUAUUAUAGAAAAGCA
env-2119  GAAUUUCAAUAAUAAAAAGAAAUACUUUUUGUUUAUUAAAAUAAAGCAGUAAUUCUAUUAGGUUUAUUAUAGAAAAGCA
env-2120  GACUUUCAAUAAUAAAUAGAAAUACUUUUUGUCUAUUAAAAUAAAGCAAUAAUUCUAUUAGGUUUAUUAUAGAAAAGCA
env-2121  GACUUUCAAUAAUAAAUAGAAAUACUUUUUGUCUAUUAAAAUAAAGCAAUAAUUCUAUUAGGUUUAUUAUAGAAAAGCA
env-2124  GACUCUCAAUAACAAAAAAAGUUUACUUUUAUCUAUUAAAAUAAAGCAGUAAUUGUAUUAGGUAUAUUAUAGAAAAGCA
env-2125  ACUUUUAUUAAAGAAAAAGGUUUAAUUUUUUCCAAUUAAAAUAAAGCGGAAAUUCUAUUAGGUAUAUUAUAGAAAAGCA
env-2126  ACUUUUAAUAACGAAAAAGGUUUAAUUUUUUCCUAUUAAAAUAAAGUGAAAAUUCUAUUCGCUAUUUUAUAGAAAAUCA
env-2127  AUUUGAUAUUAAAUAGACCUCUUAGAGAAAAAACCGCUUAAAAAAAAGCAAUUAUUACAUCCUAUUUAAUAAGAAAAACA
env-2128  GACUUUCAAUAAUAAAAAGAAAUACUUUUUGUCUAUUAAAAUAAAGCACUAAUUCUAUUAGGUAUAUUAUAGAAAAGCA
env-2129  AAUGGUUUAACCUUAGUUUUGUAUUGGUUAUUUUAAAAAAUAAAAAACAGCAAUACACACUCUAUAUUACAUAAAAUCA
env-2130  AAUGGUUUAACCUUAGUUUUGUAUUGGUUAUUUNAAAAAAUAAAAAACAGCAAUACACACUCUAUAUUACAUAAAAUCA
env-2131  AAUGGUUUAACCUUAGUUUUGUAUUGGUUAUUUUAAAAAAUAAAAAACAGCAAUACACACUCUAUAUUACAUAAAAUCA
env-2132  AUUUAAUAUUAAAUAGACCUCUAAGGGAAAAAACCGCUUAAAAAAAAGCAAUUAUUACAAUCUAUUUAAAAAGAAAAACA
env-2133  AUUUGAUAUUGAAUAGACCACUUAGAGAAAAAACUGCUUAAAAAAAAACAAUUAUUACAUAUUAAUUAAUAAGAAAAACA
env-2134  GAUUUGAUAUUAAANAGACCUCUUAGNGAAAAAACCGCUUAAAAAAAAGCAAUUAUUACAUCUAUUUAAAAAGAAAAACA
env-2135  AUUUGAUAUUAAAUAGACCUCUUAGGGAAAAAACCGCUUAAAAAAAAACAAUUAUUACAUAUUAAUUAAAAAGAAAAACA
env-2136  AUUUGAUAUUGAAUAGACCACUUAGAGAAAAAACUGCUUAAAAAAAAGCAAUUAUUACAUCUUAAUUAAUAAGAAAAACA
env-2137  AUUUGAUAUUGAAUAGACCACUUAGAGAAAAAACUGCUUAAAAAAAAACAAUUAUUACAUAUUAAUUAAUAAGAAAAACA
env-2138  AUUUAAUAUUAAAUAGACCUCUAAGGGAAAAAACCGCUUAAAAAAAAGCAAUUAUUACAUCUAUUUAAAAAGAAAAACA
env-2139  AUUGAUUUAAUAUUAAAUAGACCUCUAAGGGAAAAACCGCUUAAAAAAGCAAUUAUUACAUCUAUUUAAAAAGAAAAACA
env-2140  AACGGCUUAAGAAAAGCAUACUAAACGCUGCGGAAGGUUAAAAUUCGUAGCGUUUUUCAUUUUAUAUCAAUUCAAUUUA
env-2141  CUUAAAUUCAAAUGGUUUAAUUUGAACUUUUAGAUUGUUAAAAAGAAAAUCGAAAUAACUUGUAAAUGGUUAAUUUUUA
env-2142  CAAACAGUAUCACUACUGCCAAUCAAUAUUUUAAGAGACAAAAAGAAUACUUAAUUAGGUUGUAAGUUCCUAAUUUUAA
env-2143  UAGAUAAAAUGAAUAACUAAUAUAUGUGAUUUUAAUAUGUAAAAAUUGAAGUAAAAAAAUUGCAAGUCUCUAAUUUUUA
env-2144  AAGGUUUAAUUAAGCUCUAUCACAUUUUUAAUUUAAGAACAAAAAGACUAUUAAAAAAGUUGCAAAUGUUUAAUUUUUA
env-2145  GACGAGCGCUUUUUAGAAUUUGUAAAUCAAGAAGUUGCGAAGUUAUUUAGCUAAAAAAUUGUAUUUUAAAAAAAUAAAA
env-2146  GACGAGCGCUUUUUAGAAUUUGUAAAUCAAGAAGUUGCGAAGUUAUUUAGCUAAAAAAUUGUAUUUUAAAAAAAUAAAA
env-2147  GACGAGCGCUUUUUAGAAUUUGUAAAUCAAGAAGUUGCGAAGUUAUUUAGCUAAAAAAUUGUAUUUUAAAAAAAUAAAA
env-2148  AAAAUUAGCGUUAUCGCUAAAAUUGUUUAGAAAUUAGAAAUUAUUUGCGAUUUAAUGUAUGAGACUAAGAUAGAAUAUA
env-2149  ..................................UAGAAAUUAUUUGCGAUUUAAUGUAUGAGACUAAGAUAGAAUAUA
env-2150  UAUUAUUAAUUUCAAUUUUUAAUACUUAAAUCCUCGAUUUAUCGGGGAUUUUUUUUAUUUGUAUUUUAAAUUAAAGUUU
env-2151  ..........UUCAAUUUUUAAUACUUAAAUCCUCGAUUUAUCGGGGAUUUUUUUUAUUUGUAUUUUAAAUUAAAGUUU
env-2152  .............AUUGUACAUGGUCAAGUGUUUGAUUAGAAAUAAUUAUAUUAAUCUUUGUAACUAUCGGAUUUAUUG
Fps-4-1   UUUUAUUUUUGAUGGAAUGAUCAAAAAUAUUGCCAAGGCUAAAUAAUAAGUAAAAACAUUGCCACAUCAAAAAUAAAGU
          ...............................................................................
          ...............................................................................
          ••••••••••••••••••••••••••••••••••••••••••••••••••••••••••••••••Y••••RR••••••
```

alignment positions 80 · · · 259

```
Pgo-1-1   AUACUU.UUGCACAAU............UGAAUAG...........AAAAC.................CUAUUCU...........................................AGC.AUAAA.AAUUAUU...GA.AA..CAAUAUUGGCAUGUACUUAACAUUAGAAGAAAAAGGCAAA

          ............................<<<<<<<<<<<<<<<<<<...........>>>>>>>>>>>>>>>>>>................<<<<<<<<.......>>>>>>>>.....................................................................
          ............................2222221111..101.00...........00.101..1111222222................11011000.......00011011.....................................................................
          •UAY•U-UUGC••CY-----------◦◦◦◦◦◦◦-----------R•R◦◦-----------------◦◦◦◦◦◦◦-----------------------------------------◦UAC-AUAA•-AAUCAUU---UA-AA--YAAUAUURG•AUGUAUUUARC•ARAGAAR••AAARRRRAR
          5'                                                                                                                                                  3'
```

```
env-1     CUACUU.UUGCACGAU............UGAAUAG..........AAAAC.................CUAUUCU.....................................AGC.AUAAA.AAUUAUU...GA.AA..CAAUAUUGGCAUGUACUUAACACAAGAAGAAAAAGGCAAA
env-2     GUACUU.UUGCGCCAU............UAAAUAGA.........AAA..................UCUAUUUU.....................................AGC.AUAAA.AAUUAUU...GA.AA..UAAUAUUGGCAUGUACUUAACAUUAGAAGAAAAAGGCAAA
env-3     UUACUU.UUGCACGAU............CGAAUAG..........AAAAA.................CUAUUCU.....................................CGC.AUAAA.AAUUAUU...GA.AA..UAAUAUUGGCAUGUACCUAACAUUAGAAGAAAAAGGCAAA
Pto-1-1   GUACUU.UUGCACGAU............UGAAUAGA.........AAA..................UCUAUUUU.....................................AGC.AUAAA.AAUUAUU...GA.AA..UAAUAUUGGCAUGUACUUAACACAAGAAGAAAAAGGCGAA
Ptr-1-1   UUACUU.UUGCAGGAU............UGAAUAGA.........AAA..................UCUAUUCU.....................................AGC.AUAAA.AAUUAUU...GA.AA..UAAUAUUGGCAUGUACUUAACAUUAGAAGAAAAAGGCAAA
env-4     GUACUU.UUGCAGGAU............UGAAUAGA.........AAA..................UCUAUUUU.....................................AGC.AUAAA.AAUUAUU...GA.AA..UAAUAUUGGCAUGUACUUAACACAAGAAGAAAAAGGCAAA
env-5     UUACUU.UUGCACGAU............CGAAUAGA.........AAA..................UCUAUUCU.....................................AGC.AUAAA.AAUUAUU...GA.AA..UAAUAUUGGCAUGUACUUAACAUUAGAAGAAAAAGGCAAA
env-6     UUACUU.UUGCACGAU............UGAAUAGA.........AAA..................UCUAUUCU.....................................CGC.AUAAA.AAUUAUU...GA.AA..UAAUAUUGGCAUGUACCUAUCAUUAGAAGAAAAAGGCAAA
env-7     UUACUU.UUGCACGAU............UGAAUAGA.........AAA..................UCUAUUCU.....................................CGC.AUAAA.AAUUAUU...GA.AA..UAAUAUUGGCAUGUACCUAUCAUUAGAAGAAAAAGGCAAA
env-8     UUACUU.UUGCACGAU............UGAAUAGA.........AAA..................UCUAUUCU.....................................CGC.AUAAA.AAUUAUU...GA.AA..UAAUAUUGGCAUGUACCUAUCAUUAGAAGAAAAAGGCAAA
Mod-8-1   GUAACU.UUGCAAUU.............GAAUUUUAG........AGAA................CUAGAAUUC....................................GUAC.AUAAA.AAUCAU....UA.AA..GAAUAUUGGCAUGUAUUUAACGAAAGAAAAAAAGCUGAG
env-9     UUAUCU.UUGCGCCU.............CGGUUGCUGA.......AAGG...............UCAG.GACCA.....................................UAC.AUAAA.AAUUAUU..UAAAU..AAAUAUUCGAAUGUAUUUAGAUUCAAAAUCAAAAACAGAU
env-11    UUAUCU.UUGCGCCU.............CGGUUGCUGA.......AAGG...............UCAG.GACCA.....................................UAC.AUAAA.AAUUAUU..UAAAU..AAAUAUUCGAAUGUAUUUAGAUUCAAAAUCAAAAACAGAU
env-12    UUAUCU.UUGCGCCU.............CGGUUGCUGA.......AAGG...............UCAG.GACCA.....................................UAC.AUAAA.AAUUAUU..UAAAU..AAAUAUUCGAAUGUAUUUAGAUUCAAAAUCAAAAACAGAU
env-13    UUAUCU.UUGCGCCU.............CGGUAACUGA.......AAGG...............UCAGG.ACCA.....................................UAC.AUAAA.AAUUAUU..UAAAU..AAAUAUUCGAAUGUACUUAGAUUUAAAAUCAAAAACAGAU
env-14    UUAUCU.UUGCGCCU.............CGGUAACUGA.......AAGG...............UCAGG.ACCA.....................................UAC.AUAAA.AAUUAUU..UAAAU..AAAUAUUCGAAUGUACUUAGAUUUAAAAUCAAAAACAGAU
env-15    CUACCU.UUGCGGUCCC...........UUUAGACGC........UUA.................GCGUUGAAU.....................................GGC.AUAAC.AAUCAUU..GA.AA..CAAUAUUGGAAUGUAUUUGACUCCAGAAAAAAAGCAAGAG
env-16    UUAGUU.UUGCCACCU............UA...............AAU........................UA..................................UCAUAC.AUAAU.AAACAUU..UAAAA..CAAUUUUGGCAUGUAUUUAUCAUCUGAAAAAAAACAAGAA
env-17    GUAUAU.UUGCACCG.............CAAAAAGGA........GUG.................UCCUUUUUG....................................AUAC.AUAAC.AAUCAUU..UAUAG..AAAUAUUAGCAUGUAUUUAUCCAAAGAAAAAAAAGAAGAA
Fbr-1-1   UUAAAU.UUGCACCG.............UC.UUACA.........AAAGC................UGUAAUGA...................................CAUAC.AUAAG.AAUUAUU..AA.AU..AAAUAUUGGAAUGUAUUUAACCAAAGAAAUUAAAGAAGAA
env-18    GUACCU.UUGCACCGU............CUUACAAA.........AAG..................UUUGUAGG....................................CUAC.AUAAU.AAUUAUU..UUUAU..CAAUAUUGGCAUGUAUUUAGAAGCAGAAAAGAAAAAAGAG
env-19    GUACCU.UUGCACCGU............CUUACAAA.........AAG..................UUUGUAGG....................................CUAC.AUAAU.AAUUAUU..UUUAU..CAAUAUUGGCAUGUAUUUAGAAGCAGAAAAGAAAAAAGAG
env-20    CUACCU.UUGCACCG.............UCUAAACGAA.......AAG................UUCGU..GGC..................................GACUAC.AUAAU.AAACAUU..UA.AA..UAAUUUUAGCAUGUAUUUAGCACCUGAAAAAAAAGAAGAG
env-21    UUAUCU.UUGCAGCC.............GAUU.............UAAA.....................AAUC....................................GUAC.AUAAC.AAACAUU..AAAAA..CAAUUUUGGCAUGUAUUUAACUACAGACAAGAAAAAAGAA
env-22    UUAUAU.UUGCGCCU.............CUGUUUA..........UAACAGAU..............UAAAUUG.....................................UAC.AUAAU.AAUUAUU..UAAAA..CAAUGUUUGAAUGUAUUUAACAAAGAAAAUUAAAGAAGAC
env-23    UUAUAU.UUGCGCCU.............CUGUUUA..........UAACAGAU..............UAAAUUG.....................................UAC.AUAAU.AAUUAUU..UAAAA..CAAUGUUUGAAUGUAUUUAACAAAGAAAAUUAAAGAAGAC
env-24    UUAUAU.UUGCGCCU.............CUGUUUA..........UAACAGAU..............UAAAUUG.....................................UAC.AUAAU.AAUUAUU..UAAAA..CAAUGUUUGAAUGUAUUUAACAAAAAAAAAUUAAAGAAGAC
env-25    UUAUAU.UUGCGCCU.............CUGUUUA..........UAACAGAU..............UAAAUUG.....................................UAC.AUAAU.AAUUAUU..UAAAA..CAAUGUUUGAAUGUAUUUAACAAAAAAAAAUUAAAGAAGAC
Fba-9-1   GUAUCU.UUGCCGUCC............GAUUACC..........AAAGA.................GGUAAUU.....................................UGC.AUUAU.AAUUAUU..UACAA..AAAUAUUAGUAUGUAUUUAACACCAGAAAAAAAAGCUGAA
env-26    CUAUAU.UUGCCCUC.............AAUUCUGA.........GAA..................UCAGAGUU..................................GCCUAC.AUAAA.AAUUAUU..UAAAU..AAAUAUUGGAAUGUACCUAACAAAUAAGAUUAAACAAGAC
env-27    CUAUAU.UUGCACCG.............UCUAAUUAA........AAG.................UUAAUUAGG....................................CUAC.AUAAG.AAUCAUU..UU.AA..CAAUAUUGGCAUGUAUAUCAAUUCAGAAAUUAAAAAGAA
env-28    GUAUCU.UUGCCGGCU............CAACGGG..........AAAAC.................CCCGUUA.....................................UAC.AUAAA.AACUAUU..UA.AA..UAAUAUUGGCAUGUAUCUUACACAAGAAAAGAAAGAUGAA
env-33    GUAUCU.UUGCCGGCU............CAACGGG..........AAAAC.................CCCGUUA.....................................UAC.AUAAA.AACUAUU..UA.AA..UAAUAUUGGCAUGUAUCUUACACAAGAAAAGAAAGAUGAA
env-34    GUAUCU.UUGCCAUC.............GCU..............UUAAUU....................AGC..................................GAAUAC.AUAAC.AAUCAUU..GA.AA..UAAUAUUAGCAUGUACUUAACUAAAGAAAAAAAAGAAGAG
env-35    GUAUCU.UUGCCAUC.............GCU..............UUAAUU....................AGC..................................GAAUAC.AUAAC.AAUCAUU..GA.AA..UAAUAUUAGCAUGUACUUAACUAAAGAAAAAAAAGAAGAG
env-36    GUAUCU.UUGCCAUC.............GCU..............UUAAUU....................AGC..................................GAAUAC.AUAAC.AAUCAUU..GA.AA..UAAUAUUAGCAUGUACUUAACUAAAGAAAAAAAAGAAGAG
env-37    GUAUCU..UGCAGC..............GCU..............UUAAUU....................AGC..................................CAAUAC.AUAAU.AAUCAUU..GA.AA..UAAUAUUUGCAUGUACUUAUCAAAAGAAACAAAAGAGAAA
env-38    GUAUCU.UUGCAGC..............GCU..............UUAAUU....................AGC..................................CAAUAC.AUAAC.AAUCAUU..GA.AA..UAAUAUUAGCAUGUACUUAUCAAAAGAACAAAGAACAAAU
env-39    GUAUAU.UUGCCA...............UCGCU............UUAAUU..................AGCGA...................................AAUAC.AUAAC.AAUCAUU..GA.AA..UAAUAUUAGCAUGUACUUAACUAAAGAAAAAAAAGAAGAG
env-40    GUAUAU.UUGCCA...............UCGCU............UUAAUU..................AGCGA...................................AAUAC.AUAAC.AAUCAUU..GA.AA..UAAUAUUAGCAUGUACUUAACUAAAG..............
env-42    UUACAU.UUGCAGCC.............UU...............AAAUUUAA...................AA...................................CAUAC.AUAAU.AAACAUU..AAAAA..CAAUUUUAGCAUGUAUUUAGCAUCAGAAAAAAAACAAGAA
env-43    CUACCU.UUGCCCCG.............UC.AGUGAA........AAG.................UUCACUUGA....................................CUAC.AUAAU.AAUUAUU..UU.AA..CAAUAUUGGAAUGUAUUUAAACCCAGAAGCCAAAAAAGCA
Myu-1-1   GUAUAU.UUGCCGC..............UUGAAAG..........AAAGC.................CUUUCAAAAAAAAAUGCCCA...AUUAGUAC...UGGGC...UGUAC.AUAAC.AAUCAUU..UA.UA..CAAUAUUAGCAUGUAUUUAACAAAAGAGAAAAAAGAAGAG
env-44    UUAUAU.UUGGCCCC.............UUAAUUGCG........AUAAA...............CGCUAUUAA....................................CCGC.AUAAA.AAUUAUU..UA.AA..UAAUAUUGGAAUGUAUUUAACAAAAGAAGC..........
env-45    CUAUCU.UUGCACCG.............UCUUUGUAAA.......AAG................UUUACCCAGA....................................CUAC.AUAAU.GAUCAUU..CA.AG..CAAUAUUGGCAUGUAUUUAGAUUCAACAAAAAAGAAAGAA
env-46    CUAUCU.UUGCACCG.............UCUUUGUAAA.......AAG................UUUACCCAGA....................................CUAC.AUAAU.GAUCAUU..CA.AG..CAAUAUUGGCAUGUAUUUAGAUUCAACAAAAAAGAAAGAA
env-47    CUAUCU.UUGCACCG.............UCUUUGUAAA.......AAG................UUUACCCAGA....................................CUAC.AUAAU.GAUCAUU..CA.AG..CAAUAUUGGCAUGUAUUUAGAUUCAAC.............
env-48    CUAUCU.UUGCACCG.............UCUUUGUAAA.......AAG................UUUACCCAGG....................................CUAC.AUAAU.GAUCAUU..CA.AG..UAAUAUUGGCAUGUAUUUAGAUUCAACAAAAAAAGAAGAA
env-49    UUAUCU.UUGCAGCG.............UUCUUUCA.........CGG..................UGAAAGAC....................................GUAC.AUAAU.AAUCAUU..GA.AA..UAAUAUUUGCAUGUAUUUAAACGCAGAAAAAAAGAAAGAA
env-50    GUACAU.UUGCACAG.............AAUUUU...........UCAAUUC................AAAAAC....................................AUAC.AUAAC.AAUCAUU..UA.AA..UAAUAUUAGCAUGUAUUUAACUAAAGAAGUUAAAGAAGAA
env-51    GUAUAU.UUGCACCGC............UU...............UAAUUAGC...................AA....................................CUAC.AUAAC.AAUCAUU..GA.AA..UAAUAUUAGCAUGUAUUUAACAAAAGAAAUAAAAGAAAAA
env-52    AUAUCU.UUGCACCG.............UCG..............AAACA.....................CGG....................................CUAC.AUAAA.AAUUAUU..UAAAA..CAAUAUUCGAAUGUAUUUAGAUUCAAAAUCAAAAACAGAC
env-53    UUACUU.UUGCACUC.............UCGCAACGA........AAA.................UCGUAGCGA....................................AUAC.AUAAA.AAUUAUU..UU.AA..UAAUAUUGGCAUGUACUUAGAAGCAGAAAAGAAAAAAGAA
Fsa-1-1   UUACCU.UUGCACCA.............CGUUUUAGG........AUAAA...............CCUAU..CG..................................CGCUAC.AUAAU.AAUCAUU..UA.AA..GAAUAUUAGCAUGUACUUAACUAAAGAAGUUAAAUCAGAA
Fca-1-1   UUACCU.UUGCACCA.............CGUUUUAGG........ACAAA...............CCUA..UUG..................................CGCUAC.AUAAU.AAUCAUU..UA.AA..GGAUAUUAGCAUGUACUUAACUAAAGAAGUUAAAUCAGAA
env-54    CUAUAU.UUGCCACU.............UGAUUGU..........CUCAA.................AAAAUUG...................................AUUAC.AUAAA.AAUUAUU..UAAAG..AAAUAUUGGAAUGUACUUAACAAAGAAAUAAAAGAAAAU
env-55    CUAUAU.UUGCCACU.............UGAUUGUU.........UCA..................AAAAAUUG...................................AUUAC.AUAAA.AAUUAUU..UAAAG..AAAUAUUGGAAUGUACUUAACAAAGAAAUAAAAGAAAAU
env-56    CUAUAU.UUGCCACU.............UGAUUGUU.........UCA..................AAAAAUUG...................................AUUAC.AUAAA.AAUUAUU..UAAAG..AAAUAUUGGAAUGUACUUAACAAAGAAAUAAAAGAAAAU
env-57    CUAUAU.UUGCCACU.............UGAUUGUU.........UCA..................AAAAAUUG...................................AUUAC.AUAAA.AAUUAUU..UAAAG..AAAUAUUGGAAUGUACUUAACAAAGAAAUAAAAGAAAAU
env-59    CUACAU.UUGCAGCCAUAAAAUCA.......................................................................................UAC.AUAAC.AAACAUU..UAAAA.UAAUUUUAGCAUGUAUUUAACAACAGAAAAAAAACAAGAA
env-60    CUAUUU.UUGCAUCC.............GAAUUAUG.........AUAAA................CAUAAUUC....................................AUAC.AUAAA.AAUCAUU..UACAA..AAAUAUUAGCAUGUAUUUAGAUCCAAAGAAAAACAAGAG
env-61    CUAUUU.UUGCAUCC.............GAAUUAUG.........AUAAA................CAUAAUUC....................................AUAC.AUAAA.AAUCAUU..UACAA..AAAUAUUAGCAUGUAUUUAGAUCCAAAGAAAAACAAGAG
env-62    CUAUUU.UUGCAUCC.............GAAUUAUG.........AUAAA................CAUAAUUC....................................AUAC.AUAAA.AAUCAUU..UACAA..AAAUAUUAGCAUGUAUUUAGAUCCAAAGAAAAACAAGAG
env-63    CUAUUU.UUGCAUCC.............GAAUUAUG.........AUAAA................CAUAAUUC....................................AUAC.AUAAA.AAUCAUU..UACAA..AAAUAUUAGCAUGUAUUUAGAUCCAAAGAAAAACAAGAG
env-65    CUAUUU.UUGCACCC.............GAAUUAUG.........AUACA................CAUAAUUC....................................AUAC.AUAAA.AAUCAUU..UACAA..AAAUAUUAGCAUGUAUUUAGAUCCAAAGAAAAACAAGAG
env-66    CUAUUU.UUGCACCC.............GAAUUAUG.........AUACA................CAUAAUUC....................................AUAC.AUAAA.AAUCAUU..UACAA..AAAUAUUAGCAUGUAUUUAGAUCCAAAGAAAAACAAGAG
env-67    CUAUUU.UUGCACCC.............GAAUUAUG.........AUAGA................CAUAAUUC....................................AUAC.AUAAA.AAUCAUU..UACAA..AAAUAUUAGCAUGUAUUUAGAUCCAAAGAAAAACAAGAG
env-68    CUAUUU.UUGCACCC.............GAAUUAUG.........AUAGA................CAUAAUUC....................................AUAC.AUAAA.AAUCAUU..UACAA..AAAUAUUAGCAUGUAUUUAGAUCCAAAGAAAAACAAGAG
env-69    CUAUUU.UUGCACUC.............GAAUUAUG.........AUAGA................CAUGAUUC....................................AUAC.AUAAA.AAUCAUU..UACAA..AAAUAUUAGCAUGUAUUUAGAUCCAAAGAAAAACAAGAG
env-70    CUAUUU.UUGCACUC.............GAAUUAUG.........AUAGA................CAUGAUUC....................................AUAC.AUAAA.AAUCAUU..UACAA..AAAUAUUAGCAUGUAUUUAGAUCCAAAGAAAAACAAGAG
env-71    CUAUUU.UUGCACUC.............GAAUUAUG.........AUAGA................CAUGAUUC....................................AUAC.AUAAA.AAUCAUU..UACAA..AAAUAUUAGCAUGUAUUUAGAUCCAAAGAAAAACAAGAG
env-76    CUAUUU.UUGCACUC.............GAAUUAUG.........AUAGA................CAUGAUUC....................................AUAC.AUAAA.AAUCAUU..UACAA..AAAUAUUAGCAUGUAUUUAGAUCCAAAGAAAAACAAGAG
env-77    CUAUUU.UUGCACUC.............GAAUUAUG.........AUAGA................CAUGAUUC....................................AUAC.AUAAA.AAUCAUU..UACAA..AAAUAUUAGCAUGUAUUUAGAUCCAAAGAAAAACAAGAG
env-79    CUAUUU.UUGCACUC.............GAAUUAUG.........AUAGA................CAUGAUUC....................................AUAC.AUAAA.AAUCAUU..UACAA..AAAUAUUAGCAUGUAUUUAGAUCCAAAGAAAAACAAGAG
env-80    CUAUUU.UUGCACUC.............GAAUUAUG.........AUAGA................CAUGAUUC....................................AUACUAUAAA.AAUCAUU..UACAA..AAAUAUUAGCAUGUAUUUAGAUCCAAAGAAAACAAGAGA
env-81    CUAUUU.UUGCACCC.............GAAUUAUG.........AUAGA................CAUGAUUC....................................AUAC.AUAAA.AAUCAUU..UACAA..AAAUAUUAGCAUGUAUUUAGAUCCAAAGAAAAAGAAGCA
env-82    CUAUUU.UUGCACCC.............GAAUUAUG.........AUAGA................CAUGAUUC....................................AUAC.AUAAA.AAUCAUU..UACAA..AAAUAUUAGCAUGUAUUUAGAUCCAAAGAAAAAGAAGCA
env-83    CUAUUU.UUGCACCC.............GAAUUAUG.........AUAGA................CAUGAUUC....................................AUAC.AUAAA.AAUCAUU..UACAA..AAAUAUUAGCAUGUAUUUAGAUCCAAAGAAAAAGAAGCA
env-84    CUAUUU.UUGCACCC.............GAAUUAUG.........AUAGA................CAUGAUUC....................................AUAC.AUAAA.AAUCAUU..UACAA..AAAUAUUAGCAUGUAUUUAGAUCCAAAGAAAAAGAAGCA
env-85    CUAUUU.UUGCACCC.............GAAUUAUGA........UGG.................UCAUGAUUC....................................AUAC.AUAAA.AAUCAUU..UACAA..AAAUAUUAGCAUGUAUUUAGAUCCAAAGAAAAACAAGAG
env-86    CUAUUU.UUGCACCC.............GAAUUAUGA........UGG.................UCAUGAUUC....................................AUAC.AUAAA.AAUCAUU..UACAA..AAAUAUUAGCAUGUAUUUAGAUCCAAAGAAAAACAAGAG
env-87    CUAUUU.UUGCACCC.............GAAUUAUGA........UGG.................UCAUGAUUC....................................AUAC.AUAAA.AAUCAUU..UACAA..AAAUAUUAGCAUGUAUUUAGAUCCAAAGAAAAACAAGAG
env-89    CUAUUU.UUGCACCC.............GAAUUAUGA........UGG.................UCAUGAUUC....................................AUAC.AUAAA.AAUCAUU..UACAA..AAAUAUUAGCAUGUAUUUAGAUCCAAAGAAAAACAAGAG

          ............................<<<<<<<<<<<<<<<<<<..........>>>>>>>>>>>>>>>>>>.........<<<<<<<<........>>>>>>>>.......................................................................
          ............................2222221111..101.00..........00.101..1111222222.........11011000........00011011.......................................................................
          •UAY•U-UUGC••CY-------------∘∘∘∘∘∘∘----------R•R∘∘-----------------∘∘∘∘∘∘∘-------------------------------------∘UAC-AUAA•-AAUCAUU---UA-AA--YAAUAUURG•AUGUAUUUARC•ARAGAAR••AAARRRRAR
          5′                                                                                                                                            3′
```

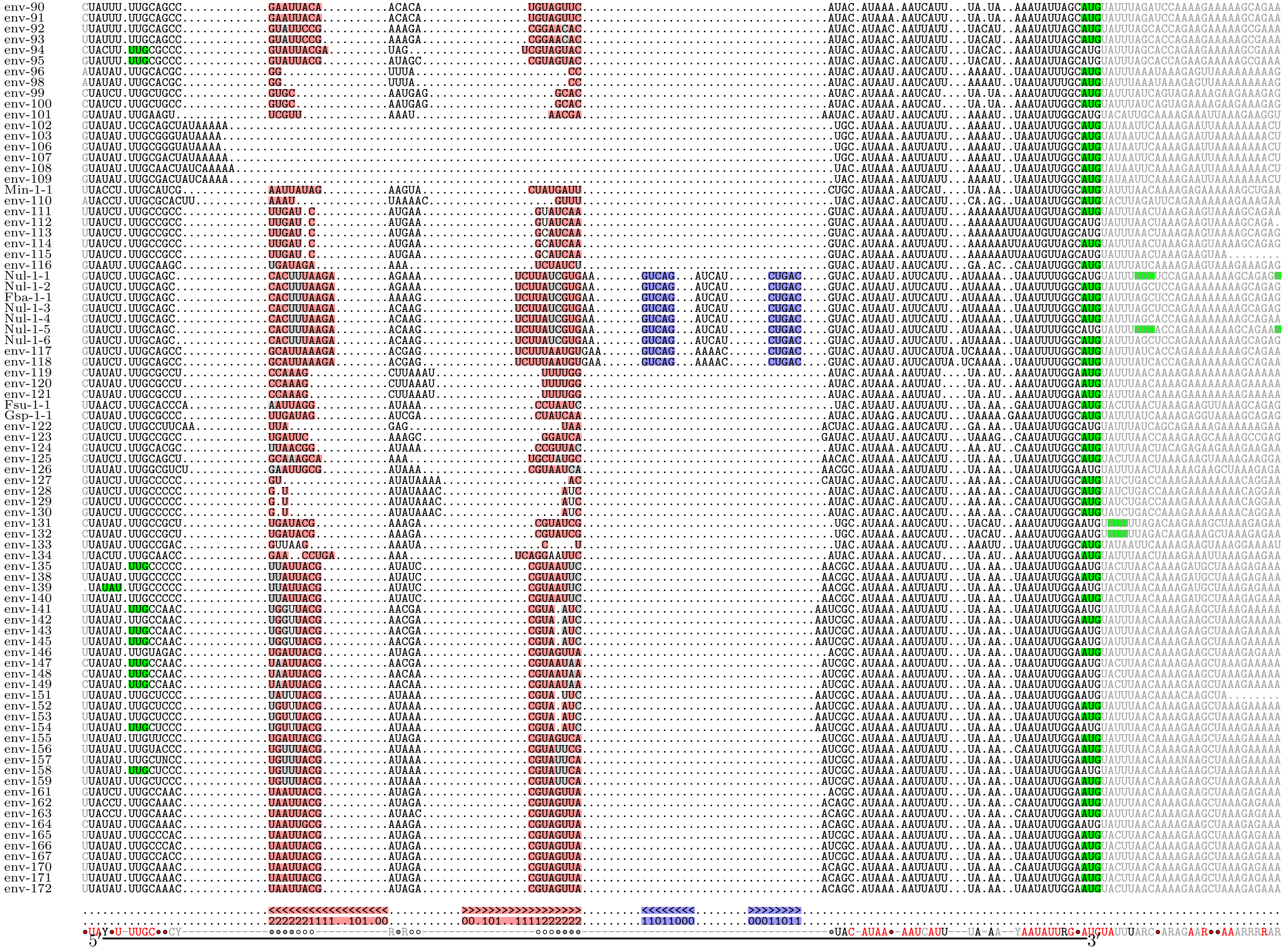

```
env-173  CUAUAU.UUGCCCAC.............UAAUUACG..........AUAGA................CGUAGUUA....................................ACAGC.AUAAA.AAUUAUU..UA.AA..UAAUAUUGGAAUGUACUUAACAAAAGAAGCUAAAGAGAAA
env-174  CUAUAU.UUGCCAAC.............UAAUUACG..........AUAGA................CGUAGUUA....................................ACAGC.AUAAA.AAUUAUU..UA.AA..UAAUAUUGGAAUGUAUUUAACAAAAGAAG...........
env-175  CUAUAU.UUGCCAAC.............UAAUUACG..........AUAGA................CGUAGUUA....................................ACAGC.AUAAA.AAUUAUU..UA.AA..UAAUAUUGGAAUGUAUUUAACAAAAGAAGCUAAAGAGAAA
env-176  CUAUAU.UUGCCAAC.............UAAUUACG..........AUAGA................CGUAGUUA....................................ACAGC.AUAAA.AAUUAUU..UA.AA..UAAUAUUGGAAUGUAUUUAACAAAAGAAGCUA........
env-177  CUAUAU.UUGCCAAC.............UAAUUACG..........AUAGA................CGUAGUUA....................................ACAGC.AUAAA.AAUUAUU..UA.AA..UAAUAUUGGAAUGUAUUUAACAAAAGAAGCUA........
env-179  CUAUAU.UUGCCAAC.............UAAUUACG..........AUAGA................CGUAGUUA....................................ACAGC.AUAAA.AAUUAUU..UA.AA..UAAUAUUGGAAUGUAUUUAACAAAAGAAGCUA........
env-180  CUAUAU.UUGCCAAC.............UAAUUACG..........AUAGA................CGUAGUUA....................................ACAGC.AUAAA.AAUUAUU..UA.AA..UAAUAUUGGAAUGUAUUUAACAAA................
env-181  CUAUAU.UUGCCAAC.............UAAUUACG..........AUAGA................CGUAGUUA....................................ACAGC.AUAAA.AAUUAUU..UA.AA..UAAUAUUGGAAUGUAUUUAACAAAAGAAGCUAAAGAGAAA
env-185  CUAUAU.UUGCCAAC.............UAAUUACG..........AUAGA................CGUAGUUA....................................ACAGC.AUAAA.AAUUAUU..UA.AA..UAAUAUUGGAAUGUAUUUAACAAA................
env-186  CUAUAU.UUGCCAAC.............UAAUUACG..........AUAGA................CGUAGUUA....................................ACAGC.AUAAA.AAUUAUU..UA.AA..UAAUAUUGGAAUGUAUUUAACAAAAGAAGCUAAAGAGAAA
env-187  CUAUAU.UUGCCAAC.............UAAUUACG..........AUAGA................CGUAGUUA....................................ACAGC.AUAAA.AAUUAUU..UA.AA..UAAUAUUGGAAUGUAUUUAACAAAAGAAGCUAAAGAGAAA
env-189  CUAUAU.UUGCGACU.............UAAUUACG..........AUAGA................CGUAGUUA.....................................ACGC.AUAAA.AAUUAUU..UA.AA..UAAUAUUGGAAUGUAUUUAACAAAAGAAGCUAAAGAGAAA
env-191  CUAUAU.UUGCAACU.............UAAUUACG..........AUAGA................CGUAGUUA.....................................ACGC.AUAAA.AAUUAUU..UA.AA..UAAUAUUGGAAUGUAUUUAACAAAAGAAGCUAAAGAGAAA
env-192  CUAUAU.UUGCAACU.............UAAUUACG..........AUAGA................CGUAGUUA.....................................ACGC.AUAAA.AAUUAUU..UA.AA..UAAUAUUGGAAUGUAUUUAACGAAAGAAGCUAAAGAGAAA
env-193  CUAUAU.UUGCAACU.............UAAUUACG..........AUAGA................CGUAGUUA.....................................ACGC.AUAAA.AAUUAUU..UA.AA..UAAUAUUGGAAUGUAUUUAACGAAAGAAGCUAAAGAGAAA
env-195  CUAUAU.UUGCUACU.............UGAUUACG..........AUAGA................CGUAGUUA.....................................GCGC.AUAAA.AAUUAUU..UA.AA..UAAUAUUGGAAUGUAUUU......................
env-196  UUAUAU.UUGACACU.............UGAUUACG..........AUAGA................CGUAAUUA....................................UUCGC.AUAAA.AAUUAUU..UA.AA..UAAUAUUGGAAUGUAUUUAACAAAAGAAGCUAAAGAGAAA
env-197  UUAUAU.UUGACACU.............UGAUUACG..........AUAGA................CGUAAUUA....................................UUCGC.AUAAA.AAUUAUU..UA.AA..UAAUAUUGGAAUGUAUUUAACAAAAGAAGCUAAAGAGAAA
env-198  UUAUAU.UUGACACU.............UGAUUACG..........AUAGA................CGUAAUUA....................................UUCGC.AUAAA.AAUUAUU..UA.AA..UAAUAUUGGAAUGUAUUUAACAAAAGAAGCUAAAGAGAAA
env-201  UUAUAU.UUGCCACU.............UGAUUACG..........AUAGA................CGUAAUUA....................................UUCGC.AUAAA.AAUUAUU..UA.AA..UAAUAUUGGAAUGUAUUUAACAAAAGAA............
env-202  UUAUAU.UUGCCACU.............UGAUUACG..........AUAGA................CGUAAUUA....................................UUCGC.AUAAA.AAUUAUU..UA.AA..UAAUAUUGGAAUGUAUUUAACAAAAGAAGCUAAAGAGAAA
env-203  GUAUCU.UUGCCCAU.............C.................AUUU........................G....................................UGUAC.AUAAU.AAUCAUU..UAAAU..AAAUAUUGGCAUGUAUUUGACAACAGAAAAAAAGAAAGAG
env-205  GUAUAU.UUGCAGAA.............UU................CAAUU.....................AA......................................UAC.AUAAU.AAUUAU...UA.AA..UAAUAUUGGAAUGUAUCUAACCAAAGAAAUUAAAAAAAG
env-206  GUAUAU.UUGCAGAA.............UU................CAAAUU.....................AA......................................UAC.AUAAU.AAUUAU...UA.AA..UAAUAUUGGCAUGUAUCUAACCAAAGAAAUUAAAAAAAG
env-207  GUAUAU.UUGCAGAA.............UU................CAAAUU.....................AA......................................UAC.AUAAU.AAUUAU...UA.AA..UAAUAUUGGCAUGUAUCUAACCAAAGAAAUUAAAAAAAG
env-210  GUAUAU.UUGCAGAA.............UU................CAAAUU.....................AA......................................UAC.AUAAU.AAUUAU...UA.AA..UAAUAUUGGCAUGUAUCUAACCAAAGAAAUUAAAAAAAG
env-211  AUAUAU.UUGCAACU.............UU................CAGAA......................AA.....................................GUAC.AUAAA.AAUUAU...UA.AA..AAAUAUUGGCAUGUAUCUAAACAAAGAAGUAAAAAAGAAG
env-212  CUAUAU.UUGCCGUU.............CAAUUCU...........UAUAAGC...............AGAAUUG....................................UUUAC.AUAAA.AAUUAUU..UAAAC..AAAUAUUGGAAUGUACUUAACAAAAAAGA...........
env-213  CUAUAU.UUGCAC...............GCAUUUU...........AGAAA.................AAAAUGC...................................AUUUAC.AUAAU.AAUCAUU..UA.AA..UAAUAUUGGCAUGUAUUUAACAGCUGAAAAGAAAAAAGAG
env-214  CUAUAU.UUGCAC...............GCAUUUU...........AGAAA.................AAAAUGC...................................AUUUAC.AUAAU.AAUCAUU..UA.AA..UAAUAUUGGCAUGUAUUUAACAGCUGAAAAGAAAAAAGAG
env-215  CUAUAU.UUGCAC...............GCAUUUU...........AGAAA.................AAAAUGC...................................AUUUAC.AUAAU.AAUCAUU..UA.AA..UAAUAUUGGCAUGUAUUUAACAGCUGAAAAGAAAAAAGAG
env-216  CUAAUU.UUGCGCCG.............UCGAAUCUAAA.......AAG...............UUUGGAG.CGA.....................................CUAC.AUAAA.AAUCAUU..UA.AA..UAAUAUUGGCAUGUAUUUAACAUCAGAAAAGAAACAAGAA
env-217  CUAAUU.UUGCGCCG.............UCGAAUCUAAA.......AAG...............UUUGGAG.CGA.....................................CUAC.AUAAA.AAUCAUU..UA.AA..UAAUAUUGGCAUGUAUUUAACAUCAGAAAAGAAACAAGAA
env-218  GUACAU.UUGUCGAUCG...........UUUG..............AAAA.....................CGAA.....................................UUAC.AUAAU.AAUCAUU..UA.AA..UAAUAUUUGCAUGUACAAUUCAAAAGAAGUAAAAGAAGAC
env-220  AUAAAU.UUGCGCCU.............CGGUAAA...........UAA...................UUUACCA......................................UAC.AUAAU.AAUCAUU..UA.AA..UAAUAUUAGCAUGUACUUAACUAAAGAAGUAAAAGAAGGU
env-221  CUAUCU.UUGCACCCG............CUACUCAA..........ACGAC................UUGAGUAC.....................................GUAC.AUAAA.AAUUAUU..UA.UA..CAAUAUUGGCAUGUAUUUAAACCCAGAAGCAAAAAAAGAA
env-222  CUAUCU.UUGCACCCG............CUACUCAA..........ACGAC................UUGAGUAC.....................................GUAC.AUAAA.AAUUAUU..UA.UA..CAAUAUUGGCAUGUAUUUAAACCCAGAAGCAAAAAAAGAA
Fli-1-1  CUAUUU.UUGCGCCAC............GUUUUAGGA.........UAG.................UCCUAUUUC....................................GUUAC.AUAAU.AAUCAUU..UA.AA..GAAUAUUUGCAUGUACUUAACUAAAGAAGUUAAAUCAGAC
Gme-1-1  CUAUCU.UUGCACCG.............UGAUUCG...........AAAAC.................CGAGUUG....................................CGUAC.AUAAC.AAUCAUU..UACAA..UAAUAUUAGCAUGUAUUUAGAUCAAAAGAAAAAGAAGCC
env-223  CUAUAU.UUGCACCU.............UGAUUCGA..........AAA..................UCGGGUCU.....................................GUAC.AUAAC.AAUCAUU..UACAA..UAAUAUUAGCAUGUAUUUAUCAAAAGAAGUAAAAAAUGAA
env-224  CUAUCU.UUGCACCC.............GCUAUUCC..........UAUGAGU..............GGAAUUGC......................................UAC.AUAAU.AAUUAUU..UAACU..AAAUAUUAGCAUGUACUUAAACACAGAAGUUAAAAAAGAA
env-225  CUAUCU.UUGCACCC.............GCUAUUCC..........UAUGAGU..............GGAAUUGC......................................UAC.AUAAU.AAUCAUU..UAACU..AAAUAUUGGCAUGUACUUAAACACAGAAGUUAAAAAAGAA
env-226  CUAUCU.UUGCAUCC.............GCUAUUCC..........UAUGAGU..............GGAAUUGC......................................UAC.AUAAU.AAUUAUU..UAACU..AAAUAUUGGCAUGUACUUAAACACAGAAGUUAAAAAAGAA
env-227  CUAUCU.UUGCACCC.............GCUAUUCC..........UAUGAGU..............GGAAUUGC......................................UAC.AUAAU.AAUUAUU..UAACU..AAAUAUUGGCAUGUACUUAAACACAGAAGUUAAAAAAGAA
env-228  CUAUCU.UUGCACCC.............GCUAUUCC..........UAUGAGU..............GGAAUUGC......................................UAC.AUAAU.AAUUAUU..UAACU..AAAUAUUGGCAUGUACUUAAACACAGAAGUUAAAAAAGAA
env-229  CUAUCU.UUGCACCC.............GCUAUUCC..........UAUGAGU..............GGAAUUGC......................................UAC.AUAAU.AAUUAUU..UAACU..AAAUAUUGGCAUGUACUUAAACACAGAAGUUAAAAAAGAA
env-230  CUAUCU.UUGCACCC.............GCUAUUCC..........UAUGAGU..............GGAAUUGC......................................UAC.AUAAU.AAUUAUU..UAACU..AAAUAUUGGCAUGUACUUAAACACAGAAGUUAAAAAAGAA
env-231  CUAUCU.UUGCACCC.............GCUAUUCC..........UAUGAGU..............GGAAUUGC......................................UAC.AUAAU.AAUUAUU..UAACU..AAAUAUUGGCAUGUACUUAAACACAGAAGUUAAAAAAGAA
env-234  CUAUCU.UUGCACCC.............GCUGUUCC..........UACGAGU..............GGAAUUGC......................................UAC.AUAAU.AAUUAUU..UAACU..AAAUAUUGGCAUGUACUUAAACACAGAAGUUAAAAAAGAA
env-235  CUAUCU.UUGCACCC.............GCUGUUCC..........UACGAGU..............GGAAUUGC......................................UAC.AUAAU.AAUUAUU..UAACU..AAAUAUUGGCAUGUACUUAAACACAGAAGUUAAAAAAGAA
env-236  CUAUCU.UUGCACCC.............GCUGUUCC..........UACGAGU..............GGAAUUGC......................................UAC.AUAAU.AAUUAUU..UAACU..AAAUAUUGGCAUGUACUUAAACACAGAAGUUAAAAAAGAA
env-237  CUAUCU.UUGCACCC.............GCUAUUCC..........UACGAGU..............GGAAUUGC......................................UAC.AUAAU.AAUUAUU..UAACU..AAAUAUUGGCAUGUACUUAAACACAGAAGUUAAAAAAGAA
env-238  CUAUAU.UUGCACCC.............GCUAUUCA..........UAUGAGU..............UGAGUGGC......................................UAC.AUAAA.AAUUAUU..UAAUC..AAAUAUUGGCAUGUAUUUAAAUCCAGAAGUAAAAAAAGGA
env-239  GUACUU.UUGCCGCU.............CGAAUCAC..........ACGAA................GUGAGUCG.....................................GUAC.AUAAA.AAUCAUU..UU.AA..UAAUAUUGGCAUGUAUUUAGCACCAGAAAAGAAAGAAGAA
env-243  GUACUU.UUGCCGCU.............CGAAUCAC..........ACGAA................GUGAGUCG.....................................GUAC.AUAAA.AAUCAUU..UU.AA..UAAUAUUGGCAUGUAUUUAGCACCAGAAAAGAAAGAAGAA
env-244  UUACAU.UUGCAACU.............UUGAUAG...........AAAAG.................CUAUCAA......................................UAC.AUAAA.AAUCAUU..GA.AA..UAAUAUUGGCAUGUAUUUAAACAAAGAAGACAAGCAAAAG
env-247  UUACAU.UUGCAACU.............UUGAUAG...........AAAAA.................CUAUCAA......................................UAC.AUAAA.AAUCAUU..GA.AA..UAAUAUUGGCAUGUAUUUAAACAAAGAAGACAAGCAAAAG
env-248  AUACAU.UUGCAACU.............UUGAUAG...........AAAAA.................CUAUCAA......................................UAC.AUAAA.AAUCAUU..GA.AA..UAAUAUUGGCAUGUAUCU......................
env-249  UUACAU.UUGCAACU.............UUGAUAG...........AAAAA.................CUAUCAA......................................UAC.AUAAA.AAUCAUU..GA.AA..UAAUAUUGGCAUGUAUUUAAACAAAGAAGACAAGCAAAAG
env-250  UUACAU.UUGCAACU.............UUGAUAG...........AAAAA.................CUAUCAA......................................UAC.AUAAA.AAUCAUU..GA.AA..UAAUAUUGGCAUGUAUUUAAACAAAGAAGACAAGCAAAAG
env-251  GUAUUU.UUGCCGCU.............CGGUAAUC..........AAAA.................GGUUAUCA...................................ACAUAC.AUAAA.AAUCAUU..UA.AA..UAAUAUUGGCAUGUAUCUAACAUCAGAAAAGAAAAAGAA
env-255  AUAUUU.UUGCCGCU.............CGGUAAUC..........AAAA.................GGUUAUCA...................................ACAUAC.AUAAA.AAUCAUU..UA.AA..UAAUAUUGGCAUGUAUUUAACACCAGAAGUAAAAAAAGAA
env-256  GUAUUU.UCGCCGCU.............CGAUAAUC..........ACAA.................GGUUAUCA...................................ACAUAC.AUAAA.AAUCAUU..UA.AA..UAAUAUUGGCAUGUAUUUAUCAUCUGAAAAAAAACAAGAA
env-257  GUAUUU.UCGCGGCU.............CGAUAAUC..........ACAA.................GGUUAUCA...................................ACAUAC.AUAAA.AAUCAUU..UA.AA..UAAUAUUGGCAUGUAUUUAUCAUCUGAGAAAAAACAAGAA
env-258  UUAUCU.UUGCGCCC.............UCAAACGG..........AUAAA................CCGUUUAG......................................UAC.AUAAA.AAUCAUU..UA.AA..GAAUAUUGGCAUGUAUUUGACUAAAGAAGCAAAAGAAGAA
env-261  UUAUCU.UUGCGCCC.............UCAAACGG..........AUAAA................CCGUUUAG......................................UAC.AUAAA.AAUCAUU..UA.AA..GAAUAUUGGCAUGUAUUUGACUAAAGAAGCAAAAGAAGAA
Nma-1-1  GUAUCU.UUGCAGCC.............GCUU.AGGGA........AUGAA..............UCCCGAAGGCACC......GUCAGA..AAA.......UCUGAC....AUAC.AUAAU.AUUCAUUAUAUA.AA..UAAUUUUGGCAUGUAUUUAGCACCAGAAAAAAAAGCUGAG
env-262  UUACUU.UUGCGCCC.............GUAUCCG...........AGAAA.................CGGCUAC......................................UAC.AUAAA.AAUCAUU..UA.AA..UAAUAUUUGCAUGUAUUUGACUAAAGAAGUUAAAGCAGAA
env-263  CUAUAU.UUGCCCGU.............CUAAUAAU..........CAA..................AUUAUUAG.....................................CUAC.AUAAU.AAUUAUU..UACAA..UAAUAUUGGCAUGUAUUUAUCAAAAGAAAAUAAAAAAAAU
env-264  CUAUAU.UUGCCCGU.............CUAAUAAU..........CAA..................AUUAUUAG.....................................CUAC.AUAAU.AAUUAUU..UACAA..UAAUAUUGGCAUGUAUUUAUCAAAAGAAAAUAAAAAAAAU
env-265  CUAUAU.UUGCCCGU.............CUAAUAAU..........CAA..................AUUAUUAG.....................................CUAC.AUAAU.AAUUAUU..UACAA..UAAUAUUGGCAUGUAUUUAUCAAAAGAAAAUAAAAAAAAU
env-266  GUAUCU.UUGCCGCU.............UCGAUAG...........AAAAA.................CUAUCGG......................................UAC.AUAAU.AAUCAUU..GA.AA..UAAUAUUGGAAUGUAUUUAACAAAAGAAGAAAAAGAACAA
env-267  GUAUCU.UUGCCGCU.............UCGAUAG...........AAAAA.................CUAUCGG......................................UAC.AUAAU.AAUCAUU..GA.AA..UAAUAUUGGAAUGUAUUUAACAAAAGAAGAAAAAGAACAA
env-268  GUAUCU.UUGCCGCU.............UCGAUAG...........AAAAA.................CUAUCGG......................................UAC.AUAAU.AAUCAUU..GA.AA..UAAUAUUGGAAUGUAUUUAACAAAAGAAGAAAAAGAACAA
env-269  GUAUCU.UUGCCGCU.............UCGAUAG...........AAAAA.................CUAUCGG......................................UAC.AUAAU.AAUCAUU..GA.AA..UAAUAUUGGAAUGUAUUUAACAAAAGAAGAAAAAGAACAA
env-270  GUAUCU.UUGCCGCU.............UCGAUAG...........AAAAA.................CUAUCGG......................................UAC.AUAAU.AAUCAUU..GA.AA..UAAUAUUGGAAUGUAUUUAACAAAAGAAGAAAAAGAACAA
env-271  GUAUCU.UUGCCGCU.............UCGAUAG...........AAAAA.................CUAUCGG......................................UAC.AUAAU.AAUCAUU..GA.AA..UAAUAUUGGAAUGUAUUUAACAAAAGAAGAAAAAGAACAA
env-272  GUAUCU.UUGCCGCU.............UCGAUAG...........AAAAA.................CUAUCGG......................................UAC.AUAAU.AAUCAUU..GA.AA..UAAUAUUGGAAUGUAUUUAACAAAAGAAGAAAAAGAACAA
Mmo-1-1  GUAUCU.UUGCCGCU.............UCGAUAG...........AAAAA.................CUAUCGG......................................UAC.AUAAU.AAUCAUU..GA.AA..UAAUAUUGGAAUGUAUUUAACAAAAGAAGAAAAAGAACAA
env-273  GUAAAU.UUGCACCACGUA.........UUUU..............AAA......................AAAA......................................UAC.AUAAC.AAUCAUU..UAAAG..CAAUAUUGGAAUGUAUCUGACAAAAGAAAAAAAACAGGAA
env-274  GUAAAU.UUGCACCACGUA.........UUUU..............AAA......................AAAA......................................UAC.AUAAC.AAUCAUU..UAAAG..CAAUAUUGGAAUGUAUCUGACAAAAGAAAAAAAACAGGAA
env-276  UUAUAU.UUGCGCCUC............UGUUUAUA..........ACAGA................UAUAAAUU.....................................AUAC.AUAAU.AAUUAUU..UAAAA..CAAUGUUUGAAUGUAUUUAACAAAGAAAAUUAAAGAAGAC
env-277  UUAUAU.UUGCGCCUC............UGUUUAUA..........ACAGA................UAUAAAUU.....................................AUAC.AUAAU.AAUUAUU..UAAAA..CAAUGUUUGAAUGUAUUUAACAAAGAAAAUUAAAGAAGAC

         ............................&lt;&lt;&lt;&lt;&lt;&lt;&lt;&lt;&lt;&lt;&lt;&lt;&lt;&lt;&lt;&lt;&lt;&lt;...........&gt;&gt;&gt;&gt;&gt;&gt;&gt;&gt;&gt;&gt;&gt;&gt;&gt;&gt;&gt;&gt;&gt;&gt;.........&lt;&lt;&lt;&lt;&lt;&lt;&lt;&lt;........&gt;&gt;&gt;&gt;&gt;&gt;&gt;&gt;.......................................................................
         ............................2222221111..101.00...........00.101..1111222222.........11011000........00011011.......................................................................
         •UAY•U-UUGC••CY-------------◦◦◦◦◦◦◦-----------R•R◦◦-----------------◦◦◦◦◦◦◦-------------------------------------◦UAC-AUAA•-AAUCAUU---UA-AA--YAAUAUURG•AUGUAUUUARC•ARAGAAR••AAARRRRAR
         5′                                                                                                                                                  3′
```

```
env-278   UUAUAU.UUGCGCCUC............UGUUUAUA..........ACAGA................UAUAAAUU.....................................AUAC.AUAAU.AAUUAUU...UAAAA..CAAUGUUUGAAUGUAUUUAACAAAGAAAAUUAAAGAAGAC
env-280   UUAUAU.UUGCCCCUC............UGUUUAUG..........ACAGA................UAUAAUUU.....................................AUAC.AUAAU.AAUUAUU...UAAAA..AAAUGUUUGCAUGUAUUUAACAAAAAAAAUAAAAGAAGAU
env-281   UUAUAU.UUGCCCCUC............UGUUUAUA..........ACAGA................UGUAAUUU.....................................GUAC.AUAAU.AAUUAUU...UA.AA..AAAUGUUUGCAUGUAUUUAACAAAAAAAAUAAAAGAAGAU
env-282   UUAUAU.UUGCGCCUC............UGUUUAUA..........ACAGA................UGUAUUUG......................................UAC.AUAAU.AAUUAUU...UAAAA..CAAUGUUUGCAUGUAUUUAACUAAAAAAAUAAAACAAGAC
env-283   UUAUAU.UUGCGCCUC............UGUUUAUA..........ACAGA................UGUAUUUG......................................UAC.AUAAU.AAUUAUU...UAAAA..CAAUGUUUGCAUGUAUUUAACUAAAAAAAUAAAACAAGAC
env-285   UUAUAU.UUGCGCCUC............UGUUUAUA..........ACAGA................UGUAUUUG......................................UAC.AUAAU.AAUUAUU...UAAAA..CAAUGUUUGCAUGUAUUUAACUAAAAAAAUAAAACAAGAC
env-286   UUAUAU.UUGCGCCUC............UGUUUAUA..........ACAGA................UGUAUUUG......................................UAC.AUAAU.AAUUAUU...UAAAA..CAAUGUUUGCAUGUAUUUAACUAAAAAAAUAAAACAAGAC
env-287   UUAUAU.UUGCGCCUC............UGUUUAUA..........ACAGA................UGUAUUUG......................................UAC.AUAAU.AAUUAUU...UAAAA..CAAUGUUUGCAUGUAUUUAACUAAAAAAAUAAAACAAGAC
env-288   UUAUAU.UUGCGCCUC............UGUUUAUA..........ACAGA................UGUAUUUG......................................UAC.AUAAU.AAUUAUU...UAAAA..CAAUGUUUGCAUGUAUUUAACUAAAAAAAUAAAACAAGAC
env-289   UUAUAU.UUGCGCCUC............UGUUUAUA..........ACAGA................UGUAUUUG......................................UAC.AUAAU.AAUUAUU...UAAAA..CAAUGUUUGCAUGUAUUUAACUAAAAAAAUAAAACAAGAC
env-290   UUAUAU.UUGCGCCUC............UGUUUAUA..........ACAGA................UGUAUUUG......................................UAC.AUAAU.AAUUAUU...UAAAA..CAAUGUUUGCAUGUAUUUAACUAAAAAAAUAAAACAAGAC
env-294   UUAUAU.UUGCGCCUC............UGUUUAUA..........ACAGA................UGUAUUUG......................................UAC.AUAAU.AAUUAUU...UAAAA..CAAUGUUUGCAUGUAUUUAACUAAAAAAAUAAAACAAGAC
env-295   UUAUAU.UUGCGCCUC............UGUUUAUA..........ACAGA................UGUAUUUG......................................UAC.AUAAU.AAUUAUU...UAAAA..CAAUGUUUGCAUGUAUUUAACUAAAAAAAUAAAACAAGAC
env-296   UUAUAU.UUGCGCCUC............UGUUUAUA..........ACAGA................UGUAUUUG......................................UAC.AUAAU.AAUUAUU...UAAAA..CAAUGUUUGCAUGUAUUUAACUAAAAAAAUAAAACAAGAC
env-297   UUAUAU.UUGCGCCUC............UGUAUAUA..........ACAGA................UAAAUUUG......................................UAC.AUAAU.AAUUAUU...UAAAA..CAAUGUUUGCAUGUAUUUAACUAAAAAAAUAAAACAAGAC
env-298   GUAUAU.UUGCAACU.............UC................AAACA......................GG.....................................AUAC.AUAAA.AACUAUU...AAAAC..UAAUAUUGGCAUGUAUAA......................
env-299   GUAUAU.UUGCAACU.............UC................AAACA......................GG.....................................AUAC.AUAAA.AACUAUU...AAAAC..UAAUAUUGGCAUGUAUAACUCAAAAGGAAUUAAAGAGGAC
env-300   GUAUAU.UUGCAACU.............UCA...............AAC.......................UGG.....................................AUAC.AUAAA.AACUAUU...AAAAC..UAAUAUUGGCAUGUAUAACUCGAAAGGAAUUAAAGAGGAC
env-301   GUAUAU.UUGCAACU.............UUA...............AAC.......................UGG.....................................AUAC.AUAAA.AACUAUU...AAAAC..UAAUAUUGGCAUGUAUAACUCGAAAGGAAUUAAAGAGGAC
env-302   GUAUAU.UUGCAACU.............U.................CAAAUAA.....................G.....................................AUAC.AUAAA.AACUAUU...AAAAC..UAAUAUUGGCAUGUAUAACUCAAAAGAAAUUAAAGAGGAU
env-303   GUAUAU.UUGCAACU.............U.................CAAACUA.....................G.....................................AUAC.AUAAA.AACUAUU...AAAAC..UAAUAUUGGCAUGUAUAACUCAAAAGAAAUUAAAGAGGAU
env-304   GUAUAU.UUGCGGCU.............UCAU..............AAU......................AUGA......................................UAC.AUAAA.AACUAUU...AACAU..UAAUAUUGGCAUGUAUAACUCAAAAGAAAUUAAACAAGAU
env-305   AUAUAU.UUGCGCCGC............UGU...............AAU.......................GCA......................................UAC.AUAAA.AAUCAUU...UAAAU..UAAUAUUGGCAUGUAUUUAACAAAAAAAAACAAAGAA....
env-306   AUAUAU.UUGCGCCGC............UGU...............AAU.......................GCA......................................UAC.AUAAA.AAUCAUU...UAAAA..UAAUAUUGGCAUGUAUUUAACAAAAAAAAACAAAGAAGAU
env-307   AUAUAU.UUGCGCCGC............UGU...............AAU.......................GCA......................................UAC.AUAAA.AAUCAUU...UAAAA..UAAUAUUGGCAUGUAUUUAACAAAAAAAAACAAAGAAGAU
env-308   CUAUCU.UUGCGCCG.............UGUUA.............GAG.....................UAACA....................................CAUAC.AUAAA.AAUCAUU...UA.AA..CAAUAUUGGCAUGUAUUUAUCAGCUGAAAAGAAAAAAGAG
Lsp-3-1   GUAUCU.UUGCCGGC.............UUAAUGAG..........AUAGA................CUCAUUAA.....................................AUAC.AUAAA.AACUAUU...UA.AA..UAAUAUUGGCAUGUAUCUUACUCAAGAAAAGAAAGAUGAA
env-309   GUAUCU.UUGCCGGC.............UCAAUGAG..........AUCAC................CUCAUUGA.....................................AUAC.AUAAA.AACUAUU...UA.AA..UAAUAUUGGCAUGUAUCUUACUCAAGAGAAAAAAGAUGAA
env-310   GUAUCU.UUGCCGGC.............UCAAUGAG..........AUCAC................CUCAUUGA.....................................AUAC.AUAAA.AACUAUU...UA.AA..UAAUAUUGGCAUGUAUCUUACUCAAGAGAAAAAAGAUGAA
env-313   GUAUCU.UUGCCGGC.............UCAAUGAG..........AUCAC................CUCAUUGA.....................................AUAC.AUAAA.AACUAUU...UA.AA..UAAUAUUGGCAUGUAUCUUACUCAAGAGAAAAAAGAUGAA
env-314   GUAUCU.UUGCCGGC.............UCAAUGAG..........AUCAC................CUCAUUGA.....................................AUAC.AUAAA.AACUAUU...UA.AA..UAAUAUUGGCAUGUAUCUUACUCAAGAGAAAAAAGAUGAA
env-315   GUAUCU.UUGCCGGC.............UCAACGAG..........AUAAC................CUCGUUGA.....................................AUAC.AUAAA.AACUAUU...UA.AA..UAAUAUUGGCAUGUAUCUUACUCAAGAGAAAAAAGAUGAA
env-316   GUAUCU.UUGCCGGC.............UCAACGAG..........AUAGC................CUCGUUGA.....................................AUAC.AUAAA.AACUAUU...UA.AA..UAAUAUUGGCAUGUAUCUUACUCAAGAAAAGAAAGAUGAA
env-317   GUAUCU.UUGCCGGC.............UCAACGAG..........AUAGC................CUCGUUGA.....................................AUAC.AUAAA.AACUAUU...UA.AA..UAAUAUUGGCAUGUAUCUUACUC.................
env-318   GUAUCU.UUGCCGGC.............UCAACGAG..........AUAGC................CUCGUUGA.....................................AUAC.AUAAA.AACUAUU...UA.AA..UAAUAUUGGCAUGUAUCUUACUCAAGAAAAGAAAGAUGAA
env-320   GUAUCU.UUGCCGGC.............UCAACGAG..........AUAGC................CUCGUUGA.....................................AUAC.AUAAA.AACUAUU...UA.AA..UAAUAUUGGCAUGUAUCUUACUCAAGAAAAGAAAGAUGAA
env-324   GUAUCU.UUGCCGGC.............UCAACGAG..........AUAGC................CUCGUUGA.....................................AUAC.AUAAA.AACUAUU...UA.AA..UAAUAUUGGCAUGUAUCUUACUCAAGAAAAGAAAGAUGAA
env-325   GUAUCU.UUGCCGGC.............UCAACGAG..........AUAGC................CUCGUUGA.....................................AUAC.AUAAA.AACUAUU...UA.AA..UAAUAUUGGCAUGUAUCUUACUCAAGAAAAGAAAGAUGAA
env-326   GUAUCU.UUGCCGGC.............UCAACGAG..........AUAGC................CUCGUUGA.....................................AUAC.AUAAA.AACUAUU...UA.AA..UAAUAUUGGCAUGUAUCUUACUCAAGAAAAGAAAGAUGAA
Fri-1-1   UUACCU.UUGCAUCCAA...........AUCAGGGA..........UAU..................UCCCUGGC......................................UAC.AUAAU.AAUCAUU...UA.AA..UAAUAUUAGCAUGUAUUUAACUAAAGAAGUUAAAGAAGAG
env-327   AUAUAU.UUGCACGCAGUUUACCA.........................................................................................UAC.AUAAU.AAUCAUU...AAAAU..UAAUAUUUGCAUGUAUUUAAAUAAAGAGUUAAAAAAAAAG
env-328   AUAUAU.UUGCACGCAGUUUACCA.........................................................................................UAC.AUAAU.AAUCAUU...AA.AU..UAAUAUUUGCAUGUAUUUAAAUAAAGAGUUAAAAAAAAAG
env-329   UUAUAU.UUGCAAUC.............C.................AAAAAC......................G....................................CGUAC.AUAAA.AAUCAUU...UAAAA..UAAUAUUGGCAUGUAUUUAACAAAAGAAAUAAAAGAAGAG
env-330   UUAUAU.UUGCAAUC.............C.................AAAAAC......................G....................................CGUAC.AUAAA.AAUCAUU...UAAAA..UAAUAUUGGCAUGUAUUUAACAAAAGAAAUAAAAGAAGAG
env-331   GUAUAU.UUGCACUC.............GAU...............UAUA......................AUU....................................AAUAC.AUAAG.AAUCAUU...AA.AA..CAAUAUUGGCAUGUAUUUAACAAAAGAAGUUAAAGAAAAA
env-332   GUAUAU.UUGCACUC.............GAU...............UAUA......................AUU....................................AGUAC.AUAAG.AAUCAUU...AA.AA..UAAUAUUGGCAUGUAUUUAACAAAAGAAGCUAAAGAAAAA
env-333   GUAUAU.UUGCACUC.............GAU...............UAUA......................AUU....................................AGUAC.AUAAG.AAUCAUU...UA.AA..UAAUAUUGGCAUGUAUUUAACAAAAGAAGUUAAAGAAAAA
env-334   GUAUAU.UUGCAAUU.............GAUA..............AAUU.....................UAUU...................................AAAUAC.AUAAU.AAUUAUU...UA.AA..UAAUAUUGGCAUGUACAUCUCAAAACAAGUAGUAGAAGAC
env-335   GUAUAU.UUGCACCC.............GAGAA.............AAAU....................UUCUC.....................................GUAC.AUAAU.AAUCAUU...GA.AA..U.AUAUUUGCAUGUAUAUUACAUC................
env-336   GUAUAU.UUGCACCC.............GAGAAA............AAU....................UUUCUC.....................................GUAC.AUAAU.AAUCAUU...GA.AA..UAAUAUUUGCAUGUAUAUUACAUCUGAAGACAAAAAGAAA
env-337   GUAUCU.UUGCAGCC.............GAGAA.............AAG.....................UUCUC.....................................AUAC.AUAAU.AAUCAUU...GA.AA..UAAUAUUUGCAUGUAUCUCACUACAGAAGAAAAAAAAGAA
env-338   GUAUAU.UUGCCAACUAAUUCAAU.........................................................................................UAC.AUAAA.AAUCAUU...AAACU..UAAUAUUGGCAUGUAUAAUUCAAAAGAAAUAAAGCAAGAU
env-339   GUAUAU.UUGCCAACUAAUUCAGU.........................................................................................UAC.AUAAA.AAUCAUU...AAACU..UAAUAUUGGCAUGUAUAAUUCAAAAGAAAUAAAGCAAGAU
env-340   GUAUAU.UUGCCAACUAAUUCAGU.........................................................................................UAC.AUAAA.AAUCAUU...AAACU..UAAUAUUGGCAUGUAUAAUUCAAAAGAAAUAAAGCAAGAU
env-341   GUAUAU.UUGCCAACUAAUUCAGU.........................................................................................UAC.AUAAA.AAUCAUU...AAACU..UAAUAUUGGCAUGUAUAAUUCAAAAGAAAUAAAGCAAGAU
env-342   GUAUAU.UUGCCAACUAAUUCAGU.........................................................................................UAC.AUAAA.AAUCAUU...AAACU..UAAUAUUGGCAUGUAUAAUUCAAAAGAAAUAAAGCAAGAU
env-343   GUAUAU.UUGCCAACUAAUUCAGU.........................................................................................UAC.AUAAA.AAUCAUU...AAACU..UAAUAUUGGCAUGUAUAAUUCAAAAGAAAUAAAGCAAGAU
env-344   GUAUAU.UUGCCAACUAAUUCAGU.........................................................................................UAC.AUAAA.AAUCAUU...AAACU..UAAUAUUGGCAUGUAUAAUUCAAAAGAAAUAAAGCAAGAU
env-345   GUAUAU.UUGCCAACAAAUUCGGU.........................................................................................UAC.AUAAA.AAUCAUU...AAGUU..UAAUAUUGGCAUGUAUAAUUCAAAAGAAAUAAAGCAAGAU
env-346   GUAUAU.UUGCCAACUAAUUCGGU.........................................................................................UAC.AUAAA.AAUCAUU...AAGUU..UAAUAUUGGCAUGUAUAAUUCAAAAGAAAUAAAGCAAGAU
env-347   GUAUAU.UUGCCAACUAAUUCAGU.........................................................................................UAC.AUAAA.AAUCAUU...AAAUU..UAAUAUUGGCAUGUAUAACUCAAAAGAAAUAAAGCAAGAU
env-352   GUAUAU.UUGCCAACUAAUUCAGU.........................................................................................UAC.AUAAA.AAUCAUU...AAAUU..UAAUAUUGGCAUGUAUAACUCAAAAGAAAUAAAGCAAGAU
env-353   GUAUAU.UUGCCAACUAAUUCAGU.........................................................................................UAC.AUAAA.AAUCAUU...AAAUU..UAAUAUUGGCAUGUAUAAUUCAAAAGAAAUAAAGCAAGAU
env-354   GUAUAU.UUGCCAACUAAUUCAGU.........................................................................................UAC.AUAAA.AAUCAUU...AAGUU..UAAUAUUGGCAUGUAUAAUUCAAAAGAAAUAAAGCAAGAU
env-355   GUAUAU.UUGCCAACUAAUUCAGU.........................................................................................UAC.AUAAA.AAUCAUU...AAGUU..UAAUAUUGGCAUGUAUAAUUCAAAAGAAAUAAAGCAAGAU
env-356   GUAUAU.UUGCCAACUAAUUCAGU.........................................................................................UAC.AUAAA.AAUCAUU...AAGUU..UAAUAUUGGCAUGUAUAAUUCAAAAGAAAUAAAGCAAGAU
env-357   GUAUCU.UCGCACCGU............AAUUCCAA..........GAG..................UUGGAGUU.....................................CUAC.AUAAU.AAUCAUU...CA.AG..UAAUAUUGGCAUGUAUUUAGAUUCAAAAGCAAAAGAAGAA
Fta-1-1   UUAUCU.UUGCAUCGU............AAUUCCAA..........AAG..................UUGGGGUU.....................................CUAC.AUAAU.AAUCAUU...CA.AG..UAAUAUUGGCAUGUAUUUAGAUUCAAAAGCAAAAGAAGAA
env-358   CUAUCU.UUGCAAAU.............CGAAAGA...........UAC...................UCUUUCG....................................UUUAC.AUAAC.AAUCAUU...UACAA..CAAUAUUAGCAUGUAUUUAGAUCAAAAAGAAAAAGAAGCA
env-359   GUAUUU.UUGCACUU.............UGAUAGA...........AAA...................UCUAUCA.....................................UUGC.AUAAU.AAUUAUU...GA.AA..CAAUAUUGGCAUGUACUUAACAAAAGAAGAAAAGG.....
env-360   GUAUUU.UUGCACUU.............UGAUAGA...........AAA...................UCUAUCA.....................................UUGC.AUAAU.AAUUAUU...GA.AA..CAAUAUUGGCAUGUACUUAACAAAAGAAGAAAAGGGCAAA
env-361   UUAUAU.UUGCGCGC.............UGAUAG............AAAGA..................CUAUCA...................................AAUUAC.AUAAU.AAUCAUU...GA.AA..UAAUAUUGGCAUGUAUUUGACAAAAGAAACAAAAAGUGAG
env-362   CUAUAU.UUGCACGC.............UGAUAGA...........AAG...................UCUAUCA...................................AAUUAC.AUAAU.AAUUAUU...GA.AA..UAAUAUUGGCAUGUAUUUGACAAAAGAAAAAAAGAGUGAG
Mod-7-1   UUACCU.UUGCAAACGU...........AAAUAUGG..........AUACA................CCAUAUUU....................................CAUGC.AUAAA.AAUCAUU...AAAAA..UAAUAUUGGCAUGUAUUUAUCAAAAGAGAAAAAAGCUGAG
Mod-6-1   UUACCU.UUGCAAACGU...........AAAUAUGG..........AUACA................CCAUAUUU....................................CAUGC.AUAAA.AAUCAUU...AAAAA..UAAUAUUGGCAUGUAUUUAUCAAAAGAGAAAAAAGCUGAG
Mod-3-1   UUACCU.UUGCAAACGU...........AAAUAUGG..........AUACA................CCAUAUUU....................................CAUGC.AUAAA.AAUCAUU...AAAAA..UAAUAUUGGCAUGUAUUUAUCAAAAGAGAAAAAAGCUGAG
Mpr-1-1   UUACCU.UUGCAAACGU...........AAAUACGG..........AUACA................CCGUAUUU....................................CAUGC.AUAAA.AAUCAUU...AAAAA..UAAUAUUGGCAUGUAUUUAUCAAAAGAGAAAAAAGCUGAG
Mod-4-1   UUACCU.UUGCAAACGU...........AAAUAUGG..........AUACA................CCGUAUUU....................................CAUGC.AUAAA.AAUCAUU...AAAAA..UAAUAUUGGCAUGUAUUUAUCAAAAGAGAAAAAAGCUGAG
Mod-1-1   UUACCU.UUGCAAACGU...........AAAUAUGG..........AAAUA................CCAUAUUA....................................UAUAC.AUAAA.AAUCAU....UA.AA..UAAUAUUAGCAUGUAUUUAUCAAAAGAGAAAAAAGCUGAG
Fin-1-1   UUACCU.UUGCACCCG............UAUUGAG...........AAACA.................CUCAAUU......................................UAC.AUAAU.AAUUAUU...UA.AG..GAAUAUUAGCAUGUACUUAUCAAAAGAGAAAAAAGCUGAA
env-363   CUAGCU.UUGCACAC.............GAAUUGCG..........AAAAC................CGCA.UUU....................................UCUAC.AUAAU.AAUCAUU...UA.AA..UAAUAUUAGCAUGUACUUAACUAAAGAAGUUAAAGAAGAA
env-364   CUAGCU.UUGCACAC.............GAAUUGCG..........AAAAC................CGCA.UUU....................................UCUAC.AUAAU.AAUCAUU...UA.AA..UAAUAUUAGCAUGUACUUAACUAAAGAAGUUAAAGAAGAA
env-366   CUAGCU.UUGCACAC.............GAAUUGCG..........AAAAC................CGCA.UUU....................................UCUAC.AUAAU.AAUCAUU...UA.AA..UAAUAUUAGCAUGUACUUAACUAAAGAAGUUAAAGAAGAA

          ............................&lt;&lt;&lt;&lt;&lt;&lt;&lt;&lt;&lt;&lt;&lt;&lt;&lt;&lt;&lt;&lt;&lt;&lt;...........&gt;&gt;&gt;&gt;&gt;&gt;&gt;&gt;&gt;&gt;&gt;&gt;&gt;&gt;&gt;&gt;&gt;&gt;.........&lt;&lt;&lt;&lt;&lt;&lt;&lt;&lt;........&gt;&gt;&gt;&gt;&gt;&gt;&gt;&gt;........................................................................
          ............................2222221111..101.00...........00.101..1111222222.........11011000........00011011........................................................................
          •UAY•U-UUGC••CY--------------◦◦◦◦◦◦◦-----------R•R◦◦-----------------◦◦◦◦◦◦◦-------------------------------------◦UAC-AUAA•-AAUCAUU---UA-A-A--YAAUAUURG•AUGUAUUUARC•ARAGAAR••AAARRRRAR
          5′                                                                                                                                                 3′
```

```
env-367   CUAGCU.UUGCACAC............GAAUUACG.........AAAAC...............CGUA.UUU...........................................UCUAC.AUAAU.AAUCAUU...UA.AA..UAAUAUUAGCAUGUACUUAACUAAAGAAGUUAAAGAAGAA
env-368   CUAGCU.UUGCACAC............GAAUUACG.........AAAAC...............CGUA.UUU...........................................UCUAC.AUAAU.AAUCAUU...UA.AA..UAAUAUUAGCAUGUACUUAACUAAAGAAGUUAAAGAAGAA
env-369   GUAUAU.UUGCAUCU............UCAAUUU.UUCCG.....AAAAU..........CGGAUUGAAUUGA..............................................AUAC.AUAAA.AAUCAUU...UAAAU..AAAUAUUGGCAUGUAUUUAACUACAGAAAAAAAGAAUGAA
env-373   AUAUAU.UUGCUUUAC...........AUU..............UAU.....................AAU........................................UUUUUAC.AUAAA.AAUCAUU...AAAAU..UAAUAUUGGCAUGUAUAAUUCAAAAGAGGUUAAGGAAAAU
env-375   AUAUAU.UUGCUUUAC...........AUU..............UAU.....................AAU........................................UUUUUAC.AUAAA.AAUCAUU...AAAAU..UAAUAUUGGCAUGUAUAAUUCAAAA...............
env-376   UUAUAU.UUGCCGCC............UGAUUAUGA........ACG..............UCAUAAUUA..............................................UUGC.AUAAA.AAUUAUU...UAAAC..AAAUAUUGGAAUGUACUUAACAAAAGAAAUUAAAGAGGAA
env-377   AUAUAU.UUGCCGCC............UGAUUAUG.........AUCGC...............CGUAAUCA...............................................AUGC.AUAAA.AAUUAUU...UAAAU..AAAUAUUGGAAUGUACUUAACAAAAGAAACAAAGAAGAA
env-378   CUAUAU.UUGCCCUU............CAAUUUUG.........AGAAA...............CAAGAUUG..............................................UUAGC.AUAAA.AAUUAUU...UA.AA..AAAUAUUGGAAUGUACCUAACAAAAAAAUUAAAGAGAGC
env-379   CUAUAU.UUGCACCU............CAAUUCUG.........AGAGC...............CAGAAUUG..............................................UUUGC.AUAAA.AAUUAUU...UAAAC..AAAUAUUGGAAUGUACUUAACAAAAAAGAUUAAAGAAAAU
env-381   CUAUAU.UUGCACCU............CAAUUCUG.........AGAGC...............CAGAAUUG..............................................UUUGC.AUAAA.AAUUAUU...UAAAC..AAAUAUUGGAAUGUACUUAACAAAAAAGAUUAAAGAAAAU
env-382   CUAUAU.UUGCCCCU............CAAUUCUG.........AGAGC...............CAGAAUUG..............................................UUUGC.AUAAA.AAUUAUU...UAAAC..AAAUAUUGGAAUGUACUUAACAAAAAAGAUUAAAGAAAAU
env-383   CUAUAU.UUGCCCCU............CAAUUCUG.........AGAGC...............CAGAAUUG..............................................UUUGC.AUAAA.AAUUAUU...UAAAC..AAAUAUUGGAAUGUACUUAACAAAAAAGAUUAAAGAAAAU
env-384   CUAUAU.UUGCCCCU............CAAUUCUG.........AGAGC...............CAGAAUUG..............................................UUUGC.AUAAA.AAUUAUU...UAAAC..AAAUAUUGGAAUGUACUUAACAAAAAAGAUUAAAGAAAAU
env-385   CUAUAU.UUGCCCCU............CAAUUCUG.........AGAGC...............CAGAAUUG..............................................UUUGC.AUAAA.AAUUAUU...UAAAC..AAAUAUUGGAAUGUACUUAACAAAAAAGAUUAAAGAAAAU
env-386   CUAUAU.UUGCCCCU............CAAUUCUG.........AGAGC...............CAGAAUUG..............................................UUUGC.AUAAA.AAUUAUU...UAAAC..AAAUAUUGGAAUGUACUUAACAAAAAAGAUUAAAGAAAAU
env-387   CUAUAU.UUGCCCCU............CAAUUCUG.........AGAGC...............CAGAAUUG..............................................UUUGC.AUAAA.AAUUAUU...UAAAC..AAAUAUUGGAAUGUACUUAACAAAAAAGAUUAAAGAAAAU
env-388   CUAUAU.UUGCCCCU............CAAUUCUG.........AGAGC...............CAGAAUUG..............................................UUUGC.AUAAA.AAUUAUU...UAAAC..AAAUAUUGGAAUGUACUUAACAAAAAAGAUUAAAGAAAAU
env-389   CUAUAU.UUGCCCCU............CAAUUCUG.........AGAGC...............CAGAAUUA..............................................UUUGC.AUAAA.AAUUAUU...UAAAA..AAAUAUUGGAAUGUACUUAACAAAAAAGAUUAAAGAAAAU
env-390   CUAUAU.UUGCCCCU............CAAUUCUGA........AAG..............UCAGAAUUG..............................................UUUGC.AUAAA.AAUUAUU...UAAAC..AAAUAUUGGAAUGUACUUAACAAAAAAGAUUAAAGAAAAU
env-391   CUAUAU.UUGCCCCU............CAGUUCUGA........AAG..............UCAGAAUUG..............................................UUUGC.AUAAA.AAUUAUU...UAAAC..AAAUAUUGGAAUGUACUUAACAAAAAAGAUUAAAGAAAAU
env-392   CUAUAU.UUGCCCCU............CAGUUCUGA........AAG..............UCAGAAUUG..............................................UUUGC.AUAAA.AAUUAUU...UAAAC..AAAUAUUGGAAUGUACUUAACAAAAAAGAUUAAAGAAAAU
env-393   CUAUAU.UUGCUCCU............CAAUUCUG.........AUCGA...............CAGAAUUG..............................................UUUAC.AUAAA.AAUUAUU...UAAAC..AAAUAUUGGAAUGUACCUAACAAAGAGAUCAAAGAGAU
env-394   CUAUAU.UUGCCCCU............CAAUUCUG.........AGCGA...............CAGAAUUG..............................................UUUAC.AUAAA.AAUUAUU...UAAAU..AAAUAUUGGAAUGUACCUAACAAAGAGAUCAAAGAGAC
env-395   CUAUAU.UUGCCCCU............CAAUUCUG.........AGCGA...............CAGAAUUG..............................................UUUAC.AUAAA.AAUUAUU...UAAAU..AAAUAUUGGAAUGUACCUAACAAAGAGAUCAAAGAGAC
env-396   GUACCU.UUGUGGGU............CGUU.............AAAA....................AACG............................................AAUUAC.AUAAU.AAUCAUU...AA.AA..UAAUAUUUGCAUGUACAAUUCAAAAGAAGUAAAAGAAGAC
env-397   GUACCU.UUGUGGGU............CGUU.............AAAA....................AACG............................................AAUUAC.AUAAU.AAUCAUU...AA.AA..UAAUAUUUGCAUGUACAAUUCAAAAGAAGUAAAAGAAGAC
env-398   GUACCU.UUGUGGGU............CGUU.............AAAA....................AACG............................................AAUUAC.AUAAU.AAUCAUU...AA.AA..UAAUAUUUGCAUGUACAAUUCAAAAGAAGUAAAAGAAGAC
env-399   GUACCU.UUGUGGGU............CGUU.............AAAA....................AACG............................................AAUUAC.AUAAU.AAUCAUU...AA.AA..UAAUAUUUGCAUGUACAAUUCAAAAGAAGUAAAAGAAGAC
env-400   GUACCU.UUGUGGGU............CGUU.............AAAA....................AACG............................................AAUUAC.AUAAU.AAUCAUU...AA.AA..UAAUAUUUGCAUGUACAAUUCAAAAGAAGUAAAAGAAGAC
env-402   GUAUAU.UUGCGGCU............UUA..............UUAG.....................UAA...............................................AUAC.AUAAA.AACUAUU...AAAAU..UAAUAUUGGCAUGUAUAACUCAAAAGAGAUUAAAGAAGAU
env-403   GUAUAU.UUGCAGCU............UUA..............UUAG.....................UAA...............................................AUAC.AUAAA.AACUAUU...AAAAU..UAAUAUUGGCAUGUAUAACUCAAAAGAGAUUAAAGAAGAU
env-404   GUAUAU.UUGCAGCU............UUA..............UUAG.....................UAA...............................................AUAC.AUAAA.AACUAUU...AAAAU..UAAUAUUGGCAUGUAUAACUCAAAAGAGAUUAAAGAAGAU
env-405   GUAUAU.UUGCGGCU............UU................UAA......................AA...............................................CUAC.AUAAA.AAUUAUU...AA.AU..UGAUAUUGGCAUGUAUAAUUCAAAAGAACUAAAAAAAACU
env-406   GUAUAU.UUGCAGCC............UU................UAA......................AA...............................................CUGC.AUAAA.AAUUAUU...AA.AU..UGAUAUUGGCAUGUAUAAUUCAAAAGAAUUAAAAAAAACU
env-407   GUAUAU.UUGCGACU............UU...............AUUGUC.....................AA...............................................AUAC.AUAAA.AACUAUU...AACAU..UAAUAUUGGCAUGUAUAAUUCAAAAGAAAUUAAAGAAGAU
env-408   GUAUAU.UUGCGACU............UU...............AUUGUC.....................AA...............................................AUAC.AUAAA.AACUAUU...AACAU..UAAUAUUGGCAUGUAUAAUUCAAA...............
env-409   GUAUAU.UUGCGACU............UU...............AUUGUC.....................AA...............................................AUAC.AUAAA.AACUAUU...AACAU..UAAUAUUGGCAUGUAUAAUUCAAAAGAAAUUAAAGAAGAU
env-410   GUAUAU.UUGCGACU............UU...............AUUGUC.....................AA...............................................AUAC.AUAAA.AACUAUU...AACAU..UAAUAUUGGCAUGUAUAAUUCAAAAGAAUUAAAGAAGAUA
env-411   GUAUAU.UUGCUGCU............UU...............AUUAUC.....................AG...............................................GUAC.AUAAA.AACUAUU...AACAU..UAAUAUUGGCAUGUAUAAUUCAAAAGAAAGUAAAGAAGAC
env-412   GUAUAU.UUGCUGCU............UU...............AUUAUC.....................AG...............................................GUAC.AUAAA.AACUAUU...AACAU..UAAUAUUGGCAUGUAUAAUUCAAAAGAAAGUAAAGAAGAC
env-413   GUAUAU.UUGCUGCU............UU...............AUUAUC.....................AG...............................................GUAC.AUAAA.AACUAUU...AACAU..UAAUAUUGGCAUGUAUAAUUCAAAAGAAAGUAAAGAAGAC
env-414   GUAUAU.UUGCUGCU............UU...............AUUAUC.....................AG...............................................GUAC.AUAAA.AACUAUU...AACAU..UAAUAUUGGCAUGUAUAAUUCAAAAGAAAGUAAAGAAGAC
env-415   GUAUAU.UUGG..CC............GCUU.............AAU...................AAGC...............................................AUAC.AUAAC.AAUCAU....UA.AA..UAAUAUUAGCAUGUAUUUAU....................
env-416   UUAUAU.UUGCAAUC............GU...............UUAA.......................AU................................................CAC.AUAAA.AAUCAUU...UAAAU..UAAUAUUGGCAUGUAUUUAUCAAAAGAAAUUAAAGAAAAU
env-417   UUAUAU.UUGCAAUC............GU...............UUAA.......................AU................................................CAC.AUAAA.AAUCAUU...UAAAU..UAAUAUUGGCAUGUAUUUAUCAAAAGAAAUUAAAGAAAAU
env-418   UUAUAU.UUGCAAUC............G................UUAAAAAU.....................U................................................UAC.AUAAU.AAUCAUU...UAAAA..UAAUAUUGGCAUGUAUUUAUCAAAAGAGAUUAAAGAAGAG
Fbe-1-1   UUACCU.UUGCACCC............AAUCGGG..........AUAGG...............CCCG..U................................................UAC.AUAAA.AAUUAUU...UAAAC..AAAUAUUGGCAUGUACUUAACUAAAGAAGUAAAAGAGCAA
env-419   GUAUCU.UUGCCCGC............UGAUAG...........ACACG................CUAUCA..............................................UAUAC.AUAAA.AAUCAUU...UAAAA..CAAUAUUGGAAUGUAUUUAACAAAAGAGAAGAAGGAAGAA
Ksp-1-1   GUAUCU.UUGCACGC............UGAUAG...........AGACG................CUAUCA...............................................AUAC.AUGAA.UUUCAUU...UAAAA..UAAUACUGGAAUGUAUUUAACAAAAGAACAAAAGCAAGAG
Dsp-2-1   GUAUCU.UUGCACGC............UGAUAG...........AGACG................CUAUCA...............................................AUAC.AUGAA.UUUCAUU...UAAAA..UAAUACUGGAAUGUAUUUAACAAAAGAAGAAAAACAAGAG
Dsp-1-1   AUAUCU.UUGCACGC............UGAUAG...........AGAAA................CUAUCA...........................................CGACAUAC.AUCGA.UCUCAUU...UAAAA..CAAUAUUGGAAUGUAUUUAUCAAAAGAAGUAAAAGAAGAA
Ddo-1-1   AUAUCU.UUGCACGC............UGAUAG...........AGAAA................CUAUCA...........................................CGACAUAC.AUCGA.UCUCAUU...UAAAA..CAAUAUUGGAAUGUAUUUAACAAAAGAAGUAAAAGAAGAA
Iha-1-1   UUAUCU.UUGCGCCC............UGAUAG...........AGCAC................CUAUCA...............................................AUAC.AUAAA.AAUCAUU...AA.AA..AAAUAUUGGCAUGUAUUUAACAAAAGAAGUAAAAGCAGAC
env-420   CUAUAU.UUGCAGCC............UUCGA............AAUAAAC..............UCGAA..............................................GUUGC.AUAAA.AAUCAUU...UACAA..UAAUAUUGGCAUGUCAUUAAGUAAAGAAAACAAAGCAGAA
env-421   UUAUAU.UUGCAGCC............UUCGAG...........AUAAA................CUCGAA..............................................AUUGC.AUAAU.AAUUAUU...UACAA..CAAUAUUGGCAUGUUAAACAAAGAAGAAAAAGCAGCAAUU
env-422   CUAUAU.UUGCAGCC............GUCGAG...........AUAAA................CUCGAA..............................................GUUGC.AUAAG.AAUCAUU...UACAA..UAAUAUUGGCAUGUUACAUAAAGAAGAAAAAGCGGCAAUU
env-423   UUAUAU.UUGCAGCC............UUCGUG...........AUAAA................CUCGAA..............................................GUUGC.AUAAG.AAUCAUU...UACAA..UAAUAUUGGCAUG..........................
env-424   CUAUAU.UUGCAGCC............UUCGAG...........AGAAA................CUCGAA..............................................GUUGC.AUAAA.AAUCAUU...UACAA..UAAUAUUGGCAUGUCAUUAAGUAAAGAAAACAAAGCAGAA
env-427   CUAUAU.UUGCAGCC............UUCGAG...........AGAAA................CUCGAA..............................................GUUGC.AUAAA.AAUCAUU...UACAA..UAAUAUUGGCAUGUCAUUAAGUAAAGAAAACAAAGCAGAA
env-428   CUAUAU.UUGCAGCC............UUCGAG...........AGAAA................CUCGAA..............................................GUUGC.AUAAA.AAUCAUU...UACAA..UAAUAUUGGCAUGUCAUUAAGUAAAGAAACCAAAGCAGAA
env-429   CUAUAU.UUGCAGCC............UUCGAG...........AGAAA................CUCGAA..............................................GUUGC.AUAAA.AAUCAUU...UACAA..UAAUAUUGGCAUGUCAUUAAGUAAAGAAAACAAAGCAGAA
env-430   UUAUAU.UUGCAACC............UUCGAG...........AUAAA................CUCUAA..............................................GUUGC.AUAAU.GAUCAUU...UACAA..AAAUAUUGGCAUGUUAAAAAAGAAGAAAAAGCAGCAAUU
env-431   UUAUAU.UUGCAGCC............UUCGAG...........AUAAA................CUCGAA..............................................GUUGC.AUAAU.GAUCAUU...UACAA..AAAUAUUGGCAUGUUAAAAAAAGAAGAAAAAGCAGCAAUU
env-432   CUAUAU.UUGCAGCC............UUCGAG...........AUAAA................CUCGAA..............................................GUUGC.AUAAU.AAUCAUU...UACAA..UAAUAUUGGCAUGUUAAAAAAAGAAGAAAAAGCAGCAAUU
env-433   CUAUAU.UUGCAGCC............UUCGAG...........AUAAA................CUCGAA..............................................GUUGC.AUAAU.AAUCAUU...UACAA..AAAUAUUGGCAUGUUAAAAAAA.................
env-434   CUAUAU.UUGCAGCC............UUCGAG...........AUAAA................CUCGAA..............................................GUUGC.AUAAU.AAUCAUU...UACUA..UAAUAUUGGCAUGUUAAACAAAGAAGAAAAAGCAGCAAUU
env-435   CUAUAU.UUGCAGCC............UUCGAG...........AUAAA................CUCGAA..............................................GUUGC.AUAAU.AAUCAUU...UACAA..UAAUAUUGGCAUGUUAAACAAAGAAGAAAAAGCAGCAAUU
env-437   CUAUAU.UUGCAGCC............UUCGAG...........AUAAA................CUCGAA..............................................GUUGC.AUAAU.AAUCAUU...UACAA..UAAUAUUGGCAUGUUAAACAAAGAAGAAAAAGCAGCAAUU
env-438   UUAUAU.UUGCAGUC............UUCGCG...........GUAAA................CUCGAA..............................................GUUGC.AUAAU.AAUUAUU...UACAA..UAAUAUUGGCAUGUUAAACAAAGAAGAAAAAGCAGCAAUU
env-439   UUAUAU.UUGCAGCC............UUCGAG...........UUUAU................UUCGAA..............................................GUAGC.AUAAG.AAUCAUU...UACAA..UAAUAUUGGCAUGUUAAACAAAGAAGAAAAAGCGGCAAUU
env-440   UUAUAU.UUGCAGCC............UUCGAG...........UUUAU................UUCGAA..............................................GUAGC.AUAAG.AAUCAUU...UACAA..UAAUAUUGGCAUGUUAAACAAAGAAGAAAAAGCGGCAAUU
env-441   UUAUAU.UUGCAGCC............UUCGAG...........UUUAU................UUCGAA..............................................GUAGC.AUAAG.AAUCAUU...UACAA..UAAUAUUGGCAUGUUAAACAAAGAAGAAAAAGCGGCAAUU
env-442   UUAUAU.UUGCAGCC............UUCGAG...........UUUAU................UUCGAA..............................................GUAGC.AUAAG.AAUCAUU...UACAA..UAAUAUUGGCAUGUUAAACAAAGAAGAAAAAGCGGCAAUU
env-444   UUAUAU.UUGCAGCC............UUCGAG...........UUUAU................UUCGAA..............................................GUAGC.AUAAG.AAUCAUU...UACAA..UAAUAUUGGCAUGUUAAACAAAGAAGAAAAAGCGGCAAUU
Kje-1-1   GUAUAU.UUGCGACU............UGAUACG..........AGAAA...............CGUAUUA...............................................AUAC.AUAAA.AAUUAUU...AA.AA..CAAUAUUGGCAUGUACUUAACUAAAGAAGAUUAAAGCUGAA
Kal-1-1   GUAUAU.UUGCGGCU............UGAUACG..........AGAAA...............CGUAUUA...............................................AUAC.AUAAA.AAUUAUU...AA.AA..CAAUAUUGGCAUGUACUUAACUAAAGAAGUUAAAGAAGAA
Kzh-1-1   GUAUAU.UUGCGGCU............UGAUACG..........AUAAA...............UGUAUCA...............................................UUGC.AUAAA.AAUUAUU...AA.AA..UAAUAUUGGCAUGUAUUUAACUAAAGAAAUUAAAGCGGAA
env-445   UUACUU.UUGCAACC............GUUAUAUA.........AUAGC..............UAUAUAAC...............................................AUAC.AUAAA.AAUCAUU...UAACA..CAAUAUUGGAAUGUAUUUAACAAAAGAAAUUAAAGAAGAA
env-446   AUACCU.UUGCAGCC............CUGA.............AAA.....................UCAG..............................................GAUAC.AUAAC.AAUCAUU...AAAAA..CAAUAUUGAAAUGUAUUUAUCAUCAGAUAAAAAGAAAGAA
env-447   AUACCU.UUGCAGCC............CUGA.............AAA.....................UCAG..............................................GAUAC.AUAAC.AAUCAUU...AAAAA..CAAUAUUGAAAUGUAUUUAUCAUCAGAUAAAAAGAAAGAA
env-449   CUACAU.UUGCAACU............GCGA.............AACG....................UCGC...............................................AUAC.AUAAA.AAUCAUU...AA.AA..UAAUAUUGGCAUGUAUUUAAGUACAGAAGAGAAAAAAAG
env-450   CUAUAU.UUGCGGCU............CAUUCCUGA........AAA..............UCAGGUGUG...............................................CUAC.AUAAA.AAUUAUU...UAAAA..AAAUAUUGGAAUGUAUUUAACGAAAAAAAUUAAAGAGGAU
env-451   UUAUAU.UUGCCGCU............CAUUCUUGA........AAA..............UCAGGAAUU...............................................UUAC.AUAAA.AAUUAUU...UAAAA..AAAUAUUGGAAUGUAUUUAACAAAAAAAAUUAAAGAGGAU

          ...........................<<<<<<<<<<<<<<<<<<..........>>>>>>>>>>>>>>>>>>.........<<<<<<<<........>>>>>>>>...........................................................................
          ...........................2222221111..101.00..........00.101..1111222222.........11011000........00011011...........................................................................
          •UAY•U-UUGC••CY------------◦◦◦◦◦◦◦-----------R•R◦◦----------------◦◦◦◦◦◦◦-----------------------------------◦UAC-AUAA•-AAUCAUU---UA-AA--YAAUAUURG•AUGUAUUURC•ARAGAAR••AAARRRRAR
          5'                                                                                                                                                    3'
```

```
env-452   UUAUAU.UUGCCGCU.............CAUUCUUGA.........AAA................UCAGGAAUU.....................................UUAC.AUAAA.AAUUAUU...UAAAA..AAAUAUUGGAAUGUAUUUAACAAAAAAAAUUAAAGAGGAU
env-453   CUAUAU.UUGCCGCU.............CAUUCCUGA.........AAA................UCAGGUAUG.....................................UUAC.AUAAA.AAUUAUU...UA.AA..AAAUAUUGGAAUGUAUUUAACAAAAAAAAUUAAAGAGGAU
env-454   CUAUAU.UUGCCGCU.............CAUUCCUGA.........AAA................UCAGGUAUG.....................................UUAC.AUAAA.AAUUAUU...UAAAA..AAAUAUUGGAAUGUAUUUAACAAAAAAAAUUAAAGAGGAU
env-455   CUAUAU.UUGCCGCU.............CAUUCCUGA.........AAA................UCAGGUAUG.....................................UUAC.AUAAA.AAUUAUU...UAAAA..AAAUAUUGGAAUGUAUUUAACAAAAAAAAUUAAAGAGGAU
env-456   GUAUAU.UUGCAGCC.............UCA...............UUUU.....................UGA....................................GGUAC.AUAAG.AAUCAUU...UA.AA..CAAUAUUGGAAUGUAUUUAUCUAGCGAAAAGAAACAAGAA
env-457   GUAUAU.UUGCAGCC.............UCA...............UUUU.....................UGA....................................GGUAC.AUAAG.AAUCAUU...UA.AA..CAAUAUUGGAAUGUAUUUAUCUAGCGAAAAGAAACAAGAA
env-458   AUAUCU.UUGCCGCU.............GAUGGA............AAA...................UCCAUU....................................UGUAC.AUAAA.AAUCAUU...GA.AA..UAAUAUUGGAAUGUAUUUAACGAAA...............
env-459   GUAUAU.UUGCCAAC.............CACU..............UUAU....................AGUG....................................UUUAC.AUAAA.AAUCAUU...AAAAU..UAAUAUUGGCAUGUAUAAUUCAAAAGAAGUAAAAGAAAAU
env-460   GUAUAU.UUGCCAAC.............CACU..............UUAU....................AGUG....................................UUUAC.AUAAA.AAUCAUU...AAAAU..UAAUAUUGGCAUGUAUAAUUCAAAAGAAGUAAAAGAAAAU
env-461   GUAUAU.UUGCCAAC.............CACU..............UUAU....................AGUG....................................UUUAC.AUAAA.AAUCAUU...AAAAU..UAAUAUUGGCAUGUAUAAUUCAAAAGAAGUAAAAGAAAAU
env-462   GUAUAU.UUGCCAAC.............CACU..............UUUAU...................AGUG....................................UUUAC.AUAAA.AAUCAUU...AAAAU..UAAUAUUGGCAUGUAUAAUUCAAAAGAAGUAAAAGAAAAU
env-463   GUAUAU.UUGCCAAC.............CACU..............UUUAU...................AGUG....................................UUUAC.AUAAA.AAUCAUU...AAAAU..UAAUAUUGGCAUGUAUAAUUCAAAAGAAGUAAAAGAAAAU
env-464   GUAUAU.UUGCCAAC.............CACU..............UUUAU...................AGUG....................................UUUAC.AUAAA.AAUCAUU...AAAAU..UAAUAUUGGCAUGUAUAAUUCAAAAGAAGUAAAAGAAAAU
env-465   GUAUAU.UUGCCAAC.............CACU..............UUUAU...................AGUG....................................UUUAC.AUAAA.AAUCAUU...AAAAU..UAAUAUUGGCAUGUAUAAUUCAAAAGAAGUAAAAGAAAAU
env-466   GUAUAU.UUGCCAAC.............CACU..............UUAAU...................AGUG....................................UUUAC.AUAAA.AAUCAUU...AAAAU..UAAUAUUGGCAUGUAUAAUUCAAAAGAAGUAAAAGAAAAU
env-467   GUAUAU.UUGCCAAC.............CACU..............UUGAU...................AGUG....................................UUUAC.AUAAA.AAUCAUU...AAAAU..UAAUAUUGGCAUGUAUAAUUCAAAAGAAGUAAAAGAAAAU
env-469   GUAUAU.UUGCCAAC.............CACU..............UUGAU...................AGUG....................................UUUAC.AUAAA.AAUCAUU...AAAAU..UAAUAUUGGCAUGUAUAAUUCAAAAGAAGUAAAAGAAAAU
env-470   GUAUAU.UUGCCGAC.............CACU..............UAUAU...................AGUG....................................UUUAC.AUAAA.AAUCAUU...AAAAU..UAAUAUUGGCAUGUACAAUUCAAAAGAAGUAAAAGAAAAG
env-471   CUACUU.UUGCAACCGU...........CUAAACCGA.........GAG................UCGGUGUGG.....................................CUAC.AUAAA.AAUUAUU...UA.UA..CAAUAUUGGCAUGUAUUUAAACCCAGAAGCAAAAAAAGAA
env-472   GUAUCU.UUGCAUGG.............CA................AUUU......................UG....................................CAUAC.AUAAA.AAUCAUU...UA.AA..UAAUAUUGGCAUGUACUUAACAAACGAAAAAAAGAAAGAG
env-473   GUAUCU.UUGCAUGG.............CA................AUUU......................UG....................................CAUAC.AUAAA.AAUCAUU...UA.AA..UAAUAUUGGCAUGUACUUAACAAACGAAAAAAAGAAAGAG
env-474   GUAUCU.UUGCAUGG.............CA................AUUU......................UG....................................CAUAC.AUAAA.AAUCAUU...UA.AA..UAAUAUUGGCAUGUACUUAACAAACGAAAAAAAGAAAAG
env-475   UUAUAU.UUGCACGC.............UGUCCGA...........AAA..................UCGGCAG....................................CAUAC.AUAAA.AAUCAUU...UAAAA..UAAUAUUGGCAUGUAUUUAAGCAAGGAAGUAAAAGCCGAC
env-476   GUAUCU.UUGCGGCG.............CUUA..............AAA.....................UAUG....................................CAUAC.AUAAC.AAUCAU....UA.AA..UAAUAUUAGCAUGUAUUUAUCUAAAGAAGUAAAAGAAAAG
env-477   GUAUAU.UUGCCCCG.............CUU...............AAC......................AAG....................................CAUAC.AUAAC.AAUCAU....UA.AA..UAAUAUUAGCAUGUAUUUAUCAAAAGAAGUAAAAGAAACG
env-478   GUAUAU.UUGCCCCG.............CUU...............AAC......................AAG....................................CAUAC.AUAAC.AAUCAU....UA.AA..UAAUAUUAGCAUGUAUUUAUCAAAAGAAGUAAAAGAAACG
env-479   GUAUAU.UUGCCCCG.............CUU...............AAC......................AAG....................................CAUAC.AUAAC.AAUCAU....UA.AA..UAAUAUUAGCAUGUAUUUAUCAAAAGAAGUAAAAGAAACG
env-480   GUAUAU.UUGCCCCG.............CUUA..............AUAA....................UAAG....................................CAUAC.AUAAC.AAUUAU....UA.AA..UAAUAUUAGCAUGUAUUUAUCAAAAAAAGUAAAAGAAACG
env-481   GUAUAU.UUGCACCG.............CUUA..............AUAA....................UAAG....................................CAUAC.AUAAC.AAUCAU....UA.AA..UAAUAUUAGCAUGUAUUUAUCAAAAGAAUUAAAGAAACG
env-482   GUAUAU.UUGCACCG.............CUUA..............AUAA....................UAAG....................................CAUAC.AUAAC.AAUCAU....UA.AA..UAAUAUUAGCAUGUAUUUGACAAAAGAAGUUAAAGAAGGG
env-484   GUAUAU.UUGCACCG.............CUUA..............AUAA....................UAAG....................................CAUAC.AUAAA.AAUCAU....UA.AA..UAAUAUUAGCAUGUAUUUGACAAAAGAAGUUAAAGAAGGG
env-486   GUAUAU.UUGCACCG.............CUUA..............AUAA....................UAAG....................................CAUAC.AUAA..AAUCAU....UA.AA..UAAUAUUAGCAUGUAUUUGACAAAAGAAGUUAAAGAAGGG
env-487   GUAUAU.UUGCCCCG.............CUUA..............AUAA....................UAAG....................................CAUAC.AUAAC.AAUCAU....UA.AA..UAAUAUUAGCAUGUAUUUAUCAAAAGAAGUAAAAGAAACG
env-488   GUAUAU.UUGCCCCG.............CUUA..............AUAA....................UAAG....................................CAUAC.AUAAC.AAUCAU....UA.AA..UAAUAUUAGCAUGUAUUUAUCAAAAGAAGUAAAAGAAACG
env-489   GUAUAU.UUGCCCCG.............CUUA..............AUAA....................UAAG....................................CAUAC.AUAAC.AAUCAU....UA.AA..UAAUAUUAGCAUGUAUUUAUCAAAAGAAGUAAAAGAAACG
env-490   GUAUAU.UUGCACCG.............CCUA..............AUAA....................UAGG....................................CAUAC.AUAAC.AAUCAU....UA.AA..UAAUAUUAGCAUGU..........................
env-491   GUAUAU.UUGCACCG.............CCUA..............AUAA....................UAGG....................................CAUAC.AUAAC.AAUCAU....UA.AA..UAAUAUUAGCAUGUAUUUGACAAAAGAAGUUAAAGAAGGG
env-492   GUAUAU.UUGCACCG.............CCUA..............AUAA....................UAGG....................................CAUAC.AUAAC.AAUCAU....UA.AA..UAAUAUUAGCAUGUAUUUGACAAAAGAAGUUAAAGAAGGG
env-494   GUAUAU.UUGCACCG.............CCUA..............AUAA....................UAGG....................................CAUAC.AUAAC.AAUCAU....UA.AA..UAAUAUUAGCAUGUAUUUGACAAAAGAAAUUAAAGAAGGG
env-496   GUAUAU.UUGCACCG.............CCUA..............AUAA....................UAGG....................................CAUAC.AUAAC.AAUCAU....UA.AA..UAAUAUUAGCAUGUAUUUGACAAAAGAAAUUAAAGAAGGG
env-498   GUAUAU.UUGCACCG.............CCUA..............AUAA....................UAGG....................................CAUAC.AUAAC.AAUCAU....UA.AA..UAAUAUUAGCAUGUAUUUGACAAAAGAAAUUAAAGAAGGG
env-500   GUAUAU.UUGCACCG.............CCUA..............AUAA....................UAGG....................................CAUAC.AUAAC.AAUCAU....UA.AA..UAAUAUUAGCAUGUAUUUGACAAAAGAAG...........
env-501   GUAUAU.UUGCACCG.............CUUA..............AUAA....................UAGG....................................CGUAC.AUAAC.AAUCAU....UA.AA..UAAUAUUAGCAUGUAUUUGACAAAAGAAGUUAAAGAAGGG
env-502   GUAUCU.UUGCAGCG.............CCUU..............AAAA....................AAGG....................................CAUAC.AUAAC.AAUCAU....UA.AA..UAAUAUUAGCAUGUAUUUAUCAAAAGAAGUAAAAGAAAAG
env-503   UUACCU.UUGCACCC.............AAUCAGGA..........AAA.................UCCUGAAU.....................................UUAC.AUAAU.AAUUAUU...UA.AA..UAAUAUUGGCAUGUACUUAACUAAAGAAGUUAAACAAGAA
env-504   UUACCU.UUGCACCC.............AAUCAGG...........AUAGA................CCUGAGG.....................................UUAC.AUAAA.AAUUAUU...UA.AA..UAAUAUUGGCAUGUACUUGACUAAAGAAGUUAAAGAACAA
env-505   UUACCU.UUGCACCC.............AAUCAGG...........AUAGA................CCUGAGG.....................................UUAC.AUAAA.AAUUAUU...UA.AA..UAAUAUUGGCAUGUACUUGACUAAAGAAGUUAAAGAACAA
env-506   CUAUCU.UUGCACGC.............UGAUUC............ACUUAAAAC.............GAAUCG....................................CAUAC.AUAAA.AAUCAUU...UAAAA..UAAUAUUGGCAUGUAUUUAACAUCAGAAAAGAAACAAGAA
env-507   CUAACU.UUGCAGCC.............CUGAA.............UUU....................UUCAG....................................GCUAC.AUAAC.AAUCAU....UA.AA..UAAUAUUGGAAUGUACUUAGAUACAGAAAAGAAAAAAGAA
env-508   CUAACU.UUGCAGCC.............CUGAA.............UUU....................UUCAG....................................GCUAC.AUAAC.AAUCAU....UA.AA..UAAUAUUGGAAUGUACUUAGAUACAGAAAAGAAAAAAGAA
env-509   CUAACU.UUGCAGCC.............CUGAA.............UUU....................UUCAG....................................GCUAC.AUAAC.AAUCAU....UA.AA..UAAUAUUGGAAUGUACUUAGAUACAGAAAAGAAAAAAGAA
env-510   CUAACU.UUGCAGCC.............CUGAA.............UUU....................UUCAG....................................GCUAC.AUAAC.AAUCAU....UA.AA..UAAUAUUGGAAUGUACUUAGAUACAGAAAAGAAAAAAGAA
env-511   CUAUCU.UUGCAAAC.............UUGAAAG...........AAGAC................CUUUCAA....................................CAUAC.AUAAC.AAUCAUU...UACAA..UAAUAUUAGCAUGUAUUUAGACAAAAAGAAAAAGAAAGCC
Fhi-1-1   CUAAUU.UUGCACCG.............UCUUACA...........AAAGA................UGUACGA.....................................CUAC.AUAAA.AAUCAU....UA.AA..UAAUAUUGGAAUGUAUUUAACUAAAGAGAAAAAGAAGAG
Fsp-9-1   GUAAUU.UUGCACCG.............UCUUACA...........AAAGA................UGUACGA.....................................CUAC.AUAAA.AAUCAU....UA.AA..UAAUAUUGGAAUGUAUUUAACUAAAGAAAAAAAGGAAGAG
Asu-1-1   UUACUU.UUGCAACC.............UUGAUUAUC.........AAAAC..............GGUAAUCGA......................................UAC.AUAAA.AAUCAUU...UACAC..AAAUAUUAGCAUGUAUUUAGACAUUAAAGAAAAGAAGAU
Aca-1-1   CUAUCU.UUGCCGCC.............UUGAUUAUC.........AAAAC..............GGUAGUCAG......................................UAC.AUAAU.AAUCAUU...UU.AA..CAAUAUUGGCAUGUAUUUAUCACAGAAGAAAAAGGAAAA
env-513   GUACUU.UUGCAGCC.............UUGAUUAUC.........AAAAC..............GGUAAUCAG......................................UAC.AUAAA.AAUCAUU...UU.AA..UAAUAUUGGCAUGUAUUUAUCCACAGAAGAAAAAGGUAAA
env-514   GUACUU.UUGCACCC.............UUGAUUAUC.........AAAAC..............GGUAGUCAG......................................UAC.AUAAA.AAUCAUU...UU.AA..UAAUAUUGGCAUGUAUUUAUCCACAGAAGAAAAAGGAAAA
env-515   GUACUU.UUGCACCC.............UUGAUUAUC.........AAAAA..............GGUAGUCAG......................................UAC.AUAAA.AAUCAUU...UU.AA..CAAUAUUGGCAUGUAUUUAUCAACAGAAGAAAAAGGAAAA
env-517   GUACUU.UUGCACCC.............UUGAUUAUC.........AAAAA..............GGUAGUCAG......................................UAC.AUAAA.AAUCAUU...UU.AA..CAAUAUUGGCAUGUAUUUAUCAACAGAAGAAAAAGGAAAA
Avl-1-1   GUACUU.UUGCACCC.............UUGAUUAUC.........AAAAA..............GGUAGUCAG......................................UAC.AUAAA.AAUCAUU...UU.AA..CAAUAUUGGCAUGUAUUUAUCAACAGAAGAAAAAGGAAAA
env-518   GUACUU.UUGCACCC.............UUGAUUAUC.........AAAAA..............GGUAGUCAG......................................UAC.AUAAA.AAUCAUU...UU.AA..CAAUAUUGGCAUGUAUUUAUCAACAGAAGAAAAAGGAAAA
env-519   GUACUU.UUGCACCC.............UUGAUUAUC.........AAAAA..............GGUAGUCAG......................................UAC.AUAAA.AAUCAUU...UU.AA..CAAUAUUGGCAUGUAUUUAUCAACAGAAGAAAAAGGAAAA
env-520   GUACUU.UUGCAACC.............UUGAUUAUCA........AAA...............UGGUAAUCAG......................................UAC.AUAAA.AAUCAUU...UU.AA..UAAUAUUGGCAUGUAUUUAUCCACAGAAGAAAAAGGUAAA
env-521   GUACUU.UUGCAACC.............UUGAUUAUCA........AAA...............UGGUAAUCAG......................................UAC.AUAAA.AAUCAUU...UU.AA..CAAUAUUGGCAUGUAUUUAUCCCAAGAAGAAAAAGAAAAA
env-522   GUACUU.UUGCAACC.............UUGAUUAUCA........AAA...............UGGUAAUCAG......................................UAC.AUAAA.AAUCAUU...UU.AA..CAAUAUUGGCAUGUAUUUAUCCCAAGAAGAAAAAGAAAAA
env-523   GUACUU.UUGCAACC.............UUGAUUAUCA........AAA...............UGGUAAUCAG......................................UAC.AUAAA.AAUCAUU...UU.AA..CAAUAUUGGCAUGUAUUUAUCCCAAGAAGAAAAAGAAAAA
env-527   GUACUU.UUGCAACC.............UUGAUUAUCA........AAA...............UGGUAAUCAG......................................UAC.AUAAA.AAUCAUU...UU.AA..CAAUAUUGGCAUGUAUUUAUCCCAAGAAGAAAAAGAAAA.
env-528   GUACUU.UUGCAGCC.............UUGAUUAUCA........AAA...............UGGUAAUCAG......................................UAC.AUAAA.AAUCAUU...UU.AA..CAAUAUUGGCAUGUAUUUAUCCCAAGAAGAAAAAGAAAAA
env-531   GUACUU.UUGCAGCC.............UUGAUUAUCA........AAA...............UGGUAAUCAG......................................UAC.AUAAA.AAUCAUU...UU.AA..CAAUAUUGGCAUGUAUUUAUCCCAAGAAGAAAAAGAAAAA
env-532   GUACUU.UUGCGGCC.............UUGAUUAUCA........AAA...............UGGUAGUCAG......................................UAC.AUAAA.AAUCAUU...UU.AA..CAAUAUUGGCAUGUAUUUAUCACAAGAAGUAAAAGAAAAA
env-533   GUACUU.UUGCAGCC.............UUGAUUAUCA........AAA...............UGGUAGUCAG......................................UAC.AUAAA.AAUCAUU...UU.AA..CAAUAUUGGCAUGUAUUUAUCACAAGAAGUAAAAGAAAAA
env-534   UUAUAU.UUGCCGCC.............UU................AUAUA.....................AA....................................CAUAC.AUAAU.AAUUAUU...UA.AA..UAAUAUUAGCAUGUAUUUAACAAAAGAAGUUAAAGAAAAG
env-536   UUAUAU.UUGCCGUC.............UU................AUUUA.....................AG....................................UAUAC.AUAAU.AAUUAUU...GA.AC..UAAUAUUGGCAUGUAUUUAGCAAAAGAAACUAAAGAAAAG
env-537   UUAUAU.UUGCCACC.............UU................AUUUA.....................AG....................................CAUAC.AUAAU.AAUUAUU...GA.AC..UAAUAUUGGCAUGUAUUUAACAAAAGAAACUAAAGAAAAG
env-539   UUAUAU.UUGCCACC.............UU................AUUUA.....................AG....................................CAUAC.AUAAU.AAUUAUU...GA.AC..UAAUAUUGGCAUGUAUUUAACAAAAGAAACUAAAGAAAAG
env-540   UUAUAU.UUGCCACC.............UU................AUUUA.....................AG....................................CAUAC.AUAAU.AAUUAUU...GA.AC..UAAUAUUGGCAUGUAUUUAACAAAAGAAACUAAAGAAAAG
Apa-1-1   GUAUAU.UUGCACGU.............UAAUAGCA..........AAA.................UGCUAUUA....................................UAUAC.AUAAA.AAUCAUU...UA.AA..UAAUAUUGGAAUGUAUUUAACGAAAGAAGUUAAAGAAGAA
Ama-1-1   UUAUAU.UUGCACGC.............UAAUAGC...........AAAGA................GCUAUUU.....................................AUAC.AUAAA.AAUCAUU...UA.AA..UAAUAUUGGAAUGUAUUUAACGAAAGAAGUUAAAGAAGAA
Aat-1-1   UUAUAU.UUGCACGC.............UAAUAGC...........AAAGA................GCUAUUG.....................................AUAC.AUAAA.AAUCAUU...UA.AA..UAAUAUUGGAAUGUAUUUAACAAAAGAAGUUAAAGAAGAA
Ame-1-1   UUAUAU.UUGCACGC.............UAAUAGC...........AAAGA................GCUAUUG.....................................AUAC.AUAAA.AAUCAUU...UA.AA..UAAUAUUGGAAUGUAUUUAACAAAAGAAGUUAAAGAAGAA
Ala-1-1   CUAUAU.UUGCACGC.............UGAUAGCA..........GAA.................UGCUAUUU.....................................AUAC.AUAAA.AAUCAUU...UAAAA..UAAUAUUGGAAUGUAUUUAACAAAAGAAACAAAAGAAGAA

          ............................<<<<<<<<<<<<<<<<<<..........>>>>>>>>>>>>>>>>>>.........<<<<<<<<........>>>>>>>>........................................................................
          ............................2222221111..101.00..........00.101..1111222222.........11011000........00011011........................................................................
          •UAY•U-UUGC••CY-------------∘∘∘∘∘∘∘-----------R•R∘∘------------------∘∘∘∘∘∘∘------------------------------------∘UAC-AUAA•-AAUCAUU---UA-AA--YAAUAUURG•AUGUAUUUARC•ARAGAAR••AAARRRRAR
          5′                                                                                                                                                  3′
```

```
Amu-1-1  GUAUAU.UUGCACGC............UCAUAGUA..........AAG..................UGCUAUGG...................................ACAUAC.AUAAA.AAUCAUU..UA.AA..UAAUAUUGGAAUGUAUUUAACAAAAGAAGUUAAAGAAGAA
Fsu-3-1  UUACCU.UUGCACCCG...........UAUUGAGA..........UCG..................UCUCAAUC......................................UGC.AUAAU.AAUCAUU..UU.AA..UAAUAUUAGCAUGUAUUUAACUAAAGAAGUAAAAGCCGAU
Fen-1-1  CUAACU.UUGCACCCG...........UAUUGAGA..........CUG..................UCUCAAUU......................................UAC.AUAAU.AAUCAUU..UU.AA..AAAUAUUAGCAUGUACUUAACUAAAGAAGUUAAAUCAGAA
Fco-1-1  UUAAAU.UUGCGUCCG...........UAUCGAGA..........AAA..................UCUCGGUC......................................UAC.AUAAU.AAUCAUU..UA.AA..UAAUAUUAGCAUGUAUUUAUCAAAGGAGAAAAAAGCUGAA
env-541  GUAAAU.UUGCACCCG...........UAUUGAGA..........CAA..................UCUCAAUC......................................UAC.AUAAU.AAUCAUU..UA.AA..UAAUAUUAGCAUGUAUUUGACUAAAGAAACAAAAGCAGAG
env-542  GUAAAU.UUGCACCCG...........UAUUGAG...........AUAAA.................CUCAAUC......................................UAC.AUAAU.AAUCAUU..UA.AA..UAAUAUUAGCAUGUAUUUGACUAAAGAAACAAAAGCUGAC
env-543  GUAAAU.UUGCACCCG...........UAUUGAG...........AUAAA.................CUCAAUC......................................UAC.AUAAU.AAUCAUU..UA.AA..UAAUAUUAGCAUGUAUUUGACUAAA...............
env-545  GUAAAU.UUGCACCCG...........UAUUGAG...........AUAAA.................CUCAAUC......................................UAC.AUAAU.AAUCAUU..UA.AA..UAAUAUUAGCAUGUAUUUGACUAAAGAAACAAAAGCUGAC
env-546  UUAUAU.UUGCGCCU............UGAUAUG...........AACGA.................CGUAUCA.....................................UUAC.AUAAU.AAUCAUU..UACAA..UAAUAUUGGCAUGUAUUUAGACAAGAAAGAAAAAGAAAAA
Fps-1-1  GUAAUU.UUGCACUC............GAAUCCGA..........GAA..................UCGGAAUU....................................CAUAC.AUAAA.AAUCAUU..AA.AC..AAAUAUUGGAAUGUAUUUAACUAAAGAGAUUAAAGAAGAA
Jma-1-1  UUAUCU.UUGCGCCC............ACUGAUAGA.........AAA.................UCUAUCAAA......................................UAC.AUAAG.AAUCAUU..UA.AA..UAAUAUUGGCAUGUAUUUAGCUAAAGAAGUAAAAGCAGAC
env-547  GUAUAU.UUGCGCCCG...........AU................AAAA.......................AU......................................UAC.AUAAU.AAUCAUU..AA.AC..AAAUAUUGGCAUGUAUUUAACAAAAGAAAAAAACAAGAA
env-548  CUAUAU.UUGCGCGG............CU................GAAAGA.....................AG....................................CGUAC.AUAAU.AAUCAUU..AA.AA..UAAUAUUUGCAUGUAUUUAACUAAAAAAAUAAAGGAAGAU
env-549  CUAUAU.UUGCGCGG............CU................GAAAGA.....................AG....................................CGUAC.AUAAU.AAUCAUU..AA.AA..UAAUAUUUGCAUGUAUUUAACUAAAAAAAUAAAGGAAGAU
env-550  CUAUAU.UUGCGCGG............CU................GAAAGA.....................AG....................................CGUAC.AUAAU.AAUCAUU..AA.AA..UAAUAUUUGCAUGUAUUUAACUAAAAAAAUAAAGGAAGAU
env-551  CUAUAU.UUGCACUU............UCAAUAGA..........AAA..................UCUGUUGA....................................AAUAC.AUAAA.AAUCAUU..UA.AA..UAAUAUUGGCAUGUACUUAACUAAAGAAGUAAAAGAAAGC
env-552  CUAUAU.UUGCACUU............UCAAUAGA..........AAA..................UCUGUUGA....................................AAUAC.AUAAA.AAUCAUU..UA.AA..UAAUAUUGGCAUGUACUUAACUAAAGAAGUAAAAGAAAGC
env-553  CUAUAU.UUGCACUU............UCAAUAGA..........AAA..................UCUGUUGA....................................AAUAC.AUAAA.AAUCAUU..UA.AA..UAAUAUUGGCAUGUACUUAACUAAAGAAGUAAAAGAAAGC
env-554  UUAUAU.UUGCAAAC............UCAACGGA..........UAU..................UCCAUUGA....................................UAUAC.AUAAU.AAUCAUU..UA.AA..UAAUAUUGGCAUGUACUUAACUAAAGAAGUAAAAGAAAGC
env-556  UUAUAU.UUGCAAAC............UCAACGGA..........UAU..................UCCAUUGA....................................UAUAC.AUAAU.AAUCAUU..UA.AA..UAAUAUUGGCAUGUACUUAACUAAAGAAGUAAAAGAAAGC
env-557  UUAUAU.UUGCAAAC............UCAACGGA..........UAU..................UCCAUUGA....................................UAUAC.AUAAU.AAUCAUU..UA.AA..UAAUAUUGGCAUGUACUUAACUAAAGAAGUAAAAGAAAGC
Ffi-1-1  UUACCU.UUGCACUC............GAAUUGGG..........AUAGA................CCC.UUUC.....................................AUAC.AUCAA.AAUCAUU..UA.AA..UAAUAUUGGAAUGUAUUUAACUAAAGAAACAAAAGCUGAA
env-558  GUAUUU.UUGCACCG............UCCUAUA...........AUCGC.................UAUAGGA....................................CAUAC.AUAAA.AAUCAUU..AA.AC..CAAUAUUGGAAUGUAUUUAACUAAAGAGAUUAAAGAAGAG
env-559  CUAUAU.UUGCCGCC............UGAAAG............AAAGA..................CUUUCG...................................GAUUAC.AUAAC.AAUUAUU..UACAC..AAAUAUUAGCAUGUACGUUACAAAAGAAGAAAAAGCAAAA
env-561  CUAUAU.UUGCCGCU............CUAUUUU...........AGAUA.................AUAGUAA..................................AUCGUAC.AUAAU.AAUCAUU..UA.AA..UAAUAUUGGAAUGUAUUUAACAAAAAAAACAAAAGAAACU
env-562  CUAUAU.UUGCCGCU............CUAUUUU...........AGAUA.................AUAGUAA..................................AUCGUAC.AUAAU.AAUCAUU..UA.AA..UAAUAUUGGAAUGUAUUUAACAAAAAAAACAAAAGAAACU
env-563  CUAUAU.UUGCCGCU............CUAUUUU...........AAGUA.................AUAGUAG..................................AUCAUAC.AUAAU.AAUCAUU..UA.AA..UAAUAUUGGAAUGUAUUUAACAAAAAAAUCAAAAGAAACU
env-564  CUAUAU.UUGCCGCU............CUAUUUU...........AAGUA.................AUAGUAG..................................AUCAUAC.AUAAU.AAUCAUU..UA.AA..UAAUAUUGGAAUGUAUUUAACAAAAAAGUCGAAAGAAACU
env-565  CUAUAU.UUGCCGCU............CUAUUUU...........AAGUA.................AUAGUAA..................................ACCAUAC.AUAAU.AAUCAUU..UA.AA..UAAUAUUGGAAUGUAUUUAACAAAAAAAUCAAAAGAAACU
env-566  CUAUAU.UUGCCGCU............CUAUUUU...........AAAUA.................AUAGUAA..................................AUCAUAC.AUAAU.AAUCAUU..UA.AA..UAAUAUUGGAAUGUAUUUAACAAAAAAAUCGAAAGAAACU
env-567  CUAUAU.UUGCCGCU............CUAUUUU...........AAAUA.................AUAGUAA..................................AUCAUAC.AUAAU.AAUCAUU..UA.AA..UAAUAUUGGAAUGUAUUUAACAAAAAAGUCGAAAGAAACU
env-569  CUAUAU.UUGCCGCU............CUAUUUU...........AAAUA.................AUAGUAA..................................AUCAUAC.AUAAU.AAUCAUU..UA.AA..UAAUAUUGGAAUGUAUUUAACAAAAAAAUCGAAAGAAACU
env-571  CUAUAU.UUGCCGCU............CUAUUUU...........AAAUA.................AUAGUAA..................................AUCAUAC.AUAAU.AAUCAUU..UA.AA..UAAUAUUGGAAUGUAUUUAACAAAAAAAUC..........
env-572  CUAUAU.UUGCCGCU............CUAUUUU...........AAAUA.................AUAGUAA..................................AUCAUAC.AUAAU.AAUCAUU..UA.AA..UAAUAUUGGAAUGUAUUUAACAAAAAAGUCGAAAGAAACU
env-573  CUAUAU.UUGCCGCU............CUAUUUU...........AAAUA.................AUAGUAA..................................AUCAUAC.AUAAU.AAUCAUU..UA.AA..UAAUAUUGGAAUGUAUUUAACAAAAAAAUCGAAAGAAACU
env-574  CUAUAU.UUGCCGCU............CUAUUUU...........AAGUA.................AUAGUAA..................................AUCAUAC.AUAAU.AAUCAUU..UA.AA..UAAUAUUGGAAUGUAUUUAACAAA................
env-575  CUAUAU.UUGCCGCU............CUAUUUU...........AAGUA.................AUAGUAA..................................AUCAUAC.AUAAU.AAUCAUU..UA.AA..UAAUAUUGGAAUGUAUUUAACAAAAAAAUCAAAAGAAACU
env-576  CUAUAU.UUGCCGCU............CUAUUUU...........AAGUA.................AUAGUAA..................................AUCAUAC.AUAAU.AAUCAUU..UA.AA..UAAUAUUGGAAUGUAUUUAACAAAAAAAUCAAAAGA....
env-577  CUAUAU.UUGCCGCU............CUAUUUU...........AAGUA.................AUAGUAA..................................AUCAUAC.AUAAU.AAUCAUU..UA.AA..UAAUAUUGGAAUGUAU........................
env-578  CUAUAU.UUGCCGCU............CUAUUUU...........AAGUA.................AUAGUAA..................................AUCAUAC.AUAAU.AAUCAUU..UA.AA..UAAUAUUGGAAUGUAUUUAACAAAAAAAUCAAAAGAAACU
env-579  CUAUAU.UUGCCGCU............CUAUUUU...........AAGUA.................AUAGUAA..................................AUCAUAC.AUAAU.AAUCAUU..UA.AA..UAAUAUUGGAAUGUAUUUAACAAAAAAAUCAAAAGAAACU
env-580  CUAUAU.UUGCCGCU............CUAUUUU...........AAGUA.................AUAGUAA..................................AUCAUAC.AUAAU.AAUCAUU..UA.AA..UAAUAUUGGAAUGUAUUUAACAAAAAAAUCAAAAGAAACU
env-581  CUAUAU.UUGCCGCU............CUAUUUU...........AAGUA.................AUAGUAA..................................AUCAUAC.AUAAU.AAUCAUU..UA.AA..UAAUAUUGGAAUGUAUUUAACAAAAAAAUCAAAAGAAACU
env-582  CUAUAU.UUGCCGCU............CUAUUUU...........AAGUA.................AUAGUAA..................................AUCAUAC.AUAAU.AAUCAUU..UA.AA..UAAUAUUGGAAUGUAUUUAACAAAAAAAUCAAAAGAAACU
env-583  CUAUAU.UUGCCGCU............CUAUUUU...........AAGUA.................AUAGUAA..................................AUCAUAC.AUAAU.AAUCAUU..UA.AA..UAAUAUUGGAAUGUAUUUAACAAAAAAAUCGAAAGAAACU
env-584  CUAUAU.UUGCCGCU............CUAUUUU...........AAGUA.................AUAGUAA..................................AUCAUAC.AUAAU.AAUCAUU..UA.AA..UAAUAUUGGAAUGUAUUUAACAAAAAAAUC..........
env-585  CUAUAU.UUGCCGCU............CUAUUUU...........AAGUA.................AUAGUAA..................................AUCAUAC.AUAAU.AAUCAUU..UA.AA..UAAUAUUGGAAUGUAUUUAACAAA................
env-586  CUAUAU.UUGCCGCU............CUAUUUU...........AAGUA.................AUAGUAA..................................AUCAUAC.AUAAU.AAUCAUU..UA.AA..UAAUAUUGGAAUGUAUUUAACAAA................
env-587  CUAUAU.UUGCCGCU............CUAUUUU...........AAGUA.................AUAGUAA..................................AUCAUAC.AUAAU.AAUCAUU..UA.AA..UAAUAUUGGAAUGUAUUUAACAAA................
env-588  CUAUAU.UUGCCGCU............CUAUUUU...........AAGUA.................AUAGUAA..................................AUCAUAC.AUAAU.AAUCAUU..UA.AA..UAAUAUUGGAAUGUAUUUAACAAAAAANUCGAAAGAAACU
env-589  CUAUAU.UUGCCGCU............CUAUUUU...........AAGUA.................AUAGUAA..................................AUCAUAC.AUAAU.AAUCAUU..UA.AA..UAAUAUUGGAAUGUAUUUAACAAAAAAAUCAAAAGAAACU
env-590  CUAUAU.UUGCCGCU............CUAUUUU...........AAGUA.................AUAGUAA..................................AUCNUAC.AUAAU.AAUCAUU..UA.AA..UAAUAUUGGAAUGUAUUUAACAAAAAAAUCAAAAGAAACU
env-591  CUAUAU.UUGCCGCU............CUAUUUU...........AAUUA.................AUAGUAA..................................AUCGUAC.AUAAU.AAUCAUU..UA.AA..UAAUAUUGGAAUGUAUUUAACAAAAAAAUCAAAAGAAACU
env-592  CUAUAU.UUGCCGCU............CUAUUUU...........AAGUA.................AUAGUAA..................................AUCGUAC.AUAAU.AAUCAUU..UA.AA..UAAUAUUGGAAUGUAUUUAACAAAAAAAUCAAAAGAAACU
env-593  CUAUAU.UUGCCGCU............CUAUUUU...........AAGUA.................AUAGUAA..................................AUCGUAC.AUAAU.AAUCAUU..UA.AA..UAAUAUUGGAAUGUAUUUAACAAAAAAAUCAAAAGAAACU
env-594  CUAUAU.UUGCCGCU............CUAUUUU...........AAGUA.................AUAGUAA..................................AUCGUAC.AUAAU.AAUCAUU..UA.AA..UAAUAUUGGAAUGUAUUUAACAAAAAAAUCAAAAGAAACU
env-596  CUAUAU.UUGCCGCU............CUAUUUU...........AAGUA.................AUAGUAA..................................AUCGUAC.AUAAU.AAUCAUU..UA.AA..UAAUAUUGGAAUGUAUUUAACAAAAAAAUCAAAAGAAACU
env-598  CUAUAU.UUGCCGCU............CUAUUUU...........AAGUA.................AUAGUAA..................................AUCGUAC.AUAAU.AAUCAUU..UA.AA..UAAUAUUGGAAUGUAUUUAACAAAAAAAUCAAAAGAAACU
env-599  CUAUAU.UUGCCGCU............CUAUUUU...........AAGUA.................AUAGUAA..................................AUCGUAC.AUAAU.AAUCAUU..UA.AA..UAAUAUUGGAAUGUAUUUAACAAAAAAAUCAAAAGAAACU
env-600  CUAUAU.UUGCCGCU............CUAUUUU...........AGUUA.................AUAGUAA..................................AUCGUAC.AUAAU.AAUCAUU..UA.AA..UAAUAUUGGAAUGUAUUUAACAAAAAAAUCAAAAGAAACU
env-601  CUAUAU.UUGCGCUU............CCA.UUUUA.........AAA.................UAAAACUGG.....................................UUAC.AUAAG.AAUUAUU..UACAA..AAAUAUUGGCAUGUAUUUAUCUAAAGAAAAGAAAAAAGAA
env-602  CUAUAU.UUGCGCCU............CUGAUUU...........AACAA.................AAAUUGG.....................................UUAC.AUAAG.AAUUAUU..UACAA..AAAUAUUGGCAUGUAUUUAUCAAAAGAAAACAAAAAAGAA
env-603  CUAUAU.UUGCGCCU............CUGAUUU...........AAUAA.................AAAUUGG.....................................UUAC.AUAAG.AAUUAUU..UACAA..AAAUAUUGGCAUGUAUUUAUCAAAAGAAAACAAAAAAGAA
env-604  CUAUAU.UUGCGCCU............CAAUUU............AAAUUA.................AAAUUG....................................GUUAC.AUAAG.AAUUAUU..UACAA..AAAUAUUGGCAUGUAUUUAUCUAAAAAAAAGAAAAAGAAA
env-605  CUAUAU.UUGCGCUU............CAAUUUUA..........AAA..................UAAAAUUG....................................GUUAC.AUAAG.AAUUAUU..UACAA..AAAUAUUGGCAUGUAUUUAUCUAAAGAAAACAAAAAAAAA
env-606  UUAUAU.UUGCCGCU............CCAAUA............AAAAAA.................UAUUGG.....................................CUAC.AUAAU.AAUCAUU..UACAA..CAAUAUUGGCAUGUAUUUAUCAAACGAAAACAAAAAAGAA
env-607  UUAUAU.UUGCCGCU............CCAAUA............AAAAAA.................UAUUGG.....................................CUAC.AUAAU.AAUCAUU..UACAA..CAAUAUUGGCAUGUAUUUAUCAAACGAAAACAAAAAAGAA
env-608  CUAUAU.UUGCGCGU............CCAAUU............AAAUUA.................AAUUGG.....................................UUAC.AUAAG.AAUUAUU..UACAA..AAAUAUUGGCAUGUAUCUAUCAAAAGAAAAUAAAAAGAAA
env-609  CUAUAU.UUGCGCCU............CCAAUA............AUUAAA.................UAUUGG.....................................UUAC.AUAAU.AAUUAUU..UACAA..AAAUAUUGGCAUGUAUUUAUCUAAGGAAAAUAAAAAGGAA
env-610  CUAUAU.UUGCGCCU............CCAAUA............AUUAAA.................UAUUGG.....................................UUAC.AUAAU.AAUUAUU..UACAA..AAAUAUUGGCAUGUAUUUAUCUAAGGAAAAUAAAAAGGAA
env-611  CUAUAU.UUGCGCCU............CCAAUA............AUUAAA.................UAUUGG.....................................UUAC.AUAAU.AAUUAUU..UACAA..AAAUAUUGGCAUGUAUUUAUCUAAGGAAAAUAAAAAGGAA
env-612  CUAUAU.UUGCGCCU............CCAAUA............AUUAAA.................UAUUGG.....................................UUAC.AUAAU.AAUUAUU..UACAA..AAAUAUUGGCAUGUAUUUAUCUAAGGAAAAUAAAAAGGAA
env-613  CUAUAU.UUGCGCCU............CCAAUA............AUUAAA.................UAUUGG.....................................UUAC.AUAAU.AAUUAUU..UACAA..AAAUAUUGGCAUGUAUUUAUCUAAGGAAAAUAAAAAGGAA
env-614  CUAUAU.UUGCGCCU............CCAAUA............AUUAAA.................UAUUGG.....................................UUAC.AUAAU.AAUUAUU..UACAA..AAAUAUUGGCAUGUAUUUAUCUAAGGAAAAUAAAAAGGAA
env-615  CUAUAU.UUGCGCCU............CCAAUA............AUUAAA.................UAUUGG.....................................UUAC.AUAAU.AAUUAUU..UACAA..AAAUAUUGGCAUGUAUUUAUCUAAGGAAAAUAAAAAGGAA
env-616  CUAUAU.UUGCGCCU............CCAAUA............AUUAAA.................UAUUGG.....................................UUAC.AUAAU.AAUUAUU..UACAA..CAAUAUUGGCAUGUAUUUAUCUAAAGAAAAUAAAAAAGAA
env-617  CUAUAU.UUGCGCCU............CCAAUA............AUUAAA.................UAUUGG.....................................UUAC.AUAAU.AAUUAUU..UACAA..UAAUAUUGGCAUGUAUUUAUCUAAAGAAAAUAAAAAAGAA
env-618  CUAUAU.UUGCGCCU............CCAAUA............AUUAAA.................UAUUGG.....................................UUAC.AUAAU.AAUUAUU..UACAA..UAAUAUUGGCAUGUAUUUAUCUAAAGAAAAUAAAAAAGAA
env-619  CUAUAU.UUGCGCGU............CCAAUA............AAUUAA.................UAUUGG.....................................UUAC.AUAAG.AAUUAUU..UACAA..AAAUAUUGGCAUGUAUUUAUCUAAAGAAAACAAAAAAGAA
env-620  CUAUAU.UUGCGCCU............CCGAUA............AAUUAA.................UAUUGG.....................................UUAC.AUAAG.AAUUAUU..UACAA..AAAUAUUGGCAUGUAUUUAUCUAAAGAAAACAAAAAAGAA
env-621  CUAUAU.UUGCGCCU............CCGAUA............AAUUAA.................UAUUGG.....................................UUAC.AUAAG.AAUUAUU..UACAA..AAAUAUUGGCAUGUAUUU......................
env-622  CUAUAU.UUGCGCCU............CCGAUA............AAUUAA.................UAUUGG.....................................UUAC.AUAAG.AAUUAUU..UACAA..AAAUAUUGGCAUGUAUUUAUCUAAAGAAAACAAAAAAGAA
env-623  CUAUAU.UUGCGCCU............CCGAUA............AAUUAA.................UAUUGG.....................................UUAC.AUAAG.AAUUAUU..UACAA..AAAUAUUGGCAUGUAUUUAUCUAAAGAAAACAAAAAAGAA

         ...........................<<<<<<<<<<<<<<<<<<...........>>>>>>>>>>>>>>>>>>.........<<<<<<<<........>>>>>>>>.......................................................................
         ...........................2222221111..101.00...........00.101..1111222222.........11011000........00011011.......................................................................
         •UAY•U-UUGC••CY------------ooooooo-----------R•Roo------------------ooooooo-----------------------------------•UAC-AUAA•-AAUCAUU---UA-AA--YAAUAUURG•AUGUAUUUARC•ARAGAAR••AAARRRRAR
          5'                                                                                                                                                     3'
```

```
env-624  CUAUAU.UUGCGCCU.............CCAUA............AAUAAA.................UAUUGG.....................................UUAC.AUAAG.AAUUAUU...UACAA..AAAUAUUGGCAUGUAUUUAUCUAAAGAAAACAAAAAAGAA
env-625  CUAUAU.UUGCGCCU.............CCAAUA............AAUAAA.................UAUUGG.....................................UUAC.AUAAG.AAUUAUU...UACAA..AAAUAUUGGCAUGUAUUUAUCUAAAGAAAACAAAAAAGAA
env-626  CUAUAU.UUGCGCCU.............CCAAUA............AAUAAA.................UAUUGG.....................................UUAC.AUAAG.AAUUAUU...UACAA..AAAUAUUGGCAUGUAUUUAUCUAAAGAAAACAAAAAAGAA
env-627  CUAUAU.UUGCGCCU.............CCAAUA............AAUAAA.................UAUUGG.....................................UUAC.AUAAG.AAUUAUU...UACAA..AAAUAUUGGCAUGUA.........................
env-628  CUAUAU.UUGCGCCU.............CCAAUA............AAUAAA.................UAUUGG.....................................UUAC.AUAAG.AAUUAUU...UACAA..AAAUAUUGGCAUGUACUUAUCUAAAGAAAACAAAAAAGAA
env-629  CUAUAU.UUGCGCCU.............CCAAUA............AAUAAA.................UAUUGG.....................................UUAC.AUAAG.AAUUAUU...UACAA..AAAUAUUGGCAUGUAUUUAUCUAAAGAAAACAAAAAAGAA
env-633  CUAUAU.UUGCGCCU.............CCAAUA............AAUAAA.................UAUUGG.....................................UUAC.AUAAG.AAUUAUU...UACAA..AAAUAUUGGCAUGUAUUUAUCAAAAGAAAACAAAAAAGAA
env-634  CUAUAU.UUGCGCCU.............CCAAUA............AAUAAA.................UAUUGG.....................................UUAC.AUAAG.AAUUAUU...UACAA..AAAUAUUGGCAUGUAUUU......................
env-635  CUAUAU.UUACGCCU.............CCAAUA............AAUUAA.................UAUUGG.....................................UUAC.AUAAG.AAUUAUU...UACAA..AAAUAUUGGCAUGUAUUUAUCUA.................
env-636  CUAUAU.UUGCGCCU.............CCAAUA............AAUUAA.................UAUUGG.....................................UUAC.AUAAG.AAUUAUU...UACAA..AAAUAUUGGCAUGUAUUUAUCUAAAGAAAACAAAAAAGAA
env-637  CUAUAU.UUGCGCCU.............CCAAUA............AAUUAA.................UAUUGG.....................................UUAC.AUAAG.AAUUAUU...UACAA..AAAUAUUGGCAUGUAUUUAUCUAAAGAAAACAAAAAAGAA
env-638  CUAUAU.UUGCGCCU.............CCAAUA............AAUUAA.................UAUUGG.....................................UUAC.AUAAG.AAUUAUU...UACAA..AAAUAUUGGCAUGUAUUUAUCUAAAGAAAACAAAAAAGAA
env-639  CUAUAU.UUGCGCCU.............CCAAUA............AAUUAA.................UAUUGG.....................................UUAC.AUAAG.AAUUAUU...UACAA..AAAUAUUGGCAUGUAUUUAUCUAAAGAAAACAAAAAAGAA
env-640  CUAUAU.UUGCGCCU.............CCAAUA............AAUUAA.................UAUUGG.....................................UUAC.AUAAG.AAUUAUU...UACAA..AAAUAUUGGCAUGUAUUUAUCUAAAAAAAAGAAAAAAGAA
env-641  UUAUAU.UUGCCGCU.............CCAAUA............AAUUAA.................UAUUGG.....................................UUAC.AUAAU.AAUUAUU...UACAA..UAAUAUUGGCAUGUAUUUAACAAAAGAAAACAAAAAAGAA
env-648  UUAUAU.UUGCCGCU.............CCAAUA............AAUUAA.................UAUUGG.....................................UUAC.AUAAU.AAUUAUU...UACAA..UAAUAUUGGCAUGUAUUUAACAAAAGAAAACAAAAAAGAA
env-649  UUAUAU.UUGCCGCU.............CCAAUA............AAUUAA.................UAUUGG.....................................UUAC.AUAAU.AAUUAUU...UACAA..UAAUAUUGGCAUGUAUUUAACAAAAGAAAACAAAAAAGAA
env-650  UUAUAU.UUGCCGCU.............CCAAUA............AAUUAA.................UAUUGG.....................................UUAC.AUAAU.AAUUAUU...UACAA..UAAUAUUGGCAUGUAUUUAACAAAAGAAAACAAAAAAGAA
env-651  UUAUAU.UUGCCGCU.............CCAAUA............AAUUAA.................UAUUGG.....................................UUAC.AUAAU.AAUUAUU...UACAA..UAAUAUUGGCAUGUAUUUAACAAAAGAAAACAAAAAAGAA
env-652  UUAUAU.UUGCCGCU.............CCAAUA............AAUUAA.................UAUUGG.....................................UUAC.AUAAU.AAUUAUU...UACAA..UAAUAUUGGCAUGUAUUUAACAAAAGAAAACAAAAAAGAA
env-653  UUAUAU.UUGCCGCU.............CCAAUA............AAUUAA.................UAUUGG.....................................UUAC.AUAAU.AAUUAUU...UACAA..UAAUAUUGGCAUGUAUUUAACAAAAGAAAACAAAAAAGAA
env-655  UUAUAU.UUGCCGCU.............CCAAUA............AAUUAA.................UAUUGG.....................................UUAC.AUAAU.AAUUAUU...UACAA..UAAUAUUGGCAUGUAUUUAACAAAAGAAAACAAAAAAGAA
env-656  UUAUAU.UUGCCGCU.............CCAAUA............AAUUAA.................UAUUGG.....................................UUAC.AUAAU.AAUUAUU...UACAA..UAAUAUUGGCAUGUAUUUAACAAAAGAAAACAAAAAAGAA
env-657  UUAUAU.UUGCCGCU.............CCAAUA............AAUUAA.................UAUUGG.....................................UUAC.AUAAU.AAUUAUU...UACAA..UAAUAUUGGCAUGUAUUUAACAAAAGAAAACAAAAAAGAA
env-658  UUAUAU.UUGCCGCU.............CCAAUA............AAUUAA.................UAUUGG.....................................UUAC.AUAAU.AAUUAUU...UACAA..UAAUAUUGGCAUGUAUUUAACAAAAGAAAACAAAAAAGAA
env-659  UUAUAU.UUGCCGCU.............CCAAUA............AAUUAA.................UAUUGG.....................................UUAC.AUAAU.AAUUAUU...UACAA..UAAUAUUGGCAUGUAUUUAACAAAAGAAAACAAAAAAGAA
env-660  UUAUAU.UUGCCGCU.............CCAAUA............AAUUAA.................UAUUGG.....................................UUAC.AUAAU.AAUUAUU...UAUAA..CAAUAUUGGCAUGUAUUUAACAAAAGAAAACAAAAAAAAA
env-661  UUAUAU.UUGCCGCU.............CCAAUA............AAUUAA.................UAUUGG.....................................UUAC.AUAAU.AAUUAUU...UAUAA..UAAUAUUGGCAUGUAUUUAACAAAAGAAAACAAAAAAGAA
env-662  UUAUAU.UUGCCGCU.............CCAAUA............AAUUAA.................UAUUGG.....................................UUAC.AUAAU.AAUUAUU...UAUAA..UAAUAUUGGCAUGUAUUUAACAAAAGAAAACAAAAAAGAA
env-663  UUAUAU.UUGCGCGU.............CCAAUAUA..........AAUU.................UAUAUUGG.....................................UUAC.AUAAA.AAUUAUU...UA.CA..AAAUAUUGGCAUGUAUUUAUCAAAAGAAAAAAAGAAAGAA
env-664  GUACUU.UUGCGCCU.............CGAUUACCA.........AAG.................UGGUAGUCU.....................................GUAC.AUAAA.AAUCAUU...UA.AA..UAAUAUUGGAAUGUAUUUAGCACCAGAAGAAAAAGAAAAA
env-669  GUACUU.UUGCGCCU.............CGAUUACCA.........GAG.................UGGUAGUCA......................................UAC.AUAAU.AAUUAUU...UA.AG..UAAUAUUGGAAUGUAUUUAGAACCAAAAGAAAAAGCAAAA
env-670  GUACUU.UUGCGCCU.............CGAUUACCA.........GAG.................UGGUAGUCA......................................UAC.AUAAU.AAUUAUU...UA.AG..UAAUAUUGGAAUGUAUUUAGAACCAAAAGAAAAAGCAAAA
env-671  GUACUU.UUGCGCCU.............CGAUUACCA.........GAG.................UGGUAGUCA......................................UAC.AUAAU.AAUUAUU...UA.AG..UAAUAUUGGAAUGUAUUUAGAACCAAAAGAAAAAGCAAAA
env-672  CUAAAU.UUGCACCC.............GUC...............CAAAGA....................GAC......................................UAC.AUAAU.AAUUAUU...UACAA..UAAUAUUGGCAUGUAUUUAGAUAAAAAGAUUAAAAAAGAU
env-673  CUAAAU.UUGCACCC.............GUUCA.............CAG.....................UGAAC......................................UAC.AUAAG.AAUUAUU...UACAA..AAAUAUUGGCAUGUACUUAGAUAAAAAAGUAAAACAAGAC
env-674  CUAAAU.UUGCACCC.............GUUCA.............UAG.....................UGAAC......................................UAC.AUAAG.AAUUAUU...UACAA..AAAUAUUGGCAUGUACUUAGAUAAAAAAGUAAAACAAGAC
env-675  CUAAAU.UUGCACCC.............GUCCA.............AAG.....................UGGAC......................................UAC.AUAAU.AAUUAUU...UACAA..AAAUAUUGGCAUGUAUCUAGAUAAAAAAGUGAAAAAAGAC
env-676  CUAAAU.UUGCACCC.............GUCC..............AAAGA....................GGGC......................................UAC.AUAAU.AAUCAUU...UACAA..UAAUAUUGGCAUGUAUUUAGAUAAAAAAGUUAAAAAAGAU
env-677  CUAAAU.UUGCACCC.............GUCC..............AAAGA....................GGGC......................................UAC.AUAAU.AAUCAUU...UACAA..UAAUAUUGGCAUGUAUUUAGAUAAAAAAGUUAAAACAGAC
env-678  CUAAAU.UUGCACCC.............GUCC..............AAAGA....................GGGC......................................UAC.AUAAU.AAUCAUU...UACAA..UAAUAUUGGCAUGUAUUUAGAUAAAAAAGUUAAAAAAGAU
env-679  CUAAAU.UUGCACCC.............GUCC..............AAAGC....................GGGC......................................UAC.AUAAU.AAUCAUU...UACAA..UAAUAUUGGCAUGUAUUUAGAUAAAAAAGUUAAAAAAGAU
env-680  CUAAAU.UUGCACCC.............GUCC..............AAAGC....................GGGC......................................UAC.AUAAU.AAUCAUU...UACAA..UAAUAUUGGCAUGUAUUUAGAUAAAAAAGUUAAAAAAGAU
env-681  CUAAAU.UUGCACCC.............GUCC..............AAAGA....................GGAC......................................UAC.AUAAU.AAUUAUU...UACAA..AAAUAUUGGCAUGUAUUUAGAUAAAAAAGUGAAAAAGGAU
env-682  CUAAAU.UUGCACCC.............GUCC..............AAAGA....................GGAC......................................UAC.AUAAU.AAUUAUU...UACAA..AAAUAUUGGCAUGUAUUUAGAUAAAAAAGUGAAAAAGGAU
env-683  CUAAAU.UUGCACCC.............GUCC..............AAAGA....................GGAC......................................UAC.AUAAU.AAUUAUU...UACAA..AAAUAUUGGCAUGUAUUUAGAUAAAAAAGUGAAAAAGGAU
env-684  CUAAAU.UUGCACCC.............GUCC..............AAAGA....................GGAC......................................UAC.AUAAU.AAUUAUU...UACAA..AAAUAUUGGCAUGUAUUUAGAUAAAAAAGUGAAAAAGGAU
env-685  CUAAAU.UUGCACCC.............GUCC..............AAAGA....................GGAC......................................UAC.AUAAU.AAUUAUU...UACAA..AAAUAUUGGCAUGUAUUUAGAUAAAAAAGUGAAAAAGGAU
env-686  CUAAAU.UUGCACCC.............GUCC..............AAAGA....................GGAC......................................UAC.AUAAU.AAUUAUU...UACAA..AAAUAUUGGCAUGUAUUUAGAUAAAAAAGUGAAAAAGGAU
env-687  CUAAAU.UUGCACCC.............GUCC..............AAAGA....................GGAC......................................UAC.AUAAU.AAUUAUU...UACAA..AAAUAUUGGCAUGUAUUUAGAUAAAAAAGUGAAAAAGGAU
env-688  CUAAAU.UUGCACCC.............GUCC..............AAAGA....................GGAC......................................UAC.AUAAU.AAUUAUU...UACAA..AAAUAUUGGCAUGUAUUUAGAUAAAAAAGUGAAAAAGGAU
env-689  CUAAAU.UUGCACCC.............GUCC..............AAAGA....................GGAC......................................UAC.AUAAU.AAUUAUU...UACAA..AAAUAUUGGCAUGUAUUUAGAUAAAAAAGUGAAAAAGGAU
env-690  CUAAAU.UUGCACCC.............GUCC..............AAAGA....................GGAC......................................UAC.AUAAU.AAUUAUU...UACAA..AAAUAUUGGCAUGUAUUUAGAUAAAAAAGUGAAAAAGGAU
env-691  CUAAAU.UUGCACCC.............GUCC..............AAAGA....................GGAC......................................UAC.AUAAU.AAUUAUU...UACAA..AAAUAUUGGCAUGUAUUUAGAUAAAAAAGUGAAAAAGGAU
env-692  CUAAAU.UUGCACCC.............GUCC..............AAAGA....................GGGC......................................UAC.AUAAU.AAUUAUU...UACAA..AAAUAUUGGCAUGUAUUUAGAUAAAAAAGUGAAAAAAGAC
env-693  CUAAAU.UUGCACCC.............GUCC..............AAAGA....................GGGC......................................UAC.AUAAU.AAUUAUU...UACAA..AAAUAUUGGCAUGUAUUUAGAUAAAAAAGUGAAAAAAGAC
env-694  GUAUAU.UUGCCGAUC............GUU...............AAUUA.....................AGC...................................UUAUAC.AUAAA.AAUCAUU...AAAAU..UAAUAUUGGCAUGUACAUUUCAAGAAAAGUUAAAGAAGAU
env-698  GUAUAU.UUGCCGAUC............GUU...............AAUUA.....................AGC...................................UUAUAC.AUAAA.AAUCAUU...AAAAU..UAAUAUUGGCAUGUACAUUUCAAGAAAAGUUAAAGAAGAU
env-699  GUAUAU.UUGCCGAUC............GUU...............AAUUA.....................AAC...................................UUAUAC.AUAAA.AAUCAUU...AAAAU..UAAUAUUGGCAUGUACAUUUCAAGAAAAGUUAAAGAAGAU
env-700  GUAUAU.UUGCCGAUC............GUU...............AAUAA.....................AGC...................................UUAUAC.AUAAA.AAUCAUU...AAAAU..UAAUAUUUGCAUGUACAUUUCAAGAAAAGUUAAAGAAGAU
env-701  GUAUAU.UUGCCGAUC............GUU...............AAUAUA....................AGC...................................UUAUAC.AUAAA.AAUCAUU...AAAAU..UAAUAUUGGCAUGUACAUUUCAAGAAAAGUUAAAGAAGAU
env-702  GUAUAU.UUGCCGAUC............CUU...............AAUU......................AAG..................................CUUAUAC.AUAAA.AAUCAUU...AAAAU..UAAUAUUGGCAUGUACAUUUCAAGAAAAGUUAAAGAAGAU
env-703  GUAUAU.UUGCAGCUC............GU................AAAAA......................AU.....................................UUAC.AUAAA.AAUCAUU...AAAAU..UAAUAUUGGCAUGUACAAUUCAAAA...............
env-704  GUAUAU.UUGCAGCUC............GU................AAAAA......................AU.....................................UUAC.AUAAA.AAUCAUU...AAAAU..UAAUAUUGGCAUGUACAA......................
env-705  GUAUAU.UUGCAGCUC............GUU...............UAAA......................AAU.....................................UUAC.AUAAA.AAUCAUU...AAAAU..UAAUAUUGGCAUGUACAAUUCAAAAGAAGUAAAAGAAACU
env-706  GUAUAU.UUGCAGCUC............GUU...............AAAA......................AAU.....................................UUAC.AUAAA.AAUCAUU...AAAAU..UAAUAUUGGCAUGUACAAUUCAAAAGAAGUAAAAGAAA..
env-707  GUAUAU.UUGCGGCUC............GU................AAAAAAA....................AC......................................UAC.AUAAA.AAUCAUU...AAAAU..UAAUAUUGGCAUGUACAAUUCAAAAGAAGUAAAAGAAACU
env-708  GUAUAU.UUGCGGCUC............GU................AAAAAAA....................AC......................................UAC.AUAAA.AAUCAUU...AAAAC..UAAUAUUGGCAUGUACAUUUCAAAAGAAGUAAAAGAAACU
env-709  AUAUAU.UUGCCGCUCAA...............................................................................................UAC.AUAAU.AAUCAUU...AAAAU..UAAUAUUGGCAUGUAUCUAACAUCAGAAGUUAAAAAAGAA
env-711  UUAUAU.UUGCGGUUCAAAAUCA..........................................................................................UAC.AUAAU.AAUCAUU...AAAAA..UAAUAUUGGAAUGUAUUUAACACCAGAAGUAAAAAAAGAA
env-712  GUAUAU.UUGCCCUU.............CGUU..............AAA......................AACG...................................AACUAC.AUAAU.AAUAAUU...AAAAU..UAAUAUUGGCAUGUAUAUUUCAAGAGAAGUAAAAGAAGGU
Fba-2-1  GUAUAU.UUGCCGUU.............CGUU..............AAAAA....................AACG...................................AAUUAC.AUAAU.AAUCAUU...AAAAU..UAAUAUUGGCAUGUACAUUUCAAAAGAAGUUAAAGAAGGU
env-713  GUAUAU.UUGCCGUU.............CGUU..............AAAAA....................AACG...................................AAUUAC.AUAAU.AAUCAUU...AAAAU..UAAUAUUGGCAUGUACAUUUCAAAAGAAGUUAAAGAAGGU
env-718  GUAUAU.UUGCCGUU.............CGUU..............AAAAA....................AACG...................................AAUUAC.AUAAU.AAUCAUU...AAAAU..UAAUAUUGGCAUGUACAUUUCAAAAGAAGUUAAAGAAGGU
env-727  GUAUAU.UUGCCGUU.............CGUU..............AAAAA....................AACG...................................AAUUAC.AUAAU.AAUCAUU...AAAAU..UAAUAUUGGCAUGUACAUUUCAAAAGAAGUUAAAGAAGGU
env-728  GUAUAU.UUGCCGUU.............CGUU..............AAAAA....................AACG...................................AAUUAC.AUAAU.AAUCAUU...AAAAU..UAAUAUUGGCAUGUACAUUUCAAAAGAAGUUAAAGAAGGU
env-729  GUAUAU.UUGCCGUU.............CGUU..............AAAAA....................AACG...................................AAUUAC.AUAAU.AAUCAUU...AAAAU..UAAUAUUGGCAUGUACAUUUCAAAAGAAGUUAAAGAAGGU
env-730  GUAUAU.UUGCCCUU.............CGU...............GAAAAA....................ACG...................................AACUAC.AUAAU.AAUCAUU...AAAAU..UAAUAUUGGCAUGUAUAUUUCAAGAGAAGUAAAAGAAGGU
env-731  GUAUAU.UUGCCGCU.............CGUUA.............AAA.....................UAACG....................................AAUAC.AUAAA.AAUCAUU...AAAAU..UAAUAUUGGCAUGUACAAUUCAAAAGAAGUUACAGAAAGU
env-734  GUAUAU.UUGCCGCU.............CGUUA.............AAA.....................UAACG....................................AAUAC.AUAAA.AAUCAUU...AAAAU..UAAUAUUGGCAUGUACAAUUCAAAAGAAGUUACAGAAAGU
env-735  GUAUAU.UUGCCGCU.............CGUUA.............AAA.....................UAACG....................................AAUAC.AUAAA.AAUCAUU...AAAAU..UAAUAUUGGCAUGUACAAUUCAAAAGAAGUUACAGAAAGU
env-736  GUAUAU.UUGCCGCU.............CGUUA.............AAA.....................UAACG....................................AAUAC.AUAAA.AAUCAUU...AAAAU..UAAUAUUGGCAUGUACAAUUCAAAAGAAGUUACAGAAAGU
env-737  GUAUCU.UUGCCGCU.............CGUUA.............AAA.....................UAACG....................................AAUAC.AUAAA.AAUCAUU...AAAAU..UAAUAUUGGCAUGUACAAUUCAAAAGA.............

         ............................<<<<<<<<<<<<<<<<<<...........>>>>>>>>>>>>>>>>>>.........<<<<<<<<........>>>>>>>>........................................................................
         ............................2222221111..101.00...........00.101..1111222222.........11011000........00011011........................................................................
         •UAY•U-UUGC••CY-------------◦◦◦◦◦◦◦-----------R•R◦◦------------------◦◦◦◦◦◦◦-------------------------------------◦UAC-AUAA•-AAUCAUU---UA-AA--YAAUAUURG•AUGUAUUUARC•ARAGAAR••AAARRRRAR
         5′                                                                                                                                            3′
```

```
env-738  GUAUCU.UUGCCGCU.............CGUUA.............AAA.....................UAACG....................................AAUAC.AUAAA.AAUCAUU...AAAAU..UAAUAUUGGCAUGUACAAUUCAAAAGAAGUCAAAGAAGAA
env-739  GUAUCU.UUGCCGCU.............CGUUA.............AAA.....................UAACG....................................AAUAC.AUAAA.AAUCAUU...AAAAU..UAAUAUUGGCAUGUACAAUUCAAAA...............
env-740  GUAUAU.UUGCCGAU.............CGUUA.............AUU.....................UAACG....................................UAUAC.AUAAA.AAUCAUU...AAAAU..UAAUAUUGGCAUGUACAUUUCAAAAAAGUAAAAGAAGAU
env-741  GUAUAU.UUGCGUUCAUUU..............................................................................................UAC.AUAAA.AAUUAUU...UA.AA..AAAUAUUGGCAUGUAUUUAAACAAAAAAGUUAAGGAGGG
env-742  CUAUAU.UUGCAGUCAUUU..............................................................................................UAC.AUAAA.AAUAAUU...GA.AA..UAAUAUUGGCAUGUAUUUAAACAAAAAAGUUAAAGAGGG
env-743  GUAUAU.UUGCACACAUUU..............................................................................................UAC.AUAAA.AAUUAUU...UA.AA..CAAUAUUGGCAUGUAUCUCAAUAAAGAAGUAAAACAAGAG
env-744  GUAUAU.UUGCACACAUUU..............................................................................................UAC.AUAAA.AAUUAUU...UA.AA..CAAUAUUGGCAUGUAUCUCAAUAAAGAAGUAAAACAAGAG
env-745  GUAUAU.UUGCACACAUUU..............................................................................................UAC.AUAAA.AAUUAUU...UA.AA..CAAUAUUGGCAUGUAUCUCAAUAAAGAAGUAAAACAAGAG
env-746  GUAUAU.UUGCACACAUUU..............................................................................................UAC.AUAAA.AAUUAUU...UA.AA..CAAUAUUGGCAUGUAUCUCAAUAAAGAAGUAAAACAAGAG
env-747  GUAUAU.UUGCACACAUUU..............................................................................................UAC.AUAAA.AAUUAUU...UA.AA..CAAUAUUGGCAUGUAUCUCAACAAAGAAGUAAAACAAGAG
env-748  UUAUAU.UUGCACUCGUUU..............................................................................................UAC.AUAAA.AAUCAUU...GA.UA..UAAUAUUGGCAUGUACUUAGACAAAAAAAUUAAAAAAGGG
env-749  UUAUAU.UUGCACUCAUUU..............................................................................................UAC.AUAAA.AAUCAUU...GA.AA..UAAUAUUGGCAUGUACUUAGACAAAAAAAUUAAAAAAGAG
env-750  UUAUAU.UUGCACUCAUUU..............................................................................................UAC.AUAAA.AAUCAUU...GA.AA..UAAUAUUGGCAUGUACUUAGACAAAAAAAUUAAAAAAGAG
env-751  UUAUAU.UUGCACUCAUUU..............................................................................................UAC.AUAAA.AAUCAUU...GA.AA..UAAUAUUGGCAUGUAUUUAGACAAAAAAAUUAAAAAAGGG
env-752  UUAUAU.UUGCACUCAUUU..............................................................................................UAC.AUAAA.AAUCAUU...GA.AA..UAAUAUUGGCAUGUAUUUAGACAAAAAAAUUAAAAAAGGG
env-753  CUAUAU.UUGCACACAUUU..............................................................................................UAC.AUAAA.AAUUAUU...GA.AC..UAAUAUUGGCAUGUACUUAAACAAAGACGUUAAAAAAGGG
env-754  CUAUAU.UUGCACACAUUU..............................................................................................UAC.AUAAA.AAUUAUU...GA.AC..UAAUAUUGGCAUGUACUUAAACAAAGACGUUAAAAAAGGG
env-755  GUAUAU.UUGCAGACAUUUU.............................................................................................UAC.AUAAA.AAUUAUU...GA.AA..UAAUAUUGGCAUGUAUUUAAACAAAAAAGUUAAAGAAGGG
env-756  UUAUAU.UUGCACACAUUU..............................................................................................UAC.AUAAA.AAUUAUU...GA.AA..UAAUAUUGGCAUGUACUUAAACAAAGACGUUAAAGAAGGG
env-758  UUAUAU.UUGCACACAUUU..............................................................................................UAC.AUAAA.AAUUAUU...GA.AA..UAAUAUUGGCAUGUACUUAAACAAAGACGUUAAAGAAGGA
env-759  GUAUAU.UUGCAC.AAAUUU.............................................................................................UAC.AUAAA.AAUUAUU...UA.AA..CAAUAUUGGCAUGUAUCUCAAUAAAGAAGUAAAACAAGAG
env-760  GUAUAU.UUGCAC.AAAUUU.............................................................................................UAC.AUAAA.AAUUAUU...UA.AA..CAAUAUUGGCAUGUAUCUCAAUAAAGAAGUAAAACAAGAG
env-761  AUAUAU.UUGCACCUUUAUUUAUCA........................................................................................UAC.AUAAA.AAUUAUU...UA.AA..UAAUAUUGGCAUGUAUUUAAACAAAGAAAUUAAAAAGAAG
env-762  AUAUAU.UUGCACCUUUAUUUAUCA........................................................................................UAC.AUAAA.AAUUAUU...UA.AA..UAAUAUUGGCAUGUAUUUAAACAAAGAAAUUAAAAAGAAG
env-763  AUAUAU.UUGCAGCUUUAUUUAUCA........................................................................................UAC.AUAAA.AAUUAUU...UA.AA..UAAUAUUGGCAUGUAUUUAAACAAAGAAAUUAAAAAGAAG
env-764  AUAUAU.UUGCAGCUUUAUUUAUCA........................................................................................UAC.AUAAA.AAUUAUU...UA.AA..UAAUAUUGGCAUGUAUUUAAACAAAGAAAUUAAAAAGAAG
env-765  AUAUAU.UUGCAGCUUUAUUUAUUA........................................................................................UAC.AUAAA.AAUUAUU...UA.AU..UAAUAUUGGCAUGUAUUUAAACAAAGAAAUUAAAAAGAAG
env-766  AUAUAU.UUGCAGCUUUAUUUAUUA........................................................................................UAC.AUAAA.AAUUAUU...UA.AU..UAAUAUUGGCAUGUAUUUAAACAAAGAAAUUAAAAAGAAG
env-767  AUAUAU.UUGCAGCUUUAUUUAUUA........................................................................................UAC.AUAAA.AAUUAUU...UA.AA..UAAUAUUGGCAUGUAUUUAAACAAAGAAAUUAAAAAGAAG
env-768  AUAUAU.UUGCAGCUUUAUUUAUUA........................................................................................UAC.AUAAA.AAUUAUU...UA.AA..UAAUAUUGGCAUGUAUUUAAACAAAGAAAUUAAAAAGAAG
env-769  AUAUAU.UUGCAGCUUUAUUUAUUA........................................................................................UAC.AUAAA.AAUUAUU...UA.AA..UAAUAUUGGCAUGUAUUUAAGCAAAGAAAUUAAAAAGAAG
env-770  AUAUAU.UUGCAGCUUUAUUUAUUA........................................................................................UAC.AUAAA.AAUUAUU...UA.AA..UAAUAUUGGCAUGUAUUUAAACAAAGAAAUUAAAAAGAAG
env-771  AUAUAU.UUGCAGCUUUAUUUAUUA........................................................................................UAC.AUAAA.AAUUAUU...UA.AA..UAAUAUUGGCAUGUAUUUAAACAAAGAAAUUAAAAAGAAG
env-772  AUAUAU.UUGCAGCUUUAUUUAUUA........................................................................................UAC.AUAAA.AAUUAUU...UA.AA..UAAUAUUGGCAUGUAUUUAAACAAAGAAAUUAAAAAGAAG
env-773  AUAUAU.UUGCAGCUUUAUUUAUUA........................................................................................UAC.AUAAA.AAUUAUU...UA.AA..UAAUAUUGGCAUGUAUUUAAACAAAGAAAUUAAAAAGAAG
env-774  AUAUAU.UUGCAGCUUUAUUUAUUA........................................................................................UAC.AUAAA.AAUUAUU...UA.AA..UAAUAUUGGCAUGUAUUUAAACAAAGAAAUUAAAAAGAAG
env-775  AUAUAU.UUGCAGCUUUAUUUAUUA........................................................................................UAC.AUAAA.AAUUAUU...UA.AA..UAAUAUUGGCAUGUAUUUAAACAAAGAAAUUAAAAAGAAG
env-776  AUAUAU.UUGCAGCUUUAUUUAUUA........................................................................................UAC.AUAAA.AAUUAUU...UA.AA..UAAUAUUGGCAUGUAUUUAAACAAAGAAAUUAAAAAGAAG
env-777  AUAUAU.UUGCACCUUAAUUUACAA........................................................................................UAC.AUAAA.AAUUAUU...AAAAA..AAAUAUUUGCAUGUAUCUAAACAAAGAAAUAAAAAAAAAG
env-778  AUAUAU.UUGCACCUUAAUUUACAA........................................................................................UAC.AUAAA.AAUUAUU...AAAAA..AAAUAUUUGCAUGUAUCUAAACAAAGAAAUAAAAAAAAAG
env-779  AUAUAU.UUGCACCUUAAUUUACAA........................................................................................UAC.AUAAA.AAUUAUU...AAAAA..AAAUAUUUGCAUGUAUCUAAACAAAGAAAUAAAAAAAAAG
env-780  AUAUAU.UUGCAGCUUCAUUUACAA........................................................................................UAC.AUAAA.AAUUAUU...AA.AA..AAAUAUUGGCAUGUAUCUAAACAAAGAAAUAAAAAAAAAG
env-781  AUAUAU.UUGCAGCUUCAUUUACAA........................................................................................UAC.AUAAA.AAUUAUU...AA.AA..AAAUAUUGGCAUGUAUCUAAACAAAGAAAUAAAAAAAAAG
env-783  AUAUAU.UUGCAGCUUCAUUUACAA........................................................................................UAC.AUAAA.AAUUAUU...AA.AA..AAAUAUUGGCAUGUAUCUAAACAAAGAAAUAAAAAAAAAG
env-784  AUAUAU.UUGCACCUUCAUUUACAA........................................................................................UAC.AUAAA.AAUUAUU...AAAAA..AAAUAUUGGCAUGUAUCUAAACAAAGAAAUAAAAAAAAAG
env-785  AUAUAU.UUGCACCUUCAUUUACAA........................................................................................UAC.AUAAA.AAUUAUU...AAAAA..AAAUAUUGGCAUGUAUCUAAACAAAGAAAUAAAAAAAAAG
env-786  AUAUAU.UUGCAGACUCAUUCAUAA........................................................................................UAC.AUAAA.AAUUAUU...AA.AA..AAAUAUUGGCAUGUAUCUUAACAAAGACAUAAAAAAGAAG
env-787  AUAUAU.UUGCAGACUCAUUCAUAA........................................................................................UAC.AUAAA.AAUUAUU...AA.AA..AAAUAUUGGCAUGUAUCUUAACAAAGACAUAAAAAAGAAG
env-788  AUAUAU.UUGCAGACUCAUUCAUAA........................................................................................UAC.AUAAA.AAUUAUU...AA.AA..AAAUAUUGGCAUGUAUCUUAACAAAGACAUAAAAAAGAAG
env-789  CUAUCU.UUGCCGCCU............GAUUACC...........AAAGC.................GGUAAUU...................................CUAUAC.AUAAA.AAUCAUU...UA.AA..UAAUAUUGGCAUGUAUCCAAGUAAAGAAGAAAAAGAAUUA
env-791  UUAUAU.UUGCACAU.............UCAAUAAU..........AAUGA................AUUAUUGA....................................UAUAC.AUAAU.AAUCAUU...UA.AA..UAAUAUUGGCAUGUACUUAACUAAAGAAGUAAAAGAAAGC
env-794  UUAUAU.UUGCACAU.............UCAAUAAU..........AAUGA................AUUAUUGA....................................UAUAC.AUAAU.AAUCAUU...UA.AA..UAAUAUUGGCAUGUACUUAACUAAAGAAGUAAAAGAAAGC
env-795  UUAUAU.UUGCACAU.............UCAAUAAU..........AAUGA................AUUAUUGA....................................UAUAC.AUAAU.AAUCAUU...UA.AA..UAAUAUUGGCAUGUACU.......................
env-796  UUAUAU.UUGCACAU.............UCAAUAAU..........AAUGA................AUUAUUGA....................................UAUAC.AUAAU.AAUCAUU...UA.AA..UAAUAUUGGCAUGUACUUAACUAAAGAAGUAAAAGAAAGC
env-797  UUAUAU.UUGCACAU.............UCAAUAAU..........AAUGA................AUUAUUGA....................................UAUAC.AUAAU.AAUCAUU...UA.AA..UAAUAUUGGCAUGUACUUAACUAAAGAAGUAAAAGAAAGC
env-798  UUAUAU.UUGCACAU.............UCAAUAAU..........AAUGA................AUUAUUGA....................................UAUAC.AUAAU.AAUCAUU...UA.AA..UAAUAUUGGCAUGUACUUAACUAAAGAAGUAAAAGAAAGC
env-799  CUAUCU.UUGCGCCCGAUA..............................................................................................UAC.AUAAU.AUUCAUU...UAAAA..UAAUUUUGGCAUGUAUUUAUCAGCAGAAAAAAAAGAAGAA
Cca-4-1  GUAUCU.UUGCCACCUAAAAAUCAG........................................................................................UAC.AUAAU.AAUCAUU...UAAAA..CAAUAUUGGAAUGUAUUUAACAAAGGAAAUUAAAGCGGAA
Cca-4-2  GUAUCU.UUGCCACCUAAAAAUCAG........................................................................................UAC.AUAAU.AAUCAUU...UAAAA..CAAUAUUGGAAUGUAUUUAACAAAGGAAAUUAAAGCGGAA
Cca-4-3  GUAUCU.UUGCCACCUAAAAAUCAG........................................................................................UAC.AUAAU.AAUCAUU...UAAAA..CAAUAUUGGAAUGUAUUUAACAAAGGAAAUUAAAGCGGAA
env-800  GUAUCU.UUGCCGAC.............U.................UAUA........................A......................................UAC.AUAAG.AAUCAUU...UC.AA..UAAUAUUGGCAUGUAUUUAUCCAAAGAGAAAAAAGCAGAA
env-801  CUAUAU.UUGCACCC.............UGUUUG............AAAAC..................CAAUCA....................................GAUAC.AUAAA.AAUCAUU...AA.AC..CAAUAUUGGCAUGUAUUUAAGCAAAGAAGCAAAGUCGGAC
env-802  GUAUCU.UUGCGCCC.............UCAUUGA...........ACG...................UCAAUGA....................................GAUAC.AUAAA.AAUCAUU...AA.UA..CAAUAUUGGAAUGUAUUUAAGUAAAGAAAAGAAGGCUGAA
env-803  GUAUAU.UUGCGCCCGAUUAAAAU.........................................................................................UAC.AUAAU.AAUCAUU...AA.AC..UAAUAUUGGCAUGUAUUUAACAAAAGAAACAAAACAAGAA
env-805  CUAUAU.UUGCCGCG.............C.................AAUAAAA.....................G.....................................CUAC.AUAAG.AAUCAUU...UU.AA..UAAUAUUGGCAUGUAUUUAACAAAAGAAAACAAAAAAGAA
env-806  CUAUAU.UUGCCGCG.............C.................AAUAAAA.....................G.....................................CUAC.AUAAG.AAUCAUU...UU.AA..UAAUAUUGGCAUGUAUUUAACAAAAGAAAACAAAAAAGAA
env-807  UUAUAU.UUGCCGCG.............CU................AUCA.......................AG.....................................CUAC.AUAAU.AAUCAUU...UU.AA..UAAUAUUGGCAUGUAUUUAACAAAAGAAAAUAAAAAAGAA
env-808  UUAUAU.UUGCCGCG.............CU................AUUA.......................AG.....................................UUAC.AUAAG.AAUCAUU...UU.AA..UAAUAUUGGCAUGUAUUUAACAAAAGAAAACAAAAAAGAA
env-809  UUAUAU.UUGCCGCG.............CU................AUUA.......................AG.....................................UUAC.AUAAG.AAUCAUU...UU.AA..UAAUAUUGGCAUGUAUUUAACAAAAGAAAACA........
Zga-1-1  GUAUCU.UUGCCGCU.............UGACCGA...........AAA...................UCGGCCA....................................UAUAC.AUAAA.AAUCAUU...UAAAA..UAAUAUUGGCAUGUAUCUAACAAAAGAAGUAAAAGCCGAA
env-810  CUACCU.UUGCACAC.............GAAUAGGA..........AAA..................UCCUAAUU...................................CACUAC.AUAAU.AAUCAUU...UA.AA..CAAUAUUAGCAUGUAUUUAACUAAAGAAGUUAAAGCGGAA
env-813  GUACUU.UUGCACGG.............UCUUAUA...........AUAGC.................UGUAAGA....................................CAUAC.AUAAC.AAUCAUU...UA.AA..UAAUAUUAGCAUGUAUUUAACUAAAGAGAUUAAAGAAGAG
env-814  GUACUU.UUGCACGG.............UCUUAUA...........AUAGC.................UGUAAGA....................................CAUAC.AUAAA.AAUCAUU...AA.AC..AAAUAUUGGAAUGUAUUUAACUAAAGAGAUUAAAGAAGAG
env-815  GUACUU.UUGCACGG.............UCUUAUA...........AUAGC.................UGUAAGA....................................CAUAC.AUAAA.AAUCAUU...AA.AC..AAAUAUUGGAAUGUAUUUAACUAAAGAGAUUAAAGAAGAG
env-816  GUACUU.UUGCACGG.............UCUUAUA...........AUAGC.................UGUAAGA....................................CAUAC.AUAAA.AAUCAUU...AA.AC..AAAUAUUGGAAUGUAUUUAACAAAAGAGAUUAAAGAAGAG
env-817  GUACUU.UUGCCCGG.............UCUUAUA...........AUAGC.................UGUAAGA....................................CAUAC.AUAAA.AAUCAUU...AA.AC..AAAUAUUGGAAUGUAUUUAACUAAAGAGAUUAAAGAAGAG
env-818  GUACUU.UUGCCCGG.............UCUUAUA...........AUAGC.................UGUAAGA....................................CAUAC.AUAAA.AAUCAUU...AA.AC..AAAUAUUGGAAUGUAUUUAACUAAAGAGAUUAAAGAAGAG
env-819  GUACUU.UUGCCCGG.............UCUUAUA...........AUAGC.................UGUAAGA....................................CAUAC.AUAAA.AAUCAUU...AA.AC..AAAUAUUGGAAUGUAUU.......................
env-820  GUACUU.UUGCCCGG.............UCUUAUA...........AUAGC.................UGUAAGA....................................CAUAC.AUAAA.AAUCAUU...AA.AC..AAAUAUUGGAAUGUAUUUAACUAAAGAGA...........
env-821  GUACUU.UUGCCCGG.............UCUUAUA...........AUAGC.................UGUAAGA....................................CAUAC.AUAAA.AAUCAUU...AA.AC..AAAUAUUGGAAUGUAUUUAACUAAAGAGAUUAAAGAAGAG
env-822  GUACUU.UUGCACGG.............UCUUAUA...........AUAGA.................UAUGAGG....................................CAUAC.AUAAA.AAUCAUU...AA.AC..AAAUAUUGGAAUGUAUUUAACUAAAGAGAUUAAAGAAGAG
env-823  GUACUU.UUGCACGG.............UCUUAUA...........AUAGA.................UAUGAGG....................................CAUAC.AUAAA.AAUCAUU...AA.AC..AAAUAUUGGAAUGUAUUUAACUAAAGAGAUUAAAGAAGAG
env-824  AUAUAU.UUGCGCCG.............C.................UGUAUU......................G....................................CAUAC.AUAAA.AAUCAUU...UAAAA..UAAUAUUGGCAUGUAUUUAACAAAAAAAAACAAAGAAGAU

         ............................<<<<<<<<<<<<<<<<<<...........>>>>>>>>>>>>>>>>>>.........<<<<<<<<........>>>>>>>>........................................................................
         ............................2222221111..101.00...........00.101..1111222222.........11011000........00011011........................................................................
         •UAY•U-UUGC••CY-------------•••••○○-----------R•R○○-----------------○○○○•••-------------------------------------•UAC-AUAA•-AAUCAUU---UA-AA--YAAUAUURG•AUGUAUUUARC•ARAGAAR••AAARRRRAR
          5′                                                                                                                                                3′
```

```
env-825  UUAUAU.UUGCCAGG.............CUAAUUAA..........AAA..................UUGAUUGG......................................CUAC.AUAAG.AAUCAUU...UAUAA..CAAUAUUGGCAUGUAUUUAACAAAAGAAAACAAAAAAGAA
env-826  UUAUAU.UUGCCGCG.............CUAAUUA...........AAAU..................UAAUUAG......................................CUAC.AUAAA.AAUCAUU...UAUAA..CAAUAUUGGCAUGUAUUUAACAAAAGAAAACAAAAAAAAA
env-827  UUAUAU.UUGCCGCG.............CUAAUUAA..........AAA..................UUAAUUAG......................................CUAC.AUAAA.AAUCAUU...UAUAA..CAAUAUUGGCAUGUAUUUAACAAAAGAAAGUAAAAAAGAA
env-828  UUAUAU.UUGCCGCG.............CUAAUUAA..........AAA..................UUAAUUAG......................................CUAC.AUAAU.AAUCAUU...UAUAA..CAAUAUUGGCAUGUAUUUAACAAAAGAAAACAAAAAAGAA
env-829  UUAUAUUUUGCCGCG.............CUAAUUAA..........AAA..................UUAAUUAG......................................CUAC.AUAAA.AAUCAUU...UAUAA..CAAUAUUGGCAUGUAUUUAACAAAAGAAAACAAAAAAGAA
env-830  UUAUAU.UUGCCGCG.............CUAAUUAA..........AAA..................UUAAUUAG......................................CUAC.AUAAG.AAUCAUU...UAUAA..CAAUAUUGGCAUGUAUUUAACAAAAGAAAACAAAAAAGAA
env-831  UUAUAU.UUGCCGCG.............CUAAUUAA..........AAA..................UUAAUUAG......................................CUAC.AUAAG.AAUCAUU...UAUAA..CAAUAUUGGCAUGUAUUUAACAAAAGAAAACAAAAAAGAA
env-832  UUAUAU.UUGCCGCG.............CUAAUUAA..........AAA..................UUAAUUAG......................................CUAC.AUAAG.AAUCAUU...UAUAA..CAAUAUUGGCAUGUAUUUAACAAAAGAAAACAAAAAAGAA
env-833  UUAUAU.UUGCCGCG.............CUAAUUAA..........AAA..................UUAAUUGG......................................CUAC.AUAAG.AAUCAUU...UAUAA..CAAUAUUGGCAUGUAUUUAACAAAAGAAAACAAAAAAGAA
env-834  UUAUAU.UUGCCGCG.............CUAAUUAA..........AAA..................UUAAUUGG......................................CUAC.AUAAG.AAUCAUU...UAUAA..CAAUAUUGGCAUGUAUUUAACAAAAGAAAACAAAAAAGAA
env-835  UUAUAU.UUGCCGCG.............CUAAUUAA..........AAA..................UUAAUUGG......................................CUAC.AUAAG.AAUCAUU...UAUAA..CAAUAUUGGCAUGUAUUUAACAAAAGAAAACAAAAAAGAA
env-836  UUAUAU.UUGCCGCG.............CUAAUUA...........AAAU..................UAAUUAG......................................CUAC.AUAAA.AAUCAUU...UAAAA..UAAUAUUGGCAUGUAUUUAACAAAAGAAAACAAAAAAGAA
env-837  UUAUAU.UUGCCGCG.............CUAAUUA...........AAAU..................UAAUUAG......................................CUAC.AUAAA.AAUCAUU...UAAAA..UAAUAUUGGCAUGUAUUUAACAAAAGAAAACAAAAAAGAA
env-838  UUAUAU.UUGCCGCG.............CUAAUUA...........AAAU..................UAAUUAG......................................CUAC.AUAAA.AAUCAUU...UAAAA..UAAUAUUGGCAUGUAUUUAACAAAAGAAAACAAAAAAGAA
env-847  UUAUAU.UUGCCGCG.............CUAAUUAA..........AAA..................UUAAUUAG......................................CUAC.AUAAA.AAUCAUU...UAAAA..UAAUAUUGGCAUGUAUUUAACAAAAGAAAACAAAAAAGAA
env-848  UUAUAU.UUGCCGCG.............CUAAUUAA..........AAA..................UUAAUUAG......................................CUAC.AUAAA.AAUCAUU...UAAAA..UAAUAUUGGCAUGUAUUUAACAAAAGAAAACAAAAAAGAA
env-849  UUAUAU.UUGCCGCG.............CUAAUUAA..........AAA..................UUAAUUAG......................................CUAC.AUAAA.AAUCAUU...UAAAA..UAAUAUUGGCAUGUAUUUAACAAAAGAAAACAAAAAAGAA
env-850  UUAUAU.UUGCCGCG.............CUAAUUAA..........AAA..................UUAGUUAG......................................CUAC.AUAAA.AAUCAUU...UAAAA..UAAUAUUGGCAUGUAUUUAACAAAAGAAAACAAAAAAGAA
env-851  UUAUAU.UUGCCGCG.............CUAAUUAA..........AAA..................UUAGUUAG......................................CUAC.AUAAA.AAUCAUU...UAAAA..UAAUAUUGGCAUGUAUUUAACAAAAGAAAACAAAAAAGAA
env-852  UUAUAU.UUGCCGCG.............CUAAUUAA..........AAA..................UUAGUUAG......................................CUAC.AUAAA.AAUCAUU...UAAAA..UAAUAUUGGCAUGUAUUUAACAAAAGAAAACAAAAAAGAA
env-853  UUAUAU.UUGCCGCG.............CUAAUUAA..........AAA..................UUAGUUAG......................................CUAC.AUAAA.AAUCAUU...UAAAA..UAAUAUUGGCAUGUAUUUAACAAAAGAAAACAAAAAAGAA
env-854  UUAUAU.UUGCCGCG.............CUAAUUAA..........AAA..................UUAGUUAG......................................CUAC.AUAAA.AAUCAUU...UAAAA..UAAUAUUGGCAUGUAUUUAA....................
env-855  UUAUAU.UUGCCGCG.............CUAAUUAA..........AAA..................UUAGUUAG......................................CUAC.AUAAA.AAUCAUU...UAAAA..UAAUAUUGGCAUGUAUUUAACA..................
env-856  UUAUAU.UUGCCGCG.............CUAAUUAA..........AAA..................UUAGUUAG.....................................GCUAC.AUAAA.AAUCAUU...UAAAA..UAAUAUUGGCAUGUAUUUAACAAAAGAAAACAAAAAAGAA
env-857  UUAUAU.UUGCCGCG.............CUAAUUAA..........AAA..................UUAGUUAG......................................CUAC.AUAAA.AAUCAUU...UAAAA..UAAUAUUGGCAUGUAUUUAACAAAAGAAAACAAAAAAGAA
env-860  UUAUAU.UUGCCGCG.............CUAAUUAA..........AAA..................UUAGUUAG......................................CUAC.AUAAA.AAUCAUU...UAAAA..UAAUAUUGGCAUGUAUUUAACAAAAGAAAACAAAAAAGAA
env-861  UUAUAU.UUGCCGCG.............CUAAUUAA..........AAA..................UUAGUUAG......................................CUAC.AUAAA.AAUCAUU...UAAAA..UAAUAUUGGCAUGUAUUUAACAAAAGAAAACAAAAAAGAA
env-862  .UAUAU.UUGCCGCG.............CUAAUUAA..........AAA..................UUAGUUAG......................................CUAC.AUAAA.AAUCAUU...UAAAA..UAAUAUUGGCAUGUAUUUAACAAAAGAAAACAAAAAAGAA
env-863  UUAUAU.UUGCCGCG.............CUAAUUAA..........AAA..................UUAGUUAG......................................CUAC.AUAAA.AAUCAUU...UAAAA..UAAUAUUGGCAUGUAUUUAACA..................
env-864  UUAUAU.UUGCCGCG.............CUAAUUAA..........AAA..................UUAGUUAG......................................CUAC.AUAAA.AAUCAUU...UAAAA..UAAUAUUGGCAUGUAUUUAACAAAAGAAAACAAAAAAGAA
env-865  UUAUAU.UUGCCGCG.............CUAAUUAA..........AAA..................UUAGUUAG......................................CUAC.AUAAA.AAUCAUU...UAAAA..UAAUAUUGGCAUGUAUUUAACAAAAGAAAACAAAAAAGAA
env-866  UUAUAU.UUGCCGGG.............CUAAUUAA..........AAU..................UUAAUUAG......................................CUAC.AUAAA.AAUCAUU...UAAAA..UAAUAUUGGCAUGUAUUUAACAAAAGAAAACAAAAAAGAA
env-868  UUAUAU.UUGCCGCG.............CUAAUUAA..........AAU..................UUAUUUAG......................................CUAC.AUAAA.AAUCAUU...UAAAA..UAAUAUUGGCAUGUAUUUAACAAAAGUAAACAAAAAAGAA
env-869  UUAUAU.UUGCCGCG.............CUAAUUAA..........AAU..................UUAAUUAG......................................CUAC.AUAAA.AAUCAUU...UAAAA..CAAUAUUGGCAUGUAUUUAACAAAAGAAAACAAAAAAGAA
env-870  UUAUAU.UUGCCGCG.............CUAAUUAA..........AAU..................UUAAUUAG......................................CUAC.AUAAA.AAUCAUU...UAAAA..UAAUAUUGGCAUGUAUUUAACAAAAGAAAACAAAAAAGAA
env-871  UUAUAU.UUGCCGCG.............CUAAUUAA..........AAU..................UUAAUUAG......................................CUAC.AUAAA.AAUCAUU...UAAAA..UAAUAUUGGCAUGUAUUUAACAAAAGAAAACAAAAAAGAA
env-875  UUAUAU.UUGCCGCG.............CUAAUUAA..........AAU..................UUAAUUAG......................................CUAC.AUAAA.AAUCAUU...UAAAA..UAAUAUUGGCAUGUAUUUAACAAAAGAAAACAAAAAAGAA
env-876  UUAUAU.UUGCCGCG.............CUAAUUAA..........AAU..................UUAAUUAG......................................CUAC.AUAAA.AAUCAUU...UAAAA..UAAUAUUGGCAUGUAUUUAACAAAAGAAAACAAAAAAGAA
env-877  UUAUAU.UUGCCGCG.............CUAAUUAA..........AAU..................UUAAUUAG......................................CUAC.AUAAA.AAUCAUU...UAAAA..UAAUAUUGGCAUGUAUUUAACAAAAGAAAACAAAAAAGAA
env-878  UUAUAU.UUGCCGCG.............CUAAUUAA..........AAU..................UUAAUUAG......................................CUAC.AUAAA.AAUCAUU...UAAAA..UAAUAUUGGCAUGUAUUUAACAAAAGAAAACAAAAAAGAA
env-879  UUAUAU.UUGCCGCG.............CUAAUUAA..........AAU..................UUAAUUAG......................................CUAC.AUAAA.AAUCAUU...UAAAA..UAAUAUUGGCAUGUAUUUAACAAAAGAAAACAAAAAAGAA
env-881  UUAUAU.UUGCCGCG.............CUAAUUAA..........AAU..................UUAAUUAG......................................CUAC.AUAAA.AAUCAUU...UAAAA..UAAUAUUGGCAUGUAUUUAACAAAAGAAAACAAAAAAGAA
env-883  UUAUAU.UUGCCGCG.............CUA...............UAA.......................UAG......................................CUAC.AUAAG.AAUCAUU...UU.AA..UAAUAUUGGCAUGUAUUUAACAAAAGAAAACAAAAAAGAA
env-884  UUAUAU.UUGCCGCG.............CUA...............UAA.......................UAG......................................CUAC.AUAAG.AAUCAUU...UU.AA..UAAUAUUGGCAUGUAUUUAACAAAAGAAAACAAAAAAGAA
env-886  UUAUAU.UUGCCGCG.............CUA...............UAA.......................UAG......................................CUAC.AUAAG.AAUCAUU...UU.AA..UAAUAUUGGCAUGUAUUUAACAAAAGAAAACAAAAAAGAA
env-887  UUAUAU.UUGCCGCG.............CUA...............UAA.......................UAG......................................CUAC.AUAAG.AAUCAUU...UU.AA..UAAUAUUGGCAUGUAUUUAACAAAAGAAAACAAAAAAGAA
env-888  UUAUAU.UUGCCGCG.............CUA...............UAA.......................UAG......................................CUAC.AUAAG.AAUCAUU...UU.AA..UAAUAUUGGCAUGUAUUUAACAAAAGAAAACAAAAAAGAA
env-889  UUAUAU.UUGCCGCG.............CUA...............UAA.......................UAG......................................CUAC.AUAAG.AAUCAUU...UU.AA..UAAUAUUGGCAUGUAUUUAACAAAAGAAAACAAAAAAGAA
env-890  UUAUAU.UUGCCGCG.............CUA...............UAA.......................UAG......................................CUAC.AUAAG.AAUCAUU...UU.AA..UAAUAUUGGCAUGUAUUUAACAAAAGAAAACAAAAAAGAA
env-891  UUAUAU.UUGCCGCG.............CUA...............UAA.......................UAG......................................CUAC.AUAAG.AAUCAUU...UU.AA..UAAUAUUGGCAUGUAUUUAACAAAAGAAAACAAAAAAGAA
env-892  UUAUAU.UUGCCGCG.............CUA...............UAA.......................UAG......................................CUAC.AUAAG.AAUCAUU...UU.AA..UAAUAUUGGCAUGUAUUUAACAAAAGAAAACAAAAAAGAA
env-894  UUAUAU.UUGCCGCG.............CUA...............UAA.......................UAG......................................CUAC.AUAAG.AAUCAUU...UU.AA..UAAUAUUGGCAUGUAUUUAACAAAAGAAAACAAAAAAGAA
env-895  UUAUAU.UUGCCGCG.............CUA...............UAA.......................UAG......................................CUAC.AUAAG.AAUCAUU...UU.AA..UAAUAUUGGCAUGUAUUUAACAAAAGAAAACAAAAAAGAA
env-896  UUAUAU.UUGCCGCG.............CUA...............UAA.......................UAG......................................CUAC.AUAAG.AAUCAUU...UU.AA..UAAUAUUGGCAUGUAUUUAACAAAAGAAAACAAAAAAGAA
env-897  UUAUAU.UUGCCGCG.............CUA...............UAA.......................UAG......................................CUAC.AUAAG.AAUCAUU...UU.AA..UAAUAUUGGCAUGUAUUUAACAAAAGAAAACAAAAAAGAA
env-898  UUAUAU.UUGCCGCG.............CUA...............UAA.......................UAG......................................CUAC.AUAAG.AAUCAUU...UU.AA..UAAUAUUGGCAUGUAUUUAACAAAAGAAAACAAAAAAGAA
env-899  UUAUAU.UUGCCGCG.............CUA...............UAA.......................UAG......................................CUAC.AUAAG.AAUCAUU...UU.AA..UAAUAUUGGCAUGUAUUUAACAAAAGAAAACAAAAAAGAA
env-900  UUAUAU.UUGCCGCG.............CUA...............UAA.......................UAG......................................CUAC.AUAAG.AAUCAUU...UU.AA..UAAUAUUGGCAUGUAUUUAACAAAAGAAAACAAAAAAGAA
env-901  UUAUAU.UUGCAUCG.............CAA...............AAA.......................UUG.....................................CAUAC.AUAAA.AAUCAUU...UAAAA..CAAUAUUGGCAUGUAUUUAAAGAAAGAAGUAAAAGAAACG
env-902  UUAUAU.UUGCAUCG.............CAA...............AAA.......................UUG.....................................CAUAC.AUAAA.AAUCAUU...UAAAA..CAAUAUUGGCAUGUAUUUAAAGAAAGAAGUAAAAGAAACG
env-903  UUAUAU.UUGCACCG.............CAA...............AAA.......................UUG.....................................CGUAC.AUAAA.AAUCAUU...UAAAA.ACAAUAUUGGCAUGUAUUUAAACAAAGAAGUAAAAGAAACG
env-905  UUAUAU.UUGCACCG.............CAA...............AAA.......................UUG.....................................CGUAC.AUAAA.AAUCAUU...UAAAA.ACAAUAUUGGCAUGUAUUUAAACAAAGAAGUAAAAGAAACG
env-906  CUAUCU.UUGCGCAC.............GAAU.UAGG.........AAAG................UCUUGAUUC......................................AUAC.AUAAU.AAUUAUU...UA.AA..UAAUAUUGGAAUGUAUUUAACGAAAGAAGUUAAAGAAGAA
env-907  CUAUCU.UUGCGCAC.............GAAU.UAGG.........AAAG................UCUUGAUUC......................................AUAC.AUAAU.AAUUAUU...UA.AA..UAAUAUUGGAAUGUAUUUAACGAAAGAAGUUAAAGAAGAA
env-908  CUAUCU.UUGCGCAC.............GAAU.UAGG.........AAAG................UCUUGAUUC......................................AUAC.AUAAU.AAUUAUU...UA.AA..UAAUAUUGGAAUGUAUUUAACGAAAGAAGUUAAAGAAGAA
Fge-1-1  UUAUCU.UUGCACCC.............GAAUUA............CUACCAUA...............UAAUUC......................................AUAC.AUAAU.AAUCAUU...UAAAG..CAAUAUUAGCAUGUACUUAACUAAAGAAGUAAAAGCAGAA
Ale-2-1  GUACUU.UUGCCGCU.............CGAUUACC..........AAAGC................GGUCGUCG......................................GUAC.AUAAA.AAUCAUU...UA.AA..UAAUAUUGGCAUGUAUUUAGCACCAGAAGAAAAAGAAAAG
env-909  UUACCU.UUGCAACC.............GAAUUAGG..........AAAAC................CCCAAUUC......................................AUAC.AUAAU.AAUCAUU...UA.AA..CAAUAUUAGCAUGUACUUAACUUCAGAAGUAAAAGCUGAG
env-910  GUACUU.UUGCCGCU.............UGAUUAUC..........AAAGA................GGUAGUCA......................................GUAC.AUAAA.AAUCAUU...UA.AA..UAAUAUUGGAAUGUAUUUAGCACCAGAAGAAAAAGCAAAA
env-911  CUAUAU.UUGCCGUU.............CAAUUCUGA.........AAA.................UCAGGGUUA......................................UUAC.AUAAA.AAUUAUU...AA.AA..UAAUAUUGGAAUGUAUCUAACAAAAAAAAUUAAAGAGAAU
env-912  CUAUAU.UUGCCGUU.............CAAUUCUGA.........AAA.................UCAGAGUUA......................................UUAC.AUAAA.AAUUAUU...AA.AA..UAAUAUUGGAAUGUAUCUAACAAAAAAAAAUUAAAGAGAU
env-913  CUAUAU.UUGCCAAU.............CAAUUCUGA.........AAA.................UCAGAGUUA......................................UUAC.AUAAA.AAUUAUU...AAUAA..UAAUAUUGGAAUGUAUCUAACAAAAAAAAAUUAAAGAGAU
env-914  CUAUAU.UUGCCACU.............CAAUUCUGA.........AAA.................UCAGAGUUA......................................UUAC.AUAAA.AAUUAUU...AAAAA..UAAUAUUGGAAUGUAUUUAACAAAAAAAAAUUAAAGAGAU
env-915  CUAUCU.UUGCCACU.............CAAUUCUGA.........AAA.................UCAGAGUUA......................................UUAC.AUAAA.AAUCAUU...AAAAA..UAAUAUUGGAAUGUAUUUAACAAAAAAAAAUUAAAGAGAU
env-916  CUAUAU.UUGCCACU.............CAAUUCUGA.........AAA.................UCAGAGUUA......................................UUAC.AUAAA.AAUUAUU...AA.AA..UAAUAUUGGAAUGUAUCUAACAAAAAAAAAUUAAAGAGAU
Man-1-1  GUAUAU.UUGCAGCC.............UGAUUGA...........AAA...................UCAUUCA.....................................GAUAC.AUAAA.AAUCAUU...AA.AC..GAAUAUUGGAAUGUAUUUAACAAAAGAAGUAAAAGCCGAA
env-917  CUAAAU.UUGCACCC.............GUUC..............AAC......................GGAA......................................CUAC.AUAAU.AAUCAUU...UACAA..CAAUAUUAGCAUGUACUUAGAUAAAGAAGUAAAAAAGAAA
env-919  CUAAAU.UUGCACCC.............GUUC..............AAC......................GGAA......................................CUAC.AUAAU.AAUCAUU...UACAA..CAAUAUUAGCAUGUACUUAGAUAAAGAAGUAAAAAAGAAG
env-920  CUAAAU.UUGCACCC.............GUUC..............AAC......................GGAA......................................CUAC.AUAAU.AAUCAUU...UACAA..UAAUAUUAGCAUGUACUUAGAUAAAGAAGUAAAAAAGAAA
env-921  CUAAAU.UUGCACCC.............GUUC..............AAC......................GGAA......................................CUAC.AUAAU.AAUCAUU...UACAA..UAAUAUUAGCAUGUACUUAGAUAAAGAAGUAAAAAAGAAA
env-922  CUAAAU.UUGCACCC.............GUUC..............AAC......................GGAA......................................CUAC.AUAAU.AAUCAUU...UACAA..UAAUAUUAGCAUGUACUUAGAUAAAGAAGUAAAAAAGAAA
env-923  CUAAAU.UUGCACCC.............GUUC..............AAC......................GGAA......................................CUAC.AUAAU.AAUCAUU...UACAA..UAAUAUUAGCAUGUACUUUAGAUAAAGAAGUAAAAAGAAA
env-924  CUAAAU.UUGCACCC.............GUUC..............AAC......................GGAA......................................CUAC.AUAAU.AAUCAUU...UACAA..UAAUAUUAGCAUGUACUUAGAUAAAGAAGUAAAAAAGAAA

         ............................<<<<<<<<<<<<<<<<<<...........>>>>>>>>>>>>>>>>>>..........<<<<<<<<........>>>>>>>>.........................................................................
         ............................2222221111..101.00...........00.101..1111222222..........11011000........00011011.........................................................................
         •UAY•U-UUGC••CY-------------∘∘∘∘∘∘∘-----------R•R∘∘-----------------∘∘∘∘∘∘∘------------------------------------•UAC-AUAA•-AAUCAUU---UA-AA--YAAUAUURG•AUGUAUUARC•ARAGAAR••AAARRRRAR
         5′                                                                                                                                                               3′
```

```
env-925   CUAAAU.UUGCACCC.............GUUC..............AAC......................GGAA.....................................CUAC.AUAAU.AAUCAUU...UACAA..UAAUAUUAGCAUGUACUUAGAUAA................
env-927   CUAAAU.UUGCACCC.............GUUC..............AAC......................GGAA.....................................CUAC.AUAAU.AAUCAUU...UACAA..UAAUAUUAGCAUGUACUUAGAUAAAGAAGUAAAAAAGAAA
env-928   CUAAAU.UUGCACCC.............GUUC..............AGA......................GGAA.....................................CUAC.AUAAU.AAUCAUU...UACAA..CAAUAUUGGCAUGUAUUUAAAUAAAGAAGUAAAAAAGAAG
env-929   CUAAAU.UUGCACCC.............GUUC..............AGA......................GGAA.....................................CUAC.AUAAU.AAUCAUU...UACAA..CAAUAUUGGCAUGUAUUUAAAUAAAGAAGUAAAAAAGAAG
env-930   CUAAAU.UUGCACCC.............GUUC..............AGA......................GGAA.....................................CUAC.AUAAU.AAUCAUU...UACAA..CAAUAUUGGCAUGUAUUUAAAUAAAGAAGUAAAAAAGAAG
env-932   AUACAU.UUGCACCC.............GUUC..............AAUG.....................GAAC.....................................AUAC.AUAAU.AAUCAUU...UACAA..UAAUAUUAGCAUGUACUUAGACAAAGAAGUUAAAAAGAAA
env-933   UUACCU.UUGCAACC.............GAAU.UAGGA........AAA................UCCUAAAUUC.....................................AUAC.AUAAA.AAUCAUU...UAAUA..UAAUAUUGGAAUGUAUUUAACGAAAGAAACUAAAGCUGAA
env-934   UUACCU.UUGCAACC.............GAA.UUAGG.........AAACA...............CCU.AUUUC.....................................AUAC.AUAAA.AAUCAUU...UAAAA..UAAUAUUGGAAUGUAUUUAACUAAAGAAACAAAAGCUGAA
env-936   CUAUCU.UUGCAGCU.............GUGUU.............AAAAGA..................AACAC.....................................GUAC.AUAAA.AAUCAUU...AAAAA..CAAUAUUGGAAUGUAUUUAGCAACUGAAAAGAAAGAAGAA
env-937   CUAUAU.UUGCGCGU.............CUAUUAU...........AAA...................AUAGUAG.................................AAAUAUAC.AUAAU.AAUCAUU...UAAAA..UAAUAUUGGAAUGUAUUUAACUAAAAAAACAAAGAAUCA
env-938   UUAUAU.UUGCAUUCA............AU................UUA........................AU....................................CAUAC.AUAAA.AAUCAUU...ACAAA..UAAUAUUGGCAUGUAUUUAUCAAAGGAAAUAAAAGAAAAG
env-939   UUAUAU.UUGCAGUCA............AU................UUA........................AU....................................CAUAC.AUAAA.AAUCAUU...AAAAA..UAAUAUUGGCAUGUAUUUAUCAAAGGAAAUAAAAGAAAAG
env-940   UUAUAU.UUGCAGUCA............AU................UUA........................AU....................................CAUAC.AUAAA.AAUCAUU...AAAAA..UAAUAUUGGCAUGUAUUUAUCAAAGGAACUAAAAGAAAAG
env-941   UUAUAU.UUGCAGUCA............AU................UUA........................AU....................................CAUAC.AUAAA.AAUCAUU...AAAAA..UAAUAUUGGCAUGUAUUUAUCAAAGGAACUAAAAGAAAAG
env-942   UUAUAU.UUGCAGUCA............AU................UUA........................AU....................................CAUAC.AUAAA.AAUCAUU...AAAAA..UAAUAUUGGCAUGUAUUUAUCAAAGGAAAUAAAAGAAAAG
env-943   UUAUAU.UUGCAGUCA............AU................UUA........................AU....................................CAUAC.AUAAA.AAUCAUU...AAAAA..UAAUAUUGGCAUGUAUUUAUCAAAGGAAAUAAAAGAAAAG
env-944   UUAUAU.UUGCAGUCA............AU................UUA........................AU....................................CAUAC.AUAAA.AAUCAUU...AAAAA..UAAUAUUGGCAUGUAUUUAUCAAAGGAACUAAAAGAAAAG
env-945   UUAUAU.UUGCAGUCA............AU................UUA........................AU....................................CAUAC.AUAAA.AAUCAUU...AAAAA..UAAUAUUGGCAUGUAUUUAUCAAAGGAAAUAAAAGAAAAG
env-946   UUAUAU.UUGCAGUCA............AU................UUA........................AU....................................CAUAC.AUAAA.AAUCAUU...AAAAA..UAAUAUUGGCAUGUAUUUAUCAAAGGAA............
Tma-1-1   GUAUAU.UUGCUGCU.............UUGAUGA...........AAAC..................UCAUCAA....................................UAUAC.AUAAU.AAUCAUU...UA.AA..UAAUAUUGGAAUGUAUUUAACUAAAGAAGUAAAAGAAGGA
env-947   CUAUAU.UUGCACCC.............GUCUA.............AAG.....................UAGAC.....................................AUAC.AUAAG.AAUCAU....UA.AA..UAAUAUUGGAAUGUAUUUAGCACCAGAAAAGAAAACAGAA
Fsp-12-1  UUAUUU.UUGCACCC.............GUCU.UACAA........AAG................UUGUAAAGAC.....................................AUAC.AUAAA.AAUCAU....UA.AA..UAAUAUUGGAAUGUAUUUAACUAAGGAAAAGAAAGAAGAG
Fde-1-1   UUAAUU.UUGCACCC.............GUC.UUACAA........AAG................UUGUAUUGAC.....................................AUAC.AUAAA.AAUCAU....UA.AA..UAAUAUUGGAAUGUAUUUAACUAAAGAAAAGAAAGAAGAG
Fba-7-1   AUAUAU.UUGCACCC.............UUGAAAG...........AAAAC.................CUUUCAA.....................................AUAC.AUAAG.AAUCAUU...UA.AA..UAAUAUUGGAAUGUAUUUAACAAAAGAAGUUAAAGAGGAA
Ale-1-1   UUAUAU.UUGCACGC.............UUGAAAG...........AAAAA.................CUUUCAA....................................GCUAC.AUAAC.AAUCAUU...UACAA..UAAUAUUAGAAUGUAUUUAUCUAAAGAAGAGAAACAAGAA
Ale-1-2   UUAUAU.UUGCACGC.............UUGAAAG...........AAAAA.................CUUUCAA....................................GCUAC.AUAAC.AAUCAUU...UACAA..UAAUAUUAGAAUGUAUUUAUCUAAAGAAGAGAAACAAGAA
Ffr-1-1   UUACUU.UUGCACCG.............UCUAACA...........AUAGA.................UGCUAGA....................................CAUAC.AUAAA.AAUCAUU...AA.AC..AAAUAUUGGAAUGUAUUUAACUAAAGAGAUUAAAGAAGAC
env-948   UUAUAU.UUGCAGCC.............GAAUUG............CCACCAUA...............CAAUUC.....................................AUAC.AUAAA.AAUCAUU...AAAUA..UAAUAUUGGCAUGUACUUGACAAAAGAAACUAAAGCUGAA
Fsp-1-1   UUAUAU.UUGCAGCC.............GAAUUG............CUACCAUA...............CAAUUC.....................................AUAC.AUAAU.AAUCAUU...UA.AA..UAAUAUUGGCAUGUACUUAACUAAUGAAGUUAAAGCUGAA
env-949   CUAUCU.UUGCACCC.............GAAUUG............CAAUCAUA...............CAAUUC.....................................AUAC.AUAAA.AAUCAUU...UAAAA..UAAUAUUGGCAUGUACUUGACAAAAGAAACUAAAGCUGAA
env-950   UUAUCU.UUGCACCC.............GAAUUG............CUACCAUA...............CAAUUC.....................................AUAC.AUAAA.AAUCAUU...UAAAA..UAAUAUUGGCAUGUACUUGACAAAAGAAACUAA.......
Fsa-2-1   CUAUCU.UUGCACCC.............GAAUUG............CAACCAUA...............CAAUUC.....................................AUAC.AUAAA.AAUCAUU...UAAAA..UAAUAUUGGCAUGUACUUGACAAAAGAAACUAAAGCUGAA
env-951   CUACCU.UUGCAACC.............GAAUUG............CAACAAUA...............CAAUUC.....................................AUAC.AUAAU.AAUCAUU...UAAAA..UAAUAUUAGCAUGUACUUGACUAAAGAAA..........
env-952   CUACCU.UUGCAUCC.............GAAUUG............CAACNAUA...............CAAUUC.....................................AUAC.AUAAU.AAUCAUU...UAAAA..UAAUAUUAGCAUGUACUUGNCUAAAGAAACUAAAGCUGAA
Fba-3-1   CUACCU.UUGCAUCC.............GAAUUG............CAACCAUA...............CAAUUC.....................................AUAC.AUAAU.AAUCAUU...UAAAA..UAAUAUUAGCAUGUACUUGACUAAAGAAACAAAAGCUGAA
env-953   UUACCU.UUGCAUCC.............GAAUUG............CAACCAUA...............CAAUUC.....................................AUAC.AUAAU.AAUCAUU...UAAAA..UAAUAUUAGCAUGUACUUGUCUAAAGAAACUAAAGCCGAA
env-954   CUACCU.UUGCAUCC.............GAAUUG............CAACAAUA...............CAAUUC.....................................AUAC.AUAAU.AAUCAUU...UAAAA..UAAUAUUAGCAUGUACUUGUCUAAAGAAACUAAAGCUGAA
env-955   CUAUAU.UUGCCCCC.............UAUUUUA...........GUUAA.................UAGAAUA...................................UCAUAC.AUAAU.AAUCAUU...UA.AA..UAAUAUUGGAAUGUAUUUAACAAAAAAAUCAAAGAAACU
env-956   CUAUAU.UUGCCCCC.............UAUUUUA...........GUUAA.................UAGAAUA...................................UCAUAC.AUAAU.AAUCAUU...UA.AA..UAAUAUUGGAAUGUAUUUAACAAAAAAAUCAAAGAAACU
env-957   CUAUAU.UUGACCCC.............UAUUUUA...........GUUAA.................UAGAAUA...................................UCAUAC.AUAAU.AAUCAUU...UA.AA..UAAUAUUGGAAUGUAUUUAACAAAAAAAUCAAAGAAACU
env-958   GUAUAU.UUGCGCCGAUGAUAUAAGCC......................................................................................UAC.AUAAU.AAUCAUU...UA.AA..UAAUAUUGGAAUGUAUUUAAAUAAAAAAACUAAAGAAGAU
env-959   GUACUU.UUGCCCCC.............GA................AAAAC.......................UC.....................................GUAC.AUAAU.AAUCAUU...AA.AU..UAAUAUUGGAAUGUAUUUAACAAAAGAAGUAAAAGCGGAU
Csp-1-1   CUACCU.UUGCGCCC.............GAA...............AAAA.......................UUC.....................................GUAC.AUAAU.AAUCAUU...AA.AU..UAAUAUUGGAAUGUAUUUAACUAAAGAAGUAAAAGCGGAA
env-960   CUACCU.UUGCGCCC.............GAA...............AAAA.......................UUC.....................................GUAC.AUAAU.AAUCAUU...AA.AU..UAAUAUUGGAAUGUAUUUAACUAAAGAAGUAAAAGCGGAA
env-961   CUACCU.UUGCGCCC.............GAA...............AAAA.......................UUC.....................................GUAC.AUAAU.AAUCAUU...AA.AU..UAAUAUUGGAAUGUAUUUAACUAAAGAAGUAAAAGCGGAA
Csp-3-1   CUACCU.UUGCGCCC.............GAA...............AAAA.......................UUC.....................................GUAC.AUAAU.AAUCAUU...AA.AU..UAAUAUUGGAAUGUAUUUAACUAAAGAAGUAAAAGCGGAA
env-962   CUACCU.UUGCGCCC.............GAA...............AAAA.......................UUC.....................................GUAC.AUAAU.AAUCAUU...AA.AU..UAAUAUUGGAAUGUAUUUAACUAAAGAAGUAAAAGCGGAA
env-965   CUACCU.UUGCGCCCGA...........AA................AAAAA.......................UU.....................................GUAC.AUAAU.AAUCAUU...AA.AU..UAAUAUUGGAAUGUAUUUAACUAAAGAAGUAAAAGCGGAA
env-966   CUACCU.UUGCGGCCG............AAA...............AAA.......................UUU.....................................GUAC.AUAAU.AAUCAUU...AA.AU..UAAUAUUGGAAUGUAUUUAACUAAAGAAGUAAAAGCGGAA
env-967   CUACCU.UUGCGCCCG............AAA...............AAA.......................UUU.....................................GUAC.AUAAU.AAUCAUU...AA.AU..UAAUAUUGGAAUGUAUUUAACUAAAGAAGUAAAAGCGGAA
env-968   CUACCU.UUGCGCCCG............AAA...............AAA.......................UUU.....................................GUAC.AUAAU.AAUCAUU...AA.AU..UAAUAUUGGAAUGUAUUUAACUAAAGAAGUAAAAGCGGAA
env-972   CUACCU.UUGCGCCCG............AAA...............AAA.......................UUU.....................................GUAC.AUAAU.AAUCAUU...AA.AU..UAAUAUUGGAAUGUAUUUAACUAAAGAAGUAAAAGCGGAA
env-973   CUACCU.UUGCGCCCG............AAA...............AAA.......................UUU.....................................GUAC.AUAAU.AAUCAUU...AA.AU..UAAUAUUGGAAUGUAUUUAACUAAAGAAGUAAAAGCGGAA
env-974   CUACCU.UUGCGCCCG............AAA...............AAA.......................UUU.....................................GUAC.AUAAU.AAUCAUU...AA.AU..UAAUAUUGGAAUGUAUUUAACUAAAGAAGUAAAAGCGGAA
env-976   CUACCU.UUGCGCCCG............AAA...............AAA.......................UUU.....................................GUAC.AUAAU.AAUCAUU...AA.AU..UAAUAUUGGAAUGUAUUUAACUAAAGAAGUAAAAGCGGAA
env-977   CUACCU.UUGCGCCCG............AAA...............AAA.......................UUU.....................................GUAC.AUAAU.AAUCAUU...AA.AU..UAAUAUUGGAAUGUAUUUAACUAAAGAAGUAAAAGCGGAA
env-978   CUACCU.UUGCGCCCG............AAA...............AAA.......................UUU.....................................GUAC.AUAAU.AAUCAUU...AA.AU..UAAUAUUGGAAUGUAUUUAACUAAAGAAGUAAAAGCGGAA
env-980   CUACCU.UUGCGCCCG............AAA...............AAA.......................UUU.....................................GUAC.AUAAU.AAUCAUU...AA.AU..UAAUAUUGGAAUGUAUUUAACUAAAGAAGUAAAAGCGGAA
env-982   CUACCU.UUGCGCCCG............AAA...............AAA.......................UUU.....................................GUAC.AUAAU.AAUCAUU...AA.AU..UAAUAUUGGAAUGUAUUUAACUAAAGAAGUAAAAGC....
env-983   CUACCU.UUGCGCCCG............AAA...............AAA.......................UUU.....................................GUAC.AUAAU.AAUCAUU...AA.AU..UAAUAUUGGAAUGUAUUUAACUAAAGAAGUAAAAGCGGAA
env-984   CUACCU.UUGCGCCCG............AAA...............AAA.......................UUU.....................................GUAC.AUAAU.AAUCAUU...AA.AU..UAAUAUUGGAAUGUAUUUAACUAAAGAAGUAAAAGCGGAA
env-985   CUACCU.UUGCGCCCG............AAA...............AAA.......................UUU.....................................GUAC.AUAAU.AAUCAUU...AA.AU..UAAUAUUGGAAUGUAUUUAACUAAAGAAGUAAAAGCGGAA
env-986   CUACCU.UUGCGCCCG............AAA...............AAA.......................UUU.....................................GUAC.AUAAU.AAUCAUU...AA.AU..UAAUAUUGGAAUGUAUUUAACUAAAGAAGUAAAAGCGGAA
env-988   CUACCU.UUGCGCCCG............AAA...............AAA.......................UUU.....................................GUAC.AUAAU.AAUCAUU...AA.AU..UAAUAUUGGAAUGUAUUUAACUAAAGAAGUAAAAGCGGAA
env-989   CUACCU.UUGCGCCCG............AAA...............AAA.......................UUU.....................................GUAC.AUAAU.AAUCAUU...AA.AU..UAAUAUUGGAAUGUAUUUAACUAAAGAAGUAAAAGCGGAA
env-990   CUACCU.UUGCGCCCG............AAA...............AAA.......................UUU.....................................GUAC.AUAAU.AAUCAUU...AA.AU..UAAUAUUGGAAUGUAUUUAACUAAAGAAGUAAAAGCGGAA
env-993   CUACCU.UUGCGCCCG............AAA...............AAA.......................UUU.....................................GUAC.AUAAU.AAUCAUU...AA.AU..UAAUAUUGGAAUGUAUUUAACUAAAGAAGUAAAAGCGGAA
env-994   CUACCU.UUGCGCCCG............AAA...............AAA.......................UUU.....................................GUAC.AUAAU.AAUCAUU...AA.AU..UAAUAUUGGAAUGUAUUUAACUAAAGAAGUAAAAGCGGAA
env-995   CUACCU.UUGCGCCCG............AAA...............AAA.......................UUU.....................................GUAC.AUAAU.AAUCAUU...AA.AU..UAAUAUUGGAAUGUAUUUAACUAAAGAAGUAAAAGCGGAA
env-999   CUACCU.UUGCGCCCG............AAA...............AAA.......................UUU.....................................GUAC.AUAAU.AAUCAUU...AA.AU..UAAUAUUGGAAUGUAUUUAACUAAAGAAGUAAAAGCGGAA
env-1001  CUACCU.UUGCGCCCG............AAA...............AAA.......................UUU.....................................GUAC.AUAAU.AAUCAUU...AA.AU..UAAUAUUGGAAUGUAUUUAACUAAAGAAGUAAAAGCGGAA
env-1002  CUACCU.UUGCGCCCG............AAA...............AAA.......................UUU.....................................GUAC.AUAAU.AAUCAUU...AA.AU..UAAUAUUGGAAUGUAUUUAACAAAAGAAGUAAAAGCGGAA
env-1003  CUACCU.UUGCGCCCG............AAA...............AAA.......................UUU.....................................GUAC.AUAAU.AAUCAUU...AA.AU..UAAUAUUGGAAUGUAUUUAACUAAAGAAGUAAAAGCGGAA
env-1004  CUACCU.UUGCGCCCG............AAA...............AAA.......................UUU.....................................GUAC.AUAAU.AAUCAUU...AA.AU..UAAUAUUGGAAUGUAUUUAACUAAAGAAGUAAAAGCGGAA
env-1005  CUACCU.UUGCGCCCG............AAA...............AAA.......................UUU.....................................GUAC.AUAAU.AAUCAUU...AA.AU..UAAUAUUGGAAUGUAUUUAACUAAAGAAGUAAAAGCGGAA
env-1006  CUACCU.UUGCGCCCG............AAA...............AAA.......................UUU.....................................GUAC.AUAAU.AAUCAUU...AA.AU..UAAUAUUGGAAUGUAUUUAACAAAAGAAGUAAAAGCGGAA
env-1007  CUACCU.UUGCGCCCG............AAA...............AAA.......................UUU.....................................GUAC.AUAAU.AAUCAUU...AA.AU..UAAUAUUGGAAUGUAUUUAACAAAAGAAGUAAAAGCGGAA
env-1008  CUACCU.UUGCGCCCG............AAA...............AAA.......................UUU.....................................GUAC.AUAAU.AAUCAUU...AA.AU..UAAUAUUGGAAUGUAUUUAACUAAAGAAGUAAAAGCGGAA
env-1009  CUACCU.UUGCGCCCG............AAA...............AAA.......................UUU.....................................GUAC.AUAAU.AAUCAUU...AA.AU..UAAUAUUGGAAUGUAUUUAACUAAAGAAGUAAAAGCGGAA
env-1010  CUACCU.UUGCGCCCG............AAA...............AAA.......................UUU.....................................GUAC.AUAAU.AAUCAUU...AA.AU..UAAUAUUGGAAUGUAUUUAACUAAAGAAGUAAAAGCGGAA
env-1011  CUACCU.UUGCGCCCG............AAA...............AAA.......................UUU.....................................GUAC.AUAAU.AAUCAUU...AA.AU..UAAUAUUGGAAUGUAUUUAACUAAAGAAGUAAAAGCGGAA
env-1012  AUAUAU.UUGCACCG.............CU................AAAU.......................GG....................................CAUAC.AUAAA.AAUCAUU...UAAAA..UAAUAUUGGCAUGUAUUUAACAAAAAAAAACAAAGAGGAU
env-1013  AUAUAU.UUGCACCG.............CU................AAAU.......................GG....................................CAUAC.AUAAA.AAUCAUU...UAAAA..UAAUAUUGGCAUGUAUUUAACAAAGAAAAAUAAAGAAGAU

          ............................<<<<<<<<<<<<<<<<<<...........>>>>>>>>>>>>>>>>>>.........<<<<<<<<........>>>>>>>>.........................................................................
          ............................2222221111..101.00...........00.101..1111222222.........11011000........00011011.........................................................................
          •UAY•U-UUGC••CY-------------○○○○○○○-----------R•R○○----------------○○○○○○○---------------------------------------○UAC-AUAA•-AAUCAUU---UA-AA--YAAUAUURG•AUGUAUUUARC•ARAGAAR••AAARRRRAR
          5′                                                                                                                                                       3′
```

```
env-1015  AUAUAU.UUGCACCG.............CU..................AAAU.......................GG....................................CAUAC.AUAAA.AAUCAUU...UAAAA..UAAUAUUGGCAUGUAUUUAACAAAAAAAAACAAAGAGGAU
env-1016  AUAUAU.UUGCACCG.............CU..................AAAU.......................GG....................................CAUAC.AUAAA.AAUCAUU...UAAAA..UAAUAUUGGCAUGUAUUUAACAAAAAAAAACAAAGAGGAU
env-1017  AUAUAU.UUGCGCCG.............CU..................GAAU.......................AG....................................CAUAC.AUAAA.AAUCAUU...UAAAA..UAAUAUUGGAAUGUAUUUAACAAAAAAAAACAAAGAAGAU
env-1018  AUAUAU.UUGCGCCG.............CU..................GAAU.......................AG....................................CAUAC.AUAAA.AAUCAUU...UAAAA..UAAUAUUGGAAUGUAUUUAACAAAAAAAAACAAAGAAGAU
env-1019  UUAUAU.UUGCGCCG.............CU..................GAAU.......................AG....................................CAUAC.AUAAA.AAUCAUU...UAAAA..UAAUAUUGGCAUGUAUUUAACAAAAAAAAACAAAGAAGAU
env-1021  UUAUAU.UUGCGCCG.............CU..................GAAU.......................AG....................................CAUAC.AUAAA.AAUCAUU...UAAAA.ACAAUAUUGGCAUGUAUUUAACAAAAAAAAACAAAGAAGAU
env-1022  AUAUAU.UUGCGCCG.............CU..................GAAU.......................AG....................................CAUAC.AUAAA.AAUCAUU...UAAAA..CAAUAUUGGCAUGUAUUUAACAAAAAAAAACAAAGAAGAU
env-1023  AUAUAU.UUGCGCCG.............CU..................GAUU.......................AG....................................CAUAC.AUAAA.AAUCAUU...UAAAA..UAAUAUUGGCAUGUAUUUAACAAAAAAAAAUAAAGAGGAU
env-1024  AUAUAU.UUGCGCCG.............CU..................GAUU.......................AG....................................CAUAC.AUAAA.AAUCAUU...UAAAA..UAAUAUUGGCAUGUAUUUAACAAAAAAAAACAAAGAAGAU
env-1025  AUAUAU.UUGCGCCG.............CU..................GAUU.......................AG....................................CAUAC.AUAAA.AAUCAUU...UAAAA..UAAUAUUGGCAUGUAUUUAACAAAAAAAAACAAAGAAGAU
env-1026  AUAUAU.UUGCGCCG.............CU..................AAAG.......................AG....................................CAUAC.AUAAA.AAUCAUU...UAAAA..UAAUAUUGGCAUGUAUUUAACAAAAAAAAACAAAGAAGAU
env-1027  AUAUAU.UUGCGCCG.............CU..................GAAG.......................AG....................................CAUAC.AUAAA.AAUCAUU...UAAAA..UAAUAUUGGCAUGUAUUUAACAAAAAAAAAUAAAGAAGAU
env-1028  GUAUAU.UUGCGCCG.............CUAA..............UAU......................UUGC.....................................AUAC.AUAAU.AAUCAUU...AA.AU..UAAUAUUGGAAUGUAUUUAACAAAGAAAAACAAAGAAGAU
env-1029  UUAUAU.UUGCACCA.............CUU...............AUAU......................AAG....................................UAUAC.AUAAU.AAUCAUU...UA.AA..UAAUAUUGGAAUGUAUUUAACAAAAAAAACUAAAGAAGAU
env-1030  UUAUAU.UUGCACCA.............CUU...............AUAU......................AAG....................................UAUAC.AUAAU.AAUCAUU...UA.AA..UAAUAUUGGAAUGUAUUUAACAAAAAAA............
env-1032  UUAUAU.UUGCACCA.............CUU...............AUAU......................AAG....................................UAUAC.AUAAU.AAUCAUU...UA.AA..UAAUAUUGGAAUGUAUUUAACAAAAAAAACUAAAGAAGAU
env-1033  UUAUAU.UUGCACCA.............CUU...............AUAU......................AAG....................................UAUAC.AUAAU.AAUCAUU...UA.AA..UAAUAUUGGAAUGUAUUUAACAAAAAAAACAAAGAAGAUU
env-1034  UUAUAU.UUGCACCA.............CUU...............AUAU......................AAG....................................UAUAC.AUAAU.AAUCAUU...UA.AA..UAAUAUUGGAAUGUAUUUAACAAAAAAAACUAAAGAAGAU
env-1035  UUAUAU.UUGCGCCA.............CU................GGUAUA.....................AG....................................UAUAC.AUAAU.AAUCAUU...UA.AA..UAAUAUUGGAAUGUAUUUAACAAAAAAA............
env-1036  GUAUAU.UUGCGCCG.............CUG...............AUAU......................CAG....................................CAUAC.AUAAU.GAUCAAU...UUAAA..UAAUAUUGGAAUGUAUUUAAACAAAAAAACAAAAGAAGAU
env-1037  GUAUAU.UUGCGCCG.............CU................GAUAG......................AG....................................CAUAC.AUAAU.AAUCAUU...UA.AA..UAAUAUUGGAAUGUAUUUAACAAAAAAAACAAAAGAAGAU
env-1038  CUAUAU.UUGCGCCG.............CU................GAUAUA.....................AG....................................CAUAC.AUAAU.AAUCAUU...UA.AA..UAAUAUUGGAAUGUAUUUAACAAAAAAAAAUAAAGAAGAU
env-1039  CUAUAU.UUGCGCCG.............CUA...............UUUA......................UAG....................................CAUAC.AUAAU.AAUCAUU...UA.AA..UAAUAUUGGAAUGUAUUUAACAA.................
env-1040  CUAUAU.UUGCGCCG.............CUA...............UUUA......................UAG....................................CAUAC.AUAAU.AAUCAUU...UA.AA..UAAUAUUGGAAUGUAUUUAACAAA................
env-1041  GUAUAU.UUGCGCCG.............CUAA..............UAU......................UUAG....................................CAUAC.AUAAU.AAUCAUU...UA.AA..UAAUAUUGGAAUGUAUUUAACA..................
env-1042  GUAUAU.UUGCGCCG.............CUAA..............UAU......................UUAG....................................CAUAC.AUAAU.AAUCAUU...UA.AA..UAAUAUUGGAAUGUAUUUAACAAAAAAAACGAAGGAAGAU
env-1044  GUAUAU.UUGCGCCG.............CUAA..............UGUA.....................UUAG....................................CAUAC.AUAAU.AAUCAUU...UA.AA..UAAUAUUGGAAUGUAUUUAACAAAAAAAACGAAAGAAGAU
env-1045  GUAUAU.UUGCGCCG.............CUG...............AUAU......................UAG....................................CAUAC.AUAAU.AAUCAUU...UA.AA..AAAUAUUGGAAUGUAUUUAACAAAAAAAACUAAAGAAGAU
env-1046  UUAUAU.UUGCGCCG.............CUG...............AUAU......................UAG....................................CAUAC.AUAAU.AAUCAUU...AA.AA..UAAUAUUGGAAUGUAUUUAACAAAAAAAACGAAAGAAGAU
env-1047  CUAUAU.UUGCGCCG.............CUG...............AAUA......................CAG....................................CAUAC.AUAAU.AAUCAUU...UA.AA..UAAUAUUGGAAUGUAUUUAACAAAAAAAAAUAAAGAAGAU
env-1048  CUAUAU.UUGCGCCG.............CUG...............AAUA......................UAG....................................CAUAC.AUAAU.AAUCAUU...UA.AA..UAAUAUUGGAAUGUAUUUAACAA.................
env-1049  CUAUAU.UUGCGCCG.............CUG...............AAUA......................UAG....................................CAUAC.AUAAU.AAUCAUU...UA.AA..UAAUAUUGGAAUGUAUUUAACAAAAAAAAAUAAAGAAGAU
env-1051  CUAUAU.UUGCGCCG.............CUG...............AAUA......................UAG....................................CAUAC.AUAAU.AAUCAUU...UA.AA..UAAUAUUGGAAUGUAUUUAACA..................
env-1053  CUAUAU.UUGCGCCG.............CUG...............AAUA......................UAG....................................CAUAC.AUAAU.AAUCAUU...UA.AA..UAAUAUUGGAAUGUAUUUAACAAAAAAAAAUAAAGAAGAU
env-1055  CUAUAU.UUGCGCCG.............CUG...............AAUA......................UAG....................................CAUAC.AUAAU.AAUCAUU...UA.AA..UAAUAUUGGAAUGUAUUUAACAAAAAAAAAUAAAGAAGAU
env-1056  GUAUAU.UUGCGCCG.............CUA...............AUAA......................UAG....................................CAUAC.AUAAU.AAUCAUU...UA.AA..UAAUAUUGGAAUGUAUUUAACAAAAAAAACAAAAGAAGAU
env-1057  CUAUAU.UUGCGCCG.............CUA...............AUAA......................UAG....................................CGUAC.AUAAU.AAUCAUU...UA.AA..UAAUAUUGGAAUGUAUUUAACAAAGAAAAACAAAGAAGAU
env-1058  UUAUAU.UUGCGCCG.............CUG...............AUAU......................CAG....................................UAUAC.AUAAU.AAUCAUU...UA.AA..UAAUAUUGGAAUGUAUUUAACAAAAAAAA...........
env-1059  UUAUAU.UUGCGCCG.............CUG...............UUAA......................CAG....................................CAUAC.AUAAU.AAUCAUU...UA.AA..CAAUAUUGGAAUGUAUUUAACAAAGAAAAAUAAAGAAGAU
env-1060  GUAUAU.UUGCGCCG.............CUG...............UUAA......................CAG....................................CAUAC.AUAAU.AAUCAUU...UA.AA..UAAUAUUGGAAUGUAUUUGACAA.................
env-1061  CUAUAU.UUGCGCCG.............CUG...............AUAA......................CAG....................................CAUAC.AUAAG.AAUCAUU...UA.AA..UAAUAUUGGAAUGUAUUUAACAAAGAAAAAGAAAGAAGAU
env-1062  UUAUAU.UUGCGCCG.............CUG...............AUAA......................CAG....................................CAUAC.AUAAG.AAUCAUU...UA.AA..UAAUAUUGGAAUGUAUUUAACAAAAAAAAAGAAAGAAGAU
env-1063  CUAUAU.UUGCGCCG.............CUG...............AUAA......................CAG....................................CAUAC.AUAAU.AAUCAUU...UA.AA..UAAUAUUGGAAUGUAUUUAACAAAAAAACAUAAAGAAGAU
env-1064  GUAUAU.UUGCGCCG.............CUG...............AUAA......................CAG....................................CAUAC.AUAAU.AAUCAUU...UA.AA..UAAUAUUGGAAUGUAUUUAACAAAAAAAACGAAAAAAGAU
env-1065  CUAUAU.UUGCGCCG.............CUG...............AUAA......................CAG....................................CAUAC.AUAAU.AAUCAUU...UA.AA..UAAUAUUGGAAUGUAUUUAACAAAAAAACACAAAGAAGAU
env-1066  CUAUAU.UUGCGCCG.............CUG...............AUAA......................CAG....................................CAUAC.AUAAU.AAUCAUU...UA.AA..UAAUAUUGGAAUGUAUUUAACAAAGAAAAACUAAAGAAGA
env-1067  CUAUAU.UUGCGCCG.............CUG...............AUAA......................CAG....................................CAUAC.AUAAU.AAUCAUU...UA.AA..UAAUAUUGGAAUGUAUUUAACAAAAAAACAUAAAGAAGAU
env-1068  CUAUAU.UUGCGCCG.............CUG...............AUAA......................CAG....................................CAUAC.AUAAU.AAUCAUU...UA.AA..UAAUAUUGGAAUGUAUUUAACAAAAAAAACUAAAGAAGAU
env-1069  CUAUAU.UUGCGCCG.............CUG...............AUAA......................CAG....................................CAUAC.AUAAU.AAUCAUU...UA.AA..UAAUAUUGGAAUGUAUUUAACAAAAAAACACAAAGAAGAU
env-1070  GUAUAU.UUGCGCCG.............CUG...............AUAA......................CAG....................................CAUAC.AUAAU.AAUCAUU...UA.AA..UAAUAUUGGAAUGUAUUUAACAAAAAAAACUAAAGAAGAU
env-1077  CUAUAU.UUGCGCCG.............CUG...............AUAA......................CAG....................................CAUAC.AUAAU.AAUCAUU...UA.AA..UAAUAUUGGAAUGUAUUUAACAAAAAAACACAAAGAAGAU
env-1078  GUAUAU.UUGCGCCG.............CUG...............AUAA......................CAG....................................CAUAC.AUAAU.AAUCAUU...UA.AA..UAAUAUUGGAAUGUAUUUAACAAAAAAAACUAAAGAAGAU
env-1079  GUAUAU.UUGCGCCG.............CUG...............AUAA......................CAG....................................CAUAC.AUAAU.AAUCAUU...UA.AA..UAAUAUUGGAAUGUAUUUAACAAAAAAACACAAAGAAGAU
env-1080  CUAUAU.UUGCGCCG.............CUG...............AUAA......................CAG....................................CAUAC.AUAAU.AAUCAUU...UA.AA..UAAUAUUGGAAUGUAUUUAACAAAGAAACACAAAGAAGAU
env-1081  CUAUAU.UUGCGCCG.............CUG...............AUAA......................CAG....................................CAUAC.AUAAU.AAUCAUU...UA.AA..UAAUAUUGGAAUGUAUUUAACAAAAAAACACAAAGAAGAU
env-1082  CUAUAU.UUGCGCCG.............CUG...............AUAA......................CAG....................................CAUAC.AUAAU.AAUCAUU...UA.AA..UAAUAUUGGAAUGUAUUUAACAAAAAAACACAAAGAAGAU
env-1085  GUAUAU.UUGCGCCG.............CUG...............AUGU......................CAG....................................CAUAC.AUAAU.AAUCAUU...UA.AA..UAAUAUUGGAAUGUAUUUAACAAAAAAAACGAAAGAAGAU
env-1086  GUAUAU.UUGCGCCG.............CUG...............AUAU......................CAG....................................CAUAC.UUAAU.AAUCAUU...UA.AA..UAAUAUUGGAAUGUAUUUAACAAAAAAAACAAAAGAAGAU
env-1087  GUAUAU.UUGCGCCG.............CUG...............AUAU......................CAG....................................CAUGC.AUAAU.AAUCAUU...UA.AA..UAAUAUUGGAAUGUAUUUAACAAAAAAAAC..........
env-1088  CUAUAU.UUGCGCCG.............CUG...............AUAU......................CAG....................................CAUAC.AUAAU.AAUGAUU...UA.AA..UAAUAUUGGAAUGUAUUUAACAAAGAAAAACAAAGAAGAU
env-1089  GUAUAU.UUGCGCCG.............CUG...............AUAU......................CAG....................................CAUAC.AUAAU.AAUCAUU...UA.AA..UAAUAUUGGAAUGUAUUUAACAAAAAAAACGAAAGAAGAU
env-1090  GUAUAU.UUGCGCCG.............CUG...............AUAU......................CAG....................................CAUAC.AUAAU.AAUCAUU...UA.AA..UAAUAUUGGAAUGUAUUUAACAAAAAAAACGAAAGAAGGU
env-1091  CUAUAU.UUGCGCCG.............CUG...............AUAU......................CAG....................................CAUAC.AUAAU.AAUCAUU...UA.AA..UAAUAUUGGAAUGUAUUUAACAAAAAAAACAAAAGAAGAU
env-1092  GUAUAU.UUGCGCCG.............CUG...............AUAU......................CAG....................................CAUAC.AUAAU.AAUCAUU...UA.AA..UAAUAUUGGAAUGUAUUUAACAAAAAAAACGAAAGAAGAU
env-1093  CUAUAU.UUGCGCCG.............CUG...............AUAU......................CAG....................................CAUAC.AUAAU.AAUCAUU...UA.AA..UAAUAUUGGAAUGUAUUUAACAAAG...............
env-1094  CUAUAU.UUGCGCCG.............CUG...............AUAU......................CAG....................................CAUAC.AUAAU.AAUCAUU...UA.AA..UAAUAUUGGAAUGUAUUUAACAA.................
env-1095  CUAUAU.UUGCGCCG.............CUG...............AUAU......................CAG....................................CAUAC.AUAAU.AAUCAUU...UA.AA..UAAUAUUGGAAUGUAUUUAACAAAGAAAAACAAAGAAGAU
env-1096  CUAUAU.UUGCGCCG.............CUG...............AUAU......................CAG....................................CAUAC.AUAAU.AAUCAUU...UA.AA..UAAUAUUGGAAUGUAUUUAACAAAGAAAAACAAAGAAGAU
env-1097  GUAUAU.UUGCGCCG.............CUG...............AUAU......................CAG....................................CAUAC.AUAAU.AAUCAUU...UA.AA..UAAUAUUGGAAUGUAUUUAACAAAAAAAACAAAAGAAGAU
env-1098  UUAUAU.UUGCGCCG.............CUG...............AUAU......................CAG....................................CAUAC.AUAAU.AAUCAUU...UA.AA..UAAUAUUGGAAUGUAUUUAACAAAAAAAACAAAAGAAGAU
env-1099  CUAUAU.UUGCGCCG.............CUG...............AUAU......................CAG....................................CAUAC.AUAAU.AAUCAUU...UA.AA..UAAUAUUGGAAUGUAUUUAACAAAGAAAAACAAAGAAGAU
env-1101  UUAUAU.UUGCGCCG.............CUG...............AUAU......................CAG....................................CAUAC.AUAAU.AAUCAUU...UA.AA..UAAUAUUGGAAUGUAUUUAACAAAAAAAACGAAAGAAGAU
env-1102  CUAUAU.UUGCGCCG.............CUG...............AUAU......................CAG....................................CAUAC.AUAAU.AAUCAUU...UA.AA..UAAUAUUGGAAUGUAUUUAACAAAAAAACACAAAGAAGAU
env-1103  GUAUAU.UUGCGCCG.............CUG...............AUAU......................CAG....................................CAUAC.AUAAU.AAUCAUU...UA.AA..UAAUAUUGGAAUGUAUUUAACAAAAAAAACAAAAGAAGAU
env-1104  CUAUAU.UUGCGCCG.............CUG...............AUAU......................CAG....................................CAUAC.AUAAU.AAUCAUU...UA.AA..UAAUAUUGGAAUGUAUUUAACAAAGAAAAACAAAGAAGAU
env-1105  CUAUAU.UUGCGCCG.............CUG...............AUAU......................CAG....................................CAUAC.AUAAU.AAUCAUU...UA.AA..UAAUAUUGGAAUGUAUUUAACAAAGAAACACAAAGAAGAU
env-1106  GUAUAU.UUGCGCCG.............CUG...............AUAU......................CAG....................................CAUAC.AUAAU.AAUCAUU...UA.AA..UAAUAUUGGAAUGUAUUUAACAAAAAAAACCAAAGAAGAU
env-1107  GUAUAU.UUGCGCCG.............CUG...............AUAU......................CAG....................................CAUAC.AUAAU.AAUCAUU...UA.AA..UAAUAUUGGAAUGUAUUUAACAAAAAAAACAAAAGAAGAU
env-1108  CUAUAU.UUGCGCCG.............CUG...............AUAU......................CAG....................................CAUAC.AUAAU.AAUCAUU...UA.AA..UAAUAUUGGAAUGUAUUUAACAAAGAAAAACAAAGAAGAU
env-1109  CUAUAU.UUGCGCCG.............CUG...............AUAU......................CAG....................................CAUAC.AUAAU.AAUCAUU...UA.AA..UAAUAUUGGAAUGUAUUUAACAAAGAAAAACAAAGAAGAU
env-1110  CUAUAU.UUGCGCCG.............CUG...............AUAU......................CAG....................................CAUACCAUAAU.AAUCAUU...UA.AA..UAAUAUUGGAAUGUAUUUAACAAAAAAAACUAAAGAAGAU
env-1111  GUAUAU.UUGCGCCG.............CUG...............AUAU......................CAG....................................CAUAC.AUAAU.AAUCAUU...UA.AA..UAAUAUUGGAAUGUAUUUAACAAAAAAAACUAAAGAAGAU
env-1112  GUAUAU.UUGCGCCG.............CUG...............UUAU......................CAG....................................CAUAC.AUAAU.AAUCAUU...UA.AA..UAAUAUUGGAAUGUAUUUGACAAAGAAAAACAAAGAAGAU

          ............................<<<<<<<<<<<<<<<<<<...........>>>>>>>>>>>>>>>>>>.........<<<<<<<<........>>>>>>>>........................................................................
          ............................2222221111..101.00...........00.101..1111222222.........11011000........00011011........................................................................
          •UAY•U-UUGC••CY-------------◦◦◦◦◦◦◦-----------R•R◦◦-----------------◦◦◦◦◦◦◦-----------------------------------◦UAC-AUAA•-AAUCAUU---UA-A--YAAUAUURG•AUGUAUUUARC•ARAGAAR••AAARRRRAR
           5′                                                                                                                                                       3′
```

```
env-1113  GUAUAU.UUGCGCCG.............CUG...............UUAU......................CAG....................................CAUAC.AUAAU.AAUCAUU...UA.AA..UAAUAUUGGAAUGUAUUUGACAAAGAAAAACAAAGAAGAU
env-1115  GUAUAU.UUGCGCCG.............CUG...............UUAU......................CAG....................................CAUAC.AUAAU.AAUCAUU...UA.AA..UAAUAUUGGAAUGUAUUUGACAAAGAAAAACAAAGAAGAU
env-1116  GUAUAU.UUGCGCCG.............CUG...............UUAU......................CAG....................................CAUAC.AUAAU.AAUCAUU...UA.AA..UAAUAUUGGAAUGUAUUUGACAAAGAAAAACAAAGAAGAU
env-1121  GUAUAU.UUGCGCCG.............CUG...............UUAU......................CAG....................................CAUAC.AUAAU.AAUCAUU...UA.AA..UAAUAUUGGAAUGUAUUUGACAAAGAAAAACAAAGAAGAU
env-1122  GUAUAU.UUGCGCCG.............CUG...............UUAU......................CAG....................................CAUAC.AUAAU.AAUCAUU...UA.AA..UAAUAUUGGAAUGUAUUUGACAAAAAAAAUAAAACAGGAU
env-1123  CUAUAU.UUGCGCCG.............CUG...............UUAU......................CAG....................................CAUAC.AUAAU.AAUCAUU...UA.AA..UAAUAUUGGAAUGUAUUUAACAAAGAAAAACAAAGAAGAU
env-1124  GUAUAU.UUGCGCCG.............CUG...............UUAU......................CAG....................................CAUAC.AUAAU.AAUCAUU...UA.AA..UAAUAUUGGAAUGUAUUUGACAAAGAAAAAAAAAGAAGAU
env-1125  GUAUAU.UUGCGCCG.............CUG...............UUAU......................CAG....................................CAUAC.AUAAU.AAUCAUU...UA.AA..UAAUAUUGGAAUGUAUUUGACAAAGAAAAACAAAGAAGAU
env-1126  GUAUAU.UUGCGCCG.............CUG...............UUAU......................CAG....................................CAUAC.AUAAU.AAUCAUU...UA.AA..UAAUAUUGGAAUGUAUUUGACAAAGAAAAACAAAGAAGAU
env-1127  GUAUAU.UUGCGCCG.............CUG...............UUAU......................CAG....................................CAUAC.AUAAU.AAUCAUU...UA.AA..UAAUAUUGGAAUGUAUUUGACAAAGAAAAACAAAGAAGAU
env-1128  GUAUAU.UUGCGCCG.............CUG...............UUAU......................CAG....................................CAUAC.AUAAU.AAUCAUU...UA.AA..UAAUAUUGGAAUGUAUUUGACAAAAAAAAUAAAACAGGAU
env-1129  CUAUCU.UUGCACCCG............UAUUGAUCA.........UGG.................UGAUCAGUC......................................UAC.AUAAU.AAUCAUU...UACAA..AAAUAUUAGCAUGUAUUUAGAUUCAGCAAAGAAGAAAGAA
env-1130  GUAUCU.UUGCAGCCG............UAUUGAUCA.........AGG.................UGAUCAGUC......................................UAC.AUAAU.AAUCAUU...UACAC..AAAUAUUAGCAUGUAUUUAGAUUCAGCAAAGAAGAAAGAA
env-1131  AUAUCU.UUGCAGCCG............UAUUGACCA.........AGG.................UGGUUAGUC......................................UAC.AUAAU.AAUCAUU...UACAC..AAAUAUUAGCAUGUACUUAGAUUCAGCAAAGAAGAAAGAG
env-1134  AUAUCU.UUGCAGCCG............UAUUGACCA.........AGG.................UGGUUAGUC......................................UAC.AUAAU.AAUCAUU...UACAC..AAAUAUUAGCAUGUACUUAGAUUCAGCAAAGAAGAAAGAG
env-1135  AUAUCU.UUGCAGCCG............UAUUGACCA.........AGG.................UGGUUAGUC......................................UAC.AUAAU.AAUCAUU...UACAC..AAAUAUUAGCAUGUACUUAGAUUCAGCAAAGAAGAAAGAG
env-1137  AUAUCU.UUGCAGCCG............UAUUGACCA.........AGG.................UGGUUAGUC......................................UAC.AUAAU.AAUCAUU...UACAC..AAAUAUUAGCAUGUACUUAGAUUCAGCAAAGAAGAAAGAG
env-1138  UUAUCU.UUGCAGCCG............UAUUGACCA.........AGG.................UGGUCAGUC......................................UAC.AUAAU.AAUCAUU...UACAC..AAAUAUUAGCAUGUAUUUAGAUU.................
env-1139  CUAUCU.UUGCAGCCG............UAUUGACCA.........AGG.................UGGUCAGUC......................................UAC.AUAAU.AAUCAUU...UACAC..AAAUAUUAGCAUGUAUUUAGAUUCAGCAAAGAAGAAAGAA
env-1140  UUAUCU.UUGCACCC.............GAAUAAGGA.........UAC.................UCUUUAUUC.....................................AUAC.AUAAU.AAUCAUU...UAAAA..UAAUAUUAAAAUGUAUUUAACUAAAGAAACUAAAGCUGAG
env-1141  UUAUAU.UUGCACCU.............GA................AUUUA......................UC.....................................GUAC.AUAAU.AAUUAUU...AA.AA..UAAUAUUGGCAUGUAUUUAAACAAAGAAGUUAAAAAAGAG
env-1142  UUAUAU.UUGCACCU.............GA................AUUUA......................UC.....................................GUAC.AUAAU.AAUUAUU...AA.AA..UAAUAUUGGCAUGUAUUUAAACAAAGAAGUUAAAAAAGAG
env-1143  UUAUAU.UUGCACCU.............GA................AUUUA......................UC.....................................GUAC.AUAAU.AAUUAUU...AA.AA..UAAUAUUGGCAUGUAUUUAAACAAAGAAGUUAAAAAAGAG
Lsp-4-1   GUAUCU.UUGCAGCC.............UUAACGG...........AAAAA.................CCGUUAU.....................................GUAC.AUAAA.AAUCAUU...UA.AA..UAAUAUUGGAAUGUAUUUAGACGCAGCAAAAAAAGAAGAA
Cca-5-1   UUAUCU.UUGCAUCC.............G.................AAU.........................U......................................UAC.AUAAU.AAUCAU....UA.AA..UAAUAUUGGAAUGUAUUUAACAUCAGAAAAGAAAGCGGAA
env-1144  UUAUCU.UUGCAUCC.............G.................AAU.........................U......................................UAC.AUAAU.AAUCAU....UA.AA..UAAUAUUGGAAUGUAUUUAACAUCAGAAAAGAAAGCAGAG
Fan-1-1   UUACUU.UUGCACGC.............AAAUACG...........AUAAA.................CGUAU.C......................................UAC.AUAAU.AAUCAU....UA.AA..UAAUAUUAGCAUGUACUUAACUAAAGAAGUUAAAGCAGAA
Fte-1-1   UUACUU.UUGCACGC.............AAAUACG...........AUAAA.................CGUAU.C......................................UAC.AUAAU.AAUCAU....UA.AA..UAAUAUUAGCAUGUACUUAACUAAAGAAGUUAAAGCAGAA
env-1145  UUAUAU.UUGCCGCU.............UGAUAG............AAAAA..................CUAUCU.....................................GUAC.AUAAU.AAUCAUU...GA.AA..UAAUAUUGGAAUGUAUUUAACUAAAGAAGAGAAACAAGGU
env-1146  UUAUAU.UUGCCGCU.............UGAUAG............ACGAA..................CUAUCU.....................................GUAC.AUAAU.AAUCAUU...GA.AA..UAAUAUUGGAAUGUAUUUAACAAAGGAAGAAAAGCAAGAU
env-1147  GUAUCU.UUGCCGCU.............CGAUAG............AAAAC..................CUAUCG.....................................GUAC.AUAAU.AAUCAUU...GA.AA..UAAUAUUGGAAUGUAUUUAACAAAAGAAGAAAAAGAACAA
env-1159  GUAUCU.UUGCCGCU.............CGAUAG............AAAAC..................CUAUCG.....................................GUAC.AUAAU.AAUCAUU...GA.AA..UAAUAUUGGAAUGUAUUUAAC...................
env-1161  GUAUCU.UUGCCGCU.............CGAUAG............AAAAC..................CUAUCG.....................................GUAC.AUAAU.AAUCAUU...GA.AA..UAAUAUUGGAAUGUAUUUAACAAAAGAAGAAAAAGAACAA
env-1162  GUAUAU.UUGCCGCU.............GAAUAG............AAAUA..................CUAUUC.....................................GUAC.AUAAU.AAUCAUU...GA.AA..UAAUAUUGGAAUGUAUUUAUCAAAAGAAGAAAAACAACAA
env-1163  UUAUAU.UUGCAAAC.............UCAAUGG...........ACAUA.................CCAUUGA....................................CAUAC.AUAAU.AAUCAUU...UA.AA..UAAUAUUGGCAUGUACUUAACUAAAGAAGUAAAAGAAAGC
env-1165  UUAUAU.UUGCAAAC.............UCAAUGG...........ACAUA.................CCAUUGA....................................CAUAC.AUAAU.AAUCAUU...UA.AA..UAAUAUUGGCAUGUACUUAACUAAAGAAGUAAAAGAAAGC
env-1167  CUAUAU.UUGCAAAC.............UCAAUG............AAAAAA.................CAUUGA....................................CAUAC.AUAAU.AAUCAUU...UA.AA..UAAUAUUGGCAUGUACUUAACUAAAGAAGUAAAAGAAAAC
env-1168  GUACAU.UUGCAACU.............UCAAU.............AGAAAAAAUC..............AUUGA....................................UGUAC.AUAAA.AAUCAAU...UUAAA..UAAUAUUGGCAUGUAUUUAACAAAAGAAGUAAAAGAAACA
env-1169  GUACAU.UUGCAACU.............UCAAU.............AGAAAAAAUC..............AUUGA....................................UGUACAAUAAA.AAUCAUU...UA.AA..UAAUAUUGGCAUGUAUUUAACAAAAGAAGUAAAAGAAACA
Psp-6-1   AUACCU.UUGCAGCU.............UCAAUGA...........AAAUA.................UCAUUGA....................................UAUAC.AUAAU.AAUCAUU...UA.AA..UAAUAUUGGAAUGUAUUUAACUAAAGAAGUAAAAGAAAGC
env-1170  AUAUAU.UUGCAUCC.............UCAAUGA...........AAACA.................UCAUUGA....................................CAUAC.AUAAA.AAUCAUU...UA.AA..CAAUAUUGGAAUGUAUUUAACUAAAGAGGUAAAAGAAAGC
env-1171  AUAUAU.UUGCAUCC.............UCAAUGA...........AAACA.................UCAUUGA....................................CAUAC.AUAAA.AAUCAUU...UA.AA..CAAUAUUGGAAUGUAUUUAACUAAAGAGGUAAAAGAAAGC
env-1172  UUAUAU.UUGCAGCU.............UCAGUGAA..........AAAU.................UUCGCUGA....................................UAUAC.AUAAU.AAUCAUU...UA.AA..UAAUAUUGGCAUGUACUUAAGUAAAGAAGAAAGAGAGCUU
env-1173  GUAUAU.UUGCAGCU.............UCAAUGAAA.........AAA.................UUUCAUUGA....................................AAUAC.AUAAU.AAUCAUU...UA.AA..GAAUAUUGGCAUGUACUUAAGUAAAGAAGGAAAAGAAGAA
env-1174  GUACAU.UUGCAACU.............UCAAAUGA..........AAAAA................UCA.UUGA....................................UGUAC.AUAAA.AAUCAUU...UA.AA..UAAUAUUGGCAUGUAUUUAACAAAAGAAGUAAAAGAAACA
Tov-1-1   GUACAU.UUGCUGCU.............UCAAUGA...........AUACA.................UCAUUGA.....................................GUAC.AUAAA.AAUCAUU...UA.AA..UAAUAUUGGAAUGUAUUUAACAAAAGAAGUAAAAGAAGGA
Pir-1-1   CUAUCU.UUGCAGCU.............UCAAUGAA..........AAA..................UUUAUUGA....................................AAUAC.AUAAA.AAUCAUU...UA.AA..UAAUAUUGGCAUGUACUUAACUAAAGAAGCAAAAGAAGUU
env-1175  GUAUAU.UUGCAGCU.............UCAAUGAA..........AAAAA................UUUAUUGA....................................UAUAC.AUAAU.AAUCAUU...UA.AA..UAAUAUUGGCAUGUACUUAAGUAAGGAAGCAAAAGAAGUU
env-1176  GUAUAU.UUGCAGCU.............UCAAUGAA..........AAAAA................UUUAUUGA....................................UAUAC.AUAAU.AAUCAUU...UA.AA..UAAUAUUGGCAUGUACUUAAGUAAGGAAGCAAAAGAAGUU
env-1177  GUAUAU.UUGCAGCU.............UCAAUGAA..........AAAAA................UUUAUUGA....................................UAUAC.AUAAU.AAUCAUU...UA.AA..UAAUAUUGGCAUGUACUUAAGUAAGGAAGCAAAAGAAGUU
env-1181  GUAUAU.UUGCAGCU.............UCAAUGAA..........AAAAA................UUUAUUGA....................................UAUAC.AUAAU.AAUCAUU...UA.AA..UAAUAUUGGCAUGUACUUA.....................
env-1182  GUAUAU.UUGCAGCU.............UCAAUGAA..........AAAAA................UUUAUUGA....................................UAUAC.AUAAU.AAUCAUU...UA.AA..UAAUAUUGGCAUGUACUUAAGUAAGGAAGCAAAAGAAGUU
env-1183  CUAUAU.UUGCAGCU.............UCAAUGA...........AAACA.................UCAUUGA....................................AAUAC.AUAAA.AAUCAUU...UA.AA..UAAUAUUGGCAUGUACUUAANUAAAGAAGUAAAGGAAGGA
env-1184  .UAUAU.UUGCAGGC.............UCAAUGA...........AAACA.................UCAUUGA....................................UAUAC.AUAAA.AAUCAUU...UA.AA..UAAUAUUGGAAUGUAUUUAACUAAAGAAGUAAAAGAAGGC
env-1185  AUAUAU.UUGCAGCU.............UCAAUGA...........AAACA.................UCAUUGA....................................UGUAC.AUAAA.AAUCAUU...UA.AA..UAAUAUUGGAAUGUAUUUAACUAAAGAAGUAAAAGAAAGC
env-1186  AUAUAU.UUGCAGCU.............UCAAUGA...........AAACA.................UCAUUGA....................................UGUAC.AUAAA.AAUCAUU...UA.AA..UAAUAUUGGAAUGUAUUUAACUAAAGAAGUAAAAGAAAGC
env-1187  CUAUAU.UUGCAGCU.............UCAAUGA...........AAACA.................UCAUUGA....................................UGUAC.AUAAA.AAUCAUU...UA.AA..UAAUAUUGGAAUGUAUUUAACUAAAGAAGUAAAAGAAAGC
env-1188  AUAUAU.UUGCAGCU.............UCAAUGA...........AAACA.................UCAUUGA....................................UGUAC.AUAAA.AAUCAUU...UA.AA..UAAUAUUGGAAUGUAUUUAACUAAAGAAGUAAAAGAAAGC
env-1190  AUAUAU.UUGCAGCU.............UCAAUGA...........AAACA.................UCAUUGA....................................UGUAC.AUAAA.AAUCAUU...UA.AA..UAAUAUUGGAAUGUAUUUAACUAAAGAAGUAAAAGAAAGC
env-1191  AUAUAU.UUGCAGCU.............UCAAUGA...........AAACA.................UCAUUGA....................................UGUAC.AUAAA.AAUCAUU...UA.AA..UAAUAUUGGAAUGUAUUUAACUAAAGAAGUAAAAGAAAGC
env-1192  AUAUAU.UUGCAGCU.............UCAAUGA...........AAACA.................UCAUUGA....................................UGUAC.AUAAA.AAUCAUU...UA.AA..UAAUAUUGGAAUGUAUUUAACUAAAGAAGUAAAAGAAAGC
env-1193  AUAUAU.UUGCAGCU.............UCAAUGA...........AAACA.................UCAUUGA....................................UGUAC.AUAAA.AAUCAUU...UA.AA..UAAUAUUGGAAUGUAUUUAACUAAAGAAGUAAAAGAAAGC
env-1194  AUAUAU.UUGCAGCU.............UCAAUGA...........AAACA.................UCAUUGA....................................UGUAC.AUAAA.AAUCAUU...UA.AA..UAAUAUUGGAAUGUAUUUAACUAAAGAAGUAAAAGAAAGC
env-1196  AUAUAU.UUGCAGCU.............UCAAUGA...........AAACA.................UCAUUGA....................................UGUAC.AUAAA.AAUCAUU...UA.AA..UAAUAUUGGAAUGUAUUUAACUAAAGAAGUAAAAGAAAGC
env-1197  AUAUAU.UUGCAGCU.............UCAAUGA...........AAACA.................UCAUUGA....................................UAUAC.AUAAA.AAUCAUU...GA.AA..UAAUAUUGGAAUGUAUUUAACUAAAGAAGUAAAAGAAAGC
env-1198  CUAUAU.UUGCGGCU.............UCAAUGA...........AAACA.................UCAUUGA....................................UAUAC.AUAAA.AAUCAUU...UA.AA..UAAUAUUGGAAUGUAUUUAACAAAAGAAGUAAAAGAAAGC
env-1199  AUAUAU.UUGCAGCU.............UCAAUGA...........AAACA.................UCAUUGA....................................UAUAC.AUAAA.AAUCAUU...UA.AA..UAAUAUUGGAAUGU..........................
env-1200  AUAUAU.UUGCAGCU.............UCAAUGA...........AAACA.................UCAUUGA....................................UAUAC.AUAAA.AAUCAUU...UA.AA..UAAUAUUGGAAUGUAUUUAACUAAAGAAGUAAAAGAAAGC
env-1203  CUAUAU.UUGCAGCU.............UCAAUGA...........AAACA.................UCAUUGA....................................UAUAC.AUAAA.AAUCAUU...UA.AA..UAAUAUUGGAAUGUAUUUAACUAAAGAAGUAAAAGA....
env-1204  AUAUAU.UUGCAGCU.............UCAAUGA...........AAACA.................UCAUUGA....................................UAUAC.AUAAA.AAUCAUU...UA.AA..UAAUAUUGGAAUGUAUUUAACUAAAGAAGUAAAAGAAAGC
env-1205  CUAUAU.UUGCACCC.............UCAAUGA...........AAACA.................UCAUUGA....................................AAUAC.AUAAA.AAUCAUU...UA.AA..UAAUAUUGGCAUGUACUUAACAAAAGAAGUAAAAGAAAAC
Psp-3-1   CUAUAU.UUGCACCU.............UCAAUGA...........AAACA.................UCAUUGA....................................ACUAC.AUAAA.AAUCAUU...UA.AA..UAAUAUUGGCAUGUACUUAACUAAAGAAGUAAAGGAAGGA
env-1208  GUAUAU.UUGCAGCU.............UCAAUGA...........AAAAA.................UCAUUGA.....................................GUAC.AUAAA.AAUCAUU...UAAAA..CAAUAUUGGCAUGUAUUUAACAAAAGAAGUAAAAGAGGAA
env-1209  GUAUAU.UUGCAGCU.............UCAAUGA...........AAAAA.................UCAUUGA.....................................AUAC.AUAAA.AAUCAUU...UAAAA..CAAUAUUGGCAUGUAUUUAACAAAAGAAGUAAAAGAGGAA
env-1210  GUAUAU.UUGCAGCU.............UCAAUGA...........AAAAA.................UCAUUGA.....................................AUAC.AUAAA.AAUCAUU...UAAAA..CAAUAUUGGCAUG...........................
env-1211  GUAUAU.UUGCAGCU.............UCAAUGA...........AAAAA.................UCAUUGA.....................................AUAC.AUAAA.AAUCAUU...UAAAA..CAAUAUUGGCAUGUAUUUAACAAAAGAAGUAAAAGAGGAA
env-1212  AUAUAU.UUGCAGCU.............UCAAUGAA..........AAC..................UUCAUUGA....................................UAUAC.AUAAA.AAUCAUU...UA.AA..UAAUAUUGGCAUGUACUUAACAAGAGAAAUUAAAGAAGGA
env-1213  GUAUAU.UUGCAGCU.............UCAAUGAA..........AAA..................UUCAUUGA....................................UGUAC.AUAAA.AAUCAUU...UAAAA..UAAUAUUGGCAUGUAUUUAACGAAAGAAGUGAAAGAAGAA
Psp-2-1   AUAUAU.UUGCAGCU.............UCAAUGA...........AACAA.................UCAUUGA....................................UAUAC.AUAAU.AAUCAUU...UA.AA..UAAUAUUGGAAUGUAUUUAACUAAAGAAGUAAAAGAAAGC
Psp-4-1   CUAUAU.UUGCACCU.............UCAAUGA...........AUACA.................UCAUUGA....................................UAUAC.AUAAA.AAUCAUU...UA.AA..UAAUAUUGGCAUGUACUUAACUAAAGAAGUAAAAGAAAGC
env-1215  GUACAU.UUGCAACU.............UCAAUGA...........AAAA..................UCAUUGA....................................UGUAC.AUAAA.AAUCAUU...UA.AA..UAAUAUUGGCAUGUAUUUAACAAAAGAAGUAAAGAAACAA
env-1216  GUACAU.UUGCAACU.............UCAAUGA...........AAAAA.................UCAUUGA....................................UGUAC.AUAAA.AAUCAUU...UA.AA..UAAUAUUGGCAUGUAUUUAACAAAAGAAGUAAAAGAAACA
env-1217  GUACAU.UUGCAACU.............UCAAUGA...........AAAAA.................UCAUUGA....................................UGUAC.AUAAA.AAUCAUU...UA.AA..UAAUAUUGGCAUGUAUUUAACAAAAGAAGUAAAAGAAACA

          ............................<<<<<<<<<<<<<<<<<<...........>>>>>>>>>>>>>>>>>>.........<<<<<<<<........>>>>>>>>........................................................................
          ............................2222221111..101.00...........00.101..1111222222.........11011000........00011011........................................................................
          •UAY•U-UUGC••CY-------------∘∘∘∘∘∘∘-----------R•R∘∘-----------------∘∘∘∘∘∘∘------------------------------------•UAC-AUAA•-AAUCAUU---UA-AA--YAAUAUURG•AUGUAUUUARC•ARAGAAR••AAARRRRAR
          5′                                                                                                                                              3′
```

```
env-1218    GUACAU.UUGCAACU.............UCAAUGA...........AAAAA.................UCAUUGA....................................UGUAC.AUAAA.AAUCAUU...UA.AA..UAAUAUUGGCAUGUAUUUAACAAAAGAAGUAAAAGAAACA
env-1219    GUACAU.UUGCAACU.............UCAAUGA...........AAAAA.................UCAUUGA....................................UGUAC.AUAAA.AAUCAUU...UA.AA..UAAUAUUGGCAUGUAUUUAACAAAAGAAGUAAAAGAAACA
env-1220    GUACAU.UUGCAACU.............UCAAUGA...........AAAAA.................UCAUUGA....................................UGUAC.AUAAA.AAUCAUU...UA.AA..UAAUAUUGGCAUGUAUUUAACAAAAGAAGUAAAAGAAACA
env-1222    GUACAU.UUGCAACU.............UCAAUGA...........AAAAA.................UCAUUGA....................................UGUAC.AUAAA.AAUCAUU...UA.AA..UAAUAUUGGCAUGUAUUUAACAAAAGAAGUAAAAGAAACA
env-1223    GUACAU.UUGCAACU.............UCAAUGA...........AAAAA.................UCAUUGA....................................UGUAC.AUAAA.AAUCAUU...UA.AA..UAAUAUUGGCAUGUAUUUAACAAAAGAAGUAAAAGAAACA
env-1226    GUACAU.UUGCAACU.............UCAAUGA...........AAAAA.................UCAUUGA....................................UGUAC.AUAAA.AAUCAUU...UA.AA..UAAUAUUGGCAUGUAUUUAACAAAAGAAGUAAAAGAAACA
env-1227    GUACAU.UUGCAACU.............UCAAUGA...........AAAAA.................UCAUUGA....................................UGUAC.AUAAA.AAUCAUU...UA.AA..UAAUAUUGGCAUGUAUUUAACAAAAGAAGUAAAAGAAACA
env-1229    GUACAU.UUGCAACU.............UCAAUGA...........AAAAA.................UCAUUGA....................................UGUAC.AUAAA.AAUCAUU...UA.AA..UAAUAUUGGCAUGUAUUUAACAAAAGAAGUAAAAGAAACA
env-1230    GUACAU.UUGCAACU.............UCAAUGA...........AAAAAA................UCAUUGA....................................UGUAC.AUAAA.AAUCAUU...UA.AA..UAAUAUUGGCAUGUAUUUAACAAAAGAAGUAAAAGAAACA
env-1231    GUACAU.UUGCAACU.............UCAAUGA...........AAAAAA................UCAUUGA....................................UGUAC.AUAA..AAUCAUU...UA.AA..UAAUAUUGGCAUGUAUUUAACAAAAGAAGUAAAAAGAAAA
env-1232    GUACAU.UUGCAACU.............UCAAUGA...........AAAAA.................UCAUUGA....................................UGUAC.AUAAA.AAUCAUU...UA.AA..UAAUAUUGGCAUGUAUUUAACAAAAGAAGUAAAAGAAACA
env-1233    GUACAU.UUGCAACU.............UCAAUGA...........AAAAA.................UCAUUGA....................................UGUAC.AUAAA.AAUCAUU...UA.AA..UAAUAUUGGCAUGUAUUUAACAAAAGAAGUAAAAGAAACA
env-1234    GUACAU.UUGCAACU.............UCAAUGA...........AAAAA.................UCAUUGA....................................UGUAC.AUAAA.AAUCAUU...UA.AA..UAAUAUUGGCAUGUAUUUAACAAAAGAAGUAAAAGAAACA
env-1235    GUACAU.UUGCAACU.............UCAAUGA...........AAAAA.................UCAUUGA....................................UGUAC.AUAAA.AAUCAUU...UA.AA..UAAUAUUGGCAUGUAUUUAACAAAAGAAGUAAAAGAAACA
env-1236    GUACAU.UUGCAACU.............UCAAUGA...........AAAAA.................UCAUUGA....................................UGUAC.AUAAA.AAUCAUU...UA.AA..UAAUAUUGGCAUGUAUUUAACAAAAGAAGUAAAAGAAACA
env-1237    GUACAU.UUGCAACU.............UCAAUGA...........AAAAA.................UCAUUGA....................................UGUAC.AUAAA.AAUCAUU...UA.AA..UAAUAUUGGCAUGUAUUUAACAAAAGAAGUAAAAGAAACA
env-1238    GUACAU.UUGCAACU.............UCAAUGA...........AAAAA.................UCAUUGA....................................UGUAC.AUAAA.AAUCAUU...UA.AA..UAAUAUUGGCAUGUAUUUAACAAAAGAAGUAAAAGAAACA
env-1239    GUACAU.UUGCAACU.............UCAAUGA...........AAAAA.................UCAUUGA....................................UGUAC.AUAAA.AAUCAUU...UA.AA..UAAUAUUGGCAUGUAUUU......................
env-1240    GUACAU.UUGCAACU.............UCAAUGA...........AAAAA.................UCAUUGA....................................UGUAC.AUAAA.AAUCAUU...UA.AA..UAAUAUUGGCAUGUAUUUAACAAAAGAAGUAAAAGAAACA
env-1241    GUACAU.UUGCAACU.............UCAAUGA...........AAAA..................UCAUUGA....................................UGUAC.AUAAA.AAUCAUU...UA.AA..UAAUAUUGGCAUGUAUUUAACAAAGAAGUAAAGAAACAAU
env-1242    GUACAU.UUGCCACU.............UCAAUGA...........AAAAA.................UCAUUGA....................................UGUAC.AUAAA.AAUCAUU...UA.AA..UAAUAUUGGCAUGUAUUUGACAAAAGAAGUAAAAGAAACA
env-1243    GUACAU.UUGCCACU.............UCAAUGA...........AAAAA.................UCAUUGA....................................UGUAC.AUAAA.AAUCAUU...UA.AA..UAAUAUUGGCAUGUAUUUGACAAAAGAAGUAAAAGAAACA
env-1245    GUACAU.UUGCCACU.............UCAAUGA...........AAAAA.................UCAUUGA....................................UGUAC.AUAAA.AAUCAUU...UA.AA..UAAUAUUGGCAUGUAUUUGACAAAAGAAGUAAAAGAAACA
env-1246    GUACAU.UUGCCACU.............UCAAUGA...........AAAAA.................UCAUUGA....................................UGUAC.AUAAA.AAUCAUU...UA.AA..UAAUAUUGGCAUGUAUUUGACAAAAGAAGUAAAAGAAACA
env-1247    GUACAU.UUGCCACU.............UCAAUGA...........AAAAA.................UCAUUGA.....................................GUAC.AUAAA.AAUCAUU...UA.AA..UAAUAUUGGCAUGUAUUUGACAAAAGAAGUAAAAGAAAGU
env-1248    GUACAU.UUGCCACU.............UCAAUGA...........AAAAA.................UCAUUGA....................................UGUAC.AUAAA.AAUCAUU...UA.AA..UAAUAUUGGCAUGUAUUUGACAAAAGAAGUAAAAGAAACA
env-1250    GUACAU.UUGCCACU.............UCAAUGA...........AAAAA.................UCAUUGA....................................UGUAC.AUAAA.AAUCAUU...UA.AA..UAAUAUUGGCAUGUAUUUAACAAAAGAAGUUAAAGAAACA
env-1252    GUACAU.UUGCCACU.............UCAAUGA...........AAAAA.................UCAUUGA....................................UGUAC.AUAAA.AAUCAUU...UA.AA..UAAUAUUGGCAUGUAUUUGACAAAAGAAGUAAAAGAAACA
env-1253    GUACAU.UUGCCACU.............UCAAUGA...........AAAAA.................UCAUUGA....................................UGUAC.AUAAA.AAUCAUU...UA.AA..UAAUAUUGGCAUGUAUUUGACAAAAGAAGUAAAAGAAACA
env-1254    GUACAU.UUGCCACU.............UCAAUGA...........AAAAA.................UCAUUGA....................................UGUAC.AUAAA.AAUCAUU...UA.AA..UAAUAUUGGCAUGUAUUUGACAAAAGAAGA..........
env-1255    GUACAU.UUGCGUCU.............UCAAUGA...........AAAAA.................UCAUUGA....................................UGUAC.AUAAA.AAUCAUU...UA.AA..UAAUAUUGGCAUG...........................
env-1256    GUACAU.UUGCCAGA.............UCAAUGA...........AAAAA.................UCAUUGA....................................UGUAC.AUAAA.AAUCAUU...UA.AA..UAAUAUUGGCAUGUAUUUGACAAAAGAAGUAAAAGAAACA
env-1257    GUACAU.UUGCCAGA.............UCAAUGA...........AAAAA.................UCAUUGA....................................UGUAC.AUAAA.AAUCAUU...UA.AA..UAAUAUUGGCAUGUAUUUGACAAAAGAAGUAAAAGAAACA
env-1259    GUACAU.UUGCCAGA.............UCAAUGA...........AAAAA.................UCAUUGA....................................UGUAC.AUAAA.AAUCAUU...UA.AA..UAAUAUUGGCAUGUAUUUGACAAAAGAAGUAAAAGAAACA
env-1260    GUACAU.UUGCCAGA.............UCAAUGA...........AAAAA.................UCAUUGA....................................UGUAC.AUAAA.AAUCAUU...UA.AA..UAAUAUUGGCAUGUAUUUGACAAAAGAAGUAAAAGAAACA
env-1262    GUACAU.UUGCCAGA.............UCAAUGA...........AAAAA.................UCAUUGA....................................UGUAC.AUAAA.AAUCAUU...UA.AA..UAAUAUUGGCAUGUAUUUGACAAAAGAAGUAAAAGAAACA
env-1263    GUACAU.UUGCCAGA.............UCAAUGA...........AAAAA.................UCAUUGA....................................UGUAC.AUAAA.AAUCAUU...UA.AA..UAAUAUUGGCAUGUAUUUGACAAAAGAAGUAAAAGAAACA
env-1264    GUACAU.UUGCCCCG.............UCAAUGA...........AAAAA.................UCAUUGA....................................UGUAC.AUAAA.AAUCAUU...UA.AA..UAAUAUUGGCAUGUAUUUGACAAAAGAAGUAAAAGAAACA
env-1268    GUACAU.UUGCCCCG.............UCAAUGA...........AAAAA.................UCAUUGA....................................UGUAC.AUAAA.AAUCAUU...UA.AA..UAAUAUUGGCAUGUAUUUGACAAAAGAAGUAAAAGAAACA
env-1269    GUACAU.UUGCCCCG.............UCAAUGA...........AAAAA.................UCAUUGA....................................UGUAC.AUAAA.AAUCAUU...UA.AA..UAAUAUUGGCAUGUAUUUGACAAAAGAAGUAAAAGAAACA
env-1270    GUACAU.UUGCCCCG.............UCAAUGA...........AAAAA.................UCAUUGA....................................UGUAC.AUAAA.AAUCAUU...UA.AA..UAAUAUUGGCAUGUAUUUGACAAAAGAAGUAAAAGAAACA
env-1271    GUACAU.UUGCCCCG.............UCAAUGA...........AAAAA.................UCAUUGA....................................UGUAC.AUAAA.AAUCAUU...UA.AA..UAAUAUUGGCAUGUAUUUGACAAAAGAAGUAAAAGAAACA
env-1272    GUACAU.UUGCCCCG.............UCAAUGA...........AAAAA.................UCAUUGA....................................UGUAC.AUAAA.AAUCAUU...UA.AA..UAAUAUUGGCAUGUAUUUGACAAAAGAAGUAAAAGAAACA
env-1273    GUACAU.UUGCCCCG.............UCAAUGA...........AAAAA.................UCAUUGA....................................UGUAC.AUAAA.AAUCAUU...UA.AA..UAAUAUUGGCAUGUAUUUGACAAAAGAAGUAAAAGAAACA
env-1274    GUACAU.UUGCCCCG.............UCAAUGA...........AAAAA.................UCAUUGA....................................UGUAC.AUAAA.AAUCAUU...UA.AA..UAAUAUUGGCAUGUAUUUGACAAAAGAAGUAAAAGAAACA
env-1275    GUACAU.UUGCCCCG.............UCAAUGA...........AAAAA.................UCAUUGA....................................UGUAC.AUAAA.AAUCAUU...UA.AA..UAAUAUUGGCAUGUAUUUGACAAAAGAAGUAAAAGAAACA
env-1277    GUACAU.UUGCCCCG.............UCAAUGA...........AAAAA.................UCAUUGA....................................UGUAC.AUAAA.AAUCAUU...UA.AA..UAAUAUUGGCAUGUAUUUGACAAAAGAAGUAAAAGAAACA
env-1278    GUACAU.UUGCCACG.............UUAAUGA...........AAAAA.................UCAUUGA....................................CGUAC.AUAAA.AAUCAUU...UA.AA..UAAUAUUGGCAUGUAUUUAACAAAAGAAGUAAAAGAAACA
Ccy-1-1     AUACAU.UUGCAACU.............UCA...............AAAAC.....................U.A.....................................GUAC.AUAAU.AAUCAU....UA.AA..UAAUAUUGGAAUGUAUUUAACAAAAGAAGUAAAAGCAGAA
Ccy-1-3     AUACAU.UUGCAACU.............UCA...............AAAAC.....................U.A.....................................GUAC.AUAAU.AAUCAU....UA.AA..UAAUAUUGGAAUGUAUUUAACAAAAGAAGUAAAAGCAGAA
Tme-1-1     GUACAU.UUGCAUCU.............UCAAUGU...........AAAU..................GCAUUGA.....................................GUAC.AUAAU.AAUCAUU...UA.AA..UAAUAUUGGCAUGUAUUUAACAAAAGAAGUUAAAGAACAA
Tsp-1-1     GUACAU.UUGCAACU.............UCAAUGU...........AAUA..................GCAUUGA.....................................GUAC.AUAAU.AAUCAUU...UA.AA..UAAUAUUGGCAUGUAUUUAACAAAAGAAGUUAAAGAACAA
env-1280    CUAUAU.UUGCAGCU.............UCAAUGAAA.........UAA.................UUUUGUUGA....................................UAUAC.AUAAU.AAUCAUU...UA.AA..UAAUAUUGGCAUGUACUUGACUAAAGAAGUAAAAGAACAA
env-1281    CUAUAU.UUGCAGCU.............UCAAUGAAA.........UAA.................UUUUGUUGA....................................UAUAC.AUAAU.AAUCAUU...UA.AA..UAAUAUUGGCAUGUACUUGACUAAAGAAGUAAAAGAACAA
env-1282    CUAUAU.UUGCAGCU.............UCAAUGAAA.........UAA.................UUUCGUUGA....................................UAUAC.AUAAU.AAUCAUU...UA.AA..UAAUAUUGGCAUGUACUUGACUAAAGAAGUAAAAGAACAA
env-1283    CUAUAU.UUGCAGCU.............UCAAUGAAA.........UAA.................UUUCGUUGA....................................UAUAC.AUAAU.AAUCAUU...UA.AA..UAAUAUUGGCAUGUACUUGACUAAAGAAGUAAAAGAACAA
env-1284    CUAUAU.UUGCAGCU.............UCAAUGAAA.........UAA.................UUUCGUUGA....................................UAUAC.AUAAU.AAUCAUU...UA.AA..UAAUAUUGGCAUGUACUUGACUAAAGAAGUAAAAGAACAA
env-1285    CUAUAU.UUGCAGCU.............UCAAUGAAA.........UAA.................UUUCGUUGA....................................UAUAC.AUAAU.AAUCAUU...UA.AA..UAAUAUUGGCAUGUACUUGACUAAAGAAGUAAAAGAACAA
env-1287    CUAUAU.UUGCAGCU.............UCAAUGAAA.........UAA.................UUUCGUUGA....................................UAUAC.AUAAU.AAUCAUU...UA.AA..UAAUAUUGGCAUGUACUUGACUAAAGAAGUAAAAGAACAA
env-1293    CUAUAU.UUGCAGCU.............UCAAUGAAA.........UAA.................UUUCGUUGA....................................UAUAC.AUAAU.AAUCAUU...UA.AA..UAAUAUUGGCAUGUACUUGACUAAAGAAGUAAAAGAACAA
env-1294    CUAUAU.UUGCAGCU.............UCAAUGAAA.........UAA.................UUUCGUUGA....................................UAUAC.AUAAU.AAUCAUU...UA.AA..UAAUAUUGGCAUGUACUUGACUAAAGAAGUAAAAGAACAA
env-1295    CUAUAU.UUGCAGCU.............UCAAUGAAA.........UAA.................UUUCGUUGA....................................UAUAC.AUAAU.AAUCAUU...UA.AA..UAAUAUUGGCAUGUACUUGACUAAAGAAGUAAAAGAACAA
env-1296    CUAUAU.UUGCAGCU.............UCAAUGAAA.........UAA.................UUUCGUUGA....................................UAUAC.AUAAU.AAUCAUU...UA.AA..UAAUAUUGGCAUGUACUUGACUAAAGAAGUAAAAGAACAA
env-1298    CUAUAU.UUGCAGCU.............UCAAUGAAA.........UAA.................UUUCGUUGA....................................UAUAC.AUAAU.AAUCAUU...UA.AA..UAAUAUUGGCAUGUACUUGACUAAAGAAGUAAAAGAACAA
env-1299    CUAUAU.UUGCAGCU.............UCAAUGAAA.........UAA.................UUUCGUUGA....................................UAUAC.AUAAU.AAUCAUU...UA.AA..UAAUAUUGGCAUGUACUUGACUAAAGAAGUAAAAGAACAA
env-1300    GUACAU.UUGCCACG.............UCAGUAA...........AAAAA.................UUACUGA....................................UGUAC.AUAAA.AAUCAUU...UA.AA..UAAUAUUGGCAUGUAUUUGACAAAAGAAGUAAAAGAAUCA
env-1301    GUACAU.UUGCCACG.............UCAGUGA...........AAAAA.................UCGCUGA....................................UGUAC.AUAAA.AAUCAUU...UA.AA..UAAUAUUGGCAUGUAUUUGACAAAAGAAGUAAAAGAAUCA
env-1302    GUACAU.UUGCCACG.............UCAGUGA...........AAAAA.................UCGCUGA....................................UGUAC.AUAAA.AAUCAUU...UA.AA..UAAUAUUGGCAUGUAUUUGACAAAAGAAGUAAAAGAAUCA
env-1303    AUAUAU.UUGCAACC.............UCAAUGA...........AUAAA.................UCAUUGA....................................UAUAC.AUAAU.AAUCAUU...UA.AA..UAAUAUUGGAAUGUAUUUAACCAAAGAAGUAAAAGAAAGA
env-1304    AUAUAU.UUGCAUGC.............UCAAUGA...........GAAAA.................UCAUUGA....................................AAUAC.AUAAA.AAUCAUU...UU.AA..UAAUAUUGGCAUGUACUUAACAAAAGAAAUUAAAGAAAAC
env-1305    UUAUCU.UUGCAGCC.............UGAUUGA...........AAA...................UCAGUCA....................................GAUAC.AUAAA.AAUGAUU...UCAAA..UAAUAUUAGCAUGUAUUUAACAAAAGAAGUAAAAGCCGAC
env-1306    UUAUCU.UUGCAGCC.............UGAUUGA...........AAA...................UCAGUCA....................................GAUAC.AUAAA.AAUGAUU...UCUAA..UAAUAUUAGCAUGUAUUUAACAAAAGAAGUAAAAGCCGAC
env-1307    UUAUCU.UUGCAGCC.............UGAUUGA...........AAA...................UCAGUCA....................................GAUAC.AUAAA.AAUGAUU...UCUAA..UAAUAUUAGCAUGUAUUUAACAAAAGAAGUAAAAGCCGAC
env-1308    CUAUAU.UUGCAGCC.............UGAUUGA...........AAA...................UCAGUCA....................................GAUAC.AUAAA.AAUCAUU...UA.AA..UAAUAUUAGCAUGUAUUUAACGAAAGAAGUUAAAGCUGAG
env-1309    UUAUAU.UUGCAGCC.............UGAUUGA...........AAA...................UCAAUCA....................................AAUAC.AUAAU.AAUCAUU...UACAA..CAAUAUUAGCAUG...........................
env-1310    UUAUAU.UUGCAGCC.............UGAUUGA...........AAA...................UCAAUCA....................................GAUAC.AUAAU.AAUCAUU...UACAA..UAAUAUUAGCAUGUACUUAACAAAAGAAGUAAAAGCCGAA
env-1311    UUAUAU.UUGCAGCC.............UGAUUGA...........AAA...................UCAAUCA....................................GAUAC.AUAAU.AAUCAUU...UACAA..UAAUAUUAGCAUGUACUUAACAAAAGAAGUAAAAGCCGAA
Tna-1-1     UUAUCU.UUGCGGCU.............UGAAAGA...........ACG...................UCUUUCA....................................UAUAC.AUAAA.AAUCAUU...UA.AA..UAAUAUUAGCAUGUAUUUAUCAAAAGAAGUAAAAGAAGAA
env-1312    UUACAU.UUGCGCCC.............UGAUUGA...........AAA...................UCAAUCA....................................CAUAC.AUCAU.AAUCAUU...UAAUA..UAAUAU.GGAAUGUAUAACUCCCUUUAUUUG.........
Cly-1-1     GUAUAU.UUGCGGCU.............UGAUUG............AAAAC..................CAAUCA....................................CAUAC.AUAAA.AAUCAUU...AA.AU..AAAUAUUGGAAUGUAUUUAACAAAAGAAGUAAAAGCCGAA
Cge-1-1     GUAUAU.UUGCGGCU.............UGAUUG............AAAAC..................CAAUCA....................................CAUAC.AUAAA.AAUCAUU...AA.AU..AAAUAUUGGAAUGUAUUUAACAAAAGAAGUAAAAGCCGAA
Cly-2-1     GUAUAU.UUGCGGCU.............UGAUUG............AAAAC..................CAAUCA....................................CAUAC.AUAAA.AAUCAUU...AA.AU..AAAUAUUGGAAUGUAUUUAACAAAAGAAGUAAAAGCCGAA
Sxi-1-1     UUAUAU.UUGCGCCC.............UGGUAG............AAAAA..................CUGCCA.....................................AUAC.AUAAA.AAUCAUU...AAAAU..UAAUAUUGGAAUGUAUUUAACUAAGGAAGCAAAAGAAGAA

            ............................<<<<<<<<<<<<<<<<<<...........>>>>>>>>>>>>>>>>>>.........<<<<<<<<........>>>>>>>>........................................................................
            ............................2222221111..101.00...........00.101..1111222222.........11011000........00011011........................................................................
            •UAY•U-UUGC••CY-------------∘∘∘∘∘∘∘-----------R•R∘∘-----------------∘∘∘∘∘∘∘------------------------------------∘UAC-AUAA•-AAUCAUU---UA-AA--YAAUAUURG•AUGUAUUUARC•ARAGAAR••AAARRRRAR
            5′                                                                                                                                                   3′
```

```
Ste-1-1   UUAUAU.UUGCGCCC.............UGGCAG............AAAAG..................CUGUCA.....................................AUAC.AUAAA.AAUCAUU...AAAAU..UAAUAUUGGAAUGUAUUUAACUAAAGACGCAAAAGAAGAA
Cal-1-1   AUAUCU.UUGCAAAC.............UGGUUGAA..........AAA..................UUCAAUCA....................................UAUAC.AUAAA.AAUCAUU...UAAAC.AUAAUAUUGGAAUGUAUUUAACUAAAGAAGGAAAAGCUGAG
env-1313  UUACCU.UUGCACAC.............GAAUUGGA..........AAAAC................UUCGAUUC.....................................AUAC.AUAAU.AAUCAUU...UA.AA..UAAUAUUAGCAUGUACUUAACUAAAGAAGUAAAGGAACAA
Mze-2-1   UUAUCU.UUGCCGCC.............UGAUUG............AAACC..................CAGUCA....................................GGUAC.AUAAA.AAUCAUU...UAAAA..CAAUAUUGGAAUGUAUUUAAGCAAAGAGGCUAAGGCCGAA
env-1314  UUACCU.UUGCAACC.............GAAUUAGG..........AAAAC................CCUGAUUC.....................................AUAC.AUAAU.AAUCAUU...UAAUA..UAAUAUUGGAAUGUAUUUAACUAAAGAAACAAAAGCUGAA
env-1316  UUACCU.UUGCAACC.............GAAUUAGG..........AAAAC................CCUGAUUC.....................................AUAC.AUAAU.AAUCAUU...UAAUA..UAAUAUUGGAAUGUAUUUAACUAAAGAAACAAAAGCUGAA
env-1317  UUACCU.UUGCACAC.............GAAUUGAGA.........GAA.................UCUCGGUUC.....................................AUAC.AUAAU.AAUCAUU...UA.AA..GAAUAUUAGCAUGUACUUAACAUCAGAAGUUAAAGCAGAA
env-1318  UUAUCU.UUGCACAC.............GAAUUGAGA.........GAA.................UCUCGAUUC.....................................AUAC.AUAAU.AAUCAUU...AAAAA..UAAUAUUGGAAUGUAUUUAAGUAAAGAAAAAAAAGCUGAA
Fgi-1-1   UUACUU.UUGCACCC.............AUCUUGCA..........ACAGA................UGUAAGAA......................................UAC.AUAAU.AAUCAUU...UA.AA..AAAUAUUGGAAUGUACUUAACUAAAGAAGUUAAAGAAGAA
Fsu-2-1   CUACUU.UUGCACCC.............GUCUUACA..........AGAGUA...............UGUUCGAC.....................................AUAC.AUAAA.AAUCAUU...UA.AA..UAAUAUUAGCAUGUAUUUAACUAAAGAAGUUAAAGCGGAA
env-1319  GUACUU.UUGCACCC.............GUCUUACA..........AUAGC................UGUAUGAC.....................................AUAC.AUAAA.AAUCAUU...UA.AA..UAAUAUUAGCAUGUAUUUAACUAAAGAGAUUAAAGAAGAG
env-1320  CUACUU.UUGCACCC.............GUCUUACA..........AUAGA................UGUAUGAC.....................................AUAC.AUAAC.AAUCAUU...UA.AA..UAAUAUUAGCAUGUAUUUAACAAAAGAGGUUAAAGAAGAA
env-1321  CUACUU.UUGCACCC.............GUCUUACA..........AUAGA................UGUAUGAC.....................................AUAC.AUAAC.AAUCAUU...UA.AA..UAAUAUUAGCAUGUAUUUAACAAAAGAGGUUAAAGAAGAA
Fjo-1-1   CUAAUU.UUGCACCC.............GUCUUACAA.........AAG.................UUGUAUGAC.....................................AUAC.AUAAA.AAUCAUU...AA.AA..CAAUAUUGGAAUGUAUUUAACUAAAGAAAAGAAAGAAGAA
Fsp-7-1   CUAAUU.UUGCACCC.............GUCUUACAA.........AAG.................UUGUAUGAC.....................................AUAC.AUAAA.AAUCAUU...UAAAA..CAAUAUUGGAAUGUAUUUAACUAAAGAAAAGAAAGAAGAA
Fsp-13-1  CUAAUU.UUGCACCC.............GUCUUACAA.........AAG.................UUGUAUGAC.....................................AUAC.AUAAA.AAUCAU....UA.AA..CAAUAUUGGAAUGUAUUUAACUAAAGAAAAGAAAGAAGAG
Fsp-6-1   GUAUUU.UUGCACCC.............GUCUUACAA.........AAG.................UUGUAUGAC.....................................AUAC.AUAAA.AAUCAU....UA.AA..UAAUAUUGGAAUGUAUUUAACUAAAGAAAAGAAAGAAGAG
Fsp-8-1   CUAUUU.UUGCACCC.............GUCUUACAA.........AAG.................UUGUAUGAC.....................................AUAC.AUAAA.AAUCAU....UA.AA..UAAUAUUGGAAUGUAUUUAACUAAAGAAAAAAAGGAAGAG
Fsp-10-1  CUAUUU.UUGCACCC.............GUCUUACAA.........AAG.................UUGUAUGAC.....................................AUAC.AUAAA.AAUCAU....UA.AA..UAAUAUUGGAAUGUAUUUAACUAAAGAAAAAAAGGAAGAG
Fhy-1-1   CUAUUU.UUGCACCC.............GUCUUACAA.........AAG.................UUGUAUGAC.....................................AUAC.AUAAA.AAUCAU....UA.AA..UAAUAUUGGAAUGUAUUUAAGUAAAGAGAAAAAAGAAGAG
env-1323  GUAUUU.UUGCACCC.............GUCUUACAA.........AAG.................UUGUAUGAC.....................................AUAC.AUAAA.AAUCAU....UA.AA..UAAUAUUGGAAUGUAUUUAACUAAAGAAAAGAAAGAAGAG
env-1324  GUAAUU.UUGCACCC.............GUCUUACAA.........AAG.................UUGUCUGAC.....................................AUAC.AUAAA.AAUCAU....UA.AA..UAAUAUUGGAAUGUAUUUAACUAAAGAAAAGAAAGAAGAG
Fsp-11-1  UUAUUU.UUGCACCC.............GUCUUACA..........AAAGA................UGUAUGAC.....................................AUAC.AUAAA.AAUCAUU...UA.AA..UAAUAUUGGAAUGUAUUUAACUAAAGAAAAGAAAGAAGAG
Fch-1-1   GUAAUU.UUGCACCC.............GUCUUACA..........AAAGC................UGUAUGAC.....................................AUAC.AUAAA.AAUCAU....UA.AA..UAAUAUUGGAAUGUAUUUAACUAAAGAAAAGAAAGAAGAG
env-1325  AUACUU.UUGCACUC.............GAAUACG...........AAAAA.................CGUAUUC.....................................AUAC.AUAAU.AAUCAUU...UA.AA..UAAUAUUGGAAUGUACUUAACUAAAGAAGUUAAAGAAGAA
env-1326  AUACUU.UUGCACUC.............GAAUACG...........AAAAA.................CGUAUUU.....................................AUAC.AUAAU.AAUCAUU...UA.AA..UAAUAUUGGAAUGUACUUAACUAAAGAAGUUAAAGAAGAA
env-1327  AUACUU.UUGCACUC.............GAAUACG...........AAAAA.................CGUAUUU.....................................AUAC.AUAAU.AAUCAUU...UA.AA..UAAUAUUGGAAUGUACUUAACUAAAGAAGUUAAAGAAGAA
env-1328  AUACUU.UUGCACUC.............GAAUACG...........AAAAA.................CGUAUUU.....................................AUAC.AUAAU.AAUCAUU...UA.AA..UAAUAUUGGAAUGUACUUAACUAAAGAAGUUAAAGAAGAA
env-1330  AUACUU.UUGCACAC.............GCAUACG...........AAAAA.................CGUAU.C......................................UAC.AUAAU.AAUCAUU...UA.AA..UAAUAUUAAAAUGUACUUAACUAAAGAAGUUAAAGAAGAA
env-1331  AUACUU.UUGCACAC.............GCAUACG...........AAAAA.................CGUAU.C......................................UAC.AUAAU.AAUCAUU...UA.AA..UAAUAUUAAAAUGUACUUAACUAAAGAAGUUAAAGAAGAA
env-1332  UUACUU.UUGCACCC.............ACUUGAU...........AAAC..................AUCAAGA......................................UAC.AUAAU.AAUCAUU...UAAAA..CAAUAUUGGCAUGUAUUUAACUAAAGAAGUUAAACAAGAA
env-1333  UUACCU.UUGCGCCC.............ACUUGAU...........AAAC..................AUCAAGU......................................UAC.AUAAU.AAUCAUU...UAAAA..CAAUAUUAGCAUGUAUUUAACUAAAGAAGUUAAACAAGAU
Gsp-4-1   GUAUCU.UUGCCGCU.............UGACAGA...........AAA...................UCUGUUU.....................................AUAC.AUAAA.AAUCAUU...AAAAU..UAAUAUUGGAAUGUAUUUAUCCAUAGAAGAAAAACAAGAU
Gli-1-1   GUAUCU.UUGCCGCU.............UGACAGA...........AAA...................UCUGUUC.....................................AUAC.AUAAA.AAUCAUU...AAAAU..UAAUAUUGGAAUGUAUUUAACUAAAGAAGAAAAGCAAGAU
env-1334  GUAUCUUUUGCCGCU.............UGACAGA...........AAA...................UCUGUUC.....................................AUAC.AUAAA.AAUCAUU...AAAAU..UAAUAUUGGAAUGUAUUUAACUAAAGAAGAAAAGCAAGAU
env-1335  GUAUAU.UUGCCGCU.............UGACAG............AAAUG..................CUGUCA.....................................AUAC.AUAAA.AAUCAUU...AAAAU..UAAUAUUGGAAUGUAUUUAACUAAAGAAGAAAAACAAGAU
env-1336  GUACAU.UUGCCGCU.............UGACAG............AAAAA..................CUGUCC.....................................AUAC.AUAAA.AAUCAUU...AAAAU..UAAUAUUGGAAUGUAUUUAACAAUUGAAGAAAAACAAGAG
env-1337  GUACAU.UUGCCGCU.............UGACAG............AAAAA..................CUGUCC.....................................AUAC.AUAAA.AAUCAUU...AAAAU..UAAUAUUGGAAUGUAUUUAACAAUUGAAGAAAAACAAGAG
env-1338  GUAUAU.UUGCGCCU.............UGACAG............AAAUA..................CUGUCU.....................................GUAC.AUAAA.AAUCAUU...AAAAU..UAAUAUUGGAAUGUAUUUAACUAAGGAAGAAAAGCAAG..
env-1339  GUAUAU.UUGCGCCU.............UGACAG............AAAUA..................CUGUCU.....................................GUAC.AUAAA.AAUCAUU...AAAAU..UAAUAUUGGAAUGUAUUUAACUAAGGAAGAAAAGCAAGAG
env-1340  GUAUAU.UUGCGCCU.............UGACAG............AAAUA..................CUGUCU.....................................GUAC.AUAAA.AAUCAUU...AAAAU..UAAUAUUGGAAUGUAUUUAACUA.................
env-1341  GUAUAU.UUGCGCCU.............UGACAG............AAAUA..................CUGUCU.....................................GUAC.AUAAA.AAUCAUU...AAAAU..UAAUAUUGGAAUGUAUUUAACUAAGGAAGAAAAGCAAGAG
Gsp-5-1   GUAUAU.UUGCGCCU.............UGACAG............AAAUA..................CUGUCU.....................................GUAC.AUAAA.AAUCAUU...AAAAU..UAAUAUUGGAAUGUAUUUAACUAAGGAAGAAAAGCAAGAG
env-1342  GUAUAU.UUGCGCCU.............UGACAG............AAAUA..................CUGUCU.....................................GUAC.AUAAA.AAUCAUU...AAAAU..UAAUAUUGGAAUGUAUUUAACUAAGGA.............
env-1343  GUAUAU.UUGCGCCU.............UGACAG............AAAUA..................CUGUCU.....................................GUAC.AUAAA.AAUCAUU...AAAAU..UAAUAUUGGAAUGUAUUUAACUAAGGAAGAAAAG......
env-1344  GUAUAU.UUGCGCCU.............UGACAG............AAAUA..................CUGUCU.....................................GUAC.AUAAA.AAUCAUU...AAAAU..UAAUAUUGGAAUGUAUUUAACUAAGGAAGAAAAGCAAGAG
env-1345  GUAUAU.UUGCGCCU.............UGACAG............AAAUA..................CUGUCU.....................................GUAC.AUAAA.AAUCAUU...AAAAU..UAAUAUUGGAAUGUAUUUAACUAAGGAAGAAAAGCAA...
env-1346  GUAUAU.UUGCGCCU.............UGACAG............AAAUA..................CUGUCU.....................................GUAC.AUAAA.AAUCAUU...AAAAU..UAAUAUUGGAAUGUAUUUAACUAAGGAAGAAAAGCAAG..
Gsp-3-1   GUAUAU.UUGCGCCU.............UGACAG............AAAUA..................CUGUCU.....................................GUAC.AUAAA.AAUCAUU...AAAAU..UAAUAUUGGAAUGUAUUUAACUAAGGAAGAAAAGCAGGAG
Gsp-2-1   GUAUAU.UUGCGCCU.............UGACAG............AAAUA..................CUGUCU.....................................GUAC.AUAAA.AAUCAUU...AAAAU..UAAUAUUGGAAUGUAUUUAACUAAGGAAGAAAAGCAAGAG
env-1347  GUAUAU.UUGCGCCU.............UGACAG............AAAUA..................CUGUCU.....................................GUAC.AUAAA.AAUCAUU...AAAAU..UAAUAUUGGAAUGUAUUUAACUAAGGAAGAAAAGCAAG..
env-1348  GUAUAU.UUGCCGCU.............UGACAG............AAAAA..................CUGUCU.....................................GUAC.AUAAA.AAUCAUU...AAAAU..UAAUAUUGGAAUGUAUUUAACAAUUGAAGAAAAACAAGAG
env-1349  GUAUAU.UUGCCGCU.............UGACAG............AAAAA..................CUGUCU.....................................GUAC.AUAAA.AAUCAUU...AAAAU..UAAUAUUGGAAUGUAUUUAACAAUUGAAGAAAAACAAGAG
env-1350  GUAUAU.UUGCCGCU.............UGACAG............AAAAA..................CUGUCU.....................................GUAC.AUAAA.AAUCAUU...AAAAU..UAAUAUUGGAAUGUAUUUAACAAUUGAAGAAAAACAAGAG
Gec-1-1   GUAUCU.UUGCCGCC.............UGACAGA...........AAA...................UCUGUCC.....................................AUAC.AUAAA.AAUCAUU...UA.AA..UAAUAUUGGAAUGUAUUUAGCAAAAGAGAAAAAAGAAGAG
Gpo-1-1   AUAUCU.UUGCCGCC.............UGACAG............AAAAG..................CUGUCA.....................................AUAC.AUAAA.AAUCAUU...UA.AA..UAAUAUUGGAAUGUAUCUAGCAAAAGAGAAAAAAGAAGAG
env-1351  AUAUCU.UUGCCGCC.............UGACAG............AAAUG..................CUGUCA.....................................AUAC.AUAAA.AAUCAUU...UA.AA..UAAUAUUGGAAUGUAUCUAGCAAAAGAAAGAAAAGAAGAG
env-1352  GUAUCU.UUGCCGCC.............UGACAG............AAAAA..................CUGUCC.....................................AUAC.AUAAA.AAUCAUU...UA.AA..UAAUAUUGGAAUGUAUUUAACUAGUGAAAAGAAAGAAGAG
env-1353  GUAUCU.UUGCCGCC.............UGACAG............AAAAA..................CUGUCC.....................................AUAC.AUAAA.AAUCAUU...UA.AA..UAAUAUUGGAAUGUAUUUAACUAGUGAAAAGAAAGAAGAG
Gfo-1-1   GUAUCU.UUGCCGCC.............UGACAG............AAAAA..................CUGUCC.....................................AUAC.AUAAA.AAUCAUU...UA.AA..UAAUAUUGGAAUGUAUUUAACUAGUGAAAAAAAAGAAGAG
env-1354  GUACAU.UUGCGGCC.............UGACAG............AAAAG..................CUGUCA.....................................AUAC.AUAAA.AAUCAUU...UA.AA..UAAUAUUGGAAUGUAUUUAACUAAAGAAAAGAAGGAAGAA
env-1355  GUACAU.UUGCGGCC.............UGACAG............AAAAG..................CUGUCA.....................................AUAC.AUAAA.AAUCAUU...UA.AA..UAAUAUUGGAAUGUAUUUAACUAAAGAAAAGAAGGAAGAA
env-1356  GUACAU.UUGCGGCC.............UGACAG............AAAAG..................CUGUCA.....................................AUAC.AUAAA.AAUCAUU...UA.AA..UAAUAUUGGAAUGUAUUUAACUAAAGAAAAGAAGGAAGAA
env-1357  GUACAU.UUGCGGCC.............UGACAG............AAAAG..................CUGUCA.....................................AUAC.AUAAA.AAUCAUU...UA.AA..UAAUAUUGGAAUGUAUUUAACUAAAGAAAAGAAGGAAGAA
env-1358  GUACAU.UUGCGGCC.............UGACAG............AAAAG..................CUGUCA.....................................AUAC.AUAAA.AAUCAUU...UA.AA..UAAUAUUGGAAUGUAUUUAACUAAAGAAAAGAAGGAAGAA
env-1359  GUACAU.UUGCGGCC.............UGACAG............AAAAG..................CUGUCA.....................................AUAC.AUAAA.AAUCAUU...UA.AA..UAAUAUUGGAAUGUAUUUAACUAAAGAAAAGAAGGAAGAA
env-1360  GUACAU.UUGCGGCC.............UGACAG............AAAAG..................CUGUCA.....................................AUAC.AUAAA.AAUCAUU...UA.AA..UAAUAUUGGAAUGUAUUUAACUAAAGAAAAGAAGGAAGAA
env-1367  GUACAU.UUGCGGCC.............UGACAG............AAAAG..................CUGUCA.....................................AUAC.AUAAA.AAUCAUU...UA.AA..UAAUAUUGGAAUGUAUUUAACUAAAGAAAAGAAGGAAGAA
env-1370  GUACAU.UUGCGGCC.............UGACAG............AAAAG..................CUGUCA.....................................AUAC.AUAAA.AAUCAUU...UA.AA..UAAUAUUGGAAUGUAUUUAACUAAAGAAAAGAAGGAAGAA
env-1371  GUACAU.UUGCGGCC.............UGACAG............AAAAG..................CUGUCA.....................................AUAC.AUAAA.AAUCAUU...UA.AA..UAAUAUUGGAAUGUAUUUAACUAAAGAAAAGAAGGAAGAA
env-1373  GUACAU.UUGCGGCC.............UGACAG............AAAAG..................CUGUCA.....................................AUAC.AUAAA.AAUCAUU...UA.AA..UAAUAUUGGAAUGUAUUUAACUAAAGAAAAGAAGGAAGAA
env-1374  GUACAU.UUGCGGCC.............UGACAG............AAAAG..................CUGUCA.....................................AUAC.AUAAA.AAUCAUU...UA.AA..UAAUAUUGGAAUGUAUUUAACUAAAGAAAAGAAGGAAGAA
env-1375  GUACAU.UUGCGGCC.............UGACAG............AAAAG..................CUGUCA.....................................AUAC.AUAAA.AAUCAUU...UA.AA..UAAUAUUGGAAUGUAUUUAACUAAAGAAAAGAAGGAAGAA
env-1376  GUACAU.UUGCGGCC.............UGACAG............AAAAG..................CUGUCA.....................................AUAC.AUAAA.AAUCAUU...UA.AA..UAAUAUUGGAAUGUAUUUAACUAAAGAAAAGAAGGAAGAA
env-1377  GUACAU.UUGCGGCC.............UGACAG............AAAAG..................CUGUCA.....................................AUAC.AUAAA.AAUCAUU...UA.AA..UAAUAUUGGAAUGUAUUUAACUAAAGAAAAGAAGGAAGAA
env-1378  GUAUCU.UUGCCGCC.............UGAUAG............AUAAA..................CUGUCA.....................................AUAC.AUAAA.AAUCAUU...AAAAU..UAAUAUUGGAAUGUAUUUAGCAAAAGAGAAGAAAGAAGAA
env-1379  GUAUCU.UUGCCGCC.............UGACGG............AAAAA..................CUGUCA.....................................AUAC.AUAAA.AAUCAUU...AAAAU..UAAUAUUGGAAUGUAUUUAGCCAAAGAAAGAAAAGAAGAA
env-1380  GUAUCU.UUGCCGCC.............UGACGG............AAAAA..................CUGUCA.....................................AUAC.AUAAA.AAUCAUU...AAAAU..UAAUAUUGGAAUGUAUUUAGCCAAAGAAAGAAAAGAAGAA
env-1381  GUAUCU.UUGCCGCC.............UGACGG............AAAAA..................CUGUCA.....................................AUAC.AUAAA.AAUCAUU...AAAAU..UAAUAUUGGAAUGUAUUUAGCCAAAGAAAGAAAAGAAGAA
env-1382  GUAUCU.UUGCCGCC.............UGACGG............AAAAA..................CUGUCA.....................................AUAC.AUAAA.AAUCAUU...AAAAU..UAAUAUUGGAAUGUAUUUAGCCAAAGAAAGAAAAGAAGAA
env-1383  GUAUCU.UUGCCGCC.............UGACGG............AAAAA..................CUGUCA.....................................AUAC.AUAAA.AAUCAUU...AAAAU..UAAUAUUGGAAUGUAUUUAGCCAAAGAAAGAAAAGAAGAA
Ssp-1-1   GUAUCU.UUGCCGAC.............UGAUGG............AAAAA..................CUGUCA.....................................AUAC.AUAAA.AAUCAUU...AAAAU..UAAUAUUGGAAUGUAUUUAGCCAAAGAAAGAAAAGAAGAA
Tse-1-1   UUAUAU.UUGCCGCU.............UGAAAG............AAAAC..................CUUUCA....................................UAUAC.AUAAA.AAUCAUU...UU.AA..AAAUAUUAGCAUGUAUUUAACUAAAGAAGUCAAAGAAGAA

          ............................<<<<<<<<<<<<<<<<<<...........>>>>>>>>>>>>>>>>>>.........<<<<<<<<........>>>>>>>>........................................................................
          ............................2222221111..101.00...........00.101..1111222222.........11011000........00011011........................................................................
          •UAY•U-UUGC••CY-------------∘∘∘∘∘∘∘-----------R•R∘∘-----------------∘∘∘∘∘∘∘-------------------------------------∘UAC-AUAA•-AAUCAUU---UA-AA--YAAUAUURG•AUGUAUUARC•ARAGAAR••AAARRRRAR
          5'                                                                                                                                                            3'
```

```
Lsp-2-1   GUAUAU.UUGCCGCU.............UGAAAG............AUAAG..................CUUUCA....................................UAUAC.AUAAC.AAUCAUU...UACAA..CAAUAUUAGAAUGUAUCUAACAAAAGAAGAAAAAGCAAAA
Jpa-1-1   CUAUAU.UUGCCGCU.............UGAAAG............AAAGA..................CUUUCA...................................UAGUAC.AUAAC.AAUCAUU...UACAA..CAAUAUUAGCAUGUAUUUAACAAAAGAAGCAAAAGAAGAA
Ssp-2-1   AUAUAU.UUGCAGCC.............UGAAAGA...........AAA...................UCUUUCA...................................ACGUAC.AUAAC.AAUCAUU...UACAA..CAAUAUUAGCAUGUAUUUAGAUCAAAAAGAAAAAGAGGCC
env-1384  GUAUAU.UUGCCGCU.............CGAAAG............AAGAG..................CUUUCG....................................UAUAC.AUAAC.AAUCAUU...UACAA..UAAUAUUAGAAUGUAUUUAGAUCCAAAAGAAAAAGAAGCC
env-1385  CUAUAU.UUGCCGCU.............CGAAAGA...........AAA...................UCUUUCA...................................GCGUAC.AUAAA.AAUCAUU...UACAA..UAAUAUUAGCAUGUAUUUAGAUCCAAAAGAAAAAGAAGCA
env-1386  CUAUAU.UUGCCGCU.............CGAAAGA...........AAA...................UCUUUCA...................................GCGUAC.AUAAA.AAUCAUU...UACAA..UAAUAUUAGCAUGUAUUUAGAUCCAAAAGAAAAAGAAGCA
env-1387  CUAUAU.UUGCAACU.............UGAAAGA...........AAA...................UCUUUCA...................................UUGUAC.AUAAC.AAUCAUU...UACAA..UAAUAUUAGCAUGUAUUUAGAUCCAAAAGAAAAAGAAGCC
env-1388  CUAUAU.UUGCAACU.............UGAAAGA...........AAA...................UCUUUCA...................................UUGUAC.AUAAC.AAUCAUU...UACAA..UAAUAUUAGCAUGUAUUUAGAUCCAAAAGAAAAAGAAGCC
env-1389  CUAUAU.UUGCAACU.............UGAAAGA...........AAA...................UCUUUCA...................................UUGUAC.AUAAC.AAUCAUU...UACAA..UAAUAUUAGCAUGUAUUUAGAUCCAAAAGAAAAAGAAGCC
env-1391  CUAUAU.UUGCAACU.............UGAAAGA...........AAA...................UCUUUCA...................................UCGUAC.AUAAC.AAUCAUU...UACAA..UAAUAUUAGCAUGUAUUUAGAUCCAAAAGAAAAAGAAGCC
env-1392  UUAUAU.UUGCCGCU.............UGAAAG............AAAAC..................CUUUCA...................................AAGUAC.AUAAC.AAUCAUU...UACAA..CAAUAUUAGAAUGUAU........................
env-1393  UUAUAU.UUGCCGCU.............UGAAAG............AAAAC..................CUUUCA...................................AAGUAC.AUAAC.AAUCAUU...UACAA..CAAUAUUAGAAUGUAUUUAUCUAAAGAAAAAAAAGAAGAA
env-1394  UUAUAU.UUGCCGCU.............UGAAAG............AAAAC..................CUUUCA...................................AAGUAC.AUAAC.AAUCAUU...UACAA..CAAUAUUAGAAUGUAUUU......................
env-1395  UUAUAU.UUGCCGCU.............UGAAAG............AAAAC..................CUUUCA...................................AAGUAC.AUAAC.AAUCAUU...UACAA..CAAUAUUAGAAUGUAUUUAUCUAAAGAAAAAAAAGAAGAA
env-1396  CUAUAU.UUGCAGCU.............UGGAAGA...........AAA...................UCUUUCA...................................AAGUAC.AUAAA.AAUCAUU...UACAA..UAAUAUUAGCAUGUAUUUAGACAAAAAAGAAAAAGAAGCC
env-1397  CUAUAU.UUGCAGCU.............UGGAAGA...........AAA...................UCUUUCA...................................AAGUAC.AUAAA.AAUCAUU...UACAA..UAAUAUUAGCAUGUAUUUAGACAAAAAAGAAAAAGAAGCC
env-1398  UUAUAU.UUGCCGAC.............UGAAAGA...........AAA...................UCUUUCA...................................AAGUAC.AUAAC.AAUCAUU...UACAA..CAAUAUUAGAAUGUAUU.......................
env-1399  UUAUAU.UUGCCGCU.............UGAAAGA...........AAA...................UCUUUCA...................................AAGUAC.AUAAC.AAUCAUU...UACAA..CAAUAUUAGAAUGUAUUUAUCUAAAGAGAAAAAAGAAGAA
env-1400  UUAUAU.UUGCCGCU.............UGAAAGA...........AAA...................UCUUUCA...................................AAGUAC.AUAAC.AAUCAUU...UACAA..CAAUAUUAGAAUGUAU........................
env-1401  UUAUAU.UUGCCGCU.............UGAAAGA...........AAA...................UCUUUCA...................................AAGUAC.AUAAC.AAUCAUU...UACAA..CAAUAUUAGAAUGUAUUUAUCUAAAGAGAAAAAAGAAGAA
env-1402  UUAUAU.UUGCCGCU.............UGAAAGA...........AAA...................UCUUUCA...................................AAGUAC.AUAAC.AAUCAUU...UACAA..CAAUAUUAGAAUGUAUUUAUCUAAAGAGAAAAAAGAAGAA
env-1403  UUAUAU.UUGCCGCU.............UGAAAGA...........AAA...................UCUUUCA...................................UAGUAC.AUAAC.AAUCAUU...UACAA..UAAUAUUAGAAUGUAUUUAACUAAAGAGAAAAAAGAAGAA
env-1404  CUAUAU.UUGCCGCU.............UGAAAGA...........AAA...................UCUUUCA...................................UAGUAC.AUAAC.AAUCAUU...UACAA..UAAUAUUAGAAUGUAUUUAACAAAAGAAGCUAAAGAGAAA
env-1405  CUAUAU.UUGCCGCU.............UGGAAGA...........AAA...................UCUUUCA...................................UAGUAC.AUAAA.AAUCAUU...UACAA..UAAUAUUAGCAUGUAUUUAGAUACAAAAGAAAAAGAAGCC
env-1406  CUAUAU.UUGCCGCU.............UGGAAGA...........AAA...................UCUUUCA...................................UAGUAC.AUAAA.AAUCAUU...UACAA..UAAUAUUAGCAUGUAUUUAGAUACAAAAGAAAAAGAAGCC
env-1411  CUAUAU.UUGCCGCU.............UGAAAGA...........AAA...................UCUUUCA...................................UAGUAC.AUAAA.AAUCAUU...UACAA..UAAUAUUAGCAUGUAUUUAGAUACAAAAGAAAAAGAAGCC
Wsp-1-1   CUAUAU.UUGCCGCU.............UGAAAGA...........AAA...................UCUUUCA...................................UAGUAC.AUAAA.AAUCAUU...UACAA..UAAUAUUAGCAUGUAUUUAGAUCAAAAAGGAAAAGAAAAA
env-1412  UUAUAU.UUGCCGCU.............UGAAAGA...........AAA...................UCUUUCA...................................UAGUAC.AUAAA.AAUCAUU...UACAA..UAAUAUUAGCAUGUAUUUAACAAAAGAAGCUAAAGAGAAA
env-1413  UUAUAU.UUGCCGCU.............UGAAAGA...........AAA...................UCUUUCA...................................UAGUAC.AUAAA.AAUCAUU...UACAA..UAAUAUUAGCAUGUAUUUAACAAAAGAAGCUAAAGAGAAA
env-1414  UUAUAU.UUGCCGCU.............UGAAAGA...........AAA...................UCUUUCA...................................UAGUAC.AUAAA.AAUCAUU...UACAA..UAAUAUUAGCAUGUAUUUAACAAAAGAAGCUAAAGAGAAA
env-1415  UUAUAU.UUGCCGCU.............UGAAAGA...........AAA...................UCUUUCA...................................UAGUAC.AUAAA.AAUCAUU...UACAA..UAAUAUUAGCAUGUAUUUAACAAAAGAAGCUAAAGAGAAA
env-1416  UUAUAU.UUGCCGCU.............UGAAAGA...........AAA...................UCUUUCA...................................UUGUAC.AUAAA.AAUCAUU...UACAA..UAAUAUUAGCAUGUAUUUAGACAAAAAAGAAAAAGAGAAA
Bar-1-1   CUAUAU.UUGCAGCU.............UGAAAG............AAAAU..................CUUUCAUCAUG....UACUCAAUUUA.....AUUGGGUA....AUAC.AUAAC.AAUCAUU...UACAA..UAAUAUUAGAAUGUAUUUAUCAAAAGAAGAAAAGCAAGAU
env-1417  CUAUAU.UUGCAGCU.............UGAAAGA...........AAA...................UCUUUCA....................................UAUAC.AUAAC.AAUCAUU...UACAA..UAAUAUUAGAAUGUAUUUAUCAAAAGAAGAAAAGCAAGAG
env-1418  CUAUAU.UUGCAGCU.............UGAAAGA...........AAA...................UCUUUCA....................................UAUAC.AUAAC.AAUCAUU...UACAA..UAAUAUUAGAAUGUAUUUAUCAAAAGAAGAAAAGCAAGAG
env-1419  CUAUAU.UUGCCGCU.............UGAAAG............AGAAA..................CUUUCA....................................UAUAC.AUAAA.AAUCAUU...UACAA..UAAUAUUAGCAUGUAUUUAGAUCAAAAAGAAAAAGAAGCC
env-1420  CUAUAU.UUGCCGCU.............UGAAAG............AGAAA..................CUUUCA....................................UAUAC.AUAAA.AAUCAUU...UACAA..UAAUAUUAGCAUGUAUUUAGAUCAAAAAGAAAAAGAAGCC
Pme-1-1   CUAUAU.UUGCCGCU.............UGAAAG............AGAAA..................CUUUCA...................................ACGUAC.AUAAA.AAUCAUU...UACAA..CAAUAUUAGCAUGUAUUUAAGUCAAAAAGAAAAAGAAGCC
env-1421  CUAUCU.UUGCAGCU.............UGAAAG............AACGA..................CUUUCA...................................ACGUAC.AUAAA.AAUCAUU...UAUAA..CAAUAUUAGCAUGUAUUUAUCAAAAGAGGAAAAAGAAGAA
Psp-5-1   GUAUCU.UUGCAGCU.............UGAAAG............AAAGA..................CUUUCA...................................ACGUAC.AUAAA.AAUCAUU...UACAA..UAAUAUUAGCAUGUAUUUAGAUCAAAAAGAAAAAGAAGCC
Pbu-1-1   GUAUAU.UUGCAGCU.............UGAAAG............AAUGA..................CUUUCA...................................ACGUAC.AUAAA.AAUCAUU...UACAA..UAAUAUUAGCAUGUAUUUAUCAAAAGAAGAAAAACAAGAA
env-1422  GUAUAU.UUGCAGCU.............UGAAAG............AAUGA..................CUUUCA...................................ACGUAC.AUAAA.AAUCAUU...UACAA..UAAUAUUAGCAUGUAUUUAACUCAAAAAGAAAAAGAGGCC
env-1425  CUAUCU.UUGCCGCU.............UGAAAG............AAAGA..................CUUUCA...................................ACGUAC.AUAAA.AAUCAUU...UAUAA..UAAUAUUAGCAUGUAUUUAGAUCAAAAAGAAAAAGAAGCC
Pja-1-1   CUAUAU.UUGCCGCU.............UGAAAG............AACGA..................CUUUCA...................................ACGUAC.AUAAA.AAUCAUU...UACAA..CAAUAUUAGCAUGUAUUUAUCAAAGACAGAAAAAGAGAAA
env-1426  CUAUAU.UUGCCGCU.............UGAAAG............AAAGA..................CUUUCA...................................ACAUAC.AUAAA.AAUCAUU...UACAA..UAAUAUUAGCAUGUAUUUAGAUCAAAAAGGAAAAGAGAAA
env-1427  CUAUAU.UUGCCGCU.............UGAAAG............AAAGA..................CUUUCA...................................ACGUAC.AUAAA.AAUCAUU...UACAA..UAAUAUUAGCAUGUAUUUAGAUCAAAAAGAAAAAGAAGAA
env-1428  CUAUAU.UUGCCGCU.............UGAAAG............AACGA..................CUUUCA...................................ACGUAC.AUAAA.AAUCAUU...UACAA..UAAUAUUAGCAUGUAUUUAGAUCAAAAAGAAAAAGAAGAA
env-1429  UUAUAU.UUGCCGCU.............UGAAAGA...........AAA...................UCUUUCA...................................ACAUAC.AUAAC.AAUCAUU...UACAA..UAAUAUUAGCAUGUAUUUAGAUCAAAAAGAAAAAGAAGCU
env-1433  UUAUAU.UUGCCGCU.............UGAAAGA...........AAA...................UCUUUCA...................................ACAUAC.AUAAC.AAUCAUU...UACAA..UAAUAUUAGCAUGUAUUUAGAUCAAAAAGAAAAAGAAGCU
env-1434  CUAUAU.UUGCGGUU.............CGAAAG............AAAAC..................CUUUCG...................................UUGUAC.AUAAC.AAUCAUU...UACAA..UAAUAUUAGCAUGUAUUUAUCAAAACAAGCAAAAGAAGAA
env-1435  CUAUAU.UUGCGGUU.............CGAAAG............AAAAC..................CUUUCG...................................UUGUAC.AUAAC.AAUCAUU...UACAA..UAAUAUUAGCAUGUAUUUAUCAAAACAAGCAAAAGAAGAA
env-1436  CCUUUU.UUGCCGCU.............CGAAAG............AAAGG..................CUUUCG....................................UGUAC.AUAAC.AAUCAUU...UACAA..UAAUAUUAGCAUGUAUUUAGAAAAGAAAGAAAAAGAGAAA
Oma-1-1   CUAUAU.UUGCGGCU.............CGAAAG............AUAAA..................CUUUCG....................................UGUAC.AUAAC.AAUCAUU...UAUAA..UAAUAUUAGAAUGUAUUUAGAUCAAAAAGAAAAAGAAGAA
env-1437  CUAUAU.UUGCGGCU.............CGAAAG............AUAAA..................CUUUCG....................................UGUAC.AUAAC.AAUCAUU...UAUAA..UAAUAUUAGAAUGUAUUUAGAUCAAAAAGAAAAAGAAGAA
env-1438  CUAUAU.UUGCGGCU.............CGAAAG............AUAAA..................CUUUCG....................................UGUAC.AUAAC.AAUCAUU...UAUAA..UAAUAUUAGAAUGUAUUUAGAUCAAAAAGAAAAAGAAGAA
env-1439  CUAUAU.UUGCGGCU.............CGAAAG............AUAAA..................CUUUCG....................................UGUAC.AUAAC.AAUCAUU...UAUAA..UAAUAUUAGAAUGUAUUUAGAUCAAAAAGAAAAAGAAGAA
Sze-1-1   UUAUAU.UUGCCGCU.............CGAAAG............AUAGC..................CUUUCG....................................GUUAC.AUAAC.AAUUAUU...UACAA..UAAUAUUAGCAUGUAUUUAGCAAAAGAAGUAAAAGAAGAA
Gsa-1-1   CUAUAU.UUGCCGCU.............CGAAAG............AUAGC..................CUUUCG....................................UUUAC.AUAAC.AAUUAUU...UACAA..UAAUAUUAGCAUGUAUUUAUCAAAAGAAGUAAAAGAAGAA
env-1441  CUAUAU.UUGCCGCU.............CGAAAG............AAAAG..................CUUUCG....................................UAUAC.AUAAA.AAUCAUU...UACAA..UAAUAUUAGCAUGUAUUUAACAAAAGAAGCUAAAGAGAAA
env-1442  CUAUAU.UUGCGCCU.............CGAAAG............AAAGG..................CUUUCG....................................UGUAC.AUAAA.AAUCAUU...UACAA..UAAUAUUAGCAUGUAUUUAGACAAAAAAGAAAAAGAAAAA
Mze-1-1   UUAUAU.UUGCCGCU.............CGAAAG............AUAAC..................CUUUCG....................................UGUAC.AUAAC.AAUCAUU...UACAA..UAAUAUUAGCAUGUAUUUAUCAAAAGAAAAAAAAGAAGAA
env-1444  UUAUAU.UUGCCGCU.............CGAAAG............AUAAC..................CUUUCG....................................UGUAC.AUAAC.AAUCAUU...UACAA..UAAUAUUAGCAUGUAUUUAUCAAAAGAAAAAAAAGAAGAA
env-1445  CUAUAU.UUGCCGCU.............CGAAAG............AAAUA..................CUUUCG....................................UGUAC.AUAAC.AAUCAUU...UACAA..UAAUAUUAGCAUGUAUUUAGAUCAAAAAGCAAAAGAGAAA
env-1447  GUAUAU.UUGCGCCU.............CGAAAG............AAAGG..................CUUUCG....................................UGUAC.AUAAC.AAUCAUU...UACAA..UAAUAUUAGAAUGUAUUUAUCAAAAGAAGAGAAACAAGAC
env-1448  UUAUAU.UUGCGCCU.............CGAAAG............AAAGG..................CUUUCG....................................UGUAC.AUAAC.AAUCAUU...UACAA..UAAUAUUAGAAUGUAUUUAUCAAAAGAAGAGAAACAAGAC
env-1449  GUAUAU.UUGCGCCU.............CGAAAG............AAAGG..................CUUUCG....................................UGUAC.AUAAC.AAUCAUU...UACAA..UAAUAUUAGAAUGUAUUUAUCAAAAGAAGAGAAACAAGAC
env-1454  CUAUAU.UUGCCGCU.............CGAAAG............AUAGG..................CUUUCG....................................UGUAC.AUAAA.AAUCAUU...UACAA..CAAUAUUAGCAUGUAUUUAGAUCAAAAAGCAAAAGAAGAA
env-1455  GUAUAU.UUGCCGCU.............CGAAAG............AGAGG..................CUUUCG....................................UGUAC.AUAAA.AAUCAUU...UACAA..UAAUAUUAGCAUGUAUUUAGAUCAAAAAGCUAAAGAGGAA
Wps-1-1   GUAUAU.UUGCCGCU.............CGAAAG............AGAGG..................CUUUCG....................................UGUAC.AUAAA.AAUCAUU...UACAA..UAAUAUUAGCAUGUAUUUAGAUCAAAAAGCAAAAGAAGAG
env-1456  GUAUAU.UUGCCGCU.............CGAAAG............AGAGG..................CUUUCG....................................UGUAC.AUAAA.AAUCAUU...UACAA..UAAUAUUAGCAUGUAUUUAGAUCAAAAAGCUAAAGAGGAA
env-1457  CUAUAU.UUGCCGCU.............CGAAAG............AUAGG..................CUUUCG....................................UGUAC.AUAAA.AAUCAUU...UACAA..UAAUAUUAGCAUGUAUUUAGAUCAAAAAGCAAAAGAAGAA
Osp-2-1   CUAUAU.UUGCAGCU.............CGAAAG............AAAGG..................CUUUCG....................................UGUAC.AUAAC.AAUCAUU...UACAA..UAAUAUUAGAAUGUAUUUAGAUCAAAAAGAAAAGGAAGAG
Osp-1-1   CUAUAU.UUGCAGCU.............CGAAAG............AAAGG..................CUUUCG....................................UGUAC.AUAAC.AAUCAUU...UACAA..UAAUAUUAGAAUGUAUUUAGAUCAAAAAGAAAAGGAAGAG
env-1458  GUAUAU.UUGCCGCU.............CGAAAG............AGAGG..................CUUUCG....................................UGUAC.AUAAC.AAUCAUU...UACAA..UAAUAUUAGAAUGUAUUUAGAUCAAAAAGGAAAAGAGGAA
env-1459  GUAUAU.UUGCCGCU.............CGAAAG............AGAGG..................CUUUCG....................................UGUAC.AUAAC.AAUCAUU...UACAA..UAAUAUUAGAAUGUAUUUAGAUCAAAAAGGAAAAGAGGAA
env-1460  CUAUAU.UUGCCGCU.............CGAAAG............AGAGG..................CUUUCG....................................UGUAC.AUAAC.AAUCAUU...UACAA..UAAUAUUAGAAUGUAUUUAGAUCCAAAAGAAAAAGAGAAA
env-1461  GUAUAU.UUGCCGCU.............CGAAAG............AGAAG..................CUUUCG....................................UGUAC.AUAAC.AAUCAUU...UACAA..UAAUAUUAGAAUGUAUUUAGACCCAAAAGAAAAAGAAGCC
env-1462  GUAUAU.UUGCCGCU.............CGAAAG............AGAAG..................CUUUCG....................................UGUAC.AUAAC.AAUCAUU...UACAA..UAAUAUUAGAAUGUAUUUAGACCCAAAAGAAAAAGAAGCC
env-1463  GUAUAU.UUGCCGCU.............CGAAAG............AGAAG..................CUUUCG....................................UGUAC.AUAAC.AAUCAUU...UACAA..UAAUAUUAGAAUGUAUUUAUCAAAAGAAGCAAAAGAAGAA
Lsp-1-1   CUAUAU.UUGCCGCU.............CGAAAG............AUAAG..................CUUUCG....................................UGUAC.AUAAC.AAUCAUU...UACAA..UAAUAUUAGAAUGUAUUUAACAAAAGAAAACAAAGAAAAG
env-1464  CUAUAU.UUGCCGCU.............CGAAAG............AAAGG..................CUUUCG....................................UGUAC.AUAAC.AAUCAUU...UACAA..UAAUAUUAGAAUGUAUUUAGAUCCAAAAGAAAAAGAGAAA
env-1465  CUAUAU.UUGCCGCU.............CGAAAG............AAAGG..................CUUUCG....................................UGUAC.AUAAC.AAUCAUU...UACAA..UAAUAUUAGAAUGUAUUUAGAUCCAAAAGAAAAAGAGAAA
env-1466  CUAUAU.UUGCCGCU.............CGAAAG............AAAGG..................CUUUCG....................................UGUAC.AUAAC.AAUCAUU...UACAA..UAAUAUUAGAAUGUAUUUAGAUCCAAAAGAAAAAGAGAAA

          ............................<<<<<<<<<<<<<<<<<<...........>>>>>>>>>>>>>>>>>>.........<<<<<<<<........>>>>>>>>........................................................................
          ............................2222221111..101.00...........00.101..1111222222.........11011000........00011011........................................................................
          •UAY•U-UUGC••CY-------------○○○○○○○-----------R•R○○-----------------○○○○○○○-------------------------------------○UAC-AUAA•-AAUCAUU---UA-AA--YAAUAUURG•AUGUAUUUARC•ARAGAAR••AAARRRRAR
           5′                                                                                                                                         3′
```

```
env-1467  GUAUAU.UUGCCGCU............CGAAAG...........AAAGG.................CUUUCG.................................UGUAC.AUAAC.AAUCAUU...UACAA..UAAUAUUAGAAUGUAUUUAUCAAAAGAAGCAAAAGAAGAA
env-1468  CUAUAU.UUGCCGCU............CGAAAG...........AAAGG.................CUUUCG.................................UGUAC.AUAAC.AAUCAUU...UACAA..UAAUAUUAGAAUGUAUUUAGAUCCAAAAGAAAAAGAGAAA
env-1469  CUAUAU.UUGCCGCU............CGAAAG...........AAAGG.................CUUUCG.................................UGUAC.AUAAC.AAUCAUU...UACAA..UAAUAUUAGAAUGUAUUUAGAUCCAAAAGAAAAAGAGAAA
env-1471  UUAUAU.UUGCCGCU............CGAAAG...........AAAGA.................CUUUCG.................................AGUAC.AUAAA.AAUCAUU...UACAA..UAAUAUUAGCAUGUAUUUAUCAAAAGAAAAGAAAGAAGAA
Fba-5-1   CUAUAU.UUGCCGCU............CGAAAG...........AAAGG.................CUUUCG.................................AGUAC.AUAAA.AAUCAUU...UACAA..UAAUAUUAGCAUGUAUUUAGAUCAAAAGGAAAAGAAAAAA
Fic-1-1   CUAUAU.UUGCCGCU............CGAAAG...........AAAGG.................CUUUCG.................................UGUAC.AUAAA.AAUCAUU...UACAA..UAAUAUUAGCAUGUAUUUAGAAAAAAAAGAAAAGGAAAAA
env-1472  GUAUAU.UUGCCGCU............CGAAAG...........AAAGG.................CUUUCG.................................UGUAC.AUAAA.AAUCAUU...UACAA..UAAUAUUAGCAUGUAUUUAGAUCAAAAGGAAAGAGAAA
Lja-1-1   UUAUAU.UUGCCGCU............CGAAAG...........AAAGG.................CUUUCG.................................UGUAC.AUAAA.AAUCAUU...UACAA..UAAUAUUAGAAUGUAUUUAACAAAAGAAGAAAAAGAGAG
env-1473  UUAUAU.UUGCCGCU............CGAAAG...........AAAAG.................CUUUCG.................................UGUAC.AUAAA.AAUCAUU...UACAA..UAAUAUUAGAAUGUAUUUAACUAAAGAAGCAAAAGAAGAA
Fba-8-1   CUAUCU.UUGCCGCU............CGAAAG...........AAAGG.................CUUUCG.................................UGUAC.AUAAC.AAUCAUU...UACAA..UAAUAUUAGCAUGUAUUUAGCAAAAGAAGCAAAGAAAAAA
env-1474  CUAUAU.UUGCCGCU............CGAAAG...........AUAGG.................CUUUCG.................................UGUAC.AUAAC.AAUCAUU...UACAA..UAAUAUUAGCAUGUAUUUAGAUCAAAAGCAAAAGAGAAA
env-1475  CUAUAU.UUGCCGCU............CGAAAG...........AUAGG.................CUUUCG.................................UGUAC.AUAAC.AAUCAUU...UACAA..UAAUAUUAGCAUGUAUUUAGAUCAAAAGUAAAGAGGAA
env-1476  CUAUAU.UUGCCGCU............CGAAAG...........AAAAG.................CUUUCG.................................UGUAC.AUAAC.AAUCAUU...UACAA..UAAUAUUAGCAUGUAUUUAGAAAAGAAAGAAAAAGAAAAA
env-1477  UUAUAU.UUGCCGCU............CGAAAG...........AAAAG.................CUUUCG.................................UGUAC.AUAAC.AAUCAUU...UACAA..UAAUAUUAGCAUGUAUUUAGAAAAGAAAGAAAAAGAGAAA
env-1478  UUAUAU.UUGCCGCU............CGAAAG...........AAAAG.................CUUUCG.................................UGUAC.AUAAC.AAUCAUU...UACAA..UAAUAUUAGCAUGUAUUUAGAAAAGAAAGAAAAAGAGAAA
env-1479  UUAUAU.UUGCCGCU............CGAAAG...........AAAAG.................CUUUCG.................................UGUAC.AUAAC.AAUCAUU...UACAA..UAAUAUUAGCAUGUAUUUAGAAAAGAAAGAAAAAGAGAAA
env-1480  CUAUAU.UUGCCGCU............CGAAAG...........AAAGG.................CUUUCG.................................UGUAC.AUAAC.AAUCAUU...UACAA..UAAUAUUAGCAUGUAUUUAUCAAAAGAAGCAAAGAAGAA
env-1481  CUAUAU.UUGCCGCU............CGAAAG...........AAAGG.................CUUUCG.................................UGUAC.AUAAC.AAUCAUU...UACAA..UAAUAUUAGCAUGUAUUUAGAAAAGAAAGAAAAAGAGAAA
env-1482  CUAUAU.UUGCCGCU............CGAAAG...........AAAGG.................CUUUCG.................................UGUAC.AUAAC.AAUCAUU...UACAA..UAAUAUUAGCAUGU...........................
env-1483  CUAUAU.UUGCCGCU............CGAAAG...........AAAGG.................CUUUCG.................................UGUAC.AUAAC.AAUCAUU...UACAA..UAAUAUUAGCAUGUAUUUAUCAAAAGAAAAAAAAGAAGA
env-1484  CUAUAU.UUGCCGCU............CGAAAG...........AAAGG.................CUUUCG.................................UGUAC.AUAAC.AAUCAUU...UACAA..UAAUAUUAGCAUGUAUUUAGAAAAGAAAGAAAAAGAGAAA
env-1485  CUAUAU.UUGCCGCU............CGAAAG...........AAAGG.................CUUUCG.................................UGUAC.AUAAC.AAUCAUU...UACAA..UAAUAUUAGCAUGUAUUUAUCAAAAGAAGCAAAAGAAGAA
env-1486  CUAUAU.UUGCCGCU............CGAAAG...........AAAGG.................CUUUCG.................................UGUAC.AUAAC.AAUCAUU...UACAA..UAAUAUUAGCAUGUAUUUAGAAAAGAAAGAAAAAGAGAAA
env-1487  CUAUAU.UUGCCGCU............CGAAAG...........AAAGG.................CUUUCG.................................UGUAC.AUAAC.AAUCAUU...UACAA..UAAUAUUAGCAUGUAUUUAUCAAAAGAAGCAAAAGAAGAA
env-1488  CUAUAU.UUGCCGCU............CGAAAG...........AAAGG.................CUUUCG.................................UGUAC.AUAAC.AAUCAUU...UACAA..UAAUAUUAGCAUGUAUUUAGAAAAGAAAGAAAAAGAGAAA
env-1489  CUAUAU.UUGCCGCU............CGAAAG...........AAAGG.................CUUUCG.................................UGUAC.AUAAC.AAUCAUU...UACAA..UAAUAUUAGCAUGUAUUUAGAAAAGAAAGAAAAAGAGAAA
env-1490  UUAUAU.UUGCCGAU............CGAAAG...........AAAGG.................CUUUCG.................................UGUAC.AUAAC.AAUCAUU...UACAA..UAAUAUUAGCAUGUAU........................
env-1491  UUAUAU.UUGCCGAU............CGAAAG...........AAAGG.................CUUUCG.................................UGUAC.AUAAC.AAUCAUU...UACAA..UAAUAUUAGCAUGUAUUUAGAUCAAAAGAAAAAGAGAAA
env-1497  UUAUAU.UUGCCGAU............CGAAAG...........AAAGG.................CUUUCG.................................UGUAC.AUAAC.AAUCAUU...UACAA..UAAUAUUAGCAUGUAUUUAGAUCAAAAGAGAAAGAGAAA
env-1498  UUAUAU.UUGCCGAU............CGAAAG...........AAAGG.................CUUUCG.................................UGUAC.AUAAC.AAUCAUU...UACAA..UAAUAUUAGCAUGUAUUUAGAUCAAAAGAGAAAGAGAAA
env-1499  UUAUAU.UUGCCGAU............CGAAAG...........AAAGG.................CUUUCG.................................UGUAC.AUAAC.AAUCAUU...UACAA..UAAUAUUAGCAUGUAUUUAGAUCAAAAGAGAAAGAGAAA
env-1500  UUAUAU.UUGCCGAU............CGAAAG...........AAAGG.................CUUUCG.................................UGUAC.AUAAC.AAUCAUU...UACAA..UAAUAUUAGCAUGUAUUUAGAUCAAAAGAGAAAGAGAAA
env-1501  UUAUAU.UUGCGGCU............CGAAAG...........AAAGG.................CUUUCG.................................UGUAC.AUAAC.AAUCAUU...UACAA..UAAUAUUAGCAUGUAUUUAGAUCAAAAGAAAAAGAGAAA
env-1502  UUAUAU.UUGCGGCU............CGAAAG...........AAAGG.................CUUUCG.................................UGUAC.AUAAC.AAUCAUU...UACAA..UAAUAUUAGCAUGUAUUUAGAUCAAAAGAAAAAGAGAAA
env-1503  UUAUAU.UUGCGGCU............CGAAAG...........AAAGG.................CUUUCG.................................UGUAC.AUAAC.AAUCAUU...UACAA..UAAUAUUAGCAUGUAUUUAGAUCAAAAGAAAAAGAGAAA
env-1504  UUAUAU.UUGCGGCU............CGAAAG...........AAAGG.................CUUUCG.................................UGUAC.AUAAC.AAUCAUU...UACAA..UAAUAUUAGCAUGUAUUUAGAUCAAAAGAAA........
env-1505  CUAUAU.UUGCCGCU............CGAAAG...........AAAGG.................CUUUCG.................................UGUAC.AUAAA.AAUUAUU...UACAA..CAAUAUUAGAAUGUAUUUAGAUCAAAAGAAAAAGAAACC
env-1506  CUAUAU.UUGCCGCU............CGAAAG...........AAGAG.................CUUUCG.................................UGUAC.AUAAA.AAUCAUU...UACAA..UAAUAUUAGCAUGUAUUUAGAUCAAAAGCAAAAGAAGAA
Pda-1-1   CUAUAU.UUGCCGCU............CGAAAG...........AAGAG.................CUUUCG.................................UAUAC.AUAAA.AAUCAUU...UACAA..CAAUAUUAGCAUGUAUUUAGAUCAAAAGAAAAAGAAGCC
env-1507  CUAUAU.UUGCCGCU............CGAAAGA..........AAA.................UCUUUCG.................................UAUAC.AUAAC.AAUCAUU...UU.AA..UAAUAUUAGCAUGUAUUUAACAAAAGAAGCUAAAGAAGAG
env-1508  CUAUAU.UUGCCGCU............CGAAAGA..........AAA.................UCUUUCG.................................UAUAC.AUAAA.AAUCAUU...UU.AA..UAAUAUUAGCAUGUAUUUAACAAAAGAAGUUAAAGAAGAA
env-1515  NUAUAU.UUGCCGCU............CGAAAGA..........AAA.................UCUUUCG.................................UAUAC.AUAAN.AAUCAUU...UUAUAA..UAAUAUUAGCAUGUAUUUAACAAAAGAAGUUAAAGAAGAA
env-1516  UUAUAU.UUGCCGCC............UGAAAG...........AAAGA.................CUUUCA................................GCAUAC.AUAAU.AAUCAUU...GA.AA..UAAUAUUGGAAUGUAUUUAACAAAAGAGACAAAAGAAGAA
env-1517  GUAUCU.UUGCCGCC............UGAAAG...........AAAAA.................CUUUCA..................................AUAC.AUAAA.AAUCAUU...UA.AA..UAAUAUUGGCAUGUAUCUAGCUAAAGAAAAAAAAGAGAG
env-1519  CUAUAU.UUGCCGCC............UGAAAG...........AAAAA.................CUUUCA.................................CAUAC.AUAAA.AAUCAUU...UA.AA..UAAUAUUGGAAUGUAUUUAACAAAAGAAGUUAAAGAGGAA
env-1520  CUAUAU.UUGCCGCC............UGAAAG...........AAAAA.................CUUUCA.................................CAUAC.AUAAA.AAUCAUU...UA.AA..UAAUAUUGGAAUGUAUUUAACAAAAGAAGUUAAAGAGGAA
env-1521  CUAUAU.UUGCCGCC............UGAAAG...........AAAAA.................CUUUCA.................................CAUAC.AUAAA.AAUCAUU...UA.AA..UAAUAUUGGAAUGUAUUUAACAAAAGAAGUUAAAGAGGAA
env-1522  GUAUAU.UUGCAGCU............UGAAAG...........AAAAA.................CUUUCAUCACU....GUCCAA..CUGUAA....UUGGGC...CAUAC.AUAAA.AAUCAUU...UAAAA..UAAUAUUGGAAUGUAUUUAACUAAAGAAGUAAAAGAAGAG
env-1524  GUAUAU.UUGCAGCU............UGAAAG...........AAAAA.................CUUUCAUCACU....GUCCAA..CUGUAA....UUGGGC...CAUAC.AUAAA.AAUCAUU...UAAAA..UAAUAUUGGAAUGUAUUUAACUAAAGAAGUAAAAGAAGAG
env-1525  UUAUAU.UUGCAGCU............UGAAAG...........AAAAA.................CUUUCA.................................UGUAC.AUAAA.AAUCAUU...UAAAA..UAAUAUUGGAAUGUAUUUAGAUCAAAAGAAAAAGAAGCC
env-1526  UUAUAU.UUGCAGCU............UGAAAG...........AAAAA.................CUUUCA.................................UGUAC.AUAAA.AAUCAUU...UAAAA..UAAUAUUGGAAUGUAUUUAGAUCAAAAGAAAAAGAAGCC
env-1527  CUAUAU.UUGCCGCC............UGAAAGA..........AAA.................UCUUUCA................................GCGUAC.AUAAA.AAUCAUU...UA.AA..UAAUAUUGGAAUGUAUUUAACAAAAGAAGUUAAAGAGGAA
env-1528  CUAUAU.UUGCCGCC............UGAAAGA..........AAA.................UCUUUCA...............................ACAUAC.AUAAA.AAUCAUU...UA.AA..UAAUAUUGGAAUGUAUCUUCAAAAGAAGCUAAAGAGGAC
env-1529  CUAUAU.UUGCCGCC............UGAAAGA..........AAA.................UCUUUCG...............................ACAUAC.AUAAA.AAUGAUU...UA.AA..UAAUAUUGGAAUGUAUCUAACAAAAGAAGUUAAAGAGGAA
env-1530  CUAUAU.UUGCCGCC............UGAAAGA..........AAA.................UCUUUCG...............................ACAUAC.AUAAA.AAUGAUU...UA.AA..UAAUAUUGGAAUGUAUCUAACAAAAGAAGUUAAAGAGGAA
1 Fag-1-1 CUAUAU.UUGCCGCU............UGGAAG...........AAAAG.................CUUCCA.................................UGUAC.AUAAA.AAUCAUU...UAAAU..AAAUAUUGGAAUGUAUUUAACAAAAGAGUCUAAAGAAGAG
env-1534  AUAC.......................UGAAAG...........AAAAC.................CUUUCA.................................UGUAC.AUAAA.AAUCAUU...UAAAU..AAAUAUUGGAAUGUAUUUAACAAAAGAAGCAAAAGAAGAA
env-1535  CUAUAU.UUGCCGCU............UGAAAG...........AAAAC.................CUUUCA.................................UGUAC.AUAAA.AAUCAUU...UAAAU..AAAUAUUGGAAUGUAUUUAACAAAAGAAGCAAAAGAAGAA
env-1536  CUAUAU.UUGCCGCU............UGAAAG...........AAAAA.................CUUUCA.................................UGUAC.AUAAA.AAUCAUU...UAAAU..AAAUAUUGGAAUGUAUUUAACAAAAGAAGCAAAAGAAGAA
env-1537  CUAUAU.UUGCCGCU............UGAAAG...........AAAAA.................CUUUCA.................................UGUAC.AUAAA.AAUCAUU...UAAAU..AAAUAUUGGAAUGUAUUUAACAAAAGAAGCAAAAGAAGAA
env-1538  UUAUAU.UUGCCGAU............UGAAAGA..........AAA.................UCUUUCA...............................UUGUAC.AUAAA.AAUCAUU...UAAAU..AAAUAUUGGAAUGUAUUUAACAAAAGAAGCUAAAGAAAAA
env-1539  UUAUAU.UUGCCGAU............UGAAAGA..........AAA.................UCUUUCA...............................UUGUAC.AUAAA.AAUCAUU...UAAAU..AAAUAUUGGAAUGUAUUUAACAAAAGAAGCUAAAGAAAAA
env-1540  UUAUAU.UUGCCGAU............UGAAAGA..........AAA.................UCUUUCA...............................UAGUAC.AUAAA.AAUCAUU...UAAAU..AAAUAUUGGAAUGUAUUUAACAAAAGAAGCUAAAGAGGAA
env-1541  UUAUAU.UUGCCGAU............UGAAAGA..........AAA.................UCUUUCA...............................UAGUAC.AUAAA.AAUCAUU...UAAAU..AAAUAUUGGAAUGUAUUUAACAAAAGAAGCUAAAGAGGAA
env-1542  UUAUAU.UUGCCGAU............UGAAAGA..........AAA.................UCUUUCA...............................UAGUAC.AUAAA.AAUCAUU...UAAAU..AAAUAUUGGAAUGUAUUUAACAAAAGAAGCUAAAGAGGAA
env-1543  UUAUAU.UUGCCGAU............UGAAAGA..........AAA.................UCUUUCA...............................UAGUAC.AUAAA.AAUCAUU...UAAAU..AAAUAUUGGAAUGUAUUUAACAAAAGAAGCUAAAGAGGAA
env-1546  UUAUAU.UUGCCGAU............UGAAAGA..........AAA.................UCUUUCA...............................UAGUAC.AUAAA.AAUCAUU...UAAAU..AAAUAUUGGAAUGUAUUUAACAAAAGAAGCUAAAGAGGAA
env-1547  UUAUAU.UUGCCGAU............UGAAAGA..........AAA.................UCUUUCA...............................UAGUAC.AUAAA.AAUCAUU...UAAAU..AAAUAUUGGAAUGUAUUUAACAAAAGAAGCUAAAGAGGAA
env-1548  UUAUAU.UUGCCGCU............UGAAAGA..........AAA.................UCUUUCA...............................UAGUAC.AUAAA.AAUCAUU...UAAAU..AAAUAUUGGAAUGUAUUUAACAAAAAAAGCUAAAGAGAAA
env-1549  CUAUAU.UUGCCGCC............UGAAAG...........AAAAG.................CUUUCA................................GUGUAC.AUAAC.AAUCAUU...UACAA..UAAUAUUAGAAUGUAUUUAU.......................
env-1550  CUAUAU.UUGCCGCU............CGAAAG...........AAGAA.................CUUUCG.................................UAUAC.AUAAA.AAUCAUU...UAAAA..UAAUAUUGGAAUGUAUUUAGACAAAAAAGAGAAAGAGAAA
env-1551  CUAUAU.UUGCCGCU............CGAAAG...........AAGAA.................CUUUCG.................................UAUAC.AUAAA.AAUCAUU...UAAAA..UAAUAUUGGAAUGUAUUUAGACAAAAAAGAGAAAGAGAAA
env-1552  CUAUAU.UUGCCGCU............CGAAAG...........AAGCA.................CUUUCG.................................UAUAC.AUAAA.AAUCAUU...AAAAA..UAAUAUUCGAAUGUAUUUAGACAAAAAAGAAAAAGAGAAA
env-1553  UUAUAU.UUGCCGCU............CGAAAG...........AAGCA.................CUUUCG.................................UAUAC.AUAAA.AAUCAUU...AAAAA..UAAUAUUCGAAUGUAUUUAGACAAAAAAGAAAAAGAGAAA
env-1554  CUAUAU.UUGCCGCU............CGAAAG...........AAGCA.................CUUUCG.................................UAUAC.AUAAA.AAUCAUU...AAAAA..UAAUAUUCGAAUGUAUUUAGACAA..............
env-1555  UUAUAU.UUGCCGAC............CGAAAG...........AAAAC.................CUUUCG................................GUGUAC.AUAAA.AAUCAUU...UACAA..UAAUAUUAGCAUGUAUUUAGACAAAGAAGAAAAAGAAGCC
env-1556  UUAUAU.UUGCCGAC............CGAAAG...........AAAAC.................CUUUCG................................GUGUAC.AUAAA.AAUCAUU...UACAA..UAAUAUUAGCAUGUAUUUAGACAAAGAAGAAAAAGAAGCC
env-1557  UUAUAU.UUGCCGAC............UGAAAG...........AAAUC.................CUUUCG................................GUGUAC.AUAAA.AAUCAUU...UACAA..UAAUAUUAGCAUGUAUUUAGACAAAAAAGAAAAAGAAGCC
env-1559  UUAUAU.UUGCCGAU............CGAAAG...........AAAAG.................CUUUCG.................................UGUAC.AUAAA.AAUCAUU...UAAAU..AAAUAUUGGAAUGUAUUUAACAAAAGAAGUUAAAGAGAAA
env-1560  UUAUAU.UUGCCGAU............CGAAAG...........AAAAG.................CUUUCG.................................UGUAC.AUAAA.AAUCAUU...UAAAU..AAAUAUUGGAAUGUAUUUAACAAAAGAAGUUAAAGAGAAA
env-1561  CUACAU.UUGCCGCUG...................................................................................................UAC.AUAAA.AAUCAUU...AA.AA..CAAUAUUGGAAUGUAUUUAACAAAAGAAGUAAAAGCAGAA
env-1562  CUACAU.UUGCCGCCG...................................................................................................UAC.AUAAA.AAUCAUU...AA.AA..CAAUAUUGGAAUGUAUUUAACAAAAGAAGUAAAAGCAGAA
env-1563  CUACAU.UUGCCGCCG...................................................................................................UAC.AUAAA.AAUCAUU...AA.AA..CAAUAUUGGAAUGUAUUUAACAAAAGAAGUAAAAGCAGAA

          ...........................<<<<<<<<<<<<<<<<<<..........>>>>>>>>>>>>>>>>>>........<<<<<<<<........>>>>>>>>..............................................................................
          ...........................2222221111..101.00..........00.101..1111222222........11011000........00011011..............................................................................
          UAY U-UUGC  CY-------------       -----------R R  ----------------       ---------------------------------- UAC-AUAA -AAUCAUU---UA-AA--YAAUAUURG AUGUAUUUARC ARAGAAR  AAARRRRAR
          5'                                                                                                                                                        3'
```

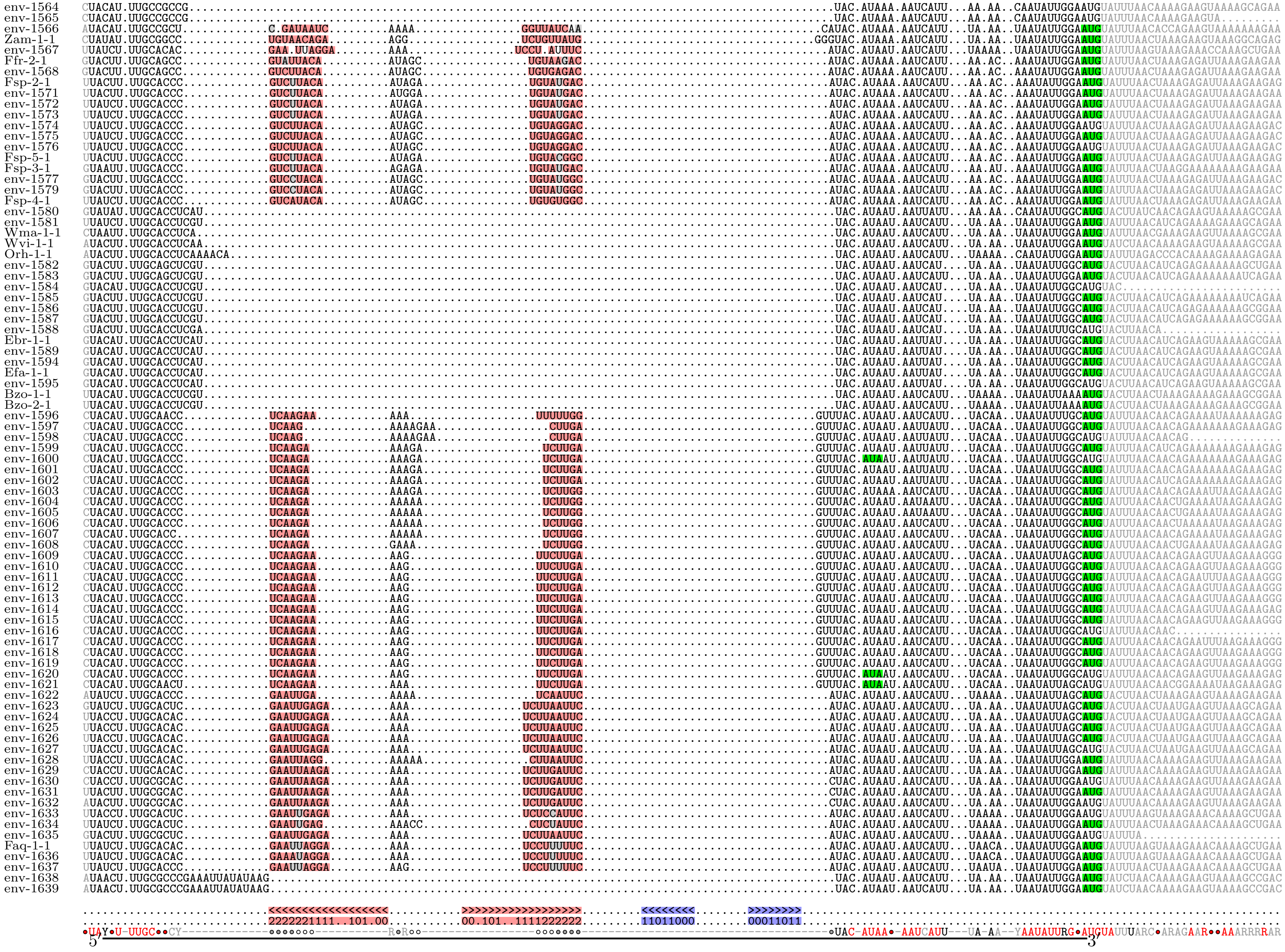
env-1564
env-1565
env-1566
Zam-1-1
env-1567
Ffr-2-1
env-1568
Fsp-2-1
env-1571
env-1572
env-1573
env-1574
env-1575
env-1576
Fsp-5-1
Fsp-3-1
env-1577
env-1579
Fsp-4-1
env-1580
env-1581
Wma-1-1
Wvi-1-1
Orh-1-1
env-1582
env-1583
env-1584
env-1585
env-1586
env-1587
env-1588
Ebr-1-1
env-1589
env-1594
Efa-1-1
env-1595
Bzo-1-1
Bzo-2-1
env-1596
env-1597
env-1598
env-1599
env-1600
env-1601
env-1602
env-1603
env-1604
env-1605
env-1606
env-1607
env-1608
env-1609
env-1610
env-1611
env-1612
env-1613
env-1614
env-1615
env-1616
env-1617
env-1618
env-1619
env-1620
env-1621
env-1622
env-1623
env-1624
env-1625
env-1626
env-1627
env-1628
env-1629
env-1630
env-1631
env-1632
env-1633
env-1634
env-1635
Faq-1-1
env-1636
env-1637
env-1638
env-1639
5'
3'

```
env-1640   AUAACU.UUGCGCCCGAAAUUAUAUAAG....................................................................................UAC.AUAAU.AAUCAUU...UA.AA..UAAUAUUGGAAUGUAUCUAACAAAAGAAGUAAAAGCCGAC
env-1641   AUAACU.UUGCGCCCGAAAUUAUAUAAG....................................................................................UAC.AUAAU.AAUCAUU...UA.AA..UAAUAUUGGAAUGUAUCUAACAAAAGAAGUAAAAGCCGAC
env-1642   AUAACU.UUGCGCCCGAAAUUAUAUAAG....................................................................................UAC.AUAAU.AAUCAUU...UA.AA..UAAUAUUGGAAUGUAUCUAACAAAAGAAGUAAAAGCCGAC
Csp-6-1    UUAUCU.UUGCGCCCGAAAAAAAAAAAU....................................................................................UAC.AUAAU.AAUCAUU...AA.AU..UAAUAUUGGAAUGUAUUUAACAAAAGAAGUAAAAGCGGAA
env-1643   UUAUCU.UUGCGCCCGAAAAAAAAAAU.....................................................................................UAC.AUAAU.AAUCAUU...AA.AU..UAAUAUUGGAAUGUAUUUAACAAAA...............
env-1644   UUAUCU.UUGCGCCCGAAAAAAAAAAU.....................................................................................UAC.AUAAU.AAUCAUU...AA.AU..UAAUAUUGGAAUGUAUUUAACAAAAGAAGUAAAAGC....
env-1645   UUAUCU.UUGCGCCCGAAAAAAAAAAU.....................................................................................UAC.AUAAU.AAUCAUU...AA.AU..UAAUAUUGGAAUGUAUUUAACAAAAGAAGUAAAAGCGGAA
Coc-2-1    UUAUCU.UUGCGCCCGAAAAAAAAAAU.....................................................................................UAC.AUAAU.AAUCAUU...AA.AU..UAAUAUUGGAAUGUAUUUAACAAAAGAAGUAAAAGCGGAA
env-1646   UUAUCU.UUGCGCCCGAAAAAAAAAAU.....................................................................................UAC.AUAAU.AAUCAUU...AA.AU..UAAUAUUGGAAUGUAUUUAACANNNNNNNNNNNNNNNNNN
Csp-9-1    UUAUCU.UUGCGCCCGAAAAAAAAAU......................................................................................UAC.AUAAU.AAUCAUU...AA.AU..UAAUAUUGGAAUGUAUUUAACAAAAGAAGUAAAAGCGGAA
env-1647   UUAUCU.UUGCGCCCGAAAAAAAAAU......................................................................................UAC.AUAAU.AAUCAUU...AA.AU..UAAUAUUGGAAUGUAUUUAACAAAAGAAGUAAAAGCGGAA
env-1649   UUAUCU.UUGCGCCCGAAAAAAAAAU......................................................................................UAC.AUAAU.AAUCAUU...AA.AU..UAAUAUUGGAAUGUAUUUAACAAAA...............
env-1650   UUAUCU.UUGCGCCCGAAAAAAAAAU......................................................................................UAC.AUAAU.AAUCAUU...AA.AU..UAAUAUUGGAAUGUAUUUAACAAAAGAAGUAAAAGCGGAA
env-1651   UUAUCU.UUGCACCCGAAAAAAAAAU......................................................................................UAC.AUAAU.AAUCAUU...AA.AU..UAAUAUUGGAAUGUAUUUAACAAAAGAAGUAAAAGCGGAA
env-1652   UUAUCU.UUGCACCCGAAAAAAAAAU......................................................................................UAC.AUAAU.AAUCAUU...AA.AU..UAAUAUUGGAAUGUAUUUAACAAAAGAAGUAAAAGCGGAA
env-1655   UUAUCU.UUGCACCCGAAAAAAAAAU......................................................................................UAC.AUAAU.AAUCAUU...AA.AU..UAAUAUUGGAAUGUAUUUAACAAAAGAAGUAAAAGCGGAA
env-1656   UUAUCU.UUGCACCCGAAAAAAAAAU......................................................................................UAC.AUAAU.AAUCAUU...AA.AU..UAAUAUUGGAAUGUAUUUAACAAAA...............
env-1657   UUAUCU.UUGCGCCCGAAAAAAAAU.......................................................................................UAC.AUAAU.AAUCAUU...AA.AU..UAAUAUUGGAAUGUAUUUAACAAAAGAAGUAAAAGCGGAA
Coc-3-1    UUAUCU.UUGCGCCCGAAAAAAAAU.......................................................................................UAC.AUAAU.AAUCAUU...AA.AU..UAAUAUUGGAAUGUAUUUAACAAAAGAAGUAAAAGCGGAA
Coc-1-1    UUAUCU.UUGCGCCCGAAAAAAAAUA......................................................................................UAC.AUAAU.AAUCAUU...AA.AU..UAAUAUUGGAAUGUAUUUAACAAAAGAAGUAAAAGCAGAA
Csp-7-1    UUAUCU.UUGCGCCCGAAAAAAAU........................................................................................UAC.AUAAU.AAUCAUU...AA.AU..UAAUAUUGGAAUGUAUUUAACAAAAGAAGUAAAAGCGGAA
Csp-10-1   UUAUCU.UUGCGCCCGAAAAAAAU........................................................................................UAC.AUAAU.AAUCAUU...AA.AU..UAAUAUUGGAAUGUAUUUAACAAAAGAAGUAAAAGCGGAA
Csp-2-1    UUAUCU.UUGCGCCCGAAAAAAAU........................................................................................UAC.AUAAU.AAUCAUU...AA.AU..UAAUAUUGGAAUGUAUUUAACAAAAGAAGUAAAAGCGGAA
env-1658   UUAUCU.UUGCACCC.............GAAUUGGG..........ACAGA................CUCAAUUC....................................AUAC.AUAAU.AAUCAUU...UA.AA..UAAUAUUGGAAUGUAUCUAAGUAAAGAAACAAAAGCAGAA
env-1659   UUACUU.UUGCAGCC.............GUUACUCA..........AUAGC................UGUGUAAC....................................AUAC.AUAAU.AAUCAUU...UA.AA..UAAUAUUGGAAUGUAUUUAACUAAAGAAAUUAAAGAAG..
env-1660   UUACUU.UUGCAGCC.............GUUACUCA..........AUAGC................UGUGUAAC....................................AUAC.AUAAU.AAUCAUU...UA.AA..UAAUAUUGGAAUGUAUUUAACUAAAGAAAUUAAAGAAGAA
env-1661   UUACUU.UUGCAGCC.............GUUACUCA..........AUAGC................UGUGUAAC....................................AUAC.AUAAU.AAUCAUU...UA.AA..UAAUAUUGGAAUGUAUUUAACUAAAGAAAUUAAAGAAGAA
env-1662   GUACUU.UUGCAGCC.............GUUACGCA..........AUAGC................UGUGUAAC....................................AUAC.AUAAU.AAUCAUU...UA.AA..UAAUAUUGGAAUGUAUUUAACUAAAGAAAUUAAAGAAGAA
env-1663   GUACUU.UUGCAGCC.............GUUACGCA..........AUAGC................UGUGUAAC....................................AUAC.AUAAU.AAUCAUU...UA.AA..UAAUAUUGGAAUGUAUUUAACUAAAGAAAUUAAAGAAGAA
env-1664   AUACUU.UUGCACCCA............CAAA..............AUA......................UUUG.....................................UAC.AUAAU.AAUCAUU...AA.AU..UAAUAUUGGAAUGUAUUUAACUAAAGAAGUAAAAGCGGAA
env-1665   AUACUU.UUGCACCCA............CAAA..............AUA......................UUUG.....................................UAC.AUAAU.AAUCAUU...AA.AU..UAAUAUUGGAAUGUAUUUAACUAAAGAAGUAAAAGCGGAA
env-1666   AUACUU.UUGCACCCA............CAAA..............AUA......................UUUG.....................................UAC.AUAAU.AAUCAUU...AA.AU..UAAUAUUGGAAUGUAUUUAACUAAAGAAGUAAAAGCGGAA
env-1668   AUACUU.UUGCACCCA............CAAA..............AUA......................UUUG.....................................UAC.AUAAU.AAUCAUU...AA.AU..UAAUAUUGGAAUGUAUUUAACUAAAGAAGUAAAAGCGGAA
env-1669   AUACUU.UUGCACCCA............CAAA..............AUA......................UUUG.....................................UAC.AUAAU.AAUCAUU...AA.AU..UAAUAUUGGAAUGUAUUUAACUAAAGAAGUAAAAGCGGAA
env-1673   AUACUU.UUGCACCCA............CAAA..............AUA......................UUUG.....................................UAC.AUAAU.AAUCAUU...AA.AU..UAAUAUUGGAAUGUAUUUAACUAAAGAAGUAAAAGCGGAA
env-1674   AUACUU.UUGCACCCA............CAAA..............AUA......................UUUG.....................................UAC.AUAAU.AAUCAUU...AA.AU..UAAUAUUGGAAUGUAUUUAACUAAAGAAGUAAAAGCGGAA
env-1676   AUACUU.UUGCACCCA............CAAA..............AUA......................UUUG.....................................UAC.AUAAU.AAUCAUU...AA.AU..UAAUAUUGGAAUGUAUUUAACUAAAGAAGUAAAAGCGGAA
env-1677   AUACUU.UUGCACCCA............CAAA..............AUA......................UUUG.....................................UAC.AUAAU.AAUCAUU...AA.AU..UAAUAUUGGAAUGUAUUUAACUAAAGAAGUAAAAGCGGAA
env-1679   AUACUU.UUGCACCCA............CAAA..............AUA......................UUUG.....................................UAC.AUAAU.AAUCAUU...AA.AU..UAAUAUUGGAAUGUAUUUAACUAAAGAAGUAAAAGCGGAA
env-1680   AUACUU.UUGCACCCA............CAAA..............AUA......................UUUG.....................................UAC.AUAAU.AAUCAUU...AA.AU..UAAUAUUGGAAUGUAUUUAACUAAAGAAGUAAAAGCGGAA
env-1681   AUACUU.UUGCACCCA............CAAA..............AUA......................UUUG.....................................UAC.AUAAU.AAUCAUU...AA.AU..UAAUAUUGGAAUGUAUUUAACUAAAGAAGUAAAAGCGGAA
env-1682   AUACUU.UUGCACCCA............CAAA..............AUA......................UUUG.....................................UAC.AUAAU.AAUCAUU...AA.AU..UAAUAUUGGAAUGUAUUUAACUAAAGAAGUAAAAGCGGAA
env-1683   AUACUU.UUGCACCCA............CAAA..............AUA......................UUUG.....................................UAC.AUAAU.AAUCAUU...AA.AU..UAAUAUUGGAAUGUAUUUAACUAAAGAAGUAAAAGCGGAA
env-1684   AUACUU.UUGCACCCA............CAAA..............AUA......................UUUG.....................................UAC.AUAAU.AAUCAUU...AA.AU..UAAUAUUGGAAUGUAUUUAACUAAAGAAGUAAAAGCGGAA
env-1685   AUACUU.UUGCACCCA............CAAA..............AUA......................UUUG.....................................UAC.AUAAU.AAUCAUU...AA.AU..UAAUAUUGGAAUGUAUUUAACUAAAGAAGUAAAAGCGGAA
env-1686   AUACUU.UUGCACCCA............CAAA..............AUA......................UUUG.....................................UAC.AUAAU.AAUCAUU...AA.AU..UAAUAUUGGAAUGUAUUUAACUAAAGAAGUAAAAGCGGAA
env-1687   AUACUU.UUGCACCCA............CAAA..............AUA......................UUUG.....................................UAC.AUAAU.AAUCAUU...AA.AU..UAAUAUUGGAAUGUAUUUAACUAAAGAAGUAAAAGCGGAA
env-1688   AUACUU.UUGCACCCA............CAAA..............AUA......................UUUG.....................................UAC.AUAAU.AAUCAUU...AA.AU..UAAUAUUGGAAUGUAUUUAACUAAAGAAGUAAAAGCGGAA
env-1689   AUACUU.UUGCACCCA............CAAA..............AUA......................UUUG.....................................UAC.AUAAU.AAUCAUU...AA.AU..UAAUAUUGGAAUGUAUUUAACUAAAGAAGUAAAAGCGGAA
env-1691   AUACUU.UUGCACCCA............CAAA..............AUA......................UUUG.....................................UAC.AUAAU.AAUCAUU...AA.AU..UAAUAUUGGAAUGUAUUUAACUAAAGAAGUAAAAGCGGAA
env-1692   AUACUU.UUGCACCCA............CAAA..............AUA......................UUUG.....................................UAC.AUAAU.AAUCAUU...AA.AU..UAAUAUUGGAAUGUAUUUAACUAAAGAAGUAAAAGCGGAA
env-1693   AUACUU.UUGCACCCA............CAAA..............AUA......................UUUG.....................................UAC.AUAAU.AAUCAUU...AA.AU..UAAUAUUGGAAUGUAUUUAACUAAAGAAGUAAAAGCGGAA
env-1694   AUACUU.UUGCACCCA............CAAA..............AUA......................UUUG.....................................UAC.AUAAU.AAUCAUU...AA.AU..UAAUAUUGGAAUGUAUUUAACUAAAGAAGUAAAAGCGGAA
env-1695   AUACUU.UUGCACCCA............CAAA..............AUA......................UUUG.....................................UAC.AUAAU.AAUCAUU...AA.AU..UAAUAUUGGAAUGUAUUUAACUAAAGAAGUAAAAGCGGAA
env-1696   AUACUU.UUGCACCCA............CAAA..............AUA......................UUUG.....................................UAC.AUAAU.AAUCAUU...AA.AU..UAAUAUUGGAAUGUAUUUAACUAAAGAAGUAAAAGCGGAA
env-1697   AUACUU.UUGCACCCA............CAAA..............AUA......................UUUG.....................................UAC.AUAAU.AAUCAUU...AA.AU..UAAUAUUGGAAUGUAUUUAACUAAAGAAGUAAAAGCGGAA
env-1698   AUACUU.UUGCACCCA............CAAA..............AUA......................UUUG.....................................UAC.AUAAU.AAUCAUU...AA.AU..UAAUAUUGGAAUGUAUUUAACUAAAGAAGUAAAAGCGGAA
env-1699   AUACUU.UUGCACCCA............CAAA..............AUA......................UUUG.....................................UAC.AUAAU.AAUCAUU...AA.AU..UAAUAUUGGAAUGUAUUUAACUAAAGAAGUAAAAGCGGAA
env-1700   AUACUU.UUGCACCCA............CAAA..............AUA......................UUUG.....................................UAC.AUAAU.AAUCAUU...AA.AU..UAAUAUUGGAAUGUAUUUAACUAAAGAAGUAAAAGCGGAA
env-1701   AUACUU.UUGCACCCA............CAAA..............AUA......................UUUG.....................................UAC.AUAAU.AAUCAUU...AA.AU..UAAUAUUGGAAUGUAUUUAACUAAAGAAGUAAAAGCGGAA
env-1702   AUACUU.UUGCACCCA............CAAA..............AUA......................UUUG.....................................UAC.AUAAU.AAUCAUU...AA.AU..UAAUAUUGGAAUGUAUUUAACUAAAGAAGUAAAAGCGGAA
env-1703   AUACUU.UUGCACCCA............CAAA..............AUA......................UUUG.....................................UAC.AUAAU.AAUCAUU...AA.AU..UAAUAUUGGAAUGUAUUUAACUAAAGAAGUAAAAGCGGAA
env-1705   AUACUU.UUGCACCCA............CAAA..............AUA......................UUUG.....................................UAC.AUAAU.AAUCAUU...AA.AU..UAAUAUUGGAAUGUAUUUAACUAAAGAAGUAAAAGCGGAA
env-1706   AUACUU.UUGCACCCA............CAAA..............AUA......................UUUG.....................................UAC.AUAAU.AAUCAUU...AA.AU..UAAUAUUGGAAUGUAUUUAACUAAAGAAGUAAAAGCGGAA
env-1707   AUACUU.UUGCACCCA............CAAA..............AUA......................UUUG.....................................UAC.AUAAU.AAUCAUU...AA.AU..UAAUAUUGGAAUGUAUUUAACUAAAGAAGUAAAAGCGGAA
env-1708   AUACUU.UUGCACCCA............CAAA..............AUA......................UUUG.....................................UAC.AUAAU.AAUCAUU...AA.AU..UAAUAUUGGAAUGUAUUUAACUAAAGAAGUAAAAGCGGAA
env-1709   AUACUU.UUGCACCCA............CAAA..............AUA......................UUUG.....................................UAC.AUAAU.AAUCAUU...AA.AU..UAAUAUUGGAAUGUAUUUAACUAAAGAAGUAAAAGCGGAA
env-1710   AUACUU.UUGCACCCAC...........AA.................AAUA......................UU...................................CGUAC.AUAAU.AAUCAUU...AA.AU..UAAUAUUGGAAUGUAUUUAACUAAAGAAGUAAAAGCGGAA
Cko-1-1    GUAAUU.UUGCAGCCGU...............................................................................................UAC.AUAAU.AAUCAUU...AA.AA..UAAUAUUGGAAUGUACUUAACAACUGACAAGAAGAAAGAA
env-1711   GUACUU.UUGCAGCCAGAAAAUG.........................................................................................UAC.AUAAU.AAUCAUU...AA.AA..UAAUAUUGGAAUGUAUUUAACAAAAGAAGUAAAAGCGGAA
env-1712   GUACUU.UUGCAGCCAGAAAAUG.........................................................................................UAC.AUAAU.AAUCAUU...AA.AA..UAAUAUUGGAAUGUAUUUAACAAAAGAAGUAAAAGCGGAA
env-1713   GUACUU.UUGCAGCCAGAAAAUG.........................................................................................UAC.AUAAU.AAUCAUU...AA.AA..UAAUAUUGGAAUGUAUUUAACAAAAGAAGUAAAAGCGGAA
env-1714   GUACUU.UUGCAGCCAGAAAAUG.........................................................................................UAC.AUAAU.AAUCAUU...AA.AA..UAAUAUUGGAAUGUAUUUAACAAAAGAAGUAAAAGCGGAA
env-1717   GUACUU.UUGCAGCCAGAAAAUG.........................................................................................UAC.AUAAU.AAUCAUU...AA.AA..UAAUAUUGGAAUGUAUUUAACAAAAGAAGUAAAAGCGGAA
env-1718   GUACUU.UUGCAGCCAGAAAAUG.........................................................................................UAC.AUAAU.AAUCAUU...AA.AA..UAAUAUUGGAAUGUAUUUAACAAAA...............
env-1720   GUACUU.UUGCAGCCAGAAAAUG.........................................................................................UAC.AUAAU.AAUCAUU...AA.AA..UAAUAUUGGAAUGUAUUUAACAAAAGAAGUAAAAGCGGAA
env-1722   GUACUU.UUGCAGCCAGAAAAUG.........................................................................................UAC.AUAAU.AAUCAUU...AA.AA..UAAUAUUGGAAUGUAUUUAACAAAAGAAGUAAAAGCGGAA
Csp-5-1    GUACUU.UUGCAGCCGGAAAAUG.........................................................................................UAC.AUAAU.AAUCAUU...AA.AA..UAAUAUUGGAAUGUAUUUAACAAAAGAAGUAAAAGCGGAA
env-1723   GUACUU.UUGCAGCCGGAAAAUG.........................................................................................UAC.AUAAU.AAUCAUU...AA.AA..UAAUAUUGGAAUGUAUUUAACAAAA...............
env-1724   UUACUU.UUGCAGCC.............GUCUUACA..........AUAGA................UGUUUGAC....................................AUAC.AUAAU.AAUCAUU...UA.AA..UAAUAUUGGAAUGUACUUAACUAAAGAAGUUAAAUCAGAA
env-1725   UUACUU.UUGCAGCC.............GUCUUACA..........AUAGA................UGUUUGAC....................................AUAC.AUAAU.AAUCAUU...UA.AA..UAAUAUUGGAAUGUACUUAACUAAAGAAGUUAAAUCAGAA
env-1726   UUACUU.UUGCAGCC.............GUCUUACA..........AUAGA................UGUUUGAC....................................AUAC.AUAAU.AAUCAUU...UA.AA..UAAUAUUGGAAUGUACUUAACUAAAGAAGUUAAAUCAGAA
env-1727   UUACUU.UUGCAGCC.............GUCUUACA..........AUAGA................UGUUUGAC....................................AUAC.AUAAU.AAUCAUU...UA.AA..UAAUAUUGGAAUGUACUUAACUAAAGAAGUUAAAUCAGAA

           ............................<<<<<<<<<<<<<<<<<<...........>>>>>>>>>>>>>>>>>>........<<<<<<<<........>>>>>>>>........................................................................
           ............................2222221111..101.00...........00.101..1111222222........11011000........00011011........................................................................
           •UAY•U-UUGC••CY-------------◦◦◦◦◦◦◦-----------R•R◦◦-----------------◦◦◦◦◦◦◦------------------------------------◦UAC-AUAA•-AAUCAUU---UA-AA--YAAUAUURG•AUGUAUUUARC•ARAGAAR••AAARRRRAR
           5′                                                                                                                                           3′
```

```
Csp-8-1    UUAACU.UUGCCGCCGAUUUAAAA........................................................................................UAC.AUAAU.AAUCAUU...UA.AA..UAAUAUUGGAAUGUAUCUAACAAAAGAAAUUAAAGCUGAA
env-1728   UUAACU.UUGCCGCCGAUUUAAAA........................................................................................UAC.AUAAU.AAUCAUU...UA.AA..UAAUAUUGGAAUGUAUCUAACAAAAGAAAUUAAAGCUGAA
env-1729   CUACCU.UUGCCCCCGAUUAAAAAAA......................................................................................UAC.AUAAU.AAUCAUU...UA.AA..CAAUAUUGGAAUGUAUCUAACAAAAGAAACAAAAGCAGAA
env-1730   CUACCU.UUGCCCCCGAUUUAAAAAA......................................................................................UAC.AUAAU.AAUCAUU...UA.AA..CAAUAUUGGAAUGUAUCUAACAAAAGAAACAAAAGCAGAA
env-1744   CUACCU.UUGCCCCCGAUUUAAAAAA......................................................................................UAC.AUAAU.AAUCAUU...UA.AA..CAAUAUUGGAAUGUAUCUAACAAAAGAAACAAAAGCAGAA
env-1745   CUACCU.UUGCCCCCGAUUUAAAAAA......................................................................................UAC.AUAAU.AAUCAUU...UA.AA..CAAUAUUGGAAUGUAUCUAACAAAAGAAACAAAAGCAGAA
env-1746   CUACCU.UUGCCCCCGAUUUAAAAAA......................................................................................UAC.AUAAU.AAUCAUU...UA.AA..CAAUAUUGGAAUGUAUCUAACAAAAGAAAC..........
env-1747   CUACCU.UUGCCCCCGAUUUAAAAAA......................................................................................UAC.AUAAU.AAUCAUU...UA.AA..CAAUAUUGGAAUGUAUCUA.....................
env-1748   CUACCU.UUGCCCCCGAUUUAAAAAA......................................................................................UAC.AUAAU.AAUCAUU...UA.AA..CAAUAUUGGAAUGUAUCUAACAAAAGAAACAAAAGCAGAA
env-1749   CUACCU.UUGCCCCCGAUUUAAAAAA......................................................................................UAC.AUAAU.AAUCAUU...UA.AA..CAAUAUUGGAAUGUAUCUAACAAAAGAAACAAAAGCAGAA
env-1750   CUACCU.UUGCCCCCGAUUUAAAAAA......................................................................................UAC.AUAAU.AAUCAUU...UA.AA..CAAUAUUGGAAUGUAUCUAACAAAAG..............
env-1752   CUACCU.UUGCCCCCGAUUUAAAAAA......................................................................................UAC.AUAAU.AAUCAUU...UA.AA..CAAUAUUGGAAUGUAUCUAACAAAAGAAACAAAAGCAGAA
env-1753   CUACCU.UUGCCCCCGAUUUAAAAAA......................................................................................UAC.AUAAU.AAUCAUU...UA.AA..CAAUAUUGGAAUGUAUCUAACAAAAGAAACAAAAGCAGAA
env-1754   CUACCU.UUGCCCCCGAUUUAAAAAA......................................................................................UAC.AUAAU.AAUCAUU...UA.AA..CAAUAUUGGAAUGUAUCUAACAAAAGAAACAAAAGCAGAA
env-1755   CUACCU.UUGCCCCCGAUUUAAAAAA......................................................................................UAC.AUAAU.AAUCAUU...UA.AA..CAAUAUUGGAAUGUAUCUAACAAAAGAAACAAAAGCAGAA
Cgi-1-1    CUACCU.UUGCCCCCGAUUUAAAAAA......................................................................................UAC.AUAAU.AAUCAUU...UAAAA..CAAUAUUGGAAUGUAUCUAACAAAAGAAACAAAAGCAGAA
env-1756   CUACCU.UUGCCCCCGAUUUAAAAAA......................................................................................UAC.AUAAU.AAUCAUU...UA.AA..CAAUAUUGGAAUGUAUCUAACAAAAG..............
env-1757   CUACCU.UUGCCCCCGAUUUAAAAAA......................................................................................UAC.AUAAU.AAUCAUU...UA.AA..CAAUAUUGGAAUGUAUCUAACAAAAG..............
env-1765   CUACCU.UUGCCCCCGAUUUAAAAAA......................................................................................UAC.AUAAU.AAUCAUU...UA.AA..CAAUAUUGGAAUGUAUCUA.....................
env-1766   CUACCU.UUGCCCCCGAUUUAAAAAA......................................................................................UAC.AUAAU.AAUCAUU...UA.AA..CAAUAUUGGAAUGUA.........................
env-1767   CUACCU.UUGCCCCCGAUUUAAAAAA......................................................................................UAC.AUAAU.AAUCAUU...UA.AA..CAAUAUUGGAAUGUAUCUAACAAAAGAAACAAAAGCAGAA
env-1768   CUACCU.UUGCCCCCGAUUUAAAAAA......................................................................................UAC.AUAAU.AAUCAUU...UA.AA..CAAUAUUGGAAUGUAUCUAACAAAAGAAACAAAAGCAGAA
env-1769   CUACCU.UUGCCCCCGAUUUAAAAAA......................................................................................UAC.AUAAU.AAUCAUU...UA.AA..CAAUAUUGGAAUGUAUCUAACAAAAGAAACAAAAGCAGAA
env-1770   CUACCU.UUGCCCCCGAUUUAAAAAA......................................................................................UAC.AUAAU.AAUCAUU...UA.AA..CAAUAUUGGAAUGUAUCUAACAAAAGAAACAAAAGCAGAA
env-1771   CUACCU.UUGCCCCCGAUUUAAAAAA......................................................................................UAC.AUAAU.AAUCAUU...UA.AA..CAAUAUUGGAAUGUAUCUAACAAAAGAAACAAAAGCAGAA
env-1772   CUACCU.UUGCCCCCGAUUUAAAAAA......................................................................................UAC.AUAAU.AAUCAUU...UA.AA..CAAUAUUGGAAUGUAUCUAACAAAAGAAACAAAAGCAGAA
env-1773   CUACCU.UUGCCCCCGAUUUAAAAAA......................................................................................UAC.AUAAU.AAUCAUU...UA.AA..CAAUAUUGGAAUGUAUCUAACAAAAGAAACAAAAGCAGAA
env-1774   CUACCU.UUGCCCCCGAUUUAAAAAA......................................................................................UAC.AUAAU.AAUCAUU...UA.AA..CAAUAUUGGAAUGUAUCUAACAAAAGAAACAAAAGCAGAA
env-1775   CUACCU.UUGCCCCCGAUUUAAAAAA......................................................................................UAC.AUAAU.AAUCAUU...UA.AA..CAAUAUUGGAAUGUAUCUAACAAAAGAAACAAAAGCAGAA
env-1779   CUACCU.UUGCCCCCGAUUUAAAAAA......................................................................................UAC.AUAAU.AAUCAUU...AUAAA..CAAUAUUGGAAUGUAUCUAACAAAAGAAACAAAAGCAGAA
env-1780   CUACCU.UUGCCCCCGAUUUAAAAAA......................................................................................UAC.AUAAU.AAUCAUU...UUAAA..CAAUAUUGGAAUGUAUCUAACAAAAGAAACAAAAGCAGAA
env-1783   CUACCU.UUGCCCCCGAUUUAAAAAA......................................................................................UAC.AUAAU.AAUCAUU...UUAAA..CAAUAUUGGAAUGUAUCUA.....................
env-1786   CUACCU.UUGCCCCCGAUUUAAAAAA......................................................................................UAC.AUAAU.AAUCAUU...UUAAA..CAAUAUUGGAAUGUAUCUAACAAAAGAAACAAAAGCAGAA
env-1795   CUACCU.UUGCCCCCGAUUUAAAAAA......................................................................................UAC.AUAAU.AAUCAUU...UUAAA..CAAUAUUGGAAUGUAUCUAACAAAAGAAACAAAAGCAGAA
env-1797   CUACCU.UUGCCCCCGAUUUAAAAAA......................................................................................UAC.AUAAU.AAUCAUU...UUAAA..CAAUAUUGGAAUGUAUCUAACAAAAGAAACAAAAGCAGAA
env-1798   CUACCU.UUGCCCCCGAUUUAAAAAA......................................................................................UAC.AUAAU.AAUCAUU...UUAAA..CAAUAUUGGAAUGUAUCUAACAAAAGAAACAAAAGCAGAA
env-1800   CUACCU.UUGCCCCCGAUUUAAAAAA......................................................................................UAC.AUAAU.AAUCAUU...UUAAA..CAAUAUUGGAAUGUAUCUAACAAAAGAAACAAAAGCAGAA
env-1801   CUACCU.UUGCCCCCGAUUUAAAAAA......................................................................................UAC.AUAAU.AAUCAUU...UUAAA..CAAUAUUGGAAUGUAUCUAACAAAAGAAACAAAAGCAGAA
env-1802   CUACCU.UUGCCCCCGAUUUAAAAAAA.....................................................................................UAC.AUAAU.AAUCAUU...UUAAA..CAAUAUUGGAAUGUAUCUAACAAAAG..............
env-1804   CUACCU.UUGCCCCCGAUUUAAAAAAA.....................................................................................UAC.AUAAU.AAUCAUU...UUAAA..CAAUAUUGGAAUGUAUCUAACAAAAGAAACAAAAGCAGAA
env-1805   CUACCU.UUGCCCCCGAUUUAAAAAA......................................................................................UAC.AUAAU.AAUCAUU...UUAAA..CAAUAUUGGAAUGUAUCUAACAAAAG..............
env-1818   CUACCU.UUGCCCCCGAUUUAAAAAA......................................................................................UAC.AUAAU.AAUCAUU...UUAAA..CAAUAUUGGAAUGUAUCUAACAAAAGAAACAAAAGCAGAA
env-1821   CUACCU.UUGCCCCCGAUUUAAAAAA......................................................................................UAC.AUAAU.AAUCAUU...UUAAA..CAAUAUUGGAAUGUAUCUAACAAAAGAAACAAAAGCAGAA
env-1823   CUACCU.UUGCCCCCGAUUUAAAAAA......................................................................................UAC.AUAAU.AAUCAUU...UUAAA..CAAUAUUGGAAUGUAUCUAACAAAAGAAACAAAAGCAGAA
env-1824   CUACCU.UUGCCCCCGAUUUAAAAAA......................................................................................UAC.AUAAU.AAUCAUU...UUAAA..CAAUAUUGGAAUGUAUCUAACAAAAGAAACAAAAGCAGAA
env-1826   CUACCU.UUGCCCCCGAUUUAAAAAA......................................................................................UAC.AUAAU.AAUCAUU...UUAAA..CAAUAUUGGAAUGUAUCUAACAAAAGAAACAAAAGCAGAA
env-1827   CUACCU.UUGCCCCCGAUUUAAAAAA......................................................................................UAC.AUAAU.AAUCAUU...UUAAA..CAAUAUUGGAAUGUAUCUAACAAAAGAAACAAAAGCAGAA
env-1828   CUACCU.UUGCCCCCGAUUUAAAAAA......................................................................................UAC.AUAAU.AAUCAUU...UUAAA..CAAUAUUGGAAUGUAUCUAACAAAAGAAACAAAAGCAGAA
env-1829   CUACCU.UUGCCCCCGAUUUAAAAAA......................................................................................UAC.AUAAU.AAUCAUU...UUAAA..CAAUAUUGGAAUGUAUCUAACAAAAGAAGUAAAAGCAGAA
env-1830   CUACCU.UUGCCCCCGAUUUAAAAAAA.....................................................................................UAC.AUAAU.AAUCAUU...UUAAA..CAAUAUUGGAAUGUAUCUAACAAAAGAAACAAAAGCAGAA
env-1832   CUACCU.UUGCCCCCGAUUUAAAAAAA.....................................................................................UAC.AUAAU.AAUCAUU...UUAAA..CAAUAUUGGAAUGUAUCUAACAAAAGAAACAAAAGCAGAA
env-1833   CUACCU.UUGCCCCCGAUUUAAAAAA......................................................................................UAC.AUAAU.AAUCAUU...UUAAA..CAAUAUUGGAAUGUAUCUAACAAAAGAAACAAAAGCAGAA
env-1834   UUACCU.UUGCCCCCGAUUUUAAAA.......................................................................................UAC.AUAAU.AAUCAU....UA.AA..UAAUAUUGGAAUGUAUCUAACAAAAGAAACUAAAGCAGAA
env-1835   UUACCU.UUGCCCCCGAUUUUAAAA.......................................................................................UAC.AUAAU.AAUCAU....UA.AA..UAAUAUUGGAAUGUAUCUAACAAAAGAAACUAAAGCAGAA
env-1836   UUACCU.UUGCCCCCGAUUUUAAAA.......................................................................................UAC.AUAAU.AAUCAU....UA.AA..UAAUAUUGGAAUGUAUCUAA....................
env-1837   UUACCU.UUGCCCCCGAUUUUAAAA.......................................................................................UAC.AUAAU.AAUCAU....UA.AA..UAAUAUUGGAAUGUAUCUAACAAAAG..............
env-1838   UUACCU.UUGCCCCCGAUUUUAAAA.......................................................................................UAC.AUAAU.AAUCAU....UA.AA..UAAUAUUGGAAUGUAUCUAACAAAAGAAACUAAAGCAGAA
env-1839   UUACCU.UUGCCCCCGAUUUUAAAA.......................................................................................UAC.AUAAU.AAUCAU....UA.AA..UAAUAUUGGAAUGUAUCUAACAAAAGAAACCAAAGCAGAA
env-1840   UUACCU.UUGCCCCCGAUUUUAAAA.......................................................................................UAC.AUAAU.AAUCAU....UA.AA..UAAUAUUGGAAUGUAUCUAACAAAAGAAACUAAAGCAGAA
env-1846   UUACCU.UUGCCCCCGAUUUUAAAA.......................................................................................UAC.AUAAU.AAUCAU....UA.AA..UAAUAUUGGAAUGUAUCUAACAAAAG..............
env-1847   UUACCU.UUGCCCCCGAUUUUAAAA.......................................................................................UAC.AUAAU.AAUCAU....UA.AA..UAAUAUUGGAAUGUAUCUAACAAAAGAAACUAAAGCAGAA
env-1848   UUACCU.UUGCCCCCGAUUUUAAAA.......................................................................................UAC.AUAAU.AAUCAU....UA.AA..UAAUAUUGGAAUGUAUCUAA....................
env-1850   UUACCU.UUGCCCCCGAUUUUAAAA.......................................................................................UAC.AUAAU.AAUCAU....UA.AA..UAAUAUUGGAAUGUAUCUAA....................
env-1851   UUACCU.UUGCCCCCGAUUUUAAAA.......................................................................................UAC.AUAAU.AAUCAU....UA.AA..UAAUAUUGGAAUGUAUCUAACAAAAGAAACUAAAGCAGAA
env-1852   UUACCU.UUGCCCCCGAUUUUAAAA.......................................................................................UAC.AUAAU.AAUCAU....UA.AA..UAAUAUUGGAAUGUAUCUAACAAAAGAAACUAAAGCAGAA
env-1853   UUACCU.UUGCCCCCGAUUUUAAAA.......................................................................................UAC.AUAAU.AAUCAU....UA.AA..UAAUAUUGGAAUGUAUCUAACAAAAGAAACUAAAGCAGAA
env-1854   UUACCU.UUGCCCCCGAUUUUAAAA.......................................................................................UAC.AUAAU.AAUCAU....UA.AA..UAAUAUUGGAAUGUAUCUAACAAAAGAAACUAAAGCAGAA
env-1855   GUAUCU.UUGCCCUCGAUUUUAAAA.......................................................................................UAC.AUAAU.AAUCAUU...UA.AA..UAAUAUUGGAAUGUAUCUAACAAAAGAAGUAAAAGCUGAA
env-1856   GUAUCU.UUGCCCCCGAUUUUAAAA.......................................................................................UAC.AUAAU.AAUCAUU...UA.AA..UAAUAUUGGAAUGUAUCUAACAAAAGAAGUAAAAGCUGAA
env-1857   GUAUCU.UUGCCCCCGAUUUUAAAA.......................................................................................UAC.AUAAU.AAUCAUU...UA.AA..UAAUAUUGGAAUGUAUCUAACAAAAGAAGUAAAAGCUGAA
env-1859   GUAUCU.UUGCCCCCGAUUUUAAAA.......................................................................................UAC.AUAAU.AAUCAUU...UA.AA..UAAUAUUGGAAUGUAUCUAACAAAAGAAGUAAAAGCUGAA
env-1860   GUAUCU.UUGCCCCCGAUUUUAAAA.......................................................................................UAC.AUAAU.AAUCAUU...UA.AA..UAAUAUUGGAAUGUAUCUAACAAAAGAAGUAAAAGCUGAA
env-1862   UUAUCU.UUGCCCCCGAUUUAAAA........................................................................................UAC.AUAAU.AAUCAU....UA.AA..UAAUAUUGGAAUGUAUCUAACAAAAGAAACCAAAGCAGAA
env-1863   CUAUCU.UUGCCCCCGAUUUAAAA........................................................................................UAC.AUAAU.AAUCAUU...AA.AA..UAAUAUUGGAAUGUAUCUAACA..................
env-1864   CUAUCU.UUGCCCCCGAUUUAAAA........................................................................................UAC.AUAAU.AAUCAUU...AA.AA..UAAUAUUGGAAUGUAUCUAACAAAAGAAACCAAAGCAGAA
env-1865   CUAUCU.UUGCAUCCGAAU.............................................................................................UAC.AUAAU.AAUCAUU...UAAAA.ACAAUAUUGGAAUGUAUAUAACAUUAGAAGAAAAAGGAAAA
Eme-2-1    GUAUCU.UUGCACCCGU...............................................................................................UAC.AUAAU.AAUCAUU...UA.AA..UAAUAUUGGAAUGUAUUUAACUAAAGAAAAGAAGGCUGAG
Cca-1-1    CUAUCU.UUGCACCCGAAU.............................................................................................UAC.AUAAU.AAUCAUU...UAAAA..UAAUAUUGGCAUGUAUUUAGCACCAGAAAAGAAAGAAGAA
Emi-1-1    GUAUAU.UUGCAGCCGAAU.............................................................................................UAC.AUAAU.AAUCAUU...UA.AA..UAAUAUUGGAAUGUAUUUAACUAAAGAAAAGAAGGCUGAG
Cte-1-1    CUAUCU.UUGCACCCGAAU.............................................................................................UAC.AUAAU.AAUCAU....UA.AA..UAAUAUUGGAAUGUAUUUAACACCAGAAAAAAAGUCAGAA
Csp-15-1   CUAUCU.UUGCACCCGAAU.............................................................................................UAC.AUAAU.AAUCAU....UA.AA..UAAUAUUGGAAUGUAUUUAACAACUGAAAAAAAGUCAGAA
Cha-2-1    CUAUCU.UUGCACCCGAAU.............................................................................................UAC.AUAAU.AAUCAUU...UA.AA..UAAUAUUGGAAUGUAUCUAACAACAGAAAAAAAGUCAGAA
Ean-3-1    GUAUAU.UUGCACCCGAAU.............................................................................................UAC.AUAAU.AAUCAUU...UA.AA..UAAUAUUGGAAUGUAUUUAACUAAAGAAAAGAAGGCUGAG

           ............................<<<<<<<<<<<<<<<<<<...........>>>>>>>>>>>>>>>>>>.........<<<<<<<<........>>>>>>>>.......................................................................
           ............................2222221111..101.00...........00.101..1111222222.........11011000........00011011.......................................................................
           •UAY•U-UUGC••CY-------------∘∘∘∘∘∘∘-----------R•R∘∘-----------------∘∘∘∘∘∘∘-----------------------------------∘UAC-AUAA•-AAUCAUU---UA-AA--YAAUAUURG•AUGUAUUUARC•ARAGAAR••AAARRRRAR
            5′                                                                                                                                                   3′
```

```
Ean-1-3    GUAUAU.UUGCACCCGAAU..............................................................................................UAC.AUAAU.AAUCAUU...UA.AA..UAAUAUUGGAAUGUAUUUAACUAAAGAAAAGAAGGCUGAG
Csp-18-1   GUACUU.UUGCACCCGU................................................................................................UAC.AUAAU.AAUCAUU...AA.AC..AAAUAUUGGAAUGUACUUAACAACUGAAAAGAAGAAAGAA
Cje-1-1    GUACUU.UUGCACCCAU................................................................................................UAC.AUAAU.AAUCAUU...AA.AA..UAAUAUUGGAAUGUACCUAACAACUGACACGAAGAAAGAG
env-1866   GUACUU.UUGCACCCGU................................................................................................UAC.AUAAU.AAUCAUU...AA.AA..UAAUAUUGGAAUGUACUUAACAACUGACAAGAAGAAAGAA
Cha-1-1    GUACUU.UUGCACCCGU................................................................................................UAC.AUAAU.AAUCAUU...AA.AA..UAAUAUUGGAAUGUACUUAACAACUGACAAGAAGAAAGAA
env-1869   GUACUU.UUGCACCCGU................................................................................................UAC.AUAAU.AAUCAUU...AA.AA..UAAUAUUGGAAUGUACUUAACAACUGACAAGAAGAAAGAA
env-1870   GUACUU.UUGCAGCCGU................................................................................................UAC.AUAAU.AAUCAU....UA.AA..UAAUAUUGGAAUGUACUUAACAUCAGACAAAAAAGCAGAA
Chi-1-1    GUACUU.UUGCAGCCGU................................................................................................UAC.AUAAU.AAUCAUU...UA.AA..UAAUAUUGGAAUGUACUUAAAUACAGAAAAAAAGCAGGAA
env-1871   GUACUU.UUGCACCCGU................................................................................................UAC.AUAAU.AAUCAU....UA.AA..UAAUAUUGGAAUGUACUUAACAACAGAAAAGAAGAAAGAA
Cso-2-1    GUACUU.UUGCACCCGU................................................................................................UAC.AUAAU.AAUCAUU...UAAAA..UAAUAUUGGAAUGUACUUAACAACUGACAAGAAGAAAGAA
Cda-1-1    GUACUU.UUGCACCCGU................................................................................................UAC.AUAAU.AAUCAU....UA.AA..CAAUAUUGGAAUGUACUUAACAACAGAAAAAAAGCAGGAA
Cpi-1-1    GUACUU.UUGCACCCGU................................................................................................UAC.AUAAU.AAUCAU....UA.AA..CAAUAUUGGAAUGUACUUAACAACAGAAAAAAAGCAGGAA
env-1872   GUACUU.UUGCACCCGU................................................................................................UAC.AUAAU.AAUCAU....UA.AA..CAAUAUUGGAAUGUACUUAACAACAGAAAAAAAAGCAGAA
Fba-4-1    GUACUU.UUGCACCCGU................................................................................................UAC.AUAAU.AAUCAU....UA.AA..CAAUAUUGGAAUGUACUUAACAACUGACAAGAAGAAAGAA
Cfo-1-1    GUACUU.UUGCACCCGU................................................................................................UAC.AUAAU.AAUCAU....UA.AA..CAAUAUUGGAAUGUACUUAACAACAGAAAAAAAAGCAGAA
env-1873   GUACUU.UUGCACCCGU................................................................................................UAC.AUAAU.AAUCAU....UA.AA..CAAUAUUGGAAUGUACUUAACAACAGAAAAAAAGCAGGAA
Clu-1-1    GUAAUU.UUGCAGCCGU................................................................................................UAC.AUAAU.AAUCAU....UA.AA..CAAUAUUGGAAUGUACUUAACAACAGAAAAAAAAGCAGAA
env-1874   GUAAUU.UUGCAGCCGU................................................................................................UAC.AUAAU.AAUCAU....UA.AA..CAAUAUUGGAAUGUACUUAACAACAGAAAAAAAGCAGGAA
Cta-1-1    GUAAUU.UUGCAGCCGU................................................................................................UAC.AUAAU.AAUCAU....UA.AA..CAAUAUUGGAAUGUACUUAACAACAGAAAAAAAAGCAGAA
Csp-29-1   GUAAUU.UUGCAGCCGU................................................................................................UAC.AUAAU.AAUCAU....UA.AA..CAAUAUUGGAAUGUACUUAACAACAGAAAAAAAAGCAGAA
Csp-24-1   GUAAUU.UUGCACCCGU................................................................................................UAC.AUAAU.AAUCAU....UA.AA..CAAUAUUGGAAUGUACUUAACAACAGACAAAAAGCAGGAA
Csp-19-1   GUAAUU.UUGCACCCGU................................................................................................UAC.AUAAU.AAUCAU....UA.AA..CAAUAUUGGAAUGUACUUAACAACAGAAAAAAAAGCAGAA
Cor-1-1    GUAAUU.UUGCACCCGU................................................................................................UAC.AUAAU.AAUCAU....UA.AA..CAAUAUUGGAAUGUACUUAACAACAGAAAAAAAGCAGGAA
Csp-16-1   GUAAUU.UUGCACCCGU................................................................................................UAC.AUAAU.AAUCAU....UA.AA..CAAUAUUGGAAUGUACUUAACAACAGAAAAAAAGCAGGAA
Can-1-1    GUAAUU.UUGCACCCGU................................................................................................UAC.AUAAU.AAUCAU....UA.AA..CAAUAUUGGAAUGUACUUAACAACAGAAAAAAAGCAGGAA
Cgl-1-1    GUAAUU.UUGCACCCGU................................................................................................UAC.AUAAU.AAUCAU....UA.AA..CAAUAUUGGAAUGUACUUAACAACAGACAAAAAGCAGGAA
Csp-26-1   GUAAUU.UUGCACCCGU................................................................................................UAC.AUAAU.AAUCAU....UA.AA..CAAUAUUGGAAUGUACUUAACAACAGACAAAAAGCAGGAA
Csp-23-1   GUAAUU.UUGCACCCGU................................................................................................UAC.AUAAU.AAUCAU....UA.AA..CAAUAUUGGAAUGUACUUAACAACAGACAAAAAGCAGGAA
Cso-1-1    GUAAUU.UUGCACCCGU................................................................................................UAC.AUAAU.AAUCAU....UA.AA..CAAUAUUGGAAUGUACUUAACAAAAGAAAAAAAGCAGGAA
Csp-17-1   GUAAUU.UUGCACCCGU................................................................................................UAC.AUAAU.AAUCAU....UA.AA..CAAUAUUGGAAUGUACUUAACAACAGAAAAAAAGCAGGAA
Csp-20-1   GUAAUU.UUGCACCCGU................................................................................................UAC.AUAAU.AAUCAU....UA.AA..CAAUAUUGGAAUGUACUUAACAACAGAAAAAAAGCAGGAA
env-1875   GUAAUU.UUGCACCCGU................................................................................................UAC.AUAAU.AAUCAU....UA.AA..CAAUAUUGGAAUGUACUUAACAACAGACAAAAAGCAGGAA
env-1876   GUAAUU.UUGCACCCGU................................................................................................UAC.AUAAU.AAUCAU....UA.AA..CAAUAUUGGAAUGUACUUAACAACAGAAAAAAAAGCAGAA
Csp-27-1   GUAAUU.UUGCACCCGU................................................................................................UAC.AUAAU.AAUCAU....UA.AA..CAAUAUUGGAAUGUACUUAACAACAGACAAAAAGCAGGAA
Csp-28-1   GUAAUU.UUGCACCCGU................................................................................................UAC.AUAAU.AAUCAU....UA.AA..CAAUAUUGGAAUGUACUUAACAACAGACAAAAAGCAGGAA
Cin-1-1    GUAAUU.UUGCACCCGU................................................................................................UAC.AUAAU.AAUCAU....UA.AA..CAAUAUUGGAAUGUACUUAACAACAGACAAAAAGCAGGAA
Csp-13-1   GUAAUU.UUGCACCCGU................................................................................................UAC.AUAAU.AAUCAU....UA.AA..CAAUAUUGGAAUGUACUUAACAACAGAAAAAAAGCAGGAA
Csp-21-1   GUAAUU.UUGCACCCGU................................................................................................UAC.AUAAU.AAUCAU....UA.AA..CAAUAUUGGAAUGUACUUAACAACAGAAAAAAAGCAGGAA
Csp-25-1   GUAAUU.UUGCACCCGU................................................................................................UAC.AUAAU.AAUCAU....UA.AA..CAAUAUUGGAAUGUACUUAACAACAGAAAAAAAAGCAGAA
Csp-22-1   GUAAUU.UUGCACCCGU................................................................................................UAC.AUAAU.AAUCAU....UA.AA..CAAUAUUGGAAUGUACUUAACAACAGACAAAAAGCAGGAA
Cga-1-1    GUAAUU.UUGCACCCGU................................................................................................UAC.AUAAU.AAUCAU....UA.AA..CAAUAUUGGAAUGUACUUAACAACAGAAAAAAAGCAGGAA
Cvr-1-1    GUAAUU.UUGCACCCGU................................................................................................UAC.AUAAU.AAUCAU....UA.AA..CAAUAUUGGAAUGUACUUAACAACAGAAAAAAAGCAGGAA
Csp-14-1   GUAAUU.UUGCACCCGU................................................................................................UAC.AUAAU.AAUCAU....UA.AA..CAAUAUUGGAAUGUACUUAACAACAGACAAAAAGCAGGAA
env-1877   GUAAUU.UUGCACCCGU................................................................................................UAC.AUAAU.AAUCAU....UA.AA..CAAUAUUGGAAUGUACUUAACAACAGAAAAAAAGCAGGAA
env-1878   CUACCU.UUGCCCCGAUUUAAAAAA........................................................................................UAC.AUAAU.AAUCAUU...UA.AA..CAAUAUUGGAAUGUAUCUAACAAAAGAAACAAAAGCAGAA
env-1879   CUACCU.UUGCCCCGAUUUAAAAAA........................................................................................UAC.AUAAU.AAUCAUU...UUAAA..CAAUAUUGGAAUGUAUCUA.......................
env-1880   CUAUCU.UUGCACCCGAAU..............................................................................................UAC.AUAAU.AAUUAUU...AA.AA..CAAUAUUGGAAUGUAUCUAACAACAGAAAAAAAAGCAGAA
env-1881   CUAUCU.UUGCACCCGAAU..............................................................................................UAC.AUAAU.AAUUAUU...AA.AA..CAAUAUUGGAAUGUAUCUAACAACAGAAAAAAAAGCAGAA
env-1883   GUACAU.UUGCACCC.............GAAUAGAGA.........AAA.................UCUUUAUUC.....................................AUAC.AUAAU.AAUCAUU...UA.AA..UAAUAUUGGAAUGUAUUUAACUAAAGAAACAAAAGCUGAA
env-1884   CUACAU.UUGCACCCUCA...............................................................................................UAC.AUAAU.AAUCAUU...UA.AA..UAAUAUUGGAAUGUAUCUAACAAAAGAAGUAAAAGCAGAA
env-1885   CUACAU.UUGCACCCUCA...............................................................................................UAC.AUAAU.AAUCAUU...UA.AA..UAAUAUUGGAAUGUAUCUAACAAAAGAAGUAAAAGCAGAA
env-1886   CUACAU.UUGCACCCUCA...............................................................................................UAC.AUAAU.AAUCAUU...UA.AA..UAAUAUUGGAAUGUAUCUAACAAAAGAAGUAAAAGCAGAA
env-1892   CUACAU.UUGCACCCUCA...............................................................................................UAC.AUAAU.AAUCAUU...UA.AA..UAAUAUUGGAAUGUAUCUAACAAAAGAAGUAAAAGCAGAA
env-1893   CUACAU.UUGCACCCUCA...............................................................................................UAC.AUAAU.AAUCAUU...UA.AA..UAAUAUUGGAAUGUAUCUAA......................
env-1898   CUACAU.UUGCACCCUCA...............................................................................................UAC.AUAAU.AAUCAUU...UA.AA..UAAUAUUGGAAUGUAUCUAA......................
env-1900   UUACAU.UUGCACCCUCA...............................................................................................UAC.AUAAU.AAUCAUU...UA.AA..UAAUAUUGGAAUGUAUCUAACAAAAGAAGUAAAAGCAGAA
env-1901   CUACAU.UUGCACCCUCA...............................................................................................UAC.AUAAU.AAUCAUU...UA.AA..UAAUAUUGGAAUGUAUCUAACAAAAGAAGUAAAAGCAGAA
env-1902   UUACAU.UUGCACCCUCA...............................................................................................UAC.AUAAU.AAUCAUU...UA.AA..UAAUAUUGGAAUGUAUUUAACAAAA.................
env-1903   CUACAU.UUGCACCCUCA...............................................................................................UAC.AUAAU.AAUCAUU...UA.AA..UAAUAUUGGAAUGUAUCUAACAAAAGAAGUAAAAGCAGAA
env-1904   CUACAU.UUGCACCCUCA...............................................................................................UAC.AUAAU.AAUCAUU...UA.AA..UAAUAUUGGAAUGUAUCUAACAAAAGAAGUAAAAGCAGAA
env-1905   CUACAU.UUGCACCCUCA...............................................................................................UAC.AUAAU.AAUCAUU...UA.AA..UAAUAUUGGAAUGUAUCUAACAAAAGAAGUAAAAGCAGAA
env-1906   CUACAU.UUGCACCCUCA...............................................................................................UAC.AUAAU.AAUCAUU...UA.AA..UAAUAUUGGAAUGUAUCUAACAAAAGAAGUAAAAGCAGAA
env-1907   CUACAU.UUGCACCCUCA...............................................................................................UAC.AUAAU.AAUCAUU...UA.AA..UAAUAUUGGAAUGUAUCUAACAAAAGAAGUAAAAGCAGAA
env-1908   .UACAU.UUGCACCCUCA...............................................................................................UAC.AUAAU.AAUCAUU...UA.AA..UAAUAUUGGAAUGUAUCUAACAAAAGAAGUAAAAGCAGAA
env-1909   CUACAU.UUGCACCCUCA...............................................................................................UAC.AUAAU.AAUCAUU...UA.AA..UAAUAUUGGAAUGUAUCUAACAAAAGAAGUAAAAGCAGAA
env-1910   CUACAU.UUGCACCCUCA...............................................................................................UAC.AUAAU.AAUCAUU...UA.AA..UAAUAUUGGAAUGUAUCUAACAAAAGAAGUAAAAGCAGAA
env-1911   CUACAU.UUGCACCCUCA...............................................................................................UAC.AUAAU.AAUCAUU...UA.AA..UAAUAUUGGAAUGUAUCUAACAAAAGAAGUAAAAGCAGAA
env-1912   CUACAU.UUGCACCCUCA...............................................................................................UAC.AUAAU.AAUCAUU...UA.AA..UAAUAUUGGAAUGUAUCUAACAAAAGAAGUAAAAGCAGAA
env-1913   UUACAU.UUGCACCCUCA...............................................................................................UAC.AUAAU.AAUCAUU...UA.AA..UAAUAUUGGAAUGUAUUUAACAAAAGAAGUAAAAGCAGAA
env-1915   CUACAU.UUGCACCCUCA...............................................................................................UAC.AUAAU.AAUCAUU...UA.AA..UAAUAUUGGAAUGUAUCUAACAAAAGAAGUAAAAGCAGAA
env-1916   CUACAU.UUGCACCCUCA...............................................................................................UAC.AUAAU.AAUCAUU...UA.AA..UAAUAUUGGAAUGUAUCUAACAAAAGAAGUAAAAGCAGAA
env-1917   CUACAU.UUGCACCCUCA...............................................................................................UAC.AUAAU.AAUCAUU...UA.AA..UAAUAUUGGAAUGUAUCUAACAAAAGAAGUAAAAGCAGAA
env-1918   CUACAU.UUGCACCCUCA...............................................................................................UAC.AUAAU.AAUCAUU...UA.AA..UAAUAUUGGAAUGUAUCUAACAAAAGAAGUAAAAGCAGAA
env-1921   CUACAU.UUGCACCCUCA...............................................................................................UAC.AUAAU.AAUCAUU...UA.AA..UAAUAUUGGAAUGUAUCUAACAAAAGAAGUAAAAGCAGAA
env-1922   CUACAU.UUGCACCCUCA...............................................................................................UAC.AUAAU.AAUCAUU...UA.AA..UAAUAUUGGAAUGUAUCUAACAAAAGAAGUAAAAGCAGAA
env-1924   CUACAU.UUGCACCCUCA...............................................................................................UAC.AUAAU.AAUCAUU...UA.AA..UAAUAUUGGAAUGUAUCUAACAAAAGAAGUAAAAGCAGAA
env-1925   CUACAU.UUGCACCCUCA...............................................................................................UAC.AUAAU.AAUCAUU...UA.AA..UAAUAUUGGAAUGUAUCUAACAAAAGAAGUAAAAGCAGAA
env-1926   UUACAU.UUGCACCCUCA...............................................................................................UAC.AUAAU.AAUCAUU...UA.AA..UAAUAUUGGAAUGUAUCUAACAAAAGAAGUAAAAGCAGAA
env-1927   CUACAU.UUGCACCCUCA...............................................................................................UAC.AUAAU.AAUCAUU...UA.AA..UAAUAUUGGAAUGUAUCUAACAAAAGAAGUAAAAGCAGAA
env-1928   CUACAU.UUGCACCCUCA...............................................................................................UAC.AUAAU.AAUCAUU...UA.AA..UAAUAUUGGAAUGUAUCUAACAAAAGAAGUAAAAGCAGAA
env-1929   CUACAU.UUGCACCCUCA...............................................................................................UAC.AUAAU.AAUCAUU...UA.AA..UAAUAUUGGAAUGUAUCUAACAAAAGAAGUAAAAGCAGAA
env-1930   CUACAU.UUGCACCCUCA...............................................................................................UAC.AUAAU.AAUCAUU...UA.AA..UAAUAUUGGAAUGUAUCUAACAAAAGAAGUAAAAGCAGAA
env-1931   CUACAU.UUGCACCCUCA...............................................................................................UAC.AUAAU.AAUCAUU...UA.AA..UAAUAUUGGAAUGUAUCUAACAAAAG..............

           ............................<<<<<<<<<<<<<<<<<<...........>>>>>>>>>>>>>>>>>>.........<<<<<<<<........>>>>>>>>........................................................................
           ............................2222221111..101.00...........00.101..1111222222.........11011000........00011011........................................................................
           •UAY•U-UUGC••CY------------◦◦◦◦◦◦◦-----------R•R◦◦----------------◦◦◦◦◦◦◦--------------------------------◦UAC-AUAA•-AAUCAUU---UA-AA--YAAUAUURG•AUGUAUUUARC•ARAGAAR••AAARRRRAR
           5′                                                                                                                                                 3′
```

```
env-1935   .UACAU.UUGCACCCUCA..........................................UAC.AUAAU.AAUCAUU...UA.AA..UAAUAUUGGAAUGUAUCUAACAAAAGAAGUAAAAGCAGAA
env-1936   CUACAU.UUGCACCCUCA..........................................UAC.AUAAU.AAUCAUU...UA.AA..UAAUAUUGGAAUGUAUCUAACAAAAGAAACAAAAGCAGAA
env-1937   CUACAU.UUGCACCCUCA..........................................UAC.AUAAU.AAUCAUU...UA.AA..UAAUAUUGGAAUGUAUCUAACAAAAGAAGUAAAAGCAGAA
env-1938   CUACAU.UUGCACCCUCA..........................................UAC.AUAAU.AAUCAUU...UA.AA..UAAUAUUGGAAUGUAUCUAACAAAAG..............
env-1939   CUACAU.UUGCACCCUCA..........................................UAC.AUAAU.AAUCAUU...UA.AA..UAAUAUUGGAAUGUAUCUAACAAAAGAAGUAAAAGCAGAA
env-1940   CUACAU.UUGCACCCUCA..........................................UAC.AUAAU.AAUCAUU...UA.AA..UAAUAUUGGAAUGUNNNNNNNNNNNNNNNNNNNNNNNNNN
env-1941   CUACAU.UUGCACCCUCA..........................................UAC.AUAAU.AAUCAUU...UA.AA..UAAUAUUGGAAUGUAUCUAACAAAAGAAGUAAAAGCAGAA
env-1942   UUACAU.UUGCACCCUCA..........................................UAC.AUAAU.AAUCAUU...UA.AA..UAAUAUUGGAAUGUAUCUAACAAAAGAAGUAAAAGCAGAA
env-1943   CUACAU.UUGCACCCUCA..........................................UAC.AUAAU.AAUCAUU...UA.AA..UAAUAUUGGAAUGUAUCUAACAAAAGAAGUAAAAGCAGAA
env-1944   CUACAU.UUGCACCCUCA..........................................UAC.AUAAU.AAUCAUU...UA.AA..UAAUAUUGGAAUGUAUUUAACAAAAGAAGUAAAAGCAGAA
env-1945   CUACAU.UUGCACCCUCA..........................................UAC.AUAAU.AAUCAUU...UA.AA..UAAUAUUGGAAUGUAUCUAACAAAAGAAGUAAAAGCAGAA
env-1946   CUACAU.UUGCACCCUCA..........................................UAC.AUAAU.AAUCAUU...UA.AA..UAAUAUUGGAAUGUAUCUAACAAAAGAAGUAAAAGCAGAA
env-1947   CUACAU.UUGCACCCUCA..........................................UAC.AUAAU.AAUCAUU...UA.AA..UAAUAUUGGAAUGUAUCUAACAAAAGAAGUAAAAGCAGAA
env-1948   CUACAU.UUGCACCCUCA..........................................UAC.AUAAU.AAUCAUU...UA.AA..UAAUAUUGGAAUGUAUCUAACAAAAGAAGUAAAAGCAGAA
env-1949   CUACAU.UUGCACCCUCA..........................................UAC.AUAAU.AAUCAUU...UA.AA..UAAUAUUGGAAUGUAUCUAACAAAAGAA............
env-1950   UUACAU.UUGCAGUCUCA..........................................UAC.AUAAU.AAUCAUU...UA.AA..UAAUAUUGGAAUGUAUCUAACAAAAGAAGUAAAAGCAGAA
env-1951   UUACAU.UUGCAGUCUCA..........................................UAC.AUAAU.AAUCAUU...UA.AA..UAAUAUUGGAAUGUAUCUAA....................
env-1952   UUACAU.UUGCAGUCUCA..........................................UAC.AUAAU.AAUCAUU...UA.AA..UAAUAUUGGAAUGUAUCUAACAAAAGAAGUAAAAGCAGAA
env-1954   UUACAU.UUGCAGUCUCA..........................................UAC.AUAAU.AAUCAUU...UA.AA..UAAUAUUGGAAUGUAUCUAACAAAAG..............
env-1955   UUACAU.UUGCAGUCUCA..........................................UAC.AUAAU.AAUCAUU...UA.AA..UAAUAUUGGAAUGUAUCUAACAAAAGAAGUAAAAGCAGAA
env-1957   UUACAU.UUGCAGCCUCA..........................................UAC.AUAAU.AAUCAUU...UA.AA..UAAUAUUGGAAUGUAUCUAACAAAAGAAGUAAAAGCAGAA
env-1958   UUACAU.UUGCAGCCUCA..........................................UAC.AUAAU.AAUCAUU...UA.AA..UAAUAUUGGAAUGUAUCUAACAAAAGAAGUAAAAGCAGAA
env-1959   UUACAU.UUGCAGCCUCA..........................................UAC.AUAAU.AAUCAUU...UA.AA..UAAUAUUGGAAUGUAUCUAACAAAAGAAGUAAAAGCAGAA
env-1961   UUACAU.UUGCAGCCUCA..........................................UAC.AUAAU.AAUCAUU...UA.AA..UAAUAUUGGAAUGUAUCUAACAAAAG..............
env-1962   UUACAU.UUGCAGCCUCA..........................................UAC.AUAAU.AAUCAUU...UA.AA..UAAUAUUGGAAUGUAUCUAACAAAAGAAGUAAAAGCAGAA
env-1963   UUACAU.UUGCAGCCUCA..........................................UAC.AUAAU.AAUCAUU...UA.AA..UAAUAUUGGAAUGUAUCUAACAAAAGAAGUAAAAGCAGAA
env-1964   UUACAU.UUGCAGCCUCA..........................................UAC.AUAAU.AAUCAUU...UA.AA..UAAUAUUGGAAUGUAUCUAACAAAAG..............
env-1965   UUACAU.UUGCAGCCUCA..........................................UAC.AUAAU.AAUCAUU...UA.AA..UAAUAUUGGAAUGUAUCUAACAAAAGAAGUAAAAGCAGAA
env-1966   UUACAU.UUGCAGCCUCA..........................................UAC.AUAAU.AAUCAUU...UA.AA..UAAUAUUGGAAUGUAUCUAACAAAAGAAGUAAAAGCAGAA
env-1967   UUACAU.UUGCAGCCUCA..........................................UAC.AUAAU.AAUCAUU...UA.AA..UAAUAUUGGAAUGUAU........................
env-1968   UUACAU.UUGCAGCCUCA..........................................UAC.AUAAU.AAUCAUU...UA.AA..UAAUAUUGGAAUGUAUCUAACAAAAGAAGUAAAAGCAGAA
env-1969   UUACAU.UUGCAGCCUCA..........................................UAC.AUAAU.AAUCAUU...UA.AA..UAAUAUUGGAAUGUAUCUAACAAAAGAAGUAAAAGCAGAA
env-1970   UUACAU.UUGCAGCCUCA..........................................UAC.AUAAU.AAUCAUU...UA.AA..UAAUAUUGGAAUGUAUCU......................
env-1971   UUACAU.UUGCAGCCUCA..........................................UAC.AUAAU.AAUCAUU...UA.AA..UAAUAUUGGAAUGUAUCUAACAAAAGAAGUAAAAGCAGAA
env-1972   UUACAU.UUGCAGCCUCA..........................................UAC.AUAAU.AAUCAUU...UA.AA..UAAUAUUGGAAUGUAU........................
env-1973   UUACAU.UUGCAGCCUCA..........................................UAC.AUAAU.AAUCAUU...UA.AA..UAAUAUUGGAAUGUAUCUAACAAAAGAAGUAAAAGCAGAA
env-1974   UUACAU.UUGCAGCCUCA..........................................UAC.AUAAU.AAUCAUU...UA.AA..UAAUAUUGGAAUGUAUCUAACAAAAGAAGUAAAAGCAGAA
env-1975   UUACAU.UUGCAGCCUCA..........................................UAC.AUAAU.AAUCAUU...UA.AA..UAAUAUUGGAAUGUAUCUAACAAAAGAAGUAAAAGCAGAA
env-1976   UUACAU.UUGCAGCCUCA..........................................UAC.AUAAU.AAUCAUU...UA.AA..UAAUAUUGGAAUGUAUCUAACAAAAGAAGUAAAAGCAGAA
env-1977   UUACAU.UUGCAGCCUCA..........................................UAC.AUAAU.AAUCAUU...UA.AA..UAAUAUUGGAAUGUAUCUAACAAAAGAAGUAAAAGCAGAA
env-1978   UUACAU.UUGCAGCCUCA..........................................UAC.AUAAU.AAUCAUU...UA.AA..UAAUAUUGGAAUGUAUCUAACAAAAGAAGUAAAAGCAGAA
env-1979   UUACAU.UUGCAGCCUCA..........................................UAC.AUAAU.AAUCAUU...UA.AA..UAAUAUUGGAAUGUAUCUAACAAAAGAAGUAAAAGCAGAA
env-1980   UUACAU.UUGCAGCCUCA..........................................UAC.AUAAU.AAUCAUU...UA.AA..UAAUAUUGGAAUGUAUCUAACAAAAGAAGUAAAAGCAGAA
env-1981   UUACAU.UUGCAGCCUCA..........................................UAC.AUAAU.AAUCAUU...UA.AA..UAAUAUUGGAAUGUAUCUAACAAAAGAAGUAAAAGCAGAA
env-1982   UUACAU.UUGCAGCCUCA..........................................UAC.AUAAU.AAUCAUU...UA.AA..UAAUAUUGGAAUGUAUUUAACAAAAGAAGUAAAAGCAGAA
env-1983   UUACAU.UUGCAGCCUCA..........................................UAC.AUAAU.AAUCAUU...UA.AA..UAAUAUUGGAAUGUAUCUAA....................
env-1984   UUACAU.UUGCAGCCUCA..........................................UAC.AUAAU.AAUCAUU...UA.AA..UAAUAUUGGAAUGUAUCUAACAAAAGAAGUAAAAGCAGAA
env-1985   UUACAU.UUGCAGCCUCA..........................................UAC.AUAAU.AAUCAUU...UA.AA..UAAUAUUGGAAUGUAUCUAACAAAAGAAGUAAAAGCAGAA
env-1986   UUACAU.UUGCAGCCUCA..........................................UAC.AUAAU.AAUCAUU...UA.AA..UAAUAUUGGAAUGUAUCUAACAAAAGAAGUAAAAGC....
env-1987   UUACAU.UUGCAGCCUCA..........................................UAC.AUAAU.AAUCAUU...UA.AA..UAAUAUUGGAAUGUAUCUAACAAAAGAAGUAAAAGCAGAA
env-1988   UUACAU.UUGCAGCCUCA..........................................UAC.AUAAU.AAUCAUU...UA.AA..UAAUAUUGGAAUGUAUCUAA....................
env-1990   UUACAU.UUGCAGCCUCA..........................................UAC.AUAAU.AAUCAUU...UA.AA..UAAUAUUGGAAUGUAUCUAACAAAAGAAGUAAAAGCAGAA
env-1991   UUACAU.UUGCAGCCUCA..........................................UAC.AUAAU.AAUCAUU...UA.AA..UAAUAUUGGAAUGUAUCUAACAAAAGAAGUAAAAGCAGAA
env-1992   UUACAU.UUGCAGCCUCA..........................................UAC.AUAAU.AAUCAUU...UA.AA..UAAUAUUGGAAUGUAUUUAACAAAAGAAGUAAAAGCAGAA
env-1993   UUACAU.UUGCAGCCUCA..........................................UAC.AUAAU.AAUCAUU...UA.AA..UAAUAUUGGAAUGUAUCUAACAAAAGAAGUAAAAGCAGAA
env-1994   UUACAU.UUGCAGCCUCA..........................................UAC.AUAAU.AAUCAUU...UA.AA..UAAUAUUGGAAUGUAUCUAACAAAAGAA............
env-1995   UUACAU.UUGCAGCCUCA..........................................UAC.AUAAU.AAUCAUU...UA.AA..UAAUAUUGGAAUGUAUCUAA....................
env-1996   UUACAU.UUGCAGCCUCA..........................................UAC.AUAAU.AAUCAUU...UA.AA..UAAUAUUGGAAUGUAUCUAACAAAAGAAGUAAAAGCAGAA
env-1997   UUACAU.UUGCAGCCUCA..........................................UAC.AUAAU.AAUCAUU...UA.AA..UAAUAUUGGAAUGUAUCUAACAAAAGAAGUAAAAGCAGAA
env-1998   UUACAU.UUGCAGCCUCA..........................................UAC.AUAAU.AAUCAUU...UA.AA..UAAUAUUGGAAUGUAUCUAACAAAAGAAGUAAAAGCAGAA
env-1999   UUACAU.UUGCAGCCUCA..........................................UAC.AUAAU.AAUCAUU...UA.AA..UAAUAUUGGAAUGUAUCUAACAAAAGAAGUAAAAGCAGAA
env-2000   UUACAU.UUGCAGCCUCA..........................................UAC.AUAAU.AAUCAUU...UA.AA..UAAUAUUGGAAUGUAUCUAACAAAAGAAGUAAAAGCAGAA
env-2001   UUACAU.UUGCAGCCUCA..........................................UAC.AUAAU.AAUCAUU...UA.AA..UAAUAUUGGAAUGUAU........................
env-2002   UUACAU.UUGCAGCCUCA..........................................UAC.AUAAU.AAUCAUU...UA.AA..UAAUAUUGGAAUGUAUCUAACAAAAG..............
env-2003   UUACAU.UUGCAGCCUCA..........................................UAC.AUAAU.AAUCAUU...UA.AA..UAAUAUUGGAAUGUAUCUAACAAAAGAAGUAAAAGCAGAA
env-2004   UUACAU.UUGCAGCCUCA..........................................UAC.AUAAU.AAUCAUU...UA.AA..UAAUAUUGGAAUGUAUCUAACAAAAGAAGUAAAAGCAGAA
env-2005   UUACAU.UUGCAGCCUCA..........................................UAC.AUAAU.AAUCAUU...UA.AA..UAAUAUUGGAAUGUAUUUAACAAAAGAAGUAAAAGCAGAA
Ran-4-1    CUAAAU.UUGCAGCCUCA..........................................UAC.AUAAU.AAUCAU....UA.AA..UAAUAUUGGAAUGUAUUUAACUAAAGAAAAAAAGGCAGAA
Ran-2-1    CUAAAU.UUGCAGCCUCA..........................................UAC.AUAAU.AAUCAU....UA.AA..UAAUAUUGGAAUGUAUUUAACUAAAGAAAAAAAGGCAGAA
Cca-2-1    AUACAU.UUGCAACCGCAUAUCAUAG..................................UAC.AUAAU.AAUCAUU...AA.AA..UAAUAUUGGAAUGUAUUUAACAAAAGAAGUAAAAUCAGAG
Cca-2-2    AUACAU.UUGCAACCGCAUAUCAUAG..................................UAC.AUAAU.AAUCAU....UA.AA..UAAUAUUGGAAUGUAUUUAACAAAAGAAGUAAAAUCAGAG
env-2006   GUACAU.UUGCACCCGCA..........................................UAC.AUAAA.AAUCAU....UA.AA..UAAUAUUGGCAUGUAUUUAACAACAGAAAAAAAACAGGAA
env-2007   UUAUCU.UUGCACGC............UGAUUG.CGCGUCCUUCGAGAGA.....CGGAGGAACUGACAAUCA.......AUAC.AUAAA.AAUCAUU..UA.AA..UAAUAUUGGAAUGUAUUUAACCAAAGAAGUAAAGGCCGAG
Mru-1-1    AUAUCU.UUGCACGC............UGAUUGCUUC..UCCU..GAGCAUGUCGA..AGGAACUGA.CAAUCA.......AUAC.AUAAA.AAUCAUU..UA.AA..UAAUAUUGGAAUGUAUUUAACAAAAGAAGUAAAGGCCGAG
env-2008   AUAUCU.UUGCACGC............UGAUUG...........AAGGA.................CAAUCA.......AUAC.AUAAA.AAUCAUU..UA.AA..UAAUAUUGGAAUGUAUUUAACAAAAGAAGCAAAAGCCGAG
env-2009   AUAUCU.UUGCACGC............UGAUUG...........AAGGA.................CAAUCA.......AUAC.AUAAA.AAUCAUU..UA.AA..UAAUAUUGGAAUGUAUUUAACAAAAGAAGCAAAAGCCGAG
env-2010   UUAUCU.UUGCACGC............UGAUUG...........AAGGC.................CAAUCA.......AUAC.AUAAA.AAUCAUU..UA.AA..UAAUAUUGGAAUGUAUUUA...................
env-2011   AUAUCU.UUGCACGC............UGAUUG...........AAAGC.................CAAUCA.......AUAC.AUAAA.AAUCAUU..UA.AA..UAAUAUUGGAAUGUAUUUAACAAAAGAAGUAAAAGCCGAG
env-2012   AUAUCU.UUGCACGC............UGAUUGA..........AGG.................UCAAUCA.......AUAC.AUAAA.AAUCAUU..UA.AA..UAAUAUUGGAAUGUAUUUAACAAAAGAAGUUAAAGCCGAG
env-2014   CUAUCU.UUGCACGC............UGAUUGA..........AGG.................UCAAUCA.......AUAC.AUAAA.AAUCAUU..UA.AA..UAAUAUUGGAAUGUAUUUAACUAAAGAAGUAAAAGCCGAU
env-2015   AUAUCU.UUGCACGC............UGAUUGA..........AGG.................UCAAUCA.......AUAC.AUAAA.AAUCAUU..UA.AA..UAAUAUUGGAAUGUAUUUAACAAAAGAAGUUAAAGCCGAG
env-2016   AUAUCU.UUGCACGC............UGAUUGA..........AGG.................UCAAUCA.......AUAC.AUAAA.AAUCAUU..UA.AA..UAAUAUUGGAAUGUAUUUAACAAAAGAAGUUAAAGCCGAG
env-2017   CUAUCU.UUGCACGC............UGAUUGA..........AGG.................UCAAUCA.......AUAC.AUAAA.AAUCAUU..UA.AA..UAAUAUUGGAAUGUAUUUAACAAAAGAAGUAAAGGCCGAG

           ...........................<<<<<<<<<<<<<<<<<<..........>>>>>>>>>>>>>>>>>>.........<<<<<<<<........>>>>>>>>.........................................
           ...........................2222221111..101.00..........00.101..1111222222.........11011000........00011011.........................................
           •UAY•U-UUGC••CY------------•••••••----------R•R••-----------------•••••••----------------------------------•UAC-AUAA•-AAUCAUU---UA-AA--YAAUAUURG•AUGUAUUUARC•ARAGAAR••AAARRRRAR
           5'                                                                                                                  3'
```

```
env-2018  AUAUCU.UUGCACGC.............UGAUUGA...........AGG...................UCAAUCA......................................AUAC.AUAAA.AAUCAUU...UA.AA..UAAUAUUGGAAUGUAUUUAACAAAAGAAGUUAAAGCCGAG
Msp-2-1   CUAUCU.UUGCGCGC.............UGAUUGA...........AGG...................UCAAUCA......................................AUAC.AUAAA.AAUCAUU...UAAAU..AAAUAUUGGAAUGUAUUUAACGAAAGAAGUAAAAGCCGAG
Cba-2-1   AUAUAU.UUGCACGC.............UGAUUGA...........AAA...................UCAAUCA.....................................UAUAC.AUAAA.AAUCAU....UA.AA..UAAUAUUGGAAUGUAUUUAACAAAAGAAGUAAAGGCUGAC
Csp-11-1  AUAUAU.UUGCACGC.............UGAUUGA...........AAA...................UCAAUCA.....................................UAUAC.AUAAA.AAUCAU....UA.AA..UAAUAUUGGAAUGUAUUUAACAAAAGAAGUAAAGGCUGAC
Cba-4-1   AUAUAU.UUGCACGC.............UGAUUGA...........AAA...................UCAAUCA.....................................UAUAC.AUAAA.AAUCAU....UA.AA..UAAUAUUGGAAUGUAUUUAACAAAAGAAGUAAAGGCUGAC
env-2019  CUAUCU.UUGCGCCC.............UGAUUGA...........GCG...................UCAAUCG.....................................GAUAC.AUAAA.AAUCAUU...UACAA..CAAUAUUGGCAUGUAUUUAACCAAAGAAACCAAGGCCGAG
env-2020  GUAUCU.UUGCGCCC.............UGAUUG............AAAAC..................CAAUCA.....................................CGUAC.AUAAA.AAUCAUU...UAAAA..UAAUAUUGGAAUGUAUUUAACAAAAGAAGUAAAAGCCGAA
Csp-12-1  GUAUCU.UUGCAUCU.............UGAUUGA...........AAA...................UCAAUCA.....................................UAUAC.AUAAA.AAUCAUU...UAAAA..UAAUAUUGGAAUGUAUUUAACACAAGAAGUAAAAGCUGAA
Zul-1-1   GUAUCU.UUGCAGCU.............UGAUUGA...........AAA...................UCAGUCA.....................................UAUAC.AUAAA.AAUCAUU...UAAAA..UAAUAUUGGAAUGUAUUUAACUAAGGAAGUUAAAGCCGAA
env-2021  GUAUAU.UUGCGCCC.............UGAUUG............AAAAA..................CAAUCA.....................................CAUAC.AUAAA.AAUCAUU...UAAAA..CAAUAUUGGAAUGUAUUUAUCAAAAGAAGUAAAAGCCGAA
env-2027  GUAUAU.UUGCGCCC.............UGAUUG............AAAAA..................CAAUCA.....................................CAUAC.AUAAA.AAUCAUU...UAAAA..CAAUAUUGGAAUGUAUUUAUCAAAAGAAGUAAAAGCCGAA
env-2028  GUAUAU.UUGCGCCC.............UGAUUG............AAAAA..................CAAUCA.....................................CAUAC.AUAAA.AAUCAUU...UAAAA..CAAUAUUGGAAUGUAUUUAUCAAAAGAAGUAAAAGCCGAA
env-2029  GUAUAU.UUGCGCCC.............UGAUUG............AAAAA..................CAAUCA.....................................CAUAC.AUAAA.AAUCAUU...AA.AC..UAAUAUUGGAAUGUAUUUAACAAAAGAAGUAAAAGCCGAA
Ace-1-1   CUAUCU.UUGCGCCC.............UGAUUG............AAAAA..................CAAUCA.....................................CAUAC.AUAAA.AAUCAUU...AA.AC..CAAUAUUGGAAUGUAUUUAAGCAAAGAAGUAAAGGCCGAC
Ala-2-1   CUAUCU.UUGCGCCC.............UGAUUG............AAAAA..................CAAUCA.....................................CAUAC.AUAAA.AAUCAUU...AA.AC..AAAUAUUGGAAUGUAUUUAAGUAAAGAAGUAAAGGCCGAA
env-2030  CUAUCU.UUGCGCCC.............UGAUUGA...........AAA...................UCAAUCA.....................................AAUAC.AUAAA.AAUCAUU...AA.AC..AAAUAUUGGAAUGUAUUU......................
env-2031  CUAUCU.UUGCGCCC.............UGAUUGA...........AAA...................UCAAUCA.....................................AAUAC.AUAAA.AAUCAUU...AA.AC..AAAUAUUGGAAUGUAUUUAACCAAGGAAAUUAAAGCCGAC
env-2032  CUAUCU.UUGCGCCC.............UGAUUGA...........AAA...................UCAAUCA.....................................AAUAC.AUAAA.AAUCAUU...AA.AC..AAAUAUUGGAAUGUAUUUAACCAAGGAAAUUAAAGCCGAC
env-2033  CUAUCU.UUGCGCCC.............UGAUUGA...........AAA...................UCAAUCA.....................................AAUAC.AUAAA.AAUCAUU...AA.AC..AAAUAUUGGAAUGUAUUUAACCAAGGAAAUUAAAGCCGAC
env-2034  CUAUCU.UUGCGCCC.............UGAUUGA...........AAA...................UCAAUCA.....................................AAUAC.AUAAA.AAUCAUU...AA.AC..AAAUAUUGGAAUGUAUUUAACCAAGGAAAUUAAAGCCGAC
env-2035  CUAUCU.UUGCGCCC.............UGAUUGA...........AAA...................UCAAUCA.....................................AAUAC.AUAAA.AAUCAUU...AA.AC..AAAUAUUGGAAUGUAUUUAACCAAGGAGAUUAAAGCUGAC
env-2036  CUAUCU.UUGCGCCC.............UGAUUGA...........AAA...................UCAAUCA.....................................AAUAC.AUAAA.AAUCAUU...AA.AC..AAAUAUUGGAAUGUAUUUAACCAAGGAGAUUAAAGCUGAC
env-2037  CUAUCU.UUGCGCCC.............UGAUUGA...........AAA...................UCAAUCA.....................................AAUAC.AUAAA.AAUCAUU...AA.AC..AAAUAUUGGAAUGUAUUUAACCAAGGAGAUUAAAGCCGAC
env-2038  CUAUCU.UUGCGCCC.............UGAUUGA...........AAA...................UCAAUCA.....................................AAUAC.AUAAA.AAUCAUU...AA.AC..AAAUAUUGGAAUGUAUUUAACCAAGGAGAUUAAAGCUGAC
env-2039  CUAUCU.UUGCGCCC.............UGAUUGA...........AAA...................UCAAUCA.....................................AAUAC.AUAAA.AAUCAUU...AA.AC..AAAUAUUGGAAUGUAUUUAACCAAGGAGAUUAAAGCUGAC
env-2040  UUAUCU.UUGCGCCC.............UGAUUGA...........AAA...................UCAAUCA.....................................AAUAC.AUAAA.AAUCAUU...AA.AC..AAAUAUUGGAAUGUAUUUAACCAAGGAGAUUAAAGCUGAC
env-2041  CUAUCU.UUGCGCCC.............UGAUUGA...........AAA...................UCAAUCA.....................................AAUAC.AUAAA.AAUCAUU...AA.AC..AAAUAUUGGAAUGUAUUUAACCAAGGAAAUUAAAGCCGAC
env-2042  UUAUCU.UUGCGCCC.............UGAUUGA...........AAA...................UCAAUCA.....................................AAUAC.AUAAA.AAUCAUU...AA.AC..AAAUAUUGGAAUGUAUUUAACCAAGGAGAUUAAAGCUGAC
env-2043  CUAUCU.UUGCGCCC.............UGAUUGA...........AAA...................UCAAUCA.....................................AAUAC.AUAAA.AAUCAUU...AA.AC..AAAUAUUGGAAUGUAUUUAACCAAGGAAAUUAAAGCCGAC
env-2044  CUAUCU.UUGCGCCC.............UGAUUGA...........AAA...................UCAAUCA.....................................AAUAC.AUAAA.AAUCAUU...AA.AC..AAAUAUUGGAAUGUAUUUAACCAAGGAAAUUAAAGCCGAC
env-2045  CUAUCU.UUGCGCCC.............UGAUUGA...........AAA...................UCAAUCA.....................................AAUAC.AUAAA.AAUCAUU...AA.AC..AAAUAUUGGAAUGUAUUUAACCAAAGAGAUUAAGGCCAAG
env-2046  CUAUCU.UUGCGCCC.............UGAUUGA...........AAA...................UCAAUCA.....................................AAUAC.AUAAA.AAUCAUU...AA.AC..AAAUAUUGGAAUGUAUUUAACCAAGGAAAUUAAAGCCGAC
env-2049  CUAUAU.UUGCGCCC.............UGAUUGA...........AAA...................UCAAUCA.....................................AAUAC.AUCAA.AAUCAUU...UAAAA..CAAUAUUGGAAUGUAUUUAACCAAAGAAGUUAAAGCCGAG
env-2050  CUAUAU.UUGCGCCC.............UGAUUGA...........AAA...................UCAAUCA.....................................AAUAC.AUCAA.AAUCAUU...UAAAA..CAAUAUUGGAAUGUAUUUAACCAAAGAAGUUAAAGCCGAG
env-2051  CUAUAU.UUGCGCCC.............UGAUUGA...........AAA...................UCAAUCA.....................................AAUAC.AUCAA.AAUCAUU...UAAAA..CAAUAUUGGAAUGUAUUUAACCAAAGAAGUUAAAGCCGAG
env-2052  CUAUAU.UUGCGCCC.............UGAUUGA...........AAA...................UCAAUCA.....................................AAUAC.AUCAA.AAUCAUU...UAAAA..CAAUAUUGGAAUGUAUUUAACUAAAGAAGUUAAAGCCGAG
env-2053  CUAUAU.UUGCGCCC.............UGAUUGA...........AAA...................UCAAUCA.....................................AAUAC.AUCAA.AAUCAUU...UAAAA..CAAUAUUGGAAUGUAUUUAACNAAAGAAGUUAAAGCCGAG
env-2054  CUAUAU.UUGCGCCC.............UGAUUGA...........AAA...................UCAAUCA.....................................AAUAC.AUCAA.AAUCAUU...UAAAA..CAAUAUUGGAAUGUAUUUAACCAAAGAAGUUAAAGCCGAG
env-2055  CUAUAU.UUGCGCCC.............UGAUUGA...........AAA...................UCAAUCA.....................................AAUAC.AUCAA.AAUCAUU...UAAAA..CAAUAUUGGAAUGUAUUUAACCAAAGAAGUUAAAGCCGAG
Ead-1-1   CUAUCU.UUGCG.CU.............CGAUUGA...........AAA...................UCAAUCA.....................................GAUAC.AUAAA.AAUCAUU...UAAAA..CAAUAUUGGAAUGUAUUUAACAAAAGAAGUUAAAGCGGAA
env-2068  CUAUCU.UUGCG.CU.............CGAUUGA...........AAA...................UCAAUCA.....................................GAUAC.AUAAA.AAUCAUU...UAAAA..UAAUAUUGGAAUGUAUUUAACAAAAGAAGUUAAGGCAGAA
env-2069  CUAUCU.UUGCG.CU.............CGAUUGA...........AAA...................UCAAUCA.....................................GAUAC.AUAAA.AAUCAUU...UAAAA..UAAUAUUGGAAUGUAUUUAACAAAAGAAGUUAAGGCAGAA
env-2070  CUAUCU.UUGCG.CU.............CGAUUGA...........AAA...................UCAAUCA.....................................GAUAC.AUAAA.AAUCAUU...UAAAA..UAAUAUUGGAAUGUAUUUAACAAAAGAAGUUAAGGCAGAA
env-2071  CUAUCU.UUGCG.CU.............CGAUUGA...........AAA...................UCAAUCA.....................................GAUAC.AUAAA.AAUCAUU...UAAAA..UAAUAUUGGAAUGUAUUUAACAAAAGAAGUUAAGUCAGAA
env-2072  CUAUCU.UUGCG.CU.............CGAUUGA...........AAA...................UCAAUCA.....................................GAUAC.AUAAA.AAUCAUU...UAAAA..UAAUAUUGGAAUGUAUUUAACAAAAGAAGUUAAGGCAGAA
env-2073  CUAUCU.UUGCCCCC.............UGAUUGA...........AAA...................UCAAUCA.....................................UGUAC.AUAAA.AAUCAUU...AA.AA..UAAUAUUGGAAUGUAUUUAACUAAGGAAGUAAAGGCCGAG
env-2074  CUAUCU.UUGCGCCC.............UGAUUGA...........AAA...................UCAAUCA.....................................UGUAC.AUAAA.AAUCAUU...AA.AA..UAAUAUUGGAAUGUAUUUAACUAAAGAAGUAAAGGCCGAG
env-2075  CUAUCU.UUGCGCCC.............UGAUUGA...........AAA...................UCAAUCA.....................................UGUAC.AUAAA.AAUCAUU...AA.AA..UAAUAUUGGAAUGUAUUUAACUAAAGAAGUAAAGGCCGAG
env-2076  CUAUCU.UUGCGGCC.............UGAUUGA...........AAA...................UCAAUCA.....................................UGUAC.AUAAA.AAUCAUU...AA.AG..UAAUAUUGGAAUGUAUUUAACUAAAGAAGUAAAGGCCGAG
env-2077  CUAUCU.UUGCGGCC.............UGGUUGA...........AAA...................UCAAUCA.....................................UGUAC.AUAAA.AAUCAUU...AA.AA..UAAUAUUGGAAUGUAUUUAACUAAGGAAGUUAAGGCCGAG
env-2078  UUAUCU.UUGCGCUC.............UGAUUGA...........AAA...................UCAAUCA.....................................AGUAC.AUAAA.AAUCAUU...AAAAA..CAAUAUUGGAAUGUAUUUAACAAAAGAGAUUAAGUCCGAG
env-2079  GUAUCU.UUGCGCCC.............UGAUUGA...........AAA...................UCAAUUA.....................................UGUAC.AUAAA.AAUCAUU...UAAAA..CAAUAUUGGAAUGUAUUUAUCAAAAGAAAUAAAAGCUGAC
Fba-6-1   UUAUCU.UUGCGCCC.............UGAUUGA...........AAA...................UCAAUCA.....................................AGUAC.AUAAA.AAUCAUU...UAAAA..CAAUAUUGGAAUGUAUCUAACAAAAGAAAUCAAAGCCGAG
env-2080  GUAUCU.UUGCGCCC.............UGAUUGA...........AAA...................UCAAUCA.....................................UGUAC.AUAAA.AAUCAUU...UAAAA..CAAUAUUGGAAUGUAUUUAACAAAAGAAGUAAAAGCCGAC
env-2081  CUAUCU.UUGCGGCC.............UGAUUGA...........AAA...................UCAAUCA.....................................GAUAC.AUAAA.AAUCAUU...UAAAA..CAAUAUUGGAAUGUAUUUAACCAAAGAAGUAAAAGCCGGA
env-2082  CUAUCU.UUGCGGCC.............UGAUUGA...........AAA...................UCAAUCA.....................................GAUAC.AUAAA.AAUCAUU...UAAAA..CAAUAUUGGAAUGUAUUUAACCAAAGAAGUAAAAGCCGGA
env-2083  GUAUCU.UUGCGGCC.............UGAUUGA...........AAA...................UCAAUCA....................................UAAUAC.AUAAA.AAUCAUU...UAAAA..UAAUAUUGGAAUGUAUCUAACAAAAGAAGUAAAGUCUGAA
Msp-1-1   GUAUCU.UUGCGCCC.............UGAUUGA...........AAA...................UCAAUCA....................................UAAUAC.AUAAA.AAUCAUU...UAAAA..UAAUAUUGGAAUGUAUCUUACAAAAGAAGUAAAAGCAGAA
env-2085  GUAUCU.UUGCGCCC.............UGAUUGA...........AAA...................UCAAUCA....................................UAAUAC.AUAAA.AAUCAUU...UAAAA..UAAUAUUGGAAUGUAUCUUACAAAAGAAGUAAAAGCAGAA
env-2086  AUAUCU.UUGCGCCC.............UGAUUGA...........AAA...................UCAAUCG.....................................GAUAC.AUAAA.AAUCAUU...UAAAA..UAAUAUUGGAAUGUAUUUAACCAAAGAAGUAAAAGCCGAC
env-2087  GUAUCU.UUGCGCCC.............UGAUUGA...........AAG...................UCAAUCG.....................................GAUAC.AUAAA.AAUCAUU...UAAAA..CAAUAUUGGAAUGUAUUUAACCAAGGAAAUAAAGUCCGAC
env-2088  GUAUCU.UUGCGCCC.............UGAUUGA...........AAG...................UCAAUCG.....................................GAUAC.AUAAA.AAUCAUU...UAAAA..CAAUAUUGGAAUGUAUUUAACCAAGGAAAUAAAGUCCGAC
Mth-1-1   CUAUCU.UUGCGCCC.............UGAUUGA...........AAA...................UCAAUCG.....................................GAUAC.AUAAA.AAUCAUU...UAAAA..CAAUAUUGGAAUGUAUUUAACGAAAGAAGUUAAGGCCGAA
env-2089  GUAUAU.UUGCGCCC.............UGAUUGA...........AAA...................UCAAUCG.....................................GAUAC.AUAAA.AAUCAUU...UAAAG..CAAUAUUGGCAUGUAUUUAAGCAAAGAAGUAAAAGCCGAC
env-2090  GUAUAU.UUGCGCCC.............UGAUUGA...........AAA...................UCAAUCG.....................................GAUAC.AUAAA.AAUCAUU...UAAAG..CAAUAUUGGCAUGUAUUUAAGCAAAGAAGUAAAAGCCG..
env-2091  GUAUAU.UUGCGCCC.............UGAUUGA...........AAA...................UCAAUCG.....................................GAUAC.AUAAA.AAUCAUU...UAAAG..CAAUAUUGGCAUGUAUUUAAGCAAAGAAGUAAAAGCCGAC
env-2092  GUAUAU.UUGCGCCC.............UGAUUGA...........AAA...................UCAAUCG.....................................GAUAC.AUAAA.AAUCAUU...UAAAG..CAAUAUUGGCAUGUAUUUAAGCAAAGAAGUAAAAGCCGAC
Rbi-1-1   CUAUCU.UUGCGGCC.............UGAUUGA...........AAA...................UCAAUCG.....................................GAUAC.AUAAA.AAUCAUU...UACAA..CAAUAUUGGCAUGUAUUUAACCAAAGAGAAAAAAGCCGAA
Mfo-1-1   UUACCU.UUGCGGCU.............UGAUUGA...........AAA...................UCAAUCA.....................................CGUAC.AUAAA.AAUCAUU...UAAAA..UAAUAUUGGAAUGUAUUUAUCAAAAGAAGUAAAAGCCGAA
env-2099  UUAUAU.UUGCUGCCG..................................................................................................UAC.AUAAA.AAUCAUU...AA.AA..UAAUAUUGGAAUGUAUUUAACAAAAGAAGUAAAAGCAGAA
env-2100  UUACCU.UUGCAACC.............GAAUUGGG..........AAACA................CCUAAUUC......................................AUAC.AUAAU.AAUCAUU...UA.AA..UAAUAUUGGAAUGUAUUUAACUAAAGAAACAAAAGCUGAA
Fse-1-1   AUAUUU.UUGCAGCC.............GUCUUACA..........AUAAC................UGUCUGAC......................................AUAC.AUAAA.AAUCAUU...UAAAA..AAAUAUUAGCAUGUAUUUAACUAAAGAAGGAAAAGCGGAA
Fda-1-1   UUACUU.UUGCUGCC.............GUCUUACA..........AUAAC................UGUAAGGC......................................AUAC.AUAAA.AAUCAUU...UAAAA..AAAUAUUAGCAUGUAUUUAACUAAAGAAGGAAAAGCGGAA
env-2102  CUAUAU.UUGCACCU.............CGAAAG............AUAAG..................CUUUCG.....................................AUUAC.AUAAC.AAUCAUU...UACAA..CAAUAUUAGAAUGUAUUUAACAAAAGAAGCAAAAGCUGAA
env-2103  CUAUAU.UUGCACCC.............GAAUUGGA..........AUAAUA...............UCCAUUUC......................................AUAC.AUAAU.AAUCAUU...UAAAA..UAAUAUUAGCAUGUACUUAACUAAAGAAGUUAAAGAAGAA
env-2105  UUACUU.UUGCACUC.............GAAUAGG...........AAAAA.................CUUAUUC.....................................GAUAC.AUAAU.AAUCAU....UA.AA..CAAUAUUAGCAUGUACUUAACUAAAGAAGUUAAAGAAGAA
Fso-1-1   UUACUU.UUGCACUC.............GAAUAAG...........ACGAA.................CUUAUUC.....................................ACUAC.AUAAU.AAUCAU....UA.AA..CAAUAUUAGCAUGUACUUAACUAAAGAAGUUAAAGAAGAA
env-2106  UUACCU.UUGCACUC.............GAAUAGG...........AAAAG.................CCUAUUC......................................CUAC.AUAAU.AAUCAUU...UA.AA..UAAUAUUAGCAUGUACUUAACUAAAGAAGUUAAAGAGCAA
env-2107  UUACCU.UUGCACUC.............GAAUAAG...........AAAAA.................CUUAUUC......................................AUAC.AUAAA.AAUCAUU...UAAAA..UAAUAUUGGCAUGUACUUAACU..................
env-2108  UUACCU.UUGCACAC.............GAAUAAG...........AAAAA.................CUUAUUC......................................AUAC.AUAAU.AAUCAUU...UA.AA..UAAUAUUAGCAUGUACUUAACUAAUGAAGUUAAAGCAGAA
env-2109  UUACCU.UUGCACUC.............GAAUAAG...........AAAAA.................CUUAUUC.....................................AUUAC.AUAAU.AAUCAUU...UAAAC..AAAUAUUGGCAUGUACUUAACUAAAGAAGUUAAAGAGCAA
env-2110  UUACCU.UUGCACUC.............GAAUAAG...........AAAAA.................CUUAUUC.....................................AUUAC.AUAAU.AAUCAUU...UAAAC..AAAUAUUGGCAUGUACUUAACUAAAGAAGUUAAAGAGCAA
Aag-1-1   GUACAU.UUGCAGCC.............UUGAAAGA..........AAA..................UCUUUCAA......................................AUGC.AUAAU.AAUCAUU...UA.AA..UAAUAUUGGAAUGUAUUUAACAAAAGAAGUUAAAGAAGAA

          ............................<<<<<<<<<<<<<<<<<<...........>>>>>>>>>>>>>>>>>>..........<<<<<<<<........>>>>>>>>........................................................................
          ............................2222221111..101.00...........00.101..1111222222..........11011000........00011011........................................................................
          •UAY•U-UUGC••CY-------------◦◦◦◦◦◦◦-----------R•R◦◦-----------------◦◦◦◦◦◦◦-------------------------------------◦UAC-AUAA•-AAUCAUU---UA-AA--YAAUAUURG•AUGUAUUUARC•ARAGAAR••AAARRRRAR
           5′                                                                                                                                                         3′
```

```
env-2111  GUAUUU.UUGCCGCC............UGAUUACC.........AUAGC...............GGUAGUCU...................................GUAC.AUAAA.AAUCAUU...UAAAU..AAAUAUUGGAAUGUAUUUAACACCAGAAAAAAAAGAAGAG
env-2112  UUAUCU.UUGCCGCU............UGAUUACCA........AAG................UGGUAUUCA...................................GUAC.AUAAA.AAUCAUU...UU.AA..UAAUAUUGGCAUGUAUUUAACUGCAGACGAGAAAAAAGAA
env-2113  UUAUCU.UUGCCGCU............UGAUUACCA........AAA................UGGUAUUCA...................................GUAC.AUAAA.AAUCAUU...UU.AA..UAAUAUUGGCAUGUAUUUAACUGCAGAAGAGAAACAAAAA
env-2114  UUAUCU.UUGCCGCU............UGAUUACCA........AAA................UGGUAUUCA...................................GUAC.AUAAA.AAUCAUU...UU.AA..UAAUAUUGGCAUGUAUUUAACUGCAGAAGAGAAACAAAAA
env-2115  UUAUCU.UUGCCGCU............UGAUUACCA........AAA................UGGUAUUCA...................................GUAC.AUAAA.AAUCAUU...UU.AA..UAAUAUUGGCAUGUAUUUAACUGCAGAAGAGAAACAAAAA
env-2116  AUAUCU.UUGCGCCC............UGAUUAUCA........UAG................UGGUAAUCU...................................GUAC.AUAAA.AAUCAUU...AA.AC..UAAUAUUGGCAUGUAUUUAUCACCUGAAGAAAAAGGAAAA
env-2117  AUAUCU.UUGCGCCC............UGAUUAUCA........UAG................UGGUAAUCU...................................GUAC.AUAAA.AAUCAUU...AA.AC..UAAUAUUGGCAUGUAUUUAUCACCUGAAGAAAAAGGAAAA
env-2118  AUAUCU.UUGCGCCC............UGAUUAUCA........UAG................UGGUAAUCU...................................GUAC.AUAAA.AAUCAUU...AA.AC..UAAUAUUGGCAUGUAUUUAUCACCUGAAGAAAAAGGAAAA
env-2119  AUAUCU.UUGCGCCC............UGAUUAUCA........UAG................UGGUAAUCU...................................GUAC.AUAAA.AAUCAUU...AA.AC..UAAUAUUGGCAUGUAUUUAUCACCUGAAGAAAAAGGAAAA
env-2120  AUAUCU.UUGCGCCC............UGAUUAUCA........UAG................UGGUAAUCU...................................GUAC.AUAAA.AAUCAUU...AA.AC..UAAUAUUGGCAUGUAUUUAUCACCAGAAGAAAAAGGAAAA
env-2121  AUAUCU.UUGCGCCC............UGAUUAUCA........UAG................UGGUAAUCU...................................GUAC.AUAAA.AAUCAUU...AA.AC..UAAUAUUGGCAUGUAUUUAUCACCUGAAGAAAAAGGAAAA
env-2124  AUAUCU.UUGCCGCC............UGAUUAUCA........UAG................UGGUAAUCU...................................GUAC.AUAAA.AAUCAUU...AA.AC..UAAUAUUGGCAUGUAUUUAUCACCAGAAGAAAAAGGAAAA
env-2125  AUAUAU.UUGCCGCC............UGAUUACCA........UAG................UGGUAAUCU...................................GUAC.AUAAA.AAUCAUU...AA.AC..UAAUAUUGGCAUGUAUUUAUCACCAGAAGAAAAAGGAAAA
env-2126  AUAUAU.UUGCCGCC............UGAUUACC.........AUAGC...............GGUAAUCU...................................GUAC.AUAAA.AAUCAUU...AA.AC..CAAUAUUGGCAUGUAUUUAUCACCAGAAGAAAAAGGA...
env-2127  AUAUAU.UUGCCGCC............UGAUUACCA........AAG................UGGUAGUCU...................................GUAC.AUAAA.AAUCAUU...UAAAA..UAAUAUUGGCAUGUAUUUAUCACCUGAGGAAAAAGGAAAA
env-2128  AUAUCU.UUGCGGCC............UGAUUAUC.........AUAGC...............GGUAAUCU...................................GUAC.AUAAA.AAUCAUU...AA.AC..UAAUAUUGGCAUGUAUUUAUCACCAGAAGAAAAAGGAAAA
env-2129  AUAUCU.UUGCCGCC............UGAUUAUC.........AUAGC...............GGUAAUCU...................................GUAC.AUAAA.AAUCAUU...UAAAA..UAAUAUUGGCAUGUAUUUAUCACCUGAAGAAAAAGGAAAA
env-2130  AUAUCU.UUGCCGCC............UGAUUAUC.........AUAGC...............GGUAAUCU...................................GUAC.AUAAA.AAUCAUU...UAAAA..UAAUAUUGGCAUGUAUUUAUCACCUGAAGAAAAAGGAAAA
env-2131  AUAUCU.UUGCCGCC............UGAUUAUC.........AUAGC...............GGUAAUCU...................................GUAC.AUAAA.AAUCAUU...UAAAA..UAAUAUUGGCAUGUAUUUAUCACCUGAAGAAAAAGGAAAA
env-2132  AUAUAU.UUGCCGCC............UGAUUACC.........AAAGC...............GGUAAUCU...................................GUAC.AUAAA.AAUCAUU...UAAAA..UAAUAUUGGACAUGUAUUUAUCACCUGAAGAAAAAGGAAAA
env-2133  AUAUAU.UUGCCGCC............UGAUUAUC.........AAAGC...............GGUAAUCU...................................GUAC.AUAAA.AAUCAUU...UAAAA..UAAUAUUGGCAUGUAUUUAUCACCUGAAGAAAAAGGAAAA
env-2134  AUAUAU.UUGCCGCC............UGAUUAUC.........AAAGC...............GGUAAUCU...................................GUAC.AUAAA.AAUCAUU...UAAAA..UAAUAUUGGCAUGUAUUUAUCACCUGAAGAAAAAGGAAAA
env-2135  AUAUAU.UUGCCGCC............UGAUUAUC.........AAAGC...............GGUAAUCU...................................GUAC.AUAAA.AAUCAUU...UAAAA..UAAUAUUGGCAUGUAUUUAUCACCUGAAGAAAAAGGAAAA
env-2136  AUAUAU.UUGCCGCC............UGAUUAUC.........AAAGC...............GGUAAUCU...................................GUAC.AUAAA.AAUCAUU...UAAAA..UAAUAUUGGCAUGUAUUUAUCACCUGAAGAAAAAGGAAAA
env-2137  AUAUAU.UUGCCGCC............UGAUUAUC.........AAAGC...............GGUAAUCU...................................GUAC.AUAAA.AAUCAUU...UAAAA..UAAUAUUGGCAUGUAUUUAUCACCUGAAGAAAAAGGAAAA
env-2138  AUAUAU.UUGCCGCC............UGAUUACC.........AAAGC...............GGUAAUCU...................................GUAC.AUAAAAAAUCAUU...UAAAA..UAAUAUUGGCAUGUAUUUAUCACCUGAAGAAAAAGGAAAA
env-2139  AUAUAU.UUGCCGCC............UGAUUACC.........AAAGC...............GGUAAUCU...................................GUAC.AUAAA.AAUCAUU...UAAAA..UAAUAUUGGCAUGUAUUUAUCACCUGAAGAAAAAGGAAAA
env-2140  CUAUUU.UUGCAGCC............UGAUUGCC.........AAAGA...............GGUAAUCG.....................................UAC.AUAAA.AAUCAUU...UA.AA..UAAUAUUUGCAUGUAUUUAGCACCAGAAGAAAAAGAAAAG
env-2141  CUAUCU.UUGCCGCU............UGAAUACG.........AAAAG...............CGUA.UCA...................................AUAC.AUAAA.AAUCAUU...UAAAU..AAAUAUUGGAAUGUAUUUAACAAAAGAAGCUAAAGAGAAA
env-2142  CUAUCU.UUGCAGCU............UGAUUACGA........AAA................UCGU.AUCA...................................AUAC.AUAAA.AAUCAUU...UACAA..UAAUAUUAGCAUGUAUUUAACAAAAGAAGCUAAAGAGAAA
env-2143  CUAUCU.UUGCAGCU............UGAAUACG.........AAAAC...............CGUAGUCA..................................AGUAC.AUAAA.AAUCAUU...UACAA..UAAUAUUAGCAUGUAUUUAACAAAAGAAGCUAAAGAGAAA
env-2144  CUAUCU.UUGCAGCU............UGAAUACG.........AAAAA...............CGUA.UCA...................................AUAC.AUAAA.AAUCAUU...UACAA..CAAUAUUAGCAUGUAUUUAGACAAGAAAGAGAAAGAGAAA
env-2145  UUAUCU.UUGCACAC............GAAUUAGG.........ACAAG...............CUUAAUUC...................................ACGC.AUAAU.AAUUAUU...UA.AA..CAAUAUUAGCAUGUACUUAACUAAAGAAGUUAAAGCAGAA
env-2146  UUAUCU.UUGCACAC............GAAUUAGG.........ACAAG...............CUUAAUUC...................................ACGC.AUAAU.AAUUAUU...UA.AA..CAAUAUUAGCAUGUACUUAACUAAAGAAGUUAAAGCAGAA
env-2147  UUAUCU.UUGCACAC............GAAUUAGG.........ACAAG...............CUUAAUUC...................................ACGC.AUAAU.AAUUAUU...UA.AA..CAAUAUUAGCAUGUACUUAACUAAAGAAGUUAAAGCAGAA
env-2148  CUAUCU.UUGCCCCG............UCGC.............AUAAG...................GCGA...................................CUAC.AUAAG.AAUCAUU...UAAAU..AAAUAUUGGCAUGUACAUCAAUUCAGAAAUUAAAAAAGAA
env-2149  CUAUCU.UUGCCCCG............UCGC.............AUAAG...................GCGA...................................CUAC.AUAAG.AAUCAUU...UAAAU..AAAUAUUGGCAUGUACAUCAAUUCA..............
env-2150  GUAUAU.UUGCAUCG............UCU..............UUA......................AGA...................................CUAC.AUAAU.AAUCAUU...UAAAU..CAAUAUUGGCAUGUAUUUAGCACCAGAAAAAAAACAGAG
env-2151  GUAUAU.UUGCAUCG............UCU..............UUA......................AGA...................................CUAC.AUAAU.AAUCAUU...UAAAU..CAAUAUUGGCAUGUAUUUAGCACCAGAAAAAAAACAGAG
env-2152  CUAUAU.UUGCAUCG............UCC..............AAAAGA...................GGG...................................CUAC.AUAAU.AAUCAUU...UAAAA..UAAUAUUUGCAUGUAUUUAGAAACAGAAAAAAAGAAAGAG
Fps-4-1   GUAAUU.UUGCACUC............GAAUCCG..........AGAAA.................CGGAAUU...................................CAUAC.AUAAA.AAUCAUU...AA.AC..AAAUAUUGGAAUGUAUUUAACUAAAGAGAUUAAAGAAGAA
          ...........................<<<<<<<<<<<<<<<<<..........>>>>>>>>>>>>>>>>>>.........<<<<<<<<.........>>>>>>>>...............................................................
          ...........................2222221111..101.00.........00.101..1111222222.........11011000.........00011011...............................................................
          •UAY•U-UUGC••CY-----------ooooooo----------R•Roo---------------ooooooo---------------------------------oUAC-AUAA•-AAUCAUU---UA-AA--YAAUAUURG•AUGUAUUUARC•ARAGAAR••AAARRRRAR
          5'                                                                                                                                              3'
```

alignment positions 260 · · · 439

```
Pgo-1-1   UUAUUUGCUAAACACGGUAAAGACGAGAAAGAUUCUGGUUCUGCCGAAGGUCAAAUCGCAUUGAUGACCAGGAGAAUUGAAUACAUCUCUCAACACCUCAAAACAAACAGAAAAGACUAUAAUUCUGAACGUUCUCUAGUAAAGCUUGUAGGUAAAAGAAGAAACCUAUUGGAUUAUCUG
env-1     UUAUUUGCUAAACAUGGGAAAAAUGAAAAAGAUUCUGGGUCUGCUGAAGGGCAAAUCGCUUUGAUGACACAAAGAAUCCAGCAUAUUUCUCAACACCUAAAAACAAACAGAAAAGAUUUUAAUUCUGAAAGAUCUCUAGUGAAAUUAGUAGGUAAACGUAGAAACUUAUUAGAUUACCUC
env-2     UUAUUUGCCAAACAUGGUAAGAACGAGAAAGACACAGGUUCUACCGAAGGGCAAAUUGCUUUGAUGACUCAAAGAAUUGAACACAUCUCUCAGCACUUAAAAACAA.............................................................................
env-3     UUAUUUGCUAAACACGGAAAGAACGAGAAAGACACUGGAUCUGCUGAAGGUCAGAUCGCUUUGAUGACUCAGAGAAUCGAGCACAUUUCUCAACACUUAAAAACAAAUAGAAAAGAUUUUAAUUCUGAAAGAUCUUUGGUCAAACUAGUUGGUAAAAGAAGAAAUCUUCUUGACUACCUC
Pto-1-1   UUAUUUGCCAAACACGGUAAGGAUGAGAAAGAUUCCGGAUCUGCAGAGGUCAAAUCGCAUUGAUGACCGGAAGAAUUGAAUUCAUUUCUCAACACCUUAAGACAAACAGAAAAGAUUUAUAUUCUGAACGUUCUCUAGUAAACCUUGUAGGUAAAAGAAGAAACCUAUUGGAUUAUCUU
Ptr-1-1   UUAUUUGCUAAACACGGUAAGAACGAGAAAGAUUCUGGUUCUGCAGAAGGCCAGAUCGCUUUGAUGACUCAGAGAAUUGAGCACAUUUCUCAACACUUAAAAACAAACAGAAAAGAUUUUAAUUCUGAAAGAUCUUUGGUUAAACUAGUUGGUAAAAGAAGAAAUCUUCUCGACUACCUC
env-4     UUAUUUGCUAAACACGGUAAGGACGAGAAAGAUUCUGGUUCUACCGAAGGUCAAAUCGCAUUGAUGACCGGAAGAAUUGAAUACAUUUCUCAACACCUUAAGACAAAUAGAAAAGAUUAUAAUUCUGAACGUUCUCUAGUAAAACUUGUAGGUAAAAGAAGAAACCUAUUGGAUUAUCUA
env-5     UUAUUUGCUAAACAC.........................................................................................................................................................................
env-6     UUAUUUGCUAAACACGGAAAGAACGAGAAAGACACUGGAUCUGCAGAAGGUCAGAUCGCUUUGAUGACUCAGAGAAUUGAGCACAUUUCUCAACACUUAAAAACAAACAGAAAAGAUUUUAAUUCUGAAAGAUCUUUGG............................................
env-7     UUAUUUGCUAAACACGGAAAGAACGAGAAAGACACUGGAUCUGCAGAGGUCAGAUCGCUUUGAUGACUCAGAGAAUUGAGCACAUUUCUCAACACUUA.....................................................................................
env-8     UUAUUUGCUAAACACGGAAAGAACGAGAAAGACACUGGAUCUGCAGAAGGUCAGAUCGCUUUGAUGACUCAGAGAAUUGAGCACAUUUCUCAACACUUAAAAACAAACAGAAAAGAUUUUAAUUCUGAAAGAUCUUUGGUCAAAUUAGUUGGUA...........................
Mod-8-1   AUCUUCCAACAACACGGAGGUGUAGCUGCAAACACAGGAUCUGCAGAGGACAAAUCGCUUUGUUCACUUACAGAAUCUCUCACUUAACUGAGCACUUGAAAAGAAAUCGUAAAGAUUACAACACGGAGCGUUCAUUAGUUCUUUUAGUUGGUAAAAGAAGAGACUUCUUGACUACUUA
env-9     AUCUUCAAAAAACACGGAGGUUCUGAAACAAACACUGGUUCUUCAGAAGGACAAGUAGCAUUAUUUUCUUACAGAAUUUCUCAUUUAACUGAGCAUUUGAAGAAAAAUCGUAAAGAUUACGCUACGCAAAGAGCACUACAAUUAUUAGUUGGUAAGCGAAGAAGCUUAUUGGAUUAUUUG
env-11    AUCUUCAAAAAACACGGAGGUUCUGAAACAAACACUGGUUCUUCAGAAGGACAAGUAGCAUUAUUUUCUUACAGAAUUUCUCAUUUAACUGAGCAUUUGAAGAAAAAUCGUAAAGAUUACGCUACGCAAAGAGCACUACAAUUAUUAGUUGGUAAGCGAAGAAGCUUAUUGGAUUAUUUG
env-12    AUCUUCAAAAAACACGGAGGUUCUGAAACAAACACUGGUUCUUCAGAAGGACAAGUAGCAUUAUUUUCUUACAGAAUUUCUCAUUUAACUGAGCAUUUGAAGAAAAAUCGUAAAGAUUACGCUACGCAAAGAGCACUACAAUUAUUAGUUGGUAAGCGAAGAAGCUUAUUGGAUUAUUUG
env-13    AUCUUCAAAAAGCAUGGAGGUUCUGAAACAAACACUGGUUCUUCAGAAGGACAAGUAGCAUUAUUUUCUUAUAGAAUUUCUCAUUUAACUGAGCACUUAAAGAAAAAAUCGUAAAGAUUACGCUACGCAAAGAGCACUACAAUUAUUAGUUGGUAAGCGAAGAAGCUUAUUGGAUUAUUUG
env-14    AUCUUCAAAAAGCAUGGAGGUUCUGAAACAAACACUGGUUCUUCAGAAGGACAAGUAGCAUUAUUUUCUUAUAGAAUUUCUCAUUUAACUGAGCACUUAAAGAAAAAAUCGUAAAGAUUACGCUACGCAAAGAGCACUACAAUUAUUAGUUGGUAAGCGAAGAAGCUUAUUGGAUUAUUUG
env-15    UUCUUCGCCAAGUACGGCAAGAACGAGAAGGACACCGGCUCUGCAGAAGCGCAGAUCGCCAUGUUCUCCUU................................................................................................................
env-16    AUUUUUAAACAGUACGGCGGAGACGCUAAAAACACUGGCUCAACUGAAGGUCAAAUCGCAUUAUUUACUUUUAGAAUUAACCAUUUAACUGAACACUUAAAGAAAAACAAAAAGAUAAAGGAACUCAAAAAGCUCUUUUAGAUUUAGUGGAAAAAGAAGAAGUUUGUUGGAUUACUAU
env-17    AUUUUCGCUAAACACGGGAAAAGUAAAAGUAAUACUGGUUCUGCCGAAGGACAGAUUGCCUUGUUUACUUACAGAAUUAAUCACUUAACUGAACACUUAAAAAAAGAAUCGUAAAGACUAUAACACAGAGCGUUCAUUAGUAAAACUCGUAGGUAAAAGAAGAGCGUUACUAGAUUAUUUA
Fbr-1-1   AUCUUCGCUAAACACGGAGGAAAUGCUGCAAACACAGGAUCUGCAGAGGGACAAAUUGCGUUAUUCACUUACAGAAUUACACAUUUAACAGAACACUUGAAAAAAAAUCGUCAUGAUUAUAAUACAGAGCGUUCACUAGUAUGUUAGUAGGAAAAAGAAGAGCGUUAUUGGACUACUUA
env-18    AUCUUCAGCAAAUUCGGGGAUCUGAGACCAACACAGGUUCUGUAGAAGGUCAAAUCGCGUUAUUCUCAUACCGCAUUAGUCAUUUAACAGAGCACUUGAAGAAAAACCGUAAGGAUUUGGUACUCAAAGAGCCCUUCAGUUAUUGGUUGGUAAACGAAGAAGCUUGUUGGAUUAUCUG
env-19    AUCUUCAGCAAAUUCGGGGGAUCUGAGACCAACACAGGUUCUGUAGAAGGUCAAAUCGCGUUAUUCUCAUACCGCAUUAGUCAUUUAACAGAGCACUUGAAGAAAAACCGUAAGGAUUUUGGUACUCAAAGAGCCCUUCAGUUAUUGGUUGGUAAACGAAGAAGCUUGUUGGAUUAUCUG
env-20    AUUUUCGCAAAAUAUGCGGGCUCUGCAAAAACACAGGUUCGGCAGAGGGGCAAAUCGCCUUGUUUUCAUACCGGAUUUCCCAUUUAACUGAACACCUGAAGAAAAAUCGUCAUGAUUUUGCAACUCAGCGUUCAUUACAGUUAUUGGUUGGUAAACGAAGAAUCUGCUGGAUUACUUGA
env-21    UUUUUCAAAUCCUACGGGAAAAACGAGAGCG.......................................................................................................................................................
env-22    UUAUUUAAAAAGCACGGCAAAAAAGCCGAAAAUACUGGAAGUGCUGAAGGACAGAUUGCACUAUUUACCUUCAGAAUAAAUCAUUUAACAGAACACCUUAAGCAAAAUAAAAAAGAUUUCAACACAGAAAGAUCUCUUGUAAAAUUAGUAGGAAAAAGAAGAGCGUUGUUGGAUUAUCUA
env-23    UUAUUUAAAAAGCACGGCAAAAAAGCCGAAAAUACUGGAAGUGCUGAAGGACAGAUUGCACUAUUUACCUUCAGAAUAAAUCAUUUAACAGAACACCUUAAGCAAAAUAAAAAAGAUUUCAACACAGAAAGAUCUCUUGUAAAAUUAGUAGGAAAAAGAAGAGCGUUGUUGGAUUAUCUA
env-24    UUAUUUAAAAAGCACGGCAAAAAAGCCAAAAAUACUGGAAGUGCUGAAGGACAGAUUGCACUAUUUACCUUCAGAAUAAAUCAUUUAACAGAACACCUUAAGCAAAAUAAAAAAGAUUUCAACACAGAAAGAUCUCUUGUAAAAUUAGUAGGAAAAAGAAGAGCGUUGUUGGAUUAUCUA
env-25    UUAUUUAA..............................................................................................................................................................................
Fba-9-1   AUUUUCGAAAAGUACGGAAACUCAGCAACUGACACAGGAUCUGCAGAAGGUCAAAUUGCAUUGUUCACUUACAGAAUUGCUCACUUAACAGAGCACUUAAAAAAGAACAGAAAAGAUUUUAACACACAACGUUCUUUUGGUAAAAAUGGUAGGAAAGCGUAGAGCUUUAUUAGACUACUUA
env-26    AUCUUUAAAAACAUGGAAAAGAUGCCAAAAAUACAGGUUCUCAGAGGGACAAAUCGCAUUAUUUACCCUCAGGAUUAAUCACCUAACUGAGCACCUUAAAAACAACCGAAAAGAUUACAACACAGAGCGAUCUCUUGUUAAGCUUGUCGGAAAGCGUCGUUCUUUACUAGACUAUCUU
env-27    AUCUUCGCUAAACACGGAGGAGCAGAAACAAAACACAGGUUCUACGAAGGGGCAAAUUGCAUUAUUUACUUUCCGUAUUUCCAUUUAACUGAGCACUUGAAAAAAAUCGUAAAGAUUUCGGAA.........................................................
env-28    AUUUUCAGCACGUACGGUAAAGACAAGAACGAUACCGGUUCUGCUGAAGGACAGAUCGCGUUAUUCACAUAUCGCAUCAACCAUUUAACAGGACAUUUAAAAAAAGAACCGUAAAGACUUUAAUACAGAGCGUUCUUUUGGUAAAGUUAGUAGGUAAGCGUAGAGCUUUGCUUGACUAUUUA

          ......................................................................................................................................................................................
          ......................................................................................................................................................................................
          AUYUUYR•RAARCACGG•RRR•••R•RR••RAYAC•GG••C•RC•GAAGG•CAAAUYGC•YURUUYAC•Y•Y•G•AUYRAYCAYYUR•C•RA•CAYYURAAAAR•AAYCR•AARGAUU•YAAYAC•GAR•G•UC•YURGU•••RYURGU•GG•AAR•G•RG••••YU•YU•GAYUAYYU•
```

```
env-33    AUUUUCAGCACGUACGGUAAAGACAAGAACGAUACCGGUUCUGCUGAAGGACAGAUCGCGUUAUUCACAUAUCGCAUCAACCAUUUAACAGGACAUUUAAAAAAGAACCGUAAAGACUUUAAUACAGAGCGUUCUUUGGUAAAGUUAGUAGGUAAGCGUAGAGCUUUGCUUGACUAUUUA
env-34    AUUUUUGCUGAACACGGUUCUUCAAAAACCAACACAGGCUCUGCCGAAGGACAAAUUGCUUUGUUCUCUCACCGCAUUAACCACCUAUCUGACCACUUGAAAAGUAACCGUAAAGAUUACAAUACGGAACGCUCUUUGGUUAAAUUGGUAGGUAAGCGCAGAAGCUUAUUGGAUUAUUUA
env-35    AUUUUUGCUGAACACGGUUCUUCAAAAACCAACACAGGCUCUGCCGAAGGACAAAUUGCUUUGUUCUCUCACCGCAUUAACCACCUAUCUAACCACUUGAAAAGUAACCGUAAAGAUUACAAUACGGAACGCUCUUUGGUUAAAUUGGUAGGUAAGCGCAGAAGCUUAUUGGAUUAUUUA
env-36    AUUUUUGCUGAACACGGUUCUUCAAAAACCAACACAGGCUCUGCCGAAGGACAAAUUGCUUUGUUCUCUCACCGCAUUAACCACCUAUCUGACCACUUGAAAAGUAACCGUAAAGAUUACAAUACGGAACGCUCUU.........................................
env-37    AUUUUCGCAGAGCACGGAAAAUCUAAAACCGACACGGGUUCUGCAGAAGGACAGAUUGCACUAUUUUCUCACCGCAUCAACCACCUUUCUGAACACUUAAAAAGUAACCGUAAAGAUUACAAUACAGAACGUUCGUUGGUAAAAUUAGUAGGAAAGCGUAGAAGCUUACUCGAUUACCUU
env-38    UUUCGCAGAGCACGGAACGUCUAAAACCGAUACCGGAUCCACCGAAGGUCAAAUUGCUUUAUUCUCUCACCGUAUCAAUCACCUCUCUGAACACUUAAAAAGUAAUCGUAAAGAUUAUAAUACAGAGCGUUCACUGGUUAAAUUAGUAGGAAAGAGAAGAAGCUUAUUAGAUUACCUUAA
env-39    AUUUUUGCUGAACACGGUUCAUCAAAAACCAACACAGGUUCUGCCGAAGGACAAAUUGCAUUGUUCUCGCACCGCAUUAACCACCUAUCUGAUCACUUGAAAAACAACCGUAAAGAUUACAAUACGGAACGCUCUCUUGUAAAAUUAGUGGGAAAACGCAGAAGUUUAUUGGAUUAUUUA
env-40    .....................................................................................................................................................................................
env-42    AUUUUUAAAGAAUUUAGCGGAGACGCUAAAAACACAGGAUCAACAGAAGGGCAAAUAGCUUUGUUUACCUAUAGAAUUAGUCAUUUAACAGAGCAUUUAAAGAAAAACAAAAAAGACUUUAGAACUCAAAAAGCUUUGUUAGAUUUAGUUGGUAAAAGAAGAAAACUAUUAGAUUAUUAC
env-43    AUCUUCGCUAAACACGGAGGUUCAGAAACAAACACAGGAUCUACAGAAGGUCAAAUCGCAUUGUUUACCUAUAGAAUCAGUCAUUUAACCGAGCAUUUAAAGAAAAAUCGUAAAGAUUUCGGUACGCAACGCUCGCUUCAAUUGUUGGUAGGUAAAAGAAGAAGUCUUUUGGAUUACUUA
Myu-1-1   AUCUUCCAAAAAUACGGAAGCAGUAAAACAAACACUGGUUCUACAGAAGGUCAGAUUGCCUUGUUUACGUUUAGAAUUAACCACCUAACAGGCCACUUGAAAAACAAUCACAAAGAUUUUAACACCGAGCGUGCUCUGGUAAAGUUGGUAGGUAAGCGUAGAUCGUUACUAGAUUAUUUA
env-44    .....................................................................................................................................................................................
env-45    AUCUUCGCUAAACACGGAGGUGCAGAAACAAACACAGGUUCAACAGAGGGACAAAUUGCAUUGUUCACAUUUCGUAUCGCUCACUUAACAGAGCAUUUGAAAAAGAACCGUAAAGACUACGGAACACAACGUUCAUUACAAUUAUUAGUUGGUAAACGUCGUAGUUUCUUAGAUUAUUUA
env-46    AUCUUCGCUAAACACGGAGGUGCAGAAACAAACACAGGUUCAACAGAGGGACAAAUUGCAUUGUUCACAUUUCGUAUCGCUCACUUAACAGAGCA........................................................................................
env-47    .....................................................................................................................................................................................
env-48    AUCUUCGC.............................................................................................................................................................................
env-49    AUCUUCAGUAAACACGGAGGUUCUGACACAAACACCGGUUCGGCUGAAGGUCAAAUCGCUUUAUUUUCUUAUCGUAUUGCUCACUUGACUGAGCAUUUGAAAAGAAUCGUAAAGAUUACGGUACACAACGCGCACUUCAAUUAUUGGUUGGUAAGAGAAGAAGUUUGUUGGAUUACCUA
env-50    AUCUUCGCGAAAACACGGAGGUGCUGCAACAAACACAGGUUCAGCAGAAGGACAAAUCGCAUUAUUCACAUUCAGAAUCUCUCACUUAACAGAACACUUG...................................................................................
env-51    AUUUUCGCAGAUCAUGGAACUUCAAAAACCGACACUGGUUCUGCCGAAGGUCAAAUUGCUUUAUUCUCGCACCGUAUCAACCACUUGUCUGAGCACUUAAAAAGUAACCGUAAGGAUUACAAUACAGAGCGUUCACUAGUGAAAUUGGUGGAGGUAAGCGCAGAAGCUUACUAGAUUACCUU
env-52    AUCUUCAAAAAGCACGGAGGUUCUGAAACAAACACUGGUUCUUCAGAAGGACAAGUAGCAUUAUUUUCUUACAGAAUUUCUCACUUAACUGAGCAUUUAAAGAAAAAUCGUAAAGAUUACGCUACACAAAGAGCUCUUCAAUUAUUGGUAGGUAAGCGUAGAAGCUUAUUAGAUUACUUG
env-53    AUUUUCAGUAAAUUCGGGGGUAGCGAGACGAACACCGGAUCCGUGGAAGGACAAAUCGCGUUGUUUUCUUACCGAAUCAAUCAUCUUACGGAGCAUUUGAAGAAAAAUCGUAAAGAUUUUGGAACGCAACGUUCCUUGCAAUUGUUGGUAGGUAAACGAAGAAGUUUGUUGGAUUACCUA
Fsa-1-1   AUCUUCGCUAAACACGGAGGAGCAGAAGCCAACACCGGAUCUGCAGAAGGUCAAAUCGCGUUAUUCACAUUCAGAAUUUCUCACUUGACUGAGCACUUGAAAAAAAAUCGUCAUGAUUAUAACACAGAGCGCUCAUUAGUUUUAUUGGUAGGUAAAAGAAGAAGUCUUCUUGACUACUUA
Fca-1-1   AUCUUCGCUAAACACGGAGGUGCAGCUGCCAACACUGGUUCUGCAGAAGGUCAAAUCGCGUUAUUCACAUUCAGAAUUUCUCACUUGACUGAGCACUUGAAAAAAAAUCGUCACGAUUACAACACAGAGCGCUCAUUAGUUUUAUUGGUAGGUAAAAGAAGAAGUCUUCUUGACUACUUA
env-54    AUAUUCAAAAAACACGGUAAAAGCUCUAAAAAUACUGGUUCAGCGGAGGGACAAAUAGCCCUAUUCACUCACCGAAUUAAUCACCUUACCGAACACCUAAAAAUCAACCUCAAGGAUUUAAUACUGAGCGUUCGCUAGUUAAAUUGGUCGGUAAAAGACGUUCCUUAUUAGACUAUUUA
env-55    AUAUUCAAAAAACACGGUAAAAGCUCUAAAAAUACUGGUUCAGCGGAAGGACAGAUAGCCCUAUUUACUCACCGAAUUAAUCACCUUACCGAGCAUCUAAAAUCUAACCGCAAAGACUUUAAUACUGAACGUUCACUGGUUAAAUUAGUUGGUAAAAGACGUUCCUUAUUAGACUAUUUA
env-56    AUAUUCAAAAAACACGGUAAAAGCUCUAAAAAUACUGGUUCAGCGGAGGGACAGAUAGCCCUAUUCACCCACCGAAUUAAUCACCUUACCGAGCAUCUAAAAUCUAACCGCAAAGACUUUAAUACUGAACGUUCACUGGUUAAAUUAGUUGGUAAAAGACGUUCCUUAUUAGACUAUUUA
env-57    AUAUUCAAAAAACACGGUAAAAGCUCUAAAAAUACUGGUUCAGCGGAGGGACAGAUAGCCCUAUUCACCCACCGAAUUAAUCACCUUACCGAGCAUCUAAAAUCUAACCGCAAAGACUUUAAUACUGAACGUUCACUGGUUAAAUUAGUUGGUAAAAGACGUUCCUUAUUAGACUAUUUA
env-59    UUCUUUAAAAAGUACGGUGGAGACGAAAAAACACUGGAUCUACAGAAGCACAAAUUGCUAUGUUUACUUACAGAAUUAGCCAUUUAACUGGUCAUUUAAAAAACAACAGAAAAGAUUUUGGAACACAAAAAGCGUUGUUGGAUUUAGUUGGAAAAGAAGAAGAUUAUUAAACUACUAU
env-60    AUUUUCGCAAAGCACGGAAA.................................................................................................................................................................
env-61    AUUUUCGCAAAGCACGGAAAAUCUGCAAAAGA.....................................................................................................................................................
env-62    AUUUUCGCAAAGCACGGAAAAUCU.............................................................................................................................................................
env-63    AUUUUCGCAAAGCACGGAAAAUCUGCAA.........................................................................................................................................................
env-65    AUUUUCGCAAAACACGGAAAAUCUGCAACAGACACUGGUAACACAGAAGGUCAAAUCGCAUUAUUCACCUACAGAAUUCAGCACCUAACGGGACACUUGAAAGCAAAUCACAAAGAUUACAACACAGAGCGUUCUUUGGUGAAAUUAGUAGGUAAGCGUCGUUCUUUAUUAGAUUAUCUA
env-66    AUUUUCGCAAAACACGGAAAAUCUGCAACAGACACUGGUAACACAGAAGGUCAAAUCGCAUUAUUCACCUACAGAAUUCAGCACCUAACGGGACACUUGAAAGCAAAUCACAAAGAUUACAACACAGAGCGUUCUUUGGUGAAAUUAGUAGGUAAGCGUCGUUCUUUAUUAGAUUAUCUA
env-67    AUUUUCGCAAAGCACGGAAAAUCUGCAAAAGACACUGGUAACACAGAAGCUCAAAUUGCAUUAUUCACUUACAGAAUUCAACACCUUACGGGGCAUUUGAAAGCAAACCACAAAGAUUACAACACAGAGCGUUCUCUAGUGAUGCUAGUAGGUAAGCGUCGUUCUUUAUUAGACUAUCUA
env-68    AUUUUCGCAAAACACGGAAAAUCUGCGACAGACACUGGUAACACAGAAGGUCAAAUUGCAUUAUUUACCUACAGAAUUGAGCACCUUACGGGGCACUUGAAAGCAAAUCACAAAGAUUACAACACAGAACGUUCUCUAGUGAAACUAGUAGGUAAGCGUCGUUCUUUAUUAGACUAUCUA
env-69    AUUUUCGCAAAGCACGGAAAAUCUGCAAAAGACACUGGUAACACAGAAGCUCAAAUUGCAUUAUUCACUUACAGAAUUCAACACCUUACGGGGCAUUUGAAAGCAAACCACAAAGAUUACAACACAGAGCGUC.........................................
env-70    AUUUUCGCAAAGCACGGAAAAUCUGCAAAAGACACUGGUAACACAGAAGCUCAAAUUGCAUUAUUCACUUACAGAAUUCAACACCUUACGGGGCAUUUGAAAGCAAAUCACAAAGAUUACAACACUGAGCGUUCUCUAGUGAUGCUUGUAGGUAAGCGUCGUUCUUUAUUAGACUAUCUA
env-71    AUUUUCGCAAAGCACGGAAAAUCUGCAAAAGACACUGGUAACACAGAAGCUCAAAUUGCAUUAUUCACUUACAGAAUUCAACACCUUACGGGGCAUUUGAAAGCAAACCACAAAGAUUACAACACAGAGCGUUCUCUAGUGAUGCUAGUAGGUAAGCGUCGUUCUUUAUUAGACUAUCUA
env-76    AUUUUCGCAAAGCACGGAAAAUCU.............................................................................................................................................................
env-77    AUUUUCGCAAAGCACGGAAAAUCUGCAAAAGA.....................................................................................................................................................
env-79    AUUUUCGCAAAGCACGGAAAAUCUGCAAAAGACACUGGUAACACAGAAGCUCAAAUUGCAUUAUUCACUUACAGAAUUCAACACCUUACGGGGCAUUUGAAAGCAAACCACAAAGAUUACAACACAGAGCGUUCUCUAGUGAUGCUA..............................
env-80    UUUUCGCAAAGCACGGAAAAUCUGCAAAAGACACUGGUAACACAGAAGCUCAAAUUGCAUUAUCACUUACAGAAUUCAACACCUUA...............................................................................................
env-81    AUUUUCGCAAAGCACGGAAAAUCUGCAACAGACACUGGUAACACAGAAGGUCAAAUCGCAUUAUUCACUUACAGAAUUCAACACCUUACGGGGCAUUUGAAAUCAAAUCACAAAGAUUACAACACAGAACGUUCUUUGGUGAAAUUAGUAGGUAAACGUCGUUCUUUAUUAGACUAUCUA
env-82    AUUUUCGCAAAGCACGGAAAAUCUGCAACAGACACUGGUAACACAGAAGGUCAAAUCGCAUUAUUCACUUACAGAAUUCAACACCUUACGGGGCAUUUGAAAUCAAAUCACAAAGAUUACAACACAGAACGUUCUUUGGUGAAAUUAGUAGGUAAACGUCGUUCUUUAUUAGACUAUCUA
env-83    AUUUUCGCAAAGCACGGAAAAUCUGCAACAGACACUGGUAACACAGAAGGUCAAAUCGCAUUAUUCACUUACAGAAUUCAACACCUUACGGGGCAUUUGAAAUCAAAUCACAAAGAUUACAACACAGAACGUUCUUUGGUGAAAUUAGUAGGUAAACGUCGUUCUUUAUUAGACUAUCUA
env-84    AUUUUCGCAAAGCACGGAAAAUCUGCAACAGACACUGGUAACACAGAAGGUCAAAUCGCAUUAUUCACUUACAGAAUUCAACACCUUACGGGGCAUUUGAAAUCAAAUCACAAAGAUUACAACACAGAACGUUCUUUGGUGAAAUUAGUAGGUAAACGUCGUUCUUUAUUAGACUAUCUA
env-85    AUUUUCGCAAAGCACGGAAAAUCUGCAA.........................................................................................................................................................
env-86    AUUUUCGCAAAGCACGGAAAAUCUGCAACAGACACUGGUAACACAGAAGGUCAAAUCGCAUUAUUCACUUACAGAAUUCAGCACCUUACGGGACACUUGAAAGCAAAUCACAAAGAUUACAACACAGAGCGUUCUUUGGUAAAAUUGGUAGGUAAGCGUCGUUCUUUAUUAGAUUAUCUA
env-87    AUUUUCGCAAAGCACGGAAAAUCUGCAACAGACACUGGUAACACAGAAGGUCAAAUCGCAUUAUUCACUUACAGAAUUCAGCACCUUACGGGACACUUGAAAGCAAAUCACAAAGAUUACAACACAGAGCGUUCUUUGGUAAAAUUGGUAGGUAAGCGUCGUUCUUUAUUAGAUUAUCUA
env-89    AUUUUCGCAAAGCACGGAAAAUCUGCAACAGACACUGGUAACACAGAAGGUCAAAUCGCAUUAUUCACUUACAGAAUUCAGCACCUUACGGGACACUUGAAAGCAAAUCACAAAGAUUACAACACAGAGCGUUCUUUGGUAAAAUUGGUAGGUAAGCGUCGUUCUUUAUUAGAUUAUCUA
env-90    AUUUUCGCAAAGCACGGAAAAUCUGCAGCAGACACUGGUAACACAGAAGGUCAAAUUGCAUUAUUCACUUACAGAAUUCAACACCUUACGGGGCAUUUGAAAGCAAACCACAAAGAUUACAACACAGAACGUUCUCUUGUGAAGCUAGUAGGUAAGCGUCGUUCUUUAUUAGAUUAUCUA
env-91    AUUUUCGCAAAGCACGGAAAAUCAGCAACAGACACUGGUAACACAGAAGGUCAAAUUGCAUUAUUCACUUACAGAAUUCAACACCUUACGGGGCAUUUGAAAGCAAACCACAAAGAUUACAACACAGAACGUUCUCUUGUGAAGCUAGUAGGUAAGCGUCGUUCUUUAUUAGAUUAUCUA
env-92    AUUUUCGCAAAGCACGGUAAAGACGCAAACGACACAGGUUCUACAGAAGGACAGAUCGCAUUGUUCACCUACCGUAUCGAGCAUUUAACAAAACACUUAAAAGCAAACCACAAGGACUACAACACAGAGCGUUCUUUGGUGAAGUUGGUAGGAAAGCGUCGUUCUUUAUUGAACUACUUG
env-93    AUUUUCGCAAAGCACGGUAAAGACGCAAACGACACAGGUUCUACAGAAGGACAGAUCGCAUUGUUCACCUACCGUAUCGAGCAUUUAACAAAACACUUAAAAGCAAACCACAAGGACUACAACACAGAGCGUUCUUUGGUGAAGUUGGUAGGAAAGCGUCGUUCUUUAUUGAACUACUUG
env-94    AUUUUCGCAAAGCACGGGAAAGACGGGAAAGACACAGGUUCUGCCGAAGGACAAAUUGCAUUAUUCACCUACCGUAUUGAGCACCUAACAGCACCUUAAAGCAAACCACAAAGAUUACAAUACAGAGCGUAGUCUUGUAAAGCUAGUAGGUAAGCGCCGUUCUUUAUUGAACUAUUUA
env-95    AUUUUCGCAAAACACGGAAAAGACGGUAAAGACACAGGUUCUGCAGAAGGACAAAUCGCGUUAUUUACCUACCGUAUCGAGCACUUAACAAAACACCUUAAAGCAAACCAUAAAGACUACAAUACAGAGCGUAGCCUUGUAAAGCUGGUAGGGAAGCGUCGUUCUUUAUUAAACUACUUA
env-96    AUUUUCAAGAAGCACGGAAGUCUGAAGCAGAUACUGGUUCUGCUGAAGGCAAAUUGCAUUAUUUUCCUACCGAAUCGAUCACUUAUCUAACCAUCUUAAGAAUAACAAAAGGAUUUCAAUACGGAACGAUCAUUAGUUAAACUAGUAGGCAAACGCCGUCAACUUUUAGAUUAUCUC
env-98    AUUUUCAAGAAGCACGGAAAGUCUGAAGCAGAUACUGGUUCUGCUGAAGGGCAAAUUGCAUUAUUUUCCCAUCGAAUCGAUCACUUAUCUAACCAUCUUAAGAAUAACAAAAGGAUUUCAAUACGGAACGAUCAUUAGUUAAACUAGUAGGCAAACGCCGUCAACUUUUAGAUUAUCUC
env-99    AUUUUCAAGAAGCAUGGAAAAUCUGAAAAUGACACCGGUUCAGCGGAAGGGCAAAUUGCGCUUUUUACUUACAGAAUCGGUCAUUUGACGGAGCACUUGAAGAAAAAUCGAAAAGAUUUUGGAACGCAACGUUCGCUUCAAUUGUUAGUCGGUAAGCGCAGAAGUUUGCUCGACUAUUUG
env-100   AUUUUCAAGAAGCAUGGAAAAUCUGAAAAUGACACCGGUUCAGCGGAAGGGCAAAUUGCGCUUUUUACUUACAGAAUCGGUCAUUUGACGGAGCACUUGAAGAAAAAUCGAAAAGAUUUUGGAACGCAACGUU.........................................
env-101   AUCUUUAAACAACACGGUAAUCUAAAAAAGACACAGGUUCUGCAGAAGGACAAAUUGCUUUAUUUACUCAUCGUAUCAAUCACUUAUCUAAUCACCUAAAAACAAACCGUAAAGAUUACAAUACGGAGCGCUCUUUGGUUCGAUUAGUUGGAAAGCGUCGUAGUUUACUAGAUUAUUUA
env-102   AUUUUUAAAAAACACGGUAAAUCUGAAAAAGACACUGGUUCAUCACAGAAGGCCAAAUUGCUCUUUUUACUCACCGUAUCGAUCACUUAUCAAAACACCUAGAGGGCAAUCGAAAUGAUUUUAAUACCGAACGCUCCUUUAGUUCGUUUAGUGGGAAAAGAAGAAGUCUACUUGAUUACUUA
env-103   AUUUUUAAAAAACACGGUAAAUCUGAAAAAGACACUGGUUCAUCAGAAGGCCAAAUUGCUCUUUUCACUCACCGUAUCGAUCACUUAUCAAAACACCUAGAGGGCAAUCGAAAUGAUUUUAAUACCGAACGCUCCUUAGUUCGUUUAGUGGGAAAAGAAGAAGUCUACUUGAUUACUUA
env-106   AUUUUUAAAAAACACGGUAAAUCUGAAAAAGACACUGGUUCAUCAGAAGGCCAAAUUGCUCUUUUCACUCACCGUAUCGAUCACUUAUCAAAACACCUAGAGGGCAAUCGAAAUGAUUUUAAUACCGAACGCUCCUUAGUUCGUUUAGUGGGAAAAGAAGA.................
env-107   GUUUUUAAAAAAUACGGUAAAUCUGAAAAAGACACUGGUUCAUCAGAAGGCCAAAUUGCUCUUUUCACUCACCGUAUCGAUCACUUAUCAAAACACCUAGAGGGCAAUCGAAAUGAUUUUAAUACCGAACGCUCCUUAGUUCGUUUAGUGGGAAAAGAAGAAGUCUACUUGAUUACUUA
env-108   AUUUUUAAGAAACAUGGUAAGUCAGAAAAAGAUACUGGUUCAUCAGAAGGCCAAAUUGCUCUUUUCACUCACCGUAUCGAUCACUUAUCAAAACACUUAGAGGACAACCGAAAUGAUUUUAAUACAGAACGCUCCUUGGUACGAUUGGUGGGUAAAAGAAGAAGUCUACUUGACUACUUA
env-109   AUUUUUAAAAAACACGGUAAAUCUGAAAAAGAUUCUGGUUCAUCACAGAAGGCCAAAUUGCUCUUUUCACUCACCGUAUCGAUCACUUAUCAAAACACUUAGAGGGUAACCGAAAUGAUUUUAAUACCGAACGCUCCUUGGUUCGUUUAGUGGGAAAAGAAGAAGUCUACUUGACUACUUA
Min-1-1   AUCUUCUCUACACAUGGAGGAGAAGCAACUAACACUGGAUCUGCAGAAGGGCAAAUCGCAUUGUUCACUUAUAGAAUAUCUCACUUAACUGAGCACUUAAAGAAAAAUCGUAAAGAUUAUAACACUGAGCGUUCUUUAGUUCUUUUAGUAGGUAAAAGAAGAAGACUUCUUGACUAUUUA
env-110   UUCUUCAAAAAGUUUGGAGGUUCUGAAGAAAACACCGGAAGUACUGAAGCACAAAUCGCUUUAUUCACAUUCCGUAUCGCUCACUUAACUGGGCAUUUGAAGUCGAAUAAAAAAGACUUUGGAACACAGCGAGCUUUGAUCAAAUUAGUAGGUAAACGUAGAAGCUUACUUGACUAUUUA
env-111   AUUUUCGUGAAACACGGAAAAACUGCAACAAACACUGGUUCUGCAGAAGGACAAAUU............................................................................................................................
env-112   .....................................................................................................................................................................................
env-113   AUUUUCGUAA...........................................................................................................................................................................
env-114   AUUUUCGUAAAACACGGAAAAACUGCAACAAACACUGGUUCUGCAGAAGGACAAAUUGCAUUAUUCACUCACCGAAUUAACCACUUAACUGGACACUUAAAAAGCAAUCGUAAAGAUUACAACACUGAGCGCUCAUUAGUAAAACUAGUAGGUAAGAGAAGAAGUUUAUUAGAUUAUUUA
env-115   .....................................................................................................................................................................................
env-116   CUUUUUGCUAAGCACGGCAAAAACGAGCAGGAUACAGGUUCUGCAGAGGGACAAAUCGCAUUAAUGACAAAGCGAAUUGAACACAUCUCUCAACACAUGAAAGAAAACAAAAAAGAUUUUAAUUCAAAAAAGUCGCUUGUAAGUCUUGUAGGUAAACGAAGAAAUUUGUUAAACUAUUUA
Nul-1-1   ■■■UUCGCAAAGUACGGAAAAGACGCAAACGACACAGGUAACACUGAUGCGCAAAUCGCAUUAUUCACUUACCGUAUUAACUAUUUAACAGGACACCUUAAAAACAACCGUCAUGACUACAAUACAGAGCGUAGUCUAGUGAAACUGGUAGGUAAAAGACGUUCUCUUCUUGAUUACAAAA
Nul-1-2   AUUUUCGCAAAGUACGGAAAAGACGCAAACGACACAGGUAACACUGAUGCGCAAAUCGCAUUAUUCACUUACCGUAUUAACUAUUUAACAGGACACCUUAAAAACAACCGUCAUGACUACAAUACAGAGCGUAGUCUAGUGAAACUGGUAGGUAAAAGACGUUCUCUUCUUGAUUACAAAA
Fba-1-1   AUUUUCGCAAAGUACGGAAAAGACGCAAACGACACAGGUAACACUGAUGCGCAAAUCGCAUUAUUCACUUACCGUAUUAACUAUUUAACAGGACACCUUAAAAACAACCGUCACGACUACAAUACAGAGCGUAGUCUAGUGAAACUGGUAGGUAAAAGACGUUCUCUUCUUGAUUACAAA
Nul-1-3   AUUUUCGCAAAGUACGGAAAAGACGCAAACGACACAGGUAACACUGAUGCGCAAAUCGCAUUAUUCACUUACCGUAUUAACUAUUUAACAGGACACCUUAAAAACAACCGUCAUGAUCUACAAUACAGAGCGUAGUCUAGUGAAACUGGUAGGUAAAAGACGUUCUCUUCUUGAUUACAAA
Nul-1-4   AUUUUCGCAAAGUACGGAAAAGACGCAAAUGACACAGGUAACACUGACUCGCAAAUCGCGUUAUUCACUUACCGUAUCAACUAUUUAACAGGACACUUAAAAAACAACCGUCACGACUACAACACAGAGCGUAGUCUAGUGAAACUGGUAGGUAAAAGACGUUCACUUCUUGAUUACAAG
Nul-1-5   ■■■UUCGCAAAGUACGGAAAAGACGCAAAUGACACAGGUAACACUGACUCGCAAAUCGCGUUAUUCACUUACCGUAUCAACUAUUUAACAGGACACUUAAAAAACAACCGUCACGACUACAACACAGAGCGUAGUCUAGUGAAAUUGGUAGGUAAAAGACGUUCACUUCUUGAUUAUAAGA
Nul-1-6   AUUUUCGCAAAGUACGGAAAAGACGCAAACGACACAGGUAACACUGAUGCGCAAAUCGCAUUAUUCACUUACCGUAUUAACUAUUUAACAGGACACCUUAAAAACAACCGUCAUGACUACAAUACAGAGCGUAGUCUAGUGAAACUGGUAGGUAAAAGACGUUCUCUUCUUGAUUACAAA
env-117   AUUUUCGCCAAGUACGGAAAAGACGCAAACGACACCGGUAACACAGACGCACAAAUCGCGUUGUUCACUUACCGCAUUAACUAUUUAACAGGACACCUUAAAAACAACCGUCACGACUACAACACUGAGCGUAGUCUAGUGAAACUGGUAGGUAAAAGACGUUCCCUUCUUGAUUAUAAA

          .....................................................................................................................................................................................
          .....................................................................................................................................................................................
          AUYUUYR•RAARCACGG•RRR•••R•RR••RAYAC•GG••C•RC•GAAGG•CAAAUYGC•YURUUYAC•Y•Y•G•AUYRAYCAYYUR•C•RA•CAYYURAAAAR•AAYCR•AARGAUU•YAAYAC•GAR•G•UC•YURGU•••RYURGU•GG•AAR•G•RG••••YU•YU•GAYUAYYU•
```

```
env-118  AUUUUCGCCAAGUACGGAAAAGACGCAAACGACACCGGUAACACAGACGCACAAAUCGCGUUGUUCACUUACCGCAUUAACUAUUUAACAGGACACCUUAAAAACAACCGUCACGACUACAACACUGAGCGUAGUCUAGUGAAACUGGUAGGUAAAAGACGUUCCCUUCUUGAUUAUAAA
env-119  AUUUUUAAAGAACAUGGAGGUUCUAAAAAAAAUACCGGAUCUGCUGAAGGCCAAAUUGCACUAUUCACAGAAAGAAUAAAUCACCUAACAAACCAUCUUAAAAAAAACCGAAAAGAUUUCAACACCGAAAGAUCAUUGGUCAUGAUGGUAGGUAAGAGAAAAAGUUUACUAAACUAUCUC
env-120  AUUUUUAAAGAACAUGGAGGUUCUAAAAAAAAUACCGGAUCUGCUGAAGGCCAAAUUGCACUAUUCACAGAAAGAAUAAAUCACCUAACAAACCAUCUUAAAAAAAACCGAAAAGAUUUCAACACCGAAAGAUCAUUGGUCAUGAUGGUAGGUAAGAGAAAAAGUUUACUAAACUAUCUC
env-121  AUUUUUAAAGAACAUGGAGGUUCUAAAAAAAAUACCGGAUCUGCUGAAGGCCAAAUUGCACUAUUCACAGAAAGAAUAAAUCACCUAACAAACCAUCUUAAAAAAAACCGAAAAGAUUUCAACACCGAAAGAUCAUUGGUCAUGAUGGUAGGUAAGAGAAAAAGUUUACUAAACUAUCUC
Fsu-1-1  AUUUUUGCAAAACAUGGCCAGGCUGCUACCAACACAGGUUCUGCCGAAGGACAAAUUGCACUAUUUACAUUCAGGAUUAACCACCUUACAGGACACCUUAAAAAAAAUCGUCAUGAUUACAAUACAGAGCGUUCUCUUGUAAAACUGGUAGGUAAAAGAAGAAGCCUACUUGAUUACCUG
Gsp-1-1  AUUUUUAAAAAAACACGGUGGAAGUGAAAACAACACUGGAUCUGCAGAAGGACAAAUCGCUUUGUUCACACACAGAAUUAAACACUUAACUGGACACUUGAAAAACAAUCAAAAAGAUUUCAACACAGAGCGUUCAUUAGUUAUGUUGGUGGGUAAAAGAAGAAGCCUGCUAGAUUAUUUG
env-122  UUUUUUAAAAA...........................................................................................................................................................................
env-123  AUUUUUAAAAAGCACGGCGGGUCUGAGGGGAACACAGGUUCUACCGAAGGCCAGAUUGCGUUGUUCACGCACCGCAUCAACCAUUUGACAACACACCUAAAGAAAAAUCACAAAGAUUUUAAUACAGAGCGUUCAUUAGUUCGCUUGGUAGGUAAAAGAAGAUCUCUUUUAGACUACCUG
env-124  AUUUUUGAAAAGUACGGAAAAGGAAAGAAUGACACUGGUUCUGCAGAAGGACAGAUUGCAUUAUUCACUUACCGUAUAAACCACCUUACAGAGCAUUUAAAAAA.........................................................................
env-125  AUCUUCGAAAAACACGGUAAAGGUAAAAACGAUACUGG...............................................................................................................................................
env-126  AUCUCUUUAAAAAGCACGGAAAAGAUGCCAAAGAUACGGGGUCUUCCGAAGGGCAAAUUGCUUUAUUCACUGAAAGAUAAACCAUUUCACUAGUCACCUGAAGAAAAACCGUAAAGAUUACAACACAGAACGUUCAUUAGUAAAGCUCGUUGGAAAACGUCGCUCAUUACUUGUAUUACU
env-127  AUUUUCGNNNNNNNNNNNNNNNNNNNNNNNNNNNNNNNNNNNNNNNNNNNNNNNNNNNNNNNNNNNNNNNNNNNNNNNNNNNNNNNNNNNNNNNNNNNNNNNNNNNNNNNNGGCAAAUCGGCUACUGACACGGGUUCGACCGAAGGGCAGAUCGCUCUUUUCACCUAUCGCAUCAACCACCUGACCGAGCAU
env-128  AUUUUCGCUCAGUACGGCAAAUCGGCUACUGACACGGGUUCGACCGAAGGGCAGAUCGCUCUUUUCACCUAUCGUAUCAACCACCUGACCGAACAUCUGAAGAAGAACAUUCACGACUUCGGUACGGAGCGCGCGUUGGUCAAACUCGUCGGUAAACGCCGCAGUCUGCUGGAGUACCUC
env-129  AUUUUCGCUCAGUACGGCAAAUCGGCUCCUGACACGGGUUCGACCGAAGGGCAGAUCGCUCUUUUCACCUAUCGUAUCAACCACCUGACCGAACAUCUGAAGAAGAACAUUCACGACUUCGGUACGGAGCGCGCGUUGGUCAAACUCGUUGGUAAACGCCGCAGUCUGCUGGAGUACCUC
env-130  AUUUUCGCUCAGUACGGCAAAUCGGCUACUGACACGGGUUCGACCGAAGGGCAGAUCGCUCUUUUCACCUAUCGUAUCAACCACCUGACCGAACAUCUGAAGAAGAAC...............................................................................
env-131  ACUCUUUAAAAAACACGGCAAGAACGAGAAUGAUACUGGUUCUGCAGAAGGACAAAUUGCAUUGUUCACUGAAAGAAUCAAUCAUUUAACAGAACACUUAAAAAAGAAUCGUAAAGAUUACAAUACUGAGCGUUCGUUGGUAAUGUUAGUAGGUAAGCGUAGAUCGCUUCUAGACUACUU
env-132  ACUCUUUAAAAAACACGGCAAGAACGAGAAUGAUACUGGUUCUGCAGAAGGACAAAUUGCAUUGUUCACUGAAAGAAUCAAUCAUUUAACAGAACACUUAAAAAAGAAUCGUAAAGAUUACAAUACUGAGCGUUCGUUGGUAAUGUUAGUAGGUAAGCGUAGAUCGCUUCUAGACUACUU
env-133  AUAUUUAAAAAACAUGGUAAAUCUAAGAAAGAUACUGGAUCUACCGAGGGACAAAUAGCACUUUUUACCCAUCGCAUUCAAUAUCUUUCAGAACACCUCAAAGAAAAUAGAAAAGAUUUCAAUACAGAACGUUCACUAGUUCGUUUGGUUGGAAAGCGAAGAAGCCUUUUAGACUAUUUA
env-134  AUCUUCGCAAAACACGGAGAAAAAACAAACACAGGUAAAGCCGAAAGUCAAAUUGCACUUUUUACUUACAGAAUCACACAUUUAACAGAACACUUGAAAAAAAAUCGUCAUGAUUAUAACACAGAGCGUUCAUUAGUAUUAUUAGUAGGUAAAAGAAGAUCUUUGUUAGAUUAUUUAAAG
env-135  CUCUUUAAAAAACACGGAAAAGAUGCCAAAGACACGGGAUCUUCAGAAGGUCAAAUAGCUUUAUUCACUGAAAGAAUAAAUCAUUUGACAAAUCACUUGAAACAAAACCGUAAAGAUUACAACACAGAACGUUCGUUAGUAAAACUAGUUGGAAAACGUCGCUCAUUACUUGACUACUUA
env-138  CUCUUUAAAAAACACGGAAAAGAUGCCAAAGACACGGGAUCUUCAGAAGGUCAAAUAGCUUUAUUCACUGAAAGAAUAAAUCAUUUGACAAAUCACUUGAAACAAAACCGUAAAGAUUACAACACAGAACGUUCGUUAGUAAAACUAGUUGGAAAACGUCGCUCAUUACUUGACUACUUA
env-139  CUCUUUAAAAAACACGGAAAAGAUGCCAAAGACACGGGAUCUUCAGAAGGUCAAAUAGCUUUAUUCACUGAAAGAAUAAAUCAUUUGACAAAUCACUUGAAACAAAACCGUAAAGAUUACAACACAGAACGUUCGUUAGUAAAACUAGUUGGAAAACGUCGCUCAUUACUUGACUACUUA
env-140  CUCUUUAAAAAACACGGAAAAGAUGCCAAAGACACGGGAUCUUCAGAAGGUCAAAUAGCUUUAUUCACUGAAAGA..........................................................................................................
env-141  CUCUUUAAAAAACACGGAAAAGAUGCCAAAGACACGGGAUCUUC.........................................................................................................................................
env-142  CUCUUUAAAAAACACGGAAAAGAUGCCAAAGACACGGGAUCUUCAGAAGGACAAAUUGCCUUAUUCACAGAAAGAAUCAACCAUUUAACUGAGCAUUUAAAAACAAAAUCGUAAAGAUUAUAAUACGGAACGCUCUUUGGUUAAACUCGUUGGGAAACGUCGCUCAUUAUUAGAUUAUCUA
env-143  CUCUUUAAAAAACACGGAAAAGAUGCCAAAGACAC..................................................................................................................................................
env-145  CUCUUUAAAAAACACGGAAAAGAUGCCAAAGACACGGGAUCUUCAGAAGGACAAAUUGCCUUAUUCACAGAAAGAAUCAACCAUUUAACUGAGCAUUUAAAAACAAAAUCGUAAAGAUUAUAAUACGGAACGCUCUUUGGUUAAACUCGUUGGGAAACGUCGCUCAUUAUUAGAUUAUCUA
env-146  CUCUUUAAAAAACACGGGAAAGAUGCCAAAGACACGGGGUCUUCUGAAGGACAAAUUGCUUUAUUCACAGAAAGAAUAAACCAUUUAACAGGUCACUUAAAACAAAAUCGUAAAGAUUACAACACAGAACGAUCACUAGUAAAACUAGUUGGUAAGCGUCGCUCAUUAUUAGAUUAUUUA
env-147  CUCUUUAAAAAACACGGAAAAGAUGCCAAAGACACGGGAUCUUCUGAAGGACAAAUUGCAUUAUUCACAGAAAGAAUCAACCAUUUAACUGAGCAUUUAAAAAUUAAAUCGUAAAGAUUAUAAUACGGAACGCUCUUUGGUUAAACUCGUUGGGAAACGUCGCUCAUUAUUAGAUUAUCUA
env-148  CUCUUUAAAAAACACGGAA..................................................................................................................................................................
env-149  CUCUUUAAAAAACACGGAAAAGAUGCCAAAGACACGGGAUCUUCUGAAGGACAAAUUGCAUUAUUUACAGAAAGAAUCAACCAUUUAACUGAGCAUUUAAAAAUUAAAUCGUAAAGAUUAUAAUACGGAACGCUCUUUGGUUAAACUCGUUG..........................
env-151  .....................................................................................................................................................................................
env-152  CUCUUUAAAAAACACGGAAAAGAUGCUAAAGAUACGGGAUCUUCCGAAGGUCAAAUUGCUUUAUUCACAGAAAGAAUAAACCACUUAACAGGGCAUUUAAAAACAAAACCGUAAAGAUUAUAACACAGAACGUUCACUAGUAAAACUAGUGGAAAACGUCGCUCAUUAUUAGACUACUUG
env-153  CUCUUUAAAAAACACGGAAAAGAUGCUAAAGAUACGGGAUCUUCCGAAGGUCAAAUUGCUUUAUUCACAGAAAGAAUAAACCACUUAACAGGGCAUUUAAAAACAAAACCGUAAAGAUUAUAACACAGAACGUUCACUAGUAAAACUAGUGGAAAACGUCGCUCAUUAUUAGACUACUUG
env-154  CUCUUUAAAAAACACGGAAAAGAUGCUA.........................................................................................................................................................
env-155  CUCUUUAAAAAACACGGAAAAGAUUCCAAAGAUACGGGAUCUUCUGAAGGUCAAAUUGCUUUAUUCACACAAAGAAUAAACCACUUAACAGGCAUUUAAAAACAAAACCGUAAAGAUUAUAACUCUGAACGUUCACUAGUAAAACUAGUCGGAAAACGUCGCUCAUUAUUAGACUACUUA
env-156  CUCUUUAAAAAACACGGAAAAGAUGCCAAAGACACGGGAUCUUCUGAAGGUCAAAUUGCUUUAUUCACAGAAAGAAUAAACCACUUAACAGGGCAUUUAAAAACAAAACCGUAAAGAUUAUAACACAGA...............................................
env-157  CUCUUUAAAAAACACGGUAAAGAUGCUAAAGACACGGGAUCUUCUGAAGGUCAAAUUGCUUUAUUCACAGAAAGAAUAAACCACUUAACAGGGCAUUUAAAAACAAAACCGUAAAGAUUAUAACACAGA...............................................
env-158  CUCUUUAAAAAACACGGAAAAGAUGCUAAAGAUACGGGAUCUUCCGAAGGUCAAAUUGCUUUAUUCACAGAAAGAAUAAACCACUUAACAGGGCAUUUAAAAACAAAACCGUAAAGAUUAUAACACAGAACGUUCACUAGUAAAACUAGUGGAAAACGUCGCUCAUUAUUAGACUACUUG
env-159  CUCUUUAAAAAACACGGAAAAGAUGCUAAAGAUACGGGAUCUUCCGAAGGUCAAAUUGCUUUAUUCACAGAAAGAAUAAACCACUUAACAGGGCAUUUAAAAACAAAACCGUAAAGAUUAUAACACAGAACGUUCACUAGUAAAACUAGUGGAAAACGUCGCUCAUUAUUAGACUACUUG
env-161  CUCUUUAAAAAACACGGAAAAGAUGCCAAAGACACGGGAUCUUCAGAAGGGCAAAUUGCAUUAUUCACAGAAAGAAUCAAUCACUUAACUGAGCAUUUAAAAACAAAAUCGUAAAGAUUAUAACACAGAACGCUCGUUAGUAAAACUAGUUGGAAAACGUCGCUCAUUAUUAGAUUAUCUUA
env-162  CUCUUUAAAAAACAUGGAAAAGAUGCCAAAGACACGGGGUCUUCAGAAGGUCAAAUUGCAUUAUUCACAGAAAGAAUCAAUCAUUUAACUGAGCAUUUAAAAACAAAAUCGUAAAGAUUAUAACACAGAGCGCUCAUUAGUAAAACUAGUUGGAAAACGUCGCUCAUUAUUAGACUACUUA
env-163  CUCUUUAAAAAACACGGAAAAGAUGCCAAAGAUACGGGAUCUUCAGAAGGACAAAUUGCUUUGUUUACAGAAAGAAUCAAUCACUUAACUGAGCAUUUAAAAACAAAAUCGUAAAGAUUUUAAUACCGAACGUGCAUUAGUAAAACUAGUUGGAAAACGUCGUGCAUUAUUAGAUUACCUA
env-164  CUCUUUAAAAAACACGG....................................................................................................................................................................
env-165  CUCUUUAAAAAACACGGGAAAGAUGCCAAAGACACUGGCUCUUCUGAAGGUCAAAUUGCCUUAUUCACUGAAAGAAUAAACCACCUCACAAGUCAUUUAAAACAAAAUCGUAAAGAUUACAACACAGAACGUUCAUUAGUAAAACUAGUUGGAAAACGUCGCUCAUUAUUAGAUUAUCUUA
env-166  CUCUUUAAAAAACACGGGAAAGAUGCCAAAGACACUGGCUCUUCUGAAGGUCAAAUUGCCUUAUUCACUGAAAGAAUAAACCACCUCACAAGUCAUUUAAAACAAAUCGUAAAGAUUACAACACAGAACGUUCAUUAGUAAAACUAGUUGGAAAACGUCGCUCAUUAUUAGAUUACUUAA
env-167  CUCUUUAAAAAACACGGAAAAGAUGCCAAAGAUACAGGAUCUUCCGAAGGACAAAUUGCCUUAUUCACAGAAAGAAUCAAUCACUUGACUGAGCAUUUAAAAACAAAAUCGUAAAGAUUAUAACACAGAACGCUCGUUAGUAAAACUAGUAGGAAAACGUCGCUCAUUAUUAGAUUAUUUA
env-170  CUCUUUAAAAAACACGGAAAAGAUGCCAAAGAUACGGGAUCUUCAGAAGGACAA...............................................................................................................................
env-171  CUCUUUAAAAAACACGGAAAAGAUGCCAAAGACACGGGAUCUUCAGAAGGACAAAUUGCCUUAUUUACAGAAAGAAUCAAUCACUUAACUGAGCAUUUAAAAACAAAAUCGUAAAGAUUAUAAUACAGAACGUGCAUUAGUAAAACUAGUAGGAAAACGUCGCUCAUUAUUAGAUUAUCUA
env-172  CUCUUUAAAAAACACGGAAAAGAUGCCAAAGAUACGGGAUCUUCAGAAGGACAAAUUGCCUUAUUUACAGAAAGAAUCAAUCACUUAACUGAGCAUUUAAAAACAAAAUCGUAAAGAUUAUAAUACAGAACGUGCAUUAGUAAAACUAGUAGGAAAACGUCGCUCAUUAUUAGAUUAUCUA
env-173  CUCUUUAAA............................................................................................................................................................................
env-174  .....................................................................................................................................................................................
env-175  CUCUUUAAAAAACACGGGAAAGAUGCCAAAGACACGGGAUCUUCAGAAGGACAGAUUGCAUUAUUCACAGAAAGAAUCAAUCACUUAACUGAGCAUUUAAAAAUUAAAUCGUAAAGAUUACAAUACAGAACGCUCGUUAGUAAAACUAGUUGGAAAACGUCGCUCAUUAUUAGACUAUUUA
env-176  .....................................................................................................................................................................................
env-177  .....................................................................................................................................................................................
env-179  .....................................................................................................................................................................................
env-180  .....................................................................................................................................................................................
env-181  CUCUUUAAAAAACACGGAAAAGAUGCCAAAGAUACGGGAUCUUCAGAAGGACAAAUUGCAUUAUUCACAGAAAGAAUCAAUCACUUAACUGAGCAUUUAAAACUAAAUCGUAAAGAUUACAACACAGAACGCUCGCUAGUUAAACUGGUUGGAAAACGUCGCUCAUUAUUAGACUAUUUA
env-185  .....................................................................................................................................................................................
env-186  CUCUUUAAAAAACACGGAAAAGAUGCCAAAGAUACGGGAUCUUCAGAAGGACAAAUUGCAUUAUUCACAGAAAGAAUCAAUCACUUAACUGAGCAUUUAAAACUAAAUCGUAAAGAUUACAACACAGAACGCUC............................................
env-187  CUCUUUAAAAAACACGGAAAAGAUGCCAAAG......................................................................................................................................................
env-189  CUCUUUAAAAAACACGGAAAAGAUGCCAAAGACACGGGAUCUUCAGAAGGGCAAAUUGCAUUAUUCACAGAAAGAAUCAAUCACUUAACUGAGCAUUUAAAAACAAAAUCGUAAAGAUUAUAACACAGAACGCUCGUUAGUAAAACUAGUUGGAAAACGUCGCUCAUUAUUAGACUACUUA
env-191  CUCUUUAAAAAACACGGAAAAGAUGCCAAAGACA...................................................................................................................................................
env-192  CUCUUUAAAAAACACGGAAAAGAUGCCAAAGACACGGGAUCUUCAGAAGGACAAAUUGCACUAUUCACAGAAAGAAUCAAUCACUUAACUGAGCAUUUAAAAACAAAAUCGUAAAGAUUACAACACAGAACGCUCAUUAGUUAAACUAGUUGGAAAACGUCGCUCAUUAUUAGACUACUUA
env-193  CUCUUUAAAAAACACGGAAAAGAUGCCAAAGACACGGGAUCUUCAGAAGGACAAAUUGCACUAUUCACAGAAAGAAUCAAUCACUUAACUGAGCAUUUAAAAACAAAAUCGUAAAGAUUACAACACAGAACGCUCAUUAGUUAAACUAGUUGGAAAACGUCGCUCAUUAUUAGACUACUUA
env-195  .....................................................................................................................................................................................
env-196  CUCUUUAAAAAACACGGAAAAGAUGCCAAAGACACGGGAUCUU..........................................................................................................................................
env-197  CUCUUUAAAAAACACGGAAAAGAUGCCAAAGACACGGGAUCUUCUGAAGGACAAAUUGCUUUAUUCACAGAAAGAAUAAACCAUUUA.............................................................................................
env-198  CUCUUUAAAAAACACGGAAAAGAUGCCAAAGACACGGGAUCUUCUGAAGGACAAAUUGCUUUAUUCACAGAAAGAAUAAACCAUUUAACAAGUCACCUGAAACAAAACCGUAAAGAUUAUAACACAGAACGUUCACUAGUUAAACUAGUUGGAAAGCGUCGCUCAUUAUUAGACUACUUA
env-201  .....................................................................................................................................................................................
env-202  CUCUUUAAAAAACACGGAAAAGAUGCCAAAGACACGGGUUCCUCAGAAGGACAAAUUGCUUUAUUCACAGAAAGAAUAAACCAUUUAACAAGUCACCUGAAACAAAACCGUAAAGAUUAUAACACAGAACGUUCACUAGUUAAACUAGUUGGAAAGCGUCGCUCAUUAUUAGACUACUUA
env-203  AUUUUCAAAAAAUACGGGAAAUCUGAAAAUGAUACCGGAUCUACAGAAGGCCAAAUUGCCUUAUUCUCACAUCGCAUCGAUCACUUGUCCAGCCAUCUCAAAAAUAACGCUAAAGAUUUUAACACAGAGCGUUCAUUGGUUCGUUUGGUAGGAAAGAGAAGAAGUUUACUUGACUAUCUC
env-205  AUUUUCAAAAGUUUUGGUAAAUCUGAUAAGGACACAGGCUCUGCUGAAAGCCAAAUAGCUUUGUUUUCUCAUAGGAUUAAUCAUUUAUCUAGUCAUUUAAAAAAAAAUAAGAAAGAUUUUAAUACGGAAAGAUCACUUGUUAGACUCGUUGGAAAAAGAAGGAAUCUUCUAGAAUAUCUA
env-206  AUUUUCAAAAGUUUUGGUAAAUCUGAUAAGGACACAGGCUCUGCUGAAAGCCAAAUAGCUUUGUUUUCUCAUAGGAUUAAUCAUUUAUCAAGUCAU....................................................................................
env-207  AUUUUCAAAAGUUUUGGUAAAUCUGAUAAGGACACAGGCUCUGCUGAAAGCCAAAUAGCUUUGUUUUCUCAUAGGAUUAAUCAUUUGUCUAGUCAUCUAAAAAAAAAAUAAGAAGGAUUUUAAUACGGAAAGAUCACUUGUUAGACUCGUUGGAAAAAGAAGGAAUCUUCUAGAAUAUCUA
env-210  AUUUUCAAAAGUUUUGGUAAAUCUGAUAAGGACACAGGCUCUGCUGAAAGCCAAAUAGCUUUGUUUUCUCAUAGGAUUAAUCAUUUGUCUAGUCAUUUAAAAAAAAAUAAGAAAGAUUUUAAUACGGAAAGAUCACUUGUUAGACUCGUUGGAAAAAGAAGGAAUCUUCUAGAAUAUCUA
env-211  AUUUUCAAGAAAUAUGGAAAAUCUGAGGCUGAUACUGGUUCUGCUGAGGGACAAAUAGCAUUAUUUUCUCACCGAAUCGAUCAUUUAUCACUUCAUCUAAAGAAAAAUCAGAAGGAUUUUAACACUGAACGAUCGUUAGUAAAACUUGUAGGUAAACGCCGUCAACUUCUUGAUUAUCUG
env-212  .....................................................................................................................................................................................
env-213  AUUUUUACUAAGUAUGGAGCUAGUGAAACAAACACAGGUUCUACAGAAGGACAAGUUGCAUUGUUUACUUUCCGUAUCAAUCACCUAACUGAGCACUUAAAAGGAAAUCGUAAAGAUUUCAACACUGAGCGUUCUCUAGUGAAAUUGGUAGGUAAGCGUCGUUCUUUACUUGAUUACUUA
env-214  AUUUUUACUAAGUAUGGGGCUAGUGAAACAAACACAGGUUCUACAGAAGGACAAGUUGCAUUGUUUACUUUCCGUAUCAAUCACCUAACUGAGCACUUAAAAGGAAAUCGUAAAGAUUUCAACACUGAGCGUUCUCUAGUGAAAUUGGUAGGUAAGCGUCGUUCUUUACUUGAUUACUUA
env-215  AUUUUUACUAAGUAUGGAGCUAGUGAAACAAACACAGGUUCUACAGAAGGACAAGUUGCNNNNNNNNNNNNNNNNNNNNNNNNNNNNNNNNNNNNNNNNNNNNNNNNNNNNNNNNNNNNNNNNNNNNNNNNNNNNNNNNNNNNNNNNNNNNNNNNAAUAAUCUUAUCUAAAAGUAUUUA
env-216  AUUUUCGAAAAGUAUGCUGGAGGCGCAGUGAACACAGGUUCAGCCGAAGGUCAAAUUGCUUUAUUUUCAUUCAGAAUUCAGAACUUAACUGAGCACUUGAAGAAAAAUCGUCACGACUACAGAACGCAGCGUUCUUUGCAGUUGUUGGUUGGUAAAAGAAGAAGUUUGCUAGACUAUUUG
env-217  AUUUUCGAAAAGUAUGCUGGAGGCGCAGUGAACACAGGUUCAGCCGAAGGUCAAAUUGCUUUAUUUUCAUUCAGAAUUCAGAACUUAACUGAGCACUUGAAGAAAAAUCGUCACGACUACAGAACGCAGCGUUCUUUGCAGUUGUUGGUUGGUAAAAGAAGAAGUUUGCUAGACUAUUUG
env-218  AUCUUUAAAAAACACGGGAAGUC..............................................................................................................................................................

         .....................................................................................................................................................................................
         .....................................................................................................................................................................................
         AUYUUYR•RAARCACGG•RRR•••R•RR••RAYAC•GG••C•RC•GAAGG•CAAAUYGC•YURUUYAC•Y•Y•G•AUYRAYCAYYUR•C•RA•CAYYURAAAAR•AAYCR•AARGAUU•YAAYAC•GAR•G•UC•YURGU•••RYURGU•GG•AAR•G•RG••••YU•YU•GAYUAYYU•
```

```
env-220    AUUUUUGCAAAACACGGAG.............................................................................................................................
env-221    AUCUUCGCUAAACACGGAGGUUCCGAUACAAACACAGGAUCUACAGAAGGUCAGGUUGCUUUGUUCACGUAUCGUAUUAGUCAUUUGACAGAGCACUUGAAGAAAAACCGUAAGGAUUUUGGAACGCAACGCUCUCUUCAAUUAUUAGUAGGAAAAGAAGAAGUUUAUUAGAUUACCUA
env-222    AUCUUCGCUAAACACGGAGGUUCCGAAAUAUUGGCAAAUAUUUAAACCCAGAAGCAAAAAAAGAAAUCUUCGCUAAACACGGAGGUUCCGAAACAAACACAGGAUCUACAGAAGGUCAGGUUGCUUUGUUCACGUAUCGUAUUAGUCAUUUGACAGAGCACUUGAAGAAAAACCGUAAGG
Fli-1-1    AUCUUCGCUAAACACGGAGGCGCAGCAGCUAACACUGGUUCUGCAGAAGGUCAAAUCGCGUUAUUCACUUUCAGAAUUUCACACUUGACUGAGCACUUGAAAAAAAAUCGUCAUGAUUACAACACUGAGCGCUCAUUGGUUUUAUUAGUAGGAAAAAGAAGAAGCCUUCUUGACUACUUA
Gme-1-1    AUCUUUGCAAAACAUGGUAAAGGAGCAAAAGACACGGGUCAGCUGCGGAAGGCAAAUUGCUUUGUUUACCUACAGAAUCAGCAUUUAACAGAACACUUAAAAAACAAUCGUAAAGAUUAUAAUACGGAGCGUUCAUUGGUCAUGUUAGUUGGUAAGCGUCGUGCAUUACUUGACUACCUA
env-223    AUCUUCGCUAAACACGGAGGAAAAGCUGAAAACACAGGUUCUGCUGAAGGGCAAAUCGCGUUAUUCACACACAGAAUUAAUCACUUAACUGAGC..................................................................
env-224    AUCUUCGCAAAACACGGAGGUUCGGAGA....................................................................................................................
env-225    AUCUUCGCUAAACACGGAGGUUCGGAGACAAACACAGGGUCUACAGAAGGUCAAGUUGCUUUGUUCACGUAUCGUAUUAGUCAUUUGACAGAGCACUUGAAGAAAAAUCGUAAGGAUUUUGGAACGCAACGCUCUCUUCAAUUAUUAGUAGGAAAAGAAGAAGUUUAUUGGAUUACUUA
env-226    AUCU............................................................................................................................................
env-227    ................................................................................................................................................
env-228    ................................................................................................................................................
env-229    AUCUUCGCUAAACACGGAGGUUCGGAGACAAACAC.............................................................................................................
env-230    AUCUUCGCUAAACACGGAGGUUCGGAGACAAACACAGGGUCUACAGAAGGUCAAGUUGCUUUGUUCACGUAUCGUAUUAGUCAUUUGACAGAGCACUUGAAGAAAAAUCGUAAGGAUUUUGGAACGCAACGCUCUCUUCAAUUAUUAGUAGGAAAAGAAGAAGUUUAUUGGAUUACUUA
env-231    AUCUUCGCUAAACACGGAGGUUCGGAGACAAACACAGGGUCUACAGAAGGUCAGGUUGCUUUGUUCACGUAUCGUAUUAGUCAUUUGACAGAGCACUUGAAGAAAAAUCGUAAGGAUUUUGGAACGCAACGCUCUCUUCAAUUAUUAGUAGGAAAAGAAGAAGUUUAUUGGAUUACUUA
env-234    AUCUUCGCUAAACACGGAGGUUCGGAGACAAACACAGGAUCUACAGAAGGUCAAGUUGCUUUGUUCACGUAUCGUAUUAGUCAUUUGACAGAGCACUUGAAGAAAAAUCGUAAGGAUUUUGGAACCCAACGCUCUCUUCAAUUAUUAGUAGGAAAAGAAGAAGUUUAUUGGAUUACUUA
env-235    AUCUUCGCUAAACACGGAGGUUCGG.......................................................................................................................
env-236    AUCUUCGCUAAACACGGAGGUUCGGAGACAAACACAGGGUCUACAGAAGGUCAAGUUGCUUUGUUCACGUAUCGUAUUAGUCAUUUGACAGAGCACUUGAAGAAAAAUCGCAAGGAUUUUGGAACACAACGCUCUCUUCAAUUAUUAGUAGGAAAAGAAGAAGUUUAUUGGAUUACUUA
env-237    AUCUUCGCUAAACACGGAGGUUCGGAGACAAACACAGGGUCUACAGAAGGUCAAGUUGCUUUGUUCACGUAUCGUAUUAGUCAUUUGACAGAGCACUUGAAGAAAAAUCGCAAGGAUUUUGGAACGCAACGCUCUCUUCAAUUAUUAGUAGGAAAAGAAGAAGUUUAUUGGAUUACUUA
env-238    AUCUUCGCAAAACACGGUGGGACAGAAACCAACACCGGUUCAACAGAAGGUCAGGUGGCAU...................................................................................
env-239    AUUUUUGCAAAACACGGGAAAUCGAAAACCGAUACCGGUUCUACUGAAGGACAGAUCGCGUUGUUCACUUACCGUAUUGAGCACCUUACAAAACACCUAAAGAACAACCAAAAGGACUUCAACACAGAGCGUUCUUUGGUACGUUUGGUAGGAAAGCGUCGUUCGCUUCUUGAUUAUUUA
env-243    AUUUUUGCAAAACACGGGAAAUCGAAAACCGAUACCGGUUCUACUGAAGGACAGAUCGCGUUGUUC..............................................................................
env-244    AUCUUUGCCAAACAUGGUAAGGACAAGAAUGACAGUGGAACAGCAGAAGCUCAGAUCGCUUUAUUUACCAGAAGGAUAGAAGCGAUAUCAAAUCACUUAAAAACGAACAAGAAAGACCACAAUUCAAAAAAAGCGUUGGUCCAACUAGUAGGUAAACGAAGAGACUUACUUAAUUAUUUG
env-247    AUUUUUGCCAAAUACGGCAAA...........................................................................................................................
env-248    ................................................................................................................................................
env-249    AUUUUUGCCAAAUACGGCAAAGACAAAAAUGACAGCGGAUCAGCUGAAGCUCAAAUCGCUUUAUUUACAAGAAGAAUUGAAGCUAUUUCAGCGCAUUUGAAAACAAACAAAAAAGAUUUUAAUUCAAAAAAAUCUUUAGUAGAUUUAGUAGGUAAACGAAGAGAUUUACUUAAUUAUUUA
env-250    AUUUUUGCCAAAUACGGCAAAGACAAAAAUGACAGCGGAUCAGCUGAAG...............................................................................................
env-251    AUUUUCAAAAAACACGGAAAGAACGAGAACGAUUCUGGUUCUGCAGAAGGACAAAUUGCAUUGUUCACACACCGUAUCGAACACUUAACACAGCACCUUAAACAAAACCACAAAGACUACAACACAGAGCGUUCUUUGGUAAAAUGGUUGGUAAGCGUCGUUCUUUGUUAAACUAUUUA
env-255    AUUUUCAAAAAACACGGUAAAGGUAAAACGAUACCGGUACUGCGGAAGGGCAAAUUGCUUUGUUCACAUACCGAAUUGAACACCUUACACAACACCUGAAAAAAAACCACAAGGAUUUCAACACUGAAAGAUCUUUGGUUAAGAUGGUAGGAAAGCGUCGUUCUUUAUUAAACUAUUUA
env-256    AUUUUUAAAAAACACGGCAAAGUGCUACUGAUACUGGUACUGCCGAGGGUCAAAUCGCGUUAUUUACCUACCGCAUUGAUCACCUGACCCAACACCUAAAACGCAACCACAAGAGCUUUAACACUGAGCGCUCUCUAGUGAAAAUGGUCGGUAAAAGACGAUCACUUUUAGACUAUUUG
env-257    AUUUUUAAAAAACACGGCAAAAGUGCUACCGAUACUGGUACUGCCGAAGGUCAAAUCGCUCUUUUCACUUACCGAAUUCAGCACUUGACUGAGCACCUUAAACGCAAUCACAAAGAUUUUAACACAGAGCGCUCUCUAGUAAAACUUGUAGGUAAAAGACGAUCACUUCUCGAUUACCUC
env-258    AUUUUGGAAAAACACGGUAACAGCAAGAACGACACAGGUUCUGCUGAAGGACAAAUCGCUUUGUUCACCUACCGUAUUCAACCACCUUGACCGAGCACCUUAAGAAAAACCGCAAGGAUUUAACACGGAGCGUUCACUUGUAAAGCUGGUAGGAAAGCGCAGGUCGCUUCUUGACUAUCUU
env-261    AUUUUUGAAAAACACGGUAACAGCAAGAACGACACAGGUUCUGCUGAAGGACAAAUCGCUUUGUUCACCUACCGUAUCAACCACUUGACCGAGCACCUUAAGAAAAACCGCAAGGAUUUUAAC....................................
Nma-1-1    GUUUUCGCAAAGUACGGGAAAUCGGCAAAUGACACAGGGAAUACCGAUUCUCAAAUCGCGUUGUUCACCUACCGUAUCAACCACUUGACAGAACACCUUAAGAGAAAUCGUCACGACUACAACACAGAGCGCAGCCUUGUGAAGCUUGUAGGAAAGCGUCGUUCCCUUCUUGAUUACAAG
env-262    AUCUUCGCAAAACACGGUGGUUCAGCAACCAACACCGGUUCAGCCGAAGGACAGAUCGCGUUGUUCACGUACCGCAUCAACCACCUG.........................................................
env-263    UUAUUUAAAAAGCACGGAAAAUCUGAAUCAGAUACCGGCUCUGCUGAAGGACAAAUAGCAUUGUUUACAGCUAGAAUCAAUCAUUUAACUGAACAUCUUAAAAAUAAUCGUAAAGAUUUUAAUACUGAAAGAUCUUUAGUUAUGAUGGUUGGUAAAAGAAAAAGCUUAUUAAACUAUCUU
env-264    UUAUUUAAAAAGCACGGAAAAUCUGAAUCAGAUACCGGCUCUGCUGAAGGACAAAUAGCAUUGUUUACAGCUAGAAUCAAUCAUUUAACUGAACAUCUUAAAAAUAAUCGUAAAGAUUUUAAUACUGAAAGAUCUUUGGUUAUGAUGGUUGGUAAAAGAAAAAGCUUAUUAAACUAUCUU
env-265    UUAUUUAAAAAGCACGGAAAAUCUGAAUCAGAUACCGGCUCUGCUGAAGGACAAAUAGCAUUGUUUACAGCUAGAAUCAAUCAUUUAACUGAACAUCUUAAAAAUAAUCGUAAAGAUUUUAAUACUGAAAGAUCUUUGGUUAUGAUGGUUGGUAAAAGAAAAAGCUUAUUAAACUAUCUU
env-266    UUGUUCGAAAAACACGGUAAAGGUAAAAACGAUACCGGUUCAGCAGAAGGUCAAAUCGCAUUAUUCACAAAAAGAAUUGCGUAUUUUCUGACCACUUAAAAACAAACAGAAAAGAUUACAACUCUGAGCGUUCACUUGUAGCGCUAGUAGGUAAAAGAAGAAGCUUACUUGAUUACUUA
env-267    UUGUUCGAAAAACACGGUAAAGGUAAAAACGAUACCGGUUCAGCAGAAGGUCAAAUCGCAUUAUUCACGAAAAGAAUUGCGUAUUUUCUGACCACUUAAAAACAAACAGAAAAGAUUACAACUCUGAGCGUUCACUUGUAGCGCUAGUAGGUAAAAGAAGAAGCUUACUUGAUUACUUA
env-268    UUGUUCGAAAAGCACGGUAAAAGUAAAAAUGAUACCGGCUCAGCACAGAAGGUCAAAUUGCGUUAUUCACUAAAAGAAUUGCUCAUAUUUCAGACCACUUAAAAAACAACAAAAAAGACUUUAACUCAGAGCGAUCUCUUGUAGCGUUGGUAGGUAAAAGAAGAAGCUUACUAGACUACUUA
env-269    UUGUUCGAAAAACACGGUAAAGGUAAAAACGAUACCGGUUCAGCAGAAGGUCAAAUCGCAUUAUUCACGAAAAGAAUUGCGUAUUUUCUGACCACUUAAAAACAAACAGAAAAGAUUACAACUCUGAGCGUUCACUAGUAGCGCUAGUAGGUAAAAGAAGAAGCUUACUUGAUUACUUA
env-270    UUGUUCGAAAAACACGGUAAAGGUAAAAACGAUACCGGUUCAGCAGAAGGUCAAAUCGCAUUAUUCACGAAAAGAAUUGCGUAUUUUCUGACCACUUAAAAACAAACAGAAAAGAUUACAACUCUGAGCGUUCACU........................
env-271    UUGUUCGAAAAACACGGUAAAGGUAAAAACGAUACCGGUUCAGCAGAAGGUCAAAUCGCAUUAUUCACGAAAAGAAUUGCGUAUUUUCUGACCACUUAAAAACAAACAGAAAAGAUUACAACUCUGAGCGUUCACUUGUAGCGCUAGUAGGUAAAAGAAGAAGCUUACUUGAUUACUUA
env-272    UUGUUCGAAAAACACGGUAAAGGUAAAAACGAUACCGGUUCAGCAGAAGGUCAAAUCGCAUUAUUCACGAAAAGAAUUGCGUAUUUUCUGACCACUUAAAAACAAACAGAAAAGAUUACAACUCUGAGCGUUCACUAGUAGCGCUAGUAGGUAAAAGAAGAAGCUUACUUGAUUACUUA
Mmo-1-1    UUGUUCGAAAAACACGGUAAAGGUAAAAACGAUACCGGUUCAGCAGAAGGUCAAAUCGCAUUAUUCACAAAAAGAAUUGCGUAUUUUCUGAUCACUUGAAAACAAACAGAAAAGAUUACAACUCUGAGCGUUCACUUGUAGCGCUAGUAGGUAAAAGAAGAAGCUUACUUGAUUACUUA
env-273    AUUUUCGCUCAGUACGGGAAAUCCGCUACCGACACAGGUUCGUCAGAGGGCAGAUAGCUCUUUUCACUUAUCGAAUCA..................................................................
env-274    AUUUUCGCUCAGUACGGGAAAUCCGCUACCGACACAGGUUCGUCAGAAGGGCAGAUAGCUCUUUUCACUUAUCGAAUCAACCAUUUGACCGAACACCUUAAAAAGAACAUCCAUGAUUUCAACACGGAACGUUCGCUCGUUAAGUUAGUCGGUAAGCGCCGUUCGCUUCUUGAAUAUCUG
env-276    UUAUUUAAAAAGCACGGCAAAAAAGCCGAAAAUACUGGAAGUACUGAAGGACAGAUUGCACUAUUUACCUUCAGAAUAAAUCAUUUAACAGAACACCUUAAGCAAAAUAAAAAAGAUUUCAACACAGAAAGAUCUCUUGUAAAAUUAGUAGGAAAAGAAGAGCGUUGUUGGAUUAUCUA
env-277    UUAUUUAAAAAGCACGGCAAAAAAGCCGAAAAUACUGGAAGUACUGAAGGACAGAUUGCACUAUUUACCUUCAGAAUAAAUCAUUUAACAGAACACCUUAAGCAAAAUAAAAAAGAUUUCAACACAGAAAGAUCUCUUGUAAAAUUAGUAGGAAAAGAAGAGCGUUGUUGGAUUAUCUA
env-278    UUAUUUAAAAAGCACGGCAAAAAAGCCGAAAAUACUGGAAGUACUGAAGGACAGAUUGCACUAUUUACCUUCAGAAUAAAUCAUUUAACAGAACACCUUAAGCAAAAUAAAAAAGAUUUCAACACAGAAAGAUCUCUUGUAAAAUUAGUAGGAAAAGAAGAGCGUUGUUGGAUUAUCUA
env-280    AUAUUUAAAAAGCACGGCAAAAAAGCCGAGACACUGGAAGUACUGAAGGACAGAUUGCUCUAUUUACCUUCAGAAUAAAUCAUUUAACAGAACAUCUUAAGAAAAACAAAAAAGAUUUCAACACGGAAAGAUCUCUUGUAAAGCUAGUAGGAAAAGAAGAGCCUUAUUAGAUUAUUUA
env-281    AUAUUUAAAAAGCACGGCAAAAAAGCCAAAAACACUGGAAGUGCUGAAGGACAGAUUGCUCUAUUUACCUUCAGAAUAAAUCAUUUAACAGAACAUCUUAAAAAAACAAAAAAGAUUUUAACACGGAAAGAUCUCUUGUUAAGCUAGUAGGAAAAGAAGAGCCUUGUUAGAUUAUUUA
env-282    AUAUUUAAAAAGCACGGGAAAAAAGCCGAAAAUACUGGAAGUGCUGAAGGUCAGAUUGCCCUUUUUACAUUUAGAAUAAAUCAUUUAACUGAACAUCUUAAGAAUAAUAAGAAAGAUUUUAAUACAGAAAGAUCUCUUGUUAAGCUAGUAGGUAAAAGAAGGGCACUAUUAGACUAUUUA
env-283    AUAUUUAAAAAGCACGGGAAAAAAGCCGAAAAUACUGGAAGUGCUGAAGGUCAGAUUGCCCUUUUUACAUUUAGAAUAAAUCAUUUAACUGAACAUCUUAAGAAUAAUAAGAAAGAUUUUAAUACAGAAAGAUCUCUUGUUAAGCUAGUAGGUAAAAGAAGGGCACUAUUAGACUAUUUA
env-285    AUAUUUAAAAAGCACGGGAAAAAAGCCGAAAAUACUGGAAGUGCUGAAGGUCAGAUUGCCCUUUUUACAUUUAGAAUAAAUCAUUUAACUGAACAUCUUAAGAAUAAUAAGAAAGAUUUUAAUACAGAAAGAUCUCUUGUUAAGCUAGUAGGUAAAAGAAGGGCACUAUUAGACUAUUUA
env-286    AUAUUUAAAAAGCA..................................................................................................................................
env-287    AUAUUUAAAAAGCACGGGAAAAAAGCCGAAAAUACUGGAAGUGCUGAAGGUCAGAUUGCCCUUUUUACAUUUAGAAUAAAUCAUUUAACUGAACAUCUUAAGAAUAAUAAGAAAGAUUUUAAUACAGAAAGAUCUCUUGUUAAGCUAGUAGGUAAAAGAAGGGCACUAUUAGACUAUUUA
env-288    AUAUUUAAAAAGCACGGGAAAAAAGCCGAAAAUACUGGAAGUGCUGAAGGUCAGAUUGCCCUUUUUACAUUUAGAAUAAAUCAUUUAACUGAACAUCUUAAGAAUAAUAAGAAAGAUUUUAAUACACAAAGAUCUCUUGUUAAACUAGUAGGUAAAAGAAGAGC...............
env-289    AUAUUUAAAAAGCACGGGAAAAAAGCCGAAAAUACUGGAAGUGCUGAAGGUCAGAUUGCCCUUUUUACAUUUAGAAUAAAUCAUUUAACUGAACAUCUUAAGAAUAAUAAGAAAGAUUUUAAUACAGAAAGAUCUCUUGUUAAGCUAGUAGGUAAAAGAAGGGCACUAUUAGACUAUUUA
env-290    AUAUUUAAAAAGCACGGGAAAAAAGCCGAAAAUACUGGAAGUGCUGAAGGUCAGAUUGCCCUUUUUACAUUUAGAAUAAAUCAUUUAACUGAACAUCUUAAGAAUAAUAAGAAAGAUUUUAAUACAGAAAGAUCUCUUGUUAAGCUAGUAGGUAAAAGAAGGGCACUAUUAGACUAUUUA
env-294    AUAUUUAAAAAGCACGGGAAAAAAGCCGAAAAUACUGGAAGUGCUGAAGGUCAGAUUGCCCUUUUUACAUUUAGAAUAAAUCAUUUAACUGAACAUCUUAAGAAUAAUAAGAAAGAUUUUAAUACAGAAAGAUCUCUUGUUAAGCUAGUAGGUAAAAGAAGGGCGCUAUUAGACUAUUUA
env-295    AUAUUUAAAAAGCACGGGAAAAAAGCCGAAAAUACUGGAAGUGCUGAAGGUCAGAUCGCCCUUUUUACACAUAGAAUAAAUCAUUUAACUGAACAUCUUAAGAAUAAUAAGAAAGAUUUUAAUACACAAAGAUCUCUUGUUAAACUAGUAGGUAAAAGAAGAGCUUUAUUAGACUAUUUA
env-296    AUAUUUAAAAAGCACGGGAAAAAAGCCGAAAAUACUGGAAGUGCUGAAGGUCAGAUUGCCCUUUUUACAUUUAGAAUAAAUCAUUUAACUGAACAUCUUAAGAAUAAUAAGAAAGAUUUUAAUACAGAAAGAUCU........................
env-297    AUAUUUAAAAAGCACGGGAAAAAAGCCGAAAAUACUGGAAGUGCUGAAGGUCAGAUCGCCCUUUUUACACAUAGAAUAAAUCAUUUAACUGAACAUCUUAAGAAUAAUAAGAAAGAUUUUAAUACACAAAGAUCUCUUGUUAAACUAGUAGGUAAAAGAAG..................
env-298    ................................................................................................................................................
env-299    AUUUUCAAAAAGUUUGGAAAGUCAAAAAAAAUACAGGAUCAACUGAAGGACAAAUAGCCCUUUUUACUCACAGAAUUCAUCACUUAUCUAAUCAUCUUAAAUCUAAUAGAAAAGAUUAUAAUACUGAAAGAUCAUUAGUUCGUUUAGUAGGAAAAAGAAGAAGUCUUCUAGACUACCUG
env-300    AUUUUCAAAAAAUUUGGAAAGUCAAAAAAAAUACAGGAUCAACUGAAGGACAAAUAGCCCUUUUUACUCACAGAAUUCAUCACUUAUCUAAUCAUCUUAAAUCUAAUAGAAAAGAUUAUAAUACUGAAAGAUCAUUAGUUCGUUUAGUAGGAAAAAGAAGAAGUCUUCUAGACUAUCUG
env-301    AUUUUCAAAAAAUUUGGAAAGUCAAAAAAAAUACA.............................................................................................................
env-302    AUUUUCAAAAAAUUUGGAAAGUCAAAAAAAAUACAGGAUCACCCGAAGGACAAAUAGCCCUUUUUACCCACAGGAUUCAUCACUUAUCUAACCAUCUUAAAUCUAAUAGAAAAGAUUAUAAUACUGAAAGAUCAUUAGUUCGUUUAGUAGGAAAACGAAGAAGUCUUCUAGACUACCUA
env-303    AUUUUCAAAAAAUUCGGAAAGUCAAAAAAAAUACUGGAUCGCCUGAAGGACAAAUAGCCCUUUUUACUCACAGGAUUCAUCACUUAUCUAAUCAUCUUAAAUCUAAUAGAAAGGAUUAUAAUACUGAAAGAUCAUUAGUUCGUUUAGUUGGAAAACGAAGAAGUCUUCUAGACUACCUA
env-304    AUUUUCAAAAAAUUUGCAAGUUCAAAAACAAACACUGGAUCACCCGAAGGCCAAAUAGCACUUUUUACGCACAGAAUACAUCACCUAUCUAAUCACCUUAAAUCUAACAGAAAAGACUUCAAUACUGAAAGAUCAUUAGUUCGUUUAGUUGGUAAACGCAGAAGUCUUCUUGAUUAUUUA
env-305    ................................................................................................................................................
env-306    UUAUUCAAAAAACACGGUAAAAGUGCAAAAGAUACCGGAAUUGCUGAAGGGCAAAUUGCUUUAUUUACGCUAAGAAUAAAUCAUUUAACUGAGCACUUAAAAAAAUAACAAAAAAGAUUUCAACACUGAAAGAGCCUUAAUAAAGCUUGUUGGAAAAAGAAGAUCGCUUCUUGAUUACUUA
env-307    UUAUUCAAAAAACACGGUAAAAGUGCAAAAGAUACCGGAAUUGCUGAAGGGCAAAUUGCUUUAUUUACGCUAAGAAUAAAUCAUUUAACUGAGCACUUAAAAAAAUAACAAAAAAGAUUUCAACACUGAAAGAGCCUUAAUAAAGCUUGUUGGAAAAAGAAGAUCGCUUCUUGAUUACUUA
env-308    AUUUUCAAGAAACACGGGAAAUCAGAGAAUGACACAGGUUCUGCAGAGGGACAGAUCGCAUUGUUCACAUACAGAAUUGGUCAUUUAACGAAUCACCUGAAAGAAAAUCGAAAAGAUUUACGGUACACAAAGAGCGUUGCAGCUUCUUGUAGGUAAAAGAAGAAGUUUGUUGGAUUAUUUA
Lsp-3-1    AUUUUUAGCAAAUACGGUAAAGGUAAAAACGAUACCGGGUCUGCAGAGGGACAAAUUGCGUUAUUCACAUACCGCAUUAGCCACCUUACUGAGCAUUUAAAAACAAACCGUAAGGAUUUUAACACAGAGCGUUCAUUAGUAAUGUUGGUAGGUAAACGCAGAGCGUUGCUUGAUUACUUA
env-309    AUUUUCAGCCAGUUCGGUAAAGACAAAAACGAUACCGGUUCUGCAGAAGGACAAAUUGCGUUGUUCACACACCGCAUCAACCACCUUACAGAGCACUUAAAAAGAAACCGCAAAGACUUUAACACUGAGCGUUCUCUAGUAAAGUUAGUAGGUAAGCGUAGAGCAUUACUUGACUAUUUA
env-310    AUUUUCAGCCAGUUCGGUAAAGACAAAAACGAUACCGGAUCUGCAGAAGGACAAAUUGCGUUGUUCACACACCGCAUCAACCACCUUACAGAGCACUUAAAAAGAAACCGCAAAGACUUUAACACUGAGCGUUCUCUAGUAAAGUUAGUAGGUAAGCGUAGAGCAUUACUUGACUAUUUA
env-313    AUUUUCAGCCAGUUCGGUAAAGACAAAAACGAUACCGGAUCUGCAGAAGGACAAAUUGCGUUGUUCACACACCGCAUCAACCACCUUACAGAGCACUUAAAAAGAAACCGCAAAGACUUUAACACUGAGCGUUCUCUAGUAAAGUUAGUAGGUAAGCGUAGAGCAUUACUUGAUUAUUUA
env-314    AUUUUCAGCCAGUUCGGUAAAGACAAAAACGAUACCGGAUCUGCAGAAGGACAAAUUGCGUUGUUCACACACCGCAUCAACCACCUUACAGAGCACUUAAAAAGAAACCGCAAAGACUUUAACACUGAGCGUUCUCUAGUAAAGUUAGUAGGUAAGCGUAGAGCAUUACUUGACUAUUUA
env-315    AUUUUCAGCCAGUUCGGUAAAGACAAAAACGAUACCGGAUCUGCAGAAGGACAAAUUGCGUUGUUCACACACCGCAUCAACCACCUUACAGAGCACUUAAAAAGAAACCGCAAAGACUUUAACACUGAGCGUUCUCUAGUAAAGUUAGUAGGUAAGCGUAGAGCAUUACUUGACUAUUUA
env-316    AUUUUCAGCAAGUUCGGUAAAGAUAAAAACGAUACCGGAUCUGCAGAAGGACAGAUCGCGUUGUUUACACACCGCAUCAACCACUUAACGGGACACUUAAAAAAGAACCGCAAAGACUUUAACACAGAGCGUUCUCUAGUAAAAUUAGUUGGUAAGCGUAGAGCAUUACUUGAUUAUUUA
env-317    ................................................................................................................................................
env-318    AUUUUCAGCAAGUUCGGUAAAGACAAAAACGAUACCGGAUCUGCGGAAGGACAGAUUGCGUUGUUUACACAUCGCAUCAACCACUUAACGGGACACUUGAAAAAGAACCGCAAAGACUUUAACACAGAGCGUUCUCUAGUAAAAUUAGUUGGUAAGCGUAGAGCAUUACUUGACUAUUUA

           ................................................................................................................................................
           ................................................................................................................................................
           AUYUUYR•RAARCACGG•RRR•••R•RR••RAYAC•GG••C•RC•GAAGG•CAAAUYGC•YURUUYAC•Y•Y•G•AUYRAYCAYYUR•C•RA•CAYYURAAAAR•AAYCR•AARGAUU•YAAYAC•GAR•G•UC•YURGU•••RYURGU•GG•AAR•G•RG••••YU•YU•GAYUAYYU•
```

env-320 AUUUUCAGCAAGUUCGGUAAAGACAAAAACGAUACCGGAUCUGCGGAAGGACAGAUUGCGUUGUUUACACAUCGCAUCAACCACUUAACGGGACACUUGAAAAAGAACCGCAAAGACUUUAACACAGAGCGUUCUCUAGUAAAAUUAGUUGGUAAGCGUAGAGCAUUACUUGACUAUUUA
env-324 AUUUUCAGCAAGUUCGGUAAAGACAAGAACGAUACCGGAUCUGCAGAAGGACAAAUUGCGUUGUUCACACACCGCAUCAACCACCUUACAGAGCACUUAAAAAGAAACCGCAAAGACUUUAACACUGAGCGUUCUCUAGUAAAGUUAGUAGGUAAGCGUAGAGCAUUACUUGACUAUUUA
env-325 AUUUUCAGCAAGUUCGGUAAAGACAAGAACGAUACCGGAUCUGCAGAAGGACAAAUUGCGUUGUUCACACACCGCAUCAACCACCUUACAGAGCACUUAAAAAGAAACCGCAAAGACUUUAACACUGAGCGUUCUCUAGUAAAGUUAGUAGGUAAGCGUAGAGCAUUACUUGACUAUUUA
env-326 AUUUUCAGCAAGUUCGGUAAAGACAAAAACGAUACCGGAUCUGCGGAAGGACAGAUUGCGUUGUUUACACAUCGCAUCAACCACUUAACGGGACACUUGAAAAAGAACCGCAAAGACUUUAACACAGAGCGUUCUCUAGUAAAA.....................................
Fri-1-1 AUUUUUGCAAAACACGGCGGAGCUGCUGUUAACACAGGUUCAUCAGAAGGCCAGAUUGCAUUAUUUACAUUCAGGAUUAACCACUUAACAGGCCACCUUAAAAAAAAUCGCCACGAUUUUAAUACAGAGCGUUCGCUUGUUAAACUGGUAGGUAAAAGAAGGAGCCUUCUUGACUACCUG
env-327 AUUUUCAAGAAGCACGGAAAGUCUGAAGCAGAUACUGGUUCUGCUGAAGGGCAAAUUGCAUUAUUUUCCCAUCGAAUCGAUCACUUAUCUAACCAUCUUAAGAAUAACAAAAAGGAUUUCAAUACGGAACGAUCAUUAGUUAAACUAGUAGGUAAACGCCGUCAACUUUUAGAUUAUCUC
env-328 AUUUUCAAGAAGCACGGAAAGUCUGAAGUAGAUACUGGUUCUUCUGAAGGACAAAUUGCAUUAUUUUCCCACCGAAUCGAUCACUUAUCUAACCAUCUUAAGAAUAAUAAAAAGGAUUUCAAUACGGAACGAUCAUUAGUUAAAUUAGUAGGCAAACGCCGUCAACUUUUAGAUUAUCUC
env-329 AUUUUCAAUAAAUACGGAAAAUCUAAAUCUAACACUGGUUCUACUGAGGGUCAAAUUGCUCUUUUUUCACACCGCAUAGAACACUUAUCAGAACAUUUAAAAAACAAUAGAAAAGAUUUUAAUACCGAAAGAUCGCUUGUUGCUUUGGUUGGUAAAAGAAGAAGUCUUUUAGAUUAUUUG
env-330 AUUUUCAAUAAAUACGGAAAAUCUAAAUCUAAUACUGGUUCUACUGAGGGUCAAAUUGCUCUUUUUUCACACCGCAUAGAUCAUUUAUCAGAACAUUUAAAAAACCAAUAGAAAAGAUUUUAAUACCGAAAGAUCACUUGUUGCUUUGGUUGGUAAAAGAAGAAGUCUUUUAGAUUAUUUG
env-331 AUUUUUAAAAAGCACGGUGGUGCUGCCUCUAAUACAGGCUCUACAGAAGGUCAAAUAGCCUUGUUUACCCACAGGAUCAAGCACUUAACUGAUCACUUAAAAUCAAACCGUAAAGAUUUUGUAACCCAACGAGCAGUUCAAUUACUAGUUGGUAAAAGAAGAAGUUUGUUGGAUUAUCUU
env-332 AUUUUUAAAAAGCACGGUGGUGCUACCUCUAAUACAGGUUCUACAGAAGGUCAAAUAGCAUUGUUUACUCACAGAAUUAAGCAUUUAACUGGCCACCUAAAAUCAAAUCGUAAAGAUUUUGUAACUCAACGAGCAGUUCAAUUACUAGUUGGUAAGAGAAGAAGUUUGUUGGAUUAUCUU
env-333 AUUUUUAAAAAGCACAGUGGUGCUUCCUCUAACACAGGUUCAACAGAAGGUCAAAUAGCGUUGUUUACUCAUAGAAUUAAGCACUUAACAGGCCACUUAAAAGAAAAUCGUAAAGAUUUUGUAACACAACGAGCAGUUCAAUUACUAGUUGGUAAGAGAAGAAGUUUGUUGGAUUAUCUU
env-334 AUUUUUAAAAAACACGGUAAAUCUGAAAAGGAUACGGGUUCAUCAGAAGGACAAAUAGCAUUAUUCACUCACCGAAUCGAACAUCUAUCAAACCACUUGAAAUCGAAUCGUAAAGAUUUUAACACGGAACGCUCACUUGUCCGAUUGGUUGGAAAGCGCCGUAGUCUGUUAGAUUAUUUA
env-335 ........................................................................................................................................................................................................................................
env-336 UUAUUUGCCAAGCACGGUAAAACGAAAAAGAUACCGGCUCAACCGAAGGACAGAUUGCAUUGAUGACCAAGCGCAUCCAACACAUGUCCGAUCACUUAAAGAAAAACACCAAAGACUUUAAUUCUGAAAGAGCUUUGGUAAGAACAGUUGGUAAACGAAGAGAUUUACUUAACUACUUAA
env-337 UUAUUUAAAAAGCACGGUAAAAACGAAAAAGAUACCGGGUCAACAGAAGGACAAAUCGCUUUAAUGACAAAGCGUAUCCAACACAUGUCUGAUCACUUUGAAGAAAAAUCCUAAAGAUUUUAAUUCAGAAAGAACACUUGUAAAGACUGUUGGUAGACGAAGAAAUCUACUAAAUUAUCUU
env-338 AUUUUCAAAAAACAUGGUAAAUCUGAAAAAGAUACAGGUUCACCGGAAGGACAAAUAGCACUCUUCACUCAUAGAAUCAACCACCUAUCAGAACACCUUAAAUCAAAUCGUAAAGACUUUAACACCGAAAGAUCUUUAGUUCGUUUAGUUGGGAGAGAAAAGCCUUCUUGACUAUCUAA
env-339 AUUUUCAAAAAACAUGGUAAAUCUGAAAAAGAUACAGGUUCACCGGAGGGACAAAUAGCGCUCUUCACUCAUAGAAUCCAACACCUAUCAGAACACCUUAAAUCAAAUCGUAAAGACUUUAAUACAGAAAGAUCUUUAGUUCGUUUAGUUGGGAAGAGAAGAAGCAUUCUUGACUACCUA
env-340 AUUUUCAAAAAACAUGGUAAAUCUGAAAAAGAUACAGGUUCACCGGAGGGACAAAUAGCGCUCUUCACUCAUAGAAUCCAACACCUAUCAGAACACCUUAAAUCAAAUCGUAAAGACUUUAAUACAGAAAGAUCUUUAGUUCGUUUAGUUGGGAAGAGAAGAAGCAUUCUGACUAUCUAA
env-341 AUUUUCAAAAAACAUGGUAAAUCUGAAAAAGAUACAG.......................................................................................................................................................................................................
env-342 AUUUUCAAAAAACAUGGUAAAUCUGAAAAAGAUACAGGUUCACCGGAGGGACAAAUAGCGCUCUUCACUCAUAGAAUCCAACACCUAUCAGAACACCUUAAAUCAAAUCGUAAAGACUUUAAUACAGAAAGAUCUUUAGUUCGUUUAGUUGGGAAGAGAAGAAGCAUUCUUGACUACCUA
env-343 AUUUUCAAAAAACAUGGUAAAUCUGA..................................................................................................................................................................................................................
env-344 AUUUUC......................................................................................................................................................................................................................................
env-345 AUAUUCAAAAAACAUGGUAAAUCUGAAAAAGAUACAGGUUCACCAGAGGGACAAAUAGCGCUCUUCACUCAUAGAAUCCAACACCUAUCAGAACACCUUAAAUCAAAUCGUAAAGACUUUAACACUGAAAGAUCUUUAGUCCGUUUAGUUGGGAAGAGAAGAAGCCUUCUUGACUAUCUA
env-346 AUUUUCAAAAAACAUGGUAAAUCUGAAAAAGAUACAGGUUCACCAGAGGGACAAAUAGCGCUCUUCACUCAUAGAAUCCAACACCUAUCAGAACACCUUAAAUCAAAUCGUAAAGAUUUUAAUACAGAAAGAUCUUUAGUUCGUUUAGUUGGGAAGAGAAGAAGCAUUCUUGACUAUCUA
env-347 AUUUUCAAAAAACAUGGUAAAUCUGAAAAAGAUACAGGUUCACCGGAGGGACAAAUAGCGCUCUUCACUCAUAGAAUCCAACACCUAUCAGAACACCUUAAAUCAAAUCGCAAAGACUUUAAUACAGAAAGAUCUUUAGUUCGUUUAGUUGGGAAGAGAAGAAGCAUACUUGACUAUCUA
env-352 AUUUUCAAAAAACAUGGUAAAUCUGAAAAAGAUACAGGU.....................................................................................................................................................................................................
env-353 AUUUUCAAAAAACAUGGUAAAUCUGAAAAAGAUACAGGUUCACCGGAGGGACAAAUAGCGCUCUUCACUCAUAGAAUCCAACACCUAUCAGAACACCUUAAAUCAAAUCGUAAAGAUUUUAAUACAGAAAGAUCUUUAGUUCGUUUAGUUGGGAAGAG.....................
env-354 AUUUUCAAAAAACAUGGUAAAUCUGAAAAAGAUACAGGUUCACCGGAGGGACAAAUAGCGCUCUUCACUCAUAGAAUCCAACACCUAUCAGAACACCUUAAAUCAAAUCGUAAAGACUUUAAUACAGAAAGAUCUUUAGUUCGUUUAGUUGGGAAGAGAAGAAGCAUUCUUGACUAUCUA
env-355 AUUUUCAAAAAACAUGGUAAAUCUGA..................................................................................................................................................................................................................
env-356 AUUUUCAAAAAACAUGGUAAAUCUGAAAAAGAUACAGGUUCACCGGAGGGACAAAUAGCGCUCUUCACUCAUAGAAUCCAACACCUAUCAGAACACCUUAAAUCAAAUCGUAAAGAUUUUAAUACAGAAAGAUCUUUAGUUCGUUUAGUUGGGAAGAGAAGAAGCAUUCUUGACUAUCUA
env-357 AUCUUCGCUAAACACGGAGGUUCAGCAACAAACACAGGUUCAACAGAAGGUCAGGUAGCAUUGUUUACUUACCGUAUUGCUCACUUAACAGGGCAUUUGAAAAAGAACCGUAAAGAUUAUGGAACACAAAGAUCUUUGCAAUUGUUGGUUGGUAAACGUCGUAGUAUGUUAGAUUACCUU
Fta-1-1 AUCUUCGCAAAACACGGAGGUUCAGCAACAAACACAGGUUCAACAGAAGGUCAAGUAGCAUUAUUUACUUACCGUAUUGCUCACUUAACAGGGCAUUUGAAAAAGAACCGUAAAGAUUAUGGAACUCAAAGAUCUUUACAAUUGUUAGUUGGUAAACGUCGUAGUAUGUUAGAUUACCUG
env-358 AUCUUCACAAAACACGGUAACGGAGCAAAAGAUACUGGUUCUGCUGAAGGACAAAUCGCUUUAUUCACGUACAGAAUUAGCCAUUUAACAGAACACUUAAAAAAGAAUCGUAAAGAUUAUAAUACUGAGCGUUCAUUAGUAAAAUUAGUAGGUAAAAGAAGAGCUUUACUUGACUAUUUA
env-359 ........................................................................................................................................................................................................................................
env-360 AUGUUUGCUCAAUUUGGCAAGAACGAACAAGACACAGGUUCUGCUGAAGGUCAAAUCGCUUUGAUGACCAAAAGAAUUGAUCACAUUUCAAAGCAUUUAAAAACAAACCGAAAAGAUUUUAAUUCUGAACGCUCGCUAGUUAAGCUUGUUGGUAAAAGAAGAAAUCUUUUAAAUUAUCUU
env-361 UUAUUUGCUAAACACGGCAAAAACGAAAAAGACACCGGUUCUGCAGAAGGGCAAAUUGCUUUAUUAACAGCAAGAAUUCUUCAUAUUUCAGAACACAUGAAACAAAACAAGAAAGAUUUUAAUUCUAAAAAAUCACUUGUAAAAUUAGUGGUAAGCGUAGAAACCUACUCAACUACUUA
env-362 UUAUUUGCUAAACACGGCAAAAACGAAAAGGACACAGGUUCUGCUGAAGGACAAAUUGCAUUAUUAACAACAAGAAUUCUUCACAUUUCAGAACACAUGAAACAAAACAAGAAAGAUUUUAAUUCUAAAAAAUCACUUGUAAAAUUAGUUGGUAAGCGAAGAAACUUACUUAACUACCUA
Mod-7-1 AUCUUCACACAAUACGGUGGAGCUGCAGCUAACACUGGAUCUGCUGAAGGACAAAUCGCUUUGUUCACAUACAGAAUCACGCAUUUAACUGAGCACUUGAAAAAAAAUCGUAAAGAUUAUAAUACAGAGCGUUCAUUAGUUCUUUUAGUAGGUAAAAGAAGAAGACUUCUUGACUACUUA
Mod-6-1 AUCUUCGCACACAACACGGUGGAGCUGCAGCUAACACUGGAUCUGCUGAAGGACAAAUCGCUUUGUUCACAUACAGAAUCACGCAUUUAACUGAGCACUUGAAAAAAAAUCGUAAAGAUUAUAAUACAGAGCGUUCAUUAGUUCUUUUAGUAGGUAAAAGAAGAAGACUUCUUGACUACUUA
Mod-3-1 AUCUUCGCACACAACACGGUGGAGCUGCAGCUAACACUGGAUCUGCUGAAGGACAAAUCGCUUUGUUCACAUACAGAAUCACGCAUUUAACUGAGCACUUGAAAAAAAAUCGUAAAGAUUAUAAUACAGAGCGUUCAUUAGUUCUUUUAGUAGGUAAAAGAAGAAGACUUCUUGACUACUUA
Mpr-1-1 AUCUUCGCACACAACACGGUGGAGCUGCAGCUAACACUGGAUCUGCUGAAGGACAAAUCGCUUUGUUCACAUACAGAAUCACGCAUUUAACUGAGCACUUGAAAAAAAAUCGUAAAGAUUAUAAUACAGAGCGUUCAUUAGUUCUUUUAGUAGGUAAAAGAAGAAGACUUCUUGACUACUUA
Mod-4-1 AUCUUCGCACACAACACGGUGGAGCUGCAGCUAACACUGGAUCUGCUGAAGGACAAAUCGCUUUGUUCACAUACAGAAUCACGCAUUUAACUGAGCACUUGAAAAAAAAUCGUAAAGAUUAUAAUACAGAACGUUCAUUAGUUCUUUUAGUAGGUAAAAGAAGAAGACUUCUUGACUACUUA
Mod-1-1 AUCUUCGCUAAACACGGAGGAAAAGCUGAAAACACUGGAUCUGCUGAAGGACAAAUCGCUUUGUUCACAUUCAGAAUUCACACUUAACUGAGCACUUAAAAAGAAAUCGUAAAGAUUACAAUACUGAGCGUUCAUUAGUUCUUUUAGUAGGUAAAAGAAGAAGACUUCUUGACUACUUA
Fin-1-1 AUUUUUGCAACUCAUGGUGGAGCUGCAACUAACACAGGUUCAACAGAAGGACAAAUCGCAUUGUUUACUUUCAGAAUCAACCACUUAACUGAGCACUUAAAAAGAAAUCGUCAUGAUUACAACACUGAGCGUUCAUUAGUGAAGUUGGUAGGAAAAAGAAGAAGUCAAUUAGACUACUUA
env-363 CUCUUCGCUAAACACGGAGGAAAAGCAGAAAACACAGGAUCUGCAGAGGCCAAAUCGCCUUGUUUACAUUCAGAAUCAACCACUUGACUGAGCACUUGAAAAAAAAUCGUCACGAUUACAACACCGAGCGUUCGUUGGUAAUGUUGGUGGAGGAAAAGACGUAGUUUGUUGGAUUACUUG
env-364 CUCUUCGCUAAACACGGAGGAAAAGCAGAAAACACAGGAUCUGCAGAAGGCCAAAUCGCCUUGUUUACAUUCAGAAUCAACCACUUGACUGAGCACUUGAAAAAAAAUCGUCACGAUUACAACACCGAGCGUUCGUUGGUAAUGUUGGUAGGAAAAAGACGUAGUUUGUUGGAUUACUUA
env-366 CUCUUCGCUAAACACGGAGGAAAAGCAGAAAACACAGGAUCUGCAGAAGGCCAAAUCGCCUUGUUUACAUUCAGAAUCAACCACUUGACUGAGCACUUGAAAAAAAAUCGUCACGAUUACAACACCGAGCGUUCGUUGGUAAUGU................................
env-367 CUCUUCGCUAAACACGGAGGAAAAGCAGAAAACACAGGAUCUGCAGAAGGCCAAAUCGCCUUGUUUACAUUCAGAAUCAACCACUUGACUGAGCACUUGAAAAAAAAUCGUCACGAUUACAACACCGAGCGUUCGUUGGUAAUGUUGGUAGGAAAAAGACGUAGUUUGUUGGAUUACUUA
env-368 CUCUUCGCUAAACACGGAGGAAAAGCAGAAAACACAGGAUCUGCAGAAGGCCAAAUCGCCUUGUUUACAUUCAGAAUCAACCACUUGACUGAGCACUUGAAAAAAAAUCGUCACGAUUACAACACCGAGCGUUCGUUGGUAAUGUUGGUAGGAAAAAGACGUAGUUUGUUGGAUUACUUA
env-369 AUCUUUAAAGAGCACGGUGGCUCUGAUAAAAACACCGGUUCUGCAACUGAAGGUCAAAUCGCGUUAUUCACAUACCGCAUCUCGCAUUUAACCAAUCACCUUAAAGAGAACAGAAAAGACAUGAAUACCAAAAGAUCGCUCGUCGGUUAGUGGUAGGUAAAAGAAGAAGAUCUUUUAGAUUACCUC
env-373 AUAUUCAAGAAGCAUGGUAAAUCUAAGAAAGAUACUGGAUCUACCGAGGGACAAAUAGCUCUUUUUACUCAUCGUAUUCAACACCUUUCAAAACACCUUGAAGAAAAUAGAAAAGAUUUCAAUACAGAACGUUCACUAGUUCGUUUGGUUGGAAAACGUCGAAGUCUUUUAGAUUAUUUA
env-375 ........................................................................................................................................................................................................................................
env-376 AUCUUUACAAAACACGGAAAAGAUGCCAAAGACACCGGUUCUGCCGAAGGUCAAAUUGCAUUAUUUACACACCGAAUCAAUCACUUAACAGAACACCUAAAGCAAAAUCGAAAGAUUUUAAUACCGAGCGUUCAUUAGUAAAACUAGUAGGAAAACGUCGUUCUCUUUUGGAUUAUUUA
env-377 AUCUUUAAAAAAUACGGAAAAGAUGCUAAAAAUACCGGUUCUGCUGAAGGCCAAAUUGCAUUAUUUACACACCGUAUCAAUCACCUCACCGAUCACUUAAAGCAAAACCGAAAAGACUUUAAUACAGAGCGUUCGUUGGUAAAGCUUGUAGGAAAGCGUCGAUCUCUAUUGGAUUAUUUA
env-378 AUCUUUAAAAAACAUGGUGGAGAUGUGAAAAAUACUGGCUCAACGGAGGGUCAAAUCGCAUUGUUCACUCACAGAAUUAAUCACCUCACUGAUCAUUUAAAGCAAAAUCGUAAAGAUUUAAUACACAGAACGUUCGCUUGUAAAGUUGGUGGUAGGUAAAACGCAGGUCUUUAUUAGAUUACCUU
env-379 CUCUUUAAAAAACACGGUAAAGAUGCAAAGGACACCGGAUCUACGGAAGGCCAAAUUGCAUUGUUUACCCACCGAAUCAAUCACCUUACAGGACACUUAAAGCAAAAUCGAAAGAUUUUAACACAGAAAGAUCGCUUGUAAAGCUUGUAGGAAAACGCCGCUCAUUGUUGGAUUACCUU
env-381 CUCUUUAAAAAACACGGUAAAGAUGCAAAGGACACCGGAUCUACGGAAGGCCAAAUUGCAUUGUUUACCCACCGAAUCAAUCACCUUACAGGACACUUAAAGCAAAAUCGAAAGAUUUUAACACAGAAAGAUCGCUUGUAAAGCUUGUAGGAAAACGCCGCUCAUUGUUGGAUUACCUU
env-382 CUCUUUAAAAAACACGGUAAAGAUGCAAAGGACACCGGAUCUACGGAAGGCCAAAUUGCAUUGUUUACCCACCGAAUCAAUCACCUUACAGGACACUUAAAGCAAAAUCGAAAGAUUUUAACACAGAAAGAUCGCUUGUAAAGCUUGUAGGAAAACGCCGCUCAUUGUUGGAUUACCUU
env-383 CUCUUUAAAAAACACGGUAAAGAUGCAAAGGACACCGGAUCUACGGAAGGCCAAAUUGCAUUGUUUACCCACCGAAUCAAUCACCUUACAGGACACUUAAAGCAAAAUCGAAAGAUUUUAACACAGAAAGAUCGCUUGUAAAGCUUGUAGGAAAACGCCGCUCAUUGUUGGAUUACCUU
env-384 CUCUUUAAAAAACACGGUAAAGAUGCAAAGGACACCGGAUCUACGGAAGGCCAAAUUGCAUUGUUUACCCACCGAAUCAAUCACCUCACAAAACACUUAAAGCAAAAUCGAAAGAUUUUAACACAGAAAGAUCGCUUGUAAAGCUUGUAGGAAAACGCCGCUCAUUGUUGGAUUACCUU
env-385 CUCUUUAAAAAACACGGUAAAGAUGCAAAGGACACCGGAUCUACGGAAGGCCAAAUUGCAUUGUUUACCCACCGAAUCAAUCACCUUACAGGACACUUAAAGCAAAAUCGAAAGAUUUUAACACAGAAAGAUCGCUUGUAAAGCUUGUAGGAAAACGCCGCUCAUUGUUGGAUUACCUU
env-386 CUCUUUAAAAAACACGGUAAAGAUGCAAAGGACACCGGAUCUACGGAAGGCCAAAUUGCAUUGUUUACCCACCGAAUCAAUCACCUUACAGGACACUUAAAGCAAAAUCGAAAGAUUUUAACACAGAAAGAUCGCUUGUAAAGCUUGUAGGAAAACGCCGCUCAUUGUUGGAUUACCUU
env-387 CUCUUUAAAAAACACGGCAAAGAUGCAAAGGACACCGGAUCUACGGAAGGCCAAAUUGCAUUGUUUACCCACCGAAUCAAUCACCUUACAGGACACUUAAAGCAAAAUCGAAAGAUUUUAACACAGAAAGAUCGCUUGUAAAGCUUGUAGGAAAACGUCGCUCAUUGUUGGAUUACCUU
env-388 CUCUUUAAAAAACACGGUAAAGAUGCAAAGGACACCGGAUCUACGGAAGGCCAAAUUGCAUUGUUUACCCACCGAAUCAAUCACCUUACAGGACACUUAAAGCAAAAUCGAAAGAUUUUAACACAGAAAGAUCGCUUGUAAAGCUUGUAGGAAAACGCCGCUCAUUGUUGGAUUACCUU
env-389 CUCUUUAAAAAACACGGGAAUGAUGCAAAGGACACCGGAUCUACCGAAGGCCAAAUUGCAUUGUUUACCCACCGAAUUAAUCACCUUACAGGACACUUAAAGCAAAAUCGAAAGAUUUUAACACAGAAAGAUCGCUUGUAAAGCUUGUAGGAAAACGCCGCUCAUUGUUGGAUUACCUU
env-390 CUCUUUAAAAAACACGGUAAAGAUGCAAAAGAUACCGGUUCUGCGGAAGGCCAAAUUGCAUUGUUCACUCACCGAAUCAAUCACCUUACAGGACACUUAAAGCAAAAUCGAAAGAUUUUAACACCGAAAGGUCGCUUGUAAAGCUUGUAGGAAAACGCCGCUCAUUACUAGAUUACCUU
env-391 CUCUUUAAAAAACACGGUAAAGAUGCAAAAGAUACCGGUUCUGCGGAAGGCCAAAUUGCAUUAUUCACUCACCGAAUCAAUCACCUUACAGGACACUUAAAGCAAAAUCGAAAGAUUUUAACACCGAAAGGUCGCUUGUAAAGCUUGUAGGAAAACGCCGCUCAUUACUAGAUUACCUU
env-392 CUCUUUAAAAAACAUGGUAGAGAUGCAAAAGAUACCGGUUCUGCGGAAGGCCAAAUUGCAUUGUUCACUCACCGAAUCAAUCACCUUACAGGACACUUAAAGCAAAAUCGAAAGAUUUUAACACCGAAAGGUCGCUUGUAAAGCUUGUAGGAAAACGCCGCUCAUUACUAGAUUACCUU
env-393 AUCUUUAAAAAACAUGGAAAAGAUGCGAAAAAUACAGGUUCCUCAGAGGGACAAAUCGCAUUGUUUACCCACAGAAUUAAUCACCUAACUGAGCACCUUAAGAACAACCGAAAAGACUACAACACAGAGCGAUCUCUUGUAAAGCUUGUAGGAAAGCGUCGUUCCUUGUUGGACUAUCUC
env-394 AUCUUUAAAAAACAUGGAAAAGAUGCGAAAAACACAGGUUCCUCAGAGGGACAAAUCGCAUUGUUUACCCACAGAAUUAAUCACCUAACUGAGCACCUUAAGAACAACCGAAAAGACUACAACACAGAGCGAUCUCUUGUAAAGCUUGUAGGAAAGCGUCGUUCCUUGCUGGACUAUCUC
env-395 AUCUUUAAAAAACAUGGAAAAGAUGCGAAAAACACAGGUUCCUCAGAGGGACAAAUCGCAUUGUUUACCCACAGAAUUAAUCACCUAACUGAGCACCUUAAGAACAACCGAAAAGACUACAAUACAGAGCGAUCUCUUGUAAAGCUUGUAGGAAAGCGUCGUUCCUUACU..........
env-396 AUCUUUAAAAAACACGGAAAGUCUAAAAAGAUACUGGAUCAGCAGAGGACAAAUCGCUUUAUUUACUCAUCGUAUCAAUCACCUAUCGGG.....................................................................................................
env-397 AUCUUUAAAAAACACGGGAAGUCUAAAAAGAUACUGGAUCAGCAGAAGGACAAAUUGCUUUAUUUACUCAUCGUAUCAAUCACCUAUCGGGUCACUUAAAAAACAAAUCGUAAAGACUACAACACGGAACGAUCUCUAGUAAAUCUAGUAGGAAAACGUCUUAGUCUAUUAGAUUAUUUG
env-398 AUCUUUAAAAAACACGGGAAGUCUAAAAAGAUACUGGAUCAGCAGAGGACAAAUUGCUUUAUUUACUCAUCGUAUCAAUCACCUAUCGGGUCACUUAAAAAACAAAUCGUAAAGACUACAACACGGAACGAUCUCUAGUAAAUCUAGUAGGAAAACGUCGUAGUCUAUUAGAUUAUUUG
env-399 AUCUUUAAAAAACACGGGAAGUCUAAAAAGAUACUGGAUCAGCAGAAGGACAAAUUGCUUUAUUUACUCAUCGUAUCAAUCACCUAUCGGGUCACUUAAAAAUAAAUCGUAAAGAUUACAAUACGGAACGAUCUCUAGUGAAUCUAGUAGGAAAACGUCGUAGUCUAUUAGAUUAUUUG
env-400 AUCUUUAAAAAACACGGGAAGUCUAAAAAGAUACUGGAUCAGCAGAAGGACAAAUUGCUUUAUUUACUCAUCGUAUCAAUCACCUAUCGGGUCACUUAAAAAACAAAUCGUAAAGACUACAACACGGAACGAUCUCUAGUAAAUCUAGUAGGAAAACGUCGUAGUCUAUUAGAUUAUUUG
env-402 AUUUUCAAAAAAUUUGGAAAUUCAAAAACAAACACUGGUUCACCUGAAGGCCAAAUAGCACUUUUUACCCACAGAAUACACCACCUAUCUAAUCACCUUAAAUCUAACAGAAAAGACUUCAACACUGAAAGAUCAUUAGUUCGUUUAGUUGGUAAACGCAGAAGUCUCCUUGAUUAUUUA
env-403 AUUUUCAAAAAAUUUGGAAAUUCAAAAACAAACACUGGAUCACCCGAAGGCCAAAUAGCACUUUUUACUUACA...................................................................................................................
env-404 AUUUUCAAAAAAUUUGGAAAUUCAAAAACAAACACUGGAUCACCCGAAGGCCAAAUAGCACUUUUUACCCACAGAAUACACCACCUAUCUAAUCACCUUAAAUCUAACAGAAAAGACUUCAACACUGAAAGAUCAUUAGUUCGUUUAGUUGGUAAACGCAGAAGUCUCCUUGAUUAUUUA
env-405 AUUUUUAAAAAACACGGUAAAUCAGAAAAUGAUACUGGUUCUACAGAAGGACAAAUCGCACUUUUCACUCACCGUAUAAACCACUUAUCAAAACACUUAGAGGGCAAUCGUAAAGAUUUUAAUACCGAACGUUCUUUAGUUCGUUUAGUAGGAAAAAGAAGGAGUUUACUAGACUACCUA
env-406 AUUUUUAAAAAACACGGUAAAUCAGAAAUGAUACUGGUUCUGCAGAAGGGCAAAUCGCACUUUUCACUCACCGUAUAGAUCACUUAUCAAAACACUUAGAGGGCAAUCGUAAAGAUUUUAAUACCGAACGCUCUUUAGUUCGUUUAGUAGGAAAAAGAAG..................
env-407 AUUUUCAAAAAAUUUGGAAAUUCAAAAACAAACACUGGUUCACCUGAAGGCCAAAUAGCUCUUUUUACUCACAGAAUACAUCACCUUUCUAAUCACCUUAAAUCUAACAAAAAGGACUUCAAUACUGAGAGAUCAUUAGUUCGUUUAGUUGGUAAACGCAGAAGUCUACUAGAUUAUUUA
env-408 ........................................................................................................................................................................................................................................

........................................................................................................................................................................................................................................
........................................................................................................................................................................................................................................
AUYUUYR•RAARCACGG•RRR•••R•RR••RAYAC•GG••C•RC•GAAGG•CAAAUYGC•YURUUYAC•Y•Y•G•AUYRAYCAYYUR•C•RA•CAYYURAAAAR•AAYCR•AARGAUU•YAAYAC•GAR•G•UC•YURGU•••RYURGU•GG•AAR•G•RG••••YU•YU•GAYUAYYU•

```
env-409  AAUUUUCAAAAAAUUUUGGAAAUUCAAAAACAAACACUGGUUCACCUGAAGGCCAAAUAGCUCUUUUUACUCACAGAAUACAUCACCUUUCUAAUCACCUUAAAUCUAACAAAAAGGACUUCAAUACUGAGAGAUCAUUAGUUCGUUUAGUUGGUAAACGCAGAAGUCUACUAGAUUAUU
env-410  UUUUCAAAAAUUUGGUAAAUUCAAAAACAAACACUGGUUCACCUGAAGGCCAAAUAGCUCUUUUUACUCACAGAAUACAUCACCUUUUCUAAUCACCUUAAAUC..........................................................................................
env-411  AUUUUCAAAAAAUUCGGAAAUUCAAAAAAAAAUACUGGAUCACCCGAAGGCCAAAUAGCGCUUUUUACACAUAGGAUUCAUCACUUAUCUAAUCAUCUUAAAUUUAAUCGAAAAGAUUUCAAUACAGAAAGAUCAUUAGUUCGUUUAGUAGGUAAGCGACGAAGUCUUCUAGAUUACUUA
env-412  AUUUUCAAAAAAUUCGGAAAUUCAAAAAAAAAUACUGGAUCACCCGAAGGCCAAAUAGCGCUUUUUACACAUAGGAUUCAUCACUUAUCUAAUCAUCUUAAAUUUAAUCGAAAAGAUUUCAAUACAGAAAGAUCAUUAGUUCGUUUAGUAGGUAAGCGACGAAGUCUUCUAGAUUACUUA
env-413  AUUUUCAAAAAAUUCGGAAAUUCAAAAAAAAAUACUGGAUCACCCGAAGGUCAAAUAGCGCUUUUUACACAUAGGAUUCAUCACUUAUCUAAUCAUCUUAAAUUUAAUCGAAAAGAUUUCAAUACAGAAAGAUCAUUAGUUCGUUUAGUAGGUAAGCGACGAAGUCUUCUAGAUUACUUA
env-414  AUUUUCAAAAAAUUCGGAAAUUCAAAAAAAAAUACUGGAUCACCCGAAGGCCAAAUAGCGCUUUUUACACAUAGGAUUCAUCACUUAUCUAAUCAUCUUAAAUUUAAUCGAAAAGAUUUCAAUACAGAAAGAUCAUUAGUUCGUUUAGUAGGUAAGCGACGAAGUCUUCUAGAUUACUUA
env-415  ..........................................................................................................................................................................................
env-416  AUAUUUAAGAAAUACGGGAAAUCUAAAUCAAAUACUGGAUCUACAGAGGGUCAAAUUGCUCUUUUCUCUCAUCGAAUAAAUCAUCUAUCUGAACAUUUGAAAAAAAACAGAAAAGACUUUAAUACUGAAAGAUCGCUGGUAGGUUUAGUUGGUAAAAGAAGAAGUCUUCUUGAUUAUUUG
env-417  AUAUUUAAGAAAUUCGGGAAAUCUAAAUCAAAUACUGGAUCUACAGAGGGUCAAAUUGCUCUUUUCUCUCAUCGAAUAAAUCAUCUAUCUGAACAUUUGAAAAAAAACAGAAAAGACUUUAAUACUGAAAGAUCGCUGGUAGGUUUAGUUGGUAAAAGAAGAAGUCUUCUUGAUUAUUUG
env-418  AUUUUUACCAAAUAUGGAAAAUCAAAAUCUAACACUGGGUCUACAGAGGUCAAAUCGCCCUCUUCUCCCAUCGAAUAAAUCAUUUAUCGGAACAUUUGAAAAAUAAUAGAAAAGAUUUUAACACUGAAAGAUCUCUUGUAGGUCUUGUGGUAAAAGAAGAAGUCUUCUUGAUUAUUUA
Fbe-1-1  AUCUUUACAAAACACGGUAAAUCUGCAACUGACACAGGAACUGCAGAAGGACAGAUUGCGUUAUUCACAUUCAGAAUUAACCACUUAACUGAGCACUUAAAAAAGAAUCGUCACGAUUUUAACACUGAGCGUUCUCUUGUAAGCUAGUAGGUAAAAGAAGAAGUCUUCUGGAUUACCUU
env-419  CUUUUUGAAAAACACGGUAAAGGAAAAACGAUACUGGCUCUGCCGAAGGUCAGAUCGCUUUAUUUACGCACCGUAUCAAUCACUUGACUGAGCACUUAAAAAACAAUCGUAAAGAUUAUAAUACAGAGCGUUCACUAGUAAAACUCGUAGGUAAGCGUCGUUCUUUAUUGGACUACCUU
Ksp-1-1  CUUUUUGCAAAACACGGUAAAGGAAAGCACGAUACAGGAUCUGCUGAAGGUCAAAUCGCAAUCUUUACACAACGUAUCUCACACUUAUCAAACCACCUGAAAACAAAUCGUAAAGAUUAUGUUACAGAGCGUUCACUAGUAAUGCUAGUAGGUAAGCGUCGUUCAUUACUUGAUUACUUA
Dsp-2-1  CUUUUUGCAAAGCACGGUAAAGGGAAAAACGAUACAGGAUCUGCUGAGGGUCAAAUCGCAAUCUUUACACAACGUAUCUCACACUUAUCUAACCACCUGAAAACAAAUCGUAAAGAUUAUGUUACAGAGCGUUCACUAGUAAUGCUAGUAGGUAAGCGUCGUUCUUUACUUGAUUACUUA
Dsp-1-1  AUUUUUGCAAAGCACGGAAAAGGGAAAAACGAUACUGGAUCUGCAGAGGGUCAAAUCGCUUUAUUCACGCACCGUAUCACACACUUAACUGAGCACUUAAAGAAAAACCGCAAGGAUUAUAACACUGAGCGUUCACUAGUAAUGCUGGUAGGUAAGCGUCGUUCUUUACUUGAUUACUUA
Ddo-1-1  AUUUUUGCAAAGCACGGAAAAGGUAAAAACGAUACAGGAUCUGCAGAGGGUCAAAUCGCUUUAUUCACGCACCGUAUCACUCACUUAACUGAGCACUUAAAGAAAAACCGUAAGGAUUUUAACACAGAGCGUUCACUAGUAAUGCUGGUAGGUAAGCGUCGUUCUUUACUUGAUUACUUA
Iha-1-1  AUUUUUGCUAAACACGGUGGAGCAGCAUCUAACACAGGUUCUGCAGAGGACAAAUUGCUCUUUUUCACACAGAAUUAGUCAUUUAACUGAGCACUUGAAAAGAAAUCGUAAAGAUUACAAUACAGAGCGUUCAUUGGUUAAAUUGGUAGGUAAAAGAAGAAGCCUACUCGAUUAUUUG
env-420  AUCUUCAAAAAGUACGGUAAGUCUGAAAAAGACACCGGAAGCACAGAAGGACAAAUCGCUUUAUUUACAAACCGAAUCAAUUUCCUAACAGGACACUUGAAAAAAAUCAUAAAGAUUAUAAUACCGAGCGUUCGUUAGUUAUGUUGGUAGGUAAAAGAAGAAGUCUUUUGGACUAUUUA
env-421  UUCAAGAAGUACGGAAAGUUUGAAAAAAACACCGGAAGUACGGAAGGACAAAU.....................................................................................................................................
env-422  UUCAAAAAGUACGGAAAGUCUGAAAAAGACACUGGAAGCACUGAAGGACAAAUUGCACUAUUCACAAAGCGUAUCAACGAUCUAACAGGACACCUGAAAAACAAUCAUAAAGAUUACAACACAGAGCGUUCAUUGGUAAUGUUGGUAGGUAAACGAAGAAGCCUUUUAGACUAUCUAAAA
env-423  ..........................................................................................................................................................................................
env-424  AUCUUCAAAAAGUACGGUAAGUCUGAAAAAGACACCGGAAGCACAGAAGGACAAAUCGCUUUAUUUACAAACCGAAUCAAUUUCCUAACAGGACACUUGAAAAAAAAUCAUAAAGAUUAUAAUACCGAGCGUUCGUUAGUUAUGUUGGUAGGUAAAAGAAGAAGUCUUUUGGACUAUUUA
env-427  AUCUUCAAAAAGUACGGUAAGUCUGAAAAAGACACCGGAAGCACAGAAGGACAAAUCGCUUUAUUUACAAACCGAAUCAAUUUCCUAACAGGACACUUGAAAAAAAAUCAUAAAGAUUAUAAUACCGAGCGUUCGUUAGUUAUGUUGGUAGGUAAAAGAAGAAGUCUUUUGGACUAUUUA
env-428  AUCUUCAAAAAGUACGGUAAGUCUGAAAAAGACACGGGCAGUACAGAAGGACAGAUUGCAUUAUUUACAAACCGAAUCAACUUCCUAACGGGACACUUGAAAAAAAAUCAUAAAGAUUACAAUACGGAGCGUUCGUUAGUUAUGUUGGUAGGUAAAAGAAGAAGUCUUUUGGAUUAUUUA
env-429  AUCUUCAAAAAGUACGGUAAGUCUGAAAAAGACACCGGAAGCACAGAAGGACAAAUCGCUUUAUUUACAAACCGAAUCAAUUUCCUAACAGGACACUUGAAAAAAAAUCAUAAAGAUUAUAACACCGAGCGUUCGUUAGUUAUGUUGGUAGGUAAAAGAAGAAGUCUUUUGGACUAUUUA
env-430  UUCAAGAAGUACGGAAAGUCUGAAAAAAACACCGGAAGCACAGAUGGACAAAUUGCGCUGUUCACGAAGCGUAUCAACGACCUAACAGUACACCUGAAAACCAAUCAUAAAGAUUAUAACACAGAGCGUUCAUUGGUUAUGUUAGUUGGUAAGCGAAGAAGUCUUUUAGAUUAUCUAAAA
env-431  UUCAAGAAGUACGGAAAGUCUGAAAAAAACACCGGAAGCACAGAUGGACAAAUUGCGCUGUUCACGAAGCGUAUCAACGACCUAACAGUACACCUGAAAACCAAUCAUAAAGAUUAUAACACAGAGCGUUCAUUGGUUAUGUUAGUUGGUAAGCGAAGAAGUCUUUUAGAUUAUCUAAAA
env-432  UUCAAGAAGUACGGAAAGUCUGAAAAAAACACCGGAAGUACAGAAGGACAAAUUGCGCUGUUCACAAAGCGUAUCAACGAGCUAACAGGACACCUGAAAACCAAUCAUAAAGAUUAUAACACAGAGCGUUCAUUGGUUAUGUUAGUUGGUAAGCGAAGAAGUCUUUUAGAUUAU......
env-433  ..........................................................................................................................................................................................
env-434  UUCAAGAAGUACGGAAAGUCUGAAAAAAACACCGGAAGUACAGAAGGACAAAUUGCGCUGUUCACAAAGCGUAUCAACGAGCUAACAGGACACCUGAAAACCAAUCAUAAAGAUUAUAACACAGAGCGUUCAUUGGUUAUGUUAGUUGGUAAGCGAAGAAGUCUUUUAGAUUAUUUAAAA
env-435  UUCAAGAAGUACGGAAAGUCUGAAAAAAACACCGGAAGUACAGAAGGACAAAUUGCGCUGUUCACAAAGCGUAUCAACGAGCUAACAGGACACCUGAAAACCAAUCAUAAAGAUUAUAACACAGAGCGUUCAUUGGUUAUGUUAGUUGGUAAGCGAAGAAGUCUUUUAGAUUAUUUAAAA
env-437  UUCAAAAAGUACGGAAAGUCUGAAAAAAACACCGGAAGUACAGAAGGACAAAUCGCGCUGUUCACGAAACGUAUCAACGACCUAACAGGACACCUGAAAGACAAUCAUAAAGAUUAUAACACAGAGCGAUCAUUGGUU.............................................
env-438  UUUAAAAAGUACGGAAAGUCUGAAAAAAACACCGGAAGUACGGAAGGGCAAAUUGCUCUGUUCACAAAGCGUAUCAACGAUCUAACAGGACACCUGAAAAACAAUCAUAAAGAUUAUAACACAGAGCGUUCAUUGGUUAUGUUAGUUGGUAAGCGAAGAAGUCUUUUAGAUUAUUUAAAA
env-439  UUCAAGAAGUACGGAAAGUCUGAAAAAAACACUGGAAGCACGGAAGGACAAAUUGCACUAUUUACAAAGCGUAUUAACGACCUAACAGGGCAUCUAAAAAACAAUCAUAAAGAUUACAAUACAGAGCGUUCAUUGGUUAUGUUAGUAGGUAAGCGAAGAAGUCUUUUAGAUUAUUUAAAA
env-440  UUCAAGAAGUACGGAAAGUCUGAAAAAAACACUGGAAGCACGGAGGGACAAAUUGCACUAUUUACAAAGCGUAUUAACGACCUAACAGGGCACCUAAAAAACAAUCAUAAAGAUUACAAUACAGAGCGUUCAUUGGUUAUGUUAGUAGGUAAGCGAAGAAGUCUUUU............
env-441  UUCAAGAAGUACGGAAAGUCUGAAAAAAACACUGGAAGCACGGAGGGACAAAUUGCACUAUUUACAAAGCGUAUUAACGACCUAACAGGGCAUCUAAAAAACAAUCAUAAAGAUUACAAUACAGAGCGUUCAUUGGUUAUGUUAGUAGGUAAGCGAAGAAGUCUUUUAGAUUAUUUAAAA
env-442  UUCAAGAAGUACGGAAAGUCUGAAAAAAACACUGGAAGCACGGAGGGACAAAUUGCACUAUUUACAAAGCGUAUUAACGACCUAACAGGGCACCUAAAAAACAAUCAUAAAGAUUACAAUACAGAGCGUUCAUUGGUUAUGUUAGUAGGUAAGCGAAGAAGUCUUUUAGAUUAUUUAAAA
env-444  UUCAAGAAGUACGGAAAGUCUGAAAAAAACACUGGAAGCACGGAGGGACAAAUUGCACUAUUUACAAAGCGUAUUAACGACCUAACAGGGCACCUAAAAAACAAUCAUAAAGAUUACAAUACAGAGCGUUCAUUGGUUAUGUUAGUAGGUAAGCGAAGAAGUCUUUUAGAUUAUUUAAAA
Kje-1-1  AUCUUCGCAAAGCACGGAAAAUCUGCUACAGAUACAGGUUCUGCAGAGGUCAGAUCGCUUUAUUUACGCACAGAAUUAAUCACUUAACGGAGCAUUUAAAGAGAAAUCGUAAAGAUUACAACACAGAACGCUCUCUAGUAAUGUUAGUAGGUAAAAGAAGAAGCUUACUAGAUUACUCA
Kal-1-1  AUCUUCGCAAAGCACGGAAAGUCUGCAACAGACACAGGUUCGUCAGAAGGUCAAAUCGCGUUAUUUACGCACAGAAUUAAUCACUUAACGGAGCACUUAAAGAAAAAUCGUAAAGAUUAUAACACAGAGCGUUCUCUAGUAAAGUUAGUAGGUAAAAGAAGAAGCUUACUUGACUACCUA
Kzh-1-1  AUUUUCGCAAAGCACGGAAAAUCUACUACUGACACAGGUUCAUCAGAAGGACAAAUUGCAUUGUUCACACACAGAAUCAAUCACUUAACAGAGCACUUAAAGAAAAAUCGUAAAGAUUAUAAUACUGAGCGUUCUCUUGUGAAAUUAGUAGGUAAAAGAAGAAGUUUACUUGAUUAUCUA
env-445  AUCUUCGCUAAACACGGAGGAAAAGCAGAAAAUACUGGUUCUGCAGAAGGUCAAAUCGC...............................................................................................................................
env-446  AUCUUUAAACAGUACGCAGGAUCAGAGAAAACA.........................................................................................................................................................
env-447  AUCUUUAAACAGUACGCAGGAUCAGAGAAAACACUGGUUCAGCAGAGGGUCAAAUCGCGCUUUUUACUCACCGUAUCAAGCAUUUAACAGAGCACAUGAAGAUCAACCGGAAAGAUACCGGUACUAAAAAAGCUCUUAUCCGCUUAGUAGGAAAAGACGUAGCUUACUUGAUUAUCUU
env-449  AUUUUCAAAAA...............................................................................................................................................................................
env-450  AUCUUUAAAAAACACGGUAAGGAAGCAAAAAACACAGGCUCUGCUGAAGGUCAAAUAGCUUUAUUUACAUAUAGAAUUAAUCACUUAACAGAUCAUUUAAAGAACAACAAAAAGAUUUUAAUACUGAGCGAUCAUUAGUAAAGCUGGUGGAAAGCGAAGAUCGUUAUUAGACUACUUG
env-451  AUCUUUAAAAAACACGGUAAGGAAGCGAAAAAUACUGGCUCUGCUGAAGGUCAAAUUGCUUUAUUUACAUACAGAAUUAAUCAUUUAACAUCUCAUUUAAAAAAUAUAAAAAAGAUUUUAAUACUGAGCGAUCAUUAGUAAAGCUAGUGGGAAAGCGAAGAUCAUUAUUAGACUACUUG
env-452  AUCUUUAAAAAACACGGUAAGGAAGCGAAAAAUACUGGCUCUGCUGAAGGUCAAAUUGCUUUAUUUACAUACAGAAUUAAUCAUUUAACAUCUCAUUUAAAAAAUAUAAAAAAGAUUUUAAUACUGAGCGAUCAUUAGUAAAGCUAGU..............................
env-453  AUCUUUAAAAAACACGGUAAGGAAGCCAAAAAUACUGGCUCUGCUGAAGGUCAAAUAGCUUUAUUUACAUACAGAAUUAAUCACUUAACAGAUCAUUUAAAGAAUAAUAAAAAGAUUUUAAUACUGAGCGAUCAUUAGUAAAGCUGU.............................
env-454  AUCUUUAAAAAACACGGUAAGGAAGCAAAAAAUACAGGGUCUGCUGAAGGUCAAAUAGCUUUAUUUACAUACAGAAUUAAUCACUUAACAGAUCAUUUAAAGAAUAAUAAAAAGAUUUUAAUACUGAGCGAUCAUUAGUAAAGCUAGUGGGAAAGCGAAGAUCUUUAUUAGACUACUUG
env-455  AUCUUUAAAAAACACGGUAAAGAAGCAAAAAAUACAGGCUCUGCUGAAGGUCAAAUAGCUUUAUUUACAUACAGAAUUAAUCACUUAACAGAUCAUUUAAAGAAUAAUAAAAAGAUUUUAAUACUGAGCGAUCAUUAGUAAAGCUGGUGGAAAGCGAAGAUCGUUAUUAGACUACUUG
env-456  UUUUUCAAAGAGUACGGUGGAGACGAAAAAAACACCGGUUCAACAGAAGCUCAAAUUGCUUUGUUCACUUACAGAAUUGAACAUUUAACACAGCAUUUGAAGAAGAAUAAGAAAGACUACAAUACACAGAGAUCUCUGAUUAGACUUGUAGGUAAACGUAAGUCACUUUUAGCCUACCUU
env-457  UUUUUCAAAGAGUACGG.........................................................................................................................................................................
env-458  ..........................................................................................................................................................................................
env-459  AUUUUUAAAGACCACGGUAAAUCAAAAAAAGACACUGGAUCUACCGAAGGUCAGAUAGCACUUUUCACUCAUCGUAUUCAACAUUUAUCUGAACACCUUAAAAAUAAUAGAAAAGAUUUUAAUACUGAGCGCUCAUUAGUACAUCUUGUUGGGAAGCGGAGGAGUCUUUUAAAUUAUUUA
env-460  AUUUUUAAAGACCACGGUAAAUCAAAAAAAGACACUGGAUCUACCGAAGGUCAGAUAGCACUUUUCACUCAUCGUAUUCAACAUUUAUCUGAACACCUUAAAAAUAAUAGAAAAGAUUUUAAUACUGAGCGCUCAUUAGUACAUCUUGUUGGGAAGCGGAGGAGUCUUUUAAAUUAUUUA
env-461  AUUUUUAAAGACCACGGUAAAUCAAAAAAAGACACUGGAUCUACCGAAGGUCAGAUAGCACUUUUCACUCAUCGUAUUCAACAUUUAUCUGAACACCUUAAAAAUAAUAGAAAAGAUUUUAAUACUGAGCGCUCAUUAGUACAUCUUGUUGGGAAGCGGAGGAGUCUUUUAAAUUAUUUA
env-462  AUUUUUAAAGACCACGGUAAAUCAAAAAAAGACACUGGAUCUACCGAAGGUCAGAUAGCACUUUUCACUCAUCGUAUUCAACAUUUAUCUGAACACCUUAAAAAUAAUAGAAAAGAUUUUAAUACUGAGCGCUCAUUAGUACAUCUAGUGGGAAAGCGGAGGAGUCUUUUAAAUUAUUUA
env-463  AUUUUUAAAGACCACGGUAAAUCAAAAAAAGACACUGGAUCUACCGAAGGUCAGAUAGCACUUUUCACUCAUCGUAUUCAACAUUUAUCUGAACACCUUAAAAAUAAUAGAAAAGAUUUUAAUACUGAGCGCUCAUUAGUACAUCUAGUGGGAAAGCGGAGGAGUCUUUUAAAUUAUUUA
env-464  AUUUUUAAAGACCACGGUAAAUCAAAAAAAGACACCGGAUCUACCGAAGGUCAGAUAGCACUUUUCACUCAUCGUAUUCACAUUUAUCUGACACUUAAAAUAUAGAAAGAUUUAUACUGAGCGCUCAUAGUACAUCUAGUGGAAGCGAGAGUCUCUUAAAUUAUUAAAUCGAAGAUAUUU
env-465  AUUUUUAAAGACCACGGUAAAUCAAAAAAAGACACUGGAUCUACCGAAGGUCAGAUAGCACUUUUCACUCAUCGUAUUCAACAUUUAUCUGAACACCUUAAAAAUAAUAGAAAAGAUUUUAAUACUGAGCGCUCAUUAGUACAUCUUGUGGGAAAGCGUAGGAGUCUUUUAAAUUAUUUA
env-466  AUUUUUAAAGACCACGGUAAAUCAAAAAAAGACACUGGUUCUACCGAAGGUCAGAUAGCACUUUUCACUCAUCGUAUUCAACAUUUAUCUGAACACCUUAAAAAUAAUAGAAAAGAUUUUAAUACUGAGCGCUCAUUAGUAAAUCUUGUGGGAAAGCGGAGAGUCUUUUAAAUUAUUUA
env-467  AUUUUUAAAGACUACGGUAAAUCAAAAAAAGACACUGGAUCUACCGAAGGCCAAAUAGCACUUUUCACUCAUCGUAUUCAACAUUUAUCGGAACAUCUUAAAAACAAUAGAAAAGAUUUUAAUACUGAGCGAUCACUAGUACAUCUAGUGGGAAAGCGAAGGAGUCUUUUAAAUUAUUUA
env-469  AUUUUUAAAGACUACGGUAAAUCAAAAAAAGACACUGGAUCUACCGAAGGCCAAAUAGCACUUUUCACUCAUCGUAUUCAACAUUUAUCGGAACAUCUUAAAAACAAUAGAAAAGAUUUUAAUACUGAGCGAUCACUAGUACAUCUAGUGGGAAAGCGAAGGAGUCUUUUAAAUUAUUUA
env-470  AUUUUUAAAGAUCACGGUAAAUCAAAAAAGAUACAGGAUCUACUGAAGGUCAAAUAGCACUUUUCACUCAUCGCAUUCAACAUUUAUCGGAACACCUUAAAAACAAUAGAAAAGAUUUUAAUACUGAGCGCUCAUUAGUUCACUUAGUGGGAAAGCGAAGGAGUCUUUUGAAUUACUUA
env-471  AUCUUCGCUAAACACGGAGGUUCCGAUACAAACACAGGAUCUACAGAAGGUCAGGUUGCUUUGUUCACGUAUCGUAUUAGUCAUUUGACAGAGCACUUGAAGAAAAACCGUAAGGAUUUUGGAACGCAACGCUCUCUUCAAUUACUAGUAGGUAAAAGAAGAAGUUUAUUAGAUUACCUA
env-472  CUUUUCAAAAAAUACGGAAAAUCUGAAAACGACACGGGAUCUACUGAAGGUCAGAUUGCUUUGUUUUCGCACCGUAUCAACCACCUUUCUGACCACUUAAAAAAACAACGCAAAAGACUUUAAUACAGAGCGUUCGCUAGUGCGUUUGGUAGGAAAAA.......................
env-473  CUUUUCAAAAAAUACGGAAAAUCUGAAAACGACACGGGAUCUACUGAAGGUCAGAUUGCUUUGUUUUCGCACCGUAUCAACCACCUUUCUGACCACUUAAAAAAACAACGCAAAAGACUUUAACACAGAGCGUUCGCUAGUGCGUUUGGUAGGAAAAAGAAGAAGCCUUCUCGACUAUCUA
env-474  AGCUUUUUCAAAAAAAUAACGGAAAAUCUGAAAACGACACGGGAUCUACUGAAGGUCAGAUUGCAUUGUUUUCGCACCGUAUCAAUCACCUUUCUGACCACUUAAAAAAACAACGCAAAAGACUUUAACACAGAGCGUUCGCUAGUGCGUUUGGUAGGAAAAAG..............
env-475  AUCUUUACAAAGCAUGGCGGGAAAGCGGAGAACACAGGUUCUACCGAAGGGCAAGUUGCUCUUUUCACUCAUCGUAUCAAUCACCUUACGGGGCAUUUGAAAAACAAUCGAAAAGAUUUUAACACCGAACGAUCACUAGUAAAAUUGGUGGGUAAGCGACGAGUCUUCUCGAUUAUUUG
env-476  AUUUUCGAAAAACACGGGAAAUCUAAAUCAGACACCGGAUCUGCUGAGGACAAAUUGCUUUGUUUUCUCACAGAAUUAACCAUUUGUCUGAACAUUUAAAAAAAAUAAUCGUAAAGACUUUAACACAGAACGUUCUCUUGUAAAACUAGUAGGUAAAAGAAGAAGUCUUUUGGACUAUUUA
env-477  AUUUUCAAAAAACACGGAAAAUCAAAAACUGACACCGGAACUGCAGAAGGACAAAUUGCUUUGUUUUCUCACAGAAUCGACCACUUGUCAGAGCAUUUAAAAAACAACAGAAAAGACUACAAUACAGAACGUUCACUUGUGAAACUAGUAGGUAAAAGAAGAAGUCUUUUAGAUUAUUUA
env-478  AUUUUCAAAAAACACGGAAAAUCAAAAACUGACACCGGAACUGCAGAGGACAAAUUGCUUUGUUUUCUCACAGAAUCGACCACUUGUCAGAGCAUUUAAAAAACAACAGAAAAGACUACAAUACAGAACGUUCACUUGUGAAACUAGUAGGUAAAAGAAGAAGUCUUUUAGAUUAUUUA
env-479  AUUUUCAAAAAACACGGAAAAUCAAAAACUGACACCGGAACUGCAGAAGGACAAAUUGCUUUGUUUUCUCACAGAAUCGACCACUUGUCAGAGCAUUUAAAAAACAACAGAAAAGACUACAAUACAGAACGUUCACUUGUGAAACUAGUAGGUAAAAGAAGAAGUCUUUUAGAUUAUUUA
env-480  AUUUUCAAAAAACACGGAAAAUCAAAAACUGACACCGGAACUGCAGAGGACAAAUUGCUUUGUUUUCUCACAGAAUCGACCACUUGUCAGAGCACUUAAAAAAUAACAGAAAAGACUAUAAUACAGAACGUUCACUUGUGAAACUAGUAGGUAAAAGAAGAAGUCUUUUAGAUUAUUUA
env-481  AUUUUCAAAAAACACGGAAAAUCAAAAACUGACACCGGAACUGCAGAAGGACAAAUUGCUUUGUUUUCUCACAGAAUCGACCACUUGUCAGAGCAUUUAAAAAACAACAGAAAAGACUACAAUACAGAACGUUCACUUGUCAAACUAGUAGGUAAAAGAAGAAGUCUUUUAGAUUAUUUA
env-482  AUUUUCGAAAAACACGGAAAAUCAAAAUCAGACACUGGUACUGCAGAAGGACAAAUCGCUUUGUUUUCUCACAGAAUCAAUCACUUAUCUGAUCACUUAAAAAGUAACCGAAAAGACUACAAUACAGAACGUUCCCUAGUAAACUAGUAGGUAAAAGAAGAAGUCUUUUAGAUUAUUUG
env-484  AUUUUCGAAAAACACGGAAAAUCAAAAUCAGACACUGGUACUGCAGAGGACAAAUCGCUUUGUUUUCUCACAGAAUCAAUCACUUAUCUGAUCACUUNAAAAGUAACCGAAAAGACUACAACACAGAACGUUCCCUAGUNAAANUAGUAGGUAAAAGAAGAAGUCUUUUAGAUUAUUUG
env-486  AUUUUCGAAAAACACGGAAAAUCAAAAUCAGACACUGGUACUGCAGAAGGACAAAUCGCUUUGUUUUCUCACAGAAUCAAUCACUUAUCUGAUCACUUAAAAAGUAACCGAAAAGACUACAACACAGAACGUUCCCUAGUUAAAUUAGUAGGUAAAAGAAGAAGUCUUUUAGAUUAUUUGA
env-487  AUUUUCAAAAAACACGGAAAAUCAAAAACUGACACCGGAACUGCAGAGGACAAAUUGCUUUGUUUUCUCACAGAAUCGACCACUUGUCAGAGCAUUUAAAAAACAACAGAAAAGACUACAAUACAGAACGUUCACUUGUGAAACUAGUAGGUAAAAGAAGAAGUCUUUUAGAUUAUUUA
env-488  AUUUUCAAAAAACACGGAAAAUCAAAAACUGACACCGGAACUGCAGAAGGACAAAUUGCUUUGUUUUCUCACAGAAUCGACCACUUGUCAGAGCAUUUAAAAAACAACAGAAAAGACUACAAUACAGAACGUUCACUUGUGAAACUAGUAGGUAAAAGAAGAAGUCUUUUAGAUUAUUUA
env-489  AUUUUCAAAAAACACGGAAAAUCAAAAACUGACACCGGAACUGCAGAAGGACAAAUUGCUUUGUUUUCUCACAGAAUCGACCACUUGUCAGAGCAUUUAAAAAACAACAGAAAAGACUACAAUACAGAACGUUCACUUGUGAAACUAGUAGGUAAAAGAAGAAGUCUUUUAGAUUAUUUA
env-490  ..........................................................................................................................................................................................
env-491  AUUUUCAAAAAACACGGGAAAUCCAAAUCAGACACCGGUACUGCAGAAGGACAAAUUGCUUUGUUUUCUCACAGAAUCAACCACUUGUCUGAUCACUUAAAAAAACAAUCAAAAAGACUAUAAUACAGAACGUUCCCUAGUAAAACUAGUAGGAAAAAGAAGAAGUCUUUUAGAUUACUUA

         ..........................................................................................................................................................................................
         ..........................................................................................................................................................................................
         AUYUUYR•RAARCACGG•RRR•••R•RR••RAYAC•GG••C•RC•GAAGG•CAAAUYGC•YURUUYAC•Y•Y•G•AUYRAYCAYYUR•C•RA•CAYYURAAAAR•AAYCR•AARGAUU•YAAYAC•GAR•G•UC•YURGU•••RYURGU•GG•AAR•G•RG••••YU•YU•GAYUAYYU•
```

| | |
|---|---|
| env-492 | AUUUUCAAAAAACACGGGAAAUCCAAAUCAGACACCGGUACUGCAGAAGGACAAAUUGCUUUGUUUUCUCACAGAAUCAACCACUUGUCUGAUCACUUAAAAAAACAAUCGAAAAGACUAUAAUACAGAACGUUCCCUAGUAAAACUAGUAGGAAAAAGAAGAAGUCUUUUAGAUUACUUA |
| env-494 | AUUUUCAAAAAACACGGGAAAUCCAAAUCAGACACCGGUACUGCAGAAGGACAAAUUGCUUUGUUUUCUCACAGAAUCAACCACUUGUCUGAUCACUUAAAAAAACAAUCGAAAAGACUAUAAUACAGAACGUUCCCUAGUAAAACUAGUAGGAAAAAGAAGAAGUCUCUUAGAUUACUUA |
| env-496 | AUUUUCAAAAAACACGGGAAAUCCAAAUCAGACACCGGUACUGCAGAAGGACAAAUUGCUUUGUUUUCUCACAGAAUCAACCACUUGUCUGAUCACUUAAAAAAACAAUC....................................................................... |
| env-498 | AUUUUCAAAAAACACGGGAAAUCCAAAUCAGACACCGGUACUGCAGAAGGACAAAUUGCUUUGUUUUCUCACAGAAUCAACCACUUGUCUGAUCACUUAAAAAAACAAUCGAAAAGACUAUAAUACAGAACGUUCCCUAGUAAAACUAGUAGGAAAAAGAAGAAGUCUUUUAGAUUACUUA |
| env-500 | ..................................................................................................................................................................................... |
| env-501 | AUUUUCGAAAAACACGGAAAAUCUAAAUCAGACACUGGUACUGCAGAAGGACAAAUUGCUUUGUUUUCUCACAGAAUCAACCACUUAUCUGAACAUUUAAAAAGUAACCGAAAAGACUACAAUACAGAACGUUCCCUAGUGAAACUAGUAGGUAAAAGAAGAAGUCUUUUAGAUUAUUUA |
| env-502 | AUUUUCGAAAAACACGGAAAAUCUAAAUCAGACACAGGAACUGCUGAGGGACAAAUCGCCUUAUUUUCUCACAGAAUAAACCAUUUGUCUGAACAUUUAAAAAAACAAUCGUAAAGACUUUAACACAGAACGCUCUCUUGUAAAGCUAGUAGGUAAAAGAAGAAGUCUUCUAGAUUAUCUA |
| env-503 | AUUUUCGAAAAGCACGGUAAAGCAGCAACCAACACAGGUUCAGCAGAAGGUCAAAUCGCGCUAUUCACAUUCAGAAUCAAUCACCUUACAGAGCACCUUAAAAAAAAAUCGUCAUGAUUACAAUAC....................................................... |
| env-504 | AUUUUCGCAAAGCACGGCAAAAACGGACAGGACACAGGUACUGCCGAAGGCCAGAUUGCGCUUUUCACAUUCAGGAUUAACCACCUUACAGAGCACCUUAAAAAAAAAUCGUCAUGAUUUUAACACAGAGCGCUCGCUUGUUAAGCUGGUAGGUAAAAGAAGGAGCCUUCUUGACUACCUC |
| env-505 | AUUUUCGCAAAGCACGGCAAAAACGGACAGGACACAGGUACUGCCGAAGGCCAGAUUGCGCUUUUCACAUUCAGGAUUAACCACCUUACAGAGCACCUUAAAAAAAAAUCGUCAUGAUUUUAACACAGAGCGCUCGCUUGUUAAGCUGGUAGGUAAAAGAAGGAGCCUUCUUGACUACCUC |
| env-506 | AUUUUCAAAAAACACGGUAAAGGCGCUAAUGACACAGGUUCUGCCGAAGGUCAAAUCGCUUUAUUUACACAUCGUAUCGAUCACUUAACCCAACACCUUAAAAACAACCACAAGGACUACAACACAGAGCGUUCUUUGGUAAAAUUGGUAGGGAAGCGUCGUUCUUUAUUAGAUUAUUUA |
| env-507 | AUUUUUAAACAGUACUCCGGUUCAGAUUCGAAUACGGGGUCAGCAGAAGGGCAAAUUGCUCUGUUUACUUACCGCAUCAAGCACCUAACGGAGCACAUGAAGCAAAAUAAAAAGGAUAUGGGUACAAAAAAAGCCCUCAUCCGUUUAGUAGGUAAAAGAAGAAGUUUACUCGACUAUCUU |
| env-508 | AUUUUUAA............................................................................................................................................................................. |
| env-509 | AUUUUUAAACAGUACUCCGGUUCAGAUUCGAAUACGGGGUCAGCAGAAGGGCAAAUUGCUCUGUUUACUUACCGCAUCAAGCACCUAACGGAGCACAUGAAGCAAAAUAAAAAGGAUAUGGGUACAAAAAAAGCCCUCAUCCGUUUAGUAGGUAAAAGAAGAAGUUUACUCGACUAUCUU |
| env-510 | AUUUUUAAACAGUACUCCGGUUCAGAUUCGAAUACGGGGUCAGCAGAAGGGCAAAUUGCUCUGUUUACUUACCGCAUCAAGCACCUAACGGAGCACAUGAAGCAAAAUAAAAAGGAUAUGGGUACAAAAAAAGCCCUCAUCCGUUUAGUAGGUAAAAGAAGAAGUUUACUCGACUAUCUU |
| env-511 | CUUUUUAAAAAACACGGUAAAGAUGCCAAGAACACCGGUUCUGCCGAGGGGCAAAUUGCAUUGUUCACACACAGGAUCAAUCAUUUGACCCAACACCUAAAAAAGAACCACAAGGAUUACGCAACCGAAAGGUCUUUGGUAAAGUUAGUGGGUAAGCGUAGGGCCUUAUUGGAUUACCUU |
| Fhi-1-1 | AUUUUCGCACAACACGGUGAAGCAACAAACACUGGAAAAGCUGAAGCUCAAAUCGCAUUGUUUACUUACAGAAUUAGUCACUUAACUGAGCACUUGAAAAAAAAUCGUCAUGAUUAUAAUACUGAGCGUUCAUUAGUACUAUUAGUAGGUAAAAGAAGAUCGUUGCUUGAUUACUUGAAA |
| Fsp-9-1 | AUUUUCGCUCAACACGGUAGUGCAACAAAUACUGGAAGCGCAGAAGGUCAAAUUGCAUUGUUCACUUACAGAAUUUCACACUUAACUGAGCACUUGAAAAAAAAUCGUCACGAUUACAACACUGAGCGUUCACUUGUACUAUUAGUAGGUAAAAGAAGAGCUUUGUUGGAUUACUUGAAA |
| Asu-1-1 | AUUUUCAAAAAACACGGAAAAUCUGUAAAAGACACAGGUUCUGCAGAAGGACAAAUUGCAUUAUUCACUUAUAGAAUAGAGCAUUUAACUAAGCACCUUAAAGCAAACCACAAGGACUACAACACUGAGCGUUCUUUGGUAAUGUUAGUAGGUAAAAGAAGAGCUCUUUUGGACUAUUUA |
| Aca-1-1 | AUUUUCGAAAAACACGGAAAGUCUAAAACUGACACUGGUUCAGCAGAAGGACAGAUUGCGCUAUUCACUUACCGCAUUUCACAUUUAACAAAGCACCUUAAAAAUAAUCAUAAAGAUUAUAACACAGAGCGUUCUUUGGUAAUGUUGGUUGGUAAGAGACGUUCAUUAUUAGAUUAUCUU |
| env-513 | AUUUUCGAAAAACACGGAAAGUCUAAAACUGACACUGGUUCAGCAGAAGGACAGAUUGCGCUAUUCACGUAUCGCAUCGAGCAUUUAACAAAACAUCUUAAAACAAACCACAAAGACUAUAACACUGAGCGUUCUUUGGUAAUGUUGGUAGGUAAAAGACGUUCGCUACUAGAUUAUCUU |
| env-514 | AUUUUCGAAAAACACGGAAAGUCUAAAACUGAUACUGGUUCAGCAGAAGGACAGAUUGCGUUAUUCACGUAUCGCAUCGAGCAUUUAACAAAAC....................................................................................... |
| env-515 | AUUUUCGAAAAACACGGAAAGUCCAAGACUGACACGGGUUCAGCAGAAGGACAGAUUGCGCUAUUCACAUACCGCAUCGAGCAUUUAACAAAACACCUAAAAGCAAAUCACAAGGAUUACAAUACAGAGCGUUCUUUGGUAAUGUUAGUAGGUAAGCGCAGAGCGCUGCUGGAUUAUCUU |
| env-517 | AUUUUCGAAAAACACGGAAAGUCCAAGACUGACACGGGUUCAGCAGAAGGACAGAUUGCGCUAUUCACAUACCGCAUCGAGCAUUUAACAAAACACCUAAAAGCAAAU......................................................................... |
| Avl-1-1 | AUUUUCGAAAAACACGGAAAGUCCAAGACUGACACUGGUUCAGCAGAAGGACAGAUUGCGCUAUUCACAUACCGCAUCGAGCAUUUAACAAAACACCUAAAAGCAAAUCACAAGGAUUACAAUACAGAGCGUUCUUUGGUAAUGUUAGUAGGUAAGCGCAGAGCGCUGCUGGAUUAUCUU |
| env-518 | AUUUUCGAAAAACACGGAAAGUCCAAGACUGACACGGGUUCAGCAGAAGGACAGAUUGCGCUAUUCACAUACCGCAUCGAGCAUUUAACAAAACACCUAAAAGCAAAUCACAAGGAUUACAAUACAGAGCGUUCUUUGGUAAUGUUAGUAGGUAAGCGCAGAGCGCUGCUGGAUUAUCUU |
| env-519 | AUUUUCGAAAAACACGGAAAGUCCAAGACUGACACUGGUUCAGCAGAAGGACAGAUUGCGCUAUUCACAUACCGCAUCGAGCAUUUAACAAAACACCUAAAAGCAAAUCACAAGGAUUACAAUACAGAGCGUUCUUUGGUAAUGUUAGUAGGUAAGCGCAGAGC................. |
| env-520 | AUUUUCGAAAAACACGGAAAGUCUAAAACUGAUACUGGUUCAGCAGAAGGACAGAUUGCGCUAUUCACAUAUCGCAUCGAGCAUUUAACAAAGCAUCUUAAAACAAACCACAAAGACUAUAACACUGAGCGUUCUUUGGUAAUGUUGGUAGGUAAAAGACGUUCGCUACUAGAUUAUCUU |
| env-521 | AUUUUCGCAAAACACGGAAAGUCCAAGACUGACACUGGUUCAGCCGAAGGACAGAUUGCGCUAUUCACGUACCGCAUCGAGCAUUUAACAAAACAUCUUAAAGCAAAUCACAAGGACUACAACACUGAGCGUUCUUUGGUAAUGUUGGUAGGUAAGCGCAGAGCAUUGUUGGAUUAUCUU |
| env-522 | AUUUUCGCAAAACACGGAAAGUCCAAGACUGACACUGGUUCAGCCGAAGGACAGAUUGCGCUAUUCACGUACCGCAUCGAGCAUUUAACAAAACAUCUUAAAGCAAAUCACAAGGACUACAACACUGAGCGUUCUUUGGUAAUGUUGGUAGGUAAGCGCAGAGCAUUGUUGGAUUAUCUU |
| env-523 | AUUUUCGCAAAACACGGAAAGUCCAAGACUGACACUGGUUCAGCUGAAGGACAGAUUGCGCUAUUCACGUACCGCAUCGAGCAUUUAACAAAACAUCUUAAAGCAAAUCACAAGGACUACAACACUGAGCGUUCUUUGGUAAUGUUGGUAGGUAAGCGCAGAGCAUUGUUGGAUUAUCUU |
| env-527 | ..................................................................................................................................................................................... |
| env-528 | AUUUUCGCAAAACACGGAAAGUCCAAGACUGACACUGGUUCGGCAGAAGGACAAAUUGCGCUAUUCACGUACCGCAUUGAGCAUUUAACCAAACACCUUAAAGCAAACCACAAGGACUACAACACUGAGCGUUCUUUGGUAAUGUUGGUAGGUAAGCGCAGAGCAUUGUUGGAUUAUCUU |
| env-531 | AUUUUCGCAAAACACGGAAAGUCCAAGACUGACACUGGUUCGGCAGAAGGACAAAUUGCGCUAUUCACGUACCGCAUUGAGCAUUUAACCAAACACCUUAAAGCAAACCACAAGGACUACAACACUGAGCGUUCUUUGGUAAUGUUGGUAGGUAAGCGCAGAGCAUUGUUGGAUUAUCUU |
| env-532 | AUUUUUGAAAAA......................................................................................................................................................................... |
| env-533 | AUUUUUGAAAAACACG..................................................................................................................................................................... |
| env-534 | AUUUUCAAGAAACACGGAUUCUCUGUCAAAAACACCGGAUCUACAGAGGGCCAAAUUGCUUUAUUUACCUACCGUAUCAAUCACCUUUCAGAGCACUUAAAAAAGAAUAAGAAAGACUUCAACACUGAACGCUCAUUAGUCCGAUUGGUUGGUAAAAGAAGACAGCAACUAGAUUAUUUA |
| env-536 | AUUU................................................................................................................................................................................. |
| env-537 | AUUUUCAAAAAACACGGAUCCUCUGCCAAAAACACAGGAUCUACAGAAGGUCAAAUUGCUUUAUUUACCUACCGUAUCAAUCACCUUUCAGAGCAUUUAAAAAAGAACAAGAAAGACUUCAACACUGAACGCUCAUUAGUACGAUUGGUGGGUAAAAGAAGACAGCAACUAGACUAUCUU |
| env-539 | AUUUUCAAAAAACA....................................................................................................................................................................... |
| env-540 | AUUUUCAAAAAACACGGAUCCUCUGCCAAAAACACAGGAUCUACAGAAGGUCAAAUUGCUUUAUUUACCUACCGUAUCAAUCACCUUUCAGAGCAUUUAAAAAAGAACAAGAAAGACUUCAACACUGAACGCUCAUUAGUACGAUUGGUGGGUAAAAGAAGACAGCAACUAGACUAUCUU |
| Apa-1-1 | AUUUUUGCAAAACACGGUAAAGGAAAGCAUGAUAGCGGUUCUGCAGAAGGUCAAAUCGCAUUAUUUACGCACAGAAUUACUCACCUUACUGAGCACUUGAAAAAAAAUCGUAAAGAUUUUAACACAGAGCGUUCAUUAGUUAAAUUGGUAGGUAAGCGUAGAGAUUUAUUAGAUUACCUU |
| Ama-1-1 | AUCUUCGCAAAGCACGGUAAAGGAAAGAAUGACAGUGGUUCUGCAGAAGGUCAAAUUGCAUUGUUUACGCACAGAAUUACUCAUCUUACUGAGCACUUAAAAAAGAAUCGUAAAGAUUUUAAUACAGAGCGUUCUUUGGUAAAGCUAGUAGGGAAACGUAGAGAUCUGUUAGACUACCUU |
| Aat-1-1 | AUCUUCGCAAAGCACGGUAAAGGAAAGAAUGACAGUGGUUCUGCAGAAGGUCAAAUUGCAUUAUUUACUCACAGAAUUACUCACCUUACUGAGCACUUAAAAAAGAAUCGUAAAGAUUUCAACACAGAGCGUUCUUUGGUGAAACUGGUAGGUAAACGUAGAGAUCUACUUGAUUACCUU |
| Ame-1-1 | AUCUUCGCAAAGCACGGUAAAGGAAAGAACGACAGUGGUUCUGCAGAAGGUCAAAUUGCAUUGUUUACUCACAGAAUUACUCAUCUUACUGAGCACUUAAAAAAGAAUCGUAAAGAUUAUAAUACAGAGCGUUCUUUGGUAAAGUUGGUAGGAAAACGUAGAGAUCUGUUAGACUAUCUU |
| Ala-1-1 | AUCUUCGCAAAACACGGAAAAGGUAAAAAUGACACCGGUUCUGCAGAAGGACAAAUUGCAUUAUUUACGCACAGAAUUACUCAUUUAACUGAGCACUUAAAAAACAACCGUAAGGAUUUUAACACUGAACGUUCGUUAGUAAAGUUAGUAGGUAAGCGUAGAGAUCUUUUAGACUAUCUU |
| Amu-1-1 | AUUUUCGCUAAACACGGUAAAGGAAAAAAUGACAGUGGUUCUGCGGAAGGUCAAAUUGCAUUAUUCACGCACAGAAUUACUCAUCUUACUGAACACUUAAAAAAGAAUCGUAAAGAUUUCAACACAGAGCGUUCUCUAGUAAAAUUAGUAGGAAAGCGUAGAGAUCUAUUAGAUUAUCUC |
| Fsu-3-1 | CUUUUCGCGAAACACGGAGGAAAAGCUGAGAACACAGGUUCUGCUGAAGGACAAAUCGCUUUGUUCACAUUCAGAAUCAAUCACCUGACCGGACACUUGAAAGCUAAUCGUCACGAUUACAACACCGAGCGUUCGUUAGUGAAAUUGGUAGGUAAAAGAAGAAGUCUUCUUGACUACUUA |
| Fen-1-1 | AUCUUCGCUAAACACGGAGGUGCUGCUGCCAACACUGGUUCUGCAGAAGGUCAAAUCGCGUUAUUCACAUUCAGAAUUUCUCACUUGACUGAGCACUUGAAAAAAAAUCGUCACGAUUUCAACACAGAGCGCUCACUAGUUUUAUUAGUAGGUAAAAGAAGAAGUCUUCUUGACUACUUA |
| Fco-1-1 | AUCUUCGCUCAACACGGAGGGAAAGCUCAAAACACAGGUUCUGCAGAAGGGCAAGUAGCUUUAUUUACUUUUAGAAUUAACCACCUAACUGAGCACUUGAAAAGAAAUCGUCAUGAUUACAACACAGAGCGCUCGUUAGUAAAACUAGUAGGUAAAAGAAGAUCUCUUCUUGACUACUUA |
| env-541 | AUCUUCGCUAAACACGGAGGAAAAGCAGAAAACACAGGUUCAGCAGAAGGACAAAUUGCGUUAUUCACAUUCAGAAUUUCUCACUUAACUGAGCACUUGAAAAAAAAUCGUCACGAUUAUAAUACUGAAAGAUCAUUAGUACUUUUAGUAGGUAAAAGAAGAAGUCUUUUAGAUUACUUA |
| env-542 | AUCUUCGCUAAACACGGAGGAAAAGCAGAAAACACAGGUUCUGCUGAAGGACAAAUUGCAUUAUUCACAUUUAGAAUUUCGCACUUAACAGAACACUUGAAAAAAAAUCGUCACGAUUACAACACAGAAAGAUCGUUGGUGCUUUUAGUAGGUAAAAGAAGAAGUCUUUUAGAUUACUUA |
| env-543 | ..................................................................................................................................................................................... |
| env-545 | AUCUUCGCUAAACACGGAGGAAAAGCAGAAAACACAGGUUCUGCAGAAGGACAAAUUGCAUUAUUCACGUUUAGAAUUUCGCACUUAACAGAACACUUGAAAAAAAAUCGUCACGAUUACAACACAGAAAGGUCGUUGGUGCUUUUAGUAG.............................. |
| env-546 | AUCUUUAAAA........................................................................................................................................................................... |
| Fps-1-1 | AUCUUCGCUAAACACGGAGAAAAAACAAACACAGGAAAAGCAGAAGCACAAAUUGCUUUAUUUACUUUCAGAAUUAGCCACUUAACAGAGCACCUGAAAAAAAAUCGUCACGAUUAUAACACUGAGCGUUCACUUGUUCUUUUAGUAGGUAAAAGAAGAUCUUUACUAGACUACUUGAAG |
| Jma-1-1 | AUAUUUAAACAAUAUGGUGGUGAUGCUAAAAACACCGGAUCUGCAGAAGGACAAAUUGCUUUGUUCACACACAGAAUUAAUCAUUUAACUGGACACUUAAAAAACAAUCGUAAAGAUUUCAAUACAGAGCGUUCAUUAGUUAUGCUAGUAGGUAAAAGAAGAAGCUUACUAGAUUAUUUG |
| env-547 | AUUUUUAAAAAACACGGUAAAUCUGAAUC........................................................................................................................................................ |
| env-548 | UUAUUCAAAAAGCAUGGUAAAAAUGCUAAAAAUACGGGUAGUGCUGAAGGUCAAAUUGCUCUAUUCACAUUCAGAAUAAAUCAUUUGACAGAUCACCUUAAGAAAAAUAAAAAAGAUUACAAUACAGAAAGAGCUCUUGUUAAACUUGUUGGUAAGAGAAGAGCACUUCUUGAUUAUCUU |
| env-549 | UUAUUCAAAAAGCAUGGUAAAAAUGCUAAAAAUACGGGUAGUGCUGAAGGCCAAAUUGCUCUAUUUACAUUCAGAAUAAAUCAUUUGACAGAUCACCUUAAGAAAAAUAAAAAAGAUUACAAUACAGAAAGAGCUCUUGUUAAACUUGUUGGUAAGAGAAGAGCACUACUUGAUUAUCUU |
| env-550 | UUAUUCAAAAAGCAUGGUAAAAAUGCAAAAAAUACGGGUAGUGCUGAAGGUCAAAUUGCUCUAUUCACAUUCAGAAUAAAUCAUUUGACAGAUCACCUUAAGAAAAAUAAAAAAGAUUACAAUACAGAAAGAGCUCUUGUUAAACUUGUUGGUAAGAGAAGAGCACUUCUUGAUUAUCUU |
| env-551 | AUCUUCGAAAAACACGGUAAAGGAAAAAACGAUACGGGUUCUUCAGAAGGGCAAAUUGCGUUAUUUACGCACAGAAUUAAUCACUUAACUGAGCAUUUAAAAAACAAUCGUAAAGAUUAUAAUACAGAACGUUCAUUAGUAAUGAUGGUAGGNAAACGUAGAAGUUUGUUAGACUAUCUA |
| env-552 | AUCUUCGAAAAACACGGUAAAGGAAAAAACGAUACGGGUUCUUCAGAAGGGCAGAUUGCGUUAUUUACGCACAGAAUUAAUCACUUAACUGAGCAUUUAAAAAACAAUCGUAAAGAUUAUAAUACAGAACGUUCAUUAGUAAUGAUGGUAGGAAAACGUAGAAGUUUGUUAGACUAUCUA |
| env-553 | AUCUUCGAAAAACACGGUAAAGGAAAAAACGAUACGGGUUCUUCAGAAGGGCAAAUUGCGUUAUUUACGCACAGAAUUAAUCACUUAACUGAGCAUUUAAAAAACAAUCGUAAAGAUUAUAAUACAGAACGUUCAUUAGUAAUGAUGGUAGGAAAACGUAGAAGUUUGUUAGACUAUCUA |
| env-554 | AUCUUCGAAAAACACGGUAAAGGAAAAAACGAUACCGGUUCUUCAGAAGGUCAAAUCGCGUUAUUUACGCACAGAAUUAACCAUUUAACAGAGCACUUAAAGAAAAAUCGUAAAGAUUAUAAUACAGAGCGUUCGUUAGUAAUGAUGGUUGGUAAGCGUAGAAGCUUGUUAGAUUACCUG |
| env-556 | AUCUUCGAAAAACACG..................................................................................................................................................................... |
| env-557 | AUCUUCGAAAAACACGGUAAAGGAAAAAACGAUACCGGUUCAUCAGAAGGUCAAAUCGCGUUAUUUACGCACAGAAUCAACCAUUUAACAGAGCACUUAAAGAAAAAUCGUAAAGAUUACAAUACAGAGCGUUCCUUAGUAAUGAUGGUUGGUAAGCGUAGAAGCUUGUUAGAUUACCUG |
| Ffi-1-1 | AUCUUCGCUAAACACGGAGGAAAAGCUGAAAACACAGGUUCUGCAGAGGGACAAAUUGCAUUGUUCACUUUCAGAAUUUCUCACUUAACGGAGCACUUGAAAAAAAAUCGUCAUGAUUACAACACCGAGCGUUCGUUAGUGUUAUUGGUAGGUAAAAGAAGAAGUCUUCUUGACUACUUA |
| env-558 | AUCUUCGCUAAACACGGAGAAAAAACAAACACUGGAAAAUCAGAAAGCCAAAUCGCUUUGUUCACUUUCAGAAUCAGUCACCUUACANAGCACUUGAAAAAAAAUCGUCAUGAUUUUAACACCGAGCGUUCUCUAGUAUUAUUAGUAGGUAAAAGAAG...................... |
| env-559 | AUAUUUGCAAAGCACGGUAAAGGUGCAAACGACACUGGUUCUGCUGAAGGACAAAUUGCCCUAUUUACAGCAAGAAUUAACCACUUAACUGAACACUUAAAAAAUAAUCGUAAAGAUUAUAACACUGAGCGUUCGUUAGUAAAAUUAGUAGGAAAGCGUCGCUCAUUACUAGAUUACUUA |
| env-561 | CUAUUCAAAAAACACGGUAAAAAUGAAAAAGAUACCGGAAGCGCUGAAGGUCAGAUCGCUUUAUUCACACUAAGAAUCAACCAUCUAACAGAGCAUUUAAAAAAUAAUCGCAAAGAUUAUAAUACUGAAAGAUCUCUUGUAAAGCUUGUAGGUAAAAGAAGAUCACUUCUAGAUUACUUA |
| env-562 | CUAUUCAAAAAACACGGUAAAAAUGAAAAAGAUACCGGAAGCGCUGAAGGUCAAAUCGCUUUAUUCACACUAAGAAUCAACCAUCUAACAGAGCAUUUAAAAAAUAAUCGUAAGGAUUAUAACACCGAAAGAUCUCUUGUUA....................................... |
| env-563 | CUAUUCAAAAAACACGGUAAAAAUGAAAAAGAUA................................................................................................................................................... |
| env-564 | CUAUUCAAAAAACACGGUAAAAAUGAAAAAGAUACCGGAAGCGCUGAAGGUCAGAUCGCUUUAUUCACACUAAGAAUUAACCAUUUAACAGAGCAUUUAAAAAAUAAUCGUAAAGAUUAUAAUACAGAAAGAUCUCUUGUUAAGCUUGUAGGUAAAAGAAGAUCACUUCUAGAUUACUUA |
| env-565 | ..................................................................................................................................................................................... |
| env-566 | CUAUUCAAAAAACACGGUAAAAAUGAAAAAGAUACCGGAAGCGCUGAAGGUCAGAUCGCUUUAUUCACACUAAGAAUUAACCAUUUAACAGAGCAUUUAAAAAAUAAUCGUAAAGAUUAUAAUACAGAAAGAUCUCUUGUUAAGCUAGUAGGUAAAAGAAGAUCACUUCUAGAUUACUUA |
| env-567 | CUAUUCAAAAAACACGGUAAAAAUGAAAAAGAUACCGGAAGCGCUGAAGGUCAGAUCGCUUUAUUCACACUAAGAAUUAACCAUUUAACAGAGCAUUUAAAAAAUAAUCGUAAAGAUUAUAAUACAGAAAGAUCUCUUGUUAAGCUAGUAGGAAAAAGAAGAUCACUUCUAGAUUACUUA |
| env-569 | CUAUUCAAAAAACACGGUAAAAAUGAAAAAGAUACCGGAAGCGCUGAAGGUCAGAUCGCUUUAUUCACACUAAGAAUUAACCAUUUAAC.............................................................................................. |
| env-571 | ..................................................................................................................................................................................... |
| env-572 | CUAUUCAAAAAACACGGUAAAAAUGAAAAAGAUACCGGAAGUGCUGAAGGUCAGAUCGCUUUAUUCACACUAAGAAUUAACCAUUUAACAGAGCAUUUAAAAAAUAAUCGUAAAGAUUAUAAUACAGAAAGAUCUCUUGUUAAGCUAGUAGGAAAAAGAAGAUCACUUCUAGAUUACUUA |
| env-573 | CUAUUCAAAAAACACGGUAAAAAUGAAAAAGAUACCGGAAGCGCUGAAGGUCAGAUCGCUUUAUUCACACUAAGAAUUAACCAUUUAACAGAGCAUUUAAAAAAUAAUCGUAAAGAUUAUAAUACAGAAAGAUCUCUUGUUAAGCUAGUAGGAAAAAGAAGAUCACUUCUAGAUUACUUA |
| env-574 | ..................................................................................................................................................................................... |
| env-575 | CUAUUCAAAAAACACGGUAAAAAUGAAAAAGAUA................................................................................................................................................... |
| env-576 | ..................................................................................................................................................................................... |
| | ..................................................................................................................................................................................... |
| | ..................................................................................................................................................................................... |
| | AUYUUYR•RAARCACGG•RRR•••R•RR••RAYAC•GG••C•RC•GAAGG•CAAAUYGC•YURUUYAC•Y•Y•G•AUYRAYCAYYUR•C•RA•CAYYURAAAAR•AAYCR•AARGAUU•YAAYAC•GAR•G•UC•YURGU•••RYURGU•GG•AAR•G•RG••••YU•YU•GAYUAYYU• |

```
env-577  ....................................................................................................................................................................................
env-578  CUAUUCAAAAAACACGGUAAGAAUGAAAAAGAUACCGGAAGCGCUGAAGGUCAGAUCGCUUUAUUCACGCUGAGAA.............................................................................................................
env-579  CUAUUCAAAAAACACGGUAAAAAUGAAAAAGAUACCGGAAGCGCUGAAGGUCAGAUCGCUUUAUUCACACUGAGAAUUAACCAUCUAACAGAGCAUUUAAAAAAAUAAUCGUAAAGAUUAUAAUACAGAAAGAUCUCUAGUUAAACUUGUAGGUAAAAGAGAUCACUUCUUGAUUACUUA
env-580  CUAUUCAAAAAACACGGUAAAAAUGAAAAAGAUACCGGAAGCGCUGAAGGUCAGAUCGCUUUAUUCACACUGAGAAUUAACCAUCUAACAGAGCAUUUAAAAAAAUAAUCGUAAAGAUUAUAAUACAGAAAGAUCUCUUGUUAAACUUGUAGGUAAAAGAGAUCACUUCUAGAUUACUUA
env-581  CUAUUCAAAAAACACGGUAAAAAUGAAAAAGAUACCGGAAGCGCUGAAGGUCAGAUCGCUUUAUUCACACUAAGAAUUAACCAUUUAACAGAGCAUUUAAAAAAAUAAUCGUAAAGAUUAUAAUACAGAAAGAUCUCUUGUUAAGCUUGUAGGUAAAAGAGAUCACUUCUAGAUUACUUA
env-582  CUAUUCAAAAAACACGGUAAAAAUGAAAAAGAUACCGGAAGCGCUGAAGGUCAGAUCGCUUUAUUCACACUAAGAAUCAAUCAUUUAACAGAGCAUUUAAAAAAAUAAUCGUAAAGAUUAUAAUACGGAAAGAUCUCUUGUUAAGCUUGUAGGUAAAAGAGAUCACUUCUAGAUUACUUA
env-583  CUAUUCAAAAAACACGGUAAAAAUGAAAAAGAUACCGGAAGCGCUGAAGGUCAGAUCGCUUUAUUCACACUAAGAAUUAACCAUUUAACAGAGCAUUUAAAAAAAUAAUCGUAAAGAUUAUAAUACAGAAAGAUCUCUUGUUAAGCUAGUAGGUAAAAGAGAUCACUUCUAGAUUACUUA
env-584  ....................................................................................................................................................................................
env-585  ....................................................................................................................................................................................
env-586  ....................................................................................................................................................................................
env-587  ....................................................................................................................................................................................
env-588  CUAUUCAAAAAACACGGUAAAAAUGAAAAAGAUACCGGAAGCGCUGAAGGUCAGAUCGCUUUAUUCACACUAAGAAUUAACCAUUUAACAGAGCAUUUAAAAAAAUAAUCGUAAAGAUUAUAAUACAGAAAGAUCUCUUGUUAAGCUAGUAGGAAAAAGAGAUCACUUCUAGAUUACUUA
env-589  CUAUUCAAAAAACACGGUAAAAAUGAAAAAGAUACCGGAAGCGCUGAAGGUCAGAUCGCUUUAUUCACACUAAGAAUCAAUCAUUUAACAGAGCAUUUAAAAAAAUAAUCGUAAAGAUUAUAAUACGGAAAGAUCUCUUGUUAAGCUUGUAGGUAAAAGAGAUCACUUCUAGAUUACUUA
env-590  CUAUUCAAAAAACACGGUAAAAAUGAAAAAGAUACCGGAAGCGCUGAAGGUCAGAUCGCUUUAUUCACACUGAGAAUUAACCAUCUAACAGAGCAUUUAAAAAAAUAAUCGUAAAGAUUAUAAUACAGAAAGAUCUCUUGUUAAACUUGUAGGUAAAAGAGAUCACUUCU..........
env-591  CUAUUCAAAAAACACGGUAAAAAUGAAAAAGAUACCGGAAGCGCUGAAGGUCAGAUCGCUUUAUUCACACUAAGAAUCAACCAUUUAACAGAGCAUUUAAAAAAAUAAUCGUAAAGAUUAUAAUACGGAAAGAUCUCUUGUUAAGCUUGUAGGUAAAAGAGAUCACUUCUAGAUUACUUA
env-592  CUAUUCAAAAAACACGGUAAAAAUGAAAAAGAUACCGGAAGCGCUGAAGGUCAGAUCGCUUUAUUCACACUGAGAAUUAACCAUCUAACAGAGCAUUUAAAAAAAUAAUCGUAAAGAUUAUAAUACAGAAAGAUCUCUAGUUAAACUUGUAGGUAAAAGAGAUCACUUCUUGAUUACUUA
env-593  CUAUUCAAAAAACACGGUAAAAAUGAAAAAGAUACCGGAAGC..........................................................................................................................................
env-594  CUAUUCAAAAAACACGGUAAAAAUGAAAAAGAUACCGGAAGCGCUGAAGGUCAGAUCGCUUUAUUCACNCUNAGAAUNAANCAUCUAACAGAGCAUUUAAAAAAAUAAUCGUAAAGAUUAUAAUACAGAAAGAUCUCUNGUUAAACUUGUAGGUAAAAGAGAUCACUUCUUGAUUACUUA
env-596  CUAUUCAAAAAACACGGUAAAAAUGAAAAAGAUA..................................................................................................................................................
env-598  CUAUUCAAAAAACACGGUAAAAAUGAAAAAGAUACCGGAAGCGCUGAAGGUCAGAUCGCUUUAUUCACACUGAGAAUUAACCAUCUAACAGAGCAUUUAAAAAAAUAAUCGUAAAGAUUAUAAUACAGAAAGAUCUCUAGUUAAACUUGUAGGUAAAAGAGAUCACUUCUUGAUUACUUA
env-599  CUAUUCAAAAAACACGGUAAAAAUGAAAAAGAUACCGGAAGCGCUGAAGGUCAGAUCGCUUUAUUCACACUNAGAAUNAANCAUNUAACAGAGCAUUUAAAAAAAUAAUCGUAAAGAUUAUAAUACAGAAAGAUCUCUAGUUAAACUUGUAGGUAAAAGAGAUCACUUCUUGAUUACUUA
env-600  UUAUUCAAAAAACACGGUAAAAAUGAAAAAGAUACCGGAAGCGCUGAAGGUCAGAUCGCUUUAUUCACACUAAGAAUCAACCAUUUAACAGAGCAUUUAAAAAAAUAAUCGUAAAGAUUAUAAUACGGAAAGAUCUCUUGUUAAACUUGUAGGUAAAAGAGAUCACUUCUUGAUUACUUA
env-601  AUAUUUAAAGAACACGGAAAAUCUGACUCAGAUACUGGUUCUGCAGAAGGUCAGAUAGCUUUAUUUACAGCUAGAAUCAACCAUUUAACUGAACAUCUAAAAAAUAAUCGUAAAGAUUUUAAUACUGAAAGAUCAUUAGUGAUGAUGGUAGGUAAAAGAAAAAGUUUGUUGAGCUAUCUU
env-602  AUAUUUAAAAAACACGGAAAAUCUGGCUCAGAUACUGGUUCUGCAGAAGGUCAAAUAGCUUUAUUUACAGCUAGAAUCAACCAUUUAACUGAACAUUUGAAAAAUAAUCGUAAAGAUUUUAAUACAGAAAGAUCAUUGGUGAUGAUGGUUGGUAAAAGAAAAAGCUUAUUAAGCUAUCUU
env-603  AUAUUUAAAAAACACGGAAAAUCUGGCUCAGAUACUGGUUCUGCAGAAGGUCAGAUAGCUUUAUUUACAGCUAGAAUCAACCAUUUAACUGAACAUUUGAAAAAUAAUCGUAAAGAUUUUAAUACAGAAAGAUCAUUGGUGAUGAUGGUUGGUAAAAGAAAAAGCUUAUUAAGCUAUCUU
env-604  UAUUCAAAGAACACGGAAAAUCUGACUCAGAUACUGGCUCUGCAGAAGGUCAGAUAGCUUUAUUUACAGCUAGAAUCAACCAUUUAACUGAACAUACUUAAAAAUAAUACGUAAAGAUUUUAACACUGAAAGAUCAUUAGUGAUGAUGGUAGUAAAAGAAAAGUUUG............
env-605  AUAUUUAAAGAACACGGAAAAUCUGACUCAGAUACUGGUUCUGCAGAAGGUCAGAUAGCUUUAUUUACAGCUAGAAUCAACCAUUUAACUGAACAUCUAAAAAAUAAUCGUAAAGAUUUUAAUACUGAAAGAUCAUUAGUGAUGAUGGUAGGUAAAAGAAAAAGUUUGUUGAGCUAUCUU
env-606  AUUUUCAAAGAACACGGAAAGUCUGAUAAAGACACCGGUUCAUCUGAAGGACAAAUUGCUUUGUUCACAAAAAGAAUUAACCAUUUAACUGAGCAUCUUAAAAACAAUCGUAAAGAUUACAAUACAGAAAGAUCUUUGGUAAUGAUGGUUGGUAAAAGAAAAAGUUUAUUAAACUAUCUU
env-607  AUUUUCAAAGAACACGGAAAGUCUGAUAAAGACACCGGUUCAUCUGAAGGACAAAUUGCUUUGUUCACAAAAAGAAUUAACCAUUUAACUGAGCAUCUUAAAAACAAUCGUAAAGAUUACAAUACAGAAAGAUCUUUGGUAAUGAUGGUUGGUAAAAGAAAAAGUUUAUUAAACUAUCUU
env-608  AUAUUUAAAGAACACGGAAAAUCUGACGUAGAUACUGGAUCUGCAGAAGGUCAAAUAGCUUUAUUUACAACUAGAAUAAAUCAUUUAACUGAACAUCUUAAAAAAAAAUCGUAAAGAUUUUAAUA..............................................................
env-609  A...................................................................................................................................................................................
env-610  AUAUUUAAAAAGCACGGUAAAUCUGAGUCAGAUACGGGCUCUGCUGAAGGACAAAUAGCAUUGUUUACAGCUAGA.........................................................................................................
env-611  AUAUUUAAAAAGCACGGUAAAUCUGAGUCAGAUACGGGCUCUGCUGAAGGACAAAUAGCAUUGUUUACAGCUAGAAUCAACCAUUUAACUAAACAUCUAAAAAAAUAAUCAUAAAGAUUUUAAUACUGAAAGGUCUUUAGUAAUGAUGGUGGGUAAAAGAAAAAGCUUAUUGAACUAUCUU
env-612  AUAUUUAAAAAGCACGGUAAAUCUGAGUCAGAUACGGGCUCUGCUGAAGGACAAAUAGCAUUGUUUACAGCUAGAAUCAACCAUUUAACUGAACAUCUAAAAAAAUAAUCGCAAAGAUUUUAAUACUGAAAGGUCUUUAGUAAUGAUGGUGGGUAAAAGAAAAAGCUUAUUGAACUAUCUU
env-613  AUAUUUAAAAAGCACGGUAAAUCUGAAUCAGAUACGGGCUCCGCUGAAGGACAAAUAGCAUUGUUUACAGCUAGAAUCAACCAUUUAACUAAACAUCUAAAAAAAUAAUCAUAAAGAUU..................................................................
env-614  AUAUUUAAAAAGCACGGUAAAUCUGAAUCAGAUACGGGCUCCGCUGAAGGACAAAUAGCAUUGUUUACAGCUAGAAUCAACCAUUUAACUAAACAUCUAAAAAAAUAAUCAUAAAGAUUUUAAUACUGAAAGGUCUUUAGUAAUGAUGGUGGGUAAAAGAAAAAGCUUAUUGAACUAUCUU
env-615  AUAUUUAAAAAGCACGGUAAAUCUGAAUCAGAUACGGGCUCCGCUGAAGGACAAAUAGCAUUGUUUACAGCUAGAAUCAACCAUUUAACUGAACAUCUAAAAAAAUAAUCGCAAAGAUUUU................................................................
env-616  AUAUUUAAAAAGCACGGUAAAUCUGAAUCAGAUACGGGCUCUGCUGAAGGACAAGUAGCAUUGUUUACAGCUAGAAUCAACCAUUUAACUAAACAUCUAAAAAAAUAAUCGUAAAGAUUUUAACACUGAAAGGUCUUUAGUAAUGAUGGUGGGUAAAAGAAAAAGCUUAUUAAACUAUCUU
env-617  AUAUUUAAAAAGCACGGUAAAUCUGAAUCAGAUACGGGCUCUGCUGAAGGACAAAUAGCAUUGUUUACAGCUAGAAUCAACCAUUUAACUGAACAUCUAAAAAAAUAAUCGUAAGGAUUUUAAUACCGAAAGGUCUUUAGUAAUGAUGGUUGGUAAAAGAAAAAGCUUAUUGAACUAUCUU
env-618  AUAUUUAAAAAGCACGGUAAAUCUGAAUCAGAUACGGGCUCUGCUGAAGGACAAAUAGCAUUGUUUACAGCUAGAAUCAACCAU..................................................................................................
env-619  AUAUUUAAAAAACACGGAAAAUCUGAAUAGAUACCGGAUCUGCAGAAGGGCAAAUAGCUUUAUUCACUGCUAGAAUAAAUCACCUAACUGAACAUCUAAAAAAAAAAUCGUAAAGAUUUUAACACUGAAAGAUCAUUAGUAAUGAUGGUAGGUAAAAGAAAAAGUUUGUUAAGCUACCUU
env-620  AUAUUUAAAGAACACGGAAAAUCUGACUCAGAUACUGGCUCUGCUGAAGGUCAGAUAGCUCUAUUUACAGCUAGGAUCAACCAUUUAACUGAACACCUUAAAAACAAUCGUAAAGAUUUUAAUACCGAAAGAUCUUUAGUAAUGAUGGUAGGUAAAAGAAAAAGUCUUUUAAGCUACUUG
env-621  ....................................................................................................................................................................................
env-622  AUAUUUAAAGAACACGGAAAGUCUGAUACAGAUACUGGUUCUGCAGAAGGUCAAAUAGCUUUAUUUACAGCUAGAAUCAACCAUUUAACUGAACACCUUAAAAACAAUCGUAAAGAUUUUAAUACCGAAAGAUCUUUAGUAAUGAUGGUAGGUAAAAGAAAAAGUCUUUUAAGCUACUUG
env-623  AUAUUUAAAGAACACGGAAAGUCUGAUACAGAUACUGGUUCUGCAGAAGGUCAAAUAGCUUUAUUUACAGCUAGAAUCAACCAUUUAACUGAACACCUUAAAAACAAUCGUAAAGAUUUUAAUACCGAAAGAUCUUUAGUAAUGAUGGUAGGCAAAAGAAAAAGUCUUUUAAGCUACUUG
env-624  AUUUUUAAAAAACACGGAAAGUCAGAUUCAGAUACUGGAUCUGCUGAGGGUCAAAUCGCUUUAUUUACAGCUAGAAUAAAUCAUUUAACUGAACAUCUUAAAAAAAAAUCGUAAAGAUUUUAAUACUGAAAGAUCUUUGGUUAUGAUGGUUGGUAAAAGAAAAAGUUUACUUAGCUACUUA
env-625  AUUUUUAAAAAACACGGAAAGUCAGAUUCAGAUACUGGAUCUGCUGAGGGUCAAAUCGCUUUAUUUACAGCUAGAAUAAAUCAUUUAACUGAACACCUUAAAAAAAAAUCGUAAAGAUUUUAAUACUGAAAGAUCUUUGGUUAUGAUGGUUGGUAAAAGAAAAAGUUUGCUUAGCUACUUA
env-626  AUUUUUAAAACACACGGAAAGUCAGAUUCAGAUACUGGAUCUGCUGAGGGUCAAAUCGCUUUAUUUACAGCUAGAAUAAAUCAUUUAACUGAACAUCUUAAAAAAAAAUCGUAAAGAUUUUAAUACUGAAAGAUCUUUGGUUAUGAUGGUUGGUAAAAGAAAAAGUUUGCUUAGCUACUUA
env-627  ....................................................................................................................................................................................
env-628  AUUUUUAAAACACACGGAAAGUCAGAUUCAGAUACUGGAUCUGCUGAGGGUCAAAUCGCUUUAUUUACAGCUAGAAUAAAUCAUUUAACUGAACAUCUUAAAAAAAAAUCGUAAAGAUUUUAAUACUGAAAGAUCUUUGGUUAUGAUGGUUGGUAAAAGAAAAAGUUUGCUUAGCUACUUA
env-629  AUUUUUAAAAAACACGGAAAGUCAGAUUCAGAUACUGGAUCUGCUGAGGGUCAAAUCGCUUUAUUUACAGCUAGAAUAAAUCAUUUAACUGAACAUCUUAAAAAAAAAUCGUAAAGAUUUUAAUACUGAAAGAUCUUUGGUUAUGAUGGUUGGUAAAAGAAAAAGUUUGCUUAGCUACUUA
env-633  AUUUUUAAAAAACACGGAAAGUCAGACUCAGAUACUGGAUCUGCUGAGGGUCAAAUCGCUUUAUUUACAGUUAGAAUAAAUCAUUUAACUGAACAUCUUAAAAAAAAAUCGUAAAGAUUUUAAUACUGAAAGAUCUUUGGUUAUGAUGGUUGGUAAAAGAAAAAGUUUACUUAGCUACUUA
env-634  ....................................................................................................................................................................................
env-635  ....................................................................................................................................................................................
env-636  AUAUUUAAAGAACACGGAAAAUCUGAUACAGAUACCGGAUCCGCAGAAGGACAAAUAGCUUUAUUCACUACUAGAAUUAAUCACUUAACUGAACAUCUAAAAAAAAAAUCGUAAAGAUUUUAACACGGAAAGAUCAUUAGUAAUGAUGGUAGGUAAAAGAAAAAGCUUAUUA.........
env-637  AUAUUUAAAGAAUACGGAAAGUCUGAUACAGAUACUGGUUCUGCAGAGGGUCAAAUAGCUUUAUUUACUGCUAGAAUCAACCAUUUAACUGAACAUCUUAAAAACAAUCGUAAAGAUUUUAAUACCGAAAGAUCUUUAGUAAUGAUGGUAGGUAAAAGAAAAAGUCUUUUAAGCUACUUG
env-638  AUAUUUAAAGAACACGGAAAAUCUGACUCAGAUACUGGUUCUGCAGAAGGUCAAAUAGCU........................................................................................................................
env-639  AUAUUUAAAGAACACGGAAAAUCUGACUCAGAUACUGGUUCUGCAGAAGGUCAGAUAGCUUUAUUUACAGCUAGAAUCAAUCAUUUAACUGAACACCUUAAAAAAUAAUCGUAAAGAUUUUAAUACUGAAAGGUCUUUAGUGAUGAUGGUAGGUAAAAGAAAAAGUUUGUUAAGCUAUUUG
env-640  AUAUUCAAAGAACACGGAAAAUCUGACUCAGAUACUGGCUCUGCAGAAGGUCAGAUAGCUUUAUUUACAGCUAGAAUCAACCAUUUAACUGAACAUCUUAAAAAAUAAUCGUAAAGAUUUUAACACUGAAAGAUCAUUAGUGAUGAUGGUAGGUAAAAGAAAAAGUUUGUUAGGCUACCUA
env-641  AUUUUCAAGGAACACGGAAAAUCUGAUACAGACACGGGUUCAUCAGAAGGACAAAUUGCUUUGUUCACUGCAAGAAUCAAUCAUUUAACCGAGCAUCUCAAAAAAAAUCGUAAAGAUUUUAAUACAGAAAGGUCGUUAGUGAUGAUGGUUGGUAAAAGAAAAAGCUUAUUAAGCUAUCUU
env-648  AUUUUCAAGGAACACGGAAAAUCUGAUACAGACACGGGUUCAUCAGAAGGACAAAUUGCUUUGUUCACUGCAAGAAUCAAUCAUUUAACCGAGCAUCUCAAAAAAAAUCGUAAAGAUUUUAAUACAGAAAGGUCAUUAGUGAUGAUGGUUGGUAAAAGAAAAAGCUU............
env-649  AUUUUCAAGGAACACGGAAAAUCUGAUACAGACACGGGUUCAUCAGAAGGACAAAUUGCUUUGUUCACUGCAAGAAUCAAUCAUUUAACCGAGCAUCUCAAAAAAAAUCGUAAAGAUUUUAAUACAGAAAGGUCAUUAGUGAUGAUGGUUGGUAAAAGAAAAAGCUUAUUUAUUAGUGAUG
env-650  AUUUUCAAGGAACACGGAAAAUCUGAUACAGACACGGGUUCAUCAGAAGGACAAAUUGCUUUGUUCACUGCAAGAAUCAAUCAUUUAACCGAGCAUCUCAAAAAAAAUCGUAAAGAUUUUAAUACAGAAAGGUCGUUAGUGAUGAUGGUUGGUAAAAGAAAAAGCUUAUUAAGCUAUCUU
env-651  AUUUUCAAGGAACACGGAAAAUCUGAUACAGACACGGGUUCAUCAGAAGGACAAAUUGCUUUGUUCACUGCAAGAAUCAAUCAUUUAACCGAGCAUCUCAAAAAAAAUCGUAAAGAUUUUAAUACAGAAAGGUCAUUAGUGAUGAUGGUUGGUAAAAGAAAAAGCUU............
env-652  AUUUUCAAGGAACACGGAAAAUCUGAUACAGACACGGGUUCAUCAGAAGGACAAAUUGCUUUGUUCACUGCA...........................................................................................................
env-653  AUUUUCAAGGAACACGGAAAAUCUGAUACAGACACGGGUUCAUCAGAAGGACAAAUUGCUUUGUUCACUGCAAGAAUCAAUCAUUUAACCGAGCAUCUCAAAAAAAAUCGUAAAGAUUUUAAUACAGAAAGGUCAUUAGUGAUGAUGGUUGGUAAAAGAAAAAGCUUAUUAAGCUAUCUU
env-655  AUUUUCAAGGAACACGGAAAAUCUGAUACAGACACGGGUUCAUCAGAAGGACAAAUUGCUUUGUUCACUGCAAGAAUCAAUCAUUUAACCGAGCAUCUCAAAAAAAAUCGUAAAGAUUUUAAUACAGAAAGGUCAUUAGUGAUGAUGGUUGGUAAAAGAAAAAGCUUAUUAAGCUAUCUU
env-656  AUUUUCAAGGAACACGGAAAAUCUGAUACAGACACGGGUUCAUCAGAAGGACAAAUUGCUUUGUUCACUGCAAGAAUCAAUCAUUUAACCGAGCAUCUCAAAAAAAAUCGUAAAGAUUUUAAUACAGAAAGGUCGUUAGUGAUGAUGGUUGGUAAAAGAAAAAGCUUAUUAAGCUAUCUU
env-657  AU..................................................................................................................................................................................
env-658  AUUUUCAAGGAACACGGAAAAUCUGAUACAGACACGGGUUCAUCAGAAGGACAAAUUGCUUUGUUCACUGCAAGAAUCAAUCAUUUAACCGAGCAUCUCAAAAAAAAUCGUAAAGAUUUUAAUACAGAAAGGUCAUUAGUGAUGAUGGUUGGUAAAAGAAAAAGCUUAUUAAGCUAUCUU
env-659  AUUUUCAAGGAACACGGAAAAUCUGAUACAGACACGGGUUCAUCAGAAGGACAAAUUGCUUUGUUCACUUCAAGAAUCAAUCAUUUAA............................................................................................
env-660  AUUUUUAAAGAACACGGAAAGUCUGAUACAGACACGGGUUCAUCUGAAGGACAAAUUGCUUUGUUCACAAAAAGAAUUAAUCAUUUAACUGAACAUCUUAAAAAUAAUCGUAAAGAUUAUAACACUGAGAGAUCCUUAGUUAUGAUGGUUGGUAAAAGAAAAAGCUUAUUAAGCUAUCUA
env-661  AUUUUCAAGGAACACGGAAAAUCUGAUACAGACACGGGUUCAUCAGAAGGACAAAUUGCUUUGUUCACUGCAAGAAUCAAUCAUUUAACCGAGCAUCUCAAAAAAAAUCGUAAAGAUUUUAAUACAGAAAGGUCAUUAGUGAUGAUGGUUGGUAAAAGAAAAAGCUUAUUAAGCUAUCUU
env-662  AUUUUCAAGGAACACGGAAAAUCUGAUACAGACACGGGUUCAUCAGAAGGACAAAUUGCUUUGUUCACUGCAAGAAUCAAUCAUUUAACCGAGCAUCUCAAAAAAAAUCGUAAAGAUUUUAAUACAGAAAGGUCAUUAGUGAUGAUGGUUGGUAAAAGAAAAAGCUUAUUAAGCUAUCUU
env-663  AUUUUUAAAGAACAUGGAAAGUCUGAUUCAGACACCGGCUCAACAGAAGGUCAGAUAGCUUUGUUUACAACCAGGAUUAAUCAUCUUACUGAUCAUUUAAAAAAUAAUCGAAAGGACUUUAAUACCGAAAGAUCCCUAGUUAUGAUGGUUGGCAAAAGAAAGAGUUUAUUGAGC......
env-664  AUCUUCGCAAAGCACGGAAAAUCAAAGACUGAUACCGGUUCUGCAGAAGGCCAGAUUGCUCUAUUUACUUAUAGAAUAGAACAUUUGACAAAACACCUUAAAACAAAUCACAAAGACUACAAUACUGAACGUUCUUUGGUUAUGUUAGUUGGUAAACGAAGAGCACUUCUUGAUUAUCUU
env-669  AUCUUUGCAAAACACGGAAAGUCGGCUACCGACACCGGUUCUGCUGAAGGGCAAAUUGCCUUAUUCACUUACCGAAUAGAGCAUUUAACAAAACACCUUAA.............................................................................
env-670  AUCUUUGCAAAACACGGAAAGUCGGCUACCGACACCGGUUCUGCUGAAGGGCAAAUUGCCUUAUUCACUUACCGAAUAGAGCAUUUAACAAAACACCUUAAAGCCAAUCAUAAAGAUUACAACACUGAAAGAUCACUGGUAAUGCUAGUAGGAAAGCGUCGAUCACUUUUAGAUUAUCUU
env-671  AUCUUUGCAAAACACGGAAAGUCGGCUACCGACACCGGUUCUGCUGAAGG..................................................................................................................................
env-672  AUUUUCAAGAAGCACGGAGGUUCUGAGAAGAUACUGGUUCGACAGAAGGGCAAAUUGCUUUGUUUACAUACAGAAUCAGUCAUCUGACAAAUCACCUUAAAAAGAUCGAAAAGAUUUUGGUACACAAAGAGCACUUCAAUUACUUGUUGGUAAAAGAAGAAGUUUAUUAGAUUACCUU
env-673  AUUUUCAAGAAACACGGUGGUUCAGAUACGAAUACUGGUUCAACAGAAGGGCAAGUUGCUCUUUUUACAUACAGAAUUAGUCAUUUGACUGAGCACUUGAAAAAGAAUCGAAAAGAUUUUGGUACACAAAGAGCACUACAAUUACUAGUUGGUAAGAGAAGAAGCUUAUUGGAUUACCUA
env-674  AUUUUCAAGAAACACGGAAAGUCUGAUACGAAUACUGGUUCAACAGAAGGGCAAGUUGCUCUUUUUACAUACCGAAUUAGUCAUUUGACUGAGCACUUGAAAAAGAAUCGAAAAGAUUUUGGUACACAAAGAGCACUACAAUUACUAGUUGGUAAGAGAAGAAGCUUAUUGGAUUACCUA
env-675  AUUUUCAAGAAACACGGGGGUUCUGAUAAAAACACCGGUUCUACAGAGGGGCAAAUUGCUCUUUUUACAUACCGAAUUAGUCACCUAACGAACCACUUG................................................................................
env-676  AUUUUCAAGAAGCACGGAGGUUCUGAUACGAACACGGGUUCCACAGAAGGGCAGAUUGCUCUAUUUACCUACCGAAUCAGUCACCUGACGAAUCACUUGAAAAAGAAUCGAAAAGAUUUUGGUACACAAAGAGCUCUUCAAUUACUUGUUGGUAAAAGAAGAAGUUUAUUGGACUACCUA

         ....................................................................................................................................................................................
         ....................................................................................................................................................................................
         AUYUUYR•RAARCACGG•RRR•••R•RR••RAYAC•GG••C•RC•GAAGG•CAAAUYGC•YURUUYAC•Y•Y•G•AUYRAYCAYYUR•C•RA•CAYYURAAAAR•AAYCR•AARGAUU•YAAYAC•GAR•G•UC•YURGU•••RYURGU•GG•AAR•G•RG••••YU•YU•GAYUAYYU•
```

```
env-677  AUUUUCAAGAAACACGGAGGUUCUGAUACGAACACUGGGUCGACAGAAGGACAAAUUGCUUUAUUCACUUACCGAAUCAGUCACCUAACGAAUCACUUGAAAAAGAAUCGAAAAGAUUUUGGUACACAAAGAGCACUUCAAUUGCUAGUUGGUAAAAGAAGGAGUUUAUUGGACUACCUA
env-678  AUUUUCAAGAAGCACGGAGGUUCUGAUACGAACACGGGUUCCACAGAAGGGCAGAUUGCUCUAUUUACCUACCGAAUCAGUCACCUGACGAAUCACUUGAAAAAGAAUCGAAAAGAUUUUGGUACACAAAGAGCUCUUCAAUUACUUGUUGGUAAAAGAAGAAGUUUAUUGGACUACCUA
env-679  AUUUUCAAGAAGCAUGGAGGUUCUGACACGAAUACGGGGUCUACAGAAGGGCAAAUUGCUUUAUUUACCUACAGAAUCAGUCACCUGACAAAUCACUUGAAAAAGAAUCGAAAAGAUUUUGGUACACAAAGAGCACUUCAAUUACUUGUUGGUAAAAGAAGAAGUUUAUUGGACUACCUA
env-680  AUUUUCAAGAAGCAUGGAGGUUCUGACACGAAUACGGGGUCUACAGAAGGGCA...........................................................................................................................................
env-681  AUUUUCAAGAAACACGGAGGUUCGGACAAAAACACCGGUUCUACAGAAGGGCAAAUCGCUCUUUUUACAUACCGAAUCGGUCACCUUACCAACCACAUGAAAAAGAAUAAAAAAGAUUUUGGUACCCAAAGAGCGCUUCAGCUACUCGUUGGUAAAAGAAGAAGUCUAUUGGACUACUUG
env-682  AUUUUCAAGAAACACGGAGGUUCGGACAAAAACACCGGUUCUACAGAAGGGCAAAUCGCUCUUUUUACAUACCGAAUCGGUCACCUUACCAACCACAUGAAAAAGAAUAAAAAAGAUUUUGGUACCCAAAGAGCGCUUCAGCUACUCGUUGGUAAAAGAAGAAGUCUAUUGGACUACUUG
env-683  AUUUUCAAGAAACACGGAGGUUCGGACAAAAACACCGGUUCUACAGAAGGGCAAAUCGCUCUUUUUACAUACCGAAUCGGUCACCUUACCAACCACAUGA.................................................................................................
env-684  AUUUUCAAGAAACACGGAGGUUCGGACAAAAACACCGGUUCUACAGAAGGGCAAAUCGCUCUUUUUACAUACCGAAUCGGUCACCUUACCAACCACAUGAAAAAGAAUAAAAAAGAUUUUGGUACCCAAAGAGCGCUUCAGCUACUCGUUGGUAAAAGAAGAAGUCUAUUGGACUACUUG
env-685  AUUUUCAAGAAACACGGAGGUUCGGACAAAAACACCGGUUCUACAGAAGGGCAAAUCGCUCUCUUUACUUACCGAAUCGGUCACCUUACGAACCACAUGAAAAAGAAUAAAAAAGAUUUUGGUACCCAAAGAGCACUUCAGCUACUCGUUGGUAAAAGAAGAAGUCUAUUGGACUACUUG
env-686  AUUUUCAAGAAACACGGAGGUUCGGACAAAAACACCGGUUCUACAGAAGGGCAAAUCGCUCUUUUUACAUACCGAAUCGGUCACCUUACCAACCACAUGAAAAAGAAUAAAAAAGAUUUUGGUACCCAAAGAGCGCUUCAGCUACUCGUUGGUAAAAGAAGAAGUCUUUUGGACUACUUG
env-687  AUUUUCAAGAAACACGGAGG................................................................................................................................................................................
env-688  AUUUUCAAGAAACACGGAGGUUCGGACAAAAACACCGGUUCUACAGAAGGGCAAAUCGCUCUUUUUACUUACCGAAUCGGUCACCUUACCAACCACAUGAAAAAGAAUAAAAAAGAUUUUGGUACCCAAAGAGCGCUUCAGCUACUCGUUGGUAAAAGAAGAAGUCUAUUGGACUACUUG
env-689  AUUUUCAAGAAACACGGAGGUUCGGACAAAAACACCGGUUCUACAGAAGGGCAAAUCGCUCUUUUUACAUACCGAAUCGGUCACCUUACCAACCACAUGAAAAAGAAUAAAAAAGAUUUUGGUACCCAAAGAGCGCUUCAGCUACUCGUUGGUAAAAGAAGAAGUCUAUUGGACUACUUG
env-690  AUUUUCAAGAAACACGGAGGUUCGGACAAAAACACCGGUUCUACAGAAGGGCAAAUCGCUCUUUUUACUUACCGAAUCGGUCACCUUACCAACCACAUGAAAAAGAAUAAAAAAGAUUUUGGUACCCAAAGAGCGCUUCAGCUACUCGUUGGUAAAAGAAGAAGUCUAUUGGACUACUUG
env-691  AUUUUCAAGAAACACGGAGGUUCGGACAAAAACACCGGUUCUACAGAGGGGCAAAUCGCUCUUUUUACUUACCGAAUCGGUCACCUUACCAACCACAUGAAAAAGAAUAAAAAAGAUUUUGGUACCCAAAGAGCGCUUCA......................................
env-692  AUUUUCAAGAAACACGGAGGUUCUGAUAAAAACACCGGUUCUACAGAAGGGCAAAUUGCUCUUUUUACAUACCGAAUUAGUCACCUAACGAACCACUUGAAAAAGAAUAGAAAAGAUUUUGGUACCCAAAGAGCACUUCAGCUACUUGUUGGUAAAAGAAGAAGCCUAUUGGACUACUUG
env-693  AUUUUCAAGAAACACGGAGGUUCUGAUAAAAACACCGGUUCUACAGAAGGGCAAAUUGCUCUUUUUACAUACCGAAUUAGUCACCUAACGAACCACUUGAAAAAGAAUAGAAAAGAUUUUGGUACCCAAAGAGCACUUCAGCUACUUGUUGGUAAAAGAAGAAGCCUAUUGGACUACUUG
env-694  AUCUUUAAAAAACAUGGUAAAUCAGAAAAAGACACUGGAUCAACUGAAGGUCAAAUUGCGCUUUUUACCCAUCGUAUUCAACAUCUAUCAGAACACCUUAAACAAAACAGAAAAGAUUUCAAUACUGAACGUUCACUAGUCCGUUUAGUUGGAAAGCGAAGAAGUCUUCUUGAUUAUUUA
env-698  AUCUUUAAAAAACAUGGGAAAUCAGAAAAAGACACUGGAUCAACUGAAGGUCAAAUUGCGCUUUUUACCCAUCGUAUUCAACAUCUAUCAGAACACCUUAAACAAAACAGAAAAGAUUUCAAUACUGAACGUUCACUAGUCCGUUUAGUUGGAAAGCGAAGAAGUCUUCUUGAUUAUUUG
env-699  AUCUUUAAAAAACAUGGGAAAUCAGAAAAAGACACUGGAUCAACUGAAGGUCAAAUUGCGCUUUUUACCCAUCGUAUUCAACAUCUAUCAGAACACCUUAAACAAAACAGAAAAGAUUUCAAUACUGAACGUUCACUAGUCCGUUUAGUUGGAAAGCGAAGAAGUCUUCUUGAUUAUUUG
env-700  AUCUUUAAAAAACAUGGUAAAUCAGAAAAAGACACUGGAUCAACUGAAGGUCAAAUUGCGCUUUUUACCCAUCGUAUUCAACAUCUAUCAGAACACCUUAAACAAAACAGAAAAGAUUUCAAUACUGAACGUUCACUAGUCCGUUUAGUUGGAAAGCGAAGAAGUCUUCUUAAUUAUUUA
env-701  AUCUUUAAAAAACAUGGGAAAUCAGAAAAAGACACUGGAUCAACUGAAGGUCAAAUUGCGCUUUUUACCCAUCGUAUUCAACAUCUAUCAGAACACCUUAAACAAAACAGAAAAGAUUUCAAUACUGAACGUUCACUAGUCCGUUUAGUUGGAAAGCGAAGAAGUCUUCUUGAUUAUUUG
env-702  AUCUUUAAAAGACAUGGUAAAUCAGAAAAAGACACUGGAUCAACUGAAGGUCAAAUUGCGCUUUUUACCCAUCGUAUUCAACAUCUAUCAGAACACCUUAAACAAAACAGAAAAGAUUUCAAUACUGAACGUUCACUAGUCCGUUUAGUUGGAAAGCGAAGAAGUCUUCUUGAUUAUUUA
env-703  ....................................................................................................................................................................................................
env-704  ....................................................................................................................................................................................................
env-705  AUUUUUAAAAAACACGGUAAAUCUGCGAAGGACACUGGGUCUGCAGAAGGACAAAUCGCUCUUCACUCACCGUAUCGAACAUGUCCAGAUGUAC.....................................................................................................
env-706  ....................................................................................................................................................................................................
env-707  AUUUUUAAAAAACACGGGAAAUCAGAAAAAGACACGGGUUCUGCUGAAGGACAAAUCGCUCUUUUUACUCACCGUAUCGAACACCUAUCUUCUCACCUAAAGAGCAAUCGUAAAGAUUACAAUACAGAACGCUCACUCGUACGUUUGGUAGGAAAACGAAGAAGUUUAUUAGAUUACCUA
env-708  AUUUUUAAAAAACACGGGAAAUCUGAAAAUGACACAGGUUCUGCAGAAGGACAAAUUGCGCUUUUCACUCACCGUAUUGAACACCUAUCUAACCACCUAAAAAGUAAUCGUAAAGAUUUUAAUACGGAACGCUCUUUAGUUCGAUUGGUAGGGAAACGAAGAAGUUUAUUAGACUACUUA
env-709  AUUUUCGCAAAACACGGUAAAGAUGAAAAAAAUACCGGUAGCGCAGAAGGACAGGUAGCUUUGUUUACCUUUAGAAUUAACCACCUAACCCAGCAUCUAAAAAGAAAUCAUAAAGAUUUCAACACAGAAAGGGCGCUAGUGGCCUUAGUGGGAAAAAGAAAAAAUUACUUGAUUACUUA
env-711  AUCUUUUCAAAGUACGGAAAAUCUGACAAAGACACAGGUUCACCCGAAGGACAAAUCGCAUUGUUUACACACCGUAUCAAUCAUUUGACCAAUCACUUAAAGAACAACAGACAUGACUAUAGCACUGAAAAAGCAUUGGUCAAGAUGGUAGGAAAAAGGCGAAGUCUUCUAAACUAUCUA
env-712  AUUUUCAAACAACACGGUAAAUCAGAAAAAGAUACAGGUUCUGCAGAAGGGCAAAUCGCUUUAUUCACACACCGUAUCAACCACCUUUCUAAUCACCUAAAAACAAACCGUAAAGAUUACAAUACGGAACGUUCUUUGGUACGCUUAGUAGGGAAACGUCGUAGUUUACUAGAUUAUUUA
Fba-2-1  AUCUUCAAACAACACGGUAAAUCUGAAAAAGAUACCGGUUCCGCAGAGGGACAAAUCGCUUUAUUUACUCAUCGUAUCAACCACCUAUCUAACCACCUAAAAACAAAUCGUAAAGAUUUUAACACGGAGCGUUCAUUGGUUAACUUAGUUGGAAAACGCCGUAGUCUACUAGAUUAUUUA
env-713  AUCUUCAAACAACACGGUAAAUCUGAAAAAGAUACCGGUUCCGCAGAGGGACAAAUCGCUUUAUUUACUCAUCGUAUCAACCACCUAUCUAACCACCUAAAAACAAAUCGUAAAGAUUUUAAUACGGAGCGUUCAUUGGUUAACUUAGUUGGAAAACGCCGUAGUCUACUAGAUUAUUUA
env-718  AUCUUCAAACAACACGGUAAAUCUGAAAAAGAUACCGGUUCCGCAGAGGGACAAAUCGCUUUAUUUACUCAUCGUAUCAACCACCUAUCUAACCACCUAAAAACAAAUCGUAAAGAUUUUAAUACGGAGCGUUCAUUGGUUAACUUAGUUGGAAAACGCCGUAGUCUACUAGAUUAUUUA
env-727  AUCUUCAAACAACACGGUAAAU..............................................................................................................................................................................
env-728  AUCUUCAAACAACACGGUAAAUCU............................................................................................................................................................................
env-729  AUCUUCAAACAACACGGUAAAUCUGAAAAAGAUACCGGUUCCGCAGAGGGACAAAUCGCUUUAUUUACUCAUCGUAUCAACCACCUAUCUAACCACCUAAAAACAAAUCGUAAAGAUUUUAAUACGGAGCGUUCAUUGGUUAACUUAGUUGGAAAACGCCGUAGUCUACUAGAUUAUUUA
env-730  AUCUUCAAACAACACGGUAAAUCAGAAAAGGAUACAGGUUCUGCUGAAGGGCAAAUCGCUUUAUUCACACACCGUAUCAACCACCUUUCUAAUCACCUAAAAACAAACCGUAAAGAUUACAAUACGGAACGUUCUUUGGUGCGCUUA.................................
env-731  AUCUUUAAAAAGCACGGAAAGUCUGAAAAAGACACAGGCUCACCUGAGGGACAAAUCGCCCUCUUUACCCACCGAAUUCAACAUUUAUCGGAUCACCUAAAAUCAAAUCGUAAAGAUUUUAACACGGAACGUUCCUUGGUUCGUUUGGUAGGAAAACGCAGAAGUCUCCUGGAUUAUUUA
env-734  AUCUUUAAAAAGCACGGAAAGUCUGAAAAAGACACAGGCUCACCUGAGGGACAAAUCGCCCUCUUUACCCACCGAAUUCAACAUUUAUCGGAUCACCUAAAAUCAAAUCGUAAAGAUUUUAACACGGAACGUUCCUUGGUUCGUUUGGUAGGAAAACGCAGAAGUCUCCUGGAUUAUUUA
env-735  AUCUUUAAAAAGCACGGAAAGUCUGAAAAAGACACAGGCUCACCUGAGGGACAAAUCGCCCUCUUUACCCACCGAAUUCAACAUUUAUCGGAUCACCUAAAAUCAAAUCGUAAAGAUUUUAACACGGAACGUUCCUUGGUUCGUUUGGUAGGAAAACGCAGAAGUCUCCUGG........
env-736  AUCUUUAAAAAGCACGGAAAGUCUGAAAAAGACACAGGCUCACCUGAGGGACAAAUCGCCCUCUUUACCCACCGAAUUCAACAUUUAUCGGAUCACCUAAAAUCAAAUCGUAAAGAUUUUAACACGGAACGUUCCUUGGUUCGUUUGGUAGGAAAACGCAG..................
env-737  ....................................................................................................................................................................................................
env-738  AUCUUCAAAAAGCACGGGAA................................................................................................................................................................................
env-739  ....................................................................................................................................................................................................
env-740  CUUUUUAAAAAACACGGUAAAUCUGAAAAAGACACUGGAUCAACUGAAGGCCAAAUUGCGCUUUUUACCCAUCGUAUUCAACAUCUAUCAGAACACCUUAAACAAAAUAGAAAGGAUUUCAAUACUGAACGUUCACUAGUUCGUUUAGUUGGAAAGCGAAGAAGUCUUCUAGAUUAUUUA
env-741  AUUUUUAAAAAACAUGGAAAAUCCCAAAAAGAUACGGGCUCUGCUGAGGGCCAGAUCGCCCUUUUCUCCUAUAGAAUUAAUCACCUUUCAAACCAUCUCAAAAAAAAUCAAAAGGAUUUCAAUACAGAGCGAUCUUUAGUUAAAUUGGUAGGAAAGCGUCGUCACCUUUUAGAUUAUUUA
env-742  AUUUUUAAAAAACAUGGAAAAUCCGAAAAAGAUACGGGCUCUGCUGAGGGCCAGAUUGCCCUUUUCUCCCAUAGAAUUAAUCACCUUUCAAACCAUCUCAAAAAAAAUCAAAAGGAUUUCAAUACAGAGCGAUCUUUAGUUAAAUUGGUAGGAAAGCGUCGUCACCUUUUAGAUUAUCUA
env-743  AUUUUCAAGAAGUACGGAAAAUCAGAGAAAAACACAGGUUCUGCAGAGGGUCAAAUUGCCCUUUUUUCACACAGAAUCGACCACCUUUCCAAACAUUUAAAAAUCAAUAAAAAAGACUUUAAUACGGAACGUUCCUUGGUGAAAUUAGUAGGAAAACGUCGUCAUCUUUUAGAUUAUCUA
env-744  AUUUUCAA............................................................................................................................................................................................
env-745  AUUUUCAAGAAGUACGGAAAAUCAGAGAAAAACACAGGUUCUGCAGAGGGUCAAAUUGCCCUUUUUUCACACAGAAUCGACCACCUUUCCAAACAUUUAAAAACCAAUAAAAAAGACUUUAAUACGGAACGUUCCUUGGUGAAAUUAGUAGGAAAACGUCGUCAUCUUUUAGAUUAUCUA
env-746  AUUUUCAAGAAGUACGGAAAAUCAGAGAAAAACACAGGUUCUGCAGAGGGUCAAA.............................................................................................................................................
env-747  AUUUUCAAGAAGUACGGAAAAUCAGAGAAAAACACAGGUUCUGCAGAGGGUCAAAUUGCCCUUUUUUCACACAGAAUCGACCACCUUUCCAAACAUUUAAAAACCAAUAAAAAAGACUUUAAUACGGAACGUUCCUUGGUGAAAUUAGUAGGAAAACGUCGUCAUCUUUUAGAUUAUCUA
env-748  AUUUUCAAAAAACACGGAAAAUCUGAAAAAGAUACUGGUUCUAUUGAAGGACAAAUUGCACUUUUUUCACUCAGAAUUGAUCACCUUUCCAACCACCUAAAAAAAAACAAAAAGGAUUUUAAUACAGAGCGUUCUCUGGUGAAAUUGGUUGGGAAACGCCGCAGUCUAUUAGAUUACCUA
env-749  AUUUUCAAAAAGCACGGAAAGUCUGAUAAAGAUACUGGUUCUAUUGAAGGACAAAUUGCACUUUUUUCACUCAGAGUUGAUCACCUCUCUAACCACCUGAAAAAAAACAAAAAAGAUUUCAAUACAGAACGUUCUCUGGUAAAAUUGGUUGGGAAACGCCGUAGCCUUUUAGAUUACCUC
env-750  AUUUUCAAAAAGCACGGAAAGUCUGAUAAAGAUACUGGUUCUAUUGAAGGACAAAUUGCACUUUUUUCACUCAGAGUUGAUCACCUCUCUAACCACCUGAAAAAAAACAAAAAAGAUUUCAAUACAGAACGUUCUCUGGUAAAAUUGGUUGGGAAACGCCGUAGCCUUUUAGAUUA....
env-751  AUUUUCAAAAAACACGGAAAAUCUGAGAAAGAUACUGGGUCUACUGAAGGACAAAUUGCACUUUUUUCGCUCAGAGUUGAUCACCUUUCCAACCACCUGAAAAAAAACAAAAAGGAUUUCAAUACAGAGCGUUCUCUGGUAAAAUUGGUUGGGAAACGUCGUAGCCUAUU..........
env-752  AUUUUCAAAAAACACGGAAAAUCUGAGAAAGAUACUGGGUCUACUGAAGGACAAAUUGCACUUUUUUCGCUCAGAGUUGAUCACCUUUCCAACCACCUGAAAAAAAACAAAAAGGAUUUCAAUACAGAGCGUUCUCUGGUAAAAUUGGUUGGGAAACGUCGUAGCCUAUUAGAUUACCUU
env-753  AUUUUCAAAAAACAUGGAAAAUCCGAAAAAGACACCGGAUCUUCUGAGGGACAAAUUGCACUUUUUUCUCACAGAAUUGACCAUCUUUCCAACCACCUUAAAAAAAACAAAAAGGAUUUCAAUACAGAGCGUUCCCUCGUUAAAUUGGUAGGGAAACGCCGUCACCUAUUGGAUUACCUU
env-754  AUUUUCAAAAAACAUGGAAAAUCCGAAAAAGACACCGGAUCUUCUGAGGGACAAAUUGCACUUUUUUCUCACAGAAUUGACCAUCUUUCCAACCACCUUAAAAAAAACAAAAAGGAUUUCAAUACAGAGCGUUCCCUCGUUAAAUUGGUAGGGAAACGCCGUCACCUAUUGGAUUACCUU
env-755  AUUUUUAAAAAACAUGGAAAAUCCGAAAAAGAUACGGGCUCUGCUGAGGGUCAGAUUGCCCUUUUCUCCCAUAGAAUUAAUCACCUUUCAAACCAUCUCAAAAAAAAUCAAAAGGAUUUCAAUACAGAGCGAUCUUUAGUUAAAUUGGUAGGAAAGCGUCGUCACCUUUUAGAUUAUCUA
env-756  AUUUUCAAAAAACAUGGAAAAUCUGAAAAAGACACAGGAUCAACUGAGGGACAAAUCGCACUUUUCUCCCACAGAAUUGACCACCUUUCCAACCACCUUAAAAAGAAUAAAAAGGAUUUUAACACAGAACGUUCCCUGGUAAAAUUGGUAGGAAAACGCCGUCAUCUUUUGGAUUACUUA
env-758  AUUUUCAAAAAACACGGAAAAUCUGAAAAAGACACCGGAUCUACUGAAGGACAAAUUGCACUUUUUUCUCACAGAAUUGAUCACCUUUCAAACCACCUUAAAAAAAACAAAAAGGAUUUCAAUACGGAACGUUCCUUGGUCAAAUUGGUAGGGAAACGCCGUCACUUAUUGGAUUACCUC
env-759  AUUUUCAAGAAGUACGGAAAAUCAGAGAAAAACACAGGUUCUGCAGAGGGUCAAAUUGCCCUUUUUUCACACAGAAUCGACCACCUUUCCAAACAUUUAAAAACCAAUAAAAAAGACUUUAAUACGGAACGUUCCUUGGUGAAAUUAGUAGGAAAACGUCGUCAUCUUUUAGAUUAUCUA
env-760  AUUUUCAAGAAGUACGGAAAAUCAGAGAAAAACACAGGUUCUGCAGAGGGUCAAAUUGCCCUUUUUUCACACAGAAUCGACCACCUUUCCAAACAUUUAAAAACCAAUAAAAAAGACUUUAAUACGGAACGUUCCUUGGUGAAGUUAGUAGGAAAACGUCGUCAUCUUUUAGAUUAUCUA
env-761  AUUUUCAAAAAACAUGGAAAGUCCGAAGC.......................................................................................................................................................................
env-762  AUUUUCAAAAAACAUGGAAAGUCCGAAGCUGACACCGGCUCUGCCGAGGGCCAAAUUGCAUUAUUUUCCCUUAGAAUUGAUCACCUAUCCUCCCAUCUUAAGAAAAACCGUAAAGAUUUCAAUACUGAAAGAUCACUAGUUAAGCUGGUAGGAAAACGUCGAAAAUUAUUAGAUUAUCUG
env-763  AUUUUCAAAAAACAUGGAAAGUCAGAAUCUGACACCGGCUCUACUGAGGGCCAAAUCGCAUUAUUUUCCCUCAGAAUUGAUCACCUAACCUCCCAUCUUAAGAAAAACCGUAAAGAU.........................................................................
env-764  AUUUUCAAAAAACAUGGAAAGUCAGAAUCUGACACCGGCUCUACUGAGGGCCAAAUCGCAUUAUUUUCCCUCAGAAUUGAUCACCUAACCUCCCAUCUUAAGAAAAACCGUAAAGAUUUCAAUACUGAAAGAUCACUAGUUAAGCUGGUAGGAAAACGUAGAAAAUUAUUGGAUUAUCUG
env-765  AUUUUCAAAAAACAUGGAAAGUCAGAAGCUGACACCGGCUCUACUGAGGGCCAAAUUGCAUUAUUUUCCCUUAGAAUUGAUCACCUUACCUCCCAUCUUAAGAAAAACCGUAAAGAUUUCAAUACUGAAAGAUCACUGGUUAAGCUGGUAGGAAAACGUAGAAAAUUAUUGGAUUAUCUU
env-766  AUUUUCAAAAAACAUGGAAAGUCAGAAGCUGACACCGGCUCUACUGAGGGCCAAAUUGCAUUAUUUUCCCUCAGAAUUGAUCACCUAACCUCACAUCUUAAGAAAAACCGUAAAGAUUUCAAUACUGAAAGAUCACUAGUUAAGCUGGUAGGAAAACGUAGAAAAUUAUUGGAUUAUCUU
env-767  AUUUUCAAAAAACAUGGAAAGUCAGAAGCUGACACCGGCUCUACUGAGGGUCAAAUUGCAUUAUUUUCCCUCAGAAUUGAUCACCUAACCUCCCAUCUUAAGAAAAACCGUAAAGAUUUCAAUACUGAAAGAUCACUAGUUAAGCUGGUAGGAAAACGUAGAAAAUUAUUGGAUUAUCUG
env-768  AUUUUCAAAAAACAUGGAAAGUCAGAAGCUGACACCGGCUCUACUGAGGGUCAAAUUGCAUUAUUUUCCCUCAGAAUUGAUCACCUAACCUCCCAUCUUAAGAAAAACCGUAAAGAUUUCAAUACUGAAAGAUCACUAGUUAAGCUGGUAGGAAAACGUAGAAAAUUAUUGGAUUAUCUU
env-769  AUUUU...............................................................................................................................................................................................
env-770  AUUUUCAAAAAACAUGGAAAGUCAGAAGCUGACACCGGCUCUACUGAGGGCCAAAUUGCAUUAUUUUCCCUCAGAAUUGAUCACCUAACCUCCCAUCUUAAGAAAAACCGUAAAGAUUUCAAUACUGAAAGAUCACUAGUUAAGCUGGUAGGAAAACGUAGAAAAUUAUUGGAUUAUCUG
env-771  AUUUUCAAAAAACAUGGAAAGUCAGAAGCUGACACCGGCUCUACUGAGGGUCAAAUUGCAUUAUUUUCCCUCAGAAUUGAUCACCUAACCUCCCAUCUUAAGAAAAACCGUAAAGAUUUCAAUACUGAAAGAUCACUAGUUAAGCUGGUAGGAAAACGUAGAAAAUUAUUGGAUUAUCUG
env-772  AUUUUCAAAAAACAUGGAAAGUCAGAAGCUGACACCGGCUCUACUGAGGGCCAAAUUGCAUUAUUUUCCCUCAGAAUUGAUCACCUAACCUCCCAUCUUAAGAAAAACCGUAAAGAUUUCAAUACUGAAAGAUCACUAGUUAAGCUGGUAGGAAAACGUAGAAAAUUAUUGGAUUAUCU.
env-773  AUUUUCAAAAAACAUGGAAAGUCAGAAUCUGACACCGGCUCUACUGAGGGCCAAGUUGCAUUAUUUUCCCUCAGAAUUGAUCACCUAACCUCCCAUCUUAAGAAAAAUCGUAAAGAUUUCAAUACUGAAAGAUCACUAGUUAAGCUGGUAGGAAAACGUAGAAAAUUAUUGGAUUAUCUA
env-774  AUUUUCAAAAAACAUGGAAAGUCAGAAGCUGACACCGGCUCUACUGAGGGCCAAAUUGCAUUAUUUUCCCUCAGAAUUGAUCAUCUAACCUCCCAUCUUAAGAAAAACCGUAAAGAUUUCAAUACUGAAAGAUCACUAGUUAAGCUGGUAGGAAAACGUAGAAAAUUAUUGGAUUAUCUU
env-775  AUUUUCAAAAAACAUGGAAAGUCAGAAGCUGACACCGGCUCUACUGAGGGUCAAAUUGCAUUAUUUUCCCUCAGAAUUGAUCACCUAACCUCCCAUCUUAAGAAAAACCGUAAAGAUUUCAAUACUGAAAGAUCACUAGUUAAGCUGGUAGGAAAACGUAG..................
env-776  AUUUUCAAAAAACAUGGAAAGUCAGAAGCUGACACCGGCUCUACUGAGGGUCAAAUUGCAUUAUUUUCCCUCAGAAUUGAUCACCUAACCUCCCAUCUUAAGAAAAACCGUAAAGAUUUCAAUACUGAAAGAUCACUAGUUAAGCUGGUAGGAAAACGUAGAAAAUUAUUGGAUUAUCUU
env-777  AUUUUCAAGAAACAUGGAAAAUCUGAGAUUGAUACUGGUUCUACAGAGGCACAAAUAGCAUUAUUUUCUCACCGAAUUGACCACUUAUCUAAUCAUCUUAAGAAUAAUCAGAAGGAUUUCAAUACGGAACGAUCACUAGUUAAACUAGUUGGUAAGCGCCGACAACUUUUAGAUUAUCUU
env-778  AUUUUCAAGAAACAUGGAAAAUCUGAGAUUGAUACUGGUUCUACAGAGGCACAAAUAGCAUUAUUUUCUCACCGAAUUGACCACUUAUCUAAUCAUCUUAAGAAUAAUCAGAAGGAUUUCAAUACGGAAAGAUCACUAGUUAAACUAGUUGGUAAGCGCCGACAACUUUUAGAUUAUCUU

         ....................................................................................................................................................................................................
         ....................................................................................................................................................................................................
         AUYUUYR•RAARCACGG•RRR•••R•RR••RAYAC•GG••C•RC•GAAGG•CAAAUYGC•YURUUYAC•Y•Y•G•AUYRAYCAYYUR•C•RA•CAYYURAAAAR•AAYCR•AARGAUU•YAAYAC•GAR•G•UC•YURGU•••RYURGU•GG•AAR•G•RG••••YU•YU•GAYUAYYU•
```

```
env-779  AUUUUCAAGAAACAUGGAAAAUCUGAAGUUGACACUGGUUCUACCGAGGGACAAAUAGCAUUAUUUUCUCACCGAAUUGACCACUUAUCUAAUCAUCUUAAGAAUAAUCAGAAGGAUUUCAAUACGGAAAGAUCACUUGUAAAACUAGUUGGUAAGCGCCGUAAACUUUUAGAUUAUCUU
env-780  AUUUUCAAGAAACAUGGAAAAUCUGAAGUUGACACUGGUUCUACCGAGGGACAAAUAGCAUUAUUUUCUCACCGAAUUGACCACUUAUCUAAUCAUCUUAAGAAUAAUCAGAAGGAUUUCAAUACAGAACGAUCACUUGUAAAACUAGUUGGUAAGCGCCGUAAACUUUUAGAUUAUCUU
env-781  AUUUUCAAGAAACAUGGAAAAUCUGAAGUUGACACUGGUUCUACCGAGGGACAAAUAGCAUUAUUUUCUCACCGAAUUGA....................................................................................................
env-783  AUUUUCAAGAAACAUGGAAAAUCUGAAGUUGACACUGGUUCUACCGAGGGACAAAUAGCAUUAUUUUCUCACCGAAUUGAUCACUUAUCUAAUCAUCUUAAGAAUAAUCAGAAGGAUUUCAAUACGGAACGAUCACUUGUAAAACUAGUUGGUAAGCGCCGUAAACUUUUAGAUUAUCUU
env-784  AUUUUCAAGAAACACGGAAAAUCUGAAGUUGACACUGGUUCUACCGAGGGACAAAUAGCAUUAUUUUCUCACCGAAUUGACCACUUAUCUAAUCAUCUUAAGAAUAAUCAGAAGGAUUUCAAUACGGAACGAUCACUUGUUAAACUAGUUGGUAAGCGCCGUAAGCUUUUAGAUUAUCUU
env-785  AUUUUCAAGAAACACGGAAAAUCUGAAGUUGACACUGGUUCUACCGAGGGACAAAUAGCAUUAUUUUCUCACCGAAUUGACCACUUAUCUAAUCAUCUUAAGAAUAAUCAGAAGGAUUUCAAUACGGAAAGAUCACUUGUAAAACUAGUUGGUAAGCGCCGUAAACUUUUAGAUUAUCUU
env-786  AUUUUCAAGAAACAUGGAAAAUCUGAAGUUGACACCGGUUCUGCCGAGGGACAAAUAGCAUUAUUUUCUCACAGAAUAGACCACUUAUCUAAUCAUCUUAAGAAUAAUCAGAAAGAUUUUAAUACGGAAAGAUCAUUAGUUAAACUUGUAGGUAAGCGCCGUCAACUUUUAGAUUAUCUU
env-787  AUUUUCAAGAAACAUGGAAAAUCUGAAGUUGACACCGGUUCUGCCGAGGGACAAAUAGCAUUAUUUUCUCACAGAAUAGACCACUUAUCUAAUCAUCUUAAGA.............................................................................
env-788  AUUUUCAAGAAACAUGGAAAAUCUGAAGUUGACACCGGUUCUGCCGAGGGACAAAUAGCAUUAUUUUCUCACAGAAUAGACCACUUAUCUAAUCAUCUUAAGAAUAAUCAGAAAGAUUUUAAUACGGAAAGAUCAUUAGUUAAACUUGUAGGUAAGCGCCGUCAACUUUUAGAUUAUCUU
env-789  AUUUCGAAAAACACGGUCAGUCUAAGACAAACACCGGUUCUGCUGAAGGGCAAAUCGCUUUGUUCACUGCACGCAUUGAGCACUUGUCUAAGCACUUAAAAAACAACCACAAAGAUUACAACACUGAAAAGUCUUUGGUGAAGUUGGUAGGUAAAAGAAGGUCACUUCUCGACUACUUA
env-791  AUUUUCGAGAAACACGGUAAAGGAAAAAACGAUACUGGUUCUUCAGAAGGUCAAAUCGCAUUAUUUACGCACAGAAUUAACCAUUUAACAGAACACUUAAAGAAAAAUCGUAAAGAUUUUAAUACAGAACGUUCUCUAGUAAUGAUGGUUGGUAAACGUAGAAGUUUAUUAGAUUAUCUU
env-794  AUUUUCGAGAAACACGGUAAAGGAAAAAACGAUACUGGUUCUUCAGAAGGUCAAAUCGCAUUAUUUACGCACAGAAUUAACCAUUUAACAGAACACUUAAAGAAAAAUCGUAAAGAUUUUAAUACAGAACGUUCUCUAGUAAUGAUGGUUGGUAAACGUAGAAGUUUAUUAGAUUAUCUU
env-795  ....................................................................................................................................................................................
env-796  AUUUCGAGAAACACGGUAAAGGAAAAAACGAUACUGGUUCUUCAGAAGGUCAAAUCGCAUUAUUUACGCACAGAAUUAACCAUUUAACAGAACACUUAAAGAAAAAUCGUAAAGAUUUUAAUACAGAACGUUCUCUAGUAAUGAUGGUUGGUAAACGUAGAAGUUUAUUAGAUUAUCUUA
env-797  AUUUUCGAGAAACACGGUAAAGGAAAAAACGAUACUGGUUCUUCAGAAGGUCAAAUCGCAUUAUUUACGCACAGAAUUAACCAUUUAACAGAACACUUAAAGAAAAAUCGUAAAGAUUUUAAUACAGAACGUUCUCUAGUAAUGAUGGUUGGUAAACGUAGAAGUUUAUUAGAUUAUCUU
env-798  AUUUUCGAGAAACACGGUAAAGGAAAAAACGAUACUGGUUCUUCAGAAGGUCAAAUCGCAUUAUUUACGCACAGAAUUAACCAUUUAACAGAACACUUAAAGAAAAAUCGUAAAGAUUUUAAUACAGAACGUUCUCUAGUAAUGAUGGUUGGUAAACGUAGAAGUUUAUUAGAUUAUCUU
env-799  AUAUUUAAAAAGCACGGAUCUAAUGAAAAAGAUACCGGAAGUUCAGAAGGACAAAUCGCUUUGUUCACUUACAGAAUUGCUCACUUAACAGAACAUCUUAAGAAUAAUCGUAAGGAUUAUG...........................................................
Cca-4-1  AUUUUCAAAAAGUACGCAGGUUCAGAAACGAACACAGGUUCAUCAGAAGGGCAAAUAGCUCUUUUUACACACAGAAUCAACCACUUAACGCGACAUUUAAAAGCAAAUCGUAAAGAUUAUAACACAGAGCGUGCGUUGGUUAAAUUGGUAGGUAAAAGAAGAAGUUUGCUUGACUACUUG
Cca-4-2  AUUUUCAAAAAGUACGCAGGUUCAGAAACGAACACAGGUUCAUCAGAAGGGCAAAUAGCUCUUUUUACACACAGAAUCAACCACUUAACGCAACACUUAAAAGCAAAUCGUAAAGAUUAUAACACAGAGCGUGCGUUGGUUAAAUUGGUAGGUAAAAGAAGAAGUUUGCUUGACUACUUG
Cca-4-3  AUUUUCAAAAAGUACGCAGGUUCAGAAACGAACACAGGUUCAUCAGAAGGGCAAAUAGCUCUUUUUACACACAGAAUCAACCACUUAACGCAACAUUUAAAAGCAAAUCGUAAAGAUUAUAACACAGAGCGUGCGUUGGUUAAAUUGGUAGGUAAAAGAAGAAGUUUGCUUGACUACUUG
env-800  UUAUUCAAGCAGUACGGUGGCUCCGACAAAAACACCGGAAGCACAGAAGGACAAAUUGCGUUGUUCACACACCGCAUUGAUCACCUUACCCAACACCUGAAAAACAACC.......................................................................
env-801  AUUUUCCAGAAACAUGGAGGUGACGCAAAAAACACAGGGAACACAGAAGGUCAAAUCGCCUUGUUCACCUUUAGAAUUAACCACCUGACUGAGCACUUAAAAAGAAACCGCAAGGAUUUCAAUACAGAGCGCUCACUGGUGAAAUUGGUAGGUAAAAGAAGAAGCCUCUUGGAUUACCUA
env-802  AUCUUCAAACAGUACGGUGGAACCGAAAAGAACACAGGUUCAACAGAAGGGCAAAUUGCUCUUUUCACCUACCGAAUCAAUUAUCUAACCGAACACUUAAAGAAAAAUCACAAGGAUUUUAACACAGAGCGUUCAUUAGUAAAACUAGUAGGUAAACGCAGAAGUUUACUCGAUUACCUC
env-803  AUUUUUAAAAAACACGGCAAAUCUGAAUCAGAUACAGGGACUUCUGAAGGACAAAUUGCAUUAUUUACAUUCAGAAUUAAUCAUUUGACCGAACAUUUAAAAAACAACAGAAAAGAUUAUAAUACCGAACGCUCUUUGGUAAAAUUAGUUGGUAAACGUCGUAGUUUAUUAGACUAUUUA
env-805  AUCUUCAAAGAACACGGUAAAUCAGAAUCUGACACUGGUUCAGCAGAAGGACAAAUUGCUCUUUUUACAAAGAGAAUCAAUCAUUUAACUGAACAUCUCAAAAAAAACAGAAAAGAUUUUAACACGGAAAGAUCCCUUGUUAUGAUGGUUGGUAAAAGAAAAAGUCUAUUAAGCUAUCUA
env-806  AUCUUCAAAGAACACGGUAAAUCAGAAUCUGACACUGGUUCAGCAGAAGGACAAAUUGCUCUUUUUACAAAGAGAAUCAAUCAUUUAACUGAACAUCUCAAAAAAAACAGAAAAGAUUUUAACACGGAAAGAUCCCUUGUUAUGAUGGUUGGUAAAAGAAAAAGUCUAUUAAGCUAUCUU
env-807  AUCUUCAAAGAACAUGGAAAAUCUGAAUCAGAUACUGGUUCUGCUGACGUCAAAUUGCAUUAUUUACUAAACGUAUCAAUCAUUUAACUGAACACUUGAAAAAAACAGAAAGGAUUUUAAUACAGAGAGAUCCUUGGUUAUGAUGGUUGGAAAAGAAAAGUUUAUUGAGUUAUCUUAAAA
env-808  AUCUUCAAAGAACAUGGAAAAUCUGAAUCAGAUACUGGUUCUGCUGAAGGUCAAAUUGCAUUAUUUACUAAACGUAUCAAUCAUUUAACUGAACACUUGAAAAAAAACAGAAAGGAUUUUAAUACUGAGAGAUCCUUGGUUAUGAUGGUUGGAAAAAGAAAAAGUUUAUUGAGUUAUCUU
env-809  ....................................................................................................................................................................................
Zga-1-1  AUCUUUAAGAAACACGGCGGAAAAGAGGAGAACACUGGUUCUACCGAAGGACAAAUCGCAUUGUUUACACACCGUAUCAAUCAUUUGACCGGCCACUUAAAAAGAAACCACAAAGAUUUCAACACUGAGCGUUCGUUAGUACGUUUGGUAGGUAAGAGAAGAAGUCUACUAGAUUACUUG
env-810  AUUUUCGCAAAACAUGGUGGAAACGCUGUUAACACAGGUUCGGCUGAAGGGCAAAUUGCUUUAUUCACAUUCAGAAUUAGCCACUUGACUGAGCACUUGAAAAAAAAUCGUCACGAUUUUAACACAGAGCGUUCACUAGUUCUUUUAGUAGGUAAAAGAAGAAGCCUUCUUGACUACUUA
env-813  AUCUUCGCUAAACACGGAGAAAAAACAAACACUGGGAAAUCAGAAGCGCAAAUUGCUUUGUUUACUUUCAGAAUUACACACUUA................................................................................................
env-814  AUCUUCGCUAAACACGGAGAAAAAACUAACACUGGUAAAUCAGAAGCGCAAAUCGCACUUUUCACUUACAGAAUUAGUCAUUUGACAGAGCAUUUGAAAAAAAAUCGUCACGAUUAUAAUACCGAGCGUUCAUUAAACACUGGGAAAUCAGAAGCGCAAAUUGCUUUG............
env-815  AUCUUCGCUAAACACGGAGAAAAAACAAACACUGGGAAAUCAGAAGCGCAAAUUGCUUUGUUUACUUUCAGAAUUACACACUUAACUGAGCAUUUGAAAAAAAAUCGUCACGAUUAUAAUACAGAACGUUCAUUAGUAUUAUUAGUAGGUAAAAGAAGAAGUUUACUUGAUUACUUGAAG
env-816  AUCUUCGCUAAACACGGAGAAAAAACAAACACUGGGAAAUCAGAAGCACAAAUUGCUUUGUUUACUUUCAGAAUUAGCCAUUUGACUGAGCAUUUGAAAAAAAAUCGUCACGAUUAUAAUACAGAGCGUUCAUUAGUAUUAUUGGUUGGUAAGAGAAGAAGUUUACUUGAUUACUUGAAG
env-817  AUCUUCGCUAAACACGGAGAAAAAACUAACACUGGUAAAUCAGAAGCGCAAAUCGCACUUUUCACUUACAGAAUUAGUCAUUUGACAGAGCAUUUGAAAAAAAAUCGUCACGAUUACAAUACCGAGCGUUCACUAGUAUUAUUAGUAGGUAAAAGAAGAAGUUUAUUAGACUACUUGAAG
env-818  AUCUUCGCUAAACACGGAGAAAAAACUAACACUGGUAAAUCAGAAGCGCAAAUCGCACUUUUCACUUACAGAAUUAGUCAUUUGACAGAGCAUUUGAAAAAAAAUCGUCACGAUUAUAAUACCGAGCGUUCAUUAGUAUUAUUAGUAGGUAAAAGAAGAAGNUUAUUAGACUACUUGAAG
env-819  ....................................................................................................................................................................................
env-820  ....................................................................................................................................................................................
env-821  AUCUUCGCUAAACACGGAGAAAAAACUAACACUGGUAAAGCGGAAGCGCAGAUUGCACUUUUUACUUACAGAAUUAGUCAUUUGACAGAGCACUUGAAAAAAAAUCGUCACGAUUACAAUACGGAGCGUUCACUAGUAUUAUUAGUAGGUAAAAGAAGAAGUUUAUUAGACUACUUGAAG
env-822  AUCUUCGCUAAACACGGAGAAAAAACAAACACUGGGAAAUCAGAAGCGCAAAUUGCUUUGUUUACUUUCAGAAUUACACACUUAACUGAGCAUUUGAAAAAAAAUCGUCACGAUUAUAAUACAGAACGUUCAUUAGUAUUAUUAGUAGGUAAAAGAAGAAGUUUACUUGAUUACUUAAAG
env-823  AUCUUCGCUAAACACGGAGAAAAAACAAACACUGGGAAAUCAGAAGCGCAAAUUGCUUUGUUUACUUUCAGAAUUACACACUUAACUGAGCAUUUGAAAAAAAAUCGUCACGAUUAUAAUACAGAGCGUUCAUUAGUAUUAUUAGUAGGUAAAAGAAGAAGUUUACUUGAUUACUUAAAG
env-824  UUAUUCAAAAAACACGGUAAAAGUGCAAAAGAUACCGGAAGAGCCGAAGGGCAAAUUGCUUUAUUUACGCUUAGAAUAAAUCAUUUAACCGAGCACUUAAAAAAUAACAAAAAAGAUUUCAACACUGAAAGAGCGUUAGUAAAGCUUGUUGGAAAAAGAAGAUCGCUACUAGAUUACUUG
env-825  AUUUUCAAAGAACACGGAAAAUCAGAAUCUGAUACUGGUUCGGCUGAAGGUCAAAUUGCAUUAUUCACAAAACGAAUAAAUCACUUAACUGAACAUCUUAAAAACAACAGAAAAGAUUUUAAUACAGAAAGAUCUUUAGUGAUGAUGGUUGGUAAAAGAAAGAGUUUAUUGAGUUAUCUA
env-826  AUCUUCAAAGAACACGGGAAAUCAGAAUCCGACACUGGUUCAGCAGAAGGUCAAAUUGCAUUAUUCACAAAACGAAUCAAUCAUUUAACUGAACAUCUUAAAAAAAAUAGAAAAGAUUUCAAUACGGAAAGAUCUUUAGUUAUGAUGGUUGGAAAAAGAAAAAGUUUGUUAAGUUAUCUU
env-827  AUCUUCAAAGAACACGGGAAAUCAGAAUCCGACACCGGUUCAGCAGAAGGUCAAAUUGCAUUAUUCACAAAACGAAUCAAUCAUUUAACUGAACAUCUUAAAAAAAACAGAAAAGAUUUUAAUACUGAAAGAUCUUUAGUAAUGAUGGUUGGAAAAAGAAAAAGUUUGUUAAGUUAUCUA
env-828  AUUUUCAAAGAACACGGAAAAUCAGAAUCUGAUACUGGUUCGGCUGAAGGUCAAAUUGCAUUAUUCACAAAACGAAUAAAUCACUUAACUGAACAUCUUAAAAACAACAGAAAAGAUUUUAAUACUGAAAGAUCUUUAGUGAUGAUGGUUGGUAAAAGAAAGAGUUUGUUGAGUUAUCUA
env-829  AUUUUCAAAGAACACGGAAAAUCAGAAUCUGAUACUGGUUCGGCUGAAGGUCAAAUUGCAUUAUUCACAAAACGAAUAAAUCACUUAACUGAACAUCUUAAAAACAACAGAAAAGAUUUUAAU.........................................................
env-830  AUUUUCAAAGAACACGGAAAAUCAGAAUCUGAUACUGGUUCGGCUGAAGGU.................................................................................................................................
env-831  AUUUUCA.............................................................................................................................................................................
env-832  AUUUUCAAAGAACACGGAAAAUCAGAAUCUGAUACUGGUUCGGCUGAAGGUCAAAUUGCAUUAUUCACAAAACGAAUAAAUCACUUAACUGAACAUCUUAAAAACAACAGAAAAGAUUUUAAUACUGAAAGAUCUUUAGUGAUGAUGGUUGGUAAAAGAAAGAGUUUAUUGAGUUAUCUA
env-833  AUUUUCAAAGAACACGGAAAAUCAGAAUCUGAUACUGGUUCGGCUGAAGGUCAAAUUGCAUUAUUCACAAAACGAAUAAAUCACUUAACUGANCAUCUUAAAAACAACAGAAAAGAUUUUAAUACAGAAAGAUCUUUAGUGAUGAUGGUUGGUAAAAGAAAGAGUUUAUUG.........
env-834  AUUUUCAAAGAACACGGAAAAUCAGAAUCUGAUACUGGUUCGGCUGAAGGUCAAAUUGCAUUAUUCACAAAACGAAUAAAUCACUUAACUGAACAUCUUAAAAACAACAGAAAAGAUUUUAAUACAGAAAGAUCUUUAG.........................................
env-835  AUUUUCAAAGAACACGGAAAAUCAGAAUCUGAUACUGGUUCNGCUGAAGGUCAAAUUGCAUUAUUCACAAAACGAAUAAAUCACUUAACUGAACAUCUUAAAAACAANAGAAAAGAUUUUAAUACAGAAAGAUCUUUAGUGAUGAUGGUUGGUAAAAGAAAGAGUUUAUUGAGUUAUCUA
env-836  AUCUUCAAAGAACAUGGUAAAUCAGAAUCUGAUACAGGUUCAGCUGAAGGUCAAAUUGCAUUAUUCACAAAACGAAUUAA....................................................................................................
env-837  AUCUUCAAAGAACAUGGUAAAUCAGAAUCUGAUACAGGUUCAGCUGAAGGUCAAAUUGCAUUAUUCACAAAACGAAUUAACCAUUUAACUGAUCAUCUUAAAAAUAACAGAAAAGAUUUUAACACAGAAAGAUCUUUAGUAAUGAUGGUUGGUAAAAGAAAAAGUUUGUUGAGCUAUCUA
env-838  AUCUUCAAAGAACAUGGUAAAUCAGAAUCUGAUACAGGUUCAGCUGAAGGUCAAAUUGCAUUAUUCACAAAACGAAUUAACCAUUUAACUGAUCAUCUUAAAAAUAACAGAAAAGAUUUUAACACAGAAAGAUCUUUAGUAAUGAUGGUUGGUAAAAGAAAAAGUUUGUUGAGCUAUCUA
env-847  AUCUUCAAAGAACAUGGUAAAUCAGAAUCUGAUACAGGUUCAGCUGAAGGUCAAAUUGCAUUAUUCACAAAACGAAUUAACCAUUUAACCGAGCAUCUUAAAAAUAACAGAAAAGAUUUUAACACAGAAAGAUCUUUAGUAAUGAUGGUUGGUAAAAGAAAAAGUUUGUUGAGCUAUCUA
env-848  AUCUUCAAAGAACAUGGUAAAUCAGAAUCUGAUACAGGUUCAGCUGAAGGUCAAAUUGCAUUAUUCACAAAACGAAUUAACCAUUUAACCGAUCAUCUUAAAAAUAACAGAAAAGAUUUUAACACAGAAAGAUCUUUAGUAAUGAUGGUUGGUAAAAGAAAAAGUUUGUUGAGCUAUCUA
env-849  AUCUUCAAAGAACAUGGUAAAUCAGAAUCUGAUACAGGUUCAGCUGAAGGUCAAAUUGCAUUAUUCACAAAACGAAUUAACCAUUUAACCGAUCAUCUUAAAAAUAACAGAAAAGAUUUUAACACAGAAAGAUCUUUAGUAAUGAUGGUUGGUAAAAGAAAAAGUUUGUUGAGUUAUCUA
env-850  AUCUUCAAAGAGCACGGUAAAUCAGAAACUGAUACAGGUUCGGCUGAAGGUCAAAUUGCAUUAUUCACAAAACGAAUUAAUCACUUAACUGAACAUCUUAAAAAUAACAGAAAAGAUUUUAACACUGAAAGAUCUUUAGUGAUGAUGGUUGGUAAAAGAAAAAGUUUGUUGAGUUAUCUU
env-851  AUCUUCAAAGAGCACGGUAAAUCAGAAACUGAUACAGGUUCGGCUGAAGGUCAAAUUGCAUUAUUCACAAAACGAAUUAAUCACUUAACUGAACAUCUUAAAAAUAACAGAAAAGAUUUUAACACUGAAAGAUCUUUAGUGAUGAUGGUUGGUAAAAGAAAAAGUUUGUUGAGUUAUCUU
env-852  AUCUUCAAAGAGCANGGUAAAUCAGANNCUGAUACAGGUUCGGCUGAAGGUCAAAUUGCAUUAUUCACAAAACGAAUUAAUCACUUAACUGAACAUCUUAAAAAUAACAGAAAAGAUUUUAACACUGAAAGAUCUUUAGUGAUGAUGGUUGGUAAAAGAAAAAGUUUGUUGAGUUAUCUU
env-853  AUCUUCAAAGAGCACGGUAAAUCAGAAACUGAUACAGGUUCGGCUGAAGGUCAAAUUGCAUUAUUCACAAAACGAAUUAAUCACUUAACUGAACAUCUUAAAAAUAACAGAAAAGAUUUUAACACUGAAAGAUCUUUAGUGAUGAUGGUUGGUAAAAGAAAAAGUUUGUUGAGUUAUCUU
env-854  ....................................................................................................................................................................................
env-855  ....................................................................................................................................................................................
env-856  AUCUUCAAAGAGCACGGUAAAUCAGAAACUGAUACAGGU.............................................................................................................................................
env-857  AUCUUCAAAGAGCACGGUAAAUCAGAAACUGAUACAGGUUCGGCUGAAGGUCAAAUUGCAUUAUUCACAAAACGAAUUAAUCACUUAACUGAACAUCUUAAAAAUAACAGAAAAGAUUUUAACACUGAAAGAUCUUUAGUGAUGAUGGUUGGUAAAAGAAAAAGUUUGUUGAGUUAUCUU
env-860  AUCUUCAAAGAGCACGGUAAAUCAGAAACUGAUACAGGUUCGGCUGAAGGUCAAAUUGCAUUAUUCACAAAACGAAUUAAUCACUUAACUGAACAUCUUAAAAAUAACAGAAAAGAUUUUAACACUGAAAGAUCUUUAGUGAUGAUGGUUGGUAAAAGAAAAAGUUUGUUGAGUUAUCUU
env-861  AUCUUCAAAGAGCACGGUAAAUCAGAAACUGAUACAGGUUCGGCUGAAGGUCAAAUUGCAUUAUUCACAAAACGAAUUAAUCACUUAACUGAACAUCUUAAAAA............................................................................
env-862  AUCUUCAAAGAGCACGGUAAAUCAGAAACUGAUACAGGUUCGGCUGAAGGUCAAAUUGCAUUAUUCACAAAACGAAUUAAUCACUUAACUGAACAUCUUAAAAAUAACAGAAAAGAUUUUAACACUGAAAGAUCUUUAGUGAUGAUGGUUGGUAAAAGAAAAAGUUUGUUGAGUUAUCUU
env-863  ....................................................................................................................................................................................
env-864  AUCUUCAAAGAGCACGGUAAAUCAGAAAUUGAUACAGGUUCGGCUGAAGGUCAAAUUGCAUUAUUCACAAAACGAAUUAAUCACUUAACUGAACAUCUUAAAAAUAACAGAAAAGAUUUUAACACUGAAAGAUCUUUAGUGAUGAUGGUUGGUAAAAGAAAAAGUUUGUUGAGUUAUCUU
env-865  AUCUUCAAAGAGCACGGUAAAUCAGAAACUGAUACAGGUUCGGCUGAAGGUCAAAUUGCAUUAUUCACAAAACGAAUUAAUCACUUAACUGAACAUCUUAAAAAUAACAGAAAAGAUUUUAACACUGAAAGAUCUUUAGUGAUGAUGGUUGGUAAAAGAAAAAGUUUGUUGAGUUAUCUU
env-866  AUCUUCAAAGAACACGGUAA................................................................................................................................................................
env-868  AUCUUCAAAGAACACGGUAAAUCAGAAUCCGAUACUGGUUCAGCUGAAGGUCAAAUUGCAUUAUUCACAAAACGAAUUAAUCAUUUAA............................................................................................
env-869  AUCUUCAAAGAACACGGUAAAUCAGAAUCCGAUACUGGUUCAGCUGAAGGUCAAAUUGCAUUAUUCACAAAACGAAUUAAUCAUUUAACAGAACAUCUUAAAAAGAAUAGAAAAGAUUAUAAUACUGAAAGAUCUUUAGUAAUGAUGGUUGGUAAAAGAAAAAGUUUGUUGAGUUAUCUA
env-870  AUCUUCAAAGAACACGGUAAAUCAGAAUCCGAUACUGGUUCAGCUGAAGGUCAAAUUGCAUUAUUCACAAAACGAAUUAAUCAUUUAACAGAACAUCUUAAAAAGAAUAGAAAAGAUUUUAAUACUGAAAGAUCUUUAGUAAUGAUGGUUGGUAAAAGAAAAAGUUUGUUGAGUUAUCUA
env-871  AUCUUCAAAGAACACGGUAAAUCAGAAUCCGAUACUGGUUCAGCUGAAGGUCAAAUUGCAUUAUUCACAAAACGAAUUAAUCAUUUAACAGAACAUCUUAAAAAGAAUAGAAAAGAUUAUAAUACUGAAAGAUCUUUAGUAAUGAUGGUUGGUAAAAGAAAAAGUUUGUUGAGUUAUCUA
env-875  AUCUU...............................................................................................................................................................................
env-876  AUCUUCAAAGAACACGGUAAAUCAGAAUC.......................................................................................................................................................
env-877  AUCUUCAAAGAACACGGUAAAUCAGAAUCCGAUACUGGUUCAGCUGAAGGUCAAAUUGCAUUAUUCACAAAACGAAUUAAUCAUUUAA............................................................................................
env-878  AUCUUCAAAGAACACGGUAAAUCAGAAUCCGAUACUGGUUCAGCUGAAGGUCAAAUUGCAUUAUUCACAAAACGAAUUAAUCAUUUAACAGAACAUCUUAAAAAGAAUAGAAAAGAUUAUAAUACUGAAAGAUCUUUAGUAAUGAUGGUUGGUAAAAGAAAAAGUUUGUUGAGUUAUCUA
env-879  AUCUUCAAAGAACACGGUAAAUCAGAAUCCGAUACUGGUUCAGCUGAAGGUCAAAUUGCAUUAUUCACAAAACGAAUUAAUCAUUUAACAAAACAUCUUAAAAAAAAUAGAAAAGAUUAUAAUACUGAAAGAUCUUUAGUAAUGAUGGUUGGUAAAAGAAAAAGUUUGUUGAGUUAUCUA

         ....................................................................................................................................................................................
         ....................................................................................................................................................................................
         AUYUUYR•RAARCACGG•RRR•••R•RR••RAYAC•GG••C•RC•GAAGG•CAAAUYGC•YURUUYAC•Y•Y•G•AUYRAYCAYYUR•C•RA•CAYYURAAAAR•AAYCR•AARGAUU•YAAYAC•GAR•G•UC•YURGU•••RYURGU•GG•AAR•G•RG••••YU•YU•GAYUAYYU•
```

| | |
|---|---|
| env-881 | AUCUUCAAAGAACACGGUAAAUCAGAAUCCGAUACUGGUUCAGCUGAAGGUCAAAUUGCAUUAUUCACAAAACGAAUUAAUCAUUUAACAGAACAUCUUAAAAAGAAUAGAAAAGAUUAUAACACUGAAAGAUCUUUAGUAAUGAUGGUUGGUAAAAGAAAAAGUUUGUUGAGUUAUCUA |
| env-883 | AUCUUCAAGGAGCACGGUAAGUCGGAAAGUGAUACUGGCUCAGCUGAAGGUCAAAUUGCAUUAUUUACAAAACGUAUCAAUCACUUAACAGAGCAUCUAAAAAACAACAGAAAAGAUUUCAAUACUGAAAGAUCACUUGUUAUGAUGGUUGGUAAAAGAAAAAGUUUAUUGAGUUAUCUU |
| env-884 | AUCUUCAAGGAGCACGGUAAGUCGGAAAGUGAUACUGGCUCAGCUGAAGGUCAAAUCGCAUUAUUUACAAAACGUAUCAAUCACUUAACAGAGCAUCUAAAAAAUAACAGAAAAGAUUUCAAUACUGAAAGAUCACUUGUUAUGAUGGUUGGUAAAAGAAAAAGUUUAUUGAGUUAUCUU |
| env-886 | AUCUUCAAGGAGCACGGUAAGUCGGAAAGUGAUACUGGCUCAGCUGAAGGUCAAAUUGCAUUAUUUACAAAACGUAUCAACCACUUAACAGAGCAUUUAAAAAAACAACAGAAAAGAUUUCAAUACUGAAAGAUCGCUUGUUAUGAUGGUUGGUAAAAGAAAAAGUUUAUUGAGUUAUCUU |
| env-887 | AUCUUCAAGGAGCACGGUAAGUCGGAAAGUGAUACUGGCUCAGCUGAAGGUCAAAUUGCAUUAUUUACAAAACGUAUCAAUCACUUAACAGAGCAUCUAAAAAACAACAGAAAAGAUUUCAAUACUGAAAGAUCACUUGUUAUGAUGGUUGGUAAAAGAAAAAGUUUAUUGAGUUAUCUU |
| env-888 | AUCUUCAAGGAGCACGGUAAGUCGGAAAGUGAUACUGGCUCAGCUGAAGGUCAAAUUGCAUUAUUUACAAAACGUAUCAAUCACUUAACAGAGCAUCUAAAAAAUAACAGAAAAGAUUUCAAUACUGAAAGAUCACUUGUUAUGAUGGUUGGUAAAAGAAAAAGUUUAUUGAGUUAUCUU |
| env-889 | AUCUUCAAGGAGCACGGUAAGUCGGAAAGUGAUACUGGCUCAGCUGAAGGUCAAAUUGCAUUAUUUACAAAACGUAUCAAUCACUUAACAGAGCAUCUAAAAAACAACAGAAAAGAUUUCAAUACUGAAAGAUCACUUGUUAUGAUGGUUGGUAAAAGAAAAAGUUUAUUGAGUUAUCUU |
| env-890 | AUCUUCAAGGAGCACGGUAAGUCGGAAAGUGAUACUGGCUCAGCUGAAGGUCAAAUUGCAUUAUUUACAAAACGUAUCAAUCACUUAACAGAGCAUCUAAAAAACAACAGAAAAGAUUUCAAUACUGAAAGAUCACUUGUUAUGAUGGUUGGUAAAAGAAAAAGUUUAUUGAGUUAUCUU |
| env-891 | AUCUUCAAGGAGCACGGCAAGUCGGAAAGUAAUACUGGGUCAGCUGAAGGUCAAAUUGCAUUAUUUACAAAACGUAUCAAUCACUUAACAGAACAUCUAAAAAACAACAGAAAAGAUUUCAAUACUGAAAGAUCGCUUGUUAUGAUGGUUGGUAAAAGAAAAAGUUUGUUGAGUUAUCUU |
| env-892 | AUCUUCAAGGAGCACGGUAAGUCGGAAAGUGAUACUGGCUCAGCUGAAGGUCAAAUUGCAUUAUUUACAAAACGUAUCAAUCACUUAACAGAGCAUCUUAAAAACAACAGAAAAGAUUUCAAUACUGAAAGAUCACUUGUUAUGAUGGUUGGUAAAAGAAAAAGUUUAUUGAGUUAUCUU |
| env-894 | AUCUUCAAGGAGCACGGUAAGUCGGAAAGUGAUACUGGCUCAGCUGAAGGUCAAAUCGCAUUAUUUACAAAACGUAUCAAUCACUUAACAGAGCAUCUAAAAAACAACAGAAAAGAUUUCAAUACUGAAAGAUCACUUGUUAUGAUGGUUGGUAAAAGAAAAAGUUUAUUGAGUUAUCUU |
| env-895 | AUCUUCAAGGAGCACGGUAAGUCGGAAAGUGAUACUGGCUCAGCUGAAGGUCAAAUUGCAUUAUUUACAAAACGUAUCAAUCACUUAACAGAGCAUCUAAAAAACAACAGAAAAGAUUUCAAUACUGAAAGAUCACUUGUUAUGAUGGUUGGUAAAAGAAAAAGUUUAUUGAGUUAUCUU |
| env-896 | AUCUUCAAGGAGCACGGUAAGUCGGAAAGUGAUACUGGCUCAGCUGAAGGUCAAAUUGCAUUAUUUACAAAACGUAUCAAUCACUUAACAGAGCAUCUAAAAAAUAACAGAAAAGAUUUCAAUACUGAAAGAUCACUUGUUAUGAUGGUUGGUAAAAGAAAAAGUUUAUUGAGUUAUCUU |
| env-897 | AUCUUCAAGGAGCACGGUAAGUCGGAAAGUGACACUGGCUCAGCUGAAGGUCAAAUUGCAUUAUUUACAAAACGUAUCAAUCACUUAACAGAGCAUCUAAAAAACAACAGAAAAGAUUUCAAUACUGAAAGAUCACUUGUUAUGAUGGUUGGUAAAAGAAAAAGUUUAUUGAGUUAUCUU |
| env-898 | AUCUU............................................................................................................................................................................... |
| env-899 | AUCUUCAA............................................................................................................................................................................ |
| env-900 | AUCUUCAANNNNNNNNNNNNNNNNNNNNNNNNNNNNNNNNNNNNNNNNNNNNNNNNNNNAAAAACAACAGAAAAGAUUUCAAUACUGAAAGAUCACUUGUUAUGAUGGUUGGUAAAAGAAAAAGUUUAUUGAGUUAUCUUAAAAAAUAAAGAUAUCAAAAGAUAUCGUGAGAUUAUCAAAA |
| env-901 | AUUUUCAAAAAACACGGAAAAUCUAAAAAAGACACCGGUACUCCU....................................................................................................................................... |
| env-902 | AUUUUCAAAAAACACGGAAAAUCUAAAAAAGACACCGGUACUCCUGAAGGACAAAUCGCAUUAUUUUCUCAUAGGAUUAAGCACCUGUCAGGUCAUCUCAAAAACAAUCGAAAGGAUUAUAAUACGGAGCGUUCUCUGGUAAAAAUAGUAGGAAAGCGAAGACAACUAUUGGAUUAUUUG |
| env-903 | AUUUUCAAAAAACACGGAAAAUCUGAAAAAGAUACCGGUACUCCUGAAGGACAAAUCGCAUUGUUUUCUCACAGGAUUAACCACUUGUCAGGUCAUCUUAAAAACAACCGAAAGGAUUAUAACACGGAGCGUUCUCUAGUAAAAUUAGUAGGAAAGCGAAGACACCUUUUGGAUUAUUUG |
| env-905 | AUUUUCAAAAAACACGGAAAAUCUGAAAAAGAUACCGGUACUCCUGAAGGACAAAUCGCAUUGUUUUCUCACAGGAUUAACCACUUGUCAGGUCAUCU.................................................................................. |
| env-906 | AUCUUCGCUAAACACGGAGAGAAAACAAACACUGGAAAAUCAGAAGCACAAAUCGCUUUGUUCACUUUCAGAAUUAGCCACCUAACAGAGCACUUGAAAAAAAAUCGCCACGAUUAUAACACCGAGCGUUCCUUAGUAUUGCUAGUAGGUAAAAGAAGAUCUUUGUUGGAUUACUUGAAG |
| env-907 | AUCUUCGCUAAACACGGAGAGAAAACAAACACUGGAAAAUCAGAAGCACAAAUCGCUUUGUUCACUUUCAGAAUUAGCCA.................................................................................................... |
| env-908 | AUCUUCGCUAAACACGGAGAGAAAACAAACACUGGAAAAUCAGAAGCACAAAUCGCUUUGUUCACUUUCAGAAUUAGCCACCUAACAGAGCACUUGAAAAAAAAUCGCCACGAUUAUAACACCGAGCGUUCCUUAGUAUUGCUAGUAGGUAAAAGAAGAUCUUUGUUGGAUUACUUGAAG |
| Fge-1-1 | AUCUUCGCUAAACACGGAGGAAAAGCAGAAAACACAGGAUCUGCAGAAGGACAAAUCGCGUUAUUCACACACAGAAUUUCACAUUUAACAGAACACUUGAAAAAAAAUCGUCAUGAUUAUAACACAGAGCGUUCACUAGUUCUUUUGGUAGGAAAAAGAAGAAGUCUUCUUGAUUACUUA |
| Ale-2-1 | AUCUUCGCAAAACACGGAAAGUCUAAAACAGACACCGGUUCAGCCGAAGGUCAGAUCGCGUUAUUCACGUAUCGCAUCGAGCAUUUAACAAAACACCUUAAAAACAACCACAAGGAUUAUAAUACAGAACGUUCUUUGGUAAAGUUAGUAGGAAAGCGUCGUUCUCUUUAGAUUACCUU |
| env-909 | AUCUUCGCUAAACACGGAGGAGACGCAACAAACACCGGCAAAUCAGAAAGCCAGAUUGCUUUGUUCACAUUUAGGAUCAAUCACUUGACCGAGCACUUGAAAAAAAAUCGUCACGAUUACAACACCGAGCGUUCACUAGUGAAACUGGUAGGUAAAAGAAGAAGCCUUCUUGACUACCUC |
| env-910 | AUUUUUAAAAAACACGGUAAAAGUGUUACAGACACAGGUUCUGCUGAAGGACAAAUUGCAUUGUUCACUUACAGAAUUGAACAUUUAACAGGUCACCUUAAGGCGAAUCACAAAGAUUACAACACUGAAAGAUCUUUGGUUAAAUUAGUAGGUAAGCGUCGUUCUUUAUUGGAUUAUUUA |
| env-911 | AUCUUUAAAAAACAUGGUAAAGAAUCAAAAAAUACUGGUUCUGCAGAAGGUCAAAUUGCUUUAUUUACUUACAGAAUUAAUCAUCUAACUGAACAUUUAAAGAAAAAUCGAAAAGAUUUUAAUACUGAACGAUCUCUUGUCAAACUUGUAGGAAAACGUAGAUCUUUACUAGAUUACUUA |
| env-912 | AUCUUUAAAAAACAUGGUAAAGAAUCAAAAAAUACUGGUUCUGCAGAAGGUCAAAUUGCUUUAUUUACCUACAGAAUUAAUCAUCUAACUGAACAUUUAAAGAAAAAUCGAAAAGAUUUUAACACUGAAAGAUCUCUUGUUAAACUCGUAGGAAAACGCAGAUCUUUACUAGAUUACUUA |
| env-913 | AUCUUUAAAAAACAUGGUAAAGAAUCAAAAAAUACUGGUUCUGCAGAAGGUCAAAUUGCUUUAUUUACCUACAGAAUUAAUCAUCUAACUGAACAUUUAAAGAAAAAUCGAAAAGAUUUUAAUACUGAACGAUCUCUUGUCAA..................................... |
| env-914 | AUCUUUAAAAAACAUGGUAAAGAAUCAAAAAACACUGGUUCUGCAGAAGGUCAAAUUGCUUUAUUUACCUAUAGGAUUAACCAUCUAACUGAACAUUUAAAGAAAAAUCGAAAAGAUUUUAAUACUGAACGAUCUCUUGUUAAACUCGUAGGAAAACGCAGAUCUUUACUAGAUUACUUA |
| env-915 | AUCUUUAAAAAACAUGGUAAAGAAUCAAAAAAUACCGGUUCUGCAGAAGGUCAAAUUGCUUUAUUUACCUAUAGGAUUAACCAUCUAACUGAACAUUUAAAGAAAAAUCGAAAAGAUUUUAAUACUGAACGAUCUCUUGUUAAACUCGUAGGAAAACGCAGAUCUUUACUAGAUUACUUA |
| env-916 | AUCUUUAAAAAACAUGGUAAAGAAUCAAAAAACACUGGUUCUGCAGAAGGUCAAAUUGCUUUAUUUACCUACAGGAUUAAUCAUCUAACUGAACAUUUAAAGAAAAAUCGAAAAGAUUUUAAUACUGAACGAUCUCUUGUUAAGCUUGUAGGAAAACGUAGAUCUUUACUAGAUUACUUA |
| Man-1-1 | AUCUUUAAGAAACACGGCGGAAAAGUGGAAAACACAGGUUCUACGGAAGGUCAAAUCGCGUUGUUCACGCACCGUAUUAAUCAUUUGACUGGUCACUUGAAAAAAAAUCACAAAGACUACAACACGGAACGAUCAUUGGUAAAGUUGGUGGGUAAAAGGAGAAGUCUUUUAGAUUAUAUG |
| env-917 | AUUUUCAAAAAGCACGGAGGUUCGGAUACAAAUACAGGAUCUACAGAAGCCCAAGUAGCUUU...................................................................................................................... |
| env-919 | AUUUUCAAAAAACACGGUGGAUCCGAUACAAACACGGGGUCAACUGAAGGUCAAAUUGCUCUAUUCACAUAUAGAAUAGGUCAUUUAACUGAACAUUUGAAAAAAAAUAGAAAAGAUUUUGGAACCCAGAGAGCCCUUCAAUUACUUGUGGGUAAAAGAAGAAGCCUUUUGGACUAUCUU |
| env-920 | AUUUUCAAAAAGCACGGAGGUUCGGAUACAAAUACAGGAUCUACAGAAGCCCAAGUAGCUUUGUUCACCUACAGAAUAAGCCACUUAACUGAGCACUUGAAGAAAAAUAGAAAAGAUUUUGGAACCCAAAGAGCCCUUCAAUUACUUGUGGGUAAGAGAAGAAGCCUUUUGGAUUAUCUG |
| env-921 | AUUUUCAAAAAGCACGGAGGUUCGGAUACAAAUACAGGAUCUACAGAAGCCCAAGUAGCUUUGUUCACCUACAGAAUAAGCCACUUAACUGAGCACUUGAAGAAAAAUAGAAAAGAUUUUGGAACCCAAAGAGCCCUUCAAUUACUUGUGGGUAAGAGAAGAAGCCUUUUGGAUUAUCUG |
| env-922 | AUUUUCAAAAAGCACGGAGGUUCGGAUACAAAUACAGGAUCUACAGAAGCCCAAGUAGCUUUGUUCACCUACAGAAUAAGCCACUUAACUGAGCACUUGAAGAAAAAUAGAAAAGAUUUUGGAACCCAAAGAGCCCUUCAAUUACUUGUGGGUAAGAGAAGAAGCCUUUUGGAUUAUCUG |
| env-923 | AUUUUCAAAAAGCACGGAGGUUCGGAUACAAAUACAGGAUCUACAGAAGCCCAAGUAGCUUUGUUCACAUACAGAAUAAGCCACUUAACUGAGCACUUGAAGAAAAAUAGAAAAGAUUUUGGAACCCAAAGAGCCCUUCAAUUACUUGUGGGUAAGAGAAGAAGCCUUUUGGAUUAUCUG |
| env-924 | AUUUUCAAAAAGCACGGAGGUUCGGAUACAAAUACAGGAUCUACAGAAGCCCAAGUAGCUUUGUUCACAUAUAGAAUAAGCCACUUAACCGAGCACUUGAAGAAAAAUAGAAAAGAUUUUGGAACCCAAAGAGCCCUUCAAUUACUUGUAGGUAAAAGAAGGAGCCUUUUGGACUAUCUA |
| env-925 | .................................................................................................................................................................................... |
| env-927 | AUUUUCAAAAAGCACGGAGGUUCGGAUACAAAUACAGGAUCUACAGAAGCCCAAGUAGCUUUGUUCACAUACAGAAUAAGCCACUUAACUGAGCACUUGAAGAAAAAUAGAAAAGAUUUUGGAACCCAAAGAGCCCUUCAAUUACUUGUGGGUAAGAGAAGAAGCCUUUUGGAUUAUCUG |
| env-928 | AUUUUCAAAAAACACGGUGGGUCCGAAACAAACACGGGAUCUAC........................................................................................................................................ |
| env-929 | AUUUUCAAAAAACACGGUGGGUCCGAAACAAACACGGGAUCUACCGAAGGUCAAAUUGCCUUAUUCACAUAUAGAAUAGGUCAUUUAACUGAUCAUUUGAAAAAAAACAGAAAAGAUUUUGGAACCCAGAGAGCCCUUCAAUUACUUGUUGGUAAGAGAAGAAGCCUUUUGGACUAUCUC |
| env-930 | AUUUUCAAAAAACACGGUGGGUCCGAAACAAACACGGGAUCUACCGAAGGUCAAAUUGCCUUAUUCACAUAUAGAAUAGGUCAUUUAACUGAUCAUUUGAAAAAAAACAGAAAAGAUUUUGGAACCCAGAGAGCCCUUCAAUUACUUGUUGGUAAGAGAAGAAGCCUUUUGGACUAUCUC |
| env-932 | AUUUUCAAAAAGCACGGAGGAUCUGAUACAAAUACAGGUUCUACAGAAGGUCAGAUUGCGUUAUUUACUUAUAGAAUAGAGCAUCUUACCAAGCAUUUGAAAAAAAACAGGAAAGAUUUUGGAACCCAAAGGGCUCUUCAAUUACUUGUAGGUAAGAGAAGAAGUUUAUUGGAUUAUCUG |
| env-933 | AUCUUCGCAAAACACGGAGGAAAAGCUGAAAACACUGGUUCUACAGAAGGACAAGUUGCCCUUUUUACUUUUAGAAUCGCUCACUUAACUGAGCAUUUGAAAAAAAACCGUCACGAUUACAACACUGAACGUUCAUUAGUACUUUUGGUAGGUAAAAGAAGAAGUCUUCUUGACUACUUA |
| env-934 | AUCUUCGCGAAACACGGAGGAAAGGCUGAAAACACUGGUUCUACAGAAGGACAAGUUGCCCUUUUUACUUUUAGAAUCGCUCACUUAACUGAGCAUUUGAAAAAAAAUCGUCAUGAUUACAACACUGAACGUUCAUUAGUACUUUUAGUAGGUAAAAGAAGAAGUCUUCUUGACUACUUA |
| env-936 | AUUUUUAAAAAGCACGGUAAGUCUGAAAAAGACACUGGUUCUGCAGAAGGACAAAUCGCUUUGUUUACAUACCGUAUCAGUCAUUUGACAGGU....................................................................................... |
| env-937 | UUAUUCAAAAAACACGGUAAAAACGAAAAAGAUACCGGAAGCGCUGAAGGUCAGAUAGCUUUAUUCACUCUAAGAAUCAAUCACUUAACAAAACAUUUAAAAUCAAAUCGUAAAGAUUUUAAUACUGAGAGAUCUCUAGUUAAACUCGUUGGCAAAAGAAGAGCACUACUAGAUUACCUU |
| env-938 | AUUUUCAAAAAGUAUGGGAAAUCGAAAUCAGAUACAGGUUCAACUGAGGGUCAAAUAGCUCUUUUCUCUCACAGAAUUAAUCAUUUGUCUGAACAUUUAAAAAAUAAUAGAAAAGAUUAUAAUACAGAAAGAUCCUUAGUUACUUUAGUUGGCAAAAGAAGGAAUCUUUUAGAUUAUUUA |
| env-939 | AUUUUCAAAAAGUAUGGGAAAUCAAAAUCAGAUACAGGUUCAACUGAGGGUCAAAUAGCUCUUUUCUCUCACAGAAUUAAUCAUUUGUCUGAACAUUUAAAAAAUAAUAGAAAAGAUUAUAAUACAGAAAGAUCUUUAGUUACUUUAGUUGGUAAAAGAAGGAAUCUUUUGGAUUAUUUA |
| env-940 | A................................................................................................................................................................................... |
| env-941 | AUUUUCAAAAAGUAUGGGAAAUCAAAAUCAGAUACAGGUUCAACUGAGGGUCAAAUAGCUCUUUUCUCUCACAGAAUUAAUCACUUGUCUGAACAUUUAAAAAAUAAUAGAAAAGAUUAUAAUACAGAAAGAUCUUUGGUUACUCUAGUUGGUAAAAGAAGGAAUCUUUUGGAUUAUUUA |
| env-942 | AUUUUCAAAAAGUAUGGGAA................................................................................................................................................................ |
| env-943 | AUUUUCAAAAAGUAUGGGAAAUCAAAAUCAGAUACAGGUUCAACUGAGGGUCAAAUAGCUCUUUUCUCUCACAGAAUUAAUCAUUUGUCUGAACAUUUAAAAAAUAAUAGAAAAGAUUAUAAUACAGAAAGAUCUUUAGUUACUUUAGUUGGUAAAAGAAGGAAUCUUUUGGAUUAUUUA |
| env-944 | AUUUUCAAAAAGUAUGGGAAAUCAAAAUCAGAUACAGGUUCAACUGAGGGUCAAAUAGCUCUUUUCUCUCAAAGAAUUAAUCACUUGUCUGAACAUUUAAAAAAUAAUAGAAAAGAUUAUAAUACAGAAAGAUCUUUGGUUACUCUAGUUGGUAAAAGAAGGAAUCUUUUGGAUUAUUUA |
| env-945 | AUUUUCAAAAAGUAUGGGAAAUCAAAAUCAGAUACAGGUUCAACUGAGGGUCAAAUAGCUCUUUUCUCUCACAGAAUUAAUCAUUUGUCUGAACAUUUAAAAAAUAAUAGAAAAGAUUAUAAUACAGAAAGAUCUUUAGUUACUUUAGUUGGUAAAAGAAGGAAUCUUUUGGAUUAUUUA |
| env-946 | .................................................................................................................................................................................... |
| Tma-1-1 | AUCUUCGAAAAGCACGGUAAAGGAAAAAAUGAUACUGGAACAUCAGAAGGUCAAAUUGCACUAUUCACUUUUAGAAUAAACCAUUUAACAGAACACUUAAAAAAUAAUCGUAAAGAUUUUAACACAGAGCGCUCUUUAGUAAAAAUGGUUGGUAAGCGUAGAAGCUUAUUAGAUUAUUUA |
| env-947 | AUCUUCGCAAAGUACGCGGGAUCUGAAACAAACACAGGAUCUGCAGAAGGUCAAAUCGCUUUAUUUUCUUACAGAAUCUCUCACUUGACAGAGCAUUUAAAGAAAAAUAGAAAAGAUUAUGGAACACAACGUUCUUUGCAAUU..................................... |
| Fsp-12-1 | AUUUUCGCUCAACACGGUGCUGCAACAAACACUGGAAGCGCAGAAGGUCAGAUUGCAUUGUUCACUUACAGAAUUUCACACUUAACUGAGCACUUGAAAAAAAAUCGUCACGAUUACAACACUGAGCGUUCACUUGUACUAUUAGUAGGUAAAAGAAGAGCUUUGUUGGAUUACUUGAAA |
| Fde-1-1 | AUUUUCGCACAACACGGUGGUGCAACAAACACUGGAAGCGCAGAAGGUCAAAUUGCAUUGUUCACUUACAGAAUUUCUCACUUAACUGAACACUUGAAAAAAAAUCGUCACGAUUACAACACUGAGCGUUCUCUUGUACUAUUAGUAGGUAAAAGAAGAGCUUUGUUGGAUUACUUGAAA |
| Fba-7-1 | AUCUUUGCUAAACACGGUAAAGGUAAAAAUGACAGUGGAACUGCAGAAGCACAAAUUGCUUUAUUUACGCACAGAAUUACACACCUAACUGAGCACUUGAAAAAAAAUCGUAAAGAUUAUAAUACUGAACGUUCAUUAGUAAAAUUAGUAGGUAAACGCAGAAGCUUACUAGAUUACUUA |
| Ale-1-1 | AUGUUUGCCAAACAUGGCAAAAACAAAAGCGAUACUGGUUCUGCAGAAGGACAAAUUGCCUUAUUUACACAAAGAAUUGAGCACUUAACUCAACACUUAAAAAAGAAUCGUAAAGAUUACAACACUGAGCGUUCUUUAGUAAUGUUAGUAGGUAAAAGAAGAAGCUUACUUGAUUAUUUA |
| Ale-1-2 | AUGUUUGCCAAACAUGGCAAAAACAAAAGCGAUACUGGUUCUGCAGAAGGACAAAUUGCCUUAUUUACACAAAGAAUUGAGCACUUAACUCAACACUUAAAAAAGAAUCGUAAAGAUUACAACACUGAGCGUUCUUUAGUAAUGUUAGUAGGUAAAAGAAGAAGCUUACUUGAUUAUUUA |
| Ffr-1-1 | AUCUUCGCACAACACGGAGAAAAAACAAACACUGGAAAAGCUGAAGCACAAAUCGCUUUAUUCACAUUCAGAAUUAGCCAUUUGACUGAGCACUUGAAAAAAAAUCGUCAUGAUUACAAUACAGAGCGUUCACUAGUAUUACUAGUAGGUAAAAGAAGAUCUUUGUUAGACUACUUGAAA |
| env-948 | AUCUUCUCUAAACACGGAGGAAAAGCUGAAAACACAGGAUCUGCUGAAGGGCAAAUCGCAUUAUUUACUUUUAGAAUUUCUCA................................................................................................. |
| Fsp-1-1 | AUCUUCGCAAAACACGGAGGAAAAGCUGAAAACACUGGAUCUGCUGAAGGGCAAAUCGCUUUGUUUACAUUUAGAAUUAAUCACUUAACAGGACACUUGAAACAAAAUCGUCAUGAUUACAACACUGAGCGUUCGUUAGUGAAACUAGUAGGUAAAAGAAGAAGUCUUCUUGACUACUUA |
| env-949 | AUCUUCGCUAAACACG.................................................................................................................................................................... |
| env-950 | .................................................................................................................................................................................... |
| Fsa-2-1 | AUUUUCGCUAAACACGGAGGAAAAGCUGAAAACACUGGAUCUGCUGAAGGGCAAAUCGCAUUAUUUACAUUUAGAAUUUCUCACUUAACUGAGCACUUGAAAAAAAAUCGUCAUGAUUAUAACACUGAGCGUUCGUUAGUACUUUUAGUAGGUAAAAGAAGAAGUCUUCUUGACUACUUA |
| env-951 | .................................................................................................................................................................................... |
| env-952 | AUNUUCGCNAAACACGGAGGAAAAGCUGAAAACACUGGAUCUGCUGAAGGGCAAAUCGCAUUAUUUACAUUUAGAAUUUCUCACUUAACUGAGCACUUGAAAAAAAAUCGUCAUGAUUUUAACACUGANCGUUCAUUAGUACUUUUAGUAGGAAAAAGAAGAAGUCUUCUUGACUAUUUA |
| Fba-3-1 | AUUUUCGCUAAACACGGAGGUAAAGCUGAAAACACUGGAUCUGCUGAAGGGCAAAUCGCAUUAUUUACAUUUAGAAUUUCUCACUUAACUGAGCACUUGAAAAAAAAUCGUCAUGAUUUUAACACAGAACGUUCAUUAGUACUUUUAGUAGGAAAAAGAAGAAGUCUUCUUGACUAUUUA |
| env-953 | AUCUUCGCAAAACACGGAGGAAAAGCUGAAAACACUGGAUCUGCUGAAGGGCAAAUCGCAUUAUUUACAUUUAGAAUUUCACACUUAACAGAGCACUUGAAAAAAAAUCGUCACGAUUACAACACAGAGCGUUCGUUAGUGCUUUUAGUAGGAAAAAGAAGAAGUCUUCUUGAUUACUUA |
| env-954 | AUCUUUGCUAAACACG.................................................................................................................................................................... |
| env-955 | UUAUUCAAAAAACACGGUAAAAAUGAAAAAGAUACCGGAAGCUCUGAAGGCCAGAUUGCUUUAUUCACGUUAAGAAUUAACCACCUAACAGAACACUUGAAAAAUAACCGUAAGGAUUAUAAUACGGAAAGAUCUCUUGUUAAACUUGUAGGUAAAAGAAGAUCCUUACUGGAUUAUCUU |
| env-956 | UUAUUCAAAAAACACGGUAAAAAUGAAAAAGAUACCGGAAGCUCUGAAGGCCAGAUUGCUUUAUUCACCUUAAGAAUUAACCACCUAACAGAACACUUGAAAAAUAAUCGUAAGGAUUAUAAUACGGAAAGAUCUCUUGUUAAACUUGUAGGUAAAAGAAGAUCCUUACUGGAUUAUCUU |
| env-957 | UUAUUCAAAAAACACGGUAAAAAUGAAAAAGAUACCGGAAGCUCUGACGCCAGAUUGCUUUAUUCACGUUAAGAAUUAACCACCUAACAGACACUUGAAAAUAACCGUAACGAUUAUAAUACGGAAAGAUCUCUUGUUAACUUGUAGUAAAGAGAUCCUUACUGAUAUCUUACUAAGUAG |
| env-958 | UUAUUCAAAAAGCACGGAAAAGAUGAUAAAGAUACGGGAAGU.......................................................................................................................................... |
| env-959 | AUCUUUAAGAAACACGCAGGUUCAGAAACCAACACUGGUUCAGCAGAAGGGCAAAUCGCUCUUUUUACCUUCAGAAUUAAUCACCUUACUGAACACCUUAAGGUAAAUCGUAAAGAUUACAAUACCGAGCGUUCAUUGGUAAAAUUGGUAGGUAAACGUCGUCGUUUGCUCGAUUAUUUG |
| | .................................................................................................................................................................................... |
| | .................................................................................................................................................................................... |
| | AUYUUYR•RAARCACGG•RRR•••R•RR••RAYAC•GG••C•RC•GAAGG•CAAAUYGC•YURUUYAC•Y•Y•G•AUYRAYCAYYUR•C•RA•CAYYURAAAAR•AAYCR•AARGAUU•YAAYAC•GAR•G•UC•YURGU•••RYURGU•GG•AAR•G•RG••••YU•YU•GAYUAYYU• |

```
Csp-1-1   AUUUUUAAAAAGUACGCAGGUUCAGAAACCAACACUGGUUCAGCAGAAGGGCAAAUCGCUCUUUUUACCCUUAGAAUCAAUCACCUUACAGAACACUUGAAAGUAAAUCGCAAAGAUUUCAAUACCGAGCGUUCAUUGGUGAAAUUGGUAGGUAAACGCAGAAGUUUGCUCGAUUAUUUG
env-960   AUUUUUAAAAAGUACGCAGGUUCAGAAACCA......................................................................................................................................................
env-961   AUUUUNNNNNNNNNNNNNNNNNNNNNNNNNNNNNNCAAUACCGAGCGUUCAUUGGUGAAAUUGGUAGGUAAACGCAGAAGUUUGCUCGAUUAUUUGAAAAAUAACGAUAUCGAGAGAUACCGUGCGAUUAUCAAAGAACUAAACAUCAGAAAAUAAGCAAUACUAAGGAGGCUUUGUGCC
Csp-3-1   AUUUUUAAAAAGUACGCAGGUUCAGAAACCAACACUGGUUCAGCAGAAGGGCAAAUCGCUCUUUUUACCCUUAGAAUCAAUCACCUUACAGAACACUUAAAGGUAAAUCGCAAAGAUUUCAAUACCGAGCGUUCAUUGGUGAAAUUGGUAGGUAAACGCAGAAGUUUGCUCGAUUAUUUG
env-962   AUUUUUAAAAAGUACGCAGGUUCAGAAACCAACACUGGUUCAGCAGAAGGGCAAAUCGCUCUUUUUACCCUUAGAAUCAAUCACCUUACAGAACACUUGAAAGUAAAUCGCAAAGAUUUCAAUACCGAGCGUUCUUUGGUAAGAUUGGUAGGUAAACGCAGAAGUUUGCUCGACUAUUUG
env-965   AUUUUUAAAAAGUACGCAGGUUCAGAAACCAACACUGGUUCAGCAGAAGGGCAAAUCGCUCUUUUUACCCUUAGAAUCAAUCACCUUACAGAACACUUGAAAGUAAAUCGCAAAGAUUUCAAUACCGAGCGUUCAUUGGUGAAAUUGGUAGGUAAACGCAGAAGUUUGCUCGAUUAUUUG
env-966   AUUUUCA..............................................................................................................................................................................
env-967   AUUUUUAAAAAGUACGCAGGUUCAGAAACCA......................................................................................................................................................
env-968   AUUUUUAAAAAGUACGCAGGUUCAGAAACCAACACUGGUUCAGCAGAAGGGCAAAUCGCUCUUUUUACCCUUAGAAUCAAUCACCUUACAGAACACUUGAAAGUAAAUCGUAAAGAUUUCAAUACCGAGCGUUCUUUGGUAAGAUUGGUAGGUAAACGCAGAAGUUUGCUCGACUAUUUG
env-972   AUUUUUAAAAAGUACGCAGGUUCAGAAACCAACACUGGUUCAGCAGAAGGGCAAAUCGCUCUUUUUACCCUUAGAAUCAAUCACCUUACAGAACACUUGAAAGUAAAUCGCAAAGAUUUCAAUACCGAGCGUUCUUUGGUAAGAUUGGUAGGUAAACGCAGAAGUUUGCUCGACUAUUUG
env-973   AUUUUUAAAAAGUACGCAGGUUCAGAAACCAACACUGGUUCAGCAGAAGGGCAAAUCGCUCUUUUUACCCUUAGAAUCAAUCACCUUACAGAACACUUGAAAGUAAAUCGCAAAGAUUUCAAUACUGAGCGUUCAUUGGUGAAAUUGGUAGGUAAACGCAGAAGUUUGCUCGAUUAUUUG
env-974   AUUUUUAAAAAGUACGCAGGUUCAGAAACCAACACUGGUUCAGCAGAAGGGCAAAUCGCUCUUUUUACCCUUAGAAUCAAUCACCUUACAGAACACUUGAAAGUAAAUCGCAAAGAUUUCAAUACCGAGCGUUCUUUGGUAAGAUUGGUAGGUAAACGCAGAAGUUUGCUCGACUAUUUG
env-976   AUUUUUAAAAAGUACGCAGGUUCAGAAACCAACACUGGUUCAGCAGAAGGGCAAAUCGCUCUUUUUACCCUUAGAAUCAAUCACCUUACAGAACACUUGAAAGUAAAUCGCAAAGAUUUCAAUACCGAGCGUUCUUUGGUAAGAUUGGUAGGUAAACGCAGAAGUUUGCUCGACUAUUUG
env-977   AUUUUUAAAAAGUACGCAGGUUCAGAAACCAACACUGGUUCAGCAGAAGGGCAAAUCGCUCUUUUUACCCUUAGAAUCAAUCACCUUACAGAACACUU...................................................................................
env-978   AUUUUUAAAAAGUACGCAGGUUCAGAAACCA......................................................................................................................................................
env-980   AUUUU................................................................................................................................................................................
env-982   .....................................................................................................................................................................................
env-983   AUUUUUAAAAAGUACGCAGGUUCAGAAAACAACACUGGUUCAGCAGAAGGGCAAAUCGCUCUUUUUACCCUUAGAAUCAAUCACCUUACAGAACACUUGAAAGUAAAUCGCAAAGAUUUCAAUACCGAGCGUUCAUUGGUGAAAUUGGUAGGUAAACGCAGAAGUUUGCUCGAUUAUUUG
env-984   AUUUUUAAAAAGUACGCAGGUUCAGAAACCAACACUGGUUCAGCAGAAGGGCAAAUCGCUCUUUUUACCCUUAGAAUCAAUCACCUUACAGAACACUUGAAAGUAAAUCGCAAAGAUUUCAAUACCGAGCGUUCAUUGGUGAAAUUGGUAGGUAAACGCAGAAGUUUGCUCGAUUAUUUG
env-985   AUUUUUAAAAAGUACGCAGGUUCAGAAACCAACACUGGUUCAGCAGAAGGGCAAAUCGCUCUUUUUACCCUUAGAAUUAAUCACCUUACAGAACACUUGAAAGUAAAUCGCAAAGAUUUCAAUACCGAGCGUUCAUUGGUGAAAUUGGUAGGUAAACGCAGAAGUUUGCUCGAUUAUUUG
env-986   AUUUUUAAAAAGUACGCAGGUUCAGAAACCAACACUGGUUCAGCAGAAGGGCAAAUCGCUCUUUUUACCCUUAGAAUCAAUCACCUUACAGAACACUUGAAAGUAAAUCGCAAAGAUUUCAAUACCGAGCGUUCAUUGGUGAAAUUGGUAGGUAAACGCAGAAGUUUGCUCGAUUAUUUG
env-988   AUUUUUAAAAAGUACGCAGGUUCAGAAACCAACACUGGUUCAGCAGAAGGGCAAAUCGCUCUUUUUACCCUUAGAAUCAAUCACCUUACAGAACACUUGAAAGUAAAUCGCAAAGAUUUCAAUACCGAGCGUUCAUUGGUGAAAUUGGUAGGUAAACGCAGAAGUUUGCUCGAUUAUUUG
env-989   AUUUUUAAAAAGUACGCAGGUUCAGAAACCAACACUGGUUCAGCAGAAGGGCAAAUCGCUCUUUUUACCCUUAGAAUUAAUCACCUUACAGAACACUUGAAAGUAAAUCGCAAAGAUUUCAAUACCGAGCGUUCAUUAGUGAAAUUGGUAGGUAAACGCAGAAGUUUGCUCAAUUAUUUG
env-990   AUUUUUAAAAAGUACGCAGGUUCAGAAACCAACACUGGUUCAGCAGAAGGGCAAAUCGCUCUUUUUACCCUUAGAAUCAAUCACCUUACAGAACACUUGAAAGUAAAUCGCAAAGAUUUCAAUACCGAGCGUUCUUUGGUAAGAUUGGUAGGUAAACGCAGAAGUUUGCUCGACUAUUUG
env-993   AUUUUUAAAAAGUACGCAGGUUCAGAAACCAACACUGGUUCAGCAGAAGGGCAAAUCGCUCUUUUUACCCUUAGAAUCAAUCACCUUACAGAACACUUGAAAGUAAAUCGCAAAGAUUUCAAUACCGAGCGUUCAUUGGUGAAAUUGGUAGGUAAACGCAGAAGUUUGCUCGAUUAUUUG
env-994   AUUUUUAAAAAGUACGCAGGUUCAGAAACCAACACAGGUUCAGCAGAAGGGCAAAUUGCACUUUUCACCCUCAGAAUCAAUCACCUUACUGAGCACUUAAAAGUGAAUCGCAAGGACUUUAAUACUGAGCGUUCGUUAGUAAAGCUCGUAGGUAAGCGUAGAAGUUUGCUAGACUAUUUG
env-995   AUUUUUAAAAAGUACGCAGGUUCAGAAACCAACACUGGUUCAGCAGAAGGGCAAAUCGCUCUUUUUACCCUUAGAAUCAAUCACCUUACAGAACACUUGAAAGUAAAUCGCAAAGAUUUCAAUACCGAGCGUUCAUUGGUGAAAUUGGUAGGUAAACGCAGAAGUUUGCUCGAUUAUUUG
env-999   AUUUUUAAAAAGUACGCAGGUUCAGAAACCAACACUGGUUCAGCAGAAGGGCAAAUCGCUCUUUUUACCCUUAGAAUCAAUCACCUUACAGAACACUUGAAAGUAAAUCGCAAAGAUUUCAAUACCGAGCGUUCUUUGGUAAGAUUGGUAGGUAAACGCAGAAGUUUGCUCGACUAUUUG
env-1001  AUUUUUAAAAAGUACGCAGGUUCAGAAACCAACACUGGUUCAGCAGAAGGGCAAAUCGCUCUUUUUACCCUUAGAAUCAAUCACCUUACAGAACACUUGAAAGUAAAUCGCAAAGAUUU..............................................................
env-1002  AUUUUUAAAAAGUACGCAGGUUCAGAGACCAACACUGGUUCAGCAGAAGGGCAAAU.............................................................................................................................
env-1003  AUUUUUAAAAAGUACGCAGGUUCAGAAACCAACACUGGUUCAGCAGAAGGGCAAAUCGCUCUUUUUACCCUUAGAAUCAAUCACCUUACAGAACACUUGAAAGUAAAUCGCAAAGAUUUCAAUACUGAGCGUUCAUUGGUGAAAUUGGUAGGUAAACGCAGAAGUUUGCUCGAUUAUUUG
env-1004  AUUUUUAAAAAGUACGCAGGUUCAGAAACCAACACUGGUUCAGCAGAAGGGCAAAUCGCUCUUUUUACUCUUAGAAUCAAUCACCUUACAGAACACUUGAAAGUAAAUCGUAAAGAUUUCAAUACCGAGCGUUCUUUGGUAAGAUUGGUAGGUAAACGCAGAAGUUUGCUCGACUAUUUG
env-1005  AUUUUUAAAAAGUACGCAGGUUCAGAAACCAACACUGGUUCAGCAGAAGGGCAAAUCGCUCUUUUUACCCUUAGAAUCAAUCACCUUACAGAACACUUGAAAGUAAAUCGCAAAGAUUUCAAUACCGAGCGUUCAUUGGUGAAAUUGGUAGGUAAACGCAGAAGUUUGCUCGA.......
env-1006  AUUUU................................................................................................................................................................................
env-1007  AUUUUUAAAAAGUACGCAGGUUCAGAGACCAACACUGGUUCAGCAGAAGGGCAAAUCGCUCUUUUUACCCUUAGAAUCAAUCACCUUACAGAACACUUGAAAGUAAAUCGCAAAGAUUUCAAUACUGAGCGUUCAUUGGUGAAAUUGGUAGGUAAACGCAGAAGUUUGCUCGAUUAUUUG
env-1008  AUUUUUAAAAAGUACGCAGGUUCAGAAACCAACACUGGUUCAGCAGAAGGGCAAAUCGCUCUUUUUACCCUUAGAAUCAAUCA..................................................................................................
env-1009  AUUUUUAAAAAGUACGCAGGUUCAGAAACCAACACUGGUUCAGCAGAAGGGCAAAUCGCUCUUUUUACCCUUAGAAUCAAUCACCUUACAGAACACUUGAAAGUAAAUCGCAAAGAUUUCAAUACCGAGCGUUCAUUGGUGAAAUUGGUAGGUAAACGCAGAAGUUUGCUCGAUUAUUUG
env-1010  AUUUUUAAAAAGUACGCAGGUUCAGAAACCA......................................................................................................................................................
env-1011  AUUUUUAAAAAGUACGCAGGUUCAGAAACCAACACUGGUUCAGCAGAAGGGCAAAUCGCUCUUUUUACCCUUAGAAUCAAUCACCUUACAGAACACUUGAAAGUAAAUCGCAAAGAUUUCAAUACUGAGCGUUCAUUGGUGAAAUUGGUAGGUAAACGCAGAAGUUUGCUCGAUUAUUUG
env-1012  UUAUUCAAAAAACACGGUAAAAGUGUAAAAGAUACCGGAAGUGCUGAAGGGCAAAUUGCUUUAUUUACACUCAGAAUAAAUCAUUUAACUGAGCACUUAAAAAAUAACAAAAAAGAUUUCAACACAGAAAGAGCUUU............................................
env-1013  UUAUUCAAAAAACACGGUAAAAGUGUAAAAGAUACCGGAAGUGCUGAAGGGCAAAUUGCUUUAUUUACACUCAGAAUAAAUCAUUUAACUGAGCACUUAAAAAAUAACAAAAAAGAUUUCAACACAGAAAGAGCUUUGGUAAAGCUUGUUGGAAAAAGAAGAUCGCUUCUAGAUUACUUA
env-1015  UUAUUCAAAAAACACGGUAAAAGUGUAAAAGAUACCGGAAGUGCUGAAGGGCAAAUUGCUUUAUUUACACUCAGAAUAAAUCAUUUAACUGAGCACUUAAAAAAUAACAAAAAAGAUUUCAACACAGAAAGAGCUUUGGUAAAGCUUGUUGGAAAAAGAAGAUCGCUUCUAGAUUACUUA
env-1016  UUAUUCAAAAAACACGGUAAAAGUGUAAAAGAUACCGGAAGUGCUGAAGGGCAAAUUGCUUUAUUUACACUCAGAAUAAAUCAUUUAACUGAGCACUUAAAAAAUAACAAAAAAGAUUUCAACACAGAAAGAGCUUUGGUAAAGCUUGUUGGAAAAAGAAGAUCGCUUCUUGAUUACUUA
env-1017  UUAUUCAAAAAACACGGAAAAGGUGCAAAAGAUACCGGAAGUGCUGAAGGUCAAAUUGCUUUAUUUACCCUUAGGAUAAAUCAUUUAACUGAGCACUUAAAAAAUAAUAAAAAAGAUUUCAAUACAGAAAGAGCAUUAGUAAAGCUGGUUGGAAAAAGAAGAUCGCUACUUGAUUAUUUA
env-1018  UUAUUCAAAAAACACGGAAAAGGUGCAAAAGAUACCGGAAGUGCUGAAGGUCAAAUUGCUUUAUUUACCCUUAGGAUAAAUCAUUUAACAGAGCACUUAAAAAAUAAUAAAAAAGAUUUUAACACAGAAAGAGCAUUAGUAAAGCUGGUUGGAAAAAGAAGAUCGCUACUUGAUUAUUUA
env-1019  UUAUUCAAAAAACAUGGUAAAAGUGCAAAGAAUACCGGAAGUGCUGAAGGGCAAAUUGCUUUAUUUACACUUAGGAUAAAUCAUUUAACUAAGCACUUAAAAAAUAACAAAAAGGAUUUCAACACUGAAAGAGCUUUGGUAAAGCUUGUAGGAAAAAGAAGAUCUCUACUUGAUUACUUA
env-1021  CUUUUCAAAAAACACGGUAAAAGUGCAAAAGACACUGGAAACGCUGAAGGUCAAAUUGCUUUAUUUACCCUUAGAAUAAAUCAUUUAACUGAGCACUUAAAAAAUAACAAAAAAGAUUUCAA...........................................................
env-1022  UUAUUCAAAAAACACGGUAAAAGUGCAAAAAAUACCGGAAGUGCUGAAGGACAAAUUGCUUUAUUUACACUAAGAAUAAAUCAUUUAACUGAGCACUUAAAAAAUAACAAAAAAGAUUUCAAUACUGAAAGAGCUUUAGUAAAGCUUGUUGGAAAAAGAAGAUCGCUUUUAGAUUACUUA
env-1023  UUAUUCAAAAAACACGGUAAAAGUGCUAAAGAUACCGGAAGCGCUGAAGGGCAAAUUGCUUUAUUUACCCUCAGAAUAAAUCAUUUAACUGAGCAUUUAAAAAAUAACAAAAAAGAUUUCAACACUGAAAGAGCGUUAGUAAAGCUUGUUGGAAAAAGAAGAUCGCUACUAGAUUACUUA
env-1024  UUAUUCAAAAAACACGGUAAAAGUGCAAAAGAUACCGGAAGUGCUGAAGGGCAAAUUGCUUUAUUUACCCUCAGAAUAAAUCAUUUAACUGAGCACUUGAAAAAUAACAAAAAAGAUUUCAACACUGAAAGAGCGUUAGUAAAGCUUGUUGGAAAAAGAAGAUCGCUACUAGAUUACUUA
env-1025  UUAUUCAAAAAACACGGUAAAAGUACAAAAGAUACCGGAAGUGCUGAAGGGCAAAUUGCUUUAUUUACCCUUAGAAUAAAUCAUUUAACUAAGCACUUGAAAAAUAACAAAAAAGAUUUCAACACUGAAAGAGCGUUAGUAAAGCUUGUUGGAAAAAGAAGAUCGCUUCUUGAUUACUUA
env-1026  UUAUUCAAUAAACACGGUAAAAGUGUAAAAGAUACCGGAAGUGCUGAAGGGCAAAUUGCUUUAUUCACACUUAGAAUAAAUCAUUUAACUGAGCAUUUAAAAAAUAACAAAAAAGAUUUCAACACAGAAAGAGCUUUGGUAAAGCUUGUUGGAAAAAGAAGAUCGCUUCUUAAUUACUUA
env-1027  UUAUUCAAAAAACACGGUAAAAGUACUAAAGAUACCGGAAGUGCUGAAGGGCAAAUUGCUUUAUUUACCCUUAGAAUAAAUCAUUUAACUGAGCACUUGAAAAAUAACAAAAAAGAUUUCAACACUGAAAGAGCAUUAGUAAAGCU...................................
env-1028  UUAUUUAAAAAGCACGGAAAAGAUGCUAAAGACACGGGAAGUGCUGAAGGUCAAAUUGCCCUUUUUACCCUAAGAAUAAAUCAUUUAACAGAGCAUCUAAAGAGAAAUAAGAAAGAUUAUAACACGGAAAGAGCACUUGUAAAGCU...................................
env-1029  UUAUUCAAAAAGCACGGAAAGGAUGCCAAAGAUACGGGAAGUGCUGAAGGUCAAAUUGCCCUUUUUACCUACAGAAUAAAUCAUUUAACAGAGCAUCUAAAAAGAAAUAAGAAGGAUUAUAACACGGAAAGAGCUCUUGUAAAGCUCGUAGGAAAAAGAAGAGCACUAUUAGAUUAUCUU
env-1030  .....................................................................................................................................................................................
env-1032  UUAUUCAAAAAGCACGGAAAGGAUGCCAAAGAUACGGGAAGUGCUGAAGGUCAAAUUGCCCUUUUUACCUACAGAAUAAAUCAUUUAACAGAGCAUCUAAAAAGAAAUAAGAAGGAUUAUAACACGGAAAGAGCUCUUGUAAAGCUCGUAGGAAAAAGAAGAGCACUAUUAGAUUAUCUU
env-1033  UAUUCAAAAAGCACGGUAAGGAUGCCAAAGAUACGGGAAGUGCUGAAGGUCAAAUUGCCCUUUUUACAUACAGAAUAAAUCAUUUAACAGAGCAUCUAAAAAGAAAUAAGAAGGAUUAUAACACGGAAAGAGCUCUUGUAAAGCUCGUAGGAAAAAGAAGAGCACUAUUAGAUUAUCUUA
env-1034  UUAUUCAAAAAGCACGGAAAGGAUGCCAAAGAUACGGGAAGUGCUGAAGGUCAAAUUGCCCUUUUUACCUACAGAAUAAAUCAUUUAACAGAGCAUCUAAAAAGAAAUAAGAAGGAUUAUAACACGGAAAGAGCUCUUGUAAAGCUCGUA...............................
env-1035  .....................................................................................................................................................................................
env-1036  UUAUCAAAAAGCACGGAAAAGAUGCUAAAGAUACGGGAAGUGCUGAAGGUCAAAUUGCCCUUUUUACCCACAGAAUAAAUCAUUUAACAGAGCAUCUUAAGAGAAAUAAGAAAGAUUAUAACACGGAAAGAGCACUUGUAAAGCUUGUAGGAAAAGAAGAGCUUUACUGGAUUAUCUUAC
env-1037  UUAUUCAAAAAGCACGGAAAAGAUGCCAAAGAUACGGGAAGUGCCGAGGGCCAAAUUGCCCUUUUUACCCAUAGAAUAAAUCAUUUAACAGAGCAUCUAAAAAGAAAUAAGAAAGAUUAUAACACGGAAAGAGCACUUGUUAAGCUCGUAGGAAAAAGAAGAUCGUUAUUAGACUAUCUU
env-1038  UUAUUCAAAAAGCAUGGAAAAGAUGCUAAAGAUACAGGAAGUGCUGAAGGUCAGAUUGCCCUUUUUACACUUAGAAUAAAUCAUUUAACUGAGCAUCUAAAAAAUAAACAAGAAGGAUUAUAACACGGAAAGAGCGCUUGUAAAGCUCGUAGGAAAAAGAAGAGCACUAUUAGACUAUCUU
env-1039  .....................................................................................................................................................................................
env-1040  .....................................................................................................................................................................................
env-1041  .....................................................................................................................................................................................
env-1042  UUAUUCAAAAAGCAUGGAAAAGAUGCCAAAGAUACGGGUAGUGCUGAGGGCCAAAUUGCCCUUUUUACCCACAGAAUUAAUCAUUUAACAGAGCAUCUAAAAAGAAAUAAGAAAGAUUAUAACACGGAAAGAGCACUUGUUAAGCUCGUAGGAAAAAGAAGGUCGUUAUUAGACUAUCUU
env-1044  UUAUUCAAAAAGCACGGAAAAGAUGCCAAAGAUACGGGUAGUGCUGAGGGCCAAAUUGCCCUUUUUACCCACAGAAUAAAUCAUUUAACAGAGCAUCUAAAAAGAAAUAAGAAAGAUUAUAACACGGAAAGAGCACUUGUUAAGCUCGUAGGAAAAAGAAGGUCGUUAUUAGACUAUCUU
env-1045  UUAUUCAAAAAGCACGGAAAAGAUGCUAAAGAUACGGGAAGUGCUGAAGGUCAAAUUGCCCUUUUUACCCACAGAAUAAAUCACCUAACUGAGCAUCUUAAAAGAAAUAAGAAAGAUUAUAACACGGAAAGAGCACUUGUAAAGCUUGUAGGAAAAAGAAGAGCUUUGCUAGAUUAUCUU
env-1046  UUAU.................................................................................................................................................................................
env-1047  UUAUUCAAAAAACACGGUAAGGAUGCUAAAAAUACUGGAAGUGCCGAAGGUCAAAUUGCCCUUUUUACUCUUAGAAUAAAUCAUUUAACAGAGCAUCUAAAAAGAAAUAAGAAAGAUUAUAACACAGAAAGAGCAUUAGUAAAGCUUGUUGGAAAAGAAGAGCACUAUUGGAUUAUCUC
env-1048  .....................................................................................................................................................................................
env-1049  UUAUUCAAAAAACACGGUAAGGAUGCUAAAAAUACUGGAAGUGCCGAAGGUCAAAUUGCCCUUUUUACUCUUAGAAUAAAUCAUUUAACAGAGCAUCUAAAAAGAAAUAAGAAAGAUUAUAACACAGAAAGAGCAUUAGUAAAGCUUGUUGGAAAAGAAGAGCACUAUUGGAUUAUCUC
env-1051  .....................................................................................................................................................................................
env-1053  UUAUUCAAAAAACACGGUAAGGAUGCUAAAAAUACUGGAAGUGCCGAAGGUCAAAUUGCCCUUUUUACUCUUAGAAUAAAUCAUUUAACAGAGCAUCUAAAAAGAAAUAAGAAAGAUUAUAACACAGAAAGAGCAUUAGUAAAGCUUGUUGGAAAAGAAGAGCACUCUUGGAUUAUCUC
env-1055  UUAUUCAAAAAACACGGUAAGGAUGCUAAAAAUACUGGAAGUGCCGAAGGUCAAAUUGCCCUUUUUACUCUUAGAAUAAAUCAUUUAACAGAGCAUCUAAAAAGAAAUAAGAAAGAUUAUAACACAGAAAGAGCAUUAGUAAAGCUUGUUGGAAAAGAAGAGCACUAUUGGAUUAUCUC
env-1056  UUAUUCAAAAAACACGGAAAAGAUGCCAAAGACACGGGAAGUGCUGAAGGGCAAAUUGCUCUUUUUACCCAUAGAAUAAAUCAUUUAACAGAGCAUCUGAAAAGAAAUAAGAAGGAUUAUAACACGGAAAGAGCACUAGUUAAGCUCGUAGGAAAAAGAAGAGCACUAUUGGAUUAUCUU
env-1057  UUAUUCAAAAAACACGGAAAAGAUGCUAAAGACACGGGAAGUGCUGAGGGUCAGAUUGCCCUUUUCACACUCAGAAUAAAUCAUUUGACCGAGCAUCUGAAAAAAAAUAAGAAAGAUUAUAAUACGGAAAGAGCGCUUGUAAAGCUCGUAGGAAAAAGAAGAGCACUAUU.........
env-1058  .....................................................................................................................................................................................
env-1059  UUAUUCAA.............................................................................................................................................................................
env-1060  .....................................................................................................................................................................................
env-1061  UUAUUCAAAAAGCACGGAAAAGAUGCUAAAGACACAGGUAGUGCAGAAGGGCAGAUUGCUCUUUUUACUGAAAGAAUAAAUCAUUUAACAGAGCAUCUUAAGAAAAAUAAGAAAGAUUAUAACACGGAAAGAGCUUUAGUAAAGCUUGUAGGAAAAAGAAGAGCACUUUUGGAUUAUCUU
env-1062  UUAUUCAAAAAGCACGGAAAAGAUGCUAAAGAUACAGGUAGUGCAGAAGGGCAGAUUGCUCUUUUUACUGAAAGAAUAAAUCAUUUAACAGAGCAUCUUAAGAAAAAUAAGAAAGAUUAUAACACGGAAAGAGCUUUAGUAAAGCUUGUAG..............................
env-1063  UUAUUCAAGAAGCACGGAAAAGAUGCUAAAGAUACCGGAAGUGCUGAGGGGCAAAUUGCCUUAUUUACUGAUAGAAUAAAUCAUUUAACAGAGCAUCUUAAAAGAAAUAAGAAAGAUUAUAACACGGAAAGAGCCUUAGUAAAGCUUGUAGGAAAAAGAAGAGCACUAUUAAACUAUCUU

          .....................................................................................................................................................................................
          .....................................................................................................................................................................................
          AUYUUYR•RAARCACGG•RRR•••R•RR••RAYAC•GG••C•RC•GAAGG•CAAAUYGC•YURUUYAC•Y•Y•G•AUYRAYCAYYUR•C•RA•CAYYURAAAAR•AAYCR•AARGAUU•YAAYAC•GAR•G•UC•YURGU•••RYURGU•GG•AAR•G•RG••••YU•YU•GAYUAYYU•
```

```
env-1064  UUAUUCAAAAAGCACGGUAAAGAUGCUAAAGAUACGGGAAGUGCUGAAGGUCAAAUUGCCCUUUUUACCCACAGAAUAAAUCAUUUAACAGAGCAUCUUAAGAGAAAUAAGAAAGAUUAUAACACGGAAAGAGCACUUGUAAAGCUUGUAGGUAAAAGAAGAGCUUUACUAGAUUAUCUU
env-1065  UUAUUCAAGAAGCACGGAAAAGAUGCUAAAGAUACAGGAAGUGCUGAAGGUCAGAUUGCCCUUUUUACACACAGAAUAAAUCAUUUAACUGAGCAUCUAAAAAGAAAUAAGAAAGAUUAUAACACGGAAAGAGCGCUUGUAAAGCUCGUAGGAAAAAGAAGAGCACUAUUAGACUAUCUU
env-1066  UUUAUUCAAAAAGCACGGUAAAGAUGCUAAAGAUACGGGAAGUGCUGAAGGUCAAAUUGCCCUUUUUACCCAUAGAAUAAAUCAUUUGACAGAGCAUCUUAAAAGAAAUAAGAAAGAUUAUAACACGGAAAGAGCCCUAGUAAAGCUUGUAGGAAAAAGAAGAUCACUAUUAGAUUAUCU
env-1067  UUAUUCAAGAAGCACGGAAAAGAUGCUAAAGAUACCGGAAGUGCUGAGGGGCAAAUUGCCUUAUUUACUGAUAGAAUAAAUCAUUUAACAGAGCAUCUUAAAAGAAAUAAGAAAGAUUAUAACACGGAAAGAGCCUUAGUAAAGCUUGUAGGAAAAAGAAGAGCACUAUUAAACUAUCUU
env-1068  UUAUUCAAAAAGCACGGUAAAGAUGCUAAAGAUACGGGAAGUGCUGAAGGUCAAAUUGCCCUUUUUACCCACAGAAUAAAUCAUUUAACAGAGCAUCUUAAGAGAAAUAAGAAAGAUUAUAACACGGAAAGAGCACUUGUAAAGCUUGUAGGUAAAAGAAGAGCUUUACUAGAUUAUCUU
env-1069  UUAUUCAAAAAGCACGGAAAAGAUGCAAAAGAUACCGGAAGUGCUGAGGGCCAAAUUGCCUUAUUUACUGAUAGAAUAAAUCAUUUAACAGAGCAUCUUAAAAGAAAUAAGAAAGAUUAUAACACGGAAAGAGCUUUAGUAAAGCUUGUAGGAAAAAGAAGAGCACUAUUAAACUAUCUU
env-1070  UUAUUCAAAAAGCACGGUAAAGAUGCUAAAGAUACGGGAAGUGCUGAAGGUCAAAUUGCCCUUUUUACCCAUAGAAUAAAUCAUUUGACAGAGCAUCUUAAAAGAAAUAAGAAAGAUUAUAACACGGAAAGAGCCCUAGUAAAGCUUGUAGGAAAAAGAAGAUCACUAUUAGAUUAUCUA
env-1077  UUAUUCAAAAAGCACGGAAAAGAUGCUAAAGAUACCGGAAGUGCUGAGGGGCAAAUUGCCUUAUUUACUGAUAGAAUAAAUCAUUUAACAGAGCAUCUUAAAAGAAAUAAGAAAGAUUAUAACACGGAAAGAGC.............................................
env-1078  UUAUUCAAAAAGCACGGUAAAGAUGCUAAAGAUACGGGAAGUGCUGAAGGUCAAAUUGCCCUUUUUACCCAUAGAAUAAAUCAUUUGACAGAGCAUCUUAAAA..........................................................................
env-1079  UUAUUCAAAAAGCACGGAAAAGAUGCUAAAGAUACCGGAAGUGCUGAGGGGCAAAUUGCCUUAUUUACUGAUAGAAUAAAUCAUUUAACAGAGCAUCUUAAAAGAAAUAAGAAAGAUUAUAACACGGAAAGAGCCUUAGUAAAGCUUGUAGGAAAAAGAAGAGCACUAUUAAACUAUCUU
env-1080  UUAUUCAAAAAGCACGGAAAAGAUGCUAAAGAUACCGGAAGUGCUGAGGGGCAAAUUGCCUUAUUUACUGAUAGAAUAAAUCAUUUAACAGAGCAUCUUAAAAGAAAUAAGAAAGAUUAUAACACGGAAAGAGCCUUAGUAAAGCUUGUAGGAAAAAGAAGAGCACUAUUAAACUAUCUU
env-1081  UUAUUCAAAAAGCACGGAAAAGAUGCUAAAGAUACCGGAAGUGCUGAGGGGCAAAUUGCCUUAUUUACUGAUAGAAUAAAUCAUUUAACAGAGCAUCUUAAAAGAAAUAAGAAAGAUUAUAACACGGAAAGAGCCUUAGUAAAGCUUGUAGGAAAAAGAAGAGCACUAUUAAACUAUCUU
env-1082  UUAUUCAAAAAGCACGGAAAAGAUGCUAAAGAUACCGGAAGUGCUGAGGGGCAAAUUGCCUUAUUUACUGAUAGAAUAAAUCAUUUAACAGAGCAUCUUAAAAGAAAUAAGAAAGAUUAUAACACGGAAAGAGCCUUAGUAAAGCUUGUAGGAAAAAGAAGAGCACUAUUAAACUAUCUU
env-1085  UUAUUCAAAAAGCACGGAAAAGAUGCUAAAGAUACGGGAAGUGCUGAAGGUCAAAUUGCCCUUUUUACCCACAGAAUAAAUCAUUUAACAGAGCAUCUAAAGAGAAAUAAGAAAGAUUAUAACACGGAAAGAGCACUUGUAAAGCUUGUAGGAAAAAGAAGAGCAUUACUAGAUUAUCUU
env-1086  UUAUUCAAAAAGCACGGAAAAGAUGCUAAAGAUACGGGAAGUGCGGAAGGUCAAAUUGCCCUUUUUACCCACAGAAUCAAUCAUUUAACAGAGCAUCUUAAGAGAAAUAAGAAAGAUUAUAACACGGAAAGAGCACUUGUAAAGCUCGUAGGAAAAAGAAGAGCUUUACUAGAUUAUCUU
env-1087  ..........................................................................................................................................................................
env-1088  UUAUUCAAAAAGCACGGAAAAGAUGCUAAAGAUACAGGAAGUGCUGAAGGUCAAAUUGCCCUUUUUACACAUAGAAUAAAUCAUUUAACUGAGCAUCUUAAAAAGUAAUAAGAAGGAUUAUAACACGGAAAGAGCACUUGUAAAGCUCGUAGGGAAAAGAAGAGCACUAUUAGACUAUCUU
env-1089  UUAUUCAAAAAGCACGGUAAAGAUGCUAAAGAUACGGGAAGUGCUGAAGGUCAAAUUGCCCUUUUUACCCACAGAAUAAAUCAUUUAACAGAGCAUCUUAAGAGAAAUAAGAAAGAUUAUAACACGGAAAGAGCACUUGUAAAGCUUGUAGGUAAAAGAAGAGCUUUACUAGAUUAUCUU
env-1090  UUAUUCAAAAAGCACGGUAAAGAUGCUAAAGAUACGGGAAGUGCUGAAGGUCAAAUUGCCCUUUUUACCCACAGAAUAAAUCAUUUAACAGAGCAUCUUAAGAGAAAUAAGAAAGAUUAUAACACGGAAAGAGCACUUGUAAAGCUUGUAGGUAAAAGAAGAGCUUUACUAGAUUAUCUU
env-1091  UUAUUCAAAAAGCACGGAAAAGAUGCUAAAGAUACGGGAAGUGCUGAAGGUCAAAUUGCCCUUUUUACCCACAGAAUAAAUCAUUUAACAGAGCAUCUUAAGAGAAAUAAGAAAGAUUAUAACACGGAAAGAGCACUUGUAAAGCUUGUAGGAAAAAGAAGAGCUUUACUAGAUUAUCUU
env-1092  UUAUUCAAAAAGCACGGAAAAGAUGCUAAAGAUACGGGAAGUGCUGAAGGUCAAAUUGCCCUUUUUACCCACAGAAUAAAUCACUUAACUGAGCAUCUUAAGAGAAAUAAGAAAGAUUAUAACACGGAAAGAGCACUUGUAAAGCUUGUAGGAAAAAGAAGAGCUUUGCUAGAUUAUCUU
env-1093  ..........................................................................................................................................................................
env-1094  ..........................................................................................................................................................................
env-1095  UUAUUCAAAAAGCACGGAAAAGAUGCUAAAGAUACAGGAAGUGCUGAAGGUCAGAUUGCCCUUUUUACACACAGAAUAAAUCAUUUAACUGAGCAUCUAAAAAGAAAUAAGAAAGAUUAUAACACGGAAAGAGCGCUUGUAAAGCUCGUAGGAAAAAGAAGAGCACUAUUAGACUAUCUU
env-1096  UUAUUCAAAAAGCACGGAAAAGAUGCUAAAGAUACAGGAAGUGCUGAAGGUCAGAUUGCCCUUUUUACACACAGAAUAAAUCAUUUAACUGAGCAUCUAAAAAGAAUAAGAAAGAUUAUAACACGGAAAGAGACG..........................................
env-1097  UUAUUCAAAAAGCACGGAAAAGAUGCUAAAGAUACGGGAAGUGCUGAAGGUCAAAUUGCCCUUUUUACCCACAGAAUAAAUCAUUUAACAGAGCAUCUUAAGAGAAAUAAGAAAGAUUAUAACACGGAAAGAGCACUUGUAAAGCUCGUAGGAAAAAGAAGAGCUUUACUAGAUUAUCUU
env-1098  UUAUUCAAAAAGCACGGAAAAGAUGCAAAAGAUACCGGAAGUGCUGAGGGCCAAAUUGCCUUAUUUACUGAAAGAAUAAAUCAUUUAACAGAGCAUCUUAAAAGAAACAAGAAAGAUUAUAACACGGAAAGAGCCUUAGUAAAGCUUGUAGGAAAAAGAAGAGCACUAUUAAACUAUCUU
env-1099  UUAUUCAAAAAGCACGGAAAAGAUGCUAAAGAUACAGGAAGUGCUGAAGGUCAGAUUGCCCUUUUUACACACAGAAUAAAUCAUUUAACUGAGCAUCUUAAAAAGAAAUAAGAAAGAUUAUAACACGGAAAGAGCGCUUGUAAAGCUCGUAUGAAAAGAAGAGCAUUAUUAGACUAUCUUA
env-1101  UUAUUCAAAAAGCACGGUAAAGAUGCUAAAGAUACGGGAAGUGCUGAAGGUCAAAUUGCCCUUUUUACCCACAGAAUAAAUCAUUUAACAGAGCAUCUUAAGAGAAAUAAGAAAGAUUAUAACACGGAAAGAGCACUUGUAAAGCUUGUAGGUAAAAGAAGAGCUUUACUAGAUUAUCUU
env-1102  UUAUUCAAGAAGCACGGAAAAGAUGCUAAAGAUACCGGAAGUGCUGAGGGGCAAAUUGCCUUAUUUACUGAAAGAAUAAAUCAUUUAACAGAGCAUCUUAAAAGAAAUAAGAAAGAUUAUAACACGGAAAGAGCCUUAGUAAAGCUUGUAGGAAAAAGAAGAGCACUAUUAAACUAUCUU
env-1103  UUAUUCAAAAAGCACGGUAAAGAUGCUAAAGAUACGGGAAGUGCUGAAGGUCAAAUUGCCCUUUUUACCCACAGAAUAAAUCAUUUAACAGAGCAUCUUAAGAGAAAUAAGAAAGAUUAUAACACGGAAAGAGCACUUGUAAAGCUUGUAGGUAAAAGAAGAGCUUUACUAGAUUAUCUU
env-1104  UUAUUCAAAAAGCACGGAAAAGAUGCUAAAGAUACAGGAAGUGCUGAAGGUCAGAUUGCCCUUUUUACACAAAGAAUAAAUCAUUUAACUGAGCAUCUAAAAAGAAAUAAGAAGGAUUAUAACACGGAAAGAGCGCUUGUAAAGCUCGUAGGAAAAAGAAGAGCACUAUUAGACUAUCUU
env-1105  UUAUUCAAAAAGCACGGAAAAGAUGCUAAAGAUACCGGAAGUGCUGAGGGGCAAAUUGCCUUAUUUACUGAUAGAAUAAAUCAUUUAACAGAGCAUCUUAAAAGAAAUAAGAAAGAUUAUAACACGGAAAGAGCCUUAGUAAAGCUUGUAGGAAAAAGAAGAGCACUAUUAAACUAUCUU
env-1106  UUAUUCAAAAAGCACGGAAAAGAUGCCAAAAACACGGGAAGUGCUGAGGGCCAAAUUGCCCUUUUUACCCAUAGAAUAAAUCAUUUAACUGAGCAUCUAAAAAGAAAUAAGAAAGAUUAUAACACGGAAAGAGCACUUGUAAAGCUCGUAGGAAAAAGAAGAUCGUUAUUAGACUAUCUU
env-1107  UUAUCAAAAAGCACGGAAAAGAUGCUAAAGACACGGGAAGUGCUGAAGGUCAAAUUGCCCUUUUUACCCACAGGAUAAAUCAUUUAACUGAGCAUCUUAAGAGAAAUAAGAAAGAUUAUAACACGGAAAGAGCACUUGUAAAGCUC.................................
env-1108  UUAUUCAAAAAGCACGGAAAAGAUGCUAAAGAUACAGGAAUCUUCCACCUAUA.......................................................................................................................
env-1109  UUAUUCAAAAAGCACGGAAAAGAUGCUAAAGAUACAGGAAGUGCUGAAGGUCAGAUUGCCCUUUUUACACACAGAAUAAAUCAUUUAACUGAGCAUCUAAAAAGAAAUAAGAAGGAUUAUAACACGGAAAGAGCGCUUGUAAAGCUCGUAGGAAAAAGAAGAGCACUAUUAGACUAUCUU
env-1110  UUAUUCAAAAAGCACGGUAAAGAUGCUAAAGAUACGGGAAGUGCUGAAGGUCAAAUUGCCCUUUUUACCCAUAGAAUAAAUCAUUUGACAGAGCAUCUUAAAAGAAAUAAGAAAGAUUAUAACACGGAAAGAGCCCUAGUAAAGCUUGUAGGAAAAAGAAGAUCACUAUUAGAUUAUCUA
env-1111  UUAUUCAAAAAGCACGGAAAAGAUGCUAAAGACACGGGAAGUGCUGAAGGUCAAAUCGCCCUUUUUACCCACAGAAUAAAUCAUUUAACAGAGCAUCUUAAAAGAAAUAAGAAAGAUUAUAACACGGAAAGAGCACUUGUAAAGCUCGUAGGAAAAAGAAGAGCUUUACUGGAUUAUCUU
env-1112  UUAUUCAAAAAGCACGGUAAAGAUGCCAAAGACACGGGAAGUGCUGAAGGACAAAUUGCCCUUUUUACCCAUAGAAUUAAUCAUUUAACUGAGCAUCUAAAAAGAAAUAAGAAAGAUUAUAACACGGAAAGAGCACUUGUAAAGCUCGUUGGAAAAAGAAGAGCACUUUUAGACUAUCUU
env-1113  UUAUUCAAAAAGCACGGUAAAGAUGCCAAAGACACGGGAAGUGCUGAAGGACAAAUUGCCCUUUUUACCCAUAGAAUUAAUCAUUUAACUGAGCAUCUAAAAAGAAAUAAGAAAGAUUAUAACACGGAAAGAGCACUUGUAAAGCUCGUUGGAAAAAGAAGAGCACUGUUAGACUAUCUG
env-1115  UUAUUCAAAAAGCACGGUAAAGAUGCCAAAGACACGGGAAGUGCUGAAGGACAAAUUGCCCUUUUUACCCAUAGAAUUAAUCAUUUAACUGAGCAUCUAAAAAGAAAUAAGAAAGAUUAUAACACGGAAAGAGCACUUGUAAAGCUCGUUGGAAAAAGAAGAGCACUGUUAGACUAUCUG
env-1116  UUAUUCAAAAAGCACGGUAAAGAUGCCAAAGACACGGGAAGUGCUGAAGGACAAAUUGCCCUUUUUACCCAUAGAAUUAAUCAUUUAACUGAGCAUCUAAAAAGAAAUAAGAAAGAUUAUAACACGGAAAGAGCACUUGUAAAGCUCGUUGGAAAAAGAAGAGCACUGUUAGACUAUCUG
env-1121  UUAUUCAAAAAGCACGGUAAAGAUGCCAAAGACACGGGAAGUGCUGAAGGACAAAUUGCCCUUUUUACCCAUAGAAUUAAUCAUUUAACUGAGCAUCUAAAAAGAAAUAAGAAAGAUUAUAACACGGAAAGAGCACUUGUAAAGCUCGUUGGAAAAAGAAGAGCACUGUUAGACUAUCUG
env-1122  UUAUUCAAAAAGCACGGAAAAGAUGCUAAAGAUACUGGAAGUGCUGAGGGUCAAAUUGCACUUUUUACUCAUAGAAUAAAUCACUUAACCGAGCAUUUAAAAAGUAAUAAAAAAGAUUACAAUACUGAAAGAGCUCUUGUAAAACUUGUUGGAAAAAGAAGAGCAUUAUUAGAUUAUCUU
env-1123  UUAUUCAAAAAGCACGGAAAAGAUGCUAAAGAUACAGGAAGUGCUGAAGGUCAGAUUGCCCUUUUUACACACAGAAUAAAUCAUUUAACUGAGCAUCUAAAAAGAAAUAAGAAAGAUUAUAACACGGAAAGAGCGCUUGUAAAGCUCGUAGGAAAAAGAAGAGCACUAUUAGACUAUCUU
env-1124  UUAUUCAAAAAGCACGGUAAAGAUGCCAAAGACACGGGAAGUGCUGAAGGACAAAUUGCCCUUUUUACCCAUAGAAUUAAUCAUUUAACUGAGCAUCUAAAAAGAAAUAAGAAAGAUUAUAACACGGAAAGAGCACUUGUAAAGCUCGUUGGAAAAAGAAGAGCACUUUUAGACUAUCUU
env-1125  UUAUUCAAAAAGCACGGUAAAGAUGCCAAAGACACGGGAAGUGCUGAAGGACAAAUUGCCCUUUUUACCCAUAGAAUUAAUCAUUUAACUGAGCAUCUAAAAAGAAAUAAGAAAGAUUAUAACACGGAAAGAGCACUUGUAAAGCUCGUUGGAAAAAGAAGAGCACUGUUAGACUAUCUG
env-1126  UUAUUCAAAAAGCACGGUAAAGAUGCCAAAGACACGGGAAGUGCUGAAGGACAAAUUGCCCUUUUUACCCAUAGAAUUAAUCAUUUAACUGAGCAUCUAAAAAGAAAUAAGAAAGAUUAUAACACGGAAAGAGCACUUGUAAAGCUCGUUGGAAAAAGAAGAGCACUGUUAGACUAUCUG
env-1127  UUAUUCAAAAAGCACGGUAAAGAUGCCAAAGACACGGGAAGUGCUGAAGGACAAAUUGCCCUUUUUACCCAUAGAAUUAAUCAUUUAACUGAGCAUCUAAAAAGAAAUAAGAAAGAUUAUAACACGGAAAGAGCACUUGUAAAGCUCGUUGGAAAAAGAAGAGCACUGUUAGACUAUCUG
env-1128  UUAUUCAAAAAGCACGGAAAAGAUGCUAAAGAUACUGGAAGUGCUGAGGGUCAAAUUGCACUUUUUACUCAUAGAAUAAAUCACUUAACCGAGCAUUUAAAAAGUAAUAAAAAAGAUUACAAUACUGAAAGAGCUCUUGUAAAACUUGUUGGAAAAAGAAGAGCAUUAUUAGAUUAUCUU
env-1129  AUCUUCGCAAAACACGG.............................................................................................................................................................
env-1130  AUCUUCGCAAAACACGGCGGUUCUGAAACCAACACUGGUUCAACAGAAGGUCAAAUCGCAUUGUUCACAUUUCGCAUCAGCCACUUGACUGAGCACCUUAAGAAGAAUCGCAAAGACUACGGUACACAGCGCUGCACUUCAACUUUUAGUUGGUAAGCGUCGUAGCCUUUUAGAUUACCU
env-1131  AUCUUCGCCAAACACGGUGGUUCAGAAAUCAACACCGGUUCUACAGAAGGUCAAAUAGCCUUAUUCACGUUUCGUAUUGCUCACCUCACUGAGCACUUGAAAAAGAACCGCAAAGACUUCGGCACCCAACGUUCACUUCAAUUGUUGGUUGGUAAACGUCGCAGUUUGUUAGACUACCUA
env-1134  AUCUUCGCCAAACACGGUGGUUCAGAAAUCAACACCGGUUCUACAGAAGGUCAAAUAGCCUUAUUUACGUUUCGUAUUGCUCACCUCACUGAGCACUUGAAAAAGAACCGCAAAGACUUCGGCACCCAACGUUCACUUCAAUUGUUGGUUGGUAAACGUCGCAGUUUGUUAGACUACCUA
env-1135  AUCUUCGCCAAACACGGUGGUUCAGAAAUCAACACCGGUUCUACAGAAGGUCAAAUAGCCUUAUUCACGUUUCGUAUUGCUCACCUCACUGAGCACUUGAAAAAGAACCGCAAAGACUUCGGCACCCAACGUUCACUUCAAUUGUUGGUUGGUAAACGUCGCAGUUUGUUAGACUACCUA
env-1137  AUCUUCGCCAAACACGGUGGUUCAGAAAUCAACACCGGUUCUACAGAAGGUCAAAUAGCCUUAUUCACGUUUCGUAUUGCUCACCUCACUGAGCACUUGAAAAAGAACCGCAAAGACUUCGGCACCCAACGUUCACUUCAAUUGUUGGUUGGUAAACGUCGCAGUUUGUUAGACUACCUA
env-1138  ..........................................................................................................................................................................
env-1139  AUCUUCGCCCAACACGGUGGUUCAGAAACCAAUACCGGUUCUACAGAAGGCCAAAUUGCCUUGUUUACUUUUCGCAUCAGUCACCUUACUGAGCACUUGAAAAAGAACCGCAAAGAUUACGGCACCCAACGCGCACUUCAGUUGUUAGUUGGAAAGCGUCGUAGUCUUCUUGAUUACUUA
env-1140  AUCUUCGCUAAACACGGAGGAAAAGCUGAAAACACAGGUUCAACAGAAGGUCAAGUAGCACUUUUCACUUUUAGAAUCUCUCACUUAACAGAGCAUUUGAAAAAAAAUCGUCACGAUUAUAAUACAGAGCGUUCGUUAGUACUUUUGGUAGGUAAAAGAAGAAGUCUUUUAGAUUACUUA
env-1141  AUUUUCAAAAAACACGGAAAAUCUGAAACUGACACUGGUUCAACUGAAGGGCAAAUUGCAUUAUUUUCUCACCGAAUCAAUCACUUAUCCAACCACCUUAAGAAAACCAAAAAGACUUUAACACGGAGCGUUCACUAGUUAAAUUAGUAGGGAAGCGCCGUCACUUAUUGGAUUAUCUU
env-1142  AUUUUCAAAAAACACGGAAAAUCUGAAACUGACACUGGUUCAACUGAGGGGCAAAUUGCAUUAUUUUCUCACCGAAUCAAUCACUUAUCCAACCACCUUAAGAAAACCAAAAAGACUUUAACACGGAGCGUUCACUAGUUAAAUUAGUAGGGAAGCGCCGUCACUUAUUGGAUUAUCUU
env-1143  AUUUUCAAAAAACACGGAAAAUCUGAAACUGACACUGGUUCAACUGAAGGGCAAAUUGCAUUAUUUUCUCACCGAAUCAAUCACUUAUCCAACCACCUUAAGAAAACCAAAAAGACUUUAACACGGAGCGUUCACUAGUUAAAUUAGUAGGGAAGCGCCGUCACUUAUUGGAUUAUCUU
Lsp-4-1   AUUUUUGAAAAGCACGGUAAAAGCAAAACGGAUACCGGUUCAGCAGAAGGACAAAUCGCUUUGUUCACACUUCGAAUCAACCACCUUACCGAGCAUUUAAAAAAUAACCGUAAAGACUUUAACACCGAGCGCUCCUUGUGAGACUGGUAGGAAAGCGCAGAUCACUUUUAGAUUAUCUU
Cca-5-1   AUCUUCGCAAAACACGGAAAAUCUGCACAAGACACAGGAACAGCUGAAGGACAAGUUGCUCUAUUCACUUUUAGAAUCAACCACUUGACUCAGCAUUGAAAGCUAAUCACAAAGACUUCAACACAGAAAGAUCUCUUGUAAAGUUAGUAGGUAAAAGAAAAGCUCUUUUAGAUUAUUUG
env-1144  AUCUUCGCAAAACACGGAAAAUCUGCACAAGACACAGGAACUGCUGAAGGACAAGUUGCUCUAUUCACUUUUAGGAUCAACCACCUUACUCAGCAUUUGAAAGCUAACCACAAAGACUUCAACACAGAAAGAUCUCUUGUGAAGUUAGUAGGUAAAAGAAAAGCUCUUUUAGAUUAUUUG
Fan-1-1   AUCUUCGCUAAACACGGAGGAACAGCAGCAAAUACUGGUAAAUCUGAAGGACAAAUCGCUUUGUUUACAUUUAGAAUUAACCACUUAACUGAACACUUGAAAAAAAAUCGCCACGAUUACAAUACAGAGCGUUCAUUAGUAAUGCUGGUAGGUAAAAGAAGAAGUUUGUUGGAUUACUUG
Fte-1-1   AUCUUCGCUAAACACGGAGGAUCAGAAGCAAACACUGGAAAUCUGAAUCUCAGAUCGCAUUGUUUACAUUUAGAAUCAACCACCUUACUGAGCACUUGAAAAAAAUCGUCACGAUUUUAACACAGAGCGUUCAUUAGUAAUGUUGGUAGGUAAAAGAAGAAGUUUGUUGGAUUACUUG
env-1145  UUAUUUGAAAAGCACGGUAAAGGAAAAAACGAUACCGGUUCAACUGAAGGGCAAAUUGCUUUAUUCACAAAAAGAAUUGCACAUAUUUCAAGUCACCUUAAGACAAACAAAAGGAUUACAAUUCAGAGCGUUCAUUAGUAAUGCUAGUAGGUAAAAGAAGAAGCUUGCUAGACUACUUA
env-1146  UUAUUUGCAAAACACGGUAAAGGGAAAAACGAUACCGGUUCAACUGAGGGGCAAAUUGCUUUAUUCACUAAAAGAAUAGCGUAUAUUUCUGGUCACUUAAAGACAAAUAAGAAAGAUUACAACUCUGAGCGAUCAUUAGUAAUGCUUGUAGGUAAAAGAAGAAGUUUACUAGAUUAUUUA
env-1147  UUGUUCGAAAAGCACGGUAAAAGUAAAAAUGAUACCGGCUCAACAGAAGGUCAAAUUGCGUUAUUCACUAAAAGAAUUGCUCAUAUUUCAGACCACUUAAAAAACAACAAAAAGACUUUAACUCAGAGCGAUCUCUUGUAGCGUUGGUAGGUAAAAGAAGAAGCUUACUAGACUACUUA
env-1159  ..........................................................................................................................................................................
env-1161  UUGUUCGAAAAACACGGUAAAGGUAAAAACGAUACCGGUUCAGCAGAAGGUCAAAUCGCAUUAUUCACGAAAAGAAUUGCGUAUAUUUCUGACCACUUAAAAACAAACAGAAAAGAUUACAACUCUGAGCGUUCAUUAGUAGCGCUGGUAGGUAAAAGAAGAAGCUUACUUGAUUACUUA
env-1162  UUAUUUGAAAAGCACGGUAGCGGGAAAAACGAUACCGGUUCUGCAGAAGGACAAAUUGCUUUAUUCACAAAAAGAAUGCGCUCACAUUUCUGACCACUUAAAAACGAACAGAAAAGAUUUAACUCUGAGCGUUCUCUAGUAAUGCUAGUAGGUA........................
env-1163  AUCUUCGAAAAACACGGUAAAGGAAAAAACGAUACCGGUUCAUCAGAAGGUCAAAUCGCGUUAUUUACGCACAGAAUCAACCAUUUAACAGAGCACUUAAAGAAAAAUCGUAAAGAUUACAAUACAGAGCGUUCCUUAGUAAUGAUGGUUGGUAAGCGUAGAAGCUUGUUAGAUUACCUG
env-1165  AUCUUCGAAAAACACGGUAAAGGAAAAAACGAUACCGGUUCAUCAGAAGGUCAAAUCGCGUUAUUUACGCACAGAAUCAACCAUUUAACAGAGCACUUAAAGAAAAAUCGUAAAGAUUACAAUACAGAGCGUUCCUUAGUAAUGAUGGUUGGUAAGCGUAGAAGCUUGUUAGAUUACCUG
env-1167  AUCUUCGAAAAACACGGUAAAGGAAAAAACGAUACCGGUUCUACAGAAGGUCAGAUCGCGUUAUUUACGCACAGAAUUAACCAUUUAACAGAGCACUUAAAGAAAAAUCGAAAAGAUUACAACACAGAGCGUUCCUUAGUAAUGAUGGUUGGUAAGCGAAGAAGUUUGUUAGAUUACUUG
env-1168  AUCUUACGAAAACACGGUAACAAGUAAAACUAACACCGGUUCUGCCGAAGGACAAAUUGCGUUAUUUACGCACAGAAUUAACCAUUUAACAGGACACUUAAAGCAAAAUCGUAAAGAUUUUAACACAGAGCGUUCUUUAGUAAAAAUGGUUGGAAAGCGUAGAAGUCUUUUAGACUAU
env-1169  AUCUUACGAAAAACACGGUAACAAGUAAAACUAAUACCGGUUCUGCCGAAGGACAAAUUGCGUUAUUUACGCACAGAAUUAAUCAUUUAACAGGACACUUAAAGCAAAAUCGUAAAGAUUUUAAUACAGAGCGUUCUUUAGUAAAAAUGGUUGGAAAGCGUAGAAGUCUUUUAGACUAUCU
Psp-6-1   AUCUUCGAGAAACACGGUAAAGGAAAAAACGAUACUGGUUCUUCAGAAGGUCAAAUUGCUUUAUUCACACAGAAUUAACCAUUUAACUGGGCACUUAAAAAAUAAUCGUAAAGAUUAUAACACAGAGCGUUCAUUAGUAAUGUUAGUUGGUAAGCGUAGAAGUUUAUUAGAUUACUA
env-1170  AUUUUCGAAAAACACGGUAAGGGUAAAAACGAUACUGGUUCUUCAGAAGGUCAAAUUGCGUUAUUCACGCACAGAAUUAACCAUUUAACAGGACAUUUAAAAAAUAAUCGUAAAGAUUAUAAUACAGAGCGUUCAUUAGUAAUGAUGGUAGGUAAACGUAGAAGUUUACUUGACUAUUUA
env-1171  AUUUUCGAAAAACACGGUAAGGGUAAAAACGAUACUGGUUCUUCAGAAGGUCAAAUUGCGUUAUUCACGCACAGAAUUAACCAUUUAACAGGACAUUUAAAAAAUAAUCGUAAAGAUUAUAAUACAGAGCGUUCAUUAGUAAUGAUGGUAGGUAAACGUAGAAGUUUACUUGACUAUUUA
env-1172  UCGAGAACACAGUAAAGUAAAACAAACCGUUCUAAGAGGGUCAAUGGCGUUAUUCGCCGGAUCACCAUUUAACAGCAUUUAAAAGAAUCGUAAGAUAU.......................................................................
env-1173  CUUUUCGAGAAACACGGUAAAGGUAAAAAUGAUACUGGUUCGACAGAAGGGCAAGUGGCGUUAUUUACGCACAGAAUUAACCAUUUAACUGAGCAUUUAAAAAGAAUCGUAAAGAUUUUAAUACAGAACGCUCAUUAGUAAUGAUGGUAGGUAAACGUAGAAGUUUAUUGGAUUAUCUU

          ..........................................................................................................................................................................
          ..........................................................................................................................................................................
          AUYUUYR•RAARCACGG•RRR•••R•RR••RAYAC•GG••C•RC•GAAGG•CAAAUYGC•YURUUYAC•Y•Y•G•AUYRAYCAYYUR•C•RA•CAYYURAAAAR•AAYCR•AARGAUU•YAAYAC•GAR•G•UC•YURGU•••RYURGU•GG•AAR•G•RG••••YU•YU•GAYUAYYU•
```

```
env-1174  AUCUUCGAAAAACACGGUAACAGUAAAACUAACACCGGUUCUGCCGAAGGACAAAUUGCGUUAUUUACGCACAGAUUAACCAUUUAACAGGGACACUUAAAGCAAAAUCGUAAAGAUUUUAACACAGAGCGUUCUUUAGAAAAAUGGUUGAAAGCGUAGAAGUCUUUUAGACUAAUCUAA
Tov-1-1   UUAUUCGAAAAGCACGGUAAAGGUAAAGCUGAUACUGGUACUUCGGAAGGACAAAUUGCAUUGUUCACAUUUAGAAUCAACCACUUAACAGAGCACUUAAAAAAAGAAUCGUAAAGAUUUUAACACUGAGCGUUCAUUAGUAAAAAUGGUAGGUAAGCGUAGAAGUUUACUUGAUUACUUA
Pir-1-1   CUUUUUGAAAAACACGGAAAAGGUAAAAACGACACCGGUUCAUCAGAAGGUCAAAUCGCGUUAUUCACGCACAGAAUUAGCCAUUUAACUGAGCAUUUAAAGAAAAAUCGUAAAGAUUUUAACACAGAGCGCUCAUUAGUAAUGAUGGUAGGUAAACGUAGAAGUUUAUUAGAUUACCUA
env-1175  CUUUUUGAAAAACACGGUAAAGGUAAAAACGAUACUGGUUCUUCAGAAGGUCAAAUCGCGUUAUUCACGCACAGAAUUAGCCAUUUAACAGAGCAUUUGAAGAAAAAUCGUAAAGAUUUUAACACAGAACGUUCGUUAGUAAUGAUGGUAGGUAAACGUAGAAGUUUAUUAGAUUAUCUA
env-1176  CUUUUUGAAAACACGGUAAAGUAAAACGAUACUGGUUCUUCAGAACUCAAUCGCGUUAUCACGCACAGAAUAGCAUUUAACAGAGCAUUUGAAGAAAUCGUAAAGAUUUACACAGACGUCGUAGUAUGAUGGUAGUAACGUAGAAGUUAUUAGAUAUCUAAGAAACUGAAUCACGAUAUC
env-1177  CUUUUUGAAAAACACGGUAAAGGUAAAAACGAUACUGGUUCUUCAGAAGGUCAAAUCGCGUUAUUCACGCACAGAAUUAGCCAUUUAACAGAGCAUUUGAAGAAAAAUCGUAAAGAUUUUAACACAGAACGUUCGUUAGUAAUGAUGGUAGGUAAACGUAGAAGUUUAUUAGAUUAUCUA
env-1181  ....................................................................................................................................................................................
env-1182  CUUUUUGAAAAACACGGUAAAGGUAAAAACGAUACUGGUUCUUCAGAAGGUCAAAUCGCGUUAUUCACGCACAGAAUUAGCCAUUUAACAGAGCAUUUGAAGAAAAAUCGUAAAGAUUUUAACACAGAACGUUCGUUAGUAAUGAUGGUAGGUAAACGUAGAAGUUUAUUAGAUUAUCUA
env-1183  AUCUUCGAAAAACACGGUAAAGGAAAAAACGAUACUGGAACUUCAGAAGGACAAAUUGCGUUAUUUACGCACAGAAUUAACCAUUAACUGGGCACUUAAAGAAAUAAGAACUGACCGAUCUUGAUCCAGUAGGUCGGCAGCCGACNNNNGCUUUUUUUUCUCUUUUUGUCUUUUUCUUU
env-1184  AUCUUUGAAAAACACGGUAAAGGAAAAAACGAUACGGGUUCUUCGGAAGGUCAAAUUGCACUAUUCACUCACAGAAUUAAUCACUUAACUGAACAUCUAAAAAGAAAUCGUAAAGAUUUUAACACUGAGCGUUCAUUAGUAAAGUUAGUAGGUAAGCGUAAAAGCUUGUUAGAUUAUCUU
env-1185  AUCUUCGAAAAACACGGUAAAGGAAAAAACGAUACAGGUUCUUCAGAAGGUCAAAUCGCGUUAUUUACGCACAGAAUUAACCAUUUAACUGGGCAUUUAAAGAAGAAUCGUAAAGAUUUUAACACUGAACGCUCAUUAGUAAUGAUGGUAGGUAAACGUAGAAGUUUACUAGAUUAUUUA
env-1186  AUCUUCGAAAAACACGGUAAAGGAAAAAACGAUACAGGUUCUUCAGAAGGUCAAAUCGCGUUAUUUACGCACAGAAUUAACCAUUUAACUGGGCAUUUAAAGAAAAAUCGUAAAGAUUUUAACACUGAACGCUCAUUAGUAAUGAUGGUAGGUAAACGUAGAAGUUUACUAGAUUAUUUA
env-1187  AUCUUCGAAAAACACGGUAAAGGAAAAAGUGAUACCGGUUCUUCAGAAGGUCAAAUUGCAUUAUUCACUCACAGAAUCAACCAUUUAACUGAGCAUCUUAAAAAAAAAUCGUAAAGAUUUCAAUACGGAACGUUCACUAGUAAUGUUAGUUGGUAAGCGUAAAAGCUUAUUAGAUUAUCUU
env-1188  AUCUUCGAAAAACACGGUAAAGGAAAAAGUGAUACCGG..............................................................................................................................................
env-1190  AUCUUCGAAAAACACGGUAAAGGAAAAAGUGAUACCGGUUCUUCAGAAGGUCAAAUUGCAUUAUUCACUCACAGAAUCAACCAUUUAACUGAGCAUCUUAAAAAAAAUCGUAAAGAUUUUAAUACUGAACGUUCAUUAGUAAUGUUAGUAGGUAAGCGUAAAAGUUUAUUAGAUUAUCUUA
env-1191  AUCUUCGAAAAACACGGUAAAGGAAAAAACGAUACAGGUUCUUCAGAAGGUCAAAUCGCGUUAUUUACGCACAGAAUUAACCAUUUAACUGGGCAUUUAAAGAAGAAUCGUAAAGAUUUUAACACUGAACGCUCAUUAGUAAUGAUGGUAGGUAAACGUAGAAGUUUACUAGAUUAUUUA
env-1192  AUCUUCGAAAAACACGGUAAAGGAAAAAGUGAUACCGGUUCUUCAGAAGGUCAAAUUGCAUUAUUCACUCACAGAAUCAACCAUUUAACUGAGCAUCUUAAAAAAAAAUCGUAAAGAUUUCAAUACUGAACGUUCAUUAGUAAUGUUAGUAGGUAAGCGUAAAAGCUUAUUAGAUUAUCUU
env-1193  AUCUUCGAAAAACACGGUAAAGGAAAAAGUGAUACCGGUUCUUCAGAAGGUCAAAUUGCAUUAUUCACUCACAGAAUCAACCAUUUAACUGAGCAUCUUAAAAAAAAAUCGUAAAGAUUUCAAUACGGAACGUUCAUUAGUAAUGUUAGUAGGUAAGCGUAAAAGCUUAUUAGAUUAUCUU
env-1194  AUC.................................................................................................................................................................................
env-1196  AUCUUCGAAAAACACGGUAAAGGAAAAAGUGAUACCGGUUCUUCAGAAGGUCAAAUUGCAUUAUUCACUCACAGAAUCAACCAUUUAACUGAGCAUCUUAAAAAAAAAUCGUAAAGAUUUCAAUACUGAACGUUCAUUAGUAAUGUUAGUAGGUAAGCGUAAAAGCUUAUUAGAUUAUCUU
env-1197  AUCUUCGAAAAACACGGUAAAGGAAAAAACGAUACUGGUUCUUCAGAAGGUCAAAUUGCGUUAUUUACGCACAGAAUCAAUCAUUUAACUGGGCAUUUAAAAAAAGAAUCGUAAAGAUUUUAAUACGGAACGCUCAUUAGUAAUGAUGGUAGGUAAACGUAGAAGUUUAUUAGAUUAUUUA
env-1198  AUUUUCGAAAAACACGGUAAAGGAAAAAACGAUACAGGUUCUUCAGAAGGUCAAAUUGCAUUAUUUACACACAGAAUUAACCACUUAACUGGGCAUUUAAAGAAAAAUCGUAAAGAUUUUAAUACAGAGCGUUCAUUAGUAAUGUUGGUUGGUAAGCGUAAAAAUUUAUUAGAUUAUCUU
env-1199  ....................................................................................................................................................................................
env-1200  AUCUUCGAGAAACACGGUAAAGGAAAAAACGAUACUGGUUCUUCAGAAGGUCAAAUUGCGUUAUUUACGCACAGAAUCAAUCAUUUAACUGGGCAUUUAAAAAAAGAAUCGUAAAGAUUUUAAUACGGAACGCUCAUUAGUAAUGAUGGUAGGUAAACGUAGAAGUUUAUUAGAUUAUUUA
env-1203  ....................................................................................................................................................................................
env-1204  AUCUUCGAGAAACACGGUAAAGGAAAAAACGAUACUGGUUCUUCAGAAGGUCAAAUUGCGUUAUUUACGCACAGAAUCAAUCAUUUAACUGGGCAUUUAAAAAAAGAAUCGUAAAGAUUUUAAUACGGAACGCUCAUUAGUAAUGAUGGUAGGUAAACGUAGAAGUUUAUUAGAUUAUUUA
env-1205  AUCUUCGAAAAACACGGUAAAGGAAAAAACGAUACGGGUUCUUCAGAAGGGCAAAUUGCGUUGUUUACGCACAGAAUUAACCAUUUAACUGAACAUUUAAAAAGAAAUCGUAAAGAUUUUAAUACAGAGCGUUCGUUAGUAAUGAUGGUAGGUAAACGUAGAAGUUUAUUAGAUUACUUA
Psp-3-1   AUCUUCGAAAAACACGGUAAAGGAAAAAACGAUACUGGAACUUCAGAAGGUCAAAUUGCGUUAUUUACGCACAGAAUUAACCAUUUAACUGGGCACUUAAAAAAGAAUCGUAAAGAUUAUAAUACAGAGCGUUCUUUAGUAAUGAUGGUAGGUAAGCGUAGAAGUUUAUUAGAUUAUUUA
env-1208  AUCUUUUCAAAACACGGUAAGUCAGCAACAGACACUGGUAACUCAGAAGGACAAAUUGCUUUGUUCACAUUUAGAAUCAACCAUUUAACAGAAC......................................................................................
env-1209  AUCUUUUCAAAACACGGUAAGUCAGCAACUGACACUGGAACAUCAGAAGGGCAAAUUGCUUUGUUCACAUUUAGAAUCAACCAUUUAACUCAACACUUAAAAAACAAUCGUAAAGAUUAUAAUACAGAGCGUUCUUUAGUAAAAUUGGUAGGUAAGCGUAGAAGCUUAUUAGAUUAUUUA
env-1210  ....................................................................................................................................................................................
env-1211  AUCUUUUCAAAACACGGUAAGUCAGCAACUGACACUGGAACAUCAGAAGGGCAAAUUGCUUUGUUCACAUUUAGAAUCAACCAUUUAACUCAACACUUAAAAAACAAUCGUAAAGAUUAUAAUACAGAGCGUUCUUUAGUAAAAUUGGUAGGUAAGCGUAGAAGCUUAUUAGAUUAUUUA
env-1212  AUCUUCGAAAAACACGGUAAAGGAAAGAAUGAUACUGGUUCUUCAGAAGGUCAAAUUGCGUUAUUUACGCACAGAAUUAACCAUUUAACAGAGCAUUUAAAACAAAAUCGUAAAGAUUUUAACACAGAGCGUUCGUUAGUAAGAAUGGUAGGUAAACGUAGAAGUUUAUUAGAUUAUUUA
env-1213  AUUUUCACAAAACACGGUAAGUCAGUUACUGACACUGGAACAUCAGAAGGACAAAUUGCAUUGUUCACAUUUAGAAUCAAC...................................................................................................
Psp-2-1   AUCUUCGAAAAACACGGUAAAGGAAAAAACGAUACUGGUUCUUCAGAAGGUCAAAUUGCAUUAUUUACACACAGAAUUAACCACUUAACGGAGCACUUAAAGAGAAAUCGUAAAGAUUUUAACACUGAGCGUUCGUUAGUAAUGUUAGUAGGUAAGCGUAAAAGUUUAUUAGAUUAUCUU
Psp-4-1   AUCUUCGAAAAACACGGUAAAGGAAAAAACGAUACUGGUUCUUCAGAAGGUCAAAUUGCGUUAUUCACGCACAGAAUUAACCACUUAACUGAGCACUUAAAAAACAAUCGUAAAGAUUUUAAUACAGAGCGUUCUUUAGUAAUGAUGGUUGGUAAGCGUAGAAGUUUAUUAGAUUAUCUA
env-1215  UCUUCGAAAAACACGGUAACAGUAAAACUAACACCGGUUCUGCCGAAGGACAAAUUGCGUUAUUUACGCACAGAAUUAACCAUUUAACAGGACACUUAAAGCAAAAUCGUAAAGAUUUUAACACAGAGCGUUCUUUAGUAAAAAANGGUUGGAAAGCGUAGAAGUCUUUUAGACUAUCUA
env-1216  AUCUUCGAAAAACACGGUAACAGUAAAACUAACACCGGUUCUGCCGAAGGACAAAUUGCGUUAUUUACGCACAGAAUUAACCAUUUAACAGGACACUUAAAACAAAAUCGUAAAGAUUUUAACACAGAACGUUCUUUAGUAAAAAUGGUUGGAAAGCGUAGAAGUCUUUUAGACUAUCUA
env-1217  AUCUUCGAAAAACACGGUAACAGUAAAACUAACACCGGUUCUGCCGAAGGACAAAUUGCGUUAUUUACGCACAGAAUUAACCAUUUAACAGGACACUUAAAGCAAAAUCGUAAAGAUUUUAACACAGAGCGUUCUUUAG..........................................
env-1218  AUCUUCGAAAAACACGGUAACAGUAAAACUAACACCGGUUCUGCCGAAGGACAAAUUGCGUUAU....................................................................................................................
env-1219  AUCUUCGAAAAACACGGUAACAGUAAAACUAACACCGGUUCUGCCGAAGGACAAAUUGCGUUAUUUACGCACAGAAUUAACCAUUUAACAGGACACUUAAAGCAAAAUCGUAAAGAUUUUAACACAGAGCGUUCUUUAGUAAAAAUGGUUGGAAAGCGUAGAAGUCUUUUAGACUAUCUA
env-1220  AUCUUCGAAAAACACGGUAACAGUAAAACUAAUACCGGUUCUGCCGAAGGACAAAUUGCGUUAUUUACGCACAGAAUUAAUCAUUUAACAGGACACUUAAAGCAAAAUCGUAAAGAUUUUAACACAGAGCGUUCUUUAGUAAAAAUGGUUGGAAAGCGUAGAAGUCUUUUAGACUAUCUA
env-1222  AUCUUCGAAAAACACGGUAACAGUAAAACUAAUACCGGUUCUGCCGAAGGACAAAUUGCGUUAUUUACGCACAGAAUUAAUCAUUUAACAGGACACUUAAAGCAAAAUCGUAAAGAUUUUAAUACAGAGCGUUCUUUAGUAAAAAUGGUUGGAAAG.........................
env-1223  AUCUUCGAAAAACACGGUAACAGUAAAACUAACACCGGUUCUGCCGAAGGACAAAUUGCGUUAUUCACGCACAGAAUUAACCAUUUAACAGGACACUUAAAACAAAAUCGUAAAGAUUUUAACACAGAACGUUCUUUAGUAAAAAUGGUUGGAAAGCGUAGAAGUCUUUUAGACUAUCUA
env-1226  AUCUUCGAAAAACACGGUAACAGUAAAACUAACACCGGUUCUGCCGAAGGACAAAUUGCGUUAUUUACGCACAGAAUUAACCAUUUAACAGGACACUUAAAGCAAAAUCGUAAAGAUUUUAACACAGAGCGUUCUCUAGUAAAAAUGGUUGGAAAGCGUAGAAGUCUUUUAGACUAUCUA
env-1227  AUCUUCGAAAAACACGGUAACAGUAAAACUAACACCGGUUCUGCCGAAGGACAAAUUGCGUUAUUUACGCACAGAAUUAACCAUUUAACAGGACACUUAAAGCAAAAUCGUAAAGAUUUUAACACAGAGCGUUCUCUAGUAAAAAUGGUUGGAAAGCGUAGAAGUCUUUUAGACUAUCUA
env-1229  AUCUUCGAAAAACACGGUAACAGUAAAACUAACACCGGUUCUGCCGAAGGACAAAUUGCGUUAUUCACGCACAGAAUUAACCAUUUAACAGGACACUUAAAACAAAAUCGUAAAGAUUUUAACACAGAACGUUCUUUAGUAAAAAUGGUUGGAAAGCGUAGAAGUCUUUUAGACUAUCUA
env-1230  AUCUUCGAAAAACACGGUAACAGUAAAACUAACACCGGUUCUGACCGAAGGACAAAUUGCGUUAUUUACGCACAGAAUUAACCAUUUAACAGGACACUUAAAGCAAAAUCGUAAAAGAUUUUAACACAGAGCGUUCUUUAAGUAAAAAUGG..............................
env-1231  AAUACUUACGAAAAACC...................................................................................................................................................................
env-1232  AUCUUCGAAAAACACGGUAACAGUAAAACUAACACCGGUUCUGCCGAAG...................................................................................................................................
env-1233  AUCUUCGAAAAACACGGUAACAGUAAAACU......................................................................................................................................................
env-1234  AUCUUCGAAAAAACACGGUAACAGUAAAACUAACACCGGUUCUGCCGAAGGACAAAUUGCGUUAUUUACGCACAGAAUUAACCAUUUAACAGGACACUUAAAAGCAAAAUCGUAAAGAUUUU............................................................
env-1235  AUCUUCGAAAAACACGGUAACAGUAAAACUAACACCGGUUCUGCCGAAGGACAAAUUGCGUUAUUUACGCACAGAAUUAACCAUUUAACAGGACACUUAAAGCAAAAUCGUAAAGAUUUUAACACAGAGCGUUCUUUAGUAAAAAUGGUUGGAAAGCGUAGAAGUCUUUUUAGACUAUCU
env-1236  AUCUUCGAAAAACACGGUAACAGUAAAACUAACACCGGUUCUGCCGAAGGACAAAUUGCGUUAUUUACGCACAGAAUUAACCAUUUAACAGGACACUUAAAACAAAAUCGUAAAGAUUUUAACACAGAACGUUCUUUA...........................................
env-1237  AUCUUCGAAAAACACGGUAANAGUAAAACUAACACCGGUUCUGCCGAAGGACAAAUUGCGUUAUUUACGCACAGAAUUAACCAUUUAACAGGNCACUUAAAGCAAAAUCGUAAAGAUUUUAACACAGAGCGUUCUNUAGUAAAAAUGGUUGGAAAGCGUAGAAGUCUUUUAGACUAUCUA
env-1238  AUCUUCGAAAAACACGGUAACAGUAAAACUAACACCGGUUCUGCCGAAGGACAAAUUGCGUUAUUUACGCACAGAAUUAACCAUUUAACAGGACACUUAAAGCAAAAUCGUAAAGAUUUUAACACAGAGCGUUCUUUAGUAAAAAUGGUUGGAAAGCGUAGAAGUCUUUUAGACUAUCUA
env-1239  ....................................................................................................................................................................................
env-1240  AUCUUCGAAAAACACGGUAACAGUAAAACUAACACCGGUUCUGCCGAAGGACAAAUUGCGUUAUUUACGCACAGAAUUAACCAUUUAACAGGACACUUAAAGCAAAAUCGUAAAGAUUUUAACACAGAGCGUUCUUUAGUAAAAAUGGUUGGAAAGCGUAGAAGUCUUUUAGACUAUCUA
env-1241  CUUCGAAAAACACGGUAACAGUAAAACUAACACCGGUUCUGCCGAAGGACAAAUUGCGUUAUUUACGCACAGAAUUAACCAUUUAACAGGACACUUAAAGCAAAAUCGUAAAGAUUUUAACACAGAGCGUUCUUUAGUAAAAAUGGUUGGAAAGCCGUAGAAGUCUUUUAGUACUAUCUA
env-1242  AUCUUCGAAAAACACGG...................................................................................................................................................................
env-1243  AUCUUCGAAAAACACGGUAA................................................................................................................................................................
env-1245  AUCUUCGAAAAACACGGUAAUAGUAAAACUAACACCGGUUCUGCCGAAGGACAAAUUGCGUUAUUUACGCACAGAAUUAAUCAUUUAACGGGGCACUUAAAGCAAAAUCGUAAAGAUUUUAAUACAGAGCGUUCUCUAGUAAAAAUGGUUGGAAAGCGUAGAAGUCUUUUAG.......
env-1246  AUCUUCGAAAAACACGGUAAUAGUAAAACUAACACCGG..............................................................................................................................................
env-1247  AUCUUCGAAAAACACGGUAAAGGAAAAAAUGAUACCGGUACAUCAG......................................................................................................................................
env-1248  AUCUUCGAAAAAC.......................................................................................................................................................................
env-1250  AUUUUUGAAAAACACGGUAACAGUAAAACUGAUACAGGUUCUUCAGAAGGACAAAUUGCGUUAUUCUCGCACAGAAUUAACCACCUAACAAGUCACUUAAAAGAAAAUCGUAAAGAUUUUAAUACUGAGCGUUCUUUAGUAAAGUUGGUAGGAAAGCGUAGAAGUCUUUUAGACUACUUA
env-1252  AUCUUCGAAAAACACGGUAAUAGUAAAACUAACACCGGUUCUGCCGAAGGACAAAUUGCGUUAUUUACGCACAGAAUUAAUCAUUUAACGGGGCACUUAAAGCAAAAUCGUAAAGAUUUUAAUACAGAGCGUUCUCUAGUAAAAAUGGUUGGAAAGCGUAGAAGUCUUUUAGACUAUCUA
env-1253  AUCUUCGAAAAACACGGUAAUAGUAAAACUAACACCGGUUCUGCCGAAGGACAAAUUGCGUUAUUUACGCACAGAAUUAAUCAUUUAACGGGGCACUUAAAGCAAAAUCGUAAAG..................................................................
env-1254  ....................................................................................................................................................................................
env-1255  ....................................................................................................................................................................................
env-1256  AUCUUCGAAAAACACGG...................................................................................................................................................................
env-1257  AUCUUCGAAAAACACGGUAA................................................................................................................................................................
env-1259  AUCUUCGAAAAACACGGUAAUAGUAAAACUAAUACUGGUUCAGCAGAAGGACAAAUUGCAUUAUUCACGCACAGAAUUAACCAUUUAACAGGGCACUUAAAACAAAAUCGUAAAGAUUUUAACACAGAGCGUUCCUUAGUGAAAAUGGUUGGAAAGCGAAGAAGUCUUUUAGACUACUUA
env-1260  AUCUUCGAAAAACACGGUAAUAGUAAAACUAACACCGGUUCUGCCGAAGGACAAAUUGCGUUAUUUACGCACAGAAUUAACCAUUUAACAGGGCACUUAAAGCAAAAUCGUAAAGAUUUUAACACAGAGCGUUCUCUAGUAAAAAUGGUUGGAAAGCGUAGAAGUCUUUUAGACUAUUUA
env-1262  AUCUUCGAAAAACACGGUAAUAGUAAAACUAACACCGGUUCUGCCGAAGGACAAAUUGCGUUAUUUACGCACAGAAUUAACCAUUUAACAGGACACUUAAAGCAAAAUCGUAAAGAUUUUAACACAGAGCGUUCUUUAGUAAAAAUGGUUGGAAAGCGUAGAAGUCUUUUAGACUAUCUA
env-1263  AUCUUCGAAAAACACGGUAANAGUAAAACUAACACCGGUUCUGCCGAAGGACAAAUUGCGUUAUUUACGCACAGAAUUAACCAUUUAACAGGNCACUUAAAGCAAAAUCGUAAAGAUUUUAACACAGAGCGUUCUNUAGUAAAAAUGGUUGGAAAGCGUAGAAGUCUUUUAGACUAUNUA
env-1264  AUCUUCGAAAAACACGGUAAUAGUAAGACUAAUACUGGUUCAGCAGAAGGACAGAUUGCAUUAUUCACGCACAGAAUUAACCAUUUAACAGGGCACUUAAAACAAAAUCGUAAAGAUUUUAAUACAGAGCGUUCUUUAGUGAAAAUGGUUGGAAAGCGAAGAAGUCUUUUAGACUACUUA
env-1268  AUCUUCGAAAAACACGGUAAUAGUAAGACUAAUACUGGUUCAGCAGAAGGACAGAUUGCAUUAUUCACGCACAGAAUUAACCAUUUAACAGGGCACUUAAAACAAAAUCGUAAAGAUUUUAAUACA.......................................................
env-1269  AUCUUCGAAAAACACGG...................................................................................................................................................................
env-1270  AUCUUCGAAAAACACGGUAAUAGUAAGACUAAUACUGGUUCAGCAGAAGGACAGAUUGCAUUAUUCACGCACAGAAUUAACCAUUUAACAGGGCACUUAAAACAAAAUCGUA.....................................................................
env-1271  AUCUUCGAAAAACACGGUAAUAGUAAAACUAAUACUGGUUCAGCAGAAGGACAAAUUGCAUUAUUCACGCACAGAAUUAACCAUUUAACAGGGCACUUAAAACAAAAUCGUAAAGAUUUUAACACAGAGCGUUCCUUAGUGAAAAUGGUUGGAAAGCGAAGAAGUCUUUUAGACUACUUA
env-1272  AUCUUCGAAAAACACGGUAAUAGUAAGACUAAUACUGGUUCAGCAGAAGGACAGAUUGCAUUAUUCACGCACAGAAUUAACCAUUUAACAGGGCACUUAAAACAAAAUCGUAAAGAUUUUAAUACAGAGCGUUCUUUAGUGAAAAUGGUUGGAAAGCGAAGAAGUCUUUUAGACUACUUA
env-1273  AUCUUCGAAAAACACGGUAAUAGUAAGACUAAUACUGGUUCAGCAGAAGGACAGAUUGCAUUAUUCACGCACAGAAUUAACCAUUUAACAGGGCACUUAAAACAAAAUCGUAAAGAUUUUAAUACAGAGCGUUCUUUAGUGAAAAUGGUUGGAAAGCGAAGAAGUCUUUUAGACUACUUA
env-1274  AUC.................................................................................................................................................................................

          ....................................................................................................................................................................................
          ....................................................................................................................................................................................
          AUYUUYR•RAARCACGG•RRR•••R•RR••RAYAC•GG••C•RC•GAAGG•CAAAUYGC•YURUUYAC•Y•Y•G•AUYRAYCAYYUR•C•RA•CAYYURAAAAR•AAYCR•AARGAUU•YAAYAC•GAR•G•UC•YURGU•••RYURGU•GG•AAR•G•RG••••YU•YU•GAYUAYYU•
```

```
env-1275   AUCUUCGAAAAACACGGUAA..........................................
env-1277   AUCUUCGAAAAACACGGUAAUAGUAAGACUAAUACUGGUUCAGCAGAAGGACAGAUUGCAUUAUUCACGCACAGAAUUAACCAUUUAACAGGGCACUUAAAACAAAAUCGUAAAGAUUUUUAAUACAGAGCGUUCUUUAGUGAAAAUGGUUGGAAAGCGAAGAAGUCUUUUUA.......
env-1278   AUCUUCGAAAAACACGGUAGUAGCAAAACUAAUACUGGUUCAGCAGAAGGACAAAUUGCAUUAUUCACGCAUAGAAUUAACCAUUUAACAGGGCAUUUAAAACAAAAUCGUAAGGAUUUUAAUACAGAGCGUUCAUUAGUUAAAAUGGUUGGAAAGCGAAGAAGUCUUUUAGACUACUUA
Ccy-1-1    AUUUUUAAAAAACACGGUGGUUCAGAAACCAACACAGGUUCAGCAGAAGGGCAAAUUGCCUUAUUCACUUUCAGAAUUAAUCACUUAACAGAACACUUGAAAAGAAAUAGAAAAGAUUUCAACACAGAGCGUUCAUUGGUGAAAUUAGUAGGUAAACGUAGAAGUUUACUUGACUAUCUG
Ccy-1-3    AUUUUUAAAAAACACGGUGGUUCAGAAACCAACACAGGUUCAGCAGAAGGGCAAAUUGCCUUAUUCACUUUCAGAAUUAAUCACUUAACAGAACACUUGAAAAGAAAUAGAAAAGAUUUCAAUACGGAGCGUUCAUUAGUGAAAUUAGUAGGUAAACGUAGAAGUUUACUUGACUAUCUG
Tme-1-1    AUCUUCGCAAAACACGGUAAAGGAGCAAAUGAUACUGGUACUGCAGAAGGUCAAAUCGCGUUAUUUACGCACAGAAUUAACCACUUAACUGAGCAUUUAAAGAAAAAUCGUAAAGAUUUUAACACAGAACGCUCGUUAGUACGUAUGGUAGGUAAGCGUAGAAGCUUAUUAGAUUAUCUA
Tsp-1-1    AUCUUCGCAAAACACGGUAAAGGAGCAAAUGAUACUGGUACUGCAGAAGGACAAAUCGCGUUAUUUACGCACAGAAUUAACCACUUAACUGAGCACUUAAAAAGAAAUCGUAAAGAUUUUAACACUGAGCGUUCAUUAGUUAAAAUGGUAGGUAAGCGUAGAAGUUUGUUAGAUUAUCUA
env-1280   CUUUUUGAGAAACAC...............................................
env-1281   CUUUUUGAGAAACACGGUAAAGGUAAAAACGAUACUGGUUCUUCAGAAGGGCAAAUUGCGUUAUUUACGCACAGAAUUAACCAUUUAACUGGACACUUAAAAAAUAAUCGUAAAGAUUAUAAUACAGAGCGUUCAUUAGUUAUGUUGGUAGGUAAACGUAGAAGUUUACUAGACUAUUUA
env-1282   CUUUUUGAGAAACACGGUAAAGGUAAAAACGAUACUGGUUCUUCAGAAGGACAAAUUGCGUUAUUUACGCACAGAAUUAACCAUUUAACUGGACACUUAAAAAAUAAUCGUAAAGAUUAUAAUACAGAGCGUUCAUUAGUUAUGUUGGUAGGUAAACGUAGAAGUUUACUAGACUAUUUA
env-1283   CUUUUUGAGAAACACGGUAAAGGUAAAAACGAACUCCAACUUAC....................
env-1284   CUUUUUGAGAAACACGGUAAAGGUAAAAACGAUAC.............................
env-1285   CUUUUUGAGAAACACGGUAAAGGUAAAAACGAUACUGGUUCUUCAGAAGGACAAAUUGCGUUAUUUACGCACAGAAUUAACCAUUUAACUGGACACUUGAAAAAUAAUCGUAAAGAUUAUAAUACAGAGCGUUCAUUAGUUAUGUUGGUAGGUAAACGUAGAAGUUUACUAGACUAUUUA
env-1287   CUUUUUGAGAAACACGGUAAAGGUAAAAACGAUACUGGUUCUUCAGAAGGACAAAUUGCGUUAUUUACGCACAGAAUUAACCAUUUAACUGGACACUUAAAAAAUAAUCGUAAAGAUUAUAAUACAGAGCGUUCAUUAGUUAUGUUGGUAGGUAAACGUAGAAGUUUACUAGACUAUUUA
env-1293   CUUUUUGAGAAACACGGUAAAGGUAAAAACGAUACUGGUUCUUCAGAAGGACAAAUUGCGUUAUUUACGCACAGAAUUAACCAUUUAACUGGACACUUAAAAAAUAAUCGUAAAGAUUAUAAUACAGAGCGUUCAUUAGUUAUGUUGGUAGGUAAACGUAGAAGUUUACUAGACUAUUUA
env-1294   CUUUUUGAGAAAAACGGUAAAGGUAAAACGAUACUGGUUCUUCAGAAGACAAAUUGC.......
env-1295   CUUUUUGAGAAACACGGUAAAGGUAAAAACGAUACUGGUUCUUCAGAAGGACAAAUUGCGUUAUUUACGCACAGAAUUAACCAUUUAACUGGACACUUAAAAAAUAAUCGUAAAGAUUAUAAUACAGAGCGUUCAUUAGUUAUGUUGGUAGGUAAACGUAGAAGUUUACUAGACUAUUUA
env-1296   CUUUUUGAGAAACACGGUAAAGGUAAAAACGAUACUGGUUCUUCAGAAGGACAAAUUGCAUUAUUUACCCACAGAAUUAACCAUUUAACUGGACACUUGAAAAAUAAUCGUAAAGAUUAUAAUACAGAGCGUUCAUUAGUUAUGUUGGUAGGUAAACGUAGAAGUUUACUAGACUAUUUA
env-1298   CUUUUUGAGAAACACGGUAAAGGUAAAAACGAUACUGGUUCUUCAGAAGGACAAAUUGCGUUAUUUACCCACAGAAUUAACCAUUUAACUGGACACUUGAAAAAUAAUCGUAAAGAUUAUAAUACAGAGCGUUCAUUAGUUAUGUUGGUAGGUAAACGUAGAAGUUUACUAGACUAUUUA
env-1299   CUUUUUGAGAAACACGGUAAAGUAAAACGAUACUGGUUCUUCAGAAGACAAAUUGCGUUAUUUACGCACAGAAUUAACCAUUUAACUGGACACUUAAAAAAAUCGUAAAGAUUUUAAUACAGAGCGUCA..........
env-1300   AUCUUCGAAAAACACGGUAAUAGUAAAACUAACACCGGUUCAGCAGAAGGACAAAUUGCAUUAUUCACGCACAGAAUUAACCACUUAACAGGACACUUAAAACAAAAUCGUAAAGAUUUUAAUACAGAGCGUUCUCUAGUAAAGAUGGUUGGAAAGCGAAGAAGUCUUUUAGACUACUUA
env-1301   AUCUUCGAAAAACACGGUAGUAGUAAAACUAAUACGGGUUCAGCAGAAGGACAAAUUGCAUUAUUCACGCACAGAAUUAACCACUUAACAGGGCACUUGAAACAAAAUCGUAAAGAUUUUAAUACAGAGCGUUCUUUAGUGAAAAUGGUUGGAAAGCGAAGAAGUCUUUUAGACUACUU
env-1302   AUCUUCGAAAAACACGGUAGUAGUAAAACUAAUACGGGUUCAGCAGAAGGACAAAUUGCAUUAUUCACGCACAGAAUUAACCACUUAACAGGGCACUUGAAACAAAAUCGUAAAGAUUUUAAUACAGAGCGUUCUUUAGUGAAAAUGGUUGGAAAGCGAAGAAGUCUUUUAGACUACUUA
env-1303   CAUCUUCGAAAAACACGGUAAAGGAAAGAACGAUACUGGGUCUUCAGAAGGUCAAAUUGCACUGUUUACACACAGAAUUAACCACCUAACUGAGCACUUAAGAAAAAAUCGUAAAGAUUUUAACACUGAGCGUUCACUUAGUAAUGUUGUAGGU..........
env-1304   AUUUUUGAAAAGCACGGUAAAGGUAAGACCGAUACUGGAUCUGCUGAGGGACAAAUCGCGUUAUUUACGCACAGAAUUAACCAUUUAACAAGUCACUUGAAAAACAAUCGUAAAGAUUAUAAUACAGAGCGUUCGUUAGUUAAAAUGGUAGGGAAGCGUAGAAGCCUUUUAGAUUACCUU
env-1305   AUCUUUAAGAAACACGGCGGAAAAGCAGAGAACACCGGUUCUACUGAAGGUCAAAUCGCAUUGUUUACACAUCGUAUCAAUCACCUGACCGGACAUCUUAAAAAAAAUCAUAAAGACUACAAUACUGAACGUUCGUUAGUAAAACUUGUAGGUAAGCGAAGAAGUCUUUUAGAUUAUAUG
env-1306   AUCUUUAAGAAACACGGCGGAAAAGCAGAGAACACCGGUUCUACUGAAGGUCAAAUCGCAUUGUUUACACAUCGUAUCAAUCACCUGACCGGACAUCUUAAAAAAAAUCAUAAAGACUACAAUACUGAACGUUCGUUAGUAAAACUUGUAGGUAAGCGAAGAAGUCUUUUAGAUUAUAUG
env-1307   AUCUUUAAGAAACACGGCGGAAAAGCAGAGAACACCGGUUC.......................
env-1308   AUUUUCACCAAACA..................................................
env-1309   ................................................................
env-1310   AUCUUUAAGAAACACGGCGGAAAAGCAGAGAACACAGGUUCUACUGAAGGGCAGAUUGCCUUAUUCACACACCGCAUCAAUCAUUUGACCGGUCACCUGAAAAAUAAUCACAAAGAUUUCAACACCGAACGUUCAUUGGUAAAGUUGGUAGGUAAAAGAAGAAGCUUACUUGAUUACAUG
env-1311   AUCUUUAAGAAACACGGCGGAAAAGCAGAGAACACAGGUUCUACUGAAGGGCAGAUUGCCUUAUUCACACACCGCAUCAAUCAUUUGACCGGUCACCUGAAAAAUAAUCACAAAGAUUUCAACACCGAACGUUCAUUGGUAAAGUUGGUAGGUAAAAGAAGAAGCUUACUUGAUUACAUG
Tna-1-1    AUCUUCGCAAAACACGGUAAAGGAAAAAACGAUACUGGUAGUGCAGAAGGACAAAUUGCGUUAUUUACGCACAGAAUUAACCAUUUAACUGAACACUUAAAACAAAAUCGUAAAGAUUUUAAUACCGAGCGUUCGUUAGUAAAAUUAGUAGGUAAGCGUCGUGCAUUACUAGAUUAUUUA
env-1312   ................................................................
Cly-1-1    AUCUUUAAGAAAUACGGUAAAAACGAAAACGAUUCUGGUACUGCAGAAGCACAAAUUGCUUUGUUUACACACAGAAUUAACCACUUAACAGGUCACUUAAGAAAUAAUCAUAAAGAUUAUAACACAGAGCGUUCUCUAGUUAAAUUGGUAGGUAAAAGAAGAAGUCUUUUAGAUUACUUA
Cge-1-1    AUCUUUAAGAAAUACGGUAAAAACGAAAACGAUUCUGGUACUGCAGAAGCACAAAUUGCUUUGUUCACACACAGAAUUAACCACUUAACAGGUCACUUAAGAAAUAAUCAUAAAGAUUAUAACACAGAGCGUUCUCUAGUUAAAUUGGUAGGUAAAAGAAGAAGUCUUUUAGAUUACUUA
Cly-2-1    AUCUUUAAGAAAUACGGUAAAAACGAAAACGAUUCUGGUACUGCAGAAGCACAAAUUGCUUUGUUCACACACAGAAUUAACCACUUAACAGGUCACUUAAGAAAUAAUCAUAAAGAUUAUAACACAGAGCGUUCUCUAGUUAAAUUGGUAGGUAAAAGAAGAAGUCUUUUAGAUUACUUA
Sxi-1-1    CUUUUUGCAAAACACGGUAAGGGUAAGAACGACACAGGAUCUGCCGAAGGUCAAAUCGCUUUAUUUACUCACCGUAUCGAACACUUGUCUCAGCACCUAAAGAAAAACCGUAAAGAUUAUAAUACCGAGCGGUCACUUGUAAUGCUGGUAGGUAAGAGAAGAAGCUUACUGGAUUAUUUG
Ste-1-1    CUUUUUGCAAAACACGGUAAAGGAAAGAACGACACAGGAUCUGCCGAAGGGCAAAUCGCCUUGUUCACUCACCGUAUUGAACAUUUGACUCAGCACUUGCAGGAUUUUAAUCGUAAAGAUUACAAUACAGAGCGUUCACUGGUAAAAUUAGUAGGUAAGAGAAGAAGCCUGCUGGAUUAUUUA
Cal-1-1    CUUUUCAAAAAACACGGUGGCGAAGAAAAAAACACUGGAUCUGCUGAAGGGCAAAUUGCAUUGUUUACGCACCGUAUUGAGCACUUAAGUCAACACUUAAGAACUAACCGUAAAGAUUACAACACGGAACGUUCUCUAGUAAUGUUAGUAGGUAAAAGAAGAAGCUUAUUAGAUUAUUUA
env-1313   AUUUUUGCAAAACACGGUAAAGGAAAGAACGACACAGGAUCUGCUGAAGGACAAAUCGCAUUAUUCACACACAGAAUUUCGCACCUUACAGAGCACUUGAAAAAGAAUCGCCACGAUUACAACACAGAGCGUUCACUCGUUUUGUUAGUAGGUAAAAGAAGGUCGCUACUUGACUACCUC
Mze-2-1    AUUUUCAAGAAACACGGUGGUUCUGAAGGAAACACAGGCUCUACUGAAGGACAAAUUGCGUUGUUCACUUACAGAAUUAACCAUUUAACACAGCACUUAAAAAGAAAUCAUAAAGAUUUCAACACAGAGCGUUCUUUGGUAAAAUUGGUAGGUAAAAGACGUAGUCUUUUAGAUUAUCUU
env-1314   AUCUUCGCUAAACACGGAGGAAAAGCUGAAAACACAGGUUCUGCUGAGGGACAAAUUGCAUUGUUCACUUUCAGAAUUUCACACUUAACAGAGCACUUGAAAAAAAAAUCGUCAUGAUUACAAUACCGAGCGUUCGUUAGUAUUGCUAGUAGGUAAAAGAAGAAGUCUUUUAGAUUACUUA
env-1316   AUCUUCGCUAAACACGGAGGAAAAGCUGAAAACACAGGUUCUGCUGAGGGACAAAUUGCAUUGUUCACUUUCAGAAUUUCACACUUAACAGAGCACUUGAAAAAAAAAUCGUCAUGAUUACAAUACCGAGCGUUCGUUAGUAUUGCUAGUAGGUAAAAGAAGAAGUCUUUUAGAUUACUUA
env-1317   AUCUUCGCUAAACACGGAGGAAACGCAACAAACACAGGUUCUGGAAAAUCAGAAAGCCAGAUCGCUUUGUUCACAUUUAGAAUCAAUCACUUAACAGAACACUUGAAAGAAAAAUCGUCACGAUUACAAUACCGAGCGUUCUUGUGAAACUAGUAGGUAAAAGAAGAUCUUUGUGGAUUACUUG
env-1318   AUCUUCGCUCAACACGGAGGAAAAGCUGAAAACACUGGUUCUGUUGAAGGACAAGUAGCAUUGUUCACAUUUAGAAUCUCUCACUUAACUGAGCACUUGAAAAAAAAAUCGUCACGAUUAUAACACUGAGCGUUCGUUAGUUCUUUUAGUAGGUAAAAGAAGAAGUCUUCUUGACUACUUG
Fgi-1-1    AUCUUCGCUAAACACGGAGGAAAAGCAGAAAACACUGGAUCUGCAGAAGGACAAAUCGCGUUAUUUACUCACAGAAUCACGCACCUUACUGAGCACUUGAAAAAAAAAUCGUCACGAUUACAACACAGAGCGUUCAUUGGUACUAUUAGUAGGUAAAAGAAGAUCUUUGUGGAUUACUUG
Fsu-2-1    AUUUUCGCUAAACACGGAGGAAAAGCAGAAAACACUGGAUCUGCAGAAGGACAAAUCGCAUUGUUCACUCACAGAAUUAGCCACCUUACUGAGCACUUGAAAAAAAAAUCGUCACGAUUAUAAUACUGAGCGUUCUUUGGUACUAUUAGUAGGUAAAAGAAGAGCUUUGUUGGAUUACUUG
env-1319   AUCUUCGCUAAACACGGAGAAAAAACAAACACUGGAAAGUCUGAAGCGCAAAUUGCUUUAUUCACAUACAGAAUUACUCACCUUACUGAGCACUUGAAAAAAAAUCGUCAUGAUUACAACACUGAGCGUUCAUUAGUAUUGUUAGUAGGUAAAAGAAGAUCAUUGUGGAUUACUUGAAA
env-1320   AUCUUCGCUAAACACGGAGGAAAAGCAGAAAACACAGGAUCUGCAGAAGGACAAAUCGCACUAUUCACACACCGAAUUACGCACCUUACUGAGCACUUGAAAAAA..........
env-1321   AUCUUCGCUAAACACGGAGGAAAAGCAGAAAACACAGGAUCUGCAGAAGGACAAAUCGCACUAUUCACACACCGAAUUACGCACCUUACUGAGCACUUGAAAAAAAAAUCGUCACGAUUAUAACACAGAGCGUUCAUUGGUACUACUAGUAGGUAAAAGAAGAUCGUUGUGGAUUACUUG
Fjo-1-1    AUCUUCGCACAACACGGUGAUGCAAAAAACACUGGUAAAGCUGAAGGUCAAAUCGCAUUGUUCACUUACAGAAUUUCUCACUUAACUGAACACUUGAAAAAAAAAUCGUCACGAUUACAACACUGAGCGUUCUUGUACUAUUAGUAGGUAAAAGAAGAGCUUUGUGGACUACUUGAAA
Fsp-7-1    AUCUUCGCACAACACGGUGGUGCAACAAACACAGGAAGUGCAGAAGGUCAAAUCGCGUUGUUCACUUACAGAAUUUCUCACUUAACUGAACACUUGAAAAAAAAAUCGUCACGAUUACAACACUGAGCGUUCUCUUGUACUAUUAGUAGGUAAAAGAAGAGCUUUGUGGACUACUUGAAA
Fsp-13-1   AUUUUCGCACAACACGGUGGUGCAACAAACACUGGAAGCGCAGAAGGUCAAAUUGCAUUGUUCACUUACAGAAUUUCACACUUAACUGAACACUUGAAAAAAAAAUCGUCACGAUUACAACACUGAGCGUUCACUUGUACUAUUAGUAGGUAAGAGAAGAGCUUUGUUGGACUACUUGAAA
Fsp-6-1    AUUUUCGCUCAACACGGUGGAGCAACAAACACUGGAAGCGCAGAAGGUCAGAUUGCAUUGUUCACUUACAGAAUUUCACACUUAACUGAGCACUUGAAAAAAAAAUCGUCACGAUUACAACACUGAGCGUUCACUGGUACUAUUAGUAGGUAAAAGAAGAGCUUUGUUGGAUUACUUGAAA
Fsp-8-1    AUUUUUGCACAACACGGUGAUGUUACAAACACUGGAAAAGCUGAAGGUCAAAUCGCAUUGUUCACUUACAGAAUUUCACACUUAACUGAACACUUGAAAAAAAAAUCGUCACGAUUAUAACACUGAGCGUUCACUUGUACUUCUAGUAGGUAAAAGAAGAGCUUUGUUAGAUUACUUGAAG
Fsp-10-1   AUUUUUGCACAACACGGUGAUGUUACAAACACUGGAAAAGCUGAAGGUCAAAUCGCAUUGUUCACUUACAGAAUUUCACACUUAACUGAACACUUGAAAAAAAAAUCGUCACGAUUAUAACACUGAGCGUUCACUUGUACUUUUAGUAGGUAAAAGAAGAGCUUUGUUAGAUUACUUGAAG
Fhy-1-1    AUUUUCGCACAACACGGUGGUGCAACAAACACUGGAAGCGCAGAAGGUCAGAUUGCAUUGUUCACGUUCAGAAUUUCACACUUAACUGAGCACUUGAAAAAAAAAUCGUCAUGAUUACAACACUGAGCGUUCACUUGUACUAUUAGUAGGUAAAAGAAGAGCUUUGUUGGAUUACUUGAAA
env-1323   AUUUUCGCUCAACACGGUGGAGCAACAAACACUGGAAGCGCAGAAGGUCAGAUUGCAUUGUUCACUUACAGAAUUUCACACUUAACUGAGCACUUGAAAAAAAAAUCGUCACGAUUACAACACUGAGCGUUCACUAGUACUAUUA..........
env-1324   AUUUUCGCACAACACGGUGGUGCAACAAACACUGGAAGCGCAGAAGGUCAAAUUGCAUUGUUCACUUACAGAAUUUCUCACUUAACUGAACACUUGAAAAAAAAAUCGUCACGAUUACAACACUGAGCGUUCUCUUGUACUAUUAGUAGGUAAAAGAAGAGCUUUGUUAGAUUACUUGAAA
Fsp-11-1   AUUUUCGCACAACACGGUGGUGCAACAAACACUGGAAGCGCAGAAGGUCAGAUUGCAUUGUUCACUUACAGAAUUUCACACUUAACUGAGCACUUGAAAAAAAAAUCGUCACGAUUACAACACAGAGCGUUCACUUGUACUAUUAGUAGGUAAAAGAAGAUCUUUGUUGGAUUACUUGAAG
Fch-1-1    AUUUUCGCACAACACGGUGGUGCAACAAACACAGGUAGUGCAGAAGGUCAAAUUGCAUUGUUCACUUACAGAAUUUCACACUUAACUGAACACUUGAAAAAAAAAUCGUCACGAUUACAACACUGAGCGUUCACUUGUACUAUUAGUAGGUAAAAGAAGAGCUUUGUUAGAUUACUUGAAA
env-1325   AUCUUCGCUAAACACGGAGGAAAAGCAGAAAACACAGGAUCUGCAGAGGGCAAAUCGCACUAUUCACUCACCGAAUCACGCACCUUACAGAGCAUUUGAAAAAAAAAUCGUCACGAUUAUAACACAGAGCGUUCAUUAGUACUGUUAGUAGGUAAAAGAAGAUCUUUGUGGAUUACUUG
env-1326   AUCUUCGCUAAACACGGAGGUAAAGCAGAAAACACAGGAUCUGCAGAAGGACAAAU..........
env-1327   AUCUUCGCUAAACACGGAGGAAAAGCAGAAAACACUGGAUCUGCAGAAGGGCAAAUCGCACUAUUCACUCACCGAAUCACGCACCUUACAGAGCAUUU..........
env-1328   AUCUUCGCUAAACACGGAGGAAAAGCAGAAAACACUGGAUCUGCAGAAGGGCAAAUCGCACUAUUCACUCACCGAAUCACGCACCUUACAGAGCAUUUGAAAAAAAAAUCGUCACGAUUAUAACACAGAGCGUUCAUUAGUACUGUUAGUAGGUAAAAGAAGAUCUUUGUGGAUUACUUG
env-1330   AUCUUCGCUAAACACGGAGAAAAAACGAACACAGGAAAACCAGAAGCGCAAAUUGCAUUGUUCACUUUCAGAAUUUCGCAUUUGACAGAACACUUAAAAAGAAAUCGUCACGAUUACAAUACCGAGCGUUCACUAGUUUUGCUAGUAGGUAAAAGAAGGGCUUUGUUGGAUUACUUGAAG
env-1331   AUCUUCGCUAAACACGGAGAAAAAACNACACAGGAAAACCAGAAGCGCAAAUUGCAUUGUUCACUUUCAGAAUUUCGCAUUUGACAGAACACUUAAAAAGAAAUCGUCACGAUUACAAUACCGAGCGUUCACUAGUUUUGCUAGUAGGUAAAAGAAGGGCUUUGUUGGAUUACUUGAAG
env-1332   UUCUUCAAAAAAUUUGGAGGUUCAGAAUCAAACACCGGUAGCACCGAAGGACAAAUUGCUUUGUUUACACAUCGCAUCAACCACCUAACAGGUCACUUGAAAAAAAAUCAAAAAGAUUUUAAUACUGAGAGAGCCUUAAUCAAAUUGGUAGGUAAACGCAGAAGUUUGUUAGAUUAUUUA
env-1333   UUCUUCAAAAAAUUUGGAGGUUCUGAAACAAACACCGGUAGCACCGAAGGACAAAUUGCUUUGUUUACACAUCGCAUCAAUCACUUGACAGGUCACUUGAAAAAAAAUCAAAAAGAUUUUAACACAGAGAGAGCCUUGAUCAAAUUGGUAGGUAAACGCAGAAGUUUGUUAGAUUAUUUA
Gsp-4-1    CUAUUCGCUAAACACGGUAAAGGUAAAAAUGAUACCGGUUCUGCUGAAGGGCAAAUCGCAUUAUUUACUCACCGUAUUUCCCACCUAUCAGAUCACCUGAGAACUAAUCGUAAAGAUUAUAACACAGAAAGAUCUCUAGUAUUACUAGUAGGUAAAAGAAGAAGACUCCUUGAUUACUUA
Gli-1-1    CUUUUCGCAAAACAUGGAAAAGGUACGAAUGAUACCGGAUCUGCAGAAGGCCAAAUUGCCCUUUUCACCCAACGUAUCUCUCAUCUUUCAGGUCACUUGAAAACCAACAGAAAAGACUAUAAUACAGAACGUUCAUUGGUAAUGCUGGUAGGUAAAAGAAGAAGUUUACUCGACUAUUUA
env-1334   CUUUUCGCAAAACAUGGAAAAGGUACGAAUGAUACCGGAUCUGC....................
env-1335   CUUUUCGCAAAACACGGUAAAGGUAAAAAUGACACCGGUAGUGCAGAAGGUCAAAUCGCGUUGUUCACGCAGCGUAUCUCACACCUUUCAGAUCACCUUAGAAAAAAUCGCAAGGAUUUCAACACCGAGCGUUCAUUGGUAAUGCUGGUAGGGAAAAGAAGAAGCUU............
env-1336   CUGUUUGCAAAACACGGAAAAGGUAAAAAUGAUACCGGUUCUGCUGAAGGUCAAAUCGCAUUAUUUACCCAACGUAUUUCACACCUUUCGGAUCAUUUAAAAACGAACCGAAAAGACUACAAUACAGAGCGUUCAUUAGUAUUAUUGGUAGGUAAAAGAAGAAGUCUUCUUGAUUAUUUG
env-1337   CUGUUUGCAAAACACGG...............................................
env-1338   ................................................................
env-1339   CUUUUUAAAAAACAUGGAAAUAGUGCGAAAAACACUGGAUCCGCAGAAGGUCAAAUCGCAUUGUUCACACAACGUAUUUCACACCUUUCUGGACAUUUAAAAACAAAUCGUAAAGAUUANAACACUGAGCGAUCUCUUGUAAGAUUGGUAGGAAAAGAAGAAAUCUACUUGAUUAUUUA
env-1340   ................................................................
env-1341   CUUUUUAAAAAACAUGGAAAUAGUGCUACAAACACUGGAUCCGCAGAAGGUCAAAUCGCAUUGUUUACACAACGUAUUUCACACCUUUCUGGACAUUUAA..........
Gsp-5-1    CUUUUUAAAAAACACGGAAAUAGUGCCAAAAACACUGGAUCCGCAGAAGGUCAAAUCGCAUUGUUCACACAACGUAUUUCACACCUUUCUGGACAUUUAAAAACAAAUCGUAAAGAUUAUAAUACUGAGCGAUCUCUUGUAAGAUUAGUAGGUAAAAGAAGAAAUCUACUUGAUUAUUUA
env-1342   ................................................................
env-1343   ................................................................
env-1344   C...............................................................

           ................................................................
           ................................................................
           AUYUUYR•RAARCACGG•RRR•••R•RR••RAYAC•GG••C•RC•GAAGG•CAAAUYGC•YURUUYAC•Y•Y•G•AUYRAYCAYYUR•C•RA•CAYYURAAAAR•AAYCR•AARGAUU•YAAYAC•GAR•G•UC•YURGU•••RYURGU•GG•AAR•G•RG••••YU•YU•GAYUAYYU•
```

| | |
|---|---|
| env-1345 | ........................................................................................................................................................................ |
| env-1346 | ........................................................................................................................................................................ |
| Gsp-3-1 | CUUUUUAAAAAACACGGAAAAAGUACUAAAGAUACUGGAUCCGCAGAAGGUCAAAUCGCAUUGUUCACACAACGUAUUUCACACCUUUCUGGUCAUUUAAAAACAAAUCGUAAAGAUUAUAAUACUGAGCGAUCUCUUGUAAGAUUGGUAGGGAAAAGAAGAAAUCUACUUGAUUAUUUA |
| Gsp-2-1 | CUUUUUAAAAAACAUGGAAAUAGUGCUACAAACACUGGAUCCGCAGAAGGUCAAAUCGCAUUGUUUACACAACGUAUUUCACACCUUUCUGGACAUUUAAAAACAAAUCGUAAAGAUUAUAACACUGAGCGAUCUCUUGUAAGAUUGGUAGGGAAAAGAAGAAAUCUACUUGAUUAUUUA |
| env-1347 | ........................................................................................................................................................................ |
| env-1348 | CUGUUUGCAAAACACGG........................................................................................................................................................ |
| env-1349 | CUGUUUGCAAAACACGGAAAAGGUAAAAAUGAUACCGGUUCUGCUGAAGGUCAAAUCGCAUUAUUUACCCAACGUAUUUCACACCUUUCGGAUCAUUUAAAAACGAACCGCAAAGAUUACAAUACAGAGCGUUCACUAGUAUUAUUGGUAGGUAAAAGAAGAAGUCUUCUUGAUUAUUUG |
| env-1350 | CUGUUUGCAAAACACGGAAAAGGUAAAAAUGAUACCGGUUCUGCUGAAGGUCAAAUCGCAUUAUUUACCCAACGUAUUUCACACCUUUCGGAUCAUUUAAAAACGAACCGUAAAGAUUACAAUACAGAGCGUUCACUAGUAUUAUUGGUAGGUAAAAGAAGAAGUCUUCUUGAUUAUUUG |
| Gec-1-1 | AUCUUCGAAAAGCACGGUAAAAGCAAAACUGAUACCGGAUCAGCUGAAGGACAGAUCGCAUUAUUUACUUACAGAAUCGCUCACCUUUCUGAGCACCUGAAGAAAAAUCGUAAGGAUUUCAAUUCAGAACGUUCUCUAGUAAUGCUAGUAGGAAAGAGAAGAAGUCUUCUUGAUUACUUA |
| Gpo-1-1 | AUCUUCGAAAAACACGGGAAAUCGAAAACCGAUACCGGUUCAGCAGAAGGACAAAUCGCAUUAUUUACUUACAGAAUUGCUCACCUUUCUGAGCACCUGAAGAAAAACCGUAAGGAUUUUAACUCAGAGCGUUCACUGGUAAAGCUUGUUGGUAAAAGAAGAAGCUUACUUGAUUACUUA |
| env-1351 | AUCUUCGAAAAACAUGGUAAAUCAAAAAAUGAUACCGGUUCAGCAGAAGGACAAAUCGCAUUAUUUACUUACAGAAUUGCUCACCUAUCUGAGCACCUGAAGAAAAACCGUAAGGAUUUUAAUUCAGAGCGUUCGCUGGUAAAGCUUGUUGGUAAAAGAAGAAGCUUACUUGAUUACUUG |
| env-1352 | AUUUUCUCAAAGCACGGUAAAGGUAAAAAUGAUACCGGUUCUGCUGAAGGACAGAUCGCAUUGUUUACCCACCGAAUUGCCCACUUAUCAGAGCACCUGAAAACAAAUCGUAAAGAUUUUAACACAGAACGUUCUUUAGUAAGAUUAGUUGGUAAAAGAAGAAGUCUGUUAGACUACUUG |
| env-1353 | AUUUUCUCAAAGCACGGUAAAGGUAAAAAUGAUACCGGUUCUGCUGAAGGACAGAUCGCAUUGUUUACCCACCGAAUUGCCCACUUAUCAGAGCACCUGAAAACAAAUCGUAAAGAUUUUAACAC.......................................... |
| Gfo-1-1 | AUCUUCUCAAAGCAUGGUAAAGGUAAAAACGAUACCGGUUCUGCUGAAGGACAGAUUGCAUUGUUUACUCACCGAAUUGCCCACUUAUCAGAACACCUGAAAACAAAUCGUAAAGAUUAUAAUACAGAACGCUCUCUGGUAAUGUUGGUUGGUAAAAGAAGAAGUCUACUAGACUAUUUG |
| env-1354 | AUCUUCGAAAAGUACGGAAACGGUAAAAACGAUUCUGGUUCAGCUGAAGGACA.......................................................................................................................... |
| env-1355 | AUCUUCGAAAAGUACGGUGAAGGAAAAUCCAACACUGGUUCCGCUGAAGGACAAAUUGCAUUAUUCACUUACAGAAUCUCUCACUUAUCUGAUCACUUAAAAAGUAAUCGUAAAGAUUUUAAUACAGAGCGUUCUCUAGUAAGAUUGGUAGGUAAAAGAAGAAGCCUUCUUGAUUACUUA |
| env-1356 | AUCUUCGAAAAGUACGGAAACGGUAAAAACGAUUCUGGUUCAGCUGAAGGACAAAUUGCAUUAUUCACUUACAGAAUUUCUCAUUUGUCUGAGCAUUUAAAAAGCAAUCGUAAAGACUUCAACACAGAGCGUUCUCUUGUUGGUCUUGUAGGAAAAAGAAGAAGACUUCUUGAUUACUUA |
| env-1357 | AUCUUCGAAAAGUACGGUGAAGGAAAAUCCAACACUGGUUCCGCUGAAGGACAAAUUGCAUUAUUCACUUACAGAAUCUCUCACUUAUCUGAUCACUUAAAAAGUAAUCGUAAAGAUUUUAACACAGAGCGUUCUCUAGUAAGAUUAGUAGGUAAAAGAAGAAGCCUUCUUGAUUACUUA |
| env-1358 | AUCUUCGAAAAGUACGGUGAAGGAAAAUCCAACACUGGUUCCGCUGAAGGACAAAUUGCAUUAUUCACUUACAGAAUCUCUCACUUAUCUGAUCACUUAAAAAGUAAUCGUAAAGAUUUUAAUACAGAGCGUUCUCUAGUAAGAUUGGUAGGUAAAAGAAGAAGCCUUCUUGAUUACUUA |
| env-1359 | AUCUUCGAAAAGUACGGUGAAGGAAAAUCCAACACUGGUUCCGCUGAAGGACAAAUUGCAUUAUUCACUUACAGAAUCUCUCACUUAUCUGAUCACUUAAAAAGUAAUCGUAAAGAUUUUAACACAGAGCGUUCUCUAGUAAGAUUAGUAGGUAAAAGAAGAAGCCUUCUUGAUUACUUA |
| env-1360 | AUCUUCGAAAAGUACGGUGAAGGAAAAUCCAACACUGGUUCCGCUGAAGGACAAAUUGCAUUAUUCACUUACAGAAUCUCUCACUUAUCUGAUCACUUAAAAAGUAAUCGUAAAGAUUUUAAUACAGAGCGUUCUCUAGUAAGAUUGGUAGGUAAAAGAAGAAGCCUUCUUGAUUACUUA |
| env-1367 | AUCUUCGAAAAGUACGGUGAAGGAAAAUCCAACACUGGUUCCGCUGAAGGACAAAUUGCAUUAUUCACUUACAGAAUCUCUCACUUAUCUGAUCACUUAAAAAGUAAUCGUAAAGAUUUUAAUACAGAGCGUUCUCUAGUAAGAUUGGUAGGUAAAAGAAGAAGCCUUCUUGAUUACUUA |
| env-1370 | AUCUUCGAAAAGUACGGUGAAGGAAAAUCCAACACUGGUUCCGCUGAAGGACAAAUUGCAUUAUUCACUUACAGAAUCUCUCACUUAUCUGAUCACUUAAAAAGUAAUCGUAAAGAUUUUAAUACAGAGCGUUCUCUAGUAAGAUUGGUAGGUAAAAGAAGAAGCCUUCUUGAUUACUUA |
| env-1371 | AUCUUCGAAAAGUACGGUGAAGGAAAAUCCAACACUGGUUCCGCUGAAGGACAAAUUGCAUUAUUCACUUACAGAAUCUCUCACUUAUCUGAUCACUUAAAAAGUAAUCGUAAAGAUUUUAACACAGAGCGUUCUCUAGUAAGAUUAGUAGGUAAAAGAAGAAGCCUUCUUGAUUACUUA |
| env-1373 | AUCUUCGAAAAGUACGGUGAAGGAAAAUCCAACACUGGUUCCGCUGAAGGACAAAUUGCAUUAUUCACUUACAGAAUCUCUCACUUAUCUGAUCACUUAAAAAGUAAUCGUAAAGAUUUUAAUACAGAGCGUUCUCUAGUAAGAUUGGUAGGUAAAAGAAGAAGCCUUCUUGAUUACUUA |
| env-1374 | AUCUUCGAAAAGUACGGUGAAGGAAAAUCCAACACUGGUUCCGCUGAAGGACAAAUUGCAUUAUUCACUUACAGAAUCUCUCACUUAUCUGAUCACUUAAAAAGUAAUCGUAAAGAUUUUAAUACAGAGCGUUCUCUAGUAAGAUUGGUAGGUAAAAGAAGAAGCCUUCUUGAUUACUUA |
| env-1375 | AUCUUCGAAAAGUACGGUGAAGGAAAAUCCAACACUGGUUCCGCUGAAGGACAAAUUGCAUUAUUCACUUACAGAAUCUCUCACUUAUCUGAUCACUUAAAAAGUAAUCGUAAAGAUUUUAAUACAGAGCGUUCUCUAGUAAGAUUGGUAGGUAAAAGAAGAAGCCUUCUUGAUUACUUA |
| env-1376 | AUCUUCGAAAAGUACGGUGAAGGAAAAUCCAACACUGGUUCCGCUGAAGGACAAAUUGCAUUAUUCACUUACAGAAUCUCUCACUUAUCUGAUCACUUAAAAAGUAAUCGUAAAGAUUUUAAUACAGAGCGUUCUCUAGUAAGAUUGGUAGGUAAAAGAAGAAGCCUUCUUGAUUACUUA |
| env-1377 | AUCUUCGAAAAGUACGGUGAAGGAAAAUCCAACACUGGUUCCGCUGAAGGACAAAUUGCAUUAUUCACUUACAGAAUCUCUCACUUAUCUGAUCACUUAAAAAGUAAUCGUAAAGAUUUUAAUACAGAGCGUUCUCUAGUAAGAUUGGUAGGUAAAAGAAGAAGCCUUCUUGAUUACUUA |
| env-1378 | AUUUUUGAAAAACAUGGUAAGAGUAAAACCGACACCGGUUCUACCGAAGGACAAAUCGCCCUUUUCACUUACCGAAUAGCUCACCUUUCAGACCAUCUAAAAACUAACCGUAAGGAUUACAAUUCCGAGCGUUCUUUGGUAAUGCUGGUUGGUAAAAGAAGGAGUUUACUCGAUUAUUUA |
| env-1379 | AUUUUUGAGAAGCACGGAAAAAGCAAAACCGACACUGGUUCUGCUGAAGGACAAAUCGCUUUGUUCACUUACCGAAUCUCUCACCUUUCUGACCACUUAAAAAAGAAUCGUAAAGACUACAACUCAGAGCGUUCUUUGGUAAGACUGGUUGGUAAAAGAAGAAGCUUACUUGAUUACUUA |
| env-1380 | AUUUUUGAGAAGCACGGAAAAAGCAAAACCGACACUGGUUCUGCUGAAGGACAAAUCGCUUUGUUCACUUACCGAAUCUCUCAUCUUUCUGACCACUUAAAAAAGAAUCGUAAAGACUACAACUCAGAGCGUUCUUUGGUAAGACUGGUUGGUAAAAGAAGAAGCUUACUUGAUUACUUA |
| env-1381 | AUUUUUGAGAAACACGGAAAAAGCAAAACCGACACUGGUUCUGCUGAAGGACAAAUCGCUUUGUUCACUUACCGAAUCUCUCACCUUUCUGACCACUUAAAAAAGAAUCGUAAAGACUACAACUCAGAGCGUUCUUUGGUAAGACUGGUUGGUAAAAGAAGAAGCUUACUUGAUUACUUA |
| env-1382 | AUUUUUGAGAAGCACGGAAAAAGCAAAACCGACACUGGUUCUGCUGAAGGACAAAUUGCUUUGUUCACUUACCGAAUCUCUCACCUUUCUGACCACUUAAAAAAGAAUCGUAAAGACUACAACUCAGAGCGUUCUUUGGUAAGACUGGUUGGUAAAAGAAGAAGCUUGCUUGAUUACUUA |
| env-1383 | AUUUUUGAGAAGCACGGAAAAAGCAAAACCGACACUGGUUCUGCUGAAGGACAAAUUGCUUUGUUCACUUACCGAAUCUCUCACCUUUCUGG......................................................................................... |
| Ssp-1-1 | AUUUUUGAGAAGCACGGAAAAAGUAAAACCGACACCGGUUCUGCUGAAGGACAAAUUGCUUUGUUCACUUACCGAAUCUCUCACCUUUCAGAUCACUUAAAAAAGAAUCGUAAAGACUACAACUCAGAGCGUUCUUUGGUAAUGUUGGUUGGUAAAAGAAGGAGCUUGCUUGAUUACUUA |
| Tse-1-1 | AUCUUCGUAAAACACGGUAAAGGGAAAAACGAUACGGGUUCUGCAGAAGGUCAAAUCGCGUUAUUUACGCACAGAAUUAAUCACUUAACAGAACACUUAAAGAAGAAUCGUAAAGAUUAUAACACCGAGCGUUCUUUAGUAAAAUUAGUAGGUAAGCGUCGCGCUUUACUAGAUUAUUUA |
| Lsp-2-1 | CUUUUUGCUAAACACGGUAAAGCUAAAAACGAUACUGGUUCUGCAGAAGGACAAAUUGCAUUAUUCACUCACAGAAUUAAGCACUUAACAGAACACUUAAAAACCAAUCGUAAAGAUUAUAAUACAGAGCGUUCUUUAGUCAUGUUAGUAGGUAAAAGAAGAAGCCUACUAGAUUACUUA |
| Jpa-1-1 | AUUUUCUCUAAGUACGGUAAAGGAAAAAACGAUACUGGUACAGCAGAAGCACAAAUUGCGUUAUUUACGCACAGAAUUAACCAUUUAACCCAACACUUAAAGAACAAUCAAAAAGAUUAUAAUACAGAGCGUUCUUUGGUAAAACUGGUAGGUAAACGUCGUGCUUUAUUAGAUUACUUA |
| Ssp-2-1 | AUCUUUAAAAAACAUGGUAAAGACGCAAAAGAUACUGGCACUGCCGAAGGACAAAUUGCUCUUUUCACUUACAGAAUCAACCAUUUAACAGGACACUUAAAAAACAAUCGUAAAGAUUACAAUACAGAGCGUUCGUUAGUAAAAUUAGUUGGUAAGCGUAGGUCUUUAUUAGACUACCUC |
| env-1384 | AUCUUUAAGAAACAUGGCAAAAGUGCAAAAGACACUGGUUCUGCAGAAGGACAGAUUGCGUUGUUCACUUACAGAAUUAAUUACUUAACAGGGCACUUAAAAAACAAUCGUAAAGAUUAUAAUACUGAGCGUUCUUUAGUAAAAUUAGUAGGUAAAAGAAGAAGUUUACUAGAUUACUUA |
| env-1385 | AUCUUUAAAAAACACGGUAAAAACGUUAAGGACACAGGAACAUCUGAAGGUCAAAUUGCGUUAUUUACUUACAGAAUUAAUCACUUAACUGAACAUUUAAAAAGCAAUCGUAAAGAUUUUAACACAGAACGUUCUUUAGUAAAACUAGUAGGAAAGCGUCGCUCACUAUUAGAUUACUUA |
| env-1386 | AUCUUUAAAAAACACGGUAAAAACGUUAAGGACACAGGAACAUCU.................................................................................................................................... |
| env-1387 | AUCUUUAAAAAGCAUGGCAAAAACGUUAAGGACACAGGUUCUGCAGAAGGACAAAUUGCUUUGUUCACUUACAG.......................................................................................................... |
| env-1388 | AUCUUUAAAAAGCAUGGCAAA............................................................................................................................................................ |
| env-1389 | AUCUUUAAAAAGCAUGGCAAAAACGUUAAGGACACAGGUUCUGCAGAAGGACAAAUUGCUUUGUUCACUUACAGAAUUAACUAUUUAACUGGUCACUUAAAAAAUAAUCGUAAAGAUUAUAAUACAGAGCGUUCAUUAGUAAAACUAGUAGGAAAGCGUCGCUCAUUACUAGAUUACUUA |
| env-1391 | AUCUUUAAAAAGCAUGGUAAAAACGCAAAGGACACAGGUUCUGCAGAAGGGCAAAUUGCUUUGUUCACUUACAGAAUUAAUUACU........................................................................................... |
| env-1392 | ........................................................................................................................................................................ |
| env-1393 | AUCUUCACAAAACACGGUAAAGGAAAAAAUGAUACCGGUUCUGCUGAAGGACAAAUUGCGUUAUUUACGUACAGAAUUAACCAUUUAACUGAGCACUUAAAAAACAAUCGUAAAGAUUUUAACUCUGAGCGUUCUCUAGUAAAAUUAGUAGGUAAGCGUAGAGCUUUAUUAGACUACUUA |
| env-1394 | ........................................................................................................................................................................ |
| env-1395 | AUCUUCACAAAACACGGUAAAGGAAAAAAUGAUACCGGUUCUGCUGAAGGACAAAUUGCGUUAUUUACGUACAGAAUUAACCAUUUAACUGAGCACUUAAAAAACAAUCGUAAAGAUUUUAACUCUGAGCGUUCUCUAGUAAAAUUAGUAGGUAAGCGUAGAGCUUUAUUAGACUACUUA |
| env-1396 | CUGUUUAAAAAACACGGUAAAAAUGGAAAAGACACUGGUUCUGCAGAAGGGCAAAUUGCUUUAUUCACACACAGAAUCAAUCAUUUAACCCAGCAUUUAAAGAACAAUCAAAAAGAUUACAAUACUGAACGUUCUUUGGUAAAAUUAGUAGGUAAGCGUAGAUCGCUUUUAGAUUACCUU |
| env-1397 | CUGUUUAAAAAACACGGCAAAAAUGGAAAAGACACUGGUUCUGCAGAAGGGCAAAUUGCUUUAUUCACACACAGAAUCAAUCAUUUAACCCAGCAUUUAAAGAACAAUCAAAAAGAUUACAAUACUGAACGUUCUUUGGUAAAAUUAGUAGGUAAGCGUAGAUCGCUUUUAGAUUACCUU |
| env-1398 | ........................................................................................................................................................................ |
| env-1399 | AUCUUCGCAAAACACGGUAAAGGAAAAAAUGAUACUGGUUCUGCUGAAGGACAAAUUGCGUUAUUUACGUACAGAAUUAAUCACUUAACUGAGCACUUAAAGAACAAUCGUAAAGAUUUUAACUCUGAGCGUUCUCUAGUAAAGUUAGUAGGUAAGCGUAGAGCUUUAUUAGACUACUUA |
| env-1400 | ........................................................................................................................................................................ |
| env-1401 | AUCUUCGCAAAACACGGUAAAGGAAAAAAUGAUACUGGUUCUGCUGAAGGACAAAUUGCGUUAUUUACGUACAGAAUUAAUCACUUAACUGAGCACUUAAAGAACAAUCGUAAAGAUUUUAACUCUGAGCGUUCUCUAGUAAAGUUAGUAGGUAAGCGUAGAGCUUUAUUAGACUACUUA |
| env-1402 | AUCUUCGCAAAACACGGUAAAGGAAAAAAUGAUACUGGUUCUGCUGAAGGACAAAUUGCGUUAUUUACGUACAGAAUUAAUCACUUAACUGAGCACUUAAAGAACAAUCGUAAAGAUUUUAACUCUGAGCGUUCUCUAGUAAAGUUAGUAGGUAAGCGUAGAGCUU.............. |
| env-1403 | AUCUUCACAAAGUACGGUAGCGGAAAGAAUGAUACCGGUUCUGCUGAAGGACAAAUUGCGUUAUUUACGCACAGAAUUAACCAUUUAACAGAGCAUUUAAAGAAUAAUCGUAAAGAUUUUAAUUCCGAGCGUUCUUUGGUAAAGUUGGUAGGUAAG......................... |
| env-1404 | AUCUUUAAAAAACACGGUAAAGAUGCAAAAAACACUGGUUCUGCAGAAGGACAGAUUGCAUUAUUUACAGCAAGAAUUGAUCACUUAACUGAACACUUAAAAAAUAAUCGUAAAGAUUAUAAUACAGAGCGUUCAUUAGUAAUGUUAGUAGGAAAACGUCGUUCGUUAUUAGACUACUUA |
| env-1405 | AUCUUUAAGAAACACGGUAAAGAUGCAAAAGACACGGGUUCUGCCGAAGGACAAAUUGCUUUGUUCACAUACAGAAUUAAUUAUUUAACGGAUCACCUUAAACAAAAUCGUAAAGAUUUUAAUACGGAACGUUCAUUAGUGAAAUUAGUAGGUAAGCGUCGUUCUUUACUAGACUAUUUA |
| env-1406 | AUCUUUAAGAAACACGGUAAAGAUGCAAAAGACACGGGUUCUGCCGAAGGACAAAUUGCUUUGUUCACAUACAGAAUUAAUUAUUUAACGGAUCACCUUAAACAAAAUCGUAAAGAUUUUAAUACGGAACGUUCAUUAGUGAAAUUAGUAGGUAAGCGUCGUUCUUUACUAGACUAUUUA |
| env-1411 | AUCUUUAAGAAACACGGUAAAGAUGCAAAAGACACGGGUUCUGCCGAAGGACAAAUUGCUUUGUUCACCUACAGAAUUAAUUACUUAACGAAUCACCUUAAACAAAAUCGCAAAGA.................................................................... |
| Wsp-1-1 | CUUUUCAAAAAACACGGUAAAAGCGCAAAAGACACUGGUUCUGCAGAAGGACAAAUCGCGUUAUUUACAGCAAGAAUUAAUCACUUAACUGAACACUUAAAAACAAAAUCGUAAAGAUUUUAAUACUGAGCGUUCGUUAGUAAAGUUAGUUGGUAAGCGUCGUGCUUUACUAGACUACUUA |
| env-1412 | CUCUUUAAAAAGCAUGGUAAAAAUGCAAAGAACACAGGUUCUGCUGAAGGUCAAAUCGCAUUAUUUACUAACAGAAUUAAUCACUUAACAGAACACUUAAAGAAUAAUCGUAAAGAUUAUAAUACAGAGCGUUCGUUAGUAAAACUAGUUGGUAAGCGUCGUGCUUUACUAGACUAUUUA |
| env-1413 | CUCUUUAAAAAGCAUGGUAAAAAU......................................................................................................................................................... |
| env-1414 | AUCUUUAAAAAGCAUGGUAAAACUGCAAAAAACACUGGCUCUACAGAAGGACAGAUUGCAUUAUUUACACACAGAAUUGAGCACUUAACUGAGCACUUAAAAAACAAUCGUAAAGAUUAUAAUACAGAGCGUUCGUUAGUAAAAUUAGUAGGUAAAAGAAGAAGCUUACUAGAUUACUUA |
| env-1415 | AUCUUUAAAAAGCAUGGUAAAACUGCAAAAAACACUGGCUCUACAGAAGGACAGAUUGCAUUAUUUACACACAGAAUUGAGCACUUAACUGAGCACUUAAAAAACAAUCGUAAAGAUUAUAAU.................................................. |
| env-1416 | AUCUUUAAAAAGCAUGGUAAAAAUGCAAAGAACACAGGUUCAGCCGAAGGUCAAAUUGCAUUAUUUACUCACAGAAUUAAUCACUUAACAGAGCACUUAAAAAAAUAAUCG........................................................... |
| Bar-1-1 | CUUUUUGCUAAACACGGUAGCGGAAAAGCCGAUACUGGAAGUGCAGAAGGACAAAUUGCAUUAUUUACGCAACGUAUCGAACACUUAACUGGACACUUAAAAAAUAAUCGUAAAGAUUAUAAUACAGAGCGUUCAUUAGUAAUGUUAGUUGGUAAGCGUCGCGCUUUACUUGAUUACUUA |
| env-1417 | CUUUUCGCAAAACACGGUAGCGGAAAAAACGAUACUGGAAGCGCAGAAGGACAAAUAGCAAUAUUUACACAACGCAUCGAGCAUUUAACACAACACUUAAAAACGAAUCGUAAAGAUUUCAAUACGGAACGUUCAUUAGUGAUGUUAGUUGGUAAGCGUCGUUCGCUACUUGAUUACUUA |
| env-1418 | CUUUUCGCAAAACACGGUAGCGGAAAAAACGAUACUGGAAGUGCAGAAGGACAAAUAGCGUUAUUUACACAACGUAUUAAUCACUUAACAGAACAC................................................................................ |
| env-1419 | AUCUUUAAAAAACACGGUAAAGAUGCAAAAGACACAGGUUCUGCAGAAGGGCAAAUUGCUUUAUUCACGUACAGAAUUAACCAUUUAACUGAGCACUUAAAAAACAAUCGUAAAGAUUAUAACACUGAGCGUUCAUUAGUAAAGUUAGUAGGUAAAAGAAGAAGCUUACUAGAUUACUUA |
| env-1420 | AUCUUUAAAAAACACGGUAAAGAUGCAAAAGACACAGGUUCUGCAGAAGGGCAAAUUGCUUUAUUCACUUACAGAAUUAACCAUUUAACUGAGCACUUAAAAAACAAUCGUAAAGAUUUUAACACUGAGCGUUCAUUAGUAAAGUUAGUAGGUAAAAGAAGAAGCUUACUAGAUUACUUA |
| Pme-1-1 | AUCUUUAAAAAACAUGGCAAAAAUGCCAAAGACACUGGAACUGCAGAAGGUCAAAUUGCAUUAUUUACUCACAGAAUUAACCACUUAACUGAGCACUUAAAAAACAAUCGUAAAGAUUAUAAUACUGAGCGUUCAUUAGUAAAGUUAGUAGGUAAGCGUCGUGCAUUACUUGAUUAUUUA |
| env-1421 | UUAUUCGCAAAACACGGUAAAGGAAAAAACGAUACGGGUUCUGCAGAGGGACAGAUCGCAAUUUUUACGCAAAGAAUUGAGCACUUAACUGCGCACUUAAAAAAGAAUCGUAAAGAUUAUAAUACCGAACGUUCAUUAGUAAUGUUGGUAGGUAAGAGAAGAAGCUUACUAGAUUACCUA |
| Psp-5-1 | AUCUUUAAAAAACAUGGCAAAAGUGCGCAAGACACUGGAACUGCAGAAGGUCAAAUCGCAUUAUUCACUUACAGAAUAAACCACUUAACUGGGCACUUAAAAAAUAAUCGUAAAGAUUAUAACACUGAGCGCUCUCUAGUAAAAUUAGUUGGUAAGAGAAGAUCAUUAUUAGAUUACUUA |
| Pbu-1-1 | UUAUUCGCAAAACACGGUAAAGGGAAAAACGAUACUGGUACUGCAGAAGGACAAAUUGCAUUAUUUACGCAAAGAAUUGAGCACUUAACUGCUCAUUUAAAAAACAAUCGUAAAGAUUACAAUACAGAACGUUCAUUAGUUAUGUUAGUAGGUAAGAGAAGAAGCUUACUAGAUUACUUA |
| env-1422 | AUCUUUAAAAAACACGGCAAAAAUGCAAAGGACACUGGAACUGCAGAAGCACAGAUUGCAUUAUUUACACAAAGAAUUGAGCACUUAACGGCUCAUUUAAAAAACAAUCGUAAAGAUUACAACACAGAACGUUCAUUAGUAAUGUUAGUUGGUAAAAGAAGAAGUUUACUUGAUUACUUA |
| env-1425 | AUCUUUAAAAAACAUGGCAAAAAUGUAAAAGACACUGGUUCUGCAGAAGGUCAAAUCGCAUUAUUCACUUACAGAAUUAAUUACUUAACUGGAC.................................................................................. |
| Pja-1-1 | CUUUUUAAAAAACACGGUAAAAACGCACAAGACACUGGUUCUGCAGAAGGACAGAUUGCGAUUUUCACACAGAGAAUUGAGCACUUAACUGCUCAUUUAAAAAGCAAUCGUAAAGAUUAUAACACAGAACGCUCAUUAGUUAUGUUAGUAGGAAAGAGAAGAAGCUUACUAGAUUACUUA |
| env-1426 | CUUUUUAAAAAACACGGUAAAAACGCACAAGACACUGGUUCUGCAGAAGGUCAAAUUGCAUUAUUUACUCAUAGAAUUGAGCACUUAACUGCUCAUUUAAAAAACAAUCGUAAAGAUUACAACACAGAGCGUUCAUUAGUAAUGCUAGUUGGUAAAAGAAGAAGUUUAUUAGAUUACUUA |
| env-1427 | AUCUUCAAAAAACACGGUAAAAACGCUAAAGACACUGGAACAGCAGAAGGUCAAAUCGCAUUAUUUACUUACAGAAUUAACCACUUAACAGAACACUUAAAAAAUAAUCGUAAAGAUUACAAUACUGAACGUUCAUUAGUAAAGUUAGUUGGUAAAAGAAGAAGCUUACUAGAUUACUUA |
| env-1428 | AUCUUCAAAAAACACGGUAAAAACGCUAAA................................................................................................................................................... |
| | ........................................................................................................................................................................ |
| | ........................................................................................................................................................................ |
| | AUYUUYR•RAARCACGG•RRR•••R•RR••RAYAC•GG••C•RC•GAAGG•CAAAUYGC•YURUUYAC•Y•Y•G•AUYRAYCAYYUR•C•RA•CAYYURAAAAR•AAYCR•AARGAUU•YAAYAC•GAR•G•UC•YURGU•••RYURGU•GG•AAR•G•RG••••YU•YU•GAYUAYYU• |

```
env-1429 AUUUUCAAAAAGCACGGUAAAAAUGUCAAGGACACAGGUACCUCAGAAGGACAAAUUGCCUUAUUUACUUACAGAAUUAACCAUUUAACUGAACAUCUCAAAAAUAAUCGUAAAGAUUUUAAUACAGAACGUUCAUUGGUUAAACUAGUAGGUAAAAGAAGAAGCUUACUAGAUUAUUUG
env-1433 AUUUUCAAAAAGCACGGUAAAAAUGUCAAGGACACAGGUACCUCAGAAGGACAAAUUGCCUUAUUUACUUACAGAAUUAACCAUUUAACUGAACAUCUCAAAAAUAUCGUAAAGAUUUUAANACAGAACGUUCAUUGGUUAAACUAGUAGGUAAAAGAAGAAGCUUACUAGAUUAUUUG
env-1434 AUGUUUGCAAAACACGGUAACGGAAAAAACGAUACUGGUUCUGCAGAAGGACAAAUUGCGUUAUUUACGCACAGAAUUAACCAUUUAACACAGCACUUAAAGAACAAUCAGAAAGAUUACAAUACUGAGCGUUCUCUGGUAAAAUUAGUAGGUAAGCGUCGUGCAUUACUAGAUUACCUA
env-1435 AUGUUUGCAAAACACGGUAACGGAAAAAACGAUACUGGUUCUGCAGAAGGACAAAUUGCGUUAUUUACGCACAGAAUUAACCAUUUAACACAGCACUUAAAGAACAAUCAGAAAGAUUACAAUACUGAGCGUUCUCUGGUAAAAUUAGUAGGUAAGCGUCGUGCAUUACUAGAUUACCUA
env-1436 CUAUUUAAAAAACACGGUAAAGAUGCAAAAGAUACUGGUUCUGCAGAAGGACAGAUUGCAUUGUUUACUCACAGAAUUGAGCACUUAACUUCGCACUUAAAAAAACAAUCGUAAAGAUUACAAUACAGAACGUUCAUUAGUUAUGUUAGUAGGUAAAAGAAGAAGCUUACUAGAUUACUUA
Oma-1-1 AUUUUUGCUAAACACGGUAAAAAUUCAAAAGACACAGGUACUGCAGAAGGGCAAAUUGCUUUAUUUACUUACCGUAUCAACCACUUAACUGAGCACUUAAAGAAAAAUCGUAAAGAUUAUAACACUGAGCGUUCUUUAGUAAAACUAGUAGGAAAAAGAAGAAGCUUACUAGAUUACUUA
env-1437 AUUUUUACUAAACACGGUAAAAAUGCAAAGGACACAGGUACUGCAGAAGGGCAAAUAGCUUUAUUUACUUACCGUAUCAACCACUUAACUGAGCACUUAAAGAAAAAUCGUAAAGAUUAUAACACUGAGCGUUCUUUAGUAAAACUAUAGGAAAAGAAGAAGCUACUAGAUUACUAACUAG
env-1438 AUUUUUNCUAAACACGGUAAAAAUNCAAANGACACAGGUACUGCAGAAGGGCAAAUNGCUUUAUUUACUUACCGUAUCAACCACUUAACUGAGCACUUAAAGAAAAAUCGUAAAGAUUAUAACACUGAGCGUUCUUUAGUAAAACUAGUAGGAAAAAGAAGAAGCUUACUAGAUUACUUA
env-1439 AUUUUUACUAAACACGGUAAAAAUGCAAAGGACACAGGUACUGCAGAAGGGCAAAUAGCUUUAUUUACUUACCGUAUCAACCACUUAACUGAGCACUUAAAGAAAAAUCGUAAAGAUUAUAACACUGAGCGUUCUUUAGUAAAACUAGUAGGAAAAAGAAGAAGCUUACUAGAUUACUUA
Sze-1-1 AUCUUCGCAAAACACGGUAAAGGAGCAAACGAUACUGGAAGUGCAGAAGGACAAAUCGCGUUAUUUACGCACAGAAUUAAUCACUUAACAGAGCACUUAAAAAAGAAUCGUAAAGAUUAUAAUACUGAGCGUUCUUUAGUAAAACUAGUAGGUAAGCGUCGUGCUUUACUAGAUUACUUA
Gsa-1-1 AUCUUCGCAAAACACGGUAACGGCAAAAAUGAUACCGGUUCAGCUGAAGGACAAAUUGCGUUAUUCACGCACAGAAUUAAUCACUUAACUGAGCACUUAAAAAAAGAAUCGUAAAGAUUUUAACACUGAGCGCUCGUUAGUAAAACUAGUAGGUAAGCGUCGUGCUUUACUUGAUUACUUA
env-1441 CUCUUUAAAAAGCACGGUAAAGAUGCAAAGAACACUGGUUCUGCAGAAGGACAAAUUGCAUUGUUUACUCACAGAAUUAAUCACUUAACAGAACACUUAA..........................................................................................
env-1442 CUUUUCAAAAAACACGGUAAAAACGCAAAGGACACAGGUUCUGCAGAAGGACAAAUUGCAAUUUUUACUCAAAGAAUUGAGCAUUUAACUGCGCACUUAAAAAAAUAAUCGUAAAGAUUACAAUACAGAACGUGCUUUAGUUAUGUUGGUAGGAAAGCGCCGUGCUUUGCUUGAUUACUUA
Mze-1-1 AUCUUCGCAAAACACGGUAAAGGAGCAAACGAUACUGGUUCUGCAGAAGGACAGAUUGCAUUAUUCACUUACAGAAUUAACCACUUAACAGAACACUUAAAAAAAGAAUCGUAAAGAUUUUAACUCAGAGCGUUCUCUUGUAAAAAUGGUAGGUAAGCGUCGUGCGCUUUUAGACUAUUUA
env-1444 AUCUUCGCAAAACACGGUAAAGGAGCAAACGAUACUGGUUCUGCAGAAGGACAAAUUGCGUUAUUCACUUACAGAAUUAACCACUUAACAGAACACUUAAAAAAAGAAUCGUAAAGAUUUUAACUCAGAGCGUUCUCUUGUAAAAAUGGUAGGUAAGCGUCGUGCGCUUUUAGACUAUUUA
env-1445 CUCUUUAAAAAACACGGUAAAGAUGCAAAAGACACUGGUUCUGCAGAAGGACAGAUUGCAUUAUUUACACACAGAAUUGAGCACUUAACAGAACACUUAAAAAGUAAUCGUAAAGAUUAUAAUACAGAGCGUUCUUUAGUAAUGUUAGUAGGUAAAAGAAGAAGCUUACUAGAUUACUUA
env-1447 AUGUUUGCAAAACAUGGUAAAGGGAAAAACGAUACCGGUUCUGCAGAAGGACAAAUUGCCUUGUUUACACAAAGAAUUAAUCACUUAACAGAGCACUUAAAAAAAUAAUCGUAAAGAUUAUAAUACUGAGCGUUCAUUAGUGAUGUUAGUAGGUAAAAGAAGAAGCUUACUAGAUUACUUA
env-1448 AUGUUUGCAAAACAUGGUAAAGGGAAAAACGAUACCGGUUCUGCAGAAGGACAAAUUGCCUUGUUUACACAAAGAAUUAAUCACUUAACAGAGCACUUAAAAAAAUAAUCGUAAAGAUUAUAAUACUGAGCGUUCAUUAGUGAUGUUAGUAGGUAAAAGAAGAAGCUUACUAGAUUACUUA
env-1449 AUGUUUGCAAAACAUGGUAAAGGGAAAAACGAUACCGGUUCUGCAGAAGGACAAAUUGCCUUGUUUACACAAAGAAUUAAUCACUUAACAGAGCACUUAAAAAAAUAAUCGUAAAGAUUAUAAUACUGAGCGUUCAUUAGUGAUGUUAGUAGGUAAAAGAAGAAGCUUACUAGAUUACUUA
env-1454 UUAUUCAAAAAACACGGUGGUAACGCAAAAAAACACUGGUUCUGCAGAAGGACAAAUCGCAAUAUUUACGCAAAGAAUUGAGCACUUAACACAACACUUAAAAAGUAAUCGUAAAGAUUAUAAUACAGAGCGUUCGUUAGUAAUGUUAGUAGGAAAAA......................
env-1455 AUCUUUAAAAAACAUGGUGGUGAUGCAAAAAACACUGGUUCUGCACAAGACAGAUUGCAUUAUUACGGCAAUAACAUCUGACCAAUUCUUGAUCCAGUAUCUCGCACUUCC............................................................
Wps-1-1 UUAUUCACUAAACAUGGUGGUAAUGCAAAGAAUACUGGUUCUGCAGAAGGACAGAUUGCAUUAUUCACACAUAGAAUAGAACACUUAACACAACACUUAAAGAAAAACCGUAAAGAUUAUAAUACGGAACGUUCUUUAGUAAUGUUAGUAGGUAAGAGAAGAAGCUUAUUAGAUUACUUA
env-1456 AUCUUUAAAAAACAUGGUGGUGAUGCAAAAAACACUGGUUCUGCAGAAGGACAGAUUGCAUUAUUUACGGCAAGAAUUAAUCACUUAACAGAACACUUAAAACAUAAUCGUAAAGAUUUUAACACAGAGCGUUCGUUAGUAAAAUUAGUAGGUAAGCGUCGUGCUUUACUAGACUACUUA
env-1457 UUAUUUAAAAAACACGGUGGUAACGCAAAAAACACUGGUUCUGCAGAAGGACAGAUUGCAUUAUUUACAAAAAGAAUUGAGCACUUAACACAACACUUAAAAAGUAAUCGUAAAGAUUAUAAUACAGAGCGUUCGUUAGUAAUGUUAGUAGGAAAAAGAAGAAGCUUACUAGAUUACUUA
Osp-2-1 AUUUUUGCUAAACAUGGUAAAAACGCUCAAGACACAGGUACUGCUGAAGGGCAAAUUGCUUUAUUCACUUACAGAAUUAACCACUUAACAGAACACUUAAAGAAAAACCGUAAAGAUUAUAACACUGAGCGUUCACUUGUAAAAUUAGUAGGUAAACGUCGUUCAUUAUUAGACUACUUA
Osp-1-1 AUUUUUGCUAAACAUGGUAAAAACGCUCAAGACACAGGUACUGCUGAAGGGCAAAUUGCUUUAUUCACUUACAGAAUUAACCACUUAACAGAACACUUAAAGAAAAACCGUAAGGAUUAUAACACUGAGCGUUCACUUGUAAAAUUAGUAGGUAAACGUCGUUCAUUAUUAGACUACUUA
env-1458 AUCUUUAAAAAACAUGGUGGUGAUGCAAAAAAUACUGGUUCUGCAGAAGGACAGAUUGCAUUAUUUACUGCUAGAAUUAAUCACUUAACAGAACACUUAAAACGAAAUCGUAAAGAUUAUAACACAGAGCGUUCAUUAGUAAAAUUGGUAGGUAAAAGAAGAAGCUUACUAGAUUACUUA
env-1459 AUCUUUAAAAAACAUGGUGGUGAUGCAAAAAAUACUGGUUCUGCAGAAGGACAGAUUGCAUUAUUUACUGCUAGAAUUAAUCACUUAACAGAACACUUAAAAAAAGAAUCGUAAAGAUUAUAACACAGAGCGUUCAUUAGUAAAAUUGGUAGGUAAAAGAAGAAGCUUACUAGAUUACUUA
env-1460 AUCUUUAAAAAACAUGGUAAAAAUGCUAAAGACACUGGAACUGCAGAAGGUCAGAUAGCAUUAUUUACAUACAGAAUCAAUUACUUAACUGGGCACUUAAAAAAAUAAUCGUAAAGAUUAUAACACUGAGCGUUCGUUAGUAAAAUUAGUAGGUAAGCGAAGAGCUUUAUUAGACUACUUA
env-1461 AUUUUUAAAAAACAUGGCAAAAACGAAAAAGACACAGGUUCUGCAGAAGGGCAAAUUGCUUUAUUCACUUACAGAAUCAAUUACUUAACUGAGCACUUAAAAAAACAAUCGUAAAGAUUUUAAUACUGAGCGUUCAUUAGUAAAGUUAGUAGGUAAGCGUCGUGCUUUAUUAGAUUACUUA
env-1462 AUUUUUAAAAAACAUGGCAAAAACGCAAAAGACACAGGUUCUGCAGAAGGGCAAAUUGCCUUAUUCACUUACAGAAUCAAUUACUUAACUGAGCACUUAAAAAAACAAUCGUAAAGAUUUUAAUACUGAGCGUUCAUUAGUAAAGUUAGUAGGUAAGCGUCGUGCUUUAUUAGAUUACUUA
env-1463 CUUUU.........................................................................................................................................................................
Lsp-1-1 CUUUUUGCAAAGCACGGUAAAAAUGCUCAAGACACUGGUUCUGCCGAAGGUCAAAUUGCAUUAUUCACAGAAAGAAUUAACCACUUAACACAGCACUUAAAAAACUAAUCGUAAAGAUUAUAACACAGAGCGUUCGUUAGUGAAGUUAGUAGGAAAGCGUCGUGCUUUAUUAGACUACUUA
env-1464 AUCUUUAAAAAACAUGGUAAAAAUGCUAAAGACACUGGAACUGCAGAAGGUCAGAUAGCAUUAUUUACAUACAGAAUCAAUUACUUAACUGGGCACUUAAAAAAAUAAUCGUAAAGAUUAUAACACUGAGCGUUCGUUAGUAAAAUUAGUAGGUAAGCGAAGAGCUUUAUUAGACUACUUA
env-1465 AUCUUUAAAAAACACGGUAAAAAUGCUAAAGACACUGGAACUGCAGAAGGUCAGAUAGCAUUAUUUACAUACAGAAUCAAUUACUUAACUGGGCACUUAAAAAAAUAAUCGUAAAGAUUAUAACACUGAGCGUUCGUUAGUAAAAUUAGUAGGUAAGCGAAGAGCUUUAUUAGACUACUUA
env-1466 AUCUUUAAAAAACAUGGUAAAAAUGCUAAAGACACUGGAACUGCAGAAGGUCAGAUAGCAUUAUUUACAUACAGAAUCAAUUACUUAACUGGGCACUUAAAAAAAUAAUCGUAAAGAUUAUAACACUGAGCGUUCGUUAGUAAAAUUAGUAGGUAAGCGAAGAGCUUUAUUAGACUACUUA
env-1467 CUCUUCGCAAAACACGGUAAAGGAAAAAACGAUACUGGUUCAGCAGAAGGACAAAUCGCAAUAUUUACUCAAAGAAUUGAGCAUUUAACUGCGCAUCUAAAAAAAUAAUCGUAAAGAUUAUAAUACAGAACGUUCGUUAGUUAUUUUAGUAGGUAAGAGAAGAAGCUUACUAGAUUACUUA
env-1468 AUCUUUAAAAAACACGGUAAAAAUGCUAAAGACACUGGAACUGCAGAAGGUCAGAUAGCAUUAUUUACAUACAGAAUCAAUUACUUAACUGGGCACUUAUGCUAAA[illegible]ACUGGAACUGCAGAAGGUCAGAUAGCAUUAUUUACAUACAGAAUCAAUUACUUAACUGGGCACUUAAAAAA
env-1469 AUCUUUAAAAAACAUGGUAAAAAUGCUAAAGACACUGGAACUGCAGAAGGUCAGAUAGCAUUAUUUACAUACAGAAUCAAUUACUUAACUGGGCACUUAAAAAAAUAAUCGUAAAGAUUAUAACACUGAGCGUUCGUUAGUAAAAUUAGUAGGUAAGCGAAGAGCUUUAUUAGACUACUUA
env-1471 AUCUUCACAAAACACGGUAAAGGAAAGAACGAUACGGGUUCUGCUGAAGGACAAAUUGCAUUAUUCACUUACAGAAUUAACCACUUAACAGAGCACUUAAAACAAAAUCGUAAAGAUUUUAACUCUGAGCGUUCUCUGGUAAAAGUUAGUAGGUAAGCGUCGUGCUUUACUAGAUUACUUA
Fba-5-1 CUUUUCAAAAAACAUGGUAAAGACGCAAAAGACACAGGUUCUGCAGAAGGGCAAAUUGCGUUAUUUACAGCAAGAAUUAACCACUUAACUGAACACUUAAAAACAAAAUCGCAAAGAUUUUAAUACUGAGCGUUCAUUAGUAAAAUUAGUAGGUAAAGCGUCGUGCUUUAUUAGACUACUUA
Fic-1-1 CUUUUCAAAAAACACGGUAAAAAUGCAAAAGACACUGGUUCUGCAGAAGGACAGAUUGCAUUGUUUACGCACAGAAUUAAUCACUUAACUGAACACUUAAAAAAACAAUCGUAAAGAUUAUAAUACAGAGCGUUCUUUAGUGAAAUUAGUAGGUAAAAGAAGAAGCUUACUUGAUUACUUA
env-1472 CUCUUUAAAAAACAUGGUG...........................................................................................................................................................
Lja-1-1 CUGUUUAAAAAACACGGUAAAGAUGCAAAAGACACUGGUUCUGCAGAAGGACAGAUUGCAAUAUUUACACAAAGAAUUGAGCACUUAUCAAAGCACUUAAAAACAAAUCGUAAAGAUUAUAAUACUGAGCGUUCGUUAGUAAUGUUAGUAGGAAAAAGAAGAAGCCUACUGGAUUACUUA
env-1473 AUCUUCGCAAAACACGGUAAAGGAAAAAACGAUACUGGUACAGCUGAAGGACAAAUUGCAUUAUUCACUGCAGAAUUAACCACUUAACAGAGCACUUAAAACAAAAUCGUAAAGAUUAUAAUACAGAGCGUUCAUUAGUAAAGUUAGUAGGUAAGCGUCGUGCCUUACUUGACUACUUA
Fba-8-1 CUUUUCGCAAAACACGGUAAAGGAAAGAACGAUACUGGUUCUGGAAGUGCAGAAGGGCAAAUUGCGUUAUUUACGCAUAGAAUUAAUCACUUAACUGAGCACUUAAAAAAAUAAUCGUAAAGAUUAUAAUACUGAGCGUUCUUUAGUGAAAUUAGUAGGAAAGCGUCGUGCUUUACUAGAUUACUUA
env-1474 CUUUUUAAAAAACACGGUAAAAACGCAAAGGAUACUGGUUCUGCAGAAGGACAGAUUGCAUUAUUUACUGAAAGAAUUAAUCACUUAACAGAACACUUAAAAAAAUAAUCGUAAAGAUUAUA..................................................
env-1475 CUCUUUAAAAAACAUGGUGGUGAUGCAAAAAACACCGGUUCUGCUGAAGGACAGAUUGCAUUAUUUACUGCAAGAAUUAAUCACUUAACAGAACACUUAAAAAAAGAAUCGUAAAGAUUAUAAUACUGAGCGUUCAUUAGUAAAGUUAGUGGUA..........................
env-1476 CUUUUUAAAAAACACGGUAAAGAUGCAAAAGACACAGGUUCUGCAGAAGGGCAAAUUGCAUUAUUUACACAAAGAAUUAAUCACCUUAACAGAACACUUAAAAAAAGAAUCGUAAAGAUUAUAAUACAGAGCGUUCAUUAGUAAAGUUAGUAGGUAAAAGAAGAAGCUUUACUUGAUUAUUUA
env-1477 CUUUUUAAAAAACACGGUAAAGAUGCAAAAGACACAGGUUCUGCAGAAGGACAAAUUGCAUUGUUUACGUCAAGAAUUAACCACUUAACAGAACACUUAAAAAAAUAAUCGUAAAGAUUACAAUACAGAGCGUUCAUUAGUAAAGUUAGUAGGUAAAAGAAGAAGCUUACUUGAUUAUUUA
env-1478 CUUUUUAAAAAACACGGUAAAGAUGCAAAAGACACAGGUUCUGCAGAAGGACAAAUUGCAUUAUUUACAUCAAGAAUUAACCACUUAACAGAACACUUAAAAAAAUAAUCGUAAAGAUUAUAAUACAGAGCGUUCAUUGGUAAAGUUAGUAGGUAAAAGAAGAAGCUUACUUGAUUAUUUA
env-1479 CUUUUUAAAAAACACGGUAAAGAUGCAAAAGACACAGGUUCUGCAGAAGGACAAAUUGCAUUAUUUACAUCAAGAAUUAACCACUUAACAGAACACUUAAAAAAAUAAUCGUAAAGAUUAUAAUACAGAGCGUUCAUUGGUAAAGUUAGUAGGUAAAAGAAGAAGCUUACUUGAUUAUUUA
env-1480 AUGUUUGCAAAACACGGUAAAGGGAAAAACGAUACCGGUUCUGCAGAAGGACAAAUUGCGUUAUUUACGCAAAGAAUUAAUCACUUAACAGAACACUUAAAAAAAUAAUCGUAAAGAUUAUAAUACAGAGCGUUCUCUAGUAAAGUUAGUAGGAAAAA......................
env-1481 CUNUUUAAAAAANCACGGUAAAGAUGCAAAAGAUACUGGUUCUGCAGAAGGACAGAUUGCAUUGUUUACUCACAGAAUUGAGCAUUUAACUGCGCACUUAAAAAAACAAUCGUAAAGAUUACAACACAGAACGUUCAUUAGUUAUGCUAGUAGGUAAAAGAAGAAGCUUACUAGAUUACUUA
env-1482 ..............................................................................................................................................................................
env-1483 AUCUUCUCAAAACACGGUAAAGGAGCAAACGAUACUGGUUCUGCAGAAGGACAAAUUGCGUUAUUCACUUACAGAAUUAACCACUUAACAGAACACUUAAAAAAAGAAUCG..........................................................
env-1484 CUUUUUAAAAAACACGGUAAAGAUGCAAAAGACACAGGUUCUGCAGAAGGACAAAUUGCAUUGUUUACGUCAAGAAUUAACCACUUAACAGAACACUUAAAAAAAUAAUCGUAAAGAUUACAAUACAGAGCGUUCAUUAGUAAAGUUAGUAGGUAAAAGAAGAAGCUUACUUGAUUAUUUA
env-1485 AUGUUUGCAAAACACGGUAAAGGGAAAAACGAUACCGGUUCUGCAGAAGGACAAAUUGCGUUAUUU...........................................................................................................
env-1486 CUAUUUAAAAAGCACGGUAAAGAUGCAAAAGAUACUGGUUCUGCAGAAGGACAGAUUGCAUUGUUUACUCACAGAAUUGAGCAUUUAACUGCGCACUUAAAAAAACAAUCGUAAAGAUUACAACACAGAACGUUCAUUAGUUAUGCUAGUAGGUAAAAGAAGAAGCUUACUAGAUUACUUA
env-1487 AUGUUUGCAAAACACGGUAAAGGGAAAAACGAUACCGGUUCUGCAGAAGGACAAAUUGCGUUAUUUACGCAAAGAAUUAAUCACUUAACAGAACACUUAAAAAAA...........................................................................
env-1488 CUUUUUAAAAAACACGGUAAAGAUGCAAAAGAUACUGGUUCUGCAGAAGGACAGAUUGCAUUAUUCACACAUAGAAUUGAGCAUUUAACUGCGCAUUUAAAAAAACAAUCGUAAAGAUUACAACACAGAACGUUCAUUAGUUAUGCUAGUAGGUAAAAGAAGAAGCUUACUAGAUUACUUA
env-1489 CUCUUUAAAAAACACG..............................................................................................................................................................
env-1490 ..............................................................................................................................................................................
env-1491 AUCUUUAAAAAACACGGUAAUAAUGCAAAAGACACUGGAUCUGCAGAGGGACAGAUUGCAUUGUUUACGCACAGAAUUAAUUACUUAACAGGACACUUAAAAAACAAUCGUAAAGAUUACAAUACUGAGCGUUCAUUAGUAAAGUUAGUAGGUAAAAGAAGAAGCUUACUUGAUUACUUA
env-1497 AUCUUUAAAAAACACGGUAGUAAUGCAAAAGACACUGGUUCUGCAGAAGGACAGAUUGCAUUGUUUACUUACAGAAUUAAUUACCUAACAGGACACUUAAAAAAAAUAAUCGUAAAGAUUACAAUACUGAGCGUUCAUUAGUAAAGUUAGUAGGUAAAAGAAGAAGCUUACUAGAUUACUUA
env-1498 AUCUUUAAAAAACACGGUAGUAAUGCAAAAGACACUGGUUCUGCAGAAGGA...................................................................................................................................
env-1499 AUCUUUAAAAAACACGGUAGUAAUGCAAAAGACACUGGUUCUGCAGAAGGACAGAUUGCAUUGUUUACUUACAGAAUUAAUUACCUAACAGGACACUUAAAAAAAAUAAUCGUAAAGAUUACAAUACUGAGCGUUCAUUAGUAAAGUUAGUAGGUAAAAGAAGAAGCUUACUAGAUUACUUA
env-1500 AUCUUUAAAAAACACGGUAGUAAUGCAAAAGACACUGGUUCUGCAGAAGGACAGAUUGCAUUGUUUACUUACAGAAUUAAUUAC.........................................................................................
env-1501 AUCUUUAAAAAACACGGUAGUAAUGCAAAAGAUACUGGUUCUGCAGAAGGACAGAUUGCAUUGUUUACGUACAGAAUUAAUUACUUAACAGGNCACUUAAAAAAAAUAAUCGUAAAGAUUAUAAUACUGAGCGUUCAUUAGUAAAGUUAGUAGGAAAAAGAAGAAGCUUACUAGAUUACUUA
env-1502 AUCUUUAAAAAACACGGUAGUAAUGCAAAAGAUACUGGUUCUGCAGAAGGACAGAUUGCAUUGUUUACGUACAGAAUUAAUUACUUAACAGGACACUUAAAAAAAAUAAUCGUAAAGAUUAUAAUACUGAGCGUUCAUUAGUAAAGUUAGUAGGAAAAAGAAGAAGCUUACUAGAUUACUUA
env-1503 AUCUUUAAAAAACACGGUAGUAAUGCAAAAGAUACUGGUUCUGCAGAAGGACAGAUUGCAUUGUUUACGUACAGAAUUAAUUACUUAACAGGACACUUAAAAAAAAUAAUCGUAAAGAUUACAAUACUGAGCGUUCAUUAGUAAAGUUAGUAGGUAAAAGAAGAAGCUUACUAGAUUACUUA
env-1504 ..............................................................................................................................................................................
env-1505 AUCUUUACAAAACAUGGUAAAGGUGCCAAAGACACUGGUUCUGCAGAAGGACAGAUUGCUUUAUUUAGUUACAGAAUUAUUCACUUAACAGAACAUUUAAAAAAAGAAUCGUAAAGAUUAUAAUACUGAGCGUUCGUUAGUGAAUUUAGUAGGUAAAAGAAGAAGCUUUACUAGAUUACUUA
env-1506 UUAUUCGCAAAACAUGGUGGUAACGCAAAAAAUACUGGUUCUGCAGAAGGACAGAUUGCAUUAUUCACUCACAGAAUUGAGCACUUAACACAACACUUAAAAAAUAAUCGUAAAGAUUAUAAUACAGAGCGUUCGUUAGUAAUGUUAGUAGGAAAAAGAAGAAGCUUACUAGAUUACUUA
Pda-1-1 AUCUUUAAAAAACAUGGCAAAAUGCACAAGACACUGGAACUGCAGAAGGUCAAAUCGCAUUAUUUACUUACAGAAUUAACCACUUAACAGAGCACUUAAAAAAACAAUCGUAAAGAUUAUAAUACUGAGCGUUCAUUAGUAAAGUUAGUAGGUAAAAGAAGAAGCUUACUAGAUUACUUA
env-1507 CUCUUCUCAAAACACGGUAAAAGUAAGAACGAUACUGGUACUG...................................................................................................................................
env-1508 AUCUUCUCAAAACACGGUAAAGGUAAGAACGAUACUGGUACUGCAGAAGGACAAAUUGCGUUAUUUACGCACAGAAUCAAUCACUUAACUGAGCACUUAAAAAAACAAUCGUAAAGAUUUUAAUACGGAGCGUUCAUUAGUAAAAUUAGUGGUAAACGCAGAGCUUUACUAGAUUAUUUA
env-1515 AUCUUCGCAAAACACGGUAAAGUUCAAACGAUACUGGUACAGCAGAAGGUCAAAUUGCGUUAUUUACGCACAGAAUCAAUCACUUAAC.....................................................................................
env-1516 AUCUUCGCUAAACACGGUAAAGGAAAAAAUGAUACCGGUUCUGCAGAAGGUCAAAUCGCAUUAUUUACUCACCGAAUUAAUCACCUUACAGAGCACCUUAAACAAAAUCGUAAAGAUUUUAAUACGGAGCGUGCACUAGUUAAGUUAGUGGGUAAAAGACGCGCCUUACUUGAUUACUUA
env-1517 AUCUUCGCAAAACACGGAAAAAGUAAAACCGAUACCGGUUCUGCAGAAGGUCAGAUUGCAUUGUUCACUUACAGAAUUGCACACCUGUCUGAGCAUUUAAAGAGCAACAGAAAAGAUUUUAACUCCGAGCGUUCACUGGUAAGACUUGUUGGUAAAAGAAGAAGUCUUUUGGAUUACCUA
env-1519 AUCUUUAAAAAACAUGGUGAAAAUGCAAAG................................................................................................................................................
env-1520 AUCUUUAAAAAACAUGGUGAAAAUGCAAAGAACACAGGUUCUGCUGAAGGGCAAAUUGCAUUAUUUACACUUAGAAUCAAUCACUUGACUGAGCACUUAAAAAAAUAAUCGUAAAGAUUUUAAUACAGAGCGUUCGUUGGUAAAAUUAGUUGGUAAACGCAGAUCAUUGCUUGAUUACUUA
env-1521 AUCUUUAAAAAACAUGGUGAAAAUGCAAAGAACACAGGUUCUGCUGAAGGGCAAAUUGCAUUAUUUACACUUAGAAUCAAUCACUUGACUGAGCACUUAAAAAAAUAAUCGUAAAGAUUUUAAUACAGAGCGUUCGUUGGUAAAAUUAGUUGGUAAACGCAGAUCAUUGCUUGAUUACUUA
env-1522 AUUUUCGCAAAACACGGUAAAGGGAAGAACGAUACUGGUACUGCAGAAGGUCAAAUUGCGUUAUUUACGCACAGAAUUAACCACUUAACCGAACACCUAAAAAACAAUCGUAAAGAUUACAACACAGAGCGUUCGUUAGUAAAACUGGUAGGUAAGCGUCGUGCAUUACUUGACUAUUUA

..............................................................................................................................................................................
..............................................................................................................................................................................
AUYUUYR•RAARCACGG•RRR•••R•RR••RAYAC•GG••C•RC•GAAGG•CAAAUYGC•YURUUYAC•Y•Y•G•AUYRAYCAYYUR•C•RA•CAYYURAAAAR•AAYCR•AARGAUU•YAAYAC•GAR•G•UC•YURGU•••RYURGU•GG•AAR•G•RG••••YU•YU•GAYUAYYU•
```

```
env-1524  AUUUUCGCAAAACACGGUAAAGGGAAGAACGAUACUGGUACUGCAGAAGGUCAAAUUGCGUUAUUUACGCACAGAAUUAACCACUUAACCGAACACCUAAAAAACAAUCGUAAAGAUUACAACACAGAGCGUUCGUUAGUAAAACUGGUAGGUAAGCGUCGUGCAUUACUUGACUAUUUA
env-1525  CUGUUUACAAAACAUGGUAAAAACGCAAAAGACACUGGUACUGCUGAGGGUCAGAUCGCUUUAUUCACUUACAGAAUUGAGCAUUUAACAACUCACCUAAAGAAAAAUCGUAA...................................................................
env-1526  CUGUUUACAAAACAUGGUAAAAACGCAAAAGACACUGGUACUGCUGAGGGUCAGAU............................................................................................................................
env-1527  AUCUUUAAAAAACAUGGUAGCAAUGCAAAGAACACAGGUUCAGCUGAAGGGCAAAUUGCAUUGUUUACACACAGAAUCAAUCACUUGACAGAACACUUAAAAAACAAUCGUAAAGAUUAUAAUACUGAGCGUUCAUUGGUAAAAUUAGUAGGUAAAAGAAGAAAUUUACUUGACUACUUA
env-1528  AUCUUUAAAAAACACGGUGAAAAUGCAAAGAACACAGGUUCAUCUGAAGGUCAAAUUGCAUUGUUUACACUUAGAAUCAAUCACUUGACAGAACACUUGAAAAAC...........................................................................
env-1529  AUCUUUAAAAAACACGGUGAAAAUGCAAAGAACACAGGCUCAGCUGAAGGUCAGAUUGCAUUGUUUACACUUAGAAUCAAUCACUUGACAGAGCACUUAAAAAACAAUCGUAAAGAUUUUAAUACAGAGCGUUCAUUGGUAAAAUUAGUUGGUAAACGCAGAUCAUUGCUUGAUUACCUU
env-1530  AUCUUUAAAAAACACGGUGAAAAUGCAAAGAACACAGGCUCAGCUGAAGGUCAGAUUGCAUUGUUUACACUUAGAAUCAAUCACUUGACAGAGCACUUAAAAAACAAUCGUAAAGAUUUUAAUACAGAGCGUUCGUUGGUAAAAUUAGUUGGUAAACGCAGAUCAUUGCUUGAUUACCUU
1 Fag-1-1  CUUUUCGCAAAGCACGGUAAAGGAAAAAACGAUACUGGUACUGCAGAAGGACAAAUUGCGUUAUUCACGCACAGAAUUAAUCACUUAACAGAACACUUAAAAAAGAAUCGUAAAGAUUAUAAUACUGAGCGUUCAUUAGUAAAAUUAGUAGGUAAGCGUAGAUCAUUACUUGACUACUUA
env-1534  CUUUUCGCAAAACACGGUAAAGGAAAAAACGAUACUGGUUCUGCAGAAGGACAAAUCGCAAUAUUUACUCAAAGAAUUGAGCAUUUAACUGCGCAUUUAAAAAACAAUCGUAAAGAUUAUAAUACAGAACGUUCGUUAGUUAUAUUAGUAGGUAAAAGAAGAAGCUUACUAGA.......
env-1535  CUUUUCGCAAAACACGGUAAAGGAAAAAACGAUACUGGUUCUGCAGAAGGACAAAUCGCAAUAUUUACUCAAAGAAUUGAGCAUUUAACUGCGCAUUUAAAAAACAAUCGUAAAGAUUAUAAUACAGAACGUUCGUUAGUUAUAUUAGUAGGUAAAAGAAGAAGCUUACUAGA.......
env-1536  CUCUUCGCAAAACACGGUAAAGGAAAAAACGAUACUGAAAGAAAACCUUUCAUGUACAUAAAAAUCAUUUAAAUAAAUAUUGGAAUGUAUUUAACAAAAGAAGCAAAAGAAGAACUUUUCGCAAAACACGGUAAAGGAAAAAACGAUACUGGUUCUGCAGAAGGACAAAUCGCAAUAUUU
env-1537  CUCUUCGCAAAACACGGUAAAGGAAAAAACGAUACUGGUUCUGCAGAAGGACAAAUCGCAAUAUUUACUCAAAGAAUUGAGCAUUUAACUGCGCAUUUAAAAAACAAUCGUAAAGAUUAUAAUACAGAACGUUCGUUAGUUAUAUUAGUAGGUAAAAGAAGAAGCUUACUAGAUUACUUA
env-1538  AUCUUUAAAAAGCAUGGUAAAGAUGCAAAGAACACAGGUUCUGCCGAAGGUCAAAUUGCAUUGUUUACACACAGAAUUAAUCACUUAACCGAACAUUUAAAAAGCAAUCGUAAAGAUUACAAUACAGAACGUUCAUUAGUGAAAUUAGUAGGUAAACG......................
env-1539  AUCUUUAAAAAGCAUGGUAAAGAUGCAAAGAACACAGGUUCUGCCGAAGGUC................................................................................................................................
env-1540  AUCUUUAAAAAGCAUGGUAAAGAUGCAAAGAACACAGGUUCUGCCGAAGGUCAAAUUGCAUUGUUUACACACAGAAUUAAUCACUUAACCGAGCAUUUAAAAAGCAAUCGUAAAGAUUACAAUACAGAACGUUCAUUAGUGAAAUUAGUAGGUAAACGUCGUUCUUUACUAGAUUACUUA
env-1541  AUCUUUACAAAACAUGGUAAAGAUGCAAAGAACACAGGUUCUGCCGAAGGUCAAAUUGCAUUGUUUACACACAGAAUUAAUCACUUAACCGAGCAUUUAAAAAGCAAUCGUAAAGAUUACAAUACAGAACGUUCAUUAGUGAAAUUAGUAGGUAAACGUCGUUCUUUACUAGAUUACUUA
env-1542  AUCUUUAAAAAACAUGGUAAAGAUGCAAAGAACACAGGUUCUGCCGAAGGUCAAAUUGCAUUGUUUACACACAGAAUUAAUCACUUAACCGAACAUUUAAAAAGCAAUCGUAAAGAUUACAAUACAGAACGUUCAUUAGUGAAAUUAGUAGGUAAACGUCGUUCUUUACUAGAUUACUUA
env-1543  AUCUUUAAAAAACAUGGUAAAGAUGCAAAGAACACAGGUUCUGCCGAAGGUCAAAUUGCAUUGUUUACACACAGAAUUAAUCACUUAACCGAGCAUUUAAAAAGCAAUCGUAAAGAUUACAAUACAGAACGUUCAUUAGUGAAAUUAGUAGGUAAACGUCGUUCUUUACUAGAUUACUUA
env-1546  AUCUUUAAAAAACAUGGUAAAGAUGCAAAGAACACA................................................................................................................................................
env-1547  AUCUUUAAAAAACAUGGUAAAGAUGCAAAGAACACAGGUUCUGCCGAAGGUCAAAUUGCAUUGUUUACACACAGAAUUAAUCACUUAACCGAACAUUUAAAAAGCAAUCGUAAAGAUUACAAUACAGAACGUUCAUUAGUGAAAUUAGUAGGUAAACGUCGUUCUUUACUAGAUUACUUA
env-1548  AUCUUUAAAAAACAUGGUAAAGAUGCAAAGAACACAGGUUCUGCCGAAGGUCAAAUUGCAUUGUUUACACACAGAAUUAAUCACUUAACCGAACAUUUAAAAAGCAAUCGUAAAGAUUAUAAUACGGAACGUUCAUUAGUGAAAUUAGUAGGUAAACGAAGAGCUUUACUAGACUAUUUA
env-1549  ....................................................................................................................................................................................
env-1550  AUCUUUAAAAAACAUGGUAAAGAUGCAAAAAACACAGGUUCAGCAGAAGGACAAAUUGCAUUGUUUACUUACAGAAUUAAUCACUUAACCGAUCACUUAAAAAAGAAUAGAAAAGAUUAUAAUAGUGAACGUUCAUUAGUUAAACUAGUUGGUAAAAGAAGAGCUUUACUUGAUUAUUUA
env-1551  AUCUUUAAAAAACAUGGUAAAGAUGCAAAAAACACAGGUUCAGCAGAAGGACAAAUUGCAUUGUUUACUUACAGAAUUAAUCACUUAACCGAUCACUUAAAAAAGAAUAGAAAAGAUUAUAAUAGUGAACGUUCAUUAGUUAAACUAGUUGGUAAAAGAAGAGCUUUACUUGAUUAUUUA
env-1552  AUCUUUAAAAAACACGGUAAAGAUGCAAAAAAUACUGGUUCUGCAGAAGGACAAAUUGCAUUGUUUACUUACAGAAUUAAUCACUUGACAAAACACUUAAAAAAAUAAUCGAAAAGAUUAUAAUUC......................................................
env-1553  AUCUUUAAAAAACACGGUAAAGAUGCAAAAAAUACUGGUUCUGCAGAAGGACAAAUUGCAUUGUUUACUUACAGAAUUAAUCACUUGACAAAACACUUAAAAAAAUAAUCGAAAAGAUUAUAAUUCAGAGCGUUCAUUAGUGAAAUUAGUUGGUAAAAGAAGAUCUUUACUAGACUAUUUA
env-1554  ....................................................................................................................................................................................
env-1555  AUCUUUAAAAAACACGGUAAAAAUGCAAAUGACACGGGCUCUGCCGAAGGUCAAAUUGCAUUAUUUACCCACAGAAUUAAUCAUUUAACUAAACACUUGAAGAACAAUCGCAAAGAUUUCAAUACUGAGCGUUCACUUGUAAAAUUGGUAGGAAAGCGUAGAUCUUUACUUGAUUAUCUU
env-1556  AUCUUUAAAAAACACGGUAAAAAUGCAAAUGACACGGGCUCUGCCGAAGGUCAAAUUGCAUUAUUUACCCACAGAAUUAAUCAUUUAACUAAACACUUGAAGAACAAUCGCAA...................................................................
env-1557  AUCUUUAAAAAACACGGUAAAAAUCCAAAUGACACGGGCUCUGCCGAAGGUCAAAUUGCAUUAUUUACUCAUAGAAUCAAUCACUUAACUGAACAUCUUAAGAAUAAUCGCAAAGAUUUUAAUACUGAGCGUUCACUUGUGAAAUUGGUAGGAAAGCGUAGAUCUUUACUUGAUUAUCUU
env-1559  AUCUUUAAAAAGCAUGGUAAAGAUGCAAAGAACACAGGUUCUGCCGAAGGGCAAAUUGCAUUGUUUACUCAUAGAAUCAAUCACUUAACAGAGCACUUAAAAAACAAUC.......................................................................
env-1560  AUCUUUAAAAAGCAUGGUAAAGAUGCAAAGAACACAGGUUCUGCCGAAGGGCAAAUUGCAUUGUUUACUCAUAGAAUCAAUCACUUAACAGAGCACUUAAAAAACAAUCGUAAAGAUUAUAAUUCUGAACGUUCAUUAGUAAUGUUAGUUGGUAAACGAAGAUCUUUAUUAGACUACUUG
env-1561  CUUUUCGCAAAGCACGGAAAAUCUGCGCAGGACACAGGGAGUGCAGAAGGGCAAAUCGCUCUUUUCACAUUUAGAAUUAACCAUUUGUCUAACCACCUAAAAGCAAACCAUAAGGAUUUCAACACUGAGAGGUCUCUCGUGAAGUUAGUAGGUAAAAGAAGAAGAUUAUUAGAUUAUCUU
env-1562  CUUUUCGCAAAGCACGGAAAAUCUGCGCAGGACACAGGGAGUGCAGAAGGGCAAAUCGCUCUUUUCACAUUUAGAAUUAACCAUUUAUCUAACCACCUAAAAGCAAACCAUAAGGAUUUCAACACUGAGAGGUCUCUUGUGAAGUUAGUAGGUAAAAGAAGAAGAUUAUUAGAUUAUCUU
env-1563  CUUUUCGCAAAGCACGGAAAAUCUGCGCAGGACACAGGGAGUGCAGAAGGGCAAAUCGCUCUUUUCACAUUUAGAAUUAACCAUUUAUCUAACCACCUAAAAGC............................................................................
env-1564  CU..................................................................................................................................................................................
env-1565  ....................................................................................................................................................................................
env-1566  AUUUUUAAAAAACACGGUAAAAGUGAAAAUGACACUGGUUCUGCAGAAGGACAAAUUGCUUUAUUCACACACCGAAUUCAACACUUAACCCAACACCUUAAGAAAAAUCAUAAGGACUUCAACACAGAGCGUUCUUUGGUUAAAAUGGUAGGUAAAAGAAGAGCAUUGCUUGAUUAUCUA
Zam-1-1   AUCUUUAAAAAGCACGGUGGAGCUGAAAACAACACUGGUUCAACAGAAGGACAAAUUGCAUUGUUCACUCACCGAAUCAGCCACUUAACUGGGCACCUUAAAAAGAAUCGUAAAGAUUUAACACAGAGCGCGCUCUAGUUAAAUUGGUAGGUAAAAGAAGAAGUUUACUAGACUAUUUA
env-1567  AUCUUCGCUAAACACGGAGGAAAAGCUGAAAACACUGGAUCUGCUGAAGGGCAAAUCGCAUUGUUCACAUUUAGAAUCACACAUUUAACNGAGCAUUUGAAAAAAAAUCGUCACGAUUAUAACACAGAGCGUUCGUUAGUUCUUUUGGUAGGAAAAAGAAGAAGUCUUUUAGAUUAUUUG
Ffr-2-1   AUCUUCGCUAAACACGGAGAAAAAACAAACACAGGUAAAUCAGAAGCUCAAAUUGCUUUGUUCACUUUCAGAAUUAGCCACCUAACAGAACACUUGAAAAAAAAUCGUCACGAUUACAACACAGAGCGUUCAUUAGUAUUGCUAGUUGGAAAAAGAAGAUCUUUGUUGGAUUACUUGAAA
env-1568  AUCUUCGCUAAACACGGAGAAAAAACAAACACAGGAAAAUCAGAAGCUCAAAUUGCUUUGUUCACUUUCAGAAUUGCACACUUAACAGAACACUUGAAAAAAAAUCGUCAUGAUUAUAACACAGAGCGUUCAUUAGUAUUGCUAGUUGGAAAAAGAAGAUCUUUGUUGGAUUACUUGAAA
Fsp-2-1   AUCUUCGCUAAACACGGAGAAAAAACAAACACUGGAAAAUCUGAAGCACAAAUCGCUUUAUUCACUUUCAGAAUUAGCCACUUAACUGAGCACUUGAAAAAAAAUCGUCAUGAUUAUAACACAGAGCGUUCAUUAGUAUUACUAGUAGGUAAAAGAAGAGCUUUGUUAGAUUACUUGAAG
env-1571  AUCUUCGCUAAACACGGAGAAAAAACAAAUACUGGAAAAUCAGAAGCUCAAAUUGCUUUGUUCACUUUCAGAAUUAGCCACUUGACUGAGCAUUUGAAAAAAAAUCGUCACGAUUAUAACACAGAGCGUUCANUAGUAUUGCUUGUAGGUAAAAGAAGAUCUUUGUUGGAUUACUUGAAG
env-1572  AUCUUCGCACAACACGGAGAAAAAACUAACACAGGUAAAGCAGAAGCUCAAAUUGCUUUAUUCAGUUUCAGAAUUAGCCACCUAACAGAACAUUUGAAAAAAAAUCGUCACGAUUAUAACACAGAGCGUUCAUUAGUAUUGCUAGUAGGAAAAAGAAGAUCU..................
env-1573  AUCUUCGCACAACACGGAGAAAAAACUAACACAGGUAAAGCAGAAGCUCAAAUUGCUUUAUUCAGUUUCAGAAUUAGCCACCUAACAGAACAUUUGAAAAAAAAUCGUCACGAUUAUAACACAGAGCGUUCAUUAGUAUUGCUAGUAGGA..............................
env-1574  AUCUUCGC............................................................................................................................................................................
env-1575  AUCUUCGCACAACACGGAGAAAAAACAAACACAGGUAAAGCAGAAGCUCAAAUUGCUUUGUUCACUUUCAGAAUUAGCCACCUAACAGAACAUUUGAAAAAAAAUCGUCACGAUUAUAACACAGAGCGUUCAUUAGUAUUGCUAGUAGGAAAAAGAAGAUCUUUGUUAGAUUACUUGAAA
env-1576  AUCUUCGCACAACACGGAGAAA..............................................................................................................................................................
Fsp-5-1   AUCUUCGCUAAACACGGAGAAAAAACAAACACUGGAAAAUCAGAAGCACAAAUCGCUUUAUUCACUUUCAGAAUUAGCCACUUAACUGAGCACUUGAAAAAAAAUCGUCAUGAUUAUAACACAGAGCGUUCAUUAGUAUUACUAGUAGGUAAAAGAAGAGCUUUGUUGGAUUACUUGAAG
Fsp-3-1   AUCUUCGCACAACACGGUGAUGCUAAAAACACUGGAAAAGCAGAAGGUCAGAUUGCAUUGUUCACUUACAGAAUUUCACACUUAACUGAACACUUGAAAAAAAAUCGUCACGAUUAUAACACUGAGCGUUCUCUUGUACUAUUAGUAGGUAAAAGAAGAGCUUUGUUGGACUACUUGAAA
env-1577  AUCUUCGCACAACACGGAGAAAAAACAAACACAGGUAAAGCAGAAGCUCAAAUUGCUUUGUUCACUUUCAGAAUUAGCCACCUAACAGAACAUUUGAAAAAAAAUCGUCACGAUUAUAACACAGAGCGUUCAUUAGUAUUGCUAGUAGGAAAAAGAAGAUCUUUGUUGGAUUACUUGAAA
env-1579  AUCUUCGCACAACACGGAGAAAAAACAAACACAGGUAAAG............................................................................................................................................
Fsp-4-1   AUCUUCGCACAACACGGAGAGAAAACAAACACAGGAAAAACAGAAGGCCAAAUUGCUUUGUUUACGUUCAGAAUUACGCACCUUACUGAACAUUUGAAAAAAAAUCGUCAUGAUUAUAACACAGAACGUUCAUUAGUAUUGUUAGUAGGAAAGAGAAGAUCUUUGUUAGAUUACUUGAAA
env-1580  AUUUUCGCAAAACAUGGGAAAUCAGCAACAGAUACAGGAUCAGCUGAAGGGCAAAUCGCAUUAUUCACUUACCGUAUCAACCAUUUAACUGGUCACCUAAGAAAAAAUAUCCACGAUUUUAACACAGAGAGAUCUUUAGUUAUGUUAGUAGG............................
env-1581  AUCUUCGCAAAACACGGAAAAUCUGCACAAGACACAGGAACUGCUGAAGGACAAGUUGCUCUAUUCACAUUUAGAAUUAACCACUUAACUCAGCACUUG.................................................................................
Wma-1-1   AUUUUCGCAAAACAUGGAAAAUCUGCUCAAGACACCGGAUCAGCAGAAGGUCAGAUCGCGUUAUUCACACACAGAAUCAACCACUUGACUCAACACUUGAAAAGAAAUCAAAAAGAUUUUAAUACCGAGAGAUCUUUGGUGAAAUUAGUAGGAAAGCGUAAAGCUUUAUUGGAUUAUUUG
Wvi-1-1   AUUUUCGCAAAACAUGGAAAAUCUGCUCAAGACACCGGAUCACCAGAAGGUCAGAUUGCAUUAUUCACGUACAGAAUUAACCACCUGACUCAACACUUGAAAAAAAAUCAUAAAGAUUACAAUACAGAGAGAUCUUUGGUGAAAUUAGUAGGAAAACGUAAAGCUUUAUUGGAUUAUUUG
Orh-1-1   AUUUUUGAAAAGUACGGAGAAAACGCUAACGAUACAGGAUCUCCAGAGAGCCAAAUCGCACUUUUCACUUACAGAAUCAACCACCUAUCUGAGCACUUAAAAGUGAACAGAAAAGAUUUCAACACUGAAAGAGCUUUGGUGAAAUUGGUAGGUAAAAGAAAAAGUUUACUAGAUUACUUA
env-1582  AUCUUCGAAAAACACGGAGGAAAAGCAGAAAACACAGGUUCUGCAGAAGGGCAGGUAGCACUUUUUACCUACCGUAUUAACCACCUUACUUCCCACCUUAAAAAGAACCACAAAGAUUACGCUACGGAGCGUUCCCUUGUGAAAAUGGUAGGUAAGAGAAAAAGAUUGUUGGAUUAUCUU
env-1583  AUCUUCGCUAAGCACGGAGGAAAUGCAGCUAACACAGGUUCUGCAGAAGGGCAAAUUGCUCUAUUUACUUUUAGAAUUAACCACCUUUCAGGUCACUUGAAAAAAAAUCACAAAGAUUAUGCUACAGAAAAAUCUCUAGUGAAAUUGGUAGGUAAAAGAAAAAGACUGUUAGAUUAUCUU
env-1584  ....................................................................................................................................................................................
env-1585  AUCUUCGCUAAACACGGAGGAAACGCUGCAAACACAGGUUCUGCAGAAGGACAAAUCGCUUUAUUCACAUUUAGAAUUAACCAUUUGUCUGGUCACUUAAAGAAAAACCACAAAGAUUAUGCUACUGAGAAAUCUCUAGUUAAAUUAGUAGGUAAAAGAAAAAGAUUGUUAGAUUAUCUU
env-1586  AUUUUCGCAAAACACGGAGGAAAAGCAGAAAACACAGGUUCUGCUGAAGGUCAGAUUGCAUUAUUCACUUUCAGAAUCAAUCAUCUUACAGGUCAUUUGAAAGCAAAUCAUAAAGAUUAUGCUACCGAAAGAUCGCUGGUGAAACUGGUAGGAAAAAGAAAAAGCCUUCUGGAUUAUCUU
env-1587  AUUUUCGCAAAACACGGAGGAAAAGCAGAAAACACAGGUUCUGCUGAAGGUCAGAUUGCAUUAUUCACUUUCAGAAUCAAUCAUCUUACAGGUCAUUUGAAAGCAAAUCAUAAAGAUUAUGCUACCGAAAGAUCGCUGGUGAAACUGGUUGGAAAAAGGAAAAGCCUUCUGGAUUAUCUU
env-1588  ....................................................................................................................................................................................
Ebr-1-1   UUAUUCGCAAAGCACGGUAAAUCUGCAACAGAUACAGGAUCUGCAGAAGGACAAAUCGCAUUAUUCACAUUUCGUAUCAACCAUUUAACUGGUCACUUAAGAAAGAAUAUUCACGAUUACAACACAGAAAGAUCAUUAGUGAUGUUAGUAGGUAAGCGUAAAGCGUUAUUAAACUACUUA
env-1589  UUAUUCGCAAAGCAUGGUAAAUCUGCAACAGAUACAGGAUCUGCAGAAGGGCAAAUCGCAUUAUUCACUUACAGAAUUAACCACUUAACUGGUCACUUGAAAAAAAAUCACAAAGAUUUUAAUACAGAAAGAUCAUUAGUUAUGUUAGUUGGUAAGCGUAAAGCGUUAUUAAACUACUUA
env-1594  UUAUUCGCAAAGCAUGGUAAAUCUGCAACAGAUACAGGAUCUGCAGAAGGGCAAAUCGCAUUAUUCACUUACAGAAUUAACCACUUAACUGGUCACUUGAAAAAAAAUCACAAAGAUUU.............................................................
Efa-1-1   UUAUUCGCAAAGCAUGGUAAAUCUGCAACAGAUACAGGAUCUGCAGAAGGGCAAAUCGCAUUAUUCACUUACAGAAUUAACCACUUAACUGGUCACUUGAAAAAAAAUCAUAAAGAUUUUAAUACAGAAAGAUCAUUAGUUAUGUUAGUUGGUAAGCGUAAAGCGUUAUUAAACUACUUA
env-1595  UUAUUCG.............................................................................................................................................................................
Bzo-1-1   AUUUUCGCAAAACACGGAGGAAAAGCUGAAAACACAGGUUCAGCAGAAGGGCAAAUUGCACUUUUUACUUACAGAAUUAACCACCUUACUCAGCACUUGAGAGCAAAUCAUAAAGAUUAUGCUACAGAGAGAGCACUGGUAAAAUUGGUAGGUAAGAGAAAAAGCCUUUUGGAUUAUCUA
Bzo-2-1   AUUUUCGCAAAACACGGAGGAAAAGCUGAAAAUACAGGUUCAGCAGAAGGGCAAAUUGCACUUUUUACUUACAGAAUUAACCACCUUACUCAGCACUUGAGAGCAAAUCAUAAAGAUUACGCUACAGAGAGAGCACUGGUAAAAUUGGUAGGUAAGAGAAAAAGCCUUUUGGAUUAUCUA
env-1596  AUUUUUAAGAAAUAUGGGGCGUCCGAAACAAAUACUGGAUCAACCGAAGGCCAAAUCGCUCUCUUUACGCAUCGAAUCAAUCAUUUAUCCAAUCACUUGAAACAAAAUCACAAAGACUUUAAUACCGAAAGAUCUCUAGUAAAGCUCGUUGGUAAACGAAGAAGUCUUCUAGAUUAUUUA
env-1597  AUUUUUAAGAAAUACGGUUCUUCCGAAACAAAUACUGGGUCGACCGAAGGCCAAAUCGCUCUCUUUACGCAUCGGAUCAAUCAUUUAUCCAAUCAUCUCAAAACAAAUCACAAAGACUUUAAUACCGAAAGAUCGCUUGUAAAGCUCGUUGGUAAACGAAGAAGUCUUCUCGAUUAUUUG
env-1598  ....................................................................................................................................................................................
env-1599  AUAUUUAAGAAAUACGGUUCUUCCGAAACAAAUACUGGAUCGACCGAAGGCCAAAUCGCUCUCUUUACUCAUCGGAUCAAUCAUUUAUCCAGUCAUCU..................................................................................
env-1600  AUUUUUAAGAAAUACGGUUCUUCCGAAACAAAUACUGGAUCGACCGAAGGCCAAAUAGCUCUCUUUACACAUCGGAUCAAUCAUUUAUCCAGUCAUCUUAGAACAAAUCACAAAGACUUUAAUACCGAAAGAUCGCUUGUAAAGCUCGUUGGUAAAAGAAGAAGUCU.............
env-1601  AUUUUUAAGAAAUACGGUUCUUCCGAAACAAAUACUGGAUCGACCGAAGGCCAAAUCGCUCUCUUUACACAUCGGAUCAAUCAUUUAUC..............................................................................................
env-1602  AUAUUUAAGAAAUACGGUUCUUCCGAAACAAAUACUGGAUCGACCGAAGGCCAAAUCGCUCUCUUUACGCAUCGGAUCAAUCAUUUAUCUAAUCAUCUUAAAACAAAUCAUAAAGACUUUAAUACCGAAAGAUCGCUUGUAAAGCUCGUUGGUAAACGAAGAAGUCUUCUCGAUUAUUUG
env-1603  AUUUUUAAAAAAUACGGUUCGUCCGAAACAGAUACCGGAUCAACCGAAGGCCAAAUCGCCCUCUUUACGCAUCGCAUCAAUCAUUUAUCCAAUCACCUGAAAAAAAAUCACAAAGACUUUAAUA.......................................................
env-1604  AUUUUUAAGAAAUACGGUUCAUCCGAAACAAAUACUGGAUCUACCGAAGGCCAAAUCGCUCUCUUUACGCAUCGAAUCAAUCACUUAUCCAAUCACCUGAAAAAAAAUCACAAAGACUUUAAUACCGAAAGAUCGCUGGUAAAGAUCGUUGGUAAACGAAGAAGUCUUCUCGAUUAUUUG
env-1605  AUUUUUAAGAAAUACGGUUCAUCCGAAACAAAUACUGGAUCUACUGAAGGCCAAAUCGCUCUCUUUACGCAUCGAAUCAAUCACUUAUCCAAUCACCUGAAAAAAAAUCACAAAGACUUUAAUACCGAAAGAUCGCUGGUAAAGCUCGUUGGUAAACGAAGAAGUCUUCUCGAUUAUUUA
env-1606  AUUUUUAAGAAAUACGGUUCAUCCGAAACAAAUACUGGAUCUACCGAAGGCCAAAUCGCUCUCUUUACACAUCGGAUCAAUCAUUUAUCGAAUCACCUAAAAAAAAAAUCACAAAGACUUUAAUACCGAAAGAUCGCUGGUAAAGCU................................

          ....................................................................................................................................................................................
          ....................................................................................................................................................................................
          AUYUUYR•RAARCACGG•RRR•••R•RR••RAYAC•GG••C•RC•GAAGG•CAAAUYGC•YURUUYAC•Y•Y•G•AUYRAYCAYYUR•C•RA•CAYYURAAAAR•AAYCR•AARGAUU•YAAYAC•GAR•G•UC•YURGU•••RYURGU•GG•AAR•G•RG••••YU•YU•GAYUAYYU•
```

```
env-1607 AUUUUUAAGAAAUACGGUUCAUCCGAAACAAAUACUGGAUCUACCGAAGGCCAAAUCGCUCUCUUUACGCAUCGGAUCAAUCAUUUAUCCAAUCACGUGAAAAAAAAUCACAAAGACUUUAAUACCGAAAGAUCGCUGGUAAAGCUCGUUGGUAAACGAAGAAGUCUUCUCGAUUAUUUG
env-1608 AUUUUUAAGAAAUACGGUUCAUCCGAAACAAAUACUGGAUCUACUGAAGGCCAAAUCGCUCUCUUUACGCAUCGGAUCAAUCAUUUAUCCAAUCACCUGAAAAAAAAUCACAAAGACUUUAAUACCGAAAGAUCGCUGGUAAAGCUCGUUGGUAAACGAAGAAGUCUUCUCGAUUAUUUG
env-1609 AUUUUUAAGAAAUACGGUUCUUCCGAAACAAAUACCGGAUCUACCGAAGGCCAAAUCGCUCUCUUUACACAUCGGAUCAAUCAUUUAUCCAAUCACCUCAAAAAAAAUCACAAAGACUUUAAUACCGAAAGAUCGCUUGUUAAGCUCGUUGGUAAACGAAGAAGUCUUCUCGAUUAUUUG
env-1610 AUUUUUAAGAAAUACGGUUCUUCCGAAACAAAUACCGGAUCUACCGAAGGCCAAAUCGCUCUCUUUACACAUCGGAUCAAUCAUUUAUCCAAUCACCUCAAAAAAAAUCACAAAGACUUUAAUACCGAAAGAUCGCUUGUUAAGCUCGUUGGUAAACGAAGAAGUCUUCUCGAUUAUUUG
env-1611 AUUUUUAAGAAAUACGGUUCUUCCGAAACAAAUACCGGAUCUACCGAAGGCCAAAUCGCUCUCUUUACACAUCGGAUCAAUCAUUUAUCCAAUCACCUCAAAAAAAAUCACAAAGACUUUAAUACCGAAAGAUCGCUUGUUAAGCUCGUUGGUAAACGAAGAAGUCUUCUCGAUUAUUUG
env-1612 AUUUUUAAGAAAUACGGUUCUUCCGAAACAAAUACCGGAUCUACAGAAGGCCAAAUCGCUCUCUUUACACAUCGGAUCAAUCAUUUAUCCAAUCACCUCAAAAAAAAUCACAAAGACU...........................................................................
env-1613 AUUUUUAAGAAAUACGGUUCUUCCGAAACAAAUACCGGAUCUACCGAAGGCCAAAUCGCUCUCUUUACACAUCGGAUCAAUCAUUUAUCCAAUCACCUCAAAAAAAAUCACAAAGACUUUAAUACCGAAAGAUCGCUUGUUAAGCUCGUUGGUAAACGAAGAAGUCUUCUCGAUUAUUUG
env-1614 AUUUUUAAGAAAUACGGUUCUUCCGAAACAAAUACCGGAUCUACCGAAGGCCAAAUCGCUCUCUUUACGCAUCGGAUCAAUCAUUUAUCCAAUCA..............................................................................................................
env-1615 AUUUUUAAGAAAUACGGUUCUUCCGAAACAAAUACCGGAUCUACCGAAGGCCAAAUCGCUCUCUUUACACAUCGGAUCAAUCACUUAUCCAAUCACCUCAAAAAAAAUCACAAAGACUUUAAUACCGAAAGAUCGCUUGUUAAGCUCGUUGGUAAACGAAGAAGUCUUCUCGAUUAUUUG
env-1616 ..............................................................................................................................................................................................................
env-1617 AUUUUUAAGAAAUACGGUUCUUCCGAAACAAAUACCGGAUCUACCGAAGGCCAAAUCGCUCUCUUUACACAUCGGAUCAAUCAUUUAUCCAAUCACCUCAAAAAAAAUCACAAAGACUUUAAUACCGAAAGAUCGCUUGUUAAGCUCGUUGGUAAACGAAGAAGUCUUCUCGAUUAUUUG
env-1618 AUUUUUAAGAAAUACGGUUCUUCCGAAACAAAUACCGGAUCUACCGAAGGCCAAAUCGCUCUCUUUACACAUCGGAUCAAUCAUUUAUC..................................................................................................................
env-1619 AUUUUUAAGAAAUACGGUUCUUCCGAAACAAAUACCGGAUCUACCGAAGGCCAAAUCGCUCUCUUUACACAUCGGAUCAAUCAUUUAUCCAAUCACCUCAAAAAAAAUCACAAAGACUUUAAUACCGAAAGAUCGCUUGUUAAGCUCGUUGGUAAACGAAGAAGUCUUCUCGAUUAUUUG
env-1620 AUUUUUAAGAAAUACGGUUCUUCCGAAACAAAUACCGGAUCUACCGAAGGCCAAAUCGCUCUCUUUACACAUCGGAUCAAUUAUUUAUCCAAUCACCUCAAAAAAAAUCACAAAGACUUUAAUACCGAAAGAUCGCUUGUUAAGCUCGUUGGUAAACGAAGAAGUCUUCUCGAUUAUUUG
env-1621 AUUUUUAAAAAAUACGGGGCGUCCGAAACAAAUACUGGAUCAACUGAAGGCCAAAUCGCUCUCUUUACGCAUCGGAUCAACCAUUUAUCCAAUCACCUAAAAAAAAAUCACAAAGACUUUAAUACCGAAAGAUCACUAGUAAAGAUCGUUGGUAAACGAAGAAGUCUUCUAGAUUAUUUA
env-1622 AUCUUCGCAAAAUUUGGCGGAGAAGCAAAGAACACAGGUUCUACAGAAGGACAAAUUGCAUUGUUCACACACAGAAUCAAUCACCUUACGGGUCACUUAAAAGCGAACCGUCACGAUUACAACACCGAAAGAGCACUUGUAAAAUUAGUAGGUAAAAGAAGAAGUCUUUUAGACUAUUUA
env-1623 AUCUUCGCUAAACACGGAGGAAAAGCAGAAAACACAGGUUCUGCUGAAGGACAAAUAGCAUUGUUCACAUUUAGAAUCUCUCACUUAACAGAGCACUUGAAAAAAAAUCGUCACGAUUACAAUACGGAGCGUUCGUUAGUACUUUUGGUAGGUAAAAGAAGAAGUCUUCUUGAUUACUUG
env-1624 AUCUUCGCUAAACACGGAGGAAAAGCAGAAAACACUGGGUCUGCAGAAGGACAAAUUGCAUUGUUCACAUUUAGAAUCUCUCACUUAACAGAGCACUUGAAAAAAAAUCGUCAUGAUUACAAUACUGAGCGUUCGUUGGUACUUUUAGUAGGUAAAAGAAGAUCUCUUCUUGACUAC...
env-1625 AUCUUCGCUAAACACGGAGGAAAAGCAGAAAACACAGGUUCUGCUGAAGGACAAAUAGCAUUGUUCACAUUUAGAAUCUCUCACUUAACAGAGCACUUGAAAAAAAAUCGUCACGAUUACAAUACGGAGCGUUCGUUAGUACUUUUGGUAGGUAAAAGAAGAAGUCUUCUUGAUUACUUG
env-1626 AUCUUCGCUAAACACGGAGGAAAAGCAGAAAACACUGGGUCUGCAGAAGGACAAAUUGCAUUGUUCACAUUUAGAAUCUCUCACUUAACAGAGCACUUGAAAAAAAAUCGUCAUGAUUACAAUACUGAGCGUUCGUUGGUACUUUUAGUAGGUAAAAGAAGAUCUCUUCUUGACUACUUG
env-1627 AUCUUCGCUAAACACGGAGGAAAAGCAGAAAACAC...........................................................................................................................................................................
env-1628 AUCUUCGCUAAACACGGAGAAAAAACAAACACAGGAAAGGCAGAAGCUCAAAUCGCUUUAUUCACUUUCAGAAUUUCUCACCUUACAGAGCACUUGAAAAAAAAUCGUCAUGAUUACAACACCGAGCGUUCAUUAGUAUUGUUGGUAGGUAAAAGAAGAAGUUUAUUAGACUACUUAAAG
env-1629 AUCUUCGCUAAACACGGAGAAAAAACAAACACAGGAAAGGCAGAAUCACAAAUCGCUUUAUUCACUUUCAGAAUUAGCCACUUGACUGAGCACUUGAAAAAAAAUCGUCACGAUUAUAACACAGAGCGUUCACUAGUAUUGCUAGUAGGUAAAAGAAGAUCUUUGUUGGAUUACUUGAAG
env-1630 AUCUUCGCUAAACAC...............................................................................................................................................................................................
env-1631 AUCUUCGCUAAACACGGAGGAGCUGCAACAAACACAGGUUCAGCAGAAGGACAAAUCGCAUUGUUUACACACAGAAUCACUCACCUAACGGAACACUUGAAAAAAAAUCGUCACGAUUAUAAUACGGAGCGUUCAUUAGUUUUGUUGGUAGGUAAAAGAAGAUCUUUACUAGAUUACUUG
env-1632 AUCUUCGCUA....................................................................................................................................................................................................
env-1633 AUCUUCG.......................................................................................................................................................................................................
env-1634 AUCUUCGCUAAACACGGAGGAAAAGCUGAAAACACAGGUUCUACAGAAGGUCAAGUAGCACUUUUCACUUUUAGAAUCUCUCACUUAACUGAGCACUUGAAAAAAAAUCGUCACGAUUAUAACACAGAGCGUUCAUUAGUUCUUUUGGUAGGAAAAAGAAGAAGUCUUCUUGAUUACUUG
env-1635 ..............................................................................................................................................................................................................
Faq-1-1  AUCUUCGCUAAACACGGAGGAAAAGCUGAAAACACAGGUUCUACAGAAGGUCAAGUAGCACUUUUCACUUUUAGAAUCUCUCACUUAACAGAGCACUUGAAAAAAAAUCGUCACGAUUAUAACACUGAGCGUUCAUUAGUUCUUUUAGUAGGUAAAAGAAGAUCUCUUCUUGACUACUUA
env-1636 AUCUUCGCUAAACACGGAGGAAAAGCUGAAAACACAGGUUCUACAGAAGGUCAAGUAGCACUUUUCACUUUUAGAAUCUCUCACUUAACAGAGCACUUGAAAAAAAAUCGUCACGAUUAUAACACAGAGCGUUCAUUAGUUCUUUUAGUAGGUAAAAGAAGAUCUCUUCUUGACUACUUA
env-1637 AUCUUUGCUAAACACGGAGGAAAAGCAGAAAACACUGGUUCUGCUGAAGGGCAAAUCGCAUUGUUCACAUUUAGAAUCUCUCACUUAACGGAGCACUUGAAAAAAAAUCGUCACGAUUACAAUACCGAGCGUUCAUUAGUUCUUUUAGUAGGUAAAAGAAGAAGUCUUCUUGACUACUUG
env-1638 AUUUUCCAAAAAUAUGCCGGUUCUGAAACUAACACAGGUUCAGCAGAAGGGCAAAUAGCUCUAUUUACACACCGUAUCAACCACUUAACAGAACACUUACGUAAAAAUCGUAAAGAUUAUAAUACAGAGCGUUCGUUAGUAAAACUUGUUGGUAAGCGUAGAAGCUUACUUGAUUAUUUG
env-1639 AUUUUCCAAAAAUAUGCCGGUUCUGAAACUAACACAGGUUCAGCAGAAGGGCAAAUAGCUCUAUUUACACACCGUAUCAACCACUUAACAGAACACUUACGUAAAAAUCGUAAAGAUUAUAAUACGGAGCGUUCGUUAGUAAAACUUGUUGGUAAGCGUAGAAGCUUACUUGAUUAUUUG
env-1640 AUUUUCCAAAAAUAUGCCGGUUCUGAAACUAACACAGGUUCAGCAGAAGGGCAAAUUGCUCUAUUUACACACCGUAUCAACCACUUAACAGAACACUUACGUAAAAAUCGUAAAGAUUAUAAUACAGAGCGUUCGUUAGUAAAACUUGUUGGUAAGCGUAGAAGCUUACUUGAUUAUUUG
env-1641 AUUUUCCAAAAAUAUGCCGGUUCUGAAACUAACACAGGUUCAGCAGAAGGGCAAAUUGCUCUAUUUACACACCGUAUCAACCACUUAACAGAACACUUACGUAAAAAUCGUAAAGAUUAUAAUACAGAGCGUUCGUUAGUAAAACUUGUUGGUAAGCGUAGAAGCUUACUUGAUUAUUUG
env-1642 AUCUUCCAAAAAUAUGCCGGUUCUGAAACUAACACAGGUUCAGCAGAAGGGCAAAUUGCUCUAUUUACACACCGUAUCAACCACUUAACAGAACACUUACGUAAAAAUCGUAAAGAUUAUAAUACAGAGCGUUCGUUAGUAAAACUUGUUGGUAAGCGUAGAAGCUUACUUGAUUAUUUG
Csp-6-1  AUUUUCAAAAAGUACGCAGGUUCAGAAACCAACACUGGUUCAGCAGAAGGGCAAAUCGCUCUUUUUACCCUUAGAAUCAAUCACCUUACAGAACACUUGAAAGUAAAUCGCAAAGAUUUUAAUACCGAGCGUUCCUUGGUGAAAUUAGUAGGUAGACGCAGAAGUUUGCUCGAUUAUUUG
env-1643 ..............................................................................................................................................................................................................
env-1644 ..............................................................................................................................................................................................................
env-1645 AUUUUCAAAAAGUACGCAGGUUCAGAAACCAACAC...........................................................................................................................................................................
Coc-2-1  AUUUUCAAAAAGUACGCAGGUUCAGAAACCAACACUGGUUCAGCAGAAGGGCAAAUCGCUCUUUUUACCCUUAGAAUCAAUCACCUUACAGAACACUUGAAAGUAAAUCGCAAAGAUUUUAAUACCGAGCGUUCUUUGGUAAGAUUGGUAGGUAAACGUAGAAGUUUGCUCGACUAUUUG
env-1646 NNNNNNNNNNNNNNNNNNNNNNNNNACUUGAAAGUAAAUCGCAAAGAUUUUAAUACCGAGCGUUCCUUGGUGAAAUUAGUAGGUAGACGCAGAAGUUUGCUCGAUUAUUUGAAGAACACAGAUAUCGAAAGAUACCGUGCGAUUAUCAAAGAGCUAAACAUCAGAAAAUAAGCAAUAG
Csp-9-1  AUUUUCAAAAAGUACGCAGGUUCAGAAACCAACACUGGUUCAGCAGAAGGGCAAAUCGCUCUUUUUACCCUUAGAAUCAAUCACCUUACAGAACACUUGAAAGUAAAUCGCAAAGAUUUUAAUACCGAGCGUUCCUUGGUGAAAUUAGUAGGUAGACGCAGAAGUUUGCUCGAUUAUUUG
env-1647 AUUUUCA.......................................................................................................................................................................................................
env-1649 ..............................................................................................................................................................................................................
env-1650 AUUUUCAAAAAGUACGCAGGUUCAGAAAACAACACUGGUUCAGCAGAAGGGCAAAUCGCUCUUUUUACCCUUAGAAUUAAUCACCUUACAGAACACUUAAAGGUAAAUCGCAAAGAUUUUAAUACCGAGCGUUCCUUGGUGAAAUUAGUAGGUAGACGCAGAAGUUUACUCGAUUAUUUG
env-1651 AUUUUCAAAAAGUACGCAGGUUCAGAAACCAACACUGGUUCAGCAGAAGGGCAAAUCGCUCUUUUUACCCUUAGAAUCAAUCACCUUACAGAACACUUGAAAGUAAAUCGCAAAGAUUU......................................................................................
env-1652 AUUUUCA.......................................................................................................................................................................................................
env-1655 AUUUUCAAAAAGUACGCAGGUUCAGAAACCAACACUGGUUCAGCAGAAGGGCAAAUCGCUCUUUUUACCCUUAGAAUCAAUCAUCUUACAGAACACUUGAAAGUAAAUCGCAAAGAUUUUAAUACCGAGCGUUCCUUGGUGAAAUUAGUAGGUAGACGCAGAAGUUUGCUCGAUUAUUUG
env-1656 ..............................................................................................................................................................................................................
env-1657 AUUUUCAAAAAGUACGCAGGUUCAGAAACUAACACUGGUUCAGCAGAAGGGCAAAUCGCUCUUUUUACCCUUAGAAUCAAUCACCUUACAGAACACUUAAAGGUAAAUCGCAAAGAUUUUAAUACCGAGCGUUCCUUGGUGAAAUUAGUAGGUAGACGCAGAAGUUUGCUCGAUUAUUUG
Coc-3-1  AUUUUCAAAAAGUACGCAGGUUCAGAAACCAACACUGGUUCAGCAGAAGGGCAAAUCGCUCUUUUUACCCUUAGAAUCAAUCACCUUACAGAACACUUGAAAGUAAAUCGCAAAGAUUUUAAUACCGAGCGUUCCUUGGUGAAAUUAGUAGGUAGACGCAGAAGUUUGCUCGAUUAUUUG
Coc-1-1  AUUUUUAAAAAGUACGCAGGUUCAGAAACCAACACAGGUUCAGCAGAAGGGCAAAUAGCUCUUUUUACCCUUAGAAUCAAUCACCUUACAGAACACUUGAAAGUAAAUCGCAAAGAUUUCAAUACCGAGCGUUCUUUGGUAAGAUUAGUAGGUAAACGCAGAAGUUUGCUCGACUAUUUG
Csp-7-1  AUUUUCAAAAAGUACGCAGGUUCAGAAACCAACACUGGUUCAGCAGAAGGGCAAAUCGCUCUUUUUACCCUUAGAAUCAAUCAUCUUACAGAACACUUGAAAGUAAAUCGCAAAGAUUUUAAUACCGAGCGUUCCUUGGUGAAAUUAGUAGGUAGACGAAGAAGUUUGCUCGAUUAUUUG
Csp-10-1 AUUUUCAAAAAGUACGCAGGUUCAGAAACCAACACUGGUUCAGCAGAAGGGCAAAUCGCUCUUUUUACCCUUAGAAUCAAUCACCUUACAGAACACUUAAAGGUAAAUCGCAAAGAUUUUAAUACCGAGCGUUCCUUGGUGAAAUUAGUAGGUAGACGCAGAAGUUUGCUCGAUUAUUUG
Csp-2-1  AUUUUCAAAAAGUACGCAGGUUCAGAAACCAACACUGGUUCAGCAGAAGGGCAAAUCGCUCUUUUUACCCUUAGAAUCAAUCACCUUACAGAACACUUGAAAGUAAAUCGCAAAGAUUUUAAUACCGAGCGUUCCUUGGUGAAAUUAGUAGGUAGACGAAGAAGUUUGCUCGAUUAUUUG
env-1658 AUCUUCGCUAAACACGGAGGAAAAGCAGAAAACACAGGAUCUGCAGAAGGGCAAAUCGCAUUGUUCACAUUCAGAAUCUCUCACUUGACUGAGCACCUGAAAAAAAAUCGUCACGAUUACAACACU.............................................................................
env-1659 ..............................................................................................................................................................................................................
env-1660 AUCUUCGCUAAACACGGAGGAAAAGCAGAAAACACUGGAUCUGCAGAAGGUCAAAUCGCUUUAUUCACACACAGAAUUAGCCAUUUAACUGAGCACUUGAAAAAAAAUCGUCACGAUUAUAACACAGAGCGUUCUCUAGUAUUAUUAGUAGGUAAAAGAAGAGCUUUGUUGGAUUACUUG
env-1661 AUCUUCGCUAAACACGGAGGAAAAGCAGAAAACACUGGAUCUGCAGAAGGUCAAAUCGCUUUAUUCACUCACAGAAUUAGCCAUUUAACUGAGCACUUGAAAAAAAAUCGUCANGAUUAUAACACAGAGCGUUCUCUAGUAUUAUUAGUAGGUAAAAGAAGAGCUUUGUUGGAUUACUUG
env-1662 AUCUUCGCUAAACACGGA............................................................................................................................................................................................
env-1663 AUCUUCGCUAAACACGGAGGAAAAGCAGAAAACACUGGGUCUGCAGAAGGUCAAAUCGCUUUAUUCACACACAGAAUUAGCCAUUUAACUGAGCACUUGAAAAAAAAUCGUCACGAUUAUAACACAGAGCGUUCACUAGUAUUAUUAGUAGGUAAAAGAAGAGCUUUGUUGGAUUACUUG
env-1664 AUUUUCAAAAAGUACGCAGGUUCAGAAACCAACACAGGUUCAGCAGAAGGGCAAAUCGCACUUUUCACCCUCAGAAUCAAUCACCUUACAGAGCACUUAAAAGUGAAUCGCAAGGAUUUUAAUACUGAGCGUUCGUUAGUAAAGCUCGUAGGUAAGCGUAGAAGUUUGCUAGACUAUUUG
env-1665 AUUUUCAAAAAGUACGCAGGUUCAGAAACCAACACAGGUUCAGCAGAAGGGCAAAUCGCACUUUUCACCCUCAGAAUCAAUCACCUUACUGAGCACUUAAAAGUGAAUCGCAAGGACUUUAACACCGAGCGUUCGUUAGUAAAGCUCGUAGGUAAGCGUAGAAGUUUGCUAGACUAUUUG
env-1666 AUUUUCAAAAAGUACGCAGGUUCAGAAACCAACACAGGUUCAGCAGAAGGGCAAAUCGCACUUUUCACCCUCAGAAUCAAUCACCUUACAGAGCACUUAAAAGUGAAUCGCAAGGACUUUAACACUGAGCGUUCAUUAGUAAAGCUCGUAGGUAAGCGUAGAAGUUUGCUAGACUAUUUG
env-1668 AUUUUCAAAAAGUACGCAGGUUCAGAAACCAACACAGGUUCAGCAGAAGGGCAAAUCGCACUUUUCACCCUUAGAAUCAAUCACCUUACAG..................................................................................................................
env-1669 AUUUUCAAAAAGUACGCAGGUUCAGAAACCAACACAGGUUCAGCAGAAGGGCAAAUCGCACUUUUCACCCUCAGAAUCAAUCACCUUACUGAGCACUUAAAAGUGAAUCGCAAGGACUUUAACACUGAGCGUUCGUUAGUAAAGCUCGUAGGUAAGCGUAGAAGUUUGCUAGACUAUUUG
env-1673 AUUUUCAAAAAGUACGCAGGUUCAGAAACCAACACAGGUUCAGCAGAAGGGCAAAUCGCACUUUUCACCCUCAGAAUCAAUCACCUUACUGAGCACUUAAAAGUGAAUCGCAAGGACUUUAACACUGAGCGUUCGUUAGUAAAGCUCGUAGGUAAGCGUAGAAGUUUGCUAGACUAUUUG
env-1674 AUUUUCAAAAAGUACGCAGGUUCAGAAACCAACACAGGUUCAGCAGAAGGGCAAAUCGCACUUUUCACCCUCAGAAUCAAUCACCUUACUGAGCACUUAAAAGUGAAUCGCAAGGACUUUAACACCGAGCGUUCGUUAGUAAAGCUCGUAGGUAAGCGUAGAAGUUUGCUAGACUAUUUG
env-1676 AUUUUCAAAAAGUACGCAGGUUCAGAAACCAACACAGGUUCAGCAGAAGGGCAAAUCGCACUUUUCACCCUUAGAAUCAAUCACCUUACAGAGCACUUAAAAGUGAAUCGCAAGGACUUUAACACUGAGCGUUCGUUAGUAAAGCUCGUAGGUAAGCGUAGAAGUUUGCUAGACUAUUUG
env-1677 AUUUUCAAAAAGUACGCAGGUUCAGAAACCA...............................................................................................................................................................................
env-1679 AUUUUCAAAAAGUACGCAGGUUCAGAAACCAACACAGGUUCAGCAGAAGGGCAAAUCGCACUUUUCACCCUCAGAAUCAACCACCUUACUGAGCACUUAAAAGUGAAUCGCAAGGACUUUAACACUGAGCGUUCGUUAGUAAAGCUCGUAGGUAAGCGCAGAAGUCUGCUAGACUAUUUG
env-1680 AUUUUCA.......................................................................................................................................................................................................
env-1681 AUUUUCAAAAAGUACGCAGGUUCAGAAACCAACACAGGUUCAGCAGAAGGGCAAAUCGCACUUUUCACCCUCAGAAUUAAUCACCUUACUGAGCACUUAAAAGUGAAUCGCAAGGACUUUAACACUGAGCGUUCGUUAGUAAAGCUCGUAGGUAAGCGUAGAAGUUUGCUAGACUAUUUG
env-1682 AUUUUCAAAAAGUACGCAGGUUCAGAAACCA...............................................................................................................................................................................
env-1683 AUUUU.........................................................................................................................................................................................................
env-1684 AUUUUCAAAAAGUACGCAGGUUCAGAAACCAACACAGGUUCAGCAGAAGGGCAAAUCGCACUUUUCACCCUCAGAAUCAAUCACCUUACUGAGCACUUAAAAGUGAAUCGCAAGGACUUUAACACUGAGCGUUCGUUAGUAAAGCUCGUAGGUAAGCGUAGAAGUUUGCUAGACUAUUUG
env-1685 AUUUUCAAAAAGUACGCAGGUUCAGAAACCA...............................................................................................................................................................................
env-1686 AUUUUCAAAAAGUACGCAGGUUCAGAAACCAACACAGGUUCAGCAGAAGGGCAAAUCGCACUUUUCACCCUUAGAAUCAAUCACCUUACUGAGCACUUAAAAGUGAAUCGCAAGGACUUUAACACCGAGCGUUCGUUAGUAAAGCUCGUAGGUAAGCGCAGAAGUCUGCUUGACUAUUUG
env-1687 AUUUUCAAAAAGUACGCAGGUUCAGAAACCAACACAGGUUCAGCAGAAGGGCAAAUCGCACUUUUCACCCUCAGAAUCAAUCACCUUACAGAGCACUUAAAAGUGAAUCGCAAGGACUUUAACACUGAGCGUUCGUUAGUAAAGCUCGUAGGUAAGCGUAGAAGUUUGCUAGACUAUUUG
env-1688 AUUUUCAAAAAGUACGCAGGUUCAGAAACCAACACAGGUUCAGCAGAAGGGCAAAUCGCACUUUUCACCCUCAGAAUCAAUCACCUUACUGAGCACUUAAAAGUGAAUCGCAAGGACUUUAACACCGAGCGUUCGUUAGUAAAGCUCGUAGGUAAGCGUAGAAGUUUGCUAGACUAUUUG
env-1689 AUUUUCAAAAAGUACGCAGGUUCAGAAACCAACACAGGUUCAGCAGAAGGGCAAAUCGCACUUUUCACCCUCAGAAUCAAUCACCUUACAGAGCACUUAAAAGUGAAUCGCAAGGACUUUAAUACUGAGCGUUCGUUAGUAAAGCUCGUAGGUAAGCGUAGAAGUUUGCUAGACUAUUUG
env-1691 AUUUUCAAAAAGUACGCAGGUUCAGAAACCAACACAGGUUCAGCAGAAGGGCAAAUCGCACUUUUCACCCUCAGAAUCAAUCACCUUACUGAGCACUUAAAAGUGAAUCGCAAGGACUUUAACACCGAGCGUUCGUUAGUAAAGCUCGUAGGUAAGCGUAGAAGUUUGCUAGACUAUUUG

         ..............................................................................................................................................................................................................
         ..............................................................................................................................................................................................................
         AUYUUYR•RAARCACGG•RRR•••R•RR••RAYAC•GG••C•RC•GAAGG•CAAAUYGC•YURUUYAC•Y•Y•G•AUYRAYCAYYUR•C•RA•CAYYURAAAAR•AAYCR•AARGAUU•YAAYAC•GAR•G•UC•YURGU•••RYURGU•GG•AAR•G•RG••••YU•YU•GAYUAYYU•
```

```
env-1692  AUUUUCAAAAAGUACGCAGGUUCAGAAACCAACACGGGUUCAGCAGAAGGGCAAAUCGCACUUUUCACCCUCAGAAUCAAUCACCUUACUGAGCACUUAAAAGUGAAUCGCAAGGACUUUAACACCGAGCGUUCGUUAGUAAAGCUCGUAGGUAAGCGUAGA..................
env-1693  AUUUUCAAAAAGUACGCAGGUUCAGAAACCAACACAGGUUCAGCAGAAGGGCAAAUCGCACUUUUCACCCUUAGAAUCAAUCACCUUACUGAGCACUUAAAAGUGAAUCGCAAGGACUUUAACACUGAGCGUUCGUUAGUAAAGCUCGUAGGUAAGCGUAGAAGUUUGCUAGACUAUUUG
env-1694  AUUUUCAAAAAGUACGCAGGUUCAGAAACCAACACAGGUUCAGCAGAAGGGCAAAUCGCACUUUUCACCCUCAGAAUCAAUCACCUUACUGAGCACUUAAAAGUGAAUCGCAAGGACUUUAAUACUGAGCGUUCGUUAGUAAAGCUCGUAGGUAAGCGUAGAAGUUUGCUAGACUAUUUG
env-1695  AUUUUCAAAAAGUACGCAGGUUCAGAAACCAACACAGGUUCAGCAGAAGGGCAAAUCGCACUUUUCACCCUCAGAAUCAAUCACCUUACAGAGCACUUAAAAGUGAAUCGCAAGGACUUUAACACCGAGCGUUCGUUAGUAAAGCUCGUAGGUAAGCGUAGAAGUUUGCUAGACUAUUUG
env-1696  AUUUUCAAAAAGUACGCAGGUUCAGAAACCAACACAGGUUCAGCAGAAGGGCAAAUCGCACUUUUCACCCUCAGAAUCAAUCACCUUACAGAGCACUUAAAAGUGAAUCGCAAGGACUUUAACACUGAGCGUUCGUUAGUAAAGCUCGUAGGUAAGCGUAGAAGUUUGCUAGACUAUUUG
env-1697  AUUUUCAAAAAGUACGCAGGUUCAGAAACCAACACAGGUUCAGCAGAAGGGCAAAUCGCACUUUUUACCCUCAGAAUCAAUCACCUUACUGAGCACUUAAAAGUGAAUCGCAAGGACUUUAACACCGAGCGUUCGUUAGUAAAGCUCGUAGGUAAGCGUAGAAGUUUGCUAGACUAUUUG
env-1698  AUUUUCAA............................................................................................................................................................................
env-1699  AUUUUCAAAAAGUACGCAGGUUCAGAAACCAACACAGGUUCAGCAGAAGGGCAAAUCGCACUUUUUACCCUCAGAAUCAAUCACCUUACAGAGCACUUAAAAGUGAAUCGCAAGGACUUUAACACCGAGCGUUCGUUAGUAAAGCUCGUAGGUAAGCGUAGAAGUUUGCUAGACUAUUUG
env-1700  AUUUUCAAAAAGUACGCAGGUUCAGAAACCAACAC.................................................................................................................................................
env-1701  AUUUUCAAAAAGUACGCAGGUUCAGAAACCAACACAGGUUCAGCAGAAGGGCAAAUCGCACUUUUCACCCUCAGAAUUAAUCACCUUACAGAGCACUUAAAAGUGAAUCGCAAGGACUUUAACACCGAGCGUUCGUUAGUAAAGCUCGUAGGUAAGCGUAGAAGUUUGCUAGACUAUUUG
env-1702  AUUUUCAAAAAGUACGCAGGUUCAGAAACCAACACAGGUUCAGCAGAAGGGCAAAUCGCACUUUUCACCCUUAGAAUUAAUCACCUUACUGAGCACUUAAAAGUGAAUCGCAAGGACUUUAACACCGAGCGUUCGUUAGUAAAGCUCGUAGGUAAGCGUAGAAGUUUGCUAGACUAUUUG
env-1703  AUUUUCAAAAAGUACGCAGGUUCAGAAACCAACACAGGUUCAGCAGAAGGGCAAAUUGCACUUUUCACCCUCAGAAUCAAUCACCUUACUGAGCACUUAAAAGUGAAUCGCAAGGACUUUAACACCGAGCGUUCGUUAGUAAAGCUCGUAGGUAAGCGUAGAAGUUUACUAGACUAUUUG
env-1705  AUUUUCAAAAAGUACGCAGGUUCAGAAACCAACACAGGUUCAGCAGAAGGGCAAAUCGCACUUUUCACCCUUAGAAUCAAUCACCUUACAGAGCACUUAAAAGUGAAUCGCAAGGACUUUAACACCGAGCGUUCGUUAGUAAAGCUCGUAGGUAAGCGUAGAAGUUUGCUAGACUAUUUG
env-1706  AUUUUCAAAAAGUACGCAGGUUCAGAAACCAACACAGGUUCAGCAGAAGGGCAAAUCGCACUUUUCACCCUCAGAAUCAAUCACCUUACUGAGCACUUAAAAGUGAAUCGCAAGGACUUUAAUACUGAGCGUUCGUUAGUAAAGCUCGUAGGUAAGCGUAGAAGUUUGCUAGACUAUUUG
env-1707  AUUUUCAAAAAGUACGCAGGUUCAGAAACCAACACAGGUUCAGCAGAAGGGCAAAUCGCACUUUUCACCCUCAGAAUCAAUCACCUUACAGAGCACUUAAAAGUGAAUCGCAAGGACUUUAACACCGAGCGUUNNNNAAAAGAGAGGAGGCUUAGUGCCUCCUUUUGCUUAAUGUUUUCA
env-1708  AUUUUCAAAAAGUACGCAGGUUCAGAAACCAACACUGGUUCAGCAGAAGGGCAAAUCGCUCUUUUUACCCUUAGAAUCAAUCACCUUACAGAGCACUUAAAAGUGAAUCGCAAGGACUUUAACACCGAGCGUUCGUUAGUAAAGCUCGUAGGUAAGCGUAGAAGUUUGCUAGACUAUUUG
env-1709  AUUUUCAAAAAGUACGCAGGUUCAGAAACCAACACAGGUUCAGCAGAAGGGCAAAUCGCACUUUUCACCCUCAGAAUCAAUCACCUUACAGAGCACUUAAAAGUGAAUCGCAAGGAUUUUAAUACUGAGCGUUCGUUAGUAAAGCUCGUAGGUAAGCGUAGAAGUUUGCUAGACUAUUUG
env-1710  AUUUUCAAAAAGUACGCAGGUUCAGAAACCAACACAGGUUCAGCAGAAGGGCAAAUCGCACUUUUCACCCUCAGAAUCAACCACCUUACUGAGCACUUAAAAGUGAAUCGCAAGGACUUUAACACUGAGCGUUCGUUAGUAAAGCUCGUAGGUAAGCGCAGAAGUCUGCUAGACUAUUUG
Cko-1-1   AUCUUCGCAAAACACGGAAAGUCUGAUGCAGACACAGGAAGCGCAGAAGGACAAGUAGCCCUUUUCACUUUCAGAAUCAAUCACUUAUCUCAACACCUGAAAAAAAACCAAAAGGAUUUCAACACUGAAAAAUCUUUGGUGAAACUGGUAGGUAAAAGAAAGAGACUUUUGGAUUACCUU
env-1711  AUUUUCAAGAAGUACGCGGGUUCAGAAAGCAAUACGGGUUCAGCAGAAGGACAGAUUGCACUUUUUACCUUUAGAAUCAAUCACCUUACAGAGCACUUGAAAGUGAAUCGUAAGGAUUUCAACACAGAGCGUUCAUUGGUAAAAUUAGUAGGUAAACGCAGAAGUUUGCU..........
env-1712  AUUUUCAAGAAGUACGCGGGUUCAGAAAGCAAUACAGGUUCAGCAGAAGGACAGAUUGCACUUUUUACCUUUAGAAUCAAUCACCUUACAGAGCACUUGAAAGUGAAUCGUAAGGAUUUUAACACGGAGCGUUCAUUGGUAAAAUUGGUAGGUAAACGCAGAAGUU..............
env-1713  AUUUUCAAGAAGUACGCGGGUUCAGAAAGCAAUACGGGUUCAGCAGAAGGACAGAUUGCACUUUUUACCUUUAGAAUCAAUCACCUUACAGAGCACUUGAAAGUGAAUCGUAAGGAUUUCAACACAGAGCGUUCAUUGGUAAAAUUAGUAGGUAAACGCAGAAGUUUGCUUGACUAUUUG
env-1714  AUUUUCAAGAAGUACGCGGGUUCAGAAAGCAAUACGGGUUCAGCAGAAGGACAGAUUGCACUUUUUACCUUUAGAAUCAAUCACCUUACAGAGCACUUGAAAGUGAAUCGUAAGGAUUUCAACACAGAGCGUUCAUUGGUAAAAUUAGUAGGUAAACGCAGAAGUUUGCUUGACUAUUUG
env-1717  AUUUUCAAGAAGUACGCGGGUUCAGAAAGCAAUACAGGUUCAGCAGAAGGACAGAUUGCACUUUUUACCUUUAGAAUCAAUCACCUUACAGAGCACUUGAAAGUGAAUCGUAAGGAUUUUAACACAGAGCGUUCAUUGGUAAAAUUAGUAGGUAAACGCAGAAGUUUGCUUGACUAUUUG
env-1718  ....................................................................................................................................................................................
env-1720  AUUUUCAAGAAGUACGCGGGUUCAGAAAGCAAUACAGGUUCAGCAGAAGGACAGAUUGCACUUUUUACCUUUAGAAUCAAUCACCUUACAGAGCACUUGAAAGUGAAUCGUAAGGAUUUUAACACAGAGCGUUCAUUGGUAAAAUUGGUAGGUAAACGCAGAAGUUUGCUUGACUAUUUG
env-1722  AUUUUCAAGAAGUACGCGGGUUCAGAAAGCAAUACAGGUUCAGCAGAAGGACAGAUUGCACUUUUUACCUUUAGAAUCAAUCACCUUACAGAGCACUUGAAAGUGAAUCGUAAGGAUUUUAACACAGAGCGUUCAUUGGUAAAAUUAGUAGGUAAACGCAGAAGUUUGCUUGACUAUUUG
Csp-5-1   AUUUUCAAGAAGUACGCGGGUUCAGAAAGCAAUACAGGUUCAGCAGAAGGACAGAUUGCACUUUUUACCUUUAGAAUCAAUCACCUUACAGAGCACUUGAAAGUGAAUCGUAAGGAUUUUAACACAGAGCGUUCAUUGGUAAAAUUGGUAGGUAAACGCAGAAGUUUGCUUGACUAUUUG
env-1723  ....................................................................................................................................................................................
env-1724  AUCUUCGCUAAUCACGGAGGAAAAGCAGAAAACACAGGAUCUGCAGAAGGGCAAAUCGCACUAUUCACUCACCGAAUCACGCACCUUACAGAGCACUUGAAAAAAAAUCGUCACGAUUAUAACACAGAGCGUUCAUUAGUACUGUUAGUAGGUAAAAGAAGAUCUUUGUUGGAUUACUUG
env-1725  AUCUUCGCUAAUCACGGAGGAAAAGCAGAAAACACAGGAUCUGCAGAAGGGCAAAUCGCACUAUUCACUCACCGAAUCACGCACCUUACAGAGCACUUGAAAAAAAAUCGUCACGAUUAUAACACAGAACGUUCAUUAGUACUGUUAGUAGGUAAAAGAAGAUCUUUGUUGGAUUACUUG
env-1726  AUCUUCGCUAAUCACGGAGGAAAAGCAGAAAACACAGGAUCUGCAGAAGGGCAAAUCGCACUAUUCACUCACCGAAUCACGCACCUUACAGAGCACUUGAAAAAAAAUCGUCACGAUUAUAACACAGAACGUUCAUUAGUACUGUUAGUAGGUAAAAGAAGAUCUUUGUUGGAUUACUUG
env-1727  AUCUUCGCUAAUCACGGAGGAAAAGCAGAAAACACAGGAUCUGCAGAAGGGCAAAUCGCACUAUUCACUCACCGAAUCACGCACCUUACAGAGCACUUGAAAAAAAAUCGUCACGAUUAUAACACAGAGCGUUCAUUAGUACUGUUAGUAGG............................
Csp-8-1   AUUUUUAAAAAGCAUGGAGGCAGUGAAACUAAUACAGGAUCUGCUGAAGGACAAAUCGCUCUAUUUACGCACCGUAUCAAUCAUUUAACAGAGCACAUGCGUAUUAAUCGUAAGGAUUAUAAUACACAGCGCUCUUUAGUAAGACUUGUAGGUAAGCGUAGGCGUUUACUAGAUUAUCUA
env-1728  AUUUUUAAAAAGCAUGGAGGCAGUGAAACUAAUACAGGAUCUGCUGAAGGACAAAUUGCUCUAUUUACGCACCGUAUCAAUCAUUUAACAGAGCACAUGCGUAUUAAUCGUAAGGAUUAUAAUACACAGCGCUCUUUAGUAAGACUUGUAGGUAAGCGUAGGCGUUUACUAGAUUAUCUA
env-1729  AUCUUCAAAAAACAUGGAGGAAGUGAAACCAAUACAGGUUCCACAGAAGGGCAAAUCGCUCUAUUUACCUUCCGUAUCAAUCACCUAACAGAGCACUUGCGUGCCAACCGUAAGGACUACAAUACAGAGCGUUCGUUAGUAAGGCUCGUAGGAAAGCGCAGAAGCCUGUUAGACUAUCUA
env-1730  AUCUUCAAAAAACAUGGAGGAAGUGAAACCAAUACAGGUUCCACAGAAGGGCAAAUCGCUCUAUUUACCUUCCGUAUCAAUCACCUAACAGAGCACUUGCGUGCCAAUCGUAAGGAUUACAAUACAGAGCGUUCGUUAGUAAGGCUCGUAGGAAAGCGCAGAAGCUUGUUAGACUAUCUA
env-1744  AUCUUCAAAAAACAUGGAGGAAGUGAAACCAAUACAGGUUCCACAGAAGGGCAAAUCGCUCUAUUUACCUUCCGUAUCAAUCACCUAACAGAGCACUUGCGUGCCAAUCGUAAAGAUUACAAUACAGAGCGUUCGUUAGUAAGGCUCGUAGGAAAGCGCAGAAGCUUGUUAGAUUAUCUA
env-1745  AUCUUCAAAAAACAUGGAGGAAGUGAAACCAAUACAGGUUCCACAGAAGGGCAAAUCGCUCUAUUUACCUUCCGUAUCAAUCACCUAACAGAGCACUUGCGUGCCAACCGUAAGGAUUACAAUACAGAGCGUUCGUUAGUAAGGCUCGUAGGAAAGCGCAGAAGCUUGUUAGAUUAUCUA
env-1746  ....................................................................................................................................................................................
env-1747  ....................................................................................................................................................................................
env-1748  AUCUUCAAAAAACAUGGAGGAAGUGAAACCAAUACAGGUUCCACAGAAGGGCAAAUCGCUCUAUUUACCUUCCGUAUCAAUCACCUAACAGAGCACUUGCGUGCCAAUCGUAAGGAUUACAAUACAGAGCGUUCGUUAGUAAGGCUCGUAGGAAAGCGCAGAAGCUUGUUAGACUAUCUA
env-1749  AUCUUCAAAAAACAUGGAGGAAGUGAAACCAAUACAGGUUCCACAGAAGGGCAAAUCGCUCUAUUUACCUUCCGUAUCAAUCACCUAACAGAGCACUUGCGUGCCAACCGUAAGGAUUACAAUACAGAGCGUUCGUUAGUAAGGCUCGUAGGAAAGCGCAGAAGCUUGUUAGACUAUCUA
env-1750  ....................................................................................................................................................................................
env-1752  AUCUUCAAAAAACAUGGAGGAAGUGAAACCAAUACAGGUUCCACAGAAGGGCAAAUCGCUCUAUUUACCUUCCGUAUCAAUCACCUAACAGAGCACUUGCGUGCCAACCGUAAGGAUUACAAUACGGAGCGUUCGUUAGUAAGGCUCGUAGGAAAGCGCAGAAGCUUGUUAGAUUAUCUA
env-1753  AUCUUCAAAAAACAUGGAGGAAGUGAAACCAAUACAGGUUCCACAGAAGGGCAAAUCGCUCUAUUUACCUUCCGUAUCAAUCACCUAACAGAGCACUUGCGUGCCAACCGUAAGGAUUACAAUACAGAGCGUUCGUUAGUAAGGCUCGUAG.............................
env-1754  AUCUUCAAAAAACAUGGAGGAAGUGAAACCAAUACAGGUUCCACAGAAGGGCAAAUCGCUCUAUUUACCUUCCGUAUCAAUCACCUAACAGAGCACUUGCGUGCCAAUCGUAAAGAUUA.............................................................
env-1755  AUCUUCAAAAAACAUGGAGGAAGUGAAACCAAUACAGGUUCCACAGAAGGGCAAAUCGCUCUAUUUACCUUCCGUAUCAAUCACCUAACAGAGCACUUGCGUGCCAACCGUAAGGAUUACAAU.........................................................
Cgi-1-1   AUCUUCAAAAAACAUGGAGGAAGUGAAACCAAUACAGGUUCCACAGAAGGGCAAAUCGCUCUAUUUACCUUCCGUAUCAAUCACCUAACAGAGCACUUGCGUGCCAACCGUAAGGAUUACAAUACAGAGCGUUCGUUAGUAAGGCUCGUAGGAAAGCGCAGAAGCUUGUUAGACUAUCUA
env-1756  ....................................................................................................................................................................................
env-1757  ....................................................................................................................................................................................
env-1765  ....................................................................................................................................................................................
env-1766  ....................................................................................................................................................................................
env-1767  AUCUUCAAAAAACAUGGAGGAAGUGAAACCAAUACAGGUUCCACAGAAGGGCAAAUCGCUCUAUUUACCUUCCGUAUCAAUCACCUAACAGAGCACUUGCGUGCCAAUCGUAAAGAUUACAAUACAGAGCGUUCGUUAGUAAGGCUCGUAGGAAAGCGCAGAAGCUUGUUAGACUAUCUA
env-1768  AUCUUCAAAAAACAUGGAGGAAGUGAAACCAAUACAGGUUCCACAGAAGGGCAAAUCGCUCUAUUUACCUUCCGUAUCAAUCACCUAACAGAGCACUUGCGUGCCAAUCGUAAGGAUUACAAUACAGAGCGUUCGUUAGUAAGGCUCGUAGGAAAGCGCAGAAGCUUGUUAGACUAUCUA
env-1769  AUCUUCAAAAAACAUGGAGGAAGUGAAACCAAUACAGGUUCCACAGAAGGGCAAAUCGCUCUAUUUACCUUCCGUAUCAAUCACCUAACAGAGCACUUGCGUGCCAAUCGUAAGGAUUACAAUACAGAGCGUUCGUUAGUAAGGCUCGUAGGAAAGCGCAGAAGCUUGUUAGA.......
env-1770  AUCUUCAAAAAACAUGGAGGAAGUGAAACCAAUACAGGUUCCACAGAAGGGCAAAUCGCUCUAUUUACCUUCCGUAUCAAUCACCUAACAGAGCACUUGCGUGCCAAUCGUAAGGAUUACAAUACAGAGCGUUCGUUAGUAAGGCUCGUAGGAAAGCGCAGAAGCUUGUUAGACUAUCUA
env-1771  AUCUUCAAAAAACAUGGAGGAAGUGAAACCAAUACAGGUUCCACAGAAGGGCAAAUCGCUCUAUUUACCUUCCGUAUCAAUCACCUAACAGAGCACUUGCGUGCCAAUCGUAAAGAUUACAAUACAGAGCGUUCGUUAGUAAGGCUCGUAGGAAAGCGCAGAAGCUUGUUAGAUUAUCUA
env-1772  AUCUUCAAAAAACAUGGAGGAAGUGAAACCAAUACAGGUUCCACAGAAGGGCAAAUCGCUCUAUUUACCUUCCGUAUCAAUCACCUAACAGAGCACUUGCGUGCCAAUCGUAAAGAUUACAAUACAGAGCGUUCGUUAGUAAGGCUCGUAGGAAAGCGCAGAAGCUUGUUAGACUAUCUA
env-1773  AUCUUCAAAAAACAUGGAGGAAGUGAAACCAAUACAGGUUCCACAGAAGGGCAAAUCGCUCUAUUUACCUUCCGUAUCAAUCACCUAACAGAGCACUUGCGUGCCAAUCGUAAAGAUUACAAUACAGAGCGUUCGUUAGUAAGGCUCGUAGGAAAGCGCAGAAGCUUGUUAGACUAUCUA
env-1774  AUCUUCAAAAAACAUGGAGGAAGUGAAACCAAUACAGGUUCCACAGAAGGGCAAAUCGCUCUAUUUACCUUCCGUAUCAAUCACCUAACAGAGCACUUGCGUGCCAAUCGUAAGGAUUACAAUACAGAGCGUUCGUUAGUAAGGCUCGUAGGAAAGCGCAGAAGCUUGUUAGACUAUCUA
env-1775  AUCUUCAAAAAACAUGGAGGAAGUGAAACCAAUACAGGUUCCACAGAAGGGCAAAUCGCUCUAUUUACCUUCCGUAUCAAUCACCUAACAGAGCACUUGCGUGCCAACCGUAAGGAUUACAAUACAGAGCGUUCGUUAGUAAGGCUCGUAGGAAAGCGCAGAAGCUUGUUAGAUUAUCUA
env-1779  AUCUUCAAAAAACAUGGAGGAAGUGAAACCAAUACAGGUUCCACAGAAGGGCAAAUCGCUCUAUUUACCUUCCGUAUCAAUCACCUAACAGAGCACUUGCGUGCCAAUCGUAAAGAUUACAAUACAGAGCGUUCGUUAGUAAGGCUCGUAGGAAAGCGCAGAAGCUUGUUAGACUAUCUA
env-1780  AUCUUCAAAAAGCAUGGAGGAAGUGAAACCAAUACAGGUUCCACAGAAGGGCAAAUCGCUCUAUUUACCUUCCGUAUCAAUCACCUAACAGAGCACUUGCGUGCCAACCGUAAGGACUACAAUACAGAGCGUUCGUUAGUAAGGCUCGUAGGAAAGCGCAGAAGCCUGUUAGACUAUCUA
env-1783  ....................................................................................................................................................................................
env-1786  AUCUUCAAAAAGCAUGGAGGAAGUGAAACCAAUACAGGUUCCACAGAAGGGCAAAUCGCUCUAUUUACCUUCCGUAUCAAUCACCUAACAGAGCACUUGCGUGCCAACCGUAAGGACUACAAUACAGAGCGUUCGUUAGUAAGGCUCGUAGGAAAGCGCAGAAGCCUGUUAGACUAUCUA
env-1795  AUCUUCAAAAAGCAUGGAGGAAGUGAAACCAAUACAGGUUCCACAGAAGGGCAAAUCGCUCUAUUUACCUUCCGUAUCAAUCACUUAACAGAGCACUUGCGUGCCAACCGUAAGGACUACAAUACGGAGCGUUCGUUAGUAAGGCUCGUAGGAAAGCGCAGAAGCCUGUUAGACUAUCUA
env-1797  AUUUUCAAAAAGCAUGGAGGAAGUGAAACCAAUACAGGUUCCACAGAAGGGCAAAUCGCUCUAUUUACCUUCCGUAUCAAUCACCUAACAGAGCACUUGCGUGCCAACCGUAAGGACUACAAUACAGAGCGUUCGUUAGUAAGGCUCGUAGGAAAGCGUAGAAGCCUGUUAGACUAUCUA
env-1798  AUCUUCAAAAAGCAUGGAGGAAGUGAAACCAAUACAGGUUCCACAGAAGGGCAAAUCGCUCUAUUUACCUUCCGUAUCAAUCACUUAACAGAGCACUUGCGUGCCAACCGUAAGGACUACAAUACAGAGCGUUCGUUAGUAAGGCUCGUAGGAAAGCGCAGAAGCCUGUUAGACUAUCUA
env-1800  AUCUUCAAAAAGCAUGGAGGAAGUGAAACCAAUACAGGUUCCACAGAAGGGCAAAUCGCUCUAUUUACCUUCCGUAUCAAUCACCUAACAGAGCACUUGCGUGCCAACCGUAAGGAC...............................................................
env-1801  AUCUUCAAAAAGCAUGGAGGAAGUGAAACCAAUACAGGUUCCACAGAAGGGCAAAUCGCUCUAUUUACCUUCCGUAUCAAUCACCUAACAGAGCACUUGCGUGCCAACCGUAAGGACUACAAUACAGAGCGUUCGUUA..........................................
env-1802  ....................................................................................................................................................................................
env-1804  AUCUUCAAAAAGCAUGGAGGAAGUGAAGCCAAUACAGGUUCCACAGAAGGGCAAAUCGCUCUAUUUACCUUCCGUAUCAAUCACCUAACAGAGCACUUGCGUGCCAACCGUAAGGACUACAAUACAGAGCGUUCAUUAGUAAGGCUCGUAGGAAAGCGCAGAAGCCUGUUAGACUAUCUA
env-1805  ....................................................................................................................................................................................
env-1818  AUCUUCAAAAAGCAUGGAGGAAGUGAAACCAAUACAGGUUCCACAGAAGGGCAAAUCGCUCUAUUUACCUUCCGUAUCAAUCACCUAACAGAGCACUUGCGUGCCAACCGUAAGGACUACAAUACAGAGCGUUCAUUAGUAAGGCUCGUAGGAAAGCGCAGAAGCCUGUUAGACUAUCUA
env-1821  AUCUUCAAAAAGCAUGGAGGAAGUGAAACCAAUACAGGUUCCACAGAAGGGCAAAUCGCUCUAUUUACCUUCCGUAUCAAUCACUUAACAGAGCACUUGCGUGCCAACCGUAAGGACUACAAUACAGAGCGUUCGUUAGUAAGGCUCGUAGGAAAGCGCAGAAGCCUGUUAGACUAUCUA
env-1823  AUCUUCAAAAAGCAUGGAGGAAGUGAAACCAAUACAGGUUCCACAGAAGGGCAAAUCGCUCUAUUUACCUUCCGUAUCAAUCACCUAACAGAGCACUUGCGUGCCAACCGUAAGGACUACAAUACAGAGCGUUCGUUAGUAAGGCUCGUAGGAAAGCGCAGAAGUCUGUUAGACUAUCUA
env-1824  AUCUUCAAAAAGCAUGGAGGAAGUGAAACCAAUACAGGUUCCACAGAAGGGCAAAUCGCUCUAUUUACCUUCCGUAUCAAUCACCUAACAGAGCACUUGCGUGCCAACCGUAAGGACUACAAUACAGAGCGUUCGUUAGUAAGGCUCGUAGGAAAGCGCAGAAGCCUGUUAGACUAUCUA
env-1826  AUCUUCAAAAAGCAUGGAGGAAGUGAAACCAAUACAGGUUCCACAGAAGGGCAAAUCGCUCUAUUUACCUUCCGUAUCAAUCACCUAACAGAGCACUUGCGUGCCAACCGUAAGGACUACAAUACAGAGCGUUCGUUAGUAAGGCUCGUAGGAAAGCGCAGAAGCCUGUUAGACUAUCUA
env-1827  AUCUUCAAAAAGCAUGGAGGAAGUGAAACCAAUACAGGUUCCACAGAAGGGCAAAUCGCUCUAUUUACCUUCCGUAUCAAUCACCUAACA..........................................................................................
env-1828  AUUUUCAAAAAGCAUGGAGGAAGUGAAACCAAUACAGGUUCCACAGAAGGGCAAAUCGCUCUAUUUACCUUCCGUAUCAAUCACUUAACAGAGCACU...................................................................................
env-1829  CUUUUCGCAAAGCAUGGAAAAUCUGCGCAGGACACAGGAAGUGCUGAAGGGCAAAUCGCUCUUUUCACAUUUAGAAUUAAUCAUUUAUCUCAACACUUGAAAUCUAACCACAAAGAUUAUGCUACUGAGAGAUCACUAGUAAAAUUGGUAGGUAAGAGAAAAAGACUUCUGGAUUAUCUU
env-1830  AUCUUCAAAAAGCAUGGAGGAAGUGAAGCCAAUACAGGUUCCACAGAAGGGCAAAUCGCUCUAUUUACCUUCCGUAUCAAUCACCUAACAGAGCACUUGCGUGCCAACCGUAAGGACUACAAUACAGAGCGUUCAUUAGUAAGGCUCGUAGGAAAGCGCAGAAGCCUGUUAGACUAUCUA
env-1832  AUCUUCAAAAAGCAUGGAGGAAGUGAAGCCAAUACAGGUUCCACAGAAGGGCAAAUCGCUCUAUUUACCUUCCGUAUCAAUCACCUAACAGAGCACUUGCGUGCCAACCGUAAGGACUACAAUACAGAGCGUUCAUUAGUAAGGCUCGUAGGAAAGCGCAGAAGCCUGUUAGACUAUCUA

          ....................................................................................................................................................................................
          ....................................................................................................................................................................................
          AUYUUYR•RAARCACGG•RRR•••R•RR••RAYAC•GG••C•RC•GAAGG•CAAAUYGC•YURUUYAC•Y•Y•G•AUYRAYCAYYUR•C•RA•CAYYURAAAAR•AAYCR•AARGAU•YAAYAC•GAR•G•UC•YURGU•••RYURGU•GG•AAR•G•RG••••YU•YU•GAYUAYYU•
```

| | |
|---|---|
| env-1833 | AUUUUCAAAAAGCAUGGAGGAAGUGAAACCAAUACAGGUUCCACAGAAGGGCAAAUCGCUCUAUUUACCUUCCGUAUCAAUCACNNNNNNNNNNNNNNNNNNNNNNNNNNNNNNNNNNNNNNNNNNNNNNNNNNNNNNNNNNNNNNNNNNNNNNNNNNCUAACAGAGCACUUGCGUGCCA |
| env-1834 | AUUUUUAAACAACACGGUGGUAGUGAGAAAAAUACUGGCUCAGCAGAGGGACAAAUAGCCCUUUUCACCUUUCGCAUCAAUCACUUAACAGAGCACUUGCGCGCCAAUCGCAAAGAUUACAAUACAGAGCGCUCCCUUGUGAGACUCGUAGGUAAGCGUAGAAGCCUACUGGAUUAUCUC |
| env-1835 | AUUUUUAAACAACACGGUGGUAGCGAGAAAAAUACUGGCUCAGCAGAGGGACAAAUAGCCCUUUUCACCUUUCGUAUCAAUCACUUAACAGAGCACUUGCGCGCCAAUCGCAAAGAUUACAAUACAGAGCGCUCCCUUGUGAGACUCGUAGGUAAGCGCAGAAGUCUACUGGAUUAUCUC |
| env-1836 | .................................................................................................................................................................................... |
| env-1837 | .................................................................................................................................................................................... |
| env-1838 | AUUUUUAAACAACACGGUGGUAGCGAGAAAAAUACUGGCUCAGCAGAGGGACAAAUAGCCCUUUUCACCUUUCGUAUCAAUCACUUAACAGAGCACUUGCGCGCCAAUCGCAAAGAUUACAAUACAGAGCGCUCCCUUGUGAGACUCGUAGGUAAGCGCAGAAGCCUACUGGAUUAUCUC |
| env-1839 | AUUUUUAAACAACACGGUGGUAGCGAGAAAAAUACUGGCUCAGCAGAGGGACAAAUAGCCCUUUUCACCUUUCGUAUCAAUCACUUAACAGAGCACUUGCGCGCCAAUCGCAAAGAUUACAAUACAGAGCGCUCCCUUGUGAGACUCGUAGGUAAGCGCAGAAGCCUACUGGAUUAUCUC |
| env-1840 | AUUUUUAAACAACACGGUGGUAGCGAGAAAAAUACUGGCUCAGCAGAGGGACAAAUAGCCCUUUUCACCUUUCGUAUCAAUCACUUAACAGAGCACUUGCGCGCCAAUCGCAAAGAUUACAAUACAGAGCGCUCCCUUGUGAGACUCGUAGGUAAGCGCAGAAGCCUACUGGAUUAUCUC |
| env-1846 | .................................................................................................................................................................................... |
| env-1847 | AUUUUUAAACAACACGGUGGUAGCGAGAAAAAUACUGGCUCAGCAGAGGGACAAAUAGCCCUUUUCACCUUUCGUAUCAAUCACUUAACAGAGCACUUGCGCGCCAAUCGCAAAGAUUACAAUACAGAGCGCUCCCUUGUGAGACUCGUAGGUAAGCGCAGAAGCCUACUGGAUUAUCUC |
| env-1848 | .................................................................................................................................................................................... |
| env-1850 | .................................................................................................................................................................................... |
| env-1851 | AUUUUUAAACAACACGGUGGUAGCGAGAAAAAUACUGGCUCAGCAGAGGGACAAAUAGCCCUUUUCACCUUUCGUAUCAAUCACUUAACAGAGCACUUGCGCGCCAAUCGCAAAGAUUACAAUACAGAGCGCUCCCUUGUGAGACUCGUAGGUAAGCGCAGAAGCCUACUGGAUUAUCUC |
| env-1852 | AUUUUUAAACAACACGGUGGUAGCGAGAAAAAUACUGGCUCAGCAGAGGGACAAAUAGCCCUUUUCACCUUUCGUAUCAAUCACUUAACAGAGCACUUGCGCGCCAAUCGCAAAGAUUACAAUACAGAGCGCUCCCUUGUGAGACUCGUAGGUAAGCGCAGAAGCCUACUGGAUUAUCUC |
| env-1853 | AUUUUUAAACAACACGGUGGUAGCGAGAAAAAUACUGGCUCAGCAGAGGGACAAAUAGCCCUUUUCACCUUUCGUAUCAAUCACUUAACAGAGCACUUGCGCGCCAAUCGCAAAGAUUACAAUACAGAGCGCUCCCUUGUGAGACUCGUAGGUAAGCGCAGAAGCCUACUGGAUUAUCUC |
| env-1854 | AUUUUUAAACAACACGGUGGUAGCGAGAAAAAUACUGGCUCAGCAGAGGGACAAAUAGCCCUUUUCACCUUUCGUAUCAAUCACUUAACAGAGCACU................................................................................... |
| env-1855 | AUUUUCAAAAAACACGGUGGUAGUGAGACCAAUACAGGGUCUGCAGAAGGUCAGAUAGCCUUAUUUACACAUCGUAUUAAUCACCUAACAGAGCACUUACGUGCAAAUCGUAAAGAUUAUAAUACACAGCGUGCUUUAGUAAGGCUUGUUGGUAAAAGAAGAAGUCUUCUAGAUUAUCUG |
| env-1856 | AUUUUCAAAAAACACGGUGGUAGUGAGACCAAUACAGGGUCAGCAGAAGGUCAGAUAGCCUUAUUUACACAUCGUAUUAAUCACCUAACAGAGCACUUACGUGCAAAUCGUAAAGAUUAUAAUACACAGCGUGCUUUAGUAAGGCUUGUUGGUAAAAGAAGAAGUCUUCUAGAUUAUCUG |
| env-1857 | AUUUUCAAAAAACACGGUGGUAGUGAGACCAAUACAGGGUCAGCAGAAGGUCAGAUAGCCUUAUUUACACAUCGUAUUAAUCACCUAACAGAGCACUUACGUGCAAAUCGUAAAGAUUAUAAUACACAGCGUGCUUUAGUAAGGCUUGUUGGUAAAAGAAGAAGUCUUCUAGAUUAUCUG |
| env-1859 | AUUUUCAAAAAACACGGUGGUAGUGAGACCAAUACAGGGUCAGCAGAAGGUCAGAUAGCCUUAUUUACACAUCGUAUUAAUCACCUAACAGAGCACUUACGUGCAAAUCGUAAAGAUUAUAAUACACAGCGUGCUUUAGUAAGGCUUGUUGGUAAAAGAAGAAGUCUUCUAGAUUAUCUG |
| env-1860 | AUUUUCAAAAAACACGGUGGUAGUGAGACCAAUACAGGGUCAGCAGAAGGUCAGAUAGCCUUAUUUACACAUCGUAUUAAUCACCUAACAGAGCACUUACGUGCAAAUCGUAAAGAUUAUAAUACACAGCGUGCUUUAGUAAGGCUUGUUGGUAAAAGAAGAAGUCUUCUAGAUUAUCUG |
| env-1862 | AUCUUCAAAAAACAUGGUGGUAGCGAGACCAACACUGGAUCCGCUGAAGGACAAAUAGCCCUUUUCACCUUCCGAAUCAACCACUUAACGGAGCACUUGCGUACCAAUCGCAAAGACUAUAACACCCAACGCUCCUUGGUGAAGCUCGUAGGUAAGCGCAGAAGCCUAC.......... |
| env-1863 | .................................................................................................................................................................................... |
| env-1864 | AUUUUUAAACAACACGGUGGCAGCGAGAAAAAUACUGGCUCAGCUGAGGGACAAAUAGCCCUUUUCACCUUUCGUAUCAAUCACUUAACAGAGCACUUGCGCACUAAUCGCAAGGAUUACAAUACAGAACGCUCUCUUGUAAGACUCGUAG............................. |
| env-1865 | AUGUAUGCUAAGCACGGUAAAGAUGCUAAAGAUACAGGAAGUGCAGAAGGUCAAAUAGCAAUGUUUACUUUCAAAAUUGCUCAC................................................................................................ |
| Eme-2-1 | AUCUUCGCAAAACAUGGAAAGUCUGCAACAGACACAGGUAGUUCAGAAGGACAAAUCGCUUUGUUCACUUUCAGAAUUAACCACCUUUCUCAACACCUAAAGAAAAACCACAAAGAUUUUGCAACUGAAAAAUCUCUUGUGAAGUUAGUAGGUAAAAGAAAAAGAUUAUUAGAUUAUCUU |
| Cca-1-1 | AUAUUUAAAGAGCACGGAUCCAGCGCAAAAGACACUGGAAGUUCUGAAGGUCAAAUCGCACUAUUUACUUAUAGAAUUGCGCAUUUAACUGAGCACUUAAAGAAAAACAGAAAAGAUUACGGAACGCAAAGAGCGCUUCAAAUCUUAGUAGGUAAACGUAGAAGUUUACUUGAUUAUUUG |
| Emi-1-1 | AUCUUCGCAAAACACGGACAAUCCGCAACUGACACUGGUAGCUCUGAAGGACAAAUCGCUUUGUUCACUUUCAGAAUCAACCACCUUUCUCAACACCUUAAAAAGAACCAUAAAGAUUUCGCUACUGAGAAAUCAUUGGUGAAAUUAGUAGGUAAAAGAAAAAGAUUGUUAGAUUAUCUU |
| Cte-1-1 | AUUUUCGCAAAGCACGGAAAAUCUGCAAAUGACACAGGAAGCGCUGAAGGACAGAUUGCUCUAUUCACUUUUAGAAUCAACCACCUUUCUCAGCAUUUGAAAGCUAACCGUCACGAUUUUGCUACAGAGAGAUCUCUAGUAAUGUUGGUAGGUAAGAGAAAAUCUCUUCUUGAUUACUUG |
| Csp-15-1 | AUUUUCGCAAAGCACGGAAAAUCUGCAAAUGACACAGGAAGCGCUGAAGGACAGAUUGCUCUAUUCACUUUUAGAAUCAACCACCUUUCUCAGCACUUGAAAGCUAACCGUCACGAUUUUGCUACUGAGAGAUCUCUAGUGAAAUUGGUAGGUAAGAGAAAAUCUCUUCUUGAUUAUUUG |
| Cha-2-1 | AUUUUCGCAAAGCACGGAAAAUCUGCAAAUGACACAGGAAGCGCUGAAGGACAGAUUGCUCUAUUCACUUUUAGAAUCAACCACCUUUCUCAGCACUUGAAAGCUAACCGUCACGAUUUUGCUACUGAGAGGUCUCUAGUGAAAUUGGUAGGUAAGAGAAAAUCUCUUCUUGAUUACUUG |
| Ean-3-1 | AUCUUCGCAAAACACGGACAAUCCGCAACUGACACUGGUAGCUCUGAAGGACAAAUCGCUUUGUUCACUUUCAGAAUUAACCACCUUUCUCAACACCUUAAAAAGAACCACAAAGAUUAUGCAACUGAAAAAUCUUUGGUGAAAUUAGUAGGUAAAAGAAAAAGAUUGUUAGAUUAUCUU |
| Ean-1-3 | AUCUUCGCAAAACACGGACAAUCCGCAACUGACACUGGUAGCUCUGAAGGACAAAUCGCUUUGUUCACUUUCAGAAUUAACCACCUUUCUCAACACCUUAAAAAGAACCACAAAGAUUAUGCAACUGAAAAAUCUUUGGUGAAAUUAGUAGGUAAAAGAAAAAGAUUGUUAGAUUAUCUU |
| Csp-18-1 | AUCUUCGCAAAACACGGAAAGUCUGAUACAGACACAGGAAGCUCAGAAGGACAAAUUGCCCUUUUUACUUUCAGAAUCAAUCAUUUAUCUCAACACCUUAAAGCAAACCGUCACGAUUACGCUACGGAAAGAUCUUUGGUAAUGUUGGUAGGUAAAAGAAAAAGUUUAUUAGACUAUCUU |
| Cje-1-1 | AUCUUCGCAAAACACGGAAAGUCUGACGCAGACACAGGAAGCGCAGAAGGACAAGUAGCCCUUUUUACUUUCAGAAUUAAUCACCUUUCAGGUCACCUGAAAAAAAAACCACAAAGAUUACGCAACUGAGAGAUCAUUGGUACUUUUGGUAGGAAAAAGAAAAGCCCUUUUGGAUUAUCUU |
| env-1866 | AUCUUCGCAAAACACGGAAAGUCUGAUGCAGACACAGGAAGCGCAGAAGGACAAGUAGCCCUUUUCACUUUCAGAAUUAAUCACUUAUCUCAGCACCUUAAAAAAAAACCAUAAGGAUUUCGCUACAGAAAAAUCUUUGGUAAAAUUGGUAGGGAAAAGAAAAAGAUUGUUAGAUUAUCUU |
| Cha-1-1 | AUCUUCGCAAAACACGGAAAGUCUGACGCAGACACAGGAAGCGCUGAAGGGCAAAUUGCAUUAUUUACCUACAGAAUCAAUCACUUAUCUCAACACUUAAAAGCUAAUCGUCACGAUUACAAUACCGAAAGAUCUUUGGUAAAAUUAGUAGGGAAAAGAAAAAGUUUAUUAGAUUAUCUU |
| env-1869 | AUCUUCGCAAAACACGGAAAGUCUGAUGCAGACACAGGAAGCGCAGAAGGACAAGUAGCCCUUUUCACUUUCAGAAUUAAUCACUUAUCUCAGCAC.................................................................................... |
| env-1870 | AUUUUCGCAAAGCACGGUAAAUCUGCACAAGACACAGGAAGCGCAGAAGGACAAAUCGCGCUUUUCACUUACAGAAUCAACCACCUUUCUGGUCACUUGAAAGCAAUCAUAAAGAUUAUGCUACAGAGAGAUCUUUGGUGAAAUUGGUAGGGAAAAGAAAAAGUUUAUUGGAUUAUCUUA |
| Chi-1-1 | AUUUUCGCAAAACAUGGAAAAUCUGCACAAGACACAGGAAGUGCUGAAGGACAAAUUGCACUUUUCACUUUCAGAAUCAACCACCUAUCUGCUCACUUAAAGAGCAAUCACAAAGAUUAUAACACAGAAAGAUCUUUGGUAAAACUGGUAGGUAAAAGAAAAAGAUUAUUAGAUUAUCUU |
| env-1871 | AUCUUCGCAAAACACGGAAAGUCUGAUGCAGACACAGGAAGCGCUGAGGGACAAGUUGCCCUUUUCACUUACAGAAUCAACCACUUAUCUCAACACCUUAAAGCAAACCGUCACGACUUCAACACGGAGCGUUCACUGGUAGCUUUGGUAGGUAAAAGAAAAGCGCUUUGGAUUAUCUU |
| Cso-2-1 | AUCUUCGCAAAACAUGGAAAGUCUGAUGCUGAUACCGGCACAGCAGAAGGACAAAUUGCCCUUUUUACUUACAGAAUCAACCACCUAUCAGGUCACCUGAAAAAAAAUCACAAAGAUUACGCAACUGAGCGUUCUUUGGUACUUUUAGUAGGUAAAAGAAAAGCCCUUUUAGAUUAUCUG |
| Cda-1-1 | AUUUUCUCUAAACACGGGAAAUCUGCACAGGACACAGGAAGUGCUGAAGGACAAGUAGCACUUUUCACUUUCAGAAUCAAUCAUUUAUCUCAGCAUUUAAAGGCUAACCGUCAUGAUUUCAACACAGAGAGAUCUUUGGUUAAAUUGGUAGGUAAAAGAAAAAGAUUACUAGAUUAUCUU |
| Cpi-1-1 | AUUUUCUCAAAACACGGGAAAUCUGCACAGGACACAGGAAGUGCUGAAGGACAAGUUGCACUUUUUACUUACAGAAUUAACCACCUUUCUGCUCAUUUAAAGAGCAAUCACAAAGAUUACGCAACUGAAAGAUCUUUGGUGAAAUUGGUAGGUAAAAGAAAAAGUUUACUAGAUUACCUU |
| env-1872 | AUUUUCGCAAAGCACGGUAAAUCUGCACAAGACACAGGAACAGCUGAAGGACAAGUAGCUCUUUUCACGUUCAGAAUUAACCACUUAUCUCAGCACUUGAAGGCAAAUCGUCACGAUUUCAACACUGAAAGAUCUUUGGUAAAAUUAGUAGGUAAGAGAAAAAGUUUAUUGGAUUAUCUU |
| Fba-4-1 | AUCUUCGCAAAACACGGAAAAUCUGACGCAGAUACCGGAAGCGCGGAAGGGCAAGUGGCUCUUUUCACGUACAGAAUCAACCAUUUAUCUUCGCACCUUAAGGCUAACCAUAAAGACUACGCAACUGAGCGUUCUCUUGUUGCUUUGGUAGGUAAAAGAAAAGCACUUUUAGAUUAUCUA |
| Cfo-1-1 | AUUUUCGCAAAGCACGGUAAAUCUGCACAAGACACAGGAAGCGCUGAAGGACAGGUUGCUCUAUUCACUUACAGAAUUAACCACUUAUCUCAGCAUUUGAAGGCUAAUCGUCACGAUUUCAACACGGAGAGAUCUUUGGUGAAAUUAGUAGGUAAGAGAAAAAGCUUAUUAGAUUAUCUU |
| env-1873 | AUUUUCGCAAAACACGGAAAAUCUGCACAGGACACAGGAACUGCUGAAGGGCAAAUUGCACUUUUCACUUUCAGAAUCAACCACCUUUCUGCUCAUUUAAAGAGCAAUCACAAAGAUUACAACACAGAAAGAUCUUUGGUGAAACUGGUAGGUAAAAGAAAAAGUUUAUUAGAUUACCUU |
| Clu-1-1 | AUUUUCGCAAAACAUGGAAAAUCUGCACAGGACACAGGAAGUGCUGAAGGACAAGUAGCUCUUUUCACUUUCAGAAUCAACCACUUAUCUCAGCAUUUAAAGGCUAACCGUCACGAUUUCAACACAGAGAGAUCUCUAGUGAAAUUGGUAGGUAAAAGAAAAAGCUUAUUGGAUUAUCUU |
| env-1874 | AUUUUCGCAAAACACGGAAAAUCUGCACAAGACACAGGAAGUGCUGAAGGGCAAGUUGCACUUUUCACUUACAGAAUCAACCACCUUUCUGCUCACUUAAAGAGCAAUCACAAAGAUUACAACACAGAAAGAUCUUUGGUGAAAUUGGUAGGUAAAAGAAAAAGAUUACUAGAUUACCUU |
| Cta-1-1 | AUUUUCGCAAAACAUGGAAAAUCUGCACAGGACACAGGGAGCGCUGAAGGACAAGUAGCUCUUUUCACUUUCAGAAUUAACCACUUAUCUCAGCACUUGAAAGCUAAUCGUCAUGAUUUCAAUACUGAAAGAUCUUUGGUGAAAUUGGUAGGUAAAAGAAAAAGUUUAUUGGAUUAUCUU |
| Csp-29-1 | AUUUUCGCAAAGCACGGUAAAUCUGCACAGGACACAGGAAGCGCUGAAGGACAGGUAGCUCUUUUCACUUUCAGAAUUAACCACUUAUCUCAGCACUUGAAAGCUAAUCGUCAUGAUUUCAAUACUGAAAGAUCUUUGGUGAAAUUGGUAGGUAAAAGAAAAAGUUUAUUGGAUUAUCUU |
| Csp-24-1 | AUUUUCGCACAACAUGGAAAAUCUGCACAGGACACAGGAAGCGCAGAAGGACAAAUCGCUCUUUUCACUUUCAGAAUCAACCACUUAUCUCAGCACUUAAAGGCUAAUCGUCACGAUUUCAACACAGAGAGAUCUCUAGUGAAAUUGGUAGGUAAGAGAAAAAGUUUAUUAGAUUACCUU |
| Csp-19-1 | AUUUUCGCAAAACAUGGAAAAUCUGCACAAGACACAGGAAGCGCUGAAGGACAAGUAGCUCUUUUUACAUUUAGAAUCAAUCACUUAUCUCAGCAUUUGAAGGCUAAUCGUCAUGACUUCAAUACUGAAAGAUCUUUGGUGAAAUUGGUAGGUAAAAGAAAAAGCUUAUUGGAUUAUCUU |
| Cor-1-1 | AUUUUCGCAAAACACGGAAAAUCUGCAACAGACACAGGAAGUGCUGAAGGACAAGUUGCUCUUUUCACUUACAGAAUCAAUCACUUAUCUCAACACUUAAAGGCUAACCGUCACGAUUUCAACACAGAGAGAUCUCUAGUGAAAUUGGUAGGUAAGAGAAAAAGUUUACUAGAUUACCUU |
| Csp-16-1 | AUUUUCUCAAAACACGGAAAAUCUGCACAAGACACAGGAAGUGCUGAAGGACAAAUUGCACUUUUUACUUUCAGAAUCAACCACUUAUCUCAGCACUUAAAAGCUAAUCGUCACGAUUUUAAUACAGAAAGAUCGUUGGUAAAGCUAGUAGGUAAGAGAAAAAGUUUACUAGAUUACCUU |
| Can-1-1 | AUUUUCUCAAAACAUGGAAAAUCUGCAACAGACACAGGAAGUGCUGAAGGACAAAUCGCUCUUUUCACUUUCAGAAUCAAUCACUUAUCUCAACACUUAAAGGCUAAUCGUCACGAUUUCAACACAGAGAGAUCUCUAGUGAAAUUGGUAGGUAAGAGAAAAAGUUUACUAGAUUACCUU |
| Cgl-1-1 | AUUUUCGCAAAACACGGAAAAUCUGCACAAGACACAGGAAGUGCUGAAGGACAAAUUGCACUUUUCACUUUCAGAAUUAACCACCUUUCUGCUCACUUGAAGAGCAAUCACAAAGAUUUCAACACUGAAAAAUCUUUGGUUAAAUUGGUAGGUAAGAGAAAAAGAUUACUAGAUUACCUU |
| Csp-26-1 | AUUUUCGCAAAACACGGGAAAUCUGCACAAGACACAGGAAGUGCUGAAGGACAAAUCGCUCUUUUCACUUUCAGAAUCAAUCACUUAUCUCAACACCUAAAGGCUAACCGUCACGAUUUCAACACAGAGAGAUCUCUAGUGAAAUUGGUAGGUAAGAGAAAAAGUUUAUUAGAUUACCUU |
| Csp-23-1 | AUUUUCGCAAAACACGGAAAAUCUGCACAAGACACAGGAAGUGCUGAAGGACAAAUUGCUCUUUUCACUUUCAGAAUCAACCACCUUUCUGCUCACUUAAAGAGUAACCACAAAGAUUUCGCAACUGAAAAAUCUUUGGUGAAAUUGGUAGGUAAAAGAAAAAGAUUACUAGAUUACCUU |
| Cso-1-1 | AUUUUCGGACAACAUGGAAAAUCUGCAACAGACACAGGAAGUGCUGAAGGACAAGUAGCUCUUUUCACUUUCAGAAUCAACCACCUAUCUCAACACUUAAAGGCUAACCGUCACGAUUUCGCAACGGAAAGAUCUUUGGUAAAAUUAGUAGGUAAAAGAAAAAGUUUAUUAGAUUACCUU |
| Csp-17-1 | AUUUUCUCAAAACAUGGAAAAUCUGCAACAGACACAGGAAGUGCUGAAGGACAAAUCGCUCUUUUCACUUUCAGAAUCAAUCACUUAUCUCAACACCUAAAGGCUAACCGUCACGAUUUCAACACAGAGAGAUCUCUAGUGAAAUUGGUAGGUAAGAGAAAAAGUUUACUAGAUUACCUU |
| Csp-20-1 | AUUUUCUCAAAACAUGGAAAAUCUGCAACAGACACAGGAAGUGCUGAAGGACAAAUCGCUCUUUUCACUUUCAGAAUCAAUCACUUAUCUCAACACCUAAAGUCUAACCGUCACGAUUUCAACACAGAGAGAUCUCUUGUGAAAUUGGUAGGUAAGAGAAAAAGUUUACUAGAUUACCUU |
| env-1875 | AUUUUCGCAAAACACGGAAAAUCUGCACAAGACACAGGAAGUGCUGAAGGACAAAUCGCUCUUUUCACUUUCAGAAUCAAUCACUUAUCUCAGCACUUAAAGGCUAACCGUCAUGAUUUCAACACAGAGAGAUCUCUAGUGAAAUUGGUAGGUAAGAGAAAAAGUUUACUAGAUUACCUU |
| env-1876 | AUUUUCGCAAAACAUGGAAAAUCUGCACAAGACACAGGAAGCGCUGAAGGACA............................................................................................................................... |
| Csp-27-1 | AUUUUCGCACAACAUGGAAAAUCUGCACAGGACACAGGAAGCGCAGAAGGACAAAUCGCUCUUUUCACUUUCAGAAUCAACCACUUAUCUCAACACCUAAAGGCUAACCGUCACGAUUUCAACACAGAGAGAUCUCUAGUGAAAUUGGUAGGUAAGAGAAAAAGUUUAUUAGAUUACCUU |
| Csp-28-1 | AUUUUCGCAAAACACGGAAAAUCUGCACAAGACACAGGAAGUGCUGAAGGACAAAUCGCUCUUUUCACUUUCAGAAUCAACCACUUAUCUCAGCACUUAAAGGCUAACCGUCAUGAUUUCAACACAGAGAGAUCUCUAGUGAAAUUGGUAGGUAAGAGAAAAAGUUUACUAGAUUACCUU |
| Cin-1-1 | AUUUUCGCACAACACGGAAAAUCUGCACAGGACACAGGAAGCGCAGAAGGACAAAUCGCUCUUUUCACUUUCAGAAUUAAUCACUUAUCUCAGCACUUAAAGGCUAACCGUCAUGAUUUCAACACAGAGAGAUCUCUAGUGAAAUUGGUAGGUAAGAGAAAAAGUUUACUAGAUUACCUU |
| Csp-13-1 | AUUUUCGCAAAACAUGGAAAAUCUGCAACAGACACUGGAAGUGCUGAAGGACAAGUUGCUCUUUUCACUUACAGAAUCAAUCACUUAUCUCAACACUUAAAGGCUAACCGUCACGAUUUCAACACAGAGAGAUCUCUAGUGAAAUUGGUAGGUAAGAGAAAAAGUUUACUAGAUUACCUU |
| Csp-21-1 | AUUUUCGCAAAACACGGAAAAUCUGCACAAGACACAGGAAGUGCUGAAGGACAAAUUGCACUUUUCACUUUCAGAAUCAACCAUCUUUCUGCUCACUUAAAGAGCAACCAUAAAGAUUUCAACACUGAAAAAUCUUUGGUUAAGCUGGUAGGUAAAAGAAAAAGAUUACUAGAUUACCUU |
| Csp-25-1 | AUUUUCGCAAAACAUGGAAAAUCUGCACAAGACACAGGAAGCGCUGAAGGACAAGUAGCUCUUUUUACUUUCAGAAUCAACCACUUAUCUCAGCACUUGAAGGCUAAUCGUCAUGACUUCAAUACUGAAAGAUCUUUGGUGAAAUUAGUAGGUAAGAGAAAAAGCUUAUUGGAUUAUCUU |
| Csp-22-1 | AUUUUCGCACAACACGGAAAAUCUGCACAGGACACAGGAAGCGCAGAAGGACAAAUCGCUCUUUUCACUUUCAGAAUUAAUCACUUAUCUCAGCACUUAAAGGCUAACCGUCAUGAUUUCAACACAGAGAGAUCUCUAGUGAAAUUGGUAGGUAAGAGAAAAAGUUUACUAGAUUACCUU |
| Cga-1-1 | AUUUUCGCAAAACACGGAAAAUCUGCACAAGACACAGGAAGUGCUGAAGGACAAAUUGCACUUUUCACUUUCAGAAUCAACCAUCUUUCUGCUCACUUAAAGAGCAACCAUAAAGAUUUCAACACUGAAAAAUCUUUGGUUAAGCUGGUAGGUAAAAGAAAAAGAUUACUAGAUUACCUU |
| Cvr-1-1 | AUUUUCUCAAAACAUGGAAAAUCUGCAACAGACACAGGAAGUGCUGAAGGACAAAUCGCUCUUUUCACUUUCAGAAUCAAUCACUUAUCUCAACACCUAAAGGCUAACCGUCACGAUUUCAACACAGAGAGAUCUCUAGUGAAAUUGGUAGGUAAGAGAAAAAGUUUACUAGAUUACCUU |
| Csp-14-1 | AUUUUCGCAAAACACGGAAAAUCUGCACAAGACACAGGAAGUGCUGAAGGACAAAUCGCUCUUUUCACUUUCAGAAUCAACCACUUAUCUCAACACCUAAAGGCUAACCGUCACGAUUUCAACACAGAGAGAUCUCUAGUGAAAUUGGUAGGUAAGAGAAAAAGUUUAUUAGAUUACCUU |
| env-1877 | AUUUUCGCAAAACAUGGAAAAUCUGCAACAGACACAGGAAGUGCUGAAGGACAAGUUGCUCUUUUCACUUACAGAAUCAAUCACUUAUCUCAACACUUAAAGGCUAACCGUCACGAUUUCAACACAGAGAGAUCUCUAGUGAAAUUGGUAGGUAAGAGAAAAAGUUUACUAGAUUACCUU |
| env-1878 | AUCUUCAAAAAACAUGGAGGAAGUGAAACCAAUACAGGUUCCACAGAAGGGCAAAUCGCUCUAUUUACCUUCCGUAUCAAUCACCUAACAGAGCACUUGCGUGCCAACCGUAAGGAUUACAAUACAGAGCGUUCGUUAGUAAGGCUCGUAGGAAAGCGCAGAAGCUUGUUAGACUAUCUA |
| env-1879 | .................................................................................................................................................................................... |
| env-1880 | AUUUUCGCAAAACACGGUAAAUCUGCACAAGACACAGGAAGCGCUGAAGGACAAGUUGCUCUAUUCACUUUUAGAAUCAACCACCUUUCUCAGCACUUAAAAGCUAACCACAAAGAUUUCGCUACAGAAAAAUCUCUUGUAAAAUUGGUAGGAAGAGAAAAGCUCUUCUUGAUUAUUUG |
| env-1881 | AUUUUCGCAAAACACGGUAAAUCUGCACAAGACACAGGAAGCGCUGAAGGACAAGUUGCUCUAUUCACUUUUAGAAUCAACCACCUUUCUCAGCACUUAAAAGCUAACCACAAAGAUUUCGCUACAGAAAAAUCUCUUGUAAAAUUGGUAGGAAGAGAAAAGCUCUUCUUGAUUAUUUG |
| env-1883 | AUCUUCGCUAAACACGGAGGAAAAGCAGAGAACACAGGUUCUACAGAAGGGCAAAUCGCUC....................................................................................................................... |
| env-1884 | CUUUUCGCAAAGCAUGGAAAAUCUGCGCAGGACACAGGAAGUGCUGAAGGGCAAAUCGCUCUUUUCACAUUUAGAAUUAAUCAUUUAUCUCAACACUUGAAAUCUAACCACAAAGAUUAUGCUACUGAGAGAUCACUAGUAAAAUUGGUAGGUAAGAGAAAAAGACUUCUGGAUUAUCUU |
| env-1885 | CUUUUCGCAAAGCAUGGAAAAUCUGCGCAGGACACAGGAAGUGCUGAAGGGCAAAUCGCUCUUUUCACAUUUAGAAUUAAUCAUUUAUCUCAACACUUGAAAUCUAACCACAAAGAUUAUGCUACUGAGAGAUCACUAGUAAAAUUGGUAGGUAAGAGAAAAAGACUUCUGGAUUAUCUU |
| env-1886 | CUUUUCGCAAAGCAUGGAAAAUCUGCGCAGGACACAGGAAGUGCUGAAGGGCAAAUCGCUCUUUUCACAUUUAGAAUUAAUCAUUUAUCUCAACACUUGAAAUCUAACCACAAAGAUUAUGCUACUGAGAGAUCACUAGUAAAAUUGGUAGGUAAGAGAAAAAGACUUCUGGAUUAUCUU |
| | .................................................................................................................................................................................... |
| | .................................................................................................................................................................................... |
| | AUYUUYR•RAARCACGG•RRR•••R•RR••RAYAC•GG••C•RC•GAAGG•CAAAUYGC•YURUUYAC•Y•Y•G•AUYRAYCAYYUR•C•RA•CAYYURAAAAR•AAYCR•AARGAUU•YAAYAC•GAR•G•UC•YURGU•••RYURGU•GG•AAR•G•RG••••YU•YU•GAYUAYYU• |

```
env-1892 CUUUUCGCAAAGCAUGGAAAAUCUGCGCAGGACACAGGAAGUGCUGAAGGGCAAAUCGCUCUUUUCACAUUUAGAAUUAAUCAUUUAUCUCAACACUUAAAAUCUAACCACAAAGAUUAUGCUACUGAGAGAUCACUAGUAAAAUUGGUAGGUAAGAGAAAAAGACUUCUGGAUUAUCUU
env-1893 ....................................................................................................................................................................................
env-1898 ....................................................................................................................................................................................
env-1900 CUUUUCGCAAAGCAUGGAAAAUCUGCACAGGACACAGGAAGUGCUGAAGGGCAAAUCGCUCUUUUCACAUUUAGAAUUAAUCAUUUAUCUCAACACUUAAAAUCUAACCACAAAGAUUAUGCUACUGAGAGAUCACUAGUAAAAUUGGUAGGUAAGAGAAAAAGACUUCUGGAUUAUCUU
env-1901 CUUUUCGCAAAACAUGGAAAAUCUGCGCAGGACACAGGAAGUGCUGAAGGGCAAAUCGCUCUUUUCACAUUUAGAAUUAAUCAUUUAUCUCAACACUUGAAAUCUAACCACAAAGAUUAUGCUACUGAGAGAUCACUAGUAAAAUUGGUAGGUAAGAGAAAAAGACUUCUGGAUUAUCUU
env-1902 ....................................................................................................................................................................................
env-1903 CUUUUCGCAAAGCAUGGGAAAUCUGCGCAGGACACAGGAAGUGCUGAAGGGCAAAUCGCUCUUUUCACAUUUAGAAUUAAUCAUUUAUCUCAACACUUGAAAUCUAACCACAAAGAUUAUGCUACUGAGAGAUCACUAGUAAAAUUGGUAGGUAAGAGAAAAAGACUUCUGGAUUAUCUU
env-1904 CUUUUCGCAAAGCAUGGAAAAUCUGCGCAGGACACAGGAAGUGCUGAAGGGCAAAUCGCUCUUUUCACAUUUAGAAUUAAUCAUUUAUCUCAACACUUGAAAUCUAACCACAAAGAUUAUGCUACUGAAAGAUCACUAGUAAAAUUGGUAGGUAAGAGAAAAAGACUUCUGGAUUAUCUU
env-1905 CUUUUCGCAAAGCAUGGAAAAUCUGCGCAGGACACAGGAAGUGCUGAAGGGCAAAUUGCUCUUUUCACAUUUAGAAUUAAUCAUUUAUCUCAACACUUGAAAUCUAACCACAAAGAUUAUGCUACUGAGAGAUCACUAGUAAAAUUGGUAGGUAAGAGAAAAAGACUUCUGGAUUAUCUU
env-1906 CUUUUCGCAAAGCAUGGAAAAUCUGCACAGGACACAGGAAGUGCUGAAGGGCAAAUCGCUCUUUUCACAUUUAGAAUUAAUCAUUUAUCUCAACACUUGAAAUCUAACCACAAAGAUUAUGCUACUGAGAGAUCACUAGUAAAAUUGGUAGGUAAGAGAAAAAGACUUCUGGAUUAUCUU
env-1907 CUUUUCGCAAAGCAUGGAAAAUCUGCGCAGGACACAGGAAGUGCUGAAGGGCAAAUCGCUCUUUUCACAUUUAGAAUUAAUCAUUUAUCUCAACACUUGAAAUCUAACCACAAAGAUUAUGCUACUGAGAGAUCACUAGUAAAAUUGGUAGGUAAGAGAAAAAGACUUCUGGAUUAUCUU
env-1908 CUUUUCGCAAAGCAUGGAAAAUCUGCGCAGGACACAGGAAGUGCUGAAGGGCAAAUCGCUCUUUUCACAUUUAGAAUUAAUCAUUUAUCUCAACACUUAAAAUCUAACCACAAAGAUUAUGCUACUGAGAGAUCACUAGUAAAAUUGGUAGGUAAGAGAAAAAGACUUCUGGAUUAUCUU
env-1909 CUUUUCGCAAAGCAUGGAAAAUCUGCGCAGGACACAGGAAGUGCUGAAGGGCAAAUCGCUCUUUUCACAUUUAGAAUUAAUCAUUUAUCUCAACACUUGAAAUCUAACCACAAAGAUUAUGCUACUGAGAGAUCACUAGUAAAAUUGGUAGGUAAGAGAAAAAGACUUCUGG........
env-1910 CUUUUCGCAAAGCAUGGAAAAUCUGCGCAGGACACAGGAAGUGCUGAAGGGCAAAUCGCUCUUUUCACAUUUAGAAUUAAUCAUUUAUCUCAACACUUGAAAUCUAACCACAAAGAUUAUGCUACUGAAAGAUCACUAGUAAAAUUGGUAGGUAAGAGAAAAAGACUUCUGGAUUAUCUU
env-1911 CUUUUCGCAAAGCAUGGAAAAUCUGCGCAGGACACAGGAAGUGCUGAAGGGCAAAUCGCUCUUUUCACAUUUAGAAUUAAUCAUUUAUCUCAACACUUGAAAUCUAACCACAAAGAUUAUGCUACUGAGAGAUCACUAGUAAAAUUGGUAGGUAAGAGAAAAAGACUUCUGGAUUAUCUU
env-1912 CUUUUCGCAAAGCAUGGAAAAUCUGCGCAGGACACAGGAAGUGCUGAAGGGCAAAUCGCUCUUUUCACAUUUAGAAUUAAUCAUUUAUCUCAACACUUGAAAUCUAACCACAAAGAUUAUGCUACUGAGAGAUCACUAGUAAAAUUGGUAGGUAAGAGAAAAAGACUUCUGGAUUAUCUU
env-1913 CUUUUCGCAAAACAUGGAAAAUCUGCACAGGACACAGGAAGUGCUGAAGGGCAAAUCGCUCUUUUCACAUUUAGAAUUAAUCAUUUAUCUCAACACUUGAAAUCUAACCACAAAGAUUAUGCUACUGAGAGAUCACUAGUAAAAUUGGUAGGUAAGAGAAAAAGACUUCUGGAUUAUCUU
env-1915 CUUUUCGCAAAGCAUGGAAAAUCUGCGCAGGACACAGGAAGUGCUGAAGGGCAAAUCGCUCUUUUCACAUUUAGAAUUAAUCAUUUAUCUCAACACUUGAAAUCUAACCACAAAGAUUAUGCUACUGAGAGAUCACUAGUAAAAUUGGUAGGUAAGAGAAAAAGACUUCUGGAUUAUCUU
env-1916 CUUUUCGCAAAGCAUGGAAAAUCUGCGCAGGACACAGGAAGUGCUGAAGGGCAAAUCGCUCUUUUCACAUUUAGAAUU......................................................................................................
env-1917 CUUUUCGCAAAGCAUGGAAAAUCUGCGCAGGACACAGGAAGUGCUGAAGGGCAAAUUGCUCUUUUCACAUUUAGAAUUAAUCAUUUAUCUCAAC......................................................................................
env-1918 CUUUUCGCAAAGCAUGGAAAAUCUGCGCAGGACACAGGAAGUGCUGAAGGGCAAAUCGCUCUUUUCACAUUUAGAAUUAAUCAUUUAUCUCAACACUUAAAAUCUAACCACAAAGAUUAUGCUACUGAGAGAUCACUAGUAAAAUUGGUAGGUAAGAGAAAAAGACUUCUGGAUUAUCUU
env-1921 CUUUUCGCAAAGCAUGGAAAAUCUGCGCAGGACACAGGAAGUGCUGAAGGGCAAAUCGCUCUUUUCACAUUUAGAAUUAAUCAUUUAUCUCAACACUUGAAAUCUAACCACAAAGAUUAUGCUACUGAGAGAUCACUAGUAAAAUUGGUAGGUAAGAGAAAAAGACUUCUGGAUUAUCUU
env-1922 CUUUUCGCAAAACAUGGAAAAUCUGCGCAGGACACAGGAAGUGCUGAAGGGCAAAUCGCUCUUUUCACAUUUAGAAUUAAUCAUUUAUCUCAACACUUGAAAUCUAACCACAAAGAUUAUGCUACUGAGAGGUCACUAGUAAAAUUAGUAGGUAAGAGAAAAAGACUUCUGGAUUAUCUU
env-1924 CUUUUCGCAAAGCAUGGAAAAUCUGCGCAGGACACAGGAAGUGCUGAAGGGCAAAUCGCUCUUUUCACAUUUAGAAUUAAUCAUUUAUCUCAACACUUGAAAUCUAACCACAAAGAUUAUGCUACUGAGAGAUCACUAGUAAAAUUGGUAGGUAAGAGAAAAAGACUUCUGGAUUAUCUU
env-1925 CUUUUCGCAAAGCAUGGAAAAUCUGCGCAGGACACAGGAAGUGCUGAAGGGCAAAUCGCUCUUUUCACAUUUAGAAUUAAUCAUUUAUCUCAACACUUGAAAUCUAACCACAAAGAUUAUGCUACUGAGAGAUCACUAGUAAAAUUGGUAGGUAAGAGAAAAAGACUUCUGGAUUAUCUU
env-1926 CUUUUCGCAAAGCAUGGAAAAUCUGCGCAGGACACAGGAAGUGCUGAAGGGCAAAUCGCUCUUUUCACAUUUAGAAUUAAUCAUUUAUCUCAACACUUAAAAUCUAACCACAAAGAUUAUGCUACUGAGAGAUCACUAGUAAAAUUGGUAGGUAAGAGAAAAAGACUUCUGGAUUAUCUU
env-1927 CUUUUCGCAAAACAUGGAAAAUCUGCGCAGGACACAGGAAGUGCUGAAGGGCAAAUCGCUCUUUUCACAUUUAGAAUUAAUCAUUUAUCUCAACACUUGAAAUCUAACCACAAAGAUUAUGCUACUGAGAGAUCACUAGUAAAAUUGGUAGGUAAGAGAAAAAGACUUCUGGAUUAUCUU
env-1928 CUUUUCGCAAAGCAUGGAAAAUCUGCGCAGGACACAGGAAGUGCUGAAGGGCAAAUCGCUCUUUUCACAUUUAGAAUUAAUCAUUUAUCUCAACACUUGAAAUCUAACCACAAAGAUUAUGCUACUGAGAGAUCACUAGUAAAAUUGGUAGGUAAGAGAAAAAGACUUCUGGAUUAUCUU
env-1929 CUUUUCGCAAAGCAUGGAAAAUCUGCGCAGGACACAGGAAGUGCUGAAGGGCAAAUCGCUCUUUUCACAUUUAGAAUUAAUCAUUUAUCUCAACACUUGAAAUCUAACCACAAAGAUUAUGCUACUGAGAGAUCACUAGUAAAAUUGGUAGGUAAGAGAAAAAGACUUCUGGAUUAUCUU
env-1930 CUUUUCGCAAAGCAUGGAAAAUCUGCGCAGGACACAGGAAAUGCUGAAGGGCAAAUCGCUCUUUUCACAUUUAGAAUUAAUCAUUUAUCUCAACACUUGAAAUCUAACCACAAAGAUUAUGCUACUGAGAGAUCACUAGUAAAAUUGGUAGGUAAGAGAAAAAGACUUCUGGAUUAUCUU
env-1931 ....................................................................................................................................................................................
env-1935 CUUUUCGCAAAGCAUGGAAAAUCUGCGCAGGACACAGGAAGUGCUGAAGGGCAAAUCGCUCUUUUCACAUUUAGAAUUAAUCAUUUAUCUCAACACUUGAAAUCUAACCACAAAGAUUAUGCUACUGAGAGAUCACUAGUAAAAUUGGUAGGUAAGAGAAAAAGACUUCUGGAUUAUCUU
env-1936 AUCUUCAAAAAACAUGGAGGAAGUGAAACCAAUACAGGUUCCACAGAAGGGCAAA.............................................................................................................................
env-1937 CUUUUCGCAAAGCAUGGAAAAUCUGCGCAGGACACAGGAAGUGCUGAAGGGCAAAUCGCUCUUUUCACAUUUAGAAUUAAUCAUUUAUCUCAACACUUGAAAUCUAACCACAAAGAUUAUGCUACUGAGAGAUCACUAGUAAAAUUGGUAGGUAAGAGAAAAAGACUU............
env-1938 ....................................................................................................................................................................................
env-1939 CUUUUCGCAAAACAUGGAAAAUCUGCACAGGACACAGGAAGUGCUGAAGGGCAAAUCGCUCUUUUCACAUUUAGAAUUAAUCAUUUAUCUCAACACUUGAAAUCUAACCACAAAGAUUAUGCUACUGAGAGAUCACUAGUAAAAUUGGUAGGUAAGAGAAAAAGACUUCUGGAUUAUCUU
env-1940 NNNNNGCAGAACUUUUCGCAAAACAUGGAAAAUCUGCGCAGGACACAGGAAGUGCUGAAGGGCAAAUCGCUCUUUUCACAUUUAGAAUUAAUCAUUUAUCUCAACACUUGAAAUCUAACCACAAAGAUUAUGCUACUGAGAGAUCACUAGUAAAAUUAGUAGGUAAGAGAAAAAGACUU
env-1941 CUUUUCGCAAAGCAUGGAAAAUCUGCGCAGGACACAGGAAGUGCUGAAGGGCAAAUCGCUCUUUUCACAUUUAGAAUUAAUCAUUUAUCUCAACACUUGAAAUCUAACCACAAAGAUUAUGCUACUGAAAGAUCACUAGUAAAAUUGGUAGGUAAGAGAAAAAGACUUCUGGAUUAUCUU
env-1942 CUUUUCGCAAAGCAUGGAAAAUCUGCGCAGGACACAGGAAGUGCUGAAGGGCAAAUCGCUCUUUUCACAUUUAGAAUUAAUCAUUUAUCUCAACACUUAAAAUCUAACCACAAAGAUUAUGCUACUGAGAGAUCACUAGUAAAAUUGGUAGGUAAGAGAAAAAGACUUCUGGAUUAUCUU
env-1943 CUUUUCGCAAAGCAUGGAAAAUCUGCGCAGGACACAGGAAGUGCUGAAGGGCAAAUCGCUCUUUUCACAUUUAGAAUUAAUCAUUUAUCUCAACACUUGAAAUCUAACCACAAAGAUUAUGCUACUGAGAGAUCACUAGUAAAAUUGGUAGGUAAGAGAAAAAGACUUCUGGAUUAUCUU
env-1944 CUUUUCGCAAAACAUGGAAAAUCUGCGCAGGACACAGGAAGUGCUGAAGGGCAAAUUGCUCUUUUCACAUUUAGAAUUAAUCAUUUAUCUCAACACUUGAAAUCUAACCACAAAGAUUAUGCUACUGAGAGAUCACUAGUAAAAUUGGUAGGUAAGAGAAAAAGACUUCUGGAUUAUCUU
env-1945 CUUUUCGCAAAGCAUGGAAAAUCUGCGCAGGACACAGGAAGUGCUGAAGGGCAAAUCGCUCUUUUCACAUUUAGAAUUAAUCAUUUAUCUCAACACUUGAAAUCUAACCACAAAGAUUAUGCUACUGAGAGAUCACUAGUAAAAUUGGUAGGUAAGAGAAAAAGACUUCUGGAUUAUCUU
env-1946 CUUUUCGCAAAACAUGGAAAAUCUGCACAGGACACAGGAAGUGCUGAAGGGCAAAUCGCUCUUUUCACAUUUAGAAUUAAUCAUUUAUC...........................................................................................
env-1947 CUUUUCGCAAAACAUGGAAAAUCUGCGCAGGACACAGGAAGUGCUGAAGGGCAAAUCGCUCUUUUCACAUUUAGAAUUAAUCAUUUAUCUCAACACUUGAAAUCUAACCACAAAGAUUAUGCUACUGAGAGAUCACUAGUAAAAUUAGUAGGUAAGAGAAAAAGACUUCUGGAUUAUCUU
env-1948 CUUUUCGCAAAGCAUGGAAAAUCUGCGCAGGACACAGGAAGUGCUGAAGGGCAAAUUGCUCUUUUCACAUUUAGAAUUAAUCAUUUAUCUCAACACUUGAAAUCUAACCACAAAGAUUAUGCUACUGAGAGAUCACUAGUAAAAUUGGUAGGUAAGAGAAAAAGACUUCUGGAUUAUCUU
env-1949 ....................................................................................................................................................................................
env-1950 CUUUUCGCAAAGCAUGGAAAAUCUGCACAGGACACAGGAAGUGCUGAAGGGCAAAUCGCUCUUUUCACAUUUAGAAUUAAUCAUUUAUCUCAACACUUGAAAUCUAACCACAAAGAUUAUGCUACUGAGAGAUCACUAGUAAAAUUGGUAGGUAAGAGAAAAAGACUUCUGGAUUAUCUU
env-1951 ....................................................................................................................................................................................
env-1952 CUUUUCGCAAAACAUGGAAAAUCUGCACAGGACACAGGAAGUGCUGAAGGGCAAAUCGCUCUUUUCACAUUUAGAAUUAAUCAUUUAUCUCAACACUUGAAAUCUAACCACAAAGAUUAUGCUACUGAGAGAUCACUAGUAAAAUUGGUAGGUAAGAGAAAAAGGCUUCUGGAUUAUCUU
env-1954 ....................................................................................................................................................................................
env-1955 CUUUUCGCAAAACAUGGAAAAUCUGCACAGGACACAGGAAGUGCUGAAGGGCAAAUCGCUCUUUUCACAUUUAGAAUUAAUCAUUUAUCUCAACACUUGAAAUCUAACCACAAAGAUUAUGCUACUGAGAGAUCACUAGUAAAAUUGGUAGGUAAGAGAAAAAGACUUCUGGAUUAUCUU
env-1957 CUUUUCGCAAAACAUGGAAAAUCUGCGCAGGACACAGGAAGUGCUGAAGGGCAAAUCGCUCUUUUCACAUUUAGAAUUAAUCAUUUAUCUCAACACUUGAAAUCUAACCACAAAGAUUAUGCUACUGAGAGAUCACUAGUAAAAUUGGUAGGUAAGAGAAAAAGACUUCUGGAUUAUCUU
env-1958 CUUUUCGCAAAACAUGGAAAAUCUGCGCAGGACACAGGAAGUGCUGAAGGGCAAAUCGCUCUUUUCACAUUUAGAAUUAAUCAUUUAUCUCAACACUUGAAAUCUAACCACAAAGAUUAUGCUACUGAGAGAUCACUAGUAAAAUUGGUAGGUAAGAGAAAAAGACUUCUGGAUUAUCUU
env-1959 CUUUUCGCAAAGCAUGGAAAAUCUGCGCAGGACACAGGAAGUGCUGAAGGGCAAAUCGCUCUUUUCACAUUUAGAAUUAAUCAUUUAUCUCAACACUUAAAAUCUAACCACAAAGAUUAUGCUACUGAGAGAUCACUAGUAAAAUUAGUAGGUAAGAGAAAAAGACUUCUGGAUUAUCUU
env-1961 ....................................................................................................................................................................................
env-1962 CUUUUCGCAAAACAUGGAAAAUCUGCACAGGACACAGGAAGUGCUGAAGGGCAAAUCGCUCUUUUCACAUUUAGAAUUAAUCAUUUAUCUCAACACUUGAAAUCUAACCACAAAGAUUAUGCUACUGAGAGAUCACUAGUAAAAUUGGUAGGUAAGAGAAAAAGACUUCUGGAUUAUCUU
env-1963 CUUUUCGCAAAGCAUGGAAAAUCUGCGCAGGACACAGGAAGUGCUGAAGGGCAAAUCGCUCUUUUCACAUUUAGAAUUAAUCAUUUAUCUCAACACUUGAAAUCUAACCACAAAGAUUAUGCUACUGAGAGAUCACUAGUAAAAUUGGUAGGUAAGAGAAAAAGACUUCUGGAUUAUCUU
env-1964 ....................................................................................................................................................................................
env-1965 CUUUUCGCAAAACAUGGAAAAUCUGCACAGGACACAGGAAGUGCUGAAGGGCAAAUCGCUCUUUUCACAUUUAGAAUUAAUCAUUUAUCUCAACACUUAAAAUCUAACCACAAAGAUUAUGCUACUGAGAGAUCACUAGUAAAAUUAGUAGGUAAGAGAAAAAGACUUCUGGAUUAUCUU
env-1966 CUUUUCGCAAAGCAUGGAAAAUCUGCGCAGGACACAGGAAGUGCUGAAGGGCAAAUCGCUCUUUUCACAUUUAGAAUUAAUCAUUUAUCUCAACACUUGAAAUCUAACCACAAAGAUUAUGCUACUGAGAGAUCACUAGUAAAAUUGGUAGGUAAGAGAAAAAGACUUCUGGAUUAUCUU
env-1967 ....................................................................................................................................................................................
env-1968 CUUUUCGCAAAGCAUGGAAAAUCUGCACAGGACACAGGAAGUGCUGAAGGGCAAAUCGCUCUUUUCACAUUUAGAAUUAAUCAUUUAUCUCAACACUUGAAAUCUAACCACAAAGAUUAUGCUACUGAGAGAUCACUAGUAAAAUUGGUAGGUAAGAGAAAAAGACUUCUGGAUUAUCUU
env-1969 CUUUUCGUAAAACAUGGAAAAUCUGCACAGGACACAGGAAGUGCUGAAGGGCAAAUCGCUCUUUUCACAUUUAGAAUUAAUCAUUUAUCUCAACACUUAAAAUCUAACCACAAAGAUUAUGCUACUGAGAGAUCACUAGUAAAAUUAGUAGGUAAGAGAAAAAGACUUCUGGAUUAUCUU
env-1970 ....................................................................................................................................................................................
env-1971 CUUUUCGCAAAACAUGGAAAAUCUGCACAGGACACAGGAAGUGCUGAAGGGCAAAUCGCUCUUUUCACAUUUAGAAUUAAUCAUUUAUCUCAACACUUGAAAUCUAACCACAAAGAUUAUGCUACUGAGAGAUCACUAGUAAAAUUGGUAGGUAAGAGAAAAAGACUUCUGGAUUAUCUU
env-1972 ....................................................................................................................................................................................
env-1973 CUUUUCGCAAAGCAUGGAAAAUCUGCACAGGACACAGGAAGUGCUGAAGGGCAAAUUGCUCUUUUCACAUUUAGAAUUAAUCAUUUAUCUCAACACUUAAAAUCUAACCACAAAGAUUAUGCUACUGAGAGAUCACUAGUAAAAUUGGUAGGUAAGAGAAAAAGACUUCUGGAUUAUCUU
env-1974 CUUUUCGCAAAACAUGGAAAAUCUGCACAGGACACAGGAAGUGCUGAAGGGCAAAUCGCUCUUUUCACAUUUAGAAUUAAUCAUUUAUCUCAACACUUGAAAUCUAACCACAAAGAUUAUGCUACUGAGAGAUCACUAGUAAAAUUGGUAGGUAAGAGAAAAAGACUUCUGGAUUAUCUU
env-1975 CUUUUCGCAAAGCAUGGAAAAUCUGCGCAGGACACAGGAAGUGCUGAAGGGCAAAUUGCUCUUUUCACAUUUAGAAUUAAUCAUUUAUCUCAACACUUGAAAUCUAACCACAAAGAUUAUGCUACUGAGAGAUCACUAGUAAAAUUGGUAGGUAAGAGAAAAAGACUUCUGGAUUAUCUU
env-1976 CUUUUCGCAAAACAUGGAAAAUCUGCACAGGACACAGGAAGUGCUGAAGGGCAAAUCGCUCUUUUCACAUUUAGAAUUAAUCAUUUAUCUCAACACUUGAAAUCUAACCACAAAGAUUAUGCUACCGAGAGAUCACUAGUAAAAUUGGUAGGUAAGAGAAAAAGACUUCUGGAUUAUCUU
env-1977 CUUUUCGCAAAACAUGGAAAAUCUGCACAGGACACAGGAAGUGCUGAAGGGCAAAUCGCUCUUUUCACAUUUAGAAUUAAUCAUUUAUCUCAACACUUAAAAUCUAACCACAAAGAUUAUGCUACUGAGAGAUCACUAGUAAAAUUGGUAGGUAAGAGAAAAAGACUUCUGGAUUAUCUU
env-1978 CUUUUCGCAAAACAUGGAAAAUCUGCACAGGACACAGGAAGUGCUGAAGGGCAAAUCGCUCUUUUCACAUUUAGAAUUAAUCAUUUAUCUCAACACUUGAAAUCUAACCACAAAGAUUAUGCUACUGAGAGAUCACUAGU........................................
env-1979 CUUUUCGCAAAGCAUGGAAAAUCUGCACAGGACACAGGAAGUGCUGAAGGGCAAAUCGCUCUUUUCACAUUUAGAAUUAAUCAUUUAUCUCAACACUUGAAAUCUAACCACAAAGAUUAUGCUACUGAGAGAUCACUAGUAAAAUUGGUAGGUAAGAGAAAAAGACUUCUGGAUUAUCUU
env-1980 CUUUUCGCAAAACAUGGAAAAUCUGCACAGGACACAGGAAGUGCUGAAGGGCAAAUCGCUCUUUUCACAUUUAGAAUUAAUCAUUUAUCUCAACACUUGAAAUCUAACCACAAAGAUUAUGCUACUGAGAGAUCACUAGUAAAAUUGGUAGGUAAGAGAAAAAGACUUCUGGAUUAUCUU
env-1981 CUUUUCGCAAAACAUGGAAAAUCUGCGCAGGACACAGGAAGUGCUGAAGGGCAAAUCGCUCUUUUCACAUUUAGAAUUAAUCAUUUAUCUCAACACUUGAAAUCUAACCACAAAGAUUAUGCUACUGAGAGAUCACUAGUAAAAUUGGUAGGUAAGAGAAAAAGACUUCUGGAUUAUCUU
env-1982 CUUUUCGCAAAACAUGGAAAAUCUGCACAGGACACAGGAAGUGCUGAAGGGCAAAUCGCUCUUUUCACAUUUAGAAUUAAUCAUUUAUCUCAACACUUAAAAUCUAACCACAAAGAUUAUGCUACUGAGAGAUCACUAGUAAAAUUAGUAGGUAAGAGAAAAAGACUUCUGGAUUAUCUU
env-1983 ....................................................................................................................................................................................
env-1984 CUUUUCGCAAAACAUGGAAAAUCUGCGCAGGACACAGGAAGUGCAGAAGGGCAAAUUGCUCUUUUCACAUUUAGAAUUAAUCAUUUAUCUCAACACUUGAAAUCUAACCACAAAGAUUAUGCUACUGAGAGAUCACUAGUAAAAUUGGUAGGUAAGAGAAAAAGACUUCUGGAUUAUCUU
env-1985 CUUUUCGCAAAACAUGGAAAAUCUGCACAGGACACAGGAAGUGCUGAAGGGCAAAUCGCUCUUUUCACAUUUAGAAUUAAUCAUUUAUCUCAACACUUGAAAUCUAACCACAAAGAUUAUGCUACUGAGAGAUCACUAGUAAAAUUAGUAGGUAAGAGAAAAAGACUUCUGGAUUAUCUU
env-1986 ....................................................................................................................................................................................
env-1987 CUUUUCGCAAAACAUGGAAAAUCUGCGCAGGACACAGGAAGUGCUGAAGGGCAAAUCGCUCUUUUCACAUUUAGAAUUAAUCAUUUAUCUCAACACUUGAAAUCUAACCACAAAGAUUAUGCUACUGAGAGAUCACUAGUAAAAUUGGUAGGUAAGAGAAAAAGACUUCUGGAUUAUCUU
env-1988 ....................................................................................................................................................................................
env-1990 CUUUUCGCAAAACAUGGAAAAUCUGCACAGGACACAGGAAGUGCAGAAGGGCAAAUUGCUCUUUUCACAUUUAGAAUUAAUCAUUUAUCUCAACACUUGAAAUCUAACCACAAAGAUUAUGCUACUGAGAGAUCACUAGUAAAAUUGGUAGGUAAGAGAAAAAGACUUCUGGAUUAUCUU
env-1991 CUUUUCGCAAAACAUGGAAAAUCUGCACAGGACACAGGAAGUGCAGAAGGGCAAAUUGCUCUUUUCACAUUUAGAAUUAAUCAUUUAUCUCAACACUUGAAAUCUAACCACAAAGAUUAUGCUACUGAGAGAUCACUAGUAAAAUUGGUAGGUAAGAGAAAAAGACUUCUGGAUUAUCUU

         ....................................................................................................................................................................................
         ....................................................................................................................................................................................
         AUYUUYR•RAARCACGG•RRR•••R•RR••RAYAC•GG••C•RC•GAAGG•CAAAUYGC•YURUUYAC•Y•Y•G•AUYRAYCAYYUR•C•RA•CAYYURAAAAR•AAYCR•AARGAUU•YAAYAC•GAR•G•UC•YURGU•••RYURGU•GG•AAR•G•RG••••YU•YU•GAYUAYYU•
```

```
env-1992  CUUUUCGCAAAACAUGGAAAAUCUGCACAGGACACAGGAAGUGCUGAAGGGCAAAUCGCUCUUUUCACAUUUAGAAUUAAUCAUUUAUCUCAACACUUGAAAUCUAACCACAAAGAUUAUGCUACUGAGAGAUCACUAGUAAAAUUGGUAGGUAAGAGAAAAAGACUUCUGGAUUAUCUU
env-1993  CUUUUCGCAAAACAUGGAAAAUCUGCACAGGACACAGGAAGUGCUGAAGGGCAAAUCGCUCUUUUCACAUUUAGAAUUAAUCAUUUAUCUCAACACUUGAAAUCUAACCACAAAGAUUAUGCUACUGAGAGAUCACUAGUAAAAUUGGUAGGUAAGAGAAAAAGACUUCUGGAUUAUCUU
env-1994  ....................................................................................................................................................................................
env-1995  ....................................................................................................................................................................................
env-1996  CUUUUCGCAAAACAUGGAAAAUCUGCACAGGACACAGGAAGUGCUGAAGGGCAAAUCGCUCUUUUCACAUUUAGAAUUAAUCAUUUAUCUCAACACUUGAAAUCUAACCACAAAGAUUAUGCUACUGAGAGAUCACUAGUAAAAUUGGUAGGUAAGAGAAAAAGACUUCUGGAUUAUCUU
env-1997  CUUUUCGCAAAACAUGGAAAAUCUGCACAGGACACAGGAAGUGCUGAAGGGCAAAUCGCUCUUUUCACAUUUAGAAUUAAUCAUUUAUCUCAACACUUGAAAUCUAACCACAAAGAUUAUGCUACUGAGAGAUCACUAGUAAAAUUGGUAGGUAAGAGAAAAAGACUUCUGGAUUAUCUU
env-1998  CUUUUCGCAAAACAUGGAAAAUCUGCACAAGACACAGGAAGUGCUGAAGGGCAAAUCGCUCUUUUCACAUUUAGAAUUAAUCAUUUAUCUCAACACUUGAAAUCUAACCAUAAAGAUUAUGCUACUGAGAGAUCACUAGUAAAAUUGGUAGGUAAGAGAAAAAGACUUCUGGAUUAUCUU
env-1999  CUUUUCGCAAAACAUGGAAAAUCUGCACAAGACACAGGAAGUGCUGAAGGGCAAAUCGCUCUUUUCACAU..............................................................................................................
env-2000  CUUUUCGCAAAACAUGGAAAAUCUGCACAAGACACAGGAAGUGCUGAAGGGCAAAUCGCUCUUUUCACAUUUAGAAUUAAUCAUUUAUCUCAACACUUGAAAUCUAACCACAAAGAUUAUGCUACUGAGAGAUCACUAGUAAAAUUGGUAGGUAAGAGAAAAAGACUUCUGGAUUAUCUU
env-2001  ....................................................................................................................................................................................
env-2002  ....................................................................................................................................................................................
env-2003  CUUUUCGCAAAACAUGGAAAAUCUGCACAGGACACAGGAAGUGCUGAAGGGCAAAUCGCUCUUUUCACAUUUAGAAUUAAUCAUUUAUCUCAACACUUGAAAUCUAACCACAAAGAUUAUGCUACUGAGAGAUCACUAGUAAAAUUGGUAGGUAAGAGAAAAAGACUUCUGGAUUAUCUU
env-2004  CUUUUCGCAAAACAUGGAAAAUCUGCGCAGGACACAGGAAGUGCUGAAGGGCAAAUCGCUCUUUUCACAUUUAGAAUUAAUCAUUUAUCUCAACACUUGAAAUCUAACCACAAAGAUUAUGCUACUGAGAGAUCACUAGUAAAAUUGGUAGGUAAGAGAAAAAGACUUCUGGAUUAUCUU
env-2005  CUUUUCGCAAAACAUGGAAAAUCUGCACAGGACACAGGAAGUGCUGAAGGGCAAAUCGCUCUUUUCACAUUUAGAAUUAAUCAUUUAUCUCAACAUUUGAAAUCUAACCACAAAGAUUAUGCUACUGAGAGAUCACUAGUAAAAUUGGUAGGUAAGAGAAAAAGACUUCUGGAUUAUCUU
Ran-4-1   AUCUUCGCAAAGCACGGAAAAUCUGCGCAGGACACAGGUAGUGCAGAAGGGCAAAUCGCUCUUUUCACUUACAGAAUUAACCACCUAUCUCAGCACUUAAAGUCUAAUCACAAAGACUUCGCUACUGAGAAAUCACUUGUUAAAAUGGUAGGUAAAAGAAAGAGAUUAUUAGAUUAUCUU
Ran-2-1   AUCUUCGCAAAGCACGGAAAAUCUGCGCAGGACACAGGUAGUGCAGAAGGGCAAAUCGCUCUUUUCACUUACAGAAUUAACCACCUAUCUCAGCACUUAAAGUCUAAUCACAAAGACUUCGCUACUGAGAAAUCACUUGUUAAAAUGGUAGGUAAAAGAAAGAGAUUAUUAGAUUAUCUU
Cca-2-1   AUUUUUAAAAAACACGCAGGUUCUGAAGCUAAUACAGGUUCUGCAGAAGGACAGAUUGCCUUAUUCACUUUCAGAAUCAAUCACUUGACUGAGCACUUGAAAAGAAAUAGAAAAGAUUUCAAUACCGAGCGUUCAUUGGUGAAAUUAGUUGGUAAACGUAGAAGUUUACUUGAUUAUUUG
Cca-2-2   AUUUUUAAAAAACACGCAGGUUCUGAAGCUAAUACAGGUUCUGCAGAAGGACAGAUUGCCUUAUUCACUUUCAGAAUCAAUCACUUGACUGAGCACUUGAAAAGAAAUAGAAAAGAUUUCAAUACCGAGCGUUCAUUGGUGAAAUUGGUUGGUAAACGUAGAAGUUUACUUGAUUAUUUG
env-2006  AUUUUUGCAAAACACGGUAAAUCACAAGCAGAUACCGGUUCUGCCGAAGGCCAAGUAGCUUUGUUCACUUACCGAAUCAAUCACCUGACCGGACAUUUAAAAAACAAUAAAAAGGAUUUUAACACCGAAAAGGCACUGGUGAGAUUGGUGGGCAAAAGAAAAAAAUGUUGGAUUAUCUG
env-2007  AUUUUCAAGAAACACGGCGGCUCAGAAGGCAACACAGGUUCUACGGAAGGACAGAUCGCUUUGUUCACACACCGUAUCAACCACUUGACAGAACACCUUAAAAGGAACCACAAAGAUUAUAACACCGAGCGUUCCUUGGUUCGUUUGGUAGGUAAAAGACGUAGUCUUUUGGAUUACCUU
Mru-1-1   AUUUUUAAAAAACACGGCGGCUCUGAGAGCAACACCGGUUCUACGGAAGGACAAAUUGCUUUGUUCACACACCGUAUCAACCACCUAACAGGACACCUUAAAAAGAACCACAAAGAUUACAACACCGAGCGUUCUUUGGUGGCAUUGGUAGGUAAAAGACGUAGUCUUUUGGAUUACCUA
env-2008  AUUUUUAAAAAGCACGGCGGCUCUGAAGGCAACACUGGUUCUACGGAAGGACAAAUUGCUUUGUUCACACACCGUAUCAACCACUUAACAGGACACCUUAAAAGGAACCACAAAGAUUACAACACCGAGCGUUCUUUGGUGGCAUUGGUAGGUAAAAGACGUAGUCUUUUGGAUUACCUA
env-2009  AUUUUUAAAAAGCACGGCGGCUCUGAAGGCAACACUGGUUCUACNGAAGGACAAAUUGCUUUGUUCACACACCGUAUCAACCACUUAACNGGACACCUUAAAAGGAACCACAAAGAUUACAACACCGAGCGUUCUUUGGUGGCAUUGGUAGGUAAAAGACGUAGUCUUUUGGAUUACCUA
env-2010  ....................................................................................................................................................................................
env-2011  AUUUUCAAAAAACACGGCGGCUCUGAGAGCAAUACAGGUUCUACGGAAGGACAAAUUGCUUUGUUCACACACCGUAUCAACCAUUUAACAGGACACCUUAAAAAGAACCACAAAGAUUAUAACACCGAGCGUUCUUUGGUGGCAUUGGUAGGUAAAAGACGUAGUCUUUUGGAUUACCUA
env-2012  AUUUUUAAGAAACACGGCGGCUCUGAGAGCAACACCGGUUCUACGGAAGGACAAAUUGCGUUGUUCACGCACCGUAUCAACCAUUUGACAGAACACCUUAAGAGAAACCACAAAGAUUACAACACCGAGCGCUCCUUGGUUGCGUUGGUAGGUAAAAGACGUAGUCUUUUGGAUUAUCUA
env-2014  AUUUUCAAAAAAUACGGCGGCUCUGAGAGCAACACCGGUUCUACGGAAGGGCAAAUCGCCUUGUUCACCUACCGUAUCAACCACUUGACAGGACACCUUAAAAAGAACCAUAAGGAUUAUAACACCGAGCGUUCUUUGGUGGCAUUGGUAGGUAAAAGACGUAGUCUUUUGGAUUACCUA
env-2015  AUUUUUAAGAAACACGGCGGCUCUGAG.........................................................................................................................................................
env-2016  AUUUUUAAGAAACACGGCGGCUCUGAGAGCAACACCGGUUCUACGGAAGGACAAAUUGCGUUGUUCACGCACCGUAUCAACCAUUUGACAGAACACCUUAAGAGAAACCACAAAGAUUACAACACCGAGCGCUCCUUGGUUGC.....................................
env-2017  AUUUUCAAGAAACACGGCGGCUCUGAGAGCAACACCGGUUCUACGGAAGGACAAAUCGCUUUGUUCACGCACCGUAUCAACCACUUGACAGGACACCUUAAAAAGAACCAUAAAGAUUACAACACUGAGCGUUCUUUGGUGGCAUUGGUAGGUAAAAGACGUAGUCUUUUGGAUUACCUG
env-2018  AUUUUUAAGAAACACGGCGGCUCUGAGAGCAACACCGGUUCUACGGAAGGACAAAUUGCGUUGUUCACGCACCGUAUCAACCAUUUGACAGAACACCUUAAGAGAAACCACAAAGAUUACAACACCGAGCGCUCCUUGGUUGCGUUGGUAGGUAAAAGACGUAGUCUUUUGGAUUAUCUA
Msp-2-1   AUUUUCAAAAAACACGGCGGCUCUGAGAGCAACACCGGUUCUACGGAAGGUCAAAUUGCAUUGUUCACACACCGUAUCAACCACUUAACAGAACACCUUAAAAGGAACCACAAGGAUUACAACACUGAACGUUCUUUGGUGGCUUUGGUAGGUAAAAGACGUAGUCUUUUGGAUUACCUA
Cba-2-1   AUUUUUAAAAAACACGGUGGUGAAGAAAAAAACACUGGUUCUGCUGAAGGGCAAAUCGCAUUGUUUACACACCGUAUCAAUCACUUAACUGAGCAUUUGAAGAAAAAUCGUAAAGAUUUCAACACAGAGCGUUCAUUAGUAAAAUUGGUAGGUAAAAGAAGAAGCUUACUUGAUUACUUA
Csp-11-1  AUUUUUAAAAAACACGGUGGUGAAGAAAAAAACACUGGUUCUGCUGAAGGGCAAAUCGCAUUGUUUACACACCGUAUCAAUCACUUAACUGAGCAUUUGAAGAAAAAUCGUAAAGAUUUCAACACAGAGCGUUCAUUAGUAAAAUUGGUAGGUAAAAGAAGAAGCUUACUUGAUUACUUA
Cba-4-1   AUUUUUAAAAAACACGGUGGUGAAGAAAAAAACACUGGUUCUGCUGAAGGGCAAAUCGCAUUGUUUACACACCGUAUCAAUCACUUAACUGAGCAUUUGAAGAAAAAUCGUAAAGAUUUCAACACAGAGCGUUCAUUAGUAAAAUUGGUAGGUAAAAGAAGAAGCUUACUUGAUUACUUA
env-2019  AUAUUCAAGAAAUACGGCGGCUCAGAUUCCAACACGGGAAGCACCGAGGGGCAGAUUGCCCUCUUCACC...............................................................................................................
env-2020  AUUUUUAAGAAGCACGGCGGCAAAGCUGAGAACACAGGUUCUACAGAAGGUCAGAUUGCUUUAUUCACACACCGUAUCAAUCACUUGACCGGUCACUUGAAAAAAAAUCAUAAAGACUACAACACUGAGCGUUCACUUGUGAAACUAGUAGGUAAAAGAAGAAGUCUUCUUGAUUAUAUG
Csp-12-1  AUUUUUAAAAAACACGGUGGCGAAGCAAAAAACACUGGAUCUGCUGAAGGGCAAAUUGCAUUGUUUACACACCGAAUUAACCACUUAACAGGGCAUUUAAAAAACAACCGUAAGGAUUAUAACACGGAGCGUUCUUUAGUAAAGCUGGUGGGUAAAAGAAGAAGUCUUUUAGAUUAUUUG
Zul-1-1   AUCUUUAAGAAACACGGCGGAAAAGCAGAGAACACUGGUUCUACUGAAGGUCAGAUUGCAUUGUUCACACACCGUAUCAAUCACUUGACCGGUCACUUGAAAAGAAAUCACAAAGACUACAACACUGAGCGUUCAUUGGUAAAACUCGUAGGUAAAAGAAGAAGCCUACUAGAUUACAUG
env-2021  AUUUUUAAAAAAUACGGCGGUAGUGCAGAAAACACUGGCUCUACAGAAGGACAAAUUGCUUUGUUCACACACAGAAUCAACCACUUGACCGGUCACUUGAAAAAGAAUCACAAAGAUUACAACACUGAGCGUUCACUUGUAAAAUUGGUGGGUAAAAGAAGAAGUCUUUUAGAUUACAUG
env-2027  AUUUUUAAAAAAUACGGCGGUAGUGCAGAAAACACUGGCUCUACAGAAGGACAAAUUGCUUUGUUCACACACAGAAUCAACCACUUGACCGGUCACUUGAAAAAGAAUCACAAAGAUUACAACACUGAGCGUUCACUUGUAAAAUUGGUGGGUAAA........................
env-2028  AUUUUUAAAAAAUACGGCGGUAGUGCAGAAAACACUGGCUCCACAGAAGGACAAAUUGCUUUGUUCACACACAGAAUCAACCACUUGACCGGUCACUUGAAAAAGAAUCACAAAGAUUACAACACUGAGCGUUCACUUGUAAAAUUGGUGGGUAAAAGAAGAAGUCUUUUAGAUUACAUG
env-2029  AUCUUUAAGAAGCACGGCAAAAAUGAAAAAGAUUCUGGAACUGCAGAAGGGCAAAUUGCUUUAUUCACACACAGAAUUAACCACCUAACUGGUCACUUAAAAAAUAAUCACAAAGAUUACAACACAGAGCGAUCGCUAGUUAAAUUAGUAGGUAAAAGAAGAAGUCUUUUAGAUUAUUUA
Ace-1-1   AUUUUUAAAAAACACGGCGGUGAUGCAAAGAACACAGGUUCUACUGAAGGACAAAUAGCUUUGUUCACUUAUAGAAUUGAGCACCUAUCCCAACACCUAAAGAAAAAUCGUAAAGAUUUUAACACUGAGCGUUCCUUGGUGAAAUUGGUAGGUAAGAGAAGAUCACUUUUAGAUUACCUA
Ala-2-1   AUUUUUAAAAAACACGGCGGUGACGCAAAGAACACAGGUUCUACUGAAGGACAAAUAGCUUUGUUCACUUACAGAAUUGAACACCUAUCUCAACACUUGAAGAAAAAUCGUAAAGAUUUUAACACUGAGCGUUCAUUGGUAAAAUUGGUAGGUAAGAGAAGAUCUCUUUUAGAUUAUUUA
env-2030  ....................................................................................................................................................................................
env-2031  AUUUUUAAAAAACACGGCGGAAAAGAAGAUAACACAGGUUCAACAGAAGGGCAAGUUGCUUUGUUCACACAUCGCAUUAACCACUUAACUGGGCACCUUAAAAAAAAAUCACAAAGAUUAUAACACCGAACGUUCGUUAGUAAGACUGGUAGGUAAAAGAAGAAGUCUUUUAGAUUACUUG
env-2032  AUUUUUAAAAAACACGGCGGAAAAGAAGAUAACACAGGUUCAACAGAAGGGCAAGUUGCUUUGUUCACACAUCGCAUUAACCACUUAACUGGGCACCUUAAAAAAAAAUCACAAAGAUUAUAACACCGAACGUUCGUUAGUUAAACUGGUAGGUAAAAGAAGAAGUCUUUUAGAUUACUUG
env-2033  AUUUUUAAAAAACACGGCGGAAAAGAAGAUAACACAGGUUCAACAGAAGGGCAAGUUGCUUUGUUCACACAUCGCAUUAACCACUUAACUGGGCACCUUAAAAAAA..........................................................................
env-2034  AUUUUUAAAAAACACGGCGGAAAAGAAGAUAACACAGGUUCAACAGAAGGGCAAGUUGCUUUGUUCACACAUCGCAUUAACCACUUAACUGGGCACCUUAAAAAAAAAUCACAAAGAUUAUAACACCGAACGUUCGUUAGUAAGACUGGUAGGUAAAAGAAGAAGUCUUUUAGAUUACUUG
env-2035  AUUUUUAAAAAAUACAGCGGAAAAGAAGAGAACACUGGUUCAACAGAAGGGCAAGUUGCUUUGUUCACACAUCGCAUUAACCACUUGACAGGGCACCUUAAAAAUAAUCAUAAAGAUUAUAACACCGAACGUUCGUUGGUAAGACUGGUAGGUAAAAGAAGAAGUCUUUUAGAUUACUUG
env-2036  AUUUUUAAAAAAUACAGCGGAAAAGAAGAGAACACAGGUUCAACAGAAGGGCAAGUUGCUUUGUUCACACAUCGCAUCAACCACUUAACAGGACACCUUAAAAGAAAUCACAAAGAUUAUAACACCGAACGUUCGUUAGUAAGACUGGUAGGUAAGAGAAGAAGUCUUUUAGAUUACUUG
env-2037  AUUUUUAAAAAACACGGCGGAAAAGAAGA.......................................................................................................................................................
env-2038  AUUUUUAAAAAAUACAGCGGAAAAGAAGAGAACACUGGUUCAACAGAAGGGCAAGUUGCUUUGUUCACACAUCGCAUUAACCACUUGACAGGGCACCUUAAAAAUAAUCAUAAAGAUUAUAACACCGAACGUUCGUUGGUAAGACUGGUAGGUAAAAGAAGAAGUCUUUUAGAUUACUUG
env-2039  AUUUUUAAAAAAUACAGCGGAAAAGAAGAGAACACAGGUUCAACAGAAGGGCAAGUUGCUUUGUUCACACAUCGCAUCAACCACUUAACAGGGCACCUUAAAAGAAAUCACAAAGAUUAUAACACCGAACGUUCGUUAGUAAGACUGGUAGGUAAGAGAA....................
env-2040  AUUUUUAAAAAAUACAGCGGAAAAGAAGAGAACACAGGUUCAACAGAAGGGCAAGUUGCUUUGUUCACACAUCGCAUCAACCACUUAACAGGGCACCUUAAAAGAAAUCACAAAGAUUAUAACACCGAACGUUCGUUAGUAA......................................
env-2041  AUUUUUAAAAAACACGGCGGAAAAGAAGAUAACACAGGUUCAACAGAAGGGCAAGUUGCUUUGUUCACACAUCGCAUUAACCACUUAACUGGGCACCUUAAAAAA...........................................................................
env-2042  AUUUUUAAAAAAUACAGCGGAAAAGAAGAGAACACAGGUUCAACAGAAGGGCAAGUUGCUUUGUUCACACAUCGCAUCAACCACUUAACAGGGCACCUUAAAAGAAAUCACAAAGAUUAUAACACCGAACGUUCGUUAGUAAGACUGGU...............................
env-2043  AUUUUUAAAAAACACGGCGGAAAAGAAGAUAACACAGGUUCAACAGAAGGGCAAGUUGCUUUGUUCACACAUCGCAUUAACCACUUAACUGGGCACCUUAAAAAAAAAUCACAAAGAUUAUAACAC......................................................
env-2044  AUUUUUAAAAAACACGGCGGAAAAGAAGAUAACACAGGUUCAACAGAAGGGCAAGUUGCUUUGUUCACACAUCGCAUUAACCACUUAACUGGGCACCUUAAAAAAAAAUCACAAAGAUUAUAACACCGAACGUUCGUUAGUAAGACUGGUAGGUAAAAGAAGAAGUCUUUUAGAUUACUUG
env-2045  AUUUUUAAGAAACACGGCGGAAAAGAAGAGAACACAGGUUCCACGGAAGGACAAAUUGCUUUGUUCACACAUCGCAUUAACCAUUUAACG..........................................................................................
env-2046  AUUUUUAAAAAACACGGCGGAAAAGAAGAUAACACAGGUUCAACAGAAGGGCAAGUUGCUUUGUUCACACAUCGCAUUAACCACUUAACUGGGCACCUUAAAAAAAAAUCACAAAGAUUAUAACACCGAACGUUCGUUAGUAAGACUGGUAGGUAAAAGAAGAAGUCUUUUAGAUUACUUG
env-2049  AUUUUUGCAAAACACGGCAAGUCUGCCACUGACACUGGCAACUCAGAAGGGCAAAUCGCCUUAUUCACACACAGAAUUAAUCACCUUACGGAGCAUUUAAAAAACAACCGUAAAGAUUACAAUACAGAACGUUCUUUAGUUCGUUUGGUAGGUAAAAGAAGGAGUUUAUUAGACUACCUG
env-2050  AUUUUUACAAAACACGGCAAAU..............................................................................................................................................................
env-2051  AUUUUUACAAAACACGGCAAAUCAGCCACUGACACUGGUAAUUCAGAAGGGCAAAUCGCCUUAUUCACCCACAGAAUUAAUCACCUUACGGAGCAUUUAAAAAAGAACCGUAAAGAUUACAAUACAGAGCGUUCUUUAGUUCGUUUGGUAGGUAAAAGAAGGAGUUUAUUAGACUACCUG
env-2052  AUUUUUACAAAACACGGCAAAUCUGCCACUGACACUGGUAAUUCAGAAGGGCAAAUCGCCUUAUUCACACACAGAAUUAAUCACCUUACGGAGCAUUUAAAAAAGAACCGUAAAGAUUA.............................................................
env-2053  AUUUUUNCAAAACACGGCAANUCUGCCACUGACACUGGNAAUUCAGAAGGGCAAAUCGCCUUAUUCACACACAGAAUUAAUCACCUUACGGAGCAUUUAAAAAAGAACCGUAAAGAUUACAAUACNGAACGUUCUUUAGUUCGUUUGGUAGGUAAAAGAAGGAGUUUAUUAGACUACCUG
env-2054  AUUUUUACAAAACACGGCAAAUCAGCCACUGACACUGGUAAUUCAGAAGGGCAAAUCGCCUUAUUCACCCACAGAAUUAAUCACCUUACGGAGCAUUUAAAAAAGAACCGUAAAGAUUACAAUACAGAGCGUUCUUUAGUUCGUUUGGUAGGUAAAAAGAAGGAGUUUUAUUAGACUACC
env-2055  AUUUUUACAAAACACGGCAAAUCAGCCACUGACACUGGUAAUUCAGAAGGGCAAAUCGCCUUAUUCACCCACAGAAUUAAUCACCUUACGGAGCAUUUAAAAAAGAACCGUAAAGAUUACAAUACAGAGCGUUCUUUAGUUCGUUUGGUAGGUAAAAGAAGGAGUUUAUUAGACUACCUG
Ead-1-1   AUCUUUAAAAAACACGGAGGUUCAGAAGGUAAUACCGGCUCCACGGAAGGUCAGAUUGCGUUGUUUACGCAUCGGAUUAAUCAUUUAACAGGACACUUAAAAAACAACCAAAAGGAUUACAAUACGGAGAGAUCCCUGGUAAGACUCGUUGGUAAAAGAAGAAGUUUGCUGGAUUAUUUG
env-2068  AUUUUUAAAAAACACGGUGGUUCAGAAGGUAACACAGGUUCCACUGAGGGUCAGAUUGCAUUAUUUACGCAUCGGAUUAAUCAUUUAACCGGACAUUUAAAAAACAACCAAAAAGAUUAUAAUACGGAGAGAUCUCUGGUAAAACUUGUGGGUAAAAGAAGAAGUUUACUGGAUUAUUUA
env-2069  AUUUUUAAAAAACACGGUGGUUCAGAAGGUAACACAGGUUCCACUGAGGGUCAGAUUGCAUUAUUUACGCAUCGGAUUAAUCAUUUAACCGGACAUUUAAAAAACAACCAAAAAGAUUAUAAUACGGAGAGAUCUCUGGUAAAACUUGUGGGUAAAAGAAGAAGUUUACUGGAUUAUUUA
env-2070  AUUUUUAAAAAACACGGUGGUUCAGAAGGUAACACAGGUUCCACUGAGGGUCAGAUUGCAUUAUUUACGCAUCGGAUUAAUCAUUUAACCGGACAUUUAAAAAACAACCAAAAAGAUUAUAAUACGGAGAGAUCUCUGGUAAAACUUGUGGGUAAAAGAAGAAGUUUACUGGAUUAUUUA
env-2071  AUCUUUAAAAAACAUGGUGGUUCAGAAGGUAACACGGGUUCCACUGAAGGUCAGAUUGCGUUGUUUACGCAUCGGAUUGAUCAUUUAACAGGACAUUUAAAAAACAACCAAAAGGAUUAUAAUACAGAGAGAUCCCUGGUAAGACUUGUUGGUAAAAGAAGAAGUUUACUGGAUUAUUUG
env-2072  AUUUUUAAAAAACACGGUGGUUCAGAAGGUAACACAGGUUCCACUGAGGGUCAGAUUGCAUUAUUUACGCAUCGGAUUAAUCAUUUAACCGGACAUUUAAAAAACAACCAAAAAGAUUAUAAUACGGAGAGAUCUCUGGUAAAACUUGUGGGUAAAAGAAGAAGUUUACUGGAUUAUUUA
env-2073  AUUUUUAAGAAACACGGCGGGGAAGAAAAAAACACAGGUUCCGCUGAAGGGCAGAUUGCCUUGUUCACCCACAGAAUCAACCACUUGACCGAGCAUUUGAAAAAGAACCGUAAGGAUUUUAACACCGAACGCUCAUUGGUAAAACUGGUAGGAAAGAGAAGAUCUCUUUUAGAUUACUUG
env-2074  AUUUUUAAGAAACACGGCGGGGAAGAAAAAAACACAGGUUCUGCUGAAGGGCAGAUUGCCUUGUUUACCCACAGAAUCAACCACUUAACAGAGCAUUUAAAAAAGAACCGUAAGGAUUUUAACACGGAGCGUUCUUUGGUAAAGUUGGUA..............................
env-2075  AUUUUUAAGAAACACGGCGGGGAAGAAAAAAACACAGGUUCUGCUGAAGGGCAGAUUGCCUUGUUUACCCACAGAAUCAACCACUUAACAGAGCAUUUAAAAAAGAACCGUAAGGAUUUUAACACGGAGCGUUCUUUGGUAAAGUUGGUAGGUAAGAGAAGAUCUCUUCUAGAUUACUUG
env-2076  AUUUUCAAAAAACACGGCGGAGAAGCAAAAAACACAGGUUCCGCUGAAGGGCAG..............................................................................................................................
env-2077  AUUUUCACGAAACACGGCGGAGAUGCAAAAAACACUGGGUCAUCUGAAGGUCAGAUUGCUCUAUUUACCCACAGAAUUAAUCACCUAACUGAACAUUUAAAGAAAAACCGUAAGGAUUUUAACACAGAACGUUCCUUGGUAAAACUGGUAGGUAAAAGAAGAUCACUUCUAGAUUAUUUA
env-2078  AUUUUUAAAAAACACGGUGGAAAAGAAGAGAACACUGGUUCUACUGAAGGUCAAGUUGCUUUGUUCACACAUCGCAUCAACCAUCUAACUG.........................................................................................
env-2079  AUAUUCAAGAAAUAUGGUGGUAAAGCAGAGAACACAGGUUCUACAGAAGGGCAAAUCGCUCUAUUCACACACCGUAUCAAUCACUUGACCGGUCACUUAAAAAAAAAAUCAUAAAGACUACAAUACCGAGCGUUCAUUAGUAAAGUUAGUAGGUAAAAGAAGAAGUCUUCUUGAUUACAUG
Fba-6-1   AUUUUCAAAAAACACGGCGGAAAAGCAGAGAACACAGGUUCUACAGAAGGUCAAGUUGCAUUGUUCACACACCGUAUCAAUCAUUUGACCGGUCACCUUAAAAACAAUCACAAAGAUUAUAAUACUGAGCGUUCAUUGGUGAAACUGGUAGGUAAAAGAAGGAGUCUUUUAGAUUACUUG
env-2080  AUUUUUAAAAAAUAUGGCGGUAAAGCAGAGAACACAGGUUCUACUGAAGGACAAAUUGCAUUGUUCACGCACCGAAUCAAUCACUUGACCGGUCACUUGAAAAAAAAUCACAAAGACUACAA..........................................................

          ....................................................................................................................................................................................
          ....................................................................................................................................................................................
          AUYUUYR•RAARCACGG•RRR•••R•RR••RAYAC•GG••C•RC•GAAGG•CAAAUYGC•YURUUYAC•Y•Y•G•AUYRAYCAYYUR•C•RA•CAYYURAAAAR•AAYCR•AARGAUU•YAAYAC•GAR•G•UC•YURGU•••RYURGU•GG•AAR•G•RG••••YU•YU•GAYUAYYU•
```

env-2081 AUUUUCAAACAGUACGGCGGAUCUGAGAUCAACACCGGAUCAACUGAGGGACAGGUUGCCCUGUUUACGCAUCGUAUCAACCACCUUACCGGGCACCUGAAAAACAACCCGAAAGAUUACAACACCGAACGCUCGCUGGUAAAACUGGUUGGGCAGCGCAGGAGCCUGCUCGAUUAUCUU
env-2082 AUUUUCAAACAGUACGGCGGAUCUGAGAUCAACACCGGAUCAACUGAGGGACAGGUUGCCCUGUUUACGCAUCGUAUCAAC..............................................................................................
env-2083 AUUUUCAAGAAACACGGCGGAAAAGCUGAGAACACUGGUUCUACUGAAGGCCAAAUCGCAUUGUUCACACACCGAAUCAAUCACUUGACCGGUCACUUGAAAAAUAAUCAUAAAGAUUACAACACAGAGCGUUCACUUGUGAAACUCGUAGGUAAAAGAAGAAGUCUUCUUGAUUAUAUG
Msp-1-1 AUUUUCAAGAAACACGGUGGUAAAGCAGAAAACACUGGUUCUACAGAAGGGCAAAUUGCCUUAUUUACACACAGAAUCAACCACUUAACUGGUCACUUGAAAAAAAAUCAUAAAGAUUACAACACUGAGCGUUCACUUGUGAAACUAGUAGGUAAAAGAAGAAGUCUUCUUGAUUACAUG
env-2085 AUUUUCAAGAAACACGGUGGUAAAGUAGAAAACACUGGUUCUACAGAAGGGCAAAUUGCCUUAUUCACACACAGAAUCAAUCACUUGACUGGUCACUUGAAAAGAAAUCACAAAGACUACAACACUGAACGUUCACUUGUGAAACUAGUAGGUAAAAGAAGAAGUCUUCUUGAUUACAUG
env-2086 AUAUUUAAGAAACAUGGCGGUAAAGCAGAGAACACAGGUUCUACAGAGGGACAAAUAGCUUUGUUCACACACCGAAUCAAUCACUUGACCGGUCACCUUAAAAACAACCACAAAGAUUACAAUACAGAGCGAUCUUUGGUAAGAUUGGUAGGUAAGCGUAGAAGCUUACUGGAUUAUCUU
env-2087 AUCUUUACAAAGCAUGGCGGCAAAGCGGAGAACACGGGCUCCACAGAAGGACAGAUCGCCCUGUUCACCCACCGGAUCAACCAUUUGACCGGACAUUUAAAAAGAAACCACAAAGACUACAACACAGAGCGGUCUUUGGUACGUUUGGUAGGUAAAAGAAGAAGCCUAUUGGAUUAUUUG
env-2088 AUCUUUACAAAGCAUGGCGGCAAAGCGGAGAACACGGGCUCCACAGAAGGACAGAUCGCCCUGUUCACCCACCGGAUCAACCAUUUGACCGGACAUUUAAAAAGAAACCACAAAGACUACAACACAGAGCGGUCUUUGGUACGUU.....................................
Mth-1-1 AUUUUUAAAAAAUACGGCGGAAAAGCGGAGAACACAGGUUCUACCGAAGGUCAGAUUGCAUUGUUUACACACCGUAUCAAUCAUUUGACAAAUCACUUAAAGAAGAAUCACAAAGAUUACAACACAGAGCGUUCGUUGGUAAAACUUGUGGGUAAAAGAAGAAGUCUUCUUGAUUAUAUG
env-2089 AUUUUUAAAAAAUACGGCGGUAGUGAAGGGAACACAGGUUCCACCGAAGGUCAAAUCGCGUUGUUUACGCACCGCAUCAACCAUUUGACCGAACAUUUAAAAAGAAACAAGAAAGACUACAA...........................................................
env-2090 ......................................................................................................................................................................................
env-2091 AUUUUUAAAAAAUACGGCGGUAGUGAAGGGAACACAGGUUCCACCGAAGGUCAAAUCGCGUUGUUUACGCACCGCAUCAACCAUUUGACCGAACAUUUAAAAAGAAACAAGAAAGACUACAACACCGAGCGUUCUUUGGUUCGUUUGGUAGGUAAG.........................
env-2092 AUUUUUAAAAAAUACGGCGGUAGUGAAGGGAACACAGGUUCCACCGAAGGUCAAAUCGCGUUGUUUACGCACCGCAUCAACCAUUUGACCGAACAUUUAAAAAGAAACAAGAAAGACUACAACACCGAGCGUUCUUUGGUUCGUUUGGUAGGUAAGAGAAGAAGUCUUCUUGACUAUAUG
Rbi-1-1 AUCUUUAAGAAAUUCGGCGGAUCGGACAGCAACACCGGCUCCACCGAGGGCCAGAUUGCGCUGUUCACGGAACGGAUCAACCACCUGACCGGGCACCUGAAGAAGAACCAGAAAGACUACAACACCGAGCGUUCCCUGGUUCGCCUGGUGGGUAAACGCCGCAGCCUGCUGGACUAUCUG
Mfo-1-1 AUUUUCAAAAAAACACGGCGGUAGUGAAACAAACACUGGUUCUACAGAAGGACAAAUUGCUUUGUUUACACACAGAAUUAAUCACUUAACUGGUCACUUGAAAAAGAACCAUAAAGAUUACAACACAGAGCGUUCAUUAGUAAAGCUCGUGGGUAAAAGAAGAAGUCUUCUUGAUUACAUG
env-2099 CU....................................................................................................................................................................................
env-2100 AUCUUCGCUAAACACGCAGGUAAAGCUGAAAACACGGGUCUGCUGAGGGACAAAUUGCAUUGUUCACUUUCAGAAUUUCACCACUUAACAGAGCAUUUGAAAAAAAAUCGUCACGAUUAUAAUACAGAGCGUUCGUUAGUAUUAUUGGUAGGUAAAAGAAGAAGUCUUUUAGAUUACCUU
Fse-1-1 AUUUUCGCUCAACACGGAGGAAAAGCAGAAAACACAGGAUCUGCAGAAGGACAAAUCGCUUUGUUCUCAUUCAGAAUCAACCACUUGACUGAACACUUGAAAAAAAAUCGUCACGAUUACAACACAGAGCGUUCUCUUGUGAAAUUAGUAGGUAAAAGAAGAUCUUUACUUGACUACUUG
Fda-1-1 AUUUUCGCUCAACACGGAGGAAAAGCAGAAAACACUGGUUCUGCAGAGGACAAAUCGCUUUGUUCUCUUUCAGAAUCAACCACUUGACUGAGCACUUGAAAAAAAAUCGUCACGAUUACAACACUGAGCGUUCACUAGUGAAAUUAGUAGGUAAAAGAAGAUCUUUACUUGACUAUUUG
env-2102 CUUUUUGCAAAACACGGUAAAGACGCUAAGAAUACUGGUUCUGCAGAAGGUCAAAUAGCAUUAUUCACAGCAAGAAUAGCACAUUUAACAGAGCACUUAAAGAAAAAUCGUAAAGAUUAUAAUACUGAGCGUUCUUUAGUAAUGUUAGUAGGAAAAAGAAGAAGCCUUACUAGAUUACUUA
env-2103 AUCUUCGCUAAACACGGAGGAAAAGCAGAAAACACAGGGUCUGCUGAAGGACAAAUCGCUUUGUUCACAUUUAGAAUCAACCACUUAACAGGACACUUAAAAAACAAUCGUCAUGAUUACAACACAGAGCGUUCGUUAGUGAAAUUAGUAGGUAAAAGAAGAAGCCUUCUUGACUACUUA
env-2105 AUCUUCGCUAAACACGGAGGAAAAGCUGAAAACACUGGAUCUGCUGAAGGGCAAAUCGCUUUGUUUACAUUUAGAAUCAACCACUUAACUGGUCACUUGAAAAAAAAUCGUCACNNNNNNNNNNNNNNNNNNNNNNNNNNNNNNNNNNNNNNNNNNNNNNNNNNNNNNNNNAU[illegible]AAUACA
Fso-1-1 AUCUUCGCUAAACACGGAGGAAAAGCUGAAAACACUGGAUCUGCUGAAGGGCAAAUCGCUUUAUUCACAUUUAGAAUCAACCACUUAACUGGUCACUUGAAAAAAAAUCGUCACGAUUAUAAUACAGAGCGUUCAUUGGUGAAAUUGGUAGGUAAAAGAAGAAGCUUACUAGAUUAUCUU
env-2106 AUUUUCGCUAAACACGGAGGAUCAGCAACAAACACUGGUUCUGCUGAAGGUCAAAUCGCAUUAUUCACGCACAGAAUCAGCCACUUGACAGAGCAUUUGAAAAAAAAACCGUCACGAUUACAACACCGAGCGUUCGCUUGUCCUAUUAGUAGGUAAAAGAAGAAGCUUGCUUGAUUAUCU
env-2107 ......................................................................................................................................................................................
env-2108 AUCUUCGCUAAACACGGAGGAAAAGCUGAAAAUACAGGUUCAGCAGAAGGUCAAAUCGCGUUGUUCACAUUCAGAAUCAGCCACUUGACAGAGCAUUUGAAAAAAAAUCGUCACGAUUACAAUACUGAGCGUUCGCUUGUCCUACUAGUAGGUAAAAGAAGAAGCUUGCUUGAUUA....
env-2109 AUCUUCGCUAAACACGGAGGAUCAGCUAAAAACACAGGUUCUGCUGAAGGACAAAUCGCGUUAUUCACGCACAGAAUUGGUCACUUGACUGAACACUUGAAGAAAAAUCGUCACGAUUACAACACGGAGCGUUCGCUUGUCCUACUAGUAGGUAAAAGAAGAAGCUUGCUUGAUUAUCUU
env-2110 AUCUUCGCUAAACACGGAGGAUCAGCUAAAAACACAGGUUCAGCUGAAGGACAAAUCGCGUUAUUCACGCACAGAAUUGGUCACUUGACUGAACACUUGAAGAAAAAUCGUCACGAUUACAACACGGAGCGUUCGCUUGUCCUACUAGUAGGUAAAAGAAGAAGCUUGCUUGAUUAUCUU
Aag-1-1 AUCUUUGCUAAACACGGUAAAGGAAAAACGAUAGUGGAACUGCAGAAGGACACAAAUUGCUUUGUUUACACACAGAAUUACACAUUUAACUGAGCACUUAAAACAAAAUCGUAAAGAUUAUAAUACUGAGCGUUCUUUAGUAAAAUUAGUAGGUAAGCGUAGAAGUUUAUUAGACUACUUA
env-2111 CUUUUUGCAAAACACGGAAAAUCAAAAACCGAUACCGGUUCUGCAGAAGGACAAAUUGCGUUGUUUACACAUCGUAUCGAGCACUUAUCAGGUCAUCUUAAAAACAACCACAAGGACUACAACACAGAACGUUCUUUAGUAAUGUUAGUAGGUAAAAGAAGAUCUCUUUUAGAUUACUUA
env-2112 UUAUUUAAAAAACACGGUAAGUCUGAAAACGACACCGGUUCUACAGAAGGGCAAAUCGCGUUGUUUACACAACGUAUUGAGCACCUUUCAAAACACCUUAAAAGCAACCAGAAAGACUUCAAUACUGAACGAUCUUUGGUGAAAUUAGUAGGUAAGCGUAGAUCACUUCUUGAUUAUCUU
env-2113 UUAUUCAAAAAACACGGGAAGUCUGAAAAAGACACCGGUUCUACCGAAGGACAAAUCGCUUUGUUUACACAGCGUAUUGAGCAUCUUUCAAAACACCUUAAAACUAACCAGAAAGAUUUCAACACAGAAAGAUCUUUGGUAAAGUUAGUAGGUAAGCGUAGAUCGCUUCUUGAUUAUCUA
env-2114 UUAUUCAAAAAACACGGGAAGUCUGAAAAAGACACCGGUUCUACCGAAGGACAAAUCGCUUUGUUUACACAGCGUAUUGAGCAUCUUUCAAAACACCUUAAAAC.....................................................................................
env-2115 UUAUUCAAAAAACACGGAAAGUCUGAAAAAGACACCGGUUCUACCGAAGGGCAAAUCGCUUUGUUUACACAGCGUAUUGAGCAUCUUUCAAAACACCUUAAAACUAACCAGAAAGAUUUCAACACAGAAAGAUCUUUGGUAAAGUUAGUAGGUAAGCGUAGAUCACUUCUUGAUUAUCUA
env-2116 AUUUUCAAAAAACACGGUAAGUCUGAAAAAGACUCUGGUUCUGCUGAAGGACAAAUCGCUUUAUUCACAUACCGUAUCGAGCAUUUAACUCACAUCUAAGAAAAAAUCACAA..............................................................................
env-2117 AUUUUCAAAAAACACGGUAAGUCUGAAAAAGACUCUGGAUCAGCUGAAGGGCAAAUUGCUUUAUUCACCUACCGUAUCGAGCAUUUAACUCAACACCUAAGAAAAAACCACAAGGAUUACAAUACAGAACGUUCUUUGGUAAUGUUAGUAGGUAAAAGACGAUCGCUUUUAAAUUAUCUC
env-2118 AUUUUCAAAAAACACGGUAAGUCUGAAAAAGACUCUGGUUCUGCUGAAGGACAAAUCGCUUUAUUCACCUACCGUAUCGAGCAUUUAACUCACAUCUAAGAAAAAAUCACAAGGACUACAACACAGAACGUUCUUUGGUUAUGUUAGUAGGUAAAAGACGAUCUCUUUUAAAUUAUCUU
env-2119 AUUUUCAAAAAACACGGUAAGUCUGAAAAAGACUCUGGUUCUGCUGAAGGACAAAUCGCUUUAUUCACCUACCGUAUCGAGCAUUUAACUCACAUCUAAGAAAAAAUCACAAGGACUACAACACAGAACGUUCUUUGGUUAUGUUAGUAGGUAAAAGACGAUCUCUUUUAAAUUAUCUU
env-2120 AUUUUCAAAAAACACGGUAAGUCUGAAAAAGACUCUGGAUCAGCUGAAGGGCAAAUUGCUUUAUUCACCUACCGUAUCGAGCAUUUAACUCAACACCUAAGAAAAAACCACAAGGAUUACAAUACAGAACGUUCUUUGGUAAUGUUAGUAGGUAAAAGACGAUCGCUUUUAAAUUAUCUC
env-2121 AUUUUCAAAAAACACGGUAAGUCUGAAAAAGACUCUGGAUCAGCUGAAGGGCAAAUUGCUCUAUUCACCUACCGUAUCGAGCAUUUAACUCAACACCUAAGAAAAAACCACAAGGAUUACAAUACAGAACGUUCUUUGGUAAUGUUAGUAGGUAAAAGACGAUCGCUUUUAAAUUAUCUA
env-2124 AUUUUCAAAAAACACGGUAAGUCUGAAAAAGACUCUGGAUCAGCUGAAGGGCAAAUUGCUUUAUUCACCUACCGUAUCGAGCAUUUAACUCAACACCUAAGAAAAAACCACAAGGAUUACAACACAGAACGUUCUUUGGUAAUGUUAGUAGGUAAAAGAAGAUCUCUUUUAAAUUAUCUU
env-2125 AUUUUCAAAAAACACGGUAAGUCUGAAAAAGACUCUGGAUCUGCUGAAGGGCAAAUCGCUUUAUUCACGUACCGUAUCGAGCAUUUAACUCAACACCUAAGAAAAAACCACAAGGACUACAACACAGAACGUUCUUUGGUAAUGUUAGUAGGUAAAAGAAGGUCUCUUUUAAAUUAUCUU
env-2126 ......................................................................................................................................................................................
env-2127 AUUUUCAAAAAACACGGCAAAUCUAAUAAAGACACUGGAUCUGCUGAAGGACAAAUCGCAUUAUUUACUUACAGAAUUGACCAUUUAACGAAGCACUUAAAGCAAAAUCAUAAGGAUUUAACACAGAGCGCUCUUUGGUAAAACUAGUAGGUAAAAGAAGAUCUCUUUUAAAUUAUCUA
env-2128 AUUUUCAAAAAACACGGUAAGUCUGAAAAAGACUCUGGAUCUGCUGAAGGGCAAAUUGCUUUAUUCACCUACCGUAUCGAGCAUUUAACUCAACACCUAAGAAAAAACCACAAGGAUUAUAACACAGAACGUUCUUUGGUAAUGUUAGUAGGUAAAAGAAGAUCUCUUUUAAAUUAUCUU
env-2129 AUUUUCAAAAAACAUGGUAAGUCUGAAAAAGACUCUGGAUCUGCUGAAGGGCAAAUUGCGUUAUUCACUUACCGAAUCAACCACUUAACUAAACACUUAAGACAAAACCACAAGGAUUUCAACACAGAACGUUCUUUGGUAAUGUGGUAGGUAAAAGACGAUCUUUAUUAAACUAUCUU
env-2130 AUUUUCAAAAAACAUGGUAAGUCUGAAAAAGACUCUGGAUCUGCUGAAGGGCAAAUCGCGUUAUUCACUUACCGAAUUAACCACUUAACUAAACACUUAAGACAAAACCACAAGGAUUUCAACACAGA...............................................................
env-2131 AUUUUCAAAAAACACGGGAAGUCUGCAAAAAGACUCUGGUUCUGCUGAAGGUCAAAUUGCUUUGUUCACAUACCGUAUCGAA...............................................................................................
env-2132 AAUUUUCAAAAAAACACGGCAAAUCAAAUAAAGACACUGGUUCUGCUGAAGGACAAAUCGCACUAUUUACUUACAGAAUUGACCAUUUAACGAAGCACUUAAAACAAAACCACAAAGAUUUUAAUACAGAGCGCUCACUGGUAAAAUUAGUAGGUAAAAGAAGAUCUCUUUUAAACUAU
env-2133 AUUUUCAAAAAACACGGCAAAUCUAAUAAAGAUACUGGAUCUGCUGAAGGACAAAUCGCAUUAUUUACCUACAGAAUUGACCAUUUAACUAAACACUUAAAACAAAAUCAUAAAGAUUUUAAUACAGAGCGUUCCCUGGUAAAACUAGUAGGUAAAAGAAGAUCUCUUUUAAACUAUCUU
env-2134 AUUUUCAAAAAACACGGCAAAUCAAAUAAAGACACUGGUUCUGCUGAAGGACAAAUCGCACUAUUUACUUACAGAAUUGACCAUUUAACGAAGCACUUAAAACAAAAUCACAAAGAUUUUAAUACAGAGCGCUCACUGGUAAAAUUAGUAGGUAAAAGAAGAUCUCUUUUAAACUAUCUA
env-2135 AUUUUCAAAAAACACGGCAAAUCAAAUAAAGACACGGGAUCUGCUGAAGGACAAAUCGCAUUAUUUACCUAUAGAAUUGACCAUUUAACGAAGCACUUAAAACAAAAUCAUAAAGAUUUUAACACAGAGCGUUCCUUGGUAAAACUAGUAGGUAAAAGAAGAUCUCUUUUAAACUAUCUU
env-2136 AUUUUCAAAAAACACGGCAAAUCUAAUAAAGACACUGGAUCUGCUGAAGGACAAAUCGCAUUAUUUACUUACAGAAUUGACCAUUUAACGAAACACUUAAAACAAAAUCAUAAAGAUUUUAAUACAGAGCGUUCCCUGGUAAAACUAGUAGGUAAAAGAAGGUCUCUUUUAAACUAUCUU
env-2137 AUUUUCAAAAAACACGGCAAAUCUAAUAAAGAUACUGGAUCUGCUGAAGGACAAAUCGCAUUAUUUACCUACAGAAUUGACCAUUUAACUAAACACUUAAAACAAAAUCAUAAAGAUUUUAAUACAGAGCGUUCCCUGGUAAAACUAGUAGGUAAAAGAAGAUCUCUUUUAAACUAUCUU
env-2138 AUU...................................................................................................................................................................................
env-2139 AUUUUCAAAAAACACGGCAAAUCAAAUAAAGACACUGGUUCUGCUGAAGGACAAAUCGCACUAUUUACUUACAGAAUUGACCAUUUAACGAAGCACUUAAAACAAAACCACAAAGAUUUUAAUACAGAGCGCUCACUGGUAAAAUUAGUAGGUAAAAGAAGAUCUCUUUUAAACUAUCUA
env-2140 AUCUUCGCAAAGCACGGUAAGUCUGGAAAAGACACNGGUUCUGCAGAAGGCCAAAUUGCGUUAUUCACAUACCGUAUCGAGCACUUAACAAAACACCUUAAAGCAACCACAAGGACUACAACACAGAACGUUCUUUGGUAAAGCUUGUAGGUAAGCGUCGUUCAUUACUUGAUUAUCUU
env-2141 AUCUUUAAAAAACAUGGUAAAGAUGCAAAGA.......................................................................................................................................................
env-2142 CUCUUUAAAAAGCACGGUAAAGAUGCAAAGAACACUGGUUCUACAGAAGGACAAAUUGCAAUAUUUACACAAAGAAUUGACCACUUAACAGAACACUUAAAAAACAAUCG................................................................................
env-2143 CUCUUUAAAAAGCAUGGUAAAGAUGCAAAAAACACAGGUUCUACAGAAGGACAAAUUGCAUUAUUUACUCACAGAAUUAAUCACUUAACUGAACACUUAAAAAACAAUCGUAAAGAUUUUAAUACUGAGCGUUCUCUAGUAAAACUAGUAGGUAAGCGUCGUGCGUUACUAGAUUAUUUA
env-2144 AUCUUUAAAAAACAUGGUAAAGAUGCAAAGAACACAGGUUCAGCUGAAGGGCAAAUUGCAUUAUUUACAUACAGAAUUAAUCACUUAACGAACACUUAAAAAAUAAUCGUAAGGAUUAUAAUUCUGAGCGUUCUUUAGUAAAGUUGGUGGUAAAACGUCGCGCGUUAUUAGACUACUUA
env-2145 AUCUUCGCUAAACACGGAGGAAAAGCAGAAAACACAGGAUCUGCUGAAGGACAAAUUGCAUUGUUCACACACAGAAUUACACACUUAACUGAGCACUUAAAAAAGAAUCGUCACGAUUACAACACUGAGCGUUCGUUAGUACUUUUAGUAGGUAAAAGAAGAAGUCUUCUUGACUACUUA
env-2146 AUCUUCGCUAAACACGGAGGAAAAGCAGAAAACACAGGAUCUGCUGAAGGACAAAUUGCAUUGUUCACACACAGAAUUACACACUUAACUGAGCACUUAAAAAAGAAUCGUCACGAUUACAACACUGAGCGUUCGUUAGUACUUUUAGUAGGUAAAAGAAGAAGUCUUCUUGACUACUUA
env-2147 AUCUUCGCUAAACACGGAGGAAAAGCAGAAAACACAGGAUCUGCUGAAGGACAAAUUGCAUUGUUCACACACAGAAUUACACACUUAACUGAGCACUUAAAAAAGAAUCGUCACGAUUACAACACUGAGCGUUCGUUAGUACUUUUAGUAGGUAAAAGAAGAAGUCUUCUUGACUACUUA
env-2148 AUCUUCGCUAAGCACGGUGGUGCAGAAACAAACACUGGAUCUGCUGAAGCACAAAUCGCUU.......................................................................................................................
env-2149 ......................................................................................................................................................................................
env-2150 AUCUUCGUUAAACACGCAGGAUCUGAAACAAACACAGGAUCUGCAGAAGGUCAAAUCGCAUUGUUUUCUUACAGAAUUUCUCAUUUAACUGAGCAUUUAAAGAAAAAUAGAAAAGACU.....................................................................
env-2151 AUCUUCGUUAAACACGCAGGAUCUGAAACAAACACAGGAUCUGCAGAAGGUCAAAUCGCAUUGUUUUCUUACAGAAUUUCUCAUUUAACUGAGCAUUUAAAGAAAAAUAGAAAAGACUAUGUUACGCAAAGGUCAUUCUUCAAUUGCUUGUUGGUAAAAGAAGAAGUUU..............
env-2152 AUUUUUGCAAAAUACGCAGGAUCUGAAACAAACACAGGAUCUGCUGAAGGUCAAAUCGCAUUGUUUUCUUACAGAAUUUCUCAUUUAACUGAGCAUUUAAAGAAAAACAGAAAGAUUACGGAACGCAAAGAUCACUUCAAUUGUUAGUAGGUAAAAGAAGAAGUCUUUUGGAUUACUUA
Fps-4-1 AUCUUCGCUAAACACGGAGAAAAAACAAACACAGGAAAAGCAGAAGCACAAAUUGCUUUAUUUACUUUCAGAAUUAGCCACUUAACAGAGCACCUGAAAAAAAAUCGUCACGAUUAUAACACUGAGCGUUCACUUGUUCUUUUAGUAGGUAAAAGAAGAUCUUUACUAGACUACUUGAAG

......................................................................................................................................................................................
......................................................................................................................................................................................
AUYUUYR•RAARCACGG•RRR•••R•RR••RAYAC•GG••C•RC•GAAGG•CAAAUYGC•YURUUYAC•Y•Y•G•AUYRAYCAYYUR•C•RA•CAYYURAAAAR•AAYCR•AARGAUU•YAAYAC•GAR•G•UC•YURGU•••RYURGU•GG•AAR•G•RG••••YU•YU•GAYUAYYU•

alignment positions 440 · · · 530

Pgo-1-1 AUGAAGAAAGAUAUCCUGAGAUAUAGAGCUAUUGUAAAAGAUUUAGGAUUAAGAAAAUAAAUUUAAAGGGGCAACAUUGCCCCUUUUUUAG
env-1 AUGAAAAA...................................................................................
env-2 ...........................................................................................
env-3 AUGAAAAAAGAUAUCAUGAGAUAUAGAGCCAUUGUUAAAGAAUUAGGAUUAAGAAAAUAAAUAUCAAAGGGGCAUUUUGCUCUUUUUUAA
Pto-1-1 AUGAAGAAACAUAUUAUGAGGUAUAGAGCUAUUGUAAAAGAUUUAGGAUUAAGAAAAUAAAUUUAAAGGGGCAAUUUUGCCCCUUUUUUAG

...........................................................................................
...........................................................................................
A•RAARAARGAYAUY••••RRUAY•G•G••AU•RU•AAAGAAYURR••U••GAAAAUAA•••••••••••••••••••••••••••••••

```
Ptr-1-1   AUGAAAAAAGACAUCAUGAGAUACAGAGCCAUCGUUAAAGAAUUAGGAUUAAGAAAAUAAAAAUUAAAGGGGCACUUUGCCCCUUUUUUAA
env-4     AUGAAGAAACAUAUCAUGAGAUAUAGAGCUAUCGUAAAAGAUUUAGGAUUAAGAAAAUAAAUUUAAAGGGGCAACAUUGCCCCUUUUUUAG
env-5     ...........................................................................................
env-6     ...........................................................................................
env-7     ...........................................................................................
env-8     ...........................................................................................
Mod-8-1   AAGAAAACUGAGAUUAACAGAUAUCGUGAGAUCAUUAAAGUAUUGGGUAUUAGAAAAUAAUCAAUAUAGGAAAGAGGUGCCUGCGCGCCUC
env-9     AAGAAGAAGGAUAUUGAAAGAUAUCGUGCAAUCGUUAAAGAAUUAGGAUUACGUCGUUAAUCUUACUUAACAUAUAAUAAAAUUGGGGGGC
env-11    AAGAAGAAGGAUAUUGAAAGAUAUCGUGCAAUCGUUAAAGAAUUAGGAUUACGUCGUUAAUCUUACUUAACAUAUAAUAAAAUUGGGGGGC
env-12    AAGAAGAAGGAUAUUGAAAGAUAUCGUGCAAUCGUUAAAGAAUUAGGAUUACGUCGUUAAUCUUACUU.......................
env-13    AAGAAAAAAGAUAUUGAGAGAUAUCGUUCAAUUGUUAAAGAGUUAGGAUUACGUCGUUAAUCUUACUUAACAUAUAACAAAAUUGGGGGGC
env-14    AAGA.......................................................................................
env-15    ...........................................................................................
env-16    AAAGAUAGAGAUAUUGUUAAAUAUCGUGAAUUGAUUAAAGAGUUAGGAAUUAGAAAAUAAUUUGAACUUACAACAAUAACAUUAAAGAUUA
env-17    ACUAAGAAAGAUAUUUUAAGAUACCGUGCCAUCAUUAAAGAAUUAGGAUUAAGAAAAUAAUUCCAAAAAGAGGCUUUCGAGCCUCUUUUUG
Fbr-1-1   AAGAAAACUGAAAUUAACAGAUACCGUGAGAUAAUCAAAGUAUUAAAUAUCAGAAAAUAAUCCUAAAGAGGUGCUUUUGCACCUCUUUUUU
env-18    AAACGAAAAGAUAUUGAAAAAUACCGUGCGUUAAUUAAGGAGUUAGGACUUCGUCGUUAAUCACAAAGUGUUAAUUUUAUUGAAGAGGGGA
env-19    AAACGAAAAGAUAUUGAAAAAUACCGUGCGUUA..........................................................
env-20    AAAAUAAGAGAUUUCAAUACGUCAUCAA...............................................................
env-21    ...........................................................................................
env-22    ACUAAGAAAGAUAUUUUAAGAUAUCGUGCAAUUGUAAAAGAGUUAGGCUUAAGAAAAUAAUAUAGA.........................
env-23    ACUAAGAAAGAUGUUUUAAGAUAUCGUGCAAUUGUAAAAGAGUUAGGCUUAAGAAAAUAACAUAUCAAAUCCUAUUAUAAUAAUUAUUCAU
env-24    ACUAAGAAAGAUAUUUUAAGAUAUCGUGCAAUUGUAAAAGAGUUA..............................................
env-25    ...........................................................................................
Fba-9-1   ACUAAGACAGACAUUUUGCGUUACCGUGCAAUUGUAAAAGAAUUAGGAUUAAGAAAAUAAUUCAGAAUAUAAAGAGGCUUAAUCGCCUCUU
env-26    AUGAAAAAAGACAUCCUUAGAUACCGUGCUAUUGUAAAAGAAUUAGGAUUGAGAAAAUAAUACCAAAACAAAGGCACUAUCCCUUUUUUUG
env-27    ...........................................................................................
env-28    ACUAAGAAAGAUAUCAACAGAUAUCGCGCUAUCGUAAAAGAAUUAGGUUUAAGAAAAUAAUUCUCAUAAAGGGGCUGUUUACAGCCCCUUU
env-33    ACUAAGAAAGAUAUCAACAGAUAUCGCGCUAUCGUAAAAGAAUUAGGUUUAAGAAAAUAAUUCUCAUAAAGGGGCUGUUUACAGCCCCUU.
env-34    AAAGCCAAAGACAUUGCUCGUUACCGUGCAAUUGUUGCUAAACUUGGUUUACGUAAAUAAACUCUCUUUAAUUUUAUCGCACUAUUAAAAA
env-35    AAAGCCAAAGACAUUGCUCGUUACCGUGCAAUUGUUGCUAAACUUGGUUUACGUAAAUAAACUCUCUUUAAUUUUAUCGCACUAUUAAAAA
env-36    ...........................................................................................
env-37    AAAGCAAAGGAUAUUGAGCGCUACCGUGCAAUCGUCAAAAAAUUAGGUUUACGUAAAUAAACCUAUACUACUUUUACUUAUUGAAAAAAGC
env-38    AGCAAAAGACAUUCAGCGCUACCGU..................................................................
env-39    AAAGCCAAAGACAUCGUUCGUUACCGUGCAAUUGUCGCAAAACUUGGUUUACGUAAAUAAACUCUUUUUUAUCUUAUCGCAUUAUUGAAAA
env-40    ...........................................................................................
env-42    AAAAACAAGGAUGUUGUAAAGUAUAGAGAGUUGAUUAAAAAACUUUCAAUAAGAAGAUAAUUGAAAAUGAGUUUUUCAAAUUCGUAUUAAA
env-43    AAAAGCAAUGACAUUGAAAAGUACCGUGCGUUAAUCAAGGAGUUAGGUCUUCGACGUUAAUCGUAAAGUACCAUAUUUUAUUGAAAGGGGG
Myu-1-1   GCGAAAAAAGAUAUCGUUAAAUAUCGUGAGCUAAUCAAGGAAUUAGGUAUUAGAAAAUAACAUUUAAGAGGCUUUGCGGCCUCUUUUUUGU
env-44    ...........................................................................................
env-45    AAGCGUAACGAUAUUGAGAAAUACCGUGCUUUAAUCAAAGAAUUAGGUUUACGUCGUUAAUACGUUGCGAGCAAUCGCACAAACGAUACAA
env-46    ...........................................................................................
env-47    ...........................................................................................
env-48    ...........................................................................................
env-49    AAGAACAAAGAUAUCGAGCGCUAUCGUGCGAUCGUUAAGGAACUUGGUUUGAGAAGAUAAUAAGUUUCGAAUUUUAUUGAGAGGGGGCAGU
env-50    ...........................................................................................
env-51    AAAGCAAAAGAUAUCGAGCGUUACCGUGCGAUCGUGAAAAAACUAGGUUUACGUAAAUAAACCACUAUUUCUUUUUAUCGGAGUAUUGAAA
env-52    AAAAAGAAAGAUAUCGAGAGAUAUCGUGCAAUUGUUAAAGAAUUAGGAUUACGUCGUUAAUCAUCUUUAACAUAUAAAUAAAAUUGGGGGG
env-53    AAAGGAAAGGAUAUUGAAAAAUACAGGGCAUUAAUCAAGGAGCUUGGGCUUCGUCGUUAAUCGCACAGGAUUGAGCAUUUUAUGGAGAGGG
Fsa-1-1   AAGAAAAAAGAUAUCAACAGAUAUCGUGCGAUUAUCAAAGAAUUAAACAUCAGAAAAUAAUCAGAAGAAGAGGUGCUCGCGCGCCUCUUUU
Fca-1-1   AAGAAAAAAGAUAUCAACAGAUAUCGUGCGAUUAUCAAAGAAUUAAACAUCAGAAAAUAAUCAGAAGAAGAGGUGCUCGCGCGCCUCUUUU
env-54    AUGAAAAAAGAUAUUUUAAGAUAUCGUGCGAUAAUCAAAGAACUUGGCCUUAGAAAGUAAAAUAAAUUUAAGCUUUCAAAGUUCAAUUGAA
env-55    AUGAAAAAAGAUAUUUUAAGAUAUCGUGCGAUAAUCAAAGAACUUGGCCUUAGAAAGUAAAAUACAUUUAAGCUUUCAAAGUUCAAUUGAA
env-56    AUGAAAAAAGAUAUUUUAAGAUAUCGUGCGAUAAUCAAAGAACUUGGCCUUAGAAAGUAAAAUACAUUUAAGCUUUCAAAGUUCAAUUGAA
env-57    AUGAAAAAAGAUAUUUUAAGAUAUCGUGCGAUAAUCAAAGAACUUGGCCUUAGAAAGUAAAAUAAAUUUAAGCUUUCAAAGUUCAAUUGAA
env-59    AUGCACAAAGAUAUUGUUAAAUAUAGAGAGUUAAUUAAAGAAUUAGGUAUCAGAAAAUAAGCUUUGAAAACUAUUUUCUAAUACAAC....
env-60    ...........................................................................................
env-61    ...........................................................................................
env-62    ...........................................................................................
env-63    ...........................................................................................
env-65    AUGAAAAAAGACAUUUUACGCUACAGAGCAAUUGUAAAAGAAUUAGGAUUACGUAAGUAAUCUAUGCAAGGGGCUCAGUGAGCCCUUACUU
env-66    AUGAAAAAAGACAUUUUACGCUACAGAGCAAUUGUAAAAGAAUUAGGAUUACGUAAGUAAUCUAUGCAAGGGGCUCAGUGAGCCCCUUACU
env-67    AUGAAAAAAGACAUUUUACGUUACAGAGCAAUUGUAAAAGAAUUAGAAUUACGUAAAUAAUCUAUACAAGGGGCUCAGUGAGCCCCUUACU
env-68    AUGAAAAAAGACAUUUUACGUUACAGAGCAAUUGUAAAAGAAUUAGGAUUACGUAAAUAAU..............................
env-69    ...........................................................................................
env-70    AUGAAAAAAGACAUUUUACGCUACAGAGCAAUUGUAAAAGAAUUAGGAUUACGUAAGUAAUCUAUGCAAGGGGCUCAGUGAGCCCUUACUU
env-71    AUGAAAAAAGACAUUUUACGUUACAGAGCAAUAGUAAAAGAAUUAGGAUUACGUAAAUAAUCCAUACUAGGGGCUCAGUGAGCCCCUUACU
env-76    ...........................................................................................
env-77    ...........................................................................................
env-79    ...........................................................................................
env-80    ...........................................................................................
env-81    AUGAAAAAAGAUGUAUUACGUUACAGAGCAAUUGUAAAA....................................................
env-82    AUGAAAAAAGAUGUAUUACGUUACAGAGCAAUUGUAAAAGAAUUAGCAUUACGUAAAUAAUCUAUCUAAGGGGCUUAACUAAGCCCCUUAA
env-83    AUGAAAAAAGAUGUAUUACGUUACAGAGCAAUUGUAAAAGA..................................................
env-84    AUGAAAAAAGAUGUAUUACGUUACAGAGCAAUUGUAAAAGAAUUAGCAUUACGUAAAUAAUCUACCUAAGGGGCUUAACUAAGCCCCUUAA
env-85    ...........................................................................................
env-86    AUGAAAAAAGACAUUUUGCGUUACAGAGCAAUUGUAAAAGAAUUAGGAUUACGUAAAUAAUCCAUACAAGGGGCUUAUAAGUCCCUUACUU
env-87    AUGAAAAAAGACAUUUUGCGUUACAGAGCAAUUGUAAAAGAAUUAGGAUUACGUAAAUAAUCCACACAAGGGGCUUAGCUAAGCCCCUUAC
env-89    AUGAAAAAAGACAUUUUGCGUUACAGAGCAAUUGUAAAAGAAUUAGGAUUACGUAAAUAAUCCAUACAAGGGGCUUACUAAGCCCCUUACU
env-90    AUGAAAAAAGAUGUAUUACGUUACAGAGCAAUUGUAAAA....................................................
env-91    AUGAAAAAAGAUGUAUUACGUUACAGAGCAAUUGUAAAAGA..................................................
env-92    AUGAAAAAAGACAUCAUGCGCUAUCGUGCCAUUGUUAAAGAAUUAGGAUUACGAAAGUAACACUAUUAAGGAGCGCCCAUAAAGCGCUCCU
env-93    AUGAAAAAAGACAUCAUGCGCUAUCGUGCCAUUGUUAAAGAAUUAGGAUUACGAAAGUAACACUAUUAAGGAGCGCCCAUAAAGCGCUCCU
env-94    AUGAAGAAAGACAUUAUGCGUUACCGUGCAAUUGUUAAAGAAUUAAAUUUACGUAAGUAAAUUAUUGAUUAUUAAAGGGCUCUUACGAGCC

          ...........................................................................................
          ...........................................................................................
          A•RAARAARGAYAUY••••RRUAY•G•G••AU•RU•AAAGAAYURRR••U••GAAAAUAA•••••••••••••••••••••••••••••••
```

```
env-95    AUGAAGAAAGACAUUAUGCGUUACCGUGCCAUCGUAGCAGAAUUGAAGUUACGUAAGUAAUUUAUUGAUUAUUCAGGGGGCUCUUAUGAGC
env-96    AAAACAAAAGAUAUUGAGCGUUAUCGAUCAAUUAUCAGUGAAUUAAGUUUAAGAAAAUAAGUUUCAAAAAUAAUUCUAUUUUCUUAAAUAG
env-98    AAAACAAAAGAUAUUGAGCGUUAUCGAUCAAUUAUCAGUGAAUUAAGUUUAAG......................................
env-99    AAGAAAAAAGACAUUGAGCGCUAUCGUUCGAUUGUCAAAGAGCUUGGAUUGAGAAGAUAGUUUUUAAUUCAACAUAAAAAAUGGUUUAAAG
env-100   ...........................................................................................
env-101   AAAGCCAAAGACAUUACUCGCUAUCGUGAGAUUGUAAAAGAA.................................................
env-102   AAGUCCAAGGAUAUAAAUCGUUACAGAGCGAUUGUAAAAAAGUUAGGGCUGCGUAAAUAAGAAAUUCUUCUUUCAUUUGAAUCUUAGUUUU
env-103   AAGUCAAAGGAUAUAAAUCGCUACAGAGCGAUUGUAAAAAAGUUAGGGCUGCGUAAAUAAGAAAUUUCUCUUUUAAUUGAAUCUUAGUUUU
env-106   ...........................................................................................
env-107   AAGUCCAAGGAUAUAAAUCGCUACAGAGCGAUUAUUAAAAAGUUAGGGCUGCGUAAAUAAGAAAUUCUUCUUCCAGUUGAAUCUUAGUUUU
env-108   AAGUCAAAGGAUAUUAAUCGCUACAGAGCGAUUGUAAAAAAAAUUAGGACUACGUAAAUAAGAAAUUCUUCUUUCAUUUGAAUCUUAGUUUU
env-109   AAGUCCAAGGAUAUUAAUCGCUACAGAGCGAUUGUAAAAAAAAUUAGGACUACGUAAAUAAGAAAUUCUUCUUUCAAUUGAAUCUUAGUUUU
Min-1-1   AAGAAAACUGAAAUUAAUAGAUAUCGUGAGAUUAUUAAAGUAUUGAAUAUUAGAAAAUAAUCAAUAGAGAAGAGGUGCCCUUGUGCGCCUC
env-110   AAGAGAAAAGACAUUAAUCGCUACCGUGCGAUUAUCGCUGAACUCGGUCUUAGAAAAUAAAGACACACAACAAAAAGGGCAAUCAUAGUAU
env-111   ...........................................................................................
env-112   ...........................................................................................
env-113   ...........................................................................................
env-114   NUAAAGUCAGAUAUUGUAAGAUAUCGUGCAAUUGUUAAAGAACUAGGAUUAAGAAAAUAAUUNAAAAGAGGCUGUAUGCCUCUUUUUUUUA
env-115   ...........................................................................................
env-116   AUGAAAAAAGACAUUUUGCGCUAUCGAGCAAUAAUCAAAGAACUUGGAUUAAGAAAGUAAGAUUUUAAAGGGGCAUUUGCCCCUUUUUUUA
Nul-1-1   UGAAGAAAGAUAUCGUUGCUUAUCGUGAGUUGAUUAAGGACUUAGGUAUUAGAAAGUAAAGAAUUGGGGGACCUUGUGUCCCCUUUUUUAU
Nul-1-2   AUGAAGAAAGAUAUCGUUGCUUAUCGUGAGUUGAUUAAGGACUUAGGUAUUAGAAAGUAAAGAAUUGGGGACCUUGUGUCCCCUUUUUUAU
Fba-1-1   AUGAAGAAAGAUAUCGUUGCUUAUCGUGAGUUGAUUAAGGACUUAGGUAUUAGAAAGUAAAGAAUUGGGGACCUUGUGUCCCCUUUUUUAU
Nul-1-3   AUGAAGAAAGAUAUCGUUGCUUAUCGUGAGUUGAUUAAGGACUUAGGUAUUAGAAAGUAAAGAAUUGGGGACCUUGUGUCCCCUUUUUUAU
Nul-1-4   AUGAAGAAAGAUAUCGUCGCUUAUCGUGAAUUGAUUAAGGACUUAGGUAUUAGAAAAUAAUAAAUUGGGGACCUUGUGUCCCCCCUUUUU
Nul-1-5   UGAAGAAAGAUAUCGUCGCUUAUCGUGAAUUGAUUAAGGACUUAGGUAUUAGAAAAUAAUAAAUUGGGGGGGACCUUGUGUCCCCUUUUUU
Nul-1-6   AUGAAGAAAGAUAUCGUUGCUUAUCGUGAGUUGAUUAAGGACUUAGGUAUUAGAAAGUAAAGAAUUGGGGGGGACCUUGUGUCCCCCCUUU
env-117   AUGAAGAAAGAUAUCGUUGCUUAUCGUGAGUUGAUUAAGGAACUAGGAAUUAGAAAGUAAAAAAAAAGGGGACUGCUUGUCCCCUUUUUUG
env-118   AUGAAGAAAGAUAUCGUUGCUUAUCGUGAGUUGAUUAAGGAACUAGGAAUUAGAAAGUAAAAAAAAAGGGGACU.................
env-119   AUGAACAAAGAUAUAAGUAGAUAUAGAGAUAUUAUUAAAAAAUUGAAACUACGAAAAUAAUUAACUACUACAAUCAUAGUUUUUCAUUGGU
env-120   AUGAACAAAGAUAUAAGUAGAUAUAGAGAUAUUAUUAAAAAAUUGAAACUACGAAAAUAAUUAAC..........................
env-121   AUGAACAAAGAUAUAAGUAGAUAUAGAAAUAUUAUUAAAAAAUUGAAACUACGAAAAUAAUUAACUACAAUAAUCAUAGUUUUUCAUUGGU
Fsu-1-1   AAAAAGAAAGAUAUCAACAGGUAUCGUGAGAUUAUCAAAGAACUAAGCAUCCGAAAAUAAUCGAGAUGAAAAGAGGUGCCCGCGCGCCUCU
Gsp-1-1   AAGAAGAAAGAUAUUACAAGAUAUCGAGCUAUCAUCAAAGAAUUAGGAUUAAGAAAGUAAAUUAUAAAGGGGAGCAAUUUGCUCCCUUUUU
env-122   ...........................................................................................
env-123   AAAAAGAAAGAUAUUGAAAAGUACCGUUCUUUGAUAAAAGAACUAGGUAUCAGAAAAUAAAAAUAAUGGGAGCGGUAUUGCUCCCAUUAUU
env-124   ...........................................................................................
env-125   ...........................................................................................
env-126   UAAUGAAGUCUG...............................................................................
env-127   CUGAAGAAGAACAUCCACGAUUUCGGUACGGAACGCGCGUUGGUCAAGCUCGUAGGUAAACGCCGCAGCCUGCUGGAGUAUCUCAAACGNN
env-128   AAACGGGUGGAUAUCGAACGUUAUCGUAAGAUCGUCGAACAACUUGGCUUGAGAAAAUAAGCGAAUCCGUCC...................
env-129   AAACGGGUAGAUAUCGAACGUUAUCGUAAGACCGUCGAACAACUUGGCUUGAGAAAAUAAGCGAAUCCGUCCGCCGGAUCGAAAAAAGAAA
env-130   ...........................................................................................
env-131   AACAAAGAAGGAUGUC...........................................................................
env-132   AACAAAGAAGGAUGUCUUAAGAUAUCGUGCCAUCGUUAAAGAAUUAGGAUUAAGAAAAUAAAAACUAAACCACGCCUAUAAGCGUGGUUUU
env-133   AAAUCAAAAGAUAUCCUUCGUUAUCGUGAAAUCGUAAAAGAAUUGGGUUUACGAAAGUAAUCUGAAAAUUCAUCUAAAAAUUGAAUUUACU
env-134   AAAACAGAGAUUAAUCGUUACCGUGAAAUUAUUAAAGAAUUGGGUAUUAGAAAAUAAUAAUCAAUACAAAAGGAGGUGCGAAAGUGCCUCC
env-135   AUGAAAUCUGAUAUUUUAAGAUACCGUGCCAUUGUAAAGGAAUUAGGAUUAAGAAAAUAAAAUAAAAACCACGCUUUUAGCGUGGUUUUUA
env-138   AUGAAAUCUGAUAUUUUAAGAUACCGUGCCAUUGUAAAGGAAUUAGGAUUAAGAAAAUAAAAUAAAAACCACGCUUUUAGCGUGGUUUUUG
env-139   AUGAAAUCUGAUAUUUUAAGAUACCGUGCCAUUGUAAAGGAAUUAGGAUUAAGAAAAUAAAAUAAAA........................
env-140   ...........................................................................................
env-141   ...........................................................................................
env-142   AUGAAGUCUGAUGUUUUAAGAUAUCGUGCAAUCGUUAAAGAAUUAGGUUUAAGAAAAUAAUAGUAAAAACCACGCUUUAUGCGUGGUUUUU
env-143   ...........................................................................................
env-145   AUGAAGUCUGAUGUUUUAAGAUAUCGUGCAAUCGUUAAAGAAUUAGGAUUAAGAAAAUAAUAGUAAAAACCACGCAUAAAGCGUGGUUUUU
env-146   AUGAAGUCAGAUAUCUUAAGAUAUCGUGCGAUUGUUAAAGAAUUAGGAUUAAGAAAAUAAUAUUAAAAACCACGCUUCAUGCGUGGUUUUU
env-147   AUGAAGUCUGACAUCUUAAGAUAUCGUGCUAUCGUUAAAGAAUUAGGAUUAAGAAAAUAAUAGUACAAACCACGCAUAAAGCGUGGUUUUU
env-148   ...........................................................................................
env-149   ...........................................................................................
env-151   ...........................................................................................
env-152   AUGAAGUCUGAUGUCUUAAGAUAUCGUGCGAUUGUGAAAGAAUUAGGAUUAAGAAAAUAAUACAAAAACCACACAUUUAGUGUGGUUUUUU
env-153   AUGAAGUCUGAUGUCUUAAGAUAUCGUGCGAUUGUGAAAGAAUUAGGAUUAAGAAAAUAAUACAAAAACCACACAUUUAGUGUGGUUUUUU
env-154   ...........................................................................................
env-155   AUGAAGUCUGAUGUCUUAAGAUAUCGUGCGAUUAUAAAAGAAUUAGG............................................
env-156   ...........................................................................................
env-157   ...........................................................................................
env-158   AUGAAGUCUGAUGUCUUAAGAUAUCGUGCGAUUGUGAAAGAAUUAGGAUUAAGAAAAUAAUACAAAAACCACACUAAAUGUGUGGUUUUUU
env-159   AUGAAGUCUGAUGUCUUAAGAUAUCGUGCGAUUGUGAAAGAAUUAGGAUUAAGAAAAUAAUACAAAAACCACUCAUUUAGUGUGGUUUUUU
env-161   AUGAAGUCAGAUAUCUUAAGAUAUCGUGCGAUUGUCAAAGAAUUAGGAUUAAGAAAAUAAGAUUACAAACCACGCUUUAUGCGUGGUUUUU
env-162   AUGAAGUCAGAUAUCUUAAGAUAUCGAGCAAUUGUCAAAGAAUUAGGAUUAAGAAAAUAAGAUUACAAACCACGCUUUUGAGCGUGGUUUU
env-163   AUGAAAACUGAUAUCUUAAGGUAUCGUGCAAUCGUUAAAGAAUUAGGAUUAAGAAAAUAAUAUUAAAAACCACGCACAAAGCGUGGUUUUU
env-164   ...........................................................................................
env-165   AUGAAGUCAGAUAUCUUAAGAUAUCGUGCGAUUGUAAAAGAAUUAGGAUUAAGAAAAUAACACUAAAACUUUUAAAGUUCAUAAAUAAUAG
env-166   UGAAGUCAGAUAUCUUAAGAUAUCGUGCGAUUGUAAAAGAAUUAGGAUUAAGAAAAUAACACUAAACUUUUUAAAGUUACAUAAAUAAUAG
env-167   AUGAAAUCAGAUAUCUUAAGAUAUCGUGCGAUCGUAAAAGAAUUAGGAUUAAGAAAAUAAUAUCAAAAGCCACGCACAAAGCGUGGUUUUU
env-170   ...........................................................................................
env-171   AUGAAAUCAGAUAUCUUAAGAUAUCGUGCGAUCGUAAAAGAAUUAGGAUUAAGAAAAUAAUAUCAAAAACCACGCUUUAUGCGUGGUUUUU
env-172   AUGAAAUCAGAUAUCUUAAGAUAUCGUGCGAUCGUAAAAGAAUUAGGAUUAAGAAAAUAAUAUCAAAAACCACGCUUUAUGCGUGGUUUUU
env-173   ...........................................................................................
env-174   ...........................................................................................
env-175   AUGAAGUCAGAUAUUUUAAGAUAUCGUGCGAUCGUAAAAGAAUUAGGAUUAAGAAAAUAAUAUUACAAACCACGCUUUAUGCGUGGUUUUU
env-176   ...........................................................................................
env-177   ...........................................................................................
env-179   ...........................................................................................

          ...........................................................................................
          ...........................................................................................
          A•RAARAARGAYAUY••••RRUAY•G•G••AU•RU•AAAGAAYURRR••U••GAAAAUAA•••••••••••••••••••••••••••••••
```

| | |
|---|---|
| env-180 | ........................................................................................... |
| env-181 | AUGAAGUCUGAUAUUUUAAGAUAUCGUGCUAUCGUAAAAGAAUUAGGAUUAAGAAAAUAAUAUUACAAACCACGCUUCAUGCGUGGUUUUU |
| env-185 | ........................................................................................... |
| env-186 | ........................................................................................... |
| env-187 | ........................................................................................... |
| env-189 | AUGAAGUCAGAUAUUUUAAGAUAUCGUGCGAUUGUAAAAGAAUUAGGAUUAAGAAAAUAAGAUUACAAACCACGCUUUAUGCGUGGUUUUU |
| env-191 | ........................................................................................... |
| env-192 | AUGAAGUCAGAUAUCUUGAGAUAUCGUGCGAUCGUCAAAGAAUUAGGAUUAAGAAAAUAAGAUUACAAACCACGCUUUAUGCGUGGUUUUU |
| env-193 | AUGAAGUCAGAUAUCUUGAGAUAUCGUGCGAUCGUCAAAGAAUUAGGAUUAAGAAAAUAAGAUUACAAACCACGCUUUAUGCGUGGUUUUU |
| env-195 | ........................................................................................... |
| env-196 | ........................................................................................... |
| env-197 | ........................................................................................... |
| env-198 | AUGAAGUCUGAUAUCUUAAGAUAUCGUGCGAUUGUUAAAGAAUUAGGAUUAAGGAAAUAAUAUAAAAAACCACGCUUUCUGCGUGGUUUUU |
| env-201 | ........................................................................................... |
| env-202 | AUGAAGUCUGAUAUCUUAAGAUAUCGUGCGAUUGUUAAAGGAUUAGGAUUAAGGAAAUAAUAUAACAAAACUACACUUUCUGCGUGGUUUU |
| env-203 | AAGGCCAAGGAUAUCGUUCGCUACCGUGAGGUUAUUAAAGAAUUAGGUAUUCGAAAAUAAUCGACCCUUUUUUAAUACUGAAUUGUAGUUU |
| env-205 | AAGUCUAAAGACAUCGAUAAAUAUCGAAAAAUUAUUAAAGAUUUAGGUCUGAGAAAAUAAUUUAUUUCUCACUAAAUCAUCUCCUCUUGGA |
| env-206 | ........................................................................................... |
| env-207 | AAGUCUAAAGACAUUGAUAAAUAUCGAAAAAUUAUUAAAGAUUUAGGUCUGAGAAAAUAAUUCAUUUCUCACUAAAUCAUCUCCUCUUGGA |
| env-210 | AAGUCUAAAGACAUUGAUAAAUAUCGAAAAAUUAUUAAAGAUUUAGGUCUGAGAAAAUAAUUUAUUUCUCACUAAAUCAUCUCCUCUUGGA |
| env-211 | AAGAUAAAAGAUAUUGAGCGUUAUCGAUCAAUCAUUAGUAAAUUAAGUUUAAGAAAAUAGGCCUUGUAAAUAUUUCUAUUUUAUUAAAUAG |
| env-212 | ........................................................................................... |
| env-213 | AAAGCAAAUGAUAUUGAAAAAUACCGUGAGCUAAUUAAAGAUUUAGGUAUUAGAAAAUAAUCUUAUCUAAAAGUAUUUAAAAAGAGGCUUC |
| env-214 | AAAGCAAAUGAUAUUGAAAAAUACCGUGAGCUAAUUAAAGAUUUAGGUAUUAGAAAAUAAUCUUAUCUAAAAGUAUUUAAAAAGAGGCUUC |
| env-215 | AAAAGAGGCUUCAUUUUGUUGCCUCUUUUUUUAUUUUUGCCAAAACUAUAAUUAUUUAAAUGCAUAAGUUAAGUACAAUCAUCAUUCUUUU |
| env-216 | AAAGCGAAGGAUAUCGAAAGAUACCGCGCAAUAGUUAAAGAAUUAGGAUUACGUCGUUAAUCUUUGUUCGAGAAAUUUAUUGGAGAGGGGA |
| env-217 | AAAGCGAAGGAUAUCGAAAGAUACCGCGCAAUAGUUAAAGAAUUAGGAUUACGUCGUUAAUCUUUGUUCGAGAAAUUUAUUGGAGAGGGGA |
| env-218 | ........................................................................................... |
| env-220 | ........................................................................................... |
| env-221 | AAAAGUAAAGACAUAGAGAAAUAUCGUGCUUUAGUGAAAGAAUUAGGAUUACGUCGUUAAUUCUUAACUGAGUACACAAAUUUUAUUGGAG |
| env-222 | AUUUUGGAACGCAACGCUCUCUUCAAUUA.............................................................. |
| Fli-1-1 | AAGAAAAAAGAUAUCAACAGAUAUCGUGCGAUUAUCAAAGAAUUAAACAUCAGAAAAUAAUCAGAAGAAAGAGGUGCCCGCGCGCCUCUUU |
| Gme-1-1 | ACUAAGAAAGAUGUCUUAAGAUAUCGUGCCAUCGUAAAAGAAUUAGGAUUAAGAAAAUAAUCCCUUCAAAGACCACGCCUUUCAGCGUGGU |
| env-223 | ........................................................................................... |
| env-224 | ........................................................................................... |
| env-225 | AAAAGUAAAGACAUAGANAAAUACCGUGCNUUNNUUAAAGAAUUAGGAUUACGUCGUUAAUUCUAAGCACAAAUUUAUUGAUUGGGGGGAC |
| env-226 | ........................................................................................... |
| env-227 | ........................................................................................... |
| env-228 | ........................................................................................... |
| env-229 | ........................................................................................... |
| env-230 | AAAAGUAAAGACAUAGAGAAAUACCGUGCAUUGGUUAAAGAAUUAGGAUUACGUCGUUAAUUCUAAGCACAAAUUUAUUGAUUGGGGGGAC |
| env-231 | AAAAGUAAAGACAUAGAGAAAUACCGUGCAUUGGUUAAAGAAUUAGGAUUACGUCGUUAAUUCUAAGCACAAAUUUAUUGAUUGGGGGGAC |
| env-234 | AAAAGUAAAGACAUAGAGAAAUACCGUGCAUUGGUUAAAGAAUUAGGAUUACGUCGUUAAUUNUAAGCACAAAUUUAUUGAUUGGGGGGAC |
| env-235 | ........................................................................................... |
| env-236 | AAAAGUAAAGACAUAGAGAAAUACCGUGCAUUGGUUAAAGAAUUAGGAUUACGUCGUUAAUUCUAAGCACAAAUUUAUUGAUUGGGGGGAC |
| env-237 | AAAAGUAAAGACAUAGAGAAAUACCGUGCAUUGGUUAAAGAAUUAGGAUUACGUCGUUAAUUCUAAGCACAAAUUUAUUGAUUGGGGGGAC |
| env-238 | ........................................................................................... |
| env-239 | AUGAAGAAGGACAUUAUGCGUUACCGCGCCAUCGUAAAAGAAUUAGGAUUGCGUAAGUAAAAAAUAAAAGGGGCUUUGUGCCCCUUUUUUA |
| env-243 | ........................................................................................... |
| env-244 | GUCAAAAAGGACAUCACAAGAUAUAGGAACAUUAUUAAAGAGCUUAAACUAAGAAAAUAGAUGAAAAGGGGCUUUUACAGCCUCUUUUUUU |
| env-247 | ........................................................................................... |
| env-248 | ........................................................................................... |
| env-249 | GUUAAAAAAGAUAUUACACGUUAUAGAAAUAUCCUUAAAGAUUUAAAACUAAGAAAAUAAAUAAAAGAGGCUUCACGCCUCUUUUUUUGAU |
| env-250 | ........................................................................................... |
| env-251 | AUGAAGAAAGAUAUUACCCGUUACCGUGCUAUUGUAAAAGAAUUAGGACUACGUAAGUAAUCUAUACCAAGGGACACUUUUGUGUCCCUUU |
| env-255 | AUGAAGAAAGACAUUACACGUUACCGUGCAAUUGUUAAAGAACUAGGAUUACGUAAGUAAUUUUGAAAAGGGGUGCCAAGGCACUCCUUU |
| env-256 | AUGAAAAAAGACAUCAUGCGCUACAGAGCAAUUGUAAAAGAACUCGGAUUGCGUAAAUAAAAACACAAAAGGGGUGCCGUUGGUGUCCCUU |
| env-257 | AUGAAAAAAGAUAUUAUGCGCUACCGUGCUAUAGUCAAAGAACUAGGCUUGCGUAAAUAAAUUAAAAAGGGAUACCAUUGGUGUCCCUUUU |
| env-258 | ACUAAAAAGGACAUUACCCGUUACCGUGCGAUAGUUAAGGAACUUGGUUUAAGAAAAUAAUUUUCUAUAAAGGGGCAUUUUUGCCCCUUUU |
| env-261 | ........................................................................................... |
| Nma-1-1 | AUGAAGAAAGACAUUGUUGCUUACCGUGAGUUGAUUAAAGAAUUAGGAAUUAGAAAGUAAAAAAUUGGGGACUUCGUGUCCCCUUUUUUAG |
| env-262 | ........................................................................................... |
| env-263 | AAAAACAAAGAUAUUACUCGUUACAGAGAAAUAAUUAAAAAUUUAAAUCUCCGUAAAUAAGUAUCCUCAUCAUAGUUUUUCAUUGGUAUUC |
| env-264 | AAAAACAAAGAUAUUACUCGUUACAGAGAAAUAAUUAAAAAUUUAAAUCUCCGUAAAUAAGUAUCCUCAUCAUAGUUUUUCAUUGGUAUUC |
| env-265 | AAAAACAAAGAUAUUACUCGUUACAGAGAAAUAAUUAAAAAUUUAAAUCUCCGUAAAUAAGUAUCCUCAUCAUAGUUUUUCAUUGGUAUUC |
| env-266 | AUGAAGAAGGAUAUUUUAAGAUAUCGUGCUAUCGUUAAGGAAUUAGGAUUAAGAAAAUAAUAUUAAAAGGGGCUUUACGCCCCUUUUUUUU |
| env-267 | AUGAAGAAAGAUAUUUUAAGAUAUCGUGCUAUCGUUAAGGAAUUAGGAUUAAG...................................... |
| env-268 | AAGAAAAAAGAUAUUAUGAGAUAUCGUGAAAUCAUUAAGGAAUUAGGUUUAAGAAAAUAAACAUUGAAAGGGGCUUUUUACGCCCCUUUUU |
| env-269 | AUGAAGAAAGAUAUUUUAAGAUAUCGUGCUAUCGUUAAGGAAUUAGGAUUAAGAAAAUAAUAUUAAAAGGGGCUUUACGCCCCUUUUUUUU |
| env-270 | ........................................................................................... |
| env-271 | AUGAAGAAAGAUAUUUUAAGAUAUCGUGCUAUCGUUAAGGAAUUAGGAUUAAGAAAAUAAUAUUAAAAGGGGCUUUACGCCCCUUUUUUUA |
| env-272 | AUGAAGAAAGAUAUUUUAAGAUAUCGUGCUAUCGUUAAGGAAUUAGGAUUAAGAAAAUAAUAUUAAAAGGGGCUUUACGCCCCUUUUUUUA |
| Mmo-1-1 | AUGAAAAAAGAUAUUUUAAGAUAUCGUGCUAUCGUUAAGGAAUUAGGAUUAAGAAAAUAAUAUUAAAAGGGGCUUUACGCCCCUUUUUUUU |
| env-273 | ........................................................................................... |
| env-274 | AAAAGGGAAGACAUCGAACGUUAUCGUAAGAUUGUCGAGCAGCUCGGACUCAGAAAGUAGGCUGUAAGCUUUUAAAAGGAAUAAAAAACCGG |
| env-276 | ACUAAGAAAGAUAUUUUAAGAUAUCGUGCAAUUGUAAAAGAGUUAGGCUUAAGAAAAUAAUAUAGAUUAAAUCCUAUUAUAAUAAUUAUUC |
| env-277 | ACUAAGAAAGAUAUUUUAAGAUAUCGUGCAAUUGUAAAAGAGUUAGGCUUAAGAAAAUAAUAUAGAUUAAAUCCUA............... |
| env-278 | ACUAAGAAAGAUAUUUUAAGAUAUCGUGCAAUUGUAAAAGAGUUAGGCUUAAGAAAAUAAUAUAUUAAAUCCUAUUAUAAUAAUUAUUCAU |
| env-280 | ACUAAAAAAGAUGUUUUAAGAUAUCGAGCAAUUGUAAAAGAGUUAGGCUUAAGAAAAUAAUUUAAGUAAUAUAUCUAUAUUCUAAUAUUAU |
| env-281 | ACUAAAAAAGAUAUUUUAAGAUAUCGAGCAAUUGUAAAAGAGUUAGGCUUAAGAAAAUAAUUUAAGUAAUAUAUCCUAUUCUAAAUAUUAU |
| env-282 | AUUAAGAAAGAUAUUUUAAGAUAUCGUGCAAUUGUAAAAGAAUUAGGCU.......................................... |
| env-283 | AUUAAGAAAGAUAUUUUAAGAUAUCGUGCAAUUGUAAAAGAAUUAGGCUUAAGAAAAUAAUAUAAAUCAAAUCCUAUUUAUAAUAAUUAUU |
| env-285 | AUUAAGAAAGAUAUUUUAAGAUAUCGUGCAAUUGUAAAAGAAUUAGGCUUAAGAAAAUAAUAUAAAUCAAAUCCUAUUUAUAAUAAUUAUU |
| env-286 | ........................................................................................... |
| | ........................................................................................... |
| | ........................................................................................... |
| | A•RAARAARGAYAUY••••RRUAY•G•G••AU•RU•AAAGAAYURRR••U••GAAAAUAA••••••••••••••••••••••••••••••• |

| | |
|---|---|
| env-287 | AUUAAGAAAGAUAUUUUAAGAUAUCGUGCAAUUGUAAAAGAAUUAGGCUUAAGAAAAUAAUAUAAAUCAAAUCCUAUUUAUAAUAAUUAUU |
| env-288 | ........................................................................................... |
| env-289 | AUUAAGAAAGAUAUUUUAAGAUAUCGUGCAAUUGUAAAAGAAUUAGGCUUAAGAAAAUAAUAUAAAUCAAAUCCUAUUUAUAAUAAUUAUU |
| env-290 | AUUAAGAAAGAUAUUUUAAGAUAUCGUGCAAUUGUAAAAGAAUUAGGCUUAAGAAAAUAAUAUAAAUCAAAUCCUAUUUAUAAUAAUUAUU |
| env-294 | AUUAAGAAAGAUAUUUUAAGAUAUCGUGCAAUUGUAAAAGAAUUAGGCUUAAGAAAAUAAUAUAAAUCAAAUCCUAUUUAUAAUAAUUAUU |
| env-295 | AUUAAGAAAGAUAUUUUAAGAUAUCGUGCAAUUGUAAAAGAGUUAGGCUUAAGAAAAUAAUAUAAAUCAAAUCCUAUUUAUAAUAAUUAUU |
| env-296 | ........................................................................................... |
| env-297 | ........................................................................................... |
| env-298 | ........................................................................................... |
| env-299 | AAAACUAAAGAUAUAGAGAGGUACAGAAAUAUUGUGAAAGAAUUAGGUUUACGUAAAUAAUUUAGAUCUUAUGUUUUUCAUUGGAUGCGAC |
| env-300 | AAAACUAAAGAUAUAGAGAGAUACAGAAAUAUUGUGAAAGAAUUAGGUUUACGUAAAUAAUUUAGAUCUUAUGUUUUUCAUUGGAUGCGAC |
| env-301 | ........................................................................................... |
| env-302 | AAAAAUAAAGAUAUAGAAAGGUAUAGAAAUAUUGUGAAAGAAUUAGGUUUACGUAAAUAAUUUAGAUCUUAUGUUUUUCAUUGGAUGCGGC |
| env-303 | AAAACUAAAGAUAUAGAAAGGUAUAGAAAUAUUGUGAAAGAACUAGGUUUACGUAAAUAAUUUAGAUCUUAUGUUUUUCAUUGGAUGCGAC |
| env-304 | AAGUCGAAGGACAUCGAAAGGUAUAGAAAUAUUGUAAAAGAAUUAGGUUUACGUAAAUAGUGAAAUUAAUCAAACAUUGUGUUUUUCAUUG |
| env-305 | ........................................................................................... |
| env-306 | ACUAAAAAAGAUAUUUUGAGAUAUAGAGCGAUUGUAAAAGAACUAGGAUUAAGAAAAUAAUUAUUUCUUGACUCUUAUUAGUUAUAGUUAU |
| env-307 | ACUAAAAAAGAUAUUUUGAGAUAUAGAGCGAUUGUAAAAGAACUAGGAUUAAGAAAAUAAUUAUUUCUUGACUC................. |
| env-308 | AAGAAGACAGACAUCGAGCGUUACCGUGCGAUCGUAAAAGAGCUUGGAUUGAGAAGAUAAUUUUUAGUUCAACAUAAAAUUUGGAUAAAAG |
| Lsp-3-1 | AUUAAGAAAGAUAUUUUAAGAUAUCGUGCUAUCGUUAAAGAAUUAGGUUUAAGAAAAUAAUUCUUAUUUUUAAAAGGGGCAGUUUGCCCCU |
| env-309 | ACUAAGAAAGA................................................................................ |
| env-310 | ACUAAGAAAGAUAUCAACAGAUAUCGUGCUAUCGUUAAAGAAUUAGGUUUAAGAAAAUAAUUCUUACGUAUAAAAGGGGCUGUUUUUCAGU |
| env-313 | ACUAAGAAAGAUAUCAACAGAUAUCGUGCUAUCGUUAAAGAAUUAGGUUUAAGAAAAUAAUUCUUACGUAUAAAAGGGGCUGUUUUUCAGU |
| env-314 | ACUAAGAAAGAUAUCAACAGAUAUCGUGCUAUCGUUAAAGAAUUAGGUUUAAGAAAAUAAUUCUUAUUUAUAAAAGGGGCUGUUUUUCAGU |
| env-315 | ACUAAGAAAGAUAUCAACAGAUAUCGUGCUAUCGUUAAAGAAUUAGGUUUAAGAAAAUAAUUCUUAUUUAUAAAAGGGGCUGUUUUUCAGU |
| env-316 | ACUAAGAAAGAUAUCAACAGAUAUCGUGCUAUCGUUAAAGAAUUAGGUUUAAGAAAAUAAUUCUUAUUUAUAAAAGGGGCUGUUUUUCAGU |
| env-317 | ........................................................................................... |
| env-318 | ACUAAGAAAGAUAUCAAUAGAUAUCGUGCUAUCGUUAAAGAAUUAGGUUUAAGAAAAUAAUUCUUAUUUAUAAAAGGGGCUGUUUUUCAGU |
| env-320 | ACUAAGAAAGAUAUCAAUAGAUAUCGUGCUAUCGUUAAAGAAUUAGGUUUAAGAAAAUAAUUCUUAUUUAUAAAAGGGGCUGUUUUUCAGU |
| env-324 | ACUAAGAAAGAUAUCAA.......................................................................... |
| env-325 | ACUAAGAAAGAUAUCAACAGAUAUCGUGCUAUCGUUAAAGAAUUAGGUUUAAGAAAAUAAUUCUUAUUUAUAAAAGGGGCUGUUUUUCAGU |
| env-326 | ........................................................................................... |
| Fri-1-1 | AAAAAGAAAGAUAUCAACAGGUAUCGUGAGAUUAUCAAAGAACUAAACAUCAGAAAAUAAUCAAGAUGAAAAGAGGUGCCUGCGCGCCUCU |
| env-327 | AAAACAAAAGAUAUUGAGCGUUAUCGAUCAAUUAUCAGUGAAUUAAGUUUAAGAAAAUAAGUUUCAAAAAUAAUUCUAUUUUCUUAAAUAG |
| env-328 | AAAAUAAAAGAUAUUGAGCGUUAUCGAUCAAUUAUCAGUGAAUUAAGUUUAAGAAAAUAAGCUUCAAAAAAUAAUUCUAUUUUCUUAAAUA |
| env-329 | AAAUUUAAAGAUAUCGAAAGGUAUCGUUCUCUAAUAAAGGAAUUAGGUAUACGAAAAUAACUCCAAGCUUAUAAAAGAGUAUAUACUUUAU |
| env-330 | AAAUUUAAAGAUAUCGAAAGGUAUCGUUCUCUAAUAAAGGAAUUAGGUAUACGAAAAUAACUCCAAGCUUAUAAAAGAGUAUAUACUUUAU |
| env-331 | AAAUCAAAGGAUAUUGAAAAAUAUAGAUCUUUGAUUAAAGAACUUGGAUUAAGACGUUAAUCCAUUUGUAAAAGUAUUGCAAAAAGAGGGG |
| env-332 | AAAUCAAAGGAUAUUGAAAAAUAUAGAUCUUUGAUUAAAGAUCUUGGAUUAAGACGUUAAUCCAAUUAAUAAAAGUAUUGCAAAAAGAGGG |
| env-333 | AAAUCAAAGGAUAUUGAAAAAUAUAGAGCUUUGAUUAAAGAGCUUGGAUUAAGACGUUAAUCCAUUAAUAAAAGUAUUGUAAAAAGAGGGG |
| env-334 | AAAGAAAAAAAUAUCACACGUUAUCGUGACAUAGUUAAAGAAUUAGGCCUACGUAAAUAAAUCUCAGAUGCAGUCUUAAAAAACUGCAUCU |
| env-335 | ........................................................................................... |
| env-336 | UGAAAAAGACAUUGAACGAUAUAGAGCUAUCGUUAAAGAUUUAGGUUUAAGAAAAUAAAUUUGCUAACCAAAAAGGGGCGUUUGCCCCUU. |
| env-337 | AUGAAGAAAGACAUUAUGCGAUACAGAGCUAUCGUUAAAGAAUUAGGAUUAAGAAAAUAAUUAUUAGGGGCUUUAAAGUCCCUUUUUUGUU |
| env-338 | UCUAAGAUAUUGACGAUAUAGAAGUAUUGUGAAAAUAAGUCUACGAAAGUAAUAAUAAGAUUUAAAGUUUAGUUUUCAUGGUGCUCACACC |
| env-339 | AAAUCUAAAGAUAUUGAACGAUAUAGAAGUAUUUUGAAAAAAUUAGGCCUACGAAAGUAAAUAAUUUAAGAUUUUAAAAGUUUUAGUUUUU |
| env-340 | AUCUUAAGCCGUAA............................................................................. |
| env-341 | ........................................................................................... |
| env-342 | AAAUCUAAAGAUAUUGAACGAUAUAGAAGUAUUUUGAAAAAAUUAGGCCUACGAAAGUAAAUAAUUUAAGAUUUUAAAAGUUUUAGUUUUU |
| env-343 | ........................................................................................... |
| env-344 | ........................................................................................... |
| env-345 | AAAUCAAAAAAUAUUGAACGAUACAGAAGUAUUGUAAAAAAAUUAGGUUUACGAAAGUAAAUAAUUAAAAAUUUUAAAAGUUAUAGUUUUU |
| env-346 | AAAUCUAAAGAUAUUGAACGAUAUAGAAGUAUUGUGAAAAAAUUAGGUCUACGAAAGUAAAUAAUUAAAGAUUUUAAAAGUUUUAGUUUUU |
| env-347 | AAAUCUAAAGAUAUUGAACGAUAUAGAAGUAUUGUGAAAAAAUUAGGCCUACGAAAGUAAAUAAUUUAAGAUUUUAAAAGUUUUAGUUUUU |
| env-352 | ........................................................................................... |
| env-353 | ........................................................................................... |
| env-354 | AAAUCCAAAGAUAUUGAACGAUAUAGAAGUAUUGUGAAAAAAUUAGGCCUACGAAAGUAAAUAAUUUAAGAUUUUAAAAGUUUUAGUUUUU |
| env-355 | ........................................................................................... |
| env-356 | AAAUCUAAAGAUAUUGAACGAUAUAGAAGUAUUGUGAAAAAAUUAGGUCUACGAAAGUAAAUAAUUAAAGAUUUUAAAAGUUUUAGUUUUU |
| env-357 | AAGAAAAAGGAUAUCGAGAAAUACCGUGCGUUGAUCAAAGAAUUAGGUUUACGUCGUUAAUCAGAAAGAGGUCACUGAGCGUAGUCGAAGU |
| Fta-1-1 | AAGAAAAAGGAUAUCGAGAAAUACCGUGCGUUGAUUAAAGAAUUAGGUUUACGUCGUUAAUAAAAAACAGGUCACUGAAUACUUGCGCUAA |
| env-358 | ACUAAGAAAGAUGUCUUAAGAUAUCGUGCAAUUGUUAAAGAACUAGGAUUAAGAAAAUAAUCUCACUCUUAAACCACGCCUACAAGCGUGG |
| env-359 | ........................................................................................... |
| env-360 | AAAAUAAAGAUAUUUUACGUUACAGAGCGAUCGUUAAAGAUUUAGGAUU.......................................... |
| env-361 | AUGAAAAAGAAUAUUACUAGAUAUAGAAGUAUCAUUAAAGAAUUAGGAUUGAGAAAAUAAUUUAUUUAAGGGGCUUUAUUGCCCCUUUUUU |
| env-362 | AUGAAAAAGGAUAUUACCCGAUACAGAAAUAUCAUUAAAGAAUUAGGAUUAAGAAAAUAAGAAUUAUAGGGGCAAUUUUGCCCCUUUUUUU |
| Mod-7-1 | AAGAAUACAGAAAUUAACAGAUAUCGUGAGAUUAUCAAAGUAUUGGGUAUUAGAAAAUAAUCAAUAUAGAGAAGAGGUGCCUUUGUGCGCC |
| Mod-6-1 | AAGAAUACAGAAAUUAACAGAUAUCGUGAGAUUAUCAAAGUAUUGGGUAUUAGAAAAUAAUCAAUAUAGAGAAGAGGUGCCUUUGUGCGCC |
| Mod-3-1 | AAGAAUACAGAAAUUAACAGAUAUCGUGAGAUUAUCAAAGUAUUGGGUAUUAGAAAAUAAUCAAUAUAGAGAAGAGGUGCCUUUGUGCGCC |
| Mpr-1-1 | AAGAAUACAGAAAUUAACAGAUAUCGUGAGAUUAUCAAAGUAUUGGGUAUUAGAAAAUAAUCAAUAUAGAGAAGAGGUGCCUUUGUGCGCC |
| Mod-4-1 | AAGAAUACAGAAAUUAACAGAUAUCGUGAGAUUAUCAAAGUAUUGGGUAUUAGAAAAUAAUCAAUAUAGAGAAGAGGUGCCUUUGUGCGCC |
| Mod-1-1 | AAGAAUACUGAAAUUAACAGAUACCGUGAGAUUAUCAAAGAAUUGAAUAUUAGAAAAUAAUCAAUAGUAAAGAGGUGCCCUUGUGCGCCUC |
| Fin-1-1 | AAGAAAAAAGAUAUCAACAAAUAUCGUGAGAUUCUGAAAACAUUAAACAUCAGAAAAUAAUUCAAGGGGGCUUUGUGCCCCUUUUUGUUUA |
| env-363 | AAGAAAACCGAGAUCAACAGAUACCGUGAGAUUAUUAAAGUAUUGAAUAUUAGAAAAUAAUCAACAAAAAAGAGGUGCCCACGCGCCUCUU |
| env-364 | AAGAAAACCGAGAUCAACAGAUACCGUGAGAUUAUUAAAGUAUUGAAUAUUAGAAAAUAAUCAACAAAAAAGAGGUGCCCACGCGCCUCUU |
| env-366 | ........................................................................................... |
| env-367 | AAGAAAACCGAGAUCAACAGAUACCGUGAGAUUAUUAAAGUAUUGAAUAUUAGAAAAUAAUCAACAAAAAAGAGGUGCCCACGCGCCUCUU |
| env-368 | AAGAAAACCGAGAUCAACAGAUACCGUGAGAUUAUUAAAGUAUUGAAUAUUAGAAAAUAAUCAACAAAAAAGAGGUGCCCACGCGCCUCUU |
| env-369 | AAAAAAACAGAUAUCGCGAGAUACCGAGCGAUUAUCAAAAAACUCAACUUACGAAAAUAAUUUUUUUAAAAGGCAGUGCUUUGGCAUUGCC |
| env-373 | AAGUCAAAAGAUAUUUUACGCUAUCGUGAAAUCGUAAAAGAAUUGGGUUUAAGAAAGUAAUUUAAAAACAUAUACAAAUAUCAAGCUGAUU |
| env-375 | ........................................................................................... |
| env-376 | AUGAAAAAAGAUAUUCUCAGAUACCGUGCGAUUGUAAAAGAGCUAGGAUUAAGAAAAUAAAUAUUCAAAAAGAGGCAUAAGCCUCUUUUUU |
| env-377 | AUUAAAAAAAGAUAUCCUUAGGUACCGUGCGAUUGUUAAAGAGUUAGGAUUAAGAAAAUAAAUACACAAAAAAGAGGC............. |
| env-378 | AUGAAAAAGGAUAUAUUAAAGUAUCGUGCUAUUGUAAAAGAGUUAGGAUUGAGAAAGUAAUAGAAGAAGCUAAUUAAAGUUUUUCAUUGGU |
| | ........................................................................................... |
| | ........................................................................................... |
| | A•RAARAARGAYAUY••••RRUAY•G•G••AU•RU•AAAGAAYURRR••U••GAAAAUAA•••••••••••••••••••••••••••••••• |

```
env-379   AUGAAGAAAGACAUCUUGAGAUACCGUGCGAUUGUAAAAGAAUUAGGAUUAAGAAAAUAAUAUUAAAAGAGAGGCCUUAUGCCUCUUUUUU
env-381   AUGAAGAAAGACAUCUUGAGAUACCGUGCGAUUGUAAAAGAAUUAGGAUUAAGAAAAUAAUACUAAAAGAGAGGCCUUAUGCCUCUUUUUU
env-382   AUGAAGAAAGACAUCUUGAGAUACCGUGCGAUUGUAAAAGAAUUAGGAUUAAGAAAAUAAUACCAAAAGAGAGGCCUUAUGCCUCUUUUUU
env-383   AUGAAGAAAGACAUCUUGAGAUACCGUGCGAUUGUAAAAGAAUUAGGAUUAAGAAAAUAAUACUAAAAGAGAGGCCUUAUGCCUCUUUUUU
env-384   AUGAAGAAAGACAUCUUGAGAUACCGUGCGAUUGUAAAAGAAUUAGGAUUAAGAAAAUAAUACCAAAAGAGAGGCCUUAUGCCUCUUUUUU
env-385   AUGA.......................................................................................
env-386   AUGAAGAAAGACAUCUUGAGAUACCGUGCGAUUGUAAAAGAAUUAGGAUUAAGAAAAUAAUACUAAAAGAGAGGCCUUAUGCCUCUUUUUU
env-387   AUGAAGAAAGACAUCUUGAGAUACCGUGCGAUUGUAAAAGAAUUAGGAUUAAGAAAAUAAUACUAAAAGAGAGGCCUAUGCCUCUUUUUUA
env-388   AUGAAGAAAGACAUCUUGAGAUACCGUGCGAUUGUAAAAGAAUUAGGAUUAAGAAAAUAAUACUAAAAGAGAGGCCUUAUGCCUCUUUUUU
env-389   AUGAAGAAAGACAUCUUGAGAUACCGUGCGAUUGUAAAAGAAUUAGGAUUAAGAAAAUAAUACCAAAAGAGAGGCCUUAUGCCUCUUUUUU
env-390   AUGAAGAAAGACAUCUUGAGAUACCGUGCGAUUGUAAAAGAAUUAGGAUUAAGAAAAUAACAUUAAAAGAGAGGCUAUGUGCCUCUUUUUU
env-391   AUGAAGAAGGAUAUUUUGAGAUACCGUGCGAUUGUAAAAGAAUUAGGAUUAAGAAAAUAACAUUAAAAGAGAGGCUAUGUGCCUCUUUUUU
env-392   AUGAAGAAGGACAUUUUGAGAUACCGUGCGAUUGUAAAAGAAUUAGGAUUAAGAAAAUAACAUUAAAAGAGAGGCUAUGUGCCUCUUUUUU
env-393   AUGAAAAAAGACAUCUUGAGAUACCGUGCGAUUGUCAAAGAAUUAGGAUUGAGAAAAUAAUACCAAAAAAGAGGUCGCAUGCCUCUUUUUG
env-394   AUGAAAAAAGACAUCUUGAGAUACCGUGCGAUUGUCAAAGAAUUAGGAUUGAGAAAAUAAUACCAAAAAAGAGGCCACAUGCCUCUUUUUG
env-395   ...........................................................................................
env-396   ...........................................................................................
env-397   AAUGACAAGCAUAUAUAGCGUUAUCGUGCUAUUGUUAUUGAAUUAGGUUUACGUAAAUAAAAUUUAAAAUAUAUACUGAAUCUUAGUUUUU
env-398   AAUGACAAGCAUAUAGAGCGUUAUCGUGCUAUUGUUAUUGAAUUAGGUUUACGUAAAUAAAAUUUAAAAUAUAUACUGAAUCUUAGUUUUU
env-399   AAAGACAAGGACAUAGUGCGUUAUCGUGCUAUUGUUAUUGAAUUAGGCUUACGUAAAUAAAAUCUAAAAUAUAUACUGAAUCUUAGUUUUU
env-400   AAUGACAAGCAUAUAGAGCGUUAUCGUGCUAUUGUUAUUGAAUUAGGUUUACGUAAAUAAAAUUUAAAAUAUAUACUGAAUCUUAGUUUUU
env-402   AAGUCAAAAGAUAU.............................................................................
env-403   ...........................................................................................
env-404   AAGUCAAAAGAUAUCGAAAGGUAUAGAAAAAUUGUAAAAGAAUUAGGUUUACGUAAAUAGUGCAAUUAACUAGACAUUAUGUUUUUCAUUG
env-405   AAAACUAAAGAUAUCAAUCGAUACAGAGCGAUAGUAAAAGAAUUAAGUUUACGUAAAUAAAAUCCUUCUUACAAUCGAAUCUUAGUUUUUC
env-406   ...........................................................................................
env-407   AAGUCGAAGGAUAUCGAAAGGUACAGAAAUAUUGUAAAAGAAUUAGGUCUACGUAAAUAGUGAUAUUAAUCAGACAUUCUGUUUUUCAUUG
env-408   ...........................................................................................
env-409   UAAAGUCGAAGGAU.............................................................................
env-410   ...........................................................................................
env-411   AAGUCCAAGGACAUAGAAAGGUAUAGAAAUAUUGUAAAAGAAUUAGGUUUACGUAAAUAGUAAAAAAAAUUAAGCCCUUAUGUUUUUCAUU
env-412   AAGUCCAAGGACAUAGAAAGGUAUAGAAAUAUUGUAAAAGAAUUAGGUUUACGUAAAUAGUAAAAACAAUUAAGACCUUAUGUUUUUCAUU
env-413   AAGUCCAAGGACAUAGAAAGGUAUAGAAAUAUUGUAAAAGAAUUAGGUUUACGUAAAUAGUAAAAAAAAUUAAGACCUUAUAUUUUUCAUU
env-414   AAGUCCAAGGACAUAGAAAGGUAUAGAAAUAUUGUAAAAGAAUUAGGUUUACGUAAAUAGUAAAAAAAAUUAAGCCCUUAUGUUUUUCAUU
env-415   ...........................................................................................
env-416   AUGUCAAAAGAUAUUGAAAGAUAUCGAACAAUUAUUAAAGAUUUAGGGAUUCGUAAAUAAAAAUCUAAUUUAUUUUCUGAAAUUAUCAAUU
env-417   AUGUCAAAAGAUAUUGAAAGAUAUCGAACAAUUAUUAAAGAUUUAGGGAUUCGUAAAUAAAAAUCUAAUUUAUUUUCUGAAAUUAUCAAUU
env-418   AUGUCAAAAGAUAUAGAAAGGUAUCGUAUUAUAAUUAAAGAUUUAGGUAUUCGUAAAUAACUGCAUAGACAUUAUAAAUAAUCUAUGAUAU
Fbe-1-1   AAGAAGAAAGAGAUUAACAGAUACCGUGAGAUUAUCAAAGAAUUAAACAUCAGAAAAUAAUCGAGAUGAAAAGAGGCGCCUGCGCGCCUCU
env-419   AUCAAAAAAGAUAUCUUGAGAUAUCGUGCGAUUAUCAAAGAAUUGAAUAUCAGAAAAUAAUUUCGAGGGGGCCGUAUGGCUCCCUUUUUAU
Ksp-1-1   AUCAAGAAAGACAUUUUACGUUACAGAGCUAUUAUUAAGGAACUAGGUAUCCGUAAAUAACAAUUUUUUAAAAAGGGAGCCUCACGGCUCC
Dsp-2-1   AAGAAAAAAGAUAUCUUACGUUACAGAGCUAUUAUUAAGGAACUAGGUAUCCGUAAAUAAUAAUUUUUUAAAAAGGGAGCUUCACGGCUCC
Dsp-1-1   AAGAAAAAAGACAUCUUACGUUACAGAGCUAUCAUUAAGGAGCUAGGUAUCCGUAAAUAAGAAUUUUUUAAAAAAGGGAGCUUCACGGCUC
Ddo-1-1   AAGAAAAAAGAUAUCUUACGUUACAGAGCUAUCAUUAAGGAGCUAGGUAUCCGUAAAUAAGAAUUUUUUAAAAAGGGAGCUUCACGGCUCC
Iha-1-1   AAGAACAAAGAUAUCAAUAGAUAUCGUGCUAUCAUCAAAGAAUUAGGAUUAAGAAAGUAAUAGAAAAGAGGCUUCGUGCCUCUUUUUUUUA
env-420   AAGAACAAAGACAUCGAACGCUACCGUUCCUUAAUUAAGGAAUUAGGUAUUCGAAAGUAAGCACAUAAAAGCAUCCGCAUCAUACCGGCUU
env-421   ...........................................................................................
env-422   GCAAAAGACAUAGAACGCUACCGUUCCUUGAUUAAGGAAUUGGGCAUUCGUAAGUAAACACUUAAAAGCAUCCGCAUCAUCGGUUUUUCAA
env-423   ...........................................................................................
env-424   AAGAACAAAGACAUCGAACGCUACCGUUCCUUAAUUAAGGAAUUAGGUAUUCGAAAGUAAGCACAUAAAAGCAUCCGCAUCAUACCGGCUU
env-427   AAGAACAAAGACAUCGAACGCUAUCGUUCCUUAAUUAAGGAAUUAGGUAUUCGAAAGUAAGCACAUAAAAGCAUCCGCAUCAUACCGGCUU
env-428   AAAGACAAAGACAUCGAACGCUACCGUUCCUUAAUUAAGGAAUUAGGCAUUCGAAAGUAAGCACAUAAAAGCAUCCGCAUCAUACCGGCUU
env-429   AAGAACAAAGACAUCGAACGCUAUCGUUCCUUAAUUAAGGAAUUAGGUAUUCGAAAGUAAGCACAUAAAAGCAUCCGCAUCAUACCGGCUU
env-430   GAUAAAGACAUCGAGCGCUACCGUUCCUUGAUAAAGGAAUUGGGCAUUCGUAAGUAAAACACUUAAAAGCAUCCGCAUCAUCGGUUUUUCA
env-431   GAUAAAGACAUCGAGCGCUACCGUUCCUUGAUAAAGGAAUUGGGCAUUCGUAAGUAAAACACUUAAAAGCAUCCGCAUCAUCGGUUUUUCA
env-432   ...........................................................................................
env-433   ...........................................................................................
env-434   GAUAAAGACAUCGAGCGCUACCGUUCCUUGAUUAAGGAAUUGGGCAUUCGUAAGUAAGCACUUAAAAGCAUCCGCAUCAUCGGUUUUUCAG
env-435   GAUAAAGACAUCGAGCGCUACCGUUCCUUGAUUAAGGAAUUGGGCAUUCGUAAGUAAGCACUUAAAAGCAUCCGCAUCAUCGGUUUUUCAG
env-437   ...........................................................................................
env-438   GACGAAGAUAUCGAUCGCUACCGUUCCUUGAUUAAGGAAUUGGGCAUUCGUAAAUAAGCAUUUAAAAGCAUCCGCAUCAUCAGUUUUUCAG
env-439   GAUAAAGAUAUCGAACGUUACCGUUCCUUAAUUAAAGAAUUGGGCAUUCGUAAGUAAGAACUUAAAAGCAUUCGCAUCAUCAGUUUUUCAA
env-440   ...........................................................................................
env-441   GAUAAAGAUAUCGAACGCUACCGUUCCUUAAUUAAAGAAUUGGGCAUUCGUAAGUAAGAACUUAAAAGUAUUCGCAUCAUCAGUUUUUCAA
env-442   GAUAAAGAUAUCGAACGCUACCGUUCCUUAAUUAAAGAAUUGGGCAUUCGUAAGUAAGAACUUAAAAGUAUUCGCAUCAUCAGUUUUUCAA
env-444   GAUAAAGAUAUCGAACGCUACCGUUCCUUAAUUAAAGAAUUGGGCAUUCGUAAGUAAGAACUUAAAAGUAUUCGCAUCAUCAGUUUUUCAA
Kje-1-1   AAGAAAACGGAUAUCACAAGAUAUCGUGCAAUCGUAAAAGAAUUAGGACUAAGAAAAUAAAAAGAAAGAGGCUUCGUGCCUCUUUUUUUAA
Kal-1-1   AAGAAAAAAGAUAUCAACAGAUAUCGUGCAAUCGUAAAAGAAUUAGGAUUAAGAAAAUAAAACUAAAGAGGCUUCGUGCCUCUUUUUUUAA
Kzh-1-1   AAGAAAAAAGAUAUCAACAGAUAUCGUGCAAUCGUAAAAGAAUUAGGAUUAAGAAAAUAAGAAGAAAGAGGCUACGCGCCUCUUUUUUUAA
env-445   ...........................................................................................
env-446   ...........................................................................................
env-447   UACGCUAAAGAUAUUGAGCGUUACAGAGCGAUAAUUAAAGAAUUGGGCUUAAGAAAGUAAUAGCCCUCAGAAAAAAGGCAAUUGACUUCAA
env-449   ...........................................................................................
env-450   AUGAAAAAGGAUAU.............................................................................
env-451   AUGAAAAAGGAUAUAUUAAGAUACCGUGCAAUUGUAAAAGAAUUAGGUUUAAGAAAGUAAUAUCAAAGAAGAGUCAAUACCUCUUCUUUUU
env-452   ...........................................................................................
env-453   ...........................................................................................
env-454   AUGAAAAAGGAUAUAUUAAGAUACCGUGCAAUUGUAAAAGAAUUAGGUUUAAGAAAGUAAUAUCAAAGAAGAGGCUAUACCUCUUCUUUUU
env-455   AUGAAAAAGGAUAUAUUAAGAUACCGUGCAAUUGUAAAAGAAUUAGGUUUAAGAAAGUAAUAUCAAAGAAGAGGCUAUACCUCUUCUUUUU
env-456   AAGGCAGGAGAUAUUGAAAAGUACAGAGCUUUGAUUAAGAAGCUAGGCUUAAGAAAAUAAAUUGACACCAAAAAAGAAAGGCAAUCCAACA
env-457   ...........................................................................................
env-458   ...........................................................................................
env-459   AAAUCGAAAGAUAUUUUAAGAUACCGUGAAAUAGUUAAAAAAUUAGGAUUACGUAAGUAAAUUUAGUUUUAAUCUAAUGCACAAAAAUUGA
env-460   AAAUCGAAAGAUAUUUUAAGAUACCGUGAAAUAGUUAAAAAAUUAGGAUUACGUAAGUAAAUUUAGUUUUAAUCUAAUGCACAAAAAUUGA

          ...........................................................................................
          ...........................................................................................
          A•RAARAARGAYAUY••••RRUAY•G•G••AU•RU•AAAGAAYURRR••U••GAAAAUAA•••••••••••••••••••••••••••••••
```

```
env-461    AAAUCGAAAGAUAUUUUAAGAUACCGUGAAAUAGUUAAAAAAUUAGGAUUACGUAAGUAA...............................
env-462    AAAUCGAAAGAUAUUUUAAGAUACCGUGAAAUAGUUAAAAAAUUAGGAUUACGUAAGUAAAUUUAGUUUUAAUCUAAUACACAGAAAUUGA
env-463    AAAUCGAAAGAUAUUUUAAGAUACCGUGAAAUAGUUAAAAAAUUAGGAUUACGUAAGUAAAUUUAGUUUUAAUCUAAUGCACAAAAAUUGA
env-464    UAGAAACGUGAAUAGUAAAAAUUAGGAUUAC............................................................
env-465    AAAUCGAAAGAUAUUUUAAGAUACCGUGAAAUAGUUAAAAAAUUAGGAUUACGUAAGUAAAUUUAGUUUUAAUCUAAUACACAAAAAUUGA
env-466    AAAUCGAAAGAUAUUUUAAGAUACCGUGAAAUAGUUAAAAAAUUAGGAUUACGUAAGUAAAUUUAGUUUUAAUCUAAUGCACAAAAAUUGA
env-467    AAAUCGAAAGAUAUUUUAAGAUACCGUGAAAUAGUUAAAAAACUUGGAUUACGUAAGUAAAUUUAGUUUUUAUUUUACACACAAAAAUUGA
env-469    AAAUCGAAAGAUAUUUUAAGAUACCGUGAAAUAGUUAAAAAAUUAGGAUUACGUAAGUAAAUUUAGUUUUAAUCUAAUGCACAAAAAUUGA
env-470    AAAGCAAAAGAUAUCUUAAGAUAUCGUGAAAUAGUUAAAAAAUUAGCAUUGCGCAAGUAAACUUAUUUUUUAUUUAAUCAUUAAAAAUUGA
env-471    AAAAGUAAAGACAUAGAGAAAUAUCGUGCUUUAGUGAAAGAAUUAGGAUUACGUCGUUAAUUCUUAACUGAGUACACAAAUUUUAUUGGAG
env-472    ...........................................................................................
env-473    AAAGCCAAAGAUAUCUUGCGUUACCGUGAGAUUAUCAAGGAAUUAGGUAUCCGUAAAUAAUUCUACUUCUUGCUACACUGAAUUGUAGUUU
env-474    ...........................................................................................
env-475    AAGAAAAAAGAUGUUGUUCGCUAUCGUGAACUAAUUAAAGAACUGAACAUCAGAAAGUAAGACAGAGCAUUUUCUGUCCAAACAAAUAAAG
env-476    AAAACAAAAGAUAUUACACGUUAUCGUGCAAUCGUAAAAGAAUUAGGUUUACGUAAAUAAACCAAAUUUCAUCCCUUAGGGAAUAUAAAAG
env-477    AAGUCAAAAGACAUCACACGUUACCGGGCUAUUGUUAAAGAAUUAGGCUUACGUAAGUAAUCGAAAAAAAUUUCCCUCGGGAAUAUUAAAA
env-478    AAGUCAAAAGACAUCACACGUUACCGGGCCAUUGUUAAAGAAUUAGGCUUACGUAAGUAAUCGAAAAUAAUUUCCCUCGGGAAUAUUAAAA
env-479    AAGUCAAAAGACAUCACACGUUACCGUGCUAUUGUUAAAGAAUUA..............................................
env-480    AAGUCAAAAGACAUCACACGUUACCGUUCG.............................................................
env-481    AAGUCAAAAGACAUCACACGUUAUCGGGCCAUUGUUAAAGAAUUAGGCUUACGUAAGUAAUCGAAAAUAAUUUCCCUCGGGAAUAUUAAAA
env-482    AAAGCAAAAGAUAUUUUACGUUACCGUGCAAUUGUUAAAGAAUUAGGUUUACGUAAAUAAUUCAAACGCAUUUCCUUCGGGAGUAUUAAAA
env-484    AAAGCAAAAGAUAUUUUACGUUACCGUGCAAUUGUUAAAGAAUUAGGUUU.........................................
env-486    AAGCAAAAGAUAUUUUAC.........................................................................
env-487    AAGUCAAAAGACAUCACACGUUACCGGGCCAUUGUUAAAGAAUUAGGCUUACGUAAGUAAUCGAAAAUAAUUUCCCUCGGGAAUAUUAAA.
env-488    AAGUCAAAAGACAUCACACGUUACCGGGCUAUUGUUAAAGAAUUAGGCUUACGUAAGUAAUCGAAAAAAAUUUCCCUCGGGAAUAUUAAAA
env-489    AAGUCAAAAGACAUCACACGUUACCGUGCUAUUGUUAAAGAAUUAGGCUUACGUAAGUAAUCGAAAAAAAUUUCCCUCGGGAAUAUUAAAA
env-490    ...........................................................................................
env-491    AAAGCAAAAGAUAUUUUACGUUACCGUGCAAUUGUAAAAGAAUUAGGUCUACGUAAAUAAUUCAAACGCAUUUCCUUCGGGAGUAUUAAAA
env-492    AAAGCAAAAGAUAUUUUACGUUACCGUGCAAUUGUAAAAGAAUUAGGUCUACGUAAAUAAUUCAAACGCAUUUCCUUCGGGAGUAUUAAAA
env-494    AAAGCAAAAGAUAUUUUACGUUACCGUGCAAUUGUAAAAGAAUUAGGUCUACGUAAAUAAUUCAAACGCAUUUCCUCCGGGAGUAUUAAAA
env-496    ...........................................................................................
env-498    AAAGCAAAAGAUAUUUUACGUUACCGUGCAAUUGUAAAAGAAUUAGGUCUACGUAAAUAAUUCAAACGCAUUUCCUUCGGGAGUAUUAAAA
env-500    ...........................................................................................
env-501    AAAGCAAAAGAUAUUUUACGUUACCGUGCAAUUGUUAAAGAAUUAGGUUUACGUAAAUAAUUCGAACGCAUUUCCUUCGGGAGUAUUAAAA
env-502    AAAUCAAAAGACAUAACACGUUACCGUGCUAUCGUAAAAGAAUUAGGUUUACGUAAAUAAAUCAAAUUUUGUCCCUUUUGGGGGAAUAAAA
env-503    ...........................................................................................
env-504    AAGAAAAAAGAGAUUAACAGGUAUCGUGAGAUUAUCAAAGAAUUGAACAUCAGAAAAUAAUCGAGAUGAGAAGAGGUGCCCGCGCGCCUCU
env-505    AAGAAAAAAGAGAUUAACAGGUAUCGUGAGAUUAUCAAAGAAUUGAACAUCAGAAAAUA................................
env-506    AUGAAGAAAGAUAUCUUGAGAUACCGUGCAAUUGUUAAAGAAUUAGGAUUACGUAAGUAAUACAUUUAAGGGGCUUUAUGCCCCUUUUUAC
env-507    UACAAAAAAGAGAUUGGCAGAUACAGAGCAAUCAUUAAAGAAUUAGGCUUAA.......................................
env-508    ...........................................................................................
env-509    UACAAAAAAGAGAUUGGCAGAUACAGAGCAAUCAUUAAAGAAUUAGGCUUAAGAAAAUAGGCCACAACAAAAAGGCAAUUUGUAACAAUUG
env-510    UACAAAAAAGAGAUUGG..........................................................................
env-511    ACCAAAAAAGAUAUUUUAAGAUAUCGUGCUAUCGUUAAGGAGCUUGGAUUAAGAAAAUAAUUUGAAUAUAAAGACCACGUCAAUAGCGUGG
Fhi-1-1    AAGAAAGAGAUCAACAGAUAUCGUGAGAUUAUCAAAGUAUUGAAUAUCAGAAAAUAAUCAAUAUAGAAAAGAGGUGCGAGAGUGCCUCUUU
Fsp-9-1    AAGAAAGAGAUCAACAGAUAUCGUGAGAUUAUCAAAGUAUUGAAUAUCAGAAAAUAAUCAAUAUAGAAAAGAGGUGCGAGAGUGCCUCUUU
Asu-1-1    ACCAAGAAAGACGUACUUCGUUACCGUGCUAUCGUUAAAGAAUUAGGAUUACGUAAGUAAUUUUUUUAAAGGGCUCUUUACGAGCCCUUUU
Aca-1-1    AUGAAAAAAGAUAUUCUACGUUACCGUGCUAUCGUUAAAGAAUUAGGAUUACGUAAGUAAUUUUUUAAAAGGGCUCUUUCGAGCCCUUUUA
env-513    AUGAAAAAAGAUAU.............................................................................
env-514    ...........................................................................................
env-515    AUGAAAAAAGAUAUUCUACGUUAUCGUGCGAUCGUUAAAGAAUUAGGAUUACGUAAAUAAAAAUUUACAAUCAAGGCGAAGUAAUCUCAAA
env-517    ...........................................................................................
Avl-1-1    AUGAAAAAAGAUAUUCUACGUUACCGUGCGAUCGUUAAAGAAUUAGGAUUACGUAAAUAAAAAUUUACAAUCAAGGCGAAGUAAUCUCAAA
env-518    AUGAAAAAAGAUAUUCUACGUUACCGUGCGAUCGUUAAAGAAUUAGGAUUACGUAAAUAAAAAUUUACAAUCAAGGCGAAGUAAUCUCAAA
env-519    ...........................................................................................
env-520    AUGAAAAAAGAUAU.............................................................................
env-521    AUGAAAAAAGACAUCCUUCGUUACCGUGCUAUCGUUAAAGAAUUAGGAUUACGUAAGUAAUAUUUUAAAGGGCUCUUACGAGCCCUUUAUU
env-522    AUGAAAAAAGACAUCCUUCGUUACCGUGCUAUCGUUAAAGAAUUAGGAUUACGUAAGUAAUAUUUUAAAGGGCUCUUACGAGCCCUUUAUU
env-523    AUGAAAAAAGACAUCCUUCGUUACCGUGCCAUCGUUAAAGAAUUAGGAUUACGUAAGUAAUAUUUUAAAGGGCUCUUACGAGCCCUUUAUU
env-527    ...........................................................................................
env-528    AUGAAGAAAGACAUCCUUCGUUACCGUGCAAUCGUUAAAGAAUUAGGAUUACGUAAGUAAUAUUUUAAAGGGCUCUUACGAGCCCUUUAUU
env-531    AUGAA......................................................................................
env-532    ...........................................................................................
env-533    ...........................................................................................
env-534    AAAUCCAAAAAUAUUGAACGAUAUCGGUCUAUUAUCAAGGAAUUGAAAAUAAGAAAAUAAAAACCUCCUUCGGGUAUUUAAACAUACUGAA
env-536    ...........................................................................................
env-537    AAAUCCAAAGAUAUUGAACGAUACCGCUCCAUUAUCAAGGAACUGAAAAUAAGAAAAUAAAAAACUCCUUCGGGAAUUCAUACUUGCUGAA
env-539    ...........................................................................................
env-540    AAAUCCAAAGAUAUUGAACGAUACCGCUCCAUUAUCAAGGAACUGAAAAUAAGAAAAUAAAAAACUCCUUCGGGAAUUCAUACUUGCUGAA
Apa-1-1    AUCAAAAAAGAUAUCUUGAGAUAUCGUGCAAUUGUAAAAGAAUUAGGAUUAAGAAAAUAAUCAAAAAAAGAGAGGCAUUUAUGUCUCUCUU
Ama-1-1    AUUAAAAAGGAUAUCAUGAGAUACCGUGCAAUUGUAAAAGAAUUAGGAUUAAGAAAGUAAUAAAAAAGAGGCUGCAUGCCUCUUUUUUUAU
Aat-1-1    AAGAAGAAAGAUAUCAUGAGAUAUCGUGCAAUUGUAAAAGAAUUAGGAUUAAGAAAGUAACAAAAAAAGAGGCUGCGUGCCUCUUUUUUUA
Ame-1-1    AUUAAAAAGGAUAUCAUGAGAUACCGUGCAAUUGUAAAAGAAUUAGGAUUAAGAAAGUAAUAAAAAGAGGCUGCGUGCCUCUUUUUUUAUA
Ala-1-1    GUUAAGAAAGAUAUUAUGAGAUAUCGUGCAAUUGUAAAAGAAUUAGGAUUAAGAAAAUAAUCAAAAAAGAGAGGCAGUACGCCUCUCUUUU
Amu-1-1    AUUAAAAAAGAUAUUUUAAGAUACCGUGCAAUAGUAAAGGAAUUAGGAUUAAGAAAGUAAUGAAAAGAGAGGCUUCCCGCCUCUUUUUUUG
Fsu-3-1    AAGAAAAAAGAUAUCAACAGAUAUCGUGCGAUUAUCAAAGAAUUAAACAUCAGAAAAUAAUCAUGAAGAAGAGGUGCCUGCGCGCCUCUUU
Fen-1-1    AAGAAAAAAGAUAUCAACAGAUAUCGUGCGAUUAUCAAAGAAUUAAACAUCAGAAAAUAAUCAUGAGAAGAGGUGCCUGCGCGCCUCUUUU
Fco-1-1    AAAAACAAGGAUAUCAAUAGAUAUCGUGCGAUUAUCAAAGAAUUAAACAUCAGAAAAUAAUCCAGAUGAAAAGAGGUGCCCGCGCGCCUCU
env-541    AAGAAGAAAGAUAUCGUAAGAUACCGUGAAAUUAUUAAAGAAUUAGGAAUUAGAAAAUAAUACUGAAGGGGCUUUGUGCCCCUUUUUGUUU
env-542    AAAAAGAAAGAUAUCUUAAGAUACC..................................................................
env-543    ...........................................................................................
env-545    ...........................................................................................
env-546    ...........................................................................................
Fps-1-1    AAAAAAGAAAUCAACCGUUACCGUGAGAUUAUCAAAGAAUUGGGUAUUAGAAAAUAAUCAAUACAAAAGAGGUGCGAAAGUGCCUCUUUUU

           ...........................................................................................
           ...........................................................................................
           A•RAARAARGAYAUY••••RRUAY•G•G••AU•RU•AAAGAAYURRR••U••GAAAAUAA•••••••••••••••••••••••••••••••
```

| | |
|---|---|
| Jma-1-1 | AAGAAGAAAGAUAUUACAAGAUAUCGUGCUAUCAUCAAAGAAUUAGGAUUAAGAAAGUAAGAUAUAAGGGAGCAAAUUUGCUCCCUUUUUU |
| env-547 | ...................................................................................... |
| env-548 | ACUAAAAAAGACAUCCUAAGAUAUCGUGCUAUUGUUAAAGAACUAGGCUUAAGAAAAUAAGAGGCUAGUCUUAUAAUUGUUCAUUGGUUAU |
| env-549 | ACUAAAAAAGACAUCCUAAGAUAUCGUGCUAUUGUUAAAGAACUAGGCCUAAGAAAAUAAGAGGCUAGUCUUAUAAUUGUUCAUUGGUUAU |
| env-550 | ACUAAAAAAGACAUCCUAAGAUAUCGUGCUAUUGUUAAAGAACUAGGCUUAAGAAAAUAAGAGGCUAGUCUUAUAAUUGUUCAUUGGUUAU |
| env-551 | AAGAAAAAAGAUAUUGCAAGAUANCGUGCAAUUAUUAAAGAAUUAGGAAUUAGAAAAUAAUUCUUACACUAAAAAAGAGGAGCUAUAAACG |
| env-552 | AAGAAAAAAGAUAUUGCAAGAUAUCGUGCAAUUAUUAAAGAAUUAGGAAUUAGAAAAUAAUUCUUACACUAAAAAAGAGGAGCUAUAAACG |
| env-553 | AAGAAAAAAGAUAUUGCAAGAUAUCGUGCAAUUAUUAAAGAAUUAGGAAUUAGAAAAUAAUUCUUACACUAAAAAAGAGGAGCUAUAAACG |
| env-554 | AAGAAAUCAGAAAUCAACAGAUAUCGUGCAAUUAUCAAAGAAUUAGGAAUUAGAAAGUAAUUCUUACAAAAAGAGGGGCUAUAAACGCACC |
| env-556 | ...................................................................................... |
| env-557 | AAGAAAUCAGAAAUCAACAGAUAUCGUGCAAUUAUCAAAGAAUUAGGAAUUAGAAAGUAAUUCUUACAAAAAGAGGGGCUAUAAACGCACC |
| Ffi-1-1 | AAGAAAAAAGAUAUCAACAGAUAUCGUGAGAUUAUCAAAGAAUUGAAUAUUAGAAAAUAAUCAAUAGGAAAGAGGUGCGAAAGUGCCUCUU |
| env-558 | ...................................................................................... |
| env-559 | ACAAAGAAAGAUGUCUUAAGAUAUCGUGCAAUAGUUAAAGAAUUAGGAUUAAGAAAAUAAUCUAACUACAAAUACCACGCCUAUCAGCGUG |
| env-561 | ACUAAAGUGGAUGUAUUGAGAUAUCGUGCAAUUGUAAAAGAAUUAGAUCUAAGAAAAUAAUUUAAAAGAGGCCAUUUGAGCCUCUUUUAUU |
| env-562 | ...................................................................................... |
| env-563 | ...................................................................................... |
| env-564 | ACUAAAGUUGAUGUCUUAAGAUAUCGUGCAAUUGUAAAAGAAUUAGAUCUAAGAAAAUAAUUUAAAAGAGGCCAAUUGUGCCUCUUUUAUU |
| env-565 | ...................................................................................... |
| env-566 | ACUAAAGUUGAUGUCUUAAGAUAUCGUGCAAUUGUAAAAGAAUUAGAUCUUAGAAAAUAAUUUAAAAGAGGCCACUAGAGCCUCUUUUAUU |
| env-567 | ACUAAAGUUGAUGUCUUAAGAUAUC............................................................. |
| env-569 | ...................................................................................... |
| env-571 | ...................................................................................... |
| env-572 | ACUAAAGUUGAUGUCUUAAGAUAUCGUGCAAUUGUAAAAGAAUUAGAUCUAAGAAAAUAAUUUAAAAGAGGCCAAUUGUGCCUCUUUUAUU |
| env-573 | ACUAAAGUUGAUGUCUUAAGAUAUCGUGCAAUUGUAAAAGAAUUAGAUCUUAGAAAAUAAUUUAAAAGAGGCCACUAGAGCCUCUUUUAUU |
| env-574 | ...................................................................................... |
| env-575 | ...................................................................................... |
| env-576 | ...................................................................................... |
| env-577 | ...................................................................................... |
| env-578 | ...................................................................................... |
| env-579 | ACUAAAGUGGAUGUCUUAAGAUAUCGUGCAAUUGUAAAAGAAUUAGAUCUUAGAAAAUAAUUUAAAAGAGGCCCUAGAGCCUCUUUUAUUA |
| env-580 | ACUAAAGUGGAUGUCUUAAGAUAUCGUGCAAUUGUAAAAGAAUUAGAUCUUAGAAAAUAAUUUAAAAGAG.................... |
| env-581 | ACUAAAGUUGAUGUCUUAAGAUAUCGUGCAAUUGUAAAAGAAUUAGAUCUAAGAAAAUAAUUUAAAAGAGGCCAACUAGAGCCUCUUUUAU |
| env-582 | ACUAAAGUUGAUGUCUUAAGAUAUCGUGCAAUUGUAAAAGAAUUAGAUCUUAGAAAAUAAUUUAAGAGAGGCCACUAGAGCCUCUUUUAUU |
| env-583 | ACUAAAGUUGAUGUCUUAAGAUAUCGUGCAAUUGUAAAAGAAUUAGAUCUUAGAAAAUAAUUUAAAAGAGGCCACUAGAGCCUCUUUUAUU |
| env-584 | ...................................................................................... |
| env-585 | ...................................................................................... |
| env-586 | ...................................................................................... |
| env-587 | ...................................................................................... |
| env-588 | ACUAAAGUUGAUGUCUUAAGAUAUCGUGCAAUUGUAAAAGAAUUAGAUCUAAGAAAAUAAUUUAAAAGAGGCCAAUUGUGCCUCUUUUAUU |
| env-589 | ACUAAAGUUGAUGUCUUAAGAUAUCGUGCAAUUGUAAAAGAAUUAGAUCUUAGAAAAUAAUCUAAAAGAGGCCACUAGAGCCUCUUUUAUU |
| env-590 | ...................................................................................... |
| env-591 | ACUAAAGUUGAUGUCUUAAGAUAUCGUGCAAUUGUAAAAGAAUUAGAUCUAAGAAAAUAAUUUAAAAGAGGCCACUAGAGCCUCUUUUAUU |
| env-592 | ACUAAAGUGGAUGUCUUGAGAUAUCGUGCAAUUGUAAAAGAAUUAGAUCUUAGAAAAUAAUUUAAAAGAGGCCCUAGAGCCUCUUUUAUUA |
| env-593 | ...................................................................................... |
| env-594 | ACUAAAGUGGAUGUCUUAAGAUAUCGUGCAAUUGUAAAAGAAUUAGAUCUUAGAAAAUAAUUUAAAAGAGGCCCUAGAGCCUCUUUUAUUA |
| env-596 | ...................................................................................... |
| env-598 | ACUAAAGUGGAUGUCUUAAGAUAUCGUGCAAUUGUAAAAGAAUUAGAUCUUAGAAAAUAAUUUAAAAGAGGCCCUAGAGCCUCUUUUAUUA |
| env-599 | ACUAAAGUGGAUGUCUUNAGAUAUCGUGCAAUUGUAAAAGAAUUAGAUCUUAGAAAAUAAUUUAAAAGAGGCCCUAGAGCCUCUUUUAUUA |
| env-600 | ACUAAAGUAGAUGUCUUAAGAUAUCGUGCAAUUGUAAAAGAAUUAGAUCUAAGAAAAUAAUUUAAAAGAGGCCACUAGAGCCUCUUUUAUU |
| env-601 | AAAAAUAAAGAUAUUACUCGUUACAGAGCAAUUAUUAAAAAACUUAAACUUAGAAAAUAAGUAUUUUCUACAUAGUUUUUCAUUGGUAUUC |
| env-602 | AAAAAUAAAGAUAUAACUCGUUACAGAGCAAUUAUUAAAAAACUUAAACUCAGGAAAUAAGUAUUUUCUACAUAGUUUUUCAUUGGUAUUC |
| env-603 | AAAAAUAAAGAUAUAACUCGUUACAGAGCAAUUAUUAAAAAACUUAAACUCAGGAAAUAAGUGUUUUCUACAUAGUUUUUCAUUGGUAUUC |
| env-604 | ...................................................................................... |
| env-605 | AAAAAUAAAGAUAUUACUCGUUACAGAGCAAUUAUUAAAAAACUUAAACUUAGAAAAUAAGUAUUUUCUACAUAGUUUUUCAUUGGUAUUC |
| env-606 | AAAAAUAAAGAUAUCAAUCGUUACAGAGAGAUUAUAAAAAAACUUAAUCUACGUAAAUAGAUAUUUUUAUCAUAGUUUUUCAUUGGUAUUC |
| env-607 | AAAAAUAAAGAUAUCAAUCGUUACAGAGAGAUUAUAAAAAAACUUAAUCUACGUAAAUAGAUAUUUUUAUCAUAGUUUUUC.......... |
| env-608 | ...................................................................................... |
| env-609 | ...................................................................................... |
| env-610 | ...................................................................................... |
| env-611 | AAAAAUCAAGAUAUUACUCGUUAUAGAGAAAUAAUUAAAAAAUUAAAUCUCCGUAAAUAAAUAUCCAUCUCAUAGUUUUUCAUUGGUAUUC |
| env-612 | AAAAAUCAAGAUAUUACUCGUUAUAGAGAAAUAAUUAAAAAAUUAAAUCUCCGUAAAUAAAUAUCCAUCUCAUAGUUUUUCAUUGGUAUUC |
| env-613 | ...................................................................................... |
| env-614 | AAAAAUAAAGAUAUUACUCGUUAUAGAGAAAUAAUUAAAAAAUUAAAUCUCCGUAAAUAAAUAUCCAUCUCAUAGUUUUUCAUUGGUAUUC |
| env-615 | ...................................................................................... |
| env-616 | AAAAACAAAGAUAUUAAUCGUUACAGAGAUAUAAUUAAAAAAUUAAAUCUCCGUAAAUAAAUAUCCAUAUCAUAGUUUUUCAUUGGAAUUC |
| env-617 | AAAAAUAAAGAUAUUACUCGUUACAGAGAAAUAAUUAAAAAAUUAAAUCUCCGUAAAUAAAUAUCCAUAUCAUAGUUUUUCAUUGGUAUUC |
| env-618 | ...................................................................................... |
| env-619 | AAAAGUAAAGAUAUUACUCGUUAUAGAGAAAUCAUUAAAAAACUUAAUCUGCGUAAAUAAAUAUUUUCUACAUAGUUUUUCAUUGGAAUUC |
| env-620 | AAAAGUAAAGAUAUUAGUCGUUACAGGGAAAUUAUAAAGAAACUUAAUCUGCGUAAAUAAAUAUUUUUUACAUAGUUUUUCAUUGGUAUUC |
| env-621 | ...................................................................................... |
| env-622 | AAAAGUAAAGAUAUUAGUCGUUACAGGGAAAUUAUAAAGAAACUUAAUCUGCGUAAAUAAAUAUUUUUUACAUAGUUUUUCAUUGGUAUUC |
| env-623 | AAAAGUAAAGAUAUUAGUCGUUACAGGGAAAUUAUAAAGAAACUUAAUCUGCGUAAAUAAAUAUUUUUUACAUAGUUUUUCAUUGGUAUUC |
| env-624 | AAAAAUAAAGAUAUUACUCGUUACAGAGAAAUUAUUAAGAAACUUAAUCUGCGUAAAUAGAUAUUUUUUACAUAGUUUUUCAUUGGUAUUC |
| env-625 | AAAAAUAAAGAUAUUACUCGUUACAGAGAAAUUAUUAAGAAACUUAAUCUGCGUAAAUAGAUAUUUUUUACAUAGUUUUUCAUUGGUAUUC |
| env-626 | AAAAAUAAAGAUAUUACUCGUUACAGAGAAAUUAUUAAGAAACUUAAUCUGCGUAAAUAGAUAUUUUUUACAUAGUUUUUCAUUGGUAUUC |
| env-627 | ...................................................................................... |
| env-628 | AAAAAUAAAGAUAUUACUCGUUACAGAGAAAUUAUUAAGAAACUUAAUCUGCGUAAAUAGAUAUUUUUUACAUAGUUUUUCAUUGGUAUUC |
| env-629 | AAAAAUAAAGAUAUUACUCGUUACAGAGAAAUUAUUAAGAAACUUAAUCUGCGUAAAUAGAUAUUUUUUACAUAGUUUUUCAUUGGUAUUC |
| env-633 | AAAAAUAAAGAUAUUACUCGUUACAGAGAAAUUAUUAAGAAACUUAAUCUGCGUAAAUAAAUAUUUUUUACAUAGUUUUUCAUUGGUAUUC |
| env-634 | ...................................................................................... |
| env-635 | ...................................................................................... |
| env-636 | ...................................................................................... |
| env-637 | AAAAGUAAAGAUAUUAGUCGUUACAGGGAAAUUAUAAAAAAACUUAAUCUUCGUAAAUAAAUAUUUUUUACAUAGUUUUUCAUUGGUAUUC |
| | ...................................................................................... |
| | ...................................................................................... |
| | A•RAARAARGAYAUY••••RRUAY•G•G••AU•RU•AAAGAAYURRR••U••GAAAAUAA••••••••••••••••••••••••••••••• |

```
env-638   ..........................................................................................
env-639   AAAAGUAAAGAUAUUACUCGAUACAGAGCAAUUAUUAAAAAGCUUAAUCUGAGAAAAUAAAUAUUUUCUUCAUAGUUUUUCAUUGGUAUUC
env-640   AAAAGUAAAGAUAUUACUCGUUAUAGAGAAAUUAUUAAAAAACUUAAACUUAGAAAAUAAAUAUUUUCUACAUAGUUUUUCAUUGGUAUUC
env-641   AAAAAUAAAGACAUUAAUCGUUACAGAGAGAUUAUUAAAAAGCUUAAUCUACGUAAAUAGAUGUUUUUAUCAUAGUUUUUCAUUGGUAUUC
env-648   ..........................................................................................
env-649   AUGGUUGGUAAAAGAAAAAGCUUAUUAAGCUAUCUUAAAAAUAAAGACAUUAAUCGUUACAGAGAGAUUAUUAAAAAGCUUAAUCUACGUA
env-650   AAAAAUAAAGACAUUAAUCGUUACAGAGAGAUUAUUAAAAAGCUUAAUCUACGUAAAUAGAUGUUUUUAUCAUAGUUUUUCAUUGGUAUUC
env-651   ..........................................................................................
env-652   ..........................................................................................
env-653   AAAAAUAAAGACAUUAAUCGUUACAGAGAGAUUAUUAAAAAGCUUAAUCUACGUAAAUAGAUGUUUUUAUCAUAGUUUUUCAUUGGUAUUC
env-655   AAAAAUAAAGACAUUAAUCGUUACAGAGAGAUUAUUAAAAAGCUUAAUCUACGUAAAUAGAUGUUUUUAUCAUAGUUUUUCAUUGGUAUUC
env-656   AAAAAUAAAGACAUUAAUCGUUACAGAGAGAUUAUUAAAAAGCUUAAUCUACGUAAAUAGAUGUUUUUAUCAUAGUUUUUCAUUGGUAUUC
env-657   ..........................................................................................
env-658   AAAAAUAAAGACAUUAAUCGUUACAGAGAGAUUAUUAAAAAGCUUAAUCUACGUAAAUAGAUGUUUUUAUCAUAGUUUUUCAUUGGUAUUC
env-659   ..........................................................................................
env-660   AAGAAUAAAGAUAUCAAUCGUUACAGAGAGAUUAUUAAAAAACUUAAUCUACGUAAAUAGAUAUUUUUAUCAUAGUUUUUC.........
env-661   AAGAAUAAAGACAUUAAUCGUUACAGGGAGAUUAUUAAAAAGCUUAAUCUACGUAAAUAGAUGUUUUUAUCAUAAUUUUUCAUUGGUAUUC
env-662   AAGAAUAAAGACAUUAAUCGUUACAGGGAGAUUAUUAAAAAGCUUAAUCUACGUAAAUAGAUGUUUUUAUCAUAAUUUUUCAUUGGUAUUC
env-663   ..........................................................................................
env-664   AUUAAGAAAGAUAUCACAAGAUAUCGUGCGAUUGUUAAAGAAUUAGGAUUACGUAAAUAAUAAUAAAGGGGGCAAAAUGCUCCCUUUAUUA
env-669   ..........................................................................................
env-670   AUGAAAAAAGAUGUUUUAC.......................................................................
env-671   ..........................................................................................
env-672   AAAAAUAAGGAUAUUGAAAAAUAUCGUAAGCUAGUAAAAGAAUUAGGACUUCGUCGCUAAUUUUAAUUGGGUUACAAAACGUAUUGAAGAG
env-673   AAGAAUAAGGAUAUUGAAAAAUAUCGUGCAUUGGUUAAAGAUUUAGGGCUUCGUCGUUAAUUUUUCAUUGCAAGCGUA............
env-674   AAGAAUAAGGAUAUUGAAAAAUAUCGUGCAUUGGUUAAAGAUUUAGGGCUUCGUCGUUAAUUUUUCAUUGCAAGCGUAAUAAAGAGGGGA
env-675   ..........................................................................................
env-676   AAGAAUAAGGAUAUUGAAAAGUAUCGUGAAUUGGUUAAAGAAUUAGGGCUUCGUCGCUAAUUUUAAUUGGUUUAUAAAAUGUAUGAAAGAG
env-677   AAGAAUAAGGAUAUUGAAAAAUAUCGUGAAUUGGUUAAAGAAUUAGGGCUUCGUCGCUAAUUUUAAUUGGUUUAUAAAAUGUAUUUAAGAG
env-678   AAGAAUAAGGAUAUUGAAAAGUAUCGUGAAUUGGUUAAAGAAUUAGGGCUUCGUCGCUAAUUUUAAUUGGUUUAUAAAAUGUAUGAAAGAG
env-679   AAGAAUAAGGAUAUUGAAAAAUAUCGUGAAUUGGUUAAAGAAUUAGGGCUUCGUCGCUAAUUUUUAAUUGGUUUAUAAAAUGUAUGAAAGA
env-680   ..........................................................................................
env-681   AAAAAUAAGGAUAUUGAGAAAUAUCGUAAACUAGUAAAAGAAUUAGGACUUCGUCGCUAAUUUUAAUUGGUUUACAAAACGUAUUGAAGAG
env-682   AAAAAUAAGGAUAUUGAGAAAUAUCGUAAACUAGUAAAAGAAUUAGGACUUCGUCGCUAAUUUUAAUUGGUUUACAAAACGUAUUGAAGAG
env-683   ..........................................................................................
env-684   AAAAAUAAGGAUAUUGAGAAAUAUCGUAAACUA.........................................................
env-685   AAAAAUAAGGAUAUUGAGAAAUAUCGUAAACUAGUAAAAGAAUUAGGACUUCGUCGCUAAUUUUAAUUGGUUUACAAAACGUAUUGAAGAG
env-686   AAAAAUAAGGAUAUUGAGAAAUAUCGUAAACUAGUAAAAGAAUUAGGACUUCGUCGCUAAUUUUAAUUGGUUUACAAAACGUAUUGAAGAG
env-687   ..........................................................................................
env-688   AAAAAUAAGGAUAUUGAGAAAUAUCGUAAACUAGUAAAAGAAUUAGGACUUCGUCGCUAAUUUUAAUUGGUUUACAAAACGUAUUGAAGAG
env-689   AAAAAUAAGGAUAUUGAGAAAUAUCGUAAACUAGUAAAAGAAUUAGGACUUCGUCGCUAAUUUUAAUUGGUUUACAAAACGUAUUGAAGAG
env-690   AAAAAUAAGGAUAUUGAGAAAUAUCGUAAACUAGUAAAAGAAUUAGGACUUCGUCGCUAAUUUUAAUUGGUUUACAAAACGUAUUGAAGAG
env-691   ..........................................................................................
env-692   AAAAAUAAGGAUAUUGAAAAAUAUCGUGAAUUGGUUAAAGAAUUAGGGCUUCGUCGCUAAUCAUAAUUGGUUUACAAGAAGUAUUAAAGAG
env-693   AAAAAUAAGGAUAUUGAAAAAUAUCGUGAAUUGGUUAAAGAAUUAGGGCUUCGUCGCUAAUCAUAAUUGGUUUACAAGAAGUAUUAAAGAG
env-694   AAAUCUAAAGACAUCUUGCGUUAUCGAGAAAUAGUAAAAGAAUUAGGUUUACGUAAGUAAUUUAAAUCACUUUAAAUACAAGCUGAAUCAU
env-698   AAAUCUAAAGACAUCUUGCGUUAUCGAGAAAUAGUAAAAGAAUUAGGUUUACGUAAGUAAUUUAAAUCACUUUAAAUACAAGCUGAAUCAU
env-699   AAAUCUAAAGACAUCUUGCGUUAUCGAGAAAUAGUAAAAGAAUUAGGUUUACGUAAGUAAUUUAAAUCACUUUAAAUACAAGCUGAAUCAU
env-700   AAAUCUAAAGACAUCUUGCGUUAUCGAGAAAUAGUAAAAGAAUUAGGUUUACGUAAGUAAUUUAAAUCACUUUAAAUACAAGCUGAAUCAU
env-701   AAAUCUAAAGACAUCUUGCGUUAUCGAGAAAUAGUAAAAGAAUUAGGUUUACGUAAGUAACUUAAAUCACUUUAAAUACAAGCUGAAUCAU
env-702   AAAUCUAAAGACAUCUUGCGUUAUCGAGAAAUAGUAAAAGAAUUAGGUUUACGUAAGUAAUUUAAAUCACUUUAAAUACAAGCUGAAUCAU
env-703   ..........................................................................................
env-704   ..........................................................................................
env-705   ..........................................................................................
env-706   ..........................................................................................
env-707   AAAGCAAAAGACAUUCUACGUUAUCGUGCAAUUGUCAAAGAAUUAGGUUUACGUAAAUAAUUCUGUAUUUCUUCGAAAACUGAAUCUUAGU
env-708   AAAGCUAAAGACAUAACACGUUAUCGUUCGAUUGUCAAAGAAUUAGGUUUACGUAAGUAAAACCUUAUUUCUUCGAAAACUGAAUCUUAGU
env-709   AAGAAAAAAGAUAUCAAUAGGUAUAGAGCAAUUAUUGCAGAAUUAGGUAUUCGUAAGUAGAAGAAAUUAUAAAGAGGCAAUUAAUAAAUUG
env-711   AAAAACAAUGAUGUCGCAAGAUAUCGAACAAUUGUAAAAGAACUUAGCUUAAGAAAAUAAGCUUACACUUAUCAAUCAAAAGGGCAAUUUU
env-712   AAAGCUAAAGACAUUACUCGUUAUCGUGAGAUUGUAAAAGAAUUAGGUUUACGUAAAUAAUUUCGUACACAUCAUUUACCUAAUCUUAGUU
Fba-2-1   AAAUCAAAAGACAUUACACGUUACCGUGCGAUUGUAAAAGAAUUAGGUUUACGUAAAUAAUCACAUUUACAGAUUUUUAGCGAAUCUUAGU
env-713   AAAUCAAAAGACAUUACACGUUACCGUGCGAUUGUAAAAGAAUUAGGUUUACGUAAAUAAUCACAUUUACAGAUUUUUAGCGAAUCUUAGU
env-718   AAAUCAAAAGACAUUACACGUUACCGUGCGAUUGUAAAAGAAUUAGGUUUACGUAAAUAAUCACAUUUACAGAUUUUUAGCGAAUCUUAGU
env-727   ..........................................................................................
env-728   ..........................................................................................
env-729   AAAUCAAAAGACAUUACACGUUACCGUGCGAUUGUAAAAGAAUUAGGUUUACGUAAAUAAUCACAUUUACAGAUUUUUAGCGAAUCUUAGU
env-730   ..........................................................................................
env-731   AAAUCCAAAGACAUUGAGCGUUAUAGAAGUAUUGUAAAAGAAUUAGGUUUACGUAAGUAAUCCAGAAAACCCUAAAGUUAUACUGAAUCUU
env-734   AAAUCCAAAGACAUUGAGCGUUAUAGAAGUAUUGUAAAAGAAUUAGGUUUACGUAAGUAAUCC...........................
env-735   ..........................................................................................
env-736   ..........................................................................................
env-737   ..........................................................................................
env-738   ..........................................................................................
env-739   ..........................................................................................
env-740   AAAUCUAAAGACAUCUUGCGUUAUAGAGAAAUAGUAAAAGAAUUAGGUUUACGUAAGUAAUUUAAAUCACUUCAAAUACAAGCUGAAUUUU
env-741   AAAGAAAAGGAUAUUGAACGUUAUCGAUCUAUCGUCAAAUUAUUGGGGUUAAGAAAAUAAUCCUAGUUAUAAUAAAAGCGUAACUAUUUAC
env-742   AAAGAAAAGGAUAUCGAACGUUAUCGAUCUAUCGUCAAAUUAUUGGGGUUAAGAAAAUAAUCUUAGUUCUAUUAAAAGCCUAACUAUUUAC
env-743   AAGUCAAAAGACAUUGAACGUUAUCGUUCUAUCGUCAAAGAACUGGGAUUGAGAAAAUAAUUCCAAUUUCUUUAAAGCACAUCUAUUUACU
env-744   ..........................................................................................
env-745   AAGUCAAAAGACAUUGAACGUUAUCGUUCUAUCGUCAAAGAACUGGGAUUGAGAAAAUAAUUCCAAUUUCUUUAAAGCACAUCUAUUUACU
env-746   ..........................................................................................
env-747   AAGUCAAAAGACAUUGAACGUUAUCGUUCUAUCGUCAAAGAACUGGGAUUGAGAAAAUAAUUCCAAUUUCUUUAAAGCACAUCUAUUUACU
env-748   AAAGCAAAAGAUAUUGAGCGUUAUCGUGCCAUCAUCAAAGAAUUGGGACUAAGAAAAUAAUCUUAAACGCAAUAAAAGCAUAUCUAUUUAC
env-749   AAAGCAAAAAAUAUAGAGCGUUAUCGUGCCAUCAUCAAA...................................................

          ..........................................................................................
          ..........................................................................................
          A•RAARAARGAYAUY••••RRUAY•G•G••AU•RU•AAAGAAYURRR••U••GAAAAUAA••••••••••••••••••••••••••••••
```

```
env-750   ...........................................................................................
env-751   ...........................................................................................
env-752   AAAGCAAAAGAUAUCGAGCGUUAUCGUGCCAUCAUCAAAGAAUUGGGACUAAGAAAAUAAUCUUAAACGCAAUAAAAGCAUAUCUAUUUAC
env-753   AAAGCAAAGGAUAUCGAGCGUUAUC..................................................................
env-754   AAAGCAAAGGAUAUCGAGCGUUAUCGUGCGAUCAUCAAAGAAUUGGGAUUAAGAAAAUAAUCUCAAUCGUAUUAAAAGCGUAUCUAUUUAC
env-755   AAAGAAAAGGAUAUCGAACGUUAUCGAUCUAUCGUCAAAUUAUUGGGGUUAAGAAAAUAAUCUUAGUUCUAUUAAAAGCGUAACUAUUUAC
env-756   AAAGCCAAAGAUAUUGAGCGAUACCGUUCGAUCAUCAAGGAAUUGGGAUUGAGAAAAUAAUCCCAUUCACAUAAAAGGUGUAUCUAUUUAC
env-758   AAAGCAAAAGAUAUCGAGCGUUAUCGUGCGAUCAUCAAAGAAUUGGGAUUAAGAAAAUAAUCUCAAACGUAUUAAAAGCGUAUCUAUUUAC
env-759   AAGUCAAAAGACAUUGAACGUUAUCGUUCUAUCGUCAAAGAACUGGGAUUGAGAAAAUAAUUCCAACUUCUUUAAAGCACAUCUAUUUACU
env-760   AAGUCAAAAGACAUUGAACGUUAUCGUUCUAUCGUCAAAGAACUGGGAUUGAGAAAAUAAUUUCAACUUCUUUAAAGCACAUCUAUUUACU
env-761   ...........................................................................................
env-762   AAGUCAAAAGAUAUUGAAAAAUAUCGACUAA............................................................
env-763   ...........................................................................................
env-764   AAAUCAAAAGACAUUGAGAGAUAUCGAUUAAUUGUAAGCGAACUAGGUUUAAGGAAAUAAUCUUCAACUGAAAAUUUUUAAUUUUCAAAAU
env-765   AAGUCAAAAGACAUUGAGAGAUAU...................................................................
env-766   AAGUCAAAAGACAUUGAGAGAUAUCGGCUAAUUGUAAGCGAACUAGGUUUAAGGAAAUAAUCUUCAUCUGAAAAUUUUUAAUUUUCAAAAU
env-767   AAGUCAAA...................................................................................
env-768   AAGUCAAAAGACAUUGAGAGAUAUCGGCUAAUUGUAAGCGAACUAGGUUUAAGGAAAUAAUCUUCAUCUGAAAAUUUU.............
env-769   ...........................................................................................
env-770   AAGUCAAAAGACAUUGAGAGAUAUCGACUAAUUUUAAGCGAACUAGGUUUAAGGAAAUAAUCUUCAUCUGAAAAUUUUUAAUUUUCAAAAU
env-771   AAGUCAAAAGACAUUGAGAGAUAUCGACUAAUUGUAAGCGAACUAGGUUUAAGGAAAUAAUCUUCAUCUGAAAAUUUUUAAUUUUCAAAAU
env-772   ...........................................................................................
env-773   AAGUCAAAAGACAUUGAGAGAUAUCGACUAAUUGUAAGCGAACUAGGUUUAAGGAAAUAAUCUUCAUCUGAAAA.................
env-774   AAGUCAAAAGACAUUGAGAGAUAUCGGCUAAUUGUAAGCGAACUAGGUUUAAGGAAAUAACCUUCACCUGAAAAUUUUUAAUUUUCAAAAU
env-775   ...........................................................................................
env-776   AAAUCAAAAGACAUUGAGAGAUAUCGGCUAAUUGUAAGCGAACUAGGUUUAAGGAAAUAACCUUCACCUGAAAAUUUUUAAUUUUCAAAAU
env-777   AAGACCAAAGACAUUGAGCGUUAUCGAUCUAUUAUCAGUGAAUUAAGUUUAAGAAAAUAAGCUUUAAAAAUAUUUCAAUUUUUAAAUAGUU
env-778   AAGACCAAAGACAUUGAGCGUUAUCGAUCUAUUAUCAGUGAAUUAAGUUUAAGAAAAUAAGCUUUAAAAAUAUUUCAAUUUUUAAAUAGUU
env-779   AAAACUAAAGAUAUUGAGCGUUAUCGUUCAAUCAUCAGUGAAUUAAGUUUAAGAAAAUAAUCUUU..........................
env-780   AAAACUAAAGAUAUUGAGCGUUAUCGUUCAAUCAUCAGUGAAUUAAGUUUAAGAAAAUAAUCUUUAAAAAUAUUUCAAUUUUUAAAUAGUU
env-781   ...........................................................................................
env-783   AAAACUAAAGAUAUUGAGCGUUAUCGUUCAAUCAUCAGUGAAUUAAGUUUAAGAAAAUAAUCUUUAAAAAUAUUUCAAUUUUUAAAUAGUU
env-784   AAAACUAAAGAUAUUGAGCGUUAUCGAUCAAUUAUCAGUGAAUUAAGUUUAAGAAAAUAAGCUUUAAAAAUAUUUCAAUUUUUAAAUAGUU
env-785   AAAACUAAAGAUAUUGAGCGUUAUCGUUCAAUCAUCAGUGAAUUAAGUUUAAGAAAAUAAUCUUU..........................
env-786   AAAAGAAAAGAUAUUGAGCGUUAUCGAUCAAUUAUCAGUGAUUUAAGUUUAAGAAAAUAAGCUUUUAAAUUUUUAAAUAGUUUUUCCUUGG
env-787   ...........................................................................................
env-788   AAAAGAAAAGAUAUUGAGCGUUAUCGAUCAAUUAUCAGUGAUUUAAGUUUAAGAAAAUAAGCUUUUAAAUUUUUAAAUAGUUUUUCCUUGG
env-789   AUGAAAAAAGACAUUUUGAGAUACCGAGCUAUUGUAAAAGAACUUGGUUUAAGAAAAUAAUUUUCAAGGGGGCUAUAUGCCCCCUUUUUUA
env-791   AAGAAAACAGAAAUCAAUAGAUAUCGUGCGAUUAUCAAAGAAUUAGGAAUUAGAAAAUAAUUCUUACAAAAAAGAGGCUCUAUAAACGUGC
env-794   AAGAAAACAGAAAUCAAUAGAUAUCGUGCGAUUAUCAAAGAAUUACGAAUUAGAAAAUAAUUCUUACAAAAAGAGGCUCUAUAAACGUGCC
env-795   ...........................................................................................
env-796   AGAAAACAGAAAUCAAUAGAUAUCGUGCGAUUAUCAAAGAAUUAGGAAUUAGAAAAUAAUUCUUACAAAAAAGAGGCUCUAUAAACGUGCC
env-797   AAGAAAACAGAAAUCAAUAGAUAUCGUGCGAUUAUCAAAGAAUUAGGAAUAGACUCAACUUUU............................
env-798   AAGAAAACAGAAAUCAAUAGAUAUCGUGC..............................................................
env-799   ...........................................................................................
Cca-4-1   AAAAACAACGACAUCGAAAGAUAUCGUAAUAUAAUCAAAGAACUUGGUAUUAGAAAAUAAUCAAAAAGGGGCUUUUUGCCUCUUUUUUUUU
Cca-4-2   AAAAACAACGACAUCGAAAGAUAUCGUAAUAUAAUCAAAGAACUUGGUAUUAGAAAAUAAUCAAAAAGGGGCUUUUUGCCUCUUUUUUUUU
Cca-4-3   AAAAACAACGACAUCGAAAGAUAUCGUAAUAUAAUCAAAGAACUUGGUAUUAGAAAAUAAUCAAAAAGGGGCUUUUUGCCUCUUUUUUUUU
env-800   ...........................................................................................
env-801   AUCAAGACAGACAUUUUGCGCUACCGCGCC.............................................................
env-802   AAGAAUAACGAUAUUGAAAAAUACCGUUCUUUAAUCAAAGAAUUAGGUAUUCGUAAAUAAGAAUAAGGCACCGCUAAAAGCGGUGCUUUUA
env-803   AAAGCUUCAGAUAUUACACGUUAUCGUGCAAUAAUUAAAGAGUUAGGUUUAAGAAAAUAAUCUAAAUUAAAAGAGGCUUAUGCCUCUUUUU
env-805   AAAAAUAAAGAUAUCAACAGAUAUCGAGAAAUAAUCAAAAAACUAAAUUUAAGAAAAUAGAUAAUUACAUAGUUUUCAUUGGUAUUCUAC
env-806   AAAAAUAAAGAUAUCAACAGAUAU...................................................................
env-807   AUAAAGAUAU.................................................................................
env-808   AAAAAUAAAGAUAUAAAUCGCUACAGAGAGAUAAUUAAAAAAUUAAACUUACGUAAAUAAAAUUUUUCCAUAGUUUUUCAUUGGUAUUCUA
env-809   ...........................................................................................
Zga-1-1   AAAAAGAAAGAUAUUGUAAAAUAUCGUGAGCUUAUCAAGGAACUAGGCAUUAGAAAAUAAGUAUAUCAUAAGGGCUGAAAUUUUCGGCCCU
env-810   AAGAAAAAAGAUAUCAACAGAUAUCGUGAGAUUAUCAAAGAAUUAAACAUCAGAAAAUAAUCACAAGAAAAGAGGUGCCCGCGCGCCUCUU
env-813   ...........................................................................................
env-814   ...........................................................................................
env-815   AAAACUGAAAUCAACAGAUAUCGUGAGAUUAUCAAAGUAUUGAAUAUUAGAAAAUAAUCAAUAAAAAAAGAGGUGCACACGCGCCUCUUUUU
env-816   AAAACUGAAAUCAACAGAUAUCGUGAGAUUAUCAAAGUAUUGAAUAUUAGAAAAUAAUCAAUAAAAAAAGAGGUGC...............
env-817   AAAACCGAAAUUAACAGAUAUCGUGAGAUUAUCAAAGUAUUGAAUAUUAGAAAAUAAUCAAUAAAAAAAGAGGUGCCCACGCGCCUCUUUUU
env-818   AAAACNGAAAUUAANAGAUAUCGUGAGAUUAUNAAAGUAUUGAAUAUUAGAAAAUAAUCAAUAAAAAAAGAGGUGCCCACGCGCCUCUUUUU
env-819   ...........................................................................................
env-820   ...........................................................................................
env-821   AAAACCGAAAUUAACAGAUAUCGUGAGAUAAUCAAAGUAUUGAAUAUUAGAAAAUAAUCAAUAAAAAAAGAGGUGCCCACGCGCCUCUUUUU
env-822   AAAACUGAAAUCAACAGAUAUCGUGAGAUUAUCAAAGUAUUGAAUAUUAGAAAAUAAUCAAUAAAAAAAGAGGUGCCCACGCGCCUCUUUUU
env-823   AAAACUGAAAUCAACAGAUAUCGUGAGAUUAUCAAAGUAUUGAAUAUUAGAAAAUAAUCAAUAAAAAAAGAGGUGCCCACGCGCCUCUUUUU
env-824   ACUAAAAAAGACAUUUUGAGAUACAGAGCGAUUGUGAAAGAACUAGGAUUAAGAAAGUAAUUGUAUCCUAACCUUUAUUACUUAUAGUUAU
env-825   AAGAAAAAAGAUAUUAAUAGAUAUCGUGAAAUAAUCAAAAAAUUAAACUUACGUAAAUAGUUUUACAUAGUUUUUCAUUGGUAUUCUACUU
env-826   AAGAAAAAAGAUAUUAGUAGAUACCGUGAAAUAAUCAAAAAAUUAAACUUACGUAAAUAGUUUUUUACACAGUUUUUCAUUGGAAUCUAC
env-827   AAGAAAAAAGAUAUUAGUAGAUACCGUGAAAUAAUCAAAAA..................................................
env-828   AAGAAAAAAGACAUUAAUAGAUACCGUGAGAUAAUCAAGAAAUUAAACUUACGUAAAUA................................
env-829   ...........................................................................................
env-830   ...........................................................................................
env-831   ...........................................................................................
env-832   AAGAAAAAAGAUAUUAAUAGAUAUCGUGAAAUAAUCAAAAAAUUAAACUUACGUAAAUAGUUUUACAUAGUUUUUCAUUGGUAUUCUACUU
env-833   ...........................................................................................
env-834   ...........................................................................................
env-835   AAGAAAAAAGAUAUUAAUAGAUANCGUGAAAUAAUCAAAAAAUUAAACUUACGUAAAUANUUUUACAUAGUUUUUCAUUGGUAUUCUACUU
env-836   ...........................................................................................
env-837   AAGAAAAAAGACAUUAAUAGAUAUCGUGAAAUAAUCAAAAAAUUAAACUUACGUAAAUAGUUUU...........................

          ...........................................................................................
          ...........................................................................................
          A•RAARAARGAYAUY••••RRUAY•G•G••AU•RU•AAAGAAYURRR••U••GAAAAUAA•••••••••••••••••••••••••••••••
```

```
env-838   AAGAAAAAAGACAUUAAUAGAUAUCGUGAAAUAAUCAAAAAAUUAAACUUACGUAAAUAGUUUUUUACACAGUUUUUCAUUGGUAUUCUAC
env-847   AAGAAAAAAGAUAUUAAUAGAUAUCGUGAAAUAAUCAAAAAAUUAAACUUACGUAAAUAGUUUUUUACACAGUUUUUCAUUGGUAUUCUAC
env-848   AAGAAAAAAGAUAUUAAUAGAUAUCGUGAAAUAAUCAAAAAAUUAAACUUACGUAAAUAGUUUUUUACACAGUUUUUCAUUGGUAUUCUAC
env-849   AUGAAAAAAGACAUUAAUAGAUAUCGUGAAAUAAUCAAGAAAUUAAACUUACGUAAAUAGUUUUUUACACAGUUUUUCAUUGGUAUUCUAC
env-850   AAGAAAAAAGAUAUUAACAGAU.....................................................................
env-851   AAGAAAAAAGAUAUUAACAGAUAUCGUGAAAUAAUCAAAAAAUUAAACUUACGUAAAUAGUUUUUU.........................
env-852   AAGAAAAAAGAUAUUAACAGAUAUCGUGAAAUAAUCAAAAAAUUAAACUUACGUAAAUAGUUUUUUACACAGUUUUUCAUUGGUAUUCUAC
env-853   AAGAAAAAAGAUAUUAACAGAUAUCGUGAAAUAAUCAAAAAAUUAAACUUACGUAAAUAGUUUUUUACACAGUUUUUCAUUGGUAUUCUAC
env-854   ...........................................................................................
env-855   ...........................................................................................
env-856   ...........................................................................................
env-857   AAGAAAAAAGAUAUUAACAGAUAUCGUGAAAUAAUCAAAAAAUUAAACUUACGUAAAUAGUUUUUUACACAGUUUUUCAUUGGUAUUCUAC
env-860   AAGAAAAAAGAUAUUAACAGAUAUCGUGAAAU...........................................................
env-861   ...........................................................................................
env-862   AAGAAAAAAGAUAUUAACAGAUAUCGUGAAA............................................................
env-863   ...........................................................................................
env-864   AAGAAAAAAGAUAUUAACAGAUAUCGUGAAAUAAUCAAAAAAUUAAACUUACGUAAAUAGUUUUUU.........................
env-865   AAGAAAAAA..................................................................................
env-866   ...........................................................................................
env-868   ...........................................................................................
env-869   AAGAAAAAAGAUAUUAAUAGAUACCGUGAAAUAAUCAAAAAAUUAAACUUACGUAAAUAGUUUUUACACAGUUUUUCAUUGGUAUUCUACU
env-870   AAGAAAAAAGAUAUUAAUAGAUACCGUGAAAUAAUCAAAAAAUUAAACUUACGUAAAUAGUUUUUUACACAGUUUUUCAUUGGUAUUCUAC
env-871   AAGAAAAAAGAUAUUAAUAGAUACCGUGAAAUAAUCAAAAAAUUAAACUUACGUAAAUAGUUUUUACACAGUUUUUCAUUGGUAUUCUACU
env-875   ...........................................................................................
env-876   ...........................................................................................
env-877   ...........................................................................................
env-878   AAGAAAAAAGAUAUUAAUAGAUACCGUGAAAUAAUCAAAAAAUUAAACUUACGUAAAUAGUUUUUACACAGUUUUUCAUUGGUAUUCUACU
env-879   AAGAAAAAAGAUAUUAAUAGAUACCGUGAAAUAAUCAAAAAAUUAAACUUACGUAAAUAGUUUUUACACAGUUUUUCAUUGGUAUUCUACU
env-881   AAGAAAAAAGAUAUUAAUAGAUACCGUGAAAUAAUCAAAAAAUUAAACUUACGUAAAUAGUUUUUACACAGUUUUUCAUUGGUAUUCUACU
env-883   AAAAUAAAGAUAUCAAAAGAUAUCGUG................................................................
env-884   AAAAUAAAGAUAUCAAAAGAUAUCGUGAGAUUAUCAAAAAACUUAACCUACGUAAAUAAGUUCUUUACAUAGUUUUUCAUUGGUAUUCUA
env-886   AA.........................................................................................
env-887   AAAAUAAAGAUAUCAAAAGAUAUCGUGAGAUUAUCAAAAAACUUAACCUACGUAAAUAAGUUCUUUACAUAGUUUUUCAUUGGUAUUCUA
env-888   AAAAUAAAGAUAUCAAAAGAUAUCGUGAGAUUAUCAAAAAACUUAACCUACGUAAAUAAGUUCUUUACAUAGUUUUUCAUUGGUAUUCUA
env-889   AAAAUAAAGAUAUCAAAAGAUAUCGUGAGAUUAUCAAAAAACUUAACCUACGUAAAUAAGUUCUUUACAUAGUUUUUCAUUGGUAUUCUA
env-890   AAAAUAAAGAUAUCAAAAGAUAUCGUGAGAUUAUCAAAAAACUUAACCUACGUAAAUAAGUUCAUUACAUAGUUUUUCAUUGGUAUUCUA
env-891   AAAAUAAAGAUAUCAAAAGAUAUCGUGAGAUUAUCAAAAAACUUAACCUACGUAAAUAAGUUCUUUACAUAGUUUUUCAUUGGU......
env-892   AAAAUAAAGAUAUCAAAAGAUAUCGUGAGAUUAUCAAAAAACUUAACCUACGUAAAUAAGUUCUUUACAUAGUUUUUCAUUGGUAUUCUA
env-894   AAAAUAAAGAUAUCAAAAGAUAUCGUGAGAUUAUCAAAAAACUUAACCUACGUAAAUAAGUUCUUUACAUAGUUUUUCAUUGGUAUUCUA
env-895   AAAAUAAAGAUAUCAAAAGAUAUCGUGAGAUUAUCAAAAAACUUAACCUACGUAAAUAAGUUCUUUACAUAGUUUUUCAUUGGUAUUCUA
env-896   AAAAUAAAGAUAUCAAAAGAUAUCGUGAGAUUAUCAAAAAACUUAACCUACGUAAAUAAGUUCUUUACAUAGUUUUUCAUUGGUAUUCUA
env-897   AAAAUAAAGAUAUCAAAAGAUAUCGUGAGAUUAUCAAAAAACUUAACCUACGUAAAUAAGUUCUUUACAUAGUUUUUCAUUGGUAUUCUA
env-898   ...........................................................................................
env-899   ...........................................................................................
env-900   AACUUAACCUACGUAAAUAAGUUCUUUACAUAGUUUUUCAUUGGUAUUCUACUUAAGACACAACACAACUUAAUUUAAUUAAUCAAAAAUA
env-901   ...........................................................................................
env-902   AAAGCGAAAGAUAUCGAAAGGUACAGAGCUAUCAUCAUAGAGCUUGGCAUAAGAAAAUAAGUUUAAGUUCUUAUUCUCAAAAGCAUUUCUU
env-903   AAAGCGAAAGAUAUCGAAAGAUACAGAGCUAUCAUCAAAGAACUUGGCAUUAGAAAAUAAGUUUAAGUUACUUAUUCUUAAAAGCAUUUCU
env-905   ...........................................................................................
env-906   AAAACAGAAAUCAACAGGUACCGUGAGAUUAUCAAAGUAUUGAAUAUCAGAAAAUAAUCAAUAGUAAAGAGGUGCGAAAGUGCCUCUUUUU
env-907   ...........................................................................................
env-908   AAAACAGAAAUCAACAGGUACCGUGAGAUUAUCAAAGUAUUGAAUAUCAGAAAAUAAUCAAUAGUAAAGAGGUGCGAAAGUGCCUCUUUUU
Fge-1-1   AUUAAGAAAGACAUUUUGAGAUAUCGUGAGAUUAUCAAAGAAUUAAACAUCAGAAAAUAAAUUAUACUAGGGGCUUUGUGCCCCUUUUAUU
Ale-2-1   AUGAAAAAGGAUAUCUUGAGAUACCGUGCAAUCGUUAAAGAAUUAGGUUUACGUAAGUAAUACAAAGAGGGGCGUUUUUCGCCCCUUUUUA
env-909   AAGAAAACGGAAAUCAACAGAUACCGUGAGAUCAUUAAAGAAUUGAAUAUCAGAAAAUAAUCAAUCGGAAAAGAGGUGCUCACGCGCCUCU
env-910   AUGAAGAAAGACAUUAUGAGAUAUCGUGCUAUAGUAAAAGAAUUAGGAUUACGUAAGUAAUUGUUUAUACUGGGGGCUUUUUGCCCCCAUA
env-911   AUGAAAAAAGACAUAUUAAGAUAUCGUGCAAUUGUUAAAAAAUUGAAUUUAAGAAAGUAAAAAAAGAGUUUAAAACUCCACUUUCUUUAAU
env-912   AUGAAAAAAGACAUAUUAAGAUAUCGUGCAAUUGUUAAAAAAUUGAAUUUAAGAAAGUAAAAAAAGAGUUUAUAACUCCAUUUUUUUAAUA
env-913   ...........................................................................................
env-914   AUGAAAAAAGACAUAUUAAGAUAUCGUGCAAUUGUUAAAAAACUGAAUUUAAGAAAGUAAUAAAAAGAGUUCAUAACUACACUUUUCUUAA
env-915   AUGAAAAAAGACAUAUUAAGAUAUCGUGCAAUUGUUAAAAAACUGAAUUUAAGAAAGUAAUAAAAAGAGUUCACAACUACACUUUUU....
env-916   AUGAAAAAAGAUAUAUUAAGAUAUCGUGCAAUUGUUAAAAAAUUGAAUUUAAGAAAGUAAAAAA...........................
Man-1-1   AUAAAGAAUGAUGUUGUAAAGUAUCGUGAGCUUAUCAAAGAAUUGGGUAUUAGAAAAUAAUUUAAAAAGGGGAAAGCUAGCUUUCCCCUUU
env-917   ...........................................................................................
env-919   AAAACGAAGGAUAUUGAAAGAUAUCGUGAGUUGAUUAAAGAGUUGAAAUUACGUCGUUAAUUUUAAUUUGCAAACGUAUUGAAGAGGGGAC
env-920   AAAAUAAGGAUAUUGAAAA........................................................................
env-921   AAAAUAAGGAUAUUGAAAAAUAUCGUGAACUUAU.........................................................
env-922   AAAAUAAGGAUAUUGAAAAAUAUCGUGAACUUAUAAAGGAAUUAAAAUUACGUCGUUAAUUUUAAUUUCGAAAUGUAUUGAAGAGGGGAC
env-923   AAAAUAAGGAUAUUGAAAAAUAUCGUGAACUUAUAAAGGAAUUAAAAUUACGUCGUUAAUUUUAAUUUCGAAAUGUAUUGAAGAGGGGAC
env-924   AAAAUAAGGAUAUUGAAAAGUAUCGUGAACUUAUUAAAGAAUUGAAAUUACGUCGUUAAUUUUAAUUUUUAAAUGUAUUGAAGAGGGGAC
env-925   ...........................................................................................
env-927   AAAAUAAGGAUAUUGAAAAAUAUCGUGAACUUAUAAAGGAAUUAAAAUUACGUCGUUAAUUUUAAUUUCGAAAUGUAUUGAAGAGGGGAC
env-928   ...........................................................................................
env-929   AAGUCUAAGGAUAUUGAAAAAUAUCGUGAGCUGAUAAAAGAAUUGAAAUUACGUCGUUAAUUUUAAUUUUUAACGUAUUGAAGAGGGGACA
env-930   AAGUCUAAGGAUAUUGAAAAAUAUCGUGAGCUGAUAAAAGAAUUGAAAUUACGUCGUUAAUUUUAAUUUUUAACGUAUUGAAGAGGGGACA
env-932   AAAAUAAGGAUAUUGAAAAAUAUCGUGAAUUAAUUAAGGAGCUGAAAUUGCGUCGUUAAUUUAGUUCUAAAACGUAUUGAAGAGGGGACA
env-933   AAGAAAAAAGAUGUUGUGAAAUAUCGUGAGUUAAUUAAAGAAUUAGGAAUUAGAAAAUAAUCAACUAAGGGGCUUUGUGCCCCUUUUUGUU
env-934   AAGAAAAAAGAUGUUGUGAAAUAUCGUGAGUUAAUUAAAGAAUUAGGAAUUAGAAAAUAAUCAACUAAGGGGCUUUGUGCCCCUUUUUGUU
env-936   ...........................................................................................
env-937   ACUAAAGUUGAUGUCUUAAGAUAUCGUGCAAUAGUUAAAGAAUUAGGUUUAAGAAAAUAACAGUAAAAGAGACUAUUUUUUAGUCUCUUUU
env-938   AAAUCUAAAGAUAUUGAGAAAUAUCGAAAAUUAAUAAAAGAGUUAGGAAUUCGAAAAUAACUGAAAAUUUUGGAAUAAUAUCUUAGUUUUU
env-939   AAAUCUAAAGAUAUUGAGAAAUAUCGAAAAUUAAUAAAAGAGUUAGGAAUUCGAAAAUAACUAAAACUUCGAAUAACACCUUAGUUUUUCA
env-940   ...........................................................................................
env-941   AAAUCUAAAGAUAUUGAGAAAUAUCGAAAAUUAAUAAAAGAGUUAGGAAUUCGAAAAUAACUAAAACUUCGAAUAAUAUCUUAGUUUUUCA

          ...........................................................................................
          ...........................................................................................
          A•RAARAARGAYAUY••••RRUAY•G•G••AU•RU•AAAGAAYURRR••U••GAAAAUAA•••••••••••••••••••••••••••••
```

```
env-942    ...........................................................................................
env-943    AAAUCUAAAGAUAUUGAGAAAUAUCGAAAAUUAAUAAAAGAGUUAGGAAUUCGAAAAUAACUGAAAAUUUUGGAAUAAUAUCUUAGUUUUU
env-944    AAAUCUAAAGAUAUUGAGAAAUAUCGAAA..............................................................
env-945    AAAUCUAAAGAUAUUGAGAAAUAUCGAAAAUUAAUAAAGGAGUUAGGAAUUCGAAAAUAACUAAAACUUCGAAUAA...............
env-946    ...........................................................................................
Tma-1-1    AAGAAAAAAGAUAUCAAUAGAUAUCGUGCGAUAAUCAAAGAAUUAGGAAUUAGAAAAUAAUUUCUACAAAAAAGAGGCUCUACAAACGUGC
env-947    ...........................................................................................
Fsp-12-1   AAGAAAGAGAUCAACAGAUAUCGUGAGAUUAUCAAAGUAUUGAAUAUCAGAAAAUAAUCAAUAUAGGAAAGAGGUGCGAAAGUGCCUCUUU
Fde-1-1    AAGAAAGAUAUCAACAGAUAUCGUGAGAUCAUUAAAGUAUUGAAUAUCAGAAAAUAAUCAAUAUAGAAAAGAGGUGCGAGAGUGCCUCUUU
Fba-7-1    AUCAAAAAAGACAUUUUAAGAUACCGUGCUAUUGUAAAAGAGCUUGGUUUAAGAAAAUAAGUCAUCAAUACUUUAAAAAGCGGAGCUUGCA
Ale-1-1    AUCAAGAAAGAUAUCUUAAGAUAUCGUGCUAUAGUAAAAGAAUUAGGAUUAAGAAAAUAAUCUCAAUAAAAAAACCACGCUUUUCAGCGUG
Ale-1-2    AUCAAGAAAGAUAUCUUAAGAUAUCGUGCUAUAGUAAAAGAAUUAGGAUUAAGAAAAUAAUCUCAAUAAAAAACCACGCUUUUCAGCGUGG
Ffr-1-1    AAGAAAGAAAUUAACAGAUAUCGUGAGAUUAUCAAAGUAUUGAAUAUCAGAAAAUAAUCAAUAUAAAAAAGAGGUGCGCAAGUGCCUCUUU
env-948    ...........................................................................................
Fsp-1-1    AAGAAAAAAGAUAUCAACAGAUAUCGUGAGAUUAUUAAAGAAUUAGGAAUUAGAAAAUAAUCAACUAAAGGGGCUUCGCGCCCCUUUUUGU
env-949    ...........................................................................................
env-950    ...........................................................................................
Fsa-2-1    AAAAAGAAAGAUAUCAACAGAUAUCGUGAGAUUAUUAAAGAAUUAGGUAUUAGAAAAUAAUCAACUAAAGGGGCUUCGUGCCCCUUUUUGU
env-951    ...........................................................................................
env-952    AAAAAGAAAGAUAUCAACAG.......................................................................
Fba-3-1    AAGAAAAAAGAUAUCAACAGAUAUCGUGAGAUUAUUAAAGAAUUAGGAAUUAGAAAAUAAUCAACUAAAGGGGCUUUGUGCCCCUUUUUGU
env-953    AAAAAGAAAGAUAUCAAUAGAUAUCGUGAGAUUAUUAAAGAAUUAGGAAUUAGAAAAUAAUCAACUUAAGGGGCUUCGUGCCCCUUUUUGU
env-954    ...........................................................................................
env-955    ACUAAGGUAGAUGUCUUAAGAUACCGUGCGAUUGUUAAAGAAUUAGAUUUAAGAAAAUAAUAUAAAAGAGGCUAAUGGCCUCUUUUAUAAA
env-956    ACUAAGGUAGAUGUCUUAAGAUACCGUGCAAUUGUUAAAGAAUUAGAUUUAAGAAAAUAAUAUAAAAGAGGCUAAUGGCCUCUUUUAUAAA
env-957    AUGUCUUAGAUACGUGCAUUGUAAAGAUAGAUUAGAAAUAUAUAAAAGAGCUAUGCUCUUUGUAAUUAUACAUAAUUAUAUAAAGCUAGUA
env-958    ...........................................................................................
env-959    AAAAAUACAGAUAUUGAAAGAUAUCGUGCUAUUAUCAAAGAACUAAAUAUCCGAAAAUAAGCAAAUACAAGGAGGCUUUGUGCCUCCUUUU
Csp-1-1    AAAAAUAACGAUAUCGAGAGAUACCGUGCGAUUAUCAAAGAGCUAAACAUCAGAAAAUAAGCAAUACUAAGGAGGCUUUGUGCCUCCUUUU
env-960    ...........................................................................................
env-961    UCCUUUUGUUUAUUACAGCAGUUUUUCAUUGGUUAAUCCCUAAAGCCCUCAACACAACAAUUAUUAAUUUAUUAAUUCACAAAAAGAAUGA
Csp-3-1    AAAAAUAACGAUAUCGAGAGAUACCGUGCGAUUAUCAAAGAACUAAACAUCAGAAAAUAAGCAAUACUAAGGAGGCUUUGUGCCUCCUUUU
env-962    AAAAACACUGAUAUUGAAAGAUAUCGUGCGAUUAUCAAAGAGCUAAACAUCAGAAAAUAAGCAAUACUAAGGAGGCUUUGUGCCUCCUUUU
env-965    AAAAACAACGAUAUCGAGAGAUACCGUGCGAUUAUCAAAGAACUAAACAUCAGAAAAUAAGCAAUACUAAGGAGGCUUUGUGCCUCCUUUU
env-966    ...........................................................................................
env-967    ...........................................................................................
env-968    AAAAACACUGAUAUUGAAAGAUAUCGUGCGAUUAUCAAAGAGCUAAACAUCAGAAAAUAAGCAAUACUAAGGAGGCUUUGUGCCUCCUUUU
env-972    AAAAACACUGAUAUUGAAAGAUAUCGUGCGAUUAUCAAAGAGCUAAACAUCAGAAAAUAAGCAAUAC........................
env-973    AAAAACAACGAUAUUGAGAGAUACCGUGCGAUUAUCAAAGAACUAAACAUCAGAAAAUAAGCAAUACUAAGGAGGCUUUGUGCCUCCUUUU
env-974    AAAAACACUGAUAUUGAAAGAUAUCGUGCGAUUAUCAAAGAGCUAAACAUCAGAAAAUAAGCAAUACUAAGGAGGCUUUGUGCCUCCUUUU
env-976    AAAAACACUGAUAUUGAAAGAUAUCGUGCGAUUAUCAAAGAGCUAAACAUCAGAAAAUAAGCAAUACU.......................
env-977    ...........................................................................................
env-978    ...........................................................................................
env-980    ...........................................................................................
env-982    ...........................................................................................
env-983    AAAAACAACGAUAUCGAGAGAUACCGUGCGAUUAUCAAAGAACUAAACAUCAGAAAAUAAGCAAUACUAAGGAGGCUUUGUGCCUCCUUUU
env-984    AAAAAUAACGAUAUCGAGAGAUACCGUGCAAUUAUCAAAGAACUAAACAUCAGAAAAUAAGCAAUACUAAGGAGGCUUUGUGCCUCCUUUU
env-985    AAAAACAACGAUAUCGAGAGAUACCGUGCGAUUAUCAAAGAACUAAACAUCAGAAAAUAAGCAAUACUAAGGAGGCUUUGUGCCUCCUUUU
env-986    AAAAACAACGAUAUCGAGAGAUACCGUGCGAUUAUCAAAGAACUAAACAUCAGAAAAUAAGCAAUACUAAGGAGGCUUUGUGCCUCCUUUU
env-988    AAAAACAACAAUAUCGAGAGAUACCGUGCGAUUAUCAAAGAACUAAACAUCAGAAAAUAAGCAAUACUAAGGAGGCUUUGUGCCUCCUUUU
env-989    AAAAACAACGAUAUCGAGAGAUACCGUGCGAUUAUCAAAGAACUAAACAUCAGAAAAUAAGCAAUACUAAGGAGGCUUUGUGCCUCCUUUU
env-990    AAAAACACUGAUAUUGAAAGAUAUCGUGCGAUUAUCAAAGAGCUAAACAUCAGAAAAUAAGCAAUACUAAGGAGGCUUUGUGCCUCCUUUU
env-993    AAAAACAACGAUAUCGAGAGAUACCGUGCGAUUAUCAAAGAACUAAACAUCAGAAAAUAAGCAAUACUAAGGAGGCUUUGUGCCUCCUUUU
env-994    AAGAAUACCGAUAUUGAAAGAUAUCGUGCAAUUAUCAAAGAGCUUAACAUUAGAAAAUAAAAAGAGAGGAGGCUUAGUGCCUCCUUUUGCU
env-995    AAAAACAACGAUAUCGAGAGAUACCGUGCGAUUAUCAAAGAACUAAACAUCAGAAAAUAAGCAAUACUAAGGAGGCUUUGUGCCUCCUUUU
env-999    AAAAACACUGAUAUUGAAAGAUAUCGUGCGAUUAUCAAAGAGCUAAACAUCAGAAAAUAAGCAAUACUAAGGAGGCUUUGUGCCUCCUUUU
env-1001   ...........................................................................................
env-1002   ...........................................................................................
env-1003   AAAAACAACGAUAUCGAGAGAUACCGUGCGAUUAUCAAAGAACUAAACAUCAGAAAAUAAGCAAUAC........................
env-1004   AAAAACACUGAUAUUGAAAGAUAUCGUGCGAUUAUCAAAGAGCUAAACAUCAGAAAAUAAGCAAUACUAAGGAGGCUUUGUGCCUCCUUUU
env-1005   ...........................................................................................
env-1006   ...........................................................................................
env-1007   AAAAACAACGAUAUCGAGAGAUACCGUGCGAUUAUCAAAGAACUAAACAUCAGAAAAUAAGCAAUACUAAGGAGGCUUUGUGCCUCCUUUU
env-1008   ...........................................................................................
env-1009   AAAAACAACGACAUCGAGAGAUACCGUGCGAUUAUCAAAGAACUAAACAUCAGAAAAUAAGCAAUACUAAGGAGGCUUUGUGCCUCCUUUU
env-1010   ...........................................................................................
env-1011   AAAAACAACGAUAUCGAGAGAUACCGUGCGAUUAUCAAAGAACUAAACAUCAGAAAAUAAGCAAUACUAAGGAGGCUUUGUGCCUCCUUUU
env-1012   ...........................................................................................
env-1013   ACUAAAAAAGAUAUUUUGAGAUAUAGAGCGAUUGUAAAAGAACUAGGAUUAAGAAAGUAAUUAAAUCCUUUCUUAUUACUUAUAGUUAUUC
env-1015   ACUAAAAAAGAUAUUUUGAGAUAUAGAGCGAUUGUAAAAGAACUAGGAUUAAGAAAGUAAUUAAAUCCUUUCUUAUUACUUAUAGUUAUUC
env-1016   ACAAAAAAAGAUAUUUUGAGAUAUAGAGCGAUUGUAAAAGAACUAGGAUUAAGAAAGUAAUUGAGUCCUUUUAUUAUUACUUGUAGUUAUU
env-1017   ACUAAAAAAGAUAUUUUGAGAUAUAGAGCGAUUGUUAAGGAACUAGGAUUAAGAAAGUAGUUAUAUACUAGUUUUGUAACUUAUAGUUAUU
env-1018   ACUAAAAAAGAUAUUUUGAGAUAUAGAGCGAUUGUUAAAGAACUAGGAUUAAGAAAGUAGUUAUAUCCUAGUUGUGUAACUUAUAGUUAUU
env-1019   ACUAAAAAAGAUAUUUUAAGAUAUAGAGCAAUUGUGAAAGAGCUAGGAUUAAGAAAGUAAUUAAGUACUUAUUUUUAUCACUUAUAGUUAU
env-1021   ...........................................................................................
env-1022   ACUAAAAAAGAUAUUUUGAGAUACAGAGCGAUUGUGAAAGAAUUAGGAUUAAGAAAAUAGUUAUAUCCUAAUAUAUAUUACUUAUAGUUAU
env-1023   ACUAAAAAAGACAUUUUGAGAUACAGAGCGAUUGUGAAAGAACUAGGAUUAAGAAAGUAAUUGUAUCCUAACCUUUAUUACUUAUAGUUAU
env-1024   ACUAAAAAAGACAUUUUGAGAUACAGAGCGAUUGUGAAAGAACUAGGAUUAAGAAAGUAAUUGUAUCCUAACUUUUAUUACUUAUAGUUAU
env-1025   ACUAAAAAAGAUAUUUUGAGAUACAGAGCGAUUGUGAAAGAACUAGGAUUAAGAAAAUAAUUGUAUCCUAACUUUUAUUACUUAUAGUUAU
env-1026   ACUAAAAAAGAUAUUUUG.........................................................................
env-1027   ...........................................................................................
env-1028   ...........................................................................................
env-1029   ACUAAAAAAGAUAUUCUUAGAUAUCGUGCAAUUGUUAAAGAGU................................................
env-1030   ...........................................................................................

           ...........................................................................................
           ...........................................................................................
           A•RAARAARGAYAUY••••RRUAY•G•G••AU•RU•AAAGAAYURRR••U••GAAAAUAA•••••••••••••••••••••••••••••••
```

```
env-1032 ACUA.......................................................................................
env-1033 CUAAAAAAAGAUAUUCUUAGAUAUCGUGCAAUUGUUAAAGA..................................................
env-1034 ...........................................................................................
env-1035 ...........................................................................................
env-1036 UAAAAAGAUAUUUUAAGAUAUCGUGCCAUCGUAAAAGAGUUAGGAUUAAGAAAAUAACUCUUUUCCAUAUUUUGUCUUAUAGUUAUUCAUU
env-1037 ACAAAAAAAGAUAUUCUUAGAUAUCGUGCUAUCGUAAAAGAGCUAGGAUUAAGAAAAUAACUCUAUUCUACAUUUUGUCUUAUAGUUAUUC
env-1038 AUCAAAAAAAGAUAUUCUUAGAUAUCGUGCGAUUAUCAAAGAGUUAGGAUUAAGAAAAUAACUCUUUUAUCCAUUUUGUCUUAAAGUUAUUC
env-1039 ...........................................................................................
env-1040 ...........................................................................................
env-1041 ...........................................................................................
env-1042 ACUAAAAAAAGAUAUUCUUAGAUAUCGUGCUAUCGUAAAAGAGCUAGGAUUAAGAAAAUAACUCUAUUCUACAUUUUGUCUUAUAGUUAUUC
env-1044 ACUAAAAAAAGAUAUUCUUAGAUAUCGUGCUAUCGUAAAAGAGCUAGGAUUAAGAAAAUAACUCUAUUCUACAUUUUGUCUUAUAGUUAUUC
env-1045 ACUAAAAAAAGAUAUUUUAAGAUAUCGUGCUAUAGUAAAAGAGUUAGGAUUAAGAAAAUAACUCUUUUCCAUAUUUUGUCUUAUAGUUAUUC
env-1046 ...........................................................................................
env-1047 ACUAAAAAAAGAUAUUUUAAGAUAUCGUGCAAUUAUAAAAGAAUUAGGGUUAAGAAAAUAAAUCCUUUUCAACAUUUUGUUUAUAGUUAUUC
env-1048 ...........................................................................................
env-1049 ACUAAAAAAAGAUAUUUUAAGAUAUCGUGCAAUUAUAAAAGAAUUAGGGUUAAGAAAAUAAAUCCUUUUCAACAUUUUGUUUAUAGUUAUUC
env-1051 ...........................................................................................
env-1053 ACUAAAAAAAGAUAUUUUAAGAUAUCGUGCAAUUAUAAAAGAAUUAGGGUUAAGAAAAUAAAUCCUUUUCAACAUUUUGUUUAUAGUUAUUC
env-1055 ACUAAAAAAAGAUAUUUUAAGAUAUCGUGCAAUUAUAA.....................................................
env-1056 ACUAAAAAAAGAUGUUCUAAGAUAUCGUGCAAUCGUAAAAGAGUUAGGAUUAAGAAAAUAACUUUUUUCUAUAUUUUGUCUUAUAGUUAUUC
env-1057 ...........................................................................................
env-1058 ...........................................................................................
env-1059 ...........................................................................................
env-1060 ...........................................................................................
env-1061 ACUAAAAAAGGAUGUCCUUAGAUAUCGUGCAAUUGUUAAAGAGUUAGGAUUAAGAAAAUAACUCAUUUCUAAAUUUUGUUUAUAGUUAUUCA
env-1062 ...........................................................................................
env-1063 ACUAAAAAAAGAUGUUUUAAGAUAUCGUGCAAUUGUAAAAGAGUUAGGAUUAAGAAAAUAACUCGUUUCUAUAUUUUGUUUAUAGUUAUUCA
env-1064 ACUAAAAAAAGAUAUUUUAAGAUAUCGUGCUAUCGUAAAAGAGUUGGGAUUAAGAAAAUAACUCUUUUCCACAUUUUGUCUUAUAGUUAUUC
env-1065 ACUAAAAAAAGAUAUUCUUAGAUAUCGUGCAAUUGUUAAAGAGUUAGGAUUAAGAAAAUAACUCUUUUAUCCAUUUUGUCUUAUAGUUAUUC
env-1066 AACAAAGAAAGAUAUCUUAAGAUAUCGUGCUAUUGUUAAAGAGUUAGGAUUAAGAAAAUAACUCUUUUCUACAUUUUGUUUUAUAGUUAUU
env-1067 ACUAAAAAAAGAUGUUUUAAGAUAUCGUGCAAUUGUAAAAGAGUUAGGAUUAAGAAAAUAACUCGUUUCUAUAUUUUGUUUAUAGUUAUUCA
env-1068 ACUAAAAAAAGAUAUUUUAAGAUAUCGUG..............................................................
env-1069 ACUAAAAAAAGAUGUUUUAAGAUAUCGUGCUAUUGUAAAAGAGUUGGGAUUAAGAAAAUAACUCGUUUCUAUAUUUUGUUUAUAGUUAUUCA
env-1070 ACAAAGAAAGAUAUCUUAAGAUAUCGUGCUAUUGUUAAAGAGUUAGGAUUAAGAAAAUAACUCUUUUCUACAUUUUGUUUUAUAGUUAUUC
env-1077 ...........................................................................................
env-1078 ...........................................................................................
env-1079 ACUAAAAAAAGAUGUUUUAAGAUAUCGUGCAAUUGUAAAAGAGUUAGGAUUAAGAAAAUAACUCGUUUCUAUAUUUUGUUUAUAGUUAUUCA
env-1080 ACUAAAAAAAGAUGUUUUAAGAUAUCGUGCAAUUGUAAAAGAGUUAGGAUUAAGAAAGUAACUCGUUUCUAUAUUUUGUUUAUAGUUAUUCA
env-1081 ACUAAAAAAAGAUGUUUUAAGAUAUCGUGCAAUUGUAAAAGAGUUAGGAUUAAGAAAAUAACUCGUUUCUAUAUUUUGUUUAUAGUUAUUCA
env-1082 ACUAAAAAAAGAUGUUUUAAGAUAUCGUGCAAUUGUAAAAGAGUUAGGAUUAAGAAAAUAACUCGUUUCUAUAUUUUGUUUAUAGUUAUUCA
env-1085 ACUAAAAAAAGACAUUCUAAGAUAUCGUGCUAUCGUAAAAGAGUUAGGAUUAAGAAAAUAACUCUUUUCCAUAUUUUGUCUUAUAGUUAUUC
env-1086 ACUAAAAAAAGAUAUUUUAAGAUAUCGUGCUAUCGUAAAAGAGUUAGGAUUAAGAAAACAACUCUUUUCCAUAUUUUGUCUUAUAGUUAUUC
env-1087 ...........................................................................................
env-1088 ACUAAAAAAGGAUAUUCUUAGAUAUCGUGCGAUUGUUAAAGAGUUAGGAUUAAGAAAAUAACUCUUUUAUCCAUUUUGUCUUAUAGUUAUUC
env-1089 ACUAAAAAAAGAUAUUUUAAGAUAUCGUGCUAUCGUAAAAGAGUUGGGAUUAAGAAAAUAACUCUUUUCCACAUUUUGUCUUAUAGUUAUUC
env-1090 ACUAAAAAAAGAUAUUUUAAGAUAUCGUGCUAUCGUAAAAGAGUUGGGAUUAAGAAAAUAACUCUUUUCCACAUUUUGUCUUAUAGUUAUUC
env-1091 ACUAAAAAAAGAUAUUUUAAGAUAUCGUGCUAUCGUAAAAGAGUUGGGAUUAAGAAAAUAACUCUUUUCCACAUUUUGUCUUAUAGUUAUUC
env-1092 ACUAAAAAAAGAUAUUUUAAGAUAUCGUGCUAUCGUAAAAGAGUUGGGAUUAAGAAAAUAACUCUUUUCCA....................
env-1093 ...........................................................................................
env-1094 ...........................................................................................
env-1095 ACUAAAAAAAGAUAUUCUUAGAUAUCGUGCAAUUGUUAAAGAGUUAGGAUUAAGAAAAUAACUCUUUUAUCCAUUUUGUCUUAUAGUUAUUC
env-1096 ...........................................................................................
env-1097 ACUAAAAAAAGAUAUUUUAAGAUAUCGUGCUAUCGUAAAAGAGUUAGGAUUAAGAAAAUAACUCUUUUCCAUAUUUUGUCUUAUAGUUAUUC
env-1098 ACUAAAAAAAGAUAUAUUAAGAUAUCGUGCAAUUGUAAAAGAGUUAGGAUUAAGAAAAUAACUCGUUUCUAUAUUUUGUUUAUAAUUAUUCA
env-1099 CUAAAAAGAUAUUCUGAGAUAUCGUGCAAUUGUUUA.......................................................
env-1101 ACUAAAAAAAGAUAUUUUAAGAUAUCGUGCUAUCGUAAAAGAGUUGGGAUUAAGAAAAUAACUCUUUUCCACAUUUUGUCUUAUAGUUAUUC
env-1102 ACUAAAAAAAGAUAUUUUAAGAUAUCGUGCAAUUGUAAAAGAGUUAGGAUUAAGAAAAUAACUCGUUUCUAUAUUUUGUUUAUAGUUAUUCA
env-1103 ACUAAAAAAAGAUAUUUUAAGAUAUCGUGCGAUCGUAAA....................................................
env-1104 AC.........................................................................................
env-1105 ACUAAAAAAAGAUAUUUUAAGAUAUCGUGCAAUUGUAAAAGAGUUAGGAUUAAGAAAGUAACUCAUUUCUAUAUUUUGUUUAUAGUUAUUCA
env-1106 ACAAAAAAAGAUAUUCUUAGAUAUCGUGCUAUCGUAAAAGAGCUAGGAUUAAGAAAAUAACUCUAUUCUACAUUUUGUCUUAUAGUUAUUC
env-1107 ...........................................................................................
env-1108 ...........................................................................................
env-1109 ACUAAAAAAAGAUAUUCUUAGAUAUCGUGCAAUUGUUAAAGAGUUAGGAUUAAGAAAAUAACUCUUUUAUCCAUUUUGUCUUAUAGUUAUUC
env-1110 ACAAAGAAAGAUAUCUUAAGAUAUCGUGCUAUUGUUAAAGAGUUAGGAUUAAGAAAAUAACUCUUUUCUACAUUU................
env-1111 ACUAAAAAAAGAUAUUUUAAGAUAUCGUGCUAUCGUAAAAGAGUUGGGAUUAAGGAAAUAACUCUUUUCCAUAUUUUGUCUUAUAGUUAUUC
env-1112 ACCAAAAAAAGAUAUUCUUAGAUAUCGUGCAAUUGUUAAAGAGUUAGGUUUAAGAAAAUAACUCAUAUAUCCAUUUUGUCUUAUAGUUAUUC
env-1113 ACAAAAAAAGAUAUUCUUAGAUAUCGUGCAAUUGUUAAAGAGUUAGGUUUAAGAAAAUAACUCAUAUAUCCAUUUUGUCUUAUAGUUAUUC
env-1115 ACAAAAAAAGAUAUUCUUAGAUAUCGUGCAAUUGUUAAAGAGUUAGGUUUAAGAAAAUAACUCAUAUAUCCAUUUUGUCUUAUAGUUAUUC
env-1116 ACAAAAAAAGAUAUUCUUAGAUAUCGUGCAAUUGUUAAAGAGUUAGGUUUAAGAAAAUAACUCAUAUAUCCAUUUUGUCUUAUAGUUAUUC
env-1121 ACAAAAAAAGAUAUUCUUAGAUAUCGUGCAAUUGUUAAAGAGUUAGGUUUAAGAAAAUAACUCAUAUAUCUAUUUUGUCUUAUAGUUAUUC
env-1122 ACAAAAAAAGAUAUUCUUAGAUAUCGUGCAAUUGUUAAAGAGUUAGGAUUAAGAAAAUAACUCAAUAAAAUUUUUGUCCUACAAUUAUUCA
env-1123 ACUAAAAAAAGAUAUUUUAAGAUAUCGUGCAAUUGUUAAAGAGUUAGGAUUAAGAAAAUAACUCUUUUAUCCAUUUUGUCUUAUAGUUAUUC
env-1124 ACCAAAAAAAGAUAUUCUUAGAUAUCGUGCAAUUGUUAAAGAGUUAGGUUUAAGAAAAUAACUCAUAUAUCCAUUUUGUCUUAUAGUUAUUC
env-1125 ACAAAAAAAGAUAUUCUUAGAUAUCGUGCAAUUGUUAAAGAGUUAGGUUUAAGAAAAUAACUCAUAUAUCCAUUUUGUCUUAUAGUUAUUC
env-1126 ACAAAAAAAGAUAUUCUUAGAUAUCGUGCAAUUGUUAAAGAGUUAGGUUUAAGAAAAUAACUCAUAUAUCCAUUUUGUCUUAUAGUUAUUC
env-1127 ACAAAAAAAGAUAUUCUUAGAUAUCGUGCAAUUGUUAAAGAGUUAGGUUUAAGAAAAUAACUCAUAUAUCCAUUUUGUCUUAUAGUUAUUC
env-1128 ACAAAAAAAGAUAUUCUUAGAUAUCGUGCAAUUGUUAAAGAGUUAGGAUUAAGAAAAUAACU.............................
env-1129 ...........................................................................................
env-1130 CAAGGCAAACGAUAUUGAGAAAUACCGUGCAUUGAUUAAAGAAUUAGGGCUUCGUCGCUAAUCUUUUAUCAAGCAUAGCAAAUUUAUUGGG
env-1131 AAGGCAAAUGACAUCGAGAAAUACCGUGCCUUGAUCAAAGAAUUAGGACUACGUCGUUAAUUCUGGGUCUGACUCGGAAAAUUUUAUGGGA
env-1134 AAGGCAAAUGACAUCGAGAAAUACCGUGCCUUGAUCAAAGAAUUAGGACUACGUCGUUAAUUCUGGGUCUGACUCGGAAAAUUUUAUGGGA
env-1135 AAGGCAAAUGACAUCGAGAAAUACCGUGCCUUGAUCAAAGAAUUAGGACUACGUCGUUAAUUCUGGGUCUGACUCGGAAAAUUUUAUGGGA

         ...........................................................................................
         ...........................................................................................
         A•RAARAARGAYAUY••••RRUAY•G•G••AU•RU•AAAGAAYURRR••U••GAAAAUAA•••••••••••••••••••••••••••••••
```

```
env-1137  AAGGCAAAUGACAUCGAGAAAUACCGUGCCUUGAUCAAAGAAUUAGGACUACGUCGUUAAUUCUGGGUCUGACUC................
env-1138  ...........................................................................................
env-1139  AAGGCGAAAGACAUCGAGAAAUACCGCGCCUUGAUCAAAGAAUUAGGACUACGUCGUUAAUUCUGAGAUCUGCUCGGAAAAUUUUAUGGGA
env-1140  AAGAAAAAAGAUGUUGUAAAAUAUCGUGAGUUAAUUAAAGAAUUAGGUAUUAGAAAAUAAUACAAAUUAUAAGAGGUGCUUUCGCGCCUCU
env-1141  AAAUCAAAAGAUAUUGAGCGUUAUCGCUCAAUCAUCAAUGAGUUGGGCUUAAGAAAAUAAGUUUCUAUAUUUCACAAAAGCAUUUCUAUUU
env-1142  AAAUCAAAAGAUAUUGAGCGUUAUCGCUCAAUCAUCAAUGAGUUGGGCUUAAGAAAAUAAGUUUCUAUAUUUUACAAAAGCAUUUCUAUUU
env-1143  AAAUCAAAAGAUAUUGAGCGUUAUCGCUCAAUCAUCAAUGAGUUGGGCUUAAGAAAAUAAGUUUCUAUAUUUUACAAAAGCAUUUCUAUUU
Lsp-4-1   ACUAAGAAAGAUGUCUUGAGAUAUCGUGCUAUUGUUAAGGAAUUGAACUUGAGAAAAUAAUAUCAAGGGGCUUUGCGGCCCCUUUUUAUUU
Cca-5-1   AAGAAAAAAGAUAUCAACAGAUAUAGAGCAAUCAUCGCUGAACUAGGAAUCAGAAAGUAAUCCUAAGCUCACGCAAAAAAUUAAAGCAACU
env-1144  AAGAAAAAAGAUAUCAACAGAUAUAGAGCAA............................................................
Fan-1-1   AAAAAAUCAGAAAUCAACAGAUAUCGUGAAAUUAUCAAAGAAUUGGGUAUUAGAAAAUAAUCAAUACGGAAAAGAGGUGCACACGCGCCUC
Fte-1-1   AAAAAGACUGAAAUCAACAGAUAUCGUGAAAUUAUCAAAGAAUUGGGUAUUAGAAAAUAAUCAAUAUGAAAAAGAGGUGCACACGCGCCUC
env-1145  AUGAAAAAAGAUAUCGUUAGAUAUCGUGAAAUCGUGAAAGAAUUAGGAUUAAGAAAAUAAUUUGAAAGGGGCUUUACGCCCCUUUUUUUAU
env-1146  AUGAAGAAAGAUGUUGUAAGAUAUCGUGAAAUCGUUAAAGAAUUAGGAUUAAGAAAAUAAUUUUCAAGGGGCUUUAUGCCCCUUUUUUUAU
env-1147  AAGAAAAAAGAUAUUAUGAGAUAUCGUGAAAUCAUUAAGGAAUUAGGUUUAAGAAAAUAAACAUUGAAAGGGGCUUUUUACGCCCCUUUUU
env-1159  ...........................................................................................
env-1161  AUGAAGAAAGAUAUUUUAAGAUAUCGUGCUAUCGUUAAGGAAUUAGGAUUAAGAAAAUAAUAUUAAAAGGGGCUUUACGCCCCU.......
env-1162  ...........................................................................................
env-1163  AAGAAAUCAGAAAUCAACAGAUAUCGUGCGAUUAUCAAAGAAUUAGGAAUUAGAAAAUAAUUCUUACAAAAAGAGGGGCUAUAAACGCACC
env-1165  AAGAAAUCAGAAAUCAACAGAUAUCGUGCGAUUAUUAAAGAAUUAGGAAUUAGAAAAUAAUUCUUACAAAAAGAGGGGCUAUAAACGCACC
env-1167  AAGAAAUCAGAAAUCAACAGAUAUCGUGCGAUUAUCAAAGAAUUAGGAAUUAGAAAAUAAUUCUUACAAAAAGAGGGGCUAUAAACGCACC
env-1168  CUAAAGAAAAAAGAUAUCACAAGAUAUCGUGCAAUAAUUAAAGAAUUAGGAAUUAGAAAAUAAUACCUAUUCAAAAGAG............
env-1169  AAAGAAA....................................................................................
Psp-6-1   AAGA.......................................................................................
env-1170  AAGAAAACCGAAAUCAAUAGAUAUCGUGCAAUCAUUAAAGAAUUAGGAAUUAGAAAAUAAUUCUUACAAAAAAAGAGGUGCUAUAAACGCG
env-1171  AAGAAAACCGAAAUCAAUAGAUAUCGUGCAAUCAUUAAAGAAUUAGGAAUUAGAAAAUAAUUCUUACAAAAAAAGAGGUGCUAUAAACGCG
env-1172  ...........................................................................................
env-1173  AAGAAAAGUGAAAUCAACAGAUAUCGUGCGAUUAUCAAGGAAUUAGGAAUUAGAAAAUAAUUCUUACAAAAAAGAGGUGCUAUAAACGCGC
env-1174  AGAAAAAAGAU................................................................................
Tov-1-1   AAGAAAACAGAAAUCAAUAGAUAUCGUUCGAUUAUUAAAGAAUUAGGAAUUAGAAAAUAAUUUCUUCAAAAAGAGGCUCUAUAAACGUGCC
Pir-1-1   AAGAAAACUGAAAUCAGCAGAUAUCGUGCGAUUAUCAAAGAUUUAGGAAUUAGAAAAUAAUUUUUAAAAAAAGAGGUGCUAUAAACGCACC
env-1175  AAGAAAACUGAAAUCAACAGAUAUCGUGCGAUUAUCAAGGAAUUAGGAAUUAGAAAAUAAUUCUUACAAAAAAGAGGUGCUAUAAACGCGC
env-1176  GUGCGAUAUCAAGGAAUAGGAAUAAAAAAAUCUUACAAAAAAGUGCUUAACGCCCUCUUUUUCCCAUUAAC....................
env-1177  AAGAAAACUGAAAUCAACAGAUAUCGUGCGAUUAUCAAGGAAUUAGGAAUUAGAAAAUAAUUCUUACAAAAAAGAGGUGCUAUAAACGCGC
env-1181  ...........................................................................................
env-1182  AAGAAAACUCUACCAAGUC........................................................................
env-1183  UCUUUUUCUGUUGUUUCUUUCUGCC..................................................................
env-1184  AAGAAAACAGAAAUCAAUAGAUAUCGUGCAAUCAUCAAAGAAUUAGGAAUUAGAAAAUAAUUCUCAGAAAAGAGGCGCUAUAAAUGUGCCU
env-1185  AAGAAAACCGAAAUCAAUAGAUAUCGUGCAAUCAUUAAAGAAUUAGGAAUUAGAAAAUAAUUCGCACAAAAAAGAGGCUCUAUAAACGUGC
env-1186  AAGAAAACCGAAAUCAAUAGAUAUCGUGCAAUCAUUAAAGAAUUAGGAAUUAGAAAAUAAUUCGCACAAAAAAGAGGCUCUAUAAACGUGC
env-1187  AAGAAAACAGAAAUUAAUAGAUAUCGUGCAAUCAUUAAAGAAUUAGGAAUUAGAAAAUAAUUCUAACAAAAAGAGGCGCUAUAAACGUGCC
env-1188  ...........................................................................................
env-1190  AGAAAACAGAAAUUAAUAGAUAUCGUGCAAUCAUUAAAGAAUU................................................
env-1191  AAGAAAACCGAAAUCAAUAGAUAUCGUGCAAUCAUUAAAGAAUUAGGAAUUAGAAAAUAAUUCGCACAAAAAAGAGGCUCUAUAAACG...
env-1192  AAGAAAACAGAAAUUAAUAGAUAUCGUGCAAUCAUUAAAGAAUUAGGAAUUAGAAAAUAAUUCUAACAAAAAGAGGCUCUAUAAACGUGCC
env-1193  AAGAAAACAGAAAUUAAUAGAUAUCGUGCAAUCAUUAAAGAAUUAGGAAUUAGAAAAUAAUUCUAACAAAAAGAGGCUCUAUAAACGUGCC
env-1194  ...........................................................................................
env-1196  AAGAAAACAGAAAUCAAUAGAUAUCGUGCAAUCAUUAAAGAAUUAGGAAUUAGAAAAUAAUUCUAACAAAAAGAGGCUCUAUAAACGUGCC
env-1197  AAGAAAACCGAAAUCAAUAGAUAUCGUGAAAUCAUUAAAGAAUUAGGAAUUAGAAAAUAAUUCUUACAAAAAAAGAGGGGCUAUAAACGCG
env-1198  AAGAAAACAGAAAUCAAUAGAUAUCGUGCAAUUAUUAAAGAAUUAGGAAUUAGAAAGUAAUUCUUACAAAAAGAGGUGCUAUAAAUGUGCC
env-1199  ...........................................................................................
env-1200  AAGAAAACCGAAAUCAAUAGAUAUCGUGAAAUCAUUAAAGAAUUAGGAAUUAGAAAAUAAUUCUUACAAAAAAAGAGGGGCUAUAAACGCG
env-1203  ...........................................................................................
env-1204  AAGAAAACCGAAAUCAAUAGAUAUCGUGAAAUCAUUAAAGAAUUAGGAAUAGAAAAUAAUCUACACUCAACGUAG................
env-1205  AAGAAAAAAGAUAUUGAAAGAUAUCGUGCAAUAAUUAAAGAAUUAGGAAUUAGAAAAUAAUUUCUUUAAAAGAGGCACUAUAAACGUGCCU
Psp-3-1   AAAAAAUCUGAUAUUACAAGAUAUCGUGCAAUUAUCGCAGAAUUAGGAAUUAGAAAAUAAUUCUUACAAAAAGAGGCUCUAUAAACGUGC
env-1208  ...........................................................................................
env-1209  AAGAAAACUGAUAUUACUAGAUAUCGUGCAAUUAUCGCAGAAUUAGGAAUUAGAAAAUAAAACCUAACAUAAAAAGAGGCUCUAUAAACGU
env-1210  ...........................................................................................
env-1211  AAGAAAACUGAUAUUACUAGAUAUCGUGCAAUUAUCGCAGAAUUAGGAAUUAGAAAAUAAAACCUAACAUAAAAAGAGGCUCUAUAAACGU
env-1212  AAGAAAACAGAUAUUGCAAGAUACCGUGCAAUUAUUAAAGAAUUAGGAAUUAGAAAAUAAUUCUCAUUUAAAAAAGAGGCUCUAUAAACGU
env-1213  ...........................................................................................
Psp-2-1   AAGAAAACGGAAAUCAAUAGAUAUCGUGCAAUUAUUAAAGAAUUAGGAAUUAGAAAAUAAUUCUUACAAAAAGAGGCUCUAUAAACGUGCC
Psp-4-1   AAGAAAAAAGAUAUUGCAAGAUAUCGUGCAAUUAUUAAAGAAUUAGGAAUUAGAAAAUAAUUUCUUACAAUAAAGAGGCUCUAUAAACGUG
env-1215  AAGAAAAAAGAUAUCACAAGAUAUCGUGCAAUAAUUAAAGAAUUAGGAAUUAG......................................
env-1216  AAGAAAAAAGAUAUCACAAGAUAUCGUGCAAUAAUUAAAGAAUUAGGAAUUAGAAAAUAAUACCUAUUCAAAAGAGGUAGCUUUAAAGCUA
env-1217  ...........................................................................................
env-1218  ...........................................................................................
env-1219  AAGAAAAAAGAUAUCACAAGAUAUCGUGCAAUAAUUAAAGAAUUAGGA...........................................
env-1220  AAGAAAAAAGAUAUCACAAGAUAUCGUGCAAUAAUUAAAGAAUUAGGAAUUAGAAAAUAAUACCUAUUCAAAAGAGGUAGCUUUAAAGCUA
env-1222  ...........................................................................................
env-1223  AAGAAAAAAGAUAUCACAAGAUAUCGUGCAAUAAUUAAAGAAUUAGGAAUUAGAAAAUAAUACCUAUUCAAAAGAGGUAGCUUUAAAGCUA
env-1226  AAGAAAAAAGAUAUCACAAGAUAUCGUGCAAUAAUUAAAGAAUUAGGAAUUAGAAAAUAAUACCUAUUCAAAAGAGGUAGCUUUAAAGCUA
env-1227  AAGAAAAAAGAUAUCACAAGAUAUCGUGCAAUAAUUAAAGAAUUAGGAAUUAGAAAAUAAUACCUAUUCAAAAGAGGUAGCUUUAAGCUAC
env-1229  AAGAAAAAAGAUAUCACAAGAUAUCGUGCAAUAAUUAAAGAAUUAGGAAUUAGAAAAUAAUACCUAUUCAAAAGAGGUAGCUUUAAAGCUA
env-1230  ...........................................................................................
env-1231  ...........................................................................................
env-1232  ...........................................................................................
env-1233  ...........................................................................................
env-1234  ...........................................................................................
env-1235  AAAGAAAAAAGAUAUCACAAGAUAUCGUGCAAUAAUUAAAGAAUUAGGAAUUAGAAAAUAAUACCUAUUCAAAAGAGGUAGCUUUAAAGCU
env-1236  ...........................................................................................
env-1237  AAGAAAAAAGAUAUCACNAGAUAUCGUGCAAUAAUUAAAGAAUUAGGAAUUAGAAAAUAAUACCUAUUCAAAAGAGGUAGCUUUAAGCUAC
env-1238  AAGAAAAAAGAUAUCACAAGAUAUCGUGCAAUAAUUAAAGAAUUAGGAAUUAGAAAAUAAUACCUAUUCAAAAGAGGUAGCUUUAAAGCUA

          ...........................................................................................
          ...........................................................................................
          A•RAARAARGAYAUY••••RRUAY•G•G••AU•RU•AAAGAAYURRR••U••GAAAAUAA••••••••••••••••••••••••••••••
```

```
env-1239   ..........................................................................................
env-1240   AAGAAAAAAGAUAUCACAAGAUAUCGUGCAAUAAUUAAAGAAUUAGGAAUUAGAAAAUAAUACCUAUUCA......................
env-1241   AAGAAAAAAGAAUAUCACAAGAUAUCGUGC............................................................
env-1242   ..........................................................................................
env-1243   ..........................................................................................
env-1245   ..........................................................................................
env-1246   ..........................................................................................
env-1247   ..........................................................................................
env-1248   ..........................................................................................
env-1250   AAGAAAAAAGAUAUCACAAGAUAUCGUGCAAUUAUCAAAGAAUUAGGAAUUAGAAAAUAAUACCUAUAUAAAAGAGGUAGCUUUAUGCUGC
env-1252   AAGAAAAAAGAUAUCACUAGAUAUCGUGCAAUAAUUAAAGAAUUAGGAAUUAGAAAAUAAUACCUAUUCAAAAGAGGUAGCUUAAGCUACC
env-1253   ..........................................................................................
env-1254   ..........................................................................................
env-1255   ..........................................................................................
env-1256   ..........................................................................................
env-1257   ..........................................................................................
env-1259   AAGAAGAAAGAUAUCGCAAGAUAUCGUGCAAUUAUCAAAGAGUUAGGAAUUAGAAAAUAAUACCUACAUAAAAGAGGUAGCUAUAUGCUAC
env-1260   AAGAAAAAAGAUAUCACUAGAUAUCGUGCAAUAAUUAAAGAAUUAGGAAUUAGAAAAUAAUACCUAUUCAAAAGAGGUAGCUUUAAUGCUA
env-1262   AAGAAAAAAGAUAUCACUAGAUAUCGUGCAAUAAUUAAAGAAUUAGGAAUUAGAAAAUAAUACCUAUUCAAAAGAGGUAGCUUUAAGCUAC
env-1263   AAGAAAAAAGAUAUCACNAGAUAUCGUGCAAUAAUUAAAGAAUUAGGAAUUAGAAAAUAAUACCUAUUCAAAAGAGGUAGCUUUAAUGCUA
env-1264   AAGAAGAAAGAUAUCGCAAGAUAUCGUGCAAUUAUCAAAGAGUUAGGAAUUAGAAAAUAAUACCUACAUAAAAGAGGUAGCUAUAUGCUAC
env-1268   ..........................................................................................
env-1269   ..........................................................................................
env-1270   ..........................................................................................
env-1271   AAGAAGAAAGAUAUCGCAAGAUAUCGUGCAAUUAUCAAAGAGUUAGGAAUUAGAAAAUAAUACCUACAUAAAAGAGGUAGCUAUAUGCUAC
env-1272   AAGAAGAAAGAUAUCGCAAGAUAUCGUGCA............................................................
env-1273   AAGAAGAAAGAUAUCGCAAGAUAUCGUGCAAUUAUCAAAGAGUUAGGAAUUAGAAAAUAAUACCUACAUAAAAGAGGUAGCUAUAUGCUAC
env-1274   ..........................................................................................
env-1275   ..........................................................................................
env-1277   ..........................................................................................
env-1278   AAGAAAAAAGAUAUCGCAAGAUAUCGUGCAAUUAUCAAAGAAUUAGGAAUUAGAAAAUAAUACCUAUACAAAAGAGGUAGCUUUAUGCUAC
Ccy-1-1    AAAACAAAUGAUAUUGAGAGAUAUCGUGCAAUUAUCAAAGAAUUAGGUAUUCGAAAAUAAUCAAAAAGGGGCUUUUUGCCUCUUUUUUUAG
Ccy-1-3    AAAACAAAUGAUAUUGAGAGAUAUCGUGCAAUUAUCAAAGAAUUAGGUAUUCGAAAAUAAUCAAAAAGGGGCUUUUUGCCUCUUUUUUUAG
Tme-1-1    AAGAAAAAAGAGAUUAAUAGAUAUCGUGCGAUUAUUAAAGAAUUAGGAAUUAGAAAAUAAUUCUACAAGAAAAGAGGCUCUAUAAACGUGC
Tsp-1-1    AAGAAAAAAGAGAUUAACAGAUAUCGUGCGAUUAUUAAAGAAUUAGGAAUUAGAAAAUAAUUCUACAAAAAAAGAGGCUCUAUAAACGUGC
env-1280   ..........................................................................................
env-1281   AAGAAAACCGAAAUCAAUAGAUAUCGUGCUAUCAUUAAAGAAUUAGGAAUUAGAAAGUAAUUCUUAUUAAAAGAGGCACUAUAAACGUGCC
env-1282   AAGAAAACCGAAAUCAAUAGAUAUCGUGCUAUCAUUAAAGAAUUAGGAGUAGAAAUAAUUUUACUAAAAGAGCACUAUAAACGUGCCUCUU
env-1283   ..........................................................................................
env-1284   ..........................................................................................
env-1285   AAGAAAACCGAAAUCAAUAGAUAUCGUGCUAUCAUUAAAGAAUUAGGAAUUAGAAAAUAAUUCUUAUUAAAAGAGGCACUAUAAACGUGCC
env-1287   AAGAAAACCGAAAUCAAUAGAUAUCGUGCUAUCAUUAAAGAAUUAGGAAUUAGAAAAUAAUUCUUACUAAAAGAGGCACUAUAAACGUGCC
env-1293   AAGAAAACCGAAAUCAAUAGAUAUCGUGCUAUCAUUAAAGAAUUAGGAAUUAGAAAAUAAUUCUUACUAAAAGAGCACUAUAAACGUGCCU
env-1294   ..........................................................................................
env-1295   AAGAAAACCGAAAUCAAUAGAUAUCGUGCUAUCAUUAAAGAAUUAGGAAUUAGAAAAUAAUUCUUAUUAAAAGAGGCACUAUAAACGUGCC
env-1296   AAGAAAACCGAAAUCAAUAGAUAUCGUGCUAUCAUUAAAGAAUUAGGAAUUAGAAAAUAAUUCUUAUUAAAAGAGGCACUAUAAACGUGCC
env-1298   AAGAAAACCGAAAUCAAUAGAUAUCGUGCUAUCAUUAAAGAAUUAGGAAUUAGAAAAUAAUUCUUAUUAAAAGAGGCACUAUAAACGUGCC
env-1299   ..........................................................................................
env-1300   AAGAAAAAAGACAUUUCAAGAUAUCGUGCAAUUAUCAAAGAGUUAGGAAUUAGAAAAUAAUACCUACACAAAAGAGGUAGCUUUUGCUACC
env-1301   AAAGAANAAAGAUAUUGCUAAGAUAUCGUGCAAUUAUCAAAA................................................
env-1302   AAGAAGAAAGAUAUUGCAAGAUAUCGUGCAAUUAUCAAAGAGCUAGGAAUUAGAAAAUAAUACCUACAUAAAAGAGGUAGCUAUAUGCUAC
env-1303   ..........................................................................................
env-1304   AAAAGAAAAGAUAUCGAGCGAUAUCGUGCAAUCAUUAAAGAAUUAGGAAUUAGAAAGUAAUUCCAUCAAAAGAGGCUUUCAUUGCCUCUUU
env-1305   AUGAGAAACGAUAUUGUAAAGUAUCGUGAGCUUAUUAAAGAAUUAGGUAUUAGAAAGUAAUAUUGAUAAGGGCUGAAAAUUUUCAGCCCUU
env-1306   AUGAGAAACGAUAUUGUAAAGUAUCGUGAGCUUAUUAAAGAAUUAGGUAUUAGAAAGUAAUAUUGAUAAGGGCUGAAAAUUUUCAGCCCUU
env-1307   ..........................................................................................
env-1308   ..........................................................................................
env-1309   ..........................................................................................
env-1310   AUAAAGAAUGAUAUUGUAAAAUAUCGUGAACUCAUCAAGGAACUAGGUAUUAGAAAGUAAUAUUUUCAGAGGGAAGCUAAGCUUCCCUCUU
env-1311   AUAAAGAAUGAUAUUGUAAAAUAUCGUGAACUCAUCAAGGAACUAGGUAUUAGAAAGUAAUAUUUUCAGAGGGAAGCUAAGCUUCCCUCUU
Tna-1-1    ACUAAGAAAGAUAUUUUAAGAUAUCGUGCCAUAGUUAAAGAAUUAGGUUUAAGAAAAUAAUUAAAAAGAGGCUUUUUAGCCUCUUUUUUGU
env-1312   ..........................................................................................
Cly-1-1    AUCAAAAAAGAUAUUGUACGUUACCGUGCUAUUAUUAAAGAAUUAGGUAUUAGAAAAUAAUACAUAAAGGGAAAGCAAUGCUUUCCCUUUU
Cge-1-1    AUCAAAAAAGAUAUUGUACGUUACCGUGCUAUUAUUAAAGAAUUAGGUAUUAGAAAAUAAUACAUAAAGGGAAAGCAAUGCUUUCCCUUUU
Cly-2-1    AUCAAAAAAGAUAUUGUACGUUACCGUGCUAUUAUUAAAGAAUUAGGUAUUAGAAAAUAAUACAUAAAGGGAAAGCAAUGCUUUCCCUUUU
Sxi-1-1    AUGAAGAAGGAUAUUUUGAGAUAUCGUGCGAUCAUUAAAGAAUUAGGAUUAAGAAAAUAAUUAAAAGGGGCUAUUAUGGCCCCUUUUUUUA
Ste-1-1    AUGAAGAAAGAUAUCUUGAGAUAUCGUGCGAUCGUUAAAGAAUUAGGAUUAAGAAAAUAAUUAAAAAGGGGCUCUUGCGGCCCCUUUUUUU
Cal-1-1    AUUAAGAAAGAUAUUGUAAGAUAUCGUGAAAUAAUUAAAGAAUUAGGUAUUAGAAAAUAAUUUUCGAAGGGGGAAAGCUUAGCUUCCCCCU
env-1313   AAGAAAAAAGAGAUCAACAGAUACCGUGAGAUUAUCAAAGAAUUGAAUAUCAGAAAAUAAUUCAAUAUUUAAAG................
Mze-2-1    AAAAAGAAAGAUAUUAAUAAGUACCGUGCCUUAAUUAAGGAUUUAGGUAUCAGAAAAUAAAUUUUAUUUGGGGAAGCUUUGCUUCCCCUU
env-1314   AAGAAAAAAGAUAUCAACAGAUAUCGUGAGAUUAUUAAAGAAUUGAAUAUUAGAAAAUAAUCAAUAGGAAAGAGGUGCGAAAGUGCCUCUU
env-1316   AAGAAAAAAGAUAUCAACAGAUAUCGUGAGAUUAUUAAAGAAUUGAAUAUU.......................................
env-1317   AAGAAAACAGAAAUCAACAGAUACCGUGAGAUUAUCAAAGAAUUGAAUAUUAGAAAAUAAUCAAUACGAAAAAGAGGUGCUCACGCGCCUC
env-1318   AAAAAGAAAGAUGUUGUAAAAUAUCGUGAAUUAAUUAAAGAAUUAGGUAUUAGAAAAUAAUACAAAUUAUAAGAGGUGCUUUUGCGCCUCU
Fgi-1-1    AAAAAGAAAGAGAUCAACAGAUAUCGUGAGAUUAUCAAAGUAUUGAAUAUCAGAAAAUAAUCAAUAUAAUAAAGAGGUGCGAGAGUGCCUC
Fsu-2-1    AAAAAGAAAGAAAUCAACAGAUACCGUGAGAUUAUCAAAGUAUUGGGUAUUAGAAAAUAAUCAAUAUACAAAAAAGAGGUGCGAAAGUGCC
env-1319   AAGACAGAGAUCAACAGAUAUCGUGAGAUUAUCAAAGUAUUGGGUAUU..........................................
env-1320   ..........................................................................................
env-1321   AAAAAGAAAGAAAUCAACAGGUAUCGUGAGAUUAUCAAAGUAUUGGGUAUUAGAAAAUAAACAAUAUAAAAAAGAGGUGCGUGAGUGCCUC
Fjo-1-1    AAGAAAGAUAUCAACAGAUAUCGUGAGAUUAUCAAAGUAUUGAAUAUCAGAAAAUAAUCAAUAUAGAAAAGAGGUGCGAAAGUGCCUCUUU
Fsp-7-1    AAGAAAGAUAUCAACAGAUAUCGUGAGAUUAUCAAAGUAUUGAAUAUCAGAAAAUAAUCAAUAUAGAAAAGAGGUGCGAGAGUGCCUCUUU
Fsp-13-1   AAGAAAGAGAUCAACAGAUAUCGUGAGAUUAUCAAAGUAUUGAAUAUCAGAAAAUAAUCAAUAUAGAAAAGAGGUGCGAGAGUGCCUCUUU
Fsp-6-1    AAGAAAGAGAUCAACAGAUAUCGUGAGAUUAUCAAAGUAUUGAAUAUCAGAAAAUAAUCAAUAUAAAAAAGAGGUGCGAGAGUGCCUCUUU
Fsp-8-1    AAAAAAGAAAUCAACAGAUAUCGUGAGAUUAUCAAAGUAUUGAAUAUCAGAAAAUAAUCAAUAUAAAAAAGAGGUGCGAGAGUGCCUCUUU

           ..........................................................................................
           ..........................................................................................
           A•RAARAARGAYAUY••••RRUAY•G•G••AU•RU•AAAGAAYURRR••U••GAAAAUAA••••••••••••••••••••••••••••••
```

```
Fsp-10-1   AAAAAAGAAAUCAACAGAUAUCGUGAGAUUAUCAAAGUAUUGAAUAUCAGAAAAUAAUCAAUAUAAAAAAGAGGUGCGAGAGUGCCUCUUU
Fhy-1-1    AAGAAAGAGAUCAACAGAUAUCGUGAGAUUAUCAAAGUAUUGAAUAUCAGAAAAUAAUCAAUAUAAAAAAGAGGUGCGAGAGUGCCUCUUU
env-1323   .........................................................................................
env-1324   AAGAAAGAAAUUAACAGAUAUCGUGAGAUUAUCAAANNNNNNNNNNNNNNNNNNNNNNNNNNNNNNNNNNNNNNNNNNNNNNNNNNNNN
Fsp-11-1   AAGAAAGAGAUCAACAGAUAUCGUGAGAUUAUCAAAGUAUUGAAUAUCAGAAAAUAAUCAAUAUAGAAAAGAGGUGCGAAAGUGCCUCUUU
Fch-1-1    AAGAAAGAGAUCAACAGAUAUCGUGAGAUUAUCAAAGUAUUGAAUAUCAGAAAAUAAUCAAUAUAGAAAAGAGGUGCGAAAGUGCCUCUUU
env-1325   AAGAAAAAAGAAAUCAACAGAUAUCGUGAGAUUAUCAAAGUA...............................................
env-1326   .........................................................................................
env-1327   .........................................................................................
env-1328   AAGAAAAAAGAAAUCAACAGAUAUCGUGAGAUUAUCAAAGUAUUGGGUAUUAGAAAAUAAACAAUAUAAAAAAGAGGUGCGUGAGUGCCUC
env-1330   AAAACAGAAAUCAACCGUUACCGUGAGAUUAUCAAAGAAUUGAAUAUUAGAAAAUAAUCAAUAUAAAAGAGGUGCGAAAGUGCCUCUUUUU
env-1331   AAAACAGAAAUCAACCGUUACCGUGAGAUUAUCAAAGAAUUGAAUAUUAGAAAAUAAUCAAUAUAAAAGAGGUGCGAAAGUGCCUCUUUUU
env-1332   AUUAAAACCGAUAUUGUAAAAUAUCGUGAAUUGAUUAAAGAACUCGGCAUUCGUAAAUAGCACCAAAAUCCAUGAGAGACGUUGCAAAGCG
env-1333   AUUAAAACCGAUAUUGUAAAAUAUCGUGAAUUGAUUAAAGAACUCGGCAUUCGUAAAUAACACCAAAAUCGAGAGACGUUGCAAAGCGUCU
Gsp-4-1    AUGAAAAAAGAUAUCAUGAGAUAUCGUGCUAUUGUUAAGGAAUUAGGAUUGAGAAAAUAAAUUUUAAAGGGGGCUCAUGCCCCCUUUUUUU
Gli-1-1    AUGAAGAAAGAUAUUAUGAGAUAUCGUGCAAUCGUUAAGGAAUUAGGAUUAAGAAAAUAAAUUAUAAAGGGGCUCUAACGCCCCUUUUUUC
env-1334   .........................................................................................
env-1335   .........................................................................................
env-1336   AUGAAAAAAGAUAUCUUGAGAUAUCGUGCUAUCGUUAAAGAAUUACAAUUAAGAAAAUAAAUUUAAGGGGCGUUUUUUCGCCCCUUUUUUA
env-1337   .........................................................................................
env-1338   .........................................................................................
env-1339   AUGAAGAAAGAUAUCUUAAGAUACCGUGCUAUCGUAAAAGAAUUAGGAUUAAGAAAAUAAUUUUCAAGGGAGCUAUAUGCUCCCUUUUUUU
env-1340   .........................................................................................
env-1341   .........................................................................................
Gsp-5-1    AUGAAGAAAGAUGUCUUAAGAUACCGUGCUAUCGUAAAAGAAUUAGGAUUAAGAAAAUAAUUUUCAGGGGAGCUAUAUGCUCCCUUUUUUU
env-1342   .........................................................................................
env-1343   .........................................................................................
env-1344   .........................................................................................
env-1345   .........................................................................................
env-1346   .........................................................................................
Gsp-3-1    AUGAAGAAAGAUAUCUUAAGAUACCGUGCUAUCGUAAAAGAAUUAGGAUUAAGAAAAUAAUUUUCAAGGGAGCUUUAUGCUCCCUUUUUUU
Gsp-2-1    AUGAAGAAAGAUAUCUUAAGAUACCGUGCUAUCGUAAAAGAAUUAGGAUUAAGAAAAUAAUUUUCAAGGGAGCUAUAUGCUCCCUUUUUUU
env-1347   .........................................................................................
env-1348   .........................................................................................
env-1349   AUGAAAAAAGAUAUCUUGAGAUAUCGUGCUAUCGUUAAAGAAUUACAAUUAAGAAAAUAAAUUUAAGGGGCGUAUUUUCGCCCCUUUUUUA
env-1350   AUGAAAAAAGAUAUCUUGAGAUAUCGUGCUAUCGUUAAAGAAUUACAAUUAAGAAAAUAAAUUUAAGGGGGCGUAUUUUCGCCCCUUUUUU
Gec-1-1    AUGAAGAAAGAUAUCAUGAGGUAUCGUGCCAUUGUUAAGGAACUAGGAUUAAGAAAAUAAUUUUCAGGGGGCUUCGUGCCCCCUUUUUUUU
Gpo-1-1    AUGAAGAAGGAUAUCAUGAGAUAUCGUGCGAUCAUCAAGGAACUUGGACUAAGAAAAUAAUUUUCAGGGGGCUUUAUGCCCCCUUUUUUUA
env-1351   AUGAAAAAGGAUAUCAUGAGAUAUCGUGCAAUCAUUAAGGAACUUGGACUAAGAAAAUAAUUUUCAGGGGGCUUUAUGCCCCCUUUUUUUA
env-1352   A........................................................................................
env-1353   .........................................................................................
Gfo-1-1    AUGAAAAAGGAUAUCAUUAGGUAUCGUGCCAUCGUGAAGGAACUAGGAUUAAGAAAAUAAUUUUCAGGGGGCUUUAUGCCCCCUUUUUUUA
env-1354   .........................................................................................
env-1355   AUGAAAAAGGAUAUCAUGAGAUAUCGUGCCAUUAUUAAGGAAUUAGGAUUAAGAAAAUAAUUUUCAGGGGGCUUUUUGCCCCCUUUUUUUA
env-1356   AUGAAAAAGGAUAUCAUGAGAUAUCGUGCAAUCAUUAAGGAAUUAGGAUUAAGAAAAUAAUUUUCAGGGGGCUUUUUGCCCCCUUUUUUUA
env-1357   AUGAAAAAGGAUAUCAUGAGAUAUCGUGCCAUUAUUAAGGAACUAGGAUUAAGAAAAUAAUUUUCAGGGGGCUUUUUGCCCCCUUUUUUUA
env-1358   AUGAAAAAGGAUAUCAUGAGAUAUCGUGCCAUUAUUAAGGAACUAGGAUUAAGAAAAUAAUUUUCAGGGGGCUUUUUGCCCCCUUUUUUUA
env-1359   AUGAAAAAGGAUAUCAUGAGAUAUCGUGCCAUUAUUAAGGAAUUAGGAUUAAGAAAAUAAUUUUCAGGGGGCUUUUUGCCCCCUUUUUUUA
env-1360   AUGAAAAAGGAUAUCAUGAGAUAUCGUGCCAUUAUUAAGGAACUAGGAUUAAGAAAAUAAUUUUCAGGGGGCUUUUUGCCCCCUUUUUUUA
env-1367   AUGAAAAAGGAUAUCAUGAGAUAUCGUGCCAUUAUUAAGGAAUUAGGAUUAAGAAAAUAAUUUUCAGGGGGCUUUUUGCCCCCUUUUUUUA
env-1370   AUGAAAAAGGAUAUCAUGAGAUAUCGUGCCAUUAUUAAGGAAUUAGGAUUAAGAAAAUAAUUUUCAGGGGGCUUUUUGCCCCCUUUUUUUA
env-1371   AUGAAAAAGGAUAUCAUGAGAUAUCGUGCCAUUAUUAAGGAAUUAGGAUUAAGAAAAUAAUUUUCAGGGGGCUUUUUGCCCCCUUUUUUUA
env-1373   AUGAAAAAGGAUAUCAUGAGAUAUCGUGCCAUUAUUAAGGAACUAGGAUUAAGAAAAUAAUUUUCAGGGGGCUUUUUGCCCCCUUUUUUUA
env-1374   AUGAAAAAGGAUAUCAUGAGAUAUCGUGCCAUUAUUAAGGAAUUAGGAUUAAGAAAAUAAUUUUCAGGGGGCUUUUUGCCCCCUUUUUUUA
env-1375   AUGAAAAAGGAUAUCAUGAGAUAUCGUGCCAUUAUUAAGGAAUUAGGAUUAAGAAAAUAAUUUUCAGGGGGCUUUUUGCCCCCUUUUUUUA
env-1376   AUGAAAAAGGAUAUCAUGAGAUAUC................................................................
env-1377   AUGAAAAAGGAUAUCAUGAGAUAUCGUGCCAUUAUUAAGGAAUUAGGAUUAAGAAAAUAAUUUUCAGGGGGCUUUUUGCCCCCUUUUUUUA
env-1378   AUGAAGAAAGAUAUUUUGAGAUAUCGUGCUAUCGUUAAAGAAUUAGGAUUAAGAAAAUAAAUUUCGAAAGGAGGCUUUACGCCCCCUUUUU
env-1379   AUGAAGAAAGAUAUUAUGAGAUAUCGUGCGAUCGUUAAAGAAUUAGGAUUAAGAAAAUAAAUUUCAAAGGGGGCUUUACGCCCCCUUUUUU
env-1380   AUGAAGAAAGAUAUUAUGAGAUAUCGUGCGAUCGUUAAAGAAUUAGGAUUAAGAAAAUAAAUUUCAAGGGGGCUUUACGCCCCCUUUUUUU
env-1381   AUGAAGAAAGAUAUU..........................................................................
env-1382   AUGAAGAAAGAUAUUAUGAGAUAUCGUGCGAUCGUUAAAGAAUUAGGAUUAAGAAAAUAAAUUUCAAAGGGGGCUAUACGCCCCCUUUUUU
env-1383   .........................................................................................
Ssp-1-1    AUGAAGAAAGAUAUUAUGAGAUAUCGUGCGAUCGUUAAAGAGUUAGGAUUAAGAAAAUAAAUUUCAAAGGGGGCUAUACGCCCCCUUUUUU
Tse-1-1    ACGAAGAAAGAUAUUUUAAGAUAUCGUGCUAUAGUUAAAGAAUUAGGUUUAAGAAAAUAAUUUAAAAGAGGCUUCGCGCCUCUUUUUUUAA
Lsp-2-1    ACUAAGAAAGAUGUCUUAAGAUAUCGUGCAAUAGUAAAGGAAUUAGGAUUAAGAAAAUAAUCUCAAUAAAAAGACCACGCCUUCCAGCGUG
Jpa-1-1    ACUAAAAAAGAUAUUUUAAGAUAUCGUGCCAUAGUUAAAGAACUAGGAUUAAGAAAAUAAAAAAACAAAAGAGGCUUGCAUAGGUAAGCCU
Ssp-2-1    AUCAAAAAAGAUAUCUUAAGAUAUCGUGCCAUCGUUAAAGAAUUAGGAUUAAGAAAAUAAUCCAAAUAUAAAGACCACGCCUUUAAGCGUG
env-1384   ACUAAGAAAGAUGUCUUAAGAUAUCGUGCAAUAGUAAAAGAAUUAGGAUUAAGAAAAUAAUCUCAAUAAAAAAACCNCGCUUAACAGCGUG
env-1385   GCGAAAUCAGAUGUAUUAAGGUAUCGUGCGAUAGUUAAGGAAUUAGGAUUAAGAAAAUAAUAAAAAAGAGGCUUCGCGCCUCUUUUUUAGU
env-1386   .........................................................................................
env-1387   .........................................................................................
env-1388   .........................................................................................
env-1389   ACUAAGAAAGAUGUCUUAAGAUAUCGUGCAAUCGUUAAAGAAUUAGGAUUAAGAAAAUAAUCGCAACAUAAAGACCACGCCUUUAGCGUGG
env-1391   .........................................................................................
env-1392   .........................................................................................
env-1393   ACUAAAAAGGAUAUCUUAAGAUACCGUGCUAUCGUUAAAGAAUUAGGAUUAAGAAAAUAAUCUCUGAUAGAAAGACCACGCUUUUAGCGUG
env-1394   .........................................................................................
env-1395   ACUAAAAAGGAUAUCUUAAGAUACCGUGCUAUCGUUAAAGAAUUAGGAUUAAGAAAAUAAUCUCUGAUAGAAAGACCACGCUUUUAGCGUG
env-1396   ACCAAAAAGGAUAUCAUGAGAUAUCGUGCCAUUGUUAAAGAACUUGGAUUAAGAAAAUAAUCAAUAAAGGCCACGCUAAAUACGUGGUCUU
env-1397   ACCAAAAAGGAUAUCAUGAGAUAUCGUGCCAUUGUUAAAGAACUUGGAUUAAGAAAAUAAUCAAUAAAGGCCACGCUAAAUACGUGGUCUU
env-1398   .........................................................................................
env-1399   ACUAAAAAGGAUAUCUUAAGAUACCGUGCAAUAGUAAAAGAAUUAGGAUUAAGAAAAUAAUCUCUAAUAGAAAGACCACGCUUUUAGCGUG
env-1400   .........................................................................................

           .........................................................................................
           .........................................................................................
           A•RAARAARGAYAUY••••RRUAY•G•G••AU•RU•AAAGAAYURRR••U••GAAAAUAA•••••••••••••••••••••••••••••••
```

```
env-1401  ACUAAAAAGGAUAUCUUAAGAUACCGUGCUAUCGUUAAAGAAUUAGGAUUAAGAAAAUAAUCUCUGAUAGAAAGACCACGCUUUUAGCGUG
env-1402  ...........................................................................................
env-1403  ...........................................................................................
env-1404  ACUAAGAAAGAUGUCUUAAGAUAUCGUGCAAUAGUGAAAGAAUUAGGAUUA........................................
env-1405  ACUAAGAAAGAUGUCUUAAGAUAUCGUGCAAUCGUUAAAGAAUUAGGAUUAAGAAAAUAAUCUCAAUAUAAAGACCACGCCUUUAGCGUGG
env-1406  ACUAAGAAAGAUGUCUUAAGAUAUCGUGCAAUCGUUAAAGAAUUAGGAUUAAGAAAAUAAUCUCAAUAUAAAGACCACGCCUUUAGCGUGG
env-1411  ...........................................................................................
Wsp-1-1   ACUAAGAAAGAUAUCUUAAGAUAUCGUGCAAUAGUUAAAGAAUUAGGAUUAAGAAAAUAAUCUCAAUAUAAAGACCACGCAAAACGCGUGG
env-1412  ACUAAAAAAGAUGUCUUGAGAUAUCGUGCAAUAGUUAAAGAAUUAGAAUUAAGAAAAUAAAAAAUGACCACGCUUUUUUAGUGUGGUUUUU
env-1413  ...........................................................................................
env-1414  ACUAAGAAGGAUGUCUUAAGAUAUCGUGCAAUAGUGAAAGAAUUAGGAUUAAGAAAAUAAAAAAAACCACGCUUUUAGUGUGGUUUUUUGU
env-1415  ...........................................................................................
env-1416  ...........................................................................................
Bar-1-1   AUUAAGAAAGAUGUUUUAAGAUAUCGUGCUAUAAUUAAAGAAUUAGGAUUAAGAAAAUAAUCACACUAUAAGAGGCUUCGCGGCCUCUUUU
env-1417  AUUAAGAAAGAUGUCUUAAGAUAUCGUGCAAUUAUUAAGGAAUUAGGAUUAAGAAAAUAAUCACAUUUAAGAGGNUUCGCGCCUCUUUUUU
env-1418  ...........................................................................................
env-1419  ACUAAGAAAGAUAUCUUA.........................................................................
env-1420  ACUAAGAAAGAUAUCUUAAGAUAUCGUGCCAUAGUAAAAGAAUUAGGAUUAAGAAAAUAAUCAAAUUAAGAGGCUUCGCGCCUCUUUUUUU
Pme-1-1   ACUAAGAAAGAUGUCCUAAGAUAUCGUGCCAUAGUUAAAGAAUUAGGAUUAAGAAAAUAAUAUUAAAGAGGCUUCGUGCCUCUUUUUUUUA
env-1421  ACUAAGAAAGAUGUCUUGAGAUAUCGUGCAAUAGUUAAAGAAUUAGGAUUAAGAAAAUAAUCUCAAUACAAAGACCACGCUUUUUAGCGUG
Psp-5-1   ACCAAGAAAGAUGUCUUAAGAUAUCGUGCAAUUGUUAAAGAAUUAGGAUUAAGAAAAUAAAAAAAAGAGGCUUUAUAGCCUCUUUUUGUUU
Pbu-1-1   ACUAAGAAAGAUGUCCUAAGAUAUCGUGCGAUAGUUAAAGAAUUAGGUUUAAGAAAAUAAUCUCAAUACAAAAACCACGCUUAUUAGCGUG
env-1422  ACUAAGAAAGAUGUCUUAAGAUAUCGUGCAAUAGUUAAGGAAUUAGGAUUAAGAAAAUAAUAUUAAAGAGGCUUCGUGCCUCUUUUUUUAU
env-1425  ...........................................................................................
Pja-1-1   ACUAAGAAAGAUGUCUUAAGAUAUCGUGCGAUAGUUAAAGAAUUAGGAUUAAGAAAAUAAUCUCAACAAAAAGACCACGCCUAACAGCGUG
env-1426  ACUAAGAAAGAUGUCUUAAGAUAUCGUGCGAUAGUUAAAGAAUUAGGAUUAAGAAAAUAAUCUCAAUCAAAAGACCACGCCUAACAGCGUG
env-1427  ACUAAGAAGGAUGUCUUAAGAUAUCGUGCGAUAGUUAAAGAAUUAGGAUUAAGAAAAUAAUAUUAAAGAGGCCUUGUGCCUCUUUUUUUUU
env-1428  ...........................................................................................
env-1429  GCAAAGAAAGAUAUCAUGAGAUAUCGUGCAAUUGUCAAGGAAUUAGGACUUAGAAAGUAAUUAAAUUAGAGGCUAUUAUAUGCCUCUUUUU
env-1433  GACAAAGAAAGAUAUCAUGAGAUAUCGUGCAAUUGUCAAGGAAUUAGGACUUAGAAAGUAAUUAAA.........................
env-1434  ACUAAGAAAGAUAUUUUAAGAUAUCGUGCCAUAGUUAAAGAAUUAGGAUUAAGAAAAUAAACAAGAUAAAAAGAGGUUUACAUUUAGUGAA
env-1435  ACUAAGAAAGAUAUUUUAAGAUAUCGUGCCAUAGUUAAAGAAUUAGGAUUAAGAAAAUAAACAAGAUAAAAAGAGGUUUACAUUUAGUGAA
env-1436  ACUAAGAAAGAUGUCUUAAGAUAUCGUGCGAUAGUAAAAGAAUUAGGAUUAAGAAAAUAAAAAAAAGAGGCUUUAUGCCUCUUUUUUUUUA
Oma-1-1   AUUAAAAAAGAUAUCUUAAGAUAUCGUGCAAUUAUUAAAGAAUUAGGAUUAAGAAAAUAAUCAUAAUAAAAAACCACGCUUUUUAGCGUGG
env-1437  AAAGAUCUGACCGAUCAA.........................................................................
env-1438  ACUAAGAAAGAUAUCUUAAGAUAUCGUGCAAUUAUUAAAGAAUUAGGAUUAAGAAAAUAAUCAUAAUAUAAAACCACGCUUUUUAGCGUGG
env-1439  ACUAAGAAAGAUAUCUUAAGAUAUCGUGCAAUUAUUAAAGAAUUAGGAUUAAGAAAAUAAUCAUAAUAUAAAACCACGCUUUUUAGCGUGG
Sze-1-1   ACGAAGAAAGAUAUCUUAAGAUAUCGUGCAAUCGUAAAAGAAUUAGGAUUAAGAAAAUAAUAAAAAGAGGCUUUAUGGCCUCUUUUUUGUU
Gsa-1-1   ACUAAGAAAGAUAUCUUAAGAUAUCGUGCGAUAGUAAAAGAAUUAGGAUUAAGAAAAUAAAAAAAAGAGGCUUUUAUAGCCUCUUUUUUGU
env-1441  ...........................................................................................
env-1442  ACUAAGAAAGAUAUUUUAAGAUAUCGUGCCAUAGUAAAAGAAUUAAAUUUAAGAAAAUAAAUUACAAUAAAAAGAGGCUUUUACAGCUUCU
Mze-1-1   ACUAAAAAAGAUAUCAUGAGAUAUCGUGCAAUAGUGAAAGAAUUAGGAUUAAGAAAAUAAUUAAAAAGAGGCUUCGCGCCUCUUUUUUUCA
env-1444  ACUAAA.....................................................................................
env-1445  ACUAAGAAAGAUGUCUUAAGAUAUCGUGCAAUAGUGAAAGAAUUAGGAUUAAGAAAAUAACAAAAAAGAGGCUUCACGCCUCUUUUUUUCA
env-1447  ACUAAGAAAGAUAUCUUAAGAUAUCGUGCCAUAGUAAAAGAAUUAGAAUUAAGAAAAUAAUUUAAAAAAGAGGCUUUUGUUAGCCUCUUUU
env-1448  ACUAAGAAAG.................................................................................
env-1449  ACUAAGAAAGAUAUCUUAAGAUAUCGUGCCAUAGUAAAAGAAUUAGAAUUAAGAAAAUAAUUUAAAAAAGAGGCUUUUGUUAGCCUCUUUU
env-1454  ...........................................................................................
env-1455  ...........................................................................................
Wps-1-1   ACUAAGAAAGAUGUCUUAAGAUAUCGUGCGAUAGUUAAAGAAUUAGGAUUAAGAAAAUAAUCUCAAUAUAAAGGCCACGCAUUUAGCGUGG
env-1456  ACUAAGAAAGAUGUCUUAAGAUAUCGUGCGAUAGUUAAAGAAUUAGGAUUAAGAAAAUAAUCUCAAUAUAAAGGCCACGCAUUUAGCGUGG
env-1457  ACUAAGAAAGAUGUCUUAAGAUAUCGUGCAAUAGUAAAAGAAUUAGGAUUAAGAAAAUAAUCUCGAUACAAAGACCACGCUUAACGGCGUG
Osp-2-1   ACUAAAAAGGAUAUCUUAAGAUAUCGUGCAAUUGUUAAAGAAUUAGGAUUAAGAAAAUAAUCUAAAUAAAAAGACCACGCUUUUCAGCGUG
Osp-1-1   ACUAAAAAGGAUAUCUUAAGAUAUCGUGCAAUUGUUAAAGAAUUAGGAUUAAGAAAAUAAUCUAAAUAAAAAGACCACGCUUUUCAGCGUG
env-1458  ...........................................................................................
env-1459  ACUAAGAAAGAUAUCUUGAGAUAUCGUGCAAUAGUUAAAGAAUUAGGAUUAAGAAAAUAAUCUCAAUAUAAAAGACCACGCCUAACAGCGU
env-1460  ACUAAGAAAGAUGUCUUAAGAUAUCGUGCAAUAGUGAAAGAAUUAAAUUUAAGAAAAUAAUAAAAAGAGGCUUCGUGCCUCUUUUUUUCUU
env-1461  ACUAAGAAAGAUAUUUUAAGAUAUCGUGCGAUAGUAAAAGAAUUAGGAUUAAGAAAAUAAUCUCAAUAAAAAGACCACGCCUAACAGCGUG
env-1462  ACUAAGAAAGAUAUUUUAAGAUAUCGUGCGAUAGUAAAAGAAUUAGGAUUAAGAAAAUAAUCUCAAUAAAAAGACCACGCCUAACAGCGUG
env-1463  ...........................................................................................
Lsp-1-1   ACUAAAAAGGAUAUCUUAAGAUAUCGUGCGAUAGUAAAAGAAUUAGGAUUAAGAAAAUAAUCUCAAUAAAAAGACCACGCCUAACAGCGUG
env-1464  ACUAAGAAAGAUGUCUUAAGAUAUCGUGCAAUAGUGAAAGAAUUAAAUUUAAGAAAAUAAUAAAAAGAGGCUUCGUGCCUCUUUUUUUCUU
env-1465  ACUAAAAAAGAUGUCUUAAGAUAUCGUGCAAUAGUGAAAGAAUUAAAUUUAAGAAAAUAAUAAAAAGAGGCUUCGUGCCUCUUUUUUUCUU
env-1466  ACUAAGAAAGAUGUCUUAAGAUAUCGUGCAAUAGUGAAAGAAUUAAAUUUAAG......................................
env-1467  ACUAAGAAAGAUGUCUUAAGAUAUCGUGCUAUAGUAAAAGAAUUAGGAUUAAGAAAAUAAUCUUAAUAUAAAGACCACGCCUAACAGCGUG
env-1468  UAAUCGUAAAGAUUAUAACACUGAGCGUUCGUUAGUAAAAUUAGUAGGUAAGCGAAGAGCUUUAUUAGACUACUUAACUAAAAAAGAUGUC
env-1469  ACUAAAAAAGAUGUCUUAAGAUAUCGUGCAAUAGUGAAAGAAUUAAAUUUAAGAAAAUAAUAAAAAGAGGCUUCGUGCCUCUUUUUUUCUU
env-1471  ACUAAAAAGGAUAUCUUAAGAUACCGUGCAAUAGUAAAAGAAUUAGGAUUAAGAAAAUAAUCAGAAAAAGAGGCUUCGCGCCUCUUUUUUU
Fba-5-1   ACUAAGAAAGAUAUCUUAAGAUAUCGUGCCAUAGUGAAAGAAUUAGGAUUAAGAAAAUAAUAAAAAGAGGCUUUUUAGCCUCUUUUUUGUU
Fic-1-1   ACUAAGAAAGAUGUUUUAAGAUAUCGUGCUAUAGUUAAAGAAUUAGGUUUAAGAAAAUAAACUUAAAAAAUAAAAGACCACGCUUUUCGCG
env-1472  ...........................................................................................
Lja-1-1   ACUAAGAAAGAUGUCUUAAGAUAUCGUGCGAUAGUAAAAGAAUUAGGAUUAAGAAAAUAAUCUCAAUGCAAAGACCACGCCUAACAGCGUG
env-1473  ACAAAGAAAGAUGUCUUAAGAUAUCGUGCAAUUAUUAAAGAAUUAGGAUUAAGAAAAUAAUCUCAAUAAAAAGACCACGCUUUUCAGCGUG
Fba-8-1   ACUAAGAAAGAUAUCUUAAGAUAUCGUGCCAUAGUAAAAGAAUUAAAUUUAAGAAAAUAAUAUUUAUAAAAAAGAGGCUAUUUCUAGCCUC
env-1474  ...........................................................................................
env-1475  ...........................................................................................
env-1476  ACGAAAAAAGAUAUCUUAAGAUAUCGUGCAAUCGUAAAAGAAUUAGGGUUAAGAAAAUAACUAAAAGAGGCUUUAUGCCUCUUUUUUUUAA
env-1477  ACGAAAAAAGAUGUCUUAAGAUAUCGUGCUAUCGUAAAAGAAUUAGGAUUAAGAAAAUAAUUAAAAGAGGCUUUAUGCCUCUUUUUUUUUA
env-1478  ACGAAAAAAGAUGUCUUAAGAUAUCGUGCUAUCGUAAAAGAAUUAGGAUUAAGAAAAUAAUUAAAAGAGGCUUUAUGCCUCUUUUUUUUUA
env-1479  ACGAAAAAAGAUGUCUUAAGAUAUCGUGCUAUCGUAAAAGAAUUAGGAUUAAGAAAAUAAUUAAAAGAGGCUUUAUGCCUCUUUUUUUUUA
env-1480  ...........................................................................................
env-1481  ACGAAGAAAGAUGUCUUAAGAUAUCGUGCAAUAGUAAAAGAAUUAGGAUUAAGAAAAUAAAAAAAAGAGGCUUUACGCCUCUUUUUUUUAAU
env-1482  ...........................................................................................
env-1483  ...........................................................................................

          ...........................................................................................
          ...........................................................................................
          A•RAARAARGAYAUY••••RRUAY•G•G••AU•RU•AAAGAAYURR••U••GAAAUAA•••••••••••••••••••••••••••••••••
```

```
env-1484   ACGAAAAAAGAUGUCUUAAGAUAUCGUGCUAUCGUAAAAGAAUUAGGAUUAAGAAAAUAAUUAAAAGAGGCUUUAUGCCUCUUUUUUUUUA
env-1485   ...........................................................................................
env-1486   ACGAAGAAAGAUGUCUUAAGAUAUCGUGCAAUAGUAAAAGAAUUAGGAUUAAGAAAAUAAAAAAAGAGGCUUUACGCCUCUUUUUUUUAAU
env-1487   ...........................................................................................
env-1488   ACGAAGAAAGAUGUCUUAAGAUAUCGUGCAAUAGUAAAAGAAUUAGGAUUAAGAAAAUAAAAAAAGAGGCUUUACGCCUCUUUUUUUUAAU
env-1489   ...........................................................................................
env-1490   ...........................................................................................
env-1491   ACAAAGAAAGAUAUCUUAAGAUAUCGUGCCAUAGUUAAAGAAUUAGGAUUAAGAAAAUAAUUUAAAAAGAGGCUUUUAUUAGCCUCUUUUU
env-1497   ACUAAGAAAGAUAUCUUAAGAUAUCGUGCCAUAGUAAAAGAAUUAGAUUUAAGAAAAUAAUUUAUAAAAAAGAGGCUUUUACUAGCCUCUU
env-1498   ...........................................................................................
env-1499   A..........................................................................................
env-1500   ...........................................................................................
env-1501   ACUAAGAAAGAUGUCUUAAGAUAUCGUGCCAUAGUAAAAGAAUUAGAUUUAAGAAAAUAAAUUAUAAAAAGAGGCUUUUACUAGCCUCUUU
env-1502   ACUAAGAAAGAUGUCUUAAGAUAUCGUGCCAUAGUAAAAGAAUUAGAUUUAAGAAAAUAAAUUAUAAAAAGAGGCUUUUACUAGCCUCUUU
env-1503   ACUAAGAAAGAUGUCUUAAGAUAUCGUGCCAUAGUAAAAGAAUUAGAUUUAAGAAAAUAAUUUAUAAAAAAGAGGCUUUUACUAGCCUCUU
env-1504   ...........................................................................................
env-1505   AUUAAGAAAGAUAUCUUAAGAUAUCGUGCGAUAGUUAAAGAAUUAGGAUUAAGAAAAUAAAAAAAAGAGGCUUUUCAAGCCUCUUUUUUGU
env-1506   AUUAAGAAA..................................................................................
Pda-1-1    ACUAAGAAAGAUAUUUUAAGAUAUCGUGCAAUAGUUAAAGAAUUGAAUUUAAGAAAAUAAUAUUAAAGAGGCUUCGUGCCUCUUUUUUUUA
env-1507   ...........................................................................................
env-1508   ACUAAAAAAGAUAUCUUAAGAUAUCGUGCCAUAGUGAAAGAAUUAGGAUUAAGAAAAUAAUAAAAAGAGGCUUCACGCCUCUUUUUUGUUU
env-1515   ...........................................................................................
env-1516   ACAAAAGUAGAUAUCUUAAGAUAUCGUGCCAUUGUUAAAGAGUUAGGCUUAAGAAAAUAAUACGAAACCACGCCAUUACGCGUGGUUUUUU
env-1517   AUGAAAAAAGACAUCAUGAGGUAUCGAGCCAUCGUAAAAGAGCUUGGACUAAGAAAAUAAAUUUCAGGGGGCUUUUUGCCCCCUUUUUUUA
env-1519   ...........................................................................................
env-1520   ACUAAAAAAGAUAUCUUAAGAUAUCG.................................................................
env-1521   ACUAAAAAAGAUAUCUUAAGAU.....................................................................
env-1522   ACUAAAAAGGAUAUCUUGAGAUAUCGUGCCAUUGUUAAAGAAUUAGGAUUAAGAAAAUAAUCACAUUAUUAAGAGGCUUCGCGCCUCUUUU
env-1524   ACUAAAAAGGAUAUCUUGAGAUAUCGUGCCAUUGUUAAAGAAUUAGGAUUAAGAAAAUAAUCACAUUAUUAAGAGG...............
env-1525   ...........................................................................................
env-1526   ...........................................................................................
env-1527   AUUAAAAACGAUAUUGUUCGAUAUCGUGCAAUUAUUAAAGAAUUAGAUUUAAGAAAAUAAUCUUUUUAAAAGAGGCAAUUAAUGCCUCUUU
env-1528   ...........................................................................................
env-1529   ACUAAAAAAGAUAUCUUAAGAUAUCGUGCCAUAGUAAAAGAAUUAGGAUUAAGAAAAUAAUCAGGAAAAAACCACGCCCGAUAGCGUGGUU
env-1530   ACUAAAAAAGAUAUCUUAAGAUAUCGUGCCAUAGUAAAAGAAUUAGGAUUAAGAAAAUAAUCAGGAAAAAACCACGCCCGAUAGCGUGGUU
1 Fag-1-1  ACUAAGAAAGAUGUCUUAAGAUAUCGUGCUAUUGUUAAAGAAUUAGGAUUAAGAAAAUAAUAAAAGAGAGGCUUUAUGCCUCUUUUUUUUA
env-1534   ...........................................................................................
env-1535   ...........................................................................................
env-1536   ACUCAAAGAAUUGAGCAUUUAACUGCGCAUUUAAAAAACAAUCGUAAAGAUUAUAAUACAGAACGUUCGUUAGUUAUAUUAGUAGGUAAAA
env-1537   ACUAAGAAAGAUGUCUUAAGAUAUCGUGCGAU...........................................................
env-1538   ...........................................................................................
env-1539   ...........................................................................................
env-1540   ACUAAGAAAGAUGUCUUAAGAUAUCGUGCCAUAGUUAAAGAAUUAGAAUUAAGAAAAUAAUCACUAAAAGAGGCUUUACGCCUCUUUUUUG
env-1541   ACUAAGAAAGAUGUCUUAAGAUAUCGUGCCAUAGUUAAAGAAUUAGAAUUAAGAAAAUAAUCACUAAAAGAGGCUUUACGCCUCUUUUUUG
env-1542   ACUAAGAAAGAUGUCCUAAGAUAUCGUGCCAUAGUUAAAGAAUUAGAAUUAAGAAAAUAAUCACUAAAAGAGGCUUUACGCCUCUUUUUUG
env-1543   ACUAAGAAAGAUGUCUUAAGAUAUCGUGCCAUAGUUAAAGAAUUAGAAUUAAGAAAAUAAUCACUAAAAGAGGCUUUACGCCUCUUUUUUG
env-1546   ...........................................................................................
env-1547   ACUAAGAAAGAUGUCUUAAGAUAUCGUGCCAUAGUUAAAGAAUUAGAAUUAAGAAAAUAAUCACAAAAAGAGGCUUUACGCCUCUUUUUUG
env-1548   ACUAAAAAGGAUGUCUUAAGAUAUCGUGCUAUAGUUAAAGAAUUAGGAUUAAGAAAAUAAUCACAAAAAGAGGCUUUACGCCUCUUUUUUG
env-1549   ...........................................................................................
env-1550   ACUAAAAAAGAUAUCUUAAGAUAUCGUGCCAUAGUGAAAGAAUUAGGAUUAAGAAAAUAAAGAAAAGAGGCUUCACAGCCUCUUUUUUGUU
env-1551   ACUAAAAAAGAUAUCUUAA........................................................................
env-1552   ...........................................................................................
env-1553   ACUAAAAAAGAUGUAUUAAGAUAUAGAGUAAUAGUAAAAGAAUUAGGAUUAAGAAAAUAAUAAAAAGAGGCUUCGCAGCCUCUUUUUUGUU
env-1554   ...........................................................................................
env-1555   ACCAAAAAAGAUAUUAUGAGAUAUCGUGCUAUAGUUAAAGAACUUGGUUUAAGAAAAUAAUCAUUACUUUAAGACCACGCUUUAUGCGUGG
env-1556   ...........................................................................................
env-1557   ACCAAAAAAGAUAUCAUGAGAUAUCGUGCUAUAGUUAAAGAACUUGGUUUAAGAAAAUAAUUAGUACUUUAAAACCACGCUUUAUGCGUGG
env-1559   ...........................................................................................
env-1560   ACUAAGAAAGAUGUCUUAAGAUAUCGUGCAAUAGUUAAAGAAUUAGGAUUAAGAAAAUAAAAAAAAGAGGCUUUGCGCCUCUUUUUUUGAA
env-1561   AAAAACAAGGACAUCACUAGAUACAGAGCUAUCAUUGCUGAACUUGGUAUUAGAAAAUAAUAAUUCGGGUAACAGAAGAAGCUUUCCAAUU
env-1562   AAAAACAAGGACAUCACUAGAUACAGAGCUAUCAUUGCUGAACUUGGUAUU........................................
env-1563   ...........................................................................................
env-1564   ...........................................................................................
env-1565   ...........................................................................................
env-1566   AUGAAAAAAGAUAUAGAGCGUUACCGUGCUAUUGUUAAAGAAUUAGGAUUACGUAAGUAAUUUUGAGAGGGCUUCUAAAAGAGGCCCUUUC
Zam-1-1    AAAAAUAAAGAUAUCGAAAGAUAUCGUUCGAUAGUUAAGGAACUAGGAUUAAGAAAGUAAAUUUAAAGAGGCUACGUGCCUCUUUUUUUUA
env-1567   AAGAAAAAAGAUAUCAACAGAUAUCGUGAGAUCAUUAAAGAAUUAGGUAUUAGAAAAUAAUACAUAUUUAAAGAGGUGCCUGCGCGCCUCU
Ffr-2-1    AAGAAAGAAAUCAACAGAUACCGUGAGAUUAUCAAAGUAUUGGGUAUUAGAAAAUAAUCAAUAUAUUAAAGAGGCGCGCAAGUGCCUCUUU
env-1568   AAGAAAGAAAUCAACAGAUAUCGUGAGAUUAUCAAAGUAUUGGGUAUCAGAAAAUAAUCAAUACAAUAAAGAGGCGCGCAAGUGCCUCUUU
Fsp-2-1    AAAACAGAAAUCAACAGAUAUCGUGAGAUUAUCAAAGUAUUGAAUAUCAGAAAAUAAUCAAUAUAAAAGAGGUGCGAAAGUGCCUCUUUUU
env-1571   AAAACAGAAAUCAACAGAUACCGUGAAAUUAUCAAAGUAUUGAAUAUCAGAAAAUAAUCAAUAUAAAAGAGGUGCGAAAGUGCCUCUUUUU
env-1572   ...........................................................................................
env-1573   ...........................................................................................
env-1574   ...........................................................................................
env-1575   AAGAAAGAAAUCAACAGAUAUCGUGAGAUUAUCAAAGUAUUGGGUAUCAGAAAAUAAUCAAUACAUUAAAGAGGCGCGAGAGUGCCUCUUU
env-1576   ...........................................................................................
Fsp-5-1    AAAACAGAAAUCAACAGAUAUCGUGAGAUUAUCAAAGUAUUGAAUAUCAGAAAAUAAUCAAUAUAAAAGAGGUGCGAAAGUGCCUCUUUUU
Fsp-3-1    AAGACAGAGAUCAACAGAUAUCGUGAGAUUAUCAAAGUAUUGAAUAUCAGAAAAUAAUCAAUAUAGAAAAGAGGUGCGAAAGUGCCUCUUU
env-1577   AAGAAAGAAAUCAACAGAUAUCGUGAGAUUAUCAAAGUAUUGGGUAUCAGAAAAUAAUCAAUAUACUAAAGAGGCGCGAGAGUGCCUCUUU
env-1579   ...........................................................................................
Fsp-4-1    AAGAAAGAAAUCAACAGAUAUCGUGAGAUUAUCAAAGUAUUGGGUAUCAGAAAAUAAUCAAUAUAUUAAAGAGGCGCGAGAGUGCCUCUUU
env-1580   ...........................................................................................
env-1581   ...........................................................................................

           ...........................................................................................
           ...........................................................................................
           A•RAARAARGAYAUY••••RRUAY•G•G••AU•RU•AAAGAAYURRR••U••GAAAAUAA•••••••••••••••••••••••••••••••
```

```
Wma-1-1    AAGAAAACGGAAAUUGACCGUUACCGUAAUAUUAUUGCCGAAUUAGGAAUUCGUAAAUAAUAGUAUAUCCAAACAAAAAAAAGGCAAUUUU
Wvi-1-1    AAGAAAACGGAAAUUGAACGUUACCGUAAUAUUAUUGCCGAAUUAGGAAUUCGUAAAUAAUAAUCUAUCCAAAAACAAGAAAAGGCAAUUU
Orh-1-1    AUCAAAAACGACAUCGAAAGAUACCGUGCAAUCAUUAAAGAAUUAGGUAUCAGAAAAUAAUUUUUUUCGCACAAAAUUUUUUGAAAGCAG
env-1582   AAGAAAAAAGAUAUUACCCGCUACAGAGCGAUUAUCGCUGAAUUGGGUCUUAGAAAAUAAGCAAACCAAUUUCGUAAAUAAAACCGCAGCU
env-1583   AAGAAAACUGAAAUUUCUCGUUA....................................................................
env-1584   ...........................................................................................
env-1585   AAGAAAACUGAAAUUUCUCGUUAUAGAGCAAUUAUUGCUGAACUAGGAAUCAGAAAAUAAUCAGAUACAAGAAAAACAAAAAAAGCAACUC
env-1586   AAAAAGAAAGAUAUCACCCGUUACAGAGCAAUCAUUGCGGAAUUAGGAUUAAGAAAAUAAUCUUUCUGUGAAAAAAUAAAGAGGCUGUCCG
env-1587   AAAAAGAAAGAUAUCACCCGUUACAGAGCAAUCAUUGCGGAAUUAGGAUUAAGAAAAUAAUCUUUCUGUGAAAAAAUAAAGAGGCUGUCCG
env-1588   ...........................................................................................
Ebr-1-1    AAGAAAAAAGAUAUCACAAGAUACCGUGCGAUUAUCGCUGAAUUAGGAUUACGUAAGUAAUUUUAAACACAAUAUAAAAGGGCAAUUACAU
env-1589   AAGAAAAAAGAUAUCACAAGAUACCGUGCGAUUAUCGCUGAAUUAGGAUUACGUAAGUAAUUUUAAACACAAUAUAAAAGGGCAAUUACGU
env-1594   ...........................................................................................
Efa-1-1    AAGAAAAAAGAUAUCACAAGAUACCGUGCGAUUAUCGCUGAAUUAGGAUUACGUAAGUAAUUUUAAACACAAUAUAAAAGGGCAAUUACGU
env-1595   ...........................................................................................
Bzo-1-1    AAGAAAAAAGAUAUUGAAAGAUAUAGAGCAAUCAUUGCUGAGCUAGGAUUAAGAAAAUAAUCUAUGCCAUGAUAAUAAUAGAAAGCAGCCC
Bzo-2-1    AAGAAAAAAGAUAUUGAAAGAUAUAGAGCAAUCAUUGCUGAGCUAGGAUUAAGAAAAUAAUCUAUGCCAUGAUAAUAAUAGAAAGCAGCCC
env-1596   AAGAAAAAGGAUAUCAAUAAAUACAGAGAGUUAAUCAAAGAACUCAAUAUCCGAAAAUAACUUAAAAAGCAUCAUGAUUUUCAUUGGUGAC
env-1597   AAGAAAAAGG.................................................................................
env-1598   ...........................................................................................
env-1599   ...........................................................................................
env-1600   ...........................................................................................
env-1601   ...........................................................................................
env-1602   AAGAAAAAGGAUAUCAAUAAAUACAGAGAGUUAAUCAAAGAACUCAACAUCCGAAAAUAACAUAAAAAGCAUCAUAGUUUUUCAUUGGUGA
env-1603   ...........................................................................................
env-1604   AAGAAAAAGG.................................................................................
env-1605   AAGAAAAAGGAUAUCAAUAAAUACAGAGAGUUAAUCAAAGAACUCAACAUCCGAAAAUAACAUAAAAAGCAUCAUGGUUUUUCAUUGGUGA
env-1606   ...........................................................................................
env-1607   AAGAACAAGGAUAUCAAUAAAUACAGAGAGUUAAUCAAAGAACUCAACAUCCGAAAAUAACAUAAAAAGCAUCAUGAUUUUUCAUUGGUGA
env-1608   AAGAAAAAGGAUAUCAAUAAAUACAGAGAGUUAAUCAAAGAGCUCAACAUCCGAAAAUAACAUAAAAAGCAUCAUGGUUUUUCAUUGGUGA
env-1609   AAGAAAAAGGAUAUCAAUAAAUACAGAGAGUUAAUCAAAGAACUCAACAUCCGAAAAUAACAUAAAAAGCAUCAUCGUUUUUCAUUGGUGA
env-1610   AAGAAAAAGGAUAUCAAUAAAUACAGAGAGUUAAUCAAAGAACUCAACAUCCGAAAAUAACAUAAAAAGCAUCAUCGUUUUUCAUUGGUGA
env-1611   AAGAAAAA...................................................................................
env-1612   ...........................................................................................
env-1613   AAGAAAAAGGAUAUCAA..........................................................................
env-1614   ...........................................................................................
env-1615   AAGAAAAAGGAUAUCAAUAAAUACAGAGAGUUAAUCAAAGAACUCAACAUCCGAAAAUAACAUAAAAAGCAUCAUCGUUUUUCAUUGGUGA
env-1616   ...........................................................................................
env-1617   AAGAAAAAGGAUAUCAAUAAAUACAGAGAGUUAAUCAAAGAACUCAACAUCCGAAAAUAACAUAAAAAGCAUCAUCGUUUUUCAUUGGUGA
env-1618   ...........................................................................................
env-1619   AAGAAA.....................................................................................
env-1620   AAGAAAAAGGAUAUCAAUAAAUACAGAGAGUUAAUCAAAGAACUCAACAUCCGAAAAUAACAUAAAAAGCAUCAUCGUUUUUCAUUGGUGA
env-1621   AAGAAAAAGGAUAUCAAUAAAUACAGAGAAUUAAUCAAAGAACUCAACAUCCGAAAAUAGCAUAAAAAGCAUCUCAGUUUUUCAUUGGUGA
env-1622   AAGAAGAAAGAUGUUGUAAAAUAUCGUGAAUUAAUUAAAGAAUUAAACAUCAGAAAAUAAUUCUAAAUAGAGGUGCGUUUGCGCCUCUUUU
env-1623   AAGAAAAAAGAGAUUAACAGAUAUCGUGAGAUCAUCAAAGAAUUGAAUAUUAGAAAAUANUCAAUAUAAAAGAGGUGCCCACGCGCCUCUU
env-1624   ...........................................................................................
env-1625   AAGAAAAAAGAGAUUAACAGAUAUCGUGAGAUCAUCAAAGAAUUGAAUAUUAGAAAAUAGUCAAUAUAAAAGAGGUGCCCACGCGCCUCUU
env-1626   AAGAAAAAAGACAUCAACAGAUAUCGUGAGAUUAUCAAAGAAUUGAAUAUUAGAAAAUAGUCAAUAUAAAAGAGGUGCCCACGCGCCUCUU
env-1627   ...........................................................................................
env-1628   AAAACCGAAAUCAACAGAUAUCGUGAGAUUAUCAAAGUAUUGAAUAUUAGAAAAUAAUCAAUAUAAAAAAGAGGUGCCCACGCGCCUCUUU
env-1629   AAAACAGAAAUCAACAGAUAUCGUGAGAUUAUCAAAGAAUUGAAUAUCAGAAAAUAAUCAAUACUAAAGAGGUGCGCAAGUGCCUCUUUUU
env-1630   ...........................................................................................
env-1631   AAGAAAACUGAAAUCAACAGAUAUCGUGAGA............................................................
env-1632   ...........................................................................................
env-1633   ...........................................................................................
env-1634   AAGAAAAAAGAUGUUGUAAAAUAUCGUGAGUUAAUUAAAGAUUUAGGUAUUAGAAAAUAAUACAUAAAAAGAGGUGCCCACGCGCCUCUUU
env-1635   ...........................................................................................
Faq-1-1    AAAAAGAAAGAUGUUGUAAAAUAUCGUGAGUUAAUUAAAGAAUUAGGUAUUAGAAAGUAAUACAUAAAAAGAGGUGCUCACGCGCCUCUUU
env-1636   AAAAAGAAAGAUGUUGUAAAAUAUCGUGAGUUAAUUAAAGAAUUAGGUAUUAGAAAGUAAUACAUAAAAAGAGGUGCUCACGCGCCUCUUU
env-1637   AAGAAAAAAGAUAUCAACAGAUAUCGUGAAAUUAUUAAAGAAUUAGGUAUUAGAAAAUAAUACAUAAUAUAAAGAGGUGCUUUUGCGCCUC
env-1638   AAGAAUAAAGAUAUCGAAAGAUAUCGUGCCAUAAUUAAGGAAUUAGGUAUCCGAAAAUAAUUUUAACAAUUUUAAGGGGCUUAUUGCCCCU
env-1639   AAGAAUAAAGAUAUCGAAAGAUAUCGUGCCAUAAUUAAGGAAUUAGGUAUCCGAAAAUAAUUUUAACAAUUUUAAGGGGCUUAUUGCCCCU
env-1640   AAGAAUAAAGAUAUCGAAAGAUAUCGUGCCAUAAUUAAGGAAUUAGGUAUCCGAAAAUAAUUUUAACAAUUUUAAGGGGCUUAUUGCCCCU
env-1641   AAGAAUAAAGAUAUCGAAAGAUAUCGUGCCAUAAUUAAGGAAUUAGGUAUCCGAAAAUAAUUUUAACAAUUUUAAGGGGCUUAUUGCCCCU
env-1642   AAGAAUAAAGAUAUCGAAAGAUAUCGUGCCAUAAUUAAGGAAUUAGGUAUCCGAAAAUAAUUUUAACAAUUUUAAGGGGCUUAUUGCCCCU
Csp-6-1    AAGAACACAGAUAUCGAAAGAUACCGUGCGAUUAUCAAAGAGCUAAACAUCAGAAAAUAAGCAAUAGUAAGGAGGCUUUGUGCCUCCUUUU
env-1643   ...........................................................................................
env-1644   ...........................................................................................
env-1645   ...........................................................................................
Coc-2-1    AAAAACACUGAUAUUGAAAGAUAUCGUGCGAUUAUCAAAGAGCUAAACAUCAGAAAAUAAGCAAUACUAAGGAGGCUUUGCGCCUCCUUUU
env-1646   UAAGGAGGCUUUGUGCCUCCUUUUG..................................................................
Csp-9-1    AAGAACACAGAUAUCGAAAGAUACCGUGCGAUUAUCAAAGAGCUAAACAUCAGAAAAUAAGCAAUAGUAAGGAGGCUUUGUGCCUCCUUUU
env-1647   ...........................................................................................
env-1649   ...........................................................................................
env-1650   AAGAACACAGAUAUCGAAAGAUACCGUGCGAUUAUCAAAGAGCUAAACAUCAGAAAAUAAGCAAUAGUAAGGAGGCUUUGUGCCUCCUUUU
env-1651   ...........................................................................................
env-1652   ...........................................................................................
env-1655   AAGGACACAGAUAUCGAAAGAUACCGUGCGAUUAUCAAAGAGCUAAACAUCAGAAAAUAAGCAAUAGUAAGGAGGCUUUGUGCCUCCUUUU
env-1656   ...........................................................................................
env-1657   AAGAACACAGAUAUCGAAAGAUACCGUGCGAUUAUCAAAGAGCUAAACAUCAGAAAAUAAGCAAUAGUAAGGAGGCUUUGUGCCUCCUUUU
Coc-3-1    AAGAACACAGAUAUCGAAAGAUACCGUGCGAUUAUCAAAGAGCUAAACAUCAGAAAAUAAGCAAUAGUAAGGAGGCUUUGUGCCUCCUUUU
Coc-1-1    AAAAACACUGAUAUUGAAAGAUAUCGAGCGAUUAUCAAAGAGCUAAACAUCAGAAAAUAAGCAAUACUAAGGAGGCUUUGUGCCUCCUUUU
Csp-7-1    AAGAACACAGAUAUCGAAAGAUACCGUGCGAUUAUCAAAGAGCUAAACAUCAGAAAAUAAGCAAUAGUAAGGAGGCUUUGUGCCUCCUUUU
Csp-10-1   AAGAACACAGAUAUCGAAAGAUACCGUGCGAUUAUCAAAGAGCUAAACAUCAGAAAAUAAGCAAUAGUAAGGAGGCUUUGUGCCUCCUUUU

           ...........................................................................................
           ...........................................................................................
           A•RAARAARGAYAUY••••RRUAY•G•G••AU•RU•AAAGAAYURRR••U••GAAAAUAA•••••••••••••••••••••••••••••••
```

```
Csp-2-1    AAGAACACAGAUAUCGAAAGAUACCGUGCGAUUAUCAAAGAGCUAAACAUCAGAAAAUAAGCAAUAGUAAGGAGGCUUUGUGCCUCCUUUU
env-1658   ...........................................................................................
env-1659   ...........................................................................................
env-1660   AAGAAAAAAGAAAUCAACAGAUAUCGUGAAAUUAUCAAAGUAUUGGGUAUUAGAAAAUAAUCAAUAUAAAAAAGAGGUGCGAAAGUGCCUC
env-1661   AAGAAAAAAGAAAUCAACAGAUAUCGUGAAAUUAUCAAAGUAUUGGGUAUUAGAAAAUAAUCAAUAUAAAAAAGAGGUGCGAAAGUGCCUC
env-1662   ...........................................................................................
env-1663   AAGAAAAAAGAAAUCAACAGAUAUCGUGAAAUUAUCAAAGUAUUGGGUAUUAGAAAAUAAUCAAUAUAAAAAAGAGGUGCGAAAGUGCCUC
env-1664   AAGAAUACCGAUAUUGAAAGAUAUCGUGCAAUUAUCAAAGAGCUUAACAUUAGAAAAUAAAAAGAGAGGAGGCUUAGUGCCUCCUUUUGCU
env-1665   AAGAAUACCGAUAUUGAAAGAUAUCGUGCAAUUAUCAAAGAGCUUAACAUUAGAAAAUAAAAAGAGAGGAGGCUUAUUGCCUCCUUUUGCU
env-1666   AAGAAUACCGAUAUUGAAAGAUAUCGUGCAAUUAUCAAAGAGCUUAACAUUAGAAAAUAAAAAGAGAGGAGGCUUAUUGCCUCCUUUUGCU
env-1668   ...........................................................................................
env-1669   AAGAAUACCGAUAUUGAAAGAUAUCGUGCAAUUAUCAAAGAGCUUAACAUUAGAAAAUAAAAAGAGAGGAGGCUUAGUGCCUCCUUUUGCU
env-1673   AAGAAUACCGAUAUUGAAAGAUAUCGUGCAAUUAUCAAAGAGCUUAACAUUAGAAAAUAAAAAGAGAGGAGGCUUAGUGCCUCCUUUUGCU
env-1674   AAGAAUACCGAUAUUGAAAGAUAUCGUGCAAUUAUCAAAGAGCUUAACAUUAGAAAAUAAAAAGAGAGGAGGCUUAGUGCCUCCUUUUGCU
env-1676   AAGAAUACCGAUAUUGAAAGAUAUCGUGCAAUUAUCAAAGAGCUUAACAUUAGAAAAUAAAAAGAGAGGAGGCUUAGUGCCUCCUUUUGCU
env-1677   ...........................................................................................
env-1679   AAGAAUACCGAUAUUGAAAGAUAUCGUGCAAUUAUCAAAGAGCUUAACAUUAGAAAAUAAAAAGAGAGGAGGCUUAGUGCCUCCUUUUGCU
env-1680   ...........................................................................................
env-1681   AAGAAUACCGAUAUUGAAAGAUAUCGUGCAAUUAUCAAAGAGCUUAACAUUAGAAAAUAAAAAGAGAGGAGGCUUAGUGCCUCCUUUUGCU
env-1682   ...........................................................................................
env-1683   ...........................................................................................
env-1684   AAGAAUACCGAUAUUGAAAGAUAUCGUGCAAUUAUCAAAGAGCUUAACAUUAGAAAAUAAAAAGAGAGGAGGCUUAGUGCCUCCUUUUGCU
env-1685   ...........................................................................................
env-1686   AAGAAUACCGAUAUUGAAAGAUAUCGUGCAAUUAUCAAAGAGCUUA.............................................
env-1687   AAGAAUACCGAUAUUGAAAGAUAU...................................................................
env-1688   AAGAAUACCGAUAUUGAAAGAUAUCGUGCAAUUAUCAAAGAGCUUAACAUUAGAAAAUAAAAAGAGAGGAGGCUUAGUGCCUCCUUUUGCU
env-1689   AAGAAUACCGAUAUUGAAAGAUAUCGUGCAAUUAUCAAAGAGCUUAACAUUAGAAAAUAAAAAGAGAGGAGGCUUAGUGCCUCCUUUUGCU
env-1691   AAGAAUACCGAUAUUGAAAGAUAUCGUGCAAUUAUCAAAGAGCUUAACAUUAGAAAAUAAAAAGAGAGGAGGCUUAGUGCCUCCUUUUGCU
env-1692   ...........................................................................................
env-1693   AAGAAUACCGAUAUUGAAAGAUAUCGUGCAAUUAUCAAAGAGCUUAACAUUAGAAAAUAAAAAGAGAGGAGGCUUAGUGCCUCCUUUUGCU
env-1694   AAGAAUACCGAUAUUGAAAGAUAUCGUGCAAUUAUCAAAGAGCUUAACAUUAGAAAAUAAAAAGAGAGGAGGCUUAGUGCCUCCUUUUGCU
env-1695   AAGAAUACCGAUAUUGAAAGAUAUCGUGCAAUUAUCAAAGAGCUUAACAUUAGAAAAUAAAAAGAGAGGAGGCUUAGUGCCUCCUUUUGCU
env-1696   AAGAAUACCGAUAUUGAAAGAUAUCGUGCAAUUAUCAAAGAGCUUAACAUUAGAAAAUAAAAAGAGAGGAGGCUUAGUGCCUCCUUUUGCU
env-1697   AAGAAUACCGAUAUUGAAAGAUAUCGUGCAAUUAUCAAAGAGCUUAACAUUAGAAAAUAAAAAGAGAGGAGGCUUAGUGCCUCCUUUUGCU
env-1698   ...........................................................................................
env-1699   AAGAAUACCGAUAUUGAAAGAUAUCGUGCAAUUAUCAAAGAGCUUAACAUUAGAAAAUAAGAAGAGAGGAGGCUUAGUGCCUCCUUUUGCU
env-1700   ...........................................................................................
env-1701   AAGAAUACCGAUAUUGAAAGAUAUCGUGCAAUUAUCAAAGAGCUUAACAUUAGAAAAUAAAAAGAGAGGAGGCUUAGUGCCUCCUUUUGCU
env-1702   AAGAAUACCGAUAUUGAAAGAUAUCGUGCAAUUAUCAAAGAGCUUAACAUUAGAAAAUAAAAAGAGAGGAGGCUUAGUGCCUCCUUUUGCU
env-1703   AAGAAUACCGAUAUUGAAAGAUAUCGUGCAAUUAUCAAAGAGCUUAACAUUAGAAAAUAAAAAGAGAGGAGGCUUAGUGCCUCCUUUUGCU
env-1705   AAGAAUACCGAUAUUGAAAGAUAUCGUGCAAUUAUCAAAGAGCUUAACAUUAGAAAAUAAAAAGAGAGGAGGCUUAGUGCCUCCUUUUGCU
env-1706   AAGAAUACCGAUAUUGAAAGAUAUCGUGCAAUUAUCAAAGAGCUUAACAUUAGAAAAUAAAAAGAGAGGAGGCUUAGUGCCUCCUUUUGCU
env-1707   UUGGUAGCCUUUUUAAACCACACAACACAACUAUUCAAUUAACAAUUUUCACUUAUCAAUUAACAAGUAUUGAAUGAUACCAGAAGUAAAA
env-1708   AAGAAUACCGAUAUUGAAAGAUAUCGUGCAAUUAUCAAAGAGCUUAACAUUAGAAAAUAAAAAGAGAGGAGGCUUAGUGCCUCCUUUUGCU
env-1709   AAGAAUACCGAUAUUGAAAGAUAUCGUGCAAUUAUCAAAGAGCUUAACAUUAGAAAAUAAAAAGAGAGGAGGCUUAGUGCCUC........
env-1710   AAGAAUACCGAUAUUGAAAGAUAUCGUGCAAUUAUCAAAGAGCUUAACAUUAGAAAAUAAAAAGAGAGGAGGCUUAGUGCCUCCUUUUGCU
Cko-1-1    AAGCAAAAAGAUAUCGCACGAUACAGAGCGAUUAUUGCGGAACUUGGUUUAAGAAAAUAAUCUUCCGAAAAAGAAAAUAAGCAACUCAGAA
env-1711   ...........................................................................................
env-1712   ...........................................................................................
env-1713   AAAAACACGGAUAUUGAAAGAUAUCGUGCAAUUA.........................................................
env-1714   AAAAACACGGAUAUUGAAAGAUAUCGUGCAAUUAUCAAAGAGUUAAACAUCAGAAAAUAGACGAAAAAGAGAGGAGGCUUCAGCGCCUCCU
env-1717   AAAAACACGGAUAUUGAAAGAUACCGUGCGAUUAUCAAAGAGUUAAACAUCAGAAAAUAGACGAAAAAGAGAGGAGGCUUCAGCGCCUCCU
env-1718   ...........................................................................................
env-1720   AAAAACACGGAUAUUGAAAGAUACCGUGCAAUUAUCAAAGAGUUAAACAUCAGAAAAUAGACGAAAAAGAGAGGAGGCUUCAGCGCCUCCU
env-1722   AAAAACACGGAUAUUGAAAGAUACCGUGCGAUUAUCAAAGAGUUAAACAUCAGAAAAUAGACGAAAAAGAGAGGAGGCUUCAGCGCCUCCU
Csp-5-1    AAAAACACGGAUAUUGAAAGAUAUCGUGCAAUUAUCAAAGAGUUAAACAUCAGAAAAUAGACGAAAAAGAGAGGAGGCUUCAGCGCCUCCU
env-1723   ...........................................................................................
env-1724   AAGAAAAAAGAAAUCAACAGAUAUCGUGAGAUUAUCAAAGUAUUGGGUAUUAGAAAAUAAACAAUAUAAAAAAGAGGUGCGUGAGUGCCUC
env-1725   AAGAAAAAAGAAAUCAACAGAUAUCGUGAGAUUAUCAAAGUA.................................................
env-1726   AAGAAAAAAGAAAUCAACAGAUAUCGUGAGAUUAUCAAAGUAUUGGGUAUUAGAAAAUAAACAAUAUAAAAAAGAGGUGCGUGAGUGCCUC
env-1727   ...........................................................................................
Csp-8-1    AAAGAAAAAGAUAUUGAAAGAUAUCGUACUAUUAUUAAGGAAUUAGGUAUCCGAAAGUAGUUUUGUUAUUGAACAAGAGGGGCUUUUUGCC
env-1728   AAAGAAAAAGAUAUUGAAAGAUAUCGUACUAUUAUUAAGGAAUUAGGUAUCCG......................................
env-1729   AAAGAAAAAGAUAUUGAGAGAUAUCGUGCUAUUAUUAAGGAAUUAGGUAUCCGAAAAUAAUUUUGUUACACAACAAAAGGGGGCUCUAUGC
env-1730   AAAGAAAAAGAUAUUGAGAGAUAUCGUGCUAUUAUUAAGGAAUUAGGUAUCCGAAAAUAAUUUUGUUACACGACAAAAGGGGGCUCUAUGC
env-1744   AAAGAAAAAGAUAUUGAGAGAUAUCGUGCUAUUAUUAAGGAAUUAGGUAUCCGAAAAUAAUUUUGUUACACAACAAAAGGGGGCUCUAUGC
env-1745   AAAGAAAAAGAUAUUGAGAGAUAUCGUGCUAUUAUUAAGGAAUUAGGUAUCCGAAAAUAAUUUUGUUACACAACAAAAGGGGGCUCUAUGC
env-1746   ...........................................................................................
env-1747   ...........................................................................................
env-1748   AAAGAAAAAGAUAUUGAGAGAUAUCGUGCUAUUAUUAAGGAAUUAGGUAUCCGAAAAUAAUUUUGUUACACGACAAAAGGGGGCUCUAUGC
env-1749   AAAGAAAAAGAUAUUGAGAGAUAUCGUGCUAUUAUUAAGGAAUUAGGUAUCCGAAAAUAAUUUUGUUACACAACAAAAGGGGGCUCUAUGC
env-1750   ...........................................................................................
env-1752   AAAGAAAAAGAUAUUGAGAGAUAUCGUGCUAUUAUUAAGGAAUUAGGUAUCCGAAAAUAAUUUUGUUACACAACAAAAGGGGGCUCUAUGC
env-1753   ...........................................................................................
env-1754   ...........................................................................................
env-1755   ...........................................................................................
Cgi-1-1    AAAGAAAAAGAUAUUGAGAGAUAUCGUGCUAUUAUUAAGGAAUUAGGUAUCCGAAAAUAAUUUUGUUACACAACAAAAGGGGGGCUCUAUG
env-1756   ...........................................................................................
env-1757   ...........................................................................................
env-1765   ...........................................................................................
env-1766   ...........................................................................................
env-1767   AAAGAAAAAGAUAUUGAGAGAUAUCGUGCUAUUAUUA......................................................
env-1768   AAAGAAAAAGAUAUUGAGAGAUAUCGUGCUAUUAUUAAGGAAUUAGGUAUCCGAAAAUAAUUUUGUUACACGACAAA..............
env-1769   ...........................................................................................

           ...........................................................................................
           ...........................................................................................
           A•RAARAARGAYAUY••••RRUAY•G•G••AU•RU•AAAGAAYURRR••U••GAAAAUAA•••••••••••••••••••••••••••••••
```

| | |
|---|---|
| env-1770 | AAAGAAAAAGAUAUUGAGAGAUAUCGUGCUAUUAUUAAGGAAUUAGGUAUCCGAAAAUAAUUUUGUUACAUGACAAAAGGGGGCUCUAUGC |
| env-1771 | AAAGAAAAAGAUAUUGAGAGAUAUCGUGCUAUUAUUAAGGAAUUAGGUAUCCGAAAAUAAUUUUGUUACACAACAAAAGGGGGCUCUAUGC |
| env-1772 | AAAGAAAAAGAUAUUGAGAGAUAUCGUGCUAUUAUUAAGGAAUUAGGUAUCCGAAAAUAAUUUUGUUACACAACAAAAGGGGGGCUCUAUG |
| env-1773 | AAAGAAAAAGAUAUUGAGAGAUAUCGUGCUAUUAUUAAGGAAUUAGGUAUCCGAAAAUAAUUUUGUUACACAACAAAAGGGGGCUCUAUGC |
| env-1774 | AAAGAAAAAGAUAUUGAGAGAUAUCGUGCUAUUAUUAAGGAAUUAGGUAUCCGAAAAUAAUUUUGUUACACGACAAAAGGGGGCUCUAUGC |
| env-1775 | AAAGAAAAAGAUAUUGAGAGAUAUCGUGCUAUUAUUAAGGAAUUAGGUAUCCGAAAAUAAUUUUGUUACACAACAAAAGGGGGCUCUAUGC |
| env-1779 | AAAGAAAAAGAUAUUGAGAGAUAUCGUGCUAUUAUUAAGGAAUUAGGUAUC........................................ |
| env-1780 | AAAGAAAAAGAUAUUGAGAGAUAUCGUGCUAUUAUUAAGGAAUUAGGUAUCCGAAAAUAAUUUUGUUACACAACAAAAGGGGGCUCUAUGC |
| env-1783 | ........................................................................................... |
| env-1786 | AAAGAAAAAGAUAUUGAGAGAUAUCGUGCUAUUAUUAAGGAAUUAGGUAUCCGAAAAUAAUUUUGUUACACAACAAAAGGGGGCUCUAUGC |
| env-1795 | AAAGAAAAAGAUAUUGAGAGAUAUCGUGCUAUUAUUAAGGAAUUAGGUAUCCGAAAAUAAUUUUGUUACACAACAAAAGGGGGCUCUAUGC |
| env-1797 | AAAGAAAAAGAUAUUGAGAGAUAUCGUGCUAUUAUUAAGGAAUUAGGUAUCCGAAAAUAAUUUUGUUACACAACAAAAGGGGGCUCUAUGC |
| env-1798 | AAAGAAAAAGAUAUUGAGAGAUAUCGUGCUAUUAUUAAGGAAUUAGGUAUCCGAAAAUAAUUUUGUUACACAACAAAAGGGGGCUCUAUGC |
| env-1800 | ........................................................................................... |
| env-1801 | ........................................................................................... |
| env-1802 | ........................................................................................... |
| env-1804 | AAAGAAAAAGAUAUUGAGAGAUAUCGUGCUAUUAUUAAGGAAUUAGGUAUCCGAAAAUAAUUUUGUUACACAACAAAAGGGGGCUCUAUGC |
| env-1805 | ........................................................................................... |
| env-1818 | AAAGAAAAAGAUAUUGAGAGAUAUCGUGCUAUUAUUAAGGAAUUAGGUAUCCGAAAAUAAUUUUGUUACACAACAAAAGGGGGCUCUAUGC |
| env-1821 | AAAGAAAAAGAUAUUGAGAGAUAUCGUGCUAUUAUUAAGGAAUUAGGUAUCCGAAAAUAAUUUUGUUACACAACAAAAGGGGGCUCUAUGC |
| env-1823 | AAAGAAAAAGAUAUUGAGAGAUAUCGUGCUAUUAUUAAGGAAUUAGGUAUCCGAAAAUAAUUUUGUUACACAACAAAAGGGGGCUCUAUGC |
| env-1824 | AAAGAAAAAGAUAUUGAGAGAUAUCGUGCUAUUAUUAAGGAAUUAGGUA.......................................... |
| env-1826 | AAAGAAAAAGAUAUUGAGAGAUAUCGUGCUAUUAUUAAGGAAUUAGGUAUCCGAAAAUAAUUUUGUUACACAACAAAAGGGGGCUCUAUGC |
| env-1827 | ........................................................................................... |
| env-1828 | ........................................................................................... |
| env-1829 | AAAAAGAAAGAUAUCAACAGAUAUAGAGCUAUCAUUGCAGAGCUUGGAAUCAGAAAAUAAUAAUUCAGAUAACAUAAAAAAGCGACUCAGU |
| env-1830 | AAAGAAAAAGAUAUUGAGAGAUAUCGUGCUAUUAUUAAGGAAUUAGGUAUCCGAAAAUAAUUUUGUUACACAACAAAAGGGGGCUCUAUGC |
| env-1832 | AAAGAAAAAGAUAUUGAGAGAUAUCGUGCUAUUAUUAAGGAAUUAGGUAUCCGAAAAUAAUUUUGUUACACAACAAAAGGGGGCUCUAUGC |
| env-1833 | ACCGUAAGGACUACAAUACAGAGCGUUCGUUAGUAAGGCUCGUAGGAAAGCGUAGAAGCCUGUUAGACUAUCUAAAAGAAAAAGAUAUUGA |
| env-1834 | AAACAAAAAGAUAUCGAAAGAUACCGUGCUAUUAUUAAGGAAU................................................ |
| env-1835 | AAACAAAAAGAUAUCGAAAGAUACCGUGCUAUUAUUAAGGAAUUAGGUAUCCGAAAAUAAUUUUGUUAUCAAACAAAAGGGGCUUUAUGCC |
| env-1836 | ........................................................................................... |
| env-1837 | ........................................................................................... |
| env-1838 | AAACAAAAAGAUAUCGAAAGAUACCGUGCUAUUAUUAAGGAAUUAGGUAUCCGAAAAUAAUUUUGUUAUCAAACAAAAGGGGCUUUAUGCC |
| env-1839 | AAACAAAAAGAUAUCGAAAGAUACCGUGCUAUUAUUAAGGAAUUAGGUA.......................................... |
| env-1840 | AAACAAAAAGAUAUCGAAAGAUACCGUGCUAUUAUUAAGGAAUUAGGUAUCCGAAAAUAAUUUUGUUAUCAAACAAAAGGGGCUUUAUGCC |
| env-1846 | ........................................................................................... |
| env-1847 | AAACAAAAAGAUAUCGAAAGAUACCGUGCUAUUAUUAAGGAAUUAGGUAUCCGAAAAUAAUUUUGUUAUCAAACAAAAGGGGCUUUAUGCC |
| env-1848 | ........................................................................................... |
| env-1850 | ........................................................................................... |
| env-1851 | AAACAAAAAGAUAUCGAAAGAUACCGUGCUAUUAUUAAGGAAUUAGGUAUCCGAAAAUAAUUUUGUUAUCAAACAAAAGGGGCUUUAUGCC |
| env-1852 | AAACAAAAAGAUAUCGAAAGAUACCGUGCUAUUAUUAAGGAAUUAGGUAUCCGAAAAUAAUUUUGUUAUCAAACAAAAGGGGCUUUAUGCC |
| env-1853 | AAACAAAAAGAUAUCGAAAGAUACCGUGCUAUUAUUAAGGAAUUAGGUA.......................................... |
| env-1854 | ........................................................................................... |
| env-1855 | AAGAACAAGGAUAUCGAGAGAUAUCGUGCUAUUAUUAAGGAAUUGGGUAUCCGAAAAUAAUUUUACUAAUGUGAAAAGAGGGGCUUUUUUG |
| env-1856 | AAGAACAAAGAUAUCGAGAGAUAUCGUGCUAUUAUUAAGGAAUUGGGUAUCCGAAAAUAAUUUUACUAAUGUGAAAAGAGGGGCUUUUUUG |
| env-1857 | AAGAACAAGGAUAUCGAGAGAUAUCGUGCUAUUAUUAAGGAAUUGGGUAUCCGAAAAUAAUUUUACUAAUGUGAAAAGAGGGGCUUUUUUG |
| env-1859 | AAGAACAAGGAUAUCGAGAGAUAUCGUGCUAUUAUUAAGGAAUUGGGUAUCCGAAAAUAAUUUUACUAAUGUGAAAAGAGGGGCUUUUUUG |
| env-1860 | AAGAACAAGGAUAUCGAGAGAUAUCGUGCUAUUAUUAAGGAAUUGGGUAUCCGAAAAUAAUUUUACUAAUGUGAAAAGAGGGGCUUUUUUG |
| env-1862 | ........................................................................................... |
| env-1863 | ........................................................................................... |
| env-1864 | ........................................................................................... |
| env-1865 | ........................................................................................... |
| Eme-2-1 | AAAAAUACUGAAAUUGAGCGUUACAGAGCAAUCAUCGCUGAAUUAGGUAUCAGAAAGUAAGAUAUAAAAGAAUUCCGGAAAGCAACUCGAA |
| Cca-1-1 | AAAAAGAUUGACAUCGAACGAUACAGAGCAAUCGUUAAGAAAUUGAGCUUAAGACGAUAGUUUUGAAUUUCAUACUGAAUUAAUUUUAAGG |
| Emi-1-1 | AAAAAUACUGAAAUUGAGCGUUACAGAGCAAUCAUCGCUGAAUUAGGUAUCAGAAAAUAAGAUACAAGAAAUUCUGUUAAAGCAACUCGAA |
| Cte-1-1 | AAGAAGAAAGAAAUCUCUAGAUACAGAGCUAUCAUCGCUGAAUUGGGAAUCAGAAAAUAAUUUCCGCUUUCAAGCAAAAAAGAAAAGCAAC |
| Csp-15-1 | AAGAAGAAAGAAAUCUCUAGAUACAGAGCUAUCAUCGCUGAAUUGGGAAUCAGAAAAUAAUUUCCGCUUUCAAGCAAAAAACAAAAGCAAC |
| Cha-2-1 | AAGAAGAAAGAAAUCUCUAGAUACAGAGCUAUCAUCGCUGAAUUGGGAAUCAGAAAAUAAUUUCCGCUUUCAAGCAAAAAAUUAAAGCAAC |
| Ean-3-1 | AAAAAUACUGAAAUUGAGCGUUACAGAGCAAUCAUCGCUGAAUUAGGUAUCAGAAAAUAAGAUAUAAGAAAUUCUGUUAAAGCAACUCGAA |
| Ean-1-3 | AAAAAUACUGAAAUUGAGCGUUACAGAGCAAUCAUCGCUGAAUUAGGUAUCAGAAAAUAAGAUACAAGAAAUUCUGUUAAAGCAACUCGAA |
| Csp-18-1 | AAGAAAACCGAAAUUGCACGUUACAGAGCAAUUAUCGCAGAAUUAGGAAUUAGAAAAUAAUUUUCGAUUACAAUUAAAAAAAAGCAACUCA |
| Cje-1-1 | AAGAAAAAAGAUGUACUUCGAUACAGAGCUAUUAUUGCUGAAUUAGGAUUAAGAAAAUAAUCUUUUUCGCAAUAACAAAAAAAGAGCAACU |
| env-1866 | AAAAGUAAAGAUAUCAGCCGUUACAGAGCGAUUAUCGCUGAUUUGGGAUUAAGAAAAUAAUCCGGAUUAAAAAUUACAAAAAAGCUGUUUC |
| Cha-1-1 | AAGAAAAAAGAAAUUGCACGUUACAGAGCAAUUAUCGCUGAAUUAGGAAUCAGAAAAUA................................ |
| env-1869 | ........................................................................................... |
| env-1870 | AAAAGAAAGAUAUCGCAAGAUACAGAGCGAUUAUCGCUGAAUUGGGAUUAA........................................ |
| Chi-1-1 | AAAAAUAAAAAUAUCGAAAGAUACAGAGCAAUUAUUGCUGAAUUAGGUUUAAGAAAGUAAUCUACAAAGAUUUUCAAAAGAGCAACUUCGA |
| env-1871 | AAGAAAAAAGAUAUCAACCGAUACAGAGCGAUUAUCGCAGAACUUGGUUUAAGAAAAUAAUCAUUUUCCGGGAAAAAUUAAAGCAACUCAA |
| Cso-2-1 | AAGAAAAAAGAUAUUACUAGAUAUAGAGCUAUCAUCGCAGAACUAGGAUUGAGAAAAUAAUCUUCCGCGAAGCGAAAAAAAAUAAAAGCAA |
| Cda-1-1 | AAAAACAAAGAUAUCAACAGAUAUAGAGCGAUUAUCGCUGAACUAGGUUUAAGAAAAUAAUCUACAAGAUUUUAAGAUAAAAGCAAUCUCG |
| Cpi-1-1 | AAAAAUAAAGAUAUCGCAAGAUACAGAGCAAUUAUUGCUGAACUAGGUUUAAGAAAAUAAUCUACAAAGAUUUUCAAAUAAAAGCAACUUC |
| env-1872 | AAGAAAAAAGAUAUUACAAGAUACAGAGCGAUUAUCGCUGAACUAGGAUUAAGAAAAUAAUCUAUCCGGUUUCCAAAAUAAAGAAGUAACU |
| Fba-4-1 | AAGAAGAAAGAUAUUGCAAGAUACCGUGCGAUCAUCGCUGAAUUAGGCUUGAGAAAAUAAUCUUCAGGCAAAAAAAAUUAAGCAACCUUUA |
| Cfo-1-1 | AAGAAAAAAGAUAUUACAAGAUACAGAGCAAUUAUUGCUGAACUUGGUCUAAGAAAAUAAUCUUUGGAUUCCAAAAGAAACAGAAGCAACU |
| env-1873 | AAAAAUAAGGAUAUCGCAAGAUACAGAGCAAUUAUUGCUGAAUUAGGUUUAAGAAAAUAAUCUACAAAGAUUUUCACAUAAAAAGCAAUCU |
| Clu-1-1 | AAGAAAAAAGAUAUUACAAGAUACAGAGCUAUCAUCGCUGAACUAGGUUUAAGAAAAUAAUCUACAGGGAUUUUCAAAUAAAAGCGACUUC |
| env-1874 | AAAAACAAAGAUAUCGCAAGAUAUAGAGCAAUUAUUGCUGAAUUAGGAUUAAGAAAAUAAUCUAUAAAGAUUUUCAACAUAAAAGCAACUU |
| Cta-1-1 | AAGAAAAAAGAUAUUACAAGAUACAGAGCUAUUAUCGCUGAACUAGGUUUAAGAAAAUAAUCUGUAAAGAUUUUCAAAUAAAAGCAACUUC |
| Csp-29-1 | AAGAAAAAAGAUAUUACAAGAUACAGAGCUAUUAUCGCUGAACUAGGUUUAAGAAAAUAAUCUGUAAAGAUUUUCAAAUAAAAGCAACUUC |
| Csp-24-1 | AAAAAGAAAGAUAUCGCAAGAUAUAGAGCUAUUAUUGCUGCAUUAGGUUUAAGAAAAUAAUCUAUAAAGAUUUUCAAAAUAAAAAGCAACU |
| Csp-19-1 | AAGAAAAAAGAUAUUACAAGAUACAGAGCUAUCAUUGCUGAACUAGGUUUAAGAAAAUAAUCUGUAAAGAUUUUCAACAAAAAAAGCAACU |
| Cor-1-1 | AAAAAUAAAGAUAUCGCAAGAUACAGAGCGAUCAUCGCUGAACUAGGUUUAAGAAAAUAAUCUACAAAGAUUGUCAAGAUAAAGCAGUUUC |
| Csp-16-1 | AAAAACAAAGAUAUCGCGAGAUAUAGAGCAAUCAUUGCUGAAUUAGGUUUAAGAAAAUAAUCUAUAAAGAUUUUCAAAAUAAAAGCAAUCU |
| | ........................................................................................... |
| | ........................................................................................... |
| | A•RAARAARGAYAUY••••RRUAY•G•G••AU•RU•AAAGAAYURRR••U••GAAAAUAA••••••••••••••••••••••••••••••• |

| | |
|---|---|
| Can-1-1 | AAAAACAAAGAUAUCGCAAGAUAUAGAGCAAUUAUUGCUGAACUAGGUUUAAGAAAAUAAUCUACAAAGAUUUUCAAAUAAAGCAGUUUCG |
| Cgl-1-1 | AAAAAUAAAGAUAUCGCAAGAUACAGAGCGAUCAUCGCUGAACUAGGUUUAAGAAAAUAAUCUAUAAAGAUUUUCAAAAACAAAAAGCAAC |
| Csp-26-1 | AAAAACAAAGAUAUCGCAAGAUAUAGAGCAAUUAUUGCUGAACUAGGUUUAAGAAAAUAAUCUAUAAAGAUUUUCAAAAUAAAAAAGCAAC |
| Csp-23-1 | AAGAAUAAAGAUAUCGCAAGAUAUAGAGCGAUCAUCGCUGAACUAGGUUUAAGAAAAUAAUCUACAAAGAUUUUCAAAAACAAGGCAACUU |
| Cso-1-1 | AAAAAUAAAGAUAUCGCAAGAUAUAGAGCAAUUAUUGCUGAACUAGGUUUAAGAAAAUAAUCUACAAAGAUUUUCAAAUAAGAAGCAGUUU |
| Csp-17-1 | AAAAACAAAGAUAUCGCAAGAUAUAGAGCAAUUAUUGCUGAACUAGGUUUAAGAAAAUAAUCUACAAAGAUUUUCAAAUAAAAGCAACUUC |
| Csp-20-1 | AAAAACAAAGAUAUCGCAAGAUAUAGAGCAAUUAUUGCUGAACUAGGUUUAAGAAAAUAAUCUACAAAGAUUUUCAAAUAAAAGCAACUUC |
| env-1875 | AAAAACAAAGAUAUCGCAAGAUAUAGAGCAAUUAUUGCUGAACUAGGUUUAAGAAAAUAAUCUAUAAAGAUUUUCAAAAUAAAAAGCAACU |
| env-1876 | ........................................................................................... |
| Csp-27-1 | AAAAAGAAAGAUAUCGCAAGAUACAGAGCUAUUAUUGCUGCAUUAGGUUUAAGAAAAUAAUCUAUAAAGAUUUUCAAAAUAAAAAGCAACU |
| Csp-28-1 | AAAAACAAAGAUAUCGCAAGAUAUAGAGCAAUUAUUGCUGAACUAGGUUUAAGAAAAUAAUCUAUAAAGAUUUUCAAAAUAAAAAGCAACU |
| Cin-1-1 | AAGAAAAAAGAUAUCGCAAGAUACAGAGCAAUUAUUGCUGCAUUAGGAUUAAGAAAAUAAUCUAUAAAGAUUUUCAAAAUAAAAGCAACUU |
| Csp-13-1 | AAAAAUAAGGAUAUCGCAAGAUACAGAGCGAUCAUCGCUGAACUAGGUUUAAGAAAAUAAUCUACAAAGAUUGUCAAGAUAAAGCAGUUUC |
| Csp-21-1 | AAAAAUAAAGAUAUCGCCAGAUAUAGAGCGAUCAUCGCUGAACUAGGUUUAAGAAAAUAAUCUAUAAAGAUUUUCAAAAACAAAGCAACUU |
| Csp-25-1 | AAGAAAAAAGAUAUUACAAGAUACAGAGCUAUCAUCGCUGAACUAGGUUUAAGAAAAUAAUCUACAGAGAUUUUCAAAUAAAAGCAACUUC |
| Csp-22-1 | AAGAAAAAAGAUAUCGCAAGAUAUAGAGCAAUUAUUGCUGCAUUAGGACUAAGAAAAUAAUCUAUAAAGAUUUUCAAAAUAAAGCAACUUC |
| Cga-1-1 | AAAAAUAAAGAUAUCGCCAGAUAUAGAGCGAUCAUCGCUGAACUAGGUUUAAGAAAAUAAUCUAUAAAGAUUUUCAAAAACAAAGCAACUU |
| Cvr-1-1 | AAAAACAAAGAUAUCGCAAGAUAUAGAGCAAUUAUUGCUGAACUAGGUUUAAGAAAAUAAUCUACAAAGAUUUUCAAAUAAAAGCAACUUC |
| Csp-14-1 | AAAAAUAAAGAUAUCGCAAGAUAUAGAGCAAUUAUUGCUGAACUAGGUUUAAGAAAAUAAUCUAUAAAGAUUUUCAAAAUAAAAAGCAACU |
| env-1877 | AAAAAUAAGGAUAUCGCAAGAUACAGAGCGAUCAUCGCUGAACUAGGUUUAAG...................................... |
| env-1878 | AAAGAAAAAGAUAUUGAGAGAUAUCGUGCUAUUAUUAAGGAAUUAGGUAUCCGAAAAUAAUUUUGUUACACAACAAAAGGGGGCUCUAUGC |
| env-1879 | ........................................................................................... |
| env-1880 | AAAAAUAAAGAUAUCAAUAGAUAUAGAGCGAUUAUCGCAGAAUUAGGAAUCAGAAAAUAAUUC............................ |
| env-1881 | AAAAAUAAAGAUAUCAAUAGAUAUAGAGCGAUUAUCGCAGAAUUAGGAAUCAGAAAAUAAUUCCAAGUUCAGGUAAAACAAAAAAGCAACU |
| env-1883 | ........................................................................................... |
| env-1884 | AAGAAGAAAGAUAUCAACAGAUAUAGAGCUAUCAUUGCAGAGCUUGGAAUCAGAAAAUAAUAAUUCAGAUAACAUAAAAAAAGCAACUCAGU |
| env-1885 | AAAAAGAAAGAUAUCAACAGAUAUAGAGCUAUCAUUGCAGAGCUUGGAAUCAGAAAAUAAUAAUUCAGAUAACAUAAAAAAAGCGACUCAGU |
| env-1886 | AAGAAGAAAGAUAUCAACAGAUAUAGAGCUAUCAUUGCAGAGCUUGGAAUCAGAAAAUAAUAAUUCAGAUAACAUAAAAAAAGCGACUCAGU |
| env-1892 | AAGAAGAAAGAUAUCAACAGAUAUAGAGCUAUCAUUGCAGAGCUUGGAAUCAGAAAAUAAUAAUUCAGAUAACAUAAAAAAAGCGACUCAGU |
| env-1893 | ........................................................................................... |
| env-1898 | ........................................................................................... |
| env-1900 | AAAAAGAAAGAUAUCAACAGAUAUAGAGCUAUCAUUGCAGAGCUUGGAAUUAGAAAAUAAUAAUUCAGAUAAUAUAAAAAAAGCAACUCAGU |
| env-1901 | AAGAAGAAAGAUAUCAACAGAUAUAGAGCUAUCAUUGCAGAGCUUGGAAU......................................... |
| env-1902 | ........................................................................................... |
| env-1903 | AAGAAGAAAGAUAUUAACAGAUAUAGAGCUAUCAUUGCAGAGCUUGGAAUCAGAAAAUAAUAAUUCAGAUAACAUAAAAAAAGCGACUCAGU |
| env-1904 | AAGAAGAAAGAUAUCAACAGAUAUAGAGCUAUCAUUGCAGAGCUUGGAAUCAGAAAAUAAUAAUUCAGAUAACAUAAAAAAAGCGACUCAGU |
| env-1905 | AAAAAGAAAGAUAUC............................................................................ |
| env-1906 | AAAAAGAAAGAUAUCAACAGAUAUAGAGCUAUCAUUGCAGAGCUUGGAAUCAGAAAAUAAUAAUUCAGAUAA................... |
| env-1907 | AAAAAGAAAGAUAUCAACAGAUAUAGAGCUAUCAUUGCAGAGCUUGGAAUCAGAAAAUAAUAAUUCAGAUAACAUAAAAAAAGCGACUCAGU |
| env-1908 | AAGAAGAAAGAUAUCAACAGAUAUAGAGCUAUCAUUGCAGAGCUUGGAAUCAGAAAAUAAUAAUUCAGAUAACAUAAAAAAAGCGACUCAGU |
| env-1909 | ........................................................................................... |
| env-1910 | AAGAAGAAAGAUAUCAACAGAUAUAGAGCUAUCAUUGCAGAGCUUGGAAUCAGAAAAUAAUAAUUCAGAUAACAUAAAAAAAGCGACUCAGU |
| env-1911 | AAAAAGAAAGAUAUCAACAGAUAUAGAGCUAUCAUUGCAGAGCUUGGAAUCAGAAAAUAAUAAUUCAGAUAACAUAAAAAAAGCGACUCAGU |
| env-1912 | AAGAAGAAAGAUAUCAACAGAUAUAGAGCUAUCAUUGCAGAGCUUGGAAUCAGAAAAU................................. |
| env-1913 | AAAAAGAAAGAUAUCAACAGAUAUAGAGCUAUCAUUGCAGAGCUUGGAAUUAGAAAAUAAUAAUUCAGAUAACAUAAAAAAAGCAACUCAGU |
| env-1915 | AAGAAGAAAGAUAUCAACAGAUAUAGAGCUAUCAUUGCAGAGCUUGGAAUCAGAAAAUAAUAAUUCAGAUAACAUAAAAAAAGCGACUCAGU |
| env-1916 | ........................................................................................... |
| env-1917 | ........................................................................................... |
| env-1918 | AAGAAGAAAGAUAUCAACAGAUAUAGAGCUAUCAUUGCAGAGCUUGGAAUCAGAAAAUAAUAAUUCAGAUAACAUAAAAAAAGCGACUCAGU |
| env-1921 | AAGAAGAAAGAUAUCAACAGAUAUAGAGCUAUCAUUGCAGAGCUUGGAAUCAGAAAAUAAUAAUUCAGAGAACAUAAAAAAAGCGACUCAGU |
| env-1922 | AAAAAGAAAGAUAUCAACAGAUAUAGAGCUAUCAUUGCAGAGCUUGGAAUCAGAAAAUAAUAAUUCAGAUAACAUAAAAAAAGCGACUCAGU |
| env-1924 | AAGAAGAAAGAUAUCAACAGAUAUAGAGCUAUCAUUGCAGAGCUUGGAAUCAGAAAAUAAUAAUUCAGAUAACAUAAAAAAAGCGACUCAGU |
| env-1925 | AA......................................................................................... |
| env-1926 | AAGAAGAAAGAUAUCAACAGAUAUAGAGCUAUCAUUGCAGAGCUUGGAAUCAGAAAAUAAUAAUUCAGAUAACAUAAAAAAAGCGACUCAGU |
| env-1927 | AAAAAGAAAGAUAUCAACAGAUAUAGAGCUAUCAUUGCAGAGCUUGGAAUCAGAAAAUAAUAAUUCAGAUAACAUAAAAAAAGCAACUCAGU |
| env-1928 | AAAAAGAAAGAUAUCAACAGAUAUAGAGCUAUCAUUGCAGAGCUUGGAAUCAGAAAAUAAUAAUUCAGAUAACAUAAAAAAAGCGACUCAGU |
| env-1929 | AAAAAGAAAGAUAUCAACAGAUAUAGAGCUAUCAUUGCAGAGCUUGGAAUCAGAAAAUAAUAAUUCGGAUAACAUAAAAAAAGCGACUCAGU |
| env-1930 | AAGAAGAAAGAUAUCAACAGAUAUAGAGCUAUCAUUGCAGAGCUUGGAAUCAGAAAAUAAUAAUUCAGAUAACAUAAAAAAAGCGACUCAGU |
| env-1931 | ........................................................................................... |
| env-1935 | AAGAAGAAAGAUAUCAACAGAUAUAGAGCUAUCAUUGCAGAGCUUGGAAUCAGAAAAUAAUAAUUCAGAUAACAUAAAAAAAGCGACUCAGU |
| env-1936 | ........................................................................................... |
| env-1937 | ........................................................................................... |
| env-1938 | ........................................................................................... |
| env-1939 | AAAAAGAAAGAUAUCAACAGAUAUAGAGCUAUCAUUGCAGAGCUUGGAAU......................................... |
| env-1940 | CUGGAUUAUCUUAAGAAGAAAGAUAUCAACAGAUAUAGAGCUAUCAUUGCAGAGCUUGGAAUCAGAAAAUAAUAAUUCAGAUAACAUAAAA |
| env-1941 | AAGAAGAAAGAUAUCAACAGAUAUAGAGCUAUCAUUGCAGAGCUUGGAAUCAGAAAAUAAUAAUUCAGAUAACAUA............... |
| env-1942 | AAGAAGAAAGAUAUCAACAGAUAUAGAGCUAUCAUUGCAGAGCUUGGAAUCAGAAAAUAAUAAUUCAGAUAACAUAAAAAAAGCGACUCAGU |
| env-1943 | AA......................................................................................... |
| env-1944 | AAGAAGAAAGAUAUCAACAGAUAUAGAGCUAUCAUUGCAGAGCUUGGAAUCAGAAAAUAAUAAUUCAGAUAACAUAAAAAAAGCAACUCAGU |
| env-1945 | AAAAAGAAAGAUAUCAACAGAUAUAGAGCUAUCAUUGCAGAGCUUGGAAUCAGAAAAUAAUAAUUCAGAUAACAUAAAAAAAGCAACUCAGU |
| env-1946 | ........................................................................................... |
| env-1947 | AAAAAGAAAGAUAUCAACAGAUAUAGAGCUAUCAUUGCAGAGCUUGGAAUCAGAAAAUAAUAAUUCAGAUAACAUAAAAAAAGCGACUCAGU |
| env-1948 | AAAAAGAAAGAUAUCAACAGAUAUAGAGCUAUCAUUGCAGAGCUUGGAAUCAGAAAAUAAUAAUUCAGAUAACAUAAAAAAAGCGACUCAGU |
| env-1949 | ........................................................................................... |
| env-1950 | AAGAAGAAAGAUAUCAACAGAUAUAGAGCUAUCAUUGCAGAGCUUGGAAUCAGAAAAUAAUAAUUCAGAUAACAUAAAAAAAGCGACUCAGU |
| env-1951 | ........................................................................................... |
| env-1952 | AAAAAGAAAGAUAUCAACAGAUAUAGAGCUAUCAUUGCAGAGCUUGGAAUCAGAAAAUAAUAAUUCAGAUAAGAUAAAAAAAGCAACUCAGU |
| env-1954 | ........................................................................................... |
| env-1955 | AAAAAGAAAGAUAUCAACAGAUAUAGAGCUAUCAUUGCAGAGCUUGGAAUUAGAAAAUAAUAAUUCAGAUAACAUAAAAAAAGCAACUCAGU |
| env-1957 | AAAAAGAAAGAUAUCAACAGAUAUAGAGCUAUCAUUGCAGAGCUUGGAAUCAGAAAAUAAUAAUUCAGAUAACAUAAAAAAAGCAACUCAAU |
| env-1958 | AAAAAGAAAGAUAUCAACAGAUAUAGAGCUAUCAUUGCAGAGCUUGGAAUCAGAAAAUAAUAAUUCAGAUAACAUAAAAAAAGCAACUCAGU |
| env-1959 | AAAAAGAAAGAUAUCAACAGAUAUAGAGCUAUCAUUGCAGAGCUUGGAAUCAGAAAAUAAUAAUUCAGAUAACAUAAAAAAGGCAACUCAAU |
| env-1961 | ........................................................................................... |
| env-1962 | AAAAAGAAAGAUAUCAACAGAUAUAGAGCUAUCAUUGCAGAGCUUGGAAUCAGAAAAUAAUAAUUCAGAUAAGAUAAAAAAGCAACUCAGUA |
| | ........................................................................................... |
| | ........................................................................................... |
| | A•RAARAARGAYAUY••••RRUAY•G•G••AU•RU•AAAGAAYURR••U••GAAAAUAA•••••••••••••••••••••••••••••••• |

```
env-1963  AAAAAGAAAGAUAUCAACAGAUAUAGAGCUAUCAUUGCAGAGCUUGGAAUCAGAAAAUAAUAAUUCAGAUAACAUAAAAAAGCAACUCAGU
env-1964  ...........................................................................................
env-1965  AAAAAGAAAGAUAUCAACAGAUAUAGAGCUAUCAUUGCAGAGCUUGGAAUCAGAAAAUAAUAAUUCAGAUAAGAUAAAAAGCAACUCAGUA
env-1966  AAAAAGAAAGAUAUCAACAGAUAUAGAGCUAUCAUUGCAGAGCUUGGAAUCAGAAAAUAAUAAUUCAGAUAACAUAAAAAAGCGACUCAGU
env-1967  ...........................................................................................
env-1968  AAAAAGAAAGAUAUCAACAGAUAUAGAGCUAUCAUUGCAGAGCUUGGAAUCAGAAAAUAAUAAUUCAGAUAACAUAAAAAAGCAACUCAAU
env-1969  AAAAAGAAAGAUAUCAACAGAUAUAGAGCUAUCAUUGCAGAGCUUGGAAUCAGAAAAUAAUAAUUCAGAUAAGAUAAAAAAGCAACUCAGU
env-1970  ...........................................................................................
env-1971  AAAAAGAAAGAUAUCAACAGAUAUAGAGCUAUCAUUGCAGAGCUUGGAAUCAGAAAAUAAUAAUUCAGAUAACAUAAAAAAGCAACUCAGU
env-1972  ...........................................................................................
env-1973  AAAAAGAAAGAUAUCAACAGAUAUAGAGCUAUCAUUGCAGAGCUUGGAAUUAGAAAAUAAUAAUUCAGAUAAUAUAAAAAAGCAACUCAGU
env-1974  AAAAAGAAAGAUAUCAACAGAUAUAGAGCUAUCAUUGCAGAGCUUGGAAU.........................................
env-1975  AAAAAGAAAGAUAUCAACAGAUAUAGAGCUAUCAUUGCAGAGCUUGGAAUCAGAAAAUAAUAAUUCAGAUAACAUAAAAAAGCAAUCAGUA
env-1976  AAAAAGAAAGAUAUCAACAGAUAUAGAGCUAUCAUUGCAGAGCUUGGAAUCAGAAAAUAAUAAUUCAGAUAACAUAAAAAAGCAACUCAGU
env-1977  AAAAAGAAAGAUAUCAACAGAUAUAGAGCUAUCAUUGCAGAGCUUGGAAUCAGAAAAUAAUAAUUCAGAUAAGAUAAAAAGGCAACUCAGU
env-1978  ...........................................................................................
env-1979  AAAAAGAAAGAUAUCAACAGAUAUAGAGCUAUCAUUGCAGAGCUUGGAAUUAGAAAAUAAUAAUUCAGAUAACAUAAAAAAGCAACUCAGU
env-1980  AAAAAGAAAGAUAUCAACAGAUAUAGAGCUAUCAUUGCAGAGCUUGGAAUCAGAAAAUAAUAAUUCAGAUAACAUAAAAAAGCAACUCAGU
env-1981  AAGAAGAAAGAUAUCAACAGAUAUAGAGCUAUCAUUGCAGAGCUUGGAAUUAGAAAAUAAUAAUUCAGAUAACAUAAAAAAGC........
env-1982  AAAAAGAAAGAUAUCAACAGAUAUAGAGCUAUCAUUGCAGAGCUUGGAAUCAGAAAAUAAUAAUUCAGAUAAGAUAAAAAGCAACUCAGUA
env-1983  ...........................................................................................
env-1984  AAAAAGAAAGAUAUCAACAGAUAUAGAGCUAUCAUUGCAGAGCUUGGAAUUAGAAAAUAAUAAUUCAGAUAAGAUAAAAAAGCAACUCAAU
env-1985  AAAAAGAAAGAUAUCAACAGAUAUAGAGCUAUCAUUGCAGAGCUUGGAAUCAGAAAAUAAUAAUUCAGAUAACAUAAAAAAGCAACUCAAU
env-1986  ...........................................................................................
env-1987  AAAAAGAAAGAUAUCAACAGAUAUAGAGCUAUCAUUGCAGAGCUUGGAAUUAGAAAAUAAUAAUUCAGAUAACAUAAAAAAGCAACUCAGU
env-1988  ...........................................................................................
env-1990  AAAAAGAAAGAUAUCAACAGAUAUAGAGCUAUCAUUGCAGAGCUUGGAAUUAGAAAAUAAUAAUUCAGAUAAGAUAAAAAAGCAACUCAAU
env-1991  AAAAAGAAAGAUAUCAACAGAUAUAGAGCUAUCAUUGCAGAGCUUGGAAUUAGAAAAUAAUAAUUCAGAUAAGAUAAAAAAGCAACUCAAU
env-1992  AAAAAGAAAGAUAUCAACAGAUAUAGAGCUAUCAUUGCAGAGCUUGGAAU.........................................
env-1993  AAAAAGAAAGAUAUCAACAGAUAUAGAG...............................................................
env-1994  ...........................................................................................
env-1995  ...........................................................................................
env-1996  AAAAAGAAAGAUAUCAACAGGUAUAGAGCUAUCAUUGCAGAGCUUGGAAUUAGAAAAUAAUAAUUCAGAUAACAUGAAAAAGCAACUCAGU
env-1997  AAAAAGAAAGAUAUCAACAGAUAUAGAGCUAUCAUUGCAGAGCUUGGGAUCAGAAAAUAAUAAUUCAGAUAACAUAAAAAAGCAACUCAGU
env-1998  AAAAAGAAAGAUAUCAACAGAUAUAGAGCUAUCAUUGCAGAGCUUGGAAUCAGAAAAUAAUAAUUCAGAUAACAUAAAAAAGCAACUCAGU
env-1999  ...........................................................................................
env-2000  AAAAAGAAAGAUAUCAACAGAUAUAGAGCUAUCAUUGCAGAGCUUGGAAUCAGAAAAUAAUAAUUCAGAUAACAUAAAAAAGCAACUCAGU
env-2001  ...........................................................................................
env-2002  ...........................................................................................
env-2003  AAAAAGAAAGAUAUCAACAGAUAUAGAGCUAUCAUUGCAGAGCUUGGAAUUAGAAAAUAAUAAUUCAGAUAAGAUAAAAAA..........
env-2004  AAAAAGAAAGAUAUCAACAGAUAUAGAGCUAUCAUUGCAGAGCUUGGAAUUAGAAAAUAAUAAUUCAGAUAACAUAAAAAAGCAACUCAGU
env-2005  AAAAAGAAAGAUAUCAACAGAUAUAGAGCUAUCAUUGCAGAGCUUGGAAUUAGAAAAUAAUAAUUCAGAUAACAUAAAAAAGCAACUCA..
Ran-4-1   AAAAACAAAGAUAUCGAAAGAUAUAGAGCAAUUAUUGCUGAACUAGGAUUAAGAAAAUAAUCAGUCCAAAAAGACAUCAGAAGCAACUCUU
Ran-2-1   AAAAACAAAGAUAUCGAAAGAUAUAGAGCAAUUAUUGCUGAACUAGGAUUAAGAAAAUAAUCAGUCCAAAAAGACAUCAGAAGCAACUCUU
Cca-2-1   AAAGCAAAAGAUAUCGAGAGAUAUCGUGCUAUCAUCAAAGAGCUAAAUAUCAGAAAGUAAUCAAAAGAGGCUUUUCGCCUCUUUUUUAUUU
Cca-2-2   AAAGCAAAAGAUAUCGAGAGAUAUCGUGCUAUCAUCAAAGAGCUAAAUAUCAGAAAGUAAUCAAAAGAGGCUUUUCGCCUCUUUUUUAUUU
env-2006  AAAUCGAAAGACAUUGAAAGAUAUCGUGCAAUCAUUGCCGAACUUGGUCUCCGUAAAUAAGAAAAAUUCAAAUCAAAAUAUAAAGGCAAUU
env-2007  AAAAAUAAGGAUAUUGAAAAAUACCGUUCUUUGAUCAAAGAAUUAGGUAUCAGAAAAUAAUAACAACGGGGAGGCUUUGGCUUCCCCUUUG
Mru-1-1   AAGAAAAAAGAUAUUGAAAAAUACCGUAUUCUAAUUAAAGAAUUAGGUAUCAGAAAAUAAAAACAAAAAGGGGAGGCUUAUGCUUCCCCUU
env-2008  AAGAAAAAAGAUAUUGUAAAAUACCGUGCCUUGAUUAAGGAACUAGGUAUCAGAAAAUAAGAACAAAAAGGGGAGGCAUUGGCUUCCCCUU
env-2009  AAGAAAAAAGAUAUUGUAAAAUACCGUGCCUUGAUUAAGGAACUAGGUAUCAGAAAAUAAGAACAAAAAGGGGAGGCAUUGGCUUCCCCUU
env-2010  ...........................................................................................
env-2011  AAGAAAAAAGAUAUUGAAAAAUACCGUUUCCUAAUUAAGGAAUUAGGUAUCAGAAAAUAAAAACAAAAAGGGGAGGCAUUGGCUUCCCCUU
env-2012  AAGAAAAAAGAUAUUGCAAAAUACCGUACUUUGAUUAAAGAACUAGGUAUUAGAAAAUAAAAACACAAGGGGAGGCAUUGGCUUCCCCUUU
env-2014  AAGAAAAAAGAUAUUGAAAAAUACCGUUCCCUAAUCAAGGAAUUAGGUAUCAGAAAAUAAAAAUACAAGGGGAGGCUCUGGCUUCCCCUUU
env-2015  ...........................................................................................
env-2016  ...........................................................................................
env-2017  AAGAAAAAAGACAUUGCAAAAUAUCGUUCCUUGAUCAAGGAAUUGAAUAUUAGAAAAUAAAAACACAAGGGGAGGCUUAGGCUUCCCCUUU
env-2018  AAGAAAAAAGAUAUUGCAAAAUACCGUAC..............................................................
Msp-2-1   AAGAAAAAAGACAUUGAAAAAUAUCGUUCCUUGAUCAAGGAAUUGAAUAUUAGAAAAUAAAAAUAAAGGGGAGGCUUUGGCUUCCCUUUUG
Cba-2-1   AUUAAGAAAGAUAUUGUAAGAUAUCGUGAAAUAAUUAAAGAAUUAGGUAUUAGAAAAUAAUUUUCGAAAGGGGAAAGCUUAGCUCUCCCCU
Csp-11-1  AUUAAGAAAGAUAUUGUAAGAUAUCGUGAAAUAAUUAAAGAAUUAGGUAUUAGAAAAUAAUUUUCGAAAGGGGAAAGCUUAGCUCUCCCCU
Cba-4-1   AUUAAGAAAGAUAUUGUAAGAUAUCGUGAAAUAAUUAAAGAAUUAGGUAUUAGAAAAUAAUUUUCGAAAGGGGAAAGCUUAGCUCUCCCCU
env-2019  ...........................................................................................
env-2020  AUUAAAAAUGAUAUUGUAAAAUACAGAGAGCUAAUUAAAGAGCUUGGUAUUAGAAAAUAAUAUUGCAAAAAGGGGAUAGUUAACUAUCCCC
Csp-12-1  AUUAAAAAAGAUAUUGUAAGAUAUCGUGCAAUUAUCAAAGAAUUAGGUAUUAGAAAAUAAUUUUUAAAGGGGAAGCUAAGCUUCCCCUUUU
Zul-1-1   AUAAAGAAUGAUAUUGUUAAGUAUCGUGAGCUUAUCAAAGAACUAGGUAUUAGAAAAUAAGUAUUACAAUAAGGGCUGAAAUUUUCGGCCC
env-2021  AUUAAAAAUGACAUUGUAAAAUACAGAGAGCUUAUUAAAGAACUAGGUAUUAGAAAAUAAUUUUUAAAAAGGGGAUAGUUAACUAUCCCCU
env-2027  ...........................................................................................
env-2028  AUUAAAAAUGACAUUGUAAAAUACAGAGAGCUUAUUAAAGAACUAGGUAUUAGAAAAUAAUUUUUAAAAAGGGGAUAGUUAACUAUCCCCU
env-2029  AUAAAAAAAGACAUUGUACGUUACCGUGCUAUUAUUAAAGAAUUAGGUAUUAGAAAAUAAUACAAUCGAGGGGAAAGCUUGCUUUCCCCUU
Ace-1-1   AUUAAAACUGAUAUCGUAAGGUAUCGUGAGAUAAUUAAAGAAUUAGGAAUCAGAAAAUAAAUUUAAAAAGGGGAAACGUUUUACAUUUCCC
Ala-2-1   AUUAAGACUGAUAUCGUUAGAUACCGUGAGAUAAUUAAAGAAUUAGGAAUUAGAAAAUAAAUUGAAAAAGGGGAAACGUUUAACUUUCCCC
env-2030  ...........................................................................................
env-2031  AAAAAGAAAGAUAUUGUAAAAUAUCGUGAGCUUAUCAAGGAACUAAAUAUUAGAAAAUAAGACAAAAAGGGGAAGCUAGGCUUCCCCUUUU
env-2032  AAAAAGAAAGAUAUUGUAAAAUAUCGUGAGCUUAUCAAGGAACUAAAUAUCAGAAAAUAAGACAAAAAGGGGAAGCUAUGCUUCCCCUUUU
env-2033  ...........................................................................................
env-2034  AAAAAGAAAGAUAUUGUAAAAUAUCGUGAGCUUAUCAAGGAACUAAAUAUCAGAAAAUAAGAC............................
env-2035  AAAAAGAAAGAUAUUGUAAAAUAUCGUGAACUUAUCAAGGAACUGAAUAUUAGAAAAUAAGACAAAAAGGGGAAGCUAGGCUUCCCCUUUU
env-2036  AAAAAGAAAGAUAUUGUAAAAUAUCGUGAGCUUAUCAAGGAACUGAAUAUUAGAAAAUAAGACAAAAAGGGGAAGCUAGGCUUCCCCUUUU
env-2037  ...........................................................................................
env-2038  AAAAAGAAAGAUAUUGUAAAAUAUCGUGAACUUAUCAAGGAACUGAAUAUUAGAAAAUAAGACAAAAAGGGGAAGCUAGGCUUCCCCUUUU
env-2039  ...........................................................................................
env-2040  ...........................................................................................

          ...........................................................................................
          ...........................................................................................
          A•RAARAARGAYAUY••••RRUAY•G•G••AU•RU•AAAGAAYURRR••U••GAAAAUAA•••••••••••••••••••••••••••••••
```

```
env-2041  ...........................................................................................
env-2042  ...........................................................................................
env-2043  ...........................................................................................
env-2044  AAAAAGAAAGAUAUUGUAAAAUAUCGUGAGCUUAUCAAGGAACUAAAUAUUAGAAAAUAAGACAAAAAGGGGAAGCUAGGCUUCCCCUUUU
env-2045  ...........................................................................................
env-2046  AAAAAGAAAGAUAUUGUAAAAUAUCGUGAGCUUAUCAAGGAACUAAAUAUUAGAAAAUAAGACAAAAAGGGGAAGCUAGGCUUCCCCUUUU
env-2049  AUUAAAGUAGACGUCUUGAGAUAUCGUGCGAUAAUCAAAGAGCUUAAUAUUCGUAAAUAAUUUAAAAGGGAAGCUUAGGCUUCCCUUUUUU
env-2050  ...........................................................................................
env-2051  AUUAAAGUAG.................................................................................
env-2052  ...........................................................................................
env-2053  AUUAAAGUAGACGUCUUGAGAUAUCGUGCGAUAAUCAAAGAGCUUAAUAUUCGUAAAUAAUUUAAAAGGGAAGCUUAGGCUUCCCUUUUUU
env-2054  UGAU.......................................................................................
env-2055  AUUAAAGUAGACGUCUUGAGAUAUCGUGCGAUAAUCAAAGAGCUUAAUAUUCGUAAAUAAUUUAAAAGGGAAGCUAAGGCUUCCCUUUUUU
Ead-1-1   AAGAAAAAGGACAUUGAAAAGUACCGUAUACUAAUAAAAGAAUUGAACAUUAGAAAAUAAUAUCCGGGGAGGCAUUGCUUCCCCUUUUUCU
env-2068  AAGAAAAAGGAUAUAGAAAAAUAUCGUAUCCUUAUU.......................................................
env-2069  AAGAAAAAGGAUAUAGAAAAAUAUCGUAUCCUUAUUAAAGAAUUGAAUAUUAGAAAAUAAUAUAAGGGGGAGGCAUUGCUUCCCCUUUUUU
env-2070  AAGAAAAAGGAUAUAGAAAAAUAUCGUAUCCUUAUUAAAGAAUUGAAUAUUAGAAAAUAAUAUAAGGGGGAGGCAUUGCUUCCCCUUUUUU
env-2071  AAGAAAAAGGACAUAGAAAAAUAUCGUGCCCUUAUUAAAGUAUUAAAUAUUAGAAAAUAAUAUAAAGGGGAGGCAUUGCUUCCCUUUUUUU
env-2072  AAGAAAAAGGAUAUAGAAAAAUAUCGUAUCCUUAUUAAAGAAUUGAAUAUUAGAAAAUAAUAUAAGGGG......................
env-2073  AUUAAGAAGGAUAUCAUUAGAUAUCGUGCCAUAAUUAAAGAAUUAGGAAUUAGAAAAUAAGUUACCAAAAGGGGAAAGUUUUACUUUCCCC
env-2074  ...........................................................................................
env-2075  AUUAAAAAAGAUAUCAUUAGAUAUCGUGCUAUAAUUAAAGAAUUAGGAAUUAGAAAAUAAUUUACCAAAAGGGGAAAGUGUUACUUUCCCC
env-2076  ...........................................................................................
env-2077  AUUAAAACUGAUAUUGUAAGAUAUCGUGAGAUAAUUAAAGAACUUGGAAUUAGAAAAUAAUUCAUAAAAAGGGGAAAAGUAAAAUUUUCCC
env-2078  ...........................................................................................
env-2079  AUUAAAAAUGAUAUUGUAAAAUACAGAGAGUUGAUUAAGGAGCUUGGUAUUAGAAAAUAAUAUUGUAAAAAGGGGUAGUAAACUACCCCC
Fba-6-1   AAGAAAAAGGAUAUUGUAAAAUAUCGUGAACUUAUCAAGGAGCUUAAUAUUAGAAAAUAAGUAUAAAAGGGGAAGCUUGGCUUCCCCUUUU
env-2080  ...........................................................................................
env-2081  AAGCGCAAGGAUAUUGAAAAAUACCGGGAGCUUAUUAAAGAAUUGAAUAUCAGAAAAUAACACAUAAUAUAAAGGGGAAGGACAUCUUCCC
env-2082  ...........................................................................................
env-2083  AUUAAAAAUGAUAUUGUAAAAUACAGAGAACUUAUCAAAGAACUUGGUAUUAGAAAAUAAUAUUACAAAAAGGGGAUAGCGAGCUAUCCCC
Msp-1-1   AUUAAAAAUGAUAUUGUAAAAUACAGAGAGCUAAUUAAAGAGCUUGGUAUUAGAAAAUAAUAUUGCAAAAAAGGGGAUAGUUAACUAUCCC
env-2085  AUUAAAAAUGAUAUUGUAAAAUACAGAGAGCUUAUUAUAGAGCUUGGUAUUAGAAAAUAAUAUUGCAAAAAAGGGGAUAGUUUACUAUCCC
env-2086  AAAAGAAAGAUAUCGUAAAAUACAGAGAGCUUAUUAAAGAGCUUGGUAUUAGAAAAUAAAUAAAAAA........................
env-2087  AAGAAAAAGGAUAUUGUUAAAUACAGAGAGCUCAUUAAAGAACUCGGUAUCAGGAAAUAAAAACAUUAAAGGGGAUGGUUUUGCCGUCCCC
env-2088  ...........................................................................................
Mth-1-1   AUCAAAAAAGAUAUUGUAAAAUACAGAGAGCUUAUAAAAGAGCUUGGUAUUAGAAAAUAAGAAUUCUAAAGGGGAUAGUUAACUAUCCCCU
env-2089  ...........................................................................................
env-2090  ...........................................................................................
env-2091  ...........................................................................................
env-2092  AUUAAGAAAGAUAUUGUAAAGUAUCGUGAAUUAAUCAAAGAAUUGAAUAUUAGAAAAUAAUUCGAUUGGGGGUCUUAUAUUUAGACCCCCU
Rbi-1-1   AAGAAAAAGGAUAUUGAGAAAUAUCGGAGCCUGAUCAAGGAACUGAAUAUUCGUAAAUAACUUAUUGCAGGGGAGCCCGUCGGGCUCCCCU
Mfo-1-1   AUUAAAAAUGAUAUUGUAAAAUACAGAGAGCUAAUUAAAGAGCUUGGUAUUAGAAAAUAAUAUUGCAAAAGGGGAUAGCUAGCUAUCCCCU
env-2099  ...........................................................................................
env-2100  AAGAAAAAAGAUAUCGUGAGAUAUCGUGAGAUUAUUAAAGAAUUAGGAAUUAGAAAAUAAUCAACUAAAGGGGCUUUGUGCCCCUUUUUGU
Fse-1-1   AAAAAGAAAGAUAUCAACAGAUAUCGUGAAAUUAUCAAAGUAUUGGGUAUCAGAAAAUAGUCAAUAUAAAAAAGAGGUGCGCAAGUACCUC
Fda-1-1   AAAAAGAAAGAUAUCAACAGAUAUCGUGAAAUCAUCAAAGUAUUGGGUAUCAGAAAAUAAUCAAUACAAAAAAGAGGUGCGCAAGUACCUC
env-2102  AUAAAGAAAGAUGUCUUAAGAUAUCGUGCAAUAGUAAAAGAAUUAGGAUUAAGAAAAUAAUCUCAAUACAAAGACCACGCCUAACAGCGUG
env-2103  AAAAAGAAAGAUAUCAACAGAUAUCGUGAGAUUAUCAAGGAAUUAAAUAUCAGAAAAUAAUCAACUUAAAGGGGCUUUGCGCCCCUUUUUG
env-2105  GAGCGUUCAUUGGUGAAAUUGGUAGGUAAAAGAAGAAGCUUACUAGAUUAUCUUAAGAAAACGGAAAUCAACAGAUACCGUGAGAUUAUCA
Fso-1-1   AAGAAAACGGAAAUCAACAGAUACCGUGAGAUUAUCAAAGAAUUGGGUAUUAGAAAAUAAUCCAUACAGAAAAGAGGUGCUCACGCGCCUC
env-2106  GAAAAGAAAGAUAUCAACAGAUAUCGUGAGAUUAUC.......................................................
env-2107  ...........................................................................................
env-2108  ...........................................................................................
env-2109  AAGAAAAAAGAUAUCAAC.........................................................................
env-2110  AAGAAAAAAGAUAUCAACAGAUAUCGUGAGAUUAUCAAAGAAUUGAAUAUUAGAAAAUAAUUCAAUAGAAAAGAGGUGCUCACGCGCCUCU
Aag-1-1   AUCAAAAAGGAUAUCUUAAGAUACCGUGCAAUUGUAAAAGAGCUUGGUUUAAGAAAAUAAGCCAACGAUAUUUAUAUAAAAAAAACGGAGC
env-2111  AUGAAGAAAGAUAUCUUAAGAUAUCGUGCCAUAGUAAAAGAAUUAGGAUUACGUAAGUAAUAAUUGAAUGGGGCUUUAUGCCCCAUUUUAU
env-2112  AUGAAAAAAGAUAUUGUAAGAUACCGUGCAAUCGUUAAAGAAUUAGGAUUACGUAAGUAAUUAUUUUAAAGGGGCUUUUUUAAGCCCCUUU
env-2113  AUGAAAAAAGAUAUUCUAAGAUACCGUGCAAUUGUUAAAGAAUUAGGAUUACGUAAGUAAUUAUUUUUAAGGGGCUUUUUUAGCCCCUUUU
env-2114  ...........................................................................................
env-2115  AUGAAAAAAGAUAUUCUAAGAUACCGUGCAAUUGUUAAAGAAUUAGGAUUACGUAAGUAAUUAUUUUUAAGGGGCUUUUUUAGCCCCUUUU
env-2116  ...........................................................................................
env-2117  AUGAAAAAAGACAUUAUGAGAUAUCGUGCAAUUGUUAAAGAAUUGAGUUUACGUAAGUAAUAGCAUUGGGAGCCUUGUGCUCCCUUUUUUA
env-2118  AUGAAAAAAGAUAUUAUGAGAUAUCGUGCAAUUGUUAAAGAAUUGAAUUUACGUAAGUAAUAGUAUUGGGAGCCUUGUGCUCCCUUUUUAA
env-2119  AUGAAAAAAGAUAUUAUGAGAUAUCGUGCAAUUGUUAAAGAAUUGAAUUUACGUAAGUAAUAGUAUUGGGAGCCUNGAUCGNUCCCUUUUU
env-2120  AUGAAAAAAGAUAUUAUGAGAUAUCGUGCAAUUGUUAAAGAAUUGAGUUUACGUAAGUAAUAGUAUUGGGAGCCUUGUGCUCCCUUUUUUA
env-2121  AUGAAAAAAGAUAUUAUGAGAUAUCGUGCAAUUGUUAAAGAAUUGAGUUUACGUAAGUAAUAGUAUUGGGAGCCUUGUGCUCCCUUUUUUA
env-2124  AUGAAAAAAGAUAUUAUGAGAUAUCGUGCAAUUGUUAAAGAAUUGAGUUUACGUAAGUAAUAGUAUUGGGAGCCUUGUGCUCCCUUUUUUA
env-2125  AUGAAAAAAGAUAUUAUGAGAUAUCGUGCAAUUGUUAAAGAAUUGAGUUUACGUAAGUAAUAGUAUUGGGAGCCUUGUGCUCCCUUUUUUA
env-2126  ...........................................................................................
env-2127  AUGAAAAAAGAUAUUUUGAGAUAUCGUGCAAUUGUUAAAGAUUUAAAUUUACGAAAGUAAUAUAUGGGAGCCUUUUGCUCCCUUUUAUUUU
env-2128  AUGAAAAAAGAUAUUAUGAGAUAUCGUGCAAUAGUUAAAGAAUUGAGUUUACGUAAGUAAUAGUAUUGGGAGCCUUGUGCUCCCUUUUUUA
env-2129  AUGAAAAAAGAUAUUUUAAGAUAUCGUGCAAUAGUUAAAGAAUUACAAUUACGUAAGUAAUUAUCUUAGGGAGCGUAAUUACGCUCCCUUU
env-2130  ...........................................................................................
env-2131  ...........................................................................................
env-2132  CUAAUGAAAAAAGAUAUUUUGAGAUACCGUGCAAUUGUGAAAGAUUUA...........................................
env-2133  AUGAAAAAAGAUAUUUU..........................................................................
env-2134  AUGAAAAAAGAUAUUUUGAGAUACCGUGCAAUUGUGAAAGAUUUAAAUUUACGUAAAUAAUAUAUGGGAGCCUUGUGCUCCCUUUUUAUUU
env-2135  AUUAAAAAAGAUAUUUUGAGAUACCGUGCAAUUGUGAAAGAUUUAAAUUUACGUAAGUAAUAUAUGGGAGCCUUG................
env-2136  AUGAAAAAAGAUAUUUUGAGAUACCGUGCAAUUGUGAAAGAUUUAAA............................................
env-2137  AUGAAAAAAGAUAUUUUGAGAUACCGUGCAAUUGUGAAAGAUUUAAAUUUACGUAAGUAAUAUAUGGGAGCUUUGUGCUCCCUUUUUAUUU
env-2138  ...........................................................................................
env-2139  AUGAAAAAAGAUAUUUUGAGAUACCGUGCAAUUGUGAAAGAUUUAAAUUUACGUAAAUAAUAUAUGGGAGCCUUAGUGCUCCCUUUUUAUU

          ...........................................................................................
          ...........................................................................................
          A•RAARAARGAYAUY••••RRUAY•G•G••AU•RU•AAAGAAYURRR••U••GAAAAUAA••••••••••••••••••••••••••••••••
```

```
env-2140   AUGAAGAAAGAUAUCUUGAGAUAUCGUGCAAUCGUAAAAGAAUUAGGAUUACGUAAGUAACCUAUAUAUAAAGGAGCGCUUAUAGGCGCUC
env-2141   .........................................................................................
env-2142   .........................................................................................
env-2143   ACUAAGAAAGAUAUCUUAAGAUAUCGUGCCAUAGUAAAAGAAUUAGGAUUNAGAAAAUAAAAAAAAGAGGCUUCGCAGCCUCUUUUUUGUU
env-2144   ACUAAAAAAGAUGUCUUAAGAUAUCGUGCCAUAGUUAAAGAAUUAGGAUUAAGAAAAUAACAAAAAAGAGGCUUUUACGCCUCUUUUUAUU
env-2145   AAGAAAAAAGAUAUCAACAGAUAUCGUGAGAUUAUUAAAGAAUUAGGAAUUAGAAAAUAAUCAACUAAAGGGGCUUUGUGCCCCUUUUUGU
env-2146   AAGAAAAAAGAUAUCAACAGAUAUCGUGAGAUUAUUAAAGAAUUAGGAAUUAGAAAAUAAUCAACUAAAGGGG..................
env-2147   AAGAAAAAAGAUAUCAACAGAUAUCGUGAGAUUAUUAAAGAAUUAGGAAUUAGAAAAUAAUCAACUAAAGGGGCUUU..............
env-2148   .........................................................................................
env-2149   .........................................................................................
env-2150   .........................................................................................
env-2151   .........................................................................................
env-2152   AAGAAAAAAGAAAUCAACCGUUACCGUGCGAUUGUUAAAGAACUUGGUUUACGUCGUUAAUCAUAUAUCUUUAUCGAAAUGGGGGGCAAUU
Fps-4-1    AAAAAAGAAAUCAACCGUUACCGUGAGAUUAUCAAAGAAUUGGGUAUUAGAAAAUAAUCAAUACAAAAGAGGUGCGAAAGUGCCUCUUUUU
           .........................................................................................
           .........................................................................................
           A•RAARAARGAYAUY••••RRUAY•G•G••AU•RU•AAAGAAYURR••U••GAAAUAA••••••••••••••••••••••••••••••
```

### 23 S16-Flavobacteria

#### 23.1 Taxa

The taxonomy of each organism containing a putative S16-Flavobacteria RNA is listed, with abbreviations identifying each hit (e.g., "Eco-1-1" and "Eco-1-2" might hypothetically represent two distinct RNAs in *E. coli*). The abbreviations will be used to identify each individual S16-Flavobacteria RNA in Sections 23.2 and 23.4:.

| abbrev. of hits | taxonomy of species |
|---|---|
| Csp-1-1 | Bacteria Bacteroidetes Flavobacteria Flavobacteriales Flavobacteriaceae *Capnocytophaga sputigena* ATCC 33612 |
| Kal-1-1 | Bacteria Bacteroidetes Flavobacteria Flavobacteriales Flavobacteriaceae *Kordia algicida* OT-1 |
| Ale-1-1 | Bacteria Bacteroidetes Flavobacteriia Flavobacteriales Flavobacteriaceae *Algibacter lectus* |
| Cca-1-1 | Bacteria Bacteroidetes Flavobacteriia Flavobacteriales Flavobacteriaceae *Capnocytophaga canimorsus* |
| Cca-2-1 | Bacteria Bacteroidetes Flavobacteriia Flavobacteriales Flavobacteriaceae *Capnocytophaga canis* |
| Cly-1-1 | Bacteria Bacteroidetes Flavobacteriia Flavobacteriales Flavobacteriaceae *Cellulophaga lytica* DSM 7489 |
| Gme-1-1 | Bacteria Bacteroidetes Flavobacteriia Flavobacteriales Flavobacteriaceae *Gelidibacter mesophilus* DSM 14095 |
| Gli-1-1 | Bacteria Bacteroidetes Flavobacteriia Flavobacteriales Flavobacteriaceae *Gillisia limnaea* DSM 15749 |
| Gsp-1-1 | Bacteria Bacteroidetes Flavobacteriia Flavobacteriales Flavobacteriaceae *Gillisia sp.* CBA3202 |
| Gsp-2-1 | Bacteria Bacteroidetes Flavobacteriia Flavobacteriales Flavobacteriaceae *Gillisia sp.* Hel_I_29 |
| Gsp-3-1 | Bacteria Bacteroidetes Flavobacteriia Flavobacteriales Flavobacteriaceae *Gillisia sp.* JM1 |
| Gpo-1-1 | Bacteria Bacteroidetes Flavobacteriia Flavobacteriales Flavobacteriaceae *Gramella portivictoriae* DSM 23547 |
| Jpa-1-1 | Bacteria Bacteroidetes Flavobacteriia Flavobacteriales Flavobacteriaceae *Jejuia pallidilutea* |
| Kje-1-1 | Bacteria Bacteroidetes Flavobacteriia Flavobacteriales Flavobacteriaceae *Kordia jejudonensis* |
| Lsp-1-1 | Bacteria Bacteroidetes Flavobacteriia Flavobacteriales Flavobacteriaceae *Lacinutrix sp.* 5H-3-7-4 |
| Lsp-2-1 | Bacteria Bacteroidetes Flavobacteriia Flavobacteriales Flavobacteriaceae *Lacinutrix sp.* Hel_I_90 |
| Myu-1-1 | Bacteria Bacteroidetes Flavobacteriia Flavobacteriales Flavobacteriaceae *Mangrovimonas yunxiaonensis* |
| Mth-1-1 | Bacteria Bacteroidetes Flavobacteriia Flavobacteriales Flavobacteriaceae *Maribacter thermophilus* |
| Osp-1-1 | Bacteria Bacteroidetes Flavobacteriia Flavobacteriales Flavobacteriaceae *Olleya sp.* VCSA23 |
| Psp-1-1 | Bacteria Bacteroidetes Flavobacteriia Flavobacteriales Flavobacteriaceae *Polaribacter sp.* Hel_I_88 |
| Pto-1-1 | Bacteria Bacteroidetes Flavobacteriia Flavobacteriales Flavobacteriaceae *Psychroflexus torquis* ATCC 700755 |
| Sxi-1-1 | Bacteria Bacteroidetes Flavobacteriia Flavobacteriales Flavobacteriaceae *Salinimicrobium xinjiangense* DSM 19287 |
| Sze-1-1 | Bacteria Bacteroidetes Flavobacteriia Flavobacteriales Flavobacteriaceae *Siansivirga zeaxanthinifaciens* CC-SAMT-1 |
| Tna-1-1 | Bacteria Bacteroidetes Flavobacteriia Flavobacteriales Flavobacteriaceae *Tamlana nanhaiensis* |
| Tme-1-1 | Bacteria Bacteroidetes Flavobacteriia Flavobacteriales Flavobacteriaceae *Tenacibaculum mesophilum* |
| Tsp-1-1 | Bacteria Bacteroidetes Flavobacteriia Flavobacteriales Flavobacteriaceae *Tenacibaculum sp.* 47A_GOM-205m |
| Wsp-1-1 | Bacteria Bacteroidetes Flavobacteriia Flavobacteriales Flavobacteriaceae *Winogradskyella sp.* PG-2 |
| env-1 to env-130 | environmental samples |

#### 23.2 Gene contexts

Each S16-Flavobacteria RNA (indicated by "RNA→") is listed. For each hit, the downstream genes predicted to reside in a regulated operon are listed. If the nearest downstream gene is encoding in the opposite strand (and therefore presumed to not be a part of a regulated operon), then that gene is still depicted. Some environmental sequences and some RefSeq entries lack gene annotations, and so no genes are available for such sequences. The direction of each gene is indicated with an arrow (→), and each predicted conserved domain in the gene is named. Conserved domains associated with more than one S16-Flavobacteria RNA are assigned a color; other domains are gray. Information about these conserved domains is given in Section 23.3. The accession of the sequence containing each S16-Flavobacteria RNA is given in the column named "Seq. accession". Accessions beginning with "NC_", "NS_", "NW_" or "NZ_" are contained in RefSeq. Other accession refer to environmental samples. Nucleotide coordinates are given for the 5′ and 3′ boundaries of each S16-Flavobacteria RNA. If the 5′ coordinate is greater than the 3′ coordinate, the RNA is present on the reverse-complement strand of the containing genomic DNA sequence. Each hit is denoted by an abbreviation (like "Eco-1-1") that refers to a taxonomy given in Section 23.1.

| abbrev. | Seq. accession | | 5′ at | 3′ at | genes |
|---|---|---|---|---|---|
| env-1 | CESM01053608.1 | - | 202 | 170 | **RNA**→ rpsP (PRK14521)Ribosomal_S16 (pfam00886)→ |
| Psp-1-1 | NZ_JHZZ01000001.1 | - | 457084 | 457051 | **RNA**→ RpsP (COG0228)rpsP (PRK00040)→ rimM (PRK00122)PRC (pfam05239)→ hypo→ hypo→ hypo→ |
| env-2 | CESM01219967.1 | + | 5701 | 5735 | **RNA**→ RpsP (COG0228)S16 (TIGR00002)→ |
| env-3 | JGI20154J14316_10001333 | - | 21676 | 21642 | **RNA**→ RpsP (COG0228)rpsP (PRK14521)→ RimM (COG0806)rimM (PRK00122)→ |
| Cca-2-1 | NZ_CDOI01000172.1 | + | 530 | 565 | **RNA**→ RpsP (COG0228)rpsP (PRK14521)→ RimM (COG0806)rimM (PRK00122)→ |
| env-4 | JCVI_SCAF_1096627433698 | + | 434 | 468 | **RNA**→ RpsP (COG0228)rpsP (PRK14521)→ |
| env-5 | KVRMV2_103388609 | - | 189 | 154 | **RNA**→ rpsP (PRK14521)Ribosomal_S16 (pfam00886)→ |
| env-6 | KVRMV2_102516834 | - | 164 | 134 | **RNA**→ rpsP (PRK14521)Ribosomal_S16 (pfam00886)→ |
| env-7 | JGI25321J39296_1021846 | + | 360 | 395 | **RNA**→ rpsP (PRK14521)Ribosomal_S16 (pfam00886)→ rimM (PRK00122)RimM (pfam01782)→ |
| env-8 | JGI12109J20283_11786692 | + | 191 | 225 | **RNA**→ RpsP (COG0228)rpsP (PRK00040)→ |
| env-9 | KGI_S1_ANT01_95m_c10031201 | + | 1995 | 2031 | **RNA**→ RpsP (COG0228)rpsP (PRK14521)→ |
| Mth-1-1 | NZ_LDAS01000001.1 | - | 2478880 | 2478844 | **RNA**→ RpsP (COG0228)rpsP (PRK14521)→ RimM (COG0806)rimM (PRK00122)→ |
| env-10 | TFO_HY__Contig_1945 | + | 10420 | 10454 | **RNA**→ RpsP (COG0228)rpsP (PRK14521)→ RimM (COG0806)rimM (PRK00122)→ PRK13759 (PRK13759)GALNS_like (cd16026)→ |
| env-11 | 2236876002_175824 | + | 203 | 238 | **RNA**→ |
| env-12 | CETA01149051.1 | + | 1248 | 1282 | **RNA**→ RpsP (COG0228)rpsP (PRK14521)→ |
| Cly-1-1 | NC_015167.1 | - | 1900499 | 1900464 | **RNA**→ rpsP (PRK14521)Ribosomal_S16 (pfam00886)→ RimM (COG0806)rimM (PRK00122)→ YjhB (COG1051)PRK05379 (PRK05379)→ PrsA (COG0462)ribP_PPkin (TIGR01251)→ PRK09198 (PRK09198)PBEF_like (cd01569)→ AdoMet_MTases (cd02440)Methyltransf_25 (pfam13649)→ PLN02526 (PLN02526)CaiA (COG1960)→ |
| env-13 | P_1C_Liq_3_UnCty_1040456 | + | 269 | 303 | **RNA**→ |
| env-14 | CEGD01011474.1 | + | 251 | 285 | **RNA**→ RpsP (COG0228)TolA (COG3064)rpsP (PRK14521)→ rimM (PRK00122)RimM (pfam01782)→ |
| env-15 | CEVP01343318.1 | - | 72 | 38 | **RNA**→ |
| env-16 | TDF_OR_ARG04_113m_c1016836 | - | 288 | 254 | **RNA**→ RpsP (COG0228)rpsP (PRK00040)→ |
| env-17 | ElkS_mat_MD6A_1002363 | - | 3313 | 3279 | **RNA**→ RpsP (COG0228)rpsP (PRK00040)→ RimM (COG0806)rimM (PRK00122)→ |
| env-18 | JGI24723J26617_10000107 | - | 52649 | 52615 | **RNA**→ RpsP (COG0228)rpsP (PRK14521)→ |
| env-19 | P_2C_Liq_1_UnCty_1011637 | - | 752 | 718 | **RNA**→ rpsP (PRK00040)Ribosomal_S16 (pfam00886)→ RimM (pfam01782)→ |
| env-20 | JGI11876J14442_10621969 | + | 123 | 157 | **RNA**→ rpsP (PRK14521)Ribosomal_S16 (pfam00886)→ |
| env-21 | P_1C_Liq_1_UnCty_1063218 | - | 143 | 109 | **RNA**→ rpsP (PRK14521)Ribosomal_S16 (pfam00886)→ |
| env-22 | ElkS_mat_CD2A_1075435 | - | 84 | 50 | **RNA**→ rpsP (PRK14521)→ |
| Pto-1-1 | NC_018721.1 | + | 2465369 | 2465403 | **RNA**→ RpsP (COG0228)S16 (TIGR00002)RR_TM4-6 (pfam06459)→ RimM (COG0806)rimM (PRK00122)→ |
| env-23 | JGI1221J11331_1000253 | - | 14608 | 14574 | **RNA**→ RpsP (COG0228)S16 (TIGR00002)→ RimM (COG0806)rimM (PRK00122)→ |
| env-24 | LV_Brine_h2_0102_1093498 | + | 2 | 36 | **RNA**→ RpsP (COG0228)S16 (TIGR00002)→ |
| env-25 | SL_9KL_010_SED_10009602 | + | 426 | 460 | **RNA**→ RpsP (COG0228)rpsP (PRK14521)→ RimM (COG0806)rimM (PRK00122)→ |
| env-26 | KGI_S1_ANT02_95m_c10124961 | + | 605 | 639 | **RNA**→ |
| env-27 | LV_Brine_h2_0102_1076447 | + | 206 | 240 | **RNA**→ RpsP (COG0228)rpsP (PRK00040)→ |
| env-28 | KGI_S1_ANT02_95m_c10037012 | + | 512 | 546 | **RNA**→ RpsP (COG0228)rpsP (PRK14521)→ rimM (PRK00122)RimM (pfam01782)→ |
| env-29 | CESN01195947.1 | - | 1665 | 1631 | **RNA**→ rpsP (PRK00040)Ribosomal_S16 (pfam00886)→ RimM (COG0806)rimM (PRK00122)→ |
| env-30 | 3300001197_1087811 | + | 340 | 374 | **RNA**→ |

| | | | | | |
|---|---|---|---|---|---|
| env-31 | JCVI_SCAF_1096628019256 | - | 205 | 171 | **RNA**→ rpsP (PRK14521)Ribosomal_S16 (pfam00886)→ |
| Cca-1-1 | NZ_CDOE01000079.1 | + | 30020 | 30055 | **RNA**→ RpsP (COG0228)rpsP (PRK14521)→ rimM (PRK00122)RimM (pfam01782)→ |
| env-32 | ElkS_mat_MD6A_1036189 | - | 417 | 383 | **RNA**→ RpsP (COG0228)Pth2 (COG1990)rpsP (PRK00040)→ |
| env-33 | ElkS_mat_MD6A_1084536 | + | 371 | 406 | **RNA**→ |
| env-34 | ElkS_mat_MD6A_1142320 | - | 275 | 241 | **RNA**→ RpsP (COG0228)rpsP (PRK00040)→ |
| env-35 | CEPZ01015035.1 | - | 63972 | 63938 | **RNA**→ RpsP (COG0228)rpsP (PRK14521)→ rimM (PRK00122)RimM (pfam01782)→ hypo→ hypo→ |
| Tsp-1-1 | NZ_JADN01000004.1 | - | 154893 | 154859 | **RNA**→ RpsP (COG0228)rpsP (PRK14521)→ rimM (PRK00122)PRC (pfam05239)→ |
| Tme-1-1 | NZ_LDOD01000012.1 | - | 29532 | 29498 | **RNA**→ RpsP (COG0228)rpsP (PRK14521)→ rimM (PRK00122)RimM (pfam01782)→ |
| env-36 | KVWG_1_1_1_newblercontig40966 | - | 246 | 212 | **RNA**→ |
| env-37 | JGI20156J14371_10166570 | - | 50 | 15 | **RNA**→ |
| env-38 | JGI20154J14316_10055217 | + | 1573 | 1608 | **RNA**→ ←hypo |
| env-39 | JGI20153J14295_10043163 | - | 740 | 705 | **RNA**→ RpsP (COG0228)rpsP (PRK14521)→ |
| env-40 | 2236876009_178640 | + | 118 | 153 | **RNA**→ rpsP (PRK14521)Ribosomal_S16 (pfam00886)→ RpsP (COG0228)rpsP (PRK14521)→ |
| env-41 | SA_S1_NOR05_45m_c10081110 | - | 672 | 638 | **RNA**→ RpsP (COG0228)rpsP (PRK14521)→ |
| env-42 | CESF01293558.1 | + | 708 | 742 | **RNA**→ RpsP (COG0228)rpsP (PRK14521)→ |
| env-43 | CERM01139738.1 | - | 74 | 40 | **RNA**→ |
| env-44 | CEOM01051344.1 | - | 585 | 551 | **RNA**→ rpsP (PRK14521)Ribosomal_S16 (pfam00886)→ |
| env-45 | CESZ01096664.1 | - | 375 | 341 | **RNA**→ rpsP (PRK00040)Ribosomal_S16 (pfam00886)→ |
| env-46 | CERK01075026.1 | + | 203 | 237 | **RNA**→ RpsP (COG0228)rpsP (PRK00040)→ |
| env-47 | CEWG01225384.1 | + | 660 | 694 | **RNA**→ RpsP (COG0228)rpsP (PRK14521)→ RimM (COG0806)rimM (PRK00122)→ |
| | | | | | AdoMet_MTases (cd02440)Methyltransf_25 (pfam13649)→ PLN02526 (PLN02526)CaiA (COG1960)→ |
| | | | | | SGNH_hydrolase_like_1 (cd01832)Lipase_GDSL_2 (pfam13472)→ |
| env-48 | CETK01073817.1 | + | 1419 | 1453 | **RNA**→ RpsP (COG0228)S16 (TIGR00002)→ |
| env-49 | CESO01307771.1 | + | 1268 | 1302 | **RNA**→ rpsP (PRK14521)→ |
| env-50 | JCVI_SCAF_1101668667308 | + | 755 | 789 | **RNA**→ rpsP (PRK00040)Ribosomal_S16 (pfam00886)→ |
| env-51 | JCVI_SCAF_1101667024969 | - | 694 | 660 | **RNA**→ RpsP (COG0228)rpsP (PRK14521)→ |
| env-52 | CEWO01363464.1 | + | 1092 | 1126 | **RNA**→ RpsP (COG0228)S16 (TIGR00002)→ |
| env-53 | JGI24723J26617_10000040 | - | 77894 | 77860 | **RNA**→ RpsP (COG0228)rpsP (PRK14521)→ RimM (COG0806)rimM (PRK00122)→ |
| env-54 | KGI_S1_ANT01_95m_c10031519 | + | 710 | 744 | **RNA**→ RpsP (COG0228)rpsP (PRK14521)→ RimM (COG0806)rimM (PRK00122)→ |
| env-55 | KGI_S2_ANT06_2345m_c1013054 | - | 98 | 64 | **RNA**→ RpsP (COG0228)rpsP (PRK14521)→ |
| env-56 | NCBI_READ_1112493730138 | + | 626 | 660 | **RNA**→ rpsP (PRK14521)Ribosomal_S16 (pfam00886)→ |
| env-57 | CETX01019456.1 | - | 113 | 79 | **RNA**→ |
| env-58 | CEOS01080067.1 | - | 1787 | 1753 | **RNA**→ RpsP (COG0228)rpsP (PRK14521)→ rimM (PRK00122)RimM (pfam01782)PRC (pfam05239)→ |
| env-59 | CEVA01364946.1 | + | 106 | 140 | **RNA**→ rpsP (PRK14521)Ribosomal_S16 (pfam00886)→ |
| env-60 | JGI1221J11331_1019162 | + | 230 | 264 | **RNA**→ rpsP (PRK14521)Ribosomal_S16 (pfam00886)→ |
| | | | | | rimM (PRK00122)RimM (pfam01782)PRC (pfam05239)→ |
| Sxi-1-1 | NZ_AUIG01000033.1 | - | 1535 | 1501 | **RNA**→ rpsP (PRK00040)Ribosomal_S16 (pfam00886)→ |
| | | | | | rimM (PRK00122)RimM (pfam01782)PRC (pfam05239)→ hypo→ hypo→ |
| env-61 | CENM01034188.1 | - | 902 | 868 | **RNA**→ rpsP (PRK14521)Ribosomal_S16 (pfam00886)→ rimM (PRK00122)RimM (pfam01782)→ |
| env-62 | CEOS01020083.1 | - | 32981 | 32947 | **RNA**→ RpsP (COG0228)rpsP (PRK14521)→ rimM (PRK00122)RimM (pfam01782)→ |
| | | | | | AdoMet_MTases (cd02440)Methyltransf_25 (pfam13649)→ PLN02526 (PLN02526)CaiA (COG1960)→ |
| env-63 | JGI20156J14371_10003640 | + | 2978 | 3012 | **RNA**→ RpsP (COG0228)rpsP (PRK00040)→ RimM (COG0806)rimM (PRK00122)→ |
| | | | | | AdoMet_MTases (cd02440)Methyltransf_25 (pfam13649)→ |
| env-64 | DelMOSum2010_c10016191 | + | 3980 | 4015 | **RNA**→ RpsP (COG0228)rpsP (PRK00040)→ |
| env-65 | KVRMV2_100000226 | + | 6455 | 6489 | **RNA**→ RpsP (COG0228)rpsP (PRK00040)→ RimM (COG0806)rimM (PRK00122)→ |
| env-66 | JGI12037J13215_1008219 | + | 266 | 300 | **RNA**→ RpsP (COG0228)rpsP (PRK14521)→ |
| env-67 | CGUI_GFCP3IQ02IOEQO | - | 70 | 36 | **RNA**→ |
| env-68 | JGI20160J14292_10003172 | - | 10084 | 10050 | **RNA**→ RpsP (COG0228)rpsP (PRK14521)→ RimM (COG0806)rimM (PRK00122)→ |
| | | | | | AdoMet_MTases (cd02440)Methyltransf_25 (pfam13649)→ PLN02526 (PLN02526)CaiA (COG1960)→ |
| | | | | | SGNH_hydrolase_like_1 (cd01832)Lipase_GDSL_2 (pfam13472)→ |
| env-69 | 2236876010_0096165 | - | 74 | 40 | **RNA**→ hypo→ |
| env-70 | CETB01226394.1 | + | 29 | 63 | **RNA**→ rpsP (PRK14521)Ribosomal_S16 (pfam00886)→ |
| env-71 | CENY01353123.1 | + | 2985 | 3019 | **RNA**→ rpsP (PRK00040)Ribosomal_S16 (pfam00886)→ |
| env-72 | CEOQ01279817.1 | - | 551 | 517 | **RNA**→ rpsP (PRK00040)Ribosomal_S16 (pfam00886)→ |

| | | | | | |
|---|---|---|---|---|---|
| env-73 | CEUC01055477.1 | + | 3449 | 3483 | **RNA**→ RpsP (COG0228)S16 (TIGR00002)→ |
| env-74 | CEOP01159386.1 | + | 4003 | 4037 | **RNA**→ rpsP (PRK14521)Ribosomal_S16 (pfam00886)→ |
| env-75 | 2236876007_0680529 | - | 113 | 79 | **RNA**→ |
| env-76 | JCVI_SCAF_1101668515375 | - | 1138 | 1104 | **RNA**→ rpsP (PRK14521)Ribosomal_S16 (pfam00886)→ |
| Gsp-3-1 | NZ_KE384218.1 | - | 412048 | 412014 | **RNA**→ RpsP (COG0228)rpsP (PRK14521)→ RimM (COG0806)rimM (PRK00122)→ |
| | | | | | AdoMet_MTases (cd02440)Methyltransf_25 (pfam13649)→ |
| | | | | | AdoMet_MTases (cd02440)Methyltransf_25 (pfam13649)→ PLN02526 (PLN02526)CaiA (COG1960)→ |
| Gsp-1-1 | NZ_AJLT01000052.1 | + | 38402 | 38436 | **RNA**→ RpsP (COG0228)rpsP (PRK14521)→ RpsP (COG0228)rpsP (PRK14521)→ |
| | | | | | rimM (PRK00122)RimM (pfam01782)→ rimM (PRK00122)RimM (pfam01782)→ |
| | | | | | AdoMet_MTases (cd02440)Methyltransf_25 (pfam13649)→ |
| | | | | | AdoMet_MTases (cd02440)Methyltransf_25 (pfam13649)→ |
| | | | | | AdoMet_MTases (cd02440)Methyltransf_25 (pfam13649)→ PLN02526 (PLN02526)CaiA (COG1960)→ |
| | | | | | PLN02526 (PLN02526)CaiA (COG1960)→ |
| env-77 | KGI_S1_ANT01_95m_c10008968 | - | 71 | 37 | **RNA**→ |
| env-78 | JGI24724J26744_10116716 | + | 457 | 491 | **RNA**→ RpsP (COG0228)rpsP (PRK00040)→ |
| env-79 | CESN01042302.1 | + | 640 | 674 | **RNA**→ ←hypo |
| Lsp-1-1 | NC_015638.1 | - | 1487340 | 1487306 | **RNA**→ RpsP (COG0228)rpsP (PRK14521)→ RpsP (COG0228)rpsP (PRK00040)→ |
| env-80 | JGI24724J26744_10001441 | + | 16124 | 16158 | **RNA**→ rpsP (PRK14521)Ribosomal_S16 (pfam00886)→ |
| Lsp-2-1 | NZ_JYNQ01000001.1 | - | 2613222 | 2613188 | **RNA**→ RpsP (COG0228)rpsP (PRK00040)→ |
| env-81 | AYRF01036350.1 | + | 43 | 77 | **RNA**→ rpsP (PRK14521)Ribosomal_S16 (pfam00886)→ |
| env-82 | KGI_S2_ANT05_2345m_c1138509 | + | 19 | 53 | **RNA**→ RpsP (COG0228)S16 (TIGR00002)→ |
| env-83 | JGI24025J20009_10003996 | + | 740 | 774 | **RNA**→ RpsP (COG0228)rpsP (PRK00040)→ |
| env-84 | KGI_S1_ANT01_95m_c10219429 | - | 306 | 272 | **RNA**→ RpsP (COG0228)S16 (TIGR00002)→ |
| Gme-1-1 | NZ_KE383968.1 | + | 18032 | 18066 | **RNA**→ RpsP (COG0228)rpsP (PRK00040)→ |
| env-85 | DelMOSum2010_c10000289 | + | 5459 | 5493 | **RNA**→ RpsP (COG0228)rpsP (PRK14521)→ RimM (COG0806)rimM (PRK00122)→ |
| | | | | | AdoMet_MTases (cd02440)Methyltransf_25 (pfam13649)→ PLN02526 (PLN02526)CaiA (COG1960)→ |
| | | | | | SGNH_hydrolase_like_1 (cd01832)Lipase_GDSL_2 (pfam13472)→ |
| env-86 | BS_KBA_SWE12_21m_c10003677 | - | 2677 | 2643 | **RNA**→ RpsP (COG0228)rpsP (PRK14521)→ rimM (PRK00122)RimM (pfam01782)→ |
| env-87 | JGI20156J14371_10011188 | + | 1622 | 1657 | **RNA**→ RpsP (COG0228)rpsP (PRK14521)→ RimM (COG0806)rimM (PRK00122)→ |
| | | | | | AdoMet_MTases (cd02440)Methyltransf_25 (pfam13649)→ PLN02526 (PLN02526)CaiA (COG1960)→ |
| env-88 | 2236876011_215066 | - | 266 | 231 | **RNA**→ RpsP (COG0228)rpsP (PRK00040)→ |
| env-89 | SA_S1_NOR05_45m_c10049152 | + | 794 | 829 | **RNA**→ RpsP (COG0228)rpsP (PRK00040)→ |
| Tna-1-1 | NZ_JTDV01000002.1 | + | 21488 | 21524 | **RNA**→ RpsP (COG0228)rpsP (PRK00040)→ |
| Sze-1-1 | NZ_CP007202.1 | + | 440776 | 440811 | **RNA**→ RpsP (COG0228)rpsP (PRK14521)→ |
| env-90 | KGI_S1_ANT01_95m_c10132422 | - | 525 | 489 | **RNA**→ RpsP (COG0228)rpsP (PRK14521)→ |
| env-91 | JGI24723J26617_10166124 | + | 450 | 486 | **RNA**→ rpsP (PRK14521)Ribosomal_S16 (pfam00886)→ |
| env-92 | JGI24025J20009_10047375 | + | 382 | 418 | **RNA**→ RpsP (COG0228)S16 (TIGR00002)→ |
| Ale-1-1 | NZ_BBNQ01000006.1 | + | 160473 | 160508 | **RNA**→ RpsP (COG0228)rpsP (PRK14521)→ |
| env-93 | KGI_S1_ANT01_95m_c10442974 | + | 162 | 197 | **RNA**→ rpsP (PRK14521)Ribosomal_S16 (pfam00886)→ |
| env-94 | CESF01207615.1 | - | 398 | 364 | **RNA**→ RpsP (COG0228)rpsP (PRK14521)→ |
| env-95 | JCVI_SCAF_1096624993332 | - | 386 | 352 | **RNA**→ RpsP (COG0228)rpsP (PRK14521)→ |
| env-96 | 2236876009_175930 | - | 360 | 326 | **RNA**→ RpsP (COG0228)rpsP (PRK00040)→ rpsP (PRK14521)→ |
| env-97 | JGI24024J18818_10507174 | + | 229 | 264 | **RNA**→ rpsP (PRK14521)→ |
| Gsp-2-1 | NZ_JQLP01000005.1 | - | 2435334 | 2435300 | **RNA**→ RpsP (COG0228)rpsP (PRK14521)→ rimM (PRK00122)RimM (pfam01782)PRC (pfam05239)→ |
| | | | | | AdoMet_MTases (cd02440)Methyltransf_25 (pfam13649)→ PLN02526 (PLN02526)CaiA (COG1960)→ |
| env-98 | CEQH01183744.1 | + | 368 | 402 | **RNA**→ rpsP (PRK14521)Ribosomal_S16 (pfam00886)→ |
| env-99 | KVRMV2_102741663 | - | 269 | 235 | **RNA**→ RpsP (COG0228)rpsP (PRK00040)→ |
| Wsp-1-1 | NZ_AP014583.1 | - | 1850820 | 1850786 | **RNA**→ RpsP (COG0228)rpsP (PRK14521)→ RimM (COG0806)rimM (PRK00122)→ |
| env-100 | KVRMV2_100400240 | - | 91 | 57 | **RNA**→ |
| Myu-1-1 | NZ_JPFK01000005.1 | - | 589182 | 589148 | **RNA**→ RpsP (COG0228)rpsP (PRK14521)→ |
| Kje-1-1 | NZ_LBMG01000002.1 | + | 77869 | 77903 | **RNA**→ RpsP (COG0228)rpsP (PRK14521)→ rimM (PRK00122)RimM (pfam01782)→ |
| | | | | | RibA (COG0807)ribA (PRK00393)→ |
| env-101 | KGI_S1_ANT01_95m_c10001015 | - | 6200 | 6166 | **RNA**→ RpsP (COG0228)rpsP (PRK14521)→ rimM (PRK00122)RimM (pfam01782)→ |
| | | | | | AdoMet_MTases (cd02440)Methyltransf_25 (pfam13649)→ PLN02526 (PLN02526)CaiA (COG1960)→ |

| | | | | | |
|---|---|---|---|---|---|
| env-102 | KGI_S1_ANT01_95m_c10178456 | + | 378 | 412 | **RNA**→ rpsP (PRK14521)Ribosomal_S16 (pfam00886)→ |
| env-103 | CERE01174386.1 | - | 139 | 105 | **RNA**→ rpsP (PRK14521)Ribosomal_S16 (pfam00886)→ |
| env-104 | 2236876002_225765 | - | 176 | 142 | **RNA**→ rpsP (PRK14521)Ribosomal_S16 (pfam00886)→ |
| env-105 | KGI_S1_ANT01_95m_c10030431 | + | 734 | 768 | **RNA**→ RpsP (COG0228)rpsP (PRK14521)→ |
| env-106 | KGI_S1_ANT01_95m_c10007706 | - | 3233 | 3199 | **RNA**→ RpsP (COG0228)rpsP (PRK14521)→ |
| env-107 | JCVI_SCAF_1101668046809 | - | 546 | 512 | **RNA**→ RpsP (COG0228)rpsP (PRK14521)→ |
| env-108 | SA_S1_NOR08_45m_c10162553 | - | 124 | 90 | **RNA**→ RpsP (COG0228)rpsP (PRK14521)→ |
| Osp-1-1 | NZ_JADS01000027.1 | + | 8919 | 8953 | **RNA**→ RpsP (COG0228)rpsP (PRK14521)→ RimM (COG0806)rimM (PRK00122)→ |
| | | | | | AdoMet_MTases (cd02440)Methyltransf_25 (pfam13649)→ |
| env-109 | SRS013252_Baylor_scaffold_30766 | - | 6014 | 5979 | **RNA**→ rpsP (PRK14521)Ribosomal_S16 (pfam00886)→ RimM (COG0806)rimM (PRK00122)→ |
| env-110 | CESG01120888.1 | - | 205 | 171 | **RNA**→ rpsP (PRK14521)Ribosomal_S16 (pfam00886)→ |
| env-111 | SRS018439_Baylor_scaffold_43479 | - | 10552 | 10517 | **RNA**→ RpsP (COG0228)rpsP (PRK14521)→ rimM (PRK00122)PRC (pfam05239)→ |
| | | | | | Transket_pyr (smart00861)TktA2 (COG3958)→ YicC (COG1561)PRK11820 (PRK11820)→ |
| | | | | | Gmk (COG0194)gmk (PRK00300)→ NMNAT (cd02165)nadD (PRK00071)→ |
| | | | | | LMWPc (smart00226)LMWPTP (cd16343)→ RecR (COG0353)recR (PRK00076)→ |
| env-112 | SRS015540_C1138365 | - | 527 | 492 | **RNA**→ RpsP (COG0228)rpsP (PRK14521)→ |
| Csp-1-1 | NZ_ABZV01000003.1 | - | 5407 | 5372 | **RNA**→ RpsP (COG0228)rpsP (PRK14521)→ RpsP (COG0228)rpsP (PRK14521)→ |
| | | | | | rimM (PRK00122)PRC (pfam05239)→ rimM (PRK00122)PRC (pfam05239)→ |
| | | | | | Transket_pyr (smart00861)TktA2 (COG3958)→ Transket_pyr (smart00861)TktA2 (COG3958)→ |
| | | | | | YicC (COG1561)PRK11820 (PRK11820)→ YicC (COG1561)PRK11820 (PRK11820)→ |
| | | | | | Gmk (COG0194)gmk (PRK00300)→ Gmk (COG0194)gmk (PRK00300)→ NMNAT (cd02165)nadD (PRK00071)→ |
| | | | | | NMNAT (cd02165)nadD (PRK00071)→ LMWPc (smart00226)LMWPTP (cd16343)→ |
| | | | | | LMWPc (smart00226)LMWPTP (cd16343)→ RecR (COG0353)recR (PRK00076)→ hypo→ |
| Kal-1-1 | NZ_DS544873.1 | + | 2949885 | 2949919 | **RNA**→ RpsP (COG0228)rpsP (PRK14521)→ rimM (PRK00122)RimM (pfam01782)→ |
| | | | | | AdoMet_MTases (cd02440)Methyltransf_25 (pfam13649)→ |
| env-113 | KGI_S1_ANT01_95m_c10275683 | + | 114 | 148 | **RNA**→ RpsP (COG0228)S16 (TIGR00002)→ |
| env-114 | JGI12097J13213_1000099 | + | 9148 | 9182 | **RNA**→ RpsP (COG0228)rpsP (PRK14521)→ rimM (PRK00122)RimM (pfam01782)→ |
| | | | | | AdoMet_MTases (cd02440)Methyltransf_25 (pfam13649)→ PLN02526 (PLN02526)CaiA (COG1960)→ |
| env-115 | KGI_S1_ANT01_95m_c10082430 | - | 337 | 303 | **RNA**→ RpsP (COG0228)rpsP (PRK00040)→ |
| env-116 | JGI24724J26744_10019130 | - | 1468 | 1434 | **RNA**→ RpsP (COG0228)rpsP (PRK00040)→ RimM (COG0806)rimM (PRK00122)→ |
| | | | | | AdoMet_MTases (cd02440)Methyltransf_25 (pfam13649)→ |
| env-117 | CEQG01049560.1 | + | 282 | 316 | **RNA**→ RpsP (COG0228)rpsP (PRK14521)→ |
| env-118 | BS_KBA_SWE12_21m_c10365013 | - | 337 | 303 | **RNA**→ RpsP (COG0228)rpsP (PRK00040)→ |
| env-119 | BS_KBA_SWE12_21m_c10000423 | + | 6152 | 6186 | **RNA**→ RpsP (COG0228)rpsP (PRK14521)→ RimM (COG0806)rimM (PRK00122)→ |
| | | | | | AdoMet_MTases (cd02440)Methyltransf_25 (pfam13649)→ |
| env-120 | P_1C_Sed_3_UnCty_1160372 | + | 82 | 116 | **RNA**→ rpsP (PRK14521)Ribosomal_S16 (pfam00886)→ |
| env-121 | SA_S1_NOR08_45m_c10086824 | + | 919 | 953 | **RNA**→ |
| env-122 | KGI_S1_ANT01_95m_c10418974 | + | 254 | 288 | **RNA**→ |
| env-123 | JCVI_SCAF_1096627435724 | + | 1230 | 1264 | **RNA**→ |
| env-124 | CERM01151318.1 | + | 572 | 606 | **RNA**→ |
| env-125 | 2236876007_0117740 | - | 261 | 227 | **RNA**→ RpsP (COG0228)rpsP (PRK00040)→ |
| env-126 | wf3_AHAI4119_b1 | - | 581 | 547 | **RNA**→ RpsP (COG0228)rpsP (PRK14521)→ |
| env-127 | CESF01282643.1 | - | 99 | 65 | **RNA**→ |
| Gpo-1-1 | NZ_AUHF01000007.1 | + | 179304 | 179338 | **RNA**→ RpsP (COG0228)rpsP (PRK14521)→ rimM (PRK00122)RimM (pfam01782)→ |
| env-128 | CEOS01080028.1 | + | 2877 | 2911 | **RNA**→ |
| Jpa-1-1 | NZ_BBNS01000030.1 | - | 35262 | 35227 | **RNA**→ RpsP (COG0228)rpsP (PRK14521)→ |
| Gli-1-1 | NZ_JH594606.1 | - | 2311466 | 2311432 | **RNA**→ RpsP (COG0228)rpsP (PRK14521)APG6 (pfam04111)→ |
| | | | | | RpsP (COG0228)rpsP (PRK14521)APG6 (pfam04111)→ rimM (PRK00122)RimM (pfam01782)→ |
| | | | | | rimM (PRK00122)RimM (pfam01782)→ AdoMet_MTases (cd02440)Methyltransf_25 (pfam13649)→ |
| | | | | | AdoMet_MTases (cd02440)Methyltransf_25 (pfam13649)→ |
| | | | | | AdoMet_MTases (cd02440)Methyltransf_25 (pfam13649)→ |
| env-129 | 3300001197_1031695 | - | 46 | 12 | **RNA**→ hypo→ |
| env-130 | CEVX01047029.1 | - | 9602 | 9568 | **RNA**→ RpsP (COG0228)S16 (TIGR00002)→ RimM (COG0806)rimM (PRK00122)→ |
| | | | | | AdoMet_MTases (cd02440)Methyltransf_25 (pfam13649)→ PLN02526 (PLN02526)CaiA (COG1960)→ |
| | | | | | SGNH_hydrolase_like_1 (cd01832)Lipase_GDSL_2 (pfam13472)→ |

#### 23.3 Conserved domains

Conserved domains found in protein-coding genes listed in Section 23.2 are shown below, with the first sentence in their description from the Conserved Domain Database (if any). Conserved domains associated with more than one S16-Flavobacteria RNA are assigned a color, while others are shown in gray. The number in parentheses after the colored domain name is the number of occurrences in Section 23.2.

cd01569 (1) pre-B-cell colony-enhancing factor (PBEF)-like.
cd01832 (4) Members of the SGNH-hydrolase superfamily, a diverse family of lipases and esterases.
cd02165 (3) Nicotinamide/nicotinate mononucleotide adenylyltransferase.
cd02440 (23) S-adenosylmethionine-dependent methyltransferases (SAM or AdoMet-MTase), class I; AdoMet-MTases are enzymes that use S-adenosyl-L-methionine (SAM or AdoMet) as a substrate for methyltransfer, creating the product S-adenosyl-L-homocysteine (AdoHcy).
cd16026 (1) galactosamine-6-sulfatase; also known as N-acetylgalactosamine-6-sulfatase (GALNS).
cd16343 (3) Low molecular weight protein tyrosine phosphatase.
COG0194 (3) Guanylate kinase [Nucleotide transport and metabolism]
COG0228 (100) Ribosomal protein S16 [Translation, ribosomal structure and biogenesis]
COG0353 (2) Recombinational DNA repair protein RecR [Replication, recombination and repair]
COG0462 (1) Phosphoribosylpyrophosphate synthetase [Nucleotide transport and metabolism, Amino acid transport and metabolism]
COG0806 (25) Ribosomal 30S subunit maturation factor RimM, required for 16S rRNA processing [Translation, ribosomal structure and biogenesis]
COG0807 (1) GTP cyclohydrolase II [Coenzyme transport and metabolism]
COG1051 (1) ADP-ribose pyrophosphatase YjhB, NUDIX family [Nucleotide transport and metabolism]
COG1561 (3) Uncharacterized conserved protein YicC, UPF0701 family [Function unknown]
COG1960 (13) Acyl-CoA dehydrogenase related to the alkylation response protein AidB [Lipid transport and metabolism]
COG1990 (1) Peptidyl-tRNA hydrolase [Translation, ribosomal structure and biogenesis]
COG3064 (1) Membrane protein involved in colicin uptake [Cell wall/membrane/envelope biogenesis]
COG3958 (3) Transketolase, C-terminal subunit [Carbohydrate transport and metabolism]
pfam00886 (35) Ribosomal protein S16.
pfam01782 (23) RimM N-terminal domain.
pfam04111 (2) Autophagy protein Apg6.
pfam05239 (9) PRC-barrel domain.
pfam06459 (1) Ryanodine Receptor TM 4-6.
pfam13472 (4) GDSL-like Lipase/Acylhydrolase family.
pfam13649 (23) Methyltransferase domain.
PLN02526 (13) acyl-coenzyme A oxidase
PRK00040 (32) 30S ribosomal protein S16; Reviewed
PRK00071 (3) nicotinic acid mononucleotide adenylyltransferase; Provisional
PRK00076 (2) recombination protein RecR; Reviewed
PRK00122 (52) 16S rRNA-processing protein RimM; Provisional
PRK00300 (3) guanylate kinase; Provisional
PRK00393 (1) GTP cyclohydrolase II; Reviewed
PRK05379 (1) bifunctional nicotinamide mononucleotide adenylyltransferase/ADP-ribose pyrophosphatase; Provisional
PRK09198 (1) putative nicotinate phosphoribosyltransferase; Provisional
PRK11820 (3) hypothetical protein; Provisional
PRK13759 (1) arylsulfatase; Provisional
PRK14521 (95) 30S ribosomal protein S16; Provisional
smart00226 (3) Low molecular weight phosphatase family.
smart00861 (3) Transketolase, pyrimidine binding domain.
TIGR00002 (12) ribosomal protein S16. [Protein synthesis, Ribosomal proteins: synthesis and modification]
TIGR01251 (1) ribose-phosphate pyrophosphokinase. [Purines, pyrimidines, nucleosides, and nucleotides, Purine ribonucleotide biosynthesis]

#### 23.4 Multiple-sequence alignment

Each S16-Flavobacteria RNA is denoted by an abbreviation (like "Eco-1-1") that refers to a taxonomy given in Section 23.1. The alignment may include sequences containing the RNA motif, as well as flanking sequence. The S16-Flavobacteria RNA itself is denoted by the line underneath marked 5′ and 3′ on either end. Nucleotides in flanking sequences (i.e., not part of the motif) are written in gray letters. Stems of predicted rho-independent transcription terminators, if any, are shaded yellow. (Note: terminator predictions have not been analyzed manually, and many are likely to be false positives. Terminator predictions are those of the RNIE software.) Nucleotides predicted to function (as DNA) as transcription-factor binding sites, if any, are shaded in green. (But note: these predictions are manually annotated, so they might be under-predicted.) Annotated start codons, if any, are shaded green. (Note: start codons are frequently misannotated, especially in environmental samples.) Nucleotides proposed to basepair as part of the consensus structure are shaded in color when they comprise Watson-Crick or G-U pairs. Otherwise they are shaded gray. Conserved stems are also indicated at the bottom of the alignment by angle brackets, where matching < and > denote base-paired columns. Below these angle brackets, the symbol "2" denotes base pairs exhibiting covariation according to the statistically well-founded R-scape method.

"1" denotes base pairs exhibiting covariation according to R2R's simplistic method. "0" denotes base pairs that are not observed to mutate and "?" denotes base pairs that have a significant frequency of non-canonical nucleotides for Watson-Crick or G-U pairs (> 5%). Below these base pair annotation is the consensus sequence: "R" = "A" or "G", "Y" = "C" or "U", red nucleotides: nucleotide identity conserved more than 97% of the time, black nucleotides: 90%, gray nucleotides: 75%, red circle (•): nucleotide is present 97% of the time, black circle (•): 90%, gray circle (•): 75%, white circle (◦): 50%. All percentages of sequences just described (e.g. 97% conserved) assume that sequences have been weighted by the GSC algorithm implemented by the Infernal software package.

alignment positions 1 · · · 47

```
env-1      UUAAUAACC..CGGGUCGAGUACCCAAAA..UAUUG.AAAUU..AUG
Psp-1-1    UUAAUAACA..AAGGUCGAGAACCUUAAAA.AAUUA.AUAAAUUAUG
env-2      AUUAAUAACC.UGGGUCGAGAACCCAAAA.UAUUGA.AAUU...AUG
env-3      AUUAUUAACC.GAGGUCGGGCACCUCAAA.AAAUAA.AAAAU..AUG
Cca-2-1    AUUAAUAACU.GAGGUCGAGAACCUCUAAAAAUUAA.ACUUUU.AUG
env-4      AUUAAUAACC.GGGGUCGUGUACCCCAAA.AUUUAA.UAUACU.AUG
env-5      AUUAAUAACC.AGGGUCGGGAACCCUUAAUAUUUAA.UGUGAU.AUG
env-6      AUUAAUAACA.AGGGUCGGGCACCCU.....AUAAA.UUA....AUG
env-7      AUUAAUAACG.AGGGUCGGGCACCCUGCAAAAUUAA.UGUGAU.AUG
env-8      AUUAAUAACA.AGGGUCGGGCACCCUACA.AAUUAA.UGUGAU.AUG
env-9      AUUAAUAACCGAGGGUCGGGCACCCUACAAAAUUAA.UGUGAU.AUG
Mth-1-1    AUUAAUAACCGAGGGUCGGGCACCCUACAUAAUUAA.UGUGAU.AUG
env-10     AUUAAUAACC.AGGGUCGGGCACCCUACA.AAUUAA.UGUGAU.AUG
env-11     AUUAAUAACCAAGGGUCGGGCACCCUACA.AAUUAA.UGUGAU.AUG
env-12     AUUAAUAACA.GGGGUCGGGCACCCCACA.AAUUAA.UGUGAU.AUG
Cly-1-1    AUUAAUAACA.GGGGUCGGGCACCCUACAAAUUUAA.UGUGAU.AUG
env-13     AUUAAUAACU.GAGGUCGAGAACCUUAAA.AUUUAA.UCUUAU.AUG
env-14     AUUAAUAACC.GAGGUCGUGUACCUCAAA.AUUUAA.ACAUU..AUG
env-15     AUUAAUAAAC.UAGGUCGAGAACCUAAAA.AUUUAA.UAAUU..AUG
env-16     AUUAAUAACU.UAGGUCGAGAACCUAAAA.AUUUAA.ACAUUU.AUG
env-17     AUUAAUAAUA.GGGGUCGAGCACCCCACA.AAUUAA.UUAAAU.AUG
env-18     AUUAAUAACC.AAGGUCGAGUACCUUAAA.AUUUAA.ACAA...AUG
env-19     AUUAAUAAUC.GGAGUCGCGAACUCCAAA.AUUUAA.UUUUU..AUG
env-20     AUUAAUAACG.GGAGUCGGGAACUCCAAA.AUUUAA.UUUUUU.AUG
env-21     AUUAAUAACG.GGAGUCGGGAACUCCAAA.AUUUAA.UUUUUUUAUG
env-22     AUUAAUAACG.GGAGUCGGGAACUCCAAA.AUUUAA.UUUUU..AUG
Pto-1-1    AUUAAUAACG.GGAGUCGAGAACUCCAAA.AUUUAA.UUUUUU.AUG
env-23     AUUAAUAACG.GGAGUCGAGAACUCCAAA.AUUUAA.UUCUUU.AUG
env-24     AUUAAUAACG.GGAGUCGAGAACUUCAAA.AUUUAA.UUUUUU.AUG
env-25     AUUAAUAACG.GGAGUCGAGCACUCCAAA.AUUUAA.UUUUU..AUG
env-26     AUUAAUAAUU.GAGGUCGAGUACCUCCCA.AAUUAA.UUUAU..AUG
env-27     AUUAAUAAUU.GAGGUCGAGAACCUCGCA.AAUUAA.UUUAU..AUG
env-28     AUUAAUAAUU.GAGGUCGAGAACCUCCCA.AAUUAA.UUUAU..AUG
env-29     AUUAAUAACC.GGGGUCGAGUACCUCACA.AAUUAA.UACUU..AUG
env-30     AUUAAUAACA.GAGGUCGGGGACCUCACA.AAUUAA.UCAUU..AUG
env-31     AUUAAUAACC.GGGGUCGAGAACCUCAAA.AUUUAA.UCGCU..AUG
Cca-1-1    AUUAAUAACC.GAGGUCGAGAACCUCAAAAAUUUAA.UCGU...AUG
env-32     AUUAAUAACA.GGGGUCGGGAACCCCAUA.AAUUAA.UUAAAA.AUG
env-33     AUUAAUAACG.GAGGUCGGGAACCUCAAAUAAUUAA.UCAUU..AUG
env-34     AUUAAUAACC.GAGGUCGGGAACCUCACU.AAUUAA.UAGUUUUAUG
env-35     AUUAAUAACC.AAGGUCGAGAACCUUAAA.AAUUAA.AAUU...AUG
Tsp-1-1    AUUAAUAACC.AAGGUCGCGAACCUUAAA.AAUUAA.AAAUU..AUG
Tme-1-1    AUUAAUAACC.AAGGUCGCGAACCUUAAA.AAUUAA.AAUU...AUG
env-36     AUUAAUAACC.UAGGUCGAGAACCUUAAA.AAUUAA.AAAUUU.AUG
env-37     AUUAAUAACC.AAGGUCGAGUACCUUAAAAAAUUAA.AAGA...AUG
env-38     AUUAAUAACC.AAGGUCGAGAACCUUAAAAAAUUAA.AAAGA..AUG
env-39     AUUAAUAACC.AAGGUCGAGAACCUUAAAAAAUUAA.AAAAA..AUG
env-40     AUUAAUAACC.AAGGUCGAGAACCUUAAAAAAUUAA.AAAA...AUG
env-41     AUUAAUAACC.AAGGUCGAGAACCUUAAA.AAUUAA.AACAA..AUG
env-42     AUUAAUAACC.AAGGUCGAGAACCUUACA.AAUUAA.UUUU...AUG
env-43     AUUAAUAACC.AAGGUCGAGAACCUUACA.AAUUAA.UUUUU..AUG
env-44     AUUAAUAACC.AAGGUCGGGAACCUUACA.AAUUAA.UUUUU..AUG
env-45     AUUAAUAACA.AAGGUCGAGAACCUUACA.AAUUAA.UUUU...AUG
env-46     AUUAAUAAUC.AAGGUCGAGAACCUUACA.AAUUAA.UUUU...AUG
env-47     AUUAAUAAUA.AAGGUCGAGAACCUUACA.AAUUAA.UUUU...AUG
env-48     AUUAAUAAUC.GAGAUCGUGUAUCUCACA.AAUUAA.UUUAU..AUG
env-49     AUUAAUAACC.GAGAUCGUGUAUCUCACA.AAUUAA.UUUAU..AUG
env-50     AUUAAUAACC.GAGAUCGUGAAUCUCACA.AAUUAA.UUUUU..AUG
env-51     AUUAAUAACG.GAGGUCGAGAACCCCAUA.AAUUAA.UUAUU..AUG
env-52     AUUAAUAACC.GAGGUCGAGAACCUCAUA.AAUUAA.UAUAU..AUG
env-53     AUUAAUAAUA.GAGGUCGAGAACCUUAUA.AAUUAA.UUUAU..AUG
env-54     AUUAAUAACA.GAGGUCGAGAACCUUAUA.AAUUAA.UUUAU..AUG
env-55     AUUAAUAACA.GAGGUCGAGAACCUCAUA.AAUUAA.UUUAU..AUG
env-56     AUUAAUAACG.GAGGUCGAGAACCCCAUA.AAUUAA.UUUAUU.AUG
env-57     AUUAAUAACA.GGGGUCGGGAACCCCAUA.AAUUAA.UUUUU..AUG
env-58     AUUAAUAAUA.GGGGUCGGGAACCCCAUA.AAUUAA.UUUUU..AUG
env-59     AUUAAUAACC.GGGGUCGGGAACCCCACA.AUUUAA.UUUUU..AUG
env-60     AUUAAUAAUC.GGGGUCGGGAACCCCACA.AUUUAA.UUUUU..AUG

           <<<<<<.....<<<<<.....>>>>>.....>>>>>.>.........
           ..102......22110.....01122......201............
           AUUAAUAAY•-RRGGUCGRG•ACCYYA•A-A•UUAA-U••••--AUG
           5′                                 3′
```

```
Sxi-1-1    AUUAAUAACC.GGGGUCGGGAACCCCACA.AAUUAA.UUUUU..AUG
env-61     AUUAAUAAUC.GGGGUCGGGAACCCCACA.AAUUAA.UUUUU..AUG
env-62     AUUAAUAAUC.GGGGUCGAGAACCCCAAA.AUUUAA.UUUAAAUAUG
env-63     AUUAAUAAUC.GGGGUCGAGAACCUCACU.AAUUAA.UAUUU..AUG
env-64     AUUAAUAACCGGCGGUCGAGAACCCCACA.AAUUAA.UUAUU..AUG
env-65     AUUAAUAAUC.GAGGUCGAGAACCUCACU.AAUUAA.UUACU..AUG
env-66     AUUAAUAAUA.GAGGUCGAGAACCUCACA.AAUUAA.UCACU..AUG
env-67     AUUAAUAACA.AAGGUCGAGAACCUUACU.AAUUAA.UUAUUU.AUG
env-68     AUUAAUAACC.GAGGUCGAGAACCUCACA.AAUUAA.UUUAUU.AUG
env-69     AUUAAUAACC.GAGGUCGAGAACCUCAAA.AAUUAA.CAAAUU.AUG
env-70     AUUAAUAACC.GGGGUCGAGUACCUCAAA.AAUUAA.UUAAUU.AUG
env-71     AUUAAUAACC.GAGGUCGUGUACCUCAAA.AAUUAA.UUAAUU.AUG
env-72     AUUAAUAACC.GGGGUCGUGUACCUCAAA.AAUUAA.UUAAUU.AUG
env-73     AUUAAUAACC.GAGGUCGAGAACCUCAAA.AAUUAA.UUAAUU.AUG
env-74     AUUAAUAAUC.GAGGUCGAGAACCUCAAA.AAUUAA.UUAAUU.AUG
env-75     AUUAAUAACC.GGGGUCGAGAACCUCAAA.AAUUAA.UUAAUAUAUG
env-76     AUUAAUAACC.GGGGUCGAGAACCUCAAA.AAUUAA.UUAAUU.AUG
Gsp-3-1    AUUAAUAAUA.GAGGUCGUGAACCUCACU.AAUUAA.UUUAAU.AUG
Gsp-1-1    AUUAAUAAUA.GAGGUCGUGAACCUCACU.AAUUAA.UUAAUU.AUG
env-77     AUUAAUAAUA.GAGGUCGUGAACCUCACU.AAUUAA.UUUAUU.AUG
env-78     AUUAAUAAUA.GAGGUCGAGAACCUCACU.AAUUAA.UUAAAU.AUG
env-79     AUUAAUAACC.GAGGUCGCGUACCUCACA.AAUUAA.UUUUU..AUG
Lsp-1-1    AUUAAUAAUC.GAGGUCGCGUACCUCACA.AAUUAA.UUUUU..AUG
env-80     AUUAAUAAUA.GAGGUCGCGUACCUCACA.AAUUAA.UUAUUU.AUG
Lsp-2-1    AUUAAUAAUA.GAGGUCGCGUACCUCACA.AAUUAA.UUUUU..AUG
env-81     AUUAAUAAUC.GAGGUCGCGAACCUCACU.AAUUAA.UUUAU..AUG
env-82     AUUAAUAAUA.GAGGUCGCGAACCUCACN.AAUUAA.UUUUU..AUG
env-83     AUUAAUAAUC.GAGGUCGCGAACCUCACU.AAUUAA.UUUUU..AUG
env-84     AUUAAUAAUA.GAGGUCGCGAACCUCACU.AAUUAA.UUUUU..AUG
Gme-1-1    AUUAAUAAUA.GAGGUCGCGAACCUCACU.AAUUAA.UUUUUU.AUG
env-85     AUUAAUAACC.GAGGUCGAGAACCUUACA.AAUUAA.UAAUU..AUG
env-86     AUUAAUAACC.GAGGUCGAGAACCUCACU.AAUUAA.UUAUU..AUG
env-87     AUUAAUAACA.GAGGUCGAGAACCUCAACUAAUUAA.UUAUU..AUG
env-88     AUUAAUAAUC.GAGGUCGAGAACCUCAAAUAAUUAA.UUAUU..AUG
env-89     AUUAAUAACC.GAGGUCGAGAACCUCAAAUAAUUAA.UUAUU..AUG
Tna-1-1    AUUAAUAACCCGAGGUCGCGAACCUCAAUUAAUUAA.UUAUU..AUG
Sze-1-1    AUUAAUAACC.GAGGUCGAGAACCUCAAUUAAUUAA.UUAUU..AUG
env-90     AUUAAUAACCCGAGGUCGAGAACCUCAAUUAAUUAA.UUAUU..AUG
env-91     AUUAAUAACCCGAGGUCGAGAACCUCAAUGAAUUAA.UUAUU..AUG
env-92     AUUAAUAACCAGAGGUCGAGAACCUCAAUGAAUUAA.UUAUU..AUG
Ale-1-1    AUUAAUAACC.GAGGUCGCGAACCUCAAAUAAUUAA.UUAUU..AUG
env-93     AUUAAUAACA.GAGGUCGAGAACCUCAAAUAAUUAA.UUAUU..AUG
env-94     AUUAAUAACC.GAGGUCGAGAACCUCACA.AAUUAA.UAAUU..AUG
env-95     AUUAAUAACC.GAGGUCGAGAACCUUACA.AAUUAA.UUAUU..AUG
env-96     AUUAAUAACC.GAGGUCGAGAACCUCACA.AAUUAA.UUAUU..AUG
env-97     AUUAAUAACAAGAGGUCGUGAACCUCAAU.AAUUAA.UUAUU..AUG
Gsp-2-1    AUUAAUAAUG.GAGGUCGAGAACCUCACA.AAUUAA.UUUUU..AUG
env-98     AUUAAUAACA.GAGGUCGUGUACCUCACU.AAUUAA.UUAUU..AUG
env-99     AUUAAUAACA.GAGGUCGUGAACCUCACA.AAUUAA.UUUUU..AUG
Wsp-1-1    AUUAAUAACA.GAGGUCGUGAACCUCACU.AAUUAA.UUAUU..AUG
env-100    AUUAAUAACA.GAGGUAGAGAACCUCACA.AAUUAA.UUUUU..AUG
Myu-1-1    AUUAAUAAUA.GAGGUCGAGAACCUCACU.AAUUAA.UUAUUU.AUG
Kje-1-1    AUUAAUAAUA.GAGGUCGAGAACCUCACA.AAUUAA.UUUUUU.AUG
env-101    AUUAAUAAUA.GAGGUCGUGAACCUCACA.AAUUAA.UUUUUU.AUG
env-102    AUUAAUAAUA.GAGGUCGUGAACCUCACA.AAUUAA.UUUUU..AUG
env-103    AUUAAUAACA.GAGGUCGAGAACCUCACA.AAUUAA.UUUUUU.AUG
env-104    AUUAAUAACA.GAGGUCGAGAACCUCACA.AAUUAA.UUUUU..AUG
env-105    AUUAAUAACA.GAGGUCGAGAACCUCACU.AAUUAA.UUAUU..AUG
env-106    AUUAAUAACA.GAGGUCGAGAACCUCACU.AAUUAA.UUAUUUUAUG
env-107    AUUAAUAACA.GAGGUCGAGAACCUCACA.AAUUAA.UUAUU..AUG
env-108    AUUAAUAACA.GAGGUCGAGAACCUCACA.AAUUAA.UUAUUUUAUG
Osp-1-1    AUUAAUAAUA.GAGGUCGAGAACCUCACA.AAUUAA.UUAUUU.AUG
env-109    AUUAAUAAUC.GAGGUCGUGAACCUCACU.AAUUAAAUUUUUU.AUG
env-110    AUUAAUAAUC.GAGGUCGUGAACCUCACU.AAUUAA.UUUUU..AUG
env-111    AUUAAUAACC.GAGGUCGAGAACCUCACU.AAUUAAAUUUUUUUAUG
env-112    AUUAAUAACC.GAGGUCGUGAACCUCACU.AAUUAAAUUUUUUUAUG
Csp-1-1    AUUAAUAACC.GAGGUCGUGAACCUCACU.AAUUAAAUUUUUU.AUG
Kal-1-1    AUUAAUAACC.GAGGUCGUGAACCUCACA.AAUUAA.UUUUUU.AUG
env-113    AUUAAUAAUC.GAGGUCGAGAACCUCACU.AAUUAA.UUUUU..AUG
env-114    AUUAAUAAUC.GAGGUCGAGAACCUCACU.AAUUAA.UUUUUU.AUG
env-115    AUUAAUAAUC.GAGGUCGAGAACCUCACA.AAUUAA.UUUUU..AUG
env-116    AUUAAUAAUC.GAGGUCGAGAACCUCACA.AAUUAA.UUUUUU.AUG
env-117    AUUAAUAACA.GAGGUCGAGAACCUCAUU.AAUUAA.UUAUU..AUG
env-118    AUUAAUAACC.GAGGUCGAGAACCUCAUU.AAUUAA.UUAUU..AUG
env-119    AUUAAUAACC.GAGGUCGAGAACCUCAUU.AAUUAA.UCAUU..AUG
env-120    AUUAAUAACA.GAGGUCGAGAACCUCACU.AAUUAA.UUUUAU.AUG
env-121    AUUAAUAACA.GAGGUCGAGAACCUCACU.AAUUAA.UUUUU..AUG
env-122    AUUAAUAACA.GAGGUCGAGAACCUCACU.AAUUAA.UUUAU..AUG
env-123    AUUAAUAACA.GAGGUCGAGCACCUCACA.AAUUAA.UUUUU..AUG
env-124    AUUAAUAACA.GAGGUCGAGUACCUCACA.AAUUAA.UUUUU..AUG
env-125    AUUAAUAACC.GAGGUCGAGAACCUCAAA.AUUUAA.UUUUU..AUG
env-126    AUUAAUAAUC.GAGGUCGAGAACCUCAAA.AUUUAA.UUUUU..AUG
env-127    AUUAAUAAUC.GGGGUCGAGAACCCCAAA.AUUUAA.UUUUU..AUG

           <<<<<<.....<<<<<.....>>>>>.....>>>>>.>.........
           ..102......22110.....01122......201...........
           AUUAAUAAY•-RRGGUCGRG•ACCYYA•A-A•UUAA-U••••--AUG
           5′                                  3′
```

```
Gpo-1-1   AUUAAUAAUC.GGGGUCGAGAACCUCAAA.AUUUAA.UUUUU..AUG
env-128   AUUAAUAACC.GGGGUCGAGAACCCCAAAA.UUUAA.UUUUU..AUG
Jpa-1-1   AUUAAUAACC.GAGGUCGGGAACCUCAAAUAAUUAA.UUUAU..AUG
Gli-1-1   AUUAAUAACC.GAGGUCGGGGACCUCACA.AAUUAA.UUUAU..AUG
env-129   AUUAAUAACA.GAGGUCGGGGACCUCACA.AAUUAA.UUUAU..AUG
env-130   AUUAAUAACC.AAGGUCGAGAACCUUAUA.AAUUAA.UAUU...AUG
          <<<<<<.....<<<<<.....>>>>>.....>>>>>.>.........
          ..102......22110.....01122......201............
          AUUAAUAAY•-RRGGUCGRG•ACCYYA•A-A•UUAA-U••••--AUG
          5′                                  3′
```
